# DAP12 deficiency alters microglia-oligodendrocyte communication and enhances resilience against tau toxicity

**DOI:** 10.1101/2023.10.26.563970

**Authors:** Hao Chen, Li Fan, Qi Guo, Man Ying Wong, Fangmin Yu, Nessa Foxe, Winston Wang, Aviram Nessim, Gillian Carling, Bangyan Liu, Chloe Lopez-Lee, Yige Huang, Sadaf Amin, Tark Patel, Sue-Ann Mok, Won-min Song, Bin Zhang, Qin Ma, Hongjun Fu, Li Gan, Wenjie Luo

## Abstract

Pathogenic tau accumulation fuels neurodegeneration in Alzheimer’s disease (AD). Enhancing aging brain’s resilience to tau pathology would lead to novel therapeutic strategies. DAP12 (DNAX-activation protein 12) is critically involved in microglial immune responses. Previous studies have showed that mice lacking DAP12 in tauopathy mice exhibit higher tau pathology but are protected from tau-induced cognitive deficits. However, the exact mechanism remains elusive. Our current study uncovers a novel resilience mechanism via microglial interaction with oligodendrocytes. Despite higher tau inclusions, Dap12 deletion curbs tau-induced brain inflammation and ameliorates myelin and synapse loss. Specifically, removal of Dap12 abolished tau-induced disease-associated clusters in microglia (MG) and intermediate oligodendrocytes (iOli), which are spatially correlated with tau pathology in AD brains. Our study highlights the critical role of interactions between microglia and oligodendrocytes in tau toxicity and DAP12 signaling as a promising target for enhancing resilience in AD.

## Introduction

The accumulation of toxic tau in the brain correlates significantly with synapse loss, impaired neuronal function, and cognitive decline in Alzheimer’s disease (AD) and other heterogeneous tauopathies. Unraveling the biological mechanisms that underlie tau-induced neurodegeneration and brain toxicity is of paramount importance in the battle against these devastating diseases. Microglia play a pivotal role in instigating tau-related neurodegeneration. In human genetic studies, microglia has been shown to have high expression of numerous AD risk genes, and in recent investigations using tau mouse models, depleting microglia effectively reduced tau seeding activity^1,2^, curbed neuroinflammation, and mitigated tau-related neurodegeneration^3,4^. This strongly supports the notion that microglia contribute to tau-driven neurodegeneration in AD. However, the precise mechanisms through which microglia mediate tau toxicity remain largely undefined.

DNAX-activation protein 12 (DAP12), also known as TYRO protein tyrosine kinase-binding protein (TYROBP), is an adaptor protein containing an immunoreceptor tyrosine-based activation motif (ITAM), variants of which have been linked to early-onset AD^5^. By binding with microglial receptors such as Triggering Receptor Expressed on Myeloid cells 2 (TREM2), a major AD risk factor, DAP12 triggers various cellular processes such as phagocytosis, proliferation, and the regulation of inflammatory cytokines^6,7,8^. Network analyses underscore DAP12’s significance as a key driver in sporadic late-onset AD, a form of AD with fewer genetic indicators, implying that DAP12 may be an important cell-level regulator of tauopathy^9^. Critically, single-cell transcriptomics studies reveal DAP12’s essential role in transitioning microglia into a disease-associated state, termed as disease-associated microglia (DAM)^10^. Deleting DAP12 in an amyloid mouse model significantly impairs the formation of microglial barriers around plaques, exacerbating dystrophic neurites bypassing plaques and increasing plaque-associated tau pathology^11^, while leaving amyloid burden unchanged^12^. In a mouse model with tau inclusions, DAP12 loss elevated tau pathology, promoting tau seeding and spreading^13^. These findings suggest that DAP12 activity constrains amyloid plaque and tau pathology and loss of DAP12 should worsen the disease. Paradoxically, inactivation of DAP12 normalizes aberrant microglial signaling associated with AD pathologies, ameliorates abnormal electrophysiological activity and improves learning deficits in both amyloid and tauopathy mouse models^13,14,15^, indicating that removal of DAP12 confers brain resilience in response to the toxicities of AD pathologies. However, the underlying mechanism remains elusive.

In this study, we analyze the effect of DAP12 deletion on microglia and related cell types via crossing homozygous P301S tau transgenic mice^16^ with Dap12-deficient mice^17^. Our results support the previous work of Haure-Mirande et al^13^ that Dap12 deletion exacerbates tau pathology while concurrently improving gliosis and synapse loss. We then investigated the mechanism behind the resilient effects of DAP12 deletion on neurodegeneration using transcriptome analysis, which unveiled a potent attenuation of tau-induced interferon signaling, related to neuroinflammatory response, due to DAP12 deficiency. Utilizing single-nuclei RNAseq analysis (SnRNAseq) on hippocampal tissues to isolate the effects on varied cell types, we uncovered that DAP12 not only plays a role in driving the formation of disease-associated microglia, but also significantly shapes the transcriptomic states of oligodendrocytes, leading to an intermediate state concurrent with brain demyelination in tauopathy. Notably, this tau-related transcriptomic state of oligodendrocytes is also found in human AD brains, highlighting the relevance of tau mouse model findings for human AD. Our discoveries unveil a novel mechanistic link between DAP12 signaling in microglia and tau-induced toxicity in oligodendrocytes, leading to demyelination in AD. This finding strongly supports the notion of targeting DAP12 as a viable therapeutic strategy for AD.

## Results

### Loss of Dap12 elevates tau inclusions but ameliorates tau-induced gliosis

Microglia process pathogenic tau via internalization and exocytosis ^1,18,19^. To investigate the role of Dap12, we exposed primary microglia from *Dap12^+/+^* or *Dap12^-/-^* mice to tau fibrils for 2 hours and quantified tau phagocytosis. Dap12 deficiency did not affect tau internalization (Supplementary Fig. 1A, B and supplementary table 1). To assess tau processing, we then remove tau fibrils from the medium, and quantified the amount of internalized tau in microglia. *Dap12^-/-^* microglia contained more intracellular tau compared to *Dap12^+/+^* counterparts, suggesting a deficiency in tau processing (Fig. 1A, B and supplementary table 1).

**Figure 1:**
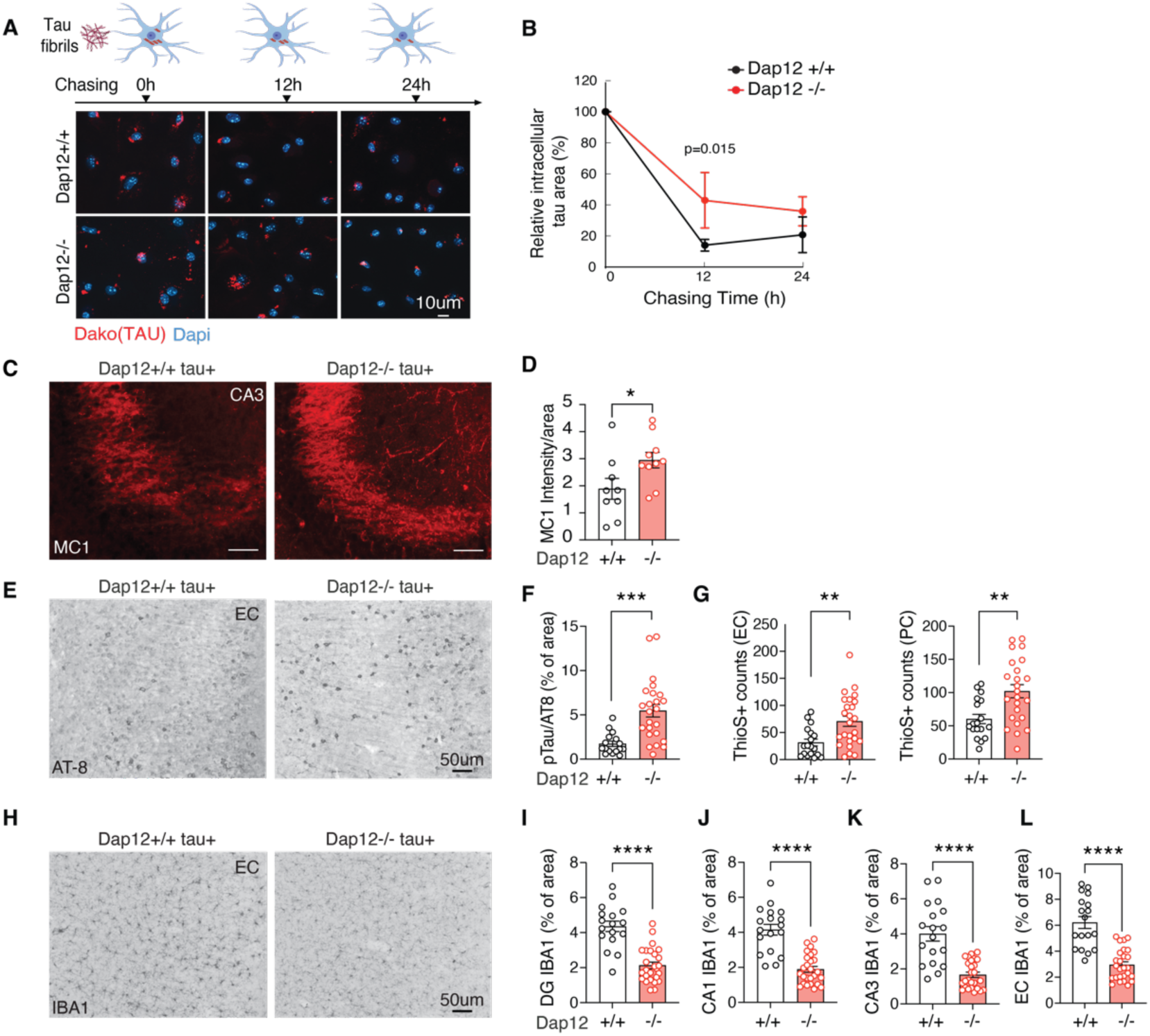
Dap12 deficiency elevates tau burden but reduces gliosis and partially rescues synapse loss in tauopathy mouse brains. A) The processing of tau fibril internalized by microglia. Tau fibrils were incubated with microglia for 2 h followed with 12 or 24 h chase in tau-free medium. B) Quantification of A. Unpaired student t-test, *p<0.05. n = 3 independent experiments. C-D) Representative images and quantification of immunohistochemical staining of MC1 in the hippocampal CA3 region of homozygous tauopathy mice. Unpaired student t-test, *p<0.05. n = 9 for *Dap12+/+ tau+*, n = 10 for *Dap12-/- tau+* mice. E-F) Representative images and quantification of immunohistochemical staining of AT-8 in the entorhinal cortex (EC) of homozygous tauopathy mice. Unpaired student t-test, ***p<0.001. n = 16 for *Dap12+/+ tau+*, n = 24 for *Dap12-/-tau+* mice. G) Quantification of Thios+ neurons in the EC or piriform cortex (PC) of homozygous tauopathy mice. Scale bar: 50 µm. Unpaired student’s t-test; **p<0.01. n = 18 *Dap12+/+ Tau+*, n = 24 (EC) or 25 (PC) *Dap12-/- Tau+* mice. H-L) Representative images of IBA1 staining and quantification of IBA1^+^ area in the hippocampal areas (CA1, CA3, dentate gyrus/DG), and entorhinal cortex (EC). Scale bar; 50 µm. Unpaired student’s t-test: ****p<0.0001. n = 18 *Dap12+/+ tau+*, n = 26 *Dap12-/- tau+*.

Homozygous P301S tau transgenic mice develop substantial tau pathology in various brain regions, including the entorhinal cortex and hippocampus, by the age of five to six months^16,20^. We examined the impact of Dap12 deletion on tau pathology in vivo. Using MC1 antibody, which recognizes conformation-specific tau relevant to AD, and AT8 antibody for phosphorylated tau at Ser202 and Thr205 epitopes, we observed a significant increase in MC1 staining within the hippocampal region upon Dap12 deletion (Fig. 1C, D and supplementary table 1). Tau inclusions detected with AT8 staining or ThioS were higher in the cortical area of *Dap12^-/-^tau^+^* mice compared to *Dap12^+/+^ tau^+^* mice (Fig. 1E-G and supplementary table 1). Tau-induced gliosis in these tau transgenic mice typically initiates as early as 2 months and expands significantly by 5-6 months of age when tau pathology peaks^16,20^. Despite elevating tau pathology, Dap12 deficiency led to a notable reduction in gliosis, as evidenced by reduced Iba1 staining (Fig. 1H-L and supplementary table 1) and GFAP staining (supplemental Fig. 1C, D and supplementary table 1). Thus, DAP12 deficiency reduces gliosis in the presence of exacerbated tau pathology, consistent with prior reports^13^.

### Dap12 mediates proinflammatory signaling in the tauopathy mouse brain

We then profiled DAP12-related signaling pathways associated with inflammation using multiplex immunoassays. Dap12 deletion reduced levels of p-AKT, p-ERK, p-JNK, p-P38, p-NFκB, and p-STAT3 in the tauopathy mouse brain (Fig. 2A-D and supplementary table 2), indicating dampened inflammatory signaling.

**Figure 2.**
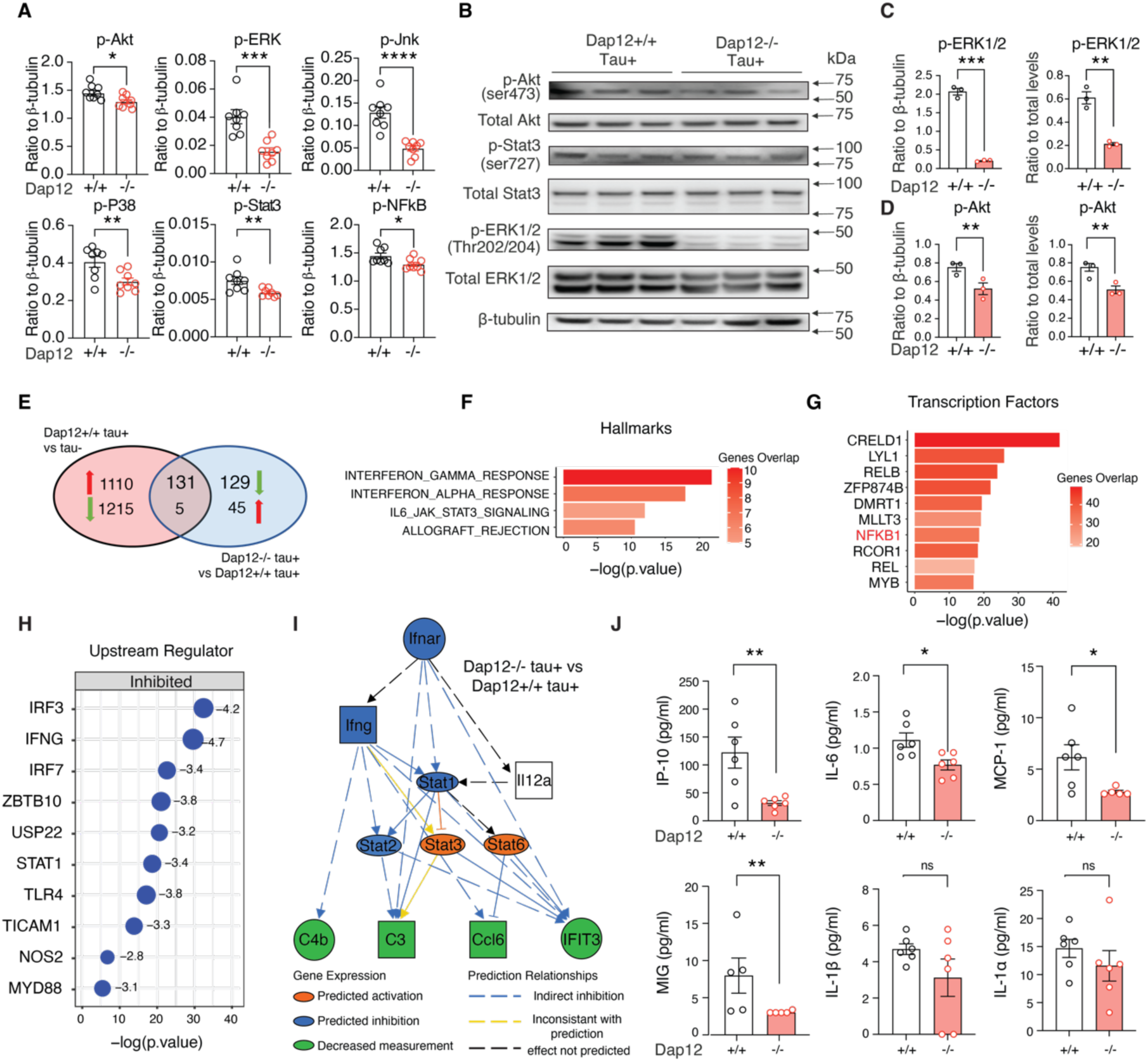
Loss of Dap12 suppresses brain-wide inflammation in mice with tauopathy revealed by transcriptomics and immune signaling analysis. A) Phosphorylation of AKT, ERK, NF-κB, JNK, P38, and STAT3 measured by cell signaling multiplex immunoassay. n = 8/genotype. B) Western blot of the phosphorylated and total AKT, STAT3, and ERK. Unpaired student’s t-test: ***p<0.001, **p<0.01, *p<0.05. *n* = 3/genotype. C-D) Quantification of (B). E) Venn diagram showing the overlapped DEGs from comparisons of frontal cortex transcriptomes between *Dap12+/+ tau+* vs *tau-* and *Dap12-/- tau+* vs *Dap12+/+ tau+* mice. n = 4/genotype. Log2FC > 0.1 or < -0.1, adjust p-value < 0.05. F-G) Hallmark pathways (F) and hallmark pathways Transcription factors (TFs) (G) predicted by GSEA for 136 DEGs (131+5) reversed by Dap12 deletion in (E). H) Upstream regulators predicted by IPA for 260 DEGs (131+129) downregulated by Dap12 deletion in tauopathy mice as in (E). I) Diagram of the interferon activation network predicted by IPA upstream regulator analysis in (H). J) The levels of IP-10, IL-6, MCP-1, MIG, IL1A, and IL1B measured by multiplex cytokine/chemokine assay. n = 6/genotype (n=5 for MCP-1 Dap12 -/-, MIG Dap12 +/+ and -/-) Unpaired student’s t-test: **p<0.01, *p<0.05, ns: not significant.

To assess the effects of Dap12 deletion on brain homeostasis at transcriptome level, we performed bulk RNA sequencing on frontal cortex tissue from *Dap12^+/+^ tau^+^*, *Dap12^-/-^tau^+^* mice (Fig. 2E, supplementary 2A, B and supplementary table 2). Ingenuity pathway analysis (IPA) of 131 differentially expressed genes (DEGs) using canonical pathways revealed that Dap12 deletion reversed tau pathology-induced cytokine storm signaling, TREM1 signaling, the complement system, and interferon (IFN) signaling (Fig. 2E, supplementary Fig. 2C, D and supplementary table 2). Gene set enrichment analysis (GSEA) also revealed Dap12 deletion reversed 131 genes enriched in tau pathology-induced interferon response and IL6-JAK-STAT3 signaling (Fig. 2F and supplementary table 2). The transcriptional factors predicted by the Gene Transcriptional Regulation Database to modulate the expression of these 131 genes include NFκB1/REL, a pivotal microglial transcriptional regulator associated with driving tau seeding and toxicity in tauopathy (Fig. 2G and supplementary table 2)^19^. Furthermore, upstream regulator analysis pinpointed cytokines IRF3, IFNG, and IRF7 as top transcriptional factors inhibited by Dap12 deletion (Fig. 2H and supplementary table 2). Mechanistic network analysis further confirmed a prominent role of Dap12 deficiency in suppressing tau pathology-activated IFN signaling (Fig. 2I and supplementary table 2).

To validate our transcriptomic findings, we extended our analysis to quantify the levels of brain cytokines and chemokines associated with proinflammatory responses. Using multiplex ELISA, we found that Dap12 deletion significantly reduced levels of CXCL10/IP-10 (Cxcl10), IL-6, MCP-1 (Ccl2), and MIG (Cxcl9) (Fig. 2J and supplementary table 2). These cytokines and chemokines are either directly or indirectly modulated by interferon signaling^21-23^ or influenced by NF-κB and ERK signaling (supplementary Fig. 2E-G). Taken together, our results demonstrated that Dap12 deletion abolished proinflammatory signaling mediated by IFN and NF-κB in the tauopathy mouse brain.

### Dap12 drives the shift of homeostatic microglia to the disease-associated state in the tauopathy mouse brain

Our results so far demonstrated that the protective effects of Dap12 deficiency are associated with the amelioration of inflammatory responses. To dissect cell type-specific mechanisms, we next performed single nuclei RNA sequencing (snRNA-Seq) to assess the effects of Dap12 deletion in response to tauopathy. Rigorous quality control steps were taken to eliminate sequencing reads derived from multiplets using DoubletFinder^24^ as well as to exclude low-quality nuclei based on thresholds for gene counts, UMI counts, and the percentage of mitochondrial genes per nucleus (Supplementary Figure 3A). Subsequent unsupervised clustering yielded 71,716 high-quality nuclei that were grouped into distinct transcriptional clusters, effectively representing the brain’s major cell types (Supplementary Figure 3B, C).

To investigate the impact of Dap12 deletion on microglial response to tauopathy, we subclustered microglia expressing microglial hallmark genes Csf1r, P2ry12, and Siglech (Supplementary Figure 3B) into four distinct subpopulations based on subcluster marker genes (Fig. 3A, supplementary Fig. 4A and supplementary table 3). MG1, cells enriched with homeostatic markers (supplementary Fig. 4A), is the predominant cluster in non-transgenic normal brains, but significantly diminishes in *Dap12^+/+^ tau^+^* brains (Fig. 3A, B). In contrast, MG2 and MG4, disease-associated cells induced by tauopathy (Fig. 3A, B), exhibit higher levels of proinflammatory genes like Apoe, Spp1, Nfkb1, and Akt3, as well as STAT3, IL-8, NF-kB, and JAK/STAT signaling (Fig. 3C-F and supplementary table 3). Markers of MG2 and MG4 are highly correlated with DAMs signatures^10^ (Supplementary Figure 4B-D). Trajectory analyses were performed to define the pseudotime trajectory from homeostatic MG1 to activated MG2 and MG4 states (Fig. 3G), as indicated by the decline of homeostatic microglial marker genes Cx3cr1 and Hexb and increase of DAM marker genes Apoe and P2rx7 (Fig. 3G). Dap12 deletion completely reversed the decline of the MG1 cell population and blocked the elevation of MG2 and MG4 cell populations (Fig. 3A, B).

**Figure 3.**
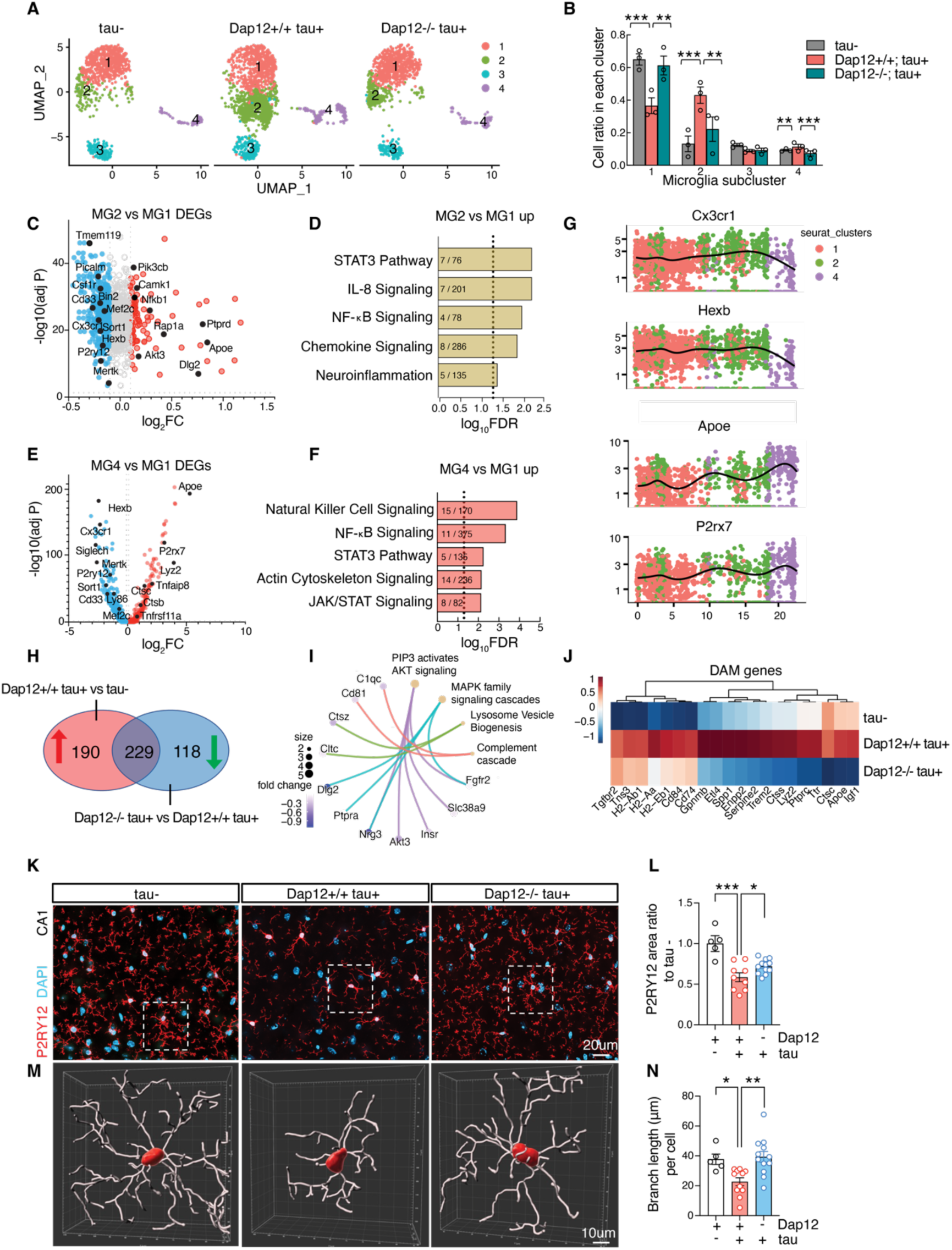
Dap12 deletion prevents loss of homeostatic microglia and blocks disease-associated microglia. A-B) UMAP (A) and cell ratios (B) of microglial subclusters (MG1-4) crossing three genotypes. One-Way ANOVA followed by Tukey test, ***p < 0.001, **p < 0.01, *p < 0.05. *n* = 3 per genotype. C) Volcano plot of DEGs (adjust p-value < 0.05, Log2FC > 0.1 or < -0.1) comparing MG2 to MG1. D) Selected IPA pathways identified for upregulated DEGs in MG2 shown in (C). E) Volcano plot of DEGs (adjust p-value < 0.05, Log2FC > 0.1 or < -0.1) comparing MG4 to MG1. F) Selected IPA pathways identified for upregulated DEGs in MG4 shown in (E). G) Expression of Cx3cr1, Hexb, Apoe, or P2ry7 in pseudotime in related to clusters in (A). H) Venn diagram of upregulated DEGs of *Dap12+/+ tau+* vs *tau-*, and downregulated DEGs of *Dap12-/- tau+* vs *Dap12+/+ tau+*. I) Selected pathways with associated genes within 229 DEGs identified in (H). J) Heatmap of DAM genes within 229 DEGs identified in (H). K) Representative images of P2RY12 staining. Scale bar: 20 µm. L) Quantification of P2RY12+ area in K across three genotypes. ***p < 0.001, *p < 0.05. n = 5 tau-, n = 11 *Dap12+/+ tau+*, n = 12 *Dap12-/- tau+* mice. M) 3D reconstructions of P2RY12 positive microglia using Imaris. Scale bar:10 µm. N) Microglial branch length crossing three genotypes. One-Way ANOVA followed by Tukey test in L and N. **p < 0.01, *p < 0.05. n = 5 *tau-*, n = 11 *Dap12+/+ tau+*, n = 12 *Dap12-/- tau+* mice.

Pseudobulk analysis of microglia revealed that 54% of the differentially expressed genes (229 genes out of 419 genes) upregulated by tau exhibited downregulation to homeostatic levels due to Dap12 deletion (Fig. 3H, and supplementary table 3). A pathway analysis of these 229 reversed genes demonstrated their involvement in processes such as the complement cascade, lysosomal vesicle biogenesis, MAPK cascade signaling, and AKT signaling (Fig. 3I). Of these 229 genes, our findings indicated that DAP12 deletion abolished the expression of signature DAM genes for both DAM1 and DAM2^10^ (Fig. 3J, supplementary Fig. 4E) while maintaining the expression of homeostatic marker genes like Cx3cr1, Csf1r, or Tmem119 (supplementary Fig. 4E).

Using immunostaining, we confirmed that Dap12 deficiency restored expression of P2ry12, a marker of homeostatic microglia (Figure 3K, L and supplementary table 3). Notably, the transition of microglial states accompanied morphological changes, transitioning from a ramified homeostatic state to a more hypertrophic activation state. Imaris analysis validated that Dap12 deficiency also halted the tau pathology-induced reduction in microglial branch length (Fig. 3M, N and supplementary table 3), along with the decrease in branch points (supplementary Fig. 4F. and supplementary table 3). Thus, Dap12 deficiency blocked the transition of microglial state from homeostatic to DAM in tauopathy, consistent with a mechanistic understanding of DAP12 mediating microglial responses to tau pathology^15,25,26^.

### Dap12 deletion reverses tau-induced gene expression alterations and reduces synapse loss in excitatory neurons

To investigate how Dap12 affects neuronal transcriptomes in tauopathy mice, we performed pseudobulk analysis of excitatory neurons (EN) and inhibitory neurons (IN) across three genotypes. Dap12 deletion significantly altered transcriptomes of EN (119 DEGs, Fig. 4A, B, and supplementary table 4), with limited effects on IN transcriptomes (merely 75 DEGs). Specifically, we identified 37 genes whose suppression by tau pathology was counteracted by Dap12 deletion. These genes were predominantly associated with protein transportation (Fig. 4C, supplementary table 4). Dap12 deletion downregulated 80 genes involved in neuronal activities (i.e. neurogenesis, neurotransmitter secretion, and synaptic signaling) out of 404 genes induced by tau pathology. (Fig. 4C, supplementary table 4).

**Figure 4.**
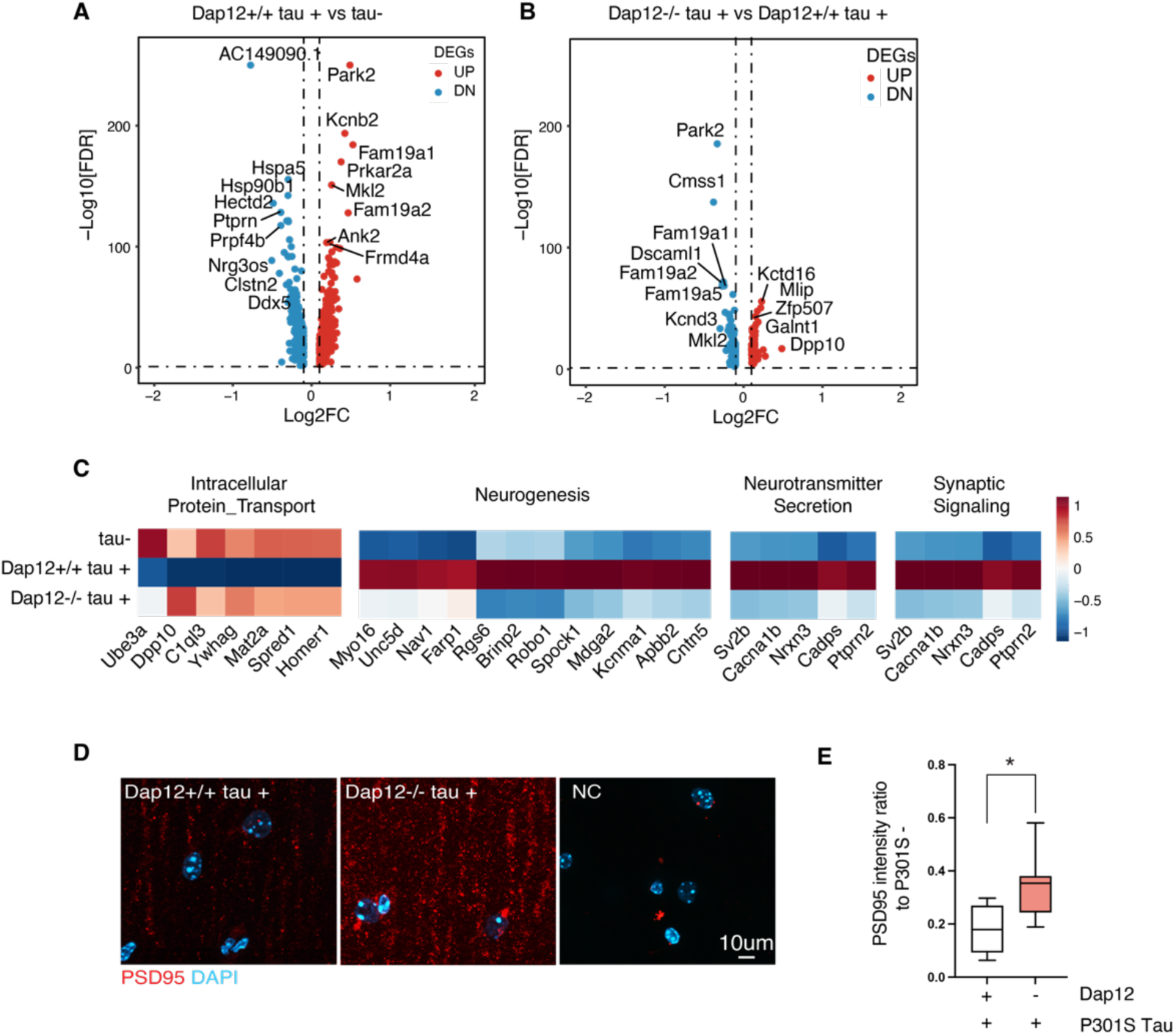
Dap12 deletion affects tau-induced excitatory neurons alterations and protects excitatory synapses. A) Volcano plot of pseudo bulk DEGs (adjust p-value < 0.05, LogFC > 0.1 or < -0.1) in comparison between *Dap12+/+ Tau+* and *Tau-* mice. B) Volcano plot of pseudo bulk DEGs (adjust p-value < 0.05, LogFC > 0.1 or < -0.1) in comparison between *Dap12-/- Tau+* and *Dap12+/+ Tau+* mice. C) Heatmap showing pathways induced by tau and reversed by Dap12 deletion. D-E) Representative images of PSD95 staining (D) and quantification (E) in the CA1 stratum radiatum region. Scale bar:10µm. Mix-model Brown-Forsythe and Welch ANOVA tests. **p<0.01, *p<0.05. n = 7 *Dap12+/+ tau+*, n = 8 *Dap12-/- tau+* mice.

To determine the effect of Dap12 on synapses, we measured tau-impaired excitatory synaptic integrity using PSD95 immunostaining^27,28^. Dap12 deletion partially rescued the tauopathy-induced decline in PSD-95 immunoreactivity at the CA1 striatum radiatum, (Fig. 4D, E and supplementary table 4), consistent with previously reported protective effects of DAP12 deletion^13^.

### Dap12 mediates tau pathology-induced transcriptomic changes in oligodendrocytes

DAP12 signaling has been implicated in brain myelination^29^. In humans, DAP12 deficiency causes Nasu–Hakola disease (NHD), an early-onset dementia characterized by myelin loss^30-32^. To assess the effects of DAP12 deletion on oligodendrocyte cells, the brain cells responsible for myelination, we subclustered OL into five distinct subpopulations based on their gene expression profiles (Fig. 5A). High-dimensional Weighted Gene Correlation Network Analysis (hdWGCNA)^33-35^ was performed across all subclusters, revealing four gene expression modules— Turquoise, Yellow, Brown, and Blue (Fig. 5B and supplementary table 5). Each module was characterized by a network of the top 10 hub genes, identified through harmonized module eigengenes (kME) (Fig. 5C, and supplementary table 5). The turquoise module, which is typically expanded in tau pathology, was diminished by Dap12 deletion (Fig. 5D, E). The turquoise module is rich with genes associated with glia-neuronal interaction, such as Neuregulin-3 (*Nrg3*), Neurexin 1 or 3 *(Nrxn1* or 3), and synaptic activities including Glutamate Ionotropic Receptor NMDA Type Subunit 2A (*Grin2a)*. Gene ontology (GO) analysis of the marker genes associated with the turquoise module revealed enrichment with membrane receptors or channels, synaptic transmission and signaling, channel and receptor activity (supplementary Fig. 5E and supplementary table 5). This cluster reduction suggests that Dap12 deletion dampens tau pathology-enhanced oligodendrocytes signaling to and from other cell types.

**Figure 5.**
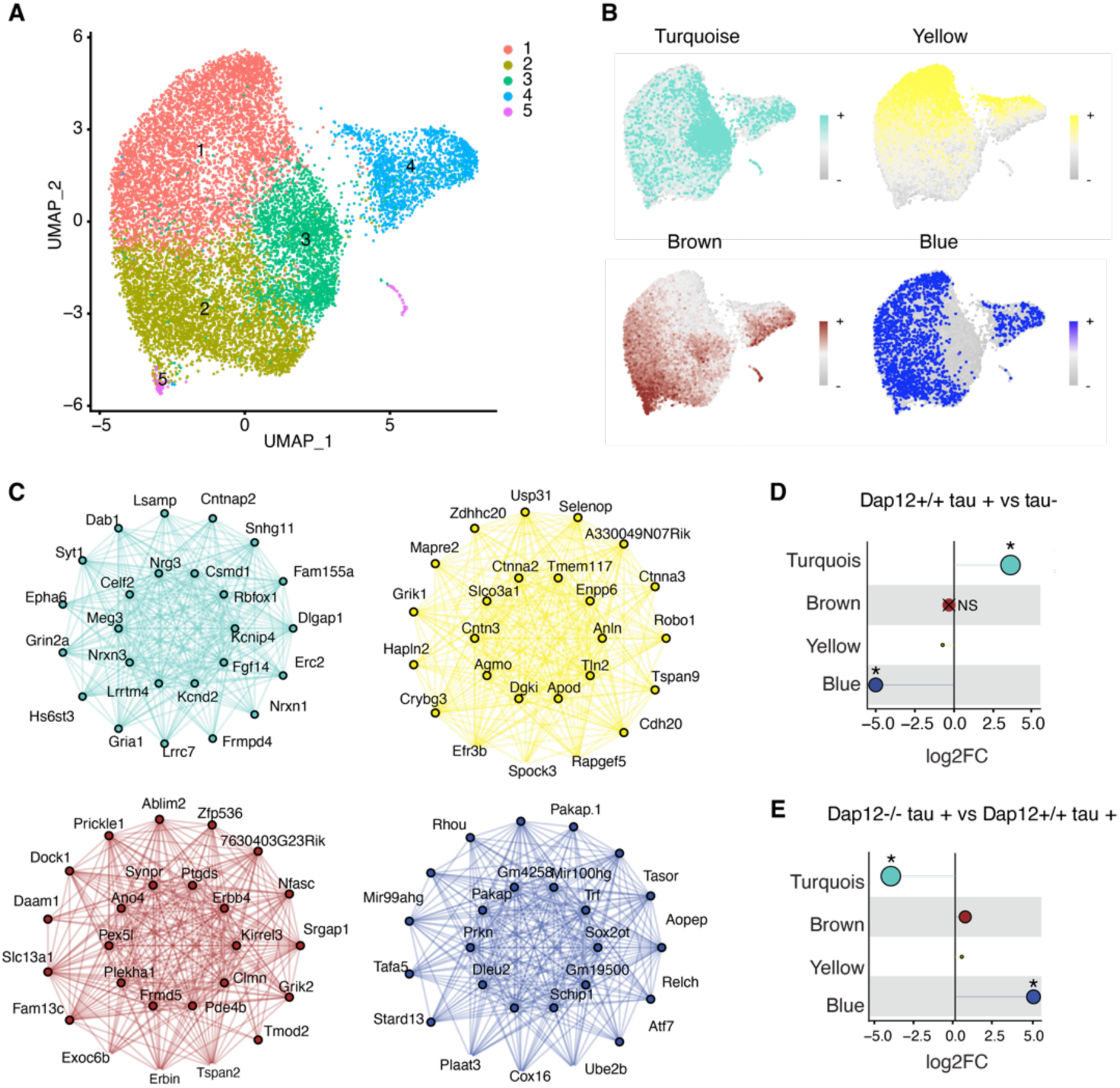
Dap12 regulates tau-induced changes of co-expression module in oligodendrocytes. A) UMAP plots of all oligodendrocytes single nuclei and subclusters. B) Feature plots showing harmonized module eigengenes (hMEs) of each co-expression module. C) Signature gene networks of each module, each node represents a gene, and each edge represents the co-expression relationship between two genes in the network. The top 10 hub genes by (eigengene-based connectivity) kME are placed in the center of the plot, while the remaining 15 genes are placed in the outer circle. D-E) Lollipop showing the fold-change for each module in comparison between *Dap12+/+ tau+* and *tau-* (D) or *Dap12-/- tau+* and *Dap12+/+ tau+* (E). The size of each dot corresponds to the number of genes in that module. *p<0.05 and “X” stands for not significant. Number of genes in different modules: Turquoise (449), Brown (150), Yellow (74), Blue (151).

In contrast, the blue module, characterized by Parkin RBR E3 Ubiquitin Protein Ligase (Prkn) and Ubiquitin Conjugating Enzyme E2 B (Ube2b), was diminished by tau pathology (Fig. 5D and supplementary table 5) and was restored by Dap12 deletion (Fig. 5E, supplementary table 5), suggesting a vulnerability of this unique OL population to tau pathology in a Dap12-dependent manner. On the other hand, the brown and yellow modules were minimally influenced by tau pathology or Dap12 deletion (Fig. 5D, E). Thus, DAP12 deletion diminishes tau pathology-induced oligodendrocyte responses, while preventing the loss of vulnerable oligodendrocytes.

### Dap12 promotes the intermediate oligodendrocyte state in the tauopathy mouse brain

We next analyzed OL1-4, which express high levels of marker genes characteristic of mature oligodendrocytes, Ptgds, Opalin, Il33, and Anln^36-38^ (see supplementary Fig. 6A), exhibiting only limited expression of previously identified disease-associated oligodendrocyte genes, such as Pan DA1/DA2 genes^38^ or DOL genes^39^ (supplementary Fig. 6B-F).

We observed that tauopathy greatly shrinked OL1 and OL2 and significantly induced OL3 (Fig. 6A, B and supplementary table 6). Strikingly, OL3 exhibits a strong resemblance to intermediate oligodendrocytes (iOli), identified in a published OL dataset obtained from the visual cortex of the human brain^40^ (Fig. 6C, D and supplementary table 6). This is evident through the expression of marker genes such as NMDA receptors (Grin2a, Grin1), adhesion molecules (Nrxn1, Nrxn3, Nrg3, Tenm2, Cntnap2), channels (Kcnip4, Dpp10), transcriptional factor (Mef2c), synaptic transmission (Dlgap1, Syn1), and axon guidance (Epha6) (Fig. 6D). OL1 exhibits a closer alignment with mature OL (Fig. 6C). Remarkably, the elimination of Dap12 completely impeded the formation of OL3 and preserved the populations of OL1 and OL2 cells (Fig. 6A, B). To further understand the effect of DAP12 deletion and tau pathology, we performed pseudobulk analysis. The dot plots demonstrate the inducibility of these iOli marker genes by tauopathy, with their upregulation fully suppressed by Dap12 deletion (Fig. 6E). The impact of tau pathology and Dap12 deletion on TEMM2 or NRG3 was further confirmed by Western blot analysis, illustrating increased levels of TEMM2 and NRG3 proteins in the cortical tissue of tauopathy-afflicted brains, which is mitigated in DAP12 loss (Fig. 6F-H and supplementary table 6).

**Figure 6.**
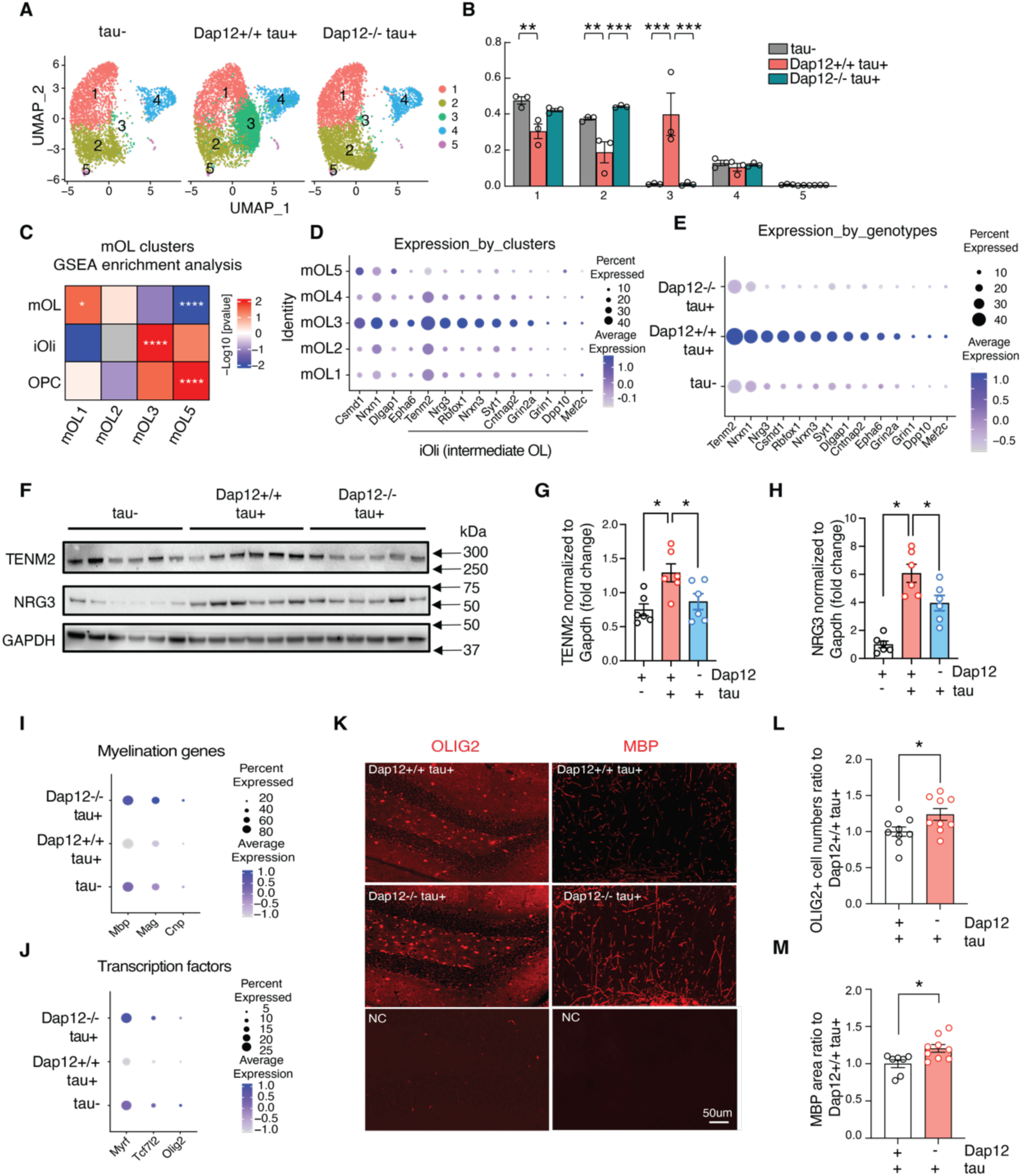
Dap12 mediates tau-induced transcriptomic changes in OPCs and oligodendrocyte. A-B) UMAP (A) and cell ratios (B) of oligodendrocyte subclusters (OL1-5) crossing three genotypes. One-Way ANOVA in B: ***p<0.001, **p<0.01, *p<0.05. C) Comparisons of the mouse oligodendrocyte clusters with iOli (intermediate or immature oligodendrocytes), mOL (mature oligodendrocytes), and OPC (oligodendrocyte precursor cells) by GSEA. Colors denote positive enrichment (+1, red) or negative enrichment (−1, blue) multiplied by the −log10P. D) Dot plot showing the expression of intermediate oligodendrocyte (iOli) marker genes across three genotypes. E) Dot plot showing the expression of intermediate oligodendrocyte (iOli) marker genes across three genotypes. F-H) Western blotting and quantification of Tenm2 or Nrg3. One-Way ANOVA, *p<0.05. *n* = 6/genotype. I-J) Dot plots of genes for myelination and transcription factors across three genotypes. K-M) Olig2 (L) and MBP (M) staining and quantification of positive cell numbers in dentate gyrus. One-Way ANOVA, ***p<0.0001, ***p<0.001, *p<0.05. *n* = 5-9/genotype.

The tau pathology-inducible OL3 was subjected to further analysis using IPA. Predominant pathways that showed upregulation in OL3 include calcium signaling, RHOGDI signaling, and the synaptogenesis signaling pathway (Supplementary Fig. 7C, and supplementary table 6). Notably, the myelination signaling pathway was downregulated in OL3 (Supplementary Fig. 7C and supplementary table 6). In an IPA upstream regulator analysis, two prominent upstream regulators suppressed in OL3 includes: Transcription Factor 7 Like 2 (Tcf7l2) and SRY-Box transcription factor 2 (Sox2) (Supplementary Fig. 7D and supplementary table 6). Both transcription factors play indispensable roles in oligodendrocyte proliferation, differentiation, and neuron myelination^41,42^, implying a compromised myelination state within these cells. Dot plots at the pseudobulk level encompassing the entire OL population highlighted a diminished expression of other myelination-related genes, including Mbp, Mag, and Cnp (Fig. 6I), along with myelination-associated transcription factors such as Myrf, Tcf7l2, and Olig2 (Fig. 6J), within the tauopathy-afflicted mouse brain. However, this downregulation was no longer observed in the tauopathy brain lacking Dap12 (Fig. 6I, J).

We observed that the number of OLIG2-positive cells in the hippocampal DG area was significantly elevated in the absence of Dap12 (Fig. 6K-M and supplementary table 6). Staining with an MBP antibody revealed the inactivation of Dap12 partially mitigated this myelin loss in hippocampal CA1 area (Fig. 6K-M and supplementary table 6).

To study the trajectory states of OL clusters from OPC, we integrated oligodendrocytes (OL) and oligodendrocyte precursor cells (OPC) and subclustered them into six subclusters (Supplementary Fig. 5A). Feature plots showcased that OPC-OL clusters 1, 2, 3, and 6 expressed OL marker genes Mbp and Plp1, whereas OPC-OL cluster 5 exhibited OPC marker genes Pdgfra and Vcan. OPC-OL Cluster 4 contained both OPC and OL subpopulations (Supplementary Fig. 5A). Pseudotime analysis of the transcriptional states revealed a trajectory transition from OPC-OL cluster 5 to cluster 2, followed by cluster 1, ultimately reaching cluster 3 (supplementary Fig. 5B and supplementary table 5). Tauopathy led to a notable decrease in OPC-OL cluster 1 and a significant induction of cluster 2; however, both these effects of tau pathology were reversed by Dap12 deletion (supplementary Fig. 5C, D and supplementary table 5). Thus, Dap12 deletion reversed the tau pathology-induced alterations in the transcriptome state of oligodendrocyte lineage. Correlation analysis of OL3 and OPC-OL Cluster 2 revealed a high similarity between these two clusters (supplementary Fig 5F), confirming a transitional trajectory state of OL3 from OPC to OL (supplementary Fig. 5B).

### Presence of Tau pathology-associated intermediate oligodendrocyte state in human AD brain

To establish the relevance of the mouse model tau pathology-inducible intermediate oligodendrocyte (iOli)-like state in relation to human AD, we conducted a re-analysis of oligodendrocyte clusters within our previously published single-nucleus RNA sequencing (snRNAseq) dataset encompassing both AD and non-AD control samples (n=8), with matched age, tau burden and clinical dementia ratings^43^. Following data integration, five distinct transcriptional clusters of human oligodendrocytes (hOL) were identified (Fig. 7A, supplementary table 7). Notably, hOL3 and hOL4 exhibited heightened cell ratio in AD brains (Fig. 7B).

**Figure 7.**
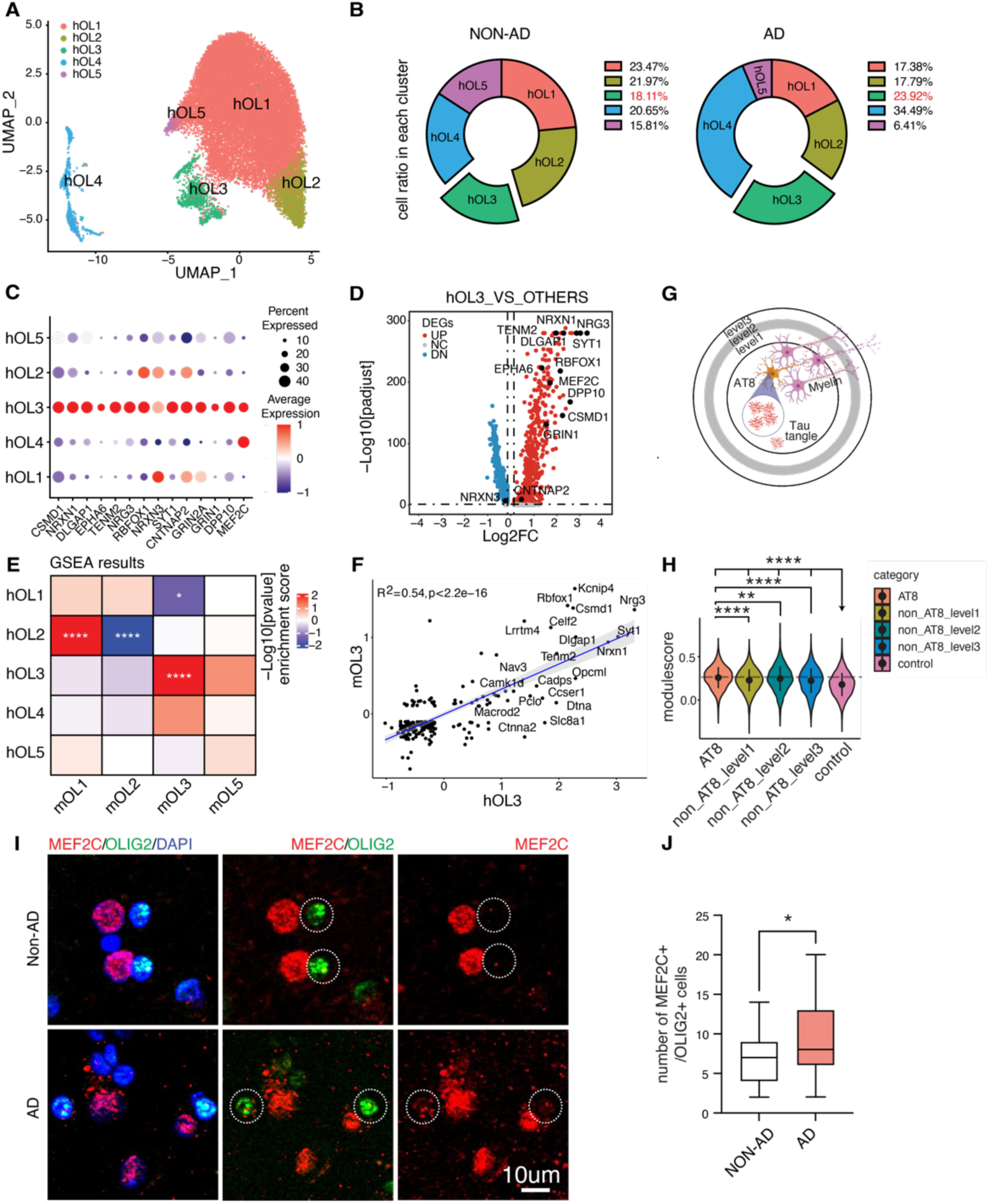
Tau-induced intermediate oligodendrocyte state in human patient. A-B) UMAP and Cell ratios of human oligodendrocyte subclusters (hOL1-5) crossing three genotypes. C) Dot plot showing the expression of iOli marker genes in human oligodendrocyte subclusters. D) Volcano plots of iOli marker genes when comparing human oligodendrocyte cluster 3 (hOL3) versus other clusters. E) Comparisons of the mouse oligodendrocyte clusters with human oligodendrocyte cluster by GSEA. Colors denote positive enrichment (+1, red) or negative enrichment (−1, blue) multiplied by the −log10P. F) Correlation scatterplot of marker genes comparing mOL3 versus hOL3. G) Diagram showing AT8+ spots and neighboring areas level1-3 based on the distance from AT8+ spots for human spatial transcriptome analysis in H. H) Violin plot showing the distribution of module scores for the iOli gene set in B across five spot groups. One-way ANOVA was used to compare mean of module scores among five groups and Wilcoxon rank sum test was used to compare mean of module scores between each pair of the five groups. ****p<0.0001, ***p<0.001, **p<0.01, *p<0.05. I-J) MEF2C,OLIG2 staining, and quantification of MEF2C+/OLIG2+ cells in grey matter region of human brain. *p<0.05.

A dot plot across the five hOL clusters revealed elevated expression of iOli marker genes within hOL3 (Fig. 7C, D and supplementary table 7). The remarkable overlap of marker genes between hOL3 and mouse OL3 was highly overlapped as indicated by Gene Set Enrichment Analysis (GSEA) (Fig. 7E and supplementary table 7). Correlation analysis demonstrated a significant similarity between OL3 in the tauopathy mouse brain and hOL3 (Fig. 7F and supplementary table 7), indicating that the intermediate oligodendrocyte transcriptomic state induced by tau in the mouse brain is conserved in the context of the human AD brain.

We next probed the spatial relationship of tau pathology in human AD brains, leveraging a published 10x Visium spatially resolved transcriptomics (SRT) dataset^44^. We focused spatial transcriptomic analysis on the middle temporal gyrus regions of three AD cases stained with AT8 for pathological tau ^44^. By comparing AT8-positive spots with neighboring spots based on their proximity to AT8-positive areas within AD cases (Fig. 7G), we observed a significant association between the iOli gene module and AT8-positive tau pathology (Fig. 7G, H and supplementary table 7).

Lastly, we investigated the levels of iOli marker genes within OL cells in human brain sections. Considering its nuclei location, we performed double staining of MEF2C, one of the iOli markers, and of OLIG2. Quantitative analysis of MEF2C and OLIG2 staining revealed a significant increase in MEF2C-expressing OLIG2-positive cells within the grey matter of AD brains compared to non-AD brains (Fig. 7I, J and supplementary table 7). Collectively, these findings reinforce the notion that the iOli-like state identified in mouse tauopathy brain is indeed conserved within the human AD brain.

## Discussion

These findings collectively emphasize the pivotal role of microglial DAP12 signaling in modulating the complex interplay between tau pathology, oligodendrocyte integrity, and neuroinflammation within the context of tauopathy. The absence of DAP12 signaling in microglia heightens resilience to brain inflammation, demyelination, and synapse loss despite increasing tau burden. Our study uncovers a novel role of DAP12 in mediating tau pathology-induced toxicity towards oligodendrocytes, precipitating an abnormal intermediate or immature state in the OL transcriptome, which is further associated with myelin loss (Fig. 8).

**Figure 8:**
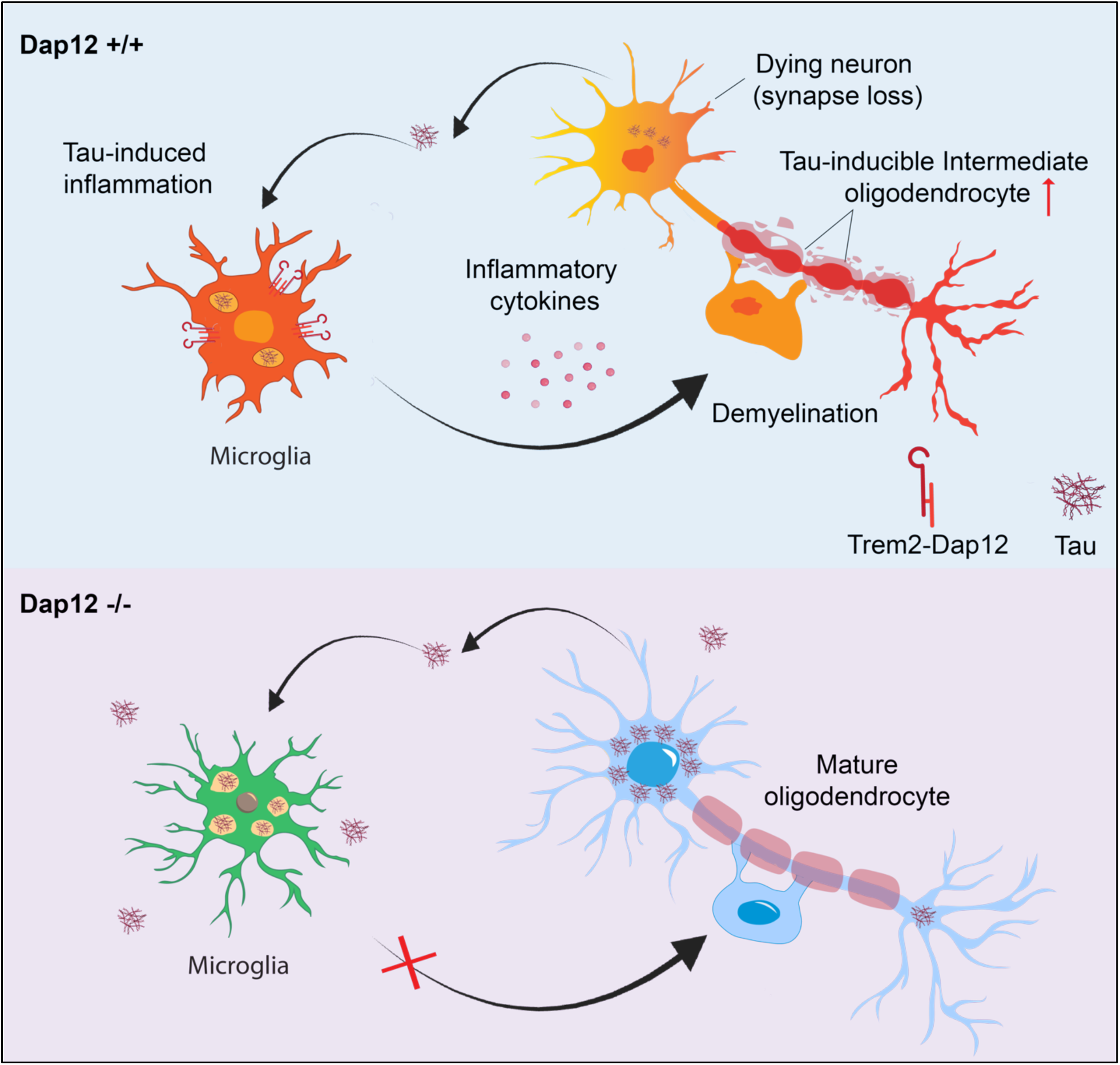
Our research identifies a novel intermediate transcriptomic state in oligodendrocytes dependent on microglial DAP12 signaling and linked to myelin loss in tauopathy mouse brains. Our findings suggest a novel mechanism whereby toxic tau released by neurons activates DAP12 in microglia, which in turn triggers oligodendrocyte toxicity. This sequence of events may contribute to white matter abnormalities and cognitive decline in AD.

The progression of AD is often accompanied by white matter abnormalities, characterized by a partial loss of myelin, axons, and oligodendrocytes^45-48^. Oligodendrocytes, being the primary cell type within white matter, play an indispensable role in forming the myelin sheath. Aberrations in oligodendrocytes have been previously reported in both AD animal models and the human brain^49-54^. Recent advancements in human brain imaging techniques have revealed compromised myelin integrity in AD brains^55-58^ yet the underlying mechanisms remaining elusive. Recent single-cell transcriptomic studies have unveiled the high heterogeneity within oligodendrocyte populations in normal brain or pathological brain, such as AD or multiple sclerosis (MS)^38,39,59-62^. To date, several disease-associated oligodendrocyte subtypes with unique molecular signatures have been identified in response to amyloid or tau pathology in brains of human AD and AD animal models ^38,39,61,63-68^. However, the driving factors for the formation of these states or their functional implications remain to be identified.

Our study uncovered a novel oligodendrocyte subpopulation responsive to brain tau pathology in brains of tauopathy mouse and human AD. Strikingly, this subpopulation possesses a distinct gene signature that sets it apart from previously reported disease-associated oligodendrocyte clusters^38,39,61,63-68^. It closely resembles the intermediate oligodendrocyte (iOli) previously identified in the visual cortex of the human brain ^40^. Importantly, our findings strongly suggest a functional linkage between this tau pathology-induced intermediate transcriptomic state and the loss of brain myelin.

In our current study, homozygous tau transgenic mice develop hippocampal tau pathology at a younger age compared to other models^16,28,38^. Memory and spine density problems start at 2.5 months when MC1+ tau and pTau tau are present, but there’s no neuronal loss, suggesting an early disease stage with synaptic connectivity issues causing memory deficits^27^. The tau pathology-inducible iOli-like oligodendrocytes identified in our study could represent an initial response of oligodendrocytes to tau toxicity, and it’s unclear if this state is temporary. Moreover, our snRNAseq data shows minimal DAP12 mRNA in oligodendrocytes, indicating that the effects of DAP12 deletion on oligodendrocytes are probably non-cell-autonomous, contrasting prior reports^29^. Future studies are needed to explore its formation and fate in later disease stages, as well as the functional implications and molecular mechanisms of these emerging cells.

Human DAP12 loss-of-function variants cause Nasu-Hakola disease (NHD), which features cerebral atrophy, myelin loss, and gliosis^30-32,69^. However, mice lacking DAP12 signaling exhibit milder phenotypes, not fully resembling human NHD^70^. This divergence suggests that DAP12-deficient mice may not be ideal for comprehensive NHD brain phenotype studies, emphasizing the need for specific insults to reveal DAP12 deficiency effects. Crossbreeding with a tauopathy model shows that DAP12 signaling is crucial for microglial responses to tauopathy, driving homeostatic microglia and oligodendrocytes into disease-associated states. This also suggests distinct mechanisms may underlie the pathogenesis of AD and NHD.

DAP12, containing an ITAM motif, can have variable effects, either positively or negatively regulating responses based on the specific receptor, ligand, or cell type involved^71^. For instance, in mice, DAP12 deletion enhances bacterial infection control^72,73^ and prevents axotomy-induced motor neuron death with reduced pro-inflammatory responses^74^. DAP12-deficient mice also exhibit heightened immune cell responsiveness, leading to increased cytokine production against infections by dendritic cells, macrophages, and NK cells^75^. In the central nervous system, Dap12 deficiency in mice provides protection in an autoimmune encephalomyelitis (EAE) mouse model immunized with myelin oligodendrocyte glycoprotein (MOG) peptide, linked to reduced IFNγ production by myelin-reactive CD4+ T cells in vivo ^17^. These findings suggest that DAP12 engages with various receptors, influencing diverse signaling networks that can either dampen or enhance immune responses to different challenges.

We speculate that TREM2 may act as a receptor partnering with DAP12 in regulating tau pathology and toxicity, leading to the positive effects of DAP12 deletion observed in our tau pathology mouse model. Studies indicate that complete loss or haploinsufficiency of microglial TREM2 worsens tau pathology, promoting tau seeding and spreading in mouse models^2,76-80^. Conversely, TREM2 deletion has a protective effect against tauopathy, reducing microglial activation and neurodegeneration^76,79^. The impact of TREM2 risk alleles like R47H on DAP12’s role in mediating tau toxicity in oligodendrocytes and myelination remains an open question, as does the exploration of other receptors interacting with DAP12 in these processes.

Earlier studies have highlighted intricate communications between oligodendrocytes and microglia^81,82^. Genetic elimination of microglia triggers demyelination, underscoring the crucial role of microglia in myelin integrity in healthy brains^83^. In diseased brains, activated microglia can exert detrimental effects on myelin-producing oligodendrocytes by releasing pro-inflammatory mediators, including chemokines and cytokines^82^. This correlation between microglial activation and oligodendrocyte impairment has been observed in the context of multiple sclerosis^84^. The release of tumor necrosis factor-alpha (TNF-α) and interleukin-1 beta (IL-1β) by microglia is associated with myelin damage^85^, while IFNγ is implicated in triggering oligodendrocyte apoptosis and hindering central nervous system remyelination^86^. Our results showed that DAP12 deletion robustly curtails the activation of interferon signaling induced by tauopathy in AD. Additionally, this deletion leads to a reduction in IP-10, as well as several cytokines and chemokines inducible by IFNγ, which are known to instigate oligodendrocyte apoptosis^87^. Collectively, these observations strongly suggest that IFNγ could be one of the downstream effectors affecting oligodendrocytes due to increased DAP12 signaling in microglia, a notion warranting further investigation.

Tau pathology is a main contributing factor in AD whose precise mechanisms remain unclear. Our study unveils a novel mechanism whereby toxic tau released by neurons activates DAP12 in microglia, which in turn triggers oligodendrocyte toxicity (Fig. 8). This sequence of events may contribute to white matter abnormalities and cognitive decline in AD. Our research identifies a novel tau-induced transcriptomic state in oligodendrocytes, an understudied cell-type in AD pathogenesis. This state is closely connected to microglial DAP12 signaling and is linked to myelin loss. Overall, our findings deepen our understanding of the complex interactions between neurons, microglia, and oligodendrocytes in tauopathy and suggest novel targets for therapeutic treatments to mitigate AD’s impact.

## Author information

Authors and Affiliations

**Helen and Robert Appel Alzheimer’s Disease Research Institute, Feil Family Brain and Mind Research Institute, Weill Cornell Medicine, New York, NY, USA**

Hao Chen, Li Fan, Man Ying Wong, Chloe Lopez, Bangyan Liu, Gloria Huang, Gillian Carling, Fangmin Yu, Nessa Foxe, Li Gan, Wenjie Luo

**Department of Biomedical Informatics, College of Medicine, Ohio State University, Columbus, OH 43210 USA**

Qi Guo, Qin Ma

**Department of Neuroscience, College of Medicine, Ohio State University, Columbus, OH 43210 USA**

Hongjun Fu

**Department of Genetics and Genomic Sciences, Mount Sinai Center for Transformative Disease Modeling, Icahn School of Medicine at Mount Sinai, New York, NY, USA**

Won-min Sung, Bin Zhang

**Department of Biochemistry, Faculty of Medicine and Dentistry, University of Alberta, Edmonton, AB Canada**

Tark Patel, Sue-Ann Mok

**Millburn High School, Millburn, New Jersey, USA**

Winston Wang

**The State University of New York at Stony Brook, Long Island, New York, USA**

Aviram Nessim

## Contributions

W.L, X.H. and L.G. conceived the project. W.L., L.G., and H.C. designed experiments. H.C., L.F., M.Y.W., Q.G., W.W., F.Y., N.F., A.N. H.F. performed experiments and/or analyses. H.C., C.L., B.L., G.H., Q.G., W.S., B.Z., Q.M., H.F. developed analytic protocols and tools. S.A., T.P., S.M. provided or prepared reagents. H.C., L.G. and W.L. wrote the manuscript. All authors read and approved the paper.

## Acknowledgements

This work was supported by the NIH R01AG064239 to W.L. and CART (Coins For Alzheimer’s Disease Trust Fund) to W.L., the NIH U54NS100717, R01AG072758, R01AG074541 (to L.G), Tau Consortium (to L.G.), JPB Foundation (to L.G.), R01 AG075092 (H.F.). We thank Dr. Xiaoyu Hu for scientific discussion and providing Dap12 knockout mice (kindly offered by Dr. Lewis L. Lanier) and Dr. Flint Beal and Dr. Michel Goedert for providing homozygous P301S tau transgenic mice. We thank Dr. Adriene Y. Tan and Dr. Jenny Xiang at the Weill Cornell medicine genomics core for performing RNA sequencing. We also thank Shuo Chen at The Ohio State University for the pilot analysis of Visium datasets and the discussion. We thank Claire Hu for editing the manuscript.

## Supplementary Information

**Supplementary Figure 1:**
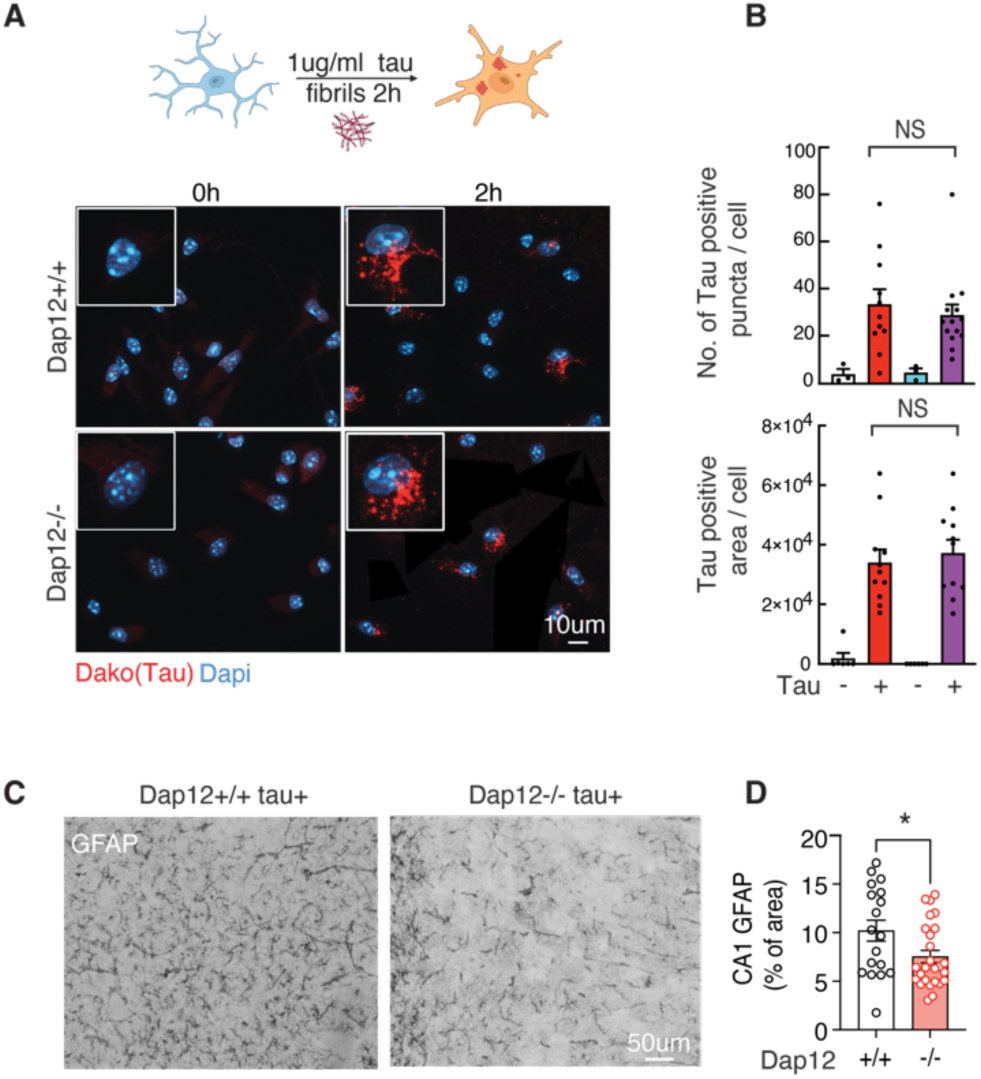
Characterization of tauopathy mouse brain with Dap12 deficiency and the effect of Dap12 deletion on microglia tau phagocytosis, related to Figure 1. A-B) Representative images (A) and quantification (B) of Tau positive puncta numbers and area in primary cultured microglia. Scale bar: 10 µm. Two-Way ANOVA with post hoc test, NS: Not significant. *n* = 11-14 from 3 independent experiments. C-D) Representative immunohistochemical staining (C) and quantification (D) of GFAP+ area in the CA1 of *Dap12^+/+^ Tau+*, and *Dap12^-/-^ Tau+* mice. Scale bar: 50 µm. Unpaired student’s t-test: *p<0.05. n = 18 *Dap12^+/+^ Tau+*, n = 26 *Dap12^-/-^ Tau+* mice.

**Supplementary Figure 2:**
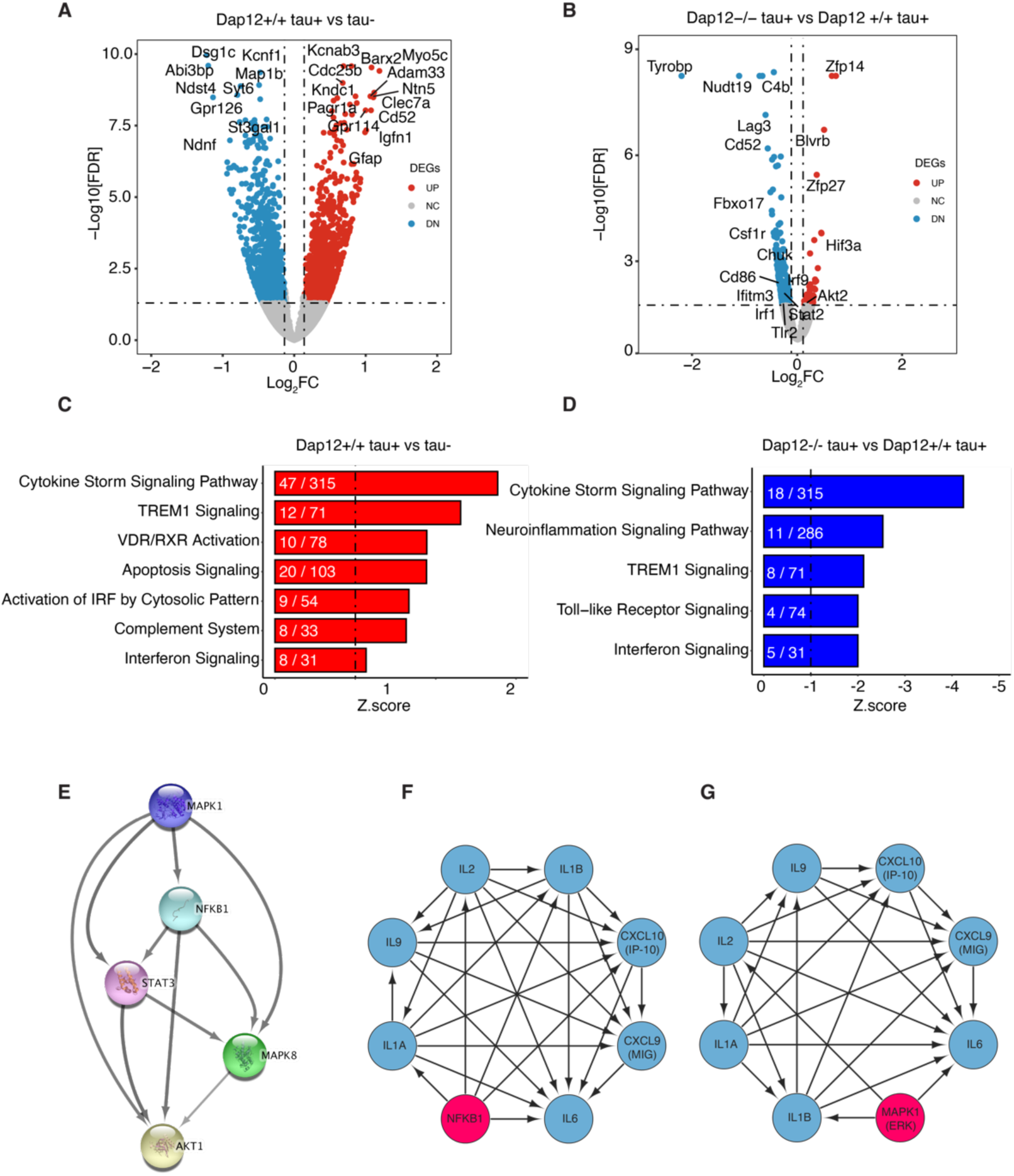
Deletion of Dap12 suppressed inflammatory signaling in tauopathy mouse brain. A-B) Volcano plot of DEGs (adjust p-value < 0.05, Log2FC > 0.1 or < -0.1) comparing *Dap12^+/+^ Tau+* versus *Tau-* mice (A) and *Dap12^-/-^ Tau+* mice versus *Dap12^+/+^ Tau+* mice. C-D) Selected top IPA canonical pathways identified from the DEGs of *Dap12^+/+^ Tau+* vs *Tau-* mice (C) or *Dap12^-/-^ Tau+* vs *Dap12^+/+^ Tau+* mice (D). IPA canonical pathways contain z score and -log10(p-value). No log2FC or adjust p-value. E) String gene network analysis showing relationships between immune regulators identified in Figure 2K. F-G) String gene network analysis showing cytokines regulated by NF-κB (F) or ERK (G).

**Supplementary Figure 3:**
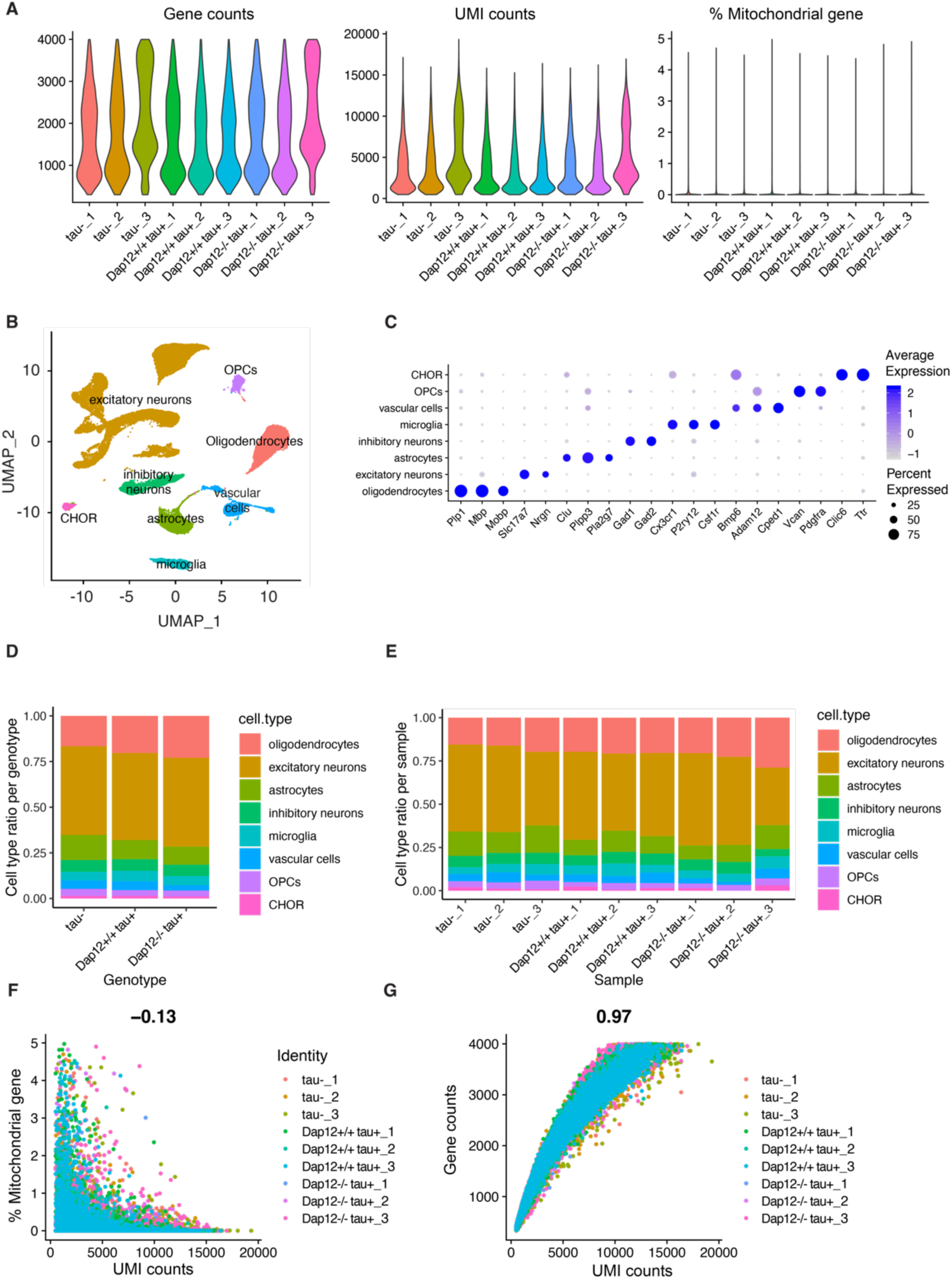
Quality control assessment of Single-Nuclei RNA-seq (related to Fig. 3-6) A) Quality control plots showing equivalent amounts of total number of genes, total number of molecules and percent mitochondrial RNA in nuclei used for downstream analyses. B) UMAP dimensional plot showing nuclei colored according to transcriptionally distinct cell clusters identified using Seurat package. C) Summary of genes used for cluster classification into different cell types. D) Proportion of each cell type detected across the different genotypes. E) Proportion of each cell type detected across the different samples. F-G) Correlation between UMI counts and percentage of mitochondrial genes per nuclei (F) and total genes detected (G) for all samples.

**Supplementary Figure 4:**
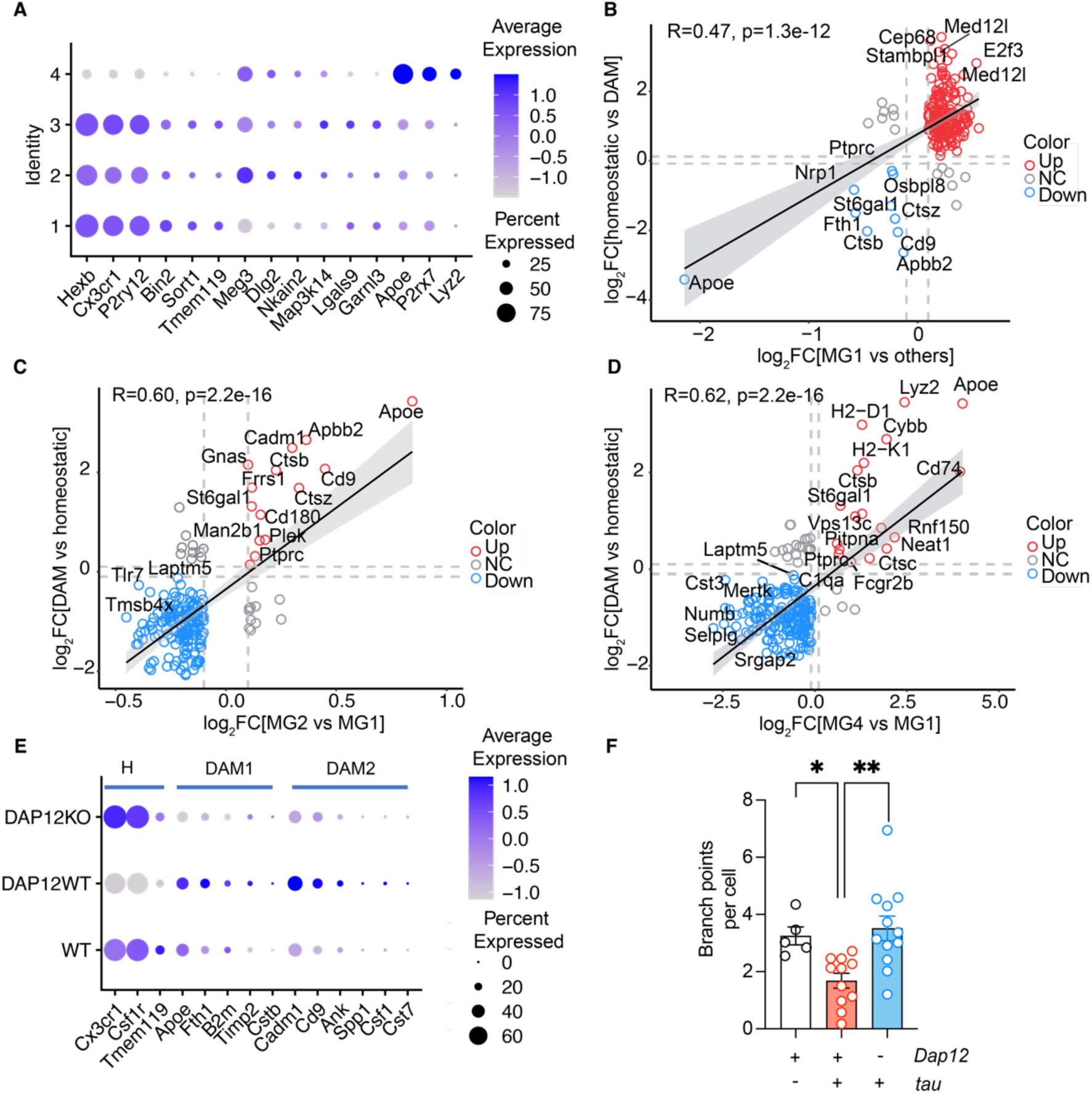
Characterization of microglia clusters (Related to Figure 3). A) Dot plot of marker genes crossing different MG clusters. B) Correlation scatterplot of marker genes comparing MG1 cluster versus other clusters. C) Correlation scatterplot of marker genes comparing MG2 versus MG1 cluster. D) Correlation scatterplot of marker genes for MG4 versus MG1 cluster. E) Dot plot of homeostatic and DAM marker genes in crossing different genotypes. F) Quantification of microglial branch points crossing three genotypes. n = 5 *tau-*, n = 11 *Dap12+/+ tau+*, n = 12 *Dap12-/- tau+* mice.

**Supplementary Figure 5.**
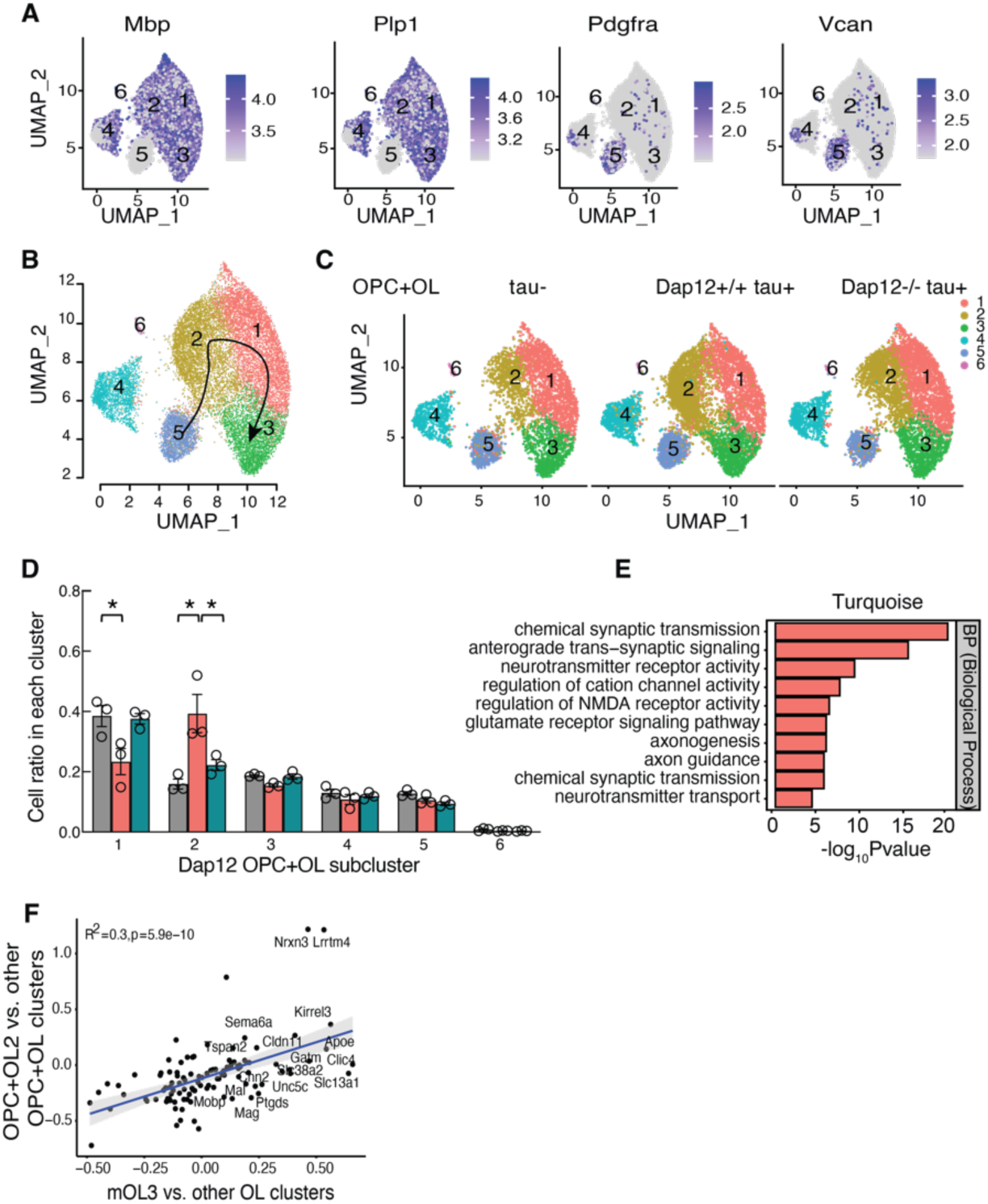
Dap12 is essential for tau-induced transcriptomic changes in oligodendrocyte lineage cells in vivo. A) Feature plots of marker genes for OPC (Pdgfra and Vcan) and OL (Mbp and Plp1) subclusters. B) Slingshot showing transition between OL lineage cell subclusters. C) UMAP of integrated OL lineage clusters with integrated OPC and OL. D) Cell ratios of OPC+OL subclusters (1,2,3,4,5 and 6) crossing three genotypes. One-Way ANOVA followed by turkey test, ***p<0.001, **p<0.01, *p<0.05. *n* = 3 per genotype. E) Bar chart of the enriched pathways for the signature gene sets of the Turquoise module identified by Gene Ontology (GO) pathways. F) Correlation scatterplot of DEGs between integrated OPC+OL2 and mouse oligodendrocyte cluster 3 (mOL3)

**Supplementary Figure 6:**
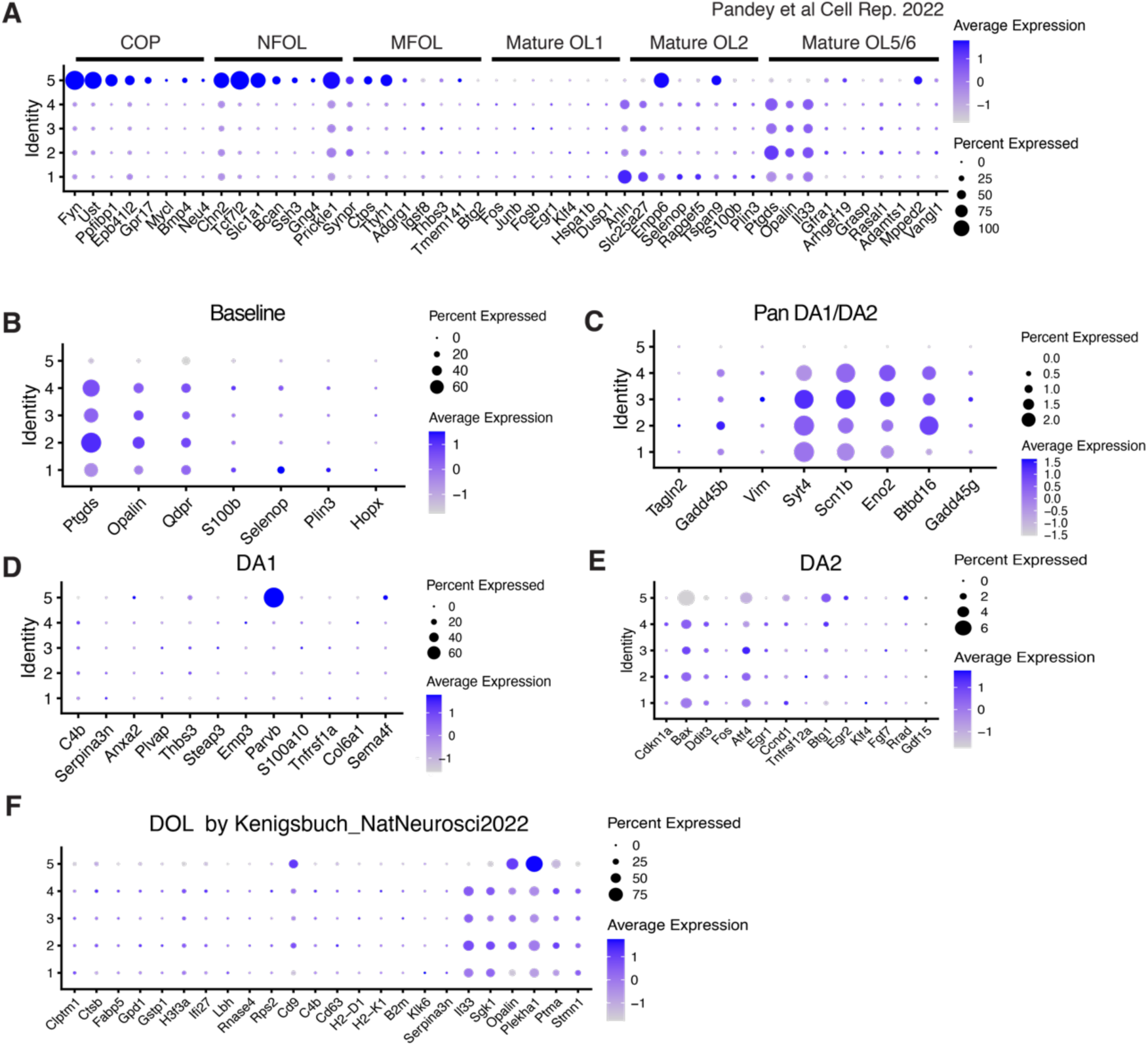
Characterization of oligodendrocyte clusters. A) Dot plot of marker genes of previous identified oligodendrocyte clusters. B-E) Dot plot of marker genes for baseline (B), pan disease associated oligodendrocyte marker genes (C), disease associated stage 1 oligodendrocyte (D), and disease associated stage 2 oligodendrocyte(E). F) Dot plot of marker genes of disease associated oligodendrocyte identified in previous report.

**Supplementary Figure 7:**
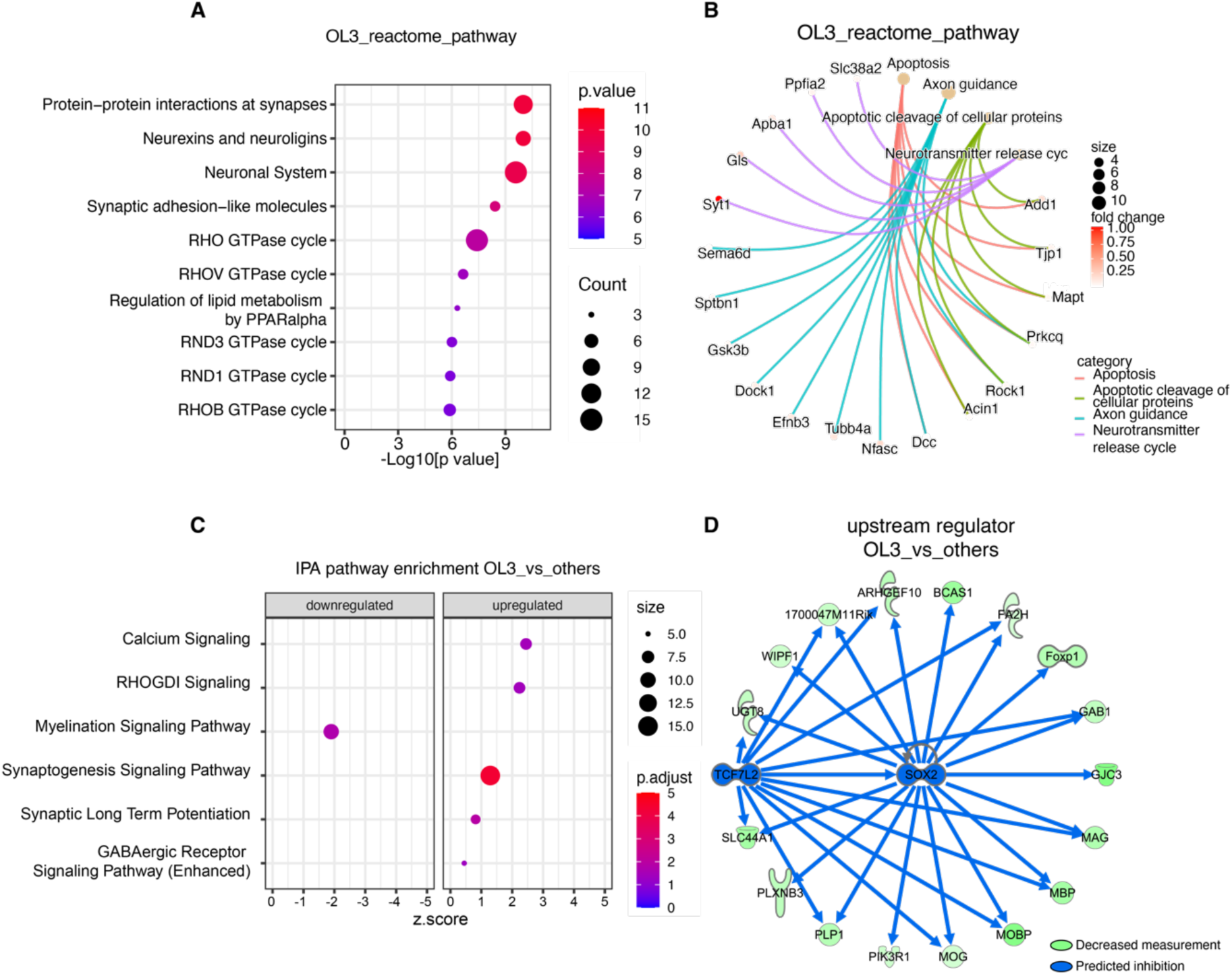
Characterization of oligodendrocyte clusters. A-B) Reactome pathways enriched in DEGs identified in OL3 (A) and selected top reactome pathways with associated genes (B). C) Selected IPA canonical pathways identified for DEGs of OL3. D) Tcf7l2 and Sox2 predicted by IPA as upstream regulators of a subset of DEGs downregulated in OL3.

## Online Methods

### Animals

Mice were housed in groups of no more than five per cage, with access to food and water ad libitum. They were kept in a pathogen-free barrier facility under controlled conditions at a temperature of 21–23 °C, humidity ranging from 30% to 70%, and a 12-hour light/12-hour dark cycle. Homozygous human *Tau P301S* transgenic mice, obtained from Dr. Michel Goedert at the MRC Laboratory of Molecular Biology, Cambridge, UK, were crossed with *Dap12^−/−^* mice, provided by Dr. Lewis Lanier at the University of California, San Francisco. This crossbreeding resulted in the generation of *P301S Dap12^+/−^* mice. Subsequent crossings of F1 litters led to the production of both *Dap12^+/+^* and *Dap12^−/−^* mice, along with their corresponding *P301S* transgenic littermates. For all analyses, female mice exhibiting accelerated tau pathology were utilized at the age of 6 months. All experimental procedures involving mice were conducted in accordance with ethical guidelines and were approved by the Institutional Animal Care and Use Committee of Weill Cornell Medicine.

### Human brain samples

The tissues used for this study were the mid-frontal cortices from brains of age-matched patients with AD and non-dementia controls. Samples were obtained from the University of Pennsylvania brain bank and Mount Sinai Hospital. All brains were donated after consent from the next-of-kin or an individual with legal authority to grant such permission. Brain tissues of University of Pennsylvania brain bank and Mount Sinai Hospital used in this study are not considered identified “human subjects” and are not subject to IRB oversight. The institutional review board has determined that clinicopathologic studies on de-identified postmortem tissue samples are exempt from human subject research according to Exemption 45 CFR 46.104(d)(2). Additional information about the donors can be found in the supplementary table 8.

### Primary microglial culture

Following established protocol^1^, primary microglia were isolated from the hippocampi and cortices of 0-3-day-old mouse pups. The isolated brain tissues were rinsed with Dulbecco’s Phosphate-Buffered Saline (DPBS), and the meninges were carefully removed. Subsequently, the brain tissues were minced, followed by treatment with 0.05% trypsin at 37 °C for 20 minutes. The trypsinization process was halted by adding 20% FBS/DMEM media, after which the digested tissues were gently triturated to generate a cell suspension. This suspension was then subjected to centrifugation at 200 x g for 15 minutes, and the pellet was resuspended in 10% FBS/DMEM. The resuspended cells were plated onto T-75 flasks coated with poly-D-lysine (PDL), facilitating the formation of mixed glial cultures. These cultures were maintained in 10% FBS/DMEM supplemented with 5ng/ml granulocyte-macrophage colony-stimulating factor (GM-CSF). By the twelfth day, when the cultures had reached confluence, microglia were isolated from the glial layer by subjecting the flasks to gentle shaking at 200 rpm for a duration of 3 hours. The microglia that floated were subsequently seeded onto plates coated with PDL at a density of 75,000 cells/cm^2^. They were then cultured in 10% FBS/DMEM devoid of GM-CSF for a 24-hour period before being employed in assays involving the phagocytosis and processing of tau fibrils.

### Tau fibril uptake and clearance by cultured microglia

Microglia were seeded into eight-well chamber slides at a density of 1 × 10^5^ cells per well. Subsequently, they were exposed to 1 μg/ml of AD-tau fibrils prepared as described previously ^2^ and incubated for a duration of 2 hours in 10% FBS/DMEM. Following the incubation period, the cells were subjected to a wash with PBS and then fixed using 4% paraformaldehyde (PFA). For the tau chasing assay, primary cultured microglia were initially exposed to AD-tau fibrils for 2 hours. Post-exposure, the medium containing tau fibrils was replaced with fresh 10% FBS/DMEM. At time intervals of 12 and 24 hours, the microglia were washed with PBS followed with fixation with 4% PFA. Immunostaining was conducted using rabbit anti-human tau antibodies (Dako, A0024, 1:500), followed by Alexa Fluor 568 conjugated goat IgG targeting rabbit (Invitrogen, A-11004, 1:200). To visualize the nuclei, DAPI staining (blue) was performed. Representative images were acquired using a ZEISS microscope at a magnification of 63×.

### Bulk RNA sequencing

Freshly perfused mouse brains were dissected to isolate the cortices. The cortices were flash-frozen and then stored at -80 °C. For RNA extraction, the cortices were thawed on ice for a duration of 30 minutes and then RNA isolation from the cortex tissue was carried out following the manufacturer’s protocol (PureLink™ RNA Mini Kit, Thermo Fisher). The isolated RNA samples were then sent to the Weill Cornell Medicine Genomics Core for assessment of RNA quality and integrity. Following successful quality control, RNA-seq libraries were prepared for sequencing using the NovaSeq platform.

### Isolation of nuclei from frozen mouse brain tissue

The protocol for isolating nuclei from frozen mouse brain tissue was adapted from previous studies with modifications ^3,4^. All procedures were done on ice or at 4 °C. In brief, mouse brain tissue was placed in 1,500 µl of nuclei PURE lysis buffer (Sigma, NUC201-1KT) and homogenized with a Dounce tissue grinder (Sigma, D8938-1SET) with 15 strokes with pestle A and 15 strokes with pestle B. The homogenized tissue was filtered through a 35 µm cell strainer and was centrifuged at 600 × g for 5 min at 4 °C and washed three times with 1 ml of PBS containing 1% BSA, 20 mM DTT, and 0.2 U µl^−1^ recombinant RNase inhibitor. Then the nuclei were centrifuged at 600 × g for 5 min at 4 °C and resuspended in 500 µl of PBS containing 0.04% BSA and 1× DAPI, followed by FACS sorting to remove cell debris. The FACS-sorted suspension of DAPI-stained nuclei was counted and diluted to a concentration of 1,000 nuclei per microliter in PBS containing 0.04% BSA.

### Droplet-based single-nuclei RNA-seq and data analysis

For droplet-based snRNA-seq, libraries were prepared with Chromium Single Cell 3’ Reagent Kits v3 (10× Genomics, PN-1000075) according to the manufacturer’s protocol. cDNA and library fragment analysis were performed using the Agilent Fragment Analyzer systems. The snRNA-seq libraries were sequenced on the NovaSeq 6000 sequencer (Illumina) with 100 cycles. Gene counts were obtained by aligning reads to the mouse genome (mm10) with Cell Ranger software (v.3.1.0) (10× Genomics). To account for unspliced nuclear transcripts, reads mapping to pre-mRNA were counted. Cell Ranger 3.1.0 default parameters were used to call cell barcodes. We further removed genes expressed in no more than three cells, cells with a unique gene count over 4,000 or less than 300, and cells with a high fraction of mitochondrial reads (>5%). Potential doublet cells were predicted and removed using DoubletFinder ^5^ for each sample. Normalization and clustering were done with the Seurat package v4.0.0. In brief, counts for all nuclei were scaled by the total library size multiplied by a scale factor (10,000), and transformed to log space. A set of 2,000 highly variable genes were identified with FindVariableFeatures function based on a variance stabilizing transformation (vst). Principal component analysis (PCA) was done on all genes, and t-SNE was run on the top 15 PCs. Cell clusters were identified with the Seurat functions FindNeighbors (using the top 15 PCs) and FindClusters (resolution = 0.1). For each cluster, we assigned a cell-type label using statistical enrichment for sets of marker genes and manual evaluation of gene expression for small sets of known marker genes. The subset() function from Seurat was used to subset each cell types. Differential gene expression analysis was done using the FindMarkers function and MAST ^6^. For pseudobulk analyses, we aggregated the expression values from all nuclei from the same cell type for genotype dependent differential expression.

### Gene network and functional enrichment analysis

Gene network and functional enrichment analysis were performed by QIAGEN’s Ingenuity® Pathway Analysis (IPA®, QIAGEN Redwood City, www.qiagen.com/ingenuity, Version 01-22-01) or by GSEA with molecular signatures database (MSigDB) ^7,8^. Significant DEGs and their log2fold change expression values and FDR were inputted into IPA for identifying canonical pathways, biological functions, and upstream regulators. Upregulated or downregulated significant DEGs were inputted into GSEA (http://www.gsea-msigdb.org/gsea/msigdb/annotate.jsp) to identify hallmark and gene ontology terms. The p-value, calculated with the Fischer’s exact test with a statistical threshold of 0.05, reflects the likelihood that the association between a set of genes in the dataset and a related biological function is significant. A positive or negative regulation z-score value indicates that a function is predicted to be activated or inhibited. Weighted gene correlation network analysis (WGCNA) was performed to identify co-expression of genes within the different oligodendrocyte clusters using weighted gene correlation network ^9-11^.

### Cell trajectory using Monocle3^12^

For oligodendrocyte trajectory analysis, the oligodendrocyte population was first isolated from the other cell types like previous report ^2^. A separate Seurat object was created for oligodendrocyte, followed by normalization with a scale factor of 10,000. FindVariableFeatures function was run again to identify the most variable gene-specific for oligodendrocyte. The oligodendrocyte Seurat object was then converted into a Monocle3 object with as.cell_data_set function. Size factor estimation of the new CDS (cell dataset) was performed using estimate_size_factors function with default parameters. Further processing of the CDS was carried out using preprocess_cds function with the num_dim parameter set to 9. UMAP was then performed to reduce the dimensionality of the data. Cell clusters were then visualized with cluster_cells function with the parameter K equals to 9. learn_graph function was used to determine the trajectory and the cluster 1 oligodendrocyte was selected as the origin of the trajectory and the gene expression along with pseudotime was generated by using plot_gene_expression function.

### Immunohistochemistry

Dulbecco’s phosphate-buffered saline (DPBS) was used for immunohistochemistry. Four brain sections per mouse that contain a series of anterior to posterior hippocampus were washed to remove cryoprotectant and then permeabilized by 0.5% Triton X-100. After blocking in 5% normal goat serum (NGS) for 1 h, brain sections were incubated with primary antibodies in the same blocking buffer overnight at 4 °C. Sections were then washed by DPBS containing 0.1% Tween-20 and incubated with Alexa-conjugated secondary antibodies for 1 h at room temperature in blocking buffer. After washing, sections were mounted on glass slides with ProLong Gold Antifade Mounting media. The primary antibodies used for immunohistochemistry were as follows: anti-IBA1 (1:500, 019-19741, Fujifilm Wako), anti-MC1 (1:500, a kind gift from P. Davies), anti-OLIG2 (1:500, ZMS1019, sigma), anti-MBP (1:800, sigma, MAB386), anti-MEF2C (1:600, abcam, ab211493), anti-GFAP (1:800, abcam, ab7260), anti-phospho-Tau (Ser202,Thr205) (AT8)(1:1000, ThermoFisher Scientific, MN1020), and anti-P2RY12 (1:600, 848002, Biolegend). The secondary antibodies used for immunohistochemistry were as follows: Goat anti-rabbit 568 (1:600, A11036, invitrogen), Goat anti-mouse 568 (1:600, A11031, invitrogen), Goat anti-rat 568 (1:600, A11077, invitrogen). Images for MC1 and IBA1 quantification were acquired on Zeiss microscope using 20x objective and analyzed with ImageJ (NIH). All images were first set the threshold manually, then the auto-measurements were performed by using the macros program in ImageJ. Regions of interest including the hippocampus and cortex were hand-traced. MC1+ areas were measured by ImageJ, whereas OLIG2+ cell numbers were counted with the Analyze Particles function. 3D structure of microglia was reconstructed using the Imaris software as described before ^13^. Experimenters performing imaging and quantification were blinded.

### Western blotting

Total brain cortex lysates were prepared in radioimmunoprecipitation assay buffer (RIPA) [1% NP-40, 0.5% sodium deoxycholate, and 0.1% sodium dodecyl (lauryl) sulfate]. Protein (20 mg) was separated by a 12% SDS-PAGE gel, then transferred to a polyvinylidene difluoride (PVDF) membrane. After blocking in TBS buffer (20 mM Tris-HCl, 150 mM sodium chloride) containing 5% (wt/vol) nonfat dry milk for 1 h at room temperature, the membranes were then probed with proper primary and secondary antibodies, which was followed by developing with Super Signal West Pico chemiluminescent substrate (34577; Thermo Scientific, Rockford, IL). Data analysis was performed by Image lab 6.1 (Bio-Rad, Hercules, CA). The following primary antibodies were used: rabbit anti-phospho-AKT (Ser473) (Cell Signaling, 4060, 1:1000), rabbit anti-total-AKT (Cell Signaling, 4691, 1:2,000), rabbit anti-phospho-STAT3 (Ser727) (Cell Signaling, 9134, 1:1,000), mouse anti-total-STAT3 (Cell Signaling, 9139, 1:3,000), rabbit anti-phospho-ERK1/2 (Thr202/204) (Cell Signaling, 4370, 1:1,000), rabbit anti-total-ERK1/2 (Cell Signaling, 9102, 1:3,000), sheep anti-TENM2 (Thermo Fisher Scientific, PA5-47638, 1:1,000), rabbit anti-NRG3 (Thermo Fisher Scientific, MA5-36144, 1:1,000), rabbit anti-GAPDH (GeneTex, GTX100118, 1:10,000), mouse anti-PSD95 (Abcam, ab2723, 1:1,000), rabbit anti-GAPDH (Cell Signaling, 2118, 1:5,000), and rabbit anti-Beta3-Tubulin (Cell Signaling, 55685, 1:5,000). The following secondary antibodies were used: HRP-goat anti-Mouse IgG (Jackson, 115-035-146, 1:2,000), HRP-goat anti-Rabbit IgG (Jackson, 111-035-144, 1:2,000).

### Multiplex bead-based immunoassay

The frontal cortex lysates were prepared by RIPA buffer with sonication at 2 °C for 5 mins at 30% amplitude with 5 secs and 2 secs pulse. Then the samples were centrifuged at 20,000 x g for 15 mins, and the resulting supernatant was collected for analysis using the MILLIPLEX MAP Mouse phosphor and total multi-pathway 9-plex Magnetic Bead Kit (Millipore, Cat.# 48-680MAG) on a MagPix System. For measuring cytokines and chemokines, the frontal cortex was homogenized in Reassembly Buffer (RAB), followed by centrifugation for 20 minutes at 50,000 x g at 4°C. The supernatant was then collected and mixed with an equal amount of RIPA buffer, followed by another centrifugation step for 20 minutes at 50,000 x g at 4°C. The resulting supernatant was used for analysis using the MILLIPLEX MAP Mouse Cytokine/Chemokine Magnetic Bead kit (Millipore, MCYTMAG-70K-PX32) on a MagPix System.

### Spatial transcriptomic analysis of intermediate OL gene set using public Visium datasets

To further validate the correlation between the expression of selected iOli gene set (CSMD1, NRXN1, DLGAP1, EPHA6, NRG3, DPP10, MEF2C, RBFOX1, TENM2, SYT1, CNTNAP2, GRIN2A, GRIN1, NRXN3, PLP1, MBP, MOBP) and AD pathological regions, we visualized the expression distribution of the gene set in six publicly available 10x Visium spatially resolved transcriptomics (SRT) datasets ^14^. These datasets included adjacent sections with pathological tau stained by AT8, consisting of three control cases and three AD cases. First, we preprocessed and integrated the six Visium datasets following the steps outlined in ^14^. After excluding noise spots as previously described ^14^, all spots were divided into five groups, including AT8+ spots group and neighboring levels 1-3 spots groups by the distance from AT8+ spots in AD cases and one control spot group in control cases. The gene module scores of the gene set were calculated using the function “AddModuleScore” by default parameters to indicate relative average expressions of gene sets in five spot groups using Seurat (v4.1.1). The module scores of the gene set can be referred to as the gene set activity. They are calculated by subtracting the aggregated expression of control feature sets from the average expression of the gene set at the single-spot level. Next, gene module scores of the gene set were visualized using the function “geom_violin” by R package ggplot2 (v3.3.5). The mean of module scores among five groups was compared using one-way analysis of variance (ANOVA) and the mean of module scores between each pair of the five groups was compared using Wilcoxon rank sum test, both performed with the function “compare_means” in R package ggpubr (v0.4.0).

To study the spatial relationship, representative spatial maps were created using a representative SRT 10x Visium dataset from an AD sample at Braak stage-IV ^14^. Two types of spatial maps were generated via the Loupe Browser (v6.4.1). First, the layer-labeled spatial map of this AD brain section showcases the layer information of each spot manually labeled as previously described ^14^. WM: white matter; noise: spots with folded tissue was excluded from gene expression analysis. Second, the log2 average expression of gene set 3 in each Visium spot of the same AD sample was computed and visualized via Loupe Browser. The colors of the spots were scaled based on the average expression of gene set 3, ranging from 1.5 to 4. This color scaling was applied to highlight spots with high expression levels of gene set.

### Data Availability

Six SRT 10x Visium datasets were downloaded from Gene Expression Omnibus (GEO: GSE220442). The AT8+ spot annotations and the cloupe files for 10x Visium datasets can be found at: https://bmbls.bmi.osumc.edu/scread/stofad-2.

### Statistics

The sample size for each experiment was determined based on previous publications^2,15^. All in vitro experiments were performed with a minimum of three biological replicates. Mean values from at least three independent experiments were used for computing statistical differences. All *in vivo* experiments were performed with a minimum of four mice per genotype. All *in vivo* data were averaged to either individual mouse (microglia number counts), individual section (MC1, AT8 tau), or individual microglia (Imaris morphology analysis), and mean values were used for computing statistical differences. Data visualization was done with Graphpad and R package ggplot2. Statistical analyses were performed with Graphpad prism 9.0 (t-test, one-way and two-way ANOVA) (Graphpad, San Diego, California). Values are reported as mean ± standard error of the mean (SEM) or standard deviation (SD). Mann–Whitney test was used when the normality test is not passed. One-way ANOVA was used to compare data with more than two groups. Two-way ANOVA was used for groups with different genotypes and/or time as factors. Tukey’s and Sidak’s post-test multiple comparisons were used to compare the statistical difference between designated groups. All P-values of enrichment analysis are calculated by right-tailed Fisher’s exact test. P < 0.05 was considered statistically significant.

