## Supplementary table 1 for "DAP12 deficiency alters microglia-oligodendrocyte communication and enhances resilience against tau toxicity"

| Figure1AB_tau_chasing | | | | | | | | | | |
|---|---|---|---|---|---|---|---|---|---|---|
| time(h) | Dap12+/+ | | | | | Dap12-/- | | | | |
| 0 | 100 | 100 | 100 | 100 | 100 | 100 | 100 | 100 | 100 | 100 |
| 12 | 13.3971973 | 17.4355586 | 16.1990283 | 13 | 21.9800545 | 57.8995805 | 32.3286868 | 61.6019308 | 48.7481391 | 20.9615194 |
| 24 | 8.01468495 | 28.2497704 | 24.7590398 | 16.44010807 | 36.9800545 | 26.38787387 | 50.0992466 | 41.8279424 | 37.0600442 | 32.0250823 |

| Figure1CD_MC1_Staining | n | 9 | 10 |
|---|---|---|---|
| | MC1 | DAP12+/+ Tau+ | DAP12-/- Tau+ |
| | | 0.464 | 1.54525 |
| | | 1.83375 | 1.723 |
| | | 4.2485 | 2.74775 |
| | | 2.84575 | 3.30425 |
| | | 1.82125 | 4.42125 |
| | | 2.30525 | 3.1345 |
| | | 1.43 | 2.906 |
| | | 0.631 | 2.8215 |
| | | 1.4245 | 2.70675 |
| | | | 4.17175 |

| Figure1EF_AT8_Staining | n | 16 | 24 | red:outlier |
|---|---|---|---|---|
| | AT8 | DAP12+/+ Tau+ | DAP12-/- Tau+ | |
| | | 2.51 | 27.29* | |
| | | 1.86 | 3.53 | |
| | | 0.63 | 13.83 | |
| | | 11.05* | 6.22 | |
| | | 0.4 | 13.64 | |
| | | 1.75 | 4.17 | |
| | | 4.66 | 7.7 | |
| | | 2.27 | 3.54 | |
| | | 6.84* | 1.37 | |
| | | 0.91 | 7.44 | |
| | | 1.21 | 3.92 | |
| | | 3.55 | 5.68 | |
| | | 1.14 | 1.98 | |
| | | 1.22 | 1.37 | |
| | | 3.05 | 6.57 | |
| | | 1.15 | 1.53 | |
| | | 0.88 | 2.92 | |
| | | 0.57 | 35.55* | |
| | | | 7.25 | |
| | | | 6.03 | |
| | | | 5.37 | |
| | | | 0.53 | |
| | | | 2.79 | |
| | | | 6.49 | |
| | | | 8.32 | |
| | | | 9.05 | |

| Figure1G_Thios_Staining | | EC_Thioflavin S | | PC_Thioflavin S | |
|---|---|---|---|---|---|
| | n | 18 | 25 | 18 | 24 |
| | | DAP12+/+ Tau+ | DAP12-/- Tau+ | DAP12+/+ Tau+ | DAP12-/- Tau+ |
| | | 78 | 193 | 113 | *363** |
| | | 4 | 34 | 101 | 124 |
| | | 49 | 50 | 45 | 121 |
| | | 56 | 86 | 26 | 53 |
| | | 20 | 133 | 48 | 170 |
| | | 69 | 76 | 82 | 62 |
| | | 7 | 68 | 88 | 181 |
| | | 24 | 11 | 45 | 43 |
| | | 88 | 44 | 108 | 102 |
| | | 44 | 39 | 61 | 74 |
| | | 37 | 27 | 91 | 44 |
| | | 9 | 125 | 15 | 110 |
| | | 3 | 8 | 53 | 38 |
| | | 11 | 5 | 49 | 15 |
| | | 6 | 34 | 31 | 62 |
| | | 25 | 24 | 30 | 88 |
| | | 26 | 42 | 54 | 93 |
| | | 11 | 107 | 45 | 145 |
| | | | 81 | | 169 |
| | | | 111 | | 90 |
| | | | 111 | | 84 |
| | | | 97 | | 133 |
| | | | 97 | | 179 |
| | | | 119 | | 120 |
| | | | 46 | | 149 |

| Figure1HIJK_IBA1_Staining | | | | | | | | | | | | |
|---|---|---|---|---|---|---|---|---|---|---|---|---|
| | n | 18 | 26 | n | 18 | 26 | n | 18 | 26 | n | 18 | 26 |
| | IBA1 EC | DAP12+/+ Tau+ | DAP12-/- Tau+ | IBA1 CA1 | DAP12+/+ Tau+ | DAP12-/- Tau+ | IBA1 CA3 | DAP12+/+ Tau+ | DAP12-/- Tau+ | IBA1 DG | DAP12+/+ Tau+ | DAP12-/- Tau+ |
| | | 4.023334 | 1.953333 | | 3.93 | 2.416667 | | 2.976667 | 2.32 | | 5.23 | 4.12 |
| | | 4.5 | 3.38 | | 4.8 | 1.966667 | | 4.343333 | 1.8 | | 9.176666 | 2.623333 |
| | | 5.06 | 2.976667 | | 5.656667 | 2.346667 | | 5.296667 | 2.203333 | | 6.61 | 3.603333 |
| | | 4.956667 | 4.506667 | | 4.41 | 3.616667 | | 3.82 | 2.856667 | | 5.336667 | 4.866667 |
| | | 6.066667 | 2.556667 | | 5.13 | 2.87 | | 4.35 | 2.68 | | 7.113333 | 5.123334 |
| | | 11.75667 | 2.766667 | | 6.81 | 2.72 | | 7.056667 | 2.59 | | 8.46 | 3.876667 |
| | | 5.213333 | 2.996667 | | 4.64 | 3.403333 | | 6.986667 | 2.54 | | 7.573333 | 5.033333 |
| | | 3.836667 | 2.963333 | | 4.12 | 2.55 | | 5.03 | 2.48 | | 7.796667 | 3.556667 |
| | | 6.626667 | 3.026667 | | 4.73 | 2.936667 | | 5.76 | 2.73 | | 8.92 | 4.093333 |
| | | 5.46 | 4.056667 | | 5.156667 | 3.186667 | | 5.846667 | 2.956667 | | 8.7 | 4.836667 |
| | | 4.563334 | 2.07 | | 5.316667 | 1.39 | | 4.476667 | 1.196667 | | 7.113333 | 2.076667 |
| | | 4.236667 | 1.68 | | 4 | 1.523333 | | 3.333333 | 1.363333 | | 5.193333 | 2.88 |
| | | 3.616667 | 0.7366667 | | 2.53 | 1.043333 | | 1.723333 | 0.7933334 | | 4.076667 | 2.093333 |
| | | 1.746667 | 1.216667 | | 2.066667 | 1.056667 | | 1.423333 | 0.6566667 | | 4.083334 | 1.94 |
| | | 3.926667 | 2.316667 | | 2.12 | 2.13 | | 2.306667 | 2.13 | | 3.276667 | 1.973333 |
| | | 2.843333 | 1.863333 | | 2.906667 | 1.376667 | | 2.14 | 0.9066667 | | 5.22 | 1.68 |
| | | 2.71 | 2.11 | | 3.726667 | 0.9766667 | | 3.066667 | 0.9466667 | | 3.813333 | 2.316667 |
| | | 4.686667 | 1.706667 | | 2.703333 | 1.633333 | | 2.133333 | 1.226667 | | 4.14 | 2.036667 |
| | | | 1.146667 | | | 0.8966667 | | | 0.79 | | | 1.63 |
| | | | 1.31 | | | 1.033333 | | | 0.73 | | | 1.36 |
| | | | 1.446667 | | | 0.7633333 | | | 0.8433333 | | | 1.663333 |
| | | | 1.196667 | | | 1.903333 | | | 1.803333 | | | 3.286667 |
| | | | 1.706667 | | | 1.736667 | | | 1.33 | | | 3.236667 |
| | | | 0.7166666 | | | 0.8566667 | | | 0.8766667 | | | 1.43 |
| | | | 1.36 | | | 1.406667 | | | 1.19 | | | 2.723333 |
| | | | 1.536667 | | | 1.476667 | | | 1.31 | | | 2.713333 |

sFigure1AB_IHC_TAKO_tau

Number of puncta

| DAP12+/+ Tau fibrils- | DAP12+/+ Tau fibrils+ 2hours | DAP12-/- Tau fibrils- | DAP12-/- Tau fibrils+ 2hours |
|---|---|---|---|
| 8 | 58 | 5 | 26 |
| 2 | 38 | 7 | 14 |
| 1 | 34 | 1 | 19 |
| | 4 | | 10 |
| | 22 | | 38 |
| | 76 | | 31 |
| | 24 | | 26 |
| | 12 | | 27 |
| | 21 | | 22 |
| | 50 | | 31 |
| | 28 | | 37 |
| | | | 22 |
| | | | 80 |
| | | | 20 |

Total tau positive area

| DAP12+/+ Tau fibrils- | DAP12+/+ Tau fibrils+ 2hours | DAP12-/- Tau fibrils- | DAP12-/- Tau fibrils+ 2hours |
|---|---|---|---|
| 769 | 22305 | 644 | 18657 |
| 153 | 6521 | 2288 | 5615 |
| 215 | 7455 | 80 | 9614 |
| | 1191 | | 3407 |
| | 3573 | | 15191 |
| | 29342 | | 11368 |
| | 4555 | | 7464 |
| | 6253 | | 9459 |
| | 10903 | | 7451 |
| | 14662 | | 9638 |
| | 17358 | | 9938 |
| | | | 6445 |
| | | | 22165 |
| | | | 9801 |

sFigure1C_IHC_GFAP

GFAP IHC

| DAP12+/+ Tau+ | DAP12-/- Tau- |
|---|---|
| 12.05 | 6.52 |
| 1.76 | 5.213333 |
| 9.663333 | 5.92 |
| 5.85 | 6.85 |
| 16.28667 | 8.596666 |
| 10.28333 | 12.03 |
| 17.17 | 13.33 |
| 13.88 | 13.44 |
| 15.53 | 10.41333 |
| 14.13333 | 10.43333 |
| 15.01 | 13.90667 |
| 13.16333 | 9.763333 |
| 8.066667 | 4.59 |
| 6.893333 | 4.763333 |
| 5.646667 | 7.113333 |
| 5.68 | 4.69 |
| 5.89 | 5.08 |
| 6.7 | 11.83333 |
| | 3.036667 |
| | 5.1 |
| | 3.453333 |
| | 6.41 |
| | 4.946667 |
| | 6.303333 |
| | 5.746666 |
| | 6.43 |
