## Supplementary table 2 for "DAP12 deficiency alters microglia-oligodendrocyte communication and enhances resilience against tau toxicity"

Figure2A_magpix_immune_regulators_measurement

| Magpix_9plex | Dap12+/+ Tau+ | | | | | | | | Dap12-/- Tau+ | | | | | | | |
|---|---|---|---|---|---|---|---|---|---|---|---|---|---|---|---|---|
| p-JNK | 0.14055 | 0.07692 | 0.13962 | 0.10661 | 0.14604 | 0.12657 | 0.19395 | 0.10052 | 0.01938 | 0.06313 | 0.04675 | 0.06509 | 0.05311 | 0.05508 | 0.02917 | 0.06113 |
| p-P38 | 0.48802 | 0.44563 | 0.44596 | 0.3618 | 0.45731 | 0.32193 | 0.4686 | 0.25766 | 0.33099 | 0.38151 | 0.2394 | 0.3658 | 0.271 | 0.2907 | 0.28173 | 0.24908 |
| p-NFkB | 0.00162 | 0.00102 | 0.0013 | 0.00135 | 0.00114 | 0.00117 | 0.00182 | 0.00179 | 0.0012 | 0.00149 | 0.00094 | 0.00111 | 0.00131 | 0.00099 | 0.00149 | 0.00144 |
| p-AKT | 1.60085 | 1.36916 | 1.34727 | 1.31984 | 1.55377 | 1.34992 | 1.68027 | 1.36803 | 1.39536 | 1.48017 | 1.24923 | 1.3614 | 1.16265 | 1.25114 | 1.22327 | 1.24361 |
| p-ERK | 0.03352 | 0.02772 | 0.03827 | 0.02601 | 0.04063 | 0.04333 | 0.0718 | 0.04184 | 0.00628 | 0.02346 | 0.02767 | 0.01616 | 0.01455 | 0.0167 | 0.00838 | 0.01239 |
| p70S6K | 0.00221 | 0.00148 | 0.00224 | 0.00184 | 0.00163 | 0.00246 | 0.00264 | 0.00206 | 0.00214 | 0.00078 | 0.00094 | 0.00401 | 0.00131 | 0.00129 | 0.00207 | 0.00253 |
| p-STAT3 | 0.00838 | 0.00648 | 0.00744 | 0.00589 | 0.00753 | 0.0079 | 0.01036 | 0.00618 | 0.00668 | 0.00661 | 0.0053 | 0.00613 | 0.00547 | 0.00555 | 0.00551 | 0.0059 |
| p-STAT5 | 0.00147016 | 0.00124986 | 0.0011812 | 0.00147221 | 0.00156272 | 0.00155461 | 0.00181769 | 0.00179179 | 0.00120313 | 0.00155561 | 0.00129534 | 0.00111458 | 0.00124748 | 0.00098583 | 0.00126335 | 0.00168462 |

Figure2BCD_western_validation_of_magpix_immune_regulators

| Pi-Akt/Total-Akt | Dap12+/+ Tau+ | Dap12-/- Tau+ | Akt/Beta-Tubu | Dap12+/+ Tau+ | Dap12-/- Tau+ |
|---|---|---|---|---|---|
| | 0.729778465 | 0.411820782 | | 0.795361805 | 0.464349695 |
| | 0.732628631 | 0.629988066 | | 0.789583665 | 0.590746983 |
| | 0.764157459 | 0.522690173 | | 0.668378133 | 0.468539655 |

| Pi-Erk/Total-Erk | Dap12+/+ Tau+ | Dap12-/- Tau+ | Erk/Beta-Tubu | Dap12+/+ Tau+ | Dap12-/- Tau+ |
|---|---|---|---|---|---|
| | 0.602110848 | 0.190121786 | | 2.182823167 | 0.184879971 |
| | 0.522866572 | 0.235587189 | | 1.890677194 | 0.224351747 |
| | 0.703570703 | 0.208230742 | | 2.125037753 | 0.1872988 |

Figure2F_Hallmarks_of_Bulkseq

Overlap Results

| | |
|---|---|
| Collection(s): | MH |
| # overlaps shown: | 7 |
| # genesets in collections: | 50 |
| # genes in comparison (n): | 129 |
| # genes in universe (N): | 42766 |

| Gene Set Name | # Genes in Gene Set (K) | # Genes in Overlap (k) | k/K | p-value | FDR q-value | Description |
|---|---|---|---|---|---|---|
| HALLMARK_INTERFERON_ GAMMA_RESPONSE | 188 | 10 | 0.0532 | 3.31E-10 | 1.66E-08 | Mouse genes annotated to HALLMARK_INTERFERON_GAMMA_RESPONSE based on orthology mappings provided by the Alliance Genome Consortium |
| HALLMARK_INTERFERON_ ALPHA_RESPONSE | 94 | 7 | 0.0745 | 1.59E-08 | 3.97E-07 | Mouse genes annotated to HALLMARK_INTERFERON_ALPHA_RESPONSE based on orthology mappings provided by the Alliance Genome Consortium |
| HALLMARK_IL6_JAK_ STAT3_SIGNALING | 85 | 5 | 0.0588 | 6.24E-06 | 1.04E-04 | Mouse genes annotated to HALLMARK_IL6_JAK_STAT3_SIGNALING based on orthology mappings provided by the Alliance Genome Consortium |
| HALLMARK_ALLOGRAFT_ REJECTION | 192 | 6 | 0.0312 | 2.72E-05 | 3.41E-04 | Mouse genes annotated to HALLMARK_ALLOGRAFT_REJECTION based on orthology mappings provided by the Alliance Genome Consortium |
| HALLMARK_EPITHELIAL_ MESENCHYMAL_TRANSITION | 194 | 4 | 0.0206 | 2.91E-03 | 2.27E-02 | Mouse genes annotated to HALLMARK_EPITHELIAL_MESENCHYMAL_TRANSITION based on orthology mappings provided by the Alliance Genome Consortium |
| HALLMARK_INFLAMMATORY_ RESPONSE | 197 | 4 | 0.0203 | 3.07E-03 | 2.27E-02 | Mouse genes annotated to HALLMARK_INFLAMMATORY_RESPONSE based on orthology mappings provided by the Alliance Genome Consortium |
| HALLMARK_E2F_TARGETS | 199 | 4 | 0.0201 | 3.18E-03 | 2.27E-02 | Mouse genes annotated to HALLMARK_E2F_TARGETS |

Figure2G_Transcription_factors_of_Bulkseq

| Gene Set Name | # Genes in Gene Set (K) | # Genes in Overlap (k) | k/K | p-value | FDR q-value | Description |
|---|---|---|---|---|---|---|
| CRELD1 | 1495 | 49 | 0.0328 | 6.06E-19 | 1.70E-16 | Genes containing one or more binding sites for UniProt:Q91XD7 (Creld1) in their promoter regions (TSS -1000,+100 bp) as identified by GTRD version 20.06 ChIP-seq harmonization. |
| LYL1 | 1658 | 41 | 0.0247 | 5.76E-12 | 8.06E-10 | Genes containing one or more binding sites for UniProt:P27792 (Lyl1) in their promoter regions (TSS -1000,+100 bp) as identified by GTRD version 20.06 ChIP-seq harmonization. |
| RELB | 1922 | 43 | 0.0224 | 4.10E-11 | 3.83E-09 | Genes containing one or more binding sites for UniProt:Q04863 (Relb) in their promoter regions (TSS -1000,+100 bp) as identified by GTRD version 20.06 ChIP-seq harmonization. |
| ZFP874B | 1966 | 42 | 0.0214 | 2.89E-10 | 2.02E-08 | Genes containing one or more binding sites for UniProt:A0A286YDG9 (Zfp874b) in their promoter regions (TSS -1000,+100 bp) as identified by GTRD version 20.06 ChIP-seq harmonization. |
| DMRT1 | 1812 | 38 | 0.021 | 3.67E-09 | 2.05E-07 | Genes containing one or more binding sites for UniProt:Q9QZ59 (Dmrt1) in their promoter regions (TSS -1000,+100 bp) as identified by GTRD version 20.06 ChIP-seq harmonization. |
| MLLT3 | 1067 | 28 | 0.0262 | 4.53988222w9 | 2.16E-07 | Genes containing one or more binding sites for UniProt:A2AM29 (Mllt3) in their promoter regions (TSS -1000,+100 bp) as identified by GTRD version 20.06 ChIP-seq harmonization. |
| NFKB1 | 1543 | 34 | 0.022 | 7.85E-09 | 3.14E-07 | Genes containing one or more binding sites for UniProt:P25799 (Nfkb1) in their promoter regions (TSS -1000,+100 bp) as identified by GTRD version 20.06 ChIP-seq harmonization. |
| RCOR1 | 1974 | 39 | 0.0198 | 1.14E-08 | 3.99E-07 | Genes containing one or more binding sites for UniProt:Q8CFE3 (Rcor1) in their promoter regions (TSS -1000,+100 bp) as identified by GTRD version 20.06 ChIP-seq harmonization. |
| REL | 560 | 19 | 0.0339 | 3.04E-08 | 9.46E-07 | Genes containing one or more binding sites for UniProt:P15307 (Rel) in their promoter regions (TSS -1000,+100 bp) as identified by GTRD version 20.06 ChIP-seq harmonization. |
| MYB | 1569 | 33 | 0.021 | 3.97E-08 | 1.11E-06 | Genes containing one or more binding sites for UniProt:P06876 (Myb) in their promoter regions (TSS -1000,+100 bp) as identified by GTRD version 20.06 ChIP-seq harmonization. |

Figure2H_IPA_updstream_regulator_of_Bulkseq

| IPA Upstream Regulator | Expr Log Ratio | Molecule Type | ted Activation | Activation z-score | p-value of overlap | Target Molecules in Dataset |
|---|---|---|---|---|---|---|
| Ifnar | | group | Inhibited | -4.15 | 3.42E-16 | APOBEC3B,CCL2,CD86,EIF2AK2,HLA-A,IFI16,IFIT1B,IFIT3,IFITM3,IRF1,IRF8,IRF9,Sp100,STAT2,TLR2,TNFSF10,UNC93B1,XAF1 |
| IRF3 | -0.119 | transcription regulator | Inhibited | -4.171 | 8.7E-15 | APOBEC3B,CD86,EIF2AK2,FCGR1A,GBP5,IFI16,IFI44,IFIT1B,IFIT3,IFITM3,IRF1,LAG3,PARP12,PARP14,PHF11,STAT2,TNFSF10,Trim30a/Trim30d |
| IFNG | 0.01 | cytokine | Inhibited | -4.659 | 1.41E-13 | BCL2L1,C4A/C4B,CCL2,Ccl6,CD14,CD86,CTSH,CTSS,CXCL16,EIF2AK2,ENDOD1,FZD2,GBP5,GBP6,HK2,HLA-A,IFI16,IFI44,IFIT1B,IFIT3,IL18BP,IRF1, |
| IRF7 | -0.265 | transcription regulator | Inhibited | -3.42 | 1.5E-10 | FCGR1A,GBP5,IFI16,IFIT1B,IFIT3,IFITM3,IRF8,PARP12,PARP14,PHF11,STAT2,Trim30a/Trim30d |
| ZBTB10 | 0.077 | transcription regulator | Inhibited | -3.801 | 7.24E-10 | CD86,CHUK,GBP5,GBP6,IFI16,IFIT1B,IFIT3,IFITM3,IRF1,LGALS9B,PARP9,STAT2,TNFRSF14,UNC93B1,XAF1 |
| USP22 | 0.019 | peptidase | Inhibited | -3.162 | 1.14E-09 | Cd33,CSF2RA,CSF2RB,CSF3R,FES,ITGAM,ITGB2,NCF2,PTPN6,PTPRC |
| STAT1 | -0.206 | transcription regulator | Inhibited | -3.434 | 7.64E-09 | APOBEC3B,BCL2L1,CCL2,CD86,CDK2,EIF2AK2,GBP5,GBP6,IFI16,IFIT1B,IFIT3,IRF1,IRF8,IRF9,ITGAX,STAT2,TNFSF10 |
| TLR4 | -0.038 | transmembrane receptor | Inhibited | -3.753 | 3.75E-08 | APOBEC3B,CCL2,CD14,CD86,IFI16,IFIT1B,IFIT3,IFITM3,IL18BP,IRF1,IRF8,ITGAM,JAG1,PLEK,RGS16,STAT2,TLR2,TNFSF10 |
| TICAM1 | -0.003 | other | Inhibited | -3.261 | 0.000000971 | APOBEC3B,CCL2,CD86,IFI16,IFIT1B,IFIT3,IL18BP,IRF1,JAG1,TLR2,TNFSF10 |
| NOS2 | -0.105 | enzyme | Inhibited | -2.8 | 0.00121 | CD14,COL4A1,CSF3R,CTSC,IFIT1B,IFIT3,TNFSF10,TYROBP |
| MYD88 | -0.129 | other | Inhibited | -3.083 | 0.00448 | APOBEC3B,CD14,CD86,IFIT1B,IRF1,IRF8,ITGAM,ITGAX,JAG1,TLR2 |

Figure2I_IPA_mechanistic_nertwork_of_Bulkseq

*Dap12+/+ tau+* vs *tau-*

Ifnar (molecules)

| Symbol | Entrez Gene Name | /Gene Symbol - mouse (Ent | ity/RPKM/FPK | Expr Log Ratio | Expr p-value | Expr False Discovery Rate (q-value) | Location | Family | Drugs | Entrez Gene ID for Human | Entrez Gene ID for Mouse | Entrez Gene ID for Rat |
|---|---|---|---|---|---|---|---|---|---|---|---|---|
| Ifnar | | | | | | | Plasma Membrane | group | | | | |
| IFNB1 | interferon beta 1 | Ifnb1 | 0 | | | | Extracellular Space | cytokine | | 3456 | 15977 | 24481 |
| IFNG | interferon gamma | Ifng | 0.29362341 | 0.010408489 | 0.789944076 | | Extracellular Space | cytokine | emapalumab | 3458 | 15978 | 25712 |
| IL12 (complex) | | | | | | | Extracellular Space | complex | /ustekinumab, ustekinumab | | | |
| IRF1 | interferon regulatory factor 1 | Irf1 | 293.14658 | 0.315925057 | 0.035553909 | 0.138398496 | Nucleus | scription regulator | | 3659 | 16362 | 24508 |
| IRF8 | interferon regulatory factor 8 | Irf8 | 253.112008 | 0.499735003 | 4.65E-04 | 0.006460974 | Nucleus | scription regulator | | 3394 | 15900 | 292060 |
| SMAD3 | SMAD family member 3 | Smad3 | 3867.65841 | 0.091008676 | 0.326952264 | 0.5613073 | Nucleus | scription regulator | | 4088 | 17127 | 25631 |
| STAT1 | signal transducer and activator of transcription 1 | Stat1 | 1256.35459 | 0.393510542 | 0.016703642 | 0.083513107 | Nucleus | scription regulator | | 6772 | 20846 | 25124 |
| STAT2 | signal transducer and activator of transcription 2 | Stat2 | 1223.06897 | 0.350726365 | 4.61E-04 | 0.006438851 | Nucleus | scription regulator | | 6773 | 20847 | 288774 |
| STAT3 | signal transducer and activator of transcription 3 | Stat3 | 2451.36959 | 0.106677692 | 0.21346546 | 0.433023546 | Nucleus | scription regul | PB-51602, TTI-1 | 6774 | 20848 | 25125 |
| STAT6 | signal transducer and activator of transcription 6 | Stat6 | 677.955564 | 0.373091623 | 1.47E-07 | 1.02E-05 | Nucleus | scription regulator | | 6778 | 20852 | 362896 |

Dap12-/- tau+ vs Dap12+/+ tau+

Ifnar (molecules)

| Symbol | Entrez Gene Name | /Gene Symbol - mouse (Entr | Expr Log Ratio | se Discovery Rate (( | Location | Family | Drugs | z Gene ID for H | z Gene ID for M | Entrez Gene ID for Rat |
|---|---|---|---|---|---|---|---|---|---|---|
| Ifnar | | | | | Plasma Membrane | group | | | | |
| IFNB1 | interferon beta 1 | Ifnb1 | | | Extracellular Space | cytokine | | 3456 | 15977 | 24481 |
| IFNG | interferon gamma | Ifng | 0.00975522 | | Extracellular Space | cytokine | emapalumab | 3458 | 15978 | 25712 |
| IL12A | interleukin 12A | Il12a | 0.03828363 | 0.923860706 | Extracellular Space | cytokine | | 3592 | 16159 | 84405 |
| IRF8 | interferon regulatory factor 8 | Irf8 | -0.49015635 | 9.41E-05 | Nucleus | transcription regulator | | 3394 | 15900 | 292060 |
| IRF9 | interferon regulatory factor 9 | Irf9 | -0.33343095 | 0.006564539 | Nucleus | transcription regulator | | 10379 | 16391 | 305896 |
| STAT1 | signal transducer and activator of transcription 1 | Stat1 | -0.2061467 | 0.263578327 | Nucleus | transcription regulator | | 6772 | 20846 | 25124 |
| STAT2 | signal transducer and activator of transcription 2 | Stat2 | -0.25731601 | 0.044114142 | Nucleus | transcription regulator | | 6773 | 20847 | 288774 |
| STAT3 | signal transducer and activator of transcription 3 | Stat3 | -0.0817989 | 0.563050456 | Nucleus | transcription regulator | CAS3/SS3, danvatirsen, OPB-31121, OPB-51602, TTI-101, | 6774 | 20848 | 25125 |
| STAT6 | signal transducer and activator of transcription 6 | Stat6 | -0.11732576 | 0.413057568 | Nucleus | transcription regulator | | 6778 | 20852 | 362896 |

*Dap12+/+ tau+* vs *tau-*

| From Molecule(s) | Relationship Type | To Molecule(s) |
|---|---|---|
| IFNB1 | activation | STAT1 |
| IFNB1 | localization | STAT1 |
| IFNB1 | phosphorylation | STAT1 |
| IFNB1 | phosphorylation | STAT2 |
| IFNB1 | phosphorylation | STAT3 |
| IFNG | activation | STAT1 |
| IFNG | activation | STAT2 |
| IFNG | activation | STAT3 |
| IFNG | localization | IRF1 |
| IFNG | localization | STAT1 |
| IFNG | modification | STAT1 |
| IFNG | phosphorylation | STAT1 |
| IFNG | phosphorylation | STAT3 |
| IFNG | regulation of binding | IRF1 |
| IFNG | regulation of binding | STAT1 |
| IFNG | regulation of binding | STAT2 |
| IFNG | translocation | STAT1 |
| IFNG | ubiquitination | IRF8 |
| IL12 (complex) | phosphorylation | STAT3 |
| IL12 (complex) | phosphorylation | STAT6 |
| Ifnar | activation | STAT1 |
| Ifnar | localization | IFNB1 |
| Ifnar | localization | IFNG |
| Ifnar | localization | IL12 (complex) |
| Ifnar | phosphorylation | STAT1 |
| STAT1 | activation | SMAD3 |
| STAT1 | activation | STAT2 |
| STAT1 | activation | STAT3 |
| STAT1 | regulation of binding | STAT6 |

Dap12-/- tau+ vs Dap12+/+ tau+

Ifnar (relationships)

| From Molecule(s) | Relationship Type | To Molecule(s) |
|---|---|---|
| IFNB1 | activation | STAT1 |
| IFNB1 | localization | IRF9 |
| IFNB1 | localization | STAT1 |
| IFNB1 | phosphorylation | STAT1 |
| IFNB1 | phosphorylation | STAT2 |
| IFNB1 | phosphorylation | STAT3 |
| IFNG | activation | STAT1 |
| IFNG | activation | STAT2 |
| IFNG | activation | STAT3 |
| IFNG | localization | STAT1 |
| IFNG | modification | STAT1 |
| IFNG | phosphorylation | STAT1 |
| IFNG | phosphorylation | STAT3 |
| IFNG | regulation of binding | STAT1 |
| IFNG | regulation of binding | STAT2 |
| IFNG | translocation | STAT1 |
| IFNG | ubiquitination | IRF8 |
| IL12A | localization | STAT1 |
| Ifnar | activation | STAT1 |
| Ifnar | localization | IFNB1 |
| Ifnar | localization | IFNG |
| Ifnar | localization | IL12A |
| Ifnar | phosphorylation | STAT1 |
| STAT1 | activation | STAT2 |
| STAT1 | activation | STAT3 |
| STAT1 | regulation of binding | STAT6 |

Figure2J_cytokine/chemokine_multiplex_measurements

| IP-10 | Dap12+/+ Tau+ | Dap12-/- Tau+ | | MCP-1 | Dap12+/+ Tau+ | Dap12-/- Tau+ |
|---|---|---|---|---|---|---|
| | 26.86 | 35.47 | | | 0.9 | 0.78 |
| | 141.3 | 44.18 | | | 0.88 | 0.59 |
| | 171.52 | 38.61 | | | 1.48 | 0.55 |
| | 109.63 | 31.25 | | | 1.02 | 0.94 |
| | 69.62 | 26.62 | | | 1.06 | 0.81 |
| | 213.88 | 14.36 | | | 1.33 | 0.94 |

| IL-6 | Dap12+/+ Tau+ | Dap12-/- Tau+ | | MIG | Dap12+/+ Tau+ | Dap12-/- Tau+ |
|---|---|---|---|---|---|---|
| | 0.9 | 0.78 | | | 3.21 | *4.29** |
| | 0.88 | 0.59 | | | 8.72 | 2.99 |
| | 1.48 | 0.55 | | | | 2.99 |
| | 1.02 | 0.94 | | | 8.41 | 2.99 |
| | 1.06 | 0.81 | | | 3.38 | 2.99 |
| | 1.33 | 0.94 | | | 16.17 | 2.99 |

| IL1a | +/+ | -/- | | IL1b | Dap12+/+ Tau+ | Dap12-/- Tau+ |
|---|---|---|---|---|---|---|
| | 8.47 | 23.3 | | | 3.7 | 4.2 |
| | 15.94 | 12.12 | | | 4.3 | 5.78 |
| | 11.87 | 7.76 | | | 5.78 | 0 |
| | 16.46 | 11.87 | | | 4.51 | 0 |
| | 15.42 | 11.13 | | | 4.71 | 5.14 |
| | 19.85 | 3.23 | | | 5.14 | 3.6 |

sFigure2CD_IPA_pathways_analysis_bulkseq

**IPA pathway analysis Dap12+/+ tau+ vs tau-**

| Ingenuity Canonical Pathways | -log(p-value) | Ratio | z-score | NO.of genes | |
|---|---|---|---|---|---|
| Phagosome Formation | 1.34 | 0.0712 | 6.782 | 46 | ,CD14,CELSR3,CLEC7A,CRHR2,FCER1G,FCGR2A,FCGR2B,FCGR3A/FCGR3B,FGR,GPR108,GPR61,GPR84,HMOX1,ITGA7,ITGAX,MAP2K3,MAP2K5,MAP2K7,MYD88,MYH3,MYH7,MYH7B,MYLK3,MYO1H,MYO5C,NAPEPLD,NPY2R,NTSR1,PIP5K1B,PLA2G4E,PLB1,PLCG1,PNPLA3,QRFP |
| Pathogen Induced Cytokine Storm Signaling Pathway | 3.83 | 0.108 | 5.488 | 34 | AIM2,C3,CASP1,CCL17,CCL3L3,CCL4,Ccl6,Ccl9,CLEC7A,COL19A1,COL23A1,COL24A1,COL26A1,COL5A1,COL5A3,COL6A2,CSF2RB,CXCL10,CXCL16,IFIH1,IL12A,IL12RB1,IRF9,JUN,MAPK13,MYD88,Naip1 (includes others),RORC,SLC2A4,STXBP2,TLR1,TLR2,TLR7,ZBP1 |
| FAK Signaling | 0 | 0.0387 | 5.303 | 32 | ADORA1,C3AR1,CAPN2,CASR,CELSR3,COL5A3,CRHR2,CSF2RB,EFNA5,FCER1G,GIT2,GPR108,GPR61,GPR84,IL12RB1,IL2RB,IL4R,ITGA7,ITGAX,JUN,MAP2K5,MAPK13,MET,NPY2R,NTSR1,PLCG1,PXN,QRFPR,RXFP2,SPARCL1,TP53,WIPF1 |
| S100 Family Signaling Pathway | 0 | 0.0464 | 5.048 | 33 | ADORA1,C3AR1,CAMK2G,CASR,CATSPER2,CCL4,CELSR3,CRHR2,FCGR2A,FCGR2B,FCGR3A/FCGR3B,GPR108,GPR61,GPR84,JUN,MAP2K3,MAP2K5,MAP2K7,MAPK13,MYD88,NCF2,NPY2R,NTSR1,PLA2G4E,PLCB4,PLCD3,PLCG1,PLCZ1,QRFPR,RXFP2,TP53,WNT2B,WNT9B |
| Cardiac Hypertrophy Signaling (Enhanced) | 0.623 | 0.063 | 4.849 | 33 | ACE,ADCY6,CAMK2G,CSF2RB,DVL2,FGF5,GNG2,HDAC10,HDAC7,IL12A,IL12RB1,IL2RB,IL4R,ITGA7,ITGAX,JUN,MAP2K3,MAP2K5,MAP2K7,MAP3K11,MAP3K3,MAPK13,NAPEPLD,PDE4C,PDE7A,PLCB4,PLCD3,PLCG1,PLCZ1,RELA,RPS6KB1,WNT2B,WNT9B |
| Superpathway of Inositol Phosphate Compounds | 1.63 | 0.0893 | 4.359 | 20 | ACP6,CDC25B,DOT1L,DUSP15,DUSP9,EYA4,INPP5B,INPP5E,INPP5K,PDCD1,PIP5K1B,PLCB4,PLCD3,PLCG1,PLCZ1,PPIP5K2,PTPN2,PTPN23,PTPRC,SSH3 |
| CREB Signaling in Neurons | 0 | 0.0343 | 4.243 | 20 | ADCY6,ADORA1,C3AR1,CAMK2G,CASR,CATSPER2,CELSR3,CRHR2,GNG2,GPR108,GPR61,GPR84,NPY2R,NTSR1,PLCB4,PLCD3,PLCG1,PLCZ1,QRFPR,RXFP2 |
| Role Of Osteoclasts In Rheumatoid Arthritis Signaling Pathway | 2.1 | 0.0906 | 3.962 | 27 | ADAM11,ADAM15,ADAM33,ADAM8,COL19A1,COL23A1,COL24A1,COL26A1,COL5A1,COL5A3,COL6A2,CSF1,CTNNA1,FCGR2A,FCGR2B,FCGR3A/FCGR3B,FOXO3,JUN,MAP2K3,MAP2K5,MAP2K7,MAPK13,Naip1 (includes others),OSCAR,PLCG1,RELA,RHOV |
| D-myo-inositol-5-phosphate Metabolism | 1.28 | 0.086 | 3.873 | 16 | ACP6,CDC25B,DOT1L,DUSP15,DUSP9,EYA4,INPP5B,PDCD1,PLCB4,PLCD3,PLCG1,PLCZ1,PTPN2,PTPN23,PTPRC,SSH3 |
| Insulin Secretion Signaling Pathway | 0.691 | 0.0684 | 3.771 | 18 | ABCC8,ADCY6,CAMK2G,CRTC2,EIF2S3,EIF4EBP3,FGR,MAPK13,NEUROD1,PLCB4,PLCD3,PLCG1,PLCZ1,RPS6KB1,SCNN1A,SLC2A4,STAT2,STAT6 |
| Hepatic Fibrosis Signaling Pathway | 0 | 0.0491 | 3.771 | 20 | COL5A3,DVL2,ITGA7,ITGAX,JUN,MAP2K3,MAP2K5,MAP2K7,MAPK13,MYD88,MYLK3,NCF2,PLCG1,PPARG,RELA,RHOV,RPS6KB1,TRPM7,WNT2B,WNT9B |
| Cardiac Hypertrophy Signaling | 0.282 | 0.0558 | 3.742 | 14 | ADCY6,GNG2,JUN,MAP2K3,MAP2K7,MAP3K11,MAP3K3,MAPK13,PLCB4,PLCD3,PLCG1,PLCZ1,RHOV,RPS6KB1 |
| of Hypercytokinemia/hyperchemokinemia in the Pathogenesis of Influ | 4.19 | 0.194 | 3.606 | 13 | CASP1,CCL4,CXCL10,EIF2AK2,IFIT2,IFIT3,IL12A,IRF9,ISG15,MYD88,Oas1b,STAT2,TLR7 |
| SNARE Signaling Pathway | 1.54 | 0.0992 | 3.606 | 13 | ADCY6,CAMK2G,CPLX3,MYH3,MYH7,MYH7B,MYO1H,MYO5C,SNAP23,SNCB,STXBP2,SYT2,SYT3 |

| | | | | | |
|---|---|---|---|---|---|
| NOD1/2 Signaling Pathway | 0.934 | 0.0798 | 3.606 | 13 | CASP1,HSPA1L,IL12A,JUN,MAP2K3,MAP2K7,MAPK13,MYD88,RELA,RNF31,TLR1,TLR2,TLR7 |
| 3-phosphoinositide Degradation | 0.883 | 0.0769 | 3.606 | 14 | ACP6,CDC25B,DOT1L,DUSP15,DUSP9,EYA4,INPP5B,INPP5E,INPP5K,PDCD1,PTPN2,PTPN23,PTPRC,SSH3 |
| Oxytocin Signaling Pathway | 0 | 0.0476 | 3.606 | 13 | GNG2,MAP2K5,MAPK13,MYH3,MYH7,MYH7B,MYO1H,MYO5C,PLA2G4E,PLCB4,PPARG,PRKAB2,RELA |
| IL-4 Signaling | 0 | 0.0348 | 3.606 | 13 | ALOX15B,COL19A1,COL23A1,COL24A1,COL26A1,COL5A1,COL5A3,COL6A2,FCER1G,IL4R,MAPK13,RPS6KB1,STAT6 |
| Neuroinflammation Signaling Pathway | 0.784 | 0.0699 | 3.578 | 20 | B2M,CASP1,CXCL10,GRINA,HLA-A,HMOX1,IL12A,JUN,KCNJ9,MAPK13,MYD88,Naip1 (includes others),NCF2,NCSTN,PLA2G4E,PLCG1,RELA,TLR1,TLR2,TLR7 |
| Pulmonary Fibrosis Idiopathic Signaling Pathway | 0.517 | 0.0629 | 3.578 | 20 | COL19A1,COL23A1,COL24A1,COL26A1,COL5A1,COL5A3,COL6A2,EIF4EBP3,EZH1,FOXO3,JUN,MAP2K3,MAPK13,RELA,RPS6KB1,RTEL1,STAT6,TP53,WNT2B,WNT9B |
| Breast Cancer Regulation by Stathmin1 | 0 | 0.0352 | 3.578 | 20 | ADORA1,ARHGEF1,ARHGEF10,C3AR1,CAMK2G,CASR,CELSR3,CRHR2,GNG2,GPR108,GPR61,GPR84,JUN,NPY2R,NTSR1,PLCB4,QRFPR,RPS6KB1,RXFP2,TP53 |
| G-Protein Coupled Receptor Signaling | 0 | 0.0522 | 3.55 | 35 | ADCY6,ADORA1,C3AR1,CAMK2G,CASR,CELSR3,CRHR2,DUSP9,FOXO3,GNG2,GPR108,GPR61,GPR84,JUN,MAP2K3,MAP2K5,MAP2K7,MAP3K11,MAP3K3,MAPK13,MYLK3,NAPEPLD,NPY2R,NTSR1,PDE4C,PDE7A,PLCB4,PLCG1,PXN,QRFPR,RELA,RGS16,RGS7,RXFP2,SIX4 |
| IL-15 Production | 1.55 | 0.103 | 3.464 | 12 | CLK1,CLK2,CLK3,CLK4,FGR,LTK,MAP2K3,MAP2K5,MAP2K7,MAP3K11,MET,RELA |
| Necroptosis Signaling Pathway | 1.03 | 0.0845 | 3.464 | 12 | CAMK2G,CAPN2,CASP1,EIF2AK2,IRF9,PLA2G4E,RNF31,STAT2,TIMM44,TP53,TRPM7,ZBP1 |
| roduction of Nitric Oxide and Reactive Oxygen Species in Macrophage | 0.556 | 0.0674 | 3.464 | 12 | CLU,IRF8,JUN,MAP2K7,MAP3K11,MAP3K3,MAPK13,NCF2,PLCG1,RELA,RHOV,TLR2 |
| 3-phosphoinositide Biosynthesis | 0.543 | 0.0663 | 3.464 | 13 | ACP6,CDC25B,DOT1L,DUSP15,DUSP9,EYA4,INPP5B,PDCD1,PIP5K1B,PTPN2,PTPN23,PTPRC,SSH3 |
| Wound Healing Signaling Pathway | 0 | 0.0502 | 3.464 | 12 | COL19A1,COL23A1,COL24A1,COL26A1,COL5A1,COL5A3,COL6A2,IL12A,JUN,MAP2K3,MAP2K5,MAP2K7 |
| Multiple Sclerosis Signaling Pathway | 0.903 | 0.0765 | 3.357 | 15 | C1QB,C1QC,C3,CAPN2,CASP1,GRINA,HLA-A,IL12A,MAPK13,PARP12,PARP3,PPARG,TLR1,TLR2,TLR7 |
| | 1.9 | 0.105 | 3.317 | 15 | C1QB,C1QC,C3,C3AR1,CASP1,CLEC7A,EIF2AK2,IFIH1,IL12A,MYD88,Oas1b,RELA,TLR1,TLR2,TLR7 |
| D-myo-inositol (1,4,5,6)-Tetrakisphosphate Biosynthesis | 0.64 | 0.0706 | 3.317 | 12 | ACP6,CDC25B,DOT1L,DUSP15,DUSP9,EYA4,INPP5B,PDCD1,PTPN2,PTPN23,PTPRC,SSH3 |
| D-myo-inositol (3,4,5,6)-tetrakisphosphate Biosynthesis | 0.64 | 0.0706 | 3.317 | 12 | ACP6,CDC25B,DOT1L,DUSP15,DUSP9,EYA4,INPP5B,PDCD1,PTPN2,PTPN23,PTPRC,SSH3 |
| Endothelin-1 Signaling | 0.83 | 0.0753 | 3.207 | 14 | ADCY6,CASP1,CASQ1,HMOX1,JUN,MAPK13,NAPEPLD,PLA2G4E,PLB1,PLCB4,PLCD3,PLCG1,PLCZ1,PNPLA3 |
| Signaling by Rho Family GTPases | 0.571 | 0.0654 | 3.207 | 17 | ARHGEF1,ARHGEF10,CYFIP1,GFAP,GNG2,ITGA7,ITGAX,JUN,MAP2K7,MAP3K11,MSN,NCF2,PIP5K1B,RELA,RHOV,SLC9A1,WIPF1 |
| TREM1 Signaling | 2.27 | 0.141 | 3.162 | 10 | CASP1,FCGR2B,ITGAX,MYD88,Naip1 (includes others),PLCG1,RELA,TLR1,TLR2,TLR7 |
| Role of MAPK Signaling in Inhibiting the Pathogenesis of Influenza | 2.18 | 0.137 | 3.162 | 10 | CXCL10,EIF2AK2,JUN,MAP2K3,MAP2K7,MAPK13,MYD88,PLA2G4E,PLB1,PNPLA3 |
| Sperm Motility | 2.06 | 0.0943 | 3.162 | 23 | CATSPER2,CLK1,CLK2,CLK3,CLK4,FGR,GNG2,LTK,MAP2K3,MAP2K5,MAP2K7,MAP3K11,MET,PDE4C,PLA2G4E,PLB1,PLCB4,PLCD3,PLCG1,PLCZ1,PNPLA3,SLC16A10,ZAN |
| Reelin Signaling in Neurons | 0.899 | 0.0815 | 3.162 | 11 | ARHGEF1,ARHGEF10,ARHGEF37,CAMK2G,FGR,GRINA,MAP2K3,MAP2K5,MAP2K7,MAP3K11,PDK2 |
| Xenobiotic Metabolism General Signaling Pathway | 0.715 | 0.0758 | 3.162 | 10 | ABCA4,GSTA3,GSTM5,HMOX1,MAP2K3,MAP2K5,MAP2K7,MAP3K11,MAP3K3,MAPK13 |
| Autophagy | 0 | 0.0478 | 3.162 | 10 | ATG13,FOXO3,JUN,MAP2K7,MYD88,PRKAB2,STX17,TP53,VPS16,VPS33B |
| Dendritic Cell Maturation | 0 | 0.0481 | 3.153 | 18 | B2M,COL5A3,FCER1G,FCGR2A,FCGR2B,FCGR3A/FCGR3B,HLA-A,IL12A,IRF8,MAPK13,MYD88,PLCB4,PLCD3,PLCG1,PLCZ1,RELA,STAT2,TLR2 |
| Thrombin Signaling | 0.65 | 0.0688 | 3.051 | 15 | ADCY6,ARHGEF1,ARHGEF10,ARHGEF37,CAMK2G,GATA2,GNG2,MAPK13,PLCB4,PLCD3,PLCG1,PLCZ1,RELA,RHOV,RPS6KB1 |
| Phospholipase C Signaling | 0 | 0.026 | 3.051 | 19 | ADCY6,ARHGEF1,ARHGEF10,FCER1G,FCGR2A,FCGR2B,GNG2,HDAC10,HDAC7,HMOX1,ITGA7,ITGAX,NAPEPLD,PLA2G4E,PLCB4,PLCD3,PLCG1,RELA,RHOV |
| PI3K Signaling in B Lymphocytes | 0 | 0.0374 | 3.051 | 15 | ATF3,ATF7,C3,CAMK2G,FCGR2B,FOXO3,IL4R,JUN,PLCB4,PLCD3,PLCG1,PLCZ1,PLEKHA4,PTPRC,RELA |
| BER (Base Excision Repair) Pathway | 3.33 | 0.209 | 3 | 9 | MAP2K3,MAP2K5,MAP2K7,MBD4,NEIL1,POLE,POLG,TP53,XRCC1 |
| Phospholipases | 2.13 | 0.143 | 3 | 9 | HMOX1,NAPEPLD,PLA2G4E,PLB1,PLCB4,PLCD3,PLCG1,PLCZ1,PNPLA3 |
| Acute Phase Response Signaling | 1.01 | 0.0809 | 3 | 14 | C1QB,C1QC,C1R,C3,C4A/C4B,HMOX1,JUN,MAP2K3,MAP2K7,MAPK13,MYD88,OSMR,RELA,SERPINA3 |
| Factors Promoting Cardiogenesis in Vertebrates | 0.366 | 0.0608 | 3 | 9 | BMP1,CAMK2G,MYH7,PLCB4,PLCD3,PLCG1,PLCZ1,WNT2B,WNT9B |
| HMGB1 Signaling | 0.313 | 0.0581 | 3 | 9 | IL12A,JUN,KAT2A,MAP2K3,MAP2K5,MAP2K7,MAPK13,RELA,RHOV |
| IL-8 Signaling | 0 | 0.0448 | 3 | 9 | GNG2,HMOX1,ITGAX,JUN,NAPEPLD,NCF2,RELA,RHOV,RPS6KB1 |
| Integrin Signaling | 0 | 0.0441 | 3 | 9 | CAPN2,ITGA7,ITGAX,MAP3K11,MYLK3,PLCG1,PXN,RHOV,WIPF1 |
| NF-κB Signaling | 0 | 0.0239 | 3 | 9 | EIF2AK2,FCER1G,MAP2K7,MAP3K3,MYD88,RELA,TLR1,TLR2,TLR7 |
| Natural Killer Cell Signaling | 2.57 | 0.112 | 2.982 | 19 | B2M,COL5A3,FCER1G,FCGR2A,FCGR3A/FCGR3B,HLA-A,HSPA1L,IL12A,IL12RB1,IL2RB,MAP2K3,MAP3K11,MAP3K3,MAPK13,MYD88,PLCG1,PXN,RELA,WIPF1 |
| GNRH Signaling | 0.643 | 0.0699 | 2.887 | 13 | ADCY6,CAMK2G,CATSPER2,GNG2,JUN,MAP2K3,MAP2K7,MAP3K11,MAP3K3,MAPK13,PLCB4,PXN,RELA |
| Adrenomedullin signaling pathway | 0.552 | 0.0667 | 2.887 | 13 | ADCY6,C3,MAP2K3,MAP2K5,MAP2K7,MAPK13,MYLK3,PLCB4,PLCD3,PLCG1,PLCZ1,PPARG,RELA |
| T Cell Receptor Signaling | 0 | 0.0297 | 2.887 | 12 | B2M,FCER1G,FOXO3,HLA-A,ICOSLG/LOC102723996,JUN,MAP2K3,MAP2K7,MAPK13,PLCG1,PTPRC,RELA |
| Role of PKR in Interferon Induction and Antiviral Response | 2.5 | 0.122 | 2.84 | 15 | ATF3,CASP1,EIF2AK2,HSPA1L,IFIH1,IRF9,JUN,MAP2K3,MAP2K7,MAPK13,MYD88,RELA,STAT2,TARBP2,TP53 |
| Toll-like Receptor Signaling | 2.63 | 0.149 | 2.828 | 11 | CD14,EIF2AK2,IL12A,JUN,MAP2K3,MAPK13,MYD88,RELA,TLR1,TLR2,TLR7 |
| Fcγ Receptor-mediated Phagocytosis in Macrophages and Monocytes | 0.87 | 0.087 | 2.828 | 8 | FCGR2A,FCGR3A/FCGR3B,FGR,HMOX1,NAPEPLD,PLCG1,PXN,RPS6KB1 |
| RAC Signaling | 0.661 | 0.0735 | 2.828 | 10 | CYFIP1,ITGA7,ITGAX,JUN,MAP2K7,MAP3K11,NCF2,PIP5K1B,RELA,RPS6KB1 |
| GP6 Signaling Pathway | 0.61 | 0.0726 | 2.828 | 9 | COL19A1,COL23A1,COL24A1,COL26A1,COL5A1,COL5A3,COL6A2,FCER1G,PLCG1 |
| Role of MAPK Signaling in Promoting the Pathogenesis of Influenza | 0.596 | 0.0734 | 2.828 | 8 | ATP6V0A1,JUN,MAP2K3,MAP2K7,MAPK13,PLA2G4E,PLB1,PNPLA3 |
| Ferroptosis Signaling Pathway | 0.387 | 0.0625 | 2.828 | 8 | ALOX15B,ALOXE3,CARS1,GLS2,HMOX1,PRKAB2,SLC11A2,TP53 |
| P2Y Purigenic Receptor Signaling Pathway | 0.361 | 0.0611 | 2.828 | 8 | ADCY6,GNG2,JUN,PLCB4,PLCD3,PLCG1,PLCZ1,RELA |
| Human Embryonic Stem Cell Pluripotency | 0 | 0.0402 | 2.828 | 8 | BMP1,DUSP9,ESRRB,MAP2K3,MAP2K5,MAP2K7,WNT2B,WNT9B |
| PKCθ Signaling in T Lymphocytes | 0 | 0.0251 | 2.828 | 9 | CAMK2G,CATSPER2,FCER1G,HLA-A,JUN,MAP3K11,MAP3K3,PLCG1,RELA |
| Role of NFAT in Cardiac Hypertrophy | 0.49 | 0.0639 | 2.714 | 14 | ADCY6,CAMK2G,CATSPER2,GNG2,HDAC10,HDAC7,MAP2K3,MAP2K7,MAPK13,PLCB4,PLCD3,PLCG1,PLCZ1,RCAN2 |
| Gustation Pathway | 0.347 | 0.0591 | 2.714 | 11 | ABCC8,ADCY6,CASR,CATSPER2,LRRC8A,P2RX6,SCN1A,SCN1B,SCNN1A,SLC2A4,TRPM4 |
| Colorectal Cancer Metastasis Signaling | 0 | 0.0492 | 2.714 | 13 | ADCY6,GNG2,JUN,MLH1,MSH2,RELA,RHOV,TLR1,TLR2,TLR7,TP53,WNT2B,WNT9B |
| HIF1α Signaling | 0.656 | 0.0697 | 2.673 | 14 | CAMK2G,HMOX1,HSPA1L,JUN,MAP2K3,MAP2K5,MAP2K7,MET,NCF2,PLCG1,RORC,RPS6KB1,SLC2A4,TP53 |
| Synaptogenesis Signaling Pathway | 0 | 0.0461 | 2.673 | 14 | ADCY6,CAMK2G,CPLX3,EFNA5,EIF4EBP3,FGR,GRINA,PLCG1,RPS6KB1,SNCB,STXBP2,SYT2,SYT3,THBS3 |
| ulation Of The Epithelial Mesenchymal Transition In Development Path | 0.725 | 0.0824 | 2.646 | 7 | BCL9,DVL2,JAG2,NCSTN,RELA,WNT2B,WNT9B |
| Salvage Pathways of Pyrimidine Ribonucleotides | 0.612 | 0.0761 | 2.646 | 7 | AK7,AK8,APOBEC3B,DMPK,EIF2AK2,MAP2K3,PRPF4B |
| HGF Signaling | 0.55 | 0.0698 | 2.646 | 9 | ITGA7,ITGAX,JUN,MAP2K7,MAP3K11,MAP3K3,MET,PLCG1,PXN |
| Apelin Cardiomyocyte Signaling Pathway | 0.542 | 0.0722 | 2.646 | 7 | MAPK13,PLCB4,PLCD3,PLCG1,PLCZ1,SLC9A1,SLC9A8 |
| GPCR-Mediated Nutrient Sensing in Enteroendocrine Cells | 0.509 | 0.069 | 2.646 | 8 | ADCY6,CASR,CATSPER2,GNG2,PLCB4,PLCD3,PLCG1,PLCZ1 |
| AMPK Signaling | 0.483 | 0.0633 | 2.646 | 15 | AK7,AK8,CRTC2,EEF2K,FOXO3,GNG2,GYS1,MAP2K3,MAPK13,PFKFB4,PPM1M,PRKAB2,RPS6KB1,SLC2A4,SMARCD2 |
| Aldosterone Signaling in Epithelial Cells | 0.348 | 0.0595 | 2.646 | 10 | DNAJB2,HSPA1L,HSPB6,PIP5K1B,PLCB4,PLCD3,PLCG1,PLCZ1,SCNN1A,SLC9A1 |
| Role Of Chondrocytes In Rheumatoid Arthritis Signaling Pathway | 0 | 0.0511 | 2.646 | 7 | CAPN2,CASP1,FOXO3,JUN,MAPK13,PRKAB2,TLR2 |
| Synaptic Long Term Depression | 0 | 0.0419 | 2.646 | 8 | CATSPER2,PLA2G4E,PLB1,PLCB4,PLCD3,PLCG1,PLCZ1,PNPLA3 |
| IL-17 Signaling | 0 | 0.0446 | 2.646 | 7 | CCL17,IL12A,JUN,MAP2K3,MAPK13,RELA,RGS16 |
| TEC Kinase Signaling | 0 | 0.026 | 2.646 | 10 | FCER1G,FGR,GNG2,ITGA7,ITGAX,PLCG1,RELA,RHOV,STAT2,STAT6 |
| lation Of The Epithelial Mesenchymal Transition By Growth Factors Pat | 0 | 0.0426 | 2.646 | 8 | FGF5,JUN,MAP2K3,MAP2K5,MAP2K7,MAPK13,MET,RELA |
| CLEAR Signaling Pathway | 1.87 | 0.0887 | 2.6 | 25 | ATP6V0A1,CLCN7,CLN3,CRTC2,CTNS,CTSA,FLCN,GBA1,GNS,HPS5,IFI30,MAPK13,MCOLN1,NPRL3,PLBD2,PLCG1,PML,PRKAB2,SCPEP1,SESN2,TLR1,TLR2,TLR7,TP53,ZKSCAN3 |
| Melatonin Signaling | 2.88 | 0.159 | 2.53 | 11 | CAMK2G,MAP2K3,MAP2K5,MAP2K7,PLCB4,PLCD3,PLCG1,PLCZ1,RORB,RORC,SLC2A4 |
| Endocannabinoid Neuronal Synapse Pathway | 0.932 | 0.0811 | 2.53 | 12 | ADCY6,CATSPER2,FAAH,GNG2,GRINA,KCNJ9,MAPK13,NAPEPLD,PLCB4,PLCD3,PLCG1,PLCZ1 |
| Role of NFAT in Regulation of the Immune Response | 0 | 0.0171 | 2.53 | 11 | FCER1G,FCGR2A,FCGR2B,FCGR3A/FCGR3B,GNG2,HLA-A,JUN,PLCB4,PLCG1,RCAN2,RELA |
| CD27 Signaling in Lymphocytes | 1.48 | 0.125 | 2.449 | 7 | JUN,MAP2K3,MAP2K5,MAP2K7,MAP3K11,MAP3K3,RELA |
| Cell Cycle Control of Chromosomal Replication | 1.15 | 0.113 | 2.449 | 6 | CDK15,CDT1,DNA2,MCM8,POLA2,POLE |
| Crosstalk between Dendritic Cells and Natural Killer Cells | 1.02 | 0.0986 | 2.449 | 7 | CAMK2G,CSF2RB,HLA-A,IL12A,IL2RB,RELA,TLR7 |
| IL-1 Signaling | 0.833 | 0.0851 | 2.449 | 8 | ADCY6,GNG2,JUN,MAP2K3,MAP2K7,MAPK13,MYD88,RELA |
| CD40 Signaling | 0.808 | 0.0909 | 2.449 | 6 | JUN,MAP2K3,MAP2K5,MAP2K7,MAPK13,RELA |
| WNT/Ca+ pathway | 0.808 | 0.0909 | 2.449 | 6 | DVL2,PLCB4,PLCD3,PLCG1,PLCZ1,RELA |
| Chemokine Signaling | 0.617 | 0.0789 | 2.449 | 6 | CAMK2G,CCL4,JUN,MAPK13,PLCB4,PLCG1 |
| RANK Signaling in Osteoclasts | 0.435 | 0.0674 | 2.449 | 6 | JUN,MAP2K7,MAP3K11,MAP3K3,MAPK13,RELA |
| DNA Methylation and Transcriptional Repression Signaling | 0.413 | 0.0648 | 2.449 | 7 | CDK15,ERCC5,GATAD2A,MBD1,MBD4,RELA,TP53 |
| Neuropathic Pain Signaling In Dorsal Horn Neurons | 0.331 | 0.0606 | 2.449 | 6 | CAMK2G,GRINA,PLCB4,PLCD3,PLCG1,PLCZ1 |
| Synaptic Long Term Potentiation | 0 | 0.0462 | 2.449 | 6 | CAMK2G,GRINA,PLCB4,PLCD3,PLCG1,PLCZ1 |
| NRF2-mediated Oxidative Stress Response | 0 | 0.0411 | 2.449 | 9 | DNAJB2,EPHX1,GSTA3,GSTM5,HMOX1,JUN,MAP2K3,MAP2K5,MAP2K7 |
| ICOS-ICOSL Signaling in T Helper Cells | 0 | 0.0293 | 2.449 | 9 | CAMK2G,FCER1G,HLA-A,ICOSLG/LOC102723996,IL2RB,PLCG1,PLEKHA4,PTPRC,RELA |
| Corticotropin Releasing Hormone Signaling | 0 | 0.0541 | 2.449 | 8 | ADCY6,CATSPER2,CRHR2,JUN,MAPK13,PLCG1,POMC,UCN3 |
| Cholecystokinin/Gastrin-mediated Signaling | 0 | 0.0531 | 2.449 | 6 | JUN,MAP2K3,MAP2K5,PLCB4,PXN,RHOV |
| Type I Diabetes Mellitus Signaling | 0 | 0.0261 | 2.449 | 8 | FCER1G,HLA-A,IL12A,MAP2K3,MAP2K7,MAPK13,MYD88,RELA |
| G Protein Signaling Mediated by Tubby | 0 | 0.0287 | 2.449 | 8 | FCER1G,GNG2,IFT140,NPY2R,PLCB4,PLCG1,TUB,WDR19 |
| Dopamine-DARPP32 Feedback in cAMP Signaling | 0 | 0.0387 | 2.449 | 7 | ADCY6,GRINA,KCNJ9,PLCB4,PLCD3,PLCG1,PLCZ1 |
| NGF Signaling | 0 | 0.0508 | 2.449 | 6 | MAP3K11,MAP3K3,PLCG1,RELA,RPS6KB1,TP53 |
| WNT/β-catenin Signaling | 0 | 0.0414 | 2.449 | 7 | BCL9,DVL2,JUN,KREMEN1,TP53,WNT2B,WNT9B |
| Th17 Activation Pathway | 0 | 0.0218 | 2.449 | 6 | FCER1G,IL12A,IL12RB1,MYD88,RELA,RORC |
| Actin Cytoskeleton Signaling | 0.801 | 0.072 | 2.333 | 17 | ARHGEF1,CD14,CYFIP1,FGF5,ITGA7,ITGAX,MSN,MYH3,MYH7,MYH7B,MYLK3,MYO1H,MYO5C,PIP5K1B,PXN,SLC9A1,SSH3 |
| Neurovascular Coupling Signaling Pathway | 0 | 0.0413 | 2.333 | 9 | ADORA1,CATSPER2,ENTPD4,GRINA,KCNJ9,PLA2G4E,PLB1,PLCB4,PNPLA3 |
| Leukocyte Extravasation Signaling | 0 | 0.0535 | 2.333 | 10 | CLDN22,CTNNA1,EDIL3,MAPK13,MSN,NCF2,PLCG1,PXN,THY1,WIPF1 |
| Glioblastoma Multiforme Signaling | 0 | 0.0536 | 2.333 | 9 | PLCB4,PLCD3,PLCG1,PLCZ1,RHOV,RPS6KB1,TP53,WNT2B,WNT9B |
| SAPK/JNK Signaling | 0 | 0.0288 | 2.333 | 9 | FCER1G,GNG2,JUN,MAP2K7,MAP3K11,MAP3K3,MAP4K2,MAP4K5,TP53 |
| Estrogen Receptor Signaling | 0 | 0.0375 | 2.324 | 15 | ADCY6,FBXO32,FOXO3,GNG2,GPS2,JUN,MED12,PLCB4,PLCD3,PLCG1,PLCZ1,PRKAB2,RELA,RPS6KB1,TP53 |
| Macrophage Classical Activation Signaling Pathway | 0.8 | 0.0764 | 2.309 | 12 | CD14,CXCL10,IL12A,IRF8,IRF9,mir-27,MYD88,NCF2,PPARG,RELA,STAT2,STAT6 |
| B Cell Receptor Signaling | 0 | 0.0339 | 2.309 | 15 | CAMK2G,CD22,FCGR2A,FCGR2B,INPP5B,INPP5K,JUN,MAP2K3,MAP2K7,MAP3K11,MAP3K3,MAPK13,PTPRC,RELA,RPS6KB1 |
| D-myo-inositol (1,4,5)-Trisphosphate Biosynthesis | 1.9 | 0.192 | 2.236 | 5 | PIP5K1B,PLCB4,PLCD3,PLCG1,PLCZ1 |
| Pyrimidine Ribonucleotides De Novo Biosynthesis | 1.67 | 0.15 | 2.236 | 6 | AK7,AK8,CTPS2,DDX3X,DHODH,ENTPD4 |
| Pyrimidine Ribonucleotides Interconversion | 1.29 | 0.135 | 2.236 | 5 | AK7,AK8,CTPS2,DDX3X,ENTPD4 |
| iNOS Signaling | 0.967 | 0.109 | 2.236 | 5 | CD14,JUN,MAPK13,MYD88,RELA |
| EGF Signaling | 0.732 | 0.0909 | 2.236 | 5 | JUN,MAP2K7,MAPK13,PLCG1,RPS6KB1 |
| Neurotrophin/TRK Signaling | 0.376 | 0.0649 | 2.236 | 5 | JUN,MAP2K3,MAP2K5,MAP2K7,PLCG1 |
| FLT3 Signaling in Hematopoietic Progenitor Cells | 0.333 | 0.0617 | 2.236 | 5 | FLT3,MAPK13,RPS6KB1,STAT2,STAT6 |
| LPS-stimulated MAPK Signaling | 0.313 | 0.0602 | 2.236 | 5 | CD14,JUN,MAP2K3,MAPK13,RELA |
| PEDF Signaling | 0.313 | 0.0602 | 2.236 | 5 | HNF1B,MAPK13,PPARG,RELA,TP53 |
| FGF Signaling | 0.286 | 0.0581 | 2.236 | 5 | FGF5,MAP2K3,MAPK13,MET,PLCG1 |
| PDGF Signaling | 0.286 | 0.0581 | 2.236 | 5 | EIF2AK2,INPP5B,INPP5K,JUN,PLCG1 |
| 14-3-3-mediated Signaling | 0 | 0.048 | 2.236 | 6 | GFAP,JUN,PLCB4,PLCD3,PLCG1,PLCZ1 |
| IL-15 Signaling | 0 | 0.0147 | 2.236 | 5 | IL2RB,MAPK13,RELA,RPS6KB1,STAT6 |
| Renin-Angiotensin Signaling | 0 | 0.05 | 2.236 | 6 | ACE,ADCY6,JUN,MAPK13,PLCG1,RELA |
| Gα12/13 Signaling | 0 | 0.0385 | 2.236 | 5 | ARHGEF1,JUN,MAP2K7,PXN,RELA |
| Regulation of IL-2 Expression in Activated and Anergic T Lymphocytes | 0 | 0.0169 | 2.236 | 5 | FCER1G,JUN,MAP2K7,PLCG1,RELA |
| ERBB Signaling | 0 | 0.0549 | 2.236 | 5 | JUN,MAP2K3,MAPK13,PLCG1,RPS6KB1 |
| IL-6 Signaling | 0 | 0.048 | 2.236 | 6 | CD14,JUN,MAP2K3,MAP2K7,MAPK13,RELA |
| Th1 Pathway | 0 | 0.055 | 2.236 | 6 | HAVCR2,HLA-A,ICOSLG/LOC102723996,IL12A,IL12RB1,NCSTN |
| White Adipose Tissue Browning Pathway | 0 | 0.0441 | 2.236 | 6 | ADCY6,CATSPER2,FCER1G,MAPK13,PPARG,PRKAB2 |
| MSP-RON Signaling In Cancer Cells Pathway | 0 | 0.0368 | 2.236 | 5 | CSF2RB,JUN,MET,RELA,ST14 |
| Ephrin Receptor Signaling | 0 | 0.04 | 2.236 | 8 | EFNA5,GNG2,GRINA,ITGA7,ITGAX,PXN,SH2D3C,WIPF1 |
| Activation of IRF by Cytosolic Pattern Recognition Receptors | 2.04 | 0.148 | 2.121 | 8 | IFIH1,IFIT2,IRF9,ISG15,JUN,RELA,STAT2,ZBP1 |
| Neuroprotective Role of THOP1 in Alzheimer's Disease | 1.09 | 0.0909 | 2.121 | 10 | ACE,ACR,C1R,ENDOU,HLA-A,HPN,HTRA2,LONP1,SERPINA3,ST14 |
| ILK Signaling | 0.314 | 0.0576 | 2.121 | 11 | DSP,FLNB,JUN,MYH3,MYH7,MYH7B,MYO1H,MYO5C,PXN,RELA,RHOV |
| Endocannabinoid Cancer Inhibition Pathway | 0.272 | 0.0559 | 2.121 | 8 | ADCY6,ATF3,CASP1,CASQ1,MAP2K3,MAP2K5,MAP2K7,PRKAB2 |
| UVA-Induced MAPK Signaling | 1.39 | 0.103 | 2 | 10 | JUN,MAPK13,PARP12,PARP3,PLCB4,PLCD3,PLCG1,PLCZ1,RPS6KB1,TP53 |
| tRNA Charging | 0.808 | 0.105 | 2 | 4 | CARS1,FARSA,SARS1,VARS1 |
| Triacylglycerol Degradation | 0.755 | 0.0926 | 2 | 5 | FAAH,NDST2,PLB1,PNPLA3,PNPLA7 |
| April Mediated Signaling | 0.7 | 0.0952 | 2 | 4 | JUN,MAP2K7,MAPK13,RELA |
| B Cell Activating Factor Signaling | 0.675 | 0.093 | 2 | 4 | JUN,MAP2K7,MAPK13,RELA |
| Spliceosomal Cycle | 0.651 | 0.0909 | 2 | 4 | DDX23,DDX46,PRPF40B,SF3B1 |
| UVB-Induced MAPK Signaling | 0.509 | 0.0784 | 2 | 4 | JUN,MAPK13,RPS6KB1,TP53 |
| PCP (Planar Cell Polarity) Pathway | 0.373 | 0.0667 | 2 | 4 | DVL2,JUN,WNT2B,WNT9B |
| VDR/RXR Activation | 0.365 | 0.0641 | 2 | 5 | CASR,CD14,CXCL10,HR,IL12A |
| Chronic Myeloid Leukemia Signaling | 0.359 | 0.0586 | 2 | 16 | DVL2,FOXO3,HDAC10,HDAC7,IRF8,JUN,PLCB4,PLCD3,PLCG1,PLCZ1,PRKAB2,RELA,SKP2,TP53,WNT2B,WNT9B |
| Thrombopoietin Signaling | 0.336 | 0.0635 | 2 | 4 | JUN,MPL,PLCG1,THPO |
| Pyridoxal 5'-phosphate Salvage Pathway | 0.314 | 0.0615 | 2 | 4 | DMPK,EIF2AK2,MAP2K3,PRPF4B |
| Unfolded protein response | 0.261 | 0.0562 | 2 | 5 | DNAJB2,EDEM1,HSPA1L,MAP2K7,PPARG |
| Immunogenic Cell Death Signaling Pathway | 0 | 0.0488 | 2 | 4 | BAK1,CASP1,CXCL10,HSPA1L |
| CCR5 Signaling in Macrophages | 0 | 0.0227 | 2 | 7 | CATSPER2,CCL4,FCER1G,GNG2,JUN,MAPK13,PLCG1 |
| fMLP Signaling in Neutrophils | 0 | 0.0312 | 2 | 4 | GNG2,NCF2,PLCB4,RELA |
| CD28 Signaling in T Helper Cells | 0 | 0.019 | 2 | 6 | FCER1G,HLA-A,JUN,PLCG1,PTPRC,RELA |
| Relaxin Signaling | 0 | 0.0464 | 2 | 7 | ADCY6,GNG2,JUN,NAPEPLD,PDE4C,PDE7A,RELA |
| Melanocyte Development and Pigmentation Signaling | 0 | 0.0515 | 2 | 5 | ADCY6,PLCG1,POMC,RPS6KB1,SH2B2 |
| ATM Signaling | 0 | 0.0408 | 2 | 4 | JUN,MAPK13,SMC2,TP53 |
| Pancreatic Adenocarcinoma Signaling | 0 | 0.0488 | 2 | 6 | HDAC10,HDAC7,HMOX1,NAPEPLD,RELA,TP53 |
| OX40 Signaling Pathway | 0 | 0.0224 | 2 | 6 | B2M,FCER1G,HLA-A,JUN,RELA,TNFRSF4 |
| Heparan Sulfate Biosynthesis | 0 | 0.0506 | 2 | 4 | B3GAT2,EXTL2,NDST2,PNPLA7 |
| Cardiac β-adrenergic Signaling | 0 | 0.0353 | 2 | 6 | ADCY6,AKAP8,GNG2,NAPEPLD,PDE4C,PDE7A |
| Xenobiotic Metabolism AHR Signaling Pathway | 0 | 0.0506 | 2 | 4 | ALDH1A1,GSTA3,GSTM5,RELA |
| Macrophage Alternative Activation Signaling Pathway | 0.721 | 0.0726 | 1.941 | 13 | CCL17,CSF1,FCER1G,FCGR2B,IL12A,IL4R,JUN,MAPK13,PPARG,RELA,RPS6KB1,STAT6,ZC3H12A |
| Myelination Signaling Pathway | 0 | 0.041 | 1.941 | 13 | BMP1,CYTH1,FGR,HDAC10,HDAC7,JUN,MYRF,NCSTN,PLCG1,RPS6KB1,WNT2B,WNT9B,XAF1 |
| Opioid Signaling Pathway | 0 | 0.0519 | 1.941 | 14 | ADCY6,CAMK2G,CATSPER2,FGR,GNG2,GRINA,KCNJ9,MAP2K3,MAP2K5,MAP2K7,POMC,RGS16,RGS7,RPS6KB1 |
| T Cell Exhaustion Signaling Pathway | 0 | 0.0315 | 1.897 | 11 | FCER1G,HAVCR2,HLA-A,IL12A,IL12RB1,IRF9,JUN,LAG3,PDCD1,PLCG1,STAT2 |
| NER (Nucleotide Excision Repair, Enhanced Pathway) | 0.999 | 0.09 | 1.89 | 9 | DDB2,DNA2,ERCC2,ERCC5,POLA2,POLE,POLK,UVSSA,XRCC1 |
| Pyroptosis Signaling Pathway | 0.761 | 0.0843 | 1.89 | 7 | AIM2,CASP1,FOXO3,MAPK13,TLR1,TLR2,TLR7 |
| IL-13 Signaling Pathway | 0.569 | 0.0737 | 1.89 | 7 | ALOX15B,FGR,IL4R,JUN,MAPK13,STAT6,TP53 |
| Pulmonary Healing Signaling Pathway | 0 | 0.0359 | 1.89 | 7 | FGR,MAPK13,MYD88,PRKAB2,TLR2,WNT2B,WNT9B |
| p70S6K Signaling | 0 | 0.0179 | 1.89 | 7 | EEF2K,IL4R,PLCB4,PLCD3,PLCG1,PLCZ1,RPS6KB1 |
| Neutrophil Extracellular Trap Signaling Pathway | 1.71 | 0.0822 | 1.826 | 30 | C1QB,C1QC,CASP1,CASQ1,CD14,CLEC7A,COL19A1,COL23A1,COL24A1,COL26A1,COL5A1,COL5A3,COL6A2,FCGR3A/FCGR3B,FGR,MAP2K3,MAP2K5,MAP2K7,MAPK13,NCF2,PLA2G4E,PLB1,PLCB4,PLCD3,PLCG1,PLCZ1,PNPLA3,TIMM44,TLR2,TLR7 |
| Sirtuin Signaling Pathway | 0.967 | 0.0743 | 1.698 | 20 | ATG13,ATG16L2,ATG4A,ATG4C,CRTC2,DOT1L,FOXO3,GABPB2,JUN,KAT2A,POLR1B,PPARG,RELA,SCNN1A,SIRT4,SIRT6,TIMM44,TP53,UCP2,WRN |
| Th2 Pathway | 1.13 | 0.0902 | 1.667 | 11 | HLA-A,ICOSLG/LOC102723996,IL12A,IL12RB1,IL2RB,IL4R,JAG2,JUN,NCSTN,STAT6,TNFRSF4 |
| Ribonucleotide Reductase Signaling Pathway | 0.274 | 0.0559 | 1.667 | 9 | FOXO3,JUN,MAPK13,MET,PARP12,PARP3,SKP2,SMARCD2,TP53 |
| Complement System | 3.47 | 0.242 | 1.633 | 8 | C1QB,C1QC,C1R,C3,C3AR1,C4A/C4B,CD46,ITGAX |
| Role of BRCA1 in DNA Damage Response | 1.13 | 0.1 | 1.633 | 8 | ATRIP,FANCG,FANCM,MLH1,MSH2,SMARCD2,TP53,UIMC1 |
| GADD45 Signaling | 0.977 | 0.102 | 1.633 | 6 | FOXO3,MAP2K3,MAP2K7,MAPK13,RELA,TP53 |
| Induction of Apoptosis by HIV1 | 0.853 | 0.0938 | 1.633 | 6 | BAK1,HTRA2,MAP2K7,Naip1 (includes others),RELA,TP53 |
| Death Receptor Signaling | 0.38 | 0.0638 | 1.633 | 6 | HTRA2,MAP2K7,Naip1 (includes others),PARP12,PARP3,RELA |
| NAD Signaling Pathway | 0 | 0.0493 | 1.633 | 7 | PARP12,PARP3,POLRMT,PRKAB2,SIRT4,SIRT6,TP53 |
| Oxytocin In Brain Signaling Pathway | 0 | 0.0321 | 1.633 | 6 | CASP1,CATSPER2,GNG2,PLA2G4E,PLB1,PNPLA3 |
| CXCR4 Signaling | 0 | 0.0366 | 1.633 | 6 | ADCY6,GNG2,JUN,PLCB4,PXN,RHOV |
| Endocannabinoid Developing Neuron Pathway | 0 | 0.0476 | 1.633 | 6 | ADCY6,GNG2,MAP2K3,MAP2K5,MAP2K7,MAPK13 |
| Apelin Endothelial Signaling Pathway | 0 | 0.0504 | 1.633 | 7 | ADCY6,GNG2,JUN,PLCB4,PRKAB2,RELA,RPS6KB1 |
| HOTAIR Regulatory Pathway | 0 | 0.0382 | 1.633 | 6 | MET,RBM38,RELA,SETDB1,WNT2B,WNT9B |
| Fc Epsilon RI Signaling | 0.545 | 0.0708 | 1.414 | 8 | FCER1G,INPP5B,INPP5K,MAP2K3,MAP2K7,MAPK13,PLA2G4E,PLCG1 |
| Role Of Osteoblasts In Rheumatoid Arthritis Signaling Pathway | 0 | 0.0343 | 1.414 | 8 | CTSA,CTSS,CTSZ,IL12A,STAT2,STAT6,WNT2B,WNT9B |
| HER-2 Signaling in Breast Cancer | 0 | 0.0411 | 1.414 | 9 | FCER1G,FGR,FOXO3,JUN,PLCG1,RAE1,RELA,RPS6KB1,TP53 |
| cAMP-mediated signaling | 0 | 0.0398 | 1.414 | 9 | ADCY6,ADORA1,AKAP8,CAMK2G,DUSP9,NAPEPLD,PDE4C,PDE7A,RGS7 |
| MIF Regulation of Innate Immunity | 1.1 | 0.119 | 1.342 | 5 | CD14,JUN,PLA2G4E,RELA,TP53 |
| PFKFB4 Signaling Pathway | 0.937 | 0.106 | 1.342 | 5 | MAP2K3,MAP2K5,MAP2K7,PFKFB4,TP53 |
| BMP signaling pathway | 0.459 | 0.069 | 1.342 | 6 | BMP1,CHRD,JUN,MAPK13,RELA,ZNF423 |
| p53 Signaling | 0.38 | 0.0638 | 1.342 | 6 | CCNG1,GNL3,JUN,PLAGL1,PML,TP53 |
| Role of JAK family kinases in IL-6-type Cytokine Signaling | 0.365 | 0.0641 | 1.342 | 5 | MAPK13,OSMR,PIAS3,POMC,SERPINA3 |
| JAK/STAT Signaling | 0.323 | 0.061 | 1.342 | 5 | JUN,PIAS3,RELA,STAT2,STAT6 |
| Amyotrophic Lateral Sclerosis Signaling | 0 | 0.0442 | 1.342 | 5 | CAPN2,CASP1,GRINA,Naip1 (includes others),TP53 |
| LPS/IL-1 Mediated Inhibition of RXR Function | 0 | 0.0519 | 1.342 | 12 | ACOX3,ALAS1,ALDH1A1,CD14,FMO2,GSTA3,GSTM5,JUN,MAP2K7,MYD88,NDST2,SLC27A2 |
| CDC42 Signaling | 0 | 0.0279 | 1.342 | 10 | B2M,EXOC1,FCER1G,HLA-A,ITGA7,ITGAX,JUN,MAP3K11,MAPK13,WIPF1 |
| Xenobiotic Metabolism PXR Signaling Pathway | 0 | 0.0292 | 1.342 | 5 | ALDH1A1,CAMK2G,GSTA3,GSTM5,NDST2 |
| Gαq Signaling | 0.37 | 0.0606 | 1.265 | 10 | GNG2,GYS1,HMOX1,NAPEPLD,PLCB4,PLCG1,RELA,RGS16,RGS7,RHOV |
| IL-33 Signaling Pathway | 0 | 0.0414 | 1.134 | 7 | CASP1,CASQ1,CCL17,JUN,MAPK13,MYD88,RPS6KB1 |
| Tumor Microenvironment Pathway | 0 | 0.0402 | 1.134 | 7 | CSF1,FGF5,FOXO3,HLA-A,JUN,RELA,SLC2A4 |
| Acute Myeloid Leukemia Signaling | 1.22 | 0.1 | 1 | 9 | CSF2RB,CSF3R,FLT3,MAP2K3,MAP2K5,MAP2K7,PML,RELA,RPS6KB1 |
| TNFR1 Signaling | 0.527 | 0.08 | 1 | 4 | JUN,MAP4K2,Naip1 (includes others),RELA |
| p38 MAPK Signaling | 0.355 | 0.0614 | 1 | 7 | CDC25B,EEF2K,MAP2K3,MAPK13,PLA2G4E,RPS6KB1,TP53 |
| ERK5 Signaling | 0 | 0.0548 | 1 | 4 | FOXO3,MAP2K5,MAP3K3,RPS6KB1 |
| RHOA Signaling | 0 | 0.0336 | 1 | 4 | ARHGEF1,MSN,MYLK3,PIP5K1B |
| NUR77 Signaling in T Lymphocytes | 0 | 0.0201 | 1 | 6 | B2M,FCER1G,HDAC7,HLA-A,MAP2K5,MAP3K3 |
| ERK/MAPK Signaling | 0 | 0.0337 | 1 | 7 | DUSP9,ITGA7,ITGAX,PLA2G4E,PLCG1,PPARG,PXN |
| MicroRNA Biogenesis Signaling Pathway | 0.421 | 0.0625 | 0.905 | 11 | DDX17,EXOSC8,HMOX1,MAPK13,PIWIL2,PNPT1,RAE1,RPS6KB1,TARBP2,TP53,ZC3H12A |
| Systemic Lupus Erythematosus In B Cell Signaling Pathway | 0 | 0.0392 | 0.894 | 20 | CD22,CD72,FCGR2A,FCGR2B,FGR,FOXO3,IFIH1,IFIT2,IFIT3,IL12A,INPP5B,INPP5K,IRF9,ISG15,JUN,MYD88,PDCD1,RELA,STAT2,TLR7 |
| Mitochondrial Dysfunction | 0 | 0.04 | 0.832 | 13 | ATP1B2,CAMK2G,CAPN2,CATSPER2,DHODH,HTRA2,MAP2K7,NCSTN,PPARG,PRKAB2,TARDBP,TP53,UCP2 |
| Interferon Signaling | 2.9 | 0.226 | 0.816 | 7 | BAK1,IFIT3,IRF9,ISG15,PTPN2,RELA,STAT2 |
| Polyamine Regulation in Colon Cancer | 1 | 0.103 | 0.816 | 6 | JUN,MAP2K3,MAP2K5,MAP2K7,PPARG,TP53 |
| PI3K/AKT Signaling | 0.525 | 0.0657 | 0.816 | 13 | CSF2RB,FOXO3,GYS1,IL12RB1,IL2RB,IL4R,INPP5B,INPP5K,ITGA7,ITGAX,RELA,RPS6KB1,TP53 |
| STAT3 Pathway | 0.328 | 0.0593 | 0.816 | 8 | CSF2RB,IL12RB1,IL2RB,IL4R,MAP3K11,MAPK13,PIAS3,PTPN2 |
| mTOR Signaling | 0 | 0.0347 | 0.816 | 7 | ATG13,EIF4EBP3,HMOX1,NAPEPLD,PRKAB2,RHOV,RPS6KB1 |
| Systemic Lupus Erythematosus In T Cell Signaling Pathway | 0 | 0.0352 | 0.775 | 15 | B2M,CASP1,CASQ1,FCER1G,HLA-A,ICOSLG/LOC102723996,JUN,MAP2K3,MAP2K5,MAP2K7,MSN,PLCG1,RHOV,RORC,RPS6KB1 |
| Senescence Pathway | 0 | 0.0528 | 0.775 | 15 | ATF3,CAPN2,CBX7,CDC25B,FOXO3,JUN,MAP2K3,MAP2K5,MAP2K7,PCGF2,PCGF6,PDK2,PML,TLR2,TP53 |
| Apoptosis Signaling | 0.682 | 0.0777 | 0.707 | 8 | BAK1,CAPN2,HTRA2,MAP2K7,Naip1 (includes others),PLCG1,RELA,TP53 |
| Sphingosine-1-phosphate Signaling | 0.533 | 0.0702 | 0.707 | 8 | ADCY6,CASP1,CASQ1,PLCB4,PLCD3,PLCG1,PLCZ1,RHOV |
| Erythropoietin Signaling Pathway | 0 | 0.0482 | 0.707 | 8 | CSF2RB,FOXO3,IL12A,JUN,PLCG1,RELA,RPS6KB1,TP53 |
| Osteoarthritis Pathway | 0 | 0.0485 | 0.707 | 11 | CASP1,CASQ1,CASR,CTNNA1,FOXO3,ITGA7,ITGAX,PPARG,PRKAB2,RELA,TLR2 |

| | | | | | |
|---|---|---|---|---|---|
| Xenobiotic Metabolism CAR Signaling Pathway | 0 | 0.0473 | 0.707 | 8 | ALDH1A1,FMO2,GSTA3,GSTM5,MAP2K3,MAP2K5,MAP2K7,NDST2 |
| IL-12 Signaling and Production in Macrophages | 1.99 | 0.0971 | 0.447 | 20 | C3,CLU,COL5A3,FCGR2A,FCGR2B,FCGR3A/FCGR3B,IFIH1,IL12A,IL12RB1,IRF8,JUN,MAP2K3,MAPK13,MYD88,PPARG,PRKAB2,RELA,STAT6,TLR2,TNIP2 |
| Aryl Hydrocarbon Receptor Signaling | 0 | 0.0541 | 0.447 | 8 | ALDH1A1,GSTA3,GSTM5,JUN,NFIC,NR2F1,RELA,TP53 |
| LXR/RXR Activation | 0 | 0.0446 | 0.447 | 5 | C3,C4A/C4B,CD14,CLU,RELA |
| CDX Gastrointestinal Cancer Signaling Pathway | 0 | 0.0469 | 0.333 | 9 | BMP1,IL12A,JUN,MAPK13,NCF2,PNN,RELA,WNT2B,WNT9B |
| Huntington's Disease Signaling | 0 | 0.0517 | 0.333 | 14 | CAPN2,CASP1,CASQ1,DNM1,GNG2,HDAC10,HDAC7,HSPA1L,JUN,MAP2K7,NEUROD1,PLCB4,TCERG1,TP53 |
| IL-10 Signaling | 0.852 | 0.0797 | 0.302 | 11 | FCGR2A,FCGR2B,HLA-A,HMOX1,IL4R,JUN,MAFB,MAP2K3,MAPK13,PRKAB2,RELA |
| Coronavirus Pathogenesis Pathway | 0.643 | 0.0699 | 0.277 | 13 | ACE,CASP1,CCDC22,HDAC10,HDAC7,IRF9,JUN,MAPK13,Oas1b,RELA,STAT2,TLR7,TP53 |
| Protein Kinase A Signaling | 0.76 | 0.0672 | 0.218 | 26 | ADCY6,AKAP8,CAMK2G,CDC25B,DUSP15,DUSP9,FLNB,GNG2,GYS1,MYLK3,NAPEPLD,PDE4C,PDE7A,PHKA2,PLCB4,PLCD3,PLCG1,PLCZ1,PTPN2,PTPN21,PTPN23,PTPN3,PTPRC,PXN,PYGM,RELA |
| Cell Cycle: G1/S Checkpoint Regulation | 0.54 | 0.0769 | 0 | 5 | GNL3,HDAC10,HDAC7,SKP2,TP53 |
| MSP-RON Signaling In Macrophages Pathway | 0.273 | 0.0566 | 0 | 6 | IL12A,JUN,NFKBIZ,RELA,SBNO2,ST14 |
| Gαi Signaling | 0 | 0.029 | 0 | 4 | ADCY6,ADORA1,GNG2,RGS7 |
| Insulin Receptor Signaling | 0 | 0.0507 | 0 | 7 | FOXO3,GYS1,INPP5B,INPP5K,RPS6KB1,SCNN1A,SLC2A4 |
| Apelin Adipocyte Signaling Pathway | 0 | 0.046 | 0 | 4 | ADCY6,MAPK13,NCF2,PRKAB2 |
| Adipogenesis pathway | 1.45 | 0.0963 | -0.277 | 13 | BSCL2,ERCC2,HDAC10,HDAC7,KAT2A,NR1D1,NR2F2,PPARG,RPS6KB1,SETDB1,SLC2A4,TP53,ZNF423 |
| Chaperone Mediated Autophagy Signaling Pathway | 0 | 0.0297 | -0.277 | 13 | ATP6V0A1,CASP1,CTSA,DDX3X,FCER1G,FLT3,GFAP,HDAC10,HSPA1L,MAPK13,PRKAB2,RELA,TP53 |
| Semaphorin Neuronal Repulsive Signaling Pathway | 0.765 | 0.0764 | -0.333 | 11 | DPYSL4,ITGA7,ITGAX,MAP2K3,MAP2K5,MAP2K7,PDE4C,PLCG1,PLXND1,SEMA6C,TP53 |
| Dilated Cardiomyopathy Signaling Pathway | 0.544 | 0.0685 | -0.333 | 10 | ADCY6,CAMK2G,CATSPER2,MAP3K3,MAPK13,MYH3,MYH7,MYH7B,MYO1H,MYO5C |
| PPARα/RXRα Activation | 0.481 | 0.0645 | -0.378 | 12 | ADCY6,JUN,MAP2K3,MAP2K7,MED12,NR2F1,PLCB4,PLCD3,PLCG1,PLCZ1,PRKAB2,RELA |
| Inhibition of ARE-Mediated mRNA Degradation Pathway | 0 | 0.0455 | -0.378 | 7 | CNOT10,CNOT3,EDC4,EXOSC8,MAPK13,PABPN1,POMC |
| Type II Diabetes Mellitus Signaling | 0 | 0.053 | -0.447 | 8 | ABCC8,CATSPER2,MAP2K7,PPARG,PRKAB2,RELA,SLC27A2,SLC2A4 |
| PTEN Signaling | 0 | 0.0537 | -0.447 | 8 | FOXO3,INPP5B,INPP5K,ITGA7,ITGAX,RELA,RPS6KB1,SIRT6 |
| CTLA4 Signaling in Cytotoxic T Lymphocytes | 0 | 0.0305 | -0.577 | 12 | AP1G2,AP1M2,B2M,CBLB,FCER1G,FGR,HLA-A,HMOX1,JUN,MAPK13,NAPEPLD,PLCG1 |
| PD-1, PD-L1 cancer immunotherapy pathway | 0.851 | 0.086 | -1.134 | 8 | B2M,CBLB,HLA-A,IL12A,IL2RB,PDCD1,PLCG1,SKP2 |
| ID1 Signaling Pathway | 0 | 0.0357 | -1.134 | 7 | ATF3,Bhlhe41,DVL2,FGR,FOXO3,MAPK13,TP53 |
| -Mediated Integration of Enteroendocrine Signaling Exemplified by an | 0.687 | 0.0833 | -1.633 | 6 | ADCY6,NPY2R,PLCB4,PLCD3,PLCG1,PLCZ1 |
| RHOGDI Signaling | 0.397 | 0.0607 | -2.236 | 13 | ARHGEF1,ARHGEF10,GNG2,ITGA7,ITGAX,MSN,MYH3,MYH7,MYH7B,MYO1H,MYO5C,PIP5K1B,RHOV |
| Antioxidant Action of Vitamin C | 2.65 | 0.131 | -3.464 | 14 | CSF2RB,HMOX1,MAPK13,NAPEPLD,PLA2G4E,PLB1,PLCB4,PLCD3,PLCG1,PLCZ1,PNPLA3,RELA,SLC23A3,SLC2A4 |

| **IPA pathway analysis Dap12-/- tau+ vs Dap12+/+ tau+** | | | | | |
|---|---|---|---|---|---|
| Ingenuity Canonical Pathways | -log(p-value) | Ratio | z-score | NO.of genes | Molecules |
| Pathogen Induced Cytokine Storm Signaling Pathway | 6.97 | 0.0603 | -4.359 | 19 | CCL2,Ccl6,COL11A1,COL11A2,COL19A1,COL4A1,COL4A2,COL5A1,COL6A2,CSF2RA,CSF2RB,CXCL16,DDX58,IRF1,IRF9,Naip1 (includes others),RYR1,TLR2,TNFSF10 |
| Wound Healing Signaling Pathway | 2.18 | 0.0377 | -3 | 9 | CHUK,COL11A1,COL11A2,COL19A1,COL4A1,COL4A2,COL5A1,COL6A2,TNFSF10 |
| Multiple Sclerosis Signaling Pathway | 2.19 | 0.0408 | -2.828 | 8 | C1QC,HLA-A,IRF1,PARP12,PARP14,PARP9,TLR2,TNFSF10 |
| Macrophage Classical Activation Signaling Pathway | 4.87 | 0.0701 | -2.714 | 11 | CD14,CD86,CHUK,IRF1,IRF8,IRF9,NCF2,PARP14,PARP9,STAT2,TNFSF10 |
| Role Of Osteoclasts In Rheumatoid Arthritis Signaling Pathway | 4.08 | 0.047 | -2.673 | 14 | AKT2,CHUK,COL11A1,COL11A2,COL19A1,COL4A1,COL4A2,COL5A1,COL6A2,CSF1R,FCGR1A,Naip1 (includes others),RHOU,TYROBP |
| Phagosome Formation | 1.15 | 0.0217 | -2.673 | 14 | AKT2,AP1G2,CD14,CX3CR1,FCGR1A,FZD2,ITGAM,ITGAX,ITGB2,LYN,PLD4,PNPLA3,TLR2,WAS |
| Role of PKR in Interferon Induction and Antiviral Response | 2.76 | 0.0569 | -2.646 | 7 | CHUK,DDX58,EIF2AK2,FCGR1A,IRF1,IRF9,STAT2 |
| Role Of Osteoblasts In Rheumatoid Arthritis Signaling Pathway | 1.34 | 0.03 | -2.646 | 7 | AKT2,CTSC,CTSH,CTSS,FZD2,STAT2,TNFSF10 |
| T Cell Exhaustion Signaling Pathway | 0.932 | 0.0229 | -2.646 | 8 | AKT2,CD86,HLA-A,IRF9,LAG3,PTPN6,STAT2,TNFRSF14 |
| Neuroinflammation Signaling Pathway | 2.62 | 0.0385 | -2.53 | 11 | AKT2,CCL2,CD86,CHUK,CSF1R,CX3CR1,HLA-A,Naip1 (includes others),NCF2,TLR2,TYROBP |
| of Hypercytokinemia/hyperchemokinemia in the Pathogenesis of Influ | 3.46 | 0.0896 | -2.449 | 6 | CCL2,DDX58,EIF2AK2,IFIT3,IRF9,STAT2 |
| GP6 Signaling Pathway | 4.21 | 0.0726 | -2.333 | 9 | AKT2,COL11A1,COL11A2,COL19A1,COL4A1,COL4A2,COL5A1,COL6A2,LYN |
| Retinoic acid Mediated Apoptosis Signaling | 3.45 | 0.114 | -2.236 | 5 | IRF1,PARP12,PARP14,PARP9,TNFSF10 |
| Necroptosis Signaling Pathway | 1.31 | 0.0352 | -2.236 | 5 | CHUK,EIF2AK2,IRF9,STAT2,TNFSF10 |
| D-myo-inositol (1,4,5,6)-Tetrakisphosphate Biosynthesis | 1.04 | 0.0294 | -2.236 | 5 | ACP6,CDC25B,DUSP11,PTPN6,PTPRC |
| D-myo-inositol (3,4,5,6)-tetrakisphosphate Biosynthesis | 1.04 | 0.0294 | -2.236 | 5 | ACP6,CDC25B,DUSP11,PTPN6,PTPRC |
| 3-phosphoinositide Degradation | 0.951 | 0.0275 | -2.236 | 5 | ACP6,CDC25B,DUSP11,PTPN6,PTPRC |
| D-myo-inositol-5-phosphate Metabolism | 0.921 | 0.0269 | -2.236 | 5 | ACP6,CDC25B,DUSP11,PTPN6,PTPRC |
| 3-phosphoinositide Biosynthesis | 0.851 | 0.0255 | -2.236 | 5 | ACP6,CDC25B,DUSP11,PTPN6,PTPRC |
| Superpathway of Inositol Phosphate Compounds | 0.688 | 0.0223 | -2.236 | 5 | ACP6,CDC25B,DUSP11,PTPN6,PTPRC |
| TREM1 Signaling | 5.19 | 0.113 | -2.121 | 8 | AKT2,CCL2,CD86,ITGAX,LAT2,Naip1 (includes others),TLR2,TYROBP |
| Caveolar-mediated Endocytosis Signaling | 5.09 | 0.11 | -2.121 | 8 | CD48,DYRK3,FLNA,FLNB,HLA-A,ITGAM,ITGAX,ITGB2 |
| Interferon Signaling | 4.19 | 0.161 | -2 | 5 | IFIT3,IFITM3,IRF1,IRF9,STAT2 |
| Crosstalk between Dendritic Cells and Natural Killer Cells | 2.5 | 0.0704 | -2 | 5 | CD86,CSF2RB,HLA-A,TNFSF10,TYROBP |
| Activation of IRF by Cytosolic Pattern Recognition Receptors | 2.17 | 0.0741 | -2 | 4 | CHUK,DDX58,IRF9,STAT2 |
| Toll-like Receptor Signaling | 1.7 | 0.0541 | -2 | 4 | CD14,CHUK,EIF2AK2,TLR2 |
| e of Pattern Recognition Receptors in Recognition of Bacteria and Viru | 1.3 | 0.035 | -2 | 5 | C1QC,DDX58,EIF2AK2,TLR2,TNFSF10 |
| Leukocyte Extravasation Signaling | 0.575 | 0.0214 | -2 | 4 | ITGAM,ITGB2,NCF2,WAS |
| Pulmonary Fibrosis Idiopathic Signaling Pathway | 1.84 | 0.0314 | -1.897 | 10 | AKT2,COL11A1,COL11A2,COL19A1,COL4A1,COL4A2,COL5A1,COL6A2,FZD2,JAG1 |
| Dendritic Cell Maturation | 1.42 | 0.0267 | -1.897 | 10 | AKT2,CD86,CHUK,COL11A2,FCGR1A,HLA-A,IRF8,STAT2,TLR2,TYROBP |
| FAK Signaling | 0 | 0.0121 | -1.897 | 10 | AKT2,COL11A2,CSF2RB,CX3CR1,ETS1,FZD2,ITGAM,ITGAX,ITGB2,WAS |
| Apoptosis Signaling | 2.69 | 0.0638 | -1.633 | 6 | CHUK,Naip1 (includes others),PARP12,PARP14,PARP9,TNFSF10 |
| Role of NFAT in Regulation of the Immune Response | 0 | 0.0093 | -1.633 | 6 | AKT2,CD86,CHUK,FCGR1A,HLA-A,LYN |
| IL-15 Production | 3.62 | 0.0684 | -1.414 | 8 | CSF1R,CSF2RA,DYRK3,FES,IRF1,LYN,RET,TIE1 |
| Fcγ Receptor-mediated Phagocytosis in Macrophages and Monocytes | 2.03 | 0.0543 | -1.342 | 5 | AKT2,FCGR1A,LYN,PLD4,WAS |
| Signaling by Rho Family GTPases | 1.52 | 0.0308 | -1.342 | 8 | ARHGEF1,IQGAP1,ITGAM,ITGAX,ITGB2,NCF2,RHOU,WAS |
| NAD Signaling Pathway | 1.31 | 0.0352 | -1.342 | 5 | AKT2,PARP12,PARP14,PARP9,RYR1 |
| HER-2 Signaling in Breast Cancer | 0.714 | 0.0228 | -1.342 | 5 | AKT2,CHUK,ETS1,ITGB2,LYN |
| IL-8 Signaling | 3.26 | 0.0498 | -1.265 | 10 | AKT2,BCL2L1,CHUK,IQGAP1,ITGAM,ITGAX,ITGB2,NCF2,PLD4,RHOU |
| roduction of Nitric Oxide and Reactive Oxygen Species in Macrophage | 2.44 | 0.0449 | -1.134 | 8 | AKT2,CHUK,IRF1,IRF8,NCF2,PTPN6,RHOU,TLR2 |
| T Cell Receptor Signaling | 0.474 | 0.0173 | -1.134 | 7 | AKT2,CD86,CHUK,HLA-A,ITGB2,PTPN6,PTPRC |
| NF-κB Activation by Viruses | 1.62 | 0.0513 | -1 | 4 | AKT2,CHUK,EIF2AK2,TNFRSF14 |
| Salvage Pathways of Pyrimidine Ribonucleotides | 1.4 | 0.0435 | -1 | 4 | AKT2,APOBEC3B,CDK2,EIF2AK2 |
| Osteoarthritis Pathway | 1.39 | 0.0308 | -1 | 7 | CHUK,FZD2,ITGAM,ITGAX,ITGB2,JAG1,TLR2 |
| MSP-RON Signaling In Cancer Cells Pathway | 0.91 | 0.0294 | -1 | 4 | AKT2,CSF2RB,ETS1,FLNA |
| TEC Kinase Signaling | 0.767 | 0.0208 | -1 | 8 | ITGAM,ITGAX,ITGB2,LYN,RHOU,STAT2,TNFSF10,WAS |
| ation Of The Epithelial Mesenchymal Transition By Growth Factors Pat | 0.57 | 0.0213 | -1 | 4 | AKT2,CHUK,ETS1,TNFSF10 |
| Pulmonary Healing Signaling Pathway | 0.536 | 0.0205 | -1 | 4 | FZD2,JAG1,LYN,TLR2 |
| Phospholipase C Signaling | 0 | 0.011 | -1 | 8 | ARHGEF1,HDAC7,ITGAM,ITGAX,ITGB2,LYN,PLD4,RHOU |
| PI3K Signaling in B Lymphocytes | 0 | 0.00998 | -1 | 4 | AKT2,CHUK,LYN,PTPRC |
| NF-κB Signaling | 0 | 0.0106 | -1 | 4 | AKT2,CHUK,EIF2AK2,TLR2 |
| IL-4 Signaling | 1.78 | 0.0294 | -0.905 | 11 | AKT2,BCL2L1,COL11A1,COL11A2,COL19A1,COL4A1,COL4A2,COL5A1,COL6A2,PRR5,PTPN6 |
| Hepatic Fibrosis Signaling Pathway | 1.22 | 0.0246 | -0.816 | 10 | AKT2,CCL2,CHUK,COL11A2,FZD2,ITGAM,ITGAX,ITGB2,NCF2,RHOU |
| Integrin Signaling | 1.17 | 0.0294 | -0.816 | 6 | AKT2,ITGAM,ITGAX,ITGB2,RHOU,WAS |
| CD28 Signaling in T Helper Cells | 0.81 | 0.0222 | -0.816 | 7 | AKT2,CD86,CHUK,HLA-A,PTPN6,PTPRC,WAS |
| CDX Gastrointestinal Cancer Signaling Pathway | 2.24 | 0.0417 | -0.707 | 8 | AKT2,BMP1,CDK2,CHUK,FZD2,NCF2,PNN,TNFSF10 |
| CTLA4 Signaling in Cytotoxic T Lymphocytes | 0.73 | 0.0204 | -0.707 | 8 | AKT2,AP1G2,BCL2L1,CD86,HLA-A,ITGB2,LYN,PLD4 |
| S100 Family Signaling Pathway | 0 | 0.0113 | -0.707 | 8 | AKT2,CATSPER2,CHUK,CX3CR1,FCGR1A,FZD2,IQGAP1,NCF2 |
| Ribonucleotide Reductase Signaling Pathway | 1.12 | 0.0311 | -0.447 | 5 | AKT2,CDK2,PARP12,PARP14,PARP9 |
| Gαq Signaling | 1.09 | 0.0303 | -0.447 | 5 | AKT2,CHUK,PLD4,RGS16,RHOU |
| Breast Cancer Regulation by Stathmin1 | 0 | 0.0088 | -0.447 | 5 | ARHGEF1,CAMK1G,CDK2,CX3CR1,FZD2 |
| G-Protein Coupled Receptor Signaling | 0 | 0.00745 | -0.447 | 5 | AKT2,CHUK,CX3CR1,FZD2,RGS16 |
| Protein Kinase A Signaling | 2.07 | 0.031 | 0 | 12 | AKAP8,CDC25B,CHUK,DUSP11,FLNA,FLNB,PHKA2,PHKG1,PTPN6,PTPRC,Ptprv,RYR1 |
| Small Cell Lung Cancer Signaling | 1.97 | 0.0526 | 0 | 5 | AKT2,BCL2L1,CDK2,CHUK,HDAC7 |
| PD-1, PD-L1 cancer immunotherapy pathway | 1.38 | 0.043 | 0 | 4 | AKT2,BCL2L1,CDK2,HLA-A |
| Opioid Signaling Pathway | 1.07 | 0.0259 | 0 | 7 | AKT2,CAMK1G,CATSPER2,LYN,RGS11,RGS16,RYR1 |
| IL-17 Signaling | 0.75 | 0.0255 | 0 | 4 | AKT2,CCL2,RGS16,TNFSF10 |
| Cardiac Hypertrophy Signaling (Enhanced) | 0.699 | 0.0191 | 0 | 10 | AKT2,CHUK,CSF2RB,FZD2,HDAC7,ITGAM,ITGAX,ITGB2,RYR1,TNFSF10 |
| Senescence Pathway | 0.682 | 0.0211 | 0 | 6 | AKT2,CDC25B,CDK2,CHUK,ETS1,TLR2 |
| Systemic Lupus Erythematosus In B Cell Signaling Pathway | 0.377 | 0.0157 | 0 | 8 | AKT2,BCL2L1,IFIT3,IRF9,LYN,PTPN6,STAT2,TNFSF10 |
| Systemic Lupus Erythematosus In T Cell Signaling Pathway | 0 | 0.00939 | 0 | 4 | AKT2,CD86,HLA-A,RHOU |
| Natural Killer Cell Signaling | 3.18 | 0.0529 | 0.333 | 9 | AKT2,CD48,COL11A2,HLA-A,LAIR1,PTPN6,TNFSF10,TYROBP,WAS |
| GM-CSF Signaling | 3.36 | 0.0857 | 0.447 | 6 | AKT2,BCL2L1,CSF2RA,CSF2RB,ETS1,LYN |
| Inhibition of ARE-Mediated mRNA Degradation Pathway | 1.19 | 0.0325 | 0.447 | 5 | AKT2,PABPN1,TIA1,TNFSF10,ZFP36 |
| HIF1α Signaling | 0.821 | 0.0249 | 0.447 | 5 | AKT2,CAMK1G,GPI,HK2,NCF2 |
| Neutrophil Extracellular Trap Signaling Pathway | 4.79 | 0.0466 | 0.728 | 17 | AKT2,C1QC,CD14,COL11A1,COL11A2,COL19A1,COL4A1,COL4A2,COL5A1,COL6A2,ITGAM,ITGB2,LAIR1,LYN,NCF2,PNPLA3,TLR2 |
| B Cell Receptor Signaling | 0.375 | 0.0158 | 0.816 | 7 | AKT2,BCL2L1,CHUK,ETS1,LYN,PTPN6,PTPRC |
| JAK/STAT Signaling | 1.55 | 0.0488 | 1 | 4 | AKT2,BCL2L1,PTPN6,STAT2 |
| Pancreatic Adenocarcinoma Signaling | 1.53 | 0.0407 | 1 | 5 | AKT2,BCL2L1,CDK2,HDAC7,PLD4 |
| Glioblastoma Multiforme Signaling | 0.68 | 0.0238 | 1 | 4 | AKT2,CDK2,FZD2,RHOU |
| ID1 Signaling Pathway | 0.532 | 0.0204 | 1 | 4 | AKT2,BCL2L1,ETS1,LYN |
| mTOR Signaling | 0.504 | 0.0198 | 1 | 4 | AKT2,PLD4,PRR5,RHOU |
| Thrombin Signaling | 0.44 | 0.0183 | 1 | 4 | AKT2,ARHGEF1,CAMK1G,RHOU |
| Erythropoietin Signaling Pathway | 1.08 | 0.0301 | 1.342 | 5 | AKT2,BCL2L1,CSF2RB,PTPN6,TNFSF10 |
| ILK Signaling | 0.886 | 0.0262 | 1.342 | 5 | AKT2,FLNA,FLNB,ITGB2,RHOU |
| Colorectal Cancer Metastasis Signaling | 0.509 | 0.0189 | 1.342 | 5 | AKT2,BCL2L1,FZD2,RHOU,TLR2 |
| Coronavirus Pathogenesis Pathway | 1.8 | 0.0376 | 1.89 | 7 | CCL2,CDK2,CHUK,DDX58,HDAC7,IRF9,STAT2 |
| Acute Myeloid Leukemia Signaling | 2.07 | 0.0556 | 2.236 | 5 | AKT2,CSF1R,CSF2RA,CSF2RB,CSF3R |
| Antioxidant Action of Vitamin C | 1.76 | 0.0467 | 2.236 | 5 | CHUK,CSF2RA,CSF2RB,PLD4,PNPLA3 |
| IL-10 Signaling | 1.88 | 0.0435 | 2.449 | 6 | AKT2,BCL2L1,BLVRB,CD86,CHUK,HLA-A |

| Figure2E_venndiagram_DEGs_of_Bulkse | | | | | | | | | | | | | | | | | | | | | | | | |
|---|---|---|---|---|---|---|---|---|---|---|---|---|---|---|---|---|---|---|---|---|---|---|---|---|
| Dap12+/+ tau+ vs tau- | | | | | | | Upregulation | Downregulation | Dap12-/- tau+ vs Dap12+/+ tau+ | | | | | | | Upregulation | Downregulation | Venn_results | | | | | | |
| gene | baseMean | log2FoldChange | lfcSE | stat | pvalue | padj | 1241 | 1220 | gene | baseMean | log2FoldChange | lfcSE | stat | pvalue | padj | 50 | 260 | | down_by_dap12 up_by_tau | 131 | Srrm1 | down_by_tau up_by_dap12 | 5 | Kcna5 |
| Xist | 9788.905385 | 8.703279608 | 0.126159893 | 68.98610486 | 0 | 0 | Xist | Vdac3 | Ambra1 | 3647.260573 | -0.000107225 | 0.042602304 | -0.002516893 | 0.997991812 | 0.999367783 | Zfp14 | Tardbp | | | | Dennd6b | | | Ppapdc2 |
| Slc25a20 | 872.0526975 | 1.445819194 | 0.063000299 | 22.94940208 | 1.49E-116 | 3.31E-113 | Slc25a20 | Cs | Stard3 | 1373.756455 | -0.000112604 | 0.061113358 | -0.001842536 | 0.998529869 | 0.999562048 | Gm1082 | Slc6a6 | | | | Ovgp1 | | | Rab3c |
| Tspan11 | 208.9624499 | 1.375178007 | 0.107852361 | 12.75056011 | 3.10E-37 | 5.33E-34 | Tspan11 | Slc30a9 | Crtam | 3.510511234 | -0.00011608 | 0.044177492 | -0.002627594 | 0.997903486 | NA | Blvrb | Luc7l2 | | | | Tlr2 | | | Tmeff1 |
| Wnt2b | 77.90124492 | 1.329179115 | 0.153024737 | 8.686040816 | 3.75E-18 | 3.06E-15 | Wnt2b | Eef1a1 | Syt7 | 15018.64047 | -0.000122317 | 0.059494473 | -0.002055936 | 0.998359601 | 0.99946048 | Kirrel2 | Usp48 | | | | 4930570G19Rik | | | Hs6st2 |
| C4b | 1933.281053 | 1.310089731 | 0.114562669 | 11.43557272 | 2.78E-30 | 4.30E-27 | C4b | Cyb5b | Fem1c | 842.9121598 | -0.000125697 | 0.074479995 | -0.001687664 | 0.99865344 | 0.99961686 | Gm6537 | Plekha5 | | | | Cd52 | | | |
| Rspo1 | 356.2679796 | 1.214775244 | 0.148470213 | 8.181945875 | 2.79E-16 | 1.88E-13 | Rspo1 | Kidins220 | Mkks | 1322.156651 | -0.000142072 | 0.063380941 | -0.002241563 | 0.998211493 | 0.999381495 | Plekhf1 | Edem1 | | | | Akap8 | | | |
| Gm765 | 60.65268662 | 1.19169213 | 0.168110463 | 7.088744567 | 1.35E-12 | 3.88E-10 | Gm765 | Eif5a | Gm11978 | 3.812917842 | -0.000152162 | 0.042243496 | -0.00360202 | 0.99712601 | NA | Zfp27 | Trub1 | | | | Rbm33 | | | |
| Ntn5 | 150.9054749 | 1.118404744 | 0.16373717 | 6.830487818 | 8.46E-12 | 2.22E-09 | Ntn5 | Dennd5a | Gm19299 | 0.224853396 | -0.000161157 | 0.009791626 | -0.016458695 | 0.986868454 | NA | Hif3a | Ddx17 | | | | Tmem181b-ps | | | |
| Clec7a | 69.16320609 | 1.107588906 | 0.163720099 | 6.76513705 | 1.33E-11 | 3.26E-09 | Clec7a | Rer1 | Magea10 | 0.224853396 | -0.000161157 | 0.009791626 | -0.016458695 | 0.986868454 | NA | Rinl | Iqgap1 | | | | Csf2rb | | | |
| Myo5c | 158.7856697 | 1.081513555 | 0.151685928 | 7.129953129 | 1.00E-12 | 2.94E-10 | Myo5c | Nap1l4 | Mirlet7c-1 | 0.224853396 | -0.000161157 | 0.009791626 | -0.016458695 | 0.986868454 | NA | Tsc22d3 | Rbm5 | | | | Rbm25 | | | |
| Cd52 | 80.47788727 | 1.07457152 | 0.162912389 | 6.596008629 | 4.22E-11 | 9.26E-09 | Cd52 | Sel1l | Tas2r143 | 0.224853396 | -0.000161157 | 0.009791626 | -0.016458695 | 0.986868454 | NA | Cbln4 | 6820431F20Rik | | | | Whamm | | | |
| Adam33 | 276.5512317 | 1.065251902 | 0.157074763 | 6.781814493 | 1.19E-11 | 2.99E-09 | Adam33 | Pdha1 | Vmn1r222 | 0.224853396 | -0.000161157 | 0.009791626 | -0.016458695 | 0.986868454 | NA | Adipor2 | Pfkfb2 | | | | Thpo | | | |
| Nr2f2 | 651.2817828 | 1.044731182 | 0.133571384 | 7.821519505 | 5.22E-15 | 2.96E-12 | Nr2f2 | Gzf1 | Klhl10 | 9.834012777 | -0.000162096 | 0.063507667 | -0.002552384 | 0.997963494 | NA | Kcna5 | Zc3h7a | | | | Pnn | | | |
| Igfn1 | 1449.509837 | 1.007539751 | 0.159582665 | 6.313591443 | 2.73E-10 | 4.59E-08 | Igfn1 | Glud1 | Phlpp2 | 2996.945819 | -0.000163591 | 0.055709574 | -0.002936497 | 0.997657018 | 0.999289749 | Pcdha10 | Heatr6 | | | | Pnpla3 | | | |
| Gpr114 | 11.3145576 | 0.998030924 | 0.151323956 | 6.595326673 | 4.24E-11 | 9.26E-09 | Gpr114 | Ubl4 | Pabpc4 | 1286.28693 | -0.000170984 | 0.064154096 | -0.002665208 | 0.997873475 | 0.999318172 | Atp4a | Rbm25 | | | | Edem1 | | | |
| Gfap | 9181.31116 | 0.990633417 | 0.157642669 | 6.284043668 | 3.30E-10 | 5.44E-08 | Gfap | Ndufb10 | Gm13889 | 524.4390432 | -0.000188979 | 0.084326027 | -0.002241049 | 0.998211903 | 0.999381495 | Ret | Srrm1 | | | | Ctss | | | |
| Entpd4 | 5040.613621 | 0.958156073 | 0.073369172 | 13.05938241 | 5.62E-39 | 1.09E-35 | Entpd4 | Micu1 | 1700009C05Rik | 1.62017637 | -0.000191074 | 0.025325175 | -0.007544828 | 0.993980155 | NA | Arrdc2 | Nxf1 | | | | Meg3 | | | |
| Scnn1a | 189.9552663 | 0.931182536 | 0.121517073 | 7.66297703 | 1.82E-14 | 8.04E-12 | Scnn1a | Btbd1 | Mapk8ip1 | 16644.26024 | -0.000209301 | 0.054429425 | -0.003845359 | 0.996931855 | 0.99913245 | Dyrk3 | Sall1 | | | | Cdc25b | | | |
| Crb1 | 33.17447847 | 0.926287801 | 0.166155692 | 5.574818343 | 2.48E-08 | 2.30E-06 | Crb1 | 15-Sep | Gm10677 | 3.695166467 | -0.000214748 | 0.039441094 | -0.005444787 | 0.99565571 | NA | Fkbp5 | Ubp1 | | | | Fam227a | | | |
| Ccl6 | 85.51648389 | 0.922088455 | 0.167185195 | 5.515371459 | 3.48E-08 | 3.08E-06 | Ccl6 | Prmt1 | Gypc | 77.59445086 | -0.000223523 | 0.093272685 | -0.002396451 | 0.998087911 | 0.999381495 | Meig1 | Myo9b | | | | Myo1f | | | |
| Sec1 | 46.14779074 | 0.906037211 | 0.167815582 | 5.399005272 | 6.70E-08 | 5.35E-06 | Sec1 | Rpl5 | Mettl10 | 495.2643526 | -0.000225672 | 0.078439725 | -0.002877011 | 0.99770448 | 0.999289749 | Ada | Tbce | | | | 2610005L07Rik | | | |
| Sytl1 | 49.0441286 | 0.877653634 | 0.165681133 | 5.297245486 | 1.18E-07 | 8.51E-06 | Sytl1 | Prune | Fau | 4848.646461 | -0.000235515 | 0.049884065 | -0.004721248 | 0.996233003 | 0.998948537 | Camk1g | Slco1a4 | | | | Itgax | | | |
| Cyp39a1 | 136.7727229 | 0.875057103 | 0.130831669 | 6.688419644 | 2.26E-11 | 5.14E-09 | Cyp39a1 | 9530068E07Rik | E130304I02Rik | 3.724907869 | -0.000246464 | 0.044546988 | -0.005532678 | 0.995585584 | NA | Prr5 | Zfp445 | | | | Srrm2 | | | |
| Miat | 16854.34769 | 0.873351165 | 0.155958359 | 5.599899691 | 2.14E-08 | 2.03E-06 | Miat | Ptp4a2 | 1700120K04Rik | 1.640092612 | -0.000247991 | 0.024836585 | -0.009984894 | 0.99203334 | NA | Klf15 | Prpf39 | | | | Ddx26b | | | |
| Thy1 | 37702.74949 | 0.870839947 | 0.079623348 | 10.93699238 | 7.67E-28 | 9.90E-25 | Thy1 | Dzank1 | Lyrm2 | 694.540003 | -0.000252561 | 0.077974908 | -0.003239009 | 0.997415649 | 0.999289749 | Ovol2 | Unc13b | | | | Tnip2 | | | |
| Clec18a | 204.5870021 | 0.870597815 | 0.155166843 | 5.610720666 | 2.01E-08 | 1.94E-06 | Clec18a | Pdgfra | Bbs4 | 2679.620795 | -0.000258814 | 0.054378791 | -0.004759468 | 0.996202508 | 0.998948537 | Rgs16 | Rbm33 | | | | Zkscan3 | | | |
| A330049N07Rik | 62.36615135 | 0.868878623 | 0.152994381 | 5.679153836 | 1.35E-08 | 1.38E-06 | A330049N07Rik | Dcaf12 | Tarm1 | 0.774374434 | -0.000259786 | 0.019922139 | -0.013040042 | 0.989595846 | NA | Tekt4 | Zfp280c | | | | Xaf1 | | | |
| Lag3 | 407.474968 | 0.862273253 | 0.158321784 | 5.446333617 | 5.14E-08 | 4.24E-06 | Lag3 | Gabrb3 | Vmn2r54 | 0.826762817 | -0.000274931 | 0.020111548 | -0.013670302 | 0.989093017 | NA | Gpr123 | Smc5 | | | | Arhgef1 | | | |
| | 446.5826992 | 0.858681124 | 0.116028675 | 7.400594041 | 1.36E-13 | 5.25E-11 | Pla2g4e | Pnpla8 | 1700011I03Rik | 3.312975103 | -0.000279252 | 0.042331044 | -0.006596863 | 0.994736503 | NA | Ppapdc2 | Slc4a7 | | | | Spag5 | | | |
| Barx2 | 133.4700717 | 0.85692625 | 0.126382557 | 6.780415499 | 1.20E-11 | 2.99E-09 | Barx2 | Pgam1 | Spc25 | 12.54616424 | -0.000281865 | 0.070408653 | -0.004003266 | 0.996805864 | NA | Plin4 | Ifi204 | | | | Parp12 | | | |
| Itgax | 89.51715568 | 0.845569665 | 0.151973521 | 5.563927566 | 2.64E-08 | 2.42E-06 | Itgax | Sec62 | Dmbx1 | 8.299835162 | -0.000296725 | 0.051619844 | -0.00574828 | 0.995413561 | NA | Rhou | Qtrtd1 | | | | Srsf11 | | | |
| Tmem215 | 450.7499935 | 0.832106122 | 0.143042301 | 5.817203131 | 5.98E-09 | 6.92E-07 | Tmem215 | Dtd1 | Ccdc68 | 3.152226755 | -0.000297175 | 0.037167116 | -0.007995653 | 0.99362046 | NA | Hs6st2 | Phka2 | | | | Naip5 | | | |
| HALLMARK_INFLAMMATORY_ RESPONSE | 24.64422152 | 0.828484152 | 0.160237074 | 5.170364943 | 2.34E-07 | 1.50E-05 | Cxcl10 | Gnb2l1 | Cacna1c | 2246.321271 | -0.00030433 | 0.066231408 | -0.004594955 | 0.996333769 | 0.998948537 | B3galt5 | Sipa1l3 | | | | Luc7l3 | | | |
| Cldn22 | 11.60959755 | 0.827013419 | 0.158006096 | 5.234060204 | 1.66E-07 | 1.13E-05 | Cldn22 | Eef1b2 | Dusp28 | 686.1498932 | -0.000332127 | 0.073693637 | -0.004506859 | 0.996404059 | 0.998948537 | Fzd2 | Itpripl2 | | | | Unc13b | | | |
| Mir27b | 19.09149771 | 0.81414511 | 0.167876699 | 4.849661183 | 1.24E-06 | 5.88E-05 | Mir27b | Cmas | Suv420h1 | 2258.553996 | -0.000336591 | 0.05051542 | -0.00666314 | 0.994683623 | 0.998727174 | Arid5b | Whamm | | | | Zc3h7a | | | |
| Endou | 142.3765054 | 0.808710506 | 0.15334902 | 5.273659449 | 1.34E-07 | 9.46E-06 | Endou | Gnb1 | Dbnl | 2681.366883 | -0.000345297 | 0.068081246 | -0.005071841 | 0.995953273 | 0.998915149 | Lfng | Srek1 | | | | Ccdc84 | | | |
| Ppl | 72.9663435 | 0.808626529 | 0.15197777 | 5.320689538 | 1.03E-07 | 7.66E-06 | Ppl | Arl2bp | Sike1 | 1089.708568 | -0.000350015 | 0.067873941 | -0.005156838 | 0.995885457 | 0.998915149 | Rhbdl3 | Phkg1 | | | | Ddb2 | | | |
| Thbs3 | 538.984351 | 0.808257379 | 0.123621781 | 6.538147008 | 6.23E-11 | 1.30E-08 | Thbs3 | Selk | Mrpl44 | 756.9796563 | -0.00035073 | 0.066150935 | -0.005301959 | 0.995769669 | 0.998915149 | Endod1 | Gigyf1 | | | | Ccl6 | | | |
| Coro6 | 1141.994843 | 0.803160271 | 0.138197986 | 5.811664096 | 6.19E-09 | 7.00E-07 | Coro6 | Rpl10a | AA415398 | 255.6004113 | -0.00035294 | 0.081476262 | -0.004331815 | 0.996543722 | 0.999019607 | Bcat1 | Csf1r | | | | Ogt | | | |
| Kcnab3 | 3234.69678 | 0.801421739 | 0.11213476 | 7.146951909 | 8.87E-13 | 2.64E-10 | Kcnab3 | Sae1 | Kcne2 | 59.05984924 | -0.000364193 | 0.025094617 | -0.014512808 | NA | NA | Rab3c | Gabpb2 | | | | Tardbp | | | |
| Slamf9 | 47.10396413 | 0.789940209 | 0.164702297 | 4.796169948 | 1.62E-06 | 7.46E-05 | Slamf9 | Naa20 | Setd5 | 5150.818357 | -0.00036514 | 0.054590511 | -0.006688713 | 0.994663219 | 0.998727174 | Ndufc2 | Ddb2 | | | | Ifi44 | | | |
| C920025E04Rik | 37.70378763 | 0.785211291 | 0.167428048 | 4.689843182 | 2.73E-06 | 0.000114804 | C920025E04Rik | Calu | Kit | 1895.80043 | -0.000365715 | 0.05345313 | -0.006841784 | 0.994541089 | 0.998727174 | Cables1 | Guf1 | | | | Capg | | | |
| Tmem145 | 1516.001754 | 0.779047173 | 0.116817463 | 6.66892737 | 2.58E-11 | 5.79E-09 | Tmem145 | Vdac1 | Creb3l4 | 12.6386198 | -0.000373165 | 0.071273245 | -0.005235701 | 0.995822534 | NA | Tmeff1 | Alkbh6 | | | | Lag3 | | | |
| Zan | 25.336803 | 0.776439059 | 0.167533641 | 4.634526263 | 3.58E-06 | 0.000140331 | Zan | Mtmr9 | Mir432 | 2.11306047 | -0.000380597 | 0.031278635 | -0.012167946 | 0.990291623 | NA | Akt2 | Safb2 | | | | Gm3002 | | | |
| Cd22 | 30.35036383 | 0.77279636 | 0.168101702 | 4.597195337 | 4.28E-06 | 0.00016222 | Cd22 | Nfyb | Cacng1 | 11.14023722 | -0.000381005 | 0.06153387 | -0.006191792 | 0.995059696 | NA | Bcl2l1 | Ccl12 | | | | Prpf39 | | | |
| Ap1m2 | 36.06865584 | 0.757293797 | 0.163898605 | 4.620501775 | 3.83E-06 | 0.000148283 | Ap1m2 | Sod2 | Arid4a | 1714.846966 | -0.000400267 | 0.072151524 | -0.005547587 | 0.995573689 | 0.998915149 | Kbtbd4 | Ogt | | | | C4b | | | |
| H2-Bl | 23.35922168 | 0.751782931 | 0.16559525 | 4.53988222w9 | 5.63E-06 | 0.000201873 | H2-Bl | Cadm1 | Fyn | 4990.959839 | -0.000401076 | 0.041912407 | -0.009569395 | 0.992364844 | 0.99829327 | Pja1 | Wsb1 | | | | Col6a2 | | | |
| Aifm3 | 3496.945545 | 0.745002989 | 0.095699294 | 7.784832698 | 6.98E-15 | 3.73E-12 | Aifm3 | Kctd20 | 1700036G14Rik | 0.23403553 | -0.000407314 | 0.009838564 | -0.041399697 | 0.966977254 | NA | Stip1 | Zfp932 | | | | Robo3 | | | |
| Col5a1 | 1179.554475 | 0.74149563 | 0.165678381 | 4.47551228 | 7.62E-06 | 0.000254543 | Col5a1 | Atpif1 | Defb41 | 0.23403553 | -0.000407314 | 0.009838564 | -0.041399697 | 0.966977254 | NA | Sccpdh | Pnn | | | | Slc38a6 | | | |
| Pnpla3 | 472.15872 | 0.741295259 | 0.133004398 | 5.573464254 | 2.50E-08 | 2.30E-06 | Pnpla3 | Marf1 | G6pc2 | 0.23403553 | -0.000407314 | 0.009838564 | -0.041399697 | 0.966977254 | NA | Gpi1 | Tmem181b-ps | | | | Ifit1 | | | |
| Fmo2 | 121.9446464 | 0.739835358 | 0.155269311 | 4.764852442 | 1.89E-06 | 8.51E-05 | Fmo2 | Ctnnb1 | Gcm2 | 0.23403553 | -0.000407314 | 0.009838564 | -0.041399697 | 0.966977254 | NA | | Uggt2 | | | | Cep70 | | | |
| C1ra | 135.4376341 | 0.736809088 | 0.160800145 | 4.58214193 | 4.60E-06 | 0.000173081 | C1ra | Cbx5 | Gm6307 | 0.23403553 | -0.000407314 | 0.009838564 | -0.041399697 | 0.966977254 | NA | | Stat2 | | | | Acp6 | | | |
| Obscn | 273.2247426 | 0.735838921 | 0.11639606 | 6.321854219 | 2.58E-10 | 4.40E-08 | Obscn | Chchd3 | H2-T3 | 0.23403553 | -0.000407314 | 0.009838564 | -0.041399697 | 0.966977254 | NA | | Pabpn1 | | | | Miat | | | |
| Gm3893 | 469.6865392 | 0.7355971 | 0.163005489 | 4.512713679 | 6.40E-06 | 0.000222847 | Gm3893 | Atp5h | Mir344f | 0.23403553 | -0.000407314 | 0.009838564 | -0.041399697 | 0.966977254 | NA | | Odf2l | | | | Irf9 | | | |
| Casr | 25.82938339 | 0.735098739 | 0.165662058 | 4.437339179 | 9.11E-06 | 0.000292771 | Casr | Rps2 | Rmnd1 | 0.23403553 | -0.000407314 | 0.009838564 | -0.041399697 | 0.966977254 | NA | | Flnb | | | | 4831440E17Rik | | | |
| 5830416P10Rik | 43.74549909 | 0.723926795 | 0.164941543 | 4.388990089 | 1.14E-05 | 0.000353593 | 5830416P10Rik | Nhp2l1 | Ces2b | 1.605141492 | -0.00045197 | 0.027333578 | -0.016535342 | 0.986807307 | NA | | Zcchc7 | | | | C1s | | | |
| Lgals3 | 25.58904438 | 0.722818856 | 0.167068041 | 4.326493872 | 1.52E-05 | 0.000445802 | Lgals3 | Ddost | BC049352 | 34.0869798 | -0.000455904 | 0.089997684 | -0.005065728 | 0.995958151 | 0.998915149 | | D4Wsu53e | | | | Snhg11 | | | |
| Thpo | 233.75172 | 0.720308059 | 0.146402317 | 4.920059153 | 8.65E-07 | 4.33E-05 | Thpo | Cyc1 | 1700030F18Rik | 0.21100899 | -0.000459554 | 0.009729044 | -0.04723532 | 0.962325678 | NA | | Szt2 | | | | Ncrna00085 | | | |
| Ncrna00085 | 411.7530876 | 0.719298456 | 0.137304392 | 5.238714101 | 1.62E-07 | 1.11E-05 | Ncrna00085 | Ndufb11 | 1700123O21Rik | 0.21100899 | -0.000459554 | 0.009729044 | -0.04723532 | 0.962325678 | NA | | Acp6 | | | | Trub1 | | | |
| Ccl9 | 67.04289552 | 0.715476926 | 0.154999199 | 4.616004021 | 3.91E-06 | 0.000150403 | Ccl9 | Srr | Btg4 | 0.21100899 | -0.000459554 | 0.009729044 | -0.04723532 | 0.962325678 | NA | | Akap8 | | | | Zcchc7 | | | |
| Chrd | 1265.389473 | 0.713978323 | 0.109066891 | 6.546242576 | 5.90E-11 | 1.25E-08 | Chrd | D5Ertd579e | Gjb4 | 0.21100899 | -0.000459554 | 0.009729044 | -0.04723532 | 0.962325678 | NA | | Trank1 | | | | Slc14a1 | | | |
| Lgals3bp | 855.1685834 | 0.709110844 | 0.13450516 | 5.271997338 | 1.35E-07 | 9.50E-06 | Lgals3bp | Myl6 | Gm19510 | 0.21100899 | -0.000459554 | 0.009729044 | -0.04723532 | 0.962325678 | NA | | Tnfrsf14 | | | | Ankrd16 | | | |
| Krt2 | 29.95118576 | 0.704135642 | 0.148487189 | 4.74206325 | 2.12E-06 | 9.29E-05 | Krt2 | Tex2 | Phyhipl | 7447.076843 | -0.000466922 | 0.058477177 | -0.007984687 | 0.993629209 | 0.998598767 | | Invs | | | | Uvssa | | | |
| Stx3 | 1104.88517 | 0.70410261 | 0.096457325 | 7.299628209 | 2.89E-13 | 9.37E-11 | Stx3 | Rps5 | Mir219-2 | 2.773660428 | -0.000467625 | 0.038040453 | -0.012292832 | 0.990191986 | NA | | Mov10 | | | | Arglu1 | | | |
| Gm5577 | 181.2691896 | 0.703416888 | 0.147188577 | 4.779018194 | 1.76E-06 | 7.98E-05 | Gm5577 | Pik3r3 | Iah1 | 330.2683267 | -0.000471838 | 0.079246122 | -0.005954088 | 0.995249353 | 0.998915149 | | Ankrd16 | | | | Tbce | | | |
| Nr4a2 | 1623.977445 | 0.699106764 | 0.086432638 | 8.088458038 | 6.04E-16 | 3.90E-13 | Nr4a2 | Arf2 | Xkr5 | 9.957613239 | -0.000477947 | 0.058384258 | -0.008186227 | 0.993468409 | NA | | Izumo4 | | | | Ifi204 | | | |
| Vwa5b2 | 2339.746938 | 0.69844737 | 0.108719406 | 6.424311883 | 1.32E-10 | 2.53E-08 | Vwa5b2 | Uqcrfs1 | Cblc | 0.955625209 | -0.000482164 | 0.023146763 | -0.020830729 | 0.983380685 | NA | | Srsf11 | | | | D4Wsu53e | | | |
| Gm6537 | 63.20351091 | 0.69558237 | 0.153572187 | 4.529351209 | 5.92E-06 | 0.000208816 | Gm6537 | Cask | Lhb | 5.764121871 | -0.000497882 | 0.044729622 | -0.011130916 | 0.991118997 | NA | | Itgam | | | | Gpatch1 | | | |
| C1ql2 | 88.92896909 | 0.693525747 | 0.16803255 | 4.127329774 | 3.67E-05 | 0.00089548 | C1ql2 | Ppp1r11 | Ms4a4b | 0.737538137 | -0.000507005 | 0.017923549 | -0.028287088 | 0.977433179 | NA | | Lgi4 | | | | Zfc3h1 | | | |
| Oas1b | 28.63052397 | 0.69301528 | 0.167831421 | 4.129234428 | 3.64E-05 | 0.0008909 | Oas1b | Cops6 | Purg | 490.1602947 | -0.000509172 | 0.075071439 | -0.006782501 | 0.994588389 | 0.998727174 | | Anxa3 | | | | Cd84 | | | |
| Arntl2 | 162.7786887 | 0.691742609 | 0.118471689 | 5.838885336 | 5.26E-09 | 6.17E-07 | Arntl2 | Ddx6 | Aox3l1 | 1.006701342 | -0.000516357 | 0.019027962 | -0.027136766 | 0.978350651 | NA | | Ifi44 | | | | Ccdc39 | | | |
| Rnf207 | 283.6688619 | 0.69161035 | 0.138242145 | 5.00289076 | 5.65E-07 | 3.10E-05 | Rnf207 | Atg12 | 9430038I01Rik | 46.22349184 | -0.000520973 | 0.094164655 | -0.005532574 | 0.995585667 | 0.998915149 | | Eml5 | | | | Luc7l2 | | | |
| Gm129 | 582.3494173 | 0.688767371 | 0.096355988 | 7.148153293 | 8.80E-13 | 2.64E-10 | Gm129 | Ran | Golt1a | 40.03716177 | -0.000523853 | 0.090997717 | -0.005756772 | 0.995406786 | 0.998915149 | | Tia1 | | | | I830012O16Rik | | | |
| Spag5 | 517.0595542 | 0.688255661 | 0.137033559 | 5.022533642 | 5.10E-07 | 2.84E-05 | Spag5 | Pgam5 | Cxxc5 | 3062.519806 | -0.000551355 | 0.069904417 | -0.007887268 | 0.993706936 | 0.998598767 | | 6720401G13Rik | | | | Apobec3 | | | |
| Ccl17 | 46.85776714 | 0.686370154 | 0.166286631 | 4.1276328 | 3.67E-05 | 0.00089548 | Ccl17 | Fau | Lrp10 | 1327.509155 | -0.000554397 | 0.081730932 | -0.006783192 | 0.994587838 | 0.998727174 | | Akap8l | | | | Stat2 | | | |
| Rnpc3 | 774.9279501 | 0.68345431 | 0.112271155 | 6.087532549 | 1.15E-09 | 1.54E-07 | Rnpc3 | Mlc1 | Atat1 | 2125.760381 | -0.000562476 | 0.054912791 | -0.010243079 | 0.991827348 | 0.99829327 | | 4930570G19Rik | | | | Med12 | | | |
| Ildr1 | 14.60061707 | 0.683200407 | 0.166648258 | 4.099655252 | 4.14E-05 | 0.000990864 | Ildr1 | Astn1 | Ide | 2117.602129 | -0.000563465 | 0.056826898 | -0.009915454 | 0.992088742 | 0.99829327 | | Sfi1 | | | | Vmn2r84 | | | |
| Mslnl | 18.16827862 | 0.683047468 | 0.166031507 | 4.113962958 | 3.89E-05 | 0.000944512 | Mslnl | 2900097C17Rik | 4933425H06Rik | 0.205932006 | -0.000576954 | 0.009704244 | -0.059453793 | 0.952590669 | NA | | Ctsh | | | | Srek1 | | | |
| Kcns1 | 673.7610618 | 0.68286652 | 0.138232449 | 4.939987123 | 7.81E-07 | 4.04E-05 | Kcns1 | Map4k4 | Cyp1a2 | 0.205932006 | -0.000576954 | 0.009704244 | -0.059453793 | 0.952590669 | NA | | Ccdc39 | | | | Catsper2 | | | |
| Ccdc108 | 321.2306714 | 0.681796538 | 0.157066532 | 4.340813587 | 1.42E-05 | 0.000424608 | Ccdc108 | Rab1 | Gm4265 | 0.205932006 | -0.000576954 | 0.009704244 | -0.059453793 | 0.952590669 | NA | | Dusp11 | | | | Rbm5 | | | |
| Zfp185 | 70.88662 | 0.680386425 | 0.165938367 | 4.100235751 | 4.13E-05 | 0.000989912 | Zfp185 | Slmo2 | Hist1h4a | 0.205932006 | -0.000576954 | 0.009704244 | -0.059453793 | 0.952590669 | NA | | Arglu1 | | | | A930013F10Rik | | | |
| Cdc25b | 817.8909763 | 0.678073689 | 0.097581754 | 6.948775353 | 3.68E-12 | 1.02E-09 | Cdc25b | Rala | Igsf23 | 0.205932006 | -0.000576954 | 0.009704244 | -0.059453793 | 0.952590669 | NA | | Tnip2 | | | | Csf3r | | | |
| Agmat | 63.00777509 | 0.677291518 | 0.153191999 | 4.421193802 | 9.82E-06 | 0.000309115 | Agmat | Ndufb3 | Lbx1 | 0.205932006 | -0.000576954 | 0.009704244 | -0.059453793 | 0.952590669 | NA | | Med12 | | | | Flnb | | | |
| Mir24-1 | 29.71920695 | 0.675921481 | 0.167765917 | 4.028955889 | 5.60E-05 | 0.001258048 | Mir24-1 | Ndufa11 | Muc5b | 0.205932006 | -0.000576954 | 0.009704244 | -0.059453793 | 0.952590669 | NA | | Flna | | | | Ncf2 | | | |
| Plekha4 | 85.21933627 | 0.6723507 | 0.144425374 | 4.655350249 | 3.23E-06 | 0.000130501 | Plekha4 | Prps1 | Olfr727 | 0.205932006 | -0.000576954 | 0.009704244 | -0.059453793 | 0.952590669 | NA | | Cx3cr1 | | | | Ap1g2 | | | |
| Slc23a3 | 46.00813392 | 0.669340157 | 0.166061419 | 4.030678308 | 5.56E-05 | 0.001250677 | Slc23a3 | Rplp2 | Snord123 | 0.205932006 | -0.000576954 | 0.009704244 | -0.059453793 | 0.952590669 | NA | | Jag1 | | | | Siglec5 | | | |
| Vsig2 | 62.16208071 | 0.665546841 | 0.152709347 | 4.358258717 | 1.31E-05 | 0.000398291 | Vsig2 | Trappc4 | Sult1d1 | 0.205932006 | -0.000576954 | 0.009704244 | -0.059453793 | 0.952590669 | NA | | Nvl | | | | C1qc | | | |
| Plxdc1 | 1523.841173 | 0.66477181 | 0.118487708 | 5.61047068 | 2.02E-08 | 1.94E-06 | Plxdc1 | Dnajc27 | Gse1 | 3019.740882 | -0.000577692 | 0.080315896 | -0.007192746 | 0.994261069 | 0.998727174 | | Fam227a | | | | Ifit3 | | | |
| BC024139 | 40.83153772 | 0.660358548 | 0.164921055 | 4.004088803 | 6.23E-05 | 0.001364151 | BC024139 | Sdhd | Gpr30 | 141.0321922 | -0.000581497 | 0.089364333 | -0.006507039 | 0.994808171 | 0.998727174 | | Col4a1 | | | | Col19a1 | | | |
| Siglec5 | 32.31699214 | 0.656799616 | 0.164795556 | 3.98554204 | 6.73E-05 | 0.001446814 | Siglec5 | Chmp5 | Gpr18 | 0.458888926 | -0.000592276 | 0.013981662 | -0.042360929 | 0.966210974 | NA | | Fmnl3 | | | | Sfi1 | | | |
| Vill | 288.3142611 | 0.656727208 | 0.106879005 | 6.144585747 | 8.02E-10 | 1.12E-07 | Vill | Rftn2 | St6galnac1 | 0.458888926 | -0.000592276 | 0.013981662 | -0.042360929 | 0.966210974 | NA | | Nckap1l | | | | Zfp280c | | | |
| Col23a1 | 646.4326187 | 0.656065592 | 0.141480978 | 4.637129323 | 3.53E-06 | 0.000139281 | Col23a1 | Hibadh | 1700065D16Rik | 0.998756267 | -0.000597607 | 0.02290445 | -0.026091309 | 0.979184509 | NA | | Dock6 | | | | Ccdc108 | | | |
| Psrc1 | 319.8846057 | 0.655551221 | 0.126468183 | 5.183526855 | 2.18E-07 | 1.42E-05 | Psrc1 | Ndufb5 | AI506816 | 1.858313118 | -0.000606898 | 0.028886122 | -0.021010023 | 0.98323766 | NA | | Slc38a6 | | | | Ttc14 | | | |
| Tdrd5 | 48.3000345 | 0.653685879 | 0.160102213 | 4.082928453 | 4.45E-05 | 0.001041174 | Tdrd5 | Dmxl2 | Bpifb3 | 1.950685126 | -0.000613196 | 0.025782673 | -0.023783274 | 0.981025482 | NA | | Zfc3h1 | | | | Col5a1 | | | |
| Tmc8 | 23.44996665 | 0.653027827 | 0.166773954 | 3.915646377 | 9.02E-05 | 0.0018119 | Tmc8 | Rwdd1 | Glp2r | 110.9693943 | -0.000621402 | 0.095061127 | -0.006536869 | 0.99478437 | 0.998727174 | | Ctss | | | | Irf8 | | | |
| Hnf1b | 16.16815233 | 0.652764166 | 0.166638127 | 3.917255792 | 8.96E-05 | 0.001802188 | Hnf1b | Apba1 | Efhb | 12.24538815 | -0.000622962 | 0.062199486 | -0.01001555 | 0.992008881 | NA | | Ccdc142 | | | | Il18bp | | | |
| Bmp1 | 2709.289137 | 0.652113949 | 0.108619921 | 6.003631224 | 1.93E-09 | 2.45E-07 | Bmp1 | Dmd | Mir494 | 0.42201798 | -0.000629897 | 0.012046709 | -0.052287898 | 0.958299296 | NA | | Atp11b | | | | Ptprc | | | |
| Vamp1 | 6675.022313 | 0.651168277 | 0.101469624 | 6.417371544 | 1.39E-10 | 2.61E-08 | Vamp1 | Slc25a5 | Gm13582 | 0.203314312 | -0.000639443 | 0.009691027 | -0.065983029 | 0.947391337 | NA | | Pion | | | | Odf2l | | | |
| Arhgef37 | 106.9466907 | 0.64857888 | 0.163450613 | 3.9680419 | 7.25E-05 | 0.001531763 | Arhgef37 | St3gal2 | Gm6904 | 0.203314312 | -0.000639443 | 0.009691027 | -0.065983029 | 0.947391337 | NA | | Col4a2 | | | | Slc11a1 | | | |
| C3ar1 | 128.7260767 | 0.647309963 | 0.129734365 | 4.989502709 | 6.05E-07 | 3.28E-05 | C3ar1 | Gcsh | Hoxd8 | 0.203314312 | -0.000639443 | 0.009691027 | -0.065983029 | 0.947391337 | NA | | Cdc25b | | | | Slamf9 | | | |
| BC061194 | 114.0649251 | 0.647240143 | 0.121346098 | 5.333835657 | 9.62E-08 | 7.23E-06 | BC061194 | Bzw2 | Zfp42 | 0.203314312 | -0.000639443 | 0.009691027 | -0.065983029 | 0.947391337 | NA | | I830077J02Rik | | | | Ly86 | | | |
| Slc2a4 | 86.66880659 | 0.647084388 | 0.168121297 | 3.848913851 | 0.000118643 | 0.002230886 | Slc2a4 | 0610009D07Rik | Ap5s1 | 600.7333367 | -0.000649907 | 0.082542392 | -0.007873616 | 0.993717828 | 0.998598767 | | Bmp1 | | | | Nvl | | | |
| Tns4 | 15.05116877 | 0.643322577 | 0.165776906 | 3.880652587 | 0.000104177 | 0.002030329 | Tns4 | 6-Mar | Pcdh8 | 1300.519623 | -0.000652918 | 0.083855261 | -0.007786253 | 0.993787531 | 0.998599729 | | Rgs11 | | | | Cep110 | | | |
| Dnahc9 | 236.1863733 | 0.642961656 | 0.123857143 | 5.191155237 | 2.09E-07 | 1.38E-05 | Dnahc9 | Rprd1a | 4930465M20Rik | 0.758715399 | -0.000653547 | 0.018261323 | -0.035788581 | 0.971450938 | NA | | Arhgef1 | | | | H2-Q4 | | | |
| E130102H24Rik | 136.2400793 | 0.641203881 | 0.148826725 | 4.308392074 | 1.64E-05 | 0.000478933 | E130102H24Rik | 2410015M20Rik | Pcnxl2 | 5259.152887 | -0.000659384 | 0.053010427 | -0.012438764 | 0.990075558 | 0.998028378 | | Tie1 | | | | Gabpb2 | | | |
| Phyhd1 | 512.6650775 | 0.641193533 | 0.149444719 | 4.290506467 | 1.78E-05 | 0.000509604 | Phyhd1 | Slc1a3 | Gpr89 | 962.9290353 | -0.000677926 | 0.058015913 | -0.011685166 | 0.990676799 | 0.998028378 | | Nsun7 | | | | Phka2 | | | |
| Lhx9 | 40.80891809 | 0.640377013 | 0.157658748 | 4.061791818 | 4.87E-05 | 0.001126148 | Lhx9 | Spcs1 | Gtf2h4 | 545.0405969 | -0.000685659 | 0.076461439 | -0.008967379 | 0.992845162 | 0.99829327 | | Tlr2 | | | | Gm5577 | | | |
| Eno3 | 254.4885681 | 0.640140949 | 0.121543379 | 5.266769408 | 1.39E-07 | 9.69E-06 | Eno3 | Nsg2 | C5ar1 | 35.3715493 | -0.000690293 | 0.090250354 | -0.007648647 | 0.993897322 | 0.99864097 | | H2-Q4 | | | | Cd14 | | | |
| Cd68 | 429.4261248 | 0.639284216 | 0.124790287 | 5.12286837 | 3.01E-07 | 1.86E-05 | Cd68 | Myh10 | Clhc1 | 6.822282124 | -0.000705404 | 0.050341925 | -0.01401225 | 0.988820208 | NA | | Ccdc84 | | | | Atp11b | | | |
| Proser2 | 107.7436 | 0.639061454 | 0.126375446 | 5.056848274 | 4.26E-07 | 2.46E-05 | Proser2 | Abcd3 | 4930479M11Rik | 0.220191124 | -0.000706246 | 0.009775455 | -0.072246912 | 0.942405412 | NA | | Pld4 | | | | Gigyf1 | | | |
| Capg | 88.0298116 | 0.638389445 | 0.14043478 | 4.54580728 | 5.47E-06 | 0.000198718 | Capg | Prmt2 | A530099J19Rik | 0.220191124 | -0.000706246 | 0.009775455 | -0.072246912 | 0.942405412 | NA | | Bin2 | | | | Bmp1 | | | |
| Flt3 | 274.0022319 | 0.638340438 | 0.115336276 | 5.534602464 | 3.12E-08 | 2.79E-06 | Flt3 | Slc38a10 | Gm11426 | 0.220191124 | -0.000706246 | 0.009775455 | -0.072246912 | 0.942405412 | NA | | Tatdn3 | | | | Hdac7 | | | |
| Sfrs18 | 5999.036191 | 0.638272853 | 0.122117262 | 5.226720985 | 1.73E-07 | 1.17E-05 | Sfrs18 | Csrnp3 | Pcdha4 | 0.220191124 | -0.000706246 | 0.009775455 | -0.072246912 | 0.942405412 | NA | | Eif2ak2 | | | | Ubp1 | | | |
| Ucn3 | 16.4104385 | 0.638030396 | 0.160218136 | 3.982260761 | 6.83E-05 | 0.001464909 | Ucn3 | Itgav | Prok1 | 0.220191124 | -0.000706246 | 0.009775455 | -0.072246912 | 0.942405412 | NA | | Parp12 | | | | Sfrs18 | | | |
| Tmc6 | 184.8515502 | 0.637606888 | 0.126603053 | 5.036267886 | 4.75E-07 | 2.68E-05 | Tmc6 | Tusc3 | Prkcdbp | 480.2667471 | -0.000727501 | 0.07996088 | -0.009098208 | 0.99274078 | 0.99829327 | | Dennd6b | | | | A230057D06Rik | | | |
| Snhg11 | 57641.62633 | 0.635492944 | 0.159991209 | 3.972049128 | 7.13E-05 | 0.001512406 | Snhg11 | Armcx2 | 2210417A02Rik | 0.430785401 | -0.000733496 | 0.013756879 | -0.053318516 | 0.957478128 | NA | | Ets1 | | | | Pabpn1 | | | |
| Cd14 | 97.32257121 | 0.6321603 | 0.130408646 | 4.847533663 | 1.25E-06 | 5.91E-05 | Cd14 | Tomm22 | Tmem88b | 3043.769463 | -0.000734643 | 0.087718965 | -0.008374963 | 0.993317824 | 0.998545698 | | 1700024P16Rik | | | | Sbsn | | | |
| Aloxe3 | 206.2570327 | 0.6321384 | 0.145349035 | 4.349106274 | 1.37E-05 | 0.000411358 | Aloxe3 | Selrc1 | Klri1 | 0.63302697 | -0.000739291 | 0.013489552 | -0.054804698 | 0.956294057 | NA | | C1qc | | | | Safb2 | | | |
| Kazald1 | 161.9342514 | 0.629186885 | 0.133488978 | 4.713399517 | 2.44E-06 | 0.00010485 | Kazald1 | Uchl1 | 4933411G11Rik | 0.24789674 | -0.00076987 | 0.010036857 | -0.076704253 | 0.938858821 | NA | | Luc7l3 | | | | Cxcl16 | | | |
| Ifit3 | 374.1335853 | 0.627249931 | 0.14617201 | 4.291176741 | 1.78E-05 | 0.000509164 | Ifit3 | Swi5 | Lrtm1 | 0.24789674 | -0.00076987 | 0.010036857 | -0.076704253 | 0.938858821 | NA | | Stx3 | | | | Stx3 | | | |
| Ifi204 | 25.57338141 | 0.626309725 | 0.15667187 | 3.997588879 | 6.40E-05 | 0.001387951 | Ifi204 | Sms | Olfr1350 | 0.24789674 | -0.00076987 | 0.010036857 | -0.076704253 | 0.938858821 | NA | | Cep70 | | | | Nxf1 | | | |
| Pvalb | 1580.452374 | 0.625764924 | 0.137677142 | 4.545162085 | 5.49E-06 | 0.000198718 | Pvalb | Mrps14 | Olfr18 | 0.24789674 | -0.00076987 | 0.010036857 | -0.076704253 | 0.938858821 | NA | | Ifitm3 | | | | Eif2ak2 | | | |
| Shmt1 | 68.83104912 | 0.62358858 | 0.158745107 | 3.928238104 | 8.56E-05 | 0.001745007 | Shmt1 | Chst10 | Ppifos | 0.24789674 | -0.00076987 | 0.010036857 | -0.076704253 | 0.938858821 | NA | | Ccdc159 | | | | Akap8l | | | |
| Gpr84 | 42.23628281 | 0.620402895 | 0.165985651 | 3.737689934 | 0.000185719 | 0.003222314 | Gpr84 | Pin1 | Prss42 | 0.24789674 | -0.00076987 | 0.010036857 | -0.076704253 | 0.938858821 | NA | | Dclre1c | | | | Uggt2 | | | |
| St14 | 36.89020823 | 0.620137979 | 0.167401967 | 3.704484417 | 0.000211821 | 0.003548499 | St14 | B230219D22Rik | Serpinb7 | 0.24789674 | -0.00076987 | 0.010036857 | -0.076704253 | 0.938858821 | NA | | Ipw | | | | Ddx17 | | | |
| Il16 | 158.1856135 | 0.619670187 | 0.163541933 | 3.789059941 | 0.000151218 | 0.002724393 | Il16 | Chchd4 | Uts2d | 0.24789674 | -0.00076987 | 0.010036857 | -0.076704253 | 0.938858821 | NA | | Ptprv | | | | Leng8 | | | |
| Gm13629 | 161.09298 | 0.615423522 | 0.132135136 | 4.657531217 | 3.20E-06 | 0.000129463 | Gm13629 | Armc1 | Mageb16-ps1 | 0.428167708 | -0.000792272 | 0.013735474 | -0.057680729 | 0.954002944 | NA | | Col6a2 | | | | Guf1 | | | |
| Syt2 | 1570.135915 | 0.615048048 | 0.111094675 | 5.5362514 | 3.09E-08 | 2.78E-06 | Syt2 | Dnajb11 | Serpinc1 | 0.428167708 | -0.000792272 | 0.013735474 | -0.057680729 | 0.954002944 | NA | | Lyn | | | | Usp48 | | | |
| 5930438M14Rik | 17.45696186 | 0.613516477 | 0.155833964 | 3.936988205 | 8.25E-05 | 0.001691032 | 5930438M14Rik | Pdxp | Cnih2 | 5558.486328 | -0.000796695 | 0.067242655 | -0.01184806 | 0.990546837 | 0.998028378 | | Lpcat2 | | | | Szt2 | | | |
| Npc1l1 | 78.63167845 | 0.611107303 | 0.149289547 | 4.093436645 | 4.25E-05 | 0.001010024 | Npc1l1 | Guk1 | Asap1 | 6955.515529 | -0.000798184 | 0.063374786 | -0.012594655 | 0.989951185 | 0.998028378 | | Rasal3 | | | | Rnf207 | | | |
| Mylk3 | 141.5205768 | 0.610234573 | 0.136596792 | 4.467415121 | 7.92E-06 | 0.000261549 | Mylk3 | Tmem242 | Ntan1 | 1448.107902 | -0.000805296 | 0.066306611 | -0.012145037 | 0.990309901 | 0.998028378 | | Liph | | | | | | | |
| Hdc | 64.38314051 | 0.607773155 | 0.153306357 | 3.964435439 | 7.36E-05 | 0.001544568 | Hdc | Smarca2 | Clca5 | 0.411864012 | -0.000821183 | 0.011963678 | -0.06863967 | 0.945276441 | NA | | Lgals9 | | | | | | | |
| Six4 | 61.13542265 | 0.607204373 | 0.154777094 | 3.923089368 | 8.74E-05 | 0.001768185 | Six4 | Dync1h1 | Smg5 | 4552.065266 | -0.000821289 | 0.050388549 | -0.016299129 | 0.986995753 | 0.997691964 | | Pnpla3 | | | | | | | |
| I830012O16Rik | 207.5166096 | 0.606921303 | 0.141195899 | 4.298434365 | 1.72E-05 | 0.000495373 | I830012O16Rik | Rps20 | 4930443O20Rik | 0.21511414 | -0.000824241 | 0.009751353 | -0.084525787 | 0.932638401 | NA | | Ccdc108 | | | | | | | |
| A930011O12Rik | 6007.734216 | 0.606534719 | 0.157869559 | 3.841999207 | 0.000122036 | 0.002280856 | A930011O12Rik | Timm22 | Fcrla | 0.21511414 | -0.000824241 | 0.009751353 | -0.084525787 | 0.932638401 | NA | | C1s | | | | | | | |
| Pparg | 162.7000464 | 0.606513618 | 0.127443221 | 4.759088908 | 1.94E-06 | 8.68E-05 | Pparg | Vta1 | Foxi2 | 0.21511414 | -0.000824241 | 0.009751353 | -0.084525787 | 0.932638401 | NA | | Csf2rb | | | | | | | |
| T2 | 28.84656258 | 0.602333471 | 0.166790932 | 3.611308271 | 0.000304656 | 0.004725069 | T2 | Higd1a | Gapt | 0.21511414 | -0.000824241 | 0.009751353 | -0.084525787 | 0.932638401 | NA | | 2610005L07Rik | | | | | | | |
| Kndc1 | 4882.897485 | 0.602118247 | 0.089168546 | 6.752585668 | 1.45E-11 | 3.46E-09 | Kndc1 | Rit1 | Gk2 | 0.21511414 | -0.000824241 | 0.009751353 | -0.084525787 | 0.932638401 | NA | | Srrm2 | | | | | | | |
| Emilin2 | 208.3914897 | 0.601685066 | 0.154238834 | 3.900995943 | 9.58E-05 | 0.001895646 | Emilin2 | Ext1 | Gm10324 | 0.21511414 | -0.000824241 | 0.009751353 | -0.084525787 | 0.932638401 | NA | | Chuk | | | | | | | |
| Cbln4 | 572.1106744 | 0.601274644 | 0.148253734 | 4.055713328 | 5.00E-05 | 0.001145662 | Cbln4 | Ep400 | Gm6567 | 0.21511414 | -0.000824241 | 0.009751353 | -0.084525787 | 0.932638401 | NA | | Ap1g2 | | | | | | | |
| Tex40 | 206.3128231 | 0.600476649 | 0.146957363 | 4.086060327 | 4.39E-05 | 0.001031585 | Tex40 | Ndufa13 | Pira2 | 0.21511414 | -0.000824241 | 0.009751353 | -0.084525787 | 0.932638401 | NA | | Ifi203 | | | | | | | |
| 5530601H04Rik | 333.8879106 | 0.599554699 | 0.097243256 | 6.165514455 | 7.03E-10 | 1.03E-07 | 5530601H04Rik | Zfp330 | Zfp600 | 0.21511414 | -0.000824241 | 0.009751353 | -0.084525787 | 0.932638401 | NA | | AF251705 | | | | | | | |
| Podn | 206.8579342 | 0.59878885 | 0.141278473 | 4.238358726 | 2.25E-05 | 0.000613905 | Podn | H3f3b | 4930455B14Rik | 0.212496447 | -0.000886988 | 0.009738512 | -0.091080455 | 0.927428663 | NA | | Irf1 | | | | | | | |
| Eif2s3x | 6066.41865 | 0.592400434 | 0.060726273 | 9.755257506 | 1.75E-22 | 1.81E-19 | Eif2s3x | Thnsl1 | 4930553E22Rik | 0.212496447 | -0.000886988 | 0.009738512 | -0.091080455 | 0.927428663 | NA | | Ddx26b | | | | | | | |
| Gsta3 | 55.33800495 | 0.590936087 | 0.162360854 | 3.639646329 | 0.000273013 | 0.004325214 | Gsta3 | Eif2s1 | Cldn18 | 0.212496447 | -0.000886988 | 0.009738512 | -0.091080455 | 0.927428663 | NA | | Slc14a1 | | | | | | | |
| A930013F10Rik | 293.0005318 | 0.590065387 | 0.160459347 | 3.677351282 | 0.000235668 | 0.003831528 | A930013F10Rik | Arl15 | Fbxw24 | 0.212496447 | -0.000886988 | 0.009738512 | -0.091080455 | 0.927428663 | NA | | Cdk2 | | | | | | | |
| Hpn | 42.1963727 | 0.589751697 | 0.16672172 | 3.537341726 | 0.000404176 | 0.005876062 | Hpn | Timm17b | Fcer1a | 0.212496447 | -0.000886988 | 0.009738512 | -0.091080455 | 0.927428663 | NA | | Vmn2r84 | | | | | | | |
| Myh7b | 303.2394482 | 0.589056212 | 0.092248831 | 6.385514096 | 1.71E-10 | 3.11E-08 | Myh7b | Csgalnact1 | Gm11747 | 0.212496447 | -0.000886988 | 0.009738512 | -0.091080455 | 0.927428663 | NA | | Csf2ra | | | | | | | |
| Pcolce2 | 167.7500345 | 0.588346204 | 0.13816828 | 4.25818576 | 2.06E-05 | 0.000572258 | Pcolce2 | Eef1e1 | Gm20611 | 0.212496447 | -0.000886988 | 0.009738512 | -0.091080455 | 0.927428663 | NA | | Lat2 | | | | | | | |
| Cst7 | 45.35413944 | 0.586135197 | 0.131382376 | 4.4612924 | 8.15E-06 | 0.000266837 | Cst7 | Stard10 | Gm6642 | 0.212496447 | -0.000886988 | 0.009738512 | -0.091080455 | 0.927428663 | NA | | Uvssa | | | | | | | |
| Cchcr1 | 200.8938411 | 0.586079184 | 0.137577279 | 4.259999808 | 2.04E-05 | 0.000569675 | Cchcr1 | Acly | Muc6 | 0.212496447 | -0.000886988 | 0.009738512 | -0.091080455 | 0.927428663 | NA | | Itgb2 | | | | | | | |
| Plcz1 | 36.57405176 | 0.584331196 | 0.168010054 | 3.477953753 | 0.000505257 | 0.006882848 | Plcz1 | Fam63b | Prss29 | 0.212496447 | -0.000886988 | 0.009738512 | -0.091080455 | 0.927428663 | NA | | Cyth4 | | | | | | | |
| Xaf1 | 204.349273 | 0.583772651 | 0.140600049 | 4.15200887 | 3.30E-05 | 0.000827609 | Xaf1 | Rasal2 | Gast | 2.16200003 | -0.000890185 | 0.032139514 | -0.027697525 | 0.977903398 | NA | | Gm3002 | | | | | | | |
| Klf14 | 19.25838698 | 0.58086797 | 0.167768834 | 3.462311544 | 0.000535557 | 0.007178129 | Klf14 | 2810407C02Rik | Mir486 | 1.485314722 | -0.000893977 | 0.025576129 | -0.034953586 | 0.972116751 | NA | | Arhgap30 | | | | | | | |
| 2010111I01Rik | 631.3388566 | 0.579226212 | 0.114069079 | 5.077854732 | 3.82E-07 | 2.25E-05 | 2010111I01Rik | Fam160a2 | Cox14 | 1395.505238 | -0.00089691 | 0.051585202 | -0.017386957 | 0.986127914 | 0.997552088 | | Hk2 | | | | | | | |
| Prrg4 | 26.58797669 | 0.57670018 | 0.166887056 | 3.455631576 | 0.000549005 | 0.007320384 | Prrg4 | Ptcd2 | Ska1 | 1.165949806 | -0.000899482 | 0.027248569 | -0.033010239 | 0.973666422 | NA | | Lcp1 | | | | | | | |
| Ankrd35 | 555.7922512 | 0.576004171 | 0.095004608 | 6.062907696 | 1.34E-09 | 1.76E-07 | Ankrd35 | Wnk1 | Lig4 | 537.6282132 | -0.000903855 | 0.083260165 | -0.010855794 | 0.991338499 | 0.998279587 | | Irf9 | | | | | | | |
| Bst2 | 133.3812876 | 0.575379262 | 0.165055871 | 3.48596665 | 0.000490362 | 0.006749647 | Bst2 | Pdzd8 | 1700017G19Rik | 0.192890664 | -0.000903865 | 0.009635283 | -0.093807822 | 0.925261818 | NA | | Lacc1 | | | | | | | |
| Ap1g2 | 260.8641313 | 0.574648173 | 0.131629733 | 4.365641102 | 1.27E-05 | 0.000388676 | Ap1g2 | Rdh14 | 1700026D11Rik | 0.192890664 | -0.000903865 | 0.009635283 | -0.093807822 | 0.925261818 | NA | | Gbp5 | | | | | | | |
| Nudt17 | 214.4373674 | 0.574134011 | 0.117377266 | 4.89135614 | 1.00E-06 | 4.88E-05 | Nudt17 | Nudt19 | 1700027A15Rik | 0.192890664 | -0.000903865 | 0.009635283 | -0.093807822 | 0.925261818 | NA | | Unc93b1 | | | | | | | |
| Krt12 | 436.6734184 | 0.573160843 | 0.104139081 | 5.503801615 | 3.72E-08 | 3.25E-06 | Krt12 | Nme2 | 2310081J21Rik | 0.192890664 | -0.000903865 | 0.009635283 | -0.093807822 | 0.925261818 | NA | | Col11a2 | | | | | | | |
| Gm3002 | 494.5042043 | 0.572045443 | 0.145621492 | 3.92830369 | 8.55E-05 | 0.001745007 | Gm3002 | Cpne5 | 4930453H23Rik | 0.192890664 | -0.000903865 | 0.009635283 | -0.093807822 | 0.925261818 | NA | | Ncrna00085 | | | | | | | |
| Atf3 | 52.25598925 | 0.571528818 | 0.168120904 | 3.399510731 | 0.000675065 | 0.008462348 | Atf3 | Jkamp | D930007P13Rik | 0.192890664 | -0.000903865 | 0.009635283 | -0.093807822 | 0.925261818 | NA | | Cd48 | | | | | | | |
| Plb1 | 48.81217099 | 0.57120599 | 0.163632616 | 3.490783215 | 0.000481607 | 0.006658136 | Plb1 | Vps29 | Fgf21 | 0.192890664 | -0.000903865 | 0.009635283 | -0.093807822 | 0.925261818 | NA | | Capg | | | | | | | |
| Kcnc3 | 5344.180163 | 0.570918041 | 0.090143566 | 6.333430868 | 2.40E-10 | 4.13E-08 | Kcnc3 | Ica1l | Gm16130 | 0.192890664 | -0.000903865 | 0.009635283 | -0.093807822 | 0.925261818 | NA | | A230057D06Rik | | | | | | | |
| Amy1 | 699.7132709 | 0.57065479 | 0.14427671 | 3.955280025 | 7.64E-05 | 0.00159628 | Amy1 | 2510009E07Rik | Hist1h3e | 0.192890664 | -0.000903865 | 0.009635283 | -0.093807822 | 0.925261818 | NA | | Sp100 | | | | | | | |
| Pdcd1 | 8.41787075 | 0.570292581 | 0.135497191 | 4.208888584 | 2.57E-05 | 0.000679696 | Pdcd1 | Chga | Mrgpra2b | 0.192890664 | -0.000903865 | 0.009635283 | -0.093807822 | 0.925261818 | NA | | Ech1 | | | | | | | |
| Il18bp | 142.584881 | 0.569456576 | 0.158984556 | 3.581835802 | 0.000341188 | 0.005167517 | Il18bp | Ndufb7 | Noxa1 | 0.192890664 | -0.000903865 | 0.009635283 | -0.093807822 | 0.925261818 | NA | | Ifit3 | | | | | | | |
| Aldob | 49.26684737 | 0.569412404 | 0.1666312 | 3.417201609 | 0.000632684 | 0.008095075 | Aldob | Mea1 | Rnf133 | 0.192890664 | -0.000903865 | 0.009635283 | -0.093807822 | 0.925261818 | NA | | Slamf9 | | | | | | | |
| Mapk13 | 24.59753223 | 0.569140682 | 0.166422005 | 3.419864339 | 0.000626524 | 0.008054403 | Mapk13 | Ppm1l | Serpinb12 | 0.192890664 | -0.000903865 | 0.009635283 | -0.093807822 | 0.925261818 | NA | | Cep110 | | | | | | | |
| Ifi30 | 113.1439711 | 0.567974349 | 0.138705738 | 4.094815075 | 4.23E-05 | 0.001005576 | Ifi30 | Zfp407 | Tac4 | 0.192890664 | -0.000903865 | 0.009635283 | -0.093807822 | 0.925261818 | NA | | Xaf1 | | | | | | | |
| Snapc4 | 1839.848641 | 0.565921866 | 0.101054208 | 5.600181114 | 2.14E-08 | 2.03E-06 | Snapc4 | Pcdhb20 | Zbtb24 | 577.5114593 | -0.000911935 | 0.06674666 | -0.013662625 | 0.989099142 | 0.998005844 | | Rnf207 | | | | | | | |
| Akr1b8 | 32.90242689 | 0.565121734 | 0.16721155 | 3.379681221 | 0.000725699 | 0.0089238 | Akr1b8 | Cnep1r1 | Ube2r2 | 3400.369608 | -0.000912615 | 0.051495165 | -0.017722341 | 0.985860358 | 0.997552088 | | Csf2rb2 | | | | | | | |
| Nxf7 | 91.38510561 | 0.564298089 | 0.161314342 | 3.498127206 | 0.000468538 | 0.006510782 | Nxf7 | Rpl15 | D7Ertd443e | 218.4410538 | -0.000913445 | 0.094484906 | -0.009667625 | 0.992286472 | 0.99829327 | | Fes | | | | | | | |
| B230312C02Rik | 25.57658329 | 0.564086595 | 0.16809139 | 3.355832779 | 0.000791264 | 0.009492476 | B230312C02Rik | Lsm6 | Iqcb1 | 995.2776678 | -0.000915244 | 0.064996278 | -0.014081479 | 0.988764977 | 0.998005844 | | Samsn1 | | | | | | | |
| Rtp4 | 88.21087806 | 0.563871015 | 0.158718159 | 3.552655975 | 0.000381363 | 0.005638202 | Rtp4 | Bloc1s1 | F5 | 171.4930781 | -0.000919092 | 0.031192135 | -0.029465489 | NA | NA | | Catsper2 | | | | | | | |
| Cpeb1 | 2576.114924 | 0.562557535 | 0.089226242 | 6.304844023 | 2.88E-10 | 4.81E-08 | Cpeb1 | Cnih3 | Cyp3a57 | 0.406628625 | -0.000921994 | 0.011919976 | -0.07734862 | 0.938346213 | NA | | Hdac7 | | | | | | | |
| Uggt2 | 1071.344552 | 0.562437283 | 0.090183341 | 6.236598431 | 4.47E-10 | 7.00E-08 | Uggt2 | Pdpk1 | Gm9999 | 0.406628625 | -0.000921994 | 0.011919976 | -0.07734862 | 0.938346213 | NA | | Thpo | | | | | | | |
| Mafb | 843.6973662 | 0.562398299 | 0.099433559 | 5.656020989 | 1.55E-08 | 1.56E-06 | Mafb | Gabra3 | Vmn2r13 | 0.406628625 | -0.000921994 | 0.011919976 | -0.07734862 | 0.938346213 | NA | | Was | | | | | | | |
| Hspb6 | 190.731772 | 0.560143428 | 0.125651625 | 4.457908359 | 8.28E-06 | 0.000269398 | Hspb6 | Rps21 | Msh4 | 0.97245336 | -0.000942409 | 0.019228272 | -0.04901162 | 0.960910036 | NA | | Hvcn1 | | | | | | | |
| Casp1 | 96.12265149 | 0.559826726 | 0.158548855 | 3.530941469 | 0.000414083 | 0.005968193 | Casp1 | Hsd17b12 | Arhgef33 | 58.43782204 | -0.000948654 | 0.092857968 | -0.010216184 | 0.991848806 | 0.99829327 | | Ifit1 | | | | | | | |
| Vmn2r84 | 21.89585925 | 0.55928314 | 0.162672249 | 3.438098043 | 0.000585816 | 0.007673777 | Vmn2r84 | Tmem70 | Gm16432 | 26.88371015 | -0.000955966 | 0.086669176 | -0.011030056 | 0.991199467 | 0.998279587 | | Plek | | | | | | | |
| Myo19 | 356.7218086 | 0.558430401 | 0.132652582 | 4.209721317 | 2.56E-05 | 0.000678356 | Myo19 | H2afv | AI462493 | 534.6690026 | -0.000960553 | 0.067411457 | -0.014249105 | 0.988631244 | 0.997974805 | | Col5a1 | | | | | | | |
| Tbc1d10c | 47.97776154 | 0.55737192 | 0.15237634 | 3.657863946 | 0.000254326 | 0.004091928 | Tbc1d10c | Ndufb8 | Agl | 2103.422878 | -0.000960751 | 0.059762761 | -0.016076085 | 0.987173692 | 0.997691964 | | Gpatch1 | | | | | | | |
| Spata32 | 28.38503249 | 0.552239221 | 0.162390024 | 3.400696708 | 0.000672144 | 0.008432544 | Spata32 | Atp5g3 | Xrcc1 | 1058.952446 | -0.000961346 | 0.061536776 | -0.015622304 | 0.987535712 | 0.997717928 | | Phf11d | | | | | | | |
| Cpne9 | 1962.2034 | 0.552008186 | 0.120911553 | 4.565388273 | 4.99E-06 | 0.000184364 | Cpne9 | Pds5b | Tpd52 | 2731.114212 | -0.000966893 | 0.057900826 | -0.01669912 | 0.986676649 | 0.997691964 | | Tmc4 | | | | | | | |
| Parp3 | 215.7890304 | 0.551963467 | 0.134055047 | 4.117438925 | 3.83E-05 | 0.000931844 | Parp3 | Rps16 | 4933402J15Rik | 1.03580242 | -0.000969439 | 0.022983628 | -0.042179551 | 0.966355564 | NA | | Ddx58 | | | | | | | |
| Rorc | 95.43351474 | 0.551602804 | 0.129582629 | 4.256765031 | 2.07E-05 | 0.000573848 | Rorc | Ralyl | Glt28d2 | 14.31486856 | -0.000984271 | 0.072994021 | -0.013484264 | 0.98924144 | NA | | 4831440E17Rik | | | | | | | |
| D4Wsu53e | 5896.814657 | 0.551128304 | 0.106799626 | 5.160395456 | 2.46E-07 | 1.56E-05 | D4Wsu53e | Serp2 | 4933404K08Rik | 0.256588971 | -0.000984451 | 0.010155606 | -0.096936706 | 0.922776659 | NA | | Spag5 | | | | | | | |
| Npff | 34.08243052 | 0.550987823 | 0.158224005 | 3.482327619 | 0.000497075 | 0.006796205 | Npff | Snx18 | C87499 | 0.256588971 | -0.000984451 | 0.010155606 | -0.096936706 | 0.922776659 | NA | | Mlxipl | | | | | | | |
| Pagr1a | 1471.115104 | 0.550577507 | 0.081945903 | 6.718792389 | 1.83E-11 | 4.24E-09 | Pagr1a | Gde1 | Sh2d1a | 0.636717407 | -0.00100889 | 0.01541517 | -0.065447861 | 0.947817418 | NA | | Parp9 | | | | | | | |
| Cerkl | 46.25713664 | 0.55052048 | 0.157939409 | 3.485643543 | 0.000490955 | 0.006749647 | Cerkl | Mzt1 | 2310039L15Rik | 20.82079808 | -0.001016019 | 0.084007027 | -0.012094452 | 0.990350258 | NA | | Ncf2 | | | | | | | |
| Esyt1 | 596.2409955 | 0.55043157 | 0.106174214 | 5.184230248 | 2.17E-07 | 1.42E-05 | Esyt1 | Fam174b | Fdxacb1 | 183.037781 | -0.001029783 | 0.085794613 | -0.012002883 | 0.990423315 | 0.998028378 | | Csf3r | | | | | | | |
| 4930570G19Rik | 287.7082891 | 0.549468517 | 0.12580114 | 4.367754674 | 1.26E-05 | 0.000386674 | 4930570G19Rik | Fxyd6 | Speer4a | 2.761868621 | -0.001031098 | 0.036931336 | -0.027919328 | 0.977726493 | NA | | Kctd15 | | | | | | | |
| Abhd14b | 557.6240189 | 0.549429402 | 0.084519451 | 6.500626688 | 8.00E-11 | 1.65E-08 | Abhd14b | Dolk | Brpf1 | 1552.828095 | -0.001033798 | 0.050951096 | -0.020290005 | 0.983812029 | 0.996711782 | | Siglec5 | | | | | | | |
| Cd84 | 164.8309008 | 0.548032146 | 0.12855625 | 4.2629755 | 2.02E-05 | 0.000563151 | Cd84 | Psmb5 | Slc10a5 | 0.41774406 | -0.001036768 | 0.013648931 | -0.075959622 | 0.939451223 | NA | | Trim30a | | | | | | | |
| Il12rb1 | 36.82283005 | 0.547840569 | 0.163880434 | 3.342928467 | 0.000828993 | 0.009842462 | Il12rb1 | Pnma2 | Plce1 | 621.2339759 | -0.001040382 | 0.081625554 | -0.012745787 | 0.989830609 | 0.998028378 | | Sfrs18 | | | | | | | |
| Crhr2 | 67.63715087 | 0.54778498 | 0.165427121 | 3.311337191 | 0.000928513 | 0.010720099 | Crhr2 | Mrps21 | Ptprcap | 5.423080402 | -0.001042194 | 0.052816627 | -0.019732311 | 0.984256915 | NA | | Zfp36 | | | | | | | |
| Stc2 | 33.38240404 | 0.547734858 | 0.162170251 | 3.37752981 | 0.0007314 | 0.008972538 | Stc2 | Cox7b | Akr1b7 | 1.471065161 | -0.001042559 | 0.026207184 | -0.039781422 | 0.968267387 | NA | | Man2b1 | | | | | | | |
| Evc | 391.0452063 | 0.546734199 | 0.1118509 | 4.888062606 | 1.02E-06 | 4.95E-05 | Evc | Nme1 | Galk2 | 935.1282756 | -0.001049982 | 0.065062557 | -0.016138042 | 0.987124264 | 0.997691964 | | Ptprc | | | | | | | |
| Col19a1 | 1105.15736 | 0.54631796 | 0.106679956 | 5.121092838 | 3.04E-07 | 1.87E-05 | Col19a1 | Usp31 | Fscn2 | 7.360289024 | -0.001073294 | 0.058618833 | -0.01830972 | 0.985391774 | NA | | Col19a1 | | | | | | | |
| Lilrb4 | 18.24260137 | 0.54491141 | 0.158174217 | 3.445007784 | 0.000571043 | 0.007510814 | Lilrb4 | Fam49a | Sema3d | 206.5692842 | -0.001079751 | 0.085597986 | -0.012614212 | 0.989935582 | 0.998028378 | | Miat | | | | | | | |
| Xylb | 299.337968 | 0.544762963 | 0.111693663 | 4.877295171 | 1.08E-06 | 5.16E-05 | Xylb | Ppdpf | Aldh8a1 | 2.991591206 | -0.001088468 | 0.039223953 | -0.027750082 | 0.977861479 | NA | | Zfp217 | | | | | | | |
| Emx1 | 445.9827917 | 0.544150679 | 0.112441129 | 4.839427388 | 1.30E-06 | 6.10E-05 | Emx1 | Prdx3 | Smarcal1 | 3782.814542 | -0.001100323 | 0.04863915 | -0.022622162 | 0.981951666 | 0.996444753 | | Aspg | | | | | | | |
| Met | 263.0066217 | 0.543404877 | 0.130774315 | 4.15528751 | 3.25E-05 | 0.000822495 | Met | Mrpl13 | Dhx16 | 1128.052173 | -0.001111174 | 0.066903028 | -0.016608731 | 0.986748759 | 0.997691964 | | Slc15a3 | | | | | | | |
| Abcc8 | 1206.904999 | 0.54237955 | 0.119633736 | 4.533667257 | 5.80E-06 | 0.000206 | Abcc8 | Myo10 | Igf2bp1 | 1.152334539 | -0.001111523 | 0.023739556 | -0.046821573 | 0.962655435 | NA | | Gbp4 | | | | | | | |
| Fcgr4 | 18.21659617 | 0.541197242 | 0.14928507 | 3.625260337 | 0.000288671 | 0.00451328 | Fcgr4 | Sephs2 | Vangl2 | 254.6860014 | -0.001123441 | 0.08824517 | -0.012730909 | 0.989842478 | 0.998028378 | | Ccl6 | | | | | | | |
| C3 | 41.06919178 | 0.541143698 | 0.16673764 | 3.245480131 | 0.001172527 | 0.012845805 | C3 | AI597468 | Foxo4 | 809.6526748 | -0.001128599 | 0.070879118 | -0.015922877 | 0.987295919 | 0.997717928 | | Ttc14 | | | | | | | |
| Acr | 39.6630239 | 0.540924191 | 0.167977053 | 3.220226707 | 0.001280893 | 0.013715376 | Acr | D630045J12Rik | Acot10 | 0.202072798 | -0.001151076 | 0.009684293 | -0.118860092 | 0.9053862 | NA | | Trim30d | | | | | | | |
| Serpina3n | 821.3954831 | 0.53937954 | 0.163707744 | 3.29477107 | 0.000985019 | 0.011255076 | Serpina3n | Slc1a2 | Gm6251 | 0.202072798 | -0.001151076 | 0.009684293 | -0.118860092 | 0.9053862 | NA | | Aif1 | | | | | | | |
| Miip | 522.0950795 | 0.538604951 | 0.113464141 | 4.746917819 | 2.07E-06 | 9.12E-05 | Miip | Scn2a1 | Pth | 0.202072798 | -0.001151076 | 0.009684293 | -0.118860092 | 0.9053862 | NA | | Tnfsf10 | | | | | | | |
| Luc7l3 | 4668.850558 | 0.538298836 | 0.107401245 | 5.012035354 | 5.39E-07 | 2.99E-05 | Luc7l3 | Casd1 | Rdh19 | 0.202072798 | -0.001151076 | 0.009684293 | -0.118860092 | 0.9053862 | NA | | Ctsc | | | | | | | |
| Mir3072 | 34.80119347 | 0.538047947 | 0.168120306 | 3.200374545 | 0.001372491 | 0.014433037 | Mir3072 | Tceb2 | Mrps18b | 833.4781751 | -0.001152745 | 0.067902576 | -0.01697645 | 0.986455403 | 0.997581728 | | Haus5 | | | | | | | |
| Hdac7 | 1242.897745 | 0.538011846 | 0.129227496 | 4.163292359 | 3.14E-05 | 0.000798084 | Hdac7 | Eif5a2 | 4930417O13Rik | 2.734422186 | -0.001159677 | 0.039576387 | -0.029302233 | 0.976623546 | NA | | Cd86 | | | | | | | |
| Col5a3 | 202.0086317 | 0.537715696 | 0.162112967 | 3.316919707 | 0.000910158 | 0.010547482 | Col5a3 | Notch1 | B230214G05Rik | 0.22897535 | -0.001163642 | 0.00981572 | -0.118548816 | 0.905632818 | NA | | Dock2 | | | | | | | |
| Syne4 | 85.68276377 | 0.537357883 | 0.160733436 | 3.3431618 | 0.000828296 | 0.009842462 | Syne4 | Snrpd2 | Ccdc64b | 0.22897535 | -0.001163642 | 0.00981572 | -0.118548816 | 0.905632818 | NA | | Meg3 | | | | | | | |
| Srpk3 | 215.437209 | 0.536711183 | 0.144951976 | 3.702682767 | 0.000213332 | 0.003556118 | Srpk3 | 1700025G04Rik | D030025P21Rik | 0.22897535 | -0.001163642 | 0.00981572 | -0.118548816 | 0.905632818 | NA | | Lair1 | | | | | | | |
| Clk1 | 2833.366666 | 0.534807707 | 0.137186122 | 3.8984097 | 9.68E-05 | 0.001913558 | Clk1 | Cplx2 | Folr4 | 0.22897535 | -0.001163642 | 0.00981572 | -0.118548816 | 0.905632818 | NA | | Gm15328 | | | | | | | |
| Catsper2 | 93.83404998 | 0.533968231 | 0.151466892 | 3.525313193 | 0.000422982 | 0.006043541 | Catsper2 | Elovl5 | Gc | 0.22897535 | -0.001163642 | 0.00981572 | -0.118548816 | 0.905632818 | NA | | Zkscan2 | | | | | | | |
| Shisa6 | 968.6956184 | 0.532741747 | 0.132302384 | 4.026698007 | 5.66E-05 | 0.001268347 | Shisa6 | Gria3 | Ifi44l | 0.22897535 | -0.001163642 | 0.00981572 | -0.118548816 | 0.905632818 | NA | | Parp14 | | | | | | | |
| Pbld2 | 41.93246654 | 0.532493738 | 0.155085353 | 3.433552731 | 0.000595726 | 0.00777606 | Pbld2 | Akap11 | Olfr46 | 0.22897535 | -0.001163642 | 0.00981572 | -0.118548816 | 0.905632818 | NA | | Emr1 | | | | | | | |
| Sh2b2 | 199.8019305 | 0.532274438 | 0.139943963 | 3.803482681 | 0.000142676 | 0.002600731 | Sh2b2 | Kcnd2 | 1700125G22Rik | 0.481442367 | -0.001165664 | 0.014155216 | -0.082348727 | 0.934369408 | NA | | Malat1 | | | | | | | |
| Ovgp1 | 285.7400495 | 0.530035867 | 0.167759261 | 3.159502877 | 0.001580385 | 0.016067252 | Ovgp1 | Gabarapl2 | Usp44 | 0.481442367 | -0.001165664 | 0.014155216 | -0.082348727 | 0.934369408 | NA | | Leng8 | | | | | | | |
| Ttc14 | 3698.981742 | 0.529844484 | 0.164163585 | 3.227539684 | 0.001248597 | 0.013490771 | Ttc14 | Ndufa8 | Mettl7b | 0.431200114 | -0.00116576 | 0.013763029 | -0.084702256 | 0.932498103 | NA | | Slc11a1 | | | | | | | |
| Meg3 | 106402.8544 | 0.529597234 | 0.16278939 | 3.253266294 | 0.001140865 | 0.012610872 | Meg3 | Reps2 | Wdr86 | 42.30540776 | -0.001167001 | 0.041231709 | -0.02830348 | 0.977420105 | 0.99502449 | | Cxcl16 | | | | | | | |
| Ptprc | 162.9082999 | 0.528853482 | 0.127446535 | 4.149610524 | 3.33E-05 | 0.000833626 | Ptprc | Zeb1 | Pwwp2a | 900.8080263 | -0.001178867 | 0.07338625 | -0.016063872 | 0.987183436 | 0.997691964 | | Itgax | | | | | | | |
| Hr | 1742.726301 | 0.528200436 | 0.103777027 | 5.089762665 | 3.59E-07 | 2.15E-05 | Hr | Gatad1 | Epgn | 2.744144728 | -0.001181049 | 0.034617385 | -0.034117224 | 0.972783674 | NA | | Ly86 | | | | | | | |
| Eqtn | 30.09263517 | 0.52756931 | 0.166071736 | 3.176755564 | 0.001489325 | 0.01539287 | Eqtn | Erh | Samd10 | 964.6014628 | -0.001195831 | 0.056201889 | -0.021277425 | 0.983024352 | 0.996711782 | | Il18bp | | | | | | | |
| Pnpla7 | 679.3112604 | 0.527251026 | 0.080060702 | 6.585640835 | 4.53E-11 | 9.75E-09 | Pnpla7 | Atp5g1 | Klre1 | 0.48193227 | -0.001205689 | 0.014158662 | -0.085155553 | 0.932137725 | NA | | Apobec3 | | | | | | | |
| Efcab12 | 208.3753231 | 0.52700164 | 0.13504409 | 3.902441338 | 9.52E-05 | 0.001886767 | Efcab12 | Celf4 | 4921509O07Rik | 1.00093551 | -0.001208107 | 0.024551784 | -0.049206494 | 0.960754736 | NA | | I830012O16Rik | | | | | | | |
| C4a | 90.02179343 | 0.526973429 | 0.163552929 | 3.222036019 | 0.001272831 | 0.013683744 | C4a | Tenm2 | 4933428C19Rik | 0.226357657 | -0.00122641 | 0.009803369 | -0.125100846 | 0.900443713 | NA | | Cd84 | | | | | | | |

| | | | | | | | | | | | | | | | | |
|---|---|---|---|---|---|---|---|---|---|---|---|---|---|---|---|---|
| Asprv1 | 40.53947689 | 0.526915855 | 0.158213556 | 3.330409021 | 0.000867185 | 0.010225679 | Asprv1 | Tmem183a | Gm5132 | 0.226357657 | -0.00122641 | 0.009803369 | -0.125100846 | 0.900443713 | NA | Cd33 |
| Gnmt | 16.5160432 | 0.526301318 | 0.16170062 | 3.254788496 | 0.001134768 | 0.012585613 | Gnmt | Dync2h1 | Krt42 | 0.226357657 | -0.00122641 | 0.009803369 | -0.125100846 | 0.900443713 | NA | A930013F10Rik |
| Irf9 | 457.3179858 | 0.526288072 | 0.116653649 | 4.511544031 | 6.44E-06 | 0.000222978 | Irf9 | Txn1 | 1700119H24Rik | 0.242744565 | -0.001261837 | 0.009967597 | -0.126593945 | 0.89926179 | NA | Naip5 |
| 4933406C10Rik | 52.29709545 | 0.525632672 | 0.166304815 | 3.160658169 | 0.001574131 | 0.016024696 | 4933406C10Rik | Tanc2 | 8030443G20Rik | 0.242744565 | -0.001261837 | 0.009967597 | -0.126593945 | 0.89926179 | NA | Slc26a10 |
| Uvssa | 459.0080038 | 0.525487171 | 0.106474448 | 4.935335936 | 8.00E-07 | 4.09E-05 | Uvssa | Spryd7 | Aifm2 | 345.3009638 | -0.001268871 | 0.08452876 | -0.015011113 | 0.988023314 | 0.997846532 | Ptpn6 |
| Cyp26b1 | 595.3830254 | 0.525322735 | 0.123661523 | 4.248069411 | 2.16E-05 | 0.000592345 | Cyp26b1 | Hsbp1 | Slc25a18 | 2317.090168 | -0.001269918 | 0.071507372 | -0.017759264 | 0.985830902 | 0.997552088 | Gm5577 |
| Mpl | 31.11447344 | 0.524515976 | 0.167727666 | 3.127188195 | 0.001764869 | 0.017394963 | Mpl | Mfap3l | Pid1 | 1712.766339 | -0.001282785 | 0.074276793 | -0.017270335 | 0.986220952 | 0.997552088 | Mpeg1 |
| Oasl2 | 242.1392103 | 0.52342252 | 0.162639367 | 3.218301516 | 0.001289522 | 0.013779208 | Oasl2 | Nab1 | Trim61 | 0.642209104 | -0.001298768 | 0.015445977 | -0.084084553 | 0.932989206 | NA | Ovgp1 |
| Pmch | 32.05889035 | 0.522935987 | 0.148988516 | 3.509908029 | 0.000448262 | 0.006323296 | Pmch | Cacybp | Nufip1 | 552.157216 | -0.001314973 | 0.069315887 | -0.018970725 | 0.98486446 | 0.997361871 | Zkscan3 |
| Adam8 | 113.0969699 | 0.522741086 | 0.153099885 | 3.414379349 | 0.000639275 | 0.008138806 | Adam8 | Ndn | 4921501E09Rik | 0.403899654 | -0.001324714 | 0.013529138 | -0.09791561 | 0.921999305 | NA | Fcgr1 |
| Cd109 | 81.48102203 | 0.522338637 | 0.167612776 | 3.116341429 | 0.001831101 | 0.017883806 | Cd109 | Phactr1 | Cd3g | 0.403899654 | -0.001324714 | 0.013529138 | -0.09791561 | 0.921999305 | NA | Snhg11 |
| P2rx6 | 86.4158704 | 0.521200156 | 0.151499789 | 3.440269835 | 0.000581134 | 0.00762413 | P2rx6 | Bsg | Dnase1 | 0.385781328 | -0.001338434 | 0.01173576 | -0.114047451 | 0.909200178 | NA | Cd14 |
| Tnip2 | 474.5824552 | 0.52102588 | 0.098551445 | 5.286841602 | 1.24E-07 | 8.93E-06 | Tnip2 | Ssbp4 | Smim5 | 13.16704612 | -0.001340532 | 0.062922382 | -0.021304534 | 0.983002727 | NA | Myo1f |
| Col6a2 | 1063.045587 | 0.520976416 | 0.106699367 | 4.882657024 | 1.05E-06 | 5.05E-05 | Col6a2 | Tcf20 | 2310040G24Rik | 38.75058803 | -0.001342782 | 0.0926922 | -0.014486465 | 0.988441878 | 0.997974805 | Cep89 |
| Mx1 | 17.65054141 | 0.520581474 | 0.163624021 | 3.181571197 | 0.001464785 | 0.015180857 | Mx1 | Rps12 | Cdh23 | 68.05795678 | -0.001344627 | 0.093263236 | -0.014417544 | 0.988496863 | 0.997974805 | Robo3 |
| Ftx | 809.9636154 | 0.520100565 | 0.145803079 | 3.567143903 | 0.000360893 | 0.005392171 | Ftx | Nedd8 | Klk1b3 | 0.424992894 | -0.001357291 | 0.012077986 | -0.112377239 | 0.910524302 | NA | Irf8 |
| Robo3 | 988.5187055 | 0.518671828 | 0.165098148 | 3.141596879 | 0.001680292 | 0.01680726 | Robo3 | Sorcs2 | 1700012L04Rik | 0.237667581 | -0.001369387 | 0.009896121 | -0.138376138 | 0.889943155 | NA | Mrps12 |
| Atp2c2 | 43.43024302 | 0.517888136 | 0.168028324 | 3.082147832 | 0.002055127 | 0.019478131 | Atp2c2 | Mrpl12 | 2310069B03Rik | 0.237667581 | -0.001369387 | 0.009896121 | -0.138376138 | 0.889943155 | NA | Gbp9 |
| Col24a1 | 404.9902976 | 0.515780321 | 0.16040753 | 3.215437084 | 0.00130246 | 0.013850598 | Col24a1 | Rbm12 | 4930564B18Rik | 0.237667581 | -0.001369387 | 0.009896121 | -0.138376138 | 0.889943155 | NA | Col11a1 |
| Clk4 | 1456.694376 | 0.514390789 | 0.118380271 | 4.345240858 | 1.39E-05 | 0.000416938 | Clk4 | Vkorc1l1 | H2-Eb2 | 0.237667581 | -0.001369387 | 0.009896121 | -0.138376138 | 0.889943155 | NA | Ryr1 |
| Ccdc11 | 11.27390938 | 0.514226207 | 0.163423258 | 3.146591337 | 0.001651857 | 0.016576338 | Ccdc11 | Ndufa2 | Rhag | 0.237667581 | -0.001369387 | 0.009896121 | -0.138376138 | 0.889943155 | NA | Klhl6 |
| Ddx26b | 1788.152824 | 0.513122808 | 0.114839497 | 4.468173577 | 7.89E-06 | 0.000261181 | Ddx26b | Pak6 | Igfbpl1 | 22.49598629 | -0.001378101 | 0.079732035 | -0.017284156 | 0.986209925 | 0.997552088 | Fbxo17 |
| Casc1 | 26.31824637 | 0.512406761 | 0.167908603 | 3.051700459 | 0.002275491 | 0.021036068 | Casc1 | Pgrmc1 | Rab17 | 0.617637615 | -0.001384567 | 0.015316353 | -0.090397957 | 0.92797098 | NA | Dmkn |
| Cep110 | 550.9013864 | 0.510986903 | 0.105475049 | 4.844623507 | 1.27E-06 | 5.97E-05 | Cep110 | Polr1d | Tmem27 | 7.579073774 | -0.001400791 | 0.036500822 | -0.03837698 | 0.969387115 | NA | Cd52 |
| 1700012B15Rik | 372.9530798 | 0.510193113 | 0.098529768 | 5.178060611 | 2.24E-07 | 1.45E-05 | 1700012B15Rik | Rps18 | 1700008I05Rik | 0.235049888 | -0.001426163 | 0.009858693 | -0.14466048 | 0.884978943 | NA | Lag3 |
| Rsad1 | 634.1711095 | 0.509872078 | 0.09648567 | 5.284433188 | 1.26E-07 | 9.00E-06 | Rsad1 | Kcnk2 | Pdc | 0.235049888 | -0.001426163 | 0.009858693 | -0.14466048 | 0.884978943 | NA | Sbsn |
| Neurod1 | 222.1712959 | 0.508871435 | 0.107637183 | 4.727654724 | 2.27E-06 | 9.94E-05 | Neurod1 | Ldoc1l | Chmp1b | 968.5667817 | -0.001430521 | 0.070211398 | -0.020374487 | 0.983744636 | 0.996711782 | C4b |
| E030003E18Rik | 239.7697447 | 0.507774735 | 0.143032662 | 3.550061423 | 0.000385141 | 0.005672413 | E030003E18Rik | Chd9 | Pnliprp1 | 0.39882267 | -0.001436782 | 0.013484198 | -0.106553045 | 0.915143571 | NA | Nudt19 |
| Cfp | 305.9397896 | 0.507097738 | 0.155441238 | 3.262311507 | 0.001105076 | 0.012318024 | Cfp | Dnalc1 | 2310008H04Rik | 124.5047635 | -0.001451834 | 0.094639739 | -0.015340635 | 0.987760425 | 0.997789147 | Tyrobp |
| Gdpd2 | 231.9702453 | 0.506007696 | 0.159293062 | 3.176583388 | 0.001490209 | 0.01539287 | Gdpd2 | Rap2b | Gm6260 | 0.927811108 | -0.001467703 | 0.020790411 | -0.070595211 | 0.943719922 | NA | |
| Tlr2 | 77.92161558 | 0.505953899 | 0.161640756 | 3.130113411 | 0.001747388 | 0.017321839 | Tlr2 | Tmsb4x | Sbno1 | 7475.514196 | -0.0014708 | 0.064063507 | -0.022958466 | 0.981683403 | 0.996444753 | |
| Boc | 889.1728922 | 0.505927061 | 0.084621444 | 5.978709843 | 2.25E-09 | 2.83E-07 | Boc | Vps13a | Inpp5j | 3132.145556 | -0.001473222 | 0.064849851 | -0.022717429 | 0.981875673 | 0.996444753 | |
| Sh3bp1 | 595.5107772 | 0.505569578 | 0.100054185 | 5.052957813 | 4.35E-07 | 2.50E-05 | Sh3bp1 | Tubb4b | Lhfpl1 | 4.541119856 | -0.001488687 | 0.04777612 | -0.031159636 | 0.97514223 | NA | |
| Rwdd3 | 128.5576472 | 0.505223793 | 0.117737359 | 4.291108594 | 1.78E-05 | 0.000509164 | Rwdd3 | Fopnl | 5430421F17Rik | 4.118278412 | -0.001489786 | 0.042767912 | -0.034834191 | 0.972211956 | NA | |
| Ssh3 | 1564.140088 | 0.50491102 | 0.121315544 | 4.161964776 | 3.16E-05 | 0.000801423 | Ssh3 | Tram1l1 | Gfral | 0.215934008 | -0.001490294 | 0.009751239 | -0.15283124 | 0.878531361 | NA | |
| Mir186 | 30.42506888 | 0.504343433 | 0.167340389 | 3.013877503 | 0.002579319 | 0.02290199 | Mir186 | Mid2 | Gimap7 | 0.215934008 | -0.001490294 | 0.009751239 | -0.15283124 | 0.878531361 | NA | |
| Nr1d1 | 5181.075221 | 0.503915447 | 0.085213962 | 5.913531513 | 3.35E-09 | 4.18E-07 | Nr1d1 | Cdh10 | Glipr1l1 | 0.215934008 | -0.001490294 | 0.009751239 | -0.15283124 | 0.878531361 | NA | |
| Wdr96 | 44.31362833 | 0.503692851 | 0.163060594 | 3.088991886 | 0.002008369 | 0.019184755 | Wdr96 | Gja1 | Gm10280 | 0.215934008 | -0.001490294 | 0.009751239 | -0.15283124 | 0.878531361 | NA | |
| Hmox1 | 233.4038846 | 0.502851508 | 0.121319761 | 4.144844199 | 3.40E-05 | 0.000847051 | Hmox1 | Sptbn1 | Hoxd4 | 0.215934008 | -0.001490294 | 0.009751239 | -0.15283124 | 0.878531361 | NA | |
| Ankrd16 | 646.0294911 | 0.502361221 | 0.089646167 | 5.603822636 | 2.10E-08 | 2.01E-06 | Ankrd16 | Grid1 | Klk4 | 0.215934008 | -0.001490294 | 0.009751239 | -0.15283124 | 0.878531361 | NA | |
| Alox8 | 246.7224836 | 0.50210292 | 0.121299727 | 4.13935739 | 3.48E-05 | 0.000857909 | Alox8 | Dgat2 | Olfr874 | 0.215934008 | -0.001490294 | 0.009751239 | -0.15283124 | 0.878531361 | NA | |
| Treml2 | 27.54744045 | 0.501679346 | 0.165546968 | 3.030435117 | 0.002442016 | 0.022066759 | Treml2 | Gpr125 | S100a2 | 0.215934008 | -0.001490294 | 0.009751239 | -0.15283124 | 0.878531361 | NA | |
| Gdap10 | 63.09027955 | 0.501216382 | 0.15520918 | 3.229295976 | 0.001240954 | 0.013431707 | Gdap10 | Mettl9 | Tm7sf3 | 1948.777735 | -0.001496195 | 0.049600663 | -0.030164818 | 0.975935607 | 0.994349486 | |
| Hook1 | 2087.80065 | 0.500913073 | 0.079815354 | 6.275898628 | 3.48E-10 | 5.61E-08 | Hook1 | Ndufb9 | Actr3 | 7768.162087 | -0.001499608 | 0.060681828 | -0.024712641 | 0.980284172 | 0.995858199 | |
| Irf8 | 253.1120082 | 0.499735003 | 0.142764546 | 3.500413913 | 0.000464536 | 0.006460974 | Irf8 | Hbegf | Olfr221 | 1.208201025 | -0.001516003 | 0.024732605 | -0.061295714 | 0.951123704 | NA | |
| Rnf112 | 13465.15019 | 0.499567263 | 0.098013838 | 5.096905413 | 3.45E-07 | 2.09E-05 | Rnf112 | Cenpv | Gm5294 | 0.445061325 | -0.001528055 | 0.01387853 | -0.110102084 | 0.912328415 | NA | |
| B3galnt2 | 683.8561278 | 0.498561888 | 0.095613863 | 5.214326399 | 1.84E-07 | 1.24E-05 | B3galnt2 | 2610507B11Rik | 1700067K01Rik | 1.205583332 | -0.001529567 | 0.024338024 | -0.062846816 | 0.949888486 | NA | |
| Naip5 | 77.38687791 | 0.498421629 | 0.16350802 | 3.048300811 | 0.002301394 | 0.021168869 | Naip5 | Atxn7l3b | Iqcf4 | 0.513177942 | -0.001540944 | 0.012718666 | -0.121156113 | 0.903567384 | NA | |
| Frem3 | 55.33041029 | 0.497691171 | 0.159946423 | 3.111611759 | 0.00186069 | 0.018120387 | Frem3 | Tulp4 | Mei4 | 28.78520641 | -0.001570498 | 0.087043126 | -0.018042758 | 0.985604743 | 0.997496292 | |
| Mirg | 554.1907585 | 0.497010713 | 0.147457804 | 3.370528383 | 0.000750242 | 0.009126188 | Mirg | Ndufa7 | Osr1 | 71.98074577 | -0.001585607 | 0.09340644 | -0.01697535 | 0.986456281 | 0.997581728 | |
| Lrrc29 | 42.95972806 | 0.49657143 | 0.165265519 | 3.004688654 | 0.002658528 | 0.023324597 | Lrrc29 | Lphn3 | Zpld1 | 5.571485153 | -0.001587218 | 0.051434481 | -0.030859028 | 0.975381965 | NA | |
| Lrrc23 | 61.69704346 | 0.496427525 | 0.166764377 | 2.976819954 | 0.00291255 | 0.024822359 | Lrrc23 | Timmdc1 | Gm5936 | 1.386410611 | -0.001604484 | 0.027665564 | -0.057995718 | 0.95375204 | NA | |
| Ccdc135 | 62.59120814 | 0.49629014 | 0.166608992 | 2.978771646 | 0.002894064 | 0.024719197 | Ccdc135 | Oxct1 | Gpr142 | 0.45890573 | -0.001615111 | 0.015498794 | -0.104208799 | 0.917003652 | NA | |
| Flnb | 3420.57541 | 0.496075006 | 0.099803472 | 4.970518523 | 6.68E-07 | 3.53E-05 | Flnb | Phactr3 | D630045J12Rik | 3009.074131 | -0.001649178 | 0.080881456 | -0.020390063 | 0.983732211 | 0.996711782 | |
| Myo1h | 66.98676742 | 0.495256245 | 0.165662142 | 2.989555959 | 0.002793833 | 0.024150338 | Myo1h | Cdc34 | Rnf114 | 2506.185198 | -0.001673202 | 0.051572494 | -0.032443688 | 0.974118223 | 0.993518395 | |
| Ace | 1023.649079 | 0.495163135 | 0.133745122 | 3.702289306 | 0.000213663 | 0.003556118 | Ace | Rpsa | Ppp1r36 | 50.68604427 | -0.001683863 | 0.093402086 | -0.018028107 | 0.985616431 | 0.997496292 | |
| Igflr1 | 213.4126547 | 0.494671856 | 0.146725447 | 3.371411489 | 0.000747841 | 0.009109311 | Igflr1 | Rgs8 | BB019430 | 0.224626239 | -0.001685212 | 0.009789551 | -0.172143964 | 0.86332435 | NA | |
| Mroh8 | 49.45192197 | 0.494563044 | 0.153448469 | 3.222991053 | 0.001268595 | 0.013659217 | Mroh8 | 2610002M06Rik | Csrp3 | 0.224626239 | -0.001685212 | 0.009789551 | -0.172143964 | 0.86332435 | NA | |
| Fermt1 | 11.79605464 | 0.494441377 | 0.154771501 | 3.194653887 | 0.001399986 | 0.014666252 | Fermt1 | Dock3 | Lgals6 | 0.224626239 | -0.001685212 | 0.009789551 | -0.172143964 | 0.86332435 | NA | |
| Espn | 31.53034311 | 0.493933235 | 0.165842309 | 2.978330665 | 0.002898231 | 0.024741154 | Espn | Lix1l | Spag11b | 0.224626239 | -0.001685212 | 0.009789551 | -0.172143964 | 0.86332435 | NA | |
| Whamm | 661.5184479 | 0.492955525 | 0.107616891 | 4.580651963 | 4.64E-06 | 0.000173896 | Whamm | Cntn1 | Tescl | 0.224626239 | -0.001685212 | 0.009789551 | -0.172143964 | 0.86332435 | NA | |
| 5830418P13Rik | 18.02715787 | 0.492797883 | 0.167719583 | 2.938225069 | 0.003300973 | 0.027132771 | 5830418P13Rik | Samd4 | 6430562O15Rik | 2.337926215 | -0.001703553 | 0.031063098 | -0.0548417 | 0.956264579 | NA | |
| Mybpc1 | 171.7075587 | 0.492514362 | 0.159582293 | 3.086271993 | 0.002026833 | 0.019325387 | Mybpc1 | Ndufs6 | 3100003L05Rik | 1.028835615 | -0.001705573 | 0.024615068 | -0.069289813 | 0.944758934 | NA | |
| Cdk15 | 18.35479181 | 0.491527816 | 0.16804288 | 2.925014231 | 0.003444405 | 0.027955792 | Cdk15 | Ppp1r12b | Efcab8 | 0.423505437 | -0.001716925 | 0.015208307 | -0.112893867 | 0.910114699 | NA | |
| Efcab4b | 31.26931778 | 0.491000666 | 0.162028958 | 3.030326626 | 0.002442894 | 0.022066759 | Efcab4b | Pard3 | Vrk3 | 997.5237429 | -0.001719095 | 0.056949349 | -0.030186388 | 0.975918405 | 0.994349486 | |
| Oscar | 72.82397407 | 0.489328862 | 0.157936155 | 3.098270079 | 0.001946539 | 0.018724864 | Oscar | Prdx4 | A630072M18Rik | 275.5621077 | -0.001729621 | 0.084379411 | -0.020498142 | 0.983645994 | 0.996711782 | |
| 4930451C15Rik | 44.73283795 | 0.489239209 | 0.158735421 | 3.082104839 | 0.002055424 | 0.019478131 | 4930451C15Rik | Pdlim5 | Cplx2 | 61181.34839 | -0.001749398 | 0.059664682 | -0.029320487 | 0.976608988 | 0.994686731 | |
| Nek5 | 13.9735857 | 0.488309147 | 0.157318604 | 3.103950425 | 0.001909553 | 0.018503201 | Nek5 | Agps | Pde6h | 5.379326757 | -0.001761937 | 0.052462549 | -0.033584666 | 0.97320835 | NA | |
| Lilra5 | 13.91655782 | 0.48700346 | 0.164229125 | 2.965390333 | 0.003022991 | 0.025445766 | Lilra5 | Mrps25 | Mir425 | 24.98596934 | -0.001774031 | 0.084434394 | -0.021010766 | 0.983237068 | 0.996711782 | |
| Arrdc1 | 272.4660883 | 0.486139135 | 0.115663437 | 4.203049354 | 2.63E-05 | 0.000692746 | Arrdc1 | Ndufa3 | Rapgef3 | 1594.753649 | -0.001774909 | 0.07792487 | -0.02277718 | 0.981828011 | 0.996444753 | |
| Fgf5 | 86.1659671 | 0.485459658 | 0.150935112 | 3.216346756 | 0.001298339 | 0.013849883 | Fgf5 | Tmem35 | Anapc4 | 2212.225623 | -0.001793342 | 0.054846123 | -0.032697696 | 0.973915662 | 0.993518395 | |
| Slc17a6 | 915.9812025 | 0.485141401 | 0.130313757 | 3.722871713 | 0.00019697 | 0.003364771 | Slc17a6 | Akap6 | Ptpn20 | 16.78955382 | -0.001811136 | 0.07339471 | -0.024676651 | 0.980312879 | NA | |
| Paxbp1 | 1499.775334 | 0.484917583 | 0.098464877 | 4.924777208 | 8.45E-07 | 4.25E-05 | Paxbp1 | Cdkl5 | Zmym4 | 2602.425746 | -0.001840826 | 0.068347522 | -0.026933324 | 0.978512914 | 0.995273517 | |
| Aldh1a1 | 1796.04932 | 0.484897496 | 0.108134416 | 4.484210615 | 7.32E-06 | 0.000246462 | Aldh1a1 | Abcb7 | Trpv1 | 5.72608507 | -0.001841908 | 0.049427506 | -0.037264831 | 0.970273847 | NA | |
| Otx1 | 194.3325165 | 0.484569328 | 0.124725842 | 3.885075618 | 0.000102298 | 0.001996229 | Otx1 | Ccm2 | 1700093K21Rik | 4.259849704 | -0.001853007 | 0.04543558 | -0.040783171 | 0.967468756 | NA | |
| Lrrc45 | 1936.627398 | 0.483971482 | 0.1350818 | 3.582803034 | 0.000339927 | 0.005158499 | Lrrc45 | Ppp1ca | Polh | 232.9251725 | -0.001863404 | 0.084399621 | -0.022078345 | 0.982385461 | 0.996546601 | |
| Vasn | 427.0462911 | 0.48298555 | 0.13936595 | 3.465592196 | 0.000529065 | 0.007115035 | Vasn | Cdk14 | Cbx8 | 270.6729617 | -0.001865893 | 0.083201169 | -0.022426286 | 0.982107912 | 0.996444753 | |
| Ifi44 | 46.20666261 | 0.482753037 | 0.158098913 | 3.053487381 | 0.002261982 | 0.020948688 | Ifi44 | Zfp827 | Pfn1 | 3182.569188 | -0.001870497 | 0.054372408 | -0.034401577 | 0.972556926 | 0.99301409 | |
| Cd72 | 34.16620951 | 0.482498999 | 0.167951457 | 2.872847956 | 0.004067898 | 0.031404095 | Cd72 | Syt5 | B230118H07Rik | 848.9419204 | -0.001883208 | 0.075201339 | -0.025042208 | 0.980021297 | 0.995718637 | |
| Dna2 | 62.42824683 | 0.482103095 | 0.151979033 | 3.172168454 | 0.001513052 | 0.015566554 | Dna2 | Lin7a | Tgfb3 | 613.8277586 | -0.001908055 | 0.08802374 | -0.02167659 | 0.982705938 | 0.996711782 | |
| Rbm38 | 73.19605725 | 0.481685913 | 0.144085832 | 3.343048421 | 0.000828634 | 0.009842462 | Rbm38 | Ube2e3 | Cbx7 | 2783.488223 | -0.00191793 | 0.048723718 | -0.039363379 | 0.968600677 | 0.992230168 | |
| Relt | 727.0285178 | 0.481383405 | 0.142614851 | 3.375408662 | 0.000737062 | 0.009013442 | Relt | Nacc2 | Adam2 | 0.684756679 | -0.001923479 | 0.0175469 | -0.109619308 | 0.912711297 | NA | |
| Dusp9 | 31.11231638 | 0.481132063 | 0.163218104 | 2.947786124 | 0.003200584 | 0.026528802 | Dusp9 | Tmem164 | Kctd11 | 95.94548898 | -0.001933297 | 0.093117186 | -0.020761981 | 0.983435526 | 0.996711782 | |
| Ccdc153 | 66.79771211 | 0.480928647 | 0.159037758 | 3.023990356 | 0.002494644 | 0.022341444 | Ccdc153 | Tmem126a | Shroom2 | 4670.04058 | -0.001960547 | 0.057952218 | -0.033830413 | 0.973012384 | 0.99304278 | |
| Isg15 | 78.27792383 | 0.479806917 | 0.16683061 | 2.876012482 | 0.004027338 | 0.031168618 | Isg15 | Rps3 | Olfr153 | 0.443599587 | -0.001966173 | 0.013869937 | -0.141757854 | 0.887271275 | NA | |
| B2m | 3100.791312 | 0.478939498 | 0.084022131 | 5.700158894 | 1.20E-08 | 1.23E-06 | B2m | Rpl36 | Ctsf | 2509.109376 | -0.001978882 | 0.06269865 | -0.031561796 | 0.97482151 | 0.99384175 | |
| Fam221a | 29.11745327 | 0.47893661 | 0.166399963 | 2.878225454 | 0.003999192 | 0.031028284 | Fam221a | Scarb1 | E130201H02Rik | 3.035542264 | -0.00198454 | 0.035833409 | -0.055382385 | 0.955833829 | NA | |
| Rassf3 | 2023.315597 | 0.478801562 | 0.096492857 | 4.962041502 | 6.98E-07 | 3.66E-05 | Rassf3 | Parvb | 1700031M16Rik | 0.46293569 | -0.001993934 | 0.014021497 | -0.142205475 | 0.886917707 | NA | |
| Gm14872 | 186.5972643 | 0.478577538 | 0.113034974 | 4.233889039 | 2.30E-05 | 0.000623245 | Gm14872 | Plcl2 | Sh2d2a | 0.46293569 | -0.001993934 | 0.014021497 | -0.142205475 | 0.886917707 | NA | |
| Pias3 | 953.0465659 | 0.478528523 | 0.091924722 | 5.205656451 | 1.93E-07 | 1.29E-05 | Pias3 | Mrpl53 | Cyp26c1 | 0.394963462 | -0.002001098 | 0.013453362 | -0.148743346 | 0.881756156 | NA | |
| Ifit1 | 136.2514187 | 0.477810762 | 0.166849983 | 2.863714784 | 0.004187048 | 0.032100014 | Ifit1 | Rap1b | Olfr570 | 0.628307206 | -0.002002362 | 0.015390481 | -0.130103937 | 0.896484195 | NA | |
| Ttc21a | 68.87641437 | 0.477126829 | 0.150794935 | 3.164077285 | 0.001555755 | 0.015900306 | Ttc21a | Mdga2 | Cr2 | 11.59745735 | -0.002002746 | 0.064276097 | -0.031158486 | 0.975143148 | NA | |
| B230216G23Rik | 53.79093156 | 0.477089856 | 0.157925875 | 3.020973319 | 0.002519636 | 0.02250152 | B230216G23Rik | 6330403K07Rik | Htr3a | 297.9637017 | -0.002004279 | 0.085753011 | -0.023372692 | 0.981352988 | 0.996444753 | |
| Morn1 | 148.9727993 | 0.477029221 | 0.124191597 | 3.841074859 | 0.000122497 | 0.002286704 | Morn1 | Mtx2 | 4921511C10Rik | 0.46301088 | -0.002020356 | 0.015532911 | -0.130069376 | 0.896511538 | NA | |
| Trpm4 | 505.6135134 | 0.477006058 | 0.116041976 | 4.11063369 | 3.95E-05 | 0.000953751 | Trpm4 | Cldn12 | 1700025N23Rik | 0.474254397 | -0.002021792 | 0.014107391 | -0.14331441 | 0.886041876 | NA | |
| Slc39a2 | 234.4910531 | 0.47695963 | 0.164503252 | 2.899393322 | 0.003738855 | 0.029707604 | Slc39a2 | 6030419C18Rik | Prph2 | 1.192608397 | -0.002022576 | 0.027661483 | -0.073118872 | 0.941711524 | NA | |
| Lime1 | 1632.314876 | 0.476643487 | 0.130300977 | 3.658019291 | 0.000254172 | 0.004091928 | Lime1 | Arhgef12 | Zfp105 | 191.0701573 | -0.002023389 | 0.085149226 | -0.023762855 | 0.981041769 | 0.996337324 | |
| Ccl3 | 19.06722801 | 0.476061748 | 0.147295898 | 3.232009549 | 0.001229229 | 0.013327979 | Ccl3 | Rnf11 | Wbp1 | 883.5808662 | -0.002029557 | 0.073836392 | -0.027487224 | 0.97807113 | 0.995129315 | |
| Ggnbp1 | 94.12601739 | 0.475929174 | 0.144261961 | 3.299062143 | 0.000970084 | 0.011141947 | Ggnbp1 | Kitl | Rap1a | 1120.474605 | -0.002040917 | 0.065194781 | -0.031304915 | 0.975026371 | 0.993940385 | |
| Sfi1 | 741.8557003 | 0.475739353 | 0.134966108 | 3.524880143 | 0.000423674 | 0.006044578 | Sfi1 | Vti1b | Gm6194 | 0.485489131 | -0.002043187 | 0.01253902 | -0.162946336 | 0.870560687 | NA | |
| Ak7 | 72.38386989 | 0.475048517 | 0.164029493 | 2.896116479 | 0.003778121 | 0.029958144 | Ak7 | Usp46 | 7630403G23Rik | 9.47790446 | -0.002053522 | 0.045563929 | -0.045069027 | 0.964052289 | NA | |
| Col26a1 | 211.9286931 | 0.475028497 | 0.144104859 | 3.296408611 | 0.000979295 | 0.01121448 | Col26a1 | Mrps36 | Os9 | 6309.895424 | -0.002054923 | 0.050206189 | -0.04092967 | 0.967351964 | 0.991795217 | |
| 4930422G04Rik | 202.0971971 | 0.474853921 | 0.101221282 | 4.691245893 | 2.72E-06 | 0.000114641 | 4930422G04Rik | Slc9a3r1 | Rpl36 | 1896.91782 | -0.002071744 | 0.061770442 | -0.033539402 | 0.973244445 | 0.993070301 | |
| Leng8 | 9341.863843 | 0.473997794 | 0.157360662 | 3.01217463 | 0.002593833 | 0.02297819 | Leng8 | Fam171a2 | S1pr2 | 124.4533986 | -0.002073455 | 0.092989789 | -0.022297661 | 0.982210515 | 0.996479252 | |
| Skp2 | 252.5865249 | 0.473736265 | 0.120315077 | 3.937463847 | 8.23E-05 | 0.00168992 | Skp2 | 3632451O06Rik | Igll1 | 0.440787404 | -0.002076767 | 0.015354875 | -0.135251329 | 0.892413165 | NA | |
| Tcea2 | 1135.620647 | 0.472711561 | 0.092883753 | 5.089281458 | 3.59E-07 | 2.15E-05 | Tcea2 | Ank2 | Otc | 0.440787404 | -0.002076767 | 0.015354875 | -0.135251329 | 0.892413165 | NA | |
| Cdan1 | 1123.913599 | 0.471901744 | 0.093743464 | 5.033969544 | 4.80E-07 | 2.69E-05 | Cdan1 | Ostc | Abra | 9.294690511 | -0.002092852 | 0.061709963 | -0.033914322 | 0.972945473 | NA | |
| Zkscan3 | 1011.359314 | 0.471207977 | 0.110462627 | 4.265768359 | 1.99E-05 | 0.000557702 | Zkscan3 | Smarcd3 | Dnase2b | 1.879966503 | -0.00209407 | 0.026407321 | -0.079298838 | 0.93679493 | NA | |
| Dcdc2b | 97.6171826 | 0.470001665 | 0.165819006 | 2.834425768 | 0.004590812 | 0.034079589 | Dcdc2b | C1d | Tmem120b | 287.0773651 | -0.002095154 | 0.080509374 | -0.026023726 | 0.979238414 | 0.995367691 | |
| Cep70 | 624.5507929 | 0.4698769 | 0.112736792 | 4.167910871 | 3.07E-05 | 0.000784665 | Cep70 | Wrb | Car6 | 1.200717788 | -0.002095213 | 0.026111744 | -0.080240261 | 0.93604617 | NA | |
| Hes3 | 9.773242983 | 0.469795598 | 0.149405714 | 3.144428588 | 0.001664115 | 0.016666969 | Hes3 | Ksr2 | Mir344g | 1.309514955 | -0.002099851 | 0.022823569 | -0.092003614 | 0.926695168 | NA | |
| Wdr62 | 69.84165674 | 0.46958471 | 0.149224588 | 3.14683202 | 0.001650498 | 0.016573434 | Wdr62 | Vps13b | B230217C12Rik | 1188.819909 | -0.0021111 | 0.058073034 | -0.0363525 | 0.971001289 | 0.993013568 | |
| Satb1 | 3895.927084 | 0.46950607 | 0.108996354 | 4.307539241 | 1.65E-05 | 0.000478982 | Satb1 | Glrx5 | Hmgb3 | 1248.657637 | -0.00211423 | 0.067614522 | -0.031268882 | 0.975055107 | 0.993940385 | |
| Wdr90 | 923.9693233 | 0.469363803 | 0.110631795 | 4.242576056 | 2.21E-05 | 0.000604892 | Wdr90 | Mrps33 | Ceacam18 | 0.437366647 | -0.002115244 | 0.015326742 | -0.138010024 | 0.890232496 | NA | |
| Slc11a1 | 141.381587 | 0.469101562 | 0.148179156 | 3.165772947 | 0.001546715 | 0.015860224 | Slc11a1 | Ptprg | Dnmt1 | 2520.133156 | -0.002119725 | 0.048896348 | -0.043351405 | 0.965421415 | 0.990993714 | |
| Wnt9b | 17.26626721 | 0.468732391 | 0.165463809 | 2.832839362 | 0.004613656 | 0.03418651 | Wnt9b | Rabl5 | Hist1h3b | 0.438854104 | -0.002125222 | 0.013832736 | -0.153637115 | 0.877895871 | NA | |
| BC051142 | 77.83953271 | 0.468026018 | 0.157832079 | 2.965341527 | 0.003023471 | 0.025445766 | BC051142 | 1110001A16Rik | Sirpb1a | 0.476780095 | -0.002127725 | 0.015638838 | -0.136053888 | 0.89177868 | NA | |
| Ccdc84 | 290.6932066 | 0.467663696 | 0.14349943 | 3.258993397 | 0.001118083 | 0.012436162 | Ccdc84 | Ptprz1 | Dtnb | 2713.301159 | -0.002127866 | 0.051167675 | -0.041586145 | 0.966828618 | 0.991586617 | |
| Arl11 | 25.4035542 | 0.467547731 | 0.166893973 | 2.801465642 | 0.005087106 | 0.036290258 | Arl11 | Rps6ka3 | Maneal | 2896.320105 | -0.002140633 | 0.049144707 | -0.043557761 | 0.965256922 | 0.990993714 | |
| E230016M11Rik | 41.37623989 | 0.467542876 | 0.160631853 | 2.910648592 | 0.003606794 | 0.028895382 | E230016M11Rik | Timm10 | Dcst1 | 1.494023757 | -0.002143558 | 0.027682237 | -0.077434438 | 0.938277945 | NA | |
| Hsf4 | 654.2707419 | 0.467490676 | 0.090715709 | 5.153359676 | 2.56E-07 | 1.62E-05 | Hsf4 | Cnn2 | Cyp2c44 | 8.971566109 | -0.002143598 | 0.062211433 | -0.034456655 | 0.972513006 | NA | |
| Tlr1 | 33.11334525 | 0.467422005 | 0.164705635 | 2.837923576 | 0.004540805 | 0.033817188 | Tlr1 | Klf16 | Cyp26b1 | 708.288211 | -0.002150776 | 0.082892071 | -0.025946701 | 0.97929985 | 0.995367691 | |
| Cdsn | 75.82981681 | 0.467340099 | 0.158737931 | 2.94409847 | 0.003238969 | 0.026736596 | Cdsn | Ranbp2 | Scube3 | 222.3713692 | -0.002164524 | 0.094834435 | -0.022824244 | 0.981790469 | 0.996444753 | |
| Fgr | 14.57654321 | 0.466914266 | 0.151354355 | 3.084908029 | 0.002036151 | 0.019331788 | Fgr | Rheb | Dusp22 | 1057.054804 | -0.002169776 | 0.056409099 | -0.038464995 | 0.969316941 | 0.992548843 | |
| Dnase1l2 | 119.7554382 | 0.466911385 | 0.167350681 | 2.790017843 | 0.005270514 | 0.037152566 | Dnase1l2 | St8sia3 | Oas1h | 0.640107034 | -0.002170223 | 0.015441567 | -0.140544229 | 0.88823001 | NA | |
| Dennd6b | 3013.996015 | 0.466728347 | 0.125203686 | 3.72775246 | 0.000193195 | 0.003311243 | Dennd6b | Coa5 | Phf3 | 3899.7618 | -0.002176205 | 0.072853552 | -0.029870949 | 0.976169975 | 0.994448773 | |
| Dsp | 39.76159768 | 0.466514801 | 0.149612224 | 3.118159652 | 0.001819842 | 0.017812149 | Dsp | Fdps | Gm12238 | 0.624360458 | -0.00217922 | 0.017242657 | -0.126385417 | 0.899426845 | NA | |
| Arrdc2 | 523.6972478 | 0.46592091 | 0.165794666 | 2.810228588 | 0.004950632 | 0.035738209 | Arrdc2 | Reep1 | Nuggc | 2.543681082 | -0.002179491 | 0.032981672 | -0.066081883 | 0.947312635 | NA | |
| Syna | 13.95154812 | 0.465681156 | 0.150933715 | 3.085335548 | 0.002033226 | 0.019331788 | Syna | Anks1b | Rimklb | 890.8495223 | -0.002185648 | 0.063175995 | -0.034596175 | 0.972401751 | 0.99301409 | |
| Gpr64 | 67.06178327 | 0.465566784 | 0.144035817 | 3.232298694 | 0.001227986 | 0.013323823 | Gpr64 | Maob | LOC171588 | 6.615725779 | -0.002186288 | 0.052984869 | -0.041262505 | 0.967086624 | NA | |
| Slc45a3 | 46.75169474 | 0.464440564 | 0.167981038 | 2.764839238 | 0.005695086 | 0.039424659 | Slc45a3 | Ndufs4 | Cnn1 | 13.66317086 | -0.002205489 | 0.062024568 | -0.035558317 | 0.971634545 | NA | |
| Apol7a | 10.63112137 | 0.464384891 | 0.162400881 | 2.859497369 | 0.004243129 | 0.032353861 | Apol7a | Usp9x | 2210404O09Rik | 177.035663 | -0.002220302 | 0.086282324 | -0.025732986 | 0.979470313 | 0.995367691 | |
| 4933439C10Rik | 776.901937 | 0.463497599 | 0.114283089 | 4.055697148 | 5.00E-05 | 0.001145662 | 4933439C10Rik | Zyg11b | Mir369 | 5.775608065 | -0.002233128 | 0.052228752 | -0.042756686 | 0.965895492 | NA | |
| Gm20751 | 29.27064596 | 0.462648674 | 0.168073602 | 2.752655193 | 0.005911412 | 0.040384223 | Gm20751 | Rps19 | Cebpe | 0.475335162 | -0.002237691 | 0.01247016 | -0.179443638 | 0.857589369 | NA | |
| Ceacam2 | 11.27049967 | 0.462617616 | 0.158145267 | 2.925270065 | 0.003441574 | 0.027955792 | Ceacam2 | Ndufs8 | Gpr171 | 0.621742765 | -0.002241714 | 0.017229099 | -0.130112108 | 0.896477731 | NA | |
| Ddx3x | 10048.43464 | 0.461210136 | 0.101412068 | 4.547882188 | 5.42E-06 | 0.000198486 | Ddx3x | Zfp428 | Csnk2a1 | 6078.6453 | -0.002244021 | 0.058198188 | -0.038558269 | 0.969242574 | 0.992548843 | |
| Car14 | 494.2670555 | 0.459803102 | 0.152395429 | 3.017171225 | 0.002551456 | 0.022693609 | Car14 | Ndrg4 | Tcp10b | 5.998320282 | -0.002256429 | 0.048854139 | -0.046187068 | 0.96316115 | NA | |
| Car4 | 1035.075922 | 0.459795566 | 0.099946695 | 4.600407896 | 4.22E-06 | 0.000160129 | Car4 | Prrc2b | Hist1h2ad | 1.924128283 | -0.002264036 | 0.024517305 | -0.09234439 | 0.926424421 | NA | |
| Mfsd7a | 35.74702236 | 0.459121206 | 0.162606118 | 2.823517422 | 0.004749984 | 0.034731599 | Mfsd7a | Triap1 | 4930556M19Rik | 11.39173576 | -0.002267331 | 0.064065721 | -0.035390697 | 0.971768203 | NA | |
| Myo1f | 165.9635705 | 0.458657398 | 0.136228283 | 3.366829462 | 0.000760377 | 0.009204125 | Myo1f | Scoc | Adamts4 | 888.6728786 | -0.002270851 | 0.087490027 | -0.025955546 | 0.979292796 | 0.995367691 | |
| Hspa1l | 14.96772533 | 0.458436397 | 0.165849496 | 2.764171189 | 0.005706759 | 0.039425264 | Hspa1l | Irf2bpl | Nsmce2 | 964.9906692 | -0.0022736 | 0.062178481 | -0.036565698 | 0.970831294 | 0.992984126 | |
| 5730480H06Rik | 21.571123 | 0.457622664 | 0.167833468 | 2.726647257 | 0.00639814 | 0.042690885 | 5730480H06Rik | Agap2 | 5430437J10Rik | 0.448676571 | -0.00228183 | 0.015419558 | -0.14798282 | 0.882356326 | NA | |
| Tacc3 | 210.6457061 | 0.45707704 | 0.11049967 | 4.136456147 | 3.53E-05 | 0.00086607 | Tacc3 | Rpl13a | Fpr1 | 0.86200232 | -0.002292304 | 0.018566172 | -0.123466685 | 0.901737554 | NA | |
| Piwil2 | 42.01146632 | 0.456232624 | 0.167892392 | 2.717410954 | 0.006579486 | 0.043453773 | Piwil2 | Nrxn3 | Tll2 | 30.89567076 | -0.002295086 | 0.088486577 | -0.025937112 | 0.979307499 | 0.995367691 | |
| Pcgf6 | 428.3916584 | 0.456061467 | 0.114091037 | 3.997347032 | 6.41E-05 | 0.001387951 | Pcgf6 | Mcee | Mir3473c | 0.469085418 | -0.002316343 | 0.015580933 | -0.148665241 | 0.88181779 | NA | |
| Il2rb | 12.36692174 | 0.455863515 | 0.162501863 | 2.805281777 | 0.005027261 | 0.036077991 | Il2rb | Xiap | 4933429O19Rik | 2.211507293 | -0.002319671 | 0.020717525 | -0.111966583 | 0.910849902 | NA | |
| Ppil6 | 80.39851164 | 0.45575248 | 0.140678881 | 3.23966524 | 0.001196701 | 0.013057525 | Ppil6 | Plekha2 | BC018242 | 7510.594424 | -0.002320214 | 0.066875031 | -0.03469478 | 0.972323124 | 0.99301409 | |
| Muc1 | 23.29115415 | 0.455382881 | 0.168012479 | 2.710411057 | 0.006719987 | 0.044107886 | Muc1 | Pdia6 | Bfar | 1537.489389 | -0.002323701 | 0.056645167 | -0.04102205 | 0.967278318 | 0.991789711 | |
| Slc50a1 | 909.4419845 | 0.454835969 | 0.084753436 | 5.366578517 | 8.02E-08 | 6.25E-06 | Slc50a1 | Panx1 | 4930528D03Rik | 2.222777668 | -0.002334451 | 0.035476399 | -0.065802934 | 0.94753472 | NA | |
| Dnaaf3 | 43.07708059 | 0.454770008 | 0.157518664 | 2.887086495 | 0.003888273 | 0.030510193 | Dnaaf3 | Mmd | Fmo6 | 0.95380211 | -0.002338591 | 0.018768855 | -0.124599552 | 0.900840583 | NA | |
| Ccdc81 | 33.80986404 | 0.454635054 | 0.1650877 | 2.753882379 | 0.005889293 | 0.040250862 | Ccdc81 | Klf7 | Snord17 | 1.198369775 | -0.002355457 | 0.027663396 | -0.085147051 | 0.932144485 | NA | |
| 1110059M19Rik | 16.77721501 | 0.453733566 | 0.125972396 | 3.601849134 | 0.000315962 | 0.004866313 | 1110059M19Rik | Mrps24 | Tmprss13 | 1.046042392 | -0.002359661 | 0.023013085 | -0.102535626 | 0.918331536 | NA | |
| Ly86 | 461.6558974 | 0.453065112 | 0.150626978 | 3.007861654 | 0.002630929 | 0.023198299 | Ly86 | Herc1 | Ccdc51 | 217.4783224 | -0.002360559 | 0.085216825 | -0.027700617 | 0.977900931 | 0.995119468 | |
| Exoc3l4 | 56.36822798 | 0.452245602 | 0.157788629 | 2.866148243 | 0.004154996 | 0.031917458 | Exoc3l4 | Neurl1b | Gm11149 | 4.221256969 | -0.002361709 | 0.032795862 | -0.072012421 | 0.942592023 | NA | |
| Kcnk4 | 445.133316 | 0.452201891 | 0.132699223 | 3.407720708 | 0.000655079 | 0.008270829 | Kcnk4 | Vstm2b | Pcbp4 | 5016.420561 | -0.002366459 | 0.05825008 | -0.040625853 | 0.967594174 | 0.991805208 | |
| Mei1 | 242.1362796 | 0.452147357 | 0.106687102 | 4.238069551 | 2.25E-05 | 0.000613905 | Mei1 | Hip1 | 1700007B14Rik | 5.201223647 | -0.002370871 | 0.045709053 | -0.051868743 | 0.95863328 | NA | |
| Usb1 | 329.0973659 | 0.451582228 | 0.106955769 | 4.222139961 | 2.42E-05 | 0.000648694 | Usb1 | Ncam2 | Adrb2 | 85.66519558 | -0.002402893 | 0.094995611 | -0.02529478 | 0.979819837 | 0.995583587 | |
| Ccdc57 | 341.7458459 | 0.451573212 | 0.125098281 | 3.609747528 | 0.000306495 | 0.004744092 | Ccdc57 | Nav3 | Hes2 | 0.455330007 | -0.00242453 | 0.013965861 | -0.173604023 | 0.86217667 | NA | |
| Zc3h12a | 38.47160976 | 0.451382516 | 0.153367731 | 2.943138774 | 0.003249027 | 0.026805341 | Zc3h12a | Bola2 | Kazald1 | 207.2002742 | -0.002438511 | 0.090718407 | -0.026880003 | 0.978555443 | 0.995273517 | |
| Pvrl4 | 237.9846808 | 0.451288363 | 0.11878786 | 3.799111822 | 0.000145216 | 0.00264081 | Pvrl4 | Hey1 | AY358078 | 0.491638859 | -0.002442439 | 0.014236904 | -0.171556874 | 0.863785914 | NA | |
| 1700112E06Rik | 8.650158005 | 0.450717204 | 0.14643425 | 3.077949358 | 0.002084303 | 0.019715626 | 1700112E06Rik | Tmem14c | Cacna1s | 49.87721707 | -0.002461839 | 0.091327765 | -0.026956081 | 0.978494763 | 0.995273517 | |
| Efhc1 | 52.1398574 | 0.450053072 | 0.167406898 | 2.688378295 | 0.00718 | 0.04643027 | Efhc1 | Rasa2 | Glce | 1883.029285 | -0.002461921 | 0.062481088 | -0.039402661 | 0.968569358 | 0.992230168 | |
| Snord2 | 25.57595321 | 0.450003461 | 0.165440366 | 2.72003424 | 0.006527516 | 0.043239925 | Snord2 | Fam162a | Klf17 | 0.825057078 | -0.002464292 | 0.018451334 | -0.13355631 | 0.893753434 | NA | |
| Myh3 | 120.1901526 | 0.449980489 | 0.150268575 | 2.994508252 | 0.002748876 | 0.023900723 | Myh3 | Gpx1 | Apol7a | 16.29096818 | -0.002486603 | 0.075757158 | -0.032823342 | 0.973815464 | NA | |
| 9830166K06Rik | 60.21086627 | 0.449650747 | 0.161015997 | 2.792584321 | 0.005228884 | 0.036943149 | 9830166K06Rik | Sar1b | Tmem165 | 848.7683778 | -0.00249157 | 0.066997623 | -0.037188932 | 0.970334363 | 0.992821601 | |
| Neil1 | 250.0495939 | 0.449641886 | 0.152062163 | 2.956961012 | 0.003106874 | 0.025894516 | Neil1 | Prex1 | Bglap | 1.582125684 | -0.002503851 | 0.024539202 | -0.102034723 | 0.918729114 | NA | |
| Apoc1 | 32.9167387 | 0.449418026 | 0.159777274 | 2.812778156 | 0.004911552 | 0.035593822 | Apoc1 | Coa3 | Mir212 | 11.84910035 | -0.00251781 | 0.066463232 | -0.037882754 | 0.969781163 | NA | |
| Rgs22 | 40.47606263 | 0.449242999 | 0.166133384 | 2.704110335 | 0.006848754 | 0.044661027 | Rgs22 | Tbc1d9 | 1810046K07Rik | 0.441471797 | -0.002521546 | 0.015362643 | -0.164134923 | 0.86962493 | NA | |
| Ccdc37 | 103.2056304 | 0.449151474 | 0.160070435 | 2.805961481 | 0.005016669 | 0.03601866 | Ccdc37 | Stxbp6 | Catsper4 | 0.991618214 | -0.002544235 | 0.021068636 | -0.12075937 | 0.903881632 | NA | |
| Icosl | 183.8017346 | 0.449083539 | 0.12699202 | 3.536313072 | 0.000405754 | 0.005884268 | Icosl | Nol4 | Abca5 | 2270.303528 | -0.002546988 | 0.071432217 | -0.035656017 | 0.971556642 | 0.99301409 | |
| Igsf6 | 43.74293926 | 0.448869066 | 0.162898542 | 2.75551309 | 0.005860016 | 0.040157049 | Igsf6 | Rnf217 | Rnf145 | 5013.386771 | -0.002553345 | 0.053099862 | -0.048085713 | 0.961647933 | 0.990572163 | |
| Tnfrsf4 | 33.97468857 | 0.448361619 | 0.167335646 | 2.679414871 | 0.007375095 | 0.047316657 | Tnfrsf4 | Pdgfrb | Acadvl | 1905.657939 | -0.002554022 | 0.072281346 | -0.035334451 | 0.971813053 | 0.99301409 | |
| 2900005J15Rik | 74.23679272 | 0.448278439 | 0.158933802 | 2.820535551 | 0.004794356 | 0.035022986 | 2900005J15Rik | Tle4 | Il9r | 0.449986343 | -0.002559688 | 0.013925377 | -0.183814623 | 0.854158884 | NA | |
| Hapln4 | 4844.91108 | 0.448028645 | 0.097185928 | 4.610015607 | 4.03E-06 | 0.000154037 | Hapln4 | Cox6b1 | Krtap7-1 | 0.608701423 | -0.002565363 | 0.017160961 | -0.149488324 | 0.881168322 | NA | |
| Gm216 | 90.47575467 | 0.447978306 | 0.153937817 | 2.910125101 | 0.003612841 | 0.028913926 | Gm216 | Cspg4 | Grpel2 | 910.0781272 | -0.002569123 | 0.057095758 | -0.044996744 | 0.964109904 | 0.990993714 | |
| Ppp1r36 | 37.06551535 | 0.447799976 | 0.156900116 | 2.854044893 | 0.004316643 | 0.032801407 | Ppp1r36 | Pde7b | Smg7 | 5717.168177 | -0.002569408 | 0.058773942 | -0.043716797 | 0.96513015 | 0.990993714 | |
| Pip5k1b | 695.5536278 | 0.447278948 | 0.115021403 | 3.888658439 | 0.0001008 | 0.001974454 | Pip5k1b | Ttc28 | Fbxo41 | 6782.836369 | -0.00257636 | 0.059528304 | -0.043279589 | 0.965478662 | 0.990993714 | |
| Casq1 | 19.50276899 | 0.447098517 | 0.168047709 | 2.660545152 | 0.007801427 | 0.04927652 | Casq1 | Fam174a | 4930433I11Rik | 0.417516903 | -0.002581817 | 0.013657868 | -0.189035154 | 0.850065265 | NA | |
| Msantd2 | 321.3338878 | 0.446856602 | 0.122915495 | 3.635478188 | 0.000277465 | 0.004355674 | Msantd2 | Neto1 | Arf3 | 61608.71545 | -0.002584133 | 0.053764687 | -0.048063752 | 0.961665435 | 0.990572163 | |
| Ankzf1 | 944.4471104 | 0.446489849 | 0.121097487 | 3.687028203 | 0.000226888 | 0.003716074 | Ankzf1 | Hmgn2 | Dtx4 | 2221.806757 | -0.002584838 | 0.053961469 | -0.047901557 | 0.961794698 | 0.990572163 | |
| Zfp57 | 301.2274226 | 0.445814719 | 0.145327214 | 3.067661626 | 0.002157408 | 0.020258711 | Zfp57 | Ephb4 | Olfr267 | 0.628998992 | -0.002602833 | 0.017273458 | -0.150683938 | 0.880225044 | NA | |
| Gm609 | 9.178286178 | 0.445636307 | 0.155687814 | 2.862371146 | 0.004204842 | 0.032204558 | Gm609 | Psd3 | Ankrd22 | 1.25525968 | -0.002609774 | 0.02438847 | -0.107008527 | 0.914782215 | NA | |
| Cd46 | 40.01352506 | 0.44495608 | 0.163594698 | 2.71986859 | 0.006530786 | 0.043239925 | Cd46 | Immp1l | 5430402E10Rik | 2.930929908 | -0.002609915 | 0.01158062 | -0.225369225 | NA | NA | |
| Hapln2 | 790.3090677 | 0.444811585 | 0.120377161 | 3.695149328 | 0.000219758 | 0.003630313 | Hapln2 | Ctxn1 | Wdpcp | 396.1461506 | -0.002617266 | 0.076520778 | -0.034203337 | 0.972715006 | 0.99301409 | |
| Abcb1b | 124.479352 | 0.444411475 | 0.123186869 | 3.607620509 | 0.000309018 | 0.004773604 | Abcb1b | Dnajb1 | Mup3 | 0.455241012 | -0.00262958 | 0.015473854 | -0.169936959 | 0.865059714 | NA | |
| Plbd2 | 3043.140017 | 0.444376299 | 0.0799921 | 5.555252352 | 2.77E-08 | 2.53E-06 | Plbd2 | Myo5a | Zfp385c | 22.64857211 | -0.002634246 | 0.078176351 | -0.033696204 | 0.973119406 | 0.993070301 | |
| 5031434O11Rik | 70.85937099 | 0.444121975 | 0.155659777 | 2.85315824 | 0.004328706 | 0.032867482 | 5031434O11Rik | Sec22a | Plac8 | 2.612746898 | -0.002641903 | 0.031553756 | -0.083727041 | 0.933273457 | NA | |
| Prpf39 | 1175.645894 | 0.443079558 | 0.111593199 | 3.970488908 | 7.17E-05 | 0.001518185 | Prpf39 | Serpini1 | Gm19303 | 0.436125133 | -0.002672891 | 0.015319177 | -0.174480061 | 0.8614882 | NA | |
| Tanc1 | 1576.170336 | 0.442933464 | 0.12597707 | 3.515984792 | 0.000438126 | 0.006199384 | Tanc1 | Pcnx | Mc5r | 11.14756877 | -0.002676085 | 0.068009102 | -0.039348929 | 0.968612197 | NA | |
| Gpat2 | 10.94092258 | 0.442765655 | 0.159509327 | 2.775797899 | 0.005506643 | 0.038519152 | Gpat2 | Snrpg | Cyp11b2 | 0.48121521 | -0.002690586 | 0.014164252 | -0.18995609 | 0.849343539 | NA | |
| Cabp7 | 277.4505003 | 0.442746738 | 0.151037819 | 2.931363416 | 0.003374777 | 0.027563943 | Cabp7 | Cycs | Olfr1033 | 2.444877895 | -0.002694957 | 0.037967195 | -0.07098119 | 0.943412726 | NA | |
| Stxbp2 | 1447.003217 | 0.442260263 | 0.109109186 | 4.053373317 | 5.05E-05 | 0.001155398 | Stxbp2 | Timm8b | Hal | 1.272710417 | -0.002696885 | 0.02619133 | -0.102968619 | 0.917987877 | NA | |
| BC048546 | 258.7041318 | 0.442013699 | 0.098706759 | 4.478048971 | 7.53E-06 | 0.000252627 | BC048546 | Ehbp1l1 | Mir344b | 0.633678068 | -0.002710549 | 0.017299642 | -0.1566824 | 0.875495159 | NA | |
| Med12 | 1713.181044 | 0.441762167 | 0.103715948 | 4.259346574 | 2.05E-05 | 0.000570317 | Med12 | Myo9a | Krt15 | 3.28384213 | -0.002714209 | 0.027389826 | -0.099095504 | 0.921062442 | NA | |
| Plcd3 | 539.3248578 | 0.441747606 | 0.098310492 | 4.493392285 | 7.01E-06 | 0.000238701 | Plcd3 | Mn1 | Trpm8 | 4.905015686 | -0.002716003 | 0.04619189 | -0.058798269 | 0.953112788 | NA | |
| Sun2 | 2983.500869 | 0.441292907 | 0.081632014 | 5.405880453 | 6.45E-08 | 5.20E-06 | Sun2 | Tm2d1 | Tigit | 0.442291665 | -0.002732824 | 0.013865234 | -0.197099031 | 0.843750042 | NA | |
| Wdr63 | 14.06777705 | 0.441078577 | 0.157117128 | 2.807323315 | 0.004995508 | 0.03594248 | Wdr63 | Atp8a1 | Bpifb1 | 6.436800035 | -0.002734233 | 0.050260888 | -0.054400817 | 0.956615828 | NA | |
| Sfxn2 | 221.7630986 | 0.439940641 | 0.117303182 | 3.750457877 | 0.000176512 | 0.003097255 | Sfxn2 | Grin3a | Zc3h12d | 2.491325265 | -0.002736191 | 0.031006717 | -0.088245093 | 0.929681878 | NA | |
| Tekt4 | 24.96881038 | 0.438625361 | 0.161052983 | 2.723484862 | 0.006459718 | 0.042939425 | Tekt4 | Tomm70a | C4bp | 0.660042781 | -0.002736506 | 0.015545619 | -0.176030685 | 0.860269836 | NA | |
| Prdm8 | 849.4752006 | 0.438233051 | 0.114953388 | 3.812267383 | 0.000137698 | 0.002515908 | Prdm8 | Atp2b4 | Gtf3a | 740.8547991 | -0.002740077 | 0.062744158 | -0.043670628 | 0.965166952 | 0.990993714 | |
| Akap8l | 4295.475447 | 0.437891996 | 0.116825733 | 3.748249508 | 0.000178073 | 0.003110557 | Akap8l | Echdc1 | Clec2e | 0.430558245 | -0.00274496 | 0.015272068 | -0.179737268 | 0.857358834 | NA | |
| Slamf8 | 15.56503004 | 0.437497472 | 0.15455723 | 2.830650319 | 0.004645348 | 0.034306491 | Slamf8 | Fem1b | Slc4a1 | 4.388138427 | -0.002757435 | 0.032685286 | -0.084363189 | 0.932767674 | NA | |
| Aim2 | 71.15896317 | 0.436468281 | 0.150967624 | 2.891138319 | 0.003838491 | 0.030256816 | Aim2 | Map2k1 | Srek1ip1 | 527.6717307 | -0.002763639 | 0.071840971 | -0.038468845 | 0.969313871 | 0.992548843 | |
| Abca4 | 167.8939466 | 0.436262523 | 0.163772426 | 2.663833796 | 0.007725572 | 0.048957062 | Abca4 | Purg | Rad54l2 | 1548.374487 | -0.002766432 | 0.059971854 | -0.046128842 | 0.963207558 | 0.990993714 | |
| Net1 | 1638.574971 | 0.43600555 | 0.124703849 | 3.49632793 | 0.000471709 | 0.006531924 | Net1 | Fasn | Gm19710 | 1.661338131 | -0.002795453 | 0.031005304 | -0.090160494 | 0.928159678 | NA | |
| Rsph4a | 168.3673849 | 0.435846851 | 0.098640488 | 4.418539065 | 9.94E-06 | 0.000311669 | Rsph4a | Il34 | 4930425O10Rik | 0.431048149 | -0.002796209 | 0.015277243 | -0.183031008 | 0.854773689 | NA | |
| Sfswap | 2048.062988 | 0.434417132 | 0.092383846 | 4.70230622 | 2.57E-06 | 0.000109196 | Sfswap | Grik2 | Snord91a | 1.473685646 | -0.002798918 | 0.029359802 | -0.095331652 | 0.924051403 | NA | |
| Eif4ebp3 | 180.6790243 | 0.43413569 | 0.127218902 | 3.412509331 | 0.000643677 | 0.008181407 | Eif4ebp3 | Tpbg | Tro | 9626.730052 | -0.002801993 | 0.058435937 | -0.047949834 | 0.961756223 | 0.990572163 | |
| Syde1 | 485.026951 | 0.434114466 | 0.09981828 | 4.349047762 | 1.37E-05 | 0.000411358 | Syde1 | Stox2 | Grxcr1 | 0.427940552 | -0.00281049 | 0.015250196 | -0.184292029 | 0.853784367 | NA | |
| Grid2ip | 111.1608945 | 0.434095213 | 0.150574921 | 2.882918417 | 0.003940095 | 0.030816672 | Grid2ip | Gda | Sfta2 | 0.427940552 | -0.00281049 | 0.015250196 | -0.184292029 | 0.853784367 | NA | |
| Ccdc176 | 158.4141381 | 0.433570523 | 0.133690279 | 3.243096853 | 0.00118238 | 0.012919462 | Ccdc176 | Chd7 | Shcbp1 | 28.9861601 | -0.00282949 | 0.086444551 | -0.032731854 | 0.973888422 | 0.993518395 | |
| Renbp | 136.9093117 | 0.43295196 | 0.111793981 | 3.87276628 | 0.000107607 | 0.002067036 | Renbp | Lrrn3 | Zfp599 | 203.6688154 | -0.002835371 | 0.093104764 | -0.030453555 | 0.975705334 | 0.994349486 | |
| Cyp27a1 | 227.0613703 | 0.43285843 | 0.13077698 | 3.309897749 | 0.000933301 | 0.010767356 | Cyp27a1 | Grik3 | Bloc1s5 | 609.9466861 | -0.002840792 | 0.071455083 | -0.03975633 | 0.968287392 | 0.992194036 | |
| Gm14446 | 131.7411296 | 0.432347412 | 0.132918517 | 3.252725209 | 0.00114304 | 0.012623135 | Gm14446 | Rpl9 | Mmab | 1529.747413 | -0.002845827 | 0.061720261 | -0.046108477 | 0.96322379 | 0.990993714 | |
| Apobec3 | 120.3943397 | 0.431709652 | 0.132312316 | 3.262807754 | 0.001103143 | 0.012305329 | Apobec3 | Garem | Cyp4f14 | 377.2152851 | -0.002858899 | 0.083074134 | -0.034413829 | 0.972547156 | 0.99301409 | |
| Cdr2l | 1028.27692 | 0.431347814 | 0.113125988 | 3.812986064 | 0.000137298 | 0.002511562 | Cdr2l | Nbea | Raph1 | 2585.685301 | -0.002859242 | 0.072935419 | -0.039202374 | 0.968729041 | 0.992230168 | |
| Il12a | 79.00272076 | 0.430821237 | 0.149071202 | 2.890036652 | 0.003851969 | 0.030341845 | Il12a | Rplp1 | Cdk5rap1 | 347.86108 | -0.002860349 | 0.094601712 | -0.030235697 | 0.97587908 | 0.994349486 | |
| Gm581 | 15.94165174 | 0.430598755 | 0.149784135 | 2.874795481 | 0.004042893 | 0.031273381 | Gm581 | Hn1 | Mir485 | 0.428430455 | -0.002861748 | 0.015255424 | -0.187588892 | 0.851198935 | NA | |
| 2810049E08Rik | 122.5407629 | 0.429492277 | 0.125402147 | 3.424919644 | 0.000614981 | 0.007960334 | 2810049E08Rik | Gp1bb | Nr1i2 | 0.428430455 | -0.002861748 | 0.015255424 | -0.187588892 | 0.851198935 | NA | |
| Snrnp70 | 5977.715205 | 0.429189101 | 0.097328823 | 4.409681425 | 1.04E-05 | 0.000323384 | Snrnp70 | Fam171a1 | Prss36 | 264.0763371 | -0.002863412 | 0.088408744 | -0.032388336 | 0.974162364 | 0.993518395 | |
| Mthfr | 1351.57992 | 0.429180644 | 0.084442751 | 5.082504277 | 3.72E-07 | 2.20E-05 | Mthfr | Uqcrq | Foxj1 | 319.8285421 | -0.00286385 | 0.094785204 | -0.030214105 | 0.9758963 | 0.994349486 | |
| Helq | 481.3091982 | 0.429020795 | 0.109245105 | 3.927139769 | 8.60E-05 | 0.001750191 | Helq | Lrp1 | Msr1 | 10.21934489 | -0.002870453 | 0.047970085 | -0.059838397 | 0.952284344 | NA | |
| Cd6 | 107.2496666 | 0.428775579 | 0.156264713 | 2.743905342 | 0.006071304 | 0.041101633 | Cd6 | Scn3b | Mypn | 178.4685264 | -0.002901569 | 0.095028919 | -0.030533533 | 0.975641551 | 0.994349486 | |
| Hdac10 | 537.4239533 | 0.428547335 | 0.113627166 | 3.771521797 | 0.000162255 | 0.002886313 | Hdac10 | Pde1b | 1700067P10Rik | 1.662250936 | -0.002905932 | 0.02903286 | -0.100091121 | 0.920271985 | NA | |
| Zc2hc1c | 121.000458 | 0.428206817 | 0.153764508 | 2.784822209 | 0.005355708 | 0.037599157 | Zc2hc1c | Htt | Fcrl6 | 1.096054655 | -0.002906255 | 0.019743624 | -0.147199649 | 0.882974438 | NA | |
| Tgm3 | 159.9422721 | 0.427301187 | 0.146006058 | 2.92659903 | 0.003426904 | 0.027877541 | Tgm3 | Cox4i1 | Gltscr1l | 2953.863421 | -0.002928629 | 0.041685746 | -0.07025492 | 0.943990763 | 0.986220499 | |
| Pomc | 104.2136796 | 0.427262683 | 0.141584063 | 3.017731465 | 0.002546745 | 0.022677737 | Pomc | Lamb1 | Cers6 | 1428.572648 | -0.002940339 | 0.086386253 | -0.03403712 | 0.97284755 | 0.99301409 | |
| G530011O06Rik | 121.7452738 | 0.426995434 | 0.138849729 | 3.075234194 | 0.002103373 | 0.019861191 | G530011O06Rik | Lsmd1 | Gli2 | 199.9130977 | -0.002946692 | 0.09154013 | -0.032190165 | 0.974320399 | 0.993609809 | |
| Tonsl | 275.5385023 | 0.426861036 | 0.108629658 | 3.929507328 | 8.51E-05 | 0.001742207 | Tonsl | Nf1 | Ms4a2 | 0.471719916 | -0.002957772 | 0.015603456 | -0.189558747 | 0.849654917 | NA | |
| C1s | 76.64378713 | 0.426434273 | 0.145497406 | 2.930871991 | 0.00338012 | 0.027584543 | C1s | Cox7c | Mtg1 | 669.296821 | -0.002972491 | 0.065717816 | -0.045231133 | 0.963923079 | 0.990993714 | |
| Cdhr3 | 79.0877983 | 0.426367305 | 0.155338658 | 2.744759808 | 0.00605552 | 0.041042969 | Cdhr3 | Thsd7a | Gm5122 | 0.710417717 | -0.002996283 | 0.019190503 | -0.156133636 | 0.875927686 | NA | |
| Tle6 | 155.1927921 | 0.425930202 | 0.141854745 | 3.002579873 | 0.002677017 | 0.023446978 | Tle6 | Fbll1 | Cntnap1 | 13343.88059 | -0.003001168 | 0.050461436 | -0.059474494 | 0.95257418 | 0.988510896 | |
| Slc38a6 | 274.2003973 | 0.425211586 | 0.134450186 | 3.162595757 | 0.001563693 | 0.015960382 | Slc38a6 | Flrt1 | Hist1h1b | 1.847495584 | -0.003018149 | 0.028906537 | -0.104410598 | 0.916843513 | NA | |
| Anxa11 | 1782.042492 | 0.424907234 | 0.120535259 | 3.525169631 | 0.000423212 | 0.006043541 | Anxa11 | Tmem100 | 4930455H04Rik | 0.439740379 | -0.003028123 | 0.015350187 | -0.197269442 | 0.843616691 | NA | |
| Klhl38 | 43.85321957 | 0.42481891 | 0.154242799 | 2.754221992 | 0.005883185 | 0.040226861 | Klhl38 | Nat14 | Pdcl | 861.4130015 | -0.003035058 | 0.074801439 | -0.040574866 | 0.967634821 | 0.991805208 | |

| | | | | | | | | | | | | | | | |
|---|---|---|---|---|---|---|---|---|---|---|---|---|---|---|---|
| Rorb | 1344.004057 | 0.424654058 | 0.103778147 | 4.091941022 | 4.28E-05 | 0.001011906 | Rorb | 1700086L19Rik | Cyp2ab1 | 0.680584311 | -0.00304767 | 0.019050048 | -0.159982247 | 0.872895059 | NA |
| Fam160a1 | 110.5692327 | 0.424522703 | 0.126933764 | 3.344442716 | 0.000824481 | 0.009818987 | Fam160a1 | Mpc2 | Tagap1 | 846.0640037 | -0.003049335 | 0.060772082 | -0.050176574 | 0.959981679 | 0.990205619 |
| Rnf39 | 141.4456213 | 0.424502989 | 0.154459052 | 2.748320559 | 0.005990141 | 0.040814094 | Rnf39 | Amotl1 | Scimp | 1.422092934 | -0.003051282 | 0.028306004 | -0.107796296 | 0.914157281 | NA |
| Cxcl16 | 73.41352993 | 0.424462181 | 0.154554809 | 2.746353765 | 0.006026174 | 0.040952059 | Cxcl16 | Nrp1 | Bola3 | 393.4482145 | -0.003062883 | 0.087215766 | -0.035118452 | 0.971985288 | 0.99301409 |
| 1500015A07Rik | 123.8775294 | 0.424236261 | 0.121731144 | 3.485026479 | 0.000492088 | 0.00675805 | 1500015A07Rik | Wnk3 | R74862 | 175.2880224 | -0.003066671 | 0.084077499 | -0.036474338 | 0.97090414 | 0.992984126 |
| Npy2r | 337.064204 | 0.423848103 | 0.146441847 | 2.894309998 | 0.003799928 | 0.030084867 | Npy2r | Tspan13 | Gm20324 | 8.18426936 | -0.003074271 | 0.057264641 | -0.05368533 | 0.957185871 | NA |
| Odf2 | 2782.072658 | 0.423824338 | 0.105568589 | 4.014682238 | 5.95E-05 | 0.001321339 | Odf2 | Kcnh7 | Hn1 | 3004.893843 | -0.003077197 | 0.050030939 | -0.061505883 | 0.95095633 | 0.988202091 |
| Crebzf | 1541.389889 | 0.422939118 | 0.094547822 | 4.473282498 | 7.70E-06 | 0.00025648 | Crebzf | Gbas | B230119M05Rik | 0.437122686 | -0.003093789 | 0.015328819 | -0.201828293 | 0.840050962 | NA |
| Jpx | 131.3863067 | 0.422480347 | 0.124085338 | 3.404756386 | 0.000662231 | 0.008335181 | Jpx | Rprml | Igfbp1 | 0.437122686 | -0.003093789 | 0.015328819 | -0.201828293 | 0.840050962 | NA |
| Rims3 | 1776.609993 | 0.422387531 | 0.102889699 | 4.105246042 | 4.04E-05 | 0.0009702 | Rims3 | Chd5 | Gns | 5561.183148 | -0.003120826 | 0.053310103 | -0.058540982 | 0.95331772 | 0.988782092 |
| Agxt2l2 | 92.79025906 | 0.422221749 | 0.136641225 | 3.090002658 | 0.002001547 | 0.019149082 | Agxt2l2 | Zfp41 | St6galnac6 | 4872.693625 | -0.003141888 | 0.04870556 | -0.064507797 | 0.948565899 | 0.987378592 |
| 4933421O10Rik | 333.1524645 | 0.421850902 | 0.092811964 | 4.545221162 | 5.49E-06 | 0.000198718 | 4933421O10Rik | Plxna4 | Mfap1b | 184.6184543 | -0.003148125 | 0.089124396 | -0.035322816 | 0.97182233 | 0.99301409 |
| Hjurp | 181.0217292 | 0.421656425 | 0.123577783 | 3.412073067 | 0.000644708 | 0.008187795 | Hjurp | BC030500 | Phlda2 | 0.459676127 | -0.003156792 | 0.014005342 | -0.22539914 | 0.821668781 | NA |
| Ncf2 | 186.8801818 | 0.421386171 | 0.123193996 | 3.420508981 | 0.000625041 | 0.008054403 | Ncf2 | Dlg2 | Tnip3 | 2.50517225 | -0.00316215 | 0.035850824 | -0.088203006 | 0.929715328 | NA |
| Cdt1 | 58.6889381 | 0.421190729 | 0.156836211 | 2.685545181 | 0.007241158 | 0.046631134 | Cdt1 | BC052040 | C8g | 32.38664308 | -0.003166471 | 0.086726251 | -0.036511098 | 0.97087483 | 0.992984126 |
| Rxfp2 | 49.83036557 | 0.421030845 | 0.157731743 | 2.669284165 | 0.007601311 | 0.048288115 | Rxfp2 | Actn2 | Il23a | 2.386181236 | -0.003185427 | 0.035016276 | -0.090969899 | 0.927516509 | NA |
| Niacr1 | 8.704669408 | 0.42011272 | 0.142905427 | 2.939795433 | 0.00328429 | 0.027038677 | Niacr1 | H3f3a | Gm5607 | 522.0120726 | -0.003186676 | 0.084279152 | -0.037810959 | 0.969838406 | 0.992732908 |
| Tlr7 | 183.9730582 | 0.418724091 | 0.136888965 | 3.058859347 | 0.002221814 | 0.020687976 | Tlr7 | Srgap1 | 6330408A02Rik | 648.1914189 | -0.00318771 | 0.070244029 | -0.045380517 | 0.96380401 | 0.990993714 |
| Esrrb | 106.1518655 | 0.418325939 | 0.132399224 | 3.159579992 | 0.001579967 | 0.016067252 | Esrrb | 4-Mar | Mesp1 | 0.922719674 | -0.003188658 | 0.020655315 | -0.154374679 | 0.877314318 | NA |
| Kdm6a | 1413.707683 | 0.417917006 | 0.083617184 | 4.997979924 | 5.79E-07 | 3.15E-05 | Kdm6a | Itgb1bp1 | Fam124b | 0.67239973 | -0.003190049 | 0.019012271 | -0.167788971 | 0.8667493 | NA |
| Cadps2 | 3889.122079 | 0.41755448 | 0.071398308 | 5.848240576 | 4.97E-09 | 5.92E-07 | Cadps2 | D430041D05Rik | Pddc1 | 1100.293617 | -0.003201581 | 0.070397309 | -0.045478736 | 0.963725724 | 0.990993714 |
| Slc26a11 | 164.1703546 | 0.417296465 | 0.108877041 | 3.832731498 | 0.000126728 | 0.002347184 | Slc26a11 | Snx10 | Gm10638 | 26.23123298 | -0.003202424 | 0.084438515 | -0.037926104 | 0.9697466 | 0.992708887 |
| Rgl2 | 1786.332434 | 0.416900757 | 0.114630883 | 3.636897378 | 0.000275942 | 0.00434933 | Rgl2 | Nhp2 | Dnttip1 | 799.0132628 | -0.003221493 | 0.0562761 | -0.057244429 | 0.954350487 | 0.988860025 |
| Kcnq4 | 214.2834339 | 0.416821847 | 0.143003149 | 2.914773916 | 0.003559463 | 0.028648293 | Kcnq4 | Ccnd2 | Gm20753 | 12.33641579 | -0.003243049 | 0.0687138 | -0.047196479 | 0.962356634 | NA |
| Pamr1 | 1324.375835 | 0.41595314 | 0.141470197 | 2.940217427 | 0.00327982 | 0.027016232 | Pamr1 | Bsn | Pop4 | 858.73021 | -0.00324363 | 0.064685724 | -0.050144449 | 0.960007279 | 0.990205619 |
| Osmr | 236.4491491 | 0.415561741 | 0.152611499 | 2.723004126 | 0.006469125 | 0.042962976 | Osmr | Tubb5 | Cenpv | 545.1090951 | -0.00326533 | 0.072295761 | -0.045166271 | 0.963974778 | 0.990993714 |
| Parp12 | 419.7776746 | 0.415409769 | 0.123159648 | 3.37293728 | 0.000743709 | 0.009066516 | Parp12 | Neurl1a | Nsa2 | 1784.717678 | -0.003266392 | 0.066618905 | -0.049031008 | 0.960894585 | 0.990548518 |
| Eya4 | 175.7902938 | 0.415135175 | 0.113460164 | 3.6588628 | 0.000253337 | 0.004084498 | Eya4 | Phlda3 | Ralgapa1 | 6907.960971 | -0.003277209 | 0.062338191 | -0.052571454 | 0.958073362 | 0.989630474 |
| Carns1 | 487.1543612 | 0.415064307 | 0.114101138 | 3.6376877 | 0.000275097 | 0.004344901 | Carns1 | Arl6 | Mir760 | 3.651487852 | -0.003287046 | 0.036317832 | -0.090507767 | 0.927883722 | NA |
| 2610005L07Rik | 599.4656951 | 0.413935451 | 0.150693211 | 2.746875238 | 0.006016602 | 0.0409224 | 2610005L07Rik | Dand5 | Smim6 | 1.367906726 | -0.003296536 | 0.025575414 | -0.128894735 | 0.897440943 | NA |
| Gm10865 | 9.873488124 | 0.413572823 | 0.154621173 | 2.674748962 | 0.007478523 | 0.047684045 | Gm10865 | Commd2 | Olfr857 | 0.458678574 | -0.003299243 | 0.015503839 | -0.212801695 | 0.831481641 | NA |
| 4930563E22Rik | 98.28359016 | 0.413304426 | 0.124492847 | 3.319905016 | 0.000900481 | 0.010498156 | 4930563E22Rik | Elmod1 | Tex29 | 0.665008488 | -0.003302769 | 0.017459755 | -0.189164662 | 0.849963763 | NA |
| Guf1 | 1073.655258 | 0.412992843 | 0.104141767 | 3.965679217 | 7.32E-05 | 0.001540028 | Guf1 | Scd1 | Wnt9b | 26.07869138 | -0.003308603 | 0.073986993 | -0.044718709 | 0.964331521 | 0.990993714 |
| Klf15 | 871.8461861 | 0.412893692 | 0.141516729 | 2.917631692 | 0.003527007 | 0.02846221 | Klf15 | Tmem150c | Ranbp1 | 1419.521914 | -0.003315314 | 0.071183209 | -0.046574377 | 0.962852454 | 0.990993714 |
| Taf1d | 416.5507127 | 0.412550898 | 0.096267806 | 4.285450297 | 1.82E-05 | 0.000518465 | Taf1d | Cox7a2 | 4930474N09Rik | 2.837501275 | -0.003340631 | 0.037804816 | -0.088365222 | 0.929586402 | NA |
| Csf2rb | 74.08500698 | 0.412294298 | 0.135251349 | 3.048356266 | 0.00230097 | 0.021168869 | Csf2rb | Btbd3 | Bid | 439.2711674 | -0.003345131 | 0.078000467 | -0.042886034 | 0.965792381 | 0.991035613 |
| Asun | 927.1327229 | 0.412202871 | 0.081087982 | 5.083402746 | 3.71E-07 | 2.20E-05 | Asun | Vhl | Tspan5 | 8940.87061 | -0.003368138 | 0.051353374 | -0.065587469 | 0.947706266 | 0.987367302 |
| Azi1 | 902.9673431 | 0.412059687 | 0.091215177 | 4.517446547 | 6.26E-06 | 0.000218909 | Azi1 | Vps13c | Kif4 | 67.40690373 | -0.00336927 | 0.095095182 | -0.035430506 | 0.97173646 | 0.99301409 |
| Nr2f1 | 1976.305543 | 0.411975361 | 0.065625128 | 6.277707565 | 3.44E-10 | 5.60E-08 | Nr2f1 | Nfya | 3632454L22Rik | 43.67757844 | -0.003379191 | 0.092596796 | -0.036493603 | 0.970888779 | 0.992984126 |
| Otx2 | 34.56064157 | 0.411885017 | 0.154112164 | 2.672631462 | 0.007525888 | 0.047926884 | Otx2 | Nhsl2 | Plekha1 | 3911.807691 | -0.00338223 | 0.064893532 | -0.052119681 | 0.958433331 | 0.989630474 |
| Dpp7 | 782.6560436 | 0.411862974 | 0.086146189 | 4.780977322 | 1.74E-06 | 7.93E-05 | Dpp7 | Sema5b | Tmem160 | 657.3014426 | -0.003390418 | 0.078221385 | -0.043343874 | 0.965427418 | 0.990993714 |
| Pisd-ps2 | 891.1261271 | 0.411808925 | 0.141550129 | 2.909279768 | 0.003622625 | 0.028942617 | Pisd-ps2 | Arxes1 | Klhdc7b | 16.61368745 | -0.00340148 | 0.075670101 | -0.044951447 | 0.96414601 | NA |
| A230057D06Rik | 222.7510123 | 0.411210893 | 0.146185577 | 2.812937507 | 0.004909119 | 0.035592836 | A230057D06Rik | Tnfrsf19 | C2cd4a | 3.283019704 | -0.003411547 | 0.042146251 | -0.080945446 | 0.935485338 | NA |
| Pole | 84.27169108 | 0.410610961 | 0.14566909 | 2.818792657 | 0.004820465 | 0.035114376 | Pole | Ubxn2b | Ticam1 | 283.0181989 | -0.003423296 | 0.076967113 | -0.044477382 | 0.96452388 | 0.990993714 |
| Efna5 | 1114.893734 | 0.410021757 | 0.088567333 | 4.629491988 | 3.67E-06 | 0.000143423 | Efna5 | Fzd3 | Abhd16a | 3019.27487 | -0.003424125 | 0.061301506 | -0.055857117 | 0.955455633 | 0.989344398 |
| Rgsl1 | 97.99706985 | 0.409837516 | 0.150586721 | 2.721604633 | 0.006496582 | 0.043126835 | Rgsl1 | Lsm4 | Bcl7a | 4411.218361 | -0.003442053 | 0.054218867 | -0.063484419 | 0.949380766 | 0.987872851 |
| H2-D1 | 646.1180434 | 0.409644746 | 0.103068222 | 3.974500954 | 7.05E-05 | 0.001501023 | H2-D1 | Rnf150 | Eml1 | 2756.897811 | -0.003444007 | 0.059672464 | -0.057715174 | 0.953975507 | 0.988860025 |
| Ttc39a | 112.1888321 | 0.40828564 | 0.141939095 | 2.876484734 | 0.004021316 | 0.031153139 | Ttc39a | Tubb2a | Cml2 | 13.95577759 | -0.003446456 | 0.070977999 | -0.048556674 | 0.961272599 | NA |
| Ctnnal1 | 644.0477964 | 0.407932335 | 0.106527333 | 3.829367767 | 0.000128473 | 0.002372538 | Ctnnal1 | Myeov2 | Krt19 | 0.640926902 | -0.003453138 | 0.01733944 | -0.199149372 | 0.842145902 | NA |
| Slc26a4 | 119.7147907 | 0.407358269 | 0.131481796 | 3.098210408 | 0.001946931 | 0.018724864 | Slc26a4 | Ttbk2 | AU022751 | 0.665083679 | -0.003456769 | 0.018977607 | -0.182149879 | 0.855465106 | NA |
| Cep164 | 638.570317 | 0.407302036 | 0.089816394 | 4.534829548 | 5.77E-06 | 0.00020534 | Cep164 | Ets1 | Stx1a | 11587.47922 | -0.003465845 | 0.070049947 | -0.049476763 | 0.960539355 | 0.990548518 |
| Srrm2 | 26989.0295 | 0.407065876 | 0.115031785 | 3.538725205 | 0.000402064 | 0.005876062 | Srrm2 | Romo1 | Pxt1 | 1.617534884 | -0.003465892 | 0.026714986 | -0.129735887 | 0.896775389 | NA |
| Tap2 | 404.9578748 | 0.406917864 | 0.120374429 | 3.380434427 | 0.000723713 | 0.008920617 | Tap2 | Tuba1a | Ndufab1 | 1399.7453 | -0.00346643 | 0.076441876 | -0.045347266 | 0.963830513 | 0.990993714 |
| Il4ra | 312.3189014 | 0.406074535 | 0.113392784 | 3.58113207 | 0.000342109 | 0.005171348 | Il4ra | Ndufa1 | Prdm12 | 8.685178865 | -0.003483265 | 0.021702956 | -0.160497274 | 0.872489369 | NA |
| Osbp2 | 4214.553581 | 0.405888975 | 0.079155281 | 5.127756096 | 2.93E-07 | 1.82E-05 | Osbp2 | Sc5d | Nlrp4f | 6.434465653 | -0.00348929 | 0.055278813 | -0.063121647 | 0.949669637 | NA |
| Kcnt1 | 6820.849648 | 0.405563177 | 0.076972785 | 5.268916491 | 1.37E-07 | 9.62E-06 | Kcnt1 | Slc4a8 | Snord47 | 2.540570966 | -0.003504851 | 0.030868233 | -0.113542326 | 0.909600608 | NA |
| Ppm1m | 632.7102671 | 0.404761785 | 0.08570283 | 4.722852007 | 2.33E-06 | 0.000100932 | Ppm1m | Dpy19l3 | Rbm6 | 2301.653597 | -0.003508267 | 0.050470329 | -0.069511485 | 0.944582491 | 0.986483951 |
| Ak8 | 73.35274009 | 0.403538322 | 0.14253715 | 2.831109795 | 0.004638679 | 0.034306301 | Ak8 | Runx1t1 | Pprc1 | 1234.624616 | -0.003513114 | 0.073205966 | -0.047989451 | 0.96172465 | 0.990572163 |
| Pgm1 | 282.1419386 | 0.402899547 | 0.095536395 | 4.217236257 | 2.47E-05 | 0.000661813 | Pgm1 | Gnb4 | A330070K13Rik | 13.90318997 | -0.003523131 | 0.069341943 | -0.050808087 | 0.959478447 | NA |
| Phka2 | 1480.198793 | 0.401821179 | 0.081290952 | 4.943000072 | 7.69E-07 | 4.00E-05 | Phka2 | Nwd1 | Ecel1 | 1206.789129 | -0.00353915 | 0.078953319 | -0.044825859 | 0.964246113 | 0.990993714 |
| Gata2 | 178.5816356 | 0.401768888 | 0.127322477 | 3.155522072 | 0.001602111 | 0.01619582 | Gata2 | Ntn3 | Gpatch11 | 1151.569811 | -0.003542001 | 0.059008694 | -0.060025066 | 0.952135671 | 0.988510896 |
| Scn1b | 14137.37845 | 0.401690107 | 0.104320172 | 3.850550677 | 0.000117853 | 0.002218721 | Scn1b | Htr1a | Cryba1 | 0.860872084 | -0.003542881 | 0.020783389 | -0.17046693 | 0.86464294 | NA |
| 4831440E17Rik | 156.4821858 | 0.400932099 | 0.147641208 | 2.71558398 | 0.006615899 | 0.043619892 | 4831440E17Rik | Sec61b | Mrps28 | 262.6467706 | -0.003555418 | 0.080156351 | -0.04435604 | 0.964620602 | 0.990993714 |
| Arglu1 | 2910.74368 | 0.40077953 | 0.107168628 | 3.739709423 | 0.000184233 | 0.003203712 | Arglu1 | Rragd | Mars | 2355.168809 | -0.003555857 | 0.054374636 | -0.065395511 | 0.947859098 | 0.987367302 |
| Zbed6 | 1317.7453 | 0.400745696 | 0.115229356 | 3.477809036 | 0.00050553 | 0.006882848 | Zbed6 | Msantd3 | 1700123L14Rik | 2.114527486 | -0.00357396 | 0.030763328 | -0.116175979 | 0.907513074 | NA |
| Nrbp2 | 5296.305868 | 0.399380579 | 0.111883609 | 3.569607577 | 0.000357516 | 0.005357214 | Nrbp2 | Fam84a | Zfp534 | 2.683628588 | -0.00359059 | 0.038093919 | -0.094256245 | 0.924905606 | NA |
| Olfml2b | 250.5628013 | 0.39911716 | 0.148895232 | 2.68052344 | 0.007350712 | 0.047199306 | Olfml2b | Hmgn1 | Tex36 | 0.800350573 | -0.003594136 | 0.018632434 | -0.192896743 | 0.847039836 | NA |
| Frmd5 | 1190.066043 | 0.398958307 | 0.098274685 | 4.059624391 | 4.92E-05 | 0.001131652 | Frmd5 | Enho | Hic1 | 118.6124997 | -0.0036069 | 0.091800109 | -0.039290803 | 0.968658539 | 0.992230168 |
| Mypn | 133.2386028 | 0.397785758 | 0.140799648 | 2.825190001 | 0.004725259 | 0.034663987 | Mypn | Med21 | Cnga2 | 20.97599604 | -0.003615722 | 0.078919358 | -0.045815398 | 0.963457386 | NA |
| Slc4a2 | 1164.158778 | 0.397675655 | 0.097579421 | 4.075404935 | 4.59E-05 | 0.001068629 | Slc4a2 | Gpr124 | Sema3b | 182.502565 | -0.003623316 | 0.057484364 | -0.063031324 | 0.949741561 | 0.987981713 |
| Fam227a | 266.1560058 | 0.39734091 | 0.132586973 | 2.996832198 | 0.002728008 | 0.023786022 | Fam227a | Map1a | D330041H03Rik | 25.39932817 | -0.003627389 | 0.085969785 | -0.042193765 | 0.966344233 | 0.99126569 |
| Qrfpr | 81.90504758 | 0.396995821 | 0.143195914 | 2.772396292 | 0.005564525 | 0.038738759 | Qrfpr | Robo2 | Clstn2 | 3200.494602 | -0.003627935 | 0.055572803 | -0.065282562 | 0.947949026 | 0.987367302 |
| Pde4c | 114.3133344 | 0.396739469 | 0.143036247 | 2.773698817 | 0.005542296 | 0.038692429 | Pde4c | Homer1 | Aqp3 | 1.912165725 | -0.003631224 | 0.032445745 | -0.111916787 | 0.910889386 | NA |
| Gm10560 | 137.6570289 | 0.396621111 | 0.138583945 | 2.861955706 | 0.004210357 | 0.032225775 | Gm10560 | Erc1 | Phrf1 | 2447.189144 | -0.003636697 | 0.05631715 | -0.064575295 | 0.948512155 | 0.987378592 |
| Akap8 | 2964.72911 | 0.396454666 | 0.10146692 | 3.907230708 | 9.34E-05 | 0.00185928 | Akap8 | Med13 | Whsc1l1 | 3737.940638 | -0.00363862 | 0.069421171 | -0.052413696 | 0.958199061 | 0.989630474 |
| Ndor1 | 1375.873407 | 0.396379179 | 0.084893467 | 4.669136417 | 3.02E-06 | 0.000123328 | Ndor1 | Cdk18 | Zbtb45 | 987.4302567 | -0.003639504 | 0.061171557 | -0.059496665 | 0.952556522 | 0.988510896 |
| Ccnl2 | 1906.477927 | 0.396049701 | 0.119327502 | 3.319014425 | 0.000903358 | 0.010507975 | Ccnl2 | Fads1 | Gkn1 | 2.912255501 | -0.003639764 | 0.03843301 | -0.094704111 | 0.924549853 | NA |
| Ccl4 | 8.703824082 | 0.395639078 | 0.129547293 | 3.054012704 | 0.002258025 | 0.020941771 | Ccl4 | Wwtr1 | Trem3 | 1.486938282 | -0.003653409 | 0.029772345 | -0.122711482 | 0.902335571 | NA |
| Slc16a10 | 102.7778088 | 0.395466299 | 0.14192466 | 2.786452332 | 0.005328845 | 0.037473304 | Slc16a10 | Ptar1 | Nup133 | 1812.331158 | -0.003665704 | 0.04639855 | -0.079004721 | 0.937028868 | 0.985461126 |
| 2810403A07Rik | 2212.842361 | 0.395099973 | 0.109819306 | 3.597727821 | 0.000321009 | 0.004932302 | 2810403A07Rik | 1110001J03Rik | Abcc3 | 154.2733847 | -0.00368985 | 0.094133527 | -0.039198039 | 0.968732497 | 0.992230168 |
| Fcgr2b | 150.5227812 | 0.393709344 | 0.136328172 | 2.887952942 | 0.003877579 | 0.030481586 | Fcgr2b | Cd97 | Msto1 | 749.0445541 | -0.003689883 | 0.067539199 | -0.054633213 | 0.956430678 | 0.989344398 |
| Ctss | 2868.987884 | 0.393089456 | 0.101346622 | 3.878663618 | 0.000105032 | 0.002036751 | Ctss | Enox2 | Hist1h1a | 0.660023499 | -0.003689921 | 0.017437486 | -0.21160853 | 0.832412454 | NA |
| Trip6 | 196.8315502 | 0.392629641 | 0.127649664 | 3.075837632 | 0.002099121 | 0.019843675 | Trip6 | Dpysl5 | Morn3 | 5.489750998 | -0.003711553 | 0.048956475 | -0.075813314 | 0.939567624 | NA |
| Nfkbiz | 170.9904054 | 0.392587452 | 0.121574444 | 3.229193898 | 0.001241397 | 0.013431707 | Nfkbiz | Chml | 9630033F20Rik | 1558.318974 | -0.00373014 | 0.055695715 | -0.066973559 | 0.946602753 | 0.98700327 |
| 1700094D03Rik | 260.8815311 | 0.392473969 | 0.135753479 | 2.89107854 | 0.003839222 | 0.030256816 | 1700094D03Rik | Kcna6 | 4933402J07Rik | 1.643947532 | -0.003732286 | 0.027764359 | -0.134427238 | 0.893064744 | NA |
| Ppip5k2 | 1173.157462 | 0.391262511 | 0.070977746 | 5.5124674 | 3.54E-08 | 3.11E-06 | Ppip5k2 | Slc32a1 | Scgn | 10.22539503 | -0.003753421 | 0.058260396 | -0.064424917 | 0.948631891 | NA |
| Atf7 | 313.9838461 | 0.3910663 | 0.094367629 | 4.14407252 | 3.41E-05 | 0.000847891 | Atf7 | Tomm7 | 4930539M17Rik | 5.21443376 | -0.003781179 | 0.045428611 | -0.083233425 | 0.933665935 | NA |
| Ntsr1 | 361.6443807 | 0.390801292 | 0.133669808 | 2.923631741 | 0.003459738 | 0.028006885 | Ntsr1 | Lrrk2 | Tub | 10736.14774 | -0.003786029 | 0.05862387 | -0.064581693 | 0.948507061 | 0.987378592 |
| Gm6548 | 132.9833753 | 0.390653733 | 0.126509694 | 3.08793516 | 0.002015524 | 0.019241242 | Gm6548 | Lzic | Cmpk2 | 1116.615678 | -0.003797865 | 0.064545255 | -0.05884034 | 0.953079277 | 0.988615469 |
| Pstpip2 | 363.3532419 | 0.390492311 | 0.129015807 | 3.026701297 | 0.002472381 | 0.022169068 | Pstpip2 | Mycbp2 | Rbl2 | 2418.098025 | -0.003798315 | 0.053823201 | -0.070570226 | 0.943739807 | 0.986029234 |
| Dgka | 438.2916731 | 0.389287384 | 0.087395892 | 4.454298448 | 8.42E-06 | 0.000273396 | Dgka | Ccdc90b | Cd5 | 5.492453844 | -0.003803212 | 0.049817997 | -0.076342124 | 0.939146913 | NA |
| Ciz1 | 1788.654606 | 0.389287067 | 0.075708734 | 5.141904338 | 2.72E-07 | 1.70E-05 | Ciz1 | Atp5k | T | 4.115627506 | -0.003803632 | 0.041451086 | -0.09176193 | 0.926887192 | NA |
| Tle2 | 1111.733298 | 0.388856397 | 0.140206334 | 2.773458135 | 0.005546398 | 0.038692429 | Tle2 | Phlda1 | Wdr16 | 50.72828986 | -0.003807327 | 0.081822835 | -0.046531354 | 0.962886744 | 0.990993714 |
| Trmt13 | 212.5896922 | 0.388451197 | 0.109934496 | 3.533478683 | 0.000410129 | 0.005927744 | Trmt13 | Cnih2 | Cd7 | 39.383808 | -0.003814839 | 0.090632815 | -0.04209114 | 0.966426043 | 0.99126569 |
| Ankrd24 | 1097.561926 | 0.388268049 | 0.142556471 | 2.723608738 | 0.006457295 | 0.042939425 | Ankrd24 | Chac2 | Vimp | 1236.997722 | -0.003815569 | 0.06423325 | -0.059401774 | 0.9526321 | 0.988510896 |
| Anxa4 | 148.4624249 | 0.386832245 | 0.12062636 | 3.206863288 | 0.001341908 | 0.014201854 | Anxa4 | Calr | Rffl | 690.8468133 | -0.003822459 | 0.07511502 | -0.050888083 | 0.959414702 | 0.990016193 |
| Odf2l | 365.1519257 | 0.386517432 | 0.114760746 | 3.368028227 | 0.000757078 | 0.009189519 | Odf2l | Arhgef19 | Ngp | 15.15045116 | -0.003825209 | 0.052400527 | -0.072999437 | 0.941806566 | NA |
| Ctps2 | 1573.638203 | 0.385517154 | 0.066985222 | 5.755256781 | 8.65E-09 | 9.39E-07 | Ctps2 | Arhgap32 | Lztfl1 | 1471.578493 | -0.003838258 | 0.069011701 | -0.055617494 | 0.955646528 | 0.989344398 |
| Plekhh3 | 409.0509653 | 0.385244042 | 0.100568286 | 3.830671236 | 0.000127794 | 0.00236282 | Plekhh3 | Itga1 | Mageh1 | 853.2570649 | -0.00383844 | 0.062042402 | -0.061868005 | 0.950667947 | 0.988202091 |
| Gpr115 | 95.86240275 | 0.385223841 | 0.13501968 | 2.853094008 | 0.004329581 | 0.032867482 | Gpr115 | St6galnac5 | 0610010F05Rik | 1691.971804 | -0.003843082 | 0.067023538 | -0.057339283 | 0.954274928 | 0.988860025 |
| Fcgr3 | 627.8433476 | 0.384514396 | 0.125639206 | 3.06046502 | 0.002209936 | 0.020614536 | Fcgr3 | Tnr | Zmat1 | 595.6323751 | -0.003852454 | 0.07264697 | -0.053029789 | 0.957708173 | 0.989630474 |
| Ifit2 | 528.9881618 | 0.383933555 | 0.112671032 | 3.407562246 | 0.00065546 | 0.008270829 | Ifit2 | Arhgap31 | Gm13498 | 0.308174694 | -0.003853093 | 0.010968742 | -0.351279401 | 0.725378747 | NA |
| Atxn7l2 | 803.2788139 | 0.383700209 | 0.123955055 | 3.095478496 | 0.001964956 | 0.018886496 | Atxn7l2 | Ndufb6 | Wfdc6a | 1.279167853 | -0.003860488 | 0.020160049 | -0.191491993 | 0.848140153 | NA |
| C77080 | 1044.137935 | 0.383297434 | 0.131411854 | 2.916764537 | 0.003536827 | 0.028526596 | C77080 | Sema6a | Megf8 | 9826.483825 | -0.003867014 | 0.063239616 | -0.061148603 | 0.951240862 | 0.988239091 |
| Pxmp2 | 119.1799182 | 0.383139418 | 0.121312385 | 3.158287723 | 0.001586988 | 0.016102684 | Pxmp2 | Brk1 | Thap2 | 483.5536159 | -0.003886391 | 0.085455731 | -0.045478417 | 0.963725978 | 0.990993714 |
| Tmem181a | 1656.620027 | 0.382756788 | 0.081951111 | 4.670550316 | 3.00E-06 | 0.000122805 | Tmem181a | Ston2 | Snrnp25 | 360.9419796 | -0.003886756 | 0.072303626 | -0.053756027 | 0.957129544 | 0.989630474 |
| 2610015P09Rik | 203.1639337 | 0.382747078 | 0.134749218 | 2.840440058 | 0.004505134 | 0.03360928 | 2610015P09Rik | Erbb4 | A630020A06 | 0.883981836 | -0.003892 | 0.020856787 | -0.186605945 | 0.851969605 | NA |
| Arid3b | 295.5686109 | 0.382067784 | 0.105966617 | 3.60554857 | 0.000311494 | 0.004807063 | Arid3b | Nfia | C430049B03Rik | 6.089537161 | -0.003922127 | 0.052675675 | -0.074458021 | 0.940645943 | NA |
| Pcdhb3 | 236.6135549 | 0.381547815 | 0.095716577 | 3.986225024 | 6.71E-05 | 0.001444662 | Pcdhb3 | Utrn | Rnf144b | 1493.494485 | -0.003936139 | 0.049698912 | -0.079199711 | 0.936873774 | 0.985461126 |
| Srsf11 | 2978.152753 | 0.381315642 | 0.087457831 | 4.359994268 | 1.30E-05 | 0.000395921 | Srsf11 | Acvr1c | Pou4f2 | 1.061619024 | -0.003942391 | 0.015187953 | -0.259573556 | 0.795192737 | NA |
| Mlip | 866.6974943 | 0.379839657 | 0.114788737 | 3.309032468 | 0.00093619 | 0.010792652 | Mlip | Uqcr10 | 1700094J05Rik | 1.091849934 | -0.003956259 | 0.020180474 | -0.196043919 | 0.84457579 | NA |
| C1qc | 1710.56872 | 0.378925295 | 0.104071683 | 3.641002863 | 0.000271578 | 0.004310233 | C1qc | Prex2 | Ankrd49 | 372.218217 | -0.003957635 | 0.073218002 | -0.054052765 | 0.956893126 | 0.989630474 |
| Pcsk4 | 157.2358002 | 0.376972912 | 0.122469657 | 3.078092317 | 0.002083304 | 0.019715626 | Pcsk4 | Syne2 | 1700001K19Rik | 0.30309771 | -0.003960746 | 0.010913552 | -0.362919938 | 0.716664692 | NA |
| Ddit4l | 1039.655694 | 0.376904493 | 0.122922457 | 3.066197184 | 0.002168003 | 0.020333557 | Ddit4l | Adra1a | Trp53tg5 | 0.30309771 | -0.003960746 | 0.010913552 | -0.362919938 | 0.716664692 | NA |
| Heph | 310.0129627 | 0.37662199 | 0.13927249 | 2.7042095 | 0.006846711 | 0.044661027 | Heph | Ddn | Gatad2a | 1424.566333 | -0.003964079 | 0.057872838 | -0.068496367 | 0.945390512 | 0.986781137 |
| Fam21 | 4914.122686 | 0.376505101 | 0.086780986 | 4.338566756 | 1.43E-05 | 0.000428145 | Fam21 | Idh1 | Tsga13 | 5.079378828 | -0.003972086 | 0.050049102 | -0.079363786 | 0.936743273 | NA |
| Sytl2 | 3140.953628 | 0.3757349 | 0.104481081 | 3.596200363 | 0.000322899 | 0.004948565 | Sytl2 | Eif2c2 | Cidea | 73.71269446 | -0.003992642 | 0.091173492 | -0.043791701 | 0.965070442 | 0.990993714 |
| Pnn | 2791.000645 | 0.375389536 | 0.088341018 | 4.249323164 | 2.14E-05 | 0.000590086 | Pnn | Setbp1 | Vmn1r65 | 1.277796275 | -0.003998214 | 0.025073419 | -0.15946026 | 0.873306265 | NA |
| Neu2 | 207.7879031 | 0.37515095 | 0.126427292 | 2.967325678 | 0.003004026 | 0.025392458 | Neu2 | Ppapdc2 | Mtnr1a | 0.751690259 | -0.003999612 | 0.014111683 | -0.283425563 | 0.776850635 | NA |
| Csf1 | 1005.125789 | 0.375019592 | 0.068799224 | 5.450927629 | 5.01E-08 | 4.17E-06 | Csf1 | Cacng4 | Parvb | 998.8332647 | -0.004005675 | 0.069500884 | -0.057634879 | 0.954039466 | 0.988860025 |
| Cc2d2a | 1088.406195 | 0.374362934 | 0.082939001 | 4.513714064 | 6.37E-06 | 0.000222297 | Cc2d2a | Churc1 | Olfr622 | 0.300480017 | -0.004016014 | 0.010882553 | -0.36903232 | 0.712103635 | NA |
| Ccdc136 | 2760.632067 | 0.373653732 | 0.097643577 | 3.826710836 | 0.000129867 | 0.002392578 | Ccdc136 | Pgm2l1 | Rp1l1 | 0.300480017 | -0.004016014 | 0.010882553 | -0.36903232 | 0.712103635 | NA |
| Ucp2 | 845.7712423 | 0.373466475 | 0.126801358 | 2.945287671 | 0.003226545 | 0.026676676 | Ucp2 | Tyro3 | Zbtb32 | 0.300480017 | -0.004016014 | 0.010882553 | -0.36903232 | 0.712103635 | NA |
| Zbp1 | 16.0414848 | 0.373138245 | 0.131747341 | 2.832226011 | 0.004622516 | 0.034225481 | Zbp1 | Spon1 | Ceacam3 | 1.105533145 | -0.004018603 | 0.023850658 | -0.16849025 | 0.866197614 | NA |
| Stat6 | 677.9555643 | 0.373091623 | 0.070977141 | 5.256503981 | 1.47E-07 | 1.02E-05 | Stat6 | Tiam2 | Prkrip1 | 576.9201059 | -0.004032275 | 0.068098463 | -0.059212426 | 0.952782912 | 0.988510896 |
| Slc14a1 | 460.2558083 | 0.372643575 | 0.12888889 | 2.891200127 | 0.003837736 | 0.030256816 | Slc14a1 | Anpep | Gpbar1 | 0.340383369 | -0.004037212 | 0.011323842 | -0.356523122 | 0.721448839 | NA |
| Camk2n2 | 2940.361701 | 0.372580667 | 0.091083597 | 4.090535266 | 4.30E-05 | 0.00101496 | Camk2n2 | Prrx1 | Nron | 0.340383369 | -0.004037212 | 0.011323842 | -0.356523122 | 0.721448839 | NA |
| Cast | 469.7537172 | 0.371928365 | 0.099962635 | 3.720673862 | 0.000198692 | 0.003386724 | Cast | Gm15800 | 2310014L17Rik | 46.37805263 | -0.004042368 | 0.093116525 | -0.043411929 | 0.965373169 | 0.990993714 |
| Cpsf4 | 587.304177 | 0.37187798 | 0.096485577 | 3.854233888 | 0.000116093 | 0.002190912 | Cpsf4 | Rgs9 | Osgep | 748.9787549 | -0.004044695 | 0.058728311 | -0.068871304 | 0.945092061 | 0.986732356 |
| Hpcal1 | 1911.672466 | 0.371098077 | 0.106309882 | 3.490720438 | 0.00048172 | 0.006658136 | Hpcal1 | Mcm5 | Atpaf1 | 2225.955664 | -0.004055748 | 0.066868836 | -0.060652296 | 0.951636124 | 0.988430624 |
| Kifc5b | 71.98163534 | 0.371066222 | 0.138547338 | 2.678263089 | 0.007400506 | 0.047350844 | Kifc5b | Chd3 | 3110007F17Rik | 58.06912255 | -0.00406121 | 0.094604299 | -0.042928386 | 0.965758621 | 0.991035613 |
| Man2c1 | 2203.38762 | 0.370688366 | 0.107201792 | 3.457856074 | 0.000544492 | 0.007285287 | Man2c1 | Prkcdbp | Defb47 | 0.917323538 | -0.004062035 | 0.019292473 | -0.210550275 | 0.833238217 | NA |
| Unc5d | 2101.251659 | 0.369527162 | 0.096597572 | 3.825429094 | 0.000130545 | 0.002399355 | Unc5d | Ehd2 | 3930402G23Rik | 11.29137687 | -0.004071743 | 0.066907755 | -0.060856074 | 0.951473832 | NA |
| 9030624G23Rik | 163.964947 | 0.368922236 | 0.111909717 | 3.296605931 | 0.000978607 | 0.01121448 | 9030624G23Rik | Unc80 | Ccne2 | 136.8195307 | -0.00407961 | 0.094969312 | -0.04295714 | 0.9657357 | 0.991035613 |
| Pcgf2 | 970.1732607 | 0.368507306 | 0.101339147 | 3.63637664 | 0.0002765 | 0.00434933 | Pcgf2 | Wfdc1 | Fst | 144.7548266 | -0.004087097 | 0.090406623 | -0.045207938 | 0.963941567 | 0.990993714 |
| Tbc1d2 | 156.2386477 | 0.368254297 | 0.134210284 | 2.743860501 | 0.006072134 | 0.041101633 | Tbc1d2 | Slc16a2 | Bag1 | 3709.71144 | -0.004089732 | 0.055693105 | -0.073433363 | 0.941461269 | 0.985461126 |
| Ccdc39 | 605.5723044 | 0.368186139 | 0.129721476 | 2.83828206 | 0.004535708 | 0.033802916 | Ccdc39 | Wdfy3 | A330009N23Rik | 498.2306525 | -0.004096805 | 0.07822868 | -0.052369608 | 0.95823419 | 0.989630474 |
| Anks6 | 444.6349305 | 0.367987208 | 0.09068614 | 4.057810898 | 4.95E-05 | 0.001138714 | Anks6 | Zswim6 | Mid1ip1 | 3375.886461 | -0.004102784 | 0.067354879 | -0.06091294 | 0.951428544 | 0.988304564 |
| 4833420G17Rik | 1280.317341 | 0.366745782 | 0.087974402 | 4.168778337 | 3.06E-05 | 0.000782975 | 4833420G17Rik | Ptprr | Chrnb4 | 18.62219421 | -0.004104223 | 0.062837896 | -0.065314455 | 0.947923633 | NA |
| Kcnh4 | 1011.259504 | 0.366222177 | 0.10525568 | 3.479357868 | 0.000502617 | 0.006855234 | Kcnh4 | Fam124a | 2210039B01Rik | 16.34856465 | -0.004107981 | 0.072051286 | -0.057014687 | 0.954533496 | NA |
| Dvl2 | 691.8969593 | 0.365570042 | 0.10401941 | 3.514440655 | 0.000440681 | 0.006229848 | Dvl2 | Birc6 | Mbd3 | 3490.526175 | -0.004108256 | 0.054747232 | -0.075040441 | 0.940182536 | 0.985461126 |
| Apol8 | 222.9614253 | 0.365031068 | 0.098706634 | 3.698141186 | 0.000217184 | 0.003602837 | Apol8 | Acyp2 | Dnajb2 | 4486.084487 | -0.004140727 | 0.051794329 | -0.079945564 | 0.936280551 | 0.985461126 |
| Slc38a7 | 1215.778497 | 0.364952929 | 0.066647459 | 5.475871585 | 4.35E-08 | 3.69E-06 | Slc38a7 | Ptn | Rbmx2 | 229.9177574 | -0.004141979 | 0.083159251 | -0.049807797 | 0.960275553 | 0.990412005 |
| Rhbdf1 | 444.8602606 | 0.364897929 | 0.101014648 | 3.61232687 | 0.000303462 | 0.004711258 | Rhbdf1 | Ncrna00086 | 0610043K17Rik | 15.21360369 | -0.004173882 | 0.067938696 | -0.061435997 | 0.951011985 | NA |
| Ptpn2 | 1044.223472 | 0.364383735 | 0.107556114 | 3.387847719 | 0.000704434 | 0.008745589 | Ptpn2 | Uqcrh | Gng7 | 12528.69263 | -0.00417406 | 0.076325128 | -0.054687886 | 0.956387121 | 0.989344398 |
| Dusp15 | 622.4026598 | 0.363932292 | 0.121204463 | 3.002631117 | 0.002676566 | 0.023446978 | Dusp15 | Zfp462 | Slc1a3 | 25730.02288 | -0.004177965 | 0.057219004 | -0.073017084 | 0.941792523 | 0.985461126 |
| Adamts17 | 194.9772686 | 0.363473049 | 0.132883709 | 2.735271702 | 0.006232881 | 0.041951456 | Adamts17 | Hspa5 | Hoxb8 | 1.835394799 | -0.004185265 | 0.026201753 | -0.159732257 | 0.87309199 | NA |
| 9330151L19Rik | 740.6043178 | 0.363002254 | 0.081927183 | 4.430791339 | 9.39E-06 | 0.000297485 | 9330151L19Rik | Ehhadh | Ccdc122 | 57.62077178 | -0.004199771 | 0.094254056 | -0.044557991 | 0.964459628 | 0.990993714 |
| Gnl3 | 885.9117574 | 0.362312976 | 0.08953426 | 4.04664066 | 5.20E-05 | 0.001183876 | Gnl3 | Nanos1 | Rps4x | 9600.652649 | -0.004207512 | 0.050243111 | -0.083743057 | 0.933260723 | 0.984471054 |
| Kcnj9 | 5660.677355 | 0.362271942 | 0.096245895 | 3.764024881 | 0.0001672 | 0.002947213 | Kcnj9 | Mageh1 | Gm16515 | 654.4632133 | -0.004210548 | 0.065090409 | -0.064687683 | 0.94842267 | 0.987378592 |
| Ccdc22 | 1247.335818 | 0.362124209 | 0.071704768 | 5.050210993 | 4.41E-07 | 2.52E-05 | Ccdc22 | Saysd1 | Gtf3c3 | 1298.514192 | -0.0042147 | 0.065032574 | -0.064809065 | 0.948326024 | 0.987378592 |
| Krba1 | 1777.096354 | 0.361598954 | 0.085273431 | 4.240464453 | 2.23E-05 | 0.000609534 | Krba1 | Mrpl54 | Spag8 | 48.39261381 | -0.00422263 | 0.092712356 | -0.045545492 | 0.963672515 | 0.990993714 |
| Rgs16 | 397.1625174 | 0.360849725 | 0.086803831 | 4.157071408 | 3.22E-05 | 0.000817437 | Rgs16 | Dclk3 | 1700030F04Rik | 0.925531857 | -0.004224836 | 0.02243494 | -0.188315024 | 0.850629709 | NA |
| Neurod2 | 2371.290604 | 0.359629948 | 0.101926648 | 3.528321143 | 0.000418204 | 0.006005245 | Neurod2 | Gfra1 | Prrc2a | 21973.51427 | -0.004234183 | 0.050502023 | -0.083841856 | 0.933182169 | 0.984471054 |
| Letm2 | 393.1244131 | 0.359541351 | 0.094616177 | 3.799998716 | 0.000144697 | 0.002634469 | Letm2 | Bmpr1b | 4922502N22Rik | 0.665186487 | -0.004249098 | 0.015590847 | -0.272537983 | 0.78520839 | NA |
| Tmem181b-ps | 8877.054085 | 0.359107604 | 0.114501339 | 3.136274265 | 0.001711091 | 0.017022514 | Tmem181b-ps | Hspa12b | 4930469G21Rik | 0.290056368 | -0.004253036 | 0.010765216 | -0.395072051 | 0.692789715 | NA |
| Car7 | 613.3628753 | 0.359092223 | 0.119930463 | 2.994170246 | 0.002751923 | 0.023913798 | Car7 | D3Bwg0562e | Ambn | 0.290056368 | -0.004253036 | 0.010765216 | -0.395072051 | 0.692789715 | NA |
| Rrnad1 | 824.4262619 | 0.358473555 | 0.10800452 | 3.31906066 | 0.000903208 | 0.010507975 | Rrnad1 | Gap43 | Mir744 | 0.290056368 | -0.004253036 | 0.010765216 | -0.395072051 | 0.692789715 | NA |
| Sirt4 | 484.8012323 | 0.358470251 | 0.126398203 | 2.836039147 | 0.004567685 | 0.033967062 | Sirt4 | Rgs20 | Pitx1 | 0.290056368 | -0.004253036 | 0.010765216 | -0.395072051 | 0.692789715 | NA |
| D930048N14Rik | 212.2627533 | 0.358134651 | 0.122139116 | 2.932186368 | 0.003365847 | 0.027505503 | D930048N14Rik | Epha7 | Pof1b | 0.290056368 | -0.004253036 | 0.010765216 | -0.395072051 | 0.692789715 | NA |
| Rbm5 | 4432.782038 | 0.357669824 | 0.066206006 | 5.402377324 | 6.58E-08 | 5.28E-06 | Rbm5 | Vps13d | 4930481A15Rik | 71.31914043 | -0.004275343 | 0.094155093 | -0.045407455 | 0.963782539 | 0.990993714 |
| Fcer1g | 321.350714 | 0.357325884 | 0.102884591 | 3.473074842 | 0.000514532 | 0.006962582 | Fcer1g | Gm561 | Abca15 | 1.86664755 | -0.004276333 | 0.032011625 | -0.133586862 | 0.893729274 | NA |
| 9430008C03Rik | 289.9068602 | 0.357255213 | 0.127117382 | 2.810435586 | 0.004947449 | 0.035738209 | 9430008C03Rik | Sh3rf1 | Zfp259 | 1524.184613 | -0.004280706 | 0.052972192 | -0.080810444 | 0.935592702 | 0.985461126 |
| Lgi3 | 4565.731009 | 0.357144893 | 0.084502183 | 4.226457606 | 2.37E-05 | 0.000639697 | Lgi3 | 1810037I17Rik | Nms | 1.115474863 | -0.004302601 | 0.023860998 | -0.18031941 | 0.856901819 | NA |
| Ctns | 694.6977618 | 0.357028161 | 0.083472835 | 4.277177855 | 1.89E-05 | 0.000533212 | Ctns | Snx7 | Apoc2 | 0.840692375 | -0.004310662 | 0.021860921 | -0.197185744 | 0.843682186 | NA |
| Parvg | 262.5668192 | 0.356994034 | 0.133330038 | 2.677521438 | 0.007416911 | 0.047350844 | Parvg | Sqle | Olfr1346 | 0.326538963 | -0.004316169 | 0.011184303 | -0.385913091 | 0.699561035 | NA |
| Arhgef1 | 1649.310176 | 0.356879914 | 0.115621561 | 3.086620783 | 0.002024457 | 0.019314614 | Arhgef1 | Cetn2 | Prmt3 | 930.193407 | -0.004318645 | 0.059490495 | -0.072593873 | 0.942129302 | 0.985461126 |
| Sec14l5 | 377.5058558 | 0.356797305 | 0.120089584 | 2.971092856 | 0.00296742 | 0.025179196 | Sec14l5 | Abcb10 | Naaa | 1451.96377 | -0.004325197 | 0.075468217 | -0.057311507 | 0.954297054 | 0.988860025 |
| Cyp4f13 | 287.6000724 | 0.356327703 | 0.127470818 | 2.795366877 | 0.005184085 | 0.03674392 | Cyp4f13 | Tmem121 | 4930451I11Rik | 2.797978986 | -0.004326416 | 0.038818551 | -0.111452283 | 0.911257702 | NA |
| Gm166 | 183.7862171 | 0.356006853 | 0.110238709 | 3.229417846 | 0.001240425 | 0.013431707 | Gm166 | Glra2 | 1700001J11Rik | 2.449557285 | -0.004328608 | 0.034469851 | -0.125576641 | 0.900067054 | NA |
| H2-Q4 | 388.8534656 | 0.35592999 | 0.115381615 | 3.084806793 | 0.002036844 | 0.019331788 | H2-Q4 | Clmp | Tsta3 | 770.5214808 | -0.004336695 | 0.06669512 | -0.065022671 | 0.948155949 | 0.987376394 |
| Cep44 | 266.9074885 | 0.355404659 | 0.104782226 | 3.391841079 | 0.000694247 | 0.008660759 | Cep44 | Ppp1r3c | Cwf19l1 | 655.4172107 | -0.004344386 | 0.060506076 | -0.071800824 | 0.942760417 | 0.985461126 |
| Ddb2 | 426.7535829 | 0.355076098 | 0.100790093 | 3.522926572 | 0.00042681 | 0.006083705 | Ddb2 | Tgfa | AA987161 | 436.5951786 | -0.004344668 | 0.086171543 | -0.050418823 | 0.959788637 | 0.990191033 |
| Rufy2 | 2253.489513 | 0.354842098 | 0.080081661 | 4.431003201 | 9.38E-06 | 0.000297485 | Rufy2 | Crip2 | Topaz1 | 0.900394332 | -0.004346429 | 0.020597226 | -0.211020121 | 0.832871571 | NA |
| Itga7 | 537.0391671 | 0.3545551 | 0.117092114 | 3.028001519 | 0.002461768 | 0.022124502 | Itga7 | Cd59a | Ptbp3 | 946.8823875 | -0.004362915 | 0.081315444 | -0.053654197 | 0.957210676 | 0.989630474 |
| Napepld | 709.4210356 | 0.353240419 | 0.090384861 | 3.908181232 | 9.30E-05 | 0.001854365 | Napepld | Ptprt | Slc2a2 | 1.690456458 | -0.004373433 | 0.02465689 | -0.177371654 | 0.859216471 | NA |
| Sh2d3c | 806.80103 | 0.353164631 | 0.081838263 | 4.315397413 | 1.59E-05 | 0.000465744 | Sh2d3c | Kcns2 | Apol11b | 0.9347476 | -0.004374767 | 0.019049056 | -0.229657928 | 0.81835759 | NA |
| Csf3r | 333.1659592 | 0.352781434 | 0.123881412 | 2.847735011 | 0.004403157 | 0.033182157 | Csf3r | Mnf1 | Zfp263 | 943.4552145 | -0.004378076 | 0.054793347 | -0.0799016 | 0.936315518 | 0.985461126 |
| Fah | 808.7980251 | 0.352217265 | 0.096741286 | 3.64081646 | 0.000271775 | 0.004310233 | Fah | Sec61g | Rnase10 | 1.582225195 | -0.004378188 | 0.029555093 | -0.148136494 | 0.882235049 | NA |
| Epb4.1l4b | 414.2891405 | 0.352062702 | 0.116893856 | 3.011815283 | 0.002596906 | 0.022992231 | Epb4.1l4b | Diras2 | Nrp | 5.007925122 | -0.00438271 | 0.047830838 | -0.091629383 | 0.926992506 | NA |
| Clcn2 | 2488.70766 | 0.35193773 | 0.079367625 | 4.434273153 | 9.24E-06 | 0.000295885 | Clcn2 | Bmpr2 | Stmn4 | 9551.004108 | -0.004392826 | 0.053554136 | -0.082025895 | 0.934626122 | 0.985294234 |
| Fam161a | 90.70667909 | 0.351657004 | 0.130943181 | 2.685569425 | 0.007240633 | 0.046631134 | Fam161a | Capn11 | Spata21 | 27.45849805 | -0.004397074 | 0.080316929 | -0.054746543 | 0.956340389 | 0.989344398 |
| Prpf4b | 3170.185032 | 0.351609355 | 0.076831034 | 4.576397556 | 4.73E-06 | 0.000176613 | Prpf4b | Gas1 | Sall4 | 7.013722063 | -0.004403226 | 0.059271082 | -0.074289615 | 0.94077994 | NA |
| Tardbp | 6429.134021 | 0.350995277 | 0.086632344 | 4.051550036 | 5.09E-05 | 0.001162723 | Tardbp | Elovl6 | Pilrb1 | 0.674595778 | -0.004423541 | 0.019030213 | -0.232448304 | 0.816189835 | NA |
| Stat2 | 1223.068967 | 0.350726365 | 0.100142316 | 3.502279359 | 0.000461296 | 0.006438851 | Stat2 | Pts | Tgs1 | 1418.371686 | -0.004425742 | 0.072102354 | -0.061381377 | 0.951055483 | 0.988202091 |
| Rtel1 | 746.5488996 | 0.350557246 | 0.130086218 | 2.694806949 | 0.007042942 | 0.045660528 | Rtel1 | Sema5a | Lhx4 | 1.064075119 | -0.004425915 | 0.02341493 | -0.189021083 | 0.850076292 | NA |
| Inpp5e | 1351.246761 | 0.350212131 | 0.065619225 | 5.337035433 | 9.45E-08 | 7.18E-06 | Inpp5e | Epha6 | 1700016H13Rik | 0.321461979 | -0.004426412 | 0.011133506 | -0.397575723 | 0.690942959 | NA |
| Safb2 | 2111.924117 | 0.350015125 | 0.097213311 | 3.600485575 | 0.000317623 | 0.004887048 | Safb2 | Shisa3 | Pars2 | 362.3155421 | -0.004426707 | 0.076559627 | -0.057820377 | 0.953891707 | 0.988860025 |
| Ogt | 11070.76687 | 0.34930878 | 0.087057659 | 4.012384256 | 6.01E-05 | 0.00132634 | Ogt | Otud1 | Ankar | 0.907937044 | -0.004440988 | 0.018611499 | -0.238615295 | 0.811403904 | NA |
| Map4k2 | 2475.690992 | 0.349150339 | 0.090162815 | 3.872442752 | 0.00010775 | 0.002067036 | Map4k2 | Mettl23 | Pgd | 1425.006829 | -0.004447787 | 0.065262836 | -0.068151917 | 0.945664702 | 0.986781137 |
| Usp54 | 3702.560475 | 0.348788941 | 0.129108445 | 2.701519193 | 0.00690235 | 0.044916004 | Usp54 | Adamts19 | Cd200r1 | 24.3849909 | -0.004457661 | 0.079948421 | -0.055756709 | 0.955535622 | 0.989344398 |
| Amt | 679.7474495 | 0.347780219 | 0.10655192 | 3.263950762 | 0.001098703 | 0.012273466 | Amt | Aldh1b1 | Zfp109 | 336.5568628 | -0.004460523 | 0.078669505 | -0.056699519 | 0.954784557 | 0.988953472 |
| Pxn | 1177.552426 | 0.347745972 | 0.102770294 | 3.383720695 | 0.000715107 | 0.008835621 | Pxn | 2010107E04Rik | Zfp493 | 153.2149501 | -0.004474885 | 0.091804397 | -0.048743685 | 0.961123562 | 0.990548518 |
| Srrm1 | 4548.41591 | 0.347485318 | 0.091854072 | 3.78301485 | 0.00015494 | 0.00277531 | Srrm1 | Atp5j2 | 4930428G15Rik | 0.318844286 | -0.004484365 | 0.01110653 | -0.403759272 | 0.686389749 | NA |
| Ppp1r13l | 149.5011414 | 0.34724911 | 0.120248328 | 2.887766635 | 0.003879876 | 0.030484179 | Ppp1r13l | Slc17a8 | Gm13944 | 0.318844286 | -0.004484365 | 0.01110653 | -0.403759272 | 0.686389749 | NA |
| Havcr2 | 188.1072072 | 0.347015228 | 0.116411771 | 2.980929039 | 0.002873754 | 0.024599967 | Havcr2 | Tma7 | Tbx6 | 39.51283578 | -0.004508295 | 0.084181395 | -0.053554528 | 0.957290085 | 0.989630474 |
| Bhlhe41 | 1655.531812 | 0.345901421 | 0.078808405 | 4.389143776 | 1.14E-05 | 0.000353593 | Bhlhe41 | Heatr3 | Ikbkg | 1993.181154 | -0.004517295 | 0.051067996 | -0.088456471 | 0.92951388 | 0.983689248 |
| Mtmr11 | 357.41333 | 0.345861613 | 0.099931997 | 3.460969704 | 0.000538233 | 0.007207768 | Mtmr11 | Kcnj4 | Tacc1 | 10441.74042 | -0.004531946 | 0.071329879 | -0.063535026 | 0.949340469 | 0.987872851 |
| Atp11b | 3451.526136 | 0.345730384 | 0.083639093 | 4.133597948 | 3.57E-05 | 0.000875529 | Atp11b | Adcy2 | Prkab1 | 607.2578945 | -0.004560789 | 0.082457057 | -0.05531108 | 0.955890635 | 0.989344398 |
| Nsun5 | 372.968735 | 0.34513212 | 0.093708652 | 3.683033685 | 0.000230475 | 0.003762882 | Nsun5 | Bcl2 | Ezh1 | 3365.78814 | -0.004568254 | 0.046612922 | -0.098004023 | 0.921929098 | 0.982822698 |
| Ypel4 | 1336.54292 | 0.344873913 | 0.07785224 | 4.429852149 | 9.43E-06 | 0.000298173 | Ypel4 | Fam101b | Msx1 | 116.7575214 | -0.004594882 | 0.072203394 | -0.063638037 | 0.949258444 | 0.987872851 |
| Fancm | 553.0535076 | 0.344632378 | 0.104006582 | 3.313563148 | 0.000921153 | 0.010651004 | Fancm | Rem2 | Tvp23b | 881.7163569 | -0.004620833 | 0.062128245 | -0.074375727 | 0.940711422 | 0.985461126 |
| Wdr67 | 291.2082765 | 0.344571742 | 0.092531309 | 3.723839482 | 0.000196216 | 0.003355592 | Wdr67 | Slc9a7 | Uts2r | 6.509905286 | -0.004650119 | 0.051467042 | -0.090351402 | 0.928007975 | NA |
| Wdr60 | 797.2392328 | 0.34450001 | 0.086051712 | 4.003406834 | 6.24E-05 | 0.001364451 | Wdr60 | Sik2 | Gm19522 | 43.85926249 | -0.004659764 | 0.088584786 | -0.052602304 | 0.958048781 | 0.989630474 |
| 2010015L04Rik | 691.6729612 | 0.344360354 | 0.071396396 | 4.82321761 | 1.41E-06 | 6.55E-05 | 2010015L04Rik | Zfp521 | Cebpg | 1461.664759 | -0.004659813 | 0.068703088 | -0.067825373 | 0.945924645 | 0.986798749 |
| Fam214a | 1950.276933 | 0.34423913 | 0.117253706 | 2.935848605 | 0.003326366 | 0.027283599 | Fam214a | 6330403M23Rik | Spata9 | 2.786785386 | -0.00467504 | 0.037317775 | -0.125276491 | 0.900304663 | NA |
| Cln3 | 541.7537533 | 0.344016268 | 0.099687509 | 3.450946587 | 0.000558624 | 0.007404262 | Cln3 | Snca | 1110032A03Rik | 1138.980879 | -0.004686032 | 0.064005411 | -0.073213052 | 0.94163658 | 0.985461126 |
| Ppox | 781.6305115 | 0.343877204 | 0.126647787 | 2.71522474 | 0.006623081 | 0.043630108 | Ppox | Cntnap3 | Tefm | 158.1871947 | -0.004701964 | 0.09212783 | -0.051037389 | 0.959295728 | 0.990016193 |
| A930017M01Rik | 477.1660834 | 0.343538779 | 0.093780295 | 3.663229868 | 0.000249055 | 0.004028033 | A930017M01Rik | Slco1c1 | Faim3 | 2.3892318 | -0.004705383 | 0.03365196 | -0.139824928 | 0.888798317 | NA |
| Fbxo27 | 483.2581327 | 0.342401876 | 0.094370067 | 3.628288988 | 0.000285306 | 0.00446518 | Fbxo27 | 4833424O15Rik | 2010308F09Rik | 10.26049401 | -0.004724382 | 0.064513363 | -0.073231066 | 0.941622245 | NA |
| Tuft1 | 90.24635769 | 0.342363264 | 0.121823713 | 2.8103171 | 0.004949271 | 0.035738209 | Tuft1 | Penk | Epg5 | 2953.966204 | -0.004725263 | 0.062072121 | -0.076125367 | 0.939319359 | 0.985461126 |
| Mettl17 | 661.8600698 | 0.342293597 | 0.114670978 | 2.98500635 | 0.002835724 | 0.024368671 | Mettl17 | Rn45s | Ccl19 | 0.510493435 | -0.004725723 | 0.012838263 | -0.368096735 | 0.712801105 | NA |
| Nt5dc3 | 6242.31945 | 0.342147904 | 0.10238873 | 3.341655918 | 0.000832802 | 0.009880121 | Nt5dc3 | Cxxc4 | Ripply1 | 0.308420637 | -0.004726334 | 0.01099736 | -0.429769866 | 0.667363055 | NA |
| Jun | 1733.054572 | 0.341968254 | 0.085701082 | 3.990244276 | 6.60E-05 | 0.001424354 | Jun | Epha5 | U90926 | 0.308420637 | -0.004726334 | 0.01099736 | -0.429769866 | 0.667363055 | NA |
| Dmpk | 531.2922133 | 0.341531986 | 0.111442362 | 3.06465136 | 0.002179239 | 0.020414223 | Dmpk | Gucy1a2 | Zyg11a | 0.308420637 | -0.004726334 | 0.01099736 | -0.429769866 | 0.667363055 | NA |
| Ltk | 1168.342884 | 0.34128215 | 0.12664514 | 2.694790725 | 0.007043285 | 0.045660528 | Ltk | Hist3h2ba | 1700016L04Rik | 1.66120297 | -0.004728728 | 0.030767658 | -0.153691511 | 0.877852978 | NA |
| 4732491K20Rik | 148.1657636 | 0.341199546 | 0.118832041 | 2.87127566 | 0.004088188 | 0.031506284 | 4732491K20Rik | Gdpd4 | Unc5b | 1589.670146 | -0.004729726 | 0.0824902 | -0.057336818 | 0.954276892 | 0.988860025 |
| Zcchc7 | 763.5175243 | 0.341078071 | 0.111068382 | 3.070883581 | 0.002134263 | 0.020096898 | Zcchc7 | Tnrc6b | 1700019G24Rik | 4.022786096 | -0.004740066 | 0.041223167 | -0.114985498 | 0.908456617 | NA |
| Mlh1 | 431.2374349 | 0.340751541 | 0.105953683 | 3.216042439 | 0.001299716 | 0.013849883 | Mlh1 | Egfr | Cym | 1.889006502 | -0.004740982 | 0.032750077 | -0.144762469 | 0.884898415 | NA |
| Gtdc1 | 2151.831098 | 0.340440626 | 0.069206793 | 4.919179345 | 8.69E-07 | 4.33E-05 | Gtdc1 | Hopx | Peg12 | 2.648344487 | -0.004742748 | 0.039787042 | -0.119203331 | 0.905114268 | NA |
| Gba | 1378.29634 | 0.339685828 | 0.077423396 | 4.387379621 | 1.15E-05 | 0.000354799 | Gba | Tet1 | Commd7 | 970.3313391 | -0.004746805 | 0.054797596 | -0.08662432 | 0.930970134 | 0.984208871 |
| Usp35 | 936.0955222 | 0.33959468 | 0.073258454 | 4.635569819 | 3.56E-06 | 0.000139979 | Usp35 | Id4 | Zfp869 | 1279.975548 | -0.004759075 | 0.055988366 | -0.085001138 | 0.932260486 | 0.984387568 |
| Gigyf1 | 2770.800586 | 0.339511098 | 0.095739763 | 3.546186948 | 0.000390849 | 0.005732886 | Gigyf1 | Nrarp | 1700071M16Rik | 0.506412023 | -0.004760294 | 0.014481862 | -0.328707328 | 0.742376915 | NA |
| Crtc2 | 995.1795021 | 0.339118138 | 0.091343487 | 3.712559597 | 0.000205174 | 0.003466698 | Crtc2 | Deptor | Ppif | 1024.156637 | -0.004763285 | 0.056234678 | -0.084703692 | 0.932496961 | 0.984387568 |
| Nphp4 | 709.628538 | 0.338924177 | 0.101211468 | 3.348673648 | 0.000811994 | 0.009692629 | Nphp4 | Hras1 | Disc1 | 35.72378749 | -0.00477442 | 0.088937342 | -0.053682961 | 0.957187759 | 0.989630474 |
| Prrt2 | 11461.04187 | 0.338893826 | 0.090389591 | 3.749257228 | 0.000177359 | 0.003101582 | Prrt2 | 1810041L15Rik | Suv39h2 | 173.4107454 | -0.004785562 | 0.088360639 | -0.054159429 | 0.956808144 | 0.989630474 |
| Pcdhga1 | 235.6751227 | 0.338581142 | 0.119396448 | 2.835772323 | 0.004571502 | 0.033971634 | Pcdhga1 | Syt17 | Pfn3 | 1.452028206 | -0.00480185 | 0.024893287 | -0.192897371 | 0.847039344 | NA |
| Nvl | 1678.933046 | 0.338191342 | 0.118745577 | 2.848033158 | 0.004399034 | 0.033182157 | Nvl | 6430573F11Rik | Antxr1 | 891.1906019 | -0.004802991 | 0.072488868 | -0.066258324 | 0.947172163 | 0.987272017 |
| Mapk1ip1 | 1139.927569 | 0.33815085 | 0.080959079 | 4.176811942 | 2.96E-05 | 0.0007596 | Mapk1ip1 | Fgf10 | 1700034I23Rik | 0.854866745 | -0.004814891 | 0.022228287 | -0.216610966 | 0.828511539 | NA |
| Vopp1 | 6048.025157 | 0.338063605 | 0.115613605 | 2.924081511 | 0.003454743 | 0.027983052 | Vopp1 | Lhfp | Pecr | 161.7224913 | -0.004815478 | 0.090061383 | -0.053468847 | 0.957358351 | 0.989630474 |
| Tprn | 705.7493628 | 0.337821789 | 0.0982861 | 3.4371268 | 0.00058792 | 0.007693612 | Tprn | Mpped1 | Tfpi2 | 0.674790267 | -0.004816177 | 0.019031211 | -0.253067301 | 0.800216206 | NA |
| Usp40 | 771.5502997 | 0.337582827 | 0.104012551 | 3.245597021 | 0.001172046 | 0.012845805 | Usp40 | 4932438A13Rik | Obfc1 | 483.7242922 | -0.004824975 | 0.070868322 | -0.068083669 | 0.945719031 | 0.986781137 |
| Rbm25 | 3887.998302 | 0.337021307 | 0.073849074 | 4.563649715 | 5.03E-06 | 0.000185456 | Rbm25 | A730017L22Rik | Nkx2-9 | 16.94863923 | -0.004826928 | 0.076961824 | -0.062718467 | 0.949990692 | NA |
| Rbm12b1 | 233.9514309 | 0.336852811 | 0.110031481 | 3.061422126 | 0.002202883 | 0.020597302 | Rbm12b1 | Dnahc5 | Spn-ps | 0.656915902 | -0.004846997 | 0.020240908 | -0.239465405 | 0.810744719 | NA |
| A230050P20Rik | 1991.438953 | 0.336448249 | 0.081271092 | 4.139826856 | 3.48E-05 | 0.000857519 | A230050P20Rik | Itpr1 | Gpr152 | 2.709964921 | -0.004882338 | 0.034797384 | -0.14030762 | 0.888416944 | NA |

| | | | | | | | | | | | | | | |
|---|---|---|---|---|---|---|---|---|---|---|---|---|---|---|
| Farsa | 1127.283223 | 0.336445413 | 0.083847012 | 4.012610665 | 6.01E-05 | 0.00132634 | Farsa | Fam210b | Lamb1 | 790.3441974 | -0.004883731 | 0.077066734 | -0.063370152 | 0.949471755 | 0.987896763 |
| Pygm | 1424.660383 | 0.336149771 | 0.083783852 | 4.012106884 | 6.02E-05 | 0.00132634 | Pygm | Gnb5 | Il2rb | 18.81958276 | -0.004885773 | 0.069244391 | -0.070558387 | 0.94374923 | NA |
| Cc2d1b | 1044.503122 | 0.33605806 | 0.111968883 | 3.001352248 | 0.002687834 | 0.023503194 | Cc2d1b | Mri1 | Fam78b | 1812.157363 | -0.004904272 | 0.051743214 | -0.094780966 | 0.924488806 | 0.983236501 |
| Cmbl | 292.6751902 | 0.335857891 | 0.103799823 | 3.235630669 | 0.001213743 | 0.01322485 | Cmbl | Rftn1 | Wnt1 | 3.788024226 | -0.004909909 | 0.038986113 | -0.125939944 | 0.899779464 | NA |
| Clasrp | 1568.212077 | 0.33559437 | 0.118531473 | 2.831268037 | 0.004636385 | 0.034305706 | Clasrp | Flrt3 | Pkdcc | 332.0798693 | -0.004913448 | 0.089227717 | -0.055066383 | 0.956085578 | 0.989344398 |
| Ccnl1 | 1004.294713 | 0.335005052 | 0.118017966 | 2.838593666 | 0.004531282 | 0.033786179 | Ccnl1 | Gpr88 | Dlgap2 | 2840.331024 | -0.004935316 | 0.066929881 | -0.073738597 | 0.941218387 | 0.985461126 |
| Mvp | 298.5631138 | 0.334685876 | 0.089969009 | 3.720012927 | 0.000199213 | 0.003388145 | Mvp | Scube1 | 1190003J15Rik | 28.26943196 | -0.004945897 | 0.073401405 | -0.067381502 | 0.946277995 | 0.986835678 |
| Pask | 159.3746101 | 0.333820147 | 0.111594357 | 2.991371205 | 0.002777277 | 0.024026312 | Pask | Rbp1 | Vmn2r-ps11 | 0.738960096 | -0.004950533 | 0.017688442 | -0.279873861 | 0.779574282 | NA |
| Masp1 | 346.0463744 | 0.333689261 | 0.085839996 | 3.887340144 | 0.000101349 | 0.001982698 | Masp1 | Ms4a15 | 4930432J09Rik | 1.322596434 | -0.004961176 | 0.022217792 | -0.223297446 | 0.823304015 | NA |
| Pabpn1 | 5170.477244 | 0.333579953 | 0.097777345 | 3.411628259 | 0.000645761 | 0.00819445 | Pabpn1 | Myl9 | Gm5424 | 0.354261383 | -0.004976809 | 0.011488129 | -0.433213175 | 0.664859909 | NA |
| Rbm33 | 3309.389709 | 0.333386399 | 0.105308453 | 3.165808527 | 0.001546526 | 0.015860224 | Rbm33 | Pcsk1 | Slfn10-ps | 5.822556287 | -0.004978509 | 0.045367045 | -0.10973844 | 0.912616814 | NA |
| Grn | 2735.47283 | 0.332777116 | 0.080333943 | 4.142422302 | 3.44E-05 | 0.000849221 | Grn | Xylt1 | Gtf2ird2 | 728.9228891 | -0.004979168 | 0.076440287 | -0.065138009 | 0.948064118 | 0.987367302 |
| Pola2 | 232.2705326 | 0.332578668 | 0.10451109 | 3.182233263 | 0.001461441 | 0.015166484 | Pola2 | Slc4a4 | Usp25 | 1541.275454 | -0.004985209 | 0.059213661 | -0.084190191 | 0.932905217 | 0.984387568 |
| C1qb | 2030.231878 | 0.332517819 | 0.124008583 | 2.681409705 | 0.00733127 | 0.047113518 | C1qb | Galnt13 | Mup4 | 0.73332599 | -0.004995669 | 0.016259118 | -0.307253409 | 0.758650496 | NA |
| 9430015G10Rik | 830.7677007 | 0.332331743 | 0.084706638 | 3.923325875 | 8.73E-05 | 0.001768185 | 9430015G10Rik | Slc13a3 | Il1rapl2 | 172.5155861 | -0.004999683 | 0.090924735 | -0.054987045 | 0.956148785 | 0.989344398 |
| Slc27a2 | 225.1095458 | 0.331923287 | 0.113817529 | 2.91627563 | 0.003542374 | 0.028541624 | Slc27a2 | Zfp704 | Rpl32 | 4250.97326 | -0.005007924 | 0.063170513 | -0.079276292 | 0.936812863 | 0.985461126 |
| Aasdh | 497.0489729 | 0.331884153 | 0.114902483 | 2.888398442 | 0.00387209 | 0.03046936 | Aasdh | Kcnj2 | Cnot1 | 5741.366166 | -0.005013341 | 0.062747031 | -0.079897659 | 0.936318651 | 0.985461126 |
| Zgpat | 1807.58806 | 0.331651004 | 0.11686739 | 2.837840425 | 0.004541988 | 0.033817188 | Zgpat | Ldha | L1cam | 7584.437566 | -0.005021605 | 0.058356125 | -0.08605104 | 0.931425843 | 0.984208871 |
| Fam19a2 | 1935.416214 | 0.331614507 | 0.122967551 | 2.696764338 | 0.00700168 | 0.045466904 | Fam19a2 | Eomes | Haus3 | 210.2873934 | -0.005023404 | 0.088739246 | -0.056608599 | 0.954856985 | 0.988953472 |
| Dcaf15 | 833.9241622 | 0.330982057 | 0.084972005 | 3.895189463 | 9.81E-05 | 0.001929318 | Dcaf15 | Mll2 | Klrd1 | 0.51821185 | -0.005049288 | 0.014732444 | -0.342732564 | 0.731799662 | NA |
| Mcoln1 | 2340.258499 | 0.330149644 | 0.073185026 | 4.511163822 | 6.45E-06 | 0.000222978 | Mcoln1 | Triqk | Scg3 | 12711.69994 | -0.005052578 | 0.064129292 | -0.078787364 | 0.937201755 | 0.985461126 |
| Pan2 | 975.5096009 | 0.32961323 | 0.093179539 | 3.537399244 | 0.000404088 | 0.005876062 | Pan2 | Luzp2 | Slc36a2 | 14.61858557 | -0.005066486 | 0.074827252 | -0.067709106 | 0.9460172 | NA |
| Dhx33 | 1744.173362 | 0.329472698 | 0.088666355 | 3.715870577 | 0.000202505 | 0.003440368 | Dhx33 | Prkca | Defb25 | 0.385490251 | -0.005082967 | 0.011791253 | -0.431079461 | 0.666410594 | NA |
| Ifih1 | 237.68493 | 0.329447824 | 0.110960888 | 2.969044597 | 0.002987272 | 0.025306068 | Ifih1 | Npy5r | Slc13a2 | 0.385490251 | -0.005082967 | 0.011791253 | -0.431079461 | 0.666410594 | NA |
| Las1l | 2084.734115 | 0.329356397 | 0.066966163 | 4.918250974 | 8.73E-07 | 4.34E-05 | Las1l | D10Bwg1379e | Vmn2r78 | 0.385490251 | -0.005082967 | 0.011791253 | -0.431079461 | 0.666410594 | NA |
| Nxf1 | 2383.938123 | 0.329312612 | 0.083159437 | 3.960014946 | 7.49E-05 | 0.001571311 | Nxf1 | Synpr | 1700027J07Rik | 0.349184399 | -0.00508888 | 0.011440739 | -0.444803437 | 0.65646181 | NA |
| Rhebl1 | 205.4009882 | 0.329018655 | 0.118082307 | 2.786350162 | 0.005330526 | 0.037473304 | Rhebl1 | Ncan | Gm4832 | 0.349184399 | -0.00508888 | 0.011440739 | -0.444803437 | 0.65646181 | NA |
| Whrn | 1215.662567 | 0.328996266 | 0.090363652 | 3.64080311 | 0.000271789 | 0.004310233 | Whrn | Vipr2 | Dmbt1 | 0.592959185 | -0.005093956 | 0.015202201 | -0.335080187 | 0.737564598 | NA |
| Pnma3 | 918.0572535 | 0.328930289 | 0.103069851 | 3.191333693 | 0.001416176 | 0.014775915 | Pnma3 | Smim18 | D630010B17Rik | 1.834930761 | -0.005097314 | 0.029084143 | -0.175260937 | 0.860874605 | NA |
| Kcns3 | 225.534418 | 0.328604135 | 0.112801713 | 2.913112973 | 0.003578451 | 0.028727732 | Kcns3 | Asb2 | 2810442I21Rik | 2.063262406 | -0.005111535 | 0.031694033 | -0.161277527 | 0.871874823 | NA |
| Cables1 | 840.2816142 | 0.328589092 | 0.066537344 | 4.938416111 | 7.88E-07 | 4.05E-05 | Cables1 | Nkain2 | Pura | 585.0067479 | -0.005138009 | 0.076222124 | -0.067408372 | 0.946256604 | 0.986835678 |
| Gm4285 | 124.083663 | 0.327332358 | 0.113805615 | 2.876240853 | 0.004024425 | 0.03116164 | Gm4285 | Dynll1 | Hephl1 | 0.346566706 | -0.00514829 | 0.011416126 | -0.450966484 | 0.652013705 | NA |
| Atp1b2 | 19777.22233 | 0.326764687 | 0.063005909 | 5.186254615 | 2.15E-07 | 1.41E-05 | Atp1b2 | Ptpn13 | Mageb18 | 0.346566706 | -0.00514829 | 0.011416126 | -0.450966484 | 0.652013705 | NA |
| Cwc27 | 617.0499789 | 0.326582686 | 0.103826399 | 3.145468676 | 0.00165821 | 0.016629319 | Cwc27 | Mina | Haao | 8.508164124 | -0.005161463 | 0.057200658 | -0.090234329 | 0.928101005 | NA |
| Bzrap1 | 14045.0598 | 0.325826249 | 0.084855594 | 3.839773333 | 0.000123148 | 0.002296094 | Bzrap1 | Lamc3 | Gm514 | 65.46510476 | -0.005165202 | 0.086904499 | -0.059435378 | 0.952605336 | 0.988510896 |
| Dot1l | 2348.723724 | 0.325487009 | 0.116396193 | 2.796371595 | 0.005167994 | 0.036680212 | Dot1l | Abcc9 | Agap3 | 10525.94679 | -0.005175689 | 0.058648834 | -0.088248801 | 0.929678931 | 0.983720582 |
| Pisd | 2971.114649 | 0.325221842 | 0.082882631 | 3.923884141 | 8.71E-05 | 0.001768185 | Pisd | Zfp536 | Zfp191 | 1394.196608 | -0.005180753 | 0.072061523 | -0.071893471 | 0.942686686 | 0.985461126 |
| Sidt1 | 2371.647412 | 0.324911083 | 0.094011485 | 3.456078639 | 0.000548095 | 0.007319044 | Sidt1 | Ccdc85c | Snord12 | 12.69521876 | -0.005195947 | 0.071299847 | -0.07287459 | 0.941905915 | NA |
| Eef2k | 1782.81568 | 0.32480299 | 0.094957472 | 3.420510093 | 0.000625038 | 0.008054403 | Eef2k | Tbx21 | Rev3l | 1797.269011 | -0.005206734 | 0.071099635 | -0.073231515 | 0.941621889 | 0.985461126 |
| Thoc1 | 919.6803635 | 0.324104407 | 0.098472957 | 3.291303709 | 0.000997242 | 0.011344541 | Thoc1 | Usmg5 | Gh | 290.9693413 | -0.005208361 | 0.034388593 | -0.151456071 | 0.879615961 | 0.973413768 |
| Abca8b | 931.8301559 | 0.324079243 | 0.090537872 | 3.579488174 | 0.000344268 | 0.005197309 | Abca8b | Rasl10b | Anp32b | 2022.265699 | -0.005209356 | 0.056385879 | -0.092387594 | 0.926390095 | 0.983236501 |
| Slc7a6 | 999.199062 | 0.323996816 | 0.09444127 | 3.430669829 | 0.000602093 | 0.007819638 | Slc7a6 | Kcnj12 | Ankrd12 | 3121.170217 | -0.005222574 | 0.075048858 | -0.069588993 | 0.944520798 | 0.986483951 |
| Tbce | 1529.441307 | 0.323969718 | 0.111236698 | 2.912435594 | 0.003586221 | 0.02876272 | Tbce | Sema3d | Slc12a2 | 3430.075009 | -0.005228085 | 0.061603538 | -0.084866639 | 0.932367414 | 0.984387568 |
| Rere | 6324.29544 | 0.323921212 | 0.109896846 | 2.947502346 | 0.003203523 | 0.026528802 | Rere | Uqcr11 | Gm136 | 4.76045503 | -0.005236 | 0.04336776 | -0.120734847 | 0.903901057 | NA |
| Cntrob | 690.0697006 | 0.322643718 | 0.108224043 | 2.981257296 | 0.002870675 | 0.024587196 | Cntrob | Prdm12 | Mrps24 | 518.7173079 | -0.005241538 | 0.072413775 | -0.072383166 | 0.942296981 | 0.985461126 |
| Acox3 | 1582.79283 | 0.322564717 | 0.072569368 | 4.44491564 | 8.79E-06 | 0.000284411 | Acox3 | Serpinf1 | Gm5797 | 0.879810723 | -0.005246963 | 0.018654851 | -0.281265322 | 0.778506906 | NA |
| Tmem108 | 727.467146 | 0.322520216 | 0.085289718 | 3.781466558 | 0.000155907 | 0.002789406 | Tmem108 | Sdk2 | Hist1h3d | 2.104481424 | -0.005248438 | 0.030618 | -0.171416764 | 0.863896074 | NA |
| Sbsn | 305.8128012 | 0.32222278 | 0.105227154 | 3.062163779 | 0.002197432 | 0.020565282 | Sbsn | Pde10a | Slc12a5 | 29614.64337 | -0.005259053 | 0.046091559 | -0.114100135 | 0.909158415 | 0.980523965 |
| Fam13a | 776.0971491 | 0.322103758 | 0.076230832 | 4.22537377 | 2.39E-05 | 0.000641668 | Fam13a | Mdn1 | Themis | 0.886846282 | -0.005277035 | 0.023562066 | -0.223963177 | 0.822785956 | NA |
| 3010026O09Rik | 345.4311167 | 0.320641882 | 0.098773127 | 3.246246147 | 0.001169377 | 0.012845805 | 3010026O09Rik | Gldc | Olfr1378 | 0.753219214 | -0.005280431 | 0.018076445 | -0.292116667 | 0.770197424 | NA |
| Adamts10 | 842.2451344 | 0.320253927 | 0.119564443 | 2.678504724 | 0.007395169 | 0.047350844 | Adamts10 | 9430020K01Rik | Anpep | 194.6729138 | -0.005288441 | 0.095048089 | -0.055639637 | 0.955628888 | 0.989344398 |
| Prr14 | 906.9558508 | 0.319805576 | 0.073391977 | 4.357500502 | 1.32E-05 | 0.000398891 | Prr14 | Agpat5 | Vsx1 | 3.067340106 | -0.005294336 | 0.031127028 | -0.170088052 | 0.86494089 | NA |
| Prkab2 | 2010.745757 | 0.319428643 | 0.055203199 | 5.786415438 | 7.19E-09 | 7.96E-07 | Prkab2 | Spryd4 | Gm3893 | 842.3859658 | -0.005305271 | 0.045080902 | -0.117683345 | 0.906318564 | 0.980118791 |
| Chfr | 1279.736261 | 0.318830516 | 0.076768597 | 4.153137207 | 3.28E-05 | 0.000825264 | Chfr | Efnb2 | Gm6583 | 2.559359672 | -0.00530846 | 0.034724626 | -0.152873067 | 0.878498376 | NA |
| Gstm1 | 6963.73249 | 0.318698504 | 0.066819478 | 4.769544923 | 1.85E-06 | 8.34E-05 | Gstm1 | Tsku | Rusc1 | 7797.96964 | -0.005311336 | 0.049742476 | -0.10677668 | 0.914966148 | 0.98170867 |
| Mfsd10 | 619.6289312 | 0.318677284 | 0.094875376 | 3.358904032 | 0.000782522 | 0.009413355 | Mfsd10 | Abl2 | Ube2d1 | 1693.881823 | -0.005315585 | 0.071660139 | -0.074177709 | 0.940868982 | 0.985461126 |
| Macrod1 | 270.6548945 | 0.317852033 | 0.102867689 | 3.089911281 | 0.002002163 | 0.019149082 | Macrod1 | Nptx2 | Cnep1r1 | 885.8621043 | -0.005319391 | 0.069591669 | -0.076437182 | 0.939071289 | 0.985461126 |
| Eif2ak2 | 313.3758622 | 0.317807974 | 0.107486007 | 2.956738128 | 0.003109121 | 0.025899308 | Eif2ak2 | Cacna1e | Uqcrq | 2238.614266 | -0.005342632 | 0.052648268 | -0.101477831 | 0.919171155 | 0.982742917 |
| Vwa5b1 | 165.1069862 | 0.317475661 | 0.114426745 | 2.774488252 | 0.005528863 | 0.03865614 | Vwa5b1 | 2810459M11Rik | Mir335 | 0.926546215 | -0.005362829 | 0.022441965 | -0.238964301 | 0.811133264 | NA |
| Gm15706 | 164.2655142 | 0.317192673 | 0.111958868 | 2.833117906 | 0.004609638 | 0.034173077 | Gm15706 | Bmp2 | Upb1 | 29.32443846 | -0.005373867 | 0.081513336 | -0.065926237 | 0.947436552 | 0.987335056 |
| Pcdhb5 | 234.8292852 | 0.316416642 | 0.109839831 | 2.880709467 | 0.003967812 | 0.030908637 | Pcdhb5 | Adam19 | Dbh | 17.48410389 | -0.005385338 | 0.074044271 | -0.072731325 | 0.942019921 | NA |
| Slc20a1 | 6211.789557 | 0.316395899 | 0.067609707 | 4.67974074 | 2.87E-06 | 0.000119315 | Slc20a1 | Lrba | Lctl | 11.45639405 | -0.005385557 | 0.06052833 | -0.088975806 | 0.929101138 | NA |
| Faah | 3515.38889 | 0.3143779 | 0.081520087 | 3.856447059 | 0.000115047 | 0.002173827 | Faah | Gadd45g | Gm14873 | 0.876617123 | -0.005396095 | 0.022296887 | -0.242011129 | 0.80877154 | NA |
| Ccdc141 | 853.3714656 | 0.31429999 | 0.094336286 | 3.331697734 | 0.00086318 | 0.010193677 | Ccdc141 | N6amt2 | Gm6568 | 0.336143057 | -0.005396283 | 0.011316387 | -0.476855656 | 0.633464911 | NA |
| Zfp280c | 753.4024207 | 0.314246635 | 0.113450273 | 2.76990637 | 0.005607241 | 0.038941547 | Zfp280c | Frzb | Nudt2 | 492.7057614 | -0.005405765 | 0.066623216 | -0.081139355 | 0.935331128 | 0.985461126 |
| Sesn2 | 426.7326696 | 0.314139299 | 0.110059658 | 2.854263808 | 0.004313669 | 0.032794894 | Sesn2 | Hivep3 | Lyrm5 | 624.8592829 | -0.005405948 | 0.070331765 | -0.076863529 | 0.938732111 | 0.985461126 |
| Smyd4 | 500.8306239 | 0.313266091 | 0.07139473 | 4.387804109 | 1.15E-05 | 0.000354799 | Smyd4 | Mpp7 | Ctdspl | 1763.560267 | -0.005439954 | 0.057550834 | -0.094524337 | 0.924692651 | 0.983236501 |
| Gm10845 | 360.462726 | 0.313211143 | 0.10826656 | 2.892962901 | 0.003816264 | 0.030167954 | Gm10845 | Tmem64 | Zfp101 | 336.5926581 | -0.005439971 | 0.080442098 | -0.067625922 | 0.94608342 | 0.986835678 |
| Atxn7 | 1165.622131 | 0.313196361 | 0.095145234 | 3.291771423 | 0.000995585 | 0.011334014 | Atxn7 | Sorl1 | 1700006F04Rik | 0.345062444 | -0.005440857 | 0.012918824 | -0.421157294 | 0.673640227 | NA |
| Cnot10 | 2009.417662 | 0.313147417 | 0.07755455 | 4.037769754 | 5.40E-05 | 0.001224134 | Cnot10 | Gm4980 | Mpv17 | 1912.263767 | -0.00544747 | 0.048602161 | -0.112082883 | 0.91075769 | 0.980847138 |
| Scrt1 | 5623.778362 | 0.313076549 | 0.0749237 | 4.17860499 | 2.93E-05 | 0.000757405 | Scrt1 | Hivep2 | Cnpy4 | 387.4161507 | -0.005450324 | 0.072481272 | -0.07519631 | 0.94005852 | 0.985461126 |
| Stx2 | 578.2145827 | 0.312874178 | 0.098696231 | 3.170072205 | 0.001524011 | 0.015659114 | Stx2 | Cntnap4 | Gm4925 | 13.03643043 | -0.005456476 | 0.072809934 | -0.074941371 | 0.94026136 | NA |
| Cecr5 | 890.9432578 | 0.312371589 | 0.094462208 | 3.306841935 | 0.000943541 | 0.010861238 | Cecr5 | Sulf1 | 1110007C09Rik | 182.3023159 | -0.005460296 | 0.090561429 | -0.060293841 | 0.951921607 | 0.988463312 |
| Timeless | 204.7374346 | 0.312166634 | 0.112923207 | 2.764415229 | 0.005702492 | 0.039425264 | Timeless | Cdh8 | Chrna10 | 25.02863998 | -0.005466026 | 0.083117418 | -0.065762702 | 0.947566751 | 0.987347864 |
| Taz | 1448.989399 | 0.311874921 | 0.098230626 | 3.17492552 | 0.00149875 | 0.015450184 | Taz | Ttc9b | Gm20757 | 0.366568861 | -0.005484773 | 0.011623202 | -0.471881406 | 0.637011441 | NA |
| Lypd1 | 2570.951793 | 0.311779271 | 0.08858678 | 3.519478525 | 0.000432396 | 0.006144506 | Lypd1 | Sez6 | Krtap4-16 | 0.366568861 | -0.005484773 | 0.011623202 | -0.471881406 | 0.637011441 | NA |
| Tspyl2 | 4051.587203 | 0.310947584 | 0.093638655 | 3.320718175 | 0.000897862 | 0.010498156 | Tspyl2 | Slc25a34 | Nrg4 | 14.91259462 | -0.005494195 | 0.071411683 | -0.076936923 | 0.938673724 | NA |
| Sdccag8 | 611.577516 | 0.310892893 | 0.08049897 | 3.862072935 | 0.000112429 | 0.002134773 | Sdccag8 | Dok6 | Dear1 | 0.526837674 | -0.005508763 | 0.01480074 | -0.372195109 | 0.709747587 | NA |
| Atad2 | 428.2355735 | 0.310825511 | 0.099693905 | 3.117798546 | 0.001822073 | 0.017822727 | Atad2 | Smpdl3a | Serpinb3c | 0.363951167 | -0.005544779 | 0.011599552 | -0.478016623 | 0.632638374 | NA |
| Mbip | 451.5009074 | 0.310688757 | 0.111249363 | 2.792723905 | 0.005226628 | 0.036943149 | Mbip | Lmo1 | Prkaa1 | 1285.024192 | -0.005547881 | 0.06035844 | -0.091915583 | 0.92676511 | 0.983431222 |
| Zfp112 | 399.7055917 | 0.310682449 | 0.083253913 | 3.731745918 | 0.000190157 | 0.003277304 | Zfp112 | Sema3c | Gm4489 | 0.309662151 | -0.005550521 | 0.012523454 | -0.443210101 | 0.657613773 | NA |
| Tenc1 | 1143.793915 | 0.310077897 | 0.099881346 | 3.104462524 | 0.001906251 | 0.018494333 | Tenc1 | Glyctk | Arhgap39 | 3216.402141 | -0.005557323 | 0.076474862 | -0.072668627 | 0.942069815 | 0.985461126 |
| Inpp5k | 1800.335274 | 0.309842966 | 0.088483931 | 3.501686254 | 0.000462324 | 0.006438851 | Inpp5k | Pkig | Cyp2d22 | 1367.153644 | -0.00555949 | 0.075777004 | -0.073366447 | 0.941514517 | 0.985461126 |
| Map7 | 3871.371567 | 0.30952292 | 0.072751631 | 4.254515168 | 2.10E-05 | 0.000578615 | Map7 | Rab40b | Ly6g6f | 17.55738223 | -0.005577303 | 0.07444087 | -0.074922588 | 0.940276305 | NA |
| Zfpm2 | 525.7586171 | 0.309449321 | 0.108514572 | 2.851684473 | 0.004348824 | 0.032932886 | Zfpm2 | Gabrg1 | Atraid | 1825.743944 | -0.005578066 | 0.058853559 | -0.094778729 | 0.924490583 | 0.983236501 |
| Ncapd3 | 1132.415469 | 0.309225511 | 0.068658795 | 4.503800432 | 6.67E-06 | 0.000228302 | Ncapd3 | Mas1 | Ctxn1 | 13355.9745 | -0.005580885 | 0.041939615 | -0.133069537 | 0.894138386 | 0.977844987 |
| Oplah | 1116.562181 | 0.308660012 | 0.076529015 | 4.033241649 | 5.50E-05 | 0.001241224 | Oplah | Dhrs13 | Sult2a5 | 1.16921559 | -0.005587928 | 0.021563915 | -0.259133299 | 0.795532393 | NA |
| Dpysl4 | 1914.72139 | 0.308419097 | 0.084639438 | 3.643917101 | 0.00026852 | 0.004275897 | Dpysl4 | Fkbp1b | Unc93a | 0.599001055 | -0.005591454 | 0.013383906 | -0.417774493 | 0.676112001 | NA |
| Rabep2 | 373.6232332 | 0.308278704 | 0.097661614 | 3.156600555 | 0.001596198 | 0.016185535 | Rabep2 | Slc8a1 | Snrpe | 620.005571 | -0.005602592 | 0.072687281 | -0.077078024 | 0.938561476 | 0.985461126 |
| Cplx3 | 699.8939804 | 0.307959893 | 0.114865188 | 2.681055061 | 0.007339044 | 0.047143925 | Cplx3 | Pcbd1 | Zfp646 | 1425.101555 | -0.005603268 | 0.059503118 | -0.094167639 | 0.924975991 | 0.983236501 |
| Kctd6 | 1089.93197 | 0.307583891 | 0.07545159 | 4.076572715 | 4.57E-05 | 0.001064877 | Kctd6 | BC056474 | Xlr4c | 1.614146089 | -0.005605904 | 0.02016957 | -0.277938707 | 0.78105941 | NA |
| Fbxo32 | 1311.789123 | 0.307430398 | 0.090675702 | 3.390438557 | 0.000697809 | 0.008684218 | Fbxo32 | Pcp4 | Zp3r | 3.036695988 | -0.005608221 | 0.038559437 | -0.145443548 | 0.884360684 | NA |
| Atg4a | 215.4847605 | 0.307263807 | 0.109932165 | 2.795030989 | 0.005189474 | 0.036748497 | Atg4a | Lix1 | C2cd3 | 1854.783901 | -0.005612616 | 0.059115851 | -0.094942664 | 0.924360368 | 0.983236501 |
| Pcdhga2 | 477.9428513 | 0.30644184 | 0.088917391 | 3.446365652 | 0.000568181 | 0.007492252 | Pcdhga2 | Cx3cl1 | Mir468 | 0.956550331 | -0.005614147 | 0.019019558 | -0.295177577 | 0.76785823 | NA |
| 4930503L19Rik | 319.053047 | 0.306301351 | 0.101508592 | 3.017491869 | 0.002548759 | 0.022682636 | 4930503L19Rik | Tesc | Lrrc10b | 1986.978125 | -0.005615853 | 0.094831925 | -0.059219017 | 0.952777663 | 0.988510896 |
| Fam57b | 2112.631604 | 0.304689943 | 0.071314729 | 4.272468652 | 1.93E-05 | 0.000543614 | Fam57b | 9830001H06Rik | Mgea5 | 10812.80011 | -0.005621 | 0.069396113 | -0.080998774 | 0.935442927 | 0.985461126 |
| Map2k3 | 503.3716609 | 0.304415422 | 0.107054515 | 2.843555207 | 0.004461328 | 0.033425443 | Map2k3 | Ctxn2 | Zdhhc17 | 5677.587165 | -0.005622685 | 0.063986541 | -0.087872928 | 0.929977673 | 0.983893349 |
| Ctsz | 1526.115012 | 0.304061443 | 0.114229987 | 2.661835572 | 0.007771583 | 0.049168195 | Ctsz | Syt9 | Snora41 | 11.92568077 | -0.005638253 | 0.058212594 | -0.096856246 | 0.922840556 | NA |
| Srrt | 4418.946013 | 0.303729867 | 0.092147405 | 3.296130466 | 0.000980265 | 0.011217299 | Srrt | Gabrb1 | 1700025E21Rik | 1.154524012 | -0.00569843 | 0.023562027 | -0.241848035 | 0.808897917 | NA |
| Atg16l2 | 576.6098939 | 0.303710352 | 0.108434572 | 2.800862744 | 0.005096619 | 0.036290258 | Atg16l2 | Tbc1d16 | 4930430F21Rik | 0.354244579 | -0.005705568 | 0.013020739 | -0.43819079 | 0.661247985 | NA |
| Sun1 | 4086.254053 | 0.303509694 | 0.066602029 | 4.55706381 | 5.19E-06 | 0.000190457 | Sun1 | Kcna3 | D030018L15Rik | 0.354244579 | -0.005705568 | 0.013020739 | -0.43819079 | 0.661247985 | NA |
| Nfic | 5115.850644 | 0.303190005 | 0.085851954 | 3.531544616 | 0.00041314 | 0.005960143 | Nfic | Akap5 | Mir5100 | 0.354244579 | -0.005705568 | 0.013020739 | -0.43819079 | 0.661247985 | NA |
| Atxn2l | 5603.951341 | 0.303033157 | 0.099477027 | 3.046262711 | 0.002317053 | 0.021280627 | Atxn2l | Adamtsl1 | Myoz1 | 0.354244579 | -0.005705568 | 0.013020739 | -0.43819079 | 0.661247985 | NA |
| Prox1 | 382.9755278 | 0.303002734 | 0.108441916 | 2.79414775 | 0.00520367 | 0.036815369 | Prox1 | Slc35f1 | Stk25 | 9744.835576 | -0.005720273 | 0.049970114 | -0.114473892 | 0.908862141 | 0.980523965 |
| Mcm8 | 190.3674274 | 0.302922841 | 0.098843651 | 3.064666652 | 0.002179127 | 0.020414223 | Mcm8 | 1700020I14Rik | Sh2d1b1 | 1.511807921 | -0.005720359 | 0.025807923 | -0.221651263 | 0.824585373 | NA |
| Pfkfb4 | 576.0966674 | 0.302913637 | 0.09545362 | 3.173411733 | 0.001506587 | 0.015520651 | Pfkfb4 | Cep112 | Pabpc1 | 5746.709121 | -0.005729407 | 0.047726002 | -0.120047908 | 0.904445197 | 0.980118791 |
| Prpf38b | 1486.065958 | 0.301509747 | 0.088277439 | 3.415479073 | 0.000636699 | 0.008131449 | Prpf38b | Chst11 | 4931440F15Rik | 24.11256228 | -0.00575234 | 0.082958043 | -0.06934035 | 0.944718709 | 0.986555281 |
| Gnl2 | 1900.318567 | 0.301496283 | 0.070129974 | 4.29910732 | 1.71E-05 | 0.000494791 | Gnl2 | Ankrd6 | Trip12 | 8214.272414 | -0.00576137 | 0.063410159 | -0.090858788 | 0.927604797 | 0.983501978 |
| Wbp2 | 15954.24471 | 0.301228942 | 0.100537949 | 2.996171545 | 0.002733926 | 0.023821531 | Wbp2 | Fat4 | Wnt7b | 1236.344943 | -0.00579019 | 0.069382447 | -0.08345324 | 0.933491157 | 0.984481307 |
| Ptpn3 | 2925.247553 | 0.30054298 | 0.105809267 | 2.8404221 | 0.004505388 | 0.03360928 | Ptpn3 | Klhdc8a | Gm9992 | 0.353527519 | -0.00579453 | 0.011502941 | -0.503743378 | 0.614441722 | NA |
| Plagl1 | 520.9508103 | 0.298715433 | 0.106742185 | 2.798475902 | 0.00513444 | 0.036475477 | Plagl1 | Kcnmb4 | Pglyrp2 | 0.353527519 | -0.00579453 | 0.011502941 | -0.503743378 | 0.614441722 | NA |
| Acp6 | 430.7721388 | 0.298687608 | 0.084503112 | 3.534634431 | 0.00040834 | 0.00590739 | Acp6 | Fam181b | Rergl | 0.353527519 | -0.00579453 | 0.011502941 | -0.503743378 | 0.614441722 | NA |
| Mdm1 | 215.0303331 | 0.298561027 | 0.112174952 | 2.661565896 | 0.007777812 | 0.049187515 | Mdm1 | Herc2 | H2-Ke6 | 610.5704531 | -0.005796709 | 0.073811342 | -0.078534122 | 0.937403189 | 0.985461126 |
| Adam11 | 6054.975929 | 0.298280293 | 0.077255317 | 3.86096784 | 0.000112939 | 0.002141828 | Adam11 | Dpysl3 | Sorcs3 | 2301.781588 | -0.005799544 | 0.065865222 | -0.088051684 | 0.929835598 | 0.983814691 |
| Zranb2 | 4255.729807 | 0.298238338 | 0.069633816 | 4.282952662 | 1.84E-05 | 0.000521593 | Zranb2 | Igsf21 | Numbl | 4094.789864 | -0.00580039 | 0.073483326 | -0.07893478 | 0.937084499 | 0.985461126 |
| Mcf2 | 260.0971539 | 0.298151199 | 0.106433974 | 2.801278446 | 0.005090058 | 0.036290258 | Mcf2 | Atp5l | Fbxo18 | 3934.543049 | -0.005805754 | 0.050853176 | -0.114166985 | 0.909105423 | 0.980523965 |
| Trub1 | 1022.216099 | 0.297550221 | 0.073765009 | 4.033758344 | 5.49E-05 | 0.001241224 | Trub1 | Mll3 | Rc3h2 | 2416.822941 | -0.005806749 | 0.076726508 | -0.075681136 | 0.939672784 | 0.985461126 |
| 4933427D14Rik | 1422.16298 | 0.297496851 | 0.074377479 | 3.999824334 | 6.34E-05 | 0.001377705 | 4933427D14Rik | Idi1 | Mef2c | 21580.62981 | -0.005828244 | 0.078174372 | -0.074554405 | 0.940569253 | 0.985461126 |
| Dhodh | 613.2768401 | 0.297421919 | 0.089532647 | 3.321938192 | 0.000893945 | 0.010481817 | Dhodh | Notch2 | Fancc | 369.8610948 | -0.005828748 | 0.078467471 | -0.074282348 | 0.940785723 | 0.985461126 |
| Sdccag3 | 3082.901601 | 0.297056165 | 0.083610307 | 3.552865389 | 0.000381059 | 0.005638202 | Sdccag3 | Cldn5 | Ggt1 | 83.3602167 | -0.005864997 | 0.09496557 | -0.061759192 | 0.950754601 | 0.988202091 |
| Ptpn21 | 408.9176946 | 0.296908519 | 0.105514815 | 2.813903605 | 0.00489439 | 0.03550266 | Ptpn21 | Glt25d2 | Ubxn4 | 2709.824582 | -0.005869872 | 0.055898946 | -0.105008628 | 0.916368963 | 0.982742917 |
| Slc25a28 | 1085.443518 | 0.296551244 | 0.084785871 | 3.497649341 | 0.000469378 | 0.006516613 | Slc25a28 | Frmd7 | Pdzd4 | 5808.908936 | -0.005871617 | 0.050720261 | -0.115764733 | 0.907839002 | 0.980520081 |
| Clk2 | 1090.389031 | 0.296430327 | 0.07725432 | 3.837071226 | 0.00012451 | 0.002318706 | Clk2 | Chsy3 | Nyx | 22.71452184 | -0.005878679 | 0.085055944 | -0.06911544 | 0.944897731 | 0.986671299 |
| Cblb | 927.0625301 | 0.296321232 | 0.06454882 | 4.590652959 | 4.42E-06 | 0.000166574 | Cblb | Cxcr7 | Pdzd11 | 1121.799801 | -0.005879344 | 0.065693538 | -0.089496534 | 0.928687308 | 0.983596265 |
| Myh7 | 541.9250231 | 0.296076649 | 0.0866067 | 3.418634455 | 0.000629362 | 0.008072299 | Myh7 | Myo18b | Arl6ip4 | 1140.053383 | -0.005880019 | 0.05847115 | -0.100562741 | 0.919897575 | 0.982742917 |
| Hook2 | 499.4074124 | 0.296071548 | 0.103699038 | 2.855104096 | 0.004302273 | 0.032740378 | Hook2 | Tekt5 | Mir99a | 1.917235761 | -0.005880338 | 0.030629899 | -0.191980317 | 0.847757623 | NA |
| Ptpn23 | 2885.085132 | 0.296010653 | 0.069040502 | 4.287492773 | 1.81E-05 | 0.000515614 | Ptpn23 | Pcdhb21 | E230019M04Rik | 1.855420341 | -0.005898507 | 0.027646657 | -0.213353374 | 0.831051346 | NA |
| Cux2 | 3319.031458 | 0.295396006 | 0.1025118 | 2.881580517 | 0.003956861 | 0.030869893 | Cux2 | Kcna4 | Myh4 | 22.32417301 | -0.005929013 | 0.077180849 | -0.076819739 | 0.938766947 | NA |
| Zswim8 | 5726.546978 | 0.295086964 | 0.069179119 | 4.265549615 | 1.99E-05 | 0.000557702 | Zswim8 | Huwe1 | Fam83a | 0.36293681 | -0.005930613 | 0.013112557 | -0.452284994 | 0.651063685 | NA |
| Zfp566 | 162.5020211 | 0.294296404 | 0.104813665 | 2.807805675 | 0.004988032 | 0.0359129 | Zfp566 | Ubr4 | Spata31d1c | 0.36293681 | -0.005930613 | 0.013112557 | -0.452284994 | 0.651063685 | NA |
| Mroh1 | 3300.376229 | 0.293868978 | 0.062650901 | 4.690578628 | 2.72E-06 | 0.000114703 | Mroh1 | Hpcal4 | Pik3r3 | 2766.769062 | -0.005937889 | 0.070579004 | -0.084131099 | 0.932952199 | 0.984387568 |
| Unc13b | 2028.390901 | 0.29371934 | 0.104841628 | 2.801552638 | 0.005085734 | 0.036290258 | Unc13b | Rab27b | Gm15179 | 0.323523361 | -0.00593921 | 0.012689672 | -0.468034922 | 0.639759617 | NA |
| Sorbs3 | 1293.567553 | 0.293297601 | 0.084119 | 3.486698609 | 0.000489022 | 0.006741022 | Sorbs3 | Slc19a3 | Rchy1 | 1256.626475 | -0.005946788 | 0.06016232 | -0.09884573 | 0.921260759 | 0.982762672 |
| Slc25a27 | 1831.970394 | 0.293018547 | 0.070120411 | 4.178791063 | 2.93E-05 | 0.000757405 | Slc25a27 | Cd93 | C730036E19Rik | 1.614645163 | -0.005953334 | 0.030353154 | -0.196135597 | 0.844504035 | NA |
| Fam5c | 1270.646754 | 0.292953368 | 0.078442276 | 3.734636266 | 0.000187987 | 0.003250786 | Fam5c | C1galt1 | A930019D19Rik | 0.703384099 | -0.005967068 | 0.015960878 | -0.373855846 | 0.708511568 | NA |
| Pnpt1 | 772.7019076 | 0.292796823 | 0.085747452 | 3.414641661 | 0.00063866 | 0.008137659 | Pnpt1 | Pcdha7 | Gsx2 | 7.42049852 | -0.005973336 | 0.053419604 | -0.111819165 | 0.910966791 | NA |
| Tmem8 | 698.9716938 | 0.292489412 | 0.100501539 | 2.910297851 | 0.003610845 | 0.028912882 | Tmem8 | Foxo1 | Angpt4 | 0.340400173 | -0.006017292 | 0.012878466 | -0.46723668 | 0.640330553 | NA |
| Fam214b | 1575.650912 | 0.292017042 | 0.076876462 | 3.798523426 | 0.000145561 | 0.002643982 | Fam214b | Aox3 | LOC100503280 | 0.340400173 | -0.006017292 | 0.012878466 | -0.46723668 | 0.640330553 | NA |
| Asl | 496.4649404 | 0.291931987 | 0.099247199 | 2.941463221 | 0.003266656 | 0.026936441 | Asl | Nrp2 | Serpina10 | 0.505990377 | -0.006033849 | 0.014670827 | -0.411282109 | 0.680865687 | NA |
| Ankrd27 | 1449.431275 | 0.291888486 | 0.066315099 | 4.401538884 | 1.07E-05 | 0.000335088 | Ankrd27 | Grm5 | Ash2l | 1558.297531 | -0.006039687 | 0.049604854 | -0.121755967 | 0.903092288 | 0.979150581 |
| Myd88 | 236.0652912 | 0.291814065 | 0.105647672 | 2.76214382 | 0.005742318 | 0.039586803 | Myd88 | Pcdh11x | Ddx21 | 1248.518501 | -0.006047691 | 0.064831856 | -0.093282707 | 0.92567897 | 0.983236501 |
| Uimc1 | 538.9098145 | 0.290979352 | 0.10297029 | 2.825857367 | 0.004715426 | 0.034613354 | Uimc1 | Emx2 | Mfap5 | 4.943196057 | -0.006062027 | 0.046823437 | -0.129465656 | 0.896989199 | NA |
| Rad52 | 455.3295744 | 0.290381844 | 0.094902277 | 3.059798485 | 0.00221486 | 0.020635619 | Rad52 | Tmem158 | Dmrta1 | 9.207022633 | -0.006072017 | 0.058602472 | -0.103613671 | 0.917475938 | NA |
| Sord | 908.8580345 | 0.290340785 | 0.097594791 | 2.9749619 | 0.002930249 | 0.024959475 | Sord | Mll1 | Med8 | 866.1745062 | -0.006078715 | 0.058309615 | -0.104248937 | 0.9169718 | 0.982742917 |
| Mbd4 | 357.6312976 | 0.290268302 | 0.084494207 | 3.435363345 | 0.00059176 | 0.007730798 | Mbd4 | Zfp365 | Mir99b | 0.582654835 | -0.006091079 | 0.015161809 | -0.401738275 | 0.687876653 | NA |
| Gabpb2 | 928.7104098 | 0.28907963 | 0.103185531 | 2.801551987 | 0.005085745 | 0.036290258 | Gabpb2 | Kcnq5 | Mir30d | 0.718908469 | -0.006101965 | 0.017945135 | -0.34003448 | 0.733830562 | NA |
| Snrnp48 | 371.5549915 | 0.288690963 | 0.094809093 | 3.04497125 | 0.002327026 | 0.021346912 | Snrnp48 | Grid2 | AA536875 | 0.334833285 | -0.006103423 | 0.012817048 | -0.476195713 | 0.633934953 | NA |
| Fyco1 | 2632.760082 | 0.288383837 | 0.091294063 | 3.158845455 | 0.001583955 | 0.016082433 | Fyco1 | Rpl41 | Guca2b | 0.334833285 | -0.006103423 | 0.012817048 | -0.476195713 | 0.633934953 | NA |
| Rnft1 | 535.2317867 | 0.288129581 | 0.084367296 | 3.415180947 | 0.000637397 | 0.008131449 | Rnft1 | Gm13889 | Gm9833 | 23.11272387 | -0.006106136 | 0.085287908 | -0.071594397 | 0.9429247 | 0.985461126 |
| Snap23 | 381.6245255 | 0.288024505 | 0.092143945 | 3.125810438 | 0.001773158 | 0.017434924 | Snap23 | Lmcd1 | Il17d | 219.5708673 | -0.006125599 | 0.089694237 | -0.068294237 | 0.945551412 | 0.986781137 |
| Slc31a2 | 346.4913312 | 0.287888504 | 0.082974544 | 3.469600301 | 0.000521233 | 0.007034835 | Slc31a2 | Lrrc7 | Ptprn | 27360.54396 | -0.006135281 | 0.06351224 | -0.096599979 | 0.923044073 | 0.98296621 |
| Scaf4 | 1297.230621 | 0.287589093 | 0.06760416 | 4.254014764 | 2.10E-05 | 0.000578879 | Scaf4 | Ndst3 | 4933438K21Rik | 0.335323189 | -0.006141311 | 0.012825248 | -0.478845398 | 0.632048618 | NA |
| Flcn | 2118.630784 | 0.287512686 | 0.095970467 | 2.995845436 | 0.002736851 | 0.023822906 | Flcn | Mlf1ip | Cd27 | 0.335323189 | -0.006141311 | 0.012825248 | -0.478845398 | 0.632048618 | NA |
| Pml | 680.1410121 | 0.287419935 | 0.073617661 | 3.904225305 | 9.45E-05 | 0.001877708 | Pml | Gpx8 | Gm14151 | 0.335323189 | -0.006141311 | 0.012825248 | -0.478845398 | 0.632048618 | NA |
| Inpp5b | 1521.8713 | 0.28716107 | 0.059963241 | 4.788951769 | 1.68E-06 | 7.69E-05 | Inpp5b | Cdh11 | Tmem55b | 2638.203295 | -0.00614162 | 0.047979537 | -0.128004996 | 0.898145019 | 0.978273929 |
| Zfc3h1 | 2436.287963 | 0.287098105 | 0.095853903 | 2.995163422 | 0.002742978 | 0.023862834 | Zfc3h1 | Tubb3 | Rasip1 | 360.4600868 | -0.006147434 | 0.079679186 | -0.07715232 | 0.938502372 | 0.985461126 |
| Ldb3 | 313.817871 | 0.286870609 | 0.10633016 | 2.697923228 | 0.006977353 | 0.04536597 | Ldb3 | Tmeff1 | Mbnl1 | 3066.142725 | -0.006156139 | 0.071835843 | -0.085697315 | 0.931707037 | 0.984289194 |
| Abcc5 | 6156.817105 | 0.286391772 | 0.063040982 | 4.542945943 | 5.55E-06 | 0.000199939 | Abcc5 | E030013I19Rik | Psg29 | 3.458559647 | -0.006160498 | 0.044809156 | -0.137483011 | 0.890649021 | NA |
| Lrrc27 | 297.6956938 | 0.28615807 | 0.091572813 | 3.124923863 | 0.001778511 | 0.017473841 | Lrrc27 | Rcn3 | Gm13003 | 16.52187612 | -0.006161357 | 0.065544912 | -0.094002062 | 0.925107519 | NA |
| Smc2 | 386.6074948 | 0.286138802 | 0.07854247 | 3.643109309 | 0.000269364 | 0.00428494 | Smc2 | Myoc | Rhd | 4.142573185 | -0.006185227 | 0.045455622 | -0.136071765 | 0.891764548 | NA |
| Fam46a | 251.8735383 | 0.286048526 | 0.101788157 | 2.810233855 | 0.004950551 | 0.035738209 | Fam46a | Chl1 | Kcnh3 | 7377.584719 | -0.00618525 | 0.069520522 | -0.088970134 | 0.929105645 | 0.983640461 |
| Top3a | 254.2554316 | 0.285927756 | 0.101152764 | 2.826692471 | 0.004703147 | 0.034552186 | Top3a | Angpt1 | Alpk3 | 24.93399999 | -0.006185438 | 0.078605066 | -0.078690063 | 0.937279149 | 0.985461126 |
| Alkbh3 | 292.483378 | 0.285695303 | 0.103363453 | 2.763987591 | 0.005709971 | 0.039425264 | Alkbh3 | Hivep1 | Tfdp2 | 1790.74725 | -0.006187503 | 0.066173538 | -0.09350419 | 0.925503021 | 0.983236501 |
| Camk2g | 11881.57758 | 0.285690176 | 0.079894857 | 3.575826859 | 0.000349123 | 0.005251754 | Camk2g | Ablim3 | Ighmbp2 | 555.7961507 | -0.006189097 | 0.067920606 | -0.091122527 | 0.927395234 | 0.983501978 |
| Ell2 | 1036.079926 | 0.285578093 | 0.077598636 | 3.680194734 | 0.000233056 | 0.003801019 | Ell2 | Gypc | Hsd11b2 | 1.075834258 | -0.006203486 | 0.024181031 | -0.256543505 | 0.797531192 | NA |
| Tox3 | 471.1725551 | 0.285390303 | 0.101587887 | 2.809294601 | 0.004965019 | 0.035795744 | Tox3 | Lrp1b | Ddx50 | 2625.077459 | -0.006207976 | 0.069799212 | -0.088940496 | 0.9291292 | 0.983640461 |
| Clk3 | 1896.046402 | 0.285251644 | 0.072964008 | 3.909484282 | 9.25E-05 | 0.001849151 | Clk3 | Adora2a | 4933440M02Rik | 0.313099713 | -0.00621683 | 0.012574609 | -0.4943955 | 0.621026892 | NA |
| Myrf | 2570.106459 | 0.284740795 | 0.099634379 | 2.85785688 | 0.004265127 | 0.032489617 | Myrf | Ptprf | Upk3b | 0.313099713 | -0.00621683 | 0.012574609 | -0.4943955 | 0.621026892 | NA |
| Mvb12a | 487.5058516 | 0.2847014 | 0.087737419 | 3.244925626 | 0.001174813 | 0.012845838 | Mvb12a | Flt1 | Twf1 | 2050.430273 | -0.006219246 | 0.049809968 | -0.124859458 | 0.900634815 | 0.979053977 |
| Ly6e | 7638.14226 | 0.284524575 | 0.097600639 | 2.915191723 | 0.003554701 | 0.028626059 | Ly6e | Vstm4 | Spic | 0.583236734 | -0.006224337 | 0.014993001 | -0.415149506 | 0.678032461 | NA |
| Pcsk7 | 1037.497521 | 0.284275344 | 0.098961071 | 2.872597694 | 0.004071122 | 0.031413328 | Pcsk7 | Trpc5 | Larp6 | 1490.169747 | -0.006224822 | 0.055190199 | -0.112788537 | 0.910198206 | 0.980590355 |
| Plxnd1 | 5240.027227 | 0.28420993 | 0.097586968 | 2.912375857 | 0.003586907 | 0.02876272 | Plxnd1 | Apcdd1 | Krtap17-1 | 0.895073387 | -0.006242043 | 0.022353177 | -0.279246347 | 0.780055778 | NA |
| Zfp940 | 643.8843164 | 0.283847092 | 0.106105275 | 2.675145907 | 0.007469673 | 0.047647229 | Zfp940 | Cyp51 | Sepw1 | 16381.45137 | -0.006245481 | 0.057626465 | -0.108378697 | 0.913695299 | 0.98170867 |
| Setd1a | 2831.862923 | 0.283765665 | 0.077139908 | 3.678584454 | 0.000234532 | 0.003821072 | Setd1a | Tmem91 | Hormad2 | 0.985901806 | -0.006248848 | 0.019473362 | -0.320892091 | 0.748292168 | NA |
| Prpf40b | 2541.661095 | 0.283651333 | 0.080610625 | 3.518783456 | 0.00043353 | 0.006145581 | Prpf40b | Grm1 | Adpgk | 625.4500734 | -0.006250188 | 0.074921927 | -0.083422678 | 0.933515457 | 0.984481307 |
| Pnpla6 | 2151.687758 | 0.283483454 | 0.06825944 | 4.153029275 | 3.28E-05 | 0.000825264 | Pnpla6 | Nid1 | F11 | 0.89196579 | -0.006259817 | 0.022346323 | -0.280127403 | 0.779379762 | NA |
| Clu | 28301.34007 | 0.283418634 | 0.074063093 | 3.826718865 | 0.000129863 | 0.002392578 | Clu | Rab3c | 2810006K23Rik | 477.253085 | -0.006261229 | 0.072146181 | -0.08678531 | 0.930842164 | 0.984208871 |
| Fnbp4 | 1797.976227 | 0.283207434 | 0.087136678 | 3.250151829 | 0.001153434 | 0.012719792 | Fnbp4 | Ccdc138 | Dhrs11 | 198.6231972 | -0.006268162 | 0.092281784 | -0.067924154 | 0.945846011 | 0.986798749 |
| Cep135 | 380.2416604 | 0.283107777 | 0.105223732 | 2.690531617 | 0.007133827 | 0.046162698 | Cep135 | Mrpl33 | Rax | 5.135484584 | -0.006293793 | 0.042380301 | -0.148507518 | 0.881942253 | NA |
| Rltpr | 1889.519764 | 0.283093426 | 0.102761228 | 2.754866137 | 0.005871615 | 0.040200975 | Rltpr | Ccr2 | Gm13154 | 0.564763665 | -0.00629936 | 0.016659354 | -0.378127504 | 0.705335877 | NA |
| Pus10 | 394.9943801 | 0.2820996 | 0.100209424 | 2.815100514 | 0.004876198 | 0.035420446 | Pus10 | Shisa2 | C1qtnf7 | 28.41915252 | -0.006340926 | 0.082184729 | -0.077154549 | 0.938500598 | 0.985461126 |
| Eml3 | 431.2331535 | 0.282017316 | 0.104668539 | 2.694384766 | 0.007051871 | 0.045697065 | Eml3 | Neto2 | Nr3c1 | 4636.656443 | -0.006342065 | 0.069869029 | -0.090770759 | 0.927674745 | 0.983501978 |
| Adcy6 | 2340.931852 | 0.28173465 | 0.079546519 | 3.541759647 | 0.000397467 | 0.005815261 | Adcy6 | Pcdhb18 | Arnt2 | 18677.36844 | -0.006362132 | 0.042012953 | -0.151432634 | 0.879634448 | 0.973413768 |
| Gns | 4767.12951 | 0.280845004 | 0.062319401 | 4.506542083 | 6.59E-06 | 0.000227381 | Gns | Gpr26 | Ephb1 | 2013.756961 | -0.006362692 | 0.068744081 | -0.092556212 | 0.926256132 | 0.983236501 |
| Scpep1 | 642.2410366 | 0.28051406 | 0.100521574 | 2.790585625 | 0.005261278 | 0.037121241 | Scpep1 | Trpc7 | Slc6a20a | 469.4821467 | -0.006365266 | 0.088383736 | -0.072018527 | 0.942587164 | 0.985461126 |
| Gm19757 | 152.1302568 | 0.280503242 | 0.105548652 | 2.657572932 | 0.007870556 | 0.049612039 | Gm19757 | Col1a2 | 1700045H11Rik | 0.097165704 | -0.006378285 | 0.008487817 | -0.751463494 | 0.452373763 | NA |
| Arhgap17 | 487.2926756 | 0.280231185 | 0.077659416 | 3.608463734 | 0.000308016 | 0.004762867 | Arhgap17 | Qpct | 4930434J06Rik | 0.097165704 | -0.006378285 | 0.008487817 | -0.751463494 | 0.452373763 | NA |
| Mitd1 | 194.336654 | 0.280161626 | 0.099339725 | 2.820237583 | 0.004798811 | 0.035023534 | Mitd1 | Rxrg | 4930487H11Rik | 0.097165704 | -0.006378285 | 0.008487817 | -0.751463494 | 0.452373763 | NA |
| Sbno2 | 741.4026484 | 0.280136684 | 0.096458033 | 2.904233843 | 0.003681531 | 0.029312249 | Sbno2 | Pbx1 | 4930503H13Rik | 0.097165704 | -0.006378285 | 0.008487817 | -0.751463494 | 0.452373763 | NA |
| Zmat4 | 2075.561661 | 0.280077682 | 0.080432483 | 3.482146405 | 0.000497412 | 0.006796205 | Zmat4 | Osbpl8 | 4930529K09Rik | 0.097165704 | -0.006378285 | 0.008487817 | -0.751463494 | 0.452373763 | NA |
| Riok1 | 492.5142375 | 0.279662668 | 0.095842684 | 2.917934424 | 0.003523584 | 0.02844941 | Riok1 | Plcxd3 | 4930542D17Rik | 0.097165704 | -0.006378285 | 0.008487817 | -0.751463494 | 0.452373763 | NA |
| Slc5a5 | 857.4069729 | 0.278953566 | 0.100320731 | 2.780617351 | 0.005425565 | 0.038020669 | Slc5a5 | Mpped2 | 4930562C15Rik | 0.097165704 | -0.006378285 | 0.008487817 | -0.751463494 | 0.452373763 | NA |
| Calcoco1 | 3201.173313 | 0.278916813 | 0.078427951 | 3.55634452 | 0.000376051 | 0.005580969 | Calcoco1 | Akap9 | 4930578E11Rik | 0.097165704 | -0.006378285 | 0.008487817 | -0.751463494 | 0.452373763 | NA |
| Celsr3 | 4967.381609 | 0.278794021 | 0.097946535 | 2.846389821 | 0.004421803 | 0.033297621 | Celsr3 | Grk5 | A230020J21Rik | 0.097165704 | -0.006378285 | 0.008487817 | -0.751463494 | 0.452373763 | NA |
| Hectd3 | 2374.141254 | 0.278637484 | 0.060405345 | 4.612795149 | 3.97E-06 | 0.000152367 | Hectd3 | Grm3 | A430088P11Rik | 0.097165704 | -0.006378285 | 0.008487817 | -0.751463494 | 0.452373763 | NA |
| Bms1 | 1511.386886 | 0.278424845 | 0.0658206 | 4.230056322 | 2.34E-05 | 0.000631746 | Bms1 | Lhx8 | A630075F10Rik | 0.097165704 | -0.006378285 | 0.008487817 | -0.751463494 | 0.452373763 | NA |
| Utp15 | 1158.978164 | 0.278360302 | 0.069710482 | 3.993091046 | 6.52E-05 | 0.001409317 | Utp15 | B230216N24Rik | A930006I01Rik | 0.097165704 | -0.006378285 | 0.008487817 | -0.751463494 | 0.452373763 | NA |
| Scn1a | 6633.89606 | 0.278343336 | 0.096943473 | 2.871192114 | 0.004089269 | 0.031506284 | Scn1a | Npffr2 | AU015836 | 0.097165704 | -0.006378285 | 0.008487817 | -0.751463494 | 0.452373763 | NA |
| Plekhm2 | 5101.002108 | 0.27827922 | 0.080065449 | 3.475646801 | 0.000509623 | 0.006914272 | Plekhm2 | Epha3 | Amy2b | 0.097165704 | -0.006378285 | 0.008487817 | -0.751463494 | 0.452373763 | NA |
| Cux1 | 3137.840507 | 0.278182169 | 0.067072418 | 4.147489786 | 3.36E-05 | 0.000840026 | Cux1 | Sv2c | Cd160 | 0.097165704 | -0.006378285 | 0.008487817 | -0.751463494 | 0.452373763 | NA |
| Fubp3 | 1172.234895 | 0.277395808 | 0.07830315 | 3.542588126 | 0.000396221 | 0.005802503 | Fubp3 | Slc6a11 | Clec4a4 | 0.097165704 | -0.006378285 | 0.008487817 | -0.751463494 | 0.452373763 | NA |
| AI854703 | 1146.145472 | 0.277139183 | 0.061717627 | 4.490438074 | 7.11E-06 | 0.000241505 | AI854703 | Sla | Cryge | 0.097165704 | -0.006378285 | 0.008487817 | -0.751463494 | 0.452373763 | NA |
| Adora1 | 10313.67738 | 0.277076233 | 0.099042012 | 2.79756266 | 0.005148978 | 0.036561991 | Adora1 | Prkar2b | Csf3 | 0.097165704 | -0.006378285 | 0.008487817 | -0.751463494 | 0.452373763 | NA |
| Fancg | 507.5489224 | 0.277050229 | 0.10044594 | 2.75820237 | 0.00581202 | 0.03995879 | Fancg | Atp5e | Entpd8 | 0.097165704 | -0.006378285 | 0.008487817 | -0.751463494 | 0.452373763 | NA |
| Rbms1 | 569.3201675 | 0.276738619 | 0.079031736 | 3.50161382 | 0.000462449 | 0.006438851 | Rbms1 | Chst15 | Foxd3 | 0.097165704 | -0.006378285 | 0.008487817 | -0.751463494 | 0.452373763 | NA |
| Dennd3 | 326.8464269 | 0.275588881 | 0.099717699 | 2.76369072 | 0.005715168 | 0.039443571 | Dennd3 | Foxo6 | Foxd4 | 0.097165704 | -0.006378285 | 0.008487817 | -0.751463494 | 0.452373763 | NA |
| Gnpda1 | 743.8797548 | 0.275482641 | 0.063318726 | 4.350729349 | 1.36E-05 | 0.000409807 | Gnpda1 | Gpr52 | Gm13242 | 0.097165704 | -0.006378285 | 0.008487817 | -0.751463494 | 0.452373763 | NA |
| Pdk2 | 4565.772657 | 0.27508552 | 0.082922403 | 3.317384809 | 0.000908644 | 0.010547482 | Pdk2 | Kctd12 | Gm14812 | 0.097165704 | -0.006378285 | 0.008487817 | -0.751463494 | 0.452373763 | NA |
| Chka | 1678.053777 | 0.275015342 | 0.096159892 | 2.859979723 | 0.004236681 | 0.032353861 | Chka | Shisa9 | Gm20199 | 0.097165704 | -0.006378285 | 0.008487817 | -0.751463494 | 0.452373763 | NA |
| Lrrc8a | 2055.320247 | 0.274907997 | 0.095482478 | 2.879146041 | 0.003987536 | 0.031002229 | Lrrc8a | Ddr2 | Gm3143 | 0.097165704 | -0.006378285 | 0.008487817 | -0.751463494 | 0.452373763 | NA |
| Per3 | 2035.965649 | 0.274901301 | 0.077834744 | 3.531858506 | 0.00041265 | 0.005958621 | Per3 | Myo16 | Gm5150 | 0.097165704 | -0.006378285 | 0.008487817 | -0.751463494 | 0.452373763 | NA |
| Polk | 494.6675454 | 0.27474836 | 0.078342053 | 3.507035486 | 0.000453129 | 0.006353644 | Polk | 4930426D05Rik | Gm5483 | 0.097165704 | -0.006378285 | 0.008487817 | -0.751463494 | 0.452373763 | NA |
| Foxo3 | 1451.09523 | 0.274144117 | 0.077859274 | 3.521020712 | 0.000429889 | 0.006116346 | Foxo3 | Alk | Gm6525 | 0.097165704 | -0.006378285 | 0.008487817 | -0.751463494 | 0.452373763 | NA |
| Lnx1 | 908.489144 | 0.273969318 | 0.079373754 | 3.451636123 | 0.000557199 | 0.007397802 | Lnx1 | AI848285 | Gm6890 | 0.097165704 | -0.006378285 | 0.008487817 | -0.751463494 | 0.452373763 | NA |
| Zfp108 | 243.448303 | 0.273240983 | 0.099526877 | 2.745398932 | 0.006043738 | 0.041016941 | Zfp108 | Lrrc9 | Gpr128 | 0.097165704 | -0.006378285 | 0.008487817 | -0.751463494 | 0.452373763 | NA |
| Sltm | 2513.626454 | 0.272462165 | 0.092528366 | 2.944633906 | 0.00323337 | 0.026704603 | Sltm | Shc3 | Gpr33 | 0.097165704 | -0.006378285 | 0.008487817 | -0.751463494 | 0.452373763 | NA |
| Zscan26 | 3193.936833 | 0.272009692 | 0.085462884 | 3.182781575 | 0.001458676 | 0.01514794 | Zscan26 | Pcsk9 | Gykl1 | 0.097165704 | -0.006378285 | 0.008487817 | -0.751463494 | 0.452373763 | NA |
| Zdhhc4 | 706.8152557 | 0.271803363 | 0.087558423 | 3.104251451 | 0.001907611 | 0.018495951 | Zdhhc4 | Alcam | H2-Ea-ps | 0.097165704 | -0.006378285 | 0.008487817 | -0.751463494 | 0.452373763 | NA |
| Nckipsd | 3224.885303 | 0.271550756 | 0.090981177 | 2.9846916 | 0.002838644 | 0.024380235 | Nckipsd | Dgkb | Habp2 | 0.097165704 | -0.006378285 | 0.008487817 | -0.751463494 | 0.452373763 | NA |
| Dcdc2a | 376.6574047 | 0.271538895 | 0.082557734 | 3.289078852 | 0.001005159 | 0.011417835 | Dcdc2a | Atp6v1c2 | Hist1h2ab | 0.097165704 | -0.006378285 | 0.008487817 | -0.751463494 | 0.452373763 | NA |
| Osbpl7 | 642.3745631 | 0.271270809 | 0.09077392 | 2.988422318 | 0.002804218 | 0.02420532 | Osbpl7 | Gng13 | Klra8 | 0.097165704 | -0.006378285 | 0.008487817 | -0.751463494 | 0.452373763 | NA |
| Zfp276 | 839.1398312 | 0.270661147 | 0.099993027 | 2.706800205 | 0.006793513 | 0.044436233 | Zfp276 | Prok2 | Krt82 | 0.097165704 | -0.006378285 | 0.008487817 | -0.751463494 | 0.452373763 | NA |
| Sel1l3 | 2790.240901 | 0.270621442 | 0.097823558 | 2.766424032 | 0.005667479 | 0.039254321 | Sel1l3 | Slc7a2 | Lce3f | 0.097165704 | -0.006378285 | 0.008487817 | -0.751463494 | 0.452373763 | NA |
| Prodh | 983.0157978 | 0.269619886 | 0.088297107 | 3.053552868 | 0.002261489 | 0.020948688 | Prodh | Tnfrsf8 | Mir195 | 0.097165704 | -0.006378285 | 0.008487817 | -0.751463494 | 0.452373763 | NA |
| Sema6c | 689.6047835 | 0.269527297 | 0.091112643 | 2.958176706 | 0.003094647 | 0.025824561 | Sema6c | Slc7a11 | Mir3074-1 | 0.097165704 | -0.006378285 | 0.008487817 | -0.751463494 | 0.452373763 | NA |
| Zfp956 | 802.9712627 | 0.269472669 | 0.079610977 | 3.384868237 | 0.000712124 | 0.008819867 | Zfp956 | Cped1 | Mir30c-2 | 0.097165704 | -0.006378285 | 0.008487817 | -0.751463494 | 0.452373763 | NA |
| Grhl1 | 514.0701098 | 0.269261792 | 0.094136075 | 2.860346477 | 0.004231784 | 0.032346946 | Grhl1 | Efcab6 | Mir5107 | 0.097165704 | -0.006378285 | 0.008487817 | -0.751463494 | 0.452373763 | NA |
| Elmod3 | 804.7182459 | 0.269060857 | 0.075817992 | 3.548773202 | 0.00038703 | 0.005694822 | Elmod3 | Trps1 | Mir654 | 0.097165704 | -0.006378285 | 0.008487817 | -0.751463494 | 0.452373763 | NA |
| Rnf31 | 1005.021949 | 0.26904605 | 0.063904177 | 4.210148134 | 2.55E-05 | 0.000678356 | Rnf31 | 9630013A20Rik | Mir711 | 0.097165704 | -0.006378285 | 0.008487817 | -0.751463494 | 0.452373763 | NA |
| Tesk2 | 243.9767993 | 0.268669483 | 0.093248934 | 2.88120701 | 0.003961554 | 0.030875408 | Tesk2 | Mgp | Mir872 | 0.097165704 | -0.006378285 | 0.008487817 | -0.751463494 | 0.452373763 | NA |
| Fbrs | 1567.645785 | 0.268057581 | 0.075947793 | 3.529497964 | 0.000416349 | 0.005984147 | Fbrs | Hey2 | Mmp10 | 0.097165704 | -0.006378285 | 0.008487817 | -0.751463494 | 0.452373763 | NA |
| Zfp335 | 1569.866418 | 0.267800344 | 0.090032125 | 2.974497638 | 0.002934687 | 0.02498354 | Zfp335 | Foxp2 | Olfr1371 | 0.097165704 | -0.006378285 | 0.008487817 | -0.751463494 | 0.452373763 | NA |
| Kctd18 | 345.9035113 | 0.267582382 | 0.09499846 | 2.816702305 | 0.004851948 | 0.035293932 | Kctd18 | Fam126a | Olfr1395 | 0.097165704 | -0.006378285 | 0.008487817 | -0.751463494 | 0.452373763 | NA |
| Ndst2 | 694.5783561 | 0.267484081 | 0.080320982 | 3.330189386 | 0.000867869 | 0.010225679 | Ndst2 | Tmem45a | Olfr150 | 0.097165704 | -0.006378285 | 0.008487817 | -0.751463494 | 0.452373763 | NA |
| Trpc2 | 511.711799 | 0.266880067 | 0.095283401 | 2.800908284 | 0.0050959 | 0.036290258 | Trpc2 | Lum | Olfr214 | 0.097165704 | -0.006378285 | 0.008487817 | -0.751463494 | 0.452373763 | NA |

| | | | | | | | | | | | | | | | |
|---|---|---|---|---|---|---|---|---|---|---|---|---|---|---|---|
| Gmppa | 1152.552036 | 0.266587613 | 0.07484588 | 3.561820807 | 0.000368292 | 0.005486838 | Gmppa | Ntrk1 | Olfr320 | 0.097165704 | -0.006378285 | 0.008487817 | -0.751463494 | 0.452373763 | NA |
| 4-Sep | 6547.743147 | 0.266501629 | 0.09947411 | 2.679105445 | 0.007381914 | 0.047340803 | 4-Sep | Ccdc80 | Olfr532 | 0.097165704 | -0.006378285 | 0.008487817 | -0.751463494 | 0.452373763 | NA |
| 2700050L05Rik | 992.9651654 | 0.266220739 | 0.076558811 | 3.477336387 | 0.000506422 | 0.00688894 | 2700050L05Rik | Aplnr | Olfr881 | 0.097165704 | -0.006378285 | 0.008487817 | -0.751463494 | 0.452373763 | NA |
| Cwf19l2 | 474.4366147 | 0.265974466 | 0.08840914 | 3.008449888 | 0.002625841 | 0.023191304 | Cwf19l2 | Cdca3 | Olfr95 | 0.097165704 | -0.006378285 | 0.008487817 | -0.751463494 | 0.452373763 | NA |
| Grina | 14385.91329 | 0.265427305 | 0.072165788 | 3.678021277 | 0.00023505 | 0.003825494 | Grina | Dcbld1 | Orm3 | 0.097165704 | -0.006378285 | 0.008487817 | -0.751463494 | 0.452373763 | NA |
| Hmha1 | 319.927829 | 0.265395447 | 0.097393904 | 2.724969805 | 0.006430737 | 0.042836559 | Hmha1 | Gng4 | Pramel3 | 0.097165704 | -0.006378285 | 0.008487817 | -0.751463494 | 0.452373763 | NA |
| Pex5l | 7514.827652 | 0.264905892 | 0.094115201 | 2.814698256 | 0.004882305 | 0.035431587 | Pex5l | Arhgap10 | Prr23a | 0.097165704 | -0.006378285 | 0.008487817 | -0.751463494 | 0.452373763 | NA |
| Szt2 | 2301.531666 | 0.26454484 | 0.091117031 | 2.903352296 | 0.003691911 | 0.029379797 | Szt2 | Zdbf2 | Prr9 | 0.097165704 | -0.006378285 | 0.008487817 | -0.751463494 | 0.452373763 | NA |
| Anks3 | 1157.292551 | 0.264533166 | 0.073178985 | 3.614878874 | 0.000300488 | 0.004669772 | Anks3 | Adra2c | Rhox4a | 0.097165704 | -0.006378285 | 0.008487817 | -0.751463494 | 0.452373763 | NA |
| Bbs2 | 1120.689392 | 0.263800713 | 0.090696917 | 2.908596238 | 0.003630554 | 0.028965914 | Bbs2 | Kcnn3 | Ros1 | 0.097165704 | -0.006378285 | 0.008487817 | -0.751463494 | 0.452373763 | NA |
| Hirip3 | 1041.647189 | 0.263528368 | 0.069536504 | 3.789784556 | 0.000150778 | 0.002722791 | Hirip3 | AI504432 | Serpina5 | 0.097165704 | -0.006378285 | 0.008487817 | -0.751463494 | 0.452373763 | NA |
| Mxra7 | 487.0823002 | 0.263473212 | 0.098190045 | 2.683298623 | 0.007289986 | 0.046926069 | Mxra7 | Fbln1 | Slc51a | 0.097165704 | -0.006378285 | 0.008487817 | -0.751463494 | 0.452373763 | NA |
| Sirt6 | 499.704828 | 0.263342774 | 0.089866252 | 2.930385644 | 0.003385416 | 0.027607174 | Sirt6 | Mei4 | Slco6d1 | 0.097165704 | -0.006378285 | 0.008487817 | -0.751463494 | 0.452373763 | NA |
| Srek1 | 1838.959289 | 0.262677687 | 0.091166617 | 2.881292462 | 0.00396048 | 0.030875408 | Srek1 | Kif27 | Spink11 | 0.097165704 | -0.006378285 | 0.008487817 | -0.751463494 | 0.452373763 | NA |
| Zfp316 | 1198.474351 | 0.262461325 | 0.078212821 | 3.355732739 | 0.00079155 | 0.009492476 | Zfp316 | Artn | Stmn1-rs1 | 0.097165704 | -0.006378285 | 0.008487817 | -0.751463494 | 0.452373763 | NA |
| Slc37a4 | 784.5343816 | 0.261652654 | 0.070185919 | 3.727993573 | 0.00019301 | 0.003311243 | Slc37a4 | Dmkn | Tas2r103 | 0.097165704 | -0.006378285 | 0.008487817 | -0.751463494 | 0.452373763 | NA |
| Chpf2 | 2194.501165 | 0.261371227 | 0.054958612 | 4.755782874 | 1.98E-06 | 8.78E-05 | Chpf2 | Egln3 | Tmprss11f | 0.097165704 | -0.006378285 | 0.008487817 | -0.751463494 | 0.452373763 | NA |
| Btbd11 | 1159.746864 | 0.261300517 | 0.080296082 | 3.254212532 | 0.001137072 | 0.012602139 | Btbd11 | Map2 | Vmn1r223 | 0.097165704 | -0.006378285 | 0.008487817 | -0.751463494 | 0.452373763 | NA |
| Lgals8 | 1017.492143 | 0.261051122 | 0.074987893 | 3.481243576 | 0.000499091 | 0.006813147 | Lgals8 | Tcfl5 | Zcchc13 | 0.097165704 | -0.006378285 | 0.008487817 | -0.751463494 | 0.452373763 | NA |
| B3gat2 | 446.4675839 | 0.260817909 | 0.089071412 | 2.928188767 | 0.00340943 | 0.027759173 | B3gat2 | Ccdc109b | Ptk2b | 25393.54815 | -0.006379224 | 0.061803477 | -0.103217887 | 0.917790044 | 0.982742917 |
| Taf1b | 675.1268158 | 0.260624888 | 0.088255243 | 2.953081076 | 0.003146193 | 0.026194045 | Taf1b | Col3a1 | Nit2 | 322.2546995 | -0.006382534 | 0.077763668 | -0.082076046 | 0.934586241 | 0.985294234 |
| Utp14a | 597.7831145 | 0.259846116 | 0.079033656 | 3.287790672 | 0.001009769 | 0.011447859 | Utp14a | Pcdhb22 | Pclo | 9072.322961 | -0.006390925 | 0.077776935 | -0.082169931 | 0.934511584 | 0.985294234 |
| Cnot3 | 1583.700684 | 0.259822933 | 0.067178437 | 3.867653738 | 0.000109888 | 0.002101972 | Cnot3 | Slc26a7 | Dscaml1 | 2779.667792 | -0.006399275 | 0.065535687 | -0.097645656 | 0.92221367 | 0.982928035 |
| Surf2 | 827.2797765 | 0.259701239 | 0.078220673 | 3.320109991 | 0.00089982 | 0.010498156 | Surf2 | Clec3b | Casc5 | 4.943653599 | -0.006406907 | 0.046585702 | -0.137529464 | 0.890612305 | NA |
| Ubp1 | 3567.846609 | 0.259582492 | 0.087962283 | 2.951065877 | 0.003166794 | 0.026337252 | Ubp1 | 2810011L19Rik | BC048943 | 9.172351895 | -0.006409401 | 0.052424647 | -0.122259313 | 0.902693654 | NA |
| Trp53 | 801.2736227 | 0.259267504 | 0.075803784 | 3.420244871 | 0.000625648 | 0.008054403 | Trp53 | Cdca7 | Pex11a | 172.9826394 | -0.006428793 | 0.086790005 | -0.074072963 | 0.940952328 | 0.985461126 |
| Apbb3 | 1424.740789 | 0.25915149 | 0.059365892 | 4.365326323 | 1.27E-05 | 0.000388676 | Apbb3 | 2610034M16Rik | Arx | 598.9354266 | -0.006437175 | 0.06834348 | -0.094188582 | 0.924959355 | 0.983236501 |
| Trex1 | 383.859503 | 0.258539066 | 0.084582579 | 3.056646766 | 0.002238279 | 0.020803774 | Trex1 | Pdgfc | Eral1 | 714.6234029 | -0.006437339 | 0.063257989 | -0.101763257 | 0.918944591 | 0.982742917 |
| Wdr19 | 906.1033872 | 0.258475462 | 0.070119223 | 3.686228287 | 0.000227602 | 0.003723832 | Wdr19 | Gm10754 | Polr1e | 254.4927396 | -0.00643974 | 0.086275362 | -0.074641702 | 0.940499794 | 0.985461126 |
| Rfxank | 641.5151331 | 0.258362765 | 0.075857229 | 3.405908271 | 0.000659443 | 0.0083136 | Rfxank | Dok4 | Syndig1 | 1759.862923 | -0.006448766 | 0.060276556 | -0.106986308 | 0.914799842 | 0.98170867 |
| Plekha6 | 5032.208944 | 0.257134812 | 0.052212076 | 4.924814984 | 8.44E-07 | 4.25E-05 | Plekha6 | Sox4 | Lrit2 | 0.321791944 | -0.006449398 | 0.012677177 | -0.508740887 | 0.610933859 | NA |
| Polrmt | 1180.968823 | 0.257033945 | 0.071465002 | 3.596640831 | 0.000322353 | 0.004945088 | Polrmt | Rgs2 | Slc30a8 | 0.533274033 | -0.006467142 | 0.016459981 | -0.392900926 | 0.694392658 | NA |
| Lrrc57 | 1263.270748 | 0.256485513 | 0.065293896 | 3.928169816 | 8.56E-05 | 0.001745007 | Lrrc57 | Csmd3 | 2410007B07Rik | 0.322281847 | -0.006487224 | 0.012685754 | -0.511378627 | 0.609085958 | NA |
| 4922501C03Rik | 415.6370696 | 0.256438059 | 0.091484063 | 2.80308996 | 0.005061555 | 0.036240172 | 4922501C03Rik | Casq2 | 7530416G11Rik | 0.322281847 | -0.006487224 | 0.012685754 | -0.511378627 | 0.609085958 | NA |
| Rhno1 | 429.9475416 | 0.256332015 | 0.093782155 | 2.733270685 | 0.006270877 | 0.042079243 | Rhno1 | Gpc6 | Spaca5 | 0.322281847 | -0.006487224 | 0.012685754 | -0.511378627 | 0.609085958 | NA |
| Polr3e | 1272.856745 | 0.256065996 | 0.067459418 | 3.795852445 | 0.000147137 | 0.002669486 | Polr3e | Fam102b | Rundc3a | 12670.92831 | -0.006497529 | 0.059288637 | -0.109591473 | 0.912733374 | 0.981664815 |
| Rffl | 562.5782036 | 0.256006693 | 0.082037893 | 3.1205908 | 0.001804886 | 0.017699311 | Rffl | Wwc1 | Il6st | 3903.252464 | -0.006498366 | 0.065238402 | -0.099609522 | 0.920654334 | 0.982762672 |
| Slc6a8 | 4564.88945 | 0.255823656 | 0.064821429 | 3.946590783 | 7.93E-05 | 0.001635816 | Slc6a8 | Tox | Crot | 1407.253973 | -0.006510436 | 0.065166226 | -0.099905062 | 0.920419697 | 0.982762672 |
| Adam15 | 4099.675672 | 0.255821041 | 0.067950603 | 3.764808986 | 0.000166676 | 0.002941327 | Adam15 | Hmgcr | Ndufaf3 | 1060.040343 | -0.00651213 | 0.060080385 | -0.108390284 | 0.913686108 | 0.98170867 |
| Arhgap15 | 418.2416294 | 0.255660185 | 0.096133849 | 2.659419012 | 0.007827555 | 0.049401276 | Arhgap15 | Ccdc3 | Rarb | 871.5478667 | -0.006518264 | 0.076321298 | -0.085405571 | 0.931938964 | 0.984387568 |
| Slc12a9 | 1121.625281 | 0.255157079 | 0.086137375 | 2.962210985 | 0.003054384 | 0.025594714 | Slc12a9 | Fam19a4 | 1700011H14Rik | 0.106347839 | -0.006533472 | 0.008543895 | -0.764694822 | 0.44445329 | NA |
| Mtcp1 | 241.7527577 | 0.254776735 | 0.094540185 | 2.69490414 | 0.007040888 | 0.045660528 | Mtcp1 | Stk32a | 1700047E10Rik | 0.106347839 | -0.006533472 | 0.008543895 | -0.764694822 | 0.44445329 | NA |
| Tmem25 | 2406.135191 | 0.254759116 | 0.085219172 | 2.989457786 | 0.002794731 | 0.024150338 | Tmem25 | Rspo2 | 1700065J11Rik | 0.106347839 | -0.006533472 | 0.008543895 | -0.764694822 | 0.44445329 | NA |
| Rcan2 | 11886.4631 | 0.254635249 | 0.077586453 | 3.28195503 | 0.0010309 | 0.011633483 | Rcan2 | Cdca5 | 1700092C02Rik | 0.106347839 | -0.006533472 | 0.008543895 | -0.764694822 | 0.44445329 | NA |
| Rhbdd3 | 559.7434643 | 0.253408777 | 0.09045072 | 2.80162257 | 0.005084632 | 0.036290258 | Rhbdd3 | Ust | 4930448F12Rik | 0.106347839 | -0.006533472 | 0.008543895 | -0.764694822 | 0.44445329 | NA |
| Lypd6b | 574.2849864 | 0.253148173 | 0.072175624 | 3.507391516 | 0.000452523 | 0.006350895 | Lypd6b | Kcnf1 | 4930563D23Rik | 0.106347839 | -0.006533472 | 0.008543895 | -0.764694822 | 0.44445329 | NA |
| Vars | 1679.577633 | 0.252971891 | 0.071339678 | 3.546019507 | 0.000391097 | 0.005732886 | Vars | St3gal1 | Aadacl3 | 0.106347839 | -0.006533472 | 0.008543895 | -0.764694822 | 0.44445329 | NA |
| Znf512b | 3330.081277 | 0.252897834 | 0.091631693 | 2.759938464 | 0.005781225 | 0.039775444 | Znf512b | Fjx1 | Ahsg | 0.106347839 | -0.006533472 | 0.008543895 | -0.764694822 | 0.44445329 | NA |
| Hrsp12 | 1043.445333 | 0.252038796 | 0.089802075 | 2.806603254 | 0.005006686 | 0.035980333 | Hrsp12 | Dst | Akr1d1 | 0.106347839 | -0.006533472 | 0.008543895 | -0.764694822 | 0.44445329 | NA |
| Ell | 865.5372805 | 0.251905219 | 0.071585962 | 3.518919238 | 0.000433309 | 0.006145581 | Ell | Steap3 | B130006D01Rik | 0.106347839 | -0.006533472 | 0.008543895 | -0.764694822 | 0.44445329 | NA |
| Rela | 820.8787464 | 0.251731491 | 0.078131853 | 3.221880457 | 0.001273523 | 0.013683744 | Rela | Pde3a | C330013F16Rik | 0.106347839 | -0.006533472 | 0.008543895 | -0.764694822 | 0.44445329 | NA |
| Exoc1 | 2049.673333 | 0.251699664 | 0.050887073 | 4.946239804 | 7.57E-07 | 3.95E-05 | Exoc1 | Aoc3 | C86695 | 0.106347839 | -0.006533472 | 0.008543895 | -0.764694822 | 0.44445329 | NA |
| Cnppd1 | 1165.702659 | 0.251658223 | 0.082404299 | 3.053945333 | 0.002258532 | 0.020941771 | Cnppd1 | Chat | Calml3 | 0.106347839 | -0.006533472 | 0.008543895 | -0.764694822 | 0.44445329 | NA |
| Zfp423 | 640.5460814 | 0.251462017 | 0.085007737 | 2.958107399 | 0.003095343 | 0.025824561 | Zfp423 | Macf1 | Cd207 | 0.106347839 | -0.006533472 | 0.008543895 | -0.764694822 | 0.44445329 | NA |
| Fam160b2 | 2431.822796 | 0.251254355 | 0.055083128 | 4.561366868 | 5.08E-06 | 0.000187038 | Fam160b2 | Hes5 | D230030E09Rik | 0.106347839 | -0.006533472 | 0.008543895 | -0.764694822 | 0.44445329 | NA |
| Acd | 1247.255523 | 0.250851148 | 0.079497429 | 3.155462404 | 0.001602439 | 0.01619582 | Acd | Acta2 | D730001G18Rik | 0.106347839 | -0.006533472 | 0.008543895 | -0.764694822 | 0.44445329 | NA |
| Rnaseh1 | 307.5830117 | 0.250480512 | 0.091588195 | 2.734855861 | 0.00624076 | 0.041968026 | Rnaseh1 | Lhfpl5 | Ear1 | 0.106347839 | -0.006533472 | 0.008543895 | -0.764694822 | 0.44445329 | NA |
| Msn | 1403.761535 | 0.250469502 | 0.055618761 | 4.503327627 | 6.69E-06 | 0.000228307 | Msn | 2810417H13Rik | Fam170a | 0.106347839 | -0.006533472 | 0.008543895 | -0.764694822 | 0.44445329 | NA |
| Katna1 | 359.0367202 | 0.25033736 | 0.091623123 | 2.732250894 | 0.006290322 | 0.042154952 | Katna1 | Gpx6 | Fam71b | 0.106347839 | -0.006533472 | 0.008543895 | -0.764694822 | 0.44445329 | NA |
| Cars | 1454.770105 | 0.250220057 | 0.04621781 | 5.413931438 | 6.17E-08 | 5.05E-06 | Cars | Scgn | Gcsam | 0.106347839 | -0.006533472 | 0.008543895 | -0.764694822 | 0.44445329 | NA |
| Smarcd2 | 559.1782418 | 0.25008658 | 0.073908746 | 3.383721059 | 0.000715106 | 0.008835621 | Smarcd2 | Mis18bp1 | Gja8 | 0.106347839 | -0.006533472 | 0.008543895 | -0.764694822 | 0.44445329 | NA |
| Sec24a | 666.7153159 | 0.249933717 | 0.074936096 | 3.335291402 | 0.000852101 | 0.010070516 | Sec24a | Tlr5 | Gm13490 | 0.106347839 | -0.006533472 | 0.008543895 | -0.764694822 | 0.44445329 | NA |
| Wdr73 | 1030.124251 | 0.249297798 | 0.071772172 | 3.473460412 | 0.000513793 | 0.006962544 | Wdr73 | Ptprb | Gm13580 | 0.106347839 | -0.006533472 | 0.008543895 | -0.764694822 | 0.44445329 | NA |
| Lrrfip1 | 1979.103503 | 0.249128055 | 0.067250966 | 3.70445317 | 0.000211847 | 0.003548499 | Lrrfip1 | Zic4 | Gm6406 | 0.106347839 | -0.006533472 | 0.008543895 | -0.764694822 | 0.44445329 | NA |
| Cables2 | 1467.331658 | 0.248956697 | 0.061931072 | 4.019899704 | 5.82E-05 | 0.001299865 | Cables2 | Pcdh10 | H2afy3 | 0.106347839 | -0.006533472 | 0.008543895 | -0.764694822 | 0.44445329 | NA |
| Zfp414 | 798.413644 | 0.248908362 | 0.085903554 | 2.897532778 | 0.003761104 | 0.029853766 | Zfp414 | Celsr1 | Hoxb2 | 0.106347839 | -0.006533472 | 0.008543895 | -0.764694822 | 0.44445329 | NA |
| C1qtnf1 | 326.247084 | 0.248296434 | 0.087234205 | 2.84631969 | 0.004422777 | 0.033297621 | C1qtnf1 | Tyrp1 | Hoxc5 | 0.106347839 | -0.006533472 | 0.008543895 | -0.764694822 | 0.44445329 | NA |
| Hps5 | 638.7026078 | 0.247686739 | 0.092061568 | 2.690446686 | 0.007135644 | 0.046162698 | Hps5 | Grm4 | Krt13 | 0.106347839 | -0.006533472 | 0.008543895 | -0.764694822 | 0.44445329 | NA |
| Timm44 | 1550.311694 | 0.247430173 | 0.076150048 | 3.249245133 | 0.001157117 | 0.012746794 | Timm44 | Lgr5 | Krt71 | 0.106347839 | -0.006533472 | 0.008543895 | -0.764694822 | 0.44445329 | NA |
| Zfp598 | 1537.487058 | 0.247382855 | 0.091273323 | 2.710352242 | 0.006721179 | 0.044107886 | Zfp598 | Adamtsl3 | Mir1967 | 0.106347839 | -0.006533472 | 0.008543895 | -0.764694822 | 0.44445329 | NA |
| Gpr137b-ps | 421.3135084 | 0.247311136 | 0.084101142 | 2.940639462 | 0.003275355 | 0.026993805 | Gpr137b-ps | Tln2 | Mir328 | 0.106347839 | -0.006533472 | 0.008543895 | -0.764694822 | 0.44445329 | NA |
| Rbmx | 2110.804637 | 0.246918016 | 0.088030362 | 2.804918768 | 0.005032926 | 0.036101925 | Rbmx | Bcl11b | Mir344e | 0.106347839 | -0.006533472 | 0.008543895 | -0.764694822 | 0.44445329 | NA |
| Sez6l2 | 15571.27814 | 0.24669124 | 0.057203884 | 4.312491053 | 1.61E-05 | 0.000471022 | Sez6l2 | Sox1 | Mir363 | 0.106347839 | -0.006533472 | 0.008543895 | -0.764694822 | 0.44445329 | NA |
| Tbc1d30 | 4777.718345 | 0.24652925 | 0.058546292 | 4.210843075 | 2.54E-05 | 0.000678356 | Tbc1d30 | Map1b | Mug2 | 0.106347839 | -0.006533472 | 0.008543895 | -0.764694822 | 0.44445329 | NA |
| Cbx7 | 2483.46923 | 0.246002687 | 0.085689616 | 2.870857627 | 0.004093599 | 0.031523964 | Cbx7 | Isoc1 | Mup11 | 0.106347839 | -0.006533472 | 0.008543895 | -0.764694822 | 0.44445329 | NA |
| Trhde | 1056.963841 | 0.245531176 | 0.087230119 | 2.814752272 | 0.004881485 | 0.035431587 | Trhde | Gnal | Myl10 | 0.106347839 | -0.006533472 | 0.008543895 | -0.764694822 | 0.44445329 | NA |
| Sf3a2 | 1214.065511 | 0.24526687 | 0.080400819 | 3.050551878 | 0.002284212 | 0.021078966 | Sf3a2 | Fstl5 | Olfr1208 | 0.106347839 | -0.006533472 | 0.008543895 | -0.764694822 | 0.44445329 | NA |
| Gatad2a | 1263.692408 | 0.245047978 | 0.082698971 | 2.963132121 | 0.003045258 | 0.025559714 | Gatad2a | Chst3 | Olfr121 | 0.106347839 | -0.006533472 | 0.008543895 | -0.764694822 | 0.44445329 | NA |
| Tmem209 | 1177.039335 | 0.244813797 | 0.077653752 | 3.152633191 | 0.00161805 | 0.016318843 | Tmem209 | Dnahc8 | Olfr1342 | 0.106347839 | -0.006533472 | 0.008543895 | -0.764694822 | 0.44445329 | NA |
| Arsa | 1495.119009 | 0.244375212 | 0.071716055 | 3.407538415 | 0.000655517 | 0.008270829 | Arsa | Khdrbs2 | Olfr1437 | 0.106347839 | -0.006533472 | 0.008543895 | -0.764694822 | 0.44445329 | NA |
| Tarbp2 | 596.6402171 | 0.244276449 | 0.090406132 | 2.701989845 | 0.006892587 | 0.04487205 | Tarbp2 | Chrna5 | Olfr460 | 0.106347839 | -0.006533472 | 0.008543895 | -0.764694822 | 0.44445329 | NA |
| Brsk2 | 8256.83612 | 0.243474526 | 0.090319628 | 2.695698942 | 0.007024112 | 0.045593461 | Brsk2 | Fzd5 | Olfr707 | 0.106347839 | -0.006533472 | 0.008543895 | -0.764694822 | 0.44445329 | NA |
| Ctdp1 | 671.3858532 | 0.242888522 | 0.085145105 | 2.852642235 | 0.00433574 | 0.032878466 | Ctdp1 | Gabrg3 | Orm1 | 0.106347839 | -0.006533472 | 0.008543895 | -0.764694822 | 0.44445329 | NA |
| Zfp740 | 2700.0881 | 0.242882392 | 0.079665021 | 3.048795925 | 0.002297605 | 0.021168869 | Zfp740 | Prc1 | Rnf186 | 0.106347839 | -0.006533472 | 0.008543895 | -0.764694822 | 0.44445329 | NA |
| Camta2 | 13166.71322 | 0.242246741 | 0.085970848 | 2.817777741 | 0.004835727 | 0.035192464 | Camta2 | Cgref1 | Rsc1a1 | 0.106347839 | -0.006533472 | 0.008543895 | -0.764694822 | 0.44445329 | NA |
| Gng2 | 7105.148466 | 0.242130475 | 0.062533033 | 3.872041123 | 0.000107928 | 0.002067036 | Gng2 | E130309F12Rik | S100z | 0.106347839 | -0.006533472 | 0.008543895 | -0.764694822 | 0.44445329 | NA |
| Tub | 9202.683926 | 0.241341825 | 0.064192826 | 3.759638582 | 0.000170159 | 0.002992558 | Tub | Adcy5 | Ssxb6 | 0.106347839 | -0.006533472 | 0.008543895 | -0.764694822 | 0.44445329 | NA |
| Rhov | 270.874117 | 0.241235283 | 0.088902058 | 2.713494937 | 0.006657759 | 0.043784088 | Rhov | Dsel | Try10 | 0.106347839 | -0.006533472 | 0.008543895 | -0.764694822 | 0.44445329 | NA |
| Stx17 | 1127.471374 | 0.241221591 | 0.07763604 | 3.107082619 | 0.001889436 | 0.018346874 | Stx17 | Lmo4 | Ubqln3 | 0.106347839 | -0.006533472 | 0.008543895 | -0.764694822 | 0.44445329 | NA |
| Atg4c | 1520.340315 | 0.240916624 | 0.071714122 | 3.359402797 | 0.000781111 | 0.009403681 | Atg4c | Gm7244 | Vmn2r2 | 0.106347839 | -0.006533472 | 0.008543895 | -0.764694822 | 0.44445329 | NA |
| Ercc2 | 1265.212997 | 0.240822727 | 0.073099686 | 3.294442686 | 0.000986171 | 0.011259931 | Ercc2 | Fam64a | Vmn2r51 | 0.106347839 | -0.006533472 | 0.008543895 | -0.764694822 | 0.44445329 | NA |
| Cstf2 | 3899.973998 | 0.240490761 | 0.068994133 | 3.485669724 | 0.000490907 | 0.006749647 | Cstf2 | Hs3st5 | Vwde | 0.106347839 | -0.006533472 | 0.008543895 | -0.764694822 | 0.44445329 | NA |
| Cyth1 | 3261.713684 | 0.240275061 | 0.066457323 | 3.615479101 | 0.000299793 | 0.004663642 | Cyth1 | Abcc12 | Nit1 | 1231.498558 | -0.006537034 | 0.063416342 | -0.10308122 | 0.91789851 | 0.982742917 |
| Kcnk9 | 560.879223 | 0.239946381 | 0.085429878 | 2.808693941 | 0.004974291 | 0.035830621 | Kcnk9 | Cdyl2 | Ssu2 | 4.906319756 | -0.006543364 | 0.048387793 | -0.135227576 | 0.892431945 | NA |
| Map2k7 | 3612.361853 | 0.239217009 | 0.06013337 | 3.978107512 | 6.95E-05 | 0.001482515 | Map2k7 | Slc22a2 | Scrt1 | 6787.10549 | -0.006545193 | 0.057341624 | -0.114143844 | 0.909123767 | 0.980523965 |
| Mccc1 | 851.0861434 | 0.23914193 | 0.090030254 | 2.656239655 | 0.007901744 | 0.049768137 | Mccc1 | Des | Krt24 | 0.784472859 | -0.006555631 | 0.018220895 | -0.359786472 | 0.719006821 | NA |
| Dgkq | 1906.872318 | 0.238881648 | 0.089903271 | 2.657096295 | 0.007881693 | 0.049662036 | Dgkq | Zic3 | Tnnc2 | 6.708040353 | -0.006568083 | 0.053469927 | -0.122836949 | 0.902236215 | NA |
| Slc9a8 | 1546.263877 | 0.238460248 | 0.074351435 | 3.20720436 | 0.001340317 | 0.014194722 | Slc9a8 | Kcnh8 | Acot8 | 286.4455354 | -0.006588127 | 0.090641519 | -0.07268332 | 0.942058122 | 0.985461126 |
| Gls2 | 932.0295382 | 0.237450278 | 0.082450957 | 2.879897153 | 0.003978049 | 0.030957254 | Gls2 | Mest | Cib1 | 387.0930004 | -0.00658843 | 0.067487848 | -0.097623946 | 0.922230909 | 0.982928035 |
| Fan1 | 408.7100891 | 0.237423189 | 0.086461258 | 2.746006629 | 0.006032554 | 0.04097694 | Fan1 | Reln | Lgals1 | 389.8960249 | -0.006597584 | 0.090388608 | -0.07299132 | 0.941813025 | 0.985461126 |
| Vps33b | 799.5557611 | 0.236931992 | 0.065139876 | 3.637280355 | 0.000275532 | 0.004347345 | Vps33b | 1700001L19Rik | Chst9 | 22.35027679 | -0.006609123 | 0.076806812 | -0.086048655 | 0.93142774 | 0.984208871 |
| Trpm7 | 1925.682531 | 0.236438089 | 0.077825514 | 3.038053689 | 0.002381115 | 0.021727326 | Trpm7 | Fxyd2 | Dip2b | 4319.887514 | -0.006616972 | 0.056550171 | -0.117010648 | 0.906851615 | 0.980201396 |
| Kdm5c | 2944.571708 | 0.236430452 | 0.072172651 | 3.275900894 | 0.001053255 | 0.011851224 | Kdm5c | AI606473 | Fndc3c1 | 0.819350582 | -0.006620822 | 0.016414351 | -0.403355682 | 0.686686585 | NA |
| Zc3h7a | 1314.597311 | 0.236248833 | 0.067355606 | 3.507485808 | 0.000452362 | 0.006350895 | Zc3h7a | Serpinh1 | Nup43 | 129.1605596 | -0.006623194 | 0.090955462 | -0.072817992 | 0.941950953 | 0.985461126 |
| Proser1 | 940.6849074 | 0.236081856 | 0.070936165 | 3.32808881 | 0.00087444 | 0.01028745 | Proser1 | Kif26b | Slc38a9 | 954.1071972 | -0.006635747 | 0.07153838 | -0.092757863 | 0.926095927 | 0.983236501 |
| Cep85l | 382.2132178 | 0.2356606 | 0.086770658 | 2.715901957 | 0.006609549 | 0.043599054 | Cep85l | Alx4 | Snord35b | 2.180108994 | -0.006639862 | 0.030244429 | -0.219539996 | 0.826229428 | NA |
| Mfsd11 | 1225.161794 | 0.235396574 | 0.086302759 | 2.727567202 | 0.006380326 | 0.042645781 | Mfsd11 | E130008D07Rik | Nck1 | 764.9875855 | -0.006644971 | 0.076545842 | -0.086810341 | 0.930822268 | 0.984208871 |
| Safb | 3661.919541 | 0.235238504 | 0.069361439 | 3.391488212 | 0.000695142 | 0.008664942 | Safb | Agbl2 | Tada2a | 1123.870879 | -0.006653798 | 0.056165662 | -0.11846737 | 0.905697348 | 0.980118791 |
| Gpatch1 | 978.0450182 | 0.235212516 | 0.065509236 | 3.590524512 | 0.000330013 | 0.005042628 | Gpatch1 | Gm19522 | Dnajc17 | 302.304977 | -0.006669491 | 0.075497524 | -0.088340532 | 0.929606025 | 0.983715095 |
| Tstd2 | 645.6827232 | 0.234716636 | 0.069733785 | 3.36589553 | 0.000762956 | 0.009220936 | Tstd2 | Olfml2a | Cyp11b1 | 0.565118408 | -0.00667245 | 0.014900289 | -0.447806749 | 0.654292672 | NA |
| Gtpbp2 | 2522.137033 | 0.234139806 | 0.07417936 | 3.156401012 | 0.001597291 | 0.016186021 | Gtpbp2 | Cntn5 | Pcsk5 | 458.7280607 | -0.006691929 | 0.079376544 | -0.084306125 | 0.932813042 | 0.984387568 |
| Mtfmt | 384.2911971 | 0.233899175 | 0.083234456 | 2.810124401 | 0.004952235 | 0.035738209 | Mtfmt | Gm5607 | Clrn1 | 13.58346615 | -0.006701599 | 0.0733312 | -0.0913881 | 0.927184217 | NA |
| Ddx17 | 16513.67643 | 0.233648534 | 0.082148343 | 2.844227318 | 0.004451927 | 0.033404246 | Ddx17 | Dcaf12l2 | Tssk1 | 4.772582658 | -0.006710543 | 0.047943312 | -0.139968291 | 0.888685044 | NA |
| Sprtn | 370.1995883 | 0.232885513 | 0.082270359 | 2.830734143 | 0.004644131 | 0.034306491 | Sprtn | Tnnc1 | Gm17830 | 0.330974078 | -0.006717897 | 0.012785567 | -0.525428213 | 0.599285545 | NA |
| Dus3l | 2188.34161 | 0.232743448 | 0.083147916 | 2.799149493 | 0.005123741 | 0.036432879 | Dus3l | Pcsk5 | Hus1b | 0.330974078 | -0.006717897 | 0.012785567 | -0.525428213 | 0.599285545 | NA |
| Plcb4 | 2109.860872 | 0.23238591 | 0.077329999 | 3.005119805 | 0.002654763 | 0.023304754 | Plcb4 | Klhl13 | Il5ra | 0.330974078 | -0.006717897 | 0.012785567 | -0.525428213 | 0.599285545 | NA |
| St5 | 707.499199 | 0.232323052 | 0.085316581 | 2.723070358 | 0.006467828 | 0.042962976 | St5 | Neb | Smr3a | 0.330974078 | -0.006717897 | 0.012785567 | -0.525428213 | 0.599285545 | NA |
| Dnm1 | 49898.31488 | 0.232188091 | 0.075826825 | 3.062083799 | 0.002198019 | 0.020565282 | Dnm1 | Mrc1 | Pole4 | 962.9182221 | -0.006737745 | 0.073981922 | -0.091072856 | 0.927434701 | 0.983501978 |
| Ttc13 | 995.6755259 | 0.232179889 | 0.06217734 | 3.734156014 | 0.000188346 | 0.003253321 | Ttc13 | P2ry1 | Gm14744 | 2.03804561 | -0.006738743 | 0.011969462 | -0.562994625 | 0.573438544 | NA |
| Xrcc1 | 933.5953986 | 0.232040905 | 0.084545205 | 2.744577955 | 0.006058876 | 0.041047761 | Xrcc1 | Apc | Adck3 | 1739.548598 | -0.006742506 | 0.058548652 | -0.115160745 | 0.908317712 | 0.980523965 |
| Wrn | 463.7214933 | 0.231922385 | 0.07392145 | 3.137416595 | 0.001704438 | 0.017004866 | Wrn | Cdhr1 | Defb14 | 0.35787663 | -0.006747201 | 0.013072305 | -0.516144702 | 0.605753348 | NA |
| Ogdhl | 4711.592687 | 0.23130974 | 0.072653394 | 3.183743083 | 0.00145384 | 0.015115049 | Ogdhl | Slc10a4 | Gfra3 | 0.35787663 | -0.006747201 | 0.013072305 | -0.516144702 | 0.605753348 | NA |
| Edc4 | 2919.170175 | 0.230998986 | 0.062864633 | 3.674545978 | 0.000238273 | 0.003869809 | Edc4 | Loxl1 | Npm3-ps1 | 0.35787663 | -0.006747201 | 0.013072305 | -0.516144702 | 0.605753348 | NA |
| Zfp207 | 3445.073966 | 0.230304199 | 0.074268152 | 3.100981933 | 0.001928801 | 0.01865818 | Zfp207 | Ranbp3l | 1110036E04Rik | 0.120209049 | -0.006749721 | 0.008635211 | -0.781650997 | 0.434419708 | NA |
| Ganc | 367.6424506 | 0.228980041 | 0.084674312 | 2.704244501 | 0.00684599 | 0.044661027 | Ganc | Kcnb2 | 1700060C20Rik | 0.120209049 | -0.006749721 | 0.008635211 | -0.781650997 | 0.434419708 | NA |
| Zfyve27 | 3183.596224 | 0.228932052 | 0.062710735 | 3.650603863 | 0.000261624 | 0.004183292 | Zfyve27 | Dcc | 1700080O16Rik | 0.120209049 | -0.006749721 | 0.008635211 | -0.781650997 | 0.434419708 | NA |
| Sugp1 | 1827.14392 | 0.228542198 | 0.068476659 | 3.337519681 | 0.000845297 | 0.009997738 | Sugp1 | Col12a1 | 1700110M21Rik | 0.120209049 | -0.006749721 | 0.008635211 | -0.781650997 | 0.434419708 | NA |
| Gys1 | 730.880899 | 0.228230043 | 0.085740874 | 2.661858129 | 0.007771062 | 0.049168195 | Gys1 | Igsf3 | 4930502E09Rik | 0.120209049 | -0.006749721 | 0.008635211 | -0.781650997 | 0.434419708 | NA |
| Wash | 1139.960495 | 0.22803636 | 0.081683285 | 2.79171387 | 0.00524297 | 0.037025785 | Wash | Bche | 4931430N09Rik | 0.120209049 | -0.006749721 | 0.008635211 | -0.781650997 | 0.434419708 | NA |
| Zc3h3 | 897.6908638 | 0.227819291 | 0.076392928 | 2.982203927 | 0.002861813 | 0.024524851 | Zc3h3 | Adcyap1 | AI314831 | 0.120209049 | -0.006749721 | 0.008635211 | -0.781650997 | 0.434419708 | NA |
| Iffo1 | 1770.609293 | 0.22770364 | 0.080847799 | 2.816448229 | 0.004855787 | 0.035305284 | Iffo1 | Sp8 | Actl10 | 0.120209049 | -0.006749721 | 0.008635211 | -0.781650997 | 0.434419708 | NA |
| Ddx55 | 766.3982168 | 0.227493134 | 0.080543267 | 2.824483568 | 0.004735688 | 0.03467463 | Ddx55 | Homer2 | Asb17 | 0.120209049 | -0.006749721 | 0.008635211 | -0.781650997 | 0.434419708 | NA |
| Kremen1 | 1244.462007 | 0.227298875 | 0.083946491 | 2.707663792 | 0.006775863 | 0.044353703 | Kremen1 | Hs3st2 | Awat1 | 0.120209049 | -0.006749721 | 0.008635211 | -0.781650997 | 0.434419708 | NA |
| Ggt7 | 5111.810648 | 0.226939944 | 0.070575087 | 3.215581472 | 0.001301805 | 0.013850598 | Ggt7 | Fam198b | Blk | 0.120209049 | -0.006749721 | 0.008635211 | -0.781650997 | 0.434419708 | NA |
| Exosc8 | 446.6885192 | 0.226851198 | 0.079861179 | 2.840569117 | 0.004503311 | 0.03360928 | Exosc8 | Draxin | Btnl10 | 0.120209049 | -0.006749721 | 0.008635211 | -0.781650997 | 0.434419708 | NA |
| Phf20l1 | 1521.505311 | 0.226698132 | 0.070068523 | 3.235377646 | 0.001214819 | 0.013227274 | Phf20l1 | Tenm1 | Casp14 | 0.120209049 | -0.006749721 | 0.008635211 | -0.781650997 | 0.434419708 | NA |
| Trim3 | 4643.726712 | 0.226311266 | 0.081676089 | 2.77083869 | 0.005591212 | 0.038865068 | Trim3 | Tenm4 | Fam71e2 | 0.120209049 | -0.006749721 | 0.008635211 | -0.781650997 | 0.434419708 | NA |
| 5031439G07Rik | 5181.112955 | 0.225677148 | 0.065687217 | 3.435632641 | 0.000591172 | 0.007729636 | 5031439G07Rik | Fbn2 | Gm10872 | 0.120209049 | -0.006749721 | 0.008635211 | -0.781650997 | 0.434419708 | NA |
| Hdgfrp2 | 2852.160487 | 0.225591066 | 0.057950236 | 3.892841224 | 9.91E-05 | 0.001945626 | Hdgfrp2 | Nusap1 | Gm13032 | 0.120209049 | -0.006749721 | 0.008635211 | -0.781650997 | 0.434419708 | NA |
| Ezh1 | 3022.571278 | 0.225525074 | 0.080884039 | 2.788251876 | 0.005299332 | 0.037321753 | Ezh1 | Tfcp2l1 | Gm19782 | 0.120209049 | -0.006749721 | 0.008635211 | -0.781650997 | 0.434419708 | NA |
| Scrib | 1988.240606 | 0.225349352 | 0.080461449 | 2.800712074 | 0.005098999 | 0.036290258 | Scrib | Pcdh12 | Gm2848 | 0.120209049 | -0.006749721 | 0.008635211 | -0.781650997 | 0.434419708 | NA |
| F3 | 1375.846653 | 0.224962826 | 0.081913394 | 2.746349731 | 0.006026249 | 0.040952059 | F3 | Itm2a | Gm6531 | 0.120209049 | -0.006749721 | 0.008635211 | -0.781650997 | 0.434419708 | NA |
| B930041F14Rik | 1093.17697 | 0.224360183 | 0.080032062 | 2.803378746 | 0.005057024 | 0.036224473 | B930041F14Rik | Gprin3 | Gm6760 | 0.120209049 | -0.006749721 | 0.008635211 | -0.781650997 | 0.434419708 | NA |
| Ik | 4229.998476 | 0.224063645 | 0.075462476 | 2.969206099 | 0.002985703 | 0.025306068 | Ik | Slc5a7 | Gm904 | 0.120209049 | -0.006749721 | 0.008635211 | -0.781650997 | 0.434419708 | NA |
| Htra2 | 905.5619878 | 0.22396659 | 0.07662938 | 2.922724792 | 0.003469831 | 0.028073919 | Htra2 | Asic4 | Gm9047 | 0.120209049 | -0.006749721 | 0.008635211 | -0.781650997 | 0.434419708 | NA |
| Tnrc6a | 3577.033996 | 0.223813904 | 0.07845248 | 2.852859521 | 0.004332777 | 0.032875635 | Tnrc6a | Zic1 | H2-M10.3 | 0.120209049 | -0.006749721 | 0.008635211 | -0.781650997 | 0.434419708 | NA |
| Ift140 | 1236.615601 | 0.223714689 | 0.064550668 | 3.465722299 | 0.000528809 | 0.007115035 | Ift140 | Gng7 | H2afb1 | 0.120209049 | -0.006749721 | 0.008635211 | -0.781650997 | 0.434419708 | NA |
| Manba | 471.8620065 | 0.223542534 | 0.074848785 | 2.986588656 | 0.00282109 | 0.024320521 | Manba | Nes | Hist1h2ae | 0.120209049 | -0.006749721 | 0.008635211 | -0.781650997 | 0.434419708 | NA |
| Spns1 | 2129.241976 | 0.222958604 | 0.076140246 | 2.928262173 | 0.003408625 | 0.027759173 | Spns1 | Zfhx3 | Hist2h2ac | 0.120209049 | -0.006749721 | 0.008635211 | -0.781650997 | 0.434419708 | NA |
| Tpra1 | 969.257986 | 0.222889506 | 0.082956033 | 2.686839 | 0.007213171 | 0.046567031 | Tpra1 | Sfrp1 | Il12b | 0.120209049 | -0.006749721 | 0.008635211 | -0.781650997 | 0.434419708 | NA |
| Mat2a | 7325.374297 | 0.222587644 | 0.073450778 | 3.030432743 | 0.002442036 | 0.022066759 | Mat2a | Pclo | Kcnj1 | 0.120209049 | -0.006749721 | 0.008635211 | -0.781650997 | 0.434419708 | NA |
| Wipf1 | 531.6603312 | 0.222518892 | 0.069193912 | 3.21587384 | 0.00130048 | 0.013849883 | Wipf1 | Mxd3 | Leap2 | 0.120209049 | -0.006749721 | 0.008635211 | -0.781650997 | 0.434419708 | NA |
| Rps6kb1 | 1808.931475 | 0.222070932 | 0.07640764 | 2.906396933 | 0.003656173 | 0.0291553 | Rps6kb1 | Lama3 | Lypd8 | 0.120209049 | -0.006749721 | 0.008635211 | -0.781650997 | 0.434419708 | NA |
| Kdsr | 1281.022261 | 0.221775382 | 0.073892244 | 3.00133505 | 0.002687986 | 0.023503194 | Kdsr | Robo1 | Mbd3l2 | 0.120209049 | -0.006749721 | 0.008635211 | -0.781650997 | 0.434419708 | NA |
| Setdb1 | 1855.798707 | 0.221693616 | 0.078988609 | 2.806652981 | 0.005005914 | 0.035980333 | Setdb1 | Ccbe1 | Mep1b | 0.120209049 | -0.006749721 | 0.008635211 | -0.781650997 | 0.434419708 | NA |
| Srsf2 | 3193.134811 | 0.221522957 | 0.078272233 | 2.830160163 | 0.004652471 | 0.034310034 | Srsf2 | Wnt7a | Mir100 | 0.120209049 | -0.006749721 | 0.008635211 | -0.781650997 | 0.434419708 | NA |
| Coro2b | 8388.601296 | 0.22104682 | 0.064835704 | 3.409337861 | 0.000651208 | 0.008237633 | Coro2b | Pcp4l1 | Mir1193 | 0.120209049 | -0.006749721 | 0.008635211 | -0.781650997 | 0.434419708 | NA |
| Serac1 | 1140.182458 | 0.220950735 | 0.064392388 | 3.431317635 | 0.000600657 | 0.007807531 | Serac1 | Tead2 | Mir192 | 0.120209049 | -0.006749721 | 0.008635211 | -0.781650997 | 0.434419708 | NA |
| Pusl1 | 564.044989 | 0.220525598 | 0.081233306 | 2.71471898 | 0.006633203 | 0.043659664 | Pusl1 | Sebox | Mir5128 | 0.120209049 | -0.006749721 | 0.008635211 | -0.781650997 | 0.434419708 | NA |
| Plod3 | 1106.202768 | 0.220450842 | 0.077470988 | 2.845592247 | 0.004432892 | 0.033344824 | Plod3 | Rhcg | Mir7b | 0.120209049 | -0.006749721 | 0.008635211 | -0.781650997 | 0.434419708 | NA |
| Paf1 | 2096.52074 | 0.220235406 | 0.080999456 | 2.718973891 | 0.006548478 | 0.043304359 | Paf1 | Cyp2f2 | Nlrp9b | 0.120209049 | -0.006749721 | 0.008635211 | -0.781650997 | 0.434419708 | NA |
| Eif2ak4 | 1107.369832 | 0.220100668 | 0.069767968 | 3.154752443 | 0.001606343 | 0.016224697 | Eif2ak4 | Gjb2 | Olfr1126 | 0.120209049 | -0.006749721 | 0.008635211 | -0.781650997 | 0.434419708 | NA |
| Tcerg1 | 2513.03226 | 0.220091098 | 0.059400563 | 3.705202246 | 0.000211222 | 0.003545694 | Tcerg1 | Phex | Olfr1211 | 0.120209049 | -0.006749721 | 0.008635211 | -0.781650997 | 0.434419708 | NA |
| Abca7 | 1145.326111 | 0.219920398 | 0.081860181 | 2.686536907 | 0.007219697 | 0.046578655 | Abca7 | Nlrp10 | Olfr1278 | 0.120209049 | -0.006749721 | 0.008635211 | -0.781650997 | 0.434419708 | NA |
| Rbm39 | 5548.235615 | 0.219635653 | 0.072526501 | 3.028350344 | 0.002458928 | 0.022123355 | Rbm39 | Adam12 | Olfr1281 | 0.120209049 | -0.006749721 | 0.008635211 | -0.781650997 | 0.434419708 | NA |
| Atp13a1 | 2066.6212 | 0.219274921 | 0.073294217 | 2.99170833 | 0.002774212 | 0.024026036 | Atp13a1 | Tmem40 | Olfr1444 | 0.120209049 | -0.006749721 | 0.008635211 | -0.781650997 | 0.434419708 | NA |
| Bnip2 | 1212.504127 | 0.218505437 | 0.070213427 | 3.112017843 | 0.001858133 | 0.01810686 | Bnip2 | Pde1c | Olfr3 | 0.120209049 | -0.006749721 | 0.008635211 | -0.781650997 | 0.434419708 | NA |
| Bak1 | 434.0666152 | 0.218256932 | 0.073438168 | 2.971982257 | 0.002958838 | 0.025129358 | Bak1 | Oprm1 | Olfr681 | 0.120209049 | -0.006749721 | 0.008635211 | -0.781650997 | 0.434419708 | NA |
| Edem1 | 1032.399806 | 0.218055878 | 0.057862968 | 3.768487576 | 0.00016424 | 0.00291159 | Edem1 | Pcdh17 | Olfr836 | 0.120209049 | -0.006749721 | 0.008635211 | -0.781650997 | 0.434419708 | NA |
| Eps15l1 | 2228.235249 | 0.217315122 | 0.060660209 | 3.582498747 | 0.000340323 | 0.00515946 | Eps15l1 | Inhba | Pinc | 0.120209049 | -0.006749721 | 0.008635211 | -0.781650997 | 0.434419708 | NA |
| Yeats2 | 1368.08169 | 0.217197597 | 0.064101941 | 3.388315421 | 0.000703233 | 0.008740529 | Yeats2 | Pdgfrl | Plac1 | 0.120209049 | -0.006749721 | 0.008635211 | -0.781650997 | 0.434419708 | NA |
| Msh2 | 1276.275588 | 0.216356276 | 0.077517671 | 2.791057498 | 0.005253614 | 0.037084052 | Msh2 | C1ql1 | Rbmxl2 | 0.120209049 | -0.006749721 | 0.008635211 | -0.781650997 | 0.434419708 | NA |
| Kcnc1 | 8453.316143 | 0.21608876 | 0.053106579 | 4.068964044 | 4.72E-05 | 0.001095311 | Kcnc1 | Ddah2 | Rptn | 0.120209049 | -0.006749721 | 0.008635211 | -0.781650997 | 0.434419708 | NA |
| Zfp385a | 2832.997442 | 0.215414447 | 0.066069187 | 3.260437353 | 0.001112405 | 0.012381903 | Zfp385a | Asxl3 | Scarna8 | 0.120209049 | -0.006749721 | 0.008635211 | -0.781650997 | 0.434419708 | NA |
| Usp36 | 1634.107729 | 0.215036903 | 0.067326259 | 3.19395294 | 0.00140339 | 0.014691975 | Usp36 | Lrp4 | Slc17a1 | 0.120209049 | -0.006749721 | 0.008635211 | -0.781650997 | 0.434419708 | NA |
| Hs1bp3 | 669.1468416 | 0.215010552 | 0.078828054 | 2.727589243 | 0.0063799 | 0.042645781 | Hs1bp3 | Fn1 | Slc22a30 | 0.120209049 | -0.006749721 | 0.008635211 | -0.781650997 | 0.434419708 | NA |
| Slc9a1 | 2588.288683 | 0.214714003 | 0.067898202 | 3.162292891 | 0.00156532 | 0.015966474 | Slc9a1 | Ankrd63 | Slc7a15 | 0.120209049 | -0.006749721 | 0.008635211 | -0.781650997 | 0.434419708 | NA |
| Rufy1 | 1369.765958 | 0.214346916 | 0.060603036 | 3.536900636 | 0.000404852 | 0.00587889 | Rufy1 | Rrm2 | Spink4 | 0.120209049 | -0.006749721 | 0.008635211 | -0.781650997 | 0.434419708 | NA |
| Jag2 | 1352.821726 | 0.213584973 | 0.071735592 | 2.977391939 | 0.002907121 | 0.024789727 | Jag2 | Sall3 | Tex19.2 | 0.120209049 | -0.006749721 | 0.008635211 | -0.781650997 | 0.434419708 | NA |
| Usp48 | 3692.080673 | 0.213378775 | 0.07539388 | 2.830186946 | 0.004652081 | 0.034310034 | Usp48 | Hdac9 | Tmem52b | 0.120209049 | -0.006749721 | 0.008635211 | -0.781650997 | 0.434419708 | NA |
| Rbfox3 | 11023.49346 | 0.213232468 | 0.074568518 | 2.859550813 | 0.004242414 | 0.032353861 | Rbfox3 | Marcksl1 | Ttc23l | 0.120209049 | -0.006749721 | 0.008635211 | -0.781650997 | 0.434419708 | NA |
| Nckap5l | 954.9283759 | 0.212273195 | 0.070586979 | 3.007257129 | 0.002636167 | 0.023198299 | Nckap5l | Emp2 | Usp17ld | 0.120209049 | -0.006749721 | 0.008635211 | -0.781650997 | 0.434419708 | NA |
| Sparcl1 | 82107.6409 | 0.212215797 | 0.054732908 | 3.877298052 | 0.000105623 | 0.002043399 | Sparcl1 | C730002L08Rik | Vmn1r232 | 0.120209049 | -0.006749721 | 0.008635211 | -0.781650997 | 0.434419708 | NA |
| Dock9 | 5818.375937 | 0.212110072 | 0.067574082 | 3.138926435 | 0.00169568 | 0.016939309 | Dock9 | Fabp7 | Vmn2r109 | 0.120209049 | -0.006749721 | 0.008635211 | -0.781650997 | 0.434419708 | NA |
| Dennd4b | 2263.169587 | 0.211943385 | 0.070554786 | 3.003954747 | 0.00266495 | 0.023367702 | Dennd4b | Th | Vmn2r37 | 0.120209049 | -0.006749721 | 0.008635211 | -0.781650997 | 0.434419708 | NA |
| Slc11a2 | 2021.013837 | 0.211377605 | 0.054981145 | 3.844547172 | 0.000120775 | 0.002260014 | Slc11a2 | Pnp2 | Rpl5 | 9360.257815 | -0.006760105 | 0.054104705 | -0.124944863 | 0.900567201 | 0.979053977 |
| Git2 | 1332.932688 | 0.211367968 | 0.072005818 | 2.935429014 | 0.003330868 | 0.027292548 | Git2 | Kntc1 | Ndufa3 | 2167.225618 | -0.006765713 | 0.066800666 | -0.101282118 | 0.919326511 | 0.982742917 |
| Otub1 | 6737.247127 | 0.21135559 | 0.074954729 | 2.819776574 | 0.00480571 | 0.035039843 | Otub1 | Wnt7b | Ces1a | 0.405340399 | -0.006772093 | 0.012119685 | -0.558768089 | 0.576320004 | NA |
| Bin1 | 8671.158807 | 0.211301844 | 0.072497091 | 2.914625151 | 0.00356116 | 0.028648293 | Bin1 | Hspg2 | Idi2 | 3.793146691 | -0.006774129 | 0.041550258 | -0.163034582 | 0.870491206 | NA |
| Jakmip3 | 1361.412843 | 0.211097364 | 0.074094036 | 2.84904665 | 0.004385045 | 0.033093954 | Jakmip3 | Gpr6 | Gm8580 | 2.328062488 | -0.006778968 | 0.034545077 | -0.196235423 | 0.844425903 | NA |
| Rgs7 | 3215.845546 | 0.210484484 | 0.069967202 | 3.008330731 | 0.002626871 | 0.023191304 | Rgs7 | Cenpf | Anxa1 | 59.9237341 | -0.00678334 | 0.091133203 | -0.07443325 | 0.940665653 | 0.985461126 |
| Pxk | 1437.086382 | 0.210355912 | 0.076335768 | 2.755666405 | 0.00585727 | 0.040157049 | Pxk | Prss35 | Serpina3b | 0.609313426 | -0.0067865 | 0.015151049 | -0.44792277 | 0.654208935 | NA |
| Snx32 | 2891.815261 | 0.210202736 | 0.063835362 | 3.292888609 | 0.000991638 | 0.011314015 | Snx32 | Tagln | Nup188 | 1042.157766 | -0.006789451 | 0.051682687 | -0.13136799 | 0.895484209 | 0.977844987 |
| Osbpl5 | 1134.127178 | 0.209682546 | 0.069256678 | 3.027614856 | 0.00246492 | 0.022127158 | Osbpl5 | Thbd | Ybey | 271.0944102 | -0.006805713 | 0.079764906 | -0.085322143 | 0.932005288 | 0.984387568 |
| Arhgef10 | 1900.040938 | 0.209320979 | 0.074228073 | 2.819970525 | 0.004802806 | 0.035035159 | Arhgef10 | Stac | Hspa2 | 1398.296944 | -0.006825549 | 0.074696048 | -0.091377649 | 0.927192521 | 0.983501978 |
| Zfp263 | 793.5984477 | 0.209028455 | 0.07725144 | 2.70581953 | 0.006813606 | 0.044544311 | Zfp263 | Tenm3 | Cdt1 | 72.46638864 | -0.006831864 | 0.095087385 | -0.071848268 | 0.942722659 | 0.985461126 |
| Cyth3 | 1471.431858 | 0.207874083 | 0.053662351 | 3.873741596 | 0.000107177 | 0.002065424 | Cyth3 | Kif2c | Scap | 4464.600711 | -0.006863951 | 0.052354999 | -0.131104017 | 0.895693023 | 0.977844987 |
| Dnajb2 | 3706.13127 | 0.207468747 | 0.066246137 | 3.131786349 | 0.001737463 | 0.017245513 | Dnajb2 | Gpr149 | 1700020G17Rik | 0.12890128 | -0.006864007 | 0.008708041 | -0.788237772 | 0.430557639 | NA |
| Slc17a5 | 784.0980704 | 0.206661443 | 0.07717882 | 2.677696326 | 0.007413039 | 0.047350844 | Slc17a5 | Drd1a | 1700027I24Rik | 0.12890128 | -0.006864007 | 0.008708041 | -0.788237772 | 0.430557639 | NA |
| Nop56 | 2725.424052 | 0.206619088 | 0.073828409 | 2.798639306 | 0.005131843 | 0.036473748 | Nop56 | Aldh1a7 | 1700097N02Rik | 0.12890128 | -0.006864007 | 0.008708041 | -0.788237772 | 0.430557639 | NA |
| Gps2 | 733.7424196 | 0.206245491 | 0.07626673 | 2.704265553 | 0.006845556 | 0.044661027 | Gps2 | Il31ra | 1700104A03Rik | 0.12890128 | -0.006864007 | 0.008708041 | -0.788237772 | 0.430557639 | NA |
| Atrip | 764.6469971 | 0.206079681 | 0.069092446 | 2.982665886 | 0.002857497 | 0.024507848 | Atrip | Cpa6 | 2310002L09Rik | 0.12890128 | -0.006864007 | 0.008708041 | -0.788237772 | 0.430557639 | NA |
| Pde7a | 829.317474 | 0.205904682 | 0.074217274 | 2.774349846 | 0.005531216 | 0.03865614 | Pde7a | Pf4 | 2900092C05Rik | 0.12890128 | -0.006864007 | 0.008708041 | -0.788237772 | 0.430557639 | NA |
| Map4k5 | 1775.033601 | 0.205323655 | 0.065572567 | 3.131243188 | 0.00174068 | 0.017266382 | Map4k5 | Klhl30 | 4933430H16Rik | 0.12890128 | -0.006864007 | 0.008708041 | -0.788237772 | 0.430557639 | NA |
| Ercc5 | 965.5107105 | 0.204674286 | 0.076989841 | 2.658458351 | 0.007849905 | 0.049502007 | Ercc5 | Htr2a | 9030619P08Rik | 0.12890128 | -0.006864007 | 0.008708041 | -0.788237772 | 0.430557639 | NA |
| Brd8 | 2330.470484 | 0.204309109 | 0.063531991 | 3.215846156 | 0.001300605 | 0.013849883 | Brd8 | Slc6a3 | Asz1 | 0.12890128 | -0.006864007 | 0.008708041 | -0.788237772 | 0.430557639 | NA |
| Ints6 | 789.9246417 | 0.203931576 | 0.074510615 | 2.736946628 | 0.006201235 | 0.041781366 | Ints6 | Jsrp1 | Atp6v1g3 | 0.12890128 | -0.006864007 | 0.008708041 | -0.788237772 | 0.430557639 | NA |
| Kri1 | 1271.519384 | 0.203730929 | 0.065743231 | 3.09888827 | 0.001942483 | 0.018716931 | Kri1 | Rarb | Bpifa6 | 0.12890128 | -0.006864007 | 0.008708041 | -0.788237772 | 0.430557639 | NA |
| Mms19 | 1492.99769 | 0.203594273 | 0.074832328 | 2.720672702 | 0.006514923 | 0.043211566 | Mms19 | Drd5 | Ccnb1ip1 | 0.12890128 | -0.006864007 | 0.008708041 | -0.788237772 | 0.430557639 | NA |
| 9930104L06Rik | 705.0685561 | 0.20319383 | 0.073288245 | 2.77252962 | 0.005562246 | 0.038738759 | 9930104L06Rik | Fat1 | Clca6 | 0.12890128 | -0.006864007 | 0.008708041 | -0.788237772 | 0.430557639 | NA |
| Luc7l2 | 4917.28795 | 0.202934333 | 0.058789306 | 3.451891982 | 0.000556671 | 0.00739713 | Luc7l2 | Zbtb20 | Cldn16 | 0.12890128 | -0.006864007 | 0.008708041 | -0.788237772 | 0.430557639 | NA |
| Khnyn | 704.1481221 | 0.202875743 | 0.071329134 | 2.8442199 | 0.004452031 | 0.033404246 | Khnyn | Slc38a5 | D16Ertd519e | 0.12890128 | -0.006864007 | 0.008708041 | -0.788237772 | 0.430557639 | NA |
| Edil3 | 4441.239623 | 0.20287005 | 0.072183055 | 2.810494094 | 0.00494655 | 0.035738209 | Edil3 | Htr1b | Evx1 | 0.12890128 | -0.006864007 | 0.008708041 | -0.788237772 | 0.430557639 | NA |
| Atg13 | 4342.891724 | 0.202598679 | 0.064018299 | 3.164699514 | 0.001552432 | 0.015897805 | Atg13 | Kif26a | Fbxw18 | 0.12890128 | -0.006864007 | 0.008708041 | -0.788237772 | 0.430557639 | NA |
| Cyb561d1 | 1229.815026 | 0.202580603 | 0.071057204 | 2.850950953 | 0.004358869 | 0.032960625 | Cyb561d1 | Hs6st2 | Fgb | 0.12890128 | -0.006864007 | 0.008708041 | -0.788237772 | 0.430557639 | NA |
| Ephx1 | 1009.826587 | 0.202508578 | 0.073520981 | 2.754432485 | 0.005879402 | 0.040218743 | Ephx1 | Islr | Gm10664 | 0.12890128 | -0.006864007 | 0.008708041 | -0.788237772 | 0.430557639 | NA |
| Atp8b2 | 3769.512524 | 0.202137081 | 0.061289643 | 3.298062623 | 0.000973544 | 0.011167224 | Atp8b2 | Tshr | Gm12603 | 0.12890128 | -0.006864007 | 0.008708041 | -0.788237772 | 0.430557639 | NA |
| Diexf | 728.5598181 | 0.201275567 | 0.067345577 | 2.988697625 | 0.002801692 | 0.024197003 | Diexf | Tll1 | Gm19583 | 0.12890128 | -0.006864007 | 0.008708041 | -0.788237772 | 0.430557639 | NA |
| Trmt1l | 1067.310916 | 0.200470394 | 0.063278687 | 3.168055537 | 0.001534622 | 0.01575708 | Trmt1l | Strip2 | Gm5065 | 0.12890128 | -0.006864007 | 0.008708041 | -0.788237772 | 0.430557639 | NA |
| Pprc1 | 1061.371014 | 0.199223196 | 0.074336381 | 2.680022809 | 0.007361714 | 0.047250374 | Pprc1 | Wnt2 | Gm5538 | 0.12890128 | -0.006864007 | 0.008708041 | -0.788237772 | 0.430557639 | NA |
| Fam189b | 1975.149822 | 0.198671218 | 0.066364695 | 2.993628144 | 0.002756817 | 0.023916083 | Fam189b | Hs3st3b1 | Gm6289 | 0.12890128 | -0.006864007 | 0.008708041 | -0.788237772 | 0.430557639 | NA |
| Gpr108 | 946.1526402 | 0.198046143 | 0.064502297 | 3.070373482 | 0.002137912 | 0.020112213 | Gpr108 | Kdr | Hamp2 | 0.12890128 | -0.006864007 | 0.008708041 | -0.788237772 | 0.430557639 | NA |
| Repin1 | 1517.461449 | 0.197788166 | 0.071470793 | 2.767398519 | 0.005650563 | 0.039197474 | Repin1 | Isl1 | Hesx1 | 0.12890128 | -0.006864007 | 0.008708041 | -0.788237772 | 0.430557639 | NA |
| Zfp286 | 583.5372339 | 0.197153628 | 0.069404051 | 2.840664574 | 0.004501964 | 0.03360928 | Zfp286 | Man1a | Hmgb4 | 0.12890128 | -0.006864007 | 0.008708041 | -0.788237772 | 0.430557639 | NA |
| Clcn7 | 2531.298654 | 0.196876359 | 0.058631858 | 3.35783934 | 0.000785543 | 0.009435036 | Clcn7 | Gbx2 | Hnf4a | 0.12890128 | -0.006864007 | 0.008708041 | -0.788237772 | 0.430557639 | NA |
| Ubl7 | 2630.996206 | 0.196757964 | 0.07346692 | 2.678184474 | 0.007402244 | 0.047350844 | Ubl7 | Myh11 | Hyal6 | 0.12890128 | -0.006864007 | 0.008708041 | -0.788237772 | 0.430557639 | NA |
| Top3b | 1329.62685 | 0.196333337 | 0.05818472 | 3.374310971 | 0.000740007 | 0.009042328 | Top3b | Dlx6as1 | Krtap5-1 | 0.12890128 | -0.006864007 | 0.008708041 | -0.788237772 | 0.430557639 | NA |
| Ccng1 | 2917.240865 | 0.19538184 | 0.064287684 | 3.039179972 | 0.002372231 | 0.021674644 | Ccng1 | Mgat4c | LOC547349 | 0.12890128 | -0.006864007 | 0.008708041 | -0.788237772 | 0.430557639 | NA |
| Map3k11 | 992.9405756 | 0.194722724 | 0.072895191 | 2.671269821 | 0.007556488 | 0.048062489 | Map3k11 | Fndc9 | Lipi | 0.12890128 | -0.006864007 | 0.008708041 | -0.788237772 | 0.430557639 | NA |
| Lonp1 | 2678.236127 | 0.194690895 | 0.065065299 | 2.992238511 | 0.002769398 | 0.023998352 | Lonp1 | Crybg3 | Mcpt-ps1 | 0.12890128 | -0.006864007 | 0.008708041 | -0.788237772 | 0.430557639 | NA |
| Sf3b1 | 10501.41869 | 0.194511355 | 0.068711812 | 2.830828473 | 0.004642761 | 0.034306491 | Sf3b1 | Zfhx4 | Mir135a-1 | 0.12890128 | -0.006864007 | 0.008708041 | -0.788237772 | 0.430557639 | NA |
| Zfp239 | 1763.672314 | 0.194291565 | 0.068892145 | 2.820228125 | 0.004798952 | 0.035023534 | Zfp239 | Kcnq1 | Mir15a | 0.12890128 | -0.006864007 | 0.008708041 | -0.788237772 | 0.430557639 | NA |
| Coq10a | 1147.109385 | 0.192805137 | 0.06648287 | 2.900072399 | 0.003730765 | 0.029658527 | Coq10a | Prss12 | Mir218-2 | 0.12890128 | -0.006864007 | 0.008708041 | -0.788237772 | 0.430557639 | NA |
| Nprl3 | 968.6154974 | 0.192647555 | 0.062355401 | 3.089508732 | 0.002004878 | 0.019163221 | Nprl3 | Fras1 | Mir300 | 0.12890128 | -0.006864007 | 0.008708041 | -0.788237772 | 0.430557639 | NA |
| Map2k5 | 1149.534504 | 0.191064593 | 0.069174379 | 2.762071676 | 0.005743587 | 0.039586803 | Map2k5 | Top2a | Mir344d-2 | 0.12890128 | -0.006864007 | 0.008708041 | -0.788237772 | 0.430557639 | NA |

| | | | | | | | | | | | | | | | |
|---|---|---|---|---|---|---|---|---|---|---|---|---|---|---|---|
| Rsrc2 | 2248.694519 | 0.188901973 | 0.066869686 | 2.824926866 | 0.004729141 | 0.034663987 | Rsrc2 | Pbk | Mir487b | 0.12890128 | -0.006864007 | 0.008708041 | -0.788237772 | 0.430557639 | NA |
| Sart1 | 2107.474655 | 0.188402257 | 0.06833298 | 2.757120451 | 0.005831287 | 0.040013267 | Sart1 | Emid1 | Olfr123 | 0.12890128 | -0.006864007 | 0.008708041 | -0.788237772 | 0.430557639 | NA |
| Map3k3 | 1139.059462 | 0.187688536 | 0.070081808 | 2.678134922 | 0.007403339 | 0.047350844 | Map3k3 | Ccnb1 | Olfr328 | 0.12890128 | -0.006864007 | 0.008708041 | -0.788237772 | 0.430557639 | NA |
| Nxpe3 | 1012.80732 | 0.187608551 | 0.062665359 | 2.993815961 | 0.002755121 | 0.023914757 | Nxpe3 | Klhl14 | Olfr741 | 0.12890128 | -0.006864007 | 0.008708041 | -0.788237772 | 0.430557639 | NA |
| Vps16 | 2089.322466 | 0.187293614 | 0.064380431 | 2.909169947 | 0.003623898 | 0.028942617 | Vps16 | Kcna5 | Olfr883 | 0.12890128 | -0.006864007 | 0.008708041 | -0.788237772 | 0.430557639 | NA |
| Cog2 | 1021.066288 | 0.187098248 | 0.063234034 | 2.958821929 | 0.003088175 | 0.025822009 | Cog2 | Slc47a1 | P2ry10 | 0.12890128 | -0.006864007 | 0.008708041 | -0.788237772 | 0.430557639 | NA |
| Flii | 2452.874925 | 0.18675676 | 0.049972148 | 3.737217002 | 0.000186068 | 0.003224768 | Flii | Ntn1 | Pla2g2a | 0.12890128 | -0.006864007 | 0.008708041 | -0.788237772 | 0.430557639 | NA |
| Vac14 | 1119.321733 | 0.186673281 | 0.069592709 | 2.682368353 | 0.007310292 | 0.047017709 | Vac14 | D8Ertd738e | Raet1a | 0.12890128 | -0.006864007 | 0.008708041 | -0.788237772 | 0.430557639 | NA |
| Hnrnpu | 13084.91441 | 0.186524227 | 0.055542552 | 3.358222146 | 0.000784455 | 0.009429288 | Hnrnpu | Syt6 | Reg4 | 0.12890128 | -0.006864007 | 0.008708041 | -0.788237772 | 0.430557639 | NA |
| Sncb | 13839.00737 | 0.185278921 | 0.067874788 | 2.729716375 | 0.006338884 | 0.042445806 | Sncb | Asgr1 | Ssxb9 | 0.12890128 | -0.006864007 | 0.008708041 | -0.788237772 | 0.430557639 | NA |
| Sgpl1 | 1627.701312 | 0.185059172 | 0.052154185 | 3.548309155 | 0.000387713 | 0.005699453 | Sgpl1 | Prkch | Sucnr1 | 0.12890128 | -0.006864007 | 0.008708041 | -0.788237772 | 0.430557639 | NA |
| BC017158 | 831.5444776 | 0.18504505 | 0.065066489 | 2.843937849 | 0.004455974 | 0.033417647 | BC017158 | Dkk2 | Sult3a1 | 0.12890128 | -0.006864007 | 0.008708041 | -0.788237772 | 0.430557639 | NA |
| Syt3 | 2067.102553 | 0.184685935 | 0.064126723 | 2.880015188 | 0.00397656 | 0.030957254 | Syt3 | Il15 | Tex24 | 0.12890128 | -0.006864007 | 0.008708041 | -0.788237772 | 0.430557639 | NA |
| Gpr61 | 803.4361203 | 0.184450127 | 0.064255099 | 2.870591284 | 0.004097049 | 0.031534861 | Gpr61 | Serpina9 | Tlr11 | 0.12890128 | -0.006864007 | 0.008708041 | -0.788237772 | 0.430557639 | NA |
| Tspan33 | 824.9971622 | 0.184255457 | 0.063123061 | 2.918987976 | 0.003511698 | 0.028383017 | Tspan33 | Pbx3 | Tprg | 0.12890128 | -0.006864007 | 0.008708041 | -0.788237772 | 0.430557639 | NA |
| Fbf1 | 2954.201891 | 0.181905349 | 0.068067721 | 2.672417204 | 0.007530696 | 0.047937797 | Fbf1 | Gpr126 | A230070E04Rik | 632.7449953 | -0.006866434 | 0.065482839 | -0.104858522 | 0.916488072 | 0.982742917 |
| Ints1 | 4194.901813 | 0.181904089 | 0.065613991 | 2.772336905 | 0.00556554 | 0.038738759 | Ints1 | St8sia2 | Rps28 | 2111.927665 | -0.006877085 | 0.081127414 | -0.084768939 | 0.932445087 | 0.984387568 |
| Ddx23 | 2744.353771 | 0.18174851 | 0.065025335 | 2.795041497 | 0.005189305 | 0.036748497 | Ddx23 | Tac1 | Slc6a19 | 12.95992685 | -0.006879733 | 0.062256769 | -0.110505781 | 0.912008265 | NA |
| Tada2a | 983.1500646 | 0.181732361 | 0.064478002 | 2.818517262 | 0.004824602 | 0.035127997 | Tada2a | Padi6 | Aldh1l1 | 3119.825443 | -0.006908468 | 0.071450423 | -0.096688966 | 0.922973403 | 0.98296621 |
| Rbbp5 | 2289.22541 | 0.181660662 | 0.058467905 | 3.107015059 | 0.001889868 | 0.018346874 | Rbbp5 | Hmcn1 | Gm15787 | 6.628577017 | -0.006908803 | 0.057708671 | -0.119718622 | 0.904706048 | NA |
| Rnh1 | 1210.727884 | 0.180503103 | 0.067880607 | 2.659126233 | 0.00783436 | 0.049424096 | Rnh1 | Meis2 | Ap1s1 | 8898.84601 | -0.006932474 | 0.073699655 | -0.094063858 | 0.92505843 | 0.983236501 |
| Ctsa | 4034.581419 | 0.180425617 | 0.058670974 | 3.075210864 | 0.002103538 | 0.019861191 | Ctsa | Slc35d3 | Pmp2 | 1.538947633 | -0.006935315 | 0.026125205 | -0.265464512 | 0.790651645 | NA |
| Polr1b | 781.07155 | 0.180254703 | 0.06468694 | 2.78657026 | 0.005326907 | 0.037473304 | Polr1b | Clec12a | Pcdhb17 | 533.7120901 | -0.006939862 | 0.067596948 | -0.102665313 | 0.918228604 | 0.982742917 |
| Arl4c | 2056.746766 | 0.180074452 | 0.056243338 | 3.201702784 | 0.001366179 | 0.01439971 | Arl4c | Fat3 | Fry | 6724.041142 | -0.006944196 | 0.06747573 | -0.102913975 | 0.918031246 | 0.982742917 |
| Mbd1 | 1067.608251 | 0.177968009 | 0.058384587 | 3.04820189 | 0.002302152 | 0.021168869 | Mbd1 | 2610028E06Rik | Ano9 | 0.397645721 | -0.006947278 | 0.012059612 | -0.576078084 | 0.564562404 | NA |
| Taok3 | 2185.693089 | 0.176669363 | 0.052239749 | 3.381895329 | 0.000719876 | 0.008887453 | Taok3 | Cbln2 | Serpina3i | 1.433623348 | -0.006962127 | 0.020665619 | -0.336894185 | 0.736196669 | NA |
| Dpf2 | 2088.632595 | 0.175884713 | 0.064207789 | 2.739304932 | 0.006156924 | 0.041559293 | Dpf2 | Aldh1a3 | Zfp472 | 192.274666 | -0.006981502 | 0.092033069 | -0.075858625 | 0.939531574 | 0.985461126 |
| Tbcd | 1839.935948 | 0.175554281 | 0.059463888 | 2.952283913 | 0.003154328 | 0.026247665 | Tbcd | Pou3f4 | Ppp1r12a | 2907.392225 | -0.006991659 | 0.065907754 | -0.106082492 | 0.915516902 | 0.982038599 |
| Prpf6 | 2552.293501 | 0.17504458 | 0.063342568 | 2.763458865 | 0.00571923 | 0.03945403 | Prpf6 | Acsm5 | Fam47e | 9.509144535 | -0.007004238 | 0.054791241 | -0.127834998 | 0.898279553 | NA |
| Tubgcp2 | 1570.010678 | 0.174888828 | 0.06282501 | 2.783745323 | 0.005373521 | 0.037690055 | Tubgcp2 | Ndnf | Dpy19l4 | 672.3623002 | -0.007005818 | 0.079137042 | -0.088527675 | 0.929457289 | 0.983689248 |
| Traf3ip1 | 659.0589943 | 0.174314584 | 0.062917765 | 2.770514549 | 0.00559678 | 0.038886327 | Traf3ip1 | Pgm5 | Arhgef37 | 121.8525847 | -0.007038963 | 0.090439205 | -0.077830881 | 0.937962582 | 0.985461126 |
| Kat2a | 3135.53649 | 0.173507368 | 0.055508747 | 3.125766245 | 0.001773425 | 0.017434924 | Kat2a | Drd3 | E130310I04Rik | 2.526221834 | -0.007063628 | 0.036376949 | -0.194178683 | 0.846035973 | NA |
| Reps1 | 1495.331577 | 0.172439414 | 0.062371376 | 2.76472038 | 0.005697161 | 0.039424659 | Reps1 | Mki67 | Tmem141 | 488.0438216 | -0.007084942 | 0.065486611 | -0.108189168 | 0.913845637 | 0.98170867 |
| Capn2 | 3133.250182 | 0.171004796 | 0.04273218 | 4.001780254 | 6.29E-05 | 0.001370002 | Capn2 | Ly6g6e | Tll1 | 91.43329482 | -0.007093892 | 0.093973428 | -0.075488276 | 0.939826225 | 0.985461126 |
| Golga1 | 1499.963615 | 0.170413094 | 0.06096032 | 2.795475718 | 0.005182339 | 0.03674392 | Golga1 | Crym | Gm17762 | 83.33225226 | -0.007099019 | 0.095053607 | -0.074684375 | 0.94046584 | 0.985461126 |
| Ddx46 | 2850.474865 | 0.169291541 | 0.05382713 | 3.145096913 | 0.001660318 | 0.016639696 | Ddx46 | Wdfy1 | Pth2 | 0.344835288 | -0.007099476 | 0.01294162 | -0.54857709 | 0.583295712 | NA |
| Ccpg1 | 3902.468409 | 0.169286814 | 0.051509306 | 3.286528709 | 0.001014304 | 0.011471264 | Ccpg1 | Ndst4 | Gm7538 | 0.55289662 | -0.007125734 | 0.016593899 | -0.429418885 | 0.667618413 | NA |
| Bcl9 | 2155.561692 | 0.169048826 | 0.061811402 | 2.73491331 | 0.006239671 | 0.041968026 | Bcl9 | Tfap2d | Sec16a | 3347.12439 | -0.007127431 | 0.063954876 | -0.111444683 | 0.911263729 | 0.980985114 |
| Sars | 6397.6773 | 0.168967269 | 0.0561883 | 3.007161085 | 0.002637 | 0.023198299 | Sars | Abi3bp | C130083M11Rik | 1.069576044 | -0.007133663 | 0.023291401 | -0.306278846 | 0.759392344 | NA |
| Fam222b | 1476.115515 | 0.168436902 | 0.056014286 | 3.00703469 | 0.002638097 | 0.023198299 | Fam222b | Dsg1c | 1810044D09Rik | 15.16966339 | -0.007146032 | 0.074301777 | -0.096175783 | 0.923380964 | NA |
| Ctnna1 | 3137.489736 | 0.167774198 | 0.04876359 | 3.440562911 | 0.000580505 | 0.007622332 | Ctnna1 | Igfbpl1 | Bola1 | 215.6643379 | -0.007173852 | 0.087548515 | -0.081941445 | 0.934693277 | 0.985294234 |
| Alas1 | 2168.222391 | 0.167597087 | 0.06113985 | 2.741208682 | 0.006121362 | 0.041362572 | Alas1 | Prss56 | Zmiz2 | 14004.48304 | -0.007181951 | 0.052213521 | -0.13754963 | 0.890596366 | 0.977182599 |
| Cherp | 2489.767726 | 0.16728869 | 0.0545513 | 3.06663065 | 0.002164862 | 0.020316395 | Cherp | Uba1y | Pfkfb1 | 30.58378175 | -0.007183979 | 0.079609386 | -0.090240348 | 0.928096222 | 0.983501978 |
| Plcg1 | 3259.412552 | 0.166452245 | 0.058904002 | 2.825822356 | 0.004715941 | 0.034613354 | Plcg1 | Nmbr | Pld1 | 470.1117957 | -0.007184942 | 0.071803407 | -0.100064078 | 0.920293454 | 0.982742917 |
| B4galnt4 | 2923.630059 | 0.166291729 | 0.060838187 | 2.733344565 | 0.006269471 | 0.042079243 | B4galnt4 | 1-Mar | Wdr5 | 1195.502164 | -0.00719166 | 0.058169754 | -0.123632296 | 0.90160642 | 0.979150581 |
| Fam98a | 1312.567607 | 0.165852453 | 0.061062501 | 2.716109743 | 0.006605402 | 0.043599054 | Fam98a | Uty | Lin28a | 0.560996453 | -0.007195803 | 0.016648492 | -0.432219488 | 0.665581897 | NA |
| Phf12 | 2268.931824 | 0.165792252 | 0.048891151 | 3.391048273 | 0.000696258 | 0.008671887 | Phf12 | Eif2s3y | 1700017J07Rik | 0.387222073 | -0.007201819 | 0.011977925 | -0.601257653 | 0.54766839 | NA |
| Gpbp1l1 | 1404.311263 | 0.164923375 | 0.061479355 | 2.682581426 | 0.007305636 | 0.04700728 | Gpbp1l1 | Kdm5d | Arhgef38 | 0.387222073 | -0.007201819 | 0.011977925 | -0.601257653 | 0.54766839 | NA |
| Riok3 | 2221.700274 | 0.163690757 | 0.056779411 | 2.882924527 | 0.003940018 | 0.030816672 | Riok3 | Ddx3y | Ints9 | 696.7570001 | -0.007203358 | 0.063616244 | -0.11323143 | 0.909847079 | 0.980590355 |
| Elp2 | 3765.13999 | 0.162250292 | 0.042315519 | 3.8342976 | 0.000125924 | 0.002336599 | Elp2 | | Lrguk | 46.90375575 | -0.007205491 | 0.093963654 | -0.076683807 | 0.938875086 | 0.985461126 |
| Tmem110 | 892.5459094 | 0.160736471 | 0.05979372 | 2.688183179 | 0.007184197 | 0.046438029 | Tmem110 | | Gpbp1l1 | 1577.878183 | -0.007233096 | 0.052281245 | -0.138349735 | 0.889964021 | 0.977182599 |
| Extl2 | 3119.696627 | 0.160724683 | 0.055972431 | 2.871497292 | 0.004085323 | 0.031506284 | Extl2 | | 4930473O22Rik | 0.527723949 | -0.007253832 | 0.016432037 | -0.44144445 | 0.658891269 | NA |
| Polg | 2669.093502 | 0.160013919 | 0.055602224 | 2.877833091 | 0.004004169 | 0.031051351 | Polg | | Gabrr3 | 2.528714734 | -0.007254736 | 0.033928524 | -0.2138241 | 0.830684231 | NA |
| Stx5a | 1904.820363 | 0.159100409 | 0.057561117 | 2.764025726 | 0.005709304 | 0.039425264 | Stx5a | | Nek8 | 128.4052748 | -0.007254759 | 0.094448352 | -0.076811918 | 0.938773169 | 0.985461126 |
| Sgsm3 | 2538.365112 | 0.158292349 | 0.057327407 | 2.76119848 | 0.005758967 | 0.039657524 | Sgsm3 | | Scarf2 | 109.8188694 | -0.007255706 | 0.094688043 | -0.076627481 | 0.938919897 | 0.985461126 |
| Lrsam1 | 1508.892021 | 0.158049562 | 0.058249042 | 2.713341841 | 0.006660836 | 0.043785739 | Lrsam1 | | 4930545H06Rik | 0.749629687 | -0.007259859 | 0.019935404 | -0.364169161 | 0.715731694 | NA |
| Rae1 | 1392.415562 | 0.156708508 | 0.056544557 | 2.771416322 | 0.005581302 | 0.038831022 | Rae1 | | Rpa3 | 153.9752133 | -0.007264665 | 0.090121293 | -0.080609864 | 0.935752221 | 0.985461126 |
| Ncstn | 2680.334264 | 0.154969648 | 0.054689357 | 2.833634472 | 0.004602194 | 0.034134224 | Ncstn | | Kcnq2 | 7577.579728 | -0.007271747 | 0.057833335 | -0.125736257 | 0.899940701 | 0.97888996 |
| Tspan9 | 1842.380723 | 0.154198765 | 0.057815768 | 2.667071103 | 0.007651548 | 0.048587331 | Tspan9 | | Slc10a3 | 129.4227104 | -0.007282371 | 0.09110017 | -0.079938057 | 0.936286521 | 0.985461126 |
| Usp8 | 2877.496576 | 0.151044523 | 0.056225142 | 2.686423129 | 0.007222156 | 0.046578655 | Usp8 | | Dis3l | 1414.884636 | -0.007284869 | 0.055750921 | -0.130668128 | 0.896037846 | 0.977899756 |
| Rnps1 | 5042.659086 | 0.149000094 | 0.054172458 | 2.750476898 | 0.005950859 | 0.040564493 | Rnps1 | | Snai1 | 17.18609486 | -0.007286666 | 0.072759469 | -0.100147317 | 0.92022737 | NA |
| Trim66 | 2087.310147 | 0.147329434 | 0.054035701 | 2.726520283 | 0.006400602 | 0.042690885 | Trim66 | | Itgav | 3789.224334 | -0.007292145 | 0.066213704 | -0.110130457 | 0.912305914 | 0.981495756 |
| Cog1 | 1949.613719 | 0.144457439 | 0.048251005 | 2.993874225 | 0.002754595 | 0.023914757 | Cog1 | | Sirt3 | 2370.750974 | -0.007304594 | 0.054383843 | -0.13431552 | 0.893153082 | 0.977844987 |
| Fbxo18 | 3453.87785 | 0.143373486 | 0.050858847 | 2.819047148 | 0.004816644 | 0.035103051 | Fbxo18 | | Rnf144a | 1475.012214 | -0.00731326 | 0.053337145 | -0.137113832 | 0.89094082 | 0.977182599 |
| Cnnm1 | 5322.957827 | 0.13845579 | 0.051127396 | 2.708054774 | 0.006767885 | 0.044320208 | Cnnm1 | | Agbl2 | 64.25407257 | -0.007314888 | 0.094005992 | -0.077812995 | 0.937976809 | 0.985461126 |
| Cyfip1 | 3412.518478 | 0.135210249 | 0.047704999 | 2.834299395 | 0.004592628 | 0.034079589 | Cyfip1 | | 4930546C10Rik | 2.127829097 | -0.007322561 | 0.034588655 | -0.211704136 | 0.832337862 | NA |
| Atp6v0a1 | 21809.71969 | 0.131881818 | 0.049324373 | 2.673765712 | 0.007500483 | 0.047804396 | Atp6v0a1 | | 1700066B19Rik | 1.782248378 | -0.007331611 | 0.027349273 | -0.268073345 | 0.788642863 | NA |
| Map7d1 | 9546.742919 | 0.129602514 | 0.044787224 | 2.893738519 | 0.00380685 | 0.030108902 | Map7d1 | | Gypa | 1.368677131 | -0.007340371 | 0.022631034 | -0.324349772 | 0.745673242 | NA |
| Sfxn3 | 5209.780514 | 0.119885872 | 0.04218895 | 2.84164156 | 0.004488192 | 0.03354561 | Sfxn3 | | Il21 | 2.000708935 | -0.00734587 | 0.030091881 | -0.244114685 | 0.807141999 | NA |
| Bscl2 | 2783.167985 | 0.119041387 | 0.043475584 | 2.738120469 | 0.006179144 | 0.041680301 | Bscl2 | | Gm9920 | 3.907953278 | -0.007350308 | 0.047948575 | -0.153295641 | 0.878165138 | NA |
| Vdac3 | 5158.700146 | -0.108268353 | 0.04065859 | -2.662865422 | 0.007747839 | 0.049078096 | | | Plekhm2 | 5995.415628 | -0.007357022 | 0.054939122 | -0.133912256 | 0.893471958 | 0.977844987 |
| Cs | 11988.39732 | -0.11238592 | 0.040034292 | -2.807241312 | 0.004996779 | 0.03594248 | | | Chpt1 | 947.4007214 | -0.007359856 | 0.063653413 | -0.115623907 | 0.907950615 | 0.980520081 |
| Slc30a9 | 6116.525227 | -0.114094066 | 0.03984955 | -2.86312055 | 0.004194909 | 0.032144374 | | | Krt20 | 3.144813577 | -0.007360235 | 0.042066225 | -0.174967791 | 0.861104942 | NA |
| Eef1a1 | 84643.77661 | -0.117467409 | 0.038991463 | -3.01264428 | 0.002589823 | 0.022955788 | | | Dmrtc2 | 0.552498712 | -0.00736466 | 0.016594498 | -0.4438013 | 0.657186248 | NA |
| Cyb5b | 7442.71167 | -0.117754099 | 0.042299577 | -2.783812697 | 0.005372405 | 0.037690055 | | | Dagla | 6605.384912 | -0.007377801 | 0.076391891 | -0.096578325 | 0.92306127 | 0.98296621 |
| Kidins220 | 15379.33307 | -0.122269131 | 0.039443024 | -3.099892401 | 0.00193591 | 0.018676826 | | | Cyp2f2 | 8.192871199 | -0.007382304 | 0.054642306 | -0.135102349 | 0.892530953 | NA |
| Eif5a | 9717.907832 | -0.122423501 | 0.045382249 | -2.697607643 | 0.00698397 | 0.045389947 | | | Il5 | 4.173938155 | -0.00738282 | 0.044299836 | -0.166655686 | 0.867640975 | NA |
| Dennd5a | 14294.85777 | -0.123288321 | 0.042713811 | -2.88638072 | 0.003897004 | 0.030556774 | | | Ntn4 | 289.8833626 | -0.007418263 | 0.082795061 | -0.089597895 | 0.928606758 | 0.983596265 |
| Rer1 | 2177.562415 | -0.123859907 | 0.045786633 | -2.705154272 | 0.006827267 | 0.044601532 | | | Nelfa | 814.8573388 | -0.007431257 | 0.064856541 | -0.114579916 | 0.908778099 | 0.980523965 |
| Nap1l4 | 4356.610403 | -0.124575023 | 0.042517077 | -2.92999969 | 0.003389623 | 0.027626946 | | | Rgs10 | 1030.717594 | -0.007443392 | 0.072668595 | -0.10242928 | 0.918415944 | 0.982742917 |
| Sel1l | 8883.508886 | -0.129959881 | 0.045143294 | -2.878830265 | 0.003991531 | 0.031015437 | | | Ap4s1 | 1209.96338 | -0.007456695 | 0.071350707 | -0.10450765 | 0.916766498 | 0.982742917 |
| Pdha1 | 9333.704875 | -0.133561824 | 0.043812584 | -3.048480854 | 0.002300016 | 0.021168869 | | | Mir3097 | 1.174215799 | -0.007472692 | 0.023969092 | -0.311763661 | 0.755220143 | NA |
| Gzf1 | 1823.659254 | -0.133658747 | 0.043984491 | -3.038769883 | 0.002375462 | 0.021688516 | | | Mir3057 | 1.389777318 | -0.007474729 | 0.027192082 | -0.274886243 | 0.783403637 | NA |
| Glud1 | 9953.459522 | -0.134578907 | 0.0451658 | -2.979664007 | 0.002885647 | 0.024666584 | | | Bpifa5 | 0.747274741 | -0.007482899 | 0.018092578 | -0.413589405 | 0.679174834 | NA |
| Ubl4 | 3072.112219 | -0.135024756 | 0.049012747 | -2.754890621 | 0.005871176 | 0.040200975 | | | Mfi2 | 40.68885353 | -0.007483106 | 0.093082359 | -0.080392307 | 0.935925245 | 0.985461126 |
| Ndufb10 | 3183.87443 | -0.137191646 | 0.048118945 | -2.851094196 | 0.004356906 | 0.032960625 | | | Gpr39 | 10.44618563 | -0.007484463 | 0.056919466 | -0.131492158 | 0.89538599 | NA |
| Micu1 | 3353.105412 | -0.138087299 | 0.046090025 | -2.996034398 | 0.002735156 | 0.023821531 | | | Fer1l4 | 0.713561805 | -0.007489503 | 0.019509524 | -0.383889587 | 0.701060284 | NA |
| Btbd1 | 4944.287773 | -0.139264955 | 0.051511667 | -2.703561416 | 0.006860077 | 0.044716042 | | | Capns2 | 4.162440638 | -0.007523118 | 0.03941591 | -0.190865013 | 0.848631352 | NA |
| 15-Sep | 4544.817021 | -0.139769007 | 0.041841941 | -3.340404518 | 0.000836564 | 0.009917161 | | | Aspa | 674.0575175 | -0.007524586 | 0.087324773 | -0.08616783 | 0.931333003 | 0.984208871 |
| Prmt1 | 2165.964715 | -0.141638883 | 0.044760607 | -3.164364659 | 0.001554219 | 0.01589782 | | | Ces2e | 0.779254277 | -0.007543179 | 0.018468508 | -0.408434651 | 0.682954598 | NA |
| Rpl5 | 9176.039278 | -0.141733471 | 0.051387301 | -2.758141983 | 0.005813094 | 0.03995879 | | | Snord16a | 6.229619863 | -0.007545925 | 0.041295423 | -0.182730306 | 0.855009635 | NA |
| Prune | 1304.968703 | -0.141848233 | 0.051819735 | -2.737339977 | 0.006193825 | 0.041761148 | | | Slc47a2 | 3.465496222 | -0.007557572 | 0.041653651 | -0.181438396 | 0.856023483 | NA |
| 9530068E07Rik | 3226.008428 | -0.146031171 | 0.048211443 | -3.028973245 | 0.002453864 | 0.02211392 | | | 3110043O21Rik | 1028.37232 | -0.007568075 | 0.068138116 | -0.111069623 | 0.911561137 | 0.980985114 |
| Ptp4a2 | 4725.170305 | -0.146884653 | 0.05325766 | -2.758000516 | 0.005815611 | 0.03995879 | | | Fam120a | 6251.138544 | -0.007576299 | 0.055525444 | -0.136447339 | 0.891467653 | 0.977391435 |
| Dzank1 | 13288.40347 | -0.14785555 | 0.053434991 | -2.767017399 | 0.005657174 | 0.039218008 | | | Hrnr | 0.425192124 | -0.007577768 | 0.0123226 | -0.61494883 | 0.538588514 | NA |
| Pdgfra | 2428.598642 | -0.148240114 | 0.055817676 | -2.655791601 | 0.007912249 | 0.049814056 | | | 4930549G23Rik | 0.577577851 | -0.007589982 | 0.016752998 | -0.453052184 | 0.650511166 | NA |
| Dcaf12 | 1994.375653 | -0.148411725 | 0.052331936 | -2.835968527 | 0.004568695 | 0.033967062 | | | Isl2 | 0.815821339 | -0.007606136 | 0.016360259 | -0.464915403 | 0.641992039 | NA |
| Gabrb3 | 11480.14202 | -0.149244718 | 0.055279254 | -2.69983232 | 0.006937443 | 0.045125418 | | | Zfp960 | 44.76405434 | -0.007616326 | 0.090374764 | -0.084274918 | 0.932837854 | 0.984387568 |
| Pnpla8 | 4217.219252 | -0.149536597 | 0.055245478 | -2.706766311 | 0.006794206 | 0.044436233 | | | Rrh | 9.490046542 | -0.007618275 | 0.042373304 | -0.179789493 | NA | NA |
| Pgam1 | 20595.79169 | -0.149886192 | 0.045911652 | -3.264665639 | 0.001095934 | 0.012251368 | | | Snora62 | 1.117309089 | -0.00763253 | 0.026723222 | -0.285614117 | 0.77517369 | NA |
| Sec62 | 5581.05613 | -0.150248517 | 0.050316642 | -2.986060128 | 0.002825971 | 0.02433885 | | | Cfi | 0.736588345 | -0.007635924 | 0.019886545 | -0.383974367 | 0.700997446 | NA |
| Dtd1 | 1597.265016 | -0.150288047 | 0.051190033 | -2.935884959 | 0.003325976 | 0.027283599 | | | Dppa1 | 1.29059397 | -0.007649285 | 0.026733247 | -0.28613379 | 0.774775652 | NA |
| Gnb2l1 | 5682.51273 | -0.150575168 | 0.056234407 | -2.677634161 | 0.007414415 | 0.047350844 | | | Reps1 | 1713.649796 | -0.007661651 | 0.062768073 | -0.12206287 | 0.902849228 | 0.979150581 |
| Eef1b2 | 4286.864517 | -0.150755237 | 0.047469155 | -3.175856752 | 0.001493947 | 0.015410929 | | | Pinlyp | 1.005770336 | -0.007672001 | 0.019540245 | -0.392625649 | 0.694595993 | NA |
| Cmas | 3069.390949 | -0.151392975 | 0.040752296 | -3.714955689 | 0.000203239 | 0.003445285 | | | Gpr143 | 3.680114979 | -0.007675304 | 0.04222875 | -0.181755419 | 0.855774672 | NA |
| Gnb1 | 25047.69788 | -0.15192344 | 0.051223002 | -2.965922227 | 0.003017768 | 0.025425392 | | | Dedd2 | 506.6341878 | -0.007702937 | 0.075938866 | -0.101436034 | 0.919204333 | 0.982742917 |
| Arl2bp | 4816.860739 | -0.152453549 | 0.048764719 | -3.126308335 | 0.001770158 | 0.017424927 | | | Ippk | 790.5313762 | -0.00770521 | 0.076045349 | -0.101323883 | 0.919293358 | 0.982742917 |
| Selk | 2487.57478 | -0.152474639 | 0.050667885 | -3.009295488 | 0.002618543 | 0.023157366 | | | Gm1123 | 2.291310793 | -0.007706697 | 0.034422473 | -0.223885639 | 0.82284629 | NA |
| Rpl10a | 4272.874724 | -0.152653455 | 0.049082012 | -3.110171094 | 0.00186979 | 0.018186144 | | | Trim43b | 1.552173921 | -0.00770736 | 0.030006151 | -0.256859329 | 0.797287368 | NA |
| Sae1 | 3527.91207 | -0.152916694 | 0.040961443 | -3.733186207 | 0.000189073 | 0.003262241 | | | 4933402C06Rik | 0.53945737 | -0.007729642 | 0.016516238 | -0.468002576 | 0.639782748 | NA |
| Naa20 | 1495.727797 | -0.153546228 | 0.052357754 | -2.932635882 | 0.003360978 | 0.027494717 | | | Nptx1 | 19297.5259 | -0.007743641 | 0.083546407 | -0.0926867 | 0.926152463 | 0.983236501 |
| Calu | 3614.309446 | -0.154439632 | 0.045408354 | -3.401128142 | 0.000671084 | 0.008426069 | | | Mmrn1 | 1.014005331 | -0.00776183 | 0.021893672 | -0.354523894 | 0.722946301 | NA |
| Vdac1 | 16919.7671 | -0.154680132 | 0.045579338 | -3.393645881 | 0.000689688 | 0.008617765 | | | Tshz1 | 1627.556463 | -0.007769153 | 0.050044713 | -0.155244232 | 0.876628779 | 0.972549751 |
| Mtmr9 | 2721.325589 | -0.155388337 | 0.056781581 | -2.736597574 | 0.006207818 | 0.041800928 | | | Eml3 | 481.1156758 | -0.007774058 | 0.080295098 | -0.096818594 | 0.922870458 | 0.98296621 |
| Nfyb | 1592.028691 | -0.155810118 | 0.049999522 | -3.116232153 | 0.00183178 | 0.017883806 | | | 5330411J11Rik | 3.104631568 | -0.007785339 | 0.041059512 | -0.189611092 | 0.849613895 | NA |
| Sod2 | 4831.109154 | -0.156084326 | 0.04612495 | -3.383945703 | 0.000714521 | 0.008835621 | | | Sipa1l1 | 22932.85322 | -0.00778943 | 0.065564313 | -0.118805938 | 0.905429104 | 0.980118791 |
| Cadm1 | 5939.79473 | -0.156448554 | 0.051115875 | -3.060664674 | 0.002208463 | 0.020613206 | | | Mir3100 | 1.134515865 | -0.007791232 | 0.025626737 | -0.304027456 | 0.761106974 | NA |
| Kctd20 | 1812.647683 | -0.157920407 | 0.05200303 | -3.036753923 | 0.002391406 | 0.021769943 | | | Sap30l | 755.7232547 | -0.007795559 | 0.069643959 | -0.111934464 | 0.91087537 | 0.980901151 |
| Atpif1 | 3785.672016 | -0.158281607 | 0.055641298 | -2.844678568 | 0.004445626 | 0.033388526 | | | Akr1c13 | 10.89633378 | -0.007802001 | 0.061475844 | -0.126911652 | 0.899010325 | NA |
| Marf1 | 6931.181184 | -0.158603522 | 0.053962942 | -2.939119231 | 0.003291464 | 0.02708335 | | | Cep350 | 2463.311594 | -0.007817362 | 0.079206391 | -0.098696097 | 0.921379568 | 0.982762672 |
| Ctnnb1 | 18229.57502 | -0.159041907 | 0.053017948 | -2.999774868 | 0.002701792 | 0.023570703 | | | Anxa13 | 0.830491695 | -0.00782862 | 0.016900116 | -0.463228759 | 0.643200404 | NA |
| Cbx5 | 8808.556698 | -0.160239879 | 0.056376797 | -2.842301949 | 0.004478905 | 0.03349786 | | | Gm12830 | 0.414768476 | -0.007839278 | 0.012248616 | -0.640013401 | 0.522163887 | NA |
| Chchd3 | 1545.551948 | -0.160376824 | 0.060058477 | -2.670344501 | 0.007577346 | 0.048155619 | | | Tspan15 | 599.8475083 | -0.007842026 | 0.079927001 | -0.098114854 | 0.921841093 | 0.982822698 |
| Atp5h | 5903.154605 | -0.16039856 | 0.056061667 | -2.861109369 | 0.004221614 | 0.032285136 | | | Sh3kbp1 | 1785.014709 | -0.00784711 | 0.056250573 | -0.139502747 | 0.889052885 | 0.977182599 |
| Rps2 | 10086.45592 | -0.16069142 | 0.056777483 | -2.83019625 | 0.004651946 | 0.034310034 | | | Bbx | 1566.452449 | -0.007850073 | 0.067449616 | -0.116384245 | 0.907348021 | 0.980388092 |
| Nhp2l1 | 3830.890707 | -0.161209585 | 0.039086195 | -4.124463514 | 3.72E-05 | 0.000905278 | | | Chrdl2 | 6.42185293 | -0.007852053 | 0.054054599 | -0.145261522 | 0.884504393 | NA |
| Ddost | 3363.055252 | -0.162000504 | 0.054605481 | -2.96674438 | 0.003009711 | 0.025407408 | | | Sp7 | 66.82767682 | -0.007868385 | 0.092385818 | -0.085168757 | 0.932127229 | 0.984387568 |
| Cyc1 | 4938.060661 | -0.162019076 | 0.050089561 | -3.234587638 | 0.001218185 | 0.013245307 | | | Gm813 | 0.523864742 | -0.007876651 | 0.016415649 | -0.479825714 | 0.631351327 | NA |
| Ndufb11 | 2399.993251 | -0.162055629 | 0.04980939 | -3.25351564 | 0.001139865 | 0.012610872 | | | Mir431 | 5.9531147 | -0.007897989 | 0.052983884 | -0.149063995 | 0.881503136 | NA |
| Srr | 4327.800748 | -0.162653845 | 0.048123093 | -3.379954112 | 0.000724979 | 0.0089238 | | | Cys1 | 291.7728167 | -0.007908789 | 0.092807091 | -0.085217504 | 0.932088475 | 0.984387568 |
| D5Ertd579e | 6460.136281 | -0.162671354 | 0.059250005 | -2.745507867 | 0.006041732 | 0.041016941 | | | Tsc22d2 | 1312.884419 | -0.007928548 | 0.076774697 | -0.103270323 | 0.917748428 | 0.982742917 |
| Myl6 | 8423.646643 | -0.163141867 | 0.059765498 | -2.729699813 | 0.006339202 | 0.042445806 | | | Gm11961 | 1.327998927 | -0.007936014 | 0.025755457 | -0.308129435 | 0.757983845 | NA |
| Tex2 | 4719.898534 | -0.164413376 | 0.047950375 | -3.428823594 | 0.000606203 | 0.007866428 | | | Csnk1g1 | 1640.976758 | -0.007964797 | 0.062182333 | -0.128087772 | 0.898079513 | 0.978273929 |
| Rps5 | 4994.792777 | -0.165104056 | 0.042729633 | -3.863923986 | 0.00011158 | 0.002126409 | | | Grasp | 1737.91751 | -0.007965294 | 0.079894104 | -0.099698147 | 0.920583972 | 0.982762672 |
| Pik3r3 | 2696.175112 | -0.165537098 | 0.060867147 | -2.719646064 | 0.006535183 | 0.043239925 | | | Klhl4 | 632.9310612 | -0.00796834 | 0.067804802 | -0.11751881 | 0.906448939 | 0.980118791 |
| Arf2 | 1918.690885 | -0.166330681 | 0.062116932 | -2.677702751 | 0.007412897 | 0.047350844 | | | Olfr564 | 0.488314838 | -0.007970634 | 0.012816821 | -0.621888534 | 0.534015165 | NA |
| Uqcrfs1 | 3208.467197 | -0.166462621 | 0.049486367 | -3.36380771 | 0.000768751 | 0.009269281 | | | 2610301G19Rik | 4163.4591 | -0.007975823 | 0.049024589 | -0.162690252 | 0.870762322 | 0.97153269 |
| Cask | 3579.618907 | -0.166598288 | 0.050783928 | -3.280531769 | 0.001036116 | 0.011675331 | | | Tinag | 1.311727898 | -0.007976504 | 0.028756156 | -0.277384236 | 0.781485084 | NA |
| Ppp1r11 | 2036.17777 | -0.166935923 | 0.062001262 | -2.692460097 | 0.007092702 | 0.045942445 | | | Cbx6 | 19377.43988 | -0.007981776 | 0.059314359 | -0.134567346 | 0.892953961 | 0.977844987 |
| Cops6 | 3183.945707 | -0.167164286 | 0.050323197 | -3.321813737 | 0.000894344 | 0.010481817 | | | Gal3st2 | 3.365335519 | -0.007982453 | 0.039101731 | -0.204145782 | 0.838239579 | NA |
| Ddx6 | 7248.125674 | -0.167303454 | 0.061298893 | -2.729306272 | 0.006346773 | 0.042474471 | | | Mum1 | 1480.728394 | -0.00798508 | 0.074341602 | -0.107410659 | 0.914463199 | 0.98170867 |
| Atg12 | 2453.663869 | -0.168047612 | 0.053698628 | -3.129458193 | 0.00175129 | 0.01733833 | | | a | 0.591928964 | -0.007985228 | 0.01684327 | -0.47409013 | 0.635435637 | NA |
| Ran | 7038.030763 | -0.16814012 | 0.061347038 | -2.740802578 | 0.006128932 | 0.041395673 | | | Prr15 | 34.01909748 | -0.007996611 | 0.090247197 | -0.088607856 | 0.929393565 | 0.983689248 |
| Pgam5 | 1901.74222 | -0.168468658 | 0.051785458 | -3.253204003 | 0.001141115 | 0.012610872 | | | Itgb1 | 2658.92756 | -0.008016257 | 0.06068189 | -0.132102948 | 0.894902864 | 0.977844987 |
| Fau | 4785.712144 | -0.16925374 | 0.059540577 | -2.842662089 | 0.004473848 | 0.03349786 | | | Tmprss15 | 0.761135951 | -0.008023822 | 0.018407624 | -0.435896648 | 0.662911717 | NA |
| Mlc1 | 6892.100298 | -0.169439283 | 0.05519258 | -3.069964886 | 0.002140839 | 0.020127528 | | | Gprc5d | 4.113543059 | -0.008028172 | 0.041416334 | -0.193840727 | 0.846300597 | NA |
| Astn1 | 13183.82223 | -0.169751503 | 0.061717155 | -2.75047517 | 0.00595089 | 0.040564493 | | | Hecw2 | 2576.911207 | -0.008056372 | 0.074948247 | -0.107492473 | 0.914398297 | 0.98170867 |
| 2900097C17Rik | 65811.20869 | -0.169824511 | 0.057641696 | -2.94620947 | 0.003216945 | 0.026625716 | | | Gm11944 | 21.89968336 | -0.008067139 | 0.081933568 | -0.098459513 | 0.92156742 | NA |
| Map4k4 | 4532.113062 | -0.169898088 | 0.058387881 | -2.90981769 | 0.003616396 | 0.028927437 | | | Tgfb2 | 476.9701734 | -0.008074777 | 0.087467991 | -0.092316935 | 0.926446233 | 0.983236501 |
| Rab1 | 6328.232708 | -0.170081229 | 0.058166132 | -2.92405944 | 0.003454988 | 0.027983052 | | | Gm6787 | 5.971818024 | -0.008085222 | 0.052109808 | -0.155157385 | 0.876697243 | NA |
| Slmo2 | 1491.282818 | -0.170175402 | 0.062659325 | -2.715883115 | 0.006609925 | 0.043599054 | | | Fam13a | 944.0073635 | -0.008098625 | 0.068880217 | -0.117575486 | 0.906404029 | 0.980118791 |
| Rala | 2160.378613 | -0.1702305 | 0.055615487 | -3.060847062 | 0.002207118 | 0.020613072 | | | Nubp1 | 550.3504674 | -0.008127778 | 0.075702417 | -0.107364846 | 0.914499542 | 0.98170867 |
| Ndufb3 | 1582.583249 | -0.171708456 | 0.054351536 | -3.159219911 | 0.001581921 | 0.016072315 | | | D930032P07Rik | 0.804933061 | -0.008134003 | 0.018321366 | -0.443962706 | 0.657069548 | NA |
| Ndufa11 | 2031.963172 | -0.172714373 | 0.056515942 | -3.05602928 | 0.002242894 | 0.020821685 | | | Snord1c | 0.804933061 | -0.008134003 | 0.018321366 | -0.443962706 | 0.657069548 | NA |
| Prps1 | 2908.057748 | -0.172800735 | 0.061667202 | -2.802149772 | 0.00507633 | 0.036290258 | | | Padi4 | 2.337343682 | -0.008140011 | 0.031153811 | -0.261284587 | 0.793873052 | NA |
| Rplp2 | 3337.463718 | -0.172917251 | 0.058263615 | -2.967842824 | 0.002998976 | 0.025363614 | | | Med31 | 276.1023753 | -0.008143542 | 0.083047204 | -0.098059196 | 0.921885288 | 0.982822698 |
| Trappc4 | 1529.869427 | -0.173523138 | 0.050739023 | -3.419914865 | 0.000626407 | 0.008054403 | | | Dcun1d5 | 1066.683768 | -0.00814592 | 0.070367599 | -0.115762373 | 0.907840872 | 0.980520081 |
| Dnajc27 | 6705.705458 | -0.173829494 | 0.041722581 | -4.16631691 | 3.10E-05 | 0.00078887 | | | Grm7 | 2323.287845 | -0.008155044 | 0.056637386 | -0.143986943 | 0.885510781 | 0.976017118 |
| Sdhd | 2890.363856 | -0.174076958 | 0.047063021 | -3.698805458 | 0.000216617 | 0.003599861 | | | Traf4 | 322.4643552 | -0.008159269 | 0.085734717 | -0.095168783 | 0.924180765 | 0.983236501 |
| Chmp5 | 2312.633652 | -0.174497479 | 0.060188833 | -2.899167021 | 0.003741555 | 0.029713817 | | | Mir136 | 0.556841831 | -0.008159937 | 0.016633924 | -0.490559937 | 0.623737728 | NA |
| Rftn2 | 1203.996563 | -0.175011445 | 0.061521999 | -2.844696972 | 0.004445369 | 0.033388526 | | | Myl7 | 0.822840093 | -0.008171344 | 0.020230544 | -0.403911229 | 0.686277999 | NA |
| Hibadh | 1929.701794 | -0.175085094 | 0.060445635 | -2.896571328 | 0.003772649 | 0.029930066 | | | Gltp | 1690.657045 | -0.008177212 | 0.060629367 | -0.134872125 | 0.892712979 | 0.977844987 |
| Ndufb5 | 3246.140297 | -0.175949887 | 0.059394782 | -2.962379527 | 0.003052712 | 0.025594549 | | | Wisp2 | 6.363977806 | -0.008186007 | 0.051078743 | -0.160262502 | 0.872674296 | NA |
| Dmxl2 | 10066.71195 | -0.176039862 | 0.050033502 | -3.518439782 | 0.000434092 | 0.006147922 | | | Hnrnpa3 | 7071.993316 | -0.008193178 | 0.062900192 | -0.1302568 | 0.896363257 | 0.97803061 |
| Rwdd1 | 1366.11154 | -0.176187197 | 0.064046844 | -2.750911436 | 0.005942971 | 0.040546187 | | | Retnlg | 2.251450064 | -0.008194992 | 0.025510104 | -0.321244934 | 0.748024783 | NA |
| Apba1 | 9000.174827 | -0.176506377 | 0.057849693 | -3.051120372 | 0.002279892 | 0.021064186 | | | Ccdc42b | 10.72285132 | -0.008217577 | 0.065540606 | -0.125381467 | 0.900221559 | NA |
| Dmd | 2511.592565 | -0.176814466 | 0.050495806 | -3.501567349 | 0.00046253 | 0.006438851 | | | Tm4sf1 | 353.308567 | -0.008218833 | 0.082126562 | -0.100075206 | 0.920284619 | 0.982742917 |
| Slc25a5 | 7442.673467 | -0.177180148 | 0.051149805 | -3.46394575 | 0.000532314 | 0.007140841 | | | Snord34 | 3.651696567 | -0.008220033 | 0.038266331 | -0.214811121 | 0.829914582 | NA |
| St3gal2 | 5037.027951 | -0.177220494 | 0.058845135 | -3.011642256 | 0.002598386 | 0.022992231 | | | 4930511A02Rik | 2.419896121 | -0.008225033 | 0.029081856 | -0.28282353 | 0.777312116 | NA |
| Gcsh | 1586.700514 | -0.177978017 | 0.05679425 | -3.133733009 | 0.001725978 | 0.017142504 | | | Tmem234 | 1293.503996 | -0.008232569 | 0.066767973 | -0.123301169 | 0.901868615 | 0.979150581 |
| Bzw2 | 1276.880679 | -0.178740258 | 0.059927956 | -2.982585595 | 0.002858247 | 0.024507848 | | | Grhl2 | 3.606149882 | -0.008235039 | 0.042447239 | -0.194006466 | 0.846170819 | NA |
| 0610009D07Rik | 1082.925922 | -0.178807012 | 0.057821125 | -3.092416688 | 0.00198534 | 0.019046966 | | | Appl1 | 4446.848321 | -0.008237799 | 0.062905618 | -0.130954899 | 0.895810985 | 0.977844987 |
| 6-Mar | 11759.12775 | -0.17935654 | 0.065420174 | -2.741609043 | 0.006113906 | 0.04133022 | | | Nxnl1 | 4.600733284 | -0.008254567 | 0.046450565 | -0.177706493 | 0.858953485 | NA |
| Rprd1a | 5401.404916 | -0.180207702 | 0.05911064 | -3.048650849 | 0.002298715 | 0.021168869 | | | Rfx7 | 1702.931677 | -0.008256396 | 0.069955864 | -0.118022933 | 0.906049487 | 0.980118791 |
| 2410015M20Rik | 1190.176276 | -0.180393732 | 0.055056287 | -3.276532826 | 0.001050901 | 0.011833328 | | | Rbck1 | 3532.506644 | -0.008280509 | 0.05353964 | -0.154661274 | 0.877088362 | 0.972549751 |
| Slc1a3 | 24809.19634 | -0.180875682 | 0.056931553 | -3.177072708 | 0.001487697 | 0.015387438 | | | Rpl31-ps12 | 1.855502249 | -0.008284371 | 0.025736869 | -0.321887299 | 0.747538076 | NA |
| Spcs1 | 2033.689478 | -0.180892589 | 0.065104782 | -2.778483901 | 0.005461322 | 0.038236657 | | | Mir1930 | 1.093290274 | -0.008286557 | 0.021383666 | -0.387518076 | 0.698372705 | NA |
| Nsg2 | 26465.299 | -0.181230185 | 0.063371631 | -2.859799891 | 0.004239084 | 0.032353861 | | | Rgs4 | 22990.81809 | -0.008287873 | 0.080388794 | -0.103097373 | 0.91788569 | 0.982742917 |
| Myh10 | 14401.10043 | -0.181637624 | 0.063287904 | -2.870021148 | 0.004104443 | 0.031576091 | | | Hist1h4c | 0.864229324 | -0.008291335 | 0.018679355 | -0.443876918 | 0.657131574 | NA |
| Abcd3 | 2862.01866 | -0.181875 | 0.063989807 | -2.842249523 | 0.004479642 | 0.03349786 | | | Krt32 | 0.995237888 | -0.008292262 | 0.021067927 | -0.393596499 | 0.693878967 | NA |
| Prmt2 | 1828.1902 | -0.183112882 | 0.062829272 | -2.91445177 | 0.003563139 | 0.028649336 | | | Slc22a6 | 332.1742669 | -0.008295693 | 0.081550993 | -0.101723994 | 0.918975756 | 0.982742917 |
| Slc38a10 | 3182.244039 | -0.183482577 | 0.060554903 | -3.030020175 | 0.002445374 | 0.022066759 | | | Enthd1 | 1.245331172 | -0.008296461 | 0.024502078 | -0.33860233 | 0.734909327 | NA |
| Csrnp3 | 4086.192875 | -0.183588217 | 0.067185468 | -2.732558445 | 0.006284452 | 0.042133838 | | | Wasf1 | 8929.277257 | -0.008301459 | 0.054312139 | -0.152847211 | 0.878518766 | 0.973035729 |
| Itgav | 3728.42287 | -0.184214363 | 0.059227433 | -3.110287841 | 0.001869051 | 0.018186144 | | | Fam129c | 74.38265268 | -0.008321688 | 0.089716481 | -0.092755396 | 0.926097887 | 0.983236501 |
| Tusc3 | 6600.70704 | -0.185119844 | 0.063710266 | -2.905651741 | 0.003664891 | 0.029209537 | | | Zkscan5 | 641.5067351 | -0.008404365 | 0.071538041 | -0.117481057 | 0.906478854 | 0.980118791 |
| Armcx2 | 3896.06452 | -0.185166701 | 0.057485056 | -3.221127613 | 0.001276873 | 0.013691257 | | | 3110009F21Rik | 1.18385265 | -0.00840583 | 0.024263842 | -0.346434427 | 0.729016256 | NA |
| Tomm22 | 2047.183508 | -0.185292339 | 0.055303652 | -3.35045394 | 0.000806792 | 0.009645401 | | | 0610038L08Rik | 0.194331409 | -0.008409681 | 0.009640818 | -0.872299489 | 0.383045016 | NA |
| Selrc1 | 493.0000477 | -0.18536028 | 0.068601535 | -2.701984426 | 0.006892699 | 0.04487205 | | | 2410021H03Rik | 0.194331409 | -0.008409681 | 0.009640818 | -0.872299489 | 0.383045016 | NA |
| Uchl1 | 17963.70546 | -0.18552021 | 0.058039683 | -3.196437364 | 0.00139136 | 0.014585749 | | | 4930550L24Rik | 0.194331409 | -0.008409681 | 0.009640818 | -0.872299489 | 0.383045016 | NA |
| Swi5 | 3323.586994 | -0.186061444 | 0.048362528 | -3.84722333 | 0.000119464 | 0.002238182 | | | A530032D15Rik | 0.194331409 | -0.008409681 | 0.009640818 | -0.872299489 | 0.383045016 | NA |
| Sms | 4538.576634 | -0.186310041 | 0.065783507 | -2.832169474 | 0.004623334 | 0.034225481 | | | A630076J17Rik | 0.194331409 | -0.008409681 | 0.009640818 | -0.872299489 | 0.383045016 | NA |
| Mrps14 | 891.8300304 | -0.187807754 | 0.067202601 | -2.794650066 | 0.005195592 | 0.036775013 | | | AA619741 | 0.194331409 | -0.008409681 | 0.009640818 | -0.872299489 | 0.383045016 | NA |
| Chst10 | 3130.053675 | -0.187883465 | 0.046815333 | -4.013289125 | 5.99E-05 | 0.001325368 | | | Ccnb3 | 0.194331409 | -0.008409681 | 0.009640818 | -0.872299489 | 0.383045016 | NA |
| Pin1 | 2505.096558 | -0.188095039 | 0.068981988 | -2.726726831 | 0.006396597 | 0.042690885 | | | Ccr4 | 0.194331409 | -0.008409681 | 0.009640818 | -0.872299489 | 0.383045016 | NA |
| B230219D22Rik | 4196.938807 | -0.1882059 | 0.066628513 | -2.824705091 | 0.004732415 | 0.034668577 | | | Gm10510 | 0.194331409 | -0.008409681 | 0.009640818 | -0.872299489 | 0.383045016 | NA |
| Chchd4 | 1146.835173 | -0.188392155 | 0.058507497 | -3.219966056 | 0.001282058 | 0.013718372 | | | Gm20745 | 0.194331409 | -0.008409681 | 0.009640818 | -0.872299489 | 0.383045016 | NA |
| Armc1 | 2980.334526 | -0.188425044 | 0.065547982 | -2.874612422 | 0.004045237 | 0.031275901 | | | Gm5771 | 0.194331409 | -0.008409681 | 0.009640818 | -0.872299489 | 0.383045016 | NA |
| Dnajb11 | 1876.55572 | -0.18874525 | 0.068044909 | -2.773833534 | 0.005540002 | 0.038692429 | | | Lrit1 | 0.194331409 | -0.008409681 | 0.009640818 | -0.872299489 | 0.383045016 | NA |
| Pdxp | 6085.790717 | -0.188768237 | 0.05921312 | -3.187946115 | 0.001432872 | 0.014919976 | | | Olfr742 | 0.194331409 | -0.008409681 | 0.009640818 | -0.872299489 | 0.383045016 | NA |
| Guk1 | 2564.350396 | -0.189284634 | 0.068257523 | -2.773095573 | 0.005552581 | 0.038718133 | | | Tas2r135 | 0.194331409 | -0.008409681 | 0.009640818 | -0.872299489 | 0.383045016 | NA |
| Tmem242 | 1239.410281 | -0.189468682 | 0.055938231 | -3.38710535 | 0.000706343 | 0.008762267 | | | Tcl1b1 | 0.194331409 | -0.008409681 | 0.009640818 | -0.872299489 | 0.383045016 | NA |
| Smarca2 | 11947.44592 | -0.189732511 | 0.070201056 | -2.702701652 | 0.006877845 | 0.044813007 | | | Ttll2 | 0.194331409 | -0.008409681 | 0.009640818 | -0.872299489 | 0.383045016 | NA |
| Dync1h1 | 25479.74989 | -0.190341877 | 0.071548914 | -2.660304202 | 0.00780701 | 0.049291695 | | | Zscan5b | 0.194331409 | -0.008409681 | 0.009640818 | -0.872299489 | 0.383045016 | NA |
| Rps20 | 3786.829768 | -0.190355044 | 0.069346441 | -2.744986493 | 0.006051339 | 0.041042969 | | | Cd209d | 2.432853711 | -0.008412486 | 0.034089346 | -0.246777567 | 0.805080373 | NA |
| Timm22 | 955.6283569 | -0.191270003 | 0.062005768 | -3.084713066 | 0.002037486 | 0.019331788 | | | Myrip | 3358.162033 | -0.008423735 | 0.057682022 | -0.146037443 | 0.88389183 | 0.975416905 |
| Vta1 | 1335.944826 | -0.1913323 | 0.070485866 | -2.714477539 | 0.00663804 | 0.043672949 | | | Wbp4 | 2302.876885 | -0.008426298 | 0.058077485 | -0.145087174 | 0.884642044 | 0.975676135 |
| Higd1a | 2675.824238 | -0.191360804 | 0.051424859 | -3.721173136 | 0.000198299 | 0.003383757 | | | Lhx3 | 7.304399762 | -0.008441256 | 0.047741086 | -0.176813233 | 0.859655094 | NA |
| Rit1 | 1031.572724 | -0.191661857 | 0.063226582 | -3.031349346 | 0.002434634 | 0.02204625 | | | Sema6b | 4549.714818 | -0.008449864 | 0.066985131 | -0.126145373 | 0.899616851 | 0.97888996 |
| Ext1 | 1673.055144 | -0.191677666 | 0.067782414 | -2.827837707 | 0.004686356 | 0.03444516 | | | Aqp11 | 459.8841116 | -0.008489679 | 0.072292617 | -0.11743494 | 0.906515397 | 0.980118791 |
| Ep400 | 4083.179035 | -0.192000293 | 0.066381874 | -2.892360242 | 0.003823593 | 0.030195078 | | | 1700030L20Rik | 36.93925134 | -0.008498034 | 0.088401483 | -0.096129988 | 0.923417335 | 0.98311054 |
| Ndufa13 | 3499.334994 | -0.1921289 | 0.055696498 | -3.449568733 | 0.000561483 | 0.007429218 | | | Cpa2 | 47.28031565 | -0.008502581 | 0.092354997 | -0.092064109 | 0.926647104 | 0.983377837 |
| Zfp330 | 793.1130917 | -0.192339612 | 0.060308318 | -3.189271711 | 0.001426317 | 0.01486171 | | | Zfp575 | 344.613131 | -0.00851241 | 0.076514494 | -0.11125225 | 0.91141632 | 0.980985114 |
| H3f3b | 5479.448395 | -0.192351253 | 0.061333172 | -3.136169954 | 0.0017117 | 0.017022514 | | | Tmem63a | 1754.561001 | -0.008518166 | 0.087004103 | -0.097905339 | 0.92200746 | 0.982834106 |
| Thnsl1 | 863.3923196 | -0.192577231 | 0.068096547 | -2.828002861 | 0.004683939 | 0.034443734 | | | Stap1 | 1.355862946 | -0.008521812 | 0.026776405 | -0.318258249 | 0.750289052 | NA |
| Eif2s1 | 1434.508531 | -0.192898108 | 0.069250112 | -2.785527743 | 0.005344067 | 0.03755146 | | | Cped1 | 184.7211006 | -0.008528604 | 0.095059336 | -0.089718742 | 0.928510722 | 0.983596265 |
| Arl15 | 1666.808741 | -0.193050821 | 0.070981634 | -2.71972917 | 0.00653354 | 0.043239925 | | | 1700066J24Rik | 3.248001682 | -0.008540746 | 0.034469605 | -0.247776158 | 0.804307603 | NA |
| Timm17b | 477.2129063 | -0.193102696 | 0.071046698 | -2.71796863 | 0.006568407 | 0.043399101 | | | Nenf | 1168.917259 | -0.008546109 | 0.060024764 | -0.142376382 | 0.886782717 | 0.976505319 |
| Csgalnact1 | 877.9778584 | -0.193183962 | 0.070988587 | -2.721338299 | 0.006501819 | 0.043143116 | | | 4931406H21Rik | 1.07426648 | -0.00854735 | 0.020193502 | -0.423272298 | 0.672096603 | NA |
| Eef1e1 | 557.9752635 | -0.193248453 | 0.065598039 | -2.945948636 | 0.003219659 | 0.026633951 | | | Snord1b | 1.248136422 | -0.008554474 | 0.022395528 | -0.381972415 | 0.702481826 | NA |
| Stard10 | 1868.215247 | -0.193401406 | 0.061796075 | -3.129671365 | 0.00175002 | 0.017336832 | | | Ccdc173 | 94.10101891 | -0.008560006 | 0.094976658 | -0.090127475 | 0.928185916 | 0.983501978 |
| Acly | 8471.035684 | -0.19397627 | 0.050188938 | -3.864920757 | 0.000111125 | 0.002123027 | | | 2310008N11Rik | 0.212695677 | -0.008577961 | 0.009736999 | -0.880965622 | 0.378336428 | NA |
| Fam63b | 3500.665236 | -0.194127117 | 0.064869168 | -2.992594505 | 0.00276617 | 0.023983794 | | | 4930425K10Rik | 0.212695677 | -0.008577961 | 0.009736999 | -0.880965622 | 0.378336428 | NA |
| Rasal2 | 4655.799076 | -0.194900065 | 0.064017893 | -3.044462353 | 0.002330966 | 0.021370407 | | | Gm6878 | 0.212695677 | -0.008577961 | 0.009736999 | -0.880965622 | 0.378336428 | NA |
| 2810407C02Rik | 5665.606864 | -0.195349578 | 0.071941356 | -2.715400262 | 0.006619571 | 0.043625536 | | | Itgb2l | 0.212695677 | -0.008577961 | 0.009736999 | -0.880965622 | 0.378336428 | NA |
| Fam160a2 | 3704.208157 | -0.195500601 | 0.051661506 | -3.784260595 | 0.000154166 | 0.002767775 | | | Kcnk16 | 0.212695677 | -0.008577961 | 0.009736999 | -0.880965622 | 0.378336428 | NA |
| Ptcd2 | 1077.061176 | -0.19563934 | 0.058001811 | -3.372986732 | 0.000743575 | 0.009066516 | | | Krtdap | 0.212695677 | -0.008577961 | 0.009736999 | -0.880965622 | 0.378336428 | NA |
| Wnk1 | 10212.70462 | -0.195668199 | 0.049876691 | -3.923038868 | 8.74E-05 | 0.001768185 | | | Lce1g | 0.212695677 | -0.008577961 | 0.009736999 | -0.880965622 | 0.378336428 | NA |
| Pdzd8 | 2967.283641 | -0.196820057 | 0.06294932 | -3.126643081 | 0.001768144 | 0.017416166 | | | Mir344d-3 | 0.212695677 | -0.008577961 | 0.009736999 | -0.880965622 | 0.378336428 | NA |
| Rdh14 | 793.9857803 | -0.196907322 | 0.065345608 | -3.013321454 | 0.00258405 | 0.022923025 | | | Mroh2b | 0.212695677 | -0.008577961 | 0.009736999 | -0.880965622 | 0.378336428 | NA |
| Nudt19 | 2127.233966 | -0.196907481 | 0.057134781 | -3.446368019 | 0.000568176 | 0.007492252 | | | Olfr437 | 0.212695677 | -0.008577961 | 0.009736999 | -0.880965622 | 0.378336428 | NA |
| Nme2 | 2158.006305 | -0.197883874 | 0.048114049 | -4.112808571 | 3.91E-05 | 0.000946283 | | | Osr2 | 0.212695677 | -0.008577961 | 0.009736999 | -0.880965622 | 0.378336428 | NA |
| Cpne5 | 5519.813838 | -0.19807441 | 0.0618166 | -3.204226834 | 0.001354257 | 0.014303247 | | | Slc36a3 | 0.212695677 | -0.008577961 | 0.009736999 | -0.880965622 | 0.378336428 | NA |
| Jkamp | 1569.531137 | -0.199050378 | 0.064183515 | -3.101269517 | 0.001926928 | 0.01865818 | | | Snord111 | 0.212695677 | -0.008577961 | 0.009736999 | -0.880965622 | 0.378336428 | NA |
| Vps29 | 2268.644399 | -0.199099184 | 0.051222525 | -3.886945893 | 0.000101513 | 0.001983415 | | | Tex16 | 0.212695677 | -0.008577961 | 0.009736999 | -0.880965622 | 0.378336428 | NA |
| Ica1l | 1544.785964 | -0.199305655 | 0.050821683 | -3.921665771 | 8.79E-05 | 0.001774122 | | | Uba1y | 0.212695677 | -0.008577961 | 0.009736999 | -0.880965622 | 0.378336428 | NA |

| | | | | | | | | | | | | | |
|---|---|---|---|---|---|---|---|---|---|---|---|---|---|
| 2510009E07Rik | 2462.912785 | -0.199470764 | 0.053777589 | -3.709180102 | 0.000207931 | 0.003505648 | Wnt8a | 0.212695677 | -0.008577961 | 0.009736999 | -0.880965622 | 0.378336428 | NA |
| Chga | 6075.46708 | -0.199622942 | 0.061472717 | -3.247342097 | 0.001164883 | 0.012809577 | Zfp957 | 0.212695677 | -0.008577961 | 0.009736999 | -0.880965622 | 0.378336428 | NA |
| Ndufb7 | 1592.123223 | -0.199745477 | 0.060163829 | -3.320025995 | 0.000900091 | 0.010498156 | Msrb2 | 694.3106236 | -0.008578242 | 0.072612342 | -0.118137523 | 0.905958692 | 0.980118791 |
| Mea1 | 2019.598271 | -0.199805989 | 0.074623723 | -2.677513025 | 0.007417097 | 0.047350844 | Mina | 435.313742 | -0.008591664 | 0.07055951 | -0.121764791 | 0.9030853 | 0.979150581 |
| Ppm1l | 2850.436383 | -0.200520314 | 0.059349513 | -3.37863453 | 0.000728468 | 0.008943645 | Reg3g | 0.528213853 | -0.008593523 | 0.017853197 | -0.481343657 | 0.630272274 | NA |
| Zfp407 | 723.649761 | -0.201442276 | 0.067761555 | -2.972810712 | 0.002950864 | 0.025079913 | Bcl2l14 | 0.203513543 | -0.008607827 | 0.009753381 | -0.88254796 | 0.377480556 | NA |
| Pcdhb20 | 424.9154934 | -0.201732851 | 0.073993086 | -2.726374331 | 0.006403433 | 0.042691391 | Ctsg | 0.203513543 | -0.008607827 | 0.009753381 | -0.88254796 | 0.377480556 | NA |
| Cnep1r1 | 886.2493507 | -0.202233766 | 0.072839997 | -2.776410951 | 0.005496269 | 0.038463952 | Gpihbp1 | 0.203513543 | -0.008607827 | 0.009753381 | -0.88254796 | 0.377480556 | NA |
| Rpl15 | 6983.480644 | -0.202484105 | 0.038270184 | -5.290910092 | 1.22E-07 | 8.77E-06 | Lrriq4 | 0.203513543 | -0.008607827 | 0.009753381 | -0.88254796 | 0.377480556 | NA |
| Lsm6 | 1233.133057 | -0.20289465 | 0.059195937 | -3.427509709 | 0.000609145 | 0.007897979 | Mir1931 | 0.203513543 | -0.008607827 | 0.009753381 | -0.88254796 | 0.377480556 | NA |
| Bloc1s1 | 978.8985218 | -0.203205424 | 0.073372609 | -2.769499779 | 0.005614244 | 0.038972715 | Ms4a3 | 0.203513543 | -0.008607827 | 0.009753381 | -0.88254796 | 0.377480556 | NA |
| Cnih3 | 2825.385613 | -0.203378977 | 0.076429609 | -2.660997231 | 0.00779096 | 0.049250563 | Nlrp5 | 0.203513543 | -0.008607827 | 0.009753381 | -0.88254796 | 0.377480556 | NA |
| Pdpk1 | 7481.442386 | -0.203620416 | 0.072554867 | -2.80643359 | 0.005009324 | 0.035982597 | Olfr329-ps | 0.203513543 | -0.008607827 | 0.009753381 | -0.88254796 | 0.377480556 | NA |
| Gabra3 | 6979.437762 | -0.204170357 | 0.066177523 | -3.085191902 | 0.002034209 | 0.019331788 | Ttc24 | 0.203513543 | -0.008607827 | 0.009753381 | -0.88254796 | 0.377480556 | NA |
| Rps21 | 4196.097987 | -0.204324777 | 0.075346081 | -2.711816922 | 0.006691554 | 0.043969015 | Dom3z | 585.7571396 | -0.008618115 | 0.075253919 | -0.114520484 | 0.90882521 | 0.980523965 |
| Hsd17b12 | 2612.578033 | -0.204361656 | 0.07006929 | -2.916565248 | 0.003539087 | 0.028529975 | Gm14743 | 4.735802286 | -0.008627364 | 0.013225891 | -0.65230873 | NA | NA |
| Tmem70 | 1094.997445 | -0.204922764 | 0.073378945 | -2.792664351 | 0.00522759 | 0.036943149 | Ubxn8 | 671.7578533 | -0.008628783 | 0.070628751 | -0.122170967 | 0.902763619 | 0.979150581 |
| H2afv | 934.5230211 | -0.205094048 | 0.066156461 | -3.100136343 | 0.001934316 | 0.018673077 | Olfr212 | 2.192704541 | -0.008638988 | 0.033051191 | -0.261382044 | 0.793797903 | NA |
| Ndufb8 | 3688.617777 | -0.205199265 | 0.070545712 | -2.908741834 | 0.003628864 | 0.028965914 | Rpph1 | 6.638601367 | -0.00864827 | 0.051847731 | -0.166801313 | 0.867526385 | NA |
| Atp5g3 | 6698.707539 | -0.205641521 | 0.053048242 | -3.876500175 | 0.00010597 | 0.002047251 | A230108P19Rik | 0.726181501 | -0.008648808 | 0.019853073 | -0.43564076 | 0.663097393 | NA |
| Pds5b | 4342.271676 | -0.205648899 | 0.071207407 | -2.88802678 | 0.003876669 | 0.030481586 | Ceacam9 | 11.85566558 | -0.008652366 | 0.068114705 | -0.127026398 | 0.898919506 | NA |
| Rps16 | 4309.207592 | -0.205712683 | 0.060236674 | -3.415073725 | 0.000637647 | 0.008131449 | Hs3st3b1 | 53.5085122 | -0.00865669 | 0.094521189 | -0.091584647 | 0.92702805 | 0.983501978 |
| Ralyl | 2226.841434 | -0.205816204 | 0.067428997 | -3.052339705 | 0.00227065 | 0.021003849 | Mmp8 | 0.872738459 | -0.008682591 | 0.01677236 | -0.517672587 | 0.604686728 | NA |
| Serp2 | 1825.538046 | -0.205860799 | 0.063267164 | -3.253833201 | 0.001138591 | 0.012609958 | Malsu1 | 215.5526816 | -0.008707943 | 0.087962601 | -0.098995969 | 0.921141471 | 0.982762672 |
| Snx18 | 957.1119867 | -0.205892181 | 0.075997035 | -2.709213344 | 0.006744296 | 0.044184405 | Mylk4 | 15.44037163 | -0.008720885 | 0.074945566 | -0.116362922 | 0.90736492 | NA |
| Gde1 | 6906.107748 | -0.20627582 | 0.058793548 | -3.508477127 | 0.00045068 | 0.006342267 | Sun3 | 2.085080895 | -0.008724846 | 0.033818974 | -0.257986712 | 0.796417163 | NA |
| Mzt1 | 1860.166132 | -0.206585931 | 0.05988008 | -3.449994211 | 0.000560599 | 0.007423858 | Becn1 | 4603.173043 | -0.008729016 | 0.041210834 | -0.211813615 | 0.832252446 | 0.960519604 |
| Fam174b | 1377.833796 | -0.206765576 | 0.076066606 | -2.718217474 | 0.006563468 | 0.043384973 | Smtnl1 | 0.456352106 | -0.008738133 | 0.012619115 | -0.692452147 | 0.488653428 | NA |
| Fxyd6 | 3992.991095 | -0.206808161 | 0.056036932 | -3.690568969 | 0.000223753 | 0.003680286 | Edar | 9.896441771 | -0.008753731 | 0.063065554 | -0.138803687 | 0.88960528 | NA |
| Dolk | 684.4458121 | -0.206886123 | 0.072527466 | -2.85252104 | 0.004337394 | 0.032878466 | Gtpbp1 | 2552.638021 | -0.00875799 | 0.048584279 | -0.180263875 | 0.856945414 | 0.967736072 |
| Psmb5 | 1781.380598 | -0.206934627 | 0.069826906 | -2.963537099 | 0.003041254 | 0.02553994 | Col14a1 | 87.66907545 | -0.00876912 | 0.094647381 | -0.092650424 | 0.926181284 | 0.983236501 |
| Pnma2 | 2969.016783 | -0.207968869 | 0.054873099 | -3.789996804 | 0.000150649 | 0.002722791 | A930016O22Rik | 32.53476756 | -0.008774967 | 0.089970205 | -0.097531924 | 0.922303983 | 0.982933797 |
| Mrps21 | 925.033669 | -0.208357331 | 0.075561317 | -2.75746031 | 0.005825228 | 0.039989406 | Ccbl2 | 245.1571875 | -0.008787435 | 0.087538612 | -0.100383531 | 0.920039844 | 0.982742917 |
| Cox7b | 3671.480743 | -0.208374788 | 0.076444561 | -2.725828834 | 0.006414025 | 0.042743615 | 0610010K14Rik | 636.0191122 | -0.008793134 | 0.075563907 | -0.116366858 | 0.907361801 | 0.980388092 |
| Nme1 | 4792.757464 | -0.208634707 | 0.062062573 | -3.361683151 | 0.00077469 | 0.009333624 | Senp3 | 2855.01886 | -0.008797076 | 0.053098649 | -0.165674195 | 0.868413355 | 0.970658528 |
| Usp31 | 5730.814556 | -0.208769454 | 0.065379466 | -3.193196044 | 0.001407074 | 0.014710663 | Dgcr14 | 1179.190837 | -0.008802453 | 0.051902703 | -0.169595274 | 0.86532844 | 0.970182416 |
| Fam49a | 10302.60065 | -0.208938773 | 0.062099451 | -3.364583265 | 0.000766593 | 0.009250467 | 1700025F24Rik | 0.240418098 | -0.008802981 | 0.009864145 | -0.892422034 | 0.372166765 | NA |
| Ppdpf | 1215.558062 | -0.209536907 | 0.060465683 | -3.465385584 | 0.000529472 | 0.007115035 | Btnl6 | 0.240418098 | -0.008802981 | 0.009864145 | -0.892422034 | 0.372166765 | NA |
| Prdx3 | 2166.488645 | -0.209842377 | 0.07492398 | -2.800737204 | 0.005098602 | 0.036290258 | Ccer1 | 0.240418098 | -0.008802981 | 0.009864145 | -0.892422034 | 0.372166765 | NA |
| Mrpl13 | 874.7496095 | -0.210444558 | 0.071804378 | -2.930804005 | 0.00338086 | 0.027584543 | Cd40lg | 0.240418098 | -0.008802981 | 0.009864145 | -0.892422034 | 0.372166765 | NA |
| Myo10 | 1760.663839 | -0.21050183 | 0.075467425 | -2.789307178 | 0.005282094 | 0.037217264 | Cyp2a4 | 0.240418098 | -0.008802981 | 0.009864145 | -0.892422034 | 0.372166765 | NA |
| Sephs2 | 1151.085914 | -0.210865158 | 0.058824576 | -3.584643934 | 0.000337538 | 0.005132306 | Ereg | 0.240418098 | -0.008802981 | 0.009864145 | -0.892422034 | 0.372166765 | NA |
| AI597468 | 1112.933861 | -0.211280938 | 0.076514875 | -2.761305399 | 0.005757081 | 0.039657524 | Gm11351 | 0.240418098 | -0.008802981 | 0.009864145 | -0.892422034 | 0.372166765 | NA |
| D630045J12Rik | 3072.036594 | -0.211491029 | 0.076855296 | -2.751808137 | 0.005926723 | 0.040455567 | Gm12794 | 0.240418098 | -0.008802981 | 0.009864145 | -0.892422034 | 0.372166765 | NA |
| Slc1a2 | 125043.0129 | -0.211759369 | 0.044956061 | -4.710363019 | 2.47E-06 | 0.00010613 | Gm20750 | 0.240418098 | -0.008802981 | 0.009864145 | -0.892422034 | 0.372166765 | NA |
| Scn2a1 | 13122.46649 | -0.211823352 | 0.070172149 | -3.018624254 | 0.002539252 | 0.022624024 | Hist1h3c | 0.240418098 | -0.008802981 | 0.009864145 | -0.892422034 | 0.372166765 | NA |
| Casd1 | 2507.605979 | -0.212233012 | 0.069921582 | -3.035300506 | 0.002402961 | 0.021823847 | Kprp | 0.240418098 | -0.008802981 | 0.009864145 | -0.892422034 | 0.372166765 | NA |
| Tceb2 | 3040.666416 | -0.212528385 | 0.056299188 | -3.774981348 | 0.00016002 | 0.002856387 | Mir199a-1 | 0.240418098 | -0.008802981 | 0.009864145 | -0.892422034 | 0.372166765 | NA |
| Eif5a2 | 2333.190621 | -0.212749616 | 0.070910817 | -3.000242079 | 0.002697651 | 0.023561109 | Mir323 | 0.240418098 | -0.008802981 | 0.009864145 | -0.892422034 | 0.372166765 | NA |
| Notch1 | 1433.46087 | -0.212755381 | 0.073014034 | -2.913897069 | 0.003569476 | 0.028685403 | Olfr1389 | 0.240418098 | -0.008802981 | 0.009864145 | -0.892422034 | 0.372166765 | NA |
| Snrpd2 | 1338.07867 | -0.212806234 | 0.066437407 | -3.203108658 | 0.001359527 | 0.014339352 | Olfr561 | 0.240418098 | -0.008802981 | 0.009864145 | -0.892422034 | 0.372166765 | NA |
| 1700025G04Rik | 4420.121962 | -0.212982072 | 0.052463671 | -4.059610584 | 4.92E-05 | 0.001131652 | Olfr609 | 0.240418098 | -0.008802981 | 0.009864145 | -0.892422034 | 0.372166765 | NA |
| Cplx2 | 59873.1095 | -0.213256739 | 0.075834503 | -2.812133415 | 0.004921409 | 0.035648576 | Olfr970 | 0.240418098 | -0.008802981 | 0.009864145 | -0.892422034 | 0.372166765 | NA |
| Elovl5 | 2724.835622 | -0.213549169 | 0.073965802 | -2.887133812 | 0.003887689 | 0.030510193 | Rxfp2 | 62.65206272 | -0.008809005 | 0.094066073 | -0.093646991 | 0.92538958 | 0.983236501 |
| Gria3 | 9985.413447 | -0.213633771 | 0.062662007 | -3.409303048 | 0.000651291 | 0.008237633 | Tjp3 | 24.45205681 | -0.008813648 | 0.061348279 | -0.14366578 | 0.885764394 | 0.976107812 |
| Akap11 | 12092.52585 | -0.214723663 | 0.077859925 | -2.757820065 | 0.005818822 | 0.039963132 | Rpl24 | 5179.205126 | -0.008836272 | 0.059674268 | -0.148075078 | 0.882283518 | 0.974530483 |
| Kcnd2 | 4019.492144 | -0.214856286 | 0.079300294 | -2.709400871 | 0.006740484 | 0.044184405 | Arl6ip6 | 261.8617282 | -0.008860525 | 0.078722509 | -0.112553889 | 0.910384244 | 0.980590355 |
| Gabarapl2 | 4780.507071 | -0.214901601 | 0.042361046 | -5.073094771 | 3.91E-07 | 2.30E-05 | Defb11 | 1.616575638 | -0.008861707 | 0.018342866 | -0.483114624 | 0.629014351 | NA |
| Ndufa8 | 2964.542542 | -0.215202682 | 0.055930349 | -3.847690655 | 0.000119236 | 0.002236622 | Dpm3 | 263.7164595 | -0.008872573 | 0.094140319 | -0.09424838 | 0.924911854 | 0.983236501 |
| Reps2 | 8345.854678 | -0.215274667 | 0.070885114 | -3.036951687 | 0.002389838 | 0.021768456 | AW046200 | 16.02101879 | -0.008876015 | 0.069443753 | -0.127815895 | 0.898294671 | NA |
| Zeb1 | 3265.258254 | -0.215305155 | 0.070923651 | -3.035731391 | 0.00239953 | 0.021823847 | Triml2 | 0.561190942 | -0.008895051 | 0.018070302 | -0.492246949 | 0.622544778 | NA |
| Gatad1 | 2382.462965 | -0.215418793 | 0.060460136 | -3.562988886 | 0.000366656 | 0.00546773 | Afmid | 141.7332787 | -0.00891647 | 0.092344775 | -0.096556302 | 0.92307876 | 0.98296621 |
| Erh | 1568.037313 | -0.215457421 | 0.04824643 | -4.465769176 | 7.98E-06 | 0.000262449 | Gpx7 | 103.0267194 | -0.008918268 | 0.093725063 | -0.095153502 | 0.924192902 | 0.983236501 |
| Atp5g1 | 4323.655338 | -0.21583791 | 0.065008111 | -3.320168934 | 0.00089963 | 0.010498156 | 3200001D21Rik | 0.257802559 | -0.008929276 | 0.009934798 | -0.898787889 | 0.368765652 | NA |
| Celf4 | 16304.22495 | -0.215866214 | 0.058419172 | -3.695126245 | 0.000219778 | 0.003630313 | 4930544D05Rik | 0.257802559 | -0.008929276 | 0.009934798 | -0.898787889 | 0.368765652 | NA |
| Tenm2 | 2511.105801 | -0.215939245 | 0.058987574 | -3.660758163 | 0.00025147 | 0.004058622 | 9530003J23Rik | 0.257802559 | -0.008929276 | 0.009934798 | -0.898787889 | 0.368765652 | NA |
| Tmem183a | 2103.510793 | -0.216436312 | 0.058608487 | -3.692917548 | 0.000221696 | 0.003650326 | Fga | 0.257802559 | -0.008929276 | 0.009934798 | -0.898787889 | 0.368765652 | NA |
| Dync2h1 | 1764.679479 | -0.216526163 | 0.059801156 | -3.620768867 | 0.000293729 | 0.004573904 | Gm1564 | 0.257802559 | -0.008929276 | 0.009934798 | -0.898787889 | 0.368765652 | NA |
| Txn1 | 1471.276216 | -0.216831216 | 0.055450046 | -3.910388399 | 9.21E-05 | 0.001844623 | Gm8179 | 0.257802559 | -0.008929276 | 0.009934798 | -0.898787889 | 0.368765652 | NA |
| Tanc2 | 8753.849108 | -0.217239088 | 0.072226312 | -3.007755512 | 0.002631848 | 0.023198299 | Hist1h4m | 0.257802559 | -0.008929276 | 0.009934798 | -0.898787889 | 0.368765652 | NA |
| Spryd7 | 1303.369018 | -0.217324671 | 0.058415596 | -3.720319313 | 0.000198971 | 0.003387755 | Krtap5-2 | 0.257802559 | -0.008929276 | 0.009934798 | -0.898787889 | 0.368765652 | NA |
| Hsbp1 | 6932.186699 | -0.217485076 | 0.040683298 | -5.345807427 | 9.00E-08 | 6.94E-06 | Lrrc72 | 0.257802559 | -0.008929276 | 0.009934798 | -0.898787889 | 0.368765652 | NA |
| Mfap3l | 2910.079189 | -0.217722191 | 0.058908813 | -3.695918839 | 0.000219093 | 0.003626738 | Prdm14 | 0.257802559 | -0.008929276 | 0.009934798 | -0.898787889 | 0.368765652 | NA |
| Nab1 | 1102.338196 | -0.218135486 | 0.076507496 | -2.851164893 | 0.004355937 | 0.032960625 | Rnu12 | 0.257802559 | -0.008929276 | 0.009934798 | -0.898787889 | 0.368765652 | NA |
| Cacybp | 2041.666175 | -0.21835868 | 0.071768822 | -3.042528402 | 0.002345997 | 0.021482786 | Art1 | 1.724394973 | -0.008937135 | 0.031488134 | -0.283825469 | 0.776544135 | NA |
| Ndn | 6907.885412 | -0.218576506 | 0.062972967 | -3.470957703 | 0.000518606 | 0.007005472 | Hist1h2bh | 1.658423627 | -0.00898891 | 0.023555224 | -0.381610047 | 0.702750631 | NA |
| Phactr1 | 25008.58176 | -0.218679675 | 0.060110038 | -3.637989277 | 0.000274775 | 0.004344247 | Cluh | 4664.25772 | -0.008994306 | 0.047780231 | -0.188243244 | 0.850685975 | 0.966301569 |
| Bsg | 13114.69301 | -0.218798481 | 0.077189634 | -2.83455782 | 0.004588915 | 0.034079589 | Dck | 1024.592299 | -0.008996815 | 0.076563364 | -0.1175081 | 0.906457426 | 0.980118791 |
| Ssbp4 | 2733.23546 | -0.218877457 | 0.073783698 | -2.966474463 | 0.003012354 | 0.025407408 | Gm16982 | 1.676041641 | -0.009015913 | 0.023177797 | -0.38898919 | 0.697284143 | NA |
| Tcf20 | 5220.32896 | -0.218901123 | 0.080375858 | -2.723468574 | 0.006460036 | 0.042939425 | Gm10390 | 0.746784837 | -0.009022343 | 0.019944113 | -0.452381246 | 0.650994355 | NA |
| Rps12 | 4654.99579 | -0.218946756 | 0.069381305 | -3.155702481 | 0.001601121 | 0.01619582 | 4933408N05Rik | 0.575450061 | -0.009029571 | 0.01816091 | -0.497198144 | 0.61904933 | NA |
| Nedd8 | 2698.066681 | -0.219274063 | 0.059122898 | -3.708784087 | 0.000208257 | 0.003507318 | Chchd2 | 4089.995581 | -0.009029571 | 0.059204314 | -0.15251542 | 0.878780429 | 0.973102951 |
| Sorcs2 | 2631.138908 | -0.219342691 | 0.075284923 | -2.913500879 | 0.003574008 | 0.028706939 | 4833428L15Rik | 1.010476088 | -0.009043075 | 0.021910881 | -0.412720726 | 0.679811238 | NA |
| Mrpl12 | 1528.721775 | -0.219477515 | 0.069962563 | -3.137070821 | 0.001706449 | 0.017013978 | E030025P04Rik | 0.217374753 | -0.009044567 | 0.009996606 | -0.904763761 | 0.365590554 | NA |
| Rbm12 | 787.1576654 | -0.21949251 | 0.081139868 | -2.705112976 | 0.006828116 | 0.044601532 | Fut4-ps1 | 0.217374753 | -0.009044567 | 0.009996606 | -0.904763761 | 0.365590554 | NA |
| Vkorc1l1 | 638.1950196 | -0.219566855 | 0.068484658 | -3.206073629 | 0.001345596 | 0.014231165 | Glyatl3 | 0.217374753 | -0.009044567 | 0.009996606 | -0.904763761 | 0.365590554 | NA |
| Ndufa2 | 1610.729781 | -0.219957898 | 0.077130134 | -2.851776436 | 0.004347567 | 0.032932886 | Gm10415 | 0.217374753 | -0.009044567 | 0.009996606 | -0.904763761 | 0.365590554 | NA |
| Pak6 | 3889.667023 | -0.220036479 | 0.065606653 | -3.35387447 | 0.000796885 | 0.009541682 | Gm5415 | 0.217374753 | -0.009044567 | 0.009996606 | -0.904763761 | 0.365590554 | NA |
| Pgrmc1 | 8558.057799 | -0.220175693 | 0.064397441 | -3.419013057 | 0.000628487 | 0.008067754 | Havcr1 | 0.217374753 | -0.009044567 | 0.009996606 | -0.904763761 | 0.365590554 | NA |
| Polr1d | 1392.615692 | -0.220204446 | 0.062242706 | -3.537835325 | 0.000403422 | 0.005876062 | Krtap11-1 | 0.217374753 | -0.009044567 | 0.009996606 | -0.904763761 | 0.365590554 | NA |
| Rps18 | 4385.517557 | -0.22023814 | 0.070397349 | -3.128500457 | 0.001757007 | 0.01736241 | Trim43c | 0.217374753 | -0.009044567 | 0.009996606 | -0.904763761 | 0.365590554 | NA |
| Kcnk2 | 2894.325898 | -0.220496965 | 0.073441897 | -3.002332084 | 0.002679197 | 0.023452816 | Ccdc114 | 115.2826705 | -0.009058687 | 0.094624184 | -0.095733313 | 0.923732383 | 0.983229764 |
| Ldoc1l | 3210.175525 | -0.220756163 | 0.074628477 | -2.958068735 | 0.003095731 | 0.025824561 | C2cd4c | 912.7639364 | -0.009075072 | 0.079510175 | -0.114137243 | 0.909128999 | 0.980523965 |
| Chd9 | 2765.180247 | -0.220845909 | 0.082681786 | -2.671034557 | 0.007561786 | 0.048076452 | Rbbp6 | 3436.528365 | -0.009076243 | 0.050187032 | -0.180848367 | 0.856486598 | 0.967598001 |
| Dnalc1 | 5506.664735 | -0.221324696 | 0.060005543 | -3.688404187 | 0.000225665 | 0.00370387 | Runx3 | 0.952960804 | -0.009110839 | 0.022921551 | -0.397479189 | 0.69101413 | NA |
| Rap2b | 2194.056811 | -0.221397533 | 0.075031754 | -2.95071781 | 0.003170364 | 0.026352802 | Pramel1 | 1.630984956 | -0.009126637 | 0.028043896 | -0.32544111 | 0.744847246 | NA |
| Tmsb4x | 16876.94959 | -0.221637628 | 0.069301923 | -3.198145429 | 0.001383145 | 0.014519275 | Patl2 | 2.566150773 | -0.009126909 | 0.039250815 | -0.232527878 | 0.816128037 | NA |
| Vps13a | 2429.544911 | -0.222028936 | 0.070217603 | -3.162012466 | 0.001566829 | 0.015971343 | 1700012P22Rik | 0.570373077 | -0.009167364 | 0.018130402 | -0.505634885 | 0.613112987 | NA |
| Tubb4b | 10125.50264 | -0.222245048 | 0.077982929 | -2.849919232 | 0.004373033 | 0.033045155 | Slc16a7 | 420.5249418 | -0.009173126 | 0.077013681 | -0.119110339 | 0.905187939 | 0.980118791 |
| Fopnl | 1211.32893 | -0.222637633 | 0.058836911 | -3.783978927 | 0.000154341 | 0.002767775 | Ccsap | 2451.185702 | -0.009176512 | 0.077341163 | -0.118649782 | 0.905552824 | 0.980118791 |
| Tram1l1 | 1079.409376 | -0.223311174 | 0.067762891 | -3.295478845 | 0.000982541 | 0.011235048 | Brca1 | 88.21459617 | -0.00918847 | 0.093363821 | -0.098415749 | 0.92160217 | 0.982762672 |
| Mid2 | 1235.345003 | -0.223400024 | 0.079097959 | -2.824346251 | 0.004737717 | 0.03467463 | 9930111J21Rik2 | 0.437549073 | -0.009196884 | 0.014203962 | -0.647487231 | 0.517316656 | NA |
| Cdh10 | 1872.274555 | -0.223563643 | 0.06134207 | -3.644540265 | 0.00026787 | 0.004269939 | Elane | 0.980505671 | -0.009203146 | 0.021499546 | -0.428062351 | 0.668605726 | NA |
| Gja1 | 6786.808247 | -0.223844228 | 0.07758358 | -2.885201062 | 0.003911638 | 0.030656 | Zfp248 | 507.7925851 | -0.00923262 | 0.073555371 | -0.125519318 | 0.900112433 | 0.97888996 |
| Sptbn1 | 40054.53704 | -0.224036527 | 0.038236364 | -5.859252989 | 4.65E-09 | 5.58E-07 | Shbg | 1.358219832 | -0.009242553 | 0.025693172 | -0.359727968 | 0.719050575 | NA |
| Grid1 | 2481.064177 | -0.22453446 | 0.057910568 | -3.877262293 | 0.000105638 | 0.002043399 | Smurf1 | 2607.484706 | -0.009246889 | 0.058074138 | -0.159225583 | 0.873491149 | 0.971912694 |
| Dgat2 | 2655.805663 | -0.224555094 | 0.060203092 | -3.729959477 | 0.000191511 | 0.003296961 | Ccr1l1 | 1.039507966 | -0.009259154 | 0.023125389 | -0.400389112 | 0.688869943 | NA |
| Gpr125 | 865.8734092 | -0.224567457 | 0.074366457 | -3.019741237 | 0.002529907 | 0.022566715 | Sppl2a | 1621.557235 | -0.009259649 | 0.06415266 | -0.144337718 | 0.885233796 | 0.975785864 |
| Mettl9 | 1995.363653 | -0.224583753 | 0.072790646 | -3.085338089 | 0.002033209 | 0.019331788 | Trpv4 | 120.8252734 | -0.009260248 | 0.058480835 | -0.158346711 | NA | NA |
| Ndufb9 | 4651.72367 | -0.224630582 | 0.075247374 | -2.985228169 | 0.002833669 | 0.024364519 | Clec4a2 | 27.31727405 | -0.009260952 | 0.085170076 | -0.108734805 | 0.913412835 | 0.98170867 |
| Hbegf | 540.6838468 | -0.224818615 | 0.079487932 | -2.828336467 | 0.00467906 | 0.034424195 | Tmsb15b2 | 1.609564543 | -0.009274967 | 0.028065118 | -0.330480242 | 0.74103712 | NA |
| Cenpv | 534.775648 | -0.225009681 | 0.077263234 | -2.912247782 | 0.003588379 | 0.02876272 | Cdc42bpb | 8489.705522 | -0.009291978 | 0.04636911 | -0.200391548 | 0.841174369 | 0.963094947 |
| 2610507B11Rik | 14041.9846 | -0.225524877 | 0.040321262 | -5.593199863 | 2.23E-08 | 2.09E-06 | Rab7l1 | 128.0619605 | -0.009303396 | 0.09390711 | -0.099070201 | 0.921082532 | 0.982762672 |
| Atxn7l3b | 8179.446081 | -0.225679697 | 0.048127415 | -4.689212912 | 2.74E-06 | 0.000114847 | 9030404E10Rik | 0.226066984 | -0.009308436 | 0.010140664 | -0.917931679 | 0.358654635 | NA |
| Tulp4 | 8180.879394 | -0.225774764 | 0.07597062 | -2.97186946 | 0.002959925 | 0.025129358 | Gm16617 | 0.226066984 | -0.009308436 | 0.010140664 | -0.917931679 | 0.358654635 | NA |
| Ndufa7 | 1711.799455 | -0.225939321 | 0.060406827 | -3.740294499 | 0.000183805 | 0.003199855 | Gm5088 | 0.226066984 | -0.009308436 | 0.010140664 | -0.917931679 | 0.358654635 | NA |
| Lphn3 | 3829.17901 | -0.225981055 | 0.075668523 | -2.986460503 | 0.002822273 | 0.024320521 | Gm6213 | 0.226066984 | -0.009308436 | 0.010140664 | -0.917931679 | 0.358654635 | NA |
| Timmdc1 | 640.732965 | -0.226316892 | 0.084250736 | -2.686230447 | 0.007226323 | 0.046578655 | Gm7616 | 0.226066984 | -0.009308436 | 0.010140664 | -0.917931679 | 0.358654635 | NA |
| Oxct1 | 11269.40151 | -0.226410841 | 0.074777728 | -3.027784426 | 0.002463538 | 0.022127158 | Gm8817 | 0.226066984 | -0.009308436 | 0.010140664 | -0.917931679 | 0.358654635 | NA |
| Phactr3 | 5455.548194 | -0.226497465 | 0.054664533 | -4.143408011 | 3.42E-05 | 0.000848284 | Lcn3 | 0.226066984 | -0.009308436 | 0.010140664 | -0.917931679 | 0.358654635 | NA |
| Cdc34 | 1222.231111 | -0.226510309 | 0.079691821 | -2.842328169 | 0.004478537 | 0.03349786 | Lgsn | 0.226066984 | -0.009308436 | 0.010140664 | -0.917931679 | 0.358654635 | NA |
| Rpsa | 12881.31304 | -0.226776535 | 0.05211697 | -4.35129927 | 1.35E-05 | 0.000409541 | Msx1as | 0.226066984 | -0.009308436 | 0.010140664 | -0.917931679 | 0.358654635 | NA |
| Rgs8 | 5912.537735 | -0.22720042 | 0.060931365 | -3.728792537 | 0.000192399 | 0.003308587 | Tespa1 | 0.226066984 | -0.009308436 | 0.010140664 | -0.917931679 | 0.358654635 | NA |
| 2610002M06Rik | 2413.129244 | -0.227289666 | 0.085230528 | -2.666763546 | 0.007658553 | 0.048611891 | Bpifb6 | 1.327498615 | -0.009313781 | 0.025366246 | -0.367172205 | 0.713490571 | NA |
| Dock3 | 14466.78316 | -0.227301123 | 0.073307809 | -3.100639985 | 0.001931029 | 0.018662475 | Zmym1 | 481.593906 | -0.009314526 | 0.086204675 | -0.108051285 | 0.913955011 | 0.98170867 |
| Lix1l | 855.7449009 | -0.227305766 | 0.078871754 | -2.881966667 | 0.003952015 | 0.030863169 | Rab9 | 952.7854024 | -0.009322824 | 0.056062301 | -0.166293997 | 0.86792559 | 0.970480832 |
| Cntn1 | 13563.94287 | -0.227370013 | 0.054740566 | -4.153592632 | 3.27E-05 | 0.000825264 | 4930455F16Rik | 0.226556888 | -0.009331758 | 0.010153338 | -0.919082739 | 0.358052297 | NA |
| Samd4 | 1406.665266 | -0.227541528 | 0.085331812 | -2.666549824 | 0.007663425 | 0.048622892 | D630045M09Rik | 0.226556888 | -0.009331758 | 0.010153338 | -0.919082739 | 0.358052297 | NA |
| Ndufs6 | 1342.603392 | -0.227597163 | 0.065461304 | -3.476819856 | 0.000507399 | 0.00689617 | Gm17677 | 0.226556888 | -0.009331758 | 0.010153338 | -0.919082739 | 0.358052297 | NA |
| Ppp1r12b | 2078.398659 | -0.227824161 | 0.076811672 | -2.966009669 | 0.00301691 | 0.025425392 | Hoxb13 | 0.226556888 | -0.009331758 | 0.010153338 | -0.919082739 | 0.358052297 | NA |
| Pard3 | 967.6525454 | -0.228014359 | 0.071295441 | -3.198161838 | 0.001383066 | 0.014519275 | Mir1983 | 0.226556888 | -0.009331758 | 0.010153338 | -0.919082739 | 0.358052297 | NA |
| Prdx4 | 417.3336089 | -0.228426108 | 0.080882368 | -2.824176817 | 0.004740223 | 0.034676586 | Spdyb | 0.226556888 | -0.009331758 | 0.010153338 | -0.919082739 | 0.358052297 | NA |
| Pdlim5 | 1232.861789 | -0.228826346 | 0.075552116 | -3.028721883 | 0.002455907 | 0.02211392 | Trim75 | 0.226556888 | -0.009331758 | 0.010153338 | -0.919082739 | 0.358052297 | NA |
| Agps | 1742.359557 | -0.229246093 | 0.073092508 | -3.136382913 | 0.001710457 | 0.017022514 | Xlr3c | 0.226556888 | -0.009331758 | 0.010153338 | -0.919082739 | 0.358052297 | NA |
| Mrps25 | 1228.800872 | -0.23041079 | 0.078504106 | -2.935015796 | 0.003335307 | 0.027312724 | Zc2hc1c | 132.7258129 | -0.009340182 | 0.092434624 | -0.10104636 | 0.919513659 | 0.982742917 |
| Ndufa3 | 2112.850974 | -0.230454898 | 0.081440128 | -2.829746264 | 0.004658493 | 0.034336846 | Atp8b4 | 3.323783005 | -0.009342042 | 0.03728193 | -0.250578278 | 0.802140178 | NA |
| Tmem35 | 2484.165498 | -0.230491158 | 0.086504925 | -2.664485971 | 0.007710608 | 0.048882228 | Aldh7a1 | 3588.296109 | -0.009376842 | 0.060711599 | -0.154448933 | 0.877255774 | 0.972549751 |
| Akap6 | 10151.29268 | -0.230590514 | 0.073098293 | -3.154526649 | 0.001607587 | 0.016226679 | Chmp4c | 3.678402829 | -0.009391255 | 0.039692718 | -0.236598941 | 0.812967942 | NA |
| Cdkl5 | 2562.332265 | -0.230651214 | 0.080684731 | -2.858672407 | 0.004254179 | 0.032422156 | Ccdc120 | 518.3985206 | -0.009391283 | 0.070593428 | -0.133033384 | 0.894166978 | 0.977844987 |
| Abcb7 | 1372.37207 | -0.230705519 | 0.076906676 | -2.999811328 | 0.002701469 | 0.023570703 | Mrps10 | 688.1369642 | -0.009391508 | 0.066877181 | -0.14042918 | 0.888320904 | 0.97703865 |
| Ccm2 | 1996.642015 | -0.230873284 | 0.064397371 | -3.585135237 | 0.000336904 | 0.005132306 | Gm10408 | 0.97901975 | -0.009404126 | 0.022986439 | -0.409116234 | 0.682454363 | NA |
| Ppp1ca | 8033.232645 | -0.231475613 | 0.061089301 | -3.789135093 | 0.000151173 | 0.002724393 | 4921507L20Rik | 0.409445549 | -0.009404997 | 0.013957946 | -0.673809565 | 0.500432393 | NA |
| Cdk14 | 8483.781205 | -0.231623159 | 0.053464618 | -4.332269974 | 1.48E-05 | 0.000438887 | Mtus1 | 2389.715737 | -0.009405923 | 0.063289836 | -0.148616649 | 0.881856135 | 0.974354774 |
| Zfp827 | 2244.603061 | -0.232856594 | 0.06542321 | -3.559235215 | 0.000371936 | 0.0055305 | 1600014K23Rik | 0.291497113 | -0.009418496 | 0.010203856 | -0.923033044 | 0.355989985 | NA |
| Syt5 | 5924.558638 | -0.232875573 | 0.057564153 | -4.045496388 | 5.22E-05 | 0.001187927 | 4930483K19Rik | 0.291497113 | -0.009418496 | 0.010203856 | -0.923033044 | 0.355989985 | NA |
| Lin7a | 2860.656039 | -0.233191432 | 0.07627387 | -3.057291192 | 0.002233472 | 0.020771557 | Aldoart1 | 0.291497113 | -0.009418496 | 0.010203856 | -0.923033044 | 0.355989985 | NA |
| Ube2e3 | 2025.595618 | -0.233193243 | 0.072121467 | -3.233340246 | 0.001223517 | 0.013284639 | Ldlrad1 | 0.291497113 | -0.009418496 | 0.010203856 | -0.923033044 | 0.355989985 | NA |
| Nacc2 | 6513.112437 | -0.233388331 | 0.07461162 | -3.128042666 | 0.001759746 | 0.017366567 | Lilra6 | 0.291497113 | -0.009418496 | 0.010203856 | -0.923033044 | 0.355989985 | NA |
| Tmem164 | 1432.32013 | -0.233571301 | 0.080665203 | -2.895564544 | 0.003784772 | 0.029995527 | Ubd | 0.291497113 | -0.009418496 | 0.010203856 | -0.923033044 | 0.355989985 | NA |
| Tmem126a | 547.3942268 | -0.233701081 | 0.070403377 | -3.319458397 | 0.000901922 | 0.010507057 | Vmn2r30 | 0.291497113 | -0.009418496 | 0.010203856 | -0.923033044 | 0.355989985 | NA |
| Rps3 | 8657.996085 | -0.233787144 | 0.060606091 | -3.857485947 | 0.000114559 | 0.002167254 | Mir874 | 0.762249383 | -0.009421132 | 0.021379696 | -0.440657906 | 0.659460675 | NA |
| Rpl36 | 1896.0831 | -0.233950509 | 0.065458725 | -3.574015673 | 0.000351548 | 0.005283106 | Trnp1 | 4671.302256 | -0.009437155 | 0.049365378 | -0.191169511 | 0.84839279 | 0.965746177 |
| Scarb1 | 1118.38109 | -0.233970993 | 0.068449074 | -3.418176164 | 0.000630423 | 0.008079216 | Mir145 | 1.595159537 | -0.009439684 | 0.028063098 | -0.336373548 | 0.736589194 | NA |
| Parvb | 1016.349827 | -0.234138962 | 0.068102163 | -3.438054726 | 0.000585909 | 0.007673777 | Aoah | 11.32795662 | -0.009440049 | 0.063731575 | -0.148122006 | 0.882246483 | NA |
| Plcl2 | 3234.985032 | -0.234156093 | 0.078438082 | -2.985234828 | 0.002833607 | 0.024364519 | Actbl2 | 0.774573665 | -0.009446498 | 0.020062243 | -0.470859496 | 0.637741074 | NA |
| Mrpl53 | 632.3209522 | -0.234259043 | 0.082809834 | -2.828879505 | 0.004671128 | 0.034398508 | Dusp13 | 9.34228095 | -0.009470546 | 0.061928507 | -0.152927086 | 0.878455775 | NA |
| Rap1b | 1750.435981 | -0.234329511 | 0.05597616 | -4.186237723 | 2.84E-05 | 0.000738546 | Lars | 2024.007247 | -0.009473059 | 0.052614413 | -0.180046844 | 0.857115793 | 0.967853235 |
| Mdga2 | 1494.131084 | -0.234339827 | 0.078938255 | -2.968647175 | 0.002991138 | 0.025311139 | 2310068J16Rik | 5.449938812 | -0.009477408 | 0.045679976 | -0.207474011 | 0.835639688 | NA |
| 6330403K07Rik | 9439.219227 | -0.23449348 | 0.051586152 | -4.545667245 | 5.48E-06 | 0.000198718 | Hsd17b10 | 840.7753128 | -0.009479198 | 0.068202303 | -0.138986483 | 0.889460831 | 0.977182599 |
| Mtx2 | 1111.998283 | -0.234640591 | 0.085408215 | -2.747283637 | 0.006009114 | 0.040889423 | Rps14 | 6558.156253 | -0.009486027 | 0.047521232 | -0.199616615 | 0.841780432 | 0.963252575 |
| Cldn12 | 1840.913707 | -0.235605854 | 0.069772714 | -3.376762073 | 0.000733445 | 0.008983392 | Bcat2 | 429.339478 | -0.009488527 | 0.086123907 | -0.110172974 | 0.912272196 | 0.981495756 |
| 6030419C18Rik | 6021.93407 | -0.235732989 | 0.063666237 | -3.702637394 | 0.00021337 | 0.003556118 | Spcs3 | 3894.241774 | -0.009494092 | 0.070930029 | -0.133851516 | 0.89351999 | 0.977844987 |
| Arhgef12 | 11573.19938 | -0.235880654 | 0.07204004 | -3.274299308 | 0.001059244 | 0.011909956 | Mapk14 | 4351.200155 | -0.009508847 | 0.056692816 | -0.16772578 | 0.866799014 | 0.97021201 |
| Rnf11 | 6279.563841 | -0.235922382 | 0.071738587 | -3.288639933 | 0.001006727 | 0.011427276 | Gsdmc | 0.423704667 | -0.009523664 | 0.014090123 | -0.675910611 | 0.499097401 | NA |
| Kitl | 1797.266147 | -0.235945227 | 0.082112454 | -2.873440193 | 0.004060279 | 0.031376544 | Slitrk1 | 3948.262583 | -0.009530065 | 0.069381086 | -0.137358257 | 0.890747624 | 0.977182599 |
| Vti1b | 4628.969292 | -0.235995426 | 0.065860405 | -3.583267148 | 0.000339323 | 0.005154387 | Sgsm3 | 2898.781809 | -0.009536674 | 0.058842254 | -0.162071868 | 0.87124926 | 0.97153269 |
| Usp46 | 4307.606762 | -0.236362944 | 0.072023566 | -3.28174451 | 0.00103167 | 0.011633698 | 2010107E04Rik | 2019.914044 | -0.009536818 | 0.060901731 | -0.156593538 | 0.875565196 | 0.972151437 |
| Mrps36 | 631.8951958 | -0.236426971 | 0.086703569 | -2.72684244 | 0.006394356 | 0.042690885 | Ascc3 | 1079.825933 | -0.009561781 | 0.077528302 | -0.123332784 | 0.901843581 | 0.979150581 |
| Slc9a3r1 | 1559.369133 | -0.236507966 | 0.055072315 | -4.294498311 | 1.75E-05 | 0.000503307 | 1810022K09Rik | 482.5878422 | -0.009570646 | 0.073983598 | -0.129361726 | 0.897071432 | 0.978039389 |
| Fam171a2 | 2705.365274 | -0.236738197 | 0.087354506 | -2.710085692 | 0.006726583 | 0.044124332 | Fbxw13 | 0.319043516 | -0.00957618 | 0.010289084 | -0.930712648 | 0.352002226 | NA |
| 3632451O06Rik | 1121.995827 | -0.237017373 | 0.086014498 | -2.755551419 | 0.005859329 | 0.040157049 | Gm13034 | 0.319043516 | -0.00957618 | 0.010289084 | -0.930712648 | 0.352002226 | NA |
| Ank2 | 20264.45172 | -0.237252924 | 0.052314343 | -4.535141036 | 5.76E-06 | 0.00020534 | Gm19424 | 0.319043516 | -0.00957618 | 0.010289084 | -0.930712648 | 0.352002226 | NA |
| Ostc | 987.4976411 | -0.237497487 | 0.059095764 | -4.018858051 | 5.85E-05 | 0.001303745 | Hist1h2bb | 0.319043516 | -0.00957618 | 0.010289084 | -0.930712648 | 0.352002226 | NA |
| Smarcd3 | 2698.468309 | -0.237531971 | 0.057792087 | -4.110112351 | 3.95E-05 | 0.000954418 | Ugt3a1 | 0.319043516 | -0.00957618 | 0.010289084 | -0.930712648 | 0.352002226 | NA |
| C1d | 879.9517057 | -0.238383592 | 0.070675629 | -3.372924964 | 0.000743742 | 0.009066516 | Psmc3ip | 71.54118031 | -0.009581876 | 0.094492017 | -0.101404076 | 0.919229701 | 0.982742917 |
| Wrb | 2033.983469 | -0.238513172 | 0.077307572 | -3.085249815 | 0.002033812 | 0.019331788 | Zscan20 | 311.6451596 | -0.009585001 | 0.082510827 | -0.116166579 | 0.907520523 | 0.980486658 |
| Ksr2 | 988.2365406 | -0.238553307 | 0.074926563 | -3.183828231 | 0.001453412 | 0.015115049 | Rtn4rl1 | 3324.918469 | -0.009587009 | 0.074192135 | -0.129218666 | 0.897184627 | 0.978039389 |
| Vps13b | 3106.92331 | -0.238990823 | 0.077269009 | -3.092971223 | 0.001981634 | 0.01902319 | Adam1b | 0.235249118 | -0.009590826 | 0.010292593 | -0.931818252 | 0.35143046 | NA |
| Glrx5 | 920.7001175 | -0.239272055 | 0.074559766 | -3.209130964 | 0.001331368 | 0.014119248 | Defb20 | 0.235249118 | -0.009590826 | 0.010292593 | -0.931818252 | 0.35143046 | NA |
| Mrps33 | 1625.72663 | -0.239272344 | 0.068839475 | -3.475801396 | 0.000509329 | 0.006914272 | Dnajc5g | 0.235249118 | -0.009590826 | 0.010292593 | -0.931818252 | 0.35143046 | NA |
| Ptprg | 2958.695308 | -0.239416329 | 0.06062175 | -3.949347041 | 7.84E-05 | 0.001629775 | Fam46d | 0.235249118 | -0.009590826 | 0.010292593 | -0.931818252 | 0.35143046 | NA |
| Rabl5 | 1335.71095 | -0.239502442 | 0.082754061 | -2.894147279 | 0.003801898 | 0.03008509 | Tmc2 | 0.235249118 | -0.009590826 | 0.010292593 | -0.931818252 | 0.35143046 | NA |
| 1110001A16Rik | 387.6976394 | -0.239689196 | 0.087245107 | -2.747308178 | 0.006008665 | 0.040889423 | Tcl1b2 | 0.769338278 | -0.009593032 | 0.02004344 | -0.478612041 | 0.632214651 | NA |
| Ptprz1 | 8916.53016 | -0.239813906 | 0.079142615 | -3.030148884 | 0.002444332 | 0.022066759 | Slc25a2 | 3.952268908 | -0.009605021 | 0.044256092 | -0.217032747 | 0.828182825 | NA |
| Rps6ka3 | 1147.772766 | -0.239865289 | 0.080903381 | -2.964836407 | 0.003028439 | 0.025459923 | Snord65 | 6.448611832 | -0.009607715 | 0.049227569 | -0.1951694 | 0.845260334 | NA |
| Timm10 | 558.5059313 | -0.239869575 | 0.083816286 | -2.861849242 | 0.004211772 | 0.032225775 | Asb2 | 119.4041281 | -0.009607974 | 0.0944975 | -0.101674377 | 0.919015141 | 0.982742917 |
| Cnn2 | 277.2405564 | -0.240031848 | 0.083765573 | -2.86551907 | 0.004163262 | 0.031965105 | Rbm4 | 461.3814006 | -0.009618824 | 0.078483601 | -0.122558388 | 0.902456808 | 0.979150581 |
| Klf16 | 2085.817085 | -0.240216806 | 0.086693292 | -2.770881127 | 0.005590483 | 0.038865068 | Thra | 20814.97132 | -0.009663449 | 0.060993289 | -0.158434624 | 0.874114332 | 0.971912694 |
| Ranbp2 | 6090.810573 | -0.240574418 | 0.07319485 | -3.286767024 | 0.001013446 | 0.011471264 | Cers2 | 2715.639997 | -0.009668049 | 0.078313367 | -0.123453365 | 0.901748101 | 0.979150581 |
| Rheb | 3197.361137 | -0.240612902 | 0.077204443 | -3.116568068 | 0.001829694 | 0.017883806 | Gtf3c1 | 6768.949666 | -0.009682639 | 0.052540805 | -0.184287994 | 0.853787532 | 0.966905493 |
| St8sia3 | 7716.409767 | -0.240626298 | 0.075397308 | -3.191444173 | 0.001415635 | 0.014775915 | Bcs1l | 778.5521498 | -0.009725164 | 0.071652728 | -0.135726358 | 0.89203761 | 0.977690693 |
| Coa5 | 4771.896106 | -0.240648456 | 0.074540909 | -3.228407839 | 0.001244814 | 0.013459276 | Hspg2 | 398.8755258 | -0.009731589 | 0.086213298 | -0.112878048 | 0.91012724 | 0.980590355 |
| Fdps | 2137.145413 | -0.240917044 | 0.083577252 | -2.882567178 | 0.00394449 | 0.030835485 | Cdhr3 | 102.0095948 | -0.009739898 | 0.093192083 | -0.104514222 | 0.916761283 | 0.982742917 |
| Reep1 | 5450.95174 | -0.241314839 | 0.063098246 | -3.824430209 | 0.000131075 | 0.002406249 | 3110015C05Rik | 0.973123212 | -0.009750883 | 0.021491986 | -0.453698569 | 0.650045797 | NA |
| Anks1b | 12133.90324 | -0.241680097 | 0.059198954 | -4.082506195 | 4.46E-05 | 0.001041174 | Cela2a | 0.423306759 | -0.009765996 | 0.014091945 | -0.693019772 | 0.488297144 | NA |
| Maob | 1130.785681 | -0.241974325 | 0.067793425 | -3.569289013 | 0.000357951 | 0.005358549 | Reg3a | 0.423306759 | -0.009765996 | 0.014091945 | -0.693019772 | 0.488297144 | NA |
| Ndufs4 | 2085.278257 | -0.242494368 | 0.061522025 | -3.941586271 | 8.09E-05 | 0.001667758 | 1700120G07Rik | 0.396404207 | -0.009769095 | 0.013847504 | -0.705476927 | 0.480513486 | NA |
| Usp9x | 11207.9479 | -0.242559842 | 0.074512608 | -3.2552859 | 0.001132783 | 0.01257259 | Map1lc3a | 5161.88873 | -0.009772253 | 0.05763983 | -0.169539938 | 0.865371961 | 0.970182416 |
| Zyg11b | 10679.49348 | -0.242698978 | 0.063472132 | -3.82370926 | 0.000131459 | 0.002410443 | Ppp1r1b | 16180.97583 | -0.009772404 | 0.086817742 | -0.112562291 | 0.910377582 | 0.980590355 |
| Rps19 | 4419.021782 | -0.243196363 | 0.068063657 | -3.57307223 | 0.000352817 | 0.005291919 | Map7d1 | 10819.92674 | -0.009784517 | 0.036785086 | -0.265991405 | 0.790245829 | 0.951607109 |
| Ndufs8 | 2008.72814 | -0.243402428 | 0.064276381 | -3.786809755 | 0.000152594 | 0.002742795 | Trpm2 | 1830.208709 | -0.009786598 | 0.064882783 | -0.150835046 | 0.88010584 | 0.973679926 |
| Zfp428 | 721.9231047 | -0.243737071 | 0.080292652 | -3.035608667 | 0.002400507 | 0.021823847 | Gjd3 | 10.78553117 | -0.009789329 | 0.058272209 | -0.167993093 | 0.866588713 | NA |
| Ndrg4 | 80486.46653 | -0.244143841 | 0.05199144 | -4.695846913 | 2.66E-06 | 0.000112397 | 1810043G02Rik | 1156.271975 | -0.009790103 | 0.060930992 | -0.160675256 | 0.872349179 | 0.97153269 |
| Prrc2b | 26769.60951 | -0.244244662 | 0.058784704 | -4.154901627 | 3.25E-05 | 0.000822541 | Grrp1 | 57.03749983 | -0.009800791 | 0.094548968 | -0.103658363 | 0.91744047 | 0.982742917 |
| Triap1 | 491.408948 | -0.244424523 | 0.083522344 | -2.926456681 | 0.003428472 | 0.027877541 | Ldb1 | 3951.470828 | -0.009820142 | 0.051986487 | -0.188897972 | 0.850172783 | 0.966260848 |
| Scoc | 4040.728539 | -0.244469075 | 0.088979296 | -2.747482698 | 0.006005468 | 0.040889423 | Pitx3 | 1.506925887 | -0.009826124 | 0.029800398 | -0.329731289 | 0.741603009 | NA |
| Irf2bpl | 2676.30094 | -0.245087378 | 0.08059155 | -3.041105136 | 0.002357115 | 0.021559115 | Derl2 | 902.739533 | -0.009842526 | 0.06196376 | -0.158843272 | 0.873792355 | 0.971912694 |
| Agap2 | 43934.69301 | -0.245474386 | 0.081628549 | -3.007212417 | 0.002636555 | 0.023198299 | Tube1 | 107.4196023 | -0.00984851 | 0.092366512 | -0.106624249 | 0.91508708 | 0.981724884 |
| Rpl13a | 8858.022542 | -0.245600197 | 0.072669462 | -3.379689202 | 0.000725678 | 0.0089238 | Nelfe | 769.0763749 | -0.009850195 | 0.066185682 | -0.14882667 | 0.881690405 | 0.97431988 |
| Nrxn3 | 8428.703657 | -0.245780785 | 0.062380551 | -3.940022681 | 8.15E-05 | 0.001676437 | Nsl1 | 68.51409678 | -0.009850857 | 0.094816183 | -0.103894262 | 0.917253261 | 0.982742917 |
| Mcee | 490.6396541 | -0.245871982 | 0.075255965 | -3.267142788 | 0.001086389 | 0.012179817 | Mir9-3 | 0.665410991 | -0.009853459 | 0.016079472 | -0.612797436 | 0.540010286 | NA |
| Xiap | 3660.133863 | -0.245992315 | 0.074158935 | -3.317096089 | 0.000909583 | 0.010547482 | Armc7 | 419.3541831 | -0.009876246 | 0.077604568 | -0.127263726 | 0.898731669 | 0.978597654 |
| Plekha2 | 1474.362606 | -0.246258442 | 0.0876375 | -2.809966516 | 0.004954665 | 0.0357391 | Duoxa2 | 0.465271493 | -0.009896549 | 0.014450844 | -0.684842264 | 0.493443462 | NA |
| Pdia6 | 3862.583827 | -0.24665017 | 0.081533212 | -3.02514966 | 0.002485101 | 0.022269612 | Ppp1r2-ps3 | 1.280040084 | -0.009925409 | 0.022991773 | -0.431693952 | 0.665963864 | NA |
| Panx1 | 577.5617808 | -0.247039215 | 0.066626155 | -3.707841404 | 0.000209033 | 0.003516574 | Pcyt2 | 2747.083534 | -0.009926236 | 0.0749709 | -0.13240118 | 0.894666981 | 0.977844987 |
| Mmd | 7643.273875 | -0.247054285 | 0.067546034 | -3.657569064 | 0.000254619 | 0.004092388 | Sstr4 | 320.2458093 | -0.009937931 | 0.089284673 | -0.111306128 | 0.911373596 | 0.980985114 |
| Klf7 | 350.4906268 | -0.247689514 | 0.086052751 | -2.878345086 | 0.003997676 | 0.031028284 | Ankrd39 | 315.1428888 | -0.009938659 | 0.075196732 | -0.132168759 | 0.894850811 | 0.977844987 |
| Mrps24 | 516.7276681 | -0.248268999 | 0.087252614 | -2.845404699 | 0.004435503 | 0.033344824 | Gpn2 | 265.8036472 | -0.009952867 | 0.079086178 | -0.125848378 | 0.899851946 | 0.97888996 |
| Herc1 | 10018.44547 | -0.248472068 | 0.074668033 | -3.327690028 | 0.000875692 | 0.010294369 | Phlpp1 | 4909.866873 | -0.009953735 | 0.063709832 | -0.156235457 | 0.875847429 | 0.972151437 |
| Neurl1b | 2294.944542 | -0.24870611 | 0.090379803 | -2.751788573 | 0.005927077 | 0.040455567 | Podnl1 | 1.355888624 | -0.009968346 | 0.025698976 | -0.387888851 | 0.698098289 | NA |
| Vstm2b | 980.1260943 | -0.249224802 | 0.066168002 | -3.766545677 | 0.000165522 | 0.002924282 | Fam193a | 2099.228464 | -0.009968367 | 0.046530589 | -0.214232558 | 0.830365708 | 0.960002081 |
| Hip1 | 2992.186094 | -0.249357203 | 0.066407034 | -3.754981785 | 0.000173354 | 0.003045293 | Pax4 | 0.766701302 | -0.009975334 | 0.020038351 | -0.497812123 | 0.61861647 | NA |
| Ncam2 | 2470.067898 | -0.249430728 | 0.083544124 | -2.985616664 | 0.002830072 | 0.024360628 | Aste1 | 211.3921407 | -0.009976502 | 0.08500585 | -0.117362535 | 0.906572771 | 0.980118791 |
| Nav3 | 2897.819276 | -0.250342557 | 0.058801309 | -4.257431713 | 2.07E-05 | 0.000573164 | Pdzrn3 | 1433.470527 | -0.009977087 | 0.084837304 | -0.11760259 | 0.906382552 | 0.980118791 |
| Bola2 | 553.6563987 | -0.250780031 | 0.079277396 | -3.163323269 | 0.00155979 | 0.015931042 | Gm5901 | 0.43199899 | -0.009981732 | 0.014172818 | -0.704287032 | 0.481254041 | NA |
| Hey1 | 1812.835411 | -0.250895648 | 0.07188967 | -3.490009733 | 0.000483003 | 0.006669918 | Asb6 | 1161.012773 | -0.009986298 | 0.073666389 | -0.135561124 | 0.892168241 | 0.977716776 |
| Tmem14c | 1014.624113 | -0.250994079 | 0.05597735 | -4.48385068 | 7.33E-06 | 0.000246462 | 9130019O22Rik | 146.647588 | -0.00999042 | 0.09268068 | -0.107793986 | 0.914159114 | 0.98170867 |
| Rasa2 | 1346.518983 | -0.25112485 | 0.093455384 | -2.687109507 | 0.007207332 | 0.046548729 | Aqp8 | 0.249110328 | -0.010000816 | 0.010509194 | -0.951625434 | 0.341286978 | NA |
| Fam162a | 708.043623 | -0.251302822 | 0.081052638 | -3.100489133 | 0.001932013 | 0.018662475 | Gm10485 | 0.249110328 | -0.010000816 | 0.010509194 | -0.951625434 | 0.341286978 | NA |
| Gpx1 | 1960.649955 | -0.251308366 | 0.067647743 | -3.714955649 | 0.000203239 | 0.003445285 | Hoxd3 | 0.249110328 | -0.010000816 | 0.010509194 | -0.951625434 | 0.341286978 | NA |
| Sar1b | 1243.417941 | -0.251488028 | 0.058417271 | -4.305028701 | 1.67E-05 | 0.00048354 | Pou1f1 | 0.249110328 | -0.010000816 | 0.010509194 | -0.951625434 | 0.341286978 | NA |
| Prex1 | 7537.498007 | -0.251518495 | 0.078509773 | -3.203658414 | 0.001356934 | 0.014321751 | Tmem245 | 1337.147633 | -0.0100075 | 0.072833227 | -0.137402953 | 0.890712297 | 0.977182599 |
| Coa3 | 1872.314449 | -0.251549445 | 0.085783011 | -2.932392359 | 0.003363615 | 0.027501768 | Stoml1 | 2452.489191 | -0.010019563 | 0.074054471 | -0.135299912 | 0.892374755 | 0.977844987 |
| Tbc1d9 | 5213.426491 | -0.251576425 | 0.05113548 | -4.919801768 | 8.66E-07 | 4.33E-05 | Apobec4 | 0.405586341 | -0.010021424 | 0.013940723 | -0.718859738 | 0.472227346 | NA |
| Stxbp6 | 2729.466581 | -0.252178537 | 0.09387931 | -2.686199295 | 0.007226997 | 0.046578655 | Cd8b1 | 0.405586341 | -0.010021424 | 0.013940723 | -0.718859738 | 0.472227346 | NA |

| | | | | | | |
|---|---|---|---|---|---|---|
| Nol4 | 3841.267403 | -0.252484477 | 0.074723268 | -3.378927099 | 0.000727693 | 0.008941218 |
| Rnf217 | 440.930607 | -0.253135429 | 0.091024327 | -2.780964572 | 0.005419765 | 0.037997213 |
| Pdgfrb | 1216.985982 | -0.253637801 | 0.069759776 | -3.635874637 | 0.000277039 | 0.00435339 |
| Tle4 | 2542.709594 | -0.253648161 | 0.077877458 | -3.257016432 | 0.001125899 | 0.012514119 |
| Cox6b1 | 5843.290998 | -0.25440655 | 0.06460162 | -3.938083146 | 8.21E-05 | 0.001687801 |
| Cspg4 | 818.8540317 | -0.254689724 | 0.090044956 | -2.828472971 | 0.004677065 | 0.034424195 |
| Pde7b | 3354.858462 | -0.254813122 | 0.090052516 | -2.829605802 | 0.004660539 | 0.034336846 |
| Ttc28 | 824.6842737 | -0.254936955 | 0.085615918 | -2.977681736 | 0.002904374 | 0.02477994 |
| Fam174a | 826.2524668 | -0.25520065 | 0.089240889 | -2.859682961 | 0.004240647 | 0.032353861 |
| Neto1 | 2877.765954 | -0.255318918 | 0.068808088 | -3.710594593 | 0.000206773 | 0.003489914 |
| Hmgn2 | 2596.161774 | -0.255446562 | 0.091156224 | -2.802294252 | 0.005074057 | 0.036290258 |
| Ephb4 | 280.9890224 | -0.255593617 | 0.087476461 | -2.921855952 | 0.003479524 | 0.028137656 |
| Psd3 | 22599.90361 | -0.255948166 | 0.084333952 | -3.034936207 | 0.002405866 | 0.021837424 |
| Immp1l | 466.715662 | -0.255978324 | 0.090016465 | -2.843683359 | 0.004459534 | 0.033425443 |
| Ctxn1 | 12894.94456 | -0.256038227 | 0.083861183 | -3.05311965 | 0.002264756 | 0.020961847 |
| Dnajb1 | 2399.650821 | -0.256133902 | 0.086340075 | -2.96657029 | 0.003011415 | 0.025407408 |
| Myo5a | 24699.48704 | -0.256323946 | 0.070488653 | -3.636385934 | 0.00027649 | 0.00434933 |
| Sec22a | 583.8708305 | -0.256368425 | 0.068155344 | -3.761530789 | 0.000168877 | 0.00297338 |
| Serpini1 | 10366.11932 | -0.256457561 | 0.079667928 | -3.219081593 | 0.001286019 | 0.01375126 |
| Pcnx | 6221.284507 | -0.25696324 | 0.086774327 | -2.961281836 | 0.003063615 | 0.025630477 |
| Snrpg | 524.2462803 | -0.257041309 | 0.089277647 | -2.87912279 | 0.00398783 | 0.031002229 |
| Cycs | 4581.692427 | -0.257106475 | 0.07550611 | -3.405108218 | 0.000661378 | 0.008331217 |
| Timm8b | 1875.278519 | -0.257265652 | 0.083685655 | -3.07419059 | 0.002110745 | 0.019904984 |
| Ehbp1l1 | 958.3001575 | -0.257320114 | 0.082407978 | -3.122514593 | 0.001793132 | 0.017606331 |
| Myo9a | 5171.231745 | -0.257835734 | 0.066035379 | -3.90450903 | 9.44E-05 | 0.001877708 |
| Mn1 | 1909.381199 | -0.257860008 | 0.079740182 | -3.233752428 | 0.001221753 | 0.013274785 |
| Tm2d1 | 637.6335583 | -0.258160068 | 0.094925485 | -2.71960757 | 0.006535943 | 0.043239925 |
| Atp8a1 | 11031.65112 | -0.258356426 | 0.059717567 | -4.32630526 | 1.52E-05 | 0.000445802 |
| Grin3a | 2151.726381 | -0.258530489 | 0.091621142 | -2.821733972 | 0.004776478 | 0.034908845 |
| Tomm70a | 8206.006034 | -0.25864699 | 0.082297445 | -3.142831334 | 0.001673222 | 0.016747355 |
| Atp2b4 | 13680.29093 | -0.259226925 | 0.075225176 | -3.446012875 | 0.000568923 | 0.007495661 |
| Echdc1 | 922.9944129 | -0.259822042 | 0.084354774 | -3.080110705 | 0.002069237 | 0.019597037 |
| Fem1b | 6727.994771 | -0.260497003 | 0.057362315 | -4.541256842 | 5.59E-06 | 0.000201026 |
| Map2k1 | 11514.50852 | -0.260840114 | 0.059022831 | -4.419308754 | 9.90E-06 | 0.000311191 |
| Purg | 511.0060247 | -0.260840317 | 0.090433025 | -2.88434802 | 0.00392225 | 0.030723633 |
| Fasn | 13324.23434 | -0.260847438 | 0.0532988 | -4.894058405 | 9.88E-07 | 4.83E-05 |
| Il34 | 2084.273178 | -0.261263587 | 0.081090389 | -3.221881061 | 0.00127352 | 0.013683744 |
| Grik2 | 1937.132698 | -0.261445858 | 0.086555275 | -3.020565269 | 0.002523033 | 0.022518362 |
| Tpbg | 587.2474673 | -0.26158311 | 0.079197772 | -3.302909966 | 0.000956871 | 0.010998338 |
| Stox2 | 6053.782691 | -0.261765874 | 0.074159431 | -3.529771863 | 0.000415918 | 0.005984147 |
| Gda | 10924.00403 | -0.261820251 | 0.093831009 | -2.790338218 | 0.005265301 | 0.037132712 |
| Chd7 | 598.8044073 | -0.261830215 | 0.088302556 | -2.965148754 | 0.003025366 | 0.025447894 |
| Lrrn3 | 2645.202435 | -0.261888786 | 0.083367887 | -3.141362879 | 0.001681635 | 0.016809844 |
| Grik3 | 5849.979912 | -0.261916832 | 0.069139414 | -3.788241998 | 0.000151717 | 0.002730202 |
| Rpl9 | 9690.250307 | -0.26200314 | 0.057817727 | -4.531536517 | 5.86E-06 | 0.000207139 |
| Garem | 1406.090467 | -0.262012659 | 0.077827482 | -3.36658277 | 0.000761057 | 0.00920517 |
| Nbea | 12146.71499 | -0.262750599 | 0.080444539 | -3.266232874 | 0.001089886 | 0.012203413 |
| Rplp1 | 11093.3193 | -0.26287478 | 0.070775833 | -3.714188445 | 0.000203857 | 0.003448207 |
| Hn1 | 2903.057988 | -0.263207859 | 0.086905359 | -3.028672347 | 0.002456309 | 0.02211392 |
| Gp1bb | 330.1562422 | -0.263473495 | 0.095208419 | -2.767334012 | 0.005651682 | 0.039197474 |
| Fam171a1 | 4272.369906 | -0.263495645 | 0.085218378 | -3.092004928 | 0.001988095 | 0.019061604 |
| Uqcrq | 2187.379794 | -0.263505744 | 0.070759396 | -3.723968239 | 0.000196116 | 0.003355592 |
| Lrp1 | 17897.30816 | -0.264025808 | 0.08267981 | -3.193352878 | 0.00140631 | 0.014710663 |
| Scn3b | 5980.761202 | -0.264448407 | 0.091769545 | -2.881657611 | 0.003955893 | 0.030869893 |
| Pde1b | 16534.44502 | -0.264751376 | 0.087267495 | -3.033791387 | 0.002415014 | 0.021882001 |
| Htt | 8429.087796 | -0.264822722 | 0.056627418 | -4.676581233 | 2.92E-06 | 0.000120521 |
| Cox4i1 | 13251.02491 | -0.265134656 | 0.061905519 | -4.282892063 | 1.84E-05 | 0.000521593 |
| Lamb1 | 819.589313 | -0.265606827 | 0.086763983 | -3.061256749 | 0.0022041 | 0.020597302 |
| Lsmd1 | 707.218225 | -0.265951943 | 0.065065234 | -4.087466144 | 4.36E-05 | 0.001026914 |
| Nf1 | 4761.694716 | -0.266125825 | 0.075926724 | -3.505035015 | 0.000456547 | 0.006390012 |
| Cox7c | 3237.854925 | -0.266505499 | 0.065933018 | -4.042064327 | 5.30E-05 | 0.001203686 |
| Thsd7a | 2015.923689 | -0.266567494 | 0.088155704 | -3.023825823 | 0.002496001 | 0.022341444 |
| Fbll1 | 840.1832984 | -0.266729676 | 0.070785028 | -3.768165136 | 0.000164452 | 0.00291202 |
| Flrt1 | 1367.906598 | -0.266965471 | 0.05924832 | -4.505874111 | 6.61E-06 | 0.000227591 |
| Tmem100 | 403.7641217 | -0.267300509 | 0.093938258 | -2.845491448 | 0.004434295 | 0.033344824 |
| Nat14 | 2137.580978 | -0.267308243 | 0.08989664 | -2.973506481 | 0.002944182 | 0.025050606 |
| 1700086L19Rik | 799.2459527 | -0.267366468 | 0.096639323 | -2.766642596 | 0.005663681 | 0.03924556 |
| Mpc2 | 1622.013928 | -0.267468917 | 0.087819195 | -3.045677171 | 0.00232157 | 0.021309478 |
| Amotl1 | 1803.631944 | -0.267545407 | 0.095518402 | -2.800982853 | 0.005094723 | 0.036290258 |
| Nrp1 | 1207.407396 | -0.267888144 | 0.095525195 | -2.804371611 | 0.005041476 | 0.036129802 |
| Wnk3 | 1689.833754 | -0.268498262 | 0.067873015 | -3.955891179 | 7.62E-05 | 0.001594353 |
| Tspan13 | 5489.225038 | -0.268533796 | 0.067844235 | -3.958093086 | 7.56E-05 | 0.001581864 |
| Kcnh7 | 1223.233201 | -0.268684675 | 0.090633627 | -2.964514207 | 0.003031612 | 0.025472778 |
| Gbas | 2749.445477 | -0.268878589 | 0.05943835 | -4.523655018 | 6.08E-06 | 0.000213063 |
| Rprml | 1218.404517 | -0.268911028 | 0.098726761 | -2.723790648 | 0.00645374 | 0.042939425 |
| Chd5 | 12626.58739 | -0.269067142 | 0.052298929 | -5.144792559 | 2.68E-07 | 1.68E-05 |
| Zfp41 | 286.9877087 | -0.269119506 | 0.094284481 | -2.854335121 | 0.004312701 | 0.032794894 |
| Plxna4 | 4845.4661 | -0.269289805 | 0.072736408 | -3.702269752 | 0.000213679 | 0.003556118 |
| BC030500 | 275.6457558 | -0.269583297 | 0.090476538 | -2.979593435 | 0.002886312 | 0.024666584 |
| Dlg2 | 18956.97038 | -0.269927555 | 0.054682573 | -4.936262913 | 7.96E-07 | 4.09E-05 |
| BC052040 | 305.307873 | -0.269987223 | 0.099613425 | -2.710349771 | 0.006721229 | 0.044107886 |
| Actn2 | 1104.620041 | -0.270039436 | 0.071646334 | -3.769061432 | 0.000163863 | 0.002908232 |
| H3f3a | 4003.980176 | -0.270058045 | 0.059104089 | -4.569193933 | 4.90E-06 | 0.000181916 |
| Srgap1 | 1201.124009 | -0.270475862 | 0.101242399 | -2.67156709 | 0.007549798 | 0.048039658 |
| 4-Mar | 1093.407799 | -0.270639012 | 0.064561249 | -4.191973002 | 2.77E-05 | 0.000723766 |
| Itgb1bp1 | 827.2080644 | -0.270676359 | 0.068571471 | -3.947361123 | 7.90E-05 | 0.001635816 |
| D430041D05Rik | 8895.851625 | -0.270948336 | 0.08348969 | -3.245290967 | 0.001173307 | 0.012845805 |
| Snx10 | 3560.434107 | -0.271057302 | 0.073967226 | -3.66455952 | 0.000247765 | 0.004011353 |
| Nhp2 | 826.8979944 | -0.271121549 | 0.074794609 | -3.624880912 | 0.000289095 | 0.004515355 |
| Ccnd2 | 2894.115089 | -0.271326709 | 0.090941152 | -2.98354158 | 0.002849333 | 0.024458487 |
| Bsn | 21922.82964 | -0.271897445 | 0.09446213 | -2.878375136 | 0.003997295 | 0.031028284 |
| Tubb5 | 17037.70134 | -0.271996295 | 0.064052323 | -4.246470398 | 2.17E-05 | 0.00059553 |
| Neurl1a | 8091.010445 | -0.272028996 | 0.087343759 | -3.114464041 | 0.001842794 | 0.017980008 |
| Phlda3 | 625.2975953 | -0.272084803 | 0.086806275 | -3.134390958 | 0.001722112 | 0.01711508 |
| Arl6 | 769.0131891 | -0.272318536 | 0.088580583 | -3.074246379 | 0.002110351 | 0.019904984 |
| Dand5 | 318.7387713 | -0.27239276 | 0.090532554 | -3.008782466 | 0.002622968 | 0.023183271 |
| Commd2 | 373.6835996 | -0.272597141 | 0.084520064 | -3.225235876 | 0.001258689 | 0.013590335 |
| Elmod1 | 10113.68959 | -0.273152044 | 0.06593415 | -4.142800745 | 3.43E-05 | 0.000849175 |
| Scd1 | 7879.33641 | -0.273542398 | 0.089668927 | -3.050581811 | 0.002283984 | 0.021078966 |
| Tmem150c | 1556.800868 | -0.273919926 | 0.072614557 | -3.772245347 | 0.000161785 | 0.002881262 |
| Cox7a2 | 4106.934733 | -0.273978707 | 0.072280504 | -3.790492456 | 0.000150349 | 0.002721919 |
| Btbd3 | 7288.471037 | -0.274244277 | 0.072549045 | -3.780122507 | 0.000156751 | 0.002801273 |
| Vhl | 2041.025207 | -0.274321206 | 0.067188994 | -4.082829508 | 4.45E-05 | 0.001041174 |
| Vps13c | 2727.000421 | -0.274972029 | 0.082233655 | -3.343789485 | 0.000826424 | 0.009834575 |
| Nfya | 1070.05673 | -0.275564693 | 0.075232807 | -3.662826158 | 0.000249448 | 0.004030182 |
| Nhsl2 | 2959.406824 | -0.275762643 | 0.102232764 | -2.697399868 | 0.00698833 | 0.045399238 |
| Sema5b | 883.0362923 | -0.275899436 | 0.093691818 | -2.944754861 | 0.003232106 | 0.026704603 |
| Arxes1 | 440.049255 | -0.276369411 | 0.0940584 | -2.938274618 | 0.003300445 | 0.027132771 |
| Tnfrsf19 | 2848.21442 | -0.276392381 | 0.085751347 | -3.223184139 | 0.00126774 | 0.013659217 |
| Ubxn2b | 1687.276205 | -0.276527401 | 0.061974402 | -4.461961597 | 8.12E-06 | 0.000266591 |
| Fzd3 | 5088.22638 | -0.277268296 | 0.081139209 | -3.417192489 | 0.000632705 | 0.008095075 |
| Lsm4 | 726.2675527 | -0.277768222 | 0.081626649 | -3.402911005 | 0.00066672 | 0.008378069 |
| Rnf150 | 4078.154573 | -0.277907678 | 0.077296743 | -3.595334903 | 0.000323975 | 0.00496014 |
| Tubb2a | 11406.40029 | -0.278208067 | 0.066191991 | -4.203047241 | 2.63E-05 | 0.000692746 |
| Myeov2 | 1382.767798 | -0.279163817 | 0.082894727 | -3.367690894 | 0.000758005 | 0.009189776 |
| Ttbk2 | 4499.553625 | -0.27924986 | 0.091303711 | -3.058472168 | 0.002224687 | 0.020702287 |
| Ets1 | 563.7972172 | -0.279800018 | 0.093075585 | -3.006159114 | 0.002645705 | 0.02323841 |
| Romo1 | 786.8249378 | -0.279890598 | 0.089226121 | -3.136868372 | 0.001707628 | 0.017014781 |
| Tuba1a | 24555.33983 | -0.280077973 | 0.085750789 | -3.266185367 | 0.001090069 | 0.012203413 |
| Ndufa1 | 1383.852938 | -0.280154 | 0.075745487 | -3.698622987 | 0.000216772 | 0.003599861 |
| Sc5d | 1706.085197 | -0.280380883 | 0.078461643 | -3.573477072 | 0.000352272 | 0.00528886 |
| Slc4a8 | 5343.381303 | -0.280504596 | 0.052758348 | -5.316781226 | 1.06E-07 | 7.79E-06 |
| Dpy19l3 | 2786.64691 | -0.280828977 | 0.066832451 | -4.201985292 | 2.65E-05 | 0.000694825 |
| Runx1t1 | 821.449698 | -0.281049249 | 0.087820928 | -3.200253689 | 0.001373067 | 0.014433037 |
| Gnb4 | 1289.234291 | -0.282679962 | 0.103197261 | -2.739219623 | 0.006158522 | 0.041559293 |
| Nwd1 | 3589.869655 | -0.283242048 | 0.064875998 | -4.365898867 | 1.27E-05 | 0.000388676 |
| Ntn3 | 382.665928 | -0.283600438 | 0.099587534 | -2.84775038 | 0.004402944 | 0.033182157 |
| Htr1a | 704.2859663 | -0.283939262 | 0.09963389 | -2.849826124 | 0.004374314 | 0.033045155 |
| Sec61b | 400.1800827 | -0.284066244 | 0.089296722 | -3.181149721 | 0.001466918 | 0.015192799 |
| Rragd | 7845.2649 | -0.284243947 | 0.061171156 | -4.64669896 | 3.37E-06 | 0.00013469 |
| Msantd3 | 375.6812436 | -0.284397132 | 0.088306674 | -3.220562136 | 0.001279395 | 0.01370881 |
| Fam84a | 2741.249843 | -0.284790976 | 0.077687553 | -3.665850775 | 0.000246518 | 0.003995339 |
| Hmgn1 | 1661.716221 | -0.284813726 | 0.083180704 | -3.424036014 | 0.000616984 | 0.007974295 |
| Enho | 1337.560229 | -0.285001367 | 0.084612906 | -3.368296655 | 0.000756342 | 0.009189519 |
| Med21 | 517.342109 | -0.285119741 | 0.08431539 | -3.381585981 | 0.000720687 | 0.008890383 |
| Gpr124 | 755.0101673 | -0.286131541 | 0.082866592 | -3.452917943 | 0.000554558 | 0.007375379 |
| Map1a | 62042.02897 | -0.28625253 | 0.071398196 | -4.009240389 | 6.09E-05 | 0.001338734 |
| Robo2 | 4828.820806 | -0.286324958 | 0.064580302 | -4.433626783 | 9.27E-06 | 0.000296018 |
| Homer1 | 6414.64479 | -0.28665955 | 0.107110646 | -2.676293721 | 0.007444137 | 0.047503899 |
| Erc1 | 2660.048639 | -0.288362174 | 0.055328716 | -5.21179953 | 1.87E-07 | 1.25E-05 |
| Med13 | 2820.827253 | -0.288401508 | 0.083728069 | -3.444502081 | 0.000572112 | 0.007518496 |
| Cdk18 | 1640.290027 | -0.288499536 | 0.100725651 | -2.86421118 | 0.004180492 | 0.032081497 |
| Fads1 | 9815.851253 | -0.288719931 | 0.088482758 | -3.26300781 | 0.001102365 | 0.012305329 |
| Wwtr1 | 667.5652034 | -0.288955195 | 0.095621116 | -3.021876402 | 0.002512131 | 0.022459872 |
| Ptar1 | 245.9913229 | -0.290153362 | 0.100678613 | -2.881976145 | 0.003951897 | 0.030863169 |
| 1110001J03Rik | 383.9704642 | -0.290169745 | 0.090193704 | -3.217184033 | 0.001294555 | 0.01382346 |
| Cd97 | 502.0610453 | -0.290193783 | 0.105721558 | -2.744887485 | 0.006053165 | 0.041042969 |
| Enox2 | 391.4511916 | -0.290667272 | 0.10600588 | -2.741991977 | 0.006106783 | 0.041300088 |
| Dpysl5 | 1463.99801 | -0.290968022 | 0.083004783 | -3.505436811 | 0.000455859 | 0.006386143 |
| Chml | 479.0348954 | -0.291022253 | 0.095916548 | -3.034119353 | 0.00241239 | 0.021882001 |
| Kcna6 | 5070.068811 | -0.291298993 | 0.099803571 | -2.918723155 | 0.003514682 | 0.028392328 |
| Slc32a1 | 4455.48704 | -0.291850313 | 0.094609159 | -3.08479977 | 0.002036892 | 0.019331788 |
| Tomm7 | 1365.140749 | -0.291889305 | 0.070329936 | -4.150285398 | 3.32E-05 | 0.000832517 |
| Lrrk2 | 3437.238212 | -0.291960367 | 0.088103951 | -3.313816938 | 0.000920317 | 0.010649289 |
| Lzic | 359.680208 | -0.292329302 | 0.076274447 | -3.832598117 | 0.000126797 | 0.002347184 |
| Mycbp2 | 15363.72493 | -0.292445904 | 0.056777002 | -5.150780981 | 2.59E-07 | 1.63E-05 |
| Ccdc90b | 411.1475481 | -0.292895391 | 0.10421604 | -2.810463639 | 0.004947018 | 0.035738209 |
| Atp5k | 2060.648747 | -0.293022495 | 0.090184517 | -3.249144162 | 0.001157528 | 0.012746794 |
| Phlda1 | 717.2971049 | -0.293340213 | 0.089662523 | -3.27160338 | 0.001069395 | 0.012015376 |
| Cnih2 | 5572.093156 | -0.293638488 | 0.075367597 | -3.896084015 | 9.78E-05 | 0.001924652 |
| Chac2 | 197.785737 | -0.293843382 | 0.09694021 | -3.031181611 | 0.002435987 | 0.02204625 |
| Calr | 12996.75824 | -0.29406075 | 0.08932617 | -3.291988775 | 0.000994816 | 0.011333586 |
| Arhgef19 | 604.0598838 | -0.294433515 | 0.108094458 | -2.723853937 | 0.006452504 | 0.042939425 |
| Arhgap32 | 20321.43569 | -0.295181582 | 0.074326588 | -3.971413036 | 7.14E-05 | 0.001514376 |
| Itga1 | 320.3860742 | -0.295300062 | 0.105415343 | -2.801300594 | 0.005089709 | 0.036290258 |
| St6galnac5 | 1318.961732 | -0.296393559 | 0.08324336 | -3.560566993 | 0.000370055 | 0.00550781 |
| Tnr | 3651.991982 | -0.296721095 | 0.088129551 | -3.366874017 | 0.000760254 | 0.009204125 |
| Arhgap31 | 1703.422183 | -0.296842791 | 0.07865307 | -3.774077606 | 0.000160601 | 0.002863462 |
| Ndufb6 | 1608.890521 | -0.296903255 | 0.054284398 | -5.46940312 | 4.52E-08 | 3.78E-06 |
| Sema6a | 1194.054276 | -0.297070256 | 0.085246547 | -3.484836253 | 0.000492438 | 0.00675805 |
| Brk1 | 5480.3681 | -0.297463365 | 0.063001046 | -4.72156232 | 2.34E-06 | 0.000101291 |
| Ston2 | 1065.013565 | -0.297793201 | 0.079589097 | -3.741633129 | 0.000182828 | 0.003186436 |
| Erbb4 | 1115.74852 | -0.298877352 | 0.091519954 | -3.265706976 | 0.001091912 | 0.012215221 |
| Nfia | 1208.783073 | -0.299005118 | 0.080940377 | -3.694140421 | 0.000220632 | 0.00363667 |
| Utrn | 3640.423923 | -0.299095429 | 0.054907726 | -5.44723756 | 5.12E-08 | 4.24E-06 |
| Acvr1c | 1099.384348 | -0.299316444 | 0.095362582 | -3.138720022 | 0.001696875 | 0.016940322 |
| Uqcr10 | 2188.451836 | -0.299482841 | 0.066999488 | -4.469927305 | 7.82E-06 | 0.000259603 |
| Prex2 | 2795.441201 | -0.300043865 | 0.063052733 | -4.758617939 | 1.95E-06 | 8.68E-05 |
| Syne2 | 1000.398004 | -0.300064439 | 0.087426316 | -3.432198137 | 0.00059871 | 0.007801861 |
| Adra1a | 424.2902146 | -0.300085241 | 0.109953862 | -2.729192358 | 0.006348966 | 0.042474471 |
| Ddn | 44170.29246 | -0.300313971 | 0.084932897 | -3.535896948 | 0.000406393 | 0.005884725 |
| Idh1 | 1481.005645 | -0.301045146 | 0.076280775 | -3.946540226 | 7.93E-05 | 0.001635816 |
| Eif2c2 | 2985.480442 | -0.301175767 | 0.101056707 | -2.980265008 | 0.002879991 | 0.024639747 |
| Setbp1 | 1681.934272 | -0.301815316 | 0.075413627 | -4.002132352 | 6.28E-05 | 0.001369891 |
| Ppapdc2 | 2006.220703 | -0.303022204 | 0.111847721 | -2.709238952 | 0.006743775 | 0.044184405 |
| Cacng4 | 490.6135266 | -0.303105591 | 0.090510684 | -3.348837687 | 0.000811513 | 0.009692629 |
| Churc1 | 671.2397872 | -0.303132204 | 0.072003783 | -4.209948281 | 2.55E-05 | 0.000678356 |
| Pgm2l1 | 13559.50154 | -0.303375422 | 0.097205401 | -3.120972896 | 0.001802546 | 0.017687556 |
| Tyro3 | 10902.4078 | -0.30356858 | 0.085064771 | -3.568675673 | 0.00035879 | 0.005365923 |
| Spon1 | 3364.181049 | -0.303576995 | 0.078045351 | -3.88975115 | 0.000100347 | 0.001968073 |
| Tiam2 | 2926.008248 | -0.303600583 | 0.087409564 | -3.473310814 | 0.00051408 | 0.006962544 |
| Anpep | 202.6453715 | -0.303631148 | 0.102274403 | -2.968789265 | 0.002989756 | 0.025311139 |
| Prrx1 | 259.6911978 | -0.303673875 | 0.100739229 | -3.014455029 | 0.002574414 | 0.02287154 |
| Gm15800 | 13692.64898 | -0.303677341 | 0.077360694 | -3.925473312 | 8.66E-05 | 0.00176004 |
| Rgs9 | 5036.503318 | -0.303951116 | 0.099785276 | -3.047783588 | 0.002305359 | 0.021185783 |
| Mcm5 | 254.5995703 | -0.30434921 | 0.113625859 | -2.678520663 | 0.007394817 | 0.047350844 |
| Chd3 | 25957.77394 | -0.304549194 | 0.061972413 | -4.914270364 | 8.91E-07 | 4.41E-05 |
| Prkcdbp | 472.2286655 | -0.304789298 | 0.101586772 | -3.000285288 | 0.002697268 | 0.023561109 |
| Ehd2 | 341.660789 | -0.304991899 | 0.08166583 | -3.734632941 | 0.000187989 | 0.003250786 |
| Unc80 | 13103.75364 | -0.305116879 | 0.067731317 | -4.504812452 | 6.64E-06 | 0.000228224 |
| Wfdc1 | 211.4390817 | -0.305144084 | 0.09943757 | -3.068700118 | 0.002149923 | 0.02020067 |
| Slc16a2 | 1908.756794 | -0.305210956 | 0.064583512 | -4.725833999 | 2.29E-06 | 0.000100023 |
| Wdfy3 | 9320.76383 | -0.306152062 | 0.095253058 | -3.214091698 | 0.001308579 | 0.013906117 |
| Zswim6 | 1342.007494 | -0.306214515 | 0.076478672 | -4.003920394 | 6.23E-05 | 0.001364151 |
| Ptprr | 773.5433013 | -0.306288759 | 0.088890716 | -3.44567771 | 0.000569629 | 0.007498586 |
| Fam124a | 1172.750571 | -0.307017094 | 0.09285312 | -3.306481179 | 0.000944757 | 0.01086716 |
| Birc6 | 5240.795754 | -0.307880226 | 0.068356539 | -4.504034756 | 6.67E-06 | 0.000228302 |
| Acyp2 | 354.3342021 | -0.308101024 | 0.085913691 | -3.58616911 | 0.000335571 | 0.005117464 |
| Ptn | 7885.585572 | -0.308298955 | 0.095306106 | -3.234828994 | 0.001217156 | 0.01324341 |
| Ncrna00086 | 942.476719 | -0.308533352 | 0.063013507 | -4.896305015 | 9.77E-07 | 4.79E-05 |
| Uqcrh | 4551.083169 | -0.308537056 | 0.071623659 | -4.307753362 | 1.65E-05 | 0.000478982 |
| Zfp462 | 2122.452167 | -0.3085816 | 0.091969098 | -3.355274843 | 0.000792862 | 0.00950085 |
| Hspa5 | 11307.71383 | -0.308810107 | 0.090935049 | -3.395941502 | 0.00068393 | 0.00855271 |
| Ehhadh | 166.7226753 | -0.309804734 | 0.102814102 | -3.013251376 | 0.002584647 | 0.022923025 |
| Nanos1 | 1163.764078 | -0.310231298 | 0.077104969 | -4.023492907 | 5.73E-05 | 0.001283882 |
| Mageh1 | 887.2726559 | -0.31049193 | 0.061589005 | -5.041353265 | 4.62E-07 | 2.62E-05 |
| Saysd1 | 197.8495034 | -0.311639463 | 0.096236384 | -3.238270692 | 0.001202566 | 0.013112291 |
| Mrpl54 | 467.531179 | -0.311910518 | 0.095653665 | -3.260831851 | 0.001110859 | 0.012373579 |
| Dclk3 | 2929.17992 | -0.313456316 | 0.095372552 | -3.286651232 | 0.001013863 | 0.011471264 |
| Gfra1 | 1802.792151 | -0.313592962 | 0.09727164 | -3.223888925 | 0.001264624 | 0.013635415 |
| Bmpr1b | 995.1696079 | -0.314259492 | 0.067679555 | -4.643344534 | 3.43E-06 | 0.000136544 |
| Hspa12b | 317.0013868 | -0.314429983 | 0.117948008 | -2.665835459 | 0.007679727 | 0.048706382 |
| D3Bwg0562e | 9157.264238 | -0.314432927 | 0.072080833 | -4.362226601 | 1.29E-05 | 0.000393448 |
| Gap43 | 8199.855136 | -0.314654271 | 0.088533242 | -3.554080516 | 0.000379303 | 0.005618472 |
| Rgs20 | 1447.0646 | -0.314922014 | 0.098659601 | -3.192005745 | 0.001412885 | 0.014761459 |
| Epha7 | 2519.553871 | -0.315456576 | 0.094767146 | -3.328754631 | 0.000872352 | 0.010270688 |
| Vps13d | 5553.789079 | -0.315506096 | 0.070727663 | -4.460858505 | 8.16E-06 | 0.000266837 |
| Gm561 | 324.0118655 | -0.316575242 | 0.087693079 | -3.610036774 | 0.000306154 | 0.004743544 |
| Sh3rf1 | 1589.979885 | -0.316720585 | 0.068587584 | -4.617753944 | 3.88E-06 | 0.000149512 |
| 1810037I17Rik | 1330.502723 | -0.316759182 | 0.062926593 | -5.033788823 | 4.81E-07 | 2.69E-05 |
| Snx7 | 565.7914671 | -0.316998941 | 0.079621006 | -3.981348086 | 6.85E-05 | 0.001468323 |
| Sqle | 2287.280488 | -0.317297592 | 0.111352659 | -2.849483754 | 0.004379024 | 0.03306462 |
| Cetn2 | 1116.672825 | -0.317396487 | 0.094241256 | -3.367914437 | 0.000757391 | 0.009189519 |
| Abcb10 | 1224.746943 | -0.317413339 | 0.07792678 | -4.073225365 | 4.64E-05 | 0.001077066 |
| Tmem121 | 383.1719074 | -0.317529577 | 0.101484475 | -3.128848775 | 0.001754926 | 0.01736241 |

| | | | | | | |
|---|---|---|---|---|---|---|
| 4931440P22Rik | 9.681013394 | -0.010026691 | 0.065477401 | -0.153132083 | 0.878294115 | NA |
| Rasal2 | 4602.60228 | -0.010037637 | 0.065119972 | -0.154140682 | 0.877498812 | 0.972570049 |
| A730020E08Rik | 58.51039697 | -0.0100455 | 0.093077105 | -0.107926645 | 0.914053881 | 0.98170867 |
| 1700125G02Rik | 0.429381296 | -0.010052491 | 0.01415197 | -0.710324479 | 0.477502943 | NA |
| Chst13 | 0.429381296 | -0.010052491 | 0.01415197 | -0.710324479 | 0.477502943 | NA |
| Fra10ac1 | 480.2927714 | -0.010059325 | 0.077939645 | -0.129065574 | 0.897305762 | 0.978039389 |
| Mterfd3 | 526.4987803 | -0.010060709 | 0.071974972 | -0.139780656 | 0.888833297 | 0.977158289 |
| Aurka | 47.22963877 | -0.010072338 | 0.09289866 | -0.108422859 | 0.913660269 | 0.98170867 |
| Gm19466 | 0.648160155 | -0.010074984 | 0.017431037 | -0.577991085 | 0.563270138 | NA |
| AU015228 | 0.360627147 | -0.010077206 | 0.010553896 | -0.954832741 | 0.339662287 | NA |
| Gdf3 | 0.360627147 | -0.010077206 | 0.010553896 | -0.954832741 | 0.339662287 | NA |
| Gm6607 | 0.360627147 | -0.010077206 | 0.010553896 | -0.954832741 | 0.339662287 | NA |
| Mir693 | 0.360627147 | -0.010077206 | 0.010553896 | -0.954832741 | 0.339662287 | NA |
| Otor | 0.360627147 | -0.010077206 | 0.010553896 | -0.954832741 | 0.339662287 | NA |
| Sftpa1 | 0.360627147 | -0.010077206 | 0.010553896 | -0.954832741 | 0.339662287 | NA |
| 4930568G15Rik | 1.578783041 | -0.010078809 | 0.030010246 | -0.335845602 | 0.7369873 | NA |
| Mcoln3 | 0.84596859 | -0.010082847 | 0.018664555 | -0.540213609 | 0.589049728 | NA |
| Pck1 | 0.800526829 | -0.010088762 | 0.014650584 | -0.68862527 | 0.491059115 | NA |
| Kdsr | 1443.447199 | -0.010097638 | 0.057056573 | -0.17697589 | 0.859527328 | 0.968543477 |
| Ncoa6 | 3654.581071 | -0.010099998 | 0.055343733 | -0.182495782 | 0.855193664 | 0.967189923 |
| Tal1 | 116.0591691 | -0.010110581 | 0.094692825 | -0.106772406 | 0.914969539 | 0.98170867 |
| Nrf1 | 662.7632218 | -0.0101121 | 0.072658173 | -0.139173605 | 0.889312966 | 0.977182599 |
| Agr2 | 0.410265417 | -0.010134484 | 0.013985708 | -0.72463146 | 0.468678154 | NA |
| Pydc4 | 0.410265417 | -0.010134484 | 0.013985708 | -0.72463146 | 0.468678154 | NA |
| Btbd6 | 2015.053841 | -0.010137435 | 0.05704709 | -0.177702924 | 0.858956289 | 0.968425881 |
| Hid1 | 5099.097141 | -0.010152643 | 0.052839306 | -0.192141864 | 0.847631083 | 0.965398237 |
| 4930594C11Rik | 0.617836846 | -0.010160213 | 0.01726803 | -0.588382891 | 0.556275319 | NA |
| 4921525O09Rik | 8.890816197 | -0.010162648 | 0.060215949 | -0.168770038 | 0.865977526 | NA |
| Snora21 | 0.776305083 | -0.010185649 | 0.021437846 | -0.475124623 | 0.634698149 | NA |
| Dytn | 3.982354247 | -0.010202063 | 0.04610121 | -0.221297065 | 0.824861135 | NA |
| Lrrc63 | 0.74940253 | -0.010206386 | 0.021329671 | -0.478506504 | 0.632289746 | NA |
| Coq2 | 2387.020376 | -0.01021391 | 0.04526949 | -0.225624582 | 0.82149342 | 0.957699646 |
| Gm14124 | 5.134910273 | -0.010219934 | 0.039528505 | -0.258545935 | 0.795985603 | NA |
| Med6 | 538.6247171 | -0.010234741 | 0.074513216 | -0.137354712 | 0.890750426 | 0.977182599 |
| Msmp | 1.001665186 | -0.010250392 | 0.023047495 | -0.444750816 | 0.656499841 | NA |
| 4930404I05Rik | 0.441671028 | -0.010255466 | 0.014264378 | -0.718956372 | 0.472167801 | NA |
| Macrod1 | 329.864913 | -0.010256288 | 0.090415443 | -0.11343514 | 0.909685582 | 0.980590355 |
| Bcl2a1d | 1.238160007 | -0.010264062 | 0.024905792 | -0.412115445 | 0.680254809 | NA |
| 4933402D24Rik | 9.614585055 | -0.010271624 | 0.061085794 | -0.168150775 | 0.866464665 | NA |
| Ppp1r3d | 333.7994168 | -0.010271957 | 0.074818231 | -0.137292165 | 0.890799863 | 0.977182599 |
| Gm10857 | 1.053336509 | -0.010290602 | 0.021669691 | -0.474884566 | 0.634869253 | NA |
| P2ry12 | 1197.301845 | -0.010292396 | 0.069914285 | -0.147214493 | 0.882962722 | 0.97494492 |
| Slc16a5 | 0.612759861 | -0.010298073 | 0.017242756 | -0.597240532 | 0.550346802 | NA |
| Mansc1 | 521.5091614 | -0.010298185 | 0.079976714 | -0.128764796 | 0.897543762 | 0.978039389 |
| Pnma5 | 6.092390332 | -0.010302884 | 0.054158512 | -0.190235726 | 0.849124417 | NA |
| Cox7a1 | 44.96097332 | -0.010305488 | 0.089793227 | -0.114769101 | 0.908628141 | 0.980523965 |
| Ms4a4c | 2.845534898 | -0.010312631 | 0.030238022 | -0.341048455 | 0.733067104 | NA |
| Scaf1 | 7342.311376 | -0.010313906 | 0.057503106 | -0.179362594 | 0.857653001 | 0.967971163 |
| Hist3h2bb-ps | 5.18475471 | -0.010319394 | 0.044200276 | -0.233469 | 0.815397239 | NA |
| Gm19434 | 1.26643968 | -0.010327594 | 0.02494206 | -0.414063379 | 0.678827691 | NA |
| Mgmt | 31.6669326 | -0.01033024 | 0.090402637 | -0.114269237 | 0.909024368 | 0.980523965 |
| B4galnt1 | 5780.755611 | -0.010345692 | 0.056802361 | -0.182134897 | 0.855476863 | 0.967189923 |
| Ifi205 | 0.418957648 | -0.010352899 | 0.014069042 | -0.73586382 | 0.461813577 | NA |
| Prr3 | 1084.5584 | -0.010355142 | 0.069517188 | -0.148958006 | 0.88158677 | 0.974279475 |
| Sema4f | 2698.173373 | -0.010356921 | 0.067412413 | -0.153635228 | 0.877897359 | 0.972792511 |
| 4930513D17Rik | 0.910332818 | -0.01035922 | 0.019241962 | -0.538366111 | 0.590324318 | NA |
| Upf3a | 1194.792755 | -0.010367729 | 0.056044123 | -0.184992266 | 0.853235102 | 0.966905493 |
| Kcnj15 | 3.032868895 | -0.010374757 | 0.039422697 | -0.263167111 | 0.792421779 | NA |
| Tceb1 | 1954.558891 | -0.010383897 | 0.073038454 | -0.142170272 | 0.886945513 | 0.976561095 |
| Gtpbp8 | 394.891646 | -0.010384563 | 0.078042598 | -0.133062754 | 0.894143751 | 0.977844987 |
| Gtf2e1 | 715.0286773 | -0.010400345 | 0.061110213 | -0.170189959 | 0.864860748 | 0.970038393 |
| 5730412P04Rik | 1.38199105 | -0.010409781 | 0.025924073 | -0.40154881 | 0.688016109 | NA |
| Acoxl | 1.027057486 | -0.010436001 | 0.023114722 | -0.451487201 | 0.651638447 | NA |
| Atf7ip | 2559.128372 | -0.010438012 | 0.063036699 | -0.165586269 | 0.868482554 | 0.970658528 |
| Wdr20a | 625.2335653 | -0.010439567 | 0.064252632 | -0.16247687 | 0.870930341 | 0.97153269 |
| Cxcl14 | 3390.507432 | -0.010448506 | 0.053760382 | -0.194353266 | 0.845899281 | 0.964996028 |
| Gm5069 | 214.5950139 | -0.010457045 | 0.091235144 | -0.114616415 | 0.908749168 | 0.980523965 |
| Ncdn | 65284.40875 | -0.010459797 | 0.065109141 | -0.160650202 | 0.872368913 | 0.97153269 |
| Coa5 | 4942.837494 | -0.010487401 | 0.056627357 | -0.185200249 | 0.853071974 | 0.966905493 |
| Alas1 | 2452.107328 | -0.010505426 | 0.053743168 | -0.195474638 | 0.845021391 | 0.964467724 |
| Hspb1 | 188.5256052 | -0.0105378 | 0.093514239 | -0.112686587 | 0.910279036 | 0.980590355 |
| Fam102a | 4740.45706 | -0.01054523 | 0.045084434 | -0.233899572 | 0.815062945 | 0.956264994 |
| 4933430M04Rik | 2.325797149 | -0.010545762 | 0.032922984 | -0.320316113 | 0.74872871 | NA |
| Spib | 0.826632487 | -0.010565712 | 0.018979127 | -0.556701679 | 0.57773127 | NA |
| Nr1h4 | 3.319054382 | -0.010568121 | 0.043245644 | -0.244374232 | 0.806940997 | NA |
| 2310039H08Rik | 68.30290355 | -0.01056905 | 0.095101607 | -0.111134297 | 0.911509852 | 0.980985114 |
| Fam89a | 29.29906413 | -0.010571839 | 0.085572759 | -0.123542108 | 0.901677832 | 0.979150581 |
| Spata25 | 2.732054541 | -0.010575298 | 0.034336273 | -0.307992015 | 0.758088408 | NA |
| Ldhal6b | 6.757425588 | -0.010588529 | 0.058082609 | -0.182301188 | 0.855346366 | NA |
| Gm4791 | 0.455532238 | -0.01060444 | 0.014387987 | -0.737034302 | 0.46110149 | NA |
| Gm3776 | 0.386703839 | -0.01061026 | 0.010827927 | -0.979897622 | 0.327136657 | NA |
| Krt10 | 222.8525473 | -0.010620803 | 0.086633456 | -0.122594705 | 0.902428047 | 0.979150581 |
| Snhg10 | 46.33150997 | -0.010624993 | 0.092785599 | -0.114511233 | 0.908832542 | 0.980523965 |
| AY512915 | 0.428629686 | -0.010629268 | 0.01416351 | -0.750468509 | 0.452972583 | NA |
| Tpm2 | 384.3063759 | -0.010630671 | 0.077462011 | -0.137237224 | 0.890843288 | 0.977182599 |
| Serpina1b | 0.468811549 | -0.010648514 | 0.014494226 | -0.734672787 | 0.462538797 | NA |
| Obox5 | 0.452914545 | -0.010676446 | 0.014368427 | -0.743049088 | 0.457451962 | NA |
| Tulp1 | 2.148672343 | -0.010719202 | 0.032776265 | -0.327041602 | 0.74363642 | NA |
| Gm16973 | 188.0957953 | -0.010736141 | 0.084771 | -0.126648746 | 0.899218414 | 0.978708077 |
| Krtcap2 | 784.6708661 | -0.010776765 | 0.063608835 | -0.169422464 | 0.865464355 | 0.970182416 |
| Mir2861 | 0.856803614 | -0.010795565 | 0.014829164 | -0.727995504 | 0.466616339 | NA |
| Ddx43 | 6.361504066 | -0.0108055 | 0.053211676 | -0.203066343 | 0.839083178 | NA |
| D830046C22Rik | 1.849948845 | -0.010826597 | 0.022191474 | -0.487871903 | 0.625640577 | NA |
| Mir24-2 | 1.008052884 | -0.010839326 | 0.02342105 | -0.462802749 | 0.643505761 | NA |
| Il10 | 0.388662817 | -0.010856415 | 0.010952562 | -0.99122153 | 0.32157742 | NA |
| Wdr53 | 331.9328779 | -0.0108569 | 0.078112128 | -0.138991225 | 0.889457084 | 0.977182599 |
| D930020B18Rik | 16.84389861 | -0.010880756 | 0.073740034 | -0.147555613 | 0.882693487 | NA |
| E030024N20Rik | 0.905180643 | -0.010883151 | 0.019238219 | -0.565704675 | 0.571594556 | NA |
| Rps10 | 4086.260649 | -0.010905049 | 0.065565312 | -0.16632345 | 0.867902413 | 0.970480832 |
| Mir337 | 1.809274879 | -0.010906935 | 0.027786292 | -0.392529336 | 0.69466714 | NA |
| Srrd | 566.4964554 | -0.01091725 | 0.06894087 | -0.158356722 | 0.874175714 | 0.971912694 |
| Slco1a5 | 8.12752002 | -0.010947728 | 0.035177864 | -0.311210702 | 0.755640448 | NA |
| Dopey2 | 4705.202021 | -0.010960985 | 0.042053686 | -0.260642666 | 0.794368084 | 0.951979493 |
| F420014N23Rik | 11.88504573 | -0.010961361 | 0.071230393 | -0.153886011 | 0.877699614 | NA |
| Abhd2 | 2695.287489 | -0.010968309 | 0.073591549 | -0.149043044 | 0.881519668 | 0.974279441 |
| Piezo1 | 378.8623929 | -0.010976521 | 0.089415016 | -0.122759261 | 0.902297735 | 0.979150581 |
| Samm50 | 3210.781337 | -0.010993845 | 0.050431355 | -0.217996227 | 0.827432051 | 0.958657209 |
| Nfyb | 1627.295796 | -0.010995623 | 0.062196043 | -0.176789746 | 0.859673544 | 0.96862798 |
| Mnf1 | 1631.707752 | -0.011006116 | 0.053657881 | -0.205116488 | 0.837481114 | 0.961831963 |
| Slc16a2 | 1878.895627 | -0.011017619 | 0.075965113 | -0.145035249 | 0.884683042 | 0.975676135 |
| 4933413G19Rik | 58.43830025 | -0.011056431 | 0.094075826 | -0.117526801 | 0.906442607 | 0.980118791 |
| Tesc | 1388.594384 | -0.01106397 | 0.079065694 | -0.139933893 | 0.888712222 | 0.977140573 |
| Wdr73 | 1211.433357 | -0.011064169 | 0.062983644 | -0.175667337 | 0.860555297 | 0.96862798 |
| Cmtm7 | 83.70242574 | -0.011082822 | 0.094644739 | -0.117099184 | 0.906781456 | 0.980201396 |
| Fam173b | 213.1752957 | -0.011098604 | 0.084029929 | -0.132079178 | 0.894921665 | 0.977844987 |
| Emilin3 | 8.02964709 | -0.011105547 | 0.050363604 | -0.220507403 | 0.825476007 | NA |
| Mpg | 458.6849364 | -0.011107418 | 0.084290006 | -0.131776213 | 0.895161302 | 0.977844987 |
| Armc3 | 39.34785784 | -0.011113994 | 0.08403649 | -0.132252 | 0.894784972 | 0.977844987 |
| Pik3r2 | 5688.676322 | -0.011120225 | 0.060921785 | -0.182532813 | 0.855164606 | 0.967189923 |
| Aknad1 | 0.450693223 | -0.011164073 | 0.014361702 | -0.77735029 | 0.436952132 | NA |
| Fuz | 526.351661 | -0.011170154 | 0.081261487 | -0.137459381 | 0.890667697 | 0.977182599 |
| Pnrc1 | 1089.528457 | -0.011175966 | 0.078674985 | -0.14205234 | 0.887038664 | 0.976589634 |
| Nkx6-1 | 3.397899967 | -0.011180986 | 0.036887788 | -0.30310806 | 0.761807512 | NA |
| A530013C23Rik | 2.257348453 | -0.011195082 | 0.029646408 | -0.37762018 | 0.705712771 | NA |
| Gm20362 | 1.793146381 | -0.011198998 | 0.032159961 | -0.348227963 | 0.727668989 | NA |
| Car4 | 1282.607159 | -0.01120698 | 0.059464793 | -0.188464115 | 0.850512844 | 0.966301569 |
| Acot12 | 10.04418092 | -0.01120991 | 0.063632511 | -0.176166392 | 0.860163223 | NA |
| 4930506C21Rik | 52.307608 | -0.011213165 | 0.094013386 | -0.119272004 | 0.905059863 | 0.980118791 |
| Ms4a8a | 1.04183065 | -0.011214262 | 0.021368766 | -0.524796909 | 0.59972438 | NA |
| 4930452A19Rik | 0.491854894 | -0.011220513 | 0.014685949 | -0.764030456 | 0.444849093 | NA |
| Mdm2 | 1799.722896 | -0.011224265 | 0.066468569 | -0.168865753 | 0.865902238 | 0.970182416 |
| Bin1 | 10429.83477 | -0.011230913 | 0.056966803 | -0.197148391 | 0.843711416 | 0.964060034 |
| Ddi2 | 529.5747245 | -0.011270205 | 0.077958222 | -0.144567234 | 0.885052569 | 0.975769681 |
| Mgll | 10334.47094 | -0.011291833 | 0.050959241 | -0.221585576 | 0.824636512 | 0.957903916 |
| Smim4 | 111.7624851 | -0.011314199 | 0.095100058 | -0.118971531 | 0.905297911 | 0.980118791 |
| Xpo6 | 4399.266636 | -0.011326281 | 0.04493665 | -0.252049967 | 0.801002442 | 0.953228631 |
| 4921530L21Rik | 1.996111418 | -0.011326426 | 0.026069353 | -0.434472855 | 0.663945103 | NA |
| Adnp2 | 1231.28136 | -0.011358981 | 0.059967203 | -0.189419882 | 0.849763743 | 0.966260848 |
| Mastl | 48.39117928 | -0.011372097 | 0.09391427 | -0.121090193 | 0.903619597 | 0.979550617 |
| Capn7 | 2008.785801 | -0.011372413 | 0.05646509 | -0.201406087 | 0.840381056 | 0.962802683 |
| Shank2 | 6273.009678 | -0.011375725 | 0.070300126 | -0.16181657 | 0.871450304 | 0.97153269 |
| Rgs7 | 3681.694182 | -0.011395466 | 0.065568649 | -0.173794431 | 0.862027021 | 0.969505811 |
| Tspyl2 | 4704.129776 | -0.011395971 | 0.075242603 | -0.151456363 | 0.879615731 | 0.973413768 |
| 1700121N20Rik | 0.459875358 | -0.011402357 | 0.014443087 | -0.789468159 | 0.429838433 | NA |
| Abca12 | 9.920506747 | -0.011415652 | 0.060759546 | -0.187882446 | 0.850968804 | NA |
| Tmem215 | 639.278744 | -0.011416071 | 0.090016092 | -0.126822556 | 0.899080843 | 0.978647308 |
| B4galnt3 | 39.66004695 | -0.011423595 | 0.08693764 | -0.13139988 | 0.895458984 | 0.977844987 |
| Hsd17b1 | 9.165575409 | -0.011424453 | 0.045702421 | -0.249974787 | 0.802606847 | NA |
| Tshz3 | 1223.705709 | -0.011431923 | 0.065221774 | -0.175277713 | 0.860861423 | 0.968844273 |
| Krt16 | 0.794999316 | -0.011434233 | 0.021519725 | -0.531337307 | 0.595185057 | NA |
| Gm1604b | 12.90969182 | -0.011447599 | 0.072846202 | -0.157147511 | 0.875128595 | NA |
| Asxl1 | 2805.421772 | -0.011461479 | 0.057266299 | -0.200143518 | 0.841368339 | 0.963098227 |
| 4933411E08Rik | 1.001886351 | -0.011463353 | 0.024688228 | -0.464324677 | 0.642415146 | NA |
| Dhh | 6.653340462 | -0.011464582 | 0.057911241 | -0.197968168 | 0.843069969 | NA |
| 8030423J24Rik | 1.830766549 | -0.011465801 | 0.030118161 | -0.380693919 | 0.703430382 | NA |
| Ppp3cc | 819.6609617 | -0.01146588 | 0.064137653 | -0.178769865 | 0.858118409 | 0.967971163 |
| Ssrp1 | 4258.250854 | -0.011468836 | 0.041920984 | -0.273582236 | 0.784405687 | 0.949699053 |
| Mettl7a3 | 4.393208858 | -0.011475014 | 0.046581039 | -0.246345182 | 0.805415037 | NA |
| Kdelc2 | 405.2574782 | -0.011478969 | 0.080968406 | -0.141770964 | 0.88726092 | 0.97661227 |
| Dnajc22 | 1.00246825 | -0.011485315 | 0.024348821 | -0.471699021 | 0.637141636 | NA |
| 4930515L03Rik | 0.700811285 | -0.011487447 | 0.017721731 | -0.648212464 | 0.516847541 | NA |
| Gpr137b | 432.2786248 | -0.011489356 | 0.071889751 | -0.159819104 | 0.873023575 | 0.971691093 |
| Npy1r | 1069.653817 | -0.011492907 | 0.066004673 | -0.174122633 | 0.861769086 | 0.969358561 |
| Exd1 | 41.40792365 | -0.011493969 | 0.090407205 | -0.127135544 | 0.89883312 | 0.978597654 |
| Prss41 | 4.238815633 | -0.01149571 | 0.047050412 | -0.244327501 | 0.806977186 | NA |
| Oxsr1 | 2324.579861 | -0.011527332 | 0.063753483 | -0.180811014 | 0.856515918 | 0.967598001 |
| Hpd | 2.333305631 | -0.011529638 | 0.031309251 | -0.368250209 | 0.712686675 | NA |
| Hbq1a | 14.55619515 | -0.011552011 | 0.074598294 | -0.154856243 | 0.876934651 | NA |
| Tmem120a | 498.7335034 | -0.011552674 | 0.071711651 | -0.161098986 | 0.872015439 | 0.97153269 |
| 2010012O05Rik | 966.3414227 | -0.011553133 | 0.070152586 | -0.164685775 | 0.869191314 | 0.970788711 |
| Gm6623 | 1.469736246 | -0.011561405 | 0.027825852 | -0.415491491 | 0.677782143 | NA |
| Neurog3 | 2.524382914 | -0.011563476 | 0.033122347 | -0.349114025 | 0.727003708 | NA |
| Mocs3 | 224.0218564 | -0.011568968 | 0.086944736 | -0.133061169 | 0.894145004 | 0.977844987 |
| Zfp229 | 97.82090343 | -0.01157357 | 0.092760472 | -0.124768341 | 0.900706951 | 0.979053977 |
| Saa2 | 2.191075973 | -0.011600951 | 0.031086471 | -0.373183277 | 0.709012042 | NA |
| Vps8 | 2868.748628 | -0.011614539 | 0.049593071 | -0.234196819 | 0.814832184 | 0.956264994 |
| Rpl39l | 0.981815442 | -0.011630113 | 0.023363963 | -0.497779989 | 0.618639122 | NA |
| Hist2h2ab | 2.007974036 | -0.011643936 | 0.033968036 | -0.342790965 | 0.731755724 | NA |
| Nkx3-1 | 63.68155376 | -0.01164997 | 0.093399375 | -0.124732848 | 0.900735051 | 0.979053977 |
| Shroom1 | 318.701116 | -0.011672915 | 0.089108027 | -0.130997344 | 0.895777409 | 0.977844987 |
| Snora74a | 3.933764526 | -0.011674549 | 0.039188884 | -0.297904598 | 0.765775979 | NA |
| Klhl20 | 931.7685458 | -0.011678111 | 0.052808465 | -0.221140882 | 0.824982739 | 0.957903916 |
| Ccnk | 1268.21783 | -0.011682621 | 0.055251887 | -0.211442935 | 0.832541657 | 0.960519604 |
| Tmem74b | 146.2096044 | -0.011684767 | 0.095049688 | -0.122933249 | 0.902159956 | 0.979150581 |
| Fbxl22 | 9.732944358 | -0.011688495 | 0.063100528 | -0.185236084 | 0.853043868 | NA |
| Mst1 | 0.657513041 | -0.01170544 | 0.017515514 | -0.668289793 | 0.503948628 | NA |
| Sec24b | 2723.814316 | -0.011706876 | 0.052312274 | -0.223788326 | 0.822922014 | 0.957903916 |
| B3gnt3 | 41.58587444 | -0.011717155 | 0.086002771 | -0.1362416 | 0.89163029 | 0.977495969 |
| Ptger3 | 64.66865513 | -0.011719386 | 0.095084676 | -0.123252097 | 0.901907472 | 0.979150581 |
| Zfp36l3 | 1.972252113 | -0.011724937 | 0.030441782 | -0.385159346 | 0.700119362 | NA |
| Ftsj3 | 1534.944843 | -0.011740926 | 0.053945973 | -0.217642309 | 0.827707817 | 0.958680788 |
| Pdlim4 | 287.0967909 | -0.011754635 | 0.085549339 | -0.137401826 | 0.890713188 | 0.977182599 |
| Dnm1 | 56731.60448 | -0.011755419 | 0.039799051 | -0.295369331 | 0.767711759 | 0.94619074 |
| 4930471I20Rik | 0.720323536 | -0.011759462 | 0.017815267 | -0.660077795 | 0.509203907 | NA |
| Plin1 | 0.883295104 | -0.011763997 | 0.018827213 | -0.624840059 | 0.532076036 | NA |
| Spatc1l | 1.945154612 | -0.0118007 | 0.032986147 | -0.357747148 | 0.720532546 | NA |
| Zfp623 | 532.4771775 | -0.011803018 | 0.070399993 | -0.167656518 | 0.866853505 | 0.97021201 |
| D17Ertd648e | 0.451410283 | -0.0118078 | 0.015842014 | -0.745347162 | 0.456061877 | NA |
| Gm5129 | 0.451410283 | -0.0118078 | 0.015842014 | -0.745347162 | 0.456061877 | NA |
| 1700024G13Rik | 7.267599535 | -0.011809329 | 0.051490071 | -0.229351575 | 0.818595671 | NA |
| Pttg1 | 1119.834693 | -0.011845281 | 0.063654462 | -0.18608721 | 0.852376372 | 0.966905493 |
| Isoc2b | 178.6466474 | -0.0118523 | 0.09163003 | -0.129349515 | 0.897081093 | 0.978039389 |
| Rspo2 | 564.3439972 | -0.011854683 | 0.0829775 | -0.142866233 | 0.886395827 | 0.976378471 |
| C330021F23Rik | 2.613490235 | -0.011855118 | 0.034443977 | -0.34418551 | 0.730706779 | NA |
| Pdzd9 | 2.025972155 | -0.011855365 | 0.033532508 | -0.353548418 | 0.723677337 | NA |
| E130308A19Rik | 368.6290194 | -0.011866381 | 0.081143903 | -0.14623873 | 0.883732932 | 0.975315649 |
| Serpinb3b | 0.62437142 | -0.011874481 | 0.01391869 | -0.853132096 | 0.393586054 | NA |
| Pnlip | 32.24304588 | -0.011876652 | 0.084038437 | -0.141324048 | 0.887613953 | 0.976852813 |
| Pax7 | 8.389591352 | -0.011877764 | 0.020108088 | -0.590695879 | 0.55472421 | NA |
| Kcnj9 | 7123.77755 | -0.011878778 | 0.057642047 | -0.206078351 | 0.836729709 | 0.961729368 |
| Rapgef6 | 1819.599547 | -0.011887543 | 0.051926464 | -0.228930333 | 0.818923064 | 0.957334483 |
| A630019I02Rik | 26.66690767 | -0.011888424 | 0.081909483 | -0.145140993 | 0.884599553 | 0.975676135 |
| Snapin | 1946.885079 | -0.011906093 | 0.061838969 | -0.192533818 | 0.84732408 | 0.965275633 |
| Slpi | 1.457270812 | -0.011916187 | 0.025955895 | -0.459093666 | 0.646166905 | NA |
| Fam109b | 23.77420839 | -0.011928856 | 0.081308296 | -0.146711431 | 0.883359796 | 0.975177867 |
| 6330416G13Rik | 658.1153801 | -0.011932574 | 0.071190115 | -0.167615597 | 0.8668857 | 0.97021201 |
| Gtf2a1l | 6.597662945 | -0.011937333 | 0.056136641 | -0.212647789 | 0.831601694 | NA |
| Fzd10 | 54.28605642 | -0.011937655 | 0.090165819 | -0.132396682 | 0.894670539 | 0.977844987 |
| Spata4 | 1.415146138 | -0.011938184 | 0.025346876 | -0.470992332 | 0.637646211 | NA |
| Itm2a | 1518.211123 | -0.011943783 | 0.060463076 | -0.197538455 | 0.843406191 | 0.963987594 |
| 4933434E20Rik | 1827.849909 | -0.011948143 | 0.04625469 | -0.258312032 | 0.796166101 | 0.952734447 |
| Ube2c | 9.170607398 | -0.01196027 | 0.057039834 | -0.209682768 | 0.833915276 | NA |
| Tmem259 | 3615.411663 | -0.011964118 | 0.062716812 | -0.190764137 | 0.848710387 | 0.965892061 |
| Bloc1s6 | 1255.94068 | -0.011972095 | 0.064472916 | -0.18569185 | 0.85268642 | 0.966905493 |
| Frmd4a | 1991.289574 | -0.011978516 | 0.054762064 | -0.218737488 | 0.826854545 | 0.958455003 |
| Gm10649 | 5.231603004 | -0.012000316 | 0.048536855 | -0.247241324 | 0.804721466 | NA |
| Mfsd3 | 197.7727406 | -0.01200572 | 0.084809056 | -0.141561775 | 0.887426162 | 0.976720142 |
| BC039771 | 6.458923014 | -0.012009958 | 0.053697512 | -0.223659485 | 0.823022274 | NA |
| Srms | 14.63049548 | -0.012012039 | 0.072448384 | -0.165801336 | 0.868313295 | NA |
| Trpc1 | 1432.356263 | -0.012021327 | 0.076068698 | -0.158032508 | 0.874431183 | 0.971912694 |
| Il1f5 | 4.376965801 | -0.01205228 | 0.044455077 | -0.271111431 | 0.78630533 | NA |
| AV039307 | 19.59031109 | -0.012076763 | 0.076660147 | -0.157536397 | 0.874822125 | NA |
| Gas2 | 275.6668239 | -0.012086819 | 0.078141613 | -0.154678386 | 0.877074871 | 0.972549751 |
| Abp1 | 2.477613783 | -0.012128187 | 0.03770605 | -0.321650957 | 0.747717136 | NA |
| Tti2 | 946.2853163 | -0.012141468 | 0.054754296 | -0.22174458 | 0.824512724 | 0.957903916 |

| | | | | | | | | | | | | | | | |
|---|---|---|---|---|---|---|---|---|---|---|---|---|---|---|---|
| Glra2 | 556.5823404 | -0.317769293 | 0.104558944 | -3.039140224 | 0.002372544 | 0.021674644 | | | D030040B21Rik | 0.300679247 | -0.012154827 | 0.011580646 | -1.049581096 | 0.293910752 | NA |
| Clmp | 1014.097593 | -0.318827512 | 0.072016174 | -4.427165394 | 9.55E-06 | 0.000301295 | | | Prame | 0.300679247 | -0.012154827 | 0.011580646 | -1.049581096 | 0.293910752 | NA |
| Ppp1r3c | 1711.517539 | -0.319668846 | 0.070525229 | -4.532687786 | 5.82E-06 | 0.000206485 | | | Rpgrip1l | 1188.773251 | -0.012171922 | 0.077160094 | -0.15774893 | 0.874654642 | 0.971956739 |
| Tgfa | 2331.659008 | -0.319697036 | 0.110520295 | -2.892654573 | 0.003820012 | 0.030182183 | | | 2300009A05Rik | 156.1417298 | -0.012182568 | 0.091421116 | -0.133257707 | 0.893989574 | 0.977844987 |
| Crip2 | 3895.34984 | -0.3197841 | 0.096295179 | -3.320873407 | 0.000897362 | 0.010498156 | | | Tmem216 | 215.990843 | -0.012200912 | 0.081148285 | -0.150353294 | 0.880485888 | 0.973779118 |
| Cd59a | 534.7187711 | -0.320208716 | 0.109050668 | -2.936329706 | 0.003321211 | 0.027270189 | | | Anks1 | 1608.176164 | -0.012202715 | 0.068570909 | -0.177957616 | 0.858756262 | 0.96835068 |
| Ptprt | 4164.382298 | -0.320666898 | 0.070842605 | -4.526469608 | 6.00E-06 | 0.000211202 | | | Tssk6 | 1.812640886 | -0.012214065 | 0.033641328 | -0.363067282 | 0.716554625 | NA |
| Kcns2 | 1728.273532 | -0.32077895 | 0.096319519 | -3.330362866 | 0.000867329 | 0.010225679 | | | Psmg4 | 129.072841 | -0.012223508 | 0.093379257 | -0.130901753 | 0.895853028 | 0.977844987 |
| Mnf1 | 1654.800376 | -0.321255486 | 0.06902765 | -4.654011609 | 3.26E-06 | 0.00013101 | | | Psd2 | 2795.734583 | -0.012246208 | 0.070668124 | -0.173291828 | 0.862422047 | 0.969759087 |
| Sec61g | 782.4474239 | -0.321353053 | 0.080999317 | -3.967355084 | 7.27E-05 | 0.001534089 | | | Gm614 | 0.674103222 | -0.012254458 | 0.017840767 | -0.68687952 | 0.492158654 | NA |
| Diras2 | 15122.23639 | -0.321489069 | 0.096877479 | -3.318511924 | 0.000904985 | 0.010519003 | | | Bgn | 793.8867094 | -0.012255526 | 0.094270818 | -0.130003386 | 0.896563748 | 0.978039389 |
| Bmpr2 | 2439.555218 | -0.322533917 | 0.089753414 | -3.593555976 | 0.000326196 | 0.004989214 | | | Acaa1a | 1447.369919 | -0.012277762 | 0.08166813 | -0.150337246 | 0.880498549 | 0.973779118 |
| Capn11 | 48.3427266 | -0.323204291 | 0.11165458 | -2.894680108 | 0.003795451 | 0.030064784 | | | F7 | 2.656525822 | -0.012278561 | 0.035379694 | -0.347051074 | 0.728552949 | NA |
| Gas1 | 235.7426758 | -0.323258158 | 0.109574237 | -2.950129214 | 0.003176411 | 0.026388904 | | | Gng8 | 46.3617978 | -0.012280204 | 0.083027019 | -0.147906113 | 0.882416864 | 0.974603662 |
| Elovl6 | 2488.470841 | -0.323325155 | 0.094816497 | -3.410009495 | 0.000649606 | 0.008229761 | | | Flii | 2811.669352 | -0.012280926 | 0.056750335 | -0.216402707 | 0.828673857 | 0.959225153 |
| Pts | 678.6571504 | -0.323508391 | 0.095355466 | -3.392657018 | 0.000692182 | 0.008641962 | | | Igsf9b | 759.8492799 | -0.012310132 | 0.093456804 | -0.131720016 | 0.895205753 | 0.977844987 |
| Sema5a | 1903.395182 | -0.323749049 | 0.093814785 | -3.450938464 | 0.000558641 | 0.007404262 | | | Fam160a1 | 133.9560094 | -0.01231321 | 0.094789776 | -0.129900192 | 0.896645392 | 0.978039389 |
| Epha6 | 822.1901054 | -0.324265701 | 0.118628544 | -2.733454283 | 0.006267382 | 0.042079243 | | | Ms4a15 | 1.341935732 | -0.012315966 | 0.026567616 | -0.463570608 | 0.642955416 | NA |
| Shisa3 | 99.94989799 | -0.324471273 | 0.121878028 | -2.662262256 | 0.007761738 | 0.049146043 | | | Hs3st4 | 2847.699485 | -0.012320887 | 0.076545558 | -0.160961482 | 0.872123738 | 0.97153269 |
| Otud1 | 634.8731627 | -0.324611732 | 0.088772631 | -3.65666455 | 0.000255518 | 0.00410251 | | | Flt3 | 358.9529399 | -0.01232323 | 0.085316475 | -0.144441383 | 0.88515194 | 0.975769681 |
| Mettl23 | 436.3612169 | -0.324932491 | 0.103862482 | -3.128487649 | 0.001757084 | 0.01736241 | | | Adig | 0.525532643 | -0.012329408 | 0.015179088 | -0.812262747 | 0.416640882 | NA |
| Adamts19 | 27.20734008 | -0.325162619 | 0.11460095 | -2.837346626 | 0.00454902 | 0.033853271 | | | Chat | 249.6982636 | -0.012334776 | 0.095064806 | -0.129751239 | 0.896763242 | 0.978039389 |
| Aldh1b1 | 370.2292102 | -0.3254819 | 0.119094786 | -2.732965154 | 0.006276697 | 0.042100064 | | | Zfa-ps | 0.712046018 | -0.012335495 | 0.018023078 | -0.684427765 | 0.493705089 | NA |
| 2010107E04Rik | 2034.57687 | -0.325830826 | 0.079358878 | -4.105789238 | 4.03E-05 | 0.000969426 | | | Cdhr2 | 33.62460414 | -0.012336489 | 0.071497433 | -0.172544498 | 0.863009481 | 0.969759087 |
| Atp5j2 | 3066.282303 | -0.326085467 | 0.069561038 | -4.687760236 | 2.76E-06 | 0.000115354 | | | Ube2o | 7756.536398 | -0.012355286 | 0.050084334 | -0.24668963 | 0.805148433 | 0.955118966 |
| Slc17a8 | 256.0028571 | -0.326233967 | 0.104302897 | -3.12775556 | 0.001761466 | 0.017372473 | | | Mir344d-1 | 0.4298712 | -0.012357999 | 0.015660003 | -0.789144071 | 0.430027806 | NA |
| Tma7 | 2265.681651 | -0.326271379 | 0.079397247 | -4.109353819 | 3.97E-05 | 0.000956068 | | | Rap2b | 2231.760971 | -0.012378335 | 0.066997476 | -0.184758237 | 0.853418666 | 0.966905493 |
| Heatr3 | 1037.872929 | -0.327157151 | 0.064976203 | -5.035030323 | 4.78E-07 | 2.69E-05 | | | Gm6377 | 0.959495378 | -0.012385604 | 0.019050066 | -0.650160672 | 0.515588442 | NA |
| Kcnj4 | 5238.358052 | -0.32720964 | 0.073359252 | -4.460373212 | 8.18E-06 | 0.000266879 | | | Calr3 | 47.605826 | -0.012410774 | 0.093598933 | -0.132595254 | 0.894513486 | 0.977844987 |
| Adcy2 | 6997.718518 | -0.327563177 | 0.072208261 | -4.536367064 | 5.72E-06 | 0.000204791 | | | Agfg1 | 1836.920975 | -0.012418527 | 0.068659733 | -0.180870605 | 0.856469142 | 0.967598001 |
| Bcl2 | 1069.470493 | -0.327564366 | 0.082453532 | -3.972714782 | 7.11E-05 | 0.001510253 | | | Mir3473d | 0.816830525 | -0.01242086 | 0.016508315 | -0.752400251 | 0.451810396 | NA |
| Fam101b | 582.9213059 | -0.328662294 | 0.082556683 | -3.981050146 | 6.86E-05 | 0.001468323 | | | Gm16863 | 0.309861382 | -0.012420872 | 0.011705926 | -1.061075556 | 0.288655566 | NA |
| Rem2 | 459.8983737 | -0.328914623 | 0.101348219 | -3.245391257 | 0.001172894 | 0.012845805 | | | Trim50 | 0.309861382 | -0.012420872 | 0.011705926 | -1.061075556 | 0.288655566 | NA |
| Slc9a7 | 796.9766947 | -0.328946606 | 0.080878241 | -4.067182966 | 4.76E-05 | 0.001102063 | | | Armc12 | 0.805688471 | -0.012423823 | 0.016697155 | -0.744068211 | 0.45683521 | NA |
| Sik2 | 1392.284125 | -0.328960483 | 0.080615205 | -4.080625771 | 4.49E-05 | 0.001048053 | | | Trmt2b | 666.3208489 | -0.01242847 | 0.070770984 | -0.175615336 | 0.860596153 | 0.96862798 |
| Zfp521 | 527.1197681 | -0.329066726 | 0.102453683 | -3.211858448 | 0.001318793 | 0.013995469 | | | Ldb3 | 405.0693128 | -0.012431752 | 0.079105637 | -0.157153802 | 0.875123636 | 0.972003598 |
| 6330403M23Rik | 1107.133131 | -0.329682636 | 0.088084085 | -3.742817305 | 0.000181969 | 0.003175023 | | | Gm3716 | 4.784573863 | -0.012438358 | 0.044010936 | -0.282619714 | 0.777468366 | NA |
| Snca | 8355.118203 | -0.329724937 | 0.086191442 | -3.825495059 | 0.00013051 | 0.002399355 | | | Pcmtd1 | 4621.669766 | -0.012438519 | 0.068072441 | -0.182724731 | 0.85501401 | 0.967189923 |
| Cntnap3 | 499.1323413 | -0.329807183 | 0.113604303 | -2.903122289 | 0.003694623 | 0.029386291 | | | Ckmt2 | 1.073482203 | -0.012440981 | 0.021662213 | -0.574317185 | 0.565753182 | NA |
| Slco1c1 | 2632.233386 | -0.329877494 | 0.061858678 | -5.332760131 | 9.67E-08 | 7.24E-06 | | | 1700110K17Rik | 1.139995443 | -0.012446879 | 0.020460107 | -0.608348677 | 0.542956243 | NA |
| 4833424O15Rik | 1496.574996 | -0.329946024 | 0.068975623 | -4.783516391 | 1.72E-06 | 7.87E-05 | | | Lsm5 | 107.6937223 | -0.012448906 | 0.094451929 | -0.131801506 | 0.895141295 | 0.977844987 |
| Penk | 13357.46749 | -0.33015191 | 0.109026064 | -3.028192515 | 0.002460213 | 0.022123355 | | | Foxp1 | 4726.978309 | -0.012466808 | 0.080966576 | -0.153974751 | 0.877629644 | 0.972570049 |
| Rn45s | 16977.48593 | -0.330302067 | 0.09646701 | -3.423989897 | 0.000617089 | 0.007974295 | | | Bccip | 1456.883299 | -0.012468214 | 0.051148144 | -0.243766689 | 0.80741152 | 0.955716697 |
| Cxxc4 | 871.6700059 | -0.331081165 | 0.096470744 | -3.431933353 | 0.000599295 | 0.007802919 | | | Pcolce | 261.0200388 | -0.012486061 | 0.073896651 | -0.168966534 | 0.865822965 | 0.970182416 |
| Epha5 | 3203.284868 | -0.331149666 | 0.093232823 | -3.551857121 | 0.000382523 | 0.005649956 | | | Sapcd1 | 6.115335559 | -0.012499312 | 0.055839043 | -0.223845379 | 0.822877618 | NA |
| Gucy1a2 | 2335.100972 | -0.33122769 | 0.113187003 | -2.926375646 | 0.003429366 | 0.027877541 | | | Gm12504 | 3.437001253 | -0.012508443 | 0.039805105 | -0.314242189 | 0.753337102 | NA |
| Hist3h2ba | 274.7343234 | -0.331726121 | 0.094726918 | -3.501920322 | 0.000461918 | 0.006438851 | | | 4930444G20Rik | 0.441181124 | -0.012513093 | 0.01575903 | -0.794026816 | 0.42717982 | NA |
| Gdpd4 | 10.93000515 | -0.331879283 | 0.121410321 | -2.733534355 | 0.006265859 | 0.042079243 | | | 1500011K16Rik | 551.2832864 | -0.012525561 | 0.065170891 | -0.192195639 | 0.847588962 | 0.965398237 |
| Tnrc6b | 3887.764246 | -0.332568813 | 0.08534825 | -3.896609611 | 9.75E-05 | 0.001922925 | | | Nlgn2 | 12292.3845 | -0.012541541 | 0.042587833 | -0.294486472 | 0.768386199 | 0.94619074 |
| Egfr | 548.6533899 | -0.332614649 | 0.076792366 | -4.331350421 | 1.48E-05 | 0.000439549 | | | Cypt12 | 0.314540458 | -0.012549443 | 0.01176596 | -1.066588938 | 0.286157498 | NA |
| Hopx | 775.0994898 | -0.332639432 | 0.107364508 | -3.098225278 | 0.001946834 | 0.018724864 | | | Prss27 | 0.314540458 | -0.012549443 | 0.01176596 | -1.066588938 | 0.286157498 | NA |
| Tet1 | 504.1623979 | -0.332654926 | 0.113721779 | -2.925164645 | 0.00344274 | 0.027955792 | | | Sept14 | 0.314540458 | -0.012549443 | 0.01176596 | -1.066588938 | 0.286157498 | NA |
| Id4 | 1572.418092 | -0.332726286 | 0.083622698 | -3.978899175 | 6.92E-05 | 0.001479626 | | | 4930509E16Rik | 1.231859255 | -0.012566017 | 0.026302142 | -0.47775641 | 0.632823589 | NA |
| Nrarp | 600.8433816 | -0.332884072 | 0.097955924 | -3.398304657 | 0.000678049 | 0.008486015 | | | 4930548H24Rik | 0.438563431 | -0.0125877 | 0.015737168 | -0.799870719 | 0.423785704 | NA |
| Deptor | 457.7023164 | -0.333184604 | 0.096653321 | -3.447213208 | 0.000566401 | 0.00748152 | | | D6Ertd527e | 2.380519081 | -0.012604681 | 0.032117682 | -0.392453017 | 0.69472352 | NA |
| Hras1 | 3978.317564 | -0.333284423 | 0.075184265 | -4.432901234 | 9.30E-06 | 0.000296317 | | | Ccng2 | 1605.189679 | -0.012606832 | 0.081731879 | -0.1542462 | 0.877415615 | 0.972570049 |
| 1810041L15Rik | 2878.681301 | -0.334149583 | 0.086290148 | -3.872395544 | 0.000107771 | 0.002067036 | | | Chst7 | 55.0833245 | -0.012610507 | 0.091420291 | -0.13793991 | 0.890287908 | 0.977182599 |
| Syt17 | 1341.397189 | -0.334364332 | 0.117097199 | -2.855442618 | 0.004297689 | 0.032721568 | | | Galnt5 | 1.457884469 | -0.012610969 | 0.026592995 | -0.474221473 | 0.635341982 | NA |
| 6430573F11Rik | 390.8860614 | -0.334494284 | 0.120093666 | -2.785278309 | 0.00534818 | 0.037563326 | | | Cldn6 | 1.00649821 | -0.012645321 | 0.024698951 | -0.511978069 | 0.608666358 | NA |
| Fgf10 | 207.8299588 | -0.33477406 | 0.105604932 | -3.17006084 | 0.00152407 | 0.015659114 | | | AI606473 | 21.79203477 | -0.012650984 | 0.077555606 | -0.163121465 | 0.870422799 | NA |
| Lhfp | 947.0737585 | -0.335311787 | 0.086044609 | -3.896952867 | 9.74E-05 | 0.001922649 | | | Has2 | 12.96911832 | -0.012688503 | 0.067380938 | -0.188309985 | 0.850633659 | NA |
| Mpped1 | 8845.1666 | -0.335800884 | 0.090686895 | -3.702860089 | 0.000213182 | 0.003556118 | | | Cdk5r2 | 10398.75056 | -0.012702989 | 0.058377986 | -0.217598957 | 0.827741597 | 0.958680788 |
| 4932438A13Rik | 5820.787091 | -0.336130938 | 0.078166329 | -4.300201164 | 1.71E-05 | 0.000493273 | | | Slc43a1 | 8.121991202 | -0.012727694 | 0.062063656 | -0.205074833 | 0.837513659 | NA |
| A730017L22Rik | 214.5836589 | -0.336162728 | 0.110801236 | -3.033925784 | 0.002413938 | 0.021882001 | | | U2af1 | 1460.892451 | -0.012746148 | 0.058755091 | -0.216936921 | 0.828257504 | 0.958819613 |
| Dnahc5 | 195.5043215 | -0.336390702 | 0.111281408 | -3.022883227 | 0.002503788 | 0.022398209 | | | Rapgef5 | 3559.277601 | -0.012749925 | 0.047089095 | -0.270761747 | 0.786574283 | 0.950484821 |
| Itpr1 | 36498.7663 | -0.336504459 | 0.095099834 | -3.538433727 | 0.000402508 | 0.005876062 | | | Gm10584 | 6.330093869 | -0.012756836 | 0.05648602 | -0.225840589 | 0.821325407 | NA |
| Fam210b | 1125.487404 | -0.336838743 | 0.095255394 | -3.536164503 | 0.000405982 | 0.005884268 | | | Brd1 | 2334.702483 | -0.012758022 | 0.057516195 | -0.22181617 | 0.824456991 | 0.957903916 |
| Gnb5 | 5733.635549 | -0.337105168 | 0.054449604 | -6.191140879 | 5.97E-10 | 8.99E-08 | | | Rps17 | 4301.080316 | -0.012758733 | 0.069705546 | -0.183037557 | 0.85476855 | 0.967189923 |
| Mri1 | 247.1534631 | -0.337303676 | 0.111878734 | -3.014904307 | 0.002570603 | 0.022850792 | | | Fam154b | 68.75033599 | -0.012772569 | 0.095095632 | -0.134312887 | 0.893155163 | 0.977844987 |
| Rftn1 | 391.7976625 | -0.337371338 | 0.098655556 | -3.419689196 | 0.000626927 | 0.008054403 | | | Sftpc | 30.26068669 | -0.012778827 | 0.082388918 | -0.155103715 | 0.876739553 | 0.972549751 |
| Flrt3 | 1201.59772 | -0.337482795 | 0.094715042 | -3.563138323 | 0.000366447 | 0.00546773 | | | Nup153 | 2393.438854 | -0.012789679 | 0.074063795 | -0.17268463 | 0.862899326 | 0.969759087 |
| Gpr88 | 8447.602446 | -0.337957101 | 0.121851955 | -2.773505777 | 0.005545586 | 0.038692429 | | | Fasn | 13137.73045 | -0.012806925 | 0.049645989 | -0.25796494 | 0.796433965 | 0.952734447 |
| Scube1 | 3362.952801 | -0.338442086 | 0.114239744 | -2.962559906 | 0.003050924 | 0.025593407 | | | Tmprss3 | 1.608423898 | -0.012810975 | 0.029469864 | -0.434714442 | 0.663769714 | NA |
| Rbp1 | 312.982462 | -0.33850729 | 0.113336939 | -2.986734009 | 0.002819749 | 0.024320521 | | | Akr1c21 | 0.506611253 | -0.012814418 | 0.015056255 | -0.851102622 | 0.394712349 | NA |
| Ms4a15 | 71.18807246 | -0.338766177 | 0.104307027 | -3.247779052 | 0.001163096 | 0.012799007 | | | Sp5 | 5.734526696 | -0.012826784 | 0.045787423 | -0.280137721 | 0.779371846 | NA |
| Myl9 | 462.6617639 | -0.338894808 | 0.114570415 | -2.957960914 | 0.003096814 | 0.025824561 | | | Ccdc33 | 47.52935367 | -0.012827295 | 0.085501788 | -0.150023703 | 0.880745915 | 0.973868856 |
| Pcsk1 | 1339.158831 | -0.339400625 | 0.110525785 | -3.07078232 | 0.002134987 | 0.020096898 | | | Tmem253 | 6.609024931 | -0.012847474 | 0.052115335 | -0.246520029 | 0.805279702 | NA |
| Xylt1 | 914.3304387 | -0.339876823 | 0.093645366 | -3.629403554 | 0.000284077 | 0.004450441 | | | Fscn3 | 0.861412953 | -0.012851954 | 0.017014756 | -0.75534162 | 0.450044032 | NA |
| Slc4a4 | 7143.742797 | -0.34000955 | 0.092199481 | -3.687759935 | 0.000226237 | 0.003709328 | | | Rad51 | 52.33237969 | -0.012867778 | 0.093977828 | -0.136923548 | 0.891091226 | 0.977273775 |
| Galnt13 | 1256.303999 | -0.340688511 | 0.096620387 | -3.526052028 | 0.000421804 | 0.006034564 | | | C330013E15Rik | 39.68580651 | -0.012884614 | 0.091583344 | -0.140687303 | 0.888116976 | 0.977036291 |
| Slc13a3 | 791.7545529 | -0.341391271 | 0.099487206 | -3.431509288 | 0.000600233 | 0.007807531 | | | Themis3 | 0.50399356 | -0.012888759 | 0.015039609 | -0.856987606 | 0.391451734 | NA |
| Zfp704 | 3463.765205 | -0.341544049 | 0.065952517 | -5.178635519 | 2.24E-07 | 1.45E-05 | | | Acap2 | 4210.641873 | -0.012907095 | 0.070741381 | -0.182454659 | 0.855225934 | 0.967189923 |
| Kcnj2 | 964.3876273 | -0.341666527 | 0.110229966 | -3.099579366 | 0.001937956 | 0.01868494 | | | Nps | 0.428139782 | -0.012907979 | 0.015649823 | -0.824800325 | 0.409484962 | NA |
| Ldha | 10348.72167 | -0.341789967 | 0.072453446 | -4.717373528 | 2.39E-06 | 0.00010311 | | | Fundc1 | 1317.063268 | -0.012909736 | 0.058102039 | -0.22219075 | 0.824165394 | 0.957903916 |
| Eomes | 106.5929677 | -0.341877523 | 0.106565971 | -3.208130325 | 0.001336009 | 0.014158776 | | | Tmem51 | 172.5953054 | -0.012914078 | 0.091445346 | -0.141221818 | 0.88769471 | 0.976867679 |
| Mll2 | 4384.654433 | -0.342139293 | 0.085793516 | -3.987938792 | 6.66E-05 | 0.001436263 | | | Cyb5r1 | 851.7955579 | -0.012922233 | 0.076128135 | -0.1697432 | 0.865212098 | 0.970182416 |
| Triqk | 384.2498901 | -0.342213695 | 0.115235638 | -2.969686302 | 0.00298104 | 0.025280915 | | | Nkg7 | 1.387827618 | -0.012923276 | 0.028843056 | -0.448055021 | 0.654113489 | NA |
| Luzp2 | 6138.995764 | -0.342319498 | 0.103493108 | -3.307655025 | 0.000940806 | 0.010837808 | | | Gm13399 | 43.88141948 | -0.012942572 | 0.092370691 | -0.140115569 | 0.88856868 | 0.977089223 |
| Prkca | 9860.561643 | -0.342682493 | 0.07636685 | -4.487319986 | 7.21E-06 | 0.00024453 | | | Oxct2b | 3.174094448 | -0.012948817 | 0.039675459 | -0.326368416 | 0.744145631 | NA |
| Npy5r | 160.6696558 | -0.34308874 | 0.105499772 | -3.252032989 | 0.001145827 | 0.012644905 | | | Mrpl4 | 1455.572116 | -0.012952896 | 0.066081201 | -0.196014845 | 0.844598547 | 0.964328285 |
| D10Bwg1379e | 2815.14733 | -0.343116092 | 0.091070613 | -3.767582975 | 0.000164836 | 0.002915485 | | | Csgalnact1 | 892.2368229 | -0.012959479 | 0.060796823 | -0.213160458 | 0.831201809 | 0.960162026 |
| Synpr | 4159.978486 | -0.343251904 | 0.110036058 | -3.119449301 | 0.001811894 | 0.017756794 | | | Gucy2d | 0.680293638 | -0.012963998 | 0.019365201 | -0.669448151 | 0.503209644 | NA |
| Ncan | 10604.43296 | -0.343589469 | 0.107443711 | -3.197855546 | 0.001384536 | 0.014524038 | | | Slc30a3 | 4867.657389 | -0.012970533 | 0.082116474 | -0.157952871 | 0.874493936 | 0.971912694 |
| Vipr2 | 77.4543463 | -0.344556224 | 0.128877676 | -2.673513635 | 0.007506122 | 0.047820667 | | | Zc3h4 | 2980.322011 | -0.012982995 | 0.055541325 | -0.233753787 | 0.815176129 | 0.956264994 |
| Smim18 | 170.5820966 | -0.344591101 | 0.109425331 | -3.149098084 | 0.001637752 | 0.016466796 | | | Mybph | 9.203483394 | -0.012995141 | 0.05722802 | -0.227076544 | 0.820364223 | NA |
| Asb2 | 130.4562002 | -0.345288696 | 0.129780489 | -2.660559376 | 0.007801097 | 0.04927652 | | | Med1 | 2082.332923 | -0.012996847 | 0.059095436 | -0.2199298 | 0.825925827 | 0.958041984 |
| Nkain2 | 2179.655224 | -0.345359911 | 0.084382008 | -4.092814571 | 4.26E-05 | 0.001011188 | | | Slc1a2 | 126585.6799 | -0.012999181 | 0.063929657 | -0.203335689 | 0.838872662 | 0.96221291 |
| Dynll1 | 7315.923822 | -0.345548646 | 0.07011187 | -4.928532769 | 8.28E-07 | 4.19E-05 | | | Ptn | 8097.002106 | -0.013003499 | 0.051488994 | -0.252549105 | 0.800616663 | 0.953228631 |
| Ptpn13 | 833.3590085 | -0.3456532 | 0.097127691 | -3.5587503 | 0.000372624 | 0.005535407 | | | Xlr4a | 5.050836268 | -0.013003671 | 0.033918884 | -0.383375562 | 0.70144132 | NA |
| Mina | 427.2017802 | -0.345781538 | 0.115019988 | -3.006273462 | 0.00264471 | 0.02323841 | | | Col9a3 | 607.5346762 | -0.013006034 | 0.069355632 | -0.187526719 | NA | NA |
| Lamc3 | 334.9395416 | -0.345974504 | 0.089536817 | -3.864047391 | 0.000111524 | 0.002126409 | | | Mirlet7a-1 | 1.214889506 | -0.013009141 | 0.026673113 | -0.487724881 | 0.625744725 | NA |
| Abcc9 | 473.2776881 | -0.346499658 | 0.088645285 | -3.908833482 | 9.27E-05 | 0.00185175 | | | 1700019G17Rik | 581.9797721 | -0.013021323 | 0.07643647 | -0.170354843 | 0.864731083 | 0.969972403 |
| Zfp536 | 615.2204977 | -0.346516081 | 0.074493238 | -4.651644781 | 3.29E-06 | 0.00013218 | | | Blm | 198.4191157 | -0.013039315 | 0.090972387 | -0.143332663 | 0.886027461 | 0.97629029 |
| Ccdc85c | 588.5678491 | -0.346848127 | 0.112194185 | -3.091498263 | 0.001991491 | 0.019082356 | | | 5330413P13Rik | 94.94763417 | -0.013042637 | 0.094505507 | -0.138009281 | 0.890233082 | 0.977182599 |
| Tbx21 | 33.81222109 | -0.347168039 | 0.117782485 | -2.947535361 | 0.003203181 | 0.026528802 | | | Dscr3 | 1426.670589 | -0.013051307 | 0.062043804 | -0.210356325 | 0.833389578 | 0.960817765 |
| Usmg5 | 3726.275804 | -0.347225825 | 0.087881505 | -3.951068253 | 7.78E-05 | 0.001620271 | | | Slc15a1 | 2.574388517 | -0.013064231 | 0.029715105 | -0.4396495 | 0.660190984 | NA |
| Rasl10b | 13699.88058 | -0.347642039 | 0.059982646 | -5.79571026 | 6.80E-09 | 7.64E-07 | | | Defb42 | 0.332904726 | -0.013064318 | 0.012003419 | -1.088383115 | 0.276426012 | NA |
| Kcnj12 | 735.6707858 | -0.348058417 | 0.076020035 | -4.578509062 | 4.68E-06 | 0.000175263 | | | Gm10639 | 0.332904726 | -0.013064318 | 0.012003419 | -1.088383115 | 0.276426012 | NA |
| Sema3d | 234.8302422 | -0.350798246 | 0.124180808 | -2.824899058 | 0.004729551 | 0.034663987 | | | Slc7a9 | 0.332904726 | -0.013064318 | 0.012003419 | -1.088383115 | 0.276426012 | NA |
| Uqcr11 | 2475.351203 | -0.351491905 | 0.08354877 | -4.20702666 | 2.59E-05 | 0.000684149 | | | Bpifb9b | 12.61121309 | -0.013067029 | 0.0198515 | -0.65823886 | 0.510384658 | NA |
| Prdm12 | 19.02797062 | -0.351805734 | 0.131179794 | -2.681859178 | 0.007321427 | 0.04706979 | | | Tph1 | 17.61168378 | -0.013073212 | 0.073617833 | -0.177582135 | 0.859051156 | NA |
| Serpinf1 | 377.8529466 | -0.352462618 | 0.127519349 | -2.763993225 | 0.005709872 | 0.039425264 | | | Efr3b | 11192.85693 | -0.013076824 | 0.047056235 | -0.277897801 | 0.781090812 | 0.948894514 |
| Sdk2 | 620.3097411 | -0.352630849 | 0.123021839 | -2.866408521 | 0.004151581 | 0.031907044 | | | Acacb | 276.8663701 | -0.013082428 | 0.094602734 | -0.138288054 | 0.890012766 | 0.977182599 |
| Pde10a | 13760.16363 | -0.353121874 | 0.109508378 | -3.224610566 | 0.001261441 | 0.013610566 | | | Fam83b | 3.387348643 | -0.013091746 | 0.043717385 | -0.299463161 | 0.764586676 | NA |
| Mdn1 | 2486.672297 | -0.35327161 | 0.119410922 | -2.958453077 | 0.003091873 | 0.025824561 | | | Cyp2c55 | 0.884519814 | -0.013110039 | 0.020843827 | -0.628965062 | 0.52937193 | NA |
| Gldc | 1325.11144 | -0.353320775 | 0.08649868 | -4.084695558 | 4.41E-05 | 0.001036094 | | | Ap3b2 | 7561.098576 | -0.013156485 | 0.055450339 | -0.237266084 | 0.812450372 | 0.956219756 |
| 9430020K01Rik | 7366.799805 | -0.353481689 | 0.072474501 | -4.877324925 | 1.08E-06 | 5.16E-05 | | | Mir541 | 12.86242223 | -0.013176257 | 0.067078133 | -0.196431482 | 0.844272457 | NA |
| Agpat5 | 1575.95567 | -0.353563213 | 0.083486386 | -4.234980475 | 2.29E-05 | 0.000621314 | | | A930001C03Rik | 1.794854418 | -0.013182037 | 0.024249313 | -0.543604535 | 0.586713638 | NA |
| Spryd4 | 278.1430381 | -0.354754786 | 0.083811892 | -4.232749998 | 2.31E-05 | 0.000625316 | | | Slco6c1 | 0.846373745 | -0.013183625 | 0.021030057 | -0.626894387 | 0.530728467 | NA |
| Efnb2 | 2587.22642 | -0.355468123 | 0.077254026 | -4.601289321 | 4.20E-06 | 0.000160129 | | | Amelx | 0.337583802 | -0.013187751 | 0.012059573 | -1.093550361 | 0.27415223 | NA |
| Tsku | 98.42037718 | -0.355821286 | 0.130009131 | -2.736894584 | 0.006202216 | 0.041781366 | | | Gm805 | 0.337583802 | -0.013187751 | 0.012059573 | -1.093550361 | 0.27415223 | NA |
| Abl2 | 3772.098611 | -0.357096436 | 0.095228881 | -3.749875369 | 0.000176922 | 0.003098526 | | | Pnpla5 | 0.337583802 | -0.013187751 | 0.012059573 | -1.093550361 | 0.27415223 | NA |
| Nptx2 | 2550.751919 | -0.357913128 | 0.124579499 | -2.87296971 | 0.004066331 | 0.031404095 | | | Pou4f3 | 0.337583802 | -0.013187751 | 0.012059573 | -1.093550361 | 0.27415223 | NA |
| Cacna1e | 7933.606378 | -0.358035221 | 0.085280561 | -4.198321569 | 2.69E-05 | 0.000704965 | | | Gm14057 | 35.85260156 | -0.01319116 | 0.084476534 | -0.156151763 | 0.875913397 | 0.972151437 |
| 2810459M11Rik | 514.48768 | -0.358523092 | 0.133399484 | -2.687589801 | 0.007196975 | 0.04650122 | | | Rufy1 | 1572.564519 | -0.013193142 | 0.058806117 | -0.224349825 | 0.822485109 | 0.95778569 |
| Bmp2 | 124.860465 | -0.358831968 | 0.12117024 | -2.961386958 | 0.003062569 | 0.025630477 | | | Acsf3 | 748.3739755 | -0.013212282 | 0.077441349 | -0.17061017 | 0.864530301 | 0.969972403 |
| Adam19 | 1045.091785 | -0.360017146 | 0.098732276 | -3.646397708 | 0.000265942 | 0.004243573 | | | Pde6g | 8.757120053 | -0.013249436 | 0.057266951 | -0.231362686 | 0.817033051 | NA |
| Lrba | 1814.378767 | -0.36073029 | 0.090968057 | -3.965461076 | 7.33E-05 | 0.001540028 | | | Gm15987 | 0.515605119 | -0.013290333 | 0.012110642 | -1.097409514 | 0.272462416 | NA |
| Gadd45g | 325.0803605 | -0.361428 | 0.122032982 | -2.961723904 | 0.00305922 | 0.025621379 | | | Gm16833 | 0.515605119 | -0.013290333 | 0.012110642 | -1.097409514 | 0.272462416 | NA |
| N6amt2 | 650.4626173 | -0.36153623 | 0.077386298 | -4.671837757 | 2.99E-06 | 0.000122685 | | | Accsl | 14.8359031 | -0.013306478 | 0.072824322 | -0.182720241 | 0.855017533 | NA |
| Frzb | 793.3125109 | -0.361718163 | 0.112287675 | -3.221352337 | 0.001275872 | 0.013689999 | | | Tdg | 1.347988832 | -0.013306906 | 0.02243323 | -0.593178316 | 0.553061823 | NA |
| Hivep3 | 2030.706399 | -0.36176914 | 0.104736569 | -3.454086225 | 0.000552161 | 0.00735613 | | | Onecut1 | 4.667185566 | -0.01333887 | 0.038686489 | -0.344665209 | 0.730346076 | NA |
| Mpp7 | 237.6069352 | -0.361769372 | 0.105895298 | -3.416293056 | 0.000634799 | 0.00811516 | | | Gm8267 | 1.730092743 | -0.013352309 | 0.029148756 | -0.458074738 | 0.646898748 | NA |
| Tmem64 | 2003.421858 | -0.362001073 | 0.067657687 | -5.350479566 | 8.77E-08 | 6.80E-06 | | | Ccpg1 | 4433.659618 | -0.01336339 | 0.052405611 | -0.254999224 | 0.798723698 | 0.953036015 |
| Sorl1 | 8861.932392 | -0.362056233 | 0.082617379 | -4.38232533 | 1.17E-05 | 0.000362409 | | | Abca4 | 217.4940995 | -0.013366194 | 0.062634164 | -0.213401005 | 0.831014197 | 0.960162026 |
| Gm4980 | 214.2852052 | -0.362969315 | 0.120706946 | -3.007029228 | 0.002638144 | 0.023198299 | | | Tbc1d19 | 1099.698519 | -0.013366841 | 0.060561422 | -0.220715436 | 0.825314012 | 0.957903916 |
| Hivep2 | 13497.52563 | -0.363061563 | 0.068329487 | -5.313395149 | 1.08E-07 | 7.90E-06 | | | Pak7 | 2535.989718 | -0.013369647 | 0.085157004 | -0.156999965 | 0.875244876 | 0.972003598 |
| Cntnap4 | 1105.743853 | -0.363064872 | 0.086915666 | -4.177208636 | 2.95E-05 | 0.000759536 | | | Acer3 | 369.1613193 | -0.01338067 | 0.070084806 | -0.190921121 | 0.848587393 | 0.96588156 |
| Sulf1 | 1077.691463 | -0.363179031 | 0.129208545 | -2.810797309 | 0.004941891 | 0.035738209 | | | H2-DMb2 | 15.73232806 | -0.01339983 | 0.070399432 | -0.190340028 | 0.849042689 | NA |
| Cdh8 | 1626.825762 | -0.36322873 | 0.081142534 | -4.476428218 | 7.59E-06 | 0.000254002 | | | Mzf1 | 277.7978555 | -0.0134076 | 0.092634461 | -0.144736631 | 0.884918816 | 0.975769681 |
| Ttc9b | 2727.648613 | -0.363386937 | 0.100327329 | -3.622013476 | 0.000292319 | 0.004556528 | | | Igsf5 | 12.94797278 | -0.013431846 | 0.07354593 | -0.182632076 | 0.855086715 | NA |
| Sez6 | 11127.77284 | -0.364118691 | 0.074159636 | -4.909930947 | 9.11E-07 | 4.50E-05 | | | Ing5 | 814.2228683 | -0.013439971 | 0.057522754 | -0.233646169 | 0.815259682 | 0.956264994 |
| Slc25a34 | 311.6324944 | -0.36492733 | 0.107799804 | -3.385231844 | 0.000711182 | 0.008815238 | | | Ankrd52 | 3743.396521 | -0.013440261 | 0.064404596 | -0.208684809 | 0.8346943 | 0.961348408 |
| Dok6 | 618.8126888 | -0.366073005 | 0.088340616 | -4.143881076 | 3.41E-05 | 0.000847891 | | | 1700061I17Rik | 0.346765936 | -0.013444599 | 0.012175679 | -1.104217574 | 0.269498768 | NA |
| Smpdl3a | 1047.420008 | -0.366096783 | 0.098882415 | -3.702344684 | 0.000213616 | 0.003556118 | | | 4833427F10Rik | 0.346765936 | -0.013444599 | 0.012175679 | -1.104217574 | 0.269498768 | NA |
| Lmo1 | 218.6309662 | -0.366243549 | 0.11983009 | -3.056357112 | 0.002240442 | 0.020811399 | | | Cabs1 | 0.346765936 | -0.013444599 | 0.012175679 | -1.104217574 | 0.269498768 | NA |
| Sema3c | 1128.107834 | -0.366573899 | 0.099952804 | -3.667469904 | 0.000244962 | 0.00397429 | | | Cst12 | 0.346765936 | -0.013444599 | 0.012175679 | -1.104217574 | 0.269498768 | NA |
| Glyctk | 254.6023416 | -0.367477908 | 0.093092854 | -3.947434097 | 7.90E-05 | 0.001635816 | | | Smg6 | 2334.176138 | -0.013449185 | 0.049666593 | -0.270789359 | 0.786553044 | 0.950484821 |
| Pkig | 2019.345202 | -0.367646146 | 0.077835234 | -4.723389713 | 2.32E-06 | 0.000100932 | | | Tbc1d23 | 1492.797589 | -0.013473195 | 0.068643628 | -0.196277428 | 0.844393027 | 0.964328285 |
| Rab40b | 1054.869768 | -0.368774995 | 0.081176314 | -4.542889153 | 5.55E-06 | 0.000199939 | | | 4930528P14Rik | 0.83948556 | -0.013491797 | 0.016864333 | -0.800019649 | 0.423699413 | NA |
| Gabrg1 | 986.6643979 | -0.369344065 | 0.110323594 | -3.347824823 | 0.000814485 | 0.009714878 | | | Gin1 | 229.621728 | -0.013494303 | 0.088335856 | -0.152761335 | 0.87858649 | 0.973036547 |
| Mas1 | 674.4205572 | -0.370121437 | 0.104321857 | -3.547880059 | 0.000388345 | 0.005703334 | | | Setmar | 269.0860962 | -0.013506422 | 0.085376548 | -0.158198266 | 0.87430057 | 0.971912694 |
| Dhrs13 | 214.7413904 | -0.3701961 | 0.121884313 | -3.037274377 | 0.00238728 | 0.021757955 | | | Rhbdl1 | 1853.68513 | -0.013513077 | 0.078401002 | -0.172358476 | 0.863155715 | 0.969773371 |
| Fkbp1b | 856.709925 | -0.370981242 | 0.096098518 | -3.860426248 | 0.000113189 | 0.002143957 | | | Efcab2 | 468.7590422 | -0.0135189 | 0.079299805 | -0.170478357 | 0.864633954 | 0.969972403 |
| Slc8a1 | 8127.002437 | -0.37164547 | 0.092166064 | -4.032346106 | 5.52E-05 | 0.001243636 | | | Mrps30 | 745.8398605 | -0.013530656 | 0.07640679 | -0.177087083 | 0.859439988 | 0.968520191 |
| Pcbd1 | 137.197122 | -0.372247651 | 0.128860721 | -2.888759649 | 0.003867646 | 0.030449849 | | | Fbxo8 | 717.0555845 | -0.013540293 | 0.066002653 | -0.205147705 | 0.837456726 | 0.961831963 |
| BC056474 | 989.7080538 | -0.372481266 | 0.069760232 | -5.339449967 | 9.32E-08 | 7.12E-06 | | | 4933427E13Rik | 0.562437198 | -0.013543243 | 0.015653755 | -0.865175341 | 0.386942558 | NA |
| Pcp4 | 8281.138623 | -0.373278771 | 0.112722367 | -3.311488063 | 0.000928012 | 0.010720099 | | | 2900092D14Rik | 622.805232 | -0.013545423 | 0.078372694 | -0.172833451 | 0.862782343 | 0.969759087 |
| Lix1 | 1952.682821 | -0.373446176 | 0.109862585 | -3.399211637 | 0.000675804 | 0.008464759 | | | Pla2g4f | 1.516743762 | -0.013570685 | 0.027219657 | -0.498561952 | 0.618088015 | NA |
| Cx3cl1 | 32935.2499 | -0.373655608 | 0.073195004 | -5.104933211 | 3.31E-07 | 2.01E-05 | | | Rab30 | 442.0431365 | -0.013574439 | 0.086898228 | -0.156210772 | 0.875866886 | 0.972151437 |
| Tesc | 1480.463594 | -0.373697521 | 0.092954967 | -4.020199594 | 5.81E-05 | 0.001299865 | | | 5730405O15Rik | 40.72667198 | -0.013592652 | 0.09095847 | -0.149437997 | 0.881208031 | 0.974259968 |
| 9830001H06Rik | 2647.79046 | -0.373976182 | 0.08940989 | -4.182716048 | 2.88E-05 | 0.000747571 | | | Cd247 | 1.982945442 | -0.013593326 | 0.0334979 | -0.405796358 | 0.684892235 | NA |
| Ctxn2 | 275.479193 | -0.37397704 | 0.101350102 | -3.689952266 | 0.000224296 | 0.003685307 | | | Uck1 | 1602.709592 | -0.013599695 | 0.058013265 | -0.23442388 | 0.814655922 | 0.956264994 |
| Syt9 | 581.687642 | -0.374321283 | 0.100781407 | -3.714189902 | 0.000203856 | 0.003448207 | | | Zfp352 | 21.84462636 | -0.013613635 | 0.077707799 | -0.175190077 | 0.860930281 | NA |
| Gabrb1 | 808.6128594 | -0.374957743 | 0.08598263 | -4.360854547 | 1.30E-05 | 0.000395144 | | | Limd2 | 2630.607001 | -0.013627483 | 0.075497488 | -0.180502467 | 0.856758117 | 0.967675019 |
| Tbc1d16 | 2909.047956 | -0.374965198 | 0.075411919 | -4.972227239 | 6.62E-07 | 3.51E-05 | | | Gm11202 | 0.354968264 | -0.013652206 | 0.012268664 | -1.112770367 | 0.265807065 | NA |
| Kcna3 | 349.8183196 | -0.375096828 | 0.115586656 | -3.245156829 | 0.00117386 | 0.012845805 | | | Olfr697 | 0.354968264 | -0.013652206 | 0.012268664 | -1.112770367 | 0.265807065 | NA |
| Akap5 | 4962.845284 | -0.375216383 | 0.08368297 | -4.483784241 | 7.33E-06 | 0.000246462 | | | Mir181c | 1.084659312 | -0.013670002 | 0.022182155 | -0.616261232 | 0.537722121 | NA |
| Adamtsl1 | 344.74579 | -0.375271588 | 0.120521011 | -3.1137441 | 0.001847296 | 0.018012592 | | | Klhl31 | 2.798314717 | -0.013675794 | 0.033655985 | -0.40634064 | 0.684492328 | NA |
| Slc35f1 | 3619.591104 | -0.375296682 | 0.069374304 | -5.409736203 | 6.31E-08 | 5.12E-06 | | | Wbp2nl | 1.975287132 | -0.013681297 | 0.028853009 | -0.47417229 | 0.635377051 | NA |
| 1700020I14Rik | 2926.67797 | -0.375547293 | 0.058990056 | -6.366281362 | 1.94E-10 | 3.49E-08 | | | Pik3ip1 | 1459.328507 | -0.013681866 | 0.068995872 | -0.198299779 | 0.842810524 | 0.963458383 |
| Cep112 | 105.3829029 | -0.376242297 | 0.124610622 | -3.019343695 | 0.00253323 | 0.022583349 | | | Bard1 | 23.9959028 | -0.013683644 | 0.082683064 | -0.165495126 | 0.868554286 | 0.970658528 |
| Chst11 | 3137.898792 | -0.377129483 | 0.0509906 | -7.39605901 | 1.40E-13 | 5.30E-11 | | | Gls2 | 1126.437479 | -0.013726612 | 0.071569171 | -0.191795047 | 0.847902751 | 0.965542582 |
| Ankrd6 | 1816.454802 | -0.377901926 | 0.091874181 | -4.113254925 | 3.90E-05 | 0.000945932 | | | Uba5 | 1886.300095 | -0.013729367 | 0.051445767 | -0.266870675 | 0.789568736 | 0.951472593 |
| Fat4 | 647.3182522 | -0.378370004 | 0.120168372 | -3.148665473 | 0.001640178 | 0.016480495 | | | AU041133 | 170.7801018 | -0.013729677 | 0.087693226 | -0.156564857 | 0.875587801 | 0.972151437 |
| Klhdc8a | 716.0204553 | -0.378634207 | 0.107015985 | -3.538108883 | 0.000403004 | 0.005876062 | | | Kptn | 755.9254643 | -0.01373009 | 0.073166195 | -0.187656201 | 0.851146167 | 0.966638801 |
| Kcnmb4 | 1936.313485 | -0.37869042 | 0.088499472 | -4.279013348 | 1.88E-05 | 0.000529798 | | | Kcna3 | 318.6664613 | -0.013750179 | 0.092085084 | -0.149320376 | 0.881300838 | 0.974259968 |
| Fam181b | 491.066758 | -0.379117375 | 0.095949726 | -3.951208537 | 7.78E-05 | 0.001620271 | | | Fam53c | 3181.347342 | -0.013754401 | 0.059082085 | -0.232801561 | 0.815915501 | 0.956264994 |
| Herc2 | 11846.88981 | -0.379328855 | 0.070680203 | -5.366833145 | 8.01E-08 | 6.25E-06 | | | S100a14 | 2.465526828 | -0.013775677 | 0.035912756 | -0.383587308 | 0.701284349 | NA |
| Dpysl3 | 636.4767929 | -0.380114796 | 0.104183706 | -3.648505209 | 0.000263771 | 0.004213259 | | | Gtf2a1 | 1339.992257 | -0.013780915 | 0.064006305 | -0.215305585 | 0.829529076 | 0.959526943 |
| Igsf21 | 1819.117811 | -0.380179591 | 0.090194782 | -4.215095202 | 2.50E-05 | 0.000666971 | | | Bag6 | 9398.78552 | -0.013785085 | 0.04970758 | -0.277323597 | 0.781531642 | 0.948894514 |
| Atp5l | 3829.251598 | -0.380180633 | 0.064842099 | -5.863175872 | 4.54E-09 | 5.50E-07 | | | Ms4a1 | 0.695642305 | -0.01378848 | 0.019450228 | -0.708910968 | 0.478379728 | NA |
| Mll3 | 3858.140982 | -0.380975137 | 0.099004723 | -3.848050104 | 0.000119062 | 0.002236051 | | | Scnm1 | 841.7269886 | -0.013813732 | 0.06774416 | -0.203910293 | 0.838423602 | 0.962077671 |
| Idi1 | 1620.958335 | -0.381210985 | 0.136776682 | -2.787105073 | 0.005318123 | 0.037437075 | | | Myo18a | 8373.099591 | -0.013849011 | 0.04477422 | -0.309307696 | 0.757087477 | 0.942139348 |
| Notch2 | 1020.963249 | -0.38136072 | 0.108151464 | -3.526172506 | 0.000421612 | 0.006034564 | | | Clgn | 334.1718788 | -0.013887579 | 0.084724198 | -0.163915146 | 0.869797944 | 0.97107716 |
| Cldn5 | 924.0491 | -0.381964078 | 0.121045932 | -3.155530075 | 0.001602067 | 0.01619582 | | | 3110039M20Rik | 119.8263547 | -0.013888853 | 0.093611862 | -0.148366378 | 0.882053633 | 0.97439733 |
| Glt25d2 | 341.1285305 | -0.382371349 | 0.109785418 | -3.482897429 | 0.000496018 | 0.006795145 | | | 3110057O12Rik | 341.3545372 | -0.013891941 | 0.078687858 | -0.176544909 | 0.859865871 | 0.96862798 |
| Frmd7 | 152.2623315 | -0.383141152 | 0.121595123 | -3.15095821 | 0.001627358 | 0.016383548 | | | Agxt2 | 0.364150398 | -0.013894694 | 0.012376423 | -1.122674507 | 0.261575759 | NA |
| Chsy3 | 503.0898584 | -0.38456634 | 0.079928622 | -4.811372056 | 1.50E-06 | 6.93E-05 | | | Terc | 0.364150398 | -0.013894694 | 0.012376423 | -1.122674507 | 0.261575759 | NA |
| Cxcr7 | 390.6885535 | -0.384626638 | 0.091040786 | -4.224772815 | 2.39E-05 | 0.000642268 | | | 4931440L10Rik | 0.548592792 | -0.013898095 | 0.015568462 | -0.89270829 | 0.37201341 | NA |
| Myo18b | 93.87978013 | -0.385053993 | 0.133524328 | -2.883774051 | 0.003929406 | 0.03076413 | | | Irf6 | 61.74919424 | -0.013900178 | 0.094535436 | -0.147036694 | 0.883103058 | 0.97499088 |
| Tekt5 | 278.2030778 | -0.385541228 | 0.112524633 | -3.426282924 | 0.000611903 | 0.007927107 | | | Trat1 | 5.257152409 | -0.013925452 | 0.052214334 | -0.266697871 | 0.789701793 | NA |
| Pcdhb21 | 92.0942294 | -0.385794202 | 0.124810754 | -3.091033338 | 0.001994612 | 0.019100448 | | | Lect1 | 28.27493804 | -0.013930692 | 0.08544606 | -0.163034925 | 0.870490937 | 0.971477516 |
| Kcna4 | 1156.017063 | -0.38607634 | 0.096110039 | -4.01702408 | 5.89E-05 | 0.001310159 | | | Mrpl36 | 723.6044324 | -0.013942889 | 0.063247544 | -0.220449486 | 0.825521109 | 0.958018665 |
| Huwe1 | 16699.80886 | -0.386175084 | 0.059640604 | -6.475036394 | 9.48E-11 | 1.93E-08 | | | Gata6 | 3.150551789 | -0.013953589 | 0.038168001 | -0.36558344 | 0.714675934 | NA |
| Ubr4 | 9260.184575 | -0.386378259 | 0.081334063 | -4.750509733 | 2.03E-06 | 8.98E-05 | | | Letmd1 | 2315.809494 | -0.013959671 | 0.055369609 | -0.252117928 | 0.800949913 | 0.953228631 |
| Hpcal4 | 50831.04813 | -0.387150414 | 0.0846597 | -4.573018979 | 4.81E-06 | 0.000179055 | | | Mmp23 | 42.24978894 | -0.013967185 | 0.093070084 | -0.15007169 | 0.880708055 | 0.973868856 |
| Rab27b | 1357.156021 | -0.388126841 | 0.089318309 | -4.345434265 | 1.39E-05 | 0.000416938 | | | 5430440P10Rik | 1.973607392 | -0.013969227 | 0.030293988 | -0.461122085 | 0.644711018 | NA |
| Slc19a3 | 164.9094588 | -0.388430463 | 0.12763295 | -3.043340012 | 0.002339678 | 0.021437595 | | | Mrpl16 | 914.4551455 | -0.013977393 | 0.068437803 | -0.204234972 | 0.838169884 | 0.962077671 |
| Cd93 | 427.9824535 | -0.388615555 | 0.121395492 | -3.20123547 | 0.001368397 | 0.014413281 | | | Plekhh1 | 2184.176064 | -0.013979222 | 0.086520771 | -0.161570706 | 0.871643927 | 0.97153269 |
| C1galt1 | 341.715848 | -0.388718125 | 0.098468531 | -3.947638115 | 7.89E-05 | 0.001635816 | | | Lcn11 | 1.23620877 | -0.0139864 | 0.024609037 | -0.568344065 | 0.569801363 | NA |
| Pcdha7 | 117.7062215 | -0.389436537 | 0.115319264 | -3.377029347 | 0.000732732 | 0.008981768 | | | Rapgef1 | 7209.017131 | -0.013996049 | 0.05442869 | -0.257144703 | 0.797067069 | 0.952885371 |
| Foxo1 | 1649.49725 | -0.389994885 | 0.07347332 | -5.307979605 | 1.11E-07 | 8.06E-06 | | | R3hdm4 | 10673.8584 | -0.013997184 | 0.061588248 | -0.227270374 | 0.820213509 | 0.957571905 |
| Aox3 | 105.9682648 | -0.390147998 | 0.13041688 | -2.991545242 | 0.002775694 | 0.024026036 | | | Cenpo | 147.6816724 | -0.014000789 | 0.090432193 | -0.154820851 | 0.876962553 | 0.972549751 |
| Nrp2 | 1666.46064 | -0.390314308 | 0.140470442 | -2.778622333 | 0.005458995 | 0.038236657 | | | Lalba | 0.556711097 | -0.014001777 | 0.015443968 | -0.90661785 | 0.364608919 | NA |
| Grm5 | 6632.334105 | -0.390362452 | 0.081951778 | -4.76331891 | 1.90E-06 | 8.55E-05 | | | Gpr37 | 3001.565326 | -0.014025389 | 0.08282177 | -0.169344232 | 0.865525886 | 0.970182416 |
| Pcdh11x | 720.7165591 | -0.390751015 | 0.117009381 | -3.339484517 | 0.00083934 | 0.009942461 | | | Vps54 | 1302.855638 | -0.014037159 | 0.062942053 | -0.223017171 | 0.823522142 | 0.957903916 |
| Emx2 | 339.3077002 | -0.393800679 | 0.113404218 | -3.472539953 | 0.000515558 | 0.006970384 | | | Fam229a | 4.049355045 | -0.014037289 | 0.044762357 | -0.313595844 | 0.753828017 | NA |
| Tmem158 | 1495.954543 | -0.39404104 | 0.119948969 | -3.285072337 | 0.001019562 | 0.011522314 | | | Spopl | 361.3453259 | -0.014042163 | 0.079936992 | -0.175665398 | 0.860556821 | 0.96862798 |
| Mll1 | 6969.910578 | -0.395232328 | 0.09221934 | -4.285785712 | 1.82E-05 | 0.000518465 | | | Gpld1 | 2151.630255 | -0.014043944 | 0.069355932 | -0.202490878 | 0.839532989 | 0.96236239 |
| Zfp365 | 23643.60402 | -0.395234793 | 0.08759501 | -4.512069749 | 6.42E-06 | 0.000222978 | | | Kynu | 4.395592164 | -0.014050821 | 0.037781319 | -0.371898636 | 0.709968321 | NA |
| Kcnq5 | 4308.845105 | -0.39538801 | 0.068030102 | -5.81195679 | 6.17E-09 | 7.00E-07 | | | Zfp558 | 137.6182573 | -0.014050992 | 0.092337245 | -0.152170361 | 0.87905257 | 0.973224531 |
| Grid2 | 425.8573006 | -0.395399531 | 0.086979373 | -4.545900005 | 5.47E-06 | 0.000198718 | | | Igfbp6 | 1288.365096 | -0.014051624 | 0.078589342 | -0.178798092 | 0.858096244 | 0.967971163 |
| Rpl41 | 7640.167928 | -0.395787114 | 0.06806047 | -5.815227438 | 6.06E-09 | 6.95E-07 | | | Slc18b1 | 867.929869 | -0.014076385 | 0.062722467 | -0.224423339 | 0.822427912 | 0.95778569 |
| Gm13889 | 575.8387015 | -0.396463769 | 0.118752699 | -3.338566402 | 0.000842119 | 0.009967755 | | | Slc17a5 | 900.8355735 | -0.014083011 | 0.06396402 | -0.220170821 | 0.82573812 | 0.958041984 |
| Lmcd1 | 230.7347163 | -0.396926575 | 0.102309739 | -3.87965583 | 0.000104604 | 0.002033551 | | | Siae | 1209.929236 | -0.01408434 | 0.064842225 | -0.217209391 | 0.828045167 | 0.958726613 |
| Lrrc7 | 2685.07043 | -0.397002339 | 0.08500096 | -4.670563019 | 3.00E-06 | 0.000122805 | | | E330020D12Rik | 25.03249191 | -0.014087636 | 0.085551829 | -0.164667851 | 0.869205423 | 0.970788711 |
| Ndst3 | 1154.764642 | -0.397976084 | 0.084522923 | -4.708498856 | 2.50E-06 | 0.000106515 | | | Baat | 5.211806029 | -0.01409674 | 0.045333931 | -0.310953403 | 0.755836046 | NA |
| Mlf1ip | 70.99086984 | -0.398573578 | 0.142581828 | -2.795402353 | 0.005183516 | 0.03674392 | | | Ddah1 | 4560.723602 | -0.014101092 | 0.054617963 | -0.25817682 | 0.796270447 | 0.952734447 |
| Gpx8 | 124.3018002 | -0.401060086 | 0.136067522 | -2.947507833 | 0.003203466 | 0.026528802 | | | Gna13 | 3009.58833 | -0.014108648 | 0.045989509 | -0.306779712 | 0.759011051 | 0.942526199 |
| Cdh11 | 2027.712629 | -0.401060434 | 0.079197557 | -5.06405058 | 4.10E-07 | 2.38E-05 | | | 4930572O03Rik | 0.928072022 | -0.014109151 | 0.020568158 | -0.685970577 | 0.492731663 | NA |
| Tubb3 | 12282.5718 | -0.401124115 | 0.083111191 | -4.826355043 | 1.39E-06 | 6.48E-05 | | | Gm5531 | 1.375280419 | -0.014112679 | 0.023962993 | -0.588936393 | 0.555903944 | NA |
| Tmeff1 | 2840.785162 | -0.401895606 | 0.066498549 | -6.043674779 | 1.51E-09 | 1.93E-07 | | | BC048403 | 526.6467308 | -0.014123441 | 0.070529579 | -0.200248484 | 0.84128625 | 0.963094947 |
| E030013I19Rik | 54.8386463 | -0.402257886 | 0.147105812 | -2.734479894 | 0.006247891 | 0.041997755 | | | Gm11166 | 1.445909434 | -0.014123651 | 0.029085234 | -0.485595233 | 0.627254177 | NA |
| Rcn3 | 152.2682099 | -0.402298342 | 0.12762043 | -3.152303613 | 0.001619877 | 0.016318843 | | | Kcnj4 | 4840.263268 | -0.014129851 | 0.071424256 | -0.197829869 | 0.843178176 | 0.963802816 |
| Myoc | 854.6466574 | -0.403915176 | 0.132811117 | -3.041275355 | 0.002355783 | 0.021559115 | | | Ypel5 | 3838.501272 | -0.014137536 | 0.058455672 | -0.241850544 | 0.808895973 | 0.955725085 |
| Chl1 | 5226.664705 | -0.404302611 | 0.085256146 | -4.742210736 | 2.11E-06 | 9.29E-05 | | | Rabggta | 1792.489877 | -0.014141796 | 0.048448818 | -0.291891448 | 0.770369622 | 0.94619074 |
| Angpt1 | 212.6973788 | -0.404486436 | 0.11377804 | -3.555048361 | 0.00037791 | 0.005603187 | | | Creld2 | 559.9558172 | -0.014142122 | 0.076417327 | -0.185064333 | 0.853178577 | 0.966905493 |
| Hivep1 | 3954.315007 | -0.405279431 | 0.089564869 | -4.524982145 | 6.04E-06 | 0.00021221 | | | Grap2 | 14.15510344 | -0.01415093 | 0.074568738 | -0.18977027 | 0.849489154 | NA |
| Ablim3 | 1535.617072 | -0.405721488 | 0.123020089 | -3.298010034 | 0.000973727 | 0.011167224 | | | Ell3 | 2.349348143 | -0.014161803 | 0.036541722 | -0.3875516 | 0.698347893 | NA |
| Gypc | 83.28495826 | -0.406910072 | 0.1300786 | -3.128186119 | 0.001758888 | 0.017366567 | | | Arap2 | 4513.038782 | -0.014170544 | 0.072065919 | -0.196633085 | 0.844114678 | 0.964266264 |
| Lrp1b | 1665.482045 | -0.40891699 | 0.107015652 | -3.821095157 | 0.00013286 | 0.002433261 | | | Gzmk | 3.16118579 | -0.014171554 | 0.042433526 | -0.333970698 | 0.73840167 | NA |
| Adora2a | 2534.098192 | -0.408970203 | 0.127281241 | -3.213122379 | 0.001313003 | 0.013943573 | | | Gm4013 | 7.653573207 | -0.014181824 | 0.052330152 | -0.271006742 | 0.786385847 | NA |
| Ptprf | 4144.378741 | -0.409466477 | 0.073773422 | -5.550325109 | 2.85E-08 | 2.58E-06 | | | Slc25a36 | 1419.034993 | -0.014238088 | 0.058466566 | -0.243525295 | 0.807598492 | 0.955716697 |
| Flt1 | 2472.320054 | -0.409549538 | 0.07180206 | -5.70386893 | 1.17E-08 | 1.21E-06 | | | Zfp804a | 930.0865896 | -0.014239176 | 0.086722114 | -0.164193136 | 0.869579104 | 0.970907461 |

| | | | | | | | | | | | | | |
|---|---|---|---|---|---|---|---|---|---|---|---|---|---|
| Vstm4 | 181.439375 | -0.410569589 | 0.129038195 | -3.181767923 | 0.001463791 | 0.015180704 | Cox17 | 1040.03762 | -0.014241719 | 0.065395731 | -0.217777507 | 0.827602471 | 0.958672527 |
| Trpc5 | 397.2779555 | -0.410827097 | 0.111469554 | -3.68555433 | 0.000228205 | 0.003729762 | Myo7b | 6.303919108 | -0.014245986 | 0.054941583 | -0.259293326 | 0.795408928 | NA |
| Apcdd1 | 971.7775461 | -0.411438765 | 0.11797788 | -3.487422939 | 0.000487699 | 0.006728776 | Dusp2 | 44.0150948 | -0.014247892 | 0.091220644 | -0.156191527 | 0.875882055 | 0.972151437 |
| Cyp51 | 2246.898954 | -0.412298282 | 0.078483852 | -5.253288076 | 1.49E-07 | 1.03E-05 | 1700016G22Rik | 0.378011608 | -0.014248474 | 0.012531915 | -1.136975044 | 0.255548718 | NA |
| Tmem91 | 538.0354061 | -0.412307367 | 0.124362831 | -3.315358492 | 0.000915257 | 0.010598645 | Ifitm5 | 1.33181971 | -0.014263215 | 0.022782701 | -0.626054607 | 0.531279124 | NA |
| Grm1 | 2078.453675 | -0.412796979 | 0.085241855 | -4.842656 | 1.28E-06 | 6.02E-05 | Wdfy1 | 948.7605502 | -0.014271831 | 0.075335532 | -0.189443559 | 0.849745187 | 0.966260848 |
| Nid1 | 433.6626812 | -0.413040271 | 0.129061037 | -3.200348309 | 0.001372616 | 0.014433037 | Acy1 | 454.7470637 | -0.014280878 | 0.083305586 | -0.171427616 | 0.863887542 | 0.969836459 |
| Rab3c | 16973.54471 | -0.413947202 | 0.083106995 | -4.980894816 | 6.33E-07 | 3.39E-05 | 6330410L21Rik | 7.002595156 | -0.014292265 | 0.050952854 | -0.280499786 | 0.77909409 | NA |
| Ccdc138 | 107.5147142 | -0.414068605 | 0.129214595 | -3.204503359 | 0.001352957 | 0.014299262 | Lmnb2 | 930.7598363 | -0.01429352 | 0.061246797 | -0.233375791 | 0.815469611 | 0.956264994 |
| Mrpl33 | 498.3041142 | -0.414337965 | 0.098859114 | -4.19119643 | 2.77E-05 | 0.000725023 | Neurl1a | 8033.44638 | -0.014299155 | 0.074195746 | -0.192722038 | 0.847176663 | 0.965275633 |
| Ccr2 | 10.19247887 | -0.414605138 | 0.144121417 | -2.876776728 | 0.004017598 | 0.031139898 | Scin | 7.860382517 | -0.0142995 | 0.058903339 | -0.24276212 | 0.808189681 | NA |
| Shisa2 | 160.5973259 | -0.415446709 | 0.133586815 | -3.109937983 | 0.001871266 | 0.018189084 | March11 | 122.416473 | -0.014309422 | 0.094321163 | -0.151709563 | 0.879416015 | 0.973413768 |
| Neto2 | 2520.131009 | -0.415486635 | 0.082960973 | -5.008217963 | 5.49E-07 | 3.04E-05 | Slc5a4b | 1.402686381 | -0.014322309 | 0.029487194 | -0.485712858 | 0.627170766 | NA |
| Pcdhb18 | 168.648665 | -0.41682465 | 0.101759428 | -4.096177217 | 4.20E-05 | 0.001001219 | Abhd14a | 965.3864621 | -0.014325282 | 0.066873972 | -0.214213113 | 0.83038087 | 0.960002081 |
| Gpr26 | 1957.299373 | -0.416980635 | 0.085134091 | -4.897927846 | 9.69E-07 | 4.76E-05 | Mpl | 44.12021185 | -0.014332196 | 0.089493991 | -0.160147018 | 0.872765264 | 0.971552474 |
| Trpc7 | 196.435079 | -0.419613083 | 0.123285402 | -3.403590979 | 0.000665063 | 0.008364026 | Trmt5 | 443.8409479 | -0.014336277 | 0.078570223 | -0.182464505 | 0.855218208 | 0.967189923 |
| Col1a2 | 891.7664996 | -0.42031474 | 0.125401943 | -3.351740251 | 0.000803053 | 0.009608114 | Zfp937 | 500.9130632 | -0.014350045 | 0.078522096 | -0.182751685 | 0.85499286 | 0.967189923 |
| Qpct | 420.8748326 | -0.421673988 | 0.10303889 | -4.092377056 | 4.27E-05 | 0.001011549 | 9230104L09Rik | 0.997185341 | -0.014359457 | 0.019780336 | -0.725946049 | 0.467871845 | NA |
| Rxrg | 969.8600865 | -0.425065021 | 0.124636469 | -3.410438572 | 0.000648585 | 0.008223547 | Il20ra | 29.0373385 | -0.014365849 | 0.080142257 | -0.179254365 | 0.857737978 | 0.967971163 |
| Pbx1 | 7473.231177 | -0.426554828 | 0.074716752 | -5.708958421 | 1.14E-08 | 1.18E-06 | Slc9a3 | 16.719281 | -0.0143692 | 0.068912284 | -0.208514345 | 0.834827383 | NA |
| Osbpl8 | 4083.251491 | -0.427470449 | 0.084718741 | -5.045760143 | 4.52E-07 | 2.57E-05 | Dmrt3 | 6.176869938 | -0.014373183 | 0.046752919 | -0.307428574 | 0.758517182 | NA |
| Plcxd3 | 345.601027 | -0.427541785 | 0.135266563 | -3.160735196 | 0.001573715 | 0.016024696 | Cxcl13 | 0.737471969 | -0.014373212 | 0.01813573 | -0.792535641 | 0.428048418 | NA |
| Mpped2 | 1304.708112 | -0.428323525 | 0.070755847 | -6.053542503 | 1.42E-09 | 1.84E-07 | 9130015A21Rik | 0.713363127 | -0.014374535 | 0.01803111 | -0.797207458 | 0.425330556 | NA |
| Akap9 | 4475.988126 | -0.428959502 | 0.109911541 | -3.902770318 | 9.51E-05 | 0.001886616 | Tbrg4 | 1354.86826 | -0.014391886 | 0.057445022 | -0.250533204 | 0.80217503 | 0.95333733 |
| Grk5 | 614.130081 | -0.429136276 | 0.070058257 | -6.125420371 | 9.04E-10 | 1.24E-07 | Gas2l2 | 16.23990206 | -0.014430126 | 0.072778813 | -0.198273724 | 0.842830909 | NA |
| Grm3 | 2977.855656 | -0.42957074 | 0.126783367 | -3.388226303 | 0.000703462 | 0.008740529 | F10 | 0.698589963 | -0.014437445 | 0.019473637 | -0.741384094 | 0.458460586 | NA |
| Lhx8 | 117.5787567 | -0.430135873 | 0.159779415 | -2.692060626 | 0.007101204 | 0.04597829 | Rell1 | 587.6470193 | -0.014447975 | 0.083874626 | -0.172256801 | 0.863235644 | 0.969788164 |
| B230216N24Rik | 134.2511896 | -0.430364302 | 0.134862419 | -3.191135871 | 0.001417146 | 0.014776085 | 2510003D18Rik | 0.959722939 | -0.01447358 | 0.019697855 | -0.734779514 | 0.462473785 | NA |
| Npffr2 | 9.440078024 | -0.430607862 | 0.147228368 | -2.924761496 | 0.003447203 | 0.027963856 | Gm15880 | 2.079917153 | -0.014494707 | 0.033243838 | -0.436011854 | 0.662828129 | NA |
| Epha3 | 224.8833475 | -0.431022572 | 0.124395574 | -3.464934954 | 0.00053036 | 0.007120792 | Dpf2 | 2410.316016 | -0.014501065 | 0.060593947 | -0.239315409 | 0.810861018 | 0.956130136 |
| Sv2c | 1345.056425 | -0.431200026 | 0.12502424 | -3.44893138 | 0.00056281 | 0.007440421 | Hbs1l | 2327.353797 | -0.01451589 | 0.04730759 | -0.306840605 | 0.7589647 | 0.942526199 |
| Slc6a11 | 8376.849193 | -0.432399271 | 0.12511868 | -3.455912985 | 0.000548432 | 0.007319044 | Fam166a | 3.554082237 | -0.014515901 | 0.038037741 | -0.381618366 | 0.702744459 | NA |
| Sla | 505.497341 | -0.432427188 | 0.08690375 | -4.975932423 | 6.49E-07 | 3.46E-05 | Dlec1 | 146.9967857 | -0.014519663 | 0.095101903 | -0.15267479 | 0.878654742 | 0.97303795 |
| Prkar2b | 2750.511227 | -0.432440352 | 0.086429695 | -5.003377067 | 5.63E-07 | 3.10E-05 | Dclre1a | 400.8582844 | -0.01453956 | 0.071746858 | -0.202650828 | 0.839407959 | 0.96231236 |
| Atp5e | 2029.569215 | -0.432453949 | 0.107616115 | -4.018486901 | 5.86E-05 | 0.001303924 | Adamts7 | 12.51700345 | -0.014548393 | 0.066507628 | -0.218747731 | 0.826846565 | NA |
| Chst15 | 2008.341138 | -0.432548801 | 0.067974309 | -6.363415923 | 1.97E-10 | 3.51E-08 | Lrrc1 | 428.9328634 | -0.014595083 | 0.086798014 | -0.168149964 | 0.866465303 | 0.97021201 |
| Foxo6 | 392.2482541 | -0.432731942 | 0.112847204 | -3.834671354 | 0.000125732 | 0.002335847 | Dnd1 | 39.77503095 | -0.014603036 | 0.080787303 | -0.180759052 | 0.856556705 | 0.967598001 |
| Gpr52 | 453.4012127 | -0.433044922 | 0.110652606 | -3.913553761 | 9.09E-05 | 0.001825314 | Tomm34 | 3184.513982 | -0.014609156 | 0.049955933 | -0.292440863 | 0.769949569 | 0.94619074 |
| Kctd12 | 3117.747668 | -0.43396618 | 0.10364205 | -4.187163237 | 2.82E-05 | 0.000736781 | Ankle1 | 8.188618525 | -0.014610343 | 0.053735216 | -0.271895112 | 0.785702669 | NA |
| Shisa9 | 2795.824996 | -0.43396713 | 0.093018132 | -4.665403627 | 3.08E-06 | 0.000125258 | Rgs9bp | 13.93581596 | -0.014632508 | 0.062892261 | -0.232659907 | 0.816025505 | NA |
| Ddr2 | 446.4424151 | -0.436666849 | 0.100952102 | -4.325485468 | 1.52E-05 | 0.000446615 | Tas2r108 | 0.706969034 | -0.014632763 | 0.019512524 | -0.749916463 | 0.453305019 | NA |
| Myo16 | 1566.656143 | -0.436991914 | 0.137263231 | -3.183605043 | 0.001454533 | 0.015115049 | Nkapl | 9.529810526 | -0.0146477 | 0.052594503 | -0.278502478 | 0.780626663 | NA |
| 4930426D05Rik | 40.55002337 | -0.437723138 | 0.163424648 | -2.678440138 | 0.007396595 | 0.047350844 | Rpp21 | 363.7612991 | -0.014675456 | 0.079291141 | -0.18508318 | 0.853163794 | 0.966905493 |
| Alk | 360.9203877 | -0.438310853 | 0.115176759 | -3.80554945 | 0.00014149 | 0.002582146 | Zfp322a | 1389.836772 | -0.014675725 | 0.072845472 | -0.201463789 | 0.840335941 | 0.962802683 |
| AI848285 | 105.4950121 | -0.438589757 | 0.149987607 | -2.924173309 | 0.003453724 | 0.027983052 | Eif1ad | 1058.222509 | -0.014679154 | 0.060316503 | -0.243368783 | 0.807719725 | 0.955716697 |
| Lrrc9 | 107.9937792 | -0.438954142 | 0.138081648 | -3.178946287 | 0.001478115 | 0.015298537 | Ptgdr | 26.59633905 | -0.014682535 | 0.083909268 | -0.174981082 | 0.861094499 | 0.968928152 |
| Shc3 | 1198.963897 | -0.439128157 | 0.088359746 | -4.969776145 | 6.70E-07 | 3.53E-05 | 1810030O07Rik | 1013.083951 | -0.014704833 | 0.060435987 | -0.24331253 | 0.807763299 | 0.955716697 |
| Pcsk9 | 238.3146009 | -0.439615476 | 0.130431555 | -3.370468726 | 0.000750404 | 0.009126188 | Hist1h2ac | 33.41247888 | -0.014709594 | 0.074045952 | -0.198654937 | 0.842532676 | 0.963292377 |
| Alcam | 6704.339778 | -0.44062504 | 0.078174827 | -5.636405673 | 1.74E-08 | 1.70E-06 | Kmo | 2.916864259 | -0.014711536 | 0.039996777 | -0.367818045 | 0.713008913 | NA |
| Dgkb | 6402.564201 | -0.440645555 | 0.110167197 | -3.999789118 | 6.34E-05 | 0.001377705 | Pdf | 1072.684113 | -0.014724817 | 0.060611829 | -0.242936366 | 0.808054694 | 0.955716697 |
| Atp6v1c2 | 64.02860352 | -0.44097743 | 0.152744845 | -2.887020045 | 0.003889095 | 0.030510193 | Rnf43 | 148.7551884 | -0.014736525 | 0.091565736 | -0.160939293 | 0.872141214 | 0.97153269 |
| Gng13 | 609.5214455 | -0.44235934 | 0.114011452 | -3.87995532 | 0.000104476 | 0.002033551 | Arsa | 1830.285942 | -0.014738956 | 0.062600851 | -0.235443386 | 0.813864616 | 0.956264994 |
| Prok2 | 11.95770798 | -0.442527696 | 0.152307242 | -2.905493458 | 0.003666745 | 0.029209537 | 4930430F08Rik | 538.053771 | -0.014744161 | 0.065607602 | -0.224732512 | 0.82218737 | 0.95778569 |
| Slc7a2 | 1535.146295 | -0.442693669 | 0.085380956 | -5.184922871 | 2.16E-07 | 1.42E-05 | Sphk2 | 1135.25874 | -0.014759156 | 0.069897854 | -0.211153211 | 0.832767719 | 0.960519604 |
| Tnfrsf8 | 59.45853693 | -0.442858344 | 0.154643175 | -2.863743216 | 0.004186673 | 0.032100014 | Mir135a-2 | 0.634600579 | -0.014761678 | 0.015937442 | -0.926226337 | 0.354328368 | NA |
| Slc7a11 | 2016.721578 | -0.442901646 | 0.082067634 | -5.396788286 | 6.78E-08 | 5.39E-06 | Gm6408 | 1.171846807 | -0.014765429 | 0.023702811 | -0.622939989 | 0.533323958 | NA |
| Cped1 | 221.5049818 | -0.443426851 | 0.12388598 | -3.579314243 | 0.000344497 | 0.005197309 | Ccdc87 | 160.9675473 | -0.014770961 | 0.092732601 | -0.159285527 | 0.873443923 | 0.971912694 |
| Efcab6 | 338.9362617 | -0.444110901 | 0.111687666 | -3.976364773 | 7.00E-05 | 0.001491362 | Cst11 | 0.91583343 | -0.014785549 | 0.02237224 | -0.66088817 | 0.508684033 | NA |
| Trps1 | 1044.161014 | -0.444463009 | 0.099518756 | -4.46612303 | 7.96E-06 | 0.000262449 | Sin3a | 1832.983482 | -0.01480855 | 0.063795035 | -0.232126991 | 0.816439382 | 0.956264994 |
| 9630013A20Rik | 48.92984847 | -0.444707705 | 0.158306451 | -2.809157183 | 0.004967139 | 0.035795744 | Sco1 | 425.3247988 | -0.014817074 | 0.068497923 | -0.216314212 | 0.828742831 | 0.959228555 |
| Mgp | 522.789843 | -0.449818358 | 0.114977346 | -3.912234649 | 9.14E-05 | 0.001832942 | Gm6602 | 0.562699945 | -0.014843974 | 0.015512614 | -0.956897035 | 0.338619228 | NA |
| Hey2 | 181.2224759 | -0.450155663 | 0.141726805 | -3.176221064 | 0.001492072 | 0.015401841 | 4930526L06Rik | 0.579264538 | -0.014871397 | 0.015800752 | -0.941182895 | 0.346611141 | NA |
| Foxp2 | 946.2517522 | -0.450656073 | 0.123285046 | -3.655399317 | 0.000256782 | 0.004114354 | Kcng2 | 175.7111684 | -0.014873423 | 0.094656003 | -0.157131328 | 0.875141348 | 0.972003598 |
| Fam126a | 875.7450863 | -0.450834309 | 0.113663263 | -3.966403014 | 7.30E-05 | 0.001538131 | Cdk11b | 3081.523234 | -0.014894228 | 0.039991842 | -0.372431662 | 0.709571484 | 0.930224827 |
| Tmem45a | 27.50803503 | -0.450852648 | 0.166069686 | -2.714840125 | 0.006630777 | 0.043659664 | Ms4a7 | 13.34998502 | -0.014896595 | 0.063448636 | -0.234781959 | 0.814377971 | NA |
| Lum | 82.67209727 | -0.45117631 | 0.139031925 | -3.245127397 | 0.001173981 | 0.012845805 | Fibp | 1282.158371 | -0.014921964 | 0.073549456 | -0.202883404 | 0.839226166 | 0.96231236 |
| Ntrk1 | 121.6366259 | -0.451497015 | 0.160994552 | -2.804424186 | 0.005040654 | 0.036129802 | Mir665 | 0.598294728 | -0.014922869 | 0.015732256 | -0.94855235 | 0.342848336 | NA |
| Ccdc80 | 303.8183343 | -0.452397036 | 0.104375668 | -4.334315143 | 1.46E-05 | 0.000435663 | Cyp4f18 | 1.396185696 | -0.014923237 | 0.02312676 | -0.645280061 | 0.518745711 | NA |
| Aplnr | 10.97508254 | -0.452603225 | 0.164309783 | -2.754572602 | 0.005876885 | 0.040218743 | Rps15a | 5791.893228 | -0.0149361 | 0.047775759 | -0.312629244 | 0.754562357 | 0.941970849 |
| Cdca3 | 23.50886033 | -0.453112299 | 0.16658306 | -2.720038265 | 0.006527436 | 0.043239925 | Ear11 | 0.576646845 | -0.014949606 | 0.015786512 | -0.946986001 | 0.343645912 | NA |
| Dcbld1 | 435.4604426 | -0.453384754 | 0.097761674 | -4.637653323 | 3.52E-06 | 0.000139281 | 5830416I19Rik | 7.346571602 | -0.014994497 | 0.058703672 | -0.255426903 | 0.798393394 | NA |
| Gng4 | 2627.038902 | -0.454575489 | 0.166667594 | -2.727437754 | 0.00638283 | 0.042645781 | Pirt | 7.287741437 | -0.014994646 | 0.051877166 | -0.289041349 | 0.772549736 | NA |
| Arhgap10 | 997.5865014 | -0.454906765 | 0.083015142 | -5.479804684 | 4.26E-08 | 3.62E-06 | Tnfrsf13b | 4.191239149 | -0.01501046 | 0.043834762 | -0.342432786 | 0.732025218 | NA |
| Zdbf2 | 1074.617536 | -0.454963452 | 0.130089915 | -3.497299928 | 0.000469993 | 0.006519313 | C8a | 1.74505306 | -0.0150248 | 0.029629655 | -0.507086562 | 0.612094083 | NA |
| Adra2c | 780.1633966 | -0.455282537 | 0.09841723 | -4.626045005 | 3.73E-06 | 0.000145462 | Gm19705 | 17.32564883 | -0.015030729 | 0.076289646 | -0.197021873 | 0.843810421 | NA |
| Kcnn3 | 1048.614526 | -0.456094211 | 0.099884443 | -4.566218681 | 4.97E-06 | 0.000184075 | Ucn | 7.337988093 | -0.015036338 | 0.050347073 | -0.298653674 | 0.765204307 | NA |
| AI504432 | 1115.839049 | -0.457149223 | 0.103519501 | -4.416068646 | 1.01E-05 | 0.000314613 | 1700020N18Rik | 1.662577843 | -0.015043638 | 0.027234797 | -0.552368281 | 0.580696058 | NA |
| Fbln1 | 304.2890215 | -0.458543731 | 0.138040043 | -3.321816781 | 0.000894334 | 0.010481817 | Gimap3 | 2.701455704 | -0.015051103 | 0.035522901 | -0.423701394 | 0.671783598 | NA |
| Mei4 | 39.15728793 | -0.458646448 | 0.163790126 | -2.800208171 | 0.005106966 | 0.036330272 | Trim34b | 1.138745056 | -0.015053915 | 0.02266037 | -0.664327836 | 0.506480509 | NA |
| Kif27 | 203.4189338 | -0.459781272 | 0.098654765 | -4.660507489 | 3.15E-06 | 0.000127939 | Top2b | 4618.694173 | -0.015064756 | 0.070052901 | -0.215048279 | 0.829729678 | 0.959659199 |
| Artn | 20.6351935 | -0.459889477 | 0.166615661 | -2.760181571 | 0.005776924 | 0.039763513 | Wdr63 | 24.662043 | -0.015094756 | 0.064776868 | -0.233026948 | 0.81574048 | 0.956264994 |
| Dmkn | 119.8857115 | -0.460134067 | 0.134827447 | -3.412762587 | 0.000643079 | 0.008180519 | 4933427G17Rik | 15.23388054 | -0.01509947 | 0.074013044 | -0.204010932 | 0.838344956 | NA |
| Egln3 | 461.6554543 | -0.460395318 | 0.090483754 | -5.088154495 | 3.62E-07 | 2.15E-05 | D430019H16Rik | 8571.480738 | -0.015104586 | 0.061246791 | -0.246618409 | 0.805203556 | 0.955118966 |
| Map2 | 26102.54727 | -0.460995134 | 0.059517995 | -7.745474847 | 9.52E-15 | 4.76E-12 | Zmym2 | 3927.327832 | -0.015114972 | 0.074860258 | -0.201909165 | 0.839987737 | 0.962655768 |
| Tcfl5 | 24.02995482 | -0.461004371 | 0.168119905 | -2.742116528 | 0.006104468 | 0.041300088 | Nmt2 | 1994.73006 | -0.015115194 | 0.062342833 | -0.242452786 | 0.808429337 | 0.955716697 |
| Ccdc109b | 64.13640588 | -0.46163123 | 0.158658664 | -2.909587278 | 0.003619063 | 0.028933832 | Cd1d1 | 33.71856723 | -0.015127163 | 0.081139341 | -0.186434385 | 0.85210413 | 0.966758161 |
| Col3a1 | 152.2011881 | -0.46220536 | 0.152597807 | -3.028912202 | 0.00245436 | 0.02211392 | Tagln2 | 436.6942107 | -0.015134026 | 0.07944656 | -0.190493153 | 0.848922708 | 0.965892061 |
| Pcdhb22 | 225.5714205 | -0.463201053 | 0.09985626 | -4.638678149 | 3.51E-06 | 0.000138949 | 1700125H03Rik | 0.620756173 | -0.015134952 | 0.015868789 | -0.953755955 | 0.340207189 | NA |
| Slc26a7 | 33.00970186 | -0.463455945 | 0.167154853 | -2.77261435 | 0.005560798 | 0.038738759 | Rab43 | 1010.413249 | -0.015138208 | 0.068154477 | -0.222116122 | 0.824223488 | 0.957903916 |
| Clec3b | 43.15523841 | -0.464923471 | 0.151398503 | -3.070859105 | 0.002134438 | 0.020096898 | Mir670 | 1.166737156 | -0.015145763 | 0.022349249 | -0.677685543 | 0.497971095 | NA |
| 2810011L19Rik | 576.5191333 | -0.464945404 | 0.132981853 | -3.496307144 | 0.000471745 | 0.006531924 | Padi3 | 0.715248492 | -0.015161004 | 0.018402352 | -0.823862271 | 0.41001782 | NA |
| Cdca7 | 149.4369424 | -0.465055668 | 0.139861983 | -3.325104207 | 0.000883855 | 0.010382448 | Rab23 | 1300.573011 | -0.015166322 | 0.068462752 | -0.221526614 | 0.824682416 | 0.957903916 |
| 2610034M16Rik | 150.2699268 | -0.466483812 | 0.130439618 | -3.576243311 | 0.000348567 | 0.005248495 | Ddrgk1 | 2009.506883 | -0.015168867 | 0.06537551 | -0.232026751 | 0.816517236 | 0.956279075 |
| Pdgfc | 331.1595035 | -0.467951917 | 0.091213411 | -5.130297329 | 2.89E-07 | 1.80E-05 | Xirp2 | 12.85751988 | -0.01517815 | 0.070795615 | -0.214393931 | 0.830239874 | NA |
| Gm10754 | 95.65462875 | -0.46856232 | 0.141257437 | -3.317080716 | 0.000909633 | 0.010547482 | Mb | 3.941334226 | -0.015187832 | 0.035767285 | -0.424629157 | 0.671107034 | NA |
| Dok4 | 686.064647 | -0.468600865 | 0.076086053 | -6.158827366 | 7.33E-10 | 1.05E-07 | Pnp2 | 30.99786526 | -0.015197088 | 0.086876156 | -0.174928186 | 0.861136062 | 0.968928152 |
| Sox4 | 436.3485372 | -0.469625086 | 0.123897127 | -3.790443718 | 0.000150378 | 0.002721919 | Epb4.2 | 0.688850708 | -0.015203593 | 0.019432969 | -0.782360772 | 0.434002581 | NA |
| Rgs2 | 2708.104062 | -0.46972086 | 0.083137382 | -5.649935656 | 1.61E-08 | 1.60E-06 | Ankrd35 | 704.4636499 | -0.015205178 | 0.072692314 | -0.20917174 | 0.834314173 | 0.961118357 |
| Csmd3 | 1312.055909 | -0.470074532 | 0.108601877 | -4.328419938 | 1.50E-05 | 0.000443226 | BC051537 | 2.72092569 | -0.015220671 | 0.032830537 | -0.46361321 | 0.642924888 | NA |
| Casq2 | 108.8846149 | -0.471048162 | 0.153932084 | -3.060103845 | 0.002212603 | 0.020626994 | Rsrc2 | 2568.191703 | -0.015228539 | 0.053687163 | -0.283653266 | 0.776676112 | 0.947430065 |
| Gpc6 | 541.3085926 | -0.471154641 | 0.092917191 | -5.070694006 | 3.96E-07 | 2.32E-05 | Tjp2 | 1500.042857 | -0.015237741 | 0.067590264 | -0.225442835 | 0.821634791 | 0.957716818 |
| Fam102b | 2977.074862 | -0.471578725 | 0.106390595 | -4.432522658 | 9.31E-06 | 0.000296317 | Krt17 | 3.364229612 | -0.015254189 | 0.042856493 | -0.355936485 | 0.721888131 | NA |
| Wwc1 | 3706.947063 | -0.472068145 | 0.07617525 | -6.197132839 | 5.75E-10 | 8.73E-08 | 4930545L23Rik | 0.397844952 | -0.015296032 | 0.012981756 | -1.17827148 | 0.238688394 | NA |
| Tox | 707.9212671 | -0.472174658 | 0.0884043 | -5.341082457 | 9.24E-08 | 7.09E-06 | Cnfn | 0.397844952 | -0.015296032 | 0.012981756 | -1.17827148 | 0.238688394 | NA |
| Hmgcr | 4416.600257 | -0.47244834 | 0.095628936 | -4.940432896 | 7.79E-07 | 4.04E-05 | Lactb | 576.9308083 | -0.015300794 | 0.067205699 | -0.227671076 | 0.81990196 | 0.957571905 |
| Ccdc3 | 807.4774521 | -0.472896806 | 0.113035946 | -4.183596657 | 2.87E-05 | 0.00074593 | Eif2c1 | 4271.622393 | -0.015310452 | 0.059349511 | -0.257970989 | 0.796429297 | 0.952734447 |
| Fam19a4 | 22.10397271 | -0.473192585 | 0.168066973 | -2.815500131 | 0.004870138 | 0.035393018 | Fbxo48 | 31.33718863 | -0.015313791 | 0.086950276 | -0.176121248 | 0.860198689 | 0.96862798 |
| Stk32a | 356.039342 | -0.474526001 | 0.109599209 | -4.329648035 | 1.49E-05 | 0.000441602 | Acd | 1482.848626 | -0.015319277 | 0.064623738 | -0.237053404 | 0.81261536 | 0.956264994 |
| Rspo2 | 614.9985749 | -0.474531863 | 0.129848705 | -3.65449825 | 0.000257685 | 0.004124564 | Fgfbp3 | 142.76027 | -0.015324846 | 0.092002369 | -0.166570125 | 0.867708302 | 0.970480832 |
| Cdca5 | 8.941967987 | -0.474538934 | 0.160964518 | -2.948096503 | 0.003197372 | 0.026528802 | A930015D03Rik | 42.77334827 | -0.015329389 | 0.092345308 | -0.16600074 | 0.868156368 | 0.970658528 |
| Ust | 1081.167932 | -0.475240707 | 0.097621669 | -4.868188722 | 1.13E-06 | 5.37E-05 | Tmem209 | 1419.646656 | -0.015342172 | 0.069791945 | -0.219827264 | 0.826005686 | 0.958041984 |
| Kcnf1 | 6721.608509 | -0.476051288 | 0.067337726 | -7.069607413 | 1.55E-12 | 4.38E-10 | Akr1c18 | 68.34866461 | -0.01534296 | 0.09470877 | -0.162001472 | 0.871304695 | 0.97153269 |
| St3gal1 | 1253.639761 | -0.476123517 | 0.070666416 | -6.737620794 | 1.61E-11 | 3.78E-09 | Wdr38 | 5.834878009 | -0.015344366 | 0.0491359 | -0.312284208 | 0.754824541 | NA |
| Fjx1 | 2017.784391 | -0.476842571 | 0.123456393 | -3.862437249 | 0.000112261 | 0.002134206 | 1700028E10Rik | 7.200830189 | -0.015352139 | 0.05938314 | -0.258526893 | 0.796000296 | NA |
| Dst | 15807.14299 | -0.477766466 | 0.052654442 | -9.073621226 | 1.15E-19 | 1.11E-16 | Pop1 | 350.915863 | -0.015356353 | 0.083125375 | -0.184737251 | 0.853435127 | 0.966905493 |
| Steap3 | 225.1031783 | -0.477927192 | 0.153256027 | -3.118488718 | 0.001817811 | 0.017803517 | 2410004N09Rik | 104.3897386 | -0.01537588 | 0.094611206 | -0.162516482 | 0.87089915 | 0.97153269 |
| Pde3a | 113.374785 | -0.478911227 | 0.15777332 | -3.035438618 | 0.002401861 | 0.021823847 | 4933422A05Rik | 1.083795384 | -0.015383075 | 0.02488722 | -0.618111435 | 0.536501885 | NA |
| Aoc3 | 43.96556511 | -0.479526232 | 0.163358724 | -2.935418565 | 0.00333098 | 0.027292548 | Tram1 | 946.5027737 | -0.015387034 | 0.060499489 | -0.254333282 | 0.799238089 | 0.953180007 |
| Chat | 302.373814 | -0.480156554 | 0.140264511 | -3.423221949 | 0.000618835 | 0.007990195 | Dleu2 | 128.8139356 | -0.015396558 | 0.092457591 | -0.166525624 | 0.86774332 | 0.970480832 |
| Macf1 | 14882.16153 | -0.480417915 | 0.077097708 | -6.231286622 | 4.63E-10 | 7.10E-08 | Camk1 | 2569.371688 | -0.015436836 | 0.057048511 | -0.270591395 | 0.786705315 | 0.950564083 |
| Hes5 | 357.7953987 | -0.480614976 | 0.137200044 | -3.503023495 | 0.000460009 | 0.006432651 | AU021092 | 94.19210239 | -0.015438801 | 0.093349686 | -0.165386753 | 0.86863958 | 0.97067917 |
| Acta2 | 546.0208198 | -0.480993306 | 0.13431229 | -3.581156325 | 0.000342077 | 0.005171348 | Pgbd5 | 7153.614185 | -0.015447756 | 0.064383854 | -0.239932147 | 0.810382859 | 0.955876688 |
| Lhfpl5 | 123.7034971 | -0.481057187 | 0.140140875 | -3.432668647 | 0.000597672 | 0.007794891 | M1ap | 1.833683077 | -0.015451212 | 0.032218954 | -0.479569002 | 0.631533892 | NA |
| 2810417H13Rik | 14.22484206 | -0.481435825 | 0.155255465 | -3.100926759 | 0.00192916 | 0.01865818 | Cdkn1c | 239.0325539 | -0.015454403 | 0.094032262 | -0.164352137 | 0.86945394 | 0.970842335 |
| Gpx6 | 40.98740255 | -0.481960137 | 0.168030922 | -2.868282411 | 0.00412707 | 0.031734401 | 2610035F20Rik | 6.052528907 | -0.015457451 | 0.051911767 | -0.297763911 | 0.765883361 | NA |
| Scgn | 60.29875569 | -0.482377882 | 0.130562632 | -3.694609063 | 0.000220225 | 0.003633836 | D16Ertd472e | 473.1518093 | -0.0154594 | 0.079210272 | -0.195169137 | 0.84526054 | 0.96466488 |
| Mis18bp1 | 10.88304371 | -0.482902521 | 0.163821547 | -2.947735075 | 0.003201113 | 0.026528802 | Sypl2 | 127.1166887 | -0.01547078 | 0.093725634 | -0.165064554 | 0.868893172 | 0.970788711 |
| Tlr5 | 20.9814671 | -0.485553471 | 0.167894193 | -2.89202064 | 0.003827728 | 0.030212337 | Lipm | 4.350341414 | -0.015473222 | 0.045029815 | -0.343621708 | 0.731130797 | NA |
| Ptprb | 2208.873738 | -0.485830172 | 0.112549699 | -4.316583484 | 1.58E-05 | 0.000464125 | Klk7 | 1.088407242 | -0.015474247 | 0.024098129 | -0.642134775 | 0.520785678 | NA |
| Zic4 | 115.8168944 | -0.486301673 | 0.165587854 | -2.936819704 | 0.003315968 | 0.027241576 | Gucy1a3 | 5725.253391 | -0.015485628 | 0.073733132 | -0.210022652 | 0.833649994 | 0.960926822 |
| Pcdh10 | 7770.362955 | -0.486566448 | 0.063181285 | -7.701116683 | 1.35E-14 | 6.33E-12 | Ppp1r3a | 2.564106344 | -0.015498568 | 0.029949929 | -0.517482617 | 0.6048193 | NA |
| Celsr1 | 381.849956 | -0.487017468 | 0.102295997 | -4.760865371 | 1.93E-06 | 8.63E-05 | Nid1 | 369.5252661 | -0.015502524 | 0.087433124 | -0.17730722 | 0.85926708 | 0.968520191 |
| Tyrp1 | 9.268890722 | -0.488700192 | 0.139250044 | -3.509515538 | 0.000448924 | 0.006323296 | Car5a | 7.286279045 | -0.015502726 | 0.049055336 | -0.316025278 | 0.751983321 | NA |
| Grm4 | 833.9925681 | -0.488740893 | 0.091609711 | -5.335033688 | 9.55E-08 | 7.22E-06 | Rad9a | 449.2376603 | -0.015527407 | 0.075831449 | -0.204762099 | 0.837758 | 0.961858836 |
| Lgr5 | 199.6824086 | -0.488976081 | 0.164818895 | -2.966747724 | 0.003009678 | 0.025407408 | Thyn1 | 1159.156575 | -0.015528792 | 0.063047671 | -0.246302394 | 0.805448157 | 0.95517511 |
| Adamtsl3 | 71.63363357 | -0.489227947 | 0.164556529 | -2.973008418 | 0.002948964 | 0.025077521 | Mnd1 | 13.68509732 | -0.015543092 | 0.072541203 | -0.2142657 | 0.830339864 | NA |
| Tln2 | 6148.558021 | -0.489390087 | 0.055067926 | -8.887026012 | 6.28E-19 | 5.40E-16 | Cenpc1 | 933.9866073 | -0.015547749 | 0.064721866 | -0.240224062 | 0.810156561 | 0.955876688 |
| Bcl11b | 4015.449145 | -0.48964874 | 0.083071657 | -5.894293619 | 3.76E-09 | 4.63E-07 | 4930447N08Rik | 5.358918605 | -0.015549127 | 0.03915464 | -0.397120928 | 0.691278288 | NA |
| Sox1 | 622.5741073 | -0.489797483 | 0.134878784 | -3.631390126 | 0.000281899 | 0.004420787 | Slc30a7 | 925.8733624 | -0.01555159 | 0.060717946 | -0.256128398 | 0.797851695 | 0.952967259 |
| Map1b | 39800.8502 | -0.491352995 | 0.070977794 | -6.922629834 | 4.43E-12 | 1.21E-09 | E330012B07Rik | 0.882475236 | -0.015560588 | 0.022277436 | -0.698490962 | 0.484870208 | NA |
| Isoc1 | 852.470721 | -0.491575813 | 0.141160478 | -3.482389819 | 0.00049696 | 0.006796205 | Gm20139 | 0.604369265 | -0.015570621 | 0.015977509 | -0.974533666 | 0.329791632 | NA |
| Gnal | 15366.84569 | -0.492662101 | 0.087409768 | -5.63623623 | 1.74E-08 | 1.70E-06 | Mylk2 | 0.846191468 | -0.015582861 | 0.018586259 | -0.838407608 | 0.401801818 | NA |
| Fstl5 | 873.7978585 | -0.49283804 | 0.125740178 | -3.919495332 | 8.87E-05 | 0.001787846 | Kcnh6 | 117.5273428 | -0.015603552 | 0.093470856 | -0.166934941 | 0.86742124 | 0.970480832 |
| Chst3 | 145.9086893 | -0.493034599 | 0.135474847 | -3.639307305 | 0.000273372 | 0.004326488 | Fam175a | 166.4856219 | -0.015618032 | 0.093689412 | -0.166700074 | 0.867606047 | 0.970480832 |
| Dnahc8 | 21.32577821 | -0.493411844 | 0.16812101 | -2.934861286 | 0.003336968 | 0.027312724 | Usp20 | 4149.550936 | -0.015630252 | 0.061765567 | -0.253057696 | 0.800223629 | 0.953228631 |
| Khdrbs2 | 187.6412855 | -0.495382893 | 0.106001766 | -4.673345669 | 2.96E-06 | 0.000122111 | Kif15 | 14.90649256 | -0.015633001 | 0.078015505 | -0.200383263 | 0.841180849 | NA |
| Chrna5 | 101.8248127 | -0.495730856 | 0.150933333 | -3.28443589 | 0.001021867 | 0.011539952 | Gm16880 | 0.411706162 | -0.015648874 | 0.013129479 | -1.191888368 | 0.23330502 | NA |
| Fzd5 | 126.8742834 | -0.497164192 | 0.134494163 | -3.696548455 | 0.000218551 | 0.003621628 | Lipf | 0.411706162 | -0.015648874 | 0.013129479 | -1.191888368 | 0.23330502 | NA |
| Gabrg3 | 247.3246116 | -0.497234577 | 0.099394789 | -5.002622211 | 5.66E-07 | 3.10E-05 | Hadha | 2494.464711 | -0.015652441 | 0.058290946 | -0.268522679 | 0.78829702 | 0.951142331 |
| Prc1 | 48.80920691 | -0.49727722 | 0.163712073 | -3.037510981 | 0.002385407 | 0.021753678 | Gm5523 | 2.312306773 | -0.015661886 | 0.030504294 | -0.513432167 | 0.607649049 | NA |
| Cgref1 | 771.3410501 | -0.49743962 | 0.061802736 | -8.048828404 | 8.36E-16 | 5.18E-13 | Slc13a4 | 588.4630619 | -0.015664938 | 0.065658601 | -0.238581658 | 0.811429989 | 0.956135608 |
| E130309F12Rik | 754.8683557 | -0.498024044 | 0.123386833 | -4.036281931 | 5.43E-05 | 0.001230118 | Upk3a | 1.885251996 | -0.015666494 | 0.029730915 | -0.526942887 | 0.59823325 | NA |
| Adcy5 | 13613.71904 | -0.498463838 | 0.082429008 | -6.047189559 | 1.47E-09 | 1.90E-07 | Skap2 | 496.2721513 | -0.01568198 | 0.066421815 | -0.236096837 | 0.813357528 | 0.956264994 |
| Dsel | 630.5473734 | -0.499066489 | 0.106674023 | -4.678425678 | 2.89E-06 | 0.000119762 | Spata20 | 5.922937722 | -0.015691818 | 0.05321143 | -0.294895634 | 0.768073607 | NA |
| Lmo4 | 6647.496987 | -0.501948986 | 0.085117901 | -5.897102487 | 3.70E-09 | 4.59E-07 | Dsg2 | 106.0236825 | -0.015694581 | 0.094743128 | -0.165654029 | 0.868429226 | 0.970658528 |
| Gm7244 | 11.75286263 | -0.50199937 | 0.162214113 | -3.09467137 | 0.001970311 | 0.018926221 | Taok2 | 6801.371059 | -0.015701304 | 0.054683402 | -0.287131073 | 0.774011961 | 0.947202842 |
| Fam64a | 10.3431049 | -0.503334519 | 0.159811692 | -3.149547516 | 0.001635235 | 0.016452164 | Uggt1 | 3063.878217 | -0.015711497 | 0.05790666 | -0.271324523 | 0.786141447 | 0.950484821 |
| Hs3st5 | 247.2097574 | -0.503862373 | 0.128444998 | -3.922787033 | 8.75E-05 | 0.001768185 | Stard13 | 605.8827332 | -0.015715423 | 0.068661689 | -0.228881978 | 0.818960648 | 0.957334483 |
| Abcc12 | 48.0976981 | -0.503892538 | 0.156422549 | -3.221354846 | 0.001275861 | 0.013689999 | BC005537 | 6808.673588 | -0.015717571 | 0.048186088 | -0.326184829 | 0.744284519 | 0.940096833 |
| Cdyl2 | 1144.80427 | -0.505191515 | 0.091465725 | -5.523287732 | 3.33E-08 | 2.96E-06 | A530053G22Rik | 1.720483642 | -0.015729018 | 0.027106167 | -0.580274527 | 0.561729502 | NA |
| Slc22a2 | 32.02283487 | -0.505432554 | 0.167308239 | -3.020966318 | 0.002519694 | 0.02250152 | Nmrk2 | 5.98511608 | -0.015747628 | 0.040310486 | -0.39065835 | 0.696049789 | NA |
| Des | 85.14853111 | -0.506622969 | 0.144284167 | -3.511285955 | 0.000445944 | 0.006298508 | 4833412C05Rik | 28.44906964 | -0.015747835 | 0.082671307 | -0.190487311 | 0.848927285 | 0.965892061 |
| Zic3 | 127.3222856 | -0.507331392 | 0.141520435 | -3.584863146 | 0.000337255 | 0.005132306 | 4930577N17Rik | 17.48180138 | -0.015757064 | 0.076215824 | -0.20674268 | 0.836210824 | NA |
| Kcnh8 | 27.42040681 | -0.508852299 | 0.168121552 | -3.026692844 | 0.002472451 | 0.022169068 | Fam161a | 114.3384175 | -0.015762087 | 0.094452781 | -0.166877953 | 0.867466081 | 0.970480832 |
| Mest | 1061.817526 | -0.510000942 | 0.093028309 | -5.482212309 | 4.20E-08 | 3.60E-06 | D830013O20Rik | 0.41620922 | -0.015769565 | 0.013179636 | -1.196509889 | 0.231497642 | NA |
| Reln | 3254.117389 | -0.510583866 | 0.10675359 | -4.782826174 | 1.73E-06 | 7.88E-05 | Spint4 | 0.41620922 | -0.015769565 | 0.013179636 | -1.196509889 | 0.231497642 | NA |
| 1700001L19Rik | 365.364897 | -0.511723503 | 0.106732134 | -4.794465192 | 1.63E-06 | 7.50E-05 | Trim55 | 0.41620922 | -0.015769565 | 0.013179636 | -1.196509889 | 0.231497642 | NA |
| Fxyd2 | 90.91497288 | -0.513027388 | 0.142391264 | -3.602941464 | 0.000314636 | 0.004850723 | Tbck | 794.6266304 | -0.015769736 | 0.066131414 | -0.238460585 | 0.811523882 | 0.956135608 |
| AI606473 | 41.40187103 | -0.513742882 | 0.162320478 | -3.164991187 | 0.001550876 | 0.015892381 | Slc35a3 | 523.7483129 | -0.01577235 | 0.076035114 | -0.207435083 | 0.835670086 | 0.961504006 |
| Serpinh1 | 598.5418462 | -0.513822388 | 0.0920663 | -5.581004015 | 2.39E-08 | 2.23E-06 | Reep2 | 5541.734573 | -0.015802088 | 0.055596527 | -0.284227966 | 0.776235685 | 0.947241001 |
| Kif26b | 346.9705902 | -0.513883098 | 0.08442379 | -6.086946563 | 1.15E-09 | 1.54E-07 | Glrp1 | 6.339215949 | -0.015812723 | 0.046536107 | -0.339794708 | 0.734011134 | NA |
| Alx4 | 94.55830759 | -0.513895215 | 0.162403315 | -3.164314823 | 0.001554485 | 0.01589782 | Olfr56 | 0.603776958 | -0.01583642 | 0.015965775 | -0.991897979 | 0.321247295 | NA |
| E130008D07Rik | 184.7770836 | -0.514051321 | 0.137886041 | -3.728088198 | 0.000192938 | 0.003311243 | Mcmbp | 1092.986833 | -0.015851695 | 0.067411739 | -0.235147403 | 0.814094328 | 0.956264994 |
| Agbl2 | 73.2121449 | -0.515412708 | 0.16243905 | -3.172960617 | 0.00150893 | 0.015534459 | Cnih4 | 1462.772037 | -0.015868084 | 0.059635335 | -0.266085264 | 0.790173544 | 0.951607109 |
| Gm19522 | 49.98368768 | -0.51541897 | 0.156610782 | -3.291082293 | 0.000998027 | 0.011345145 | Mlana | 0.407027086 | -0.015873735 | 0.013222472 | -1.200511886 | 0.229940599 | NA |
| Olfml2a | 107.5034229 | -0.516083838 | 0.145312584 | -3.551542635 | 0.00038298 | 0.005651325 | Mycs | 0.407027086 | -0.015873735 | 0.013222472 | -1.200511886 | 0.229940599 | NA |
| Cntn5 | 300.0182222 | -0.517378742 | 0.097436637 | -5.309899434 | 1.10E-07 | 8.02E-06 | Gm13497 | 1.027545051 | -0.015874426 | 0.019472628 | -0.815217453 | 0.414947849 | NA |
| Gm5607 | 604.0169666 | -0.518529593 | 0.149627345 | -3.465473462 | 0.000529299 | 0.007115035 | Tab1 | 998.4255223 | -0.015877768 | 0.071419849 | -0.222315904 | 0.824067972 | 0.957903916 |
| Dcaf12l2 | 37.56002595 | -0.519668939 | 0.164844884 | -3.15247235 | 0.001618942 | 0.016318843 | Cml1 | 315.2452292 | -0.015893163 | 0.084165461 | -0.18883237 | 0.850224201 | 0.966260848 |
| Tnnc1 | 238.9691526 | -0.520337808 | 0.150507367 | -3.457224837 | 0.000545769 | 0.007296074 | Slc17a2 | 0.593945617 | -0.015908748 | 0.015926267 | -0.998899995 | 0.317843139 | NA |
| Pcsk5 | 504.2211175 | -0.520831141 | 0.088730841 | -5.869787019 | 4.36E-09 | 5.32E-07 | Btg1 | 978.7452275 | -0.015910227 | 0.07238128 | -0.219811353 | 0.826018077 | 0.958041984 |
| Klhl13 | 2168.725409 | -0.520965248 | 0.116114226 | -4.48666167 | 7.23E-06 | 0.00024475 | Golga2 | 4877.686947 | -0.015910847 | 0.047465714 | -0.335207161 | 0.737468821 | 0.937206199 |
| Neb | 86.64729108 | -0.523143601 | 0.159120429 | -3.287721153 | 0.001010018 | 0.011447859 | Eef2k | 2025.14673 | -0.015913374 | 0.055335472 | -0.287579985 | 0.773668268 | 0.947121017 |
| Mrc1 | 208.9341952 | -0.524572285 | 0.147747236 | -3.550471063 | 0.000384542 | 0.005668982 | Zfp277 | 840.024259 | -0.015914406 | 0.072047244 | -0.220888476 | 0.825179271 | 0.957903916 |
| P2ry1 | 208.3984836 | -0.526538526 | 0.150096264 | -3.508005537 | 0.00045148 | 0.006347755 | Tex33 | 0.767812082 | -0.015966087 | 0.018915895 | -0.844056667 | 0.398637741 | NA |
| Apc | 14761.67817 | -0.526857181 | 0.071061431 | -7.414108793 | 1.22E-13 | 4.86E-11 | Kdm2b | 1071.395185 | -0.015969791 | 0.055874809 | -0.285813793 | 0.775020743 | 0.947238918 |
| Cdhr1 | 1095.455868 | -0.52730108 | 0.149507297 | -3.526925383 | 0.000420415 | 0.006030802 | Gbx1 | 13.72742537 | -0.01598929 | 0.067699697 | -0.236179643 | 0.813293275 | NA |
| Slc10a4 | 185.0474056 | -0.527329775 | 0.161383949 | -3.267547836 | 0.001084835 | 0.012171208 | Mroh2a | 0.941360207 | -0.016004608 | 0.021307679 | -0.751119257 | 0.452580887 | NA |
| Loxl1 | 85.82401008 | -0.527388471 | 0.152112538 | -3.467094027 | 0.000526118 | 0.007094577 | Ccdc125 | 11.15303906 | -0.016018143 | 0.064427827 | -0.248621497 | 0.803653579 | NA |
| Ranbp3l | 206.6090881 | -0.527575365 | 0.102062114 | -5.169159691 | 2.35E-07 | 1.50E-05 | Gpr82 | 0.817903953 | -0.016029174 | 0.018852408 | -0.850245457 | 0.395188634 | NA |
| Kcnb2 | 433.5559473 | -0.528183751 | 0.119116744 | -4.434168814 | 9.24E-06 | 0.000295885 | Pla2g7 | 6173.78166 | -0.016032404 | 0.051421933 | -0.311781441 | 0.75520663 | 0.942139348 |
| Dcc | 1013.2957 | -0.5288814 | 0.123470889 | -4.283450157 | 1.84E-05 | 0.000521593 | Mfsd7c | 125.0329891 | -0.016037589 | 0.091548082 | -0.17518214 | 0.860936518 | 0.968853723 |
| Col12a1 | 633.761478 | -0.530149493 | 0.100484782 | -5.275918211 | 1.32E-07 | 9.39E-06 | Cnnm3 | 1439.650687 | -0.016050496 | 0.060066205 | -0.267213417 | 0.789304846 | 0.951331082 |
| Igsf3 | 906.4658855 | -0.530934293 | 0.108739997 | -4.882603538 | 1.05E-06 | 5.05E-05 | Scarna13 | 4.927500618 | -0.016054689 | 0.048676901 | -0.329821509 | 0.741534834 | NA |
| Bche | 207.2537614 | -0.531509482 | 0.15877572 | -3.34754888 | 0.000815296 | 0.009717075 | 1110008L16Rik | 288.1971638 | -0.016069704 | 0.084065123 | -0.191157797 | 0.848401967 | 0.965746177 |
| Adcyap1 | 578.8085298 | -0.532217481 | 0.11357997 | -4.685839231 | 2.79E-06 | 0.000116128 | Ppig | 2515.492643 | -0.016083277 | 0.075254743 | -0.213717788 | 0.830767139 | 0.960118026 |
| Sp8 | 119.9236596 | -0.534441798 | 0.151541326 | -3.526706627 | 0.000420763 | 0.006030802 | Tnfsf9 | 11.76975247 | -0.016084405 | 0.068602273 | -0.234458772 | 0.814628836 | NA |
| Homer2 | 2865.701092 | -0.536601976 | 0.086029228 | -6.237437978 | 4.45E-10 | 7.00E-08 | Obp1a | 2.584336722 | -0.016090392 | 0.025384688 | -0.633862136 | 0.526170798 | NA |
| Hs3st2 | 1664.786729 | -0.537448585 | 0.116381354 | -4.617995626 | 3.87E-06 | 0.000149512 | Rasd2 | 8973.813095 | -0.01609549 | 0.085491936 | -0.188269105 | 0.850665703 | 0.966301569 |
| Fam198b | 254.324318 | -0.539808915 | 0.129329578 | -4.173901478 | 2.99E-05 | 0.0007681 | Treml1 | 4.460721264 | -0.01610317 | 0.043691768 | -0.368563026 | 0.712453459 | NA |
| Draxin | 33.65976406 | -0.539857461 | 0.165193998 | -3.268021039 | 0.001083023 | 0.012159682 | 4930503E14Rik | 1.497647165 | -0.016106017 | 0.029968142 | -0.537437977 | 0.590965117 | NA |
| Tenm1 | 1124.117494 | -0.54090152 | 0.109304171 | -4.948589938 | 7.48E-07 | 3.91E-05 | Whrn | 1504.147398 | -0.016106055 | 0.057145257 | -0.281844121 | 0.778063038 | 0.94784716 |
| Tenm4 | 3813.222659 | -0.541281378 | 0.092773732 | -5.834424958 | 5.40E-09 | 6.29E-07 | Tymp | 11.62648812 | -0.016107001 | 0.061360406 | -0.262498279 | 0.792937312 | NA |
| Fbn2 | 109.6071113 | -0.541846825 | 0.160485255 | -3.376302851 | 0.00073467 | 0.008991293 | Zzz3 | 2501.565554 | -0.016107693 | 0.06096176 | -0.26422618 | 0.791605639 | 0.951728598 |
| Nusap1 | 21.25068205 | -0.542459079 | 0.167309341 | -3.242252202 | 0.00118589 | 0.012948685 | Brat1 | 0.631880482 | -0.016121663 | 0.01641566 | -0.982090435 | 0.326055297 | NA |
| Tfcp2l1 | 474.7655235 | -0.54322369 | 0.085687923 | -6.339559512 | 2.30E-10 | 4.06E-08 | Sema3f | 559.2375161 | -0.016125966 | 0.078248839 | -0.206085694 | 0.836723974 | 0.961729368 |
| Pcdh12 | 16.6357907 | -0.54394977 | 0.167050933 | -3.256191151 | 0.001129177 | 0.012541555 | Tbc1d2 | 199.5177719 | -0.016131148 | 0.092832964 | -0.173765303 | 0.862049914 | 0.969505811 |
| Itm2a | 1626.313659 | -0.544095902 | 0.141268752 | -3.851495065 | 0.000117399 | 0.002212869 | Cdr1 | 0.617638168 | -0.016132876 | 0.01605715 | -1.004716048 | 0.315033599 | NA |
| Gprin3 | 396.3241914 | -0.545681997 | 0.127894143 | -4.266669179 | 1.98E-05 | 0.000556922 | Troap | 7.899980241 | -0.016137574 | 0.053224615 | -0.303197567 | 0.761739303 | NA |
| Slc5a7 | 432.889718 | -0.546010666 | 0.150739207 | -3.622220644 | 0.000292085 | 0.004556528 | Rpl30 | 8.953789965 | -0.016141321 | 0.065124522 | -0.247853194 | 0.804247996 | NA |
| Asic4 | 1140.803113 | -0.547545147 | 0.087860002 | -6.232018359 | 4.60E-10 | 7.10E-08 | Dusp8 | 7565.240847 | -0.016162111 | 0.039454896 | -0.409635124 | 0.682073627 | 0.922544285 |
| Zic1 | 538.5450475 | -0.549320527 | 0.166857702 | -3.292149659 | 0.000994247 | 0.011333586 | Gpatch2 | 755.9572319 | -0.016177158 | 0.063605474 | -0.254335943 | 0.799236034 | 0.953180007 |
| Gng7 | 14281.16362 | -0.550281163 | 0.085891828 | -6.40667657 | 1.49E-10 | 2.74E-08 | Lama1 | 167.8008718 | -0.016179034 | 0.094147626 | -0.171847504 | 0.863557418 | 0.969836459 |
| Nes | 625.753946 | -0.550390045 | 0.097642289 | -5.636799898 | 1.73E-08 | 1.70E-06 | Dnaja4 | 2535.990096 | -0.016190358 | 0.0628535 | -0.257588809 | 0.796724268 | 0.952885371 |
| Zfhx3 | 290.9259067 | -0.550950876 | 0.137251898 | -4.01415851 | 5.97E-05 | 0.001322382 | Foxp3 | 3.475135377 | -0.016200088 | 0.040946546 | -0.395639922 | 0.692370682 | NA |
| Sfrp1 | 505.7902792 | -0.550998761 | 0.096104219 | -5.73334621 | 9.85E-09 | 1.04E-06 | Hsdl2 | 1412.470834 | -0.016209401 | 0.069382147 | -0.233624958 | 0.815276151 | 0.956264994 |
| Pclo | 10455.35487 | -0.552742744 | 0.065382255 | -8.454017728 | 2.81E-17 | 1.98E-14 | Senp1 | 691.977563 | -0.016234986 | 0.072176141 | -0.22493563 | 0.822029351 | 0.95778569 |
| Mxd3 | 34.97445607 | -0.552872241 | 0.166518824 | -3.320178628 | 0.000899599 | 0.010498156 | Gpr97 | 4.195243604 | -0.016256982 | 0.040829155 | -0.398170909 | 0.690504208 | NA |
| Lama3 | 279.7090196 | -0.553979536 | 0.13261753 | -4.17727231 | 2.95E-05 | 0.000759536 | BB031773 | 32.74659619 | -0.016258463 | 0.086179572 | -0.188657964 | 0.8503609 | 0.966260848 |
| Robo1 | 1844.399059 | -0.555201819 | 0.085857011 | -6.466586857 | 1.00E-10 | 2.02E-08 | Aebp2 | 2673.855368 | -0.016267337 | 0.064414286 | -0.25254238 | 0.80062186 | 0.953228631 |
| Ccbe1 | 294.2530545 | -0.557108616 | 0.13324599 | -4.181053524 | 2.90E-05 | 0.0007518 | A630095N17Rik | 1.165960768 | -0.016273663 | 0.023724793 | -0.68593488 | 0.492754174 | NA |
| Wnt7a | 484.6670115 | -0.557153429 | 0.113010566 | -4.930100346 | 8.22E-07 | 4.18E-05 | Asb15 | 2.048915888 | -0.016276323 | 0.034117529 | -0.477066299 | 0.633314912 | NA |
| Pcp4l1 | 6945.445593 | -0.557778212 | 0.120608778 | -4.624690016 | 3.75E-06 | 0.000146048 | Pnliprp2 | 2.650133983 | -0.016278051 | 0.037094413 | -0.438827575 | 0.660786478 | NA |
| Tead2 | 98.07639631 | -0.55836474 | 0.143939994 | -3.879149396 | 0.000104822 | 0.002035235 | Gm13051 | 3.538946092 | -0.016294584 | 0.037281083 | -0.437073796 | 0.66205783 | NA |
| Sebox | 96.58272711 | -0.559710047 | 0.158231969 | -3.537275369 | 0.000404278 | 0.005876062 | Tnfsf18 | 0.902588669 | -0.016298147 | 0.022662937 | -0.719154225 | 0.4720459 | NA |
| Rhcg | 37.07514021 | -0.559916956 | 0.162121536 | -3.453686484 | 0.00055298 | 0.007360713 | Txk | 6.024010165 | -0.016309651 | 0.055945094 | -0.291529608 | 0.770646303 | NA |
| Cyp2f2 | 15.7864392 | -0.560032805 | 0.159131162 | -3.519315753 | 0.000432661 | 0.006144506 | Mir1934 | 0.657115133 | -0.016316043 | 0.02053346 | -0.794607593 | 0.4268418 | NA |
| Gjb2 | 394.7619326 | -0.56119568 | 0.122243893 | -4.590787043 | 4.42E-06 | 0.000166574 | Lce3c | 0.953969319 | -0.016324556 | 0.017471445 | -0.934356386 | 0.350120087 | NA |
| Phex | 44.42078854 | -0.565549626 | 0.168071315 | -3.364938434 | 0.000765607 | 0.009245769 | Fam163b | 7758.956276 | -0.016345973 | 0.050365725 | -0.324545175 | 0.745525024 | 0.940632024 |
| Nlrp10 | 20.16829148 | -0.565975847 | 0.160674734 | -3.522494378 | 0.000427506 | 0.006088033 | Hnf1a | 0.323722592 | -0.01635091 | 0.013414305 | -1.218915995 | 0.222876078 | NA |
| Adam12 | 210.3778322 | -0.566639059 | 0.121878369 | -4.649217627 | 3.33E-06 | 0.000133399 | Them5 | 0.323722592 | -0.01635091 | 0.013414305 | -1.218915995 | 0.222876078 | NA |
| Tmem40 | 91.26251538 | -0.569360844 | 0.163430244 | -3.483815662 | 0.00049432 | 0.006777868 | 1700007K09Rik | 4.725129896 | -0.016355126 | 0.043574854 | -0.375334037 | 0.707412056 | NA |
| Pde1c | 593.6563004 | -0.57011992 | 0.128332442 | -4.442523735 | 8.89E-06 | 0.000286993 | Map3k9 | 4708.941278 | -0.016365347 | 0.05096121 | -0.321133399 | 0.748109301 | 0.9410864 |
| Oprm1 | 65.46653988 | -0.570622301 | 0.161664629 | -3.529666979 | 0.000416083 | 0.005984147 | Tradd | 180.0105838 | -0.016369857 | 0.087639408 | -0.186786487 | 0.851828042 | 0.96669173 |
| Pcdh17 | 3009.395467 | -0.571025763 | 0.114511842 | -4.986608851 | 6.14E-07 | 3.32E-05 | Tmem243 | 213.485771 | -0.016376434 | 0.092338434 | -0.177352308 | 0.859231666 | 0.968520191 |

| | | | | | | | | | | | | | | | |
|---|---|---|---|---|---|---|---|---|---|---|---|---|---|---|---|
| Inhba | 185.1593459 | -0.571767514 | 0.109933705 | -5.201021065 | 1.98E-07 | 1.32E-05 | | | Ehf | 8.295292148 | -0.016405555 | 0.048794926 | -0.33621436 | 0.736709225 | NA |
| Pdgfrl | 34.54616712 | -0.572529831 | 0.159532252 | -3.588803046 | 0.0003322 | 0.005071036 | | | Chst2 | 6152.779704 | -0.016420508 | 0.062802243 | -0.261463718 | 0.793734925 | 0.951728598 |
| C1ql1 | 215.1760049 | -0.572670363 | 0.128824901 | -4.445339043 | 8.78E-06 | 0.000284411 | | | Nkpd1 | 26.70778424 | -0.016424195 | 0.083751783 | -0.196105618 | 0.844527499 | 0.964328285 |
| Ddah2 | 303.6091624 | -0.572737076 | 0.101963355 | -5.617087404 | 1.94E-08 | 1.89E-06 | | | Hbb-y | 1.266060517 | -0.016451359 | 0.026747618 | -0.615058846 | 0.538515859 | NA |
| Asxl3 | 467.935201 | -0.57285369 | 0.109521558 | -5.230510802 | 1.69E-07 | 1.15E-05 | | | Lrch1 | 707.4113842 | -0.016452288 | 0.069606724 | -0.236360618 | 0.813152853 | 0.956264994 |
| Lrp4 | 1783.262611 | -0.572980762 | 0.076153661 | -7.524008139 | 5.31E-14 | 2.29E-11 | | | Alx4 | 65.34837582 | -0.016482867 | 0.085490093 | -0.192804408 | 0.847112151 | 0.965275633 |
| Fn1 | 2353.942416 | -0.574702634 | 0.089283382 | -6.436837645 | 1.22E-10 | 2.37E-08 | | | Nlrp12 | 1.279829732 | -0.016486878 | 0.02676156 | -0.616065646 | 0.537851195 | NA |
| Ankrd63 | 2370.300801 | -0.576063712 | 0.09372285 | -6.146459599 | 7.92E-10 | 1.12E-07 | | | Nanog | 1.837764489 | -0.016491714 | 0.031803469 | -0.518550768 | 0.604074049 | NA |
| Rrm2 | 71.02413499 | -0.576089466 | 0.141869642 | -4.060695857 | 4.89E-05 | 0.001129761 | | | Jph2 | 50.48283032 | -0.016503107 | 0.092992992 | -0.177466134 | 0.859142264 | 0.968520191 |
| Sall3 | 314.0693231 | -0.576720517 | 0.139375943 | -4.137877051 | 3.51E-05 | 0.000862091 | | | Ccr3 | 1.122568499 | -0.016508527 | 0.025492784 | -0.647576473 | 0.517258918 | NA |
| Hdac9 | 1424.869924 | -0.57841612 | 0.091322279 | -6.333789815 | 2.39E-10 | 4.13E-08 | | | E130006D01Rik | 16.47743923 | -0.016511284 | 0.076217573 | -0.216633563 | 0.828493928 | NA |
| Marcksl1 | 1937.714658 | -0.579730554 | 0.093865646 | -6.176173917 | 6.57E-10 | 9.78E-08 | | | Kcnk5 | 27.095239 | -0.016512449 | 0.080486315 | -0.205158474 | 0.837448313 | 0.961831963 |
| Emp2 | 902.8232293 | -0.586712933 | 0.121568292 | -4.826200332 | 1.39E-06 | 6.48E-05 | | | Nhsl1 | 1087.797462 | -0.016522236 | 0.06856663 | -0.240966135 | 0.809581364 | 0.95578502 |
| C730002L08Rik | 168.9063842 | -0.587160071 | 0.12655628 | -4.639517463 | 3.49E-06 | 0.00013874 | | | Snrpg | 523.4707825 | -0.016544592 | 0.070096003 | -0.236027609 | 0.813411246 | 0.956264994 |
| Fabp7 | 865.4012846 | -0.58950708 | 0.167964267 | -3.509717219 | 0.000448583 | 0.006323296 | | | Itih4 | 4.422393775 | -0.016545823 | 0.046798174 | -0.353557025 | 0.723670885 | NA |
| Th | 283.198081 | -0.591860304 | 0.148074614 | -3.997040999 | 6.41E-05 | 0.001387951 | | | Psca | 1.500426011 | -0.016559273 | 0.027975301 | -0.591924744 | 0.553900983 | NA |
| Pnp2 | 38.38040982 | -0.592811953 | 0.164781111 | -3.597572244 | 0.000321201 | 0.004932302 | | | Tcte2 | 76.09221761 | -0.016573067 | 0.095102127 | -0.174265999 | 0.861656419 | 0.969358561 |
| Kntc1 | 23.58020688 | -0.595938928 | 0.166545025 | -3.578245152 | 0.000345909 | 0.005213534 | | | Mrps15 | 927.7097035 | -0.016578919 | 0.070530029 | -0.235061843 | 0.814160734 | 0.956264994 |
| Wnt7b | 1369.551654 | -0.597448458 | 0.109164793 | -5.472904218 | 4.43E-08 | 3.73E-06 | | | Smc6 | 1751.248001 | -0.016581085 | 0.074567475 | -0.222363504 | 0.824030921 | 0.957903916 |
| Hspg2 | 463.3904859 | -0.60344803 | 0.094049578 | -6.416275816 | 1.40E-10 | 2.61E-08 | | | Grm6 | 0.434749506 | -0.016584696 | 0.013512962 | -1.227317647 | 0.219703225 | NA |
| Gpr6 | 378.4883978 | -0.604242328 | 0.118307558 | -5.107385693 | 3.27E-07 | 1.99E-05 | | | Gpm6b | 34999.9339 | -0.016589109 | 0.05856537 | -0.283257993 | 0.776979076 | 0.947474493 |
| Cenpf | 48.7826805 | -0.604886858 | 0.163210916 | -3.706166674 | 0.00021042 | 0.003536058 | | | Il7 | 4.27378977 | -0.016595927 | 0.047313271 | -0.350766847 | 0.725763271 | NA |
| Prss35 | 131.5449012 | -0.605525326 | 0.156222717 | -3.876038888 | 0.000106171 | 0.002048579 | | | Rfc1 | 1856.858309 | -0.016601312 | 0.055671159 | -0.298203092 | 0.765548162 | 0.945308333 |
| Tagln | 312.7328641 | -0.610695238 | 0.132151033 | -4.621191565 | 3.82E-06 | 0.000148161 | | | 4930572O13Rik | 0.332414823 | -0.016609483 | 0.013519169 | -1.228587587 | 0.219226472 | NA |
| Thbd | 558.9698971 | -0.613631151 | 0.083926667 | -7.311515829 | 2.64E-13 | 8.90E-11 | | | Gm13315 | 0.332414823 | -0.016609483 | 0.013519169 | -1.228587587 | 0.219226472 | NA |
| Stac | 55.24673797 | -0.614169807 | 0.167970987 | -3.65640412 | 0.000255778 | 0.00410251 | | | Hist1h4f | 0.332414823 | -0.016609483 | 0.013519169 | -1.228587587 | 0.219226472 | NA |
| Tenm3 | 4115.8393 | -0.615308749 | 0.100536173 | -6.120272259 | 9.34E-10 | 1.27E-07 | | | Npffr2 | 3.968499126 | -0.016616684 | 0.044829519 | -0.370663885 | 0.710887893 | NA |
| Kif2c | 9.386927755 | -0.617577957 | 0.159496715 | -3.872041865 | 0.000107927 | 0.002067036 | | | Pced1b | 186.7548915 | -0.016638517 | 0.087700501 | -0.189719752 | 0.849528742 | 0.966260848 |
| Gpr149 | 173.3419624 | -0.618746005 | 0.165008278 | -3.749787655 | 0.000176984 | 0.003098526 | | | Xcl1 | 0.612959092 | -0.016639623 | 0.016318205 | -1.01969692 | 0.307872223 | NA |
| Drd1a | 2588.755088 | -0.624010223 | 0.115327163 | -5.41078273 | 6.27E-08 | 5.12E-06 | | | 4930579G18Rik | 3.399227486 | -0.016640499 | 0.040135098 | -0.414612151 | 0.678425851 | NA |
| Aldh1a7 | 35.50894822 | -0.62717093 | 0.167725019 | -3.739280728 | 0.000184548 | 0.003205583 | | | Romo1 | 790.7697913 | -0.016647236 | 0.069787633 | -0.238541348 | 0.81146125 | 0.956135608 |
| Il31ra | 12.33124961 | -0.627189031 | 0.163553465 | -3.834764553 | 0.000125685 | 0.002335847 | | | Gm13476 | 162.933881 | -0.01665316 | 0.088285524 | -0.188628432 | 0.850384048 | 0.966260848 |
| Cpa6 | 24.12097752 | -0.628285644 | 0.155224756 | -4.047586605 | 5.17E-05 | 0.00118084 | | | Pgp | 1058.514557 | -0.016654351 | 0.07615296 | -0.218696044 | 0.826886831 | 0.958455003 |
| Pf4 | 26.6843616 | -0.629235918 | 0.166919108 | -3.769705723 | 0.00016344 | 0.002904061 | | | Zfp820 | 25.81433504 | -0.016676265 | 0.082289691 | -0.202653146 | 0.839406147 | 0.96231236 |
| Klhl30 | 18.95397982 | -0.629489713 | 0.148865787 | -4.228572105 | 2.35E-05 | 0.00063482 | | | Zbtb6 | 570.8084497 | -0.01669677 | 0.073452893 | -0.22731262 | 0.820180661 | 0.957571905 |
| Htr2a | 977.2380404 | -0.63005408 | 0.124338833 | -5.067234946 | 4.04E-07 | 2.35E-05 | | | Snord66 | 1.834900853 | -0.01673042 | 0.032986509 | -0.50718977 | 0.612021672 | NA |
| Slc6a3 | 22.44684923 | -0.631175305 | 0.157404684 | -4.00988897 | 6.07E-05 | 0.001336958 | | | Nrd1 | 6773.270908 | -0.016733035 | 0.052139298 | -0.320929418 | 0.74826388 | 0.9410864 |
| Jsrp1 | 140.1323547 | -0.632165231 | 0.126419935 | -5.000518574 | 5.72E-07 | 3.12E-05 | | | Gm10549 | 1.600590109 | -0.016754276 | 0.028988823 | -0.577956425 | 0.563293539 | NA |
| Rarb | 1014.211528 | -0.633381103 | 0.102704361 | -6.167032229 | 6.96E-10 | 1.03E-07 | | | Klf3 | 657.5651006 | -0.016763517 | 0.071451008 | -0.23461554 | 0.814507147 | 0.956264994 |
| Drd5 | 112.4693383 | -0.634705805 | 0.157372468 | -4.033143866 | 5.50E-05 | 0.001241224 | | | Cenpa | 82.25759956 | -0.016766345 | 0.094991021 | -0.176504525 | 0.859897594 | 0.96862798 |
| Fat1 | 2472.149787 | -0.635631482 | 0.121384662 | -5.23650575 | 1.64E-07 | 1.12E-05 | | | Tgfb1i1 | 510.4480104 | -0.016773658 | 0.074816456 | -0.224197438 | 0.822603677 | 0.957821087 |
| Zbtb20 | 153.9481137 | -0.636141247 | 0.164648558 | -3.863630854 | 0.000111714 | 0.002126409 | | | Dnpep | 1366.595704 | -0.016787584 | 0.063814585 | -0.263068134 | 0.792498064 | 0.951728598 |
| Slc38a5 | 115.841841 | -0.640446634 | 0.159959996 | -4.003792509 | 6.23E-05 | 0.001364151 | | | Rab33a | 789.3828867 | -0.01679653 | 0.064480454 | -0.260490251 | 0.794485634 | 0.951979493 |
| Htr1b | 308.4134959 | -0.641298213 | 0.12870037 | -4.982877767 | 6.26E-07 | 3.37E-05 | | | Utf1 | 0.529654598 | -0.016801823 | 0.01704964 | -0.985465016 | 0.324395707 | NA |
| Kif26a | 392.4709979 | -0.642109763 | 0.104477397 | -6.145920392 | 7.95E-10 | 1.12E-07 | | | Nkd1 | 625.3136427 | -0.016805105 | 0.073842997 | -0.227578853 | 0.819973661 | 0.957571905 |
| Hs6st2 | 1483.031801 | -0.64377485 | 0.08678249 | -7.418257443 | 1.19E-13 | 4.84E-11 | | | Fermt1 | 16.43977992 | -0.016818313 | 0.076756259 | -0.219113243 | 0.826561835 | NA |
| Islr | 664.4305848 | -0.645081214 | 0.129633481 | -4.976192938 | 6.48E-07 | 3.46E-05 | | | Cyp46a1 | 9882.758265 | -0.016853576 | 0.067762695 | -0.248714667 | 0.803581502 | 0.953754272 |
| Tshr | 28.92724504 | -0.650990687 | 0.164948729 | -3.946624442 | 7.93E-05 | 0.001635816 | | | 4833427G06Rik | 8.604086648 | -0.016855925 | 0.062701789 | -0.268826858 | 0.788062924 | NA |
| Tll1 | 119.4395333 | -0.657391051 | 0.157577941 | -4.171846943 | 3.02E-05 | 0.00077378 | | | Ccdc30 | 426.5518498 | -0.016860233 | 0.073382726 | -0.229757518 | 0.818280198 | 0.956955008 |
| Strip2 | 2925.504315 | -0.660054149 | 0.128948252 | -5.118752218 | 3.08E-07 | 1.88E-05 | | | Tceb3 | 1572.42614 | -0.016864946 | 0.05728692 | -0.294394357 | 0.768456578 | 0.94619074 |
| Wnt2 | 209.5323649 | -0.661132857 | 0.10901458 | -6.064627822 | 1.32E-09 | 1.75E-07 | | | 1700023A16Rik | 3.649312805 | -0.016865512 | 0.039143963 | -0.430858565 | 0.666571213 | NA |
| Hs3st3b1 | 71.39058243 | -0.66506915 | 0.149847078 | -4.438319113 | 9.07E-06 | 0.000292048 | | | Hint3 | 399.416157 | -0.016866746 | 0.071741027 | -0.235105997 | 0.814126464 | 0.956264994 |
| Kdr | 638.963809 | -0.666399224 | 0.091302834 | -7.298779149 | 2.90E-13 | 9.37E-11 | | | Bmi1 | 1404.408899 | -0.016868706 | 0.072886005 | -0.231439575 | 0.816973323 | 0.956302288 |
| Isl1 | 132.1635622 | -0.667520014 | 0.162895717 | -4.097836504 | 4.17E-05 | 0.000997139 | | | Plg | 3.127638071 | -0.016887423 | 0.042057363 | -0.401533093 | 0.688027679 | NA |
| Man1a | 898.6485704 | -0.669742817 | 0.091331252 | -7.333117668 | 2.25E-13 | 7.92E-11 | | | Acvr1c | 1078.606956 | -0.016897768 | 0.089360004 | -0.189097661 | 0.850016275 | 0.966260848 |
| Gbx2 | 15.59767381 | -0.670987313 | 0.154923248 | -4.331095049 | 1.48E-05 | 0.000439549 | | | Hist1h4d | 1.970355761 | -0.016898791 | 0.03037423 | -0.556352913 | 0.577969621 | NA |
| Myh11 | 489.6863226 | -0.672791051 | 0.122388438 | -5.497178198 | 3.86E-08 | 3.36E-06 | | | AY074887 | 1.794163629 | -0.016904349 | 0.029153248 | -0.579844448 | 0.562019521 | NA |
| Dlx6as1 | 766.3631331 | -0.677375407 | 0.139112429 | -4.869265911 | 1.12E-06 | 5.36E-05 | | | Depdc1a | 1.717669478 | -0.016907996 | 0.029940213 | -0.564725317 | 0.572260612 | NA |
| Mgat4c | 187.1044456 | -0.680272279 | 0.123838817 | -5.493207205 | 3.95E-08 | 3.42E-06 | | | Gchfr | 6.732028513 | -0.016920775 | 0.057082074 | -0.296428865 | 0.766902584 | NA |
| Fndc9 | 209.234282 | -0.680379426 | 0.134433053 | -5.06110225 | 4.17E-07 | 2.41E-05 | | | Tmem71 | 21.72496814 | -0.016921574 | 0.081356066 | -0.207994008 | 0.835233646 | NA |
| Crybg3 | 395.3215998 | -0.681990337 | 0.111283829 | -6.128386699 | 8.88E-10 | 1.23E-07 | | | Cyb5d2 | 619.580471 | -0.016925601 | 0.060029986 | -0.281952447 | 0.777979973 | 0.94784716 |
| Zfhx4 | 667.4133547 | -0.682173035 | 0.109323866 | -6.239927848 | 4.38E-10 | 6.99E-08 | | | Phf11c | 13.39382483 | -0.016933556 | 0.067153371 | -0.252162417 | 0.800915527 | NA |
| Kcnq1 | 30.48005247 | -0.682977703 | 0.166955539 | -4.090775955 | 4.30E-05 | 0.00101496 | | | H1fx | 526.7986928 | -0.016941138 | 0.079008472 | -0.214421792 | 0.830218149 | 0.960002081 |
| Prss12 | 892.3969642 | -0.68309962 | 0.105701891 | -6.462510894 | 1.03E-10 | 2.05E-08 | | | 4933427E11Rik | 3.383151085 | -0.016954729 | 0.034194561 | -0.49583117 | 0.620013531 | NA |
| Fras1 | 1534.465963 | -0.685119703 | 0.11947191 | -5.734567244 | 9.78E-09 | 1.04E-06 | | | Odf1 | 1.368277527 | -0.01696788 | 0.026062873 | -0.651036417 | 0.515022979 | NA |
| Top2a | 76.86885423 | -0.688508204 | 0.16806215 | -4.09674757 | 4.19E-05 | 0.001000295 | | | Rnf222 | 1.495121871 | -0.016995912 | 0.026882847 | -0.632221436 | 0.527242192 | NA |
| Pbk | 23.22985463 | -0.696667648 | 0.168056817 | -4.145429283 | 3.39E-05 | 0.000846251 | | | Herc4 | 1376.768933 | -0.017009008 | 0.070542687 | -0.241116529 | 0.809464803 | 0.955725085 |
| Emid1 | 158.6438141 | -0.701681435 | 0.149157122 | -4.704310631 | 2.55E-06 | 0.000108426 | | | 4930543E12Rik | 0.452133968 | -0.017015856 | 0.01368602 | -1.243302033 | 0.213756555 | NA |
| Ccnb1 | 19.32549349 | -0.706826457 | 0.168121338 | -4.204263805 | 2.62E-05 | 0.000691379 | | | Adam28 | 1.873391939 | -0.017017726 | 0.034200693 | -0.497584243 | 0.618777112 | NA |
| Klhl14 | 159.6690267 | -0.732093069 | 0.133319796 | -5.491255566 | 3.99E-08 | 3.44E-06 | | | Tmem143 | 829.4186436 | -0.017020816 | 0.067603532 | -0.251774068 | 0.801215702 | 0.953228631 |
| Kcna5 | 349.5773752 | -0.73646154 | 0.144640631 | -5.091664306 | 3.55E-07 | 2.14E-05 | | | Slc22a20 | 0.346276033 | -0.017021531 | 0.013684501 | -1.243854675 | 0.213553053 | NA |
| Slc47a1 | 86.07693662 | -0.740341337 | 0.165514367 | -4.47297326 | 7.71E-06 | 0.00025648 | | | Sec31b | 15.52319493 | -0.017024803 | 0.069245635 | -0.245861032 | 0.80578981 | NA |
| Ntn1 | 632.0933595 | -0.741314527 | 0.12959358 | -5.720302872 | 1.06E-08 | 1.12E-06 | | | Trub2 | 977.4974835 | -0.017035479 | 0.05497536 | -0.309874802 | 0.756656165 | 0.942139348 |
| D8Ertd738e | 1180.125411 | -0.747207375 | 0.06646913 | -11.24141952 | 2.55E-29 | 3.60E-26 | | | Elf5 | 13.05146958 | -0.017042594 | 0.073177807 | -0.232892926 | 0.815844552 | NA |
| Syt6 | 1942.15138 | -0.750614703 | 0.108671269 | -6.907204738 | 4.94E-12 | 1.32E-09 | | | Rbfox3 | 12478.02584 | -0.017054148 | 0.050248824 | -0.339393982 | 0.734312951 | 0.936456337 |
| Asgr1 | 40.00032472 | -0.753497358 | 0.155438588 | -4.847556642 | 1.25E-06 | 5.91E-05 | | | Med22 | 1395.390669 | -0.017073154 | 0.062044172 | -0.275177401 | 0.783179949 | 0.949478217 |
| Prkch | 493.7340023 | -0.76355125 | 0.118629193 | -6.436453226 | 1.22E-10 | 2.37E-08 | | | Commd9 | 863.0233957 | -0.017093327 | 0.063223999 | -0.270361378 | 0.78688225 | 0.950698791 |
| Dkk2 | 28.60362561 | -0.766268069 | 0.162712269 | -4.709344115 | 2.49E-06 | 0.000106367 | | | Abcc12 | 37.10064146 | -0.017099338 | 0.087352322 | -0.195751384 | 0.844804764 | 0.964372024 |
| Il15 | 26.90211668 | -0.76659756 | 0.166631324 | -4.60056094 | 4.21E-06 | 0.000160129 | | | Ulk1 | 4935.197006 | -0.017120823 | 0.048038994 | -0.356394291 | 0.721545304 | 0.932513933 |
| Serpina9 | 394.2533876 | -0.771060176 | 0.134906068 | -5.715533682 | 1.09E-08 | 1.14E-06 | | | Tfec | 1.101029416 | -0.017132667 | 0.025455516 | -0.673043407 | 0.500919676 | NA |
| Pbx3 | 825.1991979 | -0.780984077 | 0.150987552 | -5.172506404 | 2.31E-07 | 1.48E-05 | | | Hsd17b2 | 5.157178834 | -0.01713949 | 0.04820764 | -0.355534728 | 0.722189033 | NA |
| Gpr126 | 192.649184 | -0.79448622 | 0.116826207 | -6.800582165 | 1.04E-11 | 2.69E-09 | | | Kcnc1 | 10143.72355 | -0.017149599 | 0.041785769 | -0.410417223 | 0.681499915 | 0.922538401 |
| St8sia2 | 229.9925071 | -0.797983371 | 0.109064861 | -7.316594566 | 2.54E-13 | 8.76E-11 | | | Akap3 | 13.62895301 | -0.01715068 | 0.06967105 | -0.246166521 | 0.805553331 | NA |
| Tac1 | 1862.400821 | -0.810545373 | 0.140841918 | -5.755000951 | 8.66E-09 | 9.39E-07 | | | BC026590 | 1250.457145 | -0.017180614 | 0.062211982 | -0.276162463 | 0.782423286 | 0.949184473 |
| Padi6 | 30.03708738 | -0.827113313 | 0.16778201 | -4.929690102 | 8.24E-07 | 4.18E-05 | | | Zfp62 | 1387.853384 | -0.017181305 | 0.073352488 | -0.234229337 | 0.814806941 | 0.956264994 |
| Hmcn1 | 198.4489543 | -0.837015838 | 0.113932474 | -7.346595828 | 2.03E-13 | 7.33E-11 | | | Rpl12 | 5248.686572 | -0.017196826 | 0.051753817 | -0.332281317 | 0.739676846 | 0.938470248 |
| Meis2 | 7112.184552 | -0.842296325 | 0.097514958 | -8.637611569 | 5.74E-18 | 4.24E-15 | | | Slc7a2 | 1350.545931 | -0.017199926 | 0.065832653 | -0.261267399 | 0.793886306 | 0.951728598 |
| Slc35d3 | 344.9785632 | -0.844357013 | 0.145888698 | -5.78767942 | 7.14E-09 | 7.95E-07 | | | Epn2 | 6187.417822 | -0.01720122 | 0.053998531 | -0.318549774 | 0.750067945 | 0.9410864 |
| Clec12a | 40.07983665 | -0.853805357 | 0.158441556 | -5.388771599 | 7.09E-08 | 5.58E-06 | | | Spaca1 | 0.794049928 | -0.017211033 | 0.018603958 | -0.925127489 | 0.354899596 | NA |
| Fat3 | 3840.366726 | -0.866225878 | 0.117425621 | -7.376804733 | 1.62E-13 | 5.98E-11 | | | Prss55 | 1.590962535 | -0.01722349 | 0.026195177 | -0.657506128 | 0.510855532 | NA |
| 2610028E06Rik | 33.5463469 | -0.876846204 | 0.164692608 | -5.3241382 | 1.01E-07 | 7.56E-06 | | | Poc1a | 353.8016106 | -0.017242201 | 0.078897055 | -0.218540483 | 0.82700802 | 0.958518989 |
| Cbln2 | 1327.489852 | -0.877116881 | 0.11218767 | -7.818300175 | 5.35E-15 | 2.96E-12 | | | Ptpn12 | 1283.228061 | -0.017274266 | 0.065389524 | -0.264174819 | 0.791645213 | 0.951728598 |
| Aldh1a3 | 52.87269128 | -0.877649708 | 0.152503366 | -5.754953008 | 8.67E-09 | 9.39E-07 | | | Scarna10 | 0.355458167 | -0.017295052 | 0.013793166 | -1.253885618 | 0.209883583 | NA |
| Pou3f4 | 172.8316529 | -0.890303679 | 0.114616852 | -7.767650806 | 8.00E-15 | 4.13E-12 | | | Wfdc12 | 0.355458167 | -0.017295052 | 0.013793166 | -1.253885618 | 0.209883583 | NA |
| Acsm5 | 16.26062706 | -0.898732704 | 0.166657553 | -5.392691101 | 6.94E-08 | 5.49E-06 | | | DQ267101 | 1.199012197 | -0.017300174 | 0.023638578 | -0.731861862 | 0.464252886 | NA |
| Ndnf | 509.431718 | -0.899595623 | 0.145982712 | -6.16234356 | 7.17E-10 | 1.04E-07 | | | Gpr27 | 236.6178855 | -0.01730156 | 0.09469948 | -0.182699633 | 0.855033704 | 0.967189923 |
| Pgm5 | 360.0563638 | -0.900829426 | 0.086790141 | -10.37939813 | 3.08E-25 | 3.41E-22 | | | Golga5 | 1261.465989 | -0.017301657 | 0.05832672 | -0.29663347 | 0.766746355 | 0.946118815 |
| Drd3 | 30.83890732 | -0.91427374 | 0.159037359 | -5.748798552 | 8.99E-09 | 9.67E-07 | | | 1810019D21Rik | 2.908809968 | -0.017308487 | 0.0314339 | -0.55063124 | 0.581886489 | NA |
| Mki67 | 78.65430023 | -0.937193313 | 0.160355298 | -5.844479863 | 5.08E-09 | 6.01E-07 | | | 2610027K06Rik | 2.058034101 | -0.017311624 | 0.031476751 | -0.549981295 | 0.582332204 | NA |
| Ly6g6e | 76.49027072 | -0.940068672 | 0.166030917 | -5.662009743 | 1.50E-08 | 1.52E-06 | | | 1700026L06Rik | 8.457015926 | -0.01733037 | 0.052469832 | -0.330292088 | 0.74117927 | NA |
| Crym | 2658.596979 | -0.997109884 | 0.111323971 | -8.956829986 | 3.34E-19 | 3.05E-16 | | | Hist1h4b | 0.589493697 | -0.017332045 | 0.017470634 | -0.992067352 | 0.321164671 | NA |
| Wdfy1 | 1568.823883 | -1.083603933 | 0.135574278 | -7.992695577 | 1.32E-15 | 7.87E-13 | | | Fryl | 3175.323234 | -0.017333999 | 0.068344709 | -0.253626062 | 0.79978446 | 0.953228631 |
| Ndst4 | 629.6451472 | -1.135988357 | 0.167953715 | -6.763698916 | 1.35E-11 | 3.26E-09 | | | Zkscan8 | 960.9636284 | -0.017348414 | 0.059677664 | -0.29070197 | 0.771279265 | 0.946250219 |
| Tfap2d | 47.43301765 | -1.166659431 | 0.155288071 | -7.512872207 | 5.78E-14 | 2.42E-11 | | | Triobp | 375.6468225 | -0.017368261 | 0.074437983 | -0.233325245 | 0.815508857 | 0.956264994 |
| Abi3bp | 698.4032325 | -1.202316504 | 0.167972945 | -7.157798552 | 8.20E-13 | 2.54E-10 | | | 4930558J18Rik | 1.241283012 | -0.017385487 | 0.022222404 | -0.782340507 | 0.434014487 | NA |
| Dsg1c | 34.39440554 | -1.223430605 | 0.168116335 | -7.277285709 | 3.41E-13 | 1.08E-10 | | | B4galnt4 | 3389.280536 | -0.017386581 | 0.059757984 | -0.290949933 | 0.771089611 | 0.946250219 |
| Igfbpl1 | 59.42862806 | -1.277622235 | 0.165657338 | -7.712439749 | 1.23E-14 | 5.98E-12 | | | Snord87 | 1.970019805 | -0.017391561 | 0.030239486 | -0.575127539 | 0.565205044 | NA |
| Prss56 | 24.32590147 | -1.293674925 | 0.168068177 | -7.697322308 | 1.39E-14 | 6.33E-12 | | | Atf5 | 621.403412 | -0.017396528 | 0.073519307 | -0.236625295 | 0.812947495 | 0.956264994 |
| Uba1y | 19.48016777 | -1.404381475 | 0.16188274 | -8.675300844 | 4.12E-18 | 3.20E-15 | | | Polr3f | 1083.670432 | -0.017397923 | 0.066478223 | -0.261708604 | 0.793546106 | 0.951728598 |
| Nmbr | 121.7881212 | -1.663346432 | 0.154595971 | -10.75931297 | 5.36E-27 | 6.38E-24 | | | Got1l1 | 29.99478459 | -0.01741178 | 0.087417493 | -0.199179581 | 0.842122271 | 0.963252575 |
| 1-Mar | 461.9146356 | -3.175418061 | 0.110532883 | -28.72826602 | 1.69E-181 | 4.37E-178 | | | Irx5 | 6.328190972 | -0.017427559 | 0.049575718 | -0.351534173 | 0.72518764 | NA |
| Uty | 237.0766996 | -4.767794529 | 0.149628733 | -31.86416421 | 8.38E-223 | 2.60E-219 | | | Dkkl1 | 728.1476348 | -0.017462029 | 0.077783704 | -0.224494692 | 0.822372397 | 0.95778569 |
| Eif2s3y | 429.3459162 | -4.920495119 | 0.153483317 | -32.05882711 | 1.65E-225 | 6.41E-222 | | | Adrbk2 | 5538.752118 | -0.017464563 | 0.060787838 | -0.287303563 | 0.773879895 | 0.947121017 |
| Kdm5d | 409.4792754 | -5.308457086 | 0.146524767 | -36.22907704 | 2.12E-287 | 1.10E-283 | | | Limk2 | 5213.806105 | -0.017475887 | 0.055862466 | -0.312837732 | 0.754403947 | 0.94196623 |
| Ddx3y | 757.8046647 | -6.167616207 | 0.138786601 | -44.43956533 | 0 | 0 | | | Lrp8 | 3850.45885 | -0.017480454 | 0.063580423 | -0.274934538 | 0.783366532 | 0.949478217 |
| Raver2 | 814.6625797 | -0.26381753 | 0.099408373 | -2.653876356 | 0.007957298 | 0.050077324 | | | Mir142 | 0.550080249 | -0.017481674 | 0.017204802 | -1.016092717 | 0.309585237 | NA |
| Irgm1 | 512.1636084 | 0.442031927 | 0.16811371 | 2.629362752 | 0.008554506 | 0.052732358 | | | Slc12a3 | 6.185878677 | -0.01748355 | 0.053953065 | -0.324051102 | 0.745899346 | NA |
| Pak3 | 4269.614667 | -0.213193997 | 0.080344689 | -2.653492112 | 0.007966363 | 0.050082274 | | | Irf2 | 1144.518555 | -0.017486822 | 0.066021798 | -0.264864372 | 0.791113945 | 0.951728598 |
| Ddx60 | 44.22567283 | 0.43813977 | 0.165593691 | 2.645872355 | 0.008148056 | 0.050926169 | | | Chchd5 | 268.4690692 | -0.017494185 | 0.079480652 | -0.220106212 | 0.825788437 | 0.958041984 |
| Tmem147 | 1766.800841 | -0.147999754 | 0.055777139 | -2.653412434 | 0.007968244 | 0.050082274 | | | Zar1 | 4.72705282 | -0.017497038 | 0.045053622 | -0.388360289 | 0.697749428 | NA |
| Purb | 18895.6317 | -0.20428154 | 0.077012755 | -2.652567597 | 0.007988213 | 0.050169994 | | | Rubie | 1.420532821 | -0.017502101 | 0.023383902 | -0.748467946 | 0.454177952 | NA |
| Acss3 | 40.66408045 | 0.437113118 | 0.166613061 | 2.623522514 | 0.008702569 | 0.053358769 | | | Nmb | 99.52427721 | -0.017516109 | 0.09478436 | -0.184799571 | 0.853386244 | 0.966905493 |
| Mcm10 | 28.48433067 | -0.435125881 | 0.164145914 | -2.650848077 | 0.008028995 | 0.050385275 | | | A130010J15Rik | 230.549164 | -0.017520356 | 0.082614843 | -0.212072738 | 0.832050288 | 0.960519604 |
| Gpnmb | 184.0650342 | 0.435719212 | 0.166499562 | 2.616939091 | 0.008872217 | 0.054017095 | | | Wnt2 | 175.1085015 | -0.017523969 | 0.089267168 | -0.196309228 | 0.844368139 | 0.964328285 |
| Zmynd15 | 15.0249667 | 0.433056287 | 0.166102801 | 2.607158247 | 0.009129716 | 0.055126973 | | | Sft2d2 | 1801.567993 | -0.017528161 | 0.055813401 | -0.314049335 | 0.753483569 | 0.94183464 |
| Gm15328 | 88.29767242 | 0.431562799 | 0.166534181 | 2.591436763 | 0.009557611 | 0.056934109 | | | Acbd4 | 1211.29055 | -0.017556799 | 0.07185668 | -0.244330785 | 0.806974643 | 0.955587482 |
| Alox12b | 252.6211388 | 0.431291711 | 0.167960582 | 2.567815059 | 0.010234174 | 0.059882286 | | | Ntsr1 | 503.2064413 | -0.017558179 | 0.08525073 | -0.205959274 | 0.836822724 | 0.961760178 |
| Otogl | 29.5151961 | -0.44502907 | 0.167974698 | -2.649381576 | 0.008063923 | 0.050516443 | | | Bmyc | 1269.620687 | -0.017580894 | 0.060823419 | -0.28904811 | 0.772544562 | 0.947121017 |
| Slc2a4rg-ps | 304.7819679 | 0.430953188 | 0.168071699 | 2.564103246 | 0.010344277 | 0.060352076 | | | Esco2 | 6.97862036 | -0.017597371 | 0.053507142 | -0.328878922 | 0.742247207 | NA |
| Tjp3 | 16.8448153 | 0.430437402 | 0.165776449 | 2.596493077 | 0.00941808 | 0.056362973 | | | Gm20735 | 1.107523121 | -0.017610646 | 0.02583458 | -0.681669513 | 0.495447949 | NA |
| Fam83f | 21.3748766 | 0.428149172 | 0.167595318 | 2.554660696 | 0.010629135 | 0.06145068 | | | Ces5a | 16.58951276 | -0.017617565 | 0.077193432 | -0.228226214 | 0.819470383 | NA |
| Epn3 | 63.74274807 | 0.426404101 | 0.167877896 | 2.539965728 | 0.011086333 | 0.063081765 | | | Smpd2 | 521.3697832 | -0.017649211 | 0.083814038 | -0.210575841 | 0.833218266 | 0.960810394 |
| Hfe2 | 38.01671704 | 0.42547106 | 0.167692105 | 2.537215806 | 0.011173805 | 0.063439698 | | | 4930429B21Rik | 427.4943965 | -0.017656307 | 0.073563665 | -0.240013967 | 0.810319429 | 0.955876688 |
| Trim43a | 19.89857973 | 0.425084875 | 0.165889881 | 2.5624521 | 0.010393592 | 0.060460709 | | | Tyw5 | 366.2367048 | -0.017672454 | 0.082983808 | -0.212962674 | 0.831356077 | 0.960263866 |
| Fcgbp | 12.15364838 | 0.424997902 | 0.161883486 | 2.62533204 | 0.008656451 | 0.053139083 | | | Cd226 | 1.020220713 | -0.017675267 | 0.022007405 | -0.803150904 | 0.421887518 | NA |
| Acot9 | 684.6268536 | -0.193904785 | 0.073365175 | -2.64300857 | 0.008217296 | 0.051276189 | | | Myl3 | 2.185320775 | -0.017710566 | 0.029749386 | -0.595325428 | 0.551625962 | NA |
| Akt2 | 4377.766625 | -0.303837945 | 0.114958666 | -2.643019067 | 0.008217041 | 0.051276189 | | | Vnn3 | 2.054711154 | -0.017719279 | 0.024278676 | -0.729828889 | 0.465494784 | NA |
| Ifitm7 | 17.85656859 | 0.424667829 | 0.167734358 | 2.531787969 | 0.011348259 | 0.064124699 | | | 5730559C18Rik | 96.96011841 | -0.017722703 | 0.095093 | -0.186372324 | 0.852152795 | 0.966758161 |
| Slc36a2 | 19.51190585 | -0.444029795 | 0.168079158 | -2.64178973 | 0.008246924 | 0.051419653 | | | Lamc1 | 1604.924255 | -0.017724064 | 0.061379118 | -0.288763746 | 0.772762177 | 0.947121017 |
| Ihh | 10.92781065 | 0.424628891 | 0.162100461 | 2.619541536 | 0.008804805 | 0.05370931 | | | Lrrc20 | 1122.299414 | -0.017735697 | 0.064310498 | -0.275782293 | 0.782715285 | 0.949296337 |
| Pdyn | 686.5699239 | -0.359056341 | 0.135960645 | -2.640884357 | 0.008268994 | 0.051499554 | | | Foxl2 | 26.68476654 | -0.017740655 | 0.084990201 | -0.208737651 | 0.834653046 | 0.961348408 |
| Tagap | 35.7175708 | 0.423186793 | 0.166926005 | 2.535175947 | 0.011239086 | 0.063685682 | | | Eif4g3 | 12517.89994 | -0.017751616 | 0.045006179 | -0.39442619 | 0.693266411 | 0.924482501 |
| Cdk3-ps | 52.34865923 | 0.419423307 | 0.166950614 | 2.512259742 | 0.011996076 | 0.066761405 | | | Sdr9c7 | 5.880920262 | -0.017765342 | 0.044503826 | -0.399186852 | 0.68975553 | NA |
| Hbq1a | 10.44393273 | 0.418286261 | 0.158764935 | 2.634626225 | 0.008422998 | 0.052216568 | | | Rgs18 | 2.147296486 | -0.017766685 | 0.03393654 | -0.523526685 | 0.600607786 | NA |
| 1700024P16Rik | 47.98445793 | 0.417937256 | 0.167998174 | 2.487748804 | 0.01285545 | 0.070009434 | | | Cryba2 | 10.37114946 | -0.017774684 | 0.062008089 | -0.286651048 | 0.774379524 | NA |
| Kank2 | 569.1855253 | -0.220487412 | 0.083593819 | -2.637604242 | 0.008349396 | 0.051912336 | | | Cacna2d4 | 108.3932638 | -0.017779778 | 0.09481285 | -0.187524981 | 0.851249039 | 0.966638801 |
| Adam21 | 15.93885895 | 0.417787618 | 0.168025432 | 2.486454657 | 0.012902301 | 0.070094058 | | | Foxn1 | 0.5396566 | -0.017826548 | 0.017137616 | -1.040199985 | 0.298246998 | NA |
| Hspe1 | 1480.166787 | -0.216229002 | 0.082033293 | -2.635868858 | 0.008392216 | 0.052136725 | | | 4732490B19Rik | 0.980038849 | -0.017832139 | 0.02191703 | -0.813620225 | 0.415862548 | NA |
| Ccdc88c | 1462.97487 | -0.173846835 | 0.065961987 | -2.635560912 | 0.008399835 | 0.05214083 | | | Ipo9 | 9290.927526 | -0.017847764 | 0.045482126 | -0.392412712 | 0.694753295 | 0.925527709 |
| Chchd1 | 622.8809436 | -0.217752002 | 0.082628978 | -2.635298278 | 0.008406338 | 0.05214083 | | | Traf2 | 427.2780109 | -0.01785964 | 0.068333475 | -0.26136005 | 0.793814862 | 0.951728598 |
| Cybrd1 | 143.0888403 | 0.416886143 | 0.162902812 | 2.559109555 | 0.010494066 | 0.060851447 | | | Hist1h4i | 21.00121644 | -0.017861032 | 0.074384344 | -0.240118164 | 0.810238653 | NA |
| Ndufb2 | 1098.622625 | -0.270415999 | 0.10260894 | -2.635403882 | 0.008403722 | 0.05214083 | | | Nr1h5 | 0.506912888 | -0.017869835 | 0.014022676 | -1.274352715 | 0.202538442 | NA |
| AA387883 | 79.88772551 | -0.396262419 | 0.150410824 | -2.634533923 | 0.008425288 | 0.052216568 | | | 2210018M11Rik | 2071.620838 | -0.01787084 | 0.057606524 | -0.310222502 | 0.75639176 | 0.942139348 |
| Glp2r | 100.010374 | 0.416774735 | 0.168041158 | 2.480194374 | 0.013131078 | 0.07088952 | | | Cables2 | 1746.602232 | -0.017878355 | 0.062042865 | -0.288161337 | 0.773223243 | 0.947121017 |
| Nsmce1 | 377.6811585 | -0.268384111 | 0.101878967 | -2.634342686 | 0.008430036 | 0.0522251 | | | Icos | 3.462577356 | -0.017891518 | 0.042765325 | -0.418365058 | 0.67568023 | NA |
| Pafah1b3 | 287.940247 | -0.233452186 | 0.088644119 | -2.633589097 | 0.008448767 | 0.052301612 | | | 2810433D01Rik | 7.44881634 | -0.017908331 | 0.051585863 | -0.347155789 | 0.728474283 | NA |
| Tnfsf13b | 35.04158362 | 0.416750175 | 0.160697549 | 2.593382269 | 0.009503707 | 0.056704891 | | | Inhbe | 9.166563898 | -0.017934669 | 0.058046494 | -0.308970762 | 0.757343768 | NA |
| Plekhf1 | 125.9128497 | 0.416564314 | 0.163426973 | 2.548932449 | 0.010805323 | 0.062098543 | | | Pla2g15 | 1230.264543 | -0.017934756 | 0.054681004 | -0.327988784 | 0.742920143 | 0.939912916 |
| Irf7 | 110.9203906 | 0.415644269 | 0.167062091 | 2.487962804 | 0.012847717 | 0.070002457 | | | Map7 | 4516.166447 | -0.017937485 | 0.061834977 | -0.290086387 | 0.771750149 | 0.946588083 |
| Npnt | 664.1860647 | -0.285122725 | 0.108323606 | -2.632138409 | 0.008484929 | 0.05246029 | | | Wdyhv1 | 680.6220696 | -0.017937668 | 0.06109274 | -0.293613734 | 0.76905308 | 0.94619074 |
| Milr1 | 13.49964511 | 0.414403684 | 0.157451209 | 2.631949844 | 0.008489639 | 0.052468478 | | | Gm711 | 13.40238324 | -0.017942683 | 0.064318541 | -0.278965955 | 0.780270952 | NA |
| Fam196b | 185.8910222 | -0.430293652 | 0.163515443 | -2.63151691 | 0.008500464 | 0.052514428 | | | Slc6a12 | 35.50689461 | -0.017949046 | 0.076525479 | -0.234549933 | 0.814558073 | 0.956264994 |
| 4930528A17Rik | 18.35362442 | 0.413291494 | 0.167244334 | 2.471183838 | 0.013466657 | 0.072123188 | | | Wnt3a | 0.575251383 | -0.017956579 | 0.01738085 | -1.033124355 | 0.301545707 | NA |
| Wisp1 | 183.6152939 | -0.373963245 | 0.14217559 | -2.63029149 | 0.008531168 | 0.052662121 | | | Dcdc2c | 140.0306626 | -0.017975735 | 0.092645344 | -0.194027398 | 0.846154429 | 0.965002688 |
| A330023F24Rik | 1931.706753 | 0.413220619 | 0.167398962 | 2.468477786 | 0.013568907 | 0.072520404 | | | Fam109a | 452.4747573 | -0.017975899 | 0.078820096 | -0.22806238 | 0.819597746 | 0.957571905 |
| Trim72 | 35.96261804 | 0.41254655 | 0.156217039 | 2.640855008 | 0.00826971 | 0.051499554 | | | Slc22a29 | 1.63290074 | -0.017977211 | 0.026923067 | -0.667725217 | 0.50430901 | NA |
| Pacrgl | 229.4907987 | -0.255296154 | 0.097096736 | -2.62929698 | 0.00855616 | 0.052732358 | | | Trpm5 | 0.95029838 | -0.017977733 | 0.021841577 | -0.823096844 | 0.410452923 | NA |
| Tcerg1l | 382.5813999 | -0.352364744 | 0.134003168 | -2.629525484 | 0.008550412 | 0.052732358 | | | Snn | 11743.64419 | -0.017994144 | 0.052175786 | -0.344875383 | 0.730188058 | 0.934182819 |
| Nanos3 | 18.31028373 | 0.412040216 | 0.166715198 | 2.471521617 | 0.013453942 | 0.072080004 | | | 9530027J09Rik | 2.367267478 | -0.018000442 | 0.033539599 | -0.536692219 | 0.591480232 | NA |
| Pdzrn4 | 280.4762648 | -0.277583845 | 0.105593137 | -2.628805732 | 0.008568529 | 0.052745648 | | | Gpr26 | 1840.511067 | -0.018002865 | 0.070235803 | -0.256320342 | 0.797703491 | 0.952967259 |
| Arhgap9 | 33.10334903 | 0.411850165 | 0.163031636 | 2.526197825 | 0.011530453 | 0.064941056 | | | Col4a4 | 34.80270465 | -0.018007197 | 0.062623064 | -0.287548954 | 0.773692024 | 0.947121017 |
| E130114P18Rik | 127.5588609 | -0.381051638 | 0.144980705 | -2.62829208 | 0.00858148 | 0.052804388 | | | 4632427E13Rik | 14.29396869 | -0.018015163 | 0.066832157 | -0.2695583 | 0.787500085 | NA |
| Ppp2r2b | 4975.855252 | -0.170584938 | 0.064912261 | -2.627930926 | 0.008590596 | 0.052839496 | | | Timm9 | 869.9663582 | -0.018023354 | 0.067240784 | -0.268041995 | 0.788666993 | 0.951272669 |
| Gskip | 1211.490148 | -0.181869351 | 0.069216394 | -2.627547312 | 0.008600288 | 0.052851067 | | | Kncn | 4.642396435 | -0.018038006 | 0.049157424 | -0.366943675 | 0.713661032 | NA |
| Stxbp3b | 32.6130831 | 0.411312499 | 0.163102092 | 2.521810081 | 0.011675272 | 0.065518534 | | | Tmco5b | 2.30639267 | -0.018041715 | 0.036543107 | -0.493710464 | 0.621510674 | NA |
| Xdh | 499.2138611 | 0.411058141 | 0.167470051 | 2.454517315 | 0.014107387 | 0.074626103 | | | Gimap1 | 58.51092558 | -0.018053346 | 0.091649568 | -0.196982342 | 0.843841356 | 0.964060034 |
| Col28a1 | 43.53800897 | 0.410365977 | 0.16137734 | 2.542897146 | 0.010993759 | 0.062791901 | | | Arhgap10 | 901.297739 | -0.018061125 | 0.059236313 | -0.304899554 | 0.760442655 | 0.943070948 |
| Cpne7 | 2053.134167 | 0.410051022 | 0.168109858 | 2.439184865 | 0.014720436 | 0.077053523 | | | Gm7102 | 31.0553998 | -0.018065105 | 0.088584403 | -0.203930988 | 0.838407429 | 0.962077671 |
| Psmb8 | 138.1169236 | 0.409618625 | 0.167460215 | 2.446065316 | 0.014442483 | 0.076009456 | | | Cd83 | 707.0422949 | -0.018075343 | 0.091776968 | -0.196948579 | 0.843867778 | 0.964060034 |
| Traf3ip3 | 26.70997236 | 0.408718251 | 0.162895611 | 2.509080804 | 0.012104579 | 0.067149427 | | | Slco1b2 | 4.555797743 | -0.018089302 | 0.042356008 | -0.427077599 | 0.669322809 | NA |
| Odf3b | 38.94316064 | 0.408559974 | 0.167591173 | 2.437837063 | 0.014775432 | 0.077237027 | | | Olfr691 | 0.755522351 | -0.018097756 | 0.020161586 | -0.897635523 | 0.369379894 | NA |
| Got1l1 | 20.09367141 | 0.4081008 | 0.16792333 | 2.430280529 | 0.01508714 | 0.078232464 | | | Lage3 | 355.6992567 | -0.018157289 | 0.086545814 | -0.209799738 | 0.833823978 | 0.961051018 |
| Mybpc2 | 9.082871711 | 0.407381604 | 0.156069127 | 2.610263872 | 0.009047241 | 0.054851779 | | | Ncapg | 10.90842384 | -0.018157478 | 0.063226818 | -0.287179998 | 0.7739745 | NA |
| Calm2 | 49721.6461 | -0.148800135 | 0.056734253 | -2.622756587 | 0.008722156 | 0.053415451 | | | Zfp90 | 549.8929987 | -0.018162549 | 0.063870288 | -0.284366171 | 0.776129781 | 0.947241001 |
| Plau | 140.7666596 | 0.407199065 | 0.155305391 | 2.621924856 | 0.008743471 | 0.053524826 | | | St6gal1 | 1171.462898 | -0.0181635 | 0.054019082 | -0.336242298 | 0.736688158 | 0.937206199 |
| Fam96a | 568.5432062 | -0.170081777 | 0.064881837 | -2.621408141 | 0.008756736 | 0.053584859 | | | Mob1a | 373.9635971 | -0.018167174 | 0.076356508 | -0.237925677 | 0.81193874 | 0.956219756 |
| Pfdn1 | 1427.856507 | -0.151237023 | 0.057696744 | -2.621240169 | 0.008761052 | 0.053590105 | | | Celsr1 | 352.2629801 | -0.018197578 | 0.085533875 | -0.21275288 | 0.831519719 | 0.960300162 |
| Scx | 43.14213191 | 0.406398603 | 0.159365241 | 2.550108171 | 0.01076895 | 0.061981468 | | | Kat2a | 3550.559765 | -0.018206707 | 0.052048041 | -0.349805795 | 0.72648445 | 0.9335794 |
| Usp6nl | 844.9485865 | -0.190985448 | 0.072880641 | -2.620523712 | 0.008779482 | 0.053660474 | | | Zfp688 | 265.4392461 | -0.018213045 | 0.085570955 | -0.212841438 | 0.831450642 | 0.960296734 |
| Mapk15 | 38.46176991 | 0.406271894 | 0.167450058 | 2.426227253 | 0.015256713 | 0.078795839 | | | 4930486I03Rik | 1.199798456 | -0.018231862 | 0.02465783 | -0.739394402 | 0.459667541 | NA |
| Pcdha8 | 21.3861232 | 0.40507413 | 0.163272321 | 2.48097245 | 0.01310245 | 0.070795299 | | | Acadm | 1317.635808 | -0.018234421 | 0.052540328 | -0.347055712 | 0.728549465 | 0.933996867 |
| Rassf6 | 22.13237468 | 0.404004639 | 0.167142443 | 2.417127765 | 0.015643525 | 0.080311722 | | | Klk13 | 0.598202733 | -0.018243386 | 0.017533333 | -1.040497311 | 0.298108913 | NA |
| Hes7 | 13.36109971 | 0.403484179 | 0.159435372 | 2.530706793 | 0.011383296 | 0.064228984 | | | Slc5a9 | 0.598202733 | -0.018243386 | 0.017533333 | -1.040497311 | 0.298108913 | NA |
| Snora34 | 19.81984439 | 0.40333706 | 0.167996416 | 2.400867056 | 0.016356278 | 0.082631847 | | | Fbxw15 | 0.581325921 | -0.018243704 | 0.017423654 | -1.047065356 | 0.295069436 | NA |
| Igf1 | 330.1659199 | -0.260893377 | 0.099615649 | -2.618999933 | 0.008818796 | 0.053773488 | | | Apof | 2.483368486 | -0.018247569 | 0.036366662 | -0.501766392 | 0.615831857 | NA |
| Enoph1 | 1422.581605 | -0.141499956 | 0.054070542 | -2.616950951 | 0.008871909 | 0.054017095 | | | Gm8801 | 0.557041062 | -0.018249557 | 0.017262776 | -1.05716233 | 0.290437509 | NA |
| Lbp | 116.5046095 | 0.403104753 | 0.168057154 | 2.398617041 | 0.016457116 | 0.08286856 | | | Usp7 | 8024.291347 | -0.01825391 | 0.051509374 | -0.354380347 | 0.723053861 | 0.932617157 |
| Sema6d | 2951.992139 | -0.214548365 | 0.081975295 | -2.617232 | 0.008864607 | 0.054017095 | | | Mir181d | 0.968508701 | -0.01825535 | 0.022237492 | -0.82092665 | 0.41168805 | NA |
| A230073K19Rik | 80.43695457 | 0.403028543 | 0.167957825 | 2.399581812 | 0.016413811 | 0.082811982 | | | Gm10548 | 57.17279584 | -0.018258133 | 0.094963495 | -0.19226475 | 0.847534829 | 0.965398237 |
| Rbm47 | 32.37786064 | 0.402868712 | 0.168094529 | 2.396679499 | 0.016544388 | 0.083172449 | | | Il2ra | 14.92368204 | -0.018279394 | 0.066377301 | -0.275386226 | 0.783019526 | NA |
| Cd36 | 46.25815205 | 0.40286397 | 0.166566815 | 2.418632851 | 0.015578956 | 0.080108712 | | | Pkib | 328.963345 | -0.018288171 | 0.084161898 | -0.217297515 | 0.827976494 | 0.958723519 |
| Ccl12 | 22.0148445 | 0.402526166 | 0.152994618 | 2.630982517 | 0.008513842 | 0.05257611 | | | Mmachc | 658.7870081 | -0.018299798 | 0.06950265 | -0.263296406 | 0.79232213 | 0.951728598 |
| Ccdc78 | 11.35947521 | 0.402335971 | 0.158822798 | 2.533238148 | 0.011301414 | 0.06390661 | | | Zxda | 48.27086049 | -0.01829987 | 0.094487641 | -0.193674751 | 0.846430563 | 0.965166054 |
| Mmrn2 | 241.7296841 | -0.302078912 | 0.115650479 | -2.611998809 | 0.009001457 | 0.054687291 | | | Tcam1 | 8.84232474 | -0.018300591 | 0.059199791 | -0.309132703 | 0.757220583 | NA |
| Rrad | 64.20442094 | 0.401675596 | 0.152156479 | 2.639884926 | 0.008293418 | 0.051605711 | | | 4930500J02Rik | 10.65566619 | -0.018308017 | 0.05247277 | -0.348905097 | 0.727160559 | NA |
| Pax6 | 334.7859712 | -0.3320233 | 0.127133991 | -2.611601329 | 0.009011928 | 0.054714267 | | | Bbs1 | 2348.631345 | -0.018313683 | 0.05895492 | -0.310638752 | 0.756075264 | 0.942139348 |
| Ccdc60 | 28.91964419 | -0.438635755 | 0.167987908 | -2.611115051 | 0.009024753 | 0.054762618 | | | Dennd4a | 3636.719824 | -0.018315799 | 0.071129762 | -0.257498389 | 0.796794059 | 0.952885371 |
| Ttc18 | 53.54381262 | 0.401007819 | 0.160910933 | 2.492110464 | 0.012698654 | 0.069499449 | | | Sorbs2 | 7309.487454 | -0.018319915 | 0.068498501 | -0.267449863 | 0.789122812 | 0.951324462 |
| 1700009P17Rik | 48.37430814 | 0.399258538 | 0.163790613 | 2.437615512 | 0.01478449 | 0.077258309 | | | Gm6225 | 18.44304911 | -0.018327863 | 0.073730029 | -0.248580716 | 0.803685127 | NA |
| Snx20 | 24.04377191 | 0.398792949 | 0.165573149 | 2.408560523 | 0.016015571 | 0.081438992 | | | Srd5a2 | 0.742151045 | -0.01833571 | 0.020389105 | -0.899289637 | 0.368498406 | NA |
| Prkcd | 636.8251155 | 0.398575485 | 0.164658697 | 2.420616061 | 0.015494232 | 0.079836261 | | | 1110002L01Rik | 525.5766389 | -0.018358144 | 0.068067413 | -0.269705324 | 0.787386964 | 0.950831248 |
| Ankrd29 | 738.9602421 | -0.234865224 | 0.090027946 | -2.608803532 | 0.009085939 | 0.055034222 | | | Stoml3 | 18.31726264 | -0.018373107 | 0.069761435 | -0.263370546 | 0.79226499 | NA |
| C1ql3 | 4383.28079 | -0.296416989 | 0.113632313 | -2.6085625 | 0.009092341 | 0.055051475 | | | Gm10714 | 9.183686047 | -0.018398892 | 0.056197379 | -0.327397685 | 0.743367117 | NA |
| Arhgef33 | 47.38334487 | 0.397435742 | 0.167782099 | 2.368761297 | 0.017847768 | 0.087510545 | | | Prosapip1 | 14260.02106 | -0.018408893 | 0.070529874 | -0.261008445 | 0.794085995 | 0.951878276 |
| Arl5a | 3846.892932 | -0.152465184 | 0.058460246 | -2.608014729 | 0.009106904 | 0.055092094 | | | Rab19 | 2.520500328 | -0.018414817 | 0.036880185 | -0.499314672 | 0.617557721 | NA |
| Timm13 | 1429.025292 | -0.203204087 | 0.077918396 | -2.607909019 | 0.009109716 | 0.055092094 | | | Polr3b | 809.7484371 | -0.018420007 | 0.060662731 | -0.303646191 | 0.761397457 | 0.943530068 |
| Fam19a3 | 16.13808345 | -0.361628583 | 0.1386794 | -2.607658979 | 0.009116373 | 0.055110838 | | | Snora20 | 0.997626583 | -0.018439638 | 0.021972079 | -0.839230454 | 0.401340001 | NA |
| Sgk1 | 3281.027366 | 0.397235072 | 0.167764677 | 2.367811148 | 0.017893667 | 0.087652379 | | | Ptk6 | 3.018710889 | -0.018440481 | 0.037591891 | -0.490544127 | 0.623748912 | NA |
| Ido1 | 138.6857691 | -0.323516708 | 0.124074147 | -2.607446555 | 0.009122031 | 0.0551171 | | | Serpinf2 | 6.813703967 | -0.018446311 | 0.046160986 | -0.399608262 | 0.68944507 | NA |
| Ybx2 | 120.0278905 | 0.396590963 | 0.160368622 | 2.472996008 | 0.013398564 | 0.071882739 | | | E130215H24Rik | 14.56544889 | -0.018453166 | 0.070740036 | -0.260858869 | 0.794201344 | NA |
| Mcin | 9.571698715 | -0.417649728 | 0.160259325 | -2.606086901 | 0.009158322 | 0.055278163 | | | Aup1 | 2123.128696 | -0.018454911 | 0.067090558 | -0.275074636 | 0.783258897 | 0.949478217 |
| Chrna2 | 40.96448014 | -0.436581027 | 0.167547535 | -2.605714408 | 0.009168287 | 0.055316761 | | | Mtmr14 | 686.9601682 | -0.01847982 | 0.0722924 | -0.255626046 | 0.798239605 | 0.952967259 |
| Edar | 13.27787921 | -0.432903459 | 0.166166133 | -2.605244837 | 0.009180863 | 0.055371074 | | | Polr2a | 3679.467281 | -0.018485577 | 0.056857602 | -0.325120586 | 0.745089809 | 0.940632024 |
| Gipc3 | 23.77989283 | 0.39649886 | 0.167739214 | 2.363781549 | 0.018089476 | 0.088276641 | | | Cntf | 3.457793133 | -0.018495716 | 0.037173783 | -0.49754731 | 0.61880315 | NA |
| Cd48 | 36.80707499 | 0.396416895 | 0.166595508 | 2.379517307 | 0.017335329 | 0.08551673 | | | Med12l | 1804.277381 | -0.018515381 | 0.060895554 | -0.304051447 | 0.761088696 | 0.943227916 |
| Mecp2 | 6988.573667 | -0.194302606 | 0.074623054 | -2.603787911 | 0.009219979 | 0.05554213 | | | 2810404M03Rik | 6.422898558 | -0.018523607 | 0.050865339 | -0.364169532 | 0.715731417 | NA |
| Gm11517 | 13.85634754 | 0.396246462 | 0.166936829 | 2.373631184 | 0.017614135 | 0.086684037 | | | Opn4 | 1.651225453 | -0.018534218 | 0.023325264 | -0.794598427 | 0.426847134 | NA |
| Asic3 | 12.41795117 | 0.395718361 | 0.16209393 | 2.441290428 | 0.01463488 | 0.076784262 | | | Rgs6 | 1591.991873 | -0.018561586 | 0.064748401 | -0.286672495 | 0.7743631 | 0.947238918 |
| Chrna6 | 14.2017664 | 0.395650315 | 0.157906615 | 2.505596837 | 0.012224492 | 0.067669266 | | | Mfsd6l | 2.19348567 | -0.018584553 | 0.033513799 | -0.554534349 | 0.579213205 | NA |
| 6720483E21Rik | 17.09136666 | 0.394455097 | 0.168046637 | 2.347295387 | 0.018910256 | 0.090767572 | | | Ifi202b | 0.517564097 | -0.018605292 | 0.014306108 | -1.300513861 | 0.193424909 | NA |
| Scml4 | 135.9778093 | 0.393618592 | 0.149496017 | 2.632970428 | 0.008464172 | 0.052352845 | | | Socs1 | 22.74104369 | -0.018610782 | 0.076172594 | -0.244323852 | 0.806980012 | 0.955587482 |
| Gmppb | 389.0998828 | -0.295911639 | 0.113697228 | -2.602628429 | 0.009251216 | 0.055600597 | | | Cd8a | 1.121830405 | -0.018640867 | 0.020496396 | -0.909470484 | 0.363101831 | NA |
| Galntl6 | 814.2043737 | -0.293602475 | 0.112835248 | -2.60204573 | 0.009266949 | 0.055673561 | | | Lzic | 351.087845 | -0.01864231 | 0.072860629 | -0.255862602 | 0.798056933 | 0.952967259 |
| Stat1 | 1256.354587 | 0.393510542 | 0.164430882 | 2.393166885 | 0.016703642 | 0.083513107 | | | Nhlrc4 | 18.70080914 | -0.018646575 | 0.070706295 | -0.26371874 | 0.791996654 | NA |
| Phf11b | 13.45256209 | 0.392874646 | 0.166435552 | 2.360521183 | 0.018249276 | 0.08874899 | | | Ttc29 | 12.7599653 | -0.01865212 | 0.065252965 | -0.285843252 | 0.774998179 | NA |

| | | | | | | | | | | | | | |
|---|---|---|---|---|---|---|---|---|---|---|---|---|---|
| 2700069I18Rik | 15.74530004 | -0.436988679 | 0.168120626 | -2.5992568 | 0.009342585 | 0.056062749 | Rnf25 | 1492.510554 | -0.018657691 | 0.068193922 | -0.273597565 | 0.784393905 | 0.949699053 |
| Mettl24 | 43.19378507 | 0.392309417 | 0.16327016 | 2.402823744 | 0.016269027 | 0.082322763 | Mirlet7e | 1.760403072 | -0.018678335 | 0.029217427 | -0.639287447 | 0.522635953 | NA |
| Crlf1 | 37.34904651 | 0.390460175 | 0.165117316 | 2.364743952 | 0.01804254 | 0.088130869 | Pdpr | 837.334499 | -0.018687972 | 0.059175626 | -0.315805233 | 0.752150346 | 0.941549114 |
| A730085K08Rik | 8.980426969 | -0.408040231 | 0.157036292 | -2.598381728 | 0.009366431 | 0.056140609 | Papolb | 4.859969328 | -0.018701188 | 0.048687638 | -0.384105475 | 0.700900274 | NA |
| Nudt21 | 834.2462704 | -0.199600703 | 0.076848224 | -2.597336558 | 0.009394982 | 0.056289966 | Zfp871 | 4066.687965 | -0.018703216 | 0.070306272 | -0.266024859 | 0.790220064 | 0.951607109 |
| Ccser2 | 4761.90317 | -0.154981226 | 0.059675538 | -2.597064586 | 0.009402424 | 0.05629968 | Tcte1 | 124.3127379 | -0.018712809 | 0.093245847 | -0.200682495 | 0.840946848 | 0.963094947 |
| Erp29 | 2311.972383 | -0.14123071 | 0.054382006 | -2.597011754 | 0.009403871 | 0.05629968 | Med15 | 2729.028774 | -0.018716281 | 0.046330313 | -0.403974848 | 0.686231215 | 0.923860706 |
| Spata18 | 21.16729931 | 0.390023094 | 0.167989837 | 2.321706489 | 0.020248745 | 0.09495708 | Pcdha8 | 28.90709352 | -0.018719367 | 0.070009993 | -0.267381366 | 0.789175545 | 0.951324462 |
| Phf11d | 26.57630504 | 0.389959622 | 0.165661044 | 2.353960918 | 0.018574564 | 0.089823437 | Gm5547 | 0.522557059 | -0.018723988 | 0.014351274 | -1.304691656 | 0.191997864 | NA |
| Hist4h4 | 30.38192396 | 0.389378455 | 0.167853673 | 2.319749381 | 0.020354439 | 0.095220916 | Ggt7 | 5862.850682 | -0.018724605 | 0.061798018 | -0.302996859 | 0.761892255 | 0.943740676 |
| Bub1 | 9.631153939 | -0.399966074 | 0.15418346 | -2.594091964 | 0.009484111 | 0.056676538 | Fam26d | 0.80879701 | -0.018745351 | 0.020679608 | -0.906465523 | 0.364689505 | NA |
| Dnali1 | 38.28962184 | 0.389283526 | 0.167736763 | 2.320800284 | 0.020297625 | 0.095098701 | Jmjd7 | 117.1452895 | -0.018753719 | 0.093631984 | -0.200291807 | 0.84125237 | 0.963094947 |
| Hprt | 4053.716439 | -0.18862834 | 0.072715897 | -2.594045419 | 0.009485396 | 0.056676538 | F630028O10Rik | 1.975428467 | -0.018754653 | 0.026539364 | -0.706673039 | 0.479769688 | NA |
| Hmgb3 | 1246.713194 | -0.24021185 | 0.092613929 | -2.593690318 | 0.009495197 | 0.056693095 | Smagp | 43.30734737 | -0.01877356 | 0.093665688 | -0.200431565 | 0.841143075 | 0.963094947 |
| Cd151 | 936.4358161 | -0.184831119 | 0.071271076 | -2.593353862 | 0.009504492 | 0.056704891 | Sez6 | 10720.64235 | -0.018786058 | 0.066088916 | -0.284254283 | 0.776215518 | 0.947241001 |
| Lrrc36 | 30.73228488 | 0.388998769 | 0.16758813 | 2.321159438 | 0.02027824 | 0.095036617 | Lrrc46 | 35.97957151 | -0.018789518 | 0.086254544 | -0.21783801 | 0.827555328 | 0.958672527 |
| Exoc3 | 4666.46096 | -0.12884067 | 0.049687981 | -2.592994685 | 0.009514424 | 0.056720464 | Ccdc41 | 1338.603656 | -0.018803411 | 0.070127498 | -0.268131778 | 0.788597886 | 0.951268301 |
| Rps24 | 8193.711025 | -0.18831099 | 0.072621383 | -2.593051553 | 0.009512851 | 0.056720464 | 1700001G11Rik | 7.729889812 | -0.018820619 | 0.059925567 | -0.314066605 | 0.753470453 | NA |
| Fap | 93.35080752 | 0.388599408 | 0.150174951 | 2.587644645 | 0.009663461 | 0.057300295 | Sart1 | 2400.448434 | -0.018823743 | 0.061536637 | -0.305894888 | 0.759684678 | 0.94272457 |
| Col16a1 | 895.8295544 | 0.388258667 | 0.167136259 | 2.323006806 | 0.020178787 | 0.094799916 | Prl | 547.633629 | -0.018836555 | 0.033462121 | -0.562921717 | 0.573488192 | 0.879485797 |
| Nip7 | 963.137193 | -0.171352023 | 0.066134946 | -2.590945246 | 0.009571272 | 0.056971681 | 0610012G03Rik | 911.0440581 | -0.018846528 | 0.058904643 | -0.319949797 | 0.749006388 | 0.9410864 |
| Rps9 | 5525.150169 | -0.156961625 | 0.060579089 | -2.591019888 | 0.009569196 | 0.056971681 | Supt5 | 7177.528473 | -0.0188638 | 0.053064331 | -0.355489258 | 0.722223091 | 0.93255988 |
| Snx31 | 9.655172613 | -0.421946834 | 0.162894287 | -2.590310816 | 0.009588931 | 0.057023545 | Chd1l | 393.6170937 | -0.018864832 | 0.071732645 | -0.262988105 | 0.792559748 | 0.951728598 |
| Tmem223 | 775.9271524 | -0.200153233 | 0.077272211 | -2.590235619 | 0.009591026 | 0.057023545 | Wdr17 | 1776.008777 | -0.018875171 | 0.073009729 | -0.258529534 | 0.795998259 | 0.952734447 |
| AW549542 | 49.97035661 | 0.388257674 | 0.164026386 | 2.367044005 | 0.017930801 | 0.087751051 | Zfp882 | 392.5939632 | -0.018876807 | 0.086282705 | -0.218778576 | 0.826822536 | 0.958455003 |
| Ms4a6c | 44.90943626 | 0.388101713 | 0.162037356 | 2.395137284 | 0.016614143 | 0.083309297 | Nhlrc1 | 691.8977754 | -0.018885148 | 0.073744576 | -0.256088636 | 0.797882396 | 0.952967259 |
| Dkk3 | 12875.11359 | -0.297464086 | 0.114898069 | -2.588938958 | 0.009627216 | 0.057150989 | Nipbl | 2325.133858 | -0.018904938 | 0.074217014 | -0.254725122 | 0.798935411 | 0.953053772 |
| Sc4mol | 2804.140156 | -0.334658407 | 0.129260044 | -2.589032126 | 0.009624611 | 0.057150989 | Mir154 | 1.947616703 | -0.018916816 | 0.034136421 | -0.554153464 | 0.579473823 | NA |
| Susd5 | 301.7701421 | -0.276897413 | 0.106952588 | -2.58897347 | 0.009626251 | 0.057150989 | Ptpn7 | 77.37257228 | -0.018930864 | 0.085968252 | -0.220207619 | 0.825709464 | 0.958041984 |
| Mme | 633.4823142 | -0.307714059 | 0.118873424 | -2.588585822 | 0.009637093 | 0.057187713 | Barhl2 | 29.80961645 | -0.018936396 | 0.084362352 | -0.224465005 | 0.822395494 | 0.95778569 |
| Gmfg | 32.90891728 | 0.387747904 | 0.16052807 | 2.415452353 | 0.015715679 | 0.080512887 | C030006K11Rik | 451.1963801 | -0.018955114 | 0.078841847 | -0.240419456 | 0.810005097 | 0.955876688 |
| Krtcap3 | 18.82308023 | 0.387596463 | 0.168072434 | 2.306127501 | 0.021103509 | 0.0974889 | Muc13 | 1.978809442 | -0.01895619 | 0.033010797 | -0.574242117 | 0.565803972 | NA |
| Bhmt2 | 12.83070366 | 0.386528659 | 0.162847833 | 2.373557272 | 0.017617661 | 0.086684037 | Hip1r | 3885.408244 | -0.018961483 | 0.059631213 | -0.317979159 | 0.750500746 | 0.9410864 |
| Rpl27a | 7433.819566 | -0.160243273 | 0.061947075 | -2.586777067 | 0.009687825 | 0.057400826 | Amz2 | 2246.322124 | -0.018964342 | 0.053107555 | -0.357093114 | 0.7210221 | 0.932513933 |
| Gpr45 | 387.6549953 | -0.191637781 | 0.074093314 | -2.586438233 | 0.009697355 | 0.057435328 | Lpar3 | 13.00001555 | -0.018972585 | 0.070427014 | -0.26939357 | 0.787626833 | NA |
| Oacyl | 63.12126134 | 0.386398094 | 0.158138969 | 2.443408447 | 0.014549261 | 0.076415676 | Alg1 | 983.4627927 | -0.018976422 | 0.070991484 | -0.267305616 | 0.789233863 | 0.951324462 |
| Jazf1 | 1124.852943 | -0.177542804 | 0.068676883 | -2.585190167 | 0.00973253 | 0.057577633 | Snord64 | 3.637874294 | -0.018982289 | 0.043217717 | -0.439224713 | 0.66049872 | NA |
| Lilrb3 | 21.61742934 | 0.386224109 | 0.168110547 | 2.297441266 | 0.02159361 | 0.099101322 | Pdcd4 | 2898.777094 | -0.018987701 | 0.063969408 | -0.296824708 | 0.766600341 | 0.94610336 |
| Trpv6 | 48.93898725 | 0.384465999 | 0.15623221 | 2.460862581 | 0.013860344 | 0.07364615 | Tcf15 | 9.759064435 | -0.019008295 | 0.063669076 | -0.298548309 | 0.765284711 | NA |
| Cfh | 1354.701137 | -0.219380876 | 0.08491732 | -2.583464433 | 0.009781355 | 0.057822326 | Mir191 | 5.066423869 | -0.019016065 | 0.038092602 | -0.49920624 | 0.617634099 | NA |
| Uba7 | 156.18167 | 0.384020309 | 0.163295871 | 2.351684135 | 0.018688637 | 0.090108701 | Mex3d | 495.7789623 | -0.01901853 | 0.075133253 | -0.253130666 | 0.800167242 | 0.953228631 |
| 6330403A02Rik | 15730.07279 | -0.271155728 | 0.105080012 | -2.580469133 | 0.009866618 | 0.058255183 | Atpbd4 | 728.2421347 | -0.019043803 | 0.074735714 | -0.254815297 | 0.79886576 | 0.953053772 |
| 1700109H08Rik | 25.25710211 | 0.382473543 | 0.167275903 | 2.286483207 | 0.022226005 | 0.101047455 | Col18a1 | 295.9233549 | -0.019059861 | 0.092711817 | -0.20558179 | 0.837117604 | 0.961831963 |
| Pnldc1 | 48.39150929 | 0.382192295 | 0.160030016 | 2.388253803 | 0.016928646 | 0.084250743 | Crcp | 1038.054103 | -0.01906477 | 0.054618312 | -0.349054554 | 0.727048354 | 0.9335794 |
| Ifi27l2a | 24.83160593 | 0.38134825 | 0.165256875 | 2.307608996 | 0.021020894 | 0.097310346 | Nfib | 2996.006185 | -0.01907945 | 0.070748368 | -0.269680419 | 0.787406126 | 0.950831248 |
| Zfand4 | 154.8127168 | 0.381100808 | 0.144506464 | 2.637257863 | 0.008357927 | 0.051944534 | U2af1l4 | 994.6607711 | -0.01909293 | 0.065297865 | -0.292397456 | 0.769982753 | 0.94619074 |
| Lrrc71 | 28.53714169 | 0.381040799 | 0.1671777 | 2.27925614 | 0.022651844 | 0.102502239 | Hmgn1 | 1651.298838 | -0.019102885 | 0.063809103 | -0.299375544 | 0.76465352 | 0.944872045 |
| Mfsd2b | 37.85891873 | 0.38097137 | 0.166143569 | 2.293025074 | 0.021846561 | 0.099761457 | Otoa | 4.725076963 | -0.019112378 | 0.046584716 | -0.410271422 | 0.681606853 | NA |
| Tmem128 | 678.8815323 | -0.180229013 | 0.069924117 | -2.57749431 | 0.009951952 | 0.058616351 | Phf5a | 742.2565555 | -0.019135023 | 0.05932727 | -0.322533353 | 0.747048675 | 0.94097069 |
| Unc13c | 2710.778719 | -0.33558413 | 0.130200541 | -2.577440354 | 0.009953506 | 0.058616351 | A730090N16Rik | 2.43617978 | -0.019170123 | 0.035018617 | -0.547426612 | 0.584085676 | NA |
| Tll2 | 33.70995509 | -0.432461503 | 0.167836353 | -2.576685532 | 0.009975266 | 0.058722175 | Cdc40 | 2072.64197 | -0.019199507 | 0.063527388 | -0.302224094 | 0.762481238 | 0.944309188 |
| Recql4 | 102.2178267 | 0.380794852 | 0.1478128 | 2.576196731 | 0.009989379 | 0.058782925 | 6430571L13Rik | 244.4606738 | -0.019199696 | 0.091778205 | -0.209196681 | 0.834294703 | 0.961118357 |
| Vmn2r1 | 18.13350838 | 0.380466106 | 0.158350002 | 2.402690882 | 0.016274938 | 0.08232579 | Gtpbp5 | 430.0149818 | -0.019200785 | 0.081776185 | -0.234796779 | 0.814366468 | 0.956264994 |
| Dapk2 | 53.41060275 | 0.380436077 | 0.150690744 | 2.52461477 | 0.011582518 | 0.065184109 | Rbm7 | 891.051306 | -0.019201919 | 0.058029742 | -0.330897881 | 0.740721623 | 0.938975678 |
| Trpv3 | 16.13461334 | 0.379824153 | 0.167691262 | 2.265020541 | 0.023511419 | 0.105133024 | Nek6 | 1264.118413 | -0.019241924 | 0.069463879 | -0.277006187 | 0.781775355 | 0.948979619 |
| Map3k15 | 88.538187 | 0.379608426 | 0.153880836 | 2.466898654 | 0.013628893 | 0.072815883 | Basp1 | 12765.52319 | -0.019262406 | 0.064000917 | -0.300970784 | 0.763436773 | 0.944624337 |
| Fezf1 | 10.38889332 | 0.379263999 | 0.151280671 | 2.507022185 | 0.012175307 | 0.067478396 | Lman1l | 2.659868646 | -0.019273043 | 0.034997015 | -0.55070535 | 0.581835677 | NA |
| 2610017I09Rik | 867.3877344 | -0.268207579 | 0.104299006 | -2.571523523 | 0.010125214 | 0.059426857 | Slamf6 | 9.123214006 | -0.019281835 | 0.060725176 | -0.317526207 | 0.750844357 | NA |
| Ptchd4 | 180.3007202 | -0.338575576 | 0.131664191 | -2.571508422 | 0.010125655 | 0.059426857 | Casp9 | 1466.136495 | -0.019298593 | 0.05236288 | -0.36855484 | 0.712459561 | 0.930507332 |
| A730056A06Rik | 137.7093341 | -0.324882338 | 0.126369686 | -2.57088823 | 0.010143806 | 0.059471971 | Sarm1 | 1501.799603 | -0.019312607 | 0.059504685 | -0.324556078 | 0.745517073 | 0.940632024 |
| Acot7 | 8096.137784 | -0.169784467 | 0.066042092 | -2.570852334 | 0.010144857 | 0.059471971 | Pag1 | 1520.402842 | -0.019313653 | 0.065200235 | -0.296220605 | 0.767061614 | 0.946118815 |
| Traip | 491.372109 | 0.379155153 | 0.143170718 | 2.648273043 | 0.008090415 | 0.050627178 | Catsper3 | 7.038414255 | -0.01931715 | 0.055064531 | -0.350809305 | 0.725731416 | NA |
| Baiap2l2 | 80.87529522 | -0.397249873 | 0.154531262 | -2.570676424 | 0.010150011 | 0.059479681 | Nckap5 | 156.1565278 | -0.019330732 | 0.093759523 | -0.206173533 | 0.836655361 | 0.961729368 |
| Hhip | 429.5842646 | -0.25608849 | 0.099637449 | -2.570203204 | 0.010163888 | 0.059538479 | 2310030G06Rik | 89.07557876 | -0.019333621 | 0.094873962 | -0.203782163 | 0.838523732 | 0.962110266 |
| Ptger1 | 43.11766507 | 0.379145039 | 0.166411614 | 2.278356837 | 0.022705326 | 0.102593826 | Dsg1a | 1.346430562 | -0.019333819 | 0.024197999 | -0.79898419 | 0.42429958 | NA |
| Zfp748 | 455.5767041 | -0.219001892 | 0.085222772 | -2.569757915 | 0.01017696 | 0.059578357 | Fggy | 487.5685305 | -0.019345682 | 0.070865226 | -0.272992599 | 0.784858905 | 0.949834246 |
| Kcng4 | 291.6605299 | 0.378820487 | 0.153462339 | 2.468491547 | 0.013568386 | 0.072520404 | Cox5b | 5438.992948 | -0.019393762 | 0.072219134 | -0.268540491 | 0.788283312 | 0.951142331 |
| Adgb | 36.98437031 | 0.378601515 | 0.167276907 | 2.263322062 | 0.023615843 | 0.1054782 | Ror1 | 54.59995757 | -0.019396141 | 0.093932503 | -0.206490192 | 0.836408026 | 0.961664 |
| Zfp389 | 15.40260289 | 0.378352219 | 0.16366456 | 2.311754118 | 0.020791237 | 0.096564577 | B9d2 | 430.515541 | -0.019402057 | 0.077449871 | -0.250511161 | 0.802192075 | 0.95333733 |
| Iqcd | 37.51066752 | 0.378204507 | 0.155502314 | 2.432147139 | 0.015009609 | 0.078013712 | Ndp | 357.5170334 | -0.019412839 | 0.083547795 | -0.232356087 | 0.816261453 | 0.956264994 |
| Dlx5 | 239.4980981 | -0.393896048 | 0.153521872 | -2.565732439 | 0.010295821 | 0.060129459 | 5830411N06Rik | 1.820817091 | -0.019413952 | 0.024664986 | -0.787105734 | 0.431219973 | NA |
| Kcnv1 | 3139.357477 | -0.286843005 | 0.111793647 | -2.565825632 | 0.010293056 | 0.060129459 | Hist2h2be | 309.7576301 | -0.019419793 | 0.080878676 | -0.24011018 | 0.810244842 | 0.955876688 |
| Hspb2 | 19.77604066 | 0.377777505 | 0.168072862 | 2.247700794 | 0.024595275 | 0.108261135 | 2310033P09Rik | 452.8840533 | -0.019423698 | 0.071635775 | -0.271145227 | 0.786279338 | 0.950484821 |
| Mycbp | 157.7033539 | -0.333435452 | 0.130042086 | -2.564058016 | 0.010345625 | 0.060352076 | Spata18 | 29.37841669 | -0.019433599 | 0.076193492 | -0.255055886 | 0.798679935 | 0.953036015 |
| Usp18 | 49.00204733 | 0.377768514 | 0.166429005 | 2.269847821 | 0.023216818 | 0.104236852 | Itpka | 5081.016021 | -0.01943968 | 0.075735597 | -0.256678243 | 0.797427169 | 0.952897915 |
| 4933408B17Rik | 25.61998967 | 0.376146819 | 0.168112913 | 2.237465355 | 0.025255941 | 0.109920099 | Necab2 | 3085.613525 | -0.019445383 | 0.074173713 | -0.262160025 | 0.793198071 | 0.951728598 |
| Lrrtm4 | 1013.18802 | -0.198608988 | 0.077473065 | -2.563587583 | 0.010359656 | 0.060365743 | BC049762 | 3.522296248 | -0.019456739 | 0.042720658 | -0.455440999 | 0.648792005 | NA |
| 4933406I18Rik | 15.8133254 | 0.375035024 | 0.167638497 | 2.237165272 | 0.02527554 | 0.109974507 | Hnrnph1 | 7883.664307 | -0.019468109 | 0.051799828 | -0.375833466 | 0.707040707 | 0.929707007 |
| Ednra | 173.6189008 | -0.274665869 | 0.107191301 | -2.562389549 | 0.010395465 | 0.060460709 | Tmem87b | 1729.103327 | -0.019495904 | 0.050664721 | -0.384802348 | 0.70038386 | 0.927053542 |
| Jrk | 431.8810855 | -0.256834836 | 0.100230364 | -2.562445414 | 0.010393792 | 0.060460709 | Klhl24 | 2900.131161 | -0.019505149 | 0.069387282 | -0.281105531 | 0.778629459 | 0.948061318 |
| Med10 | 658.123239 | -0.190768045 | 0.07444064 | -2.56268679 | 0.01038657 | 0.060460709 | Lmod1 | 49.87048321 | -0.019506672 | 0.086940778 | -0.224367347 | 0.822471476 | 0.95778569 |
| 4933407K13Rik | 118.1492937 | 0.374167069 | 0.142587059 | 2.62413063 | 0.008687046 | 0.053284676 | 0610009O20Rik | 1660.489052 | -0.019508466 | 0.057390347 | -0.339925912 | 0.733912322 | 0.936456337 |
| Trip10 | 252.6770645 | 0.374066109 | 0.146405936 | 2.554992775 | 0.010619 | 0.061415002 | Sgol2 | 45.76606423 | -0.01952491 | 0.090573302 | -0.215570256 | 0.829322743 | 0.959526943 |
| Marcks | 3298.500684 | -0.308585171 | 0.120475793 | -2.561387328 | 0.010425505 | 0.060612674 | Tigd4 | 2.814086805 | -0.019550502 | 0.038746723 | -0.504571744 | 0.61385966 | NA |
| Chp2 | 53.17426593 | -0.422928585 | 0.165147491 | -2.560914378 | 0.010439708 | 0.060672482 | Atp2c1 | 6046.212058 | -0.01956212 | 0.056703068 | -0.344992273 | 0.73010018 | 0.934152665 |
| Acad10 | 105.7457144 | 0.373817046 | 0.142924898 | 2.615478846 | 0.008910244 | 0.054181837 | Kat8 | 759.4679141 | -0.019566097 | 0.061843558 | -0.316380516 | 0.751713704 | 0.941374879 |
| 5330434G04Rik | 212.669487 | -0.288553416 | 0.112707455 | -2.560198139 | 0.01046125 | 0.060752102 | 2700069I18Rik | 11.41234745 | -0.019571758 | 0.065667788 | -0.29804198 | 0.765671123 | NA |
| Mrpl28 | 1260.636699 | -0.18211918 | 0.071146573 | -2.559774472 | 0.010474011 | 0.06080342 | Loh12cr1 | 1279.561591 | -0.019579979 | 0.06241811 | -0.313690678 | 0.753755982 | 0.94183464 |
| Flt3l | 109.2265948 | 0.373314579 | 0.143662666 | 2.598549706 | 0.009361849 | 0.056134864 | Gtsf1l | 1.933802046 | -0.019590394 | 0.022522449 | -0.869816339 | 0.384400781 | NA |
| Dnaic2 | 91.06605114 | 0.37318443 | 0.155253479 | 2.403710575 | 0.016229617 | 0.082177021 | 4930483J18Rik | 0.86768302 | -0.019590658 | 0.019361648 | -1.011827993 | 0.311620303 | NA |
| Klhl9 | 3548.340711 | -0.159257735 | 0.062228971 | -2.559221742 | 0.01049068 | 0.060851447 | Cenpj | 149.1597413 | -0.019608003 | 0.09037365 | -0.216965929 | 0.828234897 | 0.958819613 |
| Dcun1d4 | 4374.56235 | -0.14823998 | 0.057952616 | -2.557951467 | 0.010529079 | 0.061031629 | Mga | 2703.808148 | -0.019615412 | 0.074664054 | -0.262715597 | 0.792769794 | 0.951728598 |
| Apoa1 | 11.25856687 | 0.373030793 | 0.159195694 | 2.343221635 | 0.019118022 | 0.091452498 | Coq9 | 2282.230968 | -0.019622919 | 0.064003695 | -0.306590403 | 0.759155159 | 0.942526199 |
| Chrdl1 | 767.7015845 | -0.286801204 | 0.112177523 | -2.556672651 | 0.010567861 | 0.061210633 | A530088E08Rik | 1.545027225 | -0.019623354 | 0.028362643 | -0.691873277 | 0.489016914 | NA |
| Trpv4 | 92.3273928 | 0.372154975 | 0.165501429 | 2.248651129 | 0.024534701 | 0.108104516 | Dcdc2a | 446.3103645 | -0.019624713 | 0.082143092 | -0.238908861 | 0.811176254 | 0.956135608 |
| Cmpk1 | 1722.468066 | -0.229079894 | 0.0896373 | -2.555631344 | 0.010599535 | 0.061325316 | Gm4961 | 5.437016421 | -0.019630282 | 0.051282482 | -0.38278729 | 0.701877487 | NA |
| Lman2 | 3320.594971 | -0.13155135 | 0.051472644 | -2.555752706 | 0.010595839 | 0.061325316 | Ifitm10 | 349.4580706 | -0.019648562 | 0.074819915 | -0.262611397 | 0.792850115 | 0.951728598 |
| Col4a3 | 23.88964803 | 0.37118997 | 0.167432038 | 2.216959041 | 0.026625891 | 0.113961754 | Abl1 | 2854.731808 | -0.019651465 | 0.049645201 | -0.39583816 | 0.692224423 | 0.923860706 |
| Heatr8 | 244.8854986 | 0.371077819 | 0.155057308 | 2.393165615 | 0.016703699 | 0.083513107 | Acsbg2 | 1.700927331 | -0.019661064 | 0.031278689 | -0.628576989 | 0.529626029 | NA |
| Srcrb4d | 36.00718693 | 0.371011918 | 0.158549812 | 2.340033794 | 0.019281995 | 0.091924687 | Klri2 | 4.297419383 | -0.019679965 | 0.043025394 | -0.457403477 | 0.647381065 | NA |
| Gm12522 | 55.98720344 | 0.370994087 | 0.152093449 | 2.439250925 | 0.014717745 | 0.077053523 | Tspan32 | 7.120293943 | -0.019694038 | 0.05591649 | -0.352204472 | 0.724684923 | NA |
| Pdk3 | 1153.130711 | -0.1629687 | 0.063828177 | -2.553240726 | 0.01067257 | 0.061609839 | Ceacam16 | 0.724629857 | -0.01969622 | 0.016898168 | -1.165583102 | 0.243783043 | NA |
| Tgif1 | 62.43609728 | 0.370820503 | 0.158078952 | 2.345793019 | 0.018986648 | 0.091015545 | Zscan25 | 558.4562568 | -0.019708735 | 0.076289907 | -0.258340008 | 0.796144512 | 0.952734447 |
| C920021L13Rik | 20.7258669 | 0.370816472 | 0.167959409 | 2.207774338 | 0.027260008 | 0.11600752 | Ptpn22 | 36.99774802 | -0.019717198 | 0.086164811 | -0.228831206 | 0.819000111 | 0.957334483 |
| Mrpl52 | 435.2197962 | -0.238107697 | 0.093311752 | -2.551743915 | 0.010718527 | 0.06182906 | Nags | 3.325266733 | -0.019720979 | 0.037948975 | -0.51967093 | 0.603292952 | NA |
| 6430411K18Rik | 87.11345067 | 0.370725609 | 0.155249352 | 2.387936597 | 0.016943265 | 0.084250743 | Pex6 | 2347.849023 | -0.019746283 | 0.074928954 | -0.263533406 | 0.792139479 | 0.951728598 |
| Rad9b | 94.01030266 | 0.36932564 | 0.141400927 | 2.611903939 | 0.009003955 | 0.054687291 | Ampd2 | 3185.007379 | -0.019759478 | 0.067437118 | -0.293005965 | 0.769517593 | 0.94619074 |
| Tst | 770.4894314 | -0.358146629 | 0.140417724 | -2.550579935 | 0.010754386 | 0.0619667 | Bphl | 878.7874699 | -0.019823971 | 0.061786057 | -0.320848609 | 0.74832512 | 0.9410864 |
| Cd274 | 153.6026106 | 0.368607662 | 0.163725804 | 2.251371829 | 0.024361997 | 0.107754719 | Macf1 | 13108.53187 | -0.019824786 | 0.081860037 | -0.242179054 | 0.808641425 | 0.955723386 |
| Poln | 34.04675409 | 0.368445979 | 0.160787641 | 2.291506836 | 0.021934117 | 0.100013893 | Dyx1c1 | 100.8995025 | -0.019830101 | 0.095089205 | -0.208542082 | 0.834805728 | 0.961348408 |
| Srpx2 | 68.3261335 | 0.368245104 | 0.156544772 | 2.352330896 | 0.01865617 | 0.090049441 | Frs2 | 2542.218086 | -0.01986239 | 0.057311112 | -0.346571364 | 0.728913362 | 0.933996867 |
| Vat1l | 3673.592589 | -0.300640851 | 0.117900093 | -2.549962797 | 0.010773441 | 0.061984294 | Sytl5 | 154.6093884 | -0.019871889 | 0.094756609 | -0.209715072 | 0.833890061 | 0.961051018 |
| Coa6 | 315.2250508 | -0.309658564 | 0.121450604 | -2.549666721 | 0.010782594 | 0.062013925 | Kcnk2 | 2902.487803 | -0.019882805 | 0.06096545 | -0.326132347 | 0.744324224 | 0.940096833 |
| Bcl2a1b | 26.19112841 | 0.366925932 | 0.167095648 | 2.195903582 | 0.028098845 | 0.118434034 | Sstr1 | 1301.208042 | -0.019903261 | 0.073252849 | -0.271706304 | 0.785847854 | 0.950484821 |
| AI182371 | 9.735186324 | 0.366356213 | 0.161692282 | 2.2657619 | 0.023465965 | 0.104990374 | Krt7 | 4.008040747 | -0.01991817 | 0.045558321 | -0.437201585 | 0.661965161 | NA |
| Gdpd1 | 1626.402637 | -0.182492948 | 0.07160358 | -2.548656764 | 0.010813867 | 0.062124605 | Dync1li2 | 11869.66118 | -0.019918438 | 0.057102059 | -0.348821718 | 0.727223158 | 0.9335794 |
| Best1 | 41.44429403 | 0.366170272 | 0.162735601 | 2.250093213 | 0.024443029 | 0.107866787 | Arfgap1 | 4764.928585 | -0.019919348 | 0.055678151 | -0.357758787 | 0.720523835 | 0.932513933 |
| Ryr3 | 2208.2611 | -0.319152816 | 0.125308133 | -2.546944141 | 0.010867083 | 0.062360955 | Fam151a | 1.957808024 | -0.01992409 | 0.028395919 | -0.701653302 | 0.482895402 | NA |
| Snrpe | 600.7237106 | -0.190282579 | 0.07470868 | -2.546994271 | 0.010865522 | 0.062360955 | Harbi1 | 235.111235 | -0.019931153 | 0.083047475 | -0.239997096 | 0.810332508 | 0.955876688 |
| Parp10 | 110.4097956 | 0.366015921 | 0.154531118 | 2.368558029 | 0.017857579 | 0.087530948 | Fam133b | 1113.504475 | -0.019943886 | 0.068931451 | -0.289329264 | 0.77232942 | 0.947121017 |
| Mpzl1 | 716.7581133 | -0.177858949 | 0.069857709 | -2.546017494 | 0.010895973 | 0.06248046 | Rsph1 | 148.0019047 | -0.01995245 | 0.094119397 | -0.211990839 | 0.832114182 | 0.960519604 |
| Grin2b | 2290.504797 | -0.411914197 | 0.161816638 | -2.545561463 | 0.010910216 | 0.062538986 | Zfp750 | 8.85906099 | -0.019955281 | 0.06070665 | -0.328716552 | 0.742369942 | NA |
| Pnma1 | 161.9144658 | -0.323543101 | 0.127108994 | -2.545398952 | 0.010915295 | 0.062544964 | Gpr174 | 5.433178473 | -0.019970439 | 0.051013391 | -0.391474454 | 0.695446568 | NA |
| Hcst | 11.72193882 | 0.365057011 | 0.164016208 | 2.225737417 | 0.026031775 | 0.112249866 | Mvb12a | 575.5633187 | -0.019986288 | 0.076005495 | -0.262958466 | 0.792582592 | 0.951728598 |
| Gm3435 | 25.27857174 | -0.42355297 | 0.166435507 | -2.54484741 | 0.01093255 | 0.062597535 | Epha4 | 8052.688912 | -0.019999305 | 0.066855308 | -0.299143117 | 0.764830849 | 0.94492946 |
| Slc25a14 | 1897.253094 | -0.162011966 | 0.063682641 | -2.54405226 | 0.010957469 | 0.062717039 | Fam194a | 92.12351764 | -0.020000855 | 0.095100248 | -0.210313381 | 0.833423092 | 0.960817765 |
| Slc22a18 | 27.00296219 | 0.364452515 | 0.168002133 | 2.169332664 | 0.030057437 | 0.123756999 | 2810468N07Rik | 999.9604315 | -0.020015024 | 0.082159389 | -0.243612136 | 0.807531228 | 0.955716697 |
| ORF63 | 26.2326013 | -0.420781454 | 0.165422421 | -2.543678492 | 0.0109692 | 0.06273783 | Zfp827 | 2269.638638 | -0.02003775 | 0.065331342 | -0.306709598 | 0.759064424 | 0.942526199 |
| Mndal | 32.59179133 | 0.364191339 | 0.166597164 | 2.18605966 | 0.028811241 | 0.1202266 | Fam83e | 2.281906748 | -0.020056509 | 0.034857178 | -0.575391073 | 0.565026841 | NA |
| Nlrp6 | 43.20504359 | 0.363457951 | 0.167162711 | 2.174276474 | 0.029684378 | 0.122552061 | Slc25a41 | 3.406738091 | -0.020061199 | 0.042157218 | -0.475866284 | 0.634169644 | NA |
| Rab32 | 76.80530418 | 0.363455411 | 0.142960533 | 2.542347904 | 0.011011051 | 0.062813732 | Kalrn | 25732.04975 | -0.020064114 | 0.045867825 | -0.437433304 | 0.661797136 | 0.915428982 |
| Ccdc142 | 321.5963556 | 0.363215045 | 0.139434441 | 2.604916286 | 0.009189671 | 0.055394692 | Traf3ip1 | 764.5769918 | -0.020108698 | 0.062671523 | -0.320858613 | 0.748317539 | 0.9410864 |
| Rbm3 | 2276.092606 | 0.362637818 | 0.163355859 | 2.219925384 | 0.026423833 | 0.113328907 | Pkd2l1 | 55.08915914 | -0.020129136 | 0.092222725 | -0.218266548 | 0.827221437 | 0.958536901 |
| Fam171b | 9868.905014 | -0.197599745 | 0.077724557 | -2.542307747 | 0.011012316 | 0.062813732 | Obp2a | 1.181446803 | -0.020140924 | 0.02090559 | -0.963422872 | 0.335335353 | NA |
| 1700029I15Rik | 24.57306288 | 0.362550407 | 0.166994842 | 2.171027572 | 0.029929088 | 0.12336294 | Magt1 | 698.7700904 | -0.020141165 | 0.065217458 | -0.308830878 | 0.757450179 | 0.942139348 |
| Kcnip4 | 2007.893234 | -0.173751897 | 0.06834652 | -2.542220098 | 0.011015079 | 0.062813732 | Mipol1 | 108.4291163 | -0.020186405 | 0.094614637 | -0.213353938 | 0.831050906 | 0.960162026 |
| Pram1 | 15.56077495 | 0.362507314 | 0.167341322 | 2.166274946 | 0.030290183 | 0.124421022 | Gja10 | 0.735229523 | -0.020202374 | 0.017301809 | -1.167645219 | 0.242949902 | NA |
| Slc22a3 | 136.9519009 | -0.390945649 | 0.153786613 | -2.542130564 | 0.011017901 | 0.062813732 | Ddx52 | 1008.516938 | -0.020209804 | 0.06914916 | -0.29226391 | 0.77008485 | 0.94619074 |
| Socs5 | 5009.815253 | -0.128816419 | 0.050678415 | -2.541839951 | 0.011027065 | 0.062813732 | Pigx | 316.4080413 | -0.020211835 | 0.077048385 | -0.262326525 | 0.793069714 | 0.951728598 |
| Tex11 | 8.98652736 | 0.362451817 | 0.147175228 | 2.462722984 | 0.01378864 | 0.073365794 | Epb4.1l5 | 396.2296822 | -0.020212269 | 0.090438826 | -0.223491054 | 0.823153345 | 0.957903916 |
| Myh2 | 38.87337291 | 0.361700974 | 0.160702236 | 2.250752589 | 0.024401212 | 0.107805069 | Snora28 | 39.00434522 | -0.020228737 | 0.092482182 | -0.218731186 | 0.826859454 | 0.958455003 |
| Ttc25 | 61.15023481 | 0.361636371 | 0.143766143 | 2.515448792 | 0.011888095 | 0.066400194 | Sema6a | 1181.182194 | -0.020239676 | 0.080482817 | -0.251478227 | 0.801444394 | 0.953228631 |
| Tmem196 | 418.8113109 | -0.251940083 | 0.099196999 | -2.539795423 | 0.011091733 | 0.063089319 | Rab33b | 1202.553819 | -0.020248151 | 0.071969651 | -0.281342904 | 0.778447406 | 0.947998577 |
| Rpl37a | 4165.19202 | -0.184393539 | 0.072616005 | -2.539296091 | 0.011107577 | 0.063156258 | 4930452N14Rik | 0.928930868 | -0.020256816 | 0.019977316 | -1.013990908 | 0.310587091 | NA |
| Sfrp5 | 51.93560134 | 0.361439008 | 0.168108187 | 2.150038108 | 0.031552201 | 0.127809097 | Gm5803 | 2.699114828 | -0.020266166 | 0.032342936 | -0.626602539 | 0.530919804 | NA |
| Acrbp | 203.1258016 | 0.361400614 | 0.150324828 | 2.404131237 | 0.016210952 | 0.082136199 | Crb2 | 298.8308377 | -0.020268334 | 0.090748032 | -0.22334737 | 0.823265163 | 0.957903916 |
| Lpar5 | 13.08798549 | 0.360832445 | 0.165375952 | 2.181891871 | 0.029117514 | 0.120983309 | Scgb1c1 | 3.743693204 | -0.020279462 | 0.041006108 | -0.494547343 | 0.62091968 | NA |
| Vstm2l | 1890.240467 | -0.241979262 | 0.095366246 | -2.537368026 | 0.011168947 | 0.063439698 | Chka | 1860.220687 | -0.020336885 | 0.056077548 | -0.362656455 | 0.716861532 | 0.93125489 |
| Mir1249 | 35.2881246 | 0.36065409 | 0.168079106 | 2.14574017 | 0.031893714 | 0.12881459 | Dmgdh | 14.27196448 | -0.02034747 | 0.066195294 | -0.307385438 | 0.758550011 | NA |
| Pfdn5 | 3814.140752 | -0.167776788 | 0.066143121 | -2.536571966 | 0.011194373 | 0.06350993 | Frmd3 | 212.3332619 | -0.0203463 | 0.087328558 | -0.233310052 | 0.815520653 | 0.956264994 |
| Ptpmt1 | 834.8181617 | -0.173528558 | 0.068417407 | -2.536321736 | 0.011202376 | 0.06353207 | Cd19 | 1.011132555 | -0.020374785 | 0.022394471 | -0.909813195 | 0.362921035 | NA |
| Tmem200a | 575.710673 | -0.219054855 | 0.086372351 | -2.536168732 | 0.011207272 | 0.06353658 | Zic5 | 48.23536047 | -0.020377397 | 0.089427721 | -0.227864437 | 0.819751631 | 0.957571905 |
| Runx1 | 54.67550922 | 0.360086902 | 0.154347323 | 2.332964991 | 0.019649983 | 0.093049155 | Coil | 843.706678 | -0.020397952 | 0.061624627 | -0.331003254 | 0.740642028 | 0.938975678 |
| Gm2011 | 22.61763906 | 0.360078253 | 0.16811821 | 2.141815884 | 0.032208301 | 0.129592498 | C330024D21Rik | 3.579929947 | -0.020404437 | 0.045019258 | -0.45323796 | 0.650377401 | NA |
| Car9 | 29.91606227 | 0.359611238 | 0.167924643 | 2.141503668 | 0.032233443 | 0.129653418 | Zfyve21 | 493.6370105 | -0.020425632 | 0.06703928 | -0.304681555 | 0.760608699 | 0.943105713 |
| Opn3 | 167.9629247 | -0.329088965 | 0.129862169 | -2.534140365 | 0.011272357 | 0.063812166 | Loxl1 | 73.28581121 | -0.020434153 | 0.090548314 | -0.225671265 | 0.821457108 | 0.957699646 |
| Atp5o | 5375.182098 | -0.174056831 | 0.068694918 | -2.53376578 | 0.011284413 | 0.063833939 | Hey1 | 1831.846278 | -0.020437503 | 0.059190044 | -0.345286167 | 0.729879245 | 0.934125508 |
| C1galt1c1 | 635.1293776 | -0.203575367 | 0.080345012 | -2.533764849 | 0.011284443 | 0.063833939 | Utp14a | 688.4371045 | -0.020442467 | 0.070640096 | -0.289389002 | 0.77228371 | 0.947121017 |
| Pkp2 | 243.646644 | 0.358904411 | 0.15594745 | 2.301444557 | 0.021366516 | 0.098322779 | Lix1 | 1962.722114 | -0.020444191 | 0.061191886 | -0.334099697 | 0.738304329 | 0.937711697 |
| Rfx3 | 1815.425875 | -0.237778996 | 0.09387261 | -2.532996536 | 0.011309207 | 0.063927345 | Rfx8 | 9.193314498 | -0.020456302 | 0.059973281 | -0.341090263 | 0.733035632 | NA |
| Zfp54 | 44.68072237 | 0.358887679 | 0.156418853 | 2.294401676 | 0.021767437 | 0.099546833 | C920009B18Rik | 1.38569564 | -0.020457268 | 0.026528134 | -0.771153653 | 0.440615862 | NA |
| Zim1 | 138.8114445 | 0.35781858 | 0.158631694 | 2.255656301 | 0.024092164 | 0.106774598 | Baiap2 | 13106.60831 | -0.020461781 | 0.07795757 | -0.262473306 | 0.792956563 | 0.951728598 |
| Jarid2 | 1600.541071 | -0.175556257 | 0.069362069 | -2.531012393 | 0.011373383 | 0.064196429 | Nemf | 1944.073279 | -0.020463144 | 0.070331728 | -0.290951823 | 0.771088166 | 0.946250219 |
| Ywhaq | 12706.5606 | -0.216844961 | 0.085671146 | -2.531131795 | 0.011369512 | 0.064196429 | Cpt1b | 2.278823929 | -0.020470766 | 0.034138159 | -0.599644688 | 0.548743058 | NA |
| Ccdc62 | 94.66594518 | 0.357224991 | 0.15513585 | 2.302659188 | 0.021298026 | 0.09812418 | Wdr75 | 1171.699408 | -0.020471401 | 0.060503751 | -0.338349293 | 0.735099981 | 0.936456337 |
| Yaf2 | 2488.34511 | -0.143218749 | 0.056612429 | -2.529811036 | 0.011412397 | 0.064369742 | Tnrc18 | 3022.160248 | -0.02048051 | 0.06307794 | -0.324685773 | 0.745418904 | 0.940632024 |
| Prph | 24.85868427 | 0.357155118 | 0.165737679 | 2.1549422 | 0.031166358 | 0.126833011 | Cpeb2 | 2707.286743 | -0.020508642 | 0.066472616 | -0.308527684 | 0.757680839 | 0.942139348 |
| Kif18b | 9.903794219 | -0.383794341 | 0.151872233 | -2.527086972 | 0.011501302 | 0.064823997 | Anapc13 | 556.168848 | -0.020514189 | 0.075758098 | -0.270785426 | 0.78655607 | 0.950484821 |
| Acvr1 | 1144.243057 | -0.138757411 | 0.054923354 | -2.526382714 | 0.011524386 | 0.064930487 | 4931428L18Rik | 0.561390173 | -0.020524285 | 0.01864664 | -1.100696155 | 0.27102892 | NA |
| Tnfrsf10b | 35.00027664 | 0.35699303 | 0.167546508 | 2.130710062 | 0.033113036 | 0.132184168 | 4632428C04Rik | 38.03352851 | -0.020525397 | 0.091808463 | -0.223567589 | 0.823093786 | 0.957903916 |
| Ppp2r5a | 3271.737304 | -0.161666005 | 0.064014641 | -2.525453593 | 0.011554904 | 0.065055118 | Lrrc23 | 96.85106101 | -0.020533867 | 0.093662432 | -0.219232695 | 0.826468788 | 0.958429316 |
| Cap1 | 10568.90027 | -0.1504396 | 0.059591789 | -2.52450216 | 0.011586229 | 0.065184109 | Cox19 | 735.3874489 | -0.020537394 | 0.064275723 | -0.319520229 | 0.749332054 | 0.9410864 |
| Zfp692 | 1056.399207 | 0.356536924 | 0.144182675 | 2.472813909 | 0.013405393 | 0.07189448 | Hist1h2bn | 2.378402104 | -0.020538754 | 0.037856886 | -0.54253681 | 0.587448756 | NA |
| Slc22a8 | 768.4648216 | -0.358356485 | 0.141976198 | -2.524060303 | 0.011600803 | 0.065242409 | Rbm48 | 342.686583 | -0.020577444 | 0.07139053 | -0.288237723 | 0.773164775 | 0.947121017 |
| Fgf3 | 24.28948821 | -0.424131483 | 0.168065943 | -2.523601606 | 0.011615949 | 0.065248465 | Slc26a11 | 210.6901297 | -0.02063205 | 0.088673454 | -0.232674486 | 0.816014182 | 0.956264994 |
| Sstr5 | 36.12959242 | 0.356362869 | 0.16811801 | 2.11971858 | 0.034029785 | 0.134573122 | Abca3 | 5311.281086 | -0.02063771 | 0.047576905 | -0.4337758 | 0.664451257 | 0.915428982 |
| Syce2 | 34.21120355 | -0.409118455 | 0.162112881 | -2.523664082 | 0.011613885 | 0.065248465 | Fstl3 | 75.66473761 | -0.020680774 | 0.095102108 | -0.21745863 | 0.827850944 | 0.958723519 |
| Neurod4 | 13.00091171 | 0.356342214 | 0.16051976 | 2.219927409 | 0.026423696 | 0.113328907 | 1700084E18Rik | 8.641373437 | -0.020684457 | 0.055504383 | -0.372663487 | 0.709398915 | NA |
| Nop10 | 1007.761977 | -0.170212648 | 0.067481186 | -2.522371906 | 0.01165664 | 0.065437671 | Bhlha15 | 4.674872679 | -0.020702169 | 0.043266689 | -0.478478232 | 0.632309864 | NA |
| Gm7120 | 403.7331441 | 0.356298068 | 0.158454182 | 2.248587346 | 0.024538762 | 0.108104516 | LOC100503496 | 24.87964622 | -0.020705997 | 0.084731684 | -0.244371362 | 0.806943219 | 0.955587482 |
| Doc2g | 378.8774493 | -0.338442561 | 0.134223839 | -2.521478765 | 0.011686273 | 0.065556521 | Caml | 911.3940425 | -0.020736057 | 0.05815345 | -0.356574836 | 0.721410119 | 0.932513933 |
| Cald1 | 998.5525057 | -0.186006667 | 0.073778024 | -2.521166297 | 0.011696656 | 0.06559102 | Kdm2a | 4567.692233 | -0.020758925 | 0.061956802 | -0.335054824 | 0.737583731 | 0.937206199 |
| Tril | 2113.073739 | -0.314263014 | 0.124664126 | -2.520877695 | 0.011706253 | 0.065621088 | Kcnab2 | 18802.99989 | -0.020766177 | 0.046283515 | -0.448673297 | 0.653667351 | 0.914683958 |
| Amh | 28.90897381 | 0.35625165 | 0.164675532 | 2.163355082 | 0.03051388 | 0.125107714 | Gm13629 | 232.4427224 | -0.020767778 | 0.090478572 | -0.229532551 | 0.818455024 | 0.95703083 |
| Rps10 | 4001.365644 | -0.185016026 | 0.073407806 | -2.520386283 | 0.011722611 | 0.065665269 | Pet112 | 690.4927614 | -0.020805955 | 0.06597946 | -0.315339885 | 0.752503605 | 0.941691983 |
| Gm6297 | 68.78318162 | 0.356088453 | 0.146694371 | 2.427417295 | 0.015206754 | 0.07864267 | Arhgap5 | 4390.32342 | -0.020806857 | 0.079022724 | -0.263302203 | 0.792317662 | 0.951728598 |
| Vamp3 | 976.72662 | -0.172823222 | 0.068625034 | -2.51836992 | 0.011789942 | 0.065994714 | Ghrl | 3.032253809 | -0.020811307 | 0.034865386 | -0.596904532 | 0.550571121 | NA |
| Dok5 | 737.8890983 | -0.272487185 | 0.108217286 | -2.517963581 | 0.011803553 | 0.066030994 | E330023G01Rik | 0.760890561 | -0.020814195 | 0.017537465 | -1.186841736 | 0.235290053 | NA |
| Mga | 2773.137326 | -0.202494937 | 0.080421451 | -2.51792196 | 0.011804947 | 0.066030994 | 2310014F07Rik | 1.934979665 | -0.020820408 | 0.033030662 | -0.630335798 | 0.528474906 | NA |
| Scn3a | 1873.139203 | -0.262959427 | 0.104495511 | -2.516466252 | 0.011853825 | 0.066260306 | Ncr1 | 0.429580527 | -0.020828597 | 0.015124969 | -1.377100108 | 0.168481299 | NA |
| Tspan3 | 10279.70557 | -0.089707079 | 0.035648318 | -2.516446299 | 0.011854496 | 0.066260306 | Rnf44 | 7137.132753 | -0.020829145 | 0.05882061 | -0.354113044 | 0.723254167 | 0.932617157 |
| Esr2 | 15.55691602 | 0.35598228 | 0.167077388 | 2.130643073 | 0.033118558 | 0.132184168 | Acap3 | 6696.738566 | -0.020834564 | 0.065575404 | -0.31771919 | 0.750697953 | 0.9410864 |
| Rasd1 | 258.9847984 | 0.355586793 | 0.16804389 | 2.116035246 | 0.03434181 | 0.13525966 | 4930558K02Rik | 1.104512261 | -0.020840209 | 0.022228288 | -0.93755347 | 0.348473933 | NA |
| BB031773 | 39.13247164 | -0.415664011 | 0.165260255 | -2.515208581 | 0.011896198 | 0.066421511 | Far1 | 2472.868808 | -0.020856787 | 0.066624555 | -0.313049545 | 0.754243021 | 0.94196623 |
| A130049A11Rik | 59.85166435 | 0.355505057 | 0.144492452 | 2.46037113 | 0.013879341 | 0.073687366 | Gtpbp10 | 373.8056523 | -0.02087901 | 0.075248234 | -0.277468444 | 0.781420433 | 0.948894514 |
| Pde6b | 10.45540635 | 0.355273494 | 0.141529801 | 2.510238066 | 0.012064979 | 0.06702574 | Npb | 6.081205805 | -0.020903739 | 0.052581485 | -0.397549416 | 0.690962354 | NA |
| Zbtb12 | 216.6930755 | -0.235494192 | 0.093668995 | -2.514110365 | 0.011933308 | 0.066566805 | Ost4 | 961.6343622 | -0.020908776 | 0.067539594 | -0.309578056 | 0.756881846 | 0.942139348 |
| Oas1c | 42.32408205 | 0.355131174 | 0.155755809 | 2.280050909 | 0.022604669 | 0.102348552 | Rpp40 | 136.9825899 | -0.020920463 | 0.092106054 | -0.227134502 | 0.820319157 | 0.957571905 |
| Aff3 | 2436.383502 | -0.18925168 | 0.075325463 | -2.512452912 | 0.01198951 | 0.066761405 | Cobl | 5964.665241 | -0.02094495 | 0.079696475 | -0.262808984 | 0.792697811 | 0.951728598 |
| Naip1 | 9.595191009 | 0.354984398 | 0.156834619 | 2.263431381 | 0.02360911 | 0.1054782 | Ccna1 | 36.88054035 | -0.020953095 | 0.089754477 | -0.233449023 | 0.81541275 | 0.956264994 |
| Acer2 | 316.3749469 | 0.354407881 | 0.160511608 | 2.207989096 | 0.027245033 | 0.11600752 | E130012A19Rik | 2455.219556 | -0.020969289 | 0.08284526 | -0.253113927 | 0.800180177 | 0.953228631 |
| Ddah1 | 4334.129333 | -0.262785255 | 0.104606153 | -2.512139557 | 0.012000162 | 0.066761405 | 4833419F23Rik | 5.963233664 | -0.020974025 | 0.053239844 | -0.393953539 | 0.693615342 | NA |
| Itga10 | 156.0538537 | 0.354144757 | 0.150976209 | 2.345699094 | 0.018991433 | 0.091015545 | 1810007D17Rik | 1.018675763 | -0.020978868 | 0.023870444 | -0.878863764 | 0.379475148 | NA |
| Zfp503 | 213.5535354 | -0.368404696 | 0.146642708 | -2.512260597 | 0.011996047 | 0.066761405 | Madd | 12085.89484 | -0.020984635 | 0.053577484 | -0.391668921 | 0.695302857 | 0.925527709 |
| Cntnap2 | 4282.51122 | -0.19729569 | 0.078582026 | -2.510679417 | 0.012049293 | 0.066986632 | Sertad3 | 82.23851848 | -0.020991214 | 0.0948267 | -0.221363961 | 0.824809051 | 0.957903916 |
| Dlec1 | 101.2776592 | 0.353951356 | 0.159070337 | 2.225124827 | 0.02607286 | 0.112277067 | Pou3f2 | 955.504763 | -0.021032075 | 0.080322911 | -0.261844032 | 0.79344169 | 0.951728598 |
| Htr6 | 122.96441 | -0.320303293 | 0.127588227 | -2.510445523 | 0.012057893 | 0.067010397 | 4930578M01Rik | 20.50078066 | -0.021039731 | 0.079509997 | -0.264617428 | 0.791304193 | NA |
| Clcn1 | 81.47254248 | 0.353925718 | 0.157560637 | 2.246282605 | 0.024685912 | 0.10850482 | Shroom3 | 190.1372097 | -0.021049652 | 0.093955707 | -0.22403804 | 0.822727704 | 0.957837832 |
| BC028528 | 60.91635151 | -0.355981096 | 0.141838512 | -2.509763326 | 0.01208121 | 0.067067815 | Wipi1 | 728.1366484 | -0.021118907 | 0.065433529 | -0.322753598 | 0.746881857 | 0.94097069 |
| Prss16 | 32.13904646 | 0.353231679 | 0.163735951 | 2.157325116 | 0.030980341 | 0.126256066 | Thoc6 | 172.5996704 | -0.021119389 | 0.084333392 | -0.250427359 | 0.802256874 | 0.95333733 |
| Slc22a23 | 3858.830133 | -0.133431783 | 0.053178236 | -2.509142695 | 0.012102458 | 0.067149427 | Smchd1 | 1838.016417 | -0.021121229 | 0.068133699 | -0.309996801 | 0.756563389 | 0.942139348 |
| Zfhx2as | 207.2041324 | 0.35307947 | 0.161730916 | 2.183129111 | 0.029026303 | 0.120809102 | Zfp872 | 3.395008597 | -0.021136135 | 0.03594018 | -0.588092079 | 0.556470489 | NA |
| Pstpip1 | 69.545495 | 0.35288232 | 0.154861784 | 2.278692082 | 0.022685376 | 0.102564112 | Oxct2a | 1.406187698 | -0.0211515 | 0.026798132 | -0.789292675 | 0.429940967 | NA |
| Ttll8 | 51.63338781 | 0.352841496 | 0.165840956 | 2.127589615 | 0.033371124 | 0.13295248 | Olfml3 | 1071.666135 | -0.021152773 | 0.093047793 | -0.227332341 | 0.820165327 | 0.957571905 |
| Pmepa1 | 2294.364198 | -0.187419108 | 0.074759077 | -2.506974615 | 0.012176946 | 0.067478396 | Zfp51 | 208.1054058 | -0.021181373 | 0.092156883 | -0.229840385 | 0.818215802 | 0.956955008 |
| Adamts9 | 130.577071 | 0.352779046 | 0.133160035 | 2.649286226 | 0.008066198 | 0.050516443 | Mtx2 | 1128.130582 | -0.021184252 | 0.064458391 | -0.328650024 | 0.742420233 | 0.939912916 |
| Wnt10b | 49.21543297 | 0.352603055 | 0.167776231 | 2.101626986 | 0.035585964 | 0.138430561 | Gm20597 | 13.66357099 | -0.021193042 | 0.064822413 | -0.326940032 | 0.743712342 | NA |
| Atp8b5 | 14.69977951 | 0.352580741 | 0.167683754 | 2.102652964 | 0.035496116 | 0.138254608 | 4922505E12Rik | 0.443441737 | -0.021206778 | 0.015260407 | -1.389660102 | 0.164632116 | NA |
| Gad2 | 11188.00529 | -0.323058189 | 0.128948649 | -2.505324346 | 0.012233915 | 0.067697242 | Ly6i | 0.443441737 | -0.021206778 | 0.015260407 | -1.389660102 | 0.164632116 | NA |
| Nif3l1 | 740.932996 | -0.180847298 | 0.072202754 | -2.504714695 | 0.01225502 | 0.06778982 | Clic3 | 5.476162645 | -0.021216363 | 0.052712219 | -0.402494206 | 0.687320352 | NA |
| Itgal | 39.88549724 | 0.35198668 | 0.159490515 | 2.206944278 | 0.027317952 | 0.116090059 | Ccl11 | 3.20107698 | -0.021228003 | 0.039546961 | -0.536779627 | 0.591419846 | NA |
| BC021891 | 150.8479476 | 0.351776438 | 0.136361582 | 2.579732746 | 0.00988768 | 0.058317366 | Klk11 | 0.443931641 | -0.021229037 | 0.015268353 | -1.390394737 | 0.164409044 | NA |
| Plek2 | 11.25747401 | 0.35157972 | 0.162582855 | 2.162464922 | 0.030582358 | 0.125320576 | Tmem42 | 402.868127 | -0.02124098 | 0.0805264 | -0.263776593 | 0.791952072 | 0.951728598 |
| Pnp | 490.9293828 | -0.209790879 | 0.08382003 | -2.50287288 | 0.012318978 | 0.067998647 | Cald1 | 1005.736011 | -0.02125694 | 0.07221573 | -0.294353321 | 0.768487932 | 0.94619074 |
| Polr2k | 282.3981513 | -0.236112859 | 0.094336873 | -2.502869259 | 0.012319104 | 0.067998647 | Rbpms | 139.2830989 | -0.021264869 | 0.094038031 | -0.226130521 | 0.821099907 | 0.957611694 |
| Wsb1 | 2660.787774 | 0.351343111 | 0.135525846 | 2.592443584 | 0.009529681 | 0.056789568 | AV051173 | 4.227758262 | -0.021267351 | 0.042464468 | -0.500826985 | 0.616492893 | NA |
| Nras | 2028.226768 | -0.179577793 | 0.071753357 | -2.502709277 | 0.012324674 | 0.068005162 | Crlf3 | 476.6597903 | -0.021278508 | 0.074417841 | -0.28593288 | 0.77492953 | 0.947238918 |
| Star | 55.14597684 | 0.349944297 | 0.158470418 | 2.208262597 | 0.027225973 | 0.11598549 | Pde6c | 8.805477227 | -0.021305244 | 0.062035188 | -0.343438045 | 0.731268943 | NA |
| Gm16702 | 149.8892959 | 0.349793136 | 0.136442073 | 2.563675037 | 0.010357046 | 0.060365743 | Fam162a | 684.0135823 | -0.021316076 | 0.071028435 | -0.300106227 | 0.76409613 | 0.944872045 |
| Ccdc159 | 387.8843533 | 0.349554159 | 0.150146839 | 2.328082036 | 0.019907748 | 0.093909252 | Chadl | 386.8495163 | -0.021316811 | 0.079297641 | -0.268820239 | 0.788068018 | 0.951103027 |
| Adat3 | 60.72315107 | 0.349030866 | 0.152292052 | 2.291852146 | 0.021914176 | 0.099952384 | Klrb1a | 1.525437933 | -0.021326539 | 0.030707147 | -0.694513848 | 0.487360019 | NA |
| Col8a1 | 132.0863238 | 0.348900557 | 0.162472147 | 2.147448433 | 0.031757598 | 0.128439348 | Mansc4 | 7.526702359 | -0.021353213 | 0.046947747 | -0.454829358 | 0.649232008 | NA |
| Opa3 | 1763.470155 | -0.144945427 | 0.057983727 | -2.49976044 | 0.012427731 | 0.068427601 | Kctd2 | 2650.877016 | -0.021358967 | 0.05400563 | -0.395495194 | 0.692477468 | 0.924048415 |
| Tmc4 | 164.3382969 | 0.348565855 | 0.164049404 | 2.124761482 | 0.03360652 | 0.133581176 | BC017643 | 838.3609455 | -0.021365114 | 0.065650681 | -0.325436295 | 0.74485089 | 0.940516692 |
| Lamtor5 | 1115.640312 | -0.178619744 | 0.071475739 | -2.499026201 | 0.01245351 | 0.068450002 | Fam126a | 788.9090188 | -0.021384929 | 0.06006166 | -0.356049576 | 0.721803438 | 0.932513933 |

| | | | | | | | | | | | | | | |
|---|---|---|---|---|---|---|---|---|---|---|---|---|---|---|
| H2-T23 | 775.1088678 | 0.348438446 | 0.132612019 | 2.627502757 | 0.008601414 | 0.052851067 | | | Nphs2 | 12.9140783 | -0.021392246 | 0.051305744 | -0.41695616 | 0.676710474 | NA |
| Cldn1 | 67.1888557 | 0.348308357 | 0.162897936 | 2.138199943 | 0.032500519 | 0.130355434 | | | Lpin3 | 55.32720625 | -0.021394652 | 0.094150052 | -0.227239935 | 0.820237176 | 0.957571905 |
| Kdelc1 | 321.9522424 | 0.348158275 | 0.131443852 | 2.648722399 | 0.008079667 | 0.050580347 | | | Bmp7 | 357.3878193 | -0.021401386 | 0.094254302 | -0.227060048 | 0.82037705 | 0.957571905 |
| Myo1a | 39.67449104 | 0.347060107 | 0.167547343 | 2.071415168 | 0.038320015 | 0.145344016 | | | Gm17746 | 0.855988487 | -0.021403536 | 0.017872243 | -1.197585333 | 0.231078489 | NA |
| Pdzd7 | 182.1496397 | 0.347004106 | 0.150615533 | 2.303906508 | 0.021227892 | 0.097946683 | | | Rxrb | 2738.495131 | -0.021412541 | 0.063224869 | -0.338672756 | 0.734856266 | 0.936456337 |
| Srbd1 | 409.9676013 | -0.2315317 | 0.092684742 | -2.49805625 | 0.012487638 | 0.068540534 | | | Trnau1ap | 484.2052668 | -0.021420261 | 0.071512585 | -0.299531345 | 0.76453466 | 0.944872045 |
| Tshz1 | 1816.077142 | -0.341866006 | 0.136853455 | -2.498044397 | 0.012488055 | 0.068540534 | | | Otud5 | 2320.464022 | -0.021423631 | 0.048384076 | -0.442782686 | 0.657922928 | 0.915208181 |
| Nxph3 | 987.9824782 | -0.286587713 | 0.114756777 | -2.497348921 | 0.012512577 | 0.068626502 | | | Pou2af1 | 1.621032158 | -0.021424791 | 0.028260621 | -0.758114654 | 0.448382348 | NA |
| Plk5 | 810.1352229 | 0.346710775 | 0.132842479 | 2.609939058 | 0.009055835 | 0.054873333 | | | Ffar3 | 2.000452318 | -0.021429082 | 0.030185398 | -0.709915476 | 0.477756553 | NA |
| Sap30 | 122.1509836 | 0.346447399 | 0.155207329 | 2.232158759 | 0.025604472 | 0.110907378 | | | Abhd5 | 801.2237837 | -0.021448966 | 0.060736574 | -0.353147448 | 0.723977903 | 0.932714927 |
| Pcdh18 | 305.0441844 | -0.288593816 | 0.115730733 | -2.493666191 | 0.012643138 | 0.069293521 | | | Cmah | 8.722428804 | -0.021456206 | 0.061737458 | -0.347539516 | 0.728186037 | NA |
| Tert | 46.79737606 | 0.346361431 | 0.14967512 | 2.314088218 | 0.020662884 | 0.096256981 | | | Zfp518b | 726.4054561 | -0.021460236 | 0.061886239 | -0.346769114 | 0.728764782 | 0.933996867 |
| Itpr2 | 1022.74456 | -0.166243746 | 0.066684959 | -2.492972138 | 0.012667879 | 0.069378457 | | | Bcl2l10 | 1.015389671 | -0.021462643 | 0.023515085 | -0.912718081 | 0.361390832 | NA |
| Gldn | 68.72771525 | 0.34601944 | 0.15548218 | 2.225460438 | 0.026050344 | 0.112273722 | | | Dmc1 | 24.18965986 | -0.021474578 | 0.077759143 | -0.276167879 | 0.782419127 | 0.949184473 |
| Tusc5 | 27.52011876 | 0.345693761 | 0.167157105 | 2.068076982 | 0.038632785 | 0.146136809 | | | Sapcd2 | 50.50678842 | -0.021484255 | 0.093240645 | -0.230417276 | 0.817767538 | 0.956663546 |
| Matn2 | 1282.015897 | -0.282512739 | 0.113371097 | -2.491929128 | 0.012705139 | 0.069510388 | | | B4galt2 | 3335.776508 | -0.021485722 | 0.072922314 | -0.294638512 | 0.768270039 | 0.94619074 |
| Mir23b | 17.19551822 | 0.345642061 | 0.160650046 | 2.151521705 | 0.031435044 | 0.127612295 | | | Ankrd7 | 1.827834041 | -0.02148637 | 0.033001809 | -0.651066428 | 0.515003607 | NA |
| Tmem106a | 49.46717726 | 0.345014133 | 0.152697029 | 2.259468548 | 0.023854255 | 0.106157769 | | | Soga2 | 4401.90274 | -0.021494962 | 0.065508326 | -0.328125644 | 0.742816666 | 0.939912916 |
| Mrp63 | 553.5465704 | -0.246135716 | 0.098814765 | -2.490879945 | 0.012742717 | 0.069620454 | | | Znrd1 | 481.9252302 | -0.021501142 | 0.075855787 | -0.283447619 | 0.77683373 | 0.947463204 |
| Wnk4 | 258.0342013 | 0.344728973 | 0.131119628 | 2.629117995 | 0.008560665 | 0.052739143 | | | Prr12 | 3253.785862 | -0.021505107 | 0.061606275 | -0.349073315 | 0.727034269 | 0.9335794 |
| Cdc42se2 | 6584.412517 | -0.138622381 | 0.055658216 | -2.490600492 | 0.012752743 | 0.069647865 | | | Arhgap23 | 5795.798553 | -0.021521483 | 0.06121031 | -0.351598982 | 0.725139029 | 0.933334512 |
| Fanca | 45.7358904 | 0.344544539 | 0.153655542 | 2.242317685 | 0.024940848 | 0.109038796 | | | Sprn | 8159.711286 | -0.021524244 | 0.060339395 | -0.356719582 | 0.721301744 | 0.932513933 |
| Trib2 | 1610.208354 | -0.246468202 | 0.098998322 | -2.48962 | 0.012787974 | 0.069791078 | | | 1700064M15Rik | 1.425541861 | -0.021526711 | 0.025548081 | -0.842596002 | 0.399454428 | NA |
| Msi2 | 4042.303133 | -0.203190732 | 0.081646379 | -2.488668 | 0.012822264 | 0.069953577 | | | Fbxl5 | 3623.466486 | -0.021532153 | 0.062361943 | -0.345277137 | 0.729886033 | 0.934125508 |
| Slc17a9 | 15.0504936 | 0.344447085 | 0.165260549 | 2.084266854 | 0.037135901 | 0.142527531 | | | Gpatch3 | 220.0628412 | -0.021538743 | 0.085470726 | -0.252001399 | 0.801039982 | 0.953228631 |
| Lin28b | 187.4496486 | 0.344265007 | 0.138970208 | 2.47725763 | 0.01323963 | 0.071400912 | | | Ddr1 | 3414.39037 | -0.021558135 | 0.071842733 | -0.300073991 | 0.764120718 | 0.944872045 |
| Nfasc | 11960.15694 | -0.126155516 | 0.050699635 | -2.488292384 | 0.012835816 | 0.070002457 | | | Serpina9 | 292.480607 | -0.021568086 | 0.095004339 | -0.227022114 | 0.820406547 | 0.957571905 |
| Smap1 | 3609.743707 | -0.166875616 | 0.067073733 | -2.487942863 | 0.012848437 | 0.070002457 | | | Tmem221 | 8.635703972 | -0.021583116 | 0.063382361 | -0.340522433 | 0.733463132 | NA |
| Zbtb7b | 645.1483283 | 0.343790826 | 0.14807031 | 2.321807967 | 0.020243278 | 0.09495708 | | | Rhod | 18.90698801 | -0.021598223 | 0.077065869 | -0.280256658 | 0.779280601 | NA |
| Omp | 81.77302807 | -0.262601788 | 0.105562887 | -2.487633628 | 0.012859613 | 0.070009434 | | | A330035P11Rik | 23.58671429 | -0.02160417 | 0.085884863 | -0.251548051 | 0.801390417 | 0.953228631 |
| Map4k1 | 71.572548 | 0.343200707 | 0.130757842 | 2.624704575 | 0.008672418 | 0.053216016 | | | Vps37d | 628.7418342 | -0.02161355 | 0.065270537 | -0.331137921 | 0.740540309 | 0.938975678 |
| Atox1 | 818.690892 | -0.213881832 | 0.085999427 | -2.487014629 | 0.01288201 | 0.07003183 | | | Agfg2 | 1857.685967 | -0.021624019 | 0.062499694 | -0.345986004 | 0.729353233 | 0.933996867 |
| Ctnnbip1 | 1379.052416 | -0.195603699 | 0.078653769 | -2.486895436 | 0.012886327 | 0.07003183 | | | Cxcl9 | 13.8710099 | -0.021630867 | 0.01599478 | -1.352370395 | NA | NA |
| Polr2m | 6566.03823 | -0.098324381 | 0.039536789 | -2.486908617 | 0.012885849 | 0.07003183 | | | Kcnn4 | 8.696782589 | -0.021637149 | 0.064138374 | -0.337351066 | 0.735852268 | NA |
| Cd200r1 | 18.71250043 | 0.343159838 | 0.168104434 | 2.041349114 | 0.041216139 | 0.153300321 | | | AF357426 | 7.577226937 | -0.021645152 | 0.051748052 | -0.418279548 | 0.675742741 | NA |
| H2-Q5 | 15.99317857 | 0.342633203 | 0.166628699 | 2.056267642 | 0.039756725 | 0.148989976 | | | Tmem241 | 273.9296371 | -0.021646668 | 0.079051435 | -0.273830174 | 0.784215134 | 0.949699053 |
| Pip5kl1 | 188.6627441 | 0.342559891 | 0.15308141 | 2.237762846 | 0.025236525 | 0.109897333 | | | Rps2 | 10390.10425 | -0.021661621 | 0.049834336 | -0.434672613 | 0.66380008 | 0.915428982 |
| Aanat | 21.86623983 | 0.342445078 | 0.167665629 | 2.042428612 | 0.041109034 | 0.153074591 | | | Gga1 | 2976.624104 | -0.021674151 | 0.064037408 | -0.338460784 | 0.735015975 | 0.936456337 |
| Tbcc | 459.6201582 | -0.174540663 | 0.070238042 | -2.484987598 | 0.012955595 | 0.070309627 | | | Polr3a | 1521.186209 | -0.021692439 | 0.051029182 | -0.425098696 | 0.670764728 | 0.918334444 |
| 4930444P10Rik | 9.365659291 | 0.342426437 | 0.162264369 | 2.110299632 | 0.034832555 | 0.136562655 | | | Nox1 | 0.77388662 | -0.021699063 | 0.021883521 | -0.991570908 | 0.321406887 | NA |
| Liph | 52.36618567 | 0.3420732 | 0.156328277 | 2.188172268 | 0.028657056 | 0.119981547 | | | Abhd10 | 725.0112739 | -0.021699917 | 0.064873833 | -0.334494134 | 0.738006716 | 0.937497644 |
| Tatdn3 | 425.392812 | 0.341771954 | 0.145757558 | 2.344797469 | 0.019037418 | 0.09113625 | | | Mir804 | 0.461806006 | -0.021704567 | 0.015436653 | -1.406040961 | 0.159711962 | NA |
| Flrt2 | 2910.060439 | -0.193894886 | 0.078069785 | -2.483609853 | 0.013005821 | 0.070483454 | | | Snord55 | 0.68199713 | -0.021730673 | 0.018443584 | -1.178223994 | 0.238707319 | NA |
| Naip2 | 70.40954443 | 0.341650077 | 0.156998999 | 2.176129015 | 0.029545615 | 0.122139743 | | | E330033B04Rik | 48.43008222 | -0.021744764 | 0.086642809 | -0.250970214 | 0.801837138 | 0.95333733 |
| Yes1 | 420.3026607 | -0.273738464 | 0.110261944 | -2.482619608 | 0.013042028 | 0.07063026 | | | Prdm10 | 460.8632286 | -0.021749361 | 0.067952132 | -0.320068852 | 0.748916137 | 0.9410864 |
| Tmem37 | 43.53594592 | 0.341532582 | 0.163128477 | 2.093641704 | 0.03629191 | 0.140373739 | | | Afg3l1 | 1268.676015 | -0.021765519 | 0.053932653 | -0.403568486 | 0.686530064 | 0.923860706 |
| Gpr4 | 149.6672003 | -0.30580959 | 0.123237838 | -2.481458578 | 0.013084592 | 0.070737147 | | | Iba57 | 357.1305783 | -0.021766253 | 0.080762605 | -0.269509055 | 0.787537975 | 0.950858334 |
| Hrk | 2477.51507 | -0.222880156 | 0.089816716 | -2.481499714 | 0.013083082 | 0.070737147 | | | Ctse | 2.249497915 | -0.021800667 | 0.025160027 | -0.866480278 | 0.386226837 | NA |
| 1700013F07Rik | 35.26067892 | 0.341339414 | 0.15694523 | 2.174895116 | 0.029637977 | 0.122423571 | | | Phf12 | 2617.917342 | -0.021801605 | 0.047975993 | -0.454427392 | 0.64952124 | 0.913724196 |
| Zc3h7b | 9793.226436 | -0.149364354 | 0.060187444 | -2.481653031 | 0.013077454 | 0.070737147 | | | Cluap1 | 1072.128166 | -0.021807966 | 0.057877924 | -0.376792473 | 0.706327835 | 0.929442715 |
| 1110058L19Rik | 478.7329802 | -0.199866805 | 0.080561664 | -2.480917044 | 0.013104487 | 0.070795299 | | | Ucma | 19.97908135 | -0.021823872 | 0.075819938 | -0.287838173 | 0.773470616 | NA |
| 4921536K21Rik | 14.33233936 | 0.341254896 | 0.167152506 | 2.041578104 | 0.041193399 | 0.153300321 | | | Papolg | 864.9832126 | -0.021827797 | 0.072979795 | -0.299093701 | 0.764868552 | 0.94492946 |
| 1700028P14Rik | 27.78884104 | 0.341124138 | 0.165460644 | 2.061663303 | 0.039239803 | 0.147923531 | | | Ppwd1 | 285.8772863 | -0.021840246 | 0.080617776 | -0.270911043 | 0.786459451 | 0.950484821 |
| Trim7 | 49.80621069 | 0.340425498 | 0.152989635 | 2.225153993 | 0.026070902 | 0.112277067 | | | Sox30 | 5.508660513 | -0.021873506 | 0.051288326 | -0.426481177 | 0.669757261 | NA |
| Atp7b | 87.92969401 | 0.339661473 | 0.150331046 | 2.259423335 | 0.023857064 | 0.106157769 | | | Trpa1 | 11.44852968 | -0.021882047 | 0.064013724 | -0.34183368 | 0.732476062 | NA |
| Snhg3 | 141.4303023 | 0.339445503 | 0.150795213 | 2.251036322 | 0.024383237 | 0.107787125 | | | Pdilt | 1.764357048 | -0.021885491 | 0.024800471 | -0.882462722 | 0.37752663 | NA |
| BC027072 | 11.5356439 | 0.339063625 | 0.163163936 | 2.078054952 | 0.037704298 | 0.143924709 | | | Pabpc5 | 63.39373777 | -0.021896737 | 0.094623964 | -0.231407942 | 0.816997896 | 0.956302288 |
| 2810032G03Rik | 311.6239633 | -0.259296296 | 0.104683894 | -2.476945456 | 0.013251215 | 0.071413679 | | | Slc38a7 | 1455.057658 | -0.02189872 | 0.056964433 | -0.384427946 | 0.700661291 | 0.92719503 |
| Mrpl20 | 1369.550086 | -0.188633913 | 0.076154788 | -2.476980358 | 0.013249919 | 0.071413679 | | | Gm10125 | 56.78938564 | -0.021922795 | 0.094938639 | -0.230915418 | 0.817380512 | 0.956364827 |
| Calr3 | 41.82614921 | 0.337932808 | 0.160827449 | 2.101213508 | 0.035622228 | 0.138536848 | | | Lrp1b | 1589.409674 | -0.021936025 | 0.076139918 | -0.288101507 | 0.773269039 | 0.947121017 |
| Peli2 | 1083.471299 | -0.238156198 | 0.096170516 | -2.476395126 | 0.013271661 | 0.071474145 | | | B430212C06Rik | 8.203070453 | -0.021947692 | 0.054574394 | -0.40216098 | 0.687565557 | NA |
| Rwdd2a | 576.0691686 | -0.23918775 | 0.096593874 | -2.476220683 | 0.013278148 | 0.071484232 | | | Gm20755 | 1.09377793 | -0.021957341 | 0.023091384 | -0.950888892 | 0.341660781 | NA |
| Abi2 | 9088.960439 | -0.15878794 | 0.064136178 | -2.475793622 | 0.01329404 | 0.07154493 | | | Tgds | 180.535093 | -0.02196478 | 0.088887918 | -0.247106468 | 0.804825829 | 0.954826853 |
| Ctnnd1 | 3646.11893 | -0.165739105 | 0.066948813 | -2.475609316 | 0.013300904 | 0.071557014 | | | Apol9a | 3.542994518 | -0.021964887 | 0.044268704 | -0.496171893 | 0.61977314 | NA |
| C1qtnf2 | 30.89117522 | -0.39480758 | 0.159515363 | -2.475044239 | 0.013321967 | 0.071620597 | | | Zufsp | 314.2162158 | -0.021992586 | 0.087988938 | -0.249947174 | 0.802628201 | 0.953543572 |
| Agxt2l1 | 1045.036883 | 0.336873255 | 0.167333948 | 2.013179382 | 0.044095761 | 0.160530951 | | | Rsu1 | 843.5821075 | -0.021994645 | 0.062015615 | -0.354663009 | 0.722842067 | 0.93255988 |
| Zfp811 | 396.1164798 | -0.258104341 | 0.104291008 | -2.474847513 | 0.013329307 | 0.071635202 | | | Itgb3bp | 124.6024209 | -0.022009942 | 0.091251395 | -0.241201156 | 0.809399215 | 0.955725085 |
| Bud31 | 1100.227061 | -0.1565717 | 0.063292745 | -2.473770088 | 0.01336957 | 0.071801776 | | | 4930405A21Rik | 8.906832054 | -0.022014466 | 0.060592409 | -0.363320522 | 0.716365465 | NA |
| Ctrl | 19.49000184 | 0.336853736 | 0.163004335 | 2.066532375 | 0.038778239 | 0.146579661 | | | Ift88 | 812.8360044 | -0.022015298 | 0.068762942 | -0.320162254 | 0.748845336 | 0.9410864 |
| Fbxw10 | 30.80793728 | 0.336851377 | 0.165018611 | 2.041293249 | 0.041221688 | 0.153300321 | | | Atp5e | 1967.148257 | -0.022017935 | 0.069127789 | -0.318510623 | 0.750097638 | 0.9410864 |
| Ap3s1 | 1172.46874 | -0.194856077 | 0.078785741 | -2.473240403 | 0.013389404 | 0.071872933 | | | Narfl | 1035.28946 | -0.022036803 | 0.0692656 | -0.318149316 | 0.750371676 | 0.9410864 |
| Cd101 | 11.69764137 | 0.336334746 | 0.157106166 | 2.14081187 | 0.032289212 | 0.129817618 | | | Smarca1 | 1191.288121 | -0.02203758 | 0.053811816 | -0.409530507 | 0.682150384 | 0.922544285 |
| P4ha3 | 177.3235348 | 0.336193558 | 0.1428468 | 2.353525312 | 0.018596342 | 0.089872651 | | | Fgf8 | 0.475177312 | -0.022041534 | 0.015554805 | -1.417024166 | 0.156475862 | NA |
| Ank3 | 10268.14058 | -0.169225074 | 0.068438798 | -2.472648266 | 0.013411607 | 0.071902919 | | | Mettl20 | 80.3273201 | -0.022102984 | 0.094984991 | -0.232699753 | 0.815994561 | 0.956264994 |
| Parp9 | 165.8688331 | 0.33617349 | 0.134522284 | 2.499017118 | 0.01245383 | 0.068450002 | | | Gm128 | 4.982652806 | -0.022117239 | 0.042600857 | -0.519173573 | 0.603639706 | NA |
| Pdia3 | 7525.655605 | -0.19437627 | 0.078642814 | -2.471634215 | 0.013449705 | 0.072080004 | | | 4930427A07Rik | 79.52496722 | -0.022126241 | 0.095101297 | -0.232659719 | 0.816025651 | 0.956264994 |
| Cep95 | 333.0930267 | 0.336130248 | 0.139083124 | 2.416757972 | 0.015659425 | 0.080313518 | | | Pygb | 9084.037252 | -0.022129622 | 0.05013689 | -0.441384022 | 0.658935008 | 0.915208181 |
| Cd47 | 2275.66031 | -0.138062371 | 0.055894144 | -2.470068631 | 0.013508713 | 0.072298447 | | | Zfp764 | 193.5119329 | -0.02215542 | 0.090078735 | -0.245956164 | 0.805716166 | 0.95539795 |
| Sptssa | 543.9821358 | -0.186123272 | 0.075350302 | -2.470106511 | 0.013507283 | 0.072298447 | | | A930041C12Rik | 1.028348205 | -0.02216233 | 0.022595196 | -0.98084255 | 0.326670392 | NA |
| Spef2 | 33.16150609 | 0.336043686 | 0.16564792 | 2.028662266 | 0.042492702 | 0.156273895 | | | Sp4 | 808.0456685 | -0.022174164 | 0.073763656 | -0.300610964 | 0.76371117 | 0.944785254 |
| Snora16a | 17.46999578 | 0.336011578 | 0.167801377 | 2.002436351 | 0.045237822 | 0.163122567 | | | Otop1 | 1.489672776 | -0.022178896 | 0.02025692 | -1.094879983 | NA | NA |
| Gm11696 | 81.12209881 | 0.335325913 | 0.153669967 | 2.182117434 | 0.029100867 | 0.120956391 | | | D930028M14Rik | 4.488474599 | -0.022191006 | 0.044649772 | -0.49700154 | 0.619187965 | NA |
| Dhx58 | 58.61062843 | 0.335109513 | 0.16136934 | 2.076661613 | 0.037832802 | 0.144115079 | | | Fam149b | 1172.98435 | -0.022215089 | 0.055783236 | -0.398239514 | 0.690453641 | 0.923860706 |
| AI607873 | 16.91663762 | 0.334973702 | 0.156833065 | 2.135861486 | 0.032690706 | 0.130914912 | | | Fhl2 | 1410.453471 | -0.02223223 | 0.077546448 | -0.286695658 | 0.774345362 | 0.947238918 |
| Anln | 1763.645618 | 0.334644743 | 0.158902135 | 2.105980159 | 0.035206076 | 0.137470499 | | | Cuedc1 | 1150.542418 | -0.02223302 | 0.069920748 | -0.317974576 | 0.750504222 | 0.9410864 |
| Pbrm1 | 2209.209884 | -0.189171265 | 0.076694928 | -2.466542055 | 0.013642471 | 0.072838195 | | | Dppa2 | 0.677494072 | -0.022248745 | 0.018680196 | -1.191033836 | 0.233640302 | NA |
| Aurkaip1 | 1582.870454 | -0.164223904 | 0.066626798 | -2.464832599 | 0.013707728 | 0.073125764 | | | Dhx57 | 2173.04168 | -0.022263719 | 0.045841108 | -0.485671484 | 0.627200105 | 0.903033871 |
| Rgs11 | 1257.449651 | 0.33439738 | 0.164950575 | 2.027258043 | 0.04263603 | 0.156615137 | | | 5430401F13Rik | 1.77566614 | -0.022264293 | 0.029868417 | -0.745412533 | 0.45602237 | NA |
| Rxfp3 | 150.4545007 | 0.334139919 | 0.154055201 | 2.168962277 | 0.030085548 | 0.123810221 | | | Rab40c | 1730.927852 | -0.02226816 | 0.069959148 | -0.318302333 | 0.750255615 | 0.9410864 |
| Pten | 7969.889839 | -0.174139849 | 0.070655373 | -2.464637042 | 0.01371521 | 0.073125764 | | | Snord38a | 1.130209386 | -0.022288756 | 0.022649931 | -0.98405405 | 0.325088938 | NA |
| Rxfp1 | 234.9115249 | 0.334085991 | 0.15935469 | 2.096492992 | 0.036038481 | 0.139595056 | | | Tmem174 | 14.07253166 | -0.022293134 | 0.063888982 | -0.3489355 | 0.727137733 | NA |
| Inpp4b | 293.5930867 | -0.338546576 | 0.137405485 | -2.463850529 | 0.013745342 | 0.073236014 | | | 4632434I11Rik | 25.0664515 | -0.022297308 | 0.074062159 | -0.30106208 | 0.763367156 | 0.944624337 |
| Mrto4 | 694.4030856 | -0.158323657 | 0.064279903 | -2.463035076 | 0.013776644 | 0.073357204 | | | Padi2 | 1620.712642 | -0.022300426 | 0.079812938 | -0.27940866 | 0.779931226 | 0.948894514 |
| Trim30d | 68.83175796 | 0.334053504 | 0.162052288 | 2.061393316 | 0.039265532 | 0.147952372 | | | Nsun3 | 198.5255624 | -0.022303554 | 0.087102158 | -0.25606201 | 0.797902956 | 0.952967259 |
| Slc44a5 | 335.7946527 | 0.333842752 | 0.128351789 | 2.600998037 | 0.009295299 | 0.055822231 | | | Phospho1 | 1110.957084 | -0.022321515 | 0.076779231 | -0.290723351 | 0.771262911 | 0.946250219 |
| Tmem174 | 8.879555436 | 0.333825345 | 0.12963844 | 2.575049083 | 0.010022586 | 0.058942547 | | | Ascl4 | 3.919982356 | -0.02232556 | 0.03566624 | -0.625957759 | 0.531342648 | NA |
| Mrrf | 645.4882133 | -0.205969073 | 0.083646494 | -2.462375436 | 0.013802011 | 0.073411724 | | | 2810002D19Rik | 133.9927761 | -0.02233261 | 0.091299021 | -0.244609526 | 0.806758787 | 0.955587482 |
| Sdhb | 4067.832509 | -0.122950966 | 0.049939249 | -2.462010702 | 0.013816054 | 0.073461204 | | | Clec9a | 0.636822305 | -0.022335874 | 0.018447294 | -1.210794041 | 0.225974349 | NA |
| Pfdn4 | 459.6045942 | -0.195004902 | 0.079238018 | -2.461001771 | 0.013854968 | 0.07364284 | | | 1700096K18Rik | 49.56198769 | -0.022338855 | 0.094544218 | -0.236279444 | 0.813215836 | 0.956264994 |
| Slc15a3 | 103.5657395 | 0.332718787 | 0.152095787 | 2.18756084 | 0.028701607 | 0.120059603 | | | Actl7b | 0.664630415 | -0.022365955 | 0.01860811 | -1.201946623 | 0.229384209 | NA |
| Pyroxd2 | 71.89463314 | 0.332430795 | 0.163754665 | 2.030053896 | 0.042351061 | 0.156049309 | | | Snrnp200 | 6441.840876 | -0.022381205 | 0.043084905 | -0.519467419 | 0.603434828 | 0.891540503 |
| Endog | 260.3943405 | -0.263422642 | 0.107072395 | -2.460229272 | 0.013884828 | 0.073687366 | | | 4930429F24Rik | 1.254331433 | -0.022389474 | 0.027101623 | -0.826130399 | 0.408730125 | NA |
| Tmed2 | 3821.910848 | -0.123388507 | 0.050154465 | -2.460169928 | 0.013887125 | 0.073687366 | | | Wtap | 2227.672651 | -0.022403181 | 0.052670016 | -0.425349815 | 0.670581684 | 0.918256933 |
| Slc24a5 | 9.463707027 | 0.332381988 | 0.139198955 | 2.387819568 | 0.01694866 | 0.084250743 | | | 1600010M07Rik | 2.486319034 | -0.022407428 | 0.033674751 | -0.665407375 | 0.505789969 | NA |
| Bet1 | 408.0194431 | -0.242510048 | 0.098591669 | -2.459741793 | 0.013903701 | 0.07373614 | | | Skor1 | 12.57378579 | -0.022409943 | 0.06273858 | -0.357195566 | 0.720945405 | NA |
| Fam83h | 104.1186562 | 0.331585515 | 0.145851829 | 2.273440915 | 0.022999623 | 0.103506474 | | | Ep400 | 3993.024519 | -0.022411916 | 0.05880742 | -0.381106949 | 0.703123891 | 0.927855161 |
| Ppef2 | 23.68480295 | 0.331557416 | 0.168074293 | 1.972683688 | 0.048531612 | 0.17085862 | | | Eaf2 | 42.00122174 | -0.022414727 | 0.091216241 | -0.245731755 | 0.805889889 | 0.95539795 |
| Dleu7 | 149.5291586 | -0.258533973 | 0.105207965 | -2.457361215 | 0.013996188 | 0.074164478 | | | Bmp5 | 45.19765562 | -0.022415707 | 0.092803876 | -0.241538481 | 0.809137796 | 0.955725085 |
| Slc2a9 | 36.3942228 | 0.331073227 | 0.159122886 | 2.080613516 | 0.037469296 | 0.143416321 | | | Icam4 | 28.88962123 | -0.022428968 | 0.080538533 | -0.278487413 | 0.780638226 | 0.948894514 |
| D4Ertd617e | 12.44412095 | 0.330935737 | 0.160565539 | 2.061063284 | 0.039297003 | 0.148006266 | | | Deaf1 | 1617.263987 | -0.022432027 | 0.067508662 | -0.332283694 | 0.739675051 | 0.938470248 |
| Ndufa4 | 6503.007162 | -0.179506724 | 0.073112178 | -2.455223297 | 0.01407971 | 0.074530586 | | | 4833403I15Rik | 2.28468946 | -0.022433182 | 0.031418528 | -0.714011257 | 0.475220219 | NA |
| Cyld | 7809.164 | -0.186456332 | 0.075950789 | -2.454962395 | 0.014089933 | 0.074559227 | | | Nup107 | 674.7274305 | -0.022442784 | 0.065227687 | -0.344068375 | 0.730794866 | 0.934712179 |
| Col4a4 | 24.0155392 | 0.330824276 | 0.164616927 | 2.009661354 | 0.044467043 | 0.161200833 | | | Chd2 | 2742.49414 | -0.022443262 | 0.064988417 | -0.345342499 | 0.7298369 | 0.934125508 |
| 1700001O22Rik | 145.43033 | 0.330566459 | 0.126030799 | 2.622902185 | 0.00871843 | 0.053413742 | | | Gstcd | 215.2302535 | -0.022463146 | 0.080950441 | -0.277492567 | 0.781401912 | 0.948894514 |
| Ptpro | 1929.185085 | -0.410669089 | 0.167370121 | -2.453658315 | 0.014141128 | 0.074753542 | | | Otos | 2.434381313 | -0.022473201 | 0.034043819 | -0.660125731 | 0.509173148 | NA |
| Tgm7 | 9.266306776 | 0.330495245 | 0.1520202 | 2.174021912 | 0.02970349 | 0.122552061 | | | Musk | 79.78569996 | -0.022478565 | 0.092725235 | -0.24242122 | 0.808453793 | 0.955716697 |
| Rnf103 | 1638.098587 | -0.147157476 | 0.059987474 | -2.453136731 | 0.01416165 | 0.074810979 | | | Slc12a6 | 6686.683671 | -0.02252126 | 0.054157446 | -0.415847898 | 0.677521307 | 0.921073889 |
| Klhl17 | 1642.833048 | 0.330448527 | 0.127798292 | 2.585703784 | 0.009718041 | 0.057535851 | | | Neurl1b | 2359.794983 | -0.022526264 | 0.065054784 | -0.346266069 | 0.729142766 | 0.933996867 |
| Hsbp1l1 | 27.58993814 | 0.330358192 | 0.167911064 | 1.967459346 | 0.049130278 | 0.171950425 | | | Mir218-1 | 2.133570485 | -0.02252648 | 0.027781723 | -0.810838135 | 0.417458631 | NA |
| Edn1 | 113.0413102 | -0.384223666 | 0.156776199 | -2.450778041 | 0.014254782 | 0.075226021 | | | Ccrl2 | 35.01544031 | -0.022538727 | 0.090599694 | -0.248772664 | 0.803536638 | 0.953754272 |
| Apaf1 | 631.7768199 | 0.329962863 | 0.13844804 | 2.383297461 | 0.017158323 | 0.084909314 | | | Acads | 298.9708926 | -0.02253959 | 0.087077009 | -0.258846629 | 0.795753579 | 0.952700608 |
| Gli1 | 129.1466443 | 0.32973034 | 0.161687569 | 2.039305448 | 0.041419552 | 0.153824195 | | | Aspscr1 | 1235.737899 | -0.022568352 | 0.062891495 | -0.358845848 | 0.719710412 | 0.932251852 |
| Edn3 | 212.6537943 | -0.3561097 | 0.145437475 | -2.448541541 | 0.014343589 | 0.075617409 | | | Cdk1 | 13.55456425 | -0.022571659 | 0.058710266 | -0.384458468 | 0.700638673 | NA |
| Nckap5 | 153.2648546 | -0.326354035 | 0.133341381 | -2.447507532 | 0.014384812 | 0.075799835 | | | 1110038B12Rik | 217.0000935 | -0.022596112 | 0.087541332 | -0.258119357 | 0.796314793 | 0.952734447 |
| Nit2 | 310.1077565 | -0.265269897 | 0.108387201 | -2.44742825 | 0.014387977 | 0.075799835 | | | Usb1 | 412.3402773 | -0.022601642 | 0.082396833 | -0.274302312 | 0.783852308 | 0.949699053 |
| Capza1 | 1868.540963 | -0.1876753 | 0.07670898 | -2.446588393 | 0.014421543 | 0.075950845 | | | Lrp2 | 22.17130481 | -0.022602035 | 0.069947214 | -0.323129881 | 0.746596881 | NA |
| Nupr1 | 291.2407848 | 0.329040364 | 0.147013847 | 2.238158991 | 0.025210689 | 0.109877472 | | | Aoc3 | 35.31367698 | -0.022618474 | 0.083889173 | -0.269623286 | 0.787450084 | 0.950831248 |
| Lgals12 | 30.87150873 | 0.328908982 | 0.16659678 | 1.974281744 | 0.048349716 | 0.170373094 | | | Utp18 | 705.3550327 | -0.022619135 | 0.073798406 | -0.306498958 | 0.759224773 | 0.942526199 |
| Mmd2 | 5521.812297 | -0.215051694 | 0.087926605 | -2.445809142 | 0.014452749 | 0.076037652 | | | Rrp9 | 705.3567711 | -0.022624332 | 0.071796115 | -0.315119166 | 0.752671177 | 0.94179529 |
| Usp34 | 6631.766072 | -0.207727185 | 0.084956903 | -2.445088958 | 0.014481642 | 0.076161523 | | | Nek11 | 77.1371579 | -0.022654665 | 0.095101724 | -0.23821508 | 0.811714281 | 0.956204821 |
| Camkk1 | 6415.11481 | 0.328870141 | 0.129636443 | 2.536864897 | 0.011185011 | 0.063480059 | | | Efcab5 | 157.7713223 | -0.022671044 | 0.09098339 | -0.249177837 | 0.803223223 | 0.953754272 |
| Mrpl21 | 537.6861446 | -0.18141174 | 0.074201945 | -2.444838068 | 0.014491719 | 0.076165094 | | | Pde1c | 501.9408207 | -0.022674142 | 0.084895959 | -0.267081523 | 0.789406393 | 0.951374535 |
| Ltbp1 | 359.2203421 | -0.219947925 | 0.0899924 | -2.444072218 | 0.014522519 | 0.076301089 | | | Chaf1a | 225.9114935 | -0.022679593 | 0.084669257 | -0.267861009 | 0.788806308 | 0.951282725 |
| Tyrobp | 580.6805622 | 0.328746552 | 0.136359634 | 2.410878815 | 0.015914135 | 0.081184757 | | | 4930455C13Rik | 0.897270483 | -0.022684876 | 0.021503914 | -1.054918502 | 0.2914626 | NA |
| Tmtc2 | 594.9520114 | -0.202848873 | 0.083056307 | -2.442305475 | 0.014593792 | 0.07662359 | | | Chit1 | 5.977157792 | -0.022688416 | 0.048938141 | -0.463614177 | 0.642924196 | NA |
| Dcaf12l1 | 1517.828151 | -0.203775821 | 0.083456922 | -2.441688687 | 0.014618747 | 0.076728612 | | | Pm20d1 | 112.2183872 | -0.022693576 | 0.094768806 | -0.239462505 | 0.810746967 | 0.956073263 |
| 4930590J08Rik | 22.20173233 | 0.328639685 | 0.168060635 | 1.955482819 | 0.050526109 | 0.174998787 | | | E2f4 | 1165.986779 | -0.022694494 | 0.067277256 | -0.337327883 | 0.735869742 | 0.936995431 |
| Fkbp1a | 18582.69165 | -0.129539891 | 0.053064406 | -2.441182341 | 0.014639261 | 0.076784262 | | | Cyb5rl | 117.9153398 | -0.022695106 | 0.093728282 | -0.242137224 | 0.808673835 | 0.955723386 |
| Sh3rf2 | 573.6638858 | -0.31875945 | 0.130597306 | -2.440781206 | 0.014655531 | 0.076843586 | | | Birc6 | 4956.560049 | -0.022699667 | 0.072258009 | -0.314147424 | 0.753409072 | 0.94183464 |
| Fam83d | 61.14365969 | 0.328408788 | 0.163728936 | 2.005807865 | 0.044876751 | 0.16226813 | | | Pdcd5 | 965.1622359 | -0.022704725 | 0.075294982 | -0.301543673 | 0.762999953 | 0.944431816 |
| Oaf | 413.275506 | -0.208282308 | 0.085372673 | -2.439683585 | 0.014700131 | 0.077013181 | | | Ckap2 | 22.17471522 | -0.022754972 | 0.082074409 | -0.27724808 | 0.781589624 | NA |
| Tacr2 | 11.90303852 | 0.32757969 | 0.15095477 | 2.170051932 | 0.030002912 | 0.123629389 | | | Gm6498 | 0.865688855 | -0.022772076 | 0.021399474 | -1.064141848 | 0.287264446 | NA |
| Inha | 854.581333 | 0.32704529 | 0.143662473 | 2.276483785 | 0.02281707 | 0.102919267 | | | Eid3 | 4.163133451 | -0.022775517 | 0.042002773 | -0.542238411 | 0.587654277 | NA |
| Shkbp1 | 489.418943 | 0.326661472 | 0.137220152 | 2.380564863 | 0.017286118 | 0.085350894 | | | Tgm6 | 7.202611723 | -0.022794142 | 0.056382854 | -0.404274357 | 0.68601098 | NA |
| 1700037H04Rik | 2019.460639 | -0.172623529 | 0.070779241 | -2.438900548 | 0.014732022 | 0.077088129 | | | Tshb | 1.921619034 | -0.022801192 | 0.029485645 | -0.773298063 | 0.439346002 | NA |
| Upk1a | 16.28662789 | 0.326624922 | 0.16140266 | 2.023665047 | 0.043004627 | 0.15756436 | | | 5330417C22Rik | 1357.815352 | -0.022808574 | 0.057294185 | -0.398095789 | 0.690559578 | 0.923860706 |
| Plin4 | 421.809484 | 0.32650994 | 0.127321191 | 2.564458729 | 0.010333687 | 0.060327861 | | | Mrpl34 | 477.3698722 | -0.022812263 | 0.085011063 | -0.268344639 | 0.788434049 | 0.951189696 |
| Hif3a | 336.8584827 | 0.326394034 | 0.16219839 | 2.012313643 | 0.044186885 | 0.160758077 | | | Gmpr2 | 793.5347113 | -0.022814131 | 0.057174516 | -0.399026227 | 0.689873879 | 0.923860706 |
| Gm16677 | 10.84447229 | 0.326240713 | 0.150795438 | 2.163465409 | 0.030505401 | 0.125106059 | | | Cyp2d26 | 2.185797528 | -0.022825403 | 0.028371276 | -0.804525074 | 0.421093795 | NA |
| 4930432K21Rik | 109.0370228 | 0.32596401 | 0.124971087 | 2.608315396 | 0.009098907 | 0.055069716 | | | 1700092K14Rik | 2.606155627 | -0.022828962 | 0.036364105 | -0.627788356 | 0.530142595 | NA |
| Grip1 | 635.2277811 | -0.221985929 | 0.091089124 | -2.437019034 | 0.0148089 | 0.077274849 | | | Hfe | 170.5530703 | -0.022832977 | 0.095081565 | -0.240140946 | 0.810220992 | 0.955876688 |
| Il20ra | 33.82920702 | -0.408764873 | 0.167728096 | -2.437068588 | 0.014806871 | 0.077274849 | | | Ogn | 236.333343 | -0.022840713 | 0.083438961 | -0.273741582 | 0.78428322 | 0.949699053 |
| Plekhg4 | 112.0195022 | 0.325947559 | 0.155761514 | 2.092606519 | 0.036384294 | 0.140570291 | | | Mirlet7i | 1.373352305 | -0.022863272 | 0.023317233 | -0.980531089 | 0.326824031 | NA |
| Asb11 | 146.4962814 | 0.325906669 | 0.137870844 | 2.363854889 | 0.018085895 | 0.088276641 | | | Gatc | 1337.303643 | -0.022867481 | 0.059676414 | -0.383191262 | 0.701577956 | 0.92751465 |
| Mex3a | 188.1628675 | -0.294263314 | 0.120728865 | -2.437389879 | 0.014793719 | 0.077274849 | | | Mrp63 | 578.0353437 | -0.022891469 | 0.064090598 | -0.357173593 | 0.720961853 | 0.932513933 |
| Trem2 | 476.4180915 | 0.325817019 | 0.148724926 | 2.190735791 | 0.028470917 | 0.119579397 | | | Usp8 | 3229.341828 | -0.022895413 | 0.050670386 | -0.451849971 | 0.651377067 | 0.914343204 |
| Ppp2r2c | 28246.88256 | -0.172591299 | 0.070853347 | -2.435894804 | 0.014855004 | 0.077417905 | | | Ttyh1 | 20577.17498 | -0.022904502 | 0.041776624 | -0.548261195 | 0.583512568 | 0.884313345 |
| Dbndd1 | 566.9889549 | -0.224054829 | 0.092032615 | -2.434515497 | 0.014911741 | 0.077621071 | | | Cacna1e | 7234.502493 | -0.022907921 | 0.075269649 | -0.304344725 | 0.760865275 | 0.943105713 |
| Itgb1 | 2566.177853 | -0.142438265 | 0.058512103 | -2.434338487 | 0.014919036 | 0.077621071 | | | Stk31 | 2.276189835 | -0.022910186 | 0.03361188 | -0.681609772 | 0.495485734 | NA |
| Chkb | 19.72388696 | 0.325014092 | 0.168080493 | 1.933681216 | 0.053152335 | 0.180810506 | | | Gm6938 | 1.234382615 | -0.022917202 | 0.025358341 | -0.903734294 | 0.36613631 | NA |
| Ccdc171 | 128.6605904 | 0.324776658 | 0.148432207 | 2.188047091 | 0.028666172 | 0.119981547 | | | Mc2r | 1.086318744 | -0.022918591 | 0.022744646 | -1.00764775 | 0.313623597 | NA |
| Nuak2 | 44.19769673 | 0.32460354 | 0.164849962 | 1.969084719 | 0.048943363 | 0.17152872 | | | Rpl35a | 4369.42981 | -0.022923988 | 0.065441225 | -0.350298881 | 0.726114405 | 0.933575664 |
| 4833439L19Rik | 5476.223068 | -0.123407144 | 0.050699939 | -2.434068889 | 0.014930153 | 0.077652835 | | | Gm2897 | 9.877661889 | -0.022928654 | 0.059725821 | -0.383898518 | 0.701053665 | NA |
| Als2cl | 481.7774557 | 0.3241591 | 0.123672779 | 2.621103066 | 0.008764576 | 0.053590506 | | | Mpp4 | 25.25632598 | -0.022931611 | 0.084480983 | -0.271441096 | 0.786051798 | 0.950484821 |
| Mpeg1 | 1795.949944 | 0.323878968 | 0.140426767 | 2.306390549 | 0.02108882 | 0.097450096 | | | Sall3 | 261.6470168 | -0.022953943 | 0.083777356 | -0.273987442 | 0.784094272 | 0.949699053 |
| Mrfap1 | 9271.574401 | -0.140539604 | 0.057788431 | -2.431967823 | 0.015017041 | 0.07802617 | | | Ccdc59 | 614.2139827 | -0.022957061 | 0.072436099 | -0.316928459 | 0.751297887 | 0.9410864 |
| Rln1 | 14.84427357 | -0.406432031 | 0.167138247 | -2.431711707 | 0.015027663 | 0.078055184 | | | P2rx1 | 13.13994985 | -0.02296456 | 0.063198276 | -0.363373198 | 0.71632612 | NA |
| Pnmal1 | 3362.872028 | -0.179500649 | 0.073820328 | -2.431588339 | 0.015032782 | 0.078055605 | | | Csda | 843.0578766 | -0.022964593 | 0.063544185 | -0.361395658 | 0.717803693 | 0.93125489 |
| Rassf7 | 38.15156428 | 0.322784518 | 0.165191754 | 1.953998976 | 0.050701338 | 0.175451801 | | | Aak1 | 21706.13326 | -0.022985419 | 0.052734057 | -0.435874283 | 0.662927945 | 0.915428982 |
| 1810020O05Rik | 44.58619223 | 0.322496803 | 0.167396242 | 1.926547456 | 0.054036052 | 0.182682652 | | | Lins | 454.6942513 | -0.022989294 | 0.071737933 | -0.320462175 | 0.748618 | 0.9410864 |
| Ntn4 | 289.4956289 | -0.27829617 | 0.114503163 | -2.430467088 | 0.015079376 | 0.078219943 | | | Pou6f1 | 6116.836699 | -0.022994846 | 0.054939282 | -0.418550168 | 0.675544915 | 0.920229309 |
| Bag1 | 3521.01707 | -0.197817365 | 0.081400908 | -2.43016166 | 0.01509209 | 0.078232464 | | | Syndig1l | 2103.028786 | -0.023001879 | 0.095092235 | -0.241890196 | 0.808865247 | 0.955725085 |
| 1700040N02Rik | 11.50819633 | 0.321821602 | 0.163387803 | 1.969679471 | 0.048875117 | 0.171405855 | | | Ninj2 | 89.44296393 | -0.023028683 | 0.094124089 | -0.244663014 | 0.806717368 | 0.955587482 |
| Tuba8 | 541.234206 | 0.321543116 | 0.129088904 | 2.490865634 | 0.01274323 | 0.069620454 | | | Dlk2 | 1118.542873 | -0.023029665 | 0.081340829 | -0.283125525 | 0.777080616 | 0.947474493 |
| Neil2 | 71.58632919 | 0.320843557 | 0.140827823 | 2.278268241 | 0.022710601 | 0.102593826 | | | Fam154a | 1.466440785 | -0.023040127 | 0.027585891 | -0.83521417 | 0.403597133 | NA |
| Ccdc116 | 98.70811567 | 0.320592707 | 0.139754222 | 2.29397511 | 0.021791928 | 0.099600041 | | | Oraov1 | 562.7162804 | -0.023044867 | 0.067918354 | -0.339302491 | 0.734381866 | 0.936456337 |
| Ocm | 18.19864196 | 0.320262934 | 0.168069141 | 1.905542751 | 0.056709556 | 0.188149434 | | | Mir93 | 0.668067976 | -0.023056579 | 0.01863924 | -1.236991372 | 0.216090286 | NA |
| Nfkbia | 855.7529168 | 0.32023547 | 0.165412556 | 1.935980424 | 0.052870099 | 0.180155995 | | | Plekhh3 | 488.0551148 | -0.023060907 | 0.083149356 | -0.277343182 | 0.781516605 | 0.948894514 |
| A730090H04Rik | 9.780923368 | 0.319257936 | 0.160395594 | 1.990440807 | 0.046542398 | 0.166158507 | | | Dmap1 | 918.1405225 | -0.023061083 | 0.063281602 | -0.364420029 | 0.715544381 | 0.93125489 |
| Mia | 20.75224388 | 0.319169748 | 0.167793882 | 1.902153662 | 0.057151069 | 0.189047536 | | | Pskh1 | 489.7598474 | -0.023094794 | 0.079354776 | -0.291032181 | 0.771026708 | 0.946250219 |
| Fam166b | 17.81977071 | 0.319080167 | 0.16260298 | 1.962326686 | 0.049724462 | 0.173206003 | | | BC002163 | 19.3039701 | -0.02310677 | 0.077237785 | -0.299164064 | 0.764814867 | NA |
| Kctd14 | 98.58017415 | -0.377131029 | 0.155396757 | -2.426891248 | 0.01522882 | 0.078704249 | | | Mettl4 | 306.636589 | -0.023109373 | 0.079155648 | -0.291948499 | 0.770326001 | 0.94619074 |
| Taf10 | 779.6938078 | -0.213746011 | 0.088089529 | -2.426463307 | 0.015246792 | 0.078770856 | | | 6030466F02Rik | 1.590136553 | -0.02311533 | 0.027266752 | -0.847747808 | 0.396578435 | NA |
| Prkd2 | 180.3744169 | 0.319078259 | 0.124966081 | 2.55331892 | 0.010670175 | 0.061609839 | | | Zfp629 | 1611.589342 | -0.023120681 | 0.055564186 | -0.416107617 | 0.677331256 | 0.921064066 |
| Tmem170 | 105.3249497 | 0.319074137 | 0.144737442 | 2.204503087 | 0.027488983 | 0.11665689 | | | Aqp5 | 7.539287504 | -0.023144345 | 0.061584368 | -0.375815249 | 0.70705425 | NA |
| Tdrd7 | 1617.375225 | -0.132017837 | 0.054428192 | -2.42554148 | 0.015285569 | 0.078892273 | | | Gm14326 | 855.107165 | -0.023153005 | 0.074546846 | -0.310583296 | 0.756117428 | 0.942139348 |
| Slc26a2 | 209.8289778 | -0.317340961 | 0.130871528 | -2.424828112 | 0.015315636 | 0.079021136 | | | Scd4 | 17.22780539 | -0.023186887 | 0.077733055 | -0.298288634 | 0.765482878 | NA |
| Ccr1 | 12.79084354 | -0.400636525 | 0.165324645 | -2.423332134 | 0.015378859 | 0.079320919 | | | Pbx2 | 2341.268921 | -0.023193901 | 0.050310902 | -0.461011426 | 0.644790408 | 0.912353932 |
| Scd3 | 191.5562099 | 0.318535404 | 0.143398567 | 2.22132906 | 0.026328682 | 0.113124374 | | | Vipr1 | 1495.980132 | -0.023195946 | 0.074315287 | -0.312128868 | 0.754942588 | 0.942099798 |
| Sult2b1 | 247.4431425 | 0.318202091 | 0.128426785 | 2.477692565 | 0.013223503 | 0.071338773 | | | Xpc | 821.4690657 | -0.023255333 | 0.060359798 | -0.385278508 | 0.700031083 | 0.926781946 |
| LOC100503496 | 19.6772291 | 0.317584767 | 0.167997942 | 1.890408678 | 0.058703324 | 0.192512323 | | | Klrk1 | 1.238867292 | -0.0232704 | 0.025070931 | -0.928182514 | 0.353312903 | NA |
| Sdr42e1 | 339.5243965 | -0.257939479 | 0.1065796 | -2.420158057 | 0.015513762 | 0.079903737 | | | Ltf | 8.409557156 | -0.023280591 | 0.043564612 | -0.534392254 | 0.593070177 | NA |
| Ahrr | 32.7642281 | 0.317457945 | 0.162472501 | 1.953918004 | 0.050710915 | 0.175451801 | | | Ticrr | 5.776853749 | -0.023281139 | 0.043461948 | -0.535667166 | 0.5921886 | NA |
| Ccdc177 | 540.8340145 | -0.207786405 | 0.085921041 | -2.418341336 | 0.015591443 | 0.080108712 | | | Zfp84 | 896.490999 | -0.023286279 | 0.065507935 | -0.355472653 | 0.722235529 | 0.93255988 |
| Pla2g4b | 135.6899402 | 0.3172061 | 0.157652247 | 2.012062029 | 0.044213399 | 0.160808075 | | | Snord49b | 2.042164714 | -0.023298944 | 0.033897859 | -0.687327885 | 0.49187613 | NA |
| Fas | 55.81484384 | 0.317099586 | 0.164029311 | 1.933188549 | 0.053212975 | 0.180886756 | | | Crybb3 | 25.06219879 | -0.023310917 | 0.08245345 | -0.282716093 | 0.777394479 | 0.947506677 |
| Gucy1b3 | 7341.462739 | -0.179666739 | 0.074282354 | -2.418700124 | 0.015576075 | 0.080108712 | | | Palm | 8255.780365 | -0.023317549 | 0.061732497 | -0.377719191 | 0.705639209 | 0.929018955 |
| Col7a1 | 159.1235025 | 0.316953638 | 0.164336542 | 1.928686306 | 0.053769817 | 0.18214026 | | | 4930474G06Rik | 6.373690834 | -0.023324344 | 0.049254482 | -0.473547643 | 0.635822519 | NA |
| 4930520O04Rik | 20.07905756 | 0.316859162 | 0.168069205 | 1.88528983 | 0.059390718 | 0.193522564 | | | Elf2 | 1118.839471 | -0.023335296 | 0.067773278 | -0.344314103 | 0.730610079 | 0.93455812 |
| Vnn1 | 8.709869866 | -0.385208634 | 0.159289674 | -2.418290055 | 0.015593641 | 0.080108712 | | | Ift81 | 1359.555421 | -0.023375903 | 0.067397262 | -0.346837567 | 0.728713352 | 0.933996867 |
| Acadm | 1255.945702 | -0.135525956 | 0.056059465 | -2.417539242 | 0.015625849 | 0.080247565 | | | Slc9a1 | 2934.392608 | -0.02338873 | 0.055526801 | -0.421215156 | 0.673597978 | 0.919353963 |
| Insl5 | 10.69493984 | 0.316806094 | 0.156933841 | 2.018723894 | 0.043515925 | 0.158867988 | | | 1700123I01Rik | 1.677439366 | -0.023406181 | 0.028486715 | -0.821652518 | 0.41127469 | NA |
| Tob2 | 1206.785894 | 0.316793369 | 0.122299999 | 2.590297401 | 0.009589305 | 0.057023545 | | | Itprip | 48.97004031 | -0.023415407 | 0.092356323 | -0.25353334 | 0.799856101 | 0.953228631 |
| Stat4 | 83.8514627 | 0.316676075 | 0.126519264 | 2.502987019 | 0.012315006 | 0.067998647 | | | Prss32 | 1.21667189 | -0.02342308 | 0.023059031 | -1.015787668 | 0.30973051 | NA |
| Pex13 | 1161.566649 | -0.162748948 | 0.067339157 | -2.41685457 | 0.015655271 | 0.080313518 | | | Rps29 | 7567.206932 | -0.023437359 | 0.06962409 | -0.33662715 | 0.736397987 | 0.93717515 |
| Ube2v2 | 2400.334127 | -0.240008877 | 0.099322888 | -2.416450837 | 0.015672643 | 0.080354707 | | | Ribc2 | 7.833225282 | -0.023461835 | 0.047366267 | -0.49532792 | 0.620368666 | NA |
| Pkd1l3 | 54.05133579 | 0.316619364 | 0.141904367 | 2.231216503 | 0.025666791 | 0.111084152 | | | AA414768 | 311.4804665 | -0.023473008 | 0.079765548 | -0.294275021 | 0.768547758 | 0.94619074 |
| Gpr98 | 84.16520156 | -0.357341342 | 0.147941467 | -2.415423811 | 0.01571691 | 0.080512887 | | | Gm14092 | 0.933509117 | -0.023473896 | 0.02163769 | -1.084861452 | 0.277983023 | NA |
| Msrb2 | 666.4183593 | -0.177172882 | 0.073352169 | -2.415373453 | 0.015719084 | 0.080512887 | | | Gm4371 | 13.50497348 | -0.023493099 | 0.063305112 | -0.371109027 | 0.710556328 | NA |
| Slc14a2 | 22.251202 | 0.316605857 | 0.16536284 | 1.914613085 | 0.055541865 | 0.185706875 | | | Phkb | 1910.919414 | -0.023495071 | 0.061345625 | -0.382995058 | 0.701723428 | 0.92751465 |
| Mlxipl | 138.2012056 | 0.316318441 | 0.159750117 | 1.980082687 | 0.047694238 | 0.168715646 | | | Hdgfrp2 | 3262.442751 | -0.023498165 | 0.057944484 | -0.405528943 | 0.685088748 | 0.923835238 |
| Impad1 | 3170.301991 | -0.170567844 | 0.070669272 | -2.413606937 | 0.015795492 | 0.080824093 | | | Fanci | 30.295939 | -0.023506841 | 0.084390605 | -0.278548084 | 0.780591659 | 0.948894514 |
| Il17rd | 239.2957254 | -0.254104508 | 0.105306643 | -2.412996006 | 0.015821993 | 0.080932968 | | | Ppp1r9b | 37517.9857 | -0.023514106 | 0.066695542 | -0.352558879 | 0.72441917 | 0.932882346 |
| Plekho2 | 787.8535139 | -0.243894579 | 0.101103919 | -2.412315779 | 0.015851546 | 0.081057377 | | | Smad2 | 1785.587608 | -0.02354627 | 0.055888991 | -0.421304256 | 0.673532923 | 0.919353963 |
| Atp1a2 | 52691.32525 | -0.107353226 | 0.044513556 | -2.411697356 | 0.015878456 | 0.081061217 | | | Urb2 | 728.5705188 | -0.023557513 | 0.064041926 | -0.367845163 | 0.712988691 | 0.930507332 |
| Popdc2 | 14.50784079 | 0.31612689 | 0.167303401 | 1.889542518 | 0.058819171 | 0.192658904 | | | Kcnj3 | 2310.849364 | -0.023558485 | 0.05869799 | -0.4013508 | 0.688161867 | 0.923860706 |
| Irf1 | 293.1465801 | 0.315925057 | 0.15029788 | 2.101992768 | 0.035553909 | 0.138398496 | | | 5830403M04Rik | 2.249928154 | -0.02356379 | 0.034721946 | -0.678642552 | 0.497364374 | NA |
| Dpf1 | 2061.231557 | -0.224251388 | 0.092978076 | -2.411873842 | 0.015870772 | 0.081061217 | | | Apol6 | 6.750461165 | -0.023566638 | 0.047636063 | -0.494722617 | 0.620795935 | NA |
| Slc18a3 | 98.06159218 | -0.401011073 | 0.166262273 | -2.411918631 | 0.015868823 | 0.081061217 | | | Spred3 | 2143.257766 | -0.023571293 | 0.08494446 | -0.277490644 | 0.781403389 | 0.948894514 |
| Sh3gl3 | 1173.60265 | -0.181831846 | 0.075415265 | -2.411074826 | 0.015905585 | 0.081172968 | | | Slc35f3 | 1729.264494 | -0.023588422 | 0.079591557 | -0.296368399 | 0.766948756 | 0.946118815 |
| Rasal3 | 139.0768824 | 0.315412102 | 0.158420353 | 1.990982193 | 0.046482845 | 0.165984144 | | | Slc17a3 | 0.742907396 | -0.023593447 | 0.019065982 | -1.237462984 | 0.215915248 | NA |
| Lhfpl2 | 773.9357478 | -0.20110315 | 0.08342874 | -2.410478112 | 0.015931627 | 0.081184757 | | | Kctd17 | 6277.326907 | -0.023595355 | 0.054469184 | -0.433187227 | 0.664878757 | 0.915428982 |
| Nbl1 | 1679.748589 | -0.212533337 | 0.08817268 | -2.410421646 | 0.015934094 | 0.081184757 | | | Olfr1393 | 19.97134508 | -0.023606616 | 0.075556689 | -0.312435817 | 0.754709334 | NA |
| Psma5 | 2210.527955 | -0.125512509 | 0.052066677 | -2.410611083 | 0.015925821 | 0.081184757 | | | Suz12 | 1576.199654 | -0.023612504 | 0.076390398 | -0.309103032 | 0.757243153 | 0.942139348 |
| Gpr68 | 502.0228375 | 0.315392845 | 0.135486361 | 2.32785678 | 0.01991971 | 0.093909252 | | | Glrx2 | 2182.590388 | -0.023613311 | 0.048404907 | -0.487828873 | 0.625671058 | 0.902263257 |
| Adcy3 | 2639.524277 | -0.143760275 | 0.059655115 | -2.409856628 | 0.01595879 | 0.081256799 | | | Mir674 | 3.033230905 | -0.023623099 | 0.042039704 | -0.561923526 | 0.574168124 | NA |
| Tekt1 | 60.4284102 | 0.315131722 | 0.1380739 | 2.282340993 | 0.022469216 | 0.101913944 | | | Figla | 0.855438572 | -0.023631678 | 0.018269279 | -1.293520042 | 0.195831257 | NA |
| E2f3 | 1301.380637 | -0.151646768 | 0.062930803 | -2.40973831 | 0.015963966 | 0.081256799 | | | Syn1 | 29550.45756 | -0.023633002 | 0.043428409 | -0.544182996 | 0.586315552 | 0.884588669 |
| Lpcat2 | 380.1552825 | -0.352239162 | 0.14619062 | -2.409451182 | 0.015976533 | 0.081294058 | | | Clec1a | 32.50005026 | -0.02363553 | 0.085387602 | -0.276802832 | 0.781931506 | 0.948979619 |
| Slc26a10 | 188.0081752 | 0.315051767 | 0.131204755 | 2.40122218 | 0.016340412 | 0.082603047 | | | Ceacam-ps1 | 1.781110721 | -0.023651896 | 0.028042039 | -0.843444204 | 0.398980058 | NA |
| Rbms3 | 826.5907613 | -0.214626477 | 0.089114251 | -2.408441691 | 0.016020785 | 0.081438992 | | | Sarnp | 1141.920067 | -0.023656395 | 0.077343143 | -0.305862865 | 0.759709061 | 0.94272457 |
| Ccdc89 | 17.93294122 | 0.315019443 | 0.168097351 | 1.874029793 | 0.060926319 | 0.196378694 | | | Zic4 | 88.98067463 | -0.023656584 | 0.087218923 | -0.271232243 | 0.786212415 | 0.950484821 |
| Skil | 2894.546778 | -0.192410958 | 0.079900694 | -2.408126247 | 0.016034635 | 0.081482662 | | | Mfsd7a | 47.46066268 | -0.023686428 | 0.090030017 | -0.263094786 | 0.792477523 | 0.951728598 |
| Slc1a4 | 2773.012119 | -0.133413984 | 0.055407541 | -2.407866908 | 0.016046029 | 0.08151383 | | | Pnmt | 8.644115552 | -0.02370368 | 0.054839798 | -0.432235009 | 0.665570617 | NA |
| Trhr2 | 139.3885761 | 0.314770827 | 0.144331328 | 2.180890531 | 0.029191514 | 0.121258263 | | | Tpk1 | 94.61243359 | -0.023706438 | 0.095102708 | -0.249271954 | 0.803150424 | 0.953754272 |

| | | | | | | |
|---|---|---|---|---|---|---|
| Trim30a | 119.7318879 | 0.314386889 | 0.150608902 | 2.087438956 | 0.036848474 | 0.14174038 |
| Sp5 | 9.472806124 | -0.335855633 | 0.139562085 | -2.40649624 | 0.01610637 | 0.081739959 |
| Lamtor1 | 2249.70844 | -0.14891505 | 0.061909457 | -2.405368379 | 0.016156171 | 0.081931735 |
| Pcdh19 | 1546.796578 | -0.232428509 | 0.096628375 | -2.405385661 | 0.016155407 | 0.081931735 |
| Snord32a | 9.975550076 | 0.313568986 | 0.162746928 | 1.926727523 | 0.054013596 | 0.182650548 |
| Cenpa | 71.02014864 | 0.313388339 | 0.152432699 | 2.055912818 | 0.03979092 | 0.148989976 |
| AU040972 | 8.768139707 | 0.313071648 | 0.149203251 | 2.098289722 | 0.03587956 | 0.139327795 |
| Nts | 202.9411025 | -0.367821888 | 0.15300851 | -2.403930914 | 0.016219838 | 0.082154356 |
| Cdh24 | 337.5611126 | 0.312915997 | 0.129731843 | 2.412021509 | 0.015864346 | 0.081061217 |
| Lonrf2 | 7268.172454 | -0.150200575 | 0.062499791 | -2.403217246 | 0.01625153 | 0.082261091 |
| Lrrc43 | 19.75802498 | 0.312608214 | 0.160751419 | 1.944668453 | 0.051814895 | 0.177969403 |
| Ncmap | 26.711026 | 0.312396094 | 0.166162701 | 1.880061484 | 0.060099699 | 0.194686334 |
| 9530091C08Rik | 31.21152481 | 0.312162873 | 0.163777123 | 1.906022446 | 0.056647294 | 0.188063674 |
| Paqr5 | 186.9900009 | 0.311843371 | 0.153073504 | 2.037213258 | 0.041628676 | 0.154415777 |
| Pfn4 | 202.1715907 | -0.272239119 | 0.113392527 | -2.400855922 | 0.016356775 | 0.082631847 |
| Itgb2 | 293.5952065 | 0.311751747 | 0.124587808 | 2.502265281 | 0.012340142 | 0.068066273 |
| 9530082P21Rik | 77.08171998 | 0.311681198 | 0.149597658 | 2.08346309 | 0.037209033 | 0.142696449 |
| Wdr69 | 11.8005696 | 0.311543471 | 0.156781512 | 1.987118673 | 0.046909247 | 0.16704479 |
| Dmrt2 | 41.99156631 | -0.39500297 | 0.16456982 | -2.400215121 | 0.016385439 | 0.082695764 |
| Cnga2 | 18.59482687 | 0.311509509 | 0.163516642 | 1.905063023 | 0.056771879 | 0.188315885 |
| 1110018G07Rik | 7783.527919 | -0.161827239 | 0.067443496 | -2.39944915 | 0.01641976 | 0.082815028 |
| Ttc12 | 96.85539026 | 0.31045393 | 0.154141115 | 2.014089039 | 0.044000186 | 0.160220654 |
| Map3k13 | 1153.666959 | -0.255266218 | 0.106401922 | -2.399075247 | 0.016436537 | 0.082842953 |
| AB124611 | 10.95766446 | 0.310409476 | 0.164361242 | 1.888580745 | 0.058948029 | 0.192839133 |
| Zfp951 | 104.2960381 | 0.310324764 | 0.143847901 | 2.157311727 | 0.030981384 | 0.126256066 |
| Ldb2 | 1654.582872 | -0.264946849 | 0.110453564 | -2.398717055 | 0.016452622 | 0.08286856 |
| Sh3rf3 | 1447.751029 | -0.168427211 | 0.07022869 | -2.39826786 | 0.016472814 | 0.082920657 |
| Mirlet7b | 14.65435737 | 0.310266133 | 0.166683322 | 1.861410782 | 0.062686189 | 0.200012318 |
| Ebf1 | 134.4177754 | -0.281703872 | 0.117526054 | -2.396948276 | 0.016532257 | 0.083145631 |
| Fig4 | 1480.784664 | -0.159717361 | 0.06663446 | -2.396918386 | 0.016533606 | 0.083145631 |
| 9530036O11Rik | 34.24309716 | 0.310200124 | 0.158806874 | 1.95331673 | 0.050782076 | 0.175550533 |
| Tmem86b | 177.0758991 | 0.310099785 | 0.146020558 | 2.123672101 | 0.033697572 | 0.133702991 |
| Mien1 | 1600.045223 | -0.199650502 | 0.083326339 | -2.396007122 | 0.016574768 | 0.08327155 |
| 3110047P20Rik | 1871.079331 | -0.217675064 | 0.090877389 | -2.395260983 | 0.016608539 | 0.083309297 |
| Lepr | 160.3336933 | 0.310068002 | 0.136388165 | 2.273423085 | 0.023000696 | 0.103506474 |
| Itih2 | 212.6515848 | -0.396906565 | 0.165692602 | -2.395439266 | 0.016600464 | 0.083309297 |
| AW495222 | 106.1211263 | 0.309749742 | 0.1484885 | 2.086018394 | 0.036976957 | 0.142128745 |
| Snord19 | 11.2840475 | 0.309713979 | 0.143919721 | 2.151991245 | 0.031398044 | 0.127590955 |
| Zfp933 | 538.8123092 | -0.245267113 | 0.102401918 | -2.395141779 | 0.01661394 | 0.083309297 |
| Gm12216 | 19.82177914 | 0.309278009 | 0.167357985 | 1.848002704 | 0.064601952 | 0.204190665 |
| 1700007K13Rik | 55.92737795 | 0.30922307 | 0.153340161 | 2.016582401 | 0.043739111 | 0.159532436 |
| Ppm1h | 4265.55814 | -0.153782593 | 0.06422684 | -2.39436649 | 0.016649104 | 0.08337513 |
| Ryr2 | 7766.159326 | -0.236834453 | 0.098913285 | -2.394364443 | 0.016649197 | 0.08337513 |
| Ralgps2 | 1778.011528 | -0.19264432 | 0.080463925 | -2.394170059 | 0.016658023 | 0.083392379 |
| Cript | 1371.747109 | -0.178507991 | 0.074563298 | -2.39404636 | 0.016663642 | 0.083393565 |
| Xkr6 | 25.48750705 | -0.390555182 | 0.163160681 | -2.393684426 | 0.016680093 | 0.083448939 |
| Unc5a | 4732.578681 | 0.308737744 | 0.123915751 | 2.491513327 | 0.012720019 | 0.069549009 |
| Bcl2l12 | 45.417328 | 0.308718979 | 0.156147238 | 1.977101754 | 0.04803013 | 0.169632739 |
| Sub1 | 6694.016904 | -0.15812645 | 0.066085423 | -2.392758381 | 0.016722249 | 0.083578881 |
| Ralgapa1 | 6602.499185 | -0.136234445 | 0.056945068 | -2.39238358 | 0.016739338 | 0.08363731 |
| Adamts5 | 102.8277036 | -0.345467528 | 0.144438198 | -2.391801692 | 0.016765898 | 0.083743014 |
| Kdm4d | 66.92940261 | 0.308450388 | 0.152001941 | 2.029252953 | 0.042432533 | 0.15610783 |
| Calb2 | 983.2274565 | -0.383300186 | 0.160322094 | -2.390813245 | 0.016811102 | 0.08386066 |
| Pkd2l2 | 14.90525563 | 0.308358599 | 0.167305662 | 1.843085254 | 0.065316564 | 0.205426618 |
| Pcdhb17 | 511.158627 | -0.235700822 | 0.098584568 | -2.390849066 | 0.016809462 | 0.08386066 |
| Ctnnbl1 | 1049.036077 | -0.12531657 | 0.052436346 | -2.389879921 | 0.016853882 | 0.084028459 |
| 1700029J07Rik | 142.7684202 | 0.308294051 | 0.157360927 | 1.959152481 | 0.050094932 | 0.174068374 |
| Bmp4 | 122.7559221 | -0.28323722 | 0.118536011 | -2.389461376 | 0.016873098 | 0.084062773 |
| Rassf10 | 38.68127571 | -0.382789235 | 0.160199288 | -2.389456537 | 0.01687332 | 0.084062773 |
| Canx | 12922.55125 | -0.169127917 | 0.070804606 | -2.388656977 | 0.016910083 | 0.084204199 |
| Plcxd1 | 780.7515543 | -0.189170859 | 0.079197287 | -2.388602783 | 0.016912577 | 0.084204199 |
| Ptprv | 350.0405874 | 0.308234059 | 0.157023282 | 1.96298317 | 0.04964813 | 0.173082559 |
| Dchs1 | 799.0475162 | -0.28273371 | 0.118407135 | -2.387809742 | 0.016949114 | 0.084250743 |
| Fbxl17 | 3843.373087 | -0.105197285 | 0.044053897 | -2.387922326 | 0.016943922 | 0.084250743 |
| Wdr76 | 142.1958694 | 0.307757177 | 0.130486929 | 2.358528776 | 0.018347537 | 0.089087039 |
| Cdc45 | 68.87931556 | 0.307432915 | 0.159183391 | 1.931312767 | 0.053444385 | 0.181549961 |
| Rsad2 | 72.32131737 | 0.307005928 | 0.153618987 | 1.998489479 | 0.045663619 | 0.164058455 |
| Fam84b | 382.6851611 | -0.335619369 | 0.140582965 | -2.387340238 | 0.016970777 | 0.084304335 |
| Fam81a | 6024.713289 | -0.186458773 | 0.078108762 | -2.387168461 | 0.016978709 | 0.084316705 |
| Adam22 | 9401.099789 | -0.12601844 | 0.052792857 | -2.387035827 | 0.016984836 | 0.084320105 |
| Nov | 3583.438044 | -0.293309142 | 0.122918387 | -2.386210474 | 0.017023005 | 0.084482526 |
| Tex14 | 14.111638 | 0.306881355 | 0.164940217 | 1.860561121 | 0.06280618 | 0.200353911 |
| Tmem184c | 2671.823804 | -0.129623878 | 0.054338208 | -2.385501543 | 0.017055851 | 0.084591342 |
| Nhs | 237.0490173 | -0.323038881 | 0.135428146 | -2.385315683 | 0.017064471 | 0.084607013 |
| Gm16039 | 1125.569957 | -0.147624871 | 0.061902768 | -2.384786269 | 0.017089046 | 0.084701755 |
| Cdc42ep4 | 1294.033511 | -0.157829581 | 0.066201732 | -2.384070275 | 0.017122332 | 0.084804452 |
| Slc39a4 | 16.2225224 | 0.306690028 | 0.154508269 | 1.984942492 | 0.04715087 | 0.167481793 |
| Tmem50b | 3523.581493 | -0.175715821 | 0.07370171 | -2.384148509 | 0.017118692 | 0.084804452 |
| Gstm6 | 203.9989223 | 0.30629529 | 0.116215053 | 2.635590508 | 0.008399102 | 0.05214083 |
| Plat | 1229.226899 | 0.305953387 | 0.136619033 | 2.239463851 | 0.025125751 | 0.109538093 |
| Cox5b | 5388.752058 | -0.2144712 | 0.090001726 | -2.382967629 | 0.017173704 | 0.084958294 |
| Spag17 | 21.12215067 | 0.305896151 | 0.167464861 | 1.826628879 | 0.067755588 | 0.210315116 |
| Snhg7 | 123.5945182 | 0.3058899 | 0.135453124 | 2.258271279 | 0.023928752 | 0.106273139 |
| Cap2 | 9975.219708 | -0.173346537 | 0.072815455 | -2.380628359 | 0.017283139 | 0.085350894 |
| Rps8 | 7515.485747 | -0.142776088 | 0.05996433 | -2.381016993 | 0.017264915 | 0.085350894 |
| Sec11a | 632.4795626 | -0.163359317 | 0.068619099 | -2.380668345 | 0.017281263 | 0.085350894 |
| Baat1 | 11.83307264 | 0.305663517 | 0.162462501 | 1.881440432 | 0.059912031 | 0.194316101 |
| Supt4a | 1111.186566 | -0.146423761 | 0.061522721 | -2.379994863 | 0.017312879 | 0.085455799 |
| Nptx1 | 17352.25587 | 0.305360794 | 0.133587659 | 2.285845843 | 0.022263279 | 0.101179664 |
| Gcfc2 | 230.4716507 | 0.304663873 | 0.130311634 | 2.337963718 | 0.01938913 | 0.092251522 |
| Sgcd | 65.75656928 | -0.389417905 | 0.163699255 | -2.378861802 | 0.017366185 | 0.08563707 |
| Ube2j2 | 56.51516385 | 0.304627048 | 0.152433849 | 1.998421287 | 0.045671006 | 0.164058455 |
| Fam20b | 4776.139503 | -0.10263425 | 0.043157967 | -2.378106684 | 0.01740179 | 0.08575806 |
| Lmnb1 | 277.6137763 | -0.281157251 | 0.118272576 | -2.377197327 | 0.017444753 | 0.08594245 |
| Gm4532 | 32.62562595 | 0.303912691 | 0.166256364 | 1.827976293 | 0.06755311 | 0.209885442 |
| Mtm1 | 199.0857281 | 0.303855477 | 0.120949852 | 2.512243481 | 0.011996629 | 0.066761405 |
| Atp2a1 | 14.42976958 | 0.303495879 | 0.164479788 | 1.845186466 | 0.065010419 | 0.204938235 |
| Rpl27 | 2818.026145 | -0.129140066 | 0.054403625 | -2.373740087 | 0.017608941 | 0.086684037 |
| Fyb | 160.8124834 | 0.303036249 | 0.114259086 | 2.652185127 | 0.007997268 | 0.050206512 |
| Nsdhl | 937.791978 | -0.268150234 | 0.112993896 | -2.373139103 | 0.01763762 | 0.086727161 |
| Rps11 | 5109.247231 | -0.142522398 | 0.060061821 | -2.372928359 | 0.017647687 | 0.086749129 |
| 1700063D05Rik | 51.31994931 | 0.302110997 | 0.16001509 | 1.888015676 | 0.059023846 | 0.192976465 |
| Slfn8 | 55.12547367 | 0.301916286 | 0.163867331 | 1.842443418 | 0.065410316 | 0.205654918 |
| Mrps16 | 597.0168219 | -0.20335199 | 0.085823464 | -2.36942185 | 0.01781592 | 0.087492825 |
| Cd209c | 67.30200929 | 0.301880452 | 0.14830189 | 2.035580608 | 0.041792487 | 0.15472707 |
| Dock1 | 1545.824465 | -0.17984882 | 0.075923406 | -2.368819184 | 0.017844975 | 0.087510545 |
| P4ha1 | 2028.397069 | -0.246445453 | 0.104026001 | -2.369075517 | 0.017832612 | 0.087510545 |
| Pde4a | 5161.492529 | -0.143069632 | 0.060388922 | -2.369137061 | 0.017829645 | 0.087510545 |
| Srebf2 | 8693.91765 | -0.214956159 | 0.090744975 | -2.368794064 | 0.017846187 | 0.087510545 |
| Wdr52 | 157.8999337 | 0.301827207 | 0.139854776 | 2.158147303 | 0.030916381 | 0.126126654 |
| St8sia4 | 444.7314695 | -0.253243227 | 0.106933595 | -2.368228873 | 0.017873475 | 0.087581159 |
| Tcp11 | 15.6994748 | 0.301670042 | 0.165742421 | 1.820113648 | 0.0687417 | 0.21246433 |
| Zfp459 | 88.20822079 | 0.30127737 | 0.143932948 | 2.093178621 | 0.036333212 | 0.14049084 |
| Cep152 | 152.3413094 | 0.301226692 | 0.117318733 | 2.567592442 | 0.010240748 | 0.059898131 |
| Akirin2 | 1963.988884 | -0.16145039 | 0.068206361 | -2.367086986 | 0.017928718 | 0.087751051 |
| Taok1 | 5420.766461 | -0.179412309 | 0.075827577 | -2.366056216 | 0.017978714 | 0.087957751 |
| Slco5a1 | 220.8090313 | -0.391912741 | 0.165677936 | -2.365509565 | 0.018005278 | 0.088059905 |
| 4930427A07Rik | 71.19622977 | 0.30095723 | 0.154609623 | 1.946562088 | 0.051587258 | 0.177462915 |
| Prr14l | 3069.974686 | -0.179816362 | 0.076029004 | -2.365102157 | 0.018025098 | 0.088077675 |
| Slfn5 | 452.7647933 | -0.265530051 | 0.112260463 | -2.365303372 | 0.018015307 | 0.088077675 |
| Hfm1 | 63.81814348 | 0.300837231 | 0.14954987 | 2.011618143 | 0.044260205 | 0.160892133 |
| Pdia2 | 12.85152103 | 0.300466568 | 0.159839879 | 1.879797261 | 0.060135714 | 0.194734984 |
| Ccdc17 | 80.61054843 | 0.300390948 | 0.135111718 | 2.223278287 | 0.026197041 | 0.112749152 |
| Mir1188 | 15.61899507 | 0.300274796 | 0.160009958 | 1.876600679 | 0.060572844 | 0.195687166 |
| Stxbp3a | 725.6685839 | 0.300120832 | 0.11629909 | 2.58059485 | 0.009863026 | 0.058255183 |
| Fbxl12 | 377.1574979 | 0.300081804 | 0.118054697 | 2.541887879 | 0.011025554 | 0.062813732 |
| Aldh3a1 | 36.85734319 | 0.299921761 | 0.167778961 | 1.787600542 | 0.073840478 | 0.222724827 |
| Chchd2 | 3885.550337 | -0.208896162 | 0.088439196 | -2.362031453 | 0.0181751 | 0.088582888 |
| Trmt12 | 356.3919084 | -0.201327592 | 0.085243686 | -2.361788901 | 0.018186995 | 0.088612988 |
| Cckbr | 2206.5492 | -0.257810182 | 0.10919737 | -2.360955974 | 0.018227894 | 0.088728557 |
| Grin2a | 1830.157059 | -0.287698731 | 0.121853738 | -2.361016868 | 0.018224902 | 0.088728557 |
| Il1b | 13.10329417 | 0.299700174 | 0.164421222 | 1.822758461 | 0.068339982 | 0.211602653 |
| Pde4d | 3059.079742 | -0.158648385 | 0.067202141 | -2.360763859 | 0.018237339 | 0.088746651 |
| Lrrc16b | 1711.862804 | 0.29925681 | 0.125764711 | 2.379497464 | 0.017336262 | 0.08551673 |
| Suv420h2 | 422.1991259 | 0.29922621 | 0.146621651 | 2.040805078 | 0.041270206 | 0.153416643 |
| Tmcc2 | 8279.87536 | -0.166629579 | 0.070613241 | -2.359749766 | 0.018287266 | 0.088905835 |
| 4930523C07Rik | 85.01359722 | -0.316609504 | 0.134191827 | -2.359379936 | 0.018305503 | 0.088966584 |
| C030013G03Rik | 46.06686688 | 0.298595565 | 0.149756531 | 1.993873409 | 0.046165891 | 0.165337144 |
| Gm15446 | 158.1206719 | 0.298393316 | 0.13642733 | 2.187196032 | 0.028728216 | 0.12007418 |
| Gpsm3 | 71.84833671 | 0.298388646 | 0.138090008 | 2.160827206 | 0.03070869 | 0.125640464 |
| Tfr2 | 193.9693938 | 0.298066561 | 0.133848209 | 2.226899877 | 0.025953966 | 0.111982944 |
| Rasgrp1 | 23287.50574 | -0.168732706 | 0.071581867 | -2.357199009 | 0.018413376 | 0.089322744 |
| Tmem178b | 1927.931031 | -0.196697036 | 0.083445036 | -2.357204765 | 0.01841309 | 0.089322744 |
| BC030307 | 54.72657536 | 0.297645319 | 0.149009445 | 1.997492979 | 0.045771656 | 0.164277518 |
| Clptm1l | 3937.739573 | -0.120814068 | 0.051262127 | -2.356790002 | 0.018433668 | 0.089365222 |
| Bmper | 378.8123935 | -0.193070621 | 0.081956666 | -2.355764689 | 0.018484623 | 0.089556206 |
| Mrpl43 | 1035.364622 | -0.142243595 | 0.060378146 | -2.355878804 | 0.018478946 | 0.089556206 |
| Fibcd1 | 593.5034961 | 0.297614188 | 0.148963591 | 1.997898857 | 0.045727627 | 0.164195561 |
| Sft2d2 | 1819.452844 | -0.158601572 | 0.067343807 | -2.355102567 | 0.018517594 | 0.089659874 |
| Mir3060 | 12.52381966 | 0.29724734 | 0.162496035 | 1.829259035 | 0.067360812 | 0.209496054 |
| Akt3 | 6685.974434 | -0.148908785 | 0.063251394 | -2.354237204 | 0.018560763 | 0.089806328 |
| Sulf2 | 4035.195783 | -0.197211446 | 0.083771907 | -2.354147759 | 0.01856523 | 0.089806328 |
| 9330159M07Rik | 90.4850428 | 0.297178948 | 0.130200108 | 2.282478515 | 0.022461105 | 0.101906985 |
| Dlx1 | 1060.589347 | -0.39213317 | 0.166602584 | -2.353704016 | 0.018587405 | 0.089857488 |
| Xirp2 | 9.439446424 | 0.297108763 | 0.15916505 | 1.86667087 | 0.061947564 | 0.198596225 |
| Smoc2 | 629.4587014 | -0.318369396 | 0.135297912 | -2.353099114 | 0.018617671 | 0.089947675 |
| Lrrcc1 | 640.0532499 | 0.29699788 | 0.112403188 | 2.642254937 | 0.008235604 | 0.051369746 |
| Prss53 | 89.31206515 | 0.296956509 | 0.152362949 | 1.949007353 | 0.051294547 | 0.17667828 |
| C1qa | 1861.750467 | 0.296544238 | 0.123656096 | 2.398136834 | 0.016478708 | 0.082923385 |
| B3gnt2 | 1318.552831 | -0.199267049 | 0.084723758 | -2.35196187 | 0.018674689 | 0.090094901 |
| 1700020D05Rik | 33.30895522 | 0.296543604 | 0.165569081 | 1.791056651 | 0.073284205 | 0.221900619 |
| Filip1 | 747.7268209 | -0.296640352 | 0.126149484 | -2.351498729 | 0.018697953 | 0.090108701 |
| 2410137M14Rik | 12.35145875 | 0.296092344 | 0.157339932 | 1.881863941 | 0.059854491 | 0.194298238 |
| Trrap | 6414.980351 | -0.189725952 | 0.08068667 | -2.351391525 | 0.018703342 | 0.090108701 |
| Lpin3 | 42.51150674 | 0.296036156 | 0.156633875 | 1.889988073 | 0.058759555 | 0.192518619 |
| Foxj1 | 252.5669499 | 0.295971783 | 0.131080366 | 2.257941382 | 0.023949314 | 0.106293519 |
| Mmp9 | 54.49764597 | 0.295826007 | 0.16282162 | 1.816871782 | 0.069236748 | 0.213398484 |
| Ngfrap1 | 5031.795109 | -0.135800006 | 0.057797487 | -2.349583234 | 0.018794442 | 0.090463211 |
| Atp5j | 4039.508351 | -0.161324763 | 0.068669692 | -2.349286254 | 0.01880944 | 0.090479188 |
| Arsg | 1055.389691 | 0.295773048 | 0.134324704 | 2.201925933 | 0.027670541 | 0.117202668 |
| Tia1 | 2159.189722 | 0.295062542 | 0.116923094 | 2.523560847 | 0.011617295 | 0.065248465 |
| Tfap2b | 65.40873001 | -0.371337572 | 0.158161416 | -2.347839195 | 0.018882671 | 0.090762462 |
| Spns2 | 2413.318249 | 0.294936588 | 0.164424085 | 1.793755388 | 0.07285222 | 0.220938012 |
| 1700001C02Rik | 8.842111319 | 0.294694388 | 0.15172822 | 1.94225166 | 0.052106642 | 0.178615113 |
| Dgat2l6 | 24.73938272 | 0.294652895 | 0.166065327 | 1.774319175 | 0.076010371 | 0.226743296 |
| Rnf128 | 471.3599408 | -0.242232611 | 0.10318986 | -2.347445867 | 0.01890262 | 0.090767572 |
| Sycp2 | 53.01852041 | 0.294368965 | 0.157294521 | 1.871450848 | 0.061282617 | 0.197198933 |
| Pdzd4 | 5768.544316 | -0.153401151 | 0.0653716 | -2.346602348 | 0.018945462 | 0.090907708 |
| Ctnna2 | 6187.254253 | -0.144311079 | 0.061509419 | -2.346162293 | 0.018967846 | 0.090958778 |
| Gm13483 | 13.12149439 | 0.294333829 | 0.1668306 | 1.764267638 | 0.07768693 | 0.229752167 |
| 1700010I14Rik | 39.20362389 | 0.294295457 | 0.164228593 | 1.791986719 | 0.073135093 | 0.221622362 |
| Ccm2l | 137.4218963 | 0.294227869 | 0.127210273 | 2.312925381 | 0.020726742 | 0.096374032 |
| Cebpd | 66.40139963 | 0.294210054 | 0.163287342 | 1.801793399 | 0.071577917 | 0.218341833 |
| Opa1 | 7295.160246 | -0.131847867 | 0.056230324 | -2.344782268 | 0.019038194 | 0.09113625 |
| Map3k6 | 198.3903886 | 0.294120017 | 0.159310514 | 1.846205936 | 0.064862309 | 0.204600075 |
| Tcea1 | 3073.037939 | -0.136132654 | 0.058058642 | -2.344744009 | 0.019040147 | 0.09113625 |
| BC037704 | 43.3263424 | 0.294091773 | 0.15758653 | 1.866224059 | 0.062010024 | 0.198714233 |
| Trp53inp1 | 428.6488205 | 0.294006576 | 0.131091579 | 2.242757147 | 0.024912479 | 0.108976271 |
| Gpr111 | 29.67823194 | 0.293877491 | 0.16412221 | 1.790601591 | 0.073357252 | 0.221939958 |
| Gprin1 | 5672.165665 | -0.160646118 | 0.068566543 | -2.342922808 | 0.019133341 | 0.091469296 |
| Hs3st3a1 | 86.32654093 | -0.353433256 | 0.150870872 | -2.342620887 | 0.019148829 | 0.091515103 |
| Fam216b | 34.59430406 | 0.293243179 | 0.167504079 | 1.750662914 | 0.080003991 | 0.234716536 |
| Rdh9 | 23.28569845 | 0.293008104 | 0.167888759 | 1.745251476 | 0.080941104 | 0.236238712 |
| Cep350 | 2563.563985 | -0.199874439 | 0.085372085 | -2.34121538 | 0.019221075 | 0.091773148 |
| 4930581F22Rik | 90.18582821 | 0.292799766 | 0.13355759 | 2.19231094 | 0.028357062 | 0.119232798 |
| Wdr1 | 5961.029606 | -0.108109727 | 0.046184409 | -2.340827349 | 0.019241063 | 0.091814297 |
| Fgf17 | 23.5495849 | -0.392848127 | 0.167851189 | -2.340454841 | 0.019260268 | 0.091850854 |
| Cd44 | 141.6867841 | 0.292620573 | 0.131670513 | 2.22236981 | 0.026258324 | 0.112918809 |
| Tnfrsf23 | 45.52341289 | 0.291890417 | 0.157153621 | 1.857357246 | 0.063260345 | 0.201223594 |
| Adtrp | 8.362910432 | -0.350154788 | 0.149681546 | -2.339331714 | 0.019318273 | 0.092069307 |
| Trim67 | 192.0195953 | -0.345264256 | 0.147606554 | -2.339084866 | 0.019331042 | 0.092101833 |
| Lars2 | 55452.26981 | -0.20975622 | 0.089686313 | -2.338776262 | 0.019347016 | 0.092121443 |
| Stra6 | 286.7295106 | -0.278588165 | 0.119117098 | -2.338775624 | 0.019347049 | 0.092121443 |
| Rhpn1 | 332.6034345 | 0.291871561 | 0.119689548 | 2.438571844 | 0.014745427 | 0.077132225 |
| Cd209a | 27.40794034 | -0.392845576 | 0.168044844 | -2.337742513 | 0.019400609 | 0.092251522 |
| Cisd1 | 2162.321146 | -0.174341958 | 0.074573586 | -2.337851351 | 0.019394961 | 0.092251522 |
| Tpcn2 | 125.4452981 | 0.291424806 | 0.136163378 | 2.140258345 | 0.032333894 | 0.129858531 |
| Chrna1 | 20.39074904 | 0.29117839 | 0.168091697 | 1.732259213 | 0.08322742 | 0.240672947 |
| Hapln3 | 25.33553149 | -0.38670956 | 0.165480156 | -2.336893854 | 0.019444704 | 0.092416026 |
| Arpp21 | 24260.21947 | -0.173545691 | 0.074275486 | -2.336513701 | 0.019464484 | 0.092452392 |
| Exoc3l | 24.56076315 | 0.291049235 | 0.167500976 | 1.737597247 | 0.082281805 | 0.238932488 |
| Tpm2 | 380.6511522 | -0.191802801 | 0.082097233 | -2.336288247 | 0.019476224 | 0.092452392 |
| Tor3a | 313.396796 | 0.291020632 | 0.117275586 | 2.481510779 | 0.013082675 | 0.070737147 |
| Rps6kb2 | 948.0719135 | 0.2910023 | 0.124617222 | 2.335169207 | 0.019534583 | 0.092672636 |
| Gm17296 | 262.7183125 | 0.290662707 | 0.115809251 | 2.509840145 | 0.012078583 | 0.067067815 |
| Ankrd50 | 1673.840755 | -0.199697847 | 0.085521909 | -2.335048985 | 0.019540862 | 0.092674048 |
| Ildr2 | 7442.478408 | -0.160886919 | 0.068918821 | -2.334440982 | 0.019572643 | 0.092796368 |
| Gbp11 | 24.98873814 | 0.290601079 | 0.167831924 | 1.731500612 | 0.083362516 | 0.240838862 |
| Mybl1 | 132.3014691 | 0.28996759 | 0.11490611 | 2.523517601 | 0.011618724 | 0.065248465 |
| Phf7 | 154.8161466 | 0.289422653 | 0.109621966 | 2.640188488 | 0.008285993 | 0.051580222 |
| Rccd1 | 337.5748647 | 0.289164052 | 0.123188115 | 2.347337253 | 0.018908131 | 0.090767572 |
| A2m | 181.9421982 | 0.289126543 | 0.152552968 | 1.895253481 | 0.058058828 | 0.191086137 |
| Aldh5a1 | 4858.897513 | -0.166172626 | 0.071241434 | -2.332527813 | 0.019672942 | 0.093100964 |
| Tmem52 | 16.76190222 | 0.289077849 | 0.161055541 | 1.794895397 | 0.072670367 | 0.220559191 |
| 1700110C19Rik | 15.92851434 | 0.288982788 | 0.167259371 | 1.727752452 | 0.084032615 | 0.241738088 |
| Clca1 | 27.68452846 | -0.269466156 | 0.115609893 | -2.33082263 | 0.019762714 | 0.093440188 |
| Chst9 | 17.42401148 | 0.288968446 | 0.164441413 | 1.757272943 | 0.078871297 | 0.232193023 |
| Spink8 | 48.28079395 | 0.288764373 | 0.135459893 | 2.131733359 | 0.033028773 | 0.131927766 |
| Ptdss1 | 2983.413897 | -0.098019523 | 0.042088286 | -2.328902726 | 0.019864219 | 0.093834212 |
| Ifi35 | 101.166282 | 0.288747467 | 0.146234144 | 1.97455573 | 0.048318587 | 0.170302136 |
| Hps1 | 447.4882487 | 0.288530188 | 0.167850587 | 1.718970379 | 0.085619766 | 0.244578291 |
| Rilp | 49.5669599 | 0.288422388 | 0.16225641 | 1.77757161 | 0.07547424 | 0.225708912 |
| Adamts2 | 248.2203904 | 0.28839671 | 0.114424805 | 2.52040377 | 0.011722028 | 0.065665269 |
| H2afy2 | 649.8774578 | -0.174979958 | 0.0751704 | -2.32777741 | 0.019923926 | 0.093909252 |
| Rit2 | 2399.884391 | -0.174737199 | 0.075068944 | -2.327689575 | 0.019928593 | 0.093909252 |

| | | | | | | |
|---|---|---|---|---|---|---|
| Lig1 | 496.7273337 | -0.023715678 | 0.085245573 | -0.278204216 | 0.780855598 | 0.948894514 |
| Nnt | 1233.908979 | -0.023720962 | 0.093807059 | -0.252869689 | 0.800368913 | 0.953228631 |
| Col4a3bp | 3382.24449 | -0.02372457 | 0.064326801 | -0.368813145 | 0.712267006 | 0.930507332 |
| Ormdl2 | 211.8579914 | -0.023753727 | 0.08508617 | -0.279172603 | 0.780112368 | 0.948894514 |
| Maf1 | 2504.679052 | -0.023764086 | 0.052671733 | -0.451173421 | 0.651864564 | 0.914447303 |
| Impa2 | 146.4191138 | -0.023767472 | 0.09375308 | -0.253511375 | 0.799873072 | 0.953228631 |
| Iqsec2 | 9225.665809 | -0.02377568 | 0.067262241 | -0.353477375 | 0.723730588 | 0.932714927 |
| Lrrc6 | 278.1191389 | -0.023776567 | 0.082554111 | -0.288011906 | 0.773337625 | 0.947121017 |
| Daglb | 1819.046863 | -0.023777545 | 0.065515818 | -0.362928312 | 0.716658437 | 0.93125489 |
| Tgm2 | 717.8311895 | -0.023786877 | 0.073900728 | -0.321876084 | 0.747546572 | 0.9410864 |
| Mir688 | 1.225983948 | -0.023787775 | 0.027077277 | -0.878514306 | 0.379664678 | NA |
| Naaladl1 | 5.169334223 | -0.023805518 | 0.049843905 | -0.477601384 | 0.632933946 | NA |
| Scarna6 | 3.620121666 | -0.023819072 | 0.044347117 | -0.537105298 | 0.591194882 | NA |
| Stard6 | 82.47310712 | -0.023821386 | 0.095042977 | -0.250638041 | 0.802093968 | 0.95333733 |
| Fut1 | 1.713121285 | -0.023827028 | 0.030254947 | -0.787541553 | 0.430964914 | NA |
| Src | 5477.919689 | -0.023837155 | 0.062167505 | -0.383434321 | 0.70139776 | 0.92751465 |
| Serpinb6c | 5.890520457 | -0.023854682 | 0.052990272 | -0.450170969 | 0.652587167 | NA |
| Mir27a | 1.083360502 | -0.023881456 | 0.020298548 | -1.176510558 | 0.239390916 | NA |
| Arhgef19 | 614.8035819 | -0.02389123 | 0.081857198 | -0.291864746 | 0.770390039 | 0.94619074 |
| Gm3435 | 19.50219111 | -0.023923014 | 0.076916108 | -0.311027362 | 0.755779821 | NA |
| Gm13031 | 54.33205761 | -0.02393035 | 0.094512046 | -0.253198938 | 0.800114487 | 0.953228631 |
| Stk33 | 45.1613517 | -0.023957542 | 0.083780924 | -0.285954616 | 0.774912882 | 0.947238918 |
| Gm15663 | 337.4361147 | -0.023964375 | 0.083665397 | -0.286431135 | 0.774547931 | 0.947238918 |
| Arid1a | 5989.866793 | -0.023971783 | 0.056907958 | -0.421237796 | 0.673581447 | 0.919353963 |
| Gm11346 | 7.025365611 | -0.023972494 | 0.057757951 | -0.415050988 | 0.678104578 | NA |
| Mir106b | 0.705747355 | -0.024007173 | 0.016228811 | -1.479293344 | 0.139061931 | NA |
| B020018J22Rik | 1.256833248 | -0.024017592 | 0.027112915 | -0.885835849 | 0.375705989 | NA |
| Mrs2 | 977.1558393 | -0.024023021 | 0.057117597 | -0.420588785 | 0.674055384 | 0.919719003 |
| Gm4598 | 2.894554109 | -0.024030376 | 0.039047483 | -0.615414212 | 0.538281208 | NA |
| G630025P09Rik | 28.24946078 | -0.024044178 | 0.088964005 | -0.270268616 | 0.786953608 | 0.950705931 |
| Coq6 | 672.0053583 | -0.024056025 | 0.064799628 | -0.371237086 | 0.710460953 | 0.930507332 |
| Clcnkb | 1.359549035 | -0.024079298 | 0.025897425 | -0.929795066 | 0.352477201 | NA |
| Commd10 | 842.6268179 | -0.024091905 | 0.067266955 | -0.358153642 | 0.720228337 | 0.93244535 |
| Tmem208 | 605.2096735 | -0.024100187 | 0.069094544 | -0.348800146 | 0.727239354 | 0.9335794 |
| Gm14420 | 1125.63376 | -0.024114022 | 0.057549594 | -0.419012892 | 0.67520671 | 0.919907397 |
| A730085A09Rik | 5.000880027 | -0.02411975 | 0.039628942 | -0.608639758 | 0.542763246 | NA |
| Lonp1 | 2999.289013 | -0.024140904 | 0.062067492 | -0.388946013 | 0.697316083 | 0.9261158 |
| Pus7l | 117.127374 | -0.024149495 | 0.093457247 | -0.258401526 | 0.796097039 | 0.952734447 |
| Mas1 | 635.5610135 | -0.024157218 | 0.076994925 | -0.313750781 | 0.753710329 | 0.94183464 |
| Hist1h4j | 0.723986006 | -0.024157747 | 0.018977821 | -1.272946291 | 0.203037101 | NA |
| Ptcd3 | 2087.717201 | -0.024161144 | 0.050692875 | -0.476618147 | 0.633634059 | 0.906932171 |
| Slain1 | 2183.122532 | -0.024162051 | 0.06517094 | -0.370748841 | 0.710824609 | 0.930507332 |
| Gga2 | 1478.598732 | -0.024164571 | 0.054619581 | -0.442415897 | 0.658188277 | 0.915208181 |
| Samd3 | 110.3620888 | -0.024180159 | 0.095018568 | -0.254478246 | 0.799126108 | 0.953180007 |
| Zfp239 | 2066.376569 | -0.024190434 | 0.055448372 | -0.436269517 | 0.662641196 | 0.915428982 |
| Pde2a | 24756.71641 | -0.02420168 | 0.063814021 | -0.379253338 | 0.704499746 | 0.928382774 |
| Cntd1 | 20.50529426 | -0.024251746 | 0.078966799 | -0.307113198 | 0.758757212 | NA |
| Mycbpap | 99.74884166 | -0.024275537 | 0.094383741 | -0.257200409 | 0.797024068 | 0.952885371 |
| Dync2h1 | 1754.559538 | -0.024281888 | 0.066890894 | -0.363007371 | 0.716599378 | 0.93125489 |
| Onecut2 | 231.862696 | -0.024305085 | 0.09510032 | -0.255573118 | 0.798280478 | 0.952967259 |
| Islr | 521.8362234 | -0.024306679 | 0.093868595 | -0.258943678 | 0.795678697 | 0.952689478 |
| Slc6a8 | 5268.51837 | -0.024308624 | 0.05651403 | -0.430134321 | 0.667097935 | 0.916077199 |
| Tspo2 | 1.715561293 | -0.024327256 | 0.030415486 | -0.799831265 | 0.423808566 | NA |
| Clspn | 124.0718075 | -0.02433906 | 0.091592742 | -0.265731319 | 0.790446141 | 0.95168012 |
| Akap17b | 881.485694 | -0.024339618 | 0.060583334 | -0.401754345 | 0.687864825 | 0.923860706 |
| Mir434 | 5.092605892 | -0.024346155 | 0.047105327 | -0.516845037 | 0.605264338 | NA |
| F2 | 2.526762571 | -0.024369647 | 0.030042378 | -0.811175697 | 0.41726478 | NA |
| Zfp212 | 628.4210275 | -0.024392589 | 0.068902421 | -0.35401643 | 0.72332657 | 0.932617157 |
| Mir540 | 0.891101612 | -0.02439418 | 0.018295807 | -1.333320822 | 0.182426543 | NA |
| Qrfpr | 106.4185809 | -0.024409006 | 0.095085476 | -0.256705933 | 0.797405791 | 0.952897915 |
| Rps25 | 5319.221594 | -0.02440952 | 0.063904807 | -0.381966885 | 0.702485928 | 0.927855161 |
| Cd300lf | 3.804090961 | -0.024410135 | 0.0454745 | -0.536787319 | 0.591414532 | NA |
| Atp11c | 484.9555082 | -0.024435306 | 0.073620267 | -0.331910045 | 0.739957185 | 0.938661957 |
| Tdrd9 | 31.32655883 | -0.024449887 | 0.0884828 | -0.276323612 | 0.782299522 | 0.949184473 |
| Adss | 3455.360806 | -0.024460884 | 0.057353114 | -0.426496178 | 0.669746332 | 0.91780521 |
| Cyp2j12 | 164.6802121 | -0.024483208 | 0.094449199 | -0.259220912 | 0.795464797 | 0.952590407 |
| Gpr111 | 34.30868479 | -0.024494302 | 0.0866143 | -0.282797431 | 0.777332124 | 0.947506677 |
| Ccar1 | 2048.17983 | -0.024499899 | 0.070321797 | -0.348396937 | 0.727542103 | 0.9335794 |
| Chd7 | 577.6250975 | -0.024529694 | 0.090030923 | -0.272458538 | 0.785269467 | 0.95025194 |
| Asah1 | 2196.0434 | -0.024546034 | 0.049288766 | -0.498004626 | 0.618480782 | 0.898495033 |
| Ceacam15 | 1.241349419 | -0.024562624 | 0.026701544 | -0.919895281 | 0.357627485 | NA |
| Cryaa | 3.18609695 | -0.024592055 | 0.037485028 | -0.656050052 | 0.51179192 | NA |
| Zfp397 | 1535.507804 | -0.024592074 | 0.075228264 | -0.326899391 | 0.743743981 | 0.939923296 |
| Igbp1 | 1079.79087 | -0.024604335 | 0.069356773 | -0.354750284 | 0.722776677 | 0.93255988 |
| 1700056E22Rik | 1.746807055 | -0.024661831 | 0.029436414 | -0.837800107 | 0.402142979 | NA |
| Sec61a1 | 2824.68651 | -0.024669105 | 0.054511319 | -0.452550136 | 0.650872712 | 0.914280728 |
| Gm7056 | 1.476431974 | -0.024672343 | 0.029231089 | -0.84404459 | 0.398644489 | NA |
| Slc9a5 | 1923.424505 | -0.024678553 | 0.069262635 | -0.35630398 | 0.721612929 | 0.932513933 |
| Sirt1 | 564.2601079 | -0.024679167 | 0.084190286 | -0.293135565 | 0.769418533 | 0.94619074 |
| Tbl1xr1 | 2818.32923 | -0.024685373 | 0.05527163 | -0.446619225 | 0.655150012 | 0.915208181 |
| Serpinb9b | 1.032804552 | -0.024688809 | 0.020586408 | -1.199277164 | 0.230420191 | NA |
| Pcmtd2 | 2713.829262 | -0.02470164 | 0.068094703 | -0.362754202 | 0.716788506 | 0.93125489 |
| Ubtf | 6101.358076 | -0.024709141 | 0.051808028 | -0.476936531 | 0.633407318 | 0.906932171 |
| Adamts12 | 6.87992501 | -0.024714695 | 0.056122649 | -0.440369355 | 0.659669616 | NA |
| Map1s | 2081.981465 | -0.024716455 | 0.053850067 | -0.458986532 | 0.646243838 | 0.912721244 |
| Ccdc177 | 533.3644897 | -0.024739528 | 0.073792986 | -0.335255815 | 0.737432121 | 0.937206199 |
| Pkhd1 | 2.570534169 | -0.024740405 | 0.037382529 | -0.661817304 | 0.508088315 | NA |
| 4930525G20Rik | 78.35342472 | -0.024741094 | 0.095065226 | -0.260253882 | 0.794667944 | 0.952028582 |
| Snx15 | 1189.626193 | -0.024753567 | 0.058649917 | -0.422056298 | 0.672983926 | 0.919353963 |
| Slc27a6 | 4.94496703 | -0.024791847 | 0.047577267 | -0.521085987 | 0.602306872 | NA |
| Afp | 3.52164106 | -0.024795753 | 0.040887645 | -0.606436322 | 0.544225055 | NA |
| Nod2 | 51.81415763 | -0.024806535 | 0.094830468 | -0.261588239 | 0.793638913 | 0.951728598 |
| Rcbtb2 | 917.4886961 | -0.024807264 | 0.060707828 | -0.408633703 | 0.682808494 | 0.922882553 |
| Cox7b | 3745.964129 | -0.024817737 | 0.066565756 | -0.372830393 | 0.709274681 | 0.930224827 |
| Crxos1 | 5.224075751 | -0.024820368 | 0.046341044 | -0.535602259 | 0.592233468 | NA |
| Olfr110 | 0.960368879 | -0.024831325 | 0.018261248 | -1.359782479 | 0.173898768 | NA |
| Ucn2 | 1.463835896 | -0.024836302 | 0.023775593 | -1.044613358 | 0.296201703 | NA |
| Det1 | 146.1770879 | -0.024843868 | 0.093647527 | -0.265291236 | 0.790785115 | 0.951728598 |
| Mycbp2 | 14611.77692 | -0.024866331 | 0.066416717 | -0.374398676 | 0.708107728 | 0.929959854 |
| Atl3 | 1772.466941 | -0.02486708 | 0.061528041 | -0.404158483 | 0.686096181 | 0.923860706 |
| Tpr | 4880.017398 | -0.024868968 | 0.068472181 | -0.363198135 | 0.716456881 | 0.93125489 |
| Bricd5 | 15.29271671 | -0.024889418 | 0.070498003 | -0.353051389 | 0.724049915 | NA |
| Ppp2r5a | 3411.370689 | -0.024908183 | 0.064635155 | -0.38536588 | 0.699966358 | 0.926781946 |
| A630033H20Rik | 10.2089395 | -0.024919185 | 0.063585565 | -0.391900038 | 0.695132075 | NA |
| Trip10 | 270.2350669 | -0.024952163 | 0.084552335 | -0.295109092 | 0.767910544 | 0.94619074 |
| Tfcp2 | 878.7095978 | -0.024956966 | 0.062414804 | -0.39985652 | 0.689262198 | 0.923860706 |
| Gm960 | 38.4509825 | -0.024970024 | 0.092711397 | -0.269330686 | 0.78767522 | 0.950945001 |
| Rem2 | 419.141539 | -0.024978647 | 0.088749453 | -0.281451275 | 0.778364295 | 0.947976838 |
| Vtn | 2766.102544 | -0.025002945 | 0.081969298 | -0.305028166 | 0.760344701 | 0.943070948 |
| Ctdsp1 | 1224.629237 | -0.025030859 | 0.068879039 | -0.36340314 | 0.716303757 | 0.93125489 |
| Ndufc1 | 1190.510756 | -0.025061512 | 0.07696959 | -0.325602768 | 0.744724918 | 0.940439377 |
| Mir124a-2 | 6.577125017 | -0.025078641 | 0.048094198 | -0.521448369 | 0.602054464 | NA |
| Rps12 | 4592.81765 | -0.025085152 | 0.059938315 | -0.41851613 | 0.675569796 | 0.920229309 |
| Ccdc116 | 127.8115693 | -0.025090909 | 0.093036579 | -0.269688644 | 0.787399798 | 0.950831248 |
| Ints7 | 1506.187113 | -0.02509307 | 0.067086426 | -0.374040947 | 0.708373852 | 0.929959854 |
| Fam92b | 10.41981396 | -0.025152302 | 0.065877164 | -0.381806081 | 0.702605208 | NA |
| Siah1a | 860.190694 | -0.025152549 | 0.068541709 | -0.36696706 | 0.713643589 | 0.930719689 |
| Mllt4 | 4181.841489 | -0.025155128 | 0.047999419 | -0.524071513 | 0.600228801 | 0.889841393 |
| Gk5 | 81.45858571 | -0.02515867 | 0.095102734 | -0.264542023 | 0.791362288 | 0.951728598 |
| Rab44 | 2.78164657 | -0.025163898 | 0.036771772 | -0.684326493 | 0.493769021 | NA |
| Mon2 | 3992.45919 | -0.025175107 | 0.056813888 | -0.443115364 | 0.657682293 | 0.915208181 |
| Stom | 630.8424942 | -0.025190881 | 0.078528309 | -0.320787265 | 0.748371611 | 0.9410864 |
| Alg12 | 458.3531367 | -0.025204281 | 0.071549324 | -0.352264419 | 0.72463997 | 0.932965303 |
| Qk | 9641.975165 | -0.025204604 | 0.066003657 | -0.381866773 | 0.702560187 | 0.927855161 |
| Tpo | 14.35344157 | -0.025214827 | 0.070409332 | -0.358117681 | 0.720255248 | NA |
| Ppef1 | 25.22957242 | -0.025219993 | 0.084525144 | -0.298372668 | 0.765418747 | 0.945287598 |
| Fgd4 | 552.4815572 | -0.025232364 | 0.080951066 | -0.311698969 | 0.755269312 | 0.942139348 |
| Pax6os1 | 5.713588866 | -0.025239873 | 0.044467974 | -0.567596662 | 0.570308874 | NA |
| C130026I21Rik | 1.872426285 | -0.025255463 | 0.032037046 | -0.788320574 | 0.430509217 | NA |
| Arsg | 1426.317249 | -0.02526157 | 0.072932529 | -0.346369034 | 0.729065394 | 0.933996867 |
| Rps20 | 3833.126347 | -0.025263117 | 0.064071438 | -0.394296078 | 0.693362459 | 0.924525709 |
| Tuba1c | 36.94862649 | -0.025291574 | 0.076063298 | -0.332506936 | 0.739506503 | 0.938418054 |
| Atn1 | 8130.227989 | -0.025302039 | 0.054634678 | -0.463113165 | 0.643283254 | 0.911258473 |
| Klra2 | 4.23552351 | -0.025319886 | 0.0449592 | -0.563174751 | 0.573315894 | NA |
| Atp6v1e2 | 1.289684196 | -0.025353047 | 0.026514747 | -0.956186663 | 0.338977937 | NA |
| Zfp69 | 39.55926206 | -0.025363258 | 0.089460288 | -0.283514158 | 0.77678273 | 0.947463204 |
| Nufip2 | 1988.547595 | -0.025366828 | 0.066590541 | -0.380937406 | 0.703249696 | 0.927913987 |
| Ufsp1 | 292.1435533 | -0.025367899 | 0.085582246 | -0.296415435 | 0.766912839 | 0.946118815 |
| Tspan10 | 1.515838434 | -0.025374891 | 0.027554596 | -0.920895056 | 0.35710522 | NA |
| Wdr59 | 1504.457292 | -0.025383851 | 0.055387379 | -0.458296661 | 0.646739323 | 0.912750424 |
| Gm4981 | 1.121076854 | -0.025389165 | 0.024703314 | -1.027763526 | 0.304061077 | NA |
| Fbxo43 | 5.554942104 | -0.02539893 | 0.049169914 | -0.516554285 | 0.605467335 | NA |
| Endov | 1893.808257 | -0.025415663 | 0.059219599 | -0.429176555 | 0.667794743 | 0.916514214 |
| Hrg | 0.783267985 | -0.025422237 | 0.019288651 | -1.317989374 | 0.187507205 | NA |
| Pdlim5 | 1264.795765 | -0.025426282 | 0.05901741 | -0.430826808 | 0.666594305 | 0.916017403 |
| Osgin1 | 24.6887979 | -0.0254682 | 0.077925584 | -0.326827197 | 0.743798588 | 0.939923296 |
| Pigl | 272.1940361 | -0.025532692 | 0.076728361 | -0.332767336 | 0.739309917 | 0.938390909 |
| Sergef | 794.0302588 | -0.02553614 | 0.077425008 | -0.32981773 | 0.741537689 | 0.939436288 |
| Fam57b | 2522.977672 | -0.02554381 | 0.059861459 | -0.426715457 | 0.669586591 | 0.91770119 |
| Wdr76 | 165.572237 | -0.02557847 | 0.09223465 | -0.277319532 | 0.781534763 | 0.948894514 |
| Dnm3os | 1.421425898 | -0.025601519 | 0.029429333 | -0.869932001 | 0.384337566 | NA |
| Ptgir | 4.940020943 | -0.025605644 | 0.044522066 | -0.575122544 | 0.565208422 | NA |
| Pcdhga10 | 409.5123834 | -0.025620467 | 0.075473643 | -0.339462445 | 0.734261384 | 0.936456337 |
| Zfp740 | 3031.988753 | -0.025620708 | 0.047388961 | -0.540647182 | 0.58875079 | 0.885188411 |
| Slco4c1 | 66.13525866 | -0.025622794 | 0.09495121 | -0.269852209 | 0.787273955 | 0.950831248 |
| Mad2l1 | 266.1245084 | -0.025633104 | 0.08196837 | -0.31271946 | 0.75449381 | 0.94196623 |
| Zfp759 | 307.1316882 | -0.025643025 | 0.07746331 | -0.331034462 | 0.740618455 | 0.938975678 |
| Vezt | 2002.630666 | -0.02571008 | 0.056240233 | -0.457147465 | 0.647565056 | 0.912900815 |
| Rxrg | 867.438878 | -0.025711139 | 0.094060115 | -0.273347948 | 0.784585761 | 0.949699053 |
| Rbbp5 | 2570.970094 | -0.025755715 | 0.047345763 | -0.543991964 | 0.586447003 | 0.884593012 |
| Dct | 17.66044213 | -0.025755822 | 0.073279375 | -0.351474374 | 0.725232494 | NA |
| Lmo4 | 5907.638661 | -0.025761066 | 0.055872719 | -0.461066974 | 0.644750555 | 0.912353932 |
| Sprr1a | 2.480314828 | -0.025781688 | 0.03875306 | -0.665281341 | 0.505870563 | NA |
| D1Ertd622e | 2417.002069 | -0.025792079 | 0.073085346 | -0.352903559 | 0.724160742 | 0.932714927 |
| Cxcr6 | 1.574834332 | -0.025796434 | 0.025973894 | -0.993167774 | 0.3206282 | NA |
| Epyc | 3.0075103 | -0.02581491 | 0.039554563 | -0.652640502 | 0.513988068 | NA |
| Gtl3 | 1283.418068 | -0.025820429 | 0.057242002 | -0.451074879 | 0.651935582 | 0.914447303 |
| 0610039K10Rik | 1.256297036 | -0.025821506 | 0.022720445 | -1.136487687 | 0.255752516 | NA |
| Lmod3 | 1.750447079 | -0.025834907 | 0.029212861 | -0.884367588 | 0.376497814 | NA |
| P2rx4 | 577.7130213 | -0.025847294 | 0.070085188 | -0.368798239 | 0.712278118 | 0.930507332 |
| Nagpa | 424.1400848 | -0.025857334 | 0.080219676 | -0.322331573 | 0.747201517 | 0.94097069 |
| Itpkb | 1573.385111 | -0.025870752 | 0.068714442 | -0.376496572 | 0.706547763 | 0.929476902 |
| Slc44a4 | 21.47195614 | -0.025881738 | 0.081306213 | -0.318324231 | 0.750239006 | NA |
| 4933412E12Rik | 165.2499761 | -0.025887046 | 0.090848347 | -0.2849479 | 0.775684058 | 0.947241001 |
| Izumo1 | 2.447151125 | -0.025892301 | 0.038037973 | -0.680696144 | 0.496063777 | NA |
| Mtx1 | 915.3971421 | -0.025893587 | 0.055500015 | -0.46655099 | 0.640821158 | 0.910551516 |
| Oit3 | 7.775550235 | -0.02590373 | 0.041046909 | -0.631076255 | 0.527990666 | NA |
| Rabepk | 786.637916 | -0.02591679 | 0.066348544 | -0.390615804 | 0.696081242 | 0.925527709 |
| Ush1c | 6.059260386 | -0.02593677 | 0.05309171 | -0.488527686 | 0.62517612 | NA |
| Stra6 | 256.2085967 | -0.025945946 | 0.094866748 | -0.273498846 | 0.784469779 | 0.949699053 |
| Orc2 | 1314.643951 | -0.02596203 | 0.064933427 | -0.399825339 | 0.689285167 | 0.923860706 |
| Zfp819 | 5.326630336 | -0.025975504 | 0.048945907 | -0.530698192 | 0.595627939 | NA |
| Angptl6 | 264.4997176 | -0.026016232 | 0.084839089 | -0.306653838 | 0.75910687 | 0.942526199 |
| Gm21057 | 0.972280655 | -0.026018801 | 0.022277312 | -1.167950663 | 0.242826667 | NA |
| 4930583K01Rik | 1.05463012 | -0.026022832 | 0.021202773 | -1.227331551 | 0.219698002 | NA |
| Sohlh1 | 2.479969315 | -0.02604927 | 0.035841049 | -0.726799873 | 0.467348561 | NA |
| Cpd | 5308.549896 | -0.026049495 | 0.06495091 | -0.401064354 | 0.688372744 | 0.923860706 |
| Bpifb5 | 1.734144193 | -0.026084342 | 0.029466745 | -0.885212868 | 0.376041833 | NA |
| Mier1 | 1144.850262 | -0.026098609 | 0.069031373 | -0.378068804 | 0.705379482 | 0.928783753 |
| Ccdc83 | 0.780074698 | -0.026100223 | 0.01928952 | -1.353077895 | 0.176030747 | NA |
| Zfp273 | 222.3728049 | -0.026123845 | 0.08694213 | -0.30047394 | 0.763815671 | 0.944786344 |
| Fasl | 3.826065304 | -0.026127057 | 0.042344053 | -0.617018317 | 0.537222644 | NA |
| Commd8 | 1324.560879 | -0.026129044 | 0.057191974 | -0.456865576 | 0.647767668 | 0.912960943 |
| Neurl4 | 5589.489115 | -0.026133941 | 0.050914643 | -0.513289296 | 0.60774897 | 0.893916126 |
| Rpl27a | 7672.724678 | -0.026140781 | 0.062002918 | -0.421605653 | 0.67331288 | 0.919353963 |
| Il18r1 | 1.299042347 | -0.026146046 | 0.026839244 | -0.97417224 | 0.329971026 | NA |
| Pced1a | 856.4742029 | -0.026166531 | 0.066962439 | -0.390764313 | 0.695971456 | 0.925527709 |
| Atp1a4 | 4.351165372 | -0.026211824 | 0.046914804 | -0.558711153 | 0.576358867 | NA |
| 5830415F09Rik | 108.5340196 | -0.02622064 | 0.094345356 | -0.277921896 | 0.781072315 | 0.948894514 |
| Tmc3 | 14.66022472 | -0.026232522 | 0.061705137 | -0.425127035 | 0.670744071 | NA |
| Cpne4 | 2682.254625 | -0.026234179 | 0.077901297 | -0.336761772 | 0.736296493 | 0.93717515 |
| Fam86 | 163.9800653 | -0.026235511 | 0.08968582 | -0.292526852 | 0.769883832 | 0.94619074 |
| Mical3 | 4731.10071 | -0.026238927 | 0.060014173 | -0.437212177 | 0.66195748 | 0.915428982 |
| Rxfp3 | 184.5066157 | -0.026241679 | 0.092846435 | -0.282635286 | 0.777456429 | 0.947506677 |
| AU021063 | 1.630926438 | -0.026276109 | 0.026821462 | -0.979667298 | 0.327250374 | NA |
| 2410076I21Rik | 4.783323044 | -0.026290546 | 0.047257053 | -0.556330618 | 0.57798486 | NA |
| Fam184a | 1109.383772 | -0.026302864 | 0.055583111 | -0.473216838 | 0.636058486 | 0.907889736 |
| Rrm2 | 61.21839361 | -0.02631686 | 0.093586687 | -0.281203032 | 0.778554679 | 0.948049733 |
| Crhr2 | 99.91973813 | -0.026325323 | 0.093893071 | -0.280375566 | 0.779189381 | 0.948407834 |
| Klf16 | 2097.18372 | -0.026336779 | 0.071989883 | -0.365840003 | 0.714484468 | 0.931210733 |
| Ccdc63 | 10.85506877 | -0.026368126 | 0.063618665 | -0.414471536 | 0.678528808 | NA |
| 1700051A21Rik | 2.664904091 | -0.026370916 | 0.038222373 | -0.689934032 | 0.490235673 | NA |
| Npc1 | 3765.555935 | -0.026386581 | 0.066135454 | -0.398977846 | 0.689909528 | 0.923860706 |
| Bglap2 | 1.208833268 | -0.026389973 | 0.02339475 | -1.128029695 | 0.259307378 | NA |
| Slit1 | 2499.936468 | -0.026398219 | 0.083590054 | -0.315805743 | 0.752149959 | 0.941549114 |
| Ehbp1l1 | 968.1241115 | -0.026436609 | 0.071602931 | -0.369211275 | 0.711970252 | 0.930507332 |
| Mir707 | 10.40869857 | -0.026473413 | 0.058198219 | -0.454883557 | 0.649193013 | NA |
| Pts | 679.1509887 | -0.026501865 | 0.07178836 | -0.369166602 | 0.712003548 | 0.930507332 |
| Ooep | 5.386247955 | -0.026529884 | 0.044540032 | -0.595641333 | 0.551414858 | NA |
| Gm4787 | 17.8510807 | -0.026534139 | 0.077252907 | -0.343471076 | 0.731244097 | NA |
| Ipo4 | 3777.456341 | -0.026536924 | 0.057957083 | -0.457871979 | 0.647044419 | 0.912750424 |
| Fgfr1 | 2846.932721 | -0.026557975 | 0.066570546 | -0.39894483 | 0.689933856 | 0.923860706 |
| Gm8615 | 2.940016543 | -0.026596031 | 0.039251126 | -0.677586456 | 0.498033936 | NA |
| Robo4 | 385.2742879 | -0.026602019 | 0.08504654 | -0.312793666 | 0.754437428 | 0.94196623 |
| 5730409E04Rik | 4323.740723 | -0.026612069 | 0.038805775 | -0.685775994 | 0.492854377 | 0.842781259 |
| Casc3 | 2902.936162 | -0.026615674 | 0.050536501 | -0.526662388 | 0.598428059 | 0.888717024 |
| Coa6 | 323.8840699 | -0.026676602 | 0.084377073 | -0.316159368 | 0.751881546 | 0.941374879 |
| Spata19 | 6.891230852 | -0.026686635 | 0.056393246 | -0.473223947 | 0.636053414 | NA |
| Bean1 | 631.3278556 | -0.026701852 | 0.074859499 | -0.356692904 | 0.721321718 | 0.932513933 |
| S100a5 | 3.864879878 | -0.026709824 | 0.026746738 | -0.998619888 | 0.317978862 | NA |
| Ceacam1 | 129.3305535 | -0.026713865 | 0.093154383 | -0.28676981 | 0.77428858 | 0.947238918 |
| Fam98b | 1720.417333 | -0.026744814 | 0.056821806 | -0.470678698 | 0.637870199 | 0.908932006 |
| Timd4 | 3.421866688 | -0.026759223 | 0.034931234 | -0.766054339 | 0.443643971 | NA |
| 4931408D14Rik | 16.54639845 | -0.026770299 | 0.076011834 | -0.352185932 | 0.724698826 | NA |
| Fbxo30 | 310.8397207 | -0.026802095 | 0.087736903 | -0.305482577 | 0.759998637 | 0.942922806 |

| | | | | | | |
|---|---|---|---|---|---|---|
| Shcbp1l | 51.92287297 | 0.288357389 | 0.162875549 | 1.770415455 | 0.076657957 | 0.227841624 |
| Zfp619 | 206.7300945 | -0.236074103 | 0.101403426 | -2.328068323 | 0.019908476 | 0.093909252 |
| Park7 | 2892.032928 | -0.129919656 | 0.055836587 | -2.326783624 | 0.019976784 | 0.094107722 |
| Fam20c | 1566.98457 | -0.204420323 | 0.087927875 | -2.324863669 | 0.020079251 | 0.094532944 |
| Snora70 | 15.62937499 | 0.28831741 | 0.167873992 | 1.717463235 | 0.085894566 | 0.245092156 |
| Dok1 | 62.94948048 | 0.28780514 | 0.138133455 | 2.083529584 | 0.037202978 | 0.142696449 |
| Ticam2 | 32.74025237 | 0.287790444 | 0.159536005 | 1.803921588 | 0.0712436 | 0.217794507 |
| Stard4 | 836.0553654 | -0.300823348 | 0.129432835 | -2.324165639 | 0.020116618 | 0.094622611 |
| Ccnb2 | 35.20569327 | 0.287771184 | 0.162681435 | 1.76892455 | 0.076906467 | 0.228357688 |
| Tomm40l | 2502.469417 | -0.216782943 | 0.093313704 | -2.323162986 | 0.020170398 | 0.094789249 |
| Ube2n | 6979.926633 | -0.141080851 | 0.060726818 | -2.323205073 | 0.020168138 | 0.094789249 |
| 2410004P03Rik | 82.92383588 | 0.287438338 | 0.144529236 | 1.988790266 | 0.046724358 | 0.166654511 |
| Cyb5 | 1543.927305 | -0.135032844 | 0.058157211 | -2.321858998 | 0.020240529 | 0.09495708 |
| Zfp641 | 510.5076364 | 0.287424347 | 0.116854047 | 2.459686707 | 0.013905835 | 0.07373614 |
| Pigz | 319.8683362 | 0.287220481 | 0.157997659 | 1.817878077 | 0.06908277 | 0.21307269 |
| Hsd17b7 | 897.2663332 | -0.236159908 | 0.101723444 | -2.321587815 | 0.020255141 | 0.09495708 |
| Spsb1 | 725.7245026 | 0.28719305 | 0.134687542 | 2.132291126 | 0.032982921 | 0.131778592 |
| Tmem167 | 2042.608962 | -0.173801376 | 0.074853363 | -2.32189136 | 0.020238786 | 0.09495708 |
| Bub1b | 84.9209946 | 0.287069083 | 0.167284176 | 1.716056409 | 0.086151718 | 0.245554585 |
| Pld4 | 421.1596779 | 0.28663865 | 0.130932695 | 2.189206057 | 0.028581867 | 0.119878408 |
| St6galnac2 | 159.7571515 | 0.286484234 | 0.108094307 | 2.6503175 | 0.008041616 | 0.050423633 |
| Evi2b | 30.92163632 | 0.286383343 | 0.166468837 | 1.720342068 | 0.085370281 | 0.244006785 |
| Nudt5 | 440.0324851 | -0.206018146 | 0.088794644 | -2.320164113 | 0.020332001 | 0.095179354 |
| Mir425 | 20.00114247 | 0.286323532 | 0.167964694 | 1.704664982 | 0.088256924 | 0.249234687 |
| Ndst1 | 4764.696882 | -0.187275845 | 0.080722557 | -2.319993972 | 0.020341203 | 0.09518774 |
| Rgl3 | 141.7725437 | 0.285974135 | 0.117383098 | 2.436246267 | 0.014840577 | 0.077368742 |
| Wdr27 | 39.61471571 | 0.2859686 | 0.16798501 | 1.702345939 | 0.088690538 | 0.250031149 |
| G3bp2 | 10947.83141 | -0.160406708 | 0.069159788 | -2.319363781 | 0.02037532 | 0.095261076 |
| Ndufaf6 | 194.1908238 | -0.241920636 | 0.104313501 | -2.31916898 | 0.020385876 | 0.095281677 |
| Siah1b | 132.2060302 | -0.262408193 | 0.113182272 | -2.318456666 | 0.020424515 | 0.095433487 |
| Idua | 712.229041 | 0.285695428 | 0.13066562 | 2.186462118 | 0.028781813 | 0.120144436 |
| Dnajc15 | 693.4766325 | -0.208030818 | 0.089802526 | -2.316536374 | 0.020529001 | 0.095835515 |
| H2-Oa | 21.22476706 | 0.285674903 | 0.159952825 | 1.785994736 | 0.074100111 | 0.223367142 |
| Ubxn11 | 416.7485753 | 0.285569623 | 0.118000703 | 2.420067135 | 0.015517642 | 0.079903737 |
| Mctp1 | 2187.580258 | -0.264173528 | 0.114086334 | -2.315558037 | 0.020582412 | 0.096026467 |
| Col14a1 | 87.76774834 | -0.31054446 | 0.134125525 | -2.315327074 | 0.020595039 | 0.096056453 |
| Ubr1 | 2645.337175 | -0.156985098 | 0.067809236 | -2.31509906 | 0.020607511 | 0.0960857 |
| Gpr63 | 176.0202225 | -0.279605251 | 0.120782353 | -2.314951186 | 0.020615604 | 0.096094514 |
| Dhrs11 | 171.3207158 | 0.285428243 | 0.115177068 | 2.478169028 | 0.013205857 | 0.071268389 |
| Lca5l | 36.8289009 | 0.285357985 | 0.160468909 | 1.778275844 | 0.075358562 | 0.225627736 |
| Alg9 | 767.8969419 | -0.165194971 | 0.071393437 | -2.313867726 | 0.020674979 | 0.096284377 |
| Acsl4 | 2880.736316 | -0.178455681 | 0.077148044 | -2.313158853 | 0.020713907 | 0.096374032 |
| Mobp | 7684.129417 | 0.28533024 | 0.117361106 | 2.431216363 | 0.015048225 | 0.078109616 |
| Krt77 | 223.2176987 | 0.284855348 | 0.141156874 | 2.018005499 | 0.043590689 | 0.159103446 |
| Rpl10 | 8230.635221 | -0.092596678 | 0.040035419 | -2.312868956 | 0.020729845 | 0.096374032 |
| S100a16 | 1735.708879 | -0.255047108 | 0.110259799 | -2.313146873 | 0.020714565 | 0.096374032 |
| Tpgs1 | 835.5395268 | -0.247129665 | 0.106851267 | -2.312837942 | 0.020731551 | 0.096374032 |
| Vip | 1290.127885 | 0.28483478 | 0.11959138 | 2.381733362 | 0.017231369 | 0.085216353 |
| Gm5111 | 14.32056056 | 0.284464322 | 0.164834671 | 1.725755389 | 0.084391429 | 0.242410233 |
| Procr | 23.99475095 | 0.284006224 | 0.167455729 | 1.696007808 | 0.089884401 | 0.2525352 |
| Fmn2 | 3592.397969 | -0.129413846 | 0.05598414 | -2.311616227 | 0.020798842 | 0.096570947 |
| Mks1 | 187.5287445 | 0.28398343 | 0.126508756 | 2.244772926 | 0.024782714 | 0.108703579 |
| Srsf5 | 7136.155534 | 0.28367845 | 0.106915528 | 2.653295146 | 0.007971014 | 0.050082274 |
| Arpc5l | 2403.559658 | -0.128639042 | 0.055708612 | -2.309141018 | 0.020935757 | 0.097003177 |
| Commd3 | 1411.706556 | -0.112348189 | 0.048646953 | -2.309459913 | 0.020918074 | 0.097003177 |
| Dclk1 | 39858.5784 | -0.167581142 | 0.072562389 | -2.309476638 | 0.020917147 | 0.097003177 |
| Snora81 | 17.20572739 | 0.283557819 | 0.167979443 | 1.688050714 | 0.091401494 | 0.254729227 |
| Ube2c | 11.96565997 | -0.354196721 | 0.15338789 | -2.309157005 | 0.02093487 | 0.097003177 |
| Zkscan16 | 1879.419916 | 0.283218615 | 0.115836908 | 2.444977342 | 0.014486124 | 0.076161523 |
| Cxcl14 | 3211.43624 | -0.196702193 | 0.085239511 | -2.307641026 | 0.021019111 | 0.097310346 |
| Rreb1 | 2192.52444 | 0.283213521 | 0.168117156 | 1.684619981 | 0.092061914 | 0.255766056 |
| Myo15 | 17.91348268 | 0.283144409 | 0.167359873 | 1.691829737 | 0.090678442 | 0.253890779 |
| 1500004A13Rik | 1024.808388 | -0.228858893 | 0.099195208 | -2.307156746 | 0.021046083 | 0.097333324 |
| Ccdc71 | 1118.464219 | -0.123299462 | 0.053444024 | -2.307076682 | 0.021050546 | 0.097333324 |
| Cd180 | 154.7772305 | 0.282967035 | 0.160203859 | 1.766293502 | 0.077346621 | 0.229102277 |
| Ccni | 7855.03282 | -0.145028156 | 0.062868778 | -2.306839103 | 0.021063791 | 0.09736348 |
| Malt1 | 273.8656572 | 0.282831435 | 0.129189224 | 2.189280391 | 0.028576467 | 0.119878408 |
| Glis3 | 114.4998254 | 0.282770506 | 0.129487825 | 2.183761339 | 0.02897979 | 0.120734838 |
| Ccdc40 | 73.14121455 | 0.282765068 | 0.130976776 | 2.158894713 | 0.030858336 | 0.125986577 |
| Btn2a2 | 54.53343545 | 0.282058515 | 0.165466014 | 1.704631109 | 0.088263246 | 0.249234687 |
| Grcc10 | 2318.265014 | -0.159861024 | 0.069348659 | -2.305178317 | 0.021156589 | 0.097646764 |
| 4930538K18Rik | 82.48910951 | 0.281672876 | 0.141974512 | 1.983967916 | 0.047259417 | 0.167751982 |
| Cacng5 | 775.2408815 | -0.38687089 | 0.167952724 | -2.303451124 | 0.021253474 | 0.098006348 |
| Tdrd6 | 41.72990122 | 0.281605759 | 0.160058118 | 1.759396911 | 0.078510116 | 0.231393521 |
| Ccdc146 | 22.32748987 | 0.2815093 | 0.167677426 | 1.678874174 | 0.093176569 | 0.257707561 |
| Trim54 | 21.05048235 | -0.377035161 | 0.163725307 | -2.30285206 | 0.021287168 | 0.098103327 |
| Ces5a | 12.64150111 | 0.281436161 | 0.165158654 | 1.704035203 | 0.088374511 | 0.249457947 |
| Fdxr | 384.3491888 | 0.281316987 | 0.125296716 | 2.24520638 | 0.024754887 | 0.108654552 |
| Ftsjd1 | 254.8538347 | -0.224675702 | 0.097588356 | -2.302279818 | 0.021319397 | 0.098164262 |
| Gabbr1 | 31161.74343 | -0.105320341 | 0.045761985 | -2.301481059 | 0.021364455 | 0.098322779 |
| Wdr16 | 33.77385267 | 0.281232313 | 0.167471163 | 1.679287986 | 0.093095931 | 0.25758027 |
| B9d2 | 413.8175876 | -0.237292133 | 0.103112314 | -2.301297715 | 0.021374809 | 0.098331736 |
| Tnfaip6 | 311.5758205 | 0.281125414 | 0.120688801 | 2.329341351 | 0.019840989 | 0.093753062 |
| D330050I16Rik | 39.31480721 | 0.280837659 | 0.157702999 | 1.780801008 | 0.074944965 | 0.224951043 |
| Hyou1 | 5007.967909 | -0.146743993 | 0.063812629 | -2.29960737 | 0.021470474 | 0.09868393 |
| Gas2l3 | 96.30230857 | 0.280726229 | 0.146295329 | 1.918900837 | 0.054996885 | 0.184561778 |
| Tcf19 | 228.0321951 | 0.280380618 | 0.113842206 | 2.462888127 | 0.013782291 | 0.073357204 |
| Zfp563 | 561.011329 | -0.183045923 | 0.079651452 | -2.298086452 | 0.021556869 | 0.098992926 |
| Gpc1 | 3432.502557 | -0.179242687 | 0.078003263 | -2.297887047 | 0.021568219 | 0.099015698 |
| Exoc8 | 1455.031752 | -0.143907311 | 0.062640976 | -2.297335079 | 0.021599662 | 0.099101322 |
| Tmem81 | 157.2749937 | 0.280364923 | 0.112133319 | 2.500282027 | 0.012409447 | 0.068375525 |
| Cyp2a5 | 36.56313288 | -0.379531379 | 0.165232693 | -2.296950874 | 0.021621572 | 0.099172481 |
| Mirlet7c-2 | 17.75585441 | 0.28033651 | 0.157039523 | 1.785133484 | 0.074239668 | 0.223657285 |
| Sbk1 | 3810.46787 | -0.247954091 | 0.107984491 | -2.296200948 | 0.021664394 | 0.099310097 |
| Ttll6 | 12.52533153 | 0.280051347 | 0.149920671 | 1.867996889 | 0.061762504 | 0.198125931 |
| Mtor | 5602.318909 | -0.161219701 | 0.070225631 | -2.295738735 | 0.021690824 | 0.099372452 |
| Stab2 | 18.85622956 | 0.279942041 | 0.168024782 | 1.666075909 | 0.095698296 | 0.262058119 |
| Depdc7 | 43.41762778 | 0.279870471 | 0.154675101 | 1.809408689 | 0.070387534 | 0.216131 |
| Rps25 | 5419.827679 | -0.166363745 | 0.0724802 | -2.295299205 | 0.021715983 | 0.09939954 |
| Klhl5 | 1692.046546 | -0.144636657 | 0.063021699 | -2.295029463 | 0.021731436 | 0.099440894 |
| Ctla2b | 33.41014683 | 0.279741394 | 0.167808803 | 1.667024549 | 0.095509524 | 0.261638005 |
| Evc2 | 946.38831 | 0.279478095 | 0.114558123 | 2.439618306 | 0.014702788 | 0.077013181 |
| Ccna1 | 28.83746251 | 0.279397809 | 0.167574807 | 1.66730199 | 0.095454371 | 0.261533161 |
| Cyp4b1 | 25.99196954 | 0.279023365 | 0.165253764 | 1.688453911 | 0.091324129 | 0.254676254 |
| Jagn1 | 820.3731136 | -0.142870285 | 0.062288835 | -2.293674072 | 0.021809227 | 0.09964971 |
| Znhit3 | 589.3560631 | -0.151001283 | 0.065847699 | -2.293189966 | 0.021837071 | 0.099747515 |
| Per1 | 4117.479076 | 0.278894717 | 0.16264552 | 1.714739617 | 0.086392977 | 0.245880378 |
| Hebp1 | 727.5126955 | -0.296055439 | 0.129139937 | -2.292516515 | 0.021875856 | 0.099857587 |
| Tbx6 | 28.85650867 | 0.278755713 | 0.168101703 | 1.658256331 | 0.097265739 | 0.264825685 |
| Psmb9 | 60.88580777 | 0.278691384 | 0.16671232 | 1.671690392 | 0.09458539 | 0.260118216 |
| Dusp14 | 2108.113259 | -0.192906287 | 0.084169756 | -2.29187175 | 0.021913045 | 0.099952384 |
| Lad1 | 71.51435038 | 0.27868292 | 0.159457202 | 1.747697292 | 0.080516458 | 0.235510228 |
| Nav1 | 5110.355421 | -0.189417418 | 0.082665683 | -2.291367003 | 0.021942196 | 0.100021297 |
| Gm13749 | 17.29913083 | -0.372685326 | 0.162772536 | -2.289608149 | 0.022044042 | 0.100455997 |
| 2810055G20Rik | 31.86903466 | 0.278631864 | 0.165117767 | 1.687473542 | 0.091512333 | 0.254879039 |
| Spo11 | 11.35404189 | 0.278583727 | 0.159826951 | 1.743033477 | 0.081327767 | 0.237037703 |
| Wif1 | 30.54631762 | -0.384285238 | 0.167964252 | -2.287898964 | 0.022143406 | 0.100819843 |
| Tnfaip8l1 | 271.4987942 | -0.30559385 | 0.133580011 | -2.287721407 | 0.02215375 | 0.100837311 |
| Efna1 | 139.3645602 | 0.278449517 | 0.117849208 | 2.362761034 | 0.018139362 | 0.088436524 |
| Rps15 | 8574.089299 | -0.169463042 | 0.074083593 | -2.287457094 | 0.022169157 | 0.100848185 |
| Wdfy4 | 147.8910775 | -0.255176733 | 0.111566703 | -2.287212279 | 0.022183436 | 0.10088352 |
| Ppfibp2 | 493.6886026 | 0.278439172 | 0.130427023 | 2.134827312 | 0.032775119 | 0.131105179 |
| Cspp1 | 637.7998325 | 0.278385826 | 0.106944651 | 2.603083213 | 0.009238953 | 0.055586285 |
| Phf11c | 10.97839985 | 0.278360647 | 0.162316833 | 1.714921625 | 0.086359597 | 0.245830535 |
| Bmp7 | 363.1587904 | -0.183043304 | 0.080157967 | -2.283532255 | 0.022399035 | 0.101714727 |
| Svop | 8267.993609 | -0.194458064 | 0.085152909 | -2.283633846 | 0.022393059 | 0.101714727 |
| Adra1b | 822.8302902 | -0.280916644 | 0.123032054 | -2.283280118 | 0.022413873 | 0.101752287 |
| Csrp2 | 129.957248 | -0.292018185 | 0.127902764 | -2.28312646 | 0.02242292 | 0.101763541 |
| Murc | 30.43337837 | 0.27791478 | 0.168039049 | 1.653870228 | 0.098153883 | 0.266566481 |
| Rab27a | 236.2654994 | 0.277880511 | 0.135139861 | 2.056243872 | 0.039759015 | 0.148989976 |
| Unc13d | 54.85576989 | 0.277869845 | 0.15806524 | 1.757944037 | 0.078757032 | 0.231944773 |
| Gas5 | 2884.296592 | 0.277857686 | 0.104866538 | 2.649631538 | 0.00805796 | 0.050505675 |
| Slc35g1 | 89.00178292 | -0.293588764 | 0.128665947 | -2.281790722 | 0.022501699 | 0.101971723 |
| Abcb1a | 1676.413515 | -0.143784215 | 0.063036439 | -2.28096984 | 0.022550232 | 0.10216178 |
| Bbip1 | 1246.773524 | -0.176061364 | 0.077212061 | -2.280231366 | 0.02259397 | 0.102330012 |
| 2310042E22Rik | 9.810184355 | 0.277456616 | 0.158856577 | 1.746585631 | 0.080709241 | 0.235811614 |
| Gm16287 | 32.87559487 | 0.277425468 | 0.16808693 | 1.650488046 | 0.098843156 | 0.268056398 |
| Tyk2 | 736.6277638 | 0.27729153 | 0.105960995 | 2.616920778 | 0.008872693 | 0.054017095 |
| Cdh4 | 1447.533733 | -0.243666862 | 0.106917477 | -2.279018054 | 0.022665992 | 0.102536316 |
| Tomm5 | 813.1185908 | -0.191169011 | 0.083886465 | -2.278901739 | 0.022672907 | 0.10253766 |
| 1110006O24Rik | 14.94616826 | 0.277173437 | 0.167377325 | 1.655979612 | 0.097725948 | 0.265876353 |
| Chst8 | 356.3214266 | 0.276724097 | 0.114429396 | 2.418295523 | 0.015593407 | 0.080108712 |
| Pgap1 | 1539.052029 | -0.243968089 | 0.107091067 | -2.278136703 | 0.022718434 | 0.102593826 |
| Cfb | 17.33494544 | 0.276270862 | 0.145503403 | 1.898724405 | 0.057600721 | 0.190021638 |
| Qprt | 14.58791849 | -0.380900307 | 0.167195668 | -2.278170908 | 0.022716397 | 0.102593826 |
| Hunk | 815.9018934 | 0.27625373 | 0.110815616 | 2.492913356 | 0.012669976 | 0.069378457 |
| Ptpn4 | 1678.787902 | -0.199800312 | 0.087739613 | -2.277196174 | 0.022774514 | 0.102787158 |
| Dhtkd1 | 225.5715021 | 0.276196911 | 0.143871795 | 1.919743278 | 0.054890335 | 0.18437962 |
| Creb5 | 82.11932191 | 0.276062391 | 0.15364232 | 1.796786139 | 0.072369578 | 0.220034191 |
| Ppa1 | 2501.598363 | -0.130458082 | 0.057317309 | -2.276067789 | 0.022841953 | 0.103001517 |
| Fem1a | 2681.070713 | -0.152428715 | 0.066984495 | -2.275582046 | 0.022871037 | 0.103100551 |
| Sipa1l1 | 22549.3782 | -0.138845709 | 0.061018239 | -2.275478799 | 0.022877223 | 0.103100551 |
| Eif4g2 | 24694.28079 | -0.135253628 | 0.059448915 | -2.275123583 | 0.022898517 | 0.10316651 |
| Phf21b | 414.1327973 | 0.275519302 | 0.114782946 | 2.400350504 | 0.01637938 | 0.082692118 |
| Sat2 | 210.9892027 | 0.27544826 | 0.112147826 | 2.456117688 | 0.014044715 | 0.074396178 |
| Pdss1 | 185.5036119 | 0.275411798 | 0.125865161 | 2.188149568 | 0.028658709 | 0.119981547 |
| Slc3a1 | 11.91776095 | 0.275348298 | 0.146362706 | 1.881273629 | 0.059934706 | 0.194316101 |
| Mib1 | 2945.310977 | -0.183312291 | 0.080637586 | -2.273285952 | 0.023008954 | 0.103513567 |
| Phpt1 | 1172.757362 | -0.162187285 | 0.071363296 | -2.272698896 | 0.023044331 | 0.103642632 |
| Abat | 15670.01688 | -0.181085071 | 0.079702343 | -2.272016915 | 0.023085489 | 0.103767497 |
| Kpnb1 | 8956.002034 | -0.112280722 | 0.049416879 | -2.272112775 | 0.0230797 | 0.103767497 |
| Wdr86 | 27.86530411 | 0.274875208 | 0.158918415 | 1.729662413 | 0.083690608 | 0.24138166 |
| Alx3 | 33.15203347 | -0.377147353 | 0.166053571 | -2.271239035 | 0.023132512 | 0.103918566 |
| Glb1l | 367.3573928 | 0.274506489 | 0.133348241 | 2.058568511 | 0.039535592 | 0.148618256 |
| Dnaaf1 | 12.73161115 | 0.274401899 | 0.157930571 | 1.737484375 | 0.08230171 | 0.238932488 |
| Alpk2 | 16.92066073 | 0.274301548 | 0.164260664 | 1.669916218 | 0.094935941 | 0.260796651 |
| Cyp3a13 | 18.75590997 | 0.273748167 | 0.157190544 | 1.741505306 | 0.081595044 | 0.237579737 |
| Eef1g | 10085.7516 | -0.106346126 | 0.046862866 | -2.269304774 | 0.023249798 | 0.104288269 |
| Mab21l1 | 35.19397979 | 0.273704729 | 0.167458102 | 1.634466924 | 0.102160846 | 0.27423426 |
| Ahnak | 1713.340244 | -0.300875311 | 0.132615532 | -2.268778831 | 0.023281779 | 0.104316912 |
| Nrxn1 | 11631.82436 | -0.17355436 | 0.076494595 | -2.268844735 | 0.023277769 | 0.104316912 |
| Syngr3 | 6050.950194 | -0.144596391 | 0.063729781 | -2.268898266 | 0.023274513 | 0.104316912 |
| Ppp1r7 | 4112.657302 | -0.125712953 | 0.055420579 | -2.268344269 | 0.023308232 | 0.104405245 |
| E230025N22Rik | 25.68347632 | -0.374904068 | 0.165311349 | -2.267866477 | 0.023337346 | 0.104505445 |
| Ska3 | 23.86533943 | 0.273642705 | 0.167526936 | 1.633425117 | 0.102379618 | 0.27458366 |
| Zmym6 | 1472.148053 | 0.27296641 | 0.122483781 | 2.228592288 | 0.025841045 | 0.111705659 |
| Gsdmd | 107.5053849 | 0.272777818 | 0.130850735 | 2.084648724 | 0.037101198 | 0.142429625 |
| Pdia4 | 1610.538625 | -0.20287468 | 0.08956296 | -2.265162747 | 0.023502694 | 0.105124349 |
| Zwilch | 121.533348 | 0.272662856 | 0.128065239 | 2.129093408 | 0.033246533 | 0.132626618 |
| Parpbp | 17.34880898 | 0.27251942 | 0.167677757 | 1.625256834 | 0.104107845 | 0.277824139 |
| Fbxo25 | 2335.179661 | -0.143639676 | 0.063439304 | -2.264206351 | 0.023561426 | 0.105295857 |
| Otud7b | 2653.255313 | 0.272499779 | 0.124851124 | 2.182597717 | 0.029065448 | 0.120929122 |
| Lig1 | 436.0703522 | 0.272480381 | 0.10414853 | 2.616267181 | 0.008889697 | 0.054099356 |
| Zkscan2 | 757.3201273 | 0.272440915 | 0.143640118 | 1.896690973 | 0.057868736 | 0.190617208 |
| Prdx1 | 4737.77481 | -0.160236217 | 0.070819776 | -2.262591417 | 0.023660888 | 0.105618495 |
| Rgs14 | 2369.57764 | -0.207662135 | 0.091787951 | -2.262411707 | 0.023671978 | 0.105637567 |
| Atp5g2 | 3144.914635 | -0.133653854 | 0.059079112 | -2.262286087 | 0.023679733 | 0.105641748 |
| Trappc2l | 994.8086607 | -0.158576275 | 0.070114682 | -2.261670014 | 0.023717799 | 0.105781109 |
| Clmn | 4242.048565 | 0.272355866 | 0.121893879 | 2.23436868 | 0.025458825 | 0.110461784 |
| Id1 | 333.7416681 | -0.190702254 | 0.084343133 | -2.261028813 | 0.023757473 | 0.10589709 |
| Fbxl7 | 58.89310291 | -0.324580322 | 0.143573168 | -2.260731066 | 0.023775916 | 0.105948816 |
| Fgf16 | 22.28780832 | -0.348704573 | 0.154279957 | -2.260206568 | 0.023808433 | 0.106063217 |
| Golm1 | 1385.983488 | 0.272291981 | 0.128202677 | 2.123918054 | 0.033676996 | 0.133702991 |
| Snord52 | 14.4949575 | 0.272168302 | 0.167144481 | 1.628341536 | 0.103452487 | 0.276556814 |
| Tifa | 245.0144317 | 0.272155357 | 0.125102467 | 2.175459562 | 0.029595695 | 0.122281521 |
| Dlx4 | 8.926601048 | 0.271744376 | 0.144063255 | 1.886285137 | 0.059256541 | 0.193410753 |
| Arhgdig | 1680.47015 | -0.213866749 | 0.094668132 | -2.259120839 | 0.023875869 | 0.106210944 |
| Psma2 | 2660.183007 | -0.134226994 | 0.059421723 | -2.25888761 | 0.023890377 | 0.106244977 |
| Dctn3 | 2242.12878 | -0.136697296 | 0.06052717 | -2.258445188 | 0.023917918 | 0.106273139 |
| Edf1 | 2588.590742 | -0.146276707 | 0.06477479 | -2.258235125 | 0.023931004 | 0.106273139 |
| Fez1 | 7849.285884 | -0.103619154 | 0.045882335 | -2.258367063 | 0.023922784 | 0.106273139 |
| C030023E24Rik | 89.62456163 | 0.271605076 | 0.149268528 | 1.819573618 | 0.068823962 | 0.21246433 |
| Prdm5 | 93.43478864 | 0.27123949 | 0.143565721 | 1.889305391 | 0.05885092 | 0.19269572 |
| C130026L21Rik | 149.7780706 | -0.304425588 | 0.134814901 | -2.258100444 | 0.023939398 | 0.106279952 |
| Gm16119 | 10.93657159 | 0.271059605 | 0.163643924 | 1.656398838 | 0.097641076 | 0.265692103 |
| Kctd8 | 263.4639795 | -0.271206975 | 0.120134991 | -2.257518584 | 0.02397569 | 0.106380107 |
| Foxc2 | 41.5185798 | -0.379023445 | 0.167956397 | -2.256677633 | 0.024028225 | 0.106582687 |
| Ptpn6 | 215.9239118 | 0.271051454 | 0.139294015 | 1.945894479 | 0.051667416 | 0.177620357 |
| Syndig1l | 2159.093536 | -0.264516952 | 0.117266858 | -2.255683799 | 0.024090441 | 0.106774598 |
| 1700017B05Rik | 531.693281 | 0.270928801 | 0.102101613 | 2.653521263 | 0.007965675 | 0.050082274 |
| Mkks | 1273.617601 | -0.155551291 | 0.068981708 | -2.254964316 | 0.024135569 | 0.10693327 |
| Klhl40 | 240.4674714 | 0.270899519 | 0.139912968 | 1.936200221 | 0.052843183 | 0.180143517 |
| Agap1 | 5826.95639 | -0.140217458 | 0.062246391 | -2.252619889 | 0.024283126 | 0.107467225 |
| Hrasls | 1182.565824 | -0.157457203 | 0.069899548 | -2.252621196 | 0.024283044 | 0.107467225 |
| Jmjd4 | 1610.186459 | -0.113463571 | 0.050366794 | -2.252745541 | 0.024275198 | 0.107467225 |
| Homer3 | 320.4432491 | 0.270769391 | 0.138676829 | 1.952520779 | 0.050876406 | 0.175611834 |
| Pcdha12 | 300.6996188 | 0.270678478 | 0.108472907 | 2.495355622 | 0.012583095 | 0.068988845 |
| 10-Mar | 65.14516492 | 0.270613255 | 0.158182043 | 1.71077102 | 0.087123391 | 0.247103197 |
| Col2a1 | 75.45778949 | 0.270525101 | 0.155354482 | 1.741340821 | 0.081623855 | 0.237587828 |
| Gfpt1 | 3102.42671 | -0.12891471 | 0.057275311 | -2.250790223 | 0.024398827 | 0.107805069 |
| Reep4 | 102.4562888 | 0.270508231 | 0.13592741 | 1.990093314 | 0.046580657 | 0.166256785 |
| Cacna1c | 2432.963704 | -0.236236264 | 0.104964957 | -2.250620304 | 0.024409596 | 0.10781137 |
| Smpd4 | 2441.290896 | 0.270340154 | 0.14056236 | 1.923275571 | 0.054445451 | 0.183466078 |
| Rtn2 | 3534.263178 | -0.142948031 | 0.063524293 | -2.250289194 | 0.024430593 | 0.107850027 |
| Mthfd1l | 561.3525795 | 0.270311079 | 0.125661834 | 2.151099264 | 0.031468366 | 0.12763635 |
| Erp44 | 1161.354551 | -0.151181503 | 0.067195873 | -2.249862938 | 0.024457647 | 0.10788722 |
| Snrpc | 608.3236628 | -0.156663078 | 0.069634196 | -2.249800924 | 0.024461585 | 0.10788722 |
| Akr1c18 | 56.15791418 | 0.270253969 | 0.149368467 | 1.80931072 | 0.070402745 | 0.21613238 |
| Gdpd5 | 1410.55536 | -0.174961923 | 0.077802348 | -2.24880002 | 0.024525222 | 0.108104516 |
| AA474331 | 89.25430222 | 0.270157041 | 0.129005756 | 2.094147188 | 0.036246871 | 0.140261991 |
| B130034C11Rik | 46.54259718 | 0.269918802 | 0.156166636 | 1.728402484 | 0.083916091 | 0.24166991 |
| Zc3hav1 | 176.081695 | 0.269888488 | 0.128370375 | 2.102420346 | 0.03551647 | 0.138299119 |
| Cyp2j12 | 118.0317607 | 0.269874905 | 0.16461162 | 1.639464487 | 0.101116561 | 0.272280152 |
| Hck | 98.0211987 | 0.269538044 | 0.139807802 | 1.927918477 | 0.053865267 | 0.182326784 |
| 1700001L05Rik | 227.8205524 | 0.269368012 | 0.154967109 | 1.738226998 | 0.082170824 | 0.238776208 |

| | | | | | | |
|---|---|---|---|---|---|---|
| Zyx | 1847.115968 | -0.026810683 | 0.061808653 | -0.433769083 | 0.664456135 | 0.915428982 |
| Tns3 | 3399.968016 | -0.026817285 | 0.060186162 | -0.445572268 | 0.655906246 | 0.915208181 |
| 4930417O22Rik | 1.812044272 | -0.026825144 | 0.03025923 | -0.886511142 | 0.375342153 | NA |
| Nfx1 | 3823.565079 | -0.026832448 | 0.042429099 | -0.632406742 | 0.527121129 | 0.856196078 |
| Stag1 | 1550.300345 | -0.026835709 | 0.062480232 | -0.4295072 | 0.667554155 | 0.916443782 |
| Zfp518a | 452.3210531 | -0.026849945 | 0.075459231 | -0.355820556 | 0.721974954 | 0.93255988 |
| Stx17 | 1290.193638 | -0.026860097 | 0.060634396 | -0.442984482 | 0.657776959 | 0.915208181 |
| Camk2b | 38256.90206 | -0.026876927 | 0.041251707 | -0.651534891 | 0.514701262 | 0.853195313 |
| Nudt22 | 206.937743 | -0.02688526 | 0.087697618 | -0.306567737 | 0.759172414 | 0.942526199 |
| 1700039E15Rik | 1.849439134 | -0.026887322 | 0.028245025 | -0.951931255 | 0.341131847 | NA |
| Taf4b | 254.5287006 | -0.026896567 | 0.084240194 | -0.319284243 | 0.74951098 | 0.9410864 |
| Lrrc66 | 0.794511502 | -0.02691204 | 0.019758291 | -1.362063164 | 0.173177952 | NA |
| Tlcd2 | 9.851044355 | -0.026920079 | 0.056489164 | -0.476552969 | 0.633680481 | NA |
| Slc6a2 | 2.885109541 | -0.026927288 | 0.040903672 | -0.658309789 | 0.510339089 | NA |
| Map3k12 | 6354.295899 | -0.026927922 | 0.043120761 | -0.624476977 | 0.532314386 | 0.858328942 |
| Tmco5 | 59.86418556 | -0.026949921 | 0.093514335 | -0.288190264 | 0.773201101 | 0.947121017 |
| Gm10389 | 1.539676372 | -0.02695481 | 0.026934337 | -1.000760105 | 0.316942801 | NA |
| Cpsf6 | 2564.17476 | -0.026979238 | 0.065768266 | -0.410216657 | 0.681647023 | 0.922538401 |
| March9 | 662.2350963 | -0.026981206 | 0.068953404 | -0.39129621 | 0.6955783 | 0.925527709 |
| 1700120C14Rik | 16.1201867 | -0.026995396 | 0.064924501 | -0.415796744 | 0.677558742 | NA |
| Exo1 | 23.71558815 | -0.027008553 | 0.085126637 | -0.317274989 | 0.751034954 | 0.9410864 |
| Hilpda | 154.6785017 | -0.027048168 | 0.090308976 | -0.299506974 | 0.764553252 | 0.944872045 |
| Spns1 | 2465.623301 | -0.027056053 | 0.071117792 | -0.380440003 | 0.703618825 | 0.928079892 |
| Ltb4r2 | 2.557732746 | -0.027069257 | 0.033644866 | -0.804558321 | 0.421074602 | NA |
| Snora69 | 2.138970587 | -0.027084877 | 0.029800034 | -0.908887434 | 0.363409547 | NA |
| Gm3264 | 1.285375627 | -0.027095776 | 0.027235352 | -0.994875202 | 0.319796965 | NA |
| Cep68 | 1434.321039 | -0.027096348 | 0.064864284 | -0.417739109 | 0.676137875 | 0.920643418 |
| Dok2 | 24.02917382 | -0.027102098 | 0.076495302 | -0.354297548 | 0.723115905 | 0.932617157 |
| Gm10825 | 6.489731019 | -0.027112427 | 0.055760099 | -0.486233483 | 0.626801634 | NA |
| 2410137F16Rik | 1.144499055 | -0.027113908 | 0.021102536 | -1.284864929 | 0.198839488 | NA |
| Prss33 | 1.34024055 | -0.027120181 | 0.0259534 | -1.044956785 | 0.296042943 | NA |
| Slc6a5 | 13.15472404 | -0.027139727 | 0.020923007 | -1.297123625 | NA | NA |
| Zc3h13 | 2902.623559 | -0.027142092 | 0.048977831 | -0.55417098 | 0.579461837 | 0.882497656 |
| Hoxa1 | 0.807392905 | -0.027153623 | 0.019819402 | -1.370052587 | 0.170670487 | NA |
| Nkx2-2as | 33.44167404 | -0.027173011 | 0.086582711 | -0.313838763 | 0.753643502 | 0.94183464 |
| Ccdc27 | 2.030132062 | -0.027187581 | 0.029342463 | -0.926560983 | 0.35415452 | NA |
| Camp | 4.154289592 | -0.027192605 | 0.033969901 | -0.80049115 | 0.423426289 | NA |
| Pik3c2g | 8.458390493 | -0.027198586 | 0.056202011 | -0.483943295 | 0.628426113 | NA |
| Rps13 | 3524.942007 | -0.02720372 | 0.068566567 | -0.396749046 | 0.691552529 | 0.923860706 |
| Tpgs1 | 863.4208907 | -0.027208722 | 0.074961655 | -0.36296853 | 0.716628393 | 0.93125489 |
| Dmd | 2562.630837 | -0.02723794 | 0.067551214 | -0.403219107 | 0.686787045 | 0.923860706 |
| AI182371 | 11.8628424 | -0.02726412 | 0.068140527 | -0.400116079 | 0.689071022 | NA |
| Tmem116 | 78.15209665 | -0.02727244 | 0.0945119 | -0.288560914 | 0.772917409 | 0.947121017 |
| Zfp3 | 487.1218953 | -0.027273879 | 0.067473101 | -0.404218548 | 0.686052016 | 0.923860706 |
| Mical2 | 15374.50107 | -0.027284137 | 0.076600008 | -0.356189733 | 0.72169848 | 0.932513933 |
| Mlf1ip | 65.49633033 | -0.02728474 | 0.094647784 | -0.288276591 | 0.773135024 | 0.947121017 |
| Cpz | 7.936509421 | -0.027290717 | 0.041951375 | -0.650532124 | 0.515348558 | NA |
| Gm10416 | 11.58903031 | -0.027306787 | 0.068018949 | -0.401458519 | 0.688082572 | NA |
| Morc1 | 1.430629264 | -0.027343931 | 0.02631955 | -1.038920933 | 0.29884151 | NA |
| Sstr3 | 1180.740952 | -0.027352303 | 0.075329148 | -0.363103839 | 0.716527317 | 0.93125489 |
| Glis1 | 21.09973864 | -0.027416674 | 0.081809321 | -0.335128977 | 0.737527795 | NA |
| Cdh17 | 1.969219946 | -0.027445354 | 0.031102299 | -0.882422044 | 0.377548619 | NA |
| Eps8l3 | 5.085628937 | -0.027453278 | 0.044646216 | -0.614907167 | 0.538616029 | NA |
| Tmem150b | 8.431272126 | -0.027454442 | 0.044566862 | -0.616028161 | 0.537875934 | NA |
| Ghrhr | 1.167604397 | -0.027470463 | 0.019037982 | -1.442929321 | NA | NA |
| Smpdl3a | 1034.75466 | -0.027490698 | 0.078890006 | -0.3484687 | 0.727488216 | 0.9335794 |
| Gm715 | 107.3477245 | -0.027501343 | 0.092357471 | -0.297770637 | 0.765878227 | 0.945453142 |
| Slc23a3 | 68.69627719 | -0.027505131 | 0.09457426 | -0.290831042 | 0.771180543 | 0.946250219 |
| Akr1c12 | 1.349922685 | -0.027513518 | 0.023025073 | -1.194937235 | 0.232111554 | NA |
| Pou4f1 | 5.002682619 | -0.02752268 | 0.047630077 | -0.577842452 | 0.563370491 | NA |
| Ugp2 | 3276.739289 | -0.027523677 | 0.065169279 | -0.422341288 | 0.672775925 | 0.919353963 |
| Micall1 | 2803.249276 | -0.0275242 | 0.069996439 | -0.393222866 | 0.694154882 | 0.925412429 |
| Gm10440 | 2.329855727 | -0.027541761 | 0.032875755 | -0.837752973 | 0.402169455 | NA |
| Tagap | 43.57632134 | -0.027607762 | 0.084328604 | -0.327383124 | 0.743378129 | 0.939912916 |
| Txnrd2 | 522.17354 | -0.027612028 | 0.066777101 | -0.413495453 | 0.679243653 | 0.921471438 |
| Foxi1 | 3.10567469 | -0.027634719 | 0.040931961 | -0.675137904 | 0.499588154 | NA |
| Pam16 | 646.0095027 | -0.027637022 | 0.06943382 | -0.398034008 | 0.690605118 | 0.923860706 |
| Ccdc85b | 2000.741522 | -0.02763756 | 0.07139336 | -0.38711667 | 0.698669837 | 0.926409493 |
| Epb4.1 | 1351.238105 | -0.027647169 | 0.062398828 | -0.443071938 | 0.657713702 | 0.915208181 |
| 5730508B09Rik | 91.63837487 | -0.027656846 | 0.094324192 | -0.293210528 | 0.769361237 | 0.94619074 |
| Melk | 46.62873738 | -0.02766169 | 0.092102053 | -0.300337384 | 0.76391982 | 0.944801985 |
| Iqsec3 | 9870.081603 | -0.027674075 | 0.061001924 | -0.453659058 | 0.650074239 | 0.914000665 |
| Ccdc15 | 90.44954804 | -0.027678768 | 0.094399444 | -0.29320901 | 0.769362397 | 0.94619074 |
| Polr1a | 1576.26449 | -0.027681035 | 0.057428702 | -0.482006974 | 0.629800992 | 0.904807492 |
| Il4ra | 364.9141222 | -0.027686245 | 0.079788591 | -0.346995031 | 0.728595052 | 0.933996867 |
| Cep112 | 93.23294653 | -0.02769146 | 0.094902843 | -0.291787464 | 0.770449131 | 0.94619074 |
| Zfp41 | 289.6158766 | -0.027699993 | 0.079498405 | -0.348434581 | 0.727513836 | 0.9335794 |
| Tbc1d10a | 901.1617876 | -0.027700967 | 0.069091546 | -0.400931352 | 0.688470667 | 0.923860706 |
| Gm6936 | 3.216180835 | -0.027706416 | 0.04159925 | -0.666031637 | 0.505390879 | NA |
| Uevld | 697.4532374 | -0.027743215 | 0.071704231 | -0.38691182 | 0.69882149 | 0.926409493 |
| Smim8 | 349.2067987 | -0.027747684 | 0.078163329 | -0.354996186 | 0.722592449 | 0.93255988 |
| Cd69 | 0.841087458 | -0.027748874 | 0.019807242 | -1.400945865 | 0.161230262 | NA |
| Gm10731 | 0.839618383 | -0.027805479 | 0.019968184 | -1.39248916 | 0.163774322 | NA |
| Unc119b | 1267.817895 | -0.027823725 | 0.067220216 | -0.413919009 | 0.678933422 | 0.921359251 |
| Slc39a13 | 917.0454348 | -0.027827813 | 0.071300422 | -0.390289603 | 0.696322411 | 0.92555371 |
| 1700125H20Rik | 3.248894681 | -0.027841794 | 0.040645823 | -0.684985373 | 0.493353151 | NA |
| Nuak2 | 51.31285886 | -0.027850293 | 0.091532681 | -0.304266106 | 0.760925165 | 0.943105713 |
| Gzmb | 1.142123786 | -0.027855043 | 0.021442908 | -1.299032875 | 0.193932647 | NA |
| Cog1 | 2203.287363 | -0.027893606 | 0.056647027 | -0.492410755 | 0.622428997 | 0.900114632 |
| Nup88 | 2331.248676 | -0.027924867 | 0.056439879 | -0.494771906 | 0.620761138 | 0.899412757 |
| Rsbn1l | 916.3228262 | -0.02794933 | 0.085685368 | -0.326185569 | 0.744283959 | 0.940096833 |
| Lipc | 5.837605561 | -0.027953425 | 0.055749759 | -0.501408889 | 0.616083385 | NA |
| Ccdc154 | 0.9994637 | -0.027959629 | 0.022395907 | -1.248425878 | 0.211875137 | NA |
| Rdh10 | 329.5972594 | -0.027969137 | 0.081231121 | -0.344315536 | 0.730609002 | 0.93455812 |
| 4933433C11Rik | 1.021822652 | -0.028015584 | 0.022466199 | -1.247010413 | 0.21239368 | NA |
| Tomm5 | 844.1061028 | -0.028052091 | 0.06893435 | -0.40693922 | 0.684052629 | 0.923556882 |
| Klc4 | 940.4031696 | -0.028063533 | 0.074394032 | -0.377228286 | 0.706003961 | 0.929269077 |
| Diap2 | 1244.134568 | -0.028098665 | 0.080098235 | -0.350802547 | 0.725736486 | 0.933575664 |
| Trabd | 735.0852821 | -0.028126878 | 0.070370285 | -0.399698221 | 0.689378803 | 0.923860706 |
| Fam32a | 3125.426146 | -0.028148752 | 0.048016928 | -0.586225595 | 0.557723924 | 0.872673743 |
| Helt | 4.770978134 | -0.028195482 | 0.045495112 | -0.61974751 | 0.535424031 | NA |
| Gm9899 | 968.1097588 | -0.028209444 | 0.070428655 | -0.400539301 | 0.688759342 | 0.923860706 |
| 4933427I22Rik | 20.81032013 | -0.028209629 | 0.081786286 | -0.344918814 | 0.730155406 | NA |
| Gsx1 | 12.17077174 | -0.02821443 | 0.057792888 | -0.488198988 | 0.625408901 | NA |
| Colq | 48.71367921 | -0.028225655 | 0.087175459 | -0.323779826 | 0.746104731 | 0.940755508 |
| Dmxl2 | 10188.49897 | -0.028226377 | 0.060878104 | -0.463654016 | 0.642895648 | 0.911258473 |
| B4galnt2 | 6.160238756 | -0.02826212 | 0.054084621 | -0.522553725 | 0.601284851 | NA |
| Ercc6l2 | 1018.063026 | -0.028300735 | 0.055209382 | -0.512607354 | 0.608226008 | 0.894011407 |
| Gm17769 | 53.64765458 | -0.028309587 | 0.092793691 | -0.305080948 | 0.760304501 | 0.943070948 |
| Nlrx1 | 398.7624383 | -0.02831335 | 0.085618055 | -0.330693684 | 0.740875874 | 0.939089262 |
| 1700102P08Rik | 26.36959013 | -0.028357403 | 0.07888202 | -0.359491335 | 0.719227559 | 0.932229813 |
| Aqp2 | 0.892364251 | -0.028363364 | 0.023238683 | -1.220523743 | 0.222266395 | NA |
| D630037F22Rik | 568.9202018 | -0.028368093 | 0.075116188 | -0.377656184 | 0.705686021 | 0.929018955 |
| Mefv | 0.980017856 | -0.028374031 | 0.022692948 | -1.250345737 | 0.211173278 | NA |
| Dgkd | 5251.983661 | -0.02839727 | 0.05978064 | -0.475024531 | 0.634769489 | 0.907297112 |
| 5830432E09Rik | 2.993198669 | -0.028421724 | 0.037733532 | -0.753221933 | 0.451316562 | NA |
| P2ry14 | 104.6403911 | -0.028436775 | 0.094988357 | -0.299371167 | 0.764656859 | 0.944872045 |
| Lrch2 | 640.3687954 | -0.028444751 | 0.076224317 | -0.373171614 | 0.709020722 | 0.930183885 |
| Cmklr1 | 180.1897721 | -0.02845098 | 0.08971814 | -0.31711513 | 0.751156245 | 0.9410864 |
| Hoxd1 | 2.579380224 | -0.028472486 | 0.036429227 | -0.781583596 | 0.434459331 | NA |
| Exosc4 | 655.0515078 | -0.028472665 | 0.06514166 | -0.437088422 | 0.662047224 | 0.915428982 |
| Prkcd | 845.6229721 | -0.028475161 | 0.080191431 | -0.355089818 | 0.722522304 | 0.93255988 |
| Elovl7 | 860.0128634 | -0.028479973 | 0.093913815 | -0.303256482 | 0.761694408 | 0.943656543 |
| Ypel2 | 2471.946979 | -0.028499039 | 0.067176449 | -0.424241517 | 0.671389686 | 0.91883668 |
| Efhc1 | 59.87738477 | -0.028508954 | 0.09224588 | -0.30905395 | 0.757280488 | 0.942139348 |
| Fut11 | 773.763657 | -0.028515678 | 0.06267673 | -0.454964351 | 0.649134886 | 0.913724196 |
| Mapkapk5 | 1292.864931 | -0.028520728 | 0.058716913 | -0.485732759 | 0.627156654 | 0.903033871 |
| Cideb | 192.9489446 | -0.028524079 | 0.092505218 | -0.308351024 | 0.757815245 | 0.942225819 |
| Lrp1 | 16774.69883 | -0.02853424 | 0.073423053 | -0.38862781 | 0.697551492 | 0.9261158 |
| Cd3e | 2.054425479 | -0.028547691 | 0.031558639 | -0.904591977 | 0.365681587 | NA |
| Bbs5 | 601.2924875 | -0.028564342 | 0.074837684 | -0.381683942 | 0.702695812 | 0.927855161 |
| Rpl31 | 6542.748419 | -0.028591908 | 0.06314058 | -0.452829362 | 0.650671619 | 0.914175059 |
| Cilp2 | 16.9315897 | -0.028592766 | 0.074416334 | -0.384227018 | 0.700810195 | NA |
| Clmp | 974.1308965 | -0.028595851 | 0.064512733 | -0.443259029 | 0.657578387 | 0.915208181 |
| Cat | 2082.831661 | -0.028608358 | 0.053870016 | -0.531062734 | 0.595375307 | 0.887381665 |
| Kbtbd12 | 11.911247 | -0.028634534 | 0.065950993 | -0.434178974 | 0.66415848 | NA |
| Tatdn1 | 290.4920243 | -0.028649392 | 0.085237307 | -0.336113294 | 0.736785434 | 0.937206199 |
| Ldlrap1 | 159.7715177 | -0.028664089 | 0.09254054 | -0.309746286 | 0.756753901 | 0.942139348 |
| Prpf3 | 926.5945504 | -0.028684171 | 0.067342925 | -0.425941865 | 0.670150208 | 0.918098833 |
| Flcn | 2347.389585 | -0.028707301 | 0.060489351 | -0.474584381 | 0.635083241 | 0.907297112 |
| Khsrp | 4078.584727 | -0.028714447 | 0.052435084 | -0.547618966 | 0.583953563 | 0.884313345 |
| Eif2c3 | 786.2044031 | -0.028720676 | 0.086963815 | -0.330260078 | 0.741203455 | 0.939258605 |
| Mark4 | 2265.909417 | -0.028733039 | 0.061994293 | -0.463478781 | 0.643021221 | 0.911258473 |
| Mll3 | 3391.890313 | -0.028761289 | 0.078631823 | -0.365771608 | 0.714535507 | 0.931210733 |
| Hsf2bp | 14.17981283 | -0.028771157 | 0.073592706 | -0.390951203 | 0.695833305 | NA |
| Alx1 | 4.480073271 | -0.028782494 | 0.047967056 | -0.600047129 | 0.548474827 | NA |
| Pcdhgc5 | 4834.397716 | -0.028804675 | 0.077152035 | -0.373349519 | 0.708888325 | 0.930183885 |
| Myh11 | 371.4032305 | -0.028813362 | 0.089071651 | -0.323485211 | 0.746327806 | 0.940909365 |
| 2810021J22Rik | 519.9824151 | -0.028827452 | 0.068638815 | -0.419987605 | 0.674494509 | 0.919867577 |
| Mdfic | 98.87685759 | -0.028854299 | 0.089268018 | -0.323232213 | 0.746519387 | 0.94097069 |
| Scn3a | 1753.089045 | -0.028866401 | 0.066794619 | -0.432166564 | 0.665620359 | 0.915591677 |
| Mxra7 | 605.976468 | -0.028884744 | 0.078323733 | -0.368786614 | 0.712286783 | 0.930507332 |
| Pilrb2 | 2.780893249 | -0.028960047 | 0.034032944 | -0.850941571 | 0.394801811 | NA |
| Tsix | 6.039149835 | -0.028967369 | 0.053411951 | -0.542338724 | 0.587585184 | NA |
| Ace2 | 71.91481858 | -0.028968677 | 0.094292107 | -0.307222716 | 0.758673856 | 0.942526199 |
| Dnajc19 | 515.7598156 | -0.028969733 | 0.073882331 | -0.392106377 | 0.694979616 | 0.925527709 |
| 2310005G13Rik | 1.636182147 | -0.028999592 | 0.027307214 | -1.0619755 | 0.288246808 | NA |
| Chtf18 | 61.88512155 | -0.029007202 | 0.093994404 | -0.308605627 | 0.75762154 | 0.942139348 |
| Agtr1b | 3.092091181 | -0.02901575 | 0.04158378 | -0.697766043 | 0.485323518 | NA |
| Mir5119 | 4.803802496 | -0.029045965 | 0.044489868 | -0.652866966 | 0.513842047 | NA |
| Pycr1 | 38.34277971 | -0.02909239 | 0.086824662 | -0.335070582 | 0.737571844 | 0.937206199 |
| Akap1 | 2144.653175 | -0.029147633 | 0.057248135 | -0.509145553 | 0.610650204 | 0.894800514 |
| Gigyf2 | 3040.131504 | -0.029149218 | 0.054858342 | -0.531354326 | 0.595173266 | 0.887381665 |
| Cckbr | 2222.692141 | -0.029154912 | 0.074897233 | -0.389265545 | 0.697079721 | 0.925997259 |
| Mir1901 | 6.376823963 | -0.029164636 | 0.055583501 | -0.524699521 | 0.59979209 | NA |
| Calcb | 2.173775682 | -0.029169054 | 0.034427938 | -0.847249518 | 0.396856058 | NA |
| Ap1m1 | 2714.080932 | -0.029174253 | 0.06321826 | -0.461484594 | 0.644450972 | 0.91222908 |
| Gpc2 | 89.06956176 | -0.029185461 | 0.094594061 | -0.308533756 | 0.757676219 | 0.942139348 |
| Slc19a1 | 661.9542411 | -0.029207605 | 0.077546477 | -0.37664644 | 0.706436371 | 0.929476902 |
| Slc39a4 | 22.5590391 | -0.029229536 | 0.052419628 | -0.55760671 | 0.577112974 | 0.881322861 |
| Pqlc3 | 38.60128659 | -0.02925099 | 0.087741472 | -0.333377012 | 0.738849713 | 0.938076299 |
| Nop56 | 3106.47539 | -0.029255431 | 0.060797212 | -0.48119691 | 0.630376557 | 0.905249211 |
| Gm12530 | 5.800533179 | -0.029257797 | 0.054341206 | -0.538409051 | 0.590294679 | NA |
| Olfr920 | 3.984140183 | -0.029292369 | 0.042404647 | -0.690782048 | 0.489702519 | NA |
| 9430021M05Rik | 632.8507329 | -0.029303312 | 0.074798267 | -0.391764591 | 0.69523216 | 0.925527709 |
| Gm11762 | 41.7222032 | -0.029324143 | 0.092349465 | -0.317534515 | 0.750838054 | 0.9410864 |
| Osgepl1 | 482.9409453 | -0.029324206 | 0.073456354 | -0.399205841 | 0.689741539 | 0.923860706 |
| Kdm5a | 2173.544979 | -0.029337441 | 0.076375828 | -0.384119455 | 0.700889913 | 0.92732734 |
| Fbxl2 | 2273.018148 | -0.029348101 | 0.047058278 | -0.623654374 | 0.532854593 | 0.858501089 |
| Slc13a3 | 769.6376132 | -0.029348322 | 0.091852711 | -0.319515029 | 0.749335996 | 0.9410864 |
| 2310034G01Rik | 16.25403758 | -0.029369707 | 0.077291735 | -0.379985091 | 0.703956482 | NA |
| Fam135b | 1024.59786 | -0.029371435 | 0.068417188 | -0.429299061 | 0.6677056 | 0.91647846 |
| Mpst | 796.8083172 | -0.029379606 | 0.066141488 | -0.444193305 | 0.656902832 | 0.915208181 |
| Fnbp1 | 5979.701557 | -0.029386217 | 0.064326964 | -0.456825804 | 0.647796257 | 0.912960943 |
| Fbxo6 | 651.997247 | -0.029389066 | 0.071154386 | -0.413032383 | 0.679582889 | 0.921719985 |
| Rpl9 | 9497.936054 | -0.029389265 | 0.05286435 | -0.55593732 | 0.578253705 | 0.881859666 |
| Marveld1 | 456.8062013 | -0.0293992 | 0.080587465 | -0.364811073 | 0.715252439 | 0.93125489 |
| Parl | 802.1282467 | -0.029404955 | 0.064154448 | -0.458346309 | 0.646703659 | 0.912750424 |
| Pebp4 | 3.24573145 | -0.029407541 | 0.037556577 | -0.783019749 | 0.433615515 | NA |
| Akap14 | 9.170735326 | -0.029413574 | 0.059542436 | -0.493993457 | 0.6213108 | NA |
| Gm14461 | 1.417282238 | -0.029418699 | 0.023064209 | -1.275513021 | 0.202127718 | NA |
| Klk10 | 4.156257411 | -0.02943872 | 0.03908897 | -0.75312089 | 0.451377272 | NA |
| AI848285 | 57.16966845 | -0.029454214 | 0.093812766 | -0.313968078 | 0.753545284 | 0.94183464 |
| 1810013D10Rik | 623.1290827 | -0.029472811 | 0.083406979 | -0.353361452 | 0.723817481 | 0.932714927 |
| Fkbp7 | 78.86721178 | -0.029508943 | 0.094005702 | -0.313905887 | 0.753592519 | 0.94183464 |
| Mfn1 | 3182.393317 | -0.029511678 | 0.058060295 | -0.508293623 | 0.611247441 | 0.894995149 |
| Plscr1 | 76.51569174 | -0.02953966 | 0.095101063 | -0.310613355 | 0.756094573 | 0.942139348 |
| Grem2 | 1416.080935 | -0.029576198 | 0.080124145 | -0.369129653 | 0.712031088 | 0.930507332 |
| Snx5 | 3813.032458 | -0.029581069 | 0.056073018 | -0.527545522 | 0.597814813 | 0.888465109 |
| Nudt15 | 248.2148902 | -0.029582365 | 0.080829427 | -0.36598509 | 0.714376201 | 0.931210733 |
| Arih2 | 2199.04863 | -0.029630148 | 0.052041449 | -0.569356708 | 0.56911409 | 0.87745335 |
| Tmem132d | 1668.172943 | -0.029633074 | 0.081354756 | -0.364245135 | 0.715674966 | 0.93125489 |
| Fbxl13 | 13.50234993 | -0.02965555 | 0.070068172 | -0.42323853 | 0.672121238 | NA |
| 4930412C18Rik | 98.14085797 | -0.029663405 | 0.095029814 | -0.312148403 | 0.754927743 | 0.942099798 |
| Bambi-ps1 | 40.52470621 | -0.029713868 | 0.091973102 | -0.323071288 | 0.746641254 | 0.94097069 |
| St3gal6 | 506.0283028 | -0.029736553 | 0.076049911 | -0.39101365 | 0.695787146 | 0.925527709 |
| Ccl5 | 4.50238755 | -0.029765652 | 0.039131668 | -0.760653818 | 0.446863866 | NA |
| Ly6g5b | 8.822887049 | -0.02978482 | 0.056658867 | -0.525686836 | 0.599105811 | NA |
| Ascl2 | 5.693798609 | -0.029788551 | 0.05004428 | -0.595243877 | 0.551680465 | NA |
| Daam1 | 3033.21054 | -0.029800647 | 0.06640363 | -0.448780388 | 0.653590088 | 0.914683958 |
| Ints3 | 3454.674953 | -0.0298191 | 0.049592639 | -0.601280765 | 0.547652999 | 0.867553753 |
| Mpz | 5.789201278 | -0.029824851 | 0.049534407 | -0.602103728 | 0.547105093 | NA |
| 1810058I24Rik | 481.4888708 | -0.029827649 | 0.078580776 | -0.379579462 | 0.704257609 | 0.928232105 |
| Vwce | 14.49241638 | -0.02984556 | 0.06599447 | -0.452243348 | 0.651093683 | NA |
| Evi5 | 3050.968603 | -0.029859904 | 0.059848783 | -0.49892249 | 0.617833989 | 0.898138096 |
| Bbs7 | 801.190529 | -0.029885563 | 0.063540742 | -0.470337013 | 0.638114257 | 0.908932006 |
| Igsf11 | 1595.12903 | -0.02988702 | 0.070344672 | -0.424865432 | 0.670934776 | 0.918480685 |
| A830019L24Rik | 13.43938145 | -0.029897066 | 0.072020135 | -0.415120945 | 0.678053367 | NA |
| Crct1 | 0.732979872 | -0.029923963 | 0.01809103 | -1.654077343 | 0.098111799 | NA |
| Gm12360 | 3.066196279 | -0.029927609 | 0.036524543 | -0.819383543 | 0.412567621 | NA |
| Zdhhc12 | 168.0903733 | -0.02994939 | 0.090708553 | -0.330171625 | 0.741270285 | 0.939261354 |
| Madcam1 | 1.746014756 | -0.029949742 | 0.031262082 | -0.958021329 | 0.338052003 | NA |
| AF357355 | 5.026700563 | -0.029952315 | 0.040518826 | -0.739219721 | 0.459773588 | NA |
| Glb1l3 | 0.67401178 | -0.029963192 | 0.018102565 | -1.655190412 | 0.097885879 | NA |
| Atm | 1041.058208 | -0.029965932 | 0.066673282 | -0.449444377 | 0.653111125 | 0.914683958 |
| Zfp809 | 430.9711014 | -0.029968685 | 0.080330307 | -0.373068219 | 0.709097671 | 0.930183885 |
| Ror2 | 35.4003654 | -0.030025216 | 0.083644284 | -0.358963153 | 0.719622654 | 0.932251852 |
| Mtap7d3 | 2.698113209 | -0.030027285 | 0.035627552 | -0.842810787 | 0.399334274 | NA |
| Myl4 | 639.5123345 | -0.030035868 | 0.094141683 | -0.319049619 | 0.749688887 | 0.9410864 |
| Pou3f3 | 2217.521314 | -0.030046099 | 0.081193121 | -0.370057196 | 0.711339874 | 0.930507332 |
| Asb14 | 3.707845727 | -0.030058225 | 0.041684632 | -0.721086498 | 0.470856298 | NA |
| Gpank1 | 555.0422649 | -0.030063032 | 0.064480129 | -0.466237149 | 0.641045761 | 0.910692492 |
| Ankrd11 | 8942.456333 | -0.030071638 | 0.056338368 | -0.533768355 | 0.59350181 | 0.886313552 |
| Ccdc129 | 67.64882905 | -0.03008614 | 0.094941838 | -0.316890222 | 0.751326902 | 0.9410864 |
| Mitf | 334.7660166 | -0.030120816 | 0.077876267 | -0.386777861 | 0.698920669 | 0.926409493 |

| | | | | | | | | | | | | | |
|---|---|---|---|---|---|---|---|---|---|---|---|---|---|
| Atp10d | 182.1289409 | 0.268496422 | 0.101761134 | 2.638496747 | 0.00832745 | 0.051796672 | Ceacam10 | 4.579360345 | -0.030140182 | 0.045871221 | -0.657060815 | 0.511141813 | NA |
| Eps8l2 | 193.1425897 | 0.268429574 | 0.118381137 | 2.26750292 | 0.023359521 | 0.10457452 | Nae1 | 1497.610285 | -0.030161491 | 0.06649485 | -0.453591391 | 0.650122951 | 0.914000665 |
| Gpr22 | 1745.345108 | 0.268318094 | 0.124517116 | 2.154869172 | 0.031172073 | 0.126833011 | Gm13157 | 86.8986811 | -0.030175107 | 0.095094572 | -0.317316817 | 0.751003218 | 0.9410864 |
| Slc43a3 | 31.067413 | 0.268033989 | 0.168119864 | 1.594302913 | 0.110868206 | 0.289955312 | Pitpnm1 | 8973.384169 | -0.030178177 | 0.065209116 | -0.46279077 | 0.643514348 | 0.911258473 |
| Zfp709 | 233.359498 | -0.260119626 | 0.115805475 | -2.246177279 | 0.024692655 | 0.10850482 | Cysltr1 | 21.7105504 | -0.03018675 | 0.084545484 | -0.357047459 | 0.721056277 | NA |
| Fam134b | 1414.722468 | -0.179680923 | 0.080009181 | -2.245753804 | 0.024719782 | 0.108593223 | Thtpa | 1362.259897 | -0.030208492 | 0.068360548 | -0.441899496 | 0.658561935 | 0.915208181 |
| Mir17hg | 14.55705698 | 0.268030947 | 0.165968934 | 1.614946484 | 0.106322285 | 0.281828499 | A730017L22Rik | 214.4328625 | -0.030214569 | 0.082020624 | -0.368377704 | 0.71259162 | 0.930507332 |
| Elof1 | 1079.697887 | -0.15136324 | 0.067418034 | -2.24514467 | 0.024758847 | 0.108654552 | Asnsd1 | 766.9620794 | -0.030216206 | 0.06972211 | -0.433380545 | 0.664738332 | 0.915428982 |
| Fabp5 | 2531.955827 | -0.165762648 | 0.073833121 | -2.245098761 | 0.024761793 | 0.108654552 | 1700091E21Rik | 2.624174888 | -0.030227738 | 0.037125412 | -0.81420613 | 0.415526875 | NA |
| Ptpn22 | 30.47229796 | 0.267952296 | 0.162387308 | 1.650081517 | 0.098926265 | 0.268200095 | 3110039I08Rik | 30.08179067 | -0.030230698 | 0.083084785 | -0.363853596 | 0.715967338 | 0.93125489 |
| Nefh | 2990.109655 | 0.26784837 | 0.108590642 | 2.466587965 | 0.013640722 | 0.072838195 | Mir3960 | 1.394163851 | -0.030240813 | 0.019725879 | -1.533052788 | NA | NA |
| Tnfrsf12a | 138.216222 | 0.267571498 | 0.159278904 | 1.679892886 | 0.092978158 | 0.257388527 | Nek5 | 24.64126282 | -0.030241532 | 0.067065274 | -0.450926839 | 0.652042279 | 0.914508655 |
| Bend4 | 440.3532014 | -0.205521131 | 0.091568169 | -2.244460429 | 0.024802792 | 0.108716533 | Sowahc | 431.6932025 | -0.030246662 | 0.088766801 | -0.340742956 | 0.733297098 | 0.936456337 |
| Pcyt1b | 2515.354689 | -0.120359949 | 0.053625781 | -2.24444187 | 0.024803985 | 0.108716533 | Gm11837 | 27.84336635 | -0.030249387 | 0.083725887 | -0.361290735 | 0.717882119 | 0.93125489 |
| Pcdhb7 | 279.2284691 | -0.264177614 | 0.117728546 | -2.243955468 | 0.024835267 | 0.108822857 | Gimap4 | 12.90805057 | -0.03025841 | 0.065414477 | -0.462564417 | 0.643676619 | NA |
| Azi2 | 2281.498188 | -0.117453903 | 0.052353627 | -2.243472127 | 0.024866385 | 0.108893068 | Ercc5 | 1071.955071 | -0.030260413 | 0.059600043 | -0.507724686 | 0.611646434 | 0.895166661 |
| Gfod2 | 709.9540246 | -0.143782371 | 0.064091872 | -2.243379166 | 0.024872374 | 0.108893068 | Omp | 19.31650379 | -0.030318836 | 0.053830926 | -0.563223372 | 0.57328279 | NA |
| Palm3 | 93.21106216 | -0.309117475 | 0.137777732 | -2.243595323 | 0.02485845 | 0.108893068 | Rbm46 | 11.52924913 | -0.030328381 | 0.067323356 | -0.450488247 | 0.652358427 | NA |
| 6720401G13Rik | 2989.036801 | 0.267554415 | 0.138516188 | 1.931575074 | 0.053411975 | 0.181483583 | Bhmt2 | 14.77101316 | -0.030329747 | 0.0731551 | -0.414595118 | 0.678438322 | NA |
| Nnt | 1298.625809 | -0.206297799 | 0.091975259 | -2.242970566 | 0.024898713 | 0.10894681 | Fam216b | 48.56273082 | -0.030333896 | 0.088593066 | -0.342395822 | 0.732053032 | 0.935580006 |
| Ccdc113 | 93.21470943 | 0.267401669 | 0.124953793 | 2.140004421 | 0.032354409 | 0.129903918 | Gls | 15759.56626 | -0.030346187 | 0.067351717 | -0.450562934 | 0.652304586 | 0.914611624 |
| Ppp2ca | 13062.08922 | -0.132143154 | 0.05892439 | -2.242588397 | 0.024923369 | 0.108993136 | Ndst3 | 1080.902064 | -0.030348185 | 0.07323926 | -0.414370452 | 0.678602825 | 0.921316909 |
| Accsl | 12.88073023 | 0.267324761 | 0.156179386 | 1.71165202 | 0.086960816 | 0.24695214 | Ski | 7297.235713 | -0.030349895 | 0.059180471 | -0.51283633 | 0.608065814 | 0.894004456 |
| Heatr5b | 2264.040468 | -0.168058219 | 0.074952316 | -2.2422018 | 0.024948333 | 0.109040753 | Mir496 | 7.118165946 | -0.030439159 | 0.052038485 | -0.584935544 | 0.558591058 | NA |
| Ulk4 | 221.0536387 | 0.267230666 | 0.124836327 | 2.140648262 | 0.032302414 | 0.129817618 | Syvn1 | 2446.585617 | -0.030444084 | 0.059798947 | -0.50910736 | 0.610676973 | 0.894800514 |
| Mapre3 | 6850.543587 | 0.096960811 | 0.043252492 | 2.241739314 | 0.024978225 | 0.109109846 | Ubl4b | 0.697055124 | -0.030456219 | 0.018248768 | -1.668946583 | 0.095127967 | NA |
| 1700028K03Rik | 13.99096087 | 0.267176986 | 0.167584107 | 1.594285941 | 0.110872005 | 0.289955312 | Ccdc34 | 267.3767626 | -0.030456886 | 0.083865961 | -0.363161479 | 0.716484262 | 0.93125489 |
| Sult1a1 | 346.3899469 | 0.266950437 | 0.162594498 | 1.641817155 | 0.100627903 | 0.271294367 | Gm16516 | 31.98343668 | -0.030457646 | 0.089776747 | -0.339259853 | 0.734413984 | 0.936456337 |
| Cxcl12 | 2722.267082 | -0.364751399 | 0.162787794 | -2.240655701 | 0.025048386 | 0.109318646 | Mcoln1 | 2720.745436 | -0.030460053 | 0.059132282 | -0.515117153 | 0.606471157 | 0.893015715 |
| Sdc2 | 1910.222066 | -0.171081886 | 0.076356573 | -2.240565282 | 0.025054248 | 0.109318646 | Chdh | 254.3672679 | -0.030469415 | 0.092786844 | -0.328380763 | 0.742623786 | 0.939912916 |
| Nphp3 | 384.7314286 | 0.266943604 | 0.132048408 | 2.021558675 | 0.043221965 | 0.158130136 | Gm15455 | 2.783679713 | -0.030473302 | 0.035157462 | -0.86676625 | 0.386070098 | NA |
| Tmem68 | 1112.736729 | -0.217179393 | 0.096950545 | -2.240104925 | 0.025084112 | 0.109402721 | Npl | 165.7533467 | -0.030475428 | 0.088833564 | -0.343062085 | 0.731551754 | 0.935424325 |
| Mir1839 | 10.54534105 | 0.266920683 | 0.145460684 | 1.835002252 | 0.066505356 | 0.207916461 | Zfp93 | 450.2142992 | -0.030531094 | 0.070335567 | -0.434077593 | 0.664232096 | 0.915428982 |
| Rasl11a | 98.72205403 | 0.266911162 | 0.163834365 | 1.629152479 | 0.103280744 | 0.276330832 | Tmem214 | 2525.032949 | -0.030534064 | 0.060645448 | -0.503484839 | 0.614623437 | 0.89611764 |
| Mark1 | 4279.98538 | -0.102047444 | 0.045597269 | -2.238016581 | 0.025219974 | 0.109887029 | Ppapdc1b | 683.0630844 | -0.030558768 | 0.072804726 | -0.419736056 | 0.674678282 | 0.919907397 |
| Arr3 | 45.77845771 | 0.266524337 | 0.154872671 | 1.720925547 | 0.085264335 | 0.243893179 | Ces1f | 3.035197234 | -0.03057228 | 0.037204739 | -0.821730795 | 0.411230129 | NA |
| Lpxn | 59.93703947 | 0.266518226 | 0.154709572 | 1.7227003 | 0.084942736 | 0.243407204 | 2310034O05Rik | 4.645129841 | -0.030591718 | 0.045643982 | -0.67022457 | 0.502714643 | NA |
| Mutyh | 74.26758702 | 0.266479097 | 0.151050333 | 1.764174172 | 0.07770266 | 0.229752167 | Rps6kb1 | 2011.866647 | -0.030622251 | 0.068994592 | -0.443835524 | 0.657161503 | 0.915208181 |
| Rap2a | 5088.675245 | -0.171266981 | 0.076544248 | -2.237489887 | 0.02525434 | 0.109920099 | Noc3l | 680.2204322 | -0.030624409 | 0.084627836 | -0.361871591 | 0.717447991 | 0.93125489 |
| Dnajc21 | 1006.072871 | 0.266241814 | 0.101769293 | 2.616131118 | 0.00889324 | 0.05409967 | Vash1 | 2692.459352 | -0.030627485 | 0.06427556 | -0.476502817 | 0.633716201 | 0.906932171 |
| Gpc2 | 93.59860485 | -0.310421669 | 0.138833598 | -2.235926125 | 0.025356611 | 0.110296274 | Prdm16 | 575.7782272 | -0.030634072 | 0.086648855 | -0.353542727 | 0.723681602 | 0.932714927 |
| Wfdc3 | 15.57352621 | 0.265996488 | 0.16793169 | 1.583956474 | 0.113203624 | 0.294094057 | Ctnnd1 | 3668.825215 | -0.030648771 | 0.065182676 | -0.470198107 | 0.638213486 | 0.908932006 |
| 1600029O15Rik | 57.85542757 | 0.2657691 | 0.164589469 | 1.614739393 | 0.106367143 | 0.281828499 | Erp27 | 3.529799523 | -0.03067163 | 0.040359058 | -0.759968915 | 0.447273166 | NA |
| Mesdc1 | 1079.453052 | -0.22203143 | 0.099341433 | -2.235033492 | 0.02541515 | 0.110305951 | Dnahc11 | 48.48938375 | -0.030672463 | 0.090845227 | -0.337634285 | 0.735638802 | 0.936783487 |
| Pola1 | 178.6940085 | -0.223570684 | 0.100017216 | -2.235322002 | 0.025396217 | 0.110305951 | Sost | 3.510119443 | -0.030688391 | 0.038166605 | -0.804063934 | 0.421360052 | NA |
| Dnahc1 | 455.383315 | 0.265751721 | 0.112993929 | 2.351911494 | 0.018677218 | 0.090094901 | Fermt2 | 3651.786226 | -0.030690252 | 0.053837712 | -0.570051201 | 0.568642972 | 0.877419107 |
| Ticrr | 10.09579708 | -0.332275649 | 0.148639111 | -2.23545234 | 0.025387667 | 0.110305951 | Sell | 1.346184353 | -0.030696346 | 0.024073703 | -1.275098632 | 0.202274334 | NA |
| Plvap | 322.912716 | 0.265729141 | 0.101424587 | 2.619967684 | 0.00879381 | 0.053683867 | Chd6 | 4016.021013 | -0.030700326 | 0.066651475 | -0.460609858 | 0.645078538 | 0.912363974 |
| Ttll3 | 327.4116906 | 0.265522551 | 0.124815693 | 2.127317039 | 0.03339375 | 0.132974238 | Nprl3 | 1099.887991 | -0.030725748 | 0.071025111 | -0.43260401 | 0.665302476 | 0.915428982 |
| Fam76b | 522.1441007 | 0.265488141 | 0.103730059 | 2.559413766 | 0.010484886 | 0.060843757 | Lrp2bp | 19.68937987 | -0.03074927 | 0.079434605 | -0.387101693 | 0.698680924 | NA |
| Atp4a | 18.86154861 | 0.265457379 | 0.167559246 | 1.584259803 | 0.113134609 | 0.294050242 | Piwil2 | 46.37866932 | -0.030755014 | 0.094047431 | -0.327015997 | 0.743655786 | 0.939912916 |
| Snord53 | 8.176687852 | 0.265309303 | 0.157476388 | 1.684756085 | 0.092035641 | 0.255766056 | B3galt4 | 153.3745816 | -0.030756091 | 0.088631543 | -0.347010664 | 0.728583307 | 0.933996867 |
| Mycbpap | 73.12116443 | 0.265221296 | 0.159657956 | 1.661184341 | 0.09667643 | 0.263715599 | Dnlz | 1463.803995 | -0.030757449 | 0.05820221 | -0.528458446 | 0.597181181 | 0.8881595 |
| E530001F21Rik | 51.21143859 | 0.2651921 | 0.167780598 | 1.580588597 | 0.113972135 | 0.295545482 | Cotl1 | 2299.315704 | -0.030760595 | 0.069384897 | -0.44333271 | 0.6575251 | 0.915208181 |
| Cct6a | 5100.296493 | -0.101664453 | 0.04554255 | -2.232296004 | 0.025595406 | 0.11089911 | Zswim2 | 0.55897171 | -0.030791884 | 0.018347316 | -1.678277261 | 0.093292985 | NA |
| Ifi203 | 44.14980388 | 0.265172474 | 0.165922136 | 1.598174177 | 0.110004219 | 0.288637658 | Vnn1 | 5.79495185 | -0.030795319 | 0.050422108 | -0.610750339 | 0.541364875 | NA |
| Ndufaf5 | 990.5174572 | -0.151513775 | 0.067881146 | -2.232045046 | 0.025611986 | 0.110908918 | 4930467D21Rik | 2.683043962 | -0.030796305 | 0.035368233 | -0.870733483 | 0.383899692 | NA |
| Pcdhgc3 | 2693.8457 | -0.130921093 | 0.058675131 | -2.231287612 | 0.025662084 | 0.111084152 | Swt1 | 329.5878548 | -0.030825875 | 0.084402976 | -0.365222601 | 0.714945249 | 0.93125489 |
| Stap2 | 64.40375916 | 0.264974388 | 0.156915488 | 1.688643944 | 0.091287684 | 0.254676254 | Zfp945 | 566.6313989 | -0.030830273 | 0.073849265 | -0.417475698 | 0.676330496 | 0.920643418 |
| Manf | 1419.335301 | -0.211247705 | 0.094685529 | -2.231045293 | 0.025678129 | 0.111102187 | Fam19a2 | 2490.017024 | -0.030856574 | 0.084209695 | -0.366425431 | 0.714047646 | 0.931082236 |
| Igfbp2 | 1184.645477 | -0.224949541 | 0.100844007 | -2.230668406 | 0.025703102 | 0.111179191 | Grb7 | 100.1777383 | -0.03086362 | 0.094381517 | -0.327009153 | 0.743660962 | 0.939912916 |
| Dgat1 | 680.026395 | 0.264866495 | 0.103856614 | 2.550309366 | 0.010762737 | 0.061981468 | Mycn | 236.4336439 | -0.030888185 | 0.092257749 | -0.334803146 | 0.737773587 | 0.937365444 |
| Anapc16 | 1500.402122 | -0.136250073 | 0.061145443 | -2.22829481 | 0.025860862 | 0.111705659 | Mboat2 | 1302.608727 | -0.030899902 | 0.058361203 | -0.529459648 | 0.59648663 | 0.887922451 |
| Nup35 | 550.8978395 | -0.175379807 | 0.078705622 | -2.228300873 | 0.025860458 | 0.111705659 | Snora23 | 5.043238112 | -0.030904586 | 0.048832169 | -0.63287352 | 0.526816241 | NA |
| Il1r1 | 171.0612422 | 0.264798604 | 0.134890148 | 1.963068529 | 0.049638212 | 0.173082559 | L1td1 | 3.977013092 | -0.030927266 | 0.045166081 | -0.6847454 | 0.493504595 | NA |
| Zfp7 | 301.603456 | 0.264722707 | 0.121023097 | 2.18737343 | 0.028715274 | 0.120059603 | Nudt8 | 191.0709493 | -0.030931167 | 0.088062129 | -0.351242554 | 0.725406388 | 0.93340891 |
| Aff4 | 9631.926052 | -0.173643615 | 0.07793381 | -2.228090941 | 0.025874451 | 0.111708246 | Mylk | 871.0037769 | -0.030947007 | 0.080003237 | -0.386821941 | 0.698888033 | 0.926409493 |
| Emx2os | 132.7881613 | -0.335447561 | 0.15055525 | -2.228069503 | 0.025875881 | 0.111708246 | Zfp955b | 783.3153974 | -0.030985481 | 0.079344362 | -0.390518995 | 0.696152812 | 0.925527709 |
| Col15a1 | 262.4203989 | 0.264690014 | 0.162613258 | 1.627727144 | 0.103582755 | 0.276852028 | Plekha7 | 733.4470622 | -0.030987244 | 0.082193089 | -0.377005473 | 0.706169537 | 0.9294028 |
| Fgd3 | 267.2925308 | 0.264657791 | 0.105271191 | 2.514057146 | 0.011935109 | 0.066566805 | Zbed4 | 632.0216474 | -0.030991449 | 0.073927686 | -0.419213032 | 0.675060449 | 0.919907397 |
| Snrnp27 | 1025.554411 | -0.143989029 | 0.0646688 | -2.226561024 | 0.025976626 | 0.112049512 | Nipsnap3b | 340.1446934 | -0.030991516 | 0.084273455 | -0.367749434 | 0.713060077 | 0.930507332 |
| Trim21 | 92.5275991 | 0.264612676 | 0.127741932 | 2.071462914 | 0.038315557 | 0.145344016 | Clp1 | 279.6493112 | -0.031003643 | 0.083310239 | -0.372146853 | 0.709783514 | 0.930224827 |
| Ucma | 14.40894042 | 0.264126222 | 0.164358892 | 1.607009025 | 0.108052388 | 0.285049614 | C5ar2 | 45.39900486 | -0.031039252 | 0.092407139 | -0.335896694 | 0.73694877 | 0.937206199 |
| Cd33 | 258.4955267 | 0.263993573 | 0.126143108 | 2.092810121 | 0.036366108 | 0.140570291 | Rnf141 | 923.0134817 | -0.031050251 | 0.063351535 | -0.490126253 | 0.624044562 | 0.900825929 |
| Plk3 | 484.3415647 | 0.263795764 | 0.13867894 | 1.902204942 | 0.057144367 | 0.189047536 | 9130017N09Rik | 10.90361137 | -0.031053375 | 0.064161637 | -0.483986635 | 0.628395355 | NA |
| Gucy2e | 172.4471379 | 0.263571354 | 0.139527867 | 1.889023025 | 0.058888743 | 0.192738106 | Spesp1 | 1.5692159 | -0.031057606 | 0.029291975 | -1.060276958 | 0.289018619 | NA |
| Crhr1 | 894.8175787 | 0.263526267 | 0.109356162 | 2.409798069 | 0.015961352 | 0.081256799 | Nfya | 1069.335458 | -0.031077593 | 0.056978505 | -0.545426613 | 0.585460141 | 0.884512516 |
| Klhl6 | 83.04347163 | 0.263439489 | 0.158267135 | 1.66452428 | 0.096007703 | 0.262445899 | Arl15 | 1643.109601 | -0.031094739 | 0.075792232 | -0.410262875 | 0.681613123 | 0.922538401 |
| Gbp3 | 222.9413765 | 0.263317923 | 0.167556431 | 1.571517859 | 0.116062409 | 0.2986169 | Cd55 | 157.4005395 | -0.031102736 | 0.093759126 | -0.33173023 | 0.740092972 | 0.938752228 |
| Ptges | 198.4160614 | -0.268708864 | 0.120880402 | -2.222931588 | 0.026220413 | 0.112818407 | Fastkd2 | 772.7308968 | -0.031114129 | 0.066600157 | -0.46717801 | 0.640372525 | 0.910092095 |
| Mreg | 176.4369828 | 0.263304315 | 0.112224253 | 2.346233618 | 0.018964217 | 0.090958778 | Proz | 1.138838504 | -0.031126028 | 0.022498831 | -1.383450915 | 0.166526648 | NA |
| Pam16 | 631.2810988 | -0.155790479 | 0.070097818 | -2.22247258 | 0.026251385 | 0.112918809 | Gm2027 | 6.231627958 | -0.031131735 | 0.052629245 | -0.59152921 | 0.55416589 | NA |
| Tuba1c | 34.48617399 | 0.263289301 | 0.166363223 | 1.582617215 | 0.113508737 | 0.294689071 | Mir1964 | 0.57283292 | -0.031134365 | 0.018447671 | -1.687712566 | 0.091466418 | NA |
| Gm11549 | 1767.2102 | 0.26309713 | 0.130010194 | 2.023665384 | 0.043004593 | 0.15756436 | Tbx10 | 0.57283292 | -0.031134365 | 0.018447671 | -1.687712566 | 0.091466418 | NA |
| Rps27l | 242.6100206 | -0.203622587 | 0.091658639 | -2.221531865 | 0.026314959 | 0.113124374 | Gm7444 | 2.927048557 | -0.031147669 | 0.031706211 | -0.982383853 | 0.325910778 | NA |
| Tha1 | 127.5549731 | 0.263038811 | 0.145835559 | 1.803667173 | 0.071283499 | 0.217800538 | Ikbke | 56.07664125 | -0.031149713 | 0.093465611 | -0.333274587 | 0.738927021 | 0.938092458 |
| Fut7 | 8.200417491 | 0.262861688 | 0.154927185 | 1.696678915 | 0.089757381 | 0.25230422 | Sugp1 | 2063.69791 | -0.031166648 | 0.054506492 | -0.571796991 | 0.567459516 | 0.876533064 |
| Lipe | 378.8672569 | 0.262818147 | 0.122562588 | 2.144358662 | 0.03200416 | 0.129066229 | Kifc2 | 24984.93137 | -0.031176518 | 0.059443411 | -0.524473908 | 0.599948963 | 0.889725231 |
| Galnt15 | 109.1327259 | 0.262745018 | 0.164111275 | 1.601017468 | 0.10937305 | 0.287606475 | Fbxl16 | 44591.63305 | -0.03118885 | 0.067669754 | -0.460897941 | 0.644871829 | 0.912363974 |
| Gm960 | 33.73445343 | 0.262638297 | 0.16171207 | 1.624110662 | 0.104352193 | 0.278380316 | Nat8 | 59.47749196 | -0.03122531 | 0.093127585 | -0.335296038 | 0.737401782 | 0.937206199 |
| Il17ra | 923.6852512 | 0.262359945 | 0.125785646 | 2.08577015 | 0.036999449 | 0.142144671 | Gm10474 | 3.271741785 | -0.031252673 | 0.037075095 | -0.842955976 | 0.399253066 | NA |
| H2-K1 | 1086.124872 | 0.262307857 | 0.127236705 | 2.061573799 | 0.039248331 | 0.147923531 | Bspry | 13.74744501 | -0.031265828 | 0.068408507 | -0.457045908 | 0.647638049 | NA |
| Zfp458 | 161.4387853 | -0.285936566 | 0.128813397 | -2.219773497 | 0.026434147 | 0.113328907 | Bloc1s4 | 256.1819043 | -0.03128616 | 0.089306776 | -0.350322351 | 0.726096793 | 0.933575664 |
| Egflam | 178.901555 | -0.314138209 | 0.141546881 | -2.219322716 | 0.026464777 | 0.11339998 | Slc9b1 | 2.204987207 | -0.031296204 | 0.033670592 | -0.929481852 | 0.352639426 | NA |
| Scrg1 | 705.6959461 | 0.261889422 | 0.119642233 | 2.188937941 | 0.028601351 | 0.119878408 | Spink8 | 51.52842064 | -0.031307643 | 0.052559052 | -0.595666051 | 0.551398342 | 0.869755927 |
| Ube2l3 | 6215.829755 | 0.089964861 | 0.040542226 | 2.219040963 | 0.026483938 | 0.113448198 | Gm4861 | 1.818954947 | -0.03130901 | 0.024429278 | -1.281618346 | 0.199976561 | NA |
| S100a5 | 54.18756256 | -0.1853579 | 0.083554757 | -2.218400318 | 0.026527549 | 0.113572216 | Nob1 | 1018.121201 | -0.031317545 | 0.057174152 | -0.547757049 | 0.583858734 | 0.884313345 |
| Smim12 | 1177.075915 | -0.131427795 | 0.059244238 | -2.218406368 | 0.026527137 | 0.113572216 | Gm15133 | 1.124665711 | -0.031323133 | 0.022898337 | -1.367921734 | 0.171336576 | NA |
| Man2b2 | 1129.911015 | 0.261686296 | 0.101414483 | 2.580364163 | 0.009869618 | 0.058255183 | AI847159 | 12.35379395 | -0.031333189 | 0.069729291 | -0.449354767 | 0.653175756 | NA |
| Cass4 | 24.06769474 | -0.372553205 | 0.168060494 | -2.216780378 | 0.026638103 | 0.113978519 | Rps11 | 5199.604328 | -0.031335194 | 0.061316293 | -0.511041884 | 0.609321729 | 0.89440021 |
| Slc25a11 | 3768.288145 | -0.099276632 | 0.044786049 | -2.216686525 | 0.02664452 | 0.113978519 | Chst15 | 1816.326528 | -0.03136416 | 0.064896158 | -0.483297633 | 0.628884421 | 0.904383239 |
| Ptp4a3 | 2140.510833 | -0.19988915 | 0.090192397 | -2.216252781 | 0.026674195 | 0.114073965 | Galr3 | 3.259397857 | -0.031378375 | 0.040180813 | -0.780929303 | 0.434844076 | NA |
| Mns1 | 51.0060736 | 0.261629624 | 0.153497178 | 1.704458858 | 0.088295396 | 0.249280041 | Ahrr | 41.43317039 | -0.031395077 | 0.090793667 | -0.345784874 | 0.729504393 | 0.934048551 |
| Elf5 | 15.80981951 | -0.366330138 | 0.16546743 | -2.213910846 | 0.026834913 | 0.114697968 | Hrsp12 | 1293.283937 | -0.031410959 | 0.061597421 | -0.509939522 | 0.610093832 | 0.894800514 |
| Gsto2 | 24.44548743 | 0.261434582 | 0.167665235 | 1.559265291 | 0.118933603 | 0.302885807 | Ttc25 | 78.19065383 | -0.031416469 | 0.094219425 | -0.333439405 | 0.738802623 | 0.938076299 |
| Banf1 | 1171.790933 | -0.163946046 | 0.074064013 | -2.213572269 | 0.026858217 | 0.114712817 | Nck2 | 1147.882809 | -0.031417126 | 0.066011895 | -0.475931286 | 0.634123332 | 0.907220906 |
| Sdc1 | 43.28793774 | -0.358804079 | 0.16209531 | -2.213537693 | 0.026860598 | 0.114712817 | Cth | 136.9878684 | -0.031417324 | 0.092670828 | -0.33902065 | 0.734594174 | 0.936456337 |
| Asnsd1 | 732.5874788 | -0.18276482 | 0.082627522 | -2.21191216 | 0.026972735 | 0.115128252 | Slc12a1 | 5.520072124 | -0.031425853 | 0.046538048 | -0.675272266 | 0.499502801 | NA |
| Jph4 | 14605.33135 | -0.194550381 | 0.087955619 | -2.211915323 | 0.026972516 | 0.115128252 | Pet100 | 529.6934827 | -0.031428739 | 0.072439445 | -0.433862222 | 0.664388495 | 0.915428982 |
| Tmem159 | 396.377332 | -0.201170883 | 0.09096012 | -2.211638283 | 0.026991668 | 0.115177335 | C130026L21Rik | 152.7157664 | -0.031445788 | 0.092520683 | -0.339878468 | 0.733948052 | 0.936456337 |
| Mrpl46 | 697.3331217 | -0.138092039 | 0.06244189 | -2.21152882 | 0.026999238 | 0.115177918 | Ldhc | 1.080858687 | -0.031474577 | 0.023901095 | -1.316867589 | 0.187883012 | NA |
| Tmem220 | 22.64841627 | 0.261319863 | 0.167809429 | 1.557241832 | 0.11941308 | 0.303657683 | Prkag2 | 3506.670689 | -0.031476181 | 0.053954749 | -0.583381107 | 0.559636774 | 0.873713575 |
| Haus3 | 172.35982 | 0.261271121 | 0.113425639 | 2.303457336 | 0.021253125 | 0.098006348 | Lsg1 | 642.3273361 | -0.031483893 | 0.065318588 | -0.482005099 | 0.629802324 | 0.904807492 |
| Tm2d2 | 1815.132718 | -0.118269979 | 0.053537462 | -2.209106957 | 0.027167202 | 0.115798799 | Tmem98 | 426.8442742 | -0.03148561 | 0.095099025 | -0.331082368 | 0.74058227 | 0.938975678 |
| Pvrl3 | 630.7527255 | -0.260984263 | 0.118176949 | -2.208419358 | 0.027215054 | 0.115970858 | Pigq | 4841.783727 | -0.031509618 | 0.056009437 | -0.562576935 | 0.573723001 | 0.879633264 |
| 4932438H23Rik | 40.0754293 | 0.261261902 | 0.160585826 | 1.626930027 | 0.10375196 | 0.277160839 | Iqch | 4.297471241 | -0.031511144 | 0.049256686 | -0.639733333 | 0.522345981 | NA |
| 1700030J22Rik | 363.8494526 | -0.235212454 | 0.106539019 | -2.207758774 | 0.027261093 | 0.11600752 | Wbscr25 | 11.80264077 | -0.031609672 | 0.06920044 | -0.456784263 | 0.647826118 | NA |
| C230091D08Rik | 57.52774818 | 0.261137287 | 0.165799113 | 1.575022221 | 0.115251316 | 0.297319994 | 1110035M17Rik | 80.62596021 | -0.031609871 | 0.0950083 | -0.33270641 | 0.73935591 | 0.938390909 |
| Cdkl4 | 747.6635222 | 0.261075319 | 0.124950562 | 2.089428923 | 0.03666913 | 0.141256 | Lrig2 | 1503.097145 | -0.031631282 | 0.055801046 | -0.56685822 | 0.570810512 | 0.878246677 |
| Tnnt2 | 247.4143422 | 0.261053163 | 0.119874508 | 2.177720406 | 0.029426859 | 0.121935044 | Slc39a1 | 1617.050862 | -0.031664928 | 0.05189378 | -0.610187337 | 0.541737717 | 0.864471281 |
| Col4a6 | 78.52534686 | -0.306122605 | 0.138677029 | -2.207449975 | 0.027282638 | 0.116022932 | Rpl37a | 4144.273669 | -0.031671668 | 0.061285524 | -0.516788728 | 0.605303649 | 0.892291538 |
| Lgr4 | 1184.208919 | -0.161221743 | 0.07303744 | -2.207384908 | 0.02728718 | 0.116022932 | Arid4b | 2003.799241 | -0.031682999 | 0.070583051 | -0.448875451 | 0.653521506 | 0.914683958 |
| Rasgrp2 | 1904.708393 | -0.211117764 | 0.095639846 | -2.207424758 | 0.027284398 | 0.116022932 | Ddhd2 | 3352.099267 | -0.031686861 | 0.042790871 | -0.740505171 | 0.458993524 | 0.828343886 |
| Cftr | 63.01583471 | -0.325965839 | 0.147686902 | -2.207141147 | 0.0273042 | 0.116063449 | Sufu | 405.9004075 | -0.03169018 | 0.076179367 | -0.415994271 | 0.677414196 | 0.921064066 |
| Nop14 | 908.6755389 | 0.260668092 | 0.104887262 | 2.48522163 | 0.01294708 | 0.070288038 | Enoph1 | 1489.5508 | -0.031704208 | 0.058020472 | -0.546431412 | 0.584769423 | 0.884313345 |
| Pura | 635.6300238 | -0.240533321 | 0.109027408 | -2.206172977 | 0.027371891 | 0.116266709 | Rpp30 | 298.9269785 | -0.031716325 | 0.083505245 | -0.379812374 | 0.704084696 | 0.928127792 |
| Spop | 8264.730782 | -0.094208364 | 0.042702895 | -2.206135281 | 0.027374529 | 0.116266709 | Uros | 879.607677 | -0.031726094 | 0.060757628 | -0.522174661 | 0.601548727 | 0.890461307 |
| Zbtb48 | 345.5334953 | 0.260667431 | 0.105950705 | 2.460270843 | 0.01388322 | 0.073687366 | Klhl40 | 257.7856711 | -0.031727376 | 0.079193061 | -0.400633278 | 0.688690141 | 0.923860706 |
| Atp5f1 | 7229.764798 | -0.134479104 | 0.060995231 | -2.204747858 | 0.027471793 | 0.116615879 | Pcdhgb2 | 318.4397501 | -0.031747309 | 0.079712024 | -0.398275032 | 0.690427463 | 0.923860706 |
| Zfp169 | 444.1690551 | 0.260368571 | 0.098570027 | 2.641457851 | 0.008255008 | 0.051449353 | Ccnb1 | 11.94177566 | -0.03175357 | 0.069565776 | -0.456453903 | 0.648063612 | NA |
| Aldh3b2 | 65.68496793 | -0.339844636 | 0.154174978 | -2.204278798 | 0.027504743 | 0.11669181 | Racgap1 | 575.632949 | -0.031771983 | 0.079313139 | -0.400589148 | 0.688722637 | 0.923860706 |
| 2210039B01Rik | 11.43319355 | 0.259867179 | 0.162760292 | 1.596625176 | 0.110349284 | 0.289208519 | Gpr153 | 1400.0198 | -0.0318248 | 0.059183749 | -0.537728693 | 0.590764367 | 0.885693151 |
| Mcfd2 | 2533.199159 | -0.129749982 | 0.058890908 | -2.203226029 | 0.027578822 | 0.116942053 | Rassf3 | 2645.947553 | -0.031831262 | 0.060530195 | -0.525874105 | 0.598975682 | 0.889150487 |
| Snord37 | 8.138696652 | 0.259866645 | 0.151948813 | 1.710224905 | 0.087224292 | 0.247292439 | Cstf1 | 902.3240849 | -0.031837794 | 0.056421559 | -0.564284191 | 0.572560739 | 0.879277718 |
| Sec14l4 | 9.240876223 | 0.259855823 | 0.156238079 | 1.663204164 | 0.096271575 | 0.262935268 | Rasgrp2 | 1950.021513 | -0.031852374 | 0.078786903 | -0.404285143 | 0.68600305 | 0.923860706 |
| Fam163a | 331.0198716 | -0.299460911 | 0.135988058 | -2.202111835 | 0.02765741 | 0.117179083 | Igfals | 1.813150625 | -0.031858095 | 0.032102622 | -0.992382968 | 0.321010744 | NA |
| Frmd6 | 1549.798478 | 0.259564973 | 0.111803277 | 2.321622232 | 0.020253286 | 0.09495708 | Slc26a5 | 10.2863615 | -0.031877872 | 0.060116564 | -0.530267692 | 0.595926343 | NA |
| Il3ra | 50.05184338 | 0.259278391 | 0.154606426 | 1.677022085 | 0.093538163 | 0.258476958 | Kdm3a | 2218.788272 | -0.031896233 | 0.073397462 | -0.434568608 | 0.663875585 | 0.915428982 |
| Sass6 | 185.8474938 | 0.25923209 | 0.111361407 | 2.327844951 | 0.019920338 | 0.093909252 | Nol9 | 1003.416548 | -0.031903501 | 0.059615364 | -0.535155691 | 0.592542203 | 0.886226713 |
| Tbxas1 | 95.92271577 | 0.259149782 | 0.161153142 | 1.60809637 | 0.107814074 | 0.284480801 | Utp11l | 493.4128912 | -0.031911509 | 0.070309254 | -0.453873526 | 0.649919859 | 0.914000665 |
| Tuba1b | 9316.397266 | -0.197378638 | 0.089726167 | -2.199789034 | 0.027821866 | 0.117691436 | Mfsd1 | 1181.134448 | -0.031914135 | 0.057259567 | -0.557359003 | 0.57728217 | 0.881375635 |
| Zfp955a | 430.4128816 | -0.225461702 | 0.102489017 | -2.19986209 | 0.027816681 | 0.117691436 | Rai14 | 620.4672859 | -0.031917807 | 0.077537916 | -0.411641284 | 0.680602368 | 0.922327642 |
| Kcnj8 | 146.1291788 | -0.275365645 | 0.125232901 | -2.198828281 | 0.027890135 | 0.117939342 | Myl9 | 442.0100652 | -0.031919316 | 0.089969658 | -0.35477867 | 0.722755409 | 0.93255988 |
| Cobl | 5837.614543 | -0.215118876 | 0.097839358 | -2.198694689 | 0.027899639 | 0.117947341 | Prima1 | 189.2849872 | -0.031981052 | 0.093888988 | -0.34062623 | 0.73338498 | 0.936456337 |
| Ankrd17 | 6708.384731 | -0.168581064 | 0.076689881 | -2.198217837 | 0.027933586 | 0.118058643 | Amotl2 | 890.2334175 | -0.03198978 | 0.085913144 | -0.37235024 | 0.709632097 | 0.930224827 |
| Slc25a1 | 1224.820941 | -0.198390644 | 0.090271031 | -2.197722149 | 0.027968912 | 0.118143491 | Mtmr3 | 5023.602389 | -0.032004365 | 0.042051026 | -0.761084043 | 0.446606871 | 0.822210607 |
| Armc2 | 301.0133433 | 0.259087673 | 0.118822774 | 2.180454671 | 0.029223775 | 0.121359736 | Ttll6 | 17.64200936 | -0.032040031 | 0.065583494 | -0.488538033 | 0.625168793 | NA |
| Hdhd2 | 2718.824976 | -0.120245941 | 0.054723259 | -2.197346112 | 0.027995737 | 0.118192356 | Cd28 | 10.59661273 | -0.032040342 | 0.05953133 | -0.538209742 | 0.590432255 | NA |
| Ypel2 | 2401.412933 | -0.18490087 | 0.084145606 | -2.197391852 | 0.027992473 | 0.118192356 | Wdr91 | 1162.438254 | -0.032050573 | 0.059287477 | -0.540595991 | 0.588786082 | 0.885188411 |
| Sumo1 | 2861.623442 | -0.133617317 | 0.060818573 | -2.196982105 | 0.028021724 | 0.118269843 | Fam73b | 3675.49661 | -0.032054289 | 0.056398034 | -0.568358267 | 0.569791722 | 0.87770886 |
| Tmem63a | 1369.15628 | 0.258778364 | 0.122776292 | 2.10772259 | 0.035054992 | 0.137053255 | Gm5506 | 36.78957726 | -0.032057402 | 0.088860155 | -0.360762395 | 0.718277077 | 0.93125489 |
| Cradd | 174.6535522 | -0.218811317 | 0.09962127 | -2.19643171 | 0.028061058 | 0.118339149 | Abtb2 | 639.0764212 | -0.032082419 | 0.072640995 | -0.441657206 | 0.658737281 | 0.915208181 |
| Etv6 | 940.3203609 | -0.180605839 | 0.082223145 | -2.196532845 | 0.028053827 | 0.118339149 | B930041F14Rik | 1229.641557 | -0.032084742 | 0.076380649 | -0.420063749 | 0.674438884 | 0.919867577 |
| Pyhin1 | 11.48654586 | 0.258656511 | 0.15953435 | 1.621321744 | 0.104948654 | 0.279442249 | Nppa | 5.52399073 | -0.032091425 | 0.049220825 | -0.651988757 | 0.514408425 | NA |
| Rnpepl1 | 1144.618239 | 0.258594411 | 0.102994666 | 2.510755368 | 0.012047315 | 0.066986632 | Gm11413 | 7.667104193 | -0.032179158 | 0.054998896 | -0.585087343 | 0.558488989 | NA |
| Fam107b | 308.5769769 | -0.196917437 | 0.089690717 | -2.195516355 | 0.028126579 | 0.118518688 | Ptpmt1 | 868.16637 | -0.032190691 | 0.060179769 | -0.534908857 | 0.592712883 | 0.886226713 |
| Fam136a | 755.3807347 | -0.246695492 | 0.112442512 | -2.193969942 | 0.028237571 | 0.118942553 | Tfap4 | 123.1206976 | -0.032198347 | 0.095097274 | -0.33858328 | 0.73492368 | 0.936456337 |
| Gtf2a2 | 515.160977 | -0.216575347 | 0.098717007 | -2.193901063 | 0.028242523 | 0.118942553 | Gab1 | 1477.560951 | -0.03220728 | 0.08378878 | -0.38438655 | 0.700691967 | 0.92719503 |
| Batf3 | 26.4141412 | 0.258439732 | 0.1673178 | 1.54460393 | 0.122442095 | 0.308775687 | Slc44a3 | 4.415748996 | -0.032210039 | 0.037869518 | -0.850553189 | 0.395017602 | NA |
| Nomo1 | 5839.617597 | -0.134350523 | 0.061252065 | -2.193403975 | 0.028278286 | 0.119028461 | Maoa | 1364.673694 | -0.032210461 | 0.066630194 | -0.483421391 | 0.628796563 | 0.904346423 |
| E4f1 | 939.9410418 | 0.258207259 | 0.107996697 | 2.39088107 | 0.016807997 | 0.08386066 | Ggh | 403.9833674 | -0.032218808 | 0.078624543 | -0.409780543 | 0.68196694 | 0.922544285 |
| Plch2 | 3864.863156 | 0.258169869 | 0.09853474 | 2.620089832 | 0.008790661 | 0.053683867 | Kdm1a | 2591.94119 | -0.032222961 | 0.04535278 | -0.710495832 | 0.477396715 | 0.835289575 |
| Ccdc63 | 12.99156708 | 0.25812517 | 0.142475623 | 1.811714628 | 0.070030302 | 0.2152876 | Hist1h4h | 11.056691 | -0.032274588 | 0.069254564 | -0.466028309 | 0.641195237 | NA |
| Pcdhga11 | 851.8229789 | -0.163819488 | 0.074724829 | -2.192303303 | 0.028357613 | 0.119232798 | Hnrnpa1 | 117.8933061 | -0.032311795 | 0.094847284 | -0.340671805 | 0.733350667 | 0.936456337 |
| Angptl7 | 10.84652463 | -0.342442467 | 0.156287964 | -2.191099417 | 0.028444599 | 0.119533663 | Slc28a1 | 22.07906159 | -0.03233052 | 0.077871829 | -0.41517607 | 0.678013015 | NA |
| Mb21d1 | 47.71202055 | 0.257909291 | 0.168121508 | 1.534064821 | 0.125013707 | 0.312625547 | Trmt10c | 448.1149207 | -0.032331515 | 0.083169565 | -0.388742148 | 0.6974669 | 0.9261158 |
| Cpm | 558.9512007 | 0.257887833 | 0.130073407 | 1.98263302 | 0.047408436 | 0.168109493 | Opn1sw | 1.754973435 | -0.032388942 | 0.028576957 | -1.133393659 | 0.257048976 | NA |
| Bora | 62.18882426 | 0.257714923 | 0.152318692 | 1.691945483 | 0.090656369 | 0.253890779 | Ccdc106 | 1239.868657 | -0.032391944 | 0.069577728 | -0.465550469 | 0.641537304 | 0.911095592 |
| Cct3 | 4289.575189 | -0.112598699 | 0.051400846 | -2.190600115 | 0.028480743 | 0.119588246 | 4732416N19Rik | 7.744872783 | -0.032404974 | 0.060319097 | -0.53722579 | 0.591111658 | NA |
| Ssc5d | 108.8474851 | 0.25751548 | 0.153999744 | 1.672181217 | 0.094488593 | 0.260036636 | Gstz1 | 1496.802548 | -0.032409125 | 0.058896099 | -0.550276255 | 0.582129908 | 0.884020489 |
| Csmd1 | 3013.876959 | -0.165226144 | 0.075464325 | -2.189460306 | 0.028563401 | 0.119870349 | Ccl17 | 53.37243747 | -0.03241814 | 0.091857341 | -0.352918336 | 0.724149664 | 0.932714927 |
| Dffb | 224.2666979 | 0.257508973 | 0.135834506 | 1.895755222 | 0.057992419 | 0.190933818 | Lrrc27 | 363.5664686 | -0.032429989 | 0.071795102 | -0.451701974 | 0.651483696 | 0.914343204 |
| Scel | 98.31533492 | 0.257462711 | 0.13355852 | 1.927714615 | 0.053890633 | 0.182350179 | 2900041M22Rik | 27.13538567 | -0.032458543 | 0.084525769 | -0.384007665 | 0.700972766 | 0.927352496 |
| Recql5 | 903.1199365 | 0.257056064 | 0.10267852 | 2.503503793 | 0.012297036 | 0.067973084 | Gm2381 | 0.958745221 | -0.032482025 | 0.021086947 | -1.540385397 | 0.123466439 | NA |
| Rpe | 632.136526 | -0.223396439 | 0.102058703 | -2.188901414 | 0.028604006 | 0.119878408 | Grin1 | 21475.63651 | -0.032485381 | 0.043001659 | -0.755444831 | 0.449982123 | 0.823263966 |
| Masp2 | 26.44467142 | 0.256575984 | 0.167703897 | 1.529934539 | 0.126032933 | 0.313897165 | Plp2 | 261.2415118 | -0.032496203 | 0.085177757 | -0.381510434 | 0.70282453 | 0.927855161 |
| Atp13a4 | 487.8134072 | 0.256517149 | 0.104088902 | 2.464404409 | 0.013724117 | 0.073148078 | Cbr3 | 320.1270175 | -0.032505613 | 0.08746891 | -0.371624765 | 0.710172248 | 0.930478981 |
| Csf2rb2 | 50.44082239 | 0.256465118 | 0.161513195 | 1.587889569 | 0.112311313 | 0.292511595 | Klkb1 | 3.754403219 | -0.032509151 | 0.045859097 | -0.708892075 | 0.478391453 | NA |
| Klrg2 | 9.538053492 | 0.256422926 | 0.129569588 | 1.979036362 | 0.047811912 | 0.169016147 | Zfp280d | 1558.559409 | -0.032516594 | 0.073660445 | -0.441439003 | 0.658895211 | 0.915208181 |
| Retsat | 420.6220446 | 0.256372433 | 0.141432252 | 1.812687205 | 0.069880079 | 0.215083255 | Plaur | 40.02117959 | -0.032521717 | 0.093163803 | -0.349081035 | 0.727028474 | 0.9335794 |
| Yeats4 | 1002.570626 | -0.14457933 | 0.066077364 | -2.18803113 | 0.028667335 | 0.119981547 | Tmem186 | 921.8479003 | -0.032521934 | 0.063906789 | -0.508896378 | 0.610824859 | 0.894800514 |
| Gm694 | 67.13368306 | -0.350053441 | 0.160035401 | -2.187350044 | 0.02871698 | 0.120059603 | Ocln | 373.1343401 | -0.032543445 | 0.075176893 | -0.432891595 | 0.665093527 | 0.915428982 |
| Gpx4 | 8168.969268 | -0.167100753 | 0.076390596 | -2.187451892 | 0.028709552 | 0.120059603 | Zfp358 | 1129.338888 | -0.032544431 | 0.055167067 | -0.58992498 | 0.555240946 | 0.871360989 |
| Cda | 46.72362203 | 0.25572174 | 0.16214809 | 1.577087585 | 0.114775373 | 0.296753581 | Lima1 | 614.317978 | -0.032557985 | 0.07261524 | -0.448362973 | 0.65389126 | 0.914733232 |
| Vsx2 | 13.97260011 | 0.255495115 | 0.163597924 | 1.561725902 | 0.118352574 | 0.301952047 | Preb | 3055.602201 | -0.032568285 | 0.062486482 | -0.521205288 | 0.602223771 | 0.89112297 |
| Snord12 | 9.91430037 | 0.255197567 | 0.151924276 | 1.679768194 | 0.093002426 | 0.257409715 | Klhl23 | 3474.773143 | -0.03258116 | 0.072103407 | -0.451867127 | 0.651364708 | 0.914343204 |
| 4933432I09Rik | 40.61617095 | 0.25490658 | 0.164987583 | 1.545004634 | 0.122345142 | 0.308694072 | 3110082J24Rik | 35.76163169 | -0.032639121 | 0.087227682 | -0.374183062 | 0.708268125 | 0.929959854 |
| Spata6 | 133.7433974 | 0.254896927 | 0.118027923 | 2.159632412 | 0.030801137 | 0.125851079 | Sptbn2 | 47246.85343 | -0.032653947 | 0.066421137 | -0.491619811 | 0.622988136 | 0.900360726 |
| Dcp2 | 3426.32966 | -0.141981504 | 0.064937446 | -2.186434982 | 0.028783797 | 0.120144436 | Ino80d | 1560.808507 | -0.032678709 | 0.077129309 | -0.423687299 | 0.671793879 | 0.919094512 |
| Il20rb | 14.31147771 | 0.254841724 | 0.165257038 | 1.542093017 | 0.123050992 | 0.309806965 | Prpf4 | 1011.135452 | -0.032680447 | 0.05881683 | -0.555630877 | 0.578463219 | 0.881901419 |
| Col6a5 | 68.37221718 | 0.254835273 | 0.163103368 | 1.562415765 | 0.118190076 | 0.301636803 | Rho | 15.60699633 | -0.032702457 | 0.073138018 | -0.447133488 | 0.654778681 | NA |
| Ctf1 | 52.01891411 | 0.254629911 | 0.143099707 | 1.779388071 | 0.075176161 | 0.225339415 | Cyp2r1 | 8.969278955 | -0.032704843 | 0.060200251 | -0.54326755 | 0.586945603 | NA |
| Gucy2g | 321.9000507 | -0.291342177 | 0.133296228 | -2.185674568 | 0.028839423 | 0.120311798 | Fam58b | 404.789951 | -0.032708475 | 0.079886985 | -0.409434333 | 0.682220948 | 0.922544285 |
| Sybu | 3144.56938 | -0.120873286 | 0.055311866 | -2.185304803 | 0.028866505 | 0.120392365 | Dchs1 | 708.802189 | -0.032735955 | 0.081753517 | -0.400422584 | 0.688845293 | 0.923860706 |
| Slc9a3r2 | 1420.823702 | -0.161487933 | 0.073907991 | -2.184986086 | 0.028889867 | 0.120457373 | Nat6 | 524.8691185 | -0.032778222 | 0.076818116 | -0.42669912 | 0.669598492 | 0.91770119 |
| Mrpl1 | 514.1898107 | -0.205601091 | 0.094104298 | -2.184821476 | 0.028901939 | 0.120475286 | Bivm | 938.9893374 | -0.032788636 | 0.065530834 | -0.50035432 | 0.616825612 | 0.897346904 |
| 4932418E24Rik | 126.9537062 | -0.339092265 | 0.155274562 | -2.183823678 | 0.028975207 | 0.120734838 | Ppm1m | 766.2159165 | -0.032791795 | 0.081699627 | -0.401370196 | 0.688147589 | 0.923860706 |
| Vgll3 | 73.1959661 | 0.254259908 | 0.166337744 | 1.528576143 | 0.126369554 | 0.31443229 | BC049715 | 18.03857757 | -0.032805444 | 0.077408273 | -0.423797644 | 0.671713397 | NA |
| Baalc | 9374.951189 | -0.134428225 | 0.061576897 | -2.183095148 | 0.029028804 | 0.120809102 | Gm20125 | 2.036085988 | -0.032845186 | 0.03332671 | -0.98555142 | 0.324353287 | NA |
| Cyp11a1 | 102.8846507 | 0.254125712 | 0.147666278 | 1.720946145 | 0.085260597 | 0.243893179 | Oip5 | 30.54585769 | -0.032891022 | 0.084959741 | -0.387136564 | 0.69865511 | 0.926409493 |
| Vax1 | 17.44627343 | -0.36567689 | 0.167503196 | -2.183103961 | 0.029028155 | 0.120809102 | 9430076C15Rik | 1.012800456 | -0.032898707 | 0.02461216 | -1.336685049 | 0.181325464 | NA |
| Sdcbp2 | 31.37300917 | 0.254088054 | 0.167565493 | 1.516350707 | 0.129430696 | 0.318468986 | Tnni3 | 14.4201095 | -0.032903254 | 0.069563207 | -0.472997948 | 0.636214643 | NA |
| Barhl2 | 24.57405904 | 0.253802879 | 0.156455026 | 1.622209817 | 0.104758431 | 0.279301801 | Vdr | 46.35660121 | -0.032916 | 0.09109954 | -0.361319058 | 0.717860949 | 0.93125489 |
| Dip2c | 4231.51781 | -0.170467156 | 0.078116531 | -2.182216143 | 0.029093584 | 0.120956391 | Zfp282 | 722.8025257 | -0.032928505 | 0.06546199 | -0.503017173 | 0.614952198 | 0.896237146 |
| Ttc8 | 658.9703297 | 0.25338441 | 0.102426796 | 2.473809775 | 0.013368085 | 0.071801776 | Cbx2 | 132.2336821 | -0.032935791 | 0.091932932 | -0.3582589 | 0.720149573 | 0.932426935 |
| Mir374 | 17.90163355 | 0.253348668 | 0.161295332 | 1.570712957 | 0.116249338 | 0.298849717 | Gm2109 | 3.233702143 | -0.032945327 | 0.037463986 | -0.879386603 | 0.379191694 | NA |
| Rpl31 | 6469.000867 | -0.156567842 | 0.071751472 | -2.182085424 | 0.029103229 | 0.120956391 | Prrt2 | 13263.8674 | -0.032945788 | 0.042240796 | -0.779951877 | 0.435419201 | 0.815695038 |
| 1700028J19Rik | 32.1732815 | 0.253343476 | 0.167696048 | 1.510730153 | 0.130857217 | 0.320515687 | Aloxe3 | 249.3677216 | -0.032945942 | 0.089206393 | -0.369322652 | 0.711887244 | 0.930507332 |
| Pld5 | 435.0255449 | 0.253304421 | 0.105152778 | 2.408918012 | 0.015999892 | 0.081386187 | 1810062G17Rik | 3.298856266 | -0.032961664 | 0.040155667 | -0.820847132 | 0.411733348 | NA |
| Gm14378 | 106.4429904 | 0.25291999 | 0.166389607 | 1.520046804 | 0.128499213 | 0.317061653 | Cox6a1 | 7265.205531 | -0.032986973 | 0.054936781 | -0.600453324 | 0.548204159 | 0.867925189 |
| Ttpal | 3341.621795 | -0.097990887 | 0.044945792 | -2.180201564 | 0.029242524 | 0.121405054 | D630013N20Rik | 4.141915698 | -0.033000872 | 0.039808001 | -0.829000993 | 0.407103845 | NA |
| Bach2 | 436.240805 | -0.276821559 | 0.1269851 | -2.179953079 | 0.02926094 | 0.121448969 | Dync1h1 | 24765.90902 | -0.033020486 | 0.067393057 | -0.489968671 | 0.624156069 | 0.900883451 |
| Stmn3 | 12242.57949 | -0.165323721 | 0.075861906 | -2.179271922 | 0.029311474 | 0.121626131 | Itga2 | 2.495017543 | -0.033023013 | 0.035328953 | -0.934729459 | 0.349927741 | NA |
| Mdga1 | 1165.826457 | -0.322710677 | 0.148100213 | -2.179002109 | 0.029331512 | 0.12164478 | Snhg12 | 543.5105242 | -0.033025919 | 0.070292732 | -0.469834052 | 0.638473584 | 0.909172363 |
| 2210404J11Rik | 207.4924549 | 0.252900032 | 0.140364909 | 1.801732601 | 0.071587486 | 0.218341833 | Tmem18 | 1517.22293 | -0.033049524 | 0.050118922 | -0.659422079 | 0.509624768 | 0.850457637 |
| Eif3i | 2554.72046 | -0.141959719 | 0.065161779 | -2.178573389 | 0.029363375 | 0.121743679 | Vwf | 877.8542035 | -0.033063949 | 0.089875782 | -0.367884971 | 0.712959007 | 0.930507332 |
| S100a8 | 14.66361853 | -0.329496199 | 0.151277155 | -2.17809622 | 0.029398874 | 0.121858254 | Mesp2 | 23.09763834 | -0.033076919 | 0.078707507 | -0.420251129 | 0.674302008 | 0.919867577 |
| Tctex1d4 | 9.039008638 | 0.252861868 | 0.148815917 | 1.699158757 | 0.089289275 | 0.251353203 | Fut7 | 9.175731575 | -0.033113891 | 0.062565458 | -0.529267933 | 0.596619597 | NA |
| Zfp474 | 11.13058761 | 0.252684338 | 0.15826779 | 1.596561988 | 0.110363378 | 0.289208519 | Pde8b | 4126.022225 | -0.033114126 | 0.048775981 | -0.678902309 | 0.497199763 | 0.845038521 |
| Atp8b1 | 170.9281834 | 0.252528541 | 0.142569116 | 1.77127101 | 0.076515646 | 0.227636985 | Capn2 | 3597.517025 | -0.033128513 | 0.048415792 | -0.68425015 | 0.49381722 | 0.843404385 |
| C030030A07Rik | 248.1172667 | 0.251824773 | 0.097796165 | 2.574996391 | 0.010024113 | 0.058942547 | Gm13139 | 6.598793456 | -0.033140709 | 0.053116655 | -0.623923119 | 0.532678076 | NA |
| Nfatc2ip | 220.1889419 | 0.251648219 | 0.104883965 | 2.399301168 | 0.016426398 | 0.082821547 | Sptan1 | 44838.78761 | -0.033143491 | 0.051268907 | -0.646463772 | 0.517979052 | 0.853297835 |
| Mapkap1 | 1939.106199 | -0.112149329 | 0.051515321 | -2.17700922 | 0.02947988 | 0.121998198 | Ptchd2 | 1934.662846 | -0.033158501 | 0.063687583 | -0.520643112 | 0.602615411 | 0.891281205 |
| Gatsl3 | 106.3687495 | 0.251485107 | 0.119561674 | 2.103392323 | 0.035431489 | 0.138107041 | Zfp39 | 527.7564839 | -0.033173868 | 0.079099201 | -0.419395745 | 0.674926933 | 0.919907397 |
| Card9 | 53.13130101 | 0.251201328 | 0.16048511 | 1.565262516 | 0.117521368 | 0.300871791 | 4930528A17Rik | 22.41968482 | -0.033180533 | 0.082868857 | -0.400398103 | 0.688863321 | 0.923860706 |
| Ccdc173 | 70.22491207 | 0.251090976 | 0.160567696 | 1.563770191 | 0.117871547 | 0.301421803 | Zfp276 | 1033.033181 | -0.033216504 | 0.083022343 | -0.400091135 | 0.689089393 | 0.923860706 |
| 4933427I22Rik | 14.89939805 | 0.25098492 | 0.165616121 | 1.515461896 | 0.129655473 | 0.318819536 | 2310036O22Rik | 1344.550205 | -0.033235588 | 0.072828234 | -0.456355809 | 0.648134138 | 0.913259917 |
| Smad6 | 60.95485357 | -0.312260336 | 0.143523692 | -2.175671007 | 0.02957987 | 0.122248733 | Wdr43 | 1095.371847 | -0.033246156 | 0.064499911 | -0.515444997 | 0.606242096 | 0.892768925 |
| Inca1 | 79.34815103 | 0.250837208 | 0.130741411 | 1.918575043 | 0.055038137 | 0.184659741 | Sash1 | 3501.780517 | -0.033248461 | 0.063982912 | -0.519645946 | 0.603310368 | 0.891540503 |
| Tnfrsf25 | 128.2417777 | 0.250627097 | 0.15412376 | 1.62614185 | 0.103919483 | 0.277512663 | Pgbd1 | 95.79576621 | -0.03326019 | 0.093793937 | -0.354609168 | 0.722882407 | 0.93255988 |
| 3110001D03Rik | 611.0541901 | -0.164105893 | 0.075487308 | -2.173953443 | 0.029708632 | 0.122552061 | Gm4759 | 2.177775734 | -0.033298223 | 0.031549928 | -1.05541359 | 0.29123621 | NA |
| Ngp | 19.97648518 | -0.350371588 | 0.161166251 | -2.173976164 | 0.029706925 | 0.122552061 | Slc35a1 | 666.0283846 | -0.033303028 | 0.075563076 | -0.4407315 | 0.659407389 | 0.915208181 |
| Irf5 | 171.2624384 | 0.250595372 | 0.109314015 | 2.292435896 | 0.021880503 | 0.099857587 | Mbd2 | 1569.64353 | -0.03330917 | 0.05847159 | -0.56966417 | 0.568905496 | 0.87745335 |
| Pip4k2b | 7983.327125 | -0.123897001 | 0.056989656 | -2.174026141 | 0.029703172 | 0.122552061 | Cep170 | 3434.43801 | -0.033311202 | 0.069599987 | -0.478609313 | 0.632216591 | 0.906498456 |
| Pdlim7 | 1572.99402 | 0.25052761 | 0.097342551 | 2.573670061 | 0.010062618 | 0.059124082 | Tmem158 | 1436.196577 | -0.033334118 | 0.089705202 | -0.371596265 | 0.710193471 | 0.930478981 |

| | | | | | | | | | | | | | |
|---|---|---|---|---|---|---|---|---|---|---|---|---|---|
| Lair1 | 365.8334691 | 0.250499848 | 0.102798388 | 2.436807151 | 0.014817579 | 0.077274849 | Frmd8 | 1030.395003 | -0.033354576 | 0.075925938 | -0.439304102 | 0.660441202 | 0.915428982 |
| Fam193b | 3189.060558 | 0.250319872 | 0.139413497 | 1.795521073 | 0.072570718 | 0.220386262 | Rsf1 | 2432.786515 | -0.033354679 | 0.069077255 | -0.482860517 | 0.629194778 | 0.904471385 |
| Lcorl | 548.1683062 | 0.25019637 | 0.108998777 | 2.295405296 | 0.021709908 | 0.09939954 | Chi3l3 | 4.938303949 | -0.03337158 | 0.041706097 | -0.800160715 | 0.423617688 | NA |
| Crip3 | 44.58264111 | 0.250006029 | 0.168011482 | 1.488029424 | 0.136743134 | 0.329602254 | Mtfr2 | 14.09236121 | -0.033384327 | 0.070457306 | -0.473823489 | 0.635625783 | NA |
| Yap1 | 992.1040176 | -0.13301709 | 0.061299259 | -2.169962465 | 0.03000969 | 0.123629389 | 4933412O06Rik | 78.36825715 | -0.033398611 | 0.094929214 | -0.351826482 | 0.724968397 | 0.933258679 |
| Gpam | 2286.281706 | -0.121895686 | 0.056186263 | -2.169492681 | 0.030045299 | 0.123743187 | Abcg2 | 790.7512602 | -0.033416479 | 0.068057818 | -0.491001326 | 0.62342551 | 0.900570704 |
| Krt10 | 219.7425032 | -0.275184004 | 0.126857458 | -2.16923789 | 0.030064628 | 0.123756999 | Cbr2 | 43.37557645 | -0.033421108 | 0.089694569 | -0.372610157 | 0.709438612 | 0.930224827 |
| Cdnf | 57.20405843 | 0.249764279 | 0.147402062 | 1.694442229 | 0.090181281 | 0.253004419 | Clec1b | 1.078493332 | -0.033437927 | 0.023613554 | -1.416048058 | 0.156761434 | NA |
| Samd3 | 86.06833068 | 0.249731962 | 0.152281811 | 1.639932964 | 0.101019106 | 0.272073766 | Gm884 | 2.463864808 | -0.033439843 | 0.035866765 | -0.932335057 | 0.351163396 | NA |
| Efcab7 | 76.63017477 | -0.303992738 | 0.140186215 | -2.168492369 | 0.030121245 | 0.123924207 | Usp12 | 3559.360058 | -0.033441585 | 0.060286189 | -0.554713873 | 0.579090386 | 0.882452682 |
| Crocc | 2268.532449 | 0.249401255 | 0.129727136 | 1.922506443 | 0.054542064 | 0.183632059 | Rps6ka3 | 1106.720843 | -0.033477316 | 0.089293517 | -0.374913174 | 0.707725041 | 0.929959854 |
| Plek | 699.0642541 | 0.24916027 | 0.125928895 | 1.978579024 | 0.047863423 | 0.169159644 | Mgat5b | 4626.939773 | -0.033485213 | 0.076617181 | -0.43704574 | 0.662078177 | 0.915428982 |
| Zdhhc2 | 1183.227332 | -0.183945346 | 0.084855925 | -2.167737202 | 0.030178687 | 0.124061709 | Otop3 | 4.88749934 | -0.033499243 | 0.045284587 | -0.739749324 | 0.459452113 | NA |
| Gmps | 1498.636654 | -0.159203919 | 0.073448285 | -2.167564829 | 0.030191812 | 0.124082742 | Ppie | 409.9375836 | -0.033509158 | 0.071023491 | -0.471803873 | 0.637066786 | 0.908932006 |
| Col4a2 | 2738.725012 | 0.249082591 | 0.123085881 | 2.023648759 | 0.043006304 | 0.15756436 | Tnk2 | 10948.64348 | -0.03351874 | 0.061877456 | -0.54169551 | 0.588028283 | 0.885188411 |
| Tmco1 | 1072.143361 | -0.175486315 | 0.081006394 | -2.166326728 | 0.030286229 | 0.124421022 | Pelp1 | 1653.068789 | -0.033548101 | 0.052135198 | -0.643482752 | 0.519910903 | 0.853297835 |
| Psma3 | 2974.071512 | -0.104196435 | 0.048102314 | -2.166141839 | 0.03030035 | 0.124422122 | Ubap2 | 2236.191356 | -0.033550483 | 0.050146294 | -0.669052098 | 0.503462245 | 0.847807553 |
| Ucp3 | 124.0103508 | 0.249051149 | 0.135447425 | 1.8387293 | 0.065955012 | 0.206738206 | Fgfr4 | 7.884167257 | -0.033556586 | 0.06032265 | -0.556285014 | 0.57801603 | NA |
| Psph | 516.4974505 | -0.189436127 | 0.087511551 | -2.164698555 | 0.030410778 | 0.12481711 | 1700086O06Rik | 116.6824124 | -0.033608618 | 0.093674137 | -0.358782256 | 0.719757987 | 0.932251852 |
| Uggt1 | 2930.191023 | -0.100544724 | 0.046453976 | -2.164394382 | 0.030434095 | 0.12487973 | BC056474 | 945.8924814 | -0.033645551 | 0.057933824 | -0.580758327 | 0.561403345 | 0.873934985 |
| Oas2 | 35.49565973 | 0.248957196 | 0.16384446 | 1.519472775 | 0.128643536 | 0.317241369 | Pydc3 | 2.743227701 | -0.033686816 | 0.034710764 | -0.970500556 | 0.331797048 | NA |
| 4930412C18Rik | 74.90942367 | 0.248916461 | 0.154741011 | 1.608600461 | 0.107703733 | 0.284383375 | BC024582 | 2.964245369 | -0.033698285 | 0.03820352 | -0.882072779 | 0.377737451 | NA |
| Plekhn1 | 207.0382843 | 0.24882339 | 0.147084711 | 1.691701251 | 0.09070295 | 0.253890779 | Gm10220 | 2.603009291 | -0.033701148 | 0.037476506 | -0.89926066 | 0.368513837 | NA |
| Pkd1l2 | 21.00920327 | 0.248783768 | 0.164884912 | 1.508832824 | 0.13134151 | 0.321334982 | Lpp | 372.5861268 | -0.033705699 | 0.090225952 | -0.37356989 | 0.708724339 | 0.93015267 |
| Mrpl17 | 921.7257126 | -0.171142186 | 0.07914584 | -2.162364888 | 0.030590062 | 0.125320576 | Gm7694 | 153.8197865 | -0.033729768 | 0.095024423 | -0.354958932 | 0.722620358 | 0.93255988 |
| Trim68 | 176.7693508 | 0.248730674 | 0.148538422 | 1.674520784 | 0.094028293 | 0.259322601 | Mir1969 | 2.054516533 | -0.033730464 | 0.033897127 | -0.995083249 | 0.319695777 | NA |
| Xkr4 | 789.4695445 | -0.223462837 | 0.103347007 | -2.16225745 | 0.030598338 | 0.125321343 | Aurkb | 8.930552684 | -0.033755371 | 0.064453191 | -0.523719157 | 0.60047389 | NA |
| Eno4 | 119.4745534 | 0.248726777 | 0.133013739 | 1.869932976 | 0.061493126 | 0.197589069 | Abcd4 | 536.0649424 | -0.033756756 | 0.079746117 | -0.423302811 | 0.672074343 | 0.919174457 |
| Cdon | 544.6461913 | -0.177771902 | 0.082233189 | -2.161802347 | 0.030633414 | 0.125372535 | Shank1 | 17622.55666 | -0.033779313 | 0.069042432 | -0.48925439 | 0.624661607 | 0.901254917 |
| Adam1a | 188.4424594 | 0.248655352 | 0.153830811 | 1.616420992 | 0.106003323 | 0.281283693 | Scara3 | 1499.410589 | -0.033784409 | 0.077661339 | -0.435022231 | 0.663546291 | 0.915428982 |
| Cryga | 11.60957235 | 0.248513498 | 0.166326278 | 1.494132502 | 0.13514099 | 0.327189831 | Anapc5 | 8454.438082 | -0.033795162 | 0.040548947 | -0.833441172 | 0.404595962 | 0.800290023 |
| Rph3al | 27.10653739 | 0.248390953 | 0.166532494 | 1.491546465 | 0.135818081 | 0.328140551 | Relb | 196.3395466 | -0.03381557 | 0.092470092 | -0.365691974 | 0.714594936 | 0.931210733 |
| Atrnl1 | 5897.582313 | -0.188175153 | 0.087123057 | -2.159877751 | 0.030782135 | 0.125826058 | Armcx5 | 998.3691025 | -0.033817362 | 0.068949496 | -0.490465689 | 0.623804403 | 0.900733873 |
| Cpxm1 | 269.5940423 | 0.248384144 | 0.111519134 | 2.227278278 | 0.025928681 | 0.111905011 | Cubn | 33.33847989 | -0.033828842 | 0.078117654 | -0.433049892 | 0.664978524 | 0.915428982 |
| Laptm5 | 1847.709138 | 0.248314523 | 0.126262927 | 1.966646334 | 0.049223998 | 0.172129261 | Hdx | 102.7657981 | -0.033832383 | 0.094667739 | -0.357380275 | 0.720807141 | 0.932513933 |
| Hrh2 | 210.7312372 | 0.248166321 | 0.099895645 | 2.484255655 | 0.012982257 | 0.070380366 | Polb | 1348.761451 | -0.033859673 | 0.06810141 | -0.497194886 | 0.619051627 | 0.898695177 |
| Vcam1 | 1240.248454 | -0.200790008 | 0.092978474 | -2.159532187 | 0.030808903 | 0.125851079 | Gm12060 | 3.897726663 | -0.033867059 | 0.038535073 | -0.878863236 | 0.379475434 | NA |
| Mtr | 485.3805122 | -0.20434941 | 0.094638776 | -2.159256681 | 0.030830259 | 0.125905121 | Gjc2 | 569.559234 | -0.033868061 | 0.089129341 | -0.37998778 | 0.703954486 | 0.928127792 |
| Mt1 | 4270.630358 | 0.248165825 | 0.096189298 | 2.579973353 | 0.009880794 | 0.058298942 | Vwa1 | 1164.474292 | -0.03386873 | 0.060118994 | -0.563361558 | 0.573188709 | 0.879310647 |
| Cpne4 | 2625.237474 | -0.245964323 | 0.113970216 | -2.158145632 | 0.030916511 | 0.126126654 | Nup210l | 19.23016232 | -0.033870168 | 0.077430594 | -0.437426176 | 0.661802304 | NA |
| Tpt1 | 13957.01151 | -0.135086271 | 0.062593885 | -2.158138479 | 0.030917067 | 0.126126654 | Sstr5 | 43.44835017 | -0.033872864 | 0.091302788 | -0.370994848 | 0.710641369 | 0.930507332 |
| Mov10 | 344.0133928 | 0.248015679 | 0.111689924 | 2.220573434 | 0.026379867 | 0.113252883 | Gm1078 | 67.94103402 | -0.033878471 | 0.094940615 | -0.35683855 | 0.721212675 | 0.932513933 |
| Elf4 | 86.52765036 | 0.248006321 | 0.130580095 | 1.899265892 | 0.057529525 | 0.189894005 | Sh3pxd2a | 2480.92748 | -0.033879458 | 0.067338047 | -0.503125047 | 0.614876358 | 0.896216533 |
| Nrsn1 | 16341.20936 | -0.1716593 | 0.079557709 | -2.157670224 | 0.030953481 | 0.126208744 | Vip | 1440.1732 | -0.03388376 | 0.076198691 | -0.44467641 | 0.65655362 | 0.915208181 |
| C920006O11Rik | 101.3385917 | 0.247973202 | 0.143307184 | 1.730361275 | 0.083565748 | 0.241156212 | Fbxw5 | 4034.172031 | -0.033891851 | 0.062058178 | -0.546130286 | 0.584976383 | 0.884313345 |
| BC026585 | 220.8586463 | 0.247962607 | 0.122824393 | 2.01883844 | 0.043504014 | 0.158861937 | Tmem235 | 1.030691626 | -0.033962279 | 0.024659147 | -1.377268987 | 0.1684291 | NA |
| Rimklb | 837.0055581 | -0.223689156 | 0.103719395 | -2.15667624 | 0.0310309 | 0.126424601 | Acpp | 11.63108551 | -0.033972776 | 0.057496442 | -0.590867448 | 0.554609238 | NA |
| Itsn1 | 7000.462147 | -0.157981748 | 0.073276796 | -2.155958725 | 0.031086888 | 0.12661057 | Gm10818 | 2.115918774 | -0.034004131 | 0.031622919 | -1.075300223 | 0.282240336 | NA |
| Lrrc56 | 140.5814483 | 0.247937519 | 0.131995718 | 1.878375477 | 0.060329818 | 0.195227694 | Hrh2 | 248.5009117 | -0.034005615 | 0.087200599 | -0.389969969 | 0.696558754 | 0.925559134 |
| Afg3l1 | 1201.194055 | -0.171476552 | 0.079567202 | -2.155116013 | 0.031152757 | 0.126821024 | Tmem140 | 73.59117716 | -0.034016418 | 0.091649596 | -0.371157321 | 0.710520359 | 0.930507332 |
| Nktr | 4122.652561 | 0.247895388 | 0.110269269 | 2.248091329 | 0.024570367 | 0.108182229 | Plekhm1 | 1531.579563 | -0.034020323 | 0.067504978 | -0.503967621 | 0.614284131 | 0.89611764 |
| C630031E19Rik | 8.382087779 | 0.24778992 | 0.143893501 | 1.722036914 | 0.085062832 | 0.243586371 | Kcnh1 | 5240.879835 | -0.034045098 | 0.071676138 | -0.474985113 | 0.634797584 | 0.907297112 |
| Lpar3 | 15.18389111 | -0.355501921 | 0.165015973 | -2.154348543 | 0.031212849 | 0.126965576 | Slk | 4382.972673 | -0.034052738 | 0.068584523 | -0.496507617 | 0.619536315 | 0.899129234 |
| Ptgr2 | 1821.818859 | -0.16889695 | 0.078413163 | -2.153936207 | 0.031245175 | 0.127063712 | Gucy1b2 | 8.7577185 | -0.034056733 | 0.05560449 | -0.612481701 | 0.540219102 | NA |
| 1700007F19Rik | 9.385540833 | -0.331857463 | 0.154142748 | -2.152922969 | 0.031324733 | 0.127353821 | Oma1 | 262.6430264 | -0.034076334 | 0.083161838 | -0.40975927 | 0.681982547 | 0.922544285 |
| Phactr4 | 514.3173277 | 0.247274775 | 0.105016147 | 2.35463576 | 0.01854087 | 0.089744528 | Snhg9 | 5.808687506 | -0.034082696 | 0.048156451 | -0.70774933 | 0.479100937 | NA |
| Mob3c | 211.4845485 | 0.24700404 | 0.105207693 | 2.347775457 | 0.018885903 | 0.090762462 | Drd5 | 84.58536221 | -0.034087194 | 0.089328984 | -0.381591648 | 0.70276428 | 0.927855161 |
| Lin9 | 160.3153697 | 0.246873967 | 0.118154376 | 2.089418745 | 0.036670045 | 0.141256 | Mtdh | 2534.115481 | -0.034092443 | 0.073267436 | -0.465315079 | 0.641705838 | 0.911095592 |
| Hps4 | 709.2835555 | 0.246830498 | 0.112862834 | 2.186995404 | 0.02874286 | 0.120102984 | Slfn1 | 1.376489281 | -0.03413666 | 0.025432611 | -1.342239694 | 0.17951829 | NA |
| Cnpy2 | 895.8461855 | -0.175583918 | 0.081614213 | -2.151388979 | 0.03144551 | 0.127612295 | Rfng | 2652.543718 | -0.0341441 | 0.070099771 | -0.487078629 | 0.626202611 | 0.902402691 |
| Emcn | 252.8276854 | -0.284150403 | 0.132079902 | -2.151352307 | 0.031448403 | 0.127612295 | BC021785 | 31.81521089 | -0.03414717 | 0.089554961 | -0.381298471 | 0.702981788 | 0.927855161 |
| Ascc2 | 672.8611184 | 0.246690875 | 0.09503656 | 2.595747106 | 0.009438551 | 0.05646367 | 2700046G09Rik | 108.110012 | -0.034151724 | 0.095053699 | -0.359288741 | 0.719379096 | 0.932251852 |
| Tmem246 | 2757.362831 | -0.126535627 | 0.058818794 | -2.151278824 | 0.031454199 | 0.127612295 | H2-Bl | 35.3620011 | -0.034158677 | 0.087133836 | -0.392025399 | 0.695039447 | 0.925527709 |
| 1700012D14Rik | 13.98948182 | 0.246669586 | 0.159453457 | 1.546969192 | 0.121870675 | 0.307885901 | Sat2 | 241.6503722 | -0.034161431 | 0.083827912 | -0.407518569 | 0.683627159 | 0.923110101 |
| Ost4 | 907.8199939 | -0.21681163 | 0.100809282 | -2.150710986 | 0.031499019 | 0.127727243 | Rars2 | 723.8722016 | -0.03417419 | 0.064393261 | -0.530710668 | 0.595619293 | 0.887497708 |
| AI854517 | 1112.35996 | 0.246495483 | 0.132017663 | 1.867140188 | 0.061882014 | 0.198427135 | Kiss1 | 15.76579921 | -0.034185095 | 0.070581987 | -0.484331726 | 0.628150463 | NA |
| AF357359 | 192.7130186 | 0.24618781 | 0.166057054 | 1.482549541 | 0.138194132 | 0.332017349 | Etohd2 | 24.62952025 | -0.034198923 | 0.084119991 | -0.406549295 | 0.684339044 | 0.923655038 |
| Blm | 167.0256856 | 0.246108471 | 0.118493294 | 2.076982279 | 0.037803195 | 0.144089225 | Cdc42ep2 | 507.5855451 | -0.034206531 | 0.084419419 | -0.405197419 | 0.685332402 | 0.923860706 |
| Vangl2 | 256.5011432 | -0.214502586 | 0.099763719 | -2.150106151 | 0.031546819 | 0.127809097 | Anxa9 | 37.50422041 | -0.034223473 | 0.086400864 | -0.396101052 | 0.692030482 | 0.923860706 |
| Sirt7 | 1041.731851 | 0.246087994 | 0.107238261 | 2.294777915 | 0.021745855 | 0.099477495 | Treml4 | 2.573964946 | -0.034251948 | 0.033278542 | -1.029250244 | 0.303362098 | NA |
| 1810021B22Rik | 27.90805086 | 0.245973203 | 0.163236429 | 1.506852391 | 0.131848497 | 0.321812686 | Gm5622 | 2.835227714 | -0.034270465 | 0.038367772 | -0.89320967 | 0.371744901 | NA |
| Spock3 | 3533.792728 | -0.169960277 | 0.079080596 | -2.149203288 | 0.031618289 | 0.127976428 | Gnpnat1 | 357.6021379 | -0.034289294 | 0.070712917 | -0.484908492 | 0.627741258 | 0.903544344 |
| Avpr1a | 16.05992101 | -0.322575475 | 0.150124128 | -2.148725058 | 0.031656201 | 0.128096417 | Mettl23 | 435.3405592 | -0.034290267 | 0.078091319 | -0.439104721 | 0.660585658 | 0.915428982 |
| Gmfb | 6240.88514 | -0.211655877 | 0.098521695 | -2.148317452 | 0.031688545 | 0.128193818 | Clec2i | 0.753793022 | -0.034294046 | 0.019346827 | -1.772592758 | 0.076296213 | NA |
| Bcl11a | 3115.532359 | -0.12476968 | 0.058104159 | -2.147345101 | 0.031765818 | 0.128439348 | Ampd3 | 1149.829412 | -0.034306576 | 0.055308856 | -0.620272742 | 0.535078237 | 0.860420883 |
| Ncf1 | 330.0553702 | 0.245922461 | 0.104418038 | 2.355172205 | 0.018514124 | 0.089659874 | Gjc3 | 1986.131971 | -0.034331305 | 0.080460143 | -0.426687097 | 0.66960725 | 0.91770119 |
| Dlx2 | 261.0675869 | -0.330615824 | 0.15400934 | -2.146725799 | 0.031815118 | 0.128605122 | Gnat2 | 2.682671864 | -0.034362568 | 0.035977103 | -0.955123281 | 0.339515357 | NA |
| Cfdp1 | 2588.755063 | -0.124974872 | 0.058225377 | -2.146398653 | 0.031841187 | 0.128676929 | Jph4 | 15315.35718 | -0.034369647 | 0.056111899 | -0.612519751 | 0.540193934 | 0.863544778 |
| 1500032L24Rik | 2159.602029 | -0.14373654 | 0.066983896 | -2.145837272 | 0.031885964 | 0.12881459 | Avpr1b | 2.241994274 | -0.034370927 | 0.028855343 | -1.191146001 | 0.233596274 | NA |
| Evl | 4412.839041 | -0.092590995 | 0.043152707 | -2.145659028 | 0.031900192 | 0.12881459 | BC052688 | 31.22208005 | -0.03437904 | 0.086509304 | -0.397402801 | 0.69107045 | 0.923860706 |
| Ovol2 | 193.9946592 | 0.245832726 | 0.14662403 | 1.676619622 | 0.093616887 | 0.258602255 | Rmst | 68.73692086 | -0.034392366 | 0.093642233 | -0.367274091 | 0.713414578 | 0.930507332 |
| Fanci | 24.887643 | 0.24581976 | 0.167317513 | 1.469181292 | 0.141783623 | 0.337323079 | Cox18 | 343.936979 | -0.034396269 | 0.078585207 | -0.437693943 | 0.661608161 | 0.915428982 |
| Ube2b | 4780.210052 | -0.131668946 | 0.061380497 | -2.145126752 | 0.031942714 | 0.128919096 | Phf10 | 972.3977261 | -0.034402813 | 0.067823879 | -0.507237466 | 0.61198821 | 0.895245244 |
| Nrip2 | 627.5544101 | 0.245791969 | 0.124328283 | 1.976959412 | 0.048046218 | 0.16965089 | Sbf1 | 9060.890295 | -0.03440616 | 0.058312297 | -0.590032669 | 0.555168748 | 0.871360989 |
| Slc27a3 | 129.5515956 | 0.245658421 | 0.139537321 | 1.760521265 | 0.078319465 | 0.23105137 | Mst1r | 59.42694795 | -0.034455569 | 0.092016646 | -0.374449301 | 0.708070069 | 0.929959854 |
| Cdc7 | 527.7060332 | 0.245618134 | 0.142079761 | 1.728734139 | 0.083856688 | 0.24166991 | Hpse2 | 1.951739169 | -0.034458581 | 0.030467371 | -1.130999517 | 0.258055295 | NA |
| Mmp16 | 840.4209679 | -0.175923226 | 0.082047971 | -2.144150843 | 0.032020802 | 0.129099743 | Ahctf1 | 2659.381714 | -0.034474277 | 0.068062392 | -0.506509927 | 0.612498723 | 0.89535639 |
| Tcf7 | 220.7814781 | -0.248466971 | 0.115895428 | -2.143889336 | 0.032041755 | 0.129150611 | Obp2b | 2.096591503 | -0.034475855 | 0.031716796 | -1.08699049 | 0.277041012 | NA |
| Mbd6 | 1771.428608 | 0.245499738 | 0.124931154 | 1.965080209 | 0.049404955 | 0.172599858 | Prelid2 | 7.729893033 | -0.034482423 | 0.057231262 | -0.602510262 | 0.546834534 | NA |
| Slc39a8 | 150.486718 | -0.285833982 | 0.133389652 | -2.142849742 | 0.032125166 | 0.129419479 | Rsl1 | 158.2020158 | -0.034485649 | 0.093681791 | -0.368114744 | 0.712787677 | 0.930507332 |
| Ypel1 | 1040.194633 | -0.229757057 | 0.107226049 | -2.142735459 | 0.032134346 | 0.129422813 | Rps15a-ps6 | 20.72409232 | -0.034492398 | 0.071777441 | -0.480546504 | 0.630838844 | NA |
| Crk | 5730.020793 | -0.136785917 | 0.063847258 | -2.14239299 | 0.032161871 | 0.12950001 | Bicc1 | 157.9998573 | -0.034513947 | 0.095099335 | -0.362925219 | 0.716660748 | 0.93125489 |
| Dact1 | 272.5672186 | -0.269576542 | 0.125851595 | -2.142019265 | 0.032191932 | 0.129587371 | Rpl36a | 2593.246186 | -0.034534402 | 0.071351353 | -0.484004862 | 0.628382418 | 0.904028802 |
| Klhl33 | 111.2078045 | 0.245443127 | 0.134868183 | 1.819874202 | 0.068778164 | 0.21246433 | Pgam2 | 162.1047821 | -0.03458623 | 0.094146841 | -0.367364736 | 0.713346972 | 0.930507332 |
| A330070K13Rik | 8.953398796 | 0.24534387 | 0.154843783 | 1.584460581 | 0.113088945 | 0.293993308 | Ceacam2 | 17.50371015 | -0.034588647 | 0.074218175 | -0.466040125 | 0.64118678 | NA |
| Dnahc10 | 63.29001885 | 0.245303977 | 0.165140206 | 1.485428553 | 0.137430337 | 0.330592398 | Pofut1 | 1031.483556 | -0.03459256 | 0.059635699 | -0.580064642 | 0.561871026 | 0.874222208 |
| Ano3 | 5943.717289 | -0.219979756 | 0.102759521 | -2.140723842 | 0.032296315 | 0.129817618 | 3010001F23Rik | 35.27948868 | -0.034620956 | 0.08875765 | -0.390061657 | 0.696490955 | 0.92555371 |
| Acan | 206.1497196 | 0.245217216 | 0.129416421 | 1.894792128 | 0.058119947 | 0.191110028 | Ccdc78 | 18.43796605 | -0.034634141 | 0.073403972 | -0.471829252 | 0.637048669 | NA |
| Arid5a | 402.4964374 | 0.245193778 | 0.136665037 | 1.794122213 | 0.072793664 | 0.220846883 | Smyd5 | 2323.517956 | -0.034644158 | 0.04583622 | -0.75582493 | 0.449754167 | 0.823263966 |
| Khk | 266.1523313 | 0.244798417 | 0.096268524 | 2.542870787 | 0.010994588 | 0.062791901 | Nsmf | 19109.04589 | -0.034650414 | 0.067201945 | -0.515616242 | 0.606122464 | 0.892684114 |
| Tal1 | 92.897028 | 0.244637791 | 0.146192514 | 1.673394794 | 0.094249602 | 0.259655643 | Cd79a | 10.11619103 | -0.03465442 | 0.063513498 | -0.54562291 | 0.585325173 | NA |
| Stk10 | 312.8997559 | 0.244600359 | 0.114054351 | 2.144594718 | 0.031985265 | 0.129057212 | Mir377 | 15.9916664 | -0.034654606 | 0.066307374 | -0.522635776 | 0.601227739 | NA |
| Zfp661 | 332.8856708 | 0.244570203 | 0.105403217 | 2.32032959 | 0.020323055 | 0.095179354 | Mrpl24 | 1246.338287 | -0.03468994 | 0.061576902 | -0.563359622 | 0.573190027 | 0.879310647 |
| Cacna1h | 3142.803043 | -0.206384923 | 0.096465576 | -2.139467056 | 0.032397861 | 0.130044679 | Fbxl20 | 2979.165324 | -0.034699314 | 0.052127321 | -0.665664625 | 0.505625489 | 0.84841357 |
| Dad1 | 1878.900998 | -0.136342985 | 0.063737411 | -2.139135892 | 0.032424664 | 0.130084865 | Casq2 | 99.30415449 | -0.034702277 | 0.090580957 | -0.383107862 | 0.70163979 | 0.92751465 |
| Sox6 | 980.9697261 | -0.158671739 | 0.074174645 | -2.13916412 | 0.032422379 | 0.130084865 | Limch1 | 5200.64355 | -0.034711818 | 0.051375029 | -0.675655439 | 0.499259434 | 0.846241311 |
| 2810008D09Rik | 302.6760673 | 0.244508595 | 0.129040273 | 1.894823905 | 0.058115735 | 0.191110028 | Pstpip2 | 432.1183724 | -0.034720362 | 0.087521322 | -0.396707464 | 0.691583196 | 0.923860706 |
| Ankrd23 | 97.24709786 | 0.244381866 | 0.153079791 | 1.596434539 | 0.11039181 | 0.289213849 | Ptprk | 3031.743985 | -0.034740187 | 0.056806165 | -0.611556641 | 0.540831133 | 0.863847445 |
| Prss23 | 807.4464755 | -0.276371626 | 0.129334667 | -2.136871979 | 0.032608406 | 0.130652867 | Muc2 | 6.903675111 | -0.034750788 | 0.056317172 | -0.617054924 | 0.537198499 | NA |
| Slc40a1 | 463.2404825 | -0.317214027 | 0.148444877 | -2.136914607 | 0.032604938 | 0.130652867 | Pdp2 | 314.8162547 | -0.034760695 | 0.08604362 | -0.40398922 | 0.686220646 | 0.923860706 |
| Spcs2 | 2695.72003 | -0.108284025 | 0.050669884 | -2.137049014 | 0.032594005 | 0.130652867 | Best2 | 1.645174031 | -0.034810943 | 0.027350382 | -1.27277724 | 0.2030971 | NA |
| Slc9a3 | 16.98757312 | 0.244308391 | 0.159240354 | 1.534211556 | 0.124977616 | 0.312625547 | Dnaaf1 | 19.37044385 | -0.034818902 | 0.080168352 | -0.434322283 | 0.664054424 | NA |
| Ccdc33 | 35.87404782 | 0.244255818 | 0.168066274 | 1.453330354 | 0.146132053 | 0.345028899 | Lypd6 | 1328.079387 | -0.034828155 | 0.065497258 | -0.531749821 | 0.594899281 | 0.887314166 |
| Mir341 | 8.448284105 | 0.244100317 | 0.149656406 | 1.631071625 | 0.102875207 | 0.275674241 | Zdhhc16 | 1066.128811 | -0.034832319 | 0.065495211 | -0.531830015 | 0.594843732 | 0.887314166 |
| Uqcrb | 2152.724217 | -0.165155774 | 0.077338333 | -2.135496957 | 0.032720438 | 0.130966281 | Pde1b | 16336.72887 | -0.034844922 | 0.078201685 | -0.44557764 | 0.655902365 | 0.915208181 |
| A530058N18Rik | 85.55421437 | -0.284378705 | 0.13321336 | -2.134761144 | 0.032780526 | 0.131105179 | Lipo1 | 204.7496045 | -0.034848746 | 0.086318755 | -0.403721609 | 0.686417448 | 0.923860706 |
| Mbp | 60860.84191 | 0.24399672 | 0.103758657 | 2.351579401 | 0.018693899 | 0.090108701 | Phldb3 | 3.56803334 | -0.034874842 | 0.044572413 | -0.78243111 | 0.433961256 | NA |
| Flywch1 | 8319.120313 | 0.243925749 | 0.096003878 | 2.540790584 | 0.011060214 | 0.062956268 | Pgpep1l | 22.29078339 | -0.034896549 | 0.075952704 | -0.459451049 | 0.645910296 | NA |
| Tmem120b | 253.2755958 | 0.243852919 | 0.110297793 | 2.210859457 | 0.027045571 | 0.115343814 | BC030307 | 69.63131569 | -0.034898117 | 0.094991265 | -0.367382377 | 0.713333815 | 0.930507332 |
| Tmco5 | 62.76067215 | -0.317871874 | 0.148968127 | -2.133824731 | 0.032857131 | 0.131343752 | Pdzk1 | 10.24380282 | -0.034914955 | 0.0621501 | -0.561784375 | 0.574262939 | NA |
| Dbpht2 | 1590.15741 | -0.180788399 | 0.084755382 | -2.13306099 | 0.032919723 | 0.131560019 | Cst6 | 149.307521 | -0.03491965 | 0.090557365 | -0.385608062 | 0.699786961 | 0.926781946 |
| Arhgap39 | 2893.848842 | 0.243804874 | 0.110577979 | 2.204823035 | 0.027466515 | 0.116615879 | A930018P22Rik | 1.744636652 | -0.034972147 | 0.026014952 | -1.34430951 | 0.178848322 | NA |
| H2-Ob | 23.2129287 | 0.243523642 | 0.16778348 | 1.451416087 | 0.146664033 | 0.345620169 | Gm20752 | 6.029504315 | -0.03499707 | 0.046375509 | -0.75464551 | 0.450461709 | NA |
| D17Wsu104e | 908.0261084 | -0.156197351 | 0.073280264 | -2.131506376 | 0.033047448 | 0.131968339 | Kdm1b | 385.9473133 | -0.035013725 | 0.075573752 | -0.463305364 | 0.643145501 | 0.911258473 |
| Ccdc163 | 105.0446581 | 0.243481942 | 0.160955149 | 1.512731605 | 0.130347848 | 0.319760855 | A730018C14Rik | 6.521334774 | -0.035014487 | 0.05184592 | -0.675356641 | 0.499449206 | NA |
| Tspan4 | 401.8621278 | 0.243314355 | 0.10142458 | 2.398968316 | 0.016441337 | 0.082842953 | 1700113A16Rik | 40.78105288 | -0.035020374 | 0.084701634 | -0.413455713 | 0.679272764 | 0.921471438 |
| Spp1 | 470.6882354 | 0.243183104 | 0.168119289 | 1.446491389 | 0.148039426 | 0.347586431 | Sypl | 1575.14594 | -0.03504927 | 0.058863922 | -0.59542873 | 0.551556926 | 0.869755927 |
| Ms4a6b | 43.70438488 | 0.242901223 | 0.167668733 | 1.448697193 | 0.147422165 | 0.346662471 | Slfn3 | 20.78917277 | -0.035053194 | 0.079487098 | -0.440992244 | 0.659218612 | NA |
| Adamts20 | 375.589424 | 0.242882019 | 0.143886561 | 1.688010449 | 0.091409223 | 0.254729227 | Nudcd1 | 592.4015767 | -0.035069696 | 0.069784985 | -0.502539271 | 0.615288236 | 0.896636929 |
| Smc5 | 1099.575572 | 0.242633942 | 0.091709728 | 2.645672889 | 0.008152861 | 0.050935658 | Arfgap3 | 846.3549613 | -0.035077549 | 0.073585139 | -0.476693384 | 0.633580475 | 0.906932171 |
| Ldlr | 1005.89088 | -0.282710226 | 0.13282453 | -2.128448907 | 0.033299882 | 0.13273691 | Baz1a | 142.2263022 | -0.035099575 | 0.095016301 | -0.369405826 | 0.711825256 | 0.930507332 |
| Ngb | 268.9008512 | 0.242539804 | 0.15379282 | 1.577055444 | 0.114782768 | 0.296753581 | Nras | 2116.090972 | -0.035142967 | 0.064107091 | -0.548191571 | 0.583560369 | 0.884313345 |
| Mif4gd | 360.4830886 | 0.242167759 | 0.094355308 | 2.566551525 | 0.010271536 | 0.060055541 | Emilin1 | 100.6967573 | -0.035159247 | 0.094473047 | -0.372161673 | 0.709772481 | 0.930224827 |
| Cox14 | 1304.567424 | -0.162555134 | 0.076411324 | -2.127369689 | 0.033389379 | 0.132974238 | Elf3 | 3.9942423 | -0.03519462 | 0.040878944 | -0.860947398 | 0.389267015 | NA |
| Cybb | 45.33866328 | 0.241927302 | 0.168118492 | 1.439028501 | 0.150142448 | 0.350370787 | 1700001C02Rik | 12.45136468 | -0.035197635 | 0.068428136 | -0.514373724 | 0.606990727 | NA |
| Prtg | 145.3713886 | 0.24166175 | 0.131118127 | 1.843084213 | 0.065316716 | 0.205426618 | Fmo1 | 312.8572329 | -0.035234133 | 0.088141484 | -0.399745177 | 0.689344214 | 0.923860706 |
| Dph1 | 490.8284242 | 0.241461236 | 0.098402522 | 2.45381146 | 0.014135107 | 0.074747218 | Mageb3 | 4.439427723 | -0.035246154 | 0.03559266 | -0.990264669 | 0.322044771 | NA |
| Elfn1 | 1310.65856 | -0.18230414 | 0.085756048 | -2.125845871 | 0.033516095 | 0.133358597 | Bnip1 | 648.7408361 | -0.035251617 | 0.061938759 | -0.569136639 | 0.569263415 | 0.87745335 |
| Vwa1 | 1101.852367 | -0.218734173 | 0.10292484 | -2.125183504 | 0.033571304 | 0.133543975 | Ttll3 | 404.6952139 | -0.035267177 | 0.093950884 | -0.375378876 | 0.707378713 | 0.92986829 |
| Gnai1 | 5219.941118 | -0.131434769 | 0.061853726 | -2.124928878 | 0.033592548 | 0.133581176 | Tmem256 | 867.6918605 | -0.035285247 | 0.063325907 | -0.557200813 | 0.577390234 | 0.881375635 |
| Gbp5 | 135.3492143 | 0.241442967 | 0.164352476 | 1.46905585 | 0.141817641 | 0.337323079 | Ddx39 | 505.1748351 | -0.035330638 | 0.072461797 | -0.487576064 | 0.625850153 | 0.902342364 |
| Gjb6 | 2136.287412 | 0.241260456 | 0.136394472 | 1.768843358 | 0.076920019 | 0.228357688 | 2610017I09Rik | 876.3191787 | -0.035331675 | 0.085813289 | -0.41172732 | 0.680539298 | 0.922327642 |
| A330050F15Rik | 251.7232627 | 0.241221715 | 0.13813586 | 1.746264259 | 0.080765043 | 0.235930162 | Ppip5k1 | 6460.580396 | -0.035343779 | 0.043332226 | -0.815646514 | 0.414702338 | 0.805020201 |
| Gm514 | 53.02135781 | 0.241114965 | 0.168035962 | 1.434900972 | 0.151315319 | 0.352288437 | Phf11a | 3.506920076 | -0.035345644 | 0.038214001 | -0.924939641 | 0.354997305 | NA |
| Fgfr1op | 589.25957 | 0.240970209 | 0.096992609 | 2.484418254 | 0.01297633 | 0.070380366 | Gemin6 | 171.3893674 | -0.035369113 | 0.088359872 | -0.400284789 | 0.68894677 | 0.923860706 |
| Abcg1 | 3352.703806 | 0.240963463 | 0.095892992 | 2.512837034 | 0.011976465 | 0.066761405 | Cd70 | 1.700390045 | -0.035389968 | 0.029096621 | -1.216291331 | 0.223873963 | NA |
| Mir421 | 14.68276627 | 0.24082588 | 0.152666204 | 1.57746688 | 0.114688137 | 0.296657428 | Tssc4 | 1052.323024 | -0.035408163 | 0.068358483 | -0.517977592 | 0.604473904 | 0.891930651 |
| Spag8 | 34.14225429 | 0.240806585 | 0.167069472 | 1.441355995 | 0.149484136 | 0.349812294 | Otog | 7.592172514 | -0.035435292 | 0.055635784 | -0.636915471 | 0.524179905 | NA |
| Zbtb41 | 841.5755393 | -0.183872378 | 0.086569341 | -2.123989569 | 0.033671016 | 0.133702991 | Gfra4 | 1867.594379 | -0.035458554 | 0.056169457 | -0.631278206 | 0.527858634 | 0.856965327 |
| Bex1 | 1063.092331 | -0.140067901 | 0.065983478 | -2.122772329 | 0.033772935 | 0.133967703 | Gabpa | 892.6517351 | -0.035458656 | 0.076557565 | -0.463163322 | 0.643247305 | 0.911258473 |
| Scd2 | 69037.32649 | -0.124472267 | 0.058640426 | -2.122635785 | 0.033784384 | 0.133978819 | Tmem176b | 1082.688639 | -0.035459498 | 0.086294877 | -0.410910814 | 0.681137932 | 0.922451017 |
| Akap3 | 9.703246949 | 0.240559855 | 0.160163117 | 1.501967867 | 0.133105408 | 0.323960915 | Snora47 | 0.890172945 | -0.035470781 | 0.022582988 | -1.57068587 | 0.116255633 | NA |
| Lmbr1l | 316.1864538 | 0.240556361 | 0.108368893 | 2.219791616 | 0.026432916 | 0.113328907 | Fads6 | 433.448382 | -0.035471551 | 0.0808547 | -0.438707348 | 0.660873602 | 0.915428982 |
| 1700047M11Rik | 137.4973179 | 0.240445908 | 0.166978412 | 1.439982002 | 0.149872491 | 0.350139382 | Fzd5 | 100.5709769 | -0.03550725 | 0.094735425 | -0.374804354 | 0.707805976 | 0.929959854 |
| Slc25a18 | 2144.488223 | 0.240378948 | 0.114013823 | 2.108331625 | 0.035002314 | 0.136916399 | Nrde2 | 752.438431 | -0.035527972 | 0.065595534 | -0.541621806 | 0.588079067 | 0.885188411 |
| Serpine2 | 7362.563866 | -0.129019944 | 0.060819141 | -2.121370712 | 0.033890618 | 0.134237196 | Wdfy2 | 334.8950578 | -0.035531176 | 0.080494943 | -0.441408794 | 0.658917077 | 0.915208181 |
| H2-DMb1 | 66.37869106 | 0.240214581 | 0.161679727 | 1.485743363 | 0.137347016 | 0.330443271 | Pxmp2 | 144.6131378 | -0.035534928 | 0.093986556 | -0.378085223 | 0.705367285 | 0.928783753 |
| B3galt6 | 431.4777387 | -0.243382035 | 0.114776717 | -2.120482632 | 0.033965365 | 0.134455636 | Rae1 | 1604.014402 | -0.035544058 | 0.051621447 | -0.688552136 | 0.491105151 | 0.841866734 |
| Rcor3 | 316.9149482 | 0.239877984 | 0.106141597 | 2.259980926 | 0.023822435 | 0.106095086 | Pax1 | 3.185137048 | -0.035569647 | 0.037628955 | -0.945273308 | 0.344519362 | NA |
| Rps27 | 24.71073765 | 0.239877794 | 0.168121362 | 1.426813287 | 0.15363372 | 0.35488508 | Dolpp1 | 713.8268362 | -0.03558711 | 0.076575589 | -0.464731782 | 0.642123546 | 0.911258473 |
| Fst | 125.0172191 | 0.239679681 | 0.125842828 | 1.904595474 | 0.056832675 | 0.188436866 | Cirbp | 2120.298096 | -0.035591077 | 0.085789301 | -0.41486615 | 0.678239891 | 0.921316909 |
| Adamtsl4 | 303.812651 | 0.239553883 | 0.158585626 | 1.510564916 | 0.130899338 | 0.320555452 | Zfp608 | 838.0494475 | -0.035595304 | 0.080735216 | -0.44088944 | 0.659293039 | 0.915208181 |
| Mfsd7b | 409.0526651 | 0.2394743 | 0.100810099 | 2.375499108 | 0.017525235 | 0.086311503 | Brwd3 | 357.4567916 | -0.035595561 | 0.087710773 | -0.405828835 | 0.68486837 | 0.923799605 |
| Grm8 | 689.702143 | -0.218380042 | 0.103052155 | -2.119121558 | 0.034080195 | 0.134650794 | G2e3 | 464.2004146 | -0.0356026 | 0.084854779 | -0.419570948 | 0.674798915 | 0.919907397 |
| Krt9 | 560.4347328 | -0.238687063 | 0.11263742 | -2.119074298 | 0.034084188 | 0.134650794 | Ikbkap | 3529.200529 | -0.035634189 | 0.044884101 | -0.793915621 | 0.427244555 | 0.810334558 |
| Nucks1 | 4583.423902 | -0.114280163 | 0.053928729 | -2.11909618 | 0.034082339 | 0.134650794 | Sdad1 | 1005.894066 | -0.035659173 | 0.061555594 | -0.57930028 | 0.562386577 | 0.874649616 |
| Atp6v0a2 | 1955.901211 | -0.114580418 | 0.054074652 | -2.118930285 | 0.034096359 | 0.134664539 | Pyroxd1 | 521.2595987 | -0.035675011 | 0.074890292 | -0.476363622 | 0.633815346 | 0.906984703 |
| Sema3b | 139.7602293 | 0.23917753 | 0.165876814 | 1.441898505 | 0.149331008 | 0.349559546 | Lmnb1 | 284.5565653 | -0.035695069 | 0.078262257 | -0.456095571 | 0.648321255 | 0.91329006 |
| Stard6 | 68.74060725 | 0.239120501 | 0.141269276 | 1.692657504 | 0.090520679 | 0.253575736 | Akap7 | 1255.625868 | -0.035716642 | 0.062328794 | -0.573035993 | 0.566620324 | 0.875959314 |
| Tmem50a | 2850.485168 | -0.126500733 | 0.059721187 | -2.118188534 | 0.034159104 | 0.13480926 | Tmem176a | 533.289147 | -0.035723829 | 0.090360492 | -0.395347883 | 0.692586166 | 0.92408438 |
| Timm8a1 | 454.9576045 | -0.20390612 | 0.096270249 | -2.118059538 | 0.034170026 | 0.134818023 | 4931423N10Rik | 6.442202124 | -0.03576402 | 0.047208915 | -0.757569183 | 0.448708935 | NA |
| Aaas | 504.3766696 | 0.239070128 | 0.093153461 | 2.566411659 | 0.01027568 | 0.060057103 | Zfp128 | 391.7335756 | -0.03578149 | 0.078957072 | -0.453176505 | 0.650421649 | 0.914000665 |
| 4430402I18Rik | 127.6785205 | 0.238942977 | 0.135907003 | 1.758135876 | 0.078724393 | 0.231932319 | Hexa | 2478.786815 | -0.035797835 | 0.067612315 | -0.529457315 | 0.596488248 | 0.887922451 |
| Gins3 | 105.321586 | -0.264198333 | 0.124830979 | -2.116448469 | 0.034306683 | 0.135219474 | Gm6150 | 1.389506481 | -0.035811084 | 0.026946602 | -1.328964739 | 0.183859603 | NA |
| Med28 | 1866.959401 | -0.153260739 | 0.072411867 | -2.116514128 | 0.034301105 | 0.135219474 | Tmem88 | 111.0816766 | -0.035822181 | 0.094325461 | -0.379772131 | 0.704114571 | 0.928127792 |
| Col4a1 | 2146.192049 | 0.238851785 | 0.113465747 | 2.105056294 | 0.035286408 | 0.137714762 | Gdf11 | 307.5632462 | -0.035830477 | 0.081146263 | -0.441554248 | 0.658811798 | 0.915208181 |
| Cntnap5b | 582.6756277 | -0.247419356 | 0.116926732 | -2.116020449 | 0.034343068 | 0.13525966 | Bsph1 | 1.069612833 | -0.03584822 | 0.021729521 | -1.649747383 | 0.098994615 | NA |
| Gm16675 | 8.892258581 | 0.238735605 | 0.156244148 | 1.527965098 | 0.126521204 | 0.314473278 | Sec24c | 4396.030741 | -0.035860379 | 0.04371445 | -0.8203324 | 0.412026641 | 0.802795661 |
| Chrnb1 | 33.31580632 | 0.238639109 | 0.165783043 | 1.439466332 | 0.150018442 | 0.350370787 | Prim1 | 236.1473986 | -0.035860383 | 0.091413243 | -0.392288709 | 0.694844905 | 0.925527709 |
| Cenpb | 5206.01625 | -0.141645111 | 0.066959877 | -2.115372934 | 0.034398175 | 0.135407856 | Snora17 | 9.102667517 | -0.035861077 | 0.057755508 | -0.620911808 | 0.534657652 | NA |
| Coa4 | 75.58419141 | -0.318092756 | 0.150423219 | -2.114651971 | 0.034459621 | 0.135580844 | 8030442B05Rik | 2.221277528 | -0.035887038 | 0.033056713 | -1.085620283 | 0.277647022 | NA |
| Gsg2 | 8.813447853 | -0.326455306 | 0.154371431 | -2.11473913 | 0.034452187 | 0.135580844 | Ing4 | 1483.557911 | -0.035893463 | 0.057515635 | -0.624064449 | 0.53258526 | 0.858328942 |
| Mtch1 | 13618.66032 | -0.1249414 | 0.059090352 | -2.114412843 | 0.034480022 | 0.135626671 | D730005E14Rik | 5.125161974 | -0.035903773 | 0.041756564 | -0.85983541 | 0.389879777 | NA |
| Gm13845 | 77.19867594 | -0.297208446 | 0.140572827 | -2.1142667 | 0.034492495 | 0.135641298 | Kdm3b | 2794.420598 | -0.035914834 | 0.044816059 | -0.801383131 | 0.422909878 | 0.808999423 |
| Slc39a10 | 10441.43147 | -0.190042271 | 0.089899384 | -2.113944079 | 0.034520044 | 0.135715189 | Npepl1 | 517.3958374 | -0.035936381 | 0.080620122 | -0.445749518 | 0.655778191 | 0.915208181 |
| Unc79 | 3327.730961 | -0.175094186 | 0.082859111 | -2.113155489 | 0.034587462 | 0.135945746 | Ick | 1139.948388 | -0.035939521 | 0.066803181 | -0.537991168 | 0.590583146 | 0.885693151 |
| Crh | 156.4182257 | -0.298948569 | 0.141483793 | -2.112952744 | 0.034604813 | 0.135979449 | Snord83b | 5.184917167 | -0.036001299 | 0.052872168 | -0.680912114 | 0.495927103 | NA |
| Arpc5 | 2849.237723 | -0.169123045 | 0.080058778 | -2.112485973 | 0.034644788 | 0.136074988 | Snord43 | 0.917895365 | -0.036012111 | 0.022690264 | -1.58711733 | 0.112486074 | NA |
| Camkv | 21322.28554 | -0.143717619 | 0.068034471 | -2.112423559 | 0.034650136 | 0.136074988 | 4930525D18Rik | 0.945476319 | -0.036014661 | 0.022293698 | -1.61546375 | 0.106210304 | NA |
| Hsd11b1 | 405.5762954 | 0.238585833 | 0.136695646 | 1.745379903 | 0.080918762 | 0.236238712 | Creg2 | 7177.531454 | -0.036031663 | 0.070545555 | -0.510757384 | 0.609520954 | 0.89440021 |
| Shc4 | 279.8511066 | -0.224452601 | 0.106278014 | -2.111938239 | 0.034691747 | 0.136182906 | Plbd2 | 3808.60445 | -0.03604018 | 0.059397605 | -0.606761495 | 0.544009205 | 0.86604998 |
| Ccnh | 1231.281942 | -0.170079393 | 0.080563575 | -2.111120241 | 0.034761978 | 0.136424034 | Exoc3l | 26.52875741 | -0.036057052 | 0.081650105 | -0.441604481 | 0.658775441 | 0.915208181 |
| Fam159b | 12.77840498 | 0.238489955 | 0.16426088 | 1.451897465 | 0.146530118 | 0.345508696 | Dnase1l3 | 10.63385292 | -0.036075905 | 0.051289181 | -0.703382342 | 0.481817508 | NA |
| Plekhb2 | 11615.06826 | -0.091263217 | 0.043241388 | -2.110552441 | 0.034810799 | 0.136517045 | Gnb1l | 171.7942033 | -0.036086681 | 0.087615387 | -0.411876065 | 0.680430265 | 0.922298495 |
| Stard13 | 562.6615297 | 0.238439508 | 0.118576175 | 2.010855105 | 0.044340764 | 0.160962837 | Cxcr3 | 2.753435642 | -0.036088331 | 0.034822945 | -1.036337702 | 0.300044623 | NA |
| Dtx3l | 286.2162849 | 0.238376962 | 0.112222907 | 2.124138187 | 0.03365859 | 0.133702991 | Ivd | 2538.754856 | -0.036094006 | 0.062508196 | -0.577428383 | 0.563650105 | 0.875120313 |
| Zfp608 | 907.356129 | -0.22061875 | 0.104552115 | -2.110131874 | 0.034846999 | 0.136584719 | Rgp1 | 2385.926871 | -0.036097189 | 0.067804535 | -0.532371312 | 0.59446885 | 0.887214068 |
| Sptbn2 | 46970.45474 | -0.19170332 | 0.090856201 | -2.109964078 | 0.03486145 | 0.136606805 | Pxmp4 | 446.1113049 | -0.036128905 | 0.078903212 | -0.457888899 | 0.647032262 | 0.912750424 |
| Hcn3 | 659.0854656 | -0.264051746 | 0.125196686 | -2.109095332 | 0.034936354 | 0.136796529 | Gm10785 | 12.54066838 | -0.036155756 | 0.06763577 | -0.534565601 | 0.592950275 | NA |
| Ppp1r3b | 60.83814376 | -0.333822599 | 0.158277189 | -2.109101137 | 0.034935853 | 0.136796529 | Bglap3 | 11.55314369 | -0.036189215 | 0.066387172 | -0.545123614 | 0.585668503 | NA |
| Larp1b | 484.5853246 | 0.238364649 | 0.101053756 | 2.358790577 | 0.018334599 | 0.089052126 | Tspan14 | 1793.641982 | -0.036193283 | 0.061063852 | -0.592712084 | 0.553373853 | 0.870600462 |
| Zfp473 | 20.98258571 | 0.237900756 | 0.16656559 | 1.428270727 | 0.15321395 | 0.354648861 | 1700023L04Rik | 6.222151013 | -0.036212156 | 0.053547921 | -0.676256999 | 0.498877489 | NA |
| S1pr4 | 21.01762786 | -0.354352448 | 0.168057935 | -2.108513638 | 0.034986584 | 0.136916399 | Gm5741 | 3.458912933 | -0.036219313 | 0.044322836 | -0.817170482 | 0.413831009 | NA |
| Apbb2 | 2084.027992 | 0.237634991 | 0.094981039 | 2.501920325 | 0.012352172 | 0.068108382 | Ap3s1 | 1213.010322 | -0.036236481 | 0.074387865 | -0.487128927 | 0.626166968 | 0.902402691 |
| Trim39 | 1034.30367 | 0.237531354 | 0.109550277 | 2.168240567 | 0.030140388 | 0.123954281 | Gabrd | 2951.430796 | -0.036242277 | 0.068946455 | -0.525658316 | 0.599125631 | 0.889150487 |
| 2610306M01Rik | 164.644197 | -0.220220185 | 0.104476238 | -2.10784949 | 0.03504401 | 0.137044901 | Zfp605 | 603.455958 | -0.036276333 | 0.07696302 | -0.471347573 | 0.637392548 | 0.908932006 |
| Otud3 | 370.8996506 | 0.237510325 | 0.094676272 | 2.508657339 | 0.012119098 | 0.067205909 | Gm10433 | 12.71088226 | -0.036279062 | 0.067935583 | -0.534021499 | 0.59332666 | NA |
| Mrpl19 | 1162.108425 | -0.124755558 | 0.059201774 | -2.107294247 | 0.035092082 | 0.137163651 | Klhl36 | 225.3818599 | -0.036284749 | 0.082150639 | -0.441685538 | 0.658716776 | 0.915208181 |
| Gm16532 | 363.1437107 | -0.238761384 | 0.113314492 | -2.10706839 | 0.035111652 | 0.137205532 | Gm9159 | 10.53945768 | -0.036314613 | 0.060872647 | -0.596567013 | 0.5507965 | NA |
| Pcdhb16 | 298.4048613 | -0.236387646 | 0.112224897 | -2.106374362 | 0.035171847 | 0.137406099 | Cpeb4 | 2136.11034 | -0.036330476 | 0.062827815 | -0.578254641 | 0.563092214 | 0.87510382 |
| Fam96b | 467.3104517 | -0.150167119 | 0.071299882 | -2.106134185 | 0.035192698 | 0.137452903 | Arvcf | 1554.134042 | -0.036356488 | 0.064002195 | -0.568050649 | 0.570000577 | 0.8778444 |
| Olfm4 | 26.32873218 | 0.237413377 | 0.166221325 | 1.428296739 | 0.153206466 | 0.354648861 | Cdca2 | 12.33035596 | -0.036406824 | 0.066307148 | -0.549063335 | 0.582961986 | NA |
| 1810024B03Rik | 13.25266823 | 0.236900952 | 0.164011081 | 1.44442041 | 0.148620753 | 0.348423354 | Nsd1 | 6768.020628 | -0.036424779 | 0.063607261 | -0.572651271 | 0.566880837 | 0.875959314 |
| Mir3101 | 14.14223009 | 0.236800373 | 0.167687111 | 1.412156075 | 0.157904011 | 0.360746278 | 1500015L24Rik | 21.48047971 | -0.036447178 | 0.076187784 | -0.478386114 | 0.632375415 | NA |
| Ankrd13c | 2029.176599 | -0.147584055 | 0.070116083 | -2.104853098 | 0.035304097 | 0.137749103 | Zfp235 | 795.6725572 | -0.036451822 | 0.06636554 | -0.549258274 | 0.582828218 | 0.88406516 |
| Aph1b | 1152.370522 | -0.184948802 | 0.087881153 | -2.104533184 | 0.035331963 | 0.137823121 | Gripap1 | 4444.940047 | -0.036454223 | 0.047346838 | -0.76993997 | 0.441335503 | 0.818962636 |
| Six2 | 15.82095811 | -0.347471306 | 0.165118456 | -2.104375943 | 0.035345666 | 0.137841871 | Soat1 | 882.862521 | -0.036483684 | 0.067610712 | -0.539613964 | 0.589463284 | 0.885188411 |
| Pacrg | 389.5055084 | -0.204618257 | 0.097265478 | -2.103708956 | 0.035403843 | 0.138034006 | Gtpbp6 | 925.8955132 | -0.036483886 | 0.077434149 | -0.471160163 | 0.637526364 | 0.908932006 |
| 2810454H06Rik | 16.34501357 | 0.236585838 | 0.167012079 | 1.416579208 | 0.156605991 | 0.359261656 | C130030K03Rik | 150.607495 | -0.036509416 | 0.091438396 | -0.39927883 | 0.689687764 | 0.923860706 |
| D330023K18Rik | 84.87307231 | -0.282915503 | 0.13452858 | -2.10301412 | 0.035464535 | 0.138201084 | Nradd | 50.61781557 | -0.036524162 | 0.09404414 | -0.388372548 | 0.697740357 | 0.926197242 |
| Ap2b1 | 18164.42652 | -0.12870801 | 0.061205065 | -2.102898016 | 0.035474685 | 0.138205877 | Pan3 | 1383.827862 | -0.036533069 | 0.071425654 | -0.511483855 | 0.609012291 | 0.89440021 |
| Zfp36 | 211.9632348 | 0.236575276 | 0.123988723 | 1.908038651 | 0.056386221 | 0.187518375 | Rbp3 | 4.887548185 | -0.036580043 | 0.044896088 | -0.814771278 | 0.415203244 | NA |
| Stk36 | 196.1488964 | 0.236499321 | 0.122747234 | 1.926718127 | 0.054014768 | 0.182650548 | 1700030C10Rik | 47.57827614 | -0.036588376 | 0.091876827 | -0.398232905 | 0.690458513 | 0.923860706 |
| Ninl | 274.9449821 | 0.236374537 | 0.110610234 | 2.137004219 | 0.032597648 | 0.130652867 | Zc3h6 | 845.0874689 | -0.036604142 | 0.078306468 | -0.467447236 | 0.640179934 | 0.910092095 |
| Nme3 | 498.88601 | -0.169691223 | 0.080734604 | -2.101840024 | 0.035567292 | 0.138398496 | 6030407O03Rik | 12.91768032 | -0.036667695 | 0.064190533 | -0.571232126 | 0.567842304 | NA |
| Plxna2 | 7399.574262 | -0.219102504 | 0.104244035 | -2.101822941 | 0.035568789 | 0.138398496 | 4930431F12Rik | 2.225650977 | -0.036721437 | 0.029610205 | -1.240161528 | 0.214915655 | NA |
| Frrs1 | 34.09456501 | 0.235973393 | 0.162442516 | 1.452657826 | 0.146318782 | 0.345180316 | Rasgef1a | 12153.44476 | -0.036736151 | 0.072766981 | -0.504846436 | 0.613666698 | 0.895819717 |
| Rora | 422.3171381 | 0.235699476 | 0.114210417 | 2.063730101 | 0.039043314 | 0.147258302 | 6330418K02Rik | 103.1578767 | -0.036743396 | 0.095080523 | -0.386445035 | 0.699167103 | 0.926567041 |
| Snord22 | 15.79444523 | 0.235612996 | 0.165913248 | 1.420097542 | 0.155579286 | 0.357647694 | Spg20 | 1759.757005 | -0.036760438 | 0.06525257 | -0.563356164 | 0.57319238 | 0.879310647 |
| Mrps28 | 258.853129 | -0.189892783 | 0.090394247 | -2.100717583 | 0.035665765 | 0.138636568 | Nsun6 | 300.8896949 | -0.036772751 | 0.077308317 | -0.475663579 | 0.634314072 | 0.907220906 |
| Pdxk | 9376.750051 | -0.142052554 | 0.067630061 | -2.100435111 | 0.035690583 | 0.138698243 | Nudt16 | 465.4821431 | -0.036815404 | 0.074769302 | -0.492386613 | 0.622446061 | 0.900114632 |
| Col11a1 | 740.0341048 | 0.23554942 | 0.16372862 | 1.438657574 | 0.150247565 | 0.350539946 | Coro1b | 2753.538314 | -0.03682659 | 0.053913751 | -0.683064874 | 0.49456585 | 0.843609829 |
| Tmem144 | 364.3399557 | 0.235535402 | 0.100636854 | 2.340448785 | 0.01926058 | 0.091850854 | Spag17 | 28.27072378 | -0.036826822 | 0.078475384 | -0.469278646 | 0.638870477 | 0.909380945 |
| Hcls1 | 207.587284 | 0.235534442 | 0.14013538 | 1.680763574 | 0.092808847 | 0.257149547 | Ocel1 | 195.9780444 | -0.036837396 | 0.088527713 | -0.416111459 | 0.677328445 | 0.921064066 |
| Shroom2 | 4334.40958 | -0.158300015 | 0.075449051 | -2.098104792 | 0.035895889 | 0.139356279 | AI197445 | 37.85354342 | -0.036867562 | 0.089358248 | -0.412581522 | 0.679913241 | 0.921823758 |

| | | | | | | | | | | | | | | | |
|---|---|---|---|---|---|---|---|---|---|---|---|---|---|---|---|
| Lmo2 | 707.1828253 | -0.182322705 | 0.08690583 | -2.097934116 | 0.035910966 | 0.139356415 | | | Tmem51as1 | 4.350639713 | -0.036876636 | 0.04758138 | -0.775022422 | 0.43832641 | NA |
| Crebrf | 2340.210071 | 0.235486087 | 0.10212464 | 2.305869438 | 0.021117929 | 0.097526435 | | | C4bp-ps1 | 5.12714124 | -0.03688577 | 0.045277043 | -0.814668268 | 0.415262222 | NA |
| Mapk1 | 26624.26341 | -0.14331351 | 0.06832622 | -2.097489219 | 0.035950291 | 0.139462645 | | | Mir697 | 7.55552617 | -0.036890854 | 0.052039988 | -0.708894361 | 0.478390034 | NA |
| Gm608 | 3367.322114 | -0.187122987 | 0.089230745 | -2.097068532 | 0.035987509 | 0.139502243 | | | Gm4461 | 1.501906774 | -0.03689719 | 0.028622513 | -1.289096835 | 0.197364424 | NA |
| Mpdu1 | 1268.129219 | -0.125709975 | 0.059944062 | -2.097121381 | 0.035982832 | 0.139502243 | | | Plek2 | 15.16473187 | -0.036898062 | 0.074515197 | -0.495174988 | 0.620476605 | NA |
| Aspdh | 70.01622749 | 0.235485771 | 0.137741791 | 1.709617466 | 0.087336634 | 0.247497921 | | | Rasa3 | 3989.570226 | -0.036927006 | 0.048759192 | -0.757334255 | 0.448849633 | 0.823263966 |
| Adm | 67.78721592 | 0.235468475 | 0.164510115 | 1.431331287 | 0.152335291 | 0.353735457 | | | Bmp8b | 6.66869795 | -0.036931882 | 0.051793264 | -0.713063422 | 0.475806512 | NA |
| Gm6277 | 128.5533372 | 0.23542559 | 0.1287997 | 1.827842693 | 0.067573164 | 0.209899479 | | | Ivns1abp | 7768.071221 | -0.036935874 | 0.071961301 | -0.513274126 | 0.60775958 | 0.893916126 |
| Efr3a | 5894.387988 | 0.235314772 | 0.096565769 | 2.436834244 | 0.014816469 | 0.077274849 | | | Ccdc164 | 751.6754194 | -0.037014238 | 0.0639777 | -0.578549056 | 0.562893488 | 0.87510382 |
| 9330020H09Rik | 57.48230374 | 0.235052301 | 0.152351787 | 1.542826014 | 0.122872996 | 0.30949535 | | | Mef2a | 3344.526183 | -0.037026867 | 0.068573038 | -0.53996247 | 0.589222915 | 0.885188411 |
| Nrcam | 14856.3349 | -0.134758598 | 0.064323346 | -2.095018469 | 0.036169351 | 0.140031964 | | | Frat1 | 418.3901945 | -0.037032823 | 0.085521407 | -0.433024013 | 0.664997324 | 0.915428982 |
| Hnmt | 623.5794288 | -0.159824358 | 0.076291791 | -2.094908971 | 0.036179085 | 0.140034661 | | | E330017L17Rik | 2.423488993 | -0.037033105 | 0.03798264 | -0.975000818 | 0.329559854 | NA |
| Smim3 | 266.2875888 | 0.235028424 | 0.136566497 | 1.720981561 | 0.08525417 | 0.243893179 | | | Tnnt1 | 123.0218306 | -0.037052904 | 0.092834941 | -0.399126706 | 0.689799845 | 0.923860706 |
| Lemd1 | 34.69114722 | -0.348274167 | 0.166350251 | -2.093619731 | 0.036293868 | 0.140373739 | | | Capn1 | 1104.069116 | -0.037057076 | 0.059118076 | -0.626831562 | 0.530769652 | 0.858116381 |
| Olfr856-ps1 | 145.9329317 | 0.234974569 | 0.126747902 | 1.853873438 | 0.063757269 | 0.202275644 | | | Snora64 | 19.91761423 | -0.037081638 | 0.061308006 | -0.604841694 | 0.545284187 | NA |
| Fam228b | 82.57758252 | 0.234938455 | 0.153473951 | 1.530803465 | 0.125817973 | 0.313815788 | | | Mir761 | 7.789700109 | -0.037094061 | 0.057390794 | -0.646341665 | 0.51805811 | NA |
| Kcnj13 | 96.45954198 | 0.234910057 | 0.132100423 | 1.778268776 | 0.075359722 | 0.225627736 | | | Ddn | 42259.69258 | -0.037096112 | 0.074207715 | -0.499895626 | 0.617148572 | 0.897621051 |
| Dlx6 | 312.3877761 | -0.30106151 | 0.143873574 | -2.092542096 | 0.03639005 | 0.140570291 | | | Snord15b | 2.918914568 | -0.037106268 | 0.03590595 | -1.033429496 | 0.301402949 | NA |
| Il15ra | 142.4527134 | 0.234679826 | 0.116164043 | 2.020244992 | 0.043357982 | 0.158478076 | | | Ppm1j | 32.70240461 | -0.037115534 | 0.084949355 | -0.436913668 | 0.662173959 | 0.915428982 |
| Cdhr2 | 25.73751161 | 0.234604721 | 0.168067685 | 1.395894291 | 0.16274633 | 0.367900735 | | | DQ267100 | 10.80658683 | -0.037122658 | 0.063159221 | -0.587763079 | 0.556691329 | NA |
| Cnot6l | 674.8338572 | -0.201885377 | 0.09648741 | -2.092349423 | 0.03640727 | 0.140601753 | | | Trim36 | 323.929746 | -0.0371308 | 0.088528531 | -0.41942185 | 0.674907858 | 0.919907397 |
| Atrn | 10964.70067 | -0.147997934 | 0.070741493 | -2.092095154 | 0.036430005 | 0.140654496 | | | 1600029D21Rik | 7.669103177 | -0.037149915 | 0.054600131 | -0.68039974 | 0.496251386 | NA |
| Ssfa2 | 2341.970013 | 0.23448622 | 0.089395045 | 2.623033735 | 0.008715064 | 0.053413742 | | | Msl1 | 6812.601661 | -0.037154611 | 0.061362657 | -0.605492209 | 0.544852 | 0.866307897 |
| BC023829 | 1684.395711 | -0.148095199 | 0.070813887 | -2.091329905 | 0.036498501 | 0.14084876 | | | Fam57a | 227.94405 | -0.037180557 | 0.086772007 | -0.428485618 | 0.668297602 | 0.917031076 |
| Ccrn4l | 1920.709929 | -0.151046686 | 0.072229716 | -2.091198668 | 0.036510259 | 0.140859051 | | | Gm11517 | 17.14904064 | -0.037187558 | 0.077409673 | -0.480399375 | 0.630943439 | NA |
| Ralgapb | 7204.384797 | -0.139651299 | 0.066816747 | -2.090064319 | 0.036612022 | 0.141216498 | | | Dsg1b | 1.315972644 | -0.037189107 | 0.023935429 | -1.553726365 | 0.120249701 | NA |
| Arhgap28 | 36.64853367 | -0.349432309 | 0.167249526 | -2.08928729 | 0.03668187 | 0.141256 | | | Cdh15 | 85.24311831 | -0.03720383 | 0.09507817 | -0.391297288 | 0.695577503 | 0.925527709 |
| Txlnb | 93.76386259 | 0.234075461 | 0.152935189 | 1.530553317 | 0.125879827 | 0.313817527 | | | 3110002H16Rik | 1039.863742 | -0.03721687 | 0.054605921 | -0.681553736 | 0.495521177 | 0.844369474 |
| Haus6 | 222.4744739 | 0.234055149 | 0.10266909 | 2.279704126 | 0.022625242 | 0.102411775 | | | Klb | 7.817126689 | -0.037218168 | 0.058532669 | -0.635852922 | 0.52487229 | NA |
| Upf3b | 753.8853479 | 0.233964094 | 0.094529104 | 2.475048264 | 0.013321817 | 0.071620597 | | | Specc1l | 2746.471139 | -0.037220644 | 0.059841352 | -0.621988684 | 0.533949309 | 0.859470725 |
| Lpar1 | 1030.004795 | -0.251821759 | 0.120511983 | -2.08959933 | 0.036653807 | 0.141256 | | | Atp2a1 | 18.87376361 | -0.037222355 | 0.067613532 | -0.550516346 | 0.581965268 | NA |
| Paip2 | 4003.920561 | -0.13919817 | 0.066609336 | -2.089769666 | 0.036638495 | 0.141256 | | | Snora7a | 23.66617941 | -0.03722512 | 0.084269029 | -0.441741408 | 0.658676342 | 0.915208181 |
| Vcpip1 | 2361.674583 | -0.163547228 | 0.07828071 | -2.08924048 | 0.036686081 | 0.141256 | | | 1700012B09Rik | 12.24050919 | -0.037232554 | 0.058912798 | -0.631994328 | 0.527390584 | NA |
| Vars2 | 962.9283214 | 0.233949595 | 0.089555844 | 2.612331968 | 0.008992689 | 0.054661717 | | | Pqlc2 | 404.6375803 | -0.037262029 | 0.081831835 | -0.455348816 | 0.648858312 | 0.913724196 |
| Emp3 | 74.40874918 | -0.295604849 | 0.141556801 | -2.08824194 | 0.036776016 | 0.14153194 | | | Zfp119a | 108.7530164 | -0.037263523 | 0.093796184 | -0.39728187 | 0.691159615 | 0.923860706 |
| Ttll7 | 9320.98452 | -0.160881823 | 0.0770463 | -2.088118752 | 0.036787124 | 0.141539532 | | | Col1a2 | 743.2950451 | -0.037277041 | 0.093880248 | -0.397070111 | 0.69131576 | 0.923860706 |
| Sema3f | 467.4219986 | 0.233406931 | 0.100025757 | 2.333468286 | 0.019623581 | 0.092952541 | | | Gas8 | 653.1457828 | -0.037287664 | 0.078445465 | -0.475332306 | 0.634550136 | 0.907297112 |
| Ano2 | 275.5066339 | -0.277772484 | 0.133084854 | -2.087183304 | 0.036871568 | 0.141794012 | | | Pigm | 612.8545717 | -0.037300914 | 0.068142511 | -0.547395652 | 0.584106942 | 0.884313345 |
| Prox2 | 34.96773978 | 0.233303477 | 0.158924603 | 1.468013591 | 0.142100531 | 0.33788829 | | | 1600014C23Rik | 2.553059075 | -0.037311442 | 0.034846468 | -1.070738141 | 0.284287188 | NA |
| Pih1d2 | 56.93986104 | 0.233160877 | 0.147185899 | 1.584125102 | 0.113165253 | 0.294050242 | | | 2700029M09Rik | 1097.028397 | -0.037320257 | 0.076145528 | -0.490117523 | 0.624050739 | 0.900825929 |
| Lhfpl3 | 1084.297505 | -0.267995616 | 0.128481242 | -2.085873479 | 0.036990085 | 0.142143943 | | | Ubr7 | 1946.984824 | -0.037340786 | 0.060135908 | -0.620939924 | 0.534639152 | 0.860040788 |
| Impg2 | 32.8384876 | 0.233153122 | 0.167628223 | 1.390894196 | 0.164257513 | 0.369537666 | | | Rai1 | 3707.322845 | -0.037343528 | 0.054275048 | -0.688042278 | 0.491426158 | 0.841892493 |
| Ntf3 | 38.74912685 | 0.23307668 | 0.159940283 | 1.457273156 | 0.145040994 | 0.343041545 | | | 9830147E19Rik | 107.3180719 | -0.037351307 | 0.095010814 | -0.3931269 | 0.694225757 | 0.925421982 |
| Zfr | 13336.01006 | -0.119098728 | 0.057107496 | -2.085518307 | 0.037022278 | 0.142186943 | | | Zfp260 | 2647.713686 | -0.037360309 | 0.072065993 | -0.518418008 | 0.604166653 | 0.891718993 |
| Ccdc170 | 12.84278924 | 0.23294575 | 0.161698697 | 1.440616118 | 0.149693164 | 0.350068662 | | | Calcoco2 | 2.121417314 | -0.037360594 | 0.034304789 | -1.089078104 | 0.276119445 | NA |
| A330021E22Rik | 410.8324754 | 0.232607212 | 0.13930208 | 1.6698043 | 0.094958089 | 0.260796651 | | | Cda | 60.77324181 | -0.037387892 | 0.094102219 | -0.397311478 | 0.691137784 | 0.923860706 |
| Sema7a | 7349.776995 | 0.232202172 | 0.103362643 | 2.246480601 | 0.024673241 | 0.10850482 | | | Tmem63c | 2152.68242 | -0.037400686 | 0.063074279 | -0.59296256 | 0.553206209 | 0.870600462 |
| Fmo5 | 179.1854441 | 0.232115975 | 0.111784269 | 2.076463688 | 0.037851086 | 0.144115079 | | | 4930503B20Rik | 2.296046608 | -0.037451482 | 0.03391869 | -1.104154732 | 0.269526022 | NA |
| Hsph1 | 12453.36614 | -0.191940088 | 0.092120782 | -2.083569895 | 0.037199308 | 0.142696449 | | | Ier2 | 240.7875535 | -0.037454674 | 0.08268283 | -0.452992163 | 0.650554385 | 0.914098761 |
| Plxnc1 | 2773.045919 | -0.169109159 | 0.081173307 | -2.083309963 | 0.037222979 | 0.142696449 | | | Esam | 680.2306681 | -0.037456442 | 0.074335128 | -0.503886161 | 0.614341376 | 0.89611764 |
| Vma21 | 1197.871174 | -0.165462762 | 0.079424265 | -2.083277225 | 0.037225961 | 0.142696449 | | | Mreg | 199.8472193 | -0.037478296 | 0.094651317 | -0.395961693 | 0.692133288 | 0.923860706 |
| Hcfc1 | 5627.77657 | -0.098920364 | 0.047498877 | -2.082583208 | 0.037289231 | 0.142903623 | | | Zfp959 | 180.3042008 | -0.037488355 | 0.08992912 | -0.416865581 | 0.67677673 | 0.921009816 |
| Itgbl1 | 66.67599685 | -0.316598505 | 0.152035238 | -2.082402137 | 0.037305753 | 0.142931588 | | | Nxnl2 | 5.24656118 | -0.037491934 | 0.046950718 | -0.798538033 | 0.424558331 | NA |
| Car3 | 33.92960202 | 0.232086846 | 0.165955016 | 1.398492506 | 0.161965222 | 0.366938024 | | | Aif1l | 371.367335 | -0.037512726 | 0.085044902 | -0.441093175 | 0.659145544 | 0.915208181 |
| Nudt8 | 162.3650279 | 0.23195965 | 0.118611746 | 1.955621245 | 0.050509788 | 0.174998787 | | | Pdk1 | 3108.047109 | -0.037543026 | 0.078154325 | -0.480370419 | 0.630964025 | 0.905472959 |
| Mprip | 11807.21537 | -0.137034402 | 0.065859125 | -2.080720055 | 0.037459537 | 0.143414399 | | | Ttc34 | 104.3626185 | -0.037548177 | 0.093146095 | -0.403110583 | 0.686866875 | 0.923860706 |
| Rfx2 | 107.8067867 | 0.231938287 | 0.126294275 | 1.836490901 | 0.066285087 | 0.20743711 | | | Ppm1n | 1.343699343 | -0.037553304 | 0.026929102 | -1.394524954 | 0.163159139 | NA |
| Serf1 | 205.8413084 | -0.232513544 | 0.111767864 | -2.080325567 | 0.037495681 | 0.143481868 | | | Ndufb2 | 1120.611042 | -0.037575856 | 0.068051877 | -0.552164868 | 0.580835402 | 0.883755635 |
| Mid1 | 838.8949807 | -0.26148131 | 0.125702576 | -2.080158726 | 0.037510977 | 0.143504956 | | | Msh6 | 860.6782147 | -0.037612468 | 0.063594528 | -0.591441889 | 0.55422438 | 0.871360989 |
| Crim1 | 2342.157684 | -0.213936953 | 0.102854646 | -2.079993095 | 0.037526166 | 0.143527628 | | | Eif4ebp1 | 97.43826148 | -0.037620222 | 0.093526591 | -0.402240925 | 0.687506726 | 0.923860706 |
| AI480653 | 1140.265587 | -0.140316368 | 0.067466658 | -2.07978834 | 0.037544951 | 0.14353686 | | | Gm6086 | 3.976320993 | -0.037643634 | 0.044277144 | -0.850182058 | 0.395223875 | NA |
| Cgnl1 | 898.7511999 | -0.213409 | 0.102614826 | -2.079709222 | 0.037552212 | 0.14353686 | | | Gadl1 | 34.54013076 | -0.037673458 | 0.089553005 | -0.420683352 | 0.673986319 | 0.919711158 |
| Nfu1 | 1124.871718 | -0.116124526 | 0.055838122 | -2.079663893 | 0.037556372 | 0.14353686 | | | Zfp58 | 250.2950668 | -0.037689732 | 0.084885641 | -0.444005977 | 0.657038263 | 0.915208181 |
| Map3k10 | 6515.185642 | -0.160653786 | 0.077255056 | -2.07952455 | 0.037569164 | 0.143550339 | | | Kdm5b | 3260.792503 | -0.037691856 | 0.058160309 | -0.648068352 | 0.516940742 | 0.853297835 |
| Gnrh1 | 14.28045599 | -0.30038036 | 0.144462069 | -2.07930263 | 0.037589544 | 0.143592798 | | | Gm5458 | 3.15857681 | -0.037699153 | 0.037642529 | -1.001504261 | 0.316583081 | NA |
| Lyrm2 | 659.133057 | -0.19992991 | 0.096169859 | -2.078924857 | 0.037624258 | 0.14368998 | | | Hsh2d | 1.857906822 | -0.03772299 | 0.029341045 | -1.285673013 | 0.198557204 | NA |
| Zfp518b | 707.9571535 | -0.153793675 | 0.073994057 | -2.07845982 | 0.037667028 | 0.143817874 | | | Caskin2 | 1331.201844 | -0.037749846 | 0.058096916 | -0.649773663 | 0.515838434 | 0.853297835 |
| Stag3 | 35.47783023 | 0.231903759 | 0.16582158 | 1.398513752 | 0.161958846 | 0.366938024 | | | Batf | 5.354196921 | -0.037759049 | 0.045471525 | -0.830388888 | 0.406318946 | NA |
| Maats1 | 54.53201673 | 0.231707882 | 0.141806226 | 1.63397538 | 0.10226402 | 0.274407405 | | | Slc3a1 | 10.17573481 | -0.037763867 | 0.05901599 | -0.639892118 | 0.522242739 | NA |
| Ndufb4 | 1828.479802 | -0.184555102 | 0.088845212 | -2.077265584 | 0.037777054 | 0.144025017 | | | Ccno | 76.73030648 | -0.037763989 | 0.094519049 | -0.399538396 | 0.689496538 | 0.923860706 |
| Scamp1 | 9555.709007 | -0.135383924 | 0.065169459 | -2.07741367 | 0.037763396 | 0.144025017 | | | Ppp1r42 | 24.34914671 | -0.037769073 | 0.086227693 | -0.438015577 | 0.661374992 | 0.915428982 |
| Dclre1c | 575.403637 | 0.231559321 | 0.11155931 | 2.0756611 | 0.037925306 | 0.144306162 | | | Dclk3 | 2784.412726 | -0.037769252 | 0.082163182 | -0.459685852 | 0.645741726 | 0.912363974 |
| Uba52 | 9253.365395 | -0.159656524 | 0.076856047 | -2.07734499 | 0.037769729 | 0.144025017 | | | Pir | 210.8202146 | -0.037793955 | 0.083839563 | -0.450789031 | 0.652141607 | 0.914559663 |
| Pcf11 | 1185.623025 | 0.23151459 | 0.095443993 | 2.425659105 | 0.015280616 | 0.078892273 | | | Sema4d | 2216.270637 | -0.037816728 | 0.051833251 | -0.72958434 | 0.465644297 | 0.831871734 |
| Hemk1 | 377.2024423 | 0.231487924 | 0.120163041 | 1.926448617 | 0.054048382 | 0.182684474 | | | Snora78 | 4.772865748 | -0.037819981 | 0.045249887 | -0.835802773 | 0.403265866 | NA |
| Ddit4 | 1407.724289 | 0.231269922 | 0.167151693 | 1.383593052 | 0.166483097 | 0.372076569 | | | Syngr4 | 1.849443373 | -0.037866331 | 0.030452849 | -1.243441325 | 0.21370525 | NA |
| Gprasp2 | 5002.273529 | 0.231178062 | 0.100368527 | 2.303292369 | 0.021262399 | 0.098018329 | | | 4930413F20Rik | 3.823996262 | -0.037876041 | 0.0410348 | -0.923022434 | 0.355995514 | NA |
| Nphp1 | 586.6773478 | -0.14887548 | 0.071692802 | -2.076575016 | 0.037840801 | 0.144115079 | | | Myh14 | 1766.220153 | -0.037908357 | 0.073101379 | -0.518572392 | 0.604058965 | 0.891718993 |
| Pdhb | 4134.041323 | -0.105541905 | 0.050826487 | -2.076513864 | 0.03784645 | 0.144115079 | | | Prcp | 431.1652159 | -0.037909639 | 0.08688094 | -0.436340106 | 0.662589987 | 0.915428982 |
| Pqlc2 | 346.4230629 | 0.231023711 | 0.089726842 | 2.574744703 | 0.01003141 | 0.058963075 | | | Bank1 | 64.82785053 | -0.037963682 | 0.09503771 | -0.399459146 | 0.689554921 | 0.923860706 |
| Tmbim6 | 12345.85279 | -0.09837705 | 0.047394854 | -2.075690534 | 0.037922582 | 0.144306162 | | | Pds5a | 1871.246883 | -0.037970173 | 0.064853345 | -0.585477482 | 0.558226704 | 0.872714355 |
| Gal | 32.03576589 | -0.254908765 | 0.12282327 | -2.075410992 | 0.03794846 | 0.144358813 | | | 5830403L16Rik | 41.56615473 | -0.037986831 | 0.093691305 | -0.405446711 | 0.685149182 | 0.923835238 |
| Rps29 | 7473.767094 | -0.180924724 | 0.087195942 | -2.074921379 | 0.037993822 | 0.144495895 | | | Dpp4 | 165.9206796 | -0.037988248 | 0.092978262 | -0.408571284 | 0.682854308 | 0.922882553 |
| Hpx | 18.96479061 | 0.231016709 | 0.167577379 | 1.378567381 | 0.168028179 | 0.374580427 | | | Tcf4 | 9927.453659 | -0.037998864 | 0.064644715 | -0.587810829 | 0.556659274 | 0.871715461 |
| Fut4 | 14.45237673 | -0.345527258 | 0.166546253 | -2.074662454 | 0.038017829 | 0.144516254 | | | Osbpl5 | 1328.486502 | -0.038018886 | 0.064976026 | -0.585121755 | 0.558465852 | 0.872850145 |
| Psmd10 | 497.8241099 | -0.16111569 | 0.077665986 | -2.074469127 | 0.038035763 | 0.144548961 | | | St8sia2 | 172.026629 | -0.038031694 | 0.095088981 | -0.399959 | 0.689186715 | 0.923860706 |
| Snx27 | 6814.824847 | -0.1281985 | 0.061807087 | -2.074171537 | 0.038063383 | 0.144618453 | | | Fam26f | 8.113123751 | -0.038061775 | 0.061119277 | -0.622745827 | 0.533451563 | NA |
| Rplp0 | 6285.950758 | -0.139631579 | 0.067332787 | -2.073753166 | 0.038102241 | 0.1447306 | | | Mdp1 | 1535.814053 | -0.038075087 | 0.060181702 | -0.632668831 | 0.526949928 | 0.856196078 |
| Ly6d | 12.79072422 | 0.230876688 | 0.152996738 | 1.509030133 | 0.131291083 | 0.321311646 | | | Cstf3 | 975.7086267 | -0.038076973 | 0.065717946 | -0.579399918 | 0.56231936 | 0.874638721 |
| Rfx5 | 882.8432278 | 0.230846569 | 0.090699272 | 2.545186567 | 0.010921937 | 0.062559884 | | | 1700028P14Rik | 36.30497859 | -0.038105191 | 0.08840729 | -0.431018651 | 0.666454809 | 0.916017403 |
| Serpinb8 | 419.5140329 | 0.230648982 | 0.149907876 | 1.538604833 | 0.123900798 | 0.311389937 | | | Hrc | 14.59806848 | -0.038118595 | 0.073003248 | -0.522149303 | 0.601566381 | NA |
| Jph2 | 51.37896047 | -0.326585815 | 0.157641939 | -2.071693729 | 0.038294013 | 0.145316539 | | | Farp2 | 383.1558713 | -0.038122842 | 0.076015406 | -0.501514683 | 0.616008947 | 0.897147179 |
| Ms4a6d | 49.96779679 | 0.230590862 | 0.163093418 | 1.413857558 | 0.15740373 | 0.360238314 | | | Insr | 2698.427314 | -0.038139659 | 0.055773902 | -0.683826265 | 0.494084879 | 0.843468909 |
| Ttc16 | 29.41492216 | 0.230378649 | 0.168111205 | 1.370394368 | 0.170563829 | 0.377867998 | | | Tlr4 | 64.97680533 | -0.03814298 | 0.094873285 | -0.402041316 | 0.68765362 | 0.923860706 |
| Ptges2 | 1125.619704 | -0.134810802 | 0.065092311 | -2.071071065 | 0.038352155 | 0.14540359 | | | Tm4sf19 | 1.316398883 | -0.038150455 | 0.02691564 | -1.417408454 | 0.156363542 | NA |
| Rexo2 | 1257.780076 | -0.139513211 | 0.06736451 | -2.071019461 | 0.038356977 | 0.14540359 | | | Ap3d1 | 8656.017006 | -0.038154157 | 0.038042998 | -1.00292193 | 0.315898531 | 0.745993052 |
| Aspg | 119.6052189 | 0.230326552 | 0.133032896 | 1.731350362 | 0.083389294 | 0.240857981 | | | Kcna7 | 0.850958727 | -0.038172847 | 0.02039276 | -1.871882303 | 0.061222889 | NA |
| Tusc1 | 74.60636128 | -0.314279261 | 0.151818838 | -2.07009397 | 0.038443545 | 0.145669918 | | | Blzf1 | 878.427211 | -0.038209798 | 0.061104815 | -0.625315674 | 0.531763895 | 0.858328942 |
| Cdh5 | 726.7102373 | -0.171569334 | 0.08288477 | -2.069974185 | 0.038454762 | 0.145676793 | | | Car1 | 6.121786073 | -0.038213362 | 0.055333974 | -0.690594936 | 0.489820131 | NA |
| Mapk8 | 4222.53239 | -0.153275211 | 0.074059681 | -2.069617485 | 0.038488179 | 0.145767747 | | | Slc25a26 | 622.5259969 | -0.038228322 | 0.065724204 | -0.581647547 | 0.56080411 | 0.873713575 |
| Amot | 963.9198527 | -0.195909313 | 0.09467201 | -2.069347776 | 0.038513463 | 0.145792898 | | | Fbl | 1049.739853 | -0.038263438 | 0.063472302 | -0.602836776 | 0.546617279 | 0.867128614 |
| Zbtb49 | 300.237136 | 0.230282306 | 0.09941298 | 2.316420909 | 0.020535298 | 0.095835515 | | | Isca2 | 873.2575052 | -0.038292336 | 0.066346563 | -0.577156286 | 0.563833884 | 0.875120313 |
| Sh3bp2 | 166.1989452 | 0.230226356 | 0.106995197 | 2.15174477 | 0.031417462 | 0.12760348 | | | E430018J23Rik | 221.1699086 | -0.03830215 | 0.086603035 | -0.442272603 | 0.658291954 | 0.915208181 |
| Erich1 | 157.1363321 | 0.230219243 | 0.113104119 | 2.035462936 | 0.041804314 | 0.154733886 | | | Snord42a | 1.670153154 | -0.038322786 | 0.029108975 | -1.316528191 | 0.187996823 | NA |
| Cldn23 | 23.97528631 | 0.230175204 | 0.167866409 | 1.37118084 | 0.17031859 | 0.377840546 | | | Ntsr2 | 3431.572111 | -0.038331012 | 0.066663029 | -0.574996559 | 0.565293624 | 0.875364912 |
| Mfsd8 | 716.81208 | -0.23168742 | 0.11204428 | -2.067820155 | 0.038656938 | 0.146156806 | | | Zfp433 | 47.23895548 | -0.038342461 | 0.092312011 | -0.415357227 | 0.677880414 | 0.921155235 |
| Rcn1 | 987.3937432 | -0.24803602 | 0.119948955 | -2.067846437 | 0.038654465 | 0.146156806 | | | Zfp825 | 251.8944842 | -0.038372278 | 0.08992018 | -0.426737113 | 0.669570815 | 0.91770119 |
| Plcd4 | 623.9555561 | 0.230022035 | 0.127942576 | 1.797853705 | 0.072200194 | 0.219691636 | | | 1700034P13Rik | 2.301102195 | -0.038380207 | 0.032199968 | -1.191933097 | 0.233287479 | NA |
| Arl3 | 2709.021731 | -0.116939638 | 0.056617174 | -2.065444642 | 0.038880948 | 0.146932052 | | | Gm13251 | 30.20971057 | -0.03838693 | 0.083245158 | -0.461131087 | 0.64470456 | 0.912353932 |
| Klhl2 | 6899.263053 | -0.178274447 | 0.086317321 | -2.065338036 | 0.038891027 | 0.146934302 | | | Vgll4 | 457.1810566 | -0.03839284 | 0.067740343 | -0.566764767 | 0.570874011 | 0.878246677 |
| Casp3 | 265.9314145 | -0.192732773 | 0.093331764 | -2.065028708 | 0.038920284 | 0.147008991 | | | Srgn | 172.976683 | -0.03841186 | 0.088968351 | -0.431747463 | 0.665924967 | 0.915591677 |
| Inhbb | 212.5935797 | -0.243073745 | 0.117717988 | -2.064881915 | 0.038934175 | 0.147025616 | | | Rnf166 | 1172.231463 | -0.038417073 | 0.066953226 | -0.573789728 | 0.566110101 | 0.87575243 |
| Ifi27l1 | 867.8324243 | 0.229996086 | 0.099477926 | 2.312031383 | 0.020775954 | 0.096522528 | | | Srfbp1 | 423.9854066 | -0.038447324 | 0.070005393 | -0.549205166 | 0.582864659 | 0.88406516 |
| Rab11a | 3043.073394 | -0.164090131 | 0.079495678 | -2.064139033 | 0.039004536 | 0.147183702 | | | Darc | 2039.61281 | -0.038467128 | 0.065932007 | -0.583436374 | 0.559599577 | 0.873713575 |
| P2rx5 | 33.14490881 | 0.22988255 | 0.167362146 | 1.373563585 | 0.169577209 | 0.376908517 | | | Dclk2 | 3038.180846 | -0.038503007 | 0.050257032 | -0.766121782 | 0.443603843 | 0.820717992 |
| R3hdm1 | 10953.51452 | 0.229705108 | 0.102035276 | 2.251232282 | 0.024370829 | 0.107763022 | | | Scmh1 | 1895.125065 | -0.038512891 | 0.047305458 | -0.814132087 | 0.415569286 | 0.805633085 |
| Syngr2 | 319.9617977 | 0.229603547 | 0.108369534 | 2.118709372 | 0.034115036 | 0.134669647 | | | Mir3112 | 6.611335474 | -0.038517595 | 0.058243603 | -0.661318888 | 0.508407831 | NA |
| Lekr1 | 126.4758392 | 0.229553784 | 0.10725569 | 2.140248074 | 0.032334724 | 0.129858531 | | | 4930451G09Rik | 142.0340653 | -0.038518658 | 0.094391095 | -0.408075127 | 0.68321852 | 0.922993772 |
| Btd | 363.461182 | -0.198842706 | 0.096449797 | -2.06161871 | 0.039244051 | 0.147923531 | | | Mier2 | 1367.757666 | -0.038593628 | 0.067032338 | -0.575746413 | 0.564786599 | 0.875364912 |
| 5031425E22Rik | 336.4705233 | 0.229416101 | 0.107475847 | 2.134582862 | 0.032795099 | 0.131129615 | | | Hhla1 | 0.903806998 | -0.038602336 | 0.022972424 | -1.680377133 | 0.092883962 | NA |
| Rnaset2b | 10.89254366 | 0.229350487 | 0.161625987 | 1.419019866 | 0.155893224 | 0.358209939 | | | Slc29a3 | 772.4344231 | -0.038618171 | 0.062204348 | -0.620827521 | 0.534713113 | 0.860064521 |
| Crybb1 | 107.1500267 | -0.335723838 | 0.16289818 | -2.060942849 | 0.039308493 | 0.148006266 | | | Mcat | 598.5064316 | -0.038627587 | 0.071121314 | -0.543122517 | 0.58704545 | 0.884663353 |
| Foxl2os | 18.91747006 | 0.229185069 | 0.168093196 | 1.363440487 | 0.172743729 | 0.380561827 | | | Pth1r | 212.4402689 | -0.038631065 | 0.092864995 | -0.415991683 | 0.677416089 | 0.921064066 |
| Wfdc18 | 43.90132137 | -0.346057875 | 0.16790843 | -2.06099167 | 0.039303835 | 0.148006266 | | | Wasf2 | 1017.781954 | -0.038654037 | 0.06410902 | -0.602942244 | 0.546547112 | 0.867128614 |
| Uox | 20.06799221 | 0.229152507 | 0.168121552 | 1.363016847 | 0.172877201 | 0.380693484 | | | Me3 | 1976.287777 | -0.038663634 | 0.06322695 | -0.611505603 | 0.54086491 | 0.863847445 |
| Cyb5r2 | 12.54018756 | 0.229069399 | 0.164288266 | 1.394313816 | 0.163222861 | 0.368601516 | | | Slitrk2 | 1168.210587 | -0.038664636 | 0.055079544 | -0.701978153 | 0.482692788 | 0.837880119 |
| Me2 | 836.0819075 | -0.179221466 | 0.087028043 | -2.05935306 | 0.039460429 | 0.148470103 | | | Tmem30c | 8.193614122 | -0.038668885 | 0.061370921 | -0.630084807 | 0.528639099 | NA |
| Zfp580 | 394.0301461 | -0.224126701 | 0.108855639 | -2.058935155 | 0.039500451 | 0.148584604 | | | Tdrd6 | 49.85002799 | -0.038678173 | 0.094370691 | -0.409853661 | 0.6819133 | 0.922544285 |
| B3gnt3 | 34.03202724 | 0.228801256 | 0.166460379 | 1.374508801 | 0.16928378 | 0.376567454 | | | Pkhd1l1 | 22.79108359 | -0.038680031 | 0.080671027 | -0.479478592 | 0.631598194 | 0.90604339 |
| Lima1 | 625.3236354 | -0.18689693 | 0.090794696 | -2.058456469 | 0.039546336 | 0.148618256 | | | Sco2 | 121.7233621 | -0.038684439 | 0.090097605 | -0.42936146 | 0.667660196 | 0.91647846 |
| Ttpa | 100.0505957 | -0.316447154 | 0.153731424 | -2.058441567 | 0.039547765 | 0.148618256 | | | Nup160 | 596.9576961 | -0.038706587 | 0.071723313 | -0.539665348 | 0.589427841 | 0.885188411 |
| Uhmk1 | 2209.202703 | -0.157342064 | 0.076434866 | -2.058511665 | 0.039541043 | 0.148618256 | | | 1110032F04Rik | 452.4438441 | -0.038726487 | 0.086895814 | -0.445665732 | 0.655838722 | 0.915208181 |
| Zfp189 | 369.1714255 | 0.228644193 | 0.158249785 | 1.444830991 | 0.148505364 | 0.348265855 | | | Gm5134 | 3.136669825 | -0.038738966 | 0.039116532 | -0.990347653 | 0.322004222 | NA |
| Ctsc | 315.5048445 | 0.228576538 | 0.107750807 | 2.121344096 | 0.033892856 | 0.134237196 | | | Carns1 | 644.5477731 | -0.038741751 | 0.083248668 | -0.46537382 | 0.641663779 | 0.911095592 |
| Gsk3b | 8399.269487 | -0.152525168 | 0.074121588 | -2.05776985 | 0.039612232 | 0.148752282 | | | Zfp287 | 798.9211102 | -0.038742047 | 0.068616524 | -0.56461686 | 0.572334395 | 0.879277718 |
| Trmu | 448.0677832 | 0.228520784 | 0.10013765 | 2.282066587 | 0.022485409 | 0.101927716 | | | Sardh | 580.0463207 | -0.038751946 | 0.076915042 | -0.503827928 | 0.6143823 | 0.89611764 |
| 1110059E24Rik | 675.0913613 | -0.160649418 | 0.078110951 | -2.056682398 | 0.039716786 | 0.148989976 | | | Tmem201 | 3618.514701 | -0.038752915 | 0.060927141 | -0.636053395 | 0.524741621 | 0.855487854 |
| Parp16 | 135.3823216 | 0.228478349 | 0.132794051 | 1.720546571 | 0.085333136 | 0.243984426 | | | Pogz | 2912.371053 | -0.038764039 | 0.04518765 | -0.857845875 | 0.390977575 | 0.794184423 |
| Arf5 | 3231.440178 | -0.14812729 | 0.072039023 | -2.056209042 | 0.039762371 | 0.148989976 | | | Rps6ka4 | 1758.470363 | -0.038764391 | 0.063279971 | -0.612585481 | 0.540150461 | 0.863544778 |
| Pkdcc | 276.8951276 | 0.228382905 | 0.097210808 | 2.349357127 | 0.01880586 | 0.090479188 | | | Ddhd1 | 3016.050097 | -0.038782462 | 0.052982953 | -0.73198 | 0.464180775 | 0.83159896 |
| Emc6 | 927.6355634 | -0.16836515 | 0.081890454 | -2.055980186 | 0.039784426 | 0.148989976 | | | Idh3g | 4824.478471 | -0.038790864 | 0.050416355 | -0.769410331 | 0.441649757 | 0.819100918 |
| Foxs1 | 15.15181821 | -0.340001084 | 0.165303196 | -2.056833095 | 0.039702283 | 0.148989976 | | | Rps27a | 6003.498312 | -0.038798568 | 0.06066357 | -0.639569484 | 0.522452527 | 0.854248695 |
| Glipr1 | 41.71736313 | 0.228243156 | 0.159849318 | 1.42786443 | 0.153330884 | 0.354708354 | | | Smurf2 | 1426.264771 | -0.038812354 | 0.060702318 | -0.63938833 | 0.522570339 | 0.854337225 |
| Nrip3 | 5403.021662 | -0.23497779 | 0.114285153 | -2.056065749 | 0.039776179 | 0.148989976 | | | Paxip1 | 1990.035042 | -0.038818211 | 0.044211462 | -0.878012379 | 0.379937001 | 0.786275719 |
| Zbtb42 | 45.96939942 | 0.228160431 | 0.167679402 | 1.360694448 | 0.17361027 | 0.381795475 | | | Gm7457 | 2.248518919 | -0.038828142 | 0.033629707 | -1.154578686 | 0.248262999 | NA |
| Pcdhb2 | 101.9541132 | 0.228107507 | 0.116902992 | 1.951254652 | 0.051026761 | 0.176003702 | | | Trpt1 | 381.0615654 | -0.038840866 | 0.081091441 | -0.478976148 | 0.631955597 | 0.906377074 |
| Sorcs3 | 2260.869031 | -0.133706503 | 0.065032821 | -2.055984968 | 0.039783965 | 0.148989976 | | | BC024978 | 315.0953449 | -0.038846713 | 0.076950829 | -0.504825144 | 0.613681654 | 0.895819717 |
| Snx3 | 4900.082904 | -0.123860628 | 0.060250666 | -2.055755328 | 0.039806106 | 0.149010825 | | | Mecr | 485.2086906 | -0.038848034 | 0.081978248 | -0.473882222 | 0.635583897 | 0.907676507 |
| Kansl1l | 422.3079601 | 0.22806521 | 0.107149577 | 2.128475127 | 0.03329771 | 0.13273691 | | | Stk36 | 236.7802186 | -0.038852741 | 0.089486191 | -0.434175829 | 0.664160764 | 0.915428982 |
| Bbox1 | 53.76858209 | 0.228016928 | 0.167609568 | 1.360405201 | 0.173701733 | 0.381912113 | | | 3110052M02Rik | 539.4399635 | -0.038864874 | 0.078681206 | -0.493953721 | 0.621338864 | 0.899589902 |
| Alpl | 382.8306343 | -0.310857659 | 0.151407185 | -2.053123554 | 0.040060596 | 0.149854871 | | | Tcf12 | 2061.737527 | -0.03886592 | 0.05375579 | -0.723008993 | 0.469674359 | 0.833245343 |
| Shroom3 | 189.2798877 | -0.229261134 | 0.111678822 | -2.052861311 | 0.04008603 | 0.149913819 | | | Adamtsl5 | 248.1315761 | -0.038876668 | 0.090332671 | -0.430372173 | 0.666924934 | 0.916077199 |
| 5830417I10Rik | 387.1399517 | -0.190771539 | 0.092973683 | -2.051887518 | 0.040180595 | 0.150194968 | | | Ccdc107 | 605.8396544 | -0.038881751 | 0.067202844 | -0.578573 | 0.562877327 | 0.87510382 |
| Adssl1 | 796.5715147 | 0.227999521 | 0.103786935 | 2.196803679 | 0.02803447 | 0.118291417 | | | Ccdc8 | 113.7481559 | -0.038892004 | 0.093489693 | -0.416003118 | 0.677407722 | 0.921064066 |
| Ddx58 | 248.7023126 | 0.227925767 | 0.107572777 | 2.118805264 | 0.034106928 | 0.134669647 | | | Arhgap20 | 4311.892568 | -0.038900492 | 0.077891988 | -0.499415826 | 0.617486473 | 0.897948594 |
| Nedd4 | 22168.6739 | -0.110910215 | 0.0540705 | -2.051214909 | 0.040246023 | 0.150366982 | | | AA388235 | 139.7688716 | -0.038923028 | 0.091470354 | -0.425526159 | 0.670453156 | 0.918167488 |
| Klhl1 | 213.3337146 | -0.30218832 | 0.147350892 | -2.050807544 | 0.040285693 | 0.150442642 | | | Txndc17 | 919.9945139 | -0.038930218 | 0.066345731 | -0.586778041 | 0.557352787 | 0.872425278 |
| E130112N10Rik | 18.53387761 | 0.227907771 | 0.168098154 | 1.355801747 | 0.175162255 | 0.383979058 | | | Yap1 | 1000.379734 | -0.038948821 | 0.067793571 | -0.574520865 | 0.565615385 | 0.875453286 |
| Dlgap5 | 8.392274212 | -0.325903943 | 0.158924102 | -2.050689213 | 0.040297222 | 0.150449436 | | | Mrpl23 | 732.1607989 | -0.038956982 | 0.065587677 | -0.593968007 | 0.552533511 | 0.870265825 |
| Tpi1 | 22186.03448 | -0.120481373 | 0.05881293 | -2.048552457 | 0.040505898 | 0.151192093 | | | Fsd2 | 8.460259195 | -0.038994345 | 0.060878874 | -0.640523426 | 0.521832364 | NA |
| Esyt2 | 1953.413713 | -0.12015939 | 0.058661405 | -2.048355108 | 0.040525217 | 0.151227772 | | | Rnf151 | 2.090118535 | -0.039026942 | 0.032549607 | -1.198998893 | 0.230528376 | NA |
| Ssr2 | 1915.146639 | -0.108874402 | 0.053172237 | -2.047561037 | 0.04060303 | 0.15481665 | | | Slc9a8 | 1833.656561 | -0.039029053 | 0.060059912 | -0.649835343 | 0.515798587 | 0.853297835 |
| Zcchc14 | 4021.892156 | -0.158933487 | 0.077639762 | -2.047063046 | 0.040651895 | 0.151627457 | | | Enpp1 | 173.8683887 | -0.039037658 | 0.094897795 | -0.411365277 | 0.680804711 | 0.922355522 |
| Spata13 | 936.1997267 | -0.225817002 | 0.110356098 | -2.046257591 | 0.040731034 | 0.151886074 | | | Serpinb11 | 1.772136732 | -0.039043008 | 0.031031168 | -1.258186861 | 0.208324185 | NA |
| Ywhae | 23309.57805 | -0.115628404 | 0.056512293 | -2.046075262 | 0.040748966 | 0.151916383 | | | Lypd2 | 4.052908143 | -0.039056157 | 0.041832373 | -0.933634272 | 0.35049258 | NA |
| Gm17066 | 425.019724 | 0.227499013 | 0.133193927 | 1.708028424 | 0.087631068 | 0.247946635 | | | Pstk | 453.0986912 | -0.039061061 | 0.079899281 | -0.488878753 | 0.624927538 | 0.901459526 |
| Hspb1 | 228.5666377 | -0.314677189 | 0.153827395 | -2.045651155 | 0.040790705 | 0.15199884 | | | Caps2 | 8.470756141 | -0.039067226 | 0.053625267 | -0.728522732 | 0.466293658 | NA |
| Mthfsd | 859.9507372 | 0.227498736 | 0.105304651 | 2.160386398 | 0.03074277 | 0.125746693 | | | Rbpj | 1748.451627 | -0.039081443 | 0.061599495 | -0.634444211 | 0.525790965 | 0.855892499 |
| Apobr | 50.89052441 | 0.22740574 | 0.160028993 | 1.421028375 | 0.155308511 | 0.357290284 | | | Cdkn2aip | 631.2838396 | -0.039091036 | 0.069380875 | -0.563426678 | 0.573144375 | 0.879310647 |
| Rttn | 212.7957745 | 0.227401694 | 0.099009805 | 2.29675933 | 0.021632503 | 0.099193252 | | | Ears2 | 214.6948018 | -0.039109318 | 0.089120167 | -0.438838021 | 0.660778908 | 0.915428982 |
| Tm6sf2 | 43.56461501 | 0.226962542 | 0.167978105 | 1.351143599 | 0.176649438 | 0.385960345 | | | Adora2a | 2395.651432 | -0.039115145 | 0.095088911 | -0.411353382 | 0.680813432 | 0.922355522 |
| Tecta | 12.96739013 | 0.226707092 | 0.160515687 | 1.412367203 | 0.157841868 | 0.360746278 | | | Cd209c | 82.108452 | -0.039137497 | 0.094918158 | -0.412328871 | 0.68009839 | 0.921988728 |
| Gcnt4 | 608.7428908 | 0.2266443 | 0.10183281 | 2.225651048 | 0.026037564 | 0.112249866 | | | Sytl3 | 7.65893679 | -0.039143456 | 0.051615608 | -0.758364721 | 0.448232672 | NA |
| 9230112J17Rik | 28.60085269 | 0.226479897 | 0.167126628 | 1.355139512 | 0.175373112 | 0.384278178 | | | Plin2 | 264.1238248 | -0.039184136 | 0.083820772 | -0.467475254 | 0.640159893 | 0.910092095 |
| Ppp1r32 | 37.9829718 | 0.226323415 | 0.163476217 | 1.384442453 | 0.166223017 | 0.371853802 | | | BC027231 | 210.5270215 | -0.03919005 | 0.087515576 | -0.447806574 | 0.654292799 | 0.914926568 |
| Fuom | 534.6117465 | -0.149344106 | 0.073160289 | -2.041327436 | 0.041218292 | 0.153300321 | | | Gemin8 | 385.8397393 | -0.039192024 | 0.074086397 | -0.52900432 | 0.596802453 | 0.887922451 |
| Icam1 | 124.6123467 | 0.226280155 | 0.125728911 | 1.7997464 | 0.071900691 | 0.218995342 | | | Slc5a10 | 4.514881584 | -0.039255991 | 0.046164916 | -0.850342517 | 0.395134685 | NA |
| Lrrtm3 | 2125.520524 | -0.198607117 | 0.097346624 | -2.040205495 | 0.041329863 | 0.153597903 | | | Fn3krp | 1508.062582 | -0.039300587 | 0.057101311 | -0.688260679 | 0.491288638 | 0.841892493 |
| Map3k19 | 227.3353093 | -0.292635506 | 0.143440635 | -2.040115806 | 0.041338793 | 0.153597903 | | | Tmem74 | 252.6354662 | -0.039304313 | 0.087924494 | -0.447023482 | 0.654858106 | 0.915208181 |
| Mmp17 | 9402.962004 | -0.195883 | 0.096034564 | -2.03971354 | 0.041378866 | 0.153709936 | | | Scai | 1962.294568 | -0.039314022 | 0.073083216 | -0.537935032 | 0.590621902 | 0.885693151 |
| Cnr2 | 17.93677144 | 0.226260641 | 0.168116087 | 1.345859543 | 0.178347818 | 0.387849001 | | | Slc37a4 | 926.4906572 | -0.039317321 | 0.071633751 | -0.548865874 | 0.583097499 | 0.884313345 |
| 0610040J01Rik | 155.6834325 | -0.260775388 | 0.127892818 | -2.0390151 | 0.041448521 | 0.153858111 | | | Gm5176 | 14.50623841 | -0.039355265 | 0.075838563 | -0.518934738 | 0.603806252 | NA |
| Cenpw | 65.44475925 | -0.304885783 | 0.14953332 | -2.038915364 | 0.041458475 | 0.153858111 | | | Adipoq | 1.233807261 | -0.039361994 | 0.025083179 | -1.569258584 | 0.116587699 | NA |
| Ptplad2 | 45.32862608 | -0.333715575 | 0.163671011 | -2.038941249 | 0.041455891 | 0.153858111 | | | B3gnt7 | 49.60234899 | -0.039364347 | 0.088896181 | -0.442812577 | 0.657901305 | 0.915208181 |
| Zfp59 | 321.3881677 | -0.190503546 | 0.09345569 | -2.038437115 | 0.041506238 | 0.153998479 | | | Nup62-il4i1 | 3.85146023 | -0.039409261 | 0.040588525 | -0.97094587 | 0.331575234 | NA |
| Gm973 | 244.1198557 | 0.226242566 | 0.126375623 | 1.790238972 | 0.073415502 | 0.221939958 | | | Hmbs | 595.4495541 | -0.039454617 | 0.073820402 | -0.534467654 | 0.593018022 | 0.886253111 |
| Arid1a | 5937.290299 | -0.120325495 | 0.059075324 | -2.03681481 | 0.041668604 | 0.15452689 | | | Gm15910 | 9.910299235 | -0.039455712 | 0.062721331 | -0.629063687 | 0.529307362 | NA |
| Mkrn3 | 22.03490136 | -0.342168667 | 0.168032471 | -2.036324671 | 0.041717764 | 0.154598191 | | | 9330175E14Rik | 2.187960253 | -0.039510562 | 0.030382887 | -1.300421588 | 0.193456515 | NA |
| Slc6a19 | 8.716080356 | 0.226218736 | 0.14655807 | 1.543543355 | 0.122698997 | 0.30937319 | | | Mapk13 | 35.60025696 | -0.039512081 | 0.08889304 | -0.444490147 | 0.656688251 | 0.915208181 |
| Zfp599 | 198.290578 | -0.252770041 | 0.124123163 | -2.036445381 | 0.041705652 | 0.154598191 | | | Bmp4 | 111.0548631 | -0.03951826 | 0.094660042 | -0.417475626 | 0.676330549 | 0.920643418 |
| Stt3b | 3959.722342 | -0.130627974 | 0.064158709 | -2.036013125 | 0.041749037 | 0.154677089 | | | Dux | 1.160395969 | -0.039520965 | 0.025686793 | -1.538571358 | 0.123908975 | NA |
| Mdfic | 103.7097133 | -0.318846741 | 0.156612798 | -2.035891987 | 0.041761203 | 0.154685172 | | | Zfp799 | 933.4219438 | -0.039552218 | 0.057410591 | -0.688935912 | 0.490863599 | 0.841866734 |
| Serpind1 | 162.3737336 | 0.226070317 | 0.1214016 | 1.862169173 | 0.062579248 | 0.199865303 | | | Tenc1 | 1277.263668 | -0.039597119 | 0.074867478 | -0.528896119 | 0.596877514 | 0.887922451 |
| Snord88c | 9.83997425 | 0.225962295 | 0.152601018 | 1.480739107 | 0.138676109 | 0.332453601 | | | Cox11 | 547.5801396 | -0.039605971 | 0.079404021 | -0.498790496 | 0.617926984 | 0.89813962 |
| Gm15645 | 13.69571187 | 0.225813767 | 0.165280985 | 1.366241659 | 0.171863126 | 0.379485147 | | | Ihh | 15.05506724 | -0.039638108 | 0.066921815 | -0.592304734 | 0.553646546 | NA |
| Accs | 109.8823617 | 0.225687114 | 0.148330583 | 1.521514373 | 0.128130807 | 0.316541113 | | | Kank4 | 318.2608084 | -0.039664423 | 0.092068921 | -0.430812294 | 0.666604859 | 0.916017403 |
| Syt15 | 45.21824293 | -0.328966779 | 0.161692231 | -2.034524335 | 0.041898758 | 0.155009396 | | | Ammecr1 | 74.55235788 | -0.039693975 | 0.095096572 | -0.417407001 | 0.676380735 | 0.920643418 |
| Heg1 | 839.6954979 | -0.12724698 | 0.062548184 | -2.034383278 | 0.041912967 | 0.155024947 | | | Pex1 | 973.5915458 | -0.039711211 | 0.053445114 | -0.7430279 | 0.457464789 | 0.82764149 |
| Art3 | 123.7680018 | -0.249032628 | 0.122423366 | -2.034191962 | 0.041932245 | 0.155059236 | | | Tob1 | 1105.484094 | -0.03971252 | 0.074328546 | -0.534283556 | 0.593145368 | 0.886253111 |
| Slc5a3 | 1095.900571 | -0.227189116 | 0.111698213 | -2.033954795 | 0.041956153 | 0.155110628 | | | Lcorl | 617.7680595 | -0.039716187 | 0.074129529 | -0.535767429 | 0.592119295 | 0.886226713 |
| Tbkbp1 | 1793.174353 | 0.225606722 | 0.096113592 | 2.347292576 | 0.018910399 | 0.090767572 | | | Acox3 | 1843.456649 | -0.039722762 | 0.057248347 | -0.693867417 | 0.487765359 | 0.839801621 |
| Tspan31 | 2134.9313 | -0.132047695 | 0.06494222 | -2.033310438 | 0.042021169 | 0.15528611 | | | Clnk | 2.851749329 | -0.039729168 | 0.03527489 | -1.126273335 | 0.260049841 | NA |
| Raph1 | 2476.631941 | -0.131714117 | 0.064804895 | -2.032471724 | 0.042105924 | 0.155552975 | | | Zfp61 | 567.339226 | -0.03973881 | 0.07430056 | -0.534838632 | 0.592761447 | 0.886226713 |
| Tmem51 | 169.4469357 | -0.237369465 | 0.11679908 | -2.03228882 | 0.042124426 | 0.155584231 | | | Bloc1s1 | 1003.903174 | -0.0397414 | 0.061848692 | -0.642558465 | 0.520510641 | 0.8535556 |
| 4930453N24Rik | 538.6892909 | -0.151396161 | 0.074502576 | -2.032092979 | 0.042144244 | 0.155620333 | | | Gm4890 | 20.3708262 | -0.039741462 | 0.083746712 | -0.474543552 | 0.635112348 | NA |
| Aip | 2449.67361 | -0.169610819 | 0.083485661 | -2.031616171 | 0.042192528 | 0.155724401 | | | Tex14 | 18.90110364 | -0.039791498 | 0.077275137 | -0.514932738 | 0.606600023 | NA |
| Rpl22l1 | 1299.492745 | -0.248052455 | 0.122090621 | -2.031707692 | 0.042183257 | 0.155724401 | | | Eno1 | 2.300857202 | -0.039813606 | 0.032674591 | -1.218488281 | 0.223038476 | NA |
| 3425401B19Rik | 13.06232049 | 0.225545922 | 0.151035033 | 1.493335139 | 0.135349482 | 0.327438548 | | | Mir5103 | 2.300549262 | -0.039838532 | 0.036538564 | -1.090314648 | 0.275574565 | NA |
| Apopt1 | 799.6641376 | -0.138486203 | 0.068177529 | -2.031258742 | 0.042228754 | 0.155783885 | | | Ccdc64 | 1343.910024 | -0.039846843 | 0.066793967 | -0.596563501 | 0.550798846 | 0.869642174 |
| Brdt | 484.4950791 | -0.171624159 | 0.084514985 | -2.030695007 | 0.042285943 | 0.155883901 | | | Mir421 | 18.63963674 | -0.039848964 | 0.068107768 | -0.585086914 | 0.558489277 | NA |
| C1qtnf6 | 76.43563219 | -0.285823747 | 0.140748208 | -2.030745188 | 0.04228085 | 0.155883901 | | | Mybphl | 1.603244846 | -0.039857865 | 0.029034742 | -1.372764563 | 0.169825551 | NA |
| Ncam1 | 10462.49979 | -0.134933477 | 0.066446978 | -2.030693969 | 0.042286049 | 0.155883901 | | | Tagln | 232.4140614 | -0.039862593 | 0.095020036 | -0.419517764 | 0.674837775 | 0.919907397 |
| Il10ra | 283.7247094 | 0.225499409 | 0.119460867 | 1.887642484 | 0.059073963 | 0.193018131 | | | Arhgap21 | 8222.876087 | -0.039877093 | 0.052833997 | -0.754762001 | 0.450391797 | 0.823365451 |
| Tmem67 | 416.2735996 | 0.225418979 | 0.113157496 | 1.992081719 | 0.046362092 | 0.165700194 | | | Camk2n1 | 64376.91164 | -0.039879227 | 0.079465742 | -0.501841747 | 0.615778844 | 0.896906632 |
| Fry | 7183.985585 | -0.19676752 | 0.096940996 | -2.029765814 | 0.042380349 | 0.15610783 | | | Acsm3 | 5.070424228 | -0.039892822 | 0.04883841 | -0.816832936 | 0.414023907 | NA |
| Gpr27 | 243.9572567 | -0.25853388 | 0.127387643 | -2.029505171 | 0.042406863 | 0.15610783 | | | Rnaset2b | 14.40300816 | -0.039907692 | 0.067916252 | -0.587601505 | 0.5567998 | NA |
| Dagla | 5635.274089 | 0.225332447 | 0.095521164 | 2.358979269 | 0.018325279 | 0.08903477 | | | Mboat4 | 7.333695251 | -0.039912321 | 0.050243617 | -0.794375948 | 0.426976602 | NA |
| Alkbh2 | 105.0922082 | 0.224946878 | 0.114199728 | 1.969767196 | 0.048865058 | 0.171405855 | | | Gm10536 | 1.774559936 | -0.039949155 | 0.031554141 | -1.266051099 | 0.205494788 | NA |
| Lrfn5 | 1950.505291 | -0.138796829 | 0.068392277 | -2.029422563 | 0.042415269 | 0.15610783 | | | Foxd1 | 29.37742409 | -0.039962603 | 0.077034853 | -0.518760031 | 0.603928094 | 0.891718993 |
| Pm20d2 | 433.3267568 | -0.167525885 | 0.082557421 | -2.029204442 | 0.042437471 | 0.15610783 | | | Tcf7l1 | 159.4527196 | -0.039973605 | 0.094339199 | -0.423722112 | 0.671768487 | 0.919094512 |
| Smim7 | 3264.276886 | -0.12635036 | 0.06226426 | -2.029259808 | 0.042431835 | 0.15610783 | | | Cacnb3 | 12386.01636 | -0.040010587 | 0.061328828 | -0.652394443 | 0.514146748 | 0.853195313 |
| Lpcat4 | 3971.975542 | 0.224941928 | 0.091804571 | 2.450225795 | 0.014276666 | 0.075315852 | | | Dazap1 | 1337.641959 | -0.040021672 | 0.071816434 | -0.557277359 | 0.577337942 | 0.881375635 |
| Atp6v0e | 524.2076979 | 0.22488637 | 0.091498695 | 2.457809602 | 0.013978726 | 0.074097292 | | | Bai2 | 19031.54906 | -0.04003631 | 0.068881241 | -0.581236776 | 0.561080885 | 0.87387211 |
| Gm1943 | 322.2542003 | -0.173508753 | 0.085550723 | -2.028138942 | 0.042546069 | 0.156395919 | | | Pyurf | 845.8525161 | -0.040037758 | 0.071568861 | -0.559429862 | 0.575868386 | 0.880811222 |
| Epha10 | 1995.285619 | 0.224738566 | 0.088574383 | 2.537286274 | 0.011171556 | 0.063439698 | | | Foxf1 | 36.20602261 | -0.040055257 | 0.089088851 | -0.449610209 | 0.652991526 | 0.914683958 |
| Aoc2 | 162.4974398 | 0.224714249 | 0.145019243 | 1.549547807 | 0.121250089 | 0.307084755 | | | Cox7b2 | 4.835529098 | -0.040078873 | 0.048866017 | -0.820178855 | 0.412114154 | NA |
| Noxred1 | 46.60955701 | 0.224708959 | 0.164076645 | 1.369536529 | 0.170831625 | 0.378123599 | | | Traf5 | 110.0529589 | -0.040088677 | 0.092217515 | -0.434718688 | 0.663766632 | 0.915428982 |
| C230035I16Rik | 22.01888649 | 0.224682847 | 0.168116861 | 1.336468249 | 0.181396272 | 0.391933319 | | | Tnf | 2.197373991 | -0.04009571 | 0.03302635 | -1.214052122 | 0.224727821 | NA |
| Vtn | 2747.298832 | -0.145165664 | 0.071637494 | -2.026392258 | 0.042724604 | 0.156888475 | | | Man2a2 | 11686.05274 | -0.040101724 | 0.039049694 | -1.026940816 | 0.304448333 | 0.737453291 |
| Stmn1 | 21663.38994 | -0.196651353 | 0.097059708 | -2.02608638 | 0.042755934 | 0.156943955 | | | Exph5 | 1484.129813 | -0.04011695 | 0.084529163 | -0.474593017 | 0.635077085 | 0.907297112 |
| Kcnmb2 | 105.5058952 | 0.224632957 | 0.11594042 | 1.937486136 | 0.052685947 | 0.179844912 | | | Jarid2 | 1618.567028 | -0.040124334 | 0.066355971 | -0.604683097 | 0.545389581 | 0.866680785 |
| En2 | 20.38769105 | 0.224546312 | 0.167051595 | 1.344173405 | 0.178892319 | 0.388502669 | | | Lsm14b | 4152.216808 | -0.040124847 | 0.055014983 | -0.729343995 | 0.465791267 | 0.831904964 |
| Ltf | 12.61114583 | -0.292428603 | 0.144405243 | -2.025055299 | 0.042861686 | 0.157220399 | | | Csad | 766.724909 | -0.040126241 | 0.072604139 | -0.552671542 | 0.580488344 | 0.883489182 |
| Ly6g5b | 8.942715601 | 0.224501103 | 0.158318499 | 1.418034561 | 0.156180675 | 0.358551396 | | | 2610001J05Rik | 1015.115364 | -0.040141881 | 0.063052063 | -0.636646583 | 0.524355075 | 0.855487854 |
| Hars2 | 972.309758 | 0.224481144 | 0.096290101 | 2.331300324 | 0.019737529 | 0.093349596 | | | Iqgap2 | 1317.455564 | -0.040178839 | 0.092845098 | -0.43275132 | 0.665195442 | 0.915428982 |

| | | | | | | |
|---|---|---|---|---|---|---|
| Lrrc32 | 254.0460927 | 0.224420371 | 0.134456031 | 1.669098585 | 0.095097844 | 0.260971659 |
| Golga7 | 2775.652516 | -0.111159614 | 0.054922306 | -2.023942959 | 0.042976021 | 0.15756436 |
| Gm10451 | 27.12940873 | 0.224398593 | 0.167940205 | 1.336181484 | 0.181489962 | 0.391971768 |
| Zfp617 | 1263.994757 | -0.164717047 | 0.081409746 | -2.023308692 | 0.043041331 | 0.157655411 |
| Jam2 | 1956.495174 | -0.24755798 | 0.122394582 | -2.022622047 | 0.043112129 | 0.157877413 |
| Rasgrp4 | 51.39808938 | -0.339459666 | 0.167850791 | -2.022389435 | 0.043136136 | 0.157927998 |
| Postn | 120.7945585 | -0.312983343 | 0.154799778 | -2.02185912 | 0.043190908 | 0.158091172 |
| Fam120c | 3208.517417 | -0.145307309 | 0.071873827 | -2.021699899 | 0.043207364 | 0.158114054 |
| Xlr3b | 38.53565111 | 0.224356493 | 0.153076058 | 1.465653716 | 0.142742649 | 0.338950898 |
| Znrf3 | 588.5286384 | -0.163761878 | 0.081022303 | -2.021195052 | 0.043259578 | 0.158230382 |
| Dctpp1 | 253.6777673 | -0.206635155 | 0.102270197 | -2.020482612 | 0.043333352 | 0.158434995 |
| Gli3 | 513.1655299 | -0.173513374 | 0.085878266 | -2.020457354 | 0.043335969 | 0.158434995 |
| Ccbp2 | 172.4472642 | 0.224328543 | 0.152198361 | 1.473922201 | 0.140502539 | 0.335121051 |
| Kcnj5 | 56.7339992 | 0.224246153 | 0.165994229 | 1.350927411 | 0.176718687 | 0.385971149 |
| Mtss1l | 8229.557001 | -0.114818329 | 0.056848331 | -2.019730862 | 0.043411312 | 0.158598177 |
| Ap2s1 | 3728.417212 | -0.127101907 | 0.062937294 | -2.019500682 | 0.043435206 | 0.158648064 |
| Nrgn | 37700.61387 | 0.224139106 | 0.122289771 | 1.83285245 | 0.066824517 | 0.20864524 |
| Fank1 | 134.6905574 | 0.224096815 | 0.160782787 | 1.393786107 | 0.163382205 | 0.3686973 |
| Serinc4 | 14.02335306 | 0.223999096 | 0.165065329 | 1.357032982 | 0.174770731 | 0.383446291 |
| Ptchd1 | 685.1208519 | -0.229514764 | 0.113776267 | -2.017246396 | 0.043669807 | 0.159354683 |
| Ulk3 | 899.5216897 | 0.223949033 | 0.108494704 | 2.064147138 | 0.039003768 | 0.147183702 |
| Apba3 | 491.4972891 | 0.223875631 | 0.096939851 | 2.309428258 | 0.020919828 | 0.097003177 |
| Acot5 | 83.87108093 | 0.223598385 | 0.151380946 | 1.477057653 | 0.139660184 | 0.334076546 |
| Fbxo17 | 87.52423607 | 0.22350409 | 0.140921154 | 1.586022284 | 0.112734255 | 0.293317303 |
| Anapc11 | 1156.828199 | -0.162388687 | 0.080557924 | -2.015800302 | 0.043820862 | 0.159717816 |
| P2ry6 | 93.35628654 | 0.223480542 | 0.143382489 | 1.55863204 | 0.119083495 | 0.303167873 |
| Vdr | 34.58514092 | 0.223353805 | 0.16577571 | 1.347325278 | 0.177875494 | 0.387414269 |
| Lcmt2 | 264.054934 | -0.192312656 | 0.095479728 | -2.014172644 | 0.043991411 | 0.160220654 |
| 9-Sep | 2993.252897 | 0.222840286 | 0.105584629 | 2.110537195 | 0.034812111 | 0.136517045 |
| 9330175M20Rik | 8.600210793 | 0.222748957 | 0.151796501 | 1.467418259 | 0.142262311 | 0.338118002 |
| Tcte2 | 77.01315574 | -0.279749535 | 0.138988175 | -2.012757808 | 0.044140115 | 0.160654671 |
| Rho | 13.72284992 | 0.222725178 | 0.165530808 | 1.345520999 | 0.178457045 | 0.387963161 |
| Ndufc2 | 1794.110212 | -0.190512449 | 0.094674405 | -2.012290945 | 0.044189276 | 0.160758077 |
| Mir541 | 9.145460242 | 0.222626521 | 0.158791248 | 1.402007504 | 0.160913006 | 0.365623421 |
| Cggbp1 | 2189.807455 | -0.15385137 | 0.076484087 | -2.011547461 | 0.044267663 | 0.160892133 |
| Zfp648 | 10.46353473 | 0.222518667 | 0.158826 | 1.401021661 | 0.161207596 | 0.365970767 |
| Nr2e1 | 643.0924687 | -0.173989398 | 0.086489654 | -2.011678737 | 0.044253813 | 0.160892133 |
| Nlgn3 | 2936.442729 | -0.138825714 | 0.06902172 | -2.011333731 | 0.044290218 | 0.160936359 |
| Gbx1 | 22.84130351 | -0.313692828 | 0.155989785 | -2.010983144 | 0.044327237 | 0.160962837 |
| Mfge8 | 6197.098281 | -0.331623779 | 0.164914616 | -2.010881674 | 0.044337956 | 0.160962837 |
| 2810410L24Rik | 128.1307582 | 0.222185193 | 0.140655522 | 1.579640736 | 0.114189165 | 0.296009189 |
| Tsks | 31.46846595 | 0.222123795 | 0.161013955 | 1.379531328 | 0.167730994 | 0.374188394 |
| Wdhd1 | 257.8627816 | 0.221780541 | 0.110423174 | 2.008460119 | 0.04459442 | 0.161586984 |
| 2700094K13Rik | 342.5801323 | -0.158498317 | 0.07884808 | -2.010173455 | 0.044412834 | 0.161089456 |
| E230008N13Rik | 22.53036371 | 0.221755695 | 0.158553958 | 1.398613431 | 0.161928936 | 0.366938024 |
| Sncaip | 197.6563248 | -0.255667888 | 0.127178605 | -2.010305817 | 0.044398832 | 0.161089456 |
| Synj2 | 3778.113495 | 0.221607935 | 0.105106608 | 2.108411071 | 0.034995448 | 0.136916399 |
| Lingo3 | 1859.619775 | -0.159675709 | 0.079451944 | -2.009714328 | 0.044461433 | 0.161200833 |
| Afmid | 121.571033 | 0.221595043 | 0.119218283 | 1.858733728 | 0.06306489 | 0.200848388 |
| Wdr81 | 1394.502408 | 0.221594261 | 0.086869445 | 2.550888415 | 0.010744872 | 0.061955245 |
| Mgat2 | 886.3042739 | -0.137740176 | 0.068591565 | -2.008121212 | 0.044630413 | 0.161679592 |
| Ebpl | 293.036126 | -0.211397147 | 0.105288361 | -2.007792179 | 0.04466538 | 0.161768444 |
| Abhd13 | 1634.40341 | -0.138230229 | 0.068856129 | -2.007522517 | 0.044694056 | 0.161780242 |
| Snrk | 3852.552766 | -0.122770718 | 0.061154533 | -2.007548933 | 0.044691246 | 0.161780242 |
| Stard8 | 1650.700487 | -0.141928071 | 0.070700077 | -2.007466988 | 0.044699962 | 0.161780242 |
| Pknox1 | 704.1357927 | -0.181212017 | 0.090289989 | -2.00700009 | 0.044749653 | 0.161922263 |
| Cldn14 | 24.27679009 | 0.221555043 | 0.167554878 | 1.322283455 | 0.18607378 | 0.39777119 |
| Enpep | 153.8192603 | -0.215293271 | 0.107307461 | -2.006321557 | 0.044821951 | 0.16210815 |
| Nkd2 | 127.792785 | 0.221538817 | 0.143863903 | 1.539919415 | 0.123579998 | 0.31078722 |
| Rnf122 | 143.6863559 | -0.21293693 | 0.10616533 | -2.005710625 | 0.044887131 | 0.16226813 |
| Etnk2 | 288.9004254 | 0.221530414 | 0.147403446 | 1.502884904 | 0.132868726 | 0.323537333 |
| Ankrd10 | 1254.189588 | 0.221486218 | 0.102366386 | 2.163661601 | 0.03049033 | 0.125077356 |
| Cck | 6035.833451 | -0.238624809 | 0.11905924 | -2.004252741 | 0.045042994 | 0.162717683 |
| Neurod6 | 1988.816101 | -0.270906694 | 0.135173897 | -2.00413467 | 0.045055636 | 0.162725415 |
| Dnaja4 | 2447.624409 | -0.096046943 | 0.047940747 | -2.003451119 | 0.045128889 | 0.162927061 |
| Snrnp25 | 347.8560901 | -0.159980756 | 0.079853929 | -2.003417456 | 0.045132499 | 0.162927061 |
| Tchp | 403.4008906 | 0.221415473 | 0.087460698 | 2.531599658 | 0.011354355 | 0.064135753 |
| Nmt2 | 1892.492432 | -0.102410349 | 0.0511308 | -2.002909163 | 0.045187039 | 0.163047971 |
| Tmem33 | 3564.515493 | -0.178316354 | 0.089040133 | -2.002651485 | 0.04521471 | 0.163109828 |
| 2310014L17Rik | 37.63967669 | 0.221246323 | 0.157062254 | 1.408653682 | 0.158937597 | 0.362517169 |
| Adam32 | 8.848845749 | 0.221225018 | 0.154270577 | 1.43400655 | 0.151570394 | 0.35261737 |
| Ap5z1 | 861.5989474 | 0.221172604 | 0.105380299 | 2.098804101 | 0.035834173 | 0.139186432 |
| Zfp691 | 190.2602248 | 0.221099462 | 0.10031452 | 2.204062401 | 0.027519956 | 0.11672439 |
| Adam18 | 22.40908354 | -0.33421128 | 0.166998151 | -2.001287308 | 0.045361437 | 0.163448861 |
| 2700081L22Rik | 8.448350352 | -0.314548567 | 0.15718244 | -2.001168615 | 0.045374222 | 0.163456916 |
| Wdr70 | 474.513572 | 0.220964137 | 0.12516349 | 1.765404098 | 0.077495875 | 0.2293203 |
| Caln1 | 2811.706554 | -0.194167392 | 0.097115078 | -1.999353724 | 0.045570095 | 0.163895788 |
| Khsrp | 3689.281601 | 0.099440111 | 0.049731189 | 1.999552247 | 0.045548635 | 0.163895788 |
| Notch3 | 830.0802891 | -0.158251547 | 0.079146926 | -1.999465488 | 0.045558012 | 0.163895788 |
| Rdx | 2902.333307 | -0.121978458 | 0.061005865 | -1.999454603 | 0.045559189 | 0.163895788 |
| Tmem151b | 5857.611601 | -0.200889489 | 0.100472917 | -1.999439206 | 0.045560853 | 0.163895788 |
| Ush1g | 40.95785739 | -0.309837868 | 0.15493333 | -1.999814167 | 0.045520334 | 0.163895788 |
| Ppp1r2 | 5308.081901 | -0.213294204 | 0.106687012 | -1.999251836 | 0.045581113 | 0.163897368 |
| Cenpq | 126.1802555 | -0.249183453 | 0.12465119 | -1.999045914 | 0.045603386 | 0.163939412 |
| Phb | 2552.368448 | -0.098448631 | 0.049259345 | -1.998577731 | 0.045654062 | 0.164058455 |
| Cd34 | 2991.573717 | 0.220957282 | 0.105465292 | 2.095071076 | 0.036164675 | 0.140031964 |
| Tnc | 173.9900211 | -0.239212446 | 0.119705049 | -1.998348834 | 0.045678855 | 0.164058455 |
| Batf2 | 8.488925228 | 0.220847881 | 0.141526786 | 1.56046701 | 0.11864956 | 0.302391407 |
| Ehd3 | 5290.596227 | 0.220522838 | 0.093557643 | 2.357079874 | 0.018419284 | 0.089323441 |
| Emb | 907.4011932 | 0.22033939 | 0.088563728 | 2.487919103 | 0.012849296 | 0.070002457 |
| Ldhd | 562.3665884 | 0.22028979 | 0.090755228 | 2.427295863 | 0.015211845 | 0.078642751 |
| Was | 67.52605409 | 0.220237748 | 0.154974374 | 1.421123653 | 0.155280815 | 0.357279618 |
| Hspa13 | 1504.671721 | -0.179867593 | 0.090069282 | -1.99699152 | 0.045826104 | 0.164396772 |
| Arv1 | 171.6763115 | 0.220134535 | 0.125205143 | 1.758190833 | 0.078715044 | 0.231932319 |
| Tex26 | 11.558884 | 0.220034229 | 0.161475458 | 1.362648116 | 0.172993437 | 0.380811505 |
| Zbtb7c | 271.8409204 | 0.219896549 | 0.137187569 | 1.602889756 | 0.108958996 | 0.286793683 |
| Rpl19 | 9860.232707 | -0.13040246 | 0.065319733 | -1.996371599 | 0.04589349 | 0.164486174 |
| Samm50 | 2864.796395 | 0.097083249 | 0.048663974 | 1.994971675 | 0.046045969 | 0.164994507 |
| Heatr6 | 831.4745969 | 0.219784429 | 0.097697698 | 2.24963775 | 0.02447195 | 0.107902218 |
| Thap3 | 333.063253 | 0.219743868 | 0.087923999 | 2.499247875 | 0.012445722 | 0.068450002 |
| Palm | 7941.578703 | -0.137511798 | 0.068970956 | -1.993763855 | 0.046177867 | 0.165337144 |
| Tmem177 | 431.8959443 | -0.175927009 | 0.08823633 | -1.993816026 | 0.046172164 | 0.165337144 |
| Cacna2d2 | 2876.383263 | -0.240180839 | 0.120496014 | -1.993267921 | 0.046232117 | 0.165470183 |
| Rps17 | 4226.715614 | -0.154005298 | 0.077268588 | -1.993116508 | 0.046248691 | 0.165491274 |
| Htr2c | 1970.653092 | -0.316648584 | 0.158879426 | -1.993011886 | 0.046260146 | 0.165494042 |
| Polr2d | 452.104452 | -0.148986958 | 0.074774744 | -1.992476989 | 0.046318748 | 0.165635037 |
| Chgb | 18358.45937 | 0.219727192 | 0.094354167 | 2.328749211 | 0.019872355 | 0.093844034 |
| Large | 6934.451898 | -0.131842467 | 0.066175857 | -1.992304627 | 0.046337644 | 0.165656544 |
| Lrrd1 | 12.06337056 | 0.219379057 | 0.161294499 | 1.360114938 | 0.173793555 | 0.381967414 |
| Psg23 | 36.48122404 | 0.219168309 | 0.166250655 | 1.318300428 | 0.187403098 | 0.399838005 |
| Psmc3ip | 74.25716914 | -0.281242974 | 0.141199098 | -1.991818488 | 0.046390977 | 0.165732487 |
| Gcnt1 | 181.2957898 | -0.329673308 | 0.165545103 | -1.991441019 | 0.046432423 | 0.165842316 |
| Cpne6 | 6542.291153 | 0.219116175 | 0.109524089 | 2.000620846 | 0.045433265 | 0.16363157 |
| Rbpjl | 14.43624082 | 0.219034158 | 0.1640175 | 1.335431635 | 0.181735117 | 0.392282516 |
| Herc6 | 651.1058136 | 0.218920852 | 0.114953139 | 1.904435623 | 0.056853473 | 0.188465494 |
| Arhgef28 | 1517.815833 | -0.189757151 | 0.09538292 | -1.989424854 | 0.046654329 | 0.166481384 |
| Tcirg1 | 552.6248563 | 0.218779556 | 0.097667239 | 2.240050597 | 0.025087638 | 0.109402721 |
| Gm9159 | 8.588911443 | 0.218707177 | 0.148156089 | 1.476194317 | 0.139891735 | 0.334334817 |
| Dgkh | 1088.011768 | -0.173193913 | 0.087106264 | -1.988306057 | 0.046777851 | 0.166752575 |
| Capn6 | 75.64997627 | 0.218579537 | 0.164138367 | 1.331678516 | 0.182965852 | 0.393787041 |
| Tmem194 | 200.9220233 | 0.218518251 | 0.09216963 | 2.370827025 | 0.017748335 | 0.087188555 |
| Setx | 3444.651695 | -0.164354068 | 0.08267597 | -1.987930321 | 0.046819397 | 0.166839865 |
| Araf | 4917.438419 | -0.11679196 | 0.058760325 | -1.987598968 | 0.046856061 | 0.166932123 |
| F13a1 | 131.5722786 | -0.329264237 | 0.165674592 | -1.987415412 | 0.046876381 | 0.166966127 |
| Nudt22 | 183.8568566 | 0.218509077 | 0.116791215 | 1.870937618 | 0.061353728 | 0.197304828 |
| Clic1 | 364.8363775 | -0.191635675 | 0.096452227 | -1.986845522 | 0.046939518 | 0.167114177 |
| Oprl1 | 2084.905807 | 0.218369929 | 0.084669408 | 2.579088886 | 0.009906129 | 0.05838173 |
| Pld6 | 36.86469182 | -0.308005386 | 0.155039215 | -1.986628906 | 0.046963536 | 0.16712288 |
| Cd248 | 165.7146057 | -0.222926467 | 0.112226413 | -1.986399299 | 0.046989004 | 0.167136739 |
| Mmaa | 689.8980468 | 0.21822006 | 0.084882156 | 2.570859069 | 0.01014466 | 0.059471971 |
| C530008M17Rik | 1901.093845 | -0.184500995 | 0.092903508 | -1.985942174 | 0.047039745 | 0.167278818 |
| Dse | 263.7638344 | -0.20499017 | 0.10322687 | -1.985821812 | 0.047053113 | 0.16728796 |
| Synpo2 | 312.2082807 | 0.218128442 | 0.133112701 | 1.638674897 | 0.101280985 | 0.272471248 |
| Ptpn5 | 11064.15916 | -0.122204564 | 0.061558276 | -1.985184953 | 0.047123898 | 0.167424924 |
| S1pr5 | 428.3868681 | 0.218075477 | 0.156578787 | 1.39275237 | 0.163694686 | 0.368860597 |
| 4930479D17Rik | 55.94142799 | 0.218033758 | 0.148650466 | 1.466754629 | 0.142442816 | 0.33839451 |
| Fes | 160.2739849 | 0.218028581 | 0.110708068 | 1.969401011 | 0.04890706 | 0.171440268 |
| Pex11a | 179.6917316 | -0.226782288 | 0.114284452 | -1.984366942 | 0.047214949 | 0.167632542 |
| Catsperd | 41.3197543 | 0.217994027 | 0.168121363 | 1.296646799 | 0.194752742 | 0.409152405 |
| Klhl32 | 370.0551405 | -0.165605898 | 0.083479051 | -1.983801886 | 0.04727793 | 0.167779259 |
| Zbtb16 | 1489.543646 | 0.217845786 | 0.16601577 | 1.31219935 | 0.189452881 | 0.402769339 |
| Hic1 | 113.8407213 | -0.25751591 | 0.129855756 | -1.98309199 | 0.047357155 | 0.167983462 |
| Atp5c1 | 7163.682559 | -0.123835687 | 0.062461893 | -1.982579804 | 0.047414385 | 0.168109493 |
| Abtb1 | 1151.761487 | 0.217798402 | 0.094939088 | 2.294085689 | 0.021785577 | 0.099600041 |
| Mif | 3642.633674 | -0.129419009 | 0.065304194 | -1.98178709 | 0.047503075 | 0.168385414 |
| Zc3hav1l | 266.6681085 | -0.21745024 | 0.109738165 | -1.981537045 | 0.047531079 | 0.168446144 |
| Zkscan6 | 479.693555 | 0.217682586 | 0.085666722 | 2.541040237 | 0.01105232 | 0.062934453 |
| Papln | 219.0317001 | -0.289349593 | 0.14604702 | -1.981208472 | 0.0475679 | 0.168514103 |
| Cutc | 179.1464652 | 0.217602821 | 0.101840251 | 2.136707432 | 0.032621795 | 0.130672723 |
| Cdk5r1 | 11967.49081 | -0.162551194 | 0.082068347 | -1.980680744 | 0.047627088 | 0.168596639 |
| Slit2 | 1244.625576 | -0.151362628 | 0.076419795 | -1.980673041 | 0.047627952 | 0.168596639 |
| C130071C03Rik | 83.45617654 | 0.217152485 | 0.142620159 | 1.522593202 | 0.127860511 | 0.316212222 |
| Aldh6a1 | 2641.05394 | 0.216962061 | 0.085057343 | 2.550774019 | 0.010748399 | 0.061955245 |
| Rabl2 | 879.8374207 | 0.216828932 | 0.113421964 | 1.911701439 | 0.0559145 | 0.186630603 |
| Amfr | 7135.668803 | -0.088707573 | 0.044814399 | -1.979443548 | 0.04776609 | 0.168930373 |
| Elovl1 | 564.0734513 | 0.216782878 | 0.09051626 | 2.394960624 | 0.01662215 | 0.083320477 |
| Tmie | 183.1645321 | 0.216605652 | 0.121707867 | 1.779717751 | 0.075122164 | 0.22525853 |
| Gm20605 | 34.31490836 | 0.216601937 | 0.167769635 | 1.291067582 | 0.196680244 | 0.411732505 |
| Hspa4l | 5052.41623 | -0.139048345 | 0.070314058 | -1.97753263 | 0.047981456 | 0.169538125 |
| Smim15 | 1050.91118 | -0.167000402 | 0.084457456 | -1.97733167 | 0.048004152 | 0.169579647 |
| Sema4g | 1196.75279 | 0.21654403 | 0.096468761 | 2.244706229 | 0.024786998 | 0.108703579 |
| AI450353 | 53.87297458 | 0.216509014 | 0.160398901 | 1.349816065 | 0.177074991 | 0.386476955 |
| Smoc1 | 695.0822383 | 0.216390918 | 0.148853881 | 1.453713645 | 0.146025714 | 0.344896708 |
| C230081A13Rik | 1956.393236 | -0.213684673 | 0.108105962 | -1.97662246 | 0.048084321 | 0.169669432 |
| Cln8 | 1155.060086 | -0.117999897 | 0.059697483 | -1.976631016 | 0.048083354 | 0.169669432 |
| Hsp90b1 | 14339.63203 | -0.132173292 | 0.066882266 | -1.976208346 | 0.048131185 | 0.169796125 |
| Pop7 | 438.6963896 | -0.16670934 | 0.084366339 | -1.976017244 | 0.048152824 | 0.169833794 |
| Rnf7 | 2090.483091 | -0.121468392 | 0.061478871 | -1.975774595 | 0.048180312 | 0.16989207 |
| Ctxn3 | 90.45444598 | -0.329805695 | 0.166973903 | -1.975193067 | 0.048246242 | 0.170085843 |
| Cabyr | 18.83830968 | 0.216344707 | 0.167568908 | 1.291079053 | 0.196676267 | 0.411732505 |
| Pmaip1 | 88.78510312 | 0.216012613 | 0.151545205 | 1.425400511 | 0.154041461 | 0.355536778 |
| Atm | 1009.187116 | -0.122168279 | 0.061887023 | -1.974053242 | 0.048375689 | 0.170387117 |
| Pak7 | 2050.458478 | 0.215946838 | 0.121300913 | 1.780257314 | 0.07503386 | 0.225173055 |
| Yipf5 | 1333.845546 | -0.147197142 | 0.074592642 | -1.973346668 | 0.048456079 | 0.170631476 |
| Rhpn2 | 319.571945 | 0.215902796 | 0.138102452 | 1.56335237 | 0.117969737 | 0.301422015 |
| Insm1 | 157.8201947 | -0.272510667 | 0.138155226 | -1.972496256 | 0.048552983 | 0.17089503 |
| Slc22a21 | 39.18043603 | 0.215809952 | 0.165114155 | 1.30703483 | 0.191200883 | 0.405040535 |
| Rhbdl3 | 999.0210003 | -0.253956059 | 0.128769713 | -1.972172284 | 0.048589942 | 0.17094745 |
| Abhd2 | 2519.656477 | -0.174119571 | 0.088298034 | -1.97195297 | 0.048614976 | 0.170996693 |
| F420014N23Rik | 13.53092543 | 0.215801919 | 0.157747792 | 1.368018633 | 0.171306244 | 0.378651855 |
| Sra1 | 916.6028501 | -0.133549026 | 0.06774295 | -1.971408476 | 0.048677172 | 0.171098935 |
| Nfxl1 | 399.6758742 | 0.215797945 | 0.090923553 | 2.373399837 | 0.017625173 | 0.086693469 |
| Ccdc12 | 869.8145279 | -0.124794102 | 0.063318754 | -1.970886902 | 0.048736814 | 0.171242905 |
| Fam114a1 | 242.5322714 | -0.20404975 | 0.103548025 | -1.970580801 | 0.048771844 | 0.171242905 |
| G0s2 | 101.2016957 | -0.253709971 | 0.128736618 | -1.970767715 | 0.048750451 | 0.171242905 |
| Gm20063 | 142.0319636 | -0.258530634 | 0.131196045 | -1.97056728 | 0.048773392 | 0.171242905 |
| Slco4a1 | 92.25342619 | 0.215743764 | 0.15943787 | 1.353152577 | 0.176006892 | 0.385096926 |
| Zfp770 | 956.9522956 | -0.14659421 | 0.074407862 | -1.970144088 | 0.048821859 | 0.171374239 |
| Nlrp1a | 100.7171996 | 0.215419245 | 0.12489912 | 1.72474589 | 0.084573277 | 0.242797546 |
| Cish | 288.3801045 | 0.215417436 | 0.084131875 | 2.560473508 | 0.010452963 | 0.060726739 |
| Zfp595 | 113.6468897 | 0.21534906 | 0.128742975 | 1.672705321 | 0.094385321 | 0.259890913 |
| Ubxn6 | 2593.555208 | -0.11185126 | 0.056785729 | -1.969707212 | 0.048871936 | 0.171405855 |
| Tox2 | 624.4545049 | 0.215283405 | 0.094658371 | 2.274319775 | 0.022946767 | 0.103340221 |
| Tlr13 | 142.6334863 | 0.215177165 | 0.118250469 | 1.819672829 | 0.068808843 | 0.21246433 |
| Slc22a4 | 300.0352939 | 0.215163169 | 0.091750275 | 2.345095618 | 0.019022201 | 0.09113481 |
| Ppp2r2a | 4260.257295 | -0.130473113 | 0.066269238 | -1.96883375 | 0.048972185 | 0.171557863 |
| Ramp1 | 1929.127689 | -0.113369023 | 0.057582234 | -1.968819486 | 0.048973824 | 0.171557863 |
| Gm10767 | 48.58135333 | 0.215128415 | 0.155733795 | 1.381385554 | 0.167160445 | 0.373111272 |
| Ftsj2 | 263.8125868 | -0.180145443 | 0.091554665 | -1.967627121 | 0.049110957 | 0.171950425 |
| Gm13293 | 58.79498809 | 0.214951081 | 0.167182423 | 1.285727749 | 0.198538093 | 0.41450741 |
| Tro | 8701.391647 | 0.214940728 | 0.090361496 | 2.37867606 | 0.017374937 | 0.085652969 |
| Ecscr | 99.61098136 | -0.304544713 | 0.154810576 | -1.967208707 | 0.049159155 | 0.172012634 |
| Ctnnd2 | 8807.93021 | -0.113783568 | 0.057851182 | -1.966832219 | 0.049202557 | 0.17212563 |
| 1700007G11Rik | 14.94204235 | 0.214910619 | 0.163882654 | 1.311368922 | 0.189733153 | 0.403091979 |
| Rpe65 | 128.0334479 | 0.214861855 | 0.144257171 | 1.489436224 | 0.136372536 | 0.328915951 |
| Crhbp | 655.8090608 | 0.214743182 | 0.095958698 | 2.237870948 | 0.025229472 | 0.109897333 |
| As3mt | 443.246386 | -0.259577785 | 0.132059511 | -1.965612194 | 0.049343424 | 0.17246267 |
| Bai3 | 4434.438047 | -0.112880655 | 0.057435617 | -1.965342431 | 0.049374618 | 0.172532776 |
| Ppapdc1a | 148.8631456 | 0.214553355 | 0.118027043 | 1.817832162 | 0.069089789 | 0.21307269 |
| Gm14322 | 510.2207469 | 0.214519882 | 0.115791939 | 1.852632257 | 0.063935086 | 0.202703135 |
| Grk4 | 75.81620961 | 0.21447059 | 0.157238583 | 1.363981951 | 0.172573246 | 0.380349358 |
| Chmp2b | 1486.703382 | -0.13423174 | 0.068361453 | -1.963558923 | 0.049581265 | 0.17307478 |
| Ltbp3 | 1450.22034 | 0.21445592 | 0.12099558 | 1.772427721 | 0.076323584 | 0.227282425 |
| Tspan17 | 1160.232996 | 0.214448934 | 0.134951203 | 1.589085016 | 0.1120412 | 0.29211375 |
| Mblac1 | 96.29201085 | -0.243885639 | 0.12421453 | -1.963422794 | 0.049597067 | 0.17307478 |
| Nova2 | 4273.98137 | -0.10618815 | 0.054083893 | -1.963396942 | 0.049600069 | 0.17307478 |
| Spats1 | 67.66641361 | 0.214413403 | 0.150487053 | 1.424796345 | 0.154216079 | 0.355680596 |
| Ubxn10 | 70.54590193 | 0.214341644 | 0.149438465 | 1.434313744 | 0.15148275 | 0.352496758 |
| Moxd1 | 321.0978014 | 0.214130699 | 0.090586375 | 2.363828988 | 0.01808716 | 0.088276641 |
| Abi3 | 192.6927102 | 0.214108558 | 0.137216645 | 1.560368706 | 0.118672776 | 0.302391407 |
| Ugp2 | 3065.911484 | -0.191061611 | 0.097350778 | -1.962610002 | 0.049691508 | 0.173132498 |
| 0610012G03Rik | 891.0230695 | -0.127174795 | 0.064821846 | -1.961912558 | 0.049772665 | 0.173206003 |
| Clcn5 | 255.3842286 | -0.227508971 | 0.115966409 | -1.961852338 | 0.049779678 | 0.173206003 |
| Drg1 | 2567.631486 | -0.101245918 | 0.051599604 | -1.96214524 | 0.049745577 | 0.173206003 |
| Ept1 | 3299.788463 | -0.137760338 | 0.070216862 | -1.961926728 | 0.049771015 | 0.173206003 |
| Aim1l | 49.24972859 | 0.213891793 | 0.150210931 | 1.423942927 | 0.154462994 | 0.356085349 |
| Stau2 | 4178.629782 | -0.116969946 | 0.059622026 | -1.961857934 | 0.049779026 | 0.173206003 |
| Frmpd4 | 3631.074728 | -0.14079531 | 0.071804764 | -1.960807358 | 0.049901499 | 0.173512979 |
| N4bp3 | 494.4538448 | -0.204494235 | 0.104286936 | -1.960880645 | 0.049892948 | 0.173512979 |
| l7Rn6 | 850.4619408 | -0.149346161 | 0.07616521 | -1.960818605 | 0.049900187 | 0.173512979 |
| Cox6a1 | 7227.825755 | -0.156906215 | 0.080031155 | -1.960564161 | 0.049929887 | 0.173572731 |
| Mob4 | 1669.997492 | -0.150756313 | 0.076902212 | -1.960363791 | 0.049953285 | 0.173615118 |
| Mcmdc2 | 85.95875596 | 0.213788858 | 0.153580657 | 1.392029841 | 0.163913362 | 0.369138609 |
| Palb2 | 50.4393882 | 0.21378133 | 0.159413037 | 1.341052989 | 0.179903249 | 0.390176504 |
| Cep76 | 298.3568594 | 0.213710294 | 0.129824409 | 1.646148799 | 0.099733128 | 0.269632714 |

| | | | | | | |
|---|---|---|---|---|---|---|
| Utrn | 3483.326188 | -0.040217655 | 0.062801359 | -0.640394659 | 0.521916055 | 0.854113377 |
| Eif5b | 3892.350032 | -0.040246651 | 0.062592306 | -0.64299678 | 0.520226189 | 0.853297835 |
| Fkbp1b | 837.0261835 | -0.040262336 | 0.067369953 | -0.597630458 | 0.550086537 | 0.869294716 |
| Myo3b | 135.7653998 | -0.040281851 | 0.091866208 | -0.438483871 | 0.66103556 | 0.915428982 |
| Tspan4 | 447.7299201 | -0.04028548 | 0.07473701 | -0.539029861 | 0.58986625 | 0.885258542 |
| Ifitm6 | 5.926722942 | -0.040301752 | 0.047488125 | -0.848670102 | 0.396064887 | NA |
| Fbxl14 | 1140.831171 | -0.040323621 | 0.071741367 | -0.562069311 | 0.574068797 | 0.879633264 |
| D930015M05Rik | 4.88896531 | -0.040350695 | 0.047067338 | -0.857297158 | 0.391280679 | NA |
| Fam65a | 4345.699577 | -0.040398604 | 0.068652391 | -0.588451531 | 0.556229258 | 0.871546692 |
| Snora68 | 5.310437037 | -0.04043793 | 0.048064861 | -0.841320014 | 0.400168681 | NA |
| Tbc1d7 | 576.9956443 | -0.040490817 | 0.062949181 | -0.643230254 | 0.520074705 | 0.853297835 |
| Slc35f5 | 745.3700354 | -0.040496051 | 0.060213845 | -0.672537217 | 0.501241756 | 0.846601651 |
| Sgsm2 | 5861.383871 | -0.040516968 | 0.068947196 | -0.587652153 | 0.556765798 | 0.871733374 |
| Epha7 | 2474.650684 | -0.04054518 | 0.074560313 | -0.543790376 | 0.586585732 | 0.884628773 |
| Uxt | 183.5071012 | -0.040585015 | 0.08832165 | -0.45951378 | 0.645865258 | 0.912363974 |
| Fam151b | 201.978998 | -0.040607618 | 0.083763743 | -0.484787525 | 0.627827073 | 0.903578328 |
| Ccdc81 | 43.09806391 | -0.040636041 | 0.089442932 | -0.454323671 | 0.649595882 | 0.913724196 |
| Caskin1 | 7679.504247 | -0.040653241 | 0.058239483 | -0.698035749 | 0.485154838 | 0.838982004 |
| Ptprq | 4.939503204 | -0.040674249 | 0.047421273 | -0.857721572 | 0.391046226 | NA |
| Nova2 | 4478.027945 | -0.040675864 | 0.054218402 | -0.75022249 | 0.453120716 | 0.82562629 |
| Trps1 | 968.7568448 | -0.040708858 | 0.07136737 | -0.570412749 | 0.568397785 | 0.877329319 |
| Crtap | 222.9143748 | -0.040721405 | 0.089097681 | -0.457042256 | 0.647640673 | 0.912918818 |
| Ccdc96 | 81.95383742 | -0.040724289 | 0.094670429 | -0.430169053 | 0.667072672 | 0.916077199 |
| Gpr25 | 275.6256105 | -0.040779156 | 0.085525492 | -0.476807028 | 0.63349954 | 0.906932171 |
| Stx11 | 8.714227986 | -0.040781892 | 0.064067234 | -0.636548348 | 0.524419079 | NA |
| Slc22a1 | 6.326155981 | -0.040806264 | 0.055310762 | -0.737763541 | 0.460658152 | NA |
| Cers5 | 1468.316719 | -0.04081096 | 0.051120245 | -0.798332631 | 0.424677487 | 0.809188912 |
| Atg16l1 | 2835.26567 | -0.040811376 | 0.049216279 | -0.829225151 | 0.406977015 | 0.800975798 |
| Ap5m1 | 765.1691755 | -0.0408532 | 0.068975764 | -0.592283403 | 0.553660828 | 0.870774922 |
| Mtr | 450.3690184 | -0.04086802 | 0.079438964 | -0.514458113 | 0.606931739 | 0.893150688 |
| Pold4 | 311.8608596 | -0.040891124 | 0.086368558 | -0.473449192 | 0.635892742 | 0.907889736 |
| BB557941 | 9.030406223 | -0.040911704 | 0.063509201 | -0.644185459 | 0.519455179 | NA |
| Rnf111 | 1758.90836 | -0.040923717 | 0.057698806 | -0.709264547 | 0.478160325 | 0.835931746 |
| Ston2 | 1043.639745 | -0.040931075 | 0.06955218 | -0.588494491 | 0.556200431 | 0.871546692 |
| Gm765 | 118.2116051 | -0.040945479 | 0.094804435 | -0.431894129 | 0.665818362 | 0.915591677 |
| Snord68 | 7.883669935 | -0.040952396 | 0.059243998 | -0.691249701 | 0.489408635 | NA |
| C230024C17Rik | 1.505577713 | -0.040966945 | 0.027296251 | -1.500826802 | 0.133400365 | NA |
| 1700023F02Rik | 1.170791692 | -0.041007958 | 0.022998699 | -1.783055547 | 0.074577262 | NA |
| Slamf7 | 3.210977104 | -0.041012434 | 0.030343371 | -1.351611006 | 0.176499788 | NA |
| Crtc1 | 10452.1949 | -0.041020639 | 0.063956124 | -0.64138719 | 0.521271154 | 0.853911973 |
| Csmd1 | 2938.989296 | -0.04103283 | 0.076649287 | -0.535332176 | 0.592420181 | 0.886226713 |
| Ylpm1 | 4580.971564 | -0.041034953 | 0.069737714 | -0.588418376 | 0.556251506 | 0.871546692 |
| Sh2d5 | 4827.366357 | -0.041048363 | 0.069097979 | -0.594060257 | 0.552471812 | 0.870265825 |
| Pdcd11 | 1211.880023 | -0.041065159 | 0.074396676 | -0.551975716 | 0.580964991 | 0.883796098 |
| Med30 | 395.5684241 | -0.041067414 | 0.086363477 | -0.4755183 | 0.634417593 | 0.907220906 |
| Pkn2 | 1177.920086 | -0.041078256 | 0.065681694 | -0.625414077 | 0.531699326 | 0.858328942 |
| Nlrc3 | 17.73668817 | -0.041086981 | 0.071129532 | -0.577636043 | 0.563509867 | NA |
| Nrm | 87.35412634 | -0.04109002 | 0.09509655 | -0.432087391 | 0.665677899 | 0.915591677 |
| Mns1 | 60.56858392 | -0.041091798 | 0.094868119 | -0.433146548 | 0.664908308 | 0.915428982 |
| Zfp873 | 165.4441022 | -0.041106479 | 0.086046518 | -0.477723908 | 0.632846726 | 0.906637358 |
| Trim13 | 300.4118027 | -0.041108848 | 0.07770814 | -0.529015985 | 0.596794361 | 0.887922451 |
| Lgals7 | 3.58848747 | -0.041115441 | 0.043088515 | -0.954208813 | 0.339977955 | NA |
| Mroh1 | 3836.331428 | -0.041115637 | 0.065002902 | -0.632520016 | 0.527047134 | 0.856196078 |
| Pcdhb9 | 595.4219 | -0.041149414 | 0.073669129 | -0.558570663 | 0.576454766 | 0.880966011 |
| Ap4b1 | 686.7151916 | -0.041149757 | 0.065908678 | -0.624345042 | 0.53240101 | 0.858328942 |
| Krcc1 | 452.3601433 | -0.041161962 | 0.080536293 | -0.5110983 | 0.609282226 | 0.89440021 |
| Mir3093 | 1.86405858 | -0.041174963 | 0.02989543 | -1.377299541 | 0.168419657 | NA |
| Ube2t | 35.32875421 | -0.041214217 | 0.083676995 | -0.492539401 | 0.622338076 | 0.900114632 |
| Zfp760 | 380.6097328 | -0.041217283 | 0.083010902 | -0.496528554 | 0.619521547 | 0.899129234 |
| Crat | 2341.42568 | -0.041221659 | 0.053031579 | -0.777304017 | 0.436979426 | 0.816564673 |
| A330050B17Rik | 87.75817268 | -0.041225136 | 0.095098248 | -0.433500484 | 0.664651214 | 0.915428982 |
| Alg13 | 270.8259585 | -0.041248786 | 0.08916819 | -0.462595304 | 0.643654475 | 0.911368058 |
| Itgb8 | 1244.87451 | -0.041268518 | 0.068619083 | -0.601414601 | 0.547563876 | 0.867553753 |
| Gm16023 | 11.88668495 | -0.041277457 | 0.06719029 | -0.614336636 | 0.538992898 | NA |
| Coq4 | 687.3960142 | -0.041292408 | 0.070986101 | -0.581697077 | 0.560770741 | 0.873713575 |
| Sharpin | 1333.895627 | -0.041302606 | 0.06874499 | -0.600808963 | 0.547967234 | 0.867726281 |
| Gm5126 | 85.66316328 | -0.041331319 | 0.094327994 | -0.438165994 | 0.661265958 | 0.915428982 |
| Myoc | 820.4377983 | -0.041339441 | 0.09390892 | -0.440207822 | 0.659786595 | 0.915208181 |
| Zfp408 | 519.1081987 | -0.041374841 | 0.081319705 | -0.508792318 | 0.610897805 | 0.894817133 |
| Stradb | 1586.523883 | -0.041381985 | 0.060132441 | -0.688180689 | 0.491339003 | 0.841892493 |
| Zfp930 | 448.155642 | -0.041383246 | 0.087454631 | -0.473196739 | 0.636072824 | 0.907889736 |
| Zfp865 | 956.7671921 | -0.041388839 | 0.067725021 | -0.611130701 | 0.541113057 | 0.863862966 |
| 1700048O20Rik | 344.7045669 | -0.041414811 | 0.094984387 | -0.436017041 | 0.662824366 | 0.915428982 |
| Rab39 | 42.32181912 | -0.041420405 | 0.094219266 | -0.439617152 | 0.660214416 | 0.915428982 |
| Gm12070 | 163.2437605 | -0.041441266 | 0.0947927 | -0.43717782 | 0.661982394 | 0.915428982 |
| Cd248 | 162.9279379 | -0.041441272 | 0.094950064 | -0.43645333 | 0.662507853 | 0.915428982 |
| 4930505A04Rik | 4.92959076 | -0.041446561 | 0.048336196 | -0.857464283 | 0.391188346 | NA |
| B230120H23Rik | 439.9706128 | -0.04144709 | 0.078855035 | -0.525611202 | 0.599158372 | 0.889150487 |
| D430042O09Rik | 1337.313168 | -0.041451852 | 0.055114054 | -0.75211037 | 0.451984688 | 0.824608086 |
| Upf1 | 3214.246418 | -0.041452826 | 0.057076709 | -0.726265171 | 0.467676227 | 0.832353544 |
| Cyp11a1 | 133.8465451 | -0.041454216 | 0.094624419 | -0.438092156 | 0.66131948 | 0.915428982 |
| Rmi1 | 717.6004462 | -0.041468348 | 0.064466084 | -0.643258376 | 0.52005646 | 0.853297835 |
| Jam3 | 1245.852579 | -0.041483844 | 0.06954289 | -0.596521714 | 0.550826752 | 0.869642174 |
| 1700029I15Rik | 31.27177332 | -0.041508039 | 0.0882388 | -0.470405754 | 0.638065154 | 0.908932006 |
| Slc25a53 | 357.6231718 | -0.041518331 | 0.071859436 | -0.577771456 | 0.563418429 | 0.875108781 |
| 2900002K06Rik | 9.141635964 | -0.04153832 | 0.064595412 | -0.643053724 | 0.52018924 | NA |
| Hipk4 | 5.714803124 | -0.041571505 | 0.046102442 | -0.901720233 | 0.367205502 | NA |
| Snord4a | 4.97117751 | -0.041578351 | 0.050506244 | -0.823231896 | 0.410376134 | NA |
| Fam129a | 120.6896662 | -0.04159396 | 0.095102355 | -0.437359935 | 0.661850335 | 0.915428982 |
| Zfp652 | 560.4461167 | -0.04160844 | 0.071305282 | -0.58352535 | 0.559539697 | 0.873713575 |
| Ttn | 20.25288244 | -0.041618269 | 0.0805073 | -0.51695026 | 0.605190882 | NA |
| Olfr317 | 1.810214737 | -0.041621195 | 0.027238211 | -1.528044368 | 0.126501522 | NA |
| Tbcd | 2080.912493 | -0.041632431 | 0.051657192 | -0.805936789 | 0.420279297 | 0.807657282 |
| 2610019E17Rik | 71.51897576 | -0.041638954 | 0.094432592 | -0.440938381 | 0.659257607 | 0.915208181 |
| 2810417H13Rik | 8.190380104 | -0.041643582 | 0.057014565 | -0.730402523 | 0.465144177 | NA |
| Sigirr | 61.49286018 | -0.041643644 | 0.094093545 | -0.442577052 | 0.658071686 | 0.915208181 |
| Hcfc1 | 5767.180005 | -0.041684012 | 0.048561351 | -0.858378346 | 0.390683581 | 0.794106635 |
| Smg9 | 1394.519165 | -0.041714079 | 0.067623842 | -0.616854615 | 0.537330625 | 0.861996453 |
| 1700003M07Rik | 740.3104653 | -0.04172023 | 0.068119199 | -0.612459192 | 0.540233989 | 0.863544778 |
| H2-Q1 | 31.01731263 | -0.04173609 | 0.070281544 | -0.593841395 | 0.552618199 | 0.870265825 |
| Plekha6 | 5832.610833 | -0.041767132 | 0.05390851 | -0.774778085 | 0.438470801 | 0.817139233 |
| Morc4 | 163.4447336 | -0.04178008 | 0.089598577 | -0.466302941 | 0.640998674 | 0.910692492 |
| Dlgap3 | 13361.05398 | -0.041793382 | 0.074139813 | -0.563710386 | 0.572951248 | 0.879310647 |
| Cd101 | 10.97252196 | -0.041806926 | 0.06785596 | -0.616112809 | 0.537820069 | NA |
| Gldc | 1296.12851 | -0.041829227 | 0.062098748 | -0.673592123 | 0.500570662 | 0.846307849 |
| L3mbtl4 | 30.56816909 | -0.041838106 | 0.084686083 | -0.494037565 | 0.62127965 | 0.899589902 |
| Mpp3 | 2382.843616 | -0.041842612 | 0.057565179 | -0.726873652 | 0.467303359 | 0.832276679 |
| Gbp10 | 16.74372577 | -0.041842677 | 0.023842407 | -1.754968605 | NA | NA |
| 1110058L19Rik | 493.4279497 | -0.041853775 | 0.068342543 | -0.612411731 | 0.540265382 | 0.863544778 |
| Crebbp | 2204.735788 | -0.041869417 | 0.073371884 | -0.570646608 | 0.568239218 | 0.877177777 |
| C030034L19Rik | 3.85451335 | -0.041869631 | 0.043562181 | -0.961146343 | 0.336478591 | NA |
| Mfap4 | 66.18306093 | -0.041879282 | 0.087225067 | -0.480128977 | 0.631135685 | 0.905558768 |
| Sema5b | 883.0659627 | -0.041881738 | 0.071720177 | -0.583960333 | 0.559247 | 0.873695625 |
| Mrgprh | 4.081931266 | -0.041895731 | 0.047556694 | -0.880963915 | 0.378337351 | NA |
| Vps37a | 2369.623989 | -0.04189868 | 0.062720751 | -0.668019422 | 0.504121195 | 0.847807553 |
| Pin4 | 547.7474896 | -0.041911512 | 0.071111651 | -0.589376166 | 0.555608961 | 0.871360989 |
| BC021767 | 3.833913218 | -0.041936825 | 0.044150941 | -0.949851222 | 0.342187855 | NA |
| Trpm1 | 9.884618947 | -0.041937265 | 0.060046758 | -0.698410135 | 0.48492074 | NA |
| Lnx2 | 262.238318 | -0.041954787 | 0.087183693 | -0.481222867 | 0.630358111 | 0.905249211 |
| Amer1 | 511.1855869 | -0.041993177 | 0.065795559 | -0.638237254 | 0.52331925 | 0.854628731 |
| Cdh26 | 8.450385051 | -0.042011918 | 0.05916797 | -0.710044952 | 0.477676261 | NA |
| Nyap2 | 608.9440898 | -0.042011973 | 0.083060207 | -0.505801445 | 0.612996044 | 0.895428201 |
| 0610040F04Rik | 51.03505515 | -0.042034684 | 0.09495107 | -0.442698374 | 0.657983919 | 0.915208181 |
| Bcl2l15 | 11.77644069 | -0.042035922 | 0.065819065 | -0.638658752 | 0.523044952 | NA |
| Aaas | 560.5213386 | -0.042092534 | 0.064637744 | -0.651206728 | 0.514913048 | 0.853297835 |
| 4921530L18Rik | 8.357808954 | -0.042092904 | 0.062261146 | -0.676070186 | 0.498996085 | NA |
| Epha8 | 514.1184521 | -0.042097379 | 0.090000381 | -0.467746671 | 0.639965761 | 0.910092095 |
| 1190005I06Rik | 155.7561113 | -0.042101948 | 0.09038022 | -0.465831442 | 0.641336157 | 0.911015942 |
| Cyr61 | 302.8813045 | -0.042137771 | 0.072390787 | -0.582087483 | 0.560507756 | 0.873713575 |
| Fgf7 | 21.65813229 | -0.042139614 | 0.079820768 | -0.527927947 | 0.597549347 | NA |
| Atp7b | 107.4730051 | -0.042145099 | 0.094591294 | -0.445549447 | 0.655922735 | 0.915208181 |
| Trpm6 | 41.67163737 | -0.042179155 | 0.089521278 | -0.471163462 | 0.637524009 | 0.908932006 |
| Ccdc109b | 52.41279572 | -0.042179523 | 0.0944705 | -0.446483541 | 0.655247999 | 0.915208181 |
| Nup155 | 934.0509391 | -0.042182941 | 0.058854261 | -0.716735534 | 0.473537296 | 0.833983942 |
| Kif20b | 27.2371596 | -0.0421851 | 0.079912341 | -0.527892177 | 0.597574175 | 0.888198349 |
| Vps16 | 2398.202141 | -0.042207858 | 0.061393441 | -0.687497834 | 0.491769065 | 0.841892493 |
| Rin1 | 3428.076075 | -0.042216558 | 0.08013154 | -0.526840713 | 0.598304208 | 0.888717024 |
| Taf2 | 1626.599421 | -0.042242893 | 0.058299451 | -0.724584753 | 0.468706816 | 0.832446623 |
| Gm19557 | 3.52462533 | -0.042247304 | 0.037721317 | -1.119984887 | 0.262720202 | NA |
| 2810474O19Rik | 529.8974601 | -0.042249973 | 0.080655457 | -0.523832792 | 0.600394843 | 0.889841393 |
| Mdfi | 10.30369588 | -0.042278576 | 0.065250872 | -0.647938868 | 0.51702449 | NA |
| Ybx1 | 3674.499372 | -0.042292462 | 0.048684047 | -0.868712943 | 0.38500416 | 0.789016184 |
| Ntf5 | 4.827497343 | -0.04232804 | 0.047952511 | -0.882707489 | 0.377394335 | NA |
| Numb | 2059.618356 | -0.042339388 | 0.065297763 | -0.64840488 | 0.516723115 | 0.853297835 |
| Mgl2 | 14.7431898 | -0.042356849 | 0.06883287 | -0.615357875 | 0.538318405 | NA |
| Kank2 | 545.4459165 | -0.042359951 | 0.072792604 | -0.581926571 | 0.560616142 | 0.873713575 |
| Creb3l2 | 366.4912308 | -0.042364702 | 0.077343101 | -0.547750233 | 0.583863415 | 0.884313345 |
| 2810408A11Rik | 12.73600034 | -0.042416923 | 0.058926169 | -0.719831674 | 0.47162864 | NA |
| Ptpre | 1445.596094 | -0.042420429 | 0.058807324 | -0.72134602 | 0.470696649 | 0.833245343 |
| Nodal | 21.64464367 | -0.042428368 | 0.078100103 | -0.543256232 | 0.586953394 | NA |
| Syne2 | 949.7837351 | -0.042430334 | 0.061486905 | -0.690071061 | 0.490149501 | 0.84139821 |
| Tmem206 | 414.4254114 | -0.042431229 | 0.071907677 | -0.590079271 | 0.555137505 | 0.871360989 |
| Shh | 120.1114167 | -0.042472803 | 0.092843414 | -0.457467057 | 0.647335375 | 0.912750424 |
| Irg1 | 1.573195958 | -0.042473649 | 0.02701237 | -1.57237776 | 0.115862969 | NA |
| Anxa2 | 434.1638064 | -0.042489704 | 0.085701424 | -0.495787612 | 0.620044266 | 0.899216177 |
| AU023762 | 91.30270151 | -0.042498601 | 0.094605433 | -0.449219449 | 0.653273359 | 0.914683958 |
| Actr5 | 256.7364122 | -0.042537952 | 0.078160533 | -0.544238245 | 0.586277537 | 0.884588669 |
| Psors1c2 | 5.424324711 | -0.042542503 | 0.042411819 | -1.003081325 | 0.315821624 | NA |
| 1810026B05Rik | 275.8374221 | -0.042543794 | 0.086804117 | -0.49011263 | 0.624054201 | 0.900825929 |
| Sbf2 | 2876.940126 | -0.042588156 | 0.050298755 | -0.846703974 | 0.397160143 | 0.797284168 |
| Dact3 | 4727.598341 | -0.042647715 | 0.065310959 | -0.652994779 | 0.513759644 | 0.853195313 |
| Angptl3 | 5.866712144 | -0.042650094 | 0.052405969 | -0.813840391 | 0.415736393 | NA |
| Krtcap3 | 25.75287548 | -0.042665353 | 0.08574093 | -0.497607767 | 0.618760528 | 0.898575807 |
| Tectb | 3.508855062 | -0.042679724 | 0.036641991 | -1.164776358 | 0.244109531 | NA |
| Ndufa4l2 | 85.760714 | -0.042720662 | 0.094250384 | -0.45326777 | 0.650355938 | 0.914000665 |
| Gm15997 | 17.74214006 | -0.042727136 | 0.081298124 | -0.525561153 | 0.599193154 | NA |
| Hist3h2ba | 263.7124902 | -0.042730771 | 0.086304306 | -0.495117487 | 0.62051719 | 0.899412757 |
| Cgnl1 | 854.2880322 | -0.042751343 | 0.089883991 | -0.475628 | 0.634339424 | 0.907220906 |
| Atad2b | 755.7661509 | -0.042765214 | 0.074386848 | -0.574902896 | 0.565356971 | 0.875364912 |
| Pmel | 24.36044251 | -0.042765558 | 0.080835151 | -0.529046544 | 0.596773163 | 0.887922451 |
| 4930524B15Rik | 5.603620684 | -0.042768826 | 0.052068828 | -0.821390215 | 0.411424035 | NA |
| Wash | 1275.420718 | -0.042776922 | 0.063420554 | -0.674496185 | 0.499995911 | 0.846241311 |
| Acsf2 | 767.1893369 | -0.042802786 | 0.070159251 | -0.610080432 | 0.541808529 | 0.864489311 |
| Trrap | 6070.238127 | -0.042808954 | 0.071014255 | -0.602821975 | 0.546627127 | 0.867128614 |
| Cd4 | 642.3354204 | -0.042825001 | 0.094744943 | -0.452003025 | 0.651266803 | 0.914343204 |
| Scarb1 | 1134.969821 | -0.042837692 | 0.061582556 | -0.695614072 | 0.486670551 | 0.839801621 |
| Zfp386 | 636.6647695 | -0.042839845 | 0.07131279 | -0.600731571 | 0.548018787 | 0.867726281 |
| Kdm4c | 1253.125264 | -0.04285672 | 0.060451507 | -0.708943787 | 0.478359361 | 0.835936459 |
| Has2as | 2.026683379 | -0.042869551 | 0.031861522 | -1.345496027 | 0.178465104 | NA |
| Slmo1 | 1622.759108 | -0.042885196 | 0.055742897 | -0.769339211 | 0.441691965 | 0.819100918 |
| Allc | 21.18265081 | -0.042885331 | 0.080945088 | -0.529807697 | 0.596245268 | NA |
| Mlxip | 1731.936039 | -0.042889424 | 0.060741603 | -0.706096339 | 0.480128228 | 0.836914993 |
| Zfp414 | 965.4798051 | -0.042910054 | 0.085668737 | -0.50088347 | 0.616453137 | 0.897296158 |
| Tmem107 | 250.3442511 | -0.042922863 | 0.083471874 | -0.514219472 | 0.607098555 | 0.893305674 |
| 4931429L15Rik | 4.334018639 | -0.042943128 | 0.041009211 | -1.047158105 | 0.295026664 | NA |
| Gm13547 | 4.466372608 | -0.04295881 | 0.047144632 | -0.91121317 | 0.362183066 | NA |
| Pacsin2 | 3117.484608 | -0.042967505 | 0.057615311 | -0.745765388 | 0.455809151 | 0.826603827 |
| Rtp3 | 7.045666317 | -0.042971347 | 0.051422485 | -0.835652852 | 0.403350226 | NA |
| Tfeb | 325.0689475 | -0.042975074 | 0.084822067 | -0.506649691 | 0.612400636 | 0.895344807 |
| Tecrl | 10.5325273 | -0.042976136 | 0.06604609 | -0.650699171 | 0.515240698 | NA |
| Pi4k2b | 84.54333072 | -0.04298263 | 0.094941586 | -0.452727105 | 0.65074526 | 0.914190101 |
| Cnbd2 | 193.1065437 | -0.042987997 | 0.085755893 | -0.501283294 | 0.616171761 | 0.89720439 |
| BC026585 | 276.0836269 | -0.04299598 | 0.082712725 | -0.519823038 | 0.603186921 | 0.891523525 |
| Mrc2 | 359.6322112 | -0.043008502 | 0.09464073 | -0.454439664 | 0.64951241 | 0.913724196 |
| Itsn2 | 2384.277964 | -0.043012037 | 0.063745504 | -0.674746217 | 0.499837016 | 0.846241311 |
| Adamts1 | 511.678778 | -0.043022199 | 0.091962596 | -0.467822802 | 0.639911313 | 0.910092095 |
| Sirt6 | 598.9806409 | -0.043024042 | 0.084589129 | -0.50862377 | 0.611015964 | 0.89489435 |
| Lgmn | 4612.288785 | -0.043030776 | 0.060512526 | -0.711105277 | 0.477019 | 0.83524325 |
| Snord110 | 1.250600861 | -0.043052683 | 0.02640678 | -1.630364772 | 0.103024425 | NA |
| Dhx34 | 539.6156268 | -0.043067728 | 0.07721294 | -0.557778636 | 0.576995553 | 0.881236189 |
| Pln | 68.66275445 | -0.043088368 | 0.091796514 | -0.469390028 | 0.638790875 | 0.909380945 |
| 4930444M15Rik | 2.121325723 | -0.043090237 | 0.035173972 | -1.225060315 | 0.220552496 | NA |
| Omd | 26.13690973 | -0.043123343 | 0.079097071 | -0.545195193 | 0.585619277 | 0.88454878 |
| Glycam1 | 2.191804102 | -0.043127996 | 0.032741539 | -1.317225689 | 0.187762985 | NA |
| Mroh4 | 4.86843983 | -0.043130653 | 0.048650852 | -0.886534388 | 0.375329632 | NA |
| Zfp664 | 4921.272471 | -0.043142075 | 0.062610155 | -0.689058747 | 0.490786299 | 0.841866734 |
| Spem1 | 5.799838121 | -0.043144459 | 0.05480939 | -0.787172757 | 0.431180743 | NA |
| Kcnu1 | 145.7448643 | -0.043150468 | 0.092360345 | -0.467196915 | 0.640359001 | 0.910092095 |
| Wdr3 | 936.1831544 | -0.043199863 | 0.056287814 | -0.767481621 | 0.442795216 | 0.819995323 |
| Cd6 | 150.4396791 | -0.043242236 | 0.094526661 | -0.457460738 | 0.647339916 | 0.912750424 |
| Nr1h2 | 1799.39717 | -0.043269442 | 0.056729519 | -0.762732391 | 0.445623009 | 0.821773578 |
| Prokr1 | 8.034121145 | -0.043282974 | 0.05959395 | -0.726298113 | 0.467656036 | NA |
| Zfp382 | 402.6788821 | -0.043313477 | 0.078110167 | -0.554517781 | 0.57922454 | 0.882452682 |
| Bcorl1 | 1182.22536 | -0.043316296 | 0.067482196 | -0.641892202 | 0.520943178 | 0.85372574 |
| Pgls | 436.991174 | -0.043359873 | 0.080734817 | -0.537065355 | 0.591222471 | 0.8858275 |
| Tspan17 | 1499.313098 | -0.043389539 | 0.067742558 | -0.64050636 | 0.521843456 | 0.854113377 |
| Rgn | 8.798153665 | -0.043404488 | 0.060761099 | -0.714346671 | 0.47501284 | NA |
| Armc5 | 685.3417738 | -0.043411703 | 0.071364215 | -0.608311927 | 0.542980612 | 0.865503827 |
| Mir3082 | 10.21800983 | -0.043467806 | 0.055827709 | -0.778606292 | 0.436211671 | NA |
| Ccr10 | 5.212091725 | -0.043472023 | 0.046813214 | -0.928627175 | 0.353082334 | NA |
| Acaa1b | 13.06777948 | -0.043482532 | 0.065761878 | -0.661211824 | 0.50847648 | NA |
| 1700007L15Rik | 13.46687688 | -0.04349868 | 0.074089729 | -0.587108102 | 0.557131106 | NA |
| Gm2061 | 144.5439885 | -0.043503527 | 0.091010684 | -0.478004612 | 0.632646923 | 0.906637358 |
| Cd209a | 24.39922291 | -0.043544633 | 0.084947527 | -0.512606243 | 0.608226785 | 0.894011407 |
| Zfp81 | 381.5889177 | -0.04354547 | 0.090423271 | -0.481573712 | 0.630108804 | 0.905068772 |
| Tab2 | 3129.76991 | -0.043549947 | 0.061224073 | -0.711320638 | 0.476885566 | 0.835195112 |
| Tubgcp6 | 2321.103811 | -0.043558359 | 0.06184859 | -0.704274093 | 0.481262097 | 0.8371092 |
| Vsig8 | 6.936310254 | -0.043595495 | 0.058285662 | -0.747962592 | 0.454482721 | NA |
| Zzef1 | 2992.431442 | -0.043632903 | 0.060829323 | -0.717300489 | 0.473188706 | 0.833983942 |
| Enpp3 | 20.61971984 | -0.04367173 | 0.072198175 | -0.604886895 | 0.545254151 | NA |
| Snord23 | 2.158748485 | -0.043672558 | 0.030031289 | -1.454235226 | 0.145881102 | NA |
| Enthd2 | 706.8027797 | -0.043688954 | 0.072599504 | -0.601780339 | 0.547320364 | 0.867553753 |
| Ddx41 | 1530.30833 | -0.043692042 | 0.071865372 | -0.60797072 | 0.543206893 | 0.865558163 |
| Mccc2 | 703.2649048 | -0.043692317 | 0.079206561 | -0.551624972 | 0.581205323 | 0.883958169 |
| Ly6e | 9473.272843 | -0.043694421 | 0.045069094 | -0.96949856 | 0.332296498 | 0.757166892 |

| | | | | | | | | | | | | | | | |
|---|---|---|---|---|---|---|---|---|---|---|---|---|---|---|---|
| Tek | 683.292159 | -0.221948589 | 0.113341677 | -1.958225737 | 0.05020353 | 0.17432844 | | | Cyp21a1 | 1.873735045 | -0.043762589 | 0.02876664 | -1.521296512 | 0.128185445 | NA |
| Tctn1 | 414.3623719 | 0.21346218 | 0.114456849 | 1.865001373 | 0.062181212 | 0.199139251 | | | 2410006H16Rik | 382.6052973 | -0.043782841 | 0.075386509 | -0.580778199 | 0.56138995 | 0.873934985 |
| Cartpt | 282.2789716 | -0.329116406 | 0.168121218 | -1.957613742 | 0.050275354 | 0.174469598 | | | Rcc1 | 479.1378801 | -0.043802481 | 0.072871669 | -0.60109068 | 0.54777959 | 0.867631265 |
| Id3 | 856.2582333 | -0.181585752 | 0.09276432 | -1.957495646 | 0.050289223 | 0.174469598 | | | Fstl4 | 868.3593811 | -0.043817781 | 0.082767636 | -0.529407175 | 0.596523022 | 0.887922451 |
| Abhd15 | 50.80253228 | 0.213443052 | 0.168022643 | 1.270323142 | 0.203969548 | 0.421222205 | | | Ptgr1 | 36.02164019 | -0.043831879 | 0.090134753 | -0.486292771 | 0.626759604 | 0.902918401 |
| Nfs1 | 1413.552651 | -0.108994161 | 0.055701353 | -1.956759658 | 0.050375732 | 0.174730601 | | | Mtfmt | 433.2959033 | -0.043855969 | 0.076215723 | -0.575418914 | 0.565008016 | 0.875364912 |
| Map3k14 | 224.1868386 | 0.213200847 | 0.114743529 | 1.858064242 | 0.063159892 | 0.20106829 | | | Hmbox1 | 1026.298303 | -0.043875145 | 0.065576922 | -0.669063808 | 0.503454776 | 0.847807553 |
| Rbm12b2 | 278.4539419 | 0.213172543 | 0.101645474 | 2.097216284 | 0.035974434 | 0.139502243 | | | Pdia2 | 19.75583382 | -0.043875174 | 0.079000835 | -0.555376079 | 0.578637451 | NA |
| Yae1d1 | 1020.417012 | -0.124867651 | 0.06382351 | -1.956452253 | 0.050411902 | 0.174738704 | | | Rnf126 | 2154.196257 | -0.043878191 | 0.067265575 | -0.652312734 | 0.514199447 | 0.853195313 |
| Cnih | 2057.535605 | -0.105072906 | 0.053717947 | -1.956011185 | 0.050463837 | 0.174879598 | | | Mir709 | 3.225708945 | -0.043969923 | 0.042610255 | -1.031909416 | 0.302114554 | NA |
| Gm4951 | 19.11771252 | 0.21312278 | 0.123326583 | 1.728117125 | 0.083967228 | 0.24166991 | | | 2900009J06Rik | 23.55765944 | -0.043983542 | 0.083281803 | -0.528129075 | 0.597409752 | 0.888165441 |
| Gbp7 | 254.4399252 | 0.213098749 | 0.160052517 | 1.331430165 | 0.183047509 | 0.393853369 | | | Fbxo16 | 292.1191571 | -0.044007862 | 0.086354193 | -0.509620445 | 0.610317398 | 0.894800514 |
| Cpsf6 | 2306.861519 | 0.212878436 | 0.089166974 | 2.387413486 | 0.016967396 | 0.084304335 | | | Zmynd11 | 8394.373164 | -0.044018067 | 0.036487398 | -1.206390956 | 0.227666782 | 0.680415543 |
| A930016O22Rik | 25.27742302 | 0.212857275 | 0.167150813 | 1.273444448 | 0.202860373 | 0.419922328 | | | S100a11 | 155.8520556 | -0.044021861 | 0.095064316 | -0.463074504 | 0.643310965 | 0.911258473 |
| C130074G19Rik | 2663.999699 | 0.21285121 | 0.080472198 | 2.645027903 | 0.008168418 | 0.051012281 | | | Trmt6 | 1162.970233 | -0.04404283 | 0.065489362 | -0.672518853 | 0.501253442 | 0.846601651 |
| 1110002L01Rik | 492.2336206 | -0.206830442 | 0.10585816 | -1.953845054 | 0.050719544 | 0.175451801 | | | Gm3086 | 36.31725815 | -0.044045574 | 0.087764385 | -0.501861586 | 0.615764888 | 0.896906632 |
| St18 | 362.9649109 | 0.212716002 | 0.121529494 | 1.750324093 | 0.080062406 | 0.234762853 | | | Gm9573 | 2.489867221 | -0.044056017 | 0.032893698 | -1.339345212 | 0.180458317 | NA |
| Sod3 | 512.2862201 | 0.212686547 | 0.092478603 | 2.299846014 | 0.021456946 | 0.098651013 | | | Fam173a | 1698.059809 | -0.044061905 | 0.074838376 | -0.588760834 | 0.556021723 | 0.871546692 |
| Slx1b | 625.3364865 | 0.212525869 | 0.087511071 | 2.428559803 | 0.015158925 | 0.078500128 | | | Wdr93 | 16.28104093 | -0.04409977 | 0.068304594 | -0.645634028 | 0.518516394 | NA |
| Tango2 | 2402.549352 | -0.178558271 | 0.091399236 | -1.953607911 | 0.050747604 | 0.175470516 | | | A930005H10Rik | 224.4698388 | -0.044100308 | 0.080740144 | -0.546200509 | 0.584928116 | 0.884313345 |
| Xlr | 26.76001555 | 0.212467996 | 0.168077244 | 1.26410923 | 0.206190824 | 0.423815419 | | | Dbp | 3952.261596 | -0.044107895 | 0.066803654 | -0.660261705 | 0.5090859 | 0.850095619 |
| 2010204K13Rik | 430.0839923 | -0.189618986 | 0.097095325 | -1.952915707 | 0.050829584 | 0.175582561 | | | 4930512B01Rik | 1.839717732 | -0.044128896 | 0.029178661 | -1.512368766 | 0.130440076 | NA |
| AU021092 | 91.35431889 | -0.294418953 | 0.150748994 | -1.953040907 | 0.050814748 | 0.175582561 | | | Vps33b | 950.6436222 | -0.044137591 | 0.056197431 | -0.785402285 | 0.432217741 | 0.813254665 |
| Myo9b | 2369.877461 | 0.212320076 | 0.107553621 | 1.974085807 | 0.048371987 | 0.170387117 | | | Golgb1 | 4970.274831 | -0.044187162 | 0.065942261 | -0.670088678 | 0.502801262 | 0.84739426 |
| Deb1 | 782.2850529 | -0.175856968 | 0.090055582 | -1.952760319 | 0.050848003 | 0.175582561 | | | Soat2 | 7.281236281 | -0.044206721 | 0.058438124 | -0.756470568 | 0.449367112 | NA |
| Ptgdr | 35.1114014 | -0.322775333 | 0.165286493 | -1.952823407 | 0.050840524 | 0.175582561 | | | Dapl1 | 8.95994435 | -0.04425622 | 0.057810264 | -0.765542601 | 0.443948509 | NA |
| Pkp1 | 31.71495909 | 0.212302449 | 0.167313576 | 1.26888956 | 0.204480457 | 0.421576043 | | | Upk1b | 68.50374693 | -0.044292765 | 0.094543407 | -0.46849131 | 0.639433284 | 0.909767911 |
| Mff | 6370.222147 | -0.114510096 | 0.058648007 | -1.952497662 | 0.050879148 | 0.175611834 | | | Mettl2 | 594.8594167 | -0.044299675 | 0.063519468 | -0.697418853 | 0.485540706 | 0.839338843 |
| Slc2a13 | 6321.850495 | -0.143679681 | 0.073621028 | -1.951611994 | 0.050984288 | 0.175926033 | | | Nrgn | 46478.92447 | -0.044348319 | 0.051063296 | -0.868496999 | 0.385122314 | 0.789016184 |
| Zfp791 | 185.1506712 | -0.190745854 | 0.097741215 | -1.951539616 | 0.050992888 | 0.175926033 | | | Zfp369 | 665.8517258 | -0.044356569 | 0.068077737 | -0.651557628 | 0.51468659 | 0.853195313 |
| Solh | 1586.392046 | 0.212238545 | 0.084937408 | 2.498764086 | 0.012462725 | 0.068450002 | | | Znrf2 | 377.4218758 | -0.044386494 | 0.078014927 | -0.568948735 | 0.569390931 | 0.877556781 |
| Slc25a53 | 346.0892947 | -0.150067152 | 0.076913006 | -1.95112843 | 0.05104177 | 0.176016289 | | | Ubash3a | 2.263418031 | -0.044387408 | 0.027523667 | -1.612699628 | 0.106809781 | NA |
| Rps6ka1 | 603.1626318 | 0.212235267 | 0.085329401 | 2.487246658 | 0.012873611 | 0.07003183 | | | Gm19689 | 2.214672762 | -0.044439111 | 0.034353053 | -1.293600074 | 0.195803597 | NA |
| Zfp94 | 359.2682145 | 0.212097666 | 0.086511193 | 2.451678902 | 0.014219148 | 0.075063538 | | | Rdh1 | 4.068035654 | -0.044492336 | 0.04213116 | -1.056043477 | 0.290948352 | NA |
| Rem1 | 17.46079429 | -0.327883235 | 0.168113534 | -1.950367869 | 0.051132289 | 0.176210783 | | | Usp36 | 1899.463342 | -0.044506894 | 0.06012922 | -0.740187452 | 0.45918626 | 0.828381305 |
| Pnrc2 | 1169.353648 | -0.133555708 | 0.068493792 | -1.949895074 | 0.051188626 | 0.176365705 | | | 2210417K05Rik | 13.07325707 | -0.044520111 | 0.069089125 | -0.644386663 | 0.519324731 | NA |
| Adarb1 | 9992.912615 | -0.11145373 | 0.057186738 | -1.948943651 | 0.051302154 | 0.17667828 | | | Hspbap1 | 218.3290219 | -0.0445387 | 0.082812099 | -0.537828411 | 0.590695516 | 0.885693151 |
| Ctsd | 15207.27433 | 0.212034274 | 0.090115035 | 2.352928937 | 0.018626193 | 0.089960798 | | | Ttc39a | 150.325608 | -0.044547269 | 0.092638277 | -0.480873242 | 0.630606591 | 0.905249211 |
| Yars2 | 488.5549125 | -0.141642457 | 0.072681945 | -1.948798385 | 0.051319507 | 0.176698764 | | | Pemt | 65.72776903 | -0.044549122 | 0.094919191 | -0.469337357 | 0.638828517 | 0.909380945 |
| Map3k1 | 492.6648547 | -0.287225341 | 0.14739469 | -1.948681746 | 0.051333443 | 0.17670748 | | | Hlcs | 763.7866106 | -0.044565379 | 0.065377632 | -0.681660956 | 0.495453361 | 0.844369474 |
| Itpr3 | 282.3315393 | -0.204669152 | 0.105058167 | -1.948150802 | 0.051396922 | 0.176886696 | | | Nln | 1367.088097 | -0.044576616 | 0.056466234 | -0.789438451 | 0.42985579 | 0.811552535 |
| Bfsp1 | 25.10416442 | 0.211963658 | 0.166373062 | 1.27402631 | 0.202654092 | 0.419709177 | | | Slc22a8 | 777.7109672 | -0.04458361 | 0.088316448 | -0.504816613 | 0.613687646 | 0.895819717 |
| 6430550D23Rik | 26.0838998 | 0.21192096 | 0.16759192 | 1.264505833 | 0.206048528 | 0.423761947 | | | Gm11190 | 7.402993694 | -0.044589668 | 0.059692885 | -0.746984638 | 0.455072834 | NA |
| Cox16 | 287.4988112 | -0.189578446 | 0.097410692 | -1.946176982 | 0.051633484 | 0.177543098 | | | Trp73 | 50.25457952 | -0.044590176 | 0.08973926 | -0.496885937 | 0.619269489 | 0.898921615 |
| Gfra2 | 3100.363617 | -0.191536148 | 0.098415821 | -1.946192644 | 0.051631603 | 0.177543098 | | | Hnrnpa0 | 2886.35905 | -0.044593549 | 0.071060637 | -0.627542211 | 0.530303876 | 0.857915133 |
| Arhgap4 | 176.6525619 | 0.211898162 | 0.140251096 | 1.510848527 | 0.130827048 | 0.320515687 | | | Hnrnpm | 4184.749016 | -0.044646469 | 0.047240429 | -0.94509027 | 0.344612792 | 0.764478237 |
| Ppard | 954.4322572 | 0.211687087 | 0.09173922 | 2.307487308 | 0.021027669 | 0.097312635 | | | Sh3glb1 | 2315.150088 | -0.044653665 | 0.064654996 | -0.690645241 | 0.489788509 | 0.841119426 |
| Lrp2 | 21.29169247 | 0.211593368 | 0.167536841 | 1.262966204 | 0.206601326 | 0.424265201 | | | 2410127L17Rik | 307.9129223 | -0.044678069 | 0.083585391 | -0.534520068 | 0.592981769 | 0.886253111 |
| Agpat2 | 48.81788497 | 0.211587975 | 0.152582236 | 1.386714342 | 0.165528885 | 0.371158401 | | | Hmgn3 | 1160.918793 | -0.044711057 | 0.064438755 | -0.693853513 | 0.48777408 | 0.839801621 |
| Kif9 | 224.0303608 | 0.211480811 | 0.146854091 | 1.440074356 | 0.149846364 | 0.350131135 | | | Sipa1 | 594.1322035 | -0.044732645 | 0.071697065 | -0.623911805 | 0.532685507 | 0.858328942 |
| Obsl1 | 534.1506429 | 0.211319491 | 0.111303941 | 1.898580484 | 0.057619656 | 0.190029578 | | | Mir677 | 10.36118702 | -0.044735369 | 0.065022961 | -0.687993418 | 0.491456926 | NA |
| Bloc1s5 | 576.6016301 | -0.158223458 | 0.081389129 | -1.944036763 | 0.051891018 | 0.17815188 | | | Cachd1 | 622.7228952 | -0.044744641 | 0.088908683 | -0.503265143 | 0.61477787 | 0.896163393 |
| Hnrnpa0 | 2771.090071 | -0.160155718 | 0.08239676 | -1.943713767 | 0.051929977 | 0.17820666 | | | Snx33 | 493.2610494 | -0.044764412 | 0.086247606 | -0.519022081 | 0.603745343 | 0.891713763 |
| Tex9 | 376.305537 | 0.21130704 | 0.114422439 | 1.846727286 | 0.064786675 | 0.204482531 | | | D330022K07Rik | 1.990296802 | -0.044801298 | 0.031214428 | -1.43527531 | 0.151208661 | NA |
| Lxn | 1110.797032 | 0.211208684 | 0.108274387 | 1.950680035 | 0.05109512 | 0.176121866 | | | Nat1 | 57.60594234 | -0.044811222 | 0.095028392 | -0.471556138 | 0.637243641 | 0.908932006 |
| Nkd1 | 598.2562397 | -0.189394202 | 0.097478557 | -1.942931945 | 0.052024382 | 0.178451576 | | | Cxcr2 | 19.7530006 | -0.044841929 | 0.074072716 | -0.605377139 | 0.544928438 | NA |
| Dis3 | 550.9342643 | 0.210665726 | 0.088975059 | 2.367694136 | 0.017899327 | 0.087652391 | | | Acap1 | 2.194760634 | -0.044865176 | 0.032826158 | -1.366750741 | 0.171703448 | NA |
| Rps23 | 6034.698607 | -0.134123714 | 0.069048024 | -1.942469984 | 0.05208023 | 0.178564082 | | | Ltn1 | 2885.553642 | -0.044890198 | 0.063822099 | -0.703364493 | 0.481828628 | 0.837383765 |
| Leprel2 | 740.1303685 | 0.210226847 | 0.133337071 | 1.576657155 | 0.114874434 | 0.296941021 | | | Slco5a1 | 163.4932876 | -0.044902472 | 0.090509186 | -0.496109559 | 0.619817115 | 0.899216177 |
| Rtn4rl1 | 3001.220934 | 0.209983393 | 0.108346983 | 1.938064052 | 0.052615409 | 0.179774515 | | | 1810033B17Rik | 11.04505102 | -0.044916558 | 0.064816096 | -0.692984628 | 0.488319199 | NA |
| Camk2b | 36746.42362 | -0.087179452 | 0.044906992 | -1.94133357 | 0.05221783 | 0.178877528 | | | Cuzd1 | 10.4711266 | -0.044970972 | 0.06447292 | -0.697517213 | 0.485479171 | NA |
| Lcp1 | 522.7735732 | 0.20998012 | 0.106459666 | 1.972391311 | 0.048564953 | 0.170898338 | | | Shank3 | 10887.63873 | -0.044977041 | 0.066354194 | -0.677832679 | 0.497877789 | 0.845489269 |
| 1110008L16Rik | 243.4329302 | 0.209904995 | 0.093493833 | 2.245121283 | 0.024760348 | 0.108654552 | | | Xpnpep3 | 434.2429119 | -0.045010019 | 0.078181338 | -0.575713079 | 0.564809134 | 0.875364912 |
| Plekhs1 | 24.33032145 | 0.209899798 | 0.166837569 | 1.258108708 | 0.208352444 | 0.426414656 | | | Lpin2 | 4291.813414 | -0.045021207 | 0.051866486 | -0.868021158 | 0.38538275 | 0.789016184 |
| Tiprl | 1499.16675 | -0.126014516 | 0.064944409 | -1.940344343 | 0.052337856 | 0.179169849 | | | Ggact | 535.331065 | -0.045066481 | 0.079083859 | -0.569856874 | 0.568774777 | 0.877419107 |
| Zfp445 | 7620.513962 | 0.209869766 | 0.099353159 | 2.112361282 | 0.034655473 | 0.136074988 | | | Ptprz1 | 8914.845706 | -0.045212983 | 0.065764759 | -0.687495608 | 0.491770467 | 0.841892493 |
| Lzts1 | 1672.519523 | -0.201752275 | 0.104024065 | -1.93947694 | 0.05244329 | 0.179451487 | | | Unkl | 2161.799324 | -0.045217321 | 0.062320738 | -0.725558176 | 0.468109668 | 0.832353544 |
| Fam184b | 338.9119661 | -0.228070027 | 0.117616403 | -1.939100513 | 0.0524891 | 0.179568585 | | | Guca1a | 3.943673694 | -0.045240205 | 0.043790951 | -1.033094833 | 0.301559521 | NA |
| Mrpl44 | 742.2556415 | -0.12330663 | 0.063604078 | -1.938659201 | 0.05254285 | 0.179712785 | | | Znfx1 | 4274.098568 | -0.045251794 | 0.055274366 | -0.818675948 | 0.412971323 | 0.803808313 |
| Lhx5 | 11.62742051 | -0.279132071 | 0.144008915 | -1.938297159 | 0.052586979 | 0.179744684 | | | Fam114a2 | 1469.806461 | -0.045260149 | 0.052917303 | -0.855299609 | 0.392385304 | 0.794184423 |
| Sh3bgrl | 1592.247401 | -0.19222561 | 0.099171138 | -1.938322109 | 0.052583937 | 0.179744684 | | | 5430402O13Rik | 21.63667863 | -0.045267737 | 0.0821638 | -0.550945033 | 0.581671356 | NA |
| Tmem74 | 248.116035 | -0.22695138 | 0.117083633 | -1.938369812 | 0.052578121 | 0.179744684 | | | Lrit3 | 2.276302754 | -0.045270602 | 0.034288012 | -1.320304074 | 0.186733516 | NA |
| Nadsyn1 | 151.4422747 | 0.209686345 | 0.13671489 | 1.533749142 | 0.125091379 | 0.312636269 | | | Fam101a | 16.22956798 | -0.045275648 | 0.077403799 | -0.584927985 | 0.55859614 | NA |
| Prdm9 | 75.28310612 | 0.209599648 | 0.146670084 | 1.429055211 | 0.152988365 | 0.354495595 | | | Rp2h | 447.9411686 | -0.045297965 | 0.076831656 | -0.589574233 | 0.555476131 | 0.871360989 |
| H2afz | 2827.780593 | -0.107414852 | 0.055430501 | -1.937829363 | 0.052644044 | 0.179820727 | | | Nhej1 | 67.37252186 | -0.04532381 | 0.094705104 | -0.47857833 | 0.632238638 | 0.906498456 |
| Sars2 | 386.9580589 | 0.209592189 | 0.095279563 | 2.199760182 | 0.027823914 | 0.117691436 | | | Grin2c | 2122.892377 | -0.045325753 | 0.079940478 | -0.566993772 | 0.570718413 | 0.878246677 |
| Nt5e | 376.6529847 | -0.229063299 | 0.118218648 | -1.93762408 | 0.052669102 | 0.179844912 | | | Gpr21 | 126.2389897 | -0.045335794 | 0.095102259 | -0.476705753 | 0.633571665 | 0.906932171 |
| Phlpp2 | 2884.868924 | -0.103297362 | 0.053312975 | -1.937565168 | 0.052676295 | 0.179844912 | | | Gm4814 | 3.082110864 | -0.045345889 | 0.039569805 | -1.145972017 | 0.251806731 | NA |
| Adamts3 | 1327.525764 | -0.173194674 | 0.089424991 | -1.936759193 | 0.052774786 | 0.180028734 | | | Rab42 | 32.59435577 | -0.045373077 | 0.0864376 | -0.524922919 | 0.599636776 | 0.889353054 |
| Plekhg5 | 3272.728174 | 0.209370395 | 0.092815837 | 2.255761521 | 0.02408557 | 0.106774598 | | | Fbxo38 | 2236.84109 | -0.045399854 | 0.061517795 | -0.737995469 | 0.460517202 | 0.829239232 |
| Heatr2 | 288.3010222 | 0.20929798 | 0.094268439 | 2.220233866 | 0.026402897 | 0.113320354 | | | Angel1 | 582.0436381 | -0.045440344 | 0.074331925 | -0.61131666 | 0.540989963 | 0.863847445 |
| Pgk1 | 17070.29627 | -0.101468003 | 0.052391754 | -1.936717042 | 0.052779941 | 0.180028734 | | | Hook3 | 3095.875678 | -0.045470644 | 0.068248695 | -0.666249289 | 0.505251773 | 0.848183273 |
| Rcor1 | 671.7970909 | 0.209192579 | 0.08697219 | 2.40528128 | 0.016160022 | 0.081931735 | | | Il27 | 1.691845501 | -0.045473791 | 0.030034374 | -1.514058234 | 0.13001107 | NA |
| Tmem116 | 69.07292796 | 0.209097446 | 0.142826764 | 1.463993444 | 0.14319574 | 0.339776502 | | | Slc22a3 | 129.4828251 | -0.045475863 | 0.095054534 | -0.47841866 | 0.632352255 | 0.906498456 |
| Prom1 | 656.1375165 | -0.181085359 | 0.093532007 | -1.936079048 | 0.05285802 | 0.180154458 | | | Tfpi | 174.7264153 | -0.045530872 | 0.093663709 | -0.486110065 | 0.626889131 | 0.902926336 |
| Slc35b3 | 325.1914811 | 0.209037459 | 0.090323883 | 2.314309914 | 0.020650728 | 0.096229289 | | | Plagl2 | 448.9712372 | -0.045532299 | 0.076608949 | -0.594346996 | 0.552280052 | 0.870265825 |
| Chrm5 | 31.3881219 | 0.209013675 | 0.152276953 | 1.372589031 | 0.169880144 | 0.377311202 | | | Oas1c | 52.29759082 | -0.045537468 | 0.094885641 | -0.479919489 | 0.631284643 | 0.905683034 |
| Zfp804a | 993.7938261 | -0.25964602 | 0.134143586 | -1.935582819 | 0.052918816 | 0.180250711 | | | Ranbp3l | 167.6799781 | -0.045548649 | 0.092233059 | -0.493842983 | 0.621417075 | 0.899589902 |
| Gas2l2 | 10.40986047 | 0.20892815 | 0.150839005 | 1.385106919 | 0.166019777 | 0.371823021 | | | Pcdhb4 | 99.139789 | -0.045571311 | 0.093903483 | -0.485299475 | 0.627463928 | 0.903234666 |
| Eif2c3 | 775.9016469 | -0.166105969 | 0.085833733 | -1.935206162 | 0.052965002 | 0.180320752 | | | Celf2 | 14886.33288 | -0.045587228 | 0.053493006 | -0.852209128 | 0.394098031 | 0.794871981 |
| Neat1 | 5031.528199 | 0.20872049 | 0.167709764 | 1.244533924 | 0.213303122 | 0.432864253 | | | Katnbl1 | 411.9931978 | -0.04559366 | 0.080195951 | -0.568528206 | 0.569676359 | 0.87770886 |
| Tma16 | 273.6743456 | 0.208686657 | 0.147997653 | 1.410067344 | 0.158519799 | 0.361723971 | | | Tsnaxip1 | 15.30696427 | -0.045605788 | 0.06035001 | -0.75568817 | 0.449836179 | NA |
| Foxd1 | 42.46373653 | -0.324907681 | 0.168026044 | -1.933674523 | 0.053153158 | 0.180810506 | | | Tma16 | 265.1160707 | -0.045625256 | 0.085570262 | -0.533190566 | 0.593901671 | 0.886549756 |
| Wibg | 577.1176748 | -0.134563918 | 0.069590441 | -1.933655207 | 0.053155535 | 0.180810506 | | | Tfb1m | 172.4304156 | -0.045632674 | 0.088660966 | -0.514687303 | 0.606771547 | 0.893150688 |
| Cox5a | 4091.806955 | -0.176719548 | 0.091406271 | -1.933341625 | 0.053194127 | 0.180886756 | | | Plekhh2 | 513.0118713 | -0.045699392 | 0.085080199 | -0.537133105 | 0.591175675 | 0.8858275 |
| Scrt2 | 973.3405392 | 0.208517466 | 0.130676552 | 1.595676209 | 0.110561103 | 0.289412693 | | | Nup54 | 657.4756469 | -0.045745774 | 0.062993514 | -0.726198164 | 0.467717297 | 0.832353544 |
| Metap1d | 468.1833227 | 0.20837278 | 0.120539245 | 1.728671682 | 0.083867872 | 0.24166991 | | | Msrb3 | 419.2345191 | -0.045759237 | 0.081917128 | -0.558604019 | 0.576431997 | 0.880966011 |
| Dctn5 | 2976.175281 | -0.087172058 | 0.045106778 | -1.932571143 | 0.05328905 | 0.181105625 | | | Trim24 | 1215.57427 | -0.045777472 | 0.053428927 | -0.856791895 | 0.391559906 | 0.794184423 |
| Il6ra | 187.2570572 | 0.208198493 | 0.12438064 | 1.673881826 | 0.094153827 | 0.259530224 | | | Alms1-ps2 | 3.798375822 | -0.045805543 | 0.046632044 | -0.98227612 | 0.325963836 | NA |
| Wdr83 | 687.0332849 | 0.208141689 | 0.083495315 | 2.492854712 | 0.012672069 | 0.069378457 | | | Zfp628 | 527.2431936 | -0.045818706 | 0.070227788 | -0.652429858 | 0.514123908 | 0.853195313 |
| Kpna1 | 5393.446868 | -0.1336058 | 0.069185204 | -1.931132549 | 0.053466663 | 0.181549961 | | | AI427809 | 4.927230866 | -0.045824121 | 0.049038336 | -0.934455061 | 0.350069207 | NA |
| Tmprss9 | 22.33669349 | 0.207977069 | 0.16765563 | 1.240501546 | 0.214789943 | 0.43465253 | | | Ptx3 | 4.226919148 | -0.045838289 | 0.039064214 | -1.17340873 | 0.240631938 | NA |
| Cbl | 1413.689457 | -0.20003168 | 0.103598273 | -1.930839909 | 0.053502853 | 0.181633043 | | | F8 | 120.30421 | -0.045848868 | 0.093070314 | -0.492626125 | 0.622276785 | 0.900114632 |
| Kif13b | 1303.216676 | 0.207962713 | 0.099195203 | 2.096499696 | 0.036037887 | 0.139595056 | | | Asrgl1 | 6631.199879 | -0.045874391 | 0.050347579 | -0.911153854 | 0.362214314 | 0.776449324 |
| Fkbp4 | 4208.822188 | -0.104501867 | 0.054133232 | -1.930456817 | 0.053550261 | 0.181714354 | | | Gldn | 79.88109502 | -0.045889493 | 0.085580204 | -0.536216218 | 0.591809126 | 0.886226713 |
| Zar1 | 8.165889161 | -0.242975907 | 0.125872553 | -1.93033272 | 0.053565625 | 0.181726692 | | | Nol7 | 996.2078482 | -0.045894378 | 0.076277308 | -0.601677997 | 0.547388499 | 0.867553753 |
| Ppfia4 | 3262.994381 | 0.207909628 | 0.08597265 | 2.418322879 | 0.015592234 | 0.080108712 | | | Gm14405 | 11.98265158 | -0.045900639 | 0.07304358 | -0.628400726 | 0.529741462 | NA |
| Snhg12 | 510.4127891 | 0.207883298 | 0.108442691 | 1.916987633 | 0.055239503 | 0.185095341 | | | Gm13710 | 7.330188777 | -0.045916529 | 0.055526978 | -0.826922878 | 0.408280775 | NA |
| Tacr1 | 526.9605875 | -0.262203813 | 0.135880775 | -1.929660859 | 0.053648873 | 0.181889635 | | | Tmem30b | 2.858747816 | -0.045925701 | 0.035696228 | -1.286570151 | 0.198244153 | NA |
| Snhg6 | 278.3514881 | -0.226604268 | 0.117442211 | -1.929495921 | 0.053669326 | 0.181919173 | | | Stk19 | 698.5386231 | -0.045927856 | 0.079034384 | -0.58111234 | 0.561164742 | 0.87387211 |
| Abcc2 | 9.30674974 | 0.207817603 | 0.158173495 | 1.313858575 | 0.188893801 | 0.402059668 | | | Gm1715 | 4.341144704 | -0.045933168 | 0.04717292 | -0.973718978 | 0.330196092 | NA |
| P2ry13 | 554.6192313 | -0.288704034 | 0.149680247 | -1.928805163 | 0.053755055 | 0.182130071 | | | Uhrf1bp1 | 1300.239847 | -0.045944759 | 0.054719985 | -0.839633992 | 0.401113634 | 0.799277436 |
| Gmip | 348.0045897 | 0.207550487 | 0.120826402 | 1.717757743 | 0.085840812 | 0.244983891 | | | E030002O03Rik | 2.31887147 | -0.045947531 | 0.032528236 | -1.412542977 | 0.157790146 | NA |
| Stard5 | 265.7469477 | -0.210513111 | 0.109161738 | -1.928451449 | 0.053798998 | 0.182199272 | | | Ttc16 | 37.56412862 | -0.045949191 | 0.083379069 | -0.551087847 | 0.581573456 | 0.884020489 |
| Pou2f1 | 991.4750727 | -0.184918205 | 0.09589965 | -1.928246915 | 0.053824421 | 0.182245538 | | | Trerf1 | 1588.513038 | -0.045951362 | 0.07017146 | -0.654844031 | 0.512568179 | 0.852934798 |
| Gm10033 | 143.2451691 | -0.23362153 | 0.121181494 | -1.927864747 | 0.053871952 | 0.182326784 | | | Prkca | 8923.908718 | -0.045987614 | 0.071493613 | -0.643240875 | 0.520067814 | 0.853297835 |
| Ptprh | 94.63756118 | 0.207294364 | 0.144045992 | 1.439084567 | 0.150126564 | 0.350370787 | | | Hbp1 | 1074.835405 | -0.046007153 | 0.057964322 | -0.793715014 | 0.427361358 | 0.810334558 |
| I830077J02Rik | 32.14324228 | 0.207225557 | 0.161206346 | 1.285467741 | 0.198628883 | 0.41459732 | | | Actl9 | 4.933858128 | -0.046020337 | 0.042188788 | -1.090819124 | 0.27535248 | NA |
| Pawr | 75.04228703 | -0.25842569 | 0.134120272 | -1.926820506 | 0.054002003 | 0.182650548 | | | Desi2 | 994.540767 | -0.046042771 | 0.07656222 | -0.601377172 | 0.5475888 | 0.867553753 |
| Atoh8 | 44.44842071 | 0.207104189 | 0.154895359 | 1.337058717 | 0.181203471 | 0.391771527 | | | Gm5084 | 2.683582072 | -0.046054929 | 0.037810487 | -1.218046459 | 0.223206318 | NA |
| Nfkbid | 118.0953421 | 0.206978927 | 0.164062301 | 1.261587373 | 0.207097302 | 0.425155859 | | | Tfap2b | 52.86375893 | -0.046079888 | 0.084821824 | -0.543255097 | 0.586954176 | 0.884663353 |
| Pced1a | 716.7169775 | 0.206927018 | 0.078677092 | 2.63007965 | 0.008536487 | 0.052673963 | | | Ciita | 9.678908858 | -0.046084554 | 0.026622854 | -1.731014794 | NA | NA |
| A930024E05Rik | 97.08384028 | 0.206780295 | 0.140742288 | 1.469212267 | 0.141775224 | 0.337323079 | | | Dscc1 | 5.028209847 | -0.04608912 | 0.045408167 | -1.014996262 | 0.310107609 | NA |
| Hiat1 | 2059.639386 | -0.094476602 | 0.049044751 | -1.926334632 | 0.054062604 | 0.18269269 | | | Tnni2 | 9.29915725 | -0.046089515 | 0.057435268 | -0.802460162 | 0.422286824 | NA |
| Samd4b | 3285.990571 | 0.206448254 | 0.106208913 | 1.943794051 | 0.051920291 | 0.17820666 | | | Mr1 | 277.5835903 | -0.046118652 | 0.093483135 | -0.4933366 | 0.621774771 | 0.899950212 |
| Ube2a | 1086.427789 | -0.176804357 | 0.091836041 | -1.925217535 | 0.054202149 | 0.183084391 | | | Zfp454 | 200.3169338 | -0.046119828 | 0.084466347 | -0.546014241 | 0.585056148 | 0.884341895 |
| Bnip3l | 5072.891126 | -0.114040868 | 0.059242818 | -1.924973715 | 0.054232647 | 0.183107568 | | | Tomt | 3.476529954 | -0.046162558 | 0.041111398 | -1.122865188 | 0.261494755 | NA |
| Mettl7a1 | 1016.330917 | 0.206442701 | 0.115321449 | 1.790150085 | 0.073429787 | 0.221939958 | | | Guk1 | 2623.222087 | -0.046188766 | 0.054143828 | -0.853075368 | 0.39361751 | 0.794754188 |
| Fam207a | 1319.647085 | -0.116936272 | 0.060753169 | -1.924776511 | 0.054257324 | 0.183111081 | | | Thbd | 459.9607729 | -0.046199401 | 0.083325996 | -0.554441633 | 0.57927664 | 0.882452682 |
| Colec11 | 20.29710053 | 0.206367808 | 0.167685015 | 1.23068724 | 0.218439862 | 0.438862452 | | | Msl3 | 848.3414427 | -0.046249529 | 0.071730785 | -0.644765414 | 0.519079218 | 0.853297835 |
| 1500011K16Rik | 514.0058728 | -0.1718725 | 0.089312457 | -1.924395609 | 0.054305014 | 0.183192226 | | | Pus3 | 378.7075977 | -0.0462747 | 0.08175582 | -0.566011073 | 0.571386253 | 0.878389918 |
| Isyna1 | 1202.531947 | -0.138505414 | 0.071970617 | -1.924471687 | 0.054295486 | 0.183192226 | | | Ttk | 12.28877061 | -0.046332013 | 0.064725851 | -0.715819294 | 0.474102937 | NA |
| 3110039M20Rik | 123.326606 | -0.245175185 | 0.127442363 | -1.923812304 | 0.054378115 | 0.183328028 | | | Ifngr1 | 698.3689365 | -0.046366636 | 0.06178308 | -0.750474673 | 0.452968872 | 0.82547056 |
| Cdc37 | 4111.477235 | -0.098702326 | 0.051306155 | -1.923791145 | 0.054380768 | 0.183328028 | | | 4833422C13Rik | 135.6032684 | -0.046368089 | 0.094122404 | -0.492636047 | 0.622269773 | 0.900114632 |
| Foxk2 | 4801.766519 | -0.105842915 | 0.055017698 | -1.923797617 | 0.054379956 | 0.183328028 | | | Pkd1 | 6010.513083 | -0.046401024 | 0.066378376 | -0.699038259 | 0.484528122 | 0.838713365 |
| A4galt | 97.69614179 | -0.259376347 | 0.134845829 | -1.923502926 | 0.054416919 | 0.183409996 | | | Kif18a | 35.22322763 | -0.046406335 | 0.090178748 | -0.514603893 | 0.606829844 | 0.893150688 |
| Kctd21 | 781.806974 | -0.203300937 | 0.10571073 | -1.923181649 | 0.054457241 | 0.183466078 | | | Slc47a1 | 55.17437671 | -0.046420041 | 0.083581767 | -0.555384773 | 0.578631505 | 0.882065405 |
| Klk8 | 56.52797813 | 0.206321403 | 0.166942328 | 1.235884306 | 0.216501571 | 0.436443452 | | | Htra4 | 46.59015206 | -0.046423977 | 0.094141645 | -0.493129021 | 0.621921426 | 0.900011855 |
| Necap1 | 7860.732042 | -0.117872081 | 0.061298973 | -1.922904671 | 0.054492023 | 0.183543349 | | | Dhx9 | 5632.271574 | -0.046426156 | 0.062519939 | -0.742581593 | 0.457735036 | 0.827642075 |
| Cldn26 | 35.29108699 | -0.309915378 | 0.161190041 | -1.922670762 | 0.054521411 | 0.183602422 | | | D030028A08Rik | 275.0976525 | -0.046449451 | 0.079699068 | -0.582810466 | 0.560020899 | 0.873713575 |
| Fgf23 | 13.71045425 | 0.206268011 | 0.164965097 | 1.250373653 | 0.211163084 | 0.429646859 | | | Adam21 | 18.57972341 | -0.046502617 | 0.080626054 | -0.576769098 | 0.564095447 | NA |
| Dnlz | 1381.016075 | -0.164606726 | 0.085661015 | -1.92160607 | 0.054655344 | 0.18395243 | | | Zfp710 | 936.9323652 | -0.046531307 | 0.064664206 | -0.71958367 | 0.471781369 | 0.833304495 |
| Hnrnpc | 6421.726388 | -0.140460454 | 0.073101315 | -1.921449082 | 0.054675116 | 0.18395243 | | | Gm5105 | 8.235451455 | -0.046553321 | 0.052369589 | -0.888938048 | 0.374036376 | NA |
| Kcng1 | 686.3751542 | -0.223019908 | 0.116078725 | -1.921281508 | 0.054696227 | 0.18395243 | | | Mtif2 | 1010.563425 | -0.046572933 | 0.057617163 | -0.808317019 | 0.418908108 | 0.806975048 |
| Pde6d | 1136.34169 | -0.131790317 | 0.068594784 | -1.92128774 | 0.054695442 | 0.18395243 | | | Tmpo | 1088.801251 | -0.046576217 | 0.058040821 | -0.802473435 | 0.422279149 | 0.808695226 |
| Pex10 | 362.5420839 | -0.171126456 | 0.089069044 | -1.92127869 | 0.054696582 | 0.18395243 | | | Cep85 | 702.7393547 | -0.046577788 | 0.062317912 | -0.747422148 | 0.45480878 | 0.826168452 |
| Doc2a | 1540.800862 | -0.164897964 | 0.085841764 | -1.920952657 | 0.054737676 | 0.184050685 | | | Ankrd66 | 11.15524748 | -0.046635789 | 0.06780139 | -0.687829392 | 0.491560225 | NA |
| Gm14204 | 422.6636717 | -0.272268548 | 0.141792887 | -1.920184805 | 0.05483456 | 0.184315114 | | | Acad11 | 1249.1382 | -0.046653675 | 0.055890829 | -0.834728635 | 0.403870515 | 0.800260243 |
| Ptger3 | 66.47488622 | -0.259299325 | 0.135041825 | -1.92014085 | 0.054840111 | 0.184315114 | | | Slc25a43 | 7.277357917 | -0.046654282 | 0.056962918 | -0.819028995 | 0.412769871 | NA |
| Rps6ka2 | 4589.566528 | -0.167064883 | 0.087013203 | -1.919994645 | 0.054858576 | 0.184337188 | | | Mir138-1 | 3.040431364 | -0.04665914 | 0.036015218 | -1.295539582 | 0.195134156 | NA |
| Gm9833 | 16.82237226 | 0.206212781 | 0.155569105 | 1.325538137 | 0.184992729 | 0.396332597 | | | Gm10653 | 28.22248124 | -0.046666392 | 0.087940775 | -0.530657044 | 0.595656458 | 0.887497708 |
| Ndufs7 | 2212.169063 | -0.113438455 | 0.059091567 | -1.919706335 | 0.054895004 | 0.18437962 | | | Def6 | 72.95029853 | -0.046715188 | 0.095094726 | -0.49124899 | 0.623250354 | 0.900470921 |
| Olfr613 | 87.26211484 | 0.206169008 | 0.15295031 | 1.34794763 | 0.177675227 | 0.387405006 | | | Npas3 | 467.527508 | -0.046752811 | 0.07200718 | -0.649279841 | 0.516157514 | 0.853297835 |
| Npffr1 | 18.09553131 | 0.206151964 | 0.16153275 | 1.276223954 | 0.201876363 | 0.418948885 | | | Bmp8a | 5.143282247 | -0.046763005 | 0.051480628 | -0.908361212 | 0.363687412 | NA |
| Clec4a2 | 29.23073781 | -0.30901615 | 0.161012276 | -1.919208624 | 0.054957937 | 0.184471029 | | | Tchp | 464.5350482 | -0.046771015 | 0.072052892 | -0.649120582 | 0.516260439 | 0.853297835 |
| 1700034H15Rik | 102.541835 | 0.205900707 | 0.117731834 | 1.748895771 | 0.080309038 | 0.235174491 | | | Gp2 | 2.485826243 | -0.046805938 | 0.02915011 | -1.605686492 | 0.108342811 | NA |
| Fam71f2 | 17.1322985 | 0.205849466 | 0.168065478 | 1.224817066 | 0.220644153 | 0.441801565 | | | Orc6 | 607.8473079 | -0.046864588 | 0.066234865 | -0.707551642 | 0.479223732 | 0.836180652 |
| Ccdc174 | 765.345761 | 0.20562648 | 0.083977854 | 2.448579835 | 0.014342064 | 0.075617409 | | | Lrig3 | 116.7363399 | -0.046886545 | 0.094727822 | -0.494960655 | 0.620627894 | 0.899412757 |
| Klhl29 | 2881.312996 | -0.162431267 | 0.084679009 | -1.918199915 | 0.055085667 | 0.184659741 | | | Pomc | 190.1549055 | -0.046890308 | 0.057599358 | -0.814076921 | 0.415600886 | 0.805633085 |
| Lgr6 | 129.2390869 | -0.293804905 | 0.153157641 | -1.918316995 | 0.055070829 | 0.184659741 | | | Ogfod2 | 984.8879972 | -0.046891744 | 0.067310479 | -0.696648499 | 0.486022796 | 0.839772468 |
| Snora41 | 8.144679043 | 0.205510392 | 0.14856175 | 1.383333146 | 0.166562739 | 0.372076569 | | | Oc90 | 1.495963119 | -0.046919634 | 0.029465595 | -1.59235318 | 0.11130537 | NA |
| Hmg20b | 419.5021256 | 0.205502803 | 0.088847791 | 2.312975937 | 0.020723962 | 0.096374032 | | | D4Ertd617e | 17.53492791 | -0.046921335 | 0.07133023 | -0.657804347 | 0.510663861 | NA |
| Tmem179b | 143.5442264 | 0.205468242 | 0.117637869 | 1.746616493 | 0.080703884 | 0.235811614 | | | Oplah | 1347.169872 | -0.046937201 | 0.066063602 | -0.710485042 | 0.477403404 | 0.835289575 |
| Rc3h1 | 1819.729054 | -0.185674347 | 0.096865672 | -1.916822999 | 0.055260422 | 0.185125401 | | | Pomt1 | 686.8474727 | -0.046947963 | 0.072341235 | -0.64897929 | 0.516351763 | 0.853297835 |
| Sstr4 | 329.0088604 | -0.228038703 | 0.118987259 | -1.916496816 | 0.055301888 | 0.185220192 | | | Tmem82 | 20.56700936 | -0.046975255 | 0.075949242 | -0.618508543 | 0.536240168 | NA |
| Tmem169 | 636.4382066 | -0.138416192 | 0.072226726 | -1.916412382 | 0.055312626 | 0.185220192 | | | 4930473A02Rik | 7.106418008 | -0.047029369 | 0.045892125 | -1.024780798 | 0.305466629 | NA |
| Cadm2 | 6052.078997 | -0.187142657 | 0.097672415 | -1.916023647 | 0.055362086 | 0.185345756 | | | 1500032L24Rik | 2263.057393 | -0.047046921 | 0.051584884 | -0.912029215 | 0.361753338 | 0.776449324 |
| BC031361 | 243.8049379 | -0.183084504 | 0.095566112 | -1.915788982 | 0.055391961 | 0.185395636 | | | Inhba | 148.7563255 | -0.047051486 | 0.095008499 | -0.495234491 | 0.620434606 | 0.899412757 |
| Cd2bp2 | 3550.236217 | 0.099128891 | 0.051745015 | 1.915718659 | 0.055400916 | 0.185395636 | | | Pex3 | 1068.64928 | -0.047056714 | 0.069408956 | -0.677963143 | 0.497795062 | 0.845489269 |
| Nrxn2 | 13944.33774 | -0.09638522 | 0.05031907 | -1.915480945 | 0.055431197 | 0.185456913 | | | Slc25a15 | 538.7781309 | -0.047095185 | 0.068483526 | -0.687686345 | 0.491650321 | 0.841892493 |
| Arpc4 | 2862.254256 | -0.129759253 | 0.067757571 | -1.915051721 | 0.055485908 | 0.185599883 | | | Mink1 | 16581.19653 | -0.04713272 | 0.063817706 | -0.738552401 | 0.460178837 | 0.829022384 |
| Osgin1 | 29.00154326 | 0.205467253 | 0.164744461 | 1.247187622 | 0.212328711 | 0.431565137 | | | Fap | 125.5701645 | -0.047139488 | 0.095077042 | -0.495803057 | 0.620033368 | 0.899216177 |
| Gm5089 | 86.76168879 | 0.205433783 | 0.132884535 | 1.54595704 | 0.122114943 | 0.308402434 | | | Klhl34 | 4.821208803 | -0.047140724 | 0.046780648 | -1.00769711 | 0.313599893 | NA |
| Dvl3 | 1577.463177 | 0.205365568 | 0.079356245 | 2.587894226 | 0.009656463 | 0.057280718 | | | Ddx39b | 5870.487878 | -0.047202248 | 0.058203052 | -0.810992667 | 0.417369882 | 0.806153084 |
| Fam163b | 7216.941356 | -0.132125997 | 0.069058757 | -1.9132403 | 0.055717297 | 0.18621307 | | | Cnksr1 | 10.54540088 | -0.047212465 | 0.063240233 | -0.746557422 | 0.455330758 | NA |
| Rpl35a | 4398.826847 | -0.147861957 | 0.077312211 | -1.912530447 | 0.055808191 | 0.186476626 | | | Cpb1 | 3.899067489 | -0.047214967 | 0.042408905 | -1.113326722 | 0.265568134 | NA |
| Bcr | 8057.270284 | -0.170533062 | 0.089170883 | -1.91242989 | 0.055821077 | 0.186479468 | | | 2310015A10Rik | 431.3356608 | -0.047235366 | 0.077850767 | -0.606742457 | 0.544021841 | 0.86604998 |
| Retn | 19.45650277 | 0.205185317 | 0.168029025 | 1.2211302 | 0.222036727 | 0.443158192 | | | Matn1 | 13.4832559 | -0.047260567 | 0.07120238 | -0.663749816 | 0.50685045 | NA |
| Kif20a | 32.45467724 | -0.305284402 | 0.159670823 | -1.911961103 | 0.055881183 | 0.186599796 | | | 9130227L01Rik | 10.78565312 | -0.047262206 | 0.062385591 | -0.757582085 | 0.448701209 | NA |
| 1500017E21Rik | 11.82646337 | -0.301023447 | 0.157461505 | -1.911727236 | 0.055911189 | 0.186630603 | | | Fgl2 | 77.25385921 | -0.047266748 | 0.082046637 | -0.576096102 | 0.564550226 | 0.875364912 |
| Epsti1 | 34.54792512 | 0.205084006 | 0.165105882 | 1.242136276 | 0.214186286 | 0.433917667 | | | S100a8 | 9.604427418 | -0.047281306 | 0.052907521 | -0.893659447 | 0.371504129 | NA |
| Slc36a1 | 4031.638546 | -0.153157541 | 0.080132322 | -1.911307915 | 0.055965023 | 0.186759005 | | | Gpr108 | 1100.030648 | -0.047290089 | 0.064411524 | -0.734186765 | 0.462834919 | 0.831014288 |
| Bcl2l13 | 2576.132641 | -0.13033495 | 0.068212755 | -1.91071231 | 0.056041562 | 0.186826112 | | | 4930539J05Rik | 20.63338835 | -0.047354214 | 0.07877522 | -0.60113084 | 0.547752844 | NA |
| Camk1 | 2452.02424 | -0.121009745 | 0.063333263 | -1.910682291 | 0.056045422 | 0.186826112 | | | D630003M21Rik | 305.2512108 | -0.047365842 | 0.087790722 | -0.539531297 | 0.589520308 | 0.885188411 |
| Samd8 | 3560.075637 | -0.130104381 | 0.06808566 | -1.910892555 | 0.05601839 | 0.186826112 | | | Lta4h | 1758.938399 | -0.047379122 | 0.055724564 | -0.850237652 | 0.395192972 | 0.795857764 |
| Spag6 | 241.9998119 | -0.184620348 | 0.096612823 | -1.910930054 | 0.05601357 | 0.186826112 | | | Sh2b2 | 233.7494355 | -0.047385758 | 0.087640715 | -0.540682007 | 0.588726782 | 0.885188411 |
| Strn | 2962.699837 | -0.205291544 | 0.107440441 | -1.910747404 | 0.05603705 | 0.186826112 | | | Slc25a10 | 674.1428173 | -0.047398489 | 0.082168051 | -0.576848161 | 0.564042032 | 0.875251366 |
| Acat2 | 1161.34528 | -0.148267617 | 0.077615464 | -1.910284493 | 0.056096593 | 0.186952759 | | | Galnt12 | 59.06607515 | -0.047431673 | 0.093198569 | -0.508931342 | 0.61080035 | 0.894800514 |
| Mansc1 | 437.1690142 | 0.204964203 | 0.107811204 | 1.901140106 | 0.057283664 | 0.189364081 | | | Kcnk9 | 664.0534304 | -0.047461507 | 0.071573694 | -0.663113849 | 0.507257642 | 0.848958235 |
| Fbxo38 | 2060.759806 | 0.204880161 | 0.087936997 | 2.329851696 | 0.019813991 | 0.093654049 | | | 8430427H17Rik | 2891.697204 | -0.047478875 | 0.066133926 | -0.717920104 | 0.472806551 | 0.833983942 |
| Myb | 24.71277541 | 0.204879839 | 0.165679679 | 1.236602095 | 0.216234842 | 0.436413005 | | | Bbip1 | 1293.897137 | -0.047487041 | 0.060195168 | -0.788884596 | 0.430179459 | 0.811860174 |
| Trhr | 156.7015947 | 0.204829705 | 0.167503173 | 1.222840745 | 0.221389853 | 0.442484962 | | | Prpf40a | 1404.403085 | -0.047491688 | 0.063769514 | -0.744739683 | 0.456429105 | 0.82674985 |
| Gxylt1 | 695.3580599 | 0.204777153 | 0.085523828 | 2.394387121 | 0.016648167 | 0.08337513 | | | Camta2 | 15209.67056 | -0.047491778 | 0.062857789 | -0.755543252 | 0.449923091 | 0.823263966 |
| Gabra5 | 2107.397166 | -0.202989678 | 0.106350302 | -1.90868925 | 0.056302191 | 0.187399816 | | | Twist2 | 26.28527379 | -0.047491978 | 0.086327057 | -0.550140127 | 0.582223266 | 0.884020489 |
| Gtf2f2 | 415.4198925 | -0.143983024 | 0.07545138 | -1.908288811 | 0.056353898 | 0.187451116 | | | Sptssa | 556.7791981 | -0.047542485 | 0.0735561 | -0.646343202 | 0.518057115 | 0.853297835 |
| Gas8 | 579.5695564 | 0.204625127 | 0.0951529 | 2.150487549 | 0.03151667 | 0.12776538 | | | Serac1 | 1333.203496 | -0.047573625 | 0.065574069 | -0.72549448 | 0.46814873 | 0.832353544 |
| Nfrkb | 1308.327002 | 0.204422341 | 0.083569642 | 2.446131587 | 0.014439829 | 0.076009456 | | | Arl4a | 1364.464105 | -0.047577483 | 0.076867531 | -0.618954229 | 0.535946512 | 0.860988 |
| Pfkfb2 | 2073.858066 | 0.204303754 | 0.084971259 | 2.40438656 | 0.016199633 | 0.082105697 | | | Mettl8 | 459.1805188 | -0.047585562 | 0.071357486 | -0.666861521 | 0.504860591 | 0.848183273 |
| Gpr17 | 2323.309889 | -0.13287439 | 0.069681571 | -1.906879939 | 0.056536137 | 0.187855651 | | | 5031426D15Rik | 5.23184231 | -0.047589611 | 0.043611392 | -1.091219729 | 0.275176208 | NA |
| Psmb1 | 3885.826125 | -0.104059121 | 0.054568735 | -1.906936644 | 0.056528792 | 0.187855651 | | | Ldhd | 661.500099 | -0.047634407 | 0.07552541 | -0.630707034 | 0.528232099 | 0.856965327 |
| Ptrh2 | 467.5424674 | -0.139794228 | 0.073307353 | -1.906960527 | 0.056525699 | 0.187855651 | | | Klrb1b | 2.350758835 | -0.047648425 | 0.02934471 | -1.62374835 | 0.104429527 | NA |
| Lyz1 | 41.94389754 | 0.204237495 | 0.167680103 | 1.218018666 | 0.223216879 | 0.444540145 | | | March2 | 2445.142289 | -0.047649666 | 0.049357972 | -0.965389458 | 0.33434978 | 0.758238352 |
| Ppic | 51.79372059 | -0.270184318 | 0.141705413 | -1.906661935 | 0.056564379 | 0.187909196 | | | 9930013L23Rik | 891.1469304 | -0.047664722 | 0.069681046 | -0.684041432 | 0.493949004 | 0.843404385 |
| Iqck | 59.4381145 | 0.204147772 | 0.15719264 | 1.298710756 | 0.194043211 | 0.408548107 | | | Irf4 | 23.08404237 | -0.047671403 | 0.070277883 | -0.678327239 | 0.497564231 | 0.845236583 |
| Zkscan14 | 332.4571017 | 0.204141601 | 0.096402166 | 2.117603876 | 0.03420863 | 0.134921843 | | | Ebi3 | 37.06821898 | -0.047671726 | 0.09218387 | -0.517137387 | 0.605060257 | 0.892113813 |
| Flnc | 115.7884395 | 0.204079312 | 0.151164083 | 1.350051597 | 0.176999433 | 0.386420912 | | | Enpp6 | 261.8927145 | -0.047680396 | 0.090527082 | -0.526697591 | 0.598403609 | 0.888717024 |
| Npsr1 | 176.4913471 | -0.233248693 | 0.122398695 | -1.905646888 | 0.056696034 | 0.188149434 | | | Kirrel | 380.0182984 | -0.047689395 | 0.077385883 | -0.616254452 | 0.537726595 | 0.862037665 |
| Gm15708 | 11.17322191 | 0.203852718 | 0.162396039 | 1.255281347 | 0.209376644 | 0.427553024 | | | Fam184b | 340.7193542 | -0.047746855 | 0.078810509 | -0.605843756 | 0.544618511 | 0.86624804 |
| Zfp831 | 1421.431672 | -0.18033233 | 0.094632678 | -1.905603156 | 0.056701712 | 0.188149434 | | | Mirlet7b | 15.86373004 | -0.047779008 | 0.059715427 | -0.800111636 | 0.42364612 | NA |
| Ranbp9 | 3503.501605 | 0.203832846 | 0.084695413 | 2.406657426 | 0.016099264 | 0.081730666 | | | AA543186 | 9.438754294 | -0.047825678 | 0.067543822 | -0.708068878 | 0.478902486 | NA |
| Ralgapa2 | 1155.217317 | -0.130537993 | 0.068534936 | -1.90469272 | 0.056820026 | 0.188435248 | | | Mtcp1 | 284.4361514 | -0.047900078 | 0.083049743 | -0.576763716 | 0.564099084 | 0.875251366 |
| Camk2d | 4208.383815 | 0.203724484 | 0.104046394 | 1.958015798 | 0.050228159 | 0.174374881 | | | Smug1 | 728.2450196 | -0.047924425 | 0.072733803 | -0.658901679 | 0.509958908 | 0.850575622 |
| Snord69 | 11.00245022 | 0.203691037 | 0.138417999 | 1.471564675 | 0.141138471 | 0.336120422 | | | Rogdi | 4500.089056 | -0.047932935 | 0.059289213 | -0.808459635 | 0.418826034 | 0.806975048 |
| Trafd1 | 2106.066564 | 0.203573624 | 0.07989025 | 2.54816606 | 0.010829091 | 0.062189005 | | | Pbx4 | 247.1347867 | -0.047956185 | 0.089556018 | -0.535488128 | 0.592312365 | 0.886226713 |
| Zdhhc20 | 1687.488271 | 0.203318651 | 0.089733391 | 2.265808169 | 0.023463131 | 0.104990374 | | | Usp17la | 4.114706727 | -0.047993128 | 0.043051694 | -1.114779097 | 0.264945096 | NA |
| Ccdc148 | 638.1659427 | -0.196952667 | 0.103438836 | -1.904049527 | 0.056903734 | 0.188511109 | | | Hp | 3.412429754 | -0.048001677 | 0.037335477 | -1.285685382 | 0.198552885 | NA |
| Tmem176a | 435.6672337 | 0.203299998 | 0.11288579 | 1.800935242 | 0.071713087 | 0.218595825 | | | Slc29a2 | 548.3372384 | -0.048035173 | 0.074581875 | -0.644059603 | 0.519536784 | 0.853297835 |
| Exoc6 | 1321.68762 | -0.11828129 | 0.062167265 | -1.902629791 | 0.057088869 | 0.189003192 | | | Tnpo1 | 1964.34257 | -0.048070021 | 0.05721523 | -0.840161286 | 0.400817962 | 0.799169897 |
| Ids | 17630.07972 | -0.108517145 | 0.057033093 | -1.902704888 | 0.057079064 | 0.189003192 | | | Strip2 | 2330.248677 | -0.048070179 | 0.088544953 | -0.54289011 | 0.587205466 | 0.884736707 |
| Cpne2 | 875.7064033 | 0.203209377 | 0.114692508 | 1.771775515 | 0.076431829 | 0.227474981 | | | Snord58b | 5.082388009 | -0.048073682 | 0.049007184 | -0.980951735 | 0.326616543 | NA |
| Rpap1 | 1001.342642 | 0.202987301 | 0.078641981 | 2.581157015 | 0.009846978 | 0.058188056 | | | Map3k11 | 1127.767866 | -0.048139262 | 0.062285282 | -0.772883421 | 0.439591377 | 0.81760619 |
| Creg1 | 1042.278544 | 0.202608046 | 0.07935328 | 2.553240974 | 0.010672563 | 0.061609839 | | | Tcf19 | 265.9168357 | -0.048151608 | 0.083870621 | -0.574117696 | 0.565888159 | 0.875688867 |
| Cdc25a | 264.5112454 | 0.202523829 | 0.125356436 | 1.615583818 | 0.106184325 | 0.281667511 | | | 5033404E19Rik | 16.41084863 | -0.048155248 | 0.078695926 | -0.611915383 | 0.540593743 | NA |
| L1cam | 7016.747899 | -0.163857031 | 0.08617517 | -1.901441349 | 0.057244228 | 0.189315276 | | | Anapc1 | 4050.637853 | -0.048167972 | 0.043616053 | -1.104363365 | 0.269435546 | 0.711972822 |
| Ccdc94 | 231.0846861 | -0.193315818 | 0.101682774 | -1.901165847 | 0.057280293 | 0.189364081 | | | Ptprs | 19412.04024 | -0.048195339 | 0.0572571 | -0.841735599 | 0.399935967 | 0.798662339 |
| Lrrk1 | 275.9001775 | -0.185025073 | 0.097327904 | -1.901048574 | 0.057295651 | 0.189364081 | | | D14Abb1e | 1332.542042 | -0.048199023 | 0.051018966 | -0.944727537 | 0.344797995 | 0.764561711 |
| Ccdc151 | 202.892511 | 0.202473171 | 0.115975712 | 1.745823916 | 0.080841554 | 0.236064651 | | | B230208H11Rik | 4.696593207 | -0.048229179 | 0.048176383 | -1.001095893 | 0.31678045 | NA |
| Lpin1 | 1310.382718 | 0.202410454 | 0.091931818 | 2.201745348 | 0.027683302 | 0.117224672 | | | Wbscr27 | 368.8033623 | -0.048247166 | 0.07195385 | -0.67052932 | 0.502520422 | 0.84739426 |
| Iqub | 65.85057378 | 0.20233166 | 0.155705348 | 1.299452225 | 0.193788779 | 0.408226914 | | | 6530411M01Rik | 12.1557455 | -0.048325915 | 0.067596648 | -0.714915847 | 0.474661044 | NA |
| Bgn | 776.7670467 | -0.2046629 | 0.107703648 | -1.900241121 | 0.057401484 | 0.189592538 | | | Fastk | 2975.529613 | -0.048352084 | 0.07074904 | -0.683430958 | 0.494334563 | 0.843503724 |
| Pcbp3 | 2305.681298 | -0.232359288 | 0.122329151 | -1.899459662 | 0.057504065 | 0.189890876 | | | Pttg1ip | 2492.675082 | -0.048363976 | 0.045314197 | -1.067302946 | 0.285835058 | 0.722148665 |
| D430019H16Rik | 8308.975802 | -0.133791496 | 0.070443134 | -1.899283697 | 0.057527185 | 0.189894005 | | | Efcab3 | 12.93209589 | -0.048380469 | 0.059195273 | -0.817302899 | 0.41375535 | NA |
| Fuk | 711.9884873 | 0.202200007 | 0.076790511 | 2.633137919 | 0.008459999 | 0.052347931 | | | Lipt2 | 126.9123227 | -0.04838098 | 0.090335031 | -0.535572739 | 0.592253874 | 0.886226713 |
| Gas2 | 229.5652362 | 0.202196197 | 0.107716455 | 1.877115219 | 0.060502304 | 0.195540821 | | | 4930511M06Rik | 6.010115187 | -0.048382218 | 0.055683355 | -0.868881156 | 0.384912138 | NA |
| Mt3 | 5441.060187 | -0.162084143 | 0.085366212 | -1.898691995 | 0.057604985 | 0.190021638 | | | Top1mt | 552.9500387 | -0.048419894 | 0.072315854 | -0.66956126 | 0.503137517 | 0.847567618 |

| | | | | | | |
|---|---|---|---|---|---|---|
| Palmd | 637.6644949 | -0.188837182 | 0.099455749 | -1.898705546 | 0.057603202 | 0.190021638 |
| Gm15421 | 16.51977924 | 0.202191821 | 0.165075476 | 1.22484469 | 0.220633743 | 0.441801565 |
| Hmbox1 | 1038.481021 | -0.176439774 | 0.092942694 | -1.898371628 | 0.057647144 | 0.190071252 |
| Trpm3 | 1676.607019 | 0.201965484 | 0.129062226 | 1.564869059 | 0.117613615 | 0.301008482 |
| Chtf8 | 3198.402237 | -0.119842552 | 0.063137715 | -1.898113543 | 0.057681127 | 0.190110907 |
| Mgrn1 | 9530.207061 | 0.080326418 | 0.042323626 | 1.897909669 | 0.057707982 | 0.190158971 |
| Mgll | 10416.2096 | -0.179000451 | 0.094376225 | -1.896668906 | 0.05787165 | 0.190617208 |
| Myo1d | 825.1966961 | 0.20190028 | 0.131416296 | 1.536341271 | 0.124454702 | 0.311793329 |
| Slc9a5 | 1700.740138 | 0.201862697 | 0.078950813 | 2.556815941 | 0.010563509 | 0.061208308 |
| Resp18 | 2126.149205 | 0.201774641 | 0.082927871 | 2.433134212 | 0.014968751 | 0.077827461 |
| Tmem79 | 11.73558908 | 0.20150271 | 0.16544773 | 1.217923693 | 0.223252972 | 0.444554883 |
| Nfatc2 | 536.096439 | -0.200615958 | 0.105855567 | -1.89518571 | 0.058067803 | 0.191086137 |
| Opcml | 7708.301497 | -0.174376588 | 0.092013179 | -1.895126217 | 0.058075682 | 0.191086137 |
| Slc29a3 | 699.4996895 | 0.201262103 | 0.086154418 | 2.33606246 | 0.019487986 | 0.092479897 |
| Efcc1 | 227.1449971 | 0.201138806 | 0.102020206 | 1.971558516 | 0.048660027 | 0.171098935 |
| Donson | 560.7720782 | 0.201083461 | 0.093093407 | 2.160018273 | 0.030771255 | 0.125826058 |
| 0610010K14Rik | 598.1875366 | -0.196714219 | 0.103836012 | -1.894470096 | 0.05816264 | 0.191169271 |
| Tmem150a | 333.2495482 | 0.201053746 | 0.126591936 | 1.588203419 | 0.112240348 | 0.292375918 |
| Bhlhe40 | 5454.696346 | 0.200985706 | 0.157360355 | 1.277232164 | 0.201520295 | 0.418378058 |
| Slc44a4 | 16.98421301 | 0.200921069 | 0.168119469 | 1.195108875 | 0.232044495 | 0.455245894 |
| Ptplad1 | 6938.024722 | -0.127159362 | 0.067161704 | -1.893331382 | 0.058313814 | 0.191544253 |
| Cd74 | 315.5222307 | 0.200915333 | 0.134220005 | 1.49691049 | 0.13441655 | 0.325903062 |
| Fendrr | 24.17456761 | 0.200854888 | 0.16808166 | 1.194983963 | 0.232093296 | 0.455245894 |
| Gpr56 | 3082.723401 | -0.225314019 | 0.119029308 | -1.892928914 | 0.058367324 | 0.191610272 |
| Psma6 | 2735.384466 | -0.094700142 | 0.050064 | -1.891581597 | 0.05854675 | 0.192146439 |
| Acap3 | 6236.515341 | 0.200764211 | 0.110766488 | 1.81249956 | 0.069909042 | 0.215128241 |
| Hbp1 | 973.3822187 | 0.20064778 | 0.077087471 | 2.602858517 | 0.00924501 | 0.055586285 |
| Fkbp11 | 19.06464606 | -0.317754455 | 0.168096292 | -1.890312097 | 0.058716232 | 0.192512323 |
| Gm1661 | 13.62924176 | 0.200626828 | 0.167655422 | 1.19666173 | 0.231438429 | 0.454314839 |
| Rps27a | 5929.417106 | -0.12540071 | 0.06633368 | -1.890453092 | 0.058697388 | 0.192512323 |
| Sp7 | 84.27513707 | -0.292822935 | 0.154918418 | -1.8901751 | 0.058734546 | 0.192518098 |
| Acat1 | 3271.644261 | -0.098701248 | 0.052223027 | -1.889994784 | 0.058758658 | 0.192518619 |
| Grk1 | 29.27377036 | 0.200595765 | 0.164797791 | 1.217223625 | 0.223519144 | 0.444744406 |
| Dio3os | 10.14829702 | -0.293686963 | 0.155432508 | -1.889482234 | 0.058827241 | 0.192658904 |
| 5430427O19Rik | 12.51327503 | 0.200583704 | 0.159901245 | 1.2544224 | 0.209688515 | 0.427871411 |
| Pank4 | 1389.069964 | 0.200540362 | 0.076987827 | 2.604832091 | 0.009191929 | 0.055394692 |
| Irgm2 | 273.4950514 | 0.200435169 | 0.157530292 | 1.272359536 | 0.203245404 | 0.420270158 |
| Gm10814 | 30.73187582 | 0.200253042 | 0.166700476 | 1.201274571 | 0.229644711 | 0.45262882 |
| Ptrhd1 | 451.6157846 | -0.149268459 | 0.079032081 | -1.888707193 | 0.058931074 | 0.192835916 |
| Srsf6 | 3354.075592 | 0.200198755 | 0.091559821 | 2.186535003 | 0.028776487 | 0.120144436 |
| Caprin2 | 262.8574708 | 0.200117283 | 0.141236777 | 1.416892171 | 0.156514456 | 0.359104839 |
| Pou3f1 | 774.5875996 | -0.229262117 | 0.121404106 | -1.888421439 | 0.058969395 | 0.192839133 |
| Igj | 13.35680682 | 0.200064434 | 0.165025021 | 1.212327878 | 0.225386896 | 0.447194849 |
| Sparc | 10824.85432 | 0.199823053 | 0.166662501 | 1.198968283 | 0.230540278 | 0.453668953 |
| Rcbtb2 | 871.1915896 | -0.1221368 | 0.064697367 | -1.887817176 | 0.059050499 | 0.193002981 |
| Iqcc | 468.1690868 | 0.199771906 | 0.095987166 | 2.081235586 | 0.037412348 | 0.143269134 |
| Sp9 | 200.6967147 | -0.287026507 | 0.152073108 | -1.887424486 | 0.059103254 | 0.193073123 |
| Megf11 | 749.2686644 | -0.230170233 | 0.121978977 | -1.886966414 | 0.059164843 | 0.193233575 |
| Snord42b | 19.07482168 | 0.19967661 | 0.167111129 | 1.194873202 | 0.232136574 | 0.455245894 |
| 1700016K19Rik | 63.72603215 | -0.309943663 | 0.164284188 | -1.88663113 | 0.059209957 | 0.193299424 |
| Arhgef17 | 10567.95701 | 0.199501673 | 0.091234489 | 2.186691403 | 0.02876506 | 0.120144436 |
| Fam211a | 636.9511286 | -0.177412549 | 0.094061441 | -1.886134706 | 0.059276805 | 0.193436144 |
| Tbc1d15 | 1146.362106 | 0.199495369 | 0.083690137 | 2.383738121 | 0.017137793 | 0.084834813 |
| Gm11744 | 82.37011331 | 0.199433944 | 0.133202514 | 1.497223575 | 0.134335093 | 0.32583618 |
| Cib2 | 642.7847484 | -0.166206681 | 0.088153235 | -1.885429174 | 0.059371918 | 0.193522564 |
| Eps8 | 1235.879272 | -0.156062706 | 0.082779021 | -1.885292967 | 0.059390295 | 0.193522564 |
| Alox5 | 32.70542633 | 0.199405685 | 0.168099884 | 1.18623333 | 0.235530167 | 0.459475465 |
| Stmn2 | 13344.33827 | -0.1485685 | 0.078796231 | -1.885477237 | 0.059365435 | 0.193522564 |
| Susd1 | 448.3676406 | -0.171615097 | 0.091005373 | -1.88576885 | 0.05932611 | 0.193522564 |
| Gnai3 | 1200.225593 | -0.144171047 | 0.076475706 | -1.885187519 | 0.059404525 | 0.193526853 |
| Taldo1 | 2618.861743 | -0.121129659 | 0.064275643 | -1.884534361 | 0.059492732 | 0.193773467 |
| Abce1 | 3437.329459 | -0.099196014 | 0.052646285 | -1.88419778 | 0.059538228 | 0.193827647 |
| Clrn1 | 12.84448777 | 0.199239629 | 0.158978796 | 1.253246554 | 0.210115992 | 0.428360156 |
| Slc16a7 | 392.9278288 | -0.195107571 | 0.103549546 | -1.884195334 | 0.059538558 | 0.193827647 |
| Hdhd3 | 126.959801 | -0.228985312 | 0.121544052 | -1.883969709 | 0.059569073 | 0.193859109 |
| Cd302 | 481.3590355 | -0.164447525 | 0.087309408 | -1.883502932 | 0.059632243 | 0.193869931 |
| Cpq | 505.3974892 | -0.17057896 | 0.090554264 | -1.883720914 | 0.059602736 | 0.193869931 |
| S1pr1 | 5966.649379 | -0.208131518 | 0.110503065 | -1.883490912 | 0.05963387 | 0.193869931 |
| Slc39a3 | 3107.628942 | -0.122884246 | 0.065243092 | -1.883482853 | 0.059634961 | 0.193869931 |
| Vegfa | 2576.659863 | -0.181689829 | 0.096463503 | -1.883508503 | 0.059631489 | 0.193869931 |
| Qtrtd1 | 502.8900693 | 0.199208259 | 0.109631609 | 1.817069552 | 0.069206464 | 0.213347584 |
| Comt | 3370.854634 | -0.109400856 | 0.058106742 | -1.882756687 | 0.059733349 | 0.194108329 |
| B3gnt8 | 16.81483042 | -0.312297759 | 0.165885245 | -1.882613243 | 0.0597528 | 0.194118889 |
| Dclk2 | 2962.966052 | -0.125748843 | 0.066797153 | -1.882547949 | 0.059761656 | 0.194118889 |
| Trim12a | 86.87936277 | 0.199133204 | 0.13345685 | 1.492116772 | 0.135668536 | 0.327932651 |
| Enah | 2999.104855 | -0.148470405 | 0.078886075 | -1.882086342 | 0.059824294 | 0.194240906 |
| Rad51 | 42.90176671 | 0.198911849 | 0.149123663 | 1.333871802 | 0.182245874 | 0.392903171 |
| Ftsj3 | 1372.985003 | 0.198778514 | 0.077354473 | 2.569709392 | 0.010178386 | 0.059578357 |
| Foxd2 | 12.66986214 | -0.309538808 | 0.164525325 | -1.881405234 | 0.059916815 | 0.194316101 |
| Pcdhb11 | 130.2491745 | -0.231463859 | 0.123032097 | -1.881329061 | 0.05992717 | 0.194316101 |
| Popdc3 | 27.07799168 | 0.198772558 | 0.168111292 | 1.182386708 | 0.237052295 | 0.461360163 |
| Sepn1 | 1139.232577 | 0.198731793 | 0.081455034 | 2.439772996 | 0.014696494 | 0.077013181 |
| Ldb1 | 3544.473516 | 0.198727217 | 0.076435945 | 2.599918378 | 0.009324594 | 0.055976464 |
| Tmem119 | 945.1961778 | -0.245432916 | 0.130467711 | -1.881177452 | 0.059947784 | 0.194316101 |
| Trmt44 | 275.583264 | 0.198680217 | 0.113454405 | 1.751189975 | 0.079913192 | 0.234502839 |
| 4930430F08Rik | 475.2644837 | 0.198677714 | 0.08033327 | 2.473168527 | 0.013392097 | 0.071872933 |
| Gaa | 14079.33723 | 0.198588813 | 0.104824743 | 1.894484133 | 0.058160778 | 0.191169271 |
| Eln | 306.2422749 | -0.21962433 | 0.116835939 | -1.879766895 | 0.060139854 | 0.194734984 |
| Htr3a | 283.118125 | 0.198532355 | 0.127279475 | 1.559814382 | 0.118803752 | 0.302604855 |
| Bmp3 | 491.5306072 | -0.234464236 | 0.124770913 | -1.879157813 | 0.060222948 | 0.194963301 |
| Tial1 | 3053.806012 | 0.198488621 | 0.083995043 | 2.363099227 | 0.018122816 | 0.08838367 |
| Usp2 | 3083.909682 | 0.198220353 | 0.078190608 | 2.53509159 | 0.011241793 | 0.063685682 |
| Adamts1 | 560.1777684 | 0.198186585 | 0.168113011 | 1.178889038 | 0.238442363 | 0.462759891 |
| Gprin2 | 21.30903031 | -0.313286926 | 0.166864879 | -1.87748871 | 0.060451143 | 0.195497811 |
| Zkscan17 | 536.5637811 | 0.198175459 | 0.085747943 | 2.311139502 | 0.020825151 | 0.096664136 |
| Dbndd2 | 5316.625998 | 0.198133771 | 0.111797637 | 1.772253655 | 0.076352461 | 0.227282425 |
| Tmem132d | 1383.685315 | 0.198112965 | 0.103935563 | 1.906113359 | 0.0566355 | 0.188063674 |
| Ptpn14 | 626.8537632 | 0.198110501 | 0.137729439 | 1.43840346 | 0.150319612 | 0.350655234 |
| Pkmyt1 | 165.1550193 | 0.198108269 | 0.140863404 | 1.406385637 | 0.159609643 | 0.363728755 |
| Uhrf1 | 47.76490722 | -0.305476104 | 0.162778862 | -1.876632504 | 0.060568479 | 0.195687166 |
| Nalcn | 4157.041132 | -0.151598637 | 0.080798807 | -1.876248458 | 0.060621171 | 0.195720863 |
| Pappa | 35.03436232 | -0.303916027 | 0.16198067 | -1.876248734 | 0.060621133 | 0.195720863 |
| Fam50a | 804.5235151 | 0.197980397 | 0.089130942 | 2.221230832 | 0.026335331 | 0.113124374 |
| Drd2 | 1323.643414 | -0.201698693 | 0.107540767 | -1.875555642 | 0.060716322 | 0.195987229 |
| Cdc42 | 10102.09644 | -0.101926372 | 0.054354927 | -1.875200249 | 0.06076518 | 0.196104082 |
| Tmem59l | 9252.696358 | 0.197827516 | 0.075251367 | 2.628889333 | 0.008566423 | 0.052745648 |
| 1110065P20Rik | 844.9550364 | -0.160536785 | 0.085626217 | -1.874855516 | 0.060812603 | 0.19612631 |
| Catsperg1 | 33.73933621 | 0.197809486 | 0.164800273 | 1.20029829 | 0.230023514 | 0.452972079 |
| Twist1 | 14.89220231 | -0.315145412 | 0.168082165 | -1.874948551 | 0.060799802 | 0.19612631 |
| Cox6c | 5532.557521 | -0.110883571 | 0.059157006 | -1.874394575 | 0.060876061 | 0.196257529 |
| Vwa5a | 689.854735 | 0.19774887 | 0.093298451 | 2.119530046 | 0.034045697 | 0.134601693 |
| Fdft1 | 3616.44637 | -0.128260109 | 0.06845655 | -1.873598771 | 0.060985748 | 0.196529362 |
| Aif1l | 360.3120694 | -0.224849613 | 0.120021468 | -1.873411632 | 0.061011565 | 0.196571676 |
| Alox12 | 32.32659329 | -0.301456198 | 0.160927201 | -1.873245775 | 0.061034454 | 0.196604538 |
| Trak2 | 8844.268508 | -0.119625853 | 0.063874644 | -1.872822229 | 0.061092937 | 0.196752021 |
| Kif6 | 78.27570001 | 0.197734926 | 0.144700514 | 1.366511568 | 0.171778452 | 0.379406321 |
| Cyb5r4 | 1654.952277 | -0.125535886 | 0.067056729 | -1.872084843 | 0.061194867 | 0.196998392 |
| Gm9897 | 32.39055432 | 0.197634405 | 0.167348816 | 1.180972826 | 0.237613518 | 0.462046166 |
| Pan3 | 1214.579562 | 0.197435271 | 0.077290167 | 2.554468167 | 0.010635015 | 0.061461741 |
| AI464131 | 1478.414859 | -0.178861285 | 0.095596558 | -1.871001308 | 0.0613449 | 0.197304828 |
| Cdc42ep3 | 527.157063 | -0.188360737 | 0.100670429 | -1.871063229 | 0.061336318 | 0.197304828 |
| Tead3 | 82.44750613 | 0.197368793 | 0.149569636 | 1.31957794 | 0.186975972 | 0.399311607 |
| Stard3nl | 1181.515492 | -0.160391505 | 0.085745155 | -1.870560562 | 0.061406015 | 0.197431999 |
| Hmmr | 36.56939789 | -0.30710375 | 0.164219593 | -1.870079838 | 0.061472732 | 0.197564511 |
| Npr2 | 2013.40115 | 0.197299483 | 0.104335219 | 1.891015183 | 0.058622317 | 0.192353703 |
| Tnfrsf14 | 22.37885663 | 0.1972734 | 0.167354769 | 1.178773698 | 0.2384883 | 0.462759891 |
| Fam108a | 3298.831558 | -0.159243833 | 0.085170177 | -1.869713538 | 0.061523609 | 0.197596753 |
| 2700049A03Rik | 357.4740666 | 0.197214509 | 0.086908654 | 2.269216003 | 0.023255194 | 0.104288269 |
| 2810029C07Rik | 395.1301443 | 0.19715963 | 0.095331328 | 2.068151508 | 0.038625779 | 0.146136809 |
| Ccdc101 | 405.2147884 | 0.197104158 | 0.117230919 | 1.681332539 | 0.092698341 | 0.25698123 |
| Gm2694 | 42.53070387 | -0.299966686 | 0.160508271 | -1.868855014 | 0.061642988 | 0.197824454 |
| Pou6f2 | 32.79163728 | 0.197098775 | 0.165474358 | 1.1911137 | 0.233608952 | 0.457143705 |
| Slc22a7 | 8.699720755 | 0.197091147 | 0.141480683 | 1.393060468 | 0.163601506 | 0.36875934 |
| Apobec1 | 73.30956365 | 0.197077977 | 0.143525002 | 1.373126461 | 0.169713036 | 0.377102221 |
| Bfsp2 | 154.0966454 | -0.288637625 | 0.154526529 | -1.867883964 | 0.061778247 | 0.198135407 |
| Jdp2 | 1410.357959 | 0.19699364 | 0.114327036 | 1.723071345 | 0.084875624 | 0.243259881 |
| Incenp | 738.8310002 | 0.196877904 | 0.074931127 | 2.62745151 | 0.00860271 | 0.052851067 |
| Prpsap2 | 1190.020333 | -0.152197835 | 0.081539915 | -1.866543954 | 0.061965301 | 0.198611991 |
| Slc9a4 | 33.10696575 | 0.196837665 | 0.150253307 | 1.310038816 | 0.190182705 | 0.403706566 |
| Klhl3 | 749.309244 | -0.191745726 | 0.102754171 | -1.866062706 | 0.062032593 | 0.19874545 |
| Spint2 | 584.0856254 | 0.196822467 | 0.109611019 | 1.795644908 | 0.072551009 | 0.2203696 |
| Smox | 1454.352962 | 0.1967776 | 0.132523305 | 1.4848528 | 0.137582822 | 0.33085647 |
| BC039966 | 10.276408 | 0.196772817 | 0.153885058 | 1.278699956 | 0.201002735 | 0.417807403 |
| 4933413G19Rik | 42.05392704 | 0.19672884 | 0.167419207 | 1.175067327 | 0.239967781 | 0.464293306 |
| Snx29 | 314.8396535 | -0.152474186 | 0.081788666 | -1.864245898 | 0.062287181 | 0.199355004 |
| BC035044 | 33.32726087 | -0.311608985 | 0.167196997 | -1.863723571 | 0.062360534 | 0.199469076 |
| Cdk2 | 98.35195754 | 0.196692369 | 0.140963301 | 1.395344512 | 0.162911975 | 0.368167758 |
| Rps14 | 6562.15397 | -0.134561784 | 0.072197533 | -1.863800303 | 0.062349753 | 0.199469076 |
| Ube2s | 1938.556743 | -0.115161906 | 0.06179625 | -1.863574342 | 0.062381504 | 0.199492057 |
| Zfp113 | 381.5417402 | -0.173760584 | 0.093253499 | -1.863314374 | 0.062418049 | 0.199567736 |
| Ret | 241.7556416 | -0.278543101 | 0.149509734 | -1.863043247 | 0.062456182 | 0.199631622 |
| Trdn | 18.77577821 | -0.242071037 | 0.129936904 | -1.862989106 | 0.062463799 | 0.199631622 |
| Klhdc1 | 482.8569889 | -0.165187689 | 0.08867643 | -1.862813915 | 0.062488452 | 0.199669224 |
| Socs7 | 6538.312827 | -0.144533508 | 0.07760352 | -1.862460736 | 0.062538175 | 0.199786904 |
| Leprot | 985.4391701 | -0.103351429 | 0.05550781 | -1.861925876 | 0.062613539 | 0.199865303 |
| Hhatl | 370.9686276 | 0.19665292 | 0.103872035 | 1.893222949 | 0.058328227 | 0.191550985 |
| Ang | 50.78240133 | 0.196562689 | 0.14788693 | 1.329141723 | 0.183801217 | 0.394698659 |
| Tgfb1i1 | 500.8310573 | -0.199955784 | 0.107392231 | -1.861920379 | 0.062614314 | 0.199865303 |
| Nus1 | 3637.783549 | -0.139575458 | 0.074971647 | -1.861709902 | 0.062643992 | 0.199918849 |
| Kdm6b | 1164.698992 | 0.196562113 | 0.140788367 | 1.396153081 | 0.162668402 | 0.367778232 |
| Ifitm10 | 293.4467867 | 0.19639919 | 0.111623377 | 1.759480818 | 0.078495875 | 0.231393521 |
| Spata1 | 28.8230893 | 0.196293101 | 0.166751284 | 1.177160962 | 0.239131267 | 0.463246349 |
| Ddc | 344.5498891 | 0.196266797 | 0.167991214 | 1.168315845 | 0.242679386 | 0.467209584 |
| Hspb11 | 183.7228903 | -0.199980017 | 0.107539864 | -1.859589651 | 0.062943605 | 0.200668356 |
| Skp1a | 10753.18397 | -0.118571382 | 0.063767512 | -1.859432446 | 0.062965866 | 0.200698032 |
| Ppm1e | 8996.644957 | -0.158140131 | 0.085051738 | -1.859340386 | 0.062978906 | 0.200698307 |
| Snx8 | 474.7296435 | 0.196230035 | 0.089181441 | 2.200346088 | 0.02778235 | 0.117611947 |
| Slbp | 728.4719232 | 0.196212327 | 0.076966887 | 2.549308331 | 0.010793682 | 0.062054663 |
| Izumo4 | 834.1270696 | 0.196122768 | 0.139975755 | 1.401119556 | 0.161178325 | 0.365957937 |
| Matk | 5682.054505 | 0.196078418 | 0.09950268 | 1.970584291 | 0.048771445 | 0.171242905 |
| Map3k4 | 2136.478402 | 0.196021586 | 0.080519747 | 2.434453581 | 0.014914293 | 0.077621071 |
| Gm10941 | 10.77644016 | 0.195979173 | 0.153412518 | 1.277465327 | 0.201438014 | 0.418319339 |
| Spag1 | 417.2929861 | 0.195909759 | 0.114395386 | 1.712566962 | 0.086792236 | 0.246654237 |
| Crtac1 | 2539.1676 | -0.162124343 | 0.087280541 | -1.857508458 | 0.063238849 | 0.201223594 |
| Tubgcp6 | 2147.810617 | 0.195804022 | 0.089963728 | 2.176477414 | 0.029519581 | 0.122064687 |
| Cbx6 | 18585.556 | -0.127970737 | 0.068907997 | -1.857124614 | 0.063293427 | 0.201286609 |
| Zfp346 | 1483.856533 | 0.098768689 | 0.053191762 | 1.856841837 | 0.063333659 | 0.201373224 |
| Ecm1 | 303.8980988 | -0.215726306 | 0.116237515 | -1.85590948 | 0.063466462 | 0.201754074 |
| Gpr39 | 10.16146523 | 0.19567215 | 0.145289428 | 1.34677487 | 0.178052751 | 0.387578219 |
| Shmt2 | 737.9068257 | 0.195520617 | 0.084748499 | 2.30706879 | 0.021050986 | 0.097333324 |
| Mrpl23 | 705.7515705 | -0.137250239 | 0.073968223 | -1.855529758 | 0.063520614 | 0.20183165 |
| Clgn | 320.7063197 | -0.208077488 | 0.112155123 | -1.855265126 | 0.063558376 | 0.20188058 |
| Hmgb2 | 196.0277998 | -0.185478397 | 0.099982295 | -1.855112405 | 0.063580177 | 0.201908436 |
| Ncapg2 | 206.2178232 | -0.183202758 | 0.098772628 | -1.854792798 | 0.063625821 | 0.202011981 |
| Tusc2 | 3120.752287 | -0.122993171 | 0.066326371 | -1.854363045 | 0.063687238 | 0.202165554 |
| Ephx2 | 809.6284299 | -0.239168335 | 0.128991671 | -1.854137814 | 0.063719446 | 0.202226362 |
| Rara | 437.8947372 | 0.195431332 | 0.129943204 | 1.503975008 | 0.132587802 | 0.323014924 |
| Xkrx | 86.70127137 | -0.270238717 | 0.145771874 | -1.853846773 | 0.063761085 | 0.202275644 |
| Gm14169 | 344.1799889 | 0.195286282 | 0.108130094 | 1.806030826 | 0.070913525 | 0.216994211 |
| 2610203C22Rik | 12.82333829 | 0.1952206 | 0.16572931 | 1.177948549 | 0.238817119 | 0.463059624 |
| Rassf5 | 1161.603384 | 0.195203027 | 0.079601494 | 2.452253306 | 0.014196469 | 0.074969354 |
| Zfp160 | 483.885269 | 0.195104778 | 0.102315152 | 1.90690015 | 0.056533519 | 0.187855651 |
| Gtf2h5 | 1631.198143 | -0.123859066 | 0.06688244 | -1.85189216 | 0.06404131 | 0.202956853 |
| 1500012K07Rik | 10.17293774 | -0.295095903 | 0.159365173 | -1.851696307 | 0.064069445 | 0.202962988 |
| 2310003H01Rik | 449.7535824 | 0.195024619 | 0.104912976 | 1.858917991 | 0.063038764 | 0.200845046 |
| E130309D14Rik | 1810.362553 | 0.194954643 | 0.093294402 | 2.089671394 | 0.036647328 | 0.141256 |
| Dnahc11 | 70.51764961 | -0.276178785 | 0.149186393 | -1.851233079 | 0.064136029 | 0.203090871 |
| Ctps | 1646.714087 | -0.106523461 | 0.057580096 | -1.85000491 | 0.064312842 | 0.203609149 |
| Ano8 | 2026.554746 | 0.194877158 | 0.086054624 | 2.264575095 | 0.023538767 | 0.105224943 |
| Fam78a | 326.6722316 | 0.194875555 | 0.109735052 | 1.775873349 | 0.075753795 | 0.226274597 |
| Naip6 | 9.324757374 | 0.194804635 | 0.152232483 | 1.279652226 | 0.200667473 | 0.4174737 |
| Nck2 | 1144.520761 | -0.144503849 | 0.078154679 | -1.848946874 | 0.064465484 | 0.20393909 |
| Perp | 176.6429364 | -0.215854242 | 0.116779613 | -1.848389761 | 0.064545979 | 0.204138681 |
| Ankrd39 | 278.2449259 | 0.194742428 | 0.097298468 | 2.001495317 | 0.045339038 | 0.163406153 |
| Rtn4 | 26325.62958 | -0.107214358 | 0.058014493 | -1.848061616 | 0.06459343 | 0.204190665 |
| Setd4 | 226.5659771 | 0.194679247 | 0.127235392 | 1.530071506 | 0.12599903 | 0.313873335 |
| Ybx1 | 3558.259041 | -0.098014092 | 0.053044161 | -1.847782853 | 0.064633762 | 0.204249544 |
| Fancl | 178.5188047 | -0.265246507 | 0.143577607 | -1.847408605 | 0.064687943 | 0.204337408 |
| St8sia5 | 1811.642984 | -0.177987347 | 0.096343622 | -1.847422201 | 0.064685974 | 0.204337408 |
| Tm9sf3 | 6516.898438 | -0.122599143 | 0.066376491 | -1.847026573 | 0.064743289 | 0.204470551 |
| Ddx42 | 3622.450174 | 0.098768192 | 0.053479716 | 1.846834631 | 0.064771111 | 0.20447506 |
| Lpl | 965.7956696 | 0.194636179 | 0.100564908 | 1.935428408 | 0.052937746 | 0.180267569 |
| Erv3 | 17.86889303 | 0.194552991 | 0.167738635 | 1.159857959 | 0.246106642 | 0.471286158 |
| Hsd17b10 | 819.480604 | -0.130592587 | 0.070739464 | -1.846106541 | 0.064876737 | 0.204600075 |
| Kctd1 | 3439.56405 | -0.14883443 | 0.080612223 | -1.846301039 | 0.064848507 | 0.204600075 |
| Sppl2b | 1359.276204 | 0.194552752 | 0.08140818 | 2.38984279 | 0.016855586 | 0.084028459 |
| Nkiras2 | 333.6770808 | -0.17130121 | 0.092787369 | -1.846169501 | 0.064867598 | 0.204600075 |
| Iyd | 8.898279509 | 0.194469607 | 0.136253709 | 1.427261022 | 0.153504671 | 0.35484834 |
| Fbxl13 | 10.55143406 | 0.194401531 | 0.162788885 | 1.19419413 | 0.232402039 | 0.455686812 |
| Psmg4 | 131.4781166 | -0.199272336 | 0.10802976 | -1.844605934 | 0.065094883 | 0.205162758 |
| 5730508B09Rik | 89.82824416 | -0.256028774 | 0.138808451 | -1.844475404 | 0.065113887 | 0.205180917 |
| Lmtk3 | 8240.308986 | 0.194396014 | 0.08930318 | 2.176809534 | 0.029494781 | 0.122027275 |
| Nppa | 10.69718905 | -0.246384523 | 0.133597082 | -1.844235809 | 0.065148782 | 0.205207405 |
| Gm7854 | 14.17049051 | 0.194243483 | 0.156822812 | 1.238617529 | 0.215487173 | 0.435414484 |
| Gm10635 | 103.7482685 | -0.309179806 | 0.167672898 | -1.843946224 | 0.065190978 | 0.20525686 |
| Cdh23 | 70.23838618 | -0.298660992 | 0.162048793 | -1.843031271 | 0.065324445 | 0.205426618 |
| Col1a1 | 435.5905914 | -0.309012049 | 0.167651477 | -1.843181196 | 0.065302559 | 0.205426618 |
| Pcdhb8 | 127.7875134 | 0.194120905 | 0.136171632 | 1.425560535 | 0.153995235 | 0.35548304 |
| F2rl2 | 76.6447412 | 0.194101749 | 0.152534722 | 1.272508623 | 0.203192462 | 0.420270158 |
| Sesn3 | 1899.004887 | -0.183346935 | 0.099473216 | -1.843178919 | 0.065302892 | 0.205426618 |
| Sctr | 16.60404906 | 0.193946337 | 0.158495881 | 1.223667996 | 0.221077497 | 0.442129311 |
| Neu1 | 2044.273565 | 0.193882397 | 0.082734616 | 2.343425353 | 0.019107585 | 0.091430799 |
| Vps11 | 3324.454847 | 0.19387288 | 0.096405922 | 2.011005917 | 0.044324832 | 0.160962837 |
| Tbx2 | 83.4772758 | 0.193779568 | 0.137080537 | 1.413618393 | 0.157473978 | 0.360292648 |
| Mfap3 | 1203.227814 | -0.139482226 | 0.07576852 | -1.840899443 | 0.065636296 | 0.206258059 |

| | | | | | | |
|---|---|---|---|---|---|---|
| Ddx19a | 728.4905667 | -0.048442408 | 0.071060891 | -0.681702794 | 0.4954269 | 0.844369474 |
| Glmn | 472.2990274 | -0.048496788 | 0.080921303 | -0.599308047 | 0.548967483 | 0.868267067 |
| Dhrs3 | 665.0815163 | -0.048596319 | 0.080258163 | -0.605500024 | 0.544846809 | 0.866307897 |
| 4932435O22Rik | 4.514933801 | -0.04863882 | 0.047555467 | -1.022780824 | 0.306411487 | NA |
| Diexf | 830.48708 | -0.048644253 | 0.066520487 | -0.731267247 | 0.464615932 | 0.83159896 |
| Mir338 | 4.992204301 | -0.048650462 | 0.045068582 | -1.079476215 | 0.280375491 | NA |
| Cenph | 21.44171775 | -0.048653397 | 0.077666288 | -0.626441639 | 0.531025305 | NA |
| Exoc1 | 2421.699006 | -0.048656634 | 0.059200311 | -0.821898278 | 0.411134793 | 0.802622856 |
| Gpr137b-ps | 500.4556499 | -0.048676843 | 0.075673855 | -0.643245189 | 0.520065015 | 0.853297835 |
| Ccdc155 | 172.194081 | -0.048687296 | 0.091048902 | -0.534737874 | 0.592831128 | 0.886226713 |
| Mib2 | 4235.614599 | -0.048702987 | 0.07323739 | -0.665001682 | 0.506049417 | 0.848908639 |
| B4galt7 | 684.1379096 | -0.048732164 | 0.065649735 | -0.742305572 | 0.457902217 | 0.827642075 |
| Mier3 | 771.8650071 | -0.048746479 | 0.069043525 | -0.706025355 | 0.48017237 | 0.836914993 |
| D030047H15Rik | 4.749439604 | -0.048749169 | 0.047051867 | -1.036072996 | 0.300168089 | NA |
| Mir103-2 | 3.148041587 | -0.04875865 | 0.037541726 | -1.29878553 | 0.194017542 | NA |
| Papd4 | 814.6432859 | -0.048775704 | 0.066800715 | -0.730167389 | 0.465287873 | 0.831682622 |
| Mios | 963.7661168 | -0.048835134 | 0.077008981 | -0.634148554 | 0.525983878 | 0.855892499 |
| Fancb | 101.6228171 | -0.048838021 | 0.094310427 | -0.517843284 | 0.604567616 | 0.891930651 |
| Spg7 | 2078.474008 | -0.048840443 | 0.071737113 | -0.680825321 | 0.495982026 | 0.844524079 |
| Spata3 | 13.25343049 | -0.048860865 | 0.070540126 | -0.692667667 | 0.488518134 | NA |
| Phf21a | 1992.057811 | -0.048933651 | 0.053675595 | -0.911655501 | 0.361950096 | 0.776449324 |
| Flywch2 | 240.1638379 | -0.048945869 | 0.08290419 | -0.59039077 | 0.554928697 | 0.871360989 |
| Cand2 | 196.4728651 | -0.048960568 | 0.093747126 | -0.522262069 | 0.601487875 | 0.890461307 |
| Nespas | 1.69239165 | -0.048984002 | 0.024534786 | -1.996512321 | 0.045878186 | NA |
| Tbx3 | 123.9436085 | -0.048987633 | 0.094180491 | -0.520146295 | 0.602961614 | 0.891281205 |
| Cenpi | 17.90266986 | -0.049032084 | 0.078322931 | -0.626024634 | 0.531298783 | NA |
| Tmem180 | 1064.595744 | -0.049032368 | 0.066079822 | -0.74201726 | 0.458076878 | 0.827715405 |
| Arl9 | 2.454849386 | -0.049063931 | 0.032460226 | -1.511509222 | 0.130658761 | NA |
| 1110065P20Rik | 853.2581114 | -0.049067076 | 0.066080115 | -0.742539207 | 0.457760706 | 0.827642075 |
| Slc16a10 | 118.8375983 | -0.049072173 | 0.093825444 | -0.523015621 | 0.600963383 | 0.890297392 |
| 1700024F13Rik | 14.51472345 | -0.049085243 | 0.076003496 | -0.645828759 | 0.518390261 | NA |
| Ypel3 | 4972.71571 | -0.04910021 | 0.051627605 | -0.951045655 | 0.3415812 | 0.76303628 |
| St6gal2 | 515.920625 | -0.049103131 | 0.088559628 | -0.554464055 | 0.579261299 | 0.882452682 |
| Vsig10 | 99.82133378 | -0.04912083 | 0.091757199 | -0.535334894 | 0.592418301 | 0.886226713 |
| 9330158H04Rik | 21.24006886 | -0.049130843 | 0.072291235 | -0.679623796 | 0.496742698 | NA |
| Zcchc2 | 1292.460647 | -0.049134202 | 0.063630497 | -0.77218007 | 0.440007784 | 0.817961744 |
| Slc15a2 | 1109.383752 | -0.049140712 | 0.063362534 | -0.775548403 | 0.438015676 | 0.816984132 |
| Rnf8 | 1275.051034 | -0.049155588 | 0.052845692 | -0.930172104 | 0.352281983 | 0.770257962 |
| Mir453 | 8.759935517 | -0.049157301 | 0.057508055 | -0.854789832 | 0.392667508 | NA |
| Icam5 | 7735.155972 | -0.049192666 | 0.076253899 | -0.645116729 | 0.518851544 | 0.853297835 |
| Map3k2 | 1214.875371 | -0.049212822 | 0.078232787 | -0.629056225 | 0.529312247 | 0.857915133 |
| Zfp7 | 346.1904972 | -0.049240496 | 0.078125316 | -0.630275801 | 0.528514152 | 0.857228044 |
| Dab2 | 451.4014109 | -0.049253559 | 0.09483648 | -0.519352459 | 0.603514977 | 0.891540503 |
| Sf1 | 6189.680764 | -0.049267409 | 0.04889128 | -1.007693171 | 0.313601784 | 0.745844951 |
| Dnajc13 | 3378.115027 | -0.04929956 | 0.0643618 | -0.765975468 | 0.443690899 | 0.820717992 |
| 2610037D02Rik | 3.291275177 | -0.049330202 | 0.042849546 | -1.151242124 | 0.24963264 | NA |
| Rsph6a | 6.740492765 | -0.049341305 | 0.052452994 | -0.940676613 | 0.346870607 | NA |
| N4bp2l2 | 1344.319234 | -0.049347733 | 0.069619165 | -0.708823969 | 0.478433722 | 0.835936459 |
| Ccl8 | 4.761070116 | -0.049354819 | 0.044507126 | -1.108919463 | 0.267464927 | NA |
| Dus1l | 1097.469803 | -0.049382633 | 0.069315577 | -0.712431961 | 0.47619733 | 0.835195112 |
| Srgap3 | 17147.66138 | -0.049384937 | 0.059849338 | -0.825154275 | 0.409284009 | 0.801055189 |
| Mcm3ap | 2749.995317 | -0.049388265 | 0.057936401 | -0.852456558 | 0.39396074 | 0.794754188 |
| BC068281 | 329.5600589 | -0.049397855 | 0.077189689 | -0.639954068 | 0.522202461 | 0.854113377 |
| C030013G03Rik | 56.19889518 | -0.049401672 | 0.094894441 | -0.520596066 | 0.60264819 | 0.891281205 |
| Erbb2ip | 3729.267965 | -0.049404539 | 0.07349333 | -0.6722316 | 0.501436267 | 0.846601651 |
| Tmem79 | 11.98766079 | -0.049416443 | 0.07125327 | -0.693532288 | 0.48797557 | NA |
| Zfp879 | 122.6001804 | -0.049431297 | 0.093308467 | -0.529762179 | 0.596276831 | 0.887922451 |
| Ppp1r12c | 4314.09069 | -0.049461872 | 0.064033011 | -0.772443331 | 0.439851898 | 0.817776613 |
| Pygl | 215.0604082 | -0.049463978 | 0.088471909 | -0.559092466 | 0.576098616 | 0.880966011 |
| Cyp39a1 | 191.5228943 | -0.049485885 | 0.087232163 | -0.567289442 | 0.570517549 | 0.878246677 |
| Egf | 58.18632245 | -0.04949992 | 0.091523939 | -0.540841234 | 0.588617019 | 0.885188411 |
| Zfpl1 | 808.9075741 | -0.049508451 | 0.067947873 | -0.728624007 | 0.466231689 | 0.832050282 |
| Olfml2b | 304.6100365 | -0.049546896 | 0.094831769 | -0.522471498 | 0.601342087 | 0.890336882 |
| Kctd9 | 580.7946788 | -0.049553232 | 0.073388377 | -0.67521908 | 0.499536586 | 0.846241311 |
| Mctp2 | 69.19939814 | -0.049564857 | 0.090938527 | -0.545036947 | 0.585728107 | 0.884588669 |
| Hace1 | 2148.674895 | -0.049581258 | 0.070718846 | -0.701103892 | 0.483238182 | 0.838104566 |
| Dicer1 | 2764.587117 | -0.04959711 | 0.061340809 | -0.80854998 | 0.418774047 | 0.806975048 |
| Scamp2 | 1047.989014 | -0.049618979 | 0.062112708 | -0.798853902 | 0.424375131 | 0.809162684 |
| Gm16675 | 12.20743184 | -0.04961917 | 0.06786746 | -0.731118702 | 0.464706652 | NA |
| Tmod1 | 2863.068775 | -0.04962128 | 0.055176886 | -0.899312795 | 0.368486074 | 0.779452463 |
| A530064D06Rik | 2.518090489 | -0.049632277 | 0.033879238 | -1.464976196 | 0.142927412 | NA |
| 5730480H06Rik | 28.15024117 | -0.049658246 | 0.089237179 | -0.556474849 | 0.577886284 | 0.881703921 |
| Parn | 1158.35197 | -0.049680845 | 0.062422102 | -0.795885485 | 0.426098599 | 0.809519334 |
| Wdr13 | 9911.79914 | -0.049696736 | 0.057238763 | -0.868235679 | 0.385265326 | 0.789016184 |
| 3110047P20Rik | 1917.873608 | -0.049726176 | 0.077452575 | -0.642020957 | 0.520859576 | 0.85372574 |
| Apba3 | 579.6376837 | -0.049743791 | 0.079598648 | -0.624932609 | 0.532015289 | 0.858328942 |
| 6720468P15Rik | 16.7774214 | -0.049757364 | 0.077749215 | -0.639972551 | 0.522190445 | NA |
| Zfp473 | 24.81086895 | -0.049765373 | 0.086771228 | -0.573523902 | 0.56629002 | 0.875844212 |
| 9230009I02Rik | 8.788628684 | -0.049852376 | 0.062414499 | -0.7987307 | 0.424446582 | NA |
| Tstd1 | 19.84431139 | -0.049863143 | 0.080411605 | -0.620098842 | 0.535192715 | NA |
| Vsx2 | 13.5261872 | -0.049865684 | 0.073124466 | -0.681928866 | 0.495283932 | NA |
| Fam71d | 4.179820251 | -0.04987921 | 0.046037009 | -1.083458965 | 0.278604755 | NA |
| Tet1 | 494.5105663 | -0.049938576 | 0.075326078 | -0.662965302 | 0.507352777 | 0.848958235 |
| Six1 | 8.538975441 | -0.049944642 | 0.059105227 | -0.845012268 | 0.39810399 | NA |
| Foxk1 | 3315.925796 | -0.049953637 | 0.055452462 | -0.900837128 | 0.367674924 | 0.778985919 |
| Snora65 | 48.06668916 | -0.049959606 | 0.09334581 | -0.535209944 | 0.592504691 | 0.886226713 |
| Il17rb | 58.79160167 | -0.049965345 | 0.095102601 | -0.525383582 | 0.599316565 | 0.889150487 |
| Zfp958 | 269.7278944 | -0.05002133 | 0.086493522 | -0.578324576 | 0.563045005 | 0.87510382 |
| Mzb1 | 3.107203627 | -0.050025785 | 0.036291012 | -1.378462109 | 0.168060659 | NA |
| Dsc2 | 24.77883008 | -0.050026254 | 0.081823882 | -0.611389398 | 0.54094182 | 0.863847445 |
| Grm2 | 1760.554063 | -0.050036002 | 0.078576604 | -0.6367799 | 0.524268221 | 0.855487854 |
| Pla2g4c | 7.286950046 | -0.050037698 | 0.057680857 | -0.86749228 | 0.38567234 | NA |
| N4bp2 | 249.8630082 | -0.05005806 | 0.094346938 | -0.530574297 | 0.595713811 | 0.887497708 |
| Zfp60 | 1555.128762 | -0.050075884 | 0.068599594 | -0.729973476 | 0.465406398 | 0.831682622 |
| Npat | 1155.997173 | -0.050102912 | 0.068993071 | -0.72620209 | 0.467714891 | 0.832353544 |
| Snora2b | 4.81953814 | -0.050120791 | 0.043417602 | -1.154388741 | 0.248340829 | NA |
| Cad | 1015.473458 | -0.050164056 | 0.077532736 | -0.647004846 | 0.517628807 | 0.853297835 |
| Diablo | 1009.065092 | -0.050168871 | 0.063661645 | -0.788054893 | 0.430664598 | 0.812181606 |
| Myeov2 | 1353.482549 | -0.050169632 | 0.066471511 | -0.754753899 | 0.450396659 | 0.823365451 |
| Zswim5 | 887.4070209 | -0.050170068 | 0.060398312 | -0.830653479 | 0.406169414 | 0.800759397 |
| Fxyd7 | 2095.80625 | -0.05019623 | 0.069951254 | -0.717588704 | 0.473010925 | 0.833983942 |
| Gdpd4 | 3.261757732 | -0.050210324 | 0.040650531 | -1.235170194 | 0.21676717 | NA |
| Sp6 | 24.33279068 | -0.050265288 | 0.079004982 | -0.636229349 | 0.524626946 | 0.855487854 |
| Gm15104 | 4.151745343 | -0.050274045 | 0.044963296 | -1.118112983 | 0.263518742 | NA |
| Hgfac | 6.076266127 | -0.050275383 | 0.047653212 | -1.055026125 | 0.291413377 | NA |
| Plxnb1 | 4977.123788 | -0.050326862 | 0.05819132 | -0.864851697 | 0.387120192 | 0.791569245 |
| Pold3 | 1106.778103 | -0.050339005 | 0.056852446 | -0.885432526 | 0.375923396 | 0.783947144 |
| Gm11648 | 4.37029336 | -0.050343086 | 0.043734823 | -1.151098443 | 0.249691738 | NA |
| Vars2 | 1166.00833 | -0.050367696 | 0.073047848 | -0.689516488 | 0.490498301 | 0.841798006 |
| Best1 | 54.25564751 | -0.050384685 | 0.090529953 | -0.556552641 | 0.577833119 | 0.881703921 |
| Alkbh7 | 346.6968999 | -0.05039125 | 0.080217942 | -0.628179294 | 0.529886493 | 0.857915133 |
| 2310067B10Rik | 3732.224609 | -0.050416954 | 0.057525866 | -0.87642234 | 0.380800478 | 0.78650757 |
| 1600016N20Rik | 6.993231167 | -0.050435858 | 0.05121022 | -0.984878764 | 0.324683626 | NA |
| Gtf2h2 | 637.5769268 | -0.050484058 | 0.067260061 | -0.750580012 | 0.452905454 | 0.82547056 |
| Ccdc167 | 545.3180825 | -0.050486308 | 0.069132189 | -0.730286554 | 0.465215045 | 0.831682622 |
| AI429214 | 124.5127771 | -0.050490083 | 0.092554682 | -0.545516245 | 0.585398511 | 0.884512516 |
| Tepp | 4.168158204 | -0.050491564 | 0.041887921 | -1.205396754 | 0.228050174 | NA |
| Ankrd23 | 113.3906172 | -0.050503944 | 0.094430769 | -0.534825078 | 0.59277082 | 0.886226713 |
| Ankrd26 | 809.1464496 | -0.05050971 | 0.070581864 | -0.715618812 | 0.474226755 | 0.834074081 |
| Bloc1s3 | 431.5161675 | -0.050532358 | 0.071853127 | -0.703272916 | 0.481885686 | 0.837383765 |
| BC051019 | 9.741549949 | -0.050583064 | 0.056579648 | -0.894015166 | 0.371313777 | NA |
| Rbm15 | 329.4823395 | -0.05061388 | 0.080628528 | -0.627741583 | 0.53017324 | 0.857915133 |
| Mtx3 | 2514.645902 | -0.050620119 | 0.049050811 | -1.031993515 | 0.302075156 | 0.735974432 |
| Zdhhc23 | 313.2306631 | -0.050630086 | 0.075859713 | -0.667417314 | 0.50450561 | 0.847807553 |
| Ikzf2 | 127.4279702 | -0.050632742 | 0.093512912 | -0.541451884 | 0.588196154 | 0.885188411 |
| Arhgap27 | 794.1639527 | -0.050654061 | 0.069633345 | -0.727439729 | 0.466956625 | 0.832169296 |
| Mdh1b | 53.08518482 | -0.050669433 | 0.092406204 | -0.548333674 | 0.583462809 | 0.884313345 |
| Fam132a | 466.3786199 | -0.050681152 | 0.076859933 | -0.659396256 | 0.509641346 | 0.850457637 |
| Zfp846 | 450.2924265 | -0.050690832 | 0.076861585 | -0.659508024 | 0.509569595 | 0.850457637 |
| Mir22hg | 760.4971644 | -0.050694598 | 0.075210446 | -0.674036671 | 0.500288 | 0.846268338 |
| Sh3bp2 | 187.5032248 | -0.050702062 | 0.092125185 | -0.550360492 | 0.582072141 | 0.884020489 |
| D130020L05Rik | 95.17930375 | -0.050775008 | 0.093729578 | -0.541718091 | 0.588012725 | 0.885188411 |
| Xkr6 | 20.50614761 | -0.050811308 | 0.080991348 | -0.627367116 | 0.530418618 | NA |
| Sun2 | 3537.066185 | -0.050848495 | 0.063672202 | -0.798598028 | 0.424523531 | 0.809162684 |
| Josd2 | 1105.178138 | -0.05088773 | 0.081301006 | -0.625917596 | 0.531368992 | 0.858328942 |
| Layn | 157.3448347 | -0.050907435 | 0.091502022 | -0.556353113 | 0.577969485 | 0.881703921 |
| Cga | 9.106314582 | -0.050916453 | 0.048875323 | -1.041761969 | 0.297522051 | NA |
| Prpsap1 | 1557.567479 | -0.050960455 | 0.064333207 | -0.792132988 | 0.428283137 | 0.810486244 |
| 1110001J03Rik | 377.5177111 | -0.050994948 | 0.074626646 | -0.683334315 | 0.494395615 | 0.843503724 |
| Gm4980 | 191.8590147 | -0.051000843 | 0.086415906 | -0.590178883 | 0.555070728 | 0.871360989 |
| Hmgxb4 | 738.0119501 | -0.051001332 | 0.061662769 | -0.827100895 | 0.408179877 | 0.801055189 |
| Gcdh | 1025.196964 | -0.051019474 | 0.070023328 | -0.728606821 | 0.466242205 | 0.832050282 |
| Trmt61a | 731.0937212 | -0.051040389 | 0.08278078 | -0.616572945 | 0.537516444 | 0.861996453 |
| Actl6b | 3352.566048 | -0.051041672 | 0.057386294 | -0.889440115 | 0.373766594 | 0.782348609 |
| Trim28 | 5088.491881 | -0.051059813 | 0.042935522 | -1.189220746 | 0.234352813 | 0.68557298 |
| Riok1 | 569.1897864 | -0.051074982 | 0.063374414 | -0.805924334 | 0.420286479 | 0.807657282 |
| Sfxn5 | 8006.291593 | -0.051086435 | 0.0509183 | -1.003302057 | 0.315715144 | 0.745993052 |
| Fam161b | 807.9761638 | -0.05110436 | 0.059712794 | -0.855836016 | 0.392088491 | 0.794184423 |
| Rpl3l | 11.96112832 | -0.051112424 | 0.067894686 | -0.752819222 | 0.451558554 | NA |
| Cxcl12 | 2637.23013 | -0.051115855 | 0.08567403 | -0.596631847 | 0.550753204 | 0.869642174 |
| Snora30 | 5.702490234 | -0.051131577 | 0.050718575 | -1.008143009 | 0.313385813 | NA |
| Nphs1as | 10.63824656 | -0.051157412 | 0.059265672 | -0.863187919 | 0.388034155 | NA |
| Stap2 | 73.36613954 | -0.051176077 | 0.093634021 | -0.546554301 | 0.584684973 | 0.884313345 |
| Ephb6 | 5517.078892 | -0.051255672 | 0.069358488 | -0.738996386 | 0.459909192 | 0.828966488 |
| Met | 362.4192209 | -0.051274393 | 0.072800037 | -0.704318225 | 0.481234619 | 0.8371092 |
| Kif24 | 35.59236734 | -0.051281327 | 0.089159328 | -0.57516503 | 0.565179691 | 0.875364912 |
| Htr6 | 117.8693707 | -0.0512823 | 0.091698593 | -0.559248489 | 0.575992145 | 0.880907759 |
| C2 | 107.610072 | -0.051290203 | 0.092423859 | -0.554945483 | 0.57893195 | 0.882430799 |
| Slc2a10 | 80.56127861 | -0.051304815 | 0.094984976 | -0.540136101 | 0.589103176 | 0.885188411 |
| Rag1 | 8.967626615 | -0.051321704 | 0.05433263 | -0.944583457 | 0.344871576 | NA |
| Bpifc | 7.364241138 | -0.051331473 | 0.057327791 | -0.895402934 | 0.370571733 | NA |
| Sav1 | 756.9826623 | -0.051370024 | 0.072373127 | -0.709794178 | 0.47783178 | 0.835457925 |
| Hsd11b1 | 483.2881546 | -0.051383234 | 0.092618526 | -0.554783547 | 0.579042722 | 0.882452682 |
| Sh3glb2 | 6230.015403 | -0.051399537 | 0.059493504 | -0.863952075 | 0.387614218 | 0.791583702 |
| Sel1l3 | 3410.75924 | -0.051406121 | 0.06493612 | -0.791641397 | 0.428569802 | 0.810486244 |
| Eml6 | 1272.270171 | -0.051458466 | 0.069087741 | -0.744827736 | 0.456375866 | 0.82674985 |
| A330041J22Rik | 44.01756463 | -0.051475105 | 0.093031714 | -0.553307074 | 0.580053158 | 0.883120446 |
| Cbr4 | 375.8882964 | -0.051538247 | 0.072341275 | -0.712432104 | 0.476197241 | 0.835195112 |
| Hmgb2 | 201.1767463 | -0.051578391 | 0.087125323 | -0.592002287 | 0.553849057 | 0.870976659 |
| Mir3101 | 16.4571339 | -0.051582717 | 0.072637637 | -0.710137589 | 0.477618819 | NA |
| Tnfrsf13c | 12.79127943 | -0.05158357 | 0.067944263 | -0.7592042 | 0.447730414 | NA |
| Srcrb4d | 43.76959127 | -0.051597554 | 0.085127109 | -0.606123643 | 0.544432652 | 0.86624804 |
| A230072E10Rik | 23.54851746 | -0.051601081 | 0.080041784 | -0.644676797 | 0.519136656 | 0.853297835 |
| Lgals8 | 1193.10997 | -0.051601985 | 0.061020435 | -0.84565089 | 0.397747527 | 0.797651232 |
| Nlrc4 | 13.46760797 | -0.051635807 | 0.065031589 | -0.79401116 | 0.427188935 | NA |
| Wee2 | 7.266115142 | -0.051635844 | 0.05600293 | -0.922020398 | 0.356517936 | NA |
| Rbpjl | 17.68455356 | -0.051639479 | 0.076936543 | -0.671195737 | 0.502095844 | NA |
| Thap1 | 305.8990433 | -0.051644051 | 0.082720502 | -0.624319845 | 0.532417554 | 0.858328942 |
| Sun1 | 4737.759035 | -0.051658483 | 0.05521542 | -0.935580725 | 0.349489103 | 0.768268271 |
| Heatr5a | 647.5831416 | -0.051660063 | 0.077741302 | -0.664512458 | 0.506362378 | 0.848908639 |
| Mir382 | 9.292188856 | -0.05166776 | 0.059819025 | -0.863734573 | 0.387733717 | NA |
| Mir668 | 5.995332248 | -0.051676782 | 0.051500183 | -1.003429089 | 0.315653875 | NA |
| Ppp1r3e | 283.8356564 | -0.051730095 | 0.082445052 | -0.627449364 | 0.530364718 | 0.857915133 |
| Fbxl3 | 3282.596061 | -0.05173131 | 0.061922035 | -0.835426526 | 0.4034776 | 0.800224169 |
| Dnmt3a | 3698.615396 | -0.051734498 | 0.056029793 | -0.923339096 | 0.35583052 | 0.772643677 |
| Dbil5 | 14.9798487 | -0.051764629 | 0.072968529 | -0.709410336 | 0.478069876 | NA |
| Sidt2 | 4030.671608 | -0.051768282 | 0.054683062 | -0.946696832 | 0.343793285 | 0.763762832 |
| Ppcdc | 488.4705131 | -0.051779653 | 0.078077429 | -0.66318338 | 0.507213115 | 0.848958235 |
| Rhoj | 152.6372243 | -0.051781598 | 0.086401309 | -0.599314968 | 0.548962868 | 0.868267067 |
| Mro | 453.7825564 | -0.051796736 | 0.07860369 | -0.658960618 | 0.509921058 | 0.850575622 |
| Gm14308 | 2.098716026 | -0.051817746 | 0.031776178 | -1.630710439 | 0.102951433 | NA |
| Gucy2f | 46.96239142 | -0.051830031 | 0.093568818 | -0.553924179 | 0.579630737 | 0.88256982 |
| A930006K02Rik | 19.83852871 | -0.051849686 | 0.078465182 | -0.660798647 | 0.50874145 | NA |
| Epx | 12.83652883 | -0.051854212 | 0.074453734 | -0.696462217 | 0.486139411 | NA |
| Necab3 | 2929.808753 | -0.051857539 | 0.058098905 | -0.89257343 | 0.372085653 | 0.781371686 |
| Pms1 | 147.438202 | -0.051886249 | 0.089170381 | -0.581877617 | 0.560649118 | 0.873713575 |
| Wbscr17 | 2494.242365 | -0.051901885 | 0.052416104 | -0.990189684 | 0.322081414 | 0.751575039 |
| Birc3 | 82.12341621 | -0.051909469 | 0.094620877 | -0.548604812 | 0.583276682 | 0.884313345 |
| Cwc22 | 883.0338666 | -0.051948186 | 0.060645992 | -0.856580688 | 0.391676663 | 0.794184423 |
| AU040972 | 8.472841476 | -0.051950557 | 0.052213619 | -0.994961826 | 0.319754831 | NA |
| Cep55 | 8.296040583 | -0.051958825 | 0.058235223 | -0.892223344 | 0.372273232 | NA |
| Tpp1 | 3505.648535 | -0.051998752 | 0.057111903 | -0.910471368 | 0.362573975 | 0.776449324 |
| Efcab7 | 72.69426466 | -0.052026138 | 0.09493827 | -0.54799964 | 0.58369215 | 0.884313345 |
| Ttc18 | 67.77618991 | -0.052037282 | 0.093244268 | -0.558074866 | 0.576793262 | 0.881151498 |
| Fam160b1 | 1768.928673 | -0.05208029 | 0.048771111 | -1.067851214 | 0.285587632 | 0.722148665 |
| Mmgt2 | 532.439205 | -0.052081175 | 0.069361667 | -0.750863943 | 0.452734542 | 0.825461127 |
| Gm16197 | 21.06422548 | -0.052085006 | 0.076459645 | -0.681209103 | 0.495739189 | NA |
| Ctc1 | 2181.564333 | -0.052119552 | 0.053496712 | -0.974257116 | 0.329928892 | 0.756141871 |
| Apoe | 42530.03042 | -0.052163698 | 0.069070476 | -0.755224244 | 0.450114444 | 0.823263966 |
| Mmp28 | 116.8465574 | -0.052192252 | 0.090994359 | -0.57357678 | 0.566254228 | 0.875844212 |
| Pitpnm3 | 5903.629934 | -0.052231496 | 0.065871143 | -0.792934415 | 0.427816035 | 0.810334558 |
| Chp2 | 48.74438397 | -0.052243214 | 0.094295273 | -0.55403852 | 0.579552484 | 0.882543178 |
| Ptprb | 2065.623283 | -0.05224504 | 0.075220679 | -0.694556883 | 0.487333041 | 0.839801621 |
| Drd3 | 5.093873114 | -0.052266248 | 0.051017127 | -1.024484347 | 0.30560656 | NA |
| Ikzf3 | 8.304615533 | -0.052305894 | 0.059760986 | -0.875251512 | 0.381437071 | NA |
| Brd8 | 2621.370711 | -0.052316355 | 0.048707039 | -1.074102559 | 0.282776718 | 0.720920007 |
| Ppp4r4 | 1706.312515 | -0.052326845 | 0.064549071 | -0.810652177 | 0.417565444 | 0.806161037 |
| Esm1 | 10.29384499 | -0.052337094 | 0.058421115 | -0.895859214 | 0.37032796 | NA |
| Gm10790 | 71.50012643 | -0.052346976 | 0.095102567 | -0.550426534 | 0.582026853 | 0.884020489 |
| Col20a1 | 152.7110427 | -0.052384933 | 0.092438381 | -0.566701107 | 0.570917269 | 0.878246677 |
| Fxyd3 | 2.825558303 | -0.052391047 | 0.038372884 | -1.365314293 | 0.172154288 | NA |
| Zbtb44 | 942.5713906 | -0.052414994 | 0.065331561 | -0.802292083 | 0.422384021 | 0.808695226 |
| Olfm4 | 31.16336309 | -0.05242702 | 0.088747837 | -0.590741383 | 0.554693716 | 0.871360989 |
| 9530059O14Rik | 2.865997635 | -0.052453189 | 0.041366221 | -1.268019828 | 0.20479087 | NA |
| Agpat2 | 52.07817461 | -0.05246199 | 0.09271552 | -0.565838274 | 0.571503725 | 0.878389918 |
| AI662270 | 4.04762712 | -0.052474694 | 0.039196115 | -1.338772832 | 0.180644638 | NA |
| Arhgap42 | 693.0421104 | -0.052479852 | 0.078571632 | -0.667923661 | 0.504182323 | 0.847807553 |
| Kif14 | 15.47479433 | -0.052481894 | 0.073650618 | -0.712579135 | 0.476106227 | NA |
| 4930471C04Rik | 5.476811247 | -0.052486756 | 0.054256689 | -0.967378528 | 0.33335484 | NA |
| Slc4a9 | 7.77462547 | -0.052490695 | 0.059556672 | -0.881357078 | 0.378124581 | NA |
| Zfp317 | 1171.778629 | -0.052492197 | 0.051379995 | -1.021646585 | 0.3069482 | 0.739163119 |
| Ccdc13 | 200.4838213 | -0.052519302 | 0.089221813 | -0.588637462 | 0.556104498 | 0.871546692 |
| Plxdc2 | 1343.016489 | -0.052527506 | 0.063709508 | -0.824484564 | 0.409664283 | 0.801196796 |
| E2f7 | 16.08061732 | -0.052608082 | 0.076449531 | -0.688141334 | 0.491363784 | NA |
| Zfp219 | 1189.360411 | -0.052616912 | 0.069604116 | -0.75594542 | 0.44968192 | 0.823263966 |

| | | | | | | |
|---|---|---|---|---|---|---|
| Cbln1 | 249.5870599 | -0.243159523 | 0.132123754 | -1.840392179 | 0.065710681 | 0.206389882 |
| Dynlt3 | 3548.401282 | -0.123747553 | 0.067236475 | -1.840482471 | 0.065697435 | 0.206389882 |
| Chst12 | 732.352546 | 0.193701285 | 0.110565975 | 1.751906813 | 0.079789833 | 0.2341852 |
| Ncor1 | 9715.332586 | -0.12921109 | 0.070218768 | -1.84012187 | 0.065750347 | 0.206448923 |
| Llgl2 | 50.60767207 | 0.193687073 | 0.162996022 | 1.188293254 | 0.234717895 | 0.458404859 |
| Fam195a | 203.0140037 | -0.193970368 | 0.105449671 | -1.839459207 | 0.065847673 | 0.206568909 |
| Slc7a7 | 70.89198641 | 0.193664147 | 0.148972457 | 1.299999681 | 0.193601078 | 0.407978694 |
| Pole4 | 885.1868671 | -0.205743416 | 0.111848285 | -1.83948656 | 0.065843653 | 0.206568909 |
| Gtf2e1 | 681.6185281 | -0.150024125 | 0.081563938 | -1.839343822 | 0.065864631 | 0.206580283 |
| Cecr6 | 3430.962028 | -0.189540693 | 0.103065352 | -1.839034058 | 0.065910177 | 0.206681295 |
| Abca6 | 130.33335 | 0.193659521 | 0.117867329 | 1.64302968 | 0.100376793 | 0.270853017 |
| Dmbx1 | 9.447200537 | 0.193581456 | 0.132717876 | 1.458593686 | 0.144676972 | 0.342441951 |
| Caly | 4771.261857 | -0.125809417 | 0.068429204 | -1.838533983 | 0.065983759 | 0.206786481 |
| Nynrin | 615.166242 | -0.213663436 | 0.116230549 | -1.838272626 | 0.066022243 | 0.206865243 |
| Kcne1l | 147.3753399 | -0.23825585 | 0.129627903 | -1.837998176 | 0.066062674 | 0.206950075 |
| Pank3 | 2353.705901 | -0.122291843 | 0.06654915 | -1.837616913 | 0.066118875 | 0.207084262 |
| Mphosph9 | 967.7446748 | 0.193494165 | 0.077204778 | 2.506245982 | 0.01220207 | 0.067593447 |
| Sbno1 | 6932.784722 | -0.094466648 | 0.05141325 | -1.837398892 | 0.06615103 | 0.207101246 |
| Cinp | 1486.545104 | -0.121074594 | 0.065907875 | -1.837027744 | 0.0662058 | 0.207230841 |
| 1190003J15Rik | 23.2997865 | 0.193342671 | 0.165761014 | 1.166394116 | 0.243455137 | 0.468001723 |
| Mical1 | 463.3421518 | 0.193284563 | 0.095441851 | 2.025155211 | 0.042851429 | 0.157219996 |
| Col20a1 | 127.0899322 | 0.193234701 | 0.127357871 | 1.517257619 | 0.129201654 | 0.318158046 |
| Card6 | 97.69139589 | 0.193199999 | 0.120587213 | 1.60215992 | 0.109120251 | 0.287047396 |
| Mrpl49 | 1302.075244 | -0.106045471 | 0.057788752 | -1.835053846 | 0.066497712 | 0.207916461 |
| Gbp2 | 230.6271282 | 0.193073121 | 0.157300165 | 1.227418425 | 0.219665365 | 0.440386617 |
| Clic4 | 3794.803205 | -0.178113647 | 0.097103955 | -1.834257395 | 0.066615795 | 0.208219716 |
| Morf4l2 | 6786.830027 | -0.11642313 | 0.063474937 | -1.834159047 | 0.066630389 | 0.208223324 |
| Pam | 8632.802494 | -0.115128475 | 0.062778118 | -1.833894964 | 0.066669587 | 0.208303808 |
| Fbxl5 | 3218.027649 | 0.087317537 | 0.047619426 | 1.833653717 | 0.066705413 | 0.208373724 |
| 1700096K18Rik | 41.53409874 | 0.192993 | 0.155661157 | 1.239827605 | 0.215039166 | 0.434792748 |
| Fam211b | 803.2051039 | -0.200867393 | 0.10959608 | -1.832797236 | 0.06683273 | 0.20864524 |
| Gm711 | 9.963430847 | 0.192873315 | 0.156599637 | 1.231633217 | 0.218086127 | 0.438436026 |
| Zfp106 | 13024.46515 | -0.152737244 | 0.0833406 | -1.832687114 | 0.066849115 | 0.208654348 |
| Stt3a | 1343.634757 | -0.119831603 | 0.065392099 | -1.832508905 | 0.066875636 | 0.208695087 |
| Terf1 | 287.4459959 | 0.192759645 | 0.110136703 | 1.750185355 | 0.080086335 | 0.234788586 |
| Fgd6 | 987.0892307 | -0.130065645 | 0.07099844 | -1.831950742 | 0.06695876 | 0.208828306 |
| Nub1 | 4433.947982 | 0.093523485 | 0.051049279 | 1.832023614 | 0.066947903 | 0.208828306 |
| Allc | 17.23619925 | 0.192585042 | 0.168121422 | 1.145511619 | 0.251997285 | 0.477720814 |
| Ints10 | 1813.801952 | 0.192552608 | 0.092015182 | 2.092617806 | 0.036383286 | 0.140570291 |
| Bmpr1a | 2459.308416 | -0.155184909 | 0.084761963 | -1.830831928 | 0.067125633 | 0.209138259 |
| Gm6225 | 16.97815084 | 0.192524317 | 0.162210906 | 1.186876524 | 0.235276328 | 0.459056974 |
| Notum | 105.7742969 | -0.294432796 | 0.160813266 | -1.830898678 | 0.067115668 | 0.209138259 |
| Gimap9 | 12.78829912 | 0.192388452 | 0.16316376 | 1.179112639 | 0.238353326 | 0.462671816 |
| Plin5 | 21.71819375 | 0.192304778 | 0.167121885 | 1.150685793 | 0.249861523 | 0.475246064 |
| Rnd3 | 600.9682254 | -0.22134018 | 0.120922056 | -1.830436789 | 0.067184651 | 0.209220587 |
| Pfn2 | 15152.19096 | -0.101909387 | 0.055691678 | -1.829885396 | 0.067267078 | 0.209410509 |
| Acin1 | 4532.398558 | 0.192209252 | 0.079157166 | 2.42819776 | 0.015174067 | 0.078552285 |
| Bnc2 | 22.69267497 | -0.303905976 | 0.166099486 | -1.829662354 | 0.067300444 | 0.209430221 |
| Ado | 2096.056265 | -0.139079588 | 0.07603844 | -1.829069453 | 0.067389204 | 0.209496054 |
| D430036J16Rik | 118.889761 | -0.269958568 | 0.147571144 | -1.829345221 | 0.067347908 | 0.209496054 |
| Ldlrad3 | 549.1791533 | -0.150068117 | 0.082045535 | -1.829083302 | 0.06738713 | 0.209496054 |
| Ccdc42 | 55.37902446 | 0.192187296 | 0.155495842 | 1.235964215 | 0.216471866 | 0.436443452 |
| Abcb4 | 269.0683441 | 0.192117333 | 0.097189697 | 1.976725306 | 0.048072689 | 0.169669432 |
| Ptprn2 | 16650.61737 | -0.104549355 | 0.057186739 | -1.82820976 | 0.067518077 | 0.20985458 |
| Agtpbp1 | 7828.426726 | -0.132185532 | 0.072313022 | -1.827963042 | 0.067555099 | 0.209885442 |
| Ovca2 | 425.3348675 | 0.192104233 | 0.09531099 | 2.015551763 | 0.043846868 | 0.159775017 |
| Medag | 1007.076983 | 0.192068537 | 0.095229516 | 2.016901333 | 0.043705811 | 0.159448513 |
| Dleu2 | 126.1663542 | -0.23774803 | 0.130083839 | -1.827652314 | 0.067601749 | 0.209946182 |
| Ebf2 | 32.69457032 | -0.307123054 | 0.168094373 | -1.827087057 | 0.067686681 | 0.210167822 |
| Lrrc4c | 4524.979428 | -0.13881811 | 0.075998458 | -1.826591142 | 0.067761266 | 0.210315116 |
| Galns | 303.3851779 | 0.192011039 | 0.101976369 | 1.88289738 | 0.059714276 | 0.194087057 |
| Dnmt3a | 3630.299426 | -0.124700538 | 0.068285195 | -1.826172391 | 0.067824299 | 0.210426449 |
| Sssca1 | 496.8088328 | -0.146431418 | 0.080181928 | -1.826239676 | 0.067814167 | 0.210426449 |
| Il13ra1 | 284.5274408 | -0.203399808 | 0.111407287 | -1.825731627 | 0.067890697 | 0.210590281 |
| Rasl10a | 1100.205473 | -0.228133168 | 0.124973421 | -1.825453496 | 0.067932623 | 0.210678154 |
| Emc2 | 1929.570916 | -0.177658102 | 0.097332885 | -1.825262868 | 0.067961371 | 0.210725131 |
| Nfkb2 | 113.9475407 | 0.192008867 | 0.127275974 | 1.508602616 | 0.131400365 | 0.321426786 |
| Ndufc1 | 1115.936163 | -0.222307805 | 0.12187734 | -1.824029024 | 0.068147685 | 0.211218288 |
| Atp5d | 5227.551254 | -0.138125865 | 0.07573787 | -1.82373581 | 0.068192022 | 0.211313439 |
| Dgcr8 | 1538.815447 | 0.19196399 | 0.07959411 | 2.411786371 | 0.01587458 | 0.081061217 |
| Tfam | 1232.283988 | -0.104974753 | 0.057576242 | -1.82323039 | 0.068268504 | 0.211465854 |
| Pcdhga10 | 388.1040694 | -0.17971501 | 0.098583764 | -1.822967618 | 0.068308296 | 0.211546818 |
| Sec24d | 580.0889352 | 0.191826566 | 0.083437764 | 2.299037704 | 0.021502798 | 0.098803191 |
| Ube3a | 1897.642477 | -0.160274013 | 0.087960477 | -1.822113949 | 0.068437698 | 0.211862874 |
| Prlr | 140.8214869 | 0.191816148 | 0.131579888 | 1.457792303 | 0.1448978 | 0.342806638 |
| Fbxo10 | 1728.955282 | 0.191800337 | 0.098619529 | 1.944851469 | 0.051792857 | 0.177969403 |
| Hspa1a | 204.7952997 | 0.191485969 | 0.114784316 | 1.668224155 | 0.095271238 | 0.261308648 |
| Sall1 | 1305.996014 | -0.217735005 | 0.119577007 | -1.820876869 | 0.068625576 | 0.21227484 |
| Prep | 926.9609521 | -0.125408852 | 0.068881812 | -1.8206381 | 0.068661887 | 0.212344767 |
| Arhgap29 | 1000.002038 | -0.14808598 | 0.081343248 | -1.820507348 | 0.068681778 | 0.212363894 |
| Ephb1 | 1882.668525 | -0.195263892 | 0.107266795 | -1.820357292 | 0.068704612 | 0.21239211 |
| A230065H16Rik | 45.1085192 | -0.292660974 | 0.160838206 | -1.819598602 | 0.068820155 | 0.21246433 |
| Emp1 | 90.47709606 | 0.191461842 | 0.154907238 | 1.235977376 | 0.216466973 | 0.436443452 |
| Msx1 | 95.77028498 | 0.191404743 | 0.154580504 | 1.238220459 | 0.215634328 | 0.435631624 |
| Deaf1 | 1531.56352 | 0.191353811 | 0.112461907 | 1.701498905 | 0.088849343 | 0.250382744 |
| Gm4371 | 10.32010343 | 0.191327953 | 0.157354583 | 1.215903276 | 0.22402177 | 0.44545602 |
| Tcp11l1 | 474.8577324 | -0.17207038 | 0.094546939 | -1.819946601 | 0.068767137 | 0.21246433 |
| Arfgap3 | 751.124775 | 0.191290932 | 0.076338215 | 2.505834488 | 0.012216279 | 0.067647972 |
| Irf3 | 636.0657427 | 0.191265285 | 0.107529334 | 1.77872659 | 0.075284598 | 0.225566144 |
| Uck2 | 421.0909984 | -0.172919845 | 0.095100493 | -1.81828548 | 0.069020511 | 0.212986217 |
| Basp1 | 12091.76112 | -0.159252378 | 0.087591188 | -1.818132406 | 0.069043898 | 0.213015961 |
| Phf15 | 3898.301202 | 0.191199265 | 0.126720172 | 1.508830532 | 0.131342096 | 0.321334982 |
| Fance | 446.1045254 | 0.191122812 | 0.077828104 | 2.45570433 | 0.014060879 | 0.074456344 |
| Mrps12 | 940.9270518 | -0.136092749 | 0.074880723 | -1.817460396 | 0.069146647 | 0.213205602 |
| Fam160b1 | 1619.75783 | 0.191079641 | 0.074746301 | 2.556375869 | 0.01057688 | 0.061239978 |
| Ppp4r1l-ps | 276.4139006 | 0.19099051 | 0.102422813 | 1.864726263 | 0.062219784 | 0.199180439 |
| Lgals9 | 319.9079403 | 0.190961341 | 0.127881849 | 1.493263842 | 0.135368136 | 0.327438548 |
| Cby1 | 1139.546193 | -0.104396726 | 0.057473331 | -1.816437712 | 0.069303255 | 0.213518518 |
| Ryr1 | 957.7133464 | -0.261121914 | 0.143910946 | -1.814468753 | 0.069605589 | 0.214407356 |
| Tnks2 | 6422.241663 | -0.127708987 | 0.0704325 | -1.813211039 | 0.069799278 | 0.214961243 |
| Sdf2l1 | 276.0709188 | -0.243203096 | 0.134137292 | -1.813090845 | 0.069817811 | 0.214975589 |
| Tshz3 | 1065.65911 | 0.190918939 | 0.088141065 | 2.166061189 | 0.030306511 | 0.124422122 |
| Slc9b2 | 109.740784 | 0.190901907 | 0.129317393 | 1.476227612 | 0.139882799 | 0.334334817 |
| Csda | 719.5115841 | 0.190862918 | 0.079640287 | 2.396562412 | 0.016549675 | 0.083172449 |
| Tjp1 | 4623.461295 | -0.085007727 | 0.046906179 | -1.812292753 | 0.069940974 | 0.215183766 |
| Arid1b | 3767.703507 | -0.129924078 | 0.071698182 | -1.81209723 | 0.069971174 | 0.215191222 |
| Ms4a7 | 15.27995935 | -0.294099683 | 0.162295311 | -1.812126795 | 0.069966607 | 0.215191222 |
| Tspan5 | 8568.557634 | -0.134725605 | 0.074359785 | -1.811807339 | 0.070015971 | 0.215286257 |
| Tsc22d3 | 2637.300384 | 0.190729054 | 0.118424203 | 1.610558057 | 0.107276082 | 0.283640889 |
| Ebp | 562.7961153 | -0.136795433 | 0.075514733 | -1.811506533 | 0.070062479 | 0.21534379 |
| Parp8 | 721.2038107 | 0.19070326 | 0.116089452 | 1.642726843 | 0.100439462 | 0.270934267 |
| Lsm1 | 466.1104349 | -0.13064203 | 0.072147882 | -1.810753499 | 0.070179017 | 0.215616438 |
| Il17d | 208.1002727 | -0.184210522 | 0.101779137 | -1.809904536 | 0.070310593 | 0.215977859 |
| Smad9 | 704.6195877 | 0.190695394 | 0.096059326 | 1.985183547 | 0.047124054 | 0.167424924 |
| Rp2h | 433.8713756 | -0.157173745 | 0.086864952 | -1.809403463 | 0.070388345 | 0.216131 |
| Intu | 567.0976319 | 0.190679101 | 0.094195267 | 2.02429599 | 0.042939706 | 0.157469303 |
| Car12 | 939.8742721 | -0.203626327 | 0.11255609 | -1.809109814 | 0.070433945 | 0.216185329 |
| Rnf43 | 128.3420004 | 0.190669077 | 0.120815239 | 1.57818731 | 0.114522585 | 0.296434962 |
| Trim65 | 121.9407076 | 0.190633447 | 0.115157595 | 1.655413583 | 0.097840633 | 0.266049876 |
| Fam120a | 5814.374724 | -0.089348145 | 0.04941354 | -1.808171313 | 0.070579842 | 0.216504468 |
| Fam210a | 1709.962413 | -0.109942583 | 0.060816158 | -1.807785718 | 0.070639857 | 0.216645675 |
| Cst6 | 148.1844828 | -0.208011676 | 0.115079417 | -1.807548921 | 0.070676734 | 0.216715876 |
| Trmt61a | 696.1599846 | -0.212348806 | 0.117486298 | -1.807434644 | 0.070694536 | 0.216727572 |
| BC003965 | 943.7624337 | -0.158273294 | 0.087638484 | -1.80597937 | 0.070921562 | 0.216994211 |
| Gm11992 | 44.02785316 | 0.190593899 | 0.155185352 | 1.228169388 | 0.219383388 | 0.440187284 |
| Ipw | 377.3501248 | 0.190515584 | 0.167796409 | 1.135397269 | 0.256208903 | 0.482569738 |
| Gpr30 | 135.3860865 | -0.208085434 | 0.115217685 | -1.806019916 | 0.070915229 | 0.216994211 |
| Hypk | 989.0446584 | -0.107361136 | 0.059439825 | -1.806215536 | 0.070884679 | 0.216994211 |
| Insig1 | 2788.342048 | -0.206925112 | 0.114533735 | -1.806673927 | 0.070813135 | 0.216994211 |
| Gmnc | 17.54119322 | 0.190514796 | 0.167685908 | 1.136140762 | 0.255897658 | 0.482345293 |
| Ppp1r9a | 16974.30638 | -0.156462478 | 0.086618937 | -1.806331081 | 0.07086664 | 0.216994211 |
| Csrnp1 | 484.7719378 | 0.190496116 | 0.168120817 | 1.133090592 | 0.257176212 | 0.483754792 |
| Slc7a4 | 2729.214994 | -0.149560265 | 0.082812141 | -1.806018573 | 0.070915438 | 0.216994211 |
| Cyba | 175.2882944 | 0.190403832 | 0.118089047 | 1.612375038 | 0.106880353 | 0.282884215 |
| C630043F03Rik | 218.6019754 | -0.169619995 | 0.093961455 | -1.805208259 | 0.071042099 | 0.217277198 |
| Pvr | 274.7015454 | 0.190328726 | 0.100496364 | 1.89388669 | 0.058240052 | 0.191342527 |
| St3gal4 | 811.2725644 | -0.213668586 | 0.11845671 | -1.803769378 | 0.071267468 | 0.217794507 |
| Plekhm1 | 1364.389354 | 0.190249282 | 0.074209956 | 2.563662495 | 0.010357421 | 0.060365743 |
| Txnrd3 | 205.7562633 | -0.201720817 | 0.111829378 | -1.803826697 | 0.071258479 | 0.217794507 |
| Cyp19a1 | 9.177297304 | 0.190243437 | 0.14500349 | 1.311992122 | 0.189522793 | 0.40286269 |
| Tbc1d8 | 1332.128332 | -0.136452016 | 0.0756564 | -1.803575324 | 0.071297907 | 0.217801612 |
| Snord100 | 12.15908257 | 0.190221045 | 0.161273774 | 1.179491498 | 0.23820252 | 0.462671816 |
| Dhcr7 | 1275.772649 | -0.189293831 | 0.105008725 | -1.802648605 | 0.071443419 | 0.218160096 |
| Pcdha5 | 157.4569188 | -0.208283486 | 0.115552977 | -1.802493471 | 0.071467801 | 0.218191549 |
| Nanos2 | 23.6705391 | 0.189955656 | 0.167065623 | 1.137012224 | 0.255533176 | 0.48192595 |
| Cdh19 | 421.8203825 | 0.189888339 | 0.125958039 | 1.507552356 | 0.131669135 | 0.321677952 |
| Cept1 | 1186.752667 | -0.135487793 | 0.075193828 | -1.801847255 | 0.071569441 | 0.218341833 |
| Eng | 1088.065793 | -0.152174425 | 0.084451817 | -1.801908251 | 0.071559842 | 0.218341833 |
| Mettl4 | 287.6794777 | -0.202374631 | 0.112307719 | -1.801965464 | 0.071550839 | 0.218341833 |
| Mitf | 354.9048011 | -0.200906085 | 0.111514429 | -1.801615151 | 0.071605976 | 0.218355243 |
| Rab3b | 2491.016844 | -0.151044999 | 0.083870021 | -1.800941484 | 0.071712104 | 0.218595825 |
| Ank1 | 3077.521641 | 0.189734374 | 0.081627019 | 2.324406515 | 0.020103717 | 0.094590643 |
| Acaca | 4652.113352 | -0.144226353 | 0.080124318 | -1.800032204 | 0.071855553 | 0.218968649 |
| Sesn1 | 2575.33878 | 0.189647852 | 0.122065477 | 1.553656749 | 0.120266315 | 0.30507634 |
| Dag1 | 5137.596882 | -0.108576958 | 0.060324173 | -1.799891357 | 0.071877795 | 0.218968649 |
| Actn3 | 39.49687424 | 0.189579087 | 0.155419359 | 1.21979069 | 0.222544232 | 0.443624247 |
| Trp53inp2 | 7270.061936 | -0.116579594 | 0.064796686 | -1.799159803 | 0.071993406 | 0.219234637 |
| Irx6 | 10.11734141 | -0.294792748 | 0.163866147 | -1.798985042 | 0.072021047 | 0.219275712 |
| Lif | 24.43363156 | 0.189484926 | 0.167675669 | 1.130068106 | 0.258447527 | 0.484851192 |
| Efna2 | 223.3869736 | -0.176778395 | 0.098300295 | -1.798350608 | 0.072121465 | 0.219495182 |
| Dmgdh | 12.8007829 | 0.189483914 | 0.165717444 | 1.143415618 | 0.252866071 | 0.478727628 |
| 2810433D01Rik | 9.490779877 | -0.284828909 | 0.158459124 | -1.797491374 | 0.072257646 | 0.219823281 |
| Sprn | 7659.373044 | -0.164744405 | 0.091675066 | -1.797047022 | 0.072328155 | 0.219994589 |
| Rbm46 | 8.99877052 | 0.189380659 | 0.158635131 | 1.193812854 | 0.232551183 | 0.455921552 |
| Rhog | 931.1972982 | 0.189281544 | 0.107896615 | 1.75428621 | 0.079381478 | 0.233340279 |
| A330048O09Rik | 35.85012277 | -0.286253595 | 0.159379273 | -1.796052833 | 0.072486115 | 0.220302053 |
| Mtf2 | 1270.930077 | 0.189255971 | 0.086854508 | 2.178999975 | 0.02933167 | 0.12164478 |
| Hsf1 | 1426.93034 | 0.189229035 | 0.093944272 | 2.014269011 | 0.043981298 | 0.160220654 |
| Rhoa | 5673.035807 | -0.066981468 | 0.037296701 | -1.795908616 | 0.072509052 | 0.220313701 |
| Dusp10 | 244.6904732 | 0.189185488 | 0.126870012 | 1.491175768 | 0.135915354 | 0.328305122 |
| Tnfaip8l2 | 72.56482728 | 0.189134507 | 0.144912852 | 1.305160334 | 0.191838258 | 0.405907402 |
| A830052D11Rik | 38.60046104 | 0.189050091 | 0.165178702 | 1.144518567 | 0.252408642 | 0.478270699 |
| Cdk5rap1 | 346.3580878 | -0.233335046 | 0.129978187 | -1.795186184 | 0.072624041 | 0.220461773 |
| Appl2 | 3152.496046 | 0.189032993 | 0.081034841 | 2.332737266 | 0.019661939 | 0.093077325 |
| Zfp653 | 597.3371463 | 0.188896803 | 0.075637417 | 2.497398912 | 0.012510813 | 0.068626502 |
| Ddrgk1 | 1813.813338 | 0.188886509 | 0.080682467 | 2.34110972 | 0.019226516 | 0.091773148 |
| Has1 | 33.57585029 | 0.188839618 | 0.163188562 | 1.157186604 | 0.24719613 | 0.472120974 |
| Zfp90 | 491.7518715 | 0.188836482 | 0.084483367 | 2.235191228 | 0.025404797 | 0.110305951 |
| Slc28a1 | 18.79909953 | 0.188668503 | 0.168089414 | 1.122429421 | 0.261679901 | 0.488784616 |
| D330045A20Rik | 12.48394589 | -0.292375158 | 0.163074466 | -1.79289355 | 0.072989947 | 0.221269074 |
| Metrn | 1159.415383 | -0.243680497 | 0.135934137 | -1.792636518 | 0.073031063 | 0.22135041 |
| Tmbim1 | 1423.449509 | 0.188642871 | 0.084394358 | 2.23525452 | 0.025400644 | 0.110305951 |
| Dzip1l | 891.7527709 | 0.188417453 | 0.10524079 | 1.790346234 | 0.073398268 | 0.221939958 |
| Ern1 | 324.762199 | 0.18835061 | 0.101061803 | 1.863717095 | 0.062361443 | 0.199469076 |
| Fnip1 | 2124.379587 | 0.188303115 | 0.099941483 | 1.884133679 | 0.059546896 | 0.193827647 |
| Pcdha9 | 131.5197065 | -0.211547008 | 0.118110339 | -1.791096443 | 0.07327782 | 0.221900619 |
| Zfp623 | 456.5720743 | 0.188231346 | 0.085940735 | 2.190245933 | 0.028506406 | 0.119663573 |
| Zfp410 | 842.1599106 | 0.188130226 | 0.096803601 | 1.94342178 | 0.051965218 | 0.178288106 |
| Nat9 | 454.2763008 | 0.188114915 | 0.098686737 | 1.906182335 | 0.056626553 | 0.188063674 |
| Gnb2 | 9511.257982 | -0.135920683 | 0.075935172 | -1.789956872 | 0.073460845 | 0.221939958 |
| Pms2 | 631.075283 | 0.188045762 | 0.072238828 | 2.603112034 | 0.009238176 | 0.055586285 |
| Optn | 714.5885709 | 0.188042198 | 0.087467358 | 2.149855705 | 0.03156663 | 0.127834127 |
| Mlxip | 1561.133755 | 0.188017591 | 0.075681143 | 2.484338697 | 0.012979229 | 0.070380366 |
| Cirbp | 1966.46323 | 0.187980087 | 0.10998787 | 1.70909835 | 0.087432733 | 0.247630198 |
| Hspd1 | 6431.176792 | -0.111463777 | 0.062242023 | -1.79081226 | 0.073323427 | 0.221939958 |
| Msrb3 | 365.3495788 | 0.187945766 | 0.087407203 | 2.15023201 | 0.031536867 | 0.127809097 |
| Adap2 | 198.4606307 | 0.187790893 | 0.10313186 | 1.820881461 | 0.068624878 | 0.21227484 |
| Lyz2 | 362.5027814 | 0.187782031 | 0.156472787 | 1.200093858 | 0.230102891 | 0.453013238 |
| Dancr | 88.79515554 | 0.187737546 | 0.130819108 | 1.435092688 | 0.151260687 | 0.352254563 |
| Myadml2 | 245.120023 | -0.240829747 | 0.134581435 | -1.78947227 | 0.073538789 | 0.222044989 |
| Wls | 1725.69847 | -0.189765128 | 0.106048271 | -1.789422183 | 0.073546849 | 0.222044989 |
| Tsc22d4 | 2292.721168 | 0.187720992 | 0.074068948 | 2.53440877 | 0.011263725 | 0.063786608 |
| Tulp3 | 790.6567406 | 0.187642751 | 0.084322636 | 2.225295122 | 0.026061433 | 0.112277067 |
| Cyp20a1 | 346.5384567 | -0.151260007 | 0.084617228 | -1.787579328 | 0.073843903 | 0.222724827 |
| Errfi1 | 1257.646495 | 0.187593628 | 0.130609613 | 1.43629265 | 0.150919085 | 0.351663492 |
| Zfp229 | 96.24828073 | -0.245242905 | 0.137192439 | -1.787583244 | 0.073843271 | 0.222724827 |
| Pet100 | 501.1683226 | -0.191611301 | 0.107216686 | -1.78714068 | 0.073914754 | 0.222895134 |
| Glra1 | 23.17712663 | -0.300248562 | 0.168068071 | -1.786469975 | 0.074023194 | 0.223178707 |
| 2010109I03Rik | 8.233823068 | 0.187583667 | 0.149105698 | 1.258058337 | 0.20837066 | 0.426414656 |
| Casc4 | 3805.791949 | -0.102064369 | 0.057167964 | -1.785342021 | 0.074205857 | 0.2235989 |
| Srebf1 | 2762.316142 | 0.187581243 | 0.10563187 | 1.775801588 | 0.075765626 | 0.226274597 |
| Spg7 | 1905.747602 | 0.187363884 | 0.079373664 | 2.360529601 | 0.018248862 | 0.08874899 |
| H60b | 24.99285833 | -0.288623063 | 0.161694794 | -1.784986738 | 0.074263468 | 0.223685492 |
| Anxa2 | 457.2321642 | -0.163038363 | 0.091351774 | -1.784731221 | 0.074304924 | 0.22376686 |
| Upb1 | 39.7359977 | -0.297249538 | 0.166593858 | -1.784276693 | 0.074378716 | 0.223945553 |
| Smarcc1 | 3165.507702 | -0.109173765 | 0.061206659 | -1.783690967 | 0.074473895 | 0.224188562 |
| Ache | 2348.093949 | -0.156031285 | 0.087484968 | -1.783521075 | 0.07450152 | 0.224228158 |
| Cacna1g | 3791.003629 | 0.18733001 | 0.092916827 | 2.016104254 | 0.043789075 | 0.159677085 |
| Prickle3 | 59.98461639 | 0.187323594 | 0.146349025 | 1.279978422 | 0.200552725 | 0.417375946 |
| Traf3 | 2451.843187 | -0.161358378 | 0.090486781 | -1.783225978 | 0.074549525 | 0.224267761 |
| Tsen2 | 519.0931346 | -0.139650479 | 0.078339527 | -1.782631115 | 0.074646371 | 0.224489689 |
| Prkaa2 | 2169.543426 | -0.138288144 | 0.077582455 | -1.782466722 | 0.074673153 | 0.224526652 |
| Ap5b1 | 148.2980076 | -0.225022094 | 0.126251053 | -1.782338347 | 0.074694073 | 0.224545977 |
| Dll1 | 237.2300396 | -0.226735699 | 0.127262134 | -1.781643071 | 0.074807456 | 0.224843205 |
| Pif1 | 13.9718402 | -0.289827713 | 0.16268904 | -1.781482714 | 0.074833626 | 0.22487824 |
| Fmnl1 | 4952.338391 | 0.187283911 | 0.117844466 | 1.589246556 | 0.112004739 | 0.29211375 |
| Cadm4 | 8946.957143 | -0.135691322 | 0.076177669 | -1.781248019 | 0.074871942 | 0.224884427 |
| Tomm40 | 2135.34658 | -0.107544512 | 0.060380458 | -1.781114531 | 0.074893742 | 0.224884427 |
| Snora31 | 17.82126674 | 0.187180454 | 0.167394753 | 1.118197856 | 0.2634825 | 0.490790797 |
| Cenpo | 128.4151259 | 0.187101823 | 0.139077903 | 1.3453023 | 0.178527631 | 0.388007731 |
| F830016B08Rik | 10.62410873 | 0.18692409 | 0.119553507 | 1.563518254 | 0.117930746 | 0.301421803 |
| Prpf3 | 831.2739396 | 0.186887404 | 0.082753599 | 2.258359832 | 0.023923235 | 0.106273139 |
| Cmtm3 | 299.2609895 | 0.186730654 | 0.100455877 | 1.858832544 | 0.063050878 | 0.200845046 |
| Tnmd | 18.02933645 | 0.186647311 | 0.16667502 | 1.119827745 | 0.262787173 | 0.489907889 |
| Tpd52l1 | 646.8154326 | -0.156659976 | 0.08801426 | -1.77993857 | 0.075086015 | 0.225199905 |

| | | | | | | |
|---|---|---|---|---|---|---|
| Grip2 | 1598.212922 | -0.052621109 | 0.066866975 | -0.786952132 | 0.431309888 | 0.812181606 |
| Eml4 | 1599.566124 | -0.052633387 | 0.057775784 | -0.910993911 | 0.362298582 | 0.776449324 |
| Samhd1 | 829.7792567 | -0.052647584 | 0.076972621 | -0.68397806 | 0.49398902 | 0.843404385 |
| Htatip2 | 366.6569648 | -0.052665359 | 0.079704208 | -0.660760085 | 0.508766184 | 0.850050333 |
| Mirlet7f-1 | 3.533763314 | -0.052680902 | 0.042601548 | -1.236595961 | 0.21623712 | NA |
| Ccdc17 | 99.41722785 | -0.052751138 | 0.092755898 | -0.568709259 | 0.569553464 | 0.877633217 |
| AF357425 | 6.577827015 | -0.052753393 | 0.055609686 | -0.948636765 | 0.342805386 | NA |
| Sox10 | 1784.436896 | -0.05276973 | 0.081736193 | -0.645610314 | 0.518531756 | 0.853297835 |
| Cav1 | 518.6253231 | -0.052772126 | 0.079457929 | -0.664151793 | 0.506593164 | 0.848943697 |
| Znhit6 | 570.2062352 | -0.052812831 | 0.064382779 | -0.820294365 | 0.412048318 | 0.802795661 |
| Hspb7 | 13.96690146 | -0.052873545 | 0.072778392 | -0.726500605 | 0.467531937 | NA |
| Bmf | 164.2064518 | -0.052902085 | 0.094362605 | -0.560625527 | 0.575052846 | 0.88049095 |
| Zfhx2 | 2695.866948 | -0.052930806 | 0.082273159 | -0.643354485 | 0.519994109 | 0.853297835 |
| Gm20337 | 11.50094285 | -0.05295322 | 0.068025485 | -0.778432079 | 0.436314334 | NA |
| Tra2a | 1954.24342 | -0.052960989 | 0.074148696 | -0.714253811 | 0.475070248 | 0.834699977 |
| Dock9 | 6538.722432 | -0.052974986 | 0.059865237 | -0.88490396 | 0.376208432 | 0.783947144 |
| Cpn1 | 14.71494729 | -0.052976477 | 0.074292119 | -0.713083396 | 0.475794153 | NA |
| Prkd2 | 212.221959 | -0.052995191 | 0.094629794 | -0.560026492 | 0.575461368 | 0.880811222 |
| Acss1 | 1243.746607 | -0.05299673 | 0.072571092 | -0.730273288 | 0.465223153 | 0.831682622 |
| Pusl1 | 670.1031865 | -0.053001132 | 0.071300376 | -0.743349967 | 0.457269827 | 0.827391818 |
| 1110037F02Rik | 2146.560702 | -0.053016931 | 0.063517258 | -0.834685438 | 0.403894843 | 0.800260243 |
| Jmjd1c | 3512.48661 | -0.053034833 | 0.069802008 | -0.759789508 | 0.447380416 | 0.822458097 |
| Rhoc | 460.8635305 | -0.05305834 | 0.079475076 | -0.6676098 | 0.504382701 | 0.847807553 |
| Cdca5 | 3.983261408 | -0.053065982 | 0.046755425 | -1.134969511 | 0.256388093 | NA |
| 4933422H20Rik | 1.80367557 | -0.053089341 | 0.031305239 | -1.695861227 | 0.089912164 | NA |
| Serpinf1 | 346.2986498 | -0.053125676 | 0.093028668 | -0.571067795 | 0.567953688 | 0.877016613 |
| Arpp21 | 24241.97259 | -0.053142506 | 0.078456652 | -0.677348627 | 0.498184785 | 0.845625663 |
| Bod1l | 3380.943892 | -0.053150912 | 0.076586197 | -0.694001183 | 0.487681467 | 0.839801621 |
| Lrrc2 | 73.68806584 | -0.053153549 | 0.094536903 | -0.562251861 | 0.573944432 | 0.879633264 |
| Ucp3 | 153.4182366 | -0.053182634 | 0.095065051 | -0.559434127 | 0.575865476 | 0.880811222 |
| Lysmd4 | 1104.958597 | -0.053199625 | 0.062754575 | -0.847740971 | 0.396582244 | 0.796916625 |
| Sema4c | 699.4879073 | -0.053204187 | 0.070574411 | -0.753873622 | 0.450925111 | 0.823992799 |
| Zfp787 | 508.9485262 | -0.053209113 | 0.078798501 | -0.675255398 | 0.499513516 | 0.846241311 |
| Rbbp8 | 189.3168293 | -0.053214818 | 0.084868555 | -0.627026328 | 0.530641977 | 0.858116381 |
| Phf1 | 1917.197075 | -0.053221214 | 0.065071636 | -0.817886523 | 0.413421987 | 0.804056218 |
| Lat | 17.16880597 | -0.053226555 | 0.075647475 | -0.703613113 | 0.481673743 | NA |
| Gpr156 | 50.06294547 | -0.05322972 | 0.092154597 | -0.577613289 | 0.563525233 | 0.875108781 |
| Zfp541 | 34.0941933 | -0.053244235 | 0.084612366 | -0.629272501 | 0.529170671 | 0.857868166 |
| Wdr95 | 17.5805258 | -0.053317447 | 0.078648783 | -0.677918272 | 0.497823514 | NA |
| Zglp1 | 33.7786923 | -0.053415501 | 0.075754132 | -0.705116673 | 0.480737631 | 0.836914993 |
| Nfia | 1168.07113 | -0.053449235 | 0.068889731 | -0.775866505 | 0.437827812 | 0.816984132 |
| Riok3 | 2489.775814 | -0.053455971 | 0.045795728 | -1.167269824 | 0.243101421 | 0.692772033 |
| Wfs1 | 7041.56952 | -0.053458605 | 0.095080812 | -0.562243885 | 0.573949865 | 0.879633264 |
| Il1rn | 3.319060698 | -0.053512102 | 0.033006451 | -1.621261921 | 0.104961478 | NA |
| Tor2a | 794.266343 | -0.053522722 | 0.07423944 | -0.720947282 | 0.470941951 | 0.833245343 |
| S100a9 | 22.39428828 | -0.053532631 | 0.05637212 | -0.949629558 | 0.342300514 | 0.763762832 |
| Unc45b | 59.65655461 | -0.053537567 | 0.094393209 | -0.567176048 | 0.570594579 | 0.878246677 |
| Cldn19 | 18.47388086 | -0.053581869 | 0.075227143 | -0.71226776 | 0.476298984 | NA |
| Olfr558 | 12.55436411 | -0.053614462 | 0.061506826 | -0.871683109 | 0.383381277 | NA |
| Dtna | 5584.762442 | -0.053640726 | 0.052593003 | -1.01992135 | 0.307765763 | 0.740098991 |
| Phkg2 | 1959.0334 | -0.053652198 | 0.055404568 | -0.968371381 | 0.332858927 | 0.757641999 |
| A930007I19Rik | 14.63490669 | -0.053652364 | 0.071398818 | -0.751446109 | 0.452384222 | NA |
| Ccdc9 | 1583.548336 | -0.053663167 | 0.068478173 | -0.783653606 | 0.433243392 | 0.813783127 |
| Manba | 553.2931419 | -0.053681121 | 0.073167303 | -0.733676361 | 0.463146009 | 0.831088193 |
| Gm10336 | 480.9820674 | -0.053700028 | 0.075122243 | -0.714835267 | 0.47471084 | 0.834491965 |
| Zfp1 | 729.4559288 | -0.053746491 | 0.060247006 | -0.892102274 | 0.372338116 | 0.781446026 |
| Ccr9 | 11.70624472 | -0.053762435 | 0.069153477 | -0.777436472 | 0.436901302 | NA |
| Mir25 | 3.192235021 | -0.05376271 | 0.039267345 | -1.369145545 | 0.170953784 | NA |
| Fam229b | 295.43065 | -0.053777361 | 0.077052268 | -0.697933525 | 0.485218767 | 0.838982004 |
| Ankrd32 | 629.8214639 | -0.05377792 | 0.068871626 | -0.780842908 | 0.434894894 | 0.815326688 |
| Bmp2k | 759.1263795 | -0.053780234 | 0.066379783 | -0.810189972 | 0.417831 | 0.80621897 |
| Camk1d | 9941.852593 | -0.053814887 | 0.069649058 | -0.772657786 | 0.439724936 | 0.817695544 |
| Atg2a | 2667.081051 | -0.053824658 | 0.06694491 | -0.804014202 | 0.421388773 | 0.808555694 |
| Cdc42ep1 | 982.8570644 | -0.053946416 | 0.085012098 | -0.634573396 | 0.525706684 | 0.855892499 |
| Shprh | 1754.99598 | -0.053976824 | 0.073590536 | -0.733475077 | 0.463268723 | 0.831205716 |
| Prom1 | 681.4578636 | -0.05398156 | 0.065409525 | -0.825285918 | 0.409209284 | 0.801055189 |
| Cdkn2b | 22.59345146 | -0.053993666 | 0.080003435 | -0.674891855 | 0.499744476 | 0.846241311 |
| Fhl1 | 3777.8281 | -0.054002202 | 0.053606754 | -1.007376834 | 0.313753719 | 0.745844951 |
| Fbxw7 | 8953.072079 | -0.054035005 | 0.060121405 | -0.898764842 | 0.368777931 | 0.779573594 |
| BC003331 | 2164.400692 | -0.05410879 | 0.060962506 | -0.887574897 | 0.374769462 | 0.783182445 |
| Tnfrsf23 | 58.70531957 | -0.054127954 | 0.094469483 | -0.572967615 | 0.566666621 | 0.875959314 |
| B3gnt9 | 145.8148949 | -0.054144839 | 0.092091339 | -0.58794714 | 0.556567774 | 0.871666144 |
| Actn2 | 1065.725798 | -0.054161166 | 0.084225292 | -0.643051094 | 0.520190947 | 0.853297835 |
| Hcst | 12.81806759 | -0.054167096 | 0.070060651 | -0.773145776 | 0.439436113 | NA |
| Heatr1 | 1226.754739 | -0.054167136 | 0.06504793 | -0.832726515 | 0.404998987 | 0.8005027 |
| Gm11602 | 96.77181737 | -0.054196251 | 0.094647359 | -0.572612392 | 0.566907167 | 0.875959314 |
| Ncln | 2329.108165 | -0.054213563 | 0.069453612 | -0.780572264 | 0.43505411 | 0.815326688 |
| Prrc1 | 849.3640376 | -0.054229385 | 0.072743661 | -0.745486059 | 0.455977936 | 0.826649955 |
| Tubgcp4 | 960.196399 | -0.054249276 | 0.066097436 | -0.820747052 | 0.411790363 | 0.802795661 |
| Ilvbl | 985.300961 | -0.054257551 | 0.073345702 | -0.73975093 | 0.459451139 | 0.828552109 |
| Tktl1 | 3.448760418 | -0.054267798 | 0.042651351 | -1.272358244 | 0.203245863 | NA |
| Arl14ep | 1176.477717 | -0.054267816 | 0.061099011 | -0.888194662 | 0.37443605 | 0.78293624 |
| Xlr4b | 6.538480856 | -0.054269727 | 0.04925657 | -1.101776411 | 0.270558888 | NA |
| D630033O11Rik | 3.62288829 | -0.054273443 | 0.040383892 | -1.343937882 | 0.178968475 | NA |
| Snord89 | 7.849671122 | -0.054296947 | 0.055085793 | -0.985679686 | 0.324290321 | NA |
| Ift122 | 1712.386468 | -0.05432539 | 0.047520828 | -1.143191151 | 0.252959236 | 0.700378199 |
| Vwa8 | 2461.60373 | -0.054351362 | 0.066534559 | -0.816889188 | 0.413991757 | 0.804285711 |
| Samd1 | 882.4757971 | -0.054387818 | 0.063479044 | -0.856783823 | 0.391564368 | 0.794184423 |
| Cnppd1 | 1292.686382 | -0.054389867 | 0.057254268 | -0.949970515 | 0.342127234 | 0.763752913 |
| Gm16287 | 29.8090622 | -0.054397296 | 0.076736214 | -0.708886886 | 0.478394674 | 0.835936459 |
| Slc34a3 | 10.13532032 | -0.054412509 | 0.063280416 | -0.859863325 | 0.389864387 | NA |
| Pnp | 503.1314493 | -0.05443728 | 0.068310416 | -0.79691039 | 0.425503076 | 0.809519334 |
| Aplnr | 6.431248792 | -0.054446593 | 0.053446832 | -1.018705703 | 0.308342704 | NA |
| Mllt10 | 1413.895637 | -0.054454126 | 0.062233176 | -0.875001565 | 0.381573054 | 0.786663048 |
| 4930519G04Rik | 8.420917434 | -0.054460161 | 0.060531624 | -0.899697666 | 0.368281166 | NA |
| Ppfia3 | 9435.835716 | -0.054494506 | 0.056407591 | -0.966084617 | 0.334001841 | 0.758212399 |
| Sox3 | 28.58704366 | -0.054510408 | 0.08343293 | -0.653344041 | 0.513534506 | 0.853195313 |
| Tmem45a | 21.2810456 | -0.054518733 | 0.077677575 | -0.701859354 | 0.482766879 | NA |
| Mir26a-1 | 2.532465486 | -0.054523755 | 0.036049557 | -1.512466716 | 0.130415174 | NA |
| Ahr | 418.2719965 | -0.054553298 | 0.080215373 | -0.680085323 | 0.496450437 | 0.844826506 |
| Arg2 | 96.20349935 | -0.054563131 | 0.094763612 | -0.57578146 | 0.564762907 | 0.875364912 |
| 9430060I03Rik | 7.176976948 | -0.054574743 | 0.055482765 | -0.983634158 | 0.325295424 | NA |
| 1700019N19Rik | 4.53142407 | -0.054603058 | 0.042211829 | -1.293548759 | 0.195821331 | NA |
| Gpr17 | 2329.894966 | -0.05461809 | 0.077417465 | -0.70550088 | 0.480498585 | 0.836914993 |
| Snora3 | 11.79861909 | -0.054823051 | 0.068557877 | -0.79966086 | 0.423907316 | NA |
| Gm12185 | 7.932670599 | -0.05483816 | 0.048482293 | -1.131096649 | 0.258014415 | NA |
| Tsr3 | 641.8944857 | -0.054838936 | 0.07330604 | -0.748082089 | 0.454410645 | 0.826168452 |
| Mybl1 | 152.2716196 | -0.054888895 | 0.094311347 | -0.581996722 | 0.560568889 | 0.873713575 |
| Msh5 | 11.27344889 | -0.054917245 | 0.068836097 | -0.797797191 | 0.424988192 | NA |
| Nxf3 | 5.389266607 | -0.054929174 | 0.045861493 | -1.197718824 | 0.231026498 | NA |
| Plk1 | 39.92907796 | -0.054940181 | 0.090553215 | -0.606717067 | 0.544038694 | 0.86604998 |
| Kif18b | 5.320192453 | -0.054956458 | 0.044343546 | -1.239333854 | 0.215221886 | NA |
| Ggn | 36.34540413 | -0.054962432 | 0.084134179 | -0.653271153 | 0.513581486 | 0.853195313 |
| Fhit | 100.8615289 | -0.054970796 | 0.092649127 | -0.593322332 | 0.552965456 | 0.870600462 |
| Trmt10a | 245.7641711 | -0.05499004 | 0.081829519 | -0.672007366 | 0.501579007 | 0.846611278 |
| Ikbkb | 1775.762648 | -0.05499938 | 0.060792163 | -0.904711674 | 0.365618155 | 0.777735733 |
| Gm14137 | 2.972374042 | -0.055010198 | 0.037673503 | -1.460182705 | 0.144239867 | NA |
| Cdc26 | 630.5878143 | -0.055038152 | 0.064309341 | -0.855834486 | 0.392089337 | 0.794184423 |
| Lif | 26.42078716 | -0.055068131 | 0.08406664 | -0.655053313 | 0.51243343 | 0.852934798 |
| Slc22a7 | 7.143797858 | -0.055069994 | 0.047150975 | -1.167950278 | 0.242826822 | NA |
| Pcdhb15 | 129.2942163 | -0.055073204 | 0.092731851 | -0.593897389 | 0.552580746 | 0.870265825 |
| Khdc3 | 8.979998024 | -0.055085196 | 0.055162789 | -0.998593387 | 0.317991705 | NA |
| 2210408F21Rik | 369.5793372 | -0.055088522 | 0.079241628 | -0.695196743 | 0.486932013 | 0.839801621 |
| Utp23 | 409.4833124 | -0.055150232 | 0.079529194 | -0.693458955 | 0.488021576 | 0.839945271 |
| Ier5l | 356.1154228 | -0.055216671 | 0.092752688 | -0.595310735 | 0.551635781 | 0.869755927 |
| 4933405D12Rik | 1.900325692 | -0.055221468 | 0.031725147 | -1.740621364 | 0.08174997 | NA |
| Pcdhga12 | 652.5348964 | -0.055225898 | 0.073776676 | -0.748554977 | 0.454125477 | 0.826023877 |
| Nup210 | 1505.744048 | -0.055229821 | 0.063029518 | -0.876253269 | 0.380892364 | 0.786576978 |
| Tmod3 | 934.4154551 | -0.055240174 | 0.06769498 | -0.816015822 | 0.414491086 | 0.804797733 |
| Pkp2 | 253.3057477 | -0.055245386 | 0.094287866 | -0.585922537 | 0.557927571 | 0.872673743 |
| Morc3 | 961.643923 | -0.055247957 | 0.066138553 | -0.835336649 | 0.403528188 | 0.800224169 |
| Mocs2 | 1566.440986 | -0.055268338 | 0.051897099 | -1.064960075 | 0.286893997 | 0.722256881 |
| Ap3m1 | 2452.288812 | -0.05527953 | 0.054258803 | -1.018812195 | 0.308292135 | 0.74062833 |
| Fam136a | 784.2721074 | -0.05528295 | 0.071143229 | -0.777065508 | 0.437120122 | 0.816564673 |
| Jph1 | 883.4550962 | -0.055285789 | 0.067563717 | -0.81827632 | 0.413199424 | 0.803854924 |
| Srcin1 | 14489.33571 | -0.055292495 | 0.052612135 | -1.050945667 | 0.293283545 | 0.728786362 |
| Gpr35 | 13.90372398 | -0.055297735 | 0.073377568 | -0.75360545 | 0.451086171 | NA |
| Adhfe1 | 767.5047039 | -0.055305189 | 0.068799494 | -0.803860392 | 0.421477607 | 0.808555694 |
| Trim43a | 24.27252925 | -0.055329595 | 0.078177301 | -0.707745016 | 0.479103617 | 0.83617195 |
| Jakmip3 | 1593.000602 | -0.055333806 | 0.065697971 | -0.842245278 | 0.399650675 | 0.798421909 |
| Tceanc | 80.06431823 | -0.055334019 | 0.095039337 | -0.582222275 | 0.560416971 | 0.873713575 |
| Serpina3f | 3.959729982 | -0.055338406 | 0.026425496 | -2.094129373 | NA | NA |
| Sugp2 | 4230.234792 | -0.055339466 | 0.058838076 | -0.940538328 | 0.346941499 | 0.76579125 |
| Ddx4 | 110.3017683 | -0.055380464 | 0.095081892 | -0.582450167 | 0.560263499 | 0.873713575 |
| Gm7854 | 11.39097677 | -0.055429871 | 0.066594522 | -0.832348808 | 0.405212088 | NA |
| Ska2 | 291.5998259 | -0.055443842 | 0.077744445 | -0.713155033 | 0.475749828 | 0.835195112 |
| Zfx | 846.4792592 | -0.055456141 | 0.065231359 | -0.850145414 | 0.395244245 | 0.795857764 |
| Itgae | 9.535240201 | -0.055480044 | 0.064294906 | -0.862899541 | 0.388192703 | NA |
| Dctd | 315.5842074 | -0.055490995 | 0.077390123 | -0.717029412 | 0.473355948 | 0.833983942 |
| Smarcad1 | 1069.59225 | -0.055500401 | 0.074429159 | -0.745680893 | 0.455860204 | 0.826603827 |
| Sec14l5 | 489.4635947 | -0.055520967 | 0.093240739 | -0.595458248 | 0.5515372 | 0.869755927 |
| Ebf4 | 369.5028124 | -0.055553456 | 0.089604163 | -0.619987442 | 0.535266054 | 0.860477502 |
| Thap6 | 135.0167403 | -0.055578922 | 0.093070288 | -0.59717149 | 0.550392891 | 0.869565155 |
| Zfp672 | 1188.307363 | -0.055589988 | 0.052157474 | -1.065810583 | 0.286509276 | 0.722148665 |
| Xrra1 | 61.52171339 | -0.05561222 | 0.094403058 | -0.589093422 | 0.555798606 | 0.87140103 |
| Tal2 | 4.767440794 | -0.055629951 | 0.047012928 | -1.183290496 | 0.236694038 | NA |
| Tmem110 | 1019.793154 | -0.055630925 | 0.061100266 | -0.910485807 | 0.362566363 | 0.776449324 |
| Vldlr | 3836.995625 | -0.055643775 | 0.052588337 | -1.058101053 | 0.290009375 | 0.725277354 |
| 2610028H24Rik | 2.37094498 | -0.055656226 | 0.031483044 | -1.76781593 | 0.077091681 | NA |
| Smcr7 | 616.8742522 | -0.05566423 | 0.077632282 | -0.71702427 | 0.473359121 | 0.833983942 |
| Snora75 | 7.830322142 | -0.055681168 | 0.057303294 | -0.971692288 | 0.331203654 | NA |
| Tnks1bp1 | 3876.027794 | -0.055681783 | 0.073144476 | -0.761257529 | 0.446503263 | 0.822210607 |
| Ttll8 | 54.20827939 | -0.055709105 | 0.092606854 | -0.601565674 | 0.547463284 | 0.867553753 |
| Pitpnm2 | 12606.59897 | -0.055804667 | 0.074484189 | -0.749214948 | 0.453727661 | 0.825985606 |
| Rictor | 2611.119886 | -0.055828236 | 0.072885637 | -0.765970333 | 0.443693955 | 0.820717992 |
| Neurod2 | 2832.278244 | -0.055830637 | 0.066662479 | -0.83751217 | 0.402304739 | 0.79955155 |
| Prr19 | 5.274557021 | -0.055832267 | 0.04966271 | -1.124229152 | 0.260915825 | NA |
| Ccdc74a | 589.9788752 | -0.055838534 | 0.084144116 | -0.66360593 | 0.506942561 | 0.848958235 |
| Zfp513 | 873.6875304 | -0.055843618 | 0.067210665 | -0.830874357 | 0.406044611 | 0.800733644 |
| Lrrc71 | 39.11062007 | -0.055846575 | 0.081064059 | -0.688919061 | 0.490874204 | 0.841866734 |
| Rbm26 | 1520.764486 | -0.055848972 | 0.065119052 | -0.857644114 | 0.391089008 | 0.794184423 |
| Ehd2 | 320.6655307 | -0.055862612 | 0.086634146 | -0.644810561 | 0.519049957 | 0.853297835 |
| Iqcd | 45.21330901 | -0.055881269 | 0.090833346 | -0.615206543 | 0.538418327 | 0.862082844 |
| Slc22a12 | 7.750617388 | -0.05589811 | 0.054705964 | -1.021791891 | 0.306879408 | NA |
| Epha3 | 201.2201566 | -0.055911699 | 0.091444677 | -0.6114265 | 0.540917264 | 0.863847445 |
| Ndc80 | 10.09392329 | -0.055915185 | 0.066413274 | -0.841927859 | 0.399828335 | NA |
| Map2k2 | 2731.698192 | -0.056003269 | 0.065428734 | -0.855943039 | 0.392029287 | 0.794184423 |
| Lax1 | 14.63061844 | -0.056007782 | 0.073459625 | -0.762429457 | 0.445803731 | NA |
| Cpt1a | 1483.018394 | -0.056025484 | 0.062512063 | -0.896234755 | 0.370127397 | 0.780397655 |
| Gm410 | 75.96427275 | -0.056042472 | 0.094528959 | -0.59286035 | 0.553274615 | 0.870600462 |
| Scel | 121.6691407 | -0.056092184 | 0.09328381 | -0.60130674 | 0.547635701 | 0.867553753 |
| Wdr24 | 1128.922232 | -0.056110669 | 0.069650626 | -0.805601788 | 0.420472494 | 0.807801012 |
| Mnda | 5.302428134 | -0.0561172 | 0.041474303 | -1.353059517 | 0.176036617 | NA |
| Ctsl | 3528.124934 | -0.056122674 | 0.069277248 | -0.810116955 | 0.417872961 | 0.80621897 |
| Sgol1 | 6.616605979 | -0.056124701 | 0.05721969 | -0.980863427 | 0.326660095 | NA |
| Mcm9 | 222.9135382 | -0.056160596 | 0.083658757 | -0.671305643 | 0.50202584 | 0.847070201 |
| Txlna | 2348.472007 | -0.0561711 | 0.057796156 | -0.971882974 | 0.331108771 | 0.756212998 |
| Telo2 | 856.9360634 | -0.05621911 | 0.082236829 | -0.68362449 | 0.494212316 | 0.843503724 |
| Polg2 | 73.91133661 | -0.056259667 | 0.092851596 | -0.605909538 | 0.544574825 | 0.86624804 |
| Ecscr | 100.5309614 | -0.05628716 | 0.092551097 | -0.608173881 | 0.543072156 | 0.865554766 |
| 6330549D23Rik | 60.94252516 | -0.056296321 | 0.094296268 | -0.597015368 | 0.55049712 | 0.869565155 |
| Cysltr2 | 7.910038257 | -0.056336972 | 0.060235865 | -0.935272898 | 0.349647679 | NA |
| Efna1 | 161.7267738 | -0.056408217 | 0.090145558 | -0.625745939 | 0.531481596 | 0.858328942 |
| Ppan | 568.9013378 | -0.056416896 | 0.081349953 | -0.693508652 | 0.487990398 | 0.839945271 |
| Zw10 | 899.2569113 | -0.05641988 | 0.067466104 | -0.836270014 | 0.403003019 | 0.800009209 |
| Dguok | 385.4466469 | -0.056425413 | 0.071608838 | -0.787967157 | 0.430715917 | 0.812181606 |
| Slc30a6 | 390.1327636 | -0.056472329 | 0.075828549 | -0.744737039 | 0.456430703 | 0.82674985 |
| Hsf4 | 811.0129347 | -0.056502107 | 0.075750892 | -0.745893619 | 0.45573168 | 0.826603827 |
| Rnf138 | 290.6421065 | -0.056502303 | 0.089210565 | -0.63335887 | 0.526499317 | 0.85599981 |
| Ttc22 | 54.06649903 | -0.056522476 | 0.08667985 | -0.652083231 | 0.514347481 | 0.853195313 |
| 6230400D17Rik | 17.07324714 | -0.056524159 | 0.074958545 | -0.754072257 | 0.450805835 | NA |
| Igdcc4 | 2561.996501 | -0.05652679 | 0.052407618 | -1.078598724 | 0.28076665 | 0.720080692 |
| Eif3f | 4145.123622 | -0.05653543 | 0.047183587 | -1.198201198 | 0.230838699 | 0.683402059 |
| Lmf2 | 746.427105 | -0.056547686 | 0.070013523 | -0.807668045 | 0.419281704 | 0.807114502 |
| Edn1 | 109.5330764 | -0.056551311 | 0.094838694 | -0.596289428 | 0.550981893 | 0.869642174 |
| Dsp | 54.60585995 | -0.056562 | 0.042901341 | -1.318420319 | 0.187362982 | 0.641710498 |
| Wdr69 | 17.55190969 | -0.056563473 | 0.071846771 | -0.787279258 | 0.431118409 | NA |
| Rhpn1 | 394.2323382 | -0.05658835 | 0.093899092 | -0.60265066 | 0.546741112 | 0.867128614 |
| Tnfaip8l3 | 321.7505918 | -0.056593554 | 0.077413852 | -0.731052029 | 0.464747374 | 0.83159896 |
| Ctla2b | 30.58074054 | -0.056659092 | 0.084799132 | -0.668156509 | 0.504033694 | 0.847807553 |
| Cdc6 | 17.96821184 | -0.056670355 | 0.080935519 | -0.700191405 | 0.483807779 | NA |
| Iqca | 46.96585163 | -0.056747729 | 0.093973702 | -0.603868181 | 0.545931285 | 0.866989658 |
| Dcaf11 | 3879.148454 | -0.056769952 | 0.05507294 | -1.030813899 | 0.302628099 | 0.735974513 |
| Rexo1 | 2422.990928 | -0.056813793 | 0.056630827 | -1.003230847 | 0.315749493 | 0.745993052 |
| Cntln | 599.8497336 | -0.056829891 | 0.065849401 | -0.863028218 | 0.388121953 | 0.791889115 |
| Banp | 1087.501475 | -0.056844117 | 0.089729788 | -0.633503301 | 0.526405026 | 0.855990082 |
| Ddx23 | 3015.502977 | -0.056862419 | 0.055054964 | -1.032830006 | 0.301683459 | 0.735974432 |
| Ccp110 | 3366.564975 | -0.056897236 | 0.069316425 | -0.820833393 | 0.411741175 | 0.802795661 |
| Baz1b | 6061.131448 | -0.056909995 | 0.050556941 | -1.125661361 | 0.260308884 | 0.704058139 |
| Orai2 | 3712.413504 | -0.056921874 | 0.070005965 | -0.813100339 | 0.416160532 | 0.806019718 |
| Stard8 | 1635.835967 | -0.056930977 | 0.080845566 | -0.704194177 | 0.481311857 | 0.8371092 |
| Sh2d7 | 3.390627198 | -0.056938886 | 0.040839175 | -1.394222242 | 0.163250503 | NA |
| Hebp2 | 113.550386 | -0.056961517 | 0.094317577 | -0.603933207 | 0.54588805 | 0.866989658 |
| Bend3 | 279.9162906 | -0.056972557 | 0.088586547 | -0.643128774 | 0.520140545 | 0.853297835 |
| A830082K12Rik | 321.639868 | -0.056991303 | 0.087950534 | -0.647992685 | 0.516989682 | 0.853297835 |
| Gm4952 | 6.999717798 | -0.057015519 | 0.059456676 | -0.958942245 | 0.337587841 | NA |
| Crygn | 11.07150858 | -0.057074845 | 0.061144378 | -0.933443869 | 0.350590839 | NA |
| Ehmt2 | 8345.968623 | -0.057076047 | 0.06206586 | -0.919604548 | 0.357779449 | 0.773954472 |
| Parp6 | 6094.856895 | -0.057120383 | 0.046661869 | -1.224134051 | 0.220901663 | 0.672757795 |
| Atrx | 6321.334407 | -0.057147781 | 0.070700104 | -0.80831254 | 0.418910686 | 0.806975048 |
| Epb4.1l4b | 458.1813264 | -0.057148694 | 0.079349781 | -0.72021237 | 0.471394249 | 0.833245343 |
| Wdsub1 | 380.6397965 | -0.057150571 | 0.077210575 | -0.740190984 | 0.459184117 | 0.828381305 |
| Pigp | 713.0271422 | -0.057152155 | 0.072455936 | -0.788784994 | 0.430237681 | 0.811860174 |
| Serinc4 | 15.76722809 | -0.057176714 | 0.073421907 | -0.778741877 | 0.436131782 | NA |
| Pign | 767.9057362 | -0.057177981 | 0.072013104 | -0.793994125 | 0.427198851 | 0.810334558 |

| | | | | | | | | | | | | | | | |
|---|---|---|---|---|---|---|---|---|---|---|---|---|---|---|---|
| Rnf182 | 369.3571025 | 0.186645429 | 0.111352359 | 1.676169508 | 0.093704995 | 0.258661178 | | | Serping1 | 272.8311532 | -0.057206741 | 0.080864327 | -0.707441007 | 0.479292461 | 0.836200131 |
| Fnbp1 | 4949.954058 | 0.186608782 | 0.083666244 | 2.230395122 | 0.025721223 | 0.111226523 | | | 1810014B01Rik | 123.7914981 | -0.057231822 | 0.090837518 | -0.63004608 | 0.528664436 | 0.857360679 |
| Nob1 | 906.9091006 | 0.186463878 | 0.072461454 | 2.573283695 | 0.010073859 | 0.059167695 | | | Arl13b | 394.8629945 | -0.057253391 | 0.074734821 | -0.766087215 | 0.44362441 | 0.820717992 |
| Dusp19 | 341.7103473 | -0.204715733 | 0.115099763 | -1.778593874 | 0.075306369 | 0.225566144 | | | Smpd5 | 18.08520383 | -0.057278356 | 0.077894019 | -0.735337028 | 0.462134263 | NA |
| Fbxo36 | 85.7448077 | 0.186416612 | 0.130150843 | 1.432311981 | 0.152054556 | 0.353372324 | | | Pla2g4a | 153.5100082 | -0.057292142 | 0.094056264 | -0.609126275 | 0.542440743 | 0.864833084 |
| Mertk | 1828.683998 | 0.186366955 | 0.104695709 | 1.780082078 | 0.075062529 | 0.225173055 | | | Mettl21a | 419.097279 | -0.057296092 | 0.0678893 | -0.843963517 | 0.398689793 | 0.797789838 |
| Hnrpdl | 5253.598948 | 0.18635293 | 0.104113273 | 1.789905604 | 0.073469088 | 0.221939958 | | | Mtfr1l | 2532.57332 | -0.057304172 | 0.047951147 | -1.195053191 | 0.232066249 | 0.684188011 |
| Fam65c | 17.1559238 | 0.186301137 | 0.164556354 | 1.132141861 | 0.257574798 | 0.484107923 | | | Fnip1 | 2243.74396 | -0.057318692 | 0.077934597 | -0.735471712 | 0.462052262 | 0.830604784 |
| Lrrc48 | 448.9524157 | 0.186115266 | 0.099807084 | 1.864750066 | 0.062216445 | 0.199180439 | | | Cramp1l | 1956.627087 | -0.057328916 | 0.057470603 | -0.997534604 | 0.318505086 | 0.74851037 |
| Commd8 | 1249.20763 | -0.113245066 | 0.063690615 | -1.778049492 | 0.075395727 | 0.225649903 | | | Ankrd13d | 2302.250179 | -0.057336296 | 0.074717694 | -0.767372401 | 0.442860133 | 0.820011 |
| Mafk | 355.311133 | 0.186057611 | 0.134316464 | 1.38521821 | 0.165985754 | 0.371806281 | | | Zfp53 | 179.6023293 | -0.05737472 | 0.088129563 | -0.651026944 | 0.515029094 | 0.853297835 |
| Zfp708 | 125.9846947 | -0.224784619 | 0.126426412 | -1.777987806 | 0.075405858 | 0.225649903 | | | Utp15 | 1315.891573 | -0.057381963 | 0.059137809 | -0.970309244 | 0.331892371 | 0.756958483 |
| Dnm2 | 2164.425059 | 0.186049992 | 0.080133958 | 2.321737197 | 0.020247091 | 0.09495708 | | | Ces2g | 10.11002416 | -0.057388112 | 0.062439567 | -0.919098491 | 0.358044059 | NA |
| Rgs6 | 1300.23808 | 0.186043315 | 0.109500796 | 1.699013352 | 0.089316668 | 0.251384642 | | | Kin | 328.5047284 | -0.057392693 | 0.074948398 | -0.765762763 | 0.443817474 | 0.820841945 |
| 4932415G12Rik | 1363.432851 | 0.185961481 | 0.0951049 | 1.955330179 | 0.050544111 | 0.17500122 | | | Vmn1r58 | 68.0715051 | -0.057405467 | 0.094556783 | -0.607100472 | 0.543784237 | 0.86604998 |
| Dnpep | 1314.939629 | -0.110771258 | 0.062336579 | -1.77698648 | 0.075570464 | 0.225909468 | | | 6030408B16Rik | 6.045332183 | -0.057492094 | 0.046990795 | -1.223475661 | 0.221150092 | NA |
| Tm9sf2 | 4546.496132 | -0.132447828 | 0.074531538 | -1.777070913 | 0.075556573 | 0.225909468 | | | Htr1a | 653.0955023 | -0.057493404 | 0.082053543 | -0.700681562 | 0.483501765 | 0.83839786 |
| Slc10a7 | 178.2498962 | -0.180385441 | 0.101531308 | -1.776648445 | 0.0756261 | 0.226032174 | | | Mbd1 | 1214.070183 | -0.057503195 | 0.055850207 | -1.029596811 | 0.303199313 | 0.73655042 |
| 0610010B08Rik | 10.01045317 | -0.289089807 | 0.162758166 | -1.776192336 | 0.075701221 | 0.22621306 | | | Mboat1 | 150.3644128 | -0.057506099 | 0.094746386 | -0.606947677 | 0.543885636 | 0.86604998 |
| Glt25d1 | 2120.166764 | 0.185958894 | 0.071242961 | 2.61020726 | 0.009048738 | 0.054851779 | | | Podxl | 2094.447959 | -0.057515986 | 0.07543978 | -0.762409252 | 0.445815786 | 0.821773578 |
| 6330416G13Rik | 578.9097369 | 0.185940245 | 0.07531168 | 2.46894301 | 0.01355128 | 0.072501219 | | | Mfsd11 | 1339.419073 | -0.057524599 | 0.058266029 | -0.987275087 | 0.323507801 | 0.752925264 |
| Lefty1 | 30.38633122 | 0.185931189 | 0.167764966 | 1.108283771 | 0.267739282 | 0.495740014 | | | Gm13238 | 14.33041661 | -0.057530342 | 0.070784603 | -0.812752195 | 0.416360149 | NA |
| Rab28 | 1946.928689 | -0.103866167 | 0.058498182 | -1.775545216 | 0.075807907 | 0.226357237 | | | Metrn | 1228.805674 | -0.057538486 | 0.082576192 | -0.696792677 | 0.485932549 | 0.839716418 |
| Zmynd8 | 5386.935427 | 0.185803174 | 0.100604639 | 1.846864872 | 0.064766727 | 0.20447506 | | | Recql | 770.9479037 | -0.057569474 | 0.064868861 | -0.887474711 | 0.374823376 | 0.783182445 |
| Gm16938 | 53.0004067 | 0.185756986 | 0.158624865 | 1.171045825 | 0.241580358 | 0.466262791 | | | Gmnc | 21.8116485 | -0.057578067 | 0.081364981 | -0.707651696 | 0.479161581 | NA |
| Pet112 | 653.5904501 | -0.137714294 | 0.077594611 | -1.774792006 | 0.075932237 | 0.226597472 | | | Ppp6r1 | 8269.656714 | -0.057614651 | 0.057746906 | -0.997709742 | 0.318420128 | 0.74851037 |
| Rpl39 | 2422.037477 | -0.178053153 | 0.100336403 | -1.774561855 | 0.07597026 | 0.226667285 | | | Xrcc6bp1 | 141.7020967 | -0.057640108 | 0.0929515 | -0.620109493 | 0.535185703 | 0.860443555 |
| Slc12a8 | 47.85542696 | 0.185708931 | 0.15696335 | 1.183135622 | 0.236755402 | 0.460956044 | | | Dll1 | 221.9250356 | -0.057672143 | 0.085764551 | -0.67244733 | 0.50129896 | 0.846601651 |
| Mrpl50 | 1208.897597 | -0.117463597 | 0.066212628 | -1.774036185 | 0.076057165 | 0.226839214 | | | Tnfrsf25 | 174.5804649 | -0.057684086 | 0.060869403 | -0.947669653 | 0.343297655 | 0.763762832 |
| Dennd5b | 3530.395108 | -0.140820806 | 0.079382762 | -1.773946908 | 0.076071933 | 0.226839593 | | | 9330159M07Rik | 108.1508232 | -0.057691277 | 0.093871186 | -0.614579191 | 0.53883266 | 0.862393479 |
| Lax1 | 12.95677503 | 0.185686561 | 0.166579495 | 1.114702389 | 0.264977977 | 0.492392514 | | | Rdh18-ps | 4.712843211 | -0.057705005 | 0.046911444 | -1.230083757 | 0.218665742 | NA |
| Slc16a6 | 547.4101983 | 0.185623218 | 0.136087453 | 1.363999501 | 0.172567723 | 0.380349358 | | | Mterfd1 | 552.4408775 | -0.057731657 | 0.071166854 | -0.811215527 | 0.417241911 | 0.806153084 |
| Sptan1 | 44149.52124 | -0.102459586 | 0.05779254 | -1.772886015 | 0.076247596 | 0.227147116 | | | Tdp2 | 594.3210224 | -0.057736103 | 0.068693341 | -0.840490542 | 0.400633403 | 0.799169897 |
| Styk1 | 12.15494332 | -0.283381963 | 0.159842596 | -1.772881384 | 0.076248364 | 0.227147116 | | | Ropn1l | 15.60194401 | -0.057738248 | 0.072679531 | -0.794422407 | 0.426949564 | NA |
| Trp53bp1 | 4489.486908 | -0.13492119 | 0.076098767 | -1.772974727 | 0.076232895 | 0.227147116 | | | Rela | 948.9665034 | -0.057796403 | 0.065713573 | -0.879520017 | 0.379119385 | 0.785380516 |
| Entpd7 | 454.0441139 | 0.185391939 | 0.09820686 | 1.887769738 | 0.059056869 | 0.193002981 | | | Arhgef40 | 984.7553031 | -0.057807627 | 0.071051394 | -0.813603001 | 0.415872418 | 0.805784013 |
| Strada | 967.8267689 | 0.185366986 | 0.071206353 | 2.603236616 | 0.009234819 | 0.055586285 | | | Pradc1 | 507.825454 | -0.057829335 | 0.070353993 | -0.821976578 | 0.411090227 | 0.802622856 |
| Glp1r | 23.35983173 | -0.280566485 | 0.158303151 | -1.772336707 | 0.076338681 | 0.227282425 | | | Bahd1 | 2288.078718 | -0.05783148 | 0.068320997 | -0.846467282 | 0.397292118 | 0.797340699 |
| 9930012K11Rik | 63.85689864 | 0.185303988 | 0.150043506 | 1.235001721 | 0.216829864 | 0.436760518 | | | Abcb4 | 307.0692636 | -0.057853396 | 0.083140388 | -0.695851894 | 0.486521586 | 0.839801621 |
| Cxxc1 | 1519.476865 | 0.185084255 | 0.07176082 | 2.579182542 | 0.009903444 | 0.05838173 | | | Gm6249 | 2.438780768 | -0.057870585 | 0.037150811 | -1.557720663 | 0.11929948 | NA |
| Nckap1l | 701.9189656 | 0.185009587 | 0.069800477 | 2.650549035 | 0.008036106 | 0.050409485 | | | Alg6 | 332.0361137 | -0.057885269 | 0.080772947 | -0.716641794 | 0.473595149 | 0.833983942 |
| Cacna2d4 | 90.67894028 | 0.18492413 | 0.137152015 | 1.34831508 | 0.177557064 | 0.387201851 | | | Psme1 | 1175.884263 | -0.057905148 | 0.068696415 | -0.842913685 | 0.399276719 | 0.798004076 |
| Mcm2 | 199.4468085 | -0.215811905 | 0.121869952 | -1.770837695 | 0.076587696 | 0.227807594 | | | Zfp952 | 713.1664427 | -0.057923485 | 0.069029572 | -0.839111182 | 0.401406922 | 0.799277436 |
| Ak3 | 3233.742515 | -0.086519 | 0.048867002 | -1.770499449 | 0.076643976 | 0.227841624 | | | Tank | 957.9198326 | -0.057949552 | 0.069113038 | -0.83847496 | 0.401764005 | 0.799277436 |
| Cd86 | 109.4505734 | 0.184677377 | 0.14680053 | 1.258015738 | 0.208386065 | 0.426414656 | | | Atoh7 | 11.25117341 | -0.057965656 | 0.068846331 | -0.84195708 | 0.399811978 | NA |
| Jup | 2085.422935 | -0.125929905 | 0.071121502 | -1.77063055 | 0.076622158 | 0.227841624 | | | Hist1h2bc | 456.3911049 | -0.057971255 | 0.074159602 | -0.781709366 | 0.434385396 | 0.815019667 |
| Paox | 222.5881453 | 0.184660079 | 0.100386483 | 1.839491472 | 0.065842931 | 0.206568909 | | | Tmem123 | 521.6902769 | -0.058017127 | 0.069736256 | -0.831950703 | 0.405436771 | 0.800621878 |
| Arhgef7 | 5540.58052 | -0.099684972 | 0.056342827 | -1.769257568 | 0.076850901 | 0.22832749 | | | Skida1 | 328.3612429 | -0.058053773 | 0.080304164 | -0.722923574 | 0.46972684 | 0.833245343 |
| Slc25a24 | 112.6002428 | -0.228709636 | 0.129267655 | -1.769271955 | 0.076848501 | 0.22832749 | | | Tnnt2 | 272.8664888 | -0.058055455 | 0.085829735 | -0.676402592 | 0.498785072 | 0.846052342 |
| Ikbke | 44.81917586 | 0.184562125 | 0.160859772 | 1.147347923 | 0.251237855 | 0.476971641 | | | Coro7 | 3117.855941 | -0.058076612 | 0.048050469 | -1.208658588 | 0.226794038 | 0.679834424 |
| Ebf4 | 311.3494351 | 0.18448077 | 0.143436695 | 1.286147658 | 0.198391534 | 0.414381023 | | | Myh13 | 25.55260901 | -0.058083399 | 0.082003268 | -0.708305906 | 0.47875531 | 0.835936459 |
| 9530026P05Rik | 9.714312093 | 0.184464751 | 0.157293651 | 1.172741238 | 0.240899589 | 0.465428677 | | | Osbp2 | 5120.326757 | -0.05809429 | 0.065849664 | -0.882226075 | 0.377654564 | 0.784178571 |
| Tmem43 | 1551.953568 | 0.184408023 | 0.081606133 | 2.259732385 | 0.023837865 | 0.106133299 | | | March10 | 84.42266805 | -0.05809632 | 0.094791065 | -0.61288815 | 0.539950299 | 0.863544778 |
| Fxyd1 | 707.4762706 | 0.18430582 | 0.11690467 | 1.57654796 | 0.114899575 | 0.296956466 | | | Cyp27a1 | 296.6152398 | -0.058096511 | 0.092734075 | -0.626485047 | 0.530996842 | 0.858310785 |
| L3mbtl1 | 367.5758206 | 0.184303225 | 0.153373002 | 1.201666672 | 0.229492699 | 0.452510398 | | | Tram2 | 17.75232845 | -0.058098153 | 0.078768262 | -0.737583271 | 0.460767724 | NA |
| Sdhaf2 | 1483.630257 | -0.096436536 | 0.054541204 | -1.768140938 | 0.077037345 | 0.228574612 | | | Sfxn3 | 5856.743427 | -0.058099648 | 0.04531284 | -1.282189506 | 0.199776176 | 0.654328915 |
| Me1 | 2311.926913 | -0.133664183 | 0.075627099 | -1.76741123 | 0.077159384 | 0.22884906 | | | Dap | 223.1199938 | -0.058105224 | 0.087738083 | -0.662257734 | 0.507806059 | 0.849201121 |
| Zcchc17 | 1995.663777 | -0.100844692 | 0.057056029 | -1.767467748 | 0.077149926 | 0.22884906 | | | Col23a1 | 886.3005708 | -0.058126252 | 0.091736732 | -0.633620259 | 0.526328676 | 0.855977056 |
| Mycn | 226.7310954 | -0.246298758 | 0.139363113 | -1.767316709 | 0.077175203 | 0.228852172 | | | Tmem44 | 3349.109905 | -0.058149398 | 0.059110396 | -0.983742319 | 0.325242227 | 0.75293408 |
| Bzw1 | 6140.216656 | -0.118852292 | 0.067261184 | -1.767026474 | 0.077223795 | 0.228920619 | | | 1810010D01Rik | 5.158765809 | -0.058162705 | 0.050239748 | -1.157702953 | 0.246985279 | NA |
| Elavl3 | 9588.02679 | -0.084443616 | 0.047791579 | -1.766914132 | 0.07724261 | 0.228920619 | | | Mcm4 | 573.0215155 | -0.058167651 | 0.06399202 | -0.908982889 | 0.363359158 | 0.776821522 |
| 9430037G07Rik | 83.24602483 | 0.184295946 | 0.14718523 | 1.252136144 | 0.210520259 | 0.428789391 | | | Asf1a | 377.6786308 | -0.058168766 | 0.081146669 | -0.716834926 | 0.473475959 | 0.833983942 |
| Tpm1 | 7433.48525 | -0.112442107 | 0.063641549 | -1.766803437 | 0.077261153 | 0.228931785 | | | Caap1 | 180.4764777 | -0.058171279 | 0.087802776 | -0.662522093 | 0.507636681 | 0.849139846 |
| Ppp4r1 | 1371.808668 | 0.184171279 | 0.069932064 | 2.633574191 | 0.008449137 | 0.052301612 | | | Acad12 | 89.93531835 | -0.058172483 | 0.093230683 | -0.623962851 | 0.532651982 | 0.858328942 |
| Nkpd1 | 21.6648737 | 0.184141926 | 0.165740912 | 1.111022761 | 0.266558552 | 0.494085203 | | | Mob1b | 426.929653 | -0.058215206 | 0.087073419 | -0.668576088 | 0.50376593 | 0.847807553 |
| Zfyve16 | 679.8277866 | 0.184099099 | 0.11071433 | 1.662829914 | 0.096346487 | 0.263062692 | | | Zfp687 | 1696.216077 | -0.058216806 | 0.07035958 | -0.827418323 | 0.407999999 | 0.801055189 |
| Mirlet7d | 12.29551379 | 0.184081662 | 0.139163115 | 1.322776241 | 0.185909801 | 0.397582671 | | | Ccdc137 | 1012.09381 | -0.058232617 | 0.071949662 | -0.809352198 | 0.41831259 | 0.806745709 |
| Spata7 | 688.8245567 | 0.184075699 | 0.099406 | 1.851756419 | 0.064060808 | 0.202962988 | | | Bach1 | 648.7419822 | -0.05824464 | 0.059808411 | -0.973853663 | 0.330129204 | 0.756141871 |
| Them4 | 691.2040602 | -0.129697352 | 0.073449288 | -1.765808153 | 0.07742804 | 0.229195353 | | | Helz2 | 337.0376756 | -0.058247865 | 0.086664459 | -0.672107877 | 0.501515022 | 0.846601651 |
| Tctn3 | 1017.692717 | 0.184004382 | 0.096802785 | 1.90081702 | 0.057325984 | 0.189423927 | | | Klhl38 | 52.76888205 | -0.058261006 | 0.089970527 | -0.647556569 | 0.517271795 | 0.853297835 |
| Cds2 | 29423.09203 | -0.101659891 | 0.057606048 | -1.764743359 | 0.077606908 | 0.229605009 | | | Itga4 | 484.2055957 | -0.058285476 | 0.083374781 | -0.699078014 | 0.484503278 | 0.838713365 |
| Klf9 | 6826.07298 | 0.18393396 | 0.117213659 | 1.569219504 | 0.116596801 | 0.299444859 | | | Slc30a10 | 994.569136 | -0.058291713 | 0.067684873 | -0.861222156 | 0.389115699 | 0.79252589 |
| Gpr85 | 1315.320084 | -0.124291966 | 0.070456498 | -1.764095147 | 0.077715961 | 0.229752167 | | | Lrrc16a | 1015.362008 | -0.058300891 | 0.061390603 | -0.949671254 | 0.34227932 | 0.763762832 |
| 1500012F01Rik | 584.0385071 | 0.183924738 | 0.151529684 | 1.213786849 | 0.22482913 | 0.446526045 | | | Tuba8 | 601.5741162 | -0.058317801 | 0.078173197 | -0.746007624 | 0.45566281 | 0.826603827 |
| Vamp5 | 73.6921128 | -0.22991462 | 0.130326783 | -1.764139456 | 0.077708503 | 0.229752167 | | | Tyw3 | 541.7239489 | -0.058330941 | 0.066056136 | -0.88305106 | 0.377208684 | 0.783997209 |
| Nudt1 | 126.2387949 | -0.200420406 | 0.113618962 | -1.763969695 | 0.077737082 | 0.229770763 | | | Tinf2 | 433.4806691 | -0.058335598 | 0.08082476 | -0.721754053 | 0.470445703 | 0.833245343 |
| Gng10 | 982.6368485 | -0.16782708 | 0.095149605 | -1.763823196 | 0.077761651 | 0.229799841 | | | Zfp266 | 1577.661144 | -0.058348855 | 0.072829549 | -0.801170073 | 0.423033194 | 0.808999423 |
| Fh1 | 2570.023455 | -0.085548831 | 0.048513216 | -1.76341291 | 0.077830874 | 0.229960252 | | | Commd4 | 1347.17706 | -0.058363685 | 0.074472827 | -0.783691008 | 0.43322144 | 0.813783127 |
| Dhfr | 200.4083898 | 0.183910668 | 0.098386846 | 1.869260744 | 0.061586547 | 0.197725229 | | | Snora70 | 18.562294 | -0.05838352 | 0.078661672 | -0.742210516 | 0.457959798 | NA |
| Cox4i2 | 46.29549677 | -0.279826982 | 0.158719607 | -1.763027191 | 0.077895904 | 0.229976969 | | | Gltscr1 | 847.7242948 | -0.058401319 | 0.064031778 | -0.912067734 | 0.361733062 | 0.776449324 |
| Itgb8 | 1216.768137 | -0.149410214 | 0.08473923 | -1.763176442 | 0.077870736 | 0.229976969 | | | Isg20 | 18.33046176 | -0.058405162 | 0.080669691 | -0.724003786 | 0.469063411 | NA |
| Sertad4 | 843.9375329 | -0.179624895 | 0.101880514 | -1.763093727 | 0.077884683 | 0.229976969 | | | Tnfaip3 | 299.167344 | -0.058408648 | 0.092960393 | -0.628317567 | 0.529795926 | 0.857915133 |
| Gja5 | 21.89368011 | -0.292537797 | 0.166010414 | -1.762165342 | 0.078041366 | 0.230362531 | | | Tmem200b | 199.0587817 | -0.058414644 | 0.090196893 | -0.647634776 | 0.517221199 | 0.853297835 |
| Cidea | 78.41526408 | -0.241258635 | 0.136940221 | -1.761780677 | 0.078106361 | 0.230510467 | | | Gngt2 | 68.70333517 | -0.058455727 | 0.095041809 | -0.615052762 | 0.538519876 | 0.862082844 |
| Rpl32 | 4341.525042 | -0.14986941 | 0.085106487 | -1.760963421 | 0.078244594 | 0.230874452 | | | Aqp4 | 5081.744615 | -0.058543896 | 0.064872058 | -0.902451658 | 0.36681699 | 0.778320713 |
| Tfdp2 | 1685.002767 | 0.1838495 | 0.109123784 | 1.68477938 | 0.092031145 | 0.255766056 | | | Gon4l | 2428.843195 | -0.058549075 | 0.056411231 | -1.037897477 | 0.299317788 | 0.733704078 |
| Cdkn3 | 10.11112459 | -0.287411934 | 0.163270431 | -1.760342842 | 0.078349694 | 0.231096548 | | | Snx20 | 30.53145945 | -0.058553339 | 0.090968365 | -0.643667045 | 0.519791364 | 0.853297835 |
| Vamp4 | 2356.414829 | -0.107493764 | 0.061074314 | -1.760048658 | 0.078399557 | 0.231199606 | | | Chek1 | 43.12398007 | -0.058613036 | 0.093269083 | -0.628429423 | 0.529722668 | 0.857915133 |
| Bbc3 | 151.9406988 | 0.183656659 | 0.135297922 | 1.357424091 | 0.174646498 | 0.383336568 | | | 4933436C20Rik | 16.16201907 | -0.058627073 | 0.073805998 | -0.794340218 | 0.426997397 | NA |
| Ptbp1 | 1249.047071 | 0.183584784 | 0.090628159 | 2.025692513 | 0.042796304 | 0.157054936 | | | Mast3 | 14541.41295 | -0.058628535 | 0.078081755 | -0.750860873 | 0.45273639 | 0.825461127 |
| Gpsm2 | 637.8916253 | 0.183511703 | 0.100258592 | 1.830383806 | 0.067192567 | 0.209220587 | | | Adam3 | 5.039618237 | -0.058653679 | 0.047382551 | -1.237875075 | 0.215762386 | NA |
| Ldlrap1 | 141.3888867 | 0.183495992 | 0.110027514 | 1.667728235 | 0.095369688 | 0.261469936 | | | Ttll13 | 41.10665815 | -0.058703331 | 0.087269058 | -0.672670623 | 0.501156861 | 0.846601651 |
| Enkd1 | 207.8366467 | 0.183399203 | 0.110934706 | 1.653217546 | 0.098286597 | 0.266812564 | | | Fibcd1 | 687.0053464 | -0.058708253 | 0.085817744 | -0.6841039 | 0.493909559 | 0.843404385 |
| Surf4 | 2861.176253 | -0.077940001 | 0.044333037 | -1.758056868 | 0.078737834 | 0.231932319 | | | Ablim2 | 6835.209315 | -0.058715738 | 0.066876977 | -0.877966391 | 0.379961958 | 0.786275719 |
| Hap1 | 8067.595112 | 0.183209336 | 0.142093627 | 1.289356459 | 0.197274191 | 0.412495612 | | | Adam11 | 7243.000313 | -0.058727542 | 0.068695378 | -0.854898009 | 0.392607613 | 0.794184423 |
| C630020P19Rik | 134.9841611 | 0.183193383 | 0.120903562 | 1.51520253 | 0.129721122 | 0.318879751 | | | Tubgcp2 | 1804.279564 | -0.058728403 | 0.054499884 | -1.077587674 | 0.281217805 | 0.720080692 |
| Acvr2a | 1655.222413 | 0.183139655 | 0.108294347 | 1.691128492 | 0.090812265 | 0.253890779 | | | Emp2 | 807.2634246 | -0.058736023 | 0.067249189 | -0.873408647 | 0.38244038 | 0.786954445 |
| Tspan6 | 448.4341502 | -0.158168607 | 0.090031815 | -1.756807935 | 0.078950552 | 0.232382191 | | | 1190002N15Rik | 1254.789725 | -0.058741137 | 0.08492194 | -0.691707427 | 0.489121082 | 0.840836703 |
| Mrps18c | 612.3440634 | -0.165895987 | 0.094473588 | -1.756003883 | 0.079087746 | 0.232741791 | | | 2410137M14Rik | 13.90654139 | -0.058798793 | 0.069172495 | -0.850031412 | 0.395307623 | NA |
| AI593442 | 6817.382807 | 0.18306877 | 0.10432836 | 1.7547364 | 0.079304407 | 0.233202217 | | | Ufl1 | 1485.623945 | -0.058806362 | 0.062036713 | -0.947928387 | 0.343165913 | 0.763762832 |
| Ptch1 | 1756.380356 | -0.189774142 | 0.108131908 | -1.755024457 | 0.079255124 | 0.233145794 | | | Gins1 | 28.36905246 | -0.058860524 | 0.087051144 | -0.676160257 | 0.498938902 | 0.846087613 |
| Pcdhb13 | 262.2417469 | 0.182978204 | 0.110139394 | 1.661332949 | 0.096646596 | 0.263680641 | | | Sass6 | 212.4065278 | -0.05888629 | 0.088222239 | -0.667476708 | 0.504467683 | 0.847807553 |
| D19Ertd737e | 450.9877335 | -0.140929139 | 0.080313392 | -1.754740227 | 0.079303752 | 0.233202217 | | | Opalin | 685.7542955 | -0.058983646 | 0.0931684 | -0.633086391 | 0.526677229 | 0.85599981 |
| Prtn3 | 37.87172653 | -0.283378666 | 0.16150502 | -1.754612128 | 0.079325676 | 0.233220496 | | | Prox2 | 41.86354855 | -0.059004945 | 0.089357155 | -0.660327032 | 0.509043986 | 0.850095619 |
| BC005561 | 449.3696948 | 0.182940572 | 0.099559288 | 1.83750382 | 0.066135553 | 0.207094636 | | | Golga1 | 1670.501187 | -0.059014184 | 0.053139322 | -1.110555848 | 0.266759576 | 0.709441589 |
| Pnmt | 10.95260652 | -0.280500405 | 0.159915388 | -1.754055115 | 0.079421064 | 0.23341236 | | | 2810032G03Rik | 317.5005057 | -0.059034148 | 0.078423935 | -0.752756764 | 0.451596092 | 0.824520467 |
| Gm9079 | 65.59934935 | -0.247607113 | 0.141203312 | -1.753550315 | 0.079507591 | 0.233622343 | | | Zranb3 | 391.5682378 | -0.059039474 | 0.074178404 | -0.795911886 | 0.426083252 | 0.809519334 |
| Mettl13 | 443.3139483 | 0.1828547 | 0.100278692 | 1.823465143 | 0.068232972 | 0.211398053 | | | Gng13 | 556.7487848 | -0.059052946 | 0.083415026 | -0.707941349 | 0.478981681 | 0.836059588 |
| Kcnb1 | 13905.89271 | -0.159775534 | 0.091137429 | -1.753127506 | 0.079580123 | 0.233746811 | | | Klrb1c | 2.538109108 | -0.059107691 | 0.03237013 | -1.825994842 | 0.067851039 | NA |
| St8sia6 | 72.38200638 | 0.18264235 | 0.15645479 | 1.167381004 | 0.243056539 | 0.467508587 | | | Solh | 1773.604461 | -0.059131503 | 0.077424525 | -0.763730902 | 0.445027621 | 0.821493054 |
| Commd1 | 565.1336904 | -0.124615271 | 0.07109706 | -1.752748578 | 0.079645173 | 0.233849216 | | | Mgme1 | 270.7837938 | -0.059188109 | 0.078733424 | -0.751753267 | 0.452199451 | 0.82479272 |
| Adcy7 | 371.4909536 | 0.182363454 | 0.116706471 | 1.562582194 | 0.118150899 | 0.301607001 | | | Alg5 | 542.5297613 | -0.059236689 | 0.071648017 | -0.826773594 | 0.408365399 | 0.801055189 |
| Pwwp2b | 623.7011022 | 0.182338228 | 0.091704238 | 1.98832936 | 0.046775276 | 0.166752575 | | | Zcchc5 | 5.024948625 | -0.059273857 | 0.040600267 | -1.459937618 | 0.144307219 | NA |
| 4933412O06Rik | 76.45580352 | 0.182330983 | 0.155296579 | 1.174082419 | 0.240362015 | 0.464881919 | | | I730030J21Rik | 10.31155572 | -0.05929879 | 0.063914528 | -0.927782644 | 0.353520329 | NA |
| Arrb2 | 1392.625592 | 0.182312848 | 0.071682048 | 2.543354325 | 0.010979383 | 0.0627729 | | | Fbxo27 | 576.8166377 | -0.059303809 | 0.069282886 | -0.855966196 | 0.392016478 | 0.794184423 |
| Slitrk5 | 2985.110693 | -0.146142241 | 0.083481639 | -1.750591432 | 0.080016312 | 0.234716536 | | | 4930404H11Rik | 1.828556031 | -0.059310689 | 0.030418072 | -1.94985041 | 0.051193951 | NA |
| 1700037C18Rik | 117.4821038 | -0.259480027 | 0.148245984 | -1.750334279 | 0.080060649 | 0.234762853 | | | 0610040B10Rik | 40.58498063 | -0.05933121 | 0.09271901 | -0.639903406 | 0.522235399 | 0.854113377 |
| Dusp23 | 126.8272405 | 0.182292233 | 0.119117398 | 1.530357746 | 0.125928202 | 0.313838115 | | | Adra1d | 1108.215432 | -0.059351223 | 0.087797727 | -0.675999536 | 0.49904094 | 0.846161864 |
| Gck | 51.23499475 | 0.182242581 | 0.166053602 | 1.097492491 | 0.272426161 | 0.500827117 | | | Map2k3 | 576.7383564 | -0.059358124 | 0.083586958 | -0.710136189 | 0.477619687 | 0.835457925 |
| Rnf139 | 1165.440606 | -0.118316236 | 0.0676097 | -1.749989066 | 0.0801202 | 0.234843433 | | | Klhl5 | 1754.876263 | -0.059373905 | 0.048128991 | -1.233641169 | 0.217336645 | 0.670311138 |
| Dlgap3 | 13078.72048 | -0.156333211 | 0.089362666 | -1.749424217 | 0.080217718 | 0.235084797 | | | Slirp | 620.2293177 | -0.059426122 | 0.081728605 | -0.72711534 | 0.467155302 | 0.832211278 |
| Cabin1 | 4050.570807 | 0.093188177 | 0.053280633 | 1.749006577 | 0.080289883 | 0.235162843 | | | 1700018L02Rik | 23.30451592 | -0.059437258 | 0.082974854 | -0.716328566 | 0.473788492 | 0.833983942 |
| Med12l | 1782.38903 | -0.153352328 | 0.08767589 | -1.749082085 | 0.080276832 | 0.235162843 | | | Tmed1 | 408.6220989 | -0.059464754 | 0.083259943 | -0.714206043 | 0.475099781 | 0.834699977 |
| Ndufab1 | 1319.195636 | -0.17758346 | 0.101530667 | -1.749062279 | 0.080280255 | 0.235162843 | | | C130060C02Rik | 10.75026041 | -0.059476806 | 0.065059405 | -0.914192273 | 0.360615824 | NA |
| Cercam | 281.4003672 | 0.182222885 | 0.095876162 | 1.900606798 | 0.057353534 | 0.189474555 | | | Fmod | 1033.732929 | -0.059519443 | 0.080647173 | -0.738022686 | 0.460500664 | 0.829239232 |
| Csnk1g2 | 5486.518084 | -0.142913907 | 0.081735492 | -1.748492662 | 0.080378755 | 0.235245265 | | | Phldb1 | 5284.516827 | -0.059524282 | 0.078580721 | -0.757492181 | 0.448755049 | 0.823263966 |
| Dlgap2 | 2863.632138 | -0.144091853 | 0.082407279 | -1.748533065 | 0.080371765 | 0.235245265 | | | Htr1f | 104.9981316 | -0.059541821 | 0.093764057 | -0.635017545 | 0.525416972 | 0.855879197 |
| Macrod2 | 1030.462823 | -0.141116515 | 0.080703595 | -1.748577807 | 0.080364025 | 0.235245265 | | | Cep72 | 68.102927 | -0.059552058 | 0.095102743 | -0.626186546 | 0.531192591 | 0.858328942 |
| BC089491 | 32.12494045 | -0.290797486 | 0.166353773 | -1.748066672 | 0.080452483 | 0.235416577 | | | Mars2 | 373.0186572 | -0.059584486 | 0.074454325 | -0.800282398 | 0.423547199 | 0.808999423 |
| Klk6 | 164.0258607 | 0.182093511 | 0.155835553 | 1.168497871 | 0.242605998 | 0.467209584 | | | Trappc6a | 205.4926891 | -0.059588727 | 0.087763152 | -0.678972042 | 0.497155577 | 0.845038521 |
| Hectd1 | 6626.104573 | -0.13574514 | 0.077674349 | -1.747618624 | 0.080530088 | 0.235510228 | | | Pcca | 1110.982586 | -0.059647349 | 0.065781683 | -0.906747089 | 0.364540556 | 0.777237064 |
| 2310061J03Rik | 97.70773224 | 0.182050249 | 0.123263755 | 1.476916309 | 0.139698073 | 0.334076546 | | | Spef2 | 45.1903929 | -0.059680856 | 0.093148167 | -0.640708861 | 0.521711856 | 0.854113377 |
| Ubfd1 | 2511.440972 | -0.094958721 | 0.054341983 | -1.747428334 | 0.080563066 | 0.23556221 | | | Vmn2r-ps129 | 14.36537776 | -0.059686606 | 0.071850489 | -0.830705628 | 0.406139946 | NA |
| C1qtnf9 | 11.54995185 | 0.181990174 | 0.163994829 | 1.10973117 | 0.267114887 | 0.494879596 | | | Tgif2 | 46.33734617 | -0.059712795 | 0.087420564 | -0.683052043 | 0.494573958 | 0.843609829 |
| Klf2 | 336.5266341 | 0.181856655 | 0.156422681 | 1.162597734 | 0.24499275 | 0.470024477 | | | Insl5 | 15.60169665 | -0.05972865 | 0.074772198 | -0.798808271 | 0.424401594 | NA |
| Cd55 | 156.9865048 | -0.231021268 | 0.132265496 | -1.746648025 | 0.080698411 | 0.235811614 | | | Krt9 | 499.7172269 | -0.059763215 | 0.091264887 | -0.654832504 | 0.512575601 | 0.852934798 |
| Zdhhc8 | 4860.566398 | 0.181615281 | 0.081338834 | 2.23282376 | 0.025560569 | 0.110810144 | | | Snora44 | 9.864701051 | -0.059765264 | 0.062425727 | -0.957381946 | 0.338374508 | NA |
| Mroh5 | 82.68337481 | 0.181598092 | 0.135956691 | 1.335705442 | 0.18164557 | 0.392198504 | | | Fgf11 | 240.764764 | -0.059775891 | 0.093580718 | -0.638762901 | 0.522977186 | 0.854337225 |
| Emc3 | 2833.228246 | -0.086632608 | 0.049616628 | -1.746039804 | 0.080804036 | 0.235999572 | | | Brwd1 | 5062.06958 | -0.059778791 | 0.058699074 | -1.018394104 | 0.308490703 | 0.740806186 |
| Prelp | 915.86248 | 0.181563687 | 0.136950762 | 1.32575886 | 0.184919584 | 0.396245313 | | | Akr1b10 | 664.2299068 | -0.059787797 | 0.071922992 | -0.831275172 | 0.405818197 | 0.800733644 |
| BC002163 | 22.01179899 | -0.29340867 | 0.168121487 | -1.745218148 | 0.080946903 | 0.236238712 | | | Tnk1 | 5.597180968 | -0.059789215 | 0.045432675 | -1.315995914 | 0.188175414 | NA |
| Prl | 250.8284412 | 0.181551963 | 0.080983034 | 2.241851831 | 0.02497095 | 0.109108827 | | | Asb10 | 11.10414919 | -0.059800861 | 0.060107493 | -0.994898594 | 0.319785587 | NA |
| Apoc3 | 9.787055126 | 0.181510644 | 0.158112699 | 1.147982702 | 0.250975705 | 0.476604678 | | | Mterf | 67.36165212 | -0.059842744 | 0.09488466 | -0.630689342 | 0.528243669 | 0.856965327 |
| Sgce | 913.0014181 | -0.102756393 | 0.058893205 | -1.744792018 | 0.081021079 | 0.236410658 | | | Clrn2 | 8.262242196 | -0.059854385 | 0.055935718 | -1.070056615 | 0.284593826 | NA |
| Ubiad1 | 402.0264575 | -0.152012554 | 0.087153799 | -1.744187359 | 0.081126425 | 0.236673475 | | | Krt12 | 571.1490252 | -0.05987198 | 0.083551392 | -0.716588653 | 0.473627948 | 0.833983942 |
| Akr1c14 | 68.93427523 | -0.282257214 | 0.161879771 | -1.74362252 | 0.081224934 | 0.236871659 | | | Ubr3 | 8656.689115 | -0.059900516 | 0.061045618 | -0.981241862 | 0.326473485 | 0.753373722 |
| Frk | 19.72830282 | -0.292873764 | 0.167961697 | -1.743693768 | 0.081212503 | 0.236871659 | | | Gnmt | 19.04769443 | -0.059943162 | 0.07505909 | -0.798612966 | 0.424514867 | NA |
| Ap1s2 | 343.5443525 | -0.220914229 | 0.126721301 | -1.743307769 | 0.081279869 | 0.236948306 | | | Cyb5r2 | 16.1171627 | -0.05995629 | 0.075921161 | -0.789717765 | 0.429692614 | NA |
| Ptpn7 | 78.13905147 | -0.272341447 | 0.156222087 | -1.743296694 | 0.081281802 | 0.236948306 | | | Il27ra | 17.35649141 | -0.059976482 | 0.071060668 | -0.844017981 | 0.398659358 | NA |
| Eps8l1 | 84.87109809 | 0.181419469 | 0.120766791 | 1.502229775 | 0.133037778 | 0.323847184 | | | Asgr2 | 2.938424935 | -0.060001817 | 0.030350762 | -1.976946006 | 0.048047734 | NA |
| Elac1 | 1688.93885 | 0.181404783 | 0.085417852 | 2.123733846 | 0.033692405 | 0.133702991 | | | Sult5a1 | 3.848142728 | -0.060070008 | 0.04624319 | -1.299002262 | 0.193943153 | NA |
| Gmeb1 | 902.3367923 | -0.123320153 | 0.07079968 | -1.741817941 | 0.081540306 | 0.237524069 | | | Cdk4 | 1374.9262 | -0.06008846 | 0.056762682 | -1.058590926 | 0.289786121 | 0.725139224 |
| Rnf165 | 2839.671923 | -0.165894178 | 0.095242074 | -1.741816092 | 0.08154063 | 0.237524069 | | | Apol8 | 265.1206459 | -0.060115896 | 0.081405406 | -0.738475481 | 0.460225561 | 0.829022384 |
| Dmrtb1 | 18.99287888 | -0.291887645 | 0.167591286 | -1.741663616 | 0.081567323 | 0.237557161 | | | Fkbp15 | 1848.142498 | -0.06015536 | 0.052546647 | -1.144799203 | 0.252292345 | 0.69992334 |
| Tmem132c | 347.3546684 | 0.181323759 | 0.085458332 | 2.121779761 | 0.033856237 | 0.134160751 | | | Mrvi1 | 170.3799913 | -0.060186944 | 0.093525301 | -0.643536494 | 0.519876042 | 0.853297835 |
| Mid1ip1 | 3131.064246 | -0.159414625 | 0.09154162 | -1.741444227 | 0.081605741 | 0.237579737 | | | Cln5 | 565.5562391 | -0.060194833 | 0.07529882 | -0.7994127 | 0.424051149 | 0.809005384 |
| Jak3 | 226.7371253 | 0.181152076 | 0.149740015 | 1.209777336 | 0.226364345 | 0.448731981 | | | Pcdhga2 | 564.3070954 | -0.060228632 | 0.081178808 | -0.741925554 | 0.458132442 | 0.827715405 |
| Gng12 | 2557.481828 | -0.120081468 | 0.068979681 | -1.740823754 | 0.081714477 | 0.237806931 | | | Hnrnpa2b1 | 16295.16623 | -0.060255465 | 0.038602041 | -1.560939857 | 0.118537942 | 0.550297905 |
| Idh3g | 4234.613239 | 0.095159389 | 0.054666576 | 1.740723413 | 0.081732072 | 0.23781347 | | | Gm5431 | 8.342219603 | -0.060302265 | 0.053762861 | -1.121634226 | 0.262017993 | NA |
| Pptc7 | 2924.356523 | -0.174469247 | 0.100234687 | -1.740607488 | 0.081752404 | 0.237827967 | | | Slfn2 | 94.42620863 | -0.060309965 | 0.093482 | -0.645150566 | 0.518829617 | 0.853297835 |
| Ckap2l | 33.76202319 | -0.292047909 | 0.167867222 | -1.739755413 | 0.081901975 | 0.238218358 | | | Riiad1 | 53.78677097 | -0.060319901 | 0.092136111 | -0.654682512 | 0.512672187 | 0.852934798 |
| Tmsb10 | 3506.581788 | -0.222501212 | 0.127943281 | -1.739061327 | 0.082023976 | 0.238528432 | | | Lrrc25 | 23.65945545 | -0.060381464 | 0.08463316 | -0.713449248 | 0.475567808 | 0.835017283 |
| Slc25a37 | 1326.688165 | 0.181151753 | 0.106046128 | 1.708235429 | 0.087592667 | 0.247928532 | | | Neurl2 | 62.99390615 | -0.060391227 | 0.09312008 | -0.64853066 | 0.516641787 | 0.853297835 |
| Cbr3 | 331.8053387 | -0.167865728 | 0.096548428 | -1.738668673 | 0.082093059 | 0.238639749 | | | Klf6 | 2197.030923 | -0.060401041 | 0.071557252 | -0.844093919 | 0.398616926 | 0.797789838 |
| Gm4926 | 8.407138136 | -0.256981227 | 0.147815974 | -1.738521349 | 0.082118992 | 0.238670354 | | | Tspan18 | 144.0682901 | -0.060419952 | 0.093008178 | -0.64961978 | 0.515937853 | 0.853297835 |
| Ttll9 | 44.75876019 | 0.181120724 | 0.156066471 | 1.160535784 | 0.245830733 | 0.470874197 | | | Id1 | 325.1614954 | -0.060429468 | 0.080711002 | -0.748714143 | 0.454029517 | 0.825985606 |
| Bend7 | 123.5091421 | -0.266467684 | 0.153360284 | -1.737527318 | 0.082294136 | 0.238932488 | | | Tanc1 | 1777.196687 | -0.060431781 | 0.082822825 | -0.729651288 | 0.465603364 | 0.831871734 |
| Ccnt2 | 1861.946825 | 0.181050254 | 0.090068121 | 2.010148012 | 0.044415526 | 0.161089456 | | | Efcab6 | 299.9288608 | -0.0604449 | 0.087738254 | -0.688922984 | 0.490871735 | 0.841866734 |
| Tnik | 6364.252787 | 0.181047146 | 0.069106252 | 2.619837439 | 0.008797169 | 0.053683867 | | | 1700010I14Rik | 44.44877067 | -0.060544879 | 0.092062781 | -0.657647728 | 0.510764518 | 0.85162432 |
| Rabif | 1221.467124 | -0.114279482 | 0.065767776 | -1.737621187 | 0.082277584 | 0.238932488 | | | Lsr | 263.782766 | -0.060560928 | 0.093908386 | -0.644893712 | 0.518996067 | 0.853297835 |
| Slit3 | 1367.408427 | -0.193461997 | 0.111340446 | -1.737571601 | 0.082286327 | 0.238932488 | | | Nfatc4 | 68.46158207 | -0.060571124 | 0.094024154 | -0.644208122 | 0.519440485 | 0.853297835 |
| Cep85 | 634.410626 | 0.180991359 | 0.078674222 | 2.300516666 | 0.021418966 | 0.098505628 | | | Lztr1 | 7065.51499 | -0.060603877 | 0.066772532 | -0.907616884 | 0.364080671 | 0.777191869 |
| 4930577N17Rik | 13.61345547 | 0.180981535 | 0.166799489 | 1.085024517 | 0.277910797 | 0.50694018 | | | Xlr3a | 22.13421947 | -0.060621921 | 0.064972863 | -0.933034465 | 0.350802174 | NA |
| Rbm15b | 1468.056825 | -0.123364246 | 0.071014993 | -1.73715776 | 0.082359328 | 0.238965437 | | | Mmp21 | 3.995002897 | -0.060650601 | 0.044009832 | -1.378114794 | 0.168167849 | NA |
| Scube2 | 60.84118263 | -0.276252144 | 0.159067369 | -1.736699018 | 0.082440311 | 0.239155623 | | | Pabpc1l | 14.03903673 | -0.060663303 | 0.071846806 | -0.844342384 | 0.398478108 | NA |
| Vwa3a | 281.1508617 | 0.18092853 | 0.11891681 | 1.521471435 | 0.128141574 | 0.316541113 | | | Camk2a | 142857.9117 | -0.060687364 | 0.062208175 | -0.975552876 | 0.329286087 | 0.756141871 |
| 4930447N08Rik | 9.062319972 | -0.246309647 | 0.141906086 | -1.735722923 | 0.082612838 | 0.239411823 | | | Zfp773 | 89.22095658 | -0.060742661 | 0.093512442 | -0.649567687 | 0.515971511 | 0.853297835 |
| Creb1 | 2043.444031 | -0.130065769 | 0.074934277 | -1.735731292 | 0.082611358 | 0.239411823 | | | Yod1 | 382.2712303 | -0.06074837 | 0.086242946 | -0.704386536 | 0.481192089 | 0.8371092 |
| Cystm1 | 605.3853037 | -0.133335126 | 0.076814752 | -1.735801045 | 0.082599019 | 0.239411823 | | | Prkg1 | 552.6847202 | -0.060757373 | 0.081223997 | -0.748022445 | 0.454446619 | 0.826168452 |
| Fbxl22 | 11.76143722 | -0.281036788 | 0.16192038 | -1.735648023 | 0.082626089 | 0.239411823 | | | Tbxa2r | 33.72619568 | -0.060785633 | 0.081062551 | -0.749860849 | 0.453338516 | 0.82562629 |
| Wdr13 | 9471.838789 | -0.098860436 | 0.056960789 | -1.735587542 | 0.08263679 | 0.239411823 | | | Eri2 | 456.7502944 | -0.060813953 | 0.072201035 | -0.842286447 | 0.399627636 | 0.798421909 |
| Zfp712 | 238.0018675 | -0.188751773 | 0.108749362 | -1.735658666 | 0.082624206 | 0.239411823 | | | Acot6 | 144.7968604 | -0.060816807 | 0.087559083 | -0.694580223 | 0.48731841 | 0.839801621 |
| 2500004C02Rik | 228.0279746 | -0.170396755 | 0.098198953 | -1.735219684 | 0.082701901 | 0.239496535 | | | Ikzf4 | 523.2407709 | -0.060838864 | 0.07387851 | -0.823498792 | 0.410224404 | 0.80184319 |
| Gdf10 | 299.4413777 | 0.180914617 | 0.110428646 | 1.638294263 | 0.101360324 | 0.272471248 | | | Fadd | 124.0480374 | -0.060862649 | 0.094990008 | -0.640726852 | 0.521700165 | 0.854113377 |
| Mrvi1 | 167.324469 | -0.214155939 | 0.123427618 | -1.735073092 | 0.08272786 | 0.239496535 | | | Npas4 | 1134.251097 | -0.06091809 | 0.062387409 | -0.97644848 | 0.328842267 | 0.755816895 |
| Zbed3 | 367.3401818 | 0.180911127 | 0.077523207 | 2.333638314 | 0.019614669 | 0.092938739 | | | Ccdc69 | 14.75189616 | -0.060934128 | 0.072183505 | -0.844155857 | 0.398582318 | NA |
| Tmprss6 | 23.56131969 | 0.180896363 | 0.159637955 | 1.133166375 | 0.257144392 | 0.483753668 | | | Dhrs4 | 354.0008292 | -0.060948894 | 0.075559877 | -0.80663041 | 0.419879448 | 0.807657282 |
| Mapk1ip1l | 1709.227097 | -0.105331999 | 0.06075374 | -1.733753325 | 0.08296186 | 0.240084249 | | | Il13ra1 | 281.3380115 | -0.060953693 | 0.089950807 | -0.677633649 | 0.498004006 | 0.845489269 |
| Casp8 | 110.4107026 | 0.18086627 | 0.138692274 | 1.304083233 | 0.192205205 | 0.406185958 | | | Pnpla7 | 852.3864018 | -0.061041267 | 0.071231023 | -0.856947787 | 0.391473741 | 0.794184423 |
| Fgf9 | 733.2931817 | 0.180801991 | 0.069343119 | 2.607353033 | 0.009124523 | 0.0551171 | | | Ppib | 3046.794178 | -0.061054185 | 0.054469398 | -1.120889659 | 0.262334833 | 0.704810511 |
| Tdp1 | 546.7199545 | 0.180791819 | 0.094450863 | 1.91413623 | 0.055602751 | 0.185870341 | | | Gm9776 | 129.0754226 | -0.061055995 | 0.094540793 | -0.645816405 | 0.518398262 | 0.853297835 |
| Hepacam2 | 13.35046926 | 0.18071809 | 0.165186849 | 1.094022256 | 0.273945217 | 0.502189682 | | | Lyrm9 | 1951.887335 | -0.061057636 | 0.056297373 | -1.084555693 | 0.278118488 | 0.718319852 |
| Cand1 | 6275.053374 | -0.102570919 | 0.059221644 | -1.731983644 | 0.083276475 | 0.240765129 | | | Epor | 329.2007869 | -0.061067939 | 0.077554931 | -0.787415289 | 0.4310388 | 0.812181606 |
| Rpl13 | 6060.979544 | -0.145344054 | 0.083921466 | -1.731905573 | 0.083290376 | 0.240765129 | | | Adarb2 | 512.1500155 | -0.061090113 | 0.077627855 | -0.786961234 | 0.43130456 | 0.812181606 |
| Rundc3b | 1526.247285 | -0.12809498 | 0.073967848 | -1.731765676 | 0.083315292 | 0.240792227 | | | Ctf1 | 58.91079099 | -0.061102176 | 0.0945188 | -0.646455265 | 0.517984559 | 0.853297835 |
| Clpp | 1210.58312 | -0.106611847 | 0.061567876 | -1.731614828 | 0.083342164 | 0.240824971 | | | BC024139 | 62.59192886 | -0.061147657 | 0.090964551 | -0.672214139 | 0.501447381 | 0.846601651 |
| Echdc3 | 23.31948962 | 0.180587445 | 0.167500472 | 1.078130961 | 0.280975317 | 0.510715963 | | | Map2k7 | 4069.945911 | -0.061152688 | 0.051367286 | -1.19049871 | 0.233850437 | 0.685064073 |
| Gm14827 | 186.0112022 | 0.180362093 | 0.164830266 | 1.094229221 | 0.273854459 | 0.502142128 | | | Fam105a | 890.7173033 | -0.061166827 | 0.067092722 | -0.911676039 | 0.361939281 | 0.776449324 |
| Zmiz2 | 13108.27426 | 0.180321843 | 0.105016855 | 1.71707525 | 0.085965424 | 0.245204027 | | | Haus1 | 254.1824458 | -0.061203668 | 0.08449499 | -0.724346718 | 0.468852902 | 0.832586462 |
| Smu1 | 2437.84386 | -0.099688795 | 0.05758353 | -1.73120326 | 0.083415518 | 0.240857981 | | | Ift20 | 1350.604697 | -0.061227366 | 0.054941777 | -1.114404536 | 0.265105678 | 0.707370515 |
| Dnajc12 | 389.7266943 | -0.164868018 | 0.095252804 | -1.730846874 | 0.083479079 | 0.240950978 | | | Cdk10 | 2421.355817 | -0.061235067 | 0.071527172 | -0.856109155 | 0.391937405 | 0.794184423 |
| Ugcg | 2867.810196 | -0.115452817 | 0.066702757 | -1.730855248 | 0.083477585 | 0.240950978 | | | Ankmy1 | 15.58967802 | -0.061247208 | 0.071015772 | -0.862445144 | 0.388442607 | NA |
| Cdk5rap3 | 887.2964331 | 0.18031411 | 0.070933567 | 2.54201386 | 0.01102158 | 0.062813732 | | | Wnt8b | 13.53837413 | -0.061261781 | 0.071496822 | -0.85684621 | 0.391529883 | NA |
| Cep250 | 1589.249899 | 0.180283868 | 0.076463223 | 2.357785361 | 0.018384319 | 0.08923767 | | | Sin3b | 3336.217132 | -0.061271242 | 0.058377003 | -1.049578403 | 0.293911991 | 0.729288721 |
| Haus2 | 804.8281369 | -0.111951843 | 0.064721066 | -1.729758944 | 0.083673353 | 0.241376824 | | | 4930581F22Rik | 111.6509545 | -0.061275058 | 0.095080303 | -0.644455856 | 0.519279874 | 0.853297835 |
| Mapk7 | 706.7302424 | 0.180048277 | 0.10569976 | 1.703393431 | 0.088494466 | 0.249660098 | | | Prr24 | 1075.278677 | -0.061299354 | 0.056384774 | -1.087161463 | 0.276965458 | 0.7177927 |
| Fam189a2 | 454.9321733 | 0.179894717 | 0.087601767 | 2.053551242 | 0.040019145 | 0.149735967 | | | Zp1 | 3.752133087 | -0.061299669 | 0.041992126 | -1.459789597 | 0.144347909 | NA |
| Cacna1s | 51.44499538 | -0.277099445 | 0.160303836 | -1.728588988 | 0.083882682 | 0.24166991 | | | Rere | 6847.793228 | -0.061348513 | 0.058363671 | -1.051142125 | 0.293193319 | 0.728769021 |
| Dhrs3 | 584.3765511 | 0.179884572 | 0.08476699 | 2.12210641 | 0.033828803 | 0.134120644 | | | Zdhhc21 | 2053.886243 | -0.061369434 | 0.070384436 | -0.871917681 | 0.383253287 | 0.787602461 |
| Chst2 | 5698.599544 | -0.167252404 | 0.096785849 | -1.728066711 | 0.083976265 | 0.24166991 | | | Prss45 | 12.94196397 | -0.06139966 | 0.072782048 | -0.84360995 | 0.398887402 | NA |
| Gdpd3 | 18.75690575 | 0.179845756 | 0.168120948 | 1.069740316 | 0.284736214 | 0.514480389 | | | Mir128-1 | 6.764855657 | -0.061405509 | 0.054644111 | -1.123735177 | 0.261125389 | NA |
| Hist3h2a | 897.803703 | -0.164007138 | 0.094862687 | -1.728889871 | 0.083828807 | 0.24166991 | | | Fcrl1 | 16.69972909 | -0.06153515 | 0.072603922 | -0.847545814 | 0.396690962 | NA |
| 1600002H07Rik | 318.8052919 | 0.179739272 | 0.103823405 | 1.731201853 | 0.083415769 | 0.240857981 | | | Atf1 | 402.4124264 | -0.061573796 | 0.072838633 | -0.845345298 | 0.397918077 | 0.797789838 |
| Gm8234 | 17.58122535 | 0.179671455 | 0.157474551 | 1.140955504 | 0.25388844 | 0.479783814 | | | Gipc3 | 30.12514509 | -0.061592944 | 0.088970511 | -0.692284936 | 0.488758408 | 0.840500134 |
| Nudt11 | 826.3418739 | -0.13450816 | 0.077828706 | -1.728258976 | 0.083941805 | 0.24166991 | | | BC005764 | 2626.506919 | -0.061644891 | 0.070042141 | -0.880111467 | 0.378798928 | 0.785380516 |
| Megf6 | 260.4576749 | 0.17966712 | 0.123097742 | 1.459548457 | 0.144414214 | 0.341924485 | | | Mir125a | 5.686836025 | -0.061676583 | 0.046075993 | -1.338583927 | 0.180706161 | NA |
| Ptplb | 1411.394056 | 0.179649834 | 0.095824379 | 1.874782145 | 0.0608227 | 0.19612631 | | | Ncan | 10594.13281 | -0.061684972 | 0.055398473 | -1.113477838 | 0.265503262 | 0.707651445 |
| Vav2 | 364.429964 | -0.177733745 | 0.102851689 | -1.728058593 | 0.08397772 | 0.24166991 | | | Slco2a1 | 87.21819133 | -0.061688635 | 0.094384068 | -0.653591616 | 0.513374947 | 0.853195313 |
| Eif4b | 8814.46401 | 0.179647409 | 0.094905857 | 1.892901171 | 0.058371013 | 0.191610272 | | | Phyhd1 | 579.8196451 | -0.06168976 | 0.088287639 | -0.698736099 | 0.48471697 | 0.838713365 |
| Noxo1 | 58.59938679 | 0.179640091 | 0.157822823 | 1.138238992 | 0.255020701 | 0.481364938 | | | Ppm1k | 3342.615244 | -0.061698029 | 0.072219351 | -0.854314367 | 0.392930829 | 0.79461412 |
| Prrc2c | 7871.128241 | -0.150285035 | 0.086993979 | -1.727533743 | 0.084071851 | 0.241761183 | | | Otop2 | 53.48251751 | -0.061700165 | 0.081015526 | -0.76158445 | 0.446308058 | 0.822210607 |
| Zfp706 | 7503.403422 | -0.115734692 | 0.066992187 | -1.727584914 | 0.084062669 | 0.241761183 | | | Ttc4 | 2070.909051 | -0.061702583 | 0.047448046 | -1.300424109 | 0.193455651 | 0.649744561 |
| 2610019E17Rik | 67.25870507 | 0.179632898 | 0.15308875 | 1.173390586 | 0.240639211 | 0.465243815 | | | E130307A14Rik | 594.7339284 | -0.06172431 | 0.065497939 | -0.942385525 | 0.345995295 | 0.765097832 |
| Gm715 | 106.2922074 | -0.204186773 | 0.118258931 | -1.72660764 | 0.084238153 | 0.242149526 | | | Creb3l3 | 21.70365303 | -0.061724423 | 0.079225179 | -0.779101082 | 0.435920173 | NA |
| Sdc3 | 9385.650652 | -0.107325703 | 0.062164935 | -1.726466918 | 0.084263446 | 0.242177301 | | | Gm16386 | 26.48522602 | -0.061759295 | 0.083590623 | -0.738830418 | 0.460009979 | 0.829022384 |
| Tmem55b | 2363.998148 | 0.078968194 | 0.045747731 | 1.726166362 | 0.084317488 | 0.242287677 | | | Lars2 | 56155.43219 | -0.061786428 | 0.085105231 | -0.726000355 | 0.467838553 | 0.832353544 |
| Phf1 | 1700.216856 | 0.179616256 | 0.075278112 | 2.386035613 | 0.017031102 | 0.084495642 | | | Vmn2r87 | 31.99456273 | -0.061834892 | 0.084573968 | -0.731133864 | 0.464697391 | 0.83159896 |
| Sp100 | 132.5544189 | 0.179542905 | 0.144868156 | 1.239353837 | 0.215214489 | 0.435033698 | | | Tgfa | 2346.638061 | -0.061920614 | 0.070286318 | -0.880976782 | 0.378330387 | 0.785201772 |
| Ash1l | 7780.332323 | -0.184987166 | 0.107203665 | -1.725567561 | 0.084425239 | 0.242417469 | | | Srgap2 | 3935.287052 | -0.061930038 | 0.055616816 | -1.113512817 | 0.265488247 | 0.707651445 |
| Gm16701 | 85.94746744 | -0.233393204 | 0.135251452 | -1.72562439 | 0.084415008 | 0.242417469 | | | Aptx | 1455.9738 | -0.061937714 | 0.051017397 | -1.214050849 | 0.224728307 | 0.676420098 |
| Apbb1ip | 237.252086 | 0.179488293 | 0.1348604 | 1.330919184 | 0.183215604 | 0.394060885 | | | C130036L24Rik | 25.72832874 | -0.061947841 | 0.083467338 | -0.742180632 | 0.457977902 | 0.827642075 |
| Smad4 | 1619.785957 | -0.1297286 | 0.075224023 | -1.724563444 | 0.084606176 | 0.242846998 | | | Cd300a | 62.86571438 | -0.061983023 | 0.094847069 | -0.653504891 | 0.513430837 | 0.853195313 |
| Tyms | 144.0790501 | 0.179437363 | 0.131837841 | 1.361045975 | 0.173499161 | 0.381683373 | | | Zfyve27 | 3661.631997 | -0.062000996 | 0.060696742 | -1.021488046 | 0.30702327 | 0.739163119 |
| Ptprm | 1892.207669 | -0.113742451 | 0.065970605 | -1.724138365 | 0.084682868 | 0.242973448 | | | Siglec1 | 29.15343215 | -0.062002656 | 0.069171059 | -0.896367017 | 0.370056777 | 0.780397655 |
| U2af2 | 5550.764923 | -0.071339787 | 0.041377253 | -1.724130594 | 0.08468427 | 0.242973448 | | | D830015G02Rik | 12.83432344 | -0.062004419 | 0.068817872 | -0.900992981 | 0.367592052 | NA |

| | | | | | | |
|---|---|---|---|---|---|---|
| Ppp1r14c | 389.274034 | -0.146594941 | 0.085038445 | -1.723866681 | 0.084731913 | 0.243027817 |
| Gdf11 | 315.2618099 | -0.181563704 | 0.105331166 | -1.723741517 | 0.084754516 | 0.243047655 |
| Dr1 | 986.7148141 | -0.116842897 | 0.067796968 | -1.723423622 | 0.084811946 | 0.243122348 |
| Galnt12 | 55.70476269 | 0.179416384 | 0.157278959 | 1.140752616 | 0.253972884 | 0.479797818 |
| Dbc1 | 9205.968381 | 0.179412697 | 0.081470124 | 2.202190065 | 0.027651886 | 0.117179083 |
| Sema4f | 2283.982794 | 0.179410527 | 0.085560811 | 2.096877372 | 0.036004432 | 0.139532934 |
| Plscr1 | 74.9228647 | -0.240693305 | 0.139734687 | -1.722502196 | 0.084978585 | 0.243464904 |
| 2810021J22Rik | 496.7967671 | -0.127263983 | 0.07389278 | -1.722278991 | 0.085018992 | 0.243535636 |
| Gm527 | 134.3110618 | 0.179355663 | 0.11782758 | 1.522187452 | 0.127962118 | 0.316362703 |
| Fam216a | 1427.014866 | -0.114429891 | 0.066451444 | -1.722007576 | 0.085068146 | 0.243586371 |
| Arl16 | 781.8319362 | -0.118182119 | 0.068648342 | -1.721558244 | 0.085149573 | 0.243774479 |
| Zrsr1 | 3262.177632 | 0.179349552 | 0.081940475 | 2.188778518 | 0.028612942 | 0.119883429 |
| Socs3 | 134.8086578 | 0.179299625 | 0.137114558 | 1.307662931 | 0.190987662 | 0.404854183 |
| Armc5 | 600.5726449 | 0.179284029 | 0.069391078 | 2.583675501 | 0.009775372 | 0.057809012 |
| Gm410 | 61.60330962 | 0.179169572 | 0.159916445 | 1.120394914 | 0.262545511 | 0.489693048 |
| Ywhaz | 43501.72649 | -0.10130076 | 0.058865134 | -1.720895766 | 0.08526974 | 0.243893179 |
| Qser1 | 734.9057902 | -0.156089326 | 0.090712843 | -1.720697105 | 0.085305802 | 0.243951291 |
| Klhl21 | 780.5739772 | 0.178947664 | 0.08517621 | 2.100911325 | 0.035648751 | 0.138605207 |
| Iqce | 1185.993373 | 0.178945935 | 0.100102602 | 1.787625222 | 0.073836493 | 0.222724827 |
| Klhdc8b | 953.6710966 | -0.170224476 | 0.098948728 | -1.720330112 | 0.085372453 | 0.244006785 |
| Opn1mw | 11.9270485 | 0.178874493 | 0.164240407 | 1.089101618 | 0.276109077 | 0.504901927 |
| Efemp1 | 528.4960136 | -0.19802695 | 0.115179445 | -1.719290707 | 0.085561451 | 0.244481779 |
| Xrcc5 | 769.0484957 | 0.17878257 | 0.077865796 | 2.296034716 | 0.021673897 | 0.099324269 |
| Gm5431 | 8.869499416 | 0.178721378 | 0.152430469 | 1.172478047 | 0.241005181 | 0.465428677 |
| Rfk | 2072.416895 | -0.108640251 | 0.063231761 | -1.718127875 | 0.085773293 | 0.244912142 |
| Rnf167 | 2517.952094 | 0.087490967 | 0.050924017 | 1.718068848 | 0.085784058 | 0.244912142 |
| BC026590 | 1112.873473 | 0.096096911 | 0.055941145 | 1.717821653 | 0.085829151 | 0.244983891 |
| Upk1b | 52.28559402 | 0.178717541 | 0.166424071 | 1.073868345 | 0.282881693 | 0.512567997 |
| Atg9a | 6782.900769 | 0.178708864 | 0.072393574 | 2.468573596 | 0.013565275 | 0.072520404 |
| Tmprss4 | 8.581361245 | 0.178545787 | 0.145897873 | 1.223772379 | 0.221038107 | 0.442129311 |
| Mecr | 430.2632265 | 0.178319266 | 0.098714091 | 1.806421593 | 0.070852511 | 0.216994211 |
| Cnr1 | 6686.17567 | -0.149898659 | 0.087337256 | -1.716319781 | 0.08610353 | 0.24546239 |
| Supt20 | 764.9853289 | 0.178301541 | 0.118074315 | 1.510078975 | 0.131023274 | 0.32078078 |
| Prkce | 17906.87353 | -0.12173971 | 0.070926253 | -1.716426645 | 0.086083983 | 0.24546239 |
| Clip3 | 22367.47786 | 0.178267084 | 0.08733362 | 2.041219443 | 0.04122902 | 0.153300321 |
| Cav1 | 487.2728416 | -0.214382734 | 0.124954418 | -1.715687507 | 0.086219252 | 0.245701876 |
| Tdrd12 | 10.45086136 | -0.275340066 | 0.160500396 | -1.715510194 | 0.086251728 | 0.245749223 |
| Abcd2 | 1977.864815 | 0.178048968 | 0.080422716 | 2.213913886 | 0.026834703 | 0.114697968 |
| Ch25h | 14.22578733 | 0.178042774 | 0.153831023 | 1.157391859 | 0.247112299 | 0.472061549 |
| Scarf1 | 81.19007347 | -0.222305284 | 0.129599506 | -1.715325079 | 0.086285643 | 0.245799414 |
| Nudt9 | 1537.403646 | 0.17800782 | 0.074110374 | 2.401928507 | 0.016308895 | 0.082470635 |
| Tmem243 | 192.4514313 | 0.17749012 | 0.129376483 | 1.371888583 | 0.170098125 | 0.377591627 |
| Engase | 400.3477346 | 0.177444989 | 0.099202984 | 1.788706166 | 0.073662149 | 0.222349765 |
| Btbd17 | 479.3951368 | -0.263238497 | 0.153538218 | -1.714481912 | 0.086440256 | 0.24595758 |
| Slc6a13 | 430.6781508 | -0.244530653 | 0.142631814 | -1.714418729 | 0.086451851 | 0.24595758 |
| Gpd2 | 4965.184424 | -0.126180885 | 0.073613728 | -1.714094474 | 0.086511376 | 0.246081745 |
| Eif1a | 1454.699863 | -0.085229881 | 0.049728924 | -1.713889515 | 0.086549019 | 0.246143631 |
| Gpank1 | 502.0097412 | 0.177390287 | 0.083526005 | 2.123773155 | 0.033689117 | 0.133702991 |
| AI317395 | 30.83641502 | 0.177339942 | 0.165256812 | 1.07311729 | 0.283218494 | 0.512698604 |
| Kcnh6 | 112.8628633 | -0.228815708 | 0.133606253 | -1.712612263 | 0.086783896 | 0.246654237 |
| Armcx5 | 938.1143283 | 0.177207274 | 0.102225371 | 1.733496016 | 0.083007545 | 0.240171597 |
| Eva1c | 63.83711738 | 0.177202167 | 0.158713713 | 1.116489326 | 0.264212737 | 0.49160707 |
| Lphn1 | 26793.65873 | -0.112266258 | 0.065560935 | -1.712395626 | 0.086823785 | 0.246670149 |
| Syn2 | 21412.88207 | -0.116800739 | 0.068230399 | -1.711857763 | 0.086922884 | 0.246889673 |
| Anxa3 | 449.5001652 | 0.177199447 | 0.093913798 | 1.886830799 | 0.059183087 | 0.193252424 |
| Uba6 | 811.0414721 | -0.144619229 | 0.084502152 | -1.711426575 | 0.087002395 | 0.247012669 |
| Zfp935 | 241.3951015 | -0.234630026 | 0.137101214 | -1.711363586 | 0.087014015 | 0.247012669 |
| Il2ra | 11.75439319 | 0.177050731 | 0.158531544 | 1.116817048 | 0.264072558 | 0.491417273 |
| Cnksr2 | 9288.689169 | -0.12679058 | 0.07411365 | -1.71075881 | 0.087125646 | 0.247103197 |
| Cd2ap | 926.1391301 | 0.177029037 | 0.098885809 | 1.790237035 | 0.073415814 | 0.221939958 |
| Stip1 | 6665.272649 | -0.1224701 | 0.071585376 | -1.7108257 | 0.087113294 | 0.247103197 |
| Tmem198 | 2238.030794 | -0.1770627 | 0.103497711 | -1.710788562 | 0.087120152 | 0.247103197 |
| Gm5069 | 199.2472907 | -0.206926675 | 0.120973724 | -1.710509266 | 0.087171741 | 0.247188683 |
| Slit1 | 2332.836669 | 0.177013416 | 0.109810421 | 1.611991043 | 0.106963889 | 0.282977548 |
| Med11 | 249.5512667 | -0.180696107 | 0.105675843 | -1.709909299 | 0.087282647 | 0.24741261 |
| Ctsb | 25983.30086 | 0.1769532 | 0.068092287 | 2.598726034 | 0.009357042 | 0.056127761 |
| Car5a | 10.59898722 | -0.273371719 | 0.159906345 | -1.709573928 | 0.08734469 | 0.247497921 |
| Cecr2 | 180.9749866 | 0.176848234 | 0.124687922 | 1.418326908 | 0.156095344 | 0.358438532 |
| Map3k8 | 52.985293 | 0.176807026 | 0.144828555 | 1.220802251 | 0.222160901 | 0.443251547 |
| Mmadhc | 1357.367012 | -0.087956001 | 0.051460653 | -1.709189364 | 0.087415878 | 0.247630198 |
| Trim11 | 745.8333968 | 0.176780221 | 0.080242685 | 2.203069626 | 0.027589842 | 0.116956773 |
| Vps51 | 1899.31038 | 0.095968191 | 0.056163467 | 1.708729821 | 0.087501008 | 0.247729944 |
| Lrdd | 54.02416605 | 0.176713671 | 0.162946616 | 1.084488131 | 0.278148427 | 0.507194507 |
| Atp6v0b | 6473.483573 | -0.105527731 | 0.061782244 | -1.708059208 | 0.087625357 | 0.247946635 |
| 1810055G02Rik | 691.3240904 | 0.176695464 | 0.080720275 | 2.188984924 | 0.028597936 | 0.119878408 |
| Fermt3 | 173.6451502 | 0.176687656 | 0.119728251 | 1.475739053 | 0.140013958 | 0.334412122 |
| Cldn25 | 2262.204554 | -0.099565007 | 0.058338558 | -1.706675832 | 0.087882322 | 0.248566607 |
| Wipf3 | 7852.592047 | 0.176668651 | 0.13171437 | 1.341301266 | 0.179822659 | 0.390056318 |
| Tmem117 | 260.3696503 | -0.204823427 | 0.120036562 | -1.706341994 | 0.087944424 | 0.248651624 |
| Ptpn9 | 1957.263255 | -0.133314797 | 0.078142011 | -1.706057933 | 0.087997294 | 0.248755714 |
| Cox7a2l | 5000.333194 | -0.092935886 | 0.054477911 | -1.705937038 | 0.088019803 | 0.248773956 |
| E2f2 | 31.56679426 | -0.282757058 | 0.165762538 | -1.705795903 | 0.088046086 | 0.248802856 |
| Ttk | 16.13728722 | -0.285761127 | 0.167535658 | -1.705673472 | 0.088068891 | 0.248821919 |
| Gimap1 | 50.30328664 | 0.176623084 | 0.151663605 | 1.164571318 | 0.24419256 | 0.469017976 |
| Mamld1 | 1249.894525 | 0.176340991 | 0.113427233 | 1.554661844 | 0.120026626 | 0.304817269 |
| Txlng | 457.8718545 | 0.176306458 | 0.111985731 | 1.574365378 | 0.115403003 | 0.297562678 |
| Acap2 | 3723.617134 | 0.17621286 | 0.072911864 | 2.416792696 | 0.015657932 | 0.080313518 |
| Zfp574 | 1180.306197 | 0.176115369 | 0.075337848 | 2.337674542 | 0.019404138 | 0.092251522 |
| Ascl5 | 8.90921616 | 0.176095055 | 0.151945671 | 1.158934334 | 0.246482953 | 0.471424129 |
| Ltc4s | 78.68254962 | 0.176001766 | 0.141581966 | 1.243108647 | 0.2138278 | 0.433588265 |
| Sema4b | 661.954986 | 0.17598509 | 0.128374582 | 1.370871768 | 0.170414934 | 0.377840546 |
| Pmm2 | 636.2002941 | 0.175984287 | 0.067957567 | 2.58962018 | 0.009608188 | 0.057103668 |
| Mybpc3 | 26.50999208 | 0.175763283 | 0.167115523 | 1.051747197 | 0.292915551 | 0.52352446 |
| Tcof1 | 1188.179985 | 0.1757286 | 0.069892616 | 2.514265597 | 0.011928057 | 0.066566805 |
| Hvcn1 | 180.4419663 | 0.175687217 | 0.135376389 | 1.297768524 | 0.194366889 | 0.408911918 |
| Itpripl2 | 394.3116563 | -0.148583038 | 0.087288735 | -1.702201751 | 0.088717555 | 0.250061814 |
| Stmn4 | 7949.148062 | 0.175676529 | 0.082069525 | 2.140581779 | 0.032307779 | 0.129817618 |
| Smim8 | 326.08302 | -0.177342641 | 0.104232001 | -1.701422203 | 0.088863735 | 0.250382744 |
| Mchr1 | 935.5354576 | -0.185497225 | 0.109037848 | -1.701218684 | 0.088901931 | 0.250444821 |
| Tbc1d24 | 4934.773546 | 0.095244628 | 0.055990611 | 1.701082132 | 0.088927566 | 0.250471497 |
| Oat | 3123.510016 | -0.081749545 | 0.048065735 | -1.700786337 | 0.088983116 | 0.25053687 |
| Pyurf | 790.8390421 | -0.163091188 | 0.095887967 | -1.700851447 | 0.088970886 | 0.25053687 |
| Ccnk | 1079.140416 | 0.175585103 | 0.069050189 | 2.542862004 | 0.010994864 | 0.062791901 |
| Cela1 | 73.30835476 | 0.175558891 | 0.141584095 | 1.23996195 | 0.214989468 | 0.434748998 |
| Zfp879 | 106.5962502 | 0.175495497 | 0.130330671 | 1.34654027 | 0.178128342 | 0.387629289 |
| Ube2ql1 | 7009.672199 | -0.122611389 | 0.072186557 | -1.698534949 | 0.089406842 | 0.251547052 |
| Fbxl15 | 443.9386806 | -0.159324198 | 0.093817997 | -1.698226387 | 0.089465042 | 0.251665099 |
| Laptm4a | 6182.679753 | -0.070823991 | 0.041708299 | -1.698079089 | 0.089492836 | 0.251697585 |
| Hcrtr2 | 71.45472533 | -0.228224846 | 0.134426194 | -1.697770644 | 0.089551059 | 0.251815626 |
| Myt1l | 10664.48447 | -0.114453512 | 0.067430982 | -1.697343091 | 0.089631815 | 0.251996977 |
| Iqgap2 | 1292.417988 | 0.17548709 | 0.131951913 | 1.329932145 | 0.183540629 | 0.394528903 |
| Zfp51 | 189.8590095 | 0.175468017 | 0.117140407 | 1.497929033 | 0.13415169 | 0.325535831 |
| Nlgn1 | 2666.56176 | -0.139480346 | 0.082241428 | -1.695986435 | 0.089888449 | 0.2525352 |
| BC037034 | 2044.917709 | 0.175448672 | 0.078993158 | 2.221061626 | 0.026346788 | 0.11314222 |
| Khdrbs1 | 3962.977145 | -0.120122518 | 0.070834017 | -1.695830939 | 0.089917902 | 0.252572148 |
| Gulp1 | 84.21214276 | -0.238828431 | 0.140861721 | -1.695481419 | 0.089984132 | 0.25271237 |
| Fcgrt | 623.3309307 | -0.192368948 | 0.113488766 | -1.695048368 | 0.090066246 | 0.252897139 |
| Eif1 | 11852.91944 | -0.084580022 | 0.049904135 | -1.694849989 | 0.090103882 | 0.252956975 |
| Rtkn | 2199.586687 | 0.175367651 | 0.073331748 | 2.391428738 | 0.016782942 | 0.083801127 |
| Clptm1 | 7050.626363 | -0.091941924 | 0.054254783 | -1.694632594 | 0.09014514 | 0.253004419 |
| Ints9 | 616.16819 | 0.175365434 | 0.071480145 | 2.453344691 | 0.014153464 | 0.074793239 |
| Nipa1 | 1966.409344 | -0.115504272 | 0.068170053 | -1.694355037 | 0.090197838 | 0.253004419 |
| Spc24 | 64.44160373 | -0.235616644 | 0.139041758 | -1.69457469 | 0.090156132 | 0.253004419 |
| Pcdhga4 | 489.5401906 | 0.175197408 | 0.079781162 | 2.195974624 | 0.02809376 | 0.118434034 |
| Eif2d | 1019.429815 | 0.175129687 | 0.070527406 | 2.483143726 | 0.013022853 | 0.070551079 |
| Zdhhc1 | 1347.672259 | 0.175090026 | 0.094219643 | 1.858317667 | 0.063123916 | 0.20099506 |
| Tsga14 | 300.2737847 | -0.230782144 | 0.136260613 | -1.693681966 | 0.090325733 | 0.253166952 |
| Rps15a-ps4 | 18.00614132 | -0.281231652 | 0.16609747 | -1.693172398 | 0.090422657 | 0.253392774 |
| Triobp | 359.9888177 | -0.146389837 | 0.086478298 | -1.692792748 | 0.090494924 | 0.25354943 |
| Lmln | 482.4323135 | 0.175087779 | 0.097480622 | 1.796129059 | 0.072473994 | 0.220302053 |
| Nek6 | 1075.976128 | 0.174827436 | 0.08219217 | 2.127057075 | 0.033415341 | 0.133026027 |
| Bbs1 | 2222.538176 | -0.096605983 | 0.057110658 | -1.691557859 | 0.090730307 | 0.253890779 |
| Chuk | 1179.600658 | 0.174758284 | 0.128062176 | 1.364636217 | 0.172367416 | 0.38016523 |
| Clec2l | 1261.473911 | -0.104337716 | 0.06168593 | -1.691434584 | 0.090753832 | 0.253890779 |
| Gpn3 | 475.815018 | 0.174698591 | 0.086934563 | 2.009541265 | 0.044479763 | 0.161209228 |
| Cd37 | 131.2718526 | 0.174691872 | 0.15566509 | 1.122228956 | 0.261765104 | 0.488863021 |
| Lyrm1 | 243.3445357 | -0.168175872 | 0.099456247 | -1.690953319 | 0.090845719 | 0.253890779 |
| Slc45a1 | 2084.392877 | 0.174520166 | 0.084789659 | 2.058271825 | 0.039564047 | 0.148643392 |
| Nmnat1 | 130.6143512 | -0.208403461 | 0.123252558 | -1.690865199 | 0.090862551 | 0.253890779 |
| Numb | 1976.209571 | -0.113268722 | 0.06698573 | -1.690938089 | 0.090848628 | 0.253890779 |
| Pik3ip1 | 1244.090556 | 0.17450964 | 0.073843736 | 2.36322875 | 0.018116483 | 0.088380602 |
| Sema4a | 2258.716149 | -0.182332175 | 0.107809793 | -1.691239455 | 0.090791078 | 0.253890779 |
| Slc30a1 | 1495.440478 | -0.178173011 | 0.105356867 | -1.691138086 | 0.090810433 | 0.253890779 |
| Smap2 | 7926.175978 | -0.091111014 | 0.053880377 | -1.690986944 | 0.090839296 | 0.253890779 |
| Ttyh3 | 10576.59309 | -0.102344423 | 0.060543742 | -1.69042116 | 0.09094741 | 0.254082071 |
| Zfp292 | 1653.139393 | -0.144323409 | 0.085389287 | -1.690181681 | 0.090993202 | 0.254164174 |
| H2-T24 | 76.18856033 | 0.174491835 | 0.165514419 | 1.054239475 | 0.291773291 | 0.521904338 |
| Exoc4 | 2710.992845 | -0.114294124 | 0.067638051 | -1.689790317 | 0.091068077 | 0.254235817 |
| Mlh3 | 1253.078643 | -0.097877938 | 0.057921284 | -1.689844075 | 0.091057789 | 0.254235817 |
| Thrsp | 494.995604 | 0.174464957 | 0.115182467 | 1.514683282 | 0.12985263 | 0.319051166 |
| Ccnd1 | 2340.566587 | -0.165080195 | 0.097754083 | -1.68872941 | 0.091271297 | 0.254676254 |
| Itga3 | 861.3854862 | 0.174372866 | 0.089557344 | 1.947052675 | 0.05152842 | 0.177299875 |
| Fbxo9 | 2856.870341 | -0.171881828 | 0.101784533 | -1.688683173 | 0.091280162 | 0.254676254 |
| Mysm1 | 1109.88114 | 0.1743695 | 0.075001166 | 2.324890528 | 0.020077815 | 0.094532944 |
| B430306N03Rik | 12.64067679 | 0.174293585 | 0.158970597 | 1.096388817 | 0.272908656 | 0.50135721 |
| Narg2 | 300.9766093 | 0.174282357 | 0.117654115 | 1.481311191 | 0.138523668 | 0.332190947 |
| Arvcf | 1406.994098 | 0.174154163 | 0.078877689 | 2.207901433 | 0.027251145 | 0.11600752 |
| Gpr50 | 20.09085074 | -0.271576234 | 0.160882254 | -1.688043445 | 0.091402889 | 0.254729227 |
| Zfp358 | 1033.27465 | 0.17414886 | 0.086937125 | 2.003158706 | 0.045160256 | 0.162989288 |
| 4632415K11Rik | 375.4798491 | -0.146112754 | 0.086567871 | -1.687840443 | 0.091441862 | 0.254774358 |
| Dgkk | 156.548618 | -0.248187396 | 0.147064732 | -1.687606488 | 0.091486793 | 0.254853717 |
| Cox6a2 | 146.8647848 | 0.174131808 | 0.135511334 | 1.284998106 | 0.198792946 | 0.414883877 |
| Snhg5 | 241.0829528 | 0.174127334 | 0.122303385 | 1.423732741 | 0.154523852 | 0.356106268 |
| Eif4a2 | 22339.59389 | -0.095587061 | 0.056681885 | -1.686377582 | 0.091723097 | 0.255236649 |
| Smim13 | 10071.96034 | -0.154590836 | 0.091669678 | -1.686390072 | 0.091720692 | 0.255236649 |
| Aven | 293.4004143 | 0.174125292 | 0.101883496 | 1.70906279 | 0.087439319 | 0.247630198 |
| Atp11c | 418.8469482 | 0.174097972 | 0.100453873 | 1.733113586 | 0.083075482 | 0.240323286 |
| Chchd6 | 1272.479513 | -0.156476816 | 0.092827899 | -1.685665814 | 0.091860185 | 0.255572222 |
| Pabpc1l | 14.42429854 | 0.174079906 | 0.154281651 | 1.128325402 | 0.259182519 | 0.485818287 |
| Lat2 | 86.05372282 | 0.174079649 | 0.156467072 | 1.112564113 | 0.26589568 | 0.493565074 |
| Mrpl3 | 1772.663144 | -0.110929785 | 0.065846875 | -1.684662873 | 0.092053633 | 0.255766056 |
| Pdss2 | 349.9318552 | -0.149079277 | 0.08846996 | -1.685083583 | 0.091972447 | 0.255766056 |
| Slc25a36 | 1231.374073 | 0.174070295 | 0.070675618 | 2.462946892 | 0.013780033 | 0.073357204 |
| Samd12 | 271.7780415 | -0.186682655 | 0.110811357 | -1.684688816 | 0.092048625 | 0.255766056 |
| Frem1 | 147.1729021 | 0.174053444 | 0.150897817 | 1.153452362 | 0.24872476 | 0.473781833 |
| Prr5 | 225.4614639 | 0.174033376 | 0.134389745 | 1.294989998 | 0.19532368 | 0.409894454 |
| C330013E15Rik | 34.49888404 | 0.174033226 | 0.160585567 | 1.083741391 | 0.278479478 | 0.507499533 |
| 1110032F04Rik | 439.0759845 | -0.200196716 | 0.118937634 | -1.683207487 | 0.092334932 | 0.256340698 |
| Lrrc55 | 904.3925211 | -0.248289592 | 0.147503336 | -1.683281191 | 0.09232067 | 0.256340698 |
| Dnahc7a | 36.481052 | 0.173842719 | 0.167600984 | 1.037241632 | 0.29962326 | 0.531244904 |
| Mgl2 | 16.76142413 | -0.264379531 | 0.157120404 | -1.682655623 | 0.092441777 | 0.256585309 |
| Vps37a | 2157.695645 | 0.088730677 | 0.052734851 | 1.682581361 | 0.092456163 | 0.256585309 |
| 4833418N02Rik | 26.13202795 | 0.173643961 | 0.167482199 | 1.036790553 | 0.299833479 | 0.531324702 |
| Lacc1 | 213.2393604 | 0.173639152 | 0.1011673 | 1.716356494 | 0.086096814 | 0.24546239 |
| Haus5 | 221.398265 | 0.173607339 | 0.123692657 | 1.403537958 | 0.160456481 | 0.364960763 |
| Gcn1l1 | 3511.489097 | 0.097722954 | 0.058105414 | 1.681821836 | 0.092603393 | 0.256809911 |
| Fam169b | 23.80812707 | -0.282461339 | 0.167970012 | -1.68161767 | 0.092643002 | 0.256873778 |
| Sclt1 | 308.1469929 | 0.173555084 | 0.093037225 | 1.865437023 | 0.062120172 | 0.198984896 |
| Arf6 | 1314.212621 | -0.097676965 | 0.058105088 | -1.681039785 | 0.092755187 | 0.257092821 |
| Ahcyl1 | 16672.60622 | 0.064063629 | 0.038114061 | 1.680839758 | 0.092794044 | 0.257149547 |
| Nol9 | 905.9099469 | 0.173542634 | 0.073376932 | 2.365084341 | 0.018025965 | 0.088077675 |
| Sync | 56.5783377 | -0.250785839 | 0.149246423 | -1.680347405 | 0.092889743 | 0.257327673 |
| Mrgbp | 291.6283852 | 0.173511612 | 0.121420616 | 1.429012774 | 0.153000562 | 0.354495595 |
| Rnf25 | 1295.622171 | 0.17347204 | 0.071251352 | 2.434649107 | 0.014906237 | 0.077621071 |
| Wdr8 | 705.5456928 | 0.173419964 | 0.071231183 | 2.434607393 | 0.014907955 | 0.077621071 |
| Ywhag | 44860.09164 | -0.123350282 | 0.073424641 | -1.679957575 | 0.09296557 | 0.257388527 |
| Rfng | 2281.401231 | 0.173372229 | 0.074022098 | 2.342168547 | 0.019172055 | 0.091597846 |
| Clec4a1 | 12.98577273 | -0.274277118 | 0.163331061 | -1.679271022 | 0.093099236 | 0.25758027 |
| Klhdc4 | 733.686677 | 0.173227135 | 0.081109447 | 2.135720841 | 0.032702175 | 0.130927001 |
| Elf1 | 281.7239634 | 0.173203506 | 0.113456784 | 1.52660334 | 0.126859676 | 0.314691614 |
| Ddx5 | 29185.77943 | 0.173138709 | 0.068823408 | 2.515695078 | 0.011879791 | 0.066377745 |
| 3110035E14Rik | 13419.38378 | -0.204437901 | 0.121786019 | -1.678664779 | 0.093217394 | 0.25777446 |
| Acadsb | 2051.519311 | 0.092259062 | 0.054972526 | 1.678275821 | 0.093293266 | 0.257938234 |
| Pard3b | 180.126834 | -0.202975981 | 0.120951318 | -1.678162625 | 0.093315356 | 0.257953278 |
| 6820431F20Rik | 2456.605678 | 0.173129933 | 0.110301363 | 1.569608279 | 0.116506271 | 0.299311585 |
| Cacng1 | 10.73293909 | 0.173121234 | 0.146134885 | 1.184667398 | 0.236148978 | 0.460064411 |
| Smo | 614.0120718 | -0.156566513 | 0.093365655 | -1.676917636 | 0.093558589 | 0.2584873 |
| Zyx | 1682.711905 | 0.173030328 | 0.089600385 | 1.931133753 | 0.053466514 | 0.181549961 |
| Helz | 1130.598144 | -0.191131486 | 0.114020502 | -1.676290515 | 0.093681302 | 0.258661178 |
| Lpgat1 | 13165.73612 | -0.125768623 | 0.075024918 | -1.676358027 | 0.093668085 | 0.258661178 |
| Pinx1 | 152.7327591 | -0.180269092 | 0.107544482 | -1.676228183 | 0.093693506 | 0.258661178 |
| Rmdn1 | 234.5719626 | 0.172887675 | 0.136324579 | 1.268206196 | 0.204724325 | 0.421752252 |
| Gm13704 | 23.72697792 | 0.172736499 | 0.140525394 | 1.229219107 | 0.218989667 | 0.439511127 |
| Lrrc3b | 510.4011606 | -0.144039441 | 0.085983786 | -1.675193062 | 0.093896358 | 0.259077536 |
| Man2b1 | 2189.857327 | 0.172735843 | 0.087982692 | 1.963293448 | 0.049612086 | 0.17307478 |
| Gm11602 | 88.43500679 | 0.17263427 | 0.135906138 | 1.270246304 | 0.203996908 | 0.421222205 |
| Qsox1 | 1016.655878 | 0.172560494 | 0.072383138 | 2.383987453 | 0.017126186 | 0.084804452 |
| Gramd1a | 3519.014238 | 0.172498597 | 0.07904937 | 2.182162831 | 0.029097517 | 0.120956391 |
| Med24 | 4834.310498 | 0.096890942 | 0.057871725 | 1.674236274 | 0.094084173 | 0.259384373 |
| Nudt13 | 390.4019595 | 0.172487548 | 0.095358954 | 1.808823817 | 0.070478379 | 0.216278868 |
| Lcmt1 | 2147.991473 | -0.10364376 | 0.061921432 | -1.673794617 | 0.094170971 | 0.259531309 |
| 1810009A15Rik | 352.695977 | -0.156593695 | 0.093571565 | -1.673517966 | 0.094225373 | 0.259635057 |
| Map3k5 | 1871.954592 | 0.172442273 | 0.068852441 | 2.50451939 | 0.012261788 | 0.067803051 |
| Bcas1 | 3758.257839 | 0.172213289 | 0.115305108 | 1.493544319 | 0.135294762 | 0.327438548 |
| Rbp4 | 320.082795 | -0.227421118 | 0.13594891 | -1.672842519 | 0.094358302 | 0.259862697 |
| Rab20 | 11.744965 | 0.17216067 | 0.161803544 | 1.064010504 | 0.287323941 | 0.517169742 |
| Plekhj1 | 677.2831688 | -0.118613368 | 0.070916705 | -1.672573015 | 0.094411383 | 0.259916483 |
| Mboat2 | 1234.118184 | -0.105202223 | 0.062907261 | -1.672338297 | 0.094457632 | 0.25999761 |
| Cpa2 | 42.80398667 | 0.172116622 | 0.161017526 | 1.068930977 | 0.285100772 | 0.514782819 |
| Smurf2 | 1431.563926 | -0.126691951 | 0.075770054 | -1.672058341 | 0.094512818 | 0.260057113 |
| Afap1l2 | 420.5962748 | -0.141768097 | 0.084791488 | -1.671961423 | 0.094531929 | 0.260063514 |
| Tmem141 | 460.5743786 | -0.161024697 | 0.096316831 | -1.671823047 | 0.094559221 | 0.260092414 |
| Tmem39a | 285.0909019 | 0.172107576 | 0.127210349 | 1.352936909 | 0.176075787 | 0.385109859 |
| Gm2115 | 330.6411161 | 0.171887328 | 0.148714327 | 1.155822246 | 0.247753872 | 0.472687906 |
| Fam65a | 3955.520274 | 0.171814918 | 0.081051445 | 2.119825476 | 0.034020766 | 0.134573122 |

| | | | | | | |
|---|---|---|---|---|---|---|
| P2ry2 | 32.81743136 | -0.062029861 | 0.087018816 | -0.712832737 | 0.475949265 | 0.835195112 |
| Bpifb9a | 4.120746202 | -0.062046331 | 0.032382268 | -1.916058832 | 0.055357608 | NA |
| Mbtd1 | 918.9583902 | -0.062109545 | 0.064861916 | -0.957565682 | 0.338281811 | 0.760898356 |
| Pla2g2d | 17.46767024 | -0.062110456 | 0.071802305 | -0.865020369 | 0.387027609 | NA |
| Lhcgr | 19.33113077 | -0.06212514 | 0.079668894 | -0.779791667 | 0.435513512 | NA |
| Zfyve26 | 1234.320293 | -0.062133019 | 0.067131863 | -0.925536944 | 0.354686676 | 0.77200441 |
| Polr3g | 250.5924392 | -0.062156941 | 0.081766312 | -0.760177873 | 0.447148269 | 0.822458097 |
| Ssbp4 | 2722.811622 | -0.062166719 | 0.065930301 | -0.942915755 | 0.345723996 | 0.765097832 |
| Dtymk | 604.4486659 | -0.062168994 | 0.074935196 | -0.829636779 | 0.406744176 | 0.800917161 |
| Sema4g | 1399.705581 | -0.06220793 | 0.084275657 | -0.73814827 | 0.460424354 | 0.829239232 |
| Guca1b | 19.60695319 | -0.062278809 | 0.07879898 | -0.790350442 | 0.429323136 | NA |
| AW549877 | 2731.863108 | -0.062281364 | 0.063235976 | -0.984903968 | 0.324671244 | 0.75293408 |
| Sgca | 11.34513306 | -0.062290133 | 0.069110673 | -0.901309887 | 0.367423578 | NA |
| Clcc1 | 1042.476318 | -0.062299848 | 0.051680981 | -1.205469518 | 0.228022099 | 0.680552497 |
| Neu3 | 135.1721684 | -0.062307041 | 0.089032526 | -0.69982335 | 0.48403763 | 0.838713365 |
| Dcp1b | 547.2436678 | -0.062313039 | 0.072313172 | -0.861710772 | 0.388846696 | 0.79235741 |
| Elac2 | 1294.900925 | -0.062330389 | 0.060173088 | -1.035851588 | 0.300271386 | 0.733951208 |
| Osbpl7 | 786.3433384 | -0.062343693 | 0.069050405 | -0.902872218 | 0.366593717 | 0.778210102 |
| Grtp1 | 103.7713524 | -0.062371754 | 0.092743481 | -0.672519012 | 0.501253341 | 0.846601651 |
| Mtl5 | 5.816483074 | -0.06237292 | 0.047412268 | -1.315543901 | 0.188327173 | NA |
| Zscan18 | 1072.096508 | -0.062411826 | 0.066199368 | -0.942785818 | 0.345790467 | 0.765097832 |
| Suclg2 | 1088.225728 | -0.06243495 | 0.060116766 | -1.038561351 | 0.299008788 | 0.733311102 |
| Snord99 | 14.46468841 | -0.062487898 | 0.068502207 | -0.9122027 | 0.361662023 | NA |
| Cryz | 631.4912646 | -0.062502353 | 0.0634146 | -0.985614546 | 0.324322297 | 0.75293408 |
| Hhatl | 433.9208484 | -0.062520686 | 0.087783671 | -0.71221317 | 0.476332782 | 0.835195112 |
| Lemd2 | 1176.47004 | -0.062521714 | 0.068071658 | -0.918469085 | 0.358373337 | 0.774201531 |
| Sgcb | 2615.726059 | -0.062559098 | 0.044748794 | -1.398006351 | 0.16211116 | 0.613957643 |
| Tmprss9 | 24.5538342 | -0.062560329 | 0.082405579 | -0.759175888 | 0.447747348 | 0.822750604 |
| Atad5 | 223.6187343 | -0.062659276 | 0.084545679 | -0.741129254 | 0.458615073 | 0.828043599 |
| Cxcr7 | 361.5060714 | -0.062673833 | 0.075134454 | -0.834155707 | 0.404193247 | 0.800290023 |
| Eme2 | 640.3440417 | -0.062693544 | 0.083733186 | -0.748729944 | 0.454019992 | 0.825985606 |
| Gp1bb | 314.5766309 | -0.062701764 | 0.08588408 | -0.730074346 | 0.465344742 | 0.831682622 |
| Tmem62 | 1349.026158 | -0.062710947 | 0.059832813 | -1.04810295 | 0.294591177 | 0.72962173 |
| Sept2 | 2407.146423 | -0.062745728 | 0.062285602 | -1.007387359 | 0.313748663 | 0.745844951 |
| Rbm10 | 2289.094833 | -0.062760353 | 0.057884106 | -1.084241542 | 0.278257717 | 0.718445005 |
| Pom121l2 | 1.769221581 | -0.062766658 | 0.029835665 | -2.103745884 | 0.03540062 | NA |
| Fgfr1op | 648.231962 | -0.062782519 | 0.075732897 | -0.828999307 | 0.407104799 | 0.800975798 |
| 2610020H08Rik | 30.95755699 | -0.062787752 | 0.083503984 | -0.751913249 | 0.45210323 | 0.824720773 |
| 2900055J20Rik | 174.5013905 | -0.062859967 | 0.088917983 | -0.706943246 | 0.479601748 | 0.836438346 |
| 4931409K22Rik | 5.827644362 | -0.062882078 | 0.051681656 | -1.216719486 | 0.223710963 | NA |
| 4632428N05Rik | 1447.858484 | -0.062903472 | 0.06931787 | -0.907464013 | 0.364161472 | 0.777191869 |
| Zfand2b | 567.8586234 | -0.062912305 | 0.078079999 | -0.805741619 | 0.420391847 | 0.807752905 |
| Zfp953 | 53.13583901 | -0.062946589 | 0.094875983 | -0.663461784 | 0.507034847 | 0.848958235 |
| Lor | 122.7544798 | -0.063003415 | 0.095087254 | -0.662585279 | 0.507596201 | 0.849139846 |
| Tst | 774.9081238 | -0.063016547 | 0.074179981 | -0.849508811 | 0.395598229 | 0.796239418 |
| Bms1 | 1749.2367 | -0.063027494 | 0.059414101 | -1.0608171 | 0.28877303 | 0.724444298 |
| Arsk | 270.2615336 | -0.063041184 | 0.079615317 | -0.791822309 | 0.428464293 | 0.810486244 |
| Gpr12 | 728.3296246 | -0.063066389 | 0.073734504 | -0.855317193 | 0.392375572 | 0.794184423 |
| A430105I19Rik | 376.9978254 | -0.063077791 | 0.083395549 | -0.756368801 | 0.449428108 | 0.823263966 |
| Btbd19 | 269.2447229 | -0.063096915 | 0.077695051 | -0.812109834 | 0.416728611 | 0.806153084 |
| Procr | 24.55659061 | -0.063132387 | 0.086207632 | -0.732329443 | 0.463967513 | 0.83159896 |
| Hs1bp3 | 787.2071842 | -0.063168466 | 0.071311316 | -0.885812651 | 0.375718491 | 0.783947144 |
| Trim30b | 3.061838828 | -0.063169212 | 0.032441617 | -1.947165961 | 0.051514841 | NA |
| Rps6kl1 | 1510.057494 | -0.063179504 | 0.066676491 | -0.947552918 | 0.343357104 | 0.763762832 |
| Zkscan16 | 2018.188199 | -0.063197143 | 0.073628269 | -0.858327161 | 0.390711835 | 0.794106635 |
| Gar1 | 425.2484859 | -0.063206902 | 0.072228651 | -0.875094589 | 0.381522441 | 0.786663048 |
| Tsacc | 29.05707932 | -0.06325246 | 0.086173188 | -0.734015559 | 0.462939256 | 0.831014288 |
| 0610040J01Rik | 156.8869441 | -0.063283755 | 0.094136718 | -0.672253683 | 0.50142221 | 0.846601651 |
| Ptprd | 10174.14231 | -0.063302228 | 0.061640553 | -1.026957494 | 0.30444048 | 0.737453291 |
| Parp8 | 833.1669233 | -0.063312275 | 0.074935791 | -0.844886981 | 0.398173944 | 0.797789838 |
| Eya1 | 469.4696465 | -0.063317574 | 0.083444632 | -0.758797457 | 0.447973727 | 0.822871315 |
| Atad3a | 1404.870929 | -0.06337109 | 0.060408845 | -1.049036602 | 0.294161273 | 0.729447815 |
| Whsc1 | 4268.70127 | -0.06348777 | 0.059757756 | -1.062418908 | 0.288045554 | 0.724103151 |
| Spata24 | 121.0507671 | -0.063520528 | 0.093591149 | -0.678702301 | 0.497326508 | 0.845038521 |
| Rbm12b2 | 314.8561596 | -0.063521628 | 0.087952579 | -0.722225873 | 0.470155618 | 0.833245343 |
| Wdr81 | 1517.337103 | -0.063521794 | 0.066777457 | -0.951246074 | 0.341479475 | 0.76303628 |
| Zfp185 | 83.60433733 | -0.063522146 | 0.081741241 | -0.777112579 | 0.437092353 | 0.816564673 |
| 2310003H01Rik | 522.7574211 | -0.063544231 | 0.088696177 | -0.71642582 | 0.473728456 | 0.833983942 |
| Lphn2 | 4747.061589 | -0.063549873 | 0.055858886 | -1.137686009 | 0.255251618 | 0.701952734 |
| Hspb3 | 68.75556155 | -0.063553078 | 0.093509702 | -0.679641544 | 0.496731458 | 0.8449088 |
| Znf41-ps | 47.70766417 | -0.063558418 | 0.08878813 | -0.715843636 | 0.474087905 | 0.833994865 |
| Cyp27b1 | 13.11496418 | -0.063595058 | 0.068454475 | -0.929012415 | 0.352882652 | NA |
| B330016D10Rik | 65.98635061 | -0.063609085 | 0.095070838 | -0.669070412 | 0.503450563 | 0.847807553 |
| Kcne3 | 6.355423897 | -0.06362927 | 0.053819253 | -1.18227711 | 0.237095765 | NA |
| Amdhd1 | 7.286250525 | -0.063673877 | 0.055841104 | -1.140268952 | 0.254174269 | NA |
| Rorb | 1769.736373 | -0.063683045 | 0.067859271 | -0.938457546 | 0.348009324 | 0.766401371 |
| Stag3 | 39.63246908 | -0.063717409 | 0.085549158 | -0.744804625 | 0.456389838 | 0.82674985 |
| Tmsb10 | 3647.695146 | -0.063722817 | 0.083683868 | -0.761470742 | 0.446375948 | 0.822210607 |
| Pisd | 3507.028886 | -0.063725069 | 0.073285144 | -0.869549613 | 0.384546584 | 0.788972271 |
| Actl6a | 461.1349567 | -0.063738766 | 0.0656106 | -0.971470546 | 0.331314013 | 0.75639847 |
| Ccdc66 | 597.8970399 | -0.063754259 | 0.07384882 | -0.863307752 | 0.387968283 | 0.791889115 |
| Awat2 | 7.800138389 | -0.063826857 | 0.061283446 | -1.04150241 | 0.297642435 | NA |
| Pfkfb4 | 691.094326 | -0.063836193 | 0.073487282 | -0.868669948 | 0.385027683 | 0.789016184 |
| Zcchc16 | 90.69428237 | -0.063870705 | 0.094934963 | -0.672783801 | 0.501084845 | 0.846601651 |
| 1700105P06Rik | 4.320698374 | -0.063882273 | 0.049654747 | -1.286529025 | 0.198258496 | NA |
| Fhl3 | 36.32210048 | -0.063913666 | 0.09039908 | -0.70701677 | 0.479556056 | 0.836438346 |
| Simc1 | 850.6603714 | -0.063935052 | 0.061602846 | -1.037858734 | 0.299335828 | 0.733704078 |
| Gm3230 | 135.7193894 | -0.064010737 | 0.092687825 | -0.690605668 | 0.489813385 | 0.841119426 |
| Dnm2 | 2348.150913 | -0.064020232 | 0.063478602 | -1.008532491 | 0.313198898 | 0.745612333 |
| Ctcfl | 18.0840661 | -0.064037996 | 0.076517056 | -0.836911394 | 0.402642375 | NA |
| AI593442 | 8159.17285 | -0.064068589 | 0.076256305 | -0.840174324 | 0.400810653 | 0.799169897 |
| 2610528A11Rik | 11.86244139 | -0.064081006 | 0.069517696 | -0.921794159 | 0.356635955 | NA |
| Adrb3 | 50.61204653 | -0.064096399 | 0.090068517 | -0.711640439 | 0.476687459 | 0.835195112 |
| Cct8l1 | 3.099425143 | -0.064101178 | 0.039541177 | -1.621124695 | 0.104990899 | NA |
| Sec11c | 1470.986941 | -0.064109017 | 0.065934343 | -0.972316012 | 0.330893361 | 0.756212998 |
| Ido1 | 129.1848271 | -0.064115025 | 0.095098385 | -0.674196775 | 0.50018622 | 0.846268338 |
| Pacsin3 | 1255.716577 | -0.064157367 | 0.063899273 | -1.004039072 | 0.315359781 | 0.745993052 |
| Snhg7 | 136.8122798 | -0.064160319 | 0.090286891 | -0.71062718 | 0.477315296 | 0.835289575 |
| Mecom | 96.23276985 | -0.064167629 | 0.094675583 | -0.677763232 | 0.497921827 | 0.845489269 |
| Fam217a | 35.91140464 | -0.064172834 | 0.090620702 | -0.708147617 | 0.478853592 | 0.835936459 |
| 2810408I11Rik | 12.03941746 | -0.064173505 | 0.064929059 | -0.988363383 | 0.322974714 | NA |
| Lsm2 | 212.5278799 | -0.06419257 | 0.088858276 | -0.722415206 | 0.470039241 | 0.833245343 |
| Dok1 | 72.71964384 | -0.064216915 | 0.094822672 | -0.677231653 | 0.498258988 | 0.845625663 |
| Gm5111 | 16.59091814 | -0.064232597 | 0.06935217 | -0.92618006 | 0.354352413 | NA |
| Gm10658 | 13.27176259 | -0.064262471 | 0.067497667 | -0.952069521 | 0.341061725 | NA |
| Dnaic2 | 124.140422 | -0.064263537 | 0.095099559 | -0.675750107 | 0.499199317 | 0.846241311 |
| Arid3a | 290.3720964 | -0.06428957 | 0.077750214 | -0.826873229 | 0.408308919 | 0.801055189 |
| Rcor1 | 742.1190116 | -0.064294876 | 0.073817276 | -0.871000397 | 0.383753936 | 0.78787654 |
| Aoc2 | 172.6556978 | -0.064299288 | 0.095074199 | -0.676306382 | 0.498846141 | 0.846052342 |
| Cd163l1 | 2.240102337 | -0.064320487 | 0.034650467 | -1.856266088 | 0.06341564 | NA |
| Nudt6 | 95.59982406 | -0.064359386 | 0.094633859 | -0.680088359 | 0.496448515 | 0.844826506 |
| Hectd3 | 2714.305273 | -0.064374244 | 0.049987671 | -1.287802433 | 0.197814745 | 0.653535282 |
| Plekha4 | 113.3162512 | -0.064404278 | 0.094661917 | -0.68036102 | 0.496275897 | 0.8448264 |
| Mir343 | 9.5040228 | -0.064424213 | 0.061076276 | -1.054815672 | 0.291509635 | NA |
| Slc9a4 | 31.5009191 | -0.064450509 | 0.079372633 | -0.811999129 | 0.416792131 | 0.806153084 |
| Abcc2 | 10.12131111 | -0.064469553 | 0.060313522 | -1.068907114 | 0.285111526 | NA |
| Rev1 | 1412.347901 | -0.064475769 | 0.059972623 | -1.075086687 | 0.282335919 | 0.720775319 |
| Mettl21d | 424.1845371 | -0.064482486 | 0.077673978 | -0.830168448 | 0.406443552 | 0.800759397 |
| Folr2 | 14.23968427 | -0.064492222 | 0.064238926 | -1.003943025 | 0.315406077 | NA |
| Ehmt1 | 1363.054681 | -0.064560272 | 0.050920691 | -1.267859304 | 0.2048482 | 0.658461308 |
| Dynlt1b | 37.47015881 | -0.064581216 | 0.084693223 | -0.762531093 | 0.445743093 | 0.821773578 |
| Gprc5c | 186.2160479 | -0.064601211 | 0.093401053 | -0.69165399 | 0.489154647 | 0.840836703 |
| AI317395 | 35.9728698 | -0.064650366 | 0.085776198 | -0.753709864 | 0.451023457 | 0.823992799 |
| Itgal | 53.02055003 | -0.064654393 | 0.094317039 | -0.685500664 | 0.493028042 | 0.842867752 |
| Afap1 | 846.6710691 | -0.064656506 | 0.063899614 | -1.011845029 | 0.311612156 | 0.74424258 |
| Ascl1 | 155.1298305 | -0.064673708 | 0.088860843 | -0.727808849 | 0.466730607 | 0.832095317 |
| Ccbp2 | 177.8455325 | -0.0646891 | 0.094941516 | -0.68135735 | 0.495645403 | 0.844369474 |
| Tmc5 | 8.186954316 | -0.064711304 | 0.059905532 | -1.080222519 | 0.280043103 | NA |
| Col6a3 | 154.5346183 | -0.064730068 | 0.089601464 | -0.722421988 | 0.470035072 | 0.833245343 |
| Il6 | 2.857597098 | -0.06473914 | 0.034202461 | -1.892821084 | 0.058381667 | NA |
| Arsi | 19.93692438 | -0.064771238 | 0.080238738 | -0.807231513 | 0.419533113 | NA |
| Bud13 | 456.559903 | -0.064776663 | 0.068906395 | -0.940067516 | 0.34718293 | 0.765974975 |
| Krit1 | 1658.779825 | -0.064776682 | 0.068749048 | -0.942219335 | 0.346080357 | 0.76510344 |
| Abcb6 | 889.9380279 | -0.064865079 | 0.06605957 | -0.981917978 | 0.326140258 | 0.753049596 |
| Zfp949 | 976.044994 | -0.064865891 | 0.070841556 | -0.915647457 | 0.359851832 | 0.776125867 |
| Ern1 | 344.8220218 | -0.064904838 | 0.084007022 | -0.772612064 | 0.439752002 | 0.817695544 |
| Cyth1 | 3722.072819 | -0.064917131 | 0.052841545 | -1.228524467 | 0.219250151 | 0.672421385 |
| Asl | 551.5271567 | -0.064940569 | 0.065769195 | -0.987401002 | 0.323446094 | 0.752925264 |
| Zfp142 | 1875.690942 | -0.065002029 | 0.060800371 | -1.069105792 | 0.285022002 | 0.721671536 |
| Alg8 | 304.9448542 | -0.065017367 | 0.077278388 | -0.841339586 | 0.40015772 | 0.798904608 |
| Parpbp | 20.05051672 | -0.065033091 | 0.076287294 | -0.852476043 | 0.39394993 | NA |
| C330027C09Rik | 119.8976515 | -0.065040714 | 0.095019728 | -0.684496949 | 0.493661415 | 0.843404385 |
| Elf4 | 98.92336941 | -0.065045526 | 0.093662204 | -0.694469308 | 0.487387942 | 0.839801621 |
| Surf2 | 936.6297192 | -0.065073073 | 0.070167059 | -0.927402032 | 0.353717836 | 0.77113992 |
| Tmem25 | 2937.882917 | -0.06507503 | 0.059007315 | -1.102829877 | 0.270101051 | 0.71216373 |
| Fjx1 | 1902.522624 | -0.065095739 | 0.064660263 | -1.006734829 | 0.31406222 | 0.745923449 |
| Itpr3 | 272.5163924 | -0.065108496 | 0.086959348 | -0.748723362 | 0.454023959 | 0.825985606 |
| 9130206I24Rik | 4.213188275 | -0.065120183 | 0.04354508 | -1.495465937 | 0.134792883 | NA |
| Zfp120 | 557.4392678 | -0.065139868 | 0.078862053 | -0.825997622 | 0.408805441 | 0.801055189 |
| Lmo7 | 4062.487031 | -0.065142166 | 0.064505323 | -1.009872723 | 0.312556273 | 0.744930503 |
| Efcab4b | 42.4783988 | -0.065162168 | 0.092416727 | -0.705090623 | 0.480753842 | 0.836914993 |
| Trim7 | 60.20531669 | -0.065215362 | 0.094672668 | -0.688850993 | 0.490917042 | 0.841866734 |
| Tpcn1 | 2245.824144 | -0.06528756 | 0.054943403 | -1.188269321 | 0.23472732 | 0.685908078 |
| B930003M22Rik | 25.89232793 | -0.065296818 | 0.084298086 | -0.774594308 | 0.438579422 | 0.817139233 |
| Ccdc18 | 53.91926091 | -0.065297975 | 0.091481124 | -0.713786327 | 0.475359316 | 0.834927401 |
| Ttll10 | 4.728755573 | -0.065308134 | 0.050031821 | -1.305331927 | 0.191779847 | NA |
| Gimap9 | 16.85501736 | -0.06531004 | 0.075457989 | -0.865515247 | 0.386756051 | NA |
| Gja6 | 7.776498753 | -0.065326826 | 0.057117829 | -1.143720387 | 0.252739616 | NA |
| Tmem53 | 78.30153419 | -0.065377007 | 0.095090422 | -0.687524635 | 0.491752182 | 0.841892493 |
| Sart3 | 1957.660363 | -0.065377149 | 0.057408137 | -1.138813283 | 0.25478104 | 0.701619624 |
| 2500004C02Rik | 233.8569072 | -0.065378993 | 0.084439847 | -0.774267063 | 0.438772877 | 0.817234878 |
| Fance | 509.8691662 | -0.065398697 | 0.086093275 | -0.759626081 | 0.447478125 | 0.822458097 |
| Atp2a3 | 493.009536 | -0.065417988 | 0.087407901 | -0.748421913 | 0.454205709 | 0.826066374 |
| Tor1b | 1299.475602 | -0.065445994 | 0.063737202 | -1.02680997 | 0.304509954 | 0.737453291 |
| Dgkz | 21724.65729 | -0.065485088 | 0.072700494 | -0.900751623 | 0.367720394 | 0.778985919 |
| Scyl3 | 1198.371902 | -0.065491861 | 0.064116821 | -1.021445853 | 0.307043251 | 0.739163119 |
| Rdh9 | 23.50898887 | -0.065528921 | 0.083245502 | -0.787176713 | 0.431178427 | 0.812181606 |
| Tmem200c | 418.6899815 | -0.065589862 | 0.073954927 | -0.886889685 | 0.375138296 | 0.783489748 |
| Ssr4 | 1073.851778 | -0.065596187 | 0.056404755 | -1.162954912 | 0.244847796 | 0.694524329 |
| 3425401B19Rik | 18.39658629 | -0.065627318 | 0.07043311 | -0.931767995 | 0.351456438 | NA |
| Vcan | 823.1120245 | -0.065633457 | 0.076239933 | -0.860880309 | 0.389303967 | 0.792687052 |
| Dnaic1 | 143.7616719 | -0.065664343 | 0.094100362 | -0.697811806 | 0.485294895 | 0.839013763 |
| Atp9b | 3232.450001 | -0.065704809 | 0.043979019 | -1.494003526 | 0.135174698 | 0.578555714 |
| Nhlh1 | 20.90783013 | -0.065709698 | 0.075994958 | -0.864658652 | 0.38722617 | NA |
| A830052D11Rik | 43.59084611 | -0.065720266 | 0.093856447 | -0.700221116 | 0.483789226 | 0.838706564 |
| Trmt44 | 301.1491449 | -0.065755428 | 0.082216413 | -0.799784685 | 0.423835558 | 0.808999423 |
| Stk16 | 1692.271455 | -0.065783095 | 0.060237151 | -1.092068499 | 0.274802994 | 0.716529939 |
| Safb | 4065.979174 | -0.065783693 | 0.055980941 | -1.175108735 | 0.239951216 | 0.690751361 |
| Btaf1 | 1780.846278 | -0.065826032 | 0.078591695 | -0.837569827 | 0.402272345 | 0.79955155 |
| Tead4 | 27.3794714 | -0.065840071 | 0.086154378 | -0.764210391 | 0.444741876 | 0.821493054 |
| Mgat4b | 2108.838128 | -0.065947673 | 0.065706888 | -1.003664543 | 0.315540333 | 0.745993052 |
| Sept11 | 4395.747911 | -0.065950299 | 0.045362435 | -1.453852706 | 0.145987147 | 0.592514474 |
| Gm11744 | 91.79887892 | -0.066002275 | 0.095095486 | -0.694063179 | 0.487642589 | 0.839801621 |
| Snrpn | 52.49886374 | -0.06600229 | 0.087182377 | -0.757059998 | 0.449013918 | 0.823263966 |
| Ascl5 | 11.76699077 | -0.066038797 | 0.069957853 | -0.943979765 | 0.345179989 | NA |
| B930018H19Rik | 7.539884275 | -0.066081464 | 0.051009659 | -1.295469624 | 0.195158274 | NA |
| Snhg4 | 238.8027137 | -0.066085961 | 0.082338738 | -0.802610807 | 0.42219972 | 0.808695226 |
| A630089N07Rik | 74.41043591 | -0.066110232 | 0.094646333 | -0.698497555 | 0.484866087 | 0.838771558 |
| Tnnc1 | 214.1709953 | -0.066167969 | 0.084961669 | -0.778797895 | 0.436098778 | 0.816020977 |
| Rnaseh2c | 343.4393081 | -0.066194438 | 0.08389349 | -0.789029495 | 0.430094768 | 0.811860174 |
| Chd8 | 4799.843963 | -0.066196803 | 0.050342057 | -1.314940376 | 0.188529942 | 0.64225749 |
| Gba | 1572.777947 | -0.066215266 | 0.058832108 | -1.125495385 | 0.260379172 | 0.704058139 |
| 4930519F16Rik | 2.409221783 | -0.066301763 | 0.036608881 | -1.811084107 | 0.070127833 | NA |
| Fam65c | 15.81082549 | -0.066306412 | 0.065439167 | -1.013252687 | 0.31093948 | NA |
| Pih1d2 | 66.16859713 | -0.066333208 | 0.094920697 | -0.69882766 | 0.48465974 | 0.838713365 |
| Fcf1 | 454.5302668 | -0.06634193 | 0.07826177 | -0.847692685 | 0.396609141 | 0.796916625 |
| Naip1 | 14.56683364 | -0.066343633 | 0.074687592 | -0.888281862 | 0.374389154 | NA |
| Lrrc48 | 535.1049744 | -0.066360408 | 0.069509321 | -0.954697974 | 0.339730455 | 0.761679979 |
| Slc4a3 | 7575.561451 | -0.066372354 | 0.063199528 | -1.050203317 | 0.293624645 | 0.729168132 |
| Mybpc3 | 32.5271713 | -0.066387218 | 0.08818168 | -0.752845915 | 0.451542512 | 0.824520467 |
| Tbc1d8 | 1340.760951 | -0.066482471 | 0.0799728 | -0.831313539 | 0.405796528 | 0.800733644 |
| Hist1h2bg | 10.16125153 | -0.066497472 | 0.066491197 | -1.000094372 | 0.317264839 | NA |
| Ccr6 | 6.608129769 | -0.066533633 | 0.056886605 | -1.169583469 | 0.242168633 | NA |
| Zfp866 | 1189.702765 | -0.066581337 | 0.062185406 | -1.070690721 | 0.284308516 | 0.721548306 |
| Btbd8 | 73.78480812 | -0.06659898 | 0.095060086 | -0.700598779 | 0.48355344 | 0.83839786 |
| Zfp939 | 252.3858476 | -0.066656061 | 0.084155682 | -0.792056572 | 0.42832769 | 0.810486244 |
| Art3 | 123.2049317 | -0.066660097 | 0.089515233 | -0.744678811 | 0.456465912 | 0.82674985 |
| Zfp341 | 497.6247067 | -0.066667369 | 0.082711148 | -0.806026404 | 0.420227625 | 0.807657282 |
| Mzt2 | 509.863807 | -0.066695034 | 0.074020905 | -0.901029709 | 0.367572524 | 0.778899851 |
| Kansl2 | 2056.595868 | -0.066708362 | 0.05188538 | -1.285687062 | 0.198552298 | 0.65377551 |
| Pipox | 83.58929742 | -0.066713243 | 0.093681044 | -0.712131722 | 0.476383212 | 0.835195112 |
| Cgn | 226.6762952 | -0.066763044 | 0.093857471 | -0.71132371 | 0.476883662 | 0.835195112 |
| Gm15421 | 20.34535197 | -0.066814474 | 0.071944105 | -0.928699774 | 0.353044698 | NA |
| Kdm4b | 3203.538991 | -0.066878613 | 0.062062154 | -1.077607022 | 0.281209167 | 0.720080692 |
| Prex2 | 2661.798707 | -0.066915707 | 0.082441317 | -0.811676829 | 0.416977094 | 0.806153084 |
| Pdcd2l | 418.2382283 | -0.066921804 | 0.069563408 | -0.962025965 | 0.336036561 | 0.759375714 |
| Maz | 5859.916412 | -0.066969129 | 0.057343971 | -1.167849519 | 0.24286747 | 0.692772033 |
| Tlr6 | 20.98609501 | -0.067004235 | 0.080242126 | -0.835025666 | 0.403703257 | NA |
| Mau2 | 4059.539204 | -0.067008841 | 0.058377434 | -1.1478552 | 0.251028345 | 0.699465245 |
| Btg2 | 880.436352 | -0.06702477 | 0.079871836 | -0.839153983 | 0.401382906 | 0.799277436 |
| Ino80 | 1662.866576 | -0.067038237 | 0.058370408 | -1.148496975 | 0.250763461 | 0.699068333 |
| Adamts9 | 153.1298214 | -0.067052156 | 0.094055747 | -0.712898024 | 0.475908862 | 0.835195112 |
| Mxra8 | 217.4505862 | -0.067055813 | 0.095093847 | -0.705154071 | 0.48071436 | 0.836914993 |
| Gm5464 | 4.561012357 | -0.067084083 | 0.044770371 | -1.49840355 | 0.134028435 | NA |
| Pigw | 143.4846665 | -0.067129681 | 0.092231577 | -0.727838378 | 0.466712529 | 0.832095317 |
| Pitrm1 | 1766.573322 | -0.067135894 | 0.051898565 | -1.293598273 | 0.195804219 | 0.651423791 |
| Gm13375 | 412.7650055 | -0.067235636 | 0.084554496 | -0.79517518 | 0.426511607 | 0.809764424 |
| Mitd1 | 229.1314368 | -0.067261463 | 0.085152027 | -0.789898556 | 0.429587015 | 0.811442206 |
| Sgcd | 52.74460964 | -0.067272882 | 0.093575111 | -0.718918538 | 0.472191114 | 0.833441697 |
| Kbtbd13 | 11.48134195 | -0.067301812 | 0.066098482 | -1.018205101 | 0.308580496 | NA |
| Mettl3 | 904.9520238 | -0.067302671 | 0.068402823 | -0.983916569 | 0.325156536 | 0.75293408 |
| Wdr52 | 191.477926 | -0.06733733 | 0.094518751 | -0.712422981 | 0.476202889 | 0.835195112 |
| 2810428I15Rik | 789.6459049 | -0.067342029 | 0.069165511 | -0.973635965 | 0.330237322 | 0.756141871 |
| Evc2 | 1035.903142 | -0.0673504 | 0.087672165 | -0.768207337 | 0.442364015 | 0.819629727 |
| Snora81 | 21.4831359 | -0.067385695 | 0.079002737 | -0.852953933 | 0.393684851 | NA |

| | | | | | | |
|---|---|---|---|---|---|---|
| Fam98c | 739.5984597 | -0.158710717 | 0.094995146 | -1.670724499 | 0.094776108 | 0.260503993 |
| Snupn | 446.1974328 | -0.156512608 | 0.09368971 | -1.67054214 | 0.094812149 | 0.260556836 |
| Ccdc92 | 3187.824356 | 0.171530864 | 0.068611311 | 2.500037693 | 0.012418009 | 0.068398378 |
| Ugdh | 828.1515649 | 0.171460508 | 0.087283921 | 1.964399686 | 0.049483759 | 0.172797243 |
| Pou3f3 | 2247.655894 | -0.163345511 | 0.097811683 | -1.670000009 | 0.094919362 | 0.260796651 |
| Rps7 | 7261.134736 | -0.122287136 | 0.07323633 | -1.669760582 | 0.094966742 | 0.260796651 |
| Dstn | 5523.656187 | -0.105081943 | 0.062940057 | -1.669555901 | 0.095007262 | 0.260797001 |
| Ptger4 | 38.430263 | 0.171444027 | 0.166722614 | 1.028318971 | 0.30379981 | 0.535930122 |
| Ccdc9 | 1436.275166 | 0.171404608 | 0.087325271 | 1.962829374 | 0.049666003 | 0.173082559 |
| Cage1 | 136.907383 | 0.171363167 | 0.12573564 | 1.362884594 | 0.172918885 | 0.380731164 |
| Clec4a3 | 31.05358248 | 0.171241203 | 0.162321496 | 1.054950869 | 0.291447796 | 0.521743749 |
| Ptgfr | 39.5446382 | -0.265555451 | 0.159124298 | -1.668855448 | 0.095146031 | 0.261011438 |
| Ralgds | 4074.398618 | 0.171228875 | 0.074840813 | 2.287907751 | 0.022142894 | 0.100819843 |
| 1810032O08Rik | 92.29489524 | 0.171184334 | 0.133505778 | 1.282224167 | 0.199764021 | 0.416244936 |
| Ier3ip1 | 1020.957 | -0.111607613 | 0.06692402 | -1.667676473 | 0.095379968 | 0.261469936 |
| Pnoc | 286.833138 | 0.171149561 | 0.108848495 | 1.57236497 | 0.115865933 | 0.298309534 |
| Orai3 | 653.6987739 | 0.171125454 | 0.082056993 | 2.085446325 | 0.037028806 | 0.142186943 |
| 4930506M07Rik | 1237.030035 | -0.197325837 | 0.118342856 | -1.667408111 | 0.095433282 | 0.261521626 |
| Insl6 | 14.63293602 | 0.171109139 | 0.144025691 | 1.188045946 | 0.234815309 | 0.458404859 |
| Polr1a | 1509.001665 | 0.17106375 | 0.106860287 | 1.600816864 | 0.109417487 | 0.287633958 |
| Tnfrsf11b | 90.12403785 | -0.242632981 | 0.145616931 | -1.666241551 | 0.095665313 | 0.262018448 |
| Cdk2ap2 | 546.7483056 | -0.122875382 | 0.073754778 | -1.665998945 | 0.095713624 | 0.262058119 |
| Mir377 | 12.02549698 | 0.17105077 | 0.153086088 | 1.117350196 | 0.263844621 | 0.491170078 |
| Slc25a10 | 592.3006943 | 0.171000333 | 0.066985681 | 2.55278933 | 0.010686411 | 0.061666761 |
| Zscan29 | 344.3875504 | 0.170935063 | 0.11158209 | 1.531922036 | 0.125541675 | 0.313378881 |
| 1110038B12Rik | 209.7950699 | 0.17088097 | 0.128213882 | 1.332780564 | 0.182603826 | 0.393364924 |
| Srsf1 | 4473.441582 | 0.170876057 | 0.077924142 | 2.19285132 | 0.028318093 | 0.119163643 |
| Trip4 | 1613.510281 | -0.100649912 | 0.060436994 | -1.665369257 | 0.095839109 | 0.262170049 |
| Adra1d | 1013.454481 | 0.170745827 | 0.126944551 | 1.345042585 | 0.178611483 | 0.388110364 |
| Gpr21 | 128.4859317 | -0.234514251 | 0.140856456 | -1.664916592 | 0.095929398 | 0.262324406 |
| Cfl1 | 25411.21969 | -0.101448016 | 0.060943755 | -1.664617093 | 0.095989173 | 0.262441547 |
| Zmynd10 | 122.6674478 | 0.17073535 | 0.135904402 | 1.256290061 | 0.209010824 | 0.427134075 |
| 2610019F03Rik | 844.1471793 | -0.162051938 | 0.097374352 | -1.664215831 | 0.096069305 | 0.262567969 |
| 2810004N23Rik | 410.9821461 | -0.132501116 | 0.079645009 | -1.663646189 | 0.096183156 | 0.262816719 |
| 4921508A21Rik | 15.44481682 | 0.170694098 | 0.167834178 | 1.017040157 | 0.309134326 | 0.54172842 |
| Mst1r | 53.19347207 | 0.170693755 | 0.152455325 | 1.119631305 | 0.262870909 | 0.490005037 |
| Meis3 | 1271.050039 | 0.170654726 | 0.085465317 | 1.996771709 | 0.045849988 | 0.164406322 |
| Fkbp9 | 1805.118995 | -0.11539349 | 0.069397035 | -1.66280145 | 0.096352186 | 0.263062692 |
| Tmem212 | 28.36674606 | 0.170598393 | 0.167657163 | 1.017543119 | 0.308895131 | 0.541527626 |
| Kif11 | 80.91806891 | -0.252754848 | 0.152086863 | -1.661911118 | 0.096530597 | 0.26350336 |
| Mbd3 | 3324.325022 | -0.101582375 | 0.061136017 | -1.661579869 | 0.096597043 | 0.263591859 |
| Slc1a6 | 357.0271818 | 0.170575641 | 0.120269179 | 1.418282244 | 0.156108378 | 0.358438532 |
| Gdf5 | 11.03954688 | 0.170574913 | 0.159739921 | 1.067828956 | 0.285597674 | 0.515055081 |
| Atp10a | 681.2093407 | 0.170523665 | 0.123244979 | 1.383615519 | 0.166476214 | 0.372076569 |
| Nudt16 | 450.2908964 | -0.140003 | 0.084293297 | -1.660903118 | 0.096732906 | 0.263776777 |
| Sdk1 | 418.7774382 | -0.191225753 | 0.115127831 | -1.660986328 | 0.096716193 | 0.263776777 |
| Fryl | 3309.426139 | -0.146011677 | 0.087933611 | -1.660476305 | 0.096818671 | 0.263964189 |
| Galc | 1180.328706 | 0.170353831 | 0.082669759 | 2.060654735 | 0.039335991 | 0.14807382 |
| Rrp8 | 431.2569352 | 0.170286193 | 0.114237231 | 1.490636567 | 0.136056938 | 0.328429143 |
| Crb3 | 11.58058855 | 0.170278904 | 0.154515456 | 1.102018581 | 0.270453594 | 0.498730506 |
| Sri | 1756.294573 | -0.116511864 | 0.070232058 | -1.658955584 | 0.097124743 | 0.264565887 |
| 9030612E09Rik | 43.50178527 | 0.170240496 | 0.159605634 | 1.066632123 | 0.286137989 | 0.515760851 |
| Slc39a6 | 1916.927495 | -0.115552753 | 0.06967654 | -1.658416922 | 0.097233343 | 0.264815156 |
| Etfa | 1054.696266 | -0.130899609 | 0.078939437 | -1.658228306 | 0.097271394 | 0.264825685 |
| Sdc4 | 2016.916396 | 0.170193801 | 0.103076237 | 1.651144881 | 0.098708995 | 0.267892304 |
| Rmdn3 | 1731.020164 | -0.082753092 | 0.049924944 | -1.657550024 | 0.097408324 | 0.265151893 |
| Glrx3 | 298.516579 | -0.170866607 | 0.103150643 | -1.656476403 | 0.09762538 | 0.265692103 |
| Clcc1 | 950.504799 | 0.170172252 | 0.074448784 | 2.28576268 | 0.022268146 | 0.101179664 |
| Mt2 | 2130.99088 | 0.170167945 | 0.113000188 | 1.505908509 | 0.132090662 | 0.322199735 |
| Gnpat | 2228.485269 | 0.086625459 | 0.052325344 | 1.655516294 | 0.097819814 | 0.266049876 |
| Ceacam9 | 11.68099402 | 0.170083307 | 0.158454844 | 1.073386605 | 0.283097692 | 0.512599701 |
| Tvp23b | 820.6722041 | -0.126988222 | 0.076711013 | -1.65541058 | 0.097841241 | 0.266049876 |
| Mmgt1 | 1526.887222 | -0.096164132 | 0.058094999 | -1.655291065 | 0.09786547 | 0.266069065 |
| Gm17644 | 45.26630935 | 0.170073511 | 0.16404083 | 1.036775487 | 0.299840502 | 0.531324702 |
| Gstm5 | 4756.847382 | 0.170063798 | 0.074302431 | 2.288805289 | 0.022090668 | 0.100638874 |
| Acads | 280.1342732 | -0.202933092 | 0.122640205 | -1.654702801 | 0.097984799 | 0.266253327 |
| Hspa4 | 10218.49202 | -0.11926238 | 0.072083004 | -1.654514568 | 0.098023006 | 0.266310442 |
| Mrpl27 | 1108.121268 | -0.099673201 | 0.060253965 | -1.654218113 | 0.098083204 | 0.266427272 |
| Chtop | 4697.498399 | -0.113266727 | 0.068488913 | -1.653796529 | 0.098168861 | 0.266566481 |
| Gramd1c | 48.885542 | 0.170026361 | 0.165178708 | 1.029347928 | 0.303316209 | 0.535503799 |
| Abcg4 | 2641.15172 | 0.169940472 | 0.072316238 | 2.349962849 | 0.018775285 | 0.090427188 |
| Fam161b | 675.7785654 | 0.169937168 | 0.08822425 | 1.926195655 | 0.054079948 | 0.182711451 |
| Tpp1 | 2942.456635 | 0.169895307 | 0.077302525 | 2.197797636 | 0.02796353 | 0.118143491 |
| Vwa8 | 2146.162872 | 0.169843402 | 0.065252865 | 2.602849717 | 0.009245247 | 0.055586285 |
| Shank3 | 10452.37904 | -0.137840967 | 0.083498577 | -1.650818167 | 0.098775709 | 0.267979485 |
| D630041G03Rik | 177.9648893 | 0.169805667 | 0.112590795 | 1.508166514 | 0.131511915 | 0.321598108 |
| Mtch2 | 2861.503088 | -0.083989874 | 0.050889834 | -1.650425394 | 0.098855961 | 0.268056398 |
| Csad | 667.1608305 | 0.169729977 | 0.074780458 | 2.269710323 | 0.023225165 | 0.10424412 |
| Bdh2 | 43.91755434 | 0.169639674 | 0.162931808 | 1.041169777 | 0.29779676 | 0.529540878 |
| Gpx7 | 97.8630262 | -0.242367122 | 0.146918743 | -1.64966781 | 0.099010897 | 0.268382583 |
| Stambp | 1277.816659 | 0.091428978 | 0.055427489 | 1.649524088 | 0.099040312 | 0.268415358 |
| Zfp180 | 2697.99205 | -0.13231962 | 0.080231498 | -1.649222859 | 0.099101987 | 0.268535533 |
| Tmem258 | 679.886564 | -0.134641814 | 0.08164708 | -1.649070785 | 0.099133134 | 0.268572963 |
| Htr1d | 187.7952992 | -0.245978958 | 0.149196067 | -1.648695992 | 0.099209932 | 0.268690886 |
| Olfr1344 | 26.25325534 | -0.274760288 | 0.16665379 | -1.648689104 | 0.099211343 | 0.268690886 |
| Zkscan1 | 2427.397193 | -0.088917912 | 0.053955694 | -1.647980134 | 0.099356748 | 0.269037653 |
| Rnf170 | 1485.407178 | 0.169525524 | 0.068078644 | 2.490142475 | 0.01276919 | 0.069713116 |
| Rbm17 | 1618.195647 | 0.094683426 | 0.057469407 | 1.647544841 | 0.099446107 | 0.269185532 |
| Glra3 | 158.2189142 | 0.169521803 | 0.157373135 | 1.077196579 | 0.281392453 | 0.511269957 |
| Lbr | 469.8781697 | 0.16947915 | 0.100740591 | 1.682332302 | 0.092504421 | 0.256666996 |
| Itpkb | 1430.804117 | 0.169435783 | 0.083490786 | 2.029395003 | 0.042418074 | 0.15610783 |
| Alg14 | 270.3960585 | -0.160009401 | 0.097186503 | -1.646415883 | 0.099678165 | 0.26954004 |
| BC053749 | 111.9585432 | 0.169409237 | 0.123959297 | 1.36665212 | 0.171734372 | 0.37936304 |
| 3110070M22Rik | 24.49791855 | 0.169404272 | 0.167217731 | 1.01307601 | 0.311023856 | 0.54402222 |
| 2610507I01Rik | 852.8499956 | 0.16932114 | 0.074372164 | 2.276673572 | 0.022805726 | 0.102898053 |
| Wbscr17 | 2191.991231 | 0.169314681 | 0.072110622 | 2.347985305 | 0.018875266 | 0.090762462 |
| Scg3 | 11225.70952 | 0.097127957 | 0.059018601 | 1.645717726 | 0.099821888 | 0.269778533 |
| Akr1b10 | 569.4884522 | 0.169308433 | 0.085272613 | 1.98549602 | 0.047089313 | 0.167378254 |
| N4bp2l1 | 846.8445253 | 0.169234373 | 0.091100045 | 1.857676071 | 0.063215029 | 0.201202476 |
| Eif2ak3 | 1323.162797 | 0.169226157 | 0.080169849 | 2.110845393 | 0.034785603 | 0.136482181 |
| G6pdx | 1504.992627 | 0.169169513 | 0.091652446 | 1.845771937 | 0.064925327 | 0.204711643 |
| Grp | 126.130614 | -0.214133673 | 0.130271278 | -1.643751991 | 0.100227442 | 0.270638547 |
| Alkbh7 | 328.1805171 | -0.170491382 | 0.103742255 | -1.643413114 | 0.100297489 | 0.270733325 |
| Rexo4 | 1828.670651 | 0.096415039 | 0.058664801 | 1.643490423 | 0.100281505 | 0.270733325 |
| Zfp780b | 538.6399964 | -0.140652744 | 0.085600399 | -1.643131875 | 0.100355651 | 0.270843139 |
| Traf3ip2 | 27.69905862 | 0.169099557 | 0.16416393 | 1.030065233 | 0.302979385 | 0.535092054 |
| Dydc2 | 13.71324319 | 0.169039449 | 0.165675292 | 1.020305724 | 0.30758349 | 0.540141653 |
| Plekha1 | 3581.341946 | 0.169010225 | 0.09506708 | 1.777799679 | 0.075436761 | 0.225660556 |
| Ap3b2 | 6380.834835 | 0.168990645 | 0.066571804 | 2.538471761 | 0.011133779 | 0.063282013 |
| Vldlr | 3675.032005 | -0.113434124 | 0.069066998 | -1.642378077 | 0.100511675 | 0.271028176 |
| Rtbdn | 43.71684162 | 0.168882901 | 0.167348869 | 1.009166674 | 0.312894706 | 0.545944885 |
| Elf2 | 1022.852604 | 0.168832015 | 0.084540472 | 1.997055502 | 0.045819154 | 0.164396772 |
| Rpl11 | 6142.279163 | -0.108505091 | 0.066118627 | -1.641066914 | 0.100783526 | 0.271619403 |
| Ofd1 | 397.7542304 | 0.168656765 | 0.08204905 | 2.055560246 | 0.039824923 | 0.149045255 |
| Rhbdd1 | 323.3373147 | 0.168454286 | 0.104980702 | 1.604621446 | 0.108577139 | 0.286055805 |
| Prlhr | 8.540878702 | -0.23887891 | 0.145665214 | -1.639917337 | 0.101022356 | 0.272073766 |
| Fcho1 | 3799.827286 | 0.168307397 | 0.092268192 | 1.824110711 | 0.068135337 | 0.211218288 |
| Col9a3 | 458.4563731 | 0.168281886 | 0.167849686 | 1.002574922 | 0.316066001 | 0.54917695 |
| Cep290 | 643.5127787 | 0.168280855 | 0.094489245 | 1.780952466 | 0.074920216 | 0.224920332 |
| Cdkn2aip | 561.3969644 | 0.168279137 | 0.079918806 | 2.105626259 | 0.03523683 | 0.137555919 |
| Tmem208 | 566.5572256 | -0.154465583 | 0.09423794 | -1.63910186 | 0.101192048 | 0.272341426 |
| Hcn4 | 173.6593608 | -0.257635468 | 0.157206282 | -1.638836971 | 0.101247217 | 0.272442581 |
| Aifm2 | 322.1027906 | -0.191119396 | 0.116658596 | -1.638279586 | 0.101363384 | 0.272471248 |
| 1700007L15Rik | 11.58169702 | 0.168237654 | 0.166434663 | 1.010833025 | 0.312096354 | 0.545255252 |
| Slc30a3 | 4284.380181 | 0.168160943 | 0.115762378 | 1.452638982 | 0.146324017 | 0.345180316 |
| Dennd1c | 63.65636058 | 0.168123752 | 0.158762198 | 1.058965889 | 0.289615315 | 0.520046504 |
| Ppfia1 | 1585.951337 | 0.168106661 | 0.086740762 | 1.938035332 | 0.052618912 | 0.179774515 |
| Tspan7 | 44196.6614 | -0.142686644 | 0.087087 | -1.63843793 | 0.101330372 | 0.272471248 |
| Il1rl2 | 23.97345068 | 0.16805198 | 0.16763006 | 1.002516971 | 0.316093974 | 0.54917695 |
| Cct4 | 3636.356469 | -0.099049486 | 0.060477892 | -1.637780074 | 0.10146758 | 0.272656726 |
| Bin3 | 376.9463805 | 0.167952039 | 0.090663598 | 1.852474883 | 0.063957661 | 0.202733224 |
| Nt5c3l | 1087.079698 | -0.173658306 | 0.106084901 | -1.636974758 | 0.101635744 | 0.273013907 |
| Zfp324 | 449.481359 | -0.13297048 | 0.081279985 | -1.635956013 | 0.101848794 | 0.273538778 |
| Pars2 | 338.0320474 | -0.147187755 | 0.089980349 | -1.635776655 | 0.101886339 | 0.273592191 |
| St3gal6 | 434.5377962 | 0.167941318 | 0.079221096 | 2.119906512 | 0.03401393 | 0.134573122 |
| Armcx1 | 2475.894419 | 0.167935039 | 0.070115253 | 2.395128464 | 0.016614543 | 0.083309297 |
| 9230110C19Rik | 305.2241294 | 0.167892279 | 0.113156708 | 1.483714775 | 0.137884605 | 0.331476498 |
| Fnip2 | 841.8416717 | 0.167854208 | 0.083848977 | 2.001863523 | 0.045299412 | 0.163301323 |
| Rufy3 | 8947.127118 | 0.084619323 | 0.051792247 | 1.633822196 | 0.10229619 | 0.274407405 |
| Tbl1xr1 | 2726.036943 | -0.104113545 | 0.063723885 | -1.633822948 | 0.102296032 | 0.274407405 |
| Gm9899 | 906.215188 | 0.167771914 | 0.103922348 | 1.614396887 | 0.106441366 | 0.281963162 |
| H2-Q1 | 29.07651382 | 0.167735517 | 0.167737863 | 0.999986016 | 0.317317275 | 0.550314962 |
| Rnf169 | 2399.90417 | 0.09922354 | 0.060797838 | 1.632024151 | 0.102674399 | 0.275278964 |
| Fars2 | 539.8178072 | -0.119081426 | 0.072970849 | -1.631904084 | 0.102699694 | 0.27528099 |
| Limd1 | 524.0460605 | -0.120440024 | 0.073805732 | -1.631851903 | 0.102710688 | 0.27528099 |
| Ctif | 4353.583903 | 0.167721258 | 0.072305452 | 2.319621167 | 0.02036138 | 0.095224636 |
| Gpr146 | 1009.536684 | -0.107575819 | 0.065965892 | -1.630779417 | 0.102936872 | 0.275791786 |
| Rac2 | 99.62284882 | 0.167697062 | 0.13696095 | 1.224415149 | 0.220795657 | 0.44191594 |
| Atp11a | 3091.794114 | 0.167647855 | 0.082022494 | 2.043925368 | 0.040960921 | 0.152596418 |
| Ccdc67 | 10.1339444 | 0.167614756 | 0.158813212 | 1.055420729 | 0.291232946 | 0.521539907 |
| Xab2 | 1423.771143 | 0.167553701 | 0.066930969 | 2.503380771 | 0.012301312 | 0.067973084 |
| Ino80d | 1514.926483 | -0.122269419 | 0.075049053 | -1.629193377 | 0.103272089 | 0.276330832 |
| Prdm10 | 441.9958178 | -0.122675886 | 0.075290974 | -1.629357146 | 0.103237435 | 0.276330832 |
| Zfp456 | 74.49593103 | 0.167493131 | 0.13134475 | 1.275217557 | 0.202232249 | 0.419384114 |
| Zfp385b | 1428.745521 | -0.097456541 | 0.059818291 | -1.62920971 | 0.103268632 | 0.276330832 |
| Bahcc1 | 1312.242964 | -0.151152501 | 0.092785219 | -1.629057969 | 0.103300748 | 0.276336634 |
| Sp110 | 73.17541967 | 0.167399242 | 0.144361376 | 1.159584687 | 0.246217938 | 0.471382768 |
| Htr5a | 1082.182592 | 0.167303507 | 0.084483003 | 1.980321504 | 0.047667414 | 0.168697787 |
| Kat6a | 3335.655735 | -0.13386933 | 0.08221254 | -1.628332259 | 0.103454453 | 0.276556814 |
| Ap4m1 | 470.3342553 | 0.167112541 | 0.091291102 | 1.830545779 | 0.067168368 | 0.209220587 |
| BC068157 | 1419.481458 | 0.167027024 | 0.07205617 | 2.318011402 | 0.020448701 | 0.09551769 |
| Rab9b | 1127.154169 | -0.153943389 | 0.09459196 | -1.627446861 | 0.103642226 | 0.276963202 |
| Rnf4 | 3554.416021 | -0.083085133 | 0.051055538 | -1.627348114 | 0.103663185 | 0.27697144 |
| Fpgs | 218.55434 | 0.166955835 | 0.102724857 | 1.625272019 | 0.104104611 | 0.277824139 |
| Vipr1 | 1258.568698 | 0.166901229 | 0.101923714 | 1.637511258 | 0.101523688 | 0.272760192 |
| Polr3c | 705.3069206 | 0.166878633 | 0.077165186 | 2.16261557 | 0.03057076 | 0.125307765 |
| Dpy19l1 | 4659.300404 | -0.144977362 | 0.089169963 | -1.625854235 | 0.103980668 | 0.27758037 |
| S100a9 | 30.33289457 | -0.270831916 | 0.166571589 | -1.625919023 | 0.103966883 | 0.27758037 |
| Hgsnat | 2269.134365 | 0.16665925 | 0.084162435 | 1.980209453 | 0.047679998 | 0.168703789 |
| Gm13446 | 178.2292629 | 0.166637081 | 0.140975219 | 1.182031019 | 0.237193393 | 0.461518829 |
| Zfp384 | 1531.1735 | 0.166615135 | 0.071315091 | 2.336323645 | 0.01947438 | 0.092452392 |
| Morc2a | 2229.523804 | 0.166532527 | 0.074116634 | 2.246898144 | 0.024646537 | 0.10842517 |
| Rsl1 | 125.6456871 | 0.166518542 | 0.141211125 | 1.179216874 | 0.238311829 | 0.462671816 |
| Ikbkb | 1582.845805 | 0.099804399 | 0.061483754 | 1.623264568 | 0.10453286 | 0.278766289 |
| Tfe3 | 1677.973936 | 0.166504046 | 0.08118834 | 2.050836943 | 0.040282829 | 0.150442642 |
| Kank1 | 766.814029 | -0.152027644 | 0.093724301 | -1.622072861 | 0.104787748 | 0.279301801 |
| Rrp36 | 453.1182602 | -0.124515159 | 0.076761488 | -1.622104547 | 0.104780965 | 0.279301801 |
| Spats2l | 994.1113361 | 0.166492514 | 0.068828975 | 2.418930592 | 0.01556621 | 0.080108712 |
| Gnaq | 9300.135729 | -0.095338787 | 0.058794983 | -1.621546295 | 0.10490053 | 0.279410144 |
| Pdp1 | 5947.328751 | -0.145849678 | 0.089940593 | -1.62162237 | 0.10488423 | 0.279410144 |
| Phactr2 | 1506.256242 | -0.177257012 | 0.109308578 | -1.621620326 | 0.104884668 | 0.279410144 |
| Fam92a | 1392.222016 | -0.108361745 | 0.066832737 | -1.621387208 | 0.104934622 | 0.279442249 |
| Rad54b | 60.60801379 | 0.166460019 | 0.149822599 | 1.111047469 | 0.266547918 | 0.494085203 |
| Aasdhppt | 688.7303287 | -0.124549731 | 0.076829969 | -1.621108687 | 0.104994331 | 0.279485598 |
| Zfp28 | 438.2880191 | 0.16645658 | 0.101477453 | 1.640330682 | 0.100936429 | 0.271936887 |
| Agbl3 | 164.4570717 | 0.166453871 | 0.122983502 | 1.353465047 | 0.175907109 | 0.385037626 |
| Fbxo48 | 28.08582094 | 0.166413655 | 0.167389604 | 0.994169591 | 0.32014031 | 0.553376867 |
| Cul2 | 2153.166555 | -0.108986352 | 0.06726401 | -1.620277364 | 0.105172709 | 0.279798413 |
| Themis2 | 47.28465368 | 0.166413051 | 0.167745806 | 0.992054912 | 0.32117074 | 0.554073138 |
| Fbxl6 | 854.3088637 | 0.166382667 | 0.088468901 | 1.880691019 | 0.060013962 | 0.194471862 |
| Ggt5 | 104.1716105 | -0.207771206 | 0.128317794 | -1.619192477 | 0.105405856 | 0.280274298 |
| Trabd | 640.1246797 | 0.166361458 | 0.085630749 | 1.942777101 | 0.052043096 | 0.178476256 |
| Rtn4r | 2835.298067 | 0.16630805 | 0.122933866 | 1.352825349 | 0.176111432 | 0.385133456 |
| Hdac2 | 2718.900431 | -0.097569775 | 0.060276474 | -1.618704096 | 0.105510945 | 0.280429511 |
| Wdr78 | 403.0844226 | -0.195617035 | 0.120881205 | -1.618258479 | 0.105606905 | 0.280616255 |
| Cnst | 2285.990699 | -0.100642592 | 0.062200558 | -1.618033589 | 0.105655359 | 0.280696868 |
| Zzef1 | 2995.721911 | -0.111225503 | 0.06874491 | -1.617945289 | 0.105674389 | 0.280699294 |
| Tnks1bp1 | 3512.590875 | 0.166285906 | 0.091955706 | 1.808326125 | 0.070555758 | 0.216473449 |
| Tgtp2 | 27.72436825 | 0.166154913 | 0.146821277 | 1.131681429 | 0.257768392 | 0.484143855 |
| Gpatch8 | 3282.021016 | 0.166133577 | 0.096874362 | 1.714938534 | 0.086356497 | 0.245830535 |
| Nr1d2 | 4649.563805 | 0.086801627 | 0.053688617 | 1.616760326 | 0.105930026 | 0.281226693 |
| 6230400D17Rik | 13.95585025 | 0.166073984 | 0.165489678 | 1.003530773 | 0.315604837 | 0.54876192 |
| Pgpep1 | 490.9257719 | -0.117532316 | 0.072708192 | -1.616493441 | 0.10598767 | 0.281283693 |
| Srsf3 | 3632.505017 | -0.099182794 | 0.061368076 | -1.616195276 | 0.1060521 | 0.281364937 |
| Fmn1 | 922.1155731 | 0.166057032 | 0.101209058 | 1.640732911 | 0.10085287 | 0.271759021 |
| Zbtb46 | 246.7347384 | -0.182507568 | 0.112982868 | -1.61535612 | 0.106233597 | 0.281749975 |
| Cdk7 | 2304.743907 | 0.166007308 | 0.064169842 | 2.586998864 | 0.009681591 | 0.057385835 |
| Fut9 | 3221.643881 | -0.176435101 | 0.109248661 | -1.614986391 | 0.106313642 | 0.281828499 |
| C87436 | 860.8515892 | 0.166001194 | 0.06874021 | 2.414906693 | 0.015739241 | 0.080589491 |
| Kcnk10 | 131.0118844 | -0.185661629 | 0.114978134 | -1.614755981 | 0.106363549 | 0.281828499 |
| Limch1 | 4941.186566 | -0.085310077 | 0.052832888 | -1.614715397 | 0.106372342 | 0.281828499 |
| 6330549D23Rik | 53.85239455 | 0.165994379 | 0.141481009 | 1.173262622 | 0.240690507 | 0.465284929 |
| 9530077C05Rik | 247.7295275 | 0.165960679 | 0.127406149 | 1.302611227 | 0.192707523 | 0.406509239 |
| Plcg2 | 214.2150024 | 0.16588359 | 0.099024836 | 1.675171562 | 0.093900575 | 0.259077536 |
| Pdcd6 | 1886.986529 | -0.093517844 | 0.057958438 | -1.613532859 | 0.10662879 | 0.282363097 |
| 4921525O09Rik | 12.00856785 | -0.246502749 | 0.152810011 | -1.613132202 | 0.106715789 | 0.282545187 |
| Cnn1 | 20.07974851 | -0.269708087 | 0.167212491 | -1.612966143 | 0.106751864 | 0.28259241 |
| Hlx | 25.96144508 | 0.165883455 | 0.16483163 | 1.006381208 | 0.31423223 | 0.547477136 |
| Rgcc | 339.1961596 | -0.144263739 | 0.089493131 | -1.612009077 | 0.106959965 | 0.282977548 |
| Rbfox1 | 13232.77284 | 0.165871508 | 0.08308063 | 1.996512399 | 0.045878177 | 0.164469339 |
| Usp10 | 2357.068351 | -0.173119344 | 0.107396727 | -1.611961087 | 0.106970408 | 0.282977548 |
| Khdc3 | 8.317956657 | 0.165802656 | 0.147750211 | 1.122182198 | 0.26178498 | 0.488863021 |
| Zfp426 | 1048.022472 | -0.104493999 | 0.064855034 | -1.611193339 | 0.107137589 | 0.28332306 |
| Nfyc | 1406.76252 | 0.165767821 | 0.096762465 | 1.713141761 | 0.086686463 | 0.246489274 |
| Ankrd34b | 911.0774415 | -0.163504125 | 0.101528259 | -1.61042972 | 0.107304077 | 0.283666502 |
| A830082K12Rik | 269.4009045 | 0.165733523 | 0.120888835 | 1.370958059 | 0.170388031 | 0.377840546 |
| Ophn1 | 1698.860574 | 0.165645463 | 0.066268001 | 2.499629682 | 0.012432319 | 0.068428542 |
| Arhgap21 | 7951.638197 | 0.165602363 | 0.106978505 | 1.547996605 | 0.121623114 | 0.307561372 |
| Cyb561d2 | 412.6147067 | -0.147486041 | 0.091653837 | -1.609163859 | 0.107580517 | 0.28415488 |
| Tirap | 106.33414 | -0.183850693 | 0.114248167 | -1.609222256 | 0.107567751 | 0.28415488 |
| Eif2ak1 | 3114.058678 | 0.165566051 | 0.073867728 | 2.241385452 | 0.025001118 | 0.109148302 |
| Alyref | 516.9348803 | -0.124872457 | 0.077624627 | -1.608670631 | 0.107688381 | 0.284383375 |
| Chrnb4 | 21.99394608 | -0.238360466 | 0.148195175 | -1.608422586 | 0.107742658 | 0.284437681 |
| Atg5 | 799.6865395 | -0.098898112 | 0.061498129 | -1.608148302 | 0.107802703 | 0.284480801 |
| Blmh | 2098.61921 | -0.083090393 | 0.051668105 | -1.608156374 | 0.107800935 | 0.284480801 |

| | | | | | | |
|---|---|---|---|---|---|---|
| Khk | 305.7951116 | -0.067388149 | 0.085810358 | -0.785314862 | 0.432268984 | 0.813254665 |
| Myh7b | 393.5028347 | -0.067399686 | 0.074297334 | -0.907161572 | 0.364321362 | 0.777237064 |
| 4921531C22Rik | 56.20515678 | -0.067477303 | 0.094702893 | -0.712515752 | 0.476145461 | 0.835195112 |
| Zfp316 | 1341.524016 | -0.06750123 | 0.074143294 | -0.910415841 | 0.362603247 | 0.776449324 |
| Incenp | 814.251939 | -0.067521162 | 0.058466641 | -1.154866438 | 0.248145125 | 0.697476023 |
| Aph1c | 433.2641787 | -0.067536469 | 0.083740469 | -0.80649738 | 0.419956118 | 0.807657282 |
| Hdac3 | 3708.585767 | -0.067559046 | 0.051995078 | -1.299335431 | 0.19382884 | 0.649958942 |
| Celf1 | 10265.42286 | -0.067561435 | 0.061078242 | -1.106145705 | 0.268663459 | 0.710903001 |
| A330093E20Rik | 5.245324324 | -0.067581245 | 0.048689167 | -1.388013991 | 0.165132785 | NA |
| Tfe3 | 1792.472289 | -0.067638994 | 0.053894631 | -1.255022853 | 0.209470464 | 0.664226905 |
| Ntrk3 | 3308.804061 | -0.067665109 | 0.052618 | -1.28596884 | 0.198453937 | 0.653682967 |
| Aldh4a1 | 1107.694667 | -0.067666766 | 0.074262799 | -0.911179849 | 0.36220062 | 0.776449324 |
| Bahcc1 | 1277.197701 | -0.06767693 | 0.071074274 | -0.952200096 | 0.340995513 | 0.76263292 |
| Mir1949 | 3.139771678 | -0.06768633 | 0.039718068 | -1.704169738 | 0.088349381 | NA |
| Arhgef26 | 1222.559198 | -0.067686372 | 0.073152586 | -0.925276537 | 0.35482208 | 0.772141519 |
| Zfp157 | 1070.404989 | -0.067724719 | 0.056743942 | -1.193514529 | 0.232667925 | 0.684433837 |
| Odf3b | 56.49885954 | -0.067731837 | 0.080782374 | -0.838448213 | 0.401779021 | 0.799277436 |
| Gucy2e | 224.0846723 | -0.067760317 | 0.086007705 | -0.787840081 | 0.430790253 | 0.812181606 |
| Zbtb48 | 414.094628 | -0.067767584 | 0.082605039 | -0.820380756 | 0.411999083 | 0.802795661 |
| Grik4 | 1044.036517 | -0.06777808 | 0.076647211 | -0.884286309 | 0.376541677 | 0.783953619 |
| Phtf2 | 731.7276285 | -0.06777901 | 0.078536411 | -0.863026574 | 0.388122857 | 0.791889115 |
| Las1l | 2388.982548 | -0.067793623 | 0.04990513 | -1.358449975 | 0.174320946 | 0.627089168 |
| Nubpl | 180.0093613 | -0.0678467 | 0.085011084 | -0.798092397 | 0.424816873 | 0.809188912 |
| Vmac | 485.7351243 | -0.06787817 | 0.083601199 | -0.811928195 | 0.416832835 | 0.806153084 |
| Tbx4 | 10.09789491 | -0.067902674 | 0.067318953 | -1.008670965 | 0.313132461 | NA |
| Gpcpd1 | 3931.357867 | -0.067906379 | 0.062544197 | -1.085734296 | 0.277596563 | 0.71807708 |
| S100a3 | 22.71995486 | -0.067923585 | 0.082135653 | -0.82696835 | 0.408255001 | 0.801055189 |
| Zfp318 | 2633.932134 | -0.067939329 | 0.063088562 | -1.076888214 | 0.281530209 | 0.720080692 |
| Slc44a5 | 366.5880458 | -0.067941924 | 0.085122789 | -0.798163743 | 0.424775474 | 0.809188912 |
| Slc25a21 | 20.22036376 | -0.067964742 | 0.07800203 | -0.871320169 | 0.383579361 | NA |
| Nme4 | 69.47063149 | -0.067991588 | 0.094614133 | -0.718619785 | 0.472375219 | 0.833643839 |
| Trmt1l | 1206.196472 | -0.067998987 | 0.05899926 | -1.152539661 | 0.249099382 | 0.697784358 |
| 2310050B05Rik | 9.135076595 | -0.068016466 | 0.054082707 | -1.257637959 | 0.208522719 | NA |
| Gja4 | 124.2609827 | -0.068021724 | 0.093755001 | -0.72552635 | 0.468129186 | 0.832353544 |
| Rpl22 | 3478.564206 | -0.068029976 | 0.049013123 | -1.387995133 | 0.165138527 | 0.615077498 |
| Rnu73b | 5.881157751 | -0.068050395 | 0.049806521 | -1.366294888 | 0.171846425 | NA |
| 1700123M08Rik | 71.61222827 | -0.068053372 | 0.094936922 | -0.716827249 | 0.473480696 | 0.833983942 |
| Cyp1b1 | 137.2443396 | -0.068063937 | 0.094371734 | -0.721232242 | 0.470766638 | 0.833245343 |
| 2700050L05Rik | 1137.809086 | -0.068069057 | 0.076429962 | -0.890606971 | 0.373140058 | 0.782226671 |
| Nphs1 | 88.27058411 | -0.068076752 | 0.095090635 | -0.715914377 | 0.474044221 | 0.833994865 |
| Wdr19 | 1060.867365 | -0.068119586 | 0.06356156 | -1.07171041 | 0.283850124 | 0.721344803 |
| Skiv2l | 3474.396982 | -0.06812071 | 0.066821127 | -1.01944869 | 0.307990001 | 0.740254985 |
| Fbxl19 | 2625.894953 | -0.068129666 | 0.067842153 | -1.00423796 | 0.315263929 | 0.745993052 |
| Kif23 | 55.24900184 | -0.06813966 | 0.094613023 | -0.720193246 | 0.471406022 | 0.833245343 |
| Tmem138 | 297.5613943 | -0.068227448 | 0.076204079 | -0.895325412 | 0.37061316 | 0.780761257 |
| Gm5468 | 122.6306544 | -0.068252373 | 0.095102728 | -0.717669987 | 0.472960793 | 0.833983942 |
| A730043L09Rik | 8.732907499 | -0.068256709 | 0.06033899 | -1.131220616 | 0.257962247 | NA |
| Chchd7 | 433.6110298 | -0.068267576 | 0.078370834 | -0.871083958 | 0.383708313 | 0.78787654 |
| Phc3 | 2926.023858 | -0.068303419 | 0.060821416 | -1.123015929 | 0.261430729 | 0.704173028 |
| Eda2r | 20.20779828 | -0.068324287 | 0.080001276 | -0.854039972 | 0.393082842 | NA |
| Jtb | 909.6731735 | -0.068384891 | 0.06957656 | -0.982872558 | 0.325670166 | 0.75293408 |
| Tbx22 | 3.02359733 | -0.068424097 | 0.038129717 | -1.794508369 | 0.072732064 | NA |
| Hpx | 19.63998885 | -0.068481946 | 0.082033192 | -0.834807764 | 0.403825954 | NA |
| Ppfia1 | 1808.420609 | -0.068482822 | 0.066161254 | -1.03508954 | 0.300627097 | 0.734641347 |
| Sag | 76.26016113 | -0.06848431 | 0.09509585 | -0.720160867 | 0.471425955 | 0.833245343 |
| Abhd4 | 3205.8619 | -0.068509521 | 0.063290475 | -1.082461794 | 0.279047383 | 0.718516358 |
| Espn | 39.6417337 | -0.068537052 | 0.087997166 | -0.778855213 | 0.436065009 | 0.816020977 |
| Sltm | 2753.138603 | -0.068589534 | 0.057148952 | -1.200188842 | 0.230066008 | 0.683133099 |
| Ubtd1 | 322.6724989 | -0.06860764 | 0.078671845 | -0.872073612 | 0.383168221 | 0.787602461 |
| Zc3h3 | 1023.463527 | -0.068617884 | 0.072770097 | -0.942940669 | 0.345711251 | 0.765097832 |
| Nupr1 | 297.8289533 | -0.068625405 | 0.085068782 | -0.806704921 | 0.419836508 | 0.807657282 |
| Thoc2 | 1621.198615 | -0.068634119 | 0.068277877 | -1.005217533 | 0.314792115 | 0.745993052 |
| Nhlrc3 | 205.0077852 | -0.068645263 | 0.089219666 | -0.769396096 | 0.441658206 | 0.819100918 |
| Elmo3 | 73.3383848 | -0.068648602 | 0.095102609 | -0.721837209 | 0.47039457 | 0.833245343 |
| Rab26 | 1273.04184 | -0.06865635 | 0.078741784 | -0.871917631 | 0.383253315 | 0.787602461 |
| A530072M11Rik | 25.47318442 | -0.06867278 | 0.082280251 | -0.834620457 | 0.40393144 | 0.800260243 |
| Snora31 | 21.05596023 | -0.068739844 | 0.079740023 | -0.86204947 | 0.388660296 | NA |
| Glt25d2 | 320.2818892 | -0.068755246 | 0.085183597 | -0.807141848 | 0.419584764 | 0.807483875 |
| Wnt9a | 745.567634 | -0.068813764 | 0.093749475 | -0.734017598 | 0.462938013 | 0.831014288 |
| Ecm2 | 167.2268394 | -0.068823577 | 0.094128974 | -0.731162508 | 0.464679897 | 0.83159896 |
| Cryga | 13.75682525 | -0.068827411 | 0.0742905 | -0.926463147 | 0.354205341 | NA |
| Tm6sf2 | 51.37008414 | -0.068868544 | 0.073728177 | -0.934087164 | 0.350258933 | 0.768667813 |
| Ccnj | 362.7271889 | -0.068878865 | 0.079200417 | -0.869678059 | 0.384476366 | 0.788939637 |
| Calcoco1 | 3555.78984 | -0.068906698 | 0.045228041 | -1.523539326 | 0.12762383 | 0.565235453 |
| Ccna2 | 67.30770954 | -0.068914245 | 0.09491544 | -0.726059373 | 0.467802374 | 0.832353544 |
| A830009L08Rik | 165.2782873 | -0.068919046 | 0.089073296 | -0.773734087 | 0.439088058 | 0.817545617 |
| B3gat1 | 10024.52128 | -0.068930633 | 0.060297121 | -1.143182825 | 0.252962692 | 0.700378199 |
| Foxc1 | 149.3345406 | -0.068965535 | 0.094786302 | -0.727589675 | 0.466864803 | 0.832107746 |
| Mir3064 | 47.05966274 | -0.068975838 | 0.087617741 | -0.787235977 | 0.43114374 | 0.812181606 |
| Lrdd | 68.58760997 | -0.069018789 | 0.095100951 | -0.725742364 | 0.467996726 | 0.832353544 |
| 4933439K11Rik | 4.146532363 | -0.069031416 | 0.046290027 | -1.491280533 | 0.135887858 | NA |
| Ttc37 | 1020.380132 | -0.069039516 | 0.065629294 | -1.051961898 | 0.292817031 | 0.728526371 |
| Dpagt1 | 564.6142597 | -0.069056811 | 0.067682925 | -1.020298856 | 0.307586746 | 0.739991033 |
| Zfp703 | 1487.189313 | -0.069075767 | 0.048772302 | -1.416290892 | 0.156690353 | 0.607279636 |
| Palm3 | 90.48982105 | -0.069091772 | 0.095019901 | -0.727129489 | 0.467146636 | 0.832211278 |
| Clu | 32161.34513 | -0.069095906 | 0.069339983 | -0.996479998 | 0.319016981 | 0.74887535 |
| Nat9 | 539.8454868 | -0.069119434 | 0.071618018 | -0.96511235 | 0.334488542 | 0.758238352 |
| Nrn1l | 48.15236825 | -0.069139624 | 0.092472665 | -0.747676345 | 0.454655403 | 0.826168452 |
| Tmprss5 | 67.62718429 | -0.069153974 | 0.092038524 | -0.75135901 | 0.452436625 | 0.825121708 |
| Raf1 | 3018.940928 | -0.069162975 | 0.05187012 | -1.333387586 | 0.182404644 | 0.635245711 |
| Fpgs | 249.4047334 | -0.069177966 | 0.090462757 | -0.76471212 | 0.444442988 | 0.821493054 |
| Emp3 | 65.45624216 | -0.069233944 | 0.090777652 | -0.762676088 | 0.445656594 | 0.821773578 |
| Snhg3 | 147.7173678 | -0.069294668 | 0.09233076 | -0.750504686 | 0.452950802 | 0.82547056 |
| Gca | 549.105828 | -0.069302758 | 0.0851435 | -0.813952415 | 0.415672212 | 0.805633085 |
| Bcl2l12 | 55.44640986 | -0.06933191 | 0.092679559 | -0.748082002 | 0.454410697 | 0.826168452 |
| Dusp16 | 716.3156339 | -0.069398277 | 0.079381407 | -0.874238435 | 0.381988419 | 0.786663048 |
| Ercc1 | 661.1613551 | -0.06941154 | 0.072734299 | -0.954316481 | 0.339923468 | 0.761914237 |
| 8430410A17Rik | 585.3945338 | -0.069414602 | 0.068540902 | -1.012747129 | 0.311180961 | 0.743823372 |
| AI607873 | 16.16421211 | -0.069430389 | 0.065916918 | -1.053301511 | 0.292202827 | NA |
| Zfp780b | 557.9550371 | -0.069441487 | 0.086513741 | -0.802664245 | 0.422168824 | 0.808695226 |
| Lgr4 | 1183.475228 | -0.06944167 | 0.061528633 | -1.128607388 | 0.259063493 | 0.703828284 |
| 1700112E06Rik | 14.45170484 | -0.069444345 | 0.065927995 | -1.053336231 | 0.292186919 | NA |
| T2 | 35.52360724 | -0.069451214 | 0.085274747 | -0.814440575 | 0.415392602 | 0.805633085 |
| Gm16938 | 65.86533898 | -0.069473059 | 0.094803633 | -0.732810087 | 0.463674268 | 0.83159896 |
| 4932416H05Rik | 274.9223674 | -0.069495487 | 0.089048108 | -0.780426316 | 0.435139983 | 0.815382275 |
| Gria2 | 21687.52453 | -0.069516652 | 0.065828889 | -1.056020436 | 0.290958879 | 0.726946796 |
| Cd209g | 5.318352073 | -0.069521254 | 0.052711708 | -1.318895851 | 0.187203933 | NA |
| Dsn1 | 134.4841723 | -0.069555601 | 0.093242694 | -0.74596301 | 0.455689761 | 0.826603827 |
| Zfp46 | 1980.934466 | -0.069559505 | 0.060283792 | -1.153867449 | 0.248554516 | 0.697784358 |
| Cd200r4 | 9.428398792 | -0.06956079 | 0.063904957 | -1.088503822 | 0.27637275 | NA |
| Snapc2 | 1090.295093 | -0.069567387 | 0.067221126 | -1.034903626 | 0.300713921 | 0.734641347 |
| Stxbp3a | 773.6545433 | -0.0695744 | 0.066398011 | -1.047838608 | 0.294712971 | 0.729674556 |
| Wdr62 | 81.4855094 | -0.069586012 | 0.094543475 | -0.736021308 | 0.461717731 | 0.830371642 |
| Cdc23 | 2157.126353 | -0.069650671 | 0.050062234 | -1.391281729 | 0.164140012 | 0.614351408 |
| Lpar4 | 36.14557277 | -0.069689023 | 0.090497114 | -0.770069013 | 0.441258956 | 0.818962636 |
| Hist2h4 | 4.663266761 | -0.069702817 | 0.048870723 | -1.426269419 | 0.153790588 | NA |
| Psmb10 | 1144.557962 | -0.069718474 | 0.067428769 | -1.033957397 | 0.30115608 | 0.735102205 |
| Numa1 | 4603.519742 | -0.069735588 | 0.058903728 | -1.183890918 | 0.236456246 | 0.687227576 |
| Zadh2 | 1286.92833 | -0.069776046 | 0.060267379 | -1.157774688 | 0.246955995 | 0.696511265 |
| Zfp523 | 3933.452508 | -0.069803054 | 0.068414913 | -1.02029003 | 0.30759093 | 0.739991033 |
| Angpt2 | 58.78972417 | -0.0698325 | 0.086416819 | -0.808089224 | 0.419039221 | 0.806975048 |
| Prrc2c | 7530.016035 | -0.069907505 | 0.066838512 | -1.045916546 | 0.295599564 | 0.730624344 |
| Efna4 | 7.90850905 | -0.069952823 | 0.059715696 | -1.171431102 | 0.241425537 | NA |
| Gm14327 | 4.458747783 | -0.069978267 | 0.043831064 | -1.596545026 | 0.110367162 | NA |
| Vstm4 | 157.1341039 | -0.070064018 | 0.094731882 | -0.739603358 | 0.459540704 | 0.828566657 |
| Arpc1b | 557.8208511 | -0.070075611 | 0.068286693 | -1.026197172 | 0.304798654 | 0.737453291 |
| Lrriq3 | 17.7069029 | -0.07009279 | 0.080875271 | -0.866677658 | 0.38611865 | NA |
| Map4k5 | 1956.737268 | -0.070103575 | 0.061551974 | -1.138933012 | 0.254731095 | 0.701619624 |
| Hsf2 | 879.9272426 | -0.070104758 | 0.073264606 | -0.956870753 | 0.338632495 | 0.760991092 |
| Mpzl3 | 58.39818457 | -0.070104843 | 0.095029181 | -0.737719115 | 0.460685154 | 0.829336045 |
| Mll5 | 5163.908404 | -0.070106032 | 0.060556529 | -1.157695669 | 0.246988252 | 0.696511265 |
| Ccbe1 | 229.633856 | -0.070152029 | 0.094611066 | -0.741478048 | 0.458403637 | 0.827999407 |
| H2afj | 634.5104347 | -0.070176493 | 0.084395301 | -0.831521334 | 0.405679181 | 0.800733644 |
| Gm8787 | 9.866530148 | -0.070191798 | 0.06499039 | -1.080033503 | 0.280127261 | NA |
| AF357399 | 7.506786207 | -0.070229618 | 0.05839164 | -1.202734138 | 0.229079219 | NA |
| Sgk2 | 91.95481426 | -0.070280721 | 0.094999968 | -0.739797318 | 0.459422987 | 0.828552109 |
| Oprk1 | 580.0382967 | -0.070336978 | 0.078468371 | -0.896373616 | 0.370053254 | 0.780397655 |
| Luc7l | 5198.976072 | -0.070345109 | 0.053914426 | -1.304754872 | 0.19197633 | 0.647018138 |
| Cog3 | 1568.831826 | -0.070352487 | 0.056006792 | -1.256142066 | 0.209064467 | 0.663941943 |
| Galns | 347.5656745 | -0.070417876 | 0.080537612 | -0.874347694 | 0.381928933 | 0.786663048 |
| 2010001E11Rik | 12.49285768 | -0.070467465 | 0.073813875 | -0.95466422 | 0.33974753 | NA |
| 6530402F18Rik | 687.8853714 | -0.070480257 | 0.078826351 | -0.894120499 | 0.371257422 | 0.780761257 |
| Golga4 | 4711.22674 | -0.070530091 | 0.058127462 | -1.213369526 | 0.224988572 | 0.67664265 |
| Zfp606 | 507.0176629 | -0.070590296 | 0.070244485 | -1.004922962 | 0.314933947 | 0.745993052 |
| Zfp511 | 493.0675159 | -0.070595768 | 0.075457039 | -0.935575642 | 0.34949172 | 0.768268271 |
| Tmprss4 | 11.77360598 | -0.070619779 | 0.068231763 | -1.034998597 | 0.300669566 | NA |
| Mcf2l | 7751.723824 | -0.070631671 | 0.071202804 | -0.991978788 | 0.321207873 | 0.750501136 |
| Slc9b2 | 127.0227326 | -0.070644942 | 0.094925347 | -0.74421579 | 0.456745937 | 0.827003147 |
| Hps3 | 694.6830433 | -0.070683472 | 0.068346298 | -1.034196057 | 0.301044517 | 0.735062578 |
| Gtdc1 | 2572.938064 | -0.070724609 | 0.054515281 | -1.2973355 | 0.194515775 | 0.650595474 |
| Dpysl4 | 2161.92881 | -0.070775783 | 0.07011859 | -1.00937259 | 0.312795979 | 0.745030949 |
| Gata2 | 212.8275769 | -0.070789096 | 0.091251564 | -0.775757622 | 0.43789211 | 0.816984132 |
| Smarcd3 | 2632.696689 | -0.070795395 | 0.055415478 | -1.27753829 | 0.201412271 | 0.654961865 |
| Serpinb1a | 513.9535706 | -0.070813389 | 0.092577108 | -0.764912524 | 0.444323636 | 0.821464316 |
| Myt1 | 446.6396829 | -0.070820946 | 0.07822244 | -0.90537889 | 0.365264696 | 0.777735733 |
| Gm13483 | 16.75208756 | -0.070849643 | 0.077273885 | -0.916863995 | 0.359213913 | NA |
| Pfas | 1278.675842 | -0.070867957 | 0.068522378 | -1.034230856 | 0.301028252 | 0.735062578 |
| Taf6l | 360.5524529 | -0.070877124 | 0.074937225 | -0.945819978 | 0.344240414 | 0.76410657 |
| Mapk11 | 2234.308038 | -0.070884043 | 0.089049651 | -0.796005853 | 0.426028634 | 0.809519334 |
| Nnmt | 17.5621366 | -0.070895095 | 0.077903914 | -0.910032509 | 0.362805365 | NA |
| Olfml2a | 79.53306282 | -0.070903441 | 0.094703794 | -0.748686379 | 0.454046256 | 0.825985606 |
| Slc26a1 | 40.89580576 | -0.070905366 | 0.091430555 | -0.775510615 | 0.438037995 | 0.816984132 |
| Ercc6l | 5.57820661 | -0.070907094 | 0.045829641 | -1.547188507 | 0.121817796 | NA |
| Zfp457 | 9.808236318 | -0.070955993 | 0.059029107 | -1.202050937 | 0.229343794 | NA |
| Mmp19 | 26.94178065 | -0.070977574 | 0.0868746 | -0.817011809 | 0.413921679 | 0.804285711 |
| Pias3 | 1116.777526 | -0.070982361 | 0.061999784 | -1.144880775 | 0.252258549 | 0.69992334 |
| 6030468B19Rik | 3.236310119 | -0.07098463 | 0.041324612 | -1.717732537 | 0.085845412 | NA |
| Etohi1 | 564.6064396 | -0.071008198 | 0.069103111 | -1.027568755 | 0.304152728 | 0.737453291 |
| Dock1 | 1565.847599 | -0.071038864 | 0.071151979 | -0.998410231 | 0.318080475 | 0.748272127 |
| Med25 | 2756.575204 | -0.07105062 | 0.062371435 | -1.139153209 | 0.254639258 | 0.701619624 |
| Ccdc28b | 532.5706997 | -0.071094179 | 0.079837743 | -0.890483324 | 0.373206419 | 0.782226671 |
| Zfp362 | 1156.497971 | -0.071095579 | 0.060549361 | -1.174175539 | 0.240324722 | 0.6914669 |
| Msi1 | 668.948802 | -0.071154034 | 0.072860567 | -0.976578101 | 0.328778065 | 0.755788917 |
| Ctsd | 17702.75921 | -0.071274787 | 0.084656904 | -0.841925272 | 0.399829784 | 0.798662339 |
| Mettl24 | 52.73349745 | -0.071277981 | 0.094303123 | -0.755839032 | 0.449745711 | 0.823263966 |
| Zfp366 | 149.960106 | -0.071297451 | 0.094568262 | -0.753925781 | 0.450893789 | 0.823992799 |
| Fnbp1l | 4984.004963 | -0.071343255 | 0.069206982 | -1.030867883 | 0.30260278 | 0.735974513 |
| Igf2r | 1765.609842 | -0.071364675 | 0.056791461 | -1.256609248 | 0.208895164 | 0.663694478 |
| Stra13 | 275.1533547 | -0.071366775 | 0.08454668 | -0.84411091 | 0.398607431 | 0.797789838 |
| 1700011J10Rik | 303.4725756 | -0.071371787 | 0.078234656 | -0.912278396 | 0.361622184 | 0.776449324 |
| 5830416P10Rik | 65.06033746 | -0.071396841 | 0.084642804 | -0.843507521 | 0.398944661 | 0.797789838 |
| Ap5z1 | 1038.534672 | -0.071409045 | 0.079894063 | -0.893796634 | 0.37143071 | 0.780761257 |
| Wwtr1 | 594.5628202 | -0.071411483 | 0.071837419 | -0.994070837 | 0.320188382 | 0.750154748 |
| Dido1 | 3355.769063 | -0.071416196 | 0.05169285 | -1.381548827 | 0.167110276 | 0.617356019 |
| Cyp4b1 | 32.28568738 | -0.071478836 | 0.089991835 | -0.794281353 | 0.427031657 | 0.810328132 |
| Wipf1 | 619.9860346 | -0.071527794 | 0.066653925 | -1.073122023 | 0.283216371 | 0.720995619 |
| Gm10069 | 40.33905058 | -0.07153158 | 0.090825711 | -0.787569725 | 0.430948429 | 0.812181606 |
| Cd97 | 486.4019319 | -0.071557633 | 0.081095634 | -0.882385768 | 0.377568229 | 0.784178571 |
| Kctd6 | 1241.738139 | -0.07161866 | 0.063985641 | -1.119292697 | 0.26301529 | 0.70568202 |
| Cryzl1 | 1780.900909 | -0.071634382 | 0.047489485 | -1.508426159 | 0.131445492 | 0.572183763 |
| Fahd2a | 631.3133607 | -0.071636148 | 0.065678642 | -1.090706899 | 0.275401874 | 0.717061772 |
| Rpl39 | 2378.199662 | -0.071643416 | 0.078946595 | -0.907492168 | 0.364146589 | 0.777191869 |
| Raver2 | 775.1529841 | -0.07165717 | 0.080714243 | -0.887788415 | 0.374654576 | 0.783149984 |
| Igfbp7 | 711.7604195 | -0.071666484 | 0.08063461 | -0.888780687 | 0.374120957 | 0.782728075 |
| Iyd | 8.970726697 | -0.071719635 | 0.044567543 | -1.609234646 | NA | NA |
| Tcta | 1224.574759 | -0.071732127 | 0.067992888 | -1.054994559 | 0.291427813 | 0.727742894 |
| AI314180 | 5055.791805 | -0.071735923 | 0.05763962 | -1.244559273 | 0.213293799 | 0.666732456 |
| Ggcx | 677.4294235 | -0.071801252 | 0.070103745 | -1.024214211 | 0.305734108 | 0.738091018 |
| Gng11 | 349.3889454 | -0.071808501 | 0.092433822 | -0.776863914 | 0.437239063 | 0.816681836 |
| Ralgps1 | 1431.698074 | -0.071818942 | 0.062074781 | -1.156974548 | 0.247282759 | 0.696744638 |
| Car3 | 41.24078025 | -0.071830868 | 0.092215383 | -0.778946699 | 0.436011113 | 0.816020977 |
| Gramd3 | 676.0670053 | -0.071833374 | 0.073705739 | -0.974596755 | 0.329760324 | 0.756141871 |
| Zfp384 | 1736.961917 | -0.07185065 | 0.051254982 | -1.401827632 | 0.160966725 | 0.612415571 |
| Npbwr1 | 68.85813751 | -0.071891775 | 0.089449097 | -0.803717169 | 0.421560336 | 0.808573261 |
| Hkdc1 | 630.9474858 | -0.071894478 | 0.087776271 | -0.819065073 | 0.412749288 | 0.803699216 |
| Kcnj11 | 2167.87526 | -0.071933202 | 0.062369303 | -1.153343039 | 0.248769611 | 0.697784358 |
| Hmox1 | 273.4490644 | -0.071943063 | 0.080802309 | -0.890359009 | 0.373273145 | 0.782226671 |
| Lbr | 496.4391211 | -0.071947454 | 0.078188228 | -0.920182687 | 0.3574773 | 0.773877088 |
| Usp21 | 1660.797329 | -0.071989956 | 0.061031984 | -1.179544746 | 0.23818133 | 0.688933095 |
| 4930414L22Rik | 400.521918 | -0.071998637 | 0.078716987 | -0.914651849 | 0.36037443 | 0.776337125 |
| Rad50 | 810.4534724 | -0.072021941 | 0.065671623 | -1.096698054 | 0.272773407 | 0.714142044 |
| Zfp71-rs1 | 536.2460382 | -0.072045713 | 0.078084765 | -0.922660304 | 0.356184259 | 0.772920906 |
| Vps51 | 2107.419478 | -0.072045781 | 0.05916085 | -1.217794887 | 0.223301928 | 0.674214152 |
| 1700084C01Rik | 485.6556735 | -0.072048321 | 0.086772254 | -0.830315189 | 0.406360603 | 0.800759397 |
| Rimbp3 | 126.5856325 | -0.072065386 | 0.092956869 | -0.775256161 | 0.438188309 | 0.816984132 |
| Ercc4 | 1161.076942 | -0.072141394 | 0.054722335 | -1.318317188 | 0.187397489 | 0.641710498 |
| Eppk1 | 6.822632887 | -0.072149817 | 0.053135773 | -1.357838843 | 0.174514827 | NA |
| Hs3st1 | 706.8753255 | -0.072219593 | 0.076164908 | -0.948200368 | 0.34302746 | 0.763762832 |
| Rhoh | 33.26382315 | -0.072282055 | 0.091316779 | -0.791552831 | 0.42862146 | 0.810486244 |
| Rmdn2 | 232.0974923 | -0.072343481 | 0.085152 | -0.849580521 | 0.395558346 | 0.796239418 |
| Taf1b | 820.5573803 | -0.072353704 | 0.057768112 | -1.252485178 | 0.210393126 | 0.664820157 |
| Mplkip | 217.5241651 | -0.072363729 | 0.089247349 | -0.81082217 | 0.417467801 | 0.806153084 |
| Btbd7 | 661.7401532 | -0.072365061 | 0.069127816 | -1.04682985 | 0.29517806 | 0.730203765 |
| Rbm45 | 897.5321314 | -0.072398941 | 0.066019814 | -1.096624429 | 0.272805604 | 0.714142044 |
| Fam198b | 223.8975497 | -0.072410928 | 0.088080328 | -0.82210103 | 0.411019399 | 0.802622856 |
| Zfp653 | 675.9426359 | -0.072411197 | 0.068653981 | -1.054726856 | 0.291550265 | 0.727847831 |
| Mamdc4 | 66.0465452 | -0.072420564 | 0.094223687 | -0.76860253 | 0.442129303 | 0.819493462 |
| Oasl1 | 9.526263273 | -0.072440594 | 0.056789113 | -1.275607084 | 0.202094448 | NA |
| Pkp1 | 39.04563477 | -0.072474079 | 0.087414349 | -0.82908676 | 0.407055315 | 0.800975798 |
| Tnfrsf11b | 78.06196496 | -0.072561357 | 0.092959196 | -0.780572129 | 0.435054189 | 0.815326688 |
| Spef1 | 317.3489419 | -0.072671611 | 0.083139374 | -0.874093795 | 0.382067176 | 0.786663048 |
| Entpd2 | 443.8454563 | -0.07274825 | 0.082053029 | -0.886600415 | 0.37529407 | 0.783489748 |
| Gpr160 | 37.17106739 | -0.072772472 | 0.091904119 | -0.791830367 | 0.428459593 | 0.810486244 |
| Lpcat1 | 1031.936005 | -0.072785086 | 0.063134359 | -1.152860136 | 0.248967796 | 0.697784358 |
| Bin3 | 435.021654 | -0.072825189 | 0.07155998 | -1.017680402 | 0.308829864 | 0.741252908 |

| | | | | | | | | | | | | | |
|---|---|---|---|---|---|---|---|---|---|---|---|---|---|
| Lct | 81.40493008 | 0.16550512 | 0.151117999 | 1.095204549 | 0.273427033 | 0.501833505 | Vhl | 2014.393404 | -0.072840571 | 0.058902639 | -1.236626603 | 0.216225739 | 0.670049713 |
| Ppp1cb | 7532.461685 | -0.135239393 | 0.084159318 | -1.606944968 | 0.108066441 | 0.285049614 | Unk | 1033.390654 | -0.072855638 | 0.059323978 | -1.228097644 | 0.219410315 | 0.672421385 |
| Foxred1 | 1213.646557 | 0.165478151 | 0.071436956 | 2.316422189 | 0.020535228 | 0.095835515 | Bdp1 | 1651.38374 | -0.072856927 | 0.078787466 | -0.924727388 | 0.355107729 | 0.772206867 |
| Urb1 | 646.6566295 | 0.165379619 | 0.076859116 | 2.151724197 | 0.031419083 | 0.12760348 | Slc22a13b-ps | 20.0097511 | -0.07286063 | 0.072291743 | -1.007869329 | 0.313517198 | NA |
| Car8 | 462.128477 | -0.197300427 | 0.122830559 | -1.60628128 | 0.108212121 | 0.285336727 | AA465934 | 36.43230754 | -0.072866568 | 0.085814581 | -0.849116398 | 0.395816527 | 0.796347766 |
| Ism1 | 229.6560945 | 0.16534679 | 0.13581646 | 1.217428213 | 0.223441334 | 0.444711756 | Ptrf | 1005.690595 | -0.07290895 | 0.064036278 | -1.138556957 | 0.25488799 | 0.701619624 |
| Coq10b | 800.5042631 | -0.13848206 | 0.08626639 | -1.605284057 | 0.108431306 | 0.285817397 | Esyt2 | 1958.151436 | -0.07291683 | 0.059409707 | -1.227355485 | 0.21968901 | 0.672421385 |
| Iqgap3 | 37.62195037 | -0.269801793 | 0.168115361 | -1.60486104 | 0.108524389 | 0.286014099 | Slc8a2 | 13259.86438 | -0.072949183 | 0.059908339 | -1.217679944 | 0.223345622 | 0.674214152 |
| Nav2 | 2449.636448 | -0.119941298 | 0.074745086 | -1.604671353 | 0.10856615 | 0.286055805 | B430319G15Rik | 26.13514859 | -0.07294977 | 0.08529025 | -0.855311952 | 0.392378473 | 0.794184423 |
| Bcl7b | 1595.923052 | 0.165337685 | 0.076551571 | 2.159820915 | 0.030786536 | 0.125826058 | Dhrs9 | 2.156142019 | -0.072951255 | 0.034987487 | -2.085067001 | 0.037063219 | NA |
| Zxdb | 159.9154465 | -0.246801335 | 0.153920583 | -1.603432949 | 0.108839101 | 0.286697218 | Rapgefl1 | 8032.985176 | -0.072955794 | 0.063277268 | -1.152954232 | 0.248929171 | 0.697784358 |
| Zw10 | 802.3558829 | 0.165294638 | 0.083346857 | 1.983213815 | 0.047343551 | 0.167973663 | Iqub | 79.34538647 | -0.072976176 | 0.095068737 | -0.767614866 | 0.442716028 | 0.819995323 |
| Ankrd13a | 1759.749148 | 0.165266704 | 0.091597802 | 1.804264954 | 0.071189781 | 0.217685902 | Mdn1 | 2362.690612 | -0.07298355 | 0.085544991 | -0.853159831 | 0.393570676 | 0.794754188 |
| Nucb1 | 2952.363781 | 0.09829437 | 0.061318522 | 1.603012691 | 0.108931852 | 0.286793683 | Aebp1 | 1003.636885 | -0.073022193 | 0.090690206 | -0.805182786 | 0.420714208 | 0.807944816 |
| Spcs3 | 3606.289511 | -0.140460454 | 0.087636395 | -1.602763934 | 0.108986782 | 0.286793683 | Lims1 | 1076.092656 | -0.073041297 | 0.06323284 | -1.155116493 | 0.248042725 | 0.697458115 |
| Srp14 | 2178.498353 | -0.099238903 | 0.061916805 | -1.602778171 | 0.108983638 | 0.286793683 | 1810024B03Rik | 17.46279312 | -0.073048795 | 0.07412716 | -0.985452503 | 0.324401851 | NA |
| Pifo | 19.54342346 | 0.165223906 | 0.165468737 | 0.998520377 | 0.318027088 | 0.55111416 | Sept10 | 203.6225349 | -0.073110805 | 0.092740754 | -0.788335243 | 0.430500638 | 0.812136658 |
| Dmxl1 | 2575.373531 | -0.16583825 | 0.103484062 | -1.602548709 | 0.109034325 | 0.286870069 | Colec11 | 18.23717655 | -0.073169505 | 0.078167026 | -0.936066122 | 0.349239144 | NA |
| Pcdh20 | 877.5525293 | 0.165013334 | 0.125950193 | 1.310147534 | 0.190145931 | 0.403688825 | C630031E19Rik | 11.87871877 | -0.073176179 | 0.059276914 | -1.234480243 | 0.217024004 | NA |
| Dnajc25 | 91.29773315 | 0.165001571 | 0.13700588 | 1.20433934 | 0.228458447 | 0.451324134 | Med24 | 5183.182722 | -0.073186648 | 0.057146639 | -1.280681572 | 0.200305535 | 0.654475062 |
| Fzd4 | 290.9696813 | 0.164973725 | 0.114323595 | 1.443041784 | 0.149008701 | 0.349174352 | 5730507C01Rik | 40.43037769 | -0.073187674 | 0.091356272 | -0.801123693 | 0.423060041 | 0.808999423 |
| Gga3 | 3563.319896 | 0.164927639 | 0.062293414 | 2.647593498 | 0.008106694 | 0.050688101 | Mll2 | 4004.258196 | -0.073214513 | 0.076322847 | -0.959273872 | 0.337420794 | 0.760256625 |
| Ccdc64 | 1259.548578 | 0.164889955 | 0.097657463 | 1.688452159 | 0.091324465 | 0.254676254 | Fam181b | 452.6976127 | -0.073220257 | 0.081754995 | -0.895605923 | 0.370463271 | 0.780632011 |
| Cngb1 | 17.68476354 | -0.255775901 | 0.159900626 | -1.599592867 | 0.109688932 | 0.288151968 | Prr14 | 1040.055817 | -0.073305564 | 0.067357652 | -1.08830343 | 0.276461177 | 0.717475105 |
| Lifr | 1642.096512 | 0.164788054 | 0.090458205 | 1.821703788 | 0.068499943 | 0.211987375 | Adgb | 51.86077785 | -0.07338157 | 0.088312862 | -0.830927325 | 0.406014686 | 0.800733644 |
| Mall | 35.88474245 | -0.266561964 | 0.166634022 | -1.599685105 | 0.109668458 | 0.288151968 | Ccnyl1 | 841.7177033 | -0.073386019 | 0.066212718 | -1.10833721 | 0.267716211 | 0.710152314 |
| Ppara | 294.1924763 | -0.169215628 | 0.105783854 | -1.599635687 | 0.109679427 | 0.288151968 | Dalrd3 | 2531.951886 | -0.073431395 | 0.078906208 | -0.9306162 | 0.352052131 | 0.770051086 |
| Traf5 | 87.39833263 | 0.164693593 | 0.148686884 | 1.107653803 | 0.268011358 | 0.495905972 | Ccr1 | 5.885725369 | -0.073433887 | 0.04778158 | -1.53686603 | 0.12432612 | NA |
| Fam178a | 2358.483085 | 0.164673061 | 0.105144297 | 1.566162544 | 0.117310569 | 0.300737957 | Abi3bp | 275.294842 | -0.073467767 | 0.087387861 | -0.840709065 | 0.400510941 | 0.799169897 |
| Polr2c | 1701.920615 | -0.11940682 | 0.074669419 | -1.59913953 | 0.109789603 | 0.288269803 | Oxsm | 466.5000625 | -0.073530982 | 0.077497365 | -0.948819129 | 0.342712611 | 0.763762832 |
| 1810013D10Rik | 578.2923584 | -0.199145017 | 0.124550994 | -1.598903472 | 0.109842052 | 0.288358652 | BC064078 | 82.99375902 | -0.073539037 | 0.094849049 | -0.775327091 | 0.438146405 | 0.816984132 |
| Prepl | 18068.8158 | 0.16466208 | 0.065897302 | 2.498768142 | 0.012462582 | 0.068450002 | Dgkh | 1021.258487 | -0.073545476 | 0.083882615 | -0.876766614 | 0.380613416 | 0.786275719 |
| Epc2 | 1962.074664 | 0.164583883 | 0.104797696 | 1.570491431 | 0.116300826 | 0.2988829 | Fsd1l | 1140.175678 | -0.073552945 | 0.074343995 | -0.989359604 | 0.322487231 | 0.752229878 |
| Ap4e1 | 621.255981 | 0.16447983 | 0.10922014 | 1.505947804 | 0.132080573 | 0.322199735 | 2210016F16Rik | 343.1687003 | -0.073568649 | 0.083172043 | -0.884535792 | 0.37640705 | 0.783947144 |
| Ccar1 | 1945.599696 | 0.16438698 | 0.108218864 | 1.51902334 | 0.128756621 | 0.317415289 | Hist2h3c2 | 43.44213869 | -0.073574916 | 0.090234217 | -0.815377126 | 0.414856474 | 0.805104227 |
| Znhit1 | 988.1576962 | -0.136235619 | 0.085289485 | -1.597331953 | 0.110191732 | 0.289031775 | Wdr46 | 381.8981641 | -0.073576881 | 0.074080046 | -0.99320782 | 0.320608688 | 0.750352874 |
| Adi1 | 1039.209929 | 0.164347411 | 0.066084653 | 2.486922502 | 0.012885346 | 0.07003183 | Cherp | 2781.574521 | -0.073642878 | 0.057356747 | -1.283944477 | 0.199161383 | 0.65377551 |
| Fgf22 | 28.47812483 | 0.164339062 | 0.16623539 | 0.988592512 | 0.322862551 | 0.556196616 | Rpia | 377.7627157 | -0.073681788 | 0.075272031 | -0.978873391 | 0.327642541 | 0.754731296 |
| Rad23a | 2065.252559 | -0.084574152 | 0.052966097 | -1.596760131 | 0.110319186 | 0.289208519 | Foxg1 | 2732.495687 | -0.073716725 | 0.081604915 | -0.903336827 | 0.366347157 | 0.778210102 |
| Smad7 | 741.1600625 | -0.144924406 | 0.090774773 | -1.596527327 | 0.11037111 | 0.289208519 | Ahsa2 | 1164.377702 | -0.073759546 | 0.065685006 | -1.122928194 | 0.261467992 | 0.704173028 |
| Wnt5b | 72.411179 | -0.231121844 | 0.144751634 | -1.596678658 | 0.110337355 | 0.289208519 | Fam166b | 24.15292771 | -0.073772332 | 0.068390933 | -1.078685853 | 0.280727794 | 0.720080692 |
| Nfix | 3580.065298 | 0.164274252 | 0.097829132 | 1.679195642 | 0.093113921 | 0.25758027 | Chd1 | 1127.176703 | -0.073793876 | 0.075396172 | -0.978748309 | 0.327704356 | 0.754754 |
| Fam46c | 58.19601801 | 0.164145641 | 0.163663813 | 1.002944008 | 0.315887877 | 0.549010123 | Aadat | 4.590899464 | -0.073798253 | 0.049500199 | -1.490867794 | 0.135996208 | NA |
| AI197445 | 38.80726191 | -0.260256344 | 0.163075251 | -1.595927912 | 0.110504889 | 0.289412693 | Fbxo34 | 3169.287603 | -0.073807941 | 0.071786081 | -1.028165076 | 0.303872184 | 0.737396794 |
| Ccdc18 | 54.73161788 | -0.248872317 | 0.155963705 | -1.595706617 | 0.110554311 | 0.289412693 | Nacad | 2168.965158 | -0.073859605 | 0.067830139 | -1.088890665 | 0.276202103 | 0.717475105 |
| Esd | 2094.382511 | -0.113093612 | 0.070871032 | -1.59576642 | 0.110540953 | 0.289412693 | E130003G02Rik | 23.37096452 | -0.073872632 | 0.055724586 | -1.325673938 | 0.184947724 | 0.637780733 |
| Tyw3 | 494.2534413 | 0.164115723 | 0.080222406 | 2.045759179 | 0.04078007 | 0.151995768 | Tbkbp1 | 1935.898345 | -0.073946516 | 0.070941853 | -1.042353889 | 0.297247637 | 0.732415975 |
| Slc25a45 | 138.513087 | 0.164022748 | 0.131906567 | 1.243476745 | 0.213692205 | 0.43337003 | Sacs | 277.8933766 | -0.073997752 | 0.094571941 | -0.782449327 | 0.433950554 | 0.814624677 |
| Uqcrc2 | 7766.102323 | -0.091580059 | 0.057405059 | -1.595330804 | 0.110638281 | 0.289565798 | Otud3 | 427.143216 | -0.074000458 | 0.076543008 | -0.966782715 | 0.333652667 | 0.757997911 |
| Ager | 18.68590809 | 0.164020095 | 0.167942052 | 0.97664696 | 0.328743962 | 0.563509121 | Pole2 | 16.64288983 | -0.074004743 | 0.077220236 | -0.95835945 | 0.337881535 | NA |
| Gpr165 | 363.8398862 | -0.227844482 | 0.142853687 | -1.594949957 | 0.110723427 | 0.289690776 | Hdac1 | 996.6535555 | -0.074008931 | 0.064625725 | -1.145193052 | 0.252129196 | 0.69992334 |
| 2900052N01Rik | 1484.438359 | 0.163976915 | 0.085433045 | 1.919361709 | 0.054938573 | 0.184445993 | Rftn2 | 1222.204383 | -0.074018914 | 0.063200758 | -1.171171302 | 0.241529828 | 0.692393571 |
| Cwf19l1 | 569.905139 | 0.163959084 | 0.071213299 | 2.302366091 | 0.021314536 | 0.098164262 | Gm609 | 14.43189301 | -0.074038701 | 0.071368072 | -1.037420496 | 0.29953993 | NA |
| 3110021N24Rik | 14.03335348 | 0.163931952 | 0.161537922 | 1.014820232 | 0.310191528 | 0.543062998 | Tgm7 | 13.6494803 | -0.074056718 | 0.072154605 | -1.026361633 | 0.304721156 | NA |
| Smarce1 | 1991.443973 | -0.092049414 | 0.05774151 | -1.594163622 | 0.110899392 | 0.289955312 | Polq | 81.00769293 | -0.074065053 | 0.095081618 | -0.778962906 | 0.436001566 | 0.816020977 |
| Lmo3 | 3527.69097 | -0.154016004 | 0.096620537 | -1.594029694 | 0.110929384 | 0.289984795 | Pigo | 1164.23472 | -0.074081465 | 0.070574439 | -1.049692571 | 0.29385948 | 0.729288721 |
| Dusp11 | 2401.220771 | 0.163882709 | 0.117351442 | 1.396512103 | 0.162560339 | 0.367607043 | Stard5 | 258.9675069 | -0.074094864 | 0.082630882 | -0.896697 | 0.369880622 | 0.780397655 |
| Thyn1 | 1036.639465 | 0.163830892 | 0.078848561 | 2.077791789 | 0.03772854 | 0.143981775 | Zfp286 | 648.5890508 | -0.074164851 | 0.064132031 | -1.156440086 | 0.247501193 | 0.696744638 |
| Bag6 | 8447.948463 | 0.081649769 | 0.051234763 | 1.593640021 | 0.111016685 | 0.290066193 | Mir9-1 | 3.818828355 | -0.07419866 | 0.039814414 | -1.863613008 | 0.06237607 | NA |
| Ckmt1 | 7384.853641 | -0.097900492 | 0.061442461 | -1.593368674 | 0.111077509 | 0.29017618 | Mocs1 | 364.3747968 | -0.074210014 | 0.081314782 | -0.912626361 | 0.361439085 | 0.776449324 |
| Rnf14 | 14053.44576 | -0.091591823 | 0.05750755 | -1.592692149 | 0.11122927 | 0.290523652 | Piga | 371.2521065 | -0.074223656 | 0.079011448 | -0.939403815 | 0.347523455 | 0.766038372 |
| Trf | 11240.14151 | 0.163752615 | 0.167640482 | 0.976808309 | 0.328664062 | 0.563434496 | Prdx6 | 9152.52253 | -0.074232041 | 0.05246067 | -1.415003674 | 0.157067419 | 0.607833353 |
| Pold1 | 281.9869283 | 0.163744233 | 0.104089711 | 1.573106806 | 0.115694089 | 0.297966127 | Mir3070a | 5.906870762 | -0.074264786 | 0.045620204 | -1.627892433 | 0.103547696 | NA |
| Mtus2 | 2227.374698 | 0.163713903 | 0.087198015 | 1.877495747 | 0.06045018 | 0.195497811 | Id3 | 801.7745695 | -0.074294581 | 0.084913575 | -0.874943505 | 0.381604646 | 0.786663048 |
| Osbpl6 | 2933.617494 | 0.163703201 | 0.062341703 | 2.625901967 | 0.00864197 | 0.05307122 | Prss30 | 20.4621217 | -0.074376226 | 0.081195678 | -0.916012125 | 0.359660535 | NA |
| Fzd9 | 164.5389623 | -0.203302617 | 0.127786736 | -1.590952415 | 0.111620285 | 0.291299427 | 1700012B15Rik | 479.4558291 | -0.07443072 | 0.084861234 | -0.87708741 | 0.380439162 | 0.786275719 |
| Nucb2 | 596.3961076 | -0.111188098 | 0.06990953 | -1.590456957 | 0.11173184 | 0.291541451 | Slc37a2 | 185.4429621 | -0.074435552 | 0.092928768 | -0.800995789 | 0.423134084 | 0.808999423 |
| Cand2 | 189.6004404 | -0.235667324 | 0.148286274 | -1.589272676 | 0.111998845 | 0.29211375 | BC016579 | 7.121960708 | -0.074484614 | 0.058490265 | -1.273453182 | 0.202857276 | NA |
| Tacc2 | 3163.571524 | 0.163623244 | 0.074025472 | 2.210364065 | 0.027079906 | 0.115458464 | Cwc27 | 684.7848398 | -0.074494511 | 0.0758319 | -0.982363755 | 0.325920676 | 0.753033837 |
| Mmp2 | 96.40096998 | 0.163622301 | 0.161162527 | 1.015262691 | 0.309980624 | 0.542755089 | Lats1 | 1466.823874 | -0.074516391 | 0.059843833 | -1.245180786 | 0.213065304 | 0.666732456 |
| Rapgef2 | 6507.899594 | -0.175756685 | 0.110597198 | -1.589160383 | 0.112024188 | 0.29211375 | Emcn | 254.3468512 | -0.074520091 | 0.087550865 | -0.851163383 | 0.3946786 | 0.795437883 |
| Tgfbrap1 | 2723.659081 | 0.089594043 | 0.056387492 | 1.588899231 | 0.112083145 | 0.29211375 | Cxcl1 | 3.190413697 | -0.074557887 | 0.040010849 | -1.863441771 | 0.062400138 | NA |
| Ptk2b | 23606.20628 | 0.163603913 | 0.094628435 | 1.728908566 | 0.083825461 | 0.24166991 | Otof | 1692.062104 | -0.074643907 | 0.092093923 | -0.810519355 | 0.417641746 | 0.806201195 |
| Traf7 | 1966.455543 | 0.163488604 | 0.082227422 | 1.988249178 | 0.046784138 | 0.166752575 | Ddx5 | 33424.01137 | -0.074675867 | 0.051603059 | -1.447120931 | 0.147863057 | 0.594842535 |
| Rab18 | 3410.992659 | -0.122121631 | 0.076864841 | -1.588784034 | 0.112109159 | 0.29213241 | Kctd16 | 689.3346116 | -0.074683173 | 0.067043832 | -1.113945473 | 0.26530258 | 0.707506017 |
| Syt16 | 2748.663421 | -0.12009751 | 0.075609385 | -1.588394218 | 0.112197224 | 0.292312727 | D630032N06Rik | 6.672860791 | -0.074690012 | 0.05554282 | -1.344728499 | 0.178712928 | NA |
| Cdk11b | 2774.297479 | 0.163480168 | 0.062389417 | 2.620318907 | 0.008784757 | 0.053671542 | Tmem131 | 3882.273018 | -0.074723007 | 0.045981412 | -1.625069879 | 0.10414767 | 0.523645898 |
| Fcgr1 | 152.8228721 | 0.163406527 | 0.139335542 | 1.172755528 | 0.240893857 | 0.465428677 | Zfp940 | 695.0291367 | -0.074770268 | 0.065815426 | -1.136059925 | 0.255931486 | 0.701952734 |
| Lrpap1 | 7340.852705 | -0.086299383 | 0.054370204 | -1.587255091 | 0.112454882 | 0.292787085 | Sft2d1 | 806.0837213 | -0.074788821 | 0.068511524 | -1.091623959 | 0.274998421 | 0.716782175 |
| Ubl5 | 2615.886908 | -0.109453569 | 0.068955204 | -1.587314129 | 0.112441517 | 0.292787085 | D7Ertd715e | 154.4433967 | -0.074815787 | 0.093573562 | -0.799539797 | 0.42397748 | 0.808999423 |
| 0610031J06Rik | 1276.506886 | 0.163391465 | 0.064069732 | 2.550213016 | 0.010765712 | 0.061981468 | Cdhr1 | 458.8840209 | -0.074882296 | 0.094981026 | -0.788392158 | 0.430467356 | 0.812136658 |
| Kcnn4 | 10.6378526 | -0.261776997 | 0.164989843 | -1.586624922 | 0.11259762 | 0.293060225 | Ssfa2 | 2525.46506 | -0.074918525 | 0.057379673 | -1.305663159 | 0.191667132 | 0.646997681 |
| Atn1 | 7379.82213 | 0.163388492 | 0.071592029 | 2.282216247 | 0.022476576 | 0.101917493 | Acbd7 | 5.681388768 | -0.074946753 | 0.05355366 | -1.399470245 | 0.161672015 | NA |
| Mak | 141.5644726 | 0.163369212 | 0.117484212 | 1.390563114 | 0.164357948 | 0.369577538 | Syngr2 | 377.2560454 | -0.074947461 | 0.088674706 | -0.845195479 | 0.398001706 | 0.797789838 |
| C330007P06Rik | 1739.499865 | -0.128283458 | 0.080894264 | -1.585816496 | 0.112780943 | 0.293389511 | Pdpn | 347.1562361 | -0.074968266 | 0.083027414 | -0.90293389 | 0.366560983 | 0.778210102 |
| Zfp454 | 169.2438682 | 0.163172916 | 0.116183254 | 1.404444357 | 0.16018657 | 0.364614473 | Dusp12 | 474.7995372 | -0.074972147 | 0.072378644 | -1.035832426 | 0.300280327 | 0.733951208 |
| Mxd1 | 1176.025157 | 0.163074897 | 0.082936443 | 1.96626345 | 0.049268186 | 0.172238556 | Iglon5 | 1542.183123 | -0.074985635 | 0.065382581 | -1.146874802 | 0.251433368 | 0.699465245 |
| Dynlrb1 | 4134.598283 | -0.095786912 | 0.060448432 | -1.584605393 | 0.113056019 | 0.293957032 | Arhgap15 | 497.2280867 | -0.07500364 | 0.07182031 | -1.044323539 | 0.296335726 | 0.731596911 |
| 4732471J01Rik | 152.192627 | 0.163039621 | 0.116120598 | 1.404054264 | 0.160302692 | 0.364683468 | 1700012D01Rik | 11.53481532 | -0.075027263 | 0.065504573 | -1.145374421 | 0.25205409 | NA |
| Zcchc9 | 374.5193095 | 0.163005773 | 0.089999675 | 1.81118179 | 0.070112716 | 0.215455459 | Nabp1 | 359.0363227 | -0.075040618 | 0.076488811 | -0.9810666 | 0.3265599 | 0.753373722 |
| Ogn | 272.7007834 | -0.225049527 | 0.142066497 | -1.584113993 | 0.113167781 | 0.294050242 | Gm2115 | 410.5281722 | -0.075110976 | 0.093435982 | -0.803876342 | 0.421468394 | 0.808555694 |
| Astn2 | 1763.96251 | 0.163001582 | 0.065709132 | 2.480653411 | 0.013114182 | 0.070822981 | Asxl2 | 1259.222647 | -0.075116705 | 0.070881636 | -1.059748465 | 0.289259048 | 0.724444298 |
| 2-Mar | 1646.665163 | 0.162864162 | 0.087875576 | 1.853349591 | 0.063832268 | 0.202418574 | Ripk1 | 288.431728 | -0.075120614 | 0.077115664 | -0.974129118 | 0.329992434 | 0.756141871 |
| Etv1 | 2258.183266 | -0.118927222 | 0.075093216 | -1.58372791 | 0.113255651 | 0.294179891 | Otud4 | 2617.621183 | -0.07512802 | 0.069406182 | -1.082439882 | 0.279057115 | 0.718516358 |
| Bace2 | 146.2030808 | 0.162711894 | 0.135846295 | 1.197764686 | 0.231008639 | 0.453988819 | Asprv1 | 49.08449787 | -0.075182426 | 0.093929786 | -0.80041091 | 0.423472762 | 0.808999423 |
| Nckap1 | 22128.26872 | -0.106124461 | 0.067055258 | -1.582641889 | 0.11350311 | 0.294689071 | Tnrc6a | 4046.26566 | -0.075215831 | 0.060977643 | -1.23349849 | 0.21738984 | 0.670311138 |
| Ulk1 | 4568.641924 | 0.162672493 | 0.088409361 | 1.839991729 | 0.065769452 | 0.206448923 | Rnf123 | 2914.793028 | -0.075227398 | 0.06117314 | -1.229745568 | 0.218792397 | 0.671604514 |
| Gltscr1l | 2607.37818 | 0.162670186 | 0.06714602 | 2.422633316 | 0.015408471 | 0.079447204 | Fam214b | 1821.241821 | -0.075228062 | 0.071958092 | -1.045442701 | 0.29581841 | 0.73091652 |
| Ndufaf3 | 973.3157326 | -0.15393503 | 0.097339479 | -1.581424426 | 0.113781026 | 0.295297022 | Gm7008 | 17.73926759 | -0.075244522 | 0.07857882 | -0.957567469 | 0.338280909 | NA |
| Abcc6 | 57.87193829 | -0.264677566 | 0.167385734 | -1.581243274 | 0.113822424 | 0.295331486 | Barx2 | 191.7679776 | -0.075247406 | 0.087845479 | -0.85658826 | 0.391672476 | 0.794184423 |
| LOC100504703 | 25.03549314 | 0.162639557 | 0.167363491 | 0.971774402 | 0.331162793 | 0.56633955 | 2900079G21Rik | 225.9410858 | -0.075274475 | 0.088623685 | -0.849371989 | 0.395674335 | 0.796282265 |
| Pik3c3 | 2214.485819 | 0.094484951 | 0.059758343 | 1.581117316 | 0.113851216 | 0.295331486 | Pla2g2f | 5.010397569 | -0.075296748 | 0.044997721 | -1.673345815 | 0.094259238 | NA |
| Mctp2 | 78.67363428 | -0.254341321 | 0.160884778 | -1.580891152 | 0.113902928 | 0.295415461 | Mill2 | 37.6960303 | -0.075317897 | 0.084282166 | -0.893639789 | 0.37151465 | 0.780761257 |
| Ift172 | 2648.092459 | 0.162624228 | 0.070842678 | 2.295568607 | 0.021700559 | 0.099387664 | AI450353 | 65.82770912 | -0.075331469 | 0.093632695 | -0.804542356 | 0.421083818 | 0.808470525 |
| Adrbk2 | 5162.258101 | 0.162571337 | 0.098338468 | 1.653181514 | 0.098293928 | 0.266812564 | Lin7b | 1427.492799 | -0.075331602 | 0.074340651 | -1.01332987 | 0.310902625 | 0.743730778 |
| Iffo2 | 1155.182222 | 0.162503999 | 0.090768797 | 1.790306847 | 0.073404596 | 0.221939958 | Ccl2 | 5.620617041 | -0.0753318 | 0.048472925 | -1.554100555 | 0.120160433 | NA |
| Ttll12 | 698.2303168 | -0.118929229 | 0.075294976 | -1.579510815 | 0.114218938 | 0.296036839 | Gfm2 | 1099.750046 | -0.075343756 | 0.062152058 | -1.212248773 | 0.225417167 | 0.677510814 |
| Itga9 | 458.5966153 | -0.139134629 | 0.088148128 | -1.578418414 | 0.114469518 | 0.296434962 | Arntl | 1403.538022 | -0.075360035 | 0.071597012 | -1.052558382 | 0.29254344 | 0.72840007 |
| Med27 | 710.5066759 | -0.113920357 | 0.072176087 | -1.578367049 | 0.114481311 | 0.296434962 | 9530051G07Rik | 26.44202792 | -0.07539097 | 0.083887225 | -0.898718129 | 0.368802818 | 0.779573594 |
| Asb10 | 9.120888748 | 0.162434301 | 0.155095043 | 1.047321033 | 0.294951538 | 0.526313387 | Rorc | 112.9396857 | -0.075421865 | 0.094999079 | -0.793922065 | 0.427240803 | 0.810334558 |
| Mrc2 | 361.1488894 | -0.197956235 | 0.125433896 | -1.578171781 | 0.114526151 | 0.296434962 | Zfat | 300.6720454 | -0.075429023 | 0.077377623 | -0.974817007 | 0.329651039 | 0.756141871 |
| Nxpe4 | 440.0706167 | 0.162353464 | 0.097784949 | 1.660311378 | 0.096851828 | 0.264008132 | Ythdc2 | 1359.310303 | -0.075436226 | 0.067833442 | -1.112080169 | 0.266103682 | 0.70873159 |
| Ppt1 | 4526.349989 | 0.091360471 | 0.057876196 | 1.578550042 | 0.114439301 | 0.296434962 | Nlrp1c-ps | 12.07922935 | -0.075483529 | 0.064463929 | -1.170942116 | 0.241622045 | NA |
| Setdb2 | 196.7613371 | 0.162319206 | 0.124490807 | 1.303865 | 0.192279616 | 0.406270335 | E030011O05Rik | 16.68083249 | -0.075492156 | 0.069887816 | -1.080190511 | 0.280057353 | NA |
| Cep78 | 346.1385283 | 0.16220652 | 0.105139985 | 1.542767193 | 0.122887272 | 0.30949535 | Siglece | 40.75404433 | -0.075499124 | 0.09182758 | -0.822183535 | 0.410972448 | 0.802622856 |
| Kin | 303.8894087 | 0.162203401 | 0.105894822 | 1.531740623 | 0.125586454 | 0.313426464 | Vegfa | 2639.231711 | -0.075505807 | 0.081786023 | -0.923211624 | 0.355896932 | 0.772643677 |
| Tmem127 | 5716.557241 | -0.06075356 | 0.038507852 | -1.577692798 | 0.114636202 | 0.296622129 | Nup37 | 122.9410103 | -0.075538634 | 0.0904653 | -0.835001206 | 0.403717029 | 0.800224169 |
| P2ry12 | 1133.73008 | -0.264256064 | 0.167505466 | -1.577596667 | 0.114658299 | 0.296629768 | Tgm1 | 4.393657582 | -0.075539332 | 0.042679956 | -1.769901816 | 0.076743498 | NA |
| 0610040F04Rik | 44.09892936 | 0.162148361 | 0.153739248 | 1.054697243 | 0.291563813 | 0.521749609 | Hunk | 969.1343016 | -0.075556283 | 0.093192386 | -0.810755964 | 0.417505828 | 0.806153084 |
| Card11 | 8.194410806 | 0.16204799 | 0.150679384 | 1.07544898 | 0.282173763 | 0.511995964 | Arrb2 | 1543.056135 | -0.075669921 | 0.067694366 | -1.117817116 | 0.26364511 | 0.706200557 |
| Fbxo7 | 963.58793 | 0.162027488 | 0.067212974 | 2.410657916 | 0.015923776 | 0.081184757 | Upf3b | 831.0781464 | -0.075670933 | 0.065824739 | -1.149581968 | 0.250316089 | 0.698576658 |
| Naglu | 351.5502816 | 0.162015318 | 0.11751364 | 1.378693724 | 0.167989205 | 0.374561052 | Angel2 | 2386.112988 | -0.075685882 | 0.058283284 | -1.298586435 | 0.194085896 | 0.65013848 |
| Nagpa | 360.2536231 | 0.161959403 | 0.099761933 | 1.623458948 | 0.104491332 | 0.278703512 | Fah | 1013.739191 | -0.075708483 | 0.066618827 | -1.136442754 | 0.255771311 | 0.701952734 |
| Lrrn4cl | 63.12654398 | 0.161951712 | 0.160233403 | 1.010723787 | 0.312148649 | 0.545255252 | Kdm6a | 1783.92851 | -0.075725424 | 0.065858912 | -1.149812849 | 0.250220962 | 0.698461386 |
| Rab11fip5 | 4713.725086 | 0.161916675 | 0.086187446 | 1.878657298 | 0.060291302 | 0.195143813 | Tnfrsf9 | 23.47195483 | -0.075750508 | 0.085879772 | -0.88205297 | 0.377748163 | 0.784217496 |
| Scai | 1751.717248 | 0.161891181 | 0.097192618 | 1.66567363 | 0.095778437 | 0.262132229 | 1700023F06Rik | 65.74930753 | -0.075822859 | 0.093698678 | -0.809220153 | 0.418388525 | 0.80678504 |
| Hif1an | 2500.047356 | 0.161885317 | 0.063637323 | 2.543873758 | 0.01096307 | 0.062725925 | Snord49a | 8.36987764 | -0.075825735 | 0.057152766 | -1.326720306 | 0.184601221 | NA |
| Exph5 | 1296.325006 | 0.16169477 | 0.103315269 | 1.565061695 | 0.117568444 | 0.300942586 | Mirlet7d | 16.2469985 | -0.075852849 | 0.064109737 | -1.183172063 | 0.236740963 | NA |
| Zfp871 | 3792.312549 | -0.135722454 | 0.086124083 | -1.575894337 | 0.115050156 | 0.29712047 | Hnrnpu | 14763.34268 | -0.075855576 | 0.057611568 | -1.31667264 | 0.187948378 | 0.641781416 |
| Lmod1 | 63.73070514 | -0.259564969 | 0.164733561 | -1.575665379 | 0.11510294 | 0.297184629 | Ifitm2 | 239.2989946 | -0.075856733 | 0.088440842 | -0.857711572 | 0.391051748 | 0.794184423 |
| Dlg4 | 28797.77379 | -0.125702362 | 0.079788519 | -1.575444225 | 0.115153943 | 0.297227378 | Ccdc103 | 106.8740931 | -0.075864883 | 0.094849099 | -0.799848229 | 0.423798736 | 0.808999423 |
| Dysf | 236.8655917 | -0.173144072 | 0.109908734 | -1.575344065 | 0.115177048 | 0.297227378 | 1810043H04Rik | 233.9736903 | -0.075953016 | 0.084931658 | -0.894283915 | 0.371170004 | 0.780761257 |
| Taf13 | 737.2712828 | -0.112078377 | 0.071143346 | -1.575388047 | 0.115166902 | 0.297227378 | Plcg2 | 233.2673947 | -0.075969826 | 0.089892802 | -0.845115789 | 0.398046194 | 0.797789838 |
| Hmgcs2 | 424.8222214 | 0.16168046 | 0.154369815 | 1.047357994 | 0.294934498 | 0.526313387 | Dync1i2 | 5682.971103 | -0.07606266 | 0.046256011 | -1.644384318 | 0.100096842 | 0.5211685 |
| St6galnac4 | 1125.17515 | -0.11761531 | 0.074673132 | -1.575068655 | 0.115240598 | 0.297319994 | 1700001P01Rik | 6.991761568 | -0.076086929 | 0.054965728 | -1.384261267 | 0.166278469 | NA |
| Cacng7 | 2910.802726 | -0.091413999 | 0.058046497 | -1.574840937 | 0.115293165 | 0.29737844 | 5031425E22Rik | 355.1181025 | -0.076105309 | 0.089198653 | -0.853211419 | 0.393542072 | 0.794754188 |
| Mcam | 812.8395054 | -0.213092246 | 0.135335896 | -1.57454343 | 0.115361869 | 0.297506126 | Mir5133 | 9.959030639 | -0.076115897 | 0.053205542 | -1.43060091 | 0.152544627 | NA |
| Fbxw17 | 276.9335303 | 0.161529411 | 0.09813291 | 1.646026902 | 0.09975822 | 0.269653501 | F13a1 | 88.92591791 | -0.0761175 | 0.092258757 | -0.825043629 | 0.409346822 | 0.801055189 |
| Dusp3 | 8412.359065 | -0.116230133 | 0.073842351 | -1.574030777 | 0.115480334 | 0.297712529 | Snord37 | 8.287161125 | -0.076154195 | 0.060207011 | -1.26487253 | 0.205917026 | NA |
| B4galt6 | 6221.931128 | -0.099889696 | 0.063473177 | -1.573730832 | 0.115549691 | 0.297792233 | Snrnp48 | 432.7975437 | -0.076211463 | 0.075411621 | -1.01060636 | 0.312204871 | 0.744800424 |
| Vps4a | 4182.12895 | 0.089445533 | 0.056833743 | 1.573810337 | 0.115531304 | 0.297792233 | Nbl1 | 1548.662864 | -0.076274117 | 0.08193183 | -0.930946093 | 0.35188145 | 0.769793666 |
| Kctd2 | 2405.076453 | 0.098221808 | 0.062423929 | 1.57346405 | 0.115611406 | 0.297901734 | Gm16894 | 32.87984751 | -0.076279693 | 0.085102882 | -0.896323275 | 0.370080132 | 0.780397655 |
| Celsr2 | 10627.2883 | -0.12187196 | 0.077466787 | -1.573215632 | 0.115668897 | 0.297950771 | Cybrd1 | 144.2919682 | -0.07629626 | 0.095052816 | -0.80267227 | 0.422164185 | 0.808695226 |
| Morn4 | 2251.879861 | -0.112421792 | 0.071457975 | -1.573257461 | 0.115659215 | 0.297950771 | Slitrk3 | 2605.993439 | -0.076327294 | 0.067599993 | -1.129102106 | 0.258854763 | 0.703616072 |
| Zranb1 | 1454.379031 | 0.161525682 | 0.081932964 | 1.971437055 | 0.048673906 | 0.171098935 | Plscr2 | 70.57171207 | -0.07633733 | 0.094478431 | -0.80798685 | 0.419098152 | 0.806975048 |
| Wnt6 | 53.02103312 | -0.264330773 | 0.168053893 | -1.572892886 | 0.115743623 | 0.298044156 | Dnajc1 | 989.1359312 | -0.076357351 | 0.077503068 | -0.985217131 | 0.324517428 | 0.75293408 |
| Pitrm1 | 1510.592592 | 0.161486221 | 0.075940755 | 2.126476378 | 0.033463614 | 0.133183981 | Zfp451 | 798.8260081 | -0.076369727 | 0.071954681 | -1.061358708 | 0.288526915 | 0.724444298 |
| Leo1 | 837.1390371 | 0.161459566 | 0.104528223 | 1.544650437 | 0.122430839 | 0.308775687 | Agtr2 | 21.22505221 | -0.076461953 | 0.080295841 | -0.952252969 | 0.340968703 | NA |
| Ccdc157 | 1355.056535 | 0.161451596 | 0.072682685 | 2.221321284 | 0.026329208 | 0.113124374 | Gpr151 | 28.60642181 | -0.076486709 | 0.087549929 | -0.8736353 | 0.382316896 | 0.786954445 |
| Cc2d1a | 2045.275316 | 0.161427888 | 0.08053816 | 2.004365239 | 0.04503095 | 0.162712113 | E430016F16Rik | 5.605175975 | -0.076491836 | 0.04865903 | -1.571996725 | 0.115951311 | NA |
| Zfp235 | 701.6162628 | 0.161423955 | 0.071589171 | 2.254865546 | 0.02414177 | 0.10693327 | Snora61 | 25.19290436 | -0.076505729 | 0.084361673 | -0.906877798 | 0.364471423 | 0.777237064 |
| Ripk2 | 185.3302983 | 0.161142428 | 0.111867478 | 1.440476105 | 0.149732746 | 0.350068662 | Fam50a | 873.3156325 | -0.076533809 | 0.072199162 | -1.060037359 | 0.289127604 | 0.724444298 |
| 9030617O03Rik | 336.7652492 | -0.135832636 | 0.086452388 | -1.571184311 | 0.116139843 | 0.298716921 | Ccl22 | 9.872632942 | -0.0765794 | 0.065265436 | -1.173353083 | 0.240654244 | NA |
| Ogfod2 | 870.7012968 | 0.161055929 | 0.065345278 | 2.464691149 | 0.01371314 | 0.073125764 | Foxred1 | 1394.365395 | -0.076580286 | 0.059107893 | -1.29560168 | 0.19511275 | 0.65093872 |
| Golga2 | 4426.497719 | 0.161026414 | 0.068526363 | 2.34984621 | 0.018781169 | 0.090427419 | Zfp811 | 382.3456414 | -0.076584807 | 0.078880232 | -0.970899869 | 0.331598143 | 0.756643831 |
| Yipf1 | 1065.456223 | -0.100717831 | 0.064121602 | -1.570731671 | 0.116244989 | 0.298849717 | Cpt1c | 4635.541332 | -0.076622809 | 0.0708834 | -1.080969712 | 0.279710587 | 0.718934998 |
| Telo2 | 747.7107359 | 0.160999739 | 0.095350581 | 1.688502969 | 0.091314719 | 0.254676254 | Ccdc150 | 4.821910895 | -0.076630852 | 0.048106973 | -1.592926104 | 0.11117677 | NA |
| Rock1 | 1002.287694 | 0.160893504 | 0.084235043 | 1.910054277 | 0.056126225 | 0.186974787 | Tjp1 | 4685.570654 | -0.076713775 | 0.064630168 | -1.186965425 | 0.235241259 | 0.686276191 |
| Mgat5 | 1704.955291 | -0.124155965 | 0.079082218 | -1.569960576 | 0.116424283 | 0.299150555 | Galt | 1829.338374 | -0.076772084 | 0.068039476 | -1.128346202 | 0.259173738 | 0.703893404 |
| Orc4 | 1209.872764 | 0.160729129 | 0.08632358 | 1.861937712 | 0.062611871 | 0.199865303 | Fam132b | 29.80364986 | -0.076809326 | 0.081313276 | -0.944609901 | 0.34485807 | 0.764561711 |
| Fam109b | 25.30650372 | -0.25982024 | 0.16554518 | -1.569482358 | 0.116535587 | 0.299337266 | Adam15 | 4670.192678 | -0.076846272 | 0.067093896 | -1.14535414 | 0.252062487 | 0.69992334 |
| Sirt1 | 503.3344067 | 0.160635589 | 0.110694124 | 1.451166366 | 0.14673354 | 0.345620169 | Clcn7 | 2811.327118 | -0.076849436 | 0.057274834 | -1.341766199 | 0.179671815 | 0.633459544 |
| Syt14 | 226.0519141 | 0.160612625 | 0.095240412 | 1.686391547 | 0.091720409 | 0.255236649 | Enkd1 | 242.2400945 | -0.076866576 | 0.088621232 | -0.86736072 | 0.385744397 | 0.789534044 |
| Mllt11 | 10210.30031 | -0.100075246 | 0.063786391 | -1.568912192 | 0.116668401 | 0.299529444 | Pcdhb3 | 291.9673249 | -0.076878896 | 0.079713076 | -0.964445234 | 0.334822753 | 0.758238352 |
| Cep128 | 277.827191 | -0.159379782 | 0.101621663 | -1.568364229 | 0.116796154 | 0.299758094 | Nat2 | 25.54230195 | -0.07688904 | 0.084008753 | -0.915250346 | 0.36006022 | 0.776328279 |
| Ntrk3 | 3249.607935 | -0.102884896 | 0.065598882 | -1.568394041 | 0.116789201 | 0.299758094 | Pnpla6 | 2479.164265 | -0.076901918 | 0.06871544 | -1.119135933 | 0.263082152 | 0.705730626 |
| Sorcs1 | 2117.357958 | 0.160516821 | 0.080562251 | 1.992457004 | 0.046320938 | 0.165635037 | Hspa1l | 19.2298203 | -0.076910152 | 0.078225206 | -0.983188869 | 0.325514492 | NA |
| Dnajc11 | 2472.726338 | 0.160488766 | 0.075174513 | 2.134882679 | 0.032770595 | 0.131105179 | Bpifb4 | 6.177014064 | -0.07696068 | 0.049545941 | -1.553319584 | 0.120346804 | NA |
| Uprt | 132.9470319 | -0.196073262 | 0.125109547 | -1.56721263 | 0.117064999 | 0.300298857 | Cdc42ep5 | 35.338777 | -0.077042845 | 0.087596194 | -0.879522751 | 0.379117903 | 0.785380516 |
| Ggact | 498.145158 | 0.160277808 | 0.087518318 | 1.831362991 | 0.067046381 | 0.209017431 | Mir341 | 11.39807248 | -0.0770562 | 0.066883344 | -1.152098487 | 0.249280605 | NA |
| Zfp687 | 1571.2009 | 0.160230701 | 0.073900817 | 2.168185775 | 0.030144555 | 0.123954281 | Ppap2b | 11016.5852 | -0.077114184 | 0.053774496 | -1.434028948 | 0.151564002 | 0.598905575 |
| Gm13157 | 76.53630962 | 0.160198076 | 0.142109827 | 1.127283587 | 0.259622599 | 0.486370038 | Llgl1 | 2885.323746 | -0.077115214 | 0.06045633 | -1.275552347 | 0.202113808 | 0.655443359 |
| Rgs19 | 571.321884 | -0.122350848 | 0.078122115 | -1.566148686 | 0.117313813 | 0.300737957 | Rfc2 | 1170.782329 | -0.077184797 | 0.054684692 | -1.411451613 | 0.158111494 | 0.608727155 |
| Dnajc24 | 164.7118181 | -0.188778502 | 0.120544228 | -1.566051772 | 0.117336497 | 0.300746351 | Agbl5 | 599.6993829 | -0.077204935 | 0.084790632 | -0.910536142 | 0.36253983 | 0.776449324 |
| Hgf | 262.5481898 | 0.16018627 | 0.115483065 | 1.387097489 | 0.165412038 | 0.37100378 | Rps21 | 4041.956774 | -0.077257676 | 0.060190805 | -1.283546157 | 0.1993008 | 0.65377551 |
| Arid4b | 1778.595775 | 0.16017041 | 0.067274306 | 2.380855629 | 0.01727248 | 0.085350894 | Atp8b5 | 18.7828216 | -0.077274569 | 0.080068603 | -0.965104506 | 0.33449247 | NA |
| P2rx1 | 17.20879178 | -0.260036466 | 0.166071222 | -1.565812925 | 0.117392419 | 0.300771935 | Ltbr | 324.3509927 | -0.077358414 | 0.080058917 | -0.966268553 | 0.333909817 | 0.758212399 |
| Fat2 | 14.99877462 | -0.256300803 | 0.163700588 | -1.565668185 | 0.117426318 | 0.300777546 | 2310009B15Rik | 132.6620972 | -0.077480074 | 0.091452865 | -0.847213192 | 0.396876302 | 0.797044816 |
| Zrsr2 | 1371.938477 | 0.09570301 | 0.061130634 | 1.565549118 | 0.11745421 | 0.300799261 | Meis3 | 1354.697952 | -0.077487096 | 0.05857915 | -1.322776041 | 0.185909867 | 0.639632101 |
| Bmp6 | 352.8333659 | -0.159304695 | 0.101772533 | -1.56530147 | 0.117512239 | 0.300871791 | Dnahc10 | 79.35187217 | -0.077491862 | 0.087535666 | -0.885260442 | 0.37601618 | 0.783947144 |
| Gca | 458.128898 | 0.160159214 | 0.117076125 | 1.367992096 | 0.17131455 | 0.378651855 | A330069E16Rik | 25.64240387 | -0.077500896 | 0.085020205 | -0.911558573 | 0.362001139 | 0.776449324 |
| Gltscr2 | 2054.907563 | 0.16010978 | 0.094832123 | 1.688349631 | 0.091344133 | 0.254685262 | Il15 | 15.95075998 | -0.077507123 | 0.07452442 | -1.040023169 | 0.298329137 | NA |
| Qars | 2054.775251 | 0.160040046 | 0.066487306 | 2.407076729 | 0.016080791 | 0.081663643 | Mcm6 | 372.9984355 | -0.077511602 | 0.084798621 | -0.914066769 | 0.360681763 | 0.776337125 |
| Hinfp | 438.785681 | 0.160010512 | 0.089965359 | 1.778579158 | 0.075308784 | 0.225566144 | Myo1b | 1935.035603 | -0.077555016 | 0.065503813 | -1.183977112 | 0.236422123 | 0.687227576 |
| Reep5 | 16094.18526 | -0.088752353 | 0.056734078 | -1.564357018 | 0.117733749 | 0.301216431 | Map4k4 | 4434.556103 | -0.077628251 | 0.055295958 | -1.403868439 | 0.160358029 | 0.612373044 |
| Bri3bp | 5313.813318 | -0.097565894 | 0.062381253 | -1.564025869 | 0.117811493 | 0.301365573 | Rnf20 | 2781.386681 | -0.077633748 | 0.05660972 | -1.371385472 | 0.170254824 | 0.620316915 |
| Ascc3 | 1061.176633 | -0.133019535 | 0.085063624 | -1.563765198 | 0.11787272 | 0.301421803 | Kansl1l | 452.742509 | -0.077639511 | 0.074207046 | -1.046255244 | 0.295443203 | 0.730610822 |
| Hmga1 | 12.2203665 | 0.159939799 | 0.166754199 | 0.959135062 | 0.337490709 | 0.571052559 | Slc16a3 | 105.5333292 | -0.07765653 | 0.093786601 | -0.828013062 | 0.407663103 | 0.801055189 |
| Dtnb | 2502.897206 | -0.099672224 | 0.063742468 | -1.563670619 | 0.117894941 | 0.301421803 | Mirlet7c-2 | 15.04408271 | -0.077675209 | 0.069189872 | -1.122638432 | 0.261591086 | NA |
| Mag | 6216.197743 | 0.159866272 | 0.160360726 | 0.996916611 | 0.318804988 | 0.552030005 | Gcn1l1 | 3771.260464 | -0.077685008 | 0.049908371 | -1.55655265 | 0.119576733 | 0.55282356 |
| Mtss1 | 1270.404357 | -0.128332416 | 0.082075319 | -1.563593269 | 0.117913116 | 0.301421803 | Vrk1 | 618.3192573 | -0.077695362 | 0.063458087 | -1.224357147 | 0.220817528 | 0.672757795 |
| Glt8d1 | 1025.747975 | 0.15976582 | 0.138344538 | 1.154840096 | 0.248155914 | 0.473111538 | Trim41 | 2516.760384 | -0.077714702 | 0.048700699 | -1.595761527 | 0.110542046 | 0.537034702 |
| A830010M20Rik | 6754.455905 | 0.159734705 | 0.07674677 | 2.081321532 | 0.037404486 | 0.143269134 | Slc2a8 | 895.9724486 | -0.077716548 | 0.083210132 | -0.933979393 | 0.350314523 | 0.768667813 |
| Nek8 | 121.8495658 | -0.235995359 | 0.150972184 | -1.563171133 | 0.118012348 | 0.301481174 | Arhgef16 | 25.15265204 | -0.077735986 | 0.074590005 | -1.042176977 | 0.297329635 | 0.732415975 |
| Brms1l | 2485.106741 | -0.080862154 | 0.05173763 | -1.562927295 | 0.118069697 | 0.301577956 | AW495222 | 132.0647537 | -0.077740617 | 0.094703374 | -0.820885399 | 0.411711549 | 0.802795661 |
| Lonp2 | 2878.654587 | -0.082723495 | 0.052932897 | -1.562799299 | 0.118099809 | 0.30160515 | Grhl1 | 592.7663582 | -0.077751155 | 0.084244712 | -0.922920308 | 0.356048736 | 0.772857733 |
| Acad11 | 1080.329029 | 0.159731568 | 0.062719169 | 2.546774281 | 0.010872373 | 0.062368216 | Snord88c | 12.56359718 | -0.077791147 | 0.068901632 | -1.129017473 | 0.258890463 | NA |
| Rab37 | 151.6813864 | 0.159722572 | 0.167605891 | 0.952965146 | 0.34060773 | 0.574189552 | Snhg6 | 272.7049537 | -0.07780277 | 0.090742531 | -0.85740137 | 0.391223102 | 0.794184423 |
| Cldn19 | 22.123372 | -0.245925229 | 0.157387305 | -1.562548064 | 0.118158932 | 0.301607001 | Tmsb15b1 | 10.88147611 | -0.077861651 | 0.068270617 | -1.140485531 | 0.254084077 | NA |
| Nasp | 643.147386 | 0.159691161 | 0.086842 | 1.838870145 | 0.065934288 | 0.206715066 | Gm5860 | 10.1522238 | -0.077887037 | 0.058507488 | -1.331231944 | 0.183112704 | NA |
| Fam3a | 677.9260827 | 0.159665837 | 0.07035587 | 2.269403198 | 0.023243818 | 0.104288269 | Pvrl4 | 289.3050346 | -0.07790553 | 0.09013312 | -0.864338542 | 0.387401944 | 0.791583702 |
| Tjap1 | 661.8839185 | 0.159650177 | 0.121630242 | 1.312586194 | 0.189322424 | 0.402602476 | 2310042D19Rik | 39.93868593 | -0.077915209 | 0.092382448 | -0.843398398 | 0.399005667 | 0.797791648 |
| 1500009C09Rik | 3244.64739 | -0.097660925 | 0.062539 | -1.561600361 | 0.118382165 | 0.301977816 | 9930104L06Rik | 791.8893989 | -0.07795963 | 0.060587127 | -1.286735879 | 0.198186363 | 0.65354259 |
| 2610034B18Rik | 703.1394747 | -0.136979119 | 0.087739515 | -1.561202138 | 0.118476065 | 0.302167596 | Gas2l1 | 2169.210666 | -0.077979436 | 0.069916272 | -1.11532599 | 0.264710752 | 0.707370515 |
| Msr1 | 10.90050293 | 0.159640154 | 0.142636345 | 1.119210917 | 0.263050169 | 0.490280202 | F12 | 25.86089227 | -0.077995051 | 0.085954974 | -0.907394272 | 0.364198338 | 0.777191869 |
| Cox8b | 9.345137366 | 0.159576647 | 0.14070756 | 1.134101447 | 0.256751997 | 0.483132796 | Kcnq1ot1 | 177.0989626 | -0.07799914 | 0.094426427 | -0.826030826 | 0.408786605 | 0.801055189 |
| Polr3g | 222.9594598 | 0.159454538 | 0.096820367 | 1.646911122 | 0.099576315 | 0.269443841 | Usp24 | 4109.229977 | -0.078043226 | 0.056566847 | -1.379663713 | 0.16769021 | 0.617953076 |
| Pycr2 | 687.1900332 | -0.115885856 | 0.074266396 | -1.56040769 | 0.118663569 | 0.302391407 | Nfe2l3 | 162.2934901 | -0.078053382 | 0.094714319 | -0.824092735 | 0.409886868 | 0.801456003 |
| Susd4 | 3747.038484 | -0.085126346 | 0.054547722 | -1.56058481 | 0.118621745 | 0.302391407 | Nfkb1 | 785.1846131 | -0.078128014 | 0.069324225 | -1.126994417 | 0.259744842 | 0.703893404 |
| Bcl3 | 21.62825498 | 0.159389503 | 0.167939048 | 0.94909138 | 0.342574138 | 0.575828939 | Mppe1 | 611.9790515 | -0.078168216 | 0.073111638 | -1.069162427 | 0.284996486 | 0.721671536 |
| Ikzf2 | 112.2914945 | 0.159344968 | 0.116977838 | 1.362180822 | 0.173140827 | 0.381057382 | Fcgrt | 656.6161306 | -0.078210181 | 0.073651831 | -1.061890529 | 0.288285385 | 0.724378742 |
| Ppp1r9b | 37537.85477 | -0.134917029 | 0.086491455 | -1.559888539 | 0.118786223 | 0.302604855 | Mta2 | 2255.724013 | -0.078210821 | 0.053481611 | -1.462387159 | 0.143635146 | 0.589907432 |
| Gatm | 3360.566133 | 0.159336252 | 0.12228484 | 1.302992686 | 0.192577259 | 0.406446041 | Wdr78 | 422.9098256 | -0.078245523 | 0.072383862 | -1.080980213 | 0.279705915 | 0.718934998 |
| Cxcr4 | 55.35712124 | -0.259527703 | 0.166460239 | -1.559097257 | 0.118973363 | 0.302937269 | Gm17660 | 7.425864102 | -0.07825895 | 0.059256343 | -1.320684765 | 0.186606496 | NA |
| Gab3 | 53.76478252 | 0.159325663 | 0.15155395 | 1.051280179 | 0.293129928 | 0.523847186 | Gm5617 | 53.13242562 | -0.078288238 | 0.092743792 | -0.844134533 | 0.398594232 | 0.797789838 |
| Dram1 | 47.83103794 | 0.159312357 | 0.158372551 | 1.005934143 | 0.314447251 | 0.547605452 | Lrrc57 | 1424.401461 | -0.078323262 | 0.073382105 | -1.067334625 | 0.285820758 | 0.722148665 |
| S100b | 4453.939724 | 0.159251788 | 0.081321438 | 1.958300202 | 0.050194797 | 0.17432844 | Tspan12 | 480.1323501 | -0.078326975 | 0.071697837 | -1.092459385 | 0.274631233 | 0.716339252 |
| Tuba4a | 22537.3309 | -0.152656231 | 0.097999661 | -1.557722032 | 0.119299155 | 0.303576534 | Pyhin1 | 13.546221 | -0.078353468 | 0.06454254 | -1.213981781 | 0.224754681 | NA |
| Srrm3 | 1699.212928 | 0.159191173 | 0.062947262 | 2.528961047 | 0.011440072 | 0.064502358 | Kif27 | 177.2641073 | -0.078421765 | 0.089340944 | -0.877780803 | 0.380062684 | 0.786275719 |
| Icam2 | 96.25355347 | -0.189548943 | 0.121720764 | -1.557244113 | 0.119412539 | 0.303657683 | E2f8 | 17.80085121 | -0.0784627 | 0.074506293 | -1.053101642 | 0.29229441 | NA |
| Tln1 | 2213.03291 | 0.096867361 | 0.062204241 | 1.557246884 | 0.119411881 | 0.303657683 | Tnfrsf10b | 36.82755066 | -0.07848545 | 0.086752325 | -0.904707171 | 0.365620541 | 0.777735733 |
| L3mbtl2 | 1855.018214 | 0.159174355 | 0.067417335 | 2.36103006 | 0.018224253 | 0.088728557 | Gtf3c2 | 2837.630826 | -0.078527711 | 0.043963698 | -1.786194396 | 0.074067788 | 0.466974258 |
| 4933407L21Rik | 112.8923369 | -0.220969009 | 0.141950251 | -1.556665154 | 0.119550006 | 0.303955988 | Kremen1 | 1377.676778 | -0.078645074 | 0.068910164 | -1.141269582 | 0.253757756 | 0.701619624 |
| Ppp6c | 1257.078436 | -0.109697327 | 0.07048904 | -1.556232384 | 0.119652843 | 0.30416754 | 2310045N01Rik | 535.2173316 | -0.07864621 | 0.078360723 | -1.00364324 | 0.315550605 | 0.745993052 |
| Sepw1 | 15798.26678 | -0.122107733 | 0.078485957 | -1.555790833 | 0.119757839 | 0.304384507 | Reln | 2766.245626 | -0.07870614 | 0.066685145 | -1.180264967 | 0.237894847 | 0.688758314 |
| Ggct | 368.0400453 | -0.12267539 | 0.078880359 | -1.55520832 | 0.119896463 | 0.304636898 | Clec11a | 174.8118133 | -0.078758614 | 0.08847879 | -0.890141167 | 0.37339009 | 0.782325754 |
| Jmjd6 | 742.4531876 | 0.159096313 | 0.098408763 | 1.616688485 | 0.105945541 | 0.281226693 | Fgf9 | 830.2465435 | -0.078769522 | 0.066122459 | -1.191267271 | 0.233548678 | 0.685064073 |
| Gprc5b | 7573.804917 | -0.126227628 | 0.081170194 | -1.555098262 | 0.119922669 | 0.304653522 | Cic | 8212.125144 | -0.078781639 | 0.058461827 | -1.347574023 | 0.17779543 | 0.63083938 |
| Tmx2 | 8197.922598 | -0.064727991 | 0.041628482 | -1.554896747 | 0.119970662 | 0.304725482 | Gpr61 | 888.6900959 | -0.078809753 | 0.060379039 | -1.305250206 | 0.191807664 | 0.646997681 |
| Slc7a5 | 2806.160557 | 0.159015666 | 0.079598908 | 1.997711639 | 0.045747932 | 0.164230411 | Dpf3 | 114.3367124 | -0.078823383 | 0.095032874 | -0.829432805 | 0.406859545 | 0.800975798 |
| Rev3l | 1798.237886 | -0.162848394 | 0.104753954 | -1.554579937 | 0.120046145 | 0.304817269 | A330102I10Rik | 93.60835047 | -0.078824449 | 0.095100262 | -0.828856281 | 0.407185737 | 0.801026545 |
| Cd81 | 8603.855889 | 0.083752732 | 0.053885039 | 1.554285445 | 0.120116344 | 0.30494554 | Porcn | 2363.459832 | -0.078851376 | 0.058756124 | -1.342011191 | 0.179592367 | 0.633459544 |
| Fbxl3 | 3064.172815 | 0.094700665 | 0.060936722 | 1.554082033 | 0.12016485 | 0.305018707 | Myo7a | 1158.941619 | -0.078853639 | 0.065254835 | -1.208395349 | 0.226895228 | 0.679880436 |
| 2310011J03Rik | 1020.517059 | -0.129375842 | 0.083261892 | -1.553842212 | 0.120222059 | 0.305063966 | Srpk1 | 2485.967827 | -0.078893908 | 0.06230131 | -1.266328221 | 0.205395597 | 0.659018859 |
| Fgf1 | 2459.369368 | -0.130785784 | 0.084166956 | -1.553885166 | 0.120211811 | 0.305063966 | B3gntl1 | 243.39908 | -0.078906918 | 0.088289189 | -0.893732506 | 0.371465029 | 0.780761257 |

| | | | | | | | | | | | | | |
|---|---|---|---|---|---|---|---|---|---|---|---|---|---|
| Racgap1 | 579.7161744 | -0.145432995 | 0.093603371 | -1.553715358 | 0.120252328 | 0.30507634 | Ctsa | 4537.356838 | -0.078999328 | 0.068506666 | -1.153162646 | 0.248843633 | 0.697784358 |
| Grik4 | 970.7513924 | 0.158983934 | 0.105670082 | 1.504531191 | 0.132444648 | 0.322768524 | Gm16793 | 24.44454894 | -0.079008128 | 0.076437532 | -1.033630025 | 0.301309157 | 0.735352288 |
| Ephx3 | 15.32974799 | -0.249478553 | 0.160592341 | -1.553489732 | 0.120306181 | 0.30512751 | Aars2 | 1255.119799 | -0.079068843 | 0.077343068 | -1.022313255 | 0.306632662 | 0.738806192 |
| Lman2l | 1374.339392 | -0.170288391 | 0.109658798 | -1.552893107 | 0.120448674 | 0.305438912 | Ktn1 | 3754.44259 | -0.079085656 | 0.065028303 | -1.216172857 | 0.223919081 | 0.675094439 |
| Rab39 | 44.03358225 | -0.239382986 | 0.154194087 | -1.552478372 | 0.120547804 | 0.305640267 | Baz2b | 1411.674929 | -0.079099489 | 0.058544877 | -1.351091555 | 0.176666106 | 0.629083076 |
| Mpp6 | 1329.244605 | -0.122056881 | 0.078640069 | -1.552095303 | 0.120639422 | 0.305822513 | Zfp94 | 403.1129281 | -0.079112417 | 0.075080885 | -1.053695845 | 0.292022192 | 0.727847831 |
| C7 | 29.62906944 | 0.158802575 | 0.166080605 | 0.956177723 | 0.338982453 | 0.572508625 | Bbs12 | 147.5205878 | -0.079113012 | 0.090672202 | -0.872516714 | 0.382926554 | 0.787600928 |
| Pink1 | 15505.55604 | 0.158786266 | 0.112770569 | 1.408047035 | 0.159117143 | 0.36271311 | Fgfr2 | 2245.695055 | -0.079122421 | 0.07483494 | -1.05729251 | 0.29037811 | 0.725761285 |
| Zfp804b | 54.93675083 | -0.238858323 | 0.15407806 | -1.550242278 | 0.121083376 | 0.306797355 | Pkd1l2 | 19.65486971 | -0.079202927 | 0.075252442 | -1.052496445 | 0.292571841 | NA |
| Pgp | 1016.332078 | -0.155094311 | 0.100077095 | -1.549748325 | 0.121201934 | 0.307047543 | Psd | 16405.87659 | -0.079220294 | 0.073802321 | -1.073411957 | 0.283086322 | 0.720920007 |
| Rab8b | 1795.947007 | 0.158758168 | 0.090619865 | 1.751913533 | 0.079788677 | 0.2341852 | Fpr2 | 6.806976057 | -0.079244754 | 0.052852755 | -1.499349535 | 0.133782978 | NA |
| Atp1a3 | 93513.56428 | 0.080364203 | 0.051865865 | 1.549462321 | 0.121270623 | 0.307084755 | Tsfm | 468.9294897 | -0.079256248 | 0.06634492 | -1.194609132 | 0.23223978 | 0.68428297 |
| Cacng8 | 1197.939638 | -0.131248171 | 0.08470722 | -1.549433101 | 0.121277642 | 0.307084755 | Cxcr4 | 39.95362208 | -0.079286598 | 0.091845338 | -0.863262083 | 0.387993386 | 0.791889115 |
| Cdkl2 | 1519.309092 | -0.099399196 | 0.064155124 | -1.549357095 | 0.121295902 | 0.307084755 | Gtse1 | 31.31342911 | -0.079295046 | 0.082567061 | -0.960371422 | 0.336868316 | 0.760072873 |
| Tnpo3 | 3697.988227 | 0.086040326 | 0.055541382 | 1.549121096 | 0.121352612 | 0.307178137 | Htra2 | 1057.634354 | -0.079303132 | 0.072314399 | -1.096643726 | 0.272797165 | 0.714142044 |
| Pkp4 | 8915.032567 | -0.068740314 | 0.044379946 | -1.548904849 | 0.121404595 | 0.307259522 | Pld6 | 31.34537357 | -0.079337714 | 0.087580959 | -0.905878574 | 0.365000126 | 0.777634019 |
| Tmem255a | 1070.200471 | 0.158741358 | 0.16783955 | 0.945792322 | 0.344254523 | 0.576510979 | Kri1 | 1452.862903 | -0.079350935 | 0.056282991 | -1.409856395 | 0.15858209 | 0.609496529 |
| Txn2 | 3823.552154 | -0.093641409 | 0.060470449 | -1.548548262 | 0.121490351 | 0.307420027 | Frrs1 | 37.55165805 | -0.079377203 | 0.088746104 | -0.894430279 | 0.371091718 | 0.780761257 |
| D230025D16Rik | 1548.202437 | 0.158703384 | 0.070654664 | 2.2461841 | 0.024692218 | 0.10850482 | Snord104 | 33.49594113 | -0.07942785 | 0.077912749 | -1.019446119 | 0.307991221 | 0.740254985 |
| Acvrl1 | 788.8210393 | -0.133243794 | 0.086068957 | -1.548105135 | 0.121596986 | 0.307545495 | Olfr111 | 4.028216858 | -0.079471849 | 0.038192361 | -2.080831015 | 0.037449376 | NA |
| Fam117a | 197.7174087 | 0.158667461 | 0.136405892 | 1.163200936 | 0.244747987 | 0.469671205 | Sh2d3c | 955.425228 | -0.079481795 | 0.086871225 | -0.914938119 | 0.360224117 | 0.776337125 |
| Glo1 | 3207.684991 | -0.087063549 | 0.056265627 | -1.547366549 | 0.121774883 | 0.307800027 | Lrrd1 | 13.68845613 | -0.079492089 | 0.070995862 | -1.119672144 | 0.2628535 | NA |
| Gm10421 | 33.71722981 | -0.250205852 | 0.16169881 | -1.547357409 | 0.121777086 | 0.307800027 | Lpcat4 | 4758.637429 | -0.079492855 | 0.064443688 | -1.233524302 | 0.217380216 | 0.670311138 |
| Insig2 | 1755.715745 | -0.124858269 | 0.080686571 | -1.547447944 | 0.121755268 | 0.307800027 | Zfp108 | 280.6798664 | -0.079522761 | 0.078284597 | -1.015816177 | 0.309716931 | 0.742454975 |
| Eif4e | 3552.586716 | -0.082661218 | 0.053424768 | -1.547245243 | 0.12180412 | 0.307818143 | Atp6ap1l | 377.8667171 | -0.079531898 | 0.076483372 | -1.039858667 | 0.298405568 | 0.732994423 |
| Zfp451 | 719.5329128 | 0.158631688 | 0.083126501 | 1.908316678 | 0.056350299 | 0.187451116 | Impg2 | 37.02668661 | -0.079585775 | 0.087929703 | -0.905106836 | 0.365408791 | 0.777735733 |
| Tsnaxip1 | 10.97165806 | 0.158608089 | 0.146040863 | 1.086052802 | 0.277455632 | 0.506408006 | Dvl3 | 1713.447173 | -0.079606032 | 0.061406861 | -1.296370311 | 0.194847935 | 0.65082254 |
| Xrra1 | 54.1838493 | 0.158590595 | 0.148221634 | 1.069955787 | 0.28463921 | 0.514429012 | Sp1 | 1333.778939 | -0.079610406 | 0.068773022 | -1.157581901 | 0.247034699 | 0.696511265 |
| Tagap1 | 781.8964997 | 0.158589837 | 0.09360384 | 1.694266359 | 0.09021468 | 0.253004419 | Axin2 | 1141.390776 | -0.079666351 | 0.066858245 | -1.191571081 | 0.23342947 | 0.685064073 |
| Col11a2 | 906.29531 | 0.158559349 | 0.146251358 | 1.08415642 | 0.27829545 | 0.507283495 | Slc2a6 | 1010.363617 | -0.079758293 | 0.083241341 | -0.958157233 | 0.337983479 | 0.760581219 |
| Sptbn4 | 7911.858652 | 0.158553552 | 0.07402833 | 2.141795603 | 0.032209933 | 0.129592498 | Tmem69 | 514.9401212 | -0.079875738 | 0.070768384 | -1.128692422 | 0.259027607 | 0.703828284 |
| Smc1a | 3706.677148 | 0.094678476 | 0.061255582 | 1.54563017 | 0.122193911 | 0.30848017 | Rab13 | 170.738375 | -0.08000245 | 0.092056626 | -0.869056943 | 0.384815986 | 0.789016184 |
| Dpp3 | 1385.176054 | -0.103101555 | 0.06672438 | -1.545185655 | 0.122301363 | 0.308671986 | Pstpip1 | 90.82614258 | -0.08002728 | 0.0943499 | -0.848196769 | 0.396328397 | 0.796715512 |
| Gjc1 | 147.5705827 | 0.158549696 | 0.125985061 | 1.258480137 | 0.208218166 | 0.426398661 | 1500002O10Rik | 4.6156299 | -0.080044008 | 0.047158702 | -1.697332716 | 0.089633775 | NA |
| Ipo9 | 8393.62714 | 0.158542409 | 0.063393143 | 2.500939392 | 0.012386437 | 0.068273021 | Pomt2 | 1246.709531 | -0.080053822 | 0.053442529 | -1.497942249 | 0.134148256 | 0.576690607 |
| Atad3a | 1205.017581 | 0.158478003 | 0.077941827 | 2.033285726 | 0.042023664 | 0.15528611 | Flt1 | 2150.027807 | -0.08016222 | 0.081241432 | -0.986715989 | 0.32378189 | 0.752925264 |
| Anxa5 | 1897.922498 | 0.158452801 | 0.079850528 | 1.9843676 | 0.047214875 | 0.167632542 | 1110034G24Rik | 269.6397315 | -0.08019447 | 0.078013904 | -1.027950996 | 0.30397288 | 0.737453291 |
| Cd9 | 1314.85278 | 0.158436203 | 0.110519762 | 1.433555418 | 0.151699174 | 0.352863984 | Arid3b | 366.0580446 | -0.080241084 | 0.088932263 | -0.902271925 | 0.366912436 | 0.778335591 |
| Atp1a1 | 45155.12372 | 0.15842596 | 0.089484278 | 1.770433464 | 0.076654959 | 0.227841624 | Gm19757 | 176.5459794 | -0.080296818 | 0.087021105 | -0.922728088 | 0.356148925 | 0.772920906 |
| Fech | 2277.719581 | 0.090720696 | 0.058778461 | 1.543434344 | 0.122725426 | 0.309389481 | Scx | 51.44310128 | -0.08030102 | 0.090964939 | -0.882768911 | 0.377361141 | 0.783997209 |
| Entpd2 | 418.0586274 | -0.192747317 | 0.124898611 | -1.543230268 | 0.122774916 | 0.309463893 | 4930444P10Rik | 12.66172503 | -0.080319057 | 0.073104882 | -1.098682538 | 0.271906562 | NA |
| Sfpq | 5744.535786 | 0.158354228 | 0.080520576 | 1.966630597 | 0.049225813 | 0.172129261 | Cyp2j6 | 1024.468826 | -0.080323122 | 0.064752075 | -1.240471779 | 0.214800946 | 0.668817995 |
| Scn9a | 107.6080588 | -0.257127493 | 0.166643052 | -1.542983576 | 0.122834761 | 0.30946442 | Bai1 | 20664.8244 | -0.080348516 | 0.066861623 | -1.201713513 | 0.229474544 | 0.682503526 |
| Wwc2 | 1731.746708 | -0.082245775 | 0.053301072 | -1.543041675 | 0.122820665 | 0.30946442 | Clec10a | 25.99063784 | -0.08037125 | 0.07626362 | -1.053860945 | 0.291946586 | 0.727847831 |
| B130024G19Rik | 10.9025305 | 0.158352981 | 0.164678543 | 0.961588427 | 0.336256387 | 0.570266688 | Fanca | 56.77712571 | -0.080377608 | 0.094261818 | -0.852705892 | 0.393822423 | 0.794754188 |
| Sepp1 | 7957.313696 | 0.158243279 | 0.084114333 | 1.881287927 | 0.059932762 | 0.194316101 | Bnipl | 6.133403478 | -0.08038094 | 0.051260426 | -1.5680896 | 0.116860223 | NA |
| 4922501L14Rik | 244.804466 | -0.164467501 | 0.106649421 | -1.54213216 | 0.123041481 | 0.309806965 | Pcdhb16 | 294.7106433 | -0.080408148 | 0.07699779 | -1.04429163 | 0.296350484 | 0.731596911 |
| Psme4 | 2407.205705 | 0.158239066 | 0.065913981 | 2.400690453 | 0.016364173 | 0.082642272 | Edn3 | 210.3126546 | -0.080415283 | 0.086130477 | -0.933644931 | 0.35048708 | 0.768667813 |
| Asb8 | 1691.617119 | 0.092412689 | 0.059941274 | 1.541720468 | 0.123141536 | 0.309984558 | Greb1 | 66.914309 | -0.080466491 | 0.092021356 | -0.874432783 | 0.381882611 | 0.786663048 |
| Ngf | 72.43765657 | -0.219821194 | 0.142612906 | -1.541383595 | 0.123223454 | 0.310073074 | Ip6k3 | 49.89378915 | -0.080484029 | 0.092178164 | -0.873135513 | 0.382589219 | 0.787068545 |
| Pi4kb | 2173.92391 | 0.092247076 | 0.059842769 | 1.541490763 | 0.123197389 | 0.310073074 | Nhlrc2 | 455.9609038 | -0.080563548 | 0.077672583 | -1.03721989 | 0.299633391 | 0.733704078 |
| Srprb | 1526.322889 | -0.075985353 | 0.049298595 | -1.541328993 | 0.123236736 | 0.310073074 | Mybbp1a | 2811.953097 | -0.080571451 | 0.051073134 | -1.577570151 | 0.114664394 | 0.543130934 |
| Lrp3 | 3362.099759 | 0.087898537 | 0.057080072 | 1.539916349 | 0.123580745 | 0.31078722 | Zbtb1 | 474.2136807 | -0.080599558 | 0.085522996 | -0.942431414 | 0.34597181 | 0.765097832 |
| Ctsl | 3243.483941 | 0.158072311 | 0.077964814 | 2.027482697 | 0.042613072 | 0.156567925 | Alox12 | 27.82107848 | -0.080615724 | 0.085209644 | -0.946086851 | 0.34410429 | 0.763939922 |
| Mir331 | 20.14501712 | 0.15804545 | 0.166305628 | 0.950331336 | 0.341943921 | 0.575421758 | Rps27l | 235.9756613 | -0.080620875 | 0.082303873 | -0.979551415 | 0.327307597 | 0.754198947 |
| Zfp326 | 819.4951241 | 0.097594055 | 0.063391104 | 1.539554428 | 0.123669001 | 0.310958699 | Usp19 | 6148.282076 | -0.080633375 | 0.062199972 | -1.296357094 | 0.194852487 | 0.65082254 |
| Lfng | 616.7256274 | -0.166490022 | 0.10814989 | -1.539437742 | 0.123697466 | 0.310979805 | 1500015A07Rik | 149.6894128 | -0.080636256 | 0.087260918 | -0.924082141 | 0.355443551 | 0.772370994 |
| Agbl5 | 502.7610006 | 0.157967749 | 0.129271279 | 1.221986433 | 0.221712758 | 0.442910968 | Oas1g | 2.947954269 | -0.080637149 | 0.037524003 | -2.148948457 | 0.031638486 | NA |
| Wbp7 | 2845.788741 | 0.157958422 | 0.064784311 | 2.438220301 | 0.014759776 | 0.077181224 | Stx16 | 3800.761947 | -0.080651369 | 0.054302898 | -1.485212973 | 0.137487417 | 0.582012447 |
| Ecd | 1678.802068 | -0.084937738 | 0.055215224 | -1.538302887 | 0.123974574 | 0.311524822 | Fblim1 | 80.38485613 | -0.080655662 | 0.095064678 | -0.848429346 | 0.396198905 | 0.796565439 |
| Skiv2l | 3115.62374 | 0.157956795 | 0.063631803 | 2.482356104 | 0.013051677 | 0.070657821 | Cstf2 | 4481.077771 | -0.080687467 | 0.047954576 | -1.682581163 | 0.092456201 | 0.506608364 |
| Slc2a1 | 4373.888318 | 0.157931061 | 0.156637657 | 1.008257296 | 0.313330958 | 0.546413192 | Trafd1 | 2338.666339 | -0.080698808 | 0.068201807 | -1.18323564 | 0.236715772 | 0.687431687 |
| Ddx59 | 291.7719699 | 0.157890154 | 0.101325005 | 1.558254581 | 0.119172911 | 0.303345669 | Rcvrn | 8.638259164 | -0.08072021 | 0.061997807 | -1.301984926 | 0.192921539 | NA |
| Sco2 | 106.4204618 | 0.157869998 | 0.116664553 | 1.35319592 | 0.175993048 | 0.385096926 | Snord95 | 7.156057332 | -0.080724719 | 0.056431169 | -1.430498779 | 0.152573916 | NA |
| Pter | 75.1868393 | -0.236713303 | 0.153944381 | -1.537654708 | 0.124133063 | 0.311670343 | Ppfia4 | 3742.786209 | -0.080729002 | 0.06449337 | -1.251741102 | 0.210664218 | 0.6650979 |
| B330016D10Rik | 65.98139691 | 0.157783602 | 0.148072018 | 1.065586895 | 0.286610426 | 0.516185277 | Inpp5k | 1989.657461 | -0.08075188 | 0.047556336 | -1.698025672 | 0.089502917 | 0.50004591 |
| Casc3 | 2670.207624 | 0.095995838 | 0.062457073 | 1.536989069 | 0.124295986 | 0.311791886 | D130040H23Rik | 111.4770081 | -0.080782071 | 0.093223556 | -0.866541407 | 0.386193329 | 0.790230216 |
| Cd200 | 3439.861047 | -0.122262203 | 0.079543553 | -1.537047292 | 0.124281729 | 0.311791886 | Drd4 | 8.488410245 | -0.080809111 | 0.054800555 | -1.47460387 | 0.140319074 | NA |
| Gabpa | 848.6126413 | -0.128708299 | 0.083729289 | -1.537195648 | 0.124245406 | 0.311791886 | Ccdc23 | 282.9085974 | -0.08081449 | 0.077689391 | -1.040225567 | 0.298235116 | 0.732994423 |
| Ldlrad4 | 1397.217027 | -0.13889031 | 0.090376348 | -1.536799322 | 0.124342459 | 0.311791886 | Myo15 | 20.83593343 | -0.080823125 | 0.079992087 | -1.010389002 | 0.312308955 | NA |
| Mgea5 | 9858.000217 | -0.153170218 | 0.099666538 | -1.536826916 | 0.1243357 | 0.311791886 | Mlip | 1133.432614 | -0.080825396 | 0.068233987 | -1.184532806 | 0.236202218 | 0.687039131 |
| Pafah1b2 | 7673.676699 | -0.083864261 | 0.054555366 | -1.537232124 | 0.124236476 | 0.311791886 | Sox18 | 207.7209836 | -0.080856414 | 0.09434905 | -0.856992351 | 0.391449112 | 0.794184423 |
| Wscd2 | 2302.40745 | -0.123894535 | 0.080610742 | -1.536948204 | 0.124305994 | 0.311791886 | Gtpbp3 | 860.0821673 | -0.080865194 | 0.071306995 | -1.134042932 | 0.25677654 | 0.703000737 |
| 1110032A03Rik | 1002.234043 | 0.091830314 | 0.059768463 | 1.536434262 | 0.124431909 | 0.311793329 | Mustn1 | 41.43149693 | -0.080866565 | 0.085038725 | -0.950938116 | 0.341635791 | 0.76303628 |
| Gm5860 | 9.605743896 | 0.157774278 | 0.157559207 | 1.001365015 | 0.316650371 | 0.549465881 | Zfp2 | 951.7781392 | -0.080891806 | 0.068684505 | -1.177730057 | 0.23890424 | 0.689537017 |
| Uchl5 | 1438.335401 | 0.157747969 | 0.09531047 | 1.655095912 | 0.097905044 | 0.266129957 | Ostf1 | 391.6205917 | -0.080905899 | 0.08044998 | -1.005667106 | 0.314575731 | 0.745993052 |
| Derl3 | 45.61461867 | 0.157745631 | 0.166421684 | 0.947867053 | 0.34319714 | 0.575828939 | Znrd1as | 118.9089995 | -0.080943103 | 0.091169496 | -0.887830976 | 0.374631678 | 0.783149984 |
| Pspc1 | 1437.575555 | -0.077416294 | 0.050383875 | -1.53652919 | 0.124408644 | 0.311793329 | Actn1 | 4391.513086 | -0.080943133 | 0.07194556 | -1.125060849 | 0.26056325 | 0.704058139 |
| Scrn1 | 4910.807867 | -0.096246541 | 0.062648099 | -1.536304258 | 0.124463776 | 0.311793329 | Zfp346 | 1619.865653 | -0.080979365 | 0.055685469 | -1.454227933 | 0.145883123 | 0.592514474 |
| Zmat3 | 4391.916504 | -0.10303965 | 0.067088879 | -1.535867821 | 0.124570802 | 0.312010993 | Rwdd3 | 161.0728419 | -0.080998476 | 0.091565791 | -0.884593207 | 0.376376072 | 0.783947144 |
| Tbx4 | 12.68325473 | -0.254396703 | 0.165672293 | -1.535541635 | 0.124650838 | 0.312160997 | Brip1 | 34.35903367 | -0.081077052 | 0.087979042 | -0.921549606 | 0.356763555 | 0.773088648 |
| Spsb3 | 1989.036105 | 0.157587452 | 0.11067909 | 1.423823169 | 0.154497667 | 0.356106268 | Ntng2 | 735.8011549 | -0.081094485 | 0.088709715 | -0.914155622 | 0.360635079 | 0.776337125 |
| B3galt2 | 1415.88066 | -0.160744538 | 0.104725652 | -1.534910837 | 0.124805731 | 0.312304373 | Med23 | 1719.670642 | -0.08110137 | 0.052934108 | -1.532119335 | 0.125492989 | 0.562279816 |
| Hpca | 26254.09681 | 0.157561668 | 0.089741502 | 1.755727995 | 0.079134864 | 0.23283623 | Ccdc61 | 254.5449087 | -0.081112398 | 0.083030806 | -0.976895219 | 0.328621029 | 0.755786743 |
| Rps3a1 | 10357.69369 | -0.094115367 | 0.061317015 | -1.534898049 | 0.124808873 | 0.312304373 | Arid3c | 8.899705224 | -0.081117975 | 0.062238287 | -1.303345242 | 0.192456922 | NA |
| Traf6 | 866.9067289 | 0.157399416 | 0.113054751 | 1.392240613 | 0.163849548 | 0.369048539 | Gemin2 | 197.4827138 | -0.081153189 | 0.087108315 | -0.93163539 | 0.351524987 | 0.769244462 |
| Uaca | 459.1879067 | -0.146249924 | 0.095319779 | -1.53430826 | 0.124953835 | 0.312616618 | Erf | 2143.379264 | -0.081162496 | 0.059565075 | -1.36258529 | 0.173013247 | 0.624395138 |
| D630037F22Rik | 539.8813358 | -0.1095411 | 0.071410401 | -1.533965614 | 0.125038112 | 0.312625547 | Srsf7 | 2788.49791 | -0.081211339 | 0.058617185 | -1.38545272 | 0.16591408 | 0.616228056 |
| Ppan | 486.319828 | 0.157365243 | 0.09470476 | 1.661640268 | 0.096584924 | 0.263591859 | BC016423 | 1489.49667 | -0.081213411 | 0.076171382 | -1.066193217 | 0.286336308 | 0.722148665 |
| Med14 | 3442.869361 | -0.093624169 | 0.061032924 | -1.533994499 | 0.125031006 | 0.312625547 | 1700013F07Rik | 41.45600076 | -0.081232405 | 0.091573305 | -0.887075174 | 0.375038429 | 0.783489748 |
| Kcnip2 | 6803.816169 | 0.157291989 | 0.075752066 | 2.076405267 | 0.037856485 | 0.144115079 | Acvr1 | 1165.714448 | -0.081245218 | 0.055669766 | -1.459413664 | 0.144451288 | 0.589967032 |
| 2310050B05Rik | 8.43026902 | 0.157214462 | 0.119849176 | 1.311769232 | 0.189598009 | 0.402912022 | Ephx3 | 12.14611267 | -0.081255062 | 0.07101205 | -1.144243294 | 0.252522753 | NA |
| Fam110b | 1221.767599 | -0.123869039 | 0.080764727 | -1.533702191 | 0.125102934 | 0.312636269 | Rrp1b | 700.3472306 | -0.081269978 | 0.0801897 | -1.013471534 | 0.310834986 | 0.743730689 |
| Zfp369 | 589.1210235 | 0.157105459 | 0.088254425 | 1.780142558 | 0.075052633 | 0.225173055 | Klhl35 | 23.47374864 | -0.081278438 | 0.085905456 | -0.946138254 | 0.344078074 | 0.763939922 |
| Dynll2 | 28312.97122 | -0.102867753 | 0.067084072 | -1.533415476 | 0.125173518 | 0.312711785 | 4930451C15Rik | 59.47458308 | -0.081388712 | 0.09105828 | -0.893809018 | 0.371424084 | 0.780761257 |
| Mrpl36 | 659.9605518 | -0.177549201 | 0.115786299 | -1.533421503 | 0.125172034 | 0.312711785 | Cytip | 6.595034199 | -0.081414397 | 0.048915225 | -1.664397887 | 0.096032942 | NA |
| Prdm6 | 22.54000388 | -0.257689028 | 0.168071788 | -1.533208109 | 0.125224587 | 0.312788933 | Bub1b | 99.85335276 | -0.081428146 | 0.063200027 | -1.288419484 | 0.19759998 | 0.653498553 |
| Zfp639 | 855.7434068 | 0.157068435 | 0.080236416 | 1.95757041 | 0.050280442 | 0.174469598 | Mcam | 840.6263845 | -0.081439509 | 0.093385867 | -0.872075312 | 0.383167294 | 0.787602461 |
| C78339 | 902.6663009 | -0.112901925 | 0.073678058 | -1.532368368 | 0.125431559 | 0.313154459 | Sirt2 | 5310.516425 | -0.081454555 | 0.047518915 | -1.714150118 | 0.086501159 | 0.492365141 |
| Tsnax | 8337.412509 | -0.090925277 | 0.059335431 | -1.532394315 | 0.125425159 | 0.313154459 | Pcnxl3 | 3268.614451 | -0.081482427 | 0.057124211 | -1.426407919 | 0.153750629 | 0.60340295 |
| Diablo | 895.0813416 | 0.15706399 | 0.072131155 | 2.177477813 | 0.029444935 | 0.121935044 | Baat1 | 16.37996782 | -0.081598402 | 0.073250815 | -1.113958966 | 0.265296791 | NA |
| Hexdc | 1014.032459 | 0.156968226 | 0.11111271 | 1.4126937 | 0.157745805 | 0.360648295 | Tapbpl | 430.704791 | -0.081598613 | 0.092127056 | -0.885718236 | 0.375769379 | 0.783947144 |
| Tmeff2 | 2594.284649 | -0.086801265 | 0.056670593 | -1.531680911 | 0.125601195 | 0.313426464 | Cep290 | 706.0802783 | -0.081625188 | 0.081226965 | -1.004902591 | 0.314943757 | 0.745993052 |
| Mrps17 | 1171.016773 | -0.101954791 | 0.066587177 | -1.531147521 | 0.125732937 | 0.313704691 | Ift140 | 1379.124975 | -0.081680447 | 0.053272509 | -1.533257 | 0.125212545 | 0.561716315 |
| Sft2d1 | 763.1272525 | -0.140007445 | 0.091451484 | -1.530947765 | 0.125782303 | 0.313777331 | Rpl22l1 | 1316.492292 | -0.081701908 | 0.083838259 | -0.974518189 | 0.329799312 | 0.756141871 |
| Mms22l | 41.23666157 | 0.156890907 | 0.166949858 | 0.939748666 | 0.347346496 | 0.578686732 | Adamtsl4 | 406.1536915 | -0.081705415 | 0.091393256 | -0.893998302 | 0.3713228 | 0.780761257 |
| Abhd8 | 7917.9002 | -0.120941129 | 0.079018438 | -1.530543153 | 0.125882341 | 0.313817527 | Bcl2l11 | 283.4081948 | -0.081708546 | 0.083157046 | -0.982581146 | 0.325813628 | 0.753033837 |
| Gnptg | 2314.955989 | 0.095953097 | 0.06269506 | 1.53047302 | 0.125899687 | 0.313817527 | Zp2 | 7.580646502 | -0.08176445 | 0.055020829 | -1.486063575 | 0.137262306 | NA |
| Prpf4 | 929.2660522 | 0.098546855 | 0.064386548 | 1.530550376 | 0.125880554 | 0.313817527 | Add3 | 4880.199115 | -0.081769571 | 0.046129669 | -1.772602584 | 0.076294583 | 0.472435997 |
| Pstk | 400.5562997 | 0.15686529 | 0.087636121 | 1.789961581 | 0.073460088 | 0.221939958 | BC037704 | 51.69833796 | -0.081778097 | 0.087767897 | -0.931754091 | 0.351463625 | 0.769244462 |
| Agrn | 8392.062022 | 0.156854379 | 0.099443255 | 1.577325476 | 0.114720653 | 0.296692005 | Stat3 | 2544.860005 | -0.081798899 | 0.053527117 | -1.528176818 | 0.126468643 | 0.563050456 |
| AI661453 | 25.76717833 | -0.254481273 | 0.166321647 | -1.530055034 | 0.126003107 | 0.313873335 | Rpap1 | 1099.856327 | -0.081799208 | 0.073177807 | -1.117814416 | 0.263646263 | 0.706200557 |
| Bbs7 | 707.1554011 | 0.095635938 | 0.062498452 | 1.530212913 | 0.125964036 | 0.313873335 | Bcan | 10817.40836 | -0.081810105 | 0.070911671 | -1.153690284 | 0.248627169 | 0.697784358 |
| Ugt1a7c | 21.65682924 | 0.156759933 | 0.166201109 | 0.943194267 | 0.345581545 | 0.576948985 | C130060K24Rik | 39.33274837 | -0.081823282 | 0.089023431 | -0.91912075 | 0.358032417 | 0.774087808 |
| Rad50 | 756.7166771 | 0.156711187 | 0.083704664 | 1.87219183 | 0.061180069 | 0.196991685 | Ak2 | 764.8392738 | -0.081827389 | 0.074172425 | -1.103204977 | 0.26993816 | 0.71216373 |
| Magi3 | 2953.385851 | -0.095281396 | 0.062290191 | -1.529637249 | 0.126106544 | 0.313979558 | Jund | 2348.946496 | -0.081838389 | 0.087550588 | -0.934755447 | 0.349914345 | 0.768536074 |
| Pabpc4 | 1187.027774 | -0.128348213 | 0.083906512 | -1.529657365 | 0.126101562 | 0.313979558 | Map6d1 | 3223.150811 | -0.081854472 | 0.066254542 | -1.2354545 | 0.2166614 | 0.670049713 |
| Slc1a5 | 130.9747615 | 0.156570865 | 0.128359591 | 1.219783145 | 0.222547092 | 0.443624247 | Sumf2 | 528.9120216 | -0.081892414 | 0.079344021 | -1.032118271 | 0.302016716 | 0.735974432 |
| Gabra2 | 3495.688616 | -0.181129476 | 0.118433795 | -1.529373232 | 0.126171944 | 0.314041463 | Gxylt1 | 758.2814787 | -0.08190977 | 0.066969799 | -1.223085208 | 0.221297515 | 0.672757795 |
| Armc3 | 27.73129786 | 0.156524308 | 0.166960247 | 0.937494472 | 0.348504266 | 0.579680633 | Chpf2 | 2494.170674 | -0.081918904 | 0.083235827 | -0.984178408 | 0.3250278 | 0.75293408 |
| Cir1 | 1159.914216 | 0.156487456 | 0.085375064 | 1.832941 | 0.066811346 | 0.20864524 | Zfpm1 | 1365.746766 | -0.081935769 | 0.075392905 | -1.086783541 | 0.277132483 | 0.717836386 |
| 2610301G19Rik | 3788.230515 | 0.081520342 | 0.053351053 | 1.527998741 | 0.12651285 | 0.314473278 | AI414108 | 2139.00495 | -0.081964725 | 0.089112246 | -0.91979193 | 0.357681501 | 0.773905615 |
| Peg13 | 15878.86871 | 0.156370879 | 0.071812259 | 2.177495619 | 0.029443608 | 0.121935044 | Shc1 | 541.0776682 | -0.081983812 | 0.067903508 | -1.207357547 | 0.227294477 | 0.680203999 |
| Gng5 | 394.9041953 | -0.142666495 | 0.093356796 | -1.528185425 | 0.126466507 | 0.314473278 | Xist | 21529.01429 | -0.082026289 | 0.076173693 | -1.076832245 | 0.281555217 | 0.720080692 |
| Lta4h | 1574.970836 | 0.15633639 | 0.069129857 | 2.261488711 | 0.023729011 | 0.105800661 | B830017H08Rik | 94.32034792 | -0.082089521 | 0.094182467 | -0.871600879 | 0.383426151 | 0.787702475 |
| Camsap3 | 1210.491617 | 0.15632296 | 0.067707543 | 2.30879681 | 0.020954859 | 0.097062657 | Slc35f2 | 173.2446045 | -0.082139907 | 0.089153607 | -0.921330164 | 0.356878078 | 0.773088648 |
| Nr2c2 | 6569.09945 | -0.095226195 | 0.062312868 | -1.528194703 | 0.126464203 | 0.314473278 | Ang | 52.85326398 | -0.0821864 | 0.094642708 | -0.868385976 | 0.38518307 | 0.789016184 |
| Odc1 | 2486.604891 | -0.088383656 | 0.057844959 | -1.527940521 | 0.126527306 | 0.314473278 | Nupr1l | 13.38088614 | -0.082207499 | 0.073548404 | -1.117733278 | 0.263680926 | NA |
| Plekha7 | 690.5490425 | -0.17127636 | 0.112110803 | -1.527741793 | 0.126576659 | 0.314473278 | Plekhg2 | 316.7349957 | -0.082235776 | 0.078670504 | -1.045319047 | 0.295875538 | 0.730928808 |
| Prr7 | 580.9911503 | -0.220938615 | 0.144622473 | -1.527692144 | 0.126588992 | 0.314473278 | Mfng | 144.3723482 | -0.082236204 | 0.091795054 | -0.895867484 | 0.370323542 | 0.780516508 |
| Zfp931 | 124.8478526 | -0.221603171 | 0.144993508 | -1.528366157 | 0.126421653 | 0.314473278 | 1700102H20Rik | 4.676830309 | -0.082258307 | 0.048456095 | -1.697584316 | 0.089586245 | NA |
| Rnaseh2b | 272.0695826 | -0.151580484 | 0.099237444 | -1.527452519 | 0.126648525 | 0.314520316 | 4930578C19Rik | 31.89452137 | -0.082261345 | 0.080277133 | -1.024717034 | 0.305496723 | 0.73790953 |
| Zc3h15 | 3673.107715 | -0.095440584 | 0.062481449 | -1.527502734 | 0.126636048 | 0.314520316 | Fbxo32 | 1494.481048 | -0.082265369 | 0.073793388 | -1.114806786 | 0.264933228 | 0.707370515 |
| Nudt6 | 92.13492721 | 0.156262426 | 0.135015007 | 1.157370798 | 0.2471209 | 0.472061549 | Tnfsf13b | 41.06284961 | -0.082266507 | 0.090013605 | -0.913934142 | 0.360751452 | 0.776337125 |
| Slc25a44 | 4663.980415 | 0.156252599 | 0.103019078 | 1.516734584 | 0.129333709 | 0.318382028 | Haus6 | 251.6933792 | -0.082277914 | 0.084436998 | -0.974429648 | 0.329843255 | 0.756141871 |
| 9130401M01Rik | 828.9907657 | -0.103335317 | 0.067682645 | -1.526762401 | 0.126820104 | 0.314691614 | Adrb1 | 955.4151893 | -0.082297072 | 0.086818504 | -0.94792088 | 0.343169734 | 0.763762832 |
| Dhcr24 | 5045.67271 | -0.167451222 | 0.109683771 | -1.526672726 | 0.126842413 | 0.314691614 | 4933421O10Rik | 404.6307658 | -0.082311426 | 0.074716093 | -1.101655909 | 0.270611292 | 0.712294861 |
| Gba2 | 3788.971686 | 0.15620259 | 0.067641551 | 2.309269773 | 0.020928616 | 0.097003177 | Rgs13 | 11.54510417 | -0.082320633 | 0.058441723 | -1.408593529 | 0.158955393 | NA |
| Lrrfip2 | 1288.180849 | 0.156155486 | 0.079489945 | 1.964468407 | 0.049475797 | 0.172797243 | Fbf1 | 3256.886916 | -0.082347968 | 0.059042217 | -1.394730272 | 0.163097193 | 0.614351408 |
| Tomm34 | 2639.895375 | 0.156128171 | 0.088359037 | 1.766974575 | 0.077232486 | 0.228920619 | Slc38a3 | 3569.611547 | -0.082354328 | 0.078031139 | -1.055403389 | 0.291240873 | 0.727526213 |
| Tcp11l2 | 1042.101507 | 0.156114009 | 0.087051908 | 1.793343914 | 0.07291795 | 0.221094073 | Prrx2 | 19.50130942 | -0.082358588 | 0.079048777 | -1.041870489 | 0.297471728 | NA |
| Szrd1 | 1303.567361 | -0.104345628 | 0.068370839 | -1.526171529 | 0.126967152 | 0.314857403 | Ints1 | 4733.489996 | -0.082363953 | 0.062617333 | -1.315353893 | 0.188390994 | 0.64225749 |
| Ankfy1 | 2567.334077 | -0.069960587 | 0.045846851 | -1.525962757 | 0.12701914 | 0.314885764 | Ndst2 | 795.594213 | -0.082410854 | 0.063463767 | -1.298549675 | 0.194098518 | 0.65013848 |
| Pth1r | 202.2175711 | -0.201463055 | 0.132023606 | -1.525962375 | 0.127019235 | 0.314885764 | Chrna2 | 26.99694746 | -0.082432196 | 0.086882806 | -0.948774556 | 0.342735285 | 0.763762832 |
| E330020D12Rik | 19.0215704 | 0.155986143 | 0.168101072 | 0.927930687 | 0.353443525 | 0.584133757 | Slco1c1 | 2473.91219 | -0.082432704 | 0.062132841 | -1.326717129 | 0.184602273 | 0.637780733 |
| A530054K11Rik | 418.5197414 | -0.135319135 | 0.088708669 | -1.525433043 | 0.127151122 | 0.315111883 | Poc1b | 257.1290289 | -0.082444276 | 0.076575262 | -1.076643727 | 0.281639461 | 0.720136387 |
| Gpm6b | 32369.09858 | -0.106754575 | 0.06999386 | -1.52519914 | 0.127209435 | 0.315205979 | Kif17 | 1588.607466 | -0.082452308 | 0.066951474 | -1.231523421 | 0.218127162 | 0.671011258 |
| Dgki | 1495.198678 | -0.139036244 | 0.091182716 | -1.524809199 | 0.127306695 | 0.315245314 | Gnl2 | 2195.410266 | -0.08245519 | 0.053246072 | -1.548568508 | 0.121485481 | 0.555796278 |
| Dhrs7 | 839.4037071 | -0.120004039 | 0.078701004 | -1.524809495 | 0.127306621 | 0.315245314 | Pdrg1 | 1565.039365 | -0.082464054 | 0.063513859 | -1.298363145 | 0.194162577 | 0.65013848 |
| Epm2aip1 | 5274.824409 | -0.122051651 | 0.080043882 | -1.524809245 | 0.127306683 | 0.315245314 | Unc45a | 1138.512037 | -0.082470165 | 0.073445307 | -1.122878625 | 0.261489047 | 0.704173028 |
| Brd9 | 2376.18334 | 0.155971609 | 0.110831958 | 1.407280098 | 0.159344348 | 0.363177601 | E2f5 | 290.4172252 | -0.082498549 | 0.089138159 | -0.925513271 | 0.354698984 | 0.77200441 |
| Gareml | 584.3417319 | -0.151523935 | 0.099391866 | -1.524510412 | 0.127381258 | 0.315379548 | Sp8 | 45.52650057 | -0.082505369 | 0.091332098 | -0.903355674 | 0.366337157 | 0.778210102 |
| Mcm4 | 498.0310746 | 0.155942427 | 0.077792157 | 2.004603454 | 0.045005457 | 0.162657931 | Spg11 | 1229.777915 | -0.082510557 | 0.061895905 | -1.333053575 | 0.182514223 | 0.635424885 |
| Radil | 567.1796086 | 0.155900001 | 0.109565428 | 1.422894096 | 0.154766858 | 0.356467623 | Trim65 | 133.7682302 | -0.082520463 | 0.090842789 | -0.908387597 | 0.363673476 | 0.776984985 |
| D430042O09Rik | 1191.97309 | 0.155866674 | 0.060089161 | 2.593923275 | 0.009488766 | 0.056676538 | Man2b2 | 1384.31428 | -0.082623042 | 0.076211512 | -1.084128099 | 0.278308006 | 0.718447146 |
| Spsb4 | 138.2235986 | -0.201508258 | 0.132275992 | -1.523392527 | 0.127660531 | 0.315869093 | Khnyn | 807.3312474 | -0.082681853 | 0.060937173 | -1.356837679 | 0.174832792 | 0.627997315 |
| Elavl4 | 2765.658371 | -0.122990911 | 0.080754398 | -1.523024299 | 0.127752626 | 0.316046494 | Nr1d1 | 6665.987313 | -0.082704769 | 0.061795834 | -1.338355092 | 0.180780711 | 0.634469716 |
| Gtpbp1 | 2413.407322 | -0.078252911 | 0.051389613 | -1.522737885 | 0.127824296 | 0.316173314 | Nasp | 734.0167164 | -0.082713673 | 0.066754012 | -1.239081683 | 0.21531525 | 0.669593089 |
| A230056J06Rik | 34.02046686 | -0.242785555 | 0.159463805 | -1.522511992 | 0.127880843 | 0.316212222 | Apol10b | 17.35188762 | -0.082764306 | 0.073835271 | -1.120931827 | 0.262316882 | NA |
| Cdk9 | 3174.136287 | 0.155866472 | 0.070408154 | 2.213755993 | 0.026845569 | 0.114711871 | 1700003D09Rik | 30.59079242 | -0.082766535 | 0.064282566 | -1.287542475 | 0.197905275 | 0.653535282 |
| Slc19a2 | 586.5521686 | 0.155833552 | 0.069472838 | 2.243086019 | 0.024891268 | 0.108945002 | Tmem39a | 342.7355843 | -0.082788863 | 0.088098675 | -0.93972881 | 0.347356683 | 0.766038372 |
| Prkcz | 11507.05755 | -0.098739205 | 0.06488021 | -1.521869384 | 0.128041812 | 0.316509227 | Stat4 | 96.98144758 | -0.082825134 | 0.094578165 | -0.875732089 | 0.381175696 | 0.786663048 |
| Pank1 | 605.9884951 | -0.119285948 | 0.078386132 | -1.521773625 | 0.128065812 | 0.316518057 | Ghr | 391.8857752 | -0.082872575 | 0.079204641 | -1.046309587 | 0.29541812 | 0.730610822 |
| Mettl21d | 384.4012589 | 0.155807361 | 0.094705373 | 1.645179734 | 0.099932752 | 0.269936899 | Rps19-ps3 | 47.77891455 | -0.0829102 | 0.094529551 | -0.877082344 | 0.380441914 | 0.786275719 |
| Lipc | 9.028996654 | -0.242985268 | 0.159702207 | -1.521489734 | 0.128136986 | 0.316541113 | Slc22a5 | 540.4700274 | -0.08291906 | 0.080424209 | -1.031021139 | 0.302530907 | 0.735974513 |
| Rraga | 3250.114841 | -0.077485113 | 0.050929788 | -1.521410479 | 0.128156861 | 0.316541113 | 1110019D14Rik | 71.87682369 | -0.082933643 | 0.09496182 | -0.873336704 | 0.38247958 | 0.786954445 |
| Gm17762 | 68.32139159 | 0.155802121 | 0.138294379 | 1.126597641 | 0.259912637 | 0.486656967 | Il17ra | 1136.140752 | -0.08296889 | 0.084025529 | -0.987424787 | 0.323434438 | 0.752925264 |
| Bbs12 | 148.5834345 | -0.182136044 | 0.119755909 | -1.520894013 | 0.128286437 | 0.316760167 | Ccdc24 | 49.37526723 | -0.083003796 | 0.088971322 | -0.932927539 | 0.350857382 | 0.76894302 |
| Rinl | 73.11171151 | 0.15574205 | 0.136581579 | 1.14028591 | 0.254167206 | 0.480076398 | Ugt1a6a | 59.35965329 | -0.083025676 | 0.092409755 | -0.898451428 | 0.368944929 | 0.779694884 |
| 4930525G20Rik | 78.29518982 | -0.203919042 | 0.134090107 | -1.520761268 | 0.128319757 | 0.316791957 | Mir331 | 21.87748658 | -0.0830409 | 0.08217074 | -1.010589657 | 0.312212868 | NA |
| Endod1 | 4501.532508 | 0.155651974 | 0.084403491 | 1.844141424 | 0.065162532 | 0.205208999 | Arhgef15 | 1087.043917 | -0.083050634 | 0.063174252 | -1.314627897 | 0.18863499 | 0.64246468 |
| Nek10 | 53.75182265 | 0.155585334 | 0.155289427 | 1.001905517 | 0.316389228 | 0.549320338 | Srebf1 | 2843.644072 | -0.08308956 | 0.083207493 | -0.998582655 | 0.317996906 | 0.748272127 |
| Slc39a13 | 781.0906564 | 0.155554808 | 0.08033644 | 1.936292016 | 0.052831946 | 0.180143517 | Iws1 | 1501.797471 | -0.08310803 | 0.064563741 | -1.287224515 | 0.198016045 | 0.653535282 |
| Tmem9 | 1972.796631 | -0.10652992 | 0.070085451 | -1.520000482 | 0.128510855 | 0.317061653 | Sox7 | 82.36668089 | -0.08314217 | 0.095076579 | -0.874475835 | 0.381859175 | 0.786663048 |
| Lancl3 | 265.7053155 | -0.238291614 | 0.15678147 | -1.519896542 | 0.12853698 | 0.317075619 | Sctr | 13.84736701 | -0.083156885 | 0.064629426 | -1.286672188 | 0.198208571 | NA |
| Smg6 | 2145.106377 | 0.155527583 | 0.082699296 | 1.880639755 | 0.06002094 | 0.194471862 | Dnaaf2 | 197.7179573 | -0.083221284 | 0.082788749 | -1.005224567 | 0.314788728 | 0.745993052 |
| Caskin2 | 1261.914063 | -0.11045505 | 0.072697206 | -1.519385072 | 0.128665597 | 0.317241369 | Galc | 1252.709598 | -0.083239756 | 0.053352845 | -1.560174634 | 0.118718618 | 0.550960591 |
| Cyp2j6 | 913.1263962 | 0.155482873 | 0.076538436 | 2.031435207 | 0.042210866 | 0.155754979 | AI118078 | 288.6246048 | -0.083259647 | 0.084430316 | -0.986134492 | 0.32406712 | 0.752925264 |
| Naga | 731.0207856 | 0.155423364 | 0.077957042 | 1.993705251 | 0.046184275 | 0.165337144 | Mir1898 | 5.217000903 | -0.083356027 | 0.050905137 | -1.637477702 | 0.101530694 | NA |
| Lcor | 166.1143048 | -0.19860799 | 0.13077495 | -1.518700568 | 0.128837884 | 0.317565092 | B130034C11Rik | 51.49372361 | -0.083404995 | 0.092753117 | -0.899215013 | 0.368538145 | 0.779452463 |
| Cct5 | 5433.127598 | -0.097712414 | 0.064346773 | -1.518528579 | 0.128881201 | 0.317621333 | Dpm2 | 1386.119261 | -0.08350922 | 0.056839097 | -1.469221453 | 0.141772733 | 0.587554222 |
| Mboat7 | 4885.682807 | -0.088584645 | 0.058366353 | -1.517734798 | 0.129081268 | 0.317941173 | BC037034 | 2377.968677 | -0.083510259 | 0.068774894 | -1.214255 | 0.224650364 | 0.676420098 |
| Pdf | 989.3260155 | -0.105832165 | 0.06973249 | -1.51768803 | 0.129093063 | 0.317941173 | Gm3279 | 10.6191946 | -0.083521678 | 0.063049824 | -1.324693278 | 0.185272904 | NA |
| Rab33b | 1142.061642 | -0.124029826 | 0.081711226 | -1.517904349 | 0.129038513 | 0.317941173 | Gm15708 | 10.22986784 | -0.083548751 | 0.067648501 | -1.235042162 | 0.216814813 | NA |
| Ttc30a1 | 130.4108405 | -0.196261325 | 0.129311248 | -1.51774365 | 0.129079036 | 0.317941173 | LOC100504703 | 22.90848881 | -0.083552985 | 0.078777049 | -1.060625992 | 0.288859905 | 0.724444298 |
| Mylk4 | 14.31164527 | 0.155372527 | 0.16742547 | 0.928010095 | 0.353402333 | 0.584133757 | BC061194 | 150.6377188 | -0.083629254 | 0.091780766 | -0.911184966 | 0.362197924 | 0.776449324 |
| Rnf187 | 10570.70324 | -0.096314512 | 0.063496542 | -1.5168492 | 0.129304762 | 0.318361351 | Pde3b | 451.3955057 | -0.083635231 | 0.068412471 | -1.222514401 | 0.221513161 | 0.672757795 |
| Pik3ap1 | 124.1428116 | 0.155363944 | 0.142819354 | 1.087835367 | 0.276667793 | 0.505605783 | 1810021B22Rik | 30.36539337 | -0.08375215 | 0.089755313 | -0.933116353 | 0.350759897 | 0.768845369 |
| Lamc1 | 1559.00265 | -0.092684264 | 0.061120119 | -1.516428068 | 0.129411146 | 0.318468986 | Itgb1bp2 | 11.78536701 | -0.083907248 | 0.062319103 | -1.346412953 | 0.178169376 | NA |
| Pou3f2 | 900.8554901 | -0.115701213 | 0.07630001 | -1.516398408 | 0.129418641 | 0.318468986 | Mvp | 349.2382312 | -0.083928787 | 0.077137487 | -1.088041497 | 0.276576788 | 0.717475105 |
| Slc37a2 | 159.9636739 | 0.155326312 | 0.143869258 | 1.079635181 | 0.280304669 | 0.509926094 | Myom1 | 105.4540384 | -0.083949261 | 0.094788558 | -0.885647621 | 0.375807442 | 0.783947144 |
| Cd276 | 454.4088653 | -0.142511813 | 0.093995962 | -1.516148253 | 0.129481869 | 0.318544313 | Pbld2 | 49.91111262 | -0.084008265 | 0.093967805 | -0.894011141 | 0.37131593 | 0.780761257 |
| Cdc42bpa | 13614.45285 | -0.085738886 | 0.056560744 | -1.515872656 | 0.129551556 | 0.318665154 | Epha10 | 2210.258782 | -0.084086516 | 0.075039833 | -1.120558404 | 0.262475879 | 0.704885306 |
| Dnaaf2 | 184.3207733 | 0.155324067 | 0.100554839 | 1.544670232 | 0.122426049 | 0.308775687 | Bicd2 | 3613.19503 | -0.084115913 | 0.056081456 | -1.499888193 | 0.133643367 | 0.57588358 |
| Rpl17 | 8876.804727 | -0.138518006 | 0.091402391 | -1.515474639 | 0.129652248 | 0.318819536 | Naga | 830.5125108 | -0.084121295 | 0.08090655 | -1.039734051 | 0.298463476 | 0.732994423 |
| Tnfaip8 | 212.8371065 | -0.193765031 | 0.127873111 | -1.515291442 | 0.129698615 | 0.318875013 | Brpf3 | 1905.495867 | -0.084155316 | 0.050962453 | -1.651319973 | 0.098673256 | 0.520075369 |
| Dclre1a | 339.9308686 | 0.15528437 | 0.094939458 | 1.635614676 | 0.101920257 | 0.273635845 | Derl3 | 56.16644826 | -0.084216904 | 0.092930805 | -0.906232377 | 0.364812869 | 0.777589397 |
| Agk | 1558.865458 | -0.094754138 | 0.06254512 | -1.514972509 | 0.129779366 | 0.318939271 | Car11 | 6266.504902 | -0.084241665 | 0.057093831 | -1.475495045 | 0.1400795 | 0.585376897 |
| Rpl23a | 7023.904388 | -0.088383053 | 0.058340793 | -1.514944322 | 0.129786505 | 0.318939271 | Gm581 | 16.4940275 | -0.084242652 | 0.061688923 | -1.365604186 | 0.172063232 | NA |
| Mtfr2 | 14.19835155 | 0.155272669 | 0.167249112 | 0.928391588 | 0.35320448 | 0.583987858 | BC068157 | 1585.052284 | -0.084273733 | 0.072921324 | -1.155680233 | 0.247811977 | 0.696944196 |
| Gjd2 | 226.422726 | -0.170060557 | 0.112294426 | -1.514416726 | 0.12992018 | 0.319115926 | H2-Ab1 | 96.19247695 | -0.084275653 | 0.040600806 | -2.075713806 | NA | NA |
| Pigt | 3224.80853 | 0.094457248 | 0.062370961 | 1.514442718 | 0.129913592 | 0.319115926 | Gm16119 | 13.71066085 | -0.084302102 | 0.066621945 | -1.265380381 | 0.205735005 | NA |
| Fam69c | 220.0243479 | -0.163187318 | 0.10776752 | -1.514253257 | 0.129961619 | 0.319167114 | Nr2c1 | 482.3084072 | -0.084311656 | 0.066028153 | -1.276904669 | 0.201635906 | 0.655168687 |
| Eif4g3 | 11481.24289 | 0.073164337 | 0.048324587 | 1.51401887 | 0.130021054 | 0.319211885 | Paf1 | 2280.481871 | -0.08438451 | 0.061456985 | -1.373066217 | 0.169731762 | 0.619945581 |
| Rhbdl2 | 10.38032024 | -0.247437363 | 0.163429039 | -1.51403548 | 0.130016841 | 0.319211885 | Rnaseh1 | 366.0368811 | -0.084449602 | 0.074155532 | -1.138817293 | 0.254779368 | 0.701619624 |
| Bptf | 4846.532655 | -0.156658752 | 0.103479456 | -1.513911629 | 0.130048254 | 0.319228082 | Flnc | 117.9468674 | -0.08455682 | 0.094353101 | -0.896174257 | 0.370159703 | 0.780397655 |
| Lyrm4 | 677.1186309 | -0.105418864 | 0.069653629 | -1.513472665 | 0.130159639 | 0.319450887 | 0610038B21Rik | 11.0614681 | -0.084560401 | 0.061144971 | -1.3829494 | 0.166680382 | NA |
| Styx | 359.6190854 | 0.155269769 | 0.097960487 | 1.585024461 | 0.112960777 | 0.29375869 | Rce1 | 571.8295019 | -0.08467057 | 0.075246288 | -1.125245805 | 0.260484888 | 0.704058139 |
| 1700026D08Rik | 29.59193764 | 0.155263786 | 0.166871025 | 0.930441854 | 0.352142357 | 0.583040573 | Gm10640 | 5.773040383 | -0.084691721 | 0.054100782 | -1.565443547 | 0.117478944 | NA |
| Ptma | 7991.508743 | -0.081564553 | 0.053918241 | -1.512745081 | 0.130344424 | 0.319760855 | Il6ra | 193.0856311 | -0.084695901 | 0.089428635 | -0.947078093 | 0.343598987 | 0.763762832 |
| Otud4 | 2386.775338 | 0.094461244 | 0.062461442 | 1.512312896 | 0.130454282 | 0.319971291 | Wfdc18 | 40.43246144 | -0.084706504 | 0.091622453 | -0.924516874 | 0.35521727 | 0.772206867 |
| Rnf8 | 1177.6768 | 0.15525831 | 0.075681459 | 2.051470877 | 0.040221113 | 0.15031016 | Ccdc157 | 1516.276481 | -0.084730436 | 0.062121199 | -1.363953654 | 0.172582153 | 0.623924427 |
| Erbb2ip | 3637.366522 | -0.125720489 | 0.083160344 | -1.511784136 | 0.130588787 | 0.320149155 | Aldh3b1 | 198.6197626 | -0.084750688 | 0.088139763 | -0.961548852 | 0.336276275 | 0.759681053 |
| Gucy1a3 | 5356.131008 | -0.137173133 | 0.090731479 | -1.511858216 | 0.130569936 | 0.320149155 | Arhgap22 | 232.8388054 | -0.084786584 | 0.08555899 | -0.990972239 | 0.321699136 | 0.751165673 |
| Ephb2 | 704.2007425 | 0.155176506 | 0.110738238 | 1.401291094 | 0.161127043 | 0.365948755 | Ccdc93 | 1488.835031 | -0.084787112 | 0.053343727 | -1.589448604 | 0.111959149 | 0.538992394 |
| Dcun1d5 | 997.3221253 | -0.104488932 | 0.069138869 | -1.511290739 | 0.130714393 | 0.32035571 | Cxxc1 | 1722.187743 | -0.084827282 | 0.070905357 | -1.196345167 | 0.23156189 | 0.68367236 |
| Esrrg | 1189.610342 | 0.155170217 | 0.105513146 | 1.47062449 | 0.141392697 | 0.336674112 | Srsf1 | 5136.720653 | -0.084854145 | 0.052823205 | -1.606380072 | 0.108190427 | 0.533447785 |
| 4933426M11Rik | 2269.306056 | 0.155110707 | 0.104693745 | 1.481566135 | 0.138455775 | 0.332182376 | Ifnar2 | 667.0942986 | -0.084855248 | 0.069146145 | -1.227186969 | 0.219752326 | 0.672421385 |
| Fam126b | 3376.297778 | -0.141260664 | 0.093499607 | -1.510815596 | 0.13083544 | 0.320515687 | 4833417C18Rik | 10.22409467 | -0.084900679 | 0.061754403 | -1.374811755 | 0.169189813 | NA |
| Ppil1 | 545.5437127 | -0.123035981 | 0.081442501 | -1.510709765 | 0.130862413 | 0.320515687 | S1pr3 | 508.6509617 | -0.08497405 | 0.071053723 | -1.195912698 | 0.23173063 | 0.683754831 |
| Rin2 | 1353.983476 | 0.15508627 | 0.086223724 | 1.798649643 | 0.072074119 | 0.219394186 | Nr3c2 | 1335.798233 | -0.084974067 | 0.07752077 | -1.096145813 | 0.273014969 | 0.714368645 |
| Ppm1j | 45.70529242 | -0.201916204 | 0.133715639 | -1.510041801 | 0.131032758 | 0.32078078 | 4932415G12Rik | 1625.615971 | -0.084980244 | 0.068958265 | -1.232343136 | 0.217820933 | 0.670929686 |
| Stk38 | 1085.154668 | 0.155080863 | 0.115924793 | 1.337771318 | 0.180970994 | 0.391473411 | Apool | 133.6373054 | -0.085003706 | 0.092977151 | -0.914242962 | 0.360589194 | 0.776337125 |
| Nolc1 | 2673.95883 | 0.084646996 | 0.056074795 | 1.509537319 | 0.131161526 | 0.321045289 | Fam189b | 2209.379766 | -0.085060832 | 0.067272086 | -1.264429811 | 0.206075799 | 0.660203749 |
| C5ar1 | 43.91905704 | 0.154961783 | 0.165626487 | 0.935609914 | 0.349474068 | 0.580421397 | Tmem154 | 20.74783051 | -0.085089426 | 0.081497773 | -1.044070572 | 0.29645274 | NA |
| Gja4 | 108.1403361 | 0.154899605 | 0.123530132 | 1.253941872 | 0.209863134 | 0.428013611 | Tmem86a | 641.9822601 | -0.085122039 | 0.085781895 | -0.992307746 | 0.321047426 | 0.750352874 |
| 4930507D05Rik | 31.82166681 | 0.154840563 | 0.159298146 | 0.972017359 | 0.331041913 | 0.566257827 | 2610100L16Rik | 198.2218503 | -0.085212569 | 0.092057456 | -0.925645488 | 0.354630246 | 0.77200441 |
| Pcdhb9 | 527.5164641 | 0.154816404 | 0.068173983 | 2.27090156 | 0.023152938 | 0.103980179 | Hirip3 | 1187.130136 | -0.085218552 | 0.068384998 | -1.246158585 | 0.21270618 | 0.666270994 |
| Sv2b | 23288.16128 | -0.144848035 | 0.096021068 | -1.508502648 | 0.13142593 | 0.321438572 | Fam3a | 774.6212779 | -0.085259361 | 0.065676752 | -1.29816654 | 0.194230113 | 0.65013848 |
| Stk3 | 508.0199471 | 0.154728721 | 0.080711309 | 1.917063703 | 0.055229839 | 0.185095341 | C2cd4b | 23.77087474 | -0.085275718 | 0.071087353 | -1.199590577 | 0.230298388 | 0.683402059 |
| Ndufa12 | 2343.892655 | -0.090609356 | 0.060083786 | -1.508050045 | 0.131541719 | 0.32162023 | Ly75 | 69.65168538 | -0.085302093 | 0.095062427 | -0.897327111 | 0.369544394 | 0.780397655 |
| Mrpl47 | 399.6173265 | -0.122881382 | 0.08149133 | -1.507907423 | 0.131578223 | 0.321633594 | Lrrc43 | 28.61993563 | -0.085334435 | 0.080711399 | -1.057278605 | 0.290384455 | 0.725761285 |
| Ndp | 334.4153072 | -0.159681777 | 0.105899149 | -1.507866483 | 0.131588702 | 0.321633594 | Svil | 484.8306531 | -0.085367817 | 0.084138276 | -1.014613338 | 0.310290179 | 0.743039093 |
| Ppp1r16a | 2947.241788 | -0.123483577 | 0.081903221 | -1.507676691 | 0.131637294 | 0.321650881 | Nkx2-2 | 143.7980216 | -0.08545878 | 0.093025568 | -0.918659048 | 0.358273936 | 0.774201531 |
| Vipas39 | 2135.991985 | 0.089118736 | 0.059108882 | 1.50770464 | 0.131630137 | 0.321650881 | Pdcd2 | 322.0599349 | -0.085477442 | 0.077542834 | -1.102325497 | 0.270320189 | 0.71216373 |

| | | | | | | |
|---|---|---|---|---|---|---|
| Afap1l1 | 1149.790652 | 0.154661903 | 0.089959323 | 1.719242635 | 0.0855702 | 0.244481779 |
| Glcci1 | 2301.963577 | -0.100668595 | 0.066790735 | -1.507223947 | 0.131753264 | 0.321731296 |
| Med23 | 1542.533348 | 0.099308897 | 0.065887402 | 1.507251663 | 0.131746162 | 0.321731296 |
| Pgls | 414.3518463 | -0.14465356 | 0.095965359 | -1.507351838 | 0.131720497 | 0.321731296 |
| 1110015O18Rik | 10.08385814 | -0.225518337 | 0.149654078 | -1.506930779 | 0.131828401 | 0.321812686 |
| Rps4y2 | 372.0673401 | 0.154649536 | 0.101296038 | 1.526708632 | 0.12683348 | 0.314691614 |
| Il11 | 12.61849689 | 0.154623603 | 0.161054766 | 0.960068471 | 0.337020755 | 0.570594737 |
| Tmem55a | 1400.683986 | -0.105926826 | 0.070320663 | -1.50633998 | 0.13197992 | 0.322048446 |
| Vps41 | 7906.513489 | 0.073674093 | 0.048910243 | 1.506312121 | 0.131987068 | 0.322048446 |
| Npat | 1035.616379 | 0.154550374 | 0.074685617 | 2.069345896 | 0.03851364 | 0.145792898 |
| E030030I06Rik | 19.33197641 | 0.154467083 | 0.168119263 | 0.918794672 | 0.35820298 | 0.588859095 |
| Gnl3l | 9004.24465 | -0.083307793 | 0.055330907 | -1.505628529 | 0.132162561 | 0.322324368 |
| Nipa2 | 1148.482193 | -0.129508783 | 0.086033871 | -1.505323213 | 0.132241 | 0.322464913 |
| Fam65b | 2769.490572 | -0.135133989 | 0.089788822 | -1.505020175 | 0.13231889 | 0.322604072 |
| 2610203C20Rik | 668.0500346 | 0.154430251 | 0.125229994 | 1.233173027 | 0.217511217 | 0.437645554 |
| Rnf146 | 2108.828772 | 0.154422141 | 0.082043042 | 1.882208873 | 0.059807661 | 0.194227605 |
| Mtfr1l | 2321.331311 | 0.072763078 | 0.048363148 | 1.504514911 | 0.132448837 | 0.322768524 |
| Fam203a | 293.0175487 | 0.15441258 | 0.090645581 | 1.703476092 | 0.088479008 | 0.249660098 |
| Plekhh1 | 1763.704274 | 0.154315936 | 0.119860537 | 1.287462411 | 0.197933163 | 0.413646672 |
| Plekha8 | 753.9403306 | 0.154218815 | 0.07650107 | 2.015904014 | 0.043810014 | 0.159715848 |
| Ilvbl | 833.5757193 | 0.154204126 | 0.093407674 | 1.650872129 | 0.098764688 | 0.267979485 |
| Cep97 | 820.6691898 | 0.154162583 | 0.072844373 | 2.116327945 | 0.034316926 | 0.135225444 |
| Lmna | 1394.965905 | 0.154033715 | 0.081688699 | 1.885618423 | 0.059346393 | 0.193522564 |
| Frem2 | 142.4628563 | -0.191297497 | 0.127336278 | -1.502301616 | 0.133019231 | 0.323847184 |
| Pkn3 | 61.75583106 | 0.153954202 | 0.144559765 | 1.064986523 | 0.286882029 | 0.516554346 |
| Slc19a1 | 591.0200935 | 0.153935568 | 0.091620845 | 1.680136961 | 0.092930671 | 0.257349029 |
| Slc16a1 | 1289.815611 | -0.145098572 | 0.096631711 | -1.501562681 | 0.133210087 | 0.324113865 |
| Emilin1 | 105.9379437 | -0.222389495 | 0.148155603 | -1.501053557 | 0.13334171 | 0.324383177 |
| Hist1h1c | 465.5860309 | -0.210482565 | 0.140269323 | -1.500560209 | 0.13346935 | 0.324642717 |
| Optc | 28.97770028 | 0.153871157 | 0.167325786 | 0.919590224 | 0.357786937 | 0.588549826 |
| Ppp2r4 | 6962.000776 | -0.082813386 | 0.055207775 | -1.500031204 | 0.13360632 | 0.324783353 |
| Igsf9 | 214.7619593 | 0.153796077 | 0.153555974 | 1.00156362 | 0.316554399 | 0.549376037 |
| Slc9a9 | 761.2705402 | -0.147131349 | 0.09808233 | -1.50008007 | 0.133593663 | 0.324783353 |
| Ube2m | 4277.199925 | -0.092006486 | 0.061342435 | -1.499883181 | 0.133644665 | 0.324814187 |
| Ldhb | 19038.34808 | -0.088784363 | 0.059210279 | -1.499475502 | 0.133750319 | 0.325019989 |
| Cul1 | 5092.786971 | 0.086041578 | 0.057385602 | 1.499358302 | 0.133780705 | 0.325042848 |
| Mpnd | 2254.745705 | -0.116910852 | 0.077979482 | -1.499251456 | 0.133808411 | 0.32505919 |
| Slc8a3 | 1369.26922 | -0.130635457 | 0.087154654 | -1.49889249 | 0.133901525 | 0.325234398 |
| Adck2 | 813.8792359 | 0.097136582 | 0.064816945 | 1.498629452 | 0.133969788 | 0.3253492 |
| Gm15910 | 8.59686301 | 0.153783761 | 0.157944595 | 0.973656371 | 0.330227187 | 0.565426018 |
| Rpa1 | 1820.384798 | 0.153781597 | 0.059252609 | 2.595355712 | 0.009449307 | 0.056506199 |
| Chrnb3 | 63.76631455 | 0.153734267 | 0.167261609 | 0.91912465 | 0.358030377 | 0.588749683 |
| Prr12 | 2933.101879 | 0.087954193 | 0.058715362 | 1.497975815 | 0.134139534 | 0.325535831 |
| Pex26 | 727.3870079 | 0.153721993 | 0.079864129 | 1.924793965 | 0.054255139 | 0.183111081 |
| B3gat1 | 9724.129637 | -0.102930004 | 0.068734699 | -1.497496968 | 0.134263994 | 0.325757332 |
| Grpr | 27.76504019 | 0.153658733 | 0.168119026 | 0.913987766 | 0.360723274 | 0.591496075 |
| Ints3 | 3110.376072 | 0.083320882 | 0.055650755 | 1.49721028 | 0.134338551 | 0.32583618 |
| Acbd4 | 1079.654013 | 0.153632732 | 0.072172236 | 2.128695766 | 0.033279439 | 0.132723715 |
| Fdx1 | 205.9021818 | -0.150699634 | 0.100677054 | -1.496861773 | 0.134429229 | 0.325903062 |
| Inpp4a | 6774.292024 | -0.083387642 | 0.05570275 | -1.497011225 | 0.134390337 | 0.325903062 |
| Pkia | 6377.68549 | -0.146340017 | 0.097776828 | -1.496673801 | 0.134478156 | 0.325970675 |
| Vcan | 874.5373205 | -0.153176089 | 0.102364696 | -1.496376152 | 0.134555661 | 0.326107525 |
| Hspb8 | 578.6441868 | -0.130451875 | 0.087187414 | -1.496223693 | 0.134595372 | 0.32613938 |
| Ift46 | 1283.569406 | 0.099752067 | 0.066671877 | 1.496164077 | 0.134610903 | 0.32613938 |
| Nrf1 | 632.5846544 | -0.115212085 | 0.077029513 | -1.495687568 | 0.134735091 | 0.326389229 |
| Fam195b | 1900.213392 | -0.095655413 | 0.063984213 | -1.4949846 | 0.134918461 | 0.326782342 |
| Fam129a | 121.1580265 | 0.153595739 | 0.148716511 | 1.03280892 | 0.301693329 | 0.533658642 |
| Senp1 | 631.1351365 | 0.153322157 | 0.094147637 | 1.628529005 | 0.103412764 | 0.276540795 |
| Irf2bp1 | 1543.190937 | -0.085977655 | 0.057544863 | -1.494097823 | 0.135150053 | 0.327189831 |
| Wdr91 | 1052.4502 | 0.153298334 | 0.071107019 | 2.155881893 | 0.031092889 | 0.12661057 |
| Lrrc49 | 3070.245269 | 0.153221102 | 0.080540638 | 1.902407363 | 0.057117919 | 0.18901859 |
| Hmbs | 559.2749129 | -0.126716131 | 0.084852204 | -1.49337465 | 0.135339145 | 0.327438548 |
| Pls1 | 500.9237097 | 0.15319471 | 0.099022882 | 1.547063742 | 0.121847876 | 0.307878504 |
| Rybp | 2433.994646 | -0.107352797 | 0.071893486 | -1.493220078 | 0.135379588 | 0.327438548 |
| Ydjc | 418.2412156 | -0.198994206 | 0.13325569 | -1.493326151 | 0.135351833 | 0.327438548 |
| Fbxo21 | 5791.203927 | 0.153083436 | 0.071429652 | 2.143135683 | 0.032102205 | 0.129360615 |
| Nfe2l2 | 683.6596151 | 0.153081171 | 0.079329641 | 1.929684399 | 0.053645954 | 0.181889635 |
| Atxn7l1 | 914.5557991 | 0.15307298 | 0.065380346 | 2.341269046 | 0.019218312 | 0.091773148 |
| Zfp142 | 1730.06038 | 0.153060614 | 0.073908561 | 2.070945656 | 0.038363875 | 0.14540359 |
| Mrpl14 | 331.7658772 | -0.153104853 | 0.10262124 | -1.491941172 | 0.135714568 | 0.327984607 |
| Mrpl35 | 1025.362443 | -0.103626539 | 0.069460684 | -1.491873285 | 0.135732367 | 0.327984607 |
| Gm7008 | 14.04952023 | 0.153019619 | 0.163550541 | 0.935610598 | 0.349473716 | 0.580421397 |
| Fam107a | 17423.09026 | 0.152999632 | 0.127235493 | 1.202491761 | 0.229173056 | 0.452159344 |
| Nanp | 548.3087432 | -0.110251591 | 0.073938511 | -1.491125396 | 0.135928576 | 0.328305122 |
| D730039F16Rik | 15.19634192 | 0.15294994 | 0.166901735 | 0.916407125 | 0.3594534 | 0.590476142 |
| Prkd3 | 980.7356597 | -0.112365257 | 0.075379359 | -1.490663466 | 0.136049873 | 0.328429143 |
| Adamts14 | 16.96499327 | 0.152922706 | 0.159080643 | 0.961290469 | 0.336406139 | 0.570346257 |
| Snx14 | 2195.322087 | -0.082880377 | 0.05560451 | -1.490533362 | 0.136084051 | 0.328429143 |
| Ttc9 | 1342.420241 | -0.119021212 | 0.079851803 | -1.490526284 | 0.136085911 | 0.328429143 |
| Apln | 1056.745517 | -0.245863329 | 0.16496298 | -1.49041518 | 0.136115104 | 0.328448437 |
| 1700019D03Rik | 124.2965629 | -0.167290506 | 0.112297598 | -1.489706894 | 0.136301321 | 0.328796121 |
| Synpo2l | 10.68006528 | -0.239987276 | 0.161097104 | -1.489705713 | 0.136301632 | 0.328796121 |
| Hspb3 | 64.42158644 | 0.152883552 | 0.139375843 | 1.096915711 | 0.27267824 | 0.501112164 |
| Svil | 492.4578633 | -0.143345548 | 0.096266423 | -1.489050317 | 0.136474119 | 0.329109728 |
| Mesdc2 | 2017.515739 | -0.086421271 | 0.058044682 | -1.488874912 | 0.136520311 | 0.329169888 |
| A230056P14Rik | 91.80491389 | -0.211846777 | 0.142390137 | -1.487791092 | 0.136805996 | 0.329602254 |
| Ist1 | 3331.807927 | 0.152871136 | 0.067125405 | 2.277396094 | 0.022762583 | 0.102763247 |
| Gtf3c4 | 2148.770041 | -0.079960077 | 0.053741497 | -1.487864712 | 0.136786576 | 0.329602254 |
| Baiap2l1 | 27.14052769 | 0.152800143 | 0.167151059 | 0.914144034 | 0.360641167 | 0.591424031 |
| Traf2 | 386.3406078 | 0.152790484 | 0.0806681 | 1.894063251 | 0.058216615 | 0.191306093 |
| Rfc1 | 1665.777719 | 0.072801802 | 0.04895558 | 1.487099153 | 0.136988625 | 0.329990945 |
| Cdc6 | 21.14970576 | -0.247812277 | 0.166675102 | -1.486798408 | 0.137068062 | 0.330130974 |
| Atraid | 1650.536124 | 0.088651342 | 0.059633796 | 1.486595637 | 0.137121641 | 0.330157374 |
| Gmeb2 | 1016.374487 | -0.111572871 | 0.075052035 | -1.486606876 | 0.137118671 | 0.330157374 |
| Il1rapl1 | 285.9203408 | -0.154272204 | 0.103799936 | -1.48624565 | 0.137214157 | 0.330296028 |
| Cpa4 | 9.689432344 | 0.152774147 | 0.162868192 | 0.938023225 | 0.348232475 | 0.579477388 |
| Tmem238 | 30.32097445 | 0.152757839 | 0.167037014 | 0.914514908 | 0.360446348 | 0.591344072 |
| Sh3d19 | 624.1746922 | 0.152592762 | 0.078104129 | 1.953709281 | 0.050735608 | 0.175468195 |
| 11-Sep | 4113.730641 | -0.103462864 | 0.069636047 | -1.485765894 | 0.137341054 | 0.330443271 |
| Cers4 | 1819.425559 | 0.152456894 | 0.097604678 | 1.561983477 | 0.118291881 | 0.301846905 |
| Gsx1 | 13.95927557 | -0.226820968 | 0.152745941 | -1.48495578 | 0.137555538 | 0.330842209 |
| Usp16 | 989.3432752 | 0.152441886 | 0.086181486 | 1.76884727 | 0.076919366 | 0.228357688 |
| Eci1 | 414.9117712 | -0.162024724 | 0.109165183 | -1.484216126 | 0.137751593 | 0.331210922 |
| Rnasel | 572.0764952 | 0.152402547 | 0.077366927 | 1.969866881 | 0.048853629 | 0.171405855 |
| Utp14b | 273.3511431 | -0.220647913 | 0.1487208 | -1.483638557 | 0.137904835 | 0.331476498 |
| Bloc1s3 | 421.7080836 | -0.131865217 | 0.088891885 | -1.483433688 | 0.137959222 | 0.331518804 |
| Pigf | 234.4148012 | -0.175835968 | 0.11853489 | -1.483411069 | 0.137965228 | 0.331518804 |
| Nfx1 | 3496.302576 | 0.152311891 | 0.071853617 | 2.119752581 | 0.034026916 | 0.134573122 |
| Avl9 | 2671.838888 | -0.132198631 | 0.089193508 | -1.482155298 | 0.138298978 | 0.332166232 |
| Eefsec | 616.4198431 | -0.115265898 | 0.077765102 | -1.482231679 | 0.13827866 | 0.332166232 |
| Fam57a | 180.3291564 | 0.152207956 | 0.141828685 | 1.073181753 | 0.283189576 | 0.512698604 |
| Actr6 | 459.6841273 | -0.108721481 | 0.073374578 | -1.481732282 | 0.138411544 | 0.332182376 |
| Ccdc23 | 273.3756038 | -0.140625497 | 0.094893463 | -1.481930288 | 0.138358845 | 0.332182376 |
| Fndc3b | 854.3663118 | -0.155083744 | 0.104660398 | -1.481780574 | 0.13839869 | 0.332182376 |
| Kptn | 689.1783022 | -0.183688551 | 0.123952645 | -1.481925231 | 0.138360191 | 0.332182376 |
| Pcdhgc4 | 516.3070115 | -0.132662486 | 0.089535769 | -1.481670257 | 0.138428055 | 0.332182376 |
| Plekhf2 | 286.1137699 | -0.132665315 | 0.089542399 | -1.48159214 | 0.138448852 | 0.332182376 |
| Elp4 | 495.6248542 | -0.158529584 | 0.107010623 | -1.481437822 | 0.138489942 | 0.332190947 |
| Gstz1 | 1249.876322 | 0.152141651 | 0.082260737 | 1.849505086 | 0.064384914 | 0.203768935 |
| Tmco3 | 1536.837785 | -0.093056448 | 0.06281826 | -1.481359858 | 0.138510706 | 0.332190947 |
| Gpr35 | 14.15077631 | 0.152111434 | 0.166363963 | 0.914329231 | 0.360543875 | 0.591344072 |
| Enkur | 81.54132192 | 0.152097855 | 0.13497363 | 1.12687089 | 0.259797073 | 0.486583408 |
| Gatc | 1161.18818 | 0.094318978 | 0.063710032 | 1.480441539 | 0.138755451 | 0.332592352 |
| Trpc4ap | 6582.544063 | 0.081173858 | 0.054855083 | 1.479787356 | 0.138930004 | 0.33295924 |
| B3gat3 | 3981.527165 | -0.119340014 | 0.080687923 | -1.479031923 | 0.139131783 | 0.333288164 |
| Rap1a | 1040.281206 | -0.113493791 | 0.07673337 | -1.479066941 | 0.139122425 | 0.333288164 |
| Rnf149 | 826.8663355 | -0.0962524 | 0.065073302 | -1.479138096 | 0.13910341 | 0.333288164 |
| Ift88 | 736.6688389 | 0.15206118 | 0.076152116 | 1.996808326 | 0.045846009 | 0.164406322 |
| 5730577I03Rik | 60.58208864 | -0.237438038 | 0.16063704 | -1.478102674 | 0.139380299 | 0.333742489 |
| C1qtnf4 | 2514.866642 | -0.199008237 | 0.134646921 | -1.478000657 | 0.139407602 | 0.333742489 |
| Kcnn2 | 1059.721471 | 0.151992936 | 0.108951941 | 1.395045702 | 0.163002058 | 0.368263909 |
| Clcn4-2 | 8399.115107 | -0.078001525 | 0.052794622 | -1.477452091 | 0.139554492 | 0.33393935 |
| Prss22 | 50.31606294 | -0.246671059 | 0.166954919 | -1.477471054 | 0.139549412 | 0.33393935 |
| Sncg | 104.430024 | -0.222219285 | 0.150393224 | -1.477588418 | 0.139517976 | 0.33393935 |
| Pard6g | 279.7804455 | -0.150404214 | 0.101812684 | -1.477264006 | 0.139604883 | 0.334008346 |
| Bclaf1 | 4264.653653 | 0.151883325 | 0.086143827 | 1.763136508 | 0.077877469 | 0.229976969 |
| Cort | 96.80326922 | 0.151737693 | 0.160827547 | 0.943480738 | 0.345435063 | 0.576948985 |
| Wdr17 | 1682.487955 | -0.107327642 | 0.072668652 | -1.476945559 | 0.139690231 | 0.334076546 |
| Med8 | 777.4886025 | 0.091217072 | 0.061772943 | 1.476650912 | 0.139769237 | 0.334195147 |
| Acot2 | 179.2618735 | 0.151733076 | 0.100390054 | 1.511435343 | 0.13067757 | 0.320316133 |
| Eif5b | 3558.308912 | 0.151722216 | 0.072246124 | 2.100074127 | 0.035722321 | 0.13878677 |
| Nrtn | 18.37771453 | -0.242658189 | 0.164391041 | -1.476103489 | 0.139916113 | 0.334334817 |
| Ankrd32 | 556.3327726 | 0.151652874 | 0.088417496 | 1.715190772 | 0.086310256 | 0.245799414 |
| Ddx27 | 1118.556946 | 0.151578927 | 0.067326029 | 2.251416434 | 0.024359174 | 0.107754719 |
| Tmem165 | 787.9616893 | 0.151522882 | 0.099068689 | 1.529472961 | 0.126147237 | 0.314030413 |
| Mcm3ap | 2557.832358 | 0.093178265 | 0.063143369 | 1.475661919 | 0.140034674 | 0.334412122 |
| Ninj1 | 402.0048245 | -0.161575471 | 0.109493927 | -1.475656912 | 0.140036019 | 0.334412122 |
| Itpkc | 350.3820626 | 0.151513032 | 0.103917796 | 1.45800852 | 0.144838195 | 0.342771191 |
| Leprel4 | 353.9038954 | -0.139929693 | 0.094865549 | -1.475031707 | 0.140204019 | 0.334666626 |
| Tstd1 | 15.74263358 | 0.151469001 | 0.165884591 | 0.913098681 | 0.361190644 | 0.591886604 |
| Nkx2-2as | 35.44538583 | -0.246611424 | 0.167262761 | -1.474395276 | 0.140375196 | 0.335023611 |
| Grlf1 | 10859.23531 | -0.094332221 | 0.063986128 | -1.474260496 | 0.140411467 | 0.335058566 |
| A630001G21Rik | 21.30169277 | 0.151383517 | 0.168062746 | 0.900755942 | 0.367718097 | 0.597446401 |
| Nbn | 687.5640095 | 0.151197438 | 0.092280615 | 1.638452861 | 0.10132726 | 0.272471248 |
| Tmem134 | 723.3512697 | -0.125536595 | 0.0851679 | -1.473989551 | 0.140484404 | 0.335121051 |
| Htra1 | 5332.229608 | 0.151162668 | 0.160268474 | 0.943184046 | 0.345586773 | 0.576948985 |
| Aldh1l1 | 2605.606247 | 0.151131812 | 0.09463417 | 1.597011006 | 0.110263254 | 0.289170422 |
| Dhx36 | 3148.423824 | -0.094506428 | 0.064134777 | -1.473559786 | 0.140600156 | 0.335147509 |
| Pelp1 | 1467.080903 | 0.085267668 | 0.057862297 | 1.473630889 | 0.140581 | 0.335147509 |
| Ptgis | 52.57602856 | -0.2307964 | 0.156642977 | -1.473391306 | 0.140645553 | 0.335204153 |
| Ankrd2 | 13.10159108 | 0.151048264 | 0.166652118 | 0.906368707 | 0.36474073 | 0.594935559 |
| Rpa2 | 430.6330283 | 0.151017925 | 0.093570754 | 1.61394366 | 0.106539647 | 0.282175264 |
| Irf2 | 1015.662364 | 0.151006688 | 0.077246862 | 1.954858545 | 0.050599769 | 0.175154785 |
| Ltbp2 | 30.89237574 | -0.232079361 | 0.157672446 | -1.471908167 | 0.141045678 | 0.335951073 |
| Aff1 | 1053.116401 | 0.151000021 | 0.132810803 | 1.136955868 | 0.255556735 | 0.48192595 |
| Rfx1 | 1000.477133 | 0.15097262 | 0.115307082 | 1.309309163 | 0.190429647 | 0.4040697 |
| Fam19a5 | 4321.314485 | -0.078070435 | 0.053102979 | -1.470170522 | 0.141515576 | 0.336863164 |
| Fut8 | 2311.389292 | -0.088859967 | 0.060440768 | -1.470199184 | 0.141507816 | 0.336863164 |
| Zfp882 | 366.3064222 | -0.188521408 | 0.128265234 | -1.469777905 | 0.141621915 | 0.337064509 |
| Bbs5 | 524.899273 | 0.150943252 | 0.076703051 | 1.967891115 | 0.049080567 | 0.171892927 |
| Dedd2 | 448.3808762 | 0.150908013 | 0.098385787 | 1.533839567 | 0.125069126 | 0.312636269 |
| Mknk1 | 1053.224268 | 0.150876964 | 0.085160069 | 1.77168673 | 0.076446574 | 0.22747517 |
| Gm13238 | 12.61156714 | 0.15086826 | 0.163967168 | 0.920112619 | 0.357513911 | 0.588350561 |
| Bud13 | 416.7066335 | 0.150777184 | 0.088052082 | 1.71236364 | 0.086829675 | 0.246670149 |
| Myocd | 10.85512103 | -0.231769465 | 0.157887627 | -1.467939376 | 0.142120691 | 0.33788829 |
| Ncapd2 | 982.0402497 | 0.150763629 | 0.074402206 | 2.026332788 | 0.042730693 | 0.156888475 |
| Srgn | 180.6997159 | -0.157762885 | 0.107494378 | -1.467638478 | 0.14220245 | 0.338030801 |
| Patl1 | 1056.21034 | 0.150743542 | 0.087238832 | 1.727940853 | 0.083998829 | 0.241685767 |
| Tsen54 | 256.6289705 | 0.150724056 | 0.094347588 | 1.597540111 | 0.110145364 | 0.28895907 |
| Gchfr | 8.854347792 | -0.232579006 | 0.158540453 | -1.467001019 | 0.142375778 | 0.33828712 |
| Cln6 | 474.0626384 | 0.150705707 | 0.090480879 | 1.665608349 | 0.095791447 | 0.262132229 |
| Cluap1 | 949.3418951 | 0.150693935 | 0.064316562 | 2.343003568 | 0.0191292 | 0.091469296 |
| Tpgs2 | 2281.973613 | -0.088335296 | 0.060259327 | -1.465919042 | 0.142670344 | 0.338831132 |
| 1700007P06Rik | 12.41491263 | 0.150587525 | 0.165450342 | 0.910167509 | 0.362734176 | 0.593467136 |
| Emd | 1174.406291 | -0.127322958 | 0.086893099 | -1.465282745 | 0.142843792 | 0.339139093 |
| 2310042D19Rik | 37.40346073 | 0.15052373 | 0.15629227 | 0.963091329 | 0.335501693 | 0.56979757 |
| Tmx3 | 2054.857352 | -0.115862564 | 0.079094244 | -1.464867205 | 0.142957152 | 0.339304245 |
| Commd6 | 604.0371885 | -0.121875864 | 0.083250145 | -1.463971785 | 0.143201658 | 0.339776502 |
| Fgfr1op2 | 3395.188119 | 0.150465693 | 0.0759575 | 1.980919507 | 0.047600301 | 0.168575786 |
| Ythdc1 | 2309.92404 | 0.099543014 | 0.067998616 | 1.463897656 | 0.143221914 | 0.339776502 |
| Mthfd1 | 1478.002002 | 0.068227751 | 0.046622939 | 1.463394471 | 0.143359471 | 0.33999872 |
| Slc39a5 | 12.56485186 | -0.241095826 | 0.164743224 | -1.463464299 | 0.143340376 | 0.33999872 |
| Tcf23 | 19.27040081 | 0.150452214 | 0.167374533 | 0.8988955 | 0.368708325 | 0.598383449 |
| Mybbp1a | 2521.115236 | 0.089868607 | 0.061428279 | 1.462984271 | 0.143471683 | 0.340112608 |
| Prickle2 | 9523.924345 | -0.108750874 | 0.074335266 | -1.462978198 | 0.143473345 | 0.340112608 |
| Gyg | 1046.347057 | -0.117680742 | 0.080510424 | -1.461683295 | 0.143828017 | 0.340901223 |
| Spin1 | 9041.748063 | -0.089418595 | 0.061222337 | -1.460555088 | 0.144137579 | 0.341582694 |
| Atrx | 6226.999987 | -0.10433746 | 0.071443273 | -1.460423842 | 0.144173624 | 0.341588749 |
| Secisbp2 | 614.2564613 | 0.150339178 | 0.092780748 | 1.620370394 | 0.105152736 | 0.279793317 |
| Napa | 9502.803461 | 0.150302855 | 0.068556873 | 2.192381998 | 0.028351935 | 0.119232798 |
| Dcun1d1 | 1650.657132 | -0.118147406 | 0.080921933 | -1.460017102 | 0.144285374 | 0.341723874 |
| Tnks | 3521.364202 | -0.113758683 | 0.077926413 | -1.459821892 | 0.14433903 | 0.341798706 |
| Il1rap | 1271.653903 | 0.150209744 | 0.100270482 | 1.498045495 | 0.134121431 | 0.325535831 |
| Nemf | 1714.650708 | 0.150074619 | 0.067219819 | 2.232594829 | 0.025575675 | 0.11084462 |
| Nyx | 19.80501746 | 0.150071207 | 0.168019179 | 0.893179025 | 0.371761309 | 0.601409428 |
| Xpc | 746.2894489 | 0.150057225 | 0.081714803 | 1.836353012 | 0.066305464 | 0.207458979 |
| Chmp1a | 3707.007369 | -0.097240685 | 0.066703354 | -1.457808016 | 0.144893468 | 0.342806638 |
| Ntng1 | 805.7123801 | -0.209944347 | 0.144023059 | -1.457713431 | 0.144919548 | 0.342806638 |
| Slc7a3 | 325.8718432 | 0.150012315 | 0.103687426 | 1.446774416 | 0.147960114 | 0.347505535 |
| Tep1 | 268.2153584 | 0.149907832 | 0.129293417 | 1.159439022 | 0.246277279 | 0.471389617 |
| Larp7 | 651.984239 | -0.09758254 | 0.066966535 | -1.457183653 | 0.145065692 | 0.343047593 |
| Nos2 | 33.83073835 | -0.244945827 | 0.16812107 | -1.456960906 | 0.145127172 | 0.343140607 |
| Gm20753 | 10.65851557 | 0.149760445 | 0.162033675 | 0.924255068 | 0.35535353 | 0.586284589 |
| Ssna1 | 813.9602516 | -0.089558858 | 0.061505343 | -1.456115086 | 0.145360808 | 0.343588155 |
| Smg7 | 5229.999556 | 0.087490206 | 0.060099121 | 1.455765154 | 0.145457552 | 0.343764385 |
| Dpcd | 1331.07876 | -0.113416508 | 0.07791404 | -1.455662013 | 0.145486076 | 0.34377936 |
| Tcf25 | 25576.61258 | 0.074442525 | 0.051168799 | 1.454842153 | 0.145712965 | 0.34426299 |
| A930015D03Rik | 45.55080053 | 0.149636905 | 0.167637716 | 0.892620756 | 0.3720603 | 0.601694982 |
| Tbk1 | 1929.698046 | 0.149591805 | 0.06691217 | 2.235644206 | 0.025375086 | 0.110305951 |
| Acp5 | 9.529002625 | 0.149507356 | 0.152771755 | 0.978632184 | 0.327761751 | 0.562323173 |
| Edem2 | 742.886214 | 0.149447263 | 0.084631185 | 1.765865177 | 0.077418471 | 0.229195353 |
| Scn8a | 15399.91458 | -0.101711784 | 0.06998797 | -1.45327524 | 0.146147349 | 0.345028899 |
| Trmt1 | 1826.348701 | 0.149392122 | 0.071908492 | 2.077530996 | 0.037752577 | 0.144025017 |
| 5930403L14Rik | 1943.825698 | -0.165093299 | 0.113617386 | -1.4530637 | 0.146206068 | 0.345059684 |
| Trim8 | 2636.278165 | -0.115624603 | 0.079580696 | -1.452922745 | 0.146245205 | 0.345099483 |
| Map4k3 | 3000.786688 | 0.149373219 | 0.084023341 | 1.777758629 | 0.075443506 | 0.225660556 |
| Shc2 | 1861.497472 | 0.149335781 | 0.07220524 | 2.068212507 | 0.038620045 | 0.146136809 |
| Nr1h3 | 104.3251176 | 0.149233153 | 0.117530282 | 1.269742151 | 0.204176493 | 0.421351969 |
| Map3k9 | 4234.828424 | 0.149148133 | 0.063912065 | 2.333645966 | 0.019614268 | 0.092938739 |
| 5730522E02Rik | 50.74946407 | 0.149145737 | 0.153903829 | 0.969083992 | 0.332503284 | 0.567441996 |
| Matn4 | 266.4036146 | -0.160629407 | 0.110649812 | -1.451691632 | 0.146587367 | 0.345591094 |
| Dennd1a | 2590.607086 | 0.068984363 | 0.047535222 | 1.451226274 | 0.146716863 | 0.345620169 |
| Gabpb1 | 612.6494031 | -0.107061668 | 0.07376688 | -1.451351449 | 0.146682022 | 0.345620169 |
| Gtpbp10 | 350.838254 | -0.135457575 | 0.093328422 | -1.451407526 | 0.146666415 | 0.345620169 |
| Eml2 | 5288.958745 | 0.149140858 | 0.092819726 | 1.606779771 | 0.108102687 | 0.285096687 |

| | | | | | | |
|---|---|---|---|---|---|---|
| Tes | 58.90274936 | -0.085515688 | 0.093620134 | -0.913432661 | 0.361015035 | 0.776449324 |
| 1190007I07Rik | 161.9971968 | -0.085518361 | 0.089175879 | -0.958985343 | 0.337566129 | 0.760312044 |
| Noxred1 | 48.89121935 | -0.085563085 | 0.084688541 | -1.01032659 | 0.312338846 | 0.74483997 |
| Setd2 | 3289.44041 | -0.085605435 | 0.063433707 | -1.349525968 | 0.177168085 | 0.629241039 |
| Ccdc80 | 277.2201982 | -0.085621531 | 0.079617885 | -1.075405748 | 0.282193109 | 0.720537371 |
| Dgcr8 | 1663.910489 | -0.085674683 | 0.056582925 | -1.514143759 | 0.129989382 | 0.56891406 |
| Dnahc2 | 147.5302644 | -0.085693093 | 0.094606099 | -0.905788253 | 0.36504794 | 0.777634019 |
| Efcab4a | 83.4619567 | -0.08571848 | 0.095061713 | -0.901714036 | 0.367208795 | 0.778583412 |
| Slc38a8 | 16.58338342 | -0.085755487 | 0.071491949 | -1.199512507 | 0.230328724 | NA |
| Fras1 | 1254.848581 | -0.085756813 | 0.075600168 | -1.134346848 | 0.256649085 | 0.703000737 |
| Rbpms2 | 100.9020771 | -0.085776613 | 0.094392011 | -0.908727469 | 0.363494001 | 0.776830051 |
| Armc2 | 376.6844156 | -0.085791118 | 0.080199079 | -1.069726975 | 0.28474222 | 0.721641599 |
| Rps6ka1 | 711.7026847 | -0.085913427 | 0.067577877 | -1.271324746 | 0.203613142 | 0.657622144 |
| Dhx33 | 1959.898251 | -0.085917973 | 0.062273798 | -1.379680954 | 0.1676849 | 0.617953076 |
| AI854517 | 1113.309525 | -0.085940137 | 0.078303809 | -1.097521796 | 0.272413358 | 0.714142044 |
| Ttc38 | 373.6582259 | -0.085945917 | 0.079055653 | -1.087157127 | 0.276967375 | 0.7177927 |
| Wdr44 | 760.0425194 | -0.085957612 | 0.082887195 | -1.037043316 | 0.29971567 | 0.733704078 |
| Olr1 | 2.740774214 | -0.086015346 | 0.039975589 | -2.151696761 | 0.031421245 | NA |
| Ikbip | 352.222328 | -0.086024998 | 0.070311901 | -1.223477056 | 0.221149565 | 0.672757795 |
| Rbm41 | 373.2292104 | -0.086051109 | 0.085412445 | -1.00747742 | 0.313705403 | 0.745844951 |
| Pik3c2b | 1106.362243 | -0.086056356 | 0.073301082 | -1.174012082 | 0.240390186 | 0.6914669 |
| Prkd3 | 1003.976072 | -0.0860742 | 0.070093945 | -1.227983397 | 0.219453201 | 0.672421385 |
| Apol7b | 5.474585189 | -0.086081553 | 0.047481049 | -1.812966557 | 0.06983698 | NA |
| Poli | 558.0106938 | -0.086120172 | 0.069186763 | -1.24474926 | 0.213223933 | 0.666732456 |
| Kcnk6 | 249.3126616 | -0.086123412 | 0.093228176 | -0.923791665 | 0.355594796 | 0.772370994 |
| Rrbp1 | 2318.581809 | -0.086135332 | 0.064979936 | -1.32556813 | 0.184982789 | 0.637780733 |
| Golim4 | 1016.226606 | -0.086168694 | 0.060451586 | -1.42541659 | 0.154036815 | 0.60340295 |
| Cars2 | 406.9816056 | -0.086194097 | 0.078365106 | -1.09990404 | 0.271373934 | 0.713480139 |
| Dennd4c | 951.1500377 | -0.086202413 | 0.067907796 | -1.269403774 | 0.204297091 | 0.65778952 |
| Prrt1 | 1579.274773 | -0.086243023 | 0.076167742 | -1.132277538 | 0.257517771 | 0.703271881 |
| Trim11 | 843.3912074 | -0.086337843 | 0.073168495 | -1.179986593 | 0.238005547 | 0.688758314 |
| Tcerg1 | 2868.606778 | -0.08640157 | 0.056112536 | -1.539790869 | 0.123611338 | 0.558428989 |
| Baiap2l2 | 72.44005485 | -0.086440707 | 0.094333789 | -0.916328156 | 0.359494805 | 0.775701357 |
| Gjb2 | 315.8603766 | -0.08645059 | 0.095102189 | -0.909028398 | 0.363335136 | 0.776821522 |
| Sepn1 | 1329.224137 | -0.086457935 | 0.067611781 | -1.278740689 | 0.200988386 | 0.654716629 |
| Bmp3 | 492.500929 | -0.086467611 | 0.086453693 | -1.000160994 | 0.317232602 | 0.746980188 |
| Litaf | 877.2649918 | -0.086470934 | 0.086676013 | -0.997633957 | 0.318456889 | 0.74851037 |
| Kifc5b | 86.91991258 | -0.086480433 | 0.094987072 | -0.910444247 | 0.362588272 | 0.776449324 |
| Abcb1b | 153.8916806 | -0.086486875 | 0.089523279 | -0.966082521 | 0.33400289 | 0.758212399 |
| Frmd7 | 31.29076485 | -0.086490163 | 0.083437143 | -1.036590655 | 0.29992667 | 0.733704078 |
| Slc13a5 | 585.7188719 | -0.086578576 | 0.077873637 | -1.11178288 | 0.266231515 | 0.708942068 |
| Tle1 | 1452.305997 | -0.086578612 | 0.06092846 | -1.420988025 | 0.155320241 | 0.604578396 |
| 2610305D13Rik | 11.03381714 | -0.086596407 | 0.06883322 | -1.258061254 | 0.208369604 | NA |
| Zbtb37 | 152.6082528 | -0.086649259 | 0.095037101 | -0.91174139 | 0.36190487 | 0.776449324 |
| Mtbp | 94.67729466 | -0.086693689 | 0.094716409 | -0.91529746 | 0.360035492 | 0.776328279 |
| Rgs14 | 2353.830769 | -0.086700361 | 0.076316839 | -1.136058071 | 0.255932261 | 0.701952734 |
| Gpr116 | 2036.928417 | -0.086787775 | 0.068491751 | -1.267127425 | 0.205109731 | 0.658534108 |
| Ubr2 | 3919.731524 | -0.086797043 | 0.062943204 | -1.378974033 | 0.16790276 | 0.617953076 |
| Gm9897 | 40.97565767 | -0.086809885 | 0.091291107 | -0.950912831 | 0.341648628 | 0.76303628 |
| Ccdc111 | 182.7030728 | -0.086908304 | 0.088131742 | -0.986118076 | 0.324075175 | 0.752925264 |
| Vps13c | 2606.870453 | -0.086910013 | 0.079889923 | -1.087872035 | 0.276651603 | 0.717475105 |
| Zfp524 | 177.1816601 | -0.086945951 | 0.086985549 | -0.999544779 | 0.317530858 | 0.747319062 |
| Flad1 | 1266.736573 | -0.08696902 | 0.067649358 | -1.285585291 | 0.198587833 | 0.65377551 |
| Cenpf | 36.75370689 | -0.086976291 | 0.088623259 | -0.981416072 | 0.326387604 | 0.753373722 |
| Rpl13 | 6139.449868 | -0.08698393 | 0.069696971 | -1.248030274 | 0.212019971 | 0.666270994 |
| Serinc5 | 3721.50551 | -0.086992698 | 0.051082083 | -1.702998255 | 0.088568395 | 0.497517107 |
| Kcnip2 | 7927.653454 | -0.087022716 | 0.061889079 | -1.406107784 | 0.159692121 | 0.611570723 |
| 2010107G12Rik | 18.20194058 | -0.087025473 | 0.079684591 | -1.092124237 | 0.274778498 | NA |
| Ewsr1 | 7833.508998 | -0.087070545 | 0.060723115 | -1.433894561 | 0.151602354 | 0.598905575 |
| Stx5a | 2106.675557 | -0.087096152 | 0.050089401 | -1.738814007 | 0.082067484 | 0.482582908 |
| Cyp2e1 | 21.62497288 | -0.087098007 | 0.072334556 | -1.204099562 | 0.228551099 | NA |
| Atp8b2 | 4277.578005 | -0.087104266 | 0.052585094 | -1.656444059 | 0.097631925 | 0.518456434 |
| BC055402 | 10.23975924 | -0.087105698 | 0.066224617 | -1.315306941 | 0.188406766 | NA |
| Zfp747 | 239.061103 | -0.087181608 | 0.084241366 | -1.03490259 | 0.300714405 | 0.734641347 |
| Tars2 | 1321.514692 | -0.087262901 | 0.059725028 | -1.461077602 | 0.143994146 | 0.589967032 |
| Sphkap | 7496.390236 | -0.08729261 | 0.064095914 | -1.361905996 | 0.173227554 | 0.624702941 |
| Cdh5 | 688.4143168 | -0.087339952 | 0.077088911 | -1.132976857 | 0.257223972 | 0.703199912 |
| Spag4 | 99.28848097 | -0.087350433 | 0.095101024 | -0.918501503 | 0.358356372 | 0.774201531 |
| Ptprh | 98.52009466 | -0.087370633 | 0.094763409 | -0.921987019 | 0.356535347 | 0.773064269 |
| Popdc2 | 16.58293519 | -0.087379573 | 0.073261529 | -1.192707463 | 0.232983962 | NA |
| Fbn1 | 304.2643525 | -0.087401296 | 0.094172336 | -0.928099477 | 0.353355971 | 0.770624606 |
| Hdac6 | 2432.659667 | -0.087427588 | 0.06173966 | -1.416068506 | 0.156755448 | 0.607369869 |
| Degs2 | 120.4103886 | -0.087555557 | 0.091125065 | -0.960828468 | 0.336638422 | 0.75970798 |
| Bak1 | 486.581913 | -0.087644212 | 0.066535227 | -1.317260287 | 0.187751392 | 0.641781416 |
| Hsd17b14 | 8.133562054 | -0.087689952 | 0.056223863 | -1.559657196 | 0.118840912 | NA |
| Gm17644 | 54.84319258 | -0.08769475 | 0.090570872 | -0.968244513 | 0.332922269 | 0.757641999 |
| Gnptg | 2524.363774 | -0.08772776 | 0.055485742 | -1.581086537 | 0.113858253 | 0.542263927 |
| Pnkp | 807.7235735 | -0.087742348 | 0.080619322 | -1.088353831 | 0.276438935 | 0.717475105 |
| Casz1 | 84.53915357 | -0.087796731 | 0.094499922 | -0.929066705 | 0.352854518 | 0.770519168 |
| 1700034H15Rik | 119.9015299 | -0.087833332 | 0.090215221 | -0.973597705 | 0.330256326 | 0.756141871 |
| Pla2r1 | 32.16614687 | -0.087841731 | 0.088813322 | -0.98906031 | 0.322633635 | 0.752380187 |
| 1700012D14Rik | 17.19661905 | -0.08786589 | 0.070882217 | -1.239604152 | 0.215121844 | NA |
| BC021891 | 185.8166326 | -0.087869595 | 0.094708346 | -0.927791465 | 0.353515752 | 0.770815044 |
| 4930447K03Rik | 7.223210312 | -0.087887598 | 0.059908408 | -1.467032772 | 0.142367141 | NA |
| Col1a1 | 352.8872234 | -0.087889242 | 0.084279624 | -1.04282906 | 0.29702747 | 0.732415975 |
| Zc3h11a | 4861.906056 | -0.087959282 | 0.061612413 | -1.427622747 | 0.153400473 | 0.602567676 |
| Mknk1 | 1205.156163 | -0.087991175 | 0.079397779 | -1.108232203 | 0.267761546 | 0.710152314 |
| BC018473 | 9.663879306 | -0.088011287 | 0.061144996 | -1.439386577 | 0.150041025 | NA |
| Myh3 | 144.2511269 | -0.088025957 | 0.094267297 | -0.933791039 | 0.350411693 | 0.768667813 |
| Efs | 436.2932335 | -0.088027896 | 0.084580982 | -1.040752826 | 0.297990279 | 0.732916829 |
| 4933433G15Rik | 6.836029445 | -0.088073264 | 0.052278806 | -1.68468392 | 0.09204957 | NA |
| BC055324 | 56.86938224 | -0.088089704 | 0.093761284 | -0.939510431 | 0.347468739 | 0.766038372 |
| Arhgdib | 413.8444906 | -0.088161123 | 0.087657137 | -1.005749513 | 0.314536079 | 0.745993052 |
| Per3 | 2381.507038 | -0.08824352 | 0.066269182 | -1.331592108 | 0.18299426 | 0.636231359 |
| Btn2a2 | 71.13273365 | -0.088245747 | 0.094888696 | -0.929992193 | 0.352375126 | 0.770257962 |
| Adam12 | 171.3464405 | -0.088330846 | 0.092755687 | -0.952295749 | 0.340947013 | 0.76263292 |
| Nars2 | 543.75718 | -0.088356827 | 0.066176042 | -1.335178472 | 0.181817941 | 0.635245711 |
| Fbxw9 | 567.7334341 | -0.088370329 | 0.084826003 | -1.041783488 | 0.297512071 | 0.732415975 |
| Wnk4 | 295.6890543 | -0.088373592 | 0.093373806 | -0.946449508 | 0.343919364 | 0.763762832 |
| Stab1 | 786.0820661 | -0.088376124 | 0.08792937 | -1.005080832 | 0.314857929 | 0.745993052 |
| Cep128 | 282.0334221 | -0.088391499 | 0.08627934 | -1.024480464 | 0.305608394 | 0.738032839 |
| Ccnf | 246.8331458 | -0.088408621 | 0.088914538 | -0.994310075 | 0.320071932 | 0.750154748 |
| Mtag2 | 33.62318314 | -0.088419309 | 0.083806941 | -1.055035636 | 0.291409027 | 0.727742894 |
| Zfp326 | 902.1285307 | -0.088435233 | 0.060112545 | -1.471161021 | 0.141247576 | 0.586989325 |
| Rbak | 566.285696 | -0.088450644 | 0.079904764 | -1.10695083 | 0.268315187 | 0.710391551 |
| Gusb | 613.0373484 | -0.088500696 | 0.082761006 | -1.069352594 | 0.28491082 | 0.721641599 |
| Slc35g1 | 79.7921708 | -0.088572898 | 0.095082848 | -0.931533921 | 0.351577446 | 0.769244462 |
| Sec61a2 | 2412.738344 | -0.088593859 | 0.051611178 | -1.716563384 | 0.086058978 | 0.491581875 |
| Dyrk1b | 1048.075109 | -0.088611179 | 0.066850022 | -1.325522066 | 0.184998056 | 0.637780733 |
| Cfb | 25.2761475 | -0.088653298 | 0.069443654 | -1.276622016 | 0.201735725 | 0.655168687 |
| Ankrd28 | 2644.417499 | -0.088657756 | 0.05590399 | -1.585893163 | 0.112763547 | 0.540238551 |
| Nfe2l2 | 757.9884638 | -0.088745655 | 0.075312312 | -1.178368473 | 0.238649741 | 0.689537017 |
| Ptpn18 | 33.24612234 | -0.088858418 | 0.084955107 | -1.045945575 | 0.295586161 | 0.730624344 |
| Rnf38 | 2337.252657 | -0.088871195 | 0.060020927 | -1.480670143 | 0.138694494 | 0.583119021 |
| 2010002M12Rik | 35.30513387 | -0.088909842 | 0.090448201 | -0.982991826 | 0.325611462 | 0.75293408 |
| Tmem220 | 27.97544786 | -0.088929355 | 0.078104194 | -1.138598978 | 0.254870455 | 0.701619624 |
| Morn5 | 20.7103893 | -0.088944328 | 0.073252396 | -1.214217333 | 0.224664743 | NA |
| Adcy6 | 2596.996269 | -0.088966768 | 0.067894886 | -1.31036038 | 0.19007395 | 0.644946087 |
| Parp16 | 153.4539701 | -0.0890338 | 0.087821056 | -1.013809264 | 0.310673774 | 0.743589923 |
| Nmi | 121.5871984 | -0.089037239 | 0.092017301 | -0.967614109 | 0.333237128 | 0.757730862 |
| Cox20 | 381.8797593 | -0.089049835 | 0.074357659 | -1.197587931 | 0.231077477 | 0.683402059 |
| Zfp112 | 464.3141376 | -0.089103664 | 0.072577006 | -1.227712033 | 0.219555088 | 0.672421385 |
| Mlx | 1093.55316 | -0.089112528 | 0.057745025 | -1.543207012 | 0.122780557 | 0.557484892 |
| Fgd6 | 959.1096156 | -0.089163144 | 0.072813464 | -1.224541994 | 0.220747835 | 0.672757795 |
| Cklf | 49.38225594 | -0.089163948 | 0.091953373 | -0.969664794 | 0.332213604 | 0.757166892 |
| Trmu | 505.7268358 | -0.089169691 | 0.083944075 | -1.062251158 | 0.288121681 | 0.724103151 |
| Sec16b | 72.90354911 | -0.089186045 | 0.094574198 | -0.943027236 | 0.345666972 | 0.765097832 |
| Hfe2 | 55.13705992 | -0.089244582 | 0.090840586 | -0.982430719 | 0.325887699 | 0.753033837 |
| Trim17 | 147.6025751 | -0.089256824 | 0.095035551 | -0.939194048 | 0.347631124 | 0.766038372 |
| 2610008E11Rik | 1272.190184 | -0.089390761 | 0.068657383 | -1.301983231 | 0.192922118 | 0.649602848 |
| Thsd4 | 379.6541867 | -0.089430107 | 0.094836136 | -0.942996113 | 0.345682891 | 0.765097832 |
| Bco2 | 177.3347568 | -0.089473097 | 0.088084131 | -1.015768623 | 0.309739581 | 0.742454975 |
| Prrx1 | 244.7396249 | -0.089479839 | 0.090992788 | -0.983372862 | 0.325423962 | 0.75293408 |
| Pou2f1 | 974.2152843 | -0.089513946 | 0.081211026 | -1.102238822 | 0.270357858 | 0.71216373 |
| Golm1 | 1432.156782 | -0.089518055 | 0.08007776 | -1.117889105 | 0.263614359 | 0.706200557 |
| Serpind1 | 176.0020591 | -0.089522966 | 0.094964289 | -0.942701368 | 0.345833673 | 0.765097832 |
| Epha2 | 61.88170521 | -0.089545311 | 0.09494119 | -0.943166087 | 0.345595957 | 0.765097832 |
| Nr2f1 | 2411.914694 | -0.089600056 | 0.060198023 | -1.488421901 | 0.136639664 | 0.580698586 |
| Cep78 | 387.3189306 | -0.089603617 | 0.078404918 | -1.142831581 | 0.253108524 | 0.700378199 |
| Abcg3 | 23.78328439 | -0.089648094 | 0.076450996 | -1.172621653 | 0.240947562 | 0.691863635 |
| Zfp715 | 1200.454681 | -0.089658733 | 0.069713633 | -1.286100418 | 0.198408018 | 0.653679944 |
| Wrn | 535.1215393 | -0.089692055 | 0.067681069 | -1.325216282 | 0.185099427 | 0.637780733 |
| Egfr | 494.8709966 | -0.089718548 | 0.075581063 | -1.187050624 | 0.235207653 | 0.686276191 |
| Ino80b | 330.7870061 | -0.089739288 | 0.081351836 | -1.103100951 | 0.269983328 | 0.71216373 |
| Snora52 | 17.10556745 | -0.089786822 | 0.068571044 | -1.309398501 | 0.190399399 | NA |
| 9530036O11Rik | 39.84419328 | -0.089798808 | 0.093000421 | -0.965574204 | 0.334257288 | 0.758238352 |
| 3110021A11Rik | 6.469588882 | -0.089811267 | 0.054943088 | -1.634623585 | 0.10212798 | NA |
| Svep1 | 70.16405231 | -0.089815857 | 0.086980564 | -1.032596858 | 0.301792599 | 0.735974432 |
| Pbld1 | 19.6262789 | -0.08982832 | 0.069773817 | -1.287421624 | 0.197947371 | NA |
| Gga3 | 4089.798059 | -0.089855059 | 0.061432591 | -1.462661061 | 0.143560146 | 0.589907432 |
| Vsig2 | 84.53584868 | -0.089862183 | 0.095034947 | -0.945569869 | 0.344368019 | 0.764139862 |
| D2hgdh | 585.4577529 | -0.089880363 | 0.068752995 | -1.307293783 | 0.191112955 | 0.646547945 |
| Lrriq1 | 104.5457241 | -0.089973535 | 0.092716073 | -0.970420037 | 0.331837165 | 0.756958483 |
| 4922501C03Rik | 467.1714974 | -0.089975258 | 0.06791664 | -1.324789598 | 0.185240946 | 0.63778383 |
| Lrrc55 | 907.5960819 | -0.089980218 | 0.083757653 | -1.074292501 | 0.282691605 | 0.720920007 |
| 1700110C19Rik | 19.55653517 | -0.089987805 | 0.080549744 | -1.117170598 | 0.263921389 | NA |
| Ptbp1 | 1345.265501 | -0.089989701 | 0.074843448 | -1.202372457 | 0.229219255 | 0.682442899 |
| Tbc1d24 | 5339.608572 | -0.089996451 | 0.061711917 | -1.458331807 | 0.144749107 | 0.590269908 |
| Ppp1r35 | 211.0405517 | -0.090190759 | 0.086988713 | -1.036809895 | 0.299824463 | 0.733704078 |
| Tex11 | 13.47697597 | -0.090236147 | 0.070733861 | -1.275713579 | 0.202056787 | NA |
| Dvl1 | 4406.221795 | -0.090243971 | 0.067387075 | -1.339188135 | 0.180509434 | 0.634469716 |
| March1 | 71.76353904 | -0.090268186 | 0.095081442 | -0.949377553 | 0.342428623 | 0.763762832 |
| Polg | 2988.642689 | -0.09027568 | 0.060264849 | -1.497982346 | 0.134137837 | 0.576690607 |
| Ccnjl | 131.3319175 | -0.090400047 | 0.092566235 | -0.976598509 | 0.328767958 | 0.755788917 |
| Kiss1r | 234.7777751 | -0.09047862 | 0.089776122 | -1.007824998 | 0.313538483 | 0.745844951 |
| Nxt1 | 231.5933156 | -0.090568279 | 0.088591821 | -1.022309715 | 0.306634337 | 0.738806192 |
| St5 | 783.2227207 | -0.090611682 | 0.067789084 | -1.336670691 | 0.181330153 | 0.634469716 |
| Rfc4 | 137.692496 | -0.090632906 | 0.090173838 | -1.00509093 | 0.314853067 | 0.745993052 |
| Cep97 | 903.1019127 | -0.090664144 | 0.077941544 | -1.163232582 | 0.24473515 | 0.694476029 |
| Upp1 | 55.44182546 | -0.090744311 | 0.094814705 | -0.957070009 | 0.33853192 | 0.760991092 |
| Ankrd13b | 3787.191639 | -0.090754121 | 0.069856933 | -1.299142637 | 0.193894983 | 0.650015814 |
| Heatr2 | 337.5494778 | -0.090839582 | 0.080654983 | -1.126273654 | 0.260049706 | 0.704052663 |
| Rb1 | 1270.501365 | -0.09087971 | 0.073407285 | -1.238020319 | 0.215708527 | 0.669812326 |
| Syna | 18.41737594 | -0.090904718 | 0.067376685 | -1.34920142 | 0.177272279 | NA |
| Ly96 | 26.78980905 | -0.090949793 | 0.085103664 | -1.068694208 | 0.285207482 | 0.722015317 |
| BC006965 | 57.13497006 | -0.090961856 | 0.095051649 | -0.956972948 | 0.33858091 | 0.760991092 |
| Tbk1 | 2062.750641 | -0.090994751 | 0.05840961 | -1.557872954 | 0.119263368 | 0.552077654 |
| Fam214a | 2110.588997 | -0.09102321 | 0.066667414 | -1.365332844 | 0.172148461 | 0.623232235 |
| Timeless | 226.1921685 | -0.09103242 | 0.083542188 | -1.089658084 | 0.275863786 | 0.717396826 |
| Ss18 | 1127.744518 | -0.09105825 | 0.061363315 | -1.483919979 | 0.137830151 | 0.582350428 |
| Nr1h3 | 116.0648808 | -0.091073752 | 0.094094856 | -0.967892989 | 0.333097816 | 0.757730862 |
| Apbb3 | 1628.624627 | -0.091133993 | 0.063485668 | -1.435504995 | 0.151143246 | 0.598846773 |
| Ring1 | 1162.220857 | -0.091216669 | 0.084889936 | -1.074528648 | 0.282585812 | 0.720906481 |
| Rapsn | 31.53851093 | -0.091228139 | 0.090098581 | -1.012536911 | 0.311281408 | 0.743856408 |
| Carf | 448.7465277 | -0.091252369 | 0.073582703 | -1.24013341 | 0.214926053 | 0.668817995 |
| D330050G23Rik | 77.46233582 | -0.091359413 | 0.095024852 | -0.961426519 | 0.336337756 | 0.759701795 |
| Gm10865 | 13.77090466 | -0.091384382 | 0.06992159 | -1.306955144 | 0.191227946 | NA |
| Gpsm2 | 665.9245944 | -0.091401721 | 0.071399677 | -1.280141945 | 0.200495219 | 0.654475062 |
| S100a7a | 16.0240126 | -0.091402681 | 0.077096372 | -1.185563977 | 0.235794535 | NA |
| Kctd12 | 2781.719016 | -0.091417059 | 0.059911497 | -1.525868376 | 0.127042648 | 0.564004127 |
| Clic1 | 348.5701239 | -0.091427386 | 0.083302659 | -1.097532634 | 0.272408623 | 0.714142044 |
| Tusc5 | 39.83922593 | -0.09150499 | 0.090803412 | -1.007726335 | 0.313585859 | 0.745844951 |
| Tmem240 | 791.9259352 | -0.091516357 | 0.081047281 | -1.129172448 | 0.258825094 | 0.703616072 |
| Birc2 | 1439.072025 | -0.091520859 | 0.064707234 | -1.414383741 | 0.157249261 | 0.607878516 |
| E330011O21Rik | 15.51602597 | -0.091556078 | 0.071456546 | -1.281283275 | 0.200094185 | NA |
| Vamp5 | 70.30868075 | -0.091582113 | 0.095096124 | -0.963047801 | 0.335523535 | 0.759126439 |
| Pigv | 266.2513255 | -0.091606701 | 0.082733877 | -1.107245357 | 0.268187862 | 0.71037507 |
| Gm10767 | 54.45749165 | -0.091651552 | 0.090176085 | -1.016362065 | 0.309457003 | 0.742145027 |
| Smpdl3b | 176.0870777 | -0.09175992 | 0.095030558 | -0.965583309 | 0.33425273 | 0.758238352 |
| Dennd3 | 366.2206977 | -0.091792294 | 0.081195425 | -1.130510669 | 0.258261105 | 0.703616072 |
| BC037032 | 35.72962518 | -0.091849285 | 0.087287996 | -1.052255629 | 0.292682283 | 0.728496032 |
| Dok6 | 550.4432998 | -0.091863934 | 0.07853737 | -1.169684369 | 0.242128011 | 0.692772033 |
| Ctdsp2 | 5139.616083 | -0.09188341 | 0.065688066 | -1.398783918 | 0.16187779 | 0.613952162 |
| Shc2 | 2111.498626 | -0.091906508 | 0.061354001 | -1.4979709 | 0.134140811 | 0.576690607 |
| Inca1 | 87.29347905 | -0.091947788 | 0.095042828 | -0.967435314 | 0.333326463 | 0.757730862 |
| Slc43a3 | 37.65433235 | -0.092016006 | 0.089374707 | -1.029553085 | 0.303219848 | 0.73655042 |
| A230046K03Rik | 2221.019049 | -0.092037935 | 0.064181515 | -1.434025598 | 0.151564958 | 0.598905575 |
| Ryr2 | 7444.547493 | -0.092053956 | 0.075010106 | -1.227220725 | 0.219739642 | 0.672421385 |
| BC053749 | 123.9181796 | -0.092085374 | 0.093394442 | -0.985983451 | 0.324141234 | 0.752925264 |
| Notch3 | 812.5460388 | -0.092106069 | 0.077055072 | -1.195327797 | 0.231958984 | 0.684010596 |
| Nckap5l | 1074.527991 | -0.092184464 | 0.072659027 | -1.268726925 | 0.204538476 | 0.65820246 |
| Ppp1r1a | 2942.904391 | -0.092250867 | 0.055846714 | -1.651858443 | 0.098563411 | 0.520050241 |
| Gstt3 | 130.0093962 | -0.092251329 | 0.091351527 | -1.00984988 | 0.312567218 | 0.744930503 |
| Odf4 | 12.46616216 | -0.092254 | 0.0717448 | -1.285863222 | 0.198490801 | NA |
| Ccdc58 | 209.7816637 | -0.092259134 | 0.086572009 | -1.065692413 | 0.286562708 | 0.722148665 |
| Cacng8 | 1241.053453 | -0.092266095 | 0.068301295 | -1.35086888 | 0.176737439 | 0.629083076 |
| Ist1 | 3595.0028 | -0.092286867 | 0.050533362 | -1.82625622 | 0.067811676 | 0.449662797 |
| Gm10532 | 7.833005902 | -0.09237038 | 0.051968244 | -1.777438909 | 0.075496054 | NA |
| Chd4 | 8757.919544 | -0.092379825 | 0.04693876 | -1.968092582 | 0.049057386 | 0.397217164 |
| Gm5083 | 22.41448252 | -0.092383901 | 0.076781582 | -1.203203933 | 0.228897413 | 0.681764163 |
| B9d1 | 309.1032205 | -0.092454429 | 0.078294884 | -1.180848929 | 0.237662743 | 0.688529917 |
| Smc2 | 446.9914081 | -0.092484827 | 0.082150048 | -1.125803682 | 0.260248625 | 0.704058139 |
| Rnf215 | 881.0947206 | -0.09248986 | 0.074803356 | -1.236439983 | 0.216295062 | 0.670049713 |
| Tnfaip8l2 | 80.88242622 | -0.092574417 | 0.095001495 | -0.974452208 | 0.329832058 | 0.756141871 |
| Gm4285 | 144.1901653 | -0.092621103 | 0.089086802 | -1.039672556 | 0.298492055 | 0.732994423 |
| Plxnb3 | 1433.286335 | -0.092754751 | 0.083615099 | -1.109306242 | 0.267298093 | 0.709971477 |
| S100a16 | 1773.125564 | -0.092758286 | 0.06991919 | -1.32664988 | 0.184624528 | 0.637780733 |
| Uhrf1 | 37.03406341 | -0.092780497 | 0.082832653 | -1.120095686 | 0.262672989 | 0.705284258 |
| Zfp335 | 1721.091444 | -0.092800778 | 0.070498829 | -1.316344958 | 0.188058288 | 0.641854955 |
| Slx4ip | 323.9713199 | -0.09280554 | 0.078373977 | -1.18413718 | 0.236358765 | 0.687219258 |
| Zbed3 | 416.4942774 | -0.092814928 | 0.068776519 | -1.349514763 | 0.177171682 | 0.629241039 |
| Cpne3 | 678.7827198 | -0.092843808 | 0.067377505 | -1.37796448 | 0.168214255 | 0.618133132 |
| Dhdh | 545.3377747 | -0.092862016 | 0.073804268 | -1.258220141 | 0.208312153 | 0.663292927 |
| Rilp | 53.30880602 | -0.092880165 | 0.088092991 | -1.054342289 | 0.291726234 | 0.727847831 |
| Tnik | 6917.43409 | -0.092922549 | 0.0491559 | -1.890364088 | 0.058709283 | 0.424495293 |

| | | | | | | | | | | | | | |
|---|---|---|---|---|---|---|---|---|---|---|---|---|---|
| Ptges3l | 83.67605396 | -0.228968031 | 0.157773589 | -1.451244356 | 0.14671183 | 0.345620169 | Gxylt2 | 22.14229852 | -0.092954617 | 0.084385108 | -1.101552385 | 0.270656318 | NA |
| 2610035D17Rik | 508.3643898 | 0.149135239 | 0.087573995 | 1.702962613 | 0.088575065 | 0.249841992 | Nexn | 435.134289 | -0.093003433 | 0.095059843 | -0.978367204 | 0.327892743 | 0.754778534 |
| Dyrk2 | 664.1870862 | -0.151923421 | 0.104698429 | -1.451057307 | 0.146763903 | 0.345639142 | Slc14a2 | 28.96766282 | -0.093035751 | 0.074097464 | -1.255586171 | 0.209266048 | 0.664196937 |
| Gcdh | 968.8635853 | -0.132204118 | 0.091125713 | -1.450788292 | 0.146838821 | 0.345763023 | 4932443I19Rik | 20.11316877 | -0.093039522 | 0.0763991 | -1.217809147 | 0.223296508 | NA |
| Fosl2 | 1709.855283 | -0.242013956 | 0.166892044 | -1.450122785 | 0.147024282 | 0.346147125 | Tmem8 | 854.7835553 | -0.093040881 | 0.076095457 | -1.222686416 | 0.22144816 | 0.672757795 |
| Mapk6 | 3269.280817 | 0.14910326 | 0.089013752 | 1.67505871 | 0.093922713 | 0.259077536 | Nos3 | 513.7208743 | -0.093076004 | 0.08174374 | -1.138631577 | 0.254856852 | 0.701619624 |
| Gspt1 | 4663.855509 | -0.085997981 | 0.059328604 | -1.449519703 | 0.147192502 | 0.346377881 | Myo6 | 5251.353255 | -0.093102878 | 0.063008096 | -1.477633571 | 0.139505884 | 0.584163294 |
| Lrrc73 | 1082.439978 | -0.129301114 | 0.089197751 | -1.4496006 | 0.147169929 | 0.346377881 | 9330133O14Rik | 304.0635877 | -0.093128895 | 0.087382422 | -1.065762345 | 0.286531087 | 0.722148665 |
| Myo7a | 997.6894743 | 0.149078933 | 0.083975921 | 1.775258095 | 0.075855281 | 0.226455053 | Kif13b | 1485.332264 | -0.093158867 | 0.086021641 | -1.082970124 | 0.278821684 | 0.71849589 |
| Plin3 | 554.9483673 | 0.14907686 | 0.115448044 | 1.291289612 | 0.196603271 | 0.411700377 | Acin1 | 4924.250809 | -0.093162036 | 0.064239274 | -1.450234879 | 0.146993032 | 0.594031723 |
| Tinf2 | 407.9743233 | 0.149023437 | 0.105032698 | 1.418828993 | 0.155948878 | 0.358284684 | Zranb2 | 5116.067085 | -0.093188699 | 0.058689899 | -1.587814955 | 0.112328189 | 0.538992394 |
| Htatsf1 | 2875.440198 | 0.069943214 | 0.048278811 | 1.448735206 | 0.147411545 | 0.346662471 | Atp13a1 | 2347.602284 | -0.093216172 | 0.065717814 | -1.418430817 | 0.156065024 | 0.606636483 |
| Ccdc166 | 201.1955164 | 0.148985668 | 0.109729686 | 1.357751708 | 0.174542483 | 0.383216839 | Zscan29 | 368.3683436 | -0.093228816 | 0.083618694 | -1.114927911 | 0.264881315 | 0.707370515 |
| 4921531C22Rik | 57.27701201 | -0.206447803 | 0.142589981 | -1.447842277 | 0.147661166 | 0.347157599 | Usp35 | 1137.337273 | -0.093249769 | 0.073935465 | -1.261231923 | 0.2072253 | 0.661893379 |
| Camk1d | 9260.571807 | 0.148961351 | 0.084032239 | 1.772669072 | 0.076283559 | 0.227208278 | Lss | 979.3031997 | -0.093267124 | 0.069090911 | -1.349918848 | 0.177042015 | 0.629241039 |
| Hist2h2be | 290.7999626 | -0.166032633 | 0.114692421 | -1.447633869 | 0.147719474 | 0.347157599 | Gm13293 | 76.13686526 | -0.093267588 | 0.092642025 | -1.00675248 | 0.314053736 | 0.745923449 |
| Asah2 | 633.1986194 | 0.148934045 | 0.089236293 | 1.668985122 | 0.095120328 | 0.260987138 | Vmn2r1 | 17.31394169 | -0.093275018 | 0.075589434 | -1.233968998 | 0.217214457 | NA |
| Rpn1 | 5038.823333 | -0.085150767 | 0.058825246 | -1.447520803 | 0.147751114 | 0.347172546 | Lsm10 | 224.2928109 | -0.093277923 | 0.087201991 | -1.06967652 | 0.284764939 | 0.721641599 |
| Gpd1l | 5929.346827 | -0.087578666 | 0.060508477 | -1.447378457 | 0.147790956 | 0.347213506 | Scarf1 | 77.38514423 | -0.093323082 | 0.094788568 | -0.98453942 | 0.324850359 | 0.75293408 |
| Rgs13 | 15.78194508 | -0.241833556 | 0.16710352 | -1.447208025 | 0.14783867 | 0.347272946 | Trib3 | 26.9510519 | -0.093331084 | 0.083988404 | -1.111237743 | 0.266466029 | 0.709239926 |
| Snora64 | 17.77571683 | 0.148909793 | 0.164450337 | 0.905500078 | 0.365200519 | 0.595372142 | Sytl1 | 80.41864933 | -0.093370342 | 0.09498951 | -0.982954247 | 0.325629957 | 0.75293408 |
| Zfp820 | 21.99147199 | 0.148809429 | 0.167768729 | 0.886991454 | 0.375083502 | 0.604492326 | Hemk1 | 456.4835844 | -0.093383949 | 0.088813265 | -1.051463979 | 0.293045545 | 0.728526371 |
| Zfp961 | 432.8365317 | -0.13168431 | 0.09103623 | -1.446504432 | 0.14803577 | 0.347586431 | BC020402 | 148.1229454 | -0.093416582 | 0.089932678 | -1.038739023 | 0.298926127 | 0.733276458 |
| Cox20 | 361.2016516 | -0.187986171 | 0.130005917 | -1.445981648 | 0.148182349 | 0.347869291 | Fbxo5 | 43.11722563 | -0.093429998 | 0.092356945 | -1.011618543 | 0.311720477 | 0.74424258 |
| Dcun1d3 | 797.825229 | -0.120103775 | 0.083090052 | -1.445465146 | 0.148327276 | 0.348065936 | Ddr2 | 371.6227285 | -0.093442143 | 0.090985156 | -1.027004264 | 0.304418456 | 0.737453291 |
| Fbxw4 | 780.6484257 | -0.103151471 | 0.071363226 | -1.445442944 | 0.148333508 | 0.348065936 | Ift172 | 3035.974669 | -0.093579056 | 0.058260446 | -1.606219354 | 0.108225722 | 0.533447785 |
| Samhd1 | 728.2284059 | 0.148788861 | 0.082370271 | 1.80634176 | 0.070864973 | 0.216994211 | Usp2 | 3594.314808 | -0.093620026 | 0.065875875 | -1.421157993 | 0.155270834 | 0.604578396 |
| Sash1 | 3563.382899 | -0.123791592 | 0.085648192 | -1.44534973 | 0.148359676 | 0.348074624 | Ankrd2 | 12.40681504 | -0.093626422 | 0.06779752 | -1.380971197 | 0.167287817 | NA |
| Lats1 | 1494.675448 | -0.118916336 | 0.082300062 | -1.444911865 | 0.148482643 | 0.348265855 | Morc2a | 2446.305072 | -0.093673911 | 0.047272387 | -1.981577745 | 0.04752652 | 0.392147258 |
| Ak5 | 7210.188462 | 0.148731125 | 0.080152164 | 1.855609606 | 0.063509224 | 0.20183165 | Adamts14 | 12.1249835 | -0.093760723 | 0.068062282 | -1.377572432 | 0.168335337 | NA |
| Rfesd | 380.6642763 | -0.153913388 | 0.106530231 | -1.444786018 | 0.148518 | 0.348265855 | Hyls1 | 224.6006606 | -0.093790968 | 0.083190571 | -1.127423055 | 0.259563656 | 0.703893404 |
| Grhpr | 865.732597 | 0.148658271 | 0.080378741 | 1.849472496 | 0.064389616 | 0.203768935 | Trpc6 | 381.867002 | -0.093889511 | 0.082370186 | -1.139848222 | 0.25434954 | 0.701619624 |
| Plekhg3 | 678.2524021 | 0.148645176 | 0.127478741 | 1.166038937 | 0.243598703 | 0.4681697 | Gm16576 | 54.30924684 | -0.093944564 | 0.090093975 | -1.042739699 | 0.297068866 | 0.732415975 |
| Zfp92 | 146.0444978 | -0.218500309 | 0.151351565 | -1.443660717 | 0.148834437 | 0.348871522 | F730043M19Rik | 304.5455292 | -0.093988113 | 0.080727841 | -1.164258969 | 0.244319079 | 0.6939732 |
| Ceacam1 | 129.8818171 | -0.166012966 | 0.115001521 | -1.443571913 | 0.14885943 | 0.348877328 | Amt | 838.1097188 | -0.093988523 | 0.078207884 | -1.201778109 | 0.229449509 | 0.682503526 |
| Slc5a6 | 1070.879787 | 0.148637508 | 0.06356205 | 2.338463084 | 0.019363239 | 0.092170205 | Zfr2 | 2434.405785 | -0.09400011 | 0.071751426 | -1.310080019 | 0.190168767 | 0.644966498 |
| Iars | 2497.750222 | 0.148549795 | 0.064026202 | 2.320140678 | 0.020333268 | 0.095179354 | Tmsb4x | 16178.6461 | -0.094017819 | 0.063900936 | -1.471305818 | 0.141208431 | 0.586989325 |
| Gsx2 | 10.77964239 | -0.226065627 | 0.156686147 | -1.442792684 | 0.149078881 | 0.349239561 | Hnrnph3 | 1604.236155 | -0.094061501 | 0.05569478 | -1.688874626 | 0.091243459 | 0.503727415 |
| Tmem60 | 504.1477285 | -0.135518278 | 0.093931413 | -1.442736508 | 0.149094711 | 0.349239561 | Crip1 | 102.8186905 | -0.094062548 | 0.094372943 | -0.996710971 | 0.318904823 | 0.748792456 |
| Nfat5 | 3984.97482 | -0.113576619 | 0.078730469 | -1.442600565 | 0.149133024 | 0.349254395 | Dock11 | 386.3337423 | -0.094071274 | 0.086841719 | -1.083249796 | 0.278697562 | 0.71849589 |
| Mapk8ip1 | 15560.35184 | -0.089832389 | 0.062276033 | -1.442487339 | 0.149164941 | 0.349276348 | Plekhs1 | 21.03565159 | -0.09407165 | 0.080504739 | -1.168523144 | 0.242595809 | NA |
| Trim62 | 1058.313193 | -0.13624459 | 0.09448407 | -1.441984777 | 0.149306668 | 0.349555381 | Rabep2 | 439.4031515 | -0.094077608 | 0.074598198 | -1.261124405 | 0.207264029 | 0.661893379 |
| Ramp2 | 468.3035209 | 0.148535597 | 0.097707406 | 1.52020817 | 0.128458665 | 0.317033857 | Clk3 | 2222.913405 | -0.094091624 | 0.067021119 | -1.403910072 | 0.16034563 | 0.612373044 |
| Mapre2 | 13858.70105 | -0.068772258 | 0.047698897 | -1.441799749 | 0.149358874 | 0.349571962 | Sdk1 | 425.4508709 | -0.094131378 | 0.082038272 | -1.147408101 | 0.251212995 | 0.699465245 |
| Cdkl3 | 370.6494894 | 0.148509524 | 0.127960997 | 1.160584298 | 0.245810994 | 0.470874197 | Mslnl | 24.46440051 | -0.094147342 | 0.078745263 | -1.195593726 | 0.231855141 | 0.683843204 |
| Ppp1r42 | 21.6721438 | 0.148473235 | 0.166190864 | 0.893389879 | 0.371648421 | 0.601354594 | 3830408C21Rik | 109.9481873 | -0.094163946 | 0.094338197 | -0.998152911 | 0.318205216 | 0.748361825 |
| Arhgap33 | 9924.79032 | -0.194264854 | 0.134853197 | -1.440565434 | 0.149707492 | 0.350068662 | E030030I06Rik | 20.21601571 | -0.094188772 | 0.079415537 | -1.186024487 | 0.235612629 | NA |
| Scn5a | 391.0487005 | 0.148446042 | 0.165276723 | 0.898166655 | 0.369096707 | 0.598888301 | Shmt2 | 845.928887 | -0.094281973 | 0.074458243 | -1.266239555 | 0.20542733 | 0.659018859 |
| Yipf2 | 695.9186715 | 0.148374185 | 0.100187933 | 1.480958637 | 0.138617596 | 0.332364753 | Stk11ip | 1907.26664 | -0.094328612 | 0.061549097 | -1.532575084 | 0.125380584 | 0.562122953 |
| Cdc42bpg | 275.7424976 | 0.148280217 | 0.107783872 | 1.375718041 | 0.168908944 | 0.375908529 | Orai3 | 752.1311172 | -0.094329425 | 0.079000272 | -1.194039241 | 0.232462618 | 0.684327865 |
| Unc45b | 54.48967995 | 0.148276526 | 0.152512458 | 0.97222567 | 0.330938293 | 0.566143084 | Zfp598 | 1738.806411 | -0.094332971 | 0.07679147 | -1.228430332 | 0.219285467 | 0.672421385 |
| Tnfsf10 | 141.2003222 | -0.240964294 | 0.167288853 | -1.440408553 | 0.149751845 | 0.350068662 | Aox3 | 92.43776219 | -0.094423697 | 0.094906033 | -0.994917759 | 0.319776265 | 0.749688513 |
| Tcaim | 693.161967 | -0.131831403 | 0.091533351 | -1.440255399 | 0.149795155 | 0.350117081 | Fbxl6 | 983.8707619 | -0.094441817 | 0.082786371 | -1.140789424 | 0.253957563 | 0.701619624 |
| Bcam | 725.60698 | -0.156634008 | 0.108761598 | -1.440159119 | 0.149822386 | 0.350127912 | Itpr2 | 991.8711943 | -0.094455682 | 0.065716875 | -1.437312436 | 0.150629237 | 0.598511432 |
| Dopey1 | 2445.400882 | 0.148205057 | 0.082042526 | 1.806441911 | 0.07084934 | 0.216994211 | Samd12 | 274.2030091 | -0.094465884 | 0.088698534 | -1.06502193 | 0.286866006 | 0.722256881 |
| Nup43 | 121.2885924 | 0.148127595 | 0.137989845 | 1.073467366 | 0.283061473 | 0.512599701 | Lamc3 | 309.8579945 | -0.094500983 | 0.090680055 | -1.042136367 | 0.29734846 | 0.732415975 |
| C030046E11Rik | 1741.86237 | 0.148057358 | 0.072913793 | 2.030580941 | 0.042297523 | 0.15588911 | 1600002H07Rik | 373.6520725 | -0.094552055 | 0.077199632 | -1.224773396 | 0.220660611 | 0.672757795 |
| Cops4 | 2312.059205 | -0.073026959 | 0.050748647 | -1.438993206 | 0.150152448 | 0.350370787 | Osm | 4.925129957 | -0.094616335 | 0.049106265 | -1.926767072 | 0.054008665 | NA |
| Lrrc20 | 1025.380659 | 0.148014164 | 0.083006047 | 1.78317327 | 0.074558102 | 0.224267761 | Mutyh | 92.79023509 | -0.094622206 | 0.095090693 | -0.995073257 | 0.319700636 | 0.749632194 |
| Egfl7 | 907.9797413 | -0.137597532 | 0.095599777 | -1.439308083 | 0.150063254 | 0.350370787 | Creb5 | 102.8163048 | -0.094625705 | 0.095046689 | -0.995570769 | 0.319458743 | 0.749427924 |
| Pdia5 | 133.9333575 | 0.147946442 | 0.150529521 | 0.982840051 | 0.325686166 | 0.559970267 | Snx32 | 3313.166463 | -0.09464011 | 0.067094724 | -1.410544739 | 0.158378896 | 0.609272203 |
| Jph1 | 777.9091553 | 0.147878178 | 0.096311941 | 1.535408559 | 0.124683503 | 0.312192338 | Ccdc134 | 350.5550437 | -0.094656916 | 0.090281053 | -1.048469347 | 0.294422418 | 0.729576944 |
| Cpt1c | 4011.529703 | 0.147851538 | 0.083576399 | 1.769058491 | 0.076884114 | 0.228357688 | Gm15645 | 14.50455922 | -0.094696632 | 0.070761038 | -1.338259508 | 0.180811857 | NA |
| Zcchc12 | 3004.18388 | -0.225782076 | 0.15689959 | -1.439022729 | 0.150144083 | 0.350370787 | Nfatc3 | 711.0180161 | -0.094710087 | 0.071724704 | -1.320466752 | 0.18667923 | 0.640912902 |
| Carf | 403.2810094 | 0.147827258 | 0.079590189 | 1.857355274 | 0.063260625 | 0.201223594 | Cd40 | 15.69028893 | -0.094722538 | 0.073106789 | -1.295673614 | 0.195087955 | NA |
| Nedd9 | 420.164579 | 0.147797831 | 0.129566014 | 1.1407145 | 0.253988751 | 0.479797818 | Axl | 2448.928071 | -0.094732715 | 0.063383215 | -1.49460255 | 0.1350182 | 0.578547014 |
| Cilp2 | 19.06767113 | -0.241646164 | 0.168054717 | -1.4379017 | 0.150461947 | 0.350776018 | Gtf2h3 | 533.8571523 | -0.094803513 | 0.071093384 | -1.333506827 | 0.182365536 | 0.635245711 |
| Nenf | 1089.415352 | -0.11842915 | 0.082354944 | -1.438033271 | 0.150424614 | 0.350776018 | Tnfrsf1a | 459.7421971 | -0.094809714 | 0.084705052 | -1.119292318 | 0.263015452 | 0.70568202 |
| Slc16a9 | 123.8605851 | -0.212416251 | 0.14772177 | -1.437948186 | 0.150448756 | 0.350776018 | Zfp169 | 519.6256822 | -0.094859822 | 0.072677692 | -1.305212365 | 0.191820545 | 0.646997681 |
| Ubl3 | 6292.624756 | -0.104921536 | 0.072964442 | -1.437981752 | 0.150439232 | 0.350776018 | Slc2a4 | 90.62613659 | -0.094911234 | 0.095092035 | -0.998098669 | 0.318231515 | 0.748361825 |
| 1700003E16Rik | 895.7964678 | -0.13872458 | 0.096503376 | -1.43751013 | 0.150573097 | 0.350929536 | Radil | 671.7074251 | -0.094954686 | 0.077119126 | -1.231272854 | 0.218220831 | 0.671157272 |
| Vash2 | 113.3097375 | -0.232149435 | 0.161490255 | -1.437544543 | 0.150563326 | 0.350929536 | Mir27b | 28.15194852 | -0.095089308 | 0.077109506 | -1.233172313 | 0.217511483 | 0.670311138 |
| Cdc37l1 | 2382.826196 | 0.147611459 | 0.080621439 | 1.830920667 | 0.067112385 | 0.209138259 | Prickle3 | 66.22636594 | -0.095102796 | 0.094449822 | -1.006913446 | 0.31397637 | 0.745923449 |
| Slc30a10 | 921.632983 | -0.160788357 | 0.111950719 | -1.436242295 | 0.150933408 | 0.351663492 | A430078G23Rik | 232.1620452 | -0.095113736 | 0.084146357 | -1.130336945 | 0.258334273 | 0.703616072 |
| Orc2 | 1186.304747 | 0.094284747 | 0.065659263 | 1.435970225 | 0.151010814 | 0.351790942 | Amn | 99.21245803 | -0.095117246 | 0.094643306 | -1.005007649 | 0.314893167 | 0.745993052 |
| Nudt15 | 242.2040111 | -0.132176537 | 0.092066527 | -1.435663343 | 0.151098161 | 0.351941507 | Ell | 973.6806343 | -0.095136416 | 0.060511472 | -1.572204629 | 0.115903102 | 0.54588186 |
| Mocos | 98.88467813 | 0.147606879 | 0.135097164 | 1.09259791 | 0.274570381 | 0.503038132 | Orc4 | 1376.088701 | -0.095230609 | 0.07922041 | -1.202096905 | 0.229325985 | 0.682503526 |
| Trmt10b | 401.3786115 | 0.147487503 | 0.088238264 | 1.671468775 | 0.094629121 | 0.260192299 | Mmrn2 | 225.8445753 | -0.095272201 | 0.086490261 | -1.101536746 | 0.270663121 | 0.712294861 |
| Nlrc4 | 13.23053909 | 0.147484398 | 0.165242752 | 0.892531721 | 0.372107998 | 0.601694982 | 3110001D03Rik | 626.2366294 | -0.095287302 | 0.063054576 | -1.511187742 | 0.130740625 | 0.570140593 |
| Celf6 | 885.9140165 | -0.135732105 | 0.094630247 | -1.43434166 | 0.151474787 | 0.352496758 | Rbm39 | 6462.848051 | -0.095289421 | 0.058671809 | -1.624109141 | 0.104352517 | 0.524144076 |
| Eif4a1 | 10203.51352 | -0.076146183 | 0.053090502 | -1.434271286 | 0.151494861 | 0.352496758 | Tmem236 | 9.695423514 | -0.095319519 | 0.061741574 | -1.543846613 | 0.122625496 | NA |
| Adcy9 | 2872.887658 | 0.147459496 | 0.132195289 | 1.115467103 | 0.264650308 | 0.491960633 | Xylb | 357.1806506 | -0.09532876 | 0.076906417 | -1.239542339 | 0.215144719 | 0.669349367 |
| Mier3 | 698.1506846 | 0.147371713 | 0.067684932 | 2.177319343 | 0.029456749 | 0.121935044 | Plekho1 | 1431.114809 | -0.095436356 | 0.068810108 | -1.386952579 | 0.165456223 | 0.615678595 |
| Mustn1 | 43.87914477 | 0.147318982 | 0.161959102 | 0.909606066 | 0.363030298 | 0.593603784 | Fubp1 | 3120.832801 | -0.095467068 | 0.060517833 | -1.577503089 | 0.114679812 | 0.543130934 |
| Tut1 | 878.7096007 | 0.147284227 | 0.075320561 | 1.9554319 | 0.050532114 | 0.174998787 | Map3k15 | 95.18094176 | -0.095501591 | 0.093199903 | -1.024696247 | 0.305506534 | 0.73790953 |
| Btg1 | 1044.495043 | -0.167947795 | 0.117182623 | -1.433214162 | 0.151796644 | 0.353037706 | Abca6 | 147.6934762 | -0.095555369 | 0.092126165 | -1.037222905 | 0.299631986 | 0.733704078 |
| Sh3tc1 | 109.1683346 | 0.147275554 | 0.129773829 | 1.13486328 | 0.256432607 | 0.482883668 | Dhx35 | 844.1011976 | -0.095559318 | 0.076809282 | -1.24411159 | 0.213458496 | 0.667103725 |
| Ankrd13d | 1975.047324 | 0.147265751 | 0.084873429 | 1.73512196 | 0.082719205 | 0.239496535 | Tecta | 12.76454034 | -0.09557382 | 0.071331509 | -1.339854175 | 0.180292759 | NA |
| Tmtc1 | 4978.041604 | -0.104850221 | 0.073182979 | -1.432713226 | 0.151939808 | 0.353158624 | Gpsm3 | 84.68939132 | -0.09562078 | 0.095062125 | -1.005876739 | 0.314474867 | 0.745993052 |
| Zxda | 40.77736756 | 0.147185747 | 0.156868626 | 0.938273959 | 0.348103639 | 0.57938745 | Pknox1 | 707.8692117 | -0.095654149 | 0.069850555 | -1.369411445 | 0.170870699 | 0.620885919 |
| Vps54 | 1169.758751 | 0.147177211 | 0.070367653 | 2.091546391 | 0.036479112 | 0.14080901 | Slc39a5 | 9.35167071 | -0.095747204 | 0.062369306 | -1.535165457 | 0.124743191 | NA |
| Xbp1 | 3317.519262 | -0.125886384 | 0.087902989 | -1.432105838 | 0.152113534 | 0.353456373 | Ptger1 | 59.81588585 | -0.095760664 | 0.094214002 | -1.016416475 | 0.309431103 | 0.742145027 |
| E2f8 | 20.41412082 | -0.240016853 | 0.167613213 | -1.431968565 | 0.152152818 | 0.353494641 | Mir3070b | 5.700176491 | -0.095823678 | 0.053601496 | -1.78770528 | 0.073823569 | NA |
| 1810014B01Rik | 116.553151 | -0.206643687 | 0.144347865 | -1.431567326 | 0.152267686 | 0.353708475 | Pkd1l3 | 64.09867124 | -0.095905809 | 0.095001336 | -1.009520635 | 0.31272501 | 0.745030949 |
| Fn3k | 1116.376025 | 0.147083721 | 0.063069432 | 2.332092049 | 0.019695849 | 0.093180912 | Lap3 | 2010.651727 | -0.095964832 | 0.054244285 | -1.769123349 | 0.076873292 | 0.472560915 |
| Cops5 | 2306.220243 | -0.085536922 | 0.059765544 | -1.431207958 | 0.152370623 | 0.353735457 | Sncaip | 176.0570471 | -0.095968277 | 0.094961111 | -1.010606097 | 0.312204997 | 0.744800424 |
| Esf1 | 1048.481956 | 0.147034838 | 0.075152906 | 1.956475746 | 0.050409137 | 0.174738704 | Phc1 | 2332.564676 | -0.095974298 | 0.057885353 | -1.658006617 | 0.097316131 | 0.518456434 |
| Slc35d1 | 338.501674 | 0.147017706 | 0.092122712 | 1.595889901 | 0.110513377 | 0.289412693 | Chrdl1 | 756.5385378 | -0.095995078 | 0.082200452 | -1.167816918 | 0.242880623 | 0.692772033 |
| Zdhhc12 | 162.1212063 | -0.183670499 | 0.128358489 | -1.430918203 | 0.152453659 | 0.353875206 | Dyrk4 | 8.664852141 | -0.096055291 | 0.058815483 | -1.633163352 | 0.102434645 | NA |
| Lemd3 | 1441.201453 | -0.115755275 | 0.080919845 | -1.430493034 | 0.152575564 | 0.353946068 | Mir409 | 10.07523633 | -0.096074052 | 0.058894653 | -1.631286485 | 0.102829884 | NA |
| Mgat3 | 8345.692197 | -0.096636428 | 0.067550496 | -1.430580572 | 0.152550459 | 0.353946068 | Fam71e1 | 93.25075236 | -0.09608359 | 0.095088545 | -1.01046441 | 0.312272842 | 0.744839952 |
| Psmd14 | 2563.940223 | -0.081487629 | 0.056961884 | -1.430564165 | 0.152555164 | 0.353946068 | Col4a6 | 69.39427782 | -0.096110891 | 0.094700955 | -1.014888295 | 0.310159078 | 0.743030104 |
| Nup214 | 1897.38163 | 0.14690944 | 0.069234808 | 2.121901458 | 0.033846014 | 0.134154551 | Pcsk9 | 217.4469274 | -0.096194947 | 0.090121554 | -1.067391127 | 0.285795254 | 0.722148665 |
| Sfrp2 | 58.36713029 | -0.221246506 | 0.154680019 | -1.430349625 | 0.152616699 | 0.35397807 | Sf3b1 | 11806.16552 | -0.096243176 | 0.05611212 | -1.715194067 | 0.086309652 | 0.491660395 |
| Tbrg1 | 1728.696609 | 0.086098284 | 0.060196566 | 1.430285651 | 0.152635051 | 0.35397807 | Zmym5 | 1473.952487 | -0.096267261 | 0.06124279 | -1.571895423 | 0.115974806 | 0.54588186 |
| Dis3l2 | 1045.927317 | 0.086570294 | 0.060534104 | 1.430107804 | 0.152686081 | 0.35404342 | Enpep | 150.8106724 | -0.096274607 | 0.08898863 | -1.081875371 | 0.27930791 | 0.718729265 |
| Gria1 | 17469.31063 | -0.131456836 | 0.091941576 | -1.429786637 | 0.152778266 | 0.354204167 | Ccdc88a | 2761.361918 | -0.096303684 | 0.072472579 | -1.328829258 | 0.183904307 | 0.637355821 |
| Tank | 932.2985497 | -0.097084446 | 0.067907005 | -1.429667614 | 0.15281244 | 0.354230392 | Plcd4 | 782.5803795 | -0.096312094 | 0.076532561 | -1.258445985 | 0.20823051 | 0.663292927 |
| Naa40 | 1360.918461 | 0.146853873 | 0.117219154 | 1.252814643 | 0.210273171 | 0.428462109 | Sfpq | 6225.734069 | -0.096316464 | 0.072908598 | -1.321057682 | 0.186482132 | 0.640826269 |
| Zfp457 | 8.839519458 | 0.14682617 | 0.155377233 | 0.944965795 | 0.344676339 | 0.576722182 | Pgm5 | 249.7812861 | -0.096339098 | 0.091401797 | -1.054017548 | 0.291874883 | 0.727847831 |
| Prss57 | 33.65445095 | -0.235352326 | 0.164702885 | -1.428950836 | 0.153018365 | 0.354495595 | Ap1m2 | 49.37254147 | -0.096420787 | 0.09392684 | -1.026552018 | 0.304631458 | 0.737453291 |
| Slc52a3 | 90.45263649 | -0.195684671 | 0.136940824 | -1.42897249 | 0.15301214 | 0.354495595 | Ypel1 | 1056.045679 | -0.096438362 | 0.067653119 | -1.425482867 | 0.154017669 | 0.60340295 |
| Atp5b | 41779.31123 | -0.072270017 | 0.0505784 | -1.428871143 | 0.153041273 | 0.354495662 | Lama3 | 237.7956647 | -0.096444232 | 0.085917174 | -1.122525652 | 0.261639007 | 0.704173028 |
| Jund | 2336.870245 | -0.181910866 | 0.127331192 | -1.42864339 | 0.153106756 | 0.354594332 | Zfand1 | 597.6578116 | -0.096507979 | 0.067191476 | -1.436312824 | 0.150913347 | 0.598846773 |
| 2410004B18Rik | 517.4208917 | -0.122705553 | 0.085904559 | -1.428393966 | 0.153178495 | 0.354648861 | Jpx | 169.9052701 | -0.096521219 | 0.087417549 | -1.104140071 | 0.269532381 | 0.711972822 |
| Mrpl34 | 449.7831171 | -0.133962439 | 0.093795255 | -1.428243241 | 0.153221858 | 0.354648861 | Edc4 | 3337.042521 | -0.096591232 | 0.054808568 | -1.762338191 | 0.078012175 | 0.475337606 |
| 4933436C20Rik | 11.90254202 | 0.146792999 | 0.151779216 | 0.967148221 | 0.333469943 | 0.56815299 | Cabp7 | 322.1994472 | -0.096594499 | 0.087979237 | -1.09792381 | 0.27223776 | 0.714142044 |
| Arhgap18 | 352.6553974 | 0.146746503 | 0.103627578 | 1.416095082 | 0.156747668 | 0.359427019 | Cyp3a13 | 20.35255979 | -0.096602542 | 0.069770534 | -1.384575066 | 0.16618244 | NA |
| Bbs9 | 699.0515345 | -0.107331834 | 0.075170994 | -1.427835769 | 0.153339135 | 0.354708354 | Iqgap3 | 34.88402574 | -0.096658642 | 0.089111462 | -1.084693713 | 0.278057333 | 0.718319852 |
| Spag4 | 80.56865009 | 0.146682589 | 0.157973889 | 0.928524265 | 0.353135687 | 0.583966184 | Rfx3 | 1781.699283 | -0.096688803 | 0.075525422 | -1.280215321 | 0.200469419 | 0.654475062 |
| Kcnma1 | 4028.386 | -0.101429813 | 0.071032635 | -1.427932558 | 0.153311271 | 0.354708354 | Lpar5 | 16.66724293 | -0.096736459 | 0.071097424 | -1.360618339 | 0.173634333 | NA |
| Pycrl | 524.9837824 | -0.133753861 | 0.093670607 | -1.427917094 | 0.153315723 | 0.354708354 | Rian | 24082.87652 | -0.096778479 | 0.049029859 | -1.973868189 | 0.048396733 | 0.394969953 |
| Emc1 | 3557.474732 | 0.070708584 | 0.049526699 | 1.427686178 | 0.153382206 | 0.354719364 | Kctd14 | 92.35004801 | -0.096787291 | 0.094698555 | -1.022056691 | 0.30675407 | 0.738956819 |
| Ppargc1b | 349.2025862 | -0.132000942 | 0.092459635 | -1.427660217 | 0.153389682 | 0.354719364 | Gpr157 | 145.3458164 | -0.096823221 | 0.091226763 | -1.061346665 | 0.288532386 | 0.724444298 |
| Tbx3 | 124.6664972 | -0.189561433 | 0.132802981 | -1.427388381 | 0.153467978 | 0.354847464 | 4931406P16Rik | 1826.150435 | -0.096917192 | 0.055252175 | -1.754088279 | 0.079415382 | 0.477013957 |
| Afg3l2 | 3261.522888 | 0.090954493 | 0.063729872 | 1.427187755 | 0.153525783 | 0.35484834 | 1810062O18Rik | 35.17006265 | -0.096936753 | 0.089646759 | -1.081319101 | 0.279555194 | 0.718934998 |
| Snord104 | 31.13770234 | 0.146642835 | 0.168074926 | 0.872484896 | 0.382943904 | 0.611369871 | Otx1 | 229.877529 | -0.096942426 | 0.088016844 | -1.10140766 | 0.270719273 | 0.712294861 |
| Ccdc77 | 158.9555165 | 0.14660309 | 0.114757387 | 1.277504602 | 0.201424157 | 0.418319339 | Hic2 | 470.2948362 | -0.09694329 | 0.077606422 | -1.249165815 | 0.211604432 | 0.666270994 |
| Hdac5 | 6247.872518 | -0.111143889 | 0.07788429 | -1.427038607 | 0.153568767 | 0.354868676 | Cercam | 312.8030461 | -0.097051993 | 0.089953349 | -1.07891473 | 0.280625742 | 0.720080692 |
| F2rl1 | 15.39855513 | 0.14658492 | 0.164082769 | 0.893359623 | 0.371664618 | 0.601354594 | Nek9 | 3330.294899 | -0.097081969 | 0.04309936 | -2.252515327 | 0.024289725 | 0.292321913 |
| Hipk2 | 1520.108415 | -0.180208917 | 0.126305015 | -1.426775624 | 0.15364458 | 0.35488508 | 5430405H02Rik | 177.5997933 | -0.097082332 | 0.09050032 | -1.072729162 | 0.283392652 | 0.721034821 |
| Stk11ip | 1685.380121 | 0.146529722 | 0.06575817 | 2.228312043 | 0.025859714 | 0.111705659 | Hpca | 28728.24186 | -0.097108851 | 0.061625998 | -1.575777338 | 0.115077127 | 0.543394118 |
| AU022754 | 77.12945137 | -0.222875108 | 0.156239527 | -1.426496305 | 0.153725133 | 0.354965306 | D130043K22Rik | 1874.79223 | -0.097130924 | 0.068957648 | -1.408559122 | 0.158965573 | 0.610397546 |
| Vbp1 | 1351.727283 | -0.105861679 | 0.074209034 | -1.426533574 | 0.153714383 | 0.354965306 | Gm20605 | 38.62469015 | -0.09717178 | 0.08023374 | -1.211108696 | 0.225853749 | 0.678248958 |
| Ifngr2 | 1606.269655 | 0.069614594 | 0.048821288 | 1.425906549 | 0.153895319 | 0.35530533 | Abcb1a | 1676.852 | -0.09717863 | 0.069312317 | -1.402039839 | 0.16090335 | 0.612415571 |
| Parva | 3232.852688 | 0.14645348 | 0.07120938 | 2.05665997 | 0.039718945 | 0.148989976 | 1600002K03Rik | 80.36374405 | -0.0972319 | 0.094802719 | -1.025623533 | 0.305069071 | 0.737466021 |
| Gclc | 2495.130115 | 0.146402846 | 0.087907954 | 1.665410679 | 0.09583085 | 0.262170049 | Mki67 | 47.92295484 | -0.097247371 | 0.090858662 | -1.070314801 | 0.284477635 | 0.721641599 |
| Galnt3 | 37.57596139 | 0.146370274 | 0.158958793 | 0.920806402 | 0.357151512 | 0.588003988 | Nfkbid | 136.4785951 | -0.097253075 | 0.095069438 | -1.022968862 | 0.306322568 | 0.738584943 |
| Dbf4 | 56.78960586 | 0.146320648 | 0.147410479 | 0.992606825 | 0.320901597 | 0.553865361 | Tssk4 | 13.40366221 | -0.097257496 | 0.065285588 | -1.489723836 | 0.136296864 | NA |
| Dach1 | 508.1055138 | -0.174540137 | 0.122502549 | -1.424787793 | 0.154218552 | 0.355680596 | Snx11 | 1366.228973 | -0.097258958 | 0.064207144 | -1.514768469 | 0.129831048 | 0.568885619 |
| Itgb3 | 216.678942 | 0.146269701 | 0.11527878 | 1.268834565 | 0.204500074 | 0.421576043 | Kat6b | 2640.872198 | -0.097311219 | 0.052378435 | -1.857848935 | 0.06319047 | 0.440342223 |
| Mcm6 | 325.8838534 | 0.146185996 | 0.110829777 | 1.319013718 | 0.187164526 | 0.399549073 | Capn9 | 7.405676436 | -0.097319943 | 0.059192527 | -1.64412551 | 0.10015028 | NA |
| Ift57 | 1459.063714 | -0.092687881 | 0.065072563 | -1.4243773 | 0.154337282 | 0.35590145 | Cpsf7 | 3788.570747 | -0.097331634 | 0.045162012 | -2.155166041 | 0.031148843 | 0.323885538 |
| Ngfr | 192.2685964 | -0.222118924 | 0.155956617 | -1.424235333 | 0.15437836 | 0.355943201 | Sh3tc2 | 157.3458656 | -0.0973977 | 0.090353177 | -1.077966525 | 0.281048695 | 0.720080692 |
| Nat10 | 1415.120491 | 0.146184636 | 0.085231185 | 1.715154326 | 0.086316936 | 0.245799414 | Pml | 774.6111754 | -0.097407339 | 0.064951166 | -1.499701153 | 0.133691832 | 0.575921574 |
| Dync1i1 | 5145.170262 | -0.102702121 | 0.072138818 | -1.423673464 | 0.154541019 | 0.356106268 | Col6a6 | 20.99624879 | -0.097412115 | 0.08131629 | -1.197940966 | 0.23094 | NA |
| Rrp1b | 614.049503 | 0.146168428 | 0.116222404 | 1.257661372 | 0.208514248 | 0.426610294 | Ms4a6d | 48.68311452 | -0.097431454 | 0.091238256 | -1.067879399 | 0.285574916 | 0.722148665 |
| Htatip2 | 324.7246756 | 0.146147526 | 0.111107898 | 1.315365767 | 0.188387005 | 0.401605428 | Zswim6 | 1254.49152 | -0.097459772 | 0.074081515 | -1.315574766 | 0.188316808 | 0.64225749 |
| Dlst | 4916.537688 | -0.057721991 | 0.040547844 | -1.423552653 | 0.15457601 | 0.356133933 | Npas1 | 312.4928033 | -0.097505307 | 0.086448894 | -1.127895356 | 0.259364115 | 0.703893404 |
| Extl1 | 2960.89417 | 0.146080358 | 0.098804281 | 1.478482072 | 0.139278792 | 0.333588748 | Klf2 | 388.53213 | -0.097518159 | 0.089741783 | -1.086652793 | 0.277190284 | 0.717858097 |
| Mast3 | 13134.78879 | 0.146073519 | 0.091327988 | 1.599438712 | 0.109723156 | 0.288193013 | Fndc1 | 238.0907312 | -0.0975294 | 0.091367405 | -1.067441941 | 0.285772318 | 0.722148665 |
| Pik3c2b | 1102.846641 | -0.119117517 | 0.083724736 | -1.422727887 | 0.154815053 | 0.356525629 | Frem2 | 137.7366006 | -0.097596819 | 0.093177503 | -1.047428995 | 0.294901765 | 0.73000898 |
| Smyd3 | 1021.534103 | -0.118353719 | 0.083197298 | -1.422566855 | 0.154861758 | 0.356580187 | Ppef2 | 26.65041848 | -0.097597985 | 0.083511868 | -1.168672036 | 0.242535793 | 0.692772033 |
| Ccnc | 742.0571057 | -0.102364722 | 0.07199035 | -1.42192283 | 0.155048655 | 0.356862624 | Gbp11 | 33.12723373 | -0.097653491 | 0.088189655 | -1.107312313 | 0.268158922 | 0.71037507 |
| Npm3 | 142.6160312 | -0.169277743 | 0.11904664 | -1.421944732 | 0.155042296 | 0.356862624 | Fbxw17 | 310.215332 | -0.097656262 | 0.079741092 | -1.224666723 | 0.220700817 | 0.672757795 |
| Llph | 871.8248981 | 0.145967383 | 0.076992615 | 1.89586212 | 0.057978279 | 0.190927833 | Cep57 | 563.4902292 | -0.097657912 | 0.066373673 | -1.471335055 | 0.141200528 | 0.586989325 |
| Calca | 16.71029913 | -0.238994131 | 0.168109598 | -1.421656671 | 0.155125944 | 0.356976293 | Slc25a27 | 2135.055923 | -0.097668428 | 0.054843295 | -1.780863597 | 0.074934737 | 0.468589657 |
| Il2rg | 45.1150865 | 0.145869968 | 0.154662416 | 0.943150715 | 0.345603818 | 0.576948985 | P2rx6 | 120.1929254 | -0.097681424 | 0.094794196 | -1.030457854 | 0.302795127 | 0.736257451 |
| Cited1 | 63.63005366 | 0.145856988 | 0.154170425 | 0.946076318 | 0.344109662 | 0.576410728 | Pdlim3 | 73.72600403 | -0.097742223 | 0.094776618 | -1.031290472 | 0.302404625 | 0.735974513 |
| Farp2 | 372.6309053 | 0.14575656 | 0.123644295 | 1.178837732 | 0.238462796 | 0.462759891 | H2-Eb1 | 106.1703908 | -0.097810206 | 0.050594459 | -1.933219733 | NA | NA |
| Agfg1 | 1768.209748 | -0.098075735 | 0.069047085 | -1.420418195 | 0.155485969 | 0.357592341 | Pms2 | 703.3010103 | -0.097819632 | 0.05715205 | -1.711568201 | 0.086976273 | 0.493908263 |
| Kansl2 | 1930.837496 | 0.145748618 | 0.075548345 | 1.92920995 | 0.053704803 | 0.181999612 | Ddit4l | 1349.198628 | -0.097872342 | 0.083535759 | -1.171622104 | 0.241348809 | 0.692393571 |
| Ubtd2 | 1496.755276 | -0.134246159 | 0.09452707 | -1.420187454 | 0.155553115 | 0.357640595 | Col13a1 | 58.2443769 | -0.097950245 | 0.091521781 | -1.07023971 | 0.284511424 | 0.721641599 |
| Slc2a8 | 779.3806528 | 0.145726729 | 0.092327669 | 1.578364646 | 0.114481862 | 0.296434962 | Tmem238 | 34.78583793 | -0.098092502 | 0.087207291 | -1.124819956 | 0.260665336 | 0.704058139 |
| Plcd1 | 438.6798023 | 0.145707908 | 0.121947172 | 1.194844509 | 0.232147787 | 0.455245894 | Fam71f2 | 20.72549595 | -0.098224612 | 0.082750119 | -1.18700267 | 0.235226567 | NA |
| Hyls1 | 198.7642157 | 0.145699319 | 0.098850149 | 1.473941321 | 0.140497391 | 0.335121051 | Slc22a15 | 346.7887761 | -0.098343191 | 0.081691032 | -1.203843167 | 0.228650202 | 0.68173159 |
| Elk1 | 3070.005193 | 0.145642306 | 0.083061832 | 1.753420338 | 0.079529882 | 0.233643534 | H2-M5 | 20.93562416 | -0.098352983 | 0.077533302 | -1.26852565 | 0.204610297 | NA |
| Kif5b | 10623.88286 | 0.145622524 | 0.059915567 | 2.430462242 | 0.015079577 | 0.078219943 | Hspa1b | 183.8677698 | -0.098397757 | 0.091283126 | -1.077940274 | 0.281060411 | 0.720080692 |
| Hdx | 87.05584079 | 0.145602817 | 0.146476581 | 0.994034789 | 0.320205931 | 0.553376867 | Spo11 | 11.69815952 | -0.098424354 | 0.064292833 | -1.530875993 | 0.125800043 | NA |
| Schip1 | 154.5538226 | -0.19375933 | 0.136612992 | -1.418308221 | 0.156100798 | 0.358438532 | Obscn | 366.1199401 | -0.098492413 | 0.08746457 | -1.126083552 | 0.260130156 | 0.704052663 |
| Zbtb24 | 498.7825236 | 0.145553783 | 0.085289258 | 1.706589857 | 0.087898312 | 0.248566607 | Strc | 17.72364808 | -0.098494282 | 0.075119026 | -1.311176237 | 0.189798229 | NA |
| Tada1 | 1166.557699 | -0.083836383 | 0.059105813 | -1.418411795 | 0.156070574 | 0.358438532 | Cldn22 | 20.96914417 | -0.098515231 | 0.07792961 | -1.264156598 | 0.206173825 | NA |
| Ubn1 | 2809.11202 | 0.14539162 | 0.102251212 | 1.421906085 | 0.155053516 | 0.356862624 | Glt25d1 | 2440.226471 | -0.098533996 | 0.055639334 | -1.770941316 | 0.076570461 | 0.472560915 |
| Dus1l | 983.1271972 | 0.145325867 | 0.076385574 | 1.902530272 | 0.057101865 | 0.189005832 | Ranbp17 | 268.2204097 | -0.098566476 | 0.085437629 | -1.153665865 | 0.248637184 | 0.697784358 |
| Pcdhgb4 | 166.9155259 | -0.182473915 | 0.128707324 | -1.417743055 | 0.156265795 | 0.358640531 | Mss51 | 15.61185022 | -0.098617079 | 0.073729635 | -1.337550068 | 0.181043151 | NA |
| Sstr1 | 1258.008508 | -0.130769916 | 0.092271078 | -1.417236243 | 0.156413869 | 0.358927204 | Lin37 | 737.8399561 | -0.098641319 | 0.067213082 | -1.467591077 | 0.142215333 | 0.58805008 |
| Snai2 | 18.28562654 | 0.145309436 | 0.167077263 | 0.869714007 | 0.384456716 | 0.612647573 | Cabyr | 17.34649615 | -0.098698559 | 0.074065822 | -1.332579 | 0.182670001 | NA |
| Kcnk3 | 1013.639463 | 0.145276712 | 0.093994802 | 1.545582407 | 0.122205453 | 0.30848017 | Vwa7 | 44.55976635 | -0.098861536 | 0.080263128 | -1.231717963 | 0.218054457 | 0.670929686 |
| Pak1 | 20968.67747 | 0.145174681 | 0.072198447 | 2.010772914 | 0.044349448 | 0.160962837 | Nfatc2 | 519.05603 | -0.098864866 | 0.080929662 | -1.221614717 | 0.22185336 | 0.672987912 |
| Prmt10 | 715.1386915 | 0.14515952 | 0.071582348 | 2.027867534 | 0.042573769 | 0.156460622 | Erv3 | 17.79198645 | -0.09895105 | 0.079828617 | -1.239543581 | 0.21514426 | NA |
| Azin1 | 5825.330562 | -0.121568448 | 0.085846733 | -1.416110358 | 0.156743196 | 0.359427019 | Taf15 | 1805.090109 | -0.098991933 | 0.069282379 | -1.428818339 | 0.153056453 | 0.601704476 |
| Ptk7 | 462.7486134 | 0.145133658 | 0.087154885 | 1.665238366 | 0.09586521 | 0.262195156 | Kcng3 | 59.66639143 | -0.099002411 | 0.088219034 | -1.122234131 | 0.261762904 | 0.704272634 |
| Chmp3 | 4393.253408 | -0.085436907 | 0.060344664 | -1.415815449 | 0.156829545 | 0.359508355 | Fam160b2 | 2731.568352 | -0.099009612 | 0.061176625 | -1.618422262 | 0.105571628 | 0.527531291 |
| AW209491 | 779.6424673 | -0.117685921 | 0.083141775 | -1.415484824 | 0.156926395 | 0.359528273 | Nav2 | 2474.555096 | -0.099039954 | 0.069209717 | -1.43101227 | 0.152426699 | 0.600557969 |
| Dcx | 1183.550688 | -0.231098195 | 0.163265092 | -1.415478301 | 0.156928306 | 0.359528273 | Prox1 | 446.3904098 | -0.099064405 | 0.091796654 | -1.07917228 | 0.280510934 | 0.720080692 |
| Dusp8 | 7035.265199 | -0.063386108 | 0.044776523 | -1.415610325 | 0.156889627 | 0.359528273 | Kif20a | 27.86900993 | -0.099103924 | 0.087757649 | -1.129291005 | 0.258775094 | 0.703616072 |
| Mtus1 | 2083.952227 | 0.098869869 | 0.06984955 | 1.415468931 | 0.156931051 | 0.359528273 | Rfx5 | 1003.066723 | -0.099120493 | 0.07145871 | -1.387101625 | 0.165410777 | 0.615678595 |
| Ric8b | 2758.372174 | -0.119486561 | 0.08443528 | -1.415126017 | 0.157031551 | 0.359652159 | Ubr5 | 7058.405845 | -0.099180749 | 0.056869337 | -1.744011003 | 0.081157171 | 0.479898997 |
| Tnni3 | 17.08130532 | -0.228857072 | 0.161717781 | -1.415163321 | 0.157020616 | 0.359652159 | Xrcc6 | 636.2151619 | -0.099199913 | 0.072441777 | -1.369374377 | 0.170882279 | 0.620885919 |
| Slfn9 | 22.35631945 | 0.145115473 | 0.167414662 | 0.866802649 | 0.386050151 | 0.613704655 | Cep85l | 435.2991762 | -0.09923892 | 0.087062791 | -1.139854571 | 0.254346894 | 0.701619624 |
| Foxp1 | 4573.472792 | -0.115210126 | 0.081447426 | -1.414533663 | 0.157205271 | 0.35990366 | Rpl38 | 3409.281554 | -0.09925095 | 0.076260734 | -1.301468592 | 0.193098108 | 0.649744561 |
| Pex7 | 763.4412908 | -0.096575062 | 0.068274378 | -1.414513973 | 0.157211048 | 0.35990366 | Casp8 | 129.3604155 | -0.099260994 | 0.094578704 | -1.049506811 | 0.293944922 | 0.729288721 |
| Cml3 | 173.098191 | -0.219454751 | 0.155177499 | -1.414217611 | 0.157298019 | 0.360049564 | Ifrd2 | 309.08056 | -0.099378308 | 0.081268668 | -1.22283668 | 0.221391388 | 0.672757795 |
| Tamm41 | 264.6218114 | 0.145091927 | 0.130591188 | 1.111039183 | 0.266551484 | 0.494085203 | Zdhhc24 | 964.5183211 | -0.099379422 | 0.05830289 | -1.704536806 | 0.088280846 | 0.496728133 |
| Cdkn1c | 212.2369083 | 0.145007707 | 0.128239723 | 1.130754989 | 0.25815823 | 0.484543139 | Pilra | 11.21593858 | -0.099393609 | 0.066072007 | -1.50432255 | 0.132498336 | NA |
| Mcf2l | 6946.176185 | 0.145005634 | 0.068365758 | 2.121027221 | 0.033919512 | 0.134308437 | Zfp821 | 1293.281238 | -0.099395805 | 0.059132204 | -1.680908166 | 0.092780754 | 0.506868248 |
| Atp2b1 | 27834.39602 | -0.137207053 | 0.097092836 | -1.413153222 | 0.157610677 | 0.360498942 | Ranbp10 | 1886.47753 | -0.099421611 | 0.056321755 | -1.765243483 | 0.077522854 | 0.474745772 |
| Sirt2 | 4875.095061 | 0.144992453 | 0.056581755 | 2.562530146 | 0.010391257 | 0.060460709 | Erich1 | 172.4693353 | -0.099567435 | 0.083719036 | -1.189304601 | 0.234319825 | 0.68557298 |
| Polr2a | 3517.095035 | 0.144973271 | 0.107187714 | 1.352517615 | 0.176209788 | 0.385239798 | March7 | 1401.699691 | -0.099591698 | 0.065074738 | -1.530420259 | 0.125912738 | 0.562787717 |
| Lhpp | 476.225887 | -0.191757496 | 0.135726912 | -1.412818532 | 0.157709088 | 0.360617563 | Ammecr1l | 1705.929883 | -0.099629989 | 0.059813102 | -1.66568838 | 0.095775498 | 0.515046226 |
| Rgp1 | 2087.672047 | 0.14488676 | 0.067141648 | 2.157926782 | 0.030933525 | 0.126160577 | Zbtb49 | 351.0730015 | -0.099649996 | 0.079813506 | -1.248535506 | 0.211835014 | 0.666270994 |
| App | 63331.49167 | 0.099149455 | 0.070190208 | 1.412582444 | 0.157778534 | 0.360669903 | Pik3cd | 1269.98839 | -0.099751897 | 0.06806808 | -1.465472457 | 0.142792062 | 0.589284353 |
| Zfp768 | 401.0374563 | 0.144876286 | 0.081338913 | 1.781143625 | 0.074888991 | 0.224884427 | Dzip1l | 919.2976628 | -0.099804747 | 0.074822072 | -1.33389445 | 0.182238451 | 0.635245711 |
| Slc35f4 | 456.5015815 | 0.144828864 | 0.096551734 | 1.50001307 | 0.133611017 | 0.324783353 | Map3k1 | 441.4800993 | -0.099825567 | 0.074300203 | -1.343543672 | 0.179095995 | 0.632845581 |
| Rpf2 | 514.9392298 | -0.115015853 | 0.081447193 | -1.412152454 | 0.157905077 | 0.360746278 | A930012L18Rik | 93.92366578 | -0.09991955 | 0.094445006 | -1.057965415 | 0.29007121 | 0.725277354 |
| Flna | 4240.357641 | 0.144791459 | 0.087876462 | 1.647670548 | 0.099420295 | 0.269162685 | Mrps18c | 610.0785649 | -0.099928515 | 0.080962031 | -1.234263928 | 0.217104573 | 0.670278645 |
| Klhl23 | 3329.147418 | -0.09481816 | 0.067153722 | -1.411956888 | 0.157962656 | 0.360824619 | 2610507I01Rik | 971.5975824 | -0.100059239 | 0.057536266 | -1.739063836 | 0.082023535 | 0.482582908 |
| Ltbr | 323.9983843 | -0.123376612 | 0.087397173 | -1.411677382 | 0.158044976 | 0.360959443 | Mob3c | 241.1274825 | -0.100059318 | 0.082158662 | -1.217879106 | 0.223269917 | 0.674214152 |
| Stk30 | 285.980693 | -0.119130434 | 0.084401178 | -1.411478333 | 0.15810362 | 0.36104016 | Srsf9 | 1176.312039 | -0.100074037 | 0.068852997 | -1.453444894 | 0.146100269 | 0.592807964 |
| Trmt2a | 1179.037258 | 0.144732149 | 0.091471504 | 1.582264887 | 0.113589113 | 0.294848336 | Tsks | 36.56542411 | -0.100163986 | 0.092496788 | -1.082891504 | 0.278856583 | 0.71849589 |
| Cd163 | 71.23336289 | -0.233354748 | 0.165425621 | -1.410632443 | 0.15835302 | 0.361503123 | Adat3 | 78.0716222 | -0.100203663 | 0.095090502 | -1.053771514 | 0.291987538 | 0.727847831 |
| Csk | 1112.344283 | -0.103553549 | 0.07343015 | -1.41023203 | 0.158471181 | 0.361719576 | Loxl3 | 185.8043309 | -0.100234281 | 0.08861459 | -1.131126164 | 0.258001993 | 0.703616072 |
| Efr3b | 10398.89226 | 0.094345797 | 0.066908431 | 1.410073378 | 0.158518017 | 0.361723971 | Gm166 | 222.8934967 | -0.100292924 | 0.08638713 | -1.160970661 | 0.245653831 | 0.69504627 |
| Camta1 | 10685.64734 | 0.144657653 | 0.080836112 | 1.789517691 | 0.073531481 | 0.222044989 | Npff | 41.96432999 | -0.100311834 | 0.090454475 | -1.108975918 | 0.267440571 | 0.710152314 |
| A230046K03Rik | 2006.60272 | 0.099979061 | 0.07091329 | 1.40987763 | 0.158575819 | 0.36174525 | Slc11a2 | 2260.657203 | -0.100330979 | 0.05147385 | -1.949164078 | 0.051275833 | 0.403266245 |
| Mmgt2 | 496.9204622 | -0.12606898 | 0.089416025 | -1.40991484 | 0.15856483 | 0.36174525 | Gipr | 56.4302575 | -0.100340436 | 0.091941821 | -1.091347065 | 0.275120195 | 0.716917118 |
| Ctc1 | 1948.230254 | 0.144651782 | 0.059588366 | 2.427517172 | 0.015202567 | 0.07864267 | Gm14403 | 291.6731994 | -0.100351302 | 0.089232873 | -1.124600145 | 0.260758512 | 0.704173028 |
| 1810043H04Rik | 220.3387365 | -0.193863359 | 0.137638625 | -1.408495319 | 0.158984452 | 0.362540932 | H2-Aa | 122.9707825 | -0.100363112 | 0.045501462 | -2.205711827 | NA | NA |
| Bard1 | 25.13931704 | -0.234291336 | 0.166345715 | -1.408460305 | 0.158994813 | 0.362540932 | Elk1 | 3440.205577 | -0.100376481 | 0.073509395 | -1.365491864 | 0.172098509 | 0.623232235 |
| Cep68 | 1356.355669 | -0.08176954 | 0.058069819 | -1.40812459 | 0.159094181 | 0.36271311 | Tbp | 670.3561402 | -0.100437213 | 0.071224235 | -1.410155022 | 0.158493914 | 0.609391897 |
| Cd7 | 45.25950252 | 0.144649621 | 0.167113255 | 0.865578382 | 0.386721415 | 0.614237786 | Agap2 | 43160.91705 | -0.100454356 | 0.06742721 | -1.489819252 | 0.136271768 | 0.579855848 |
| Cryab | 3015.513412 | 0.144635962 | 0.114747925 | 1.260466903 | 0.20750098 | 0.425492348 | Ska3 | 29.27804344 | -0.100534732 | 0.081853819 | -1.228222854 | 0.219363322 | 0.672421385 |
| Ino80 | 1455.924769 | 0.144627437 | 0.064688895 | 2.235738239 | 0.025368923 | 0.110305951 | Lrrc29 | 53.1732765 | -0.100564406 | 0.084232724 | -1.193887617 | 0.232521932 | 0.684327865 |
| Polr2i | 438.1155141 | -0.133113728 | 0.094665223 | -1.406152374 | 0.159678883 | 0.363833031 | C030034I22Rik | 90.82420222 | -0.100574169 | 0.094380396 | -1.065625635 | 0.286592906 | 0.722148665 |
| Cd320 | 385.3375548 | -0.122186781 | 0.086935839 | -1.405482285 | 0.159877914 | 0.364165372 | Mir3068 | 3.229482682 | -0.100660554 | 0.041236343 | -2.441064059 | 0.014644057 | NA |
| Nebl | 1307.061628 | -0.096571991 | 0.068713779 | -1.405423941 | 0.159895252 | 0.364165372 | Polr1b | 883.297398 | -0.100803701 | 0.054736667 | -1.841611977 | 0.065531928 | 0.445445383 |
| Rnf5 | 1994.740429 | -0.079465369 | 0.056539879 | -1.405474703 | 0.159880167 | 0.364165372 | Mfap3l | 2861.347111 | -0.100824386 | 0.063913999 | -1.57750082 | 0.114680334 | 0.543130934 |
| Hist1h2bc | 422.8362005 | -0.188747764 | 0.134325048 | -1.405156872 | 0.159974636 | 0.364292622 | Rasa2 | 1288.727484 | -0.100846222 | 0.069486168 | -1.451313616 | 0.146692551 | 0.593534375 |
| Epb4.1l1 | 24999.08826 | -0.082391116 | 0.058639913 | -1.40503477 | 0.160010939 | 0.364321747 | Fam120aos | 500.6851342 | -0.100924071 | 0.073494985 | -1.373210299 | 0.169686979 | 0.619945581 |
| Arfgef1 | 5351.578562 | -0.106825742 | 0.076050509 | -1.404668344 | 0.160119923 | 0.364516322 | Cybasc3 | 596.6680083 | -0.100955135 | 0.071581582 | -1.410350711 | 0.158436152 | 0.609331093 |
| Pld2 | 909.8046348 | 0.144597792 | 0.068285633 | 2.117543411 | 0.034213755 | 0.134921843 | Scrib | 2246.425418 | -0.100985667 | 0.078585844 | -1.285036361 | 0.198779578 | 0.65377551 |
| Rars2 | 648.1360739 | 0.14452757 | 0.074707664 | 1.93457488 | 0.053042485 | 0.180544873 | Batf2 | 10.06878781 | -0.101019292 | 0.055953342 | -1.805420169 | 0.071008957 | NA |
| Asb5 | 19.78645864 | -0.236037344 | 0.168114622 | -1.40402626 | 0.16031103 | 0.364683468 | Arv1 | 204.5755036 | -0.101046436 | 0.085923054 | -1.176010757 | 0.239590578 | 0.690203878 |
| Pnkd | 7470.073027 | 0.144463812 | 0.078981473 | 1.829084811 | 0.067386904 | 0.209496054 | Cyfip1 | 3776.551507 | -0.101067799 | 0.043151059 | -2.342185824 | 0.019171167 | 0.263564471 |
| Timm17a | 1668.377509 | -0.084906435 | 0.060473078 | -1.404036938 | 0.16030785 | 0.364683468 | Pcsk4 | 196.4044227 | -0.101072638 | 0.090044766 | -1.122471003 | 0.26166223 | 0.704173028 |

| | | | | | | |
|---|---|---|---|---|---|---|
| Mapkbp1 | 3164.165961 | 0.14445887 | 0.062144386 | 2.324568316 | 0.020095055 | 0.094578608 |
| Nudt18 | 1324.545848 | -0.123193357 | 0.087783729 | -1.403373476 | 0.160505498 | 0.365018668 |
| Sp4 | 779.3606622 | -0.120323806 | 0.085764523 | -1.402955466 | 0.160630119 | 0.365248469 |
| Polr3k | 1265.501501 | -0.095838885 | 0.068329028 | -1.402608636 | 0.160733575 | 0.365430083 |
| Ppp2r5c | 7965.336575 | -0.059699519 | 0.042565835 | -1.402521968 | 0.160759435 | 0.365435254 |
| Rasgef1a | 12532.56497 | -0.127286908 | 0.090765019 | -1.402378458 | 0.160802263 | 0.365478987 |
| Tmem135 | 2305.754448 | -0.078522756 | 0.056000469 | -1.402180323 | 0.160861406 | 0.365559786 |
| Dbp | 3109.427239 | 0.144451221 | 0.136101782 | 1.061347022 | 0.288532224 | 0.518626383 |
| Cyp2d22 | 1298.653052 | 0.144444394 | 0.107059726 | 1.349194502 | 0.1772745 | 0.386748959 |
| Ddx31 | 273.2381838 | 0.144423656 | 0.100106296 | 1.442703014 | 0.14910415 | 0.349239561 |
| 1500015O10Rik | 66.23948984 | 0.144417364 | 0.138910222 | 1.039645332 | 0.298504708 | 0.530150383 |
| Cox17 | 956.0233468 | -0.125414939 | 0.089505332 | -1.401200756 | 0.161154048 | 0.365956445 |
| Kctd9 | 533.0305741 | 0.144373564 | 0.095472383 | 1.512202371 | 0.130482388 | 0.319989573 |
| Zfr2 | 2026.180466 | 0.144306212 | 0.114179904 | 1.263849485 | 0.206284056 | 0.423838371 |
| Gh | 126.4188396 | 0.144199548 | 0.08076689 | 1.785379467 | 0.074199787 | 0.2235989 |
| Degs2 | 101.5245377 | 0.144082072 | 0.135588743 | 1.062640368 | 0.287945073 | 0.517926743 |
| Polq | 79.96117344 | -0.206492021 | 0.147440937 | -1.400506707 | 0.161361636 | 0.366197278 |
| Rnf115 | 1852.633198 | 0.143977512 | 0.076534688 | 1.881205963 | 0.059943907 | 0.194316101 |
| Arl10 | 367.9783038 | -0.132141159 | 0.094399518 | -1.399807558 | 0.161570954 | 0.366527139 |
| Comtd1 | 246.4065079 | -0.179501759 | 0.128282521 | -1.399269035 | 0.161732322 | 0.366839495 |
| 1110004E09Rik | 782.3391334 | 0.099118544 | 0.070845624 | 1.399077855 | 0.161789638 | 0.366915785 |
| Mll5 | 4892.70503 | 0.143957177 | 0.084440236 | 1.704840997 | 0.088224083 | 0.24921494 |
| Abcf3 | 2781.846147 | 0.143925132 | 0.08374923 | 1.718524842 | 0.085700927 | 0.244765008 |
| Foxj2 | 1738.892885 | 0.098123249 | 0.070150844 | 1.39874651 | 0.161889012 | 0.366938024 |
| Hnrnpd | 5094.106278 | -0.104498594 | 0.074701067 | -1.398890232 | 0.161845902 | 0.366938024 |
| Hk2 | 215.2843771 | 0.143885874 | 0.09908935 | 1.452082127 | 0.146478771 | 0.345440194 |
| Rev1 | 1288.111845 | 0.143858078 | 0.077704025 | 1.851359415 | 0.064117864 | 0.203074853 |
| Usp39 | 1208.387749 | -0.081742877 | 0.05844729 | -1.39857428 | 0.161940684 | 0.366938024 |
| 9330158H04Rik | 26.52249887 | -0.234941157 | 0.168045641 | -1.398079448 | 0.162089211 | 0.367061759 |
| Mrs2 | 882.881563 | 0.086908284 | 0.062162873 | 1.39807379 | 0.16209091 | 0.367061759 |
| Arnt | 1349.111187 | 0.143855015 | 0.092055907 | 1.562691842 | 0.118125094 | 0.301607001 |
| Grasp | 1802.454415 | -0.143154617 | 0.102417439 | -1.397756266 | 0.162186272 | 0.367224038 |
| Acpp | 15.58084136 | -0.226050936 | 0.161749778 | -1.397534751 | 0.162252824 | 0.367321049 |
| B3galnt1 | 1454.943431 | -0.078826843 | 0.056413803 | -1.397297088 | 0.162324251 | 0.367321739 |
| Gpc4 | 333.8838493 | -0.171828741 | 0.122965997 | -1.397367927 | 0.162302959 | 0.367321739 |
| Lpar4 | 37.38889357 | -0.223084359 | 0.159653923 | -1.39729958 | 0.162323502 | 0.367321739 |
| Kcnc2 | 4303.805015 | 0.143732081 | 0.07215472 | 1.991998312 | 0.046371243 | 0.165700194 |
| Gpr62 | 312.5098788 | 0.143680124 | 0.131568513 | 1.092055548 | 0.274808686 | 0.503236708 |
| Casp4 | 14.13253228 | 0.143676469 | 0.164166378 | 0.875188151 | 0.381471539 | 0.610149688 |
| Kcnq1ot1 | 182.7966094 | -0.21344022 | 0.152841209 | -1.396483455 | 0.162568959 | 0.367607043 |
| Fhod1 | 175.437892 | 0.143669254 | 0.102565229 | 1.400759838 | 0.161285902 | 0.366094897 |
| Eny2 | 1756.549699 | 0.143589292 | 0.078036003 | 1.840039042 | 0.065762506 | 0.206448923 |
| Wnt4 | 796.9972816 | 0.143473726 | 0.122053206 | 1.175501494 | 0.23979414 | 0.464110431 |
| Ccdc155 | 147.7129948 | 0.143392039 | 0.11676308 | 1.22805975 | 0.219424539 | 0.440212846 |
| Kcnip3 | 6085.073312 | -0.100640169 | 0.072122571 | -1.395404627 | 0.162893857 | 0.368167758 |
| Cmss1 | 302.5169231 | -0.132819225 | 0.09519833 | -1.395184402 | 0.162960239 | 0.368223122 |
| Sharpin | 1175.889754 | 0.143380368 | 0.059224009 | 2.420983834 | 0.015478566 | 0.079782068 |
| Map6d1 | 2694.846636 | 0.143352753 | 0.105703899 | 1.356172802 | 0.175044194 | 0.383828862 |
| Rnf180 | 399.624126 | -0.117957858 | 0.084589219 | -1.394478628 | 0.163173119 | 0.368542901 |
| Timp4 | 863.0914008 | 0.14334641 | 0.090955526 | 1.576005497 | 0.115024536 | 0.29712047 |
| Abcd1 | 551.4438371 | 0.143331549 | 0.095678136 | 1.498059585 | 0.13411777 | 0.325535831 |
| Cdk19 | 4218.700239 | 0.095913999 | 0.068812422 | 1.393847156 | 0.163363765 | 0.3686973 |
| Ccdc114 | 94.50101332 | 0.143312188 | 0.149574278 | 0.958133914 | 0.337995236 | 0.571651369 |
| Prrg3 | 2127.449149 | 0.088679498 | 0.063625208 | 1.393779316 | 0.163384256 | 0.3686973 |
| Gm16516 | 25.61308369 | 0.14329964 | 0.167651181 | 0.854748767 | 0.392690247 | 0.618824677 |
| Flt4 | 180.1858191 | 0.14325256 | 0.131719539 | 1.087557405 | 0.276790544 | 0.505605783 |
| Kcnd3 | 4122.20825 | -0.132737705 | 0.095267389 | -1.393317336 | 0.16352385 | 0.368744952 |
| Lyplal1 | 144.9522689 | -0.207083883 | 0.148593697 | -1.393624947 | 0.163430891 | 0.368744952 |
| Ptdss2 | 1620.171482 | -0.102560619 | 0.073602719 | -1.393435197 | 0.163488229 | 0.368744952 |
| C2 | 102.5879124 | 0.143228444 | 0.16017618 | 0.89419316 | 0.371218551 | 0.600821083 |
| AI118078 | 300.9837202 | -0.136111217 | 0.097699208 | -1.393166018 | 0.163569593 | 0.36875934 |
| Rbm15 | 312.457539 | 0.143164203 | 0.10002607 | 1.431268894 | 0.152353165 | 0.353735457 |
| Gfm2 | 972.5204655 | 0.143153271 | 0.065772347 | 2.176496298 | 0.02951817 | 0.122064687 |
| Nbr1 | 7676.570057 | 0.053720885 | 0.03857135 | 1.392766547 | 0.163690398 | 0.368860597 |
| Zcchc8 | 1010.072386 | 0.143142743 | 0.07939945 | 1.802817818 | 0.071416831 | 0.218121896 |
| Gm1673 | 253.5963533 | -0.225490559 | 0.161956523 | -1.392290688 | 0.16383439 | 0.369048539 |
| Nedd4l | 7230.270747 | -0.095936832 | 0.068900673 | -1.392393247 | 0.163803348 | 0.369048539 |
| Txnip | 1117.997253 | 0.143118035 | 0.091082161 | 1.571306974 | 0.116111362 | 0.29869325 |
| Krt222 | 1562.047825 | 0.143072473 | 0.090488277 | 1.581116124 | 0.113851489 | 0.295331486 |
| Coro1b | 2467.884385 | 0.069552689 | 0.049983781 | 1.39150516 | 0.164072296 | 0.369298607 |
| Filip1l | 186.7167389 | -0.228137115 | 0.163952785 | -1.391480569 | 0.164079748 | 0.369298607 |
| Slc8a2 | 12625.56724 | -0.133388359 | 0.095856843 | -1.391537156 | 0.164062601 | 0.369298607 |
| Sugt1 | 1997.242317 | -0.096936579 | 0.069661183 | -1.391543692 | 0.16406062 | 0.369298607 |
| Ctbp2 | 701.0141151 | -0.138173664 | 0.099313377 | -1.391289562 | 0.164137638 | 0.369375245 |
| Hus1 | 641.202604 | -0.140188815 | 0.100768039 | -1.391203166 | 0.164163828 | 0.369380532 |
| Max | 1876.150222 | 0.142999426 | 0.085967707 | 1.663408634 | 0.096230666 | 0.262869876 |
| 4632428C04Rik | 36.35364196 | -0.228562098 | 0.164420962 | -1.390103152 | 0.164497556 | 0.369577538 |
| Dpysl2 | 5681.660567 | -0.096695993 | 0.069563572 | -1.390037764 | 0.16451741 | 0.369577538 |
| Elovl2 | 661.7143353 | -0.122351221 | 0.087980197 | -1.390667731 | 0.164326207 | 0.369577538 |
| Emc9 | 515.1193963 | -0.105578575 | 0.075923052 | -1.390599717 | 0.164346842 | 0.369577538 |
| Gpatch2l | 1548.932771 | -0.08547773 | 0.06148641 | -1.390188994 | 0.164471495 | 0.369577538 |
| Hint1 | 3231.433454 | -0.111751659 | 0.080384925 | -1.390206672 | 0.164466128 | 0.369577538 |
| Zfp52 | 162.640089 | 0.142788035 | 0.11811528 | 1.208887073 | 0.226706233 | 0.449122411 |
| Mmp24 | 1916.687596 | -0.117524462 | 0.084551724 | -1.389971216 | 0.164537618 | 0.369577538 |
| Nek4 | 665.784405 | 0.096865281 | 0.069664837 | 1.390447249 | 0.164393107 | 0.369577538 |
| Nrde2 | 720.4543202 | -0.093501569 | 0.067265054 | -1.390046722 | 0.16451469 | 0.369577538 |
| Zdhhc17 | 5080.160321 | 0.069936259 | 0.050314457 | 1.38998339 | 0.164533921 | 0.369577538 |
| Rph3a | 20642.39654 | -0.132955792 | 0.095735167 | -1.38878738 | 0.164897414 | 0.370332009 |
| Ccdc91 | 1405.914582 | -0.097874138 | 0.07049673 | -1.388350041 | 0.165030481 | 0.370577142 |
| Mtmr14 | 614.9868866 | 0.142755699 | 0.087147712 | 1.638088883 | 0.101403154 | 0.27253087 |
| Mecom | 96.69695863 | -0.198714663 | 0.143166187 | -1.38799997 | 0.165137054 | 0.370698637 |
| Trim2 | 11261.98334 | -0.111038645 | 0.08000268 | -1.387936563 | 0.165156363 | 0.370698637 |
| Fam150b | 26.64825955 | 0.142731935 | 0.167821284 | 0.8504996 | 0.395047383 | 0.620462661 |
| Dnajc9 | 938.4275376 | -0.105057774 | 0.075710361 | -1.38762744 | 0.165250521 | 0.370802545 |
| Klhl11 | 1025.841382 | -0.108924759 | 0.078515662 | -1.387299758 | 0.165350377 | 0.370972884 |
| Phospho1 | 1071.2511 | 0.142722744 | 0.124544657 | 1.145956376 | 0.251813203 | 0.477551256 |
| Trpm1 | 8.455845851 | 0.142682192 | 0.150611474 | 0.947352733 | 0.343459068 | 0.575863522 |
| Zhx2 | 667.7196952 | -0.144526892 | 0.104204271 | -1.386957477 | 0.16545473 | 0.371045822 |
| Ppfibp1 | 1329.126568 | 0.142649563 | 0.085444233 | 1.669504869 | 0.095017366 | 0.260797001 |
| Xpot | 4002.586384 | -0.090459767 | 0.065237061 | -1.386631554 | 0.165554141 | 0.371161318 |
| Ier3 | 339.5199949 | 0.142514121 | 0.155687725 | 0.91538444 | 0.359989844 | 0.59091881 |
| Sla2 | 41.12153579 | -0.228573039 | 0.164924845 | -1.385922417 | 0.165770595 | 0.371539071 |
| Slc7a6os | 650.8039841 | 0.092988338 | 0.067113782 | 1.385532692 | 0.165889643 | 0.371752117 |
| Cntn3 | 1043.372782 | 0.142504997 | 0.073402502 | 1.941418788 | 0.052207501 | 0.178877528 |
| Kcnk6 | 219.6721598 | 0.1424867 | 0.118641824 | 1.200982041 | 0.229758168 | 0.452794843 |
| Rnf144a | 1383.926541 | -0.090642256 | 0.065435371 | -1.385218047 | 0.165985804 | 0.371806281 |
| Anapc4 | 1962.50138 | 0.142325252 | 0.061139315 | 2.32788432 | 0.019918247 | 0.093909252 |
| Hrct1 | 29.17826643 | -0.222688112 | 0.160803935 | -1.38484243 | 0.166100654 | 0.371823021 |
| Lpcat1 | 948.3618389 | 0.083014858 | 0.059949018 | 1.384757611 | 0.166126596 | 0.371823021 |
| Rbm42 | 1548.172434 | -0.073385741 | 0.052994099 | -1.384790807 | 0.166116443 | 0.371823021 |
| Tacr3 | 129.9070936 | -0.232397046 | 0.167815691 | -1.384835023 | 0.166102919 | 0.371823021 |
| Tmed3 | 834.059861 | -0.106948719 | 0.077234753 | -1.384722735 | 0.166137265 | 0.371823021 |
| BC031181 | 2854.241449 | -0.07421025 | 0.053596587 | -1.384607769 | 0.166172435 | 0.371848022 |
| Arih1 | 4049.527506 | -0.094562557 | 0.068303544 | -1.384445827 | 0.166221985 | 0.371853802 |
| Aatf | 462.1363089 | 0.142297026 | 0.076501609 | 1.86005273 | 0.062878066 | 0.200541942 |
| Casp9 | 1314.619231 | 0.081238744 | 0.05868998 | 1.384201242 | 0.166296843 | 0.371915182 |
| Cd38 | 102.9231255 | -0.212982149 | 0.153875768 | -1.384117542 | 0.166322466 | 0.371915182 |
| Zp2 | 8.626441853 | 0.142252359 | 0.153601453 | 0.926113366 | 0.354387069 | 0.585380817 |
| Adss | 2971.132898 | 0.096359592 | 0.06965065 | 1.383470101 | 0.166520769 | 0.372076569 |
| Pmp22 | 872.1830916 | 0.142176861 | 0.106964869 | 1.329192116 | 0.183784595 | 0.394698659 |
| Clybl | 587.4424075 | -0.11553569 | 0.083507466 | -1.383537254 | 0.166500193 | 0.372076569 |
| Btbd9 | 3734.743756 | 0.142172099 | 0.092998591 | 1.528755414 | 0.126325089 | 0.314372138 |
| Mrpl15 | 930.9247488 | -0.11188929 | 0.080871307 | -1.383547452 | 0.166497068 | 0.372076569 |
| Scap | 4005.705034 | 0.081665954 | 0.059034814 | 1.383352448 | 0.166556824 | 0.372076569 |
| Zfp810 | 600.5183561 | 0.142164028 | 0.097874521 | 1.452513134 | 0.14635898 | 0.345210235 |
| Smarca4 | 6525.420915 | -0.077188855 | 0.055825631 | -1.382677719 | 0.166763707 | 0.372471799 |
| Rars | 1621.483231 | 0.142120403 | 0.096152826 | 1.478067882 | 0.13938961 | 0.333742489 |
| Lemd2 | 1048.888766 | 0.091916142 | 0.066522563 | 1.381728818 | 0.167054982 | 0.373002426 |
| Irf4 | 20.36222075 | 0.141962996 | 0.166743425 | 0.851385873 | 0.394555037 | 0.620018297 |
| Siglec1 | 31.19796171 | 0.14194772 | 0.165288808 | 0.858786032 | 0.390458575 | 0.617448986 |
| Tmem170b | 3198.897884 | -0.093781587 | 0.067894955 | -1.38127475 | 0.167194499 | 0.373111272 |
| Tmem19 | 1423.586912 | 0.079487051 | 0.057545001 | 1.38130246 | 0.167185982 | 0.373111272 |
| Gps1 | 4715.618787 | -0.065009768 | 0.047068042 | -1.381186998 | 0.167221472 | 0.373117725 |
| Ap1ar | 3579.98083 | -0.07798429 | 0.056532031 | -1.379470881 | 0.167749618 | 0.374188394 |
| Slmo1 | 1523.658433 | 0.141895766 | 0.083756934 | 1.694137539 | 0.09023915 | 0.253015816 |
| St6gal2 | 497.1264103 | -0.145860059 | 0.105756379 | -1.379208141 | 0.167830589 | 0.374315121 |
| Eftud2 | 3969.339645 | 0.141832944 | 0.076265243 | 1.859732421 | 0.062923393 | 0.200645204 |
| Rnft2 | 4379.39951 | -0.079391978 | 0.057584771 | -1.378697474 | 0.167988049 | 0.374561052 |
| Zfp488 | 509.1308458 | 0.14181463 | 0.091423332 | 1.551186401 | 0.12085702 | 0.306324009 |
| Slc22a6 | 374.2911531 | -0.186981169 | 0.135640169 | -1.378508819 | 0.168046247 | 0.374580427 |
| B3gnt5 | 28.33625497 | -0.221153686 | 0.160450192 | -1.378332326 | 0.168100707 | 0.374647922 |
| Tfb1m | 163.3486628 | -0.170963891 | 0.124099046 | -1.377640652 | 0.168314263 | 0.375069926 |
| Pigg | 749.5945454 | -0.120599042 | 0.087563106 | -1.377281455 | 0.168425247 | 0.375263269 |
| Mtmr2 | 2142.206137 | 0.141810762 | 0.077392807 | 1.832350668 | 0.066899193 | 0.208726559 |
| Gm16576 | 46.51514197 | 0.14178154 | 0.159188837 | 0.890650015 | 0.373116958 | 0.602699272 |
| Stxbp1 | 41661.2787 | 0.091586124 | 0.066545034 | 1.3763029 | 0.168727875 | 0.375775435 |
| Dhrs7b | 571.7599068 | -0.111686063 | 0.081170038 | -1.375951846 | 0.168836542 | 0.37587621 |
| Eps15 | 8095.577494 | -0.090764922 | 0.065966481 | -1.37592487 | 0.168844895 | 0.37587621 |
| Minos1 | 2198.352063 | -0.0850644 | 0.06182358 | -1.375921613 | 0.168845903 | 0.37587621 |
| Nde1 | 796.1804172 | 0.141740845 | 0.081688234 | 1.735143945 | 0.082715312 | 0.239496535 |
| Slc39a1 | 1543.001741 | -0.070130135 | 0.050976717 | -1.375728757 | 0.168905625 | 0.375908529 |
| Ano7 | 16.97265511 | -0.231076957 | 0.168006896 | -1.375401624 | 0.169006966 | 0.376018657 |
| Srf | 2337.869843 | -0.113856634 | 0.082779233 | -1.375425094 | 0.168999694 | 0.376018657 |
| Nars2 | 495.2251346 | 0.141739182 | 0.072040901 | 1.967482101 | 0.049127657 | 0.171950425 |
| Gon4l | 2228.141765 | 0.141601194 | 0.060936012 | 2.323768655 | 0.020137897 | 0.094693951 |
| Tmem205 | 534.7581224 | -0.139785305 | 0.101708393 | -1.374373355 | 0.169325804 | 0.376567454 |
| Slc38a1 | 8583.691473 | -0.110439417 | 0.080369337 | -1.37414866 | 0.169395535 | 0.376666823 |
| Vegfc | 127.3999181 | -0.153731329 | 0.11188721 | -1.373984826 | 0.169446393 | 0.376725845 |
| Hdac11 | 7948.267176 | -0.075074663 | 0.054652412 | -1.373675184 | 0.169542544 | 0.376885536 |
| Tyw5 | 339.5131366 | 0.141452358 | 0.094062671 | 1.503809703 | 0.132630372 | 0.323058479 |
| Arl13b | 355.1801469 | 0.141426344 | 0.092025564 | 1.536815845 | 0.124338412 | 0.311791886 |
| Gpld1 | 1918.831183 | 0.141416037 | 0.078989325 | 1.790318348 | 0.073402748 | 0.221939958 |
| Chtf18 | 57.01780333 | 0.141373421 | 0.151982159 | 0.930197477 | 0.352268848 | 0.583187684 |
| Asb13 | 2061.758141 | 0.141351626 | 0.080052176 | 1.765743696 | 0.077438858 | 0.229195353 |
| Vps33a | 5485.893068 | 0.141343311 | 0.082306721 | 1.717275445 | 0.085928856 | 0.245144854 |
| Cdk4 | 1293.096878 | -0.10558869 | 0.07695372 | -1.372106385 | 0.170030322 | 0.377590628 |
| Atp6v1f | 3001.833846 | -0.085126405 | 0.062049819 | -1.371904161 | 0.170093275 | 0.377591627 |
| E030019B13Rik | 20.85045332 | -0.223218191 | 0.162710881 | -1.371870091 | 0.170103883 | 0.377591627 |
| Iqsec1 | 13203.63182 | 0.141324368 | 0.064496855 | 2.191182315 | 0.028438602 | 0.119533663 |
| Bcor | 1410.767925 | -0.116337148 | 0.084817524 | -1.371616875 | 0.170182738 | 0.377712554 |
| Wipf2 | 4199.261539 | 0.141215323 | 0.075310627 | 1.87510487 | 0.060778298 | 0.19610557 |
| Rfwd2 | 2973.851651 | -0.098098026 | 0.071536892 | -1.371292818 | 0.170283694 | 0.377828376 |
| Zfp259 | 1358.522468 | 0.141211477 | 0.058161207 | 2.427932367 | 0.015185175 | 0.078583533 |
| Camkk2 | 12136.47269 | -0.141647393 | 0.103324689 | -1.370895903 | 0.170407409 | 0.377840546 |
| Prss45 | 11.33269832 | 0.141207951 | 0.163460679 | 0.863864949 | 0.387662083 | 0.614866324 |
| Lbh | 1887.180088 | -0.095842544 | 0.069920926 | -1.370727617 | 0.170459882 | 0.377840546 |
| Ncaph | 49.54932276 | -0.210297314 | 0.153393095 | -1.370969889 | 0.170384343 | 0.377840546 |
| BC029214 | 944.8818816 | 0.1411845 | 0.074873057 | 1.885651601 | 0.059341919 | 0.193522564 |
| Srp54b | 10.47395693 | -0.223428424 | 0.162996895 | -1.370752641 | 0.170452079 | 0.377840546 |
| Chac1 | 203.7055503 | -0.154680633 | 0.112871053 | -1.370418982 | 0.17055615 | 0.377867998 |
| Lpp | 383.3123454 | -0.146847869 | 0.107156072 | -1.370411079 | 0.170558616 | 0.377867998 |
| Sidt2 | 3600.827955 | 0.141182687 | 0.056806689 | 2.48531801 | 0.012943575 | 0.070288038 |
| Med25 | 2499.322148 | 0.14116631 | 0.071054483 | 1.986733328 | 0.046951956 | 0.167120058 |
| Lsp1 | 274.13026 | -0.145816798 | 0.10641808 | -1.370225793 | 0.170616429 | 0.37791722 |
| Gpr158 | 8615.76531 | -0.095816336 | 0.069948457 | -1.369813431 | 0.170745149 | 0.378069092 |
| Magohb | 89.40205552 | -0.183085333 | 0.133661209 | -1.369771641 | 0.170758197 | 0.378069092 |
| Oxtr | 474.6347513 | -0.180326277 | 0.131637179 | -1.369873453 | 0.170726408 | 0.378069092 |
| Sipa1l2 | 2613.878189 | -0.096324992 | 0.070328715 | -1.369639577 | 0.170799439 | 0.378106373 |
| Pias1 | 1956.232371 | 0.141143949 | 0.079310323 | 1.779641572 | 0.075134638 | 0.22525853 |
| Fam108c | 915.7159766 | -0.08784519 | 0.064149906 | -1.369373643 | 0.170882509 | 0.37812819 |
| Tmem132e | 769.3993453 | 0.141009457 | 0.119681057 | 1.178210319 | 0.23871277 | 0.462963531 |
| Sh3bp4 | 875.1662784 | -0.107251703 | 0.078326563 | -1.369289023 | 0.170908948 | 0.378132692 |
| Spg20 | 1581.478148 | 0.084408152 | 0.061655729 | 1.369023663 | 0.170991878 | 0.378262159 |
| Mvd | 609.7741835 | -0.172704459 | 0.126168001 | -1.368845176 | 0.171047676 | 0.378277575 |
| Vangl1 | 134.3235673 | -0.173816725 | 0.126978811 | -1.368864017 | 0.171041786 | 0.378277575 |
| Prpf19 | 7750.999931 | 0.071758262 | 0.052440212 | 1.368382394 | 0.171192412 | 0.378543633 |
| Adar | 8679.581518 | 0.083966096 | 0.061379097 | 1.367991712 | 0.17131467 | 0.378651855 |
| Gm14393 | 23.2362761 | 0.140867971 | 0.167441477 | 0.841296756 | 0.400181707 | 0.625294007 |
| Itprip | 51.84479614 | 0.140808702 | 0.16415036 | 0.857803187 | 0.39100115 | 0.617802551 |
| Espnl | 26.28826548 | -0.229198159 | 0.167566659 | -1.367802883 | 0.171373785 | 0.378728488 |
| Cntnap5c | 228.6506983 | -0.185973181 | 0.135998381 | -1.367466139 | 0.171479244 | 0.378907502 |
| Clec16a | 4319.508769 | 0.140805275 | 0.080562823 | 1.747769874 | 0.080503884 | 0.235510228 |
| Stac2 | 4617.538443 | 0.140794788 | 0.097098875 | 1.450014616 | 0.147054444 | 0.346165535 |
| E430025E21Rik | 2080.747214 | 0.140794663 | 0.071715712 | 1.963233147 | 0.049619089 | 0.17307478 |
| Ankhd1 | 2892.331898 | 0.140780244 | 0.072364822 | 1.945423774 | 0.051723996 | 0.177775419 |
| Bcl2l15 | 10.4603284 | 0.140700492 | 0.160181273 | 0.878382907 | 0.379735957 | 0.608347533 |
| Pdzd3 | 39.55240884 | 0.140668742 | 0.165955898 | 0.84762725 | 0.396645593 | 0.621775276 |
| Ccnyl1 | 810.2592657 | -0.095670884 | 0.070051662 | -1.365718964 | 0.17202719 | 0.379739176 |
| Zfp763 | 678.794139 | -0.123934897 | 0.090762773 | -1.365481601 | 0.172101732 | 0.379849606 |
| Dock2 | 238.472003 | 0.14060824 | 0.126308601 | 1.113211917 | 0.265617426 | 0.493225838 |
| Dap | 213.9628448 | -0.160372106 | 0.117501098 | -1.364856233 | 0.17229824 | 0.38012088 |
| Zfp873 | 157.6616045 | -0.155239689 | 0.113736462 | -1.364906961 | 0.172282294 | 0.38012088 |
| Armc6 | 850.4440859 | -0.105454244 | 0.077272274 | -1.364709992 | 0.172344218 | 0.38016523 |
| Ccnjl | 118.9656999 | 0.140604968 | 0.11389019 | 1.234566104 | 0.216992031 | 0.436916767 |
| Slc29a4 | 380.7054243 | 0.140603717 | 0.145941266 | 0.963426728 | 0.335333418 | 0.569636661 |
| Golga7b | 4394.643091 | -0.125494353 | 0.09200596 | -1.363980698 | 0.172573641 | 0.380349358 |
| Slc12a2 | 2931.716607 | 0.140533033 | 0.07516581 | 1.86964036 | 0.061533777 | 0.197596753 |
| Kremen2 | 36.69721408 | -0.225373703 | 0.165224448 | -1.36404573 | 0.172553173 | 0.380349358 |
| Rbm26 | 1372.176365 | 0.140448401 | 0.064021074 | 2.193783889 | 0.02825095 | 0.11894571 |
| Nudt3 | 8353.918462 | -0.099964681 | 0.073305733 | -1.363667979 | 0.172672086 | 0.380512204 |
| Parp14 | 215.4161879 | 0.140406133 | 0.131492811 | 1.067785627 | 0.285617223 | 0.515055081 |
| Med15 | 2489.456488 | 0.140354159 | 0.068200372 | 2.057967635 | 0.03959324 | 0.14871701 |
| Dact3 | 4563.828074 | -0.11960407 | 0.087737931 | -1.363196828 | 0.172820487 | 0.380676801 |
| Emg1 | 832.3568103 | -0.102178072 | 0.074962769 | -1.363050933 | 0.172866459 | 0.380693484 |
| Mmp15 | 852.9693779 | 0.140286041 | 0.122715838 | 1.143177954 | 0.252964713 | 0.478855867 |
| Rerg | 507.1543271 | 0.140277186 | 0.130287153 | 1.076677035 | 0.281624575 | 0.511426532 |
| Rasa4 | 261.0113403 | 0.140240175 | 0.15623989 | 0.897595198 | 0.3694014 | 0.599257177 |
| Rgs9bp | 13.98943723 | 0.140177943 | 0.157224007 | 0.891581038 | 0.372617538 | 0.602302923 |
| Gltp | 1383.841908 | 0.140166491 | 0.098852132 | 1.417940996 | 0.156207992 | 0.358560982 |
| Setd7 | 8764.265114 | -0.072880207 | 0.053512057 | -1.361939924 | 0.173216845 | 0.381170544 |
| Rnf123 | 2673.948487 | 0.074048245 | 0.054383903 | 1.361583859 | 0.173329252 | 0.381363736 |
| Rxra | 1716.936304 | 0.140151991 | 0.102575526 | 1.366329735 | 0.171835492 | 0.379478209 |
| Ireb2 | 2439.486669 | -0.103261426 | 0.075876745 | -1.360910065 | 0.173542112 | 0.381723664 |
| Ubr2 | 3684.569965 | 0.09565825 | 0.070303302 | 1.360650885 | 0.173624043 | 0.381795475 |
| Gbp9 | 157.8052779 | 0.140137899 | 0.153794212 | 0.911203985 | 0.362187905 | 0.592893756 |

| | | | | | | |
|---|---|---|---|---|---|---|
| Zfp800 | 439.6752478 | -0.101074461 | 0.090708564 | -1.114276934 | 0.265160399 | 0.707386586 |
| Ankrd44 | 462.7419247 | -0.101099316 | 0.073932087 | -1.367461948 | 0.171480557 | 0.622106016 |
| Gbp8 | 10.71689967 | -0.101103492 | 0.040639282 | -2.487826749 | NA | NA |
| Atrip | 847.8544455 | -0.101144115 | 0.060633584 | -1.668120363 | 0.095291836 | 0.51433043 |
| Pax6 | 300.9743099 | -0.101156578 | 0.082962353 | -1.21930701 | 0.222727689 | 0.67374894 |
| Ripk2 | 196.4209525 | -0.101161928 | 0.086487313 | -1.169673613 | 0.242132341 | 0.692772033 |
| 2810454H06Rik | 19.41361911 | -0.101271173 | 0.079269669 | -1.277552613 | 0.201407218 | NA |
| Dock4 | 4960.376975 | -0.10128626 | 0.057393931 | -1.764755572 | 0.077604854 | 0.475047667 |
| C230004F18Rik | 179.8341295 | -0.101302456 | 0.086375815 | -1.172810423 | 0.240871838 | 0.691863635 |
| 1700040N02Rik | 14.21524246 | -0.101309729 | 0.067643338 | -1.497704455 | 0.134210054 | NA |
| Nr2e1 | 649.6091836 | -0.10133453 | 0.079941226 | -1.2676129 | 0.204936223 | 0.658461308 |
| Slfn5 | 447.7216576 | -0.10133814 | 0.089486796 | -1.132436795 | 0.257450843 | 0.703271881 |
| Mocos | 112.3910636 | -0.101361208 | 0.091964209 | -1.102181055 | 0.270382966 | 0.71216373 |
| Mus81 | 759.9069314 | -0.101435175 | 0.068398272 | -1.483007862 | 0.138072322 | 0.582865025 |
| Cpsf4 | 695.2247647 | -0.101551196 | 0.07585875 | -1.338687966 | 0.180672276 | 0.634469716 |
| Lactb2 | 342.2572352 | -0.101631305 | 0.085473399 | -1.18904017 | 0.23442386 | 0.68557298 |
| Myzap | 27.77159268 | -0.101644537 | 0.085587579 | -1.187608516 | 0.234987681 | 0.686276191 |
| Prkab2 | 2316.790048 | -0.101663501 | 0.074915998 | -1.357033256 | 0.174770644 | 0.627929353 |
| Fam203a | 327.2135359 | -0.101699724 | 0.076602762 | -1.327624772 | 0.184302097 | 0.637780733 |
| 3010026O09Rik | 383.8398411 | -0.101710452 | 0.078887464 | -1.289310705 | 0.197290091 | 0.653301923 |
| Runx2 | 335.0311196 | -0.101729375 | 0.083214476 | -1.222496121 | 0.22152007 | 0.672757795 |
| Zc3h12a | 48.03006646 | -0.101796829 | 0.09329148 | -1.091169616 | 0.275198255 | 0.716917118 |
| Slc4a11 | 157.559984 | -0.101851011 | 0.094286279 | -1.080231526 | 0.280039093 | 0.719336492 |
| Galnt4 | 186.0708219 | -0.101878485 | 0.083797636 | -1.21576801 | 0.224073308 | 0.675148076 |
| Kif22 | 47.15587626 | -0.101994333 | 0.092568457 | -1.101826005 | 0.270537322 | 0.712294861 |
| Shisa7 | 8120.467184 | -0.102026746 | 0.057256662 | -1.781919221 | 0.074762406 | 0.468388672 |
| Kctd12b | 176.3092853 | -0.102082585 | 0.087045322 | -1.172752108 | 0.240895229 | 0.691863635 |
| Adamts17 | 237.0262788 | -0.102127573 | 0.087215902 | -1.170974218 | 0.241609141 | 0.692393571 |
| Cenpt | 572.5364358 | -0.102151464 | 0.072862003 | -1.4019854 | 0.160919607 | 0.612415571 |
| Alms1 | 526.7761204 | -0.102169913 | 0.081950189 | -1.24673187 | 0.212495829 | 0.666270994 |
| Rfx2 | 121.3348945 | -0.102199274 | 0.094451174 | -1.082032862 | 0.279237926 | 0.718676491 |
| Slc27a1 | 2890.273291 | -0.102216848 | 0.078939298 | -1.294879119 | 0.195361933 | 0.651003086 |
| Hydin | 87.07739964 | -0.102227848 | 0.09493641 | -1.076803397 | 0.281568107 | 0.720080692 |
| AW112010 | 34.69728357 | -0.102305753 | 0.089164796 | -1.147378315 | 0.2512253 | 0.699465245 |
| Dcaf15 | 984.6894673 | -0.102388029 | 0.082682786 | -1.238323394 | 0.215596173 | 0.669812326 |
| Kctd18 | 414.5962677 | -0.102419939 | 0.070027566 | -1.462566039 | 0.143586162 | 0.589907432 |
| Myom3 | 128.3771191 | -0.102477138 | 0.089514253 | -1.144813632 | 0.252286367 | 0.69992334 |
| Tsen15 | 586.1426554 | -0.102643358 | 0.0645869 | -1.589228746 | 0.112008759 | 0.538992394 |
| Tsc2 | 5898.314783 | -0.102669638 | 0.056933491 | -1.803325886 | 0.07133705 | 0.459174089 |
| Sec14l3 | 112.1952406 | -0.102674473 | 0.093843113 | -1.094107709 | 0.273907742 | 0.716079153 |
| Trim46 | 3712.71765 | -0.102678967 | 0.069396347 | -1.47960189 | 0.138979522 | 0.583135914 |
| Necap2 | 1062.723274 | -0.102762372 | 0.069240223 | -1.484142707 | 0.137771065 | 0.582270147 |
| Smc4 | 280.6869833 | -0.102821876 | 0.080801455 | -1.272525044 | 0.203186632 | 0.657054544 |
| Efemp1 | 521.472493 | -0.102852301 | 0.091264885 | -1.126964671 | 0.259757419 | 0.703893404 |
| 4930507D05Rik | 35.55420516 | -0.102877951 | 0.092692865 | -1.109879931 | 0.267050769 | 0.70969255 |
| Prdm6 | 14.69580203 | -0.10294984 | 0.069073719 | -1.490434301 | 0.136110079 | NA |
| Zcchc9 | 409.8157982 | -0.102950177 | 0.081681335 | -1.260388019 | 0.207529422 | 0.66254338 |
| 9330020H09Rik | 60.76014836 | -0.102950552 | 0.093844819 | -1.097029684 | 0.272628416 | 0.714142044 |
| Cutc | 196.0940845 | -0.102966229 | 0.08442672 | -1.219592908 | 0.222619236 | 0.67374894 |
| Gm20063 | 127.866124 | -0.10299544 | 0.094331655 | -1.091843881 | 0.274901728 | 0.71665874 |
| Tmtc2 | 572.3756769 | -0.103049718 | 0.080119825 | -1.286195007 | 0.198375013 | 0.653679944 |
| Trip13 | 22.18870218 | -0.103057785 | 0.080876373 | -1.274263186 | 0.202570159 | NA |
| Klhl41 | 17.48058856 | -0.103141201 | 0.077728503 | -1.326941806 | 0.184527934 | NA |
| Slc1a5 | 144.8432452 | -0.103153761 | 0.089671501 | -1.150351678 | 0.249999054 | 0.698461386 |
| Rbm3 | 2305.54484 | -0.103199298 | 0.091109486 | -1.132695421 | 0.257342181 | 0.703271881 |
| Ulk4 | 268.5801607 | -0.103254101 | 0.080111906 | -1.288873345 | 0.197442122 | 0.653417817 |
| Trdmt1 | 175.3824903 | -0.103270123 | 0.086027335 | -1.200433819 | 0.229970901 | 0.683133099 |
| Slc12a7 | 234.8767107 | -0.103290955 | 0.092992021 | -1.110750733 | 0.266675658 | 0.709441589 |
| Pkd2 | 1510.992194 | -0.103349625 | 0.055800934 | -1.852112827 | 0.064009623 | 0.441368101 |
| 2310015D24Rik | 7.381563182 | -0.103422586 | 0.059309778 | -1.743769558 | 0.081199281 | NA |
| Iigp1 | 128.6605486 | -0.103450512 | 0.043809871 | -2.361351698 | NA | NA |
| Dok3 | 225.0450658 | -0.103450732 | 0.094306609 | -1.096961635 | 0.272658163 | 0.714142044 |
| 1500012F01Rik | 532.9781181 | -0.103481163 | 0.080798375 | -1.280733215 | 0.200287389 | 0.654475062 |
| BC005561 | 496.1957447 | -0.10355297 | 0.089035942 | -1.163046823 | 0.244810506 | 0.694524329 |
| Myo10 | 1681.414871 | -0.10355452 | 0.057613386 | -1.797403836 | 0.072271532 | 0.461579135 |
| Drp2 | 4677.431162 | -0.103591024 | 0.046115451 | -2.246340925 | 0.024682179 | 0.294362343 |
| Gm4841 | 10.56741595 | -0.103605973 | 0.038381214 | -2.699392812 | NA | NA |
| Tysnd1 | 558.5025793 | -0.103612829 | 0.076283527 | -1.358259553 | 0.17438134 | 0.627151112 |
| AU019823 | 690.9884575 | -0.103624274 | 0.078645336 | -1.317614989 | 0.187632565 | 0.641776649 |
| Plcd3 | 650.9711437 | -0.103667838 | 0.079432796 | -1.305101209 | 0.191858387 | 0.646997681 |
| Atf3 | 58.51936373 | -0.103745412 | 0.085137688 | -1.218560369 | 0.223011099 | 0.674048714 |
| Rtbdn | 51.52634375 | -0.103849045 | 0.092502123 | -1.12266662 | 0.26157911 | 0.704173028 |
| Pcdha12 | 346.8053768 | -0.103865603 | 0.084897972 | -1.223416776 | 0.22117232 | 0.672757795 |
| Cul9 | 3585.09837 | -0.103958484 | 0.062405137 | -1.6658642 | 0.095740466 | 0.515046226 |
| Tmem81 | 189.6696858 | -0.103964874 | 0.084402454 | -1.231775486 | 0.218032963 | 0.670929686 |
| Cdh9 | 770.2065615 | -0.103981754 | 0.05921598 | -1.755974553 | 0.079092754 | 0.477013957 |
| 4930590J08Rik | 24.96602838 | -0.103986081 | 0.074797543 | -1.390233907 | 0.16445786 | 0.614688762 |
| Slc16a4 | 94.85891166 | -0.104015653 | 0.092463014 | -1.124943354 | 0.260613039 | 0.704058139 |
| Hscb | 149.9744566 | -0.10403563 | 0.091825823 | -1.132967037 | 0.257228096 | 0.703199912 |
| Ints6 | 898.4791176 | -0.104118112 | 0.07127159 | -1.460864174 | 0.14405272 | 0.589967032 |
| Zeb2 | 5261.271556 | -0.10417793 | 0.070452252 | -1.47870262 | 0.139219812 | 0.583573764 |
| Cd34 | 3461.669379 | -0.104182637 | 0.072892364 | -1.429266821 | 0.152927559 | 0.601523346 |
| Mapkbp1 | 3426.56809 | -0.104276283 | 0.051401103 | -2.02867792 | 0.042491106 | 0.373397344 |
| Katna1 | 410.951674 | -0.104285324 | 0.078029393 | -1.336487703 | 0.181389917 | 0.634469716 |
| 1600029O15Rik | 61.65260869 | -0.104290825 | 0.089924839 | -1.159755483 | 0.246148374 | 0.695662109 |
| Gm973 | 297.4700787 | -0.104305604 | 0.089458613 | -1.165964915 | 0.243628631 | 0.692824881 |
| Dnahc9 | 284.6487361 | -0.104312961 | 0.09220771 | -1.131282417 | 0.257936243 | 0.703616072 |
| Nfkb2 | 122.3311822 | -0.104326336 | 0.093077732 | -1.120851715 | 0.262350987 | 0.704810511 |
| Abcc1 | 1127.263707 | -0.104333591 | 0.062987917 | -1.656406428 | 0.09763954 | 0.518456434 |
| Frmd4b | 936.6902775 | -0.104346475 | 0.070739782 | -1.47507487 | 0.140192416 | 0.585518985 |
| Pgm1 | 330.7713981 | -0.104378984 | 0.077136256 | -1.353176704 | 0.175999186 | 0.62871374 |
| Ifnar1 | 2193.148729 | -0.104391444 | 0.054164601 | -1.92730016 | 0.053942234 | 0.409506392 |
| Dysf | 223.9631151 | -0.104415668 | 0.089656376 | -1.164620661 | 0.244172577 | 0.693692714 |
| Bag3 | 395.8046842 | -0.104418173 | 0.082552771 | -1.264865765 | 0.205919451 | 0.660160219 |
| Ccdc14 | 153.8660788 | -0.104460801 | 0.088304054 | -1.182967219 | 0.236822139 | 0.687465707 |
| Shisa6 | 1175.071858 | -0.104469347 | 0.091764507 | -1.138450487 | 0.254932423 | 0.701619624 |
| Cdkl3 | 368.2099831 | -0.104532705 | 0.088237576 | -1.184673363 | 0.236146618 | 0.687039131 |
| 0610007N19Rik | 74.14745248 | -0.104534314 | 0.094972832 | -1.100675974 | 0.271037707 | 0.712854196 |
| Stk17b | 297.7872634 | -0.104605441 | 0.089444433 | -1.169501973 | 0.242201447 | 0.692772033 |
| Sytl2 | 3757.830791 | -0.104618017 | 0.063919975 | -1.636703033 | 0.101692535 | 0.522606076 |
| Rab36 | 1169.495628 | -0.104630236 | 0.070239088 | -1.489629766 | 0.136321611 | 0.579855848 |
| Zfp738 | 615.8333753 | -0.104660109 | 0.086057732 | -1.216161611 | 0.223923364 | 0.675094439 |
| Trim6 | 9.202660641 | -0.104673659 | 0.063399972 | -1.65100482 | 0.098737591 | NA |
| Ap4m1 | 529.2497043 | -0.104675982 | 0.068460517 | -1.528997818 | 0.126264985 | 0.562787717 |
| Rgsl1 | 125.5817812 | -0.104684186 | 0.093920842 | -1.114600167 | 0.265021799 | 0.707370515 |
| Fxn | 144.8573165 | -0.104706487 | 0.091878762 | -1.139615779 | 0.254446409 | 0.701619624 |
| Dbn1 | 9725.131013 | -0.104735428 | 0.061285123 | -1.708986183 | 0.087453509 | 0.495597906 |
| Baz2a | 3446.236635 | -0.104798084 | 0.057043737 | -1.837153201 | 0.066187282 | 0.446279967 |
| Slc22a21 | 48.4837803 | -0.104820171 | 0.091241692 | -1.148818798 | 0.250630706 | 0.699068333 |
| Ggct | 377.9822669 | -0.104826067 | 0.069727892 | -1.503359195 | 0.132746443 | 0.574746597 |
| Kdm4d | 77.18909466 | -0.104859783 | 0.093882304 | -1.116928106 | 0.264025066 | 0.706954491 |
| 5830454E08Rik | 25.7830862 | -0.104893632 | 0.084864592 | -1.236011745 | 0.216454198 | 0.670049713 |
| Narg2 | 355.3566288 | -0.104985983 | 0.07203219 | -1.457487033 | 0.144981988 | 0.590580022 |
| Tmem40 | 67.11100441 | -0.105021298 | 0.086337618 | -1.216402543 | 0.223831616 | 0.675094439 |
| Snx18 | 945.6355821 | -0.105022502 | 0.059215827 | -1.7735546 | 0.076136853 | 0.472435997 |
| Tprkb | 2552.806821 | -0.105047613 | 0.046987036 | -2.235672283 | 0.025373246 | 0.298582503 |
| Sgpl1 | 1804.667408 | -0.105052343 | 0.050180112 | -2.09350555 | 0.036304049 | 0.345126059 |
| Nudcd2 | 405.8377132 | -0.105067378 | 0.075597311 | -1.38982957 | 0.164580636 | 0.614688762 |
| 3000002C10Rik | 141.0389828 | -0.105069487 | 0.091632401 | -1.146641208 | 0.251529938 | 0.699465245 |
| Pbxip1 | 2896.771158 | -0.105087811 | 0.07674229 | -1.369359853 | 0.170886817 | 0.620885919 |
| Dus4l | 198.0370166 | -0.105124013 | 0.089981584 | -1.168283644 | 0.242692371 | 0.692772033 |
| Plcb3 | 814.5461466 | -0.105225714 | 0.068581351 | -1.534319652 | 0.124951033 | 0.560889589 |
| Cacna1a | 6389.351803 | -0.105253193 | 0.046587702 | -2.25924843 | 0.023867936 | 0.291530239 |
| Palb2 | 56.55460306 | -0.105286734 | 0.094135881 | -1.118454872 | 0.26337277 | 0.706200557 |
| Gm10635 | 89.77542047 | -0.105310504 | 0.0877417 | -1.200233224 | 0.230048775 | 0.683133099 |
| Eif2d | 1148.357056 | -0.105323787 | 0.057366545 | -1.835979246 | 0.066360726 | 0.446279967 |
| Nos2 | 29.41007555 | -0.105363845 | 0.078501793 | -1.342183928 | 0.179536366 | 0.633459544 |
| Adprm | 447.5005488 | -0.105367409 | 0.073380155 | -1.435911503 | 0.151027525 | 0.598846773 |
| Taf1c | 575.708282 | -0.105417975 | 0.084335877 | -1.249977808 | 0.211307654 | 0.666270994 |
| S100a13 | 454.5866654 | -0.105445143 | 0.076667733 | -1.37535231 | 0.169022247 | 0.619378697 |
| 2610524H06Rik | 376.6804476 | -0.105455621 | 0.077554404 | -1.359763158 | 0.173904885 | 0.626678827 |
| Sirt5 | 395.1571157 | -0.105493848 | 0.08496485 | -1.241617537 | 0.214377708 | 0.668471732 |
| Kdm8 | 225.0498728 | -0.105590225 | 0.084752148 | -1.245870785 | 0.212811837 | 0.666286814 |
| Rexo4 | 1994.453235 | -0.105605614 | 0.057394338 | -1.840000566 | 0.065768154 | 0.445799444 |
| Cd109 | 85.86945781 | -0.105626849 | 0.089016605 | -1.186597141 | 0.235386564 | 0.686276191 |
| Rpusd3 | 268.4154605 | -0.105632664 | 0.075560169 | -1.397994016 | 0.162114864 | 0.613957643 |
| 2610044O15Rik8 | 293.8725639 | -0.105685118 | 0.084023808 | -1.257799666 | 0.208464216 | 0.663405095 |
| Gm13704 | 15.73702858 | -0.105689095 | 0.058834845 | -1.796369057 | 0.072435842 | NA |
| Zfp595 | 131.4045542 | -0.105776314 | 0.094140765 | -1.123597355 | 0.261183879 | 0.704173028 |
| Slc26a6 | 282.4218142 | -0.105780567 | 0.082765715 | -1.278072294 | 0.201223937 | 0.654716629 |
| Pld2 | 993.6330256 | -0.105834568 | 0.057578759 | -1.838083519 | 0.066050099 | 0.446279967 |
| Il15ra | 166.5050854 | -0.105853106 | 0.087088569 | -1.21546498 | 0.224188798 | 0.675355967 |
| Slx1b | 707.9200385 | -0.105892788 | 0.070091038 | -1.510789264 | 0.130842151 | 0.570291305 |
| Kif7 | 103.1764909 | -0.105945377 | 0.094900563 | -1.11638302 | 0.264258219 | 0.707187709 |
| 2310002F09Rik | 9.331062062 | -0.105987334 | 0.059679851 | -1.77593162 | 0.075744189 | NA |
| Jun | 1971.857171 | -0.10606326 | 0.07298274 | -1.453264967 | 0.1461502 | 0.592844962 |
| Rpain | 355.9660153 | -0.106117974 | 0.075796942 | -1.400029757 | 0.161504408 | 0.613176432 |
| Rbmx | 2489.05226 | -0.106166501 | 0.059010529 | -1.799111142 | 0.072001102 | 0.460419542 |
| Mmp2 | 112.341225 | -0.106209254 | 0.092885874 | -1.143438173 | 0.252856711 | 0.700378199 |
| Tpra1 | 1137.534381 | -0.106212687 | 0.061436588 | -1.72881814 | 0.083841649 | 0.486764103 |
| Cyb561d1 | 1413.748332 | -0.106221271 | 0.070583618 | -1.504899779 | 0.132349845 | 0.573884732 |
| Rbm12b1 | 270.4289666 | -0.106225559 | 0.09399273 | -1.130146549 | 0.258414478 | 0.703616072 |
| Cldn23 | 26.66720276 | -0.106312722 | 0.084184542 | -1.262853246 | 0.206641925 | 0.661001099 |
| Dus2l | 509.7713556 | -0.106335569 | 0.068813047 | -1.545282083 | 0.122278047 | 0.556819993 |
| Tcf3 | 886.1030749 | -0.106355691 | 0.072222602 | -1.472609523 | 0.140856356 | 0.586773565 |
| Timm44 | 1702.970899 | -0.106364794 | 0.054635639 | -1.946802407 | 0.051558428 | 0.404394022 |
| Il1rl2 | 25.36087056 | -0.106444011 | 0.083215026 | -1.27914411 | 0.200846313 | 0.654716629 |
| 1700052K11Rik | 258.3134951 | -0.106461186 | 0.084743692 | -1.256272688 | 0.209017121 | 0.663936737 |
| Abcd1 | 635.2655377 | -0.106508684 | 0.075783414 | -1.405435286 | 0.15989188 | 0.61201303 |
| Ltbp1 | 344.0631741 | -0.106580427 | 0.082567758 | -1.290823797 | 0.196764784 | 0.652856385 |
| Dis3 | 610.474263 | -0.106585888 | 0.075356611 | -1.414419879 | 0.157238657 | 0.607878516 |
| Ccdc22 | 1487.104486 | -0.106617855 | 0.06136652 | -1.737394517 | 0.082317558 | 0.483019453 |
| Zfp618 | 75.99833589 | -0.106623797 | 0.095067244 | -1.121561888 | 0.262048764 | 0.704389405 |
| Nat10 | 1616.7261 | -0.10667009 | 0.062246519 | -1.71367158 | 0.086589059 | 0.492672415 |
| Slc22a4 | 335.8526383 | -0.106677328 | 0.079400973 | -1.343526701 | 0.179101486 | 0.632845581 |
| Unc13a | 16404.80586 | -0.10670484 | 0.060757184 | -1.756250581 | 0.079045632 | 0.477013957 |
| Zcwpw1 | 291.4654278 | -0.106746355 | 0.076476657 | -1.395803093 | 0.162773799 | 0.614351408 |
| Pafah2 | 334.1567681 | -0.106764106 | 0.084356163 | -1.265634921 | 0.205643818 | 0.659550987 |
| Gpx2-ps1 | 9.033993797 | -0.106805607 | 0.058067935 | -1.839321615 | 0.065867896 | NA |
| Usp16 | 1083.14042 | -0.106815652 | 0.068834414 | -1.55177688 | 0.12071562 | 0.554612895 |
| Ctu2 | 567.3627597 | -0.106915263 | 0.068997311 | -1.549556947 | 0.121247893 | 0.555423178 |
| Cdca8 | 45.5225276 | -0.106948395 | 0.091655719 | -1.166849122 | 0.243271306 | 0.692772033 |
| Nes | 525.7631733 | -0.106990093 | 0.0904315 | -1.18310647 | 0.236766954 | 0.687442889 |
| Begain | 4073.300766 | -0.10708 | 0.072419366 | -1.478610021 | 0.139244573 | 0.583573764 |
| Fam69c | 221.2046751 | -0.107123782 | 0.089160806 | -1.201467174 | 0.229570032 | 0.682539238 |
| Eps15l1 | 2490.98124 | -0.107191816 | 0.045486403 | -2.35656831 | 0.018444675 | 0.258617131 |
| Plcg1 | 3640.326508 | -0.107222894 | 0.051265418 | -2.091524818 | 0.036481044 | 0.345903163 |
| Capn6 | 94.09951572 | -0.10726415 | 0.094597785 | -1.133897054 | 0.256837733 | 0.703000737 |
| Inpp5e | 1573.982648 | -0.107267748 | 0.064330854 | -1.667438585 | 0.095427227 | 0.514541907 |
| Cd63 | 1624.442451 | -0.107274544 | 0.083717506 | -1.281387231 | 0.200057687 | 0.654380963 |
| Lamb2 | 1312.192823 | -0.107279383 | 0.075056471 | -1.42931556 | 0.152913556 | 0.601523346 |
| Tmem203 | 257.8051516 | -0.107316366 | 0.085631747 | -1.253231076 | 0.210121624 | 0.664608511 |
| Cnn2 | 253.1925632 | -0.1073873 | 0.088581704 | -1.212296615 | 0.225398859 | 0.677510814 |
| Muc1 | 25.78984972 | -0.107398676 | 0.087928048 | -1.221438203 | 0.22192015 | 0.672987912 |
| Retn | 20.75461809 | -0.107435057 | 0.070593286 | -1.521887748 | 0.12803721 | NA |
| 5033411D12Rik | 150.5386032 | -0.10746013 | 0.089620916 | -1.199051913 | 0.23050776 | 0.683402059 |
| Wdr89 | 90.58502795 | -0.107460545 | 0.095058579 | -1.130466556 | 0.258279683 | 0.703616072 |
| 6430550D23Rik | 30.84052153 | -0.107461 | 0.087575429 | -1.227067935 | 0.219797059 | 0.672421385 |
| Zcchc11 | 1449.433703 | -0.107482947 | 0.059207014 | -1.815375224 | 0.069466266 | 0.455055507 |
| Anxa4 | 187.2217748 | -0.107488152 | 0.087422464 | -1.229525537 | 0.218874829 | 0.671604514 |
| Ccdc110 | 30.46062274 | -0.107516788 | 0.083797375 | -1.283056758 | 0.199472192 | 0.65377551 |
| Trim47 | 118.268768 | -0.107525363 | 0.092289597 | -1.165086492 | 0.243983984 | 0.69367782 |
| Parp2 | 663.4165181 | -0.10765778 | 0.073838423 | -1.458018409 | 0.144835469 | 0.590269908 |
| Ppp1r18 | 373.3267814 | -0.107711931 | 0.073376856 | -1.467927854 | 0.142123821 | 0.587991041 |
| Lst1 | 51.74836349 | -0.107736886 | 0.094486774 | -1.140232459 | 0.254189468 | 0.701619624 |
| Dgat1 | 823.8501774 | -0.107758144 | 0.079616805 | -1.35345978 | 0.175908791 | 0.628658305 |
| Adh1 | 12.84448624 | -0.107770894 | 0.061092108 | -1.764072265 | 0.077719813 | NA |
| 2700049A03Rik | 397.040383 | -0.107796402 | 0.074483714 | -1.447247948 | 0.147827492 | 0.594842535 |
| Ugt1a7c | 22.51773913 | -0.107890504 | 0.079131798 | -1.363427943 | 0.172747679 | 0.624056899 |
| Cdca4 | 272.8290627 | -0.107918543 | 0.078226745 | -1.379560701 | 0.167721944 | 0.617953076 |
| Ccdc141 | 939.3841763 | -0.107920946 | 0.073621397 | -1.465891034 | 0.142677975 | 0.589124578 |
| Socs3 | 144.3490188 | -0.107989661 | 0.095093941 | -1.135610324 | 0.256119686 | 0.702218644 |
| Bmpr1b | 953.1869614 | -0.108021465 | 0.054305437 | -1.989146389 | 0.046685047 | 0.390412779 |
| 4933406C10Rik | 58.94514443 | -0.108056262 | 0.094901768 | -1.138611676 | 0.254865156 | 0.701619624 |
| F830016B08Rik | 11.40428454 | -0.108075122 | 0.043200765 | -2.501694628 | 0.012360048 | NA |
| Athl1 | 297.5027925 | -0.108136373 | 0.091283297 | -1.18462387 | 0.236166195 | 0.687039131 |
| Sfrp4 | 57.91777798 | -0.108136879 | 0.093502354 | -1.156515038 | 0.247470552 | 0.696744638 |
| Plk5 | 1052.280773 | -0.108169174 | 0.087218233 | -1.240212856 | 0.214896675 | 0.668817995 |
| Casp1 | 137.9201493 | -0.10820977 | 0.091905089 | -1.177407809 | 0.239032774 | 0.689537017 |
| Psme4 | 2665.152143 | -0.108215835 | 0.063962668 | -1.691859301 | 0.090672804 | 0.502789698 |
| Zdhhc1 | 1561.675942 | -0.108240626 | 0.072577619 | -1.491377463 | 0.135862422 | 0.579435507 |
| Rhbdf1 | 499.1221713 | -0.108276341 | 0.088358033 | -1.225427246 | 0.220414286 | 0.672757795 |
| Ccdc102a | 50.34937698 | -0.108283765 | 0.094269682 | -1.148659486 | 0.250696418 | 0.699068333 |
| Psme2 | 813.5823533 | -0.108299211 | 0.061444687 | -1.76254801 | 0.077976752 | 0.475337606 |
| Ddx27 | 1226.073768 | -0.108301904 | 0.062781057 | -1.725072956 | 0.084514326 | 0.488499377 |
| Rpusd4 | 316.7488129 | -0.108388971 | 0.073527201 | -1.474134327 | 0.140445428 | 0.585785508 |
| Dimt1 | 505.5144749 | -0.108437942 | 0.069405302 | -1.562387011 | 0.118196845 | 0.549241002 |
| Il20rb | 12.44698962 | -0.108455034 | 0.073034685 | -1.484979831 | 0.137549167 | NA |
| Zfp783 | 303.1728064 | -0.108467757 | 0.087459647 | -1.240203466 | 0.214900147 | 0.668817995 |
| Slc29a1 | 925.1203206 | -0.108527659 | 0.071645456 | -1.51478775 | 0.129826164 | 0.568885619 |
| Repin1 | 1734.687305 | -0.108550112 | 0.061453496 | -1.766378136 | 0.077332431 | 0.473845905 |
| Car9 | 39.76128406 | -0.108566589 | 0.087783911 | -1.236748135 | 0.216180602 | 0.670049713 |
| Pllp | 1971.773029 | -0.108619454 | 0.077372888 | -1.403843864 | 0.160365349 | 0.612373044 |
| Fsbp | 9.941548812 | -0.108627341 | 0.067564001 | -1.607769517 | 0.107885667 | NA |
| Cflar | 912.8302355 | -0.10865066 | 0.062288896 | -1.744302232 | 0.081106403 | 0.479898997 |
| Ubn2 | 1896.573159 | -0.108663311 | 0.080407938 | -1.351400305 | 0.176567236 | 0.629083076 |
| Slc35b3 | 357.4900336 | -0.108675674 | 0.082480044 | -1.317599618 | 0.187637714 | 0.641776649 |
| Gem | 62.74031448 | -0.108677604 | 0.094322828 | -1.152187717 | 0.249243944 | 0.697863827 |
| Lin9 | 173.5007878 | -0.108815477 | 0.090734818 | -1.199269253 | 0.230423267 | 0.683402059 |
| Maf | 534.6474511 | -0.108909778 | 0.071790454 | -1.517050976 | 0.129253814 | 0.568262986 |
| Tubb6 | 43.63800768 | -0.108923222 | 0.091293917 | -1.193104921 | 0.232828285 | 0.684628272 |
| Casc1 | 34.02248517 | -0.108961289 | 0.081978521 | -1.329144367 | 0.183800345 | 0.637355821 |
| Hnrnpl | 5843.999863 | -0.109009175 | 0.055673598 | -1.958004873 | 0.050229441 | 0.402001575 |
| 2700097O09Rik | 157.470258 | -0.109012343 | 0.087371075 | -1.247693739 | 0.212143235 | 0.666270994 |
| Erbb2 | 136.9700901 | -0.109058975 | 0.091424786 | -1.192881938 | 0.232915614 | 0.68474645 |

| | | | | | | |
|---|---|---|---|---|---|---|
| Aagab | 2213.748289 | 0.140096055 | 0.06996328 | 2.002422625 | 0.045239297 | 0.163122567 |
| Cd24a | 309.6917897 | -0.177075851 | 0.130194034 | -1.360091902 | 0.173800843 | 0.381967414 |
| Gm1653 | 16.33567347 | 0.139972077 | 0.167936053 | 0.833484382 | 0.404571603 | 0.629423879 |
| Tmem107 | 234.058013 | -0.160382631 | 0.117917526 | -1.360125481 | 0.173790219 | 0.381967414 |
| Chic2 | 192.6727476 | -0.134574688 | 0.099016352 | -1.359115798 | 0.174109898 | 0.38253811 |
| Sstr3 | 1169.601754 | -0.120391355 | 0.088578509 | -1.359148595 | 0.174099507 | 0.38253811 |
| Dpy30 | 564.7521342 | -0.1102446 | 0.081129917 | -1.358864929 | 0.174189394 | 0.38265851 |
| D030056L22Rik | 741.7312544 | -0.09137547 | 0.067256814 | -1.358605387 | 0.174271667 | 0.382784975 |
| Arhgap5 | 4139.438643 | -0.108665516 | 0.080001135 | -1.358299684 | 0.17436861 | 0.382943622 |
| Dok7 | 407.7664095 | 0.139904721 | 0.094964667 | 1.47322921 | 0.140689241 | 0.335256706 |
| Ankra2 | 1058.327639 | 0.139904036 | 0.074754908 | 1.871503027 | 0.061275391 | 0.197198933 |
| Dpagt1 | 500.9827251 | 0.139901283 | 0.072933301 | 1.918208579 | 0.055084569 | 0.184659741 |
| Galnt11 | 1491.336784 | -0.075585634 | 0.055681825 | -1.357456126 | 0.174636325 | 0.383336568 |
| Cdk1 | 15.43189986 | -0.220141621 | 0.162198593 | -1.357235081 | 0.174706527 | 0.383414014 |
| Icam5 | 7458.385829 | -0.103776131 | 0.076469162 | -1.35709778 | 0.174750144 | 0.383446291 |
| Dock6 | 797.1480952 | 0.139895853 | 0.118414946 | 1.181403684 | 0.237442395 | 0.461829333 |
| Rcan3 | 2007.475451 | -0.125496669 | 0.092490259 | -1.356863631 | 0.174824545 | 0.383510052 |
| Igdcc3 | 54.00682994 | 0.139854926 | 0.160131499 | 0.873375491 | 0.382458445 | 0.611034353 |
| Mkx | 506.7013427 | 0.139831736 | 0.099847639 | 1.400451101 | 0.161378277 | 0.366197278 |
| Trpv2 | 741.5159218 | 0.139758326 | 0.097542685 | 1.432791461 | 0.151917442 | 0.353158624 |
| Snx13 | 3649.808047 | -0.096444185 | 0.071123555 | -1.356009064 | 0.175096284 | 0.383888755 |
| Fam135a | 1157.582834 | 0.139736327 | 0.086795917 | 1.609941244 | 0.107410684 | 0.283899887 |
| Rdh12 | 73.72730301 | 0.139705925 | 0.130480182 | 1.070706089 | 0.284301604 | 0.513998723 |
| Daam2 | 4173.112571 | 0.139629735 | 0.083356724 | 1.675086637 | 0.093917234 | 0.259077536 |
| 0610009L18Rik | 66.79084824 | 0.13958861 | 0.160011987 | 0.872363458 | 0.383010129 | 0.611369871 |
| Rac3 | 550.7597499 | -0.152224285 | 0.112392611 | -1.354397622 | 0.175609556 | 0.384741863 |
| Dtwd2 | 117.6136887 | -0.174414785 | 0.128800331 | -1.354148572 | 0.175688983 | 0.384861458 |
| Atxn2 | 3684.107935 | 0.139564059 | 0.057821622 | 2.413700166 | 0.015791451 | 0.080824093 |
| Ccna2 | 66.73768245 | -0.192048347 | 0.141886266 | -1.353537255 | 0.175884056 | 0.385037626 |
| D930015E06Rik | 719.5345221 | -0.107954308 | 0.07974628 | -1.353722181 | 0.175825029 | 0.385037626 |
| Prr24 | 976.3999939 | 0.074226323 | 0.054843146 | 1.353429346 | 0.175918507 | 0.385037626 |
| Suco | 1543.067705 | -0.10598483 | 0.078308284 | -1.353430627 | 0.175918098 | 0.385037626 |
| Ubqln4 | 6413.965212 | -0.095234619 | 0.07036055 | -1.353522934 | 0.175888628 | 0.385037626 |
| Diras1 | 4931.947634 | -0.07747065 | 0.05725702 | -1.353033217 | 0.176045019 | 0.385096926 |
| Npr3 | 471.1122436 | 0.139552352 | 0.147070521 | 0.948880517 | 0.342681385 | 0.575828939 |
| Rell2 | 4389.244603 | 0.139530728 | 0.067585666 | 2.064501793 | 0.038970164 | 0.147125663 |
| Btk | 53.61841418 | 0.139379883 | 0.160587761 | 0.8679359 | 0.385429425 | 0.613251541 |
| Defb1 | 17.68320313 | 0.139318926 | 0.164699955 | 0.845895347 | 0.397611129 | 0.622847723 |
| Pou6f1 | 5530.442943 | 0.139318 | 0.076606063 | 1.818628886 | 0.068968068 | 0.212866782 |
| Fblim1 | 68.90810906 | 0.139212243 | 0.155602641 | 0.89466504 | 0.370966172 | 0.600555851 |
| Uckl1 | 689.9562338 | 0.139210228 | 0.104732425 | 1.329198937 | 0.183782346 | 0.394698659 |
| Vrk3 | 911.1796389 | -0.107930815 | 0.079843475 | -1.351780031 | 0.176445694 | 0.385701126 |
| Lrrc24 | 1002.111593 | -0.116837961 | 0.08645026 | -1.351505034 | 0.176533709 | 0.385784666 |
| Qtrt1 | 345.2620872 | 0.139174809 | 0.115529866 | 1.204665199 | 0.228332576 | 0.451219768 |
| C330018D20Rik | 333.3785293 | -0.161444008 | 0.119491277 | -1.35109451 | 0.17666516 | 0.385960345 |
| Grpel1 | 1513.643788 | -0.079988668 | 0.059206105 | -1.351020613 | 0.17668883 | 0.385960345 |
| Glrx | 1262.084873 | 0.139164225 | 0.082299508 | 1.690948449 | 0.090846649 | 0.253890779 |
| Hid1 | 4494.510704 | 0.139152894 | 0.063738422 | 2.183186985 | 0.029022043 | 0.120809102 |
| Atxn1 | 3258.477935 | -0.079362919 | 0.058774436 | -1.350296568 | 0.176920873 | 0.386303833 |
| Tbcb | 1891.596574 | -0.101041664 | 0.074825962 | -1.350355687 | 0.176901918 | 0.386303833 |
| Tbp | 604.1238734 | 0.138932645 | 0.072424466 | 1.91831094 | 0.055071596 | 0.184659741 |
| Cpsf1 | 2493.481069 | 0.138837219 | 0.084521261 | 1.642630711 | 0.100459363 | 0.270934267 |
| Egr4 | 824.0449137 | -0.221430232 | 0.164041265 | -1.349844697 | 0.177065804 | 0.386476955 |
| A730020M07Rik | 17.56249834 | -0.22273685 | 0.165022462 | -1.34973656 | 0.177100501 | 0.386478191 |
| Pid1 | 1648.06395 | -0.124333604 | 0.092144028 | -1.349339795 | 0.177227849 | 0.386701633 |
| Slc25a13 | 130.6656785 | 0.138834643 | 0.1386204 | 1.001545534 | 0.316563138 | 0.549376037 |
| Sike1 | 999.1619133 | -0.118689482 | 0.087991536 | -1.348873853 | 0.177377488 | 0.386919161 |
| Tie1 | 497.5000591 | -0.175389708 | 0.130040805 | -1.348728255 | 0.177424266 | 0.386966721 |
| Gm16793 | 21.22041269 | 0.138792206 | 0.168119772 | 0.825555522 | 0.409056275 | 0.632908847 |
| Snora61 | 22.18609337 | 0.138771248 | 0.167888433 | 0.826568247 | 0.408481821 | 0.632395817 |
| 9430038I01Rik | 44.23061993 | -0.207523968 | 0.154031408 | -1.34728346 | 0.177888957 | 0.387414269 |
| Bex4 | 368.9564351 | -0.145236703 | 0.10778515 | -1.347464873 | 0.177830559 | 0.387414269 |
| Bicc1 | 172.4875477 | -0.18384948 | 0.136448355 | -1.347392422 | 0.17785388 | 0.387414269 |
| G2e3 | 403.0692082 | 0.138689858 | 0.095210553 | 1.456664769 | 0.145208939 | 0.343281554 |
| Kif15 | 14.96683085 | -0.224300927 | 0.166489813 | -1.347235143 | 0.177904513 | 0.387414269 |
| Scd4 | 19.08053275 | 0.138607703 | 0.162929985 | 0.850719424 | 0.394925231 | 0.62033369 |
| Sgip1 | 5698.137183 | -0.091304618 | 0.067766883 | -1.347333879 | 0.177872726 | 0.387414269 |
| Tmem140 | 73.32942144 | -0.208387297 | 0.154664876 | -1.34734726 | 0.177868418 | 0.387414269 |
| Armc9 | 1077.86333 | 0.138544526 | 0.061659327 | 2.246935404 | 0.024644156 | 0.10842517 |
| Ints7 | 1312.884188 | 0.138506812 | 0.057007558 | 2.429621933 | 0.01511458 | 0.078296657 |
| Arhgap6 | 512.0272363 | 0.138490896 | 0.167326705 | 0.827667619 | 0.407858762 | 0.631999566 |
| Pgpep1l | 23.31495532 | -0.226220414 | 0.167970721 | -1.3467848 | 0.178049551 | 0.387578219 |
| Psmc6 | 2439.432814 | -0.089000215 | 0.066088088 | -1.346690729 | 0.178079859 | 0.387578219 |
| Qrich1 | 3312.669495 | 0.138448378 | 0.066767334 | 2.073594514 | 0.038116985 | 0.14475112 |
| Rbbp4 | 4690.467943 | -0.06586425 | 0.048919548 | -1.346378948 | 0.178180337 | 0.387687984 |
| Gm6682 | 25.81745905 | 0.138428811 | 0.166283737 | 0.832485571 | 0.405134919 | 0.62986703 |
| Ctbp1 | 7139.353195 | -0.072209501 | 0.053653801 | -1.345841287 | 0.178353707 | 0.387849001 |
| Dazl | 116.084771 | -0.20606381 | 0.153111792 | -1.345838928 | 0.178354468 | 0.387849001 |
| Athl1 | 246.9337955 | 0.138415607 | 0.142742931 | 0.969684495 | 0.332203781 | 0.567180759 |
| Csdc2 | 4816.610841 | -0.136751135 | 0.101626101 | -1.345630044 | 0.178421857 | 0.387941097 |
| Nr2c1 | 430.9400013 | 0.138415155 | 0.086847146 | 1.593778972 | 0.110985549 | 0.290033749 |
| Cdh2 | 4638.305819 | 0.098573414 | 0.073266161 | 1.345415302 | 0.178491156 | 0.387982881 |
| Tbc1d2b | 966.2958825 | 0.138335165 | 0.083385052 | 1.658992366 | 0.097117331 | 0.264565887 |
| Pnkp | 701.4580796 | 0.138203856 | 0.100655391 | 1.37303978 | 0.16973998 | 0.377108009 |
| Xpo1 | 2634.424829 | -0.128843851 | 0.095794622 | -1.345000876 | 0.178624952 | 0.388110364 |
| Cacna2d1 | 7371.178065 | -0.119239768 | 0.088676852 | -1.344654954 | 0.178736688 | 0.388276877 |
| Dpp4 | 155.537685 | -0.171906064 | 0.127848417 | -1.344608466 | 0.178751708 | 0.388276877 |
| 2310008H09Rik | 1116.031243 | -0.117656257 | 0.087533281 | -1.344131694 | 0.178905805 | 0.388502669 |
| Sptssb | 83.84425516 | 0.138018383 | 0.140649494 | 0.981293137 | 0.326448206 | 0.560801819 |
| Itga5 | 333.6946138 | 0.137994352 | 0.126553083 | 1.090406879 | 0.275533953 | 0.504326411 |
| Il7r | 11.16661731 | -0.221007069 | 0.164542145 | -1.343163905 | 0.179218907 | 0.389108174 |
| Gpr19 | 471.8088751 | 0.137898167 | 0.128066439 | 1.076770527 | 0.281582796 | 0.511426532 |
| Nsmaf | 1254.379289 | 0.137897823 | 0.097648214 | 1.412189912 | 0.15789405 | 0.360746278 |
| Zc4h2 | 664.8062963 | -0.105607931 | 0.078677658 | -1.342286162 | 0.179503229 | 0.389527034 |
| Mdfi | 11.8349771 | -0.206073475 | 0.1535892 | -1.341718526 | 0.179687278 | 0.389871822 |
| Lst1 | 51.58919593 | -0.225045561 | 0.167752602 | -1.341532461 | 0.179747638 | 0.38994818 |
| Mybph | 11.78802157 | 0.13783049 | 0.146132091 | 0.943191118 | 0.345583156 | 0.576948985 |
| Dhx37 | 669.6491214 | 0.137770273 | 0.07194479 | 1.914944405 | 0.055499594 | 0.185605592 |
| 1700030L20Rik | 35.83407705 | -0.219105968 | 0.163398908 | -1.340926762 | 0.179944232 | 0.390210768 |
| Ckap4 | 808.0486773 | -0.08678407 | 0.064738126 | -1.340540345 | 0.180069737 | 0.390321947 |
| Drosha | 4820.658573 | 0.055293143 | 0.041247034 | 1.340536218 | 0.180071078 | 0.390321947 |
| Pdpn | 328.1016569 | -0.159511444 | 0.118977976 | -1.340680436 | 0.180024229 | 0.390321947 |
| 3110045C21Rik | 21.71625581 | -0.22426974 | 0.167346274 | -1.340153771 | 0.180195357 | 0.390529473 |
| Fam219b | 763.8556073 | -0.089389102 | 0.066726386 | -1.339636499 | 0.180363551 | 0.390529473 |
| Kifap3 | 11935.4299 | -0.080254815 | 0.059908579 | -1.33962141 | 0.18036846 | 0.390529473 |
| Lrrc14 | 749.0038916 | -0.13591727 | 0.101449753 | -1.339749642 | 0.180326753 | 0.390529473 |
| Pcnxl3 | 3024.164392 | 0.088050392 | 0.065719239 | 1.339796289 | 0.180311583 | 0.390529473 |
| Prpsap1 | 1435.487483 | 0.09663917 | 0.072134362 | 1.339710615 | 0.180339445 | 0.390529473 |
| Serf2 | 4269.68939 | -0.113814525 | 0.084934639 | -1.340024829 | 0.180237273 | 0.390529473 |
| She | 254.2414285 | -0.142873381 | 0.106635208 | -1.339833098 | 0.180299613 | 0.390529473 |
| Arhgef25 | 6533.416218 | 0.137763466 | 0.087985022 | 1.565760419 | 0.117404716 | 0.300771935 |
| Tsc22d1 | 15341.75225 | -0.075361984 | 0.056296044 | -1.338672839 | 0.180677202 | 0.391088652 |
| Poli | 486.3938117 | 0.137688196 | 0.081164658 | 1.696405779 | 0.08980906 | 0.252403696 |
| Scaf1 | 6558.796832 | 0.091350133 | 0.068262305 | 1.338222219 | 0.180824008 | 0.391297092 |
| Hlf | 8878.864707 | 0.137686221 | 0.094111109 | 1.463017727 | 0.143462528 | 0.340112608 |
| Dnahc6 | 87.7134062 | 0.137655266 | 0.163599466 | 0.841416355 | 0.400114726 | 0.625252402 |
| Theg | 13.58280173 | -0.220309945 | 0.164670957 | -1.337879791 | 0.180935626 | 0.391473411 |
| Aspn | 37.84927768 | -0.221005457 | 0.165246545 | -1.337428612 | 0.18108277 | 0.391638252 |
| 5830403L16Rik | 34.29917644 | 0.137508677 | 0.158883116 | 0.865470672 | 0.386780506 | 0.614237786 |
| Swt1 | 291.0552203 | 0.137377087 | 0.104493676 | 1.31469283 | 0.188613157 | 0.401747886 |
| Nosip | 1563.795502 | 0.095095645 | 0.071128821 | 1.33694955 | 0.181239105 | 0.391771527 |
| Pgs1 | 2970.032842 | -0.093886322 | 0.070217222 | -1.337083971 | 0.181195229 | 0.391771527 |
| Tmem123 | 489.6015526 | -0.124389356 | 0.093046439 | -1.336852405 | 0.181270819 | 0.391771527 |
| Wfikkn2 | 74.01261014 | 0.137370412 | 0.161563604 | 0.850255929 | 0.395182813 | 0.620486675 |
| Pcdha1 | 45.92473869 | -0.206383544 | 0.154424011 | -1.336473144 | 0.181394673 | 0.391933319 |
| Gm3219 | 22.69986946 | -0.224512413 | 0.168010752 | -1.336297894 | 0.181451925 | 0.391944252 |
| Tars | 1134.249308 | 0.137324071 | 0.081256751 | 1.690001992 | 0.091027573 | 0.254214352 |
| Brap | 2872.796086 | 0.137302087 | 0.101954195 | 1.346703654 | 0.178075695 | 0.387578219 |
| Def8 | 2359.153676 | -0.072795148 | 0.054495022 | -1.335812798 | 0.181610469 | 0.392177366 |
| Gadl1 | 27.09782797 | 0.137294218 | 0.167249951 | 0.820892429 | 0.411707544 | 0.635358236 |
| 9930014A18Rik | 91.23071809 | -0.199675205 | 0.149511735 | -1.335515266 | 0.181707762 | 0.39227812 |
| Birc2 | 1324.401187 | 0.137264127 | 0.055693094 | 2.464652563 | 0.013714616 | 0.073125764 |
| Tceb1 | 1815.195052 | -0.091137036 | 0.068250504 | -1.335331329 | 0.181767929 | 0.39229869 |
| Impg1 | 58.84893574 | -0.219461318 | 0.164361149 | -1.335238408 | 0.18179833 | 0.392309655 |
| Sdf4 | 11115.16578 | 0.137190851 | 0.051816848 | 2.647610906 | 0.008106277 | 0.050688101 |
| Ddx47 | 1581.43699 | 0.088057925 | 0.065964167 | 1.334935759 | 0.181897373 | 0.392412937 |
| Sh3tc2 | 141.6138503 | 0.137190373 | 0.123836112 | 1.107838186 | 0.267931705 | 0.495872824 |
| Rpl36al | 1612.902668 | -0.095241728 | 0.071355903 | -1.334742095 | 0.181960771 | 0.392441563 |
| Etfdh | 1128.961133 | -0.078895326 | 0.059148799 | -1.333844931 | 0.182254682 | 0.392903171 |
| Rapsn | 26.77104769 | 0.137175592 | 0.166554128 | 0.823609681 | 0.410161373 | 0.63423556 |
| Spire2 | 598.3976047 | -0.096325466 | 0.072211083 | -1.33394295 | 0.182222554 | 0.392903171 |
| Tmem5 | 759.4405109 | -0.096149382 | 0.072087931 | -1.333779182 | 0.182276236 | 0.392903171 |
| Ppp4c | 1027.909594 | -0.077130189 | 0.057844715 | -1.333400791 | 0.182400313 | 0.393115934 |
| Spint1 | 165.7080776 | 0.137172023 | 0.121835763 | 1.125876508 | 0.260217794 | 0.487051764 |
| Fen1 | 542.8136946 | 0.137131071 | 0.108365535 | 1.265449129 | 0.205710373 | 0.423436651 |
| Gpr137c | 802.2638139 | -0.136980974 | 0.102769861 | -1.332890521 | 0.182567734 | 0.393364924 |
| Igsf5 | 12.88947234 | -0.210519652 | 0.157969266 | -1.33266209 | 0.182642719 | 0.393364924 |
| Ivns1abp | 6966.512743 | 0.071344646 | 0.053535444 | 1.332661899 | 0.182642782 | 0.393364924 |
| Rnf166 | 1070.203555 | 0.080183375 | 0.060172376 | 1.332561239 | 0.182675832 | 0.393381424 |
| Sytl4 | 44.75014824 | -0.214658849 | 0.161142389 | -1.332106656 | 0.182825144 | 0.393648245 |
| Sppl2a | 1492.445383 | -0.099935304 | 0.075027914 | -1.331974965 | 0.182868415 | 0.393686706 |
| Rnpep | 1496.059625 | -0.09373483 | 0.070383444 | -1.331773842 | 0.182934516 | 0.393774297 |
| Nup133 | 1622.495315 | 0.137090064 | 0.061771366 | 2.219314106 | 0.026465363 | 0.11339998 |
| Rgs10 | 939.629422 | -0.130877379 | 0.098291964 | -1.331516571 | 0.183019096 | 0.393846927 |
| Lsg1 | 599.9024546 | 0.137036087 | 0.094851767 | 1.444739415 | 0.148531094 | 0.348265855 |
| Orai2 | 3580.196339 | 0.137029286 | 0.102654425 | 1.334859999 | 0.181922172 | 0.392412937 |
| Epb4.1l5 | 370.4985634 | -0.157486859 | 0.118329531 | -1.330917623 | 0.183216118 | 0.394060885 |
| Zik1 | 680.9335632 | -0.098188408 | 0.073775667 | -1.330905054 | 0.183220254 | 0.394060885 |
| Ap2m1 | 20663.73222 | -0.078708536 | 0.059146917 | -1.33072931 | 0.183278095 | 0.394130577 |
| Dynlrb2 | 31.81926436 | 0.13701994 | 0.166598914 | 0.822453982 | 0.410818567 | 0.634618433 |
| 0610007N19Rik | 77.7980029 | -0.188936775 | 0.142081187 | -1.329780383 | 0.183590641 | 0.394528903 |
| Mau2 | 3754.47143 | 0.137005892 | 0.065306184 | 2.09790076 | 0.035913913 | 0.139356415 |
| Ndufv2 | 4053.525859 | -0.102235739 | 0.076878238 | -1.329839772 | 0.183571069 | 0.394528903 |
| Rabgap1l | 4028.004855 | -0.121230487 | 0.091151624 | -1.329987139 | 0.183522509 | 0.394528903 |
| 5830418K08Rik | 1081.683019 | 0.136893089 | 0.081829492 | 1.6729065 | 0.094345704 | 0.259862697 |
| Chchd5 | 248.2440744 | -0.129615613 | 0.097528675 | -1.329000043 | 0.183847956 | 0.394698659 |
| Chrna3 | 33.68938477 | -0.216218444 | 0.162678288 | -1.329116788 | 0.183809442 | 0.394698659 |
| Cwh43 | 113.4586499 | -0.172133652 | 0.12948668 | -1.329354117 | 0.183731168 | 0.394698659 |
| Kif1b | 39493.97554 | -0.120191802 | 0.090432898 | -1.329071656 | 0.18382433 | 0.394698659 |
| Cpt2 | 228.4015257 | 0.136843735 | 0.107352711 | 1.274711491 | 0.202411378 | 0.419384114 |
| Rmst | 59.05506884 | 0.136783083 | 0.157378188 | 0.869136217 | 0.38477263 | 0.61273055 |
| Map1s | 1873.206608 | 0.083711899 | 0.062995323 | 1.328858956 | 0.183894507 | 0.394743903 |
| Dis3l | 1213.094454 | 0.097767446 | 0.073609094 | 1.328197924 | 0.18411273 | 0.395156762 |
| Fth1 | 49965.45387 | 0.136759225 | 0.10782577 | 1.268335254 | 0.204678253 | 0.42171341 |
| Tubgcp4 | 872.2312489 | 0.093710664 | 0.070566033 | 1.327985442 | 0.184182916 | 0.395198741 |
| Aoah | 12.6493359 | -0.217203121 | 0.163610264 | -1.327564151 | 0.184322134 | 0.395333215 |
| 2310061I04Rik | 1496.157731 | 0.136723546 | 0.070637586 | 1.935563683 | 0.052921162 | 0.180250711 |
| Zmat5 | 458.8952039 | -0.129606617 | 0.097617738 | -1.327695356 | 0.184278768 | 0.395333215 |
| Atp2c1 | 5547.687789 | -0.087195973 | 0.065705137 | -1.327079998 | 0.184482221 | 0.395621804 |
| Lepre1 | 309.5186252 | 0.136692973 | 0.100671144 | 1.357816823 | 0.174521816 | 0.383216839 |
| Rnf181 | 2419.354298 | 0.082715308 | 0.062344612 | 1.326743485 | 0.184593551 | 0.395751 |
| Arl4a | 1342.237679 | -0.118568096 | 0.089433628 | -1.325766369 | 0.184917096 | 0.396245313 |
| Dcakd | 962.3464338 | -0.118113922 | 0.089084135 | -1.32586933 | 0.184882983 | 0.396245313 |
| 3000002C10Rik | 119.6177392 | 0.136646771 | 0.142789826 | 0.956978342 | 0.338578187 | 0.57220009 |
| Zfp97 | 54.7323301 | -0.213877605 | 0.161327174 | -1.325738247 | 0.184926414 | 0.396245313 |
| Angel2 | 2216.015571 | 0.136629356 | 0.066402439 | 2.057595449 | 0.039628984 | 0.148779132 |
| 6430571L13Rik | 256.821184 | -0.212414272 | 0.160285676 | -1.325223049 | 0.185097183 | 0.396391949 |
| Commd9 | 788.7430815 | 0.094793024 | 0.071523943 | 1.325332753 | 0.18506081 | 0.396391949 |
| Sec24c | 4083.199817 | 0.074141207 | 0.055945725 | 1.325234539 | 0.185093373 | 0.396391949 |
| 1810033B17Rik | 10.64209597 | 0.136616931 | 0.163451521 | 0.835825387 | 0.403253142 | 0.628129505 |
| Trim28 | 4772.818636 | 0.077349696 | 0.058385304 | 1.324814481 | 0.185232691 | 0.396605993 |
| Dpp10 | 6783.735366 | -0.092696971 | 0.070002523 | -1.324194705 | 0.185438389 | 0.396849532 |
| Glce | 1795.413439 | -0.089308678 | 0.067443848 | -1.324193108 | 0.185438919 | 0.396849532 |
| Tor1b | 1150.444994 | 0.099728257 | 0.075305772 | 1.324310936 | 0.1853998 | 0.396849532 |
| Rab31 | 3468.811264 | 0.136574205 | 0.074972035 | 1.821668644 | 0.068505278 | 0.211987375 |
| Ptprj | 6858.683197 | -0.11356251 | 0.08580147 | -1.323549691 | 0.185652643 | 0.39719719 |
| Sestd1 | 2856.895091 | 0.136523196 | 0.052807495 | 2.585299627 | 0.009729441 | 0.057577633 |
| Aars2 | 1073.679402 | 0.136499551 | 0.097966497 | 1.393328902 | 0.163520354 | 0.368744952 |
| Smg9 | 1223.485021 | 0.136496286 | 0.086889367 | 1.570920488 | 0.116201119 | 0.298824919 |
| Coro1c | 3806.55163 | -0.082203249 | 0.062156245 | -1.32252598 | 0.185993064 | 0.397705843 |
| D930016D06Rik | 436.7948959 | 0.136494909 | 0.100100938 | 1.363572728 | 0.17270208 | 0.38052418 |
| Pcdhb19 | 250.5854634 | -0.12992888 | 0.098261254 | -1.32227988 | 0.18607497 | 0.39777119 |
| F8a | 451.8620611 | -0.139710665 | 0.105725915 | -1.321442002 | 0.186354029 | 0.398312777 |
| Brd4 | 3795.098712 | 0.093128873 | 0.070507963 | 1.320827736 | 0.186558809 | 0.398530566 |
| H2-K2 | 50.65935898 | 0.136458513 | 0.16134673 | 0.845746999 | 0.397693898 | 0.622914402 |
| Siah3 | 75.07763029 | 0.136417096 | 0.14643869 | 0.931564578 | 0.351561596 | 0.582452457 |
| Syde2 | 137.069172 | -0.157867574 | 0.119515777 | -1.320893167 | 0.186536989 | 0.398530566 |
| A830009L08Rik | 138.8637268 | 0.136399395 | 0.148494535 | 0.918548251 | 0.35833191 | 0.588946072 |
| Pcdhga8 | 458.6026719 | 0.136381813 | 0.105149724 | 1.297024928 | 0.19462261 | 0.409106291 |
| BC055324 | 60.0320784 | -0.190652576 | 0.144510545 | -1.319298718 | 0.187069266 | 0.39941383 |
| Dgcr14 | 1093.894549 | 0.136336481 | 0.085934634 | 1.586513787 | 0.112622807 | 0.293076549 |
| Taf8 | 655.0113974 | 0.136326653 | 0.093429783 | 1.459134858 | 0.144527994 | 0.342141594 |
| Tgoln1 | 3730.802909 | -0.075923824 | 0.05757173 | -1.318769201 | 0.187246284 | 0.399668538 |
| Eif1ax | 1648.085098 | -0.085270625 | 0.064678507 | -1.318376523 | 0.187377636 | 0.399838005 |
| Polr2h | 510.4137418 | -0.112481351 | 0.085319622 | -1.318352674 | 0.187385616 | 0.399838005 |
| Pilra | 11.09924532 | 0.136306715 | 0.159990486 | 0.851967625 | 0.394232061 | 0.619921228 |
| Nkain3 | 31.52659367 | -0.221507462 | 0.168077162 | -1.317891493 | 0.187539974 | 0.400074949 |
| Ppp1r35 | 206.0481219 | -0.1457966 | 0.110667251 | -1.317432201 | 0.187693793 | 0.400347967 |
| B3gnt1 | 4041.51153 | -0.072835989 | 0.055311524 | -1.31683209 | 0.187894914 | 0.400721788 |
| Kxd1 | 894.1259985 | 0.075306267 | 0.057206426 | 1.316395238 | 0.18804142 | 0.400979048 |
| Zfp11 | 289.8121812 | -0.128867531 | 0.097912324 | -1.316152297 | 0.188122931 | 0.40109766 |
| Ercc1 | 596.3682081 | 0.13629815 | 0.079998832 | 1.703751761 | 0.088427474 | 0.249561981 |
| Sema3a | 943.2010615 | -0.126413608 | 0.096112824 | -1.315262656 | 0.188421644 | 0.401624014 |
| Oxld1 | 120.2555498 | -0.16486331 | 0.125376371 | -1.314947216 | 0.188527643 | 0.401684172 |
| Zfp114 | 81.24335085 | 0.136211409 | 0.134427242 | 1.013272358 | 0.310930087 | 0.543987214 |
| Brms1 | 832.8858325 | 0.136166995 | 0.076354849 | 1.783344427 | 0.074530254 | 0.224267761 |
| Selenbp1 | 339.79918 | 0.136134152 | 0.089156498 | 1.526912287 | 0.126782824 | 0.314691614 |
| Slc29a2 | 482.166667 | 0.136083072 | 0.075861634 | 1.793832589 | 0.072839894 | 0.220938012 |
| Umps | 671.4531499 | -0.102194163 | 0.07773625 | -1.314626873 | 0.188635334 | 0.401747886 |
| Tmem179 | 5003.319942 | -0.083623322 | 0.063622059 | -1.314376243 | 0.188719621 | 0.401872157 |
| Gcc2 | 3401.31645 | -0.090278049 | 0.068692938 | -1.314226053 | 0.188770143 | 0.401924502 |
| Trmt11 | 226.374449 | 0.135918447 | 0.086031371 | 1.579870743 | 0.114136471 | 0.295922101 |
| Gm10406 | 53.61151871 | 0.13589837 | 0.154331107 | 0.880563692 | 0.378554017 | 0.607597359 |
| Aurkb | 11.5885159 | -0.216811631 | 0.165031491 | -1.313759151 | 0.188927268 | 0.402059668 |
| Sdf2 | 1512.917033 | -0.083802442 | 0.063789749 | -1.313728983 | 0.188937424 | 0.402059668 |

| | | | | | | |
|---|---|---|---|---|---|---|
| Gmppa | 1349.816661 | -0.109174872 | 0.068915954 | -1.584174131 | 0.113154099 | 0.540327972 |
| Mamstr | 92.64262701 | -0.109197653 | 0.095101948 | -1.148216788 | 0.250879081 | 0.699207507 |
| Tgm3 | 180.2894973 | -0.109219513 | 0.09112506 | -1.198567257 | 0.230696256 | 0.683402059 |
| Evi2a | 708.0564637 | -0.109285873 | 0.084765623 | -1.289271151 | 0.197303837 | 0.653301923 |
| Ppp4r1 | 1522.880075 | -0.10928906 | 0.054258444 | -2.014231362 | 0.043985248 | 0.379863091 |
| Zbtb43 | 1426.348108 | -0.109313656 | 0.057045705 | -1.91624691 | 0.055333675 | 0.413057568 |
| Gm12216 | 25.78671874 | -0.109329937 | 0.08505814 | -1.285355362 | 0.198668132 | 0.65377551 |
| Pmm2 | 714.8707174 | -0.109347177 | 0.070097259 | -1.55993513 | 0.118775212 | 0.550992847 |
| Dnmbp | 643.3614615 | -0.109384169 | 0.072185223 | -1.515326317 | 0.129689787 | 0.568885619 |
| Pcsk7 | 1115.953367 | -0.109460057 | 0.05978744 | -1.830820255 | 0.067127376 | 0.447688159 |
| Myo1g | 25.41201949 | -0.109463575 | 0.082877851 | -1.320782009 | 0.18657406 | 0.640826269 |
| Sclt1 | 346.5906517 | -0.109468034 | 0.076387548 | -1.433061238 | 0.151840338 | 0.599519637 |
| Afap1l1 | 1308.383929 | -0.109549184 | 0.062379866 | -1.756162546 | 0.079060658 | 0.477013957 |
| Gramd2 | 184.8347472 | -0.109587732 | 0.093946235 | -1.166494137 | 0.243414718 | 0.692772033 |
| Setdb1 | 2097.389607 | -0.109627849 | 0.062514675 | -1.75363383 | 0.07949327 | 0.477013957 |
| Lrrc17 | 205.9323374 | -0.109656695 | 0.053009291 | -2.068631608 | 0.038580669 | 0.355824003 |
| Frem1 | 172.8655989 | -0.10971856 | 0.092968883 | -1.180164343 | 0.237934858 | 0.688758314 |
| Lamc2 | 696.5336057 | -0.109727574 | 0.08188094 | -1.340086879 | 0.180217101 | 0.63422 |
| Prdm15 | 826.3184645 | -0.109732534 | 0.072761936 | -1.508103544 | 0.131528028 | 0.572371522 |
| Il1b | 14.5472279 | -0.109734552 | 0.075912974 | -1.445530946 | 0.148308807 | NA |
| Serpinb9 | 412.0131733 | -0.109788529 | 0.070999977 | -1.546317816 | 0.122027831 | 0.55651046 |
| Tlr13 | 153.0650109 | -0.109805736 | 0.091776931 | -1.196441583 | 0.231524282 | 0.68367236 |
| Col9a1 | 24.96099362 | -0.109821498 | 0.084620564 | -1.297810979 | 0.194352296 | 0.650198399 |
| Rrp15 | 278.3243043 | -0.109836366 | 0.083706407 | -1.312162 | 0.18946548 | 0.644077615 |
| Ms4a6c | 35.41701924 | -0.109922837 | 0.08566259 | -1.283207027 | 0.199419555 | 0.65377551 |
| Ttc5 | 1429.725689 | -0.10995756 | 0.05918987 | -1.857709105 | 0.063210335 | 0.440342223 |
| Top3b | 1493.740954 | -0.109988257 | 0.061691679 | -1.782870218 | 0.074607432 | 0.468343802 |
| Zfp207 | 3801.569487 | -0.110057643 | 0.051705566 | -2.128545361 | 0.033291893 | 0.334669924 |
| Dopey1 | 2566.895081 | -0.110069163 | 0.059998131 | -1.834543203 | 0.066573401 | 0.447108619 |
| Fam98c | 740.0335082 | -0.110217736 | 0.083025756 | -1.327512588 | 0.184339179 | 0.637780733 |
| Fam181a | 75.04079135 | -0.110218263 | 0.092754635 | -1.188277694 | 0.234724023 | 0.685908078 |
| P2ry13 | 563.7015259 | -0.110253332 | 0.081332442 | -1.355588615 | 0.175230096 | 0.628026741 |
| Kdr | 510.7818089 | -0.110325838 | 0.093800555 | -1.176174687 | 0.239525077 | 0.690203878 |
| Cacna1g | 4397.059948 | -0.1103436 | 0.067202205 | -1.641963991 | 0.100597467 | 0.521512778 |
| Tex26 | 10.85960582 | -0.110364361 | 0.059445169 | -1.856574083 | 0.063371774 | NA |
| Tead3 | 83.96183618 | -0.110381342 | 0.094936926 | -1.162680813 | 0.244959029 | 0.694568584 |
| Scn7a | 215.7009389 | -0.110459492 | 0.094190413 | -1.172725422 | 0.240905933 | 0.691863635 |
| Apoa1 | 13.39905657 | -0.110550187 | 0.068133144 | -1.622561083 | 0.104683266 | NA |
| Smyd4 | 573.6050314 | -0.11055147 | 0.06654435 | -1.661320155 | 0.096649164 | 0.516802454 |
| Insc | 180.1991168 | -0.110563209 | 0.095039265 | -1.163342431 | 0.244690597 | 0.694476029 |
| Fgd5 | 388.7214099 | -0.110592019 | 0.087006626 | -1.271075834 | 0.203701671 | 0.657622144 |
| Cdk5rap2 | 625.3186453 | -0.110638938 | 0.07989606 | -1.384785911 | 0.16611794 | 0.61635484 |
| Rhbdd3 | 659.5876952 | -0.110686861 | 0.076700802 | -1.443099137 | 0.148992546 | 0.596874167 |
| Myef2 | 2318.968179 | -0.11075334 | 0.047777335 | -2.318114647 | 0.020443091 | 0.269900834 |
| Lrrc36 | 36.94045088 | -0.110767798 | 0.083473462 | -1.326982204 | 0.18451457 | 0.637780733 |
| Map3k3 | 1235.001309 | -0.110778154 | 0.058825464 | -1.883166674 | 0.059677784 | 0.428234899 |
| Mcm3 | 143.257507 | -0.110789669 | 0.092455929 | -1.198297077 | 0.230801384 | 0.683402059 |
| Snord22 | 15.68998674 | -0.11080956 | 0.071420925 | -1.551499924 | 0.120781926 | NA |
| Arr3 | 52.72600707 | -0.110854317 | 0.089722313 | -1.235526742 | 0.21663453 | 0.670049713 |
| Polrmt | 1343.863219 | -0.110983572 | 0.074501266 | -1.489687057 | 0.136306539 | 0.579855848 |
| Spice1 | 595.9757425 | -0.110994011 | 0.070117754 | -1.582965853 | 0.113429247 | 0.540575213 |
| Entpd1 | 741.7345441 | -0.111052353 | 0.073764848 | -1.505491532 | 0.132197752 | 0.573705114 |
| Ermap | 46.8852731 | -0.111167848 | 0.094086223 | -1.181552884 | 0.237383158 | 0.688415946 |
| Gps2 | 814.1890042 | -0.111176727 | 0.070060942 | -1.586857439 | 0.112544936 | 0.5393691 |
| Gpatch8 | 3367.737096 | -0.11126373 | 0.044673256 | -2.490611628 | 0.012752343 | 0.215303986 |
| Tcf7 | 207.9404799 | -0.111290569 | 0.093039945 | -1.196159013 | 0.231634512 | 0.6837478 |
| Ccdc77 | 176.9098655 | -0.11129336 | 0.08651241 | -1.286443874 | 0.198288195 | 0.653581647 |
| Cdk9 | 3500.095966 | -0.111336 | 0.062486579 | -1.781758602 | 0.074788606 | 0.468388672 |
| Cep63 | 1066.140014 | -0.111386484 | 0.054095039 | -2.059088686 | 0.039485744 | 0.359835305 |
| Nadsyn1 | 180.5197248 | -0.111415871 | 0.08886882 | -1.25371161 | 0.209946846 | 0.664564805 |
| Nupl2 | 358.6505436 | -0.111480715 | 0.077615805 | -1.436314619 | 0.150912836 | 0.598846773 |
| Tfr2 | 227.407892 | -0.111499048 | 0.090680073 | -1.229587098 | 0.218851764 | 0.671604514 |
| Mir7-1 | 6.711561705 | -0.111510616 | 0.054792025 | -2.03516145 | 0.041834631 | NA |
| Fendrr | 27.27575016 | -0.111546696 | 0.072547145 | -1.537575268 | 0.124152498 | 0.559205951 |
| Mms22l | 50.12708291 | -0.111552052 | 0.093402077 | -1.194320892 | 0.232352468 | 0.684327865 |
| Irak3 | 64.55234941 | -0.111553624 | 0.093292161 | -1.195744881 | 0.231796131 | 0.683808003 |
| Zfp867 | 221.6743728 | -0.111719603 | 0.091593068 | -1.219738628 | 0.222563973 | 0.67374894 |
| Nup205 | 1429.457373 | -0.111771206 | 0.058653227 | -1.905627563 | 0.056698543 | 0.417225452 |
| Ddx55 | 883.7208682 | -0.111831123 | 0.070183402 | -1.593412696 | 0.111067639 | 0.53778951 |
| Cdk6 | 22.03415341 | -0.11183879 | 0.085472474 | -1.308477276 | 0.190711477 | NA |
| Amh | 31.78051295 | -0.111855509 | 0.082848307 | -1.35012425 | 0.176976131 | 0.629241039 |
| Pus7 | 719.80552 | -0.111891352 | 0.070424015 | -1.58882381 | 0.112100176 | 0.538992394 |
| Inf2 | 9258.228609 | -0.111901545 | 0.07423139 | -1.507469348 | 0.131690395 | 0.572734933 |
| Cog2 | 1136.517522 | -0.111934426 | 0.069995156 | -1.599173897 | 0.109781969 | 0.535311472 |
| H2-M3 | 122.443436 | -0.111942966 | 0.09164827 | -1.221441119 | 0.221919046 | 0.672987912 |
| Emx2os | 126.53938 | -0.111996006 | 0.094189039 | -1.189055613 | 0.234417784 | 0.68557298 |
| Zdhhc4 | 796.508067 | -0.111997337 | 0.069296703 | -1.616200084 | 0.106051061 | 0.52850889 |
| Cntnap5c | 227.1770104 | -0.112023521 | 0.090658042 | -1.235671083 | 0.21658085 | 0.670049713 |
| Vps9d1 | 2375.520406 | -0.11206848 | 0.071309032 | -1.571588866 | 0.11604593 | 0.54588186 |
| Ddit3 | 909.2465648 | -0.112149475 | 0.060028232 | -1.868278827 | 0.061723216 | 0.435133477 |
| Gm10432 | 120.1451003 | -0.112253131 | 0.090077994 | -1.246177086 | 0.21269939 | 0.666270994 |
| Mrc1 | 149.5262205 | -0.112311447 | 0.095094715 | -1.181048252 | 0.237583555 | 0.688529917 |
| Zscan26 | 3710.197062 | -0.112340685 | 0.056310236 | -1.995031349 | 0.046039461 | 0.387222051 |
| Mstn | 10.98835454 | -0.112347657 | 0.065579403 | -1.713154613 | 0.086684099 | NA |
| Nup93 | 1009.620214 | -0.112463335 | 0.064670039 | -1.739033048 | 0.08202895 | 0.482582908 |
| C030046E11Rik | 1858.451244 | -0.11248603 | 0.060579268 | -1.856840383 | 0.063333866 | 0.440607157 |
| Tbx1 | 42.98428412 | -0.112493403 | 0.090045776 | -1.249291284 | 0.211558555 | 0.666270994 |
| Mb21d1 | 44.28449536 | -0.112497638 | 0.090217491 | -1.246960392 | 0.212412021 | 0.666270994 |
| Snord32a | 10.48661788 | -0.112497914 | 0.06206127 | -1.812691145 | 0.069879471 | NA |
| Ecm1 | 285.5404961 | -0.112501127 | 0.089938721 | -1.250864211 | 0.210984023 | 0.665621429 |
| Ccdc151 | 231.7858622 | -0.112520586 | 0.090318429 | -1.245820902 | 0.212830154 | 0.666286814 |
| Accs | 134.5138766 | -0.112529868 | 0.094687213 | -1.188437848 | 0.234660953 | 0.685908078 |
| 9430041J12Rik | 118.6500358 | -0.11255983 | 0.093315969 | -1.206222588 | 0.227731677 | 0.680415543 |
| Cd93 | 344.719405 | -0.112587946 | 0.093227914 | -1.207663458 | 0.227176739 | 0.680203999 |
| Ttc13 | 1128.073662 | -0.112722356 | 0.055038895 | -2.048049063 | 0.040555192 | 0.364545002 |
| Gpam | 2266.059379 | -0.112836702 | 0.052564729 | -2.146623868 | 0.031823238 | 0.326301135 |
| Abca17 | 35.67291926 | -0.112899243 | 0.090336605 | -1.249761855 | 0.211386554 | 0.666270994 |
| 4930402H24Rik | 4271.764783 | -0.112968074 | 0.058508987 | -1.930781588 | 0.053510068 | 0.407961168 |
| Faah | 4164.460774 | -0.113117576 | 0.061084824 | -1.851811428 | 0.064052906 | 0.441381647 |
| P4ha3 | 229.5619378 | -0.113122726 | 0.087517172 | -1.292577485 | 0.196157231 | 0.651733746 |
| 9330179D12Rik | 90.07962867 | -0.113171126 | 0.09495144 | -1.19188425 | 0.233306634 | 0.685064073 |
| Secisbp2 | 677.0310162 | -0.113179072 | 0.074140624 | -1.526545983 | 0.126873948 | 0.563718247 |
| Rcor3 | 341.6986057 | -0.11318398 | 0.081916074 | -1.381706603 | 0.167061806 | 0.617356019 |
| Decr1 | 819.737038 | -0.113188545 | 0.062769343 | -1.803245651 | 0.071349644 | 0.459174089 |
| Ddx46 | 3123.737039 | -0.113190753 | 0.051961961 | -2.178338761 | 0.029380825 | 0.317771905 |
| Wfdc3 | 17.37265845 | -0.113235999 | 0.075646286 | -1.496914198 | 0.134415585 | NA |
| Apobr | 54.23075335 | -0.11323986 | 0.090301143 | -1.254024667 | 0.20983304 | 0.664421221 |
| 1700063D05Rik | 65.10438178 | -0.113514336 | 0.092501293 | -1.227164856 | 0.219760636 | 0.672421385 |
| Doc2g | 157.3495688 | -0.113565019 | 0.095054011 | -1.194741989 | 0.232187852 | 0.684268763 |
| Kcns2 | 1598.376963 | -0.113600757 | 0.084171646 | -1.349632111 | 0.177134019 | 0.629241039 |
| Mafa | 12.6026177 | -0.113653216 | 0.069811116 | -1.628010296 | 0.103522703 | NA |
| Ptpla | 204.9237983 | -0.113663141 | 0.082987047 | -1.369649174 | 0.170796441 | 0.620885919 |
| Inpp5b | 1746.904207 | -0.113689005 | 0.055592848 | -2.045029331 | 0.040851966 | 0.365377968 |
| Sox2ot | 1372.800111 | -0.113695033 | 0.084115049 | -1.35166102 | 0.17648378 | 0.629083076 |
| Gng10 | 1012.641527 | -0.113782177 | 0.067442551 | -1.687097752 | 0.091584557 | 0.504530343 |
| Lcat | 751.3325746 | -0.11382146 | 0.080359012 | -1.416411884 | 0.156654947 | 0.607279636 |
| Gdpd3 | 17.07200613 | -0.113899964 | 0.074226232 | -1.534497461 | 0.124907317 | NA |
| Ddx31 | 301.9873232 | -0.113983203 | 0.074282248 | -1.534460858 | 0.124916316 | 0.560889589 |
| Ltbp3 | 1739.230394 | -0.114012334 | 0.077054249 | -1.47963721 | 0.138970091 | 0.583135914 |
| Zfp697 | 749.5312194 | -0.114027427 | 0.075880245 | -1.502728764 | 0.132909002 | 0.575107585 |
| Cnot3 | 1771.112429 | -0.114034322 | 0.059443092 | -1.918378018 | 0.055063096 | 0.412292029 |
| Tet2 | 1638.690964 | -0.114368851 | 0.066550749 | -1.718520877 | 0.08570165 | 0.490505187 |
| Swap70 | 596.138351 | -0.114401261 | 0.073779835 | -1.550576258 | 0.121003266 | 0.555002664 |
| Lrch3 | 1714.790976 | -0.114429176 | 0.069090236 | -1.656227907 | 0.097675674 | 0.518456434 |
| Tmem181a | 2008.952096 | -0.114449134 | 0.05386982 | -2.124550133 | 0.033624169 | 0.335456507 |
| Ano6 | 1254.364598 | -0.114470302 | 0.057743321 | -1.982399012 | 0.0474346 | 0.391951501 |
| Wls | 1761.751678 | -0.114507175 | 0.069193324 | -1.654887621 | 0.097947296 | 0.518695741 |
| Tdrd5 | 68.67454168 | -0.114536656 | 0.094354046 | -1.213902971 | 0.224784778 | 0.676449905 |
| Ptpn13 | 763.3909734 | -0.114553202 | 0.075289901 | -1.521494923 | 0.128135684 | 0.566072313 |
| Fam98a | 1455.342868 | -0.114613362 | 0.062224643 | -1.841928785 | 0.065485568 | 0.445445383 |
| Lrwd1 | 479.747393 | -0.114623295 | 0.081045741 | -1.414303747 | 0.157272737 | 0.607878516 |
| Tnfrsf19 | 2793.089312 | -0.114648986 | 0.051272343 | -2.236078559 | 0.025346626 | 0.298582503 |
| Map3k8 | 55.64481751 | -0.114661734 | 0.094683431 | -1.211001042 | 0.225895005 | 0.678248958 |
| Psd4 | 75.42475833 | -0.114747964 | 0.094683529 | -1.211910507 | 0.22554664 | 0.67775972 |
| Phf16 | 303.4101277 | -0.114766884 | 0.079846239 | -1.437348652 | 0.150618951 | 0.598511432 |
| C130071C03Rik | 84.37537473 | -0.114802356 | 0.095006508 | -1.20836307 | 0.226907638 | 0.679880436 |
| Arhgef25 | 7295.418901 | -0.114823028 | 0.069899077 | -1.642697349 | 0.100445568 | 0.5211685 |
| 1810012K16Rik | 6.28149351 | -0.114940486 | 0.05461072 | -2.10472386 | 0.035315352 | NA |
| Mtm1 | 233.977424 | -0.11496418 | 0.089019688 | -1.291446678 | 0.196548834 | 0.652437924 |
| Zfp26 | 881.5879932 | -0.114970273 | 0.071370028 | -1.610904125 | 0.107200621 | 0.532011007 |
| Esyt1 | 711.3280434 | -0.114974864 | 0.075632489 | -1.52017823 | 0.128466187 | 0.566659164 |
| Mmp25 | 12.11157119 | -0.115029031 | 0.067921911 | -1.693548207 | 0.090351167 | NA |
| Scpep1 | 710.6691955 | -0.115036545 | 0.074038677 | -1.553735833 | 0.120247442 | 0.554014004 |
| ORF63 | 18.62215235 | -0.115071945 | 0.075300163 | -1.528176567 | 0.126468705 | NA |
| Ttc9b | 2568.618704 | -0.115100243 | 0.063144211 | -1.822815438 | 0.068331349 | 0.45117326 |
| Trim56 | 161.6113384 | -0.115116629 | 0.093526371 | -1.230846736 | 0.218380192 | 0.671344538 |
| Sox21 | 331.4291973 | -0.115137726 | 0.09157869 | -1.257254565 | 0.208661471 | 0.663532513 |
| Erlin1 | 1188.737533 | -0.115145616 | 0.068365236 | -1.684271456 | 0.092129219 | 0.505579538 |
| 4930479D17Rik | 64.00689176 | -0.115187111 | 0.095067916 | -1.211629698 | 0.225654161 | 0.677942574 |
| Dzip1 | 2983.436817 | -0.11521455 | 0.070041912 | -1.644937237 | 0.099982755 | 0.521115714 |
| Psg23 | 41.42622043 | -0.115277384 | 0.093086706 | -1.238387187 | 0.215572529 | 0.669812326 |
| Lrrfip1 | 2215.43964 | -0.115284695 | 0.049766957 | -2.316490733 | 0.02053149 | 0.269900834 |
| Ccnb2 | 35.40722651 | -0.115364113 | 0.089242078 | -1.292709835 | 0.196111435 | 0.651730657 |
| Pram1 | 17.99966471 | -0.115382124 | 0.075737239 | -1.523453 | 0.127645411 | NA |
| 6720483E21Rik | 20.83510442 | -0.115411326 | 0.07399413 | -1.559736248 | 0.118822222 | NA |
| P2rx7 | 521.1487907 | -0.115524399 | 0.066740333 | -1.730953288 | 0.083460096 | 0.48610319 |
| 1700003F12Rik | 3.910003615 | -0.115568536 | 0.042638944 | -2.710398632 | 0.006720239 | NA |
| Fbln2 | 321.7883772 | -0.115588252 | 0.092235496 | -1.253186212 | 0.210137946 | 0.664608511 |
| Focad | 1670.183517 | -0.115620802 | 0.057553791 | -2.0089172 | 0.044545915 | 0.380589751 |
| Inmt | 61.00851825 | -0.11570219 | 0.09222598 | -1.254550939 | 0.209641823 | 0.664226905 |
| Aplf | 282.6614399 | -0.11572743 | 0.079452637 | -1.456558692 | 0.145238237 | 0.591055907 |
| Pif1 | 9.454403623 | -0.115770382 | 0.059256078 | -1.953730069 | 0.050733148 | NA |
| Tpx2 | 143.7183305 | -0.115815697 | 0.088522409 | -1.308320662 | 0.19076457 | 0.646232776 |
| Npc1l1 | 101.9992682 | -0.11592048 | 0.095041682 | -1.219680428 | 0.222586044 | 0.67374894 |
| Mospd2 | 367.9263573 | -0.116072697 | 0.072567076 | -1.599522867 | 0.109704472 | 0.535311472 |
| Pbk | 12.14453337 | -0.116272209 | 0.070149819 | -1.657484091 | 0.097421643 | NA |
| Lyz2 | 401.8701488 | -0.116318097 | 0.084565082 | -1.375486118 | 0.168980787 | 0.619378697 |
| Htr1d | 168.7756339 | -0.116467622 | 0.091650511 | -1.270779845 | 0.203806981 | 0.657622144 |
| Tsen54 | 277.0314471 | -0.116488519 | 0.090782778 | -1.283156577 | 0.199437225 | 0.65377551 |
| Dennd4b | 2585.307737 | -0.116496621 | 0.063901312 | -1.823070882 | 0.068292656 | 0.451155091 |
| Akip1 | 96.5449448 | -0.116525778 | 0.093477175 | -1.246569317 | 0.212555458 | 0.666270994 |
| Notch2 | 868.8237973 | -0.116527395 | 0.075479472 | -1.543828971 | 0.122629771 | 0.557245639 |
| Strada | 1074.319836 | -0.11653366 | 0.05699914 | -2.044481012 | 0.040906051 | 0.365437455 |
| Rnpepl1 | 1378.572454 | -0.116646749 | 0.071012144 | -1.642631011 | 0.100459301 | 0.5211685 |
| Lxn | 1349.490411 | -0.116657334 | 0.062802245 | -1.857534439 | 0.063235156 | 0.440342223 |
| Gm15760 | 12.39661844 | -0.116669836 | 0.070590843 | -1.652761624 | 0.098379387 | NA |
| Mir3060 | 16.29616498 | -0.116685762 | 0.079317274 | -1.471126731 | 0.141256847 | NA |
| Abca8b | 1072.347326 | -0.11674994 | 0.071271647 | -1.638097966 | 0.101401259 | 0.522606076 |
| Gm5077 | 7.921354853 | -0.116788683 | 0.061016275 | -1.914057892 | 0.055612759 | NA |
| Plxnb2 | 2899.084593 | -0.116824775 | 0.061740155 | -1.892200872 | 0.058464222 | 0.423355579 |
| Hlx | 25.60525893 | -0.116839891 | 0.086000161 | -1.358600833 | 0.174273111 | 0.627072384 |
| Trim39 | 1191.768146 | -0.116851263 | 0.081435232 | -1.434898146 | 0.151316124 | 0.598905575 |
| Plscr4 | 145.3188051 | -0.116857986 | 0.092081019 | -1.269077904 | 0.20441328 | 0.657945338 |
| C130074G19Rik | 3012.863975 | -0.11700246 | 0.058448245 | -2.001813044 | 0.045304843 | 0.383869599 |
| Zfand4 | 165.314276 | -0.117015456 | 0.086341547 | -1.355262447 | 0.175333955 | 0.628089033 |
| Phip | 1901.960678 | -0.117054687 | 0.075632798 | -1.547670982 | 0.121701532 | 0.555997647 |
| Ido2 | 82.65260953 | -0.117063039 | 0.095010718 | -1.232103512 | 0.21791042 | 0.670929686 |
| Lilrb4 | 20.31163142 | -0.117255691 | 0.056560889 | -2.073087817 | 0.038164108 | NA |
| Ssbp1 | 505.1876082 | -0.117260081 | 0.071476676 | -1.640536296 | 0.100893708 | 0.521542547 |
| Rfc5 | 353.7908623 | -0.117289138 | 0.073476763 | -1.596275237 | 0.110427356 | 0.536657001 |
| Tmco6 | 359.8733908 | -0.117312954 | 0.075668223 | -1.550359566 | 0.121055238 | 0.555065778 |
| Stat6 | 783.5728508 | -0.117325761 | 0.061247479 | -1.915601461 | 0.055415843 | 0.413057568 |
| Atp2c2 | 50.60192321 | -0.117333812 | 0.088515954 | -1.325566828 | 0.18498322 | 0.637780733 |
| Gtpbp2 | 2882.20412 | -0.117354657 | 0.072028955 | -1.629270569 | 0.103255754 | 0.522975271 |
| Asic3 | 12.07581403 | -0.117363325 | 0.064142454 | -1.829729258 | 0.067290434 | NA |
| Olfr464 | 18.3386413 | -0.117413161 | 0.077702511 | -1.511060068 | 0.130773147 | NA |
| Mapk12 | 240.2295634 | -0.117468117 | 0.090743273 | -1.294510472 | 0.195489155 | 0.651003086 |
| Zfp607 | 41.00724343 | -0.117579022 | 0.092579849 | -1.270028242 | 0.20407457 | 0.65778952 |
| 4732471J01Rik | 160.2173552 | -0.117652936 | 0.091290001 | -1.288782286 | 0.197473786 | 0.653417817 |
| Frk | 15.55729792 | -0.117683809 | 0.074772711 | -1.573887154 | 0.11551354 | NA |
| Raver1 | 631.1233367 | -0.117723976 | 0.058928609 | -1.997738912 | 0.045744973 | 0.385435894 |
| A230072C01Rik | 339.5441913 | -0.117831004 | 0.073759085 | -1.597511735 | 0.110151684 | 0.535496439 |
| Gm14322 | 618.2081494 | -0.117832258 | 0.067998237 | -1.732872263 | 0.083118375 | 0.485085383 |
| Tmem191c | 2802.905502 | -0.117928382 | 0.072719212 | -1.621694993 | 0.104868671 | 0.525646073 |
| Abcc10 | 488.1744212 | -0.118077024 | 0.089659046 | -1.316956066 | 0.187853351 | 0.641781416 |
| Nfat5 | 3803.664495 | -0.118131355 | 0.066057761 | -1.788303943 | 0.073726984 | 0.465836525 |
| Fcho2 | 1069.710424 | -0.118334281 | 0.068610633 | -1.724722169 | 0.084577554 | 0.488499377 |
| Ankrd10 | 1426.40361 | -0.118360084 | 0.073496048 | -1.610427857 | 0.107304483 | 0.532178904 |
| Prpf38b | 1688.819877 | -0.118538392 | 0.064550382 | -1.83637011 | 0.066302937 | 0.446279967 |
| Prr22 | 35.59219749 | -0.118551528 | 0.087794371 | -1.350331769 | 0.176909586 | 0.629241039 |
| Fbxo4 | 213.3445042 | -0.118600191 | 0.085719509 | -1.38358459 | 0.166485689 | 0.616902656 |
| 4931406C07Rik | 1114.244132 | -0.118662178 | 0.063236476 | -1.87648309 | 0.060588975 | 0.431188114 |
| Snora43 | 11.65684916 | -0.118742816 | 0.068288133 | -1.738849935 | 0.082061162 | NA |
| Snord100 | 13.99743965 | -0.118758372 | 0.070479967 | -1.684994703 | 0.091989593 | NA |
| AU022793 | 7.69148397 | -0.118792465 | 0.056352926 | -2.108008806 | 0.035030228 | NA |
| Gas1 | 213.5148891 | -0.118897689 | 0.093042931 | -1.277879876 | 0.201291785 | 0.654716629 |
| Irf3 | 736.2685615 | -0.118937138 | 0.083961867 | -1.416561384 | 0.156611205 | 0.607279636 |
| Srsf2 | 3560.401533 | -0.118940644 | 0.066610634 | -1.785610433 | 0.074162356 | 0.467117664 |
| Snhg1 | 270.7960553 | -0.119049616 | 0.086810886 | -1.371367361 | 0.170260467 | 0.620316915 |
| Cox8a | 7784.657486 | -0.119133918 | 0.068000383 | -1.751959525 | 0.079780768 | 0.478486967 |
| Thsd1 | 192.0053297 | -0.119176689 | 0.085194573 | -1.39887654 | 0.161850009 | 0.613952162 |
| Airn | 16.28575355 | -0.119249801 | 0.078098848 | -1.526908588 | 0.126783744 | NA |
| Pabpc4l | 10.41113065 | -0.119262048 | 0.059486137 | -2.004871282 | 0.04497681 | NA |
| Hsf1 | 1654.797284 | -0.119388651 | 0.070368651 | -1.696617033 | 0.089769087 | 0.50087668 |
| Ppp1r13l | 166.3876679 | -0.119404997 | 0.092311871 | -1.293495572 | 0.195839715 | 0.651423791 |
| Gbp6 | 147.0475794 | -0.119422295 | 0.050026777 | -2.387167463 | NA | NA |
| Rhobtb1 | 249.0277662 | -0.119588919 | 0.089647223 | -1.333994691 | 0.182205596 | 0.635245711 |
| Gm1141 | 12.15427643 | -0.119620557 | 0.070896283 | -1.687261334 | 0.091553112 | NA |
| Plec | 8355.872823 | -0.11966508 | 0.067949347 | -1.761092407 | 0.078222764 | 0.475440315 |
| Rasl11b | 2450.804693 | -0.119677166 | 0.069469757 | -1.722723242 | 0.084938585 | 0.488499377 |
| Trpv6 | 56.584789 | -0.119700939 | 0.094794534 | -1.262740947 | 0.206682293 | 0.661001099 |
| Plekhg5 | 3689.655429 | -0.119706892 | 0.076168423 | -1.571607852 | 0.116041524 | 0.54588186 |
| Nxpe4 | 514.7222274 | -0.11975148 | 0.064253681 | -1.863729496 | 0.062359701 | 0.436873272 |
| Adcy4 | 180.5294153 | -0.119779708 | 0.094720896 | -1.264554209 | 0.206031177 | 0.660203749 |
| BC035044 | 29.66246364 | -0.119874311 | 0.088628264 | -1.352551724 | 0.176198884 | 0.629015726 |

| | | | | | | |
|---|---|---|---|---|---|---|
| Cacna1b | 3865.04407 | -0.085557818 | 0.065129789 | -1.313651078 | 0.188963651 | 0.40206026 |
| Atp9a | 16676.76021 | -0.097948922 | 0.074569274 | -1.313529254 | 0.18900467 | 0.402092318 |
| Tnrc18 | 3087.901567 | -0.111751949 | 0.085101992 | -1.313153143 | 0.18913135 | 0.40230658 |
| Stoml1 | 2060.60092 | 0.135855039 | 0.093042842 | 1.460134232 | 0.144253186 | 0.341699873 |
| Sypl | 1365.43255 | 0.135849065 | 0.065820912 | 2.063919499 | 0.03902535 | 0.147226386 |
| Emc7 | 2345.047201 | -0.077721244 | 0.059219195 | -1.312433309 | 0.189373974 | 0.402656835 |
| Apoh | 14.97679845 | 0.135816569 | 0.159468663 | 0.851681872 | 0.394390685 | 0.619996882 |
| Znhit6 | 499.0651278 | 0.135743936 | 0.076858202 | 1.766160694 | 0.077368893 | 0.229119578 |
| Tmem130 | 8035.237347 | -0.126646244 | 0.096539694 | -1.311856688 | 0.189568494 | 0.40290456 |
| Ermn | 2191.097111 | 0.135725789 | 0.129128193 | 1.051093383 | 0.293215703 | 0.523879624 |
| Gusb | 548.0881935 | 0.135696257 | 0.101391709 | 1.338336815 | 0.180786666 | 0.391270932 |
| Lrrc14b | 8.642557725 | 0.135688498 | 0.14634983 | 0.927151729 | 0.353847762 | 0.584552427 |
| Antxr1 | 782.4282343 | 0.097535626 | 0.074390122 | 1.311136791 | 0.189811553 | 0.403199918 |
| Tubgcp3 | 1577.693832 | 0.076413214 | 0.058292768 | 1.310852387 | 0.189907639 | 0.403348727 |
| Ncs1 | 14756.34968 | -0.080910095 | 0.06173254 | -1.310655531 | 0.189974169 | 0.403434727 |
| Oxsr1 | 2054.519657 | 0.087559878 | 0.066825734 | 1.310271845 | 0.190103888 | 0.403654878 |
| Ints2 | 802.8297114 | 0.135553645 | 0.107299276 | 1.26332302 | 0.206473117 | 0.42411434 |
| Id2 | 3466.397025 | 0.135528975 | 0.072472047 | 1.870086193 | 0.061471849 | 0.197564511 |
| Tcap | 119.6426229 | 0.135511762 | 0.153902293 | 0.880505154 | 0.378585714 | 0.607597359 |
| Bmp5 | 50.88284066 | -0.214228035 | 0.163618766 | -1.30931213 | 0.190428643 | 0.4040697 |
| Tapt1 | 2127.679978 | 0.135447573 | 0.087579673 | 1.546564043 | 0.121968406 | 0.308082569 |
| Ccdc51 | 206.1003606 | -0.123672822 | 0.094474575 | -1.309059313 | 0.19051426 | 0.404152589 |
| Hps3 | 617.1261928 | 0.135431363 | 0.080389381 | 1.684692211 | 0.09204797 | 0.255766056 |
| Tbc1d8b | 257.385404 | -0.164354109 | 0.125560569 | -1.308962764 | 0.190546964 | 0.404152589 |
| Lgmn | 4190.948676 | 0.080528371 | 0.061547783 | 1.308387842 | 0.190741794 | 0.404510452 |
| Ntm | 5099.340452 | -0.083027659 | 0.063465949 | -1.308223698 | 0.190797447 | 0.404573099 |
| Rngtt | 858.4208902 | -0.112664908 | 0.086161479 | -1.307601832 | 0.191008395 | 0.404854183 |
| Jakmip2 | 2276.774044 | 0.135418468 | 0.078161305 | 1.732551268 | 0.083175457 | 0.240567581 |
| Agbl4 | 328.9752428 | 0.135408223 | 0.085039479 | 1.592298357 | 0.111317682 | 0.29060761 |
| Cdc73 | 676.3146958 | -0.09884915 | 0.075609285 | -1.307367878 | 0.191087801 | 0.40491802 |
| Slc25a23 | 17586.02334 | 0.082417937 | 0.063041545 | 1.3073591 | 0.191090781 | 0.40491802 |
| Fstl1 | 1562.563221 | -0.112876819 | 0.08634555 | -1.307268512 | 0.191121534 | 0.404927807 |
| Zfp110 | 653.3694407 | 0.135341731 | 0.079664612 | 1.69889399 | 0.08933916 | 0.251402277 |
| Msl1 | 6471.004428 | -0.072188088 | 0.055237356 | -1.30687081 | 0.191256591 | 0.405103161 |
| Grb14 | 749.9004811 | -0.103092093 | 0.078908183 | -1.306481644 | 0.191388817 | 0.40532782 |
| Ky | 146.1822601 | -0.195279351 | 0.149506235 | -1.306161918 | 0.1914975 | 0.405408899 |
| Vmn2r87 | 27.01138101 | 0.135315572 | 0.167989675 | 0.805499338 | 0.420531588 | 0.643209914 |
| Tspan12 | 450.6292005 | -0.135085611 | 0.103423379 | -1.306141914 | 0.191504302 | 0.405408899 |
| Zfp37 | 729.4334311 | -0.104673873 | 0.080155513 | -1.305884895 | 0.191591704 | 0.405535773 |
| Tbc1d5 | 1853.950946 | 0.135300785 | 0.057037671 | 2.372130237 | 0.017685855 | 0.08690918 |
| Bpifc | 9.204371208 | 0.135297007 | 0.15058815 | 0.898457196 | 0.368941855 | 0.598699739 |
| Nap1l1 | 9465.601092 | -0.069659588 | 0.053376498 | -1.30506103 | 0.191872067 | 0.405907402 |
| Porcn | 2063.71414 | 0.135296561 | 0.076292113 | 1.773401684 | 0.07616217 | 0.227064973 |
| Cacna1i | 5033.471203 | -0.083145549 | 0.063728303 | -1.304687946 | 0.191999128 | 0.406065313 |
| Dph3 | 1157.944532 | -0.074819093 | 0.057344577 | -1.304728309 | 0.191985378 | 0.406065313 |
| Atp5s | 451.7756526 | -0.12810893 | 0.09819807 | -1.304597236 | 0.19203003 | 0.40607524 |
| Blcap | 3417.506164 | -0.080511059 | 0.061728327 | -1.304280596 | 0.192137929 | 0.406161583 |
| Mocs1 | 327.7152867 | 0.135201017 | 0.090969933 | 1.486216509 | 0.137221862 | 0.330296028 |
| Prpf38a | 639.8827561 | -0.097414343 | 0.074690124 | -1.304246637 | 0.192149503 | 0.406161583 |
| Cacfd1 | 1335.107569 | 0.13518056 | 0.060536811 | 2.233030765 | 0.025546916 | 0.110781951 |
| Ccdc55 | 620.2060811 | 0.083291594 | 0.063871032 | 1.304059001 | 0.192213466 | 0.406185958 |
| Shd | 745.5049687 | 0.135173764 | 0.094847275 | 1.425172877 | 0.154107235 | 0.35558265 |
| Sucla2 | 5340.981539 | -0.075243867 | 0.057719154 | -1.303620396 | 0.192363043 | 0.40639119 |
| Adam10 | 3317.692324 | -0.08437057 | 0.064746482 | -1.303091178 | 0.192543635 | 0.406446041 |
| Afap1 | 792.0776995 | -0.109039208 | 0.083694407 | -1.302825502 | 0.192634342 | 0.406446041 |
| B3galt1 | 1751.761484 | -0.115393646 | 0.088575217 | -1.302775753 | 0.192651331 | 0.406446041 |
| Cr2 | 12.8264472 | -0.213480543 | 0.163797304 | -1.303321468 | 0.192465035 | 0.406446041 |
| Dnmbp | 592.9266138 | 0.098756201 | 0.075783548 | 1.303135096 | 0.192528644 | 0.406446041 |
| Rhob | 8018.836568 | 0.135126611 | 0.070765167 | 1.909507406 | 0.056196667 | 0.187088776 |
| Fam131c | 94.12592257 | -0.188968487 | 0.144990423 | -1.303317023 | 0.192466552 | 0.406446041 |
| Fkbp15 | 1623.275172 | 0.091037932 | 0.069873566 | 1.302895182 | 0.192610549 | 0.406446041 |
| Hs6st3 | 431.527433 | 0.1350973 | 0.096382781 | 1.401674639 | 0.161012427 | 0.365745564 |
| Limk2 | 4605.693885 | 0.085382953 | 0.065532954 | 1.302901034 | 0.192608551 | 0.406446041 |
| Tmem180 | 961.2820099 | 0.135053364 | 0.06798697 | 1.986459533 | 0.046982322 | 0.167136739 |
| Gp9 | 8.848061451 | -0.180180833 | 0.138408278 | -1.301806766 | 0.19298245 | 0.407033772 |
| Iqcg | 154.975466 | -0.188553202 | 0.144856992 | -1.301650684 | 0.193035825 | 0.407090932 |
| Taf2 | 1495.806924 | 0.085717065 | 0.065867778 | 1.301350491 | 0.193138512 | 0.407252056 |
| Clspn | 132.096537 | -0.192607051 | 0.148087084 | -1.300633693 | 0.193383868 | 0.407700176 |
| Nxn | 397.5199592 | -0.135967565 | 0.104544121 | -1.300575905 | 0.193403659 | 0.407700176 |
| Usp12 | 3336.914348 | 0.098842553 | 0.076026026 | 1.300114697 | 0.193561661 | 0.407977742 |
| Kdm5b | 3030.021834 | 0.078681034 | 0.060529325 | 1.299882898 | 0.193641107 | 0.407978694 |
| Rcbtb1 | 1814.298909 | 0.092387666 | 0.071072977 | 1.299898646 | 0.193635709 | 0.407978694 |
| Dusp5 | 165.9994193 | 0.135039658 | 0.157448441 | 0.85767542 | 0.391071716 | 0.617851042 |
| B430319G15Rik | 28.02414583 | -0.217945626 | 0.167729748 | -1.299385644 | 0.193811616 | 0.408226914 |
| Ilf3 | 3439.728647 | 0.1349301 | 0.07965155 | 1.694004685 | 0.090264393 | 0.2530408 |
| Maf | 490.6469589 | 0.134913201 | 0.09067481 | 1.487879613 | 0.136782645 | 0.329602254 |
| Polr2l | 934.1371711 | -0.104719563 | 0.080626167 | -1.298828495 | 0.194002793 | 0.408518521 |
| Ece2 | 1158.70352 | 0.134852348 | 0.12864854 | 1.048222912 | 0.294535916 | 0.525753397 |
| St7l | 796.9060952 | -0.108187131 | 0.083323744 | -1.298394987 | 0.194151641 | 0.408720859 |
| Ttc17 | 1792.053849 | 0.093986649 | 0.072398836 | 1.298179007 | 0.19422583 | 0.408821493 |
| Arap2 | 4196.060432 | -0.110667063 | 0.085286404 | -1.29759326 | 0.19442714 | 0.408911918 |
| Asxl2 | 1273.822881 | -0.108473499 | 0.083581406 | -1.297818555 | 0.194349692 | 0.408911918 |
| Cdh22 | 1699.455756 | -0.123988721 | 0.095544641 | -1.297704614 | 0.194388858 | 0.408911918 |
| Zfp783 | 247.6483314 | 0.134817471 | 0.147533049 | 0.913812006 | 0.360815637 | 0.591584918 |
| Ifitm1 | 67.59696309 | -0.203179558 | 0.156545126 | -1.297897695 | 0.194322492 | 0.408911918 |
| Ube2d2a | 5396.318419 | -0.082513862 | 0.06358908 | -1.297610568 | 0.194421189 | 0.408911918 |
| Dnajc7 | 6010.056568 | 0.098723095 | 0.07611547 | 1.297017471 | 0.194625175 | 0.409106291 |
| Itfg3 | 693.6113591 | 0.134801563 | 0.080051588 | 1.683933656 | 0.092194491 | 0.256088463 |
| Pggt1b | 280.8867488 | -0.12251653 | 0.094456791 | -1.297064286 | 0.194609069 | 0.409106291 |
| Synm | 1851.600765 | 0.089034732 | 0.068641814 | 1.297091768 | 0.194599614 | 0.409106291 |
| Mocs3 | 219.2515407 | -0.144037339 | 0.111059376 | -1.296939931 | 0.194651856 | 0.409106871 |
| Ikbip | 352.9410744 | -0.10841349 | 0.083599654 | -1.296817447 | 0.194694006 | 0.409126645 |
| Psmb7 | 4245.350658 | -0.087118326 | 0.067181579 | -1.296759136 | 0.194714075 | 0.409126645 |
| Cep192 | 381.4003644 | 0.134796892 | 0.11115772 | 1.212663341 | 0.225258561 | 0.447054713 |
| Gm3604 | 36.40572569 | 0.134794924 | 0.164888338 | 0.817492163 | 0.413647226 | 0.637388731 |
| Higd2a | 1985.032624 | -0.089682602 | 0.069205811 | -1.295882526 | 0.19501596 | 0.409594319 |
| Wdr4 | 839.2100333 | -0.096508108 | 0.074482941 | -1.29570754 | 0.195076263 | 0.409628147 |
| Zfp87 | 597.1264293 | -0.09490578 | 0.073247721 | -1.295682357 | 0.195084942 | 0.409628147 |
| Fads2 | 3501.915819 | -0.07563383 | 0.058378265 | -1.295582015 | 0.195119529 | 0.409645254 |
| Helb | 207.5760586 | 0.134792746 | 0.107634378 | 1.252320571 | 0.210453076 | 0.428708909 |
| Klhdc7a | 705.7717208 | -0.155929196 | 0.120415065 | -1.294930963 | 0.195344046 | 0.409894454 |
| Zfp420 | 499.0216027 | 0.134614754 | 0.086555879 | 1.555235249 | 0.119890052 | 0.304636898 |
| Eed | 450.5232675 | 0.134574595 | 0.089615694 | 1.501685567 | 0.133178333 | 0.324087497 |
| Mtfr1 | 278.6830154 | 0.134542907 | 0.108467845 | 1.240394399 | 0.214829552 | 0.43465253 |
| Pcdhgc5 | 4531.126476 | -0.122934663 | 0.094948576 | -1.29474993 | 0.19540651 | 0.409914495 |
| Nudt4 | 5753.555696 | 0.134523403 | 0.075075622 | 1.791838677 | 0.073158811 | 0.221650884 |
| Gxylt2 | 25.41262791 | -0.217490247 | 0.168109592 | -1.29374085 | 0.19575495 | 0.410534272 |
| Rpl35 | 3466.266528 | -0.096046598 | 0.074277722 | -1.293074088 | 0.195985436 | 0.410962017 |
| Tfip11 | 2126.658619 | 0.098477116 | 0.076163679 | 1.292966895 | 0.196022509 | 0.410984135 |
| Aldh1a2 | 355.7754064 | -0.202655323 | 0.156764741 | -1.29273535 | 0.196102607 | 0.411040827 |
| Sugp2 | 3683.208703 | 0.134519115 | 0.06188212 | 2.173796165 | 0.029720447 | 0.122568166 |
| Prdx5 | 5059.420878 | -0.080202781 | 0.062052736 | -1.292493882 | 0.196186163 | 0.411160343 |
| Gm19345 | 43.84532391 | -0.202971539 | 0.157056264 | -1.292349209 | 0.196236237 | 0.411209665 |
| Mapre1 | 3769.6599 | -0.084571326 | 0.065444733 | -1.29225564 | 0.196268629 | 0.411221925 |
| Fkbp8 | 11013.73812 | -0.089289457 | 0.069102951 | -1.292122188 | 0.196314833 | 0.41124651 |
| Setd6 | 410.2096246 | 0.134518999 | 0.104379318 | 1.288751459 | 0.197484506 | 0.412820418 |
| Dapl1 | 11.42358066 | -0.198415594 | 0.153607502 | -1.291705106 | 0.196459289 | 0.411454477 |
| Tet2 | 1530.998853 | 0.134516705 | 0.096171996 | 1.398709714 | 0.16190005 | 0.366938024 |
| Gtf2f1 | 2311.363784 | 0.134456263 | 0.0816542 | 1.646654588 | 0.099629063 | 0.269539498 |
| Rnf126 | 1931.667215 | 0.134453649 | 0.064885173 | 2.072178312 | 0.038248815 | 0.145180584 |
| Trim6 | 8.56547513 | 0.134421179 | 0.159627654 | 0.84209205 | 0.399736431 | 0.624787275 |
| Nfkb1 | 716.9560893 | 0.134400113 | 0.080430958 | 1.670999785 | 0.09472172 | 0.260400698 |
| Ppp1r8 | 1308.095862 | -0.097877789 | 0.075827436 | -1.290796492 | 0.196774255 | 0.411780159 |
| E130311K13Rik | 282.5187032 | -0.148282433 | 0.11490703 | -1.290455713 | 0.196892479 | 0.411971921 |
| C330005M16Rik | 69.68397893 | 0.134363448 | 0.156454484 | 0.858802153 | 0.390449679 | 0.617448986 |
| Fyn | 4726.147293 | -0.059523954 | 0.046146034 | -1.289904011 | 0.197083988 | 0.412261281 |
| Ssh2 | 1948.613432 | -0.113956487 | 0.088368658 | -1.289557751 | 0.197204253 | 0.412457167 |
| Crip1 | 112.0889711 | -0.171616633 | 0.133103052 | -1.289351599 | 0.19727588 | 0.412495612 |
| D130043K22Rik | 1687.534086 | 0.134314272 | 0.07290444 | 1.842333228 | 0.065426422 | 0.205663824 |
| 2900055J20Rik | 164.6071537 | -0.15861663 | 0.123060462 | -1.288932513 | 0.197421549 | 0.4127445 |
| Amph | 9332.729008 | 0.134261476 | 0.068743149 | 1.953088823 | 0.050809071 | 0.175582561 |
| Cntn4 | 2070.344764 | -0.10217319 | 0.079286561 | -1.288657103 | 0.197517322 | 0.412833318 |
| Gtf2h1 | 1673.021898 | 0.134257438 | 0.076841711 | 1.747194796 | 0.080603554 | 0.235636126 |
| Ipo11 | 2272.361253 | -0.107871645 | 0.083813544 | -1.287043106 | 0.198079264 | 0.41386462 |
| Tmed8 | 2812.514556 | -0.079366542 | 0.061667396 | -1.287009792 | 0.198090875 | 0.41386462 |
| Qk | 9090.366197 | -0.112469478 | 0.087440123 | -1.286245644 | 0.198357346 | 0.414365474 |
| Rnf144b | 1281.670203 | 0.134215975 | 0.07277155 | 1.84434679 | 0.065132617 | 0.205198203 |
| Ccdc69 | 11.51478732 | 0.134188344 | 0.157926657 | 0.849687739 | 0.395498717 | 0.620711462 |
| Iscu | 2074.212315 | -0.089436219 | 0.069546778 | -1.285986512 | 0.198447769 | 0.414386757 |
| Unk | 917.6351036 | 0.134163005 | 0.080841102 | 1.659589021 | 0.096997159 | 0.264311287 |
| Ttc7 | 467.9756467 | -0.112536031 | 0.087531195 | -1.28566771 | 0.198559055 | 0.41450741 |
| Exd2 | 3314.991285 | 0.134143404 | 0.120428171 | 1.113887242 | 0.265327563 | 0.492805714 |
| Fbxl18 | 613.091389 | 0.134106923 | 0.111000221 | 1.208168073 | 0.22698262 | 0.449440091 |
| Morc3 | 872.4706819 | 0.084124134 | 0.065477374 | 1.28478174 | 0.198868565 | 0.414985797 |
| Mtx3 | 2105.61415 | 0.134088473 | 0.092509845 | 1.449450849 | 0.147211717 | 0.346377881 |
| Rgs5 | 1547.971665 | -0.140044318 | 0.109021951 | -1.284551567 | 0.198949033 | 0.415041916 |
| Mettl16 | 879.8547229 | -0.109620721 | 0.085352507 | -1.284329244 | 0.199026779 | 0.415148211 |
| Cog7 | 2199.239511 | 0.069281782 | 0.053968473 | 1.283745454 | 0.199231035 | 0.415518327 |
| Stk33 | 43.39287773 | -0.215726171 | 0.168087477 | -1.283416081 | 0.199346343 | 0.415702858 |
| E130006D01Rik | 19.09981269 | -0.215445406 | 0.167902931 | -1.283154525 | 0.199437944 | 0.41583791 |
| Abca8a | 447.9011592 | 0.134078213 | 0.154811242 | 0.866075431 | 0.386448798 | 0.613972588 |
| Tsc2 | 5349.248144 | 0.134022019 | 0.05515311 | 2.429999313 | 0.015098851 | 0.078241339 |
| Sh3pxd2b | 1077.548197 | -0.109214287 | 0.085156744 | -1.282508951 | 0.199664167 | 0.416141593 |
| Prosc | 2558.399287 | 0.134012246 | 0.077759074 | 1.723429041 | 0.084810967 | 0.243122348 |
| Glb1 | 971.8747349 | 0.099501057 | 0.077600957 | 1.282214301 | 0.199767481 | 0.416244936 |
| Aifm1 | 876.0443065 | -0.077196622 | 0.060219316 | -1.281924583 | 0.199869104 | 0.416400684 |
| Enpp5 | 9272.519054 | 0.067606666 | 0.052751562 | 1.281605014 | 0.19998124 | 0.416578292 |
| Foxg1 | 2800.768316 | -0.107087226 | 0.083583241 | -1.281204526 | 0.200121837 | 0.416815128 |
| Gtf2ird1 | 1126.404695 | 0.13396454 | 0.094117888 | 1.423369593 | 0.154629042 | 0.356203149 |
| Atad1 | 4031.429452 | -0.074869021 | 0.058454089 | -1.280817501 | 0.200257775 | 0.416986154 |
| Glg1 | 5521.770853 | -0.102845885 | 0.080316805 | -1.280502694 | 0.200368398 | 0.417104388 |
| Otop2 | 58.20368632 | -0.204551563 | 0.159735184 | -1.280566736 | 0.20034589 | 0.417104388 |
| Kras | 3234.429241 | -0.11733977 | 0.091654766 | -1.280236419 | 0.200462001 | 0.417243182 |
| Leprel1 | 93.75609248 | 0.133951951 | 0.12353972 | 1.084282457 | 0.278239581 | 0.507283495 |
| Cybasc3 | 504.6778091 | 0.133931874 | 0.111293367 | 1.203412906 | 0.228816576 | 0.451686079 |
| Pbxip1 | 2533.337237 | 0.133857129 | 0.092600149 | 1.445539017 | 0.148306542 | 0.348065936 |
| Ccdc24 | 40.61955369 | 0.133742099 | 0.168121551 | 0.795508359 | 0.42631785 | 0.648329765 |
| Cdc23 | 1918.103671 | 0.079867868 | 0.062437342 | 1.27916829 | 0.2008378 | 0.417688708 |
| Plekhd1 | 153.0634034 | 0.133579546 | 0.126563186 | 1.055437609 | 0.29122523 | 0.521539907 |
| Dnase2a | 153.3761961 | 0.133574201 | 0.137382443 | 0.972279996 | 0.330911273 | 0.566143084 |
| Eef1a2 | 34140.58874 | 0.089059679 | 0.06964435 | 1.278778238 | 0.200975159 | 0.417806134 |
| Stoml3 | 20.31063833 | -0.178587542 | 0.139652688 | -1.27879774 | 0.20096829 | 0.417806134 |
| Vsig10 | 87.83310894 | 0.133484428 | 0.129441718 | 1.031231901 | 0.302432084 | 0.534369109 |
| Sec14l2 | 1913.44112 | 0.133470488 | 0.098483397 | 1.355258776 | 0.175335124 | 0.384249281 |
| Ccdc25 | 1653.855243 | 0.072847449 | 0.056991177 | 1.27822328 | 0.20117071 | 0.418011556 |
| Cpsf3 | 1763.54589 | 0.085101742 | 0.066575803 | 1.278268359 | 0.20115482 | 0.418011556 |
| Sptb | 9426.575295 | -0.091293915 | 0.071444313 | -1.277833195 | 0.201308247 | 0.418218019 |
| Ankrd9 | 327.5863409 | 0.133416553 | 0.107636007 | 1.239516003 | 0.215154466 | 0.434969115 |
| Ube2cbp | 73.61089701 | 0.133374218 | 0.151551325 | 0.880059731 | 0.378826952 | 0.607752012 |
| Rab39b | 1598.446589 | -0.111965255 | 0.087637013 | -1.277602354 | 0.20138967 | 0.418319339 |
| Arf1 | 15095.00588 | -0.076736423 | 0.060073147 | -1.277383105 | 0.201467026 | 0.41832352 |
| Tubg2 | 1955.438157 | 0.13333591 | 0.092624341 | 1.439534233 | 0.149999218 | 0.350370787 |
| Arhgap20 | 4233.195755 | -0.097718924 | 0.076549329 | -1.27654841 | 0.201761725 | 0.41876707 |
| Arpc1a | 5809.927702 | -0.092114265 | 0.072155494 | -1.276607779 | 0.201740754 | 0.41876707 |
| Parn | 1017.867542 | 0.133238188 | 0.064679518 | 2.059974975 | 0.039400933 | 0.148282259 |
| Ulk2 | 4997.595689 | -0.076275723 | 0.059795105 | -1.27561817 | 0.202090528 | 0.419337169 |
| C77370 | 1487.71064 | -0.122235719 | 0.095833808 | -1.275496842 | 0.202133441 | 0.41937005 |
| Arrdc3 | 1899.947523 | 0.075327392 | 0.059097159 | 1.274636415 | 0.202437962 | 0.419384114 |
| C030037D09Rik | 151.5551914 | -0.181272058 | 0.142190757 | -1.274851205 | 0.202361913 | 0.419384114 |
| Osbpl1a | 11853.92705 | 0.13323612 | 0.101709386 | 1.309968777 | 0.190206398 | 0.403706566 |
| Fxr1 | 2301.335885 | 0.092615726 | 0.072660363 | 1.27463892 | 0.202437075 | 0.419384114 |
| Gne | 1566.792001 | -0.07248684 | 0.056867861 | -1.274653887 | 0.202431775 | 0.419384114 |
| Mre11a | 368.2836338 | -0.123552668 | 0.09690961 | -1.274926888 | 0.202335121 | 0.419384114 |
| Pglyrp1 | 164.9321696 | -0.178398158 | 0.139956176 | -1.274671564 | 0.202425516 | 0.419384114 |
| Abcc10 | 438.592471 | 0.133228317 | 0.131428281 | 1.013695957 | 0.310727854 | 0.543817617 |
| Tmem198b | 380.6174795 | -0.164902179 | 0.129310361 | -1.275243358 | 0.202223119 | 0.419384114 |
| Usp27x | 661.2433355 | -0.118277192 | 0.092790968 | -1.27466276 | 0.202428633 | 0.419384114 |
| Zbtb38 | 1411.78459 | 0.133209557 | 0.108250924 | 1.230562771 | 0.218486436 | 0.438899111 |
| Gpr135 | 192.5791984 | -0.137371498 | 0.107811635 | -1.274180637 | 0.202599406 | 0.419662459 |
| Actr3 | 7471.781012 | -0.076258311 | 0.059859073 | -1.273964107 | 0.202676137 | 0.419709177 |
| Gm996 | 4853.639863 | 0.132823927 | 0.081998841 | 1.619826885 | 0.105269469 | 0.280007753 |
| Acn9 | 164.4981979 | -0.145937114 | 0.114564768 | -1.273839387 | 0.202720343 | 0.419744621 |
| Tcn2 | 868.1682202 | 0.132807141 | 0.102786461 | 1.292068431 | 0.196333447 | 0.41124651 |
| Plch1 | 496.244255 | -0.098423661 | 0.07728804 | -1.273465615 | 0.202852867 | 0.419922328 |
| Prkd1 | 451.9781371 | -0.112244895 | 0.088148391 | -1.273362952 | 0.202889278 | 0.419926058 |
| Ehbp1 | 2293.058515 | 0.098047567 | 0.077019895 | 1.273016109 | 0.203012326 | 0.420066199 |
| Grin1 | 19513.86456 | 0.132746074 | 0.063612526 | 2.08679143 | 0.036906992 | 0.141895021 |
| H13 | 2476.754898 | 0.070909506 | 0.055705176 | 1.27294285 | 0.203038322 | 0.420066199 |
| Stk32b | 143.7311265 | 0.13274166 | 0.163132014 | 0.813706993 | 0.415812827 | 0.638957051 |
| Cnksr1 | 10.07086803 | 0.132673402 | 0.154867723 | 0.856688526 | 0.391617046 | 0.618208305 |
| Lgi2 | 1298.765978 | -0.123069176 | 0.096723787 | -1.272377567 | 0.203239001 | 0.420270158 |
| Zxdc | 1113.026313 | 0.132666312 | 0.073926414 | 1.794572532 | 0.072721833 | 0.220672166 |
| Ambra1 | 3315.641175 | 0.069992812 | 0.055017953 | 1.272181305 | 0.203308708 | 0.420344959 |
| Slc16a4 | 95.11597242 | -0.21380562 | 0.168087687 | -1.271988591 | 0.203377173 | 0.420430409 |
| Uap1l1 | 527.777398 | 0.132658943 | 0.096348718 | 1.376862558 | 0.168554745 | 0.375497803 |
| Abtb2 | 574.5916698 | 0.132612553 | 0.067777061 | 1.956599353 | 0.050394591 | 0.174738704 |
| Dnajb3 | 24.78591691 | -0.213673033 | 0.168121366 | -1.270945142 | 0.203748165 | 0.420956615 |
| Mcu | 1099.339641 | -0.095702856 | 0.075295136 | -1.271036363 | 0.203715712 | 0.420956615 |
| Nbas | 2118.092973 | -0.085298373 | 0.067117006 | -1.270890624 | 0.203767563 | 0.420956615 |
| Supt16 | 2902.551955 | 0.081918121 | 0.064476081 | 1.270519553 | 0.203899623 | 0.421173277 |
| Gm19461 | 26.14918571 | 0.132504846 | 0.166086295 | 0.797807226 | 0.424982367 | 0.647312752 |
| Abca5 | 2050.874975 | 0.132493436 | 0.070091918 | 1.890281222 | 0.058720359 | 0.192512323 |
| Kifc3 | 1983.704375 | -0.078573337 | 0.061857861 | -1.270223952 | 0.204004868 | 0.421222205 |
| Cml1 | 288.073289 | -0.154952733 | 0.122013177 | -1.269967207 | 0.204096311 | 0.421255955 |
| Kirrel3 | 2011.439999 | -0.106681158 | 0.084004282 | -1.269949053 | 0.204102779 | 0.421255955 |
| Ttc4 | 1829.630066 | 0.132493186 | 0.069378311 | 1.909720547 | 0.056169204 | 0.18707774 |
| Hsf2bp | 12.81656627 | 0.132442267 | 0.166284782 | 0.796478577 | 0.425753923 | 0.647900885 |
| Cplx1 | 29668.84106 | 0.088273913 | 0.069535855 | 1.269473322 | 0.2042723 | 0.421381309 |
| Dcps | 448.422341 | -0.109212699 | 0.086024141 | -1.269558725 | 0.20424186 | 0.421381309 |
| Coq6 | 573.516756 | 0.13237303 | 0.075991674 | 1.741941229 | 0.081518729 | 0.237524069 |
| Adcyap1r1 | 9518.920507 | -0.099753813 | 0.078594244 | -1.269225438 | 0.204360671 | 0.421507486 |
| Invs | 538.7378875 | -0.132359116 | 0.139603295 | 0.948108826 | 0.343074055 | 0.575828939 |
| Mdm4 | 1509.62579 | 0.132809125 | 0.081852058 | 1.616929777 | 0.10589344 | 0.281226693 |
| Msantd1 | 73.83391776 | -0.176489791 | 0.139096804 | -1.268817078 | 0.204502745 | 0.421255955 |
| Gm10658 | 12.95065956 | -0.1323364 | 0.164225065 | -0.8058234 | 0.420344683 | 0.643196272 |

| | | | | | | |
|---|---|---|---|---|---|---|
| Mri1 | 225.6595815 | -0.119894348 | 0.09227451 | -1.299322505 | 0.193833274 | 0.649958942 |
| Lilra5 | 18.63702593 | -0.119953531 | 0.075795612 | -1.582592029 | 0.113514481 | NA |
| Naip6 | 10.87268139 | -0.119973797 | 0.064734543 | -1.853319596 | 0.063836564 | NA |
| Mphosph9 | 1078.799594 | -0.120028606 | 0.060547444 | -1.982389325 | 0.047435683 | 0.391951501 |
| Dus3l | 2510.923503 | -0.120075568 | 0.078447231 | -1.530654005 | 0.125854927 | 0.562787717 |
| Nptxr | 105.2976328 | -0.120112212 | 0.049950061 | -2.404645943 | 0.016188141 | 0.241922774 |
| Cst3 | 30741.28026 | -0.120139695 | 0.059780917 | -2.00966631 | 0.044466518 | 0.380589751 |
| Nr2f6 | 633.5407631 | -0.120146701 | 0.080099475 | -1.499968644 | 0.133622525 | 0.57588358 |
| Ofd1 | 438.2630768 | -0.120202347 | 0.074298627 | -1.617827301 | 0.105699821 | 0.527979239 |
| Mybpc2 | 9.587488879 | -0.120206882 | 0.064351649 | -1.86796894 | 0.0617664 | NA |
| Rasgef1c | 1674.1498 | -0.120238891 | 0.08436644 | -1.425198109 | 0.154099943 | 0.60340295 |
| Ppp1r14b | 569.7083115 | -0.120254051 | 0.08294271 | -1.449844725 | 0.147101824 | 0.594031723 |
| 2810029C07Rik | 412.8439442 | -0.120281271 | 0.086467667 | -1.391054886 | 0.164208784 | 0.614450488 |
| Speg | 6329.527422 | -0.120308207 | 0.070585843 | -1.704424036 | 0.088301897 | 0.496728133 |
| Bnip2 | 1334.38692 | -0.120331909 | 0.056982072 | -2.111750318 | 0.034707871 | 0.33922827 |
| Clec4a3 | 33.6772112 | -0.120337867 | 0.088212238 | -1.364185627 | 0.172509151 | 0.623815764 |
| Sprtn | 419.4713878 | -0.120340465 | 0.077165767 | -1.559505834 | 0.118876705 | 0.55116598 |
| Cyp2u1 | 218.6056096 | -0.120393822 | 0.084054783 | -1.432325652 | 0.152050645 | 0.600023818 |
| 5930438M14Rik | 20.46830921 | -0.120416576 | 0.057361388 | -2.099261896 | 0.03579382 | NA |
| 5430416O09Rik | 33.44224007 | -0.120458452 | 0.091182999 | -1.321062629 | 0.186480483 | 0.640826269 |
| Noxo1 | 68.95237976 | -0.120464875 | 0.094766103 | -1.271181061 | 0.203664243 | 0.657622144 |
| Pcdh15 | 687.7594912 | -0.120486164 | 0.073045293 | -1.649471987 | 0.099050978 | 0.520271327 |
| Stac3 | 66.10831339 | -0.120494373 | 0.086578657 | -1.391732997 | 0.164003266 | 0.614351408 |
| Cux2 | 3935.01911 | -0.120526864 | 0.081906609 | -1.471515737 | 0.141151695 | 0.586989325 |
| Klhdc4 | 829.8275319 | -0.120568198 | 0.077152578 | -1.562724165 | 0.118117488 | 0.549241002 |
| 2700081O15Rik | 1567.491693 | -0.12063883 | 0.062405641 | -1.933139834 | 0.053218974 | 0.407317206 |
| Trmt1 | 1980.034791 | -0.12076996 | 0.069266929 | -1.74354431 | 0.081238581 | 0.479898997 |
| Tnfsf8 | 3.625124784 | -0.120789172 | 0.043304684 | -2.789286558 | 0.00528243 | NA |
| Lilrb3 | 24.10685186 | -0.120850375 | 0.079272114 | -1.524500476 | 0.127383738 | 0.564894808 |
| Ppfibp2 | 502.9475231 | -0.120901614 | 0.08277932 | -1.460529204 | 0.144144687 | 0.589967032 |
| Fam135a | 1285.901534 | -0.120943326 | 0.063958064 | -1.89097853 | 0.05862721 | 0.424112977 |
| Tsga10 | 351.9140887 | -0.121094331 | 0.07742438 | -1.5640336 | 0.117809678 | 0.548846498 |
| Shmt1 | 76.58329287 | -0.121136068 | 0.094974511 | -1.27545872 | 0.202146926 | 0.655443359 |
| Nanos2 | 18.81761408 | -0.121144678 | 0.073499986 | -1.648227232 | 0.099306051 | NA |
| Fam167b | 10.99419472 | -0.121181218 | 0.062902004 | -1.926508085 | 0.054040963 | NA |
| H2-K2 | 53.63784969 | -0.121192322 | 0.093508062 | -1.296062815 | 0.194953845 | 0.650861767 |
| Kank3 | 762.9556119 | -0.121194313 | 0.073266478 | -1.654157764 | 0.098095462 | 0.519101888 |
| Rimbp2 | 4806.382694 | -0.12122277 | 0.070296142 | -1.724458364 | 0.084625129 | 0.488499377 |
| Ccl27a | 1774.101833 | -0.121234836 | 0.070093478 | -1.729616504 | 0.083698815 | 0.486580734 |
| Trmt2a | 1349.754728 | -0.121248561 | 0.075093122 | -1.614642704 | 0.106388092 | 0.529607068 |
| Phf21b | 482.1711151 | -0.121251479 | 0.090382158 | -1.3415422 | 0.179744478 | 0.633459544 |
| Fus | 8774.019803 | -0.121272886 | 0.049929868 | -2.428864553 | 0.01514619 | 0.233312354 |
| Zswim8 | 6371.789537 | -0.121346354 | 0.057560931 | -2.108137462 | 0.035019101 | 0.34025917 |
| Lyl1 | 69.29809234 | -0.121420602 | 0.094615414 | -1.283306775 | 0.19938462 | 0.65377551 |
| 9130024F11Rik | 358.1781239 | -0.121420867 | 0.093775726 | -1.294800604 | 0.195389023 | 0.651003086 |
| Slc50a1 | 1072.214212 | -0.121444293 | 0.073323671 | -1.656276777 | 0.097665781 | 0.518456434 |
| Tmem106a | 58.76656118 | -0.121559145 | 0.093007123 | -1.306987479 | 0.191216964 | 0.646682644 |
| Shroom4 | 198.2018885 | -0.12155926 | 0.090045043 | -1.349982811 | 0.177021497 | 0.629241039 |
| C230035I16Rik | 21.02954182 | -0.121583286 | 0.074697805 | -1.627668783 | 0.103595136 | NA |
| Cblb | 1042.034924 | -0.121637557 | 0.063589778 | -1.912847647 | 0.055767559 | 0.413057568 |
| Donson | 639.320635 | -0.121644167 | 0.079414989 | -1.531753252 | 0.125583336 | 0.562511112 |
| Ncf1 | 365.2881546 | -0.121644782 | 0.090967586 | -1.337232165 | 0.181146866 | 0.634469716 |
| E530001F21Rik | 58.4658805 | -0.121675763 | 0.092784368 | -1.311382142 | 0.189728689 | 0.644323531 |
| Ttc23 | 130.3513877 | -0.121688666 | 0.093178457 | -1.30597425 | 0.191561315 | 0.646997681 |
| Niacr1 | 9.219611668 | -0.121749002 | 0.045080666 | -2.700692177 | 0.006919535 | NA |
| Plch2 | 4545.689559 | -0.121776057 | 0.080894531 | -1.505368233 | 0.132229432 | 0.573705114 |
| Cdc7 | 620.492941 | -0.121787888 | 0.082451857 | -1.477078788 | 0.139654519 | 0.584617158 |
| Zdhhc8 | 5578.868938 | -0.121838732 | 0.067706418 | -1.799515261 | 0.071937212 | 0.460419542 |
| Rhbdf2 | 63.95651894 | -0.121841224 | 0.095061461 | -1.281709991 | 0.199944399 | 0.654380963 |
| Srrt | 4831.873867 | -0.121879323 | 0.062749699 | -1.942309294 | 0.052099669 | 0.406272098 |
| Plxnd1 | 6209.809292 | -0.121955724 | 0.076164353 | -1.601217886 | 0.109328669 | 0.535243338 |
| Helb | 227.6619354 | -0.121973237 | 0.08231909 | -1.481712643 | 0.138416771 | 0.583109099 |
| Snord96a | 57.4566841 | -0.122033851 | 0.095079491 | -1.283492889 | 0.199319449 | 0.65377551 |
| Zfp28 | 473.5198544 | -0.122131282 | 0.068740839 | -1.776691758 | 0.075618969 | 0.470626027 |
| Gm4951 | 23.85016393 | -0.122136414 | 0.051687794 | -2.36296435 | 0.018129413 | 0.256569934 |
| Hnrpdl | 6252.706877 | -0.122161204 | 0.061446 | -1.988106714 | 0.046799889 | 0.390923052 |
| C7 | 32.80899545 | -0.122208141 | 0.086498732 | -1.412831596 | 0.157705246 | 0.608141173 |
| Trex1 | 435.5875693 | -0.122256781 | 0.07434803 | -1.644384938 | 0.100096714 | 0.5211685 |
| Agbl3 | 192.6135161 | -0.122257979 | 0.084138945 | -1.453048626 | 0.146210253 | 0.592922988 |
| Aldh1l2 | 162.6740741 | -0.122262116 | 0.091223446 | -1.340248819 | 0.180164465 | 0.63422 |
| Slc35d2 | 27.07756562 | -0.122263765 | 0.082649695 | -1.479300863 | 0.139059922 | 0.583304774 |
| B430306N03Rik | 8.761835432 | -0.122281211 | 0.064968467 | -1.882162474 | 0.059813959 | NA |
| Notch1 | 1327.081381 | -0.122298046 | 0.065031706 | -1.880591066 | 0.060027568 | 0.429169608 |
| Fhl4 | 33.77905248 | -0.122344751 | 0.09054421 | -1.351215627 | 0.17662637 | 0.629083076 |
| Cdnf | 68.14441381 | -0.122376323 | 0.095088851 | -1.286968157 | 0.198105387 | 0.65354259 |
| Gm872 | 37.45083222 | -0.122386996 | 0.091734106 | -1.334149324 | 0.182154924 | 0.635245711 |
| Fam21 | 5475.245017 | -0.122405522 | 0.048863973 | -2.505025999 | 0.012244239 | 0.211851282 |
| Vcam1 | 1265.873557 | -0.122413477 | 0.063705681 | -1.921547257 | 0.054662751 | 0.412055587 |
| Igtp | 411.4049186 | -0.122454346 | 0.059946366 | -2.042731751 | NA | NA |
| Plod3 | 1221.492183 | -0.122546974 | 0.069856158 | -1.754275907 | 0.079383242 | 0.477013957 |
| Lrrc42 | 693.7852614 | -0.122565896 | 0.066833675 | -1.833894306 | 0.066669685 | 0.447318173 |
| Rep15 | 22.65732805 | -0.12261113 | 0.084025277 | -1.459217212 | 0.144505333 | 0.589967032 |
| C3 | 46.09510738 | -0.122625716 | 0.074731935 | -1.640874354 | 0.1008235 | 0.521542547 |
| 1700040L02Rik | 73.09868875 | -0.122630295 | 0.094888211 | -1.292365972 | 0.196230435 | 0.651827876 |
| Pank4 | 1580.248206 | -0.122699354 | 0.067454122 | -1.819004531 | 0.068910739 | 0.453262505 |
| Dcx | 1140.929455 | -0.122732692 | 0.069362773 | -1.769431749 | 0.076821851 | 0.472560915 |
| F9 | 9.353768716 | -0.122744506 | 0.067105532 | -1.82912649 | 0.067380661 | NA |
| Tonsl | 340.7670827 | -0.122751927 | 0.083447274 | -1.47101182 | 0.14128792 | 0.586989325 |
| Trit1 | 533.7245096 | -0.122754686 | 0.071238459 | -1.723151897 | 0.08486106 | 0.488499377 |
| Gm17066 | 522.2092127 | -0.122808679 | 0.080130963 | -1.532599563 | 0.125374549 | 0.562122953 |
| Gsto2 | 30.5830771 | -0.122859787 | 0.088401222 | -1.389797382 | 0.164590413 | 0.614688762 |
| Gm12942 | 92.09976788 | -0.122910345 | 0.094864696 | -1.295638421 | 0.195100086 | 0.65093872 |
| 1300002E11Rik | 274.1472159 | -0.123005117 | 0.077473955 | -1.58769636 | 0.112355017 | 0.538992394 |
| Zfp174 | 366.2062469 | -0.123091456 | 0.078780279 | -1.562465343 | 0.118178404 | 0.549241002 |
| Gins3 | 92.42081169 | -0.123161932 | 0.094774623 | -1.29952437 | 0.193764036 | 0.649958942 |
| Spata13 | 857.7231946 | -0.123180644 | 0.084250508 | -1.462075977 | 0.143720391 | 0.589907432 |
| Son | 13919.88995 | -0.123258256 | 0.048476248 | -2.542652567 | 0.011001456 | 0.199361888 |
| Lsp1 | 267.1561058 | -0.123258706 | 0.084734088 | -1.454653129 | 0.145765315 | 0.59244179 |
| Strbp | 7467.188516 | -0.123297211 | 0.059186895 | -2.083184319 | 0.037234426 | 0.348836693 |
| Dbf4 | 59.50202385 | -0.123354757 | 0.094245202 | -1.308870421 | 0.190578247 | 0.645902852 |
| Lyz1 | 50.1612296 | -0.123363901 | 0.08931846 | -1.381169148 | 0.167226959 | 0.617511666 |
| Lct | 84.89152746 | -0.12370897 | 0.058798914 | -2.103932898 | 0.035384301 | 0.342065168 |
| Sppl2b | 1536.446298 | -0.123781246 | 0.077887045 | -1.589240499 | 0.112006106 | 0.538992394 |
| Ncaph | 45.18572361 | -0.123799453 | 0.091194889 | -1.357526228 | 0.174614065 | 0.627521998 |
| Pear1 | 282.6197591 | -0.123881207 | 0.086391639 | -1.433949037 | 0.151586807 | 0.598905575 |
| 9130023H24Rik | 154.5902409 | -0.123996171 | 0.090587771 | -1.368795917 | 0.171063078 | 0.621020522 |
| Zfp661 | 400.365089 | -0.124083289 | 0.068933138 | -1.800052802 | 0.071852301 | 0.460196881 |
| Bcl6b | 83.06419659 | -0.124116656 | 0.08956671 | -1.385745404 | 0.165824659 | 0.616168889 |
| Bmx | 61.72282659 | -0.124171428 | 0.092706367 | -1.339405621 | 0.18043866 | 0.634469716 |
| Zfp956 | 905.1544437 | -0.124217833 | 0.071095778 | -1.747190016 | 0.080604383 | 0.479417114 |
| Kcp | 68.67068517 | -0.124242935 | 0.092330894 | -1.345626906 | 0.17842287 | 0.631985031 |
| Tspo | 103.8516472 | -0.124272269 | 0.094985141 | -1.308333784 | 0.190760121 | 0.646232776 |
| Mir1943 | 7.513756175 | -0.124400412 | 0.055112449 | -2.25721075 | 0.023994909 | NA |
| Gas2l3 | 106.7412231 | -0.124414185 | 0.095020565 | -1.309339564 | 0.190419353 | 0.645514942 |
| H2-DMb1 | 76.61832183 | -0.12441811 | 0.090813447 | -1.370040609 | 0.170674225 | 0.620885919 |
| 0610037L13Rik | 1098.277923 | -0.124491304 | 0.072852982 | -1.708801765 | 0.087487676 | 0.495597906 |
| C1rl | 20.83768123 | -0.124544367 | 0.069779699 | -1.784822348 | 0.074290137 | NA |
| 9630013A20Rik | 38.18985316 | -0.124609162 | 0.088325897 | -1.410788528 | 0.158306978 | 0.609156864 |
| E530011L22Rik | 308.3559851 | -0.124643873 | 0.090648149 | -1.375029432 | 0.16912232 | 0.619512712 |
| Tacc3 | 249.398851 | -0.124654753 | 0.083533024 | -1.492281094 | 0.135625471 | 0.579179191 |
| Zfp467 | 956.3990433 | -0.124681331 | 0.065711209 | -1.897413441 | 0.057773393 | 0.420449052 |
| Tlr3 | 627.1140405 | -0.124752058 | 0.075607244 | -1.650001386 | 0.098942653 | 0.520271327 |
| Itga11 | 98.90851397 | -0.124811007 | 0.094710089 | -1.317821665 | 0.187563354 | 0.641776649 |
| Prr11 | 28.0682471 | -0.124837432 | 0.088071176 | -1.417460715 | 0.156348272 | 0.60719411 |
| Fli1 | 154.3597814 | -0.124846336 | 0.089343699 | -1.397371461 | 0.162301897 | 0.613957643 |
| Cxcl5 | 7.958610589 | -0.12485085 | 0.060600599 | -2.060224682 | 0.039377066 | NA |
| Ccdc90a | 1054.148433 | -0.125020631 | 0.060091938 | -2.080489261 | 0.03748068 | 0.350065015 |
| Cpne9 | 2618.68375 | -0.125028018 | 0.064126936 | -1.949695788 | 0.051212389 | 0.403266245 |
| Crlf1 | 44.16998074 | -0.125029561 | 0.070261082 | -1.779499513 | 0.075157905 | 0.468962081 |
| Nrbp2 | 6763.313593 | -0.125042784 | 0.075213022 | -1.662515086 | 0.096409541 | 0.51670168 |
| Cybb | 43.24797018 | -0.125071134 | 0.074026224 | -1.689551721 | 0.091113749 | 0.503727415 |
| Zfyve28 | 2261.633213 | -0.125102989 | 0.061656568 | -2.029029391 | 0.042455297 | 0.373308499 |
| Abhd1 | 131.4923319 | -0.125142222 | 0.093113962 | -1.343968398 | 0.178958606 | 0.632648507 |
| Mdm1 | 243.7982616 | -0.125169247 | 0.079404461 | -1.576350322 | 0.11494509 | 0.543342787 |
| Spdl1 | 20.0072745 | -0.12517456 | 0.081099606 | -1.543466928 | 0.122717526 | NA |
| Letm2 | 465.6033568 | -0.12519144 | 0.071384638 | -1.753758837 | 0.079471839 | 0.477013957 |
| Zfp133-ps | 100.8699413 | -0.125216867 | 0.093098758 | -1.344989667 | 0.178628571 | 0.632097108 |
| Dmrta2 | 35.9385545 | -0.12522607 | 0.085855606 | -1.458566032 | 0.144684588 | 0.590269908 |
| Cyp4v3 | 379.85215 | -0.125271902 | 0.088064535 | -1.422501143 | 0.15488082 | 0.604489543 |
| Nrg2 | 253.7092912 | -0.125315292 | 0.07876586 | -1.590984871 | 0.111612981 | 0.538992394 |
| Hjurp | 231.0132728 | -0.125343205 | 0.091672406 | -1.367294819 | 0.171532915 | 0.622106016 |
| Rassf9 | 44.66290788 | -0.125362545 | 0.090048346 | -1.392169319 | 0.163871131 | 0.614351408 |
| Ctps2 | 1850.946946 | -0.125422053 | 0.060515876 | -2.072547932 | 0.038214371 | 0.354019103 |
| Rpa2 | 469.9877859 | -0.125443929 | 0.080647779 | -1.555454243 | 0.119837924 | 0.553678653 |
| Bbs2 | 1323.211703 | -0.125449099 | 0.060129603 | -2.086311786 | 0.03695039 | 0.348173348 |
| Prdm5 | 112.232596 | -0.125451286 | 0.093517999 | -1.341466737 | 0.179768963 | 0.633459544 |
| Hsd3b7 | 279.9291625 | -0.125506694 | 0.08478879 | -1.480227453 | 0.138812556 | 0.583135914 |
| Card6 | 109.4846776 | -0.125524174 | 0.09083713 | -1.381859751 | 0.167014769 | 0.617356019 |
| Ints10 | 2119.56311 | -0.125546892 | 0.051326575 | -2.446040702 | 0.014443469 | 0.230050259 |
| Plk4 | 218.0248584 | -0.125559925 | 0.086283579 | -1.455200716 | 0.145613703 | 0.591991224 |
| Hyi | 151.1822505 | -0.125665832 | 0.092793038 | -1.354259267 | 0.175653677 | 0.628459435 |
| Slc7a7 | 77.2852782 | -0.12571896 | 0.094263774 | -1.333693263 | 0.182304404 | 0.635245711 |
| Slc44a2 | 2077.922932 | -0.125731094 | 0.062252804 | -2.019685647 | 0.043416005 | 0.37703831 |
| D930048N14Rik | 240.2578163 | -0.125862136 | 0.094146109 | -1.336880918 | 0.181261511 | 0.634469716 |
| Cacng6 | 15.49176918 | -0.125872883 | 0.07576098 | -1.661447405 | 0.096623624 | NA |
| Rnf31 | 1147.52283 | -0.125945857 | 0.064186088 | -1.962198667 | 0.049739359 | 0.399757904 |
| Kcnd1 | 297.2103252 | -0.125975657 | 0.081607834 | -1.543671121 | 0.122668026 | 0.557245639 |
| Homer3 | 360.0002061 | -0.126056649 | 0.095096568 | -1.325564643 | 0.184983944 | 0.637780733 |
| Kif11 | 79.24705467 | -0.126059685 | 0.095093649 | -1.325637265 | 0.184959877 | 0.637780733 |
| Fcrls | 1297.364406 | -0.126111201 | 0.093005308 | -1.355957035 | 0.175112839 | 0.628026741 |
| Setd1a | 3095.27776 | -0.126177286 | 0.057045897 | -2.211855575 | 0.026976646 | 0.306595176 |
| 2610204G22Rik | 20.50866306 | -0.126199145 | 0.082965058 | -1.521111995 | 0.128231735 | NA |
| Col28a1 | 52.06209842 | -0.126202637 | 0.09245555 | -1.365008775 | 0.172250292 | 0.623232235 |
| Ttc32 | 48.95293652 | -0.126214002 | 0.094076139 | -1.34161546 | 0.179720711 | 0.633459544 |
| C030018K13Rik | 77.24511084 | -0.126243639 | 0.09506819 | -1.327927241 | 0.184202144 | 0.637683592 |
| Rest | 116.2152373 | -0.126268277 | 0.092394988 | -1.36661392 | 0.171746352 | 0.622296709 |
| Gm10451 | 30.7481403 | -0.126289938 | 0.075753473 | -1.667117463 | 0.09549105 | 0.514694989 |
| Ctrl | 25.50937916 | -0.126331573 | 0.081985382 | -1.54090363 | 0.123340242 | 0.558143414 |
| Gabrr1 | 5.897516715 | -0.126424481 | 0.051867842 | -2.437434756 | 0.014791883 | NA |
| Kcnj2 | 868.5950087 | -0.126487136 | 0.089921681 | -1.406636692 | 0.159535147 | 0.611570723 |
| Irak1 | 2519.299127 | -0.126488126 | 0.058228683 | -2.172264919 | 0.029835686 | 0.318437312 |
| Hes7 | 15.3203102 | -0.126554047 | 0.063462002 | -1.994170401 | 0.046133436 | NA |
| Erap1 | 468.4370834 | -0.126569855 | 0.084243451 | -1.502429601 | 0.132986196 | 0.575193212 |
| Cabin1 | 4309.508141 | -0.12657935 | 0.055137376 | -2.295708632 | 0.021692547 | 0.276408712 |
| Slc38a5 | 86.31757808 | -0.126607453 | 0.094609622 | -1.338209053 | 0.180828299 | 0.634469716 |
| Ube2l6 | 217.2373141 | -0.12662334 | 0.090896204 | -1.393054213 | 0.163603398 | 0.614351408 |
| Sox2 | 907.4834152 | -0.126634751 | 0.085516087 | -1.480829582 | 0.138651991 | 0.583109099 |
| Ncoa3 | 2277.492465 | -0.1267001 | 0.066423624 | -1.907455385 | 0.056461643 | 0.416586327 |
| Fbrsl1 | 1719.305775 | -0.126832319 | 0.067838576 | -1.869619426 | 0.061536686 | 0.435077842 |
| Sec24a | 766.4769622 | -0.1269223 | 0.064915288 | -1.955198901 | 0.050559598 | 0.403264376 |
| Tbx15 | 42.44829937 | -0.126922691 | 0.086841899 | -1.461537487 | 0.143867996 | 0.589967032 |
| Arhgef6 | 655.0655295 | -0.126934738 | 0.077720685 | -1.633216927 | 0.102423381 | 0.522975271 |
| Aim2 | 86.43046516 | -0.12695093 | 0.094356242 | -1.345442835 | 0.17848227 | 0.632041311 |
| Mir1839 | 14.16468634 | -0.126981756 | 0.069049953 | -1.838983969 | 0.065917544 | NA |
| Cdh24 | 405.6839497 | -0.126997203 | 0.090293793 | -1.406488735 | 0.159579047 | 0.611570723 |
| Emx1 | 578.9228127 | -0.127068217 | 0.084504127 | -1.503692436 | 0.132660578 | 0.574717432 |
| Plxna3 | 1635.821575 | -0.127068508 | 0.080205851 | -1.584279779 | 0.113130065 | 0.540327972 |
| Cdc42bpg | 283.9781604 | -0.127089959 | 0.084449701 | -1.504918987 | 0.132344906 | 0.573884732 |
| Znhit2 | 570.150694 | -0.127105031 | 0.080987049 | -1.569448848 | 0.11654339 | 0.546452317 |
| Snord53 | 8.567590711 | -0.127130182 | 0.062937497 | -2.0199434 | 0.043389259 | NA |
| Terf1 | 322.3265696 | -0.127185676 | 0.078248776 | -1.625401479 | 0.104077041 | 0.523645898 |
| AB124611 | 12.63636353 | -0.127352342 | 0.072012793 | -1.768468311 | 0.076982645 | NA |
| Myh9 | 3925.928972 | -0.127386544 | 0.06985629 | -1.823551516 | 0.068219902 | 0.451155091 |
| Ddx51 | 864.4577872 | -0.127393924 | 0.069336845 | -1.837319316 | 0.06616277 | 0.446279967 |
| Tmem179b | 167.0778251 | -0.127417049 | 0.085684388 | -1.487050935 | 0.137001359 | 0.581385258 |
| Fermt3 | 194.6057532 | -0.12743925 | 0.093991771 | -1.355855399 | 0.175145181 | 0.628026741 |
| Cnksr2 | 9262.92383 | -0.127474203 | 0.080206873 | -1.589317707 | 0.111988683 | 0.538992394 |
| Zfp442 | 97.62701692 | -0.127504904 | 0.094017467 | -1.356183142 | 0.175040904 | 0.628026741 |
| Mks1 | 226.5437538 | -0.127579832 | 0.088573776 | -1.440379284 | 0.149760121 | 0.598098567 |
| Mak | 161.9345077 | -0.127709228 | 0.087227883 | -1.46408721 | 0.143170121 | 0.589481061 |
| Card14 | 8.506194928 | -0.127714738 | 0.06181714 | -2.06600852 | 0.038827675 | NA |
| Dcaf17 | 621.4321147 | -0.127753211 | 0.066229284 | -1.928953521 | 0.053736633 | 0.408679751 |
| Wdr83 | 781.056369 | -0.127786742 | 0.073172406 | -1.746378842 | 0.080745144 | 0.479417114 |
| 4933407K13Rik | 140.8161676 | -0.127925876 | 0.094120449 | -1.359171963 | 0.174092103 | 0.626964546 |
| Ankrd24 | 1477.511836 | -0.127954592 | 0.076046598 | -1.682581407 | 0.092456153 | 0.506608364 |
| Tmem198b | 399.249995 | -0.127971555 | 0.076716526 | -1.668109348 | 0.095294022 | 0.51433043 |
| Gm14325 | 864.9481125 | -0.12797352 | 0.066208181 | -1.932895864 | 0.053249027 | 0.407317206 |
| Gm12522 | 69.08467253 | -0.127979569 | 0.095020459 | -1.346863308 | 0.178024261 | 0.631188775 |
| Cage1 | 147.550461 | -0.127980203 | 0.09169271 | -1.395751121 | 0.162789454 | 0.614351408 |
| Utp20 | 1149.436534 | -0.128035138 | 0.064768483 | -1.97681238 | 0.048062842 | 0.393773739 |
| Tgfb1 | 227.2592222 | -0.128095808 | 0.093358763 | -1.372081245 | 0.170038147 | 0.620316915 |
| Prdm8 | 997.1153801 | -0.128129384 | 0.080594498 | -1.589803113 | 0.111879192 | 0.538992394 |
| Fignl1 | 30.27093336 | -0.128171182 | 0.090729126 | -1.41267957 | 0.157749961 | 0.608141173 |
| Plekha8 | 829.4381955 | -0.128189767 | 0.07139432 | -1.795517739 | 0.072571249 | 0.462655961 |
| Zfp354a | 323.4616906 | -0.128202619 | 0.084228458 | -1.522081988 | 0.127988539 | 0.565782566 |
| Gm996 | 5479.062405 | -0.128276865 | 0.063735747 | -2.012636108 | 0.044152925 | 0.380406522 |
| Pltp | 3082.505687 | -0.128302797 | 0.068739257 | -1.866514162 | 0.061969464 | 0.436339526 |
| Sox13 | 398.9577887 | -0.12847674 | 0.080779416 | -1.590463837 | 0.111730291 | 0.538992394 |
| Dock7 | 1896.403227 | -0.128485938 | 0.063847969 | -2.012373143 | 0.044180617 | 0.380419472 |
| Slc25a37 | 1558.05316 | -0.128737633 | 0.065720621 | -1.958862101 | 0.050128938 | 0.401582548 |
| Inppl1 | 1724.954143 | -0.128860975 | 0.055393121 | -2.326299246 | 0.020002592 | 0.267548485 |
| Txlnb | 117.3307999 | -0.128871047 | 0.093694096 | -1.375444683 | 0.168993624 | 0.619378697 |
| Ppp4r1l-ps | 318.4820442 | -0.128874852 | 0.079094578 | -1.629376569 | 0.103233326 | 0.522975271 |
| Fubp3 | 1318.454189 | -0.128982712 | 0.071307317 | -1.808828577 | 0.070477639 | 0.456017009 |
| Fan1 | 442.7553385 | -0.128983493 | 0.068286826 | -1.888848852 | 0.058912085 | 0.425326512 |
| Siglech | 701.2074 | -0.128998761 | 0.08201777 | -1.572814781 | 0.115761712 | 0.545619664 |
| Abtb1 | 1327.6168 | -0.129042181 | 0.073580325 | -1.753759332 | 0.079471754 | 0.477013957 |
| Mkx | 556.2513477 | -0.129066444 | 0.08624681 | -1.496477891 | 0.134529165 | 0.577862888 |
| 5530601H04Rik | 427.3242011 | -0.129088771 | 0.069635728 | -1.85377212 | 0.063771769 | 0.441368101 |
| Murc | 27.06517411 | -0.129161225 | 0.087324236 | -1.479099398 | 0.139113751 | 0.58336211 |
| 4732491K20Rik | 167.2815483 | -0.129208945 | 0.092121358 | -1.402594877 | 0.16073768 | 0.612387888 |
| Tns4 | 20.98582626 | -0.129304849 | 0.07546326 | -1.713480823 | 0.086624118 | NA |
| Srrm3 | 1831.392349 | -0.129308999 | 0.064934986 | -1.991360989 | 0.046441215 | 0.388809057 |
| Krt77 | 263.3887187 | -0.12932242 | 0.093586567 | -1.378310518 | 0.167664474 | 0.6180824 |
| Rab3il1 | 418.9360023 | -0.129383648 | 0.080350282 | -1.610245092 | 0.107344361 | 0.532178904 |

| | | | | | | |
|---|---|---|---|---|---|---|
| Dzip3 | 2387.637588 | -0.128883847 | 0.101583897 | -1.268742884 | 0.204532782 | 0.421581871 |
| Atf6 | 4761.193318 | -0.115320782 | 0.090913231 | -1.268470837 | 0.204629859 | 0.42171341 |
| Aptx | 1258.035436 | 0.132327986 | 0.068055542 | 1.944411618 | 0.051845834 | 0.178036204 |
| Syn3 | 2071.169055 | -0.137879317 | 0.108707557 | -1.268350803 | 0.204672702 | 0.42171341 |
| Myom1 | 95.16259355 | 0.132270392 | 0.145438372 | 0.909460072 | 0.363107325 | 0.593603784 |
| Rpp25 | 467.210933 | 0.132260126 | 0.11296528 | 1.170803325 | 0.241677841 | 0.466262791 |
| Faf2 | 2264.661499 | -0.055683753 | 0.043926347 | -1.267661829 | 0.204918742 | 0.42204054 |
| Slitrk2 | 1109.666162 | -0.073120682 | 0.057733676 | -1.266517003 | 0.205328046 | 0.422827319 |
| Cpsf3l | 967.1027995 | -0.075646284 | 0.05973531 | -1.266357936 | 0.205384964 | 0.422843318 |
| Fam120b | 3165.000336 | 0.066631076 | 0.052616936 | 1.266342752 | 0.205390397 | 0.422843318 |
| Sart3 | 1745.175727 | 0.132209025 | 0.055189928 | 2.395528132 | 0.016596441 | 0.083309297 |
| Gm16596 | 39.19155249 | -0.203875027 | 0.161116961 | -1.265385255 | 0.205733258 | 0.423436651 |
| Irs1 | 757.552852 | -0.098829405 | 0.078125649 | -1.265005869 | 0.205869224 | 0.423660215 |
| Hmga2-ps1 | 38.21044611 | -0.212635318 | 0.168116438 | -1.264809798 | 0.205939518 | 0.423692324 |
| Ipo7 | 4321.870839 | -0.087652807 | 0.069297535 | -1.264876261 | 0.205915688 | 0.423692324 |
| 4931428F04Rik | 1185.169472 | -0.107303625 | 0.084863466 | -1.264426611 | 0.206076946 | 0.423761947 |
| Zfp383 | 472.3349313 | 0.132145655 | 0.103850768 | 1.272457177 | 0.20321073 | 0.420270158 |
| Gpi1 | 23538.58571 | -0.055072837 | 0.043555426 | -1.264431141 | 0.206075322 | 0.423761947 |
| Tpm4 | 1212.215722 | -0.108274213 | 0.085632175 | -1.264410408 | 0.206082759 | 0.423761947 |
| Lsr | 288.1094879 | -0.167886114 | 0.132789541 | -1.264302237 | 0.206121567 | 0.423777155 |
| Slc35b4 | 3194.562976 | -0.07940213 | 0.062806349 | -1.264237326 | 0.206144857 | 0.423777155 |
| Mthfd2 | 208.7466335 | 0.132143255 | 0.106556524 | 1.240123555 | 0.214929698 | 0.43470211 |
| Agpat1 | 4616.445789 | -0.108097188 | 0.085522482 | -1.263962245 | 0.206243578 | 0.423831546 |
| Xrcc6bp1 | 142.3074847 | -0.155929783 | 0.12336852 | -1.263934941 | 0.206253379 | 0.423831546 |
| Faim2 | 25017.38475 | 0.132112843 | 0.062644419 | 2.108932373 | 0.034950419 | 0.136817028 |
| Ctgf | 1357.163445 | 0.132085927 | 0.098022581 | 1.347505102 | 0.177817612 | 0.387414269 |
| Wwox | 683.3330216 | 0.132084398 | 0.101666792 | 1.299189196 | 0.193879008 | 0.408313355 |
| Bod1 | 2584.826831 | -0.086825154 | 0.068742407 | -1.263050832 | 0.206570912 | 0.424258976 |
| Neu4 | 343.6359573 | 0.132082998 | 0.099552039 | 1.326773408 | 0.18458365 | 0.395751 |
| Thoc5 | 1164.862807 | 0.066659386 | 0.052829585 | 1.261781374 | 0.207027466 | 0.42508396 |
| Gnpda2 | 634.7257135 | 0.09406153 | 0.074561371 | 1.261531653 | 0.207117363 | 0.425155859 |
| Gch1 | 55.1187699 | 0.132064223 | 0.151347778 | 0.872587789 | 0.382887799 | 0.611369871 |
| Eapp | 736.0195562 | -0.084250125 | 0.066797729 | -1.261272304 | 0.207210756 | 0.425234895 |
| Kcnj3 | 2077.376718 | 0.132006438 | 0.064624825 | 2.042658362 | 0.041086269 | 0.1530266 |
| Nos3 | 517.8592264 | -0.110573619 | 0.087673846 | -1.26119275 | 0.20723941 | 0.425237375 |
| Pdk4 | 511.773732 | 0.131811772 | 0.13066764 | 1.008756047 | 0.313091645 | 0.546226996 |
| Gm13375 | 356.0449078 | 0.131658119 | 0.097129962 | 1.355484092 | 0.175263374 | 0.384146373 |
| Pi16 | 32.79571635 | -0.205552426 | 0.163050861 | -1.260664458 | 0.207429764 | 0.425492348 |
| Rgl1 | 5243.726395 | -0.104392695 | 0.082818236 | -1.260503733 | 0.207487702 | 0.425492348 |
| Ube4b | 5068.093557 | 0.097362036 | 0.077232749 | 1.260631491 | 0.207441647 | 0.425492348 |
| Asah1 | 2023.310133 | 0.131641504 | 0.072622415 | 1.812684209 | 0.069880541 | 0.215083255 |
| Ociad1 | 10450.6343 | -0.058569014 | 0.046478447 | -1.260132755 | 0.207621476 | 0.425682886 |
| A630007B06Rik | 992.004553 | 0.131611308 | 0.080740132 | 1.6300606 | 0.10308869 | 0.276018401 |
| Tdrkh | 1042.085744 | -0.067035608 | 0.053219805 | -1.25959889 | 0.207814096 | 0.425908943 |
| Flot1 | 3744.698931 | -0.102507331 | 0.081400877 | -1.259290257 | 0.207925511 | 0.426024579 |
| Sp140 | 33.34216167 | 0.131497798 | 0.161836522 | 0.812534753 | 0.416484853 | 0.639292284 |
| Syt11 | 35077.79624 | -0.06758898 | 0.053690513 | -1.258862638 | 0.208079951 | 0.426284644 |
| Zfyve26 | 1151.840258 | 0.085220255 | 0.067703345 | 1.258730336 | 0.20812775 | 0.426326198 |
| Rnf208 | 3597.718028 | 0.131494715 | 0.07990859 | 1.64556419 | 0.099853517 | 0.26981695 |
| Slc38a4 | 44.41413628 | -0.197880728 | 0.157228619 | -1.258554128 | 0.208191425 | 0.426398661 |
| Snw1 | 3082.408186 | 0.131491818 | 0.094043975 | 1.39819503 | 0.162054509 | 0.367061759 |
| Ifrd2 | 270.9328065 | 0.131450574 | 0.115731709 | 1.135821593 | 0.256031238 | 0.482479689 |
| Cltb | 4910.369744 | -0.058133552 | 0.046199614 | -1.258312505 | 0.20827876 | 0.426414656 |
| Gna12 | 1950.243551 | -0.12803034 | 0.10177278 | -1.258001801 | 0.208391105 | 0.426414656 |
| Imp3 | 709.8767354 | -0.107503157 | 0.085439298 | -1.258240176 | 0.208304909 | 0.426414656 |
| Npl | 156.6260435 | 0.131391557 | 0.123189201 | 1.066583398 | 0.28616 | 0.515760851 |
| 1700084E18Rik | 8.614635622 | 0.131192908 | 0.149892756 | 0.875245152 | 0.38144053 | 0.610149688 |
| Zfp395 | 423.8942693 | -0.132157407 | 0.105088921 | -1.257576966 | 0.208544788 | 0.426616444 |
| Mrap | 10.06648932 | -0.205790046 | 0.163675422 | -1.257305732 | 0.208642949 | 0.426760905 |
| Olfr558 | 15.58488801 | -0.197917308 | 0.157433288 | -1.257150315 | 0.208699211 | 0.426819637 |
| Spink10 | 36.6025272 | -0.207644648 | 0.165201445 | -1.256917867 | 0.208783378 | 0.426935418 |
| Bank1 | 64.63687392 | -0.185609946 | 0.147724058 | -1.256463898 | 0.208947827 | 0.427134075 |
| Snapc3 | 568.6621205 | -0.09223849 | 0.073414525 | -1.256406547 | 0.208968609 | 0.427134075 |
| Spin4 | 22.28352849 | -0.209865459 | 0.167045761 | -1.256335136 | 0.208994488 | 0.427134075 |
| Tmem231 | 522.1618862 | -0.093820457 | 0.074681807 | -1.256269253 | 0.209018365 | 0.427134075 |
| Myo1e | 864.1682578 | 0.13113825 | 0.106606044 | 1.230120218 | 0.21865209 | 0.439118015 |
| Trappc11 | 3099.56442 | 0.061281344 | 0.048783749 | 1.256183553 | 0.209049428 | 0.427141217 |
| C030039L03Rik | 633.0159649 | 0.086995986 | 0.069286524 | 1.255597504 | 0.209261937 | 0.427406334 |
| Lrrc4 | 4343.673094 | 0.131126749 | 0.067707493 | 1.936665263 | 0.052786275 | 0.180028734 |
| Zfp367 | 759.1093129 | 0.131086963 | 0.086225268 | 1.520284782 | 0.128439417 | 0.317033857 |
| Ikzf4 | 477.0708976 | 0.130970968 | 0.08543456 | 1.532997507 | 0.125276469 | 0.312868087 |
| Nt5c2 | 2825.381946 | 0.094679547 | 0.075426998 | 1.255247445 | 0.209388947 | 0.427553024 |
| Zfp597 | 840.7139711 | -0.102618347 | 0.0817669 | -1.255010844 | 0.209474824 | 0.427672014 |
| Fktn | 1284.588412 | -0.126957351 | 0.101172434 | -1.254861088 | 0.209529192 | 0.427726653 |
| Sf3b2 | 9717.684122 | 0.130929835 | 0.066147934 | 1.979348829 | 0.047776746 | 0.168930373 |
| Micalcl | 98.89648915 | 0.130908284 | 0.144717128 | 0.904580444 | 0.3656877 | 0.595790244 |
| Neo1 | 4708.386084 | -0.063468694 | 0.05059726 | -1.254389936 | 0.209700309 | 0.427871411 |
| Prr3 | 946.8654901 | 0.091850157 | 0.073228389 | 1.254297116 | 0.209734032 | 0.427871411 |
| Slc38a3 | 3287.266443 | -0.164296544 | 0.130987973 | -1.254287242 | 0.209737619 | 0.427871411 |
| Zbtb8os | 337.9869594 | 0.130888671 | 0.089201143 | 1.467342976 | 0.142282779 | 0.338118002 |
| Nup107 | 615.7182374 | 0.097946962 | 0.078108889 | 1.253979708 | 0.209849381 | 0.428013611 |
| Kif4 | 68.30568263 | -0.180228937 | 0.143777624 | -1.253525631 | 0.210014477 | 0.4282659 |
| Ajap1 | 1980.420023 | -0.117282236 | 0.093573885 | -1.253365043 | 0.210072887 | 0.428328637 |
| Asna1 | 3079.53666 | 0.130863614 | 0.076473274 | 1.711233319 | 0.08703805 | 0.247035638 |
| Tbc1d17 | 1920.666227 | 0.130834234 | 0.069677688 | 1.877706317 | 0.060421352 | 0.195483072 |
| Kirrel | 390.3754495 | -0.143370963 | 0.114428008 | -1.252935938 | 0.210229022 | 0.428462109 |
| Fam108b | 787.2235606 | 0.130812463 | 0.080101386 | 1.633086141 | 0.102450881 | 0.274727233 |
| Trappc13 | 1792.133527 | -0.089112334 | 0.071124937 | -1.252898604 | 0.21024261 | 0.428462109 |
| Leng1 | 254.1338035 | 0.130700576 | 0.093572679 | 1.396781385 | 0.162479322 | 0.367565281 |
| Phkg2 | 1809.566854 | 0.076746989 | 0.061283207 | 1.252333104 | 0.210448511 | 0.428708909 |
| Krt73 | 47.42333646 | 0.130639722 | 0.153894396 | 0.848892006 | 0.395941388 | 0.620922658 |
| Pcdha6 | 123.2704618 | -0.193646882 | 0.154666365 | -1.252029691 | 0.210559045 | 0.428812019 |
| Doc2b | 2215.449168 | -0.148250556 | 0.118416551 | -1.251941178 | 0.210591299 | 0.428821341 |
| Gpn2 | 241.0598863 | 0.130629136 | 0.099357321 | 1.314740923 | 0.188596988 | 0.401747886 |
| Fam212b | 6107.305511 | -0.112119505 | 0.089597422 | -1.251369764 | 0.210799603 | 0.42913271 |
| Lypd6 | 1257.910282 | -0.085003515 | 0.067932838 | -1.251287554 | 0.210829585 | 0.42913736 |
| Etohd2 | 27.88126795 | -0.204181158 | 0.16326906 | -1.250580843 | 0.211087444 | 0.429605786 |
| Slc25a19 | 1428.756418 | 0.080687932 | 0.064524735 | 1.250496152 | 0.21111836 | 0.429612276 |
| Galnt6 | 655.1739531 | 0.130607894 | 0.138293314 | 0.94442667 | 0.344951658 | 0.576948985 |
| Creb3l3 | 22.83512179 | 0.130541618 | 0.168121354 | 0.77647256 | 0.437470016 | 0.658137725 |
| Dapp1 | 235.504319 | -0.17849274 | 0.142781336 | -1.25011255 | 0.211258436 | 0.429728004 |
| Umodl1 | 13.08750624 | -0.203356792 | 0.162683275 | -1.250016584 | 0.211293489 | 0.429742888 |
| Phf14 | 1462.85246 | -0.080866535 | 0.064781102 | -1.2483044 | 0.211919604 | 0.430959751 |
| Ap3s2 | 3882.244403 | 0.072734056 | 0.058291712 | 1.247759808 | 0.212119032 | 0.431308698 |
| Trib3 | 35.81015496 | -0.204669034 | 0.164054018 | -1.247570996 | 0.212188206 | 0.431392739 |
| C030018K13Rik | 76.49972424 | -0.198822402 | 0.159388231 | -1.24740955 | 0.212247367 | 0.431456403 |
| Siah2 | 583.0987224 | 0.130488291 | 0.088431319 | 1.475589106 | 0.140054231 | 0.334412122 |
| Serpinb1b | 59.42686677 | -0.205956575 | 0.165179681 | -1.246863864 | 0.212447419 | 0.431749778 |
| Elp6 | 312.8409599 | -0.145801768 | 0.117002258 | -1.246144909 | 0.2127112 | 0.432229158 |
| Rnf111 | 1595.775972 | 0.130464744 | 0.067085643 | 1.944749112 | 0.051805181 | 0.177969403 |
| Clstn2 | 2801.826474 | 0.080000989 | 0.064214323 | 1.245843384 | 0.212821899 | 0.432340697 |
| Cdc40 | 1903.615357 | 0.078558197 | 0.063068053 | 1.245609989 | 0.212907613 | 0.432401437 |
| Garnl3 | 3559.918321 | 0.130368086 | 0.102691254 | 1.269514994 | 0.204257446 | 0.421381309 |
| Ten1 | 565.9142909 | -0.103845946 | 0.083381426 | -1.245432602 | 0.212972776 | 0.432477088 |
| Slc37a1 | 246.2057962 | 0.130362402 | 0.108755509 | 1.198674012 | 0.230654727 | 0.453668953 |
| Ier2 | 255.8904783 | -0.202737524 | 0.162824149 | -1.24513179 | 0.21308331 | 0.432588157 |
| Cox8a | 7639.345278 | -0.123370387 | 0.099095184 | -1.244968536 | 0.213143316 | 0.432653287 |
| Tpd52 | 2445.70238 | 0.081903683 | 0.065793131 | 1.244866773 | 0.213180726 | 0.43267254 |
| Zfp541 | 28.74802507 | 0.130324505 | 0.166460514 | 0.782915432 | 0.433676775 | 0.65465588 |
| Dync1i2 | 4919.938148 | 0.130294326 | 0.068755784 | 1.895030773 | 0.058088325 | 0.191087156 |
| Osgep | 675.7741031 | -0.119036735 | 0.095679157 | -1.244123991 | 0.213453933 | 0.433023546 |
| Cdk10 | 2071.963481 | 0.130276903 | 0.096030656 | 1.356617863 | 0.174902663 | 0.383627104 |
| C230052I12Rik | 179.7640373 | -0.141211494 | 0.113539763 | -1.243718413 | 0.213603217 | 0.433246268 |
| Rusc2 | 6042.144061 | 0.130188654 | 0.0814055 | 1.59926115 | 0.109762588 | 0.288247719 |
| Lin54 | 515.7277537 | 0.130181905 | 0.118288548 | 1.100545289 | 0.271094608 | 0.499267783 |
| Atp13a5 | 564.2392137 | -0.132571034 | 0.106667065 | -1.242848802 | 0.213923555 | 0.433676986 |
| Smim14 | 3888.006172 | -0.061701528 | 0.049645672 | -1.242838008 | 0.213927533 | 0.433676986 |
| Adrm1 | 2276.747012 | 0.072019286 | 0.057951738 | 1.242745925 | 0.213961475 | 0.433689049 |
| Pttg1ip | 2370.702691 | -0.081941526 | 0.065942333 | -1.242624009 | 0.214006418 | 0.433723407 |
| Camk4 | 11080.12719 | -0.121622918 | 0.097905609 | -1.242246693 | 0.214145557 | 0.433891886 |
| Depdc1b | 12.48791962 | 0.130122024 | 0.149678264 | 0.869344826 | 0.384658551 | 0.61266645 |
| Fam102a | 4165.251602 | 0.13011946 | 0.051637862 | 2.519846009 | 0.011740619 | 0.065742372 |
| Pcbp2 | 7795.673722 | 0.075324138 | 0.060662337 | 1.241695285 | 0.21434901 | 0.434190556 |
| Tmed7 | 2889.924406 | -0.091853232 | 0.073983295 | -1.241540176 | 0.214406266 | 0.434249763 |
| Exosc4 | 623.9155933 | -0.084361682 | 0.067971964 | -1.241124682 | 0.214559693 | 0.434503709 |
| 9630033F20Rik | 1406.572891 | 0.074935295 | 0.06041053 | 1.240434313 | 0.214814796 | 0.43465253 |
| Aspscr1 | 1086.399461 | 0.086500306 | 0.06971483 | 1.24077339 | 0.214689474 | 0.43465253 |
| Mrpl45 | 1242.918877 | 0.084511176 | 0.068112608 | 1.24075672 | 0.214695634 | 0.43465253 |
| Spice1 | 510.8293749 | 0.130110711 | 0.110239126 | 1.180258911 | 0.237897255 | 0.462365789 |
| Phyhipl | 6731.696796 | -0.102745193 | 0.082819921 | -1.240585493 | 0.214758914 | 0.43465253 |
| Slc38a8 | 15.38911121 | 0.13007309 | 0.166568723 | 0.780897447 | 0.434862813 | 0.655998874 |
| Zfp449 | 211.9592111 | -0.134791601 | 0.108658302 | -1.240509001 | 0.214787187 | 0.43465253 |
| Sumo2 | 4477.961626 | -0.062426532 | 0.050331084 | -1.240317651 | 0.214857926 | 0.434653181 |
| Timp2 | 15280.10831 | 0.130035084 | 0.10195732 | 1.275387431 | 0.202172145 | 0.419384114 |
| Nup85 | 1158.636761 | 0.130013632 | 0.078052784 | 1.665714219 | 0.095770349 | 0.262132229 |
| Slc44a1 | 3219.778995 | 0.1299815 | 0.084507553 | 1.538105121 | 0.124022914 | 0.311545238 |
| Irf2bp2 | 1280.408169 | 0.129911239 | 0.094524144 | 1.374370973 | 0.169326543 | 0.376567454 |
| Naaa | 1279.892325 | 0.129878576 | 0.070245734 | 1.848917624 | 0.064469709 | 0.20393909 |
| Efcab3 | 11.08370414 | 0.129832478 | 0.153883846 | 0.843704402 | 0.398834607 | 0.624070229 |
| Mex3b | 548.8003879 | -0.131001924 | 0.105732947 | -1.23898867 | 0.215349694 | 0.435250216 |
| 0610030E20Rik | 785.6958151 | 0.129796666 | 0.100930644 | 1.285998599 | 0.198443551 | 0.414386757 |
| Ppap2b | 10110.45147 | -0.137295345 | 0.110844773 | -1.23862715 | 0.215483609 | 0.435414484 |
| Ccdc130 | 425.550829 | 0.129787316 | 0.102268198 | 1.269087741 | 0.204409772 | 0.421552643 |
| Ammecr1l | 1534.285692 | 0.129763724 | 0.063108719 | 2.056193293 | 0.039763889 | 0.148989976 |
| Zfp553 | 890.5760157 | 0.085422109 | 0.06900057 | 1.237991349 | 0.215719268 | 0.43571299 |
| Lipg | 256.3727519 | -0.160221485 | 0.129505703 | -1.237177057 | 0.216021357 | 0.436266281 |
| Uchl3 | 737.7666646 | -0.090484332 | 0.07314251 | -1.237096338 | 0.216051319 | 0.436269926 |
| Simc1 | 759.4926414 | 0.12974995 | 0.066828451 | 1.941537597 | 0.052193104 | 0.17887192 |
| Icmt | 3484.665416 | 0.084248124 | 0.068116002 | 1.236833065 | 0.216149064 | 0.436353563 |
| Zfp442 | 98.51912176 | -0.163704021 | 0.132370367 | -1.236712001 | 0.216194022 | 0.436387464 |
| 4931408D14Rik | 14.52117776 | 0.129607582 | 0.164883063 | 0.786057583 | 0.431833753 | 0.653295873 |
| Abhd10 | 642.2947172 | 0.087592032 | 0.070862927 | 1.236076965 | 0.216429956 | 0.436443452 |
| Pdrg1 | 1435.278895 | 0.129591425 | 0.064153633 | 2.020016937 | 0.043381631 | 0.158527119 |
| Polr3a | 1337.685989 | 0.129585721 | 0.05727145 | 2.262658279 | 0.023656763 | 0.105618495 |
| Fxyd5 | 428.8217881 | 0.129575573 | 0.104135066 | 1.244302983 | 0.213388074 | 0.432979939 |
| Zfp40 | 552.4256265 | 0.129559105 | 0.144726263 | 0.895201068 | 0.370679614 | 0.600513377 |
| 5430416N02Rik | 131.9862758 | 0.129475274 | 0.129003086 | 1.003660282 | 0.315542388 | 0.54876192 |
| Ppia | 29828.72814 | -0.091297336 | 0.073871607 | -1.235892114 | 0.216498669 | 0.436443452 |
| Sf1 | 5565.280103 | 0.06920738 | 0.055998496 | 1.235879268 | 0.216503445 | 0.436443452 |
| Zdhhc16 | 966.5522336 | 0.077503447 | 0.06268591 | 1.236377477 | 0.216318284 | 0.436443452 |
| Arpc3 | 3858.624131 | -0.088391259 | 0.071548099 | -1.235410308 | 0.216677838 | 0.436738185 |
| Zfp82 | 192.3624234 | 0.12942952 | 0.113245427 | 1.142911667 | 0.253075268 | 0.478948113 |
| Baz2a | 3263.369709 | 0.085138761 | 0.068930565 | 1.235138016 | 0.216779143 | 0.436760518 |
| Dopey2 | 4222.116807 | 0.055606204 | 0.045024237 | 1.235028219 | 0.216820002 | 0.436760518 |
| Sowahc | 386.5069707 | 0.129421395 | 0.105275324 | 1.229361163 | 0.218936425 | 0.439461195 |
| Usp32 | 7624.303373 | -0.10061915 | 0.08146394 | -1.235137282 | 0.216779416 | 0.436760518 |
| Rab3d | 511.2456451 | 0.129293818 | 0.102970457 | 1.255639939 | 0.209246544 | 0.427406334 |
| Fv1 | 203.345902 | 0.129291939 | 0.113110379 | 1.143059901 | 0.253013722 | 0.47889013 |
| Tm4sf1 | 318.1300214 | -0.150932427 | 0.122241213 | -1.234709826 | 0.216938518 | 0.436916767 |
| Atp6v1g1 | 2501.726398 | -0.069089332 | 0.055982643 | -1.234120567 | 0.217157981 | 0.437194095 |
| Tmem17 | 164.4543393 | -0.158231722 | 0.128253417 | -1.233742739 | 0.217298783 | 0.437420727 |
| Stxbp5l | 5824.395622 | -0.087711069 | 0.071112267 | -1.233416866 | 0.217420276 | 0.437608437 |
| Tnxb | 131.5199224 | 0.129261628 | 0.143746332 | 0.899234273 | 0.368527888 | 0.598153269 |
| Gprc5c | 188.5419426 | -0.139526353 | 0.113147192 | -1.233140218 | 0.217523455 | 0.437645554 |
| Shank2 | 6175.420076 | -0.102569106 | 0.083175533 | -1.233164403 | 0.217514433 | 0.437645554 |
| Cpsf7 | 3436.261118 | 0.129095883 | 0.059289211 | 2.177392482 | 0.029451296 | 0.121935044 |
| Prss30 | 21.39475252 | -0.205389491 | 0.166643416 | -1.232508888 | 0.217759049 | 0.438005803 |
| Dnajc8 | 2814.115513 | 0.070215213 | 0.056979998 | 1.232278262 | 0.217845157 | 0.438122127 |
| Lyrm5 | 565.1137524 | -0.142865011 | 0.11594493 | -1.232179887 | 0.217881895 | 0.438139141 |
| Zfp955b | 727.2684129 | -0.089808124 | 0.07290575 | -1.231838694 | 0.218009346 | 0.438338542 |
| Itk | 17.23646412 | 0.12899577 | 0.162821014 | 0.792255048 | 0.428211976 | 0.649761665 |
| Pappa2 | 173.1223308 | -0.19923811 | 0.161852373 | -1.230986647 | 0.218327859 | 0.438694312 |
| Pdcd10 | 1146.373835 | -0.111034291 | 0.090185337 | -1.231178972 | 0.218255934 | 0.438694312 |
| Upf3a | 1063.602098 | 0.128938115 | 0.058854048 | 2.190811349 | 0.028465447 | 0.119579397 |
| Wasf1 | 8418.016047 | -0.071531343 | 0.05810213 | -1.231131164 | 0.218273812 | 0.438694312 |
| Tnfrsf11a | 230.7027707 | 0.128924245 | 0.126319623 | 1.020619294 | 0.307434845 | 0.54000629 |
| Alkbh5 | 2966.234666 | 0.128891541 | 0.065058234 | 1.981171826 | 0.047572008 | 0.168514103 |
| Ttyh2 | 1507.013485 | 0.128866215 | 0.123593992 | 1.042657596 | 0.297106904 | 0.529027456 |
| Zc3hc1 | 860.113706 | -0.073450271 | 0.059709139 | -1.230134486 | 0.218646748 | 0.439118015 |
| 0610038B21Rik | 13.94850875 | -0.203016803 | 0.16505963 | -1.229960366 | 0.218711947 | 0.439129313 |
| 5033411D12Rik | 156.1623094 | -0.132587892 | 0.107799082 | -1.229953817 | 0.2187144 | 0.439129313 |
| Morn2 | 245.3325895 | -0.141555895 | 0.115110234 | -1.229742051 | 0.218793715 | 0.439231642 |
| Lrig2 | 1348.127048 | 0.128804825 | 0.062773343 | 2.051903212 | 0.04017907 | 0.150194968 |
| Nars | 7719.223909 | 0.128784099 | 0.078561565 | 1.63927615 | 0.101155761 | 0.272308699 |
| Sigmar1 | 1249.228201 | -0.083793938 | 0.068181177 | -1.228989309 | 0.219075815 | 0.439627078 |
| Scn7a | 192.6546179 | 0.128783902 | 0.123089241 | 1.046264486 | 0.295438937 | 0.527008738 |
| Hdac3 | 3300.136138 | 0.128777774 | 0.057453682 | 2.241419016 | 0.024998946 | 0.109148302 |
| Psme1 | 1074.220716 | 0.081853715 | 0.066660053 | 1.227927543 | 0.219474169 | 0.440255408 |
| Paqr6 | 220.7617775 | 0.128704093 | 0.132472425 | 0.971553839 | 0.331272557 | 0.566339732 |
| 4930441O14Rik | 138.8910197 | 0.128658148 | 0.109860547 | 1.171104202 | 0.241556895 | 0.466262791 |
| Dock11 | 422.6409196 | -0.166404045 | 0.135571978 | -1.227422119 | 0.219663977 | 0.440386617 |
| Ankrd54 | 470.1476557 | 0.128640573 | 0.103299468 | 1.245316897 | 0.213015287 | 0.43250673 |
| Ppp2r3a | 1818.965405 | 0.093344963 | 0.076052524 | 1.227374959 | 0.219681694 | 0.440386617 |
| Abca3 | 4752.108171 | 0.068026998 | 0.055431317 | 1.227230408 | 0.219736003 | 0.440438504 |
| Zswim5 | 889.6096737 | -0.112497115 | 0.09171662 | -1.226572838 | 0.219983183 | 0.440876917 |
| Zfp703 | 1364.58744 | 0.063844223 | 0.052073152 | 1.226048757 | 0.220180328 | 0.44121495 |
| Pcdhb15 | 126.4936235 | -0.145722863 | 0.118880652 | -1.225791249 | 0.220277241 | 0.441352072 |
| Chsy1 | 564.985191 | -0.118547924 | 0.096741335 | -1.225411289 | 0.220420296 | 0.441581596 |
| Mcph1 | 529.1154347 | -0.081137281 | 0.066227824 | -1.225123769 | 0.220528591 | 0.441741433 |
| Nup88 | 2085.678942 | 0.128589284 | 0.065920222 | 1.950680377 | 0.051095079 | 0.176121866 |
| Pcdhga7 | 690.94397 | 0.128500994 | 0.0926378 | 1.387133474 | 0.165401067 | 0.37100378 |
| Ppm1a | 6535.341505 | -0.079083409 | 0.064561494 | -1.224931525 | 0.220601021 | 0.441801565 |
| A230070E04Rik | 556.1036033 | 0.128453368 | 0.083604159 | 1.536447108 | 0.124428761 | 0.311793329 |
| Nkiras1 | 3047.50552 | 0.070592546 | 0.057656554 | 1.224362893 | 0.220815361 | 0.44191594 |
| Slc4a11 | 171.4043077 | 0.128383246 | 0.1572166 | 0.816601085 | 0.414156435 | 0.63764026 |
| Osbpl2 | 2536.114067 | 0.128350707 | 0.067255087 | 1.908416336 | 0.056337427 | 0.187451116 |
| Angptl6 | 239.6731691 | -0.167431002 | 0.136840253 | -1.223550807 | 0.221121727 | 0.442129311 |
| Dhx16 | 1012.827522 | 0.088513761 | 0.07232198 | 1.223884652 | 0.220995744 | 0.442129311 |
| Ctsf | 2235.774225 | 0.12834959 | 0.071470109 | 1.795849938 | 0.072518386 | 0.220313701 |
| Nmnat2 | 7365.105149 | 0.079196775 | 0.064720231 | 1.223678808 | 0.221073417 | 0.442129311 |
| Nrm | 85.00248411 | -0.177921108 | 0.145408767 | -1.223592719 | 0.221105908 | 0.442129311 |
| Upf1 | 2918.368698 | 0.128247191 | 0.060357382 | 2.12479709 | 0.033603548 | 0.133581176 |
| Nup54 | 583.0974803 | 0.128133089 | 0.078592877 | 1.630339729 | 0.103029715 | 0.275945101 |
| Mpzl2 | 77.66480809 | -0.2053997 | 0.167893936 | -1.223389632 | 0.221182567 | 0.442136847 |
| Zdhhc24 | 828.7894492 | 0.128101346 | 0.091902123 | 1.393888864 | 0.163351168 | 0.3686973 |

| | | | | | | |
|---|---|---|---|---|---|---|
| Zfp580 | 386.0110556 | -0.129404114 | 0.083332751 | -1.552860219 | 0.120456532 | 0.554583368 |
| Rttn | 235.9238054 | -0.129419946 | 0.084118676 | -1.53853998 | 0.123916641 | 0.559183948 |
| Myd88 | 260.7387984 | -0.129439174 | 0.084846287 | -1.525572647 | 0.127116328 | 0.564158811 |
| Gm3414 | 64.50861906 | -0.129562687 | 0.095102165 | -1.362352657 | 0.173086617 | 0.624504768 |
| Tert | 53.62264683 | -0.129583874 | 0.094070026 | -1.37752565 | 0.16834979 | 0.618474722 |
| Ccdc130 | 492.7954894 | -0.129591052 | 0.066205199 | -1.957415042 | 0.050298691 | 0.402334136 |
| Nid2 | 167.5688479 | -0.129644439 | 0.093918075 | -1.38039923 | 0.167463757 | 0.617953076 |
| Sema6c | 800.0252612 | -0.129841427 | 0.078914488 | -1.64534334 | 0.099899027 | 0.521115714 |
| Zbp1 | 17.60772859 | -0.129992317 | 0.047875659 | -2.715206864 | 0.006623438 | NA |
| Rad54b | 59.82860688 | -0.130006811 | 0.092011226 | -1.412945093 | 0.157671869 | 0.608141173 |
| Cacna1d | 2261.665325 | -0.130019829 | 0.053515515 | -2.429572626 | 0.015116636 | 0.233312354 |
| Mthfr | 1618.505898 | -0.130244556 | 0.071254919 | -1.827867576 | 0.067569428 | 0.448803776 |
| Rlbp1 | 836.9629526 | -0.130349758 | 0.08602124 | -1.515320612 | 0.129691231 | 0.568885619 |
| Slc38a11 | 84.89212454 | -0.130442433 | 0.094756093 | -1.37661262 | 0.168632047 | 0.619198462 |
| Mmaa | 794.8851144 | -0.130738712 | 0.070591859 | -1.852036683 | 0.064020556 | 0.441368101 |
| Lef1 | 244.7501972 | -0.130780139 | 0.084419818 | -1.549163959 | 0.121342311 | 0.555673148 |
| Epb4.1l2 | 4107.371655 | -0.130797889 | 0.048076984 | -2.720592673 | 0.0065165 | 0.155688625 |
| Gmcl1 | 1064.419704 | -0.130813664 | 0.061868555 | -2.114380479 | 0.034482784 | 0.33913129 |
| Wdr27 | 44.6007339 | -0.130831053 | 0.085061692 | -1.538072551 | 0.124030876 | 0.559205951 |
| Nedd1 | 147.1069924 | -0.130937584 | 0.088032378 | -1.487379831 | 0.136914521 | 0.581186537 |
| Arap3 | 539.8077453 | -0.130938325 | 0.08301527 | -1.57728 | 0.114731112 | 0.543130934 |
| Rad51ap2 | 34.49530744 | -0.130979568 | 0.091303801 | -1.434546722 | 0.151416306 | 0.598905575 |
| Col17a1 | 35.55108943 | -0.130983976 | 0.080683314 | -1.623433265 | 0.104496819 | 0.524427522 |
| Alkbh1 | 396.3504379 | -0.131032841 | 0.074540807 | -1.757867221 | 0.078770104 | 0.476674706 |
| Diap3 | 35.79586107 | -0.131181238 | 0.087426207 | -1.500479584 | 0.133490218 | 0.57588358 |
| Jag2 | 1517.115497 | -0.131227556 | 0.073741337 | -1.779565722 | 0.07514706 | 0.468962081 |
| Prtg | 168.2658957 | -0.131230111 | 0.091115217 | -1.440265582 | 0.149792275 | 0.598098567 |
| Shcbp1l | 61.97514073 | -0.131263198 | 0.094020134 | -1.396117954 | 0.162678978 | 0.614351408 |
| Rbm38 | 82.18855437 | -0.131280497 | 0.094508448 | -1.389087437 | 0.164806163 | 0.615071433 |
| Cyp4f13 | 367.8737408 | -0.131341276 | 0.083684807 | -1.569475755 | 0.116537124 | 0.546452317 |
| Cass4 | 17.83253685 | -0.131366541 | 0.079434961 | -1.653762263 | 0.098175826 | NA |
| Mfsd7b | 465.8565763 | -0.131371299 | 0.068874398 | -1.907403962 | 0.056468297 | 0.416586327 |
| Snord45b | 7.342379586 | -0.13141492 | 0.054860817 | -2.395424029 | 0.016601154 | NA |
| Nlrc5 | 83.70274765 | -0.131428115 | 0.053211463 | -2.469921088 | NA | NA |
| Slc25a35 | 217.7082853 | -0.131496139 | 0.087463487 | -1.503440394 | 0.132725517 | 0.574746597 |
| Nav1 | 4801.826613 | -0.131560628 | 0.068101788 | -1.931823412 | 0.053381305 | 0.407791741 |
| Cd74 | 332.7995493 | -0.13157187 | 0.050312084 | -2.615114682 | NA | NA |
| H2-Q5 | 20.2984589 | -0.131607808 | 0.077639326 | -1.695117867 | 0.090053064 | NA |
| Bfsp2 | 148.9153738 | -0.131630761 | 0.094666503 | -1.390468194 | 0.164386751 | 0.614641427 |
| Rock1 | 1115.86303 | -0.131715824 | 0.07338775 | -1.794793042 | 0.07268668 | 0.463090663 |
| Plekhg4 | 123.8804362 | -0.131731198 | 0.0948561 | -1.38874778 | 0.164909459 | 0.615071433 |
| Smg1 | 6914.010593 | -0.131832554 | 0.07000893 | -1.883081984 | 0.059689258 | 0.428234899 |
| Lama2 | 673.162143 | -0.131905234 | 0.080049452 | -1.647796849 | 0.099394366 | 0.520411863 |
| Gnl3 | 1014.18192 | -0.131952671 | 0.075758558 | -1.74175268 | 0.08155173 | 0.481161832 |
| Sec1 | 75.34318251 | -0.132146211 | 0.090509836 | -1.460020451 | 0.144284453 | 0.589967032 |
| Ptgs1 | 818.9620055 | -0.132155223 | 0.082576415 | -1.600399144 | 0.109510065 | 0.535243338 |
| Zfp560 | 277.9704453 | -0.13215672 | 0.090618636 | -1.458383455 | 0.144734878 | 0.590269908 |
| BC051142 | 89.41708228 | -0.132274599 | 0.095001151 | -1.392347337 | 0.163817244 | 0.614351408 |
| Ganc | 420.5319029 | -0.132282745 | 0.067261633 | -1.966689477 | 0.049219021 | 0.397725048 |
| Fank1 | 169.4674352 | -0.132398502 | 0.094841735 | -1.395994091 | 0.162716275 | 0.614351408 |
| Reck | 542.3347409 | -0.132469954 | 0.081464363 | -1.62610925 | 0.103926417 | 0.523645898 |
| Lrmp | 67.03359217 | -0.132497984 | 0.095101448 | -1.393227834 | 0.163550905 | 0.614351408 |
| Card9 | 64.37166191 | -0.132554426 | 0.095100511 | -1.393835063 | 0.163367417 | 0.614351408 |
| Pla2g4b | 168.9457738 | -0.132556652 | 0.095039694 | -1.394750412 | 0.163091117 | 0.614351408 |
| 1600020E01Rik | 174.1720981 | -0.132597005 | 0.089533101 | -1.480983045 | 0.138611092 | 0.583109099 |
| 1810020O05Rik | 42.49780954 | -0.132662328 | 0.093160982 | -1.424011688 | 0.154443089 | 0.60421231 |
| Scube1 | 3220.3861 | -0.132793 | 0.077337529 | -1.717057704 | 0.085968629 | 0.491258972 |
| Mfap2 | 116.0131626 | -0.132934899 | 0.094023445 | -1.413848429 | 0.157406411 | 0.608107852 |
| Cpsf1 | 2820.870974 | -0.132971281 | 0.066674932 | -1.994321953 | 0.046116882 | 0.38744582 |
| Crip3 | 52.05594756 | -0.132982235 | 0.083068339 | -1.600877499 | 0.109404054 | 0.535243338 |
| Fbxl12 | 458.8441898 | -0.133020592 | 0.086401641 | -1.539560947 | 0.123667411 | 0.558428989 |
| Snord69 | 15.83057544 | -0.133074881 | 0.076323115 | -1.74357244 | 0.081233672 | NA |
| Igf2bp2 | 36.73827311 | -0.133091455 | 0.090535941 | -1.470039995 | 0.141550922 | 0.587141261 |
| Tm7sf2 | 650.3721785 | -0.133178671 | 0.082393517 | -1.61637317 | 0.106013655 | 0.52850889 |
| Micall2 | 87.66224489 | -0.133190882 | 0.095046776 | -1.401319308 | 0.16111861 | 0.612724387 |
| Spats1 | 81.99153171 | -0.133225876 | 0.094987778 | -1.402558083 | 0.160748659 | 0.612387888 |
| Brca2 | 201.6242448 | -0.133333153 | 0.085162208 | -1.565637576 | 0.117433488 | 0.548468184 |
| Evc | 470.111337 | -0.133365239 | 0.081680187 | -1.63277342 | 0.102516659 | 0.522975271 |
| Mir374 | 22.22510651 | -0.133373361 | 0.075124661 | -1.775360576 | 0.075838369 | NA |
| Rnft1 | 595.2602143 | -0.133384529 | 0.076178982 | -1.750936099 | 0.079956918 | 0.478751136 |
| Gm5089 | 91.72100402 | -0.133483891 | 0.094215391 | -1.416794957 | 0.156542885 | 0.60719411 |
| Gm4532 | 39.1722044 | -0.133490409 | 0.090835493 | -1.46958423 | 0.141674394 | 0.587317993 |
| Mllt6 | 8366.595943 | -0.133540823 | 0.052439183 | -2.546584704 | 0.010878281 | 0.198764662 |
| Als2cl | 589.9148434 | -0.133578314 | 0.085521889 | -1.561919599 | 0.118306931 | 0.549576744 |
| Zfp40 | 675.4366362 | -0.13358551 | 0.073990476 | -1.805441953 | 0.071005551 | 0.458300422 |
| Celf5 | 11722.22512 | -0.133613092 | 0.060600465 | -2.204819579 | 0.027466758 | 0.308173879 |
| E130008D07Rik | 156.9718806 | -0.133620483 | 0.088333581 | -1.512680463 | 0.130360845 | 0.569619261 |
| Ddx20 | 527.7872978 | -0.133699396 | 0.078348486 | -1.706470708 | 0.087920476 | 0.495973914 |
| Gm8234 | 19.98504479 | -0.133703062 | 0.070236716 | -1.903606398 | 0.056961465 | NA |
| Boc | 1067.790138 | -0.133781296 | 0.072664688 | -1.841077147 | 0.065610254 | 0.4455608 |
| Lrrc16b | 2062.692951 | -0.133789706 | 0.088452824 | -1.512554379 | 0.13039289 | 0.569619261 |
| Prss22 | 46.28756175 | -0.133836031 | 0.091607738 | -1.460968616 | 0.144024054 | 0.589967032 |
| Acad10 | 133.1651685 | -0.133984852 | 0.095083927 | -1.409121991 | 0.158799099 | 0.609919544 |
| Aspn | 31.53221453 | -0.133998935 | 0.088117003 | -1.520693277 | 0.128336827 | 0.566288196 |
| Chek2 | 50.07923011 | -0.134029111 | 0.094171235 | -1.423248934 | 0.154664005 | 0.604402956 |
| Sp9 | 170.9697843 | -0.134037426 | 0.091275831 | -1.468487599 | 0.141971822 | 0.587991041 |
| Cenpe | 68.50988644 | -0.134045866 | 0.095096952 | -1.409570579 | 0.158666519 | 0.609571505 |
| Rad54l | 22.67863056 | -0.13417457 | 0.079311399 | -1.691743833 | 0.090694827 | 0.502789698 |
| Arhgap18 | 384.8337566 | -0.134199091 | 0.070657336 | -1.899294523 | 0.057525763 | 0.419820382 |
| 4933407L21Rik | 108.7815758 | -0.134226465 | 0.093358518 | -1.437752732 | 0.150504225 | 0.598511432 |
| Brd9 | 2820.315329 | -0.134242731 | 0.052462971 | -2.558809179 | 0.010503138 | 0.196333074 |
| Zmynd10 | 141.5679697 | -0.134254289 | 0.094892538 | -1.41480344 | 0.157126135 | 0.607833353 |
| Vcl | 1860.450452 | -0.134306138 | 0.06191888 | -2.169065998 | 0.030077674 | 0.319144111 |
| 5031434O11Rik | 94.5497811 | -0.134343276 | 0.094916202 | -1.415388241 | 0.156954696 | 0.607833353 |
| Mthfsd | 1025.394485 | -0.134399189 | 0.06639981 | -2.024089969 | 0.042960896 | 0.375481329 |
| S100a4 | 38.77029213 | -0.134451995 | 0.086877545 | -1.547603521 | 0.121717783 | 0.555997647 |
| 5430427O19Rik | 15.23594099 | -0.134607458 | 0.07525997 | -1.788566444 | 0.073684666 | NA |
| 4930563E22Rik | 116.1757101 | -0.134695474 | 0.09204063 | -1.463434933 | 0.143348406 | 0.589907432 |
| Mex3b | 558.2294407 | -0.134721043 | 0.080941413 | -1.664426624 | 0.096027203 | 0.515513544 |
| Dll4 | 130.1596886 | -0.134726886 | 0.094741679 | -1.42204453 | 0.155013324 | 0.604489543 |
| Slc22a18 | 28.66644486 | -0.134900214 | 0.081312015 | -1.659044031 | 0.09710692 | 0.517908272 |
| Sox17 | 194.9755461 | -0.135049548 | 0.094944691 | -1.422402311 | 0.154909493 | 0.604489543 |
| Usp49 | 720.1265382 | -0.135076883 | 0.062467302 | -2.162361415 | 0.030590329 | 0.320602541 |
| 9330151L19Rik | 849.1006789 | -0.135156554 | 0.068475807 | -1.973785493 | 0.048406139 | 0.394969953 |
| Zcchc8 | 1090.380134 | -0.135166486 | 0.057124078 | -2.366191117 | 0.017972164 | 0.255694081 |
| Tmem194 | 222.7605689 | -0.135175366 | 0.084383371 | -1.601919483 | 0.109173416 | 0.535010276 |
| Eogt | 1201.405251 | -0.135243942 | 0.060784805 | -2.224962999 | 0.026083722 | 0.302212096 |
| Pparg | 202.6920262 | -0.135415148 | 0.09325011 | -1.452171453 | 0.146453938 | 0.593251254 |
| Dact2 | 2835.294794 | -0.135445312 | 0.086869273 | -1.559185511 | 0.118952479 | 0.551341324 |
| Insrr | 22.40174239 | -0.135489157 | 0.076021891 | -1.78223871 | 0.074710313 | 0.468388672 |
| Il1r2 | 12.3452988 | -0.135565719 | 0.072334003 | -1.874163101 | 0.060907948 | NA |
| Scml2 | 37.62780447 | -0.135728575 | 0.089676681 | -1.513532537 | 0.130144443 | 0.569421137 |
| Ifitm7 | 20.43010166 | -0.13574414 | 0.080082025 | -1.695063792 | 0.09006332 | NA |
| Ccdc163 | 132.9788488 | -0.13591962 | 0.094423902 | -1.439462014 | 0.150019665 | 0.598234686 |
| 2610316D01Rik | 180.5611967 | -0.135928449 | 0.092418402 | -1.470794203 | 0.141346781 | 0.586989325 |
| Vim | 1457.748291 | -0.135949608 | 0.064695448 | -2.101378253 | 0.035607775 | 0.342728236 |
| Ube2cbp | 70.60453136 | -0.135949819 | 0.095095807 | -1.429608976 | 0.152829278 | 0.601462501 |
| 1700047M11Rik | 186.2978794 | -0.136051109 | 0.094249455 | -1.443521449 | 0.148873635 | 0.596726936 |
| Plscr3 | 645.1219281 | -0.136082143 | 0.075057826 | -1.813030692 | 0.069827088 | 0.455201066 |
| Casp7 | 110.3607678 | -0.136254384 | 0.094943956 | -1.435103302 | 0.151257663 | 0.598846773 |
| Gm20257 | 29.28663572 | -0.136383199 | 0.084022815 | -1.62316864 | 0.10455336 | 0.524427522 |
| Kif9 | 281.1062971 | -0.136393217 | 0.085200392 | -1.600851994 | 0.109409704 | 0.535243338 |
| Gm14169 | 395.8632721 | -0.136432163 | 0.087186601 | -1.564829476 | 0.117622898 | 0.548537627 |
| Tex40 | 276.8690588 | -0.136476594 | 0.092074127 | -1.482246945 | 0.1382746 | 0.582904025 |
| Tbx2 | 87.22981441 | -0.136659532 | 0.094962407 | -1.439090855 | 0.150124783 | 0.598439241 |
| Card10 | 320.6189843 | -0.137001899 | 0.083211534 | -1.646429194 | 0.099675427 | 0.521115714 |
| Fbrs | 1756.702395 | -0.137042144 | 0.066233801 | -2.06906657 | 0.03853984 | 0.355824003 |
| Cep250 | 1718.800348 | -0.137062594 | 0.071040331 | -1.92936312 | 0.053685799 | 0.408507027 |
| Ccl25 | 112.4924001 | -0.13713112 | 0.094587821 | -1.449775655 | 0.147121091 | 0.594031723 |
| Sepsecs | 254.8427767 | -0.137145374 | 0.084168428 | -1.629415875 | 0.10322501 | 0.522975271 |
| Antxr2 | 76.6736182 | -0.13722202 | 0.094883015 | -1.446223235 | 0.148114599 | 0.595162562 |
| Sema4a | 2325.548078 | -0.137351478 | 0.057435958 | -2.391384798 | 0.016784951 | 0.246336261 |
| Phf11b | 16.35940056 | -0.137402535 | 0.07712298 | -1.781603034 | 0.074813989 | NA |
| Ctsk | 130.4838285 | -0.137434246 | 0.092090912 | -1.492375776 | 0.135600662 | 0.579179191 |
| Sdccag3 | 3648.232078 | -0.137490963 | 0.06137627 | -2.240132285 | 0.025082336 | 0.29718272 |
| Cdon | 520.0464829 | -0.137535377 | 0.070741857 | -1.944186677 | 0.051872944 | 0.405765419 |
| Pigb | 310.1499431 | -0.137577972 | 0.076038427 | -1.809321637 | 0.070401049 | 0.456017009 |
| Fv1 | 216.0727659 | -0.137595696 | 0.085066871 | -1.617500377 | 0.105770314 | 0.527979239 |
| Zc3h8 | 260.7687989 | -0.137608523 | 0.078186306 | -1.760007996 | 0.078406451 | 0.475744405 |
| Itga10 | 180.6534112 | -0.137634763 | 0.094931481 | -1.449832671 | 0.147105186 | 0.594031723 |
| Cdkn2c | 62.15281764 | -0.137667831 | 0.095062031 | -1.448189452 | 0.147564074 | 0.594737171 |
| Ciz1 | 2131.239677 | -0.137754703 | 0.063915003 | -2.155279599 | 0.031139961 | 0.323885538 |
| C1qtnf4 | 2556.842732 | -0.137871074 | 0.07990721 | -1.725389651 | 0.084457275 | 0.488499377 |
| Lmo2 | 689.6803646 | -0.137898338 | 0.0711952 | -1.936905003 | 0.052756957 | 0.406864177 |
| 1700008J07Rik | 67.80167708 | -0.137925053 | 0.093653897 | -1.472710237 | 0.140829185 | 0.586773565 |
| Sestd1 | 3113.668229 | -0.137959543 | 0.054659145 | -2.52399744 | 0.011602877 | 0.204295026 |
| Nfkbiz | 192.3359947 | -0.138005906 | 0.090410673 | -1.526433784 | 0.126901869 | 0.563718247 |
| Nfam1 | 147.1800003 | -0.138015065 | 0.089552924 | -1.541156432 | 0.123278718 | 0.558038847 |
| Rnf32 | 305.0838004 | -0.138088112 | 0.078287387 | -1.763861544 | 0.077755293 | 0.475337606 |
| Fzd8 | 122.8274174 | -0.138094545 | 0.093137534 | -1.482694882 | 0.138155495 | 0.582904025 |
| 2510002D24Rik | 444.338675 | -0.138102934 | 0.070365763 | -1.962643871 | 0.049687569 | 0.399757904 |
| Fignl2 | 48.83703541 | -0.138171034 | 0.092004602 | -1.501783945 | 0.133152916 | 0.575193212 |
| Ctso | 1475.274626 | -0.13819333 | 0.05644072 | -2.448468568 | 0.014346495 | 0.229550543 |
| Susd3 | 50.77266966 | -0.138404264 | 0.094477103 | -1.464950333 | 0.142934469 | 0.589284353 |
| Stc2 | 38.73799743 | -0.138523185 | 0.091943513 | -1.506611837 | 0.131910182 | 0.572833273 |
| Uckl1 | 794.0249261 | -0.138555962 | 0.065380675 | -2.119218899 | 0.034071972 | 0.336970372 |
| Adck5 | 528.3807269 | -0.138634712 | 0.084782454 | -1.635181631 | 0.102010978 | 0.522723164 |
| Naalad2 | 76.04894275 | -0.138748129 | 0.094892856 | -1.462155689 | 0.143698551 | 0.589907432 |
| Itga8 | 231.2052956 | -0.138978482 | 0.086375784 | -1.608998208 | 0.107616734 | 0.532801865 |
| Pcdh20 | 938.0747184 | -0.138986357 | 0.087416846 | -1.589926463 | 0.111851382 | 0.538992394 |
| Exosc8 | 498.4980204 | -0.139080642 | 0.064224352 | -2.165543717 | 0.030346071 | 0.32021294 |
| Nr6a1 | 505.6576918 | -0.139124816 | 0.069890424 | -1.990613434 | 0.046523402 | 0.389285104 |
| Slc25a42 | 1547.53034 | -0.139127037 | 0.063935659 | -2.176047577 | 0.029551703 | 0.317771905 |
| Six5 | 88.28833281 | -0.139150465 | 0.094264885 | -1.476164376 | 0.13989977 | 0.58530647 |
| Chfr | 1441.505566 | -0.139191251 | 0.071857829 | -1.937036674 | 0.052740861 | 0.406864177 |
| Clcn2 | 2886.270063 | -0.139206499 | 0.063637235 | -2.18750074 | 0.028705989 | 0.315418457 |
| Gcnt4 | 691.4381233 | -0.139253782 | 0.083002592 | -1.677704018 | 0.093404895 | 0.50854554 |
| Has1 | 37.24989401 | -0.139377452 | 0.090127492 | -1.54644769 | 0.121996484 | 0.55651046 |
| Pdzd3 | 44.77838466 | -0.1393791 | 0.09248712 | -1.507010921 | 0.131807858 | 0.572833273 |
| Iffo1 | 2039.342718 | -0.139383015 | 0.07123081 | -1.956779869 | 0.050373355 | 0.402601282 |
| Gli1 | 137.8874235 | -0.139385868 | 0.089937454 | -1.549808918 | 0.121187386 | 0.555321126 |
| Trim68 | 222.8768647 | -0.139425763 | 0.09182722 | -1.518348945 | 0.128926455 | 0.567618221 |
| H2-Ob | 29.49696169 | -0.139454639 | 0.087463817 | -1.594426627 | 0.110840512 | 0.537406303 |
| Grn | 3077.845787 | -0.139476644 | 0.077204957 | -1.806576275 | 0.070828371 | 0.457674788 |
| Map4k2 | 2956.528494 | -0.139495929 | 0.07344386 | -1.899354546 | 0.057517876 | 0.419820382 |
| Kdelr3 | 16.03923588 | -0.139623216 | 0.071637938 | -1.949012218 | 0.051293966 | NA |
| AA474331 | 98.41892866 | -0.139625462 | 0.094272033 | -1.481091029 | 0.138582318 | 0.583109099 |
| 4833420G17Rik | 1452.447021 | -0.139671635 | 0.060914261 | -2.29292177 | 0.021852509 | 0.277716141 |
| BC030867 | 43.18538234 | -0.139754675 | 0.090537232 | -1.543615508 | 0.122681506 | 0.557245639 |
| Sp140 | 35.20064515 | -0.13986008 | 0.08870293 | -1.576724479 | 0.114858935 | 0.543130934 |
| Obsl1 | 623.4127347 | -0.139906738 | 0.085566205 | -1.635070036 | 0.102034367 | 0.522723164 |
| Slc31a2 | 386.0321703 | -0.139934865 | 0.085546772 | -1.635770264 | 0.101887678 | 0.522606076 |
| Chrnb1 | 31.52406175 | -0.139935805 | 0.084185302 | -1.662235593 | 0.096465546 | 0.51670168 |
| Rnf182 | 387.859798 | -0.139957239 | 0.084229715 | -1.661613583 | 0.096590278 | 0.516802454 |
| Wbp7 | 3108.802392 | -0.140075879 | 0.057387019 | -2.440898332 | 0.014650779 | 0.232079838 |
| Cdk3-ps | 68.55457882 | -0.140113611 | 0.093668408 | -1.495847038 | 0.134693521 | 0.577947814 |
| Rcor2 | 502.3597743 | -0.140118107 | 0.081413189 | -1.721073815 | 0.08523743 | 0.489390873 |
| Traip | 611.0538986 | -0.140185701 | 0.094450968 | -1.484216657 | 0.137751452 | 0.582270147 |
| Mlh1 | 522.1232483 | -0.14030257 | 0.071106122 | -1.973143336 | 0.048479234 | 0.395027926 |
| Sdccag8 | 688.4419746 | -0.140381314 | 0.066239952 | -2.119284637 | 0.034066419 | 0.336970372 |
| Tm6sf1 | 565.7723281 | -0.140382079 | 0.066949025 | -2.096850232 | 0.036006835 | 0.344080613 |
| Nlrp1b | 27.28018392 | -0.140394226 | 0.087359714 | -1.607082037 | 0.108036373 | 0.533425002 |
| Dtx3l | 327.908652 | -0.140425799 | 0.085450375 | -1.643360829 | 0.1003083 | 0.5211685 |
| Clec7a | 82.32206052 | -0.140446556 | 0.043794146 | -3.206971002 | NA | NA |
| Lox | 15.78146708 | -0.140451422 | 0.069467848 | -2.021819089 | 0.043195045 | NA |
| Dancr | 98.46820895 | -0.140488848 | 0.092993157 | -1.51074393 | 0.130853705 | 0.570291305 |
| Cd72 | 47.248882 | -0.140494521 | 0.087477903 | -1.60605726 | 0.108261329 | 0.533447785 |
| Nsmaf | 1411.451093 | -0.140609142 | 0.064099364 | -2.193612126 | 0.028263306 | 0.312827238 |
| C030039L03Rik | 662.6159213 | -0.140622176 | 0.084669636 | -1.660833593 | 0.096746872 | 0.516802454 |
| Mx2 | 42.49774316 | -0.14066445 | 0.089905369 | -1.564583417 | 0.11768062 | 0.548537627 |
| Pik3cg | 200.8150597 | -0.140670351 | 0.085176355 | -1.65151879 | 0.098632687 | 0.520050241 |
| Gm12250 | 22.8306853 | -0.140796375 | 0.053330369 | -2.640078797 | NA | NA |
| Capn12 | 31.51583232 | -0.140831416 | 0.087824781 | -1.603549869 | 0.108813308 | 0.534621387 |
| Erg | 174.8738988 | -0.140836061 | 0.091341435 | -1.541863895 | 0.123106671 | 0.557484892 |
| Pkn1 | 1791.118127 | -0.140853634 | 0.071145612 | -1.979793678 | 0.047726717 | 0.392678115 |
| Snord52 | 14.614972 | -0.140878376 | 0.075579747 | -1.863969946 | 0.062325925 | NA |
| Abca13 | 9.487931622 | -0.140900299 | 0.063635289 | -2.214184959 | 0.026816058 | NA |
| Klf8 | 218.2008194 | -0.140950074 | 0.084685414 | -1.664396117 | 0.096033295 | 0.515513544 |
| Srsf6 | 3817.196327 | -0.140970032 | 0.062657584 | -2.249847857 | 0.024458605 | 0.293140012 |
| Atp13a4 | 558.1329455 | -0.141022249 | 0.075593555 | -1.865532689 | 0.062106775 | 0.436531911 |
| Phf19 | 42.09001923 | -0.141092188 | 0.093474187 | -1.509424073 | 0.131190445 | 0.571416013 |
| Hspb2 | 23.59026371 | -0.141107796 | 0.082897078 | -1.702204705 | 0.088717001 | 0.497760972 |
| Nuf2 | 85.55900615 | -0.141177779 | 0.095027521 | -1.485651493 | 0.137371327 | 0.582012447 |
| Themis2 | 55.50492914 | -0.141255727 | 0.091550582 | -1.542925489 | 0.122848856 | 0.557484892 |
| Cep135 | 428.4065888 | -0.141515315 | 0.078592907 | -1.800611799 | 0.071764088 | 0.460037575 |
| Dnaaf3 | 52.94093693 | -0.141623415 | 0.093251042 | -1.518732787 | 0.12882977 | 0.567429121 |
| Pkmyt1 | 196.2881275 | -0.141645282 | 0.088176025 | -1.606392234 | 0.108187756 | 0.533447785 |
| Arfrp1 | 1299.37266 | -0.14165392 | 0.054307864 | -2.608350082 | 0.009097985 | 0.182034899 |
| Ptpn23 | 3290.752167 | -0.141695308 | 0.066777528 | -2.121901067 | 0.033846047 | 0.336326295 |
| Plgrkt | 381.377328 | -0.141874715 | 0.072337261 | -1.961295106 | 0.049844608 | 0.400244762 |
| Arhgap19 | 221.4718431 | -0.141877367 | 0.088649419 | -1.600431991 | 0.109502783 | 0.535243338 |
| Dtx3 | 8256.649771 | -0.141922427 | 0.068065035 | -2.085100338 | 0.037060194 | 0.348202098 |
| Tmem86b | 203.0136207 | -0.141940839 | 0.095096533 | -1.492597413 | 0.135542601 | 0.579179191 |
| Plcd1 | 508.4397483 | -0.142150026 | 0.079839133 | -1.780455521 | 0.075001442 | 0.468589657 |
| Zfp182 | 456.125923 | -0.142193708 | 0.081531163 | -1.744041208 | 0.081151905 | 0.479898997 |
| Isg15 | 83.97133234 | -0.142212549 | 0.059414461 | -2.393567949 | 0.01668539 | 0.245968033 |
| Ttf1 | 634.5481669 | -0.142370839 | 0.073296892 | -1.942385762 | 0.052090418 | 0.406272098 |
| Zfp160 | 557.5127804 | -0.142373319 | 0.067903938 | -2.096687218 | 0.036021273 | 0.344080613 |
| Cpne6 | 7355.165087 | -0.142396689 | 0.083681123 | -1.701658452 | 0.088819413 | 0.49789071 |
| Cers4 | 2097.65404 | -0.142417984 | 0.06098316 | -2.33536578 | 0.019524321 | 0.263578327 |
| Trim59 | 639.2742277 | -0.142436097 | 0.091856881 | -1.55063067 | 0.120990218 | 0.555002664 |
| Vars | 1939.050705 | -0.142499417 | 0.063377512 | -2.248422376 | 0.02454927 | 0.293500158 |
| Sat1 | 1244.529383 | -0.142653148 | 0.067706772 | -2.106925837 | 0.035124009 | 0.341050367 |
| Gtf2ird1 | 1231.874313 | -0.14288297 | 0.056693276 | -2.520280704 | 0.011726128 | 0.205471748 |
| Vasp | 539.6331441 | -0.142964191 | 0.075859502 | -1.884591732 | 0.059484979 | 0.428234899 |

| | | | | | | | | | | | | | | | |
|---|---|---|---|---|---|---|---|---|---|---|---|---|---|---|---|
| Nrg2 | 238.634586 | 0.12804285 | 0.087677448 | 1.460385238 | 0.144184228 | 0.341588749 | | | Ppip5k2 | 1359.53103 | -0.143015884 | 0.056603591 | -2.526622101 | 0.011516535 | 0.203514823 |
| Unc119b | 1244.430031 | -0.091942969 | 0.075191918 | -1.222777273 | 0.221413832 | 0.442484962 | | | Clk2 | 1253.894316 | -0.14303058 | 0.057374615 | -2.492924422 | 0.012669581 | 0.215249514 |
| Lrp6 | 2236.124704 | -0.092163079 | 0.075406546 | -1.222215891 | 0.221625996 | 0.442851842 | | | Gsta3 | 69.05903904 | -0.14303379 | 0.094611143 | -1.511807027 | 0.130582961 | 0.569624053 |
| Baz1b | 5566.544064 | 0.127988935 | 0.058905203 | 2.17279508 | 0.029795744 | 0.122845996 | | | Unc13d | 65.15363377 | -0.143108673 | 0.093729086 | -1.526833116 | 0.126802514 | 0.563718247 |
| Rspo3 | 366.9279618 | -0.155057063 | 0.126887156 | -1.222007557 | 0.221704769 | 0.442910968 | | | Npr2 | 2354.322058 | -0.14313291 | 0.067665648 | -2.115296515 | 0.034404683 | 0.338653871 |
| Necap2 | 935.8443304 | 0.127938652 | 0.087983381 | 1.454122935 | 0.145912226 | 0.344681206 | | | Sptbn4 | 8726.290236 | -0.143153842 | 0.06606437 | -2.166884232 | 0.030243683 | 0.320203887 |
| 3300002I08Rik | 24.77765357 | -0.201265728 | 0.164732052 | -1.221776365 | 0.221792209 | 0.442929929 | | | Sycp2 | 64.43173072 | -0.143265516 | 0.093711756 | -1.528789149 | 0.126316723 | 0.56284562 |
| Capza2 | 6975.491148 | -0.065476787 | 0.053593299 | -1.221734592 | 0.221808011 | 0.442929929 | | | Snora34 | 23.80395049 | -0.14331663 | 0.075669816 | -1.893973546 | 0.058228521 | 0.42291375 |
| Socs6 | 417.0393565 | -0.120474573 | 0.098629754 | -1.221483049 | 0.22190318 | 0.44306287 | | | Slc25a28 | 1174.424189 | -0.143636418 | 0.059542113 | -2.412350014 | 0.015850057 | 0.239084042 |
| Nup98 | 1105.824405 | 0.093701023 | 0.076720268 | 1.221333372 | 0.221959823 | 0.443118863 | | | R3hdm1 | 13183.71669 | -0.143698739 | 0.054397832 | -2.641626195 | 0.008250906 | 0.174077875 |
| Agl | 1917.02454 | -0.079990613 | 0.065502916 | -1.22117637 | 0.222019249 | 0.443158192 | | | C2cd5 | 2440.098642 | -0.143740531 | 0.052940306 | -2.715143579 | 0.006624704 | 0.156472283 |
| Zscan18 | 940.5353363 | 0.127934049 | 0.083070255 | 1.54007051 | 0.123543167 | 0.31078722 | | | 2310061J03Rik | 102.693532 | -0.143747034 | 0.092944481 | -1.546590319 | 0.121962066 | 0.55651046 |
| Adprh | 3094.872523 | -0.0533568 | 0.043707137 | -1.22078001 | 0.222169324 | 0.443251547 | | | Fxyd1 | 822.7900187 | -0.143922108 | 0.089784625 | -1.602970528 | 0.108941161 | 0.534621387 |
| Rnf6 | 2495.987197 | 0.127909492 | 0.092258258 | 1.386428641 | 0.165616056 | 0.371246408 | | | Eif2c4 | 704.8700998 | -0.143939125 | 0.073668616 | -1.953873076 | 0.050716229 | 0.403266245 |
| Nfib | 2903.103431 | -0.083600642 | 0.068480574 | -1.22079355 | 0.222164197 | 0.443251547 | | | Uaca | 420.0117618 | -0.144049897 | 0.088522307 | -1.627272293 | 0.10367928 | 0.523645898 |
| Slc5a11 | 33.77533449 | -0.205024479 | 0.167974594 | -1.220568383 | 0.222249483 | 0.443297309 | | | Eif2ak4 | 1228.65505 | -0.144079684 | 0.063907741 | -2.254495024 | 0.024165043 | 0.291788379 |
| Ssr1 | 5415.770971 | -0.065470127 | 0.05363633 | -1.220630249 | 0.222226048 | 0.443297309 | | | Ppfibp1 | 1394.768536 | -0.144279683 | 0.05299278 | -2.722629057 | 0.006476473 | 0.154987231 |
| Tmem8b | 4246.408514 | -0.08740621 | 0.071638475 | -1.220101495 | 0.222426402 | 0.443593084 | | | Nin | 2215.790977 | -0.14441699 | 0.080627008 | -1.791173862 | 0.073265399 | 0.46453653 |
| Aamdc | 575.091094 | -0.087465392 | 0.071702283 | -1.219841097 | 0.222525119 | 0.443624247 | | | Lgals4 | 8.303348144 | -0.144458785 | 0.060942159 | -2.370424461 | 0.017767674 | NA |
| Tex10 | 775.4214835 | 0.127786691 | 0.073866904 | 1.729958675 | 0.083637659 | 0.241318787 | | | Gm2011 | 27.39982895 | -0.144552309 | 0.083208982 | -1.737220017 | 0.082348343 | 0.483019453 |
| Mppe1 | 555.175926 | 0.127773723 | 0.073545466 | 1.737343297 | 0.082326593 | 0.238959955 | | | Syt15 | 39.74547114 | -0.144732191 | 0.092211486 | -1.569567934 | 0.116515663 | 0.546452317 |
| Wbp11 | 3524.039969 | 0.08552911 | 0.070119726 | 1.219758189 | 0.222556556 | 0.443624247 | | | Leprel1 | 101.5267641 | -0.14474289 | 0.092160521 | -1.570551993 | 0.116286749 | 0.546307021 |
| A430107P09Rik | 20.59531828 | -0.204439548 | 0.167655397 | -1.219403318 | 0.222691151 | 0.443835438 | | | Dqx1 | 29.97147434 | -0.144758474 | 0.087370294 | -1.656838584 | 0.097552114 | 0.518456434 |
| Shf | 1052.943489 | 0.127751401 | 0.071974595 | 1.7749513 | 0.075905929 | 0.226562601 | | | C530005A16Rik | 289.1343129 | -0.144844434 | 0.08143203 | -1.778715756 | 0.075286375 | 0.469360463 |
| Ubtf | 5537.519114 | 0.050944081 | 0.041801766 | 1.218706455 | 0.222955627 | 0.444248262 | | | Cst7 | 58.57180908 | -0.14485178 | 0.044716672 | -3.239323782 | 0.001198135 | 0.054729892 |
| Rbpj | 1607.941894 | 0.081825204 | 0.067152466 | 1.218498864 | 0.223034456 | 0.44429106 | | | Mms19 | 1677.284714 | -0.144872721 | 0.061656359 | -2.349680112 | 0.018789551 | 0.261433927 |
| Samd10 | 844.1962258 | 0.076043774 | 0.062407258 | 1.218508493 | 0.223030799 | 0.44429106 | | | Glis2 | 506.0661707 | -0.144890555 | 0.069873572 | -2.073610257 | 0.038115522 | 0.353328699 |
| Trpc1 | 1313.426421 | 0.087967359 | 0.072208966 | 1.218233193 | 0.223135369 | 0.444434942 | | | 1700095B22Rik | 42.22440262 | -0.144901058 | 0.088913581 | -1.629684198 | 0.103168259 | 0.522975271 |
| Alg13 | 236.2906821 | 0.127746007 | 0.101381144 | 1.260056868 | 0.207648848 | 0.425682886 | | | Matk | 6568.965917 | -0.144901734 | 0.075763558 | -1.912551861 | 0.055805447 | 0.413057568 |
| Arih2 | 2070.082701 | 0.127742952 | 0.084075107 | 1.519390889 | 0.128664134 | 0.317241369 | | | Mbd4 | 401.739327 | -0.144918905 | 0.086671186 | -1.672054018 | 0.094513671 | 0.511705397 |
| Btla | 14.16077839 | -0.202362112 | 0.166180286 | -1.217726345 | 0.223327982 | 0.444647103 | | | Zfp389 | 19.70221992 | -0.144989231 | 0.081100891 | -1.787763721 | 0.073814136 | NA |
| AW554918 | 596.6801396 | 0.098631829 | 0.081002051 | 1.217646065 | 0.223358501 | 0.444650728 | | | 2810403A07Rik | 2375.268501 | -0.145080008 | 0.068959979 | -2.10382907 | 0.03539336 | 0.342065168 |
| Cd99l2 | 3997.885305 | -0.100250235 | 0.08234684 | -1.217414467 | 0.223446561 | 0.444711756 | | | Aspdh | 79.22646283 | -0.145297188 | 0.094990817 | -1.529591938 | 0.126117766 | 0.562787717 |
| Brpf1 | 1408.262384 | 0.12771233 | 0.071701339 | 1.781170775 | 0.074884556 | 0.224884427 | | | Recql5 | 1077.131336 | -0.145304501 | 0.071323284 | -2.037265999 | 0.041623393 | 0.368446929 |
| Dock5 | 616.5031507 | -0.123316932 | 0.101310274 | -1.217220388 | 0.223520375 | 0.444744406 | | | Icosl | 203.3414158 | -0.145354936 | 0.092817213 | -1.566034264 | 0.117340596 | 0.548468184 |
| S100a6 | 436.3462323 | 0.127637418 | 0.115849218 | 1.101754677 | 0.270568339 | 0.498730506 | | | Lca5l | 42.14975927 | -0.145413719 | 0.09289971 | -1.565276348 | 0.117518127 | 0.548537627 |
| Foxm1 | 76.86541793 | -0.184648117 | 0.151731069 | -1.216943362 | 0.223625766 | 0.444896972 | | | Cdca7 | 128.187617 | -0.145448997 | 0.09098178 | -1.598660714 | 0.109896011 | 0.535492107 |
| Smc6 | 1622.652002 | 0.095936059 | 0.07886773 | 1.216417136 | 0.223826059 | 0.44523828 | | | Gas5 | 3082.567876 | -0.14555467 | 0.07199144 | -2.021833012 | 0.043193606 | 0.37570678 |
| Gltpd1 | 798.2591467 | 0.091876733 | 0.075541058 | 1.216248946 | 0.223890104 | 0.445308506 | | | She | 245.7796465 | -0.145619768 | 0.091050762 | -1.599325079 | 0.10974839 | 0.535311472 |
| Lrtm2 | 3457.775125 | 0.061726027 | 0.050760056 | 1.216035437 | 0.223971423 | 0.44541307 | | | Agxt2l2 | 113.9138907 | -0.145626229 | 0.093081838 | -1.564496715 | 0.117700964 | 0.548537627 |
| A930007I19Rik | 13.25595126 | 0.127599396 | 0.163631984 | 0.779794957 | 0.435511575 | 0.656402368 | | | Spag1 | 489.2889648 | -0.145671783 | 0.072903651 | -1.998141105 | 0.045701364 | 0.385357198 |
| Depdc5 | 2723.774465 | -0.071128335 | 0.058531911 | -1.215206096 | 0.224287496 | 0.445927174 | | | Zfp691 | 214.8860211 | -0.145707136 | 0.083012503 | -1.755243247 | 0.079217709 | 0.477013957 |
| Gpr75 | 446.1384369 | -0.108720181 | 0.089477831 | -1.215051594 | 0.224346414 | 0.445987085 | | | Urb1 | 709.3889534 | -0.145764405 | 0.065097863 | -2.239158077 | 0.025145633 | 0.29768986 |
| Slc7a1 | 2120.111597 | -0.102840137 | 0.084658529 | -1.214764052 | 0.224456095 | 0.446147882 | | | Ifi35 | 117.0458995 | -0.145984566 | 0.095100828 | -1.535050427 | 0.124771442 | 0.560776598 |
| Sox11 | 473.1803678 | -0.154346177 | 0.127097274 | -1.21439408 | 0.224597275 | 0.446371239 | | | Prss53 | 108.4733162 | -0.145991491 | 0.094934751 | -1.53780875 | 0.124095383 | 0.559205951 |
| Snhg1 | 246.8679815 | 0.127594805 | 0.104200454 | 1.224512948 | 0.220758785 | 0.44191594 | | | Cpxm1 | 284.5811399 | -0.146047555 | 0.088064983 | -1.658406672 | 0.097235411 | 0.518327184 |
| 4930452G13Rik | 12.46373813 | -0.194670667 | 0.16037503 | -1.213846488 | 0.224806351 | 0.446526045 | | | Greb1l | 458.1728604 | -0.146096824 | 0.0737448 | -1.981113573 | 0.047578539 | 0.392147258 |
| Aldh2 | 2521.000786 | -0.087702473 | 0.072258205 | -1.213737229 | 0.224848083 | 0.446526045 | | | 9230116N13Rik | 12.64693172 | -0.146153257 | 0.068845248 | -2.122924386 | 0.033760189 | NA |
| Aup1 | 1851.796753 | 0.092134572 | 0.075897273 | 1.213937846 | 0.22477146 | 0.446526045 | | | Casp2 | 509.3200995 | -0.146154241 | 0.071133985 | -2.054633109 | 0.039914455 | 0.361071695 |
| Dcaf13 | 734.6630133 | 0.07902049 | 0.065087868 | 1.214058648 | 0.224725329 | 0.446526045 | | | Afap1l2 | 412.699764 | -0.146246841 | 0.072295496 | -2.022903913 | 0.043083055 | 0.375514735 |
| Elfn2 | 4314.869525 | -0.091738796 | 0.075576235 | -1.213857717 | 0.224802062 | 0.446526045 | | | Plekho2 | 755.2281215 | -0.146297029 | 0.079471939 | -1.840863968 | 0.065641496 | 0.445564659 |
| Bcl7a | 4015.134442 | 0.078181447 | 0.064438207 | 1.213277816 | 0.225023621 | 0.446817377 | | | Tap1 | 208.1511572 | -0.146568135 | 0.068789136 | -2.130687242 | NA | NA |
| Eif4h | 12957.06104 | -0.060242298 | 0.049660783 | -1.213075877 | 0.225100812 | 0.446913375 | | | Pnldc1 | 53.75440173 | -0.146649156 | 0.089986537 | -1.629678861 | 0.103169387 | 0.522975271 |
| Stat5b | 1518.699839 | -0.094645898 | 0.078028556 | -1.21296488 | 0.225143248 | 0.446940357 | | | Zfp784 | 198.2798639 | -0.146651296 | 0.088068931 | -1.665187638 | 0.095875327 | 0.515046226 |
| Cdk5r2 | 9618.630456 | -0.114806707 | 0.094666765 | -1.212745643 | 0.225227083 | 0.447049504 | | | Ccnl1 | 1185.351787 | -0.146701415 | 0.06944887 | -2.112365751 | 0.03465509 | 0.33922827 |
| 0610009O20Rik | 1413.035145 | 0.127585189 | 0.076072933 | 1.677143021 | 0.093514518 | 0.258457713 | | | Cd274 | 182.7608905 | -0.146814934 | 0.070024972 | -2.096608261 | 0.036028267 | 0.344080613 |
| Chrna4 | 1802.433821 | -0.098084619 | 0.080902752 | -1.212376788 | 0.225368182 | 0.447194849 | | | Stk10 | 377.0270149 | -0.146852444 | 0.074468014 | -1.972020423 | 0.048607275 | 0.395027926 |
| Src | 5026.393582 | 0.127580317 | 0.083883724 | 1.520918614 | 0.128280262 | 0.316760167 | | | Dusp15 | 786.4301022 | -0.147022987 | 0.077899068 | -1.887352322 | 0.059112953 | 0.42635291 |
| Ublcp1 | 1739.040398 | 0.065800303 | 0.054291124 | 1.21199007 | 0.225516182 | 0.447394074 | | | Kif21b | 4919.488407 | -0.147054036 | 0.063705129 | -2.308354726 | 0.020979415 | 0.273070772 |
| Scn4b | 15817.88756 | -0.131104229 | 0.108200814 | -1.211675076 | 0.225636783 | 0.447576024 | | | Cr1l | 623.300326 | -0.147200572 | 0.06917514 | -2.127940351 | 0.033342029 | 0.334860976 |
| Ube2w | 1905.665933 | -0.095060138 | 0.078465643 | -1.211487393 | 0.225708663 | 0.447661294 | | | Cmtm6 | 1212.866882 | -0.147202535 | 0.062641504 | -2.349920189 | 0.018777437 | 0.261433927 |
| Lims2 | 410.3786252 | -0.135358477 | 0.111757704 | -1.211178048 | 0.225827174 | 0.447839016 | | | Proser1 | 1061.334287 | -0.147291778 | 0.063714517 | -2.311745974 | 0.020791686 | 0.271356725 |
| Hspa8 | 79645.73242 | -0.061707348 | 0.050991474 | -1.210150298 | 0.226221226 | 0.44856305 | | | Cln3 | 620.3859892 | -0.147367554 | 0.075545385 | -1.950715511 | 0.051090897 | 0.403266245 |
| Marveld3 | 19.68125727 | -0.203164071 | 0.167921583 | -1.209874679 | 0.226326985 | 0.448715329 | | | Fancf | 79.06984216 | -0.147436665 | 0.095024285 | -1.551568253 | 0.120765564 | 0.554612895 |
| Efhd1 | 787.2098693 | 0.127554614 | 0.109621389 | 1.163592391 | 0.244589236 | 0.469482858 | | | Gpr124 | 706.0051939 | -0.147451657 | 0.069819293 | -2.111904187 | 0.034694668 | 0.33922827 |
| Ubxn8 | 610.2856804 | -0.109426809 | 0.090465819 | -1.209592865 | 0.226435157 | 0.448757525 | | | Poln | 35.07360966 | -0.14769218 | 0.091941056 | -1.606378976 | 0.108190667 | 0.533447785 |
| Xk | 853.179271 | -0.080561556 | 0.066601265 | -1.209609987 | 0.226428584 | 0.448757525 | | | 9430083A17Rik | 107.2306028 | -0.147758808 | 0.095064119 | -1.554306807 | 0.120111251 | 0.553708672 |
| Gorab | 335.2005745 | -0.098535439 | 0.081493187 | -1.209124866 | 0.226614877 | 0.449017805 | | | Masp2 | 29.31607488 | -0.147773041 | 0.08350131 | -1.769709241 | 0.07677559 | 0.472560915 |
| Tgfbr1 | 1218.935316 | -0.109708693 | 0.090735836 | -1.209099945 | 0.226624451 | 0.449017805 | | | Gm10863 | 13.7541752 | -0.147845903 | 0.071055276 | -2.080716738 | 0.037459841 | NA |
| Sntb2 | 422.3698309 | 0.127446 | 0.112243188 | 1.135445296 | 0.25618879 | 0.482569738 | | | Pdss1 | 216.7109522 | -0.147897347 | 0.08368676 | -1.767272946 | 0.077182528 | 0.4735552 |
| Gucy2f | 49.22275876 | -0.192234545 | 0.159039874 | -1.208719174 | 0.226770753 | 0.449192795 | | | Irak4 | 71.2646764 | -0.147917662 | 0.095027258 | -1.556581402 | 0.119569902 | 0.55282356 |
| Mvk | 620.8295266 | -0.137366392 | 0.113681344 | -1.208345961 | 0.226914216 | 0.449416926 | | | Trim21 | 103.8013417 | -0.1479201 | 0.094472434 | -1.565748806 | 0.117407435 | 0.548468184 |
| Rps6 | 8462.512855 | -0.087572861 | 0.072477655 | -1.208273925 | 0.226941915 | 0.449416926 | | | Slc16a1 | 1317.211597 | -0.147938982 | 0.066336733 | -2.230121632 | 0.025739369 | 0.300614489 |
| Nat8l | 18840.71131 | 0.127437016 | 0.068751495 | 1.853589027 | 0.063797979 | 0.202351255 | | | Nfrkb | 1409.710376 | -0.148193526 | 0.063089791 | -2.34893036 | 0.018827428 | 0.261710254 |
| 4932443I19Rik | 15.12654626 | 0.127288671 | 0.149758044 | 0.849962167 | 0.39534612 | 0.620554431 | | | Pamr1 | 1809.134718 | -0.14824878 | 0.062107571 | -2.386967905 | 0.016987974 | 0.248007351 |
| Ncald | 20174.25527 | -0.111054383 | 0.09194722 | -1.207805769 | 0.227121982 | 0.449601123 | | | Grk4 | 87.88060051 | -0.148554815 | 0.094868678 | -1.565899489 | 0.11737215 | 0.548468184 |
| Ggh | 369.8772891 | -0.139542036 | 0.115597017 | -1.207142194 | 0.227377388 | 0.449991729 | | | Tle6 | 201.826859 | -0.14860294 | 0.090915628 | -1.634514797 | 0.102150802 | 0.522723164 |
| Msl2 | 2028.619259 | -0.072591261 | 0.060131089 | -1.207216811 | 0.227348658 | 0.449991729 | | | Mir3072 | 47.22242229 | -0.148648439 | 0.089259676 | -1.665348174 | 0.095843313 | 0.515046226 |
| Cops3 | 2205.103575 | -0.063366566 | 0.052521768 | -1.206481982 | 0.227631702 | 0.450437496 | | | Pole | 88.9540909 | -0.148866133 | 0.094837753 | -1.569692739 | 0.116486611 | 0.546452317 |
| Rpgrip1 | 26.68339034 | -0.200379139 | 0.166111191 | -1.20629524 | 0.227703672 | 0.450522372 | | | 9430037G07Rik | 99.76322476 | -0.148982312 | 0.092862711 | -1.604328685 | 0.108641621 | 0.534621387 |
| Slc25a3 | 11865.39342 | -0.065791962 | 0.054547713 | -1.206136029 | 0.227765045 | 0.450586262 | | | Ccl4 | 11.02313557 | -0.148997266 | 0.049825302 | -2.990393602 | 0.002786182 | NA |
| Bcas3 | 2529.635635 | 0.075234738 | 0.062400248 | 1.205680119 | 0.227940854 | 0.450876495 | | | Ccdc88b | 163.6501073 | -0.149009705 | 0.092674604 | -1.607880682 | 0.107861314 | 0.533015565 |
| Clns1a | 2222.744848 | -0.057411706 | 0.047630614 | -1.205353049 | 0.228067039 | 0.450971516 | | | B230209K01Rik | 66.34035792 | -0.149199405 | 0.094958595 | -1.571204859 | 0.116135072 | 0.546124329 |
| Lrch3 | 1585.957394 | 0.099208661 | 0.082308347 | 1.205329281 | 0.22807621 | 0.450971516 | | | Taf4a | 942.6251733 | -0.149210514 | 0.066916872 | -2.22978912 | 0.025761446 | 0.300614489 |
| Micall2 | 90.45157461 | -0.179288958 | 0.148743301 | -1.205358205 | 0.228065049 | 0.450971516 | | | Ascc2 | 758.425544 | -0.149225281 | 0.072879639 | -2.047557901 | 0.040603338 | 0.364752064 |
| 2310035C23Rik | 2608.006354 | 0.062100991 | 0.051525885 | 1.205238711 | 0.228111162 | 0.450983073 | | | Foxf2 | 168.6593079 | -0.149283069 | 0.094649595 | -1.577218254 | 0.114745314 | 0.543130934 |
| Lars | 1772.813595 | 0.07815956 | 0.064865644 | 1.204945414 | 0.228224375 | 0.451149333 | | | Arrdc4 | 332.8258418 | -0.149305397 | 0.074595441 | -2.001535163 | 0.045334748 | 0.383869599 |
| Elac2 | 1140.562648 | 0.09398 | 0.078015858 | 1.204626885 | 0.228347373 | 0.451219768 | | | Pola2 | 251.8079128 | -0.149321793 | 0.081568682 | -1.830626531 | 0.067156306 | 0.447688159 |
| Larp6 | 1330.007546 | 0.127269969 | 0.075247005 | 1.691362592 | 0.090767572 | 0.253890779 | | | Tex9 | 407.8750126 | -0.149368311 | 0.085011496 | -1.757036617 | 0.078911568 | 0.476674706 |
| Strap | 3966.3969 | -0.072851921 | 0.060475017 | -1.204661438 | 0.228334028 | 0.451219768 | | | Gt(ROSA)26Sor | 106.5920242 | -0.149479829 | 0.094899114 | -1.575144625 | 0.115223065 | 0.543478516 |
| Shb | 443.6533571 | -0.127203599 | 0.105613249 | -1.204428426 | 0.228424031 | 0.451313686 | | | Timm21 | 361.947193 | -0.149575938 | 0.075656354 | -1.977043972 | 0.04803666 | 0.393773739 |
| Mboat1 | 120.3048774 | 0.12726768 | 0.155951279 | 0.816073329 | 0.414458197 | 0.63764026 | | | Snord19 | 14.63033988 | -0.149635686 | 0.066095228 | -2.263940833 | 0.023577754 | NA |
| Pgam2 | 164.7044779 | -0.17089323 | 0.141914944 | -1.204194742 | 0.228514318 | 0.451376956 | | | Tubd1 | 195.6050602 | -0.149637511 | 0.091446013 | -1.636348118 | 0.101766751 | 0.522606076 |
| Enpp1 | 162.6575017 | -0.164654593 | 0.136784207 | -1.203754417 | 0.228684513 | 0.451625538 | | | Gm10941 | 12.52539075 | -0.149826866 | 0.068769957 | -2.178667436 | 0.029356383 | NA |
| Prdm15 | 772.8845985 | 0.127231357 | 0.095211216 | 1.336306397 | 0.181449146 | 0.391944252 | | | Igj | 15.31356732 | -0.149842657 | 0.077031207 | -1.945220159 | 0.051748487 | NA |
| Abhd4 | 2784.680551 | 0.127218042 | 0.076288653 | 1.66758799 | 0.095397544 | 0.261469936 | | | Tmem194b | 213.3653175 | -0.149974051 | 0.086248581 | -1.738858178 | 0.082059712 | 0.482582908 |
| Osgepl1 | 440.428318 | 0.127172156 | 0.094095274 | 1.351525433 | 0.176527178 | 0.385784666 | | | Cd53 | 281.9478423 | -0.150075402 | 0.085801905 | -1.749091717 | 0.080275167 | 0.479231997 |
| Gabrb2 | 11914.94012 | -0.093715257 | 0.077870368 | -1.20347777 | 0.228791489 | 0.451686079 | | | Fam83f | 28.82903363 | -0.150140443 | 0.088681187 | -1.69303601 | 0.090448613 | 0.502238744 |
| Med18 | 103.857058 | -0.143978704 | 0.119668454 | -1.203146681 | 0.228919564 | 0.451831812 | | | Ano8 | 2342.560518 | -0.150148299 | 0.067461111 | -2.225701525 | 0.026034181 | 0.3021678 |
| Gtpbp3 | 741.5823206 | 0.127157333 | 0.093016334 | 1.367043057 | 0.17161181 | 0.379146355 | | | Cttnbp2 | 4874.898029 | -0.150163549 | 0.056905982 | -2.638800751 | 0.008319986 | 0.174157793 |
| Thoc2 | 1498.405905 | 0.098999443 | 0.082309917 | 1.202764458 | 0.229067482 | 0.452040467 | | | Pde4c | 145.1428472 | -0.1501958 | 0.090818619 | -1.653799646 | 0.098168228 | 0.519297771 |
| Nelfa | 732.0148985 | 0.127137232 | 0.071889813 | 1.768501369 | 0.076977124 | 0.228483439 | | | Usp43 | 125.7584053 | -0.150346744 | 0.091980055 | -1.634558101 | 0.102141717 | 0.522723164 |
| Lamtor2 | 921.8769241 | -0.075354664 | 0.062679885 | -1.202214451 | 0.229280452 | 0.452292337 | | | Tarbp2 | 686.8695132 | -0.150408732 | 0.083003739 | -1.812071761 | 0.069975109 | 0.455201066 |
| Tmem185b | 497.831753 | -0.095026376 | 0.079045905 | -1.202166967 | 0.229298845 | 0.452292337 | | | Itga6 | 910.3856358 | -0.150559472 | 0.075120812 | -2.004231158 | 0.045045304 | 0.383004357 |
| 8430419L09Rik | 12703.0853 | 0.073092282 | 0.060810598 | 1.201966169 | 0.229376636 | 0.452388187 | | | Rhebl1 | 239.1888389 | -0.150572392 | 0.091310842 | -1.649008911 | 0.099145809 | 0.520271327 |
| Prr5l | 477.5146725 | 0.127081614 | 0.146135895 | 0.869612589 | 0.384512156 | 0.61266645 | | | Prr7 | 574.3429618 | -0.150723753 | 0.092594752 | -1.627778575 | 0.103571845 | 0.523462117 |
| Lsm5 | 105.4282647 | -0.142255748 | 0.118383133 | -1.201655543 | 0.229497012 | 0.452510398 | | | Agmat | 85.96677476 | -0.150758521 | 0.094935697 | -1.588006675 | 0.11228483 | 0.538992394 |
| Cul7 | 1397.301564 | -0.076178277 | 0.063401464 | -1.201522368 | 0.229548635 | 0.452554594 | | | 2410089E03Rik | 2455.684771 | -0.150807668 | 0.066009359 | -2.284640678 | 0.022333906 | 0.281750998 |
| Map1lc3a | 4712.989746 | -0.1186927 | 0.098796922 | -1.201380546 | 0.229603619 | 0.452605404 | | | Trpm7 | 2099.75254 | -0.15087907 | 0.065389601 | -2.307386311 | 0.021033294 | 0.273267769 |
| Ranbp10 | 1736.692805 | 0.12688284 | 0.062364441 | 2.034538227 | 0.041897358 | 0.155009396 | | | Rbm28 | 2102.872684 | -0.150913078 | 0.065448414 | -2.305832472 | 0.021119995 | 0.273675398 |
| Ezh2 | 161.820538 | 0.126836654 | 0.133932214 | 0.94702126 | 0.343627946 | 0.576022005 | | | Tial1 | 3403.502488 | -0.150979011 | 0.058744905 | -2.570078383 | 0.010167551 | 0.193316546 |
| Zfp781 | 95.51744913 | -0.16668869 | 0.138813608 | -1.200809441 | 0.229825129 | 0.452869203 | | | Intu | 639.5592727 | -0.150981829 | 0.068143146 | -2.215656867 | 0.026715011 | 0.305080386 |
| 6430584L05Rik | 182.651583 | -0.143646055 | 0.119669254 | -1.200358909 | 0.22999998 | 0.452972079 | | | Irgm2 | 303.3758548 | -0.151037486 | 0.068145501 | -2.216397031 | NA | NA |
| Brpf3 | 1802.637395 | 0.12672318 | 0.073981456 | 1.712904647 | 0.086730083 | 0.246568057 | | | Ophn1 | 1889.627723 | -0.151183031 | 0.046843678 | -3.22739456 | 0.001249231 | 0.056707267 |
| Map3k2 | 1202.154553 | -0.112070549 | 0.093367082 | -1.200321862 | 0.230014363 | 0.452972079 | | | Abca1 | 1691.996749 | -0.151281039 | 0.075613595 | -2.000712167 | 0.045423418 | 0.384063191 |
| Mlec | 6412.875781 | -0.078078137 | 0.065040992 | -1.20044505 | 0.229966541 | 0.452972079 | | | Ccdc88c | 1421.006506 | -0.151516005 | 0.066276594 | -2.286116345 | 0.022247453 | 0.281258923 |
| Ppp2r5e | 3608.369154 | -0.073089693 | 0.060881857 | -1.200516811 | 0.229938688 | 0.452972079 | | | Dgkq | 2219.894605 | -0.151635508 | 0.060240757 | -2.517158092 | 0.011830573 | 0.20655157 |
| Rpl3 | 17829.27268 | -0.070670019 | 0.058882734 | -1.200182372 | 0.23006852 | 0.453003132 | | | Il16 | 179.9539773 | -0.15181687 | 0.079381249 | -1.912502905 | 0.05581172 | 0.413057568 |
| Atg16l1 | 2597.244093 | 0.126692179 | 0.056973266 | 2.223712773 | 0.026167775 | 0.11265449 | | | Bora | 66.56986819 | -0.151829945 | 0.094973918 | -1.598648849 | 0.109898649 | 0.535492107 |
| Csde1 | 8449.548095 | -0.064218265 | 0.053526218 | -1.199753451 | 0.230235108 | 0.453215952 | | | Iqcc | 535.0023727 | -0.151845838 | 0.072833037 | -2.084848354 | 0.037083068 | 0.348202098 |
| Chaf1b | 19.67289757 | -0.201608843 | 0.16806522 | -1.199586942 | 0.2302998 | 0.45328571 | | | Ryr3 | 2044.482804 | -0.151871149 | 0.085831563 | -1.769409102 | 0.076825628 | 0.472560915 |
| 3110009E18Rik | 32.94612746 | -0.198195807 | 0.165361456 | -1.198561088 | 0.230698656 | 0.453668953 | | | Gm527 | 151.1447623 | -0.151938554 | 0.092147564 | -1.648861314 | 0.09917605 | 0.520271327 |
| Akip1 | 85.8667189 | 0.126675906 | 0.134298615 | 0.943240599 | 0.345557852 | 0.576948985 | | | Ninl | 318.5706394 | -0.151976554 | 0.084928192 | -1.789471201 | 0.073538961 | 0.465458367 |
| Fis1 | 3738.827597 | -0.093193917 | 0.077745271 | -1.198708491 | 0.230641315 | 0.453668953 | | | Mir1249 | 47.23218661 | -0.151998604 | 0.090268044 | -1.683858401 | 0.092209037 | 0.505826464 |
| Gdap1 | 4542.465689 | -0.076140457 | 0.063527034 | -1.19855204 | 0.230702177 | 0.453668953 | | | 2210404J11Rik | 259.2478093 | -0.152038415 | 0.082081566 | -1.852284523 | 0.063984977 | 0.441368101 |
| Gyk | 824.5423227 | -0.086866861 | 0.072494289 | -1.198257987 | 0.230816597 | 0.453668953 | | | Gm13826 | 65.35142094 | -0.152057592 | 0.088093264 | -1.726097833 | 0.084329814 | 0.488499377 |
| Klf3 | 652.7116837 | -0.099625875 | 0.08310645 | -1.198774279 | 0.230615726 | 0.453668953 | | | Il17rc | 124.1346834 | -0.152081663 | 0.095089689 | -1.599349662 | 0.109742931 | 0.535311472 |
| Psmf1 | 1353.005288 | 0.126587136 | 0.073297256 | 1.727037854 | 0.084160865 | 0.241972249 | | | Spata1 | 34.03759494 | -0.152127292 | 0.08444717 | -1.801449267 | 0.071632097 | 0.459800193 |
| Rhobtb2 | 5472.627938 | -0.084579865 | 0.070584066 | -1.19828553 | 0.230805878 | 0.453668953 | | | 2610015P09Rik | 258.4250176 | -0.1521334 | 0.089064685 | -1.70812258 | 0.0876136 | 0.495597906 |
| Gas6 | 3456.335157 | 0.126577536 | 0.07524262 | 1.682258483 | 0.092518728 | 0.256666996 | | | Cstad | 60.85532511 | -0.152172716 | 0.094735319 | -1.606293385 | 0.108209463 | 0.533447785 |
| Clec11a | 147.4760546 | 0.126518184 | 0.134909562 | 0.937799973 | 0.348347215 | 0.579556372 | | | Wnt10b | 45.20657458 | -0.152228869 | 0.089398634 | -1.70280978 | 0.088603672 | 0.497517107 |
| Zfp518a | 412.8097296 | -0.151926464 | 0.126765421 | -1.198485066 | 0.230728233 | 0.453668953 | | | Slc7a6os | 695.002622 | -0.152338855 | 0.060397291 | -2.522279588 | 0.011659699 | 0.205046966 |
| Gm15760 | 10.72703723 | 0.126436118 | 0.154631334 | 0.817661691 | 0.413550391 | 0.637376878 | | | Ctns | 794.2379829 | -0.152447081 | 0.064104877 | -2.378088661 | 0.017402641 | 0.252286191 |
| 1110038F14Rik | 460.1336451 | -0.115895004 | 0.09677964 | -1.197514317 | 0.23110615 | 0.45406327 | | | Zan | 40.55497771 | -0.152462212 | 0.090104141 | -1.692066648 | 0.090633267 | 0.502789698 |
| Sod1 | 5637.486818 | -0.080963646 | 0.067611636 | -1.197480957 | 0.231119146 | 0.45406327 | | | Alkbh2 | 115.5286466 | -0.152521816 | 0.091382762 | -1.669043628 | 0.095108734 | 0.513969296 |
| Fhad1 | 283.7044634 | 0.126426184 | 0.154869599 | 0.816339579 | 0.414305943 | 0.63764026 | | | Nfxl1 | 442.8028178 | -0.152549933 | 0.068758391 | -2.218637324 | 0.026511408 | 0.303950052 |
| Fignl2 | 54.35269759 | -0.18281599 | 0.152686377 | -1.197330068 | 0.231177929 | 0.45409113 | | | Nudt13 | 438.5675273 | -0.152633051 | 0.07811181 | -1.954032961 | 0.050697319 | 0.403266245 |
| Diap2 | 1206.214312 | -0.118666661 | 0.099119184 | -1.197211842 | 0.231223995 | 0.454124044 | | | Itga1 | 292.2322211 | -0.152635367 | 0.085194198 | -1.791616935 | 0.073194349 | 0.464395167 |
| Dexi | 1148.134135 | 0.126328838 | 0.084216932 | 1.500040845 | 0.133603822 | 0.324783353 | | | Ptar1 | 225.1959051 | -0.152639885 | 0.089272371 | -1.709822232 | 0.087298751 | 0.495352208 |
| Rab33a | 727.5078049 | 0.092328226 | 0.077139639 | 1.196897303 | 0.231346586 | 0.454249651 | | | Tep1 | 312.0351233 | -0.152658431 | 0.089993531 | -1.696326722 | 0.089824023 | 0.50087668 |
| Ctdsp2 | 4886.588314 | 0.090730267 | 0.075810439 | 1.196804403 | 0.231382802 | 0.454263194 | | | Gmip | 404.1568533 | -0.152674213 | 0.082313757 | -1.854783671 | 0.063627125 | 0.440996088 |
| Pla1a | 57.50642296 | 0.12631771 | 0.155580023 | 0.811914712 | 0.416840572 | 0.639521519 | | | Ppil6 | 96.63396439 | -0.152691387 | 0.094302155 | -1.619171767 | 0.105410311 | 0.526906461 |
| Vps39 | 3660.325859 | 0.126201465 | 0.050958627 | 2.476547594 | 0.013265994 | 0.071468465 | | | Adamts6 | 152.558716 | -0.152949887 | 0.090690086 | -1.686511654 | 0.091697292 | 0.504543511 |
| Fscn1 | 7521.526885 | -0.091902387 | 0.076856102 | -1.19577216 | 0.231785483 | 0.454880845 | | | Snora16a | 19.09954908 | -0.153064886 | 0.074860548 | -2.044666924 | 0.040887707 | NA |
| Mfsd2a | 910.5516322 | 0.126152676 | 0.167983027 | 0.75098466 | 0.452661888 | 0.671466535 | | | Dnase1l2 | 164.6030468 | -0.153087076 | 0.093016334 | -1.645808528 | 0.099803186 | 0.521115714 |
| Fam133b | 1070.402896 | 0.126122316 | 0.106177381 | 1.187845421 | 0.234894316 | 0.458484824 | | | Dgka | 506.0286943 | -0.153130676 | 0.066844651 | -2.290844124 | 0.021972431 | 0.278517602 |
| Six1 | 11.93115248 | -0.188837153 | 0.158037491 | -1.194888323 | 0.232130666 | 0.455245894 | | | Cela1 | 83.13800065 | -0.153281814 | 0.094332358 | -1.624912348 | 0.104181237 | 0.523645898 |
| Fbxo22 | 1358.376155 | 0.126111582 | 0.075055755 | 1.680238677 | 0.092910887 | 0.257340236 | | | Otogl | 15.62130939 | -0.153559298 | 0.071065358 | -2.160817885 | 0.03070941 | NA |
| Shh | 104.7925646 | 0.126097751 | 0.148029244 | 0.851840635 | 0.394302549 | 0.619921228 | | | Knstrn | 39.21201519 | -0.153638271 | 0.091060894 | -1.68720363 | 0.091564204 | 0.504530343 |
| Tax1bp1 | 6135.004462 | -0.073115983 | 0.061192157 | -1.194858722 | 0.232142233 | 0.455245894 | | | Hars2 | 1117.019929 | -0.153734278 | 0.06384024 | -2.408109328 | 0.016035378 | 0.240747652 |
| Klhdc3 | 6645.267689 | 0.1260457 | 0.084536955 | 1.491013005 | 0.13595808 | 0.328325202 | | | Mif4gd | 387.4049863 | -0.15374118 | 0.079642336 | -1.930395173 | 0.053557893 | 0.407961168 |
| Nod1 | 246.3475725 | 0.126030284 | 0.122759496 | 1.026643866 | 0.30458819 | 0.537031958 | | | Wdfy4 | 137.6597233 | -0.153787609 | 0.088180835 | -1.744002631 | 0.081158631 | 0.479898997 |
| Olig1 | 2955.154384 | -0.129746246 | 0.10875886 | -1.192971743 | 0.23288044 | 0.456378333 | | | Ncapd3 | 1276.259089 | -0.15384376 | 0.056196321 | -2.737612641 | 0.006188692 | 0.150581145 |
| Phrf1 | 2266.401425 | 0.080018319 | 0.067072739 | 1.193008081 | 0.232866208 | 0.456378333 | | | Tulp3 | 857.8146805 | -0.153879674 | 0.073438229 | -2.095362006 | 0.036138824 | 0.344457062 |
| Shq1 | 133.1838679 | -0.141146944 | 0.118327804 | -1.192846806 | 0.232929375 | 0.456378333 | | | Ntn5 | 250.090193 | -0.15389603 | 0.089520947 | -1.719106358 | 0.085595007 | 0.490281182 |
| Trappc6b | 2116.113181 | -0.065113913 | 0.054587229 | -1.192841514 | 0.232931448 | 0.456378333 | | | 2010111I01Rik | 728.4696742 | -0.153911905 | 0.091678233 | -1.678827143 | 0.093185737 | 0.507923459 |
| Ube2z | 4246.787922 | -0.073573614 | 0.061671275 | -1.192996487 | 0.232870749 | 0.456378333 | | | Rsad1 | 808.3126686 | -0.154058923 | 0.067502639 | -2.282265195 | 0.022473688 | 0.281750998 |
| Slu7 | 2534.832178 | 0.076301914 | 0.06397157 | 1.192747253 | 0.232968374 | 0.456392967 | | | Qtrt1 | 384.1259543 | -0.154122416 | 0.088394308 | -1.743578511 | 0.081232613 | 0.479898997 |
| Pik3r4 | 2085.168807 | 0.087262973 | 0.073177564 | 1.192482616 | 0.233072063 | 0.456500657 | | | Polm | 286.0286224 | -0.154229627 | 0.08747146 | -1.763199413 | 0.077866863 | 0.475337606 |
| Slc46a1 | 333.2542117 | -0.15136577 | 0.126936087 | -1.192456569 | 0.233082271 | 0.456500657 | | | Itgb5 | 2252.775838 | -0.154246573 | 0.074835261 | -2.061148327 | 0.039288892 | 0.359615903 |
| Ube2o | 6890.490458 | 0.126023472 | 0.054828193 | 2.298515867 | 0.021532446 | 0.098910085 | | | Trp53 | 878.1366384 | -0.15426613 | 0.064059587 | -2.408166184 | 0.016032881 | 0.240747652 |
| Sh3kbp1 | 1657.543605 | -0.094327066 | 0.079121233 | -1.192183979 | 0.233189114 | 0.456594481 | | | Fcgr4 | 21.22068383 | -0.154311858 | 0.054986533 | -2.806357314 | 0.00501051 | NA |
| Tom1 | 1595.942335 | 0.082552377 | 0.069258935 | 1.191938293 | 0.233285442 | 0.456725377 | | | Mapk1ip1 | 1277.521033 | -0.154339354 | 0.059707766 | -2.584912575 | 0.009740369 | 0.18840027 |
| Mlst8 | 1685.50605 | 0.074354405 | 0.062393869 | 1.191694102 | 0.233381212 | 0.456855148 | | | Memo1 | 713.8941971 | -0.154628059 | 0.062731791 | -2.464907457 | 0.013704864 | 0.224208584 |
| Sar1a | 5238.054964 | -0.062128179 | 0.052139267 | -1.191581371 | 0.233425433 | 0.456883989 | | | Slc35d1 | 369.1764293 | -0.154649186 | 0.076439689 | -2.023153006 | 0.043057375 | 0.375514735 |
| Bysl | 1094.647966 | -0.08658585 | 0.072702485 | -1.190961357 | 0.233668755 | 0.457143705 | | | Ggnbp1 | 110.105889 | -0.154692215 | 0.095041599 | -1.627626391 | 0.10360413 | 0.523462117 |
| Zc3h14 | 3252.519103 | 0.125990473 | 0.070838012 | 1.778571538 | 0.075310034 | 0.225566144 | | | Slfn9 | 22.44223661 | -0.154697308 | 0.082990696 | -1.864031948 | 0.062317218 | 0.436873272 |
| Mast1 | 3557.106992 | 0.125912761 | 0.066673989 | 1.888483987 | 0.058961006 | 0.192839133 | | | Higd1b | 77.64062183 | -0.15478375 | 0.094368456 | -1.640206444 | 0.10096225 | 0.521542547 |
| Wdr43 | 985.7000102 | 0.074050944 | 0.062176786 | 1.190974129 | 0.233663741 | 0.457143705 | | | Fxyd5 | 453.6522966 | -0.154882053 | 0.087825099 | -1.763528358 | 0.077811419 | 0.475337606 |
| Adk | 1916.002114 | -0.079358351 | 0.066655862 | -1.190568221 | 0.233823133 | 0.457315783 | | | Gpr65 | 8.840092391 | -0.15494588 | 0.063350591 | -2.445847434 | 0.014451214 | NA |
| Wnt16 | 19.30065258 | -0.198401661 | 0.166640175 | -1.190599211 | 0.233810961 | 0.457315783 | | | Pdgfrb | 1131.761926 | -0.154959584 | 0.07533371 | -2.056975356 | 0.039688596 | 0.36080087 |
| Smad5 | 1586.461262 | -0.088665355 | 0.074484747 | -1.190382704 | 0.233896008 | 0.457400574 | | | Fcho1 | 4313.482876 | -0.15507052 | 0.076770005 | -2.019936307 | 0.043389995 | 0.37703831 |
| Cib1 | 343.5618478 | 0.090044217 | 0.075659069 | 1.190131181 | 0.233994836 | 0.457478359 | | | Gm1673 | 244.4016644 | -0.155153506 | 0.094833106 | -1.63606901 | 0.101825145 | 0.522606076 |
| Slc16a11 | 468.338856 | 0.125892858 | 0.132297298 | 0.951590543 | 0.341304679 | 0.574801598 | | | Grk1 | 31.27034136 | -0.155220194 | 0.089274885 | -1.738677409 | 0.082091585 | 0.482582908 |
| Pcif1 | 1743.081628 | 0.097296818 | 0.081774898 | 1.189812761 | 0.234119993 | 0.4576653 | | | Dpp7 | 932.6501703 | -0.155240428 | 0.085573517 | -1.814117649 | 0.069659615 | 0.455184691 |
| D630003M21Rik | 285.4808509 | -0.157184957 | 0.132135502 | -1.189573995 | 0.234213872 | 0.45779106 | | | 6430584L05Rik | 180.838553 | -0.155270755 | 0.090239816 | -1.720645736 | 0.085315129 | 0.489643446 |
| Coq2 | 2160.851229 | -0.096675322 | 0.081285115 | -1.189336102 | 0.234307434 | 0.457916169 | | | Colec12 | 600.8469558 | -0.155301385 | 0.08404201 | -1.847901836 | 0.064616545 | 0.443022969 |
| Csnk2b | 2714.220313 | 0.125851187 | 0.079025593 | 1.592537079 | 0.111264079 | 0.290535356 | | | Ect2 | 20.38607892 | -0.155305497 | 0.080825759 | -1.92148516 | 0.054670572 | NA |
| Socs4 | 204.4773262 | -0.12864508 | 0.108217463 | -1.188764521 | 0.234532343 | 0.458182338 | | | Tom1l1 | 752.4558003 | -0.155475895 | 0.060915056 | -2.552339338 | 0.010700225 | 0.19724805 |
| Ubald1 | 3177.275875 | 0.077270078 | 0.064997052 | 1.188824342 | 0.234508797 | 0.458182338 | | | Bves | 29.7111861 | -0.155551489 | 0.089263055 | -1.742618931 | 0.081400201 | 0.480521762 |
| Acvr2b | 133.268294 | -0.140943413 | 0.118579926 | -1.188594206 | 0.234599389 | 0.458255539 | | | Dmpk | 625.5666504 | -0.1557769 | 0.082054221 | -1.898462962 | 0.057635122 | 0.419863484 |
| Gria2 | 20032.8255 | 0.061784645 | 0.052001328 | 1.188135912 | 0.234779868 | 0.458404859 | | | Casp6 | 95.60868786 | -0.155851837 | 0.092921087 | -1.677249401 | 0.093493722 | 0.50864787 |
| Auts2 | 2725.837832 | 0.125829202 | 0.111936431 | 1.124113039 | 0.260965075 | 0.488037763 | | | Loxl2 | 103.3420568 | -0.155871196 | 0.095008494 | -1.640602746 | 0.100879005 | 0.521542547 |
| Lect1 | 22.94122483 | 0.125817592 | 0.167171214 | 0.752627135 | 0.451674007 | 0.670329221 | | | Fam111a | 108.8155152 | -0.155882019 | 0.093580459 | -1.665753942 | 0.095762433 | 0.515046226 |
| 4930550C14Rik | 8.489261048 | 0.125781799 | 0.140159774 | 0.89741725 | 0.369496311 | 0.599285653 | | | Dcdc2b | 135.7061619 | -0.155897984 | 0.094713683 | -1.645992199 | 0.099765365 | 0.521115714 |
| Zfp287 | 714.1340679 | 0.095930248 | 0.080747702 | 1.188024488 | 0.234823762 | 0.458404859 | | | Arrdc1 | 342.0227425 | -0.155909672 | 0.085828643 | -1.816522626 | 0.06929024 | 0.454875446 |
| Sqstm1 | 16117.37628 | 0.125654935 | 0.070758565 | 1.775826507 | 0.075761517 | 0.226274597 | | | Gimap8 | 57.84724191 | -0.15592735 | 0.093745349 | -1.663307576 | 0.096250883 | 0.516109386 |
| Kif16b | 696.4921236 | 0.094634023 | 0.079704785 | 1.187306663 | 0.235106681 | 0.458829708 | | | Gli3 | 499.551502 | -0.155986738 | 0.078824969 | -1.978900092 | 0.047827256 | 0.392951764 |
| Steap2 | 629.7891626 | -0.145559034 | 0.122602157 | -1.187246926 | 0.235130236 | 0.458829708 | | | Plekhn1 | 250.2026367 | -0.155994906 | 0.089312248 | -1.7466239 | 0.080702599 | 0.479417114 |
| Itga8 | 220.6725581 | 0.12558126 | 0.100240055 | 1.252805182 | 0.210276615 | 0.428462109 | | | B230216G23Rik | 70.26702126 | -0.156068 | 0.09416964 | -1.65730697 | 0.097457429 | 0.518456434 |
| Cacng2 | 1890.182188 | 0.12557837 | 0.084020071 | 1.49462347 | 0.135012737 | 0.326959574 | | | Arhgap17 | 553.3169907 | -0.156086245 | 0.068650558 | -2.273634025 | 0.022988 | 0.28516281 |
| Cpeb2 | 2477.554734 | 0.096610582 | 0.081446623 | 1.186182783 | 0.235550124 | 0.459475465 | | | Setd6 | 448.704934 | -0.156098684 | 0.07198354 | -2.168533021 | 0.030118155 | 0.31934038 |
| Pmel | 21.29431325 | 0.125550465 | 0.168014935 | 0.747257771 | 0.454907978 | 0.673065719 | | | Adcy7 | 425.6547598 | -0.156104876 | 0.089144789 | -1.751138544 | 0.079922048 | 0.478751136 |
| 7-Mar | 1263.614658 | 0.093157831 | 0.078579933 | 1.185516799 | 0.235813177 | 0.459699215 | | | 4932418E24Rik | 111.4204912 | -0.156130365 | 0.095025476 | -1.643036914 | 0.100375296 | 0.5211685 |
| Phf17 | 1525.406485 | 0.091268276 | 0.076984626 | 1.185538993 | 0.235804407 | 0.459699215 | | | Ptges | 166.419141 | -0.156187181 | 0.090510981 | -1.725615832 | 0.084416549 | 0.488499377 |
| Slc30a5 | 1679.665966 | 0.075529165 | 0.063705157 | 1.185605197 | 0.235778249 | 0.459699215 | | | Dph1 | 576.7672237 | -0.156393054 | 0.075255678 | -2.078156203 | 0.037694974 | 0.35144878 |
| Son | 12725.10692 | 0.125524688 | 0.062547786 | 2.006860588 | 0.044764509 | 0.161938199 | | | Vwa5a | 794.5075276 | -0.1564211 | 0.071822047 | -2.177898086 | 0.029413625 | 0.317771905 |
| Ankrd13b | 3414.500451 | 0.09081033 | 0.076619735 | 1.185208091 | 0.235935182 | 0.459821346 | | | Hmg20b | 469.5491222 | -0.156566204 | 0.08070536 | -1.93997281 | 0.052382994 | 0.406355487 |
| Tspan8 | 14.84534253 | -0.189344769 | 0.159747411 | -1.185275974 | 0.23590835 | 0.459821346 | | | Mfsd10 | 723.1081004 | -0.156640755 | 0.090352 | -1.733672245 | 0.082976254 | 0.484450588 |
| Dlk1 | 581.2297118 | -0.172693054 | 0.14575703 | -1.184800855 | 0.236096195 | 0.460019421 | | | 1810032O08Rik | 101.6062169 | -0.15672345 | 0.094631037 | -1.656152726 | 0.097690894 | 0.518456434 |
| Lamp2 | 2349.474284 | 0.080911833 | 0.068290644 | 1.184815786 | 0.23609029 | 0.460019421 | | | 9030025P20Rik | 66.65325334 | -0.156725071 | 0.094830495 | -1.652686417 | 0.0983947 | 0.520050241 |
| Pitpnm1 | 8211.257429 | 0.125473512 | 0.077144411 | 1.626475721 | 0.103848494 | 0.277370896 | | | 3110056K07Rik | 215.2790016 | -0.156852166 | 0.092461746 | -1.696400655 | 0.08981003 | 0.50087668 |
| Asb3 | 516.8455345 | -0.105046194 | 0.088703376 | -1.18424122 | 0.236317589 | 0.46027715 | | | Lgals12 | 37.76544752 | -0.156972822 | 0.091407741 | -1.717281495 | 0.085927751 | 0.491218621 |
| Lipt1 | 150.8861907 | -0.154135068 | 0.130153688 | -1.184254325 | 0.236312404 | 0.46027715 | | | Hinfp | 478.2912574 | -0.157020166 | 0.077539507 | -2.025034355 | 0.042863837 | 0.375084392 |
| Sox21 | 384.6218319 | -0.158048435 | 0.133481651 | -1.184046152 | 0.236394794 | 0.460369651 | | | Supt20 | 880.2275896 | -0.157152886 | 0.073026785 | -2.151989649 | 0.03139817 | 0.32443523 |
| Slc7a10 | 1016.099328 | -0.171482637 | 0.144889057 | -1.183544434 | 0.236593447 | 0.460698614 | | | Crhr1 | 1077.973133 | -0.157173479 | 0.064923845 | -2.420889864 | 0.015482567 | 0.236284305 |
| Adora3 | 42.69240062 | 0.125459053 | 0.164643884 | 0.762002513 | 0.446058505 | 0.665501249 | | | Zbtb34 | 1007.114384 | -0.157198388 | 0.07815269 | -2.011426443 | 0.044280433 | 0.380589751 |
| Ttf1 | 557.4432912 | 0.125439729 | 0.110366211 | 1.136577288 | 0.255715039 | 0.482059717 | | | Dusp27 | 26.55567317 | -0.157206405 | 0.085559708 | -1.837388286 | 0.066152595 | 0.446279967 |
| Shc1 | 526.6805184 | 0.125435722 | 0.10216022 | 1.227833325 | 0.219509543 | 0.440269367 | | | Abca7 | 1240.612167 | -0.157426086 | 0.083369887 | -1.888284755 | 0.058987733 | 0.425661106 |
| Slco2a1 | 81.00811144 | 0.125315225 | 0.149102428 | 0.840464016 | 0.400648269 | 0.625581405 | | | Thbs2 | 379.766097 | -0.157472065 | 0.093340796 | -1.687065799 | 0.091590701 | 0.504530343 |
| Htr5b | 38.5811311 | -0.198705342 | 0.168092112 | -1.18212175 | 0.237157395 | 0.461506742 | | | Ccdc37 | 133.6101788 | -0.15755923 | 0.094921482 | -1.659890122 | 0.096936559 | 0.517493736 |
| Sox12 | 795.4184923 | 0.125155363 | 0.102301455 | 1.223397675 | 0.221179531 | 0.442136847 | | | Zfp810 | 652.4624069 | -0.157573516 | 0.073253802 | -2.151062633 | 0.031471256 | 0.32443523 |
| Ednrb | 1909.673665 | -0.114989725 | 0.097290606 | -1.181920122 | 0.237237397 | 0.461546488 | | | Col3a1 | 103.163387 | -0.157576846 | 0.08423615 | -1.87065583 | 0.061392801 | 0.434384717 |
| Fntb | 1190.957545 | 0.066074075 | 0.055919264 | 1.181597733 | 0.237365354 | 0.46173745 | | | Cd2ap | 1016.826608 | -0.157584101 | 0.063949462 | -2.464197452 | 0.013732044 | 0.224208584 |
| Rbm43 | 620.6710339 | 0.12511251 | 0.065176047 | 1.919608769 | 0.054907336 | 0.18438107 | | | Elf1 | 286.8425927 | -0.157700068 | 0.082382744 | -1.914236642 | 0.055589926 | 0.413057568 |
| Pag1 | 1448.479325 | -0.08484273 | 0.071827087 | -1.181208007 | 0.237520101 | 0.461922486 | | | Lekr1 | 139.8236272 | -0.157808001 | 0.089343151 | -1.766313364 | 0.077343291 | 0.473845905 |
| Myrip | 3075.0621 | 0.125095231 | 0.096766529 | 1.292753103 | 0.196096465 | 0.411040827 | | | Klf11 | 371.8072583 | -0.157917177 | 0.082573521 | -1.912443299 | 0.055819359 | 0.413057568 |
| Ctdnep1 | 2512.328089 | -0.065135092 | 0.055175636 | -1.180504599 | 0.237799582 | 0.462349947 | | | Igsf6 | 37.00315562 | -0.157980018 | 0.079539278 | -1.986188745 | 0.04701237 | 0.39102736 |
| Ccdc167 | 504.1807522 | -0.117868865 | 0.099859232 | -1.180350211 | 0.237860956 | 0.462353236 | | | Gramd1a | 3939.658061 | -0.15807222 | 0.06435325 | -2.456320695 | 0.014036783 | 0.227385586 |
| Col6a3 | 150.8114551 | -0.17553962 | 0.14871421 | -1.180382292 | 0.237848202 | 0.462353236 | | | 1700049G17Rik | 244.6542665 | -0.158080089 | 0.078516242 | -2.013342536 | 0.044078606 | 0.380217243 |
| Plekha3 | 1432.571969 | 0.12508276 | 0.083123676 | 1.504778973 | 0.132380912 | 0.322704506 | | | Inpp5d | 792.7526384 | -0.158089222 | 0.074407738 | -2.124634159 | 0.033617151 | 0.335456507 |
| Gm1821 | 1452.26643 | -0.095591784 | 0.081008187 | -1.180026204 | 0.237989793 | 0.462487627 | | | 9530091C08Rik | 35.25207141 | -0.158218281 | 0.090568562 | -1.746944815 | 0.080646911 | 0.479417114 |
| Parm1 | 1084.843639 | -0.076407754 | 0.064766358 | -1.179744497 | 0.238101851 | 0.462589351 | | | Etv4 | 344.1712377 | -0.158227325 | 0.089082864 | -1.776181383 | 0.075703026 | 0.470947389 |
| Rcn2 | 3859.375806 | -0.070334184 | 0.059614369 | -1.179819303 | 0.238072091 | 0.462589351 | | | Galnt3 | 40.54175999 | -0.158288585 | 0.092981285 | -1.702370372 | 0.08868596 | 0.497760972 |
| Fam181a | 75.399425 | -0.187033798 | 0.158614571 | -1.179171605 | 0.23832985 | 0.462671816 | | | Slc20a1 | 7246.849784 | -0.15831571 | 0.051731308 | -3.060346189 | 0.002210813 | 0.079261025 |
| G630025P09Rik | 32.73196021 | -0.192935826 | 0.163599291 | -1.179319447 | 0.238270997 | 0.462671816 | | | Ttll4 | 500.3312232 | -0.158390607 | 0.070245219 | -2.254824008 | 0.024144378 | 0.29178139 |
| Cyb5r3 | 2756.10856 | 0.125015232 | 0.081021813 | 1.542982407 | 0.122835045 | 0.30946442 | | | Naglu | 402.8857193 | -0.158487023 | 0.087318831 | -1.81503831 | 0.069518022 | 0.455055507 |
| Nek9 | 3048.216903 | 0.066993418 | 0.056813346 | 1.179184528 | 0.238324705 | 0.462671816 | | | BC065397 | 113.2797228 | -0.158555583 | 0.095058173 | -1.667984754 | 0.095318753 | 0.51433043 |
| Rab23 | 1149.407251 | 0.124970323 | 0.06120776 | 2.041739847 | 0.041177344 | 0.153292111 | | | Etnk2 | 357.5770794 | -0.158869392 | 0.0771554 | -2.059083248 | 0.039486264 | 0.359835305 |
| Phf16 | 278.2449715 | 0.124862116 | 0.09510458 | 1.312892771 | 0.189219083 | 0.402437952 | | | Akr1c14 | 64.30038214 | -0.158908283 | 0.094938249 | -1.673806757 | 0.094168584 | 0.510979772 |
| Zfp821 | 1157.101426 | 0.124839769 | 0.064664783 | 1.930568113 | 0.053536484 | 0.181707402 | | | Stat5a | 79.6012095 | -0.15896782 | 0.095046476 | -1.672527234 | 0.094420402 | 0.511705397 |
| AW112010 | 30.2035809 | 0.124817586 | 0.16530918 | 0.755055383 | 0.450215752 | 0.668713036 | | | Hps4 | 842.5544044 | -0.158993805 | 0.078176986 | -2.033767405 | 0.041975052 | 0.370320688 |
| Oraov1 | 496.1631857 | 0.124784681 | 0.07612328 | 1.639244678 | 0.101162312 | 0.272308699 | | | Slc16a6 | 659.2616309 | -0.159305952 | 0.085016998 | -1.873812956 | 0.06095621 | 0.432354503 |
| Rasl11b | 2141.188047 | 0.124771132 | 0.107382438 | 1.161932391 | 0.245262929 | 0.470309878 | | | Gm20743 | 46.4972765 | -0.15933675 | 0.089486801 | -1.780561471 | 0.074984119 | 0.468589657 |
| Usp20 | 3754.427122 | 0.08707076 | 0.073873707 | 1.178643433 | 0.238540189 | 0.462802616 | | | B230206H07Rik | 185.4442793 | -0.159405304 | 0.09507539 | -1.676620031 | 0.093616807 | 0.508936281 |

| | | | | | | | | | | | | | | | | | |
|---|---|---|---|---|---|---|---|---|---|---|---|---|---|---|---|---|---|
| Cryba4 | 18.35147458 | -0.197779779 | 0.167820757 | -1.178517975 | 0.238590171 | 0.462841631 | | | 9430015G10Rik | 974.8670684 | -0.159464625 | 0.068151134 | -2.339867516 | 0.019290582 | 0.263564471 | | |
| Mettl22 | 541.5173748 | 0.094214976 | 0.079954347 | 1.178359642 | 0.23865326 | 0.46290606 | | | 9530082P21Rik | 80.94987343 | -0.159483895 | 0.09507304 | -1.677488122 | 0.09344707 | 0.508584542 | | |
| Amdhd2 | 388.4523223 | 0.12473543 | 0.118045834 | 1.056669481 | 0.290662458 | 0.521060295 | | | Cnr2 | 17.00650197 | -0.159607055 | 0.069391169 | -2.300106165 | 0.021442206 | NA | | |
| Ralbp1 | 3194.750823 | 0.1247201 | 0.086957319 | 1.434267998 | 0.151495799 | 0.352496758 | | | Olfr78 | 43.23795534 | -0.159643775 | 0.093253751 | -1.71192873 | 0.086909803 | 0.493723819 | | |
| Polr2e | 1580.044169 | -0.080243287 | 0.068121937 | -1.177936081 | 0.238822089 | 0.463059624 | | | Scaf4 | 1443.686162 | -0.159769026 | 0.061708876 | -2.589076901 | 0.00962336 | 0.187266026 | | |
| Ivd | 2185.100833 | 0.124718805 | 0.083233722 | 1.49841677 | 0.134025002 | 0.32543228 | | | Oas1a | 23.62114571 | -0.159783266 | 0.075157391 | -2.12598207 | 0.033504752 | 0.335456507 | | |
| Rasl12 | 136.2705087 | -0.151465126 | 0.128618399 | -1.177631866 | 0.2389434 | 0.463178912 | | | Iqce | 1346.916099 | -0.159973966 | 0.074862617 | -2.136900546 | 0.032606082 | 0.329798129 | | |
| Crabp1 | 69.84479758 | -0.193283719 | 0.164165422 | -1.177371681 | 0.239047188 | 0.463206244 | | | Ifi30 | 126.8666609 | -0.160009454 | 0.093680053 | -1.708041883 | 0.087628571 | 0.495597906 | | |
| Dgcr6 | 1403.777269 | -0.080740776 | 0.068574328 | -1.177419855 | 0.239027969 | 0.463206244 | | | Adamts20 | 471.1894728 | -0.160213598 | 0.076169654 | -2.103378301 | 0.035432714 | 0.342120019 | | |
| Slmap | 5048.549344 | -0.091023048 | 0.077309809 | -1.177380336 | 0.239043735 | 0.463206244 | | | Msantd2 | 400.5723454 | -0.160218269 | 0.083356298 | -1.922089552 | 0.05459449 | 0.411968607 | | |
| Anapc7 | 1392.99074 | 0.124698562 | 0.06927925 | 1.799940992 | 0.071869956 | 0.218968649 | | | Aifm3 | 4608.448095 | -0.160236135 | 0.081950266 | -1.955285122 | 0.050549426 | 0.403264376 | | |
| Snx22 | 257.2681705 | 0.124685319 | 0.110387925 | 1.129519547 | 0.258678728 | 0.485226177 | | | Pyroxd2 | 90.603692 | -0.160267896 | 0.091789776 | -1.746032113 | 0.080805372 | 0.479484817 | | |
| Zfp949 | 874.7461313 | 0.124626456 | 0.068110456 | 1.829769789 | 0.06728437 | 0.209422264 | | | Hexdc | 1159.918226 | -0.160544962 | 0.076437832 | -2.100333795 | 0.035699488 | 0.342970085 | | |
| Atxn7l3 | 11988.02855 | -0.089381357 | 0.075944689 | -1.176927029 | 0.239224633 | 0.463318308 | | | Btk | 62.94161719 | -0.160588537 | 0.095093383 | -1.688745656 | 0.091268182 | 0.503727415 | | |
| Myom3 | 120.814855 | -0.178628916 | 0.151804406 | -1.176704425 | 0.239313501 | 0.463432494 | | | 2810008D09Rik | 347.0601091 | -0.160615413 | 0.082746344 | -1.941057525 | 0.0522513 | 0.406355487 | | |
| Smg5 | 4241.972481 | 0.124602918 | 0.088061193 | 1.414958319 | 0.157080717 | 0.359711592 | | | Mdc1 | 1113.266169 | -0.160661846 | 0.059643625 | -2.69369687 | 0.00706644 | 0.160386105 | | |
| Cmtm4 | 7870.075732 | -0.078249953 | 0.066547039 | -1.175859273 | 0.239651116 | 0.46397031 | | | Paxbp1 | 1825.068172 | -0.160722067 | 0.067564787 | -2.378784504 | 0.017369827 | 0.252062047 | | |
| Gramd1b | 3737.145377 | 0.124554444 | 0.086467483 | 1.440477274 | 0.149732415 | 0.350068662 | | | Ncf4 | 36.28782518 | -0.160778111 | 0.087737208 | -1.832496323 | 0.066877509 | 0.447688159 | | |
| Casz1 | 95.01282322 | -0.173323462 | 0.147452426 | -1.17545344 | 0.239813354 | 0.464110431 | | | Irgm1 | 578.2573813 | -0.160911169 | 0.076286826 | -2.109291708 | 0.03491941 | 0.340200772 | | |
| 2610305D13Rik | 9.24982342 | 0.124452936 | 0.154722605 | 0.804361687 | 0.421188121 | 0.64363258 | | | Gramd1c | 52.60213821 | -0.160958736 | 0.092723486 | -1.735900403 | 0.082581446 | 0.483895963 | | |
| Ppp1r1b | 17095.2017 | -0.139073304 | 0.11833323 | -1.175268382 | 0.23988736 | 0.464195673 | | | Trhr2 | 171.1160795 | -0.161025525 | 0.092706697 | -1.736935198 | 0.08239861 | 0.483019453 | | |
| Mepce | 1608.981537 | 0.124422084 | 0.058812329 | 2.115578255 | 0.034380693 | 0.135373432 | | | Ypel4 | 1531.213365 | -0.161052736 | 0.064372407 | -2.501890842 | 0.012353201 | 0.21214894 | | |
| Emc8 | 2625.726958 | 0.124404203 | 0.089802052 | 1.385315819 | 0.165955919 | 0.371806281 | | | Psmb9 | 70.11715032 | -0.161087123 | 0.075935666 | -2.121363139 | NA | NA | | |
| Ighmbp2 | 489.2183083 | 0.124361818 | 0.092404376 | 1.345843386 | 0.17835303 | 0.387849001 | | | D930015E06Rik | 695.2503505 | -0.161150941 | 0.061115515 | -2.63682537 | 0.00836859 | 0.174157793 | | |
| Snhg4 | 221.1965992 | 0.124302035 | 0.105825998 | 1.174588823 | 0.240159257 | 0.464547756 | | | Yeats2 | 1529.285586 | -0.161181141 | 0.059829592 | -2.694003657 | 0.007059939 | 0.160386105 | | |
| Gm11974 | 58.68638944 | 0.124187915 | 0.144537052 | 0.859211622 | 0.390223773 | 0.617203668 | | | Foxl2os | 19.71469933 | -0.161191697 | 0.080091791 | -2.012587008 | 0.044158095 | NA | | |
| 2700089E24Rik | 3410.690197 | -0.117271687 | 0.099927115 | -1.173572234 | 0.240566408 | 0.465161104 | | | Ttc8 | 761.5016844 | -0.161267057 | 0.066803788 | -2.414040605 | 0.015776704 | 0.238721253 | | |
| BC018473 | 10.49417964 | 0.124179718 | 0.164214747 | 0.756203209 | 0.449527369 | 0.668359759 | | | A330023F24Rik | 2444.630706 | -0.161319236 | 0.083858992 | -1.923696332 | 0.054392658 | 0.410949742 | | |
| Fbxo47 | 24.33760891 | 0.124159702 | 0.167961339 | 0.739215953 | 0.459775875 | 0.676957488 | | | Vwa3a | 317.3566456 | -0.161345641 | 0.088747473 | -1.818030822 | 0.069059422 | 0.45371197 | | |
| Plcb3 | 815.8222209 | -0.078992239 | 0.067339014 | -1.173053092 | 0.240774516 | 0.465389265 | | | Top2a | 47.2468573 | -0.161495531 | 0.091985401 | -1.755664804 | 0.07914566 | 0.477013957 | | |
| A930003A15Rik | 9.119688138 | -0.184541146 | 0.157328307 | -1.172968485 | 0.240808444 | 0.465396786 | | | Fam180a | 74.91564944 | -0.1615336 | 0.091919793 | -1.757332064 | 0.078861226 | 0.476674706 | | |
| Nrap | 46.11569408 | 0.124094617 | 0.158206571 | 0.784383454 | 0.432815144 | 0.654004155 | | | Col5a2 | 201.7357889 | -0.161587286 | 0.095101204 | -1.699108728 | 0.089298699 | 0.499459783 | | |
| Enthd2 | 613.7564151 | 0.124023421 | 0.079042039 | 1.56908176 | 0.116628889 | 0.299477629 | | | Pdia5 | 156.6531153 | -0.161728084 | 0.092972625 | -1.73952369 | 0.081942689 | 0.482582908 | | |
| Dusp1 | 1384.910223 | 0.124014716 | 0.164254548 | 0.755015419 | 0.450239731 | 0.668713036 | | | Lefty1 | 32.74894581 | -0.161816336 | 0.078717645 | -2.055655185 | 0.039815765 | 0.36080087 | | |
| Lrig3 | 108.3421125 | 0.124008179 | 0.12565297 | 0.986910053 | 0.323686736 | 0.557368558 | | | Prdm9 | 83.83594222 | -0.161831701 | 0.095102717 | -1.701651707 | 0.088820678 | 0.49789071 | | |
| Sympk | 5660.307043 | 0.09579863 | 0.081705839 | 1.172482059 | 0.241003571 | 0.465428677 | | | Mir143hg | 33.55061287 | -0.161892757 | 0.08970392 | -1.804745637 | 0.071114495 | 0.458300422 | | |
| Tmem30a | 11302.24965 | -0.101654496 | 0.086688873 | -1.172636029 | 0.240941795 | 0.465428677 | | | Bcl2a1a | 6.520767286 | -0.161997023 | 0.056169436 | -2.884077784 | 0.003925618 | NA | | |
| Tspan15 | 539.3444342 | -0.137528043 | 0.117340253 | -1.172044881 | 0.241179038 | 0.465706382 | | | Zmym3 | 6877.608952 | -0.162001467 | 0.060387247 | -2.682709916 | 0.00730283 | 0.16270079 | | |
| 2810006K23Rik | 428.7555294 | -0.132474073 | 0.113139789 | -1.170888457 | 0.241643616 | 0.466262791 | | | Cep44 | 320.4946813 | -0.162174758 | 0.07179019 | -2.259010032 | 0.023882761 | 0.291530239 | | |
| Lingo4 | 33.75730959 | 0.12399894 | 0.156141324 | 0.794145568 | 0.427110692 | 0.648980393 | | | Appl2 | 3560.226216 | -0.162180966 | 0.059776462 | -2.713124213 | 0.006665212 | 0.156678744 | | |
| Bad | 708.7982409 | -0.098966411 | 0.084514178 | -1.171003647 | 0.241597311 | 0.466262791 | | | Fuk | 774.5914343 | -0.162273696 | 0.079110038 | -2.051240263 | 0.040243555 | 0.363317511 | | |
| Zfp808 | 37.07228873 | 0.12399859 | 0.165161424 | 0.750772108 | 0.452789818 | 0.671470659 | | | Gm14393 | 22.48566219 | -0.162432295 | 0.081862234 | -1.984215288 | 0.047231845 | 0.391827402 | | |
| Phb2 | 2192.673428 | -0.067962653 | 0.0580464 | -1.170833216 | 0.241665823 | 0.466262791 | | | Gpr114 | 19.48681293 | -0.162459497 | 0.068537634 | -2.370369197 | 0.017770331 | NA | | |
| Prokr2 | 126.0538647 | -0.194174876 | 0.165797958 | -1.171153607 | 0.241537039 | 0.466262791 | | | Wdhd1 | 289.806395 | -0.162522036 | 0.084182558 | -1.930590368 | 0.05353373 | 0.407961168 | | |
| Cnga4 | 53.81050859 | 0.123995865 | 0.161048584 | 0.769928315 | 0.441342417 | 0.661314189 | | | Mei1 | 291.6338544 | -0.162555165 | 0.087973346 | -1.847777457 | 0.064634543 | 0.443022969 | | |
| Ppil4 | 1502.633859 | 0.12394462 | 0.071808997 | 1.726031918 | 0.084341671 | 0.242312228 | | | E130311K13Rik | 284.5516993 | -0.162685915 | 0.086543368 | -1.879819557 | 0.060132674 | 0.42944308 | | |
| Tmem57 | 2468.477217 | 0.058424917 | 0.04994407 | 1.169806875 | 0.242078696 | 0.466919871 | | | Zfp951 | 127.4613959 | -0.162794846 | 0.091231559 | -1.784413721 | 0.074356463 | 0.467576616 | | |
| C130060K24Rik | 44.99221469 | -0.195763888 | 0.167375701 | -1.169607578 | 0.242158926 | 0.467016481 | | | Cep152 | 166.8543505 | -0.162803365 | 0.089734631 | -1.814275756 | 0.069635282 | 0.455184691 | | |
| Pdcl | 794.1859622 | -0.093180061 | 0.079692334 | -1.169247491 | 0.242303932 | 0.467063589 | | | Gm10516 | 205.9208208 | -0.162847034 | 0.084987639 | -1.916126105 | 0.055349046 | 0.413057568 | | |
| Sec24b | 2498.778609 | 0.066155039 | 0.056573647 | 1.169361397 | 0.242258056 | 0.467063589 | | | Sirt7 | 1218.627001 | -0.162864329 | 0.082128796 | -1.983035635 | 0.047363449 | 0.391951501 | | |
| Zbtb10 | 289.8792558 | -0.121747407 | 0.104116308 | -1.169340412 | 0.242266507 | 0.467063589 | | | Ankhd1 | 3049.379164 | -0.162893842 | 0.063297656 | -2.573457714 | 0.010068795 | 0.19194136 | | |
| Zfp146 | 1258.687465 | 0.070317535 | 0.060136579 | 1.16929722 | 0.242283903 | 0.467063589 | | | Lmtk3 | 9300.216162 | -0.162985824 | 0.072049948 | -2.262122717 | 0.023689822 | 0.290395241 | | |
| Cdca7l | 48.72390096 | -0.196087381 | 0.16776164 | -1.168845157 | 0.242466023 | 0.467201662 | | | F2rl1 | 15.75543931 | -0.163090073 | 0.072836955 | -2.239111648 | 0.025148653 | NA | | |
| Raf1 | 2729.586524 | 0.123934005 | 0.047465536 | 2.6110314 | 0.00902696 | 0.054762618 | | | Aldob | 66.26202399 | -0.163161071 | 0.094860524 | -1.720010218 | 0.085430584 | 0.489918936 | | |
| Zc3h11a | 4325.937266 | 0.123912424 | 0.079258634 | 1.563393394 | 0.117960093 | 0.301422015 | | | Zic3 | 99.6178413 | -0.163198659 | 0.09264503 | -1.76154792 | 0.07814571 | 0.475440315 | | |
| A930003O13Rik | 12.99113037 | -0.193225481 | 0.165385458 | -1.168334164 | 0.242672 | 0.467209584 | | | Zfp446 | 495.4911271 | -0.163226276 | 0.074841187 | -2.18096856 | 0.029185742 | 0.317415456 | | |
| Ebna1bp2 | 1368.056494 | 0.123876422 | 0.062276592 | 1.989132979 | 0.046686527 | 0.166557921 | | | Rarg | 437.2887718 | -0.163451689 | 0.075951259 | -2.152060303 | 0.031392605 | 0.32443523 | | |
| Gucy2c | 15.44328467 | -0.195584567 | 0.167350618 | -1.168711349 | 0.242519948 | 0.467209584 | | | Pvrl3 | 603.7438764 | -0.16368315 | 0.090200794 | -1.814653098 | 0.069577237 | 0.455055806 | | |
| Hddc2 | 311.4537738 | -0.105868288 | 0.090601982 | -1.168498591 | 0.242605707 | 0.467209584 | | | 4933408B17Rik | 30.30399302 | -0.163723281 | 0.085242256 | -1.920682175 | 0.054771788 | 0.412110852 | | |
| Mier2 | 1251.130113 | 0.123846682 | 0.068983394 | 1.795311527 | 0.07260408 | 0.220444368 | | | Pros1 | 612.7332853 | -0.163731236 | 0.070815121 | -2.312094266 | 0.020772489 | 0.271349978 | | |
| Mrpl40 | 672.5870013 | -0.087740996 | 0.075099342 | -1.168332423 | 0.242672702 | 0.467209584 | | | Renbp | 156.4308354 | -0.163768563 | 0.09501965 | -1.723523114 | 0.084793969 | 0.488499377 | | |
| Dctd | 252.8615026 | 0.123822886 | 0.141581649 | 0.874568747 | 0.381808599 | 0.610501468 | | | Zfp114 | 85.77669782 | -0.163816695 | 0.094475267 | -1.733963822 | 0.082924502 | 0.484423387 | | |
| Gab2 | 1169.921858 | -0.101827838 | 0.087170095 | -1.168151055 | 0.242745839 | 0.467217893 | | | Ifi27l1 | 1011.166257 | -0.163845378 | 0.08089376 | -2.025439018 | 0.042822304 | 0.374984137 | | |
| Rint1 | 676.9439882 | -0.076850861 | 0.065784926 | -1.168213842 | 0.242720518 | 0.467217893 | | | Atp10d | 202.8343792 | -0.16388484 | 0.084409558 | -1.94154363 | 0.052192373 | 0.406355487 | | |
| Arfip2 | 3655.586091 | 0.097591109 | 0.083556206 | 1.167969604 | 0.242819026 | 0.467300706 | | | Mysm1 | 1233.874185 | -0.163924491 | 0.060966073 | -2.688782158 | 0.00717132 | 0.161546928 | | |
| Crat | 2102.288094 | 0.123808695 | 0.059672431 | 2.074805606 | 0.038004555 | 0.144501245 | | | Rgs9 | 4569.625627 | -0.163973127 | 0.081600145 | -2.009471017 | 0.044487205 | 0.380589751 | | |
| Kcnab2 | 17102.65301 | 0.123800552 | 0.081231042 | 1.5240547 | 0.127495048 | 0.315610843 | | | Klk8 | 74.87076071 | -0.164054437 | 0.093913465 | -1.746868097 | 0.080660221 | 0.479417114 | | |
| Tnfsf9 | 13.65695359 | -0.194468548 | 0.166555097 | -1.167592895 | 0.242971018 | 0.46741904 | | | Cep192 | 412.8996416 | -0.164056633 | 0.075803948 | -2.16422282 | 0.030447253 | 0.320602541 | | |
| Mospd1 | 332.9368397 | 0.123758518 | 0.094827101 | 1.30509651 | 0.191859987 | 0.405907402 | | | Recql4 | 120.6063395 | -0.164208937 | 0.095099232 | -1.726711498 | 0.08421949 | 0.488372178 | | |
| Alkbh1 | 359.029531 | 0.123735856 | 0.079084076 | 1.564611508 | 0.117674029 | 0.301113361 | | | 2010015L04Rik | 776.2323578 | -0.164275102 | 0.060577391 | -2.711822011 | 0.006691451 | 0.156789377 | | |
| Sf3b5 | 757.3952892 | -0.087072743 | 0.074617446 | -1.166922055 | 0.243241848 | 0.467765818 | | | Pnpt1 | 865.8025638 | -0.164388555 | 0.065989159 | -2.491144878 | 0.012733219 | 0.215303986 | | |
| Grb2 | 6526.952233 | -0.078692652 | 0.067441833 | -1.166822547 | 0.24328204 | 0.467785049 | | | Gm13446 | 193.7996208 | -0.164552982 | 0.088696245 | -1.855241795 | 0.063561706 | 0.440996088 | | |
| Rabepk | 723.4305857 | 0.123670379 | 0.077575771 | 1.594188202 | 0.110893888 | 0.289955312 | | | Cd209f | 16.9247893 | -0.164645657 | 0.077759552 | -2.117368896 | 0.034228552 | NA | | |
| Madd | 11031.96419 | 0.074169884 | 0.063585217 | 1.166464278 | 0.243426783 | 0.468001723 | | | Prdm1 | 56.44363688 | -0.164703245 | 0.094225255 | -1.747973462 | 0.080468623 | 0.479417114 | | |
| Zfp629 | 1427.430551 | 0.123633409 | 0.075087975 | 1.64651409 | 0.099657962 | 0.26954004 | | | Tmco4 | 169.7375145 | -0.164732862 | 0.092927533 | -1.772702415 | 0.076278031 | 0.472435997 | | |
| Zfyve21 | 471.801476 | -0.086166219 | 0.073897185 | -1.16602843 | 0.243602951 | 0.4681697 | | | Il3ra | 55.70510742 | -0.164814073 | 0.08909291 | -1.84991233 | 0.064326186 | 0.442634858 | | |
| 9630028B13Rik | 16.62289437 | -0.189331641 | 0.16246711 | -1.165353658 | 0.243875868 | 0.468636077 | | | Cwh43 | 106.1098031 | -0.164844756 | 0.09377314 | -1.757910163 | 0.078762796 | 0.476674706 | | |
| Syt1 | 65383.76035 | -0.059910657 | 0.051424523 | -1.165021141 | 0.244010435 | 0.468836518 | | | Gm16998 | 16.51142448 | -0.164956755 | 0.076914055 | -2.144689346 | 0.031977693 | NA | | |
| Akap1 | 1959.286914 | 0.071614173 | 0.061498376 | 1.164488848 | 0.24422596 | 0.469017976 | | | Tek | 684.1709782 | -0.165094295 | 0.074179278 | -2.225612055 | 0.026040178 | 0.3021678 | | |
| Apeh | 727.0379699 | -0.102376649 | 0.087901441 | -1.164675431 | 0.244150398 | 0.469017976 | | | Tgif1 | 66.66692358 | -0.165120953 | 0.09338657 | -1.768144539 | 0.077036743 | 0.472965229 | | |
| Epha1 | 11.23721149 | 0.123632624 | 0.163166482 | 0.757708461 | 0.448625533 | 0.667723727 | | | Ms4a6b | 34.46474964 | -0.165131654 | 0.089482092 | -1.845415653 | 0.064977098 | 0.4443509 | | |
| Maz | 5522.533108 | -0.094324219 | 0.080995387 | -1.164562855 | 0.244195987 | 0.469017976 | | | Crlf2 | 411.0540021 | -0.16534472 | 0.085693066 | -1.929499413 | 0.053668893 | 0.408507027 | | |
| Cbr2 | 42.24324239 | -0.193941001 | 0.166559258 | -1.164396402 | 0.244263405 | 0.469022123 | | | Tardbp | 7760.601967 | -0.165372985 | 0.046783129 | -3.534885097 | 0.000407953 | 0.02634142 | | |
| Gm10336 | 491.6472984 | -0.103847051 | 0.089190084 | -1.164334051 | 0.244288662 | 0.469022123 | | | Eva1b | 50.42556125 | -0.165473128 | 0.094613224 | -1.748942919 | 0.080300887 | 0.479231997 | | Slc6a6 |
| Mpp4 | 18.80791091 | 0.123630176 | 0.165813797 | 0.745596433 | 0.455911238 | 0.674115317 | | | Calcrl | 206.963729 | -0.165633811 | 0.088630227 | -1.868818526 | 0.061648066 | 0.435129208 | | Luc7l2 |
| Timm23 | 2172.598336 | -0.083190362 | 0.071493403 | -1.163608926 | 0.244582532 | 0.469482858 | | | Uimc1 | 599.1247188 | -0.165737801 | 0.074546776 | -2.22327256 | 0.026197427 | 0.302373406 | | Usp48 |
| Tnfrsf9 | 20.46195897 | 0.12358276 | 0.167949146 | 0.735834406 | 0.46183148 | 0.678212469 | | | Rnasel | 627.7691771 | -0.165766579 | 0.065559843 | -2.528477363 | 0.011455847 | 0.203432321 | | Plekha5 |
| Rrp7a | 3716.488022 | -0.071703732 | 0.061640871 | -1.163249825 | 0.244728157 | 0.469671205 | | | Gm11696 | 94.98962498 | -0.16576693 | 0.094128169 | -1.761076752 | 0.078225413 | 0.475440315 | | Edem1 |
| Mir22hg | 720.719731 | 0.123541393 | 0.100899286 | 1.224403055 | 0.220800217 | 0.44191594 | | | Ifi47 | 41.93980667 | -0.165874551 | 0.07972402 | -2.080609485 | NA | NA | | Trub1 |
| Ap3m1 | 2073.641946 | 0.123489365 | 0.089376984 | 1.381668529 | 0.167073502 | 0.373002426 | | | Mafb | 1002.130043 | -0.165944027 | 0.069163264 | -2.399308788 | 0.016426056 | 0.243474381 | | Ddx17 |
| Suclg1 | 3147.773873 | -0.058896127 | 0.050664015 | -1.162484404 | 0.245038756 | 0.470054536 | | | Lama5 | 654.8648405 | -0.165955281 | 0.076838861 | -2.159783202 | 0.030789457 | 0.321052104 | | Iqgap1 |
| Cox19 | 688.9984811 | -0.073636046 | 0.063350785 | -1.16235412 | 0.245091651 | 0.470097802 | | | Rabl2 | 1043.933166 | -0.165962035 | 0.066567523 | -2.493138212 | 0.012661955 | 0.215249514 | | Rbm5 |
| Bdnf | 853.5535749 | -0.153944457 | 0.132472256 | -1.162088288 | 0.245199604 | 0.470246648 | | | Adam17 | 729.775038 | -0.166043116 | 0.062905317 | -2.639572051 | 0.008301078 | 0.174157793 | | 6820431F20Rik |
| Top1mt | 474.0558515 | 0.123480843 | 0.098336196 | 1.255700829 | 0.209224459 | 0.427406334 | | | Zfp383 | 537.2766942 | -0.166118319 | 0.06667203 | -2.491574331 | 0.012717835 | 0.215303986 | | Pfkfb2 |
| Cog3 | 1428.913701 | 0.123468493 | 0.053556404 | 2.305391767 | 0.021144642 | 0.097620705 | | | Mir344-2 | 6.431452148 | -0.166122192 | 0.054362664 | -3.05581406 | 0.002244504 | NA | | Zc3h7a |
| Zfp36l1 | 911.5156877 | -0.119133462 | 0.10254412 | -1.1617776 | 0.245325815 | 0.47037115 | | | Cxcl10 | 27.56363512 | -0.166164706 | 0.059788998 | -2.779185345 | 0.005449542 | 0.138877279 | | Heatr6 |
| Syt4 | 7771.551305 | -0.08661102 | 0.074560249 | -1.161624606 | 0.245387983 | 0.470375034 | | | Foxm1 | 66.7735972 | -0.166236935 | 0.094536184 | -1.758447702 | 0.078671363 | 0.476674706 | | Rbm25 |
| Card10 | 316.0706401 | -0.14464655 | 0.124613179 | -1.160764466 | 0.245737697 | 0.470819915 | | | Wdr67 | 328.5304268 | -0.1662371 | 0.076560046 | -2.171329661 | 0.029906262 | 0.3187222 | | Srrm1 |
| Hipk3 | 3908.148318 | -0.094460439 | 0.081368003 | -1.160903987 | 0.245680947 | 0.470819915 | | | Atxn7 | 1333.903391 | -0.166265507 | 0.073382117 | -2.265749677 | 0.023466714 | 0.288824644 | | Nxf1 |
| Rpa3 | 143.4034611 | -0.141182894 | 0.121630246 | -1.160754824 | 0.245741619 | 0.470819915 | | | Magix | 20.85908421 | -0.166336202 | 0.082424033 | -2.018054644 | 0.043585571 | NA | | Sall1 |
| Tk1 | 13.75165579 | -0.192608955 | 0.165910083 | -1.160923744 | 0.245672912 | 0.470819915 | | | Gm11974 | 58.37907529 | -0.166364099 | 0.095062213 | -1.750054974 | 0.080108828 | 0.478984869 | | Ubp1 |
| Ocrl | 3556.615506 | 0.123412132 | 0.097905623 | 1.260521389 | 0.207481337 | 0.425492348 | | | Sox12 | 898.0973168 | -0.166631444 | 0.077285518 | -2.156050047 | 0.031079757 | 0.323630506 | | Myo9b |
| Gm5086 | 35.38221242 | 0.123315359 | 0.166216615 | 0.7418955 | 0.458150652 | 0.6759631 | | | Rfxank | 710.6413315 | -0.166764079 | 0.062278688 | -2.677707015 | 0.007412803 | 0.163894025 | | Tbce |
| Actn4 | 6690.69616 | -0.080022951 | 0.068979684 | -1.160094481 | 0.246010341 | 0.471159978 | | | Lmcd1 | 204.7086064 | -0.166788719 | 0.087710786 | -1.901575926 | 0.057226618 | 0.419207468 | | Slco1a4 |
| Rtl1 | 37.96039997 | 0.123306471 | 0.15286794 | 0.806620875 | 0.419884943 | 0.642730443 | | | Espl1 | 53.35582345 | -0.16682186 | 0.092431539 | -1.804815348 | 0.071103582 | 0.458300422 | | Zfp445 |
| Tapbp | 1210.054025 | 0.123261947 | 0.074684142 | 1.650443373 | 0.098852286 | 0.268056398 | | | Plod2 | 174.527182 | -0.166920452 | 0.091233443 | -1.829597192 | 0.067310194 | 0.448302557 | | Prpf39 |
| Vps37d | 546.3852641 | 0.09037688 | 0.077938288 | 1.159595407 | 0.246213572 | 0.471382768 | | | Mir23b | 19.35736512 | -0.166922683 | 0.071239623 | -2.343116766 | 0.019123448 | NA | | Unc13b |
| Cops8 | 3551.534459 | -0.049235661 | 0.042465529 | -1.159426539 | 0.246282364 | 0.471389617 | | | Syne4 | 109.7673514 | -0.167040894 | 0.091539059 | -1.824804579 | 0.068030525 | 0.450415406 | | Rbm33 |
| Endov | 1698.480515 | 0.12325798 | 0.071620656 | 1.720983675 | 0.085253786 | 0.243893179 | | | Dgat2l6 | 26.56374978 | -0.16705551 | 0.084842779 | -1.969000922 | 0.048952985 | 0.396592896 | | Zfp280c |
| 4931406P16Rik | 1701.835815 | 0.068318355 | 0.0589329 | 1.159256628 | 0.246351595 | 0.471424129 | | | Nek1 | 1612.923193 | -0.167701029 | 0.060225806 | -2.784537706 | 0.005360409 | 0.13805904 | | Smc5 |
| Setd8 | 1535.172038 | 0.123200978 | 0.086116558 | 1.430630534 | 0.152536132 | 0.353946068 | | | Traf3ip3 | 30.89429908 | -0.167756124 | 0.087132629 | -1.925296243 | 0.054192307 | 0.410426203 | | Slc4a7 |
| Kcnh1 | 5097.029546 | -0.095831035 | 0.082678484 | -1.159080698 | 0.246423293 | 0.471424129 | | | Ptpn2 | 1278.009567 | -0.167783785 | 0.070959773 | -2.364491558 | 0.018054839 | 0.256272607 | | Ifi204 |
| Lnpep | 1836.85406 | -0.13570367 | 0.117064221 | -1.159224126 | 0.24636484 | 0.471424129 | | | Rasgef1b | 2730.00515 | -0.167797743 | 0.079120336 | -2.120791596 | 0.033939344 | 0.336326295 | | Qtrtd1 |
| Thoc7 | 1126.694672 | -0.080013991 | 0.069038606 | -1.158974597 | 0.24646654 | 0.471424129 | | | Hmga2-ps1 | 34.24473153 | -0.167823475 | 0.086005326 | -1.951314918 | 0.051019596 | 0.403266245 | | Phka2 |
| Tppp | 25867.86801 | -0.107620489 | 0.092859908 | -1.158955366 | 0.246474379 | 0.471424129 | | | Cldn14 | 28.41128285 | -0.167995926 | 0.088038846 | -1.908202271 | 0.056365078 | 0.416282454 | | Sipa1l3 |
| Pcmtd1 | 3956.216254 | 0.094319669 | 0.081402641 | 1.158680701 | 0.24658636 | 0.4715055 | | | A730036I17Rik | 30.53031476 | -0.168104869 | 0.086963097 | -1.933059819 | 0.053228829 | 0.407317206 | | Itpripl2 |
| Slc25a16 | 1602.049887 | -0.084887435 | 0.073259387 | -1.158724339 | 0.246568567 | 0.4715055 | | | Tgtp2 | 31.30998525 | -0.168143785 | 0.067630749 | -2.486203207 | NA | NA | | Whamm |
| Gucy1b2 | 9.715393783 | -0.169977947 | 0.146724758 | -1.158481699 | 0.246667516 | 0.471602479 | | | 0610009L18Rik | 76.91325961 | -0.168203042 | 0.095060272 | -1.769435721 | 0.076821189 | 0.472560915 | | Srek1 |
| Atp5sl | 1481.649108 | 0.06204501 | 0.053565229 | 1.158307574 | 0.246738542 | 0.471680069 | | | Mir24-1 | 36.55807357 | -0.168218426 | 0.082628269 | -2.035837081 | 0.041766718 | 0.369265575 | | Phkg1 |
| Nsg1 | 11377.81936 | -0.086285513 | 0.074503254 | -1.158144224 | 0.246805185 | 0.471749265 | | | Psrc1 | 414.1945429 | -0.168403619 | 0.084102864 | -2.002352964 | 0.045246783 | 0.383869599 | | Gigyf1 |
| Lck | 55.52871928 | -0.165752077 | 0.143174321 | -1.157694175 | 0.246988862 | 0.472042115 | | | Postn | 84.73479375 | -0.168422423 | 0.091331169 | -1.844084824 | 0.065170779 | 0.444691459 | | Csf1r |
| Bai1 | 18410.74252 | 0.12316402 | 0.069455657 | 1.773275577 | 0.076183054 | 0.227083539 | | | Slc6a6 | 7760.754252 | -0.168449042 | 0.051165641 | -3.292229685 | 0.000993964 | 0.048083105 | | |
| Tsr3 | 558.8368277 | 0.123100011 | 0.08674683 | 1.419072155 | 0.155877981 | 0.358209939 | | | Acat3 | 168.0958657 | -0.168663194 | 0.092728769 | -1.818887456 | 0.068928602 | 0.453262505 | | Ddb2 |
| Trpt1 | 325.6503713 | 0.123070271 | 0.095801459 | 1.28463879 | 0.198918537 | 0.415034179 | | | Tmem136 | 923.6650443 | -0.168707113 | 0.066015968 | -2.555550076 | 0.01060201 | 0.196685574 | | Guf1 |
| C030006K11Rik | 394.6467051 | 0.12303956 | 0.093825615 | 1.311364272 | 0.189734723 | 0.403091979 | | | St14 | 40.55400058 | -0.168808072 | 0.088259997 | -1.912622681 | 0.055796374 | 0.413057568 | | Alkbh6 |
| Ifltd1 | 19.59386864 | 0.122988064 | 0.161979713 | 0.759280661 | 0.447684684 | 0.667028223 | | | Tstd2 | 723.0584299 | -0.16885398 | 0.062081676 | -2.719868262 | 0.006530793 | 0.155695086 | | Safb2 |
| Ggnbp2 | 3504.600686 | 0.122975054 | 0.049358214 | 2.49148105 | 0.012721175 | 0.069549009 | | | Metap1d | 534.6563802 | -0.168855541 | 0.078489882 | -2.151303281 | 0.031452269 | 0.32443523 | | Ccl12 |
| D330041H03Rik | 22.82113479 | 0.122943872 | 0.168107561 | 0.731340525 | 0.464571183 | 0.680539464 | | | A230077H06Rik | 184.0051021 | -0.168958979 | 0.088942646 | -1.899639665 | 0.057480423 | 0.419820382 | | Ogt |
| Tsn | 3023.995351 | -0.056324541 | 0.048694958 | -1.156681165 | 0.247402648 | 0.472366805 | | | Hps5 | 728.0910038 | -0.168975413 | 0.06927512 | -2.439193378 | 0.014720089 | 0.232209004 | | Wsb1 |
| 11-Mar | 120.2965932 | -0.153835644 | 0.133016632 | -1.156514349 | 0.247470834 | 0.472380571 | | | 5031414D18Rik | 72.34616054 | -0.169120375 | 0.095070475 | -1.778894816 | 0.075257008 | 0.469360463 | | Zfp932 |
| Slc35c1 | 681.2339309 | 0.122829095 | 0.085257512 | 1.440683552 | 0.149674104 | 0.350068662 | | | Gm13845 | 70.93188522 | -0.169139234 | 0.094761669 | -1.784890827 | 0.074279027 | 0.467418832 | | Pnn |
| Numbl | 3628.403801 | 0.09705439 | 0.083936631 | 1.156281694 | 0.247565954 | 0.47244573 | | | Arhgef39 | 32.50489761 | -0.169145719 | 0.089239988 | -1.895402756 | 0.058039064 | 0.42195968 | | Tmem181b-ps |
| Rapgefl1 | 7116.634556 | 0.088491813 | 0.076530262 | 1.156298317 | 0.247559156 | 0.47244573 | | | Gm10814 | 29.00067005 | -0.169218681 | 0.088380362 | -1.914663821 | 0.05553539 | 0.413057568 | | Uggt2 |
| Naa35 | 1716.203603 | 0.069132924 | 0.059800895 | 1.156051658 | 0.247660028 | 0.472567053 | | | Rab34 | 570.2978219 | -0.169308763 | 0.08369161 | -2.023007591 | 0.043072365 | 0.375514735 | | Stat2 |
| Pdlim4 | 267.9419049 | 0.122820664 | 0.103816382 | 1.183056674 | 0.236786687 | 0.460959031 | | | Cd44 | 157.9527791 | -0.16931619 | 0.093376405 | -1.813265254 | 0.06979092 | 0.455201066 | | Pabpn1 |
| Xrcc6 | 574.7283053 | 0.093835643 | 0.081222484 | 1.155291462 | 0.247971091 | 0.473044088 | | | Cxadr | 979.969776 | -0.169323181 | 0.066029525 | -2.564355603 | 0.010336758 | 0.194329551 | | Odf2l |
| Ankle1 | 8.668018369 | -0.176978086 | 0.153234712 | -1.154947757 | 0.248111821 | 0.473111538 | | | Exd2 | 3909.343377 | -0.169342144 | 0.062358355 | -2.715628809 | 0.006615004 | 0.156472283 | | Flnb |
| Cstad | 48.16732965 | 0.122803126 | 0.167857338 | 0.731592239 | 0.464417486 | 0.680386485 | | | Plekha2 | 1354.776382 | -0.16960698 | 0.063259733 | -2.681120691 | 0.007337605 | 0.163225184 | | Zcchc7 |
| Gpc3 | 191.8428026 | -0.159597052 | 0.138179396 | -1.154998907 | 0.248090874 | 0.473111538 | | | E4f1 | 1087.674524 | -0.169716811 | 0.077459518 | -2.191038825 | 0.028448983 | 0.313543192 | | D4Wsu53e |
| Ifnar1 | 2105.389605 | -0.068349525 | 0.059185674 | -1.154832259 | 0.248159124 | 0.473111538 | | | Sntg2 | 116.0599844 | -0.169941239 | 0.091071519 | -1.866019597 | 0.062038624 | 0.436531911 | | Szt2 |
| Ppat | 611.5321949 | 0.122774278 | 0.075144969 | 1.63383231 | 0.102294066 | 0.274407405 | | | Anks3 | 1304.288553 | -0.17019608 | 0.073748442 | -2.307792205 | 0.021010697 | 0.273233107 | | Acp6 |
| Ick | 1013.036525 | 0.079945519 | 0.069248372 | 1.15447507 | 0.248305454 | 0.473241848 | | | A230050P20Rik | 2277.398145 | -0.170216975 | 0.069864487 | -2.436387675 | 0.014834776 | 0.232209004 | | Akap8 |
| Pygo1 | 452.7040918 | -0.113799418 | 0.098575279 | -1.154441753 | 0.248319106 | 0.473241848 | | | Smoc2 | 551.9565571 | -0.170333154 | 0.094703931 | -1.798585892 | 0.072084211 | 0.460666627 | | Trank1 |
| Slc18a2 | 34.82142139 | -0.183146735 | 0.15864251 | -1.154461907 | 0.248310847 | 0.473241848 | | | Maml2 | 412.1795482 | -0.170842661 | 0.081247721 | -2.10273787 | 0.03548869 | 0.342120019 | | Tnfrsf14 |
| Ccsap | 2296.26145 | 0.122726303 | 0.091741538 | 1.337739758 | 0.180981286 | 0.391431411 | | | Slc9a2 | 223.4621993 | -0.170855459 | 0.093059789 | -1.835975141 | 0.066361333 | 0.446279967 | | Invs |
| Zfp799 | 854.1699229 | -0.126407821 | 0.109538427 | -1.154004349 | 0.248498386 | 0.473467043 | | | Tor3a | 360.6142234 | -0.170865135 | 0.089748805 | -1.903815142 | 0.056934264 | 0.418112798 | | Mov10 |
| Mcts1 | 947.6168792 | -0.124216104 | 0.10768249 | -1.15354042 | 0.248688637 | 0.47377127 | | | Morn1 | 172.5101294 | -0.170870786 | 0.088294875 | -1.935228814 | 0.052962223 | 0.406864177 | | Ankrd16 |
| Iars2 | 2138.386259 | 0.122715897 | 0.086985622 | 1.410760693 | 0.158315188 | 0.361470015 | | | Nmrk1 | 194.6561968 | -0.170985061 | 0.088088772 | -1.941053971 | 0.052251731 | 0.406355487 | | Izumo4 |
| Ggn | 39.02249532 | -0.192168034 | 0.166640768 | -1.153187405 | 0.248833472 | 0.473930647 | | | Mylip | 581.7375014 | -0.171020024 | 0.072318803 | -2.364807168 | 0.018039461 | 0.256272607 | | Srsf11 |
| Bloc1s4 | 237.9935582 | -0.146209342 | 0.126866168 | -1.152469129 | 0.249128348 | 0.474379245 | | | Cebpb | 68.19415658 | -0.17118996 | 0.095020681 | -1.801607376 | 0.0716072 | 0.459800193 | | Itgam |
| Trim46 | 3440.888662 | 0.081833145 | 0.071007084 | 1.152464526 | 0.249130239 | 0.474379245 | | | C920021L13Rik | 21.98077923 | -0.171321102 | 0.083930242 | -2.041232069 | 0.041227766 | NA | | Lgi4 |
| Ppfia3 | 8313.470176 | 0.122683488 | 0.085128575 | 1.441155201 | 0.149540842 | 0.349892147 | | | A130077B15Rik | 377.0621112 | -0.171451767 | 0.079638033 | -2.152888023 | 0.03132748 | 0.32443523 | | Anxa3 |
| Brca1 | 92.39825393 | -0.16600758 | 0.144109758 | -1.151952389 | 0.249340638 | 0.474623391 | | | Fam222a | 416.2719678 | -0.171593015 | 0.086936991 | -1.973762985 | 0.0484087 | 0.394969953 | | Ifi44 |
| Ewsr1 | 6927.860543 | 0.090705271 | 0.07874718 | 1.151854209 | 0.249380988 | 0.474623391 | | | Cobll1 | 563.4614645 | -0.171663504 | 0.089377325 | -1.920660564 | 0.054774514 | 0.412110852 | | Eml5 |
| Minpp1 | 2334.537241 | -0.06057523 | 0.052587959 | -1.151884019 | 0.249368736 | 0.474623391 | | | Phf20l1 | 1698.364099 | -0.171665463 | 0.066681248 | -2.574418858 | 0.010040864 | 0.191660425 | | Tia1 |
| Ss18 | 1029.461351 | 0.07040148 | 0.061143965 | 1.151405202 | 0.249565575 | 0.474916361 | | | 1700001L05Rik | 276.5574763 | -0.171729398 | 0.09468535 | -1.813684998 | 0.069726236 | 0.455201066 | | 6720401G13Rik |
| Smarcal1 | 3495.249201 | -0.060823952 | 0.052835071 | -1.15120414 | 0.249648263 | 0.475015373 | | | Evi2b | 30.26041139 | -0.17189824 | 0.087669953 | -1.960742912 | 0.04990902 | 0.40054057 | | Akap8l |
| Magee2 | 270.467799 | -0.131115157 | 0.113903428 | -1.151108084 | 0.249687773 | 0.475032214 | | | Cc2d1b | 1272.445497 | -0.171898972 | 0.08305081 | -2.069804879 | 0.03847062 | 0.355486152 | | 4930570G19Rik |
| Zdhhc21 | 1928.335284 | -0.070014963 | 0.060840926 | -1.150787273 | 0.249819762 | 0.475224971 | | | Prpf4b | 3593.285223 | -0.171989029 | 0.061687928 | -2.788049996 | 0.005302636 | 0.137476727 | | Sfi1 |
| Ccdc134 | 312.4128003 | 0.122674435 | 0.123531581 | 0.99306132 | 0.320680072 | 0.553706097 | | | Srsf5 | 8446.686054 | -0.17202158 | 0.065338677 | -2.632768015 | 0.008469217 | 0.175232365 | | Ctsh |
| Zadh2 | 1225.306559 | -0.098995801 | 0.086039531 | -1.150585078 | 0.249902975 | 0.475266563 | | | Celf3 | 5991.565073 | -0.172116958 | 0.073186507 | -2.351758057 | 0.018684924 | 0.260978081 | | Ccdc39 |
| Txlna | 2141.476462 | 0.122646427 | 0.068252895 | 1.79694103 | 0.072344982 | 0.220002581 | | | Foxn2 | 626.6196937 | -0.172159177 | 0.07646825 | -2.251381157 | 0.024361407 | 0.292457679 | | Dusp11 |
| 1700052N19Rik | 905.2895477 | 0.076584929 | 0.066592683 | 1.150050205 | 0.250123194 | 0.47556863 | | | Ppl | 96.16915691 | -0.172177869 | 0.09510162 | -1.810461991 | 0.070224173 | 0.456017009 | | Arglu1 |
| Zfp1 | 667.1702377 | 0.122645909 | 0.083085796 | 1.476135698 | 0.139907467 | 0.334334817 | | | Mtf2 | 1398.251006 | -0.172207721 | 0.073329916 | -2.348396537 | 0.018854437 | 0.261805126 | | Tnip2 |
| Trappc5 | 1176.839821 | -0.080309941 | 0.069882246 | -1.149218094 | 0.250466062 | 0.476103688 | | | 4933427D14Rik | 1608.487505 | -0.17224216 | 0.06415496 | -2.684503547 | 0.007263761 | 0.162353883 | | Med12 |
| 3010001F23Rik | 35.96528432 | -0.184299034 | 0.1603938 | -1.149040887 | 0.250539122 | 0.476184145 | | | Sox4 | 334.798716 | -0.172263152 | 0.089085769 | -1.933677547 | 0.053152786 | 0.407317206 | | Flna |
| Nhlrc4 | 14.10132117 | 0.122518592 | 0.160332545 | 0.76415298 | 0.444776083 | 0.664220593 | | | Fbxw10 | 33.6379734 | -0.172297767 | 0.09006359 | -1.913067958 | 0.055739353 | 0.413057568 | | Cx3cr1 |
| Sowaha | 3377.491107 | -0.117706522 | 0.102485035 | -1.148523992 | 0.250752315 | 0.476448049 | | | Top3a | 274.3791095 | -0.172466347 | 0.079504512 | -2.169264897 | 0.030062579 | 0.319144111 | | Jag1 |
| Gm16973 | 162.0726494 | 0.122511424 | 0.117659486 | 1.041237124 | 0.29776551 | 0.529540878 | | | Notum | 98.52384054 | -0.172698949 | 0.094543517 | -1.826660938 | 0.067750765 | 0.449587762 | | Nvl |
| Anapc13 | 507.7503024 | -0.1121543 | 0.097661479 | -1.148398535 | 0.250804078 | 0.476453946 | | | Slc39a8 | 138.9324356 | -0.172761026 | 0.094360706 | -1.830857708 | 0.067121784 | 0.447688159 | | Fam227a |
| Zfp710 | 916.9824609 | -0.07848273 | 0.068345727 | -1.148319494 | 0.250836694 | 0.47645749 | | | Crocc | 2712.846777 | -0.172892936 | 0.086395935 | -2.001169796 | 0.045374095 | 0.383869599 | | Col4a1 |
| Ube2l6 | 203.3456761 | 0.122410855 | 0.128873927 | 0.94984965 | 0.342188654 | 0.575602106 | | | Luc7l2 | 5384.002953 | -0.172945471 | 0.045537164 | -3.797897258 | 0.000145929 | 0.01286942 | | |
| Trim71 | 11.68374045 | 0.122390898 | 0.161241846 | 0.759051702 | 0.44782163 | 0.667103964 | | | Zfp329 | 1325.573161 | -0.172993577 | 0.070551137 | -2.452031009 | 0.014205242 | 0.228510903 | | Nckap1l |
| 2310045N01Rik | 490.8040076 | 0.085105901 | 0.074146475 | 1.147807779 | 0.251047925 | 0.476683401 | | | Ncapd2 | 1073.102104 | -0.173035328 | 0.058853772 | -2.940088992 | 0.00328118 | 0.099891275 | | Dock6 |
| Ecsit | 1479.201988 | -0.082695151 | 0.072083289 | -1.147216676 | 0.251292081 | 0.476971641 | | | Agrn | 9509.708624 | -0.173205641 | 0.071931315 | -2.407930973 | 0.016043214 | 0.240747652 | | Slc38a6 |
| Gng11 | 324.7404262 | -0.174077344 | 0.151729929 | -1.147284158 | 0.251264199 | 0.476971641 | | | Fancm | 612.4370763 | -0.173328744 | 0.075322964 | -2.301140766 | 0.021383676 | 0.275127789 | | Zfc3h1 |
| 4921507P07Rik | 37.4614982 | 0.122283928 | 0.161807994 | 0.755734777 | 0.449808229 | 0.668456617 | | | Havcr2 | 222.6576837 | -0.173431045 | 0.089455355 | -1.938744151 | 0.0525325 | 0.406864177 | | Ctss |
| Immp2l | 89.72395888 | -0.156208972 | 0.136181776 | -1.147062233 | 0.251355901 | 0.477034338 | | | Ezh2 | 151.6984262 | -0.173458884 | 0.091713977 | -1.89130261 | 0.058583961 | 0.424011266 | | Ccdc142 |
| Gm10532 | 9.209810764 | 0.122255733 | 0.149165061 | 0.819600329 | 0.412443986 | 0.636114585 | | | BC018507 | 6497.022868 | -0.173471729 | 0.055052633 | -3.151016053 | 0.001627035 | 0.065650881 | | Atp11b |
| Gstt3 | 129.3197066 | -0.154900131 | 0.13508442 | -1.146691319 | 0.251509219 | 0.477149975 | | | Zfp641 | 611.074131 | -0.173528769 | 0.067933105 | -2.554406549 | 0.010636898 | 0.197081095 | | Pion |
| Kif19a | 111.5012093 | -0.14367928 | 0.125294149 | -1.14673575 | 0.25149085 | 0.477149975 | | | Arap1 | 736.0495003 | -0.17356493 | 0.07113579 | -2.439910079 | 0.014690918 | 0.232209004 | | Col4a2 |
| Cep104 | 1444.81921 | 0.075935405 | 0.066226425 | 1.146602815 | 0.251545813 | 0.477160972 | | | Alpk1 | 217.4308053 | -0.173600071 | 0.085749467 | -2.024503204 | 0.042918403 | 0.375335772 | | Cdc25b |
| Bccip | 1333.87916 | 0.07227216 | 0.06305434 | 1.146188519 | 0.251717157 | 0.477427548 | | | Ifit2 | 608.3127271 | -0.17382122 | 0.086428359 | -2.011159568 | 0.044308605 | 0.380589751 | | I830077J02Rik |
| Epb4.9 | 14779.41777 | 0.122222227 | 0.078942987 | 1.54823413 | 0.121565937 | 0.307517164 | | | Abcc5 | 6775.532219 | -0.173838668 | 0.061263329 | -2.837564838 | 0.004545911 | 0.125301532 | | Bmp1 |
| Taf5l | 824.6266211 | -0.073183665 | 0.063868401 | -1.145850903 | 0.251856849 | 0.477575574 | | | Depdc1b | 14.49786466 | -0.173950211 | 0.076031968 | -2.287856215 | 0.022145896 | NA | | Rgs11 |
| Pecam1 | 954.9595997 | -0.120227574 | 0.104942717 | -1.145649526 | 0.251940196 | 0.477675159 | | | 9830166K06Rik | 69.52807151 | -0.174057317 | 0.094191632 | -1.847906375 | 0.064615888 | 0.443022969 | | Arhgef1 |
| Snx1 | 2660.74592 | 0.122212276 | 0.087663329 | 1.394109454 | 0.163284555 | 0.368687102 | | | Slc40a1 | 453.1770367 | -0.174174254 | 0.074228738 | -2.346453105 | 0.018953051 | 0.262451878 | | Tie1 |
| Sfxn1 | 2279.90105 | -0.070289246 | 0.061364278 | -1.145442404 | 0.252025941 | 0.477720814 | | | Gpr115 | 107.5706986 | -0.174229943 | 0.094503236 | -1.843639971 | 0.065235627 | 0.444691459 | | Nsun7 |
| Ndfip1 | 10755.5344 | -0.068880355 | 0.060139514 | -1.14534274 | 0.252067208 | 0.477740589 | | | Cspp1 | 726.1325055 | -0.174302496 | 0.077178582 | -2.258430924 | 0.023918807 | 0.29172307 | | Tlr2 |
| Actg1 | 43908.84406 | -0.099907634 | 0.087249905 | -1.145074411 | 0.252178334 | 0.477892749 | | | Pold1 | 307.674381 | -0.1744531 | 0.08222362 | -2.121690825 | 0.03386371 | 0.336326295 | | H2-Q4 |
| Rnf38 | 2064.121112 | 0.122146061 | 0.064145432 | 1.904205133 | 0.056883473 | 0.188507836 | | | Gfap | 10952.7927 | -0.174500776 | 0.080752632 | -2.160929896 | 0.030700755 | 0.321052104 | | Ccdc84 |
| AU041133 | 154.4654543 | -0.153002028 | 0.133711161 | -1.144272675 | 0.252510572 | 0.478384902 | | | Car7 | 742.2795394 | -0.174589156 | 0.085827984 | -2.034175191 | 0.041933935 | 0.370320688 | | Pld4 |
| Cpt1a | 1382.985871 | -0.090422331 | 0.079030154 | -1.144149747 | 0.25256154 | 0.478384902 | | | Rasl10a | 1057.893851 | -0.174626539 | 0.091515391 | -1.908165794 | 0.056369791 | 0.416282454 | | Bin2 |
| Sarm1 | 1331.449138 | 0.073969485 | 0.064647085 | 1.144204494 | 0.25253884 | 0.478384902 | | | Cc2d2a | 1272.458544 | -0.174705103 | 0.068531021 | -2.549284988 | 0.010794405 | 0.198480414 | | Tatdn3 |
| Gsg1l | 4597.972467 | -0.114300142 | 0.099944061 | -1.143641158 | 0.252772485 | 0.478665461 | | | Gm14378 | 132.3152922 | -0.17472303 | 0.093504516 | -1.868605263 | 0.061677753 | 0.435129208 | | Eif2ak2 |
| Ptges3 | 2927.3967 | -0.08819692 | 0.077121483 | -1.143610263 | 0.252785303 | 0.478665461 | | | Prpf40b | 2943.474344 | -0.174898222 | 0.069824204 | -2.504836616 | 0.012250797 | 0.211851282 | | Parp12 |
| Ubqln1 | 6977.531444 | 0.06713441 | 0.058706032 | 1.1435692 | 0.252802341 | 0.478665461 | | | Cfh | 1271.577537 | -0.174987772 | 0.077386322 | -2.261223523 | 0.023745419 | 0.290831331 | | Dennd6b |
| Tbata | 96.78027954 | 0.122142168 | 0.149608558 | 0.816412296 | 0.414264367 | 0.63764026 | | | D930016D06Rik | 481.674099 | -0.175071852 | 0.075059344 | -2.332445797 | 0.019677251 | 0.264659029 | | Ets1 |
| Zfp963 | 148.2951088 | 0.122142118 | 0.114401099 | 1.067665596 | 0.285671383 | 0.515092797 | | | Ano1 | 177.2527774 | -0.175183475 | 0.094567052 | -1.852478957 | 0.063957076 | 0.441368101 | | 1700024P16Rik |
| Jmjd7 | 99.31871252 | 0.122120622 | 0.140009805 | 0.872229071 | 0.383083423 | 0.611369871 | | | Usp48 | 4073.196016 | -0.175246408 | 0.052636098 | -3.329395869 | 0.000870346 | 0.044373556 | | |
| Nhlrc2 | 414.3914617 | 0.12197177 | 0.083258078 | 1.464984205 | 0.142925227 | 0.339280445 | | | Cecr5 | 1046.903996 | -0.175361223 | 0.073783112 | -2.37671221 | 0.017467711 | 0.252725064 | | Luc7l3 |
| 4933427G17Rik | 16.66359955 | -0.190279604 | 0.166525582 | -1.142644877 | 0.253186065 | 0.478982282 | | | Lca5 | 385.7175269 | -0.175438018 | 0.077708755 | -2.257635159 | 0.023968415 | 0.29172307 | | Stx3 |
| Abcg2 | 736.6336593 | 0.086621723 | 0.075803142 | 1.142719436 | 0.253155098 | 0.478982282 | | | Tha1 | 154.1998129 | -0.175444662 | 0.091402861 | -1.919465748 | 0.054925417 | 0.412110852 | | Cep70 |
| Phldb2 | 328.8468264 | -0.113021291 | 0.09890888 | -1.142680931 | 0.25317109 | 0.478982282 | | | Mroh8 | 55.47332217 | -0.17557584 | 0.094646891 | -1.855061876 | 0.063587391 | 0.440996088 | | Ifitm3 |
| Dennd2a | 1110.962134 | -0.102234355 | 0.089486118 | -1.142460499 | 0.253262656 | 0.479068685 | | | Klhl17 | 2045.542088 | -0.175578646 | 0.081635909 | -2.150752623 | 0.03149573 | 0.32443523 | | Ccdc159 |
| Dnajc5 | 15041.48259 | -0.052234202 | 0.045725942 | -1.142331901 | 0.253316086 | 0.479111259 | | | Pip5kl1 | 238.3424525 | -0.175612453 | 0.092789724 | -1.892585136 | 0.058413062 | 0.423355579 | | Dclre1c |
| Fgfrl1 | 654.4530236 | -0.133754374 | 0.117126867 | -1.141961506 | 0.253470021 | 0.479343891 | | | Prkdc | 1161.016933 | -0.175647703 | 0.079593372 | -2.206813179 | 0.027327114 | 0.308173879 | | Ipw |
| Hip1r | 3465.386484 | 0.064761806 | 0.056732106 | 1.14153715 | 0.253646462 | 0.479619023 | | | F11r | 435.0343667 | -0.175715734 | 0.087853046 | -2.00010975 | 0.045488414 | 0.384389008 | | Ptprv |
| Nodal | 22.12132244 | -0.191523343 | 0.167796873 | -1.141399953 | 0.253703524 | 0.479668384 | | | Slco2b1 | 1063.35345 | -0.175751111 | 0.05778493 | -3.041469623 | 0.002354263 | 0.081813462 | | Col6a2 |
| Bach1 | 649.5594566 | -0.094071212 | 0.082431554 | -1.141203909 | 0.253785078 | 0.479754037 | | | Msn | 1551.033671 | -0.17593495 | 0.065436309 | -2.688644175 | 0.007174285 | 0.161546928 | | Lyn |
| Ypel3 | 4549.843231 | 0.07965225 | 0.06980046 | 1.141142193 | 0.253810755 | 0.479754037 | | | Pdzd7 | 227.2644544 | -0.175949819 | 0.093406704 | -1.883695824 | 0.059606132 | 0.428234899 | | Lpcat2 |
| 2610001J05Rik | 947.0249733 | -0.079296447 | 0.069499555 | -1.140963383 | 0.253885161 | 0.479783814 | | | Mdm4 | 1629.448363 | -0.176013431 | 0.070288672 | -2.504150773 | 0.012274571 | 0.212010018 | | Rasal3 |
| Stac3 | 60.22126559 | 0.121871188 | 0.167220999 | 0.72880313 | 0.466122097 | 0.681389166 | | | Lig3 | 1374.209589 | -0.176025727 | 0.067129256 | -2.622190934 | 0.008736647 | 0.178996522 | | Liph |
| Smek2 | 1686.972935 | 0.121858325 | 0.105364257 | 1.156543295 | 0.247459001 | 0.472380571 | | | Csf1 | 1154.344337 | -0.176144659 | 0.067451887 | -2.611411885 | 0.009016922 | 0.181364305 | | Lgals9 |
| Kdm4b | 2988.166303 | 0.07199206 | 0.063103848 | 1.140850553 | 0.253932119 | 0.479797818 | | | Dffb | 267.0915877 | -0.176227027 | 0.087260011 | -2.01956229 | 0.04342881 | 0.37703831 | | Pnpla3 |
| Pacsin3 | 1072.615313 | 0.12181277 | 0.092058916 | 1.323204481 | 0.185787386 | 0.397332949 | | | Herc6 | 747.8590428 | -0.176539827 | 0.085467786 | -2.065571547 | 0.038868953 | 0.357169642 | | Ccdc108 |
| Strn4 | 9488.093494 | 0.121808582 | 0.090055023 | 1.352601747 | 0.176182894 | 0.38523536 | | | Lpxn | 67.92121673 | -0.176552206 | 0.093298964 | -1.892327624 | 0.058447343 | 0.423355579 | | C1s |
| Lrp8 | 3608.491016 | -0.089172568 | 0.078243372 | -1.139682063 | 0.254418783 | 0.480493006 | | | Lepr | 164.811167 | -0.176619549 | 0.094689723 | -1.865245174 | 0.062147046 | 0.436531911 | | Csf2rb |
| Ercc4 | 1094.972193 | 0.076494507 | 0.067168899 | 1.138838176 | 0.254770656 | 0.481098908 | | | Hook2 | 562.3414342 | -0.176754714 | 0.077588544 | -2.278103256 | 0.022720426 | 0.283293487 | | 2610005L07Rik |
| Tceal3 | 2795.574988 | 0.074586157 | 0.06550911 | 1.138561592 | 0.254886056 | 0.481258171 | | | Trpc2 | 578.9649251 | -0.176872937 | 0.078436236 | -2.254990123 | 0.024133949 | 0.29178139 | | Srrm2 |
| Alg10b | 1176.518509 | -0.112026424 | 0.098423957 | -1.138202809 | 0.255035806 | 0.481364938 | | | Col27a1 | 407.6247667 | -0.176896375 | 0.093943098 | -1.883016192 | 0.059698174 | 0.428234899 | | Chuk |
| Fkrp | 2062.800204 | 0.08064009 | 0.07084207 | 1.138307933 | 0.254991922 | 0.481364938 | | | Fancd2 | 197.18293 | -0.176939578 | 0.089744676 | -1.971588474 | 0.048656604 | 0.395027926 | | Ap1g2 |
| Wtip | 178.795008 | 0.121683882 | 0.114189306 | 1.065632905 | 0.286589619 | 0.516185277 | | | Gm14872 | 213.4711444 | -0.176980321 | 0.087522845 | -2.022104296 | 0.043165578 | 0.375695964 | | Ifi203 |
| Znfx1 | 4094.91522 | 0.098846913 | 0.086854668 | 1.138072534 | 0.255090196 | 0.481408952 | | | Fnbp4 | 2009.572349 | -0.17706392 | 0.058573211 | -3.022950555 | 0.002503231 | 0.084486004 | | AF251705 |
| Cbll1 | 672.6888457 | 0.08797691 | 0.077314181 | 1.137914283 | 0.255156277 | 0.481475015 | | | Gm2382 | 105.5620084 | -0.177183158 | 0.094067355 | -1.883577554 | 0.05962214 | 0.428234899 | | Irf1 |
| Eef2 | 42445.04113 | -0.06412787 | 0.056384322 | -1.137335125 | 0.255398216 | 0.481872864 | | | Fgfrl1 | 675.2921841 | -0.177195469 | 0.073621342 | -2.406849201 | 0.016090813 | 0.24095394 | | Ddx26b |
| Plk4 | 195.1377744 | 0.121641335 | 0.102031055 | 1.192199133 | 0.233183173 | 0.456594481 | | | Itga5 | 328.2394456 | -0.177195664 | 0.084676071 | -2.092629743 | 0.036382219 | 0.345417071 | | Slc14a1 |
| Emc10 | 6161.475952 | -0.099550422 | 0.087565207 | -1.136871875 | 0.255591851 | 0.48192595 | | | Lin28b | 225.3611139 | -0.177334495 | 0.090972787 | -1.949313643 | 0.05125798 | 0.403266245 | | Cdk2 |
| Gm10516 | 212.0865773 | -0.108475642 | 0.095420131 | -1.136821349 | 0.255612977 | 0.48192595 | | | Bst2 | 159.5387085 | -0.177382465 | 0.07648006 | -2.319329578 | 0.020377173 | 0.269758295 | | Vmn2r84 |
| Mtmr12 | 2122.165134 | 0.121606481 | 0.091596959 | 1.327625749 | 0.184301774 | 0.395333215 | | | Park2 | 246.7325046 | -0.177544472 | 0.090437364 | -1.963176105 | 0.049625715 | 0.399665044 | | Csf2ra |
| Piezo2 | 37.05556738 | -0.189285302 | 0.166455599 | -1.137151911 | 0.255474786 | 0.48192595 | | | Adamts18 | 86.40353207 | -0.177599042 | 0.094577785 | -1.877809263 | 0.060407263 | 0.430557358 | | Lat2 |
| Slc25a33 | 1043.06481 | -0.072826313 | 0.064058181 | -1.136877636 | 0.255589442 | 0.48192595 | | | Cyyr1 | 123.9333307 | -0.177611537 | 0.094840576 | -1.87273786 | 0.061104593 | 0.432978201 | | Uvssa |
| Rnf135 | 141.6339528 | 0.121583577 | 0.108293637 | 1.122721281 | 0.261555887 | 0.488729729 | | | Rasgrp4 | 38.86759462 | -0.177846419 | 0.081281397 | -2.188033493 | 0.028667163 | 0.315230284 | | Itgb2 |
| Pgbd1 | 79.64814995 | 0.121570066 | 0.142138141 | 0.855295173 | 0.392387759 | 0.618634473 | | | Trim12c | 149.7194165 | -0.177867633 | 0.091631208 | -1.941125051 | 0.052243111 | 0.406355487 | | Cyth4 |
| Ganab | 5082.637687 | -0.078504461 | 0.069108657 | -1.135956975 | 0.255974571 | 0.482431579 | | | Nphp4 | 818.3632044 | -0.177928051 | 0.085045496 | -2.092151379 | 0.036424976 | 0.34559713 | | Gm3002 |

| | | | | | | | | | | | | | | | | | |
|---|---|---|---|---|---|---|---|---|---|---|---|---|---|---|---|---|---|
| Cebpa | 315.3716242 | 0.12156839 | 0.11011169 | 1.104046178 | 0.269573107 | 0.497867046 | | | 4930538K18Rik | 86.58954393 | -0.177977477 | 0.095002042 | -1.873406847 | 0.061012225 | 0.432534692 | | Arhgap30 |
| 9930021J03Rik | 2807.385656 | -0.092825997 | 0.081754717 | -1.135420684 | 0.256199097 | 0.482569738 | | | Tor4a | 378.2787119 | -0.178085317 | 0.092082154 | -1.93398296 | 0.053115223 | 0.407317206 | | Hk2 |
| Map3k12 | 5585.219302 | 0.121523388 | 0.049863868 | 2.437103125 | 0.014805456 | 0.077274849 | | | Mir1188 | 18.60809112 | -0.178101307 | 0.073681221 | -2.417187252 | 0.015640969 | NA | | Lcp1 |
| BC021785 | 26.81849828 | 0.121501822 | 0.167709338 | 0.724478576 | 0.468771975 | 0.68339791 | | | Ube2j2 | 67.59203655 | -0.17813991 | 0.095067527 | -1.873825013 | 0.060954548 | 0.432354503 | | Irf9 |
| Ccdc120 | 456.0041186 | 0.121493063 | 0.076005343 | 1.598480551 | 0.10993607 | 0.288556577 | | | Ssc5d | 126.3808835 | -0.178155835 | 0.094931336 | -1.876681015 | 0.060561826 | 0.431188114 | | Lacc1 |
| Tmem38a | 4020.161649 | -0.083054319 | 0.073153984 | -1.135335553 | 0.256234751 | 0.482569738 | | | Shpk | 190.9802717 | -0.178206894 | 0.091997479 | -1.937084539 | 0.05273501 | 0.406864177 | | Gbp5 |
| Dll4 | 135.2778033 | 0.121448688 | 0.140147787 | 0.866575854 | 0.386174448 | 0.613806617 | | | Lhfpl2 | 756.91459 | -0.178209575 | 0.064364077 | -2.768773894 | 0.005626767 | 0.141672083 | | Unc93b1 |
| Ogfod1 | 5916.947022 | -0.061404644 | 0.054114132 | -1.134724738 | 0.256490669 | 0.482934309 | | | Aspm | 26.31198451 | -0.178335563 | 0.08751914 | -2.037675004 | 0.041582446 | 0.368308905 | | Col11a2 |
| 5330426P16Rik | 172.8342734 | -0.150525249 | 0.132688959 | -1.134421809 | 0.256617655 | 0.483114696 | | | Acr | 41.00105014 | -0.178570582 | 0.091434261 | -1.952994214 | 0.050820281 | 0.403266245 | | Ncrna00085 |
| Hbs1l | 2076.925141 | 0.121395091 | 0.0543149 | 2.23502375 | 0.02541579 | 0.110305951 | | | Plag1 | 148.1946885 | -0.178708728 | 0.09208682 | -1.940654788 | 0.052300164 | 0.406355487 | | Cd48 |
| Pphln1 | 1538.654301 | -0.080552032 | 0.071027104 | -1.134102725 | 0.256751461 | 0.483132796 | | | Olfr613 | 97.49314752 | -0.178868851 | 0.094924113 | -1.884335238 | 0.059519644 | 0.428234899 | | Capg |
| Rbm18 | 1655.317741 | -0.077305264 | 0.068160059 | -1.13417249 | 0.256722201 | 0.483132796 | | | Rac2 | 106.7236335 | -0.178945465 | 0.095100531 | -1.88164528 | 0.059884194 | 0.428722425 | | A230057D06Rik |
| Timp3 | 5575.209021 | -0.081477336 | 0.071829614 | -1.134313997 | 0.25666286 | 0.483132796 | | | Plekha5 | 2347.283512 | -0.179025446 | 0.053010161 | -3.377191124 | 0.000732301 | 0.040141174 | | |
| Cuedc2 | 3532.513372 | 0.086588125 | 0.076407451 | 1.133241906 | 0.25711268 | 0.483752747 | | | Tcea2 | 1381.156586 | -0.179025682 | 0.073938051 | -2.421292963 | 0.015465408 | 0.236284305 | | Ech1 |
| Tmem138 | 256.0541739 | 0.121394932 | 0.102385467 | 1.185665657 | 0.235754362 | 0.459699215 | | | B230378P21Rik | 11.50972309 | -0.179244634 | 0.068678426 | -2.609911792 | 0.009056557 | NA | | Ifit3 |
| Scg5 | 7026.97994 | 0.121394655 | 0.099567463 | 1.219220123 | 0.222760656 | 0.443916863 | | | Anks6 | 524.2733561 | -0.179462019 | 0.076429153 | -2.348083326 | 0.018870299 | 0.261805126 | | Slamf9 |
| 4933430I17Rik | 15.77635331 | -0.189906023 | 0.167626914 | -1.132908898 | 0.257252513 | 0.483780852 | | | Gm10845 | 430.7382562 | -0.179748508 | 0.073436817 | -2.447662019 | 0.014378646 | 0.229590624 | | Cep110 |
| Snx5 | 3450.071886 | 0.060665061 | 0.053545577 | 1.132961206 | 0.257230545 | 0.483780852 | | | C4a | 103.8722857 | -0.179770441 | 0.093853833 | -1.915429922 | 0.055437698 | 0.413057568 | | Xaf1 |
| Med30 | 385.2275989 | -0.118197179 | 0.104350426 | -1.132694742 | 0.257342467 | 0.483891284 | | | Cnot10 | 2323.337586 | -0.179810469 | 0.059344496 | -3.029943476 | 0.002445995 | 0.083252314 | | Rnf207 |
| A330009N23Rik | 450.3140024 | -0.137357242 | 0.121329597 | -1.132100047 | 0.257592375 | 0.484107923 | | | Gm15706 | 181.4353073 | -0.180020502 | 0.090642465 | -1.986050386 | 0.047027729 | 0.39102736 | | Csf2rb2 |
| Ftl1 | 9881.961791 | 0.121371189 | 0.079512841 | 1.526435068 | 0.12690155 | 0.314745096 | | | Tjap1 | 740.5990857 | -0.180064665 | 0.088825999 | -2.02716173 | 0.042645876 | 0.37389754 | | Fes |
| Pcdhb12 | 315.8682183 | -0.112264333 | 0.099146818 | -1.132303941 | 0.257506674 | 0.484107923 | | | Rgs22 | 51.25571681 | -0.180110324 | 0.092640235 | -1.944191134 | 0.051872407 | 0.405765419 | | Samsn1 |
| Rnf40 | 2641.962262 | 0.072885979 | 0.06438413 | 1.132048835 | 0.257613904 | 0.484107923 | | | Catsperd | 50.7968235 | -0.180130735 | 0.092157921 | -1.954587669 | 0.050631758 | 0.403266245 | | Catsper2 |
| Vmn2r29 | 188.2571239 | -0.154452637 | 0.136413425 | -1.132239269 | 0.257533855 | 0.484107923 | | | Polk | 555.5426304 | -0.18013709 | 0.069912253 | -2.576616863 | 0.009977247 | 0.19094795 | | Hdac7 |
| Bnip3 | 3723.073268 | -0.088530915 | 0.078238104 | -1.131557521 | 0.257820508 | 0.484143855 | | | Arl11 | 29.36215852 | -0.180417586 | 0.08381269 | -2.152628505 | 0.031347886 | 0.32443523 | | Thpo |
| Efha2 | 3098.935447 | 0.096587954 | 0.085348994 | 1.131682397 | 0.257767985 | 0.484143855 | | | Arhgef5 | 164.3021736 | -0.180638692 | 0.088836394 | -2.033386134 | 0.042013527 | 0.370320688 | | Was |
| Mob1a | 349.0685411 | -0.098130281 | 0.086720036 | -1.131575653 | 0.257812881 | 0.484143855 | | | St8sia6 | 75.50765712 | -0.180692984 | 0.094814314 | -1.905756387 | 0.05668182 | 0.417225452 | | Hvcn1 |
| Naalad2 | 64.46796623 | 0.121366929 | 0.158355062 | 0.766422795 | 0.443424774 | 0.663360379 | | | Acrbp | 244.8180833 | -0.180935236 | 0.094892918 | -1.906730667 | 0.056555474 | 0.416806094 | | Ifit1 |
| Tmem104 | 784.650637 | 0.070070818 | 0.061916861 | 1.131692016 | 0.257763939 | 0.484143855 | | | Cdan1 | 1295.455413 | -0.181282286 | 0.069232635 | -2.618451337 | 0.008832989 | 0.179451744 | | Plek |
| Rbm10 | 1987.045772 | 0.121322071 | 0.075297426 | 1.611237966 | 0.107127866 | 0.28332306 | | | Zfp36l2 | 879.1711633 | -0.1812952 | 0.091264379 | -1.98648369 | 0.046979642 | 0.39102736 | | Col5a1 |
| Rapgef3 | 1414.817631 | 0.121258742 | 0.082374569 | 1.47204098 | 0.141009812 | 0.335925279 | | | Sfxn4 | 526.9561023 | -0.181303428 | 0.070241475 | -2.581144938 | 0.009847323 | 0.189711158 | | Gpatch1 |
| Pdcl3 | 689.5905013 | -0.093360019 | 0.082536205 | -1.131140186 | 0.257996092 | 0.48433973 | | | Mapkapk3 | 140.267034 | -0.181310884 | 0.092541255 | -1.959243834 | 0.050084238 | 0.401503113 | | Phf11d |
| Phtf2 | 669.6101778 | 0.088740637 | 0.07845277 | 1.131134537 | 0.25799847 | 0.48433973 | | | Lgals3bp | 1014.970777 | -0.181401396 | 0.088702739 | -2.045048416 | 0.040850085 | 0.365377968 | | Tmc4 |
| Fgfr3 | 4680.997222 | 0.121244705 | 0.098027728 | 1.236840911 | 0.216146151 | 0.436353563 | | | Tnfrsf22 | 38.27127116 | -0.181476104 | 0.092471087 | -1.962517267 | 0.049702292 | 0.399757904 | | Ddx58 |
| Mpv17l2 | 380.9715886 | -0.099885252 | 0.088350273 | -1.130559631 | 0.258240487 | 0.48463882 | | | Nfatc2ip | 239.3431904 | -0.181783216 | 0.087784963 | -2.070778526 | 0.038379498 | 0.354869883 | | 4831440E17Rik |
| Rasgrf1 | 14672.72925 | 0.121215387 | 0.105387552 | 1.15018695 | 0.25006688 | 0.475519912 | | | Cacna1h | 2892.532099 | -0.181977547 | 0.058625166 | -3.104085807 | 0.00190868 | 0.071827671 | | Spag5 |
| Mblac2 | 834.3715977 | -0.084865749 | 0.075085179 | -1.130259662 | 0.258366826 | 0.48475849 | | | Arhgef2 | 6865.59606 | -0.182005592 | 0.058159253 | -3.129434849 | 0.001751429 | 0.068703774 | | Mlxipl |
| Idh2 | 1264.942591 | 0.121204499 | 0.106741638 | 1.135494085 | 0.256168359 | 0.482569738 | | | Nanos3 | 23.51254187 | -0.182045073 | 0.083716259 | -2.174548578 | 0.029663961 | 0.317771905 | | Parp9 |
| Pdlim2 | 309.9829665 | 0.121168021 | 0.138812731 | 0.87288839 | 0.382723915 | 0.611332406 | | | Mapk7 | 811.9406836 | -0.182134111 | 0.080101913 | -2.273779803 | 0.022979229 | 0.28516281 | | Ncf2 |
| Gkn3 | 100.8579886 | -0.171493567 | 0.151848205 | -1.129375001 | 0.258739674 | 0.485281746 | | | Gal3st4 | 182.0294937 | -0.182191483 | 0.089387466 | -2.038221815 | 0.041527755 | 0.368048911 | | Csf3r |
| Nudt2 | 442.3119106 | -0.114785199 | 0.101655614 | -1.129157501 | 0.258831398 | 0.485395024 | | | Tctn3 | 1169.159403 | -0.182199115 | 0.05824609 | -3.128091783 | 0.001759452 | 0.068703774 | | Kctd15 |
| Bop1 | 1883.774401 | -0.086229302 | 0.07637473 | -1.129029232 | 0.258885503 | 0.485437732 | | | Elk3 | 355.247573 | -0.182201671 | 0.076008303 | -2.397128511 | 0.016524127 | 0.24467836 | | Siglec5 |
| Zfp652 | 571.2856494 | -0.10969761 | 0.097170997 | -1.128913086 | 0.2589345 | 0.485470855 | | | Il21r | 81.86647701 | -0.182375338 | 0.095091411 | -1.91789495 | 0.055124333 | 0.412537896 | | Trim30a |
| Pdhx | 1677.161823 | -0.076196458 | 0.067518278 | -1.128530817 | 0.25909581 | 0.485714516 | | | Fdxr | 457.8959162 | -0.182503572 | 0.087511781 | -2.085474322 | 0.037026267 | 0.348173348 | | Sfrs18 |
| Csf2ra | 690.9675447 | 0.121148525 | 0.165455095 | 0.732213932 | 0.464038002 | 0.68007991 | | | Dera | 63.95677675 | -0.182541809 | 0.093849714 | -1.945043841 | 0.051769702 | 0.405394443 | | Zfp36 |
| Zswim1 | 1017.338473 | 0.066693435 | 0.059115374 | 1.128191037 | 0.259239249 | 0.485865842 | | | Col2a1 | 86.5619747 | -0.182737948 | 0.094755492 | -1.928520924 | 0.053790364 | 0.408874323 | | Man2b1 |
| Srfbp1 | 382.6947095 | 0.097060207 | 0.086059732 | 1.12782372 | 0.259394374 | 0.486097777 | | | Ushbp1 | 213.0383837 | -0.182765132 | 0.089623337 | -2.03925828 | 0.041424257 | 0.368029822 | | Ptprc |
| Atg2a | 2466.860051 | 0.077779849 | 0.068979488 | 1.127579399 | 0.259497591 | 0.486232395 | | | Rasa4 | 300.9072449 | -0.18302096 | 0.094769364 | -1.931224948 | 0.05345524 | 0.407906655 | | Col19a1 |
| Fn3krp | 1287.518685 | 0.121115781 | 0.074791854 | 1.619371289 | 0.105367401 | 0.280220135 | | | Eno3 | 330.4129066 | -0.183102641 | 0.087284217 | -2.097774907 | 0.035925034 | 0.344080613 | | Miat |
| Morc2b | 47.51827889 | 0.120970678 | 0.150542705 | 0.803563866 | 0.421648898 | 0.644028789 | | | Cdca7l | 41.84915241 | -0.183152794 | 0.091878968 | -1.993413696 | 0.046216165 | 0.387831322 | | Zfp217 |
| Psap | 77343.38931 | 0.12094028 | 0.075943417 | 1.59250512 | 0.111271254 | 0.290535356 | | | Epsti1 | 34.6570421 | -0.183380489 | 0.085507139 | -2.144621972 | 0.031983084 | 0.327404003 | | Aspg |
| Ankfn1 | 93.16936004 | 0.120924567 | 0.164285981 | 0.736061385 | 0.461693341 | 0.678212469 | | | Fam178a | 2663.883213 | -0.18342426 | 0.069442481 | -2.641384012 | 0.008256807 | 0.174077875 | | Slc15a3 |
| Osbpl11 | 841.690223 | 0.120839024 | 0.068414278 | 1.766283702 | 0.077348265 | 0.229102277 | | | Il10ra | 324.1009558 | -0.18347318 | 0.082955089 | -2.211717 | 0.026986225 | 0.306595176 | | Gbp4 |
| Sirt5 | 342.3834422 | 0.120818887 | 0.103362244 | 1.1688493 | 0.242464353 | 0.467201662 | | | Zmynd15 | 18.15214063 | -0.183571877 | 0.075121242 | -2.443674675 | 0.01453853 | NA | | Ccl6 |
| Dmrta2 | 40.61036895 | -0.185180845 | 0.164400288 | -1.126402192 | 0.259995319 | 0.486752958 | | | Traf7 | 2170.505058 | -0.183836023 | 0.069113441 | -2.659917099 | 0.007815989 | 0.169708597 | | Ttc14 |
| Sbspon | 13.20048409 | -0.187951554 | 0.166932654 | -1.125912458 | 0.260202576 | 0.487051764 | | | Chst14 | 75.04760106 | -0.183936519 | 0.094973105 | -1.936722174 | 0.052779314 | 0.406864177 | | Trim30d |
| Ddhd2 | 2908.00737 | 0.120799049 | 0.05280702 | 2.287556609 | 0.022163355 | 0.100848185 | | | 4933439C10Rik | 940.3748484 | -0.18400155 | 0.086590805 | -2.124954835 | 0.033590382 | 0.335456507 | | Aif1 |
| Ccdc43 | 670.2885991 | 0.07074529 | 0.062843459 | 1.125738325 | 0.260276297 | 0.48709443 | | | Tctn1 | 471.5544748 | -0.184190968 | 0.08506823 | -2.165214541 | 0.030371259 | 0.32021294 | | Tnfsf10 |
| Hdgfrp3 | 4426.881292 | -0.081755507 | 0.072628039 | -1.125674161 | 0.260303465 | 0.48709443 | | | C1qa | 2069.417702 | -0.184425404 | 0.094324013 | -1.955232794 | 0.050555599 | 0.403264376 | | Ctsc |
| Gipc1 | 2230.610072 | 0.120777915 | 0.087027541 | 1.387812568 | 0.165194127 | 0.370729693 | | | Slc7a6 | 1059.729648 | -0.184648787 | 0.065196301 | -2.8321973 | 0.004622931 | 0.126226882 | | Haus5 |
| Amn1 | 307.1942675 | -0.110212315 | 0.09795196 | -1.125167021 | 0.260518265 | 0.487378653 | | | Reep4 | 113.2158005 | -0.184763729 | 0.095100004 | -1.942836176 | 0.052035955 | 0.406272098 | | Cd86 |
| Scn2b | 13903.76542 | -0.092365518 | 0.082113698 | -1.124849082 | 0.260652992 | 0.487571828 | | | Glis3 | 121.7348104 | -0.184881195 | 0.092031247 | -2.008895751 | 0.044548174 | 0.380589751 | | Dock2 |
| Slc9a6 | 5570.718228 | -0.084104167 | 0.074810655 | -1.124227121 | 0.260916687 | 0.488006174 | | | D630041G03Rik | 191.1480397 | -0.184944936 | 0.092081723 | -2.008486915 | 0.044591575 | 0.380589751 | | Meg3 |
| Chd1 | 1049.629912 | 0.120765995 | 0.077101695 | 1.56632088 | 0.117273515 | 0.300734168 | | | Gpr34 | 424.3215779 | -0.184990173 | 0.089064971 | -2.077025021 | 0.03779925 | 0.351969169 | | Lair1 |
| Wnt5a | 316.6419987 | -0.139372185 | 0.124002517 | -1.12394642 | 0.261035757 | 0.488111033 | | | Slc16a11 | 525.3114085 | -0.184998524 | 0.087993275 | -2.102416618 | 0.035516797 | 0.342120019 | | Gm15328 |
| Brd7 | 1621.080479 | 0.068170222 | 0.060663046 | 1.123752043 | 0.261118231 | 0.488147427 | | | Tmem80 | 922.3945185 | -0.185069221 | 0.063185874 | -2.928965128 | 0.003400926 | 0.102281258 | | Zkscan2 |
| Tsta3 | 671.0610766 | 0.078678205 | 0.070013705 | 1.123754351 | 0.261117252 | 0.488147427 | | | Casp12 | 41.12608815 | -0.185132062 | 0.092198327 | -2.007976367 | 0.044645803 | 0.380589751 | | Parp14 |
| Bpifa1 | 10.11102975 | -0.112744252 | 0.100358001 | -1.123420657 | 0.261258881 | 0.488351441 | | | Echdc2 | 374.106329 | -0.185271252 | 0.084375593 | -2.19579199 | 0.028106835 | 0.312140588 | | Emr1 |
| Preb | 2677.564496 | 0.078397715 | 0.069812122 | 1.122981407 | 0.261445391 | 0.48864112 | | | Ovca2 | 483.4350975 | -0.185313076 | 0.070841013 | -2.615900973 | 0.008899236 | 0.18004221 | | Malat1 |
| Haus4 | 74.64047351 | -0.150406572 | 0.133947861 | -1.122874015 | 0.261491005 | 0.488667426 | | | Mpzl1 | 674.6399346 | -0.185393591 | 0.070561952 | -2.627387513 | 0.008604328 | 0.176784265 | | Leng8 |
| Bcl2l1 | 4176.566229 | 0.120729868 | 0.112387303 | 1.074230495 | 0.282719388 | 0.512513653 | | | Ndnf | 311.3397539 | -0.185450973 | 0.090634348 | -2.046144498 | 0.040742156 | 0.365321333 | | Slc11a1 |
| 1190002N15Rik | 1195.240707 | -0.136789066 | 0.12184543 | -1.122644204 | 0.261588634 | 0.488731978 | | | Fzd6 | 421.1218243 | -0.1854673 | 0.078573863 | -2.360419787 | 0.018254266 | 0.257188614 | | Cxcl16 |
| Sacm1l | 2973.871318 | 0.067751503 | 0.060360038 | 1.122456254 | 0.261668498 | 0.488784616 | | | Mir17hg | 13.80863866 | -0.18559634 | 0.07316871 | -2.536553412 | 0.011194966 | NA | | Itgax |
| Mob2 | 885.9952601 | 0.120699643 | 0.086111345 | 1.401669465 | 0.161013973 | 0.365745564 | | | 2900076A07Rik | 130.4427097 | -0.185708514 | 0.09437127 | -1.967850101 | 0.049085288 | 0.397221665 | | Ly86 |
| Rad18 | 178.6788782 | 0.12069146 | 0.139647286 | 0.86425926 | 0.387445485 | 0.614790257 | | | Scand1 | 236.375585 | -0.185835448 | 0.093324354 | -1.991285671 | 0.04644949 | 0.388890657 | | Il18bp |
| Igfbp3 | 441.5004479 | 0.120660584 | 0.135731142 | 0.888967577 | 0.374020505 | 0.603339272 | | | Cpeb1 | 2996.80889 | -0.18604047 | 0.067729146 | -2.746830291 | 0.006017426 | 0.147900395 | | Apobec3 |
| Vps18 | 2248.408116 | 0.093538536 | 0.083380591 | 1.121826255 | 0.261936321 | 0.48908669 | | | Zgpat | 2154.25622 | -0.186106601 | 0.078413068 | -2.373413075 | 0.017624541 | 0.253730508 | | I830012O16Rik |
| Mipep | 1108.213878 | -0.078357325 | 0.069859288 | -1.12164505 | 0.262013389 | 0.489171641 | | | Ptafr | 35.39846019 | -0.186153429 | 0.089725166 | -2.074706983 | 0.0380137 | 0.352889542 | | Cd84 |
| Mllt3 | 1287.262065 | -0.087853443 | 0.078339818 | -1.121440484 | 0.262100412 | 0.489275155 | | | Tmem173 | 185.8008231 | -0.186258639 | 0.092079701 | -2.022798041 | 0.043093974 | 0.375514735 | | Cd33 |
| Amdhd1 | 9.052811219 | -0.170854008 | 0.152403742 | -1.121061763 | 0.262261573 | 0.489476425 | | | Trpv3 | 18.14646162 | -0.186273546 | 0.076706801 | -2.4283837 | 0.015166288 | NA | | A930013F10Rik |
| Cacng3 | 4089.641714 | -0.103018956 | 0.091908159 | -1.120890215 | 0.262334596 | 0.489476425 | | | Krba1 | 2082.149685 | -0.186643225 | 0.069263316 | -2.694690878 | 0.007045396 | 0.160386105 | | Naip5 |
| Efcab1 | 121.4431669 | -0.147215089 | 0.131327401 | -1.120977708 | 0.262297351 | 0.489476425 | | | 5430435G22Rik | 68.52469412 | -0.186664775 | 0.094073469 | -1.984244624 | 0.047228576 | 0.391827402 | | Slc26a10 |
| Snx4 | 1383.410419 | 0.064555749 | 0.057592864 | 1.120898412 | 0.262331107 | 0.489476425 | | | Kcnh4 | 1189.025269 | -0.186717762 | 0.087289133 | -2.139072251 | 0.032429817 | 0.328963357 | | Ptpn6 |
| Cadm3 | 21401.14211 | 0.083737941 | 0.074714884 | 1.120766518 | 0.262387259 | 0.489515737 | | | 5930403L14Rik | 1978.88434 | -0.186726343 | 0.073891648 | -2.527029089 | 0.011503197 | 0.203514823 | | Gm5577 |
| Mrpl18 | 965.0895375 | -0.061743776 | 0.055105089 | -1.120473195 | 0.262512169 | 0.489689808 | | | Ybx2 | 156.4934039 | -0.186773117 | 0.094733441 | -1.971564784 | 0.048659311 | 0.395027926 | | Mpeg1 |
| Cyp46a1 | 8551.30337 | 0.120612562 | 0.07843031 | 1.537830998 | 0.124089942 | 0.311612571 | | | Chrd | 1632.355855 | -0.186779928 | 0.08423581 | -2.217345897 | 0.026599464 | 0.304478969 | | Ovgp1 |
| Chia | 78.98573644 | -0.148750307 | 0.132788987 | -1.120200632 | 0.262628275 | 0.489788455 | | | Tlr7 | 190.5483933 | -0.186789164 | 0.086228637 | -2.166208007 | 0.030295296 | 0.32021294 | | Zkscan3 |
| Gsk3a | 9488.231549 | 0.084298352 | 0.075258199 | 1.120121849 | 0.262661841 | 0.489792101 | | | Ptpn21 | 451.7187563 | -0.186798755 | 0.084489091 | -2.210921586 | 0.027041268 | 0.306595176 | | Fcgr1 |
| Lphn2 | 4421.044804 | 0.072231882 | 0.064499489 | 1.119883012 | 0.262763618 | 0.489907889 | | | Phf7 | 171.5324099 | -0.186896193 | 0.0857066 | -2.180651112 | 0.029209232 | 0.317415456 | | Snhg11 |
| Tmco6 | 316.6737769 | 0.120555092 | 0.100677213 | 1.197441697 | 0.23113444 | 0.45406327 | | | Sp110 | 80.06631555 | -0.186930409 | 0.095012131 | -1.967437289 | 0.049132819 | 0.397384927 | | Cd14 |
| Acat3 | 148.3868612 | 0.120445676 | 0.144591962 | 0.833003955 | 0.404842499 | 0.629676416 | | | A930024E05Rik | 102.1592212 | -0.186998332 | 0.094857215 | -1.971366454 | 0.048681975 | 0.395027926 | | Myo1f |
| Haus1 | 227.4770436 | 0.120435151 | 0.104057336 | 1.157392223 | 0.24711215 | 0.472061549 | | | Mtmr11 | 396.5878766 | -0.187095526 | 0.081336383 | -2.300268579 | 0.021433009 | 0.275274875 | | Cep89 |
| Zfp248 | 460.5362595 | 0.097612204 | 0.08722164 | 1.119128288 | 0.263085414 | 0.490286913 | | | Ccdc171 | 159.946862 | -0.187177438 | 0.091613756 | -2.043114988 | 0.041041057 | 0.366193113 | | Robo3 |
| St6galnac6 | 4817.27972 | -0.089157997 | 0.079709098 | -1.118542282 | 0.263335459 | 0.490693878 | | | Dtl | 411.1343753 | -0.187625457 | 0.086268365 | -2.174904523 | 0.029637272 | 0.317771905 | | Irf8 |
| Hcfc1r1 | 2217.179695 | -0.08278541 | 0.074017499 | -1.118457279 | 0.263371742 | 0.490702474 | | | Cntrob | 818.9833046 | -0.187717423 | 0.07814536 | -2.402156989 | 0.016298712 | 0.242552129 | | Mrps12 |
| S100a3 | 22.16049593 | 0.120343571 | 0.167778202 | 0.717277746 | 0.473202736 | 0.687592909 | | | Myo1a | 52.3338673 | -0.187757371 | 0.091385085 | -2.054573454 | 0.039920222 | 0.361071695 | | Gbp9 |
| Wdr77 | 2141.318358 | 0.12022915 | 0.05906093 | 2.035679934 | 0.041782505 | 0.15472707 | | | 9430008C03Rik | 354.7296345 | -0.187910958 | 0.079479562 | -2.364267672 | 0.018065755 | 0.256272607 | | Col11a1 |
| Ppp1r3d | 312.6578132 | -0.104775384 | 0.093742794 | -1.117690011 | 0.26369941 | 0.491076763 | | | Pde7a | 897.9388442 | -0.188028052 | 0.06054037 | -3.105829267 | 0.001897462 | 0.071591009 | | Ryr1 |
| Prpf40a | 1280.301304 | 0.089220735 | 0.07982561 | 1.117695632 | 0.263697009 | 0.491076763 | | | Tmem67 | 485.8695036 | -0.188069427 | 0.068081614 | -2.762411424 | 0.005737613 | 0.143203716 | | Klhl6 |
| Podxl2 | 5107.318845 | 0.072570579 | 0.06493731 | 1.117548281 | 0.263759968 | 0.491130507 | | | Pctp | 83.15387197 | -0.18807174 | 0.09483414 | -1.983164918 | 0.047349011 | 0.391951501 | | Fbxo17 |
| Mfsd12 | 560.829234 | 0.12022505 | 0.077640879 | 1.548476163 | 0.121507696 | 0.307420027 | | | Azi1 | 1036.396064 | -0.188090803 | 0.0771956 | -2.436548251 | 0.014828191 | 0.232209004 | | Dmkn |
| Stox1 | 154.9708997 | -0.127097962 | 0.113745088 | -1.117392979 | 0.263826336 | 0.491170078 | | | Rufy2 | 2620.854008 | -0.188261586 | 0.059019057 | -3.189844044 | 0.001423496 | 0.061125282 | | Cd52 |
| 2900011O08Rik | 10355.8693 | 0.120134992 | 0.056452802 | 2.128060753 | 0.033332047 | 0.13283095 | | | Egflam | 167.262533 | -0.188388906 | 0.093165193 | -2.022095382 | 0.043166499 | 0.375695964 | | Lag3 |
| Zfp3 | 467.284277 | -0.094603159 | 0.084700484 | -1.116914025 | 0.264031087 | 0.491399119 | | | Alkbh3 | 320.7247417 | -0.188582762 | 0.083464511 | -2.259436494 | 0.023856246 | 0.291530239 | | Sbsn |
| Alpk3 | 20.60317969 | 0.120113303 | 0.166452617 | 0.721606577 | 0.470536394 | 0.685260917 | | | Usp40 | 867.2412304 | -0.188617972 | 0.067230753 | -2.805531166 | 0.005023372 | 0.133393847 | | C4b |
| Lrch1 | 634.4789373 | 0.120086498 | 0.093081725 | 1.290118957 | 0.197009359 | 0.412160816 | | | Caprin2 | 309.0361386 | -0.188744374 | 0.081566107 | -2.314004928 | 0.020667452 | 0.27070821 | | Nudt19 |
| Mfsd7c | 113.0162254 | -0.161083228 | 0.14428716 | -1.116407229 | 0.26424786 | 0.49160707 | | | Zfp36l1 | 889.3321464 | -0.188768985 | 0.084607184 | -2.231122428 | 0.025673021 | 0.300614489 | | Tyrobp |
| Npepps | 6607.557112 | -0.058274172 | 0.052200343 | -1.116356103 | 0.264269736 | 0.49160707 | | | Fhod1 | 186.0566558 | -0.188779025 | 0.085179628 | -2.216246188 | 0.026674646 | 0.305080386 | | |
| Fem1c | 788.8668946 | -0.107202481 | 0.09605959 | -1.11599977 | 0.264422233 | 0.491723661 | | | Neu2 | 249.0108219 | -0.188927268 | 0.085061826 | -2.221058233 | 0.026347017 | 0.303114414 | | |
| Slitrk6 | 56.31074322 | -0.187617554 | 0.168118019 | -1.115987185 | 0.26442762 | 0.491723661 | | | 5430416N02Rik | 122.9885316 | -0.188977874 | 0.09173214 | -2.060105357 | 0.03938847 | 0.359829336 | | |
| Swsap1 | 475.8911343 | -0.09404897 | 0.084268479 | -1.116063465 | 0.264394969 | 0.491723661 | | | Rfx1 | 1109.95299 | -0.188993009 | 0.083613336 | -2.260321348 | 0.023801314 | 0.291270334 | | |
| Arhgef18 | 2285.955562 | 0.067397499 | 0.060414393 | 1.11558679 | 0.264599049 | 0.491924366 | | | Prrt4 | 159.4053116 | -0.189061828 | 0.089346511 | -2.116051609 | 0.034340419 | 0.338648282 | | |
| Podxl | 1895.966885 | -0.135633836 | 0.121577389 | -1.115617279 | 0.264585993 | 0.491924366 | | | 9030624G23Rik | 189.6497607 | -0.18936728 | 0.088192862 | -2.147195084 | 0.031777754 | 0.326301135 | | |
| Lmnb2 | 839.6030421 | 0.120079388 | 0.079842137 | 1.50396009 | 0.132591643 | 0.323014924 | | | Cd37 | 154.8703809 | -0.189476018 | 0.094826197 | -1.998140016 | 0.045701482 | 0.385357198 | | |
| Camkmt | 261.4541303 | -0.109159626 | 0.097887642 | -1.115152274 | 0.264785174 | 0.492152289 | | | C030023E24Rik | 106.5632592 | -0.189511746 | 0.094410795 | -2.00731014 | 0.04471665 | 0.380855808 | | |
| Map7d2 | 6884.898809 | 0.119973454 | 0.097047 | 1.236240734 | 0.216369094 | 0.436443452 | | | Gm11545 | 6.98125267 | -0.189605479 | 0.055632132 | -3.408200849 | 0.000653927 | NA | | |
| Tmem229a | 2038.721082 | -0.113266667 | 0.101608526 | -1.114735852 | 0.264963633 | 0.492392514 | | | Psmb8 | 153.3258806 | -0.189638234 | 0.076515766 | -2.478420386 | 0.013196556 | 0.219077909 | | |
| Gm16897 | 14.19940257 | -0.1867849 | 0.167619212 | -1.114340636 | 0.26513308 | 0.49262165 | | | Atg4a | 247.3572108 | -0.189698188 | 0.084349097 | -2.248965256 | 0.024514707 | 0.293328698 | | |
| Amacr | 326.3761898 | -0.096915872 | 0.086989553 | -1.114109326 | 0.265232288 | 0.492706257 | | | Dnahc7a | 42.24349653 | -0.189703865 | 0.09250732 | -2.05069032 | 0.040297114 | 0.363536189 | | |
| Sytl5 | 163.2693188 | -0.16108249 | 0.1445871 | -1.114086183 | 0.265242216 | 0.492706257 | | | Tnfrsf1b | 251.3004539 | -0.189802528 | 0.081090012 | -2.340640028 | 0.019250718 | 0.263564471 | | |
| Wnt8b | 11.34611189 | 0.119943681 | 0.154192859 | 0.77788091 | 0.436639223 | 0.657079267 | | | Ctnnal1 | 770.8452213 | -0.189853606 | 0.080680131 | -2.353164328 | 0.018614406 | 0.260494084 | | |
| Cnrip1 | 2428.538817 | -0.075154155 | 0.067508455 | -1.113255445 | 0.265598736 | 0.493225838 | | | Atf7 | 345.4430936 | -0.190178972 | 0.080492437 | -2.362693689 | 0.018142658 | 0.256569934 | | |
| Gars | 4880.853975 | 0.119938573 | 0.062305067 | 1.925021166 | 0.05422671 | 0.183107568 | | | 2810049E08Rik | 143.0393204 | -0.190236205 | 0.093555734 | -2.033399728 | 0.042012155 | 0.370320688 | | |
| Wdr41 | 1325.15404 | -0.078507369 | 0.070534112 | -1.113041143 | 0.26569076 | 0.493302892 | | | 8430429K09Rik | 197.3866758 | -0.190307523 | 0.08741109 | -2.17715537 | 0.029468977 | 0.317771905 | | |
| S100a1 | 1377.735024 | -0.073964897 | 0.066461404 | -1.112900005 | 0.265751378 | 0.49335632 | | | Tgfbr1 | 1246.408669 | -0.190365136 | 0.071422819 | -2.665326559 | 0.007691359 | 0.167597786 | | |
| Supt5 | 6515.283452 | 0.119931084 | 0.061800536 | 1.940615608 | 0.052304919 | 0.179136256 | | | Zfp57 | 385.51964 | -0.19055593 | 0.089184008 | -2.136660306 | 0.032625631 | 0.329798129 | | |
| Zcchc18 | 11679.72647 | -0.048454404 | 0.043556842 | -1.112440717 | 0.265948706 | 0.493604367 | | | Trip6 | 231.3317201 | -0.190577776 | 0.092433944 | -2.061772635 | 0.039229388 | 0.359524343 | | |
| Ankmy2 | 1846.131722 | -0.080118702 | 0.072039054 | -1.112156491 | 0.266070871 | 0.493660492 | | | Rad9b | 105.3313116 | -0.190773212 | 0.093049988 | -2.050222852 | 0.040342689 | 0.363536189 | | |
| Ap2a1 | 11902.26648 | 0.072868458 | 0.065520469 | 1.11214798 | 0.26607453 | 0.493660492 | | | Ppp1r14a | 248.8467534 | -0.191184615 | 0.094834759 | -2.01597617 | 0.043802468 | 0.378860465 | | |
| Samd1 | 829.0063508 | -0.085603399 | 0.076963339 | -1.112262024 | 0.266025507 | 0.493660492 | | | Chi3l1 | 386.9040664 | -0.191388899 | 0.084555102 | -2.263481375 | 0.023606031 | 0.289811141 | | |
| Sox3 | 28.574023 | -0.185351813 | 0.166709713 | -1.111823719 | 0.266213951 | 0.493860029 | | | Gbp2 | 238.2975561 | -0.191456037 | 0.068225498 | -2.806242116 | 0.005012582 | 0.133393847 | | |
| 1110046J04Rik | 90.59358953 | -0.165135582 | 0.148623257 | -1.111101895 | 0.266524492 | 0.494085203 | | | Dnase2a | 169.9712535 | -0.191508411 | 0.09369194 | -2.044022256 | 0.040951349 | 0.365617264 | | |
| Adamts8 | 135.329917 | -0.164131523 | 0.14771232 | -1.111156623 | 0.266500939 | 0.494085203 | | | Mical1 | 506.4218499 | -0.191568802 | 0.082393572 | -2.325045495 | 0.020069528 | 0.267704278 | | |
| Katnb1 | 1820.322883 | 0.066123759 | 0.059507466 | 1.111184257 | 0.266489046 | 0.494085203 | | | Zc3hav1 | 203.5515442 | -0.191661044 | 0.090301182 | -2.122464411 | 0.033798758 | 0.336326295 | | |
| Noc4l | 791.8126584 | 0.119901501 | 0.087092578 | 1.376713203 | 0.168600935 | 0.375546706 | | | Chkb | 22.29064703 | -0.191897996 | 0.085437011 | -2.246075722 | 0.024699158 | NA | | |
| Med20 | 737.7315826 | 0.119846274 | 0.061882708 | 1.93666823 | 0.052785912 | 0.180028734 | | | Edem1 | 1114.190989 | -0.191954292 | 0.053558505 | -3.584011423 | 0.000338357 | 0.023743854 | | |
| Mapk11 | 1900.398859 | 0.119783395 | 0.125934426 | 0.951156874 | 0.341524747 | 0.575028625 | | | Eif4ebp3 | 202.8462281 | -0.191982509 | 0.088248612 | -2.175473408 | 0.029594659 | 0.317771905 | | |
| Wnt11 | 10.45994348 | 0.119767023 | 0.160798235 | 0.74482797 | 0.456375724 | 0.674528805 | | | Kdelc1 | 386.811152 | -0.191987741 | 0.078194877 | -2.455247069 | 0.014078779 | 0.227385586 | | |
| Pcbd2 | 157.8378927 | -0.123167001 | 0.11094656 | -1.110147088 | 0.266935649 | 0.494724993 | | | Fam228b | 97.43256362 | -0.192029574 | 0.094197736 | -2.038579513 | 0.041492012 | 0.368048911 | | |
| Acsl1 | 2334.86195 | 0.068754708 | 0.061946597 | 1.109902909 | 0.267040867 | 0.494860805 | | | Mccc1 | 968.5752052 | -0.19205416 | 0.071556882 | -2.683936947 | 0.007276083 | 0.162353883 | | |
| Rab34 | 525.7814607 | 0.119759827 | 0.128377575 | 0.932871854 | 0.350886136 | 0.581831099 | | | Dnahc7b | 189.5309022 | -0.192082839 | 0.089322752 | -2.150435743 | 0.031520764 | 0.32443523 | | |
| Dusp16 | 655.4525216 | 0.092147429 | 0.083032394 | 1.109776842 | 0.267095201 | 0.494879596 | | | E130102H24Rik | 161.193477 | -0.192134138 | 0.09162258 | -2.097017319 | 0.035992042 | 0.344080613 | | |
| Avpi1 | 647.841116 | -0.092833328 | 0.083676194 | -1.109435354 | 0.267242418 | 0.495056674 | | | H2-T23 | 868.6070342 | -0.192146795 | 0.093267873 | -2.060160579 | 0.039383192 | 0.359829336 | | |
| 1110035M17Rik | 76.58679424 | -0.155990279 | 0.140623842 | -1.109273339 | 0.267312283 | 0.495119458 | | | Sash3 | 97.90768692 | -0.192199689 | 0.093527037 | -2.055017409 | 0.039877323 | 0.361071695 | | |
| Ap1g1 | 4965.994085 | -0.065520009 | 0.059069152 | -1.109208559 | 0.267340221 | 0.495119458 | | | Nphp3 | 455.3317195 | -0.192237062 | 0.079560222 | -2.416245933 | 0.015681466 | 0.237621413 | | |
| Suds3 | 3546.117545 | -0.07562864 | 0.068232454 | -1.108396886 | 0.267690449 | 0.495708835 | | | Slc25a45 | 158.554909 | -0.192352994 | 0.087299987 | -2.203356507 | 0.027569631 | 0.308295968 | | |
| Zranb3 | 347.7690402 | 0.119756723 | 0.087452356 | 1.369393905 | 0.170876178 | 0.37812819 | | | B3galnt2 | 820.925279 | -0.192447515 | 0.067941975 | -2.832527548 | 0.004618159 | 0.126226882 | | |
| Ndnl2 | 586.6742595 | 0.086054037 | 0.077657709 | 1.108119685 | 0.267810131 | 0.495811945 | | | Cd46 | 49.92900458 | -0.192763363 | 0.092505626 | -2.083801503 | 0.037178227 | 0.348662607 | | |
| Grem2 | 1303.29523 | -0.132216185 | 0.119329631 | -1.107991235 | 0.267865601 | 0.495855392 | | | Lrrk1 | 249.1810065 | -0.192846504 | 0.084818705 | -2.273631779 | 0.022988135 | 0.28516281 | | |
| Tti2 | 833.010732 | 0.11973991 | 0.064070076 | 1.868889776 | 0.061638151 | 0.197824454 | | | Naprt1 | 190.6936525 | -0.193114533 | 0.092588392 | -2.085731568 | 0.037002945 | 0.348173348 | | |
| Kansl1 | 4233.579039 | 0.119722708 | 0.062697792 | 1.90952031 | 0.056195004 | 0.187088776 | | | Fgd3 | 300.4001706 | -0.193121483 | 0.08099075 | -2.38448814 | 0.017102899 | 0.249184268 | | |
| Syk | 185.1038212 | -0.142219626 | 0.128399755 | -1.107631599 | 0.268020951 | 0.495905972 | | | Mfsd2b | 37.32123703 | -0.193300458 | 0.089364665 | -2.163052454 | 0.030537145 | 0.320602541 | | |
| Nid2 | 161.3037206 | 0.119703353 | 0.133666224 | 0.895539271 | 0.370498882 | 0.60040892 | | | Sh3bp1 | 682.4073089 | -0.193312919 | 0.08352818 | -2.314343706 | 0.020648876 | 0.27070821 | | |
| Fxr2 | 5047.86653 | 0.091854274 | 0.082940933 | 1.107466131 | 0.268092448 | 0.495979031 | | | Rgl3 | 159.4592501 | -0.193335533 | 0.090940475 | -2.125956937 | 0.033506845 | 0.335456507 | | |
| 1110059G10Rik | 375.1338284 | -0.0971328 | 0.087716709 | -1.107346601 | 0.268144103 | 0.49601537 | | | 2700038G22Rik | 28.72045214 | -0.193733145 | 0.086945511 | -2.228213315 | 0.025866294 | 0.301310169 | | |
| Aldoa | 59874.47854 | -0.079585955 | 0.07187845 | -1.107229706 | 0.268194627 | 0.496049606 | | | C920006O11Rik | 113.6099251 | -0.19374627 | 0.095045886 | -2.038449821 | 0.041504968 | 0.368048911 | | |
| Rrn3 | 2580.314146 | 0.068634152 | 0.062002175 | 1.106963606 | 0.268309663 | 0.496203142 | | | Tln1 | 2302.005325 | -0.193798896 | 0.063672066 | -3.043703586 | 0.002336853 | 0.081484567 | | |
| Tmco4 | 151.9737644 | 0.119699121 | 0.132993857 | 0.900034961 | 0.368101646 | 0.597837201 | | | Man2c1 | 2667.685955 | -0.193865243 | 0.079637801 | -2.434336986 | 0.014919098 | 0.232332035 | | |
| Adamtsl2 | 238.6595451 | -0.160683087 | 0.145342161 | -1.105550417 | 0.268921161 | 0.497156003 | | | Trim72 | 37.78313842 | -0.193915917 | 0.090730297 | -2.137278535 | 0.032575344 | 0.329798129 | | |
| Pin4 | 507.7052617 | 0.098275655 | 0.088891388 | 1.105570034 | 0.268912666 | 0.497156003 | | | Itga7 | 621.2869759 | -0.193979389 | 0.088073775 | -2.202464797 | 0.027632493 | 0.308761227 | | |
| Eif3m | 2033.687958 | -0.065618295 | 0.059363232 | -1.105369317 | 0.268999593 | 0.497241672 | | | Tmc6 | 243.9814104 | -0.194317328 | 0.088922461 | -2.185244599 | 0.028870917 | 0.316746301 | | |
| Pard6a | 707.8057943 | -0.104503968 | 0.094582962 | -1.104892113 | 0.26920634 | 0.49756448 | | | Ctsz | 1774.930645 | -0.194652183 | 0.088829286 | -2.191306409 | 0.028429627 | 0.313543192 | | |
| Aqp11 | 405.1963468 | 0.09917172 | 0.089777223 | 1.104642328 | 0.269314602 | 0.497705206 | | | Gsdmd | 118.1768303 | -0.194665834 | 0.094129471 | -2.068064677 | 0.038633942 | 0.356089237 | | |
| Zfp191 | 1291.402695 | 0.119689087 | 0.091526526 | 1.30769835 | 0.190975643 | 0.404854183 | | | H2-Oa | 25.35050634 | -0.194900368 | 0.072875339 | -2.674435133 | 0.007485526 | 0.165250372 | | |
| Eif3c | 7152.185581 | 0.068612571 | 0.062140883 | 1.104145419 | 0.269530062 | 0.497865854 | | | Fam46c | 64.67680988 | -0.195274431 | 0.092869658 | -2.102672019 | 0.03549445 | 0.342120019 | | |
| Twf1 | 1965.123656 | -0.064450038 | 0.058368987 | -1.104182914 | 0.2695138 | 0.497865854 | | | Hgf | 279.7099871 | -0.195464788 | 0.084928177 | -2.301530463 | 0.021361666 | 0.275088257 | | |
| Zfp300 | 105.8851236 | -0.152714818 | 0.138300398 | -1.104225445 | 0.269495354 | 0.497865854 | | | Oasl2 | 282.4421304 | -0.195525909 | 0.06753608 | -2.895132626 | 0.003789984 | 0.11032726 | | |
| Npdc1 | 5687.219208 | 0.119685509 | 0.071100989 | 1.683317083 | 0.092313725 | 0.256340698 | | | Rnpc3 | 999.3496069 | -0.195531265 | 0.066295649 | -2.949383059 | 0.00318409 | 0.098702162 | | |
| Tbc1d23 | 1364.600189 | 0.085488382 | 0.077436823 | 1.10397584 | 0.269603618 | 0.497867046 | | | Dhx58 | 69.97524385 | -0.195533688 | 0.094259027 | -2.074429303 | 0.038039458 | 0.352889542 | | |
| Pip5k1c | 20366.55346 | 0.119673638 | 0.089387303 | 1.338821437 | 0.18062881 | 0.391038534 | | | Mir682 | 56.29074 | -0.195577067 | 0.087162128 | -2.243830802 | 0.02484329 | 0.294831395 | | |
| Anxa1 | 77.00415017 | -0.18361089 | 0.16639775 | -1.103445746 | 0.26983364 | 0.498070338 | | | Crebzf | 1853.913958 | -0.195696465 | 0.062465796 | -3.132857958 | 0.001731132 | 0.068332671 | | |
| Gm20187 | 10.5949802 | -0.180475033 | 0.163547985 | -1.10349897 | 0.269810539 | 0.498070338 | | | Trub1 | 1163.165969 | -0.195856 | 0.057861913 | -3.384886345 | 0.000712077 | 0.039479523 | | |
| Smurf1 | 2478.401444 | -0.065454268 | 0.059310108 | -1.103593802 | 0.269769381 | 0.498070338 | | | Ddx17 | 18592.86926 | -0.195972766 | 0.049620264 | -3.949450264 | 7.83E-05 | 0.008820422 | | |
| BC025920 | 150.3095689 | -0.14055945 | 0.127423645 | -1.103087653 | 0.269989102 | 0.498297933 | | | Hook1 | 2559.032066 | -0.196058391 | 0.062298437 | -3.147083625 | 0.001649078 | 0.066172668 | | |
| Stx18 | 831.1552124 | 0.096305984 | 0.087319215 | 1.102918581 | 0.270062524 | 0.498374077 | | | Ccnt2 | 2045.763996 | -0.196134327 | 0.066325712 | -2.957138658 | 0.003105085 | 0.097207883 | | |
| Ddx18 | 805.7829513 | 0.075858226 | 0.068789473 | 1.102759227 | 0.270131739 | 0.498442439 | | | Cyba | 182.3276758 | -0.196192517 | 0.094565827 | -2.07466611 | 0.03801749 | 0.352889542 | | |
| Pcgf5 | 318.7060356 | -0.098864155 | 0.089659499 | -1.102662355 | 0.27017382 | 0.498460726 | | | Iqgap1 | 1261.324977 | -0.196294171 | 0.054644859 | -3.592180014 | 0.000327923 | 0.023585654 | | |
| 5730408K05Rik | 12.83389096 | -0.182556238 | 0.165573449 | -1.102569524 | 0.270214151 | 0.498475778 | | | Atr | 697.4807259 | -0.196796662 | 0.062287866 | -3.159470278 | 0.001580562 | 0.06480716 | | |
| Cerk | 3508.060188 | 0.095362659 | 0.086532261 | 1.102047468 | 0.270441036 | 0.498730506 | | | Rbm5 | 5107.812644 | -0.196961597 | 0.047636466 | -4.13468114 | 3.55E-05 | 0.00537837 | | |
| Ero1lb | 839.4966842 | 0.119662704 | 0.06258112 | 1.912121494 | 0.055860612 | 0.186571315 | | | Gfpt2 | 339.3242941 | -0.19697219 | 0.092667403 | -2.125582299 | 0.033538055 | 0.335456507 | | |
| Fgd1 | 632.3751565 | -0.070030327 | 0.063544369 | -1.102069754 | 0.270431347 | 0.498730506 | | | 6820431F20Rik | 2800.15088 | -0.196984419 | 0.047896515 | -4.112708809 | 3.91E-05 | 0.005796222 | | |
| Fmo1 | 297.3660851 | -0.13978359 | 0.126872158 | -1.101767263 | 0.270562866 | 0.498730506 | | | AU022754 | 70.13948773 | -0.196986281 | 0.094656093 | -2.081073428 | 0.037427186 | 0.350065015 | | |
| Klhl22 | 3293.423932 | 0.063908167 | 0.058005917 | 1.101752553 | 0.270569263 | 0.498730506 | | | Oscar | 82.91914014 | -0.197397415 | 0.094225738 | -2.094941571 | 0.036176187 | 0.344587075 | | |
| Mrpl30 | 1306.778361 | -0.069730832 | 0.063299493 | -1.101601746 | 0.270634849 | 0.498730506 | | | 5830418K08Rik | 1149.619542 | -0.197426924 | 0.062369217 | -3.165454598 | 0.001548408 | 0.064263366 | | |
| Ogfr | 1346.09774 | -0.068912409 | 0.062557479 | -1.101585461 | 0.270641932 | 0.498730506 | | | 1700020D05Rik | 41.95066779 | -0.197997201 | 0.090268262 | -2.193430965 | 0.028276343 | 0.312827238 | | |
| Srsf10 | 1858.310009 | 0.119610515 | 0.081087826 | 1.475073631 | 0.140192749 | 0.334666626 | | | Ercc2 | 1409.594554 | -0.198021355 | 0.067289434 | -2.942829853 | 0.003252271 | 0.099878415 | | |
| Wdr11 | 2164.423184 | 0.065952152 | 0.059863866 | 1.101702191 | 0.270591164 | 0.498730506 | | | Hpn | 51.73890666 | -0.198226083 | 0.088320964 | -2.244383149 | 0.02480776 | 0.294650465 | | |
| Wdr48 | 3446.581606 | 0.053418248 | 0.048502999 | 1.101339078 | 0.270749109 | 0.498868677 | | | Runx1 | 57.97645615 | -0.19856418 | 0.093441092 | -2.12501991 | 0.033584952 | 0.335456507 | | |
| Prx | 24.93524271 | -0.182984224 | 0.166169565 | -1.101189764 | 0.270814076 | 0.498929048 | | | Nbeal2 | 664.2823081 | -0.198671111 | 0.085752902 | -2.316785875 | 0.020515399 | 0.269900834 | | |
| Cttnbp2 | 4671.253769 | 0.069106949 | 0.062764996 | 1.101042846 | 0.27087801 | 0.498987503 | | | 4933406I18Rik | 15.92333845 | -0.198767054 | 0.073029783 | -2.721725934 | 0.006494198 | NA | | |
| Rab11fip2 | 2653.533019 | -0.073010552 | 0.066319134 | -1.10089724 | 0.270941384 | 0.499044912 | | | Nlrp1a | 106.0997897 | -0.198832715 | 0.094164918 | -2.111537056 | 0.034726176 | 0.33922827 | | |
| Ssrp1 | 3885.961094 | 0.119589714 | 0.063996387 | 1.868694764 | 0.061665293 | 0.197855053 | | | Nxf7 | 108.8456193 | -0.19905512 | 0.095045865 | -2.094305936 | 0.036232736 | 0.344899555 | | |
| Dnajc28 | 502.7541736 | 0.086385741 | 0.078502738 | 1.10041692 | 0.27115051 | 0.499311386 | | | Glipr1 | 42.65719365 | -0.199103503 | 0.089294779 | -2.229732869 | 0.025765182 | 0.300614489 | | |
| Sgcz | 85.58715352 | -0.149697745 | 0.136088897 | -1.0999997 | 0.271332252 | 0.499586681 | | | 0610030E20Rik | 875.410351 | -0.199238924 | 0.064257057 | -3.100654375 | 0.001930935 | 0.072477432 | | |
| Aktip | 4162.402379 | -0.060104083 | 0.054669387 | -1.09941024 | 0.271589166 | 0.499940898 | | | Cep164 | 730.9754028 | -0.199489357 | 0.076619562 | -2.60363479 | 0.009224099 | 0.183045438 | | |
| Polr2g | 1924.032436 | -0.070315137 | 0.063955187 | -1.099443844 | 0.271574515 | 0.499940898 | | | Adam8 | 147.43239 | -0.199509586 | 0.094973774 | -2.100680827 | 0.035668993 | 0.342903904 | | |
| Galnt1 | 2247.339256 | 0.078670354 | 0.071593412 | 1.098849066 | 0.271833906 | 0.500331972 | | | Vill | 370.5896287 | -0.199560017 | 0.085311435 | -2.339194242 | 0.019325383 | 0.263564471 | | |
| Sgsm1 | 5754.829528 | -0.090003293 | 0.081913342 | -1.09876231 | 0.271871756 | 0.500342201 | | | Cwf19l2 | 532.4078865 | -0.199737246 | 0.067727718 | -2.949121156 | 0.00318679 | 0.098702162 | | |
| 2900092D14Rik | 588.4398588 | -0.106264159 | 0.096749808 | -1.098339737 | 0.272056166 | 0.500535371 | | | Zmym6 | 1757.103646 | -0.199791303 | 0.0687376 | -2.906579541 | 0.00365404 | 0.107446524 | | |
| Mpi | 1944.212642 | 0.096604429 | 0.087957613 | 1.098306623 | 0.272070621 | 0.500535371 | | | Blnk | 366.0139861 | -0.199827127 | 0.080356628 | -2.486753502 | 0.012891469 | 0.216344675 | | |
| Smyd5 | 2002.07832 | 0.085350483 | 0.077711468 | 1.098299719 | 0.272073634 | 0.500535371 | | | C1qb | 2328.58751 | -0.200067074 | 0.095027151 | -2.105367476 | 0.035259333 | 0.341907254 | | |
| Htr4 | 145.1613738 | -0.183959957 | 0.167580892 | -1.097738258 | 0.272318799 | 0.500748572 | | | Apbb1ip | 271.3074809 | -0.200174686 | 0.091097069 | -2.197377906 | 0.027993468 | 0.311357668 | | |
| Syt13 | 10224.19455 | 0.089716882 | 0.08172365 | 1.097808058 | 0.272288312 | 0.500748572 | | | Mir186 | 41.80427735 | -0.200285779 | 0.090854852 | -2.204458808 | 0.027492094 | 0.308173879 | | |
| Wdr5b | 164.8584227 | -0.136647072 | 0.12447684 | -1.097771056 | 0.272304473 | 0.500748572 | | | Lepre1 | 327.4922715 | -0.200291233 | 0.088226044 | -2.270205308 | 0.023195129 | 0.286020752 | | |
| Xrcc4 | 270.6936457 | -0.128763548 | 0.117276423 | -1.097949146 | 0.272226696 | 0.500748572 | | | Aqp9 | 61.27155413 | -0.200352556 | 0.093903168 | -2.133608057 | 0.032874878 | 0.331395195 | | |
| Sardh | 530.9826734 | -0.116092465 | 0.105767201 | -1.09762255 | 0.272369342 | 0.500782079 | | | Rad51ap1 | 61.3464051 | -0.200676562 | 0.095100648 | -2.110149265 | 0.034845501 | 0.339936701 | | |
| Sec16b | 61.88444107 | 0.119578043 | 0.155389342 | 0.769538253 | 0.441573845 | 0.661485561 | | | Arnt | 1453.547339 | -0.200785946 | 0.069281152 | -2.898132345 | 0.003753921 | 0.10971723 | | |
| Heyl | 281.9341015 | -0.150458294 | 0.137106475 | -1.097382849 | 0.272474068 | 0.50085576 | | | Gcnt1 | 145.7735044 | -0.201100668 | 0.092210796 | -2.180879865 | 0.029192303 | 0.317415456 | | |
| Fam3c | 2355.629652 | -0.061344814 | 0.055907156 | -1.097262275 | 0.272526757 | 0.500893187 | | | Ssh3 | 1986.830878 | -0.201229729 | 0.086114174 | -2.336778259 | 0.019450717 | 0.263564471 | | |
| Zfp46 | 1822.538192 | 0.119573355 | 0.061850261 | 1.933271639 | 0.053202744 | 0.180886756 | | | Ints2 | 908.0333464 | -0.201256431 | 0.062540961 | -3.217993904 | 0.001290906 | 0.057875602 | | |
| Asb1 | 2493.072167 | -0.07374827 | 0.067250011 | -1.096628373 | 0.272803879 | 0.501283599 | | | Pycard | 88.71190982 | -0.201401005 | 0.09504503 | -2.119006164 | 0.034089946 | 0.336970372 | | |
| Nol10 | 875.3819833 | -0.075808387 | 0.069138312 | -1.096474358 | 0.272871239 | 0.501347916 | | | Ccdc3 | 664.3007976 | -0.20146998 | 0.087701873 | -2.297214097 | 0.021606559 | 0.276282466 | | |
| Rhoc | 430.3330208 | 0.119532542 | 0.095936035 | 1.245960828 | 0.212778777 | 0.432309778 | | | Pald1 | 564.9280957 | -0.201630709 | 0.07065961 | -2.853549702 | 0.004323376 | 0.120309131 | | |
| Mtap | 392.9806395 | -0.09166859 | 0.083627837 | -1.096149249 | 0.273013465 | 0.501430848 | | | Nbeal1 | 611.8424663 | -0.201808357 | 0.08657455 | -2.331035584 | 0.019751483 | 0.2654117 | | |
| Ncf4 | 33.15740409 | 0.119514289 | 0.167644714 | 0.712902222 | 0.475906264 | 0.689955989 | | | A630001G21Rik | 21.48783318 | -0.201825658 | 0.082844528 | -2.436197825 | 0.014842565 | NA | | |
| Ppp1r18 | 346.2834634 | 0.119506387 | 0.084533079 | 1.413723336 | 0.157443151 | 0.360275318 | | | E230016M11Rik | 52.22805279 | -0.202077046 | 0.094816156 | -2.131251191 | 0.033068454 | 0.332884518 | | |
| Nphs2 | 17.67623178 | -0.181882943 | 0.165958746 | -1.095952747 | 0.273099454 | 0.501471293 | | | Hfm1 | 74.02917977 | -0.202136161 | 0.094799093 | -2.132258378 | 0.032985612 | 0.332280856 | | |
| Cyp2u1 | 218.4817147 | -0.099519639 | 0.090841596 | -1.095529399 | 0.273284773 | 0.501649393 | | | Arhgap11a | 147.2337024 | -0.202581169 | 0.091662783 | -2.210070032 | 0.027100302 | 0.306826961 | | |
| Klhdc5 | 1314.3435 | 0.07608696 | 0.069447037 | 1.095611328 | 0.273248902 | 0.501649393 | | | Gbp3 | 254.1353809 | -0.202593069 | 0.082219058 | -2.46406457 | 0.013737136 | 0.224208584 | | |
| Tmem101 | 625.9193563 | -0.098090398 | 0.08953877 | -1.095507546 | 0.273294341 | 0.501649393 | | | Rad52 | 521.4349738 | -0.202614151 | 0.065514993 | -3.092637899 | 0.00198386 | 0.073891169 | | |
| Dusp2 | 36.02515702 | 0.11949356 | 0.162428413 | 0.735669074 | 0.461932115 | 0.678212469 | | | Sbno2 | 816.18807 | -0.202640475 | 0.083161253 | -2.436717445 | 0.014821255 | 0.232209004 | | |
| Mir568 | 19.02698099 | -0.182741848 | 0.166879308 | -1.095053963 | 0.273492996 | 0.501895118 | | | Kcnt1 | 8027.833192 | -0.202669937 | 0.063443511 | -3.194494342 | 0.00140076 | 0.06073864 | | |
| Zfp2 | 844.7152888 | 0.090957446 | 0.083072166 | 1.094920842 | 0.273551317 | 0.501942694 | | | Znf512b | 3709.018787 | -0.202743279 | 0.079674016 | -2.544659974 | 0.010938419 | 0.19886293 | | |
| Nono | 4933.752965 | -0.056561904 | 0.051665443 | -1.09477247 | 0.27361633 | 0.502002537 | | | Aasdh | 595.634955 | -0.202825386 | 0.077854368 | -2.605189552 | 0.009182344 | 0.182762906 | | |
| Cenpc1 | 849.4423517 | 0.087356127 | 0.079811698 | 1.094527854 | 0.273723538 | 0.50213977 | | | Abi3 | 222.9738028 | -0.202860336 | 0.092318581 | -2.197394438 | 0.027992288 | 0.311357668 | | |
| Galk1 | 308.8816817 | 0.11948931 | 0.090457523 | 1.320943871 | 0.18652008 | 0.398530566 | | | Tnfaip2 | 39.63919234 | -0.202898185 | 0.090667822 | -2.237819115 | 0.025232853 | 0.298479176 | | |
| Lig3 | 1254.993708 | 0.096351953 | 0.088047335 | 1.094319923 | 0.273814691 | 0.502142128 | | | Igsf9 | 254.2666241 | -0.20303859 | 0.091636075 | -2.21570588 | 0.026711652 | 0.305080386 | | |
| Acadvl | 1676.708446 | 0.119474593 | 0.063242951 | 1.889136923 | 0.058873484 | 0.192728874 | | | Gm6297 | 74.88183417 | -0.203119147 | 0.093374978 | -2.175305966 | 0.029607196 | 0.317771905 | | |
| Sorbs2 | 6514.191753 | 0.078590386 | 0.071814735 | 1.094349042 | 0.273801924 | 0.502142128 | | | Trim34a | 68.54565458 | -0.203427675 | 0.094185312 | -2.159866232 | 0.030783027 | 0.321052104 | | |
| Aes | 14483.80129 | -0.063478937 | 0.058017151 | -1.094140884 | 0.273893194 | 0.502153727 | | | Trim12a | 95.65072675 | -0.203610996 | 0.09420459 | -2.161370238 | 0.030666751 | 0.320940366 | | |
| Adra2a | 656.2493234 | 0.119401664 | 0.135703407 | 0.879872261 | 0.378928513 | 0.607752012 | | | Ajuba | 82.2640887 | -0.203623133 | 0.095089928 | -2.141374357 | 0.032243861 | 0.327711599 | | |
| Spc25 | 12.98815244 | -0.18065646 | 0.165142775 | -1.093941046 | 0.273980835 | 0.502195559 | | | Cecr2 | 180.2323834 | -0.203792905 | 0.093951898 | -2.169119624 | 0.030073603 | 0.319144111 | | |
| Chchd10 | 3756.904024 | -0.099135315 | 0.090639121 | -1.0937365 | 0.274070561 | 0.5023006 | | | Pfkfb2 | 2328.368369 | -0.203812067 | 0.052785865 | -3.861110703 | 0.000112873 | 0.011329455 | | |
| Nxph2 | 68.82851891 | -0.180334122 | 0.165004149 | -1.092906593 | 0.274434813 | 0.502902752 | | | Trim25 | 370.2814394 | -0.203871851 | 0.08365804 | -2.436966614 | 0.014811047 | 0.232209004 | | |
| Trmt2b | 605.6187536 | -0.110109235 | 0.100755124 | -1.09284006 | 0.274464029 | 0.502902752 | | | Naa40 | 1537.459359 | -0.203925423 | 0.068020934 | -2.997980325 | 0.002717752 | 0.087924428 | | |

| | | | | | | |
|---|---|---|---|---|---|---|
| 3632454L22Rik | 35.34004446 | 0.119364743 | 0.164287142 | 0.726561688 | 0.467494505 | 0.682177421 |
| Dnajc19 | 484.5306663 | -0.083942636 | 0.076848557 | -1.09231245 | 0.27469579 | 0.503089429 |
| Ttll13 | 40.55526167 | 0.11930872 | 0.168052823 | 0.709947727 | 0.477736552 | 0.691328116 |
| Ung | 76.78285397 | -0.158001594 | 0.144646958 | -1.09232573 | 0.274689955 | 0.503089429 |
| Tbc1d1 | 971.3181185 | 0.119246107 | 0.075308443 | 1.583436106 | 0.113322099 | 0.294303151 |
| Tmem175 | 1279.101553 | 0.088743909 | 0.081268697 | 1.091981444 | 0.274841257 | 0.503236876 |
| Gm13476 | 145.465822 | -0.16251424 | 0.148875361 | -1.091612736 | 0.275003356 | 0.503474181 |
| Utp18 | 648.7310528 | -0.087174542 | 0.079899517 | -1.091052176 | 0.275249925 | 0.50386606 |
| Man1c1 | 1950.140761 | 0.119225183 | 0.071411095 | 1.669561049 | 0.095006242 | 0.260797001 |
| 2410004I01Rik | 13.18370108 | -0.180510185 | 0.165568702 | -1.090243401 | 0.27560594 | 0.504398586 |
| Dram2 | 569.6489118 | -0.086758711 | 0.07962072 | -1.089649923 | 0.275867383 | 0.504817436 |
| Acad8 | 858.0057154 | 0.069773288 | 0.064060046 | 1.089185722 | 0.276071994 | 0.504893705 |
| Sepsecs | 234.3843779 | 0.11914594 | 0.103718805 | 1.148739997 | 0.250663208 | 0.476361553 |
| Lman1 | 1450.340784 | 0.060465312 | 0.055507316 | 1.089321498 | 0.276012136 | 0.504893705 |
| Mnt | 1910.083417 | 0.092700821 | 0.085105458 | 1.089246485 | 0.276045205 | 0.504893705 |
| Ppp6r3 | 3447.019568 | 0.068649341 | 0.063020237 | 1.089322173 | 0.276011838 | 0.504893705 |
| Snph | 15311.15964 | 0.118994422 | 0.10103474 | 1.17775749 | 0.2388933 | 0.463139739 |
| Atg14 | 932.5302857 | 0.064215073 | 0.058982259 | 1.088718426 | 0.276278074 | 0.505132102 |
| Erap1 | 437.660611 | 0.097180674 | 0.089275927 | 1.088542881 | 0.276355517 | 0.505132102 |
| Gtf2e2 | 518.3262086 | -0.073648188 | 0.067658808 | -1.088523289 | 0.276364161 | 0.505132102 |
| Wdr59 | 1439.094584 | -0.063329483 | 0.058179408 | -1.08852058 | 0.276365356 | 0.505132102 |
| Brf2 | 490.5940917 | -0.083204466 | 0.076492368 | -1.087748595 | 0.276706109 | 0.505605783 |
| Mrps27 | 1001.614593 | 0.118987386 | 0.060416389 | 1.969455432 | 0.048900816 | 0.171440268 |
| E2f4 | 1058.617321 | 0.118910218 | 0.087880022 | 1.353097265 | 0.176024559 | 0.385096926 |
| Lamp5 | 7298.585867 | 0.118890329 | 0.109316653 | 1.087577466 | 0.276781683 | 0.505605783 |
| Pnmal2 | 15463.46883 | -0.091234812 | 0.083900024 | -1.087422963 | 0.276849927 | 0.505605783 |
| Ptrh1 | 88.98114447 | -0.153960877 | 0.141571317 | -1.08751462 | 0.276809441 | 0.505605783 |
| Tpp2 | 2240.997529 | 0.078796419 | 0.072462065 | 1.087416142 | 0.276852941 | 0.505605783 |
| Snap91 | 12515.20191 | -0.065517121 | 0.060262355 | -1.087198141 | 0.276949252 | 0.505722064 |
| Scyl3 | 1096.174775 | 0.076139637 | 0.070044793 | 1.087013526 | 0.277030832 | 0.50581142 |
| Atp10b | 92.45460193 | -0.182558434 | 0.167984281 | -1.086759031 | 0.277143318 | 0.505957178 |
| Eltd1 | 898.5104858 | -0.112099706 | 0.103184864 | -1.086396794 | 0.277303479 | 0.506189928 |
| Pigo | 1041.196464 | 0.118884433 | 0.068543919 | 1.734427151 | 0.082842321 | 0.239783098 |
| BC017643 | 741.5838149 | 0.118854939 | 0.077346125 | 1.536663142 | 0.124375821 | 0.311793329 |
| Pigu | 816.8536786 | 0.063377251 | 0.058362133 | 1.08593102 | 0.277509512 | 0.50641482 |
| 2610018G03Rik | 206.9364362 | -0.114659825 | 0.105636213 | -1.085421572 | 0.277734982 | 0.506699118 |
| Agap3 | 9651.161794 | 0.087592008 | 0.080700447 | 1.085396819 | 0.277745941 | 0.506699118 |
| Smpdl3b | 174.321638 | 0.118836436 | 0.152863605 | 0.777401763 | 0.436921772 | 0.657440608 |
| Nipsnap3b | 315.3507926 | -0.122460509 | 0.1128939 | -1.084739829 | 0.278036902 | 0.507083953 |
| Plekhh2 | 516.391218 | -0.113074927 | 0.10424546 | -1.084698819 | 0.278055071 | 0.507083953 |
| Phc1 | 2125.808898 | 0.118793078 | 0.062578728 | 1.898298076 | 0.057656827 | 0.190071252 |
| Yars | 3649.680323 | 0.118662748 | 0.060452426 | 1.962911276 | 0.049656484 | 0.173082559 |
| Rbm28 | 1861.026815 | 0.118623264 | 0.100776285 | 1.177095026 | 0.23915758 | 0.463246349 |
| Pmpcb | 1645.025308 | -0.07844346 | 0.072351353 | -1.08420171 | 0.278275373 | 0.507283495 |
| Zcchc11 | 1331.295381 | 0.118621235 | 0.077853425 | 1.52364824 | 0.127596606 | 0.31576135 |
| Gtf2ird2 | 647.7119966 | 0.118619442 | 0.074215449 | 1.598311995 | 0.109973559 | 0.288606085 |
| Grap | 152.2924145 | -0.128699355 | 0.118789294 | -1.083425542 | 0.278619584 | 0.50765721 |
| Tfb2m | 872.882933 | -0.083069551 | 0.076674965 | -1.083398617 | 0.278631529 | 0.50765721 |
| Acsl5 | 3392.512396 | 0.086671357 | 0.080038967 | 1.082864512 | 0.278868566 | 0.507917251 |
| Klf5 | 218.619607 | -0.133305647 | 0.123105674 | -1.082855426 | 0.278872599 | 0.507917251 |
| Rexo1 | 2200.499592 | 0.118562951 | 0.05515246 | 2.149730991 | 0.0315765 | 0.12784068 |
| Emc4 | 3093.165791 | -0.05790194 | 0.053561252 | -1.081041575 | 0.27967862 | 0.509325405 |
| Eif1ad | 968.5425612 | 0.084505396 | 0.07820587 | 1.080550561 | 0.279897084 | 0.50948372 |
| Npr1 | 195.307958 | -0.174366882 | 0.161368347 | -1.080551948 | 0.279896467 | 0.50948372 |
| Rbm6 | 2030.424896 | 0.06928201 | 0.064116775 | 1.080559805 | 0.279892971 | 0.50948372 |
| Twsg1 | 1225.273733 | -0.091452686 | 0.084622128 | -1.080718348 | 0.279822419 | 0.50948372 |
| Mmp25 | 15.38811468 | -0.181530644 | 0.168026388 | -1.080369853 | 0.279977515 | 0.509510408 |
| Tmx4 | 10364.73909 | -0.074817805 | 0.069251771 | -1.080373884 | 0.279975721 | 0.509510408 |
| Rft1 | 413.7652269 | 0.08890548 | 0.082297779 | 1.080290148 | 0.280012996 | 0.509515133 |
| Eif5 | 8667.705849 | 0.07843978 | 0.072624612 | 1.08007159 | 0.280110302 | 0.509632341 |
| Ecel1 | 987.3731519 | 0.118540051 | 0.167159457 | 0.709143555 | 0.478235397 | 0.691726964 |
| Gspt2 | 1181.382337 | 0.078553371 | 0.072777137 | 1.079368802 | 0.280423353 | 0.510082112 |
| Vti1a | 2643.52934 | -0.078428135 | 0.072698809 | -1.078809077 | 0.280672848 | 0.510416091 |
| Zfp800 | 391.3196483 | 0.118538292 | 0.10308418 | 1.1499174 | 0.250177894 | 0.475614268 |
| Fgf13 | 3923.910022 | -0.102961604 | 0.095451917 | -1.078675073 | 0.280732601 | 0.510439319 |
| Igf2r | 1793.093347 | -0.086967674 | 0.080627702 | -1.078632676 | 0.280751509 | 0.510439319 |
| Inf2 | 8709.124272 | -0.107652313 | 0.099824856 | -1.078411911 | 0.280849974 | 0.51055843 |
| Zmynd12 | 12.40994413 | 0.118485914 | 0.164353153 | 0.720922673 | 0.470957092 | 0.685615821 |
| Zfx | 816.6267901 | -0.076564884 | 0.071020332 | -1.078069919 | 0.281002555 | 0.510715963 |
| Armc8 | 3112.98283 | -0.058833522 | 0.054593944 | -1.07765656 | 0.281187051 | 0.510991341 |
| Adck5 | 489.1439116 | 0.118437759 | 0.082649705 | 1.433008854 | 0.151855307 | 0.353068146 |
| Limd2 | 2551.599673 | -0.094195325 | 0.087447408 | -1.077165437 | 0.281406363 | 0.511269957 |
| Psmb4 | 3436.261866 | -0.067113778 | 0.062311002 | -1.07707749 | 0.281445649 | 0.511281379 |
| Fbxo31 | 5644.01074 | 0.118411356 | 0.107204289 | 1.104539353 | 0.269359243 | 0.497728337 |
| Rbm22 | 1253.948939 | -0.061812634 | 0.057408118 | -1.076722872 | 0.281604091 | 0.511426532 |
| Ppp1r15a | 626.2953724 | 0.118407297 | 0.12645566 | 0.936354265 | 0.349090817 | 0.58015801 |
| Ndrg1 | 4481.54286 | 0.09146947 | 0.084991009 | 1.076225259 | 0.281826526 | 0.511733293 |
| Copz2 | 145.1021692 | -0.122908114 | 0.114236767 | -1.07590679 | 0.281968945 | 0.511811954 |
| Dhx9 | 5403.381649 | -0.074143485 | 0.068911978 | -1.075915793 | 0.281964918 | 0.511811954 |
| Psma1 | 2499.518693 | -0.067229873 | 0.062482731 | -1.075975264 | 0.281938319 | 0.511811954 |
| Glt1d1 | 589.2324156 | 0.118366582 | 0.073303573 | 1.614745058 | 0.106365916 | 0.281828499 |
| Appl1 | 4111.574042 | -0.076600982 | 0.071227972 | -1.075433992 | 0.28218047 | 0.511995964 |
| Zfp318 | 2481.8788 | 0.118364233 | 0.075290806 | 1.572094126 | 0.115928724 | 0.298421606 |
| Smg1 | 6906.342286 | -0.090164405 | 0.083843856 | -1.075384765 | 0.2822025 | 0.511995964 |
| Bbs10 | 188.2486504 | -0.112686953 | 0.104818364 | -1.075068804 | 0.282343925 | 0.512132612 |
| Foxk1 | 3050.404742 | 0.061187283 | 0.056910959 | 1.075140607 | 0.282311782 | 0.512132612 |
| 5730409E04Rik | 3866.729646 | 0.074627373 | 0.069428595 | 1.074879499 | 0.282428681 | 0.512226383 |
| Prune2 | 4033.555022 | 0.118349033 | 0.084712822 | 1.397061626 | 0.16239504 | 0.367428264 |
| Def6 | 74.48293941 | -0.15722533 | 0.146349723 | -1.074312458 | 0.282682663 | 0.512513653 |
| Cyfip2 | 54647.31487 | 0.118318457 | 0.081647875 | 1.449130872 | 0.147301038 | 0.346535421 |
| Tfpt | 602.2887793 | 0.089195786 | 0.083024959 | 1.074324964 | 0.28267706 | 0.512513653 |
| Cd63 | 1499.283891 | 0.087795291 | 0.081746637 | 1.073992698 | 0.282825954 | 0.51252694 |
| Mta2 | 2121.537522 | 0.11827454 | 0.07125716 | 1.659826747 | 0.096949311 | 0.264227375 |
| Pcnp | 5172.49143 | 0.099841482 | 0.092950561 | 1.07413533 | 0.282762032 | 0.51252694 |
| Wdr95 | 15.3164765 | 0.118252638 | 0.168060151 | 0.703632821 | 0.481661467 | 0.694413582 |
| Cds1 | 4328.36351 | 0.080602494 | 0.075080916 | 1.073541696 | 0.283028141 | 0.512599701 |
| Il1r2 | 12.56616994 | 0.118245786 | 0.166554253 | 0.70995357 | 0.477732929 | 0.691328116 |
| Col27a1 | 360.1577219 | 0.118233397 | 0.138768386 | 0.852019687 | 0.394203165 | 0.619921228 |
| Sfn | 63.38660011 | -0.146903496 | 0.136824116 | -1.07366669 | 0.282972096 | 0.512599701 |
| Usp29 | 2474.751376 | -0.136975551 | 0.127597801 | -1.073494603 | 0.283049259 | 0.512599701 |
| Vps4b | 1655.086822 | 0.068669568 | 0.063971128 | 1.073446259 | 0.283070939 | 0.512599701 |
| Aars | 7112.702703 | 0.118203671 | 0.073730469 | 1.60318621 | 0.108893549 | 0.286743143 |
| Smim5 | 9.752686901 | 0.118202927 | 0.157166069 | 0.752089354 | 0.451997326 | 0.670702205 |
| Klc4 | 828.0229109 | 0.087899926 | 0.081959518 | 1.072479781 | 0.283504591 | 0.513156557 |
| 2410089E03Rik | 2325.516928 | 0.097997917 | 0.091396274 | 1.072230995 | 0.283616292 | 0.513238826 |
| Trim37 | 16383.24485 | -0.075344184 | 0.070268336 | -1.072235206 | 0.283614401 | 0.513238826 |
| Trp53bp2 | 1246.973607 | 0.07091751 | 0.066175887 | 1.071651822 | 0.283876448 | 0.513649618 |
| Pear1 | 268.8977448 | 0.092255535 | 0.086110539 | 1.071361721 | 0.284006818 | 0.513825506 |
| Ddx52 | 918.7518901 | 0.074924604 | 0.069943116 | 1.071221996 | 0.284069624 | 0.51387913 |
| Cisd2 | 1464.34161 | -0.06102846 | 0.056981551 | -1.071021395 | 0.28415981 | 0.513960079 |
| Trim32 | 7948.22561 | 0.070500183 | 0.065828044 | 1.070974902 | 0.284180715 | 0.513960079 |
| Wasl | 4406.545273 | 0.09722989 | 0.090792994 | 1.070896393 | 0.284216018 | 0.513963933 |
| Ufsp1 | 292.5084045 | -0.111328534 | 0.103967036 | -1.070806081 | 0.284256632 | 0.513977391 |
| Rapgef5 | 3251.446077 | 0.118201077 | 0.075881472 | 1.557706682 | 0.119302796 | 0.303576534 |
| Eml4 | 1497.282309 | 0.05923744 | 0.05534727 | 1.070286582 | 0.284490332 | 0.514279922 |
| Megf8 | 9102.421395 | -0.077508602 | 0.072429779 | -1.070120643 | 0.284565008 | 0.514354904 |
| Jmy | 2409.68851 | 0.118163003 | 0.083426924 | 1.416365341 | 0.156668566 | 0.359352 |
| Add1 | 18180.0682 | -0.055951844 | 0.052314653 | -1.069525293 | 0.284833038 | 0.514480389 |
| Spag16 | 23.95808064 | 0.1181322 | 0.164656134 | 0.717447915 | 0.473097764 | 0.687592909 |
| Hgs | 3150.125042 | 0.063695475 | 0.059548653 | 1.069637539 | 0.284782491 | 0.514480389 |
| Lamtor4 | 845.064462 | -0.100972698 | 0.094409015 | -1.069523905 | 0.284833663 | 0.514480389 |
| R3hcc1l | 552.8045638 | 0.08412809 | 0.07863886 | 1.069803026 | 0.28470798 | 0.514480389 |
| 0610007P14Rik | 1072.632876 | -0.072347914 | 0.06765099 | -1.069428755 | 0.284876516 | 0.514497813 |
| Mdm2 | 1693.121607 | 0.118123318 | 0.095625827 | 1.23526585 | 0.216731579 | 0.436760518 |
| Rgs7bp | 11765.7414 | -0.089389445 | 0.083619493 | -1.069002473 | 0.285068555 | 0.514782819 |
| Elp5 | 2080.900377 | -0.063857343 | 0.059752801 | -1.068692045 | 0.285208457 | 0.51491725 |
| Creb3l2 | 334.3391402 | 0.083752094 | 0.078377192 | 1.068577381 | 0.285260145 | 0.514950564 |
| Arf3 | 58150.79269 | -0.073849472 | 0.069143882 | -1.068055042 | 0.285495684 | 0.515019946 |
| Cldn10 | 1020.082093 | -0.093145657 | 0.087199015 | -1.068196214 | 0.285432012 | 0.515019946 |
| Dohh | 1667.43535 | -0.095377636 | 0.089304841 | -1.068000727 | 0.285520184 | 0.515019946 |
| Gins4 | 488.1354333 | -0.0843749 | 0.078985556 | -1.06823202 | 0.285415865 | 0.515019946 |
| Gm5113 | 839.0118194 | -0.07231284 | 0.067687511 | -1.068333557 | 0.285370077 | 0.515019946 |
| Ly75 | 67.43123078 | -0.158075087 | 0.148013683 | -1.067976174 | 0.285531259 | 0.515019946 |
| Prkaa1 | 1167.651232 | 0.074210211 | 0.069468757 | 1.068253042 | 0.285406384 | 0.515019946 |
| Arhgap19 | 223.9301013 | 0.118099783 | 0.108419901 | 1.089281412 | 0.276029807 | 0.504893705 |
| Eml5 | 2808.250243 | 0.118050904 | 0.134175169 | 0.879826757 | 0.378953167 | 0.607752012 |
| Rnd1 | 861.147822 | 0.118046492 | 0.111614517 | 1.057626695 | 0.290225667 | 0.520518172 |
| Sirt3 | 2202.687747 | -0.078015538 | 0.073106792 | -1.067144871 | 0.285906422 | 0.51545661 |
| Oscp1 | 1192.673446 | 0.118042985 | 0.057968518 | 2.036329178 | 0.041717312 | 0.154598191 |
| Gpr137b | 358.3258256 | 0.117982374 | 0.102216349 | 1.154241721 | 0.248401082 | 0.473339856 |
| Susd2 | 614.9995738 | -0.135482097 | 0.127028346 | -1.066550113 | 0.286175038 | 0.515760851 |
| Lss | 908.2744817 | 0.074338016 | 0.06973654 | 1.065983715 | 0.286431004 | 0.516162127 |
| Fam73b | 3269.264083 | 0.117966836 | 0.054569298 | 2.161780341 | 0.030635112 | 0.125372535 |
| Fahd1 | 435.695269 | -0.098830746 | 0.092728105 | -1.065812213 | 0.286508539 | 0.516185277 |
| Gramd3 | 644.2934578 | -0.091051577 | 0.085438945 | -1.065691724 | 0.28656302 | 0.516185277 |
| Wdr20a | 576.673366 | 0.117950773 | 0.084971136 | 1.388127533 | 0.165098214 | 0.370675515 |
| Zic2 | 428.9986134 | -0.132173997 | 0.124020694 | -1.065741472 | 0.286540525 | 0.516185277 |
| Six3 | 432.6716726 | -0.162211465 | 0.152250053 | -1.065427974 | 0.286682303 | 0.516254719 |
| Acacb | 231.3087356 | 0.117915214 | 0.12968594 | 0.909236682 | 0.363225205 | 0.593603784 |
| Ormdl2 | 193.0846536 | -0.139136802 | 0.130760902 | -1.064055075 | 0.287303751 | 0.517169742 |
| Daxx | 921.2361264 | 0.117905857 | 0.073630905 | 1.601309362 | 0.109308417 | 0.287493567 |
| Tacc1 | 9975.023693 | -0.068847593 | 0.064705356 | -1.064016921 | 0.287321034 | 0.517169742 |
| Rps13 | 3462.102145 | -0.090835222 | 0.085384112 | -1.063842205 | 0.287400188 | 0.517246895 |
| Arrb1 | 17019.29764 | -0.063594399 | 0.05980773 | -1.063314041 | 0.28763956 | 0.517531236 |
| G3bp1 | 2210.560185 | -0.062266945 | 0.058559585 | -1.06330918 | 0.287641763 | 0.517531236 |
| Pes1 | 1671.896537 | 0.060019125 | 0.056447546 | 1.063272518 | 0.287658384 | 0.517531236 |
| Syvn1 | 2158.563423 | 0.117850297 | 0.054763852 | 2.151972382 | 0.03139953 | 0.127590955 |
| Pdcd4 | 2624.658169 | 0.117849781 | 0.06897043 | 1.70870009 | 0.087506517 | 0.247729944 |
| Mpdz | 2635.595138 | -0.060536848 | 0.056972507 | -1.062562481 | 0.287980409 | 0.517930175 |
| Lmo7 | 4185.847381 | -0.102067498 | 0.096093763 | -1.062165693 | 0.288160471 | 0.518193865 |
| Wdr45b | 2729.671626 | -0.081151493 | 0.076411538 | -1.062031918 | 0.288221195 | 0.518242915 |
| Cbx1 | 675.7151607 | -0.091285221 | 0.08598073 | -1.061693947 | 0.288374648 | 0.518458668 |
| 2900026A02Rik | 2590.265197 | 0.117818439 | 0.09424521 | 1.250126551 | 0.211253322 | 0.429728004 |
| Ect2 | 27.01609526 | -0.175421427 | 0.165282786 | -1.061341176 | 0.288534879 | 0.518626383 |
| Ppp2r5b | 5571.73657 | 0.063500555 | 0.059836661 | 1.061231589 | 0.288584667 | 0.518655704 |
| Gm266 | 69.11805724 | -0.16209379 | 0.152888495 | -1.060209213 | 0.289049431 | 0.519370507 |
| Wbscr22 | 794.8289508 | 0.083165553 | 0.078438473 | 1.060264806 | 0.289024146 | 0.519370507 |
| Atp2b3 | 4270.213891 | 0.076759111 | 0.072407092 | 1.06010487 | 0.289096892 | 0.519395553 |
| Chi3l1 | 377.5241923 | -0.136415359 | 0.128703923 | -1.059916093 | 0.289182774 | 0.519489611 |
| Cntln | 573.6743929 | -0.086474006 | 0.081634893 | -1.059277507 | 0.289473415 | 0.519937033 |
| Mkl2 | 12851.93544 | -0.09397182 | 0.088717825 | -1.059221417 | 0.289498953 | 0.519937033 |
| Alox5ap | 113.7988001 | -0.136805406 | 0.129224252 | -1.058666651 | 0.289751621 | 0.520046504 |
| Brip1 | 36.23683618 | -0.171314781 | 0.161810941 | -1.058734229 | 0.289720835 | 0.520046504 |
| Mr1 | 243.285977 | 0.117763123 | 0.115010956 | 1.023929609 | 0.305868524 | 0.538414782 |
| Fam13c | 2549.43466 | -0.067773954 | 0.064028414 | -1.058498089 | 0.289828422 | 0.520046504 |
| Fam196a | 472.6069251 | -0.12340581 | 0.116585074 | -1.058504367 | 0.289825561 | 0.520046504 |
| Fam69b | 1769.539417 | -0.070297 | 0.066407562 | -1.058569206 | 0.289796017 | 0.520046504 |
| Tet3 | 2418.117423 | -0.081618318 | 0.077089285 | -1.058750478 | 0.289713432 | 0.520046504 |
| Zfp266 | 1438.904755 | 0.07433462 | 0.070205908 | 1.058808617 | 0.289686948 | 0.520046504 |
| U2surp | 2567.306888 | -0.067178764 | 0.063487241 | -1.058145909 | 0.289988927 | 0.520274251 |
| Chn2 | 1879.531631 | -0.122778395 | 0.116039912 | -1.058070389 | 0.290023353 | 0.520275771 |
| Ywhah | 32788.2713 | -0.068562843 | 0.064818046 | -1.057773988 | 0.290158494 | 0.520457943 |
| Gm17821 | 356.3756072 | 0.117752483 | 0.100420059 | 1.172599223 | 0.240956561 | 0.465428677 |
| Fgf11 | 234.6615145 | -0.118671515 | 0.112260806 | -1.057105499 | 0.290463442 | 0.520884325 |
| BC018242 | 6584.968432 | 0.079277584 | 0.075007528 | 1.056928365 | 0.290544282 | 0.520968997 |
| Zfp12 | 1427.247254 | 0.059241508 | 0.05605576 | 1.056831766 | 0.290588374 | 0.520987765 |
| 3110002H16Rik | 933.6123716 | 0.117692194 | 0.060981798 | 1.929956114 | 0.053612276 | 0.18184514 |
| 1700024F13Rik | 14.55627125 | -0.176283658 | 0.166882212 | -1.056335819 | 0.290814817 | 0.52115253 |
| Asb4 | 43.67168318 | -0.170024741 | 0.160955337 | -1.056347336 | 0.290809557 | 0.52115253 |
| Stxbp5 | 5076.964502 | -0.080214205 | 0.075930928 | -1.05641017 | 0.290780861 | 0.52115253 |
| D14Abb1e | 1266.402198 | -0.085404898 | 0.080868853 | -1.056091378 | 0.290926469 | 0.521292322 |
| Erc2 | 6930.720012 | -0.079955428 | 0.075717528 | -1.055969876 | 0.290981978 | 0.521331495 |
| Zfp334 | 897.2841765 | -0.060316864 | 0.057131245 | -1.055759665 | 0.29107803 | 0.521443288 |
| Itsn2 | 2250.135015 | 0.080215847 | 0.075986424 | 1.055660247 | 0.291123465 | 0.52146439 |
| Fgf7 | 18.96677089 | 0.11768871 | 0.168056051 | 0.700294391 | 0.483743474 | 0.69596223 |
| Mad1l1 | 561.4880272 | 0.117642246 | 0.087998537 | 1.336865937 | 0.181266401 | 0.391771527 |
| Col10a1 | 14.92005335 | -0.174587196 | 0.16546099 | -1.055156239 | 0.291353874 | 0.521696167 |
| Arl6ip4 | 1037.535121 | 0.064236963 | 0.060887737 | 1.055006578 | 0.291422316 | 0.521743749 |
| Zfp568 | 174.7873816 | 0.117591698 | 0.113904544 | 1.032370561 | 0.301898558 | 0.53382703 |
| Arhgef6 | 605.9003969 | 0.117585814 | 0.118266086 | 0.994247954 | 0.320102167 | 0.553376867 |
| 1700123O20Rik | 548.3879099 | -0.090792832 | 0.086081968 | -1.054725324 | 0.291550966 | 0.521749609 |
| Ccne2 | 125.365141 | 0.117554557 | 0.152987356 | 0.768393938 | 0.44225318 | 0.661991187 |
| Pafah1b1 | 24073.3691 | -0.062463313 | 0.059226622 | -1.054649259 | 0.291585766 | 0.521749609 |
| Nedd1 | 137.7732087 | 0.117502826 | 0.113594164 | 1.034408999 | 0.300944999 | 0.53259187 |
| Pcdhga5 | 446.0402153 | 0.098277424 | 0.093211924 | 1.054343908 | 0.291725493 | 0.521904338 |
| Tspyl3 | 1421.838961 | -0.070037599 | 0.066429928 | -1.05430792 | 0.291741964 | 0.521904338 |
| Rpl4 | 16421.58382 | 0.070496549 | 0.066916596 | 1.053498723 | 0.292112479 | 0.522450739 |
| Gsr | 943.8385654 | -0.082127873 | 0.07797622 | -1.053242545 | 0.292229844 | 0.522600324 |
| Unc13a | 15275.00453 | -0.10925496 | 0.103758155 | -1.052977091 | 0.292351492 | 0.522757532 |
| Fam219a | 6435.63558 | 0.053096704 | 0.050434657 | 1.052782112 | 0.292440865 | 0.522857001 |
| Ift122 | 1520.209681 | 0.066857082 | 0.063534825 | 1.052290336 | 0.292666364 | 0.523199798 |
| Cndp2 | 1841.144209 | 0.053166481 | 0.05053673 | 1.052036422 | 0.292782839 | 0.523347637 |
| 1700018L02Rik | 23.15032039 | 0.11739939 | 0.165487116 | 0.709417105 | 0.478065676 | 0.691610606 |
| Dact2 | 2524.153756 | 0.117386355 | 0.116935399 | 1.003856458 | 0.315447807 | 0.54876192 |
| Zc3h18 | 1661.076001 | 0.117374738 | 0.060486506 | 1.940511141 | 0.052317602 | 0.179140092 |
| Hspa2 | 1295.38218 | 0.094765814 | 0.090154215 | 1.051152337 | 0.29318863 | 0.523879624 |
| Nop2 | 907.9924419 | 0.082503027 | 0.078498328 | 1.051016353 | 0.293251079 | 0.523882419 |
| Prdm1 | 59.88455048 | -0.176371884 | 0.167951493 | -1.050135852 | 0.293655657 | 0.524423785 |
| Rab8a | 1261.907444 | 0.064183041 | 0.061118371 | 1.05014319 | 0.293652284 | 0.524423785 |
| Timm10b | 508.8226847 | -0.10536525 | 0.100325681 | -1.050232091 | 0.293611419 | 0.524423785 |
| Lurap1 | 524.0651879 | -0.101821913 | 0.097028869 | -1.04939812 | 0.293994923 | 0.524969153 |
| Ccne1 | 617.4708187 | -0.072640062 | 0.069250758 | -1.048942496 | 0.294204585 | 0.525282996 |
| Ufc1 | 1879.275163 | -0.061967013 | 0.059089303 | -1.048701033 | 0.294315739 | 0.525420908 |
| Lzts2 | 1495.878527 | 0.117368463 | 0.089859153 | 1.306138102 | 0.191505598 | 0.405408899 |
| Nprl2 | 1148.888695 | 0.079113261 | 0.075522796 | 1.047541472 | 0.294849915 | 0.526253264 |
| Cdk5rap2 | 535.7926005 | 0.117344401 | 0.114345344 | 1.026228065 | 0.304784096 | 0.537102144 |
| Tbr1 | 4183.424161 | 0.117330753 | 0.100897903 | 1.162866125 | 0.244883823 | 0.469873679 |
| Pfkm | 17444.5524 | 0.117216345 | 0.066603625 | 1.759909392 | 0.078423171 | 0.231225234 |
| Myo1b | 1689.92804 | 0.117191249 | 0.075242773 | 1.557508372 | 0.119349834 | 0.30364636 |
| Snx15 | 1058.0902 | 0.069191646 | 0.066132672 | 1.046255117 | 0.295443262 | 0.527008738 |
| Igdcc4 | 2304.655976 | -0.120627086 | 0.115306538 | -1.046142638 | 0.295495182 | 0.527040675 |
| Narfl | 921.6653927 | 0.074019404 | 0.070770677 | 1.045904981 | 0.295604905 | 0.52717569 |
| Pcdh9 | 4733.884826 | 0.085954252 | 0.082215981 | 1.045468917 | 0.295806299 | 0.52747414 |
| Gucd1 | 553.0228871 | -0.079256255 | 0.07581715 | -1.045360517 | 0.295856377 | 0.527502729 |
| Brwd3 | 358.2704106 | -0.10505866 | 0.100551059 | -1.044828979 | 0.296102019 | 0.527758505 |

| | | | | | | |
|---|---|---|---|---|---|---|
| Apcdd1 | 878.0516347 | -0.203944135 | 0.065868827 | -3.096216269 | 0.001960073 | 0.073192871 |
| Zc3h7a | 1472.437608 | -0.204007136 | 0.05372102 | -3.797529085 | 0.000146146 | 0.01286942 |
| Ndor1 | 1598.914242 | -0.2040246 | 0.075729522 | -2.694122362 | 0.007057425 | 0.160386105 |
| Glb1l | 436.6549673 | -0.204156431 | 0.08497814 | -2.402458215 | 0.016285295 | 0.242552129 |
| Mybl2 | 37.75923922 | -0.204187476 | 0.087797248 | -2.325670574 | 0.020036131 | 0.267640225 |
| Fancg | 595.5571862 | -0.204286709 | 0.066773759 | -3.059386078 | 0.002217911 | 0.079261025 |
| Gm6548 | 154.3783314 | -0.204521328 | 0.092335743 | -2.214974635 | 0.026761806 | 0.305374696 |
| Rgs5 | 1569.179504 | -0.204543106 | 0.067060863 | -3.050111433 | 0.002287565 | 0.080849551 |
| Zfp488 | 532.6969622 | -0.204584343 | 0.080363828 | -2.545726699 | 0.010905053 | 0.198767619 |
| Taz | 1709.933222 | -0.204629588 | 0.070077807 | -2.920034106 | 0.003499931 | 0.104182436 |
| A130049A11Rik | 66.55608356 | -0.204629672 | 0.095077629 | -2.152237846 | 0.031378626 | 0.32443523 |
| Map4k1 | 78.6395891 | -0.204791479 | 0.095102502 | -2.153376363 | 0.031289112 | 0.32443523 |
| Pan2 | 1140.4154 | -0.204813361 | 0.07207762 | -2.841566629 | 0.004489247 | 0.123974909 |
| Heatr6 | 928.7800134 | -0.204833853 | 0.060675849 | -3.375871227 | 0.000735824 | 0.040182602 |
| Tapbp | 1258.598886 | -0.204879884 | 0.079944171 | -2.562787014 | 0.010383572 | 0.194873092 |
| Adora3 | 45.92865487 | -0.205179531 | 0.092269164 | -2.223706403 | 0.026168204 | 0.302373406 |
| Ulk3 | 1039.091676 | -0.205267437 | 0.080576031 | -2.547499963 | 0.010849786 | 0.198744008 |
| Ticam2 | 32.49029428 | -0.205288106 | 0.087486942 | -2.346499967 | 0.018950668 | 0.262451878 |
| Zfp771 | 392.0822371 | -0.205354875 | 0.094841276 | -2.165247908 | 0.030368705 | 0.32021294 |
| Idua | 867.2895184 | -0.205404769 | 0.079315844 | -2.589706638 | 0.009605775 | 0.187266026 |
| Sfswap | 2411.193226 | -0.205535532 | 0.06817074 | -3.015010996 | 0.002569699 | 0.085222496 |
| Hdac10 | 655.8335272 | -0.205600465 | 0.0827466 | -2.48469986 | 0.01296607 | 0.216488663 |
| Tap2 | 456.2824726 | -0.205898702 | 0.091370938 | -2.253437546 | 0.024231574 | 0.291915574 |
| Itga2b | 95.86429326 | -0.205915028 | 0.094470227 | -2.179681722 | 0.029281062 | 0.317415456 |
| Akap13 | 1848.860655 | -0.205917302 | 0.070519881 | -2.919989345 | 0.003500434 | 0.104182436 |
| Leprel2 | 834.301798 | -0.205965696 | 0.08673652 | -2.374613318 | 0.017567343 | 0.253409359 |
| Slc12a9 | 1238.569545 | -0.206061651 | 0.082615753 | -2.494217442 | 0.012623519 | 0.215222104 |
| Stat1 | 1443.514868 | -0.206146703 | 0.088245182 | -2.336067527 | 0.019487722 | 0.263578327 |
| Oacyl | 78.12565905 | -0.206358305 | 0.095079444 | -2.170377697 | 0.029978245 | 0.318947844 |
| Asun | 1070.292666 | -0.206382875 | 0.063423835 | -3.254027059 | 0.001137814 | 0.052636599 |
| Adam1a | 223.7027506 | -0.206436075 | 0.090902445 | -2.270962841 | 0.023149228 | 0.286019362 |
| Skp2 | 302.0220928 | -0.206651106 | 0.088176248 | -2.343614188 | 0.019097915 | 0.263453293 |
| Gm5086 | 39.34251437 | -0.206708185 | 0.093072403 | -2.220939589 | 0.026355053 | 0.303114414 |
| Tbxas1 | 110.8418046 | -0.206744234 | 0.093493596 | -2.211319741 | 0.027013703 | 0.306595176 |
| Ddx11 | 80.37712385 | -0.206775333 | 0.095013372 | -2.17627612 | 0.02953462 | 0.317771905 |
| Pla2g4e | 570.7372835 | -0.206775558 | 0.085093489 | -2.429980951 | 0.015099616 | 0.233312354 |
| Zfp579 | 433.8169117 | -0.206851424 | 0.085534319 | -2.418344214 | 0.01559132 | 0.237151327 |
| B2m | 3685.558993 | -0.206919632 | 0.07795223 | -2.654441462 | 0.007943982 | 0.170584808 |
| Igfn1 | 2250.28956 | -0.207100529 | 0.095095223 | -2.177822624 | 0.029419245 | 0.317771905 |
| Foxq1 | 141.4822234 | -0.207101289 | 0.093956845 | -2.204217144 | 0.027509077 | 0.308173879 |
| Abhd14b | 652.3893308 | -0.207141517 | 0.088606062 | -2.337780431 | 0.019398641 | 0.263564471 |
| Sft2d3 | 177.8859836 | -0.207180876 | 0.090903982 | -2.279117716 | 0.022660069 | 0.28302679 |
| Hist4h4 | 36.84747879 | -0.207323973 | 0.089975268 | -2.304232905 | 0.021209573 | 0.274101653 |
| Zfp420 | 504.7053987 | -0.207373398 | 0.072777876 | -2.849401635 | 0.004380155 | 0.121423909 |
| Rnf112 | 16509.88398 | -0.207493422 | 0.082655375 | -2.510343973 | 0.012061361 | 0.209573362 |
| Tpcn2 | 141.3275075 | -0.207497647 | 0.090870425 | -2.283445336 | 0.022404149 | 0.281750998 |
| Fam76b | 584.9143322 | -0.207549324 | 0.081795224 | -2.537425937 | 0.011167099 | 0.20051086 |
| Nod1 | 274.9060099 | -0.207558606 | 0.083319968 | -2.491102811 | 0.012734726 | 0.215303986 |
| Rbm25 | 4392.493867 | -0.207559611 | 0.057890793 | -3.585364806 | 0.000336607 | 0.02373572 |
| Spata5l1 | 62.96821866 | -0.207869769 | 0.09490502 | -2.190292679 | 0.028503017 | 0.313900554 |
| Ubxn11 | 484.5464336 | -0.208141168 | 0.086130196 | -2.416587652 | 0.015666754 | 0.237621413 |
| Abcc8 | 1566.672775 | -0.208419439 | 0.067467385 | -3.089188038 | 0.002007044 | 0.074383385 |
| Stk38 | 1200.190107 | -0.208648721 | 0.072500331 | -2.877900245 | 0.004003317 | 0.113864147 |
| Clec18a | 293.4421732 | -0.208777789 | 0.095079862 | -2.195815028 | 0.028105186 | 0.312140588 |
| Xlr | 29.22583788 | -0.208780581 | 0.083736809 | -2.493295158 | 0.012656359 | 0.215249514 |
| Ccm2l | 152.6454866 | -0.208976771 | 0.094182414 | -2.218851289 | 0.026496843 | 0.303950052 |
| Srrm1 | 5284.812112 | -0.208995695 | 0.055042756 | -3.796970048 | 0.000146475 | 0.01286942 |
| 4933432I09Rik | 53.89153596 | -0.209149134 | 0.094061966 | -2.223525012 | 0.026180419 | 0.302373406 |
| Lcp2 | 142.8045705 | -0.209352504 | 0.094947441 | -2.204930464 | 0.027458974 | 0.308173879 |
| Cerkl | 59.36915581 | -0.209478529 | 0.09505277 | -2.203813 | 0.027537498 | 0.308173879 |
| Lime1 | 2046.85709 | -0.209687809 | 0.08784367 | -2.387056553 | 0.016983878 | 0.248007351 |
| Gm14827 | 235.0710097 | -0.209707123 | 0.095038917 | -2.206539499 | 0.027346248 | 0.308173879 |
| Parp4 | 398.7614629 | -0.209758748 | 0.080953372 | -2.591105737 | 0.009566809 | 0.186784235 |
| Tec | 86.88546946 | -0.209818665 | 0.094767406 | -2.214038281 | 0.026826146 | 0.305868596 |
| Nxf1 | 2696.109198 | -0.209845469 | 0.059352393 | -3.535585639 | 0.000406872 | 0.02634142 |
| Thoc1 | 1075.572384 | -0.209876517 | 0.069501866 | -3.019724931 | 0.002530043 | 0.084486004 |
| Snhg5 | 263.0220603 | -0.210020118 | 0.087878462 | -2.389892956 | 0.016853284 | 0.24653656 |
| Rasl12 | 130.3664543 | -0.210248178 | 0.091790027 | -2.290534019 | 0.021990379 | 0.278517602 |
| Inha | 1047.304125 | -0.210517446 | 0.087621675 | -2.402572729 | 0.016280197 | 0.242552129 |
| Adap2 | 213.3738463 | -0.211128765 | 0.090158892 | -2.341740901 | 0.019194035 | 0.263564471 |
| Sall1 | 1219.244421 | -0.211152762 | 0.050629631 | -4.170537234 | 3.04E-05 | 0.004695953 |
| Hck | 108.2362102 | -0.211216858 | 0.094785262 | -2.228372371 | 0.025855694 | 0.301310169 |
| Atg16l2 | 655.6892323 | -0.211363081 | 0.090150983 | -2.344545487 | 0.019050287 | 0.263085246 |
| Ubp1 | 4011.714008 | -0.211473527 | 0.057138288 | -3.701082681 | 0.000214682 | 0.017421584 |
| Rasgrp3 | 857.4523741 | -0.211589253 | 0.078504108 | -2.695263455 | 0.0070333 | 0.160386105 |
| Ap4e1 | 689.8798976 | -0.211593884 | 0.071952186 | -2.940756848 | 0.003274114 | 0.099891275 |
| Slc26a4 | 132.5080571 | -0.211701075 | 0.095080894 | -2.226536439 | 0.025978271 | 0.3021678 |
| Snap23 | 432.6682099 | -0.211708879 | 0.074955205 | -2.824472029 | 0.004735858 | 0.127298346 |
| Rmdn1 | 273.8044857 | -0.211715956 | 0.083401439 | -2.538516829 | 0.011132345 | 0.20051086 |
| Usp28 | 1248.877818 | -0.211795436 | 0.072923478 | -2.904351821 | 0.003680144 | 0.107995487 |
| 2610203C20Rik | 752.5465455 | -0.21195988 | 0.068614359 | -3.089147578 | 0.002007317 | 0.074383385 |
| Gcfc2 | 279.010927 | -0.212427034 | 0.078982186 | -2.689556267 | 0.007154708 | 0.161546928 |
| Heatr8 | 314.0527982 | -0.212493203 | 0.088785317 | -2.393337203 | 0.016695889 | 0.245968033 |
| A230073K19Rik | 115.0779278 | -0.212610316 | 0.095073615 | -2.236270457 | 0.02533406 | 0.298582503 |
| Ddx60 | 56.81493954 | -0.212843369 | 0.081682256 | -2.605747927 | 0.00916739 | 0.182762906 |
| Myo9b | 2648.749946 | -0.212887472 | 0.062999827 | -3.379175499 | 0.000727036 | 0.040003493 |
| S100a6 | 453.5323448 | -0.213122552 | 0.078752879 | -2.706219175 | 0.006805411 | 0.158676414 |
| Miip | 630.3988449 | -0.213315214 | 0.083452552 | -2.556125698 | 0.010584488 | 0.196611595 |
| Zkscan7 | 136.0297037 | -0.213781003 | 0.092062916 | -2.322118532 | 0.020226554 | 0.269057623 |
| Cldn5 | 726.5244789 | -0.213929361 | 0.082635423 | -2.588833619 | 0.009630161 | 0.187266026 |
| Wdr60 | 883.1989925 | -0.214012145 | 0.072675127 | -2.94477843 | 0.00323186 | 0.099461865 |
| Rreb1 | 2620.314921 | -0.214050945 | 0.094991501 | -2.253369427 | 0.024235866 | 0.291915574 |
| Tifab | 165.4027612 | -0.214071999 | 0.094526665 | -2.264673135 | 0.023532745 | 0.28920191 |
| Lrrcc1 | 737.6794115 | -0.214109647 | 0.072837982 | -2.939532922 | 0.003287073 | 0.099891275 |
| Zfyve16 | 778.9209442 | -0.21418955 | 0.0669428 | -3.199590529 | 0.00137623 | 0.06021419 |
| Ifi27l2a | 25.45252315 | -0.214504388 | 0.077159384 | -2.780016837 | 0.005435608 | 0.138877279 |
| Tbce | 1843.727523 | -0.214668308 | 0.060591413 | -3.542883334 | 0.000395778 | 0.02634142 |
| Parp3 | 230.876434 | -0.215076579 | 0.093671431 | -2.296074461 | 0.021671624 | 0.276408712 |
| Tbata | 106.2597532 | -0.215486178 | 0.094384554 | -2.283066132 | 0.022426473 | 0.281750998 |
| Glt8d1 | 1211.216073 | -0.215528426 | 0.066418139 | -3.245023541 | 0.001174409 | 0.05383536 |
| Mx1 | 18.93959695 | -0.215532059 | 0.066524842 | -3.239873314 | 0.001195828 | NA |
| Col5a3 | 202.3335351 | -0.215704428 | 0.091966694 | -2.345462448 | 0.019003493 | 0.262899749 |
| 1700109H08Rik | 25.57412999 | -0.215850034 | 0.085487602 | -2.524927935 | 0.011572202 | 0.204002187 |
| Naip2 | 76.808045 | -0.21617455 | 0.093117606 | -2.321521784 | 0.0202587 | 0.269238676 |
| Gm216 | 116.3069484 | -0.216234035 | 0.092669111 | -2.333399251 | 0.019627201 | 0.264230505 |
| Snord2 | 32.56535568 | -0.216252389 | 0.081910236 | -2.64011433 | 0.008287806 | 0.174157793 |
| Trmt13 | 250.2192641 | -0.21640079 | 0.083978594 | -2.576856552 | 0.009970332 | 0.19094795 |
| Gm20751 | 31.65224638 | -0.216431225 | 0.088963716 | -2.43280333 | 0.014982436 | 0.232764567 |
| Tmc8 | 27.27931103 | -0.216489737 | 0.082100941 | -2.636872773 | 0.008367421 | 0.174157793 |
| Laptm5 | 2213.804705 | -0.216616819 | 0.080395578 | -2.694387218 | 0.007051819 | 0.160386105 |
| Slco1a4 | 2061.883328 | -0.216785386 | 0.061041863 | -3.551421517 | 0.000383156 | 0.026130177 |
| Atp13a5 | 563.6572922 | -0.216855842 | 0.077966119 | -2.781411291 | 0.005412312 | 0.138877279 |
| Clec2d | 152.6503099 | -0.217298189 | 0.09394674 | -2.312993391 | 0.020723003 | 0.270947196 |
| Il10rb | 391.1982218 | -0.21736085 | 0.075158591 | -2.89202933 | 0.003827622 | 0.111200078 |
| Hhex | 75.35210837 | -0.217404428 | 0.094898108 | -2.29092479 | 0.021967764 | 0.278517602 |
| Slc24a6 | 241.0781078 | -0.21743053 | 0.087707184 | -2.479050418 | 0.013173268 | 0.219077909 |
| C1ra | 173.9585403 | -0.21780143 | 0.093181638 | -2.337385717 | 0.019419137 | 0.263564471 |
| Gna15 | 79.21182506 | -0.217858408 | 0.09404611 | -2.316506321 | 0.02053064 | 0.269900834 |
| Abca9 | 445.012748 | -0.217885842 | 0.072988358 | -2.985213662 | 0.002833803 | 0.090867064 |
| 4930447C04Rik | 81.6116268 | -0.217900481 | 0.094659705 | -2.301934931 | 0.021338842 | 0.275079973 |
| Itgb3 | 238.0905467 | -0.217909036 | 0.082667501 | -2.63596981 | 0.00838972 | 0.174347735 |
| A330021E22Rik | 489.5540278 | -0.21804977 | 0.083701057 | -2.605101752 | 0.009184698 | 0.182762906 |
| Aox1 | 292.4837882 | -0.21810412 | 0.093629246 | -2.329444377 | 0.019835536 | 0.266048941 |
| Tmem150a | 378.3251875 | -0.218160439 | 0.087617152 | -2.489928436 | 0.012776882 | 0.215303986 |
| D330050I16Rik | 40.57469832 | -0.218291029 | 0.091175188 | -2.394193356 | 0.016656965 | 0.24589337 |
| Rsad2 | 73.10873353 | -0.218402862 | 0.091957162 | -2.375050053 | 0.017546571 | 0.253361317 |
| Eya4 | 201.5054874 | -0.218537445 | 0.08977602 | -2.434251876 | 0.014922607 | 0.232332035 |
| 4930470H14Rik | 73.68447443 | -0.218562036 | 0.095050196 | -2.299438046 | 0.021480078 | 0.275635696 |
| Hpgd | 189.6485636 | -0.218651132 | 0.091049799 | -2.401445516 | 0.016330441 | 0.242552129 |
| Vegfc | 117.5595011 | -0.218659147 | 0.093265125 | -2.34448995 | 0.019053124 | 0.263085246 |
| Gm17821 | 372.011346 | -0.218669063 | 0.077169599 | -2.833616691 | 0.00460245 | 0.126226882 |
| Pagr1a | 1737.824094 | -0.218892181 | 0.072471399 | -3.020394029 | 0.00252446 | 0.084486004 |
| Odf2 | 3371.3231 | -0.219014523 | 0.078582317 | -2.787071334 | 0.005318677 | 0.137476727 |
| Nktr | 4630.989869 | -0.219016462 | 0.070473288 | -3.107794031 | 0.001884894 | 0.071301999 |
| Fam228a | 63.44013144 | -0.219266963 | 0.094628922 | -2.317124164 | 0.020496969 | 0.269900834 |
| Clcn1 | 95.75726186 | -0.219363563 | 0.095083235 | -2.307068768 | 0.021050987 | 0.273267769 |
| Lmbr1l | 353.0906235 | -0.219369007 | 0.083876481 | -2.615381629 | 0.008912781 | 0.180065445 |
| Lpl | 1062.026025 | -0.219505509 | 0.07977234 | -2.751649361 | 0.005929597 | 0.146485255 |
| Tbx18 | 104.8411077 | -0.219614059 | 0.095034799 | -2.310880435 | 0.02083946 | 0.271736083 |
| Col4a5 | 188.6446047 | -0.219649005 | 0.094893524 | -2.314689087 | 0.020629953 | 0.270705243 |
| Crtc2 | 1115.700987 | -0.220071178 | 0.077814343 | -2.828156979 | 0.004681684 | 0.126877139 |
| Kif19a | 106.3604411 | -0.220166405 | 0.092979643 | -2.367899012 | 0.017889418 | 0.255043174 |
| Zwilch | 136.9667513 | -0.220188629 | 0.090578334 | -2.430919393 | 0.015060565 | 0.233312354 |
| H2-D1 | 733.6634636 | -0.220461659 | 0.084451037 | -2.610526374 | 0.0090403 | 0.181380385 |
| Myct1 | 29.15773614 | -0.220564833 | 0.08817646 | -2.501402693 | 0.012370243 | 0.21214894 |
| Fyco1 | 2996.189938 | -0.220700433 | 0.068752123 | -3.210088967 | 0.001326939 | 0.05912612 |
| Amy1 | 848.0352953 | -0.22071235 | 0.074791008 | -2.951054638 | 0.003166909 | 0.098506464 |
| Zfp445 | 8652.042808 | -0.220750124 | 0.065931414 | -3.348178208 | 0.000813447 | 0.042689184 |
| Apobec1 | 73.84144659 | -0.220751979 | 0.094729663 | -2.330336367 | 0.01978838 | 0.265661746 |
| Ccnl2 | 2226.655765 | -0.220970763 | 0.073607622 | -3.002009257 | 0.00268204 | 0.087506957 |
| Arhgap29 | 940.6322137 | -0.221138646 | 0.073151326 | -3.023029913 | 0.002502575 | 0.084486004 |
| Mbip | 526.3336678 | -0.22133038 | 0.073266985 | -3.020874668 | 0.002520457 | 0.084486004 |
| Tlr12 | 40.80684099 | -0.221782701 | 0.087628681 | -2.530937334 | 0.011375817 | 0.202439555 |
| Asap3 | 246.0337934 | -0.221788457 | 0.084271537 | -2.63183117 | 0.008492605 | 0.175232365 |
| Pxdn | 3387.253934 | -0.221808487 | 0.08348592 | -2.656837058 | 0.007887756 | 0.169996349 |
| Bcl2a1b | 23.82138782 | -0.222674486 | 0.078181603 | -2.848169887 | 0.004397144 | 0.121662707 |
| Mettl17 | 771.604056 | -0.222824142 | 0.080132291 | -2.780703508 | 0.005424125 | 0.138877279 |
| Tnfrsf11a | 258.29117 | -0.222948261 | 0.085672857 | -2.602320835 | 0.009259518 | 0.183393127 |
| Hk3 | 85.34104637 | -0.223004719 | 0.092121728 | -2.420761358 | 0.015488041 | 0.236284305 |
| Papln | 199.4567752 | -0.223389494 | 0.091626801 | -2.438036609 | 0.014767278 | 0.232209004 |
| Ggta1 | 203.2759328 | -0.223510586 | 0.093444024 | -2.391919516 | 0.016760517 | 0.246336261 |
| Gm266 | 58.03992489 | -0.224010238 | 0.092696088 | -2.416609401 | 0.015665818 | 0.237621413 |
| 1700094D03Rik | 293.0949867 | -0.224022859 | 0.090682756 | -2.470401976 | 0.01349613 | 0.222778167 |
| Rbms1 | 633.3415844 | -0.224053444 | 0.070660383 | -3.170849579 | 0.001519938 | 0.06344432 |
| Cnga4 | 57.01044436 | -0.224082894 | 0.093707467 | -2.39130244 | 0.016788717 | 0.246336261 |
| Pkn3 | 64.49379296 | -0.224212154 | 0.095102726 | -2.357578619 | 0.01839456 | 0.258413322 |
| Zfp692 | 1310.332432 | -0.224322312 | 0.087924338 | -2.551310805 | 0.010731857 | 0.197580431 |
| Ccr5 | 217.0915648 | -0.22437717 | 0.082533072 | -2.718633441 | 0.006555221 | 0.155844743 |
| Adamts3 | 1227.458387 | -0.22446114 | 0.072436209 | -3.098742234 | 0.00194344 | 0.072758797 |
| Clec14a | 166.5784525 | -0.224624513 | 0.093767344 | -2.395551639 | 0.016595376 | 0.245483134 |
| Dennd2c | 34.16499045 | -0.224632803 | 0.086447732 | -2.598481163 | 0.009363718 | 0.184100285 |
| Zfhx2as | 248.3904996 | -0.22464211 | 0.094485814 | -2.377522091 | 0.017429399 | 0.252422183 |
| Milr1 | 14.08035286 | -0.224805488 | 0.068367105 | -3.288211318 | 0.001008261 | NA |
| E230029C05Rik | 28.90041993 | -0.224838886 | 0.083229653 | -2.70142767 | 0.00690425 | 0.159953955 |
| Pus10 | 439.3096895 | -0.224879158 | 0.074304079 | -3.02647125 | 0.002474263 | 0.083974653 |
| AW011738 | 159.228046 | -0.225488905 | 0.086742234 | -2.59952845 | 0.009335194 | 0.183993254 |
| Taf1d | 477.5356907 | -0.226145137 | 0.071602686 | -3.158333137 | 0.001586741 | 0.06480716 |
| Plb1 | 62.90552893 | -0.226440754 | 0.094328247 | -2.400561469 | 0.016369941 | 0.242890464 |
| A330076H08Rik | 85.12221156 | -0.226481197 | 0.094708306 | -2.391355154 | 0.016786306 | 0.246336261 |
| Cchcr1 | 269.7157 | -0.226643635 | 0.084401506 | -2.68530319 | 0.007246404 | 0.162353883 |
| C230091D08Rik | 65.39358164 | -0.226808865 | 0.094730105 | -2.394263833 | 0.016653765 | 0.24589337 |
| Rrnad1 | 958.8813731 | -0.226971187 | 0.075593771 | -3.002511763 | 0.002677616 | 0.087506957 |
| Rccd1 | 394.8716509 | -0.227191 | 0.080446616 | -2.824121288 | 0.004741044 | 0.127298346 |
| Sh3tc1 | 115.1227563 | -0.227493422 | 0.093442124 | -2.434591726 | 0.014908601 | 0.232332035 |
| 1700029J07Rik | 177.8657883 | -0.227584126 | 0.090025486 | -2.527996644 | 0.011471545 | 0.203462345 |
| Snapc4 | 2223.079437 | -0.227826905 | 0.081963832 | -2.779602908 | 0.005442541 | 0.138877279 |
| Pcsk1n | 8130.877654 | -0.228102642 | 0.088801726 | -2.568673516 | 0.010208859 | 0.193453318 |
| Vav1 | 196.5613509 | -0.228107416 | 0.092537279 | -2.465032672 | 0.013700076 | 0.224208584 |
| H2-K1 | 1224.405121 | -0.228149208 | 0.090272838 | -2.527329525 | 0.011493361 | 0.203514823 |
| Fgfbp1 | 52.8466015 | -0.22818963 | 0.089964507 | -2.536440613 | 0.011198573 | 0.200580121 |
| Rab32 | 83.58630381 | -0.228217975 | 0.094595877 | -2.412557305 | 0.015841047 | 0.239084042 |
| Prpf39 | 1383.500281 | -0.228811843 | 0.066289218 | -3.451720361 | 0.000557025 | 0.033435292 |
| Igflr1 | 263.4690097 | -0.228817644 | 0.095083634 | -2.406488203 | 0.016106724 | 0.24095394 |
| Tcirg1 | 612.4305361 | -0.228952355 | 0.078386895 | -2.920798882 | 0.003491351 | 0.104182436 |
| Ifih1 | 268.2169271 | -0.228959124 | 0.088215807 | -2.595443281 | 0.009446899 | 0.185189824 |
| Unc13b | 2407.33301 | -0.229044653 | 0.064330538 | -3.560434257 | 0.000370242 | 0.025610168 |
| Irf5 | 184.8896405 | -0.229066669 | 0.08868376 | -2.582960741 | 0.009795647 | 0.189217509 |
| Mndal | 41.64685387 | -0.229408151 | 0.087674682 | -2.616583785 | 0.008881456 | 0.179953183 |
| Rgl2 | 2142.26731 | -0.22949865 | 0.085084381 | -2.69730646 | 0.006990291 | 0.160386105 |
| 2810055G20Rik | 34.75497903 | -0.229524586 | 0.088746132 | -2.58630522 | 0.009701098 | 0.188141728 |
| Zfp82 | 208.193946 | -0.229887231 | 0.086084476 | -2.670484174 | 0.007574194 | 0.166701129 |
| Styx | 385.9951812 | -0.229913172 | 0.084028003 | -2.736149432 | 0.006216279 | 0.150747361 |
| Dvl2 | 811.5811287 | -0.230042792 | 0.081640031 | -2.817769535 | 0.004835851 | 0.129604368 |
| Neat1 | 6240.970233 | -0.230145223 | 0.094890537 | -2.425375908 | 0.015292543 | 0.234571782 |
| Rbm33 | 3862.600328 | -0.23033986 | 0.061806934 | -3.726764053 | 0.000193954 | 0.015917366 |
| Hexb | 5099.463387 | -0.230381288 | 0.076977465 | -2.992840675 | 0.00276394 | 0.089022155 |
| Slc2a4rg-ps | 404.0839001 | -0.230431902 | 0.094749589 | -2.43200953 | 0.015015312 | 0.233026095 |
| Zfp280c | 893.3975499 | -0.230462156 | 0.066058 | -3.488784945 | 0.000485221 | 0.030512231 |
| Col24a1 | 608.4075453 | -0.230514783 | 0.092224835 | -2.499487067 | 0.012437324 | 0.213047838 |
| Phldb2 | 303.21987 | -0.230633725 | 0.084150317 | -2.740735066 | 0.006130191 | 0.149407989 |
| Smc5 | 1223.408165 | -0.230696232 | 0.056803996 | -4.061267632 | 4.88E-05 | 0.006564539 |
| Dapp1 | 235.0321283 | -0.230705498 | 0.085547338 | -2.69681681 | 0.007000577 | 0.160386105 |
| Col16a1 | 1204.200657 | -0.230809436 | 0.090246639 | -2.557540512 | 0.010541528 | 0.196333074 |
| Casp4 | 12.76275709 | -0.230922902 | 0.069880512 | -3.304539355 | 0.000951326 | NA |
| P2ry6 | 98.37981341 | -0.23094679 | 0.094774059 | -2.436814368 | 0.014817284 | 0.232209004 |
| Atp8b1 | 199.4570568 | -0.231089534 | 0.092572155 | -2.496317955 | 0.012549006 | 0.214455135 |
| Gm10560 | 161.6435471 | -0.231129145 | 0.09377366 | -2.464755508 | 0.013710677 | 0.224208584 |
| Slc4a7 | 722.2505516 | -0.23125239 | 0.067387404 | -3.431685702 | 0.000599842 | 0.03496515 |
| Eqtn | 32.25223171 | -0.231265721 | 0.0889608 | -2.599636256 | 0.009332262 | 0.183993254 |
| Nlrp3 | 53.12146834 | -0.231327022 | 0.087224623 | -2.652083936 | NA | NA |
| Ppox | 944.2365591 | -0.231481345 | 0.080170224 | -2.887373052 | 0.003884733 | 0.111963564 |
| Adamts10 | 965.1935216 | -0.231492443 | 0.087613676 | -2.642195282 | 0.008237055 | 0.174077875 |
| Bzrap1 | 15593.29937 | -0.231570457 | 0.07534128 | -3.073619889 | 0.002114787 | 0.077094131 |
| Tmem145 | 2051.42566 | -0.231667039 | 0.076010716 | -3.047820758 | 0.002305074 | 0.081073853 |
| Pik3r6 | 168.7819858 | -0.231730528 | 0.093205898 | -2.486221723 | 0.01291075 | 0.216344675 |
| 9930012K11Rik | 68.57147636 | -0.231893753 | 0.094427273 | -2.455792123 | 0.014057445 | 0.227385586 |
| Ccl3 | 22.66162113 | -0.231960324 | 0.064867887 | -3.575888399 | NA | NA |
| Rbl1 | 89.85241024 | -0.232066072 | 0.093286412 | -2.487672827 | 0.012858196 | 0.216178423 |
| Stxbp3b | 36.01320272 | -0.232075072 | 0.091667046 | -2.531717581 | 0.011350537 | 0.202439555 |
| Ltbp4 | 2859.080522 | -0.232265592 | 0.073011068 | -3.181238086 | 0.001466471 | 0.062104815 |
| A930011O12Rik | 7787.879579 | -0.232293334 | 0.094908402 | -2.447552907 | 0.014383001 | 0.229590624 |
| Ifi204 | 27.12459308 | -0.232358995 | 0.063935451 | -3.63427474 | 0.000278764 | 0.02098094 |
| Gm684 | 52.85048096 | -0.232673302 | 0.094782745 | -2.454806528 | 0.014096043 | 0.227385586 |
| Sirt4 | 570.7969222 | -0.232741475 | 0.093040159 | -2.501516322 | 0.012366274 | 0.21214894 |
| Wtip | 183.7298073 | -0.232780864 | 0.087837241 | -2.650138613 | 0.008045875 | 0.172381098 |

| | | | | | | | | | | | | | |
|---|---|---|---|---|---|---|---|---|---|---|---|---|---|
| Gtpbp4 | 1212.11697 | 0.070424227 | 0.067393473 | 1.044971033 | 0.296036358 | 0.527758505 | 2810410L24Rik | 141.1012059 | -0.232805697 | 0.093443784 | -2.491398427 | 0.012724134 | 0.215303986 |
| Ndc80 | 12.07436441 | -0.170522977 | 0.163199897 | -1.04487184 | 0.296082206 | 0.527758505 | Serpina3n | 778.4488747 | -0.232819204 | 0.09237846 | -2.520275861 | 0.011726289 | 0.205471748 |
| Dnajc16 | 2820.594457 | 0.117174119 | 0.075806998 | 1.545689993 | 0.122179455 | 0.30848017 | 2900005J15Rik | 91.97407777 | -0.233283553 | 0.094608595 | -2.465775467 | 0.013671701 | 0.224208584 |
| Mon1b | 1763.245246 | -0.050022077 | 0.047917145 | -1.04392858 | 0.296518434 | 0.528336622 | Mrgbp | 320.6562358 | -0.233356902 | 0.084231147 | -2.770434798 | 0.005598151 | 0.141670274 |
| Plin2 | 237.0035293 | -0.139031432 | 0.133183804 | -1.043906451 | 0.296528673 | 0.528336622 | Qtrtd1 | 567.4114928 | -0.233788206 | 0.067689781 | -3.453818309 | 0.00055271 | 0.033313941 |
| Cdk17 | 8567.236864 | -0.080647638 | 0.077297899 | -1.043335453 | 0.296792957 | 0.528685913 | Xlr3b | 43.83685867 | -0.233887546 | 0.07586126 | -3.083095969 | 0.002048591 | 0.075719668 |
| Lamp1 | 12406.18079 | 0.050288243 | 0.048197183 | 1.043385508 | 0.296769783 | 0.528685913 | Phka2 | 1662.411297 | -0.23390189 | 0.06667709 | -3.507979896 | 0.000451523 | 0.028766779 |
| D830030K20Rik | 97.66554383 | -0.129967623 | 0.12461573 | -1.042947174 | 0.29697276 | 0.528945389 | Fyb | 175.049246 | -0.234258898 | 0.088666717 | -2.64201616 | 0.008241412 | 0.174077875 |
| AI462493 | 511.8579732 | -0.071408784 | 0.068503842 | -1.042405539 | 0.2972237 | 0.529027456 | Sipa1l3 | 2410.47067 | -0.234303755 | 0.067992038 | -3.446046934 | 0.000568852 | 0.03358999 |
| Gm6402 | 10.93467629 | -0.171182337 | 0.164196714 | -1.042544229 | 0.297159431 | 0.529027456 | Samd9l | 273.8234423 | -0.2343287 | 0.09333303 | -2.510672813 | 0.012050133 | 0.209573362 |
| Mthfd2l | 145.3220107 | -0.148201758 | 0.142147397 | -1.042592141 | 0.297137231 | 0.529027456 | Slc7a11 | 1693.821151 | -0.23467916 | 0.083081881 | -2.824673169 | 0.004732887 | 0.127298346 |
| Pramef8 | 712.8073667 | -0.072916584 | 0.06994759 | -1.042445979 | 0.297204959 | 0.529027456 | 4930503L19Rik | 353.4793158 | -0.23480943 | 0.07396462 | -3.174618207 | 0.001500338 | 0.063017784 |
| F8 | 110.2658355 | 0.117167003 | 0.126079358 | 0.929311545 | 0.352727654 | 0.583635441 | Hcls1 | 241.0245947 | -0.234810947 | 0.089046319 | -2.636952877 | 0.008365445 | 0.174157793 |
| Vstm2a | 2910.134783 | -0.089237156 | 0.08558649 | -1.042654689 | 0.297108251 | 0.529027456 | Cd180 | 176.9553587 | -0.234864181 | 0.076414186 | -3.073567803 | 0.002115156 | 0.077094131 |
| Cpe | 76847.68497 | 0.085522537 | 0.082083873 | 1.04189208 | 0.297461716 | 0.529268699 | Trpm4 | 614.9359421 | -0.234904406 | 0.083123554 | -2.825966822 | 0.004713815 | 0.127272995 |
| Aldh16a1 | 230.2489788 | 0.11708456 | 0.097810565 | 1.197054323 | 0.231285382 | 0.454187035 | Trem2 | 583.7764962 | -0.236312408 | 0.094239058 | -2.507584566 | 0.012155949 | 0.210964535 |
| Gpsm1 | 2230.170285 | -0.138518816 | 0.13293744 | -1.041984981 | 0.297418642 | 0.529268699 | D11Wsu47e | 324.6479892 | -0.236410927 | 0.077004315 | -3.070099723 | 0.002139873 | 0.077515702 |
| Gm14405 | 13.1899039 | -0.173267246 | 0.16631882 | -1.041777746 | 0.297514734 | 0.529302249 | Lpar6 | 139.6021702 | -0.236635116 | 0.089155853 | -2.654173649 | 0.00795029 | 0.170584808 |
| Klc2 | 10359.76807 | 0.070510219 | 0.06769405 | 1.041601424 | 0.297596508 | 0.529386946 | Ccdc57 | 409.207547 | -0.236815458 | 0.079224019 | -2.989187633 | 0.002797203 | 0.089894182 |
| D6Ertd474e | 19.21335687 | -0.174936944 | 0.167980232 | -1.041413873 | 0.297683507 | 0.529480915 | Tgfbr2 | 1031.626482 | -0.236876175 | 0.078666201 | -3.011155652 | 0.002602554 | 0.085919773 |
| Clasp1 | 5144.145952 | 0.117042384 | 0.088349388 | 1.324767352 | 0.185248326 | 0.396605993 | Col12a1 | 509.5265694 | -0.236981174 | 0.085609382 | -2.768168268 | 0.005637234 | 0.141672083 |
| Ranbp17 | 253.1732783 | 0.1169966 | 0.106920192 | 1.094242334 | 0.273848709 | 0.502142128 | Dock8 | 268.5886384 | -0.237042793 | 0.077887418 | -3.043402878 | 0.002339189 | 0.081484567 |
| Has2 | 12.76859917 | -0.170023531 | 0.163308257 | -1.041120233 | 0.29781975 | 0.529540878 | Parp10 | 124.8363507 | -0.237077712 | 0.09346547 | -2.536527253 | 0.011195803 | 0.200580121 |
| Fam105b | 547.5202546 | 0.116969845 | 0.087736652 | 1.333192493 | 0.182468642 | 0.393208504 | H2-DMa | 518.8129695 | -0.237110889 | 0.089126727 | -2.660379179 | 0.007805273 | 0.169708597 |
| Galt | 1601.633693 | 0.09190827 | 0.088333384 | 1.040470379 | 0.298121419 | 0.529836392 | Atxn2l | 6339.308554 | -0.23720114 | 0.073402409 | -3.231517122 | 0.001231349 | 0.056070791 |
| Pex19 | 3357.952168 | -0.059436821 | 0.057117371 | -1.040608485 | 0.298057292 | 0.529836392 | 6430411K18Rik | 104.4736144 | -0.237247611 | 0.095093457 | -2.494888909 | 0.012599657 | 0.215067705 |
| Tmem167b | 2140.070685 | -0.058784323 | 0.056501432 | -1.040404129 | 0.298152184 | 0.529836392 | Rps6kb2 | 1092.403585 | -0.237334345 | 0.088037432 | -2.695834491 | 0.007021254 | 0.160386105 |
| Zfp947 | 60.76940665 | -0.156419527 | 0.150346447 | -1.040393906 | 0.298156932 | 0.529836392 | Ltk | 1366.09663 | -0.237681481 | 0.089055827 | -2.668904311 | 0.007609913 | 0.166981257 |
| Ccdc127 | 3301.711676 | -0.054274336 | 0.052185848 | -1.040020192 | 0.29833052 | 0.529906191 | Col22a1 | 103.2584293 | -0.237727008 | 0.094415773 | -2.517873862 | 0.011806559 | 0.206380365 |
| Lrfn3 | 951.3312787 | -0.081109977 | 0.077984177 | -1.04008248 | 0.298301583 | 0.529906191 | Col6a4 | 215.9328858 | -0.237808469 | 0.094383739 | -2.519591536 | 0.011749109 | 0.205623558 |
| Pmf1 | 90.89429907 | -0.133170712 | 0.128046936 | -1.040014825 | 0.298333013 | 0.529906191 | Itpripl2 | 384.0352659 | -0.238466899 | 0.06899824 | -3.456130169 | 0.000547991 | 0.033305907 |
| Maneal | 2588.990214 | 0.116968119 | 0.051491776 | 2.271588388 | 0.023111383 | 0.103853761 | Tle2 | 1388.643769 | -0.238534768 | 0.081703758 | -2.919507924 | 0.003505845 | 0.104182436 |
| Fam71e1 | 90.28755069 | 0.116949653 | 0.131725119 | 0.887831064 | 0.374631631 | 0.604136395 | Whamm | 762.3049476 | -0.238621139 | 0.067150767 | -3.553513218 | 0.000380122 | 0.026045536 |
| Brd1 | 2186.468735 | 0.07476211 | 0.071923259 | 1.039470559 | 0.298585944 | 0.530212546 | Il2rg | 45.16385225 | -0.238677774 | 0.09319052 | -2.561180825 | 0.010431705 | 0.195271829 |
| Slc2a3 | 7099.238062 | -0.087717485 | 0.084390573 | -1.039422788 | 0.29860815 | 0.530212546 | Akna | 261.7962092 | -0.238710088 | 0.088590193 | -2.694543053 | 0.007048522 | 0.160386105 |
| E2f7 | 18.63773243 | -0.174131358 | 0.167631201 | -1.038776531 | 0.298908679 | 0.530536947 | Gpr183 | 59.33522384 | -0.238754698 | 0.093507973 | -2.553308459 | 0.010670495 | 0.197200523 |
| Eri1 | 551.4673448 | 0.09985909 | 0.096142793 | 1.038653929 | 0.298965715 | 0.530536947 | Gpr84 | 47.82370797 | -0.239594799 | 0.079412813 | -3.017079862 | 0.002552226 | 0.08483668 |
| Fpgt | 885.2394797 | -0.079644988 | 0.076675203 | -1.038732022 | 0.298929384 | 0.530536947 | Arhgap4 | 202.3418634 | -0.240171575 | 0.091620522 | -2.621373121 | 0.008757635 | 0.17917382 |
| Sh3bgrl2 | 1284.155747 | -0.124403938 | 0.119761229 | -1.038766379 | 0.298913401 | 0.530536947 | Sfpi1 | 113.9206449 | -0.240239354 | 0.093105946 | -2.580279383 | 0.009872041 | 0.189935457 |
| Tbl3 | 1042.503587 | 0.081852281 | 0.078804693 | 1.038672675 | 0.298956993 | 0.530536947 | Ankzf1 | 1170.269729 | -0.240339319 | 0.076624123 | -3.136601249 | 0.001709184 | 0.067650159 |
| Vps36 | 523.2545972 | -0.082839167 | 0.079761321 | -1.038588198 | 0.298996297 | 0.530536947 | Tmem170 | 107.0557786 | -0.240591487 | 0.09373756 | -2.566649783 | 0.010268627 | 0.193928045 |
| Kif1a | 64998.7361 | -0.067076298 | 0.064636638 | -1.03774423 | 0.299389147 | 0.531173188 | Suv420h2 | 505.0905168 | -0.240709549 | 0.093023035 | -2.587633784 | 0.009663766 | 0.18766827 |
| 2310022B05Rik | 2743.952165 | 0.087462104 | 0.084313831 | 1.037339938 | 0.299577459 | 0.531244904 | Helq | 560.64088 | -0.240768188 | 0.07929622 | -3.036313543 | 0.002394902 | 0.082632645 |
| Cnot6 | 1373.409647 | -0.093430199 | 0.090053105 | -1.037501142 | 0.299502364 | 0.531244904 | Sox9 | 803.0536186 | -0.240774594 | 0.083755721 | -2.874724135 | 0.004043806 | 0.11423992 |
| Lancl1 | 7738.949675 | 0.116943511 | 0.069023971 | 1.694244898 | 0.090218756 | 0.253004419 | Catsperg1 | 39.27333461 | -0.240884247 | 0.087134483 | -2.764511124 | 0.005700816 | 0.142775962 |
| Naa50 | 2059.631266 | -0.08100948 | 0.078089274 | -1.037395734 | 0.299551466 | 0.531244904 | Malt1 | 316.9152143 | -0.241608789 | 0.075787264 | -3.187986675 | 0.001432671 | 0.061208778 |
| Phf5a | 689.710471 | -0.070722291 | 0.068188749 | -1.037154828 | 0.299663706 | 0.531244904 | Ikzf1 | 214.4058847 | -0.241613854 | 0.085395678 | -2.829345218 | 0.004664336 | 0.12664325 |
| Soga3 | 2329.752946 | -0.077486309 | 0.07469334 | -1.037392473 | 0.299552985 | 0.531244904 | 9930111J21Rik1 | 92.70673649 | -0.241698628 | 0.094897786 | -2.546936425 | 0.010867323 | 0.198764662 |
| Zcchc3 | 344.5151295 | -0.082685514 | 0.079724374 | -1.037142225 | 0.299669579 | 0.531244904 | Ftx | 1032.960593 | -0.242628039 | 0.084300563 | -2.878130721 | 0.004000393 | 0.113864147 |
| Dalrd3 | 2156.714379 | 0.11693443 | 0.110868521 | 1.054712633 | 0.291556772 | 0.521749609 | Col26a1 | 272.5388456 | -0.242653813 | 0.091269888 | -2.658640405 | 0.007845665 | 0.169996349 |
| Rom1 | 170.1285862 | 0.116924707 | 0.123978269 | 0.943106467 | 0.345626448 | 0.576948985 | Wdr90 | 1070.92114 | -0.242981421 | 0.091170334 | -2.665136889 | 0.007695699 | 0.167597786 |
| Mrpl41 | 1250.811805 | 0.097612001 | 0.094150194 | 1.036768985 | 0.299843533 | 0.531324702 | Srek1 | 1992.847244 | -0.243360762 | 0.063785791 | -3.815281689 | 0.000136028 | 0.012666269 |
| Nnmt | 19.29785546 | -0.174289747 | 0.168111428 | -1.036751334 | 0.299851761 | 0.531324702 | Setd4 | 246.8814842 | -0.243466157 | 0.088719137 | -2.744234966 | 0.006065211 | 0.148321973 |
| Rragc | 3104.88049 | 0.084810199 | 0.08182099 | 1.036533512 | 0.299953314 | 0.53144387 | Phkg1 | 631.6367996 | -0.243482631 | 0.068825881 | -3.53766096 | 0.000403688 | 0.02634142 |
| Rpl12 | 5007.443237 | -0.063586093 | 0.061356658 | -1.036335659 | 0.300045576 | 0.531546553 | Podn | 269.4063983 | -0.243540338 | 0.092563765 | -2.631054791 | 0.008512031 | 0.175384063 |
| Spa17 | 64.94272403 | -0.160552586 | 0.154945048 | -1.036190491 | 0.300113282 | 0.531605715 | Mbd6 | 1973.830687 | -0.243737233 | 0.088210659 | -2.763126748 | 0.005725053 | 0.143136179 |
| Aim1 | 58.24524131 | -0.172799704 | 0.166797706 | -1.035983697 | 0.300209748 | 0.531655028 | Vwa5b2 | 2981.755551 | -0.244183351 | 0.083924301 | -2.909566704 | 0.003619301 | 0.106640918 |
| Zfp964 | 72.86458967 | -0.158597476 | 0.153088317 | -1.035986802 | 0.3002083 | 0.531655028 | Rxfp1 | 308.2707309 | -0.244209684 | 0.083650935 | -2.919389763 | 0.003507174 | 0.104182436 |
| Exosc10 | 1608.586723 | 0.116924178 | 0.067819146 | 1.724058538 | 0.084697276 | 0.242973448 | Slc39a2 | 289.17254 | -0.244350696 | 0.09450791 | -2.585505239 | 0.00972364 | 0.18832745 |
| Fam43a | 494.6485382 | 0.11692086 | 0.108693652 | 1.075691707 | 0.282065157 | 0.511926619 | Neil1 | 324.7003463 | -0.244706815 | 0.087366242 | -2.800930997 | 0.005095541 | 0.134333636 |
| Gm2762 | 14.25800011 | 0.116720984 | 0.167226569 | 0.6979811 | 0.485189014 | 0.697149058 | Dna2 | 68.11805203 | -0.244929532 | 0.095088774 | -2.5757986 | 0.010000888 | 0.191148551 |
| Ano6 | 1219.189938 | 0.095973522 | 0.092740257 | 1.034863668 | 0.300732584 | 0.53233756 | Icam1 | 131.7586345 | -0.245009191 | 0.092213383 | -2.656980821 | 0.007884393 | 0.169996349 |
| Kbtbd12 | 12.05431378 | -0.168943012 | 0.163295806 | -1.034582675 | 0.300863848 | 0.532509078 | Gm17296 | 287.682353 | -0.245020883 | 0.08536563 | -2.870252141 | 0.004101446 | 0.115138402 |
| Phf13 | 673.1897173 | 0.116667244 | 0.097354899 | 1.198370545 | 0.230772794 | 0.453668953 | Npr1 | 157.8236296 | -0.245299625 | 0.094423366 | -2.597869954 | 0.009380401 | 0.184134741 |
| Peli3 | 555.9250354 | 0.072759787 | 0.070396504 | 1.033571028 | 0.301336749 | 0.533163366 | Gigyf1 | 3159.268209 | -0.246002024 | 0.073369346 | -3.352926508 | 0.000799619 | 0.042391499 |
| Sdha | 16296.74015 | -0.038967777 | 0.037700589 | -1.033611879 | 0.301317644 | 0.533163366 | Capn3 | 227.9967817 | -0.246069506 | 0.089393354 | -2.752659946 | 0.005911326 | 0.146477633 |
| Ticam1 | 247.8635776 | 0.096963592 | 0.093826039 | 1.033440109 | 0.301397985 | 0.533210821 | B930059L03Rik | 34.65876254 | -0.246100398 | 0.091413168 | -2.692176653 | 0.007098733 | 0.160867708 |
| Msh3 | 933.6717271 | 0.116664109 | 0.103143382 | 1.131086713 | 0.258018596 | 0.48433973 | Csf1r | 5157.77762 | -0.246104525 | 0.064800532 | -3.797878143 | 0.00014594 | 0.01286942 |
| Narf | 2217.128149 | 0.116637657 | 0.082192767 | 1.419074471 | 0.155877306 | 0.358209939 | Tbc1d10c | 54.36077467 | -0.246124185 | 0.0929983 | -2.646544991 | 0.008131869 | 0.173201649 |
| Prkcq | 449.4749496 | 0.116619672 | 0.10856105 | 1.074231246 | 0.282719051 | 0.512513653 | Rtp4 | 98.62329338 | -0.246128213 | 0.091067416 | -2.702703384 | 0.006877809 | 0.15963226 |
| Rbfa | 854.4184493 | 0.076721498 | 0.074318928 | 1.032327834 | 0.301918566 | 0.53382703 | Sned1 | 271.1566463 | -0.246544825 | 0.090979715 | -2.70988785 | 0.006730596 | 0.157437424 |
| Atp9b | 2880.378434 | 0.116586739 | 0.046405772 | 2.512332701 | 0.011993596 | 0.066761405 | Fgr | 14.5826347 | -0.246762036 | 0.056092026 | -4.399235571 | 1.09E-05 | NA |
| Cdc42bpb | 7875.405969 | 0.077269899 | 0.074867645 | 1.03208667 | 0.302031518 | 0.533904888 | AF357359 | 237.315513 | -0.247114819 | 0.09510274 | -2.598398532 | 0.009365972 | 0.184100285 |
| Cox15 | 1121.951279 | -0.060251962 | 0.058375778 | -1.032139754 | 0.302006653 | 0.533904888 | Fcer1g | 369.1126506 | -0.247133576 | 0.085676106 | -2.88450989 | 0.003920234 | 0.112763021 |
| Ablim2 | 6340.608985 | 0.116557933 | 0.078094013 | 1.492533525 | 0.135559335 | 0.327770965 | Pxn | 1335.464942 | -0.247198013 | 0.077870352 | -3.174481773 | 0.001501043 | 0.063017784 |
| 2610204G22Rik | 21.89452361 | -0.173355629 | 0.168020589 | -1.031752297 | 0.302188172 | 0.534025881 | Tyms | 156.8724868 | -0.247260953 | 0.088662792 | -2.788779234 | 0.005290712 | 0.137476727 |
| B230217C12Rik | 1065.703605 | 0.078573397 | 0.076157685 | 1.031719874 | 0.302203364 | 0.534025881 | Celsr3 | 5384.476084 | -0.247297383 | 0.079090577 | -3.126761648 | 0.001767432 | 0.068830324 |
| Cebpb | 70.74134214 | 0.116455164 | 0.157362168 | 0.740045499 | 0.459272387 | 0.676917731 | Gabpb2 | 1054.44523 | -0.247381649 | 0.067933478 | -3.641527797 | 0.000271025 | 0.020563301 |
| Tnfsf12 | 146.4199015 | -0.12575955 | 0.122040992 | -1.030469751 | 0.302789545 | 0.534939705 | Coro6 | 1507.263176 | -0.247682902 | 0.087969602 | -2.815551008 | 0.004869367 | 0.130262283 |
| Pthlh | 98.13622443 | -0.145384087 | 0.141098824 | -1.030370655 | 0.302836043 | 0.534960854 | Plau | 174.3059733 | -0.247775111 | 0.09496083 | -2.609234894 | 0.009074493 | 0.181815287 |
| Mpp2 | 7689.336452 | -0.079835437 | 0.077493839 | -1.03021657 | 0.302908353 | 0.53502759 | Zbed6 | 1549.130725 | -0.247852223 | 0.084165251 | -2.94482842 | 0.003231338 | 0.099461865 |
| Usp19 | 5607.861882 | 0.116414079 | 0.069027883 | 1.686479052 | 0.091703566 | 0.255236649 | Ddb2 | 496.9790336 | -0.248035062 | 0.070189406 | -3.533796284 | 0.000409637 | 0.02634142 |
| Tmem63b | 5132.135135 | 0.116352341 | 0.046563764 | 2.498774383 | 0.012462362 | 0.068450002 | Pcgf6 | 512.1137082 | -0.248115472 | 0.084178058 | -2.947507659 | 0.003203468 | 0.099007614 |
| Mb21d2 | 1650.162609 | 0.116339475 | 0.06579615 | 1.768180589 | 0.077030718 | 0.228574612 | Cd68 | 508.8635483 | -0.24895297 | 0.093337216 | -2.667242284 | 0.007647652 | 0.1673039 |
| Rusc1 | 6695.074574 | 0.077030951 | 0.074833847 | 1.029359766 | 0.303310648 | 0.535503799 | Tyk2 | 837.052247 | -0.249351693 | 0.078840646 | -3.162730199 | 0.001562971 | 0.064431734 |
| A930011G23Rik | 22.28760303 | -0.1724977 | 0.167626678 | -1.029058749 | 0.303452069 | 0.535633715 | Rras2 | 417.6854411 | -0.249427801 | 0.083701049 | -2.979984181 | 0.002882633 | 0.092025302 |
| Jmjd1c | 3415.37995 | -0.065656916 | 0.06380379 | -1.029044135 | 0.303458936 | 0.535633715 | Lama4 | 382.7683066 | -0.249453808 | 0.093552621 | -2.6664545 | 0.007665598 | 0.16744433 |
| 2810428I15Rik | 750.6016863 | -0.094722807 | 0.092056787 | -1.028960605 | 0.303498187 | 0.535641977 | L3mbtl1 | 424.6778311 | -0.24953684 | 0.092969964 | -2.684058699 | 0.007273433 | 0.162353883 |
| Zmpste24 | 1253.062325 | -0.069347227 | 0.067401166 | -1.028872803 | 0.30353945 | 0.535653786 | Guf1 | 1262.572621 | -0.249742998 | 0.064373343 | -3.879602734 | 0.000104627 | 0.010778828 |
| Pck2 | 826.6243246 | 0.0852042 | 0.082832129 | 1.028637085 | 0.303650245 | 0.53578828 | Parvg | 313.3323539 | -0.249760824 | 0.09184095 | -2.719493018 | 0.006538208 | 0.155695086 |
| Wdr46 | 333.1147059 | 0.11630262 | 0.090086718 | 1.29100741 | 0.196701108 | 0.411732505 | Pdlim7 | 1719.583252 | -0.249811815 | 0.079144106 | -3.156417161 | 0.001597202 | 0.06480716 |
| Txndc16 | 2946.69625 | -0.05471531 | 0.053205285 | -1.028381108 | 0.303770591 | 0.535930122 | Fam193b | 3745.83994 | -0.250332017 | 0.083403077 | -3.001472207 | 0.002686776 | 0.087506957 |
| 1810011H11Rik | 23.24774067 | -0.171906427 | 0.167345425 | -1.02725501 | 0.304300401 | 0.536752096 | Clk4 | 1725.260698 | -0.25033265 | 0.087095754 | -2.874223346 | 0.004050224 | 0.11423992 |
| Slc35f6 | 1440.670803 | 0.065485305 | 0.063757818 | 1.027094504 | 0.304375966 | 0.536824271 | Nsun5 | 412.8039845 | -0.250481611 | 0.084743907 | -2.955747743 | 0.003119121 | 0.09743731 |
| Ror1 | 47.42496023 | 0.116256765 | 0.148436816 | 0.783207081 | 0.433505517 | 0.654541515 | Cd9 | 1411.09227 | -0.250557981 | 0.086699028 | -2.88997451 | 0.003852731 | 0.111261959 |
| Pycr1 | 35.7755793 | -0.172566614 | 0.168091405 | -1.026623667 | 0.304597705 | 0.537031958 | Hmha1 | 351.1610005 | -0.250629841 | 0.08228995 | -3.045691981 | 0.002321455 | 0.081452798 |
| Trmt61b | 278.5082268 | -0.086996514 | 0.084736018 | -1.026676915 | 0.304572623 | 0.537031958 | Ccl9 | 83.63069379 | -0.250637718 | 0.092380745 | -2.713094788 | 0.006665804 | 0.156678744 |
| Ccdc164 | 665.6362659 | 0.074260724 | 0.07236679 | 1.026171315 | 0.30481084 | 0.537102144 | Snrnp70 | 6880.890335 | -0.250763748 | 0.076864113 | -3.26242949 | 0.001104616 | 0.051593759 |
| 9430021M05Rik | 571.9625612 | 0.11622468 | 0.066943221 | 1.736168037 | 0.082534127 | 0.239382959 | Alkbh6 | 956.9097166 | -0.250987433 | 0.056456655 | -4.445666037 | 8.76E-06 | 0.001844591 |
| Kcnj11 | 1972.834453 | 0.084349306 | 0.082175516 | 1.026453015 | 0.3046781 | 0.537102144 | Pask | 174.7327919 | -0.251374594 | 0.090209788 | -2.786555641 | 0.005327147 | 0.137476727 |
| Loh12cr1 | 1183.854724 | 0.082881373 | 0.080753826 | 1.026346088 | 0.304728481 | 0.537102144 | Clk1 | 3345.062499 | -0.251571627 | 0.088913607 | -2.829393997 | 0.004663625 | 0.12664325 |
| Tspyl5 | 2036.013803 | -0.067637533 | 0.065911633 | -1.026185048 | 0.304804368 | 0.537102144 | Pcgf2 | 1083.254423 | -0.251729642 | 0.081930619 | -3.072473331 | 0.002122928 | 0.077094131 |
| Jam3 | 1129.693405 | 0.071629019 | 0.069846394 | 1.025522074 | 0.305116916 | 0.537580338 | 2810459M11Rik | 465.0791344 | -0.251800456 | 0.080991576 | -3.108970954 | 0.001877402 | 0.071204015 |
| Igsf10 | 178.454035 | -0.14291475 | 0.139432401 | -1.02497518 | 0.305374899 | 0.537895292 | Thbs3 | 663.4589144 | -0.251884613 | 0.090940505 | -2.769773642 | 0.005609526 | 0.141672083 |
| Prmt8 | 3775.69855 | -0.078096094 | 0.076202553 | -1.02484879 | 0.305434541 | 0.537895292 | Mcmdc2 | 96.01463404 | -0.252422979 | 0.095072004 | -2.655071625 | 0.007929157 | 0.170584808 |
| Rab11b | 8795.318171 | 0.052832795 | 0.051551451 | 1.024855621 | 0.305431317 | 0.537895292 | Oas2 | 39.36332655 | -0.252492448 | 0.080118767 | -3.151476949 | 0.00162447 | 0.065650881 |
| Zeb2 | 5077.629658 | -0.061970956 | 0.060455842 | -1.025061498 | 0.305334171 | 0.537895292 | Safb2 | 2395.916686 | -0.252803115 | 0.073936122 | -3.419209812 | 0.000628033 | 0.035775704 |
| Cdc20 | 43.03935707 | -0.160043097 | 0.156196562 | -1.024626247 | 0.305539575 | 0.538019113 | Il1a | 38.04261118 | -0.252822232 | 0.089833652 | -2.814337675 | 0.004887786 | 0.130514657 |
| Lasp1 | 6573.675095 | -0.090442132 | 0.088299833 | -1.024261642 | 0.30571171 | 0.53826105 | Bcl3 | 16.53613191 | -0.25293191 | 0.074647712 | -3.3883411 | 0.000703168 | NA |
| Kdelr1 | 2049.578652 | -0.052421481 | 0.051186536 | -1.024126359 | 0.305775596 | 0.53831236 | Pot1b | 278.3682234 | -0.252954631 | 0.083357733 | -3.034567044 | 0.002408812 | 0.082915651 |
| 2410018M08Rik | 202.234848 | 0.116202357 | 0.10391667 | 1.118226334 | 0.26347034 | 0.490790797 | Treml2 | 27.30688501 | -0.253273196 | 0.087896865 | -2.881481551 | 0.003958104 | 0.113403194 |
| Cacna2d3 | 5020.281322 | -0.069244524 | 0.067661122 | -1.023401946 | 0.30611784 | 0.538792436 | Rtel1 | 898.9632671 | -0.253986297 | 0.082936714 | -3.062410884 | 0.002195618 | 0.07914033 |
| Coro7 | 2969.545098 | -0.053459663 | 0.052270785 | -1.022744594 | 0.306428621 | 0.539175836 | Shkbp1 | 577.3892483 | -0.254112675 | 0.090623457 | -2.804049687 | 0.005046513 | 0.133434166 |
| Cstf2t | 3262.027784 | -0.055004182 | 0.053776915 | -1.022821441 | 0.306392278 | 0.539175836 | Spata32 | 31.85378847 | -0.254335599 | 0.087987827 | -2.890577113 | 0.003845352 | 0.111261959 |
| Smc4 | 260.7716712 | 0.093878977 | 0.091793395 | 1.022720389 | 0.306440068 | 0.539175836 | Ccl12 | 24.89050196 | -0.254399684 | 0.066972986 | -3.798541749 | 0.00014555 | 0.01286942 |
| Sntg1 | 851.5365817 | 0.087164691 | 0.085254996 | 1.022399804 | 0.306591713 | 0.539381402 | Ogt | 12777.40456 | -0.254510356 | 0.059828164 | -4.254022501 | 2.10E-05 | 0.003426899 |
| Cdo1 | 1087.267559 | -0.129061249 | 0.12627095 | -1.022097713 | 0.306734656 | 0.539510359 | Wsb1 | 3248.782231 | -0.254632596 | 0.065064585 | -3.913535991 | 9.10E-05 | 0.009786687 |
| Ttc19 | 3180.604789 | -0.068336332 | 0.066858899 | -1.022097781 | 0.306734623 | 0.539510359 | Sfxn2 | 252.7220463 | -0.254849121 | 0.084710553 | -3.00846957 | 0.002625671 | 0.086290716 |
| Ctage5 | 1853.612535 | 0.081111865 | 0.079400244 | 1.021556865 | 0.306990682 | 0.539605236 | Stx2 | 647.6704684 | -0.255113444 | 0.078320777 | -3.257289496 | 0.001124817 | 0.052201549 |
| Focad | 1570.081249 | 0.061097212 | 0.05981159 | 1.021494532 | 0.307020198 | 0.539605236 | Dot1l | 2620.972567 | -0.255160151 | 0.082020821 | -3.110919243 | 0.001865059 | 0.070956043 |
| Rock2 | 10323.25194 | -0.093841596 | 0.091835502 | -1.021844423 | 0.30685454 | 0.539605236 | Efcab12 | 264.9584202 | -0.255274517 | 0.084250693 | -3.02993966 | 0.002446026 | 0.083252314 |
| Sbds | 2180.472258 | 0.063441463 | 0.062100151 | 1.021599174 | 0.306970649 | 0.539605236 | Zfp932 | 1563.847263 | -0.255446771 | 0.073008618 | -3.498857761 | 0.000467256 | 0.029510249 |
| Sec13 | 1589.475292 | -0.057353256 | 0.056139614 | -1.02161829 | 0.306961597 | 0.539605236 | Fcgr3 | 628.8817501 | -0.256009567 | 0.08067982 | -3.173154918 | 0.00150792 | 0.063124068 |
| Sntg2 | 123.3358075 | -0.129321219 | 0.1266032 | -1.021468799 | 0.307032384 | 0.539605236 | Pnn | 3176.347877 | -0.256077572 | 0.063766327 | -4.015874594 | 5.92E-05 | 0.007229522 |
| Usp13 | 1088.398101 | 0.116160892 | 0.072177716 | 1.609373343 | 0.107534731 | 0.284130818 | Pisd-ps2 | 1040.580608 | -0.256181035 | 0.091937041 | -2.78648337 | 0.005328335 | 0.137476727 |
| Senp8 | 246.4874988 | -0.133538199 | 0.130760775 | -1.021240496 | 0.30714051 | 0.539734044 | Adamts16 | 174.0048673 | -0.256181527 | 0.093028484 | -2.753796655 | 0.005890836 | 0.14630719 |
| Myadm | 2615.153929 | -0.08102286 | 0.079348378 | -1.02110292 | 0.30720568 | 0.539787344 | Tmem181b-ps | 10365.76073 | -0.256282078 | 0.067519387 | -3.795681358 | 0.000147239 | 0.01286942 |
| Mfsd4 | 2043.72753 | 0.116119753 | 0.073580506 | 1.578132026 | 0.114535282 | 0.296434962 | Atxn7l2 | 934.6593169 | -0.256599871 | 0.081744045 | -3.139064989 | 0.001694879 | 0.067373669 |
| Tfcp2 | 800.3342327 | 0.071489512 | 0.070041223 | 1.020677668 | 0.307407178 | 0.54000629 | Dennd1c | 68.41112044 | -0.256670652 | 0.095077007 | -2.699608043 | 0.006942121 | 0.160386105 |
| Kdm8 | 197.4295471 | 0.116074232 | 0.113698756 | 1.020892717 | 0.30730527 | 0.539901106 | Uggt2 | 1319.686242 | -0.256895803 | 0.063564951 | -4.041469393 | 5.31E-05 | 0.006889126 |
| A530072M11Rik | 26.08005159 | 0.116062129 | 0.166381569 | 0.697566021 | 0.485448638 | 0.697149058 | Mirg | 688.173883 | -0.257060315 | 0.086890539 | -2.958438495 | 0.003092019 | 0.097207883 |
| Eid2b | 880.905984 | -0.090801327 | 0.089007419 | -1.020154591 | 0.307655149 | 0.540141653 | Spint1 | 162.9265907 | -0.257305744 | 0.094365027 | -2.726706617 | 0.006396989 | 0.154049193 |
| Frs3 | 661.4379201 | -0.08015175 | 0.078576006 | -1.020053761 | 0.307702965 | 0.540141653 | Stat2 | 1401.135806 | -0.257316014 | 0.077159072 | -3.33487697 | 0.000853371 | 0.044114142 |
| Gak | 5243.514958 | 0.082640413 | 0.081020392 | 1.019995223 | 0.307730726 | 0.540141653 | Gdap10 | 76.15709017 | -0.257356167 | 0.09501322 | -2.708635371 | 0.006756054 | 0.157778854 |
| Olfm3 | 950.2529625 | -0.098492834 | 0.096573435 | -1.019875017 | 0.30778774 | 0.540141653 | Fgd2 | 232.4174123 | -0.257720814 | 0.082697332 | -3.116434441 | 0.001830524 | 0.070354982 |
| Tmem222 | 2504.715506 | -0.072026698 | 0.070607211 | -1.020103991 | 0.307679144 | 0.540141653 | Oas1b | 30.76809102 | -0.258322771 | 0.082972391 | -3.113358166 | 0.001849714 | 0.070715853 |
| Uqcc | 2140.230045 | -0.051676576 | 0.0506635 | -1.019996175 | 0.307730275 | 0.540141653 | Pabpn1 | 5687.946795 | -0.259533746 | 0.072217126 | -3.593797747 | 0.000325893 | 0.023585654 |
| Uxs1 | 1469.772436 | -0.092311 | 0.090512643 | -1.019868564 | 0.3077908 | 0.540141653 | Odf2l | 439.5629013 | -0.260288303 | 0.067704988 | -3.844447971 | 0.000120824 | 0.011688418 |
| Mxd4 | 1037.146477 | -0.081546294 | 0.079998291 | -1.01935046 | 0.308036616 | 0.540511815 | Flnb | 3768.40108 | -0.260646137 | 0.078331368 | -3.327480991 | 0.00087635 | 0.044373556 |
| Cbx3 | 3375.00603 | -0.070724001 | 0.069406183 | -1.018987043 | 0.308209118 | 0.540692037 | Fcgr2b | 159.8117014 | -0.261007002 | 0.094946995 | -2.7489759 | 0.005978178 | 0.147434662 |
| Tbc1d14 | 4057.6187 | 0.06526454 | 0.064047359 | 1.019004383 | 0.308200886 | 0.540692037 | Myo19 | 446.3690144 | -0.261027587 | 0.090725692 | -2.877107702 | 0.004013386 | 0.113864147 |
| Braf | 5522.824927 | -0.099331544 | 0.097512324 | -1.018656302 | 0.308366165 | 0.540783855 | Jak3 | 263.8473183 | -0.261065993 | 0.088977701 | -2.934060885 | 0.003345586 | 0.101245788 |
| Plcb1 | 11559.21202 | -0.080304932 | 0.078830956 | -1.018697942 | 0.30834639 | 0.540783855 | Hps1 | 583.6512228 | -0.261323616 | 0.087558891 | -2.984546889 | 0.002839987 | 0.090867064 |
| Sec22c | 2109.11264 | -0.06611504 | 0.064901112 | -1.018704265 | 0.308343387 | 0.540783855 | Zcchc7 | 841.2395217 | -0.261415532 | 0.064952317 | -4.024729909 | 5.70E-05 | 0.007179599 |
| 9930111J21Rik1 | 101.6560598 | -0.146997172 | 0.144329255 | -1.018484939 | 0.308447554 | 0.54086537 | Ccdc176 | 185.7948878 | -0.261700869 | 0.090811882 | -2.881791065 | 0.003954218 | 0.113403194 |
| Bnipl | 8.470775996 | -0.157050817 | 0.154311912 | -1.017749145 | 0.308797186 | 0.541417177 | Pik3ap1 | 130.5533675 | -0.262719738 | 0.093866894 | -2.798854051 | 0.005128431 | 0.134955782 |
| Surf1 | 1918.790619 | 0.116056145 | 0.07049086 | 1.646399904 | 0.099681453 | 0.26954004 | Clasrp | 1776.435166 | -0.263060674 | 0.084087934 | -3.128399781 | 0.001757609 | 0.068703774 |
| Tefm | 149.5189205 | -0.117759211 | 0.115749721 | -1.01736064 | 0.308981899 | 0.541618457 | Clec5a | 105.5540405 | -0.263355223 | 0.094441514 | -2.788553589 | 0.005294399 | 0.137476727 |
| Tk2 | 1575.91435 | 0.116025761 | 0.050179674 | 2.312206355 | 0.020766314 | 0.096506681 | C3ar1 | 141.5368275 | -0.263384808 | 0.094192736 | -2.796232696 | 0.005170216 | 0.135809327 |
| Mapk8ip3 | 21408.68188 | 0.067402232 | 0.066278643 | 1.016952492 | 0.30917603 | 0.54172842 | Osmr | 216.6755115 | -0.263527621 | 0.094414246 | -2.791184944 | 0.005251546 | 0.137448561 |
| Epb4.1 | 1302.421956 | 0.116025431 | 0.102969113 | 1.126798394 | 0.25982773 | 0.486583408 | D4Wsu53e | 7279.580763 | -0.263531719 | 0.076534137 | -3.443322529 | 0.000574614 | 0.033792862 |
| Nkain4 | 756.8624741 | -0.078117337 | 0.076822023 | -1.01686123 | 0.309219449 | 0.54172842 | Szt2 | 2505.216463 | -0.263936532 | 0.069257419 | -3.810949592 | 0.000138434 | 0.012752174 |
| Prdx6 | 7950.481737 | 0.082979582 | 0.081590947 | 1.017019466 | 0.309144169 | 0.54172842 | Acp6 | 469.0758991 | -0.263966463 | 0.069479215 | -3.799214803 | 0.000145155 | 0.01286942 |
| Ssu72 | 1939.082096 | -0.063124309 | 0.062112097 | -1.016296531 | 0.3094882 | 0.542076664 | Arntl2 | 199.7486263 | -0.264138105 | 0.087891637 | -3.005270072 | 0.002653451 | 0.087006847 |
| Ubxn7 | 1711.513803 | -0.06587598 | 0.064817321 | -1.016332952 | 0.309470862 | 0.542076664 | Aanat | 24.11580607 | -0.264234753 | 0.083400104 | -3.168278457 | 0.001533446 | 0.063824735 |
| Mal2 | 5768.464621 | -0.105939136 | 0.104271622 | -1.015992018 | 0.309633188 | 0.542269313 | Arhgap9 | 34.16223775 | -0.264267597 | 0.090825932 | -2.90960513 | 0.003618857 | 0.106640918 |
| 2210408I21Rik | 382.71195 | -0.108460451 | 0.106765543 | -1.015875045 | 0.309688894 | 0.542305575 | Akap8 | 3390.384067 | -0.264554828 | 0.066471048 | -3.980000851 | 6.89E-05 | 0.008130208 |
| Dars2 | 774.7369334 | 0.116000611 | 0.068848229 | 1.684874286 | 0.092012829 | 0.255766056 | Lrrc45 | 2405.997923 | -0.264621232 | 0.08667261 | -3.053112518 | 0.00226481 | 0.080316218 |
| Gadd45a | 184.921943 | 0.115939255 | 0.149673258 | 0.774615698 | 0.438566779 | 0.658955942 | Trank1 | 19038.44546 | -0.26506235 | 0.065147328 | -4.068660319 | 4.73E-05 | 0.006479723 |
| Mrpl38 | 1275.735991 | 0.115894706 | 0.07585595 | 1.527826181 | 0.1265557 | 0.314473278 | Irf7 | 125.0842672 | -0.265219485 | 0.088456546 | -2.998302536 | 0.00271488 | 0.087924428 |
| Smarca1 | 1076.946111 | -0.102600523 | 0.101201852 | -1.013820609 | 0.310668359 | 0.543774916 | Tnfrsf14 | 22.64225463 | -0.265237028 | 0.080729996 | -3.285482967 | 0.001018077 | 0.048487168 |
| AI414108 | 1894.537704 | 0.115877987 | 0.100101058 | 1.157610007 | 0.247023224 | 0.472049559 | 4930422G04Rik | 225.466564 | -0.265571608 | 0.08639964 | -3.073758284 | 0.002113806 | 0.077094131 |
| Zbtb5 | 468.1316776 | -0.083617425 | 0.082501528 | -1.013525777 | 0.31080909 | 0.543898356 | Invs | 617.0892343 | -0.26642356 | 0.074301179 | -3.585724509 | 0.000336144 | 0.02373572 |
| Ccdc93 | 1386.479814 | 0.063966861 | 0.063127539 | 1.013295651 | 0.310918964 | 0.543987214 | Pou2f2 | 228.4269991 | -0.266457316 | 0.087145769 | -3.05760472 | 0.002231137 | 0.079435028 |
| Acot4 | 42.29219627 | 0.11581185 | 0.162013436 | 0.714828679 | 0.474714911 | 0.688950247 | Tlr1 | 35.53224131 | -0.266601472 | 0.087566284 | -3.04456762 | 0.00233015 | 0.081484567 |
| Syne3 | 92.54021392 | 0.115653041 | 0.153780622 | 0.752065112 | 0.452011903 | 0.670702205 | Bbc3 | 153.8244273 | -0.266823266 | 0.0942349 | -2.831469715 | 0.004633462 | 0.126277056 |
| Cflar | 972.3233144 | -0.098995561 | 0.097724179 | -1.013009902 | 0.311055431 | 0.54402222 | Mov10 | 381.4679661 | -0.266895199 | 0.080212208 | -3.327363801 | 0.000876718 | 0.044373556 |
| Pfkl | 6657.9081 | -0.098207655 | 0.096945497 | -1.013019251 | 0.311050966 | 0.54402222 | Ankrd16 | 744.2242263 | -0.266965577 | 0.071210013 | -3.748989321 | 0.000177549 | 0.014737552 |
| Pm20d1 | 103.7336959 | -0.153249659 | 0.151300735 | -1.012881128 | 0.311116944 | 0.544068389 | Neurl3 | 42.74228278 | -0.267719191 | 0.091175373 | -2.936310341 | 0.003321419 | 0.100724271 |
| Ank | 8933.881206 | 0.076336318 | 0.07537437 | 1.012762268 | 0.311173727 | 0.544106278 | Stxbp2 | 1700.902879 | -0.268101622 | 0.083196342 | -3.222516938 | 0.001270697 | 0.057501987 |
| Nrbf2 | 722.0221238 | 0.069228841 | 0.068383119 | 1.012367398 | 0.31136242 | 0.544374784 | Izumo4 | 970.2414993 | -0.2683417 | 0.081593475 | -3.288764232 | 0.001006283 | 0.048083105 |
| Tmed4 | 2310.752443 | 0.065589141 | 0.064835846 | 1.011618493 | 0.3117205 | 0.544939347 | Atad2 | 475.1802548 | -0.269217618 | 0.083757321 | -3.214257749 | 0.001307822 | 0.058453616 |
| Eif1b | 2800.356998 | 0.082030262 | 0.081127931 | 1.01112233 | 0.311957884 | 0.545231298 | Srsf11 | 3374.870484 | -0.269779166 | 0.061629399 | -4.377442769 | 1.20E-05 | 0.002295106 |
| Fam63a | 909.364996 | 0.067744407 | 0.066996064 | 1.011169952 | 0.311935094 | 0.545231298 | Cep95 | 391.5742724 | -0.270339831 | 0.086106839 | -3.139586044 | 0.001691867 | 0.067373669 |
| Gtf3c2 | 2578.218854 | 0.115607465 | 0.051417213 | 2.248419506 | 0.024549453 | 0.108120869 | Slfn8 | 59.03905568 | -0.270452452 | 0.091412909 | -2.95858054 | 0.003090595 | 0.097207883 |
| 9930013L23Rik | 907.141211 | -0.098904596 | 0.097846185 | -1.010817096 | 0.312103979 | 0.545255252 | Itgam | 1199.284719 | -0.271064873 | 0.06026827 | -4.497638204 | 6.87E-06 | 0.001609867 |
| Copb1 | 2535.3313 | -0.057278923 | 0.056675186 | -1.010652586 | 0.312182738 | 0.545255252 | A330049N07Rik | 85.79137884 | -0.271157842 | 0.095051751 | -2.852739037 | 0.00433442 | 0.120385822 |
| Ddi2 | 494.9362502 | 0.11559289 | 0.093047499 | 1.242299799 | 0.214125969 | 0.433891886 | Zfp566 | 166.8703862 | -0.271485823 | 0.088352925 | -3.07274288 | 0.002121012 | 0.077094131 |
| Scmh1 | 1788.742641 | -0.04470194 | 0.044229338 | -1.010685262 | 0.312167093 | 0.545255252 | Smpd4 | 2883.067784 | -0.272007553 | 0.085289335 | -3.189232885 | 0.001426509 | 0.061125282 |
| Tab1 | 933.2343497 | -0.066752835 | 0.066038861 | -1.010811434 | 0.31210669 | 0.545255252 | Usp18 | 62.80808088 | -0.272928106 | 0.090447931 | -3.017516311 | 0.002548553 | 0.08483668 |
| 1700021F05Rik | 476.2414204 | -0.079031827 | 0.078209086 | -1.01051977 | 0.312246332 | 0.545304855 | Lgi4 | 1540.32414 | -0.272967291 | 0.067361369 | -4.052282431 | 5.07E-05 | 0.006697857 |
| Fam115a | 8457.355551 | -0.056667904 | 0.056087778 | -1.010343186 | 0.312330898 | 0.545391066 | Col6a1 | 4211.869628 | -0.273388479 | 0.094993557 | -2.877968653 | 0.004002449 | 0.113864147 |
| Arf4 | 3042.880835 | -0.069512884 | 0.068830013 | -1.009921116 | 0.312533085 | 0.545436767 | Anxa3 | 446.8128622 | -0.273900667 | 0.08284667 | -3.306115594 | 0.000945991 | 0.046581215 |
| Zkscan5 | 610.4804891 | 0.115587839 | 0.1044567 | 1.106562233 | 0.268483244 | 0.496464898 | Ifi44 | 51.24543977 | -0.274185061 | 0.068349867 | -4.011493685 | 6.03E-05 | 0.007303648 |
| Mylip | 589.5415802 | -0.074689961 | 0.073937799 | -1.0101729 | 0.31241246 | 0.545436767 | Olfr856-ps1 | 153.0915155 | -0.274381881 | 0.09080229 | -3.021750678 | 0.002513175 | 0.084486004 |
| Orc5 | 834.8111864 | 0.0814945 | 0.080689935 | 1.00997108 | 0.312509146 | 0.545436767 | Eml5 | 3035.789718 | -0.274741017 | 0.074902651 | -3.667974551 | 0.00024448 | 0.019300594 |
| Tubb4a | 28586.72965 | 0.079976394 | 0.079186134 | 1.009979774 | 0.31250498 | 0.545436767 | Tia1 | 2489.824 | -0.274761543 | 0.070198388 | -3.914071987 | 9.08E-05 | 0.009786687 |
| Aldoc | 30197.82806 | 0.055072745 | 0.054570676 | 1.009200343 | 0.312878562 | 0.545944885 | Cfp | 385.9824354 | -0.275665398 | 0.094098264 | -2.929548183 | 0.003394552 | 0.102281258 |
| Npc1 | 3406.395122 | 0.115573966 | 0.073236572 | 1.578090873 | 0.114544735 | 0.296434962 | 6720401G13Rik | 3516.251139 | -0.275941129 | 0.071802193 | -3.843073834 | 0.000121503 | 0.011688418 |
| Pcdhga6 | 774.3867739 | 0.115518954 | 0.105755336 | 1.092322694 | 0.274691289 | 0.503089429 | Akap8l | 5134.106955 | -0.276626135 | 0.075979628 | -3.640793469 | 0.000271799 | 0.020563301 |
| Stard7 | 4586.21541 | -0.046003591 | 0.04560974 | -1.008635233 | 0.313149604 | 0.546266602 | 4930570G19Rik | 315.11887 | -0.276880882 | 0.083584766 | -3.312575914 | 0.00092441 | 0.046144277 |
| Gm9962 | 65.13246285 | -0.155943688 | 0.154667504 | -1.008251143 | 0.313333911 | 0.546413192 | Myo5c | 227.4913003 | -0.277984134 | 0.095088898 | -2.923413136 | 0.003462168 | 0.103907963 |
| Vps8 | 2478.922151 | 0.115517912 | 0.061564686 | 1.876366478 | 0.060604974 | 0.195720863 | Sfi1 | 933.6173876 | -0.278193583 | 0.080048553 | -3.47531059 | 0.000510262 | 0.031560853 |
| Slc6a15 | 2374.12819 | 0.058244586 | 0.057769948 | 1.008216005 | 0.313350776 | 0.546413192 | Ctsh | 461.9397725 | -0.278565507 | 0.084172087 | -3.309476057 | 0.000934708 | 0.046498502 |
| Phtf1 | 1949.7097 | 0.115509282 | 0.049098763 | 2.352590448 | 0.018643155 | 0.09001466 | Ccdc39 | 737.3523386 | -0.279633732 | 0.071224576 | -3.926084913 | 8.63E-05 | 0.00950129 |
| Crot | 1317.053435 | -0.065255041 | 0.064762357 | -1.007607565 | 0.313642896 | 0.54681929 | Hpgds | 256.6455066 | -0.279889498 | 0.086924544 | -3.219913316 | 0.001282294 | 0.057667484 |
| Csrnp2 | 1194.565587 | 0.081863065 | 0.081270598 | 1.00729006 | 0.313795405 | 0.546931265 | Gm15446 | 180.683477 | -0.27990179 | 0.088666435 | -3.156795357 | 0.001595133 | 0.06480716 |
| Nkain1 | 2343.763054 | 0.094310489 | 0.093625649 | 1.007314662 | 0.313783586 | 0.546931265 | Srpk3 | 271.4553314 | -0.280244112 | 0.094063005 | -2.979323394 | 0.002888857 | 0.092025302 |
| Psmg3 | 298.9814814 | -0.103600751 | 0.102854706 | -1.007253393 | 0.313813021 | 0.546931265 | Dusp11 | 2594.484577 | -0.28060646 | 0.072892853 | -3.849574394 | 0.000118323 | 0.011600059 |
| Hspa14 | 966.3662692 | -0.055907426 | 0.055509579 | -1.007167173 | 0.313854445 | 0.546941938 | Arglu1 | 3327.755096 | -0.281284147 | 0.073094927 | -3.848203396 | 0.000118987 | 0.011600059 |
| Smad1 | 768.6985072 | 0.091320141 | 0.090718922 | 1.006627281 | 0.314113919 | 0.547332554 | Tnip2 | 571.8799493 | -0.281547767 | 0.071191261 | -3.954807996 | 7.66E-05 | 0.008715881 |
| Drp2 | 4164.108918 | 0.115476915 | 0.060452807 | 1.910199381 | 0.056107547 | 0.186952759 | Med12 | 1864.223624 | -0.281606892 | 0.069435385 | -4.055668342 | 5.00E-05 | 0.006662124 |
| Clta | 4951.280947 | -0.046246537 | 0.045963563 | -1.006156484 | 0.314340301 | 0.547542285 | Dhtkd1 | 247.3277377 | -0.28166477 | 0.090381742 | -3.116390147 | 0.001830799 | 0.070354982 |
| Ppt2 | 968.5547507 | 0.076943899 | 0.076472667 | 1.006162093 | 0.314337603 | 0.547542285 | Flna | 4352.157067 | -0.28175527 | 0.072073649 | -3.909268804 | 9.26E-05 | 0.009815749 |
| Ptprd | 8954.94578 | 0.115464711 | 0.063555474 | 1.816754762 | 0.069254672 | 0.213411276 | Pdcd1 | 10.8440444 | -0.283258051 | 0.062031425 | -4.566363738 | 4.96E-06 | NA |
| Zfp30 | 421.0433743 | 0.115404778 | 0.092199979 | 1.251679011 | 0.210686851 | 0.428959537 | Cx3cr1 | 2682.254466 | -0.284030971 | 0.066532321 | -4.269067559 | 1.96E-05 | 0.003240153 |
| Kifc2 | 23131.26605 | 0.086171913 | 0.085688188 | 1.005645173 | 0.314586286 | 0.547727297 | Plcb2 | 175.4814559 | -0.284286122 | 0.08859203 | -3.208935642 | 0.001332273 | 0.05918226 |
| Zar1l | 15.67516203 | -0.167345046 | 0.166406219 | -1.005641779 | 0.314587919 | 0.547727297 | Jag1 | 337.4349971 | -0.284412137 | 0.084281954 | -3.374531847 | 0.000739414 | 0.040227423 |
| Mettl5 | 197.4552752 | -0.110134903 | 0.109571325 | -1.005143482 | 0.314827765 | 0.548083303 | Slc2a9 | 34.33935671 | -0.284638589 | 0.091083415 | -3.125032033 | 0.001777857 | 0.069051205 |
| Ctdspl | 1551.811491 | 0.115394139 | 0.049389485 | 2.336411051 | 0.019469829 | 0.092452392 | Nvl | 1987.591374 | -0.284658405 | 0.065486481 | -4.346827031 | 1.38E-05 | 0.00249049 |
| Ly6h | 4382.958666 | -0.075728958 | 0.075393505 | -1.004449359 | 0.315162069 | 0.548542023 | Fam227a | 302.2116472 | -0.285192236 | 0.08483518 | -3.361721364 | 0.000774582 | 0.041703834 |
| Erlec1 | 1838.985182 | 0.115390291 | 0.061464274 | 1.877355463 | 0.060469391 | 0.195516016 | Col4a1 | 2174.644013 | -0.285208609 | 0.083399161 | -3.4198019 | 0.000626668 | 0.035775704 |
| Bax | 962.4061326 | -0.065063283 | 0.064794499 | -1.004148268 | 0.315307153 | 0.548643115 | Fmnl3 | 666.2656188 | -0.286021516 | 0.070118875 | -4.07909451 | 4.52E-05 | 0.006438643 |
| Krcc1 | 413.4035114 | -0.105323343 | 0.104892406 | -1.004108372 | 0.315326381 | 0.548643115 | Nckap1l | 737.8115118 | -0.286273932 | 0.05915232 | -4.839606122 | 1.30E-06 | 0.000429497 |
| Folh1 | 262.0021354 | 0.115378132 | 0.107423125 | 1.074053015 | 0.282798921 | 0.51252694 | Dock6 | 857.7042041 | -0.286578924 | 0.083091833 | -3.448942123 | 0.000562787 | 0.03358999 |
| Kalrn | 22849.18386 | 0.115375824 | 0.072123691 | 1.599693835 | 0.10966652 | 0.288151968 | Gbp7 | 262.0714339 | -0.287393345 | 0.093643402 | -3.069018638 | 0.002147632 | 0.077603237 |
| Tmem80 | 811.6868597 | 0.115297272 | 0.110164902 | 1.046588073 | 0.295289605 | 0.526855958 | Flt3l | 125.3329142 | -0.287470513 | 0.09178436 | -3.132020688 | 0.001736076 | 0.06834213 |
| Fam131b | 7869.792651 | -0.081399406 | 0.081116533 | -1.003487239 | 0.315625831 | 0.54876192 | Slc38a6 | 331.5049198 | -0.287498677 | 0.085666341 | -3.35602846 | 0.000790704 | 0.04208229 |
| Gngt2 | 64.09989949 | -0.157316928 | 0.156774004 | -1.003463095 | 0.315637475 | 0.54876192 | Zfc3h1 | 2588.245007 | -0.287546226 | 0.065033209 | -4.421529076 | 9.80E-06 | 0.001993526 |
| Mob3a | 455.0014859 | -0.076877388 | 0.07658975 | -1.003755569 | 0.315496445 | 0.54876192 | Arl5c | 73.40569053 | -0.288340422 | 0.095096497 | -3.032082463 | 0.002428728 | 0.083206862 |
| Ppp1r14b | 556.3094291 | -0.092210354 | 0.091899819 | -1.00337906 | 0.315678004 | 0.54876192 | Tnfrsf4 | 39.93369917 | -0.288490375 | 0.09052693 | -3.186790675 | 0.001438609 | 0.061282191 |
| Tor4a | 386.1379459 | -0.141134381 | 0.140651526 | -1.003432987 | 0.315651995 | 0.54876192 | Ctss | 3301.869585 | -0.289898852 | 0.081751095 | -3.546115818 | 0.000390954 | 0.02634142 |
| Slc7a8 | 3396.663295 | 0.11507678 | 0.072140829 | 1.595168532 | 0.110674554 | 0.289611811 | Tlr9 | 59.26938402 | -0.290225602 | 0.094319535 | -3.077046578 | 0.002090626 | 0.076882121 |
| Nup37 | 116.8913676 | -0.122477188 | 0.122118625 | -1.002936187 | 0.315891651 | 0.549010123 | Ccdc142 | 374.4284979 | -0.290483565 | 0.087374878 | -3.3245662 | 0.000885562 | 0.044665537 |
| Foxq1 | 140.5985014 | 0.115071749 | 0.125978491 | 0.913423777 | 0.361019705 | 0.591756282 | Atp11b | 3806.078508 | -0.290789677 | 0.059449809 | -4.891347509 | 1.00E-06 | 0.000376211 |
| Pot1b | 256.6620085 | 0.115011607 | 0.129566037 | 0.887667865 | 0.374719436 | 0.604215105 | Pion | 276.7235449 | -0.290858433 | 0.080502663 | -3.613028705 | 0.000302641 | 0.022091286 |
| Ushbp1 | 189.5071984 | 0.1150115 | 0.129669909 | 0.88695597 | 0.375102606 | 0.604492326 | AW549542 | 51.78402886 | -0.291047747 | 0.093561209 | -3.110773697 | 0.001865979 | 0.070956043 |
| Chordc1 | 1450.491861 | -0.088049555 | 0.087860721 | -1.002149235 | 0.316271521 | 0.549267267 | Col4a2 | 2917.356857 | -0.291063732 | 0.086697712 | -3.357225074 | 0.00078729 | 0.04208229 |
| Cited4 | 208.874593 | -0.113351887 | 0.113091824 | -1.002299579 | 0.316198926 | 0.549267267 | Cdc25b | 986.7786974 | -0.291871679 | 0.080056564 | -3.645818217 | 0.000266542 | 0.020485684 |
| Eif4ebp2 | 861.2109892 | 0.074729236 | 0.074576935 | 1.002042203 | 0.31632321 | 0.549267267 | I830077J02Rik | 30.73612512 | -0.292172572 | 0.087421865 | -3.342099497 | 0.000831473 | 0.043290214 |
| Ccp110 | 2983.730692 | 0.114993342 | 0.069821268 | 1.646967252 | 0.099564777 | 0.269443841 | Bmp1 | 3285.541053 | -0.292467475 | 0.08419283 | -3.473781269 | 0.000513179 | 0.031586614 |
| Stk16 | 1555.575337 | -0.090116962 | 0.089932484 | -1.00205129 | 0.316318822 | 0.549267267 | Rgs11 | 1593.992311 | -0.292876982 | 0.086404672 | -3.389596592 | 0.000699955 | 0.039105972 |
| Usp49 | 642.2855886 | 0.114959993 | 0.092288603 | 1.245657525 | 0.212890153 | 0.432401437 | Arhgef1 | 1855.373335 | -0.293716826 | 0.083002193 | -3.538663461 | 0.000402158 | 0.02634142 |
| Col25a1 | 2026.797708 | -0.113904606 | 0.113724126 | -1.001586998 | 0.316543103 | 0.549376037 | C330005M16Rik | 68.79526308 | -0.29468707 | 0.093831237 | -3.140607332 | 0.001685979 | 0.067373669 |
| Tppp3 | 1088.163581 | 0.114943019 | 0.085584086 | 1.343041971 | 0.179258384 | 0.389108174 | Tie1 | 491.7339399 | -0.294713485 | 0.064564513 | -4.564635782 | 5.00E-06 | 0.001213916 |

| | | | | | | |
|---|---|---|---|---|---|---|
| Rasa1 | 1842.721249 | -0.086068588 | 0.085922159 | -1.001704204 | 0.316486476 | 0.549376037 |
| Trpa1 | 11.4919667 | 0.114936191 | 0.153160181 | 0.750431285 | 0.452994995 | 0.671518227 |
| Rapgef1 | 7243.845863 | 0.114884867 | 0.122768703 | 0.935783017 | 0.349384917 | 0.580421397 |
| Rab22a | 1161.229521 | -0.066705367 | 0.066619767 | -1.001284911 | 0.316689086 | 0.549471522 |
| Nt5c3 | 1563.310431 | -0.062223811 | 0.062179799 | -1.000707824 | 0.316968084 | 0.549877862 |
| Pitpna | 13582.28152 | 0.074948357 | 0.074899395 | 1.000653697 | 0.31699426 | 0.549877862 |
| Galk2 | 826.3542025 | 0.074700651 | 0.074684486 | 1.000216448 | 0.317205771 | 0.550183165 |
| Trmt5 | 374.2889622 | 0.114788924 | 0.104717842 | 1.096173501 | 0.273002854 | 0.501430848 |
| Pde8b | 3689.303714 | 0.056620484 | 0.056656368 | 0.999366636 | 0.317617116 | 0.550773318 |
| Ddx19a | 704.1560766 | 0.095096287 | 0.095181652 | 0.999103141 | 0.31774473 | 0.550877097 |
| Spata5 | 478.7337254 | -0.07163996 | 0.071704764 | -0.999096243 | 0.317748071 | 0.550877097 |
| Usp30 | 1564.160514 | 0.114781631 | 0.059177243 | 1.939624524 | 0.052425338 | 0.179429687 |
| Arhgef15 | 1007.840861 | -0.098504274 | 0.098630497 | -0.99872024 | 0.317930233 | 0.55103906 |
| Zfp202 | 185.6069599 | -0.123680448 | 0.123843527 | -0.998683191 | 0.317948186 | 0.55103906 |
| Greb1l | 431.8401566 | 0.114688336 | 0.079162973 | 1.448762361 | 0.147403958 | 0.346662471 |
| Klhdc7b | 13.44270856 | 0.114654004 | 0.15954511 | 0.718630633 | 0.472368534 | 0.68686651 |
| Supt7l | 692.2647744 | -0.069431107 | 0.069588232 | -0.997742075 | 0.318404445 | 0.551644691 |
| 2810013P06Rik | 590.3418468 | -0.076898951 | 0.077084131 | -0.997597685 | 0.318474484 | 0.551704345 |
| Pim1 | 119.8554952 | -0.150106651 | 0.150487835 | -0.997467018 | 0.318537875 | 0.55175247 |
| 1810058I24Rik | 421.7928355 | 0.082251594 | 0.082497137 | 0.99702362 | 0.318753045 | 0.552030005 |
| Gng3 | 6920.720767 | 0.065998038 | 0.066199203 | 0.996961219 | 0.318783335 | 0.552030005 |
| Ripk4 | 25.50695218 | 0.114645106 | 0.16790055 | 0.682815546 | 0.494723405 | 0.704526143 |
| Pla2g12a | 515.1491366 | -0.074536233 | 0.074773849 | -0.996822213 | 0.318850814 | 0.552047661 |
| Hspbp1 | 2226.094474 | -0.077901295 | 0.078159696 | -0.996693938 | 0.318913093 | 0.552068788 |
| Ormdl3 | 2134.65925 | 0.074199137 | 0.074448517 | 0.996650305 | 0.318934279 | 0.552068788 |
| Prkx | 455.7821389 | 0.093961629 | 0.094294745 | 0.99646729 | 0.319023152 | 0.552160939 |
| Bag2 | 309.4004141 | -0.081618744 | 0.081926334 | -0.996245522 | 0.319130866 | 0.552285674 |
| AW146154 | 141.5832176 | -0.109802699 | 0.110238199 | -0.996049462 | 0.319226113 | 0.552388809 |
| Snrpf | 443.1899895 | -0.085946329 | 0.086311917 | -0.99576434 | 0.31936466 | 0.55256684 |
| Degs1 | 2986.684062 | 0.052246758 | 0.052498355 | 0.995207539 | 0.319635336 | 0.552973413 |
| Pla2g5 | 94.28904912 | -0.147990685 | 0.148733189 | -0.995007815 | 0.319732463 | 0.55307969 |
| Sgol1 | 8.790862732 | -0.158526088 | 0.159366072 | -0.994729219 | 0.319867979 | 0.553252341 |
| Abl1 | 2637.536238 | 0.114638911 | 0.071500094 | 1.603339304 | 0.108859763 | 0.286702902 |
| Mynn | 560.4372921 | 0.114578841 | 0.081595181 | 1.404235393 | 0.160248766 | 0.364683468 |
| Ptpdc1 | 1189.789808 | 0.114566277 | 0.090205814 | 1.270054244 | 0.204065309 | 0.421255955 |
| Msc | 41.99876292 | -0.160239097 | 0.161187728 | -0.99411475 | 0.320167005 | 0.553376867 |
| Rps6ka4 | 1611.330204 | 0.067187067 | 0.067597787 | 0.993924055 | 0.320259842 | 0.553376867 |
| Snx19 | 3923.275107 | -0.05514018 | 0.055475553 | -0.993954592 | 0.320244975 | 0.553376867 |
| Tarsl2 | 1813.202553 | -0.062830788 | 0.063202139 | -0.994124394 | 0.32016231 | 0.553376867 |
| Unc45a | 1106.358599 | -0.085413042 | 0.085934394 | -0.993933134 | 0.320255422 | 0.553376867 |
| Zfp26 | 809.9906615 | 0.074063048 | 0.074516044 | 0.993920825 | 0.320261415 | 0.553376867 |
| Tsr2 | 3326.378483 | -0.046446497 | 0.046742711 | -0.993662877 | 0.320387022 | 0.553532172 |
| Pomp | 2825.773596 | -0.04697457 | 0.047280567 | -0.993528044 | 0.320452691 | 0.553583899 |
| A830082N09Rik | 943.8772894 | -0.084824373 | 0.08538533 | -0.993430292 | 0.320500306 | 0.553584696 |
| Ppap2a | 743.5537633 | 0.09742333 | 0.098072531 | 0.993380399 | 0.32052461 | 0.553584696 |
| Ndfip2 | 3669.301209 | -0.076149415 | 0.076667635 | -0.993240691 | 0.320592673 | 0.553640534 |
| Trim34a | 63.19343896 | 0.114493511 | 0.16069376 | 0.712495065 | 0.476158266 | 0.689955989 |
| Xylt2 | 965.3961535 | 0.076848562 | 0.07738904 | 0.993016094 | 0.320702111 | 0.553706097 |
| Alkbh8 | 584.0131489 | -0.070418267 | 0.070924838 | -0.992857648 | 0.320779332 | 0.553777712 |
| Grsf1 | 3935.872399 | 0.076262765 | 0.07682419 | 0.992692085 | 0.320860033 | 0.55385532 |
| Dll3 | 34.18248516 | 0.114463377 | 0.160212944 | 0.714445252 | 0.474951899 | 0.68916508 |
| Cntn6 | 765.6208954 | -0.122141397 | 0.123061623 | -0.992522232 | 0.32094284 | 0.553874845 |
| Adra2b | 46.40447594 | -0.166374692 | 0.167716848 | -0.991997488 | 0.321198751 | 0.554073138 |
| Fip1l1 | 2102.694995 | -0.054873898 | 0.055309061 | -0.992132156 | 0.321133062 | 0.554073138 |
| Nudt10 | 440.4827162 | -0.090983846 | 0.091718204 | -0.991993323 | 0.321200783 | 0.554073138 |
| Trim41 | 2269.13392 | 0.114405839 | 0.053350139 | 2.144433751 | 0.031998148 | 0.129066229 |
| Gpr123 | 2680.995365 | 0.083363915 | 0.084104034 | 0.991199953 | 0.321587954 | 0.554679256 |
| Chadl | 351.7282444 | -0.107505992 | 0.108514956 | -0.990702077 | 0.321831077 | 0.554975037 |
| Pogk | 3354.731617 | -0.056328934 | 0.056854623 | -0.990753798 | 0.321805814 | 0.554975037 |
| Nsmf | 18516.86405 | -0.083431518 | 0.084246676 | -0.990324155 | 0.322015704 | 0.555231617 |
| Elk3 | 358.178704 | -0.089342733 | 0.090247608 | -0.989973417 | 0.322187113 | 0.555465352 |
| 1700120C14Rik | 16.09144385 | 0.114354192 | 0.16209683 | 0.705468404 | 0.480518788 | 0.693475978 |
| Bcap29 | 1298.491283 | -0.091975779 | 0.092960745 | -0.989404493 | 0.322465277 | 0.555799581 |
| Il1rl1 | 11.62216241 | -0.159309111 | 0.161034844 | -0.989283482 | 0.322524463 | 0.555799581 |
| Limk1 | 2455.64453 | -0.072090263 | 0.072866682 | -0.98934467 | 0.322494536 | 0.555799581 |
| Lpar6 | 140.1215552 | -0.141376609 | 0.142960461 | -0.988921045 | 0.322701773 | 0.556002754 |
| Zfp622 | 1083.438995 | 0.083250714 | 0.084185528 | 0.988895786 | 0.322714132 | 0.556002754 |
| Zufsp | 274.5067301 | 0.114339985 | 0.095580928 | 1.196263601 | 0.231593708 | 0.454562062 |
| Shfm1 | 1038.680663 | -0.062438497 | 0.063179023 | -0.988278927 | 0.323016063 | 0.556399208 |
| Ankrd26 | 765.9500307 | 0.085813637 | 0.086919995 | 0.987271532 | 0.323509543 | 0.557187289 |
| Lyl1 | 71.17327148 | -0.148380005 | 0.150323365 | -0.987072134 | 0.323607277 | 0.557293671 |
| Igfbp4 | 2956.996421 | 0.114333663 | 0.119798985 | 0.954379229 | 0.339891716 | 0.57366909 |
| Exosc5 | 317.1288025 | -0.091673044 | 0.092930867 | -0.986464962 | 0.323905001 | 0.557682418 |
| Gmpr | 1499.908432 | -0.058413494 | 0.059221719 | -0.986352548 | 0.323960142 | 0.557715382 |
| Rabgef1 | 1485.687142 | 0.083891134 | 0.085081662 | 0.986007227 | 0.324129566 | 0.557945062 |
| Sertad1 | 380.1337679 | -0.142143477 | 0.144231772 | -0.985521258 | 0.324368095 | 0.558293631 |
| Slc22a5 | 462.5372488 | 0.11428417 | 0.113788457 | 1.004356448 | 0.315206835 | 0.548558317 |
| Parp4 | 390.0265235 | 0.092112743 | 0.093504183 | 0.985118955 | 0.324565644 | 0.558509561 |
| Klhl28 | 484.1084474 | 0.088199877 | 0.089549443 | 0.984929377 | 0.324658762 | 0.558546456 |
| Ubap2l | 5235.096056 | 0.114252964 | 0.069465681 | 1.644739718 | 0.100023498 | 0.270134928 |
| Nefm | 8992.632089 | 0.09130271 | 0.092777131 | 0.984107929 | 0.325062449 | 0.559178148 |
| Mtrr | 475.9707847 | 0.097027502 | 0.098631512 | 0.983737353 | 0.325244669 | 0.559429496 |
| Vegfb | 1866.206416 | -0.096105436 | 0.097704639 | -0.983632269 | 0.325296353 | 0.559456287 |
| Aak1 | 20381.28435 | 0.062103128 | 0.063197021 | 0.982690741 | 0.325759669 | 0.559970267 |
| Cmc1 | 306.8782707 | -0.100486729 | 0.102265053 | -0.982610643 | 0.325799104 | 0.559970267 |
| Hecw2 | 2452.726486 | -0.086744005 | 0.088277071 | -0.982633468 | 0.325787866 | 0.559970267 |
| 2010002M12Rik | 33.77311532 | 0.114169956 | 0.167606065 | 0.681180335 | 0.495757389 | 0.705449141 |
| Smpd2 | 462.145808 | 0.082124492 | 0.083580094 | 0.982584343 | 0.325812053 | 0.559970267 |
| Tbc1d22a | 1737.311906 | 0.084584179 | 0.086052867 | 0.982932727 | 0.325640549 | 0.559970267 |
| Sall2 | 2990.642919 | -0.072206844 | 0.073533545 | -0.981957885 | 0.326120597 | 0.56043839 |
| St7 | 660.3346173 | -0.076847013 | 0.078277946 | -0.981719854 | 0.326237881 | 0.560577768 |
| Nup188 | 944.641777 | 0.059704531 | 0.060827825 | 0.981533214 | 0.326329864 | 0.560673643 |
| Nelfe | 688.2859197 | -0.097270607 | 0.099130794 | -0.981235025 | 0.326476856 | 0.560801819 |
| Hepacam | 3685.666239 | 0.114038877 | 0.07043509 | 1.619063394 | 0.105433624 | 0.280300029 |
| 1110007C09Rik | 155.6381591 | 0.114037742 | 0.114176945 | 0.998780809 | 0.317900885 | 0.55103906 |
| Sgsm2 | 5583.705884 | -0.085509939 | 0.087192864 | -0.980698815 | 0.326741288 | 0.561007261 |
| Tbc1d7 | 512.1636642 | 0.083292329 | 0.084920268 | 0.98082979 | 0.326676685 | 0.561007261 |
| Vapa | 4059.633244 | -0.069371227 | 0.070721514 | -0.980906987 | 0.326638612 | 0.561007261 |
| Smad3 | 3867.658412 | 0.091008676 | 0.092840304 | 0.980271205 | 0.326952264 | 0.5613073 |
| Ncl | 10173.18772 | 0.059429508 | 0.060631904 | 0.980168912 | 0.327002747 | 0.561331771 |
| Plk1 | 39.20613737 | -0.160407184 | 0.163686481 | -0.979965988 | 0.327102908 | 0.561441504 |
| Dars | 1779.59311 | -0.080032708 | 0.081716107 | -0.979399423 | 0.327382662 | 0.561859434 |
| Pomgnt1 | 2992.520243 | -0.066354142 | 0.067757572 | -0.97928748 | 0.327437955 | 0.561892089 |
| Dio2 | 3857.314582 | -0.163490064 | 0.167035254 | -0.978775794 | 0.327690772 | 0.562263658 |
| Rnf223 | 34.54662009 | 0.114001415 | 0.16746958 | 0.680729089 | 0.496042926 | 0.705679763 |
| 1110012L19Rik | 303.9756137 | -0.087166783 | 0.089105219 | -0.978245541 | 0.327952897 | 0.562588816 |
| Herc3 | 7350.090016 | 0.082855261 | 0.084715982 | 0.978035775 | 0.32805663 | 0.562704465 |
| Brd2 | 8207.414125 | 0.073183656 | 0.074839012 | 0.977881102 | 0.328133133 | 0.562773385 |
| Rcc1 | 452.0523291 | 0.093637627 | 0.095789261 | 0.977537845 | 0.328302951 | 0.563002317 |
| Bri3 | 929.1652977 | -0.078051438 | 0.079888679 | -0.977002486 | 0.328567922 | 0.563394354 |
| Rab4a | 1447.861186 | 0.099692449 | 0.102055185 | 0.976848447 | 0.328644187 | 0.563434496 |
| B4galnt1 | 5117.175979 | 0.113999828 | 0.078443606 | 1.453271138 | 0.146148488 | 0.345028899 |
| Phyhip | 23217.32159 | 0.113835271 | 0.087275141 | 1.304326395 | 0.192122319 | 0.406161583 |
| Gm13031 | 52.40378175 | -0.147575768 | 0.151130701 | -0.976477752 | 0.328827768 | 0.563590424 |
| Gm15997 | 18.51521607 | -0.163773338 | 0.167770295 | -0.976176016 | 0.328977246 | 0.563784255 |
| Mospd3 | 454.8439959 | 0.086188777 | 0.088299353 | 0.97609749 | 0.329016155 | 0.563788576 |
| Cysltr2 | 11.12522589 | -0.154290943 | 0.158088402 | -0.975978888 | 0.329074927 | 0.563826929 |
| Zfp61 | 526.4502326 | 0.088913977 | 0.09113482 | 0.975631229 | 0.329247243 | 0.564059794 |
| Tbca | 1058.238075 | -0.084332333 | 0.086461752 | -0.975371549 | 0.329375991 | 0.564217977 |
| Cps1 | 18.15517449 | -0.163413052 | 0.167609236 | -0.974964483 | 0.329577878 | 0.564501398 |
| Alg1 | 893.8590091 | -0.081416518 | 0.083595768 | -0.9739311 | 0.330090751 | 0.565317353 |
| Irf6 | 58.86127556 | -0.148229066 | 0.152211167 | -0.973838314 | 0.330136826 | 0.565333774 |
| Acox1 | 9186.537025 | 0.11379284 | 0.074669215 | 1.523959242 | 0.127518893 | 0.315619445 |
| Acsl3 | 6925.88872 | 0.054268077 | 0.055759997 | 0.973243906 | 0.330432094 | 0.56571435 |
| Slc25a12 | 6321.78113 | 0.081454692 | 0.083707409 | 0.973088191 | 0.330509473 | 0.565744467 |
| Zfp260 | 2441.661549 | 0.063790847 | 0.065561797 | 0.972988079 | 0.330559227 | 0.565744467 |
| Zscan21 | 641.3482048 | 0.075605983 | 0.077703557 | 0.973005427 | 0.330550605 | 0.565744467 |
| Ncoa2 | 4805.521027 | -0.079476351 | 0.081719388 | -0.972551966 | 0.330776026 | 0.566052987 |
| Prr13 | 1190.916385 | 0.073538236 | 0.075625809 | 0.972396028 | 0.330853567 | 0.566123156 |
| Fcho2 | 997.492919 | 0.11376431 | 0.070282633 | 1.618668878 | 0.105518526 | 0.280429511 |
| Zfat | 281.5159272 | 0.113736674 | 0.098291778 | 1.157133142 | 0.247217968 | 0.472120974 |
| Kat2b | 1097.528834 | 0.113679373 | 0.110074597 | 1.032748483 | 0.301721618 | 0.533658642 |
| Gpcpd1 | 3351.000397 | 0.113677014 | 0.090247283 | 1.259617021 | 0.207807552 | 0.425908943 |
| Rpusd3 | 254.1102956 | 0.113628644 | 0.097258875 | 1.168311312 | 0.242681214 | 0.467209584 |
| Taf4a | 888.4713477 | 0.11362369 | 0.072300046 | 1.571557637 | 0.116053178 | 0.2986169 |
| Fkbp7 | 73.7795458 | 0.11358606 | 0.142915887 | 0.794775597 | 0.426744048 | 0.648609949 |
| Zfp870 | 450.9737637 | -0.079162468 | 0.081478728 | -0.971572217 | 0.33126341 | 0.566339732 |
| Riok2 | 806.2805603 | 0.066655769 | 0.068622996 | 0.971332834 | 0.331382563 | 0.566465295 |
| 8-Mar | 2597.930665 | -0.068114055 | 0.070130099 | -0.971252789 | 0.331422412 | 0.566470916 |
| Ak4 | 2110.117776 | -0.086760008 | 0.089375159 | -0.97073963 | 0.331677952 | 0.566732816 |
| Cdk16 | 11345.36541 | 0.068118343 | 0.070176592 | 0.970670442 | 0.331712415 | 0.566732816 |
| Cenpk | 20.61194501 | -0.162615038 | 0.167520993 | -0.970714389 | 0.331690525 | 0.566732816 |
| Habp4 | 8761.111421 | 0.059828579 | 0.061637561 | 0.9706513 | 0.331721951 | 0.566732816 |
| 2-Mar | 2226.807536 | 0.053725044 | 0.055353777 | 0.970575937 | 0.331759493 | 0.566734464 |
| Pebp1 | 14166.94793 | -0.053667235 | 0.055320877 | -0.970108169 | 0.331992577 | 0.567070113 |
| 8430429K09Rik | 207.4729391 | -0.088628472 | 0.091390969 | -0.969772751 | 0.332159778 | 0.567168129 |
| Mapkapk5 | 1177.370908 | 0.06873533 | 0.07086761 | 0.969911773 | 0.332090471 | 0.567168129 |
| Trappc2 | 871.6540559 | -0.07851808 | 0.080962647 | -0.969806232 | 0.332143086 | 0.567168129 |
| Gm872 | 34.22522344 | 0.113574674 | 0.159504313 | 0.71204767 | 0.476435257 | 0.690211698 |
| Tgm2 | 711.1601974 | -0.08493551 | 0.087611348 | -0.969457857 | 0.332316797 | 0.567311201 |
| 1700013G23Rik | 16.04302459 | -0.162031731 | 0.167167941 | -0.969275145 | 0.332407927 | 0.567404255 |
| Serp1 | 1327.144052 | -0.063963584 | 0.06599801 | -0.969174434 | 0.332458165 | 0.567427495 |
| Cdc14b | 436.2300373 | 0.11357372 | 0.115271443 | 0.985271956 | 0.324490504 | 0.558442282 |
| 4930402H24Rik | 3950.366378 | -0.105681531 | 0.109062515 | -0.968999582 | 0.332545398 | 0.567451365 |
| Thtpa | 1248.771011 | 0.053658528 | 0.055379604 | 0.968922207 | 0.332584004 | 0.567454748 |
| Pcdhac2 | 1395.671182 | -0.076511209 | 0.078977835 | -0.968768128 | 0.332660892 | 0.567523438 |
| A230077H06Rik | 179.8350525 | -0.130696424 | 0.135025926 | -0.967935772 | 0.333076448 | 0.567976249 |
| Cenpj | 138.8635373 | -0.10538385 | 0.108882305 | -0.967869392 | 0.333109602 | 0.567976249 |
| Mrpl42 | 1060.984099 | -0.075646349 | 0.07815639 | -0.967884391 | 0.333102111 | 0.567976249 |
| Rimbp2 | 4717.511023 | -0.066892202 | 0.069098032 | -0.968076804 | 0.333006014 | 0.567976249 |
| Zfp637 | 943.4473277 | -0.067274999 | 0.069491538 | -0.968103471 | 0.332992697 | 0.567976249 |
| Ergic1 | 5651.866506 | 0.051060161 | 0.052762678 | 0.967732558 | 0.333177954 | 0.568030283 |
| Tmem126b | 486.4551292 | 0.083845527 | 0.086652729 | 0.967603994 | 0.333242182 | 0.568077277 |
| Dock7 | 1723.705442 | 0.082603308 | 0.085385162 | 0.967419936 | 0.333334148 | 0.5681402 |
| Slc37a3 | 2308.314995 | -0.04881401 | 0.050459841 | -0.967383349 | 0.333352431 | 0.5681402 |
| Map2k6 | 407.4579139 | 0.113521286 | 0.120093927 | 0.945270825 | 0.344520629 | 0.576722182 |
| Lrrc16a | 943.3036367 | 0.089885163 | 0.092936892 | 0.967163424 | 0.333462344 | 0.56815299 |
| Mrpl10 | 1835.139608 | -0.05203485 | 0.053799042 | -0.967207745 | 0.333440191 | 0.56815299 |
| Cisd3 | 616.9010725 | -0.074531986 | 0.077076045 | -0.966992881 | 0.333547592 | 0.568222803 |
| Snx12 | 1715.34453 | -0.051862648 | 0.053643024 | -0.966810665 | 0.333638692 | 0.568284318 |
| Tagln3 | 8194.455318 | -0.070310652 | 0.072727087 | -0.966773936 | 0.333657057 | 0.568284318 |
| Gga2 | 1352.494945 | 0.049248976 | 0.050948575 | 0.966640891 | 0.333723585 | 0.568335154 |
| Alg11 | 1359.126618 | -0.064948477 | 0.067197353 | -0.966533256 | 0.333777414 | 0.568364354 |
| Exoc7 | 2813.596921 | 0.047492834 | 0.049142827 | 0.966424534 | 0.333831792 | 0.568394483 |
| AA414768 | 269.4824003 | 0.093797578 | 0.097067412 | 0.966313783 | 0.333887191 | 0.568426343 |
| BC029722 | 2080.928881 | -0.078525213 | 0.081280564 | -0.966100739 | 0.333993775 | 0.568482868 |
| Slc45a4 | 1874.210921 | 0.076614925 | 0.079301858 | 0.966117656 | 0.333985311 | 0.568482868 |
| 6720456H20Rik | 1048.617676 | -0.072813577 | 0.07537793 | -0.965980053 | 0.334054163 | 0.568523198 |
| Mllt10 | 1337.155133 | 0.075851891 | 0.078548311 | 0.965671826 | 0.334208422 | 0.56872326 |
| Lhfpl4 | 6285.666257 | -0.052984995 | 0.054889947 | -0.965295068 | 0.334397042 | 0.568981744 |
| Ccdc47 | 4314.204223 | 0.056049966 | 0.058072549 | 0.965171435 | 0.334458952 | 0.569024596 |
| Nxt2 | 996.5639337 | -0.075297978 | 0.078040552 | -0.964857061 | 0.334616411 | 0.569229982 |
| Ccdc132 | 2729.82367 | 0.06632758 | 0.068753778 | 0.964711783 | 0.334689192 | 0.569238517 |
| Ensa | 11811.77905 | -0.050964785 | 0.05283105 | -0.964674837 | 0.334707703 | 0.569238517 |
| Mef2c | 18167.07783 | 0.099942273 | 0.103607164 | 0.964627049 | 0.334731646 | 0.569238517 |
| Ccdc65 | 243.6260114 | -0.101425111 | 0.105162211 | -0.964463468 | 0.334813616 | 0.569315426 |
| Corin | 8.551559676 | -0.147118153 | 0.152558428 | -0.964339729 | 0.334875629 | 0.569354094 |
| Erlin2 | 2205.048776 | 0.050799989 | 0.052685657 | 0.964209085 | 0.33494111 | 0.569354094 |
| Serinc1 | 32227.4425 | -0.065388027 | 0.067815964 | -0.964198141 | 0.334946596 | 0.569354094 |
| Nr6a1 | 472.9886945 | 0.080398316 | 0.083405429 | 0.96394584 | 0.33507308 | 0.569506615 |
| Gramd4 | 2327.811112 | -0.071029662 | 0.0736987 | -0.963784461 | 0.335153999 | 0.569519199 |
| Scg2 | 11742.93462 | -0.153365967 | 0.15912377 | -0.963815567 | 0.335138401 | 0.569519199 |
| Pex2 | 783.2432322 | -0.060464712 | 0.062745263 | -0.963653811 | 0.335219518 | 0.569568069 |
| Igfbp5 | 7577.156347 | -0.09547549 | 0.099086393 | -0.963558035 | 0.335267555 | 0.569587226 |
| Thbs1 | 125.4159506 | 0.11352071 | 0.167809857 | 0.676484158 | 0.498733301 | 0.707181699 |
| Pcyt1a | 1869.743387 | 0.092989451 | 0.096544757 | 0.963174529 | 0.335459945 | 0.569789124 |
| Smarcc2 | 12777.15184 | 0.113452898 | 0.065634059 | 1.728567447 | 0.08388654 | 0.24166991 |
| Trim27 | 896.0382314 | 0.067958681 | 0.070582218 | 0.962830065 | 0.33563281 | 0.569957778 |
| Gm15663 | 295.8298831 | -0.139180781 | 0.144610731 | -0.962451268 | 0.335822971 | 0.569968355 |
| L2hgdh | 802.0516853 | 0.069257158 | 0.071956589 | 0.962485287 | 0.33580589 | 0.569968355 |
| Malsu1 | 205.3364872 | -0.098952201 | 0.102810773 | -0.962469187 | 0.335813974 | 0.569968355 |
| Pld1 | 432.5780714 | 0.080363347 | 0.083475132 | 0.962722005 | 0.335687051 | 0.569968355 |
| Dvl1 | 4085.95285 | 0.113373245 | 0.067406494 | 1.681933568 | 0.092581723 | 0.256795777 |
| Dld | 3960.59639 | -0.062111774 | 0.064544536 | -0.962308784 | 0.335894518 | 0.570014191 |
| Impdh2 | 1686.970016 | 0.09571097 | 0.099473278 | 0.962177706 | 0.335960346 | 0.570014191 |
| Cdc14a | 138.9501469 | 0.113355112 | 0.115580503 | 0.980745966 | 0.32671803 | 0.561007261 |
| Utp20 | 1045.108786 | 0.113320828 | 0.073621071 | 1.539244502 | 0.123744617 | 0.311047874 |
| Fgf18 | 45.00914696 | 0.113317653 | 0.158322067 | 0.71574137 | 0.474151061 | 0.688454365 |
| Rpl38 | 3323.726093 | -0.094302322 | 0.098055556 | -0.961723384 | 0.336188573 | 0.570214854 |
| Pip4k2c | 6537.314096 | 0.11331618 | 0.08937559 | 1.26786497 | 0.204846176 | 0.421947174 |
| Ift27 | 585.8502549 | 0.071208877 | 0.074064354 | 0.961446 | 0.336327965 | 0.570325652 |
| Dzip1 | 2590.845607 | 0.113283595 | 0.108864986 | 1.040587976 | 0.298066814 | 0.529836392 |
| Mvb12b | 5594.942764 | -0.051352611 | 0.053421333 | -0.961275354 | 0.336413737 | 0.570346257 |
| Igbp1 | 1072.545944 | 0.113278166 | 0.1196852 | 0.946467616 | 0.343910132 | 0.576370318 |
| Sox13 | 392.3321883 | -0.100144755 | 0.104209122 | -0.960997975 | 0.336553186 | 0.570457834 |
| Ogdh | 12661.32192 | -0.049389638 | 0.051401346 | -0.960862736 | 0.336621189 | 0.570503566 |
| Stag1 | 1436.256735 | 0.081873755 | 0.085214339 | 0.960797874 | 0.336653808 | 0.570503566 |
| Psenen | 1262.275784 | 0.062207624 | 0.064755004 | 0.960661274 | 0.336722509 | 0.570557585 |
| Rbm48 | 305.5668213 | 0.113267718 | 0.082585217 | 1.37152534 | 0.17021125 | 0.377721728 |
| Cyp2r1 | 10.86477611 | -0.151934272 | 0.158254866 | -0.960060669 | 0.337024682 | 0.570594737 |
| Efcab4a | 84.15184665 | -0.13566493 | 0.141312907 | -0.96003212 | 0.33703905 | 0.570594737 |
| Slc35c2 | 1829.521077 | 0.113195907 | 0.072198513 | 1.567842647 | 0.116917859 | 0.300020754 |
| Nol3 | 900.6758856 | 0.08656701 | 0.090128553 | 0.960483746 | 0.336811808 | 0.570594737 |
| Pla2g4a | 145.9192873 | -0.137989811 | 0.143729098 | -0.960068719 | 0.337020631 | 0.570594737 |
| Rnf19a | 1455.798487 | 0.071992704 | 0.074979427 | 0.960166096 | 0.336971627 | 0.570594737 |
| Rpp25l | 367.1585145 | -0.082163591 | 0.085565671 | -0.960240133 | 0.336934372 | 0.570594737 |
| Chpf | 3037.308032 | -0.070365586 | 0.073317779 | -0.959734286 | 0.337188963 | 0.570700002 |
| Ddx20 | 492.3717722 | 0.078589549 | 0.081887575 | 0.959724949 | 0.337193663 | 0.570700002 |
| Lmf2 | 663.5639468 | 0.078025354 | 0.081302744 | 0.959689065 | 0.337211728 | 0.570700002 |
| Vps53 | 2842.578568 | 0.11310721 | 0.075803166 | 1.492117225 | 0.135668417 | 0.327932651 |
| Tigd3 | 126.1530219 | -0.112466886 | 0.117259394 | -0.959129009 | 0.337493758 | 0.571052559 |
| Cog5 | 947.7284861 | -0.066155161 | 0.069015923 | -0.958549253 | 0.337785868 | 0.571422015 |
| Rnf215 | 781.404933 | 0.087210461 | 0.090981664 | 0.958549859 | 0.337785562 | 0.571422015 |
| Psd | 16358.7105 | -0.089139071 | 0.093021492 | -0.958263178 | 0.337930067 | 0.571603543 |
| Wdr6 | 6999.013636 | 0.113026846 | 0.1047678 | 1.078831912 | 0.280662666 | 0.510416091 |

| | | | | | | |
|---|---|---|---|---|---|---|
| Nsun7 | 165.5883214 | -0.295494148 | 0.085242172 | -3.466525329 | 0.000527232 | 0.032276094 |
| Tlr2 | 78.64670622 | -0.296160878 | 0.088249133 | -3.35596359 | 0.00079089 | 0.04208229 |
| H2-Q4 | 430.3689413 | -0.296610199 | 0.089305981 | -3.321280326 | 0.000896055 | 0.045038385 |
| Ccdc84 | 327.9165278 | -0.296832579 | 0.088933919 | -3.337675701 | 0.000844823 | 0.043828205 |
| Pld4 | 485.704191 | -0.296851407 | 0.086669351 | -3.425102463 | 0.000614568 | 0.035388683 |
| Col7a1 | 192.0368488 | -0.297006704 | 0.093072926 | -3.191118146 | 0.001417233 | 0.061125282 |
| Uba7 | 186.6273189 | -0.298326125 | 0.094341055 | -3.162208916 | 0.001565772 | 0.064431734 |
| Adam33 | 409.5614273 | -0.298441983 | 0.095087563 | -3.138601654 | 0.00169756 | 0.067373669 |
| Bin2 | 266.5983599 | -0.298765057 | 0.085271592 | -3.503688025 | 0.000458863 | 0.029106716 |
| Tatdn3 | 518.9302249 | -0.299045015 | 0.077038312 | -3.881770096 | 0.000103699 | 0.010759501 |
| Eif2ak2 | 335.0410949 | -0.299821311 | 0.086796367 | -3.454307154 | 0.000551709 | 0.033313941 |
| Parp12 | 508.2499547 | -0.300342063 | 0.076640717 | -3.918831601 | 8.90E-05 | 0.00971814 |
| Dennd6b | 3614.624161 | -0.300625141 | 0.079633521 | -3.775107969 | 0.000159938 | 0.013586344 |
| Ccdc62 | 109.6592516 | -0.301092509 | 0.094617053 | -3.182222432 | 0.001461495 | 0.062075089 |
| Ets1 | 514.946524 | -0.302135645 | 0.065017138 | -4.647015448 | 3.37E-06 | 0.00094076 |
| 1700024P16Rik | 48.93689332 | -0.302282052 | 0.091639104 | -3.298614225 | 0.000971633 | 0.047521706 |
| C1qc | 1930.48976 | -0.302331526 | 0.088040873 | -3.433990569 | 0.000594765 | 0.034836914 |
| Luc7l3 | 5691.921881 | -0.302664572 | 0.061898811 | -4.889667019 | 1.01E-06 | 0.000376211 |
| Col15a1 | 316.2576924 | -0.303153237 | 0.093424014 | -3.244917699 | 0.001174846 | 0.05383536 |
| Stx3 | 1309.629746 | -0.303428617 | 0.062512832 | -4.853861329 | 1.21E-06 | 0.000409027 |
| Cep70 | 745.9681042 | -0.303987948 | 0.066486698 | -4.572161942 | 4.83E-06 | 0.001213916 |
| Ifitm3 | 328.5259502 | -0.304029192 | 0.088220548 | -3.446240101 | 0.000568445 | 0.03358999 |
| Ccdc159 | 477.4810088 | -0.304130775 | 0.086507082 | -3.515674867 | 0.000438638 | 0.028068954 |
| Dclre1c | 630.0777169 | -0.304277804 | 0.078587266 | -3.871846149 | 0.000108014 | 0.01097212 |
| Slamf8 | 15.18104501 | -0.306207076 | 0.061812233 | -4.953826501 | 7.28E-07 | NA |
| Ipw | 443.7409427 | -0.306254519 | 0.092276002 | -3.318896709 | 0.000903739 | 0.045267951 |
| H2-T24 | 85.66416614 | -0.306313864 | 0.095095396 | -3.221121913 | 0.001276898 | 0.057603177 |
| Ptprv | 393.0429352 | -0.306467015 | 0.086174378 | -3.556358897 | 0.00037603 | 0.02588728 |
| Col6a2 | 1235.455651 | -0.308965186 | 0.087361085 | -3.536645488 | 0.000405243 | 0.02634142 |
| Lyn | 409.3667518 | -0.309025703 | 0.080562771 | -3.835837576 | 0.000125137 | 0.011880663 |
| Lpcat2 | 339.7725945 | -0.309336154 | 0.082230872 | -3.761800709 | 0.000168694 | 0.014083077 |
| Rasal3 | 164.7337673 | -0.309976524 | 0.094654557 | -3.274818797 | 0.001057298 | 0.049703256 |
| Liph | 54.73469408 | -0.310966335 | 0.092015688 | -3.379492591 | 0.000726198 | 0.040003493 |
| Lgals9 | 344.8743039 | -0.311089128 | 0.089248873 | -3.485636499 | 0.000490968 | 0.030740501 |
| Pnpla3 | 611.3469588 | -0.311155997 | 0.088054556 | -3.53367289 | 0.000409828 | 0.02634142 |
| Ccdc108 | 434.2583441 | -0.311297369 | 0.092958646 | -3.348772617 | 0.000811704 | 0.042689184 |
| C1s | 86.01757752 | -0.311753565 | 0.095101918 | -3.278099672 | 0.001045085 | 0.049288645 |
| Csf2rb | 75.07014121 | -0.312024936 | 0.094823906 | -3.290572477 | 0.000999837 | 0.048083105 |
| 2610005L07Rik | 718.1818116 | -0.31304165 | 0.092412203 | -3.38744927 | 0.000705458 | 0.039262366 |
| Srrm2 | 30891.1576 | -0.31526377 | 0.073469175 | -4.29110261 | 1.78E-05 | 0.003038099 |
| Chuk | 1305.646425 | -0.315493879 | 0.073627753 | -4.284985866 | 1.83E-05 | 0.003051308 |
| Ap1g2 | 284.3237942 | -0.315949893 | 0.091164437 | -3.4657143 | 0.000528825 | 0.032276094 |
| Ifi203 | 49.80148407 | -0.315950438 | 0.087422065 | -3.614081175 | 0.000301415 | 0.022091286 |
| AF251705 | 110.3470426 | -0.31629093 | 0.095025624 | -3.328480424 | 0.000873211 | 0.044373556 |
| Irf1 | 323.049198 | -0.317631309 | 0.094981787 | -3.344128565 | 0.000825415 | 0.043129406 |
| Ddx26b | 2180.945356 | -0.318737945 | 0.063189679 | -5.044145639 | 4.56E-07 | 0.000189067 |
| Slc14a1 | 550.7871664 | -0.319536827 | 0.078982568 | -4.045662644 | 5.22E-05 | 0.006827917 |
| Cdk2 | 89.54156948 | -0.321140693 | 0.093782643 | -3.424308407 | 0.000616366 | 0.035388683 |
| Vmn2r84 | 31.66865262 | -0.321825427 | 0.090955033 | -3.538291591 | 0.000402725 | 0.02634142 |
| Csf2ra | 781.1309724 | -0.322208984 | 0.085477125 | -3.769534635 | 0.000163552 | 0.013732714 |
| Lat2 | 91.19534975 | -0.322318082 | 0.095077718 | -3.390048569 | 0.000698802 | 0.039105972 |
| Uvssa | 535.0820942 | -0.323783699 | 0.072206866 | -4.484112367 | 7.32E-06 | 0.001661821 |
| Itgb2 | 311.0803894 | -0.324428717 | 0.09412839 | -3.446661698 | 0.000567559 | 0.03358999 |
| Cyth4 | 504.1714374 | -0.32451554 | 0.079685113 | -4.072473864 | 4.65E-05 | 0.006479723 |
| Gm3002 | 637.324772 | -0.325282282 | 0.070617518 | -4.606254843 | 4.10E-06 | 0.001102862 |
| Arhgap30 | 168.4926081 | -0.328316008 | 0.086161679 | -3.810464365 | 0.000138706 | 0.012752174 |
| Hk2 | 209.8444779 | -0.333115118 | 0.090229585 | -3.69186136 | 0.000222619 | 0.017965338 |
| Lcp1 | 560.9722753 | -0.333411231 | 0.076839651 | -4.339051845 | 1.43E-05 | 0.002504401 |
| Irf9 | 529.277702 | -0.333430951 | 0.082095206 | -4.061515473 | 4.88E-05 | 0.006564539 |
| Lacc1 | 219.6643466 | -0.334321224 | 0.082157253 | -4.06928435 | 4.72E-05 | 0.006479723 |
| Gbp5 | 158.3434278 | -0.335275918 | 0.094567751 | -3.545351501 | 0.00039209 | 0.02634142 |
| Unc93b1 | 693.6997405 | -0.335496374 | 0.081674012 | -4.107749389 | 4.00E-05 | 0.00580362 |
| Col11a2 | 1033.387715 | -0.336296843 | 0.077446904 | -4.342289068 | 1.41E-05 | 0.002497867 |
| Ncrna00085 | 430.7779093 | -0.336751892 | 0.090659588 | -3.714465272 | 0.000203634 | 0.016617893 |
| Cd48 | 39.5085305 | -0.337126237 | 0.088594869 | -3.805256899 | 0.000141657 | 0.01286942 |
| Capg | 96.58287031 | -0.337347481 | 0.09508756 | -3.547756201 | 0.000388528 | 0.02634142 |
| A230057D06Rik | 249.4336797 | -0.33773499 | 0.09402078 | -3.592131351 | 0.000327984 | 0.023585654 |
| Sp100 | 142.738279 | -0.337955746 | 0.092532764 | -3.652281983 | 0.00025992 | 0.020082986 |
| Ech1 | 1234.130815 | -0.339138322 | 0.058101156 | -5.837032264 | 5.31E-09 | 4.06E-06 |
| Ifit3 | 441.0825018 | -0.339259463 | 0.094842987 | -3.577064296 | 0.000347475 | 0.024266432 |
| Slamf9 | 57.17082679 | -0.339481901 | 0.089502322 | -3.792995463 | 0.000148841 | 0.01286942 |
| Cep110 | 638.3545329 | -0.340343939 | 0.078622842 | -4.32881754 | 1.50E-05 | 0.002592408 |
| Xaf1 | 257.4136361 | -0.340503169 | 0.087444061 | -3.893954192 | 9.86E-05 | 0.010306488 |
| Rnf207 | 354.4363388 | -0.340600587 | 0.087447842 | -3.894899831 | 9.82E-05 | 0.010306488 |
| Csf2rb2 | 52.90440529 | -0.341395936 | 0.093285274 | -3.659698047 | 0.000252513 | 0.019720421 |
| Fes | 165.0930382 | -0.341573725 | 0.086753147 | -3.937306446 | 8.24E-05 | 0.0092074 |
| Samsn1 | 56.79513362 | -0.341778385 | 0.093524197 | -3.654438046 | 0.000257746 | 0.02002148 |
| Catsper2 | 114.7450725 | -0.344185125 | 0.093564345 | -3.678592798 | 0.000234524 | 0.018615852 |
| Hdac7 | 1532.395537 | -0.344945448 | 0.085480715 | -4.035359895 | 5.45E-05 | 0.007008302 |
| Thpo | 289.4104594 | -0.345231241 | 0.093726273 | -3.68339879 | 0.000230145 | 0.018470065 |
| Was | 68.80934543 | -0.345541392 | 0.093920514 | -3.679083253 | 0.000234074 | 0.018615852 |
| Hvcn1 | 178.7917752 | -0.345656852 | 0.094871248 | -3.643431046 | 0.000269028 | 0.020563301 |
| Ifit1 | 155.5557556 | -0.347333644 | 0.09170584 | -3.787475755 | 0.000152186 | 0.013080751 |
| Plek | 762.7550869 | -0.347521244 | 0.086415786 | -4.02150188 | 5.78E-05 | 0.007179599 |
| Col5a1 | 1668.118593 | -0.348921463 | 0.088736904 | -3.932089677 | 8.42E-05 | 0.009337738 |
| Gpatch1 | 977.1291577 | -0.349113434 | 0.053813997 | -6.487409449 | 8.73E-11 | 1.15E-07 |
| Phf11d | 27.76999338 | -0.349630303 | 0.079833323 | -4.379503334 | 1.19E-05 | 0.002295106 |
| Tmc4 | 184.108921 | -0.353452527 | 0.0943836 | -3.744851088 | 0.000180501 | 0.01489747 |
| Ddx58 | 254.1809065 | -0.354680512 | 0.088186844 | -4.021920898 | 5.77E-05 | 0.007179599 |
| 4831440E17Rik | 178.1454702 | -0.354931177 | 0.0891007 | -3.983483582 | 6.79E-05 | 0.008086025 |
| Spag5 | 649.0770472 | -0.355277428 | 0.081755056 | -4.345632488 | 1.39E-05 | 0.00249049 |
| Mlxipl | 161.7954127 | -0.355793663 | 0.094065818 | -3.782390573 | 0.000155329 | 0.013272439 |
| Parp9 | 177.7816606 | -0.356530445 | 0.09314229 | -3.82780416 | 0.000129292 | 0.012195385 |
| Ncf2 | 204.4404366 | -0.356569866 | 0.092070442 | -3.872794117 | 0.000107595 | 0.01097212 |
| Csf3r | 361.4791217 | -0.357082924 | 0.092533519 | -3.858957567 | 0.000113872 | 0.011329455 |
| Kctd15 | 410.9900555 | -0.358287523 | 0.068369146 | -5.240485549 | 1.60E-07 | 8.88E-05 |
| Siglec5 | 36.62049432 | -0.358877236 | 0.078164886 | -4.591284561 | 4.41E-06 | 0.001148619 |
| Trim30a | 123.7714625 | -0.360269648 | 0.094956649 | -3.794043421 | 0.000148214 | 0.01286942 |
| Sfrs18 | 7556.932928 | -0.360726385 | 0.066508163 | -5.423791155 | 5.83E-08 | 3.85E-05 |
| Zfp36 | 200.9219435 | -0.360842276 | 0.085579177 | -4.216472851 | 2.48E-05 | 0.003918122 |
| Man2b1 | 2146.94953 | -0.363986954 | 0.083669608 | -4.350288765 | 1.36E-05 | 0.00249049 |
| Ptprc | 183.2461383 | -0.365003777 | 0.09183601 | -3.974516923 | 7.05E-05 | 0.008130208 |
| Col19a1 | 1298.402088 | -0.370690028 | 0.078703858 | -4.709934679 | 2.48E-06 | 0.000719898 |
| Miat | 23357.0276 | -0.371041218 | 0.08262541 | -4.490642987 | 7.10E-06 | 0.001637253 |
| Zfp217 | 175.1211143 | -0.371529204 | 0.084978552 | -4.37203501 | 1.23E-05 | 0.002322156 |
| Aspg | 121.0110147 | -0.372163274 | 0.092630705 | -4.01770961 | 5.88E-05 | 0.007229522 |
| Slc15a3 | 106.1324819 | -0.372499786 | 0.094204417 | -3.95416474 | 7.68E-05 | 0.008715881 |
| Gbp4 | 131.6410243 | -0.373560215 | 0.09091976 | -4.108680185 | 3.98E-05 | 0.00580362 |
| Ccl6 | 107.509393 | -0.37402255 | 0.08847579 | -4.227399932 | 2.36E-05 | 0.003773683 |
| Ttc14 | 4985.536852 | -0.374806833 | 0.072152436 | -5.194652478 | 2.05E-07 | 9.61E-05 |
| Trim30d | 71.96198326 | -0.374869549 | 0.094314341 | -3.974682378 | 7.05E-05 | 0.008130208 |
| Aif1 | 230.4358008 | -0.375054553 | 0.08089695 | -4.636201389 | 3.55E-06 | 0.000972612 |
| Tnfsf10 | 121.9224566 | -0.379715936 | 0.094908284 | -4.000872443 | 6.31E-05 | 0.007576254 |
| Ctsc | 306.293372 | -0.381779572 | 0.092539278 | -4.125594912 | 3.70E-05 | 0.005537515 |
| Haus5 | 232.0451918 | -0.38194108 | 0.083681997 | -4.564196526 | 5.01E-06 | 0.001213916 |
| Cd86 | 111.5448965 | -0.382234781 | 0.09496493 | -4.025009864 | 5.70E-05 | 0.007179599 |
| Dock2 | 244.9161645 | -0.389378357 | 0.086909818 | -4.480257417 | 7.46E-06 | 0.001666089 |
| Meg3 | 133078.2528 | -0.392067205 | 0.088766427 | -4.41684114 | 1.00E-05 | 0.001993526 |
| Lair1 | 380.3287927 | -0.393355672 | 0.082910601 | -4.744335041 | 2.09E-06 | 0.000620149 |
| Gm15328 | 107.6535738 | -0.394093443 | 0.091894991 | -4.288519323 | 1.80E-05 | 0.003038099 |
| Zkscan2 | 845.8572196 | -0.394282699 | 0.073151842 | -5.389921667 | 7.05E-08 | 4.45E-05 |
| Parp14 | 203.9879468 | -0.395434677 | 0.094562093 | -4.181746232 | 2.89E-05 | 0.004518347 |
| Emr1 | 327.1843711 | -0.396968152 | 0.082254537 | -4.826094277 | 1.39E-06 | 0.000449456 |
| Malat1 | 8083.783777 | -0.398032462 | 0.087083035 | -4.570723376 | 4.86E-06 | 0.001213916 |
| Leng8 | 11174.74125 | -0.399388559 | 0.08962834 | -4.456052155 | 8.35E-06 | 0.001811202 |
| Slc11a1 | 135.5536509 | -0.402194973 | 0.095095973 | -4.229358601 | 2.34E-05 | 0.003773683 |
| Cxcl16 | 75.5319207 | -0.403461318 | 0.092389725 | -4.366950101 | 1.26E-05 | 0.002346387 |
| Itgax | 102.6554611 | -0.408936713 | 0.064577544 | -6.332490946 | 2.41E-10 | 2.50E-07 |
| Ly86 | 561.0912527 | -0.41091316 | 0.089519564 | -4.590205097 | 4.43E-06 | 0.001148619 |
| Il18bp | 169.3893851 | -0.412845496 | 0.093472144 | -4.416775724 | 1.00E-05 | 0.001993526 |
| Apobec3 | 130.5387185 | -0.413054696 | 0.092788386 | -4.45157754 | 8.52E-06 | 0.001820914 |
| I830012O16Rik | 248.0010337 | -0.41313565 | 0.092716452 | -4.455904418 | 8.35E-06 | 0.001811202 |
| Cd84 | 177.1092639 | -0.413512276 | 0.094014101 | -4.398406983 | 1.09E-05 | 0.00214059 |
| Cd33 | 276.1730083 | -0.417140194 | 0.085656825 | -4.869900256 | 1.12E-06 | 0.000395584 |
| A930013F10Rik | 374.1246032 | -0.426274783 | 0.089117795 | -4.783273426 | 1.72E-06 | 0.000544609 |
| Naip5 | 88.86037488 | -0.429307414 | 0.095077715 | -4.515331654 | 6.32E-06 | 0.001505412 |
| Slc26a10 | 206.126767 | -0.429691495 | 0.084031248 | -5.11347276 | 3.16E-07 | 0.000139225 |
| Ptpn6 | 220.8980729 | -0.434744984 | 0.093432562 | -4.653035028 | 3.27E-06 | 0.000931614 |
| Gm5577 | 221.6962085 | -0.435279408 | 0.088809494 | -4.901271125 | 9.52E-07 | 0.000373823 |
| Cd22 | 29.40777017 | -0.439350672 | 0.07937693 | -5.534991971 | NA | NA |
| Mpeg1 | 1997.379678 | -0.441723497 | 0.088732926 | -4.978123854 | 6.42E-07 | 0.000259061 |
| Ovgp1 | 266.9610844 | -0.445222446 | 0.093692943 | -4.751931506 | 2.01E-06 | 0.000609737 |
| Zkscan3 | 1163.650182 | -0.446284155 | 0.070718832 | -6.310683368 | 2.78E-10 | 2.69E-07 |
| Fcgr1 | 152.2107014 | -0.449813292 | 0.092651199 | -4.854910614 | 1.20E-06 | 0.000409027 |
| Snhg11 | 73090.53635 | -0.450347664 | 0.085099513 | -5.292012267 | 1.21E-07 | 7.32E-05 |
| Cd14 | 98.2637337 | -0.461192413 | 0.094420031 | -4.884476376 | 1.04E-06 | 0.000376601 |
| Ly9 | 47.20549678 | -0.461462823 | 0.081138404 | -5.687353973 | NA | NA |
| Myo1f | 177.6744227 | -0.481449215 | 0.093620655 | -5.142553384 | 2.71E-07 | 0.00012303 |
| Cep89 | 462.1990602 | -0.487892183 | 0.075416872 | -6.469270999 | 9.85E-11 | 1.19E-07 |
| Robo3 | 1385.509849 | -0.488242926 | 0.093267151 | -5.234886258 | 1.65E-07 | 8.88E-05 |
| Irf8 | 278.0190289 | -0.490156352 | 0.094108921 | -5.208394155 | 1.90E-07 | 9.41E-05 |
| Mrps12 | 803.7912892 | -0.492843662 | 0.065027142 | -7.579045437 | 3.48E-14 | 7.22E-11 |
| Gbp9 | 153.547225 | -0.496320855 | 0.094760206 | -5.237650647 | 1.63E-07 | 8.88E-05 |
| Col11a1 | 849.3852423 | -0.512153256 | 0.085824774 | -5.967429158 | 2.41E-09 | 2.06E-06 |
| Ryr1 | 814.1892978 | -0.527323356 | 0.082111434 | -6.42204536 | 1.34E-10 | 1.50E-07 |
| Klhl6 | 80.66227421 | -0.529154941 | 0.095063864 | -5.566310035 | 2.60E-08 | 1.80E-05 |
| Fbxo17 | 72.42112456 | -0.534556476 | 0.094960969 | -5.629223046 | 1.81E-08 | 1.31E-05 |
| Dmkn | 74.88580842 | -0.562152356 | 0.094948919 | -5.920576671 | 3.21E-09 | 2.59E-06 |
| Cd52 | 94.61086518 | -0.618505511 | 0.093598114 | -6.608098035 | 3.89E-11 | 5.65E-08 |
| Lag3 | 504.3588427 | -0.664146063 | 0.094071643 | -7.060002775 | 1.66E-12 | 3.02E-09 |
| Sbsn | 264.8129095 | -0.729972413 | 0.084046097 | -8.685381472 | 3.77E-18 | 1.10E-14 |
| C4b | 2177.430204 | -0.787387232 | 0.085483244 | -9.211012536 | 3.23E-20 | 1.17E-16 |
| Nudt19 | 1406.974187 | -1.211110834 | 0.061162364 | -19.80156993 | 2.89E-87 | 2.10E-83 |
| Ints4 | 1463.528862 | -1.71E-05 | 0.060368444 | -0.000283798 | 0.999773562 | 0.999917241 |
| Tyrobp | 359.6387953 | -2.40291985 | 0.095023142 | -25.28773319 | 4.36E-141 | 6.33E-137 |
| Sh2d4b | 13.63584079 | -2.47E-05 | 0.071502681 | -0.000345895 | 0.999724016 | NA |
| Parp11 | 852.6964715 | -2.49E-05 | 0.062260012 | -0.000400189 | 0.999680696 | 0.999917241 |
| Fastkd1 | 518.4055211 | -2.70E-05 | 0.071806223 | -0.000376444 | 0.999699641 | 0.999917241 |
| 4933408J17Rik | 1.265031602 | -2.96E-05 | 0.02561675 | -0.001154799 | 0.999078604 | NA |
| Aatk | 13313.88148 | -4.29E-05 | 0.05549952 | -0.000773613 | 0.999382746 | 0.999917241 |
| Snx7 | 548.7133995 | -4.72E-06 | 0.07231238 | -6.53E-05 | 0.999947932 | 0.999947932 |
| Slc35g3 | 4.371261137 | -5.49E-06 | 0.039279626 | -0.0001397 | 0.999888535 | NA |
| Gm15070 | 0.980770624 | -5.95E-05 | 0.02135653 | -0.002786411 | 0.997776768 | NA |
| 2310010M20Rik | 12.40052891 | -6.51E-05 | 0.073277699 | -0.000888355 | 0.999291195 | NA |
| Wnt11 | 10.78524007 | -6.64E-05 | 0.066281379 | -0.001001723 | 0.99920074 | NA |
| Mir320 | 1.307155672 | -6.71E-05 | 0.023047514 | -0.002910815 | 0.997677509 | NA |
| Zfp663 | 0.805095652 | -7.04E-05 | 0.020036685 | -0.00351476 | 0.997195633 | NA |
| 4930552P12Rik | 1.178231564 | -7.62E-05 | 0.022195824 | -0.003430973 | 0.997262485 | NA |
| Cyp2g1 | 4.907342637 | -9.31E-05 | 0.013876166 | -0.006706724 | 0.994648848 | NA |
| Foxo6 | 358.5640331 | -9.39E-05 | 0.089938774 | -0.00104349 | 0.999167415 | 0.999786862 |
| 1110038F14Rik | 505.4804255 | 0.000102869 | 0.074990073 | 0.001371773 | 0.998905484 | 0.999662491 |
| 4930402F06Rik | 1.028572868 | 0.000144485 | 0.021801544 | 0.006627271 | 0.994712241 | NA |
| Nepn | 0.754372279 | 0.00014583 | 0.019841392 | 0.007349779 | 0.994135778 | NA |
| Stx6 | 3433.393598 | 0.00015798 | 0.042756454 | 0.003694892 | 0.99705191 | 0.99913245 |
| Stxbp5 | 5162.033456 | 0.00019104 | 0.059697411 | 0.003200134 | 0.997446667 | 0.999289749 |
| Grip1 | 630.1448681 | 0.00021136 | 0.073402658 | 0.00287946 | 0.997702527 | 0.999289749 |
| Mbl2 | 0.795913517 | 0.00021625 | 0.020004648 | 0.010809997 | 0.991375038 | NA |
| Pax3 | 0.795913517 | 0.00021625 | 0.020004648 | 0.010809997 | 0.991375038 | NA |
| Nek4 | 755.7873638 | 0.000225473 | 0.069988865 | 0.00322156 | 0.997429572 | 0.999289749 |
| Xrcc5 | 931.1260125 | 0.000227722 | 0.060755275 | 0.003748192 | 0.997009383 | 0.99913245 |
| Spsb3 | 2099.193824 | 0.000234595 | 0.063861567 | 0.003673497 | 0.99706898 | 0.99913245 |
| Cyp2a5 | 32.93540652 | 0.000243205 | 0.087263907 | 0.002787 | 0.997776299 | 0.999289749 |
| Nle1 | 359.2750216 | 0.000247531 | 0.086198647 | 0.002871631 | 0.997708773 | 0.999289749 |
| Ugt2b5 | 0.778565143 | 0.000269352 | 0.019660249 | 0.013700353 | 0.989069042 | NA |
| Cntn3 | 1226.946852 | 0.000281354 | 0.072625364 | 0.003874039 | 0.996908971 | 0.99913245 |
| Sesn2 | 488.8658519 | 0.000287631 | 0.074838353 | 0.003843361 | 0.996933449 | 0.99913245 |
| Sh3pxd2b | 1100.229319 | 0.000290424 | 0.064438625 | 0.004506989 | 0.996403955 | 0.998948537 |
| Mir493 | 0.834263005 | 0.000290882 | 0.020138672 | 0.014443942 | 0.988475803 | NA |
| Cdhr5 | 1.733664362 | 0.00032513 | 0.031398081 | 0.01035508 | 0.991737989 | NA |
| Marcksl1-ps4 | 0.915593775 | 0.000329591 | 0.019199722 | 0.017166451 | 0.986303827 | NA |
| Gpr141 | 0.57694017 | 0.000332632 | 0.015006459 | 0.022165933 | 0.982315593 | NA |
| Slfn14-ps | 0.57694017 | 0.000332632 | 0.015006459 | 0.022165933 | 0.982315593 | NA |
| Cxcl11 | 0.938461913 | 0.000340361 | 0.021590522 | 0.015764361 | 0.98742238 | NA |
| Rbp7 | 2.274294225 | 0.000358956 | 0.035314828 | 0.010164456 | 0.991890077 | NA |
| Muc15 | 6.353283851 | 0.000373539 | 0.054456382 | 0.006859425 | 0.994527013 | NA |
| BC051070 | 1.022498331 | 0.000419229 | 0.023303932 | 0.017989646 | 0.985647113 | NA |
| Tmem144 | 454.2430676 | 0.000419811 | 0.075902801 | 0.005530906 | 0.995586998 | 0.998915149 |
| Mat2a | 8545.739586 | 0.000419904 | 0.069733854 | 0.006021516 | 0.995195555 | 0.998915149 |
| Rnf40 | 2864.912118 | 0.000421682 | 0.046606726 | 0.009047672 | 0.992781101 | 0.99829327 |
| Odf3 | 4.861169343 | 0.000438262 | 0.0488068 | 0.008979534 | 0.992835465 | NA |
| Gm10373 | 1.147038825 | 0.000442871 | 0.020101282 | 0.022031975 | 0.98242245 | NA |
| Six3os1 | 202.6254942 | 0.00044587 | 0.094305547 | 0.004727929 | 0.996227673 | 0.998948537 |
| Rnh1 | 1420.347374 | 0.000449431 | 0.071560133 | 0.006280469 | 0.994988943 | 0.998839626 |
| Anapc10 | 308.6105478 | 0.000460099 | 0.087182523 | 0.005277423 | 0.995789245 | 0.998915149 |
| Mef2b | 5.651830588 | 0.000462772 | 0.042875932 | 0.010793271 | 0.991388383 | NA |
| Rnase13 | 1.801764817 | 0.000492263 | 0.031628601 | 0.015563857 | 0.98758234 | NA |
| Olfr316 | 42.49757096 | 0.000496071 | 0.091300609 | 0.005433381 | 0.995664811 | 0.998915149 |
| Ehhadh | 157.6520547 | 0.000499014 | 0.090891037 | 0.005490239 | 0.995619445 | 0.998915149 |
| Rps15 | 8610.970614 | 0.000508963 | 0.056907243 | 0.008943733 | 0.992864028 | 0.99829327 |
| Fchsd2 | 1973.863046 | 0.000524471 | 0.056786232 | 0.009235884 | 0.992630936 | 0.99829327 |
| Ptpn4 | 1688.941045 | 0.000537971 | 0.081295975 | 0.006617439 | 0.994720086 | 0.998727174 |
| BB123696 | 0.735831993 | 0.000546488 | 0.017914998 | 0.030504504 | 0.975664702 | NA |
| Minos1 | 2281.166637 | 0.000548473 | 0.055315912 | 0.009915282 | 0.99208888 | 0.99829327 |
| 4930461G14Rik | 0.694002734 | 0.000549709 | 0.017735642 | 0.030994587 | 0.975273857 | NA |
| Vps4b | 1837.229103 | 0.000566594 | 0.062320979 | 0.009091546 | 0.992746096 | 0.99829327 |
| Cacna1b | 4033.031622 | 0.000569997 | 0.061695684 | 0.009238854 | 0.992628566 | 0.99829327 |
| Snai2 | 24.08880296 | 0.000578675 | 0.082445096 | 0.007018919 | 0.994399759 | 0.998727174 |
| Prdm4 | 1595.927161 | 0.000580489 | 0.052768894 | 0.011000595 | 0.991222972 | 0.998279587 |
| Lin7c | 2562.375208 | 0.000582383 | 0.070825254 | 0.008222814 | 0.993439217 | 0.998595119 |
| 4933405L10Rik | 10.60579032 | 0.0005913 | 0.067065538 | 0.008816756 | 0.992965338 | NA |
| Gm10012 | 0.745256552 | 0.000594825 | 0.017959077 | 0.033121142 | 0.973577983 | NA |
| Batf3 | 37.74946985 | 0.00059902 | 0.084904061 | 0.007055263 | 0.994370761 | 0.998727174 |
| 4930467K11Rik | 2.337336422 | 0.000600468 | 0.028924239 | 0.020760043 | 0.983437072 | NA |
| Slc15a4 | 886.3789451 | 0.000603503 | 0.072098021 | 0.008370595 | 0.993321309 | 0.998545698 |
| Arfgap2 | 3135.34028 | 0.000604339 | 0.045264749 | 0.013351216 | 0.989347588 | 0.998028378 |

| | | | | | | |
|---|---|---|---|---|---|---|
| Oxr1 | 10782.37388 | -0.076594708 | 0.079955689 | -0.957964449 | 0.338080685 | 0.57173348 |
| Epb4.1l3 | 7817.170012 | 0.055234037 | 0.057676194 | 0.957657445 | 0.338235522 | 0.571901923 |
| Tes | 66.70952328 | -0.143071681 | 0.149403305 | -0.95762059 | 0.338254112 | 0.571901923 |
| Nap1l2 | 3288.145398 | 0.113018474 | 0.096069222 | 1.1764275 | 0.239424088 | 0.463588705 |
| Hnrnpm | 3771.98518 | 0.059706888 | 0.06237964 | 0.957153462 | 0.338489803 | 0.57220009 |
| Ndufaf1 | 692.6295926 | -0.068762983 | 0.071852196 | -0.957006002 | 0.338564225 | 0.57220009 |
| Tmem59 | 6563.297528 | -0.049101735 | 0.05130597 | -0.957037457 | 0.338548349 | 0.57220009 |
| Lclat1 | 1691.523974 | -0.06346229 | 0.066323991 | -0.956852701 | 0.338641608 | 0.572226272 |
| Nudc | 4225.465817 | 0.060714794 | 0.06345601 | 0.956801326 | 0.338667543 | 0.572226272 |
| Cox18 | 301.3237464 | 0.089520702 | 0.093584584 | 0.956575302 | 0.33878166 | 0.57223188 |
| Zdhhc14 | 1697.470456 | -0.07787312 | 0.081396314 | -0.956715563 | 0.338710841 | 0.57223188 |
| Zfp9 | 1044.595998 | 0.08677288 | 0.090708368 | 0.956613831 | 0.338762205 | 0.57223188 |
| Pla2g6 | 799.279865 | 0.113012479 | 0.082938064 | 1.362612936 | 0.17300453 | 0.380811505 |
| Mfhas1 | 2211.643343 | 0.075916797 | 0.079432194 | 0.955743428 | 0.339201876 | 0.572754344 |
| Prkci | 2568.762703 | 0.06427433 | 0.067246366 | 0.95580376 | 0.339171388 | 0.572754344 |
| Sirpa | 21604.14535 | -0.05384758 | 0.056365447 | -0.955329608 | 0.339411039 | 0.573045074 |
| Wdr3 | 815.922808 | 0.076336509 | 0.079914479 | 0.955227517 | 0.339462653 | 0.57306977 |
| Ipmk | 1410.877791 | -0.071711367 | 0.075082653 | -0.955099003 | 0.339527633 | 0.573117022 |
| Nsun7 | 160.3040673 | 0.113009568 | 0.14611338 | 0.7734375 | 0.439263504 | 0.659682924 |
| A730043L09Rik | 10.87153933 | -0.14310545 | 0.149999483 | -0.954039624 | 0.340063585 | 0.573834153 |
| Egr2 | 457.2895258 | -0.143054743 | 0.14994123 | -0.954072093 | 0.340047151 | 0.573834153 |
| Xpo4 | 326.9004302 | -0.094752395 | 0.09932931 | -0.953921804 | 0.340123225 | 0.573872292 |
| Anapc15 | 363.5676395 | -0.078851116 | 0.082678491 | -0.953707736 | 0.340231603 | 0.573900218 |
| Slc2a6 | 854.0496946 | 0.112991328 | 0.114720333 | 0.98492852 | 0.324659183 | 0.558546456 |
| Nfkbib | 706.8351407 | 0.094851478 | 0.099446043 | 0.953798408 | 0.340185695 | 0.573900218 |
| Rdh13 | 816.7595936 | 0.062689058 | 0.065752123 | 0.953414974 | 0.340379858 | 0.574055243 |
| Anxa6 | 4378.250549 | 0.079779906 | 0.083715165 | 0.95299228 | 0.340593982 | 0.574189552 |
| Fam78b | 1655.30227 | -0.059016924 | 0.061929041 | -0.952976546 | 0.340601954 | 0.574189552 |
| Pus3 | 350.3693 | 0.112894565 | 0.097569278 | 1.157070833 | 0.247243422 | 0.472120974 |
| Wscd1 | 2875.445517 | -0.112550338 | 0.118078885 | -0.953179206 | 0.34049928 | 0.574189552 |
| Zfp551 | 215.5689118 | -0.089116822 | 0.093541884 | -0.95269432 | 0.340744972 | 0.57435842 |
| Iqcb1 | 888.7408228 | 0.064793681 | 0.068018705 | 0.952586221 | 0.340799761 | 0.574388284 |
| Rab30 | 418.3964392 | -0.091184417 | 0.095739223 | -0.952424864 | 0.340881555 | 0.574463651 |
| Arpp19 | 12780.46044 | -0.091533975 | 0.096117456 | -0.952313752 | 0.340937885 | 0.574496095 |
| Zmiz1 | 6541.841961 | -0.081869958 | 0.085979187 | -0.952206694 | 0.340992167 | 0.574525079 |
| 2410002F23Rik | 2062.920712 | 0.058637391 | 0.061617055 | 0.951642216 | 0.341278464 | 0.574801598 |
| Lrrtm1 | 1675.315308 | -0.091196779 | 0.09582835 | -0.951668054 | 0.341265356 | 0.574801598 |
| Ddx50 | 2405.674646 | 0.112862055 | 0.082173782 | 1.373455771 | 0.169610702 | 0.376928889 |
| Zc3h4 | 2795.186763 | -0.049352145 | 0.051860112 | -0.951639768 | 0.341279706 | 0.574801598 |
| AA987161 | 397.7206078 | -0.101460087 | 0.106632199 | -0.951495776 | 0.341352762 | 0.574820095 |
| Dmwd | 4789.244944 | -0.056659352 | 0.059572099 | -0.951105501 | 0.341550822 | 0.575028625 |
| Btbd8 | 61.39246431 | 0.112776229 | 0.156380276 | 0.721166578 | 0.470807032 | 0.685526187 |
| Fto | 8641.253163 | 0.052531707 | 0.055241559 | 0.950945415 | 0.341632086 | 0.575055208 |
| Rsph9 | 316.0894234 | -0.096273299 | 0.1012414 | -0.950928169 | 0.341640841 | 0.575055208 |
| Mast2 | 6044.896164 | -0.064680904 | 0.068032641 | -0.950733408 | 0.341739725 | 0.575159167 |
| Zscan25 | 504.5870473 | 0.11261997 | 0.081506498 | 1.38172996 | 0.167054632 | 0.373002426 |
| Rest | 112.5168288 | 0.11260827 | 0.117031691 | 0.962203219 | 0.335947532 | 0.570014191 |
| Gpkow | 1637.78344 | 0.070801316 | 0.07453651 | 0.949887728 | 0.342169303 | 0.575602106 |
| Rtn4ip1 | 523.6757993 | -0.102606799 | 0.10801362 | -0.949943161 | 0.342141134 | 0.575602106 |
| Csnk1e | 3999.008011 | 0.112287099 | 0.063885205 | 1.757638535 | 0.078809032 | 0.232053809 |
| Commd7 | 884.1599229 | -0.067432206 | 0.070998159 | -0.949773996 | 0.342227102 | 0.575604289 |
| Ergic2 | 1695.388389 | -0.064762091 | 0.068198448 | -0.949612386 | 0.342309243 | 0.575617474 |
| Zfp932 | 1375.390122 | 0.112269897 | 0.134011239 | 0.837764786 | 0.402162819 | 0.626935378 |
| Acvr1b | 4792.300795 | 0.082927805 | 0.087399488 | 0.948836275 | 0.342703889 | 0.575828939 |
| Alg2 | 5429.982202 | -0.042606362 | 0.044920771 | -0.948477964 | 0.342886186 | 0.575828939 |
| Alg5 | 514.8841768 | 0.090130678 | 0.095091499 | 0.947831082 | 0.343215454 | 0.575828939 |
| Apoe | 40943.05504 | 0.07785142 | 0.082112978 | 0.948101282 | 0.343077896 | 0.575828939 |
| Arhgdia | 10904.11064 | -0.064519901 | 0.068058187 | -0.948010866 | 0.343123923 | 0.575828939 |
| Cabp1 | 3864.496703 | 0.112253423 | 0.119603718 | 0.938544595 | 0.347964611 | 0.579228723 |
| Cdh6 | 408.2508943 | 0.083402727 | 0.087950648 | 0.94829008 | 0.342981799 | 0.575828939 |
| Myc | 126.9688261 | 0.112205255 | 0.13124631 | 0.854921216 | 0.392594765 | 0.618743087 |
| Eif2b4 | 1096.711984 | 0.069478071 | 0.073287425 | 0.94802172 | 0.343118397 | 0.575828939 |
| Grin2c | 1938.439969 | -0.096050451 | 0.101195371 | -0.949158545 | 0.342539982 | 0.575828939 |
| Csnk1d | 6090.134862 | 0.112177904 | 0.082705268 | 1.356357418 | 0.174985475 | 0.383754415 |
| Kcnj10 | 9747.172351 | -0.071039328 | 0.074887283 | -0.948616713 | 0.342815588 | 0.575828939 |
| Lgalsl | 2861.428381 | -0.051697191 | 0.054511094 | -0.948379266 | 0.342936411 | 0.575828939 |
| Lypla1 | 708.5421395 | -0.082748583 | 0.087248435 | -0.948424847 | 0.342913215 | 0.575828939 |
| Slc3a2 | 4225.439469 | 0.112148057 | 0.080504925 | 1.393058341 | 0.163602149 | 0.36875934 |
| Pmvk | 1107.991716 | -0.073743898 | 0.077726974 | -0.948755543 | 0.342744957 | 0.575828939 |
| Ppwd1 | 249.1012572 | 0.086297592 | 0.091043398 | 0.947873143 | 0.343194039 | 0.575828939 |
| Rabggta | 1571.602124 | 0.059837741 | 0.063106648 | 0.94820027 | 0.34302751 | 0.575828939 |
| Tbc1d4 | 603.7358776 | 0.095120212 | 0.100325572 | 0.948115325 | 0.343070747 | 0.575828939 |
| Wdr33 | 2321.336549 | 0.063525239 | 0.067013131 | 0.947952115 | 0.343153832 | 0.575828939 |
| Zfpl1 | 713.7963941 | 0.070635119 | 0.074510916 | 0.947983501 | 0.343137854 | 0.575828939 |
| AU023762 | 83.87971847 | -0.135232141 | 0.142714597 | -0.947570491 | 0.343348154 | 0.575863522 |
| Ankrd28 | 2538.424663 | -0.057586953 | 0.060783718 | -0.947407545 | 0.343431148 | 0.575863522 |
| D1Ertd622e | 2134.236505 | 0.075898622 | 0.08011081 | 0.947420477 | 0.343424561 | 0.575863522 |
| Stk35 | 1475.626354 | 0.062217685 | 0.065655056 | 0.947644989 | 0.343310215 | 0.575863522 |
| Fam217b | 1447.552844 | 0.112132806 | 0.069148096 | 1.621632585 | 0.104882041 | 0.279410144 |
| Utp11l | 452.4697294 | 0.075470387 | 0.07965204 | 0.947501003 | 0.343383546 | 0.575863522 |
| 9330117O12Rik | 76.88595692 | 0.11212559 | 0.168005001 | 0.667394356 | 0.50452027 | 0.710734511 |
| Man1a2 | 5643.189334 | -0.054662834 | 0.057719054 | -0.947050074 | 0.343613263 | 0.576022005 |
| Taf6 | 1280.89925 | 0.060821124 | 0.064231066 | 0.946911327 | 0.343683965 | 0.576053587 |
| Eif3f | 3817.035764 | 0.112108131 | 0.059719971 | 1.877230145 | 0.060486558 | 0.195530717 |
| Rfxap | 783.1271041 | 0.071302838 | 0.07534299 | 0.946376538 | 0.343956568 | 0.576385795 |
| Arxes2 | 544.5424958 | -0.096986269 | 0.102491033 | -0.946290281 | 0.344000549 | 0.576397157 |
| Ap3b1 | 2026.488278 | 0.051859681 | 0.05481674 | 0.946055547 | 0.344120255 | 0.576410728 |
| Ddx24 | 8024.439951 | 0.112092565 | 0.070544774 | 1.58895632 | 0.112070255 | 0.29211375 |
| Syn1 | 25587.49247 | 0.071229605 | 0.07528933 | 0.946078348 | 0.344108626 | 0.576410728 |
| Nup153 | 2207.825796 | 0.067535235 | 0.071403533 | 0.945824841 | 0.344237933 | 0.576510979 |
| Ppp6r2 | 3635.610539 | 0.1120764 | 0.069633407 | 1.609520563 | 0.107502562 | 0.284094269 |
| Lrp10 | 1160.211294 | 0.07431359 | 0.078580213 | 0.945703595 | 0.344299788 | 0.57652447 |
| 9530052E02Rik | 12.81494132 | -0.15757016 | 0.16672746 | -0.945076236 | 0.344619957 | 0.576722182 |
| Arx | 599.5358166 | -0.089944953 | 0.09518283 | -0.944970358 | 0.344674009 | 0.576722182 |
| Cst3 | 29140.80583 | -0.078609023 | 0.083193939 | -0.94488882 | 0.34471564 | 0.576722182 |
| Mfap2 | 108.5860249 | 0.112047521 | 0.117490919 | 0.953669631 | 0.340250897 | 0.573900218 |
| Pgd | 1322.551735 | 0.070111783 | 0.074191763 | 0.945007643 | 0.344654974 | 0.576722182 |
| Ppm1b | 4134.029688 | -0.072974648 | 0.077225078 | -0.944960497 | 0.344679044 | 0.576722182 |
| Slc27a4 | 3908.810715 | -0.061761106 | 0.065333258 | -0.945324131 | 0.344493422 | 0.576722182 |
| Mtg1 | 577.7272388 | 0.112033781 | 0.072844418 | 1.537987169 | 0.124051752 | 0.311567166 |
| Ginm1 | 402.0289567 | 0.077387696 | 0.081908439 | 0.944807369 | 0.344757229 | 0.576729487 |
| AI467606 | 66.90449025 | -0.141277021 | 0.149673796 | -0.943899498 | 0.345221009 | 0.576948985 |
| Sypl2 | 106.7184176 | 0.112012242 | 0.140343435 | 0.79812955 | 0.424795315 | 0.647112242 |
| Mtmr1 | 1823.012354 | 0.111903554 | 0.093962177 | 1.190942542 | 0.233676142 | 0.457143705 |
| Dbi | 3219.290115 | -0.081454249 | 0.086313455 | -0.943702798 | 0.345321544 | 0.576948985 |
| Denr | 2073.786632 | 0.054314256 | 0.057594773 | 0.943041408 | 0.345659723 | 0.576948985 |
| Eif6 | 1247.762082 | 0.066326854 | 0.070310189 | 0.943346266 | 0.345503818 | 0.576948985 |
| Fignl1 | 31.85959655 | -0.154849762 | 0.16420584 | -0.943022257 | 0.345669519 | 0.576948985 |
| Mta3 | 2074.317999 | 0.111886047 | 0.075914742 | 1.473838205 | 0.140525159 | 0.335123413 |
| Hectd2 | 599.4413907 | -0.096675158 | 0.102511497 | -0.943066492 | 0.345646893 | 0.576948985 |
| Tmc7 | 710.6562234 | 0.11188011 | 0.085081438 | 1.314976722 | 0.188517726 | 0.401684172 |
| Dcp1a | 1118.767705 | 0.111789935 | 0.068106215 | 1.641405785 | 0.10071321 | 0.271477118 |
| Lrguk | 44.74658474 | -0.137989968 | 0.146252031 | -0.943508042 | 0.345421104 | 0.576948985 |
| Mapk9 | 10646.0154 | 0.048227689 | 0.051136339 | 0.943119709 | 0.345619676 | 0.576948985 |
| Med17 | 1016.252218 | 0.07463759 | 0.07908583 | 0.943754277 | 0.345295231 | 0.576948985 |
| Wrap53 | 425.7405504 | 0.111778163 | 0.107010039 | 1.044557722 | 0.296227428 | 0.527921299 |
| Nudt14 | 159.7638798 | -0.097003265 | 0.102827598 | -0.943358272 | 0.345497679 | 0.576948985 |
| Rnaseh2c | 324.2460017 | -0.112256357 | 0.118918903 | -0.94397404 | 0.345182914 | 0.576948985 |
| Tsr1 | 1422.760314 | 0.111765865 | 0.055099628 | 2.028432274 | 0.042516149 | 0.156323021 |
| Setd5 | 4738.151442 | 0.057926541 | 0.061360644 | 0.944034122 | 0.345152212 | 0.576948985 |
| Syngr1 | 26888.13004 | -0.065002566 | 0.068930173 | -0.943020502 | 0.345670416 | 0.576948985 |
| Ftsjd2 | 4483.906161 | 0.111759339 | 0.052094083 | 2.14533651 | 0.031925951 | 0.128885016 |
| Mapk4 | 4008.792027 | 0.086076525 | 0.091286516 | 0.942927044 | 0.345718221 | 0.576966622 |
| Cdc123 | 2186.89042 | -0.041046658 | 0.043539859 | -0.942737499 | 0.345815188 | 0.577066292 |
| Gm14295 | 1694.566296 | -0.1345732 | 0.142844308 | -0.942097043 | 0.346142958 | 0.577551044 |
| Fchsd2 | 1900.741819 | -0.08244709 | 0.087534574 | -0.941880287 | 0.346253934 | 0.577674001 |
| Rab3gap2 | 4312.17338 | 0.06972247 | 0.074034526 | 0.941756149 | 0.346317501 | 0.577717846 |
| D17Wsu92e | 12150.4624 | -0.073340457 | 0.077893463 | -0.941548294 | 0.346423953 | 0.577815952 |
| Fam115c | 30.53282782 | -0.156690759 | 0.166427981 | -0.941492879 | 0.346452337 | 0.577815952 |
| Fam131a | 10788.97297 | -0.100897889 | 0.107175947 | -0.941422886 | 0.34648819 | 0.577815952 |
| Ccdc160 | 74.65780813 | -0.142444032 | 0.151346401 | -0.941178852 | 0.346613212 | 0.577962237 |
| Cyp4v3 | 352.2845021 | 0.094479369 | 0.100392941 | 0.941095737 | 0.3466558 | 0.57797105 |
| Cyp1b1 | 176.1955793 | -0.145863784 | 0.155028118 | -0.940885989 | 0.346763289 | 0.578088056 |
| Haus7 | 184.251154 | -0.10380642 | 0.110353837 | -0.940668873 | 0.346874575 | 0.578166078 |
| Lancl2 | 3771.136896 | -0.069516636 | 0.073908557 | -0.940576292 | 0.346922036 | 0.578166078 |
| Vps26a | 2073.219914 | -0.061399443 | 0.06527626 | -0.940609077 | 0.346905229 | 0.578166078 |
| Gpm6a | 34413.24414 | -0.081525031 | 0.086735917 | -0.939922398 | 0.347257367 | 0.578662686 |
| Mfsd1 | 1041.927889 | 0.060548989 | 0.064427664 | 0.939798001 | 0.347321184 | 0.578686732 |
| Kansl3 | 3727.441082 | 0.11167621 | 0.066193858 | 1.687108342 | 0.091582521 | 0.255028682 |
| Rlbp1 | 768.8120905 | -0.139955276 | 0.148952384 | -0.939597424 | 0.347424098 | 0.578753787 |
| Nos1 | 1237.92031 | -0.118587493 | 0.126230879 | -0.939449158 | 0.347500184 | 0.578818303 |
| 4930578M01Rik | 19.03879293 | -0.151617445 | 0.161457616 | -0.939054153 | 0.347702941 | 0.579093774 |
| Foxp4 | 1421.150567 | 0.06521685 | 0.0694586 | 0.938931248 | 0.347766044 | 0.579099155 |
| Psmd8 | 3259.088566 | -0.057892524 | 0.061659798 | -0.938902269 | 0.347780924 | 0.579099155 |
| Asb18 | 12.39940403 | -0.155193671 | 0.165319444 | -0.938750264 | 0.34785898 | 0.579166886 |
| Wdyhv1 | 574.3404132 | 0.111624668 | 0.087260041 | 1.279218608 | 0.200820085 | 0.417688708 |
| Zbtb7a | 5091.879057 | 0.066954358 | 0.071339423 | 0.938532372 | 0.347970889 | 0.579228723 |
| Acy3 | 99.41741866 | 0.111594653 | 0.147385308 | 0.757162669 | 0.448952412 | 0.667953589 |
| Gal3st1 | 351.8062454 | -0.115409653 | 0.123016758 | -0.93816204 | 0.348161143 | 0.579420918 |
| Mars | 2027.289153 | 0.111587879 | 0.060917405 | 1.831789755 | 0.06698275 | 0.208861085 |
| AI837181 | 1954.47698 | 0.111522637 | 0.095300282 | 1.170223583 | 0.241911005 | 0.46665452 |
| Efhc2 | 71.84877043 | -0.139692285 | 0.14895977 | -0.937785315 | 0.34835475 | 0.579556372 |
| Zbtbd6 | 24.99524599 | -0.157418688 | 0.167895635 | -0.937598454 | 0.348450806 | 0.579653939 |
| Npy | 2427.223343 | 0.111458192 | 0.078869323 | 1.413200814 | 0.157596687 | 0.360498942 |
| Pcnxl4 | 1700.789845 | -0.054882366 | 0.058555237 | -0.937275104 | 0.348617066 | 0.579806013 |
| Zfp105 | 169.2982824 | 0.097163648 | 0.103687264 | 0.937083728 | 0.348715491 | 0.579907462 |
| Cpxm2 | 95.17164078 | -0.14541902 | 0.155210673 | -0.93691379 | 0.348802905 | 0.579990579 |
| Rai14 | 557.9948775 | 0.08854415 | 0.094524977 | 0.936727548 | 0.348898722 | 0.580087649 |
| Cdh20 | 859.2066795 | 0.066956818 | 0.07150201 | 0.936432669 | 0.349050464 | 0.58015801 |
| Ldoc1 | 49.41917001 | -0.147985206 | 0.158042266 | -0.936364741 | 0.349085425 | 0.58015801 |
| Mpp5 | 1461.62597 | 0.064194039 | 0.068555212 | 0.936384516 | 0.349075247 | 0.58015801 |
| Smyd2 | 1888.516259 | 0.111397738 | 0.058502589 | 1.90415055 | 0.05689058 | 0.188507836 |
| Slc25a32 | 283.5621273 | 0.111358399 | 0.092512007 | 1.203718338 | 0.228698462 | 0.451625538 |
| Chrnb2 | 2884.325367 | 0.05968216 | 0.063776433 | 0.935802722 | 0.34937477 | 0.580421397 |
| Dscaml1 | 2670.205175 | -0.070897893 | 0.075773579 | -0.935654539 | 0.349451084 | 0.580421397 |
| Dspp | 9.467365553 | -0.152568173 | 0.163035384 | -0.935797915 | 0.349377245 | 0.580421397 |
| Mbtd1 | 856.3540546 | 0.111341611 | 0.084140436 | 1.323283027 | 0.185741274 | 0.397331948 |
| Gna14 | 11.44649048 | 0.111340537 | 0.159662535 | 0.697349172 | 0.485584302 | 0.697252604 |
| Rasgef1b | 2657.181794 | -0.099782422 | 0.106668371 | -0.935445264 | 0.349558879 | 0.580500029 |
| Itm2c | 31448.61079 | -0.061524295 | 0.065784924 | -0.935233953 | 0.349667744 | 0.580618587 |
| Ddx39b | 5243.881011 | 0.062976922 | 0.067344147 | 0.935150634 | 0.349710675 | 0.580627647 |
| Nfatc3 | 658.7756281 | 0.071883518 | 0.076878143 | 0.935031917 | 0.349771851 | 0.580666995 |
| Fbxo30 | 296.0676914 | -0.10254503 | 0.109705753 | -0.934727911 | 0.349928539 | 0.580864879 |
| Mrpl48 | 1717.277959 | -0.061875357 | 0.066203354 | -0.934625704 | 0.349981228 | 0.580890106 |
| Mras | 9924.456202 | 0.083695954 | 0.089561407 | 0.93450914 | 0.350041323 | 0.58092762 |
| Btg2 | 1063.098737 | -0.153347691 | 0.164138438 | -0.934258257 | 0.350170691 | 0.581080078 |
| Enox1 | 1149.524669 | -0.09404502 | 0.100700678 | -0.933906519 | 0.350352116 | 0.581256632 |
| Pdzd11 | 1014.442888 | -0.075956426 | 0.081327208 | -0.933960814 | 0.350324107 | 0.581256632 |
| Zfp788 | 1151.886933 | 0.111331736 | 0.070164821 | 1.586717305 | 0.112576685 | 0.293054967 |
| 2210016L21Rik | 3756.668437 | 0.047652239 | 0.051055673 | 0.93333878 | 0.350645078 | 0.581493615 |
| Fubp1 | 2712.921755 | 0.089737273 | 0.096142189 | 0.933380802 | 0.350623389 | 0.581493615 |
| Vwf | 898.2523373 | -0.088861773 | 0.095199262 | -0.933429225 | 0.350598397 | 0.581493615 |
| Ralgps1 | 1313.652456 | 0.11130089 | 0.068673167 | 1.620733317 | 0.105074845 | 0.279634086 |
| Psmb6 | 2584.905107 | -0.070262929 | 0.075330075 | -0.932734086 | 0.350957281 | 0.5818475 |
| Spata2 | 2941.008464 | 0.068397382 | 0.0733321 | 0.932707265 | 0.350971133 | 0.5818475 |
| Rundc3a | 11918.53311 | -0.071072622 | 0.076228514 | -0.932362683 | 0.351149124 | 0.582080295 |
| Fam53c | 2991.643276 | 0.071990715 | 0.077229664 | 0.932164036 | 0.351251759 | 0.582188141 |
| 2810025M15Rik | 400.8737746 | -0.092732004 | 0.099542953 | -0.931577791 | 0.351554765 | 0.582452457 |
| A630019I02Rik | 24.69201386 | -0.155532108 | 0.166953218 | -0.931590956 | 0.351547959 | 0.582452457 |
| AU019823 | 610.2404072 | 0.111273904 | 0.104674243 | 1.063049527 | 0.287759492 | 0.517653032 |
| St8sia1 | 2322.547516 | -0.082102702 | 0.088118668 | -0.931728821 | 0.351476688 | 0.582452457 |
| Grm7 | 2091.015162 | -0.076380724 | 0.082016532 | -0.931284486 | 0.351706425 | 0.582567816 |
| Srp9 | 2172.704684 | -0.066741084 | 0.0716644 | -0.9313004 | 0.351698195 | 0.582567816 |
| H2-Q2 | 14.8448339 | -0.144805222 | 0.155527595 | -0.931058069 | 0.351823527 | 0.582637209 |
| Thop1 | 1521.106972 | -0.072733641 | 0.07811403 | -0.93112136 | 0.351790791 | 0.582637209 |
| AI413582 | 826.7788496 | -0.076297709 | 0.081987257 | -0.930604474 | 0.352058199 | 0.582963529 |
| Nmi | 110.0686612 | 0.111154731 | 0.120540156 | 0.9221386 | 0.356456285 | 0.587394165 |
| Tdrd9 | 27.93414474 | 0.111086267 | 0.165351082 | 0.67182062 | 0.5016979 | 0.709201588 |
| Mmp11 | 63.58565418 | -0.139471409 | 0.149954218 | -0.930093267 | 0.352322796 | 0.58321468 |
| Ebag9 | 628.66907 | -0.08193653 | 0.088120579 | -0.929822873 | 0.352462801 | 0.583347711 |
| Psmd4 | 3069.981823 | 0.06427493 | 0.069128242 | 0.929792634 | 0.352478461 | 0.583347711 |
| Aatk | 11601.78781 | 0.06194257 | 0.066641413 | 0.92949064 | 0.352634874 | 0.583544242 |
| 1-Sep | 21.21572057 | 0.11107061 | 0.167907928 | 0.661497117 | 0.508293563 | 0.713450923 |
| Caprin1 | 8137.635469 | -0.059269218 | 0.06380535 | -0.928906719 | 0.352937431 | 0.583857842 |
| Rab40c | 1603.503978 | 0.062076016 | 0.066826882 | 0.928907851 | 0.352936844 | 0.583857842 |
| Fam92b | 10.78926911 | -0.149749933 | 0.161241592 | -0.928730184 | 0.353028934 | 0.583928053 |
| Ltv1 | 582.8716848 | -0.070607947 | 0.076030485 | -0.928679424 | 0.353055247 | 0.583928053 |
| Bhlha9 | 38.28866341 | -0.155742895 | 0.167737907 | -0.928489558 | 0.353153682 | 0.583966184 |
| Lctl | 9.927637418 | 0.111048717 | 0.160671414 | 0.691154166 | 0.489468664 | 0.700390421 |
| Msto1 | 658.1647394 | 0.111022208 | 0.072764263 | 1.525779326 | 0.127064831 | 0.314948407 |
| Usp7 | 7421.263275 | 0.110988449 | 0.071127665 | 1.560411826 | 0.118662592 | 0.302391407 |
| Gdi1 | 25701.98993 | 0.042550766 | 0.045855452 | 0.927932528 | 0.35344257 | 0.584133757 |
| Wnk2 | 5566.95994 | 0.110896016 | 0.096558895 | 1.148480585 | 0.250770223 | 0.476448049 |
| Uevld | 639.2699104 | 0.071819674 | 0.077381435 | 0.928125387 | 0.353342532 | 0.584133757 |
| Ggt1 | 75.74652659 | -0.134274831 | 0.144753131 | -0.927612624 | 0.353608547 | 0.584344158 |
| Klhl4 | 532.9334684 | 0.092632412 | 0.099869897 | 0.927530871 | 0.353650971 | 0.58435194 |
| Zc3h8 | 246.4634479 | 0.082308285 | 0.088761907 | 0.92729289 | 0.353774485 | 0.584493695 |
| Cyp2j9 | 1061.31183 | 0.110886798 | 0.089944053 | 1.232841912 | 0.217634751 | 0.437812624 |
| 4933431E20Rik | 5094.994154 | 0.048455625 | 0.052329659 | 0.925968665 | 0.354462265 | 0.585380817 |
| Heatr5a | 605.1863091 | -0.077715469 | 0.08393108 | -0.925943867 | 0.354475152 | 0.585380817 |
| Tctn2 | 877.0459494 | -0.062434727 | 0.067431728 | -0.925895412 | 0.354500336 | 0.585380817 |
| Os9 | 5637.186447 | 0.110874366 | 0.052202793 | 2.12391636 | 0.033677138 | 0.133702991 |
| Cdk20 | 332.7509843 | 0.080240359 | 0.086678301 | 0.925726023 | 0.354588381 | 0.585463807 |
| Pcdhgb2 | 290.8558481 | -0.105199565 | 0.113697419 | -0.925259044 | 0.354831177 | 0.585802265 |
| Gm3279 | 12.06548888 | -0.150350391 | 0.162510563 | -0.925173036 | 0.354875907 | 0.58581369 |
| Klc3 | 78.96828778 | -0.12615745 | 0.136375254 | -0.925075819 | 0.35492647 | 0.585834742 |
| Vash1 | 2451.662182 | -0.0795175 | 0.085977914 | -0.924859605 | 0.355038941 | 0.585957963 |
| Stub1 | 3850.149428 | -0.049531576 | 0.053568833 | -0.924634218 | 0.355156208 | 0.58608907 |
| Galnt4 | 180.4409276 | -0.100999958 | 0.109285024 | -0.924188454 | 0.355388206 | 0.586284589 |
| AI316807 | 737.9161315 | 0.110845092 | 0.078288219 | 1.415859156 | 0.156816745 | 0.359508355 |
| Tmem42 | 362.3844539 | -0.11214618 | 0.121326656 | -0.924332573 | 0.355313188 | 0.586284589 |
| S100pbp | 680.6379026 | -0.078409147 | 0.084872822 | -0.923842816 | 0.355568159 | 0.58651901 |
| Capn3 | 225.8964674 | -0.131089671 | 0.141949226 | -0.923496906 | 0.355748312 | 0.586748225 |
| Gm20748 | 26.3654268 | -0.154096999 | 0.166874479 | -0.923430591 | 0.355782856 | 0.586748225 |
| Ccdc32 | 1552.422371 | -0.069814802 | 0.075648837 | -0.922880035 | 0.356069726 | 0.586971416 |

| | | | | | | |
|---|---|---|---|---|---|---|
| Ei24 | 4491.384074 | 0.000608116 | 0.059977265 | 0.010139112 | 0.991910298 | 0.99829327 |
| Ftcd | 2.8407767 | 0.000610462 | 0.036979689 | 0.016508025 | 0.9868291 | NA |
| 4921525B02Rik | 0.729403027 | 0.000619083 | 0.017892828 | 0.034599533 | 0.972399074 | NA |
| Gpr22 | 1945.48966 | 0.000624278 | 0.090218136 | 0.006919646 | 0.994478966 | 0.998727174 |
| D5Ertd605e | 1.902378653 | 0.000644155 | 0.029180455 | 0.022074873 | 0.98238823 | NA |
| Ppp2r3a | 2003.602511 | 0.000650379 | 0.074067708 | 0.008780865 | 0.992993973 | 0.998354821 |
| Vpreb3 | 1.561088071 | 0.000656472 | 0.029290921 | 0.022412133 | 0.982119202 | NA |
| Gm10556 | 0.591637739 | 0.00066069 | 0.015095854 | 0.04376635 | 0.96509065 | NA |
| Ppm1f | 4349.489281 | 0.000660905 | 0.06272323 | 0.010536854 | 0.991592963 | 0.99829327 |
| Taar3 | 0.55367566 | 0.000661977 | 0.014865518 | 0.044531053 | 0.9644811 | NA |
| Epcam | 5.809712952 | 0.000676377 | 0.052559941 | 0.012868681 | 0.989732561 | NA |
| Med13l | 5432.756733 | 0.000685494 | 0.071496512 | 0.009587798 | 0.992350161 | 0.99829327 |
| Trim38 | 1.904837943 | 0.000688206 | 0.030573761 | 0.022509706 | 0.98204137 | NA |
| 2610206C17Rik | 1.520360396 | 0.00070517 | 0.02869449 | 0.024575117 | 0.980393867 | NA |
| Tsr1 | 1584.168318 | 0.00071454 | 0.055112536 | 0.012965104 | 0.989655634 | 0.998028378 |
| Bbs10 | 200.7019849 | 0.000726717 | 0.091031 | 0.007983181 | 0.993630411 | 0.998598767 |
| Pwp2 | 1428.84286 | 0.000730462 | 0.064954158 | 0.011245815 | 0.991027327 | 0.998279587 |
| Mir129-1 | 6.227782697 | 0.00074305 | 0.051087886 | 0.01454455 | 0.988395537 | NA |
| 1500012K07Rik | 9.296915535 | 0.000743661 | 0.063614321 | 0.011690156 | 0.990672818 | NA |
| Gpr15 | 0.800262629 | 0.000744586 | 0.019741165 | 0.037717413 | 0.969912993 | NA |
| Qrich2 | 1.982759282 | 0.000745287 | 0.030543403 | 0.024400912 | 0.980532821 | NA |
| Prex1 | 7686.81451 | 0.000768219 | 0.070350318 | 0.01091991 | 0.991287345 | 0.998279587 |
| Nfe2 | 1.736274808 | 0.0007712 | 0.028053535 | 0.027490298 | 0.978068678 | NA |
| Slc24a4 | 1122.568292 | 0.000773367 | 0.076581316 | 0.010098641 | 0.991942588 | 0.99829327 |
| Aldh9a1 | 1533.266815 | 0.000787937 | 0.071996045 | 0.010944174 | 0.991267986 | 0.998279587 |
| Zbtbd6 | 27.58780106 | 0.000793403 | 0.083831194 | 0.009464291 | 0.992448701 | 0.99829327 |
| Adam10 | 3561.168864 | 0.000795099 | 0.06634238 | 0.011984786 | 0.990437753 | 0.998028378 |
| Gmds | 314.5227905 | 0.000800109 | 0.080185906 | 0.009978178 | 0.992038698 | 0.99829327 |
| Nup98 | 1198.677622 | 0.000815946 | 0.063996819 | 0.012749786 | 0.989827418 | 0.998028378 |
| Gm20110 | 0.561481615 | 0.000827798 | 0.014913642 | 0.055506106 | 0.955735265 | NA |
| Olfm1 | 38352.88984 | 0.000841136 | 0.053175128 | 0.015818225 | 0.987379409 | 0.997717928 |
| Cdk2ap1 | 3152.297831 | 0.000843294 | 0.061692906 | 0.013669218 | 0.989093881 | 0.998005844 |
| Zc3h18 | 1839.110378 | 0.000849208 | 0.045798592 | 0.01854224 | 0.985206281 | 0.997361871 |
| Mir700 | 0.570173846 | 0.000854631 | 0.01658092 | 0.051543047 | 0.958892801 | NA |
| Adat1 | 307.1992107 | 0.000865876 | 0.083222246 | 0.010404379 | 0.991698656 | 0.99829327 |
| 4930412O13Rik | 1.602984943 | 0.000866859 | 0.021207599 | 0.040874936 | 0.967395599 | NA |
| Lgr6 | 87.35356704 | 0.000876321 | 0.0950801 | 0.00921666 | 0.992646274 | 0.99829327 |
| Apol7d | 0.752792827 | 0.000899544 | 0.019838776 | 0.045342704 | 0.963834149 | NA |
| Ankk1 | 0.756170104 | 0.000906821 | 0.019572768 | 0.04633074 | 0.963046639 | NA |
| Tmem211 | 1.510697141 | 0.000914641 | 0.025768148 | 0.035495034 | 0.971685006 | NA |
| Rnf169 | 2731.328973 | 0.000915695 | 0.058077408 | 0.015766808 | 0.987420428 | 0.997717928 |
| Spats2l | 1155.315904 | 0.000916192 | 0.066782931 | 0.013718949 | 0.989054206 | 0.998005844 |
| Ccdc79 | 41.30852495 | 0.000923779 | 0.088733892 | 0.010410666 | 0.99169364 | 0.99829327 |
| Tubb2a-ps2 | 0.524155415 | 0.00093284 | 0.0162881 | 0.057271236 | 0.954329133 | NA |
| Smek1 | 1629.277246 | 0.000938023 | 0.065192756 | 0.014388448 | 0.988520075 | 0.997974805 |
| Ager | 19.84716907 | 0.000938345 | 0.083017075 | 0.01130304 | 0.990981671 | NA |
| Npc2 | 2502.742232 | 0.000951485 | 0.073691119 | 0.012911804 | 0.989698157 | 0.998028378 |
| Optn | 811.1451742 | 0.000956118 | 0.070262783 | 0.013607745 | 0.989142925 | 0.998005844 |
| Gm4566 | 0.765868818 | 0.000972758 | 0.016300351 | 0.059677118 | 0.952412796 | NA |
| 4933417E11Rik | 0.564827182 | 0.000975918 | 0.01797547 | 0.05429165 | 0.956702802 | NA |
| Pde6a | 3.357753918 | 0.000983371 | 0.040103694 | 0.024520716 | 0.980437259 | NA |
| Txlng | 509.8051626 | 0.000985945 | 0.070369213 | 0.014011029 | 0.988821182 | 0.998005844 |
| Scn4a | 1.554855131 | 0.000990897 | 0.029717643 | 0.033343738 | 0.973400475 | NA |
| Dag1 | 5225.080223 | 0.000999949 | 0.043486182 | 0.022994649 | 0.981654541 | 0.996444753 |
| Gpaa1 | 1479.46611 | 0.001003797 | 0.06197594 | 0.016196562 | 0.987077578 | 0.997691964 |
| Slfnl1 | 2.250463993 | 0.001029035 | 0.029188088 | 0.035255292 | 0.971876173 | NA |
| Strn4 | 10329.78007 | 0.001033531 | 0.068429759 | 0.015103536 | 0.98794958 | 0.99784144 |
| Nt5c1b | 0.72527368 | 0.001034131 | 0.015868836 | 0.06516743 | 0.948040693 | NA |
| 9430076G02Rik | 0.758139492 | 0.001039152 | 0.019857698 | 0.05232991 | 0.958265821 | NA |
| Plcxd1 | 790.6206687 | 0.001039554 | 0.0682175 | 0.015238819 | 0.987841653 | 0.997801811 |
| Tubgcp5 | 1336.507417 | 0.001046143 | 0.067674961 | 0.01545835 | 0.987666512 | 0.997763666 |
| Ube2u | 1.825540927 | 0.001051316 | 0.028640562 | 0.036707239 | 0.970718437 | NA |
| Gm7609 | 5.832558768 | 0.001055052 | 0.049803428 | 0.02118432 | 0.983098622 | NA |
| Dpcr1 | 0.778605831 | 0.001072146 | 0.016072664 | 0.066706164 | 0.946815627 | NA |
| Cldn20 | 8.758740003 | 0.001078603 | 0.060398489 | 0.017858104 | 0.985752052 | NA |
| Sdc1 | 38.32032578 | 0.00108236 | 0.083407493 | 0.012976771 | 0.989646325 | 0.998028378 |
| Tmem91 | 525.8558413 | 0.001089335 | 0.091914186 | 0.011851657 | 0.990543967 | 0.998028378 |
| Cenpm | 71.83074623 | 0.001100267 | 0.0940724 | 0.011695958 | 0.990668188 | 0.998028378 |
| Chst3 | 136.6735208 | 0.001103107 | 0.093385411 | 0.011812417 | 0.990575274 | 0.998028378 |
| Olfr1396 | 0.545002712 | 0.001106466 | 0.014647159 | 0.075541362 | 0.93978399 | NA |
| Yipf7 | 0.773030112 | 0.001110629 | 0.016122295 | 0.068887766 | 0.945078957 | NA |
| Klk6 | 260.1341053 | 0.001112344 | 0.095102023 | 0.011696322 | 0.990667898 | 0.998028378 |
| Dnajc15 | 722.1665811 | 0.001119356 | 0.065827297 | 0.017004441 | 0.986433073 | 0.997581728 |
| Atp13a2 | 6698.055681 | 0.001123407 | 0.068012146 | 0.016517744 | 0.986821347 | 0.997691964 |
| Spata31d1d | 1.486257525 | 0.001143285 | 0.026994352 | 0.042352758 | 0.966217488 | NA |
| Casp3 | 274.112759 | 0.001146314 | 0.079424756 | 0.014432704 | 0.988484768 | 0.997974805 |
| A530016L24Rik | 0.544512808 | 0.001147186 | 0.014645709 | 0.078329182 | 0.937566205 | NA |
| Gm19345 | 47.51595982 | 0.001150024 | 0.092636067 | 0.012414427 | 0.990094975 | 0.998028378 |
| Zfp572 | 0.531141502 | 0.001150477 | 0.012869483 | 0.089395739 | 0.92876741 | NA |
| Arfip2 | 3887.100798 | 0.001151469 | 0.043994689 | 0.026172915 | 0.979119419 | 0.995367691 |
| Ankra2 | 1161.372107 | 0.001151792 | 0.052253379 | 0.022042433 | 0.982414107 | 0.996546601 |
| Car15 | 213.5527915 | 0.001158504 | 0.091358673 | 0.012680835 | 0.989882429 | 0.998028378 |
| Stox1 | 168.6806984 | 0.001160514 | 0.08756108 | 0.013253765 | 0.989425335 | 0.998028378 |
| Mir1968 | 3.172473764 | 0.001164641 | 0.040622606 | 0.028669774 | 0.977127963 | NA |
| D730045A05Rik | 0.781166032 | 0.001165104 | 0.021332411 | 0.054616588 | 0.956443923 | NA |
| Nisch | 18828.5405 | 0.001175277 | 0.051228311 | 0.022941947 | 0.981696581 | 0.996444753 |
| Slc34a1 | 2.340456278 | 0.001180963 | 0.037225477 | 0.031724588 | 0.974691687 | NA |
| Noc4l | 859.9697702 | 0.001185118 | 0.063139138 | 0.018769947 | 0.985024628 | 0.997361871 |
| Vstm5 | 562.772975 | 0.00119869 | 0.065252496 | 0.018370018 | 0.98534367 | 0.997361871 |
| Snord85 | 0.517988882 | 0.001211529 | 0.014637749 | 0.082767464 | 0.934036441 | NA |
| Kcnj13 | 155.011273 | 0.001212253 | 0.03065523 | 0.039544746 | NA | NA |
| Bdkrb2 | 24.44729177 | 0.001217086 | 0.082361918 | 0.014777292 | 0.988209856 | 0.997965543 |
| Tnrc6c | 4388.364021 | 0.001245818 | 0.058468525 | 0.021307498 | 0.983000363 | 0.996711782 |
| 3110099E03Rik | 6.644073915 | 0.001247742 | 0.048867907 | 0.025532947 | 0.979629869 | NA |
| Best3 | 9.729391705 | 0.001250117 | 0.054461827 | 0.022954 | 0.981686966 | NA |
| 1110006O24Rik | 17.05743458 | 0.001258064 | 0.073144453 | 0.017199724 | 0.986277282 | NA |
| Acox2 | 8.518887032 | 0.001260426 | 0.0414733 | 0.030391261 | 0.975755014 | NA |
| Defb30 | 0.770412418 | 0.001268072 | 0.018053435 | 0.070239947 | 0.944002679 | NA |
| Slc30a1 | 1622.57602 | 0.001268572 | 0.072206209 | 0.017568736 | 0.985982898 | 0.997552088 |
| Tbc1d16 | 2821.498093 | 0.001272566 | 0.061495519 | 0.02069364 | 0.983490042 | 0.996711782 |
| Asphd2 | 3533.728155 | 0.001281532 | 0.069159268 | 0.018530154 | 0.985215922 | 0.997361871 |
| 2010016I18Rik | 0.558660649 | 0.00129212 | 0.016512228 | 0.078252323 | 0.937627342 | NA |
| Ccl7 | 2.706960573 | 0.001302788 | 0.035114455 | 0.03710118 | 0.970404331 | NA |
| Lrrc51 | 109.1215858 | 0.001303182 | 0.094194738 | 0.013834974 | 0.98896164 | 0.998005844 |
| Actn3 | 46.09859252 | 0.001306229 | 0.091532727 | 0.014270622 | 0.988614078 | 0.997974805 |
| 9330162B11Rik | 0.558930329 | 0.001313289 | 0.016513518 | 0.079528128 | 0.936612559 | NA |
| A230028O05Rik | 7.763369906 | 0.001320757 | 0.052895274 | 0.024969283 | 0.980079464 | NA |
| Rrm1 | 864.704614 | 0.001331827 | 0.068233751 | 0.019518598 | 0.984427401 | 0.997057065 |
| Prmt6 | 508.7583402 | 0.00134408 | 0.072307452 | 0.018588405 | 0.985169452 | 0.997361871 |
| Ctsw | 4.950156864 | 0.001349123 | 0.045264855 | 0.02980508 | 0.976222508 | NA |
| Rnf146 | 2310.574701 | 0.001358185 | 0.060444669 | 0.022469887 | 0.982073133 | 0.996444753 |
| Foxl1 | 0.765234524 | 0.001361471 | 0.016023983 | 0.084964595 | 0.932289538 | NA |
| Dgke | 3291.801127 | 0.001373158 | 0.067130327 | 0.020455114 | 0.983680318 | 0.996711782 |
| A3galt2 | 7.479995603 | 0.001382958 | 0.055494209 | 0.024920765 | 0.980118165 | NA |
| 1700055N04Rik | 0.769254926 | 0.001386439 | 0.021287703 | 0.065128641 | 0.948071576 | NA |
| C86187 | 1.91680046 | 0.001408178 | 0.029844152 | 0.04718438 | 0.962366277 | NA |
| Dab1 | 3881.395759 | 0.001422264 | 0.066282129 | 0.021457726 | 0.982880526 | 0.996711782 |
| Timm10b | 557.6468016 | 0.001434888 | 0.076489151 | 0.018759362 | 0.985033073 | 0.997361871 |
| Adam4 | 5.892466854 | 0.001434946 | 0.055055722 | 0.026063526 | 0.979206669 | NA |
| Bnc2 | 15.57552781 | 0.001450085 | 0.069451875 | 0.020878987 | 0.983342189 | NA |
| Rgr | 0.525396929 | 0.001468801 | 0.016298428 | 0.090119166 | 0.928192519 | NA |
| Mir1907 | 0.531833288 | 0.001476713 | 0.01473175 | 0.100240141 | 0.920153678 | NA |
| Dach2 | 24.33081698 | 0.001481793 | 0.08067811 | 0.018366734 | 0.98534629 | 0.997361871 |
| Mgst1 | 221.1813861 | 0.00148352 | 0.095090148 | 0.015601196 | 0.987552552 | 0.997717928 |
| Atp5k | 2026.847485 | 0.001490537 | 0.070906975 | 0.021021027 | 0.983228882 | 0.996711782 |
| Atp7a | 432.5885375 | 0.001492784 | 0.085781059 | 0.017402254 | 0.986115711 | 0.997552088 |
| Sympk | 6114.783238 | 0.001503295 | 0.05801296 | 0.025913096 | 0.979326654 | 0.995367691 |
| Arpp19 | 14002.75836 | 0.00150965 | 0.081672076 | 0.018484281 | 0.985252517 | 0.997361871 |
| Fsd1 | 1599.295283 | 0.001519525 | 0.062534749 | 0.024298888 | 0.9806142 | 0.996042366 |
| Ankdd1b | 11.30595212 | 0.001533268 | 0.061286485 | 0.025018037 | 0.980040576 | NA |
| Fam170b | 0.554225534 | 0.001535207 | 0.013011358 | 0.117989752 | 0.906075778 | NA |
| Brms1 | 945.9424076 | 0.001539789 | 0.067283096 | 0.022885228 | 0.981741824 | 0.996444753 |
| Emr4 | 1.538712315 | 0.001548459 | 0.028532765 | 0.054269509 | 0.956720442 | NA |
| Mamdc2 | 56.10741145 | 0.001557189 | 0.094928655 | 0.016403783 | 0.986912262 | 0.997691964 |
| Prr5l | 617.0124351 | 0.001566889 | 0.089261773 | 0.01755386 | 0.985994765 | 0.997552088 |
| 9230102O04Rik | 0.491929936 | 0.001568997 | 0.014457993 | 0.108521073 | 0.913582365 | NA |
| Tnpo2 | 9274.185961 | 0.001581401 | 0.043727386 | 0.036165006 | 0.971150789 | 0.99301409 |
| Kel | 0.829413177 | 0.001591566 | 0.018260046 | 0.087161124 | 0.93054344 | NA |
| Gm6034 | 1.685180203 | 0.001593278 | 0.031439332 | 0.050677864 | 0.959582216 | NA |
| Bola2 | 544.9824701 | 0.001594525 | 0.081121278 | 0.019656065 | 0.984317739 | 0.997015514 |
| Kis2 | 0.341198496 | 0.001595255 | 0.011244564 | 0.141868977 | 0.887183499 | NA |
| Aurkc | 5.238413299 | 0.001601457 | 0.047885835 | 0.033443226 | 0.973321139 | NA |
| Gsc | 0.536376888 | 0.001602396 | 0.014760442 | 0.108560138 | 0.913551379 | NA |
| Man1c1 | 2247.063226 | 0.001609764 | 0.066558671 | 0.024185634 | 0.980704537 | 0.996064474 |
| Arpc4 | 3055.208674 | 0.001609816 | 0.046633857 | 0.034520319 | 0.97246224 | 0.99301409 |
| Aldoart2 | 0.494547629 | 0.001624579 | 0.014477242 | 0.112216067 | 0.91065209 | NA |
| Olfr288 | 10.8552333 | 0.001626305 | 0.052641866 | 0.030893759 | 0.975354267 | NA |
| 1700019B03Rik | 5.966871673 | 0.001627147 | 0.047847388 | 0.03400702 | 0.972871553 | NA |
| Muc20 | 0.552207486 | 0.001653453 | 0.01647335 | 0.100371376 | 0.920049493 | NA |
| Slamf1 | 11.13686847 | 0.001664651 | 0.064723013 | 0.025719612 | 0.979480981 | NA |
| Hist1h1d | 3.194053724 | 0.001671119 | 0.029803011 | 0.056072152 | 0.955284328 | NA |
| Xrn1 | 742.4843136 | 0.001677824 | 0.072609484 | 0.023107505 | 0.981564519 | 0.996444753 |
| Edn2 | 1.743607036 | 0.001680442 | 0.031648648 | 0.053096805 | 0.957654777 | NA |
| Cnr1 | 6837.044198 | 0.001691235 | 0.078226045 | 0.021619843 | 0.982751205 | 0.996711782 |
| Speer8-ps1 | 0.346433883 | 0.001696603 | 0.01129932 | 0.150150844 | 0.880645607 | NA |
| Frmd5 | 1412.093086 | 0.001705947 | 0.061651628 | 0.027670745 | 0.977924757 | 0.995119468 |
| Zfp639 | 969.6446165 | 0.001706889 | 0.056845168 | 0.030026977 | 0.976045538 | 0.994391744 |
| Piwil1 | 7.404292387 | 0.001709463 | 0.045133779 | 0.037875468 | 0.969786973 | NA |
| Scn8a | 15640.99245 | 0.001712054 | 0.057885524 | 0.029576547 | 0.97640477 | 0.994548467 |
| Oaz3 | 2.700948297 | 0.001721038 | 0.033987006 | 0.050638115 | 0.959613891 | NA |
| Pcid2 | 1242.308043 | 0.001721527 | 0.050718158 | 0.033943003 | 0.972922601 | 0.993020918 |
| Gm8909 | 0.297797758 | 0.001727946 | 0.0107702 | 0.160437665 | 0.872536322 | NA |
| Als2cr12 | 1.38811635 | 0.001728901 | 0.027147931 | 0.063684452 | 0.949221485 | NA |
| Zcchc6 | 2754.316261 | 0.00173126 | 0.053236945 | 0.032519895 | 0.974057451 | 0.993518395 |
| Psg17 | 1.326564174 | 0.001732005 | 0.025687876 | 0.067425011 | 0.946243358 | NA |
| Zfp874a | 573.0016137 | 0.001756888 | 0.081802417 | 0.021477208 | 0.982864985 | 0.996711782 |
| Slc17a9 | 21.07770029 | 0.001757407 | 0.076493649 | 0.022974547 | 0.981670576 | NA |
| Il18rap | 1.756437712 | 0.001768343 | 0.029573094 | 0.059795682 | 0.952318365 | NA |
| Zfp936 | 0.332506265 | 0.001799388 | 0.011154591 | 0.161313707 | 0.871846328 | NA |
| Zfp78 | 146.4260065 | 0.001806742 | 0.089940602 | 0.020088165 | 0.983973041 | 0.996744384 |
| Fcnb | 0.55890461 | 0.001811485 | 0.014735798 | 0.122930899 | 0.902161817 | NA |
| Ppp2r1b | 661.0791764 | 0.001825147 | 0.078266145 | 0.023319745 | 0.981395222 | 0.996444753 |
| BC048644 | 4.505176405 | 0.001829991 | 0.04635945 | 0.039473952 | 0.968512521 | NA |
| Gm5177 | 2.247010132 | 0.001831766 | 0.033297021 | 0.055012894 | 0.956128191 | NA |
| Gm5480 | 3.566236658 | 0.001832503 | 0.027742889 | 0.066053086 | 0.947335561 | NA |
| Epha1 | 12.27091342 | 0.001840445 | 0.070927767 | 0.025948163 | 0.979298685 | NA |
| Ppt2 | 1109.02625 | 0.001856792 | 0.072084401 | 0.025758589 | 0.979449892 | 0.995367691 |
| Hltf | 1063.162478 | 0.001868246 | 0.06453233 | 0.028950542 | 0.976904036 | 0.99477799 |
| Egfl8 | 60.67257021 | 0.00187708 | 0.093491136 | 0.020077631 | 0.983981445 | 0.996744384 |
| Cdk5 | 4402.63096 | 0.001877195 | 0.053852295 | 0.034858214 | 0.972192801 | 0.99301409 |
| Ptplad2 | 45.96307611 | 0.001890737 | 0.091855232 | 0.020583884 | 0.983577597 | 0.996711782 |
| Cyp24a1 | 2.725777577 | 0.001900859 | 0.033731373 | 0.056352838 | 0.955060727 | NA |
| Igf2as | 10.66763695 | 0.001901012 | 0.058978247 | 0.032232427 | 0.974286697 | NA |
| Gpr110 | 0.337741652 | 0.001901809 | 0.011212843 | 0.169609875 | 0.865316956 | NA |
| Vmn2r18 | 0.337741652 | 0.001901809 | 0.011212843 | 0.169609875 | 0.865316956 | NA |
| Gsg1l | 4885.554324 | 0.001907747 | 0.084920204 | 0.022465177 | 0.98207689 | 0.996444753 |
| Rdh14 | 801.3052661 | 0.001912591 | 0.06051245 | 0.031606573 | 0.974785801 | 0.99384175 |
| Smc1b | 5.73928136 | 0.001927952 | 0.045623455 | 0.042257905 | 0.966293102 | NA |
| Mogat2 | 0.556118146 | 0.001947387 | 0.017922159 | 0.108658033 | 0.913473729 | NA |
| Tbr1 | 4839.668549 | 0.001960619 | 0.071720501 | 0.02733694 | 0.978190994 | 0.995181565 |
| Olfr33 | 0.333392541 | 0.001963252 | 0.012690293 | 0.154705017 | 0.877053874 | NA |
| 2010106C02Rik | 0.288615623 | 0.001973745 | 0.010664833 | 0.185070371 | 0.853173841 | NA |
| 4930471G03Rik | 0.288615623 | 0.001973745 | 0.010664833 | 0.185070371 | 0.853173841 | NA |
| Gm14139 | 0.288615623 | 0.001973745 | 0.010664833 | 0.185070371 | 0.853173841 | NA |
| Gm1553 | 0.288615623 | 0.001973745 | 0.010664833 | 0.185070371 | 0.853173841 | NA |
| Snord33 | 0.288615623 | 0.001973745 | 0.010664833 | 0.185070371 | 0.853173841 | NA |
| Nfix | 4158.861074 | 0.00197828 | 0.058935972 | 0.033566601 | 0.973222755 | 0.993070301 |
| Larp7 | 699.2378742 | 0.0019844 | 0.061170986 | 0.032440223 | 0.974120986 | 0.993518395 |
| C730002L08Rik | 149.2555482 | 0.001985854 | 0.094873302 | 0.020931639 | 0.983300188 | 0.996711782 |
| Bhlhe23 | 2.888036141 | 0.001986538 | 0.037310053 | 0.053244045 | 0.957537462 | NA |
| S100a1 | 1457.540538 | 0.001990807 | 0.072169265 | 0.02758525 | 0.977992946 | 0.995119468 |
| Lrrc9 | 97.99483767 | 0.001992389 | 0.093857302 | 0.021227851 | 0.983063897 | 0.996711782 |
| Tmem89 | 1.303856917 | 0.002002478 | 0.021846622 | 0.091660754 | 0.92696758 | NA |
| BC048507 | 0.571777182 | 0.002005396 | 0.016593835 | 0.120851694 | 0.903808346 | NA |
| Nf1 | 4622.201162 | 0.002011939 | 0.069781976 | 0.028831785 | 0.97699875 | 0.9948047 |
| Tnfsf14 | 0.555628243 | 0.002012196 | 0.017919596 | 0.112290247 | 0.910593275 | NA |
| Vmn2r28 | 1.074723478 | 0.002014275 | 0.023785503 | 0.084685005 | 0.932511818 | NA |
| Clec4a1 | 8.570775218 | 0.002020631 | 0.05860509 | 0.034478768 | 0.972495373 | NA |
| Dlx3 | 2.450194858 | 0.002025962 | 0.0316455 | 0.064020526 | 0.948953883 | NA |
| A630066F11Rik | 29.46303411 | 0.002029824 | 0.082472442 | 0.024612149 | 0.980364329 | 0.995858199 |
| Rs1 | 10.24727284 | 0.002029869 | 0.064934606 | 0.031260201 | 0.97506203 | NA |
| Ankrd42 | 577.2417736 | 0.002044172 | 0.073035684 | 0.027988681 | 0.977671179 | 0.995119468 |
| Ocstamp | 0.542256936 | 0.002045183 | 0.016410744 | 0.124624663 | 0.900820702 | NA |
| Hif1an | 2936.210625 | 0.002051857 | 0.060002767 | 0.034196037 | 0.972720827 | 0.99301409 |
| Ppp6r3 | 3753.013353 | 0.002056289 | 0.055364713 | 0.037140781 | 0.970372756 | 0.992821601 |
| Krt85 | 0.551209933 | 0.002058297 | 0.016467673 | 0.124990179 | 0.900531326 | NA |
| Gm1647 | 0.760714661 | 0.002064289 | 0.016085452 | 0.12833268 | 0.897885704 | NA |
| Mira | 1.511422223 | 0.002071933 | 0.029518052 | 0.070192076 | 0.944040781 | NA |
| Gpr173 | 529.7028759 | 0.002074518 | 0.073775225 | 0.028119437 | 0.977566892 | 0.995104181 |
| Trim42 | 0.338469525 | 0.002075488 | 0.01274613 | 0.162832788 | 0.870650092 | NA |
| Gstm3 | 1.40269462 | 0.002080778 | 0.022727347 | 0.091553919 | 0.927052465 | NA |
| Fbxl15 | 461.8507642 | 0.002083779 | 0.08088819 | 0.025761227 | 0.979447788 | 0.995367691 |
| Gtf2h5 | 1750.806132 | 0.002089739 | 0.044218335 | 0.047259554 | 0.962306363 | 0.990818134 |
| Tuba3a | 0.34789562 | 0.002094788 | 0.011319137 | 0.18506609 | 0.853177198 | NA |
| 1700007J10Rik | 5.936779414 | 0.002104248 | 0.051220152 | 0.041082433 | 0.967230179 | NA |
| Npas2 | 2252.450966 | 0.00210511 | 0.067761186 | 0.031066598 | 0.975216428 | 0.994035073 |
| 2310042E22Rik | 12.3423468 | 0.002108324 | 0.070596801 | 0.029864303 | 0.976175275 | NA |
| Ccin | 0.322082616 | 0.002134195 | 0.012566521 | 0.169831848 | 0.86514238 | NA |
| Gm15545 | 13.93130976 | 0.002135763 | 0.071232876 | 0.029982831 | 0.976080746 | NA |
| Tbc1d20 | 1526.55144 | 0.002137887 | 0.05728799 | 0.037318235 | 0.970231266 | 0.992821601 |
| Alkbh5 | 3295.265184 | 0.002139171 | 0.059672961 | 0.035848243 | 0.971403366 | 0.99301409 |
| 4930500L23Rik | 0.318645055 | 0.002148855 | 0.011013013 | 0.195119666 | 0.845299267 | NA |
| Triqk | 392.9499376 | 0.002155522 | 0.078110167 | 0.027595918 | 0.977984437 | 0.995119468 |
| Atg10 | 409.4699528 | 0.002160422 | 0.075264644 | 0.028704339 | 0.977100395 | 0.994838462 |
| Ythdc1 | 2610.998642 | 0.002161895 | 0.076042743 | 0.028429999 | 0.977319198 | 0.994991497 |

| | | | | | | | | | | | | | | | |
|---|---|---|---|---|---|---|---|---|---|---|---|---|---|---|---|
| Cirh1a | 1136.86539 | 0.068885336 | 0.074634826 | 0.922965044 | 0.356025422 | 0.586971416 | | | Cd79b | 9.467615619 | 0.002163183 | 0.065006829 | 0.033276249 | 0.973454294 | NA |
| Cnksr3 | 186.5974524 | 0.096782176 | 0.104864924 | 0.922922291 | 0.356047703 | 0.586971416 | | | A430071A18Rik | 0.520471911 | 0.002165992 | 0.01465982 | 0.147750226 | 0.882539892 | NA |
| Ncoa3 | 2191.942367 | 0.062385889 | 0.067588219 | 0.92302903 | 0.355992077 | 0.586971416 | | | Dppa5a | 2.26022915 | 0.00216735 | 0.031004749 | 0.069903818 | 0.944270214 | NA |
| Ccbl2 | 217.0903522 | 0.08856334 | 0.095997015 | 0.922563475 | 0.356234737 | 0.58718096 | | | Camk2g | 14345.41197 | 0.002177107 | 0.046319278 | 0.047002171 | 0.962511497 | 0.990888873 |
| Luzp1 | 5035.997611 | 0.088388029 | 0.095823335 | 0.922406101 | 0.356316789 | 0.587253731 | | | Tecpr2 | 4888.496435 | 0.002191045 | 0.051988403 | 0.042144882 | 0.966383201 | 0.99126569 |
| Eya3 | 1534.958712 | 0.110832499 | 0.053069258 | 2.088450124 | 0.03675725 | 0.141494865 | | | 9830132P13Rik | 0.343816189 | 0.002206381 | 0.01280494 | 0.172307012 | 0.863196172 | NA |
| Zfp366 | 144.0507112 | -0.131325731 | 0.142420685 | -0.922097311 | 0.356477819 | 0.587394165 | | | Hoxc13 | 0.343816189 | 0.002206381 | 0.01280494 | 0.172307012 | 0.863196172 | NA |
| Traf4 | 320.2069641 | -0.091164213 | 0.098902747 | -0.921756126 | 0.356655797 | 0.587624939 | | | Npy6r | 0.343816189 | 0.002206381 | 0.01280494 | 0.172307012 | 0.863196172 | NA |
| Iqsec3 | 9183.362736 | 0.074415191 | 0.080759698 | 0.921439679 | 0.35682092 | 0.587709508 | | | Nrcam | 15978.63759 | 0.00221429 | 0.052244177 | 0.042383486 | 0.966192993 | 0.99126569 |
| Nrg1 | 208.6563094 | -0.097912105 | 0.106244268 | -0.921575413 | 0.356750088 | 0.587709508 | | | Dcaf12l2 | 31.4662004 | 0.002215622 | 0.086615241 | 0.025580049 | 0.9795923 | 0.995422018 |
| Sh3glb1 | 2151.9357 | -0.053545071 | 0.058109941 | -0.921444264 | 0.356818528 | 0.587709508 | | | 4933409K07Rik | 1.306977674 | 0.002230354 | 0.026492953 | 0.084186697 | 0.932907995 | NA |
| Acsl6 | 6394.044766 | -0.045500494 | 0.049398309 | -0.92109415 | 0.357001274 | 0.587944063 | | | Stxbp5l | 5984.349757 | 0.00223159 | 0.069918258 | 0.031917129 | 0.974538138 | 0.993692335 |
| Lonrf3 | 1019.693794 | 0.110772883 | 0.133161514 | 0.831868607 | 0.405483113 | 0.630019591 | | | Gjb5 | 0.527194754 | 0.002237244 | 0.016312492 | 0.137149113 | 0.890912933 | NA |
| Sec61a2 | 2158.715885 | 0.066429803 | 0.072141486 | 0.920826657 | 0.357140935 | 0.588003988 | | | Csn1s1 | 0.323880442 | 0.002250219 | 0.011069789 | 0.203275733 | 0.838919521 | NA |
| Yipf6 | 1598.590955 | 0.058888556 | 0.063943784 | 0.920942625 | 0.357080383 | 0.588003988 | | | 0610005C13Rik | 0.546936012 | 0.002256051 | 0.017864335 | 0.12628797 | 0.899503978 | NA |
| 5430402O13Rik | 17.68293405 | 0.110711856 | 0.167711237 | 0.660133795 | 0.509167973 | 0.714070292 | | | Hcn2 | 5489.81836 | 0.002269036 | 0.061297935 | 0.037016515 | 0.970471838 | 0.992821601 |
| Copg1 | 13598.53206 | 0.042672584 | 0.046362187 | 0.92041785 | 0.357354444 | 0.588213084 | | | Sbk2 | 5.76904807 | 0.002286204 | 0.048458294 | 0.047178806 | 0.962370719 | NA |
| Adal | 570.5848755 | 0.075274805 | 0.081791672 | 0.920323595 | 0.357403682 | 0.58823164 | | | Exosc5 | 346.9434651 | 0.002302214 | 0.07780422 | 0.029589838 | 0.97639417 | 0.994548467 |
| Actn1 | 4256.179937 | 0.110701743 | 0.087053284 | 1.27165499 | 0.203495729 | 0.420609802 | | | Tipin | 397.2249815 | 0.002303285 | 0.087241235 | 0.026401337 | 0.978937227 | 0.995367691 |
| Haghl | 2481.200639 | 0.096041249 | 0.104403726 | 0.9199025 | 0.357623712 | 0.588468755 | | | Bcl9l | 1924.052684 | 0.002305779 | 0.056974013 | 0.040470716 | 0.967717853 | 0.99182033 |
| 2810429I04Rik | 23.29190464 | -0.153982532 | 0.167420514 | -0.91973515 | 0.357711179 | 0.588549826 | | | Pspn | 0.329777294 | 0.002306066 | 0.01265364 | 0.182245283 | 0.855390238 | NA |
| Nqo2 | 358.4824109 | -0.110164933 | 0.119795151 | -0.919610954 | 0.3577761 | 0.588549826 | | | Mad2l2 | 300.6812423 | 0.002306967 | 0.083054629 | 0.027776503 | 0.977840407 | 0.995119468 |
| Ethe1 | 384.7066135 | 0.110691863 | 0.094029477 | 1.177203856 | 0.23911415 | 0.463246349 | | | 1300002K09Rik | 5.912161705 | 0.002313489 | 0.040708866 | 0.056830109 | 0.954680529 | NA |
| Rfx7 | 1605.954663 | -0.061505566 | 0.066896671 | -0.919411473 | 0.357880391 | 0.588578577 | | | Fam25c | 0.348893173 | 0.002317888 | 0.012859101 | 0.180252738 | 0.856954158 | NA |
| Tubb2b | 3384.666494 | -0.118855402 | 0.129266822 | -0.919457912 | 0.35785611 | 0.588578577 | | | Slc6a20b | 2.037485204 | 0.002318312 | 0.028362908 | 0.081737451 | 0.934855497 | NA |
| Crebl2 | 1367.477755 | 0.110686369 | 0.092361411 | 1.198404911 | 0.230759422 | 0.453668953 | | | Il23r | 0.528166588 | 0.002319522 | 0.014711837 | 0.157663642 | 0.874721852 | NA |
| Gas7 | 24337.08852 | 0.068422491 | 0.074447757 | 0.919067194 | 0.358060427 | 0.588749683 | | | Ctla2a | 177.506956 | 0.002326876 | 0.094424806 | 0.024642631 | 0.980340015 | 0.995858199 |
| Bbs4 | 2443.151434 | 0.110659374 | 0.069121161 | 1.600947851 | 0.10938847 | 0.287606475 | | | Ccdc50 | 2397.839496 | 0.002361027 | 0.060222071 | 0.039205346 | 0.968726672 | 0.992230168 |
| Gpr81 | 53.61712825 | -0.154126373 | 0.167735598 | -0.918865016 | 0.358166181 | 0.588859095 | | | Tyms-ps | 0.35231393 | 0.002365814 | 0.012893442 | 0.183489743 | 0.854413766 | NA |
| Apba2 | 7754.8332 | 0.110598623 | 0.082349645 | 1.343037039 | 0.179259981 | 0.389108174 | | | Actn4 | 7174.926895 | 0.00236631 | 0.058318437 | 0.040575681 | 0.967634172 | 0.991805208 |
| Fbxo28 | 2208.429998 | 0.064980338 | 0.070738633 | 0.918597592 | 0.358306092 | 0.588946072 | | | Tmtc1 | 5225.992893 | 0.00236658 | 0.062027938 | 0.038153449 | 0.969565336 | 0.992593281 |
| Zfp775 | 307.9812816 | -0.092527817 | 0.100776443 | -0.918149261 | 0.358540728 | 0.589226776 | | | 1700113H08Rik | 0.530784281 | 0.002372216 | 0.014729279 | 0.161054453 | 0.872050513 | NA |
| Crbn | 2892.93264 | 0.073460316 | 0.080064453 | 0.917514735 | 0.358872975 | 0.589710243 | | | Mir135b | 0.351510867 | 0.00237641 | 0.012886796 | 0.184406597 | 0.853694495 | NA |
| Zfp69 | 38.9184144 | -0.152338069 | 0.166133303 | -0.916962861 | 0.359162102 | 0.590122757 | | | Tmed6 | 0.351510867 | 0.00237641 | 0.012886796 | 0.184406597 | 0.853694495 | NA |
| B4galt2 | 3045.527199 | -0.095734772 | 0.104466084 | -0.916419648 | 0.359446834 | 0.590476142 | | | AU019990 | 0.384276662 | 0.002386924 | 0.011665248 | 0.20461835 | 0.837870318 | NA |
| B630019K06Rik | 1047.945508 | 0.11058091 | 0.085387249 | 1.295051786 | 0.195302366 | 0.409894454 | | | Gm1968 | 0.384276662 | 0.002386924 | 0.011665248 | 0.20461835 | 0.837870318 | NA |
| Arhgap22 | 216.9443347 | -0.118740956 | 0.129622745 | -0.916050309 | 0.359640509 | 0.590660962 | | | Iqcf3 | 0.574236473 | 0.002392129 | 0.01803507 | 0.132637662 | 0.894479945 | NA |
| Rsf1 | 2426.264364 | -0.080291427 | 0.087649882 | -0.916047174 | 0.359642153 | 0.590660962 | | | Plscr5 | 0.574236473 | 0.002392129 | 0.01803507 | 0.132637662 | 0.894479945 | NA |
| Caap1 | 173.9304366 | -0.097581525 | 0.106535495 | -0.91595318 | 0.359691452 | 0.590679317 | | | Pard6g | 277.9627062 | 0.002392216 | 0.088892581 | 0.026911311 | 0.978530472 | 0.995273517 |
| Zmym4 | 2469.541407 | -0.057525416 | 0.062811719 | -0.915838912 | 0.359751391 | 0.590715139 | | | 1600002D24Rik | 0.309462921 | 0.002395369 | 0.010912734 | 0.219502217 | 0.826258853 | NA |
| Zfp385c | 25.43576974 | -0.153876458 | 0.168038004 | -0.915724148 | 0.359811596 | 0.59075139 | | | 4933436H12Rik | 0.309462921 | 0.002395369 | 0.010912734 | 0.219502217 | 0.826258853 | NA |
| Klf12 | 1512.472478 | -0.077771916 | 0.084944807 | -0.915558216 | 0.359898655 | 0.590831719 | | | Bpi | 0.309462921 | 0.002395369 | 0.010912734 | 0.219502217 | 0.826258853 | NA |
| Pqlc1 | 599.4602038 | 0.110514817 | 0.094548533 | 1.168868653 | 0.242456555 | 0.467201662 | | | Olfr1140 | 0.309462921 | 0.002395369 | 0.010912734 | 0.219502217 | 0.826258853 | NA |
| Smug1 | 682.6406575 | -0.061793375 | 0.067511608 | -0.915300002 | 0.360034158 | 0.590928946 | | | Snora15 | 0.309462921 | 0.002395369 | 0.010912734 | 0.219502217 | 0.826258853 | NA |
| Stab1 | 809.6821873 | -0.100242142 | 0.109570368 | -0.914865436 | 0.360262277 | 0.591240729 | | | Trem1 | 0.309462921 | 0.002395369 | 0.010912734 | 0.219502217 | 0.826258853 | NA |
| Mmp14 | 463.3730223 | -0.131912334 | 0.144208782 | -0.914731625 | 0.360332538 | 0.591293406 | | | Mir1895 | 0.587590975 | 0.002397912 | 0.016690574 | 0.143668666 | 0.885762116 | NA |
| 2310010J17Rik | 40.57674791 | -0.145823676 | 0.159490498 | -0.914309493 | 0.360554244 | 0.591344072 | | | Tcp10a | 0.913990035 | 0.002413784 | 0.018995048 | 0.12707439 | 0.898881521 | NA |
| Aspm | 29.66348298 | -0.153139769 | 0.167488881 | -0.914327977 | 0.360544534 | 0.591344072 | | | Flrt2 | 2998.242836 | 0.002413858 | 0.082798894 | 0.029153263 | 0.976742356 | 0.99470885 |
| Klhdc9 | 159.1412293 | 0.110494336 | 0.128871002 | 0.857402626 | 0.391222408 | 0.617995175 | | | Gm15350 | 0.534316316 | 0.002428288 | 0.014753208 | 0.164593889 | 0.869263642 | NA |
| Ide | 1990.478111 | -0.053357272 | 0.058353284 | -0.91438335 | 0.360515447 | 0.591344072 | | | Cep57l1 | 176.988499 | 0.002434671 | 0.091129727 | 0.026716538 | 0.978685822 | 0.995336432 |
| Cry2 | 7162.290936 | 0.11048175 | 0.09461877 | 1.167651525 | 0.242947358 | 0.46741904 | | | Fndc8 | 0.335123958 | 0.00243719 | 0.012713899 | 0.191694901 | 0.8479812 | NA |
| Gm889 | 16.39239648 | 0.11042917 | 0.166306464 | 0.664010089 | 0.506683854 | 0.712391981 | | | Hoxa7 | 0.52184809 | 0.002439561 | 0.017699155 | 0.137834905 | 0.890370897 | NA |
| Mef2d | 4954.697462 | 0.110365112 | 0.089393119 | 1.234604115 | 0.216977877 | 0.436916767 | | | 4930449E01Rik | 0.33403441 | 0.002443448 | 0.011179364 | 0.218567683 | 0.826986829 | NA |
| Gpatch4 | 639.7236164 | 0.110290565 | 0.063428231 | 1.738824544 | 0.08206563 | 0.23860478 | | | 8430436N08Rik | 0.33403441 | 0.002443448 | 0.011179364 | 0.218567683 | 0.826986829 | NA |
| 6530411M01Rik | 14.86124717 | -0.151176225 | 0.165505709 | -0.913420005 | 0.361021688 | 0.591756282 | | | Defb28 | 0.33403441 | 0.002443448 | 0.011179364 | 0.218567683 | 0.826986829 | NA |
| Ttc37 | 924.1078246 | 0.110218869 | 0.074071817 | 1.488000064 | 0.136750877 | 0.329602254 | | | Gm7904 | 0.33403441 | 0.002443448 | 0.011179364 | 0.218567683 | 0.826986829 | NA |
| Tns1 | 2632.262513 | -0.069266211 | 0.075839812 | -0.913322553 | 0.361072924 | 0.591756282 | | | Mir137 | 1.258010048 | 0.002453443 | 0.027074076 | 0.090619636 | 0.927794829 | NA |
| Tti1 | 1024.096521 | 0.047527835 | 0.052034688 | 0.913387528 | 0.361038763 | 0.591756282 | | | Inpp1 | 812.3228262 | 0.002460059 | 0.061228741 | 0.040178175 | 0.967951078 | 0.99191939 |
| Sspo | 103.1973181 | 0.110180227 | 0.140297654 | 0.78533193 | 0.43225898 | 0.65353441 | | | Pira6 | 1.327112512 | 0.002464795 | 0.025083838 | 0.098262265 | 0.921724041 | NA |
| Airn | 21.33529681 | -0.151527921 | 0.166012011 | -0.912752762 | 0.361372588 | 0.592122131 | | | 5730408K05Rik | 11.00904932 | 0.002465882 | 0.062056927 | 0.039735805 | 0.968303756 | NA |
| Fmr1 | 1838.848056 | -0.056009467 | 0.061392221 | -0.91232189 | 0.361599294 | 0.592430947 | | | 1810010H24Rik | 98.41960669 | 0.002478065 | 0.094309779 | 0.026275805 | 0.979037353 | 0.995367691 |
| A730020E08Rik | 53.90710479 | -0.140813726 | 0.154364607 | -0.912215103 | 0.361655495 | 0.592460376 | | | Elp2 | 4317.915401 | 0.002489283 | 0.040556274 | 0.0613785 | 0.951057774 | 0.988202091 |
| 4931406C07Rik | 1016.011891 | 0.064761047 | 0.071021666 | 0.911849166 | 0.361848124 | 0.592600427 | | | 1190003K10Rik | 0.314698307 | 0.002498242 | 0.010973885 | 0.227653386 | 0.819915714 | NA |
| A930004D18Rik | 103.8126059 | -0.123724394 | 0.135694267 | -0.911787925 | 0.361880368 | 0.592600427 | | | Krt76 | 0.314698307 | 0.002498242 | 0.010973885 | 0.227653386 | 0.819915714 | NA |
| Blvrb | 220.9626668 | -0.091489807 | 0.100345579 | -0.911747267 | 0.361901776 | 0.592600427 | | | Sit1 | 0.572749016 | 0.00251544 | 0.016601993 | 0.151514331 | 0.879570007 | NA |
| Gle1 | 2315.111185 | 0.074962899 | 0.082223923 | 0.911692074 | 0.361930837 | 0.592600427 | | | Nusap1 | 13.37826019 | 0.002517834 | 0.067156408 | 0.037492095 | 0.970092643 | NA |
| Tbpl1 | 2235.077881 | 0.044395098 | 0.048695417 | 0.911689446 | 0.361932221 | 0.592600427 | | | Mir324 | 0.576364262 | 0.002519484 | 0.018048746 | 0.13959328 | 0.88898135 | NA |
| Gart | 1385.748606 | 0.110114012 | 0.084122107 | 1.308978301 | 0.190541701 | 0.404152589 | | | Slc45a2 | 2.705179238 | 0.002528722 | 0.03742745 | 0.067563289 | 0.94613328 | NA |
| Gclm | 1129.860608 | 0.074474674 | 0.081725717 | 0.911275865 | 0.36215004 | 0.592893756 | | | Abca16 | 0.308221406 | 0.002531337 | 0.012410715 | 0.203963844 | 0.838381753 | NA |
| Dguok | 352.7933601 | -0.101127529 | 0.111032355 | -0.910793329 | 0.362404278 | 0.593183458 | | | Klra9 | 0.308221406 | 0.002531337 | 0.012410715 | 0.203963844 | 0.838381753 | NA |
| Gatad2b | 1097.479438 | 0.092663101 | 0.101752089 | 0.910675165 | 0.362466553 | 0.593183458 | | | Prss46 | 0.308221406 | 0.002531337 | 0.012410715 | 0.203963844 | 0.838381753 | NA |
| Rad1 | 326.3528614 | -0.086148899 | 0.094601531 | -0.910650159 | 0.362479733 | 0.593183458 | | | Asxl3 | 409.8959446 | 0.002535251 | 0.087057611 | 0.029121536 | 0.97676766 | 0.99470885 |
| Birc3 | 92.61354309 | -0.133320801 | 0.146440463 | -0.910409585 | 0.362606545 | 0.593328314 | | | Ipp | 424.7521678 | 0.002588021 | 0.080775652 | 0.032039617 | 0.974440458 | 0.993662484 |
| Dscam | 2964.854063 | 0.11010571 | 0.113316144 | 0.971668344 | 0.33121557 | 0.566339732 | | | Cyp17a1 | 0.375584431 | 0.002594219 | 0.011588524 | 0.223861007 | 0.822865458 | NA |
| Pithd1 | 2665.824443 | 0.071728979 | 0.0788141 | 0.910103383 | 0.36276799 | 0.593467136 | | | Mir873 | 0.3436217 | 0.002597312 | 0.012804432 | 0.202844744 | 0.839256384 | NA |
| Echs1 | 1664.194968 | -0.060295196 | 0.066262589 | -0.909943254 | 0.362852437 | 0.593542616 | | | Hemgn | 0.342818636 | 0.00260767 | 0.012797704 | 0.203760804 | 0.838540424 | NA |
| Dcaf11 | 3429.380127 | 0.110007429 | 0.061406978 | 1.791448343 | 0.073221377 | 0.221797071 | | | Cul4a | 2972.301681 | 0.002613346 | 0.053493412 | 0.048853606 | 0.961035962 | 0.990548518 |
| Atg7 | 1010.549692 | 0.061669283 | 0.067830907 | 0.909162001 | 0.363264619 | 0.593603784 | | | Mir149 | 4.708733545 | 0.002622699 | 0.041373653 | 0.063390563 | 0.949455502 | NA |
| Cnnm2 | 1152.05273 | -0.050804349 | 0.055872316 | -0.909293772 | 0.363195077 | 0.593603784 | | | Gm15941 | 0.523892668 | 0.002624619 | 0.016292816 | 0.161090554 | 0.87202208 | NA |
| Ddx41 | 1342.247922 | 0.076534291 | 0.084175072 | 0.909227514 | 0.363230044 | 0.593603784 | | | Rpe65 | 224.7034138 | 0.002636544 | 0.07619778 | 0.034601323 | 0.972397647 | 0.99301409 |
| Fbxl19 | 2495.799893 | -0.065705216 | 0.072250742 | -0.909405409 | 0.363136168 | 0.593603784 | | | Scaf11 | 3013.903696 | 0.002648715 | 0.064229221 | 0.041238473 | 0.967105783 | 0.991752813 |
| Gm10509 | 153.3196779 | -0.116114288 | 0.127717947 | -0.909146216 | 0.36327295 | 0.593603784 | | | Gm7244 | 7.385878191 | 0.002660123 | 0.049733173 | 0.053487897 | 0.957343173 | NA |
| Hebp2 | 109.604699 | -0.117870089 | 0.129641778 | -0.909198336 | 0.363245442 | 0.593603784 | | | Gm19897 | 2.301024175 | 0.002665436 | 0.030858259 | 0.08637676 | 0.93116692 | NA |
| Zglp1 | 29.73814788 | 0.10997057 | 0.16811999 | 0.654119535 | 0.513034796 | 0.717285791 | | | Gm14164 | 1.766862212 | 0.00267896 | 0.02393186 | 0.111941137 | 0.910870079 | NA |
| 2310040G24Rik | 30.67612911 | 0.109945273 | 0.165112844 | 0.665879589 | 0.505488068 | 0.711356233 | | | Il1rl1 | 12.5048087 | 0.002679342 | 0.067767759 | 0.039537121 | 0.968462158 | NA |
| Olfr1393 | 20.7944011 | -0.152406229 | 0.167552542 | -0.909602605 | 0.363032124 | 0.593603784 | | | 2010106E10Rik | 0.324852276 | 0.002692527 | 0.011086532 | 0.242864675 | 0.808110231 | NA |
| Ak1 | 3584.317141 | -0.055009774 | 0.060535171 | -0.908724188 | 0.363495733 | 0.593901493 | | | Gm19276 | 0.548733837 | 0.002696881 | 0.016454365 | 0.163900656 | 0.869809351 | NA |
| Pth2r | 48.31474261 | -0.138261536 | 0.152160504 | -0.908655874 | 0.363531803 | 0.593901493 | | | Tnfrsf17 | 6.656281683 | 0.002702763 | 0.053299638 | 0.050708839 | 0.959557533 | NA |
| Tecr | 10991.7111 | 0.05347133 | 0.058857614 | 0.908486191 | 0.363621406 | 0.593985247 | | | Astl | 0.315916084 | 0.002704152 | 0.012501427 | 0.216307503 | 0.828748061 | NA |
| Npc2 | 2178.494441 | 0.068632222 | 0.075577532 | 0.908103502 | 0.363823541 | 0.594252787 | | | Gm12295 | 2.113712748 | 0.002710044 | 0.03197718 | 0.084749305 | 0.932460697 | NA |
| Pex12 | 281.2537421 | -0.087386963 | 0.096251337 | -0.907903887 | 0.363929004 | 0.594362389 | | | Zfml | 4750.223563 | 0.002721591 | 0.076862826 | 0.03540842 | 0.97175407 | 0.99301409 |
| Ss18l1 | 2050.965675 | 0.109917807 | 0.056113772 | 1.958838329 | 0.050131723 | 0.174157157 | | | Aph1a | 2358.991537 | 0.002730489 | 0.049993714 | 0.054616645 | 0.956443878 | 0.989344398 |
| Capn10 | 871.8828527 | -0.07436083 | 0.081927857 | -0.907637922 | 0.364069552 | 0.594378872 | | | Slc5a12 | 1.498691249 | 0.002735365 | 0.02736856 | 0.099945506 | 0.920387588 | NA |
| Mcm9 | 195.634284 | 0.092140776 | 0.101537224 | 0.907458091 | 0.364164602 | 0.594378872 | | | Gm15401 | 1.323608543 | 0.002737488 | 0.024679972 | 0.110919427 | 0.91168024 | NA |
| Mtmr7 | 3213.639047 | 0.066922746 | 0.073748196 | 0.907449264 | 0.364169268 | 0.594378872 | | | Elmo2 | 7909.753245 | 0.002743715 | 0.057863201 | 0.047417265 | 0.962180669 | 0.990758942 |
| Prorsd1 | 55.42666473 | -0.149665218 | 0.164867284 | -0.9077921 | 0.363988073 | 0.594378872 | | | Slc24a5 | 16.70125483 | 0.002758826 | 0.045509511 | 0.060620867 | 0.951661154 | NA |
| Txndc12 | 852.2420588 | -0.067591535 | 0.074484617 | -0.907456295 | 0.364165552 | 0.594378872 | | | Arhgap8 | 6.446520908 | 0.002779014 | 0.053625985 | 0.051822156 | 0.958670401 | NA |
| Bivm | 855.8328053 | 0.059796794 | 0.065902725 | 0.907349336 | 0.364222092 | 0.594402475 | | | Cyp4x1 | 378.5281887 | 0.002780849 | 0.0793602 | 0.035040856 | 0.972047163 | 0.99301409 |
| Pygo2 | 2096.871229 | -0.059134358 | 0.065181957 | -0.907219745 | 0.364290604 | 0.594451672 | | | Ccdc101 | 431.8118579 | 0.00278178 | 0.07443659 | 0.03737114 | 0.970189084 | 0.992821601 |
| Caskin1 | 6826.030657 | 0.109912428 | 0.062705288 | 1.752841443 | 0.079629227 | 0.233846711 | | | Gcm1 | 0.531587345 | 0.00279592 | 0.016344661 | 0.171060127 | 0.864176488 | NA |
| Pcdhga12 | 572.1809851 | 0.109887872 | 0.0889085 | 1.235965874 | 0.216471249 | 0.436443452 | | | Sowahd | 0.618148956 | 0.0028 | 0.015137723 | 0.184968392 | 0.853253827 | NA |
| Slc38a9 | 880.5416938 | 0.070559978 | 0.077848658 | 0.906373721 | 0.364738077 | 0.594935559 | | | Nampt | 2200.187764 | 0.002805698 | 0.062837179 | 0.044650286 | 0.96438606 | 0.990993714 |
| Sptlc1 | 1147.111639 | -0.051133321 | 0.056413362 | -0.906404432 | 0.364721828 | 0.594935559 | | | Dsg3 | 0.299039272 | 0.002817595 | 0.012303739 | 0.229003175 | 0.818866449 | NA |
| Car11 | 5670.378617 | -0.106740073 | 0.117798015 | -0.906127939 | 0.364868139 | 0.595018097 | | | Gtf2i | 10596.72743 | 0.002825407 | 0.03474899 | 0.081309028 | 0.935196194 | 0.985461126 |
| Tgfbr2 | 1000.487006 | 0.08021665 | 0.08852127 | 0.906185034 | 0.364837923 | 0.595018097 | | | Gm12171 | 0.321262748 | 0.002835746 | 0.012564188 | 0.225700674 | 0.821434234 | NA |
| Bsdc1 | 2566.512623 | 0.068556669 | 0.075709406 | 0.905523798 | 0.365187958 | 0.595372142 | | | Gm5108 | 0.321262748 | 0.002835746 | 0.012564188 | 0.225700674 | 0.821434234 | NA |
| 3-Mar | 47.59453514 | -0.138299334 | 0.15271968 | -0.905576376 | 0.365160118 | 0.595372142 | | | Pias4 | 650.2752329 | 0.002837681 | 0.073378974 | 0.038671584 | 0.969152229 | 0.992548843 |
| Dym | 2259.376154 | 0.109843648 | 0.069939332 | 1.57055615 | 0.116285782 | 0.2988829 | | | Defb6 | 0.354045348 | 0.002838436 | 0.012915673 | 0.219766771 | 0.8260528 | NA |
| Epb4.1l2 | 3876.282409 | -0.06819043 | 0.075340447 | -0.905097225 | 0.365413882 | 0.595469362 | | | Lin54 | 546.1683159 | 0.002844538 | 0.083483449 | 0.034073076 | 0.972818878 | 0.99301409 |
| Med19 | 1206.742111 | 0.057873618 | 0.063939368 | 0.905132781 | 0.365395048 | 0.595469362 | | | Fam45a | 1135.876909 | 0.002855465 | 0.056465572 | 0.050570012 | 0.95966816 | 0.990191033 |
| Papd7 | 1928.753028 | 0.068268442 | 0.075421503 | 0.905158859 | 0.365381234 | 0.595469362 | | | Iqcj | 1.351053393 | 0.002867826 | 0.027135261 | 0.105686339 | 0.915831219 | NA |
| Srpk1 | 2292.899991 | 0.063970652 | 0.070661278 | 0.905314101 | 0.365299008 | 0.595469362 | | | Cel | 0.301656965 | 0.002877498 | 0.012335801 | 0.233264001 | 0.815556411 | NA |
| 2310015A10Rik | 384.8017204 | 0.096937008 | 0.107141887 | 0.904753595 | 0.365595941 | 0.595703387 | | | Mir134 | 5.481220332 | 0.002892601 | 0.051775597 | 0.055868034 | 0.955446937 | NA |
| Map2k4 | 8593.68337 | 0.109823127 | 0.083144404 | 1.320872153 | 0.186543996 | 0.398530566 | | | Mfsd9 | 260.3968407 | 0.002896368 | 0.083534705 | 0.034672626 | 0.972340789 | 0.99301409 |
| 2410127L17Rik | 267.3160764 | 0.088952929 | 0.098375373 | 0.904219481 | 0.365879033 | 0.596039294 | | | Abca14 | 0.356663042 | 0.002896668 | 0.012942952 | 0.223802694 | 0.822910834 | NA |
| Ccdc13 | 182.055429 | -0.134674973 | 0.148975801 | -0.904005701 | 0.365992379 | 0.596098593 | | | Prkci | 2820.941124 | 0.002904372 | 0.065583403 | 0.044285167 | 0.964677095 | 0.990993714 |
| Prkcb | 52543.29474 | -0.069594897 | 0.076984314 | -0.904013985 | 0.365987986 | 0.596098593 | | | Ppil4 | 1682.786445 | 0.002918873 | 0.063295601 | 0.046114948 | 0.963218632 | 0.990993714 |
| Bmp2k | 671.2787017 | 0.069716519 | 0.077146446 | 0.903690606 | 0.366159482 | 0.596153796 | | | Hoga1 | 34.56806819 | 0.002935201 | 0.08901292 | 0.032974996 | 0.973694527 | 0.993459767 |
| Med29 | 601.0314982 | -0.060564036 | 0.067010778 | -0.903795442 | 0.366103879 | 0.596153796 | | | Grik5 | 9775.990764 | 0.002937667 | 0.067861202 | 0.04328934 | 0.965470888 | 0.990993714 |
| Tdp2 | 538.5785242 | 0.069825332 | 0.07726536 | 0.903708104 | 0.366150201 | 0.596153796 | | | Rpl41 | 7104.953221 | 0.002939476 | 0.069715817 | 0.042163684 | 0.966368213 | 0.99126569 |
| Xpr1 | 6145.866821 | -0.067020007 | 0.07416576 | -0.903651586 | 0.366180178 | 0.596153796 | | | Traf1 | 34.71493907 | 0.002951072 | 0.078312675 | 0.037683196 | 0.969940274 | 0.992740432 |
| Dlg1 | 7741.701007 | -0.065809517 | 0.072848302 | -0.903377496 | 0.366325579 | 0.596239482 | | | Rras | 336.7052014 | 0.002956458 | 0.079828341 | 0.037035193 | 0.970456945 | 0.992821601 |
| Lingo2 | 821.3108854 | -0.092988944 | 0.102931094 | -0.90340965 | 0.36630852 | 0.596239482 | | | Dydc1 | 1.328186792 | 0.002968129 | 0.024678338 | 0.120272665 | 0.904267157 | NA |
| Tmem97 | 544.2609843 | -0.103154892 | 0.114202596 | -0.90326223 | 0.366386737 | 0.596239482 | | | Ankle2 | 1755.497957 | 0.002975141 | 0.057030818 | 0.052167257 | 0.958395422 | 0.989630474 |
| Ttc21b | 502.6656256 | 0.083581849 | 0.092529023 | 0.903304136 | 0.366364502 | 0.596239482 | | | Pcp4l1 | 5781.254648 | 0.002977709 | 0.082321231 | 0.036171822 | 0.971145354 | 0.99301409 |
| 2310039H08Rik | 70.39366605 | -0.12901247 | 0.142933085 | -0.902607464 | 0.366734263 | 0.596573703 | | | Atp1b3 | 4049.763391 | 0.002989 | 0.056711656 | 0.052705212 | 0.957966786 | 0.989630474 |
| Ap5s1 | 556.7820422 | -0.080052954 | 0.08868468 | -0.902669482 | 0.366701338 | 0.596573703 | | | 4930452G13Rik | 13.10987468 | 0.002990671 | 0.063911907 | 0.046793636 | 0.962677702 | NA |
| Rcl1 | 557.3551244 | -0.059187019 | 0.065575 | -0.902585116 | 0.366746129 | 0.596573703 | | | 9230115E21Rik | 0.30673395 | 0.00299119 | 0.012397037 | 0.241282618 | 0.809336081 | NA |
| Zfp64 | 455.2570908 | -0.080909675 | 0.089628167 | -0.902725984 | 0.366671342 | 0.596573703 | | | Gm597 | 0.30673395 | 0.00299119 | 0.012397037 | 0.241282618 | 0.809336081 | NA |
| Tor1aip1 | 1753.537879 | 0.067464274 | 0.074764475 | 0.902357352 | 0.366867069 | 0.596707785 | | | Dnmt3l | 3.785301734 | 0.002991422 | 0.045801342 | 0.065312975 | 0.947924812 | NA |
| Mrpl32 | 430.5742038 | -0.083868737 | 0.092971166 | -0.902094061 | 0.367006904 | 0.596872569 | | | Nrk | 0.32976049 | 0.002997204 | 0.01265821 | 0.236779445 | 0.812827898 | NA |
| Decr1 | 749.6465442 | -0.112999003 | 0.125314254 | -0.901725058 | 0.367202939 | 0.596940754 | | | Tmem65 | 4215.340058 | 0.003002058 | 0.065129657 | 0.04609356 | 0.963235679 | 0.990993714 |
| Fahd2a | 554.0878826 | 0.083683649 | 0.092799163 | 0.901771594 | 0.367178212 | 0.596940754 | | | Syce3 | 0.328957426 | 0.003007117 | 0.01265135 | 0.237691372 | 0.812120476 | NA |
| Rnf26 | 1333.693709 | 0.10980276 | 0.07459282 | 1.472025813 | 0.141013167 | 0.335925279 | | | Hoxc4 | 1.088186466 | 0.003009428 | 0.021894933 | 0.137448623 | 0.8906762 | NA |
| Ubr7 | 1801.222714 | 0.050009306 | 0.055458305 | 0.901746028 | 0.367191797 | 0.596940754 | | | Tcp11l2 | 1157.458083 | 0.003023454 | 0.061796176 | 0.048926239 | 0.960978078 | 0.990548518 |
| Gm12070 | 155.6847635 | -0.136545648 | 0.151443889 | -0.90162534 | 0.367255925 | 0.596964258 | | | Rlf | 1638.363162 | 0.003030469 | 0.058799085 | 0.05153939 | 0.958895715 | 0.989769328 |
| Cyp4x1 | 316.0346238 | 0.095073216 | 0.105487699 | 0.901273013 | 0.367443179 | 0.597143341 | | | 1700034J05Rik | 0.565037534 | 0.003036034 | 0.01655741 | 0.183364044 | 0.854512386 | NA |
| Oxt | 42.17968518 | -0.089592175 | 0.099399926 | -0.901330404 | 0.367412673 | 0.597143341 | | | Psmb11 | 8.45064709 | 0.003037646 | 0.063984614 | 0.047474631 | 0.962134949 | NA |
| Nadk | 2655.002254 | -0.056516961 | 0.062718214 | -0.901125154 | 0.36752178 | 0.597208439 | | | Npdc1 | 6559.044262 | 0.003042776 | 0.056946493 | 0.053432196 | 0.957387553 | 0.989630474 |
| Sowahb | 845.8800677 | 0.109789711 | 0.128357529 | 0.855342975 | 0.392361303 | 0.618634473 | | | Rpl17 | 8761.145761 | 0.003067414 | 0.072161099 | 0.042507864 | 0.966093843 | 0.99126569 |
| Ano1 | 176.7209534 | -0.137324404 | 0.152463291 | -0.900704714 | 0.367745342 | 0.597446401 | | | Ssh1 | 534.160192 | 0.003077229 | 0.070656073 | 0.043552228 | 0.965261332 | 0.990993714 |
| Ptprs | 17729.73001 | 0.053152758 | 0.059053808 | 0.900073329 | 0.368081228 | 0.597837201 | | | Szrd1 | 1386.992041 | 0.003089078 | 0.063430165 | 0.048700449 | 0.961158018 | 0.990548518 |
| Sntb1 | 73.64341992 | -0.146285423 | 0.162526967 | -0.900068622 | 0.368083733 | 0.597837201 | | | Ctrb1 | 0.567655227 | 0.003094332 | 0.016573489 | 0.186703693 | 0.85189296 | NA |
| Mta1 | 2384.853693 | 0.109763848 | 0.065820406 | 1.667626417 | 0.095389911 | 0.261469936 | | | 9030204H09Rik | 0.544628687 | 0.003098637 | 0.016430094 | 0.188595218 | 0.850410081 | NA |
| Abhd16a | 2655.929107 | 0.058327757 | 0.064814695 | 0.899915634 | 0.368165149 | 0.597863573 | | | Olfr286 | 0.544628687 | 0.003098637 | 0.016430094 | 0.188595218 | 0.850410081 | NA |
| Kif1c | 4476.555011 | 0.07240266 | 0.080459966 | 0.89985944 | 0.368195057 | 0.597863573 | | | Cndp2 | 2066.915182 | 0.003104478 | 0.05142857 | 0.060364861 | 0.951865044 | 0.988463312 |
| Cbwd1 | 396.4672691 | -0.085496613 | 0.095021807 | -0.899757811 | 0.368249151 | 0.597887471 | | | Olig3 | 3.202049419 | 0.003111048 | 0.033650293 | 0.09245234 | 0.926338656 | NA |
| Rarg | 416.4014242 | 0.084606254 | 0.094039675 | 0.899686798 | 0.368286951 | 0.597887471 | | | Trim15 | 2.092455193 | 0.003120232 | 0.030622331 | 0.101894012 | 0.918840803 | NA |
| Syt10 | 286.742948 | -0.142103008 | 0.15799951 | -0.899388917 | 0.368445541 | 0.598082264 | | | Zfp862 | 604.5282903 | 0.003122694 | 0.065462207 | 0.047702238 | 0.961953551 | 0.990595298 |
| Kctd17 | 5578.965209 | 0.109748318 | 0.071349199 | 1.538185702 | 0.124003216 | 0.311545238 | | | Nelfb | 2317.280311 | 0.003126221 | 0.050706772 | 0.06165292 | 0.950839233 | 0.988202091 |
| Sf3b4 | 1633.095674 | 0.109744666 | 0.076252944 | 1.439218743 | 0.150088556 | 0.350370787 | | | Tecpr1 | 4367.512512 | 0.003128573 | 0.060121305 | 0.052037671 | 0.958498677 | 0.989630474 |
| Rad9a | 396.4081893 | 0.109718297 | 0.07281298 | 1.506850801 | 0.131848905 | 0.321812686 | | | Mir181b-1 | 0.913601056 | 0.003133902 | 0.019001915 | 0.164925567 | 0.869002569 | NA |
| BC030867 | 41.06606576 | 0.109691075 | 0.163135234 | 0.672393528 | 0.501333202 | 0.708921843 | | | Socs4 | 209.4577036 | 0.003140828 | 0.091274517 | 0.034410782 | 0.972549586 | 0.99301409 |
| Adarb2 | 496.8110099 | -0.116396069 | 0.129603438 | -0.898093991 | 0.369135443 | 0.598888434 | | | Pf4 | 18.28679974 | 0.00314085 | 0.073507772 | 0.042728136 | 0.965918251 | NA |
| 3110052M02Rik | 451.5443453 | 0.109638583 | 0.120813853 | 0.907500091 | 0.364142402 | 0.594378872 | | | Zfp395 | 437.3462991 | 0.003154718 | 0.088567895 | 0.035619204 | 0.971585996 | 0.99301409 |
| Krit1 | 1476.235996 | 0.109521455 | 0.07172104 | 1.52704779 | 0.126749128 | 0.314669282 | | | Gm15008 | 0.685014685 | 0.003158108 | 0.015709015 | 0.201037942 | 0.840668906 | NA |
| Atp6v0a4 | 12.77901486 | -0.147225179 | 0.164044251 | -0.897472347 | 0.369466923 | 0.599285653 | | | 4921507P07Rik | 38.04780287 | 0.003170456 | 0.092466308 | 0.034287688 | 0.972647743 | 0.99301409 |
| F9 | 11.29878812 | -0.134282892 | 0.149774014 | -0.896570031 | 0.369948396 | 0.599956086 | | | Ptchd4 | 172.9067352 | 0.003199189 | 0.092096676 | 0.034737293 | 0.972289224 | 0.99301409 |
| Cpox | 1274.346763 | 0.062806156 | 0.070073576 | 0.896288718 | 0.370098583 | 0.600106315 | | | Slc4a5 | 123.835318 | 0.003207275 | 0.024559389 | 0.130592627 | NA | NA |
| Poc5 | 1261.143326 | 0.077195975 | 0.086132053 | 0.896251428 | 0.370118494 | 0.600106315 | | | Serpina3g | 16.21186766 | 0.00322837 | 0.045958476 | 0.070245365 | 0.943998367 | NA |
| Slc13a5 | 555.2860016 | -0.087283054 | 0.097404004 | -0.896093072 | 0.370203057 | 0.600180618 | | | Prrg1 | 146.4025335 | 0.003230713 | 0.093631979 | 0.034504371 | 0.972474957 | 0.99301409 |
| Snx30 | 2416.01707 | 0.074818313 | 0.083511867 | 0.895900379 | 0.370305972 | 0.600284654 | | | 1600029I14Rik | 3.540352257 | 0.003243402 | 0.041457369 | 0.078234639 | 0.937641409 | NA |
| Gpr12 | 653.1292314 | 0.080412412 | 0.089780054 | 0.895660096 | 0.370434329 | 0.60040892 | | | Cars | 1726.344533 | 0.003244607 | 0.050423305 | 0.064347375 | 0.948693632 | 0.987440792 |
| Ipo4 | 3301.817512 | 0.10951018 | 0.057521633 | 1.903808611 | 0.056935115 | 0.188574747 | | | Spdef | 8.416982083 | 0.003263416 | 0.057490557 | 0.056764381 | 0.954732888 | NA |
| Reck | 502.4898129 | 0.08066125 | 0.09006616 | 0.895577764 | 0.370478316 | 0.60040892 | | | Zfp668 | 519.8427311 | 0.003281609 | 0.070935913 | 0.0462616 | 0.963101745 | 0.990993714 |
| Arnt2 | 17196.76819 | 0.058151351 | 0.064947903 | 0.895353788 | 0.370597996 | 0.600443935 | | | Krt14 | 1.247129162 | 0.003284032 | 0.022501368 | 0.1459481 | 0.88396236 | NA |
| Atp6v1c1 | 9366.141147 | 0.052128815 | 0.058217608 | 0.895413202 | 0.370566246 | 0.600443935 | | | Mrps33 | 1682.549137 | 0.003289946 | 0.053579921 | 0.0614026 | 0.951038582 | 0.988202091 |
| Stim2 | 2855.945161 | 0.109504737 | 0.088440376 | 1.238175839 | 0.215650868 | 0.435631624 | | | 1110028F11Rik | 0.342801831 | 0.003297835 | 0.012802206 | 0.257598965 | 0.796716429 | NA |
| Actr10 | 4676.209296 | -0.04056411 | 0.045323177 | -0.894997061 | 0.370788659 | 0.60051934 | | | Cabp2 | 0.342801831 | 0.003297835 | 0.012802206 | 0.257598965 | 0.796716429 | NA |
| Ccdc117 | 712.7152667 | 0.064237742 | 0.071776764 | 0.894965694 | 0.370805427 | 0.60051934 | | | Mov10l1 | 0.342801831 | 0.003297835 | 0.012802206 | 0.257598965 | 0.796716429 | NA |
| Chd6 | 3971.662892 | -0.070233742 | 0.078481859 | -0.894904151 | 0.370838327 | 0.60051934 | | | Isyna1 | 1270.942009 | 0.003307838 | 0.064601391 | 0.051203825 | 0.959163104 | 0.989967546 |
| Rbm4 | 439.9062325 | -0.078136384 | 0.087298221 | -0.895051276 | 0.370759679 | 0.60051934 | | | Dhrs13 | 213.0520859 | 0.00332054 | 0.08670241 | 0.03829813 | 0.969449981 | 0.992567417 |
| Cygb | 2361.323845 | 0.057958251 | 0.064783555 | 0.8946445 | 0.370977156 | 0.600555851 | | | Mrpl52 | 446.5201876 | 0.003345184 | 0.077042691 | 0.043419881 | 0.96536683 | 0.990993714 |
| Dyrk3 | 309.0748985 | 0.109438214 | 0.150353011 | 0.727875111 | 0.46669004 | 0.681613104 | | | Smpd3 | 4814.58865 | 0.003354589 | 0.071270531 | 0.047068392 | 0.962458719 | 0.990888873 |
| Pde4dip | 10132.57593 | -0.062375303 | 0.069715383 | -0.894713623 | 0.370940194 | 0.600555851 | | | Il19 | 0.863694892 | 0.00336837 | 0.018492259 | 0.18215029 | 0.855464783 | NA |
| Rabl6 | 6200.363873 | -0.045985589 | 0.05141323 | -0.89443104 | 0.371091311 | 0.600677891 | | | Sla2 | 35.06453242 | 0.003368712 | 0.083534898 | 0.040327004 | 0.967832425 | 0.991867773 |
| A330050B17Rik | 73.29345392 | 0.109410183 | 0.145874763 | 0.750028176 | 0.453237735 | 0.671684884 | | | Osbpl6 | 3314.779031 | 0.003374461 | 0.065251368 | 0.051714788 | 0.958755954 | 0.989758296 |
| Dcp1b | 482.0873202 | 0.10932046 | 0.09082758 | 1.203604229 | 0.228742584 | 0.451655103 | | | Card11 | 12.35078047 | 0.003375365 | 0.058768397 | 0.057435044 | 0.954198648 | NA |
| Pwp2 | 1220.571941 | 0.10931674 | 0.077362405 | 1.413047325 | 0.15764181 | 0.36051693 | | | Pla2g6 | 949.9523014 | 0.003385252 | 0.068704302 | 0.04927278 | 0.960701912 | 0.990548518 |
| Tax1bp3 | 285.4937813 | -0.092249312 | 0.103255132 | -0.893411401 | 0.3716369 | 0.601354594 | | | Kxd1 | 1020.935424 | 0.003406474 | 0.059940372 | 0.056831041 | 0.954679787 | 0.98892381 |
| Dusp26 | 2819.367714 | -0.054465463 | 0.060981226 | -0.893151327 | 0.371776139 | 0.601409428 | | | Btnl2 | 0.347878816 | 0.003408867 | 0.012856413 | 0.26514917 | 0.79089455 | NA |
| Chmp1b | 855.8538076 | 0.109283123 | 0.058897991 | 1.855464359 | 0.063529944 | 0.20183165 | | | Mir1929 | 0.347878816 | 0.003408867 | 0.012856413 | 0.26514917 | 0.79089455 | NA |
| Msl3 | 780.5531065 | 0.109167641 | 0.074311219 | 1.469060023 | 0.14181651 | 0.337323079 | | | Frg1 | 811.0419143 | 0.003429219 | 0.078786557 | 0.043525433 | 0.965282691 | 0.990993714 |
| Mbd2 | 1497.050441 | 0.109147411 | 0.09186569 | 1.188119425 | 0.234786362 | 0.458404859 | | | Rltpr | 2153.53289 | 0.003429693 | 0.06010503 | 0.057061668 | 0.954496071 | 0.988860025 |
| 5-Sep | 24702.30582 | -0.083668077 | 0.093735768 | -0.892594985 | 0.372074106 | 0.601694982 | | | Slc35f4 | 503.3612859 | 0.003433045 | 0.077646092 | 0.044214007 | 0.964733817 | 0.990993714 |
| Usp22 | 11181.6756 | -0.048126999 | 0.053919492 | -0.892571443 | 0.372086718 | 0.601694982 | | | Slc28a2 | 4.19160987 | 0.003452925 | 0.045962436 | 0.075124933 | 0.940115311 | NA |
| Rab7 | 4673.944339 | 0.078383223 | 0.087848204 | 0.89225755 | 0.372254902 | 0.601869712 | | | 4932702P03Rik | 0.789727694 | 0.003457874 | 0.018259154 | 0.189377531 | 0.849796934 | NA |
| Acss2 | 1948.905593 | -0.102484707 | 0.11495243 | -0.891540151 | 0.372639461 | 0.602302923 | | | Syt3 | 2434.435512 | 0.00346221 | 0.063324775 | 0.054673856 | 0.956398298 | 0.989344398 |
| Ddr1 | 2983.888881 | 0.071921656 | 0.080661945 | 0.891642958 | 0.372584337 | 0.602302923 | | | Pigyl | 489.616155 | 0.003464435 | 0.070371537 | 0.049230629 | 0.960735503 | 0.990548518 |
| Lpin2 | 3929.11538 | 0.109076289 | 0.064057444 | 1.702788648 | 0.088607628 | 0.249888349 | | | Reg3b | 1.322442219 | 0.00347238 | 0.028716645 | 0.120918731 | 0.903755406 | NA |
| Ikzf1 | 206.7245943 | 0.09423357 | 0.105720341 | 0.891347575 | 0.372742733 | 0.602407 | | | Cldn15 | 4.06319975 | 0.003484218 | 0.044448173 | 0.078388319 | 0.937519165 | NA |
| Secisbp2l | 6270.940956 | -0.064474644 | 0.072361495 | -0.891007631 | 0.372925077 | 0.602638834 | | | Brd2 | 8777.746503 | 0.003485864 | 0.044899139 | 0.077637659 | 0.938116285 | 0.985461126 |
| Akap10 | 1212.387809 | -0.056643615 | 0.063607538 | -0.890517328 | 0.373188168 | 0.602699272 | | | Il1rap | 1460.17887 | 0.003488028 | 0.077524236 | 0.044992745 | 0.964113092 | 0.990993714 |
| Nrbp1 | 2988.748294 | 0.108995408 | 0.059395942 | 1.835064896 | 0.066496074 | 0.207916461 | | | Six2 | 13.25143872 | 0.003496319 | 0.053393461 | 0.065482159 | 0.947790111 | NA |
| Golga5 | 1130.748679 | 0.051055695 | 0.057338214 | 0.890430504 | 0.373234769 | 0.602699272 | | | Fam188b | 295.4877911 | 0.003501515 | 0.074970005 | 0.046705552 | 0.962747905 | 0.990993714 |
| Mbnl1 | 2822.82628 | 0.068525618 | 0.076935126 | 0.890693512 | 0.373093616 | 0.602699272 | | | Jmjd4 | 1709.149775 | 0.003514803 | 0.056837411 | 0.061839606 | 0.950690563 | 0.988202091 |
| Mpp1 | 1515.943759 | 0.056942792 | 0.063938867 | 0.890581808 | 0.373153562 | 0.602699272 | | | Klhl29 | 2811.520509 | 0.003537619 | 0.072387909 | 0.048870301 | 0.961022657 | 0.990548518 |
| Snrnp35 | 318.0252707 | -0.08381657 | 0.094122713 | -0.890503123 | 0.373195792 | 0.602699272 | | | 2310015B20Rik | 3.427190836 | 0.003538267 | 0.040332668 | 0.087727064 | 0.930093608 | NA |
| Strbp | 6754.389725 | 0.066677563 | 0.074877361 | 0.89049029 | 0.37320268 | 0.602699272 | | | 2310007B03Rik | 2.633311402 | 0.003545015 | 0.03317441 | 0.106859941 | 0.914900094 | NA |
| H1fx | 474.0105955 | -0.112432984 | 0.126278849 | -0.89035484 | 0.373275383 | 0.602702041 | | | Ttc12 | 119.1031302 | 0.003573795 | 0.094971193 | 0.037630309 | 0.969982442 | 0.992740432 |
| Uhrf1bp1 | 1194.434209 | 0.05728906 | 0.064362468 | 0.890100421 | 0.373411967 | 0.602859749 | | | Tmem184b | 3836.107586 | 0.003584062 | 0.059681438 | 0.060053214 | 0.952113253 | 0.988510896 |
| Bambi | 123.8550893 | 0.108877305 | 0.149757988 | 0.727021681 | 0.467212674 | 0.681990939 | | | D030025E07Rik | 0.333619697 | 0.003587481 | 0.012706056 | 0.282344153 | 0.777679633 | NA |
| Nkx3-1 | 58.5219506 | -0.139556198 | 0.156841524 | -0.889791135 | 0.373578047 | 0.603002215 | | | March8 | 2873.753103 | 0.003588005 | 0.060849673 | 0.058965074 | 0.952979926 | 0.988615469 |
| Inpp5f | 4186.828688 | -0.071817665 | 0.080726659 | -0.88964 | 0.373659221 | 0.603070413 | | | Seh1l | 1604.97342 | 0.003588803 | 0.053625392 | 0.066923579 | 0.946642542 | 0.98700327 |
| Cnbp | 7709.209246 | -0.039814388 | 0.044764279 | -0.889423185 | 0.373775689 | 0.603120383 | | | Dram1 | 54.93842303 | 0.003593368 | 0.0938931 | 0.038270839 | 0.969471741 | 0.992567417 |
| Ikzf3 | 9.384554634 | -0.136430991 | 0.15340326 | -0.889361748 | 0.373808696 | 0.603120383 | | | Zbtb40 | 674.7613357 | 0.003600587 | 0.087697668 | 0.041056817 | 0.967250601 | 0.991789711 |
| Mink1 | 15871.55482 | -0.060108929 | 0.067591852 | -0.889292528 | 0.373845886 | 0.603120383 | | | Nub1 | 4901.333861 | 0.00361161 | 0.037290057 | 0.096851833 | 0.922844061 | 0.98296621 |
| Txndc17 | 847.6222636 | -0.069317453 | 0.077942788 | -0.88933762 | 0.373821659 | 0.603120383 | | | Abhd3 | 2111.800821 | 0.00361696 | 0.045193962 | 0.080031936 | 0.936211856 | 0.985461126 |
| Tnfrsf1a | 444.604296 | 0.108865091 | 0.097967965 | 1.11123153 | 0.266468703 | 0.494085203 | | | Kcna4 | 1113.829681 | 0.003619613 | 0.068398944 | 0.052919134 | 0.957796339 | 0.989630474 |
| Apc2 | 6697.520261 | -0.050476854 | 0.056794634 | -0.888760972 | 0.374131555 | 0.603455581 | | | Gfi1 | 0.945824685 | 0.003624364 | 0.019089691 | 0.189859773 | 0.849419016 | NA |
| A630089N07Rik | 75.64373217 | -0.131501867 | 0.14804331 | -0.888266192 | 0.374397581 | 0.603821809 | | | Rfc3 | 219.8446472 | 0.00362635 | 0.082429543 | 0.04399333 | 0.964909721 | 0.990993714 |
| Arhgef39 | 30.90244382 | 0.108800081 | 0.164895704 | 0.659811497 | 0.509374804 | 0.714231061 | | | Gm16291 | 11.91751746 | 0.00363134 | 0.067675986 | 0.053657738 | 0.957207854 | NA |
| Dip2a | 3955.500496 | 0.108694583 | 0.047793404 | 2.274259092 | 0.022950413 | 0.103340221 | | | Ebna1bp2 | 1525.109608 | 0.003631762 | 0.048051171 | 0.075581139 | 0.939752342 | 0.985461126 |
| Fundc2 | 1532.45724 | -0.054619256 | 0.061544791 | -0.887471618 | 0.37482504 | 0.60431583 | | | 4933433G19Rik | 2.263605847 | 0.003651751 | 0.034142085 | 0.106957483 | 0.91482271 | NA |
| St13 | 3932.308687 | -0.054693001 | 0.061632387 | -0.887406826 | 0.37485991 | 0.60431583 | | | Slc22a2 | 25.43934457 | 0.003661205 | 0.069640499 | 0.052572932 | 0.958072184 | 0.989630474 |
| Srd5a3 | 648.8377223 | 0.074867009 | 0.084377246 | 0.887289072 | 0.374923288 | 0.60435512 | | | Nt5m | 1235.092451 | 0.003664635 | 0.051618955 | 0.070993982 | 0.943402545 | 0.98574776 |
| Ift74 | 745.7418799 | 0.067470071 | 0.076072884 | 0.886913544 | 0.375125449 | 0.604492326 | | | Mc1r | 2.042746324 | 0.003665331 | 0.033072602 | 0.110826801 | 0.911753692 | NA |
| Cers2 | 2319.593313 | 0.108687398 | 0.084977949 | 1.279007079 | 0.200894563 | 0.417750686 | | | Atf7ip2 | 0.854022854 | 0.003678692 | 0.018472123 | 0.199148324 | 0.842146721 | NA |
| Cuta | 1148.770803 | 0.108676465 | 0.078512359 | 1.384195639 | 0.166298558 | 0.371915182 | | | Serpinb10 | 0.67286561 | 0.003679241 | 0.015659951 | 0.234945894 | 0.814250728 | NA |
| Ppp6r1 | 7548.466109 | 0.057916824 | 0.06530892 | 0.886813376 | 0.375179385 | 0.604516367 | | | 3110021N24Rik | 15.14203202 | 0.003686303 | 0.072144827 | 0.051095871 | 0.959249126 | NA |
| Tmem38b | 326.8767344 | 0.094572367 | 0.10667503 | 0.886546424 | 0.37532315 | 0.604685129 | | | Tat | 2.810170396 | 0.003693689 | 0.034626414 | 0.106672586 | 0.915048732 | NA |
| Cltc | 27104.65603 | -0.061640891 | 0.069572857 | -0.885990507 | 0.375622642 | 0.60487861 | | | Tmem95 | 1.699291406 | 0.003697434 | 0.025337035 | 0.145930015 | 0.883976637 | NA |
| Fbxo4 | 201.5458163 | 0.09099916 | 0.102693332 | 0.886125306 | 0.375550008 | 0.60487861 | | | Tbx5 | 0.338696681 | 0.003698955 | 0.012761698 | 0.289848151 | 0.771932409 | NA |
| Golga4 | 4496.747278 | 0.06180334 | 0.069751229 | 0.886053774 | 0.375588551 | 0.60487861 | | | Alyref | 547.0065176 | 0.003705158 | 0.071087126 | 0.052121358 | 0.958431995 | 0.989630474 |
| Nr1h2 | 1600.857567 | 0.064447198 | 0.072742688 | 0.885961193 | 0.37563844 | 0.60487861 | | | Loxl4 | 5.853579617 | 0.003715358 | 0.045422118 | 0.081796231 | 0.934808753 | NA |
| Zfp646 | 1325.193226 | 0.070336248 | 0.079382428 | 0.886043046 | 0.375594332 | 0.60487861 | | | Atg13 | 4909.815825 | 0.003720117 | 0.038403552 | 0.096869099 | 0.922830349 | 0.98296621 |
| Zdhhc23 | 298.45051 | -0.079088798 | 0.089278689 | -0.885864237 | 0.37569069 | 0.60489988 | | | Zfp113 | 405.7193929 | 0.00372388 | 0.071216007 | 0.052289933 | 0.958297675 | 0.989630474 |
| Gsto1 | 846.2183455 | 0.075825399 | 0.085615582 | 0.885649519 | 0.375806419 | 0.605023343 | | | Ap3b1 | 2269.20709 | 0.003724915 | 0.047920653 | 0.077730884 | 0.938042126 | 0.985461126 |
| Lrrc39 | 10.27489823 | -0.137085745 | 0.154835176 | -0.88536564 | 0.375959458 | 0.60520684 | | | Ltb4r1 | 4.873109598 | 0.003732845 | 0.05016034 | 0.074418258 | 0.940677581 | NA |
| Bod1l | 3375.267634 | -0.07048511 | 0.079624759 | -0.885215997 | 0.376040146 | 0.605210971 | | | 4930563E18Rik | 2.611025974 | 0.003735457 | 0.032369464 | 0.115400645 | 0.908127568 | NA |
| Zbtb34 | 912.7039682 | 0.082867169 | 0.093611005 | 0.885228924 | 0.376033176 | 0.605210971 | | | Cbfa2t3 | 1426.230555 | 0.003763593 | 0.092604969 | 0.040641371 | 0.967581802 | 0.991805208 |
| Ngef | 15915.5029 | -0.075964828 | 0.085822763 | -0.885136131 | 0.376083215 | 0.605217421 | | | Qrsl1 | 528.5759043 | 0.003764217 | 0.065187891 | 0.057744117 | 0.953952452 | 0.988860025 |
| Cpne8 | 1122.290982 | -0.078305448 | 0.088487018 | -0.88493713 | 0.376190541 | 0.605327265 | | | Lrrc10 | 1.044785728 | 0.003765961 | 0.023733084 | 0.158679811 | 0.873921145 | NA |
| Dapk3 | 943.9414897 | -0.061916606 | 0.07000619 | -0.884444735 | 0.376456183 | 0.605377488 | | | E130018N17Rik | 1.953163287 | 0.003775447 | 0.022866994 | 0.165104632 | 0.868861628 | NA |
| Gabrd | 2705.939155 | -0.076987306 | 0.08704121 | -0.884492597 | 0.376430357 | 0.605377488 | | | Snord8 | 0.848365821 | 0.003777015 | 0.021905278 | 0.172424865 | 0.863103525 | NA |

| | | | | | | |
|---|---|---|---|---|---|---|
| Gamt | 277.0015097 | -0.141197049 | 0.159604474 | -0.884668493 | 0.376335453 | 0.605377488 |
| Morf4l1 | 11811.04303 | -0.067005255 | 0.07575947 | -0.884447249 | 0.376454826 | 0.605377488 |
| Rab6a | 18501.34492 | 0.056206814 | 0.063536573 | 0.88463717 | 0.376352353 | 0.605377488 |
| Zfp689 | 262.1371146 | -0.078221841 | 0.08842833 | -0.88457897 | 0.376383753 | 0.605377488 |
| D10Wsu102e | 1076.971978 | -0.075043932 | 0.084857146 | -0.884356075 | 0.376504027 | 0.605391593 |
| Nfe2l1 | 16143.58044 | -0.055122777 | 0.062349903 | -0.884087617 | 0.376648917 | 0.605561723 |
| Rspry1 | 913.1829943 | -0.062414152 | 0.070660873 | -0.883291545 | 0.377078771 | 0.606189923 |
| Polh | 209.9739348 | 0.085744076 | 0.097086162 | 0.883175049 | 0.3771417 | 0.606228188 |
| Aarsd1 | 1730.625939 | -0.058259771 | 0.06599927 | -0.882733562 | 0.377380244 | 0.60642289 |
| Cep57 | 532.5306237 | 0.076288111 | 0.086410227 | 0.882859746 | 0.377312055 | 0.60642289 |
| Lrrtm2 | 2554.357053 | -0.070070412 | 0.079374903 | -0.882777927 | 0.377356269 | 0.60642289 |
| C130036L24Rik | 30.66970351 | -0.145672941 | 0.165131492 | -0.88216329 | 0.37768851 | 0.606792387 |
| Sap130 | 1802.951407 | 0.108596305 | 0.068222115 | 1.591805026 | 0.111428521 | 0.290847961 |
| Bace1 | 3957.113346 | -0.044797396 | 0.050789824 | -0.882015193 | 0.377768591 | 0.606858119 |
| Brix1 | 681.9424542 | 0.054364027 | 0.061652989 | 0.881774396 | 0.37789882 | 0.607004387 |
| Mrpl51 | 1156.618859 | -0.071771688 | 0.081411834 | -0.881587906 | 0.377999698 | 0.607065669 |
| Zfp516 | 810.4667674 | -0.10131241 | 0.11492414 | -0.881559004 | 0.378015333 | 0.607065669 |
| Maml3 | 481.9819355 | -0.084458181 | 0.095815576 | -0.881466082 | 0.378065605 | 0.607083478 |
| Kcnt2 | 806.4586586 | 0.091755723 | 0.104114543 | 0.881295931 | 0.378157668 | 0.607105471 |
| Tsga10 | 324.3073191 | -0.122063375 | 0.138500777 | -0.881319065 | 0.37814515 | 0.607105471 |
| Snn | 10928.64642 | 0.058813638 | 0.066770081 | 0.880838202 | 0.3784054 | 0.607440247 |
| Bola1 | 204.0374117 | -0.098879515 | 0.112306913 | -0.880440143 | 0.378620918 | 0.607597359 |
| Hsf2 | 813.9028862 | 0.108538789 | 0.076052899 | 1.427148608 | 0.153537064 | 0.35484834 |
| Gm6981 | 33.44942253 | 0.108505841 | 0.160037032 | 0.678004578 | 0.497768791 | 0.706525251 |
| Tmod2 | 22212.14452 | -0.090804394 | 0.103151978 | -0.880297164 | 0.378698349 | 0.60765868 |
| Rab4b | 1880.342455 | 0.108504434 | 0.068732833 | 1.578640497 | 0.11441854 | 0.296434962 |
| Atxn10 | 9654.259973 | -0.048584118 | 0.055221832 | -0.879799088 | 0.378968159 | 0.607752012 |
| Nat8 | 48.48660238 | 0.108489245 | 0.160105549 | 0.677610774 | 0.498018513 | 0.706685488 |
| Eral1 | 636.2589448 | 0.058320565 | 0.066297255 | 0.879682949 | 0.379031089 | 0.607752012 |
| Sp2 | 665.6809862 | 0.056653326 | 0.064397857 | 0.879739294 | 0.379000558 | 0.607752012 |
| Gadd45gip1 | 873.3169231 | 0.108437587 | 0.071652131 | 1.51338957 | 0.130180733 | 0.319452055 |
| Utp3 | 914.6769507 | 0.070332181 | 0.079933458 | 0.879884123 | 0.378922086 | 0.607752012 |
| AA415398 | 228.7321846 | 0.086903696 | 0.098810876 | 0.879495243 | 0.379132812 | 0.607852213 |
| Gabbr2 | 10647.79382 | 0.076475222 | 0.086975368 | 0.879274484 | 0.379252467 | 0.607968717 |
| Kiss1r | 215.6941594 | 0.098593473 | 0.112137892 | 0.879216396 | 0.379283956 | 0.607968717 |
| Gmcl1 | 986.1843835 | 0.053697237 | 0.061089424 | 0.878994009 | 0.379404524 | 0.60803617 |
| Lrrc33 | 276.0205107 | 0.099017076 | 0.112644801 | 0.879020385 | 0.379390223 | 0.60803617 |
| Zfp592 | 1554.329327 | 0.085111292 | 0.096869444 | 0.87861857 | 0.379608124 | 0.608299541 |
| Myef2 | 2068.851141 | 0.108391306 | 0.070944305 | 1.52783659 | 0.126553115 | 0.314473278 |
| Ly6g6d | 8.26898452 | -0.129573537 | 0.14751989 | -0.878346215 | 0.379755863 | 0.608347533 |
| Tceb3 | 1485.396308 | 0.070418846 | 0.080166352 | 0.878409017 | 0.379721793 | 0.608347533 |
| Pgf | 80.08302212 | -0.117978236 | 0.134402292 | -0.877799281 | 0.380052655 | 0.608760037 |
| Fam110a | 174.4860926 | -0.088387736 | 0.100739871 | -0.877385834 | 0.380277105 | 0.609056591 |
| Ptrf | 1017.436632 | -0.074373572 | 0.084787463 | -0.877176529 | 0.380390762 | 0.609175656 |
| AW046200 | 15.71622264 | -0.145930205 | 0.166447391 | -0.876734714 | 0.380630747 | 0.609245124 |
| Cdpf1 | 278.2672707 | -0.092724063 | 0.105750186 | -0.876821745 | 0.380583466 | 0.609245124 |
| Etf1 | 3280.720858 | -0.066075972 | 0.075356396 | -0.876846232 | 0.380570164 | 0.609245124 |
| Rpgrip1l | 1074.141273 | -0.071885334 | 0.08199131 | -0.876743318 | 0.380626073 | 0.609245124 |
| Taok2 | 6324.153893 | 0.059262427 | 0.067580996 | 0.876909636 | 0.380535722 | 0.609245124 |
| Slx4ip | 303.5110228 | -0.090421441 | 0.103144397 | -0.876649081 | 0.380677272 | 0.609256652 |
| Qrsl1 | 480.0168613 | -0.073711066 | 0.084127017 | -0.876187794 | 0.380927951 | 0.609594885 |
| Magi1 | 4281.700778 | -0.044734357 | 0.05107209 | -0.875906132 | 0.381081066 | 0.609769958 |
| Tkt | 5370.194231 | -0.058845324 | 0.067187164 | -0.875841763 | 0.381116063 | 0.609769958 |
| Dbh | 16.74623483 | -0.145841352 | 0.16660807 | -0.875355871 | 0.381380303 | 0.610129728 |
| Unc93b1 | 637.3990027 | 0.108366776 | 0.136854019 | 0.791842119 | 0.42845274 | 0.650037121 |
| Dnase1l1 | 65.63983326 | 0.108225055 | 0.140409763 | 0.77078013 | 0.440837267 | 0.661093969 |
| Ufm1 | 2214.470129 | -0.047873268 | 0.054727896 | -0.874750758 | 0.381709535 | 0.610467335 |
| Rps6ka5 | 1005.819026 | 0.108152893 | 0.13380709 | 0.808274758 | 0.41893243 | 0.641714194 |
| Etv3 | 1301.764456 | 0.078333424 | 0.089568262 | 0.874566757 | 0.381809682 | 0.610501468 |
| Atcay | 10536.10617 | -0.04597697 | 0.052590166 | -0.874250335 | 0.38198194 | 0.610650864 |
| Nmd3 | 704.6826246 | 0.063522349 | 0.072654728 | 0.874304409 | 0.381952499 | 0.610650864 |
| Cd209f | 19.35221151 | -0.130029265 | 0.148773703 | -0.874007045 | 0.382114417 | 0.610799626 |
| D2hgdh | 548.5688286 | -0.077502461 | 0.088688564 | -0.873872099 | 0.382187911 | 0.610799825 |
| Stau1 | 2670.191467 | 0.056858007 | 0.065065198 | 0.873862048 | 0.382193385 | 0.610799825 |
| Asphd1 | 533.4901059 | -0.089285269 | 0.102186892 | -0.873744836 | 0.382257228 | 0.610838644 |
| Lrrc59 | 4083.792501 | -0.052956511 | 0.060613672 | -0.873672698 | 0.382296523 | 0.610838644 |
| Ctu2 | 504.8393859 | 0.108150054 | 0.084317656 | 1.282650149 | 0.199614672 | 0.416113389 |
| Pyroxd1 | 449.4111143 | 0.088397327 | 0.101243752 | 0.873113901 | 0.382600998 | 0.611199078 |
| Mum1 | 1249.928111 | 0.108082499 | 0.081846751 | 1.320547213 | 0.186652384 | 0.398675494 |
| Arid4a | 1556.25375 | 0.108063078 | 0.066351003 | 1.628657789 | 0.103385483 | 0.276515567 |
| Ccnd3 | 821.2100373 | 0.10805621 | 0.089448568 | 1.20802615 | 0.227037204 | 0.449490728 |
| Ino80c | 938.1483269 | 0.061354546 | 0.070322682 | 0.872471654 | 0.382951126 | 0.611369871 |
| Ngdn | 540.4091065 | 0.108032057 | 0.095836225 | 1.127257013 | 0.259633831 | 0.486370038 |
| Rian | 20955.46053 | 0.083914877 | 0.096211241 | 0.872194103 | 0.383102496 | 0.611369871 |
| Sh3pxd2a | 2468.916073 | -0.06881927 | 0.078890837 | -0.872335398 | 0.383025432 | 0.611369871 |
| Mark4 | 2096.709039 | 0.108026824 | 0.074338385 | 1.453176904 | 0.146174643 | 0.345038074 |
| Tmub2 | 2377.634671 | 0.055590164 | 0.063726138 | 0.872329087 | 0.383028874 | 0.611369871 |
| Zfp329 | 1179.10925 | 0.093573542 | 0.107259802 | 0.872400853 | 0.382989736 | 0.611369871 |
| Kiss1 | 15.11924878 | -0.140421768 | 0.161038633 | -0.87197566 | 0.383221656 | 0.611434079 |
| Zfp697 | 697.7158205 | 0.076718047 | 0.087976003 | 0.872033786 | 0.383189947 | 0.611434079 |
| 2210404O07Rik | 16.92486679 | -0.145834294 | 0.167363484 | -0.871362681 | 0.383556156 | 0.611876648 |
| Ablim1 | 6036.168743 | 0.107970883 | 0.048328802 | 2.234089769 | 0.025477167 | 0.110510422 |
| Abcb6 | 808.9765916 | 0.058087327 | 0.066672864 | 0.871228922 | 0.383629171 | 0.611880995 |
| Pde8a | 707.938274 | 0.0926361 | 0.106334914 | 0.871172944 | 0.38365973 | 0.611880995 |
| Prrx2 | 24.40641086 | -0.146042356 | 0.167655623 | -0.871085345 | 0.383707555 | 0.611894284 |
| Med22 | 1276.131055 | 0.051127493 | 0.058740023 | 0.870403012 | 0.384080202 | 0.612425506 |
| Commd4 | 1210.491808 | 0.064859904 | 0.074530122 | 0.870250876 | 0.384163318 | 0.612431984 |
| Tmcc3 | 2080.475811 | 0.069556178 | 0.079924338 | 0.870275314 | 0.384149966 | 0.612431984 |
| Fbxo16 | 251.9236949 | 0.089498209 | 0.102851977 | 0.870165181 | 0.384210141 | 0.612443614 |
| Mpv17 | 1718.988999 | -0.068905707 | 0.079209807 | -0.869913835 | 0.384347494 | 0.612599535 |
| Abca9 | 496.966352 | -0.102663233 | 0.118025564 | -0.869838952 | 0.384388421 | 0.612601749 |
| Rilpl1 | 2138.008236 | 0.107907165 | 0.119665344 | 0.901741147 | 0.36719439 | 0.596940754 |
| Ahsa2 | 1026.268876 | 0.076520764 | 0.088014664 | 0.869409254 | 0.384623323 | 0.61266645 |
| Cgrrf1 | 611.1345087 | 0.084452569 | 0.097135434 | 0.869431114 | 0.384611371 | 0.61266645 |
| Gm16845 | 24.44215899 | 0.107867837 | 0.168017887 | 0.642002104 | 0.520871817 | 0.723931462 |
| Luc7l | 4650.219192 | 0.107774756 | 0.051980552 | 2.073366916 | 0.038138145 | 0.144795988 |
| Spr | 689.7928637 | -0.100698521 | 0.115834538 | -0.869330706 | 0.384666272 | 0.61266645 |
| Copa | 10607.43666 | 0.037655959 | 0.043332944 | 0.868991457 | 0.384851803 | 0.61273055 |
| Ppp2r3c | 1019.619137 | -0.062624013 | 0.072067121 | -0.868967873 | 0.384864703 | 0.61273055 |
| Rhobtb1 | 234.4168945 | -0.118532138 | 0.136392604 | -0.869051063 | 0.384819202 | 0.61273055 |
| Arfgap1 | 4322.646027 | 0.10776752 | 0.066113357 | 1.630041566 | 0.103092712 | 0.276018401 |
| Rab5b | 7255.267845 | -0.046255411 | 0.053242829 | -0.868763215 | 0.384976657 | 0.612814857 |
| Zfp467 | 861.7523536 | 0.08513891 | 0.098004278 | 0.868726467 | 0.384996761 | 0.612814857 |
| Slc35e4 | 878.0769059 | 0.071013162 | 0.081755883 | 0.868600023 | 0.385065942 | 0.612862014 |
| Sema6b | 4179.248385 | -0.061644257 | 0.070976594 | -0.868515285 | 0.385112308 | 0.612872854 |
| 1810026J23Rik | 2909.346808 | 0.075320861 | 0.086731862 | 0.86843357 | 0.385157024 | 0.612881065 |
| Clip2 | 6694.133926 | 0.107741688 | 0.063100312 | 1.707466804 | 0.087735323 | 0.248196292 |
| Sin3b | 2985.410322 | 0.107666575 | 0.058256184 | 1.848157002 | 0.064579634 | 0.204190665 |
| Zfp58 | 214.3843029 | 0.095143361 | 0.109641542 | 0.867767447 | 0.385521654 | 0.613272331 |
| Arcn1 | 5231.268893 | -0.059444408 | 0.068516604 | -0.867591276 | 0.385618124 | 0.613299857 |
| Trappc3 | 2099.628001 | 0.079071342 | 0.091134366 | 0.86763475 | 0.385594317 | 0.613299857 |
| Gprasp1 | 24211.94543 | 0.062994794 | 0.07262107 | 0.867445137 | 0.38569816 | 0.613364188 |
| Impa2 | 124.6967514 | 0.107657748 | 0.131404435 | 0.819285501 | 0.412623543 | 0.636328175 |
| Txnl4b | 642.5932217 | -0.066037365 | 0.076178746 | -0.866873875 | 0.386011119 | 0.613704655 |
| Upp2 | 259.1219636 | 0.095849956 | 0.110573779 | 0.866841647 | 0.38602878 | 0.613704655 |
| Vps25 | 1406.679059 | -0.049372304 | 0.056961568 | -0.866765169 | 0.38607069 | 0.613704655 |
| Zdhhc6 | 946.0737395 | 0.107580214 | 0.070086733 | 1.534958311 | 0.124794069 | 0.312304373 |
| Grpel2 | 843.9911768 | 0.062438892 | 0.072076825 | 0.866282499 | 0.386335261 | 0.61393626 |
| Pla2r1 | 30.86322092 | -0.144740997 | 0.167074593 | -0.866325596 | 0.386311633 | 0.61393626 |
| Myo1c | 920.1079587 | 0.107456264 | 0.083975456 | 1.279615122 | 0.200680529 | 0.4174737 |
| Rbl2 | 2333.894507 | 0.089041429 | 0.102816353 | 0.866024 | 0.386477001 | 0.613972588 |
| Adck3 | 1583.360229 | 0.107443513 | 0.082910091 | 1.295903928 | 0.195008586 | 0.409594319 |
| Tmem106c | 957.9491531 | 0.107431398 | 0.077015698 | 1.394928574 | 0.163037379 | 0.368290007 |
| Slc25a35 | 191.5654539 | 0.107422148 | 0.124039686 | 0.866030471 | 0.386473452 | 0.613972588 |
| Hells | 60.93598857 | -0.12655782 | 0.146236826 | -0.865430568 | 0.38680251 | 0.614237786 |
| Rnf157 | 15890.60018 | -0.06125963 | 0.070784461 | -0.865438952 | 0.386797909 | 0.614237786 |
| Nelfb | 2027.964669 | 0.059670656 | 0.068968536 | 0.86518664 | 0.386936357 | 0.614387366 |
| Copb2 | 3983.521931 | 0.060407696 | 0.069842753 | 0.86491001 | 0.387088183 | 0.614565459 |
| Itfg1 | 9927.09633 | -0.053174404 | 0.061496088 | -0.86467946 | 0.387214746 | 0.61470341 |
| Ahsa1 | 4011.031162 | -0.058227673 | 0.067349411 | -0.864560981 | 0.387279796 | 0.614743691 |
| 2810002D19Rik | 122.2878211 | -0.118402661 | 0.137006591 | -0.864211421 | 0.387471759 | 0.614790257 |
| Cmc2 | 412.1291523 | -0.077400188 | 0.089553308 | -0.864291777 | 0.387427626 | 0.614790257 |
| Fam180a | 90.37512287 | -0.134907198 | 0.156076651 | -0.864365026 | 0.387387399 | 0.614790257 |
| Ifitm3 | 342.900116 | 0.090888873 | 0.105177645 | 0.864146298 | 0.387507529 | 0.614790257 |
| Mfn1 | 2932.826717 | 0.107384882 | 0.078361667 | 1.370375169 | 0.170569819 | 0.377867998 |
| Slc25a29 | 425.0970198 | 0.074833869 | 0.086607514 | 0.864057474 | 0.387556319 | 0.61480471 |
| Dhrs1 | 2171.286006 | 0.053096087 | 0.061465024 | 0.8638423 | 0.387674527 | 0.614866324 |
| Smek1 | 1522.498237 | 0.107267806 | 0.09308837 | 1.152322314 | 0.249188651 | 0.474432165 |
| Awat2 | 8.462637929 | -0.129504818 | 0.149995647 | -0.86339051 | 0.387922795 | 0.615197113 |
| Slc27a5 | 11.64357807 | 0.107210503 | 0.16352373 | 0.655626574 | 0.512064423 | 0.716446286 |
| 2700060E02Rik | 1891.665054 | -0.060312545 | 0.069916485 | -0.862636982 | 0.388337091 | 0.615635694 |
| Gm11190 | 9.155661789 | -0.13920346 | 0.161376897 | -0.862598445 | 0.388358286 | 0.615635694 |
| Zfp236 | 2156.341849 | -0.077706654 | 0.090079313 | -0.862647055 | 0.388331551 | 0.615635694 |
| Spef1 | 310.2043035 | -0.084012702 | 0.097469334 | -0.861939845 | 0.388720621 | 0.616047658 |
| Taf1c | 523.8634455 | 0.080157641 | 0.093007887 | 0.861837031 | 0.388777204 | 0.616047658 |
| Txndc5 | 2372.095655 | -0.044666186 | 0.051822527 | -0.86190676 | 0.388738829 | 0.616047658 |
| Vps35 | 6580.227258 | -0.061725001 | 0.071604252 | -0.862029824 | 0.388671107 | 0.616047658 |
| Pa2g4 | 2612.572635 | 0.046666893 | 0.054154072 | 0.861743006 | 0.388828954 | 0.616066655 |
| 9430016H08Rik | 378.1038584 | -0.082347612 | 0.095585238 | -0.861509724 | 0.388957368 | 0.616109828 |
| Adc | 1058.489422 | 0.076524202 | 0.088829092 | 0.861476795 | 0.388975496 | 0.616109828 |
| Pard6b | 208.2692062 | -0.086792102 | 0.100734482 | -0.861592777 | 0.388911647 | 0.616109828 |
| 2310008H04Rik | 113.8882889 | -0.111470888 | 0.12942089 | -0.861305219 | 0.389069962 | 0.616133482 |
| Rcc2 | 4119.707011 | -0.048891691 | 0.056760775 | -0.861364041 | 0.389037574 | 0.616133482 |
| Eea1 | 1946.249702 | -0.067612702 | 0.078510764 | -0.861190222 | 0.389133284 | 0.616170783 |
| Hes6 | 343.663794 | -0.07639193 | 0.088721425 | -0.861031368 | 0.389220767 | 0.616246328 |
| Rbbp6 | 3230.57207 | 0.062822498 | 0.072972368 | 0.860908037 | 0.389288695 | 0.6162909 |
| Lmbrd1 | 7694.251998 | 0.052068133 | 0.060505463 | 0.860552588 | 0.389484508 | 0.616537901 |
| Ube2e1 | 1937.880494 | -0.043270408 | 0.050298037 | -0.86028025 | 0.389634577 | 0.616712446 |
| Ddx39 | 452.8116589 | 0.07669283 | 0.089169079 | 0.860083238 | 0.38974316 | 0.616770739 |
| Wdr45 | 1383.571032 | 0.057390541 | 0.066727835 | 0.860068979 | 0.38975102 | 0.616770739 |
| BC048943 | 9.662921329 | -0.110708632 | 0.128757081 | -0.85982558 | 0.389885196 | 0.616888543 |
| Gm11110 | 21.1354512 | -0.14367875 | 0.167109218 | -0.859789493 | 0.389905092 | 0.616888543 |
| Aph1a | 2104.661621 | 0.042365146 | 0.049283879 | 0.859614684 | 0.390001478 | 0.616915049 |
| Mmab | 1334.902532 | 0.063382972 | 0.073732672 | 0.85963211 | 0.389991869 | 0.616915049 |
| Dctn1 | 16711.8334 | 0.107091467 | 0.06471608 | 1.654789146 | 0.097967276 | 0.266252409 |
| Chek1 | 36.6499643 | 0.107089065 | 0.159235192 | 0.672521344 | 0.501251857 | 0.708899185 |
| Prrt1 | 1410.196401 | 0.107085247 | 0.080496114 | 1.330315729 | 0.183414267 | 0.394368672 |
| Pros1 | 576.2951601 | 0.067880141 | 0.079064285 | 0.858543668 | 0.390592329 | 0.617597463 |
| 1700021K19Rik | 3764.971208 | -0.047498268 | 0.055363641 | -0.85793253 | 0.390929721 | 0.617802551 |
| Ap5m1 | 665.0013434 | 0.074198356 | 0.086472295 | 0.858059282 | 0.39085973 | 0.617802551 |
| Foxred2 | 883.3243643 | 0.064174369 | 0.07479815 | 0.857967331 | 0.390910503 | 0.617802551 |
| Aox1 | 271.6158382 | 0.107032281 | 0.125755789 | 0.851112154 | 0.394707055 | 0.620116721 |
| Mrps11 | 571.4761288 | -0.067440192 | 0.078611285 | -0.857894534 | 0.390950703 | 0.617802551 |
| Piga | 338.5768355 | -0.110443131 | 0.128746547 | -0.857833737 | 0.390984278 | 0.617802551 |
| Pik3cb | 1515.474815 | 0.064791901 | 0.07553095 | 0.857819226 | 0.390992292 | 0.617802551 |
| Ppp1r3f | 1244.198598 | 0.107002623 | 0.079701502 | 1.342542122 | 0.179420282 | 0.389401577 |
| Dnajc4 | 337.6439497 | 0.106998797 | 0.108104196 | 0.989774685 | 0.322284261 | 0.555571021 |
| Rpl8 | 7851.347703 | -0.064669852 | 0.075428536 | -0.857365865 | 0.391242718 | 0.617995175 |
| Etv5 | 5349.696101 | -0.099395333 | 0.115941991 | -0.857285028 | 0.391287381 | 0.618002719 |
| Bola3 | 365.3355468 | -0.093106366 | 0.108616554 | -0.857202353 | 0.391333063 | 0.618011871 |
| Utp6 | 1740.195392 | -0.053579324 | 0.06251254 | -0.857097208 | 0.391391164 | 0.618040633 |
| Abhd12 | 11366.5171 | 0.060819173 | 0.070981994 | 0.856825364 | 0.391541406 | 0.618204152 |
| Ppp4r4 | 1618.613171 | -0.047565304 | 0.055517298 | -0.856765464 | 0.391574516 | 0.618204152 |
| Pom121 | 3552.348237 | 0.106968153 | 0.075071505 | 1.424883551 | 0.154190865 | 0.355680596 |
| Bbx | 1503.858441 | 0.085222315 | 0.099520875 | 0.856326019 | 0.391817473 | 0.618286373 |
| Mettl18 | 149.908694 | -0.10139854 | 0.118402483 | -0.856388631 | 0.391782851 | 0.618286373 |
| Prelid1 | 3542.928675 | -0.05069804 | 0.059205215 | -0.856310383 | 0.39182612 | 0.618286373 |
| Ptpn20 | 18.26636562 | -0.142736729 | 0.166668255 | -0.85641221 | 0.391769813 | 0.618286373 |
| Ahctf1 | 2607.237059 | 0.08826043 | 0.103149688 | 0.855653878 | 0.392189259 | 0.618521669 |
| Fam26f | 8.347921985 | -0.129605396 | 0.151415843 | -0.855956642 | 0.392021762 | 0.618521669 |
| Higd1b | 77.09183923 | -0.121247743 | 0.141709488 | -0.8556078 | 0.392214754 | 0.618521669 |
| Pcdhb14 | 289.7421541 | 0.095575863 | 0.111694448 | 0.855690365 | 0.392169071 | 0.618521669 |
| Slc12a4 | 841.7785675 | 0.073880706 | 0.086343887 | 0.855656474 | 0.392187823 | 0.618521669 |
| Slc25a42 | 1429.537997 | 0.067754017 | 0.079183773 | 0.855655318 | 0.392188462 | 0.618521669 |
| Slc25a30 | 211.4833031 | 0.10695524 | 0.111272528 | 0.961200767 | 0.336451231 | 0.570347414 |
| Rab36 | 1043.391537 | 0.082919509 | 0.096952162 | 0.855262095 | 0.392406067 | 0.618634473 |
| Sh3bgr | 32.17154788 | 0.106870918 | 0.166358685 | 0.642412617 | 0.52060531 | 0.723755825 |
| Dom3z | 547.7972238 | 0.081507961 | 0.095318417 | 0.855112406 | 0.392488922 | 0.618702142 |
| Gla | 537.0982149 | 0.085558961 | 0.100075507 | 0.854944071 | 0.392582111 | 0.618743087 |
| Rnf20 | 2546.767764 | 0.106864631 | 0.058723358 | 1.819797692 | 0.068789819 | 0.21246433 |
| Frrs1l | 8030.202146 | -0.068674304 | 0.080357634 | -0.854608344 | 0.392768007 | 0.618824677 |
| Lrrc38 | 89.10596748 | 0.106804936 | 0.138379928 | 0.7718239 | 0.440218734 | 0.660477299 |
| Pxdn | 3137.70331 | 0.09477035 | 0.110897595 | 0.854575341 | 0.392786284 | 0.618824677 |
| Ripply3 | 8.639553801 | -0.128456112 | 0.15032208 | -0.854539212 | 0.392806293 | 0.618824677 |
| Aebp2 | 2516.979197 | 0.06253734 | 0.073196471 | 0.854376441 | 0.392896445 | 0.618903774 |
| Fert2 | 589.2462759 | -0.066331956 | 0.077646125 | -0.854285461 | 0.392946841 | 0.618920235 |
| Uap1 | 2014.437567 | -0.073329089 | 0.085866515 | -0.853989344 | 0.393110893 | 0.619115693 |
| Fzd10 | 48.87105983 | -0.141747422 | 0.165999552 | -0.853902436 | 0.39315905 | 0.619128602 |
| Bloc1s6 | 1166.614612 | -0.060155429 | 0.070480656 | -0.853502676 | 0.393380604 | 0.61923566 |
| Nom1 | 1118.685903 | 0.062730093 | 0.073481179 | 0.85368925 | 0.393277192 | 0.61923566 |
| Ralb | 642.3797652 | -0.067539251 | 0.079132909 | -0.853491322 | 0.393386898 | 0.61923566 |
| Wdr44 | 740.0976696 | -0.063655628 | 0.074574883 | -0.85357999 | 0.39333775 | 0.61923566 |
| Med1 | 1988.570684 | -0.054005313 | 0.063316165 | -0.852946684 | 0.393688871 | 0.619648047 |
| Lrwd1 | 446.5857992 | 0.071174853 | 0.083459639 | 0.852805673 | 0.393767077 | 0.619708186 |
| Frmd3 | 198.1408387 | -0.117557271 | 0.137870222 | -0.852666145 | 0.39384447 | 0.619762733 |
| Slco4c1 | 68.47457103 | -0.118970093 | 0.139538163 | -0.852598964 | 0.393881737 | 0.619762733 |
| Topors | 956.07908 | 0.082413743 | 0.096681063 | 0.852429011 | 0.393976024 | 0.619848143 |
| Zfp74 | 680.9881335 | 0.079248055 | 0.092976038 | 0.852349242 | 0.394020283 | 0.619854835 |
| 2810408M09Rik | 471.2845247 | -0.074984786 | 0.087990609 | -0.852190782 | 0.394108212 | 0.619921228 |
| Msh5 | 9.581838008 | 0.106791547 | 0.161154335 | 0.662666304 | 0.507544295 | 0.713213433 |
| Itih5 | 2035.547038 | -0.087442503 | 0.10263119 | -0.852007098 | 0.394210153 | 0.619921228 |
| Cab39l | 1263.50508 | 0.10678985 | 0.105735694 | 1.00996973 | 0.312509793 | 0.545436767 |
| Prrc1 | 757.2402699 | 0.072219284 | 0.084773437 | 0.851909349 | 0.394264407 | 0.619921228 |
| Mxi1 | 3276.322784 | 0.106781367 | 0.098510289 | 1.083961558 | 0.278381844 | 0.507381283 |
| Stat3 | 2451.369591 | 0.106677692 | 0.085747384 | 1.244092665 | 0.21346546 | 0.433023546 |
| Uvrag | 1592.244874 | 0.106659805 | 0.094358677 | 1.130365621 | 0.258322194 | 0.484733447 |
| Rnf216 | 2284.092463 | 0.083201185 | 0.097722176 | 0.851405363 | 0.394544214 | 0.620018297 |
| Timm21 | 324.7746903 | 0.087322868 | 0.102548537 | 0.851527187 | 0.394476568 | 0.620018297 |
| Pttg1 | 970.4192634 | 0.106642203 | 0.075085556 | 1.420275865 | 0.155527385 | 0.357634506 |
| Nat6 | 476.8784799 | 0.106609435 | 0.079128066 | 1.347302423 | 0.177882852 | 0.387414269 |
| Zfp945 | 483.7079954 | 0.089731996 | 0.105422217 | 0.851167794 | 0.394676151 | 0.620116721 |
| Gbgt1 | 32.00290705 | -0.13689319 | 0.160899734 | -0.850798109 | 0.394881513 | 0.620327908 |
| Sec31a | 4436.886512 | 0.106588293 | 0.066873077 | 1.593889474 | 0.110960793 | 0.290017969 |
| Pim3 | 1371.05235 | 0.106568519 | 0.102935776 | 1.035291351 | 0.300532868 | 0.532105617 |
| Raver1 | 589.9479063 | 0.055628798 | 0.065420493 | 0.850326802 | 0.395143419 | 0.620486675 |

| | | | | | | |
|---|---|---|---|---|---|---|
| Med16 | 2452.075047 | 0.003783559 | 0.065808269 | 0.057493674 | 0.954151945 | 0.988860025 |
| Olfr324 | 2.253404505 | 0.003791103 | 0.029930015 | 0.126665593 | 0.899205079 | NA |
| Ppt1 | 5056.973455 | 0.003794609 | 0.055766539 | 0.068044549 | 0.945750171 | 0.986781137 |
| Rbm11 | 248.6423534 | 0.003796371 | 0.091418475 | 0.041527395 | 0.966875453 | 0.991586617 |
| Foxb2 | 1.134372229 | 0.003799083 | 0.021823219 | 0.174084445 | 0.861799097 | NA |
| Gm11128 | 7.7388057 | 0.003807592 | 0.056090014 | 0.067883605 | 0.94587829 | NA |
| Pdcd7 | 898.1852476 | 0.003817655 | 0.055860202 | 0.068343027 | 0.945512573 | 0.986781137 |
| Wnk1 | 10331.97098 | 0.003817663 | 0.072883715 | 0.05238019 | 0.958225759 | 0.989630474 |
| 1700020L24Rik | 8.261759608 | 0.003828332 | 0.053485076 | 0.071577582 | 0.942938081 | NA |
| Ikzf5 | 1002.176369 | 0.003832939 | 0.056231761 | 0.068163243 | 0.945655687 | 0.986781137 |
| Jph3 | 11191.90093 | 0.003840381 | 0.056200933 | 0.068333048 | 0.945520517 | 0.986781137 |
| A430107P09Rik | 17.62972331 | 0.00384521 | 0.075169533 | 0.05115383 | 0.959202942 | NA |
| Tprgl | 5720.650099 | 0.003851861 | 0.049747123 | 0.077428824 | 0.938282412 | 0.985461126 |
| Cul5 | 2679.123418 | 0.003865545 | 0.070782633 | 0.054611494 | 0.956447982 | 0.989344398 |
| BC049635 | 2.650237588 | 0.003876572 | 0.034096556 | 0.113693946 | 0.909480412 | NA |
| Serpinb6e | 0.727020108 | 0.003878178 | 0.015898645 | 0.243931366 | 0.807283976 | NA |
| Lrrc69 | 2.651490013 | 0.003887929 | 0.03560724 | 0.109189282 | 0.913052361 | NA |
| Gm7120 | 451.8071838 | 0.003895733 | 0.062917628 | 0.061917987 | 0.950628144 | 0.988202091 |
| Cidec | 0.674591037 | 0.003897011 | 0.017566016 | 0.221849471 | 0.824431066 | NA |
| 4833423E24Rik | 3.375788921 | 0.003917605 | 0.035669071 | 0.10983198 | 0.912542628 | NA |
| Ftl1 | 10727.26742 | 0.003922234 | 0.05091001 | 0.077042488 | 0.938589745 | 0.985461126 |
| Psph | 507.969375 | 0.003926114 | 0.078214838 | 0.050196535 | 0.959965773 | 0.990205619 |
| Gnpda2 | 723.0288657 | 0.003926694 | 0.067904198 | 0.057826977 | 0.95388645 | 0.988860025 |
| Gm19619 | 11.12975644 | 0.003944879 | 0.062565019 | 0.063052463 | 0.949724728 | NA |
| 9130008F23Rik | 3.572183108 | 0.003948231 | 0.032140775 | 0.122841808 | 0.902232367 | NA |
| Pdcd1lg2 | 5.153699133 | 0.003955074 | 0.049178834 | 0.080422286 | 0.935901403 | NA |
| Steap1 | 21.20695591 | 0.003961434 | 0.025793684 | 0.15358154 | NA | NA |
| Sh3d19 | 740.1844444 | 0.003970509 | 0.074385202 | 0.053377667 | 0.957430999 | 0.989630474 |
| Amdhd2 | 468.3782464 | 0.003974503 | 0.09237458 | 0.043025942 | 0.965680854 | 0.991035613 |
| Wiz | 2055.749603 | 0.003977114 | 0.053850405 | 0.073854864 | 0.941125871 | 0.985461126 |
| Mir30b | 0.631190298 | 0.003977747 | 0.017121533 | 0.232324222 | 0.816286201 | NA |
| Nme5 | 311.1559055 | 0.003990603 | 0.076807837 | 0.051955681 | 0.958564007 | 0.989630474 |
| Cldn9 | 4.861988429 | 0.003991384 | 0.037368014 | 0.106812839 | 0.914937461 | NA |
| Gm9926 | 0.685637272 | 0.004005732 | 0.015918529 | 0.251639616 | 0.801319635 | NA |
| 2200002D01Rik | 1.306873329 | 0.004020911 | 0.026692904 | 0.15063594 | 0.880262909 | NA |
| C030007H22Rik | 1.111433229 | 0.004027645 | 0.025558067 | 0.157588005 | 0.874781456 | NA |
| Vmn2r5 | 0.641969186 | 0.004030547 | 0.015274634 | 0.263871909 | 0.791878622 | NA |
| Snora5c | 6.630578333 | 0.004031076 | 0.049025875 | 0.082223431 | 0.934469041 | NA |
| Il17b | 8.761059008 | 0.004035385 | 0.063475457 | 0.063573943 | 0.94930948 | NA |
| Hrasls5 | 0.730710545 | 0.004038705 | 0.0161183 | 0.250566428 | 0.802149341 | NA |
| 1110020A21Rik | 24.36474314 | 0.004041747 | 0.080081882 | 0.050470184 | 0.959747709 | 0.990191033 |
| Hibch | 566.0517258 | 0.004045771 | 0.06758447 | 0.059862439 | 0.952265195 | 0.988510896 |
| Olfr520 | 0.393522717 | 0.004048562 | 0.01185164 | 0.341603504 | 0.7326493 | NA |
| Olfr58 | 0.393522717 | 0.004048562 | 0.01185164 | 0.341603504 | 0.7326493 | NA |
| Scgb3a2 | 0.393522717 | 0.004048562 | 0.01185164 | 0.341603504 | 0.7326493 | NA |
| Igbp1b | 10.62473705 | 0.004064954 | 0.055722643 | 0.072949761 | 0.941846095 | NA |
| Rpap2 | 727.8441722 | 0.004078378 | 0.068402763 | 0.059623001 | 0.952455899 | 0.988510896 |
| Spata31d1a | 2.514100685 | 0.004106867 | 0.032353118 | 0.126938833 | 0.898988811 | NA |
| Ripk3 | 10.38469083 | 0.004128188 | 0.066432975 | 0.062140646 | 0.950450828 | NA |
| Col8a2 | 168.5281401 | 0.004130109 | 0.046322759 | 0.089159383 | 0.928955245 | 0.983640461 |
| Rsph3a | 176.9169997 | 0.004148745 | 0.088967505 | 0.046632141 | 0.962806415 | 0.990993714 |
| Mir140 | 1.58693622 | 0.004157333 | 0.027991677 | 0.148520344 | 0.881932132 | NA |
| Dhx37 | 758.1138608 | 0.004187086 | 0.068553645 | 0.061077509 | 0.951297481 | 0.988239091 |
| Htt | 8226.465366 | 0.004187436 | 0.066455245 | 0.063011377 | 0.949757445 | 0.987981713 |
| Olfr859 | 0.690783455 | 0.004199012 | 0.017647409 | 0.237939289 | 0.811928182 | NA |
| Tk2 | 1802.166996 | 0.004227135 | 0.057078409 | 0.07405839 | 0.940963924 | 0.985461126 |
| Cul7 | 1506.536895 | 0.004232673 | 0.058662909 | 0.072152459 | 0.942480578 | 0.985461126 |
| Mylpf | 20.55906142 | 0.004237316 | 0.078713567 | 0.053832087 | 0.957068945 | NA |
| Heyl | 280.6482672 | 0.004247861 | 0.094592296 | 0.04490705 | 0.964181397 | 0.990993714 |
| Gm1045 | 1.053020722 | 0.004253238 | 0.019986898 | 0.212801278 | 0.831481967 | NA |
| Xpo4 | 357.536599 | 0.004265824 | 0.083689417 | 0.050972088 | 0.959347762 | 0.990016193 |
| Cd244 | 2.377071555 | 0.004268436 | 0.028957203 | 0.147404977 | 0.882812377 | NA |
| Vax2os | 1.610696021 | 0.004269914 | 0.02673953 | 0.159685449 | 0.873128864 | NA |
| Rpl23a | 7434.902101 | 0.00427529 | 0.068476453 | 0.062434451 | 0.950216861 | 0.988202091 |
| Ptchd1 | 722.3166706 | 0.0042783 | 0.081479747 | 0.052507521 | 0.958124303 | 0.989630474 |
| Ruvbl2 | 933.1314018 | 0.004291177 | 0.064896375 | 0.066123517 | 0.947279488 | 0.98731304 |
| Krtap10-10 | 0.384340583 | 0.004296611 | 0.011777417 | 0.364817799 | 0.715247418 | NA |
| Mir5112 | 1.845075032 | 0.00431641 | 0.027904215 | 0.15468667 | 0.877068339 | NA |
| Scarna17 | 1.791971035 | 0.004323814 | 0.026362135 | 0.164016097 | 0.869718472 | NA |
| Muc5ac | 0.874635119 | 0.004329325 | 0.018907316 | 0.228976201 | 0.818887413 | NA |
| Uba6 | 852.204835 | 0.004329896 | 0.08984945 | 0.048190566 | 0.961564369 | 0.990572163 |
| Ppyr1 | 1.131070143 | 0.004332709 | 0.022321336 | 0.194106169 | 0.846092751 | NA |
| Brap | 3024.454239 | 0.004351243 | 0.056058413 | 0.077619801 | 0.938130491 | 0.985461126 |
| Anp32a | 7296.919244 | 0.004354077 | 0.048749509 | 0.089315294 | 0.92883134 | 0.983611872 |
| Dand5 | 303.4825889 | 0.004355527 | 0.075746586 | 0.057501294 | 0.954145875 | 0.988860025 |
| Pank2 | 2160.976897 | 0.004356745 | 0.060042916 | 0.07256051 | 0.942155852 | 0.985461126 |
| Vmn2r27 | 1.293232342 | 0.004358403 | 0.026681005 | 0.16335227 | 0.870241081 | NA |
| Smc3 | 2429.404374 | 0.004362012 | 0.060623482 | 0.071952522 | 0.942639692 | 0.985461126 |
| Fxr1 | 2563.692165 | 0.004364001 | 0.057988243 | 0.075256649 | 0.940010513 | 0.985461126 |
| Gm5512 | 1.105671852 | 0.004382802 | 0.025554723 | 0.171506549 | 0.863825481 | NA |
| Fn3k | 1288.393926 | 0.004385147 | 0.062856717 | 0.069764173 | 0.944381364 | 0.986415739 |
| Aqp1 | 129.0974644 | 0.004399669 | 0.026739894 | 0.16453578 | NA | NA |
| Ccdc166 | 234.1979389 | 0.004406085 | 0.090918557 | 0.048461892 | 0.961348134 | 0.990572163 |
| 1700009J07Rik | 1.072889252 | 0.004412421 | 0.025493864 | 0.173077747 | 0.862590316 | NA |
| Fhod3 | 3041.551438 | 0.004417256 | 0.070973628 | 0.062237986 | 0.950373313 | 0.988202091 |
| Acot5 | 94.24578925 | 0.004425566 | 0.092659866 | 0.047761409 | 0.961906392 | 0.990595298 |
| Atp6v0a2 | 2101.520501 | 0.00443487 | 0.047173091 | 0.094012707 | 0.925099062 | 0.983236501 |
| Gm9054 | 3.103595811 | 0.004450542 | 0.040584106 | 0.109662193 | 0.912677285 | NA |
| Il18 | 624.1089526 | 0.004462595 | 0.070689329 | 0.063129686 | 0.949663235 | 0.987981713 |
| Tgm5 | 1.242644486 | 0.004465632 | 0.022685494 | 0.196849667 | 0.843945183 | NA |
| Btbd16 | 35.73183486 | 0.00446568 | 0.09276136 | 0.048141598 | 0.961603395 | 0.990572163 |
| Eid2 | 1166.125216 | 0.004470526 | 0.059415403 | 0.075241871 | 0.940022271 | 0.985461126 |
| Efha1 | 941.162954 | 0.00447172 | 0.056550504 | 0.07907481 | 0.93697312 | 0.985461126 |
| Syde1 | 593.7899244 | 0.004474244 | 0.075564404 | 0.059211002 | 0.952784047 | 0.988510896 |
| Rab11fip2 | 2866.811064 | 0.004490416 | 0.073320698 | 0.061243499 | 0.951165288 | 0.988239091 |
| Kera | 0.86531832 | 0.004504411 | 0.016856406 | 0.267222496 | 0.789297856 | NA |
| Igf2bp3 | 0.613315933 | 0.004508259 | 0.017031874 | 0.264695416 | 0.791244109 | NA |
| Mael | 3.052938119 | 0.004516387 | 0.037226717 | 0.121321117 | 0.903436695 | NA |
| Gpr52 | 416.5177017 | 0.004527568 | 0.094375863 | 0.047973792 | 0.961737129 | 0.990572163 |
| Gm11110 | 21.02024724 | 0.004535205 | 0.081470337 | 0.055666946 | 0.955607132 | NA |
| Fam221a | 39.44116953 | 0.004536975 | 0.088044701 | 0.051530354 | 0.958902915 | 0.989769328 |
| Mir3058 | 1.091337543 | 0.004548824 | 0.026658841 | 0.17063098 | 0.864513936 | NA |
| F730035M05Rik | 0.882616281 | 0.004551823 | 0.020568865 | 0.22129676 | 0.824861373 | NA |
| Hmgcl | 801.7499349 | 0.004555191 | 0.06158994 | 0.073959989 | 0.941042222 | 0.985461126 |
| Dlx6as2 | 1.100519677 | 0.004567304 | 0.027626782 | 0.165321607 | 0.868690853 | NA |
| Cspg5 | 11569.35997 | 0.00456745 | 0.04373406 | 0.104436911 | 0.916822632 | 0.982742917 |
| Hcn3 | 688.2607413 | 0.004576112 | 0.080167215 | 0.057082092 | 0.954479802 | 0.988860025 |
| Edem2 | 888.4636587 | 0.004576827 | 0.069620076 | 0.065740052 | 0.947584784 | 0.987347864 |
| Btnl5 | 0.638578749 | 0.00457849 | 0.015269228 | 0.299850802 | 0.764290963 | NA |
| Uspl1 | 1735.579214 | 0.004584288 | 0.063965104 | 0.071668575 | 0.942865665 | 0.985461126 |
| Ppbp | 0.999363745 | 0.004586468 | 0.016842027 | 0.272322788 | 0.785373836 | NA |
| Myoz3 | 1.221922479 | 0.004589767 | 0.02281876 | 0.201140068 | 0.840589052 | NA |
| Krt8 | 23.34132622 | 0.004595956 | 0.040143256 | 0.11448887 | 0.908850269 | 0.980523965 |
| Star | 68.46508068 | 0.004605161 | 0.093223514 | 0.049399131 | 0.960601221 | 0.990548518 |
| Rdh13 | 904.8739876 | 0.004606826 | 0.054822328 | 0.084031934 | 0.933031042 | 0.984399253 |
| 2410012M07Rik | 0.424793663 | 0.004608296 | 0.012133828 | 0.379789083 | 0.704101987 | NA |
| Klhl26 | 2234.087364 | 0.004615163 | 0.056664483 | 0.081447196 | 0.935086317 | 0.985461126 |
| Gtf2h1 | 1834.280726 | 0.004617189 | 0.059593496 | 0.077478071 | 0.938243236 | 0.985461126 |
| Mapk15 | 53.84247939 | 0.004636521 | 0.084720539 | 0.054727231 | 0.956355774 | 0.989344398 |
| Cdc45 | 75.62735604 | 0.004644886 | 0.09510183 | 0.048841186 | 0.96104586 | 0.990548518 |
| Kcnc3 | 6827.119155 | 0.004647081 | 0.063727699 | 0.072920891 | 0.941869069 | 0.985461126 |
| Tmbim1 | 1668.979178 | 0.004668856 | 0.075598961 | 0.061758198 | 0.950755393 | 0.988202091 |
| Pon3 | 110.0013073 | 0.004683087 | 0.070252601 | 0.066660687 | 0.946851832 | 0.987150629 |
| Snip1 | 555.3148445 | 0.004688571 | 0.064931052 | 0.072208454 | 0.942436017 | 0.985461126 |
| Mkl1 | 3139.587201 | 0.004699493 | 0.07121769 | 0.065987722 | 0.947387601 | 0.987335056 |
| Gm17019 | 1.772317822 | 0.004723347 | 0.030263093 | 0.156076142 | 0.875973004 | NA |
| Gnat1 | 6.869559469 | 0.004726689 | 0.056310059 | 0.083940411 | 0.93310381 | NA |
| Hmgxb3 | 2623.005012 | 0.004729103 | 0.055049631 | 0.085906168 | 0.931541008 | 0.984221522 |
| Piwil4 | 1.063707118 | 0.004731631 | 0.025478887 | 0.185707906 | 0.852673828 | NA |
| Trip11 | 1172.032843 | 0.004733796 | 0.084284352 | 0.056164585 | 0.955210693 | 0.989191597 |
| Gm14858 | 0.936484148 | 0.004737043 | 0.021032606 | 0.225223784 | 0.821805188 | NA |
| Kif1c | 4801.455113 | 0.004742167 | 0.061562001 | 0.077030751 | 0.938599082 | 0.985461126 |
| Rnase12 | 0.432646743 | 0.004743167 | 0.012200629 | 0.388764151 | 0.697450622 | NA |
| 9230102K24Rik | 1.866512249 | 0.004745971 | 0.031595826 | 0.150208811 | 0.880599875 | NA |
| Omg | 5472.185215 | 0.004748015 | 0.070775941 | 0.067085159 | 0.946513908 | 0.98700327 |
| Dnajb13 | 15.90774647 | 0.004754711 | 0.071209039 | 0.066771178 | 0.946763869 | NA |
| Tex10 | 876.8678412 | 0.004767681 | 0.077416288 | 0.061584989 | 0.950893331 | 0.988202091 |
| Sod3 | 627.7371786 | 0.004778801 | 0.077795515 | 0.06142772 | 0.951018577 | 0.988202091 |
| Prkcsh | 3163.912858 | 0.004780787 | 0.056604143 | 0.084460019 | 0.93269069 | 0.984387568 |
| Psapl1 | 0.655475157 | 0.004786924 | 0.018994562 | 0.252015493 | 0.801029088 | NA |
| Meox1 | 22.00125762 | 0.004807455 | 0.082262321 | 0.058440551 | 0.953397715 | NA |
| Kcne1 | 0.910229902 | 0.004822271 | 0.020960929 | 0.230060008 | 0.81804514 | NA |
| 4921508D12Rik | 1.059196812 | 0.004823055 | 0.023760647 | 0.202985012 | 0.839146747 | NA |
| Rhot2 | 3512.82576 | 0.004824605 | 0.072648743 | 0.066410031 | 0.947051384 | 0.987216965 |
| Dlx5 | 195.0280668 | 0.00482907 | 0.08541176 | 0.0565387 | 0.954912667 | 0.988953472 |
| Dcaf10 | 979.813922 | 0.004830121 | 0.074708621 | 0.064652799 | 0.948450445 | 0.987378592 |
| 1700052N19Rik | 1001.534053 | 0.004839092 | 0.063903584 | 0.075724897 | 0.939637968 | 0.985461126 |
| 5830417I10Rik | 385.1354286 | 0.004839486 | 0.07999002 | 0.060501123 | 0.951756522 | 0.988433205 |
| Hoxa4 | 0.415611529 | 0.00486001 | 0.012067553 | 0.402733672 | 0.68714416 | NA |
| Prr15l | 0.415611529 | 0.00486001 | 0.012067553 | 0.402733672 | 0.68714416 | NA |
| Hes3 | 19.36480714 | 0.004861658 | 0.082011468 | 0.059280222 | 0.952728914 | NA |
| Rpl14 | 7111.608304 | 0.004875933 | 0.066683372 | 0.073120669 | 0.941710095 | 0.985461126 |
| Col4a3 | 32.70271373 | 0.004880889 | 0.067240805 | 0.072588198 | 0.942133819 | 0.985461126 |
| H60c | 0.631545537 | 0.004897666 | 0.01713309 | 0.28586006 | 0.774985305 | NA |
| Nol6 | 6319.547752 | 0.004898854 | 0.048887912 | 0.100205832 | 0.920180916 | 0.982742917 |
| Olfr1419 | 0.599454723 | 0.004918595 | 0.016959374 | 0.290022216 | 0.771799242 | NA |
| Gpr63 | 179.8116388 | 0.004925609 | 0.091445022 | 0.053864155 | 0.957043395 | 0.989630474 |
| Ibsp | 5.979585314 | 0.004935102 | 0.046249379 | 0.10670635 | 0.915021945 | NA |
| 5730403I07Rik | 2.007448435 | 0.004950507 | 0.031008424 | 0.159650377 | 0.873156493 | NA |
| Muc19 | 1.319024518 | 0.004953063 | 0.020648356 | 0.239876867 | 0.810425715 | NA |
| Clasp2 | 11216.05616 | 0.004956594 | 0.054804207 | 0.090441845 | 0.927936106 | 0.983501978 |
| Gfer | 436.7557836 | 0.00496093 | 0.069282697 | 0.071604172 | 0.94291692 | 0.985461126 |
| Dstyk | 3392.921403 | 0.00497503 | 0.059445494 | 0.083690619 | 0.933302416 | 0.984471054 |
| H6pd | 459.8175166 | 0.004981044 | 0.087396253 | 0.056993798 | 0.954550136 | 0.988860025 |
| 1300015D01Rik | 0.09572496 | 0.004982553 | 0.00848 | 0.587565219 | 0.556824162 | NA |
| 1700010J16Rik | 0.09572496 | 0.004982553 | 0.00848 | 0.587565219 | 0.556824162 | NA |
| 1700018B24Rik | 0.09572496 | 0.004982553 | 0.00848 | 0.587565219 | 0.556824162 | NA |
| 1700034G24Rik | 0.09572496 | 0.004982553 | 0.00848 | 0.587565219 | 0.556824162 | NA |
| 1700071K01Rik | 0.09572496 | 0.004982553 | 0.00848 | 0.587565219 | 0.556824162 | NA |
| 2310057J18Rik | 0.09572496 | 0.004982553 | 0.00848 | 0.587565219 | 0.556824162 | NA |
| 4921506M07Rik | 0.09572496 | 0.004982553 | 0.00848 | 0.587565219 | 0.556824162 | NA |
| 4930407I10Rik | 0.09572496 | 0.004982553 | 0.00848 | 0.587565219 | 0.556824162 | NA |
| 4930441J16Rik | 0.09572496 | 0.004982553 | 0.00848 | 0.587565219 | 0.556824162 | NA |
| 4930502A04Rik | 0.09572496 | 0.004982553 | 0.00848 | 0.587565219 | 0.556824162 | NA |
| 4930524O08Rik | 0.09572496 | 0.004982553 | 0.00848 | 0.587565219 | 0.556824162 | NA |
| 9230112D13Rik | 0.09572496 | 0.004982553 | 0.00848 | 0.587565219 | 0.556824162 | NA |
| 9430007A20Rik | 0.09572496 | 0.004982553 | 0.00848 | 0.587565219 | 0.556824162 | NA |
| A1bg | 0.09572496 | 0.004982553 | 0.00848 | 0.587565219 | 0.556824162 | NA |
| Abo | 0.09572496 | 0.004982553 | 0.00848 | 0.587565219 | 0.556824162 | NA |
| Adam34 | 0.09572496 | 0.004982553 | 0.00848 | 0.587565219 | 0.556824162 | NA |
| Akr1b3 | 0.09572496 | 0.004982553 | 0.00848 | 0.587565219 | 0.556824162 | NA |
| Cldn13 | 0.09572496 | 0.004982553 | 0.00848 | 0.587565219 | 0.556824162 | NA |
| Crisp1 | 0.09572496 | 0.004982553 | 0.00848 | 0.587565219 | 0.556824162 | NA |
| Crisp2 | 0.09572496 | 0.004982553 | 0.00848 | 0.587565219 | 0.556824162 | NA |
| Cxcl17 | 0.09572496 | 0.004982553 | 0.00848 | 0.587565219 | 0.556824162 | NA |
| Eppin | 0.09572496 | 0.004982553 | 0.00848 | 0.587565219 | 0.556824162 | NA |
| Fam3b | 0.09572496 | 0.004982553 | 0.00848 | 0.587565219 | 0.556824162 | NA |
| Fshb | 0.09572496 | 0.004982553 | 0.00848 | 0.587565219 | 0.556824162 | NA |
| Fxyd4 | 0.09572496 | 0.004982553 | 0.00848 | 0.587565219 | 0.556824162 | NA |
| Galntl5 | 0.09572496 | 0.004982553 | 0.00848 | 0.587565219 | 0.556824162 | NA |
| Gm10466 | 0.09572496 | 0.004982553 | 0.00848 | 0.587565219 | 0.556824162 | NA |
| Gm5434 | 0.09572496 | 0.004982553 | 0.00848 | 0.587565219 | 0.556824162 | NA |
| Gm5801 | 0.09572496 | 0.004982553 | 0.00848 | 0.587565219 | 0.556824162 | NA |
| Gm9758 | 0.09572496 | 0.004982553 | 0.00848 | 0.587565219 | 0.556824162 | NA |
| Gml | 0.09572496 | 0.004982553 | 0.00848 | 0.587565219 | 0.556824162 | NA |
| H2-M9 | 0.09572496 | 0.004982553 | 0.00848 | 0.587565219 | 0.556824162 | NA |
| Hoxa10 | 0.09572496 | 0.004982553 | 0.00848 | 0.587565219 | 0.556824162 | NA |
| Ifne | 0.09572496 | 0.004982553 | 0.00848 | 0.587565219 | 0.556824162 | NA |
| Ifnz | 0.09572496 | 0.004982553 | 0.00848 | 0.587565219 | 0.556824162 | NA |
| Il13 | 0.09572496 | 0.004982553 | 0.00848 | 0.587565219 | 0.556824162 | NA |
| Klk1 | 0.09572496 | 0.004982553 | 0.00848 | 0.587565219 | 0.556824162 | NA |
| Krt27 | 0.09572496 | 0.004982553 | 0.00848 | 0.587565219 | 0.556824162 | NA |
| Mir210 | 0.09572496 | 0.004982553 | 0.00848 | 0.587565219 | 0.556824162 | NA |
| Mir3971 | 0.09572496 | 0.004982553 | 0.00848 | 0.587565219 | 0.556824162 | NA |
| Olfr12 | 0.09572496 | 0.004982553 | 0.00848 | 0.587565219 | 0.556824162 | NA |
| Olfr1340 | 0.09572496 | 0.004982553 | 0.00848 | 0.587565219 | 0.556824162 | NA |
| Olfr1428 | 0.09572496 | 0.004982553 | 0.00848 | 0.587565219 | 0.556824162 | NA |
| Olfr523 | 0.09572496 | 0.004982553 | 0.00848 | 0.587565219 | 0.556824162 | NA |
| Olfr545 | 0.09572496 | 0.004982553 | 0.00848 | 0.587565219 | 0.556824162 | NA |
| Olfr875 | 0.09572496 | 0.004982553 | 0.00848 | 0.587565219 | 0.556824162 | NA |
| Oosp1 | 0.09572496 | 0.004982553 | 0.00848 | 0.587565219 | 0.556824162 | NA |
| Scgb2b2 | 0.09572496 | 0.004982553 | 0.00848 | 0.587565219 | 0.556824162 | NA |
| Serpinb3d | 0.09572496 | 0.004982553 | 0.00848 | 0.587565219 | 0.556824162 | NA |
| Spaca3 | 0.09572496 | 0.004982553 | 0.00848 | 0.587565219 | 0.556824162 | NA |
| Sult1b1 | 0.09572496 | 0.004982553 | 0.00848 | 0.587565219 | 0.556824162 | NA |
| Tmprss11d | 0.09572496 | 0.004982553 | 0.00848 | 0.587565219 | 0.556824162 | NA |
| Tssk5 | 0.09572496 | 0.004982553 | 0.00848 | 0.587565219 | 0.556824162 | NA |
| Vmn1r53 | 0.09572496 | 0.004982553 | 0.00848 | 0.587565219 | 0.556824162 | NA |
| Vmn2r77 | 0.09572496 | 0.004982553 | 0.00848 | 0.587565219 | 0.556824162 | NA |
| Wfdc9 | 0.09572496 | 0.004982553 | 0.00848 | 0.587565219 | 0.556824162 | NA |
| A430035B10Rik | 84.03437181 | 0.00498551 | 0.09504514 | 0.052454127 | 0.958166846 | 0.989630474 |
| Klk14 | 33.55137992 | 0.004990871 | 0.090554333 | 0.055114656 | 0.95604712 | 0.989344398 |
| Pfpl | 0.423464609 | 0.004995804 | 0.012136222 | 0.411644061 | 0.680600332 | NA |
| Sh3bgrl3 | 3806.247994 | 0.005006953 | 0.049308891 | 0.101542602 | 0.919119741 | 0.982742917 |
| Capn8 | 2.165906263 | 0.005017737 | 0.031919457 | 0.157199952 | 0.875087266 | NA |
| Fam13c | 2682.423928 | 0.005025606 | 0.06193548 | 0.081142598 | 0.935328549 | 0.985461126 |
| Ksr1 | 1612.142272 | 0.005037922 | 0.068974107 | 0.073040768 | 0.941773676 | 0.985461126 |
| Cpe | 81611.37246 | 0.005043518 | 0.052164685 | 0.09668453 | 0.922976926 | 0.98296621 |
| Nop9 | 471.2768741 | 0.005063799 | 0.070878498 | 0.071443374 | 0.94304489 | 0.985515833 |
| Echs1 | 1836.231642 | 0.005072298 | 0.058664992 | 0.086462097 | 0.931099086 | 0.984208871 |
| Pias1 | 2157.218428 | 0.005073971 | 0.063663127 | 0.079700311 | 0.936475612 | 0.985461126 |
| Gnai2 | 7232.486432 | 0.005095205 | 0.055930911 | 0.091098189 | 0.927414571 | 0.983501978 |
| 1700058G18Rik | 4.316964816 | 0.005101346 | 0.040680614 | 0.125399935 | 0.90020694 | NA |
| Homez | 768.9738387 | 0.005103519 | 0.064225974 | 0.07946192 | 0.936665219 | 0.985461126 |
| Slc6a14 | 0.434194484 | 0.005120533 | 0.013963086 | 0.36671932 | 0.713828394 | NA |

| | | | | | | | | | | | | | |
|---|---|---|---|---|---|---|---|---|---|---|---|---|---|
| Sap18 | 1114.806114 | -0.047962821 | 0.05640521 | -0.850326074 | 0.395143824 | 0.620486675 | C1ql4 | 0.655896803 | 0.005121079 | 0.017486665 | 0.292856243 | 0.769632037 | NA |
| Adcy8 | 929.920884 | 0.106548636 | 0.103454933 | 1.029903873 | 0.303055133 | 0.535164831 | Ctnnd2 | 9236.343425 | 0.005125373 | 0.037909653 | 0.135199694 | 0.892453989 | 0.977844987 |
| Zfand1 | 534.8712866 | 0.070431921 | 0.082846162 | 0.850153096 | 0.395239975 | 0.620513545 | Mettl15 | 150.4015073 | 0.005125444 | 0.092903394 | 0.055169613 | 0.956003337 | 0.989344398 |
| Sergef | 686.8141075 | 0.106543246 | 0.091712879 | 1.161704302 | 0.245355598 | 0.47037115 | Gm8369 | 1.146326815 | 0.005126568 | 0.026410094 | 0.194113968 | 0.846086645 | NA |
| E130317F20Rik | 148.7940329 | -0.100128734 | 0.11779587 | -0.850019054 | 0.395314493 | 0.620554431 | 2310001H17Rik | 6.814855266 | 0.005127499 | 0.053545858 | 0.09575902 | 0.923711965 | NA |
| Pi4k2a | 3613.738128 | 0.106515953 | 0.069845588 | 1.525020487 | 0.127253988 | 0.315245314 | Smarcc2 | 13988.81522 | 0.005131902 | 0.048083718 | 0.106728485 | 0.915004384 | 0.98170867 |
| Tmem62 | 1254.595658 | 0.044556896 | 0.052442202 | 0.849638167 | 0.395526285 | 0.620711462 | 1700003G18Rik | 0.106148608 | 0.005155541 | 0.008542008 | 0.60355139 | 0.546141939 | NA |
| 4933434E20Rik | 1666.659053 | -0.050859555 | 0.059880895 | -0.849345271 | 0.395689197 | 0.620841359 | 1700018F24Rik | 0.106148608 | 0.005155541 | 0.008542008 | 0.60355139 | 0.546141939 | NA |
| Klhl35 | 25.30494885 | -0.140898127 | 0.165882397 | -0.849385648 | 0.395666737 | 0.620841359 | 1700019O17Rik | 0.106148608 | 0.005155541 | 0.008542008 | 0.60355139 | 0.546141939 | NA |
| Tyw1 | 583.3612702 | 0.056095471 | 0.066053221 | 0.849246556 | 0.395744112 | 0.620864649 | 1700042G15Rik | 0.106148608 | 0.005155541 | 0.008542008 | 0.60355139 | 0.546141939 | NA |
| Efnb3 | 4529.005573 | 0.065755681 | 0.077455606 | 0.848946689 | 0.395910957 | 0.620922658 | 2200002J24Rik | 0.106148608 | 0.005155541 | 0.008542008 | 0.60355139 | 0.546141939 | NA |
| Rbak | 515.5859663 | 0.106485555 | 0.091179276 | 1.167870155 | 0.242859144 | 0.467319869 | 4921511I17Rik | 0.106148608 | 0.005155541 | 0.008542008 | 0.60355139 | 0.546141939 | NA |
| Rps28 | 2071.510394 | -0.086876321 | 0.102339671 | -0.848901702 | 0.395935992 | 0.620922658 | 4930428E07Rik | 0.106148608 | 0.005155541 | 0.008542008 | 0.60355139 | 0.546141939 | NA |
| Zfp747 | 217.5707374 | 0.089366503 | 0.105259309 | 0.849012822 | 0.395874157 | 0.620922658 | 4930500F04Rik | 0.106148608 | 0.005155541 | 0.008542008 | 0.60355139 | 0.546141939 | NA |
| Ikzf5 | 949.026983 | 0.076638024 | 0.090303191 | 0.848674593 | 0.396062388 | 0.620986707 | 4933401D09Rik | 0.106148608 | 0.005155541 | 0.008542008 | 0.60355139 | 0.546141939 | NA |
| Rsph3b | 103.4851568 | -0.116320305 | 0.137054727 | -0.848714288 | 0.396040294 | 0.620986707 | 4933402N03Rik | 0.106148608 | 0.005155541 | 0.008542008 | 0.60355139 | 0.546141939 | NA |
| Sh3gl1 | 1180.694777 | 0.072512235 | 0.085490766 | 0.848187913 | 0.396333328 | 0.621348637 | 4933426D04Rik | 0.106148608 | 0.005155541 | 0.008542008 | 0.60355139 | 0.546141939 | NA |
| Epm2a | 457.3143976 | 0.106464173 | 0.0944866 | 1.126764778 | 0.259841946 | 0.486583408 | 5033403H07Rik | 0.106148608 | 0.005155541 | 0.008542008 | 0.60355139 | 0.546141939 | NA |
| Asphd2 | 3393.145856 | 0.087989758 | 0.103820386 | 0.847519072 | 0.396705861 | 0.62180684 | 5430421N21Rik | 0.106148608 | 0.005155541 | 0.008542008 | 0.60355139 | 0.546141939 | NA |
| Orc6 | 562.4572421 | -0.066079333 | 0.078028652 | -0.846859858 | 0.397073239 | 0.622256779 | AA792892 | 0.106148608 | 0.005155541 | 0.008542008 | 0.60355139 | 0.546141939 | NA |
| Snx33 | 427.2596767 | 0.091359905 | 0.107876303 | 0.846895026 | 0.397053635 | 0.622256779 | Agr3 | 0.106148608 | 0.005155541 | 0.008542008 | 0.60355139 | 0.546141939 | NA |
| Gopc | 2257.562715 | 0.106438685 | 0.082204767 | 1.294799436 | 0.195389427 | 0.409914495 | B930092H01Rik | 0.106148608 | 0.005155541 | 0.008542008 | 0.60355139 | 0.546141939 | NA |
| Oaz2 | 6821.680643 | -0.047594173 | 0.056258329 | -0.845993364 | 0.397556446 | 0.622825033 | C9 | 0.106148608 | 0.005155541 | 0.008542008 | 0.60355139 | 0.546141939 | NA |
| Ubac1 | 1496.873644 | -0.050837793 | 0.060087591 | -0.846061438 | 0.397518471 | 0.622825033 | Ccdc54 | 0.106148608 | 0.005155541 | 0.008542008 | 0.60355139 | 0.546141939 | NA |
| Rab11fip3 | 5705.762383 | 0.10643118 | 0.061266444 | 1.737185535 | 0.082354427 | 0.238965437 | Clca4 | 0.106148608 | 0.005155541 | 0.008542008 | 0.60355139 | 0.546141939 | NA |
| Foxn2 | 583.394082 | 0.106398223 | 0.084827741 | 1.254285698 | 0.20973818 | 0.427871411 | Clrn3 | 0.106148608 | 0.005155541 | 0.008542008 | 0.60355139 | 0.546141939 | NA |
| Mphosph8 | 2039.5236 | 0.058566974 | 0.069267007 | 0.845524824 | 0.397817879 | 0.622982638 | Defb10 | 0.106148608 | 0.005155541 | 0.008542008 | 0.60355139 | 0.546141939 | NA |
| Srxn1 | 4706.465379 | 0.106359472 | 0.10410044 | 1.021700496 | 0.306922676 | 0.539605236 | E330017A01Rik | 0.106148608 | 0.005155541 | 0.008542008 | 0.60355139 | 0.546141939 | NA |
| Camk2n1 | 59769.82772 | -0.08461625 | 0.100116797 | -0.845175362 | 0.398012936 | 0.623225107 | Fbxw19 | 0.106148608 | 0.005155541 | 0.008542008 | 0.60355139 | 0.546141939 | NA |
| Pcsk6 | 822.8987079 | -0.096593811 | 0.114344959 | -0.844757937 | 0.398246004 | 0.62352704 | Foxa2 | 0.106148608 | 0.005155541 | 0.008542008 | 0.60355139 | 0.546141939 | NA |
| Baz1a | 149.1198714 | -0.116228252 | 0.137677505 | -0.844206554 | 0.398553993 | 0.623934497 | Fscb | 0.106148608 | 0.005155541 | 0.008542008 | 0.60355139 | 0.546141939 | NA |
| Dmrtc1a | 174.2173123 | -0.113584734 | 0.134566984 | -0.844075792 | 0.398627054 | 0.623934497 | Galp | 0.106148608 | 0.005155541 | 0.008542008 | 0.60355139 | 0.546141939 | NA |
| Mrpl9 | 1326.883885 | -0.046736221 | 0.055365494 | -0.84413987 | 0.398591251 | 0.623934497 | Gm14501 | 0.106148608 | 0.005155541 | 0.008542008 | 0.60355139 | 0.546141939 | NA |
| Dstyk | 3294.429537 | -0.058456905 | 0.06926609 | -0.843946948 | 0.398699052 | 0.623984153 | Gm16497 | 0.106148608 | 0.005155541 | 0.008542008 | 0.60355139 | 0.546141939 | NA |
| Syndig1 | 1617.480047 | -0.067242728 | 0.07968393 | -0.843868118 | 0.398743106 | 0.62399007 | Gm5420 | 0.106148608 | 0.005155541 | 0.008542008 | 0.60355139 | 0.546141939 | NA |
| Llgl1 | 2549.183849 | 0.106352478 | 0.069288177 | 1.534929647 | 0.12480111 | 0.312304373 | Gm5779 | 0.106148608 | 0.005155541 | 0.008542008 | 0.60355139 | 0.546141939 | NA |
| 6330418K02Rik | 93.2653855 | -0.120863674 | 0.143315191 | -0.84334168 | 0.399037378 | 0.624324461 | Gm5916 | 0.106148608 | 0.005155541 | 0.008542008 | 0.60355139 | 0.546141939 | NA |
| Gabra4 | 4613.988801 | -0.075777038 | 0.089862689 | -0.84325362 | 0.399086616 | 0.624338452 | Gsdma2 | 0.106148608 | 0.005155541 | 0.008542008 | 0.60355139 | 0.546141939 | NA |
| Pitpnb | 3472.50823 | 0.078174353 | 0.092718012 | 0.84314095 | 0.399149619 | 0.624370251 | Gsdma3 | 0.106148608 | 0.005155541 | 0.008542008 | 0.60355139 | 0.546141939 | NA |
| Rps6ka6 | 151.7500459 | -0.110047466 | 0.130531339 | -0.843073144 | 0.399187538 | 0.624370251 | Hand1 | 0.106148608 | 0.005155541 | 0.008542008 | 0.60355139 | 0.546141939 | NA |
| Cfl2 | 2136.518648 | -0.060204536 | 0.071426804 | -0.842884351 | 0.399293126 | 0.624385379 | Hdhd1a | 0.106148608 | 0.005155541 | 0.008542008 | 0.60355139 | 0.546141939 | NA |
| Gdnf | 44.73546183 | -0.134004782 | 0.158974385 | -0.84293317 | 0.399265821 | 0.624385379 | Hoxc6 | 0.106148608 | 0.005155541 | 0.008542008 | 0.60355139 | 0.546141939 | NA |
| Plekho1 | 1263.699703 | 0.075412156 | 0.089473902 | 0.842839694 | 0.399318105 | 0.624385379 | Khdc1b | 0.106148608 | 0.005155541 | 0.008542008 | 0.60355139 | 0.546141939 | NA |
| Begain | 4014.077453 | -0.074929023 | 0.088930364 | -0.842558384 | 0.399475474 | 0.624521618 | Klk1b4 | 0.106148608 | 0.005155541 | 0.008542008 | 0.60355139 | 0.546141939 | NA |
| Hist2h3c1 | 57.38393019 | -0.1351642 | 0.160424699 | -0.842539839 | 0.39948585 | 0.624521618 | Lce3d | 0.106148608 | 0.005155541 | 0.008542008 | 0.60355139 | 0.546141939 | NA |
| Ribc1 | 11.80555578 | -0.139888957 | 0.166097453 | -0.842210128 | 0.399670346 | 0.624747007 | Lcn8 | 0.106148608 | 0.005155541 | 0.008542008 | 0.60355139 | 0.546141939 | NA |
| Epha8 | 413.0871082 | 0.106259388 | 0.145337583 | 0.7311212 | 0.464705126 | 0.680542649 | Mepe | 0.106148608 | 0.005155541 | 0.008542008 | 0.60355139 | 0.546141939 | NA |
| Taf6l | 344.2571011 | 0.082435939 | 0.097909393 | 0.841961495 | 0.399809507 | 0.624838461 | Mir133a-1 | 0.106148608 | 0.005155541 | 0.008542008 | 0.60355139 | 0.546141939 | NA |
| Ccdc41 | 1213.0545 | 0.106259065 | 0.068515376 | 1.550879104 | 0.120930659 | 0.306460522 | Mrgprb1 | 0.106148608 | 0.005155541 | 0.008542008 | 0.60355139 | 0.546141939 | NA |
| Pdk1 | 2690.528541 | 0.106245568 | 0.095099883 | 1.117199785 | 0.263908913 | 0.491230742 | Olfr1131 | 0.106148608 | 0.005155541 | 0.008542008 | 0.60355139 | 0.546141939 | NA |
| Mtpn | 12001.84505 | -0.072605269 | 0.086313068 | -0.841185125 | 0.400244232 | 0.625328641 | Olfr594 | 0.106148608 | 0.005155541 | 0.008542008 | 0.60355139 | 0.546141939 | NA |
| Nxph4 | 182.2140989 | 0.106233213 | 0.131958544 | 0.805049902 | 0.420790884 | 0.643479466 | Olfr921 | 0.106148608 | 0.005155541 | 0.008542008 | 0.60355139 | 0.546141939 | NA |
| Gmnn | 119.5645859 | -0.101598785 | 0.120803952 | -0.841022034 | 0.400335591 | 0.625348059 | Pdcl2 | 0.106148608 | 0.005155541 | 0.008542008 | 0.60355139 | 0.546141939 | NA |
| Hnrnph1 | 6858.105057 | 0.070253469 | 0.083554821 | 0.840806882 | 0.400456131 | 0.625348059 | Prf1 | 0.106148608 | 0.005155541 | 0.008542008 | 0.60355139 | 0.546141939 | NA |
| Nmt1 | 4366.552331 | 0.079519104 | 0.094553606 | 0.840994942 | 0.400350768 | 0.625348059 | Prm2 | 0.106148608 | 0.005155541 | 0.008542008 | 0.60355139 | 0.546141939 | NA |
| Ppp2cb | 4182.77292 | -0.054445237 | 0.064750159 | -0.840851029 | 0.400431395 | 0.625348059 | Proc | 0.106148608 | 0.005155541 | 0.008542008 | 0.60355139 | 0.546141939 | NA |
| Vamp8 | 237.828524 | 0.106224776 | 0.111857594 | 0.94964296 | 0.342293702 | 0.575617474 | Prp2 | 0.106148608 | 0.005155541 | 0.008542008 | 0.60355139 | 0.546141939 | NA |
| Nsmce2 | 859.5167393 | -0.077299928 | 0.092049401 | -0.839765684 | 0.401039777 | 0.626129616 | Serpina11 | 0.106148608 | 0.005155541 | 0.008542008 | 0.60355139 | 0.546141939 | NA |
| P4htm | 2748.021378 | 0.0622595 | 0.074164906 | 0.839473865 | 0.401203449 | 0.626195854 | Slc22a26 | 0.106148608 | 0.005155541 | 0.008542008 | 0.60355139 | 0.546141939 | NA |
| Tgif2 | 50.43011266 | -0.134506822 | 0.160217595 | -0.839525908 | 0.401174257 | 0.626195854 | Speer5-ps1 | 0.106148608 | 0.005155541 | 0.008542008 | 0.60355139 | 0.546141939 | NA |
| Ttc32 | 49.19572702 | -0.138188 | 0.164604902 | -0.839513276 | 0.401181342 | 0.626195854 | Spink12 | 0.106148608 | 0.005155541 | 0.008542008 | 0.60355139 | 0.546141939 | NA |
| Atf1 | 359.4638159 | 0.070719189 | 0.084265513 | 0.839242372 | 0.401333314 | 0.626335452 | Spink5 | 0.106148608 | 0.005155541 | 0.008542008 | 0.60355139 | 0.546141939 | NA |
| Far2 | 492.2033448 | -0.087224227 | 0.103945577 | -0.83913361 | 0.401394338 | 0.626367597 | Tlx2 | 0.106148608 | 0.005155541 | 0.008542008 | 0.60355139 | 0.546141939 | NA |
| Fam58b | 389.2620919 | -0.069551039 | 0.082896428 | -0.839011289 | 0.401462975 | 0.626411615 | Tmem207 | 0.106148608 | 0.005155541 | 0.008542008 | 0.60355139 | 0.546141939 | NA |
| Alg3 | 403.5236729 | 0.068596976 | 0.081785218 | 0.838745398 | 0.401612197 | 0.626453747 | Tmem233 | 0.106148608 | 0.005155541 | 0.008542008 | 0.60355139 | 0.546141939 | NA |
| Appbp2 | 1715.530223 | -0.061445164 | 0.07324987 | -0.838843322 | 0.401557236 | 0.626453747 | Vmn1r208 | 0.106148608 | 0.005155541 | 0.008542008 | 0.60355139 | 0.546141939 | NA |
| Foxf2 | 165.5578192 | 0.091897445 | 0.109574559 | 0.83867501 | 0.401651705 | 0.626453747 | Vmn2r73 | 0.106148608 | 0.005155541 | 0.008542008 | 0.60355139 | 0.546141939 | NA |
| Dyrk1b | 945.5265789 | 0.106179739 | 0.06696748 | 1.585541809 | 0.112843286 | 0.293502412 | Zdhhc11 | 0.106148608 | 0.005155541 | 0.008542008 | 0.60355139 | 0.546141939 | NA |
| Nuak1 | 3820.564498 | 0.088958331 | 0.106084138 | 0.838563921 | 0.401714064 | 0.626466868 | Nadk | 2918.481874 | 0.005169808 | 0.045891978 | 0.112651664 | 0.910306724 | 0.980590355 |
| Twistnb | 317.1388932 | -0.089103327 | 0.106263125 | -0.838515968 | 0.401740984 | 0.626466868 | Efcab1 | 134.4967677 | 0.005173529 | 0.090364389 | 0.057251858 | 0.954344569 | 0.988860025 |
| Hlcs | 674.2645561 | 0.106140198 | 0.073319248 | 1.447644387 | 0.147716531 | 0.347157599 | Mir16-1 | 0.674396547 | 0.005179867 | 0.019092166 | 0.271308522 | 0.786153753 | NA |
| 2310069G16Rik | 78.5563848 | -0.127001029 | 0.151529161 | -0.838129292 | 0.401958093 | 0.626679281 | Atp1a1 | 52519.46437 | 0.005181679 | 0.066268284 | 0.078192437 | 0.937674978 | 0.985461126 |
| Mx2 | 41.99859868 | 0.106071765 | 0.168065037 | 0.631135224 | 0.527952112 | 0.729981578 | Cmtm1 | 0.90694462 | 0.005189266 | 0.022406007 | 0.231601534 | 0.816847516 | NA |
| Frs2 | 2386.398179 | 0.057258711 | 0.068357235 | 0.837639367 | 0.402233276 | 0.626982131 | 1700008F21Rik | 0.108766301 | 0.005191714 | 0.008560034 | 0.606506234 | 0.544178644 | NA |
| Rbx1 | 3255.55558 | -0.04023311 | 0.04805151 | -0.837291289 | 0.402428855 | 0.627223888 | 1700010B08Rik | 0.108766301 | 0.005191714 | 0.008560034 | 0.606506234 | 0.544178644 | NA |
| Prr11 | 32.43841625 | -0.137223594 | 0.163922488 | -0.837124884 | 0.402522374 | 0.627306545 | 1700039E22Rik | 0.108766301 | 0.005191714 | 0.008560034 | 0.606506234 | 0.544178644 | NA |
| Rnasek | 3555.760641 | 0.052844543 | 0.063153225 | 0.83676713 | 0.402723477 | 0.627556829 | 1700042B14Rik | 0.108766301 | 0.005191714 | 0.008560034 | 0.606506234 | 0.544178644 | NA |
| Pald1 | 544.845052 | 0.075867341 | 0.090701416 | 0.836451564 | 0.402900915 | 0.62777019 | 1810007C17Rik | 0.108766301 | 0.005191714 | 0.008560034 | 0.606506234 | 0.544178644 | NA |
| Jak1 | 7445.739883 | -0.045586735 | 0.054525185 | -0.836067509 | 0.403116926 | 0.628043604 | 4921517D22Rik | 0.108766301 | 0.005191714 | 0.008560034 | 0.606506234 | 0.544178644 | NA |
| Sh3yl1 | 271.9032342 | 0.086129093 | 0.103029299 | 0.835966995 | 0.403173471 | 0.628068546 | 4930405P13Rik | 0.108766301 | 0.005191714 | 0.008560034 | 0.606506234 | 0.544178644 | NA |
| Arrdc4 | 309.222921 | 0.106063639 | 0.097673791 | 1.085896616 | 0.277524734 | 0.50641482 | 4930448C13Rik | 0.108766301 | 0.005191714 | 0.008560034 | 0.606506234 | 0.544178644 | NA |
| Lrig1 | 1592.887045 | 0.092431273 | 0.110600655 | 0.835720852 | 0.403311962 | 0.628157975 | 4930448I18Rik | 0.108766301 | 0.005191714 | 0.008560034 | 0.606506234 | 0.544178644 | NA |
| Arhgap42 | 617.572874 | 0.078470543 | 0.093936623 | 0.835356222 | 0.403517171 | 0.628351261 | 4930480G23Rik | 0.108766301 | 0.005191714 | 0.008560034 | 0.606506234 | 0.544178644 | NA |
| Suv39h2 | 159.0044346 | -0.092478045 | 0.110703183 | -0.835369343 | 0.403509785 | 0.628351261 | 4930527F14Rik | 0.108766301 | 0.005191714 | 0.008560034 | 0.606506234 | 0.544178644 | NA |
| Fbxo8 | 669.6041852 | -0.05545176 | 0.066413896 | -0.834942138 | 0.403750288 | 0.628651086 | 4930556C24Rik | 0.108766301 | 0.005191714 | 0.008560034 | 0.606506234 | 0.544178644 | NA |
| Ccnf | 236.3197071 | 0.081991426 | 0.098298192 | 0.834109199 | 0.404219452 | 0.629203086 | 9130221F21Rik | 0.108766301 | 0.005191714 | 0.008560034 | 0.606506234 | 0.544178644 | NA |
| Fundc1 | 1211.453258 | -0.056799443 | 0.068096972 | -0.834096447 | 0.404226637 | 0.629203086 | Acer1 | 0.108766301 | 0.005191714 | 0.008560034 | 0.606506234 | 0.544178644 | NA |
| Nlrp1c-ps | 12.858969 | -0.129481885 | 0.155222695 | -0.834168516 | 0.40418603 | 0.629203086 | Actc1 | 0.108766301 | 0.005191714 | 0.008560034 | 0.606506234 | 0.544178644 | NA |
| Ciapin1 | 2196.596086 | 0.058225964 | 0.069826236 | 0.833869439 | 0.40435456 | 0.629275769 | Akr1cl | 0.108766301 | 0.005191714 | 0.008560034 | 0.606506234 | 0.544178644 | NA |
| Terf2 | 1601.221744 | 0.062927132 | 0.075462416 | 0.833887048 | 0.404344637 | 0.629275769 | Arl14epl | 0.108766301 | 0.005191714 | 0.008560034 | 0.606506234 | 0.544178644 | NA |
| Supt3 | 326.066491 | -0.073586553 | 0.088258128 | -0.833765161 | 0.404413331 | 0.629304022 | Atp1b4 | 0.108766301 | 0.005191714 | 0.008560034 | 0.606506234 | 0.544178644 | NA |
| Rnf32 | 289.0511987 | -0.093731207 | 0.112436647 | -0.833635733 | 0.404486283 | 0.629354336 | C330024C12Rik | 0.108766301 | 0.005191714 | 0.008560034 | 0.606506234 | 0.544178644 | NA |
| Ring1 | 1000.153844 | 0.105992612 | 0.111538302 | 0.950279951 | 0.341970023 | 0.575421758 | C87198 | 0.108766301 | 0.005191714 | 0.008560034 | 0.606506234 | 0.544178644 | NA |
| Prkar2a | 2727.27992 | 0.083240697 | 0.099901491 | 0.83322777 | 0.404716283 | 0.629585753 | Cfd | 0.108766301 | 0.005191714 | 0.008560034 | 0.606506234 | 0.544178644 | NA |
| Arhgap23 | 5121.246177 | 0.10596346 | 0.074263036 | 1.426866786 | 0.153618296 | 0.35488508 | Clec4b1 | 0.108766301 | 0.005191714 | 0.008560034 | 0.606506234 | 0.544178644 | NA |
| Adhfe1 | 683.4762905 | -0.102481473 | 0.123030262 | -0.832977766 | 0.404857269 | 0.629676416 | Clps | 0.108766301 | 0.005191714 | 0.008560034 | 0.606506234 | 0.544178644 | NA |
| Pcyox1l | 549.8555373 | -0.088794779 | 0.106608117 | -0.832908235 | 0.404896485 | 0.629676416 | Cpa1 | 0.108766301 | 0.005191714 | 0.008560034 | 0.606506234 | 0.544178644 | NA |
| Wdr25 | 276.6885318 | -0.074841217 | 0.089868059 | -0.832789953 | 0.404963202 | 0.629716966 | Cpb2 | 0.108766301 | 0.005191714 | 0.008560034 | 0.606506234 | 0.544178644 | NA |
| Fgf14 | 1626.235692 | -0.071056444 | 0.085356198 | -0.832469646 | 0.405143904 | 0.62986703 | Crygb | 0.108766301 | 0.005191714 | 0.008560034 | 0.606506234 | 0.544178644 | NA |
| Gm16907 | 9.2819162 | -0.130223704 | 0.156455542 | -0.832336792 | 0.405218869 | 0.62986703 | Csn1s2b | 0.108766301 | 0.005191714 | 0.008560034 | 0.606506234 | 0.544178644 | NA |
| Snx24 | 504.7597046 | 0.105920389 | 0.072110141 | 1.468869543 | 0.141868177 | 0.337391487 | Cyp2c70 | 0.108766301 | 0.005191714 | 0.008560034 | 0.606506234 | 0.544178644 | NA |
| Gpr155 | 2826.171408 | 0.066526168 | 0.079927569 | 0.832330684 | 0.405222316 | 0.62986703 | D730048I06Rik | 0.108766301 | 0.005191714 | 0.008560034 | 0.606506234 | 0.544178644 | NA |
| Tra2b | 1924.557411 | -0.042586383 | 0.051171671 | -0.832225762 | 0.405281525 | 0.629895872 | Fbxo39 | 0.108766301 | 0.005191714 | 0.008560034 | 0.606506234 | 0.544178644 | NA |
| Wdsub1 | 350.394992 | 0.071328449 | 0.085725849 | 0.832052984 | 0.405379038 | 0.629984235 | Galr2 | 0.108766301 | 0.005191714 | 0.008560034 | 0.606506234 | 0.544178644 | NA |
| Kdm2a | 4287.285582 | 0.05439588 | 0.065389213 | 0.831878496 | 0.405477531 | 0.630019591 | Gm12839 | 0.108766301 | 0.005191714 | 0.008560034 | 0.606506234 | 0.544178644 | NA |
| Actr1b | 9938.384078 | 0.105916773 | 0.065206301 | 1.624333421 | 0.104304668 | 0.278301451 | Gm14318 | 0.108766301 | 0.005191714 | 0.008560034 | 0.606506234 | 0.544178644 | NA |
| Anxa7 | 3630.638085 | 0.04954536 | 0.059572917 | 0.831675904 | 0.405591905 | 0.63006226 | Gm7714 | 0.108766301 | 0.005191714 | 0.008560034 | 0.606506234 | 0.544178644 | NA |
| Ptcra | 8.433840029 | -0.133940647 | 0.161047886 | -0.831682119 | 0.405588396 | 0.63006226 | Gm9994 | 0.108766301 | 0.005191714 | 0.008560034 | 0.606506234 | 0.544178644 | NA |
| Has3 | 91.8190172 | -0.113709155 | 0.136746673 | -0.831531421 | 0.405673486 | 0.630125813 | Gp6 | 0.108766301 | 0.005191714 | 0.008560034 | 0.606506234 | 0.544178644 | NA |
| A030009H04Rik | 4192.115389 | -0.048413281 | 0.058262174 | -0.830955626 | 0.405998697 | 0.630504542 | H2-M11 | 0.108766301 | 0.005191714 | 0.008560034 | 0.606506234 | 0.544178644 | NA |
| Zcchc4 | 198.293236 | 0.085829196 | 0.103287244 | 0.830975751 | 0.405987328 | 0.630504542 | Hamp | 0.108766301 | 0.005191714 | 0.008560034 | 0.606506234 | 0.544178644 | NA |
| Crispld1 | 205.1665269 | -0.10369086 | 0.124798171 | -0.830868433 | 0.406047958 | 0.630517846 | Itih1 | 0.108766301 | 0.005191714 | 0.008560034 | 0.606506234 | 0.544178644 | NA |
| Pygb | 8233.664118 | -0.060086542 | 0.072340785 | -0.830603964 | 0.406197395 | 0.630686686 | Kpna7 | 0.108766301 | 0.005191714 | 0.008560034 | 0.606506234 | 0.544178644 | NA |
| Ccny | 4798.18846 | -0.058392179 | 0.070352312 | -0.829996595 | 0.406540709 | 0.631050988 | Krt74 | 0.108766301 | 0.005191714 | 0.008560034 | 0.606506234 | 0.544178644 | NA |
| Ibtk | 1130.100054 | -0.062412295 | 0.075198008 | -0.829972712 | 0.406554212 | 0.631050988 | Krt84 | 0.108766301 | 0.005191714 | 0.008560034 | 0.606506234 | 0.544178644 | NA |
| Pkhd1l1 | 22.88857945 | -0.136599592 | 0.164557592 | -0.830102039 | 0.406481094 | 0.631050988 | Lce3a | 0.108766301 | 0.005191714 | 0.008560034 | 0.606506234 | 0.544178644 | NA |
| Ciao1 | 1976.322657 | 0.054739074 | 0.065996602 | 0.829422606 | 0.406865314 | 0.631280911 | Lipn | 0.108766301 | 0.005191714 | 0.008560034 | 0.606506234 | 0.544178644 | NA |
| Mtx1 | 837.147316 | -0.055769515 | 0.067236752 | -0.829449865 | 0.406849895 | 0.631280911 | Ly6g6c | 0.108766301 | 0.005191714 | 0.008560034 | 0.606506234 | 0.544178644 | NA |
| Sec23ip | 1721.098748 | 0.055115871 | 0.066447725 | 0.829462126 | 0.40684296 | 0.631280911 | Mcpt8 | 0.108766301 | 0.005191714 | 0.008560034 | 0.606506234 | 0.544178644 | NA |
| Serinc5 | 3207.766111 | 0.081253667 | 0.097950585 | 0.829537335 | 0.40680042 | 0.631280911 | Mettl7a2 | 0.108766301 | 0.005191714 | 0.008560034 | 0.606506234 | 0.544178644 | NA |
| Cdk2ap1 | 2855.375536 | 0.069799049 | 0.084194564 | 0.829020855 | 0.407092606 | 0.631443872 | Mir1251 | 0.108766301 | 0.005191714 | 0.008560034 | 0.606506234 | 0.544178644 | NA |
| Fbxw9 | 509.4931913 | 0.091271662 | 0.110091067 | 0.829056018 | 0.40707271 | 0.631443872 | Mir26b | 0.108766301 | 0.005191714 | 0.008560034 | 0.606506234 | 0.544178644 | NA |
| Slirp | 588.2824161 | -0.078975431 | 0.095248099 | -0.829154927 | 0.407016746 | 0.631443872 | Mir30e | 0.108766301 | 0.005191714 | 0.008560034 | 0.606506234 | 0.544178644 | NA |
| Chid1 | 1987.555034 | -0.088688777 | 0.107017514 | -0.828731428 | 0.407256398 | 0.631561428 | Mir484 | 0.108766301 | 0.005191714 | 0.008560034 | 0.606506234 | 0.544178644 | NA |
| Hnrnph2 | 4573.004573 | 0.048013597 | 0.057942722 | 0.828638963 | 0.407308734 | 0.631561428 | Mir708 | 0.108766301 | 0.005191714 | 0.008560034 | 0.606506234 | 0.544178644 | NA |
| Tmem63c | 1999.589695 | 0.058400106 | 0.070468729 | 0.8287379 | 0.407252735 | 0.631561428 | Mup6 | 0.108766301 | 0.005191714 | 0.008560034 | 0.606506234 | 0.544178644 | NA |
| Urod | 1326.414156 | 0.06802593 | 0.082097544 | 0.828598846 | 0.407331441 | 0.631561428 | Olfr1079 | 0.108766301 | 0.005191714 | 0.008560034 | 0.606506234 | 0.544178644 | NA |
| Fgl2 | 88.05299186 | -0.137535334 | 0.166032383 | -0.828364513 | 0.407464098 | 0.631703895 | Olfr1335 | 0.108766301 | 0.005191714 | 0.008560034 | 0.606506234 | 0.544178644 | NA |
| Zscan22 | 383.1638721 | -0.067642695 | 0.081696674 | -0.827973675 | 0.407685409 | 0.631983764 | Olfr1508 | 0.108766301 | 0.005191714 | 0.008560034 | 0.606506234 | 0.544178644 | NA |
| 2610008E11Rik | 1137.069822 | 0.105900283 | 0.086670217 | 1.221876297 | 0.22175441 | 0.442929929 | Olfr411 | 0.108766301 | 0.005191714 | 0.008560034 | 0.606506234 | 0.544178644 | NA |
| Phf8 | 1338.729295 | -0.068989141 | 0.083349794 | -0.827706198 | 0.407836908 | 0.631999566 | Olfr62 | 0.108766301 | 0.005191714 | 0.008560034 | 0.606506234 | 0.544178644 | NA |
| Taf1 | 3466.955789 | -0.060961701 | 0.073653559 | -0.827681669 | 0.407850803 | 0.631999566 | Olfr661 | 0.108766301 | 0.005191714 | 0.008560034 | 0.606506234 | 0.544178644 | NA |
| Tph2 | 37.55128812 | -0.138969695 | 0.167881074 | -0.827786553 | 0.407791391 | 0.631999566 | Rfx6 | 0.108766301 | 0.005191714 | 0.008560034 | 0.606506234 | 0.544178644 | NA |
| Gm17769 | 50.18222238 | -0.123454455 | 0.149194745 | -0.827471871 | 0.407969659 | 0.63206613 | Rhox4g | 0.108766301 | 0.005191714 | 0.008560034 | 0.606506234 | 0.544178644 | NA |
| Prkra | 457.9782439 | -0.075604401 | 0.0913706 | -0.827447782 | 0.407983307 | 0.63206613 | Rln3 | 0.108766301 | 0.005191714 | 0.008560034 | 0.606506234 | 0.544178644 | NA |
| 4933404O12Rik | 655.7883706 | 0.063711143 | 0.077029018 | 0.827105733 | 0.408177135 | 0.632239981 | Scp2d1 | 0.108766301 | 0.005191714 | 0.008560034 | 0.606506234 | 0.544178644 | NA |
| St3gal3 | 1930.834518 | 0.050444638 | 0.060984619 | 0.827169836 | 0.408140806 | 0.632239981 | Sectm1a | 0.108766301 | 0.005191714 | 0.008560034 | 0.606506234 | 0.544178644 | NA |
| Tmed10 | 5675.602002 | -0.054685228 | 0.066124616 | -0.827002575 | 0.408235602 | 0.632267335 | Serpinb9e | 0.108766301 | 0.005191714 | 0.008560034 | 0.606506234 | 0.544178644 | NA |
| 2410016O06Rik | 483.3300907 | -0.064512124 | 0.078022959 | -0.826835142 | 0.408330509 | 0.632351115 | Serpine3 | 0.108766301 | 0.005191714 | 0.008560034 | 0.606506234 | 0.544178644 | NA |
| Naa16 | 187.9179767 | 0.091631645 | 0.110848847 | 0.826635982 | 0.408443416 | 0.632395817 | Sohlh2 | 0.108766301 | 0.005191714 | 0.008560034 | 0.606506234 | 0.544178644 | NA |
| Cul4a | 2699.803422 | 0.105825932 | 0.065281228 | 1.621077536 | 0.105001011 | 0.279485598 | Speer9-ps1 | 0.108766301 | 0.005191714 | 0.008560034 | 0.606506234 | 0.544178644 | NA |
| Tars2 | 1138.734632 | 0.081558311 | 0.09865561 | 0.826697146 | 0.40840874 | 0.632395817 | Sval2 | 0.108766301 | 0.005191714 | 0.008560034 | 0.606506234 | 0.544178644 | NA |
| Cox10 | 1018.529404 | -0.055061133 | 0.066623721 | -0.826449379 | 0.408549223 | 0.632436972 | Tmem139 | 0.108766301 | 0.005191714 | 0.008560034 | 0.606506234 | 0.544178644 | NA |
| 2510003E04Rik | 6279.516197 | 0.054624626 | 0.066103435 | 0.826350795 | 0.408605127 | 0.632460323 | Ugt2a2 | 0.108766301 | 0.005191714 | 0.008560034 | 0.606506234 | 0.544178644 | NA |
| 2610020C07Rik | 21.68067674 | -0.13764763 | 0.166595625 | -0.826237962 | 0.408669117 | 0.632496185 | Uty | 0.108766301 | 0.005191714 | 0.008560034 | 0.606506234 | 0.544178644 | NA |
| Srl | 333.4644312 | 0.077880545 | 0.094311529 | 0.825779685 | 0.40892908 | 0.632835314 | Vmn1r5 | 0.108766301 | 0.005191714 | 0.008560034 | 0.606506234 | 0.544178644 | NA |
| Tfeb | 283.0313521 | 0.105804094 | 0.105282874 | 1.004950666 | 0.314920606 | 0.548183335 | Wdr20b | 0.108766301 | 0.005191714 | 0.008560034 | 0.606506234 | 0.544178644 | NA |
| Plgrkt | 338.2347216 | 0.09266329 | 0.11224466 | 0.825547423 | 0.409060871 | 0.632908847 | Zdhhc19 | 0.108766301 | 0.005191714 | 0.008560034 | 0.606506234 | 0.544178644 | NA |
| Rwdd2b | 340.5223163 | -0.077898276 | 0.09436725 | -0.825479984 | 0.409099142 | 0.632908847 | Apbb2 | 2301.215755 | 0.005209595 | 0.048660239 | 0.107060603 | 0.914740902 | 0.98170867 |
| Arhgef2 | 6543.23415 | 0.052414169 | 0.063518036 | 0.825185602 | 0.409266226 | 0.633104124 | Gm8363 | 0.700284741 | 0.005212619 | 0.017699251 | 0.294510731 | 0.768367665 | NA |
| Eepd1 | 809.3980271 | -0.05504483 | 0.066767997 | -0.824419368 | 0.409701314 | 0.633650644 | 4930419G24Rik | 1.563857722 | 0.005226039 | 0.027113248 | 0.192748538 | 0.847155909 | NA |
| Cwc22 | 788.6281913 | 0.105751507 | 0.067488547 | 1.566954852 | 0.117125245 | 0.300403666 | Tmub2 | 2659.444652 | 0.005245385 | 0.054532515 | 0.096188208 | 0.923371096 | 0.98311054 |
| H2-Aa | 119.954637 | 0.095744281 | 0.116245891 | 0.823635833 | 0.410146509 | 0.63423556 | Nkx6-3 | 2.878226646 | 0.005255419 | 0.040495821 | 0.12977683 | 0.896742994 | NA |
| AW549877 | 2398.956076 | 0.105631639 | 0.077385802 | 1.365000239 | 0.172252975 | 0.38012088 | 4921508A21Rik | 19.14415816 | 0.005255678 | 0.080042149 | 0.065661381 | 0.947647419 | NA |
| Hira | 2749.59249 | 0.090775602 | 0.110247198 | 0.823382393 | 0.410290572 | 0.634372031 | Prom2 | 23.62464428 | 0.005257777 | 0.077585713 | 0.06776733 | 0.945970851 | 0.986798749 |
| Cmtm6 | 1148.603129 | -0.076090166 | 0.092424224 | -0.823270811 | 0.410354008 | 0.634393404 | 1700095B10Rik | 0.113843286 | 0.005259477 | 0.00859368 | 0.612016898 | 0.540526578 | NA |
| G6pc3 | 1571.324411 | 0.079214362 | 0.096225718 | 0.823214044 | 0.410386284 | 0.634393404 | 2410088K16Rik | 0.113843286 | 0.005259477 | 0.00859368 | 0.612016898 | 0.540526578 | NA |
| Rai2 | 379.299967 | 0.105614415 | 0.104098542 | 1.014561902 | 0.310314708 | 0.543217273 | 4930401C15Rik | 0.113843286 | 0.005259477 | 0.00859368 | 0.612016898 | 0.540526578 | NA |
| Ints12 | 465.0077612 | -0.075047122 | 0.091204124 | -0.822847899 | 0.410594494 | 0.634502692 | 4930511E03Rik | 0.113843286 | 0.005259477 | 0.00859368 | 0.612016898 | 0.540526578 | NA |
| Prkdc | 1137.202951 | -0.084613713 | 0.102836098 | -0.822801668 | 0.410620788 | 0.634502692 | 4933406F09Rik | 0.113843286 | 0.005259477 | 0.00859368 | 0.612016898 | 0.540526578 | NA |
| Soga2 | 3993.213776 | 0.060975996 | 0.074099546 | 0.82289297 | 0.410568861 | 0.634502692 | Acsm2 | 0.113843286 | 0.005259477 | 0.00859368 | 0.612016898 | 0.540526578 | NA |
| Palm2 | 2545.371537 | 0.072461144 | 0.08808389 | 0.822637871 | 0.410713956 | 0.63458337 | Ampd1 | 0.113843286 | 0.005259477 | 0.00859368 | 0.612016898 | 0.540526578 | NA |
| Xpo6 | 4180.380516 | 0.105595049 | 0.088724359 | 1.190147224 | 0.233988532 | 0.457478359 | Apol7c | 0.113843286 | 0.005259477 | 0.00859368 | 0.612016898 | 0.540526578 | NA |
| Zdhhc13 | 801.9356219 | -0.070044432 | 0.085160933 | -0.822494886 | 0.410795296 | 0.634618433 | C6 | 0.113843286 | 0.005259477 | 0.00859368 | 0.612016898 | 0.540526578 | NA |
| Parp2 | 605.9660255 | 0.076559146 | 0.093112825 | 0.822219137 | 0.410952189 | 0.634761561 | Cyp4f40 | 0.113843286 | 0.005259477 | 0.00859368 | 0.612016898 | 0.540526578 | NA |
| Cbfb | 631.8414927 | -0.080026391 | 0.097342158 | -0.822114409 | 0.411011785 | 0.634790331 | Fmo9 | 0.113843286 | 0.005259477 | 0.00859368 | 0.612016898 | 0.540526578 | NA |
| Nae1 | 1356.568928 | 0.05116393 | 0.062259943 | 0.821779271 | 0.411202534 | 0.635021634 | Gm11186 | 0.113843286 | 0.005259477 | 0.00859368 | 0.612016898 | 0.540526578 | NA |
| Wdr92 | 561.8195381 | 0.095752398 | 0.116530929 | 0.821690848 | 0.411252869 | 0.635036073 | Gm17359 | 0.113843286 | 0.005259477 | 0.00859368 | 0.612016898 | 0.540526578 | NA |
| Kctd13 | 4420.710679 | 0.066249705 | 0.080633958 | 0.821610479 | 0.411298623 | 0.635043435 | Gm20098 | 0.113843286 | 0.005259477 | 0.00859368 | 0.612016898 | 0.540526578 | NA |
| Kank3 | 734.7644958 | -0.093258285 | 0.113529925 | -0.821442322 | 0.411394365 | 0.635127969 | Gm3925 | 0.113843286 | 0.005259477 | 0.00859368 | 0.612016898 | 0.540526578 | NA |
| Brwd1 | 4632.877631 | -0.06796798 | 0.082766808 | -0.821198522 | 0.411533199 | 0.635152443 | Gsta1 | 0.113843286 | 0.005259477 | 0.00859368 | 0.612016898 | 0.540526578 | NA |
| Eif2b2 | 1077.268774 | 0.070060379 | 0.085313779 | 0.821208248 | 0.411527659 | 0.635152443 | Hist1h2af | 0.113843286 | 0.005259477 | 0.00859368 | 0.612016898 | 0.540526578 | NA |
| Il1f9 | 50.38023512 | -0.121062945 | 0.147398691 | -0.82132985 | 0.411458409 | 0.635152443 | Hist1h2ai | 0.113843286 | 0.005259477 | 0.00859368 | 0.612016898 | 0.540526578 | NA |
| Gtse1 | 27.99827502 | 0.105577266 | 0.167262851 | 0.631205704 | 0.527906033 | 0.729981578 | Hist1h2ak | 0.113843286 | 0.005259477 | 0.00859368 | 0.612016898 | 0.540526578 | NA |
| Gm7102 | 28.89618421 | -0.134903998 | 0.164499078 | -0.820089693 | 0.412164978 | 0.635937479 | Klrc1 | 0.113843286 | 0.005259477 | 0.00859368 | 0.612016898 | 0.540526578 | NA |
| Uba1 | 14571.01628 | 0.059017348 | 0.071962359 | 0.820114145 | 0.412151039 | 0.635937479 | Krt33b | 0.113843286 | 0.005259477 | 0.00859368 | 0.612016898 | 0.540526578 | NA |
| Med16 | 2238.272062 | -0.064206655 | 0.078299744 | -0.820011051 | 0.412209807 | 0.635943319 | Mir1940 | 0.113843286 | 0.005259477 | 0.00859368 | 0.612016898 | 0.540526578 | NA |
| Ergic3 | 3016.973837 | -0.046143681 | 0.05629643 | -0.819655547 | 0.412412498 | 0.636114585 | Mir30c-1 | 0.113843286 | 0.005259477 | 0.00859368 | 0.612016898 | 0.540526578 | NA |
| Zmat1 | 497.4053865 | 0.105533172 | 0.104678352 | 1.008166158 | 0.313374701 | 0.546413192 | Mir381 | 0.113843286 | 0.005259477 | 0.00859368 | 0.612016898 | 0.540526578 | NA |
| Htra4 | 49.21213292 | -0.123666267 | 0.150878911 | -0.819639185 | 0.412421828 | 0.636114585 | Mir667 | 0.113843286 | 0.005259477 | 0.00859368 | 0.612016898 | 0.540526578 | NA |
| Satb2 | 2145.809708 | 0.105521973 | 0.124791485 | 0.845586329 | 0.397783555 | 0.622982638 | Olfr1034 | 0.113843286 | 0.005259477 | 0.00859368 | 0.612016898 | 0.540526578 | NA |
| Lpcat3 | 610.468335 | -0.060381652 | 0.073733801 | -0.818914136 | 0.412835404 | 0.636528187 | Olfr20 | 0.113843286 | 0.005259477 | 0.00859368 | 0.612016898 | 0.540526578 | NA |
| Myg1 | 842.258642 | -0.052160826 | 0.063691047 | -0.818966384 | 0.412805593 | 0.636528187 | Olfr459 | 0.113843286 | 0.005259477 | 0.00859368 | 0.612016898 | 0.540526578 | NA |
| Nup93 | 1033.578815 | 0.090402395 | 0.110434653 | 0.818605326 | 0.413011628 | 0.636706509 | Olfr53 | 0.113843286 | 0.005259477 | 0.00859368 | 0.612016898 | 0.540526578 | NA |
| Paics | 3663.105181 | -0.042766144 | 0.052245107 | -0.818567446 | 0.413033247 | 0.636706509 | Tnfsf4 | 0.113843286 | 0.005259477 | 0.00859368 | 0.612016898 | 0.540526578 | NA |
| Bcl2l11 | 256.2857164 | 0.083086951 | 0.101556976 | 0.818131399 | 0.413282161 | 0.637026841 | Vmn1r4 | 0.113843286 | 0.005259477 | 0.00859368 | 0.612016898 | 0.540526578 | NA |
| Amer3 | 1517.423964 | 0.105473052 | 0.078997433 | 1.335145306 | 0.181828794 | 0.392320753 | Vmn2r111 | 0.113843286 | 0.005259477 | 0.00859368 | 0.612016898 | 0.540526578 | NA |
| Cops7a | 4926.393974 | -0.044115589 | 0.053976321 | -0.817313737 | 0.413749158 | 0.637388731 | Stim2 | 3107.196888 | 0.005261708 | 0.07200341 | 0.07307582 | 0.941745783 | 0.985461126 |
| Derl1 | 2554.152324 | -0.042074598 | 0.051475243 | -0.817375411 | 0.413713923 | 0.637388731 | Zbtb17 | 1407.887688 | 0.005270355 | 0.075243399 | 0.070044088 | 0.944158569 | 0.986253946 |
| Gm10778 | 11.63371234 | -0.136058959 | 0.166496877 | -0.817186254 | 0.413821997 | 0.637388731 | Apob | 2.070968059 | 0.005275562 | 0.032069302 | 0.164505054 | 0.869333568 | NA |
| Tmem203 | 238.0716132 | 0.105369233 | 0.097295534 | 1.082981188 | 0.278816773 | 0.507917251 | Kcnmb3 | 0.503272123 | 0.005296638 | 0.012724059 | 0.416269575 | 0.677212754 | NA |
| Mettl14 | 764.7438829 | -0.056641088 | 0.069315905 | -0.817144166 | 0.413846046 | 0.637388731 | Acta2 | 488.1541435 | 0.00530304 | 0.090144649 | 0.058828114 | 0.953089015 | 0.988615469 |
| Mrps10 | 622.5145387 | -0.067654191 | 0.082762877 | -0.817446107 | 0.413673536 | 0.637388731 | G730013B05Rik | 7.247731364 | 0.005360199 | 0.055871145 | 0.095938591 | 0.923569345 | NA |
| Zbtb9 | 677.2038759 | -0.054749862 | 0.066989883 | -0.817285527 | 0.413765276 | 0.637388731 | Akt1s1 | 1318.408652 | 0.005370194 | 0.071154218 | 0.075472607 | 0.939838692 | 0.985461126 |
| Tmem39b | 269.517906 | 0.082622632 | 0.101141043 | 0.816905077 | 0.413982676 | 0.63753579 | Slc25a40 | 416.674286 | 0.005374549 | 0.074491146 | 0.072150173 | 0.942482398 | 0.985461126 |
| Eya2 | 101.8802245 | -0.123682022 | 0.151554111 | -0.816091496 | 0.414447807 | 0.63764026 | Ydjc | 463.3549999 | 0.005378254 | 0.082295612 | 0.065352862 | 0.947893055 | 0.987367302 |
| Dhx35 | 735.5665357 | 0.105318057 | 0.101715399 | 1.035418996 | 0.300473278 | 0.532060911 | Thsd7a | 1988.324923 | 0.005381008 | 0.071879684 | 0.074861323 | 0.94032505 | 0.985461126 |
| Ercc6l2 | 869.0277835 | 0.105301385 | 0.086048953 | 1.223738129 | 0.221051031 | 0.442129311 | Gm6300 | 0.926794364 | 0.005383581 | 0.021006997 | 0.256275627 | 0.797738016 | NA |
| Mcl1 | 2514.764332 | 0.065630005 | 0.080406092 | 0.816231754 | 0.414367599 | 0.63764026 | 4933406M09Rik | 0.44803889 | 0.005388685 | 0.014082118 | 0.382661551 | 0.701970726 | NA |
| Nceh1 | 5554.875065 | -0.050848198 | 0.062308884 | -0.816066586 | 0.414462054 | 0.63764026 | Tas2r137 | 0.44803889 | 0.005388685 | 0.014082118 | 0.382661551 | 0.701970726 | NA |
| Nploc4 | 3088.217232 | 0.053360686 | 0.065380278 | 0.816158751 | 0.414409345 | 0.63764026 | Capn13 | 0.680543483 | 0.005393226 | 0.017379742 | 0.310316794 | 0.756320062 | NA |
| Gripap1 | 4069.472909 | 0.105299179 | 0.058314888 | 1.805699769 | 0.070965249 | 0.217085008 | Stard10 | 1909.920579 | 0.005393357 | 0.057923168 | 0.093112256 | 0.925814381 | 0.983236501 |
| Xpo5 | 2047.955227 | 0.10528917 | 0.049420804 | 2.130462515 | 0.033133447 | 0.132209537 | Ucn3 | 28.53939441 | 0.005394244 | 0.058407951 | 0.092354614 | 0.926416298 | 0.983236501 |
| Tbx18 | 125.2194596 | -0.113794024 | 0.139347283 | -0.81662176 | 0.414144616 | 0.63764026 | Acsm4 | 2.126352761 | 0.005404393 | 0.032911826 | 0.164208256 | 0.869567202 | NA |
| Znrf2 | 369.8387647 | -0.071461407 | 0.087525121 | -0.816467388 | 0.414232868 | 0.63764026 | Gm11437 | 0.428231225 | 0.005418156 | 0.013917417 | 0.389307593 | 0.69704862 | NA |
| Hdlbp | 9242.996845 | -0.03802918 | 0.046613774 | -0.815835688 | 0.414594119 | 0.637716795 | Pde3a | 90.33810263 | 0.005419377 | 0.093965558 | 0.05767408 | 0.95400824 | 0.988860025 |
| Svip | 1902.590772 | 0.045412122 | 0.055662273 | 0.815851016 | 0.414585351 | 0.637716795 | 1700022I11Rik | 0.127687691 | 0.005422984 | 0.008697249 | 0.623528683 | 0.532937159 | NA |
| Rsbn1l | 845.8081726 | 0.06603205 | 0.08095079 | 0.815706064 | 0.41466827 | 0.637767538 | 1700025C18Rik | 0.127687691 | 0.005422984 | 0.008697249 | 0.623528683 | 0.532937159 | NA |
| Scfd1 | 1128.86844 | -0.05175325 | 0.063462438 | -0.815494203 | 0.414789481 | 0.637890642 | 1700060C16Rik | 0.127687691 | 0.005422984 | 0.008697249 | 0.623528683 | 0.532937159 | NA |
| Dhx57 | 1935.355047 | 0.043256257 | 0.053070945 | 0.81506475 | 0.415035248 | 0.638205253 | 1700122O11Rik | 0.127687691 | 0.005422984 | 0.008697249 | 0.623528683 | 0.532937159 | NA |
| Banp | 997.344282 | 0.080640342 | 0.098977226 | 0.814736329 | 0.415223254 | 0.63843099 | 2310016D03Rik | 0.127687691 | 0.005422984 | 0.008697249 | 0.623528683 | 0.532937159 | NA |
| Apip | 317.582073 | -0.08141154 | 0.099955383 | -0.814478789 | 0.415370718 | 0.638594355 | 2310030A07Rik | 0.127687691 | 0.005422984 | 0.008697249 | 0.623528683 | 0.532937159 | NA |
| Usp42 | 995.7528294 | 0.069133817 | 0.084889449 | 0.814398229 | 0.415416853 | 0.638601917 | 4930448H16Rik | 0.127687691 | 0.005422984 | 0.008697249 | 0.623528683 | 0.532937159 | NA |
| Zcchc16 | 86.20426533 | -0.121439751 | 0.149150197 | -0.814211138 | 0.415524006 | 0.638703269 | 4930596M17Rik | 0.127687691 | 0.005422984 | 0.008697249 | 0.623528683 | 0.532937159 | NA |
| Zfp532 | 2241.153255 | -0.046819757 | 0.057524664 | -0.813907536 | 0.415697923 | 0.638907214 | 4933400F21Rik | 0.127687691 | 0.005422984 | 0.008697249 | 0.623528683 | 0.532937159 | NA |
| Gtf2h3 | 481.9449639 | 0.070706097 | 0.08688252 | 0.813812689 | 0.415752265 | 0.638927355 | 9130409I23Rik | 0.127687691 | 0.005422984 | 0.008697249 | 0.623528683 | 0.532937159 | NA |
| Runx2 | 326.2508076 | 0.105241605 | 0.116038296 | 0.906955795 | 0.364430174 | 0.594616798 | Abcg8 | 0.127687691 | 0.005422984 | 0.008697249 | 0.623528683 | 0.532937159 | NA |
| 2210408F21Rik | 354.4407285 | -0.079536041 | 0.097766334 | -0.813531998 | 0.415913108 | 0.638959676 | Adad1 | 0.127687691 | 0.005422984 | 0.008697249 | 0.623528683 | 0.532937159 | NA |
| Fkbp10 | 265.8208026 | -0.073662097 | 0.090558929 | -0.813416169 | 0.415979492 | 0.638959676 | Adam26a | 0.127687691 | 0.005422984 | 0.008697249 | 0.623528683 | 0.532937159 | NA |
| Pnrc1 | 1025.794994 | 0.070751225 | 0.086960708 | 0.813599927 | 0.41587418 | 0.638959676 | Bc1 | 0.127687691 | 0.005422984 | 0.008697249 | 0.623528683 | 0.532937159 | NA |
| Tssc1 | 1490.908257 | -0.069686911 | 0.085670156 | -0.813432751 | 0.415969988 | 0.638959676 | Calm5 | 0.127687691 | 0.005422984 | 0.008697249 | 0.623528683 | 0.532937159 | NA |
| Cmya5 | 385.2793425 | 0.089467106 | 0.110013354 | 0.813238599 | 0.416081273 | 0.63898932 | Ccdc144b | 0.127687691 | 0.005422984 | 0.008697249 | 0.623528683 | 0.532937159 | NA |
| Zfp213 | 283.6166565 | -0.073856891 | 0.090816385 | -0.813255124 | 0.4160718 | 0.63898932 | Cd300c | 0.127687691 | 0.005422984 | 0.008697249 | 0.623528683 | 0.532937159 | NA |
| Tecpr2 | 4500.166658 | 0.055940123 | 0.068813611 | 0.81292236 | 0.416262574 | 0.639204393 | Crybb2 | 0.127687691 | 0.005422984 | 0.008697249 | 0.623528683 | 0.532937159 | NA |
| Srsf7 | 2473.840235 | 0.072041725 | 0.088636545 | 0.81277677 | 0.416346057 | 0.63926923 | Ctsll3 | 0.127687691 | 0.005422984 | 0.008697249 | 0.623528683 | 0.532937159 | NA |
| Egfl8 | 54.13830042 | -0.132163884 | 0.162650436 | -0.812563967 | 0.416468098 | 0.639292284 | Cyp2w1 | 0.127687691 | 0.005422984 | 0.008697249 | 0.623528683 | 0.532937159 | NA |
| Zbtb8b | 220.0219526 | 0.105224029 | 0.101159197 | 1.040182517 | 0.298255112 | 0.529906191 | Gm12718 | 0.127687691 | 0.005422984 | 0.008697249 | 0.623528683 | 0.532937159 | NA |
| Zfp382 | 378.0466522 | 0.06456215 | 0.079451344 | 0.812599854 | 0.416447515 | 0.639292284 | Gm20756 | 0.127687691 | 0.005422984 | 0.008697249 | 0.623528683 | 0.532937159 | NA |
| Gnl1 | 7600.382524 | -0.056992666 | 0.070154369 | -0.812389407 | 0.416568223 | 0.639356913 | Gm2516 | 0.127687691 | 0.005422984 | 0.008697249 | 0.623528683 | 0.532937159 | NA |
| Magt1 | 620.8881539 | -0.088780679 | 0.109296549 | -0.812291696 | 0.416624274 | 0.639379607 | Gm4745 | 0.127687691 | 0.005422984 | 0.008697249 | 0.623528683 | 0.532937159 | NA |
| Phf21a | 1924.937784 | 0.067762939 | 0.083440021 | 0.812115558 | 0.416725327 | 0.639471349 | Gpr31b | 0.127687691 | 0.005422984 | 0.008697249 | 0.623528683 | 0.532937159 | NA |
| Entpd5 | 1731.106209 | 0.048740507 | 0.060025355 | 0.811998648 | 0.416792407 | 0.639510949 | Gzmg | 0.127687691 | 0.005422984 | 0.008697249 | 0.623528683 | 0.532937159 | NA |
| Lenep | 202.3775702 | 0.105221215 | 0.120760339 | 0.871322621 | 0.383578023 | 0.611876648 | Hormad1 | 0.127687691 | 0.005422984 | 0.008697249 | 0.623528683 | 0.532937159 | NA |
| Prmt6 | 482.223881 | -0.075456355 | 0.093079667 | -0.810664217 | 0.417558528 | 0.640559587 | Krt34 | 0.127687691 | 0.005422984 | 0.008697249 | 0.623528683 | 0.532937159 | NA |
| Chpt1 | 857.727491 | -0.067225171 | 0.082964885 | -0.810284629 | 0.417776607 | 0.640787605 | Mir181a-1 | 0.127687691 | 0.005422984 | 0.008697249 | 0.623528683 | 0.532937159 | NA |

| | | | | | | | | | | | | | | | |
|---|---|---|---|---|---|---|---|---|---|---|---|---|---|---|---|
| Copg2 | 3456.629089 | -0.047989215 | 0.059226821 | -0.810261533 | 0.417789879 | 0.640787605 | | | Mir190 | 0.127687691 | 0.005422984 | 0.008697249 | 0.623528683 | 0.532937159 | NA |
| Mettl10 | 452.7251927 | -0.076607204 | 0.094567489 | -0.810079704 | 0.417894368 | 0.640820996 | | | Mir224 | 0.127687691 | 0.005422984 | 0.008697249 | 0.623528683 | 0.532937159 | NA |
| Thoc6 | 160.2669248 | -0.085218915 | 0.105189004 | -0.810150418 | 0.41785373 | 0.640820996 | | | Mir412 | 0.127687691 | 0.005422984 | 0.008697249 | 0.623528683 | 0.532937159 | NA |
| Sp1 | 1282.835051 | -0.066554821 | 0.082185005 | -0.80981709 | 0.418045309 | 0.640989017 | | | Mptx2 | 0.127687691 | 0.005422984 | 0.008697249 | 0.623528683 | 0.532937159 | NA |
| Elmo2 | 7295.211283 | 0.047224127 | 0.058338907 | 0.809479112 | 0.418239613 | 0.641099272 | | | Mrgpra1 | 0.127687691 | 0.005422984 | 0.008697249 | 0.623528683 | 0.532937159 | NA |
| F11r | 391.0655531 | 0.092592149 | 0.114385279 | 0.809476095 | 0.418241348 | 0.641099272 | | | Nxpe5 | 0.127687691 | 0.005422984 | 0.008697249 | 0.623528683 | 0.532937159 | NA |
| Mex3d | 465.4396971 | -0.061132512 | 0.075511521 | -0.809578598 | 0.418182412 | 0.641099272 | | | Olfr10 | 0.127687691 | 0.005422984 | 0.008697249 | 0.623528683 | 0.532937159 | NA |
| Pdlim3 | 78.02237828 | -0.123899631 | 0.153112846 | -0.809204676 | 0.418397426 | 0.641275073 | | | Olfr1030 | 0.127687691 | 0.005422984 | 0.008697249 | 0.623528683 | 0.532937159 | NA |
| Gm5820 | 31.40843979 | -0.135961341 | 0.168115768 | -0.808736401 | 0.418666785 | 0.641528526 | | | Olfr279 | 0.127687691 | 0.005422984 | 0.008697249 | 0.623528683 | 0.532937159 | NA |
| Magee1 | 6631.988667 | -0.046235952 | 0.057168679 | -0.8087637 | 0.41865108 | 0.641528526 | | | Olfr54 | 0.127687691 | 0.005422984 | 0.008697249 | 0.623528683 | 0.532937159 | NA |
| Tbxa2r | 34.20073439 | -0.133964905 | 0.165654379 | -0.808701257 | 0.418687005 | 0.641528526 | | | Olfr876 | 0.127687691 | 0.005422984 | 0.008697249 | 0.623528683 | 0.532937159 | NA |
| Fbxo46 | 409.3302056 | 0.067358406 | 0.083309328 | 0.808533785 | 0.418783365 | 0.641612722 | | | Pax9 | 0.127687691 | 0.005422984 | 0.008697249 | 0.623528683 | 0.532937159 | NA |
| H2-Eb1 | 98.66044544 | 0.105210809 | 0.127616839 | 0.824427324 | 0.409696795 | 0.633650644 | | | Prss44 | 0.127687691 | 0.005422984 | 0.008697249 | 0.623528683 | 0.532937159 | NA |
| Trerf1 | 1518.214119 | 0.068124435 | 0.084280453 | 0.808306462 | 0.418914184 | 0.641714194 | | | Ssxb2 | 0.127687691 | 0.005422984 | 0.008697249 | 0.623528683 | 0.532937159 | NA |
| Plcl1 | 1722.303482 | 0.073141667 | 0.090515872 | 0.808053505 | 0.419059782 | 0.641845815 | | | Tchhl1 | 0.127687691 | 0.005422984 | 0.008697249 | 0.623528683 | 0.532937159 | NA |
| Dus2l | 446.2408935 | 0.0823519 | 0.101973594 | 0.807580642 | 0.419332034 | 0.642189209 | | | Tlx3 | 0.127687691 | 0.005422984 | 0.008697249 | 0.623528683 | 0.532937159 | NA |
| Mrpl22 | 431.9054088 | -0.087065873 | 0.107828437 | -0.807448166 | 0.419408326 | 0.642189209 | | | Tmem181c-ps | 0.127687691 | 0.005422984 | 0.008697249 | 0.623528683 | 0.532937159 | NA |
| P2rx4 | 513.6189763 | 0.063700618 | 0.07888877 | 0.807473836 | 0.419393542 | 0.642189209 | | | Tmem213 | 0.127687691 | 0.005422984 | 0.008697249 | 0.623528683 | 0.532937159 | NA |
| Htr7 | 386.6767317 | 0.08832782 | 0.109467193 | 0.806888507 | 0.419730721 | 0.642619347 | | | Tmem239 | 0.127687691 | 0.005422984 | 0.008697249 | 0.623528683 | 0.532937159 | NA |
| Lmf1 | 543.9081029 | -0.067134682 | 0.08322977 | -0.806618609 | 0.419886249 | 0.642730443 | | | Trpv5 | 0.127687691 | 0.005422984 | 0.008697249 | 0.623528683 | 0.532937159 | NA |
| Il33 | 2018.053977 | 0.105190685 | 0.165524752 | 0.635498216 | 0.52510353 | 0.727398667 | | | Usp26 | 0.127687691 | 0.005422984 | 0.008697249 | 0.623528683 | 0.532937159 | NA |
| AI314180 | 4877.04855 | 0.056697489 | 0.070362072 | 0.805796181 | 0.420360381 | 0.643196272 | | | Wfdc8 | 0.127687691 | 0.005422984 | 0.008697249 | 0.623528683 | 0.532937159 | NA |
| Clock | 3229.71826 | -0.061895595 | 0.076829729 | -0.805620372 | 0.420461775 | 0.643196272 | | | Zg16 | 0.127687691 | 0.005422984 | 0.008697249 | 0.623528683 | 0.532937159 | NA |
| Gm19705 | 15.87710644 | 0.10515029 | 0.168118011 | 0.625455238 | 0.531672318 | 0.732808502 | | | Klf5 | 206.8216571 | 0.005424464 | 0.0816473 | 0.066437763 | 0.947029306 | 0.987216965 |
| Mbtps1 | 3831.798326 | 0.052606958 | 0.065271794 | 0.805967715 | 0.420261465 | 0.643196272 | | | Sorcs2 | 2725.897011 | 0.005426841 | 0.062035358 | 0.087479801 | 0.930290141 | 0.983937278 |
| Mrps34 | 931.8995168 | -0.058027684 | 0.072010255 | -0.805825282 | 0.420343598 | 0.643196272 | | | Efhc2 | 76.50053619 | 0.005443254 | 0.09481267 | 0.057410622 | 0.954218102 | 0.988860025 |
| Trim44 | 14729.38232 | -0.038948926 | 0.048348518 | -0.805586769 | 0.420481157 | 0.643196272 | | | Gm6927 | 0.452312811 | 0.005493258 | 0.014119057 | 0.389066898 | 0.69722666 | NA |
| Uba3 | 2563.463054 | -0.048192957 | 0.059816968 | -0.805673693 | 0.420431022 | 0.643196272 | | | Zkscan17 | 617.4866582 | 0.005497014 | 0.077082056 | 0.071313802 | 0.943148011 | 0.985552694 |
| Zfp651 | 3709.737553 | 0.104963691 | 0.101642191 | 1.032678359 | 0.301754444 | 0.533658642 | | | Tubgcp3 | 1801.822961 | 0.005507814 | 0.053771336 | 0.102430289 | 0.918415143 | 0.982742917 |
| Hps6 | 257.6418368 | 0.080173908 | 0.099567886 | 0.805218549 | 0.420693574 | 0.64339416 | | | BC107364 | 1.925270583 | 0.005508544 | 0.034236316 | 0.160897672 | 0.872173996 | NA |
| L3mbtl3 | 837.2152563 | 0.104824823 | 0.076353182 | 1.372893979 | 0.169785309 | 0.377154634 | | | 3230401D17Rik | 719.853454 | 0.005517918 | 0.068837504 | 0.080158606 | 0.936111112 | 0.985461126 |
| Gabra1 | 12773.10968 | 0.062913023 | 0.078175479 | 0.804766701 | 0.420954321 | 0.64353885 | | | Eme1 | 2.806630389 | 0.005522007 | 0.040220876 | 0.137292062 | 0.890799945 | NA |
| Ndufaf4 | 922.3848322 | -0.070473345 | 0.087563488 | -0.804825689 | 0.420920275 | 0.64353885 | | | Nox4 | 5.141287849 | 0.00552216 | 0.04377513 | 0.12614833 | 0.899614511 | NA |
| Txnl1 | 2334.867916 | -0.043560453 | 0.054126248 | -0.8047935 | 0.420938853 | 0.64353885 | | | Sox14 | 4.945473281 | 0.005538662 | 0.028079531 | 0.19724911 | 0.843632601 | NA |
| B4galt4 | 734.0594434 | -0.066234702 | 0.082319884 | -0.804601493 | 0.421049681 | 0.643621128 | | | Lmx1a | 11.2885041 | 0.005546883 | 0.041702637 | 0.133010362 | NA | NA |
| Tubd1 | 195.0895305 | 0.104811518 | 0.132228502 | 0.792654509 | 0.427979141 | 0.64966286 | | | Gnpat | 2533.631011 | 0.00556386 | 0.050060074 | 0.111143664 | 0.911502425 | 0.980985114 |
| Ankrd33b | 4640.540299 | -0.097355853 | 0.121043369 | -0.804305552 | 0.421220532 | 0.64363258 | | | Xirp1 | 5.012100265 | 0.005571967 | 0.051104964 | 0.109029857 | 0.913178809 | NA |
| BC005537 | 6027.313899 | 0.046033743 | 0.057221606 | 0.804481846 | 0.42111875 | 0.64363258 | | | Tax1bp3 | 284.4462723 | 0.005573278 | 0.079446958 | 0.070150929 | 0.944073531 | 0.986236039 |
| Lrrc3 | 645.2624931 | -0.078646845 | 0.097791636 | -0.804228753 | 0.421264876 | 0.64363258 | | | Ict1 | 1031.989255 | 0.005580709 | 0.061812583 | 0.090284349 | 0.928061257 | 0.983501978 |
| Zfp85-rs1 | 139.5306057 | -0.097269325 | 0.120945321 | -0.804242151 | 0.42125714 | 0.64363258 | | | Rnps1 | 5681.723714 | 0.005585377 | 0.043764061 | 0.127624736 | 0.898445955 | 0.978396127 |
| Prnp | 34417.04835 | 0.047120746 | 0.058601764 | 0.804084083 | 0.421348417 | 0.643696743 | | | Larp4b | 2789.322717 | 0.00558885 | 0.051959481 | 0.107561701 | 0.914343379 | 0.98170867 |
| Bap1 | 6285.644644 | -0.051718513 | 0.064333318 | -0.80391489 | 0.42144613 | 0.643782543 | | | Hn1l | 226.3204416 | 0.005597346 | 0.094133098 | 0.059462043 | 0.952584097 | 0.988510896 |
| Usp25 | 1434.09782 | 0.104753182 | 0.080923967 | 1.29446425 | 0.19550511 | 0.410065815 | | | Rrp12 | 1764.319122 | 0.005604459 | 0.058911652 | 0.095133284 | 0.924208961 | 0.983236501 |
| Rbpms | 128.2056268 | 0.097072423 | 0.120815602 | 0.803475884 | 0.42169973 | 0.644042939 | | | Cdk5rap3 | 1017.936141 | 0.005610085 | 0.065005327 | 0.086301934 | 0.931226401 | 0.984208871 |
| Zfyve28 | 2122.073797 | 0.053829886 | 0.067029907 | 0.803072666 | 0.421932736 | 0.644335285 | | | Tpst2 | 795.155637 | 0.005623575 | 0.072391114 | 0.07768322 | 0.938080043 | 0.985461126 |
| Fbxw5 | 3493.271695 | 0.063704111 | 0.079349226 | 0.802832163 | 0.422071749 | 0.644484053 | | | Ngrn | 1167.507569 | 0.005630861 | 0.051789334 | 0.108726265 | 0.913419609 | 0.98170867 |
| 5730455P16Rik | 3327.241694 | 0.057846722 | 0.072079829 | 0.80253689 | 0.422242458 | 0.644554152 | | | Mir34a | 0.406564059 | 0.005656565 | 0.013731356 | 0.411945108 | 0.680379657 | NA |
| Rpl18 | 3073.162152 | -0.062750073 | 0.078186534 | -0.802568799 | 0.422224008 | 0.644554152 | | | 2310002D06Rik | 0.489410913 | 0.005659299 | 0.012644381 | 0.447574209 | 0.65446052 | NA |
| Thg1l | 179.2911929 | 0.081577827 | 0.101641623 | 0.802602564 | 0.422204486 | 0.644554152 | | | Fcer2a | 0.489410913 | 0.005659299 | 0.012644381 | 0.447574209 | 0.65446052 | NA |
| Ube2k | 4131.808243 | -0.054454788 | 0.067863847 | -0.802412329 | 0.422314483 | 0.644600592 | | | Cabp4 | 1.908898629 | 0.005669748 | 0.031541692 | 0.179754094 | 0.857345624 | NA |
| Iba57 | 329.8193672 | -0.079272346 | 0.098806316 | -0.8023004 | 0.422379211 | 0.644635884 | | | AI747448 | 0.635802653 | 0.005682843 | 0.017163521 | 0.331100049 | 0.740568915 | NA |
| Sh3bp5 | 1959.931409 | 0.067448681 | 0.084092219 | 0.802079933 | 0.422506722 | 0.64476698 | | | 1700092M07Rik | 2.524417425 | 0.005689869 | 0.034775064 | 0.163619216 | 0.870030917 | NA |
| Kif5a | 139939.9325 | -0.067553511 | 0.084250127 | -0.801820885 | 0.422656576 | 0.644932144 | | | 1810011H11Rik | 23.44506111 | 0.00570311 | 0.076920446 | 0.074142964 | 0.940896629 | 0.985461126 |
| Dpm3 | 256.2929338 | -0.106714645 | 0.133132575 | -0.801566748 | 0.422803619 | 0.645029467 | | | Mir9-2 | 50.11727891 | 0.005709561 | 0.091483138 | 0.062411072 | 0.950235478 | 0.988202091 |
| Tex15 | 46.90644742 | -0.132946039 | 0.165853088 | -0.801589166 | 0.422790647 | 0.645029467 | | | Zkscan14 | 381.2526345 | 0.005716535 | 0.07487543 | 0.076347276 | 0.939142815 | 0.985461126 |
| Eif4enif1 | 2399.66444 | 0.080625066 | 0.100622826 | 0.801260207 | 0.422981023 | 0.645236582 | | | Creg1 | 1178.401181 | 0.005717961 | 0.067517314 | 0.08468881 | 0.932508792 | 0.984387568 |
| Cdc42ep2 | 426.3963463 | 0.095793329 | 0.119581952 | 0.801068454 | 0.423092017 | 0.645342362 | | | Atp11a | 3442.988939 | 0.00572105 | 0.060732185 | 0.094201289 | 0.924949261 | 0.983236501 |
| Il18 | 553.4087036 | -0.082596047 | 0.103159599 | -0.800662741 | 0.423326918 | 0.645637097 | | | Gpr37l1 | 8218.292541 | 0.00572122 | 0.057769522 | 0.09903527 | 0.921110266 | 0.982762672 |
| Mum1l1 | 133.1896097 | -0.104859296 | 0.130986338 | -0.800536125 | 0.423400242 | 0.645685369 | | | Prc1 | 43.57620233 | 0.005738881 | 0.089532996 | 0.064097943 | 0.94889224 | 0.987576748 |
| Hsdl2 | 1290.016573 | -0.073503449 | 0.091868666 | -0.800092699 | 0.423657091 | 0.64601348 | | | Sntb2 | 448.8835523 | 0.005743132 | 0.085022993 | 0.067547992 | 0.946145457 | 0.986835678 |
| Mzt2 | 459.3766003 | 0.1047469 | 0.076769907 | 1.364426561 | 0.172433353 | 0.380256529 | | | Tma7 | 2248.119689 | 0.005753703 | 0.06663261 | 0.086349652 | 0.93118847 | 0.984208871 |
| Tmem11 | 750.4501946 | -0.07513653 | 0.093973825 | -0.799547431 | 0.423973055 | 0.646368053 | | | Mroh5 | 91.62659939 | 0.005760071 | 0.09506686 | 0.060589682 | 0.951685991 | 0.988430624 |
| Apol10b | 17.58885436 | -0.13170982 | 0.164808377 | -0.799169448 | 0.424192165 | 0.646517445 | | | Naa25 | 1537.312407 | 0.005771626 | 0.073308017 | 0.078731168 | 0.937246454 | 0.985461126 |
| H2-DMb2 | 15.99011988 | -0.12976181 | 0.162372258 | -0.799162441 | 0.424196227 | 0.646517445 | | | Snord88a | 2.941289526 | 0.005778317 | 0.034668309 | 0.166674323 | 0.86762631 | NA |
| Ilkap | 1198.325067 | 0.053771961 | 0.067276972 | 0.799262494 | 0.424138222 | 0.646517445 | | | Cux1 | 3630.50548 | 0.005785422 | 0.055472613 | 0.104293301 | 0.916936594 | 0.982742917 |
| Eogt | 1142.148878 | -0.068373353 | 0.085572009 | -0.799015404 | 0.42428148 | 0.646583776 | | | Gm1720 | 0.611119886 | 0.005801565 | 0.017257341 | 0.336179547 | 0.736735475 | NA |
| Maged2 | 1599.491205 | -0.059231082 | 0.074184333 | -0.798431138 | 0.42462034 | 0.64703654 | | | Fam60a | 92.26203855 | 0.005810283 | 0.093353875 | 0.062239332 | 0.950372241 | 0.988202091 |
| Ccdc124 | 1960.003261 | -0.058966952 | 0.073871254 | -0.798239484 | 0.424731528 | 0.647078692 | | | Camkv | 22608.97461 | 0.005813321 | 0.062105695 | 0.093603675 | 0.92542399 | 0.983236501 |
| Fam135b | 1055.031826 | -0.083288967 | 0.104337544 | -0.798264592 | 0.424716961 | 0.647078692 | | | Cdk17 | 8819.62247 | 0.005818872 | 0.078640174 | 0.073993634 | 0.941015451 | 0.985461126 |
| Disc1 | 29.15871968 | 0.104730789 | 0.165628822 | 0.632322249 | 0.527176328 | 0.729452594 | | | Mir1938 | 0.648422349 | 0.00582903 | 0.018963286 | 0.307384999 | 0.758550345 | NA |
| Cdkn2c | 66.11176658 | -0.112761109 | 0.141360132 | -0.797686785 | 0.425052274 | 0.647312752 | | | Cdc42se1 | 2699.247703 | 0.005829529 | 0.047527156 | 0.122656798 | 0.902378876 | 0.979150581 |
| Msx2 | 9.676149083 | 0.10471414 | 0.156235091 | 0.67023445 | 0.502708346 | 0.709765183 | | | Gje1 | 0.87416202 | 0.005836135 | 0.023537049 | 0.247955253 | 0.804169029 | NA |
| Gm5141 | 103.6764198 | -0.124672156 | 0.156292105 | -0.797686843 | 0.425052241 | 0.647312752 | | | 1700019A02Rik | 1.655031696 | 0.005849449 | 0.031027712 | 0.188523383 | 0.850466387 | NA |
| Dtx2 | 265.1188358 | 0.094533246 | 0.118566889 | 0.797298865 | 0.425277479 | 0.647528426 | | | Synj2bp | 2114.156788 | 0.005858282 | 0.06927537 | 0.084565147 | 0.932607108 | 0.984387568 |
| Rpusd2 | 306.1194442 | -0.073517021 | 0.092206941 | -0.797304627 | 0.425274134 | 0.647528426 | | | Mir1192 | 0.670958985 | 0.005859154 | 0.019078877 | 0.307101606 | 0.758766035 | NA |
| Trappc9 | 3359.285744 | 0.063489554 | 0.079652655 | 0.797080193 | 0.425404459 | 0.64765812 | | | Ttl | 4101.575723 | 0.005870258 | 0.059889017 | 0.098018949 | 0.921917247 | 0.982822698 |
| 1110057K04Rik | 1560.198489 | -0.054022346 | 0.067810312 | -0.796668594 | 0.425643528 | 0.647831123 | | | Nbr1 | 8592.516243 | 0.005876576 | 0.048100829 | 0.122172026 | 0.90276278 | 0.979150581 |
| Sptlc2 | 943.6327482 | 0.051833211 | 0.065050784 | 0.796811474 | 0.42556053 | 0.647831123 | | | Gm20554 | 1.409755441 | 0.005881581 | 0.028970487 | 0.203019765 | 0.839119584 | NA |
| Zfp277 | 733.694385 | 0.06246178 | 0.07839822 | 0.796724463 | 0.425611073 | 0.647831123 | | | R3hdml | 0.394764232 | 0.005894938 | 0.013628608 | 0.432541452 | 0.665347932 | NA |
| Slc44a2 | 1917.026584 | 0.104708628 | 0.066807996 | 1.567306818 | 0.117042993 | 0.300292122 | | | 4933415F23Rik | 0.451584937 | 0.005896716 | 0.014119674 | 0.417624101 | 0.676221973 | NA |
| Tceal6 | 2052.673695 | 0.055291191 | 0.069422419 | 0.796445748 | 0.425772997 | 0.647900885 | | | Zbtb12 | 216.3888816 | 0.005900978 | 0.085527792 | 0.068994862 | 0.944993709 | 0.986700591 |
| Csnk2a1 | 5618.717578 | -0.048768771 | 0.061245492 | -0.796283452 | 0.425867302 | 0.647980749 | | | Cdkl2 | 1634.048465 | 0.005936877 | 0.069143499 | 0.085863127 | 0.931575224 | 0.984221522 |
| Gimap5 | 17.9763272 | -0.133553809 | 0.167742046 | -0.796185644 | 0.42592414 | 0.648003597 | | | Kng2 | 0.397381925 | 0.005949938 | 0.013653982 | 0.435765744 | 0.663006701 | NA |
| Nufip2 | 1794.484977 | 0.052566288 | 0.066034326 | 0.796044891 | 0.426005943 | 0.648064417 | | | Akr1e1 | 1041.958752 | 0.005951185 | 0.059477302 | 0.100058078 | 0.920298217 | 0.982742917 |
| Ipo8 | 2268.817707 | -0.061660104 | 0.077472856 | -0.795892999 | 0.426094231 | 0.648135089 | | | 1700011L22Rik | 0.732155478 | 0.00595239 | 0.01761161 | 0.337981037 | 0.735377478 | NA |
| Efcab10 | 38.59291579 | 0.104703964 | 0.15951797 | 0.656377232 | 0.511581436 | 0.715959495 | | | Ccr7 | 3.375673998 | 0.00598645 | 0.043167396 | 0.138679886 | 0.889703113 | NA |
| Cep63 | 933.4702578 | 0.079610361 | 0.100081295 | 0.795456946 | 0.426347746 | 0.648329765 | | | Dpm1 | 1407.069953 | 0.005992416 | 0.065417396 | 0.091602798 | 0.927013629 | 0.983501978 |
| Itga6 | 963.6195681 | -0.071565686 | 0.089962746 | -0.79550357 | 0.426320635 | 0.648329765 | | | Srp54b | 8.459657433 | 0.006004486 | 0.061787351 | 0.097179864 | 0.922583559 | NA |
| Kif17 | 1533.067433 | 0.064231363 | 0.080757917 | 0.795356859 | 0.426405948 | 0.648354637 | | | Gzma | 0.763547837 | 0.006009736 | 0.014197578 | 0.423293089 | 0.672081436 | NA |
| Cdc42se1 | 2556.89788 | -0.044148109 | 0.0555128 | -0.795278009 | 0.426451803 | 0.648360733 | | | Daxx | 1021.624849 | 0.006020841 | 0.062865578 | 0.095773252 | 0.923700662 | 0.983229764 |
| Dnaja1 | 10538.99716 | -0.048988992 | 0.061610565 | -0.795139479 | 0.426532372 | 0.648419601 | | | 5430403N17Rik | 1.438699779 | 0.006033482 | 0.026060551 | 0.231517809 | 0.816912552 | NA |
| Dennd4a | 3518.47007 | -0.054108514 | 0.068065839 | -0.794943761 | 0.426646217 | 0.648529038 | | | H3f3a | 4044.162289 | 0.006038027 | 0.05417953 | 0.111444795 | 0.91126364 | 0.980985114 |
| Adpgk | 554.7744291 | 0.055645117 | 0.070025884 | 0.794636411 | 0.426825032 | 0.648609949 | | | Tas1r2 | 0.489247677 | 0.00603939 | 0.012723247 | 0.474673616 | 0.635019626 | NA |
| Sh3d21 | 49.80100149 | 0.104703548 | 0.151238276 | 0.692308527 | 0.488743597 | 0.699675994 | | | Car2 | 7066.430313 | 0.006046662 | 0.076341143 | 0.079205803 | 0.936868929 | 0.985461126 |
| Ttc9c | 1293.742898 | 0.044050928 | 0.055430888 | 0.794700018 | 0.426788022 | 0.648609949 | | | Kctd4 | 758.0290171 | 0.006047437 | 0.077462679 | 0.078069041 | 0.937773133 | 0.985461126 |
| Ufd1l | 2795.565205 | 0.104695017 | 0.063627904 | 1.645426131 | 0.099881964 | 0.269846758 | | | Ngef | 17321.9276 | 0.006050192 | 0.058728196 | 0.10302023 | 0.917946915 | 0.982742917 |
| Tmem47 | 2920.286654 | -0.086118744 | 0.108472167 | -0.793924804 | 0.427239209 | 0.649112013 | | | Tmem54 | 1.897081997 | 0.006051059 | 0.029889437 | 0.202448064 | 0.839566457 | NA |
| Fam179b | 2538.359323 | -0.060327629 | 0.076003387 | -0.79374921 | 0.427341446 | 0.649140036 | | | Mir1896 | 0.856955244 | 0.006059301 | 0.022250806 | 0.272318259 | 0.785377318 | NA |
| Rpp14 | 1191.101055 | -0.052073907 | 0.065600625 | -0.793801997 | 0.42731071 | 0.649140036 | | | Zp3 | 0.892355537 | 0.006073508 | 0.02236226 | 0.27159635 | 0.785932406 | NA |
| Dirc2 | 2869.404216 | -0.041533648 | 0.052353233 | -0.793334933 | 0.427582709 | 0.649203686 | | | Tnfsf15 | 1.180454581 | 0.006077134 | 0.022048579 | 0.275624738 | 0.782836307 | NA |
| Kif2a | 3069.963507 | -0.060785011 | 0.076618908 | -0.793342173 | 0.427578492 | 0.649203686 | | | Ppil2 | 2620.639036 | 0.006086753 | 0.043733192 | 0.139179256 | 0.8893085 | 0.977182599 |
| Phka1 | 1142.013112 | 0.069813433 | 0.08800188 | 0.793317523 | 0.42759285 | 0.649203686 | | | Npm2 | 11.20811721 | 0.006090745 | 0.068124994 | 0.089405442 | 0.928759699 | NA |
| Prkrip1 | 521.9163775 | 0.056387579 | 0.071067721 | 0.79343447 | 0.427524734 | 0.649203686 | | | Cbfa2t2 | 2806.846059 | 0.006096645 | 0.060206457 | 0.10126232 | 0.919342227 | 0.982742917 |
| Vps9d1 | 2161.700972 | 0.062708275 | 0.079034942 | 0.793424688 | 0.427530432 | 0.649203686 | | | AB099516 | 0.427411356 | 0.006098352 | 0.013923256 | 0.437997556 | 0.661388055 | NA |
| Nit1 | 1045.479204 | 0.071456721 | 0.090111623 | 0.792980076 | 0.427789431 | 0.649438511 | | | Mir124a-3 | 0.427411356 | 0.006098352 | 0.013923256 | 0.437997556 | 0.661388055 | NA |
| Csrp1 | 9394.908719 | 0.104675533 | 0.087032138 | 1.202722745 | 0.229083629 | 0.452040467 | | | Olfr574 | 0.427411356 | 0.006098352 | 0.013923256 | 0.437997556 | 0.661388055 | NA |
| Dpm2 | 1251.352067 | 0.056625348 | 0.071448467 | 0.792534114 | 0.428049309 | 0.649685981 | | | Sftpb | 0.427411356 | 0.006098352 | 0.013923256 | 0.437997556 | 0.661388055 | NA |
| Mrps18a | 1242.767098 | -0.049163002 | 0.062041407 | -0.79242242 | 0.428114411 | 0.649685981 | | | Unc79 | 3242.497643 | 0.006101568 | 0.068775877 | 0.088716682 | 0.929307075 | 0.983689248 |
| Ube2e2 | 2183.232836 | 0.06496676 | 0.081986032 | 0.792412545 | 0.428120167 | 0.649685981 | | | Krt33a | 0.715278666 | 0.006101765 | 0.017545608 | 0.347765964 | 0.728015953 | NA |
| Slc23a2 | 7989.154743 | 0.104660058 | 0.056781008 | 1.843222983 | 0.065296461 | 0.205426618 | | | Setdb2 | 218.4210756 | 0.006114409 | 0.089045123 | 0.068666409 | 0.945255157 | 0.986781137 |
| Nppc | 83.93647574 | -0.10949529 | 0.138286584 | -0.791799802 | 0.428477418 | 0.650037121 | | | Arhgap28 | 26.82185867 | 0.006133232 | 0.0854277 | 0.071794421 | 0.942765513 | 0.985461126 |
| Pfkfb1 | 25.14722184 | 0.104574802 | 0.167918419 | 0.622771479 | 0.533434703 | 0.734343607 | | | 9430018G01Rik | 1.620084179 | 0.006150537 | 0.025393649 | 0.242207692 | 0.808619235 | NA |
| Farp1 | 3038.489655 | 0.058853598 | 0.074361428 | 0.791453299 | 0.428679518 | 0.650280052 | | | Zfp345 | 0.43002905 | 0.006151786 | 0.01394651 | 0.441098571 | 0.659141638 | NA |
| Zfyve1 | 2075.49585 | 0.076057328 | 0.096110057 | 0.791356606 | 0.428735925 | 0.65030195 | | | Gpnmb | 230.4196012 | 0.006155362 | 0.071482956 | 0.086109499 | 0.931379372 | 0.984208871 |
| Pja1 | 7300.473163 | -0.06706629 | 0.084777432 | -0.791086588 | 0.428893465 | 0.650413561 | | | Dnahc1 | 581.1062147 | 0.006158006 | 0.094565421 | 0.065119009 | 0.948079246 | 0.987367302 |
| Zc2hc1a | 1988.105021 | -0.059568786 | 0.075298831 | -0.791098423 | 0.428886559 | 0.650413561 | | | Sec14l4 | 13.83826438 | 0.006159514 | 0.067301508 | 0.09152119 | 0.92707847 | NA |
| 4930506C21Rik | 46.48612643 | -0.131335392 | 0.166116048 | -0.790624347 | 0.429163235 | 0.650758971 | | | Nrtn | 15.67546915 | 0.006161585 | 0.069009598 | 0.089285918 | 0.928854685 | NA |
| Slc17a7 | 51375.26665 | -0.07474471 | 0.094620719 | -0.789940199 | 0.429562693 | 0.651267729 | | | Mpc2 | 1677.973958 | 0.006162668 | 0.067601628 | 0.091161538 | 0.927364236 | 0.983501978 |
| Smcr7l | 1167.255954 | -0.039675623 | 0.050228301 | -0.78990574 | 0.429582819 | 0.651267729 | | | Tmem87a | 1307.516738 | 0.006164399 | 0.064101608 | 0.096166065 | 0.923388682 | 0.98311054 |
| 2700029M09Rik | 1019.983567 | -0.070194314 | 0.088920537 | -0.789404969 | 0.429875353 | 0.651519976 | | | Treh | 1.629630927 | 0.006165321 | 0.027326436 | 0.225617458 | 0.821498962 | NA |
| Ext2 | 1448.120252 | 0.104562501 | 0.080242763 | 1.303077037 | 0.192548462 | 0.406446041 | | | Mir3104 | 1.491974438 | 0.006170832 | 0.027516495 | 0.224259391 | 0.822555472 | NA |
| Paxip1 | 1873.197795 | 0.045914075 | 0.058156277 | 0.789494747 | 0.429822899 | 0.651519976 | | | Gna14 | 12.45963215 | 0.00617551 | 0.067068473 | 0.092077688 | 0.926636315 | NA |
| Cysltr1 | 21.95226128 | -0.130321431 | 0.165119077 | -0.789257263 | 0.42996166 | 0.651548009 | | | Tsc1 | 4187.291018 | 0.006178602 | 0.057415292 | 0.10761248 | 0.914303097 | 0.98170867 |
| Gsta4 | 1193.945956 | -0.058510922 | 0.074143536 | -0.789157422 | 0.430020004 | 0.651548009 | | | Tmem39b | 314.4369258 | 0.006184631 | 0.076878291 | 0.080447046 | 0.935881711 | 0.985461126 |
| Tbc1d10b | 3139.957104 | 0.043708827 | 0.055382308 | 0.789220024 | 0.42998342 | 0.651548009 | | | Chd9 | 2653.391874 | 0.006187895 | 0.073346201 | 0.084365577 | 0.932765775 | 0.984387568 |
| Yme1l1 | 2602.31824 | -0.043091299 | 0.05462341 | -0.788879695 | 0.430182324 | 0.651730217 | | | Slc25a31 | 2.145967408 | 0.006190344 | 0.025527208 | 0.242499862 | 0.808392864 | NA |
| Rpl29 | 5053.820781 | -0.056816545 | 0.072055588 | -0.788509901 | 0.43039851 | 0.651993988 | | | Gulp1 | 87.09385818 | 0.006199346 | 0.094709564 | 0.065456384 | 0.947810633 | 0.987367302 |
| Micall1 | 2490.690102 | 0.054575173 | 0.069279211 | 0.78775685 | 0.430838945 | 0.652275301 | | | Gabrp | 0.432488341 | 0.006200366 | 0.013968126 | 0.443893945 | 0.657119263 | NA |
| Nmb | 100.1704317 | -0.132460987 | 0.168089738 | -0.78803732 | 0.430674877 | 0.652275301 | | | Mir684-1 | 0.432488341 | 0.006200366 | 0.013968126 | 0.443893945 | 0.657119263 | NA |
| Prim1 | 205.4980727 | 0.091218618 | 0.115805437 | 0.787688555 | 0.430878902 | 0.652275301 | | | Ppp1r16a | 3154.976095 | 0.006210162 | 0.069696605 | 0.089102792 | 0.929000218 | 0.983640461 |
| Prkg1 | 510.5347522 | -0.089047253 | 0.113010037 | -0.787958796 | 0.430720808 | 0.652275301 | | | Rasl10b | 12932.17956 | 0.006226667 | 0.055867901 | 0.111453386 | 0.911256828 | 0.980985114 |
| Slc39a11 | 599.8407931 | -0.067915647 | 0.08618405 | -0.788030344 | 0.430678957 | 0.652275301 | | | Cldn4 | 0.410423267 | 0.006227516 | 0.013778152 | 0.451984871 | 0.651279881 | NA |
| Spsb2 | 81.55843434 | -0.116622515 | 0.148032108 | -0.787819053 | 0.430802555 | 0.652275301 | | | Spink2 | 0.410423267 | 0.006227516 | 0.013778152 | 0.451984871 | 0.651279881 | NA |
| Tigd5 | 212.1538662 | -0.081588426 | 0.103568937 | -0.787769273 | 0.430831677 | 0.652275301 | | | Syce1 | 7.089492757 | 0.006236594 | 0.05294523 | 0.117793313 | 0.906231428 | NA |
| Cep19 | 2233.142856 | -0.049470528 | 0.06286045 | -0.786989717 | 0.431287886 | 0.652830647 | | | Gng12 | 2639.408872 | 0.006247656 | 0.051166505 | 0.12210442 | 0.902816321 | 0.979150581 |
| Bhlhe22 | 906.2403092 | 0.087320296 | 0.110980089 | 0.786810473 | 0.431392822 | 0.652925699 | | | Pdgfrl | 29.14374952 | 0.006253245 | 0.087138635 | 0.071762026 | 0.942791294 | 0.985461126 |
| Ptpra | 6865.32804 | -0.03645477 | 0.04634141 | -0.786656473 | 0.431482991 | 0.652998385 | | | Car5b | 179.658932 | 0.006263756 | 0.085964293 | 0.072864627 | 0.941913842 | 0.985461126 |
| Helz2 | 301.8274867 | 0.075529993 | 0.09603831 | 0.786456918 | 0.431599849 | 0.653111443 | | | Golga7b | 4796.407064 | 0.00626782 | 0.058540566 | 0.107067975 | 0.914735053 | 0.98170867 |
| 2810408A11Rik | 11.68559419 | 0.10448961 | 0.164794356 | 0.63406061 | 0.526041268 | 0.728341121 | | | Uts2 | 0.563966486 | 0.006272637 | 0.013195796 | 0.475351175 | 0.634536689 | NA |
| Pcdhb6 | 103.2351802 | -0.11320932 | 0.144006785 | -0.786138796 | 0.431786178 | 0.653295873 | | | Spata31d1b | 2.465217617 | 0.006296083 | 0.025642611 | 0.245532054 | 0.806044492 | NA |
| Slc6a6 | 7755.263573 | 0.055035665 | 0.070017001 | 0.786032887 | 0.43184822 | 0.653295873 | | | Sptlc1 | 1249.912726 | 0.006313532 | 0.055221822 | 0.114330381 | 0.9089759 | 0.980523965 |
| Pum2 | 8430.12264 | -0.060077191 | 0.076452931 | -0.785806246 | 0.431981007 | 0.653320983 | | | Med10 | 681.8334371 | 0.006327614 | 0.066355398 | 0.095359444 | 0.924029328 | 0.983236501 |
| Spin2 | 174.1517619 | 0.099657057 | 0.126824252 | 0.78578865 | 0.431991317 | 0.653320983 | | | Fmr1nb | 0.880555709 | 0.006335746 | 0.022329044 | 0.283744598 | 0.776606115 | NA |
| Tram2 | 19.64504898 | -0.131018596 | 0.166708862 | -0.785912605 | 0.431918689 | 0.653320983 | | | 2810405F15Rik | 2.827417771 | 0.006336786 | 0.036380315 | 0.174181733 | 0.861722641 | NA |
| Zmym1 | 444.954953 | 0.068228287 | 0.086849302 | 0.785593951 | 0.432105409 | 0.653429749 | | | 4930471M09Rik | 0.614955356 | 0.006348657 | 0.018788327 | 0.337904331 | 0.735435284 | NA |
| AI836003 | 893.8624824 | -0.080851139 | 0.102933748 | -0.785467749 | 0.432179372 | 0.653477818 | | | Pax8 | 1.20695477 | 0.006354024 | 0.024421069 | 0.260186149 | 0.794720188 | NA |
| 3110057O12Rik | 290.8721778 | 0.104434009 | 0.108770248 | 0.960133955 | 0.336987801 | 0.570594737 | | | Foxo3 | 1690.966981 | 0.006357208 | 0.051616979 | 0.123161186 | 0.901979461 | 0.979150581 |
| Susd3 | 48.65137964 | -0.129526879 | 0.165001749 | -0.785003067 | 0.432451771 | 0.653762097 | | | Wbscr22 | 885.0188712 | 0.006364866 | 0.070958968 | 0.089697841 | 0.928527332 | 0.983596265 |
| Epc1 | 1619.454179 | 0.06630153 | 0.084487951 | 0.784745386 | 0.432602867 | 0.653926714 | | | Mgrn1 | 10738.52821 | 0.006367107 | 0.037176283 | 0.171267983 | 0.864013054 | 0.969836459 |
| Ccrl2 | 34.80790341 | -0.126734792 | 0.161526355 | -0.784607516 | 0.432683722 | 0.653973357 | | | Gm10768 | 1.528836905 | 0.006379446 | 0.025455863 | 0.250608107 | 0.802117113 | NA |
| D10Wsu52e | 4349.760596 | 0.055816602 | 0.071144841 | 0.784548834 | 0.43271814 | 0.653973357 | | | Snord42b | 21.8074519 | 0.006387179 | 0.079365539 | 0.080477988 | 0.935857102 | NA |
| Atpaf2 | 724.2690409 | 0.104382188 | 0.061308887 | 1.702562109 | 0.088650046 | 0.249962478 | | | Hapln1 | 668.0623773 | 0.006398872 | 0.086857879 | 0.073670601 | 0.941272492 | 0.985461126 |
| Wasf2 | 897.983756 | 0.058546246 | 0.074641092 | 0.784370167 | 0.432822938 | 0.654004155 | | | AI987944 | 316.6186107 | 0.006405623 | 0.076635839 | 0.08358522 | 0.933386218 | 0.984481307 |
| Mrpl55 | 633.8563917 | -0.065369722 | 0.083363764 | -0.784150322 | 0.432951911 | 0.654110921 | | | Mcu | 1151.10804 | 0.006414119 | 0.063742909 | 0.100624824 | 0.91984829 | 0.982742917 |
| Sox9 | 828.6672561 | -0.081481609 | 0.1039166 | -0.784105803 | 0.432978031 | 0.654110921 | | | Me1 | 2524.02341 | 0.006415352 | 0.049719551 | 0.129030771 | 0.8973333 | 0.978039389 |
| Vstm5 | 495.9183251 | 0.057746874 | 0.073658684 | 0.783979175 | 0.43305233 | 0.654159384 | | | Kcnk12 | 222.3240153 | 0.006424287 | 0.086959673 | 0.073876621 | 0.941108559 | 0.985461126 |
| Ramp3 | 393.2424967 | -0.114803431 | 0.146459159 | -0.783859692 | 0.433122444 | 0.654201515 | | | Gpt | 563.9166865 | 0.006424698 | 0.08894633 | 0.072231182 | 0.942417931 | 0.985461126 |
| Ube2q1 | 3872.088485 | 0.057671907 | 0.073599467 | 0.783591371 | 0.433279921 | 0.654375581 | | | Exoc6 | 1394.072023 | 0.0064285 | 0.070994169 | 0.090549685 | 0.927850413 | 0.983501978 |
| Hnrnpab | 3930.071135 | -0.052326905 | 0.066812671 | -0.783188335 | 0.433516524 | 0.654541515 | | | Herpud2 | 1698.247301 | 0.006438304 | 0.059169911 | 0.108810445 | 0.913352839 | 0.98170867 |
| Bcl6b | 98.51193222 | 0.10437519 | 0.166538093 | 0.626734628 | 0.530833201 | 0.732282217 | | | Ildr1 | 23.81969779 | 0.00644169 | 0.082146646 | 0.078416959 | 0.937496384 | 0.985461126 |
| Scnm1 | 736.2852655 | 0.058016064 | 0.074072753 | 0.783230838 | 0.433491569 | 0.654541515 | | | Gm12359 | 13.00483454 | 0.006441871 | 0.065791086 | 0.097914039 | 0.922000552 | NA |
| Ubac2 | 1229.491544 | -0.058301729 | 0.07445232 | -0.783074715 | 0.433583238 | 0.654578456 | | | Speer2 | 0.67765611 | 0.006443162 | 0.019112619 | 0.337115603 | 0.736029756 | NA |
| Klhl20 | 839.4055186 | 0.104363365 | 0.064000947 | 1.630653443 | 0.102963466 | 0.275815342 | | | Wdr4 | 906.7833943 | 0.006446595 | 0.065033145 | 0.099127831 | 0.921036775 | 0.982762672 |
| St6gal1 | 1107.905847 | -0.042227757 | 0.053973572 | -0.782378403 | 0.433992222 | 0.655068241 | | | Dse | 253.579465 | 0.006454117 | 0.088045182 | 0.073304605 | 0.941563727 | 0.985461126 |
| Srpr | 2922.297067 | -0.035239211 | 0.045053436 | -0.782164784 | 0.434117738 | 0.655094028 | | | 2310043L19Rik | 0.875876633 | 0.006455705 | 0.022313054 | 0.289324131 | 0.772333348 | NA |
| U2af1 | 1358.081605 | 0.057173761 | 0.073094946 | 0.782184872 | 0.434105934 | 0.655094028 | | | Wdr72 | 12.7707595 | 0.006463565 | 0.025519205 | 0.253282391 | 0.800050003 | NA |
| Zfp764 | 175.0975746 | 0.087250291 | 0.111554224 | 0.782133453 | 0.434136148 | 0.655094028 | | | Cd300ld | 3.823994927 | 0.006492052 | 0.043486795 | 0.149287884 | 0.881326475 | NA |
| Spdya | 34.34806141 | -0.126264595 | 0.161565335 | -0.781507959 | 0.434503797 | 0.655584948 | | | Arhgap12 | 1678.59176 | 0.006498855 | 0.072678645 | 0.089419044 | 0.928748889 | 0.983596265 |
| Dnahc2 | 149.726623 | -0.110772701 | 0.141762964 | -0.781393803 | 0.434570915 | 0.655622371 | | | Rhbg | 0.645700311 | 0.006500517 | 0.01745038 | 0.372514354 | 0.709509927 | NA |
| Tmem189 | 1457.05935 | 0.10433157 | 0.079338654 | 1.315015626 | 0.188504651 | 0.401684172 | | | Osgin2 | 665.5196933 | 0.006508385 | 0.078691513 | 0.082707584 | 0.934084054 | 0.984937933 |
| Ano10 | 890.6945432 | 0.066152424 | 0.084763946 | 0.780431148 | 0.43513714 | 0.656157163 | | | 1810013L24Rik | 1841.550695 | 0.006536595 | 0.064799957 | 0.100873443 | 0.919650925 | 0.982742917 |
| Arhgdib | 394.2675028 | -0.079195653 | 0.101473033 | -0.780460101 | 0.435120104 | 0.656157163 | | | Oas1d | 0.658320007 | 0.006548537 | 0.019019851 | 0.344300113 | 0.730620599 | NA |
| BC003331 | 1954.609424 | 0.047425587 | 0.060753944 | 0.780617429 | 0.435027537 | 0.656157163 | | | Gabbr2 | 11965.88807 | 0.006572715 | 0.054502604 | 0.120594518 | 0.904012211 | 0.979829979 |
| Srms | 14.25514479 | -0.130523576 | 0.167231492 | -0.780496391 | 0.435098751 | 0.656157163 | | | Huwe1 | 15499.57196 | 0.006572973 | 0.072594329 | 0.090543888 | 0.92785502 | 0.983501978 |
| Slc25a4 | 18279.35085 | -0.047108355 | 0.060374689 | -0.780266618 | 0.435233957 | 0.656239289 | | | Cit | 4152.018881 | 0.006578912 | 0.046167787 | 0.142500044 | 0.886685045 | 0.976496358 |
| Cdip1 | 10104.43992 | 0.058256377 | 0.074682953 | 0.780049185 | 0.435361925 | 0.65630935 | | | Ly6g6e | 45.40872896 | 0.006580107 | 0.087230784 | 0.075433312 | 0.939869956 | 0.985461126 |
| Tufm | 2094.984566 | -0.05967158 | 0.076497738 | -0.780043721 | 0.435365141 | 0.65630935 | | | Sult6b1 | 1.667044668 | 0.00658011 | 0.03058002 | 0.215176763 | 0.829629507 | NA |
| D16Ertd472e | 400.9617831 | 0.086368931 | 0.110747438 | 0.779872953 | 0.43546566 | 0.656397017 | | | Optc | 36.41011206 | 0.0065849 | 0.082056652 | 0.080248216 | 0.936039843 | 0.985461126 |
| Klhl41 | 16.45577994 | 0.104322408 | 0.167182891 | 0.624001695 | 0.532626472 | 0.733426462 | | | Gm12409 | 0.428541593 | 0.006603575 | 0.013943279 | 0.473602695 | 0.635783253 | NA |
| Ccdc110 | 30.61906568 | -0.130671212 | 0.167590718 | -0.779704348 | 0.435564919 | 0.656418914 | | | Selp | 0.428541593 | 0.006603575 | 0.013943279 | 0.473602695 | 0.635783253 | NA |
| Rap1gds1 | 16174.92772 | -0.053171994 | 0.068202448 | -0.779620016 | 0.435614571 | 0.656429893 | | | Fgfr1op2 | 3752.481763 | 0.006606606 | 0.0696291 | 0.094882831 | 0.924407894 | 0.983236501 |
| 6330408A02Rik | 572.5937471 | 0.053489501 | 0.068656029 | 0.779094012 | 0.435924337 | 0.656577397 | | | Tfdp1 | 1738.840059 | 0.00662393 | 0.056206929 | 0.117848999 | 0.906187305 | 0.980118791 |
| Ankrd66 | 12.44894353 | -0.129830402 | 0.166626417 | -0.779170581 | 0.435879238 | 0.656577397 | | | Gm10421 | 34.4608198 | 0.006643539 | 0.091624051 | 0.072508681 | 0.942197097 | 0.985461126 |
| Frmpd3 | 152.5595989 | -0.085306761 | 0.109486309 | -0.779154596 | 0.435888653 | 0.656577397 | | | Npvf | 5.076898678 | 0.006652883 | 0.011661433 | 0.57050308 | NA | NA |
| Nhlrc1 | 621.0874187 | 0.068127157 | 0.087420688 | 0.779302461 | 0.435801565 | 0.656577397 | | | Synb | 4.064533027 | 0.006654302 | 0.041456955 | 0.160511122 | 0.872478461 | NA |
| Zfp707 | 321.7717877 | 0.070815065 | 0.090890419 | 0.779125739 | 0.435905649 | 0.656577397 | | | 4930429D17Rik | 0.191449919 | 0.006655081 | 0.009624056 | 0.691504752 | 0.489248395 | NA |
| A730017C20Rik | 1430.689383 | -0.065004238 | 0.083457209 | -0.778893032 | 0.436042729 | 0.656626826 | | | 4930520P13Rik | 0.191449919 | 0.006655081 | 0.009624056 | 0.691504752 | 0.489248395 | NA |
| Adcy1 | 36973.19334 | -0.087305181 | 0.112109302 | -0.77875055 | 0.436126673 | 0.656626826 | | | 9830107B12Rik | 0.191449919 | 0.006655081 | 0.009624056 | 0.691504752 | 0.489248395 | NA |
| Tmem53 | 74.84516962 | -0.11475881 | 0.147327917 | -0.778934586 | 0.436018249 | 0.656626826 | | | Ces1e | 0.191449919 | 0.006655081 | 0.009624056 | 0.691504752 | 0.489248395 | NA |
| Tom1l1 | 695.8877736 | 0.057251463 | 0.073513892 | 0.778784276 | 0.436106802 | 0.656626826 | | | Cldn7 | 0.191449919 | 0.006655081 | 0.009624056 | 0.691504752 | 0.489248395 | NA |
| Nsd1 | 6955.261315 | -0.071992636 | 0.092491807 | -0.778367703 | 0.436352273 | 0.656902654 | | | Eif4e1b | 0.191449919 | 0.006655081 | 0.009624056 | 0.691504752 | 0.489248395 | NA |
| Mdk | 350.3763455 | -0.083545203 | 0.107350508 | -0.778246927 | 0.436423457 | 0.656945986 | | | Gdf2 | 0.191449919 | 0.006655081 | 0.009624056 | 0.691504752 | 0.489248395 | NA |
| B3galt4 | 144.5814249 | -0.081656723 | 0.104940662 | -0.778122815 | 0.436496614 | 0.65699228 | | | Gltpd2 | 0.191449919 | 0.006655081 | 0.009624056 | 0.691504752 | 0.489248395 | NA |
| Tmem201 | 3131.757534 | 0.067500583 | 0.086763551 | 0.777983173 | 0.436578933 | 0.657052355 | | | Gm20758 | 0.191449919 | 0.006655081 | 0.009624056 | 0.691504752 | 0.489248395 | NA |
| Cndp1 | 13.34192625 | 0.104297666 | 0.157441752 | 0.6624524 | 0.507681331 | 0.713276618 | | | Gm5494 | 0.191449919 | 0.006655081 | 0.009624056 | 0.691504752 | 0.489248395 | NA |
| Gm5148 | 76.73333438 | 0.104289789 | 0.141582277 | 0.736602008 | 0.461364413 | 0.677935394 | | | Gm5938 | 0.191449919 | 0.006655081 | 0.009624056 | 0.691504752 | 0.489248395 | NA |
| Zbtb14 | 1126.30684 | -0.056167151 | 0.072264611 | -0.777242841 | 0.437015511 | 0.657517803 | | | Gphb5 | 0.191449919 | 0.006655081 | 0.009624056 | 0.691504752 | 0.489248395 | NA |
| Iqca | 35.78274144 | 0.104248378 | 0.165635489 | 0.629384306 | 0.52909749 | 0.730714258 | | | Krtap13-1 | 0.191449919 | 0.006655081 | 0.009624056 | 0.691504752 | 0.489248395 | NA |
| Hnrnpul2 | 10937.83103 | -0.043864272 | 0.056500588 | -0.776350719 | 0.437541934 | 0.658182012 | | | Mir376c | 0.191449919 | 0.006655081 | 0.009624056 | 0.691504752 | 0.489248395 | NA |
| Aco2 | 20870.41951 | -0.034722332 | 0.044752525 | -0.775874251 | 0.437823237 | 0.658477309 | | | Mirlet7g | 0.191449919 | 0.006655081 | 0.009624056 | 0.691504752 | 0.489248395 | NA |
| Serping1 | 332.0367562 | -0.116937051 | 0.150708535 | -0.775915248 | 0.437799029 | 0.658477309 | | | Nppb | 0.191449919 | 0.006655081 | 0.009624056 | 0.691504752 | 0.489248395 | NA |
| Dos | 10310.52196 | -0.049510041 | 0.063841568 | -0.775514179 | 0.43803589 | 0.658642811 | | | Psg22 | 0.191449919 | 0.006655081 | 0.009624056 | 0.691504752 | 0.489248395 | NA |
| Fkbp3 | 3518.309412 | -0.052677295 | 0.067932272 | -0.775438451 | 0.438080622 | 0.658642811 | | | Rnf148 | 0.191449919 | 0.006655081 | 0.009624056 | 0.691504752 | 0.489248395 | NA |
| Paqr3 | 335.4104287 | -0.064266667 | 0.08286089 | -0.775597113 | 0.437986905 | 0.658642811 | | | Scarna9 | 0.191449919 | 0.006655081 | 0.009624056 | 0.691504752 | 0.489248395 | NA |
| Sez6l | 9236.349388 | 0.071095884 | 0.091689298 | 0.775400029 | 0.438103318 | 0.658642811 | | | Scgb1b2 | 0.191449919 | 0.006655081 | 0.009624056 | 0.691504752 | 0.489248395 | NA |
| AI846148 | 793.8136019 | -0.049554184 | 0.063955196 | -0.774826559 | 0.438442153 | 0.658832579 | | | Skint4 | 0.191449919 | 0.006655081 | 0.009624056 | 0.691504752 | 0.489248395 | NA |
| Cacnb4 | 8155.658781 | 0.073065968 | 0.094286952 | 0.774931907 | 0.438379897 | 0.658832579 | | | Tmprss11bnl | 0.191449919 | 0.006655081 | 0.009624056 | 0.691504752 | 0.489248395 | NA |
| Cyp2s1 | 225.8167933 | -0.095843307 | 0.123695624 | -0.774831829 | 0.438439039 | 0.658832579 | | | Vmn1r15 | 0.191449919 | 0.006655081 | 0.009624056 | 0.691504752 | 0.489248395 | NA |
| Cyp7b1 | 381.1451626 | -0.060815763 | 0.078479716 | -0.774923323 | 0.438384969 | 0.658832579 | | | Vmn1r31 | 0.191449919 | 0.006655081 | 0.009624056 | 0.691504752 | 0.489248395 | NA |
| Evi2a | 597.570977 | 0.104166726 | 0.131952949 | 0.789423249 | 0.429864672 | 0.651519976 | | | Vmn2r15 | 0.191449919 | 0.006655081 | 0.009624056 | 0.691504752 | 0.489248395 | NA |
| Ppp3ca | 31805.9687 | 0.104160973 | 0.07895287 | 1.319280381 | 0.187075394 | 0.39941383 | | | Chrna1 | 25.81485492 | 0.00666283 | 0.084653986 | 0.078706629 | 0.937265972 | 0.985461126 |
| Zfp438 | 223.6393599 | 0.074643294 | 0.09637907 | 0.774476184 | 0.438649247 | 0.659015944 | | | Arfip1 | 696.2051525 | 0.006663469 | 0.071261276 | 0.093507567 | 0.925500338 | 0.983236501 |
| Brcc3 | 925.9086605 | -0.050050252 | 0.064643106 | -0.774255063 | 0.438779972 | 0.659148428 | | | Nedd4l | 7436.105284 | 0.006670361 | 0.073075192 | 0.091280787 | 0.927269484 | 0.983501978 |
| Fiz1 | 802.9746057 | 0.04558231 | 0.058918852 | 0.773645595 | 0.439140401 | 0.659561979 | | | Fkbp6 | 2.363767144 | 0.006692871 | 0.028916438 | 0.231455589 | 0.816960883 | NA |
| Zfp958 | 246.6996955 | -0.088303562 | 0.11413254 | -0.773693126 | 0.439112286 | 0.659561979 | | | AI463170 | 1.011354418 | 0.006695231 | 0.021094363 | 0.317394335 | 0.750944406 | NA |
| Abca13 | 8.498165381 | 0.10410297 | 0.139622664 | 0.745602232 | 0.455907734 | 0.674115317 | | | Myb | 32.44200348 | 0.006703093 | 0.086548166 | 0.077449278 | 0.93826614 | 0.985461126 |
| Ndrg2 | 28502.24942 | 0.051030819 | 0.065995233 | 0.773250082 | 0.439374392 | 0.659781311 | | | Trim60 | 0.433618577 | 0.006705181 | 0.013987982 | 0.479352965 | 0.631687548 | NA |
| Zfp715 | 1083.492043 | 0.069127197 | 0.089406016 | 0.773182836 | 0.439414183 | 0.659781311 | | | Haus2 | 866.1103992 | 0.006708299 | 0.06545236 | 0.102491322 | 0.918366701 | 0.982742917 |
| Pik3r2 | 5246.572417 | 0.060691791 | 0.078522438 | 0.772922913 | 0.439568003 | 0.659948318 | | | Mtss1l | 8681.038097 | 0.006728412 | 0.059464938 | 0.113149232 | 0.909912244 | 0.980590355 |
| Exoc6b | 4653.121809 | 0.058608959 | 0.075904386 | 0.772141922 | 0.440030375 | 0.660415671 | | | 4933405O20Rik | 0.201873568 | 0.006733818 | 0.009680116 | 0.695634043 | 0.486658041 | NA |
| Jhdm1d | 2455.537424 | -0.065413001 | 0.084698574 | -0.772303443 | 0.439934727 | 0.660415671 | | | Clec4f | 0.201873568 | 0.006733818 | 0.009680116 | 0.695634043 | 0.486658041 | NA |
| Ptp4a1 | 371.9907172 | -0.072159339 | 0.093457433 | -0.772109145 | 0.440049786 | 0.660415671 | | | Cyp2c65 | 0.201873568 | 0.006733818 | 0.009680116 | 0.695634043 | 0.486658041 | NA |
| Rps4x | 8955.883965 | -0.03460931 | 0.044819992 | -0.772184651 | 0.440005071 | 0.660415671 | | | Hist2h2bb | 0.201873568 | 0.006733818 | 0.009680116 | 0.695634043 | 0.486658041 | NA |
| 9430091E24Rik | 46.92200696 | -0.123507922 | 0.15999401 | -0.771953413 | 0.440142021 | 0.660426154 | | | Olfr1029 | 0.201873568 | 0.006733818 | 0.009680116 | 0.695634043 | 0.486658041 | NA |
| Ankrd12 | 2918.008475 | 0.065542548 | 0.084901195 | 0.771986161 | 0.440122624 | 0.660426154 | | | Pou5f1 | 0.201873568 | 0.006733818 | 0.009680116 | 0.695634043 | 0.486658041 | NA |
| Rfc3 | 185.327006 | 0.104096933 | 0.111499729 | 0.933607048 | 0.350506628 | 0.581450717 | | | Serpinb9c | 0.201873568 | 0.006733818 | 0.009680116 | 0.695634043 | 0.486658041 | NA |
| Gpr137 | 2816.511147 | 0.054472775 | 0.070602939 | 0.771536927 | 0.440388743 | 0.660668396 | | | Slc25a23 | 20296.77918 | 0.0067353 | 0.049117964 | 0.137124993 | 0.890931998 | 0.977182599 |
| Iqgap1 | 1226.191355 | 0.055852667 | 0.072409819 | 0.771341063 | 0.440504799 | 0.660778522 | | | Gm7092 | 1.453518001 | 0.00673688 | 0.029816456 | 0.225945015 | 0.821244186 | NA |
| Katnal2 | 124.0388856 | 0.090947834 | 0.117945272 | 0.771101988 | 0.440646482 | 0.660927067 | | | Il12rb2 | 10.15642188 | 0.006749883 | 0.051288011 | 0.131607426 | 0.895294812 | NA |
| Cachd1 | 654.7657027 | -0.092980389 | 0.120648432 | -0.77067217 | 0.440901272 | 0.661093969 | | | 1110059M19Rik | 40.01463741 | 0.00675245 | 0.030425847 | 0.221931384 | 0.824367299 | 0.957903916 |
| Slc35f3 | 1609.412935 | 0.104049156 | 0.078343078 | 1.328121875 | 0.184137848 | 0.395156762 | | | 4930511M11Rik | 0.204491261 | 0.006752517 | 0.00969339 | 0.696610518 | 0.486046572 | NA |
| Ears2 | 194.2392617 | -0.09875068 | 0.128148274 | -0.770597033 | 0.44094582 | 0.661093969 | | | Akp3 | 0.204491261 | 0.006752517 | 0.00969339 | 0.696610518 | 0.486046572 | NA |
| Tcea3 | 14.20852046 | -0.129183733 | 0.167650373 | -0.770554403 | 0.440971097 | 0.661093969 | | | Fshr | 0.204491261 | 0.006752517 | 0.00969339 | 0.696610518 | 0.486046572 | NA |
| E530011L22Rik | 279.2476365 | 0.103979005 | 0.103712678 | 1.002567927 | 0.316069377 | 0.54917695 | | | Gm11570 | 0.204491261 | 0.006752517 | 0.00969339 | 0.696610518 | 0.486046572 | NA |
| Angel1 | 503.8195606 | 0.08044321 | 0.104421005 | 0.770373835 | 0.44107817 | 0.661140563 | | | Olfr5 | 0.204491261 | 0.006752517 | 0.00969339 | 0.696610518 | 0.486046572 | NA |

| | | | | | | |
|---|---|---|---|---|---|---|
| Tlr4 | 62.25508253 | -0.11202485 | 0.145419194 | -0.770358071 | 0.441087518 | 0.661140563 |
| Kat6b | 2646.855897 | -0.059763926 | 0.077594231 | -0.770210944 | 0.441174773 | 0.661207384 |
| Efna3 | 1252.100074 | 0.065556317 | 0.08512484 | 0.770119711 | 0.441228885 | 0.661224523 |
| Glyr1 | 8968.868449 | 0.103972099 | 0.069978273 | 1.485776865 | 0.137338151 | 0.330443271 |
| Klf10 | 1024.191863 | -0.127876066 | 0.166099791 | -0.769874937 | 0.441374082 | 0.661314189 |
| Kctd3 | 3248.118637 | 0.055856145 | 0.072569907 | 0.769687429 | 0.441485329 | 0.66141691 |
| Stard3 | 1229.24049 | 0.103924769 | 0.069126475 | 1.503400382 | 0.132735828 | 0.323264527 |
| Lurap1l | 305.4903035 | 0.099801273 | 0.129716091 | 0.769382365 | 0.441666354 | 0.661560179 |
| 2510002D24Rik | 404.5670678 | 0.075080422 | 0.097635212 | 0.768989185 | 0.441899731 | 0.661742575 |
| Ccdc112 | 315.5923076 | 0.066937756 | 0.08704957 | 0.768961353 | 0.441916253 | 0.661742575 |
| Cpeb3 | 2802.3721 | -0.069076952 | 0.089830784 | -0.768967485 | 0.441912613 | 0.661742575 |
| Rps26 | 2652.39605 | -0.042582526 | 0.055394184 | -0.768718359 | 0.442060523 | 0.661830684 |
| Scaf11 | 2813.169685 | 0.061890751 | 0.080504731 | 0.768784022 | 0.442021536 | 0.661830684 |
| Rap2c | 994.1624724 | -0.070376406 | 0.091571882 | -0.768537291 | 0.442168045 | 0.661927699 |
| Gtl3 | 1165.137151 | 0.103882366 | 0.065921201 | 1.575856693 | 0.115058833 | 0.29712047 |
| 4930502E18Rik | 8.639294525 | -0.121133309 | 0.157733343 | -0.767962601 | 0.442509404 | 0.66224676 |
| Scara3 | 1302.814859 | 0.076207022 | 0.099229964 | 0.767983975 | 0.442496705 | 0.66224676 |
| Mlph | 115.2625895 | 0.092306639 | 0.120259713 | 0.767560781 | 0.44274817 | 0.662540095 |
| Akap7 | 1098.413956 | 0.071226414 | 0.092820322 | 0.767357976 | 0.442868707 | 0.66265647 |
| Zcchc2 | 1202.706098 | 0.04760825 | 0.062070649 | 0.767000992 | 0.443080925 | 0.66290999 |
| Herpud2 | 1524.596479 | 0.103876005 | 0.056597761 | 1.835337701 | 0.066455668 | 0.207886962 |
| Nlk | 4774.215292 | -0.059811614 | 0.07806211 | -0.76620544 | 0.443554072 | 0.663489747 |
| Ankdd1b | 11.78846257 | -0.12746447 | 0.166414656 | -0.765944981 | 0.44370904 | 0.663529376 |
| Msh6 | 792.8022391 | 0.052456082 | 0.068477865 | 0.766029754 | 0.443658598 | 0.663529376 |
| Tollip | 6820.188828 | 0.068751756 | 0.089759586 | 0.765954464 | 0.443703397 | 0.663529376 |
| Ttll11 | 1774.884556 | 0.060693172 | 0.079247897 | 0.765864764 | 0.443756773 | 0.663536716 |
| 1110037F02Rik | 2068.876311 | -0.046992772 | 0.061366415 | -0.765773451 | 0.443811114 | 0.663553932 |
| Cdk8 | 829.1003027 | 0.054992681 | 0.071828868 | 0.765606962 | 0.4439102 | 0.66363804 |
| Kif22 | 46.70694851 | -0.117198916 | 0.153152451 | -0.765243493 | 0.444126565 | 0.663645311 |
| Ppm1k | 3113.409394 | -0.059636659 | 0.07792741 | -0.765284761 | 0.444101996 | 0.663645311 |
| Rab14 | 8204.765543 | -0.053056971 | 0.069321286 | -0.765377767 | 0.444046628 | 0.663645311 |
| Rprd1b | 1217.576775 | 0.04117444 | 0.053805986 | 0.765239024 | 0.444129226 | 0.663645311 |
| Zfp655 | 1418.393397 | 0.048859164 | 0.063825626 | 0.765510135 | 0.443967834 | 0.663645311 |
| Dyrk1a | 3588.291365 | -0.054429787 | 0.0712147 | -0.764305511 | 0.444685202 | 0.664220593 |
| Itgb3bp | 108.7609379 | -0.114894686 | 0.15038239 | -0.764016892 | 0.444857176 | 0.664220593 |
| L3hypdh | 48.98996932 | -0.11802213 | 0.154424861 | -0.764268973 | 0.444706971 | 0.664220593 |
| Cntn2 | 7031.799008 | 0.103871886 | 0.072876432 | 1.425315196 | 0.154066109 | 0.355540706 |
| Pdcd2 | 300.8540736 | 0.066163023 | 0.086594747 | 0.764053547 | 0.444835333 | 0.664220593 |
| Rtp3 | 8.874265687 | -0.11230421 | 0.146943157 | -0.76426975 | 0.444706508 | 0.664220593 |
| Tm7sf3 | 1722.170353 | 0.045383658 | 0.05939317 | 0.764122502 | 0.444794244 | 0.664220593 |
| Tmem186 | 808.1796464 | 0.063198211 | 0.082713958 | 0.764057393 | 0.444833041 | 0.664220593 |
| Txndc15 | 1971.664308 | -0.042780911 | 0.056013904 | -0.763755219 | 0.445013128 | 0.66438942 |
| Gm17455 | 8.208940609 | -0.116341437 | 0.152407732 | -0.76335653 | 0.445250797 | 0.66461617 |
| Lrrc25 | 25.75659007 | -0.125630192 | 0.164564862 | -0.76340836 | 0.445219896 | 0.66461617 |
| Klhl36 | 207.8852861 | -0.078726075 | 0.103178651 | -0.763007411 | 0.445458976 | 0.664862863 |
| Atp6v1d | 8725.989307 | 0.059161482 | 0.077569036 | 0.762694558 | 0.445645577 | 0.66501325 |
| Brsk1 | 15401.49642 | -0.063853145 | 0.083718374 | -0.762713627 | 0.445634202 | 0.66501325 |
| Zfp60 | 1434.194338 | -0.063850623 | 0.083776573 | -0.762153673 | 0.445968292 | 0.665430732 |
| 3110056K07Rik | 195.460663 | 0.103851405 | 0.123514592 | 0.840802718 | 0.400458464 | 0.625348059 |
| Cdh1 | 80.88761612 | -0.127928053 | 0.168080335 | -0.761112556 | 0.446589841 | 0.666037444 |
| Oprk1 | 535.5925476 | -0.08456735 | 0.111088689 | -0.761259772 | 0.446501923 | 0.666037444 |
| Pdxdc1 | 3953.247432 | 0.032008219 | 0.042049 | 0.761212376 | 0.446530228 | 0.666037444 |
| Thada | 582.6089293 | -0.061694294 | 0.081052339 | -0.761166123 | 0.44655785 | 0.666037444 |
| Gtf3c3 | 1175.367613 | 0.04796521 | 0.063025958 | 0.761038971 | 0.44663379 | 0.666038878 |
| Fndc5 | 3195.470871 | 0.058381555 | 0.076756133 | 0.760610946 | 0.44688948 | 0.666356039 |
| Carkd | 1176.044982 | -0.045815806 | 0.060253987 | -0.760377998 | 0.447028671 | 0.666499445 |
| Yif1b | 844.0457741 | -0.070199465 | 0.09235439 | -0.760109675 | 0.44718903 | 0.66667438 |
| Cetn4 | 216.8574237 | -0.076117077 | 0.100168132 | -0.759893147 | 0.447318458 | 0.666726681 |
| Grrp1 | 58.2147173 | -0.111688185 | 0.146990046 | -0.759835021 | 0.447353206 | 0.666726681 |
| Zer1 | 7648.197387 | 0.0398468 | 0.052433906 | 0.759943374 | 0.447288433 | 0.666726681 |
| Chrm4 | 1915.472243 | 0.059641921 | 0.078506447 | 0.759707295 | 0.447429567 | 0.666776351 |
| Cd28 | 11.68156491 | -0.121501462 | 0.159992915 | -0.759417764 | 0.44760269 | 0.666970195 |
| Gm4890 | 15.07178263 | 0.103848043 | 0.162716186 | 0.638215814 | 0.523333205 | 0.726072113 |
| 3110062M04Rik | 145.0410871 | -0.090153871 | 0.118761517 | -0.759116871 | 0.447782647 | 0.667103964 |
| B4galt1 | 267.8086848 | 0.103779404 | 0.116617621 | 0.889911863 | 0.373513214 | 0.602960381 |
| Zbtb18 | 9549.740872 | 0.078456807 | 0.10337229 | 0.758973293 | 0.447868533 | 0.667109696 |
| Ampd3 | 1129.76831 | 0.083751489 | 0.110392502 | 0.758670086 | 0.448049935 | 0.667315745 |
| Stx7 | 3523.779352 | -0.052156992 | 0.068759657 | -0.758540604 | 0.448127415 | 0.66736699 |
| Clasp2 | 9975.679647 | -0.05800507 | 0.076482189 | -0.758412781 | 0.448203909 | 0.667416757 |
| Cramp1l | 1824.153236 | 0.047939105 | 0.063228798 | 0.758184667 | 0.448340439 | 0.667491761 |
| Gpr173 | 499.4226631 | -0.065867235 | 0.08687481 | -0.758185656 | 0.448339847 | 0.667491761 |
| Ptpru | 1744.754612 | 0.070821972 | 0.093457677 | 0.757797268 | 0.448572358 | 0.667719362 |
| Rhoj | 142.3456911 | 0.09351735 | 0.123408762 | 0.757785333 | 0.448579504 | 0.667719362 |
| Trit1 | 493.5479999 | 0.103775716 | 0.091701479 | 1.131668946 | 0.257773642 | 0.484143855 |
| Kpna2 | 1186.229407 | -0.043412785 | 0.057311756 | -0.757484814 | 0.448759461 | 0.667794764 |
| Tmppe | 929.6836041 | 0.052540025 | 0.069357552 | 0.757524206 | 0.448735869 | 0.667794764 |
| Spred3 | 2075.53102 | -0.08130203 | 0.107362776 | -0.757264608 | 0.44889135 | 0.667926877 |
| Dennd6a | 1039.359057 | 0.10373584 | 0.056351837 | 1.840859939 | 0.065642086 | 0.206258059 |
| Dpm1 | 1262.101188 | 0.048755103 | 0.064409736 | 0.756952385 | 0.449078389 | 0.668076867 |
| Actr2 | 11301.45998 | -0.054340828 | 0.071839025 | -0.756424914 | 0.449394475 | 0.668226295 |
| Dph5 | 296.1303938 | -0.076515831 | 0.101150342 | -0.75645647 | 0.449375562 | 0.668226295 |
| Pip5k1a | 2200.123925 | 0.083740687 | 0.110677713 | 0.756617426 | 0.449279098 | 0.668226295 |
| Samd5 | 242.1208187 | -0.090258587 | 0.119313603 | -0.756481948 | 0.449360292 | 0.668226295 |
| Wdr85 | 775.5408446 | 0.061094838 | 0.080765117 | 0.756450809 | 0.449378955 | 0.668226295 |
| Crygn | 8.818391556 | 0.103731865 | 0.144156683 | 0.719577222 | 0.47178534 | 0.686434601 |
| Hspa9 | 9117.42648 | 0.051317977 | 0.067869481 | 0.756127448 | 0.449572787 | 0.668363151 |
| Bmyc | 1153.972216 | -0.066156315 | 0.087522135 | -0.755880955 | 0.449720574 | 0.668454582 |
| Vimp | 1148.735055 | 0.052088414 | 0.068904326 | 0.755952749 | 0.449677526 | 0.668454582 |
| Dnajb4 | 2550.780622 | 0.103682318 | 0.099508785 | 1.041941344 | 0.297438874 | 0.529268699 |
| Lrrc41 | 1858.501887 | 0.054462768 | 0.072061048 | 0.755786506 | 0.449777208 | 0.668456617 |
| Sumf2 | 482.3769159 | -0.089844066 | 0.118897915 | -0.755640382 | 0.449864837 | 0.668476627 |
| Omg | 4826.191376 | 0.057510484 | 0.076124164 | 0.755482634 | 0.449959448 | 0.668553096 |
| 5430417L22Rik | 1495.597843 | -0.065741334 | 0.087069608 | -0.755043417 | 0.450222932 | 0.668713036 |
| Mpp3 | 2159.33731 | 0.103671353 | 0.06644176 | 1.56033423 | 0.118680918 | 0.302391407 |
| Atf7ip | 2403.281245 | 0.103620935 | 0.087132578 | 1.189232975 | 0.234348002 | 0.45793769 |
| Rsbn1 | 979.6591244 | 0.056258342 | 0.074504275 | 0.7551022 | 0.450187663 | 0.668713036 |
| Rbm4b | 376.333024 | -0.062033528 | 0.082177635 | -0.754871178 | 0.450326281 | 0.668777474 |
| Wbp1 | 773.3275554 | -0.087834444 | 0.116453158 | -0.754246996 | 0.450700923 | 0.669269704 |
| Igbp1b | 9.97958867 | -0.115465599 | 0.153127055 | -0.754050932 | 0.45081864 | 0.669316213 |
| Nkx2-1 | 60.93497913 | -0.126681231 | 0.167991989 | -0.754090909 | 0.450794636 | 0.669316213 |
| Kcna1 | 10162.81788 | 0.076712958 | 0.101755747 | 0.753893124 | 0.4509134 | 0.669392758 |
| Ndufa9 | 3540.181492 | -0.045633947 | 0.060539072 | -0.753793305 | 0.450973345 | 0.669417609 |
| Saal1 | 363.4020811 | 0.056375369 | 0.074858639 | 0.753091026 | 0.451395217 | 0.669979643 |
| Anks1 | 1390.870326 | 0.103593776 | 0.072189185 | 1.435031807 | 0.151278034 | 0.352254563 |
| Sertm1 | 1156.884099 | 0.103529779 | 0.089635098 | 1.155013844 | 0.248084757 | 0.473111538 |
| Nsl1 | 56.10229366 | 0.103511294 | 0.153448585 | 0.674566624 | 0.499951144 | 0.70810314 |
| 2310009A05Rik | 116.6805004 | -0.095010814 | 0.12646792 | -0.751264145 | 0.452493703 | 0.671352814 |
| Fam19a1 | 1263.120234 | -0.070788238 | 0.094239633 | -0.751151454 | 0.452561513 | 0.67138913 |
| Mbnl2 | 7482.261466 | 0.060247556 | 0.080239273 | 0.750848728 | 0.4527437 | 0.671466535 |
| Gdap1l1 | 2605.936575 | 0.103509692 | 0.081312106 | 1.272992397 | 0.20302074 | 0.420066199 |
| Thumpd1 | 2025.798939 | 0.04503672 | 0.059976705 | 0.750903534 | 0.452710713 | 0.671466535 |
| Zbtb43 | 1305.362414 | 0.103506203 | 0.064468169 | 1.605539664 | 0.108375092 | 0.285717827 |
| Cenpi | 18.30035922 | -0.125941809 | 0.167801896 | -0.750538653 | 0.452930353 | 0.671518227 |
| Inpp5a | 2412.675289 | 0.040350143 | 0.05377446 | 0.750358867 | 0.453038598 | 0.671518227 |
| Khdrbs3 | 4830.722772 | -0.064503003 | 0.085937966 | -0.750576329 | 0.452907671 | 0.671518227 |
| Rif1 | 1092.440875 | -0.070684316 | 0.094198679 | -0.750374817 | 0.453028994 | 0.671518227 |
| Lrrn4 | 18.82943768 | 0.103496811 | 0.168114409 | 0.615633195 | 0.538136638 | 0.737397186 |
| Zfp825 | 218.0749704 | 0.076267176 | 0.101663014 | 0.7501959 | 0.453136728 | 0.671599432 |
| Ppap2c | 103.2276227 | 0.103494484 | 0.141714307 | 0.730303708 | 0.465204562 | 0.680878063 |
| 2900056M20Rik | 777.2718262 | 0.065001817 | 0.08667997 | 0.749905856 | 0.453311407 | 0.67169764 |
| Gpatch2 | 698.3224981 | 0.050207165 | 0.066954499 | 0.74986993 | 0.453333047 | 0.67169764 |
| Cd40 | 19.0224353 | -0.125966282 | 0.168112277 | -0.749298526 | 0.453677296 | 0.672143433 |
| Shisa5 | 2081.008554 | 0.064666672 | 0.08632419 | 0.749114142 | 0.453788412 | 0.672243777 |
| Cd164 | 2162.98476 | -0.05063605 | 0.067606563 | -0.748981279 | 0.453868489 | 0.672298123 |
| Apoo | 189.4165721 | -0.118959363 | 0.158872461 | -0.748772708 | 0.453994212 | 0.672303015 |
| Phyh | 2629.588045 | -0.059667464 | 0.079685842 | -0.748783757 | 0.453987551 | 0.672303015 |
| Pla2g7 | 5060.012468 | 0.083706747 | 0.111804599 | 0.748687871 | 0.454045356 | 0.672303015 |
| Synj2bp | 1954.494792 | -0.044544469 | 0.059491433 | -0.748754343 | 0.454005283 | 0.672303015 |
| Smchd1 | 1717.056779 | -0.043576185 | 0.058233061 | -0.748306631 | 0.454275226 | 0.672579106 |
| Phf23 | 1702.402482 | 0.078905177 | 0.105482405 | 0.748041125 | 0.454435352 | 0.672687623 |
| Ssb | 3407.762385 | 0.058324016 | 0.077964434 | 0.748084898 | 0.454408951 | 0.672687623 |
| Bcl7c | 288.9295897 | -0.072079842 | 0.096387633 | -0.747812145 | 0.454573475 | 0.672827802 |
| Armcx6 | 272.0169892 | 0.083674559 | 0.111906206 | 0.747720458 | 0.454628789 | 0.672845396 |
| Arhgef3 | 3335.436297 | 0.067389954 | 0.090161329 | 0.747437452 | 0.454799545 | 0.67296955 |
| Snap25 | 101385.2305 | 0.06834872 | 0.091442009 | 0.747454265 | 0.4547894 | 0.67296955 |
| Stk19 | 633.4944735 | 0.103478736 | 0.084059226 | 1.231021746 | 0.218314731 | 0.438694312 |
| Scamp2 | 913.8273747 | 0.103469888 | 0.072826477 | 1.420772949 | 0.155382777 | 0.357408069 |
| Rpain | 344.4601638 | 0.077947033 | 0.104338965 | 0.747055838 | 0.455029856 | 0.67311749 |
| Ubox5 | 714.7321449 | -0.070323952 | 0.094226718 | -0.746327086 | 0.455469853 | 0.673704048 |
| Pcid2 | 1122.09288 | 0.103467358 | 0.066969826 | 1.544984713 | 0.122349961 | 0.308694072 |
| Loxhd1 | 10.00185357 | -0.114773383 | 0.153938649 | -0.745578737 | 0.455921932 | 0.674115317 |
| Srrm4 | 1968.051841 | 0.103446415 | 0.073891493 | 1.399977341 | 0.161520104 | 0.36646544 |
| Zfp628 | 505.0823896 | 0.063133329 | 0.084673597 | 0.745608216 | 0.455904118 | 0.674115317 |
| Zfp217 | 187.4399117 | 0.078960336 | 0.105937077 | 0.745351284 | 0.456059386 | 0.67425421 |
| Zfp937 | 450.9879097 | -0.072335934 | 0.097091689 | -0.745027043 | 0.456255372 | 0.674479604 |
| Skap1 | 61.13403916 | -0.114820048 | 0.154130273 | -0.744954549 | 0.456299197 | 0.674480037 |
| Phf10 | 905.2166847 | 0.103371344 | 0.080696597 | 1.280987649 | 0.200198004 | 0.416917726 |
| Atp1b1 | 54268.86191 | -0.044498021 | 0.059821785 | -0.743843089 | 0.456971409 | 0.675299957 |
| Tmem200b | 179.0441938 | 0.10336955 | 0.121415676 | 0.851369059 | 0.394564374 | 0.620018297 |
| Ccdc97 | 2153.188899 | 0.04180549 | 0.056239593 | 0.743346234 | 0.457272087 | 0.675531438 |
| Gphn | 2820.944486 | -0.039081406 | 0.052573252 | -0.743370537 | 0.457257377 | 0.675531438 |
| Kat7 | 3929.506192 | 0.049144558 | 0.06610273 | 0.743457318 | 0.457204853 | 0.675531438 |
| Bcl2l2 | 6352.74434 | -0.046105544 | 0.062051534 | -0.743020203 | 0.457469449 | 0.675565349 |
| Dcaf7 | 10961.57824 | -0.0435692 | 0.0586285 | -0.743140269 | 0.457396762 | 0.675565349 |
| Fcf1 | 422.8307612 | 0.06619056 | 0.089079235 | 0.743052633 | 0.457449815 | 0.675565349 |
| Zfp352 | 21.927997 | -0.124639382 | 0.167721434 | -0.743133295 | 0.457400984 | 0.675565349 |
| Slx4 | 1209.719743 | 0.057207138 | 0.077013504 | 0.742819563 | 0.45759093 | 0.675680345 |
| Cav2 | 910.798954 | 0.080290919 | 0.108103857 | 0.742720208 | 0.457651093 | 0.675704787 |
| Flad1 | 1082.015187 | 0.079695057 | 0.107361303 | 0.742307091 | 0.457901297 | 0.675945378 |
| Prkg2 | 505.8075079 | -0.076940357 | 0.103649913 | -0.742309901 | 0.457899595 | 0.675945378 |
| Atg10 | 361.1437581 | -0.079848031 | 0.107640304 | -0.741804212 | 0.458205968 | 0.6759631 |
| Decr2 | 1270.197979 | 0.057210203 | 0.07712867 | 0.741750161 | 0.458238722 | 0.6759631 |
| Gcat | 340.9674043 | 0.08025262 | 0.108195089 | 0.741739953 | 0.458244908 | 0.6759631 |
| Tubgcp5 | 1217.154202 | 0.10336607 | 0.079822963 | 1.294941531 | 0.1953404 | 0.409894454 |
| Lactb | 531.7833558 | -0.056547828 | 0.076209459 | -0.742005376 | 0.458084079 | 0.6759631 |
| Metrnl | 219.0008296 | 0.095062555 | 0.128166532 | 0.741711219 | 0.458262321 | 0.6759631 |
| Mrps7 | 1350.563149 | -0.041824081 | 0.056387685 | -0.74172368 | 0.458254769 | 0.6759631 |
| Pikfyve | 2244.818641 | -0.060045446 | 0.080906107 | -0.742162098 | 0.45798913 | 0.6759631 |
| Efs | 402.1326853 | -0.098791713 | 0.133216138 | -0.741589674 | 0.458335982 | 0.676007397 |
| Arpc1b | 525.8277498 | 0.052373333 | 0.070631158 | 0.741504665 | 0.458387505 | 0.676019036 |
| Pitpnc1 | 3427.416826 | -0.058160377 | 0.078452961 | -0.741340753 | 0.458486858 | 0.676101207 |
| Rasgef1c | 1581.274398 | -0.083748599 | 0.112983195 | -0.741248281 | 0.458542914 | 0.67611952 |
| Pias2 | 2380.599884 | 0.066396883 | 0.089606764 | 0.740980707 | 0.458705139 | 0.676294359 |
| Fkbp2 | 2010.592874 | -0.05400009 | 0.072931761 | -0.74041939 | 0.459045557 | 0.676731861 |
| Fam228a | 57.61225714 | 0.103365316 | 0.167353321 | 0.617647231 | 0.536807903 | 0.736292279 |
| Etnk1 | 14726.91397 | -0.047221265 | 0.063812906 | -0.739995533 | 0.459302705 | 0.676917731 |
| Shoc2 | 2364.938354 | -0.05475215 | 0.073981004 | -0.740083898 | 0.459249088 | 0.676917731 |
| 2610318N02Rik | 28.38692282 | -0.122458467 | 0.165578887 | -0.739577791 | 0.459556222 | 0.676957488 |
| Ddx21 | 1211.273294 | 0.076382116 | 0.103377935 | 0.738862857 | 0.459990279 | 0.676957488 |
| Tnni1 | 96.48379536 | 0.103365278 | 0.161728064 | 0.639130127 | 0.522738282 | 0.72574435 |
| Lsm3 | 452.4564055 | -0.062188959 | 0.084157764 | -0.738956881 | 0.459933181 | 0.676957488 |
| Mrps5 | 1195.807599 | 0.051608911 | 0.069804248 | 0.739337681 | 0.459701973 | 0.676957488 |
| Nab2 | 1701.092071 | 0.082601356 | 0.111752768 | 0.73914371 | 0.459819738 | 0.676957488 |
| Nek2 | 66.84815621 | -0.112700102 | 0.152361038 | -0.73969109 | 0.459487456 | 0.676957488 |
| Pim2 | 1687.687445 | -0.045660911 | 0.061790684 | -0.738961095 | 0.459930622 | 0.676957488 |
| Plekhg2 | 284.984652 | 0.075911792 | 0.102613046 | 0.739786945 | 0.459429282 | 0.676957488 |
| Psmd2 | 9006.134837 | -0.037239039 | 0.050374004 | -0.739251122 | 0.459754524 | 0.676957488 |
| Ptpn1 | 2025.146636 | -0.065223546 | 0.088213893 | -0.739379525 | 0.459676572 | 0.676957488 |
| Sbf1 | 8349.287794 | 0.050025657 | 0.067644835 | 0.73953402 | 0.45958279 | 0.676957488 |
| Sf3a3 | 1109.444489 | 0.057067045 | 0.077242938 | 0.738799515 | 0.460028746 | 0.676957488 |
| Slitrk4 | 1112.109099 | -0.081867402 | 0.110708613 | -0.739485394 | 0.459612306 | 0.676957488 |
| Tbl2 | 1084.360477 | 0.050285862 | 0.067992254 | 0.739582214 | 0.459553537 | 0.676957488 |
| Trpc3 | 671.9431308 | 0.059761631 | 0.080876905 | 0.738920847 | 0.459955063 | 0.676957488 |
| Bcorl1 | 1127.038631 | 0.055551297 | 0.075208399 | 0.738631561 | 0.460130754 | 0.677038414 |
| R3hdm4 | 10168.74753 | -0.059185012 | 0.080135134 | -0.738565083 | 0.460171133 | 0.677038414 |
| Tceal1 | 559.1655831 | -0.065258208 | 0.088380471 | -0.738378135 | 0.460284698 | 0.677076911 |
| Zfp777 | 1191.012896 | 0.058237068 | 0.078870105 | 0.738392173 | 0.46027617 | 0.677076911 |
| Vsnl1 | 19691.90344 | 0.064898598 | 0.087905375 | 0.738277925 | 0.460345578 | 0.677102183 |
| Fchsd1 | 331.6514419 | -0.073932132 | 0.100188007 | -0.737933957 | 0.460554583 | 0.677345297 |
| Rbm19 | 578.012566 | 0.051828692 | 0.070244664 | 0.737831024 | 0.460617138 | 0.677373001 |
| Unkl | 1988.315132 | 0.040756215 | 0.055268593 | 0.737420887 | 0.460866437 | 0.677675295 |
| Hadhb | 2779.176812 | -0.057138833 | 0.077507098 | -0.73720774 | 0.460996027 | 0.677767216 |
| Prg4 | 262.5092609 | -0.110654087 | 0.150105756 | -0.737174172 | 0.461016438 | 0.677767216 |
| Smpd1 | 4579.140266 | -0.048544782 | 0.065868456 | -0.736995902 | 0.461124841 | 0.677862267 |
| 2700099C18Rik | 10.48386147 | -0.121922001 | 0.165465166 | -0.736843915 | 0.461217274 | 0.677933825 |
| 1500011B03Rik | 3003.95319 | -0.039778395 | 0.054002921 | -0.736597101 | 0.461367399 | 0.677935394 |
| Cenpt | 505.7952431 | 0.103357442 | 0.088541899 | 1.167328057 | 0.243077912 | 0.467508587 |
| Nefl | 10522.66853 | -0.060512368 | 0.082156004 | -0.736554423 | 0.46139336 | 0.677935394 |
| Ythdf1 | 2327.926066 | 0.059228292 | 0.080407091 | 0.736605329 | 0.461362394 | 0.677935394 |
| Pik3ca | 3204.848271 | -0.051476876 | 0.069904372 | -0.736389925 | 0.461493435 | 0.678018137 |
| Akr7a5 | 586.1360266 | 0.057297725 | 0.07788128 | 0.735705999 | 0.461909638 | 0.678212469 |
| Mical3 | 4324.217106 | 0.103347557 | 0.070017195 | 1.476031106 | 0.139935542 | 0.334334817 |
| Cars2 | 371.1748432 | 0.063476957 | 0.086255909 | 0.735914309 | 0.461782849 | 0.678212469 |
| Odf4 | 13.32431388 | 0.103264094 | 0.165675721 | 0.623290449 | 0.533093673 | 0.733939343 |
| Eri2 | 403.9451679 | 0.069333564 | 0.09424411 | 0.735680605 | 0.461925096 | 0.678212469 |
| Slc30a4 | 3010.914904 | 0.065745828 | 0.089360781 | 0.735734709 | 0.461892163 | 0.678212469 |
| Mrap2 | 343.041319 | 0.103259108 | 0.118985573 | 0.867828808 | 0.385488057 | 0.613272331 |
| Esam | 641.119318 | 0.067162091 | 0.091327059 | 0.73540188 | 0.462094777 | 0.678387008 |
| Hs3st4 | 2485.035822 | 0.073739161 | 0.100297383 | 0.73520524 | 0.462214509 | 0.678498493 |
| Arl14ep | 1061.27528 | 0.056590178 | 0.076993712 | 0.734997398 | 0.462341079 | 0.678564691 |
| Zbtb33 | 935.0810253 | -0.077115075 | 0.104920274 | -0.734987355 | 0.462347195 | 0.678564691 |
| Srsf12 | 594.249664 | -0.050376903 | 0.068567932 | -0.734700629 | 0.462521837 | 0.67875671 |
| Irak1 | 2316.033811 | 0.058814162 | 0.080068061 | 0.73455209 | 0.462612325 | 0.678825208 |
| Myh9 | 3764.109509 | -0.053397277 | 0.072770944 | -0.733771942 | 0.463087743 | 0.679458475 |
| Cadps | 11494.73447 | -0.038891329 | 0.053030915 | -0.733370881 | 0.463332254 | 0.679675919 |
| Evi5l | 1447.812306 | 0.053133093 | 0.072467563 | 0.733198284 | 0.463437502 | 0.679675919 |
| Il10rb | 381.9501679 | -0.081815771 | 0.111557029 | -0.733398613 | 0.463315345 | 0.679675919 |
| Prosapip1 | 12939.83964 | 0.057616422 | 0.078585444 | 0.733169134 | 0.463455278 | 0.679675919 |
| Zfp78 | 129.718445 | -0.098165885 | 0.133881337 | -0.73323054 | 0.463417832 | 0.679675919 |
| Hadha | 2204.849255 | 0.046136431 | 0.06296267 | 0.732758489 | 0.463705744 | 0.679978875 |
| Megf9 | 3180.721121 | -0.05774376 | 0.078828923 | -0.732519966 | 0.463851261 | 0.680013495 |
| Nrg3 | 1589.835353 | 0.065452548 | 0.089363305 | 0.732432042 | 0.463904908 | 0.680013495 |
| Tiam1 | 4978.981321 | 0.089128124 | 0.121675568 | 0.732506333 | 0.463859579 | 0.680013495 |
| Wasf3 | 2699.427375 | -0.034991156 | 0.047770299 | -0.732487689 | 0.463870954 | 0.680013495 |
| Pknox2 | 2901.230965 | 0.051108112 | 0.069791442 | 0.732297695 | 0.463986886 | 0.680069322 |

| | | | | | | |
|---|---|---|---|---|---|---|
| Plac1l | 0.204491261 | 0.006752517 | 0.00969339 | 0.696610518 | 0.486046572 | NA |
| Tmem72 | 56.84427618 | 0.006767992 | 0.026787527 | 0.252654609 | NA | NA |
| Gm853 | 0.751133396 | 0.006773783 | 0.016160089 | 0.419167436 | 0.675093769 | NA |
| Med28 | 2010.38411 | 0.006784363 | 0.053814319 | 0.126069839 | 0.899676642 | 0.97888996 |
| Kctd19 | 0.430848918 | 0.00680113 | 0.015451208 | 0.440168173 | 0.659815309 | NA |
| Brdt | 509.7457967 | 0.006802456 | 0.079994098 | 0.085036978 | 0.932231993 | 0.984387568 |
| 1700016D06Rik | 0.212297216 | 0.006810501 | 0.009734391 | 0.699633035 | 0.484156506 | NA |
| 1700085C21Rik | 0.212297216 | 0.006810501 | 0.009734391 | 0.699633035 | 0.484156506 | NA |
| 2810442N19Rik | 0.212297216 | 0.006810501 | 0.009734391 | 0.699633035 | 0.484156506 | NA |
| Aqp12 | 0.212297216 | 0.006810501 | 0.009734391 | 0.699633035 | 0.484156506 | NA |
| Chrna9 | 0.212297216 | 0.006810501 | 0.009734391 | 0.699633035 | 0.484156506 | NA |
| Cpa3 | 0.212297216 | 0.006810501 | 0.009734391 | 0.699633035 | 0.484156506 | NA |
| E330021D16Rik | 0.212297216 | 0.006810501 | 0.009734391 | 0.699633035 | 0.484156506 | NA |
| Gm3258 | 0.212297216 | 0.006810501 | 0.009734391 | 0.699633035 | 0.484156506 | NA |
| Gpha2 | 0.212297216 | 0.006810501 | 0.009734391 | 0.699633035 | 0.484156506 | NA |
| Hoxb9 | 0.212297216 | 0.006810501 | 0.009734391 | 0.699633035 | 0.484156506 | NA |
| Il3 | 0.212297216 | 0.006810501 | 0.009734391 | 0.699633035 | 0.484156506 | NA |
| Sprr2g | 0.212297216 | 0.006810501 | 0.009734391 | 0.699633035 | 0.484156506 | NA |
| Tbpl2 | 0.212297216 | 0.006810501 | 0.009734391 | 0.699633035 | 0.484156506 | NA |
| Tpte | 0.212297216 | 0.006810501 | 0.009734391 | 0.699633035 | 0.484156506 | NA |
| Vmn1r207-ps | 0.212297216 | 0.006810501 | 0.009734391 | 0.699633035 | 0.484156506 | NA |
| Rbm22 | 1321.373989 | 0.006811994 | 0.051149464 | 0.13317821 | 0.894052443 | 0.977844987 |
| 4930599N23Rik | 4.57635825 | 0.006834991 | 0.041141258 | 0.166134718 | 0.868050933 | NA |
| Lhx2 | 1633.697123 | 0.00684462 | 0.072378358 | 0.094567212 | 0.924658594 | 0.983236501 |
| 1700030M09Rik | 0.217532603 | 0.006846852 | 0.009760003 | 0.701521557 | 0.482977586 | NA |
| 2410017I17Rik | 0.217532603 | 0.006846852 | 0.009760003 | 0.701521557 | 0.482977586 | NA |
| 4930456L15Rik | 0.217532603 | 0.006846852 | 0.009760003 | 0.701521557 | 0.482977586 | NA |
| 4933432I03Rik | 0.217532603 | 0.006846852 | 0.009760003 | 0.701521557 | 0.482977586 | NA |
| 4933436E23Rik | 0.217532603 | 0.006846852 | 0.009760003 | 0.701521557 | 0.482977586 | NA |
| 8430431K14Rik | 0.217532603 | 0.006846852 | 0.009760003 | 0.701521557 | 0.482977586 | NA |
| Bpifa2 | 0.217532603 | 0.006846852 | 0.009760003 | 0.701521557 | 0.482977586 | NA |
| Cdcp2 | 0.217532603 | 0.006846852 | 0.009760003 | 0.701521557 | 0.482977586 | NA |
| Gm1587 | 0.217532603 | 0.006846852 | 0.009760003 | 0.701521557 | 0.482977586 | NA |
| Gm17727 | 0.217532603 | 0.006846852 | 0.009760003 | 0.701521557 | 0.482977586 | NA |
| Gm6880 | 0.217532603 | 0.006846852 | 0.009760003 | 0.701521557 | 0.482977586 | NA |
| Kcnk18 | 0.217532603 | 0.006846852 | 0.009760003 | 0.701521557 | 0.482977586 | NA |
| Lect2 | 0.217532603 | 0.006846852 | 0.009760003 | 0.701521557 | 0.482977586 | NA |
| Mir669a-2 | 0.217532603 | 0.006846852 | 0.009760003 | 0.701521557 | 0.482977586 | NA |
| Oas1f | 0.217532603 | 0.006846852 | 0.009760003 | 0.701521557 | 0.482977586 | NA |
| Olfr235 | 0.217532603 | 0.006846852 | 0.009760003 | 0.701521557 | 0.482977586 | NA |
| Olfr577 | 0.217532603 | 0.006846852 | 0.009760003 | 0.701521557 | 0.482977586 | NA |
| Otol1 | 0.217532603 | 0.006846852 | 0.009760003 | 0.701521557 | 0.482977586 | NA |
| Prss34 | 0.217532603 | 0.006846852 | 0.009760003 | 0.701521557 | 0.482977586 | NA |
| Pyy | 0.217532603 | 0.006846852 | 0.009760003 | 0.701521557 | 0.482977586 | NA |
| Serpina7 | 0.217532603 | 0.006846852 | 0.009760003 | 0.701521557 | 0.482977586 | NA |
| Tmem210 | 0.217532603 | 0.006846852 | 0.009760003 | 0.701521557 | 0.482977586 | NA |
| Trim31 | 0.217532603 | 0.006846852 | 0.009760003 | 0.701521557 | 0.482977586 | NA |
| Ifi27l2b | 2.267211278 | 0.006852129 | 0.029870713 | 0.229392869 | 0.818563578 | NA |
| Cndp1 | 16.30639992 | 0.006856943 | 0.056921365 | 0.120463424 | 0.904116052 | NA |
| 1700001K23Rik | 0.209568245 | 0.006861974 | 0.009771272 | 0.702260104 | 0.482516969 | NA |
| Cfc1 | 0.209568245 | 0.006861974 | 0.009771272 | 0.702260104 | 0.482516969 | NA |
| Gm11240 | 0.209568245 | 0.006861974 | 0.009771272 | 0.702260104 | 0.482516969 | NA |
| Mir592 | 0.209568245 | 0.006861974 | 0.009771272 | 0.702260104 | 0.482516969 | NA |
| Olfr701 | 0.209568245 | 0.006861974 | 0.009771272 | 0.702260104 | 0.482516969 | NA |
| Pin1-ps1 | 0.209568245 | 0.006861974 | 0.009771272 | 0.702260104 | 0.482516969 | NA |
| Pip | 0.209568245 | 0.006861974 | 0.009771272 | 0.702260104 | 0.482516969 | NA |
| Tmem161b | 650.7549454 | 0.006867192 | 0.066907342 | 0.102637353 | 0.918250796 | 0.982742917 |
| Rpl34 | 7.955198875 | 0.006877368 | 0.056657486 | 0.121384982 | 0.903386111 | NA |
| Rplp1 | 11216.18111 | 0.006880235 | 0.064344472 | 0.106928151 | 0.91484598 | 0.98170867 |
| Gm13546 | 1.402126726 | 0.006905625 | 0.025625135 | 0.269486384 | 0.787555419 | NA |
| 1700020N01Rik | 1.304871827 | 0.006912419 | 0.024506452 | 0.282065257 | 0.777893473 | NA |
| Stmnd1 | 10.55987753 | 0.006913859 | 0.067755017 | 0.102042016 | 0.918723325 | NA |
| 4930519D14Rik | 0.227686571 | 0.006914604 | 0.009807553 | 0.705028462 | 0.480792524 | NA |
| Adamdec1 | 0.227686571 | 0.006914604 | 0.009807553 | 0.705028462 | 0.480792524 | NA |
| Apoc4 | 0.227686571 | 0.006914604 | 0.009807553 | 0.705028462 | 0.480792524 | NA |
| BC094916 | 0.227686571 | 0.006914604 | 0.009807553 | 0.705028462 | 0.480792524 | NA |
| Cyct | 0.227686571 | 0.006914604 | 0.009807553 | 0.705028462 | 0.480792524 | NA |
| Gm13178 | 0.227686571 | 0.006914604 | 0.009807553 | 0.705028462 | 0.480792524 | NA |
| Gm6416 | 0.227686571 | 0.006914604 | 0.009807553 | 0.705028462 | 0.480792524 | NA |
| LOC100048884 | 0.227686571 | 0.006914604 | 0.009807553 | 0.705028462 | 0.480792524 | NA |
| Tubal3 | 0.227686571 | 0.006914604 | 0.009807553 | 0.705028462 | 0.480792524 | NA |
| Wfdc5 | 0.227686571 | 0.006914604 | 0.009807553 | 0.705028462 | 0.480792524 | NA |
| Dsc1 | 3.090523456 | 0.006935 | 0.037329119 | 0.185779901 | 0.852617367 | NA |
| AU040320 | 3672.260992 | 0.006936987 | 0.052787702 | 0.131412932 | 0.895448659 | 0.977844987 |
| N28178 | 12763.90353 | 0.006945664 | 0.063677606 | 0.109075465 | 0.913142635 | 0.98170867 |
| Sfmbt1 | 1727.169171 | 0.006955969 | 0.069510728 | 0.100070443 | 0.920288401 | 0.982742917 |
| Lsm3 | 492.8204973 | 0.006957204 | 0.070531422 | 0.098639774 | 0.921424289 | 0.982762672 |
| 1700073E17Rik | 0.448266046 | 0.006968612 | 0.01411232 | 0.493796306 | 0.621450042 | NA |
| Urm1 | 1422.852831 | 0.006974201 | 0.050824194 | 0.137222072 | 0.890855265 | 0.977182599 |
| Rap1b | 1768.888707 | 0.006975385 | 0.068687292 | 0.101552779 | 0.919111663 | 0.982742917 |
| 1700065I17Rik | 0.21491491 | 0.00699378 | 0.009864233 | 0.70900394 | 0.478322032 | NA |
| 1700069L16Rik | 0.21491491 | 0.00699378 | 0.009864233 | 0.70900394 | 0.478322032 | NA |
| Clec4e | 0.21491491 | 0.00699378 | 0.009864233 | 0.70900394 | 0.478322032 | NA |
| Pldi | 0.21491491 | 0.00699378 | 0.009864233 | 0.70900394 | 0.478322032 | NA |
| Ankrd1 | 1.191895526 | 0.007013499 | 0.025625479 | 0.273692402 | 0.784321017 | NA |
| Lrrc41 | 2012.319612 | 0.007044588 | 0.053223267 | 0.132359186 | 0.894700196 | 0.977844987 |
| Babam1 | 2575.35959 | 0.007049367 | 0.051367016 | 0.137235291 | 0.890844817 | 0.977182599 |
| Mir370 | 2.217855882 | 0.007050126 | 0.033205627 | 0.212317193 | 0.831859583 | NA |
| Gmnn | 119.1149744 | 0.00705265 | 0.094347095 | 0.074752164 | 0.940411903 | 0.985461126 |
| Il17f | 0.681515317 | 0.007058648 | 0.01913564 | 0.368874397 | 0.712221348 | NA |
| Tspan7 | 48434.60493 | 0.007067029 | 0.045711469 | 0.154600777 | 0.877136058 | 0.972549751 |
| 4930432M17Rik | 0.255375383 | 0.007083044 | 0.009924717 | 0.713677164 | 0.47542683 | NA |
| Acnat2 | 0.255375383 | 0.007083044 | 0.009924717 | 0.713677164 | 0.47542683 | NA |
| Ankrd60 | 0.255375383 | 0.007083044 | 0.009924717 | 0.713677164 | 0.47542683 | NA |
| Fabp12 | 0.255375383 | 0.007083044 | 0.009924717 | 0.713677164 | 0.47542683 | NA |
| Il4 | 0.255375383 | 0.007083044 | 0.009924717 | 0.713677164 | 0.47542683 | NA |
| Klk1b21 | 0.255375383 | 0.007083044 | 0.009924717 | 0.713677164 | 0.47542683 | NA |
| LOC100038947 | 0.255375383 | 0.007083044 | 0.009924717 | 0.713677164 | 0.47542683 | NA |
| Lamp3 | 0.255375383 | 0.007083044 | 0.009924717 | 0.713677164 | 0.47542683 | NA |
| Mir704 | 0.255375383 | 0.007083044 | 0.009924717 | 0.713677164 | 0.47542683 | NA |
| Olfr166 | 0.255375383 | 0.007083044 | 0.009924717 | 0.713677164 | 0.47542683 | NA |
| Olfr19 | 0.255375383 | 0.007083044 | 0.009924717 | 0.713677164 | 0.47542683 | NA |
| Prss3 | 0.255375383 | 0.007083044 | 0.009924717 | 0.713677164 | 0.47542683 | NA |
| Psg19 | 0.255375383 | 0.007083044 | 0.009924717 | 0.713677164 | 0.47542683 | NA |
| Myh6 | 1.149044516 | 0.007096665 | 0.024361244 | 0.291309621 | 0.77081453 | NA |
| Spats2 | 1059.499553 | 0.007105341 | 0.060131585 | 0.118163215 | 0.905938336 | 0.980118791 |
| Phactr1 | 24972.06156 | 0.007107007 | 0.055286995 | 0.128547534 | 0.897715683 | 0.978040802 |
| 3110001I22Rik | 57.75753143 | 0.007111191 | 0.090619492 | 0.07847308 | 0.937451744 | 0.985461126 |
| Adam7 | 0.219991894 | 0.00711449 | 0.009948618 | 0.715123456 | 0.474532761 | NA |
| Hoxa5 | 0.219991894 | 0.00711449 | 0.009948618 | 0.715123456 | 0.474532761 | NA |
| Naip7 | 0.219991894 | 0.00711449 | 0.009948618 | 0.715123456 | 0.474532761 | NA |
| Tmigd1 | 0.219991894 | 0.00711449 | 0.009948618 | 0.715123456 | 0.474532761 | NA |
| Csn2 | 0.53223091 | 0.007114963 | 0.013051168 | 0.545159131 | 0.585644077 | NA |
| Dhrs7c | 0.466157216 | 0.00712082 | 0.015745046 | 0.452257782 | 0.651083286 | NA |
| Rps5 | 5123.989623 | 0.007129269 | 0.047177021 | 0.151117401 | 0.879883106 | 0.973507655 |
| Usp13 | 1245.577465 | 0.007143791 | 0.064706346 | 0.110403253 | 0.912089574 | 0.981408381 |
| Snta1 | 1217.256665 | 0.0071643 | 0.065946894 | 0.108637417 | 0.913490082 | 0.98170867 |
| Arsb | 5989.766881 | 0.007165477 | 0.04481509 | 0.159889818 | 0.87296787 | 0.971691093 |
| 7420461P10Rik | 0.222609587 | 0.007176514 | 0.009991691 | 0.718248158 | 0.472604289 | NA |
| Gm4956 | 0.222609587 | 0.007176514 | 0.009991691 | 0.718248158 | 0.472604289 | NA |
| Gm6756 | 0.222609587 | 0.007176514 | 0.009991691 | 0.718248158 | 0.472604289 | NA |
| Lipk | 0.222609587 | 0.007176514 | 0.009991691 | 0.718248158 | 0.472604289 | NA |
| 1700027H10Rik | 0.223412651 | 0.00718832 | 0.009999938 | 0.718836445 | 0.472241699 | NA |
| Ddx3y | 0.223412651 | 0.00718832 | 0.009999938 | 0.718836445 | 0.472241699 | NA |
| Gm6792 | 0.223412651 | 0.00718832 | 0.009999938 | 0.718836445 | 0.472241699 | NA |
| Hdgfl1 | 0.223412651 | 0.00718832 | 0.009999938 | 0.718836445 | 0.472241699 | NA |
| Hist1h2bp | 0.223412651 | 0.00718832 | 0.009999938 | 0.718836445 | 0.472241699 | NA |
| Olfr700 | 0.223412651 | 0.00718832 | 0.009999938 | 0.718836445 | 0.472241699 | NA |
| Saa3 | 0.223412651 | 0.00718832 | 0.009999938 | 0.718836445 | 0.472241699 | NA |
| Serpinb1c | 0.223412651 | 0.00718832 | 0.009999938 | 0.718836445 | 0.472241699 | NA |
| Wwp1 | 2254.396443 | 0.007204003 | 0.068946854 | 0.104486313 | 0.916783429 | 0.982742917 |
| Peg3as | 3.422907728 | 0.007212211 | 0.040001724 | 0.1802975 | 0.856919018 | NA |
| Pter | 78.19600695 | 0.007225921 | 0.095087243 | 0.07599254 | 0.939425033 | 0.985461126 |
| Celsr2 | 10760.62116 | 0.00723003 | 0.061132083 | 0.118268993 | 0.905854525 | 0.980118791 |
| Dgcr6 | 1511.894041 | 0.007252086 | 0.059951374 | 0.120966134 | 0.903717859 | 0.979584033 |
| Cer1 | 5.52947549 | 0.007260547 | 0.053067471 | 0.136817282 | 0.891175223 | NA |
| Kcnc2 | 5076.347687 | 0.007264834 | 0.053391047 | 0.13606839 | 0.891767216 | 0.977572303 |
| Pick1 | 1828.081261 | 0.007268991 | 0.055051976 | 0.132038695 | 0.894953686 | 0.977844987 |
| D230025D16Rik | 1735.228881 | 0.007270134 | 0.058145163 | 0.125034202 | 0.900496473 | 0.979053977 |
| Pggt1b | 299.3668635 | 0.007277204 | 0.080331347 | 0.090589844 | 0.927818502 | 0.983501978 |
| Zdhhc13 | 849.6433241 | 0.007279583 | 0.081248517 | 0.089596498 | 0.928607867 | 0.983596265 |
| 1700030A11Rik | 1.480134068 | 0.007287589 | 0.029825425 | 0.244341491 | 0.806966352 | NA |
| Sis | 4.516395738 | 0.007314142 | 0.049196089 | 0.148673234 | 0.881811483 | NA |
| Ppp1r3f | 1352.533368 | 0.007316787 | 0.062870186 | 0.116379277 | 0.907351959 | 0.980388092 |
| Rapgef4 | 25921.22962 | 0.007320674 | 0.043667512 | 0.167645788 | 0.866861947 | 0.97021201 |
| Cct6b | 5.993991038 | 0.007333421 | 0.050362292 | 0.145613321 | 0.884226651 | NA |
| Srgap1 | 1164.257432 | 0.007336394 | 0.083784101 | 0.087563084 | 0.930223944 | 0.983937278 |
| Baiap2l1 | 33.82250138 | 0.007356385 | 0.078479045 | 0.093736938 | 0.925318127 | 0.983236501 |
| B3gnt8 | 13.88971931 | 0.007357303 | 0.072435897 | 0.101569848 | 0.919098114 | NA |
| Gzf1 | 1909.848379 | 0.007365403 | 0.054794129 | 0.134419571 | 0.893070807 | 0.977844987 |
| Cdk13 | 2274.56798 | 0.007366187 | 0.074118492 | 0.099383936 | 0.920833437 | 0.982762672 |
| Dbx2 | 565.1526707 | 0.007366268 | 0.071444041 | 0.103105417 | 0.917879305 | 0.982742917 |
| Lrrc15 | 1.192965478 | 0.007366515 | 0.024241181 | 0.303884334 | 0.761216013 | NA |
| Rtn4rl2 | 894.8026959 | 0.007377317 | 0.076258434 | 0.096741001 | 0.922932078 | 0.98296621 |
| Yif1a | 520.0652572 | 0.007392506 | 0.081575402 | 0.090621758 | 0.927793143 | 0.983501978 |
| Gm9705 | 0.479198558 | 0.007413413 | 0.015849836 | 0.467728085 | 0.639979055 | NA |
| Kcnmb4 | 1843.557377 | 0.007416636 | 0.067053701 | 0.110607412 | 0.911927669 | 0.981306862 |
| Zfp236 | 2148.315324 | 0.00741783 | 0.059190073 | 0.125322201 | 0.900268477 | 0.978986368 |
| Cd300lb | 2.725536848 | 0.007427612 | 0.033339599 | 0.222786495 | 0.823701679 | NA |
| 1700057H15Rik | 0.233836299 | 0.007433728 | 0.010168377 | 0.731063374 | 0.464740444 | NA |
| Gpr119 | 0.233836299 | 0.007433728 | 0.010168377 | 0.731063374 | 0.464740444 | NA |
| Gzmc | 0.233836299 | 0.007433728 | 0.010168377 | 0.731063374 | 0.464740444 | NA |
| Il1f8 | 0.233836299 | 0.007433728 | 0.010168377 | 0.731063374 | 0.464740444 | NA |
| Klrb1 | 0.233836299 | 0.007433728 | 0.010168377 | 0.731063374 | 0.464740444 | NA |
| Lmod2 | 0.233836299 | 0.007433728 | 0.010168377 | 0.731063374 | 0.464740444 | NA |
| 5530401A14Rik | 0.763355183 | 0.007438959 | 0.018228935 | 0.408085198 | 0.683211127 | NA |
| Btf3 | 3570.803107 | 0.007466997 | 0.060152167 | 0.124135132 | 0.901208282 | 0.979150581 |
| Hc | 10.69454291 | 0.007469388 | 0.062085269 | 0.120308542 | 0.904238738 | NA |
| 0610009B14Rik | 0.725539079 | 0.007470458 | 0.018074999 | 0.413303381 | 0.679384353 | NA |
| Pacrg | 417.2048372 | 0.007472835 | 0.074122027 | 0.100818004 | 0.919694935 | 0.982742917 |
| Wscd2 | 2396.084525 | 0.007476899 | 0.075630741 | 0.098860581 | 0.921248967 | 0.982762672 |
| Tnrc6b | 3555.091005 | 0.007478805 | 0.07796345 | 0.095927062 | 0.923578502 | 0.983210063 |
| 4931419H13Rik | 0.236453993 | 0.007494012 | 0.010209324 | 0.734036108 | 0.462926732 | NA |
| Gm4251 | 0.236453993 | 0.007494012 | 0.010209324 | 0.734036108 | 0.462926732 | NA |
| Rnu11 | 0.236453993 | 0.007494012 | 0.010209324 | 0.734036108 | 0.462926732 | NA |
| Kif12 | 0.46281165 | 0.007498411 | 0.015721198 | 0.476961785 | 0.633389334 | NA |
| Rhbdd1 | 389.8765516 | 0.007520044 | 0.080582128 | 0.093321487 | 0.925648163 | 0.983236501 |
| Ccdc121 | 25.92657945 | 0.007525101 | 0.079147934 | 0.095076405 | 0.924254139 | 0.983236501 |
| Wdr64 | 0.627996698 | 0.007548502 | 0.020153638 | 0.374547846 | 0.707996767 | NA |
| Fate1 | 3.345602043 | 0.007556179 | 0.043583714 | 0.173371621 | 0.86235933 | NA |
| Wdr96 | 64.28653905 | 0.007568484 | 0.086326498 | 0.087672784 | 0.930136751 | 0.983918326 |
| 9130019P16Rik | 7.618092065 | 0.007571221 | 0.05387756 | 0.140526428 | 0.888244073 | NA |
| Cck | 6296.987685 | 0.007572683 | 0.063025081 | 0.120153477 | 0.904361571 | 0.980118791 |
| Gapvd1 | 3888.919729 | 0.007600749 | 0.058350134 | 0.130261041 | 0.896359902 | 0.97803061 |
| Olfr690 | 1.72318098 | 0.00760414 | 0.031853317 | 0.238723644 | 0.811319881 | NA |
| Ryk | 1223.159318 | 0.007606354 | 0.05517914 | 0.137848352 | 0.89036027 | 0.977182599 |
| 2610016A17Rik | 0.241530977 | 0.007609484 | 0.010287283 | 0.739698176 | 0.459483155 | NA |
| Apoa5 | 0.241530977 | 0.007609484 | 0.010287283 | 0.739698176 | 0.459483155 | NA |
| Hoxb6 | 0.318445824 | 0.007617188 | 0.010286907 | 0.740474071 | 0.459012389 | NA |
| Slc7a13 | 0.318445824 | 0.007617188 | 0.010286907 | 0.740474071 | 0.459012389 | NA |
| Wbscr28 | 0.318445824 | 0.007617188 | 0.010286907 | 0.740474071 | 0.459012389 | NA |
| Metap1 | 1699.784029 | 0.007619137 | 0.059157182 | 0.128794793 | 0.897520026 | 0.978039389 |
| 4933407I05Rik | 3.219468406 | 0.007636467 | 0.03362392 | 0.227114115 | 0.820335009 | NA |
| Gm15408 | 9.047503379 | 0.007638671 | 0.05509069 | 0.138656295 | 0.889721756 | NA |
| Thbs1 | 107.9440572 | 0.007644589 | 0.085404352 | 0.089510527 | 0.928676188 | 0.983596265 |
| Bnc1 | 0.776462932 | 0.007647644 | 0.017943095 | 0.426216577 | 0.66995004 | NA |
| Snord15a | 0.435106034 | 0.007650782 | 0.015495196 | 0.493751846 | 0.621481445 | NA |
| 4921531P14Rik | 0.326298904 | 0.007651111 | 0.01030945 | 0.742145475 | 0.4579992 | NA |
| Cmtm2a | 0.326298904 | 0.007651111 | 0.01030945 | 0.742145475 | 0.4579992 | NA |
| Gm12633 | 0.326298904 | 0.007651111 | 0.01030945 | 0.742145475 | 0.4579992 | NA |
| Olfr281 | 0.326298904 | 0.007651111 | 0.01030945 | 0.742145475 | 0.4579992 | NA |
| Uroc1 | 0.326298904 | 0.007651111 | 0.01030945 | 0.742145475 | 0.4579992 | NA |
| Trim69 | 1.167652745 | 0.00765697 | 0.024777526 | 0.309028834 | 0.757299593 | NA |
| Atp6v0c | 2.952785157 | 0.00765883 | 0.040992193 | 0.186836314 | 0.851788974 | NA |
| Fgf3 | 19.48548822 | 0.007677766 | 0.081207259 | 0.094545316 | 0.924675987 | NA |
| Tex35 | 1.012063008 | 0.00768424 | 0.023417817 | 0.32813646 | 0.742808488 | NA |
| Atp6v0e | 615.6580127 | 0.007684309 | 0.074964818 | 0.102505532 | 0.918355422 | 0.982742917 |
| Ece2 | 1433.2077 | 0.007686031 | 0.086726886 | 0.08862339 | 0.929381219 | 0.983689248 |
| Mtpap | 1000.318225 | 0.007686237 | 0.06757547 | 0.113743011 | 0.909441516 | 0.980590355 |
| C1rb | 0.794369964 | 0.007687447 | 0.01663057 | 0.46224797 | 0.643903507 | NA |
| Tmem175 | 1518.97997 | 0.007691297 | 0.064402125 | 0.119426141 | 0.904937752 | 0.980118791 |
| Ercc6 | 1376.60222 | 0.007698657 | 0.070086369 | 0.109845283 | 0.912532078 | 0.981593673 |
| Stxbp4 | 1019.758238 | 0.007703322 | 0.064402692 | 0.119611797 | 0.904790675 | 0.980118791 |
| 4931431B13Rik | 0.341529857 | 0.007714019 | 0.010351116 | 0.745235492 | 0.45612937 | NA |
| Msln | 48.42781794 | 0.007722706 | 0.07998045 | 0.096557418 | 0.923077874 | 0.98296621 |
| 6720489N17Rik | 105.2280976 | 0.007722815 | 0.09446417 | 0.08175391 | 0.934842408 | 0.98537993 |
| Trap1a | 0.415500251 | 0.007731216 | 0.01532529 | 0.504474365 | 0.613928072 | NA |
| Dek | 1925.911067 | 0.007731627 | 0.072827243 | 0.106163935 | 0.915452285 | 0.982038599 |
| Tmem167 | 2140.395816 | 0.007731883 | 0.06571768 | 0.117653012 | 0.906342599 | 0.980118791 |
| Gnpda1 | 889.1186941 | 0.007731911 | 0.056008569 | 0.138048715 | 0.890201917 | 0.977182599 |
| Mrpl33 | 485.7496905 | 0.007735138 | 0.078259854 | 0.098839154 | 0.92126598 | 0.982762672 |
| 4930592A05Rik | 0.7778633 | 0.007737886 | 0.016062456 | 0.481737381 | 0.629992518 | NA |
| Cops7b | 883.3743325 | 0.007742258 | 0.061309411 | 0.126281725 | 0.899508922 | 0.97888996 |
| Nup214 | 2103.680493 | 0.007745647 | 0.048317126 | 0.16030853 | 0.87263804 | 0.97153269 |
| Leprot | 1059.170495 | 0.007749247 | 0.0539538 | 0.143627459 | 0.885794656 | 0.976107812 |
| Jakmip2 | 2492.858387 | 0.007764399 | 0.05336961 | 0.145483522 | 0.884329125 | 0.975676135 |
| Tyr | 3.876374708 | 0.007774271 | 0.042367657 | 0.183495406 | 0.854409323 | NA |
| Skint3 | 0.768228894 | 0.007792226 | 0.01825705 | 0.426806423 | 0.669520328 | NA |
| R3hcc1 | 737.7819684 | 0.007799994 | 0.076293872 | 0.102236175 | 0.918569215 | 0.982742917 |
| Nrep | 2687.971185 | 0.007808109 | 0.075805985 | 0.103001227 | 0.917961997 | 0.982742917 |
| Krt79 | 0.443603775 | 0.007811616 | 0.015566053 | 0.501836644 | 0.615782435 | NA |
| Cmtm3 | 350.6673158 | 0.007817811 | 0.08447934 | 0.092541104 | 0.926268135 | 0.983236501 |
| Henmt1 | 55.47743679 | 0.007821937 | 0.093999654 | 0.083212399 | 0.933682654 | 0.984586151 |
| Fam19a3 | 8.242350781 | 0.007823726 | 0.057757042 | 0.135459249 | 0.892248782 | NA |
| Chrm3 | 2096.367786 | 0.007832424 | 0.057292407 | 0.136709641 | 0.891260309 | 0.977385418 |

| | | | | | | | | | | | | | |
|---|---|---|---|---|---|---|---|---|---|---|---|---|---|
| Smtn | 320.1119419 | 0.103250563 | 0.107326162 | 0.962026045 | 0.336036521 | 0.57008101 | Dmrtb1 | 18.30721047 | 0.007836386 | 0.081310417 | 0.096376163 | 0.923221823 | NA |
| Snf8 | 1733.778739 | -0.042455545 | 0.057996896 | -0.73203133 | 0.464149445 | 0.680178899 | Kcna10 | 4.924735604 | 0.007840441 | 0.04611698 | 0.170012013 | 0.865000689 | NA |
| Slc41a1 | 3068.903233 | 0.051733333 | 0.070690564 | 0.731827979 | 0.464273569 | 0.680296452 | Gm16062 | 37.78092223 | 0.00784701 | 0.084457392 | 0.092910874 | 0.925974367 | 0.983236501 |
| Ppp1r13b | 3713.408321 | 0.103247166 | 0.103424328 | 0.998287034 | 0.318140193 | 0.551248507 | 4933400B14Rik | 0.516150229 | 0.007847584 | 0.014803592 | 0.530113477 | 0.596033255 | NA |
| Gm10433 | 12.8841283 | -0.111841828 | 0.152876373 | -0.731583473 | 0.464422839 | 0.680386485 | Nr2e3 | 0.516150229 | 0.007847584 | 0.014803592 | 0.530113477 | 0.596033255 | NA |
| Mkl1 | 2989.255276 | 0.10324546 | 0.101530474 | 1.016891343 | 0.309205122 | 0.54172842 | Cadm3 | 24273.37534 | 0.007858111 | 0.063936597 | 0.122904736 | 0.902182536 | 0.979150581 |
| Ankib1 | 2277.824565 | -0.052207134 | 0.071398724 | -0.73120541 | 0.464653696 | 0.680542649 | Prg4 | 161.8328851 | 0.007858443 | 0.059650578 | 0.131741272 | 0.89518894 | 0.977844987 |
| Cse1l | 2648.822125 | 0.048719927 | 0.066632248 | 0.731176393 | 0.464671418 | 0.680542649 | Mrpl9 | 1432.886224 | 0.007859572 | 0.056040908 | 0.140247041 | 0.888464806 | 0.977089223 |
| Trappc12 | 3000.529438 | 0.103227884 | 0.048940575 | 2.109249507 | 0.034923051 | 0.136796529 | Tfap2c | 5.078949204 | 0.007861233 | 0.051074612 | 0.153916653 | 0.877675453 | NA |
| Pabpc1 | 5359.08856 | 0.044606864 | 0.061021802 | 0.730998793 | 0.46477989 | 0.68058781 | Alb | 1.271591302 | 0.007871162 | 0.024499714 | 0.321275662 | 0.748001499 | NA |
| 5830415F09Rik | 100.8563872 | -0.091222234 | 0.124819481 | -0.730833306 | 0.464880977 | 0.680671505 | Cdc14a | 149.4671059 | 0.007883504 | 0.087016748 | 0.090597554 | 0.927812376 | 0.983501978 |
| Grip2 | 1430.857763 | 0.089209857 | 0.122095481 | 0.730656504 | 0.464988989 | 0.680694055 | Cd2 | 1.20151611 | 0.00788723 | 0.026597148 | 0.296544185 | 0.766814529 | NA |
| Immt | 5287.129796 | -0.041106375 | 0.056264447 | -0.730592365 | 0.465028177 | 0.680694055 | Slc12a4 | 991.6542787 | 0.007902469 | 0.077331981 | 0.102188881 | 0.918606753 | 0.982742917 |
| Vps45 | 1448.058916 | -0.047946261 | 0.065623976 | -0.730621095 | 0.465010623 | 0.680694055 | Fgd1 | 669.4661294 | 0.007907112 | 0.061369097 | 0.128845173 | 0.897480161 | 0.978039389 |
| Nicn1 | 4159.589224 | 0.040519226 | 0.055487332 | 0.730242818 | 0.465241774 | 0.680878063 | 6330403M23Rik | 1092.568648 | 0.007909661 | 0.079500096 | 0.099492473 | 0.920747264 | 0.982762672 |
| Tnrc6c | 3895.408319 | 0.10319807 | 0.069126956 | 1.492877401 | 0.135469282 | 0.327604347 | Rasal1 | 3960.031883 | 0.007909949 | 0.078059172 | 0.101332737 | 0.91928633 | 0.982742917 |
| Dhrs4 | 322.937256 | 0.057516431 | 0.078778691 | 0.730101383 | 0.465328216 | 0.680940251 | Nlrp4c | 0.617098604 | 0.007916525 | 0.013556327 | 0.583972741 | 0.559238652 | NA |
| Eef1d | 2105.631821 | 0.103196851 | 0.06382977 | 1.616751094 | 0.10593202 | 0.281226693 | 1700108F19Rik | 0.727270497 | 0.007919961 | 0.01808879 | 0.437838084 | 0.661503661 | NA |
| Elp3 | 3419.329212 | 0.042784055 | 0.058642446 | 0.729574873 | 0.465650086 | 0.68114179 | Zfp52 | 178.7123271 | 0.007940431 | 0.091752266 | 0.086542068 | 0.931035516 | 0.984208871 |
| Rnf13 | 3773.583413 | 0.057811096 | 0.079243706 | 0.729535493 | 0.465674165 | 0.68114179 | Gm3985 | 0.521227213 | 0.007940868 | 0.014841135 | 0.535057972 | 0.592609771 | NA |
| Rnf44 | 6588.656931 | -0.040808442 | 0.055939022 | -0.72951655 | 0.465685748 | 0.68114179 | Plac8l1 | 1.941924685 | 0.007950048 | 0.032467053 | 0.244865105 | 0.806560881 | NA |
| Zdhhc7 | 1305.510387 | -0.053237024 | 0.072955615 | -0.729717979 | 0.465562589 | 0.68114179 | Tpsb2 | 0.767280798 | 0.007950164 | 0.017918475 | 0.443685291 | 0.657270131 | NA |
| Rhbdl1 | 1714.675452 | -0.061271049 | 0.084014226 | -0.72929373 | 0.465822007 | 0.681148186 | A430092G05Rik | 0.426726963 | 0.007965777 | 0.015424583 | 0.516433844 | 0.605551433 | NA |
| Zfp341 | 462.1009314 | 0.056351689 | 0.077262937 | 0.729349562 | 0.465787862 | 0.681148186 | Il22 | 1.243066132 | 0.00796886 | 0.026648659 | 0.299034193 | 0.764913957 | NA |
| Zfpm1 | 1283.79896 | 0.072961232 | 0.100043325 | 0.729296358 | 0.465820399 | 0.681148186 | Timm8a2 | 0.4259239 | 0.007974079 | 0.015419736 | 0.517134634 | 0.605062178 | NA |
| Cdh15 | 78.20387337 | -0.108845964 | 0.149288768 | -0.729096808 | 0.465942447 | 0.681260005 | Arrb1 | 17918.51732 | 0.007979091 | 0.050306062 | 0.158610929 | 0.873975417 | 0.971912694 |
| Dlk2 | 925.6786684 | 0.090260243 | 0.123851126 | 0.728780155 | 0.466136153 | 0.681389166 | Zfp954 | 460.6428495 | 0.00798566 | 0.07811896 | 0.102224354 | 0.918578598 | 0.982742917 |
| Ccdc14 | 134.7202439 | 0.103188345 | 0.142848518 | 0.722362028 | 0.470071926 | 0.684906378 | 4930502E18Rik | 8.998824987 | 0.007986158 | 0.060118875 | 0.132839453 | 0.894320352 | NA |
| Tmem29 | 477.7873258 | -0.056812756 | 0.077960604 | -0.728736733 | 0.466162719 | 0.681389166 | Ulk2 | 5447.263166 | 0.008018975 | 0.05993212 | 0.13380095 | 0.893559976 | 0.977844987 |
| 1110051M20Rik | 2904.480626 | -0.0388002 | 0.053267995 | -0.728396108 | 0.466371145 | 0.681565225 | Kifc1 | 46.64749451 | 0.008023606 | 0.092766243 | 0.086492731 | 0.931074734 | 0.984208871 |
| Myo6 | 4803.10504 | -0.069764313 | 0.095773355 | -0.728431337 | 0.466349586 | 0.681565225 | Oxnad1 | 650.8212551 | 0.008039545 | 0.06282251 | 0.127972359 | 0.898170848 | 0.978273929 |
| Aig1 | 835.0992092 | -0.045030032 | 0.061866126 | -0.727862486 | 0.466697769 | 0.681613104 | Gm13212 | 6.238532706 | 0.008040558 | 0.040190636 | 0.200060474 | 0.841433286 | NA |
| Cdc16 | 1792.995564 | -0.035800389 | 0.04917973 | -0.727950101 | 0.466644132 | 0.681613104 | Hmga2 | 3.565496224 | 0.008040956 | 0.041960686 | 0.191630711 | 0.848031485 | NA |
| Dhx34 | 454.1688839 | 0.08413967 | 0.115572933 | 0.728022282 | 0.466599946 | 0.681613104 | 1300018J18Rik | 927.7246237 | 0.008046168 | 0.06233614 | 0.129077091 | 0.897296649 | 0.978039389 |
| Nol6 | 5653.385621 | 0.103159026 | 0.059682512 | 1.728463196 | 0.083905214 | 0.24166991 | Timd2 | 0.454027424 | 0.00804978 | 0.015654406 | 0.514218194 | 0.607099449 | NA |
| Lactb2 | 307.9668814 | 0.076263622 | 0.104732147 | 0.728177776 | 0.466504769 | 0.681613104 | 5430419D17Rik | 1.011123841 | 0.008059336 | 0.021908678 | 0.367860438 | 0.712977301 | NA |
| Tmem214 | 2208.099155 | 0.103150636 | 0.065619695 | 1.571946299 | 0.115963006 | 0.298460268 | Btla | 13.64167669 | 0.008068049 | 0.068125702 | 0.11842885 | 0.905727867 | NA |
| Tcf7l1 | 157.1007233 | 0.093602624 | 0.128603389 | 0.727839485 | 0.466711851 | 0.681613104 | Crip2 | 3881.458183 | 0.008076797 | 0.064599101 | 0.125029562 | 0.900500147 | 0.979053977 |
| Inpp5j | 2931.417406 | -0.091602945 | 0.125902789 | -0.727568828 | 0.466877568 | 0.681756583 | Amigo1 | 5808.236522 | 0.008077248 | 0.041825471 | 0.193117917 | 0.846866621 | 0.965275633 |
| Klc1 | 23672.97381 | 0.039734665 | 0.054619878 | 0.727476265 | 0.46693425 | 0.681756583 | Cdh16 | 0.383063074 | 0.008085644 | 0.010595513 | 0.763119678 | 0.445392025 | NA |
| Psmd12 | 2452.817284 | 0.060606443 | 0.083312011 | 0.72746345 | 0.466942097 | 0.681756583 | Cuta | 1314.525452 | 0.008091688 | 0.064285813 | 0.125870507 | 0.899834429 | 0.97888996 |
| Fam118b | 550.5490566 | -0.061677035 | 0.084798606 | -0.727335478 | 0.46702047 | 0.681806761 | Cyp7b1 | 420.1060753 | 0.00810052 | 0.07106096 | 0.113993954 | 0.909242586 | 0.980523965 |
| 1500009L16Rik | 358.4435041 | -0.067344787 | 0.09263563 | -0.726985791 | 0.46723466 | 0.681990939 | Reep3 | 1608.853786 | 0.008128013 | 0.063531506 | 0.127936723 | 0.898199049 | 0.978273929 |
| Strc | 17.33315745 | 0.103104234 | 0.16739349 | 0.615939329 | 0.537934563 | 0.737262992 | Bfsp1 | 31.90639337 | 0.008129287 | 0.082040867 | 0.099088268 | 0.921068187 | 0.982762672 |
| Polb | 1241.988059 | -0.04979657 | 0.068522612 | -0.726717333 | 0.467399133 | 0.682158898 | Pex12 | 300.6649488 | 0.008143282 | 0.08132187 | 0.100136429 | 0.920236014 | 0.982742917 |
| Zc3h13 | 2705.157401 | 0.037294999 | 0.05132427 | 0.726654254 | 0.467437784 | 0.682158898 | Cstl1 | 8.062861158 | 0.008189063 | 0.048681865 | 0.168215883 | 0.866413446 | NA |
| 2410004N09Rik | 101.5600217 | 0.103079469 | 0.141208688 | 0.729979652 | 0.465402622 | 0.680984817 | Prkcg | 12670.70839 | 0.008190158 | 0.065056784 | 0.125892456 | 0.899817054 | 0.97888996 |
| Dock10 | 2100.99105 | -0.08423246 | 0.116015112 | -0.726047316 | 0.467809765 | 0.682372648 | Sh3rf1 | 1514.720633 | 0.008195116 | 0.054982101 | 0.149050617 | 0.881513693 | 0.974279441 |
| Rbm8a | 1615.786224 | 0.044516832 | 0.061307572 | 0.726122902 | 0.46776343 | 0.682372648 | Hepacam2 | 17.986423 | 0.008203844 | 0.074965217 | 0.109435335 | 0.912857209 | NA |
| Thap1 | 288.0708085 | -0.081668496 | 0.112485799 | -0.726033838 | 0.467818027 | 0.682372648 | 2300005B03Rik | 0.382899838 | 0.008207275 | 0.010673784 | 0.768918921 | 0.441941444 | NA |
| Usp5 | 8651.93891 | -0.04236851 | 0.058360105 | -0.725984131 | 0.467848499 | 0.682372648 | Stau1 | 2871.69108 | 0.008211007 | 0.046757805 | 0.175607188 | 0.860602555 | 0.96862798 |
| Zfp711 | 300.478785 | -0.071162502 | 0.09802167 | -0.725987442 | 0.467846469 | 0.682372648 | Atp6v1b1 | 1.454548221 | 0.008212972 | 0.0294045 | 0.27931002 | 0.780006917 | NA |
| Tymp | 11.21297542 | -0.119265847 | 0.164492406 | -0.725053821 | 0.468419013 | 0.683140454 | Lpin1 | 1465.621247 | 0.008216264 | 0.070465128 | 0.116600423 | 0.907176703 | 0.980388092 |
| Park2 | 228.0408165 | 0.088954348 | 0.122716919 | 0.724874355 | 0.468529115 | 0.683236716 | Aim1l | 58.45100518 | 0.008225476 | 0.094996773 | 0.086586904 | 0.930999876 | 0.984208871 |
| Nek7 | 1648.913311 | 0.053136986 | 0.073321231 | 0.724714862 | 0.468626976 | 0.683315111 | Zfp574 | 1373.21872 | 0.008228913 | 0.067649046 | 0.121641224 | 0.903183164 | 0.979150581 |
| Lhx2 | 1566.516755 | -0.063930465 | 0.088229184 | -0.724595445 | 0.468700254 | 0.68335765 | Tbc1d1 | 1104.140064 | 0.008235774 | 0.072694691 | 0.113292649 | 0.909798545 | 0.980590355 |
| Lyzl4 | 14.53070523 | 0.10292519 | 0.16746813 | 0.614595685 | 0.538821765 | 0.737826286 | Il17c | 1.497785723 | 0.008260378 | 0.027485344 | 0.300537536 | 0.763767169 | NA |
| Dlat | 3606.037603 | -0.047748792 | 0.065931103 | -0.724222564 | 0.468929108 | 0.683562668 | Olfml1 | 574.790076 | 0.008267003 | 0.088626845 | 0.093278766 | 0.9256821 | 0.983236501 |
| Zscan20 | 270.9988051 | 0.102881159 | 0.111759597 | 0.920557712 | 0.357281389 | 0.588155316 | Rgs20 | 1474.378709 | 0.008276162 | 0.078076972 | 0.106000044 | 0.915582317 | 0.982038599 |
| Lins | 420.0686119 | -0.05970722 | 0.082579558 | -0.723026634 | 0.469663521 | 0.68450443 | Rbbp9 | 1198.542945 | 0.008281986 | 0.063345834 | 0.130742397 | 0.895979092 | 0.977899756 |
| Egln1 | 3715.397893 | -0.059583472 | 0.082465758 | -0.722523792 | 0.469972503 | 0.684890328 | Nup85 | 1270.562155 | 0.00828312 | 0.054712021 | 0.151394884 | 0.879664225 | 0.973413768 |
| Gjb1 | 446.1865323 | 0.102863419 | 0.153643049 | 0.669496079 | 0.503179081 | 0.70997253 | Rap1gap2 | 13380.93028 | 0.008294306 | 0.054182173 | 0.153081824 | 0.878333749 | 0.973035729 |
| Krr1 | 1590.232229 | 0.043731286 | 0.060535178 | 0.722411117 | 0.470041754 | 0.684906378 | Fetub | 4.099065369 | 0.008302698 | 0.037700231 | 0.220229366 | 0.825692527 | NA |
| Dnajc30 | 1132.198269 | -0.060017836 | 0.083095121 | -0.722278693 | 0.47012315 | 0.684916604 | Wdr85 | 890.5520037 | 0.008303501 | 0.057151596 | 0.145289057 | 0.884482654 | 0.975676135 |
| H2afx | 768.7425469 | -0.064265939 | 0.088992513 | -0.722149953 | 0.470202289 | 0.684967495 | Exoc3l4 | 79.95728357 | 0.00832306 | 0.09490929 | 0.08769489 | 0.930119181 | 0.983918326 |
| Fam173a | 1457.52927 | 0.10274437 | 0.086866217 | 1.182788576 | 0.236892949 | 0.461107959 | Cfhr2 | 0.725209114 | 0.008325867 | 0.018339658 | 0.453981602 | 0.649842069 | NA |
| Tab2 | 3032.939389 | 0.059756374 | 0.082801707 | 0.721680462 | 0.470490957 | 0.685260917 | Tcof1 | 1333.523405 | 0.008334114 | 0.060649719 | 0.137413894 | 0.890703649 | 0.977182599 |
| Tmcc1 | 3685.313977 | 0.055469553 | 0.076865526 | 0.721644103 | 0.470513317 | 0.685260917 | Gata3 | 8.562112109 | 0.008363513 | 0.034729304 | 0.240820069 | 0.809694575 | NA |
| Ppil2 | 2415.929794 | -0.035149189 | 0.048718387 | -0.721476854 | 0.470616176 | 0.685312691 | Rassf2 | 3369.776528 | 0.008364836 | 0.061652858 | 0.135676364 | 0.892077134 | 0.977690693 |
| B830017H08Rik | 84.04199776 | 0.102704983 | 0.137460085 | 0.747162225 | 0.454965643 | 0.673086764 | Atxn1l | 2520.162697 | 0.008366612 | 0.059379391 | 0.140900935 | 0.887948203 | 0.976924608 |
| Mterfd3 | 472.2392003 | -0.075013815 | 0.104044329 | -0.720979369 | 0.470922208 | 0.685615821 | Tbx21 | 6.367014925 | 0.008371383 | 0.051116596 | 0.163770352 | 0.869911932 | NA |
| Cebpg | 1279.096436 | 0.102691778 | 0.065582622 | 1.565838243 | 0.117386491 | 0.300771935 | Cldn2 | 72.39906469 | 0.008371427 | 0.026899615 | 0.311209928 | NA | NA |
| Apitd1 | 73.80665607 | -0.105737795 | 0.146697666 | -0.720787169 | 0.471040471 | 0.685672779 | Olfr877 | 0.444845289 | 0.008375254 | 0.015582252 | 0.53748674 | 0.590931442 | NA |
| Arid2 | 1986.007884 | -0.06297668 | 0.08744718 | -0.720168224 | 0.471421426 | 0.686115494 | Npy | 2767.435272 | 0.008378647 | 0.063373511 | 0.132210546 | 0.89481776 | 0.977844987 |
| Trmt10c | 416.7274191 | 0.057323339 | 0.07959926 | 0.720149146 | 0.471433171 | 0.686115494 | Vwa9 | 863.0864855 | 0.008383974 | 0.056867928 | 0.14742886 | 0.882793527 | 0.974871429 |
| Stx16 | 3368.826147 | 0.058306989 | 0.080995416 | 0.719880111 | 0.471598815 | 0.686292105 | F830045P16Rik | 1.441765289 | 0.008394374 | 0.025891817 | 0.324209545 | 0.745779396 | NA |
| Tfrc | 4034.068564 | 0.079625012 | 0.110636826 | 0.719697185 | 0.47171146 | 0.686391563 | Sh3yl1 | 290.9273296 | 0.00840106 | 0.078297284 | 0.10729695 | 0.914553404 | 0.98170867 |
| Dusp12 | 399.4322907 | 0.102665589 | 0.116375743 | 0.882190622 | 0.377673733 | 0.606792387 | Fbp2 | 1.191606103 | 0.008401619 | 0.026187809 | 0.320821755 | 0.748345472 | NA |
| Cd59b | 19.22995161 | -0.120901374 | 0.16812038 | -0.719135746 | 0.472057285 | 0.686650062 | Zim3 | 3.605831694 | 0.008451167 | 0.033691005 | 0.250843429 | 0.801935163 | NA |
| Mapt | 21616.63965 | -0.040552319 | 0.05638272 | -0.7192331 | 0.471997308 | 0.686650062 | Tshr | 20.99114888 | 0.00846706 | 0.082463876 | 0.102675989 | 0.918220131 | NA |
| Snx16 | 904.1328057 | -0.058741135 | 0.081684634 | -0.719120986 | 0.472066378 | 0.686650062 | Mark3 | 2814.894508 | 0.008478203 | 0.05594269 | 0.15551575 | 0.879540629 | 0.973413768 |
| Gemin2 | 182.5589399 | 0.0689541 | 0.095912781 | 0.718925039 | 0.472187108 | 0.686761199 | Lingo3 | 1955.788584 | 0.008484132 | 0.079432217 | 0.106809715 | 0.91493994 | 0.98170867 |
| Cdc26 | 589.3188005 | -0.05185574 | 0.072141746 | -0.7188035 | 0.472262001 | 0.686805655 | Mcm8 | 228.7785018 | 0.00850913 | 0.082266262 | 0.103434012 | 0.917618519 | 0.982742917 |
| Col18a1 | 256.657546 | 0.102625264 | 0.121273815 | 0.84622772 | 0.397425721 | 0.622746169 | Gnb3 | 10.72234035 | 0.008524645 | 0.057743883 | 0.14762854 | 0.88263593 | NA |
| Ostm1 | 2232.774623 | 0.053825168 | 0.074903684 | 0.718591733 | 0.472392509 | 0.68686651 | 5031410I06Rik | 2.236617333 | 0.008547142 | 0.036830533 | 0.232066752 | 0.816486168 | NA |
| 2300009A05Rik | 146.2069163 | 0.094152297 | 0.131111084 | 0.718110887 | 0.472688918 | 0.687213221 | 4932438A13Rik | 5373.247914 | 0.008557022 | 0.071599608 | 0.119512135 | 0.904869627 | 0.980118791 |
| Gale | 388.8984167 | 0.080758509 | 0.112467474 | 0.718061015 | 0.472719666 | 0.687213221 | Dupd1 | 2.199804773 | 0.008563339 | 0.035047064 | 0.244338288 | 0.806968832 | NA |
| Ip6k2 | 2476.663663 | 0.07219634 | 0.100632098 | 0.717428551 | 0.473109708 | 0.687592909 | Bace2 | 172.1222896 | 0.008572157 | 0.095097086 | 0.09014111 | 0.928175081 | 0.983501978 |
| Ptgs2 | 560.6470203 | -0.12051165 | 0.168000174 | -0.717330506 | 0.473170188 | 0.687592909 | Rassf7 | 43.50245613 | 0.008575165 | 0.090027972 | 0.095250011 | 0.924116248 | 0.983236501 |
| Clip1 | 4673.497504 | 0.102541485 | 0.091106108 | 1.125517128 | 0.260369963 | 0.487160031 | Msx2 | 13.81086082 | 0.008615976 | 0.065825245 | 0.130891666 | 0.895861008 | NA |
| Gm20743 | 38.55297641 | 0.102459361 | 0.164586712 | 0.622525111 | 0.533596637 | 0.734388578 | Kif1a | 67083.24583 | 0.008619855 | 0.053562616 | 0.160930429 | 0.872148195 | 0.97153269 |
| Ythdf3 | 2010.662846 | -0.045490972 | 0.063421518 | -0.71727978 | 0.473201481 | 0.687592909 | Syt8 | 1.193591353 | 0.008633123 | 0.022336315 | 0.386506137 | 0.699121858 | NA |
| Kcnc4 | 2745.638006 | 0.051485883 | 0.071803341 | 0.717040208 | 0.473349288 | 0.68774136 | Zfp963 | 161.8802626 | 0.008651763 | 0.087098677 | 0.09933289 | 0.920873965 | 0.982762672 |
| Peg3 | 10592.90237 | 0.081924489 | 0.114303101 | 0.716730238 | 0.473540564 | 0.687954759 | Fat3 | 2981.612239 | 0.00865395 | 0.083071646 | 0.104174535 | 0.917030842 | 0.982742917 |
| AI987944 | 282.1878581 | -0.073946081 | 0.103200717 | -0.716526813 | 0.473666117 | 0.687973655 | Zfp768 | 469.1098115 | 0.008661803 | 0.075960087 | 0.114030979 | 0.909213236 | 0.980523965 |
| Rims2 | 4866.566337 | 0.048810077 | 0.068129504 | 0.716430817 | 0.473725372 | 0.687973655 | Lnx1 | 1127.340581 | 0.008667072 | 0.070387692 | 0.12313335 | 0.902001503 | 0.979150581 |
| Slc15a2 | 1044.977058 | -0.049503713 | 0.069098595 | -0.716421405 | 0.473731182 | 0.687973655 | Olfr878 | 0.650179346 | 0.008671157 | 0.013817771 | 0.62753663 | 0.530307533 | NA |
| Uba5 | 1692.407454 | 0.037968933 | 0.052984196 | 0.716608647 | 0.473615607 | 0.687973655 | Olfr1507 | 2.123502352 | 0.008676733 | 0.02497936 | 0.347356102 | 0.728323808 | NA |
| Lrrc4b | 5068.034591 | -0.046636981 | 0.065132406 | -0.716033446 | 0.473970697 | 0.68825698 | B230319C09Rik | 15.10490354 | 0.008678559 | 0.071992087 | 0.120548788 | 0.904048434 | NA |
| Mdc1 | 1059.907789 | 0.102434736 | 0.093466529 | 1.095951002 | 0.273100218 | 0.501471293 | Rps3a1 | 10782.13131 | 0.008690448 | 0.062582959 | 0.138862852 | 0.889558527 | 0.977182599 |
| BC030336 | 1961.022127 | 0.051140469 | 0.071461876 | 0.715632893 | 0.474218057 | 0.688487123 | Snord71 | 2.224623015 | 0.008693449 | 0.03581007 | 0.242765468 | 0.808187088 | NA |
| Rab5c | 5226.872411 | -0.035298514 | 0.049340645 | -0.715404395 | 0.474359198 | 0.688627509 | Gm608 | 3227.970725 | 0.008706273 | 0.074025526 | 0.117611774 | 0.906375275 | 0.980118791 |
| Nabp2 | 1609.101212 | 0.060531375 | 0.084641233 | 0.715152335 | 0.474514918 | 0.688789033 | Mblac1 | 95.79886949 | 0.008712127 | 0.09308267 | 0.093595585 | 0.925430417 | 0.983236501 |
| Crb2 | 272.1248721 | 0.076689554 | 0.107255774 | 0.715015625 | 0.474599388 | 0.688847112 | 1700013H16Rik | 0.859859974 | 0.008712569 | 0.018592332 | 0.468610895 | 0.639347788 | NA |
| D3Ertd751e | 525.1926616 | 0.102338956 | 0.124337611 | 0.823073201 | 0.410466367 | 0.6344539 | Serpina1e | 0.515330361 | 0.008716785 | 0.014992042 | 0.581427443 | 0.560952406 | NA |
| Plk1s1 | 541.5822641 | -0.07519067 | 0.105219509 | -0.714607693 | 0.47485149 | 0.689083917 | Dtx2 | 289.2863771 | 0.008719997 | 0.086184718 | 0.101177992 | 0.919409167 | 0.982742917 |
| Phf20 | 3747.939206 | 0.102331872 | 0.078389824 | 1.305422906 | 0.191748883 | 0.405813029 | Clca3 | 1.376225769 | 0.008732599 | 0.019717934 | 0.442875971 | 0.657855449 | NA |
| Rbm34 | 692.3277105 | -0.053044479 | 0.074301038 | -0.713913033 | 0.475280958 | 0.689577971 | Pramef8 | 756.7395568 | 0.008733526 | 0.066516088 | 0.131299456 | 0.895538422 | 0.977844987 |
| Galnt10 | 565.4176672 | -0.065374953 | 0.091587708 | -0.713796145 | 0.475353244 | 0.689618273 | Hmg20a | 4135.277945 | 0.008738914 | 0.045017565 | 0.194122319 | 0.846080106 | 0.964996028 |
| Cdca2 | 13.31195699 | -0.117315753 | 0.164404742 | -0.713578891 | 0.475487615 | 0.689748628 | Cdh1 | 66.62953834 | 0.008746521 | 0.076823897 | 0.11385157 | 0.909355457 | 0.980572888 |
| Ccdc158 | 12.74728167 | -0.116014375 | 0.162599527 | -0.713497614 | 0.475537889 | 0.689756979 | Zfp97 | 58.34418934 | 0.00875001 | 0.094291185 | 0.09279775 | 0.926064238 | 0.983236501 |
| C130030K03Rik | 134.254635 | -0.107602224 | 0.15092261 | -0.712962916 | 0.475868705 | 0.689955989 | Ccdc174 | 858.9483871 | 0.008751137 | 0.068452324 | 0.12784281 | 0.89827337 | 0.978281525 |
| Lama4 | 411.0630674 | -0.076162468 | 0.10682618 | -0.712956952 | 0.475872395 | 0.689955989 | Bckdha | 1501.392653 | 0.008754562 | 0.07185693 | 0.121833225 | 0.9030311 | 0.979150581 |
| Lsm14b | 3829.813136 | -0.042172921 | 0.059189626 | -0.712505282 | 0.476151942 | 0.689955989 | Engase | 479.9984845 | 0.008773571 | 0.081853197 | 0.107186664 | 0.914640895 | 0.98170867 |
| Ica1 | 1696.847826 | 0.102245203 | 0.061064526 | 1.6743797 | 0.094056 | 0.25935285 | Ly6k | 1.231991385 | 0.008777328 | 0.026240301 | 0.334497982 | 0.738003814 | NA |
| Papd5 | 1534.384359 | 0.054410077 | 0.076374413 | 0.712412369 | 0.476209458 | 0.689955989 | Gsc2 | 0.754611645 | 0.008789005 | 0.017889249 | 0.491300966 | 0.623213597 | NA |
| Psat1 | 4389.474557 | -0.047235787 | 0.066265022 | -0.712831387 | 0.475950101 | 0.689955989 | Tdp1 | 609.7884405 | 0.008798705 | 0.070007958 | 0.125681499 | 0.899984047 | 0.97888996 |
| Sapcd2 | 45.01868081 | -0.115406939 | 0.161854864 | -0.713027316 | 0.475828854 | 0.689955989 | Lrrc38 | 101.4123467 | 0.008804483 | 0.094703703 | 0.092968728 | 0.925928405 | 0.983236501 |
| Slc43a2 | 3017.572934 | 0.046669979 | 0.065489065 | 0.712637735 | 0.476069955 | 0.689955989 | Cdk19 | 4792.631404 | 0.008809017 | 0.068481114 | 0.128634261 | 0.897647056 | 0.978039389 |
| Tmem2 | 634.9983919 | -0.079793489 | 0.111959827 | -0.712697504 | 0.476032961 | 0.689955989 | Smarca5-ps | 6.232461676 | 0.008815931 | 0.052731681 | 0.167184708 | 0.867224716 | NA |
| Tmem65 | 3643.430786 | 0.050415881 | 0.070733921 | 0.712753938 | 0.475998034 | 0.689955989 | Rab3d | 576.8000482 | 0.008817727 | 0.078453498 | 0.112394319 | 0.91051076 | 0.980653911 |
| Fancf | 70.59874138 | 0.102181794 | 0.148607493 | 0.687595167 | 0.491707753 | 0.702100612 | Armc8 | 3360.073184 | 0.008824769 | 0.055025323 | 0.160376513 | 0.87258449 | 0.97153269 |
| Wipi1 | 641.2028401 | 0.052365154 | 0.073502445 | 0.712427375 | 0.476200169 | 0.689955989 | Aym1 | 0.424594432 | 0.00882659 | 0.011066623 | 0.797586571 | 0.425110447 | NA |
| Darc | 1851.653767 | 0.102161572 | 0.080858896 | 1.263454942 | 0.20642573 | 0.424073224 | Zfp786 | 200.4972397 | 0.00882831 | 0.086907244 | 0.101583129 | 0.919087571 | 0.982742917 |
| Kcnj14 | 82.77219175 | -0.103599745 | 0.145508648 | -0.711983419 | 0.476475044 | 0.690211698 | Msh2 | 1437.400382 | 0.008857229 | 0.051469616 | 0.172086568 | 0.863369473 | 0.969788506 |
| Twist2 | 23.73766272 | -0.119608093 | 0.168015856 | -0.711885746 | 0.47653553 | 0.690234785 | Rgs3 | 1318.231437 | 0.008866277 | 0.070043868 | 0.126581773 | 0.899271424 | 0.978708077 |
| Ube2g2 | 1821.928755 | -0.068630167 | 0.096429599 | -0.711712664 | 0.476642724 | 0.690325515 | Rbm47 | 39.65496104 | 0.008878486 | 0.069001722 | 0.128670502 | 0.897618378 | 0.978039389 |
| Sac3d1 | 370.7179773 | 0.064471001 | 0.090693604 | 0.710866018 | 0.477167266 | 0.69102062 | Ucp1 | 2.143707167 | 0.008880283 | 0.030271085 | 0.293358592 | 0.769248072 | NA |
| Car13 | 64.19912326 | -0.115349012 | 0.162449018 | -0.710062848 | 0.477665164 | 0.691328116 | Pi15 | 9.752901974 | 0.008913145 | 0.064172361 | 0.138893829 | 0.889534048 | NA |
| Csnk1g3 | 2281.688109 | -0.056135588 | 0.079047251 | -0.710152313 | 0.477609689 | 0.691328116 | Lgi3 | 5428.266753 | 0.008916563 | 0.060584984 | 0.14717447 | 0.882994311 | 0.97494492 |
| Glmn | 424.3374238 | 0.066670834 | 0.093862552 | 0.710302805 | 0.477516381 | 0.691328116 | G6pc | 0.988647175 | 0.008918441 | 0.02107785 | 0.423119072 | 0.672208388 | NA |
| Ccdc121 | 18.33301211 | 0.102064876 | 0.163376046 | 0.624723628 | 0.532152463 | 0.732989631 | Cdc25c | 18.29826254 | 0.008921078 | 0.068681223 | 0.129891082 | 0.8966526 | NA |
| Ktn1 | 3390.570224 | 0.049522787 | 0.069751938 | 0.70998438 | 0.477713822 | 0.691328116 | Mia | 31.20169135 | 0.008941016 | 0.070444326 | 0.126923147 | 0.899001226 | 0.978633978 |
| Slc27a1 | 2519.537115 | 0.07720077 | 0.108693638 | 0.710260249 | 0.477542765 | 0.691328116 | Klrb1f | 0.538795351 | 0.008949191 | 0.014986482 | 0.597150872 | 0.550406656 | NA |
| Tmem132a | 9397.38563 | -0.075713894 | 0.106642327 | -0.709979761 | 0.477716687 | 0.691328116 | 2610307P16Rik | 4.58676699 | 0.008959841 | 0.046470642 | 0.192806475 | 0.847110532 | NA |
| Tspan14 | 1617.400829 | 0.102041859 | 0.081026414 | 1.25936535 | 0.207898399 | 0.426024579 | Pglyrp3 | 0.435065206 | 0.008983603 | 0.011163971 | 0.804695949 | 0.420995158 | NA |
| Fuca1 | 1083.766197 | 0.062386907 | 0.087893339 | 0.709802448 | 0.477826651 | 0.691393923 | Myo3a | 1.785826984 | 0.008985135 | 0.029152807 | 0.308208241 | 0.757923882 | NA |
| Kcnh3 | 6891.977575 | -0.054160363 | 0.076332228 | -0.709534688 | 0.477992734 | 0.691569653 | Mast2 | 6274.169317 | 0.008988245 | 0.04943752 | 0.181810188 | 0.855731689 | 0.967192539 |
| Dync1li2 | 10732.73287 | 0.102004893 | 0.04718036 | 2.162020238 | 0.030616616 | 0.125363069 | 1700109G14Rik | 0.76626644 | 0.008989742 | 0.0179365 | 0.501198226 | 0.616231622 | NA |
| Creld2 | 515.6167596 | -0.07910887 | 0.111533086 | -0.709286129 | 0.478146935 | 0.69166358 | Fam83g | 5.166910194 | 0.008996133 | 0.052117871 | 0.172611296 | 0.862956972 | NA |
| 1600020E01Rik | 169.7082311 | 0.101997638 | 0.118159353 | 0.863221027 | 0.388015955 | 0.615281875 | Slc38a4 | 46.00036715 | 0.009005751 | 0.089483022 | 0.100642007 | 0.919834649 | 0.982742917 |
| Hist1h2ac | 32.25540862 | -0.119030996 | 0.16795128 | -0.708723361 | 0.478496165 | 0.692032764 | Mrpl40 | 730.5285936 | 0.009026143 | 0.0681087 | 0.132525544 | 0.89456862 | 0.977844987 |
| Tada3 | 1206.407539 | 0.041124718 | 0.058031749 | 0.708658949 | 0.478536145 | 0.692032764 | 2010109A12Rik | 1.611207363 | 0.009034915 | 0.026053124 | 0.346788158 | 0.728750475 | NA |
| Xpnpep3 | 416.3586283 | -0.064177772 | 0.090572281 | -0.708580724 | 0.478584702 | 0.692038392 | Myh10 | 14808.47377 | 0.009035045 | 0.05316821 | 0.16993321 | 0.865062663 | 0.970182416 |
| Pcsk2 | 18273.46689 | -0.067614199 | 0.095473807 | -0.708196322 | 0.47882335 | 0.692318867 | Gkn3 | 108.2477305 | 0.009035608 | 0.091809592 | 0.098416817 | 0.921601322 | 0.982762672 |
| Kit | 1756.975653 | -0.037005736 | 0.052263377 | -0.708062478 | 0.478906459 | 0.692349819 | Ptk2 | 5983.627454 | 0.009046562 | 0.073162255 | 0.123650673 | 0.901591869 | 0.979150581 |
| Tradd | 157.1876433 | 0.077675068 | 0.109707771 | 0.708017924 | 0.478934126 | 0.692349819 | Rnf170 | 1665.722527 | 0.009051128 | 0.052683585 | 0.171801675 | 0.863593449 | 0.969836459 |
| Plekhb1 | 9509.201287 | 0.063923067 | 0.090312997 | 0.707794771 | 0.479072714 | 0.692473415 | Ube3b | 6738.534981 | 0.009053607 | 0.047951341 | 0.188808216 | 0.850243132 | 0.966260848 |
| Tac2 | 322.0488343 | 0.091156004 | 0.128799385 | 0.707736333 | 0.47910901 | 0.692473415 | 2300002M23Rik | 1.063647385 | 0.009082249 | 0.021230031 | 0.427801958 | 0.668795311 | NA |
| Zfp664 | 4553.089956 | 0.044048798 | 0.062254606 | 0.707558861 | 0.479219248 | 0.69256814 | Fam101b | 569.9483183 | 0.009084913 | 0.070614887 | 0.12865436 | 0.89763115 | 0.978039389 |
| Catsper3 | 8.428020913 | 0.099828967 | 0.141133844 | 0.707335423 | 0.479358057 | 0.692574948 | Nf2 | 4734.377592 | 0.009109088 | 0.040177458 | 0.226721364 | 0.820640414 | 0.957571905 |
| Cnnm4 | 597.8059973 | -0.059675592 | 0.084362673 | -0.707369628 | 0.479336806 | 0.692574948 | Slfn4 | 0.97778493 | 0.009111403 | 0.023344793 | 0.390297004 | 0.696316938 | NA |
| Mtmr6 | 6202.629554 | -0.049642609 | 0.070178453 | -0.707376791 | 0.479332356 | 0.692574948 | Vps13d | 5244.282046 | 0.009114382 | 0.071757336 | 0.127016725 | 0.898927161 | 0.978626692 |
| Bpnt1 | 1892.304297 | -0.050712623 | 0.071761695 | -0.706680954 | 0.479764768 | 0.692981569 | Gemin5 | 1303.06476 | 0.009139039 | 0.064214938 | 0.142319517 | 0.886827631 | 0.976505319 |
| Clpx | 1179.969015 | -0.060961967 | 0.08626694 | -0.706666627 | 0.479773673 | 0.692981569 | 9230112J17Rik | 37.72316541 | 0.009145876 | 0.090651474 | 0.100890536 | 0.919637356 | 0.982742917 |
| Dhdds | 2893.595931 | 0.045489735 | 0.064366396 | 0.706731119 | 0.479733587 | 0.692981569 | Ung | 77.58844354 | 0.009150932 | 0.094018251 | 0.097331444 | 0.922463186 | 0.98296621 |
| Cdh9 | 719.6300131 | -0.063904928 | 0.090460398 | -0.706440934 | 0.479913973 | 0.693055 | Atp1a2 | 55752.70747 | 0.009176472 | 0.057164898 | 0.160526341 | 0.872466474 | 0.97153269 |
| Rchy1 | 1179.818977 | -0.044490008 | 0.062975287 | -0.706467719 | 0.479897321 | 0.693055 | Ccdc73 | 1.47116782 | 0.009191803 | 0.025596129 | 0.359109117 | 0.719513461 | NA |
| Eif3h | 4053.105411 | -0.044186516 | 0.062604796 | -0.705800818 | 0.480312013 | 0.693397062 | Gja3 | 4.366468414 | 0.009194554 | 0.040069079 | 0.229467571 | 0.818505523 | NA |
| Kcnip1 | 1054.569209 | 0.076238188 | 0.108019298 | 0.705783031 | 0.480323077 | 0.693397062 | Fmnl1 | 5825.951168 | 0.009201342 | 0.090400746 | 0.10178392 | 0.918928189 | 0.982742917 |
| Map6 | 5059.765005 | -0.037078283 | 0.052538592 | -0.705734229 | 0.48035343 | 0.693397062 | Mir138-2 | 7.804399111 | 0.009208706 | 0.058904858 | 0.156331859 | 0.875771445 | NA |
| Tmem150b | 12.82447305 | 0.084361304 | 0.119498655 | 0.705960282 | 0.480212838 | 0.693397062 | Cadm2 | 6497.704025 | 0.009212843 | 0.079684481 | 0.115616527 | 0.907956464 | 0.980520081 |
| Zfand6 | 1080.919527 | -0.061682762 | 0.087406468 | -0.705700193 | 0.480374601 | 0.693397062 | Dnajc8 | 3113.662759 | 0.009219959 | 0.058947529 | 0.156409588 | 0.87571018 | 0.972151437 |
| 1300002E11Rik | 259.1585931 | -0.07089978 | 0.100491036 | -0.705533378 | 0.480478368 | 0.693475978 | Ust | 969.0802248 | 0.009228447 | 0.070891999 | 0.130176148 | 0.896427065 | 0.97803061 |
| Hnrnph3 | 1421.159744 | 0.101986743 | 0.080857996 | 1.261306833 | 0.20719832 | 0.425234895 | Ap2a2 | 10859.00859 | 0.009236763 | 0.053980738 | 0.1711122 | 0.864135543 | 0.969899003 |
| Lgi1 | 6475.521042 | -0.045120347 | 0.06398726 | -0.705145783 | 0.480719517 | 0.693670584 | Atpaf2 | 814.5768439 | 0.009267623 | 0.057216789 | 0.161973833 | 0.87132646 | 0.97153269 |
| Zfp930 | 401.8907281 | -0.078235825 | 0.11095584 | -0.705107767 | 0.480743173 | 0.693670584 | Prrxl1 | 0.962850976 | 0.009282282 | 0.022961647 | 0.404251589 | 0.686027721 | NA |
| Otud7a | 1776.685688 | 0.060068642 | 0.08522696 | 0.704807989 | 0.480929736 | 0.693875159 | Klhl22 | 3640.274215 | 0.009284898 | 0.054207694 | 0.171283774 | 0.864000638 | 0.969836459 |
| Antxr2 | 73.72692293 | 0.097955184 | 0.139066271 | 0.70437773 | 0.481197571 | 0.694196943 | 4921530D09Rik | 0.297598527 | 0.009286915 | 0.011356721 | 0.817746142 | 0.413502157 | NA |
| 1190005I06Rik | 151.472035 | 0.092081462 | 0.130751293 | 0.704248955 | 0.481277749 | 0.69424797 | Ptf1a | 0.297598527 | 0.009286915 | 0.011356721 | 0.817746142 | 0.413502157 | NA |
| Aplf | 273.6885648 | 0.062610636 | 0.08893046 | 0.704040388 | 0.481407622 | 0.694370666 | Pzp | 0.297598527 | 0.009286915 | 0.011356721 | 0.817746142 | 0.413502157 | NA |
| Mcmbp | 1005.053302 | 0.054622661 | 0.077605196 | 0.703853143 | 0.481524235 | 0.694413582 | Xkr9 | 0.297598527 | 0.009286915 | 0.011356721 | 0.817746142 | 0.413502157 | NA |
| Reep2 | 4828.411119 | 0.051287785 | 0.07288527 | 0.703678327 | 0.48163312 | 0.694413582 | Brsk2 | 9400.409769 | 0.009290225 | 0.045229015 | 0.205404099 | 0.837256418 | 0.961831963 |
| Stim1 | 3813.326368 | 0.041985817 | 0.059652206 | 0.703843503 | 0.481530239 | 0.694413582 | Gm11468 | 1.224102218 | 0.009292407 | 0.026631189 | 0.348929473 | 0.727142258 | NA |
| Vapb | 6504.891629 | -0.034653772 | 0.049242104 | -0.703742722 | 0.481593009 | 0.694413582 | Tmem223 | 784.9414655 | 0.00929992 | 0.080119897 | 0.116075039 | 0.907593071 | 0.980492113 |
| Mib2 | 3720.793943 | 0.101961977 | 0.077596897 | 1.313995541 | 0.188847705 | 0.402034396 | Ugt2a1 | 1.387563536 | 0.009312046 | 0.020803057 | 0.447628713 | 0.654421177 | NA |
| Gemin4 | 437.4848065 | 0.068324306 | 0.097173784 | 0.703114602 | 0.481984333 | 0.694749768 | Cnnm1 | 6011.652752 | 0.009313255 | 0.042110254 | 0.221163586 | 0.824965061 | 0.957903916 |
| Pak1ip1 | 1443.290302 | -0.042326044 | 0.060196169 | -0.703135184 | 0.481971507 | 0.694749768 | Fign | 162.4140934 | 0.009319645 | 0.090843943 | 0.102589613 | 0.918288687 | 0.982742917 |
| Aacs | 1640.434787 | -0.057814811 | 0.082261171 | -0.702820172 | 0.482167824 | 0.694820338 | Ckap2l | 33.46441539 | 0.009322954 | 0.085952257 | 0.108466652 | 0.913625532 | 0.98170867 |
| Foxc1 | 167.2301909 | -0.094577921 | 0.134543966 | -0.702951785 | 0.482085797 | 0.694820338 | Eif2s3y | 0.300216221 | 0.009343858 | 0.011391267 | 0.82026504 | 0.412065031 | NA |
| Rtca | 1769.877754 | 0.072777431 | 0.103541631 | 0.702880862 | 0.482129999 | 0.694820338 | Irx4 | 0.300216221 | 0.009343858 | 0.011391267 | 0.82026504 | 0.412065031 | NA |
| Zmym3 | 6315.213497 | 0.06587453 | 0.093748289 | 0.702674479 | 0.482258636 | 0.694886572 | Mir1956 | 0.300216221 | 0.009343858 | 0.011391267 | 0.82026504 | 0.412065031 | NA |
| Akirin1 | 2001.685598 | -0.044917674 | 0.06395737 | -0.702306456 | 0.482488068 | 0.694990786 | Ccdc172 | 1.646965382 | 0.009347868 | 0.028317237 | 0.330112291 | 0.741315117 | NA |
| Creb3 | 1435.465168 | 0.033927909 | 0.048317568 | 0.702185787 | 0.482563308 | 0.694990786 | Gm2447 | 0.587881648 | 0.009349919 | 0.015479875 | 0.604004852 | 0.545840416 | NA |
| Csnk1a1 | 6427.778591 | -0.040858372 | 0.058198261 | -0.702054862 | 0.48264495 | 0.694990786 | Trp53i13 | 140.8800767 | 0.009357369 | 0.095047086 | 0.098449822 | 0.921575115 | 0.982762672 |
| Ppm1d | 547.7392764 | 0.069359584 | 0.098741798 | 0.702433879 | 0.482408624 | 0.694990786 | Fdx1l | 25.54956931 | 0.009359543 | 0.072183277 | 0.129663599 | 0.896832584 | 0.978039389 |
| Sh2b3 | 653.7765032 | 0.058749826 | 0.083674983 | 0.702119361 | 0.482604729 | 0.694990786 | Man1b1 | 4020.703192 | 0.009360557 | 0.039083804 | 0.239499633 | 0.810718181 | 0.956073263 |
| Thumpd3 | 806.4924676 | -0.054572024 | 0.077717467 | -0.702184797 | 0.482563926 | 0.694990786 | Pip5k1c | 22788.76989 | 0.009360798 | 0.058980368 | 0.158710403 | 0.873897041 | 0.971912694 |
| Zc3h12b | 484.9744567 | -0.079882287 | 0.113756612 | -0.70222104 | 0.482541326 | 0.694990786 | A330032B11Rik | 10.25026493 | 0.009364235 | 0.065706611 | 0.142515879 | 0.886672538 | NA |
| 4931440F15Rik | 24.0031026 | -0.117951197 | 0.168059145 | -0.701843373 | 0.482776847 | 0.695105921 | Gm4827 | 1.402403799 | 0.009373086 | 0.029388142 | 0.318941094 | 0.749771182 | NA |
| Ercc3 | 1259.498238 | 0.063714091 | 0.09079821 | 0.701710865 | 0.482859496 | 0.695105921 | Rmnd5b | 2019.534382 | 0.009376656 | 0.06612711 | 0.141797452 | 0.887239997 | 0.97661227 |
| Nbeal2 | 670.8633228 | 0.084120824 | 0.119867452 | 0.701782027 | 0.482815109 | 0.695105921 | Akr7a5 | 677.3000956 | 0.009380711 | 0.068045781 | 0.137858815 | 0.890352001 | 0.977182599 |
| 4930444A02Rik | 1420.942895 | -0.046749129 | 0.066678531 | -0.701112161 | 0.483233022 | 0.69544979 | Eif4b | 9544.972267 | 0.009388764 | 0.051864464 | 0.181024999 | 0.856347954 | 0.967598001 |
| Pgr15l | 17.05242208 | -0.102982754 | 0.146865479 | -0.701204633 | 0.483175319 | 0.69544979 | Foxn4 | 0.551567013 | 0.009395243 | 0.015255111 | 0.615875079 | 0.537976971 | NA |
| Pik3c2a | 1028.897237 | -0.05259154 | 0.075005773 | -0.701166556 | 0.483199079 | 0.69544979 | 4921515E04Rik | 0.749695855 | 0.009400388 | 0.019766315 | 0.475576138 | 0.634376379 | NA |
| Daglb | 1608.36277 | 0.054273313 | 0.077434744 | 0.700890974 | 0.483371058 | 0.695583837 | Agtpbp1 | 8496.356879 | 0.009408327 | 0.060732557 | 0.15491406 | 0.876889069 | 0.972549751 |
| Fastk | 2643.718488 | 0.051343948 | 0.073319793 | 0.700274047 | 0.483756176 | 0.69596223 | Pnma3 | 1182.28157 | 0.009409418 | 0.086724517 | 0.108497783 | 0.913600839 | 0.98170867 |
| Smtnl2 | 45.80297574 | 0.101950226 | 0.16664359 | 0.611786063 | 0.540679312 | 0.739475705 | E130218I03Rik | 0.526995524 | 0.009423773 | 0.015094121 | 0.624334017 | 0.532408248 | NA |
| Hnrnpk | 12322.00153 | -0.038748741 | 0.055335274 | -0.700253889 | 0.483768763 | 0.69596223 | Tmem219 | 371.9407676 | 0.009438764 | 0.082101885 | 0.114964042 | 0.908473624 | 0.980523965 |
| Vps37c | 843.2561517 | 0.061424213 | 0.087745904 | 0.700023708 | 0.483912499 | 0.696039756 | Fmr1 | 1989.725751 | 0.009451458 | 0.076704124 | 0.123219686 | 0.901933137 | 0.979150581 |
| Zswim4 | 902.2578683 | -0.084714351 | 0.121005288 | -0.700088006 | 0.483872346 | 0.696039756 | Smarcd2 | 663.4778402 | 0.009457863 | 0.070189925 | 0.134746733 | 0.892812122 | 0.977844987 |
| Gadd45b | 404.817298 | -0.09219148 | 0.131802381 | -0.69946749 | 0.484259923 | 0.696474821 | Ppp1r14c | 409.9627382 | 0.0094748 | 0.076540961 | 0.12378732 | 0.901483672 | 0.979150581 |
| Bcl10 | 458.7087436 | -0.057500963 | 0.082254637 | -0.699060443 | 0.484514258 | 0.696517343 | Slc16a8 | 20.07484796 | 0.009475019 | 0.023380498 | 0.405253083 | NA | NA |
| Dab2ip | 6414.175407 | 0.062690208 | 0.08967392 | 0.699090748 | 0.48449532 | 0.696517343 | Krt6a | 0.54329922 | 0.009491492 | 0.015032275 | 0.631407564 | 0.527774072 | NA |
| Naf1 | 391.4548438 | -0.089807015 | 0.128427459 | -0.699282032 | 0.484375794 | 0.696517343 | Hmx2 | 1.815946383 | 0.009494477 | 0.011463217 | 0.828255908 | 0.407525587 | NA |
| Pef1 | 2440.609711 | -0.049438666 | 0.070714605 | -0.699129485 | 0.484471113 | 0.696517343 | Dpf1 | 1987.90373 | 0.009510729 | 0.080920283 | 0.117532077 | 0.906438427 | 0.980118791 |
| Upf2 | 2512.143887 | 0.063583264 | 0.090941814 | 0.699164232 | 0.484449401 | 0.696517343 | Otud7a | 1901.645031 | 0.009512281 | 0.061385464 | 0.154959837 | 0.87685298 | 0.972549751 |
| Epha2 | 74.17867061 | -0.108005771 | 0.154520012 | -0.698975939 | 0.484567068 | 0.696528634 | Ptgr2 | 1948.812509 | 0.009514134 | 0.053290613 | 0.178533011 | 0.858304399 | 0.967971163 |
| D17H6S53E | 556.4153014 | -0.055632623 | 0.079602002 | -0.698884726 | 0.484624073 | 0.696545955 | Hapln4 | 6144.371188 | 0.009527091 | 0.056553852 | 0.168460506 | 0.866221012 | 0.97021201 |
| N4bp1 | 2970.323635 | 0.065103133 | 0.093189346 | 0.698611329 | 0.484794961 | 0.69666232 | B4galt4 | 823.5341468 | 0.009530872 | 0.057927262 | 0.164531714 | 0.869312583 | 0.970833749 |
| Sik3 | 6740.653434 | -0.068955878 | 0.098701557 | -0.698630092 | 0.484783233 | 0.69666232 | Taf8 | 722.8615797 | 0.009542711 | 0.072160004 | 0.132243773 | 0.894791479 | 0.977844987 |
| Efcab2 | 416.6612204 | 0.101926046 | 0.098777455 | 1.031875612 | 0.302130392 | 0.534018742 | Foxn3 | 569.6087511 | 0.009547444 | 0.076695083 | 0.124485738 | 0.900930692 | 0.979150581 |
| Ccnt1 | 703.144811 | 0.068234163 | 0.097769573 | 0.697907957 | 0.485234758 | 0.697149058 | Pxk | 1636.585095 | 0.009564743 | 0.046679434 | 0.204902714 | 0.837648135 | 0.961858836 |
| Pkn1 | 1638.706657 | 0.101884524 | 0.064652082 | 1.575889283 | 0.115051321 | 0.29712047 | Gja5 | 19.88664392 | 0.009572995 | 0.074344704 | 0.128764985 | 0.897543613 | NA |
| Npas1 | 284.2592587 | 0.07976764 | 0.114349625 | 0.697576755 | 0.485441923 | 0.697149058 | 3830403N18Rik | 3.073693115 | 0.009575343 | 0.036050011 | 0.265612753 | 0.790537463 | NA |
| Ppif | 931.9832712 | 0.042582346 | 0.061038101 | 0.697635495 | 0.485405178 | 0.697149058 | Gm4975 | 1.006182018 | 0.009586606 | 0.021578614 | 0.444264197 | 0.656851583 | NA |
| Ranbp3 | 3018.122364 | 0.050144317 | 0.071884526 | 0.697567611 | 0.485447643 | 0.697149058 | Cav2 | 1089.292713 | 0.009602426 | 0.078752094 | 0.121932328 | 0.902952613 | 0.979150581 |
| Snai1 | 21.82421618 | -0.117208414 | 0.16797509 | -0.697772596 | 0.48531942 | 0.697149058 | Hist1h3g | 0.603271003 | 0.009617147 | 0.015576612 | 0.617409403 | 0.536964721 | NA |
| Mapk10 | 17337.19708 | 0.101869732 | 0.071168756 | 1.431382774 | 0.152320543 | 0.353735457 | 4930468A15Rik | 0.313257562 | 0.00962481 | 0.011560138 | 0.832586027 | 0.405078242 | NA |
| St6galnac3 | 318.4121481 | 0.080480323 | 0.115427831 | 0.697234991 | 0.485655744 | 0.697252604 | Olfr560 | 0.313257562 | 0.00962481 | 0.011560138 | 0.832586027 | 0.405078242 | NA |
| Ube2d3 | 4246.032617 | -0.063907349 | 0.091654293 | -0.6972652 | 0.485636843 | 0.697252604 | Skint5 | 0.313257562 | 0.00962481 | 0.011560138 | 0.832586027 | 0.405078242 | NA |
| Mis18a | 168.1543223 | -0.074169279 | 0.106417466 | -0.696965278 | 0.485824524 | 0.697365682 | Bcar3 | 634.2313261 | 0.009625732 | 0.077546409 | 0.12412866 | 0.901213406 | 0.979150581 |
| Napg | 6044.882863 | -0.043624666 | 0.062591771 | -0.696971269 | 0.485820775 | 0.697365682 | Gpr62 | 389.3153734 | 0.009638654 | 0.088647573 | 0.108730041 | 0.913416614 | 0.98170867 |
| Fads3 | 1236.608619 | 0.057728777 | 0.082854077 | 0.696752398 | 0.485957761 | 0.697492316 | Ppcs | 186.2137886 | 0.009649568 | 0.08773808 | 0.109981524 | 0.912424028 | 0.981550131 |
| Eci2 | 1086.257467 | 0.050724468 | 0.072812347 | 0.696646517 | 0.486024037 | 0.697522826 | Kcnk4 | 599.8267932 | 0.009669677 | 0.077832095 | 0.124237654 | 0.901127109 | 0.979150581 |
| Enpp6 | 230.9769829 | -0.104901648 | 0.150628848 | -0.696424687 | 0.486162907 | 0.697592895 | Usp38 | 1338.183907 | 0.009672355 | 0.069881986 | 0.138409846 | 0.889916516 | 0.977182599 |
| Tnfaip3 | 307.4276896 | -0.082494378 | 0.118446115 | -0.696471789 | 0.486133419 | 0.697592895 | Gjb3 | 3.431121622 | 0.009673946 | 0.036895206 | 0.26220061 | 0.793166783 | NA |
| Atp13a3 | 2790.094682 | -0.064770141 | 0.09304768 | -0.696096245 | 0.486368558 | 0.697779778 | 1700029H14Rik | 0.761934134 | 0.009675331 | 0.01981617 | 0.48825433 | 0.625369706 | NA |
| Rnf141 | 837.0160583 | -0.059977527 | 0.086165591 | -0.696072833 | 0.486383219 | 0.697779778 | A930001A20Rik | 5.182807737 | 0.009695238 | 0.050579981 | 0.191681337 | 0.847991826 | NA |
| Gpbp1 | 3300.873221 | -0.05100778 | 0.073308753 | -0.695793853 | 0.486557939 | 0.697836592 | Tmprss11a | 12.47182721 | 0.009701584 | 0.024998313 | 0.388089536 | NA | NA |
| Pde9a | 627.0343803 | 0.067054014 | 0.096357776 | 0.695885855 | 0.486500316 | 0.697836592 | Npw | 2.562940129 | 0.009739901 | 0.033424345 | 0.291401406 | 0.77074434 | NA |
| Tas1r3 | 12.49878659 | -0.109900186 | 0.157948378 | -0.695798133 | 0.486555259 | 0.697836592 | Xpo5 | 2303.253858 | 0.009747183 | 0.048074076 | 0.202753405 | 0.839327779 | 0.96231236 |
| Lyrm7 | 393.5479695 | -0.089515291 | 0.128672785 | -0.695681619 | 0.486628239 | 0.69787282 | Thada | 625.2692457 | 0.009747701 | 0.072188837 | 0.135030585 | 0.892587692 | 0.977844987 |
| Exoc5 | 1869.578726 | -0.067715709 | 0.097463149 | -0.694782695 | 0.487191495 | 0.698562527 | Ifng | 0.751621763 | 0.009755221 | 0.020053804 | 0.486452406 | 0.626646442 | NA |
| Notch4 | 328.8568353 | -0.100929053 | 0.14528474 | -0.694698238 | 0.487244432 | 0.698562527 | Jmjd6 | 821.3639332 | 0.009768072 | 0.077799953 | 0.125553696 | 0.900085218 | 0.97888996 |

| | | | | | | |
|---|---|---|---|---|---|---|
| Rell1 | 513.9085803 | 0.080469465 | 0.115831296 | 0.694712641 | 0.487235405 | 0.698562527 |
| Fbxl2 | 1952.357578 | 0.060646918 | 0.087354716 | 0.694260376 | 0.487518936 | 0.698826754 |
| Glrb | 7288.754177 | -0.058509877 | 0.084273369 | -0.694286675 | 0.487502446 | 0.698826754 |
| AK129341 | 687.0759947 | 0.064817492 | 0.093400547 | 0.693973372 | 0.487698908 | 0.698856463 |
| Atp13a2 | 5925.12541 | 0.047645333 | 0.068671286 | 0.693817386 | 0.487796738 | 0.698856463 |
| BC024978 | 276.3213452 | 0.06492577 | 0.093561544 | 0.693936493 | 0.487722037 | 0.698856463 |
| Slitrk3 | 2429.371237 | 0.044089803 | 0.063539899 | 0.693891613 | 0.487750184 | 0.698856463 |
| Tsfm | 423.6609091 | 0.056486028 | 0.081415929 | 0.693795778 | 0.487810291 | 0.698856463 |
| Usp21 | 1565.290165 | -0.04647983 | 0.066992702 | -0.693804384 | 0.487804893 | 0.698856463 |
| Dck | 935.9648853 | 0.050999497 | 0.073531545 | 0.693573034 | 0.48795001 | 0.698991998 |
| Ppib | 2751.892343 | 0.04420312 | 0.063742938 | 0.69345909 | 0.488021491 | 0.699029766 |
| Rragb | 2309.045396 | -0.049595229 | 0.071548161 | -0.693172656 | 0.488201206 | 0.699222544 |
| Tmem199 | 1258.391153 | 0.053003911 | 0.076478879 | 0.693052928 | 0.488276336 | 0.69926551 |
| Rnf145 | 4630.530505 | 0.046978566 | 0.067806398 | 0.692833826 | 0.488413841 | 0.699364482 |
| Tnfaip8l3 | 320.5580317 | -0.072750017 | 0.105008841 | -0.692798966 | 0.488435721 | 0.699364482 |
| 5730507C01Rik | 39.0862782 | -0.11520335 | 0.166377183 | -0.692422774 | 0.488671868 | 0.699637952 |
| Dsn1 | 129.2233824 | 0.101847916 | 0.136925263 | 0.743821217 | 0.456984642 | 0.675299957 |
| Crem | 497.1539186 | -0.082040803 | 0.118564366 | -0.6919516 | 0.488967725 | 0.699932181 |
| Prdx2 | 6338.249413 | 0.035806443 | 0.051783127 | 0.691469305 | 0.489270664 | 0.700301124 |
| D030028A08Rik | 252.3541778 | -0.074210178 | 0.107343975 | -0.691330631 | 0.489357786 | 0.700319482 |
| Gtf2h4 | 468.2559844 | 0.063502997 | 0.091859594 | 0.691305002 | 0.489373889 | 0.700319482 |
| B4galnt3 | 32.25578941 | 0.101835785 | 0.168079941 | 0.605877089 | 0.544596374 | 0.74232218 |
| Nr2f6 | 635.2188479 | -0.076082784 | 0.110100853 | -0.691028107 | 0.489547878 | 0.700413803 |
| Slc35f2 | 158.9181391 | 0.073327293 | 0.106120058 | 0.690984288 | 0.489575415 | 0.700413803 |
| Psma4 | 2700.316828 | -0.038353297 | 0.0555307 | -0.690668343 | 0.489773988 | 0.700633198 |
| Tbcel | 2140.46758 | 0.043120133 | 0.062451705 | 0.690455646 | 0.489907693 | 0.700759767 |
| Irak1bp1 | 419.1004121 | -0.061779613 | 0.089496193 | -0.690304367 | 0.490002802 | 0.700802864 |
| Pcyox1 | 5603.39692 | 0.052684659 | 0.076333349 | 0.690191904 | 0.490073514 | 0.700802864 |
| Zfp101 | 281.3517477 | 0.077901472 | 0.112858109 | 0.690260295 | 0.490030512 | 0.700802864 |
| Acta1 | 269.2933722 | 0.08280491 | 0.120001403 | 0.690032844 | 0.490173533 | 0.700872964 |
| Mrps9 | 875.5995501 | -0.039461226 | 0.057192663 | -0.689970076 | 0.490213006 | 0.700872964 |
| Mob3b | 205.0528134 | -0.098712707 | 0.143087605 | -0.689876015 | 0.490272161 | 0.700892864 |
| Ptprn | 25116.23741 | 0.044602008 | 0.064702232 | 0.68934265 | 0.490607664 | 0.701243095 |
| Rdm1 | 165.1227109 | 0.084698256 | 0.122856433 | 0.689408399 | 0.4905663 | 0.701243095 |
| Pea15a | 21177.32046 | -0.046897261 | 0.06806366 | -0.689020569 | 0.490810324 | 0.701468052 |
| Sgk3 | 538.5891274 | 0.097071043 | 0.140918206 | 0.688846713 | 0.490919736 | 0.701559711 |
| Fnbp1l | 4561.594576 | -0.05922331 | 0.085988295 | -0.688736884 | 0.490988861 | 0.701593785 |
| Ppig | 2264.101357 | 0.05486297 | 0.079679897 | 0.688542171 | 0.491111423 | 0.701612191 |
| Stam | 2173.23903 | -0.041816449 | 0.060729529 | -0.688568646 | 0.491094757 | 0.701612191 |
| Tm9sf1 | 1132.977918 | 0.040464371 | 0.05877173 | 0.688500604 | 0.49113759 | 0.701612191 |
| Ccdc90a | 997.2245169 | 0.040494006 | 0.058841747 | 0.68818498 | 0.491336302 | 0.70183135 |
| Mrpl2 | 834.0894578 | 0.0505041 | 0.073395579 | 0.688108208 | 0.491384642 | 0.701835697 |
| Tatdn2 | 1675.130667 | -0.052452271 | 0.076257602 | -0.687830062 | 0.491559803 | 0.702021162 |
| BC048403 | 460.5734932 | 0.059184455 | 0.086075918 | 0.687584363 | 0.491714558 | 0.702100612 |
| 3230401D17Rik | 649.7001163 | 0.101794543 | 0.086584944 | 1.175661015 | 0.239730363 | 0.464065748 |
| Inadl | 461.9018188 | -0.104565899 | 0.152106035 | -0.68745398 | 0.491796692 | 0.702100612 |
| Nol8 | 1052.822855 | 0.045714702 | 0.066494396 | 0.687497056 | 0.491769555 | 0.702100612 |
| Tmem237 | 742.8737308 | 0.048734861 | 0.070922479 | 0.687156763 | 0.491983948 | 0.702303232 |
| Tbrg4 | 1245.328305 | 0.041587837 | 0.060566366 | 0.686649037 | 0.492303919 | 0.702695248 |
| D7Ertd443e | 198.8571453 | 0.084820917 | 0.123550416 | 0.686528779 | 0.492379723 | 0.702738709 |
| Hyal1 | 154.1855313 | 0.080312305 | 0.117016871 | 0.68633099 | 0.492504411 | 0.702817162 |
| Nxpe2 | 10.76358979 | -0.10942783 | 0.159463315 | -0.686225732 | 0.492570774 | 0.702817162 |
| Rtkn2 | 105.9123102 | 0.098882032 | 0.144093319 | 0.686236063 | 0.49256426 | 0.702817162 |
| Ncln | 2021.093847 | 0.059565151 | 0.086823689 | 0.686047221 | 0.492683331 | 0.702913033 |
| Ddx49 | 883.1983317 | 0.045061839 | 0.065736913 | 0.685487604 | 0.49303628 | 0.703173645 |
| Fgfr4 | 8.546220118 | -0.109542516 | 0.15978629 | -0.685556414 | 0.492992875 | 0.703173645 |
| Hdac6 | 2138.40591 | 0.06155411 | 0.089785404 | 0.685569224 | 0.492984795 | 0.703173645 |
| Lcn2 | 17.81525375 | 0.069090439 | 0.100792833 | 0.685469765 | 0.493047533 | 0.703173645 |
| Dus4l | 175.6141132 | 0.07865039 | 0.114777916 | 0.685239747 | 0.493192646 | 0.703315863 |
| Cd83 | 638.0142853 | -0.067562283 | 0.098641547 | -0.684927237 | 0.493389838 | 0.703465993 |
| Etaa1 | 367.5817337 | -0.058561911 | 0.08550453 | -0.684898342 | 0.493408072 | 0.703465993 |
| Rpl37 | 4322.367656 | -0.046435719 | 0.067810643 | -0.68478511 | 0.493479533 | 0.703465993 |
| Wdr54 | 517.7601756 | -0.053855166 | 0.078643876 | -0.684797964 | 0.493471421 | 0.703465993 |
| Rassf4 | 506.7051678 | -0.088216266 | 0.128843645 | -0.684676887 | 0.493547837 | 0.703498637 |
| Ift52 | 1136.6903 | -0.052778744 | 0.077094813 | -0.684595271 | 0.493599352 | 0.703507346 |
| Gpr116 | 2065.947494 | -0.055398465 | 0.080957373 | -0.684291778 | 0.493790938 | 0.703710274 |
| Kdm1a | 2434.845811 | -0.030473402 | 0.044537052 | -0.684225839 | 0.493832568 | 0.703710274 |
| Nradd | 51.27723724 | -0.104907012 | 0.153342592 | -0.68413485 | 0.493890017 | 0.703727416 |
| Meig1 | 26.73356607 | -0.111119066 | 0.162610033 | -0.683346925 | 0.494387649 | 0.7043717 |
| Copz1 | 3264.539672 | -0.045101272 | 0.066014221 | -0.683205391 | 0.494477067 | 0.70443432 |
| Msantd4 | 2134.200277 | -0.050370667 | 0.073753104 | -0.682963349 | 0.494630003 | 0.704483316 |
| Pygl | 193.5745049 | 0.071474643 | 0.104647835 | 0.683001645 | 0.494605803 | 0.704483316 |
| Tm4sf5 | 9.785699211 | -0.107873906 | 0.157956311 | -0.682935084 | 0.494647863 | 0.704483316 |
| Zc3h10 | 568.368523 | 0.10176061 | 0.092181003 | 1.103921697 | 0.269627106 | 0.497867046 |
| Cdr2 | 644.3913054 | 0.061940385 | 0.090774699 | 0.682352965 | 0.49501579 | 0.704812961 |
| Numa1 | 4410.540145 | 0.043664244 | 0.063985739 | 0.682405872 | 0.494982344 | 0.704812961 |
| Oxa1l | 1644.934595 | 0.03698911 | 0.054240441 | 0.681947068 | 0.495272422 | 0.705048778 |
| Tmem86a | 608.4299339 | -0.061299132 | 0.089883856 | -0.681981561 | 0.49525061 | 0.705048778 |
| Dbnl | 2402.110858 | -0.056470416 | 0.082826629 | -0.681790596 | 0.495371371 | 0.705124853 |
| Trp73 | 51.39660746 | -0.114338225 | 0.167753708 | -0.681583892 | 0.495502103 | 0.70524615 |
| Usp28 | 1193.144654 | 0.060929284 | 0.089411641 | 0.681446882 | 0.495588767 | 0.705304708 |
| Anxa9 | 27.88387546 | 0.101730878 | 0.166905576 | 0.609511561 | 0.542185412 | 0.740777876 |
| Smim1 | 457.2804021 | -0.058583579 | 0.086007814 | -0.681142519 | 0.495781315 | 0.705449141 |
| Cbs | 1067.763785 | 0.085121357 | 0.124996249 | 0.680991289 | 0.495877002 | 0.705520502 |
| Ephb6 | 5020.532607 | -0.087830612 | 0.129049082 | -0.680598503 | 0.496125575 | 0.705679763 |
| Atf4 | 3968.653618 | 0.101730351 | 0.088710115 | 1.146772838 | 0.251475518 | 0.477149975 |
| Zfp575 | 295.2863084 | 0.065083914 | 0.095625086 | 0.680615477 | 0.496114831 | 0.705679763 |
| Ccdc149 | 2551.604648 | 0.087211145 | 0.12815581 | 0.68050871 | 0.496182409 | 0.705695818 |
| Zfp934 | 60.96080511 | -0.11133794 | 0.163630886 | -0.680421299 | 0.496237738 | 0.705709731 |
| Nf2 | 4600.747214 | -0.04833596 | 0.071065644 | -0.680159316 | 0.49640359 | 0.705880802 |
| Ar | 340.6826411 | -0.08499094 | 0.125002524 | -0.679913794 | 0.496559047 | 0.705999105 |
| Entpd6 | 2035.345873 | -0.039947942 | 0.058756997 | -0.679883995 | 0.496577917 | 0.705999105 |
| Acbd5 | 2582.128235 | -0.039197761 | 0.057689726 | -0.679458262 | 0.496847545 | 0.706089882 |
| Cnot2 | 2212.564471 | 0.032589682 | 0.047972717 | 0.679337837 | 0.496923827 | 0.706089882 |
| Frat1 | 403.8305114 | -0.059811163 | 0.088045056 | -0.679324491 | 0.496932282 | 0.706089882 |
| Hyal3 | 25.56481971 | -0.111406994 | 0.164045013 | -0.679124537 | 0.497058956 | 0.706089882 |
| Nagk | 1130.060745 | -0.048418546 | 0.071339694 | -0.678704137 | 0.497325344 | 0.706089882 |
| Pik3cg | 191.7182701 | -0.084855957 | 0.124937484 | -0.679187335 | 0.497019171 | 0.706089882 |
| Reep3 | 1469.207403 | -0.046861229 | 0.068977802 | -0.679366799 | 0.496905481 | 0.706089882 |
| Rnf214 | 2482.895318 | 0.05063386 | 0.074559122 | 0.679110193 | 0.497068044 | 0.706089882 |
| Robo4 | 345.5569146 | 0.071926513 | 0.105839741 | 0.679579448 | 0.496770787 | 0.706089882 |
| Rpp21 | 329.9904098 | 0.057542889 | 0.084765608 | 0.678847124 | 0.497234732 | 0.706089882 |
| Sbf2 | 2792.065342 | 0.059826722 | 0.088076698 | 0.679257096 | 0.496974976 | 0.706089882 |
| Srm | 1518.895173 | -0.047491712 | 0.069937121 | -0.679063022 | 0.497097931 | 0.706089882 |
| Tle3 | 1706.450691 | -0.058459421 | 0.086107871 | -0.67890914 | 0.497195434 | 0.706089882 |
| Tmem256 | 824.4771174 | -0.052432146 | 0.077237034 | -0.678847222 | 0.49723467 | 0.706089882 |
| Ttc26 | 418.1469905 | 0.053874198 | 0.079370336 | 0.678769935 | 0.497283646 | 0.706089882 |
| Cd164l2 | 105.1785953 | 0.082834674 | 0.122101788 | 0.67840672 | 0.49751385 | 0.706229334 |
| Slc15a4 | 847.300736 | 0.101729605 | 0.10569365 | 0.962494953 | 0.335801037 | 0.569968355 |
| Ppp2r1b | 598.8823197 | 0.101666537 | 0.086202461 | 1.179392507 | 0.238241917 | 0.462671816 |
| Acp2 | 3657.289588 | 0.036795179 | 0.054282895 | 0.677841132 | 0.497872428 | 0.706607622 |
| Myct1 | 31.35777722 | -0.113901694 | 0.168081502 | -0.677657521 | 0.497988866 | 0.706685488 |
| Pls3 | 6501.460095 | 0.101576466 | 0.079194255 | 1.282624169 | 0.199623778 | 0.416113389 |
| 2900008C10Rik | 147.6573109 | 0.082562855 | 0.12191556 | 0.677213436 | 0.498270544 | 0.706913635 |
| Hax1 | 1454.551465 | -0.058423862 | 0.086265361 | -0.677257497 | 0.498242593 | 0.706913635 |
| Atr | 625.7483048 | 0.076173788 | 0.112508689 | 0.677048043 | 0.498375472 | 0.706925704 |
| Rhoq | 1524.833155 | -0.059773239 | 0.088302772 | -0.676912372 | 0.498461554 | 0.706925704 |
| Smcr7 | 549.8537263 | 0.056730942 | 0.083789488 | 0.677065148 | 0.49836462 | 0.706925704 |
| Tom1l2 | 15169.7077 | 0.04228695 | 0.062469258 | 0.676924163 | 0.498454072 | 0.706925704 |
| Ifngr1 | 649.8619592 | 0.047555694 | 0.070305793 | 0.67641218 | 0.498778986 | 0.707181699 |
| Pcolce | 269.2047136 | -0.100607071 | 0.148725856 | -0.676459854 | 0.498748727 | 0.707181699 |
| 1810030O07Rik | 874.0631097 | 0.101554153 | 0.070862073 | 1.433124211 | 0.151822344 | 0.353044483 |
| Ndufaf2 | 422.8828843 | -0.057665789 | 0.085278547 | -0.67620511 | 0.498910428 | 0.707303338 |
| Gabarapl1 | 23987.59184 | -0.037713735 | 0.055779365 | -0.676123429 | 0.498962283 | 0.707312133 |
| Slain1 | 1858.446771 | 0.06541958 | 0.096789479 | 0.675895566 | 0.499106953 | 0.707452487 |
| BC022687 | 480.588446 | -0.061284354 | 0.090688275 | -0.675769328 | 0.499187111 | 0.707501382 |
| Aprt | 662.7637505 | -0.048744716 | 0.072191317 | -0.675215765 | 0.499538692 | 0.707924612 |
| Birc5 | 29.62744576 | -0.109176504 | 0.161740225 | -0.675011453 | 0.499668488 | 0.707924612 |
| Mfap1b | 167.2659605 | 0.076534282 | 0.113379472 | 0.675027682 | 0.499658178 | 0.707924612 |
| Pcdhb4 | 83.5469653 | 0.099891261 | 0.147983499 | 0.67501621 | 0.499665466 | 0.707924612 |
| Fam184a | 1074.633142 | -0.085720288 | 0.127016958 | -0.674872781 | 0.499756595 | 0.707984701 |
| 1700012B09Rik | 9.700104813 | 0.101539892 | 0.140347463 | 0.723489328 | 0.469379308 | 0.684154563 |
| Gm2061 | 137.8211397 | -0.078017227 | 0.115662389 | -0.674525461 | 0.499977304 | 0.70810314 |
| Gpaa1 | 1297.802725 | 0.101538642 | 0.056708289 | 1.790543205 | 0.073366628 | 0.221939958 |
| Aggf1 | 2507.456949 | -0.035177523 | 0.052159208 | -0.674425942 | 0.500040554 | 0.708127991 |
| Fam134c | 3055.710573 | -0.047224347 | 0.070030126 | -0.674343311 | 0.500093075 | 0.708137644 |
| Chrna10 | 25.70431038 | -0.10825703 | 0.160562564 | -0.674235806 | 0.500161409 | 0.708169685 |
| Fyttd1 | 2470.862867 | -0.046237429 | 0.06862353 | -0.673783891 | 0.500448718 | 0.708447001 |
| Ksr1 | 1630.779762 | 0.08417143 | 0.124918287 | 0.673811911 | 0.500430901 | 0.708447001 |
| B230378P21Rik | 10.94163064 | 0.098094216 | 0.145649491 | 0.673495084 | 0.500632376 | 0.708642246 |
| Lrfn2 | 1027.49568 | -0.078419834 | 0.116457544 | -0.673377017 | 0.500707467 | 0.708683794 |
| Mavs | 351.3194906 | 0.060035676 | 0.089168381 | 0.673284357 | 0.500766403 | 0.708702471 |
| 1810026B05Rik | 257.3006401 | 0.062690012 | 0.093173516 | 0.672830802 | 0.501054939 | 0.708899185 |
| Alad | 1011.09613 | 0.038651743 | 0.057448059 | 0.672811992 | 0.501066908 | 0.708899185 |
| Nthl1 | 85.69078635 | 0.101410065 | 0.139324711 | 0.727868476 | 0.466694102 | 0.681613104 |
| Cmpk2 | 1041.00115 | -0.042202774 | 0.062755932 | -0.672490595 | 0.501271426 | 0.708899185 |
| Hic2 | 417.8648384 | 0.082403223 | 0.122502479 | 0.672665758 | 0.501159957 | 0.708899185 |
| Pacsin2 | 2888.70102 | -0.073203915 | 0.108802882 | -0.672812273 | 0.501066729 | 0.708899185 |
| Pigb | 283.8421601 | 0.066363003 | 0.098658107 | 0.672656364 | 0.501165934 | 0.708899185 |
| Tpmt | 365.5804214 | -0.054787744 | 0.081458999 | -0.672580609 | 0.501214141 | 0.708899185 |
| Fbxl20 | 2819.183261 | 0.101348656 | 0.079185778 | 1.279884582 | 0.200585731 | 0.417388573 |
| Elavl2 | 2462.61656 | 0.049074234 | 0.073038406 | 0.67189629 | 0.501649722 | 0.709201588 |
| Gfod1 | 6825.624607 | 0.089710547 | 0.133538547 | 0.671795142 | 0.501714122 | 0.709201588 |
| Supv3l1 | 815.9103371 | 0.0495101 | 0.073683246 | 0.671931577 | 0.501627257 | 0.709201588 |
| Mast4 | 2757.671133 | 0.10128957 | 0.072530955 | 1.396501267 | 0.162563599 | 0.367607043 |
| Cul3 | 4498.466325 | -0.048274243 | 0.071878405 | -0.671609822 | 0.501832124 | 0.709238979 |
| Mgat4a | 1910.784035 | 0.045821082 | 0.068221692 | 0.671649749 | 0.501806699 | 0.709238979 |
| Tpo | 14.87062312 | -0.111443229 | 0.165993689 | -0.671370274 | 0.501984677 | 0.709389874 |
| Calm1 | 122655.312 | -0.043596818 | 0.065009068 | -0.670626717 | 0.502458358 | 0.709756076 |
| Dlgap4 | 11175.95011 | -0.049099026 | 0.073222667 | -0.670544089 | 0.502511011 | 0.709756076 |
| E2f5 | 288.1378865 | -0.076314653 | 0.113836521 | -0.670388135 | 0.502610396 | 0.709756076 |
| Figf | 87.47033351 | -0.090089064 | 0.134326303 | -0.670673296 | 0.502428678 | 0.709756076 |
| Impact | 9458.615061 | -0.049250785 | 0.073458779 | -0.670454715 | 0.502567965 | 0.709756076 |
| Kpna6 | 5357.157729 | -0.032088532 | 0.047870723 | -0.670316443 | 0.502656088 | 0.709756076 |
| Mrps23 | 1172.19521 | -0.053830078 | 0.080300039 | -0.670361792 | 0.502627185 | 0.709756076 |
| Mtmr3 | 4802.642707 | 0.047189878 | 0.070367437 | 0.670620955 | 0.50246203 | 0.709756076 |
| Parp11 | 772.7999301 | -0.057095685 | 0.085143087 | -0.670585092 | 0.502484882 | 0.709756076 |
| Sox8 | 1588.200315 | 0.101264828 | 0.101043554 | 1.002189884 | 0.316251893 | 0.549267267 |
| Pop1 | 312.5982026 | 0.062983107 | 0.093992537 | 0.670086252 | 0.502802808 | 0.709833869 |
| Dnajc2 | 1060.392244 | -0.044849501 | 0.066939773 | -0.669997799 | 0.502859193 | 0.709848792 |
| 9530051G07Rik | 25.90577971 | -0.110119645 | 0.164416206 | -0.669761504 | 0.503009837 | 0.709867422 |
| Dpp9 | 2852.17505 | -0.04951136 | 0.073919533 | -0.669800773 | 0.502984801 | 0.709867422 |
| Nyap2 | 594.4297271 | -0.068749988 | 0.102632721 | -0.669864226 | 0.502944346 | 0.709867422 |
| Ces2g | 10.95154869 | -0.10416784 | 0.155624706 | -0.669352848 | 0.503270423 | 0.70997253 |
| Wdr93 | 12.57260587 | 0.101190881 | 0.156306952 | 0.647385671 | 0.517382368 | 0.721003535 |
| Mrps18b | 757.3434769 | 0.04784847 | 0.071493438 | 0.669270797 | 0.503322752 | 0.70997253 |
| Nlgn2 | 11191.40095 | 0.031066762 | 0.046426362 | 0.669162088 | 0.503392088 | 0.70997253 |
| Ogfrl1 | 4331.60472 | -0.043732319 | 0.065355203 | -0.669148245 | 0.503400917 | 0.70997253 |
| Osbp | 3134.146657 | -0.044573331 | 0.066582804 | -0.669442075 | 0.50321352 | 0.70997253 |
| Spg21 | 1194.089071 | 0.043940716 | 0.065667278 | 0.669141727 | 0.503405074 | 0.70997253 |
| Fbxl16 | 43855.31174 | -0.06015995 | 0.089933785 | -0.668936041 | 0.503536279 | 0.710079244 |
| Sox18 | 206.1547186 | -0.102445223 | 0.153159478 | -0.668879422 | 0.503572398 | 0.710079244 |
| Klhl26 | 2033.278825 | 0.051605882 | 0.077165225 | 0.668771229 | 0.503641423 | 0.710111949 |
| Rtf1 | 4327.60451 | 0.062015958 | 0.092756708 | 0.668587304 | 0.503758774 | 0.710129757 |
| Slco3a1 | 1453.743363 | 0.059859682 | 0.089538465 | 0.668535936 | 0.503791551 | 0.710129757 |
| Zbtb44 | 887.5538306 | -0.04605729 | 0.068891649 | -0.668546782 | 0.50378463 | 0.710129757 |
| Peli1 | 1350.465465 | -0.051217819 | 0.076647272 | -0.668227555 | 0.503988349 | 0.710277922 |
| Psmc4 | 3201.606307 | 0.047085127 | 0.070461701 | 0.668237152 | 0.503982224 | 0.710277922 |
| A930005H10Rik | 206.330726 | 0.067641342 | 0.101313174 | 0.667646071 | 0.504359542 | 0.710709078 |
| Zfp658 | 260.8800028 | 0.078024101 | 0.116871726 | 0.667604598 | 0.504386022 | 0.710709078 |
| Rab35 | 3391.477458 | 0.101148139 | 0.064186142 | 1.575856342 | 0.115058914 | 0.29712047 |
| Cit | 3880.426694 | 0.046068181 | 0.069037182 | 0.667295216 | 0.504583581 | 0.710734511 |
| F2r | 194.0571893 | 0.069849632 | 0.104664324 | 0.667368112 | 0.504537029 | 0.710734511 |
| Flot2 | 4542.176595 | -0.055776444 | 0.083592173 | -0.667244809 | 0.504615773 | 0.710734511 |
| Gnas | 45382.66248 | 0.026489301 | 0.039705443 | 0.667145341 | 0.504679301 | 0.710734511 |
| Plag1 | 136.1812689 | 0.087822319 | 0.131632266 | 0.667179268 | 0.504657632 | 0.710734511 |
| Serpina3g | 10.70017179 | 0.078740231 | 0.118094834 | 0.666754235 | 0.50492913 | 0.711021715 |
| Dynlt1b | 37.24422471 | -0.111925765 | 0.167931873 | -0.666495068 | 0.505094715 | 0.711125626 |
| Pex11b | 1510.497518 | 0.045639187 | 0.06847025 | 0.666555 | 0.505056421 | 0.711125626 |
| Dhps | 1024.364 | 0.047786738 | 0.071708958 | 0.66639844 | 0.505156459 | 0.711147936 |
| 4930452B06Rik | 518.6956291 | -0.053557077 | 0.080405419 | -0.666087902 | 0.505354917 | 0.711233452 |
| Ing4 | 1314.742555 | 0.047052411 | 0.070632683 | 0.666156365 | 0.505311161 | 0.711233452 |
| Psmd5 | 1672.3725 | 0.050233529 | 0.075405133 | 0.666181821 | 0.505294891 | 0.711233452 |
| Nipal4 | 192.2892279 | 0.101108096 | 0.163095055 | 0.619933545 | 0.53530154 | 0.735675877 |
| Ccl25 | 119.5070715 | 0.095292591 | 0.143154451 | 0.66566279 | 0.505626662 | 0.71142204 |
| Ndufa6 | 1943.327302 | -0.051208084 | 0.076921318 | -0.665720313 | 0.505589887 | 0.71142204 |
| Stradb | 1445.509465 | 0.0417672 | 0.062764797 | 0.665455822 | 0.505758991 | 0.711479009 |
| Tnnt1 | 108.0620907 | 0.098918485 | 0.148642494 | 0.665479184 | 0.505744053 | 0.711479009 |
| Pdcd7 | 835.9637297 | -0.045995329 | 0.069149419 | -0.66515857 | 0.505949076 | 0.711681796 |
| Rarres2 | 168.1464343 | 0.095739281 | 0.143971481 | 0.664987816 | 0.506058286 | 0.711706189 |
| Slc2a10 | 80.26386372 | -0.098244808 | 0.147736752 | -0.664999109 | 0.506051063 | 0.711706189 |
| 4931414P19Rik | 145.8254662 | 0.081946227 | 0.123267401 | 0.664784251 | 0.506188497 | 0.711824703 |
| Col9a2 | 243.6309242 | -0.074043938 | 0.11146294 | -0.664291988 | 0.506503448 | 0.712202961 |
| Ube2g1 | 1505.342069 | 0.101097312 | 0.072558802 | 1.393315623 | 0.163524368 | 0.368744952 |
| Cend1 | 10170.52643 | -0.03501981 | 0.05279074 | -0.663370316 | 0.507093412 | 0.712838367 |
| Mon2 | 3861.898415 | 0.057654487 | 0.086909648 | 0.663384203 | 0.507084521 | 0.712838367 |
| Pofut2 | 972.0560114 | 0.050187487 | 0.075663496 | 0.66329855 | 0.507139365 | 0.712838367 |
| 9330182L06Rik | 1354.735189 | -0.051986092 | 0.078416796 | -0.662945889 | 0.50736521 | 0.713085955 |
| Pak4 | 331.9297388 | -0.059578832 | 0.08987879 | -0.662879776 | 0.507407555 | 0.713085955 |
| Kat8 | 708.1168675 | 0.101092728 | 0.088054156 | 1.148074469 | 0.250937823 | 0.476591153 |
| Ift81 | 1175.685554 | 0.101090735 | 0.076361656 | 1.323841583 | 0.185555663 | 0.397044529 |
| Rxrb | 2497.312601 | 0.037240519 | 0.056215979 | 0.662454332 | 0.507680094 | 0.713276618 |
| 2700062C07Rik | 427.7599329 | -0.061498215 | 0.092919301 | -0.661845435 | 0.508070284 | 0.713389644 |
| Aen | 879.2665494 | 0.049901505 | 0.07537438 | 0.662048632 | 0.507940055 | 0.713389644 |
| Car10 | 5023.263574 | 0.072979808 | 0.110258216 | 0.661899051 | 0.50803592 | 0.713389644 |
| Celf2 | 14229.09489 | -0.037470604 | 0.056587338 | -0.662172939 | 0.507860395 | 0.713389644 |
| Crkl | 2092.211695 | 0.068307821 | 0.103180086 | 0.662025237 | 0.507955048 | 0.713389644 |
| Gm5176 | 14.39025908 | -0.110402601 | 0.166746547 | -0.662098272 | 0.507908243 | 0.713389644 |
| Papolg | 783.0237049 | 0.046287627 | 0.069946801 | 0.661754738 | 0.508128418 | 0.713389644 |
| Tnfaip1 | 2356.170738 | -0.038717988 | 0.058508298 | -0.661752079 | 0.508130122 | 0.713389644 |
| Bambi-ps1 | 38.26029841 | -0.104440495 | 0.157896574 | -0.661448767 | 0.50832456 | 0.713450923 |
| Fastkd5 | 353.1191091 | 0.089120076 | 0.134745273 | 0.661396675 | 0.508357957 | 0.713450923 |
| Maoa | 1284.047643 | -0.052655869 | 0.079606573 | -0.661451275 | 0.508322952 | 0.713450923 |
| Gm12942 | 78.55149213 | 0.101071267 | 0.156918825 | 0.644099052 | 0.519511205 | 0.722688688 |
| Gls | 14595.87558 | -0.03950478 | 0.059746728 | -0.661204075 | 0.508481448 | 0.713494979 |

| | | | | | | |
|---|---|---|---|---|---|---|
| Kl | 662.9426602 | 0.009784804 | 0.036444849 | 0.268482492 | NA | NA |
| Wfdc15a | 0.767122395 | 0.009785254 | 0.019837694 | 0.493265713 | 0.621824852 | NA |
| Prkaa2 | 2244.601114 | 0.009785632 | 0.074747237 | 0.130916307 | 0.895841514 | 0.977844987 |
| Oas3 | 5.989800439 | 0.009786289 | 0.040681958 | 0.240556006 | 0.809899251 | NA |
| Adck1 | 1489.944413 | 0.009786329 | 0.062140995 | 0.157485869 | 0.874861944 | 0.971956739 |
| 4930590L20Rik | 0.321063518 | 0.009790192 | 0.011658371 | 0.839756417 | 0.401044974 | NA |
| Gm19589 | 0.321063518 | 0.009790192 | 0.011658371 | 0.839756417 | 0.401044974 | NA |
| Shisa2 | 131.9794624 | 0.00979915 | 0.093381006 | 0.104937296 | 0.916425564 | 0.982742917 |
| En2 | 30.5336832 | 0.009804766 | 0.086374366 | 0.113514768 | 0.909622456 | 0.980590355 |
| Gm4926 | 6.073344593 | 0.009814713 | 0.053620726 | 0.183039536 | 0.854766998 | NA |
| Zfp398 | 589.8777824 | 0.00981965 | 0.07579223 | 0.129560109 | 0.896914466 | 0.978039389 |
| Myo9a | 5087.245522 | 0.009825498 | 0.073672704 | 0.133366871 | 0.893903244 | 0.977844987 |
| Herc2 | 10721.02451 | 0.009831001 | 0.068815567 | 0.142860126 | 0.88640065 | 0.976378471 |
| Ggnbp2 | 3949.430413 | 0.009841772 | 0.060364877 | 0.163038046 | 0.870488479 | 0.971477516 |
| Csf2 | 0.323681211 | 0.009850234 | 0.011693836 | 0.842344081 | 0.399595384 | NA |
| Mir32 | 0.323681211 | 0.009850234 | 0.011693836 | 0.842344081 | 0.399595384 | NA |
| Cdrt4 | 2.250340267 | 0.009875503 | 0.030790805 | 0.320728952 | 0.748415805 | NA |
| Gm17384 | 8.315182849 | 0.009880061 | 0.060805062 | 0.162487483 | 0.870921985 | NA |
| Rps19 | 4343.316174 | 0.009888513 | 0.053943546 | 0.183312262 | 0.854553014 | 0.967189923 |
| Mgst2 | 3.205764872 | 0.009913451 | 0.037896284 | 0.261594275 | 0.793634258 | NA |
| Cd200r2 | 1.823617806 | 0.009919328 | 0.028055543 | 0.353560382 | 0.723668369 | NA |
| Nckipsd | 3707.596194 | 0.009935036 | 0.055477649 | 0.179081785 | 0.857873484 | 0.967971163 |
| Pbp2 | 4.41781871 | 0.009939376 | 0.048136606 | 0.20648269 | 0.836413885 | NA |
| Haghl | 3028.070549 | 0.009940646 | 0.068472887 | 0.145176379 | 0.884571616 | 0.975676135 |
| Gdf15 | 2.014736869 | 0.009945118 | 0.034712543 | 0.286499276 | 0.774495748 | NA |
| Pcdhb11 | 135.1990926 | 0.009949391 | 0.091322304 | 0.108948091 | 0.913243663 | 0.98170867 |
| Pabpn1l | 0.762737197 | 0.009951876 | 0.020103111 | 0.495041587 | 0.620570765 | NA |
| Prpf8 | 11075.16149 | 0.009954849 | 0.045468633 | 0.218938825 | 0.826697702 | 0.958455003 |
| Aipl1 | 1.061152941 | 0.009967512 | 0.022070598 | 0.451619469 | 0.651543142 | NA |
| Ankrd34c | 355.928771 | 0.00996903 | 0.085399274 | 0.11673436 | 0.907070561 | 0.980365102 |
| Nol8 | 1145.609368 | 0.009989474 | 0.074996853 | 0.13319857 | 0.894036342 | 0.977844987 |
| Fbxw4 | 842.9562814 | 0.009999102 | 0.063113158 | 0.158431331 | 0.874116927 | 0.971912694 |
| Phtf1 | 2268.797143 | 0.010023252 | 0.051238838 | 0.195618253 | 0.844908972 | 0.964415192 |
| Gm11201 | 2.149941643 | 0.010026705 | 0.031938409 | 0.313938766 | 0.753567546 | NA |
| Prss57 | 35.92737339 | 0.010049089 | 0.091629298 | 0.109671132 | 0.912670195 | 0.981664815 |
| Cd3d | 0.589409793 | 0.01004926 | 0.015506378 | 0.648072683 | 0.516937941 | NA |
| Dmtf1 | 2678.774545 | 0.010050058 | 0.053742736 | 0.187003097 | 0.851658205 | 0.96665003 |
| Gstp2 | 11.1338592 | 0.010055667 | 0.061385389 | 0.163812066 | 0.869879093 | NA |
| Rarres1 | 25.44891848 | 0.0100697 | 0.084968304 | 0.118511252 | 0.90566258 | 0.980118791 |
| Tcf21 | 0.743239487 | 0.010073237 | 0.014350729 | 0.701932086 | 0.482721518 | NA |
| Eci3 | 3.102882784 | 0.010094702 | 0.032333167 | 0.312208893 | 0.754881774 | NA |
| Arhgef28 | 1464.236884 | 0.01009546 | 0.076343497 | 0.132237321 | 0.894796582 | 0.977844987 |
| Chrna7 | 535.4292273 | 0.01009728 | 0.0693235 | 0.145654508 | 0.884194135 | 0.975602279 |
| Cd320 | 414.2262987 | 0.010132427 | 0.072644547 | 0.139479525 | 0.889071234 | 0.977182599 |
| Ncor1 | 9706.989055 | 0.010141257 | 0.056727595 | 0.17877114 | 0.858117407 | 0.967971163 |
| Dydc2 | 15.28233037 | 0.01015336 | 0.068766804 | 0.147649147 | 0.882619667 | NA |
| Plekhg3 | 861.44023 | 0.010170008 | 0.08686738 | 0.117075108 | 0.906800535 | 0.980201396 |
| Sycp1 | 1.193554861 | 0.010174038 | 0.019965074 | 0.509591806 | 0.610337466 | NA |
| Mepce | 1794.514287 | 0.01018091 | 0.052446556 | 0.1941197 | 0.846082157 | 0.964996028 |
| 4933401H06Rik | 0.57007369 | 0.010186007 | 0.0153948 | 0.661652462 | 0.508193978 | NA |
| Mir23a | 0.57007369 | 0.010186007 | 0.0153948 | 0.661652462 | 0.508193978 | NA |
| Tbx19 | 1.58698693 | 0.010204118 | 0.024789043 | 0.411638227 | 0.680604608 | NA |
| Slc5a1 | 2.53745628 | 0.010223741 | 0.037525083 | 0.272450863 | 0.785275368 | NA |
| Tnks | 3590.50213 | 0.010242184 | 0.062552719 | 0.163736825 | 0.869938327 | 0.971097167 |
| Xkrx | 83.54524752 | 0.010246182 | 0.095082819 | 0.107760609 | 0.91418559 | 0.98170867 |
| Lrrc39 | 10.78513989 | 0.010250803 | 0.060115546 | 0.170518342 | 0.864602511 | NA |
| Klc1 | 25785.81948 | 0.010251012 | 0.04159502 | 0.246448066 | 0.805335402 | 0.955131256 |
| Nipal4 | 256.3882201 | 0.010252575 | 0.095083229 | 0.107827377 | 0.914132626 | 0.98170867 |
| Pla2g2c | 8.447316024 | 0.010258892 | 0.0621111 | 0.165170023 | 0.86881016 | NA |
| Rp1 | 2.692349903 | 0.010260807 | 0.037052172 | 0.276928609 | 0.781834925 | NA |
| Slc5a4a | 1.34506611 | 0.010263184 | 0.024870828 | 0.412659526 | 0.679856082 | NA |
| Gcg | 1.149091228 | 0.010264067 | 0.020292456 | 0.505807056 | 0.612992104 | NA |
| St8sia4 | 461.2523997 | 0.010279669 | 0.076247335 | 0.134820047 | 0.892754156 | 0.977844987 |
| Ferd3l | 1.323332537 | 0.010279872 | 0.024854647 | 0.413599603 | 0.679167364 | NA |
| A430090L17Rik | 0.345220294 | 0.010284943 | 0.011947302 | 0.860859035 | 0.389315686 | NA |
| Prap1 | 0.345220294 | 0.010284943 | 0.011947302 | 0.860859035 | 0.389315686 | NA |
| Zfp462 | 1967.967147 | 0.010291304 | 0.064862265 | 0.158663964 | 0.873933631 | 0.971912694 |
| Eid2b | 908.5997358 | 0.010305585 | 0.073132205 | 0.14091719 | 0.887935362 | 0.976924608 |
| Llgl2 | 52.53909829 | 0.010311356 | 0.092744479 | 0.111180268 | 0.911473399 | 0.980985114 |
| AU023871 | 1.603182284 | 0.010315931 | 0.023549184 | 0.438058952 | 0.66134355 | NA |
| Ovol1 | 0.909119122 | 0.010356616 | 0.019331906 | 0.535726593 | 0.592147522 | NA |
| Lifr | 1824.003833 | 0.010370994 | 0.057373797 | 0.180761864 | 0.856554498 | 0.967598001 |
| Hoxb7 | 0.530743041 | 0.010381478 | 0.011994275 | 0.865536077 | 0.386744623 | NA |
| Prdx6b | 0.351100342 | 0.010390521 | 0.012008062 | 0.865295413 | 0.386876668 | NA |
| Rbm44 | 1.034513641 | 0.010419048 | 0.024758888 | 0.420820519 | 0.673886147 | NA |
| Acp5 | 11.66981932 | 0.010425735 | 0.070523457 | 0.147833572 | 0.882474113 | NA |
| Cd96 | 2.226441281 | 0.010430022 | 0.032108821 | 0.324833552 | 0.745307049 | NA |
| Alpl | 405.439408 | 0.010443129 | 0.092398625 | 0.113022562 | 0.910012668 | 0.980590355 |
| Rspo4 | 7.316038993 | 0.010457059 | 0.056687158 | 0.184469621 | 0.853645057 | NA |
| Tmem217 | 2.99097361 | 0.010497043 | 0.03331293 | 0.315104174 | 0.752682559 | NA |
| Hspa1a | 242.6632594 | 0.01050472 | 0.089288204 | 0.117649586 | 0.906345314 | 0.980118791 |
| Zeb1 | 3401.171368 | 0.01050901 | 0.052757899 | 0.199193116 | 0.842111684 | 0.963252575 |
| Kbtbd3 | 272.5478923 | 0.010510035 | 0.087700139 | 0.119840575 | 0.904609439 | 0.980118791 |
| Gm16596 | 41.20961124 | 0.010518396 | 0.092908203 | 0.113212781 | 0.909861863 | 0.980590355 |
| Ablim1 | 6891.982326 | 0.010527374 | 0.045345692 | 0.232158184 | 0.816415155 | 0.956264994 |
| BC030336 | 2164.28034 | 0.010543416 | 0.070055859 | 0.150500134 | 0.880370045 | 0.973779118 |
| 2310001K24Rik | 0.543831507 | 0.01054504 | 0.012087653 | 0.872381087 | 0.383000515 | NA |
| Nkx2-4 | 0.543831507 | 0.01054504 | 0.012087653 | 0.872381087 | 0.383000515 | NA |
| Dip2a | 4555.945589 | 0.010547869 | 0.052678249 | 0.20023196 | 0.841299173 | 0.963094947 |
| Meox2 | 0.898895954 | 0.010550889 | 0.018914547 | 0.55781875 | 0.576968158 | NA |
| Mettl21c | 1.547359131 | 0.010581372 | 0.028819908 | 0.367154948 | 0.713503443 | NA |
| Lrrc30 | 0.361523991 | 0.010595551 | 0.012125053 | 0.87385609 | 0.38219663 | NA |
| Mir340 | 0.361523991 | 0.010595551 | 0.012125053 | 0.87385609 | 0.38219663 | NA |
| Ppp1r14d | 0.361523991 | 0.010595551 | 0.012125053 | 0.87385609 | 0.38219663 | NA |
| Dut | 772.5311393 | 0.010596537 | 0.059373126 | 0.178473619 | 0.858351038 | 0.967971163 |
| 4930562F07Rik | 0.974347369 | 0.010596759 | 0.024278897 | 0.436459642 | 0.662503274 | NA |
| Spns3 | 8.623028829 | 0.010600972 | 0.05708681 | 0.185699152 | 0.852680693 | NA |
| Gm6981 | 35.64135011 | 0.010617293 | 0.092548978 | 0.114720801 | 0.908666426 | 0.980523965 |
| Hmcn1 | 152.7869976 | 0.010619963 | 0.09377621 | 0.113247949 | 0.909833983 | 0.980590355 |
| Cldnd2 | 0.855261623 | 0.010636678 | 0.018895719 | 0.562914698 | 0.573492971 | NA |
| Psmg1 | 363.501582 | 0.010638004 | 0.074460492 | 0.142867767 | 0.886394616 | 0.976378471 |
| Als2cr11 | 0.548112961 | 0.010638047 | 0.01683431 | 0.631926545 | 0.527434877 | NA |
| Endou | 192.6774268 | 0.010645944 | 0.088867907 | 0.119795147 | 0.904645427 | 0.980118791 |
| Ear2 | 0.364141684 | 0.010646089 | 0.012153714 | 0.875953584 | 0.381055267 | NA |
| Hist1h3f | 0.364141684 | 0.010646089 | 0.012153714 | 0.875953584 | 0.381055267 | NA |
| Mir330 | 0.364141684 | 0.010646089 | 0.012153714 | 0.875953584 | 0.381055267 | NA |
| Cecr6 | 3681.137511 | 0.010650407 | 0.075818707 | 0.140472028 | 0.888287052 | 0.97703865 |
| Siglec15 | 0.927348726 | 0.010662967 | 0.019387971 | 0.549978491 | 0.582334127 | NA |
| Hnrpll | 1890.33587 | 0.010663947 | 0.064655885 | 0.164933899 | 0.86899601 | 0.970788711 |
| Tfrc | 4169.980125 | 0.010676711 | 0.075946505 | 0.140581999 | 0.88820017 | 0.97703865 |
| Thy1 | 56123.68496 | 0.01068857 | 0.040797055 | 0.261993677 | 0.793326317 | 0.951728598 |
| Xcr1 | 2.937892694 | 0.010711131 | 0.038040411 | 0.281572439 | 0.778271376 | NA |
| Chmp6 | 1049.256316 | 0.010712906 | 0.066849336 | 0.160254487 | 0.872680609 | 0.97153269 |
| Rhox8 | 0.553189945 | 0.010739126 | 0.016869237 | 0.636610063 | 0.524378869 | NA |
| Irx2 | 9.65513574 | 0.010758271 | 0.037839111 | 0.284316171 | 0.776168095 | NA |
| Cdk2ap2 | 576.5200048 | 0.010767573 | 0.06361231 | 0.169268694 | 0.865585299 | 0.970182416 |
| 6430531B16Rik | 22.34841792 | 0.010768918 | 0.084555986 | 0.12735843 | 0.898656716 | NA |
| Hist1h2bj | 0.677809771 | 0.010773931 | 0.015946382 | 0.675634828 | 0.499272523 | NA |
| Hnrnpr | 2770.350116 | 0.010779896 | 0.056593677 | 0.190478811 | 0.848933946 | 0.965892061 |
| S1pr1 | 6522.570883 | 0.010798151 | 0.058546315 | 0.184437754 | 0.853670054 | 0.966905493 |
| Kif5b | 12321.13885 | 0.010800002 | 0.064089508 | 0.16851436 | 0.866178647 | 0.97021201 |
| Gnrh1 | 10.58063559 | 0.010822861 | 0.061820726 | 0.175068489 | 0.861025818 | NA |
| Rabggtb | 1464.600151 | 0.010823657 | 0.047988577 | 0.225546533 | 0.82155413 | 0.957699646 |
| Lphn1 | 28496.32144 | 0.010827798 | 0.056200931 | 0.192662261 | 0.847223481 | 0.965275633 |
| Adamtsl2 | 271.6674647 | 0.01083101 | 0.090631257 | 0.119506343 | 0.904874215 | 0.980118791 |
| Dcaf13 | 823.2454982 | 0.010840405 | 0.062886081 | 0.172381627 | 0.863137515 | 0.969773371 |
| Tns1 | 2737.700327 | 0.010846961 | 0.06301736 | 0.17212655 | 0.86333804 | 0.969788506 |
| Gm15800 | 12714.29872 | 0.010850832 | 0.074076252 | 0.146481929 | 0.883540955 | 0.975177867 |
| Adat2 | 211.0826864 | 0.010868765 | 0.090738977 | 0.11978056 | 0.904656982 | 0.980118791 |
| Med17 | 1100.66529 | 0.010870139 | 0.0576222 | 0.188644987 | 0.850371072 | 0.966260848 |
| Apon | 0.750337725 | 0.010877907 | 0.020059281 | 0.542288018 | 0.587620108 | NA |
| 3110062M04Rik | 155.911508 | 0.010883254 | 0.089446417 | 0.121673454 | 0.903157638 | 0.979150581 |
| B430010I23Rik | 0.523136316 | 0.010887252 | 0.016666502 | 0.653241539 | 0.513600575 | NA |
| Svopl | 2.202772154 | 0.010895173 | 0.027047144 | 0.402821577 | 0.687079487 | NA |
| D030045P18Rik | 1.073374414 | 0.010896263 | 0.022110149 | 0.492817225 | 0.622141739 | NA |
| 1500015O10Rik | 108.9475325 | 0.010925627 | 0.032989451 | 0.331185487 | NA | NA |
| Rab28 | 2084.5362 | 0.010928042 | 0.05833935 | 0.187318538 | 0.851410889 | 0.96665003 |
| Gm14047 | 2.629237912 | 0.010928706 | 0.036379093 | 0.300411719 | 0.763863126 | NA |
| Adprhl2 | 708.6101081 | 0.010931818 | 0.074988153 | 0.145780604 | 0.884094588 | 0.975566544 |
| Vegfb | 2093.009567 | 0.010936232 | 0.066230757 | 0.165123163 | 0.868847042 | 0.970788711 |
| Slc9a9 | 838.132309 | 0.010953175 | 0.065625145 | 0.166905158 | 0.867444674 | 0.970480832 |
| Pax5 | 1.391655813 | 0.010954638 | 0.020934907 | 0.523271386 | 0.60078541 | NA |
| Zbtb9 | 746.450629 | 0.010957387 | 0.059459898 | 0.184281968 | 0.853792259 | 0.966905493 |
| 2410124H12Rik | 0.855649974 | 0.010958384 | 0.01880268 | 0.582809716 | 0.560021404 | NA |
| Rnf39 | 185.4180772 | 0.010977124 | 0.095093196 | 0.115435434 | 0.908099995 | 0.980523965 |
| Gsdmc4 | 2.289432896 | 0.01097799 | 0.035003422 | 0.313626184 | 0.75380497 | NA |
| Mos | 4.28031773 | 0.010978106 | 0.047633535 | 0.230470112 | 0.817726485 | NA |
| Clasp1 | 5500.032111 | 0.010981888 | 0.052959047 | 0.20736566 | 0.835724299 | 0.961504006 |
| Dusp18 | 1734.69552 | 0.010985375 | 0.075774693 | 0.144974198 | 0.884731244 | 0.975676135 |
| 1700021F07Rik | 0.756487453 | 0.010985648 | 0.020086768 | 0.546909675 | 0.584440788 | NA |
| Wdr45 | 1514.940947 | 0.010997547 | 0.054038626 | 0.20351271 | 0.838734312 | 0.962130192 |
| Glyat | 0.793813653 | 0.011003582 | 0.020234877 | 0.543792903 | 0.586583993 | NA |
| Zfp677 | 75.08966222 | 0.011019113 | 0.095015141 | 0.115972179 | 0.907674591 | 0.980507258 |
| Dcp1a | 1255.323369 | 0.011026629 | 0.063706024 | 0.173086127 | 0.862583729 | 0.969759087 |
| F2rl3 | 6.835461511 | 0.011038114 | 0.053761691 | 0.205315598 | 0.837325558 | NA |
| Veph1 | 5.046815941 | 0.011045982 | 0.046327896 | 0.238430459 | 0.811547245 | NA |
| Ctbp2 | 733.2917133 | 0.011049755 | 0.0773146 | 0.142919383 | 0.88635385 | 0.976378471 |
| Mospd4 | 1.004578682 | 0.011056193 | 0.02469512 | 0.447707611 | 0.654364228 | NA |
| 2700070H01Rik | 14.14887958 | 0.011069538 | 0.071546069 | 0.154719026 | 0.87704283 | NA |
| Zfyve19 | 625.0546276 | 0.011070737 | 0.068796613 | 0.160919805 | 0.872156563 | 0.97153269 |
| Uck2 | 448.3182204 | 0.011073343 | 0.071525778 | 0.15481611 | 0.87696629 | 0.972549751 |
| Erlec1 | 2081.462225 | 0.011087235 | 0.064586137 | 0.171665864 | 0.863700224 | 0.969836459 |
| Trpv2 | 894.4255517 | 0.011095594 | 0.066618822 | 0.166553437 | 0.867721433 | 0.970480832 |
| Cd209e | 0.798890638 | 0.011101446 | 0.020255231 | 0.54807797 | 0.583638366 | NA |
| Vps52 | 3489.843625 | 0.011114846 | 0.05157284 | 0.21551743 | 0.829363924 | 0.959526943 |
| Map3k5 | 2173.506186 | 0.011118494 | 0.057991734 | 0.191725501 | 0.84795723 | 0.965542582 |
| Kcnq1 | 22.32249441 | 0.011126706 | 0.080932716 | 0.137480944 | 0.890650654 | NA |
| Prkcb | 55081.02425 | 0.011129493 | 0.061185359 | 0.181897979 | 0.855662791 | 0.967189923 |
| Crls1 | 888.2216621 | 0.01113035 | 0.063282323 | 0.175884026 | 0.860385055 | 0.96862798 |
| Mob3b | 240.2037888 | 0.011142405 | 0.087619322 | 0.127168358 | 0.898807148 | 0.978597654 |
| Rn4.5s | 0.535907978 | 0.011145937 | 0.016758314 | 0.665098932 | 0.505987218 | NA |
| Polr3gl | 703.5166762 | 0.011164137 | 0.070863626 | 0.157543969 | 0.874816158 | 0.971956739 |
| Nelfcd | 1829.640954 | 0.011170174 | 0.052720285 | 0.211876208 | 0.832203612 | 0.960519604 |
| Leng1 | 299.2239346 | 0.011179598 | 0.083600433 | 0.13372655 | 0.89361881 | 0.977844987 |
| Mettl21e | 0.960856407 | 0.011200871 | 0.01948884 | 0.574732545 | 0.565472193 | NA |
| S100b | 5442.853195 | 0.01120866 | 0.058453331 | 0.191754003 | 0.847934902 | 0.965542582 |
| Tbx20 | 1.078307322 | 0.011209801 | 0.020426187 | 0.548795565 | 0.583145754 | NA |
| Zpbp2 | 1.880278259 | 0.011219324 | 0.028004264 | 0.400629148 | 0.688693183 | NA |
| Zfp568 | 195.2849814 | 0.011238271 | 0.085813604 | 0.130961414 | 0.895805832 | 0.977844987 |
| Herc3 | 8361.88624 | 0.011241141 | 0.06957412 | 0.161570721 | 0.871643916 | 0.97153269 |
| Ldlrad3 | 582.5760565 | 0.011246243 | 0.066591396 | 0.168884327 | 0.865887627 | 0.970182416 |
| Kansl3 | 4135.203094 | 0.011248162 | 0.046615019 | 0.241299104 | 0.809323305 | 0.955725085 |
| Glcci1 | 2487.342573 | 0.011258021 | 0.067344239 | 0.167171253 | 0.867235303 | 0.970480832 |
| Speer4e | 1.330708063 | 0.011258228 | 0.027298755 | 0.412408094 | 0.680040332 | NA |
| Dnahc17 | 15.48493669 | 0.011275512 | 0.076055667 | 0.148253411 | 0.882142782 | NA |
| Tfap2e | 0.516571875 | 0.011276658 | 0.016626545 | 0.678232192 | 0.497624484 | NA |
| Abcb9 | 2392.384255 | 0.011283205 | 0.068467606 | 0.164796255 | 0.869104352 | 0.970788711 |
| Mettl9 | 2079.787327 | 0.011283787 | 0.059185659 | 0.190650692 | 0.848799272 | 0.965892061 |
| Serpinb2 | 1.563712549 | 0.011284524 | 0.023275465 | 0.484824867 | 0.627800582 | NA |
| Itfg3 | 808.9742712 | 0.011286059 | 0.074043348 | 0.152425024 | 0.878851721 | 0.973107714 |
| Gatm | 4266.368844 | 0.011294674 | 0.078631763 | 0.143640093 | 0.885784679 | 0.976107812 |
| Rps16 | 4338.508106 | 0.011295764 | 0.060942022 | 0.185352635 | 0.852952457 | 0.966905493 |
| Cdc20 | 39.91358251 | 0.011305038 | 0.091648479 | 0.123352164 | 0.901828235 | 0.979150581 |
| 4930415F15Rik | 5.804862899 | 0.011307453 | 0.049943353 | 0.22640556 | 0.820886004 | NA |
| Abr | 26431.83324 | 0.011337625 | 0.052735725 | 0.214989452 | 0.829775543 | 0.959659199 |
| Rnf113a2 | 308.1940034 | 0.011337702 | 0.078476285 | 0.144472974 | 0.885126996 | 0.975769681 |
| Tctex1d4 | 14.12539754 | 0.011352794 | 0.070103697 | 0.161942864 | 0.871350848 | NA |
| Ccdc38 | 3.356248116 | 0.011358316 | 0.037294148 | 0.30456027 | 0.760701083 | NA |
| Chchd10 | 4162.673959 | 0.011360866 | 0.057994419 | 0.19589585 | 0.844691686 | 0.964328285 |
| Tmprss2 | 3.008733408 | 0.011382404 | 0.040765003 | 0.279219988 | 0.780076005 | NA |
| Gdpd5 | 1500.242776 | 0.011385488 | 0.068636671 | 0.165880539 | 0.868250963 | 0.970658528 |
| Clec3a | 1.614509012 | 0.011409717 | 0.025907234 | 0.440406625 | 0.659642627 | NA |
| Tm4sf4 | 1.419526267 | 0.011410875 | 0.02255864 | 0.505831692 | 0.612974808 | NA |
| 9330117O12Rik | 109.0685205 | 0.011418709 | 0.093139374 | 0.122598086 | 0.90242537 | 0.979150581 |
| 1700091H14Rik | 9.518185756 | 0.011421288 | 0.063942093 | 0.178619232 | 0.858236692 | NA |
| Itm2b | 20904.76061 | 0.011436976 | 0.048153891 | 0.23750887 | 0.812262039 | 0.956219756 |
| A230065H16Rik | 39.22237975 | 0.011440471 | 0.081622666 | 0.140162923 | 0.888531266 | 0.977089223 |
| Pigh | 428.4001969 | 0.011440956 | 0.069443815 | 0.164751254 | 0.869139774 | 0.970788711 |
| Gp5 | 31.52605263 | 0.011449656 | 0.087899367 | 0.130258688 | 0.896361763 | 0.97803061 |
| Vezf1 | 1685.423193 | 0.011461667 | 0.074396996 | 0.154060879 | 0.877561734 | 0.972570049 |
| Tekt3 | 4.135558479 | 0.011471325 | 0.045528076 | 0.251961558 | 0.801070777 | NA |
| Zfp961 | 458.4466749 | 0.01147151 | 0.073028343 | 0.157082984 | 0.875179448 | 0.972003598 |
| Tmc7 | 838.5860981 | 0.011472304 | 0.07030303 | 0.163183641 | 0.870373846 | 0.971477516 |
| Bckdk | 1630.444517 | 0.011473976 | 0.070521231 | 0.16270244 | 0.870752726 | 0.97153269 |
| Gpr132 | 1.569175784 | 0.011477053 | 0.025336733 | 0.45298077 | 0.650562589 | NA |
| Hist1h3i | 0.916997476 | 0.011493331 | 0.018993099 | 0.605131956 | 0.545091323 | NA |
| Gm833 | 1.855722136 | 0.011528072 | 0.028549613 | 0.403790829 | 0.686366541 | NA |
| 1700054M17Rik | 0.703269883 | 0.011528433 | 0.016082734 | 0.716820454 | 0.473484889 | NA |
| Marveld3 | 18.21266796 | 0.011543008 | 0.080127201 | 0.14405854 | 0.885454243 | NA |
| Ttf2 | 110.9686642 | 0.011555854 | 0.094640293 | 0.122102901 | 0.902817525 | 0.979150581 |
| Cbs | 1064.927887 | 0.011568713 | 0.081151281 | 0.14255737 | 0.886639767 | 0.976496358 |
| Abcd2 | 2390.834851 | 0.011607071 | 0.066078407 | 0.17565604 | 0.860564173 | 0.96862798 |
| Qser1 | 771.3344142 | 0.011616152 | 0.070187648 | 0.165501373 | 0.868549369 | 0.970658528 |
| D330023K18Rik | 88.31312862 | 0.011618272 | 0.094083379 | 0.1234891 | 0.901719805 | 0.979150581 |
| 1700030K09Rik | 360.389532 | 0.011627265 | 0.079341804 | 0.146546515 | 0.883489973 | 0.975177867 |
| Papss1 | 2887.319488 | 0.011647366 | 0.04254743 | 0.273750168 | 0.784276621 | 0.949699053 |
| Cib3 | 0.534690201 | 0.011652295 | 0.016758269 | 0.695316166 | 0.486857185 | NA |
| Map1a | 59203.09631 | 0.011653337 | 0.064617602 | 0.180343067 | 0.856883247 | 0.967736072 |
| Tex21 | 0.710575567 | 0.011654619 | 0.016389593 | 0.711098775 | 0.477023029 | NA |
| Htr3b | 1.303480001 | 0.011660637 | 0.025973996 | 0.448935021 | 0.653478532 | NA |
| Ccrl1 | 9.099144152 | 0.011677893 | 0.043026854 | 0.27140941 | 0.786076165 | NA |
| Acnat1 | 6.584851945 | 0.011678917 | 0.040897183 | 0.28556777 | 0.775209191 | NA |
| Pcm1 | 4428.357916 | 0.011681623 | 0.07491289 | 0.155936087 | 0.8760834 | 0.972265831 |
| A1cf | 0.644943959 | 0.011695753 | 0.016073783 | 0.727629186 | 0.46684061 | NA |
| Brd4 | 4110.533811 | 0.011729756 | 0.051673486 | 0.226997578 | 0.820425626 | 0.957571905 |

| | | | | | | |
|---|---|---|---|---|---|---|
| Ipcef1 | 4322.602595 | -0.061627562 | 0.093203725 | -0.661213512 | 0.508475397 | 0.713494979 |
| Ints8 | 1020.09852 | 0.055765045 | 0.084351286 | 0.661104859 | 0.508545069 | 0.713519633 |
| Fbxl21 | 196.2633894 | -0.076841121 | 0.116252726 | -0.660983386 | 0.508622968 | 0.713564313 |
| Fam111a | 104.6795112 | 0.083906883 | 0.126958901 | 0.660897992 | 0.508677734 | 0.713576534 |
| D630023F18Rik | 135.7331276 | 0.081452907 | 0.123349451 | 0.660342681 | 0.509033947 | 0.713970788 |
| Pla2g16 | 1646.521191 | 0.052159074 | 0.078991058 | 0.660316191 | 0.509050942 | 0.713970788 |
| Ccdc125 | 9.840496829 | 0.101039374 | 0.155404129 | 0.650171746 | 0.515581289 | 0.71929855 |
| Pot1a | 673.4261343 | 0.039972889 | 0.060566312 | 0.659985522 | 0.50926312 | 0.714139088 |
| Pcmtd2 | 2460.232626 | 0.101034274 | 0.054037084 | 1.869721048 | 0.061522565 | 0.197596753 |
| Acaa2 | 775.6814445 | -0.045611337 | 0.069142775 | -0.659668873 | 0.509466346 | 0.714294776 |
| Scand1 | 256.0297069 | -0.110544716 | 0.167669101 | -0.659302847 | 0.509701315 | 0.714559552 |
| Srd5a1 | 381.0906811 | 0.058189768 | 0.088290038 | 0.659075131 | 0.509847525 | 0.71469986 |
| Gm20300 | 1470.080242 | -0.057975235 | 0.087978549 | -0.658970122 | 0.509914955 | 0.714729719 |
| Sdr39u1 | 1880.914238 | 0.044169919 | 0.067043251 | 0.658827225 | 0.510006722 | 0.714793682 |
| Atp8a2 | 1186.789217 | -0.045066529 | 0.068412071 | -0.658751138 | 0.510055589 | 0.714797512 |
| Acot11 | 2217.641878 | -0.044911922 | 0.068212935 | -0.65840771 | 0.510276182 | 0.714904317 |
| Glrx2 | 2004.262127 | -0.036522736 | 0.055464243 | -0.65849156 | 0.510222318 | 0.714904317 |
| Lrp11 | 7320.484085 | 0.041665119 | 0.063274997 | 0.658476824 | 0.510231784 | 0.714904317 |
| Slamf1 | 10.01821087 | -0.102334892 | 0.155442613 | -0.658345162 | 0.510316364 | 0.714904317 |
| Asap2 | 2096.4364 | -0.055209727 | 0.083904847 | -0.658004019 | 0.510535548 | 0.715146713 |
| BC005624 | 2630.260416 | 0.035681332 | 0.054246569 | 0.657762002 | 0.510691074 | 0.715235244 |
| Cdc5l | 2395.855423 | 0.059046842 | 0.089768533 | 0.657767708 | 0.510687407 | 0.715235244 |
| Grin2d | 423.2078632 | -0.072146485 | 0.109738563 | -0.657439676 | 0.510898247 | 0.715460723 |
| Sgtb | 5588.182931 | -0.044001447 | 0.066966308 | -0.657068438 | 0.511136912 | 0.715730259 |
| Plcxd2 | 4866.534976 | 0.04618678 | 0.070314334 | 0.656861508 | 0.511269971 | 0.715851882 |
| Nupl2 | 324.9585881 | 0.062066563 | 0.094500389 | 0.656786324 | 0.511318319 | 0.715854887 |
| Stk40 | 1501.653346 | 0.068828464 | 0.104815264 | 0.656664514 | 0.511396657 | 0.715899874 |
| 9230114K14Rik | 74.72735619 | -0.097603388 | 0.148707796 | -0.656343451 | 0.511603166 | 0.715959495 |
| Srgap2 | 3636.92434 | 0.101019028 | 0.056817456 | 1.777957612 | 0.075410817 | 0.225649903 |
| Ficd | 569.7072464 | -0.047824657 | 0.0728484 | -0.656495647 | 0.511505267 | 0.715959495 |
| Frmd4a | 1934.257791 | -0.051296985 | 0.078159576 | -0.656310936 | 0.511624082 | 0.715959495 |
| Hn1l | 213.5734729 | -0.059117318 | 0.090159155 | -0.65569956 | 0.512017452 | 0.716445263 |
| Psmc1 | 5287.499535 | 0.101010002 | 0.085173785 | 1.185928304 | 0.235650614 | 0.459613623 |
| Dnajc17 | 281.3286903 | -0.053439491 | 0.081521282 | -0.655528107 | 0.512127796 | 0.716470255 |
| Josd1 | 4038.679012 | -0.051677143 | 0.078875202 | -0.655176047 | 0.512354415 | 0.71672258 |
| Ankrd37 | 131.6074743 | -0.078180279 | 0.1195031 | -0.654211304 | 0.51297568 | 0.717275477 |
| Mkln1 | 2343.437338 | 0.062439143 | 0.095435254 | 0.654256582 | 0.512946513 | 0.717275477 |
| Plxnb2 | 2763.911546 | 0.043572347 | 0.066603725 | 0.65420285 | 0.512981126 | 0.717275477 |
| Rpusd1 | 2230.095845 | 0.052057969 | 0.079569104 | 0.654248525 | 0.512951703 | 0.717275477 |
| Zfp583 | 185.2716755 | 0.072050924 | 0.110092048 | 0.654460749 | 0.512815007 | 0.717275477 |
| Itpripl1 | 79.86457257 | 0.101002903 | 0.130320645 | 0.775033788 | 0.438319694 | 0.658832579 |
| Dctn4 | 3796.728018 | 0.038959984 | 0.059587899 | 0.653823762 | 0.513225356 | 0.717391525 |
| Inmt | 57.25964956 | -0.108668331 | 0.166184529 | -0.653901613 | 0.513175195 | 0.717391525 |
| Slc6a1 | 19579.76945 | -0.042153669 | 0.064476194 | -0.653786562 | 0.513249326 | 0.717391525 |
| Myo18a | 8327.400638 | -0.05511082 | 0.084327908 | -0.653530034 | 0.513414634 | 0.717557851 |
| Rrp1 | 10899.49526 | 0.037032424 | 0.056672661 | 0.653444243 | 0.513469924 | 0.717570398 |
| Slc25a51 | 3600.783044 | -0.039178021 | 0.059998946 | -0.652978476 | 0.513770155 | 0.717925214 |
| 4930470H14Rik | 74.55849228 | 0.088963175 | 0.13630062 | 0.652698245 | 0.513950834 | 0.718048172 |
| Tlcd1 | 534.5844104 | -0.067796244 | 0.103861458 | -0.65275652 | 0.513913259 | 0.718048172 |
| 4930414L22Rik | 377.4400261 | -0.064185528 | 0.098506115 | -0.65158927 | 0.514666171 | 0.718669124 |
| Abhd5 | 737.1301244 | -0.043647516 | 0.067002206 | -0.651434018 | 0.514766357 | 0.718669124 |
| Crcp | 968.8392653 | 0.041772177 | 0.064100178 | 0.651670218 | 0.514613939 | 0.718669124 |
| Dsc2 | 23.8512984 | -0.105055756 | 0.161267211 | -0.651439034 | 0.51476312 | 0.718669124 |
| Nipbl | 2271.018137 | -0.056247006 | 0.086324626 | -0.651575435 | 0.514675099 | 0.718669124 |
| Sec23a | 4109.50894 | -0.042092547 | 0.064610338 | -0.651483154 | 0.514734648 | 0.718669124 |
| Stx4a | 1582.050392 | -0.052893746 | 0.081147853 | -0.651819418 | 0.514517673 | 0.718669124 |
| Tapbpl | 381.2379022 | 0.085170422 | 0.130660001 | 0.651847711 | 0.514499419 | 0.718669124 |
| Ammecr1 | 63.94162102 | 0.090486458 | 0.13900143 | 0.65097502 | 0.515062612 | 0.718950077 |
| Ccdc36 | 27.00021538 | -0.107688336 | 0.165413621 | -0.651024599 | 0.515030608 | 0.718950077 |
| Fhod3 | 2739.679919 | 0.060192422 | 0.092484962 | 0.650834689 | 0.515153205 | 0.718950077 |
| Pcgf1 | 377.5119576 | 0.054443897 | 0.083650978 | 0.650845907 | 0.515145963 | 0.718950077 |
| Ercc6l | 9.12442639 | -0.103193516 | 0.15862621 | -0.650545178 | 0.515340129 | 0.719146173 |
| Tube1 | 91.00867004 | 0.088141395 | 0.135519221 | 0.65039774 | 0.515435336 | 0.719214256 |
| Hmg20a | 3633.93658 | 0.100931846 | 0.04485336 | 2.250262763 | 0.02443227 | 0.107850027 |
| Tmem200c | 400.2623587 | -0.055236186 | 0.084957778 | -0.650160435 | 0.515588595 | 0.71929855 |
| Sh3gl2 | 13216.14166 | 0.046136721 | 0.070986896 | 0.649932925 | 0.515735549 | 0.719438786 |
| Cnot7 | 3059.532693 | -0.040767707 | 0.062740284 | -0.649785182 | 0.515830992 | 0.719507147 |
| Acer3 | 338.6673427 | -0.069602883 | 0.107159734 | -0.649524604 | 0.515999349 | 0.719677191 |
| Mark2 | 4848.52171 | -0.04201597 | 0.064702656 | -0.649370098 | 0.516099187 | 0.719751647 |
| Dkc1 | 961.5832104 | -0.044313762 | 0.068256075 | -0.649228107 | 0.516190947 | 0.719814826 |
| Itpk1 | 2501.417543 | 0.057094067 | 0.087993865 | 0.64884145 | 0.516440863 | 0.720098518 |
| Fam129b | 1496.534332 | 0.054625308 | 0.084235839 | 0.648480606 | 0.516674151 | 0.720358642 |
| Tsen34 | 1580.711717 | -0.037735319 | 0.058196778 | -0.648409072 | 0.516720404 | 0.720358642 |
| Mcc | 653.3971365 | 0.063938071 | 0.098619825 | 0.648328779 | 0.516772324 | 0.720366207 |
| Dusp22 | 981.4855818 | -0.038997365 | 0.060158375 | -0.648244992 | 0.516826506 | 0.720376924 |
| Gpd1 | 4107.537257 | 0.086872363 | 0.134154185 | 0.647556114 | 0.51727209 | 0.720933144 |
| Ppid | 2146.842905 | -0.050151962 | 0.077474605 | -0.647334211 | 0.517415664 | 0.721003535 |
| Adamts16 | 161.0495322 | 0.100905745 | 0.14956402 | 0.674665905 | 0.499888051 | 0.70810314 |
| Tnip1 | 1096.68632 | 0.042473135 | 0.065673503 | 0.646731681 | 0.517805615 | 0.721482033 |
| Ogg1 | 187.7883143 | 0.069595619 | 0.107628123 | 0.646630428 | 0.51787116 | 0.721508475 |
| Sdad1 | 921.966244 | 0.041511823 | 0.064214799 | 0.646452582 | 0.517986296 | 0.721586379 |
| Synj1 | 17435.27485 | -0.045899033 | 0.071007147 | -0.646400185 | 0.51802022 | 0.721586379 |
| D15Ertd621e | 3147.90537 | -0.051168053 | 0.079185238 | -0.646181717 | 0.518161679 | 0.721718541 |
| Lnx2 | 238.7593756 | -0.078324106 | 0.121242407 | -0.646012465 | 0.518271283 | 0.721806315 |
| Dmap1 | 830.3518416 | 0.044788833 | 0.069347897 | 0.645857125 | 0.518371888 | 0.721881542 |
| Ncapg | 11.2407673 | -0.10423978 | 0.161429527 | -0.645729329 | 0.518454663 | 0.721931927 |
| Tssc4 | 951.2982165 | 0.041669878 | 0.064571401 | 0.645330247 | 0.518713195 | 0.722227017 |
| Lrrc2 | 62.76978077 | 0.092531546 | 0.143437642 | 0.645099465 | 0.518862731 | 0.722370307 |
| Mrps35 | 496.4656099 | 0.049927645 | 0.077407233 | 0.644999734 | 0.518927358 | 0.722395372 |
| Nek1 | 1385.700247 | 0.083251595 | 0.129122065 | 0.644751111 | 0.519088488 | 0.72255476 |
| 1700003D09Rik | 43.82252613 | 0.085030662 | 0.1319666 | 0.644334714 | 0.51935841 | 0.722636233 |
| 2410131K14Rik | 816.0703731 | 0.053641848 | 0.083255872 | 0.644301077 | 0.519380218 | 0.722636233 |
| 9330159F19Rik | 3103.338697 | 0.060516752 | 0.093896452 | 0.644505203 | 0.519247885 | 0.722636233 |
| Dbr1 | 401.35588 | 0.070488629 | 0.109393819 | 0.644356596 | 0.519344224 | 0.722636233 |
| Hist1h4i | 19.73407781 | -0.108068347 | 0.167665943 | -0.644545608 | 0.519221693 | 0.722636233 |
| Ythdc2 | 1294.903099 | 0.046270542 | 0.071824026 | 0.644220948 | 0.519432169 | 0.722643623 |
| Cnnm3 | 1326.818294 | 0.10089703 | 0.068471206 | 1.473568754 | 0.140597739 | 0.335147509 |
| BC006965 | 54.26947882 | 0.093213468 | 0.144762801 | 0.643904835 | 0.519637146 | 0.722789818 |
| Brf1 | 1415.820743 | 0.058267616 | 0.090499717 | 0.643843069 | 0.519677202 | 0.722789818 |
| Pcdhgb8 | 28.37376782 | -0.107206568 | 0.166621689 | -0.643413045 | 0.519956121 | 0.723112839 |
| Gm2518 | 9.716638256 | -0.102856743 | 0.160009862 | -0.64281502 | 0.520344136 | 0.723587511 |
| Nipal3 | 5842.832626 | -0.037013593 | 0.057596829 | -0.642632487 | 0.520462598 | 0.723687292 |
| Prrt4 | 146.166614 | 0.085590584 | 0.133222897 | 0.642461514 | 0.52057357 | 0.723755825 |
| Ccdc61 | 233.1276546 | 0.100886208 | 0.097459323 | 1.035162211 | 0.300593163 | 0.532151562 |
| Cacnb2 | 2039.924815 | 0.055614241 | 0.086622801 | 0.642027743 | 0.52085517 | 0.723931462 |
| Ip6k3 | 43.50025501 | 0.100827371 | 0.158175094 | 0.63744151 | 0.523837297 | 0.726488998 |
| Pcdhgb7 | 296.2653701 | -0.076019045 | 0.118403731 | -0.64203251 | 0.520852075 | 0.723931462 |
| Myof | 236.5276257 | -0.090006244 | 0.140356229 | -0.641270038 | 0.521347252 | 0.724527251 |
| Nkx2-2 | 131.7747054 | -0.094859034 | 0.147988809 | -0.640987889 | 0.521530551 | 0.724716983 |
| Tmf1 | 1538.944418 | 0.065097691 | 0.101581824 | 0.640839951 | 0.521626674 | 0.724785551 |
| Capn1 | 985.3789761 | 0.045722564 | 0.071415128 | 0.640236384 | 0.522018932 | 0.725200514 |
| Tlk2 | 2219.191218 | 0.05143114 | 0.08032691 | 0.640272854 | 0.521995226 | 0.725200514 |
| Ccdc106 | 1203.046213 | -0.047897149 | 0.074842159 | -0.639975511 | 0.52218852 | 0.725306045 |
| Ubxn2a | 1999.45366 | -0.049599362 | 0.077500924 | -0.639984135 | 0.522182914 | 0.725306045 |
| Rab9 | 906.0775379 | -0.040372734 | 0.063120025 | -0.639618478 | 0.522420666 | 0.725563446 |
| Mrpl39 | 836.5745324 | 0.052268282 | 0.081740295 | 0.639443275 | 0.522534605 | 0.725656643 |
| Gm10548 | 53.90553573 | -0.100998338 | 0.158018753 | -0.639154129 | 0.52272267 | 0.72574435 |
| Mtdh | 2391.944094 | 0.051508884 | 0.08057624 | 0.639256489 | 0.522656089 | 0.72574435 |
| Melk | 50.1864003 | 0.100809221 | 0.16713666 | 0.603154453 | 0.546405945 | 0.743600379 |
| Rnf168 | 1398.027547 | -0.04373162 | 0.068449387 | -0.638889871 | 0.522894578 | 0.725896299 |
| Tecpr1 | 3991.744248 | 0.047856251 | 0.074920387 | 0.638761395 | 0.522978166 | 0.725947294 |
| Pddc1 | 997.235046 | 0.040880322 | 0.064018346 | 0.638571983 | 0.523101412 | 0.726053327 |
| Bub3 | 2277.784628 | -0.036113814 | 0.056578208 | -0.638299011 | 0.523279056 | 0.726072113 |
| Prcp | 359.6245277 | 0.100797836 | 0.120186228 | 0.838680419 | 0.401648669 | 0.626453747 |
| Lztfl1 | 1260.164141 | 0.05368292 | 0.084126796 | 0.638119158 | 0.523396117 | 0.726072113 |
| Mapkapk3 | 127.0741021 | 0.0883659 | 0.138446679 | 0.638266664 | 0.523300108 | 0.726072113 |
| Snx9 | 397.0628377 | 0.057217226 | 0.089661311 | 0.638148439 | 0.523377057 | 0.726072113 |
| Zfp455 | 59.92333224 | -0.097254234 | 0.152405338 | -0.638128791 | 0.523389846 | 0.726072113 |
| Layn | 166.8733655 | -0.088601945 | 0.138906352 | -0.637853808 | 0.523568849 | 0.72624671 |
| Pabpc4l | 12.60977279 | -0.093729619 | 0.146998562 | -0.637622693 | 0.52371932 | 0.726390398 |
| Ncdn | 59910.32366 | 0.100755164 | 0.079235471 | 1.271591651 | 0.203518245 | 0.420609802 |
| Cdh3 | 13.24534967 | -0.094464389 | 0.148239638 | -0.63724109 | 0.523967817 | 0.726543599 |
| Kars | 2775.272535 | 0.046829847 | 0.073488895 | 0.637237044 | 0.523970451 | 0.726543599 |
| Egr3 | 2768.242867 | -0.106301971 | 0.166948183 | -0.636736314 | 0.524296615 | 0.7267399 |
| Fbxo6 | 613.6449152 | -0.049924584 | 0.078384947 | -0.636915448 | 0.52417992 | 0.7267399 |
| Supt6 | 7694.061734 | 0.056849634 | 0.089273494 | 0.63680306 | 0.524253133 | 0.7267399 |
| Tef | 18092.39366 | 0.045345165 | 0.071215501 | 0.636731674 | 0.524299639 | 0.7267399 |
| Tgm4 | 13.83775446 | -0.105044406 | 0.165006746 | -0.636606735 | 0.524381037 | 0.726787708 |
| Golgb1 | 4833.234087 | -0.041135464 | 0.064624803 | -0.636527499 | 0.524432664 | 0.726794248 |
| Igf1r | 3133.508271 | -0.063507302 | 0.09978715 | -0.636427654 | 0.524497721 | 0.726819399 |
| 7-Sep | 12745.39314 | -0.05304618 | 0.083373158 | -0.636250101 | 0.524613423 | 0.726900682 |
| Tmem234 | 1187.011318 | -0.048532045 | 0.076285019 | -0.636193655 | 0.524650208 | 0.726900682 |
| Capns1 | 4823.071507 | -0.040283336 | 0.063337598 | -0.63600985 | 0.524770002 | 0.727001647 |
| Ttll5 | 1373.173005 | 0.100745602 | 0.065969067 | 1.527164264 | 0.12672017 | 0.314647807 |
| Golga3 | 2238.857407 | 0.050772148 | 0.079939138 | 0.635135044 | 0.525340343 | 0.727661655 |
| Exosc6 | 231.0221759 | 0.063950315 | 0.100700333 | 0.635055647 | 0.525392122 | 0.727668324 |
| Dimt1 | 452.2432509 | 0.100711638 | 0.104720061 | 0.961722493 | 0.336189021 | 0.570214854 |
| Erg | 191.4667898 | -0.079169666 | 0.124868795 | -0.634022822 | 0.526065928 | 0.728341121 |
| Tbc1d22b | 2004.284419 | -0.039371482 | 0.062084872 | -0.634155807 | 0.525979145 | 0.728341121 |
| Zfp952 | 649.7016516 | -0.058069169 | 0.091560194 | -0.634218504 | 0.525938233 | 0.728341121 |
| Fbxo34 | 2960.242935 | 0.044421057 | 0.070095725 | 0.633719099 | 0.52626363 | 0.728549739 |
| Dsg2 | 94.47644739 | 0.080893946 | 0.127773901 | 0.63310227 | 0.52666686 | 0.728977696 |
| Fam173b | 187.0006998 | 0.066947971 | 0.105739806 | 0.633138777 | 0.526643021 | 0.728977696 |
| Aebp1 | 950.3378089 | 0.076083088 | 0.120233019 | 0.632796954 | 0.526866246 | 0.729188532 |
| Scyl2 | 1251.100999 | 0.100677447 | 0.073711666 | 1.365827868 | 0.171992997 | 0.379717797 |
| Mdh1 | 26459.33545 | -0.034885547 | 0.055173452 | -0.632288648 | 0.52719828 | 0.729452594 |
| Zfp282 | 681.6733305 | -0.043995786 | 0.069570118 | -0.632394871 | 0.527128885 | 0.729452594 |
| Atpaf1 | 2076.735692 | 0.050074687 | 0.079266253 | 0.631727692 | 0.527564829 | 0.729883917 |
| Clic6 | 341.2041643 | 0.097109486 | 0.153735152 | 0.631667414 | 0.527604226 | 0.729883917 |
| Ankle2 | 1616.249255 | -0.030098133 | 0.04767808 | -0.631278217 | 0.527858627 | 0.729981578 |
| Cx3cr1 | 2564.477999 | -0.102867546 | 0.162997696 | -0.63109816 | 0.527976344 | 0.729981578 |
| Fam89b | 774.0120373 | -0.054735119 | 0.086741355 | -0.631015267 | 0.528030542 | 0.729981578 |
| Fbxw2 | 2766.81002 | 0.045963242 | 0.072843884 | 0.63098286 | 0.528051731 | 0.729981578 |
| Snx11 | 1261.113865 | 0.100661921 | 0.061437226 | 1.63845159 | 0.101327525 | 0.272471248 |
| Ltb | 12.30251148 | -0.104778905 | 0.165967823 | -0.631320596 | 0.527830923 | 0.729981578 |
| Clstn1 | 46959.18858 | 0.100653816 | 0.08575787 | 1.173697705 | 0.24051613 | 0.465121931 |
| Tgfb2 | 442.117993 | -0.061199233 | 0.096973648 | -0.631091373 | 0.527980782 | 0.729981578 |
| Eif2b3 | 522.7548729 | 0.055382848 | 0.087823562 | 0.630614908 | 0.528292349 | 0.730249054 |
| Vps28 | 2761.78234 | -0.033859525 | 0.053715083 | -0.63035414 | 0.528462908 | 0.730419652 |
| Scube3 | 199.9302187 | -0.086657406 | 0.137547331 | -0.630018814 | 0.528682275 | 0.730657672 |
| Fam183b | 61.43152185 | 0.095710287 | 0.151940714 | 0.629918635 | 0.52874782 | 0.730663399 |
| Med6 | 500.2857781 | -0.043768209 | 0.069495823 | -0.629796262 | 0.528827892 | 0.730663399 |
| Nelfcd | 1648.764371 | 0.033798656 | 0.053660373 | 0.629862483 | 0.528784561 | 0.730663399 |
| Slc12a7 | 226.2823927 | 0.060929185 | 0.09675972 | 0.629695759 | 0.528893658 | 0.730689107 |
| Agtrap | 449.8254462 | -0.072665158 | 0.115425684 | -0.629540632 | 0.528995176 | 0.730714258 |
| Xkr8 | 250.7770511 | 0.100633531 | 0.130577934 | 0.770677927 | 0.440897859 | 0.661093969 |
| Rnmt | 2482.068486 | 0.038223503 | 0.060732024 | 0.629379696 | 0.529100507 | 0.730714258 |
| Zfp512 | 1802.820692 | 0.047356844 | 0.075233379 | 0.629465861 | 0.529044112 | 0.730714258 |
| 2700097O09Rik | 142.7715771 | 0.082579155 | 0.131329005 | 0.628796016 | 0.529482609 | 0.731176786 |
| Recql | 678.4482361 | 0.05051108 | 0.080362617 | 0.628539506 | 0.529650575 | 0.731343554 |
| Cdyl | 295.4605896 | 0.0566197 | 0.090110153 | 0.628338737 | 0.529782061 | 0.731394747 |
| Mrps31 | 593.567952 | -0.053230227 | 0.084710477 | -0.628378314 | 0.52975614 | 0.731394747 |
| Mtmr4 | 4564.498698 | -0.032761556 | 0.052155775 | -0.628148198 | 0.529906861 | 0.731501863 |
| Sft2d3 | 171.5549895 | 0.073949223 | 0.11775251 | 0.628005495 | 0.53000034 | 0.731565726 |
| Myo3b | 126.3486871 | -0.099161342 | 0.157929823 | -0.627882309 | 0.530081041 | 0.731611941 |
| Ncbp1 | 2316.705374 | 0.048992197 | 0.07804169 | 0.627769554 | 0.530154914 | 0.731648725 |
| Carhsp1 | 2368.750631 | -0.052322053 | 0.083361223 | -0.627654573 | 0.53023025 | 0.731687522 |
| H2-M5 | 17.88093326 | 0.100565126 | 0.162587423 | 0.618529554 | 0.536226323 | 0.736158233 |
| Ddt | 708.1271216 | 0.043160492 | 0.068857649 | 0.626807524 | 0.530785411 | 0.732282217 |
| Mapkapk2 | 1275.648947 | 0.041659968 | 0.066474222 | 0.626708614 | 0.530850256 | 0.732282217 |
| Rpgr | 647.8147627 | 0.041930537 | 0.066899644 | 0.626767718 | 0.530811507 | 0.732282217 |
| Slc4a3 | 6871.914398 | -0.059824271 | 0.095496172 | -0.626457269 | 0.531015056 | 0.732379142 |
| Usp33 | 5577.351477 | 0.03282913 | 0.052402143 | 0.626484492 | 0.530997206 | 0.732379142 |
| Thrap3 | 5381.623998 | 0.043353046 | 0.069220579 | 0.626302856 | 0.531116313 | 0.732453596 |
| Bicd1 | 856.095204 | -0.05028039 | 0.080299082 | -0.626163944 | 0.531207414 | 0.732514033 |
| Trak1 | 7825.333289 | -0.04780542 | 0.076364062 | -0.626019864 | 0.531301912 | 0.732579143 |
| Anapc5 | 7395.660668 | 0.044228291 | 0.070669562 | 0.625846408 | 0.531415689 | 0.732628482 |
| Msrb1 | 619.2041207 | 0.06036402 | 0.096455707 | 0.62582114 | 0.531432264 | 0.732628482 |
| Dhx40 | 1122.204085 | -0.049268844 | 0.078778998 | -0.625405831 | 0.531704736 | 0.732808502 |
| Trmt6 | 1044.242803 | 0.100409858 | 0.063191838 | 1.588968783 | 0.112067441 | 0.29211375 |
| Slco1a4 | 1929.963197 | 0.055663767 | 0.088986499 | 0.625530481 | 0.531622949 | 0.732808502 |
| Kif21b | 4825.106593 | -0.043278303 | 0.069211968 | -0.62530086 | 0.531773616 | 0.732838246 |
| Rpl7l1 | 1511.464988 | -0.05115991 | 0.081843643 | -0.625093267 | 0.531909847 | 0.732960794 |
| Abca2 | 13134.97386 | 0.100405064 | 0.080965264 | 1.240100496 | 0.214938226 | 0.43470211 |
| Dpf3 | 109.7959633 | -0.09229753 | 0.147700148 | -0.624898016 | 0.532037995 | 0.732989631 |
| Krt1 | 97.58030842 | 0.077008609 | 0.123272748 | 0.624701004 | 0.532167314 | 0.732989631 |
| Lin37 | 699.3363979 | 0.052433496 | 0.083930314 | 0.624726558 | 0.53215054 | 0.732989631 |
| Ppp1r37 | 5928.409961 | -0.044169327 | 0.070680581 | -0.6249146 | 0.532027109 | 0.732989631 |
| Clec1a | 31.45513116 | -0.104845159 | 0.167860038 | -0.624598684 | 0.532234483 | 0.733016986 |
| 1700052K11Rik | 236.121215 | -0.079151835 | 0.12684526 | -0.624003096 | 0.532625552 | 0.733426462 |
| Pcx | 3334.009733 | 0.100340958 | 0.060315891 | 1.66359074 | 0.096194244 | 0.262816719 |
| Atp5a1 | 32801.1374 | -0.044386378 | 0.071151088 | -0.623832735 | 0.532737439 | 0.733514075 |
| Cers5 | 1343.337097 | 0.100336246 | 0.061080942 | 1.642676789 | 0.100449824 | 0.270934267 |
| Zfp946 | 470.3382727 | 0.100323317 | 0.099727133 | 1.005978144 | 0.314426084 | 0.547605452 |
| A330102I10Rik | 90.9404528 | -0.078080844 | 0.125446355 | -0.622424174 | 0.533662989 | 0.734388578 |
| Aplp2 | 23520.06434 | 0.03047712 | 0.048970144 | 0.622361238 | 0.533704362 | 0.734388578 |
| Rpap3 | 919.3708231 | 0.100317204 | 0.08171714 | 1.227615203 | 0.219591452 | 0.440376644 |
| Nol12 | 601.0562568 | -0.047457393 | 0.076232766 | -0.622532739 | 0.533591623 | 0.734388578 |
| Ttc33 | 1370.817082 | -0.039638386 | 0.063683826 | -0.622424692 | 0.533662648 | 0.734388578 |
| Ndufv3 | 1645.812697 | -0.047548695 | 0.07641216 | -0.622266074 | 0.533766926 | 0.734409444 |
| Apool | 125.0352762 | -0.081257525 | 0.130656141 | -0.621918907 | 0.533995192 | 0.734639672 |
| Rpl7a | 7655.240134 | -0.039408786 | 0.06337169 | -0.621867368 | 0.534029084 | 0.734639672 |
| Edc3 | 1210.040075 | 0.052171063 | 0.083924873 | 0.621640057 | 0.534178575 | 0.734780082 |
| Creld1 | 2625.371495 | -0.046709211 | 0.075166271 | -0.62141185 | 0.534328677 | 0.734914706 |
| Nup62 | 1045.564111 | 0.041574386 | 0.066917855 | 0.621274935 | 0.534418742 | 0.734914706 |
| Senp5 | 3054.03657 | 0.045383872 | 0.073047173 | 0.621295385 | 0.53440529 | 0.734914706 |
| Adamtsl5 | 207.6198469 | 0.083627526 | 0.134747096 | 0.620625813 | 0.534845852 | 0.735428175 |
| Tceal5 | 2163.737861 | 0.046243593 | 0.074518748 | 0.620563199 | 0.53488706 | 0.735428175 |
| Cyhr1 | 5301.302516 | -0.034284229 | 0.05526523 | -0.620358022 | 0.535022103 | 0.735548577 |
| Bahd1 | 2142.065436 | 0.029429057 | 0.047471668 | 0.619928861 | 0.535304624 | 0.735675877 |
| Kbtbd11 | 6051.746415 | -0.035748122 | 0.0576477 | -0.620113583 | 0.53518301 | 0.735675877 |
| Ankrd44 | 412.8962872 | 0.100296663 | 0.103192648 | 0.971936129 | 0.331082325 | 0.566264437 |
| Pdp2 | 277.6550284 | 0.062889291 | 0.101441988 | 0.619953258 | 0.535288561 | 0.735675877 |
| Skiv2l2 | 2637.808367 | 0.036750094 | 0.059306006 | 0.619669009 | 0.535475723 | 0.735845752 |
| Ercc8 | 402.0540899 | -0.052930016 | 0.08545054 | -0.619422839 | 0.535637839 | 0.735872732 |

| | | | | | | |
|---|---|---|---|---|---|---|
| Slmap | 5241.713447 | 0.011735295 | 0.076205092 | 0.153996203 | 0.877612729 | 0.972570049 |
| Ccdc71l | 599.8711559 | 0.011746295 | 0.074860602 | 0.156908904 | 0.875316642 | 0.972008986 |
| Celf4 | 16709.70869 | 0.011750364 | 0.053030372 | 0.221578013 | 0.8246424 | 0.957903916 |
| Cdh11 | 1915.482287 | 0.011751222 | 0.076097828 | 0.154422562 | 0.877276565 | 0.972549751 |
| Kcnv2 | 0.636891649 | 0.011752306 | 0.012754461 | 0.921427092 | 0.356827489 | NA |
| 1700066B17Rik | 0.539767185 | 0.01175365 | 0.016794049 | 0.699869941 | 0.484008531 | NA |
| Ednrb | 2032.682548 | 0.011779932 | 0.077028702 | 0.152929119 | 0.878454173 | 0.973035729 |
| Rbms3 | 842.0536132 | 0.011786506 | 0.074214315 | 0.158817154 | 0.873812934 | 0.971912694 |
| 4933407E24Rik | 1.496508618 | 0.011786572 | 0.028396439 | 0.415072197 | 0.678089052 | NA |
| Acmsd | 0.723721657 | 0.011808887 | 0.021304402 | 0.554293299 | 0.579378135 | NA |
| Tctex1d1 | 10.45028329 | 0.01181661 | 0.066727215 | 0.177088318 | 0.859439018 | NA |
| Efcab11 | 8.751759098 | 0.011822572 | 0.058618302 | 0.20168738 | 0.840161129 | NA |
| Lrrc8e | 0.865135084 | 0.011826126 | 0.019225254 | 0.615134942 | 0.538465608 | NA |
| Osbpl2 | 2807.725604 | 0.011827432 | 0.047005682 | 0.251617058 | 0.801337072 | 0.953228631 |
| Myog | 0.570488403 | 0.011828053 | 0.016993193 | 0.696046506 | 0.486399706 | NA |
| Serpinh1 | 554.2349355 | 0.011829414 | 0.064814317 | 0.182512361 | 0.855180655 | 0.967189923 |
| Gm15915 | 1.165605285 | 0.011836953 | 0.022130187 | 0.534878117 | 0.592734141 | NA |
| Gm5640 | 2.472465543 | 0.011871704 | 0.032619103 | 0.363949429 | 0.715895773 | NA |
| Zfp407 | 740.275358 | 0.011881215 | 0.067470768 | 0.176094262 | 0.860219889 | 0.96862798 |
| Ttr | 10626.60361 | 0.011898093 | 0.024014963 | 0.495444974 | NA | NA |
| Zcrb1 | 1160.669351 | 0.011912671 | 0.076816815 | 0.155078955 | 0.876759073 | 0.972549751 |
| Wisp3 | 5.556606437 | 0.011921931 | 0.045597129 | 0.261462323 | 0.793736001 | NA |
| Tdo2 | 15.73434334 | 0.01192803 | 0.072632047 | 0.164225448 | 0.869553668 | NA |
| Vsig1 | 0.414170784 | 0.011937777 | 0.012862702 | 0.928092477 | 0.353359601 | NA |
| Cd300e | 7.623722105 | 0.011948184 | 0.051577745 | 0.231653872 | 0.816806862 | NA |
| Slc22a14 | 1.375413303 | 0.011951595 | 0.024913936 | 0.479715258 | 0.631429877 | NA |
| Tlr8 | 5.224068675 | 0.011958436 | 0.04382162 | 0.272888964 | 0.78493857 | NA |
| Tpm1 | 7665.884669 | 0.011960576 | 0.056612569 | 0.211270678 | 0.832676062 | 0.960519604 |
| Pck2 | 950.8260782 | 0.011960925 | 0.067904834 | 0.17614247 | 0.860182016 | 0.96862798 |
| Slc26a7 | 23.44703058 | 0.011963986 | 0.069842648 | 0.171299152 | 0.863988547 | 0.969836459 |
| Znrf1 | 6462.519745 | 0.011964105 | 0.070923202 | 0.168690988 | 0.866039707 | 0.970182416 |
| Srl | 390.3556637 | 0.011966363 | 0.073971428 | 0.161770068 | 0.871486925 | 0.97153269 |
| Sowaha | 3352.180346 | 0.011969214 | 0.080230786 | 0.149184805 | 0.881407809 | 0.974279441 |
| Pde5a | 572.2291674 | 0.011975298 | 0.075058307 | 0.159546611 | 0.873238238 | 0.971855551 |
| Arl13a | 0.403747135 | 0.011983694 | 0.012887768 | 0.929850236 | 0.352448632 | NA |
| Brsk1 | 16089.54361 | 0.011989383 | 0.059168419 | 0.202631461 | 0.839423098 | 0.96231236 |
| Thoc7 | 1221.891325 | 0.011990278 | 0.063305622 | 0.189403056 | 0.84977693 | 0.966260848 |
| Qsox1 | 1167.708755 | 0.012007536 | 0.065887617 | 0.182242689 | 0.855392273 | 0.967189923 |
| Cpxm2 | 109.6520991 | 0.012009001 | 0.085157053 | 0.141021801 | 0.88785272 | 0.976924608 |
| Sln | 8.029300378 | 0.012013516 | 0.053197554 | 0.225828345 | 0.821334931 | NA |
| Dram2 | 611.6657998 | 0.012024621 | 0.071245281 | 0.168777787 | 0.865971431 | 0.970182416 |
| Gng3 | 7853.265932 | 0.012039248 | 0.049253118 | 0.244436262 | 0.80689296 | 0.955587482 |
| Ilkap | 1318.278851 | 0.012040626 | 0.060438755 | 0.199220288 | 0.84209043 | 0.963252575 |
| Mettl13 | 501.4273588 | 0.01204822 | 0.076236975 | 0.158036438 | 0.874428086 | 0.971912694 |
| Rps18 | 4312.937453 | 0.012058638 | 0.055968778 | 0.215452951 | 0.82941419 | 0.959526943 |
| Prmt10 | 838.6071938 | 0.0120602 | 0.06579285 | 0.18330564 | 0.854558209 | 0.967189923 |
| Tmem92 | 0.557032348 | 0.012066325 | 0.016910504 | 0.713540265 | 0.475511507 | NA |
| Dars2 | 891.88776 | 0.01206886 | 0.066021992 | 0.182800602 | 0.854954477 | 0.967189923 |
| Tor1aip1 | 1899.511469 | 0.012073087 | 0.066677816 | 0.18106603 | 0.856315748 | 0.967598001 |
| Skint7 | 0.408982522 | 0.012084672 | 0.01294149 | 0.933792934 | 0.350410715 | NA |
| Ugt1a1 | 0.408982522 | 0.012084672 | 0.01294149 | 0.933792934 | 0.350410715 | NA |
| Gpr75 | 456.9022827 | 0.012108831 | 0.076571102 | 0.158138389 | 0.874347751 | 0.971912694 |
| Cdc27 | 2660.407192 | 0.012119333 | 0.062550119 | 0.193753959 | 0.846368539 | 0.965166054 |
| Deb1 | 790.5072205 | 0.012138258 | 0.075373159 | 0.161042186 | 0.872060174 | 0.97153269 |
| 2310065F04Rik | 3.374394011 | 0.01216075 | 0.042150758 | 0.288506085 | 0.772959373 | NA |
| Epo | 0.561306268 | 0.012168314 | 0.01694005 | 0.718316268 | 0.472562302 | NA |
| 4930414N06Rik | 0.427212126 | 0.012175077 | 0.012988784 | 0.937353083 | 0.348576966 | NA |
| Mir1947 | 0.427212126 | 0.012175077 | 0.012988784 | 0.937353083 | 0.348576966 | NA |
| Gm19990 | 1.254337168 | 0.012183772 | 0.025428408 | 0.479140188 | 0.631838902 | NA |
| Duox2 | 1.901967724 | 0.012208829 | 0.027857642 | 0.438257796 | 0.661199416 | NA |
| Cox5a | 4376.584947 | 0.012211699 | 0.072286336 | 0.168935099 | 0.865847691 | 0.970182416 |
| Idh2 | 1334.678393 | 0.012212989 | 0.07285747 | 0.167628503 | 0.866875546 | 0.97021201 |
| A2m | 207.2851278 | 0.012219474 | 0.054802349 | 0.22297355 | 0.823556093 | 0.957903916 |
| BC053393 | 2.691327572 | 0.012226567 | 0.037463864 | 0.326356282 | 0.74415481 | NA |
| Slc22a17 | 24948.50851 | 0.012226825 | 0.055640668 | 0.219746187 | 0.826068831 | 0.958041984 |
| Snapc3 | 611.681576 | 0.012231621 | 0.077606492 | 0.157610796 | 0.874763496 | 0.971956739 |
| Lias | 1124.591008 | 0.012255552 | 0.058463766 | 0.20962646 | 0.833959226 | 0.96105448 |
| Svip | 2125.12793 | 0.012255808 | 0.053749567 | 0.228016869 | 0.819633126 | 0.957571905 |
| Lcn12 | 0.432447513 | 0.012268512 | 0.013038109 | 0.940973227 | 0.346718579 | NA |
| Mir762 | 0.310639869 | 0.012274667 | 0.013050502 | 0.940551333 | 0.346934832 | NA |
| Ofcc1 | 0.310639869 | 0.012274667 | 0.013050502 | 0.940551333 | 0.346934832 | NA |
| Gale | 465.6589584 | 0.012276343 | 0.085965306 | 0.142805785 | 0.886443568 | 0.976378471 |
| Hbq1b | 4.426962402 | 0.012277141 | 0.047341033 | 0.259334038 | 0.795377519 | NA |
| Mir1b | 0.41913649 | 0.012277319 | 0.013043348 | 0.941270488 | 0.346566263 | NA |
| Ap2a1 | 13082.83682 | 0.012289117 | 0.055692255 | 0.220661147 | 0.825356286 | 0.957903916 |
| Wdr31 | 180.9244038 | 0.012289686 | 0.08910486 | 0.137923858 | 0.890300595 | 0.977182599 |
| Igfbp5 | 8495.664644 | 0.012294113 | 0.07630069 | 0.161127155 | 0.871993253 | 0.97153269 |
| Hccs | 848.2960044 | 0.012303456 | 0.063149555 | 0.194830442 | 0.845525689 | 0.964815881 |
| Ptms | 15928.3821 | 0.012314562 | 0.053259926 | 0.231216276 | 0.817146786 | 0.956302288 |
| Npy2r | 413.8489335 | 0.012326388 | 0.095068368 | 0.129658141 | 0.896836901 | 0.978039389 |
| Pgpep1 | 513.6042383 | 0.012327073 | 0.07120191 | 0.173128406 | 0.862550497 | 0.969759087 |
| Snx12 | 1844.879756 | 0.012327859 | 0.046635973 | 0.264342272 | 0.79151619 | 0.951728598 |
| Gm13298 | 0.839767923 | 0.012337958 | 0.019158515 | 0.643993464 | 0.519579673 | NA |
| Moap1 | 3.697028797 | 0.012338647 | 0.038958977 | 0.31670871 | 0.751464639 | NA |
| Krt78 | 1.146559101 | 0.012358345 | 0.024150256 | 0.511727298 | 0.608841878 | NA |
| Ufm1 | 2417.973957 | 0.012366585 | 0.059396007 | 0.208205654 | 0.835068394 | 0.961493618 |
| 4933416I08Rik | 1.102862949 | 0.012381148 | 0.024085222 | 0.514055805 | 0.607212975 | NA |
| Tmod4 | 7.516796618 | 0.012386477 | 0.05410268 | 0.228943869 | 0.818912544 | NA |
| Fgf20 | 0.315716853 | 0.012394842 | 0.01311372 | 0.945181266 | 0.344566342 | NA |
| Gprc6a | 0.315716853 | 0.012394842 | 0.01311372 | 0.945181266 | 0.344566342 | NA |
| Guca2a | 0.315716853 | 0.012394842 | 0.01311372 | 0.945181266 | 0.344566342 | NA |
| Olfr683 | 0.315716853 | 0.012394842 | 0.01311372 | 0.945181266 | 0.344566342 | NA |
| Slc6a18 | 0.315716853 | 0.012394842 | 0.01311372 | 0.945181266 | 0.344566342 | NA |
| 2010010A06Rik | 0.44014219 | 0.012404015 | 0.013109263 | 0.946202269 | 0.344045429 | NA |
| Capza2 | 7552.446365 | 0.012413687 | 0.069118645 | 0.17959969 | 0.857466848 | 0.967971163 |
| 4933400C23Rik | 1.526865223 | 0.012416311 | 0.028383554 | 0.437447366 | 0.66178694 | NA |
| Dhx29 | 1188.454684 | 0.012421913 | 0.066235978 | 0.187540272 | 0.851237051 | 0.966638801 |
| Paox | 260.7971857 | 0.012427306 | 0.091458613 | 0.135879011 | 0.891916929 | 0.97766264 |
| Ehd4 | 717.9992949 | 0.012451938 | 0.081352448 | 0.153061631 | 0.878349674 | 0.973035729 |
| Qars | 2318.228202 | 0.012455027 | 0.051500309 | 0.241843728 | 0.808901254 | 0.955725085 |
| Ccr2 | 6.074415662 | 0.012455119 | 0.046719645 | 0.266592756 | 0.789782734 | NA |
| Hsbp1l1 | 40.9693029 | 0.012456182 | 0.092767032 | 0.134273803 | 0.893186068 | 0.977844987 |
| Meiob | 0.318334547 | 0.012456712 | 0.013146139 | 0.947556725 | 0.343355165 | NA |
| Psg16 | 108.7579398 | 0.012469194 | 0.094581888 | 0.131834906 | 0.895114877 | 0.977844987 |
| Rab11fip1 | 164.6209819 | 0.012472512 | 0.091298343 | 0.136612691 | 0.891336944 | 0.977391435 |
| 5730488B01Rik | 0.429829819 | 0.012479621 | 0.013149382 | 0.949065186 | 0.342587459 | NA |
| Camkk2 | 13453.56029 | 0.012481506 | 0.060822335 | 0.205212549 | 0.837406065 | 0.961831963 |
| A930003O13Rik | 11.43787874 | 0.012491118 | 0.062786532 | 0.198945827 | 0.842305122 | NA |
| Hus1 | 712.7593272 | 0.01250885 | 0.073855967 | 0.169368171 | 0.865507058 | 0.970182416 |
| Gm13544 | 2.086943085 | 0.012514494 | 0.028090252 | 0.445510215 | 0.655951079 | NA |
| Actr2 | 12059.73247 | 0.012518461 | 0.06690285 | 0.187114014 | 0.851571241 | 0.96665003 |
| Irs3 | 10.91409598 | 0.012519612 | 0.057346432 | 0.218315444 | 0.827183342 | NA |
| 5730420D15Rik | 3.190337572 | 0.01252276 | 0.040082952 | 0.312421105 | 0.754720513 | NA |
| Ccdc42 | 64.57823312 | 0.012525894 | 0.089543462 | 0.139886188 | 0.888749914 | 0.977140573 |
| Gpr162 | 8079.021073 | 0.012528093 | 0.068024659 | 0.184169875 | 0.853880192 | 0.966905493 |
| Ftsj1 | 858.0402353 | 0.012530929 | 0.059142455 | 0.211877051 | 0.832202955 | 0.960519604 |
| Vmn2r57 | 27.56865001 | 0.012539787 | 0.082909937 | 0.151245886 | 0.879781755 | 0.973469666 |
| Fam174b | 1409.823147 | 0.012540782 | 0.070512861 | 0.177850992 | 0.85884 | 0.968369932 |
| Tyw1 | 653.1093517 | 0.012545993 | 0.060552806 | 0.207190941 | 0.835860742 | 0.961504006 |
| Srrm4 | 2162.002461 | 0.012554842 | 0.070260064 | 0.17869101 | 0.858180328 | 0.967971163 |
| Gm9871 | 2.197402966 | 0.012564886 | 0.03424113 | 0.366953025 | 0.713654058 | NA |
| Gba2 | 4240.566934 | 0.012586651 | 0.063845956 | 0.197140934 | 0.843717252 | 0.964060034 |
| Cwc15 | 1903.716914 | 0.012611185 | 0.075045941 | 0.168046193 | 0.866546939 | 0.97021201 |
| Dpy19l2 | 10.85011542 | 0.012616879 | 0.063192242 | 0.199658675 | 0.841747535 | NA |
| Hhipl1 | 192.2454467 | 0.012629709 | 0.084973827 | 0.148630579 | 0.881845143 | 0.974354774 |
| Nlrp2 | 3.004186556 | 0.012633036 | 0.039039172 | 0.323598984 | 0.746241657 | NA |
| Psmg3 | 331.7637159 | 0.012635072 | 0.079440105 | 0.159051552 | 0.873628258 | 0.971912694 |
| Sostdc1 | 94.29098429 | 0.01265434 | 0.038535334 | 0.328382775 | NA | NA |
| Mast1 | 4013.927499 | 0.012659342 | 0.061272864 | 0.206606004 | 0.836317571 | 0.961664 |
| Ifnk | 0.439983788 | 0.012665238 | 0.013245954 | 0.956159023 | 0.338991899 | NA |
| Xlr5a | 0.439983788 | 0.012665238 | 0.013245954 | 0.956159023 | 0.338991899 | NA |
| Prelp | 1208.443865 | 0.012665709 | 0.094565739 | 0.133935492 | 0.893453585 | 0.977844987 |
| Slitrk4 | 1127.917675 | 0.012668549 | 0.081342747 | 0.155742817 | 0.876235746 | 0.972360614 |
| 4933424G05Rik | 0.574923518 | 0.012679746 | 0.018411045 | 0.688703208 | 0.491010057 | NA |
| Ppp1r37 | 6378.701395 | 0.012687204 | 0.062023212 | 0.204555741 | 0.837919238 | 0.96195486 |
| Usf2 | 3627.870295 | 0.012688143 | 0.052201926 | 0.243058896 | 0.807959773 | 0.955716697 |
| Extl1 | 3390.34685 | 0.012691402 | 0.065129015 | 0.194865562 | 0.845498195 | 0.964815881 |
| Wbscr16 | 503.4276578 | 0.012691486 | 0.077518255 | 0.163722545 | 0.869949568 | 0.971097167 |
| Trim63 | 4.982828573 | 0.012702236 | 0.051852238 | 0.244969871 | 0.80647976 | NA |
| F630206G17Rik | 0.328758195 | 0.012702659 | 0.013274184 | 0.956944651 | 0.338595193 | NA |
| Decr2 | 1485.158105 | 0.012702843 | 0.059876576 | 0.212150461 | 0.831989654 | 0.960519604 |
| Syap1 | 942.2355852 | 0.01272579 | 0.066515031 | 0.191322021 | 0.84827331 | 0.965746177 |
| Zfp429 | 69.69011215 | 0.012734863 | 0.095018434 | 0.13402518 | 0.893382663 | 0.977844987 |
| 3110082I17Rik | 299.9144535 | 0.012740418 | 0.079939968 | 0.159374825 | 0.873373572 | 0.971912694 |
| Slc5a2 | 0.768725731 | 0.012751829 | 0.02149898 | 0.593136471 | 0.553089824 | NA |
| Nwd1 | 3572.357827 | 0.012756737 | 0.056294516 | 0.226607093 | 0.820729276 | 0.957583411 |
| 4930503O07Rik | 0.332178952 | 0.012772104 | 0.013310193 | 0.959573144 | 0.337270091 | NA |
| Gp1ba | 0.332178952 | 0.012772104 | 0.013310193 | 0.959573144 | 0.337270091 | NA |
| Rad1 | 361.353771 | 0.012773151 | 0.072564681 | 0.176024357 | 0.860274807 | 0.96862798 |
| Pgf | 83.36128123 | 0.012774468 | 0.093570581 | 0.136522269 | 0.891408421 | 0.977391435 |
| Lman1 | 1656.006792 | 0.012775806 | 0.049697747 | 0.257070132 | 0.797124634 | 0.952885371 |
| BC051665 | 1.560338262 | 0.012777045 | 0.022055202 | 0.579321152 | 0.562372496 | NA |
| Neurod1 | 282.2763105 | 0.012781217 | 0.08401987 | 0.152121356 | 0.87909122 | 0.973224531 |
| Elovl3 | 0.845218932 | 0.012829351 | 0.02079288 | 0.617006941 | 0.537230147 | NA |
| Syce1l | 6.117300296 | 0.012830361 | 0.052027579 | 0.246606918 | 0.80521245 | NA |
| En1 | 3.673177089 | 0.01284413 | 0.030320668 | 0.423609726 | 0.671850461 | NA |
| I730028E13Rik | 1.635074717 | 0.012851582 | 0.028794256 | 0.4463245 | 0.655362861 | NA |
| Gm12657 | 4.44738203 | 0.012873631 | 0.040363361 | 0.318943492 | 0.749769363 | NA |
| Krt86 | 0.337255936 | 0.012887313 | 0.013369576 | 0.96392829 | 0.335081879 | NA |
| Pogk | 3681.671189 | 0.012894028 | 0.062136153 | 0.207512497 | 0.835609634 | 0.961504006 |
| Wdr34 | 1137.36903 | 0.012895864 | 0.070671945 | 0.182475015 | 0.85520996 | 0.967189923 |
| Trim71 | 13.87673465 | 0.012898198 | 0.069051953 | 0.186789768 | 0.85182547 | NA |
| Lrp6 | 2312.212666 | 0.012903827 | 0.061952213 | 0.208286782 | 0.835005052 | 0.961493618 |
| Gm16065 | 1.077925408 | 0.012923613 | 0.023517295 | 0.549536544 | 0.582637293 | NA |
| Tmem45b | 1.661960464 | 0.012947481 | 0.028413376 | 0.455682595 | 0.64861824 | NA |
| Bclaf1 | 4679.09344 | 0.012969395 | 0.056429641 | 0.229833017 | 0.818221528 | 0.956955008 |
| Skor2 | 0.800420478 | 0.012971779 | 0.016636645 | 0.779711218 | 0.435560874 | NA |
| Plvap | 377.1127685 | 0.012973021 | 0.084558072 | 0.15342144 | 0.878065938 | 0.972905097 |
| Cp | 764.3784287 | 0.012979009 | 0.092961673 | 0.139616775 | 0.888962786 | 0.977182599 |
| Trdn | 11.93532316 | 0.012986349 | 0.065549433 | 0.198115353 | 0.842954813 | NA |
| 4930455D15Rik | 0.342602601 | 0.013011701 | 0.013433327 | 0.968613427 | 0.332738102 | NA |
| 4933433H22Rik | 0.342602601 | 0.013011701 | 0.013433327 | 0.968613427 | 0.332738102 | NA |
| Serpina1d | 0.342602601 | 0.013011701 | 0.013433327 | 0.968613427 | 0.332738102 | NA |
| Gpc6 | 517.8641111 | 0.013015837 | 0.070851873 | 0.183704908 | 0.854244958 | 0.967189923 |
| Raly | 2980.844934 | 0.013017363 | 0.051321729 | 0.253642332 | 0.799771889 | 0.953228631 |
| Fam219b | 830.3210016 | 0.013028966 | 0.065515119 | 0.198869601 | 0.84236475 | 0.963252575 |
| A930003A15Rik | 9.866450946 | 0.013029732 | 0.066777575 | 0.195121369 | 0.845297934 | NA |
| Rps19bp1 | 512.1755314 | 0.013086325 | 0.068413813 | 0.191281905 | 0.848304737 | 0.965746177 |
| Etv3 | 1394.00133 | 0.013108576 | 0.064811792 | 0.202256025 | 0.839716576 | 0.962420939 |
| A530065N20Rik | 1.025780582 | 0.013109745 | 0.01944062 | 0.674348119 | 0.500090019 | NA |
| Snd1 | 3344.794664 | 0.013110647 | 0.046059164 | 0.284647963 | 0.775913861 | 0.947241001 |
| Sdsl | 91.01492069 | 0.013125192 | 0.095048221 | 0.138089826 | 0.890169426 | 0.977182599 |
| Unc119 | 880.7050804 | 0.013151766 | 0.069530404 | 0.189151289 | 0.849974244 | 0.966260848 |
| Eef1b2 | 4513.658928 | 0.01315449 | 0.047402547 | 0.277505979 | 0.781391615 | 0.948894514 |
| Cpne1 | 1669.331385 | 0.013155312 | 0.046452957 | 0.283196439 | 0.777026258 | 0.947474493 |
| Sorcs1 | 2350.759028 | 0.013163446 | 0.062274658 | 0.211377255 | 0.832592904 | 0.960519604 |
| Akr1c19 | 0.576654935 | 0.013166549 | 0.018421288 | 0.714746351 | 0.474765791 | NA |
| Myh8 | 19.65741406 | 0.013169485 | 0.073502204 | 0.179171292 | 0.857803205 | NA |
| Cma1 | 0.892192393 | 0.013190494 | 0.020530658 | 0.642477894 | 0.520562938 | NA |
| Pot1a | 761.8136483 | 0.013195198 | 0.073924182 | 0.178496374 | 0.858333169 | 0.967971163 |
| Sptbn1 | 40931.29879 | 0.013199127 | 0.052201732 | 0.252848446 | 0.800385329 | 0.953228631 |
| Zfp868 | 494.6570626 | 0.013202539 | 0.078482317 | 0.16822311 | 0.866407761 | 0.97021201 |
| Tlcd1 | 589.0941234 | 0.013217468 | 0.081866524 | 0.161451437 | 0.871737857 | 0.97153269 |
| Sumf1 | 958.1317009 | 0.013235605 | 0.07040353 | 0.187996326 | 0.850879531 | 0.966445857 |
| Gcgr | 0.868302965 | 0.013245089 | 0.020874523 | 0.634509763 | 0.525748197 | NA |
| Commd1 | 602.4585136 | 0.013260428 | 0.067645608 | 0.196027918 | 0.844588314 | 0.964328285 |
| Prss38 | 1.921803007 | 0.013265411 | 0.032120455 | 0.41298951 | 0.6796143 | NA |
| Lrat | 6.48853038 | 0.013274535 | 0.046646702 | 0.284576063 | 0.775968952 | NA |
| Dak | 479.2199325 | 0.013296118 | 0.077853064 | 0.170784769 | 0.864393006 | 0.969963136 |
| Acp2 | 4016.718641 | 0.013302684 | 0.052082551 | 0.255415371 | 0.7984023 | 0.952967259 |
| Hoxb5 | 0.521648859 | 0.013313797 | 0.018053322 | 0.737470742 | 0.460836129 | NA |
| E030044B06Rik | 0.657073752 | 0.013341105 | 0.017806147 | 0.74924154 | 0.453711636 | NA |
| Mcrs1 | 2815.126758 | 0.01336017 | 0.069297162 | 0.192795339 | 0.847119254 | 0.965275633 |
| P4htm | 2996.02623 | 0.013367918 | 0.058679462 | 0.227812551 | 0.819791969 | 0.957571905 |
| Pcp4 | 8286.438994 | 0.013373688 | 0.084969963 | 0.15739312 | 0.874935035 | 0.971956739 |
| R3hcc1l | 626.4440671 | 0.013386962 | 0.080135151 | 0.167054807 | 0.867326925 | 0.970480832 |
| Naa35 | 1880.207888 | 0.013390068 | 0.052366794 | 0.255697693 | 0.798184277 | 0.952967259 |
| Wdr48 | 3806.363821 | 0.013397576 | 0.044423217 | 0.301589504 | 0.76296501 | 0.944431816 |
| Snord35a | 4.997169502 | 0.013402147 | 0.045552082 | 0.294215897 | 0.768592934 | NA |
| Slc7a10 | 1166.429157 | 0.013418364 | 0.081607085 | 0.164426462 | 0.869395433 | 0.970842335 |
| Prg2 | 0.840452317 | 0.013437508 | 0.020369723 | 0.659680463 | 0.509458907 | NA |
| 4931403E22Rik | 24.28644521 | 0.013442659 | 0.064278461 | 0.209131631 | 0.834345483 | 0.961118357 |
| Nbea | 11600.90382 | 0.013451697 | 0.067819399 | 0.198345865 | 0.842774469 | 0.963458383 |
| Dazl | 130.4178549 | 0.013464608 | 0.094951912 | 0.141804492 | 0.887234436 | 0.97661227 |
| Insl3 | 3.262886568 | 0.013464911 | 0.038105169 | 0.353361795 | 0.723817224 | NA |
| Zfp428 | 733.8074719 | 0.013466682 | 0.069313374 | 0.194286917 | 0.845951229 | 0.964996028 |
| Rab38 | 12.27904393 | 0.013501342 | 0.070223024 | 0.192263749 | 0.847535613 | NA |
| Thap4 | 797.9404761 | 0.013521694 | 0.065266901 | 0.207175359 | 0.835872911 | 0.961504006 |
| 1110008F13Rik | 1327.275663 | 0.013538139 | 0.057960003 | 0.233577263 | 0.815313181 | 0.956264994 |
| Rprd2 | 2354.08606 | 0.013541269 | 0.060684908 | 0.223140643 | 0.823426047 | 0.957903916 |
| Wfdc17 | 5.58392691 | 0.013546603 | 0.054031175 | 0.250718271 | 0.802031934 | NA |
| Vps39 | 4167.094697 | 0.013546611 | 0.049069029 | 0.276072524 | 0.782492363 | 0.949184473 |
| Aacs | 1801.540069 | 0.013552608 | 0.065844945 | 0.20582609 | 0.836926761 | 0.961803649 |
| Rsl24d1 | 1374.753259 | 0.013563238 | 0.064085336 | 0.211643385 | 0.832385259 | 0.960519604 |
| Msc | 46.55535208 | 0.013568484 | 0.091446427 | 0.148376312 | 0.882045793 | 0.97439733 |
| Arid2 | 2013.26159 | 0.013575355 | 0.064254131 | 0.21127599 | 0.832671918 | 0.960519604 |
| 4930520O04Rik | 26.40960403 | 0.013597435 | 0.084772173 | 0.160399746 | 0.87256619 | 0.97153269 |
| Rngtt | 931.8290965 | 0.013608706 | 0.073327301 | 0.185588528 | 0.85276745 | 0.966905493 |
| Ube3c | 4742.603854 | 0.013609404 | 0.06132873 | 0.221909111 | 0.824384638 | 0.957903916 |
| Gyltl1b | 12.73194804 | 0.013610425 | 0.044315487 | 0.307125698 | 0.758747698 | NA |
| H2-M2 | 1.035380625 | 0.01361292 | 0.024758831 | 0.549820791 | 0.582442297 | NA |
| Ppp1r21 | 3123.027819 | 0.013621798 | 0.043442477 | 0.313559409 | 0.753855693 | 0.94183464 |
| Olfr550 | 3.7527994 | 0.01363282 | 0.037026424 | 0.368191634 | 0.712730348 | NA |
| Ankfy1 | 2756.717284 | 0.01363551 | 0.05030553 | 0.271053893 | 0.786349583 | 0.950484821 |
| Kif6 | 100.3460042 | 0.013637431 | 0.094264128 | 0.144672545 | 0.884969417 | 0.975769681 |

| | | | | | | | | | | | | | |
|---|---|---|---|---|---|---|---|---|---|---|---|---|---|
| Ghitm | 15912.52682 | -0.037622113 | 0.060728998 | -0.619508215 | 0.535581611 | 0.735872732 | Frzb | 804.6025923 | 0.013669381 | 0.08680788 | 0.157467051 | 0.874876774 | 0.971956739 |
| Gm5595 | 130.8863771 | -0.082146449 | 0.132613413 | -0.619442988 | 0.535624569 | 0.735872732 | Ctsb | 29907.83668 | 0.013669619 | 0.052312921 | 0.261304827 | 0.793857445 | 0.951728598 |
| Plp2 | 252.1191319 | -0.074619661 | 0.120494753 | -0.619277263 | 0.53573372 | 0.735939202 | Gm10334 | 16.67583771 | 0.01368737 | 0.074774821 | 0.183047851 | 0.854760473 | NA |
| Card14 | 12.82305078 | -0.102694603 | 0.165891783 | -0.619045751 | 0.53588622 | 0.73608343 | Pdxk-ps | 2.898917696 | 0.013700424 | 0.041107974 | 0.333278989 | 0.738923698 | NA |
| Dpyd | 226.7685054 | 0.080379705 | 0.129957377 | 0.618508213 | 0.536240386 | 0.736158233 | Rgs1 | 7.654425543 | 0.013706567 | 0.061864114 | 0.221559261 | 0.824656999 | NA |
| Lrrc46 | 31.38138717 | 0.100290693 | 0.167246033 | 0.599659622 | 0.548733103 | 0.745142043 | Cox4i2 | 46.08836447 | 0.013713474 | 0.09282012 | 0.147742469 | 0.882546014 | 0.974672197 |
| Lamb2 | 1323.052436 | -0.046695182 | 0.075474616 | -0.618687239 | 0.536122418 | 0.736158233 | Rnpep | 1623.769919 | 0.013735907 | 0.058289014 | 0.235651737 | 0.813702924 | 0.956264994 |
| Mro | 428.4483635 | -0.05254895 | 0.084949701 | -0.618588987 | 0.536187159 | 0.736158233 | Gfra2 | 3117.68483 | 0.013743786 | 0.064255695 | 0.213892101 | 0.8306312 | 0.960113696 |
| Naa15 | 2153.295549 | -0.043399391 | 0.07017351 | -0.618458319 | 0.536273265 | 0.736158233 | 2410004P03Rik | 105.2778305 | 0.013765499 | 0.093886557 | 0.146618428 | 0.883433208 | 0.975177867 |
| Nrg4 | 15.28238527 | 0.097821348 | 0.158100612 | 0.618728457 | 0.53609526 | 0.736158233 | Eya3 | 1773.960511 | 0.013769783 | 0.046223302 | 0.297897005 | 0.765781774 | 0.945453142 |
| Trim17 | 126.1370945 | 0.094115192 | 0.152121909 | 0.618682693 | 0.536125413 | 0.736158233 | Sf3a2 | 1385.496062 | 0.013783669 | 0.056081647 | 0.245778614 | 0.805853613 | 0.95539795 |
| Agpat9 | 208.0184076 | 0.072858375 | 0.117843968 | 0.618261389 | 0.536403049 | 0.736259913 | E230008N13Rik | 35.43349274 | 0.013785332 | 0.077807341 | 0.177172638 | 0.859372787 | 0.968520191 |
| Lamb3 | 83.49485168 | 0.087757245 | 0.141961031 | 0.618178415 | 0.536457737 | 0.736259913 | Ankmy2 | 2044.826234 | 0.013797188 | 0.057997721 | 0.237891905 | 0.811964934 | 0.956219756 |
| Tcf7l2 | 186.6188788 | -0.099109552 | 0.160337812 | -0.618129627 | 0.536489894 | 0.736259913 | Palmd | 636.2300338 | 0.013802035 | 0.085268287 | 0.161865986 | 0.871411389 | 0.97153269 |
| Ehd1 | 1491.853617 | -0.052175095 | 0.084444918 | -0.617859503 | 0.536667956 | 0.73626355 | Dhrs1 | 2454.266828 | 0.013809179 | 0.05814503 | 0.237495428 | 0.812272466 | 0.956219756 |
| Hist1h2be | 58.81570444 | -0.103846947 | 0.168101004 | -0.617765179 | 0.53673014 | 0.73626355 | Gm10791 | 9.762593256 | 0.013818162 | 0.058325239 | 0.236915649 | 0.81272223 | NA |
| Vgf | 8548.986705 | 0.067707776 | 0.109574586 | 0.617914957 | 0.536631399 | 0.73626355 | Cdh7 | 704.2156165 | 0.013821288 | 0.078412046 | 0.176264854 | 0.860085872 | 0.96862798 |
| Zfyve20 | 2757.05948 | 0.028945175 | 0.04684972 | 0.61783026 | 0.536687234 | 0.73626355 | Mrps27 | 1139.056753 | 0.013853496 | 0.061077325 | 0.226818972 | 0.820564511 | 0.957571905 |
| Zkscan4 | 201.4303729 | -0.067147047 | 0.108669605 | -0.617900905 | 0.536640663 | 0.73626355 | Nfic | 6148.513303 | 0.013875945 | 0.054471163 | 0.254739289 | 0.798924469 | 0.953053772 |
| Atl2 | 2255.425341 | -0.03755591 | 0.060810497 | -0.617589261 | 0.536846126 | 0.736292279 | Cpvl | 1.058559131 | 0.013882805 | 0.024015921 | 0.578066723 | 0.563219073 | NA |
| Amhr2 | 28.53907677 | 0.100265793 | 0.16528411 | 0.606626936 | 0.54409852 | 0.742035249 | Gm5475 | 1.341305976 | 0.013902756 | 0.026904964 | 0.516735705 | 0.605340668 | NA |
| Ubn2 | 1819.359699 | -0.070445448 | 0.114093678 | -0.617435157 | 0.536947739 | 0.73636646 | Sirpa | 23465.59815 | 0.013904594 | 0.047235093 | 0.294369996 | 0.768475192 | 0.94619074 |
| Nuf2 | 81.0874181 | -0.099348095 | 0.160925455 | -0.617354755 | 0.537000758 | 0.736373993 | Ndufa4 | 6566.228299 | 0.013907763 | 0.062982756 | 0.220818593 | 0.825233686 | 0.957903916 |
| Sgms2 | 149.0550072 | 0.084245542 | 0.136566766 | 0.616881723 | 0.537312743 | 0.736698119 | P2ry4 | 1.112289045 | 0.013917685 | 0.024908246 | 0.558758126 | 0.576326804 | NA |
| Slc25a21 | 19.9167664 | -0.103663506 | 0.168052424 | -0.616852194 | 0.537332222 | 0.736698119 | Dennd5a | 15005.83809 | 0.013932876 | 0.043457777 | 0.320607188 | 0.74850809 | 0.9410864 |
| Slitrk1 | 3530.036347 | 0.040679851 | 0.065956403 | 0.616768794 | 0.537387238 | 0.736708358 | Smpx | 2.231234673 | 0.013938537 | 0.032640933 | 0.427026311 | 0.669360164 | NA |
| Scrn3 | 1159.564495 | -0.050132954 | 0.081332398 | -0.61639587 | 0.537633279 | 0.736980449 | Gprin3 | 354.6743416 | 0.013941848 | 0.092762288 | 0.150296513 | 0.880530684 | 0.973779118 |
| Csnk2a2 | 1366.077064 | 0.047843406 | 0.077672166 | 0.615965911 | 0.537917019 | 0.737262992 | Rassf1 | 494.3528427 | 0.013955707 | 0.073724985 | 0.189294131 | 0.849862295 | 0.966260848 |
| Eif3g | 1941.83215 | 0.100211182 | 0.069224612 | 1.447623599 | 0.147722347 | 0.347157599 | Sh3gl2 | 14822.09849 | 0.013955833 | 0.058165247 | 0.239934226 | 0.810381247 | 0.955876688 |
| Actb | 35544.54328 | -0.031999904 | 0.051985318 | -0.615556571 | 0.538187221 | 0.737397186 | Nmnat3 | 125.745363 | 0.013975328 | 0.093581248 | 0.149338982 | 0.881286157 | 0.974259968 |
| Dgkg | 3940.089015 | 0.072282542 | 0.117397913 | 0.61570551 | 0.5380889 | 0.737397186 | Oca2 | 27.14640978 | 0.01398712 | 0.035920952 | 0.389386098 | NA | NA |
| Dok3 | 195.4260009 | 0.096471435 | 0.156736028 | 0.61550261 | 0.538222846 | 0.737397186 | Bhlhe41 | 2044.277858 | 0.014018187 | 0.072636474 | 0.192991014 | 0.846966006 | 0.965275633 |
| Agtr2 | 20.24657334 | 0.100166497 | 0.166417765 | 0.601897865 | 0.547242126 | 0.744128707 | Abhd11 | 900.5424293 | 0.014019949 | 0.067687423 | 0.207127832 | 0.835910027 | 0.961504006 |
| Trp53i11 | 3894.626471 | 0.061101017 | 0.099285958 | 0.61540441 | 0.53828768 | 0.737420805 | Adam25 | 0.520319392 | 0.014023599 | 0.013929525 | 1.0067535 | 0.314053246 | NA |
| Zfp174 | 351.5371279 | -0.060609416 | 0.09850853 | -0.615270737 | 0.538375939 | 0.73747651 | Snf8 | 1869.125067 | 0.01402412 | 0.04943263 | 0.283701668 | 0.776639017 | 0.947430065 |
| Tbx15 | 55.83725186 | -0.100110884 | 0.162743339 | -0.615145818 | 0.538458426 | 0.737524298 | Pcdh12 | 12.56836415 | 0.014033629 | 0.068196954 | 0.205780875 | 0.836962081 | NA |
| Rnase4 | 565.5582172 | -0.046372382 | 0.075431134 | -0.614764477 | 0.538710272 | 0.737738815 | 4933411K16Rik | 1.923758284 | 0.01403653 | 0.029014759 | 0.483772056 | 0.628547649 | NA |
| Surf6 | 658.4674457 | 0.038865903 | 0.063220081 | 0.614771481 | 0.538705646 | 0.737738815 | Selenbp2 | 0.93937571 | 0.014045516 | 0.021103851 | 0.665542828 | 0.50570336 | NA |
| Lcp2 | 132.6095193 | 0.100106601 | 0.147561636 | 0.678405336 | 0.497514727 | 0.706229334 | Phf14 | 1530.776321 | 0.014046966 | 0.05769546 | 0.243467448 | 0.807643299 | 0.955716697 |
| Pde4b | 5451.238946 | -0.049282552 | 0.08020949 | -0.614422954 | 0.538935872 | 0.73791732 | Ilk | 2853.277738 | 0.014063297 | 0.05060426 | 0.277907371 | 0.781083465 | 0.948894514 |
| Ppie | 367.9067388 | -0.058542812 | 0.095339549 | -0.614045401 | 0.539185326 | 0.738193642 | 1700018A04Rik | 2.040357079 | 0.014071801 | 0.029442062 | 0.477948901 | 0.632686575 | NA |
| Nhsl1 | 1021.149801 | 0.047931214 | 0.078070729 | 0.613946035 | 0.539250988 | 0.738218308 | Nobox | 1.318592192 | 0.014090647 | 0.023115311 | 0.60958069 | 0.542139606 | NA |
| 9130024F11Rik | 331.9156161 | 0.085349611 | 0.1391079 | 0.613549704 | 0.539512928 | 0.738511645 | Dapk2 | 71.35436094 | 0.014093413 | 0.095024904 | 0.148312832 | 0.882095889 | 0.97439733 |
| Dusp4 | 353.8899367 | -0.095902776 | 0.156397259 | -0.613199851 | 0.539744203 | 0.738762958 | Garnl3 | 3961.693254 | 0.014100456 | 0.050567772 | 0.278842732 | 0.780365519 | 0.948894514 |
| Mtif3 | 610.4618012 | -0.044842761 | 0.073146366 | -0.613055218 | 0.539839829 | 0.738828576 | Folr1 | 129.7458546 | 0.014104686 | 0.027343802 | 0.515827529 | NA | NA |
| Fam172a | 1597.050508 | 0.042696262 | 0.069689995 | 0.612659845 | 0.540101279 | 0.73899057 | Ubqln4 | 6796.688491 | 0.014107151 | 0.056238808 | 0.250843704 | 0.80193495 | 0.95333733 |
| Gm4636 | 294.9924339 | -0.063781232 | 0.104089152 | -0.6127558 | 0.54003782 | 0.73899057 | Samd5 | 271.5793756 | 0.014109989 | 0.086625443 | 0.162885042 | 0.870608948 | 0.97153269 |
| Mlkl | 9.256713253 | -0.097027364 | 0.15835337 | -0.61272687 | 0.540056953 | 0.73899057 | Ero1lb | 973.709698 | 0.014119295 | 0.062387662 | 0.226315507 | 0.820956039 | 0.957611694 |
| Zmynd11 | 7853.757492 | 0.028440169 | 0.046432953 | 0.612499673 | 0.540207214 | 0.739070249 | Smlr1 | 1.895823843 | 0.014119324 | 0.02713959 | 0.520248253 | 0.602890558 | NA |
| Ppp3cc | 756.8253611 | -0.055801956 | 0.091184399 | -0.611968237 | 0.540558773 | 0.739420643 | Gm10248 | 0.72587044 | 0.014124629 | 0.016792333 | 0.84113557 | 0.40027199 | NA |
| Zfp251 | 2040.689552 | 0.056006586 | 0.091508369 | 0.612037854 | 0.540512713 | 0.739420643 | Gm20594 | 1.939887592 | 0.014143438 | 0.030784643 | 0.459431605 | 0.645924257 | NA |
| Adck1 | 1341.360054 | -0.04188061 | 0.068458864 | -0.611763142 | 0.540694479 | 0.739475705 | Magi3 | 3046.552019 | 0.014147284 | 0.060558142 | 0.233614902 | 0.815283958 | 0.956264994 |
| Dgcr2 | 5620.361633 | 0.100082575 | 0.06824226 | 1.466577677 | 0.142490976 | 0.338457027 | Gm5918 | 153.8066863 | 0.014147932 | 0.091537723 | 0.154558488 | 0.877169398 | 0.972549751 |
| S100a7a | 13.59255941 | 0.098574394 | 0.161317442 | 0.6110585 | 0.541160853 | 0.740048213 | Dpy19l1 | 5085.751615 | 0.014160996 | 0.058877538 | 0.240516104 | 0.809930181 | 0.955876688 |
| Gm16861 | 30.33648144 | -0.101760058 | 0.166561241 | -0.610946808 | 0.541234795 | 0.74008401 | Farp1 | 3259.15115 | 0.014181107 | 0.045092644 | 0.314488247 | 0.753150242 | 0.94183464 |
| Chrac1 | 344.6026352 | 0.050613122 | 0.082889082 | 0.610612644 | 0.54145605 | 0.740125279 | 2810471M01Rik | 0.530790165 | 0.01418647 | 0.014009315 | 1.012645456 | 0.31122954 | NA |
| Entpd1 | 689.9804184 | -0.063964918 | 0.104739082 | -0.610707267 | 0.541393394 | 0.740125279 | Smarcc1 | 3255.724166 | 0.014186912 | 0.049586426 | 0.286104756 | 0.774797889 | 0.947238918 |
| Kdm2b | 1016.216389 | -0.038597526 | 0.06321073 | -0.610616676 | 0.54145338 | 0.740125279 | Dennd1a | 2903.319527 | 0.014189772 | 0.057995028 | 0.244672218 | 0.806710241 | 0.955587482 |
| Pdzrn3 | 1294.359135 | 0.057466319 | 0.094090078 | 0.610758545 | 0.541359441 | 0.740125279 | Etfa | 1151.581558 | 0.014200598 | 0.056532167 | 0.251195007 | 0.801663345 | 0.953332931 |
| Mex3c | 1538.122192 | -0.049977328 | 0.081866973 | -0.61046997 | 0.54155053 | 0.740189125 | Kank1 | 751.3449037 | 0.014219352 | 0.070519072 | 0.201638386 | 0.840199434 | 0.962746468 |
| Plk2 | 8538.54075 | 0.05065636 | 0.083068933 | 0.609811135 | 0.541986925 | 0.740720245 | Hoxa2 | 0.533360734 | 0.014226237 | 0.014028712 | 1.014079985 | 0.310544588 | NA |
| Abhd1 | 112.777406 | 0.091925764 | 0.150822475 | 0.609496457 | 0.54219542 | 0.740777876 | Lactbl1 | 1.097861156 | 0.014238479 | 0.025411597 | 0.560314216 | 0.575265133 | NA |
| Actl6b | 2984.150117 | -0.062738147 | 0.103029548 | -0.608933535 | 0.542568495 | 0.740777876 | 5031434C07Rik | 0.685151557 | 0.01425129 | 0.017964736 | 0.793292487 | 0.427607433 | NA |
| Zfp182 | 402.7108213 | 0.100081773 | 0.125100801 | 0.80000905 | 0.423705554 | 0.646023799 | Zfp456 | 89.841218 | 0.014272464 | 0.094955563 | 0.150306776 | 0.880522587 | 0.973779118 |
| Cbx2 | 120.2436227 | 0.076487277 | 0.125549033 | 0.609222348 | 0.542377069 | 0.740777876 | Spata16 | 2.526488474 | 0.014273947 | 0.035459775 | 0.402539134 | 0.687287294 | NA |
| Dnajc3 | 2453.633561 | -0.050748459 | 0.083272256 | -0.609428174 | 0.542240668 | 0.740777876 | Espnl | 28.12643965 | 0.014283103 | 0.081706296 | 0.17481031 | 0.861228687 | 0.968957316 |
| Gosr1 | 2080.864821 | 0.031482748 | 0.051692449 | 0.609039595 | 0.542498194 | 0.740777876 | Dnajc5b | 0.689569867 | 0.014294014 | 0.018233 | 0.783963904 | 0.43306129 | NA |
| Hrh1 | 675.265137 | -0.06981984 | 0.114667816 | -0.608887854 | 0.542598775 | 0.740777876 | Spp2 | 3.047594687 | 0.014295179 | 0.041202307 | 0.346950945 | 0.728628172 | NA |
| Polr2j | 723.8734926 | 0.047070878 | 0.077307094 | 0.60888175 | 0.542602822 | 0.740777876 | Polr2k | 292.9442166 | 0.014302784 | 0.082437005 | 0.173499558 | 0.862258775 | 0.969665632 |
| Tmed5 | 678.4442641 | -0.052612589 | 0.086313607 | -0.609551506 | 0.542158944 | 0.740777876 | Gm14085 | 0.538437718 | 0.014303766 | 0.014066474 | 1.016869365 | 0.309215579 | NA |
| Trim16 | 83.23713713 | -0.092699429 | 0.152170806 | -0.609180117 | 0.542405058 | 0.740777876 | Foxk2 | 5019.084325 | 0.014313877 | 0.043412724 | 0.329716173 | 0.741614432 | 0.939451578 |
| Zfp472 | 168.3668712 | -0.080046087 | 0.131456198 | -0.608918317 | 0.542578582 | 0.740777876 | Fgf15 | 1.309825175 | 0.014339109 | 0.027309739 | 0.525054802 | 0.599545095 | NA |
| Zfp787 | 472.5939517 | 0.05261417 | 0.086355443 | 0.609274509 | 0.5423425 | 0.740777876 | Qrfp | 0.877224048 | 0.014342364 | 0.017047752 | 0.841305314 | 0.400176914 | NA |
| Kif3a | 7972.371217 | -0.03141031 | 0.051606339 | -0.608652174 | 0.542755014 | 0.740883918 | Alox15 | 3.143714617 | 0.014344273 | 0.036312199 | 0.395026274 | 0.692823498 | NA |
| Vmn1r58 | 66.96891914 | -0.088299379 | 0.145081219 | -0.608620325 | 0.54277613 | 0.740883918 | Mprip | 11860.43447 | 0.014344505 | 0.046393894 | 0.309189492 | 0.757177386 | 0.942139348 |
| Stxbp4 | 923.3177638 | -0.042030996 | 0.069123392 | -0.60805748 | 0.543149352 | 0.741328053 | Tcp10c | 0.692298838 | 0.014348034 | 0.01824747 | 0.786302671 | 0.431690188 | NA |
| Cdc42ep5 | 38.70031211 | -0.099393421 | 0.163540657 | -0.6077597 | 0.543346861 | 0.741401697 | Bub1 | 7.457294122 | 0.014351645 | 0.058355437 | 0.245935012 | 0.80573254 | NA |
| Fam122a | 363.3067163 | -0.060098042 | 0.098882124 | -0.607774584 | 0.543336988 | 0.741401697 | Gckr | 4.520779335 | 0.01435232 | 0.044122299 | 0.325284948 | 0.744965422 | NA |
| Snora3 | 12.30488376 | -0.100958196 | 0.166077369 | -0.607898575 | 0.543254745 | 0.741401697 | Rd3 | 19.33013986 | 0.014361153 | 0.067347956 | 0.213238143 | 0.831141218 | NA |
| 2310036O22Rik | 1320.703145 | -0.058001615 | 0.095529829 | -0.607157109 | 0.543746653 | 0.741881881 | Pde4dip | 10423.1031 | 0.014400842 | 0.055868905 | 0.257761312 | 0.796591124 | 0.952839482 |
| Arl8b | 8450.704777 | -0.039697046 | 0.065404586 | -0.606945915 | 0.543886806 | 0.741925327 | Mab21l3 | 4.987563243 | 0.014407843 | 0.050984043 | 0.282595146 | 0.777487201 | NA |
| Ccdc28b | 479.6287133 | 0.060785236 | 0.100156768 | 0.606900933 | 0.543916659 | 0.741925327 | Mpdu1 | 1355.891654 | 0.014408659 | 0.06256648 | 0.230293588 | 0.817863643 | 0.956698927 |
| Dctn6 | 1851.989778 | 0.041590867 | 0.068530845 | 0.606892659 | 0.54392215 | 0.741925327 | Dnajc4 | 409.9737754 | 0.014410841 | 0.070560093 | 0.204235001 | 0.838169861 | 0.962077671 |
| Polr2f | 719.9464122 | -0.045252234 | 0.074574504 | -0.606805691 | 0.543979871 | 0.741938743 | Rab24 | 1944.884119 | 0.014419587 | 0.060434062 | 0.238600325 | 0.811415513 | 0.956135608 |
| Plscr3 | 556.5837572 | 0.100077574 | 0.119369532 | 0.838384569 | 0.401814753 | 0.626518847 | Gm6654 | 1.087101106 | 0.014429051 | 0.024071169 | 0.599432927 | 0.548884225 | NA |
| Pgbd5 | 6570.503865 | 0.045862223 | 0.075632281 | 0.606384238 | 0.544259632 | 0.742189645 | Aqp6 | 16.20689406 | 0.014430917 | 0.072804265 | 0.198215271 | 0.84287664 | NA |
| Aph1c | 379.025436 | 0.066473668 | 0.109699259 | 0.605962779 | 0.54453947 | 0.742315495 | Dhodh | 721.5631406 | 0.014435492 | 0.07114265 | 0.202909111 | 0.839206073 | 0.96231236 |
| Ddo | 249.0116207 | -0.094477976 | 0.155915412 | -0.60595662 | 0.54454356 | 0.742315495 | Mir132 | 10.9902922 | 0.014454638 | 0.055090002 | 0.262382233 | 0.793026769 | NA |
| Myl2 | 9.41690481 | -0.0975707 | 0.161009831 | -0.605992191 | 0.544519939 | 0.742315495 | Clca2 | 2.910918961 | 0.014455499 | 0.039051464 | 0.370165348 | 0.711259294 | NA |
| Slc30a7 | 823.8962026 | 0.040480681 | 0.066797378 | 0.606022005 | 0.544500141 | 0.742315495 | 1700123K08Rik | 1.287534481 | 0.014489069 | 0.028465434 | 0.509005728 | 0.610748209 | NA |
| Sfmbt1 | 1615.190641 | 0.100071764 | 0.090332051 | 1.107821236 | 0.267939027 | 0.495872824 | Ctage5 | 2015.471967 | 0.014489571 | 0.061804667 | 0.234441368 | 0.814642346 | 0.956264994 |
| Gpr89 | 869.4841826 | -0.043166581 | 0.071268234 | -0.605691751 | 0.544719463 | 0.742359342 | Adtrp | 6.474368897 | 0.014491051 | 0.052287518 | 0.277141682 | 0.781671317 | NA |
| Wdr89 | 84.88160923 | -0.087912768 | 0.145128984 | -0.605756106 | 0.544676721 | 0.742359342 | Ggt6 | 3.887703818 | 0.014496156 | 0.045958374 | 0.315419257 | 0.752443348 | NA |
| Zbtb17 | 1311.034451 | 0.051251892 | 0.084628607 | 0.60560954 | 0.544774066 | 0.742368459 | Rbm17 | 1805.559771 | 0.014496819 | 0.052639257 | 0.27539938 | 0.78300942 | 0.949438153 |
| 6-Sep | 3378.162243 | 0.051355496 | 0.08481822 | 0.605477167 | 0.544861991 | 0.742422979 | Pi4ka | 17009.24952 | 0.014499675 | 0.043059221 | 0.336737977 | 0.736314432 | 0.93717515 |
| Cyp2c44 | 8.441753471 | -0.09161044 | 0.151342894 | -0.605317091 | 0.544968328 | 0.742435638 | Eomes | 8.104965014 | 0.014502887 | 0.051941095 | 0.279217957 | 0.780077563 | NA |
| Fsd2 | 8.105248128 | -0.079721489 | 0.131695987 | -0.605344857 | 0.544949882 | 0.742435638 | 4933404O12Rik | 765.9110929 | 0.014508782 | 0.059340162 | 0.244501895 | 0.806842134 | 0.955587482 |
| Grm2 | 1633.962397 | -0.063800397 | 0.105419103 | -0.605207171 | 0.545041351 | 0.742435638 | Pax2 | 9.250346322 | 0.014511666 | 0.057553512 | 0.252142139 | 0.8009312 | NA |
| Kif3b | 4106.107783 | -0.037845023 | 0.062548289 | -0.60505289 | 0.545143855 | 0.742435638 | Aff3 | 2412.274394 | 0.014520328 | 0.061603719 | 0.235705383 | 0.813661293 | 0.956264994 |
| Plod1 | 784.105156 | -0.056254742 | 0.09297837 | -0.605030416 | 0.545158787 | 0.742435638 | Zfp54 | 58.31056292 | 0.014521673 | 0.094921659 | 0.152985871 | 0.878409418 | 0.973035729 |
| Ttf2 | 92.69814457 | 0.084375594 | 0.139443005 | 0.605090189 | 0.545119073 | 0.742435638 | Dock10 | 2166.314931 | 0.014530933 | 0.081418997 | 0.178471043 | 0.85835306 | 0.967971163 |
| Tasp1 | 703.8749663 | -0.05229266 | 0.0864533 | -0.604865979 | 0.54526805 | 0.742519174 | Cyp4a12b | 3.071682626 | 0.014534493 | 0.031929235 | 0.455209564 | 0.648958481 | NA |
| Usp38 | 1217.293996 | 0.041083323 | 0.067945779 | 0.604648646 | 0.545412477 | 0.742650577 | Trim66 | 2364.697612 | 0.014536664 | 0.064121625 | 0.226704544 | 0.820653493 | 0.957571905 |
| Ctso | 1352.317329 | 0.046407716 | 0.076794087 | 0.604313662 | 0.545635125 | 0.742888456 | Polr2l | 1011.955389 | 0.01454828 | 0.069258014 | 0.210059162 | 0.833621498 | 0.960926822 |
| Arntl | 1384.342415 | 0.058080458 | 0.096190312 | 0.603807775 | 0.54597145 | 0.743215748 | Lrrc4b | 5580.999704 | 0.014549319 | 0.049953218 | 0.291258898 | 0.77085332 | 0.946250219 |
| Tmem161b | 613.731862 | -0.049715447 | 0.082326754 | -0.603879594 | 0.545923696 | 0.743215748 | Entpd4 | 7503.919256 | 0.01455047 | 0.065586967 | 0.221850021 | 0.824430638 | 0.957903916 |
| 2610528A11Rik | 12.6770639 | -0.10077751 | 0.167102881 | -0.603086612 | 0.546451072 | 0.743600379 | Rab11fip3 | 6297.570274 | 0.014560776 | 0.042555794 | 0.342157305 | 0.732232514 | 0.935722272 |
| Bcap31 | 1976.59574 | -0.045356381 | 0.075196393 | -0.60317229 | 0.54639408 | 0.743600379 | Ahcyl1 | 18950.55947 | 0.014573868 | 0.044522777 | 0.327335109 | 0.743414441 | 0.939912916 |
| Bcar1 | 1856.665221 | 0.100055665 | 0.077506127 | 1.290938775 | 0.196724909 | 0.411732505 | Ube2g1 | 1710.930153 | 0.014610692 | 0.084549948 | 0.172805456 | 0.862804348 | 0.969759087 |
| Rai1 | 3606.17286 | -0.039825104 | 0.066042524 | -0.603022132 | 0.546493966 | 0.743600379 | Gm4792 | 0.707957873 | 0.014628458 | 0.018328843 | 0.798111384 | 0.424805856 | NA |
| Ssbp2 | 3609.698034 | -0.045588724 | 0.075598241 | -0.603039478 | 0.546482427 | 0.743600379 | Mir568 | 18.79264089 | 0.014639767 | 0.075857655 | 0.192989975 | 0.846966819 | NA |
| Zfp606 | 451.3699727 | 0.057880202 | 0.096001543 | 0.60290908 | 0.546569175 | 0.743637408 | Tmco2 | 1.041925876 | 0.014642674 | 0.020406992 | 0.717532226 | 0.47304576 | NA |
| Slc16a12 | 127.8309968 | 0.076836147 | 0.12748981 | 0.602684612 | 0.54671852 | 0.743775288 | Egr4 | 797.8118027 | 0.014656575 | 0.080555888 | 0.181942934 | 0.85562751 | 0.967189923 |
| Nmral1 | 372.1136646 | -0.064388803 | 0.106898597 | -0.602335343 | 0.546950939 | 0.74402615 | Hoxb3 | 0.710417164 | 0.01467032 | 0.018341392 | 0.799847712 | 0.423799035 | NA |
| Agfg2 | 1700.58031 | 0.100002654 | 0.078237611 | 1.27819157 | 0.201181888 | 0.418011556 | Rragd | 7845.945252 | 0.014670612 | 0.070819017 | 0.207156389 | 0.835887725 | 0.961504006 |
| B4galt7 | 629.3408993 | 0.044530974 | 0.073989494 | 0.601855361 | 0.54727042 | 0.744128707 | Otub2 | 1513.909759 | 0.014675746 | 0.087220236 | 0.168260795 | 0.866378115 | 0.97021201 |
| B930059L03Rik | 35.36801585 | 0.099861392 | 0.165896823 | 0.601948789 | 0.547208227 | 0.744128707 | Gm15471 | 1.946779206 | 0.014683263 | 0.031192625 | 0.470728686 | 0.637834496 | NA |
| Chrna7 | 487.3587347 | 0.046675316 | 0.077562701 | 0.601775272 | 0.547323738 | 0.744128707 | Kif2a | 3281.134747 | 0.014701648 | 0.072672843 | 0.202299056 | 0.839682938 | 0.962420939 |
| Fos | 1299.876561 | -0.093032632 | 0.154567873 | -0.601888548 | 0.547248328 | 0.744128707 | Abcb11 | 1.083999946 | 0.014727821 | 0.02538691 | 0.580134442 | 0.561823959 | NA |
| Msra | 1102.766488 | 0.05869602 | 0.097559244 | 0.601644882 | 0.547410546 | 0.744128707 | Ncoa1 | 6547.411819 | 0.014735786 | 0.055719026 | 0.264465969 | 0.791420885 | 0.951728598 |
| Rab38 | 10.2809071 | 0.095869667 | 0.159341146 | 0.601662967 | 0.547398506 | 0.744128707 | Gnaq | 9758.125776 | 0.014738592 | 0.064559715 | 0.228293949 | 0.819417729 | 0.957571905 |
| Tnk2 | 10854.0892 | 0.06281706 | 0.104339423 | 0.602045307 | 0.547143979 | 0.744128707 | Clta | 5409.761339 | 0.014759352 | 0.037883201 | 0.389601491 | 0.696831249 | 0.925836525 |
| Ccdc122 | 55.59397627 | -0.093686299 | 0.155748372 | -0.601523454 | 0.547491395 | 0.74417332 | Gm4894 | 0.643456503 | 0.014761549 | 0.017755533 | 0.83137742 | 0.405760451 | NA |
| Sema4d | 1978.342213 | 0.037356054 | 0.062125614 | 0.601298752 | 0.547641021 | 0.744311401 | Vmn2r55 | 0.538596121 | 0.014768721 | 0.014291854 | 1.033366314 | 0.301432504 | NA |
| Tbc1d25 | 544.2520363 | 0.049692047 | 0.082659895 | 0.601162714 | 0.547731616 | 0.744369236 | A930011G23Rik | 23.05016019 | 0.014771393 | 0.079057578 | 0.18684348 | 0.851783355 | 0.96669173 |
| Dda1 | 2029.744976 | -0.042381477 | 0.070512288 | -0.601050941 | 0.547806058 | 0.74440511 | Cacna1f | 10.87853332 | 0.014782067 | 0.052134439 | 0.283537468 | 0.776764864 | NA |
| Etohi1 | 519.2063723 | -0.062697066 | 0.104417053 | -0.60044853 | 0.548207354 | 0.744814712 | Mfsd4 | 2358.489945 | 0.01478252 | 0.062784986 | 0.235446729 | 0.813862021 | 0.956264994 |
| Folr2 | 15.07447431 | -0.098120864 | 0.16343073 | -0.600381976 | 0.548251697 | 0.744814712 | Ppp1r1c | 1.449683927 | 0.014788065 | 0.025487935 | 0.580198614 | 0.561780688 | NA |
| Lats2 | 1000.532933 | 0.060076734 | 0.100058014 | 0.60041901 | 0.548227022 | 0.744814712 | 4930413G21Rik | 3.653778029 | 0.014793605 | 0.044879364 | 0.329630458 | 0.741679206 | NA |
| Usp14 | 5274.207639 | -0.053022814 | 0.088329773 | -0.600282464 | 0.548318003 | 0.744839483 | 6030498E09Rik | 0.833729473 | 0.014817327 | 0.021091913 | 0.702512249 | 0.482359766 | NA |
| Pde3b | 412.9722291 | 0.046245963 | 0.077062522 | 0.600109648 | 0.548433163 | 0.744930607 | Cdc73 | 696.6687525 | 0.014821681 | 0.079249982 | 0.187024408 | 0.851641496 | 0.96665003 |
| Ttc22 | 51.11747396 | -0.095444786 | 0.159067385 | -0.60002738 | 0.548487988 | 0.744939769 | 2700099C18Rik | 10.42749203 | 0.014830007 | 0.066519908 | 0.222940881 | 0.823581519 | NA |
| Gtf2a1 | 1230.187129 | -0.044010712 | 0.073364303 | -0.599892727 | 0.54857773 | 0.744996349 | Gm10471 | 3.388938125 | 0.014850207 | 0.038789773 | 0.3828382 | 0.701839736 | NA |
| Wdr74 | 836.342893 | 0.100002114 | 0.085137574 | 1.174594364 | 0.240157039 | 0.464547756 | Clcf1 | 17.25641078 | 0.014871755 | 0.078665457 | 0.189050643 | 0.850053125 | NA |
| Fdxacb1 | 163.5125176 | -0.070184499 | 0.117100473 | -0.599352826 | 0.548937628 | 0.745354448 | Dcaf8 | 6930.124131 | 0.014896877 | 0.04047613 | 0.368041039 | 0.712842634 | 0.930507332 |
| Hsdl1 | 2041.973916 | -0.039696724 | 0.066252678 | -0.599171616 | 0.549058448 | 0.745387856 | 2610203C22Rik | 18.46878777 | 0.014913641 | 0.070407481 | 0.211818983 | 0.832248258 | NA |
| Rab11fip4 | 5956.184506 | -0.042676082 | 0.071221942 | -0.59919852 | 0.54904051 | 0.745387856 | Hnrnpd | 5590.233005 | 0.014930889 | 0.055726291 | 0.267932578 | 0.788751217 | 0.951282725 |
| Ccdc71l | 520.1760712 | 0.052844221 | 0.088209005 | 0.599079662 | 0.549119763 | 0.745405783 | Rbm19 | 648.9577913 | 0.014933123 | 0.064526976 | 0.231424501 | 0.816985033 | 0.956302288 |
| 3830408C21Rik | 106.2112079 | 0.079723742 | 0.133123966 | 0.598868443 | 0.549260617 | 0.745531669 | Grifin | 2.926006549 | 0.014952193 | 0.038340902 | 0.38998021 | 0.696551181 | NA |
| Relb | 197.8866786 | -0.07631247 | 0.127466896 | -0.598684618 | 0.549383217 | 0.745632757 | Tmem38a | 4352.647314 | 0.014962083 | 0.047943184 | 0.312079452 | 0.754980142 | 0.942099798 |
| Col17a1 | 41.47010534 | -0.098689271 | 0.16487637 | -0.598565285 | 0.549462811 | 0.745675466 | Igsf21 | 1807.409356 | 0.014966766 | 0.06934942 | 0.215816747 | 0.829130594 | 0.959448021 |
| Slc26a1 | 39.79662281 | -0.097293799 | 0.162580686 | -0.598433933 | 0.54955043 | 0.745729056 | Hnrnpul1 | 3696.431959 | 0.014967928 | 0.048358589 | 0.309519536 | 0.756926353 | 0.942139348 |
| Mul1 | 1909.589671 | -0.036172095 | 0.060453377 | -0.598346977 | 0.549608437 | 0.745742458 | Gclm | 1237.698375 | 0.014979312 | 0.064514414 | 0.2321855 | 0.816393939 | 0.956264994 |
| Cntd1 | 20.69288458 | -0.099938 | 0.167131648 | -0.597959758 | 0.549866785 | 0.745962347 | Scn4b | 16662.21722 | 0.014993196 | 0.094652074 | 0.15840325 | 0.874139053 | 0.971912694 |
| Prkar1a | 25550.26677 | -0.035787418 | 0.059846173 | -0.597990088 | 0.549846547 | 0.745962347 | Olfr536 | 0.900494549 | 0.014994034 | 0.017152694 | 0.87415038 | 0.382036364 | NA |
| Atp6v1e1 | 12725.87037 | 0.044054776 | 0.073690262 | 0.597837149 | 0.5499486 | 0.746008021 | Cd177 | 1.205319222 | 0.014999449 | 0.022814877 | 0.657441613 | 0.510897002 | NA |
| Ankrd55 | 134.7097154 | 0.085103344 | 0.142537748 | 0.597058291 | 0.550468464 | 0.746061754 | Ubr1 | 2769.435415 | 0.015016405 | 0.061970762 | 0.242314358 | 0.808536589 | 0.955716697 |
| Arhgef26 | 1175.244826 | -0.050408777 | 0.084430311 | -0.597045968 | 0.55047669 | 0.746061754 | Lyzl6 | 0.861411351 | 0.015022576 | 0.022281356 | 0.674221794 | 0.500170316 | NA |
| Ccr9 | 11.45335156 | -0.097094636 | 0.162572172 | -0.597240199 | 0.550347024 | 0.746061754 | Fbxo22 | 1494.902252 | 0.015049719 | 0.052228848 | 0.288149542 | 0.773232271 | 0.947121017 |
| Fam114a2 | 1393.450406 | -0.039003024 | 0.06533342 | -0.596984274 | 0.55051788 | 0.746061754 | Tcn2 | 1030.356807 | 0.015063383 | 0.092992243 | 0.161985372 | 0.871317373 | 0.97153269 |
| Fbxl14 | 1060.2336 | 0.04885723 | 0.081839256 | 0.596990156 | 0.550513953 | 0.746061754 | Gtf3c6 | 601.7845856 | 0.015067956 | 0.063394582 | 0.237685237 | 0.812125235 | 0.956219756 |
| Nrn1l | 44.28824163 | -0.100285174 | 0.167882692 | -0.597352668 | 0.550271948 | 0.746061754 | Cdcp1 | 16.5790013 | 0.015083166 | 0.074184868 | 0.20331863 | 0.838885995 | NA |
| Rabggtb | 1352.468121 | 0.03658569 | 0.061232725 | 0.597485892 | 0.550183024 | 0.746061754 | Ncam1 | 11331.0157 | 0.01511266 | 0.061701999 | 0.244929837 | 0.806510759 | 0.955587482 |
| Six5 | 87.77942271 | -0.086543129 | 0.144958192 | -0.597021303 | 0.550493158 | 0.746061754 | Ccdc104 | 3216.753499 | 0.015122459 | 0.064651072 | 0.233908862 | 0.815055734 | 0.956264994 |
| Speg | 6024.046489 | -0.051439 | 0.086140592 | -0.597151692 | 0.550406108 | 0.746061754 | 1700003L19Rik | 1.16541587 | 0.01514449 | 0.02261133 | 0.669774404 | 0.503001612 | NA |
| Tipin | 378.0803007 | -0.06377995 | 0.106728494 | -0.597590651 | 0.550113104 | 0.746061754 | Gm6313 | 30.06735782 | 0.015146857 | 0.088896776 | 0.170387024 | 0.864705776 | 0.969972403 |
| Tns3 | 3232.820089 | -0.03276958 | 0.05487283 | -0.59719137 | 0.55037962 | 0.746061754 | 4921504E06Rik | 1.058845616 | 0.01514748 | 0.02570377 | 0.589309641 | 0.555653579 | NA |
| Hars | 3643.222555 | -0.040185484 | 0.067326228 | -0.596877112 | 0.55058943 | 0.74609346 | Fez2 | 1602.223408 | 0.015149615 | 0.051889222 | 0.291960718 | 0.770316658 | 0.94619074 |
| Junb | 1830.703675 | -0.100327165 | 0.168114651 | -0.596778237 | 0.55065545 | 0.746117669 | N6amt1 | 801.0526072 | 0.015150715 | 0.064693437 | 0.234192463 | 0.814835566 | 0.956264994 |
| Tmpo | 1013.597682 | 0.039055684 | 0.06545632 | 0.596667886 | 0.550729137 | 0.746152261 | Anxa6 | 5159.863783 | 0.015162972 | 0.058219297 | 0.260445814 | 0.794519908 | 0.951979493 |
| Tmem9b | 3357.727927 | -0.036735866 | 0.061577893 | -0.596575563 | 0.55079079 | 0.746170543 | 1700015F17Rik | 3.303416737 | 0.015171408 | 0.041406526 | 0.366401373 | 0.714065595 | NA |
| Tmtc4 | 416.2612799 | -0.060452728 | 0.101355193 | -0.596444308 | 0.550878448 | 0.746224049 | Pitx2 | 3.032052365 | 0.015172418 | 0.030316381 | 0.500469303 | 0.616744666 | NA |
| Tbck | 739.5901732 | -0.038673106 | 0.0648698 | -0.596165029 | 0.551064986 | 0.746411477 | Opa1 | 7734.422422 | 0.015193588 | 0.05834891 | 0.260391975 | 0.794561433 | 0.951979493 |
| Lyrm9 | 1764.184818 | 0.046501224 | 0.078027172 | 0.595961921 | 0.551200666 | 0.746529992 | Comtd1 | 274.5551962 | 0.015210586 | 0.078986052 | 0.192573069 | 0.847293338 | 0.965275633 |
| Erbb3 | 417.7410406 | -0.062136822 | 0.104313599 | -0.595673265 | 0.551393522 | 0.746628481 | Vav2 | 354.0151408 | 0.015220832 | 0.080310068 | 0.18952583 | 0.849680713 | 0.966260848 |
| Mkrn1 | 4765.783074 | -0.04183727 | 0.070233789 | -0.595685783 | 0.551385158 | 0.746628481 | Slc10a4 | 135.9234815 | 0.015237598 | 0.095039205 | 0.160329598 | 0.872621444 | 0.97153269 |
| Sphk2 | 1043.828299 | 0.039970846 | 0.067106083 | 0.595636707 | 0.551417949 | 0.746628481 | Pde8a | 839.9144272 | 0.015238117 | 0.088869988 | 0.171465277 | 0.863857931 | 0.969836459 |
| Zcwpw1 | 268.0337954 | -0.073548984 | 0.123519537 | -0.595444132 | 0.551546633 | 0.746737464 | Ipo13 | 4453.326033 | 0.015259204 | 0.057075939 | 0.267349146 | 0.78920035 | 0.951324462 |
| Rnase1 | 9.796111194 | 0.077105906 | 0.129518354 | 0.595328026 | 0.551624226 | 0.746777261 | Pdzk1ip1 | 7.921058025 | 0.01526092 | 0.054487529 | 0.280080968 | 0.779415386 | NA |
| Cct7 | 6515.985863 | -0.036061263 | 0.060609536 | -0.594976712 | 0.551859038 | 0.747009187 | Acvr2b | 138.3148488 | 0.015275977 | 0.088420739 | 0.172764639 | 0.862836433 | 0.969759087 |
| Zfp688 | 249.4758524 | -0.054709629 | 0.09196017 | -0.594927448 | 0.551891969 | 0.747009187 | Dock3 | 13962.45755 | 0.015282393 | 0.059457221 | 0.257031732 | 0.797154277 | 0.952885371 |
| Cdv3 | 3605.447501 | -0.028746537 | 0.048324498 | -0.594643178 | 0.552082012 | 0.74713588 | Asic5 | 0.606475725 | 0.015283342 | 0.014534743 | 1.051504152 | 0.293027103 | NA |
| Iglon5 | 1361.487352 | 0.056889551 | 0.095660213 | 0.59470441 | 0.552041073 | 0.74713588 | Trappc1 | 1173.285533 | 0.015295346 | 0.066645072 | 0.22950453 | 0.8184768 | 0.95703083 |
| B230206H07Rik | 138.9740127 | 0.086590284 | 0.145663526 | 0.594454122 | 0.552208419 | 0.747209042 | Atxn2 | 4169.331033 | 0.015297683 | 0.056438446 | 0.271050751 | 0.786351999 | 0.950484821 |
| Serpinb6b | 50.78160011 | -0.090729714 | 0.152636198 | -0.594418071 | 0.552232525 | 0.747209042 | Nosip | 1825.258263 | 0.015308625 | 0.067633247 | 0.226347627 | 0.820931058 | 0.957611694 |
| 2700046A07Rik | 25.95309645 | -0.099638578 | 0.167877997 | -0.593517793 | 0.552834679 | 0.74757312 | Cks2 | 34.14137276 | 0.015318561 | 0.090746108 | 0.168806806 | 0.865948604 | 0.970182416 |
| A730036I17Rik | 32.10432879 | -0.099794289 | 0.168096155 | -0.593673832 | 0.552730289 | 0.74757312 | 2900060B14Rik | 0.574512993 | 0.015323365 | 0.014554802 | 1.052804801 | 0.292430464 | NA |
| Acsbg1 | 6036.166675 | -0.055681372 | 0.093811902 | -0.59354273 | 0.552817995 | 0.74757312 | Crabp1 | 67.97053399 | 0.015323823 | 0.094472819 | 0.162203513 | 0.871145594 | 0.97153269 |
| Dtl | 367.2746342 | 0.07661291 | 0.129084268 | 0.593510818 | 0.552839345 | 0.74757312 | 9630028H03Rik | 0.682786739 | 0.015335166 | 0.018209764 | 0.842139717 | 0.399709752 | NA |
| Gm101 | 9.413431964 | 0.086565102 | 0.145776355 | 0.593821281 | 0.552631654 | 0.74757312 | 0610012H03Rik | 0.406364829 | 0.015342854 | 0.014574102 | 1.052747816 | 0.292456588 | NA |
| Hmgxb3 | 2440.229766 | 0.041451358 | 0.069831814 | 0.593588443 | 0.552787413 | 0.74757312 | Marc1 | 5.298693257 | 0.015354016 | 0.046089503 | 0.333134764 | 0.739032559 | NA |
| Oas1a | 24.05345707 | 0.099548704 | 0.167659242 | 0.59375614 | 0.552675228 | 0.74757312 | Rfxap | 864.8323688 | 0.015359262 | 0.065178791 | 0.235648152 | 0.813705706 | 0.956264994 |
| Esco1 | 775.1239776 | 0.052547844 | 0.088552354 | 0.593409908 | 0.55290686 | 0.74759917 | Tvp23a | 1782.15453 | 0.015381024 | 0.061735429 | 0.249144199 | 0.803249241 | 0.953754272 |
| Dnttip1 | 728.2437001 | 0.039966107 | 0.067424443 | 0.592753976 | 0.553345813 | 0.747996861 | BC030476 | 15.05141573 | 0.015385511 | 0.06671533 | 0.23061433 | 0.817614434 | NA |
| Drg2 | 1969.198591 | 0.028360797 | 0.047841489 | 0.592807579 | 0.553309935 | 0.747996861 | Krt83 | 1.898568809 | 0.015387488 | 0.02957821 | 0.520230543 | 0.6029029 | NA |
| Myh8 | 20.35493054 | -0.099456305 | 0.167757163 | -0.592858769 | 0.553275673 | 0.747996861 | Actr1b | 11157.37239 | 0.015392221 | 0.051232391 | 0.300439244 | 0.763842133 | 0.944786344 |
| Aco1 | 1314.751263 | -0.036587154 | 0.061748588 | -0.592518068 | 0.553503725 | 0.748145051 | Dfnb59 | 2.522778796 | 0.015406484 | 0.035078059 | 0.439205734 | 0.66051247 | NA |
| Ppa2 | 1168.133258 | -0.04091596 | 0.069087834 | -0.592231049 | 0.553695881 | 0.748339496 | 1110046J04Rik | 99.0606268 | 0.01541074 | 0.094903143 | 0.162383872 | 0.87100357 | 0.97153269 |
| Chm | 1773.805331 | -0.052045066 | 0.087912514 | -0.592009753 | 0.553844058 | 0.748409195 | Ncstn | 3035.677189 | 0.015429697 | 0.055033844 | 0.280367419 | 0.779195631 | 0.948407834 |
| Hmgb1 | 6512.796655 | -0.034830853 | 0.058830453 | -0.592054819 | 0.55381388 | 0.748409195 | Tmem171 | 6.384764122 | 0.015431312 | 0.047551529 | 0.324517685 | 0.745546135 | NA |
| 2610002J02Rik | 666.0513271 | -0.047657641 | 0.080535374 | -0.591760349 | 0.554011079 | 0.748569604 | Kcns3 | 290.3425046 | 0.015432367 | 0.085790942 | 0.179883413 | 0.857244098 | 0.967922873 |
| Ano4 | 825.0102154 | -0.05640346 | 0.095327501 | -0.591680888 | 0.554064297 | 0.748576231 | Slc6a7 | 3672.330954 | 0.0154511 | 0.06912067 | 0.22353805 | 0.823116773 | 0.957903916 |
| Epg5 | 2833.413321 | -0.037738014 | 0.063789735 | -0.591600102 | 0.554118406 | 0.748584059 | Enpp2 | 17413.19297 | 0.015452406 | 0.059617709 | 0.259191533 | NA | NA |
| Gabrq | 51.31806127 | -0.09524499 | 0.161088787 | -0.591257728 | 0.554347748 | 0.748771246 | Wfdc10 | 0.619517067 | 0.015462141 | 0.014618466 | 1.05771298 | 0.290186316 | NA |
| H2afy | 2227.298024 | 0.032958697 | 0.055757799 | 0.5911047 | 0.554450271 | 0.748771246 | Nox3 | 1.159403599 | 0.01547524 | 0.02452605 | 0.630971576 | 0.52805911 | NA |
| Prdm2 | 3529.744522 | -0.048321468 | 0.081730676 | -0.591228045 | 0.554367634 | 0.748771246 | Lenep | 250.0339852 | 0.015484671 | 0.080406545 | 0.192579738 | 0.847288115 | 0.965275633 |
| Rel | 58.45766292 | -0.087103075 | 0.147352973 | -0.591118546 | 0.554440994 | 0.748771246 | Olfr1377 | 0.908957928 | 0.0154869 | 0.022437179 | 0.690233842 | 0.490047144 | NA |
| Bex2 | 5446.217425 | -0.034963757 | 0.059170432 | -0.590899128 | 0.55458801 | 0.748803416 | Gm16853 | 8.67318582 | 0.015487115 | 0.056905294 | 0.272155956 | 0.785502106 | NA |
| Cic | 7663.632863 | 0.038278682 | 0.064777095 | 0.590929279 | 0.554567807 | 0.748803416 | Zdhhc18 | 1264.110425 | 0.015491727 | 0.05844747 | 0.265053845 | 0.790967982 | 0.951728598 |
| F630042J09Rik | 24.52632518 | 0.099316258 | 0.168089691 | 0.590852763 | 0.554619079 | 0.748803416 | Usp3 | 646.0074536 | 0.015495536 | 0.074611465 | 0.207683044 | 0.835476458 | 0.961504006 |
| Atf5 | 569.5793618 | -0.056606894 | 0.095954574 | -0.589934294 | 0.555234702 | 0.749154089 | 1700028K03Rik | 18.19629399 | 0.015498886 | 0.079567935 | 0.194788091 | 0.845558846 | NA |
| Atp6ap2 | 8360.627703 | -0.044212537 | 0.074911027 | -0.590200653 | 0.555056134 | 0.749154089 | Fam150a | 1.619597959 | 0.015505517 | 0.026779002 | 0.57901774 | 0.562577203 | NA |
| Ccdc107 | 562.3874758 | -0.045010429 | 0.076295817 | -0.589946219 | 0.555226706 | 0.749154089 | Lmtk2 | 10525.96123 | 0.015528062 | 0.059258308 | 0.262040249 | 0.793290412 | 0.951728598 |
| Cmtm8 | 33.45298972 | -0.097387998 | 0.165130103 | -0.589765257 | 0.555348038 | 0.749154089 | Fut10 | 469.0794231 | 0.015530917 | 0.068678823 | 0.226138365 | 0.821093806 | 0.957611694 |
| Dcn | 492.094384 | 0.084358289 | 0.142937005 | 0.590178096 | 0.555071256 | 0.749154089 | Tmem102 | 4.029851536 | 0.015569832 | 0.036902533 | 0.421917707 | 0.673085085 | NA |
| Dnajc13 | 3324.396462 | 0.055776136 | 0.094503867 | 0.590199515 | 0.555056897 | 0.749154089 | Prss43 | 8.025508264 | 0.015593337 | 0.057200993 | 0.272606051 | 0.78515606 | NA |
| Fbxl8 | 26.49390958 | 0.096619156 | 0.163831194 | 0.589748223 | 0.55535946 | 0.749154089 | Abhd12b | 14.96552665 | 0.015599063 | 0.044521332 | 0.350372788 | 0.726058946 | NA |
| Itgb5 | 2081.987797 | 0.049714232 | 0.084259873 | 0.59001076 | 0.555183436 | 0.749154089 | 1700019D03Rik | 126.9518453 | 0.015607751 | 0.091350082 | 0.170856451 | 0.86433664 | 0.969963136 |
| Lat | 15.19571426 | 0.098919616 | 0.167753671 | 0.589671843 | 0.555410676 | 0.749154089 | Serpina3m | 0.419406171 | 0.015621438 | 0.014704373 | 1.062366861 | 0.288069172 | NA |
| Mki67ip | 842.9502212 | -0.04214926 | 0.071456085 | -0.589862428 | 0.555282885 | 0.749154089 | Trappc11 | 3444.219492 | 0.015630309 | 0.043802882 | 0.356832891 | 0.721216911 | 0.932513933 |
| Scara5 | 20.67301429 | 0.098549616 | 0.167115306 | 0.589710294 | 0.555384892 | 0.749154089 | Gm6402 | 10.61500856 | 0.015641492 | 0.066978127 | 0.233531338 | 0.815348838 | NA |
| Akap17b | 812.6572692 | -0.052830968 | 0.089607865 | -0.58957959 | 0.555472538 | 0.749172311 | Defb12 | 0.59690524 | 0.01565877 | 0.014711274 | 1.064406159 | 0.287144745 | NA |
| Plekha5 | 2291.425778 | 0.039020661 | 0.066204494 | 0.58939596 | 0.555595686 | 0.749273179 | Atg9b | 232.9466722 | 0.015662917 | 0.094189561 | 0.166291428 | 0.867927612 | 0.970480832 |
| B3gntl1 | 205.6849733 | 0.078149374 | 0.132624414 | 0.589253303 | 0.555691365 | 0.749276814 | Adamts5 | 94.76126503 | 0.015667869 | 0.092891768 | 0.168668006 | 0.866057786 | 0.970182416 |
| Enc1 | 36674.0018 | -0.056774489 | 0.096362588 | -0.589175637 | 0.555743459 | 0.749276814 | Art5 | 0.871835 | 0.015683202 | 0.022638509 | 0.692766541 | 0.488456073 | NA |
| Hilpda | 144.705607 | -0.068081984 | 0.115546649 | -0.589216428 | 0.555716098 | 0.749276814 | Krt31 | 0.865465048 | 0.015685948 | 0.022620427 | 0.693441753 | 0.488032367 | NA |
| Lims1 | 993.3352613 | -0.043439393 | 0.073745253 | -0.589046635 | 0.555829991 | 0.749298616 | Efcab14 | 2737.738613 | 0.01569856 | 0.062690385 | 0.250414164 | 0.802267077 | 0.95333733 |
| Zbtb4 | 7437.542528 | -0.030650191 | 0.052037027 | -0.589007339 | 0.555856351 | 0.749298616 | Sfrp1 | 442.8883934 | 0.015705784 | 0.091874183 | 0.17094883 | 0.864264001 | 0.969963136 |
| A930009A15Rik | 11.08351095 | 0.094855232 | 0.161188962 | 0.588472252 | 0.556215353 | 0.749717328 | Slc52a2 | 558.3052121 | 0.015712909 | 0.084703277 | 0.185505329 | 0.852832701 | 0.966905493 |
| Aldh4a1 | 1012.303573 | -0.054437956 | 0.09259229 | -0.587931851 | 0.556578037 | 0.750140927 | Fam53a | 655.547041 | 0.015721673 | 0.067999442 | 0.231202968 | 0.817157124 | 0.956302288 |
| R3hcc1 | 656.1629 | 0.044806101 | 0.07622955 | 0.587778643 | 0.556680881 | 0.750184817 | Sct | 1.70644656 | 0.015723817 | 0.027623222 | 0.569224568 | 0.56920375 | NA |
| Ube2v1 | 8181.743499 | -0.032435491 | 0.055186888 | -0.587739085 | 0.556707437 | 0.750184817 | Polr2i | 462.4219065 | 0.015726382 | 0.075719231 | 0.20769337 | 0.835468395 | 0.961504006 |
| Sos1 | 2607.209177 | -0.049906049 | 0.084932574 | -0.587596101 | 0.556803429 | 0.75024892 | Pacsin1 | 21579.59504 | 0.01573293 | 0.053310868 | 0.295116751 | 0.767904694 | 0.94619074 |
| Atl3 | 1635.074579 | 0.040952703 | 0.069713175 | 0.587445673 | 0.556904427 | 0.750319756 | Acbd6 | 1389.018181 | 0.015735434 | 0.054078501 | 0.290973923 | 0.771071264 | 0.946250219 |
| Atat1 | 1838.465362 | 0.04205974 | 0.071654181 | 0.586982359 | 0.557215555 | 0.750608399 | Cabp1 | 4141.621742 | 0.015739405 | 0.058663122 | 0.268301515 | 0.78846724 | 0.951189696 |
| Dnajc14 | 2559.125205 | -0.036750028 | 0.062607638 | -0.58698952 | 0.557210745 | 0.750608399 | Phox2a | 1.091611411 | 0.015742439 | 0.025771604 | 0.610844347 | 0.541302631 | NA |
| Fam105a | 838.8125251 | 0.044857206 | 0.076473643 | 0.586570806 | 0.557491994 | 0.750915497 | Plekhg6 | 11.07522905 | 0.015744312 | 0.061499021 | 0.256009153 | 0.79794377 | NA |
| E130309D02Rik | 733.5535901 | 0.045698277 | 0.077988885 | 0.58595885 | 0.557903168 | 0.751338695 | Rdh5 | 151.8046017 | 0.015752533 | 0.048898197 | 0.322149575 | NA | NA |
| Thbs2 | 391.9664599 | -0.074719881 | 0.127512237 | -0.58598204 | 0.557887584 | 0.751338695 | Id2 | 3941.778486 | 0.015756644 | 0.067868995 | 0.232162626 | 0.816411705 | 0.956264994 |
| Myl12b | 7018.32707 | 0.041512508 | 0.07088437 | 0.58563698 | 0.558119492 | 0.751456036 | Usp45 | 1891.477701 | 0.015768531 | 0.081479655 | 0.193527219 | 0.846546091 | 0.965222019 |
| Ostf1 | 357.7107995 | 0.051262081 | 0.087537493 | 0.585601425 | 0.558143391 | 0.751456036 | Hnf4g | 1.074416698 | 0.015779568 | 0.019962554 | 0.790458365 | 0.429260128 | NA |
| Pfas | 1182.207835 | -0.054379696 | 0.092882373 | -0.585468415 | 0.558232798 | 0.751456036 | Olfr1347 | 0.427100848 | 0.01578097 | 0.014778443 | 1.067837146 | 0.285593979 | NA |
| Wtap | 2039.566126 | -0.036744336 | 0.06274556 | -0.585608536 | 0.558138611 | 0.751456036 | Pcdhgc3 | 2765.790565 | 0.015781478 | 0.063456083 | 0.248699216 | 0.803593456 | 0.953754272 |
| Zc3h12c | 621.7316892 | -0.069746806 | 0.119126938 | -0.585483078 | 0.558222942 | 0.751456036 | Apom | 1.152755946 | 0.015782884 | 0.02414648 | 0.653630839 | 0.513349671 | NA |
| Ankrd45 | 3106.934529 | -0.033943296 | 0.057999316 | -0.585236144 | 0.558388945 | 0.751590158 | Hs3st6 | 8.415898699 | 0.015788293 | 0.058633545 | 0.269270661 | 0.787721408 | NA |

| | | | | | | | | | | | | |
|---|---|---|---|---|---|---|---|---|---|---|---|---|---|
| Rfx4 | 709.3999139 | -0.064414327 | 0.11008689 | -0.585122596 | 0.558465287 | 0.751590158 | Itgb6 | 1.451383634 | 0.015801159 | 0.026279577 | 0.601271422 | 0.54765922 | NA |
| Synrg | 4110.187106 | -0.051789787 | 0.088513853 | -0.585103748 | 0.558477959 | 0.751590158 | A830080D01Rik | 261.3394527 | 0.015824817 | 0.081459979 | 0.194264931 | 0.845968443 | 0.964996028 |
| Slc16a3 | 97.11809984 | -0.08073135 | 0.138036849 | -0.584853615 | 0.55864615 | 0.751751212 | Hnf1b | 27.00949256 | 0.015832501 | 0.08082543 | 0.195885148 | 0.844700062 | 0.964328285 |
| Rab3a | 19382.83934 | 0.034549133 | 0.059092631 | 0.5846606 | 0.558775951 | 0.751795293 | Dynll2 | 31018.2382 | 0.015853038 | 0.049132848 | 0.322656597 | 0.746955326 | 0.94097069 |
| Slc38a11 | 91.18270251 | -0.081811439 | 0.139925795 | -0.58467732 | 0.558764707 | 0.751795293 | Nr5a2 | 2.530492686 | 0.015859628 | 0.025460078 | 0.622921412 | 0.533336166 | NA |
| Lama5 | 633.9109276 | 0.056431028 | 0.096544123 | 0.584510234 | 0.558877082 | 0.751866068 | Muc4 | 1.156918588 | 0.015864769 | 0.024152048 | 0.656870552 | 0.511264155 | NA |
| Leprotl1 | 2710.314875 | 0.032328651 | 0.055321311 | 0.584379695 | 0.558964885 | 0.751918903 | Bok | 988.0539052 | 0.015870035 | 0.085338347 | 0.185966045 | 0.852471389 | 0.966905493 |
| Abca1 | 1951.831964 | 0.096349053 | 0.164919817 | 0.584217563 | 0.559073946 | 0.752000323 | Nkx1-2 | 2.45143039 | 0.015871076 | 0.031772074 | 0.499529119 | 0.617406678 | NA |
| Slc12a6 | 6120.359942 | -0.028780124 | 0.049275824 | -0.584061754 | 0.559178765 | 0.752076023 | Tuft1 | 110.8998611 | 0.015872445 | 0.094669249 | 0.167662101 | 0.866849113 | 0.97021201 |
| Fhdc1 | 232.9268354 | -0.068196278 | 0.116796872 | -0.583887878 | 0.559295749 | 0.75216807 | Fam102b | 2675.364313 | 0.01587367 | 0.072125913 | 0.220082755 | 0.825806705 | 0.958041984 |
| Nlrc3 | 17.10053237 | -0.097851741 | 0.16762087 | -0.583768246 | 0.559376245 | 0.752211034 | Golt1b | 1481.261687 | 0.015881486 | 0.067748776 | 0.234417307 | 0.814661024 | 0.956264994 |
| Tifab | 151.6708886 | 0.087675122 | 0.150236193 | 0.583581893 | 0.559501645 | 0.75231437 | Tgfbrap1 | 3014.852674 | 0.015882841 | 0.039852907 | 0.398536566 | 0.69023471 | 0.923860706 |
| Pus7 | 644.553115 | 0.058586493 | 0.100406888 | 0.583490779 | 0.559562963 | 0.75233153 | Arhgap25 | 461.894994 | 0.015889625 | 0.083725008 | 0.189783494 | 0.84947879 | 0.966260848 |
| Ccr5 | 200.4138042 | -0.093915922 | 0.161038603 | -0.583188879 | 0.559766157 | 0.752539422 | Gm15850 | 1.523124282 | 0.015897864 | 0.027378103 | 0.580678075 | 0.561457441 | NA |
| Rictor | 2536.0841 | 0.049219016 | 0.084412376 | 0.583078193 | 0.559840663 | 0.752574288 | Tbc1d15 | 1280.815689 | 0.015912965 | 0.066160949 | 0.24051899 | 0.809927943 | 0.955876688 |
| Manbal | 1336.123048 | -0.038859504 | 0.066689317 | -0.582694582 | 0.560098921 | 0.752835852 | 1700011E24Rik | 0.940453988 | 0.015915848 | 0.02152624 | 0.739369626 | 0.459682581 | NA |
| Rbbp9 | 1076.451062 | -0.039697608 | 0.068133459 | -0.582644831 | 0.560132419 | 0.752835852 | Lum | 77.3735883 | 0.015916977 | 0.092051975 | 0.172912938 | 0.862719862 | 0.969759087 |
| Lingo1 | 12552.16738 | -0.06746305 | 0.115869174 | -0.582234663 | 0.560408628 | 0.753141754 | Gm12709 | 1.004606608 | 0.015917376 | 0.02006621 | 0.793242745 | 0.427636407 | NA |
| Crabp2 | 96.09040249 | -0.094996009 | 0.163180928 | -0.582151419 | 0.560464693 | 0.753151774 | Scamp4 | 2000.724114 | 0.01592097 | 0.06975534 | 0.228240166 | 0.819459537 | 0.957571905 |
| Greb1 | 70.58939524 | -0.087010581 | 0.149513922 | -0.581956382 | 0.560596062 | 0.753262976 | Osbp | 3279.439995 | 0.015938266 | 0.0523313 | 0.304564689 | 0.760697717 | 0.943105713 |
| Bspry | 12.1063203 | 0.096968537 | 0.166675725 | 0.581779601 | 0.560715146 | 0.753357655 | Spatc1 | 0.880929594 | 0.015945064 | 0.022351986 | 0.713362293 | 0.4756216 | NA |
| Ikbkap | 3197.34942 | 0.02864111 | 0.049254961 | 0.581486812 | 0.560912404 | 0.753557339 | 2310002J15Rik | 1.420358485 | 0.015948623 | 0.028441099 | 0.560759747 | 0.574961331 | NA |
| Npbwr1 | 59.09688859 | 0.096588096 | 0.166174801 | 0.58124394 | 0.561076058 | 0.753711846 | Cep104 | 1581.591695 | 0.015951116 | 0.065822533 | 0.242335182 | 0.808520454 | 0.955716697 |
| Cpped1 | 1590.52956 | -0.033208067 | 0.057161546 | -0.58095117 | 0.561273364 | 0.75391153 | 1700042O10Rik | 0.435709867 | 0.015952828 | 0.014857873 | 1.073695266 | 0.282959284 | NA |
| Dnaic1 | 123.4087539 | 0.081263076 | 0.139936702 | 0.580713103 | 0.561433829 | 0.754061698 | Ccdc152 | 0.435709867 | 0.015952828 | 0.014857873 | 1.073695266 | 0.282959284 | NA |
| Il17rc | 113.4405912 | 0.081734455 | 0.14081777 | 0.580427135 | 0.561626611 | 0.75425524 | Armc4 | 35.21768091 | 0.015954211 | 0.087061475 | 0.183252251 | 0.854600098 | 0.967189923 |
| 4933403G14Rik | 152.3025018 | -0.08478462 | 0.146094399 | -0.580341344 | 0.561684452 | 0.754267542 | Spag6 | 254.2905091 | 0.015959854 | 0.087740263 | 0.181898866 | 0.855662094 | 0.967189923 |
| Nr3c1 | 4234.877718 | -0.038412334 | 0.066199567 | -0.580250535 | 0.561745679 | 0.754284388 | 5031439G07Rik | 6193.871752 | 0.015962069 | 0.065511971 | 0.24365118 | 0.807500986 | 0.955716697 |
| Tmem255b | 42.76691906 | -0.095978633 | 0.165465179 | -0.580053356 | 0.561878637 | 0.754332172 | Mlh3 | 1339.302655 | 0.015964263 | 0.061023409 | 0.261608837 | 0.79362303 | 0.951728598 |
| Zfp867 | 222.1062431 | -0.072803195 | 0.125508187 | -0.580067296 | 0.561869237 | 0.754332172 | Ndufaf4 | 1006.791975 | 0.015966959 | 0.079314278 | 0.20131255 | 0.84045419 | 0.962810534 |
| Drd4 | 8.807071055 | 0.090545838 | 0.156173169 | 0.579778453 | 0.56206403 | 0.754515689 | Sele | 1.450016684 | 0.015971174 | 0.02801433 | 0.570107316 | 0.568604913 | NA |
| Ppp1r3g | 276.4768181 | 0.078448295 | 0.135393772 | 0.579408442 | 0.562313609 | 0.75478533 | Efcab9 | 7.328702053 | 0.015975259 | 0.053916425 | 0.296296704 | 0.767003502 | NA |
| Gm9767 | 11.64910974 | 0.095987575 | 0.165743099 | 0.579134669 | 0.562498308 | 0.754939222 | Megf9 | 3332.210653 | 0.015990017 | 0.073981598 | 0.216135061 | 0.828882471 | 0.959237314 |
| Vav1 | 186.4588382 | 0.074227743 | 0.128179079 | 0.57909406 | 0.562525708 | 0.754939222 | Lrp4 | 1614.337485 | 0.015999866 | 0.073264561 | 0.218384792 | 0.827129314 | 0.958524924 |
| Desi2 | 909.5545877 | -0.049879534 | 0.086177469 | -0.578800176 | 0.562724012 | 0.755139949 | Zcchc14 | 4062.821087 | 0.016002295 | 0.057032185 | 0.280583588 | 0.779029805 | 0.948407834 |
| Tmem66 | 8924.509462 | -0.028586529 | 0.049396767 | -0.578712553 | 0.562783144 | 0.755153896 | Slc27a2 | 274.154173 | 0.016004609 | 0.092747098 | 0.172561835 | 0.862995853 | 0.969759087 |
| Coq3 | 557.8929939 | 0.045680271 | 0.078966911 | 0.578473574 | 0.562944434 | 0.755201161 | Trim58 | 3.416377328 | 0.01601585 | 0.037334452 | 0.428983135 | 0.667935498 | NA |
| Fbn1 | 292.2813088 | 0.057767443 | 0.09986701 | 0.578443706 | 0.562964594 | 0.755201161 | Atf6b | 2405.003685 | 0.016051426 | 0.066153295 | 0.242639855 | 0.808284404 | 0.955716697 |
| Hnrnpa2b1 | 14024.76091 | 0.055694257 | 0.096262925 | 0.578563942 | 0.562883441 | 0.755201161 | Slc25a3 | 12847.5027 | 0.01606536 | 0.050443473 | 0.31848244 | 0.750119012 | 0.9410864 |
| Nampt | 2108.800118 | -0.042527922 | 0.073562535 | -0.578119303 | 0.563183576 | 0.755298738 | Gapdhs | 0.649109394 | 0.016067045 | 0.019502245 | 0.823856189 | 0.410021276 | NA |
| Scaper | 1512.215008 | -0.033351944 | 0.05768812 | -0.578142322 | 0.563168036 | 0.755298738 | Ccne1 | 647.5109777 | 0.016068722 | 0.064548141 | 0.248941669 | 0.803405902 | 0.953754272 |
| Sec23b | 1692.585276 | 0.041181269 | 0.071230234 | 0.57814311 | 0.563167503 | 0.755298738 | C77370 | 1527.262193 | 0.016071121 | 0.071521467 | 0.224703467 | 0.822209967 | 0.95778569 |
| A730046J19Rik | 24.51861197 | 0.085028044 | 0.147133686 | 0.577896509 | 0.563333993 | 0.7553697 | Timm50 | 870.6171966 | 0.01608689 | 0.066305919 | 0.242616191 | 0.808302738 | 0.955716697 |
| Rassf1 | 464.6100751 | 0.056587443 | 0.09791216 | 0.577940918 | 0.563304008 | 0.7553697 | Scyl2 | 1387.02377 | 0.01609026 | 0.0579027 | 0.277884448 | 0.781101063 | 0.948894514 |
| Gm20337 | 10.49401531 | 0.094580657 | 0.163767109 | 0.577531456 | 0.563580495 | 0.755589345 | Tpp2 | 2504.815207 | 0.016091286 | 0.076499088 | 0.210346118 | 0.833397543 | 0.960817765 |
| Snap29 | 1227.038063 | -0.047134949 | 0.081617618 | -0.577509487 | 0.563595331 | 0.755589345 | 5031425F14Rik | 3.620920833 | 0.01610045 | 0.041694021 | 0.386157296 | 0.699380178 | NA |
| Mesp2 | 21.38800553 | -0.09704882 | 0.168121542 | -0.577253924 | 0.563767934 | 0.755755353 | Gm5796 | 5.296437332 | 0.016113576 | 0.051800653 | 0.311068968 | 0.755748192 | NA |
| D8Ertd82e | 1533.56423 | -0.057099154 | 0.098954216 | -0.577025984 | 0.563921903 | 0.755826487 | Sec31a | 4877.130781 | 0.016113934 | 0.042764274 | 0.376808326 | 0.706316053 | 0.929442715 |
| Dnajc18 | 4726.297645 | 0.030132738 | 0.052214188 | 0.577098655 | 0.563872813 | 0.755826487 | Map2k6 | 431.1362792 | 0.016125099 | 0.079232565 | 0.203516054 | 0.838731699 | 0.962130192 |
| Gar1 | 386.7581865 | 0.046771109 | 0.081064914 | 0.576958718 | 0.563967343 | 0.755826487 | Adh7 | 4.741622799 | 0.016125915 | 0.033767068 | 0.477563372 | 0.632961006 | NA |
| 9230116N13Rik | 14.0430113 | 0.093568654 | 0.162203867 | 0.576858341 | 0.564035155 | 0.755839226 | 2310008H09Rik | 1135.296047 | 0.016128501 | 0.057241922 | 0.281760298 | 0.778127315 | 0.94784716 |
| Usf2 | 3409.190482 | -0.032190213 | 0.055808253 | -0.57680023 | 0.564074414 | 0.755839226 | 5830418P13Rik | 27.21353515 | 0.0161307 | 0.08702583 | 0.185355311 | 0.852950358 | 0.966905493 |
| 9430041J12Rik | 120.8946012 | 0.076742297 | 0.133138171 | 0.576410934 | 0.564337456 | 0.755931229 | Arhgef11 | 4328.282219 | 0.01613671 | 0.048743247 | 0.331055292 | 0.740602721 | 0.938975678 |
| Actr3b | 3441.665919 | 0.041823317 | 0.072536684 | 0.576581603 | 0.56422213 | 0.755931229 | Perp | 166.6477878 | 0.016148965 | 0.09479891 | 0.170349686 | 0.864735138 | 0.969972403 |
| Naa25 | 1432.854971 | -0.03016889 | 0.052334246 | -0.576465559 | 0.564300543 | 0.755931229 | Fam96a | 589.0532365 | 0.01615694 | 0.061842805 | 0.261258208 | 0.793893392 | 0.951728598 |
| Rnf138 | 265.9028744 | -0.07478353 | 0.129745941 | -0.576384357 | 0.564355416 | 0.755931229 | AF529169 | 155.478479 | 0.01617348 | 0.094382528 | 0.171360954 | 0.863939955 | 0.969836459 |
| Sstr2 | 570.4334335 | 0.071888166 | 0.124732738 | 0.576337592 | 0.564387018 | 0.755931229 | Gsdma | 1.055462159 | 0.016177923 | 0.020223103 | 0.799972354 | 0.423726815 | NA |
| Cdcp1 | 12.24717585 | 0.095193628 | 0.165307473 | 0.575857983 | 0.564711178 | 0.756103957 | Apoc1 | 35.58683875 | 0.0162101 | 0.080806749 | 0.200603293 | 0.841008783 | 0.963094947 |
| Fhit | 102.0758147 | -0.072298019 | 0.125519746 | -0.575989211 | 0.564622474 | 0.756103957 | Pcdh1 | 5096.426384 | 0.016216043 | 0.078877351 | 0.205585547 | 0.837114668 | 0.961831963 |
| Gstcd | 192.5794838 | 0.05660972 | 0.098281269 | 0.575997042 | 0.564617181 | 0.756103957 | Kidins220 | 16056.68149 | 0.016217058 | 0.047900411 | 0.338557806 | 0.734942873 | 0.936456337 |
| Plscr2 | 67.11826278 | 0.084112497 | 0.146046879 | 0.575928069 | 0.564663802 | 0.756103957 | Diap1 | 2403.909993 | 0.016217754 | 0.07266844 | 0.223174661 | 0.823399572 | 0.957903916 |
| Rbm45 | 798.6069207 | 0.052797391 | 0.091700828 | 0.575756969 | 0.564779463 | 0.756130044 | Tnfsf11 | 1.635285069 | 0.016246304 | 0.02977686 | 0.545601636 | 0.585339799 | NA |
| Acot8 | 257.4529484 | -0.059792871 | 0.103980432 | -0.575039651 | 0.565264481 | 0.756568262 | Dhrs7b | 625.1057176 | 0.016256013 | 0.063145964 | 0.257435502 | 0.796842599 | 0.952885371 |
| Cttn | 4453.828542 | 0.036685013 | 0.063789112 | 0.575098354 | 0.565224781 | 0.756568262 | Bloc1s2a | 909.6889597 | 0.016257337 | 0.05723897 | 0.284025677 | 0.776390703 | 0.947241001 |
| Letmd1 | 2074.641289 | 0.03397266 | 0.059084529 | 0.574984023 | 0.565302102 | 0.756568262 | Csnka2ip | 1.260580072 | 0.016266914 | 0.024837602 | 0.654930932 | 0.512512224 | NA |
| Ppfia2 | 3551.241972 | -0.036899425 | 0.064172115 | -0.575007148 | 0.565286463 | 0.756568262 | Dnajc2 | 1148.418212 | 0.016270545 | 0.067004005 | 0.242829432 | 0.808137535 | 0.955716697 |
| Fam35a | 213.2553049 | 0.065684066 | 0.114265865 | 0.574835414 | 0.565402613 | 0.756637424 | Angpt1 | 189.2961842 | 0.016271425 | 0.090062666 | 0.180667814 | 0.856628324 | 0.967603657 |
| Acsf3 | 629.9471688 | 0.061135083 | 0.106421963 | 0.57445927 | 0.565657055 | 0.756694044 | Uhrf2 | 1907.653354 | 0.016291615 | 0.071054964 | 0.229281873 | 0.818649841 | 0.957156117 |
| Capn7 | 1771.112866 | 0.040199315 | 0.070001026 | 0.574267515 | 0.565786788 | 0.756694044 | Mak16 | 1182.9102 | 0.01629908 | 0.062666549 | 0.260092196 | 0.794792657 | 0.952099467 |
| Hnrnpa1 | 104.0818431 | -0.085935867 | 0.149615764 | -0.574377091 | 0.565712652 | 0.756694044 | Prlhr | 7.949361486 | 0.016300045 | 0.046117233 | 0.353448033 | 0.723752581 | NA |
| Mir143hg | 32.34100031 | 0.09324339 | 0.162349678 | 0.574336771 | 0.565739931 | 0.756694044 | Ep300 | 3234.413906 | 0.016308639 | 0.069783187 | 0.233704418 | 0.815214458 | 0.956264994 |
| Ndufa5 | 2094.784946 | -0.046256126 | 0.080488749 | -0.574690587 | 0.565500575 | 0.756694044 | Ngdn | 585.8456838 | 0.016309889 | 0.066687367 | 0.244572391 | 0.806787544 | 0.955587482 |
| Nqo1 | 176.1937164 | 0.07684816 | 0.133779771 | 0.57443782 | 0.565671566 | 0.756694044 | 2210010C04Rik | 0.453828193 | 0.01631689 | 0.015024447 | 1.086022658 | 0.277468968 | NA |
| Rad51d | 1002.918748 | 0.042216804 | 0.073484898 | 0.574496332 | 0.565631982 | 0.756694044 | Arhgef12 | 11106.69345 | 0.016335824 | 0.060809349 | 0.268640005 | 0.788206724 | 0.951142331 |
| Zfp790 | 700.9034284 | 0.041881542 | 0.072997137 | 0.573742259 | 0.566142227 | 0.757104062 | Cdh10 | 1898.271721 | 0.016336503 | 0.058694266 | 0.278332183 | 0.780757373 | 0.948894514 |
| Adipor1 | 2909.099371 | -0.026157489 | 0.045618531 | -0.57339612 | 0.566376517 | 0.757352011 | Cpne5 | 5524.402414 | 0.016364517 | 0.055563935 | 0.294516882 | 0.768362965 | 0.94619074 |
| Cml2 | 10.05381525 | 0.088875638 | 0.155194272 | 0.572673442 | 0.566865823 | 0.757895294 | Tmem182 | 2.06983075 | 0.016375579 | 0.035835058 | 0.456970906 | 0.647691957 | NA |
| Fign | 148.8357352 | -0.087328793 | 0.15253438 | -0.572518753 | 0.566970585 | 0.757895294 | Gm16548 | 2.31402558 | 0.01638315 | 0.028277733 | 0.579365733 | 0.562342421 | NA |
| Hspbap1 | 199.383466 | 0.05535839 | 0.096702769 | 0.5724592 | 0.567010919 | 0.757895294 | Ly6a | 935.7630607 | 0.016383482 | 0.078730367 | 0.208096102 | 0.835153931 | 0.961504006 |
| Irak2 | 809.1641783 | -0.057672593 | 0.100731504 | -0.572537791 | 0.566957691 | 0.757895294 | Mfsd8 | 776.047092 | 0.01638879 | 0.076662599 | 0.213778173 | 0.830720047 | 0.960118026 |
| Taf11 | 1117.580581 | -0.036889806 | 0.064443671 | -0.572434894 | 0.567027382 | 0.757895294 | Supt16 | 3192.354189 | 0.016416469 | 0.05482991 | 0.299407188 | 0.764629379 | 0.944872045 |
| Colq | 36.44070418 | 0.096152554 | 0.168032009 | 0.572227602 | 0.56716779 | 0.758017574 | 1110059E24Rik | 717.1752098 | 0.016425437 | 0.065183123 | 0.251989102 | 0.801049487 | 0.953228631 |
| Gpr37l1 | 7063.508031 | -0.064193749 | 0.112199939 | -0.572137111 | 0.567229089 | 0.758034113 | 4933434I20Rik | 0.461522871 | 0.016466862 | 0.01509252 | 1.091061118 | 0.27524599 | NA |
| Ghdc | 267.4533465 | 0.056710465 | 0.099202223 | 0.571665259 | 0.567548775 | 0.758395923 | Nol3 | 1079.207368 | 0.016483735 | 0.062105292 | 0.265415953 | 0.790689048 | 0.951728598 |
| Nt5m | 1145.553573 | -0.032159557 | 0.056314135 | -0.571074323 | 0.567949263 | 0.758865633 | Lmo1 | 206.3346478 | 0.016498201 | 0.092154373 | 0.179027872 | 0.857915817 | 0.967971163 |
| Wdr7 | 9534.380612 | -0.043259893 | 0.075766263 | -0.57096511 | 0.568023294 | 0.758899104 | Eny2 | 1948.409617 | 0.016505042 | 0.056370642 | 0.292795005 | 0.769678846 | 0.94619074 |
| Pcdha3 | 94.09224325 | -0.074009865 | 0.129702684 | -0.570611675 | 0.568262903 | 0.759153769 | C030016D13Rik | 17.49752904 | 0.016515051 | 0.079569546 | 0.207554918 | 0.835576508 | NA |
| Mad2l2 | 269.0808264 | 0.05663899 | 0.09929174 | 0.570430025 | 0.56838607 | 0.759252846 | Erich2 | 16.79315303 | 0.016525277 | 0.075479723 | 0.218936637 | 0.826699406 | NA |
| Sh3bp5l | 1956.098145 | -0.040165405 | 0.070434758 | -0.570249777 | 0.568508299 | 0.759350654 | Zc3h12c | 601.2991478 | 0.016532321 | 0.079893719 | 0.206928927 | 0.836065365 | 0.961574465 |
| Fcrlb | 12.36515853 | -0.094546337 | 0.165889165 | -0.569936782 | 0.568720577 | 0.759568711 | Psmd10 | 523.6470725 | 0.016537511 | 0.069772979 | 0.237018852 | 0.812642165 | 0.956264994 |
| 1700088E04Rik | 121.0465836 | 0.071914765 | 0.126306046 | 0.569369142 | 0.569105654 | 0.759886504 | Gm889 | 20.21979035 | 0.016538916 | 0.075928415 | 0.217822483 | 0.827567426 | NA |
| Anp32a | 6641.390151 | -0.025714061 | 0.045160286 | -0.569395445 | 0.569087807 | 0.759886504 | Snord72 | 2.134308492 | 0.016542977 | 0.02923591 | 0.565844435 | 0.571499537 | NA |
| Prkcsh | 2831.081554 | 0.036551268 | 0.064195256 | 0.569376466 | 0.569100684 | 0.759886504 | Arhgap33 | 11187.42028 | 0.016543925 | 0.073889166 | 0.223901906 | 0.822833633 | 0.957884384 |
| Ccdc88b | 147.9719682 | 0.075765006 | 0.133100886 | 0.569229914 | 0.569200123 | 0.759947152 | Tmprss7 | 92.37218828 | 0.016549867 | 0.095058606 | 0.174101722 | 0.86178552 | 0.969358561 |
| 9830147E19Rik | 96.11558511 | -0.087596297 | 0.153973211 | -0.568906089 | 0.569419873 | 0.760138156 | Rad51b | 9.388619497 | 0.016553264 | 0.062526486 | 0.26474004 | 0.79120973 | NA |
| Cacnb1 | 4101.574959 | 0.053322107 | 0.093745823 | 0.568794477 | 0.569495623 | 0.760138156 | Rps8 | 7734.12279 | 0.016578894 | 0.056214404 | 0.29492252 | 0.768053068 | 0.94619074 |
| Eid1 | 4575.92487 | -0.047596371 | 0.083720798 | -0.568513111 | 0.569686606 | 0.760138156 | 1700063O14Rik | 0.892156306 | 0.016600935 | 0.022710916 | 0.730967216 | 0.464799178 | NA |
| Lypla2 | 2466.515044 | 0.046223839 | 0.081287317 | 0.568647614 | 0.569595305 | 0.760138156 | Nphp1 | 609.2267704 | 0.016605815 | 0.077560779 | 0.214100675 | 0.830468549 | 0.960002081 |
| Siae | 1028.712101 | 0.054619302 | 0.096062436 | 0.56858127 | 0.569640338 | 0.760138156 | Gm3696 | 2.099391675 | 0.0166104 | 0.030993333 | 0.535934633 | 0.592003728 | NA |
| Ttc34 | 84.66276334 | 0.085025607 | 0.14954365 | 0.56856715 | 0.569649924 | 0.760138156 | Ccm2 | 1998.720623 | 0.016614684 | 0.063999141 | 0.259607926 | 0.795166222 | 0.952311364 |
| Zfp397 | 1336.586802 | 0.050606015 | 0.088993002 | 0.568651624 | 0.569592584 | 0.760138156 | Pi4kb | 2397.418482 | 0.016615417 | 0.041664642 | 0.398789396 | 0.690048391 | 0.923860706 |
| Vmac | 445.9078806 | 0.050070105 | 0.088142392 | 0.568059295 | 0.569994706 | 0.760483766 | Olfr898 | 0.678454432 | 0.01662445 | 0.019661798 | 0.845520308 | 0.397820399 | NA |
| Tmem154 | 25.49268815 | 0.095338829 | 0.168040858 | 0.567355048 | 0.570472984 | 0.761056347 | G630090E17Rik | 1.088190654 | 0.016635694 | 0.026516168 | 0.627379265 | 0.530410656 | NA |
| Ermap | 51.51710048 | -0.083312079 | 0.146875805 | -0.567228064 | 0.570559243 | 0.76110589 | Prnp | 38453.64813 | 0.01663924 | 0.047752176 | 0.348449879 | 0.727502349 | 0.9335794 |
| Capn5 | 2350.0524 | 0.04317872 | 0.076158635 | 0.56695764 | 0.570742961 | 0.761260281 | Aldh1a7 | 27.90760048 | 0.016653385 | 0.083714195 | 0.19893143 | 0.842316383 | 0.963252575 |
| Rslcan18 | 27.96306873 | 0.093307324 | 0.164588417 | 0.566913065 | 0.570773247 | 0.761260281 | Lgr5 | 126.4892576 | 0.016655528 | 0.094573177 | 0.176112599 | 0.860205483 | 0.96862798 |
| Cks2 | 32.62427738 | -0.089742683 | 0.158386998 | -0.566603849 | 0.570983359 | 0.761290531 | Il7r | 11.8053372 | 0.016666651 | 0.070773818 | 0.235491768 | 0.813827068 | NA |
| Crlf2 | 362.7397428 | 0.07295355 | 0.128843157 | 0.566219828 | 0.571244353 | 0.761290531 | Cyp26a1 | 5.824360881 | 0.016671904 | 0.043714732 | 0.381379526 | 0.702921651 | NA |
| D6Wsu163e | 1271.217306 | 0.03930445 | 0.069371637 | 0.566578096 | 0.57100086 | 0.761290531 | Syk | 205.6052481 | 0.016673757 | 0.091606341 | 0.182015313 | 0.855570709 | 0.967189923 |
| Galr1 | 21.30730012 | -0.092683429 | 0.163560561 | -0.566661233 | 0.570944364 | 0.761290531 | Ptk7 | 526.0911573 | 0.0166849 | 0.077915095 | 0.214142077 | 0.830436264 | 0.960002081 |
| Gas2l1 | 2141.536354 | -0.045255056 | 0.079933801 | -0.566156684 | 0.571287273 | 0.761290531 | Mcts1 | 1062.70383 | 0.016685798 | 0.071177319 | 0.234425778 | 0.814654449 | 0.956264994 |
| Lipt2 | 121.3747502 | -0.064072131 | 0.113078795 | -0.566614905 | 0.570975846 | 0.761290531 | Rln1 | 12.29038666 | 0.016703557 | 0.063356921 | 0.263642183 | 0.792055651 | NA |
| Pion | 258.504343 | 0.079091862 | 0.139669728 | 0.566277765 | 0.571204973 | 0.761290531 | Zmynd12 | 16.02131086 | 0.016705455 | 0.071255094 | 0.23444576 | 0.814638937 | NA |
| Tlk1 | 4819.497231 | -0.046456874 | 0.082022307 | -0.566393173 | 0.571126534 | 0.761290531 | Rft1 | 475.0497361 | 0.016708792 | 0.07559755 | 0.221022928 | 0.82507458 | 0.957903916 |
| Ugt8a | 1655.267426 | -0.093366209 | 0.164834559 | -0.56642375 | 0.571105754 | 0.761290531 | Dntt | 1.069428219 | 0.016723725 | 0.021850706 | 0.765363154 | 0.444055328 | NA |
| Zbed4 | 596.1574726 | -0.046984692 | 0.082976179 | -0.566243138 | 0.571228509 | 0.761290531 | Med11 | 271.3910865 | 0.016726362 | 0.08392379 | 0.199304174 | 0.842024814 | 0.963252575 |
| Lcat | 678.0830077 | -0.075892238 | 0.134071956 | -0.566056022 | 0.571355698 | 0.761316235 | Tceal5 | 2332.687375 | 0.016731417 | 0.067205055 | 0.248960688 | 0.803391191 | 0.953754272 |
| A330040F15Rik | 9.76531492 | -0.088919466 | 0.157163395 | -0.565777203 | 0.571545246 | 0.761503314 | Zfp11 | 310.2885203 | 0.016733944 | 0.084176773 | 0.198795265 | 0.8424229 | 0.963252575 |
| Tdgf1 | 14.06972081 | 0.092015333 | 0.162699809 | 0.5655528 | 0.571697822 | 0.761616356 | 4930539E08Rik | 500.2937884 | 0.016740685 | 0.081058525 | 0.206525906 | 0.836380131 | 0.961664 |
| Tfap4 | 104.3016255 | 0.078262189 | 0.13839276 | 0.56550783 | 0.5717284 | 0.761616356 | 4930426L09Rik | 27.07507493 | 0.016745176 | 0.081784503 | 0.204747536 | 0.837769379 | 0.961858836 |
| Foxl2 | 27.22668218 | -0.094949901 | 0.167942601 | -0.565371148 | 0.571821344 | 0.761674683 | 4930415O20Rik | 1.535240012 | 0.016759822 | 0.026225732 | 0.639060211 | 0.522783763 | NA |
| Necab1 | 4211.967138 | -0.054060909 | 0.095644574 | -0.56522714 | 0.571919279 | 0.761739646 | Gm14005 | 6.409563778 | 0.016764845 | 0.057667659 | 0.290714843 | 0.771269419 | NA |
| Mrgpre | 490.6500124 | -0.056769992 | 0.100461634 | -0.565091265 | 0.572011689 | 0.761797242 | Vmn1r200 | 0.770647357 | 0.016765901 | 0.01897813 | 0.88343275 | 0.377002503 | NA |
| Cthrc1 | 194.5271856 | -0.059476988 | 0.105271528 | -0.564986464 | 0.572082971 | 0.761826691 | 4933425L06Rik | 0.47798497 | 0.016777915 | 0.015232716 | 1.101439536 | 0.270705406 | NA |
| Clpb | 2039.223153 | -0.027308365 | 0.048364263 | -0.564639328 | 0.572319109 | 0.761944067 | Kat6a | 3308.261731 | 0.016779702 | 0.064033954 | 0.262043818 | 0.79328766 | 0.951728598 |
| Efnb1 | 601.8756222 | -0.05801971 | 0.102781538 | -0.564495449 | 0.572416996 | 0.761944067 | Gjd4 | 0.905989442 | 0.016780833 | 0.021304741 | 0.787657226 | 0.430897232 | NA |
| Mettl21a | 383.3669285 | -0.057191105 | 0.101262814 | -0.564778943 | 0.572224131 | 0.761944067 | Sphk1 | 137.3583932 | 0.016791466 | 0.094786406 | 0.177150573 | 0.859390119 | 0.968520191 |
| Mmp23 | 35.8829567 | 0.087738363 | 0.155423845 | 0.564510309 | 0.572406886 | 0.761944067 | Chd3 | 24837.14271 | 0.016801042 | 0.051818418 | 0.324229145 | 0.745764558 | 0.940689126 |
| Sys1 | 874.9445353 | -0.052523263 | 0.093021647 | -0.564634843 | 0.572322161 | 0.761944067 | Ccdc158 | 14.2416676 | 0.01680694 | 0.073697431 | 0.228053261 | 0.819604835 | NA |
| Smc3 | 2271.429075 | 0.045807268 | 0.081159065 | 0.564413453 | 0.572472785 | 0.761952868 | Gm10754 | 80.68761794 | 0.016825309 | 0.091743232 | 0.183395644 | 0.854487594 | 0.967189923 |
| Cln5 | 522.3420494 | 0.044750993 | 0.079312333 | 0.5642375 | 0.57259251 | 0.762018442 | Syt14 | 261.208803 | 0.016830304 | 0.087354176 | 0.192667423 | 0.847219438 | 0.965275633 |
| Rilpl2 | 354.7589868 | 0.055963092 | 0.099190783 | 0.564196491 | 0.572620416 | 0.762018442 | Vaultrc5 | 1.009678759 | 0.016837257 | 0.020337984 | 0.827872494 | 0.407742714 | NA |
| Ddx10 | 707.8152486 | -0.03934421 | 0.06975832 | -0.564007415 | 0.572749086 | 0.762058766 | 4930593C16Rik | 0.667302911 | 0.016849683 | 0.019600365 | 0.859661714 | 0.389975545 | NA |
| Ptgfrn | 1921.118534 | -0.050798468 | 0.090062797 | -0.564033871 | 0.572731081 | 0.762058766 | Prkx | 505.9890009 | 0.016853072 | 0.082103474 | 0.205266243 | 0.837364117 | 0.961831963 |
| Asxl1 | 2618.141823 | 0.032641512 | 0.057958435 | 0.563188292 | 0.573306674 | 0.762342397 | BC051628 | 3.610209457 | 0.016856013 | 0.03492237 | 0.482670945 | 0.629329397 | NA |
| Gab1 | 1391.512485 | 0.051137226 | 0.090787413 | 0.563263391 | 0.573255543 | 0.762342397 | Pdgfc | 303.49703 | 0.016859281 | 0.084514812 | 0.199483153 | 0.841884821 | 0.963252575 |
| Glul | 66094.55951 | -0.051948118 | 0.09218573 | -0.56351583 | 0.573083684 | 0.762342397 | Pgc | 1.981941967 | 0.016860403 | 0.03198977 | 0.527056093 | 0.598154636 | NA |
| Ranbp6 | 1750.719123 | -0.038645892 | 0.068581408 | -0.563503913 | 0.573091796 | 0.762342397 | Slc23a2 | 8989.443049 | 0.01686683 | 0.052000286 | 0.324360337 | 0.745665244 | 0.940687916 |
| Sppl3 | 3118.161838 | 0.031712853 | 0.05630317 | 0.563251636 | 0.573263546 | 0.762342397 | Ltc4s | 96.25697335 | 0.016868683 | 0.090105259 | 0.187210857 | 0.851495312 | 0.96665003 |
| Trim56 | 168.5985045 | -0.069081865 | 0.122622083 | -0.56337214 | 0.573181504 | 0.762342397 | Gnao1 | 32710.19495 | 0.0168729 | 0.042473193 | 0.397259993 | 0.691175746 | 0.923860706 |
| Zdhhc22 | 595.2748481 | -0.061170585 | 0.108613274 | -0.563196213 | 0.573301281 | 0.762342397 | Zfp781 | 102.2134697 | 0.01688249 | 0.09417399 | 0.17926914 | 0.857726377 | 0.967971163 |
| Ash2l | 1419.037037 | -0.034304855 | 0.06093622 | -0.562963288 | 0.573459883 | 0.762480685 | 9330182L06Rik | 1482.501909 | 0.016891767 | 0.06209884 | 0.272014211 | 0.785611093 | 0.950484821 |
| Rabl3 | 295.0727364 | -0.058000561 | 0.103098755 | -0.562572859 | 0.573725778 | 0.762659199 | Tmem5 | 802.5904747 | 0.016898391 | 0.068271534 | 0.247517376 | 0.804507846 | 0.954605536 |
| Tmem232 | 86.92880545 | -0.087757309 | 0.15599337 | -0.562570757 | 0.573727209 | 0.762659199 | Gm1821 | 1591.852893 | 0.016906251 | 0.064515778 | 0.262048319 | 0.79328419 | 0.951728598 |
| Zfp354c | 1804.092385 | 0.050099332 | 0.08905767 | 0.562549319 | 0.573741811 | 0.762659199 | Tgfbi | 183.6781946 | 0.016918637 | 0.094566305 | 0.178907663 | 0.858010206 | 0.967971163 |
| Crmp1 | 7134.079123 | -0.030066659 | 0.053464533 | -0.56236644 | 0.57386638 | 0.762720771 | 1700101E01Rik | 10.5973199 | 0.016923981 | 0.058188517 | 0.29084743 | 0.771168009 | NA |
| Usp1 | 978.2084561 | 0.046926027 | 0.083448262 | 0.562336778 | 0.573886585 | 0.762720771 | 1500004A13Rik | 1091.179059 | 0.016934509 | 0.076729011 | 0.220705425 | 0.825321807 | 0.957903916 |
| Oard1 | 388.1861107 | 0.045811898 | 0.081482262 | 0.562231548 | 0.573958269 | 0.762750616 | Synpr | 4144.656626 | 0.016938336 | 0.076290494 | 0.222024195 | 0.824295048 | 0.957903916 |
| Psmd11 | 2044.35448 | 0.033792442 | 0.060115637 | 0.562123987 | 0.574031546 | 0.762779925 | Eif3k | 2846.714229 | 0.016951641 | 0.04678387 | 0.362339435 | 0.717098393 | 0.93125489 |
| Zfp760 | 346.1673176 | -0.067859959 | 0.120735518 | -0.56205465 | 0.574078785 | 0.762779925 | Htra3 | 183.7194423 | 0.016955404 | 0.090892283 | 0.186543933 | 0.85201823 | 0.966756508 |
| Apmap | 2245.332474 | 0.042210736 | 0.075169045 | 0.561544133 | 0.574426653 | 0.762944234 | Cpsf3 | 1961.611931 | 0.016964162 | 0.05125804 | 0.330956126 | 0.740677626 | 0.938975678 |
| C1qtnf7 | 28.82564837 | -0.089720617 | 0.159804576 | -0.5614396 | 0.574497895 | 0.762944234 | Gdf6 | 3.393105803 | 0.016968048 | 0.037451212 | 0.453070736 | 0.650497807 | NA |
| Glb1l2 | 22.6372275 | 0.094101933 | 0.16751737 | 0.561744333 | 0.574290224 | 0.762944234 | Uhrf1bp1l | 4259.305814 | 0.016985934 | 0.061089558 | 0.278049709 | 0.7809742 | 0.948894514 |
| Magoh | 428.1393623 | -0.056083974 | 0.099885751 | -0.561481227 | 0.574469525 | 0.762944234 | Oprd1 | 1066.366838 | 0.016993208 | 0.080671525 | 0.210646918 | 0.833162799 | 0.960810394 |
| Man1b1 | 3687.942776 | 0.0277558 | 0.049412646 | 0.561714512 | 0.574310546 | 0.762944234 | Znhit3 | 620.8518741 | 0.016993302 | 0.06161144 | 0.275814072 | 0.782690875 | 0.949296337 |
| Spred1 | 4532.063589 | -0.064632023 | 0.115106533 | -0.561497434 | 0.574458479 | 0.762944234 | Pno1 | 833.6434115 | 0.016997215 | 0.075726662 | 0.224454828 | 0.822403412 | 0.95778569 |
| Ptch2 | 74.83602779 | -0.087948297 | 0.156668534 | -0.56136542 | 0.574548453 | 0.762945983 | 5730522E02Rik | 63.70918848 | 0.01700987 | 0.095099239 | 0.178864414 | 0.858044165 | 0.967971163 |
| 5830454E08Rik | 25.96236218 | 0.093895773 | 0.167286423 | 0.561287466 | 0.574601584 | 0.762951148 | Senp2 | 4588.905566 | 0.017026048 | 0.056996377 | 0.298721576 | 0.765152493 | 0.945119482 |
| Lrp12 | 2286.212438 | 0.031896463 | 0.056861927 | 0.560945874 | 0.574834436 | 0.763194923 | Vmn2r86 | 2.729100944 | 0.017027777 | 0.030279369 | 0.562355745 | 0.573873665 | NA |
| Adprm | 419.3402751 | 0.047267874 | 0.084288369 | 0.560787617 | 0.574942329 | 0.763202562 | Calml4 | 53.76396739 | 0.017030959 | 0.036445233 | 0.467302796 | 0.640283256 | 0.910092095 |
| Lin7b | 1252.878162 | -0.07660111 | 0.136626815 | -0.560659416 | 0.575029739 | 0.763202562 | Mrpl11 | 1340.784471 | 0.017033048 | 0.056597209 | 0.300952083 | 0.763451034 | 0.944624337 |
| Nfkbil1 | 275.7508445 | -0.060013639 | 0.107033864 | -0.560697679 | 0.575003649 | 0.763202562 | Nos1 | 1267.095665 | 0.017039756 | 0.091962871 | 0.185289514 | 0.853001962 | 0.966905493 |
| Pi4ka | 16689.23417 | -0.045810195 | 0.081711991 | -0.560630022 | 0.575049781 | 0.763202562 | Lypd3 | 1.255514117 | 0.017041454 | 0.023067644 | 0.738760054 | 0.460052712 | NA |
| Pkd2l1 | 47.52111772 | 0.091870842 | 0.163886448 | 0.560576201 | 0.575086479 | 0.763202562 | Gle1 | 2481.239133 | 0.017058486 | 0.043884641 | 0.388711989 | 0.697489212 | 0.9261158 |
| Ap3d1 | 8086.209003 | -0.023396579 | 0.041763015 | -0.560222456 | 0.575327712 | 0.763230445 | Gm10046 | 2.985689659 | 0.017059974 | 0.040640957 | 0.419772942 | 0.674651333 | NA |
| Arhgap12 | 1533.532502 | -0.045165298 | 0.080592657 | -0.560414557 | 0.575196705 | 0.763230445 | Mt3 | 5885.203373 | 0.017062466 | 0.059234321 | 0.288050337 | 0.773308207 | 0.947121017 |
| Baiap2 | 12995.6329 | -0.049554778 | 0.088448175 | -0.560269087 | 0.57529591 | 0.763230445 | Fam185a | 708.0033206 | 0.017062966 | 0.070397804 | 0.24237924 | 0.808486319 | 0.955716697 |
| Ndrg3 | 23188.8046 | -0.032353624 | 0.057751553 | -0.560220855 | 0.575328804 | 0.763230445 | Adprhl1 | 4.816383266 | 0.017072047 | 0.049369027 | 0.34580481 | 0.72948941 | NA |
| Psg16 | 94.31802111 | 0.069387578 | 0.123871175 | 0.560159198 | 0.575370855 | 0.763230445 | Nrxn2 | 14980.64799 | 0.017073236 | 0.047807757 | 0.357122714 | 0.720999941 | 0.932513933 |
| Spry1 | 312.6673071 | 0.053826642 | 0.096099784 | 0.560111997 | 0.575403048 | 0.763230445 | Gm16294 | 6.976440534 | 0.017077552 | 0.056303555 | 0.303312147 | 0.76165199 | NA |
| Rab7l1 | 114.9442478 | -0.0766331 | 0.136863597 | -0.559923176 | 0.57553184 | 0.763335929 | Zmat2 | 7864.083048 | 0.017115214 | 0.057098766 | 0.299747526 | 0.764369744 | 0.944872045 |
| Dock4 | 4803.356088 | 0.034963136 | 0.062460396 | 0.559764878 | 0.575639823 | 0.763413799 | Rtdr1 | 8.886900332 | 0.017129259 | 0.058368746 | 0.29346628 | 0.76916577 | NA |
| Mettl20 | 75.15535246 | 0.082776521 | 0.14790545 | 0.559658356 | 0.575712493 | 0.763444827 | Tcp11 | 18.89066489 | 0.017135644 | 0.072546188 | 0.236203229 | 0.813274974 | NA |
| Cys1 | 262.3306415 | 0.064286837 | 0.114950688 | 0.559255783 | 0.575987167 | 0.76363536 | Gsg1 | 3.354542912 | 0.017137408 | 0.035942766 | 0.47679714 | 0.633506582 | NA |
| Glis2 | 481.2954931 | 0.046764649 | 0.083623124 | 0.559231071 | 0.576004031 | 0.76363536 | Cd2bp2 | 4015.197645 | 0.017144661 | 0.055101824 | 0.311145076 | 0.755690335 | 0.942139348 |
| Wdr18 | 2175.271159 | 0.032421459 | 0.057972875 | 0.559252217 | 0.575989601 | 0.76363536 | Wnk2 | 5935.243561 | 0.017150373 | 0.058016716 | 0.295610886 | 0.767527259 | 0.94619074 |
| Ttll4 | 506.9156926 | 0.055167771 | 0.098672322 | 0.559100771 | 0.576092949 | 0.763687898 | Zbtb39 | 588.903224 | 0.017151329 | 0.069233705 | 0.247730919 | 0.804342607 | 0.954487437 |
| Tmem18 | 1388.671869 | 0.030419555 | 0.054475334 | 0.558409706 | 0.576564646 | 0.764247808 | Pigg | 831.9668946 | 0.01715248 | 0.077573764 | 0.221111867 | 0.82500533 | 0.957903916 |
| Kif18a | 30.68812692 | 0.091822594 | 0.164519381 | 0.558126305 | 0.576758139 | 0.764373502 | Wnt6 | 40.34068802 | 0.017158509 | 0.086123722 | 0.199230929 | 0.842082107 | 0.963252575 |
| Zfp862 | 526.3511311 | 0.047363122 | 0.084855057 | 0.558164992 | 0.576731724 | 0.764373502 | Tmem63b | 5918.504264 | 0.017197481 | 0.048655124 | 0.353456734 | 0.723746059 | 0.932714927 |
| Qsox2 | 809.0852883 | -0.041931095 | 0.075146308 | -0.557992753 | 0.576849333 | 0.764428974 | Fam195a | 203.9150226 | 0.017198988 | 0.087675283 | 0.1961669 | 0.844479535 | 0.964328285 |
| 4930473A02Rik | 10.30533971 | 0.071634048 | 0.128406007 | 0.557871471 | 0.576932154 | 0.764473343 | Cpped1 | 1702.520757 | 0.01720926 | 0.054002297 | 0.318676449 | 0.749971876 | 0.9410864 |
| Nod2 | 47.73521017 | 0.086383119 | 0.154864579 | 0.557797786 | 0.576982475 | 0.764474642 | Fopnl | 1258.917805 | 0.017211326 | 0.049651294 | 0.346644052 | 0.728858746 | 0.933996867 |
| 5330417C22Rik | 1266.269681 | 0.035132721 | 0.063014822 | 0.55753107 | 0.577164637 | 0.764560121 | Lhpp | 536.3084099 | 0.017212664 | 0.073204703 | 0.235130578 | 0.814107386 | 0.956264994 |
| Hscb | 141.5269484 | 0.066514615 | 0.119296817 | 0.557555658 | 0.577147843 | 0.764560121 | Ddx18 | 900.3755974 | 0.017219261 | 0.058896623 | 0.292364148 | 0.770008217 | 0.94619074 |
| Sufu | 384.1833299 | 0.052978514 | 0.095031012 | 0.557486579 | 0.577195026 | 0.764560121 | 4430402I18Rik | 144.9287642 | 0.017220221 | 0.093248588 | 0.18467005 | 0.853487839 | 0.966905493 |
| Mfsd5 | 1241.565059 | 0.038387495 | 0.068921507 | 0.556974111 | 0.577545117 | 0.764827696 | Dkk1 | 0.73024527 | 0.017238578 | 0.018800214 | 0.9169352 | 0.359176597 | NA |
| Nat1 | 50.10764092 | -0.09064873 | 0.162720382 | -0.557082822 | 0.577470844 | 0.764827696 | Ano5 | 60.85278973 | 0.017244724 | 0.093148672 | 0.185131184 | 0.853126143 | 0.966905493 |
| Rsph1 | 116.4601944 | 0.088476213 | 0.158840237 | 0.557013857 | 0.577517962 | 0.764827696 | Pkp3 | 2.691068741 | 0.017269753 | 0.031702493 | 0.544744328 | 0.585929373 | NA |
| 4930524L23Rik | 15.53073396 | -0.093567705 | 0.168044674 | -0.556802559 | 0.577662335 | 0.764852181 | Mxd1 | 1301.175897 | 0.017275388 | 0.070305236 | 0.24571979 | 0.805899152 | 0.95539795 |
| Med26 | 352.0258037 | -0.049796488 | 0.08942817 | -0.55683224 | 0.577642054 | 0.764852181 | Prop1 | 0.733507625 | 0.017281421 | 0.018816707 | 0.918408339 | 0.358405126 | NA |
| Ptbp3 | 879.5328262 | -0.045944812 | 0.08255374 | -0.556544276 | 0.577838835 | 0.7650205 | Kcnk15 | 1.461269618 | 0.017285834 | 0.022974247 | 0.752400444 | 0.45181028 | NA |
| Ankrd42 | 513.0221254 | 0.048434334 | 0.087071724 | 0.556257901 | 0.578034562 | 0.765057436 | E230025N22Rik | 23.23881003 | 0.017304168 | 0.077962979 | 0.221953654 | 0.824349962 | 0.957903916 |
| BC064078 | 74.75305515 | 0.084642994 | 0.152229031 | 0.556023994 | 0.578194452 | 0.765057436 | Dyrk1a | 3745.7956 | 0.017317477 | 0.059424895 | 0.291417879 | 0.770731742 | 0.946250219 |
| Coch | 1537.875167 | 0.072871161 | 0.13107666 | 0.555943071 | 0.578249773 | 0.765057436 | A930009A15Rik | 14.81798236 | 0.017318997 | 0.072416736 | 0.239157376 | 0.810983553 | NA |
| Fam188b | 269.1670418 | -0.047395866 | 0.085190976 | -0.556348432 | 0.577972684 | 0.765057436 | Ascl3 | 3.112209689 | 0.017321224 | 0.035499665 | 0.487926403 | 0.625601972 | NA |
| Itga4 | 439.0183097 | 0.052010405 | 0.09352615 | 0.556105486 | 0.578138745 | 0.765057436 | Eif4e3 | 620.1677231 | 0.017322181 | 0.072983905 | 0.237342486 | 0.812391105 | 0.956219756 |
| Pelo | 443.9514889 | -0.049211581 | 0.088521892 | -0.555925544 | 0.578261755 | 0.765057436 | Ttll9 | 58.01424054 | 0.017323909 | 0.095046583 | 0.182267559 | 0.855372756 | 0.967189923 |
| Ppme1 | 9416.025652 | -0.042093702 | 0.075656451 | -0.556379548 | 0.577951417 | 0.765057436 | E2f1 | 693.4371017 | 0.017327187 | 0.07264492 | 0.238518909 | 0.811478651 | 0.956135608 |
| Pvt1 | 314.0901327 | -0.056931969 | 0.102406552 | -0.555940686 | 0.578251403 | 0.765057436 | Abcc6 | 62.82198544 | 0.017335097 | 0.093958226 | 0.184497918 | 0.85362286 | 0.966905493 |
| Rps15a-ps6 | 19.82120477 | -0.089262934 | 0.160600384 | -0.555807726 | 0.578342303 | 0.765098672 | Cntnap5a | 1096.894068 | 0.017354236 | 0.084725899 | 0.204827996 | 0.837706513 | 0.961858836 |
| Hapln1 | 576.4980755 | -0.072127569 | 0.12980944 | -0.55564194 | 0.578455655 | 0.76515473 | Gnb2l1 | 5805.721191 | 0.017362831 | 0.049615042 | 0.349950954 | 0.726375506 | 0.9335794 |
| Nrip1 | 1018.936234 | -0.061810276 | 0.111249337 | -0.555601295 | 0.578483446 | 0.76515473 | Acvr1b | 5235.688607 | 0.017366407 | 0.072679082 | 0.238946427 | 0.811147124 | 0.956135608 |
| Sf3a1 | 3348.285125 | 0.030487301 | 0.054971567 | 0.55460127 | 0.57916742 | 0.765970429 | 4931429P17Rik | 0.738051225 | 0.017371565 | 0.018838057 | 0.92215266 | 0.356448952 | NA |
| Slc6a20a | 499.8538132 | 0.080291089 | 0.144784695 | 0.554555089 | 0.579199015 | 0.765970429 | Spata17 | 16.63550244 | 0.01737976 | 0.072291943 | 0.240410751 | 0.810011845 | NA |
| Trnt1 | 1140.047267 | 0.03383133 | 0.061023039 | 0.554402583 | 0.579303359 | 0.766043035 | Tacr2 | 21.16480622 | 0.017385554 | 0.080591492 | 0.215724438 | 0.829202551 | NA |
| Poglut1 | 1981.256679 | 0.034368849 | 0.062009936 | 0.554247457 | 0.579409504 | 0.766052637 | Mad2l1bp | 317.8831195 | 0.017397887 | 0.081528975 | 0.21339514 | 0.831018771 | 0.960162026 |
| Ylpm1 | 4415.793214 | -0.034087651 | 0.061498156 | -0.554287364 | 0.579382197 | 0.766052637 | Lrr1 | 1.050075312 | 0.017401608 | 0.02168547 | 0.802454719 | 0.422289971 | NA |
| 1810044D09Rik | 13.82045169 | -0.089220727 | 0.161096983 | -0.553832385 | 0.579693562 | 0.766362804 | Hpse | 59.38766555 | 0.017414308 | 0.094315727 | 0.184638431 | 0.85351264 | 0.966905493 |
| Alg12 | 387.0680473 | 0.059346128 | 0.107233796 | 0.553427463 | 0.579970738 | 0.766607924 | Rtca | 1834.818347 | 0.017430498 | 0.059683282 | 0.292049916 | 0.77024846 | 0.94619074 |
| Rab24 | 1704.021134 | 0.038338867 | 0.06927664 | 0.553416953 | 0.579977933 | 0.766607924 | Olfr1392 | 2.685136898 | 0.017431317 | 0.038609219 | 0.45148069 | 0.651643139 | NA |
| Dusp6 | 2135.10789 | -0.087438275 | 0.158105377 | -0.553037962 | 0.580237416 | 0.766885483 | Ccdc112 | 358.8377235 | 0.017451137 | 0.075472771 | 0.23122428 | 0.817140568 | 0.956302288 |
| Plxna3 | 1390.807285 | 0.077597581 | 0.140349202 | 0.552889363 | 0.580339171 | 0.766954548 | Mir345 | 2.597795488 | 0.017476034 | 0.032652646 | 0.535210347 | 0.592504412 | NA |
| Use1 | 1638.54185 | -0.03784995 | 0.068476115 | -0.552746746 | 0.580436839 | 0.767018199 | 3830406C13Rik | 989.4279286 | 0.017490055 | 0.068136844 | 0.256690126 | 0.797417994 | 0.952897915 |
| Car5b | 153.5490642 | 0.067668483 | 0.122471997 | 0.552522089 | 0.580590705 | 0.767107029 | Ptch2 | 85.12928962 | 0.017510676 | 0.095051163 | 0.184223687 | 0.853837978 | 0.966905493 |
| Carm1 | 3215.166508 | 0.038379391 | 0.069464456 | 0.552504021 | 0.58060308 | 0.767107029 | Ric8 | 2420.674927 | 0.017513574 | 0.056959037 | 0.307476648 | 0.758480595 | 0.942526199 |
| Atp6v1b2 | 31166.81289 | -0.036898066 | 0.066793327 | -0.552421444 | 0.580659642 | 0.767116345 | Kirrel3 | 2169.599311 | 0.017519032 | 0.070424162 | 0.248764515 | 0.803542941 | 0.953754272 |
| Dyx1c1 | 90.0565935 | 0.073325528 | 0.132765546 | 0.552293348 | 0.580747387 | 0.767166853 | Spast | 2121.908136 | 0.017520624 | 0.066334835 | 0.264124034 | 0.791684345 | 0.951728598 |
| AI597479 | 1296.250587 | -0.03802538 | 0.068890231 | -0.551970564 | 0.580968521 | 0.767185714 | Cmtm5 | 1503.613946 | 0.017524013 | 0.077807509 | 0.225222646 | 0.821806073 | 0.95778569 |
| Arhgap24 | 244.9311633 | -0.067948597 | 0.123072058 | -0.552104177 | 0.58087698 | 0.767185714 | Pkd2l2 | 20.05317315 | 0.01753536 | 0.070724143 | 0.247940219 | 0.804180661 | NA |
| Mllt6 | 7928.291013 | 0.033608209 | 0.060879355 | 0.552046071 | 0.580916789 | 0.767185714 | 2010003O02Rik | 55.07511054 | 0.01754618 | 0.093206364 | 0.188250873 | 0.850679994 | 0.966301569 |
| Rmi1 | 650.6745029 | 0.044246799 | 0.080170151 | 0.551911133 | 0.581009241 | 0.767185714 | C630043F03Rik | 234.3894136 | 0.01755942 | 0.082819695 | 0.212019858 | 0.832091542 | 0.960519604 |
| Snx17 | 2932.025867 | -0.025987725 | 0.04708678 | -0.551911278 | 0.581009141 | 0.767185714 | Gm10636 | 3.494092062 | 0.017561886 | 0.041227828 | 0.425971646 | 0.670128507 | NA |
| Dera | 65.37567884 | 0.082173201 | 0.148931672 | 0.551751012 | 0.581118955 | 0.767239587 | Fam179a | 162.2638716 | 0.017572612 | 0.090389211 | 0.194410498 | 0.845854471 | 0.964996028 |
| Dnmt1 | 2374.21126 | 0.040708004 | 0.073785542 | 0.551707054 | 0.581149076 | 0.767239587 | Fam209 | 1.629426551 | 0.017575569 | 0.02248076 | 0.781804952 | 0.43432921 | NA |
| Zfp133-ps | 94.31778235 | -0.080237165 | 0.145478297 | -0.551540449 | 0.581263246 | 0.767324933 | Zbtb25 | 386.479464 | 0.017578991 | 0.075726746 | 0.232137157 | 0.816431486 | 0.956264994 |
| 4930429B21Rik | 394.6208255 | -0.059014274 | 0.107063093 | -0.551210252 | 0.581489553 | 0.767558284 | Ppp6r2 | 3995.113466 | 0.017582318 | 0.050634835 | 0.347237579 | 0.728412841 | 0.933996867 |
| Gpr162 | 7192.702998 | -0.044067649 | 0.079997304 | -0.550864182 | 0.581726783 | 0.767675221 | 2610507B11Rik | 14264.75371 | 0.01761103 | 0.040562576 | 0.434169424 | 0.664165414 | 0.915428982 |
| Slc1a1 | 5266.366623 | -0.036285107 | 0.065863416 | -0.550914438 | 0.58169233 | 0.767675221 | Acot4 | 48.10058109 | 0.017616125 | 0.093227637 | 0.188958188 | 0.850125587 | 0.966260848 |
| Ufl1 | 1370.167696 | 0.040723804 | 0.073916261 | 0.550945135 | 0.581671286 | 0.767675221 | Scamp3 | 1740.9067 | 0.01762514 | 0.060592885 | 0.290878043 | 0.771144595 | 0.946250219 |
| Capzb | 7379.898525 | 0.041903888 | 0.076094683 | 0.55068089 | 0.581852447 | 0.767761516 | Ppm1e | 8982.750348 | 0.017643951 | 0.055464545 | 0.31811225 | 0.750399792 | 0.9410864 |
| Gpr151 | 27.81127556 | 0.0899005 | 0.163270141 | 0.550624254 | 0.581891279 | 0.767761516 | Rnf135 | 163.6868104 | 0.017657596 | 0.087997182 | 0.200660929 | 0.840963712 | 0.963094947 |
| Cyr61 | 316.1565312 | -0.089105878 | 0.162037976 | -0.549907371 | 0.582382908 | 0.768194642 | Gm15612 | 14.00544087 | 0.017677703 | 0.054813238 | 0.322507907 | 0.747067949 | NA |
| Dhx29 | 1111.169425 | -0.038505297 | 0.070027604 | -0.549858841 | 0.582416196 | 0.768194642 | Lace1 | 752.5503694 | 0.017691327 | 0.059721782 | 0.296229061 | 0.767055157 | 0.946118815 |
| Exd1 | 34.73727159 | 0.087742865 | 0.159537175 | 0.549983822 | 0.58233047 | 0.768194642 | Bche | 190.9669227 | 0.017699164 | 0.094819966 | 0.186660726 | 0.851926651 | 0.96672811 |

| | | | | | | |
|---|---|---|---|---|---|---|
| Matn1 | 13.23215288 | 0.091795009 | 0.166987504 | 0.549711848 | 0.582517029 | 0.768194642 |
| Slc20a2 | 2222.290289 | 0.034621587 | 0.062973108 | 0.549783673 | 0.582467758 | 0.768194642 |
| Stc1 | 158.6219822 | 0.08183276 | 0.148845355 | 0.549783769 | 0.582467693 | 0.768194642 |
| Zfp93 | 415.5877512 | 0.045160486 | 0.08216654 | 0.54962136 | 0.582579105 | 0.768211119 |
| Pwwp2a | 831.1122896 | -0.033258183 | 0.060545393 | -0.549309894 | 0.582792798 | 0.768379032 |
| Spock2 | 29177.39221 | -0.028078876 | 0.05111838 | -0.549291198 | 0.582805627 | 0.768379032 |
| Akr1a1 | 10758.47722 | 0.035694782 | 0.065006293 | 0.549097337 | 0.582938652 | 0.768489022 |
| L3mbtl4 | 32.50067235 | -0.089098696 | 0.162297464 | -0.548983907 | 0.583016494 | 0.768526251 |
| 1700023F06Rik | 60.33701765 | 0.087006132 | 0.158663745 | 0.548368072 | 0.583439195 | 0.769018025 |
| Wdr24 | 1070.551066 | 0.040267833 | 0.073465385 | 0.548119819 | 0.583609633 | 0.769177241 |
| Fam5b | 4795.485344 | -0.046105215 | 0.084146621 | -0.547915226 | 0.583750114 | 0.76923649 |
| Tvp23a | 1624.162441 | -0.037747112 | 0.068892939 | -0.547909738 | 0.583753882 | 0.76923649 |
| Pgap3 | 404.8244089 | 0.051978025 | 0.094878736 | 0.547836401 | 0.583804242 | 0.769237428 |
| 1110019D14Rik | 66.25876234 | -0.083052663 | 0.151832207 | -0.547002932 | 0.584376718 | 0.769926264 |
| Psma7 | 2228.059734 | -0.043684699 | 0.079876391 | -0.546903771 | 0.584444845 | 0.769950551 |
| 3110039I08Rik | 24.38128652 | 0.091646113 | 0.167685971 | 0.546534167 | 0.584698808 | 0.770154155 |
| Gstt1 | 308.5353495 | -0.055963769 | 0.102388832 | -0.546580795 | 0.584666766 | 0.770154155 |
| A830018L16Rik | 2572.598838 | -0.043413223 | 0.079495834 | -0.546106894 | 0.584992461 | 0.770330198 |
| Dlg5 | 1781.896664 | 0.064298479 | 0.117756654 | 0.546028417 | 0.585046403 | 0.770330198 |
| Morn5 | 18.0922029 | 0.09001505 | 0.164934911 | 0.545761046 | 0.585230203 | 0.770330198 |
| Nfam1 | 153.1128724 | 0.075680768 | 0.138667365 | 0.545772022 | 0.585222657 | 0.770330198 |
| Sag | 79.64616371 | -0.074184569 | 0.135913318 | -0.545822663 | 0.585187843 | 0.770330198 |
| Scarb2 | 3127.367233 | -0.030378247 | 0.055658303 | -0.545799013 | 0.585204102 | 0.770330198 |
| Tnn | 17.07874531 | 0.089641927 | 0.16422241 | 0.545856847 | 0.585164343 | 0.770330198 |
| Trim9 | 12174.33499 | -0.040158729 | 0.073527527 | -0.546172712 | 0.584947222 | 0.770330198 |
| Oip5 | 25.49654365 | 0.091485345 | 0.167677769 | 0.545602112 | 0.585339472 | 0.770408578 |
| Arhgef5 | 176.5287083 | 0.070205621 | 0.128746005 | 0.545303294 | 0.585544939 | 0.770474879 |
| Cep170 | 3260.298136 | -0.036848226 | 0.067568943 | -0.545342654 | 0.585517873 | 0.770474879 |
| Evi5 | 2818.827892 | 0.031607306 | 0.057969574 | 0.545239576 | 0.585588755 | 0.770474879 |
| Megf10 | 1315.816933 | -0.055617682 | 0.101977773 | -0.545390241 | 0.585485151 | 0.770474879 |
| Gm10863 | 13.13959842 | 0.086681568 | 0.159092479 | 0.544850196 | 0.585856552 | 0.770761775 |
| Htra3 | 153.424567 | 0.068451039 | 0.1256957 | 0.544577414 | 0.586044192 | 0.770943175 |
| 5430405H02Rik | 162.3550858 | 0.061463667 | 0.11296623 | 0.544088854 | 0.58638033 | 0.771031552 |
| Arhgap44 | 6359.016289 | 0.050628972 | 0.09300847 | 0.544347967 | 0.586202044 | 0.771031552 |
| C330006A16Rik | 2868.919883 | -0.035613193 | 0.06544319 | -0.544184851 | 0.586314276 | 0.771031552 |
| Dtwd1 | 227.5989053 | -0.054349048 | 0.099884728 | -0.544117696 | 0.586360484 | 0.771031552 |
| Gpt | 472.2157925 | 0.068410455 | 0.125743924 | 0.544045805 | 0.586409953 | 0.771031552 |
| Katnbl1 | 392.328904 | -0.046582134 | 0.085597426 | -0.544200168 | 0.586303736 | 0.771031552 |
| Wars2 | 316.8494004 | -0.044798139 | 0.082355991 | -0.543957249 | 0.586470892 | 0.771046245 |
| Eno2 | 41775.84434 | -0.026118498 | 0.048026571 | -0.543834328 | 0.586555484 | 0.771090368 |
| Wdfy2 | 312.6339017 | 0.049538485 | 0.09110294 | 0.543763852 | 0.586603987 | 0.771090368 |
| 4933417O13Rik | 9.174195133 | 0.085140678 | 0.156617588 | 0.543621437 | 0.586702005 | 0.771145473 |
| Mamdc4 | 57.05258338 | 0.090454828 | 0.166412371 | 0.543558319 | 0.586745449 | 0.771145473 |
| Rbm7 | 838.1205129 | 0.037814397 | 0.069587693 | 0.543406386 | 0.58685003 | 0.771170181 |
| Siva1 | 171.1826869 | -0.065988866 | 0.121440041 | -0.543386392 | 0.586863793 | 0.771170181 |
| Etfb | 839.5823339 | -0.050697988 | 0.093335066 | -0.543182643 | 0.587004056 | 0.771289081 |
| Thnsl2 | 163.7217201 | 0.066002719 | 0.121540918 | 0.543049375 | 0.587095808 | 0.771344225 |
| Gm3230 | 126.6873184 | -0.076088414 | 0.140140668 | -0.542943137 | 0.587168954 | 0.771374917 |
| Zcchc10 | 222.358831 | -0.064346037 | 0.118533635 | -0.542850453 | 0.587232772 | 0.771393351 |
| Fam193a | 1954.879038 | 0.032421206 | 0.059774861 | 0.542388656 | 0.587550793 | 0.771745675 |
| Exo1 | 22.50942625 | -0.090635404 | 0.16720242 | -0.542069927 | 0.587770335 | 0.771945244 |
| Mss51 | 15.03948817 | 0.090493928 | 0.166992334 | 0.541904682 | 0.587884171 | 0.771945244 |
| Sec61a1 | 2555.351737 | -0.040791116 | 0.075264127 | -0.541972889 | 0.587837182 | 0.771945244 |
| Srp68 | 2779.255889 | 0.045355091 | 0.083699701 | 0.541878776 | 0.587902019 | 0.771945244 |
| Cbfa2t3 | 1293.621034 | 0.059280562 | 0.109460961 | 0.541568081 | 0.588116086 | 0.772160887 |
| Nup210 | 1339.036602 | 0.045332907 | 0.083749101 | 0.541294254 | 0.58830478 | 0.772274883 |
| Pex5 | 2129.756942 | 0.034057109 | 0.062929785 | 0.541192199 | 0.588375114 | 0.772274883 |
| Sned1 | 256.7051707 | 0.066000328 | 0.121948523 | 0.541214655 | 0.588359638 | 0.772274883 |
| Stk32c | 6147.26102 | 0.046303171 | 0.085574114 | 0.541088522 | 0.58844657 | 0.772274883 |
| Tceal7 | 17.06878168 | 0.090665906 | 0.167564556 | 0.541080457 | 0.588452128 | 0.772274883 |
| Klhl7 | 2902.61606 | -0.042744422 | 0.079017569 | -0.540948338 | 0.588543192 | 0.772328975 |
| Hint2 | 365.7362738 | -0.046319722 | 0.085678218 | -0.540624247 | 0.588766602 | 0.772556718 |
| Zfp446 | 426.1447276 | 0.074587078 | 0.138009457 | 0.540449036 | 0.588887399 | 0.772649789 |
| Bcas2 | 1531.147994 | -0.03650223 | 0.067562929 | -0.540270094 | 0.589010779 | 0.772708749 |
| Dnajb9 | 1266.195632 | 0.04245037 | 0.078595605 | 0.540111243 | 0.589120317 | 0.772708749 |
| Tcf3 | 820.5181599 | 0.04065741 | 0.075278317 | 0.540094559 | 0.589131822 | 0.772708749 |
| Ttc27 | 738.984848 | -0.03884873 | 0.071916861 | -0.540189462 | 0.58906638 | 0.772708749 |
| Gm13003 | 11.74540247 | 0.079124047 | 0.146622325 | 0.539645288 | 0.589441678 | 0.772918869 |
| Med31 | 254.704196 | -0.053928676 | 0.099931124 | -0.539658456 | 0.589432595 | 0.772918869 |
| Zfp35 | 888.5747094 | 0.04563404 | 0.084543301 | 0.53977121 | 0.589354824 | 0.772918869 |
| Clp1 | 258.3360283 | 0.060736835 | 0.112585026 | 0.539475252 | 0.589558969 | 0.772972905 |
| Sfxn4 | 448.4389421 | 0.076798213 | 0.142366315 | 0.539440903 | 0.589582664 | 0.772972905 |
| Kazn | 4001.846865 | -0.050839996 | 0.094328086 | -0.538969863 | 0.589907649 | 0.773333541 |
| B9d1 | 283.9950284 | -0.059873947 | 0.111215744 | -0.538358554 | 0.590329534 | 0.773821134 |
| Ebi3 | 36.36160618 | -0.083975887 | 0.15612682 | -0.537869707 | 0.590667003 | 0.774197999 |
| Pign | 682.7833016 | 0.046930789 | 0.087271511 | 0.537756117 | 0.590745432 | 0.7742353 |
| Rab6b | 43874.09773 | -0.030336851 | 0.056443111 | -0.537476596 | 0.590938447 | 0.774422761 |
| Skor1 | 14.50772559 | -0.087232606 | 0.162397116 | -0.537156126 | 0.591159774 | 0.774647289 |
| Lefty2 | 13.17076397 | 0.089327036 | 0.1664186 | 0.536761131 | 0.591432624 | 0.774873759 |
| Tmem191c | 2468.554151 | 0.057219425 | 0.106594058 | 0.536797512 | 0.59140749 | 0.774873759 |
| Fgfr1 | 2835.623602 | -0.047563585 | 0.088686811 | -0.53630956 | 0.591744625 | 0.775206951 |
| Serinc3 | 12117.60224 | 0.035948704 | 0.067037431 | 0.536248234 | 0.591787002 | 0.775206951 |
| Zfp362 | 1039.939701 | 0.037782068 | 0.070483892 | 0.536038328 | 0.591932061 | 0.775331419 |
| Gtpbp5 | 362.7761543 | 0.061191779 | 0.114211438 | 0.535776277 | 0.59211318 | 0.775372009 |
| Lsamp | 2075.803701 | 0.026264858 | 0.04901191 | 0.535887255 | 0.592036473 | 0.775372009 |
| Slc24a3 | 4927.338196 | -0.030894416 | 0.057657563 | -0.535825916 | 0.592078869 | 0.775372009 |
| A330069E16Rik | 25.34016021 | 0.087315723 | 0.163135436 | 0.535234555 | 0.592487674 | 0.7754411 |
| Calb1 | 4230.795365 | 0.052287493 | 0.097771552 | 0.534792499 | 0.59279335 | 0.7754411 |
| Col6a6 | 22.64783505 | -0.088277921 | 0.164976005 | -0.53509552 | 0.592583808 | 0.7754411 |
| Hk3 | 75.10203008 | 0.089639666 | 0.167502847 | 0.535153087 | 0.592544003 | 0.7754411 |
| Kcnk13 | 333.8588311 | -0.065918972 | 0.123208601 | -0.535019241 | 0.592636552 | 0.7754411 |
| Mmel1 | 32.01723255 | 0.089821862 | 0.167989748 | 0.534686568 | 0.592866611 | 0.7754411 |
| Nxph1 | 1242.408831 | 0.036095747 | 0.067490555 | 0.53482664 | 0.59276974 | 0.7754411 |
| Pold4 | 280.5835191 | -0.063054564 | 0.117916091 | -0.534740958 | 0.592828995 | 0.7754411 |
| Rlim | 2133.296295 | -0.045069941 | 0.084231059 | -0.535075088 | 0.592597936 | 0.7754411 |
| Tmub1 | 610.1568111 | 0.042684019 | 0.079816238 | 0.534778641 | 0.592802934 | 0.7754411 |
| Topbp1 | 949.0796511 | -0.038875688 | 0.072630162 | -0.535255425 | 0.592473245 | 0.7754411 |
| Trappc8 | 2578.254857 | -0.030042254 | 0.056117041 | -0.535349936 | 0.592407902 | 0.7754411 |
| Zbtb39 | 538.7564335 | 0.038390297 | 0.071768007 | 0.534922154 | 0.592703688 | 0.7754411 |
| Zbtb8a | 221.2639498 | -0.059531116 | 0.111204198 | -0.535331559 | 0.592420607 | 0.7754411 |
| Fbxl4 | 704.5238708 | 0.038152817 | 0.071365891 | 0.534608574 | 0.592920553 | 0.775446193 |
| D10Jhu81e | 2127.894553 | -0.038433928 | 0.07197716 | -0.533973948 | 0.593359558 | 0.775954845 |
| Atpbd4 | 673.1346387 | -0.03762146 | 0.070480675 | -0.533784055 | 0.593490947 | 0.776046641 |
| Cdipt | 5101.79367 | -0.036329299 | 0.068067101 | -0.533727721 | 0.593529927 | 0.776046641 |
| 4732415M23Rik | 25.93944297 | -0.088772466 | 0.166451122 | -0.533324525 | 0.593808953 | 0.776345956 |
| Papd4 | 742.9325855 | 0.039605544 | 0.074288652 | 0.533130466 | 0.593943271 | 0.776400816 |
| Sec63 | 3951.845278 | -0.037222037 | 0.069819365 | -0.533119106 | 0.593951134 | 0.776400816 |
| Rnf121 | 854.5150217 | -0.0315951 | 0.059274079 | -0.533034018 | 0.594010032 | 0.776412303 |
| Ate1 | 2963.607927 | -0.030964589 | 0.058115128 | -0.532814606 | 0.594161922 | 0.776414339 |
| Cxx1b | 1781.601242 | -0.028567546 | 0.053609856 | -0.532878618 | 0.594117607 | 0.776414339 |
| Rapgef6 | 1702.602406 | 0.04857852 | 0.091154832 | 0.532923147 | 0.594086781 | 0.776414339 |
| Armcx3 | 3691.361337 | -0.034622052 | 0.065010731 | -0.532559033 | 0.594338867 | 0.776516842 |
| Asap1 | 6369.518621 | -0.036852966 | 0.0692001 | -0.532556533 | 0.594340598 | 0.776516842 |
| Fxn | 137.9790933 | -0.067819046 | 0.127383135 | -0.532402078 | 0.594447546 | 0.776591086 |
| Ripk1 | 283.2885721 | 0.05701631 | 0.107118324 | 0.532274111 | 0.59453616 | 0.776641368 |
| 2700081O15Rik | 1492.068242 | -0.034457289 | 0.064774606 | -0.531956751 | 0.59475595 | 0.776753967 |
| 9130023H24Rik | 142.9165213 | -0.071457838 | 0.134336292 | -0.531932489 | 0.594772755 | 0.776753967 |
| Mical2 | 13542.45555 | 0.056516875 | 0.106225916 | 0.532044124 | 0.594695435 | 0.776753967 |
| 6720468P15Rik | 14.28620424 | 0.089048715 | 0.167519094 | 0.531573525 | 0.595021405 | 0.777013203 |
| Pomt1 | 581.9629841 | 0.060501188 | 0.113912629 | 0.531119227 | 0.595336161 | 0.777358712 |
| Wapal | 1700.363611 | -0.03553505 | 0.066926028 | -0.53096009 | 0.595446436 | 0.777437185 |
| Utp23 | 378.6408259 | 0.046944919 | 0.088462092 | 0.530678369 | 0.595641678 | 0.777626572 |
| Irs4 | 133.4081737 | -0.062918382 | 0.118659514 | -0.530243042 | 0.595943432 | 0.777954969 |
| Btf3 | 3333.605446 | -0.030492641 | 0.057554073 | -0.529808563 | 0.596244668 | 0.778217074 |
| Sphkap | 7731.660266 | -0.048917952 | 0.09232868 | -0.529824013 | 0.596233954 | 0.778217074 |
| Elk4 | 370.1737255 | -0.054002226 | 0.101971525 | -0.529581427 | 0.596402175 | 0.778291527 |
| Fzd2 | 471.4328076 | -0.044463106 | 0.083952739 | -0.529620673 | 0.596374958 | 0.778291527 |
| Fam103a1 | 2256.902235 | 0.046273345 | 0.087410981 | 0.529376791 | 0.596544095 | 0.778411168 |
| Glt8d2 | 570.2109351 | -0.061646914 | 0.116495469 | -0.529178641 | 0.596681532 | 0.778495955 |
| Wdr35 | 1907.025135 | -0.033072314 | 0.06250222 | -0.529138231 | 0.596709562 | 0.778495955 |
| Prmt5 | 2242.055961 | 0.043861185 | 0.082967139 | 0.528657325 | 0.597043186 | 0.778865633 |
| Mocs2 | 1454.159588 | -0.029765777 | 0.056318893 | -0.528522057 | 0.597137042 | 0.778922489 |
| 2210417K05Rik | 12.93306885 | 0.087928029 | 0.166441081 | 0.528283213 | 0.597302781 | 0.778986324 |
| A130077B15Rik | 365.3042928 | 0.050521924 | 0.095656161 | 0.528161735 | 0.597387085 | 0.778986324 |
| Gfer | 405.7905869 | -0.0418071 | 0.079141329 | -0.528258753 | 0.597319756 | 0.778986324 |
| Trpm6 | 44.37186436 | -0.084973052 | 0.160863945 | -0.528229317 | 0.597340184 | 0.778986324 |
| Ccdc8 | 104.4766446 | -0.071152161 | 0.134735766 | -0.528086658 | 0.597439191 | 0.778988708 |
| Pbx2 | 2157.282221 | 0.02903111 | 0.054998514 | 0.52785263 | 0.597601625 | 0.779069379 |
| Steap4 | 21.08238514 | -0.088726547 | 0.168067567 | -0.52792189 | 0.597553551 | 0.779069379 |
| Imp4 | 679.1388081 | -0.04017686 | 0.076141348 | -0.52766153 | 0.597734279 | 0.779176755 |
| Tpst1 | 1207.874192 | 0.030600785 | 0.058009549 | 0.527512896 | 0.597837463 | 0.779245702 |
| Ccdc88a | 2676.535623 | -0.039200295 | 0.074356931 | -0.527190871 | 0.598061047 | 0.779433947 |
| Tfdp1 | 1637.834743 | 0.039221455 | 0.074401428 | 0.527159982 | 0.598082496 | 0.779433947 |
| Mcts2 | 196.4677665 | -0.060324287 | 0.11454019 | -0.52666481 | 0.598426377 | 0.779816509 |
| Abhd11 | 772.606822 | 0.044935977 | 0.085418332 | 0.526069472 | 0.598839938 | 0.780027407 |
| Atad5 | 214.4275116 | -0.054586302 | 0.103756875 | -0.526098169 | 0.598820001 | 0.780027407 |
| Nap1l3 | 1631.661027 | -0.040644714 | 0.077239806 | -0.526214597 | 0.598739113 | 0.780027407 |
| Ssx2ip | 3474.949982 | 0.048989644 | 0.093118218 | 0.526101608 | 0.598817612 | 0.780027407 |
| Wdpcp | 348.7030177 | -0.054240656 | 0.10304961 | -0.526354792 | 0.59864172 | 0.780027407 |
| 4930405A21Rik | 9.256650748 | -0.0841704 | 0.160031708 | -0.525960766 | 0.598915467 | 0.780060209 |
| Urb2 | 685.4355988 | -0.032119228 | 0.061091086 | -0.525759648 | 0.599055214 | 0.78017664 |
| Neu3 | 117.3537397 | 0.069355613 | 0.13193603 | 0.525676061 | 0.599113299 | 0.780186709 |
| Gm10785 | 11.98684131 | 0.087050489 | 0.165702633 | 0.525341615 | 0.599345733 | 0.780423799 |
| Gm3086 | 32.84045887 | 0.083363092 | 0.158762537 | 0.525080377 | 0.599527317 | 0.780594643 |
| Adrb3 | 43.32108901 | 0.084132134 | 0.160329151 | 0.524746332 | 0.599759544 | 0.78083139 |
| Cobll1 | 552.2965673 | 0.061109291 | 0.116488241 | 0.524596219 | 0.599863916 | 0.780901656 |
| Amigo3 | 51.85024728 | -0.085252003 | 0.162591809 | -0.524331475 | 0.600048008 | 0.780910185 |
| Rrp15 | 245.6297048 | 0.065456789 | 0.124833679 | 0.524352001 | 0.600033735 | 0.780910185 |
| Senp3 | 2653.709835 | -0.026994279 | 0.051472964 | -0.524436062 | 0.59997528 | 0.780910185 |
| Thsd1 | 184.5954078 | -0.057543246 | 0.109753174 | -0.524296874 | 0.600072071 | 0.780910185 |
| Tm6sf1 | 519.364006 | 0.053925418 | 0.102929354 | 0.523907082 | 0.600343169 | 0.781197368 |
| Rgag1 | 104.1534605 | -0.08292019 | 0.158330717 | -0.523715119 | 0.600476698 | 0.781304013 |
| Ric3 | 3710.007343 | -0.026190142 | 0.05001514 | -0.52364428 | 0.600525977 | 0.781304013 |
| Cdkn2b | 23.094942 | -0.087617513 | 0.16742217 | -0.523332804 | 0.600742676 | 0.781520321 |
| Gm5617 | 44.61026715 | 0.085582857 | 0.163584754 | 0.523171354 | 0.600855014 | 0.781537017 |
| Sult4a1 | 17284.15835 | -0.040430953 | 0.07728081 | -0.523169373 | 0.600856392 | 0.781537017 |
| Mut | 1543.040115 | -0.050041275 | 0.09571147 | -0.522834671 | 0.60108931 | 0.781774344 |
| Nufip1 | 496.2177907 | 0.038621918 | 0.073895179 | 0.522658159 | 0.60121216 | 0.781868492 |
| Furin | 1472.765093 | 0.055827783 | 0.106862562 | 0.522426022 | 0.601373742 | 0.781953027 |
| Sspn | 492.1177506 | -0.060872372 | 0.116520041 | -0.522419762 | 0.601378099 | 0.781953027 |
| Aimp2 | 675.4891072 | -0.045706985 | 0.087513236 | -0.522286532 | 0.601470845 | 0.782007995 |
| B3gnt4 | 17.86325094 | -0.085253889 | 0.163372331 | -0.52183799 | 0.601783136 | 0.782282734 |
| Stk11 | 2100.619902 | 0.040654048 | 0.07789851 | 0.521884798 | 0.601750544 | 0.782282734 |
| Aurka | 42.59070592 | 0.085744342 | 0.164336306 | 0.521761404 | 0.601836466 | 0.782286426 |
| Ilf2 | 3089.11293 | 0.032060685 | 0.061476002 | 0.521515454 | 0.602007743 | 0.782377786 |
| Knstrn | 39.41195593 | -0.086390073 | 0.165643342 | -0.521542681 | 0.601988781 | 0.782377786 |
| Mier1 | 1056.509272 | 0.042878802 | 0.082267926 | 0.521209229 | 0.602221026 | 0.782589329 |
| Tmem71 | 20.98049719 | -0.087535125 | 0.168114371 | -0.520687934 | 0.602584181 | 0.78292992 |
| Trim45 | 375.9160394 | -0.049450828 | 0.094967195 | -0.520714838 | 0.602565436 | 0.78292992 |
| Wnt9a | 643.1987464 | 0.071522894 | 0.137392731 | 0.520572618 | 0.602664528 | 0.782968657 |
| Tmprss5 | 49.64822643 | 0.082641805 | 0.158781515 | 0.520474975 | 0.602732565 | 0.782991395 |
| Rpf1 | 341.7337062 | 0.051209556 | 0.098431972 | 0.520253283 | 0.602887052 | 0.783126424 |
| Creg2 | 6770.976562 | -0.040110544 | 0.077120243 | -0.520103958 | 0.60299112 | 0.783168941 |
| Napb | 23209.18757 | -0.037183007 | 0.071502906 | -0.52002092 | 0.603048994 | 0.783168941 |
| Zfp771 | 388.78494 | -0.079758962 | 0.15338594 | -0.51998874 | 0.603071423 | 0.783168941 |
| Hrh3 | 4025.517459 | 0.040434484 | 0.077798945 | 0.519730498 | 0.603251427 | 0.783302166 |
| Rogdi | 4085.288766 | 0.031133932 | 0.059913725 | 0.519646067 | 0.603310284 | 0.783302166 |
| Stk17b | 291.9778228 | 0.055418498 | 0.106651155 | 0.519623985 | 0.603325677 | 0.783302166 |
| A430078G23Rik | 208.6479762 | 0.053769645 | 0.103562485 | 0.519200025 | 0.603621262 | 0.783430843 |
| Dixdc1 | 3703.791155 | 0.038070504 | 0.073318918 | 0.519245304 | 0.60358969 | 0.783430843 |
| Dtx3 | 7443.176167 | 0.050161574 | 0.096593961 | 0.519303413 | 0.603549174 | 0.783430843 |
| Mc4r | 83.67093784 | -0.087156133 | 0.167868876 | -0.519191734 | 0.603627043 | 0.783430843 |
| Pias4 | 587.5839783 | 0.037531851 | 0.072306225 | 0.519068047 | 0.60371329 | 0.783477151 |
| Anapc10 | 288.2411336 | -0.047797355 | 0.092175298 | -0.518548422 | 0.604075685 | 0.783867561 |
| Prdm11 | 108.9645971 | 0.066483355 | 0.128224551 | 0.51849162 | 0.604115306 | 0.783867561 |
| Itgb1bp2 | 11.37530105 | 0.079877374 | 0.154197566 | 0.518019682 | 0.604444537 | 0.784170038 |
| Usp43 | 118.5062943 | 0.07683897 | 0.14833424 | 0.518012364 | 0.604449643 | 0.784170038 |
| Dap3 | 2463.786631 | -0.025541785 | 0.049316711 | -0.5179134 | 0.604518692 | 0.784193957 |
| 2310034G01Rik | 16.34958121 | -0.087034524 | 0.168117353 | -0.517701012 | 0.604666892 | 0.784264815 |
| Zbtb6 | 532.0229142 | -0.04869831 | 0.094068471 | -0.517690038 | 0.60467455 | 0.784264815 |
| Lamc2 | 635.4194175 | 0.059462563 | 0.11508302 | 0.516692757 | 0.605370653 | 0.785101941 |
| Abcc4 | 634.1941226 | 0.045533751 | 0.088145393 | 0.516575505 | 0.605452518 | 0.785142394 |
| Plec | 8670.419617 | -0.044566177 | 0.086341011 | -0.516164648 | 0.605739419 | 0.785251552 |
| Rpl24 | 5012.913876 | -0.039639601 | 0.076792423 | -0.516191568 | 0.605720619 | 0.785251552 |
| Slc24a2 | 27643.09709 | 0.050717881 | 0.098242443 | 0.516252238 | 0.60567825 | 0.785251552 |
| Zfp944 | 301.0423839 | 0.05280435 | 0.102280549 | 0.516269714 | 0.605666046 | 0.785251552 |
| Fbxo5 | 40.72755384 | -0.083500919 | 0.16186612 | -0.515864089 | 0.605949337 | 0.785457963 |
| Sox2 | 952.4562778 | -0.056606599 | 0.109798114 | -0.515551651 | 0.606167586 | 0.785675136 |
| Slc36a4 | 1124.526106 | 0.061428518 | 0.119244774 | 0.515146421 | 0.606450706 | 0.785976348 |
| Nle1 | 304.3146394 | 0.055231098 | 0.107286068 | 0.514802146 | 0.606691286 | 0.786222381 |
| 2610316D01Rik | 168.1640833 | -0.07674214 | 0.149271994 | -0.514109435 | 0.607175481 | 0.786522522 |
| Shpk | 178.074681 | 0.065502483 | 0.127390848 | 0.514185155 | 0.607122545 | 0.786522522 |
| Tgds | 158.1064716 | -0.070150925 | 0.136418302 | -0.514233969 | 0.607088421 | 0.786522522 |
| Zap70 | 47.65134935 | -0.079189652 | 0.1540332 | -0.514107683 | 0.607176706 | 0.786522522 |
| Zbtb11 | 2059.691094 | 0.029901383 | 0.058135869 | 0.514336221 | 0.607016942 | 0.786522522 |
| Cwc15 | 1775.219599 | 0.047267965 | 0.091977772 | 0.513906395 | 0.607317436 | 0.786538225 |
| E130003G02Rik | 33.35411529 | 0.059060679 | 0.114944721 | 0.513818112 | 0.607379164 | 0.786538225 |
| Otud5 | 2167.729162 | 0.030670134 | 0.059692299 | 0.513803864 | 0.607389126 | 0.786538225 |
| Pop5 | 1331.511493 | 0.055676971 | 0.108375896 | 0.513739426 | 0.607434184 | 0.786538225 |
| Psmg1 | 324.5091036 | 0.042605189 | 0.082933471 | 0.513727319 | 0.607442649 | 0.786538225 |
| Gm3558 | 28.71803363 | 0.085765921 | 0.166988897 | 0.513602538 | 0.607529905 | 0.786585472 |
| 1700102P08Rik | 26.9093646 | -0.086242428 | 0.168024785 | -0.513272065 | 0.607761022 | 0.786629774 |
| Hadh | 784.6530571 | -0.04178259 | 0.081405773 | -0.513263229 | 0.607767202 | 0.786629774 |
| Nkx2-9 | 12.68229921 | 0.083809601 | 0.16328148 | 0.513282958 | 0.607753403 | 0.786629774 |
| Pcdh15 | 634.1396526 | -0.063917332 | 0.124531094 | -0.513264034 | 0.607766639 | 0.786629774 |
| Msl3l2 | 386.9255868 | -0.048114702 | 0.093792369 | -0.512991645 | 0.607957165 | 0.786809916 |
| Il21r | 77.53425147 | 0.073605775 | 0.143525142 | 0.512842379 | 0.608061583 | 0.786879326 |
| Per2 | 1888.81075 | -0.081215589 | 0.158435347 | -0.512610285 | 0.608223957 | 0.787023718 |
| Icam4 | 26.22460807 | -0.086024895 | 0.167868897 | -0.512452853 | 0.608334109 | 0.787100517 |
| Chrm3 | 1850.603948 | 0.047306267 | 0.092374584 | 0.512113447 | 0.608571614 | 0.787210601 |
| Rab43 | 963.3199815 | 0.044507317 | 0.086900483 | 0.512164202 | 0.608536094 | 0.787210601 |
| Zpbp | 51.09197502 | -0.078690559 | 0.153642353 | -0.512167105 | 0.608534063 | 0.787210601 |
| Zfp111 | 425.2946643 | -0.043600299 | 0.085150902 | -0.512035667 | 0.608626048 | 0.787215292 |
| Nhlrc3 | 193.7832775 | 0.056429067 | 0.110262646 | 0.51176957 | 0.60881229 | 0.787324732 |
| Polr1e | 229.7882535 | 0.052726878 | 0.10301539 | 0.511834958 | 0.608766522 | 0.787324732 |

| | | | | | | |
|---|---|---|---|---|---|---|
| 1110017D15Rik | 69.71163517 | 0.01773877 | 0.094800718 | 0.187116407 | 0.851569365 | 0.96665003 |
| Trim33 | 2764.988841 | 0.017743659 | 0.068276324 | 0.259880111 | 0.794956253 | 0.952171746 |
| Nipal1 | 0.731486784 | 0.017778231 | 0.018817935 | 0.944749304 | 0.34478688 | NA |
| Iars | 2810.873002 | 0.01778415 | 0.054147795 | 0.328437199 | 0.742581121 | 0.939912916 |
| Oscp1 | 1356.59232 | 0.017800225 | 0.067298087 | 0.264498227 | 0.791396031 | 0.951728598 |
| Atox1 | 861.5300554 | 0.01781063 | 0.063174569 | 0.28192721 | 0.777999325 | 0.94784716 |
| Sap130 | 1986.377376 | 0.017813387 | 0.05526803 | 0.322309064 | 0.747218567 | 0.94097069 |
| Them4 | 750.1553198 | 0.01783179 | 0.065683102 | 0.271482147 | 0.786020229 | 0.950484821 |
| Siglecg | 3.809185785 | 0.017832034 | 0.041891038 | 0.425676588 | 0.670343524 | NA |
| Tm9sf1 | 1280.748852 | 0.017838652 | 0.062059613 | 0.287443821 | 0.773772512 | 0.947121017 |
| Tg | 7.946859883 | 0.017841692 | 0.055723046 | 0.320185145 | 0.748827983 | NA |
| Klhl2 | 6725.650091 | 0.01785 | 0.066902616 | 0.266805707 | 0.78961876 | 0.951472593 |
| Bcr | 7894.237924 | 0.017886841 | 0.067270289 | 0.265895113 | 0.790319989 | 0.951607109 |
| Mll1 | 6452.190213 | 0.017890035 | 0.088075452 | 0.203121692 | 0.839039917 | 0.96231236 |
| Cpne2 | 969.2240376 | 0.017904591 | 0.078608606 | 0.227768841 | 0.819825951 | 0.957571905 |
| 2610301B20Rik | 1554.303034 | 0.017913345 | 0.058181531 | 0.307887138 | 0.758168213 | 0.942510233 |
| Zpbp | 52.15200596 | 0.017917892 | 0.094106506 | 0.190400147 | 0.848995583 | 0.965892061 |
| 2310010J17Rik | 43.58973866 | 0.017921721 | 0.092887215 | 0.192940661 | 0.84700544 | 0.965275633 |
| 1700029M20Rik | 0.771558907 | 0.017923007 | 0.018997678 | 0.943431449 | 0.345460263 | NA |
| Ppp1r3c | 1687.902341 | 0.017926366 | 0.067854813 | 0.26418709 | 0.791635758 | 0.951728598 |
| 2700062C07Rik | 459.4380699 | 0.01792906 | 0.074977375 | 0.239126265 | 0.811007676 | 0.956135608 |
| Armcx6 | 306.0821621 | 0.017942531 | 0.085754517 | 0.209231318 | 0.834267665 | 0.961118357 |
| Tk1 | 14.33553245 | 0.017992558 | 0.070755601 | 0.254291641 | 0.799270257 | NA |
| Fat1 | 1929.096596 | 0.018001827 | 0.085240787 | 0.211187951 | 0.832740612 | 0.960519604 |
| Coprs | 703.6916941 | 0.018002806 | 0.071312183 | 0.252450633 | 0.800692768 | 0.953228631 |
| Kpna4 | 2943.215879 | 0.018011832 | 0.073903507 | 0.243720943 | 0.807446952 | 0.955716697 |
| Napa | 11036.88616 | 0.018035695 | 0.041642697 | 0.433105837 | 0.664937882 | 0.915428982 |
| Ctla4 | 3.128608248 | 0.018038707 | 0.040754198 | 0.44262205 | 0.658039133 | NA |
| Serf2 | 4756.04364 | 0.018043435 | 0.047695092 | 0.378308005 | 0.705201799 | 0.928783753 |
| Irx3 | 11.42235705 | 0.018050638 | 0.055143374 | 0.327340103 | 0.743410664 | NA |
| Hdac5 | 6733.568788 | 0.018057279 | 0.071726149 | 0.251753088 | 0.80123192 | 0.953228631 |
| Ckap5 | 8228.99655 | 0.018061055 | 0.061758779 | 0.292445151 | 0.769946291 | 0.94619074 |
| Akap11 | 11900.41975 | 0.018068437 | 0.072477707 | 0.249296476 | 0.803131457 | 0.953754272 |
| Ap1ar | 3803.455109 | 0.018068793 | 0.055112124 | 0.32785513 | 0.743021201 | 0.939912916 |
| Zscan10 | 3.183903651 | 0.018101507 | 0.02893822 | 0.625522482 | 0.531628198 | NA |
| Serpinb5 | 3.883766842 | 0.018107732 | 0.040985065 | 0.44181294 | 0.658624574 | NA |
| Foxj3 | 3650.045617 | 0.018152816 | 0.05357685 | 0.338818276 | 0.734746632 | 0.936456337 |
| Tbc1d10b | 3497.007504 | 0.018161021 | 0.043402811 | 0.418429608 | 0.675633043 | 0.920229309 |
| Naa10 | 820.5820656 | 0.018166989 | 0.064968728 | 0.279626672 | 0.779763942 | 0.948806418 |
| Adcy10 | 8.954459603 | 0.018184038 | 0.060561399 | 0.300257898 | 0.763980445 | NA |
| Cers3 | 8.631064728 | 0.018185617 | 0.057920048 | 0.313977925 | 0.753537805 | NA |
| Nudt7 | 169.3910412 | 0.018193096 | 0.086955105 | 0.209224007 | 0.834273372 | 0.961118357 |
| Serhl | 194.4272864 | 0.018194741 | 0.091443957 | 0.198971503 | 0.842285036 | 0.963252575 |
| Slc25a24 | 112.104759 | 0.018194761 | 0.091153281 | 0.19960621 | 0.841788571 | 0.963252575 |
| Zranb1 | 1620.476177 | 0.018196751 | 0.055585011 | 0.327367942 | 0.74338961 | 0.939912916 |
| Zar1l | 16.45912498 | 0.018198269 | 0.076623959 | 0.23750103 | 0.81226812 | NA |
| 4930546K05Rik | 1.171869033 | 0.018223872 | 0.025151016 | 0.72457797 | 0.468710978 | NA |
| Grin2d | 471.5495049 | 0.018225054 | 0.086099885 | 0.211673384 | 0.832361854 | 0.960519604 |
| Mlc1 | 7288.055738 | 0.018234728 | 0.05166509 | 0.352940996 | 0.724132676 | 0.932714927 |
| Mcph1 | 567.6510999 | 0.018246648 | 0.063823188 | 0.285893714 | 0.774959529 | 0.947238918 |
| Ablim3 | 1428.205356 | 0.01825735 | 0.062323303 | 0.292945807 | 0.769563575 | 0.94619074 |
| Mir26a-2 | 0.728599411 | 0.018258336 | 0.019085533 | 0.956658429 | 0.338739687 | NA |
| Gm10190 | 6.175972794 | 0.018276562 | 0.051278783 | 0.35641567 | 0.721529296 | NA |
| 4930483O08Rik | 2.29962735 | 0.01829988 | 0.03182715 | 0.57497701 | 0.565306846 | NA |
| Nmu | 0.763449516 | 0.018300029 | 0.018972495 | 0.964555746 | 0.334767374 | NA |
| Use1 | 1731.002809 | 0.018312539 | 0.064300801 | 0.284794878 | 0.775801296 | 0.947241001 |
| Nfyc | 1525.24212 | 0.018317298 | 0.056379805 | 0.324891112 | 0.745263484 | 0.940632024 |
| Ccnt1 | 740.4188523 | 0.018327872 | 0.075236475 | 0.243603545 | 0.807537883 | 0.955716697 |
| Anapc11 | 1225.968668 | 0.018335323 | 0.065381419 | 0.280436298 | 0.779142792 | 0.948407834 |
| Ftsjd2 | 5102.46938 | 0.018339791 | 0.047486638 | 0.386209498 | 0.69934152 | 0.92671364 |
| Npm3 | 153.9901569 | 0.018344399 | 0.088324506 | 0.207693194 | 0.835468532 | 0.961504006 |
| Rab35 | 3760.079226 | 0.018361533 | 0.041858071 | 0.438661714 | 0.660906672 | 0.915428982 |
| Cd300lh | 0.724429695 | 0.018363154 | 0.015914042 | 1.153896246 | 0.248542708 | NA |
| Fam168a | 16128.76744 | 0.018376162 | 0.052870744 | 0.347567689 | 0.728164875 | 0.933979954 |
| Il1r1 | 192.7285406 | 0.018376417 | 0.091118685 | 0.20167561 | 0.840170331 | 0.962746468 |
| Prkrir | 3194.677027 | 0.018395961 | 0.060328363 | 0.304930556 | 0.760419043 | 0.943070948 |
| Olfr283 | 0.985661481 | 0.018397009 | 0.022362224 | 0.822682456 | 0.410688595 | NA |
| 4930550C14Rik | 13.4583258 | 0.018409434 | 0.0742334 | 0.247993948 | 0.80413909 | NA |
| Coro1a | 3361.562305 | 0.01841059 | 0.067609854 | 0.272306314 | 0.785386502 | 0.950314396 |
| Acaa2 | 857.0997139 | 0.018481375 | 0.075470191 | 0.244883105 | 0.806546944 | 0.955587482 |
| Prkag3 | 4.307395596 | 0.018482915 | 0.042102218 | 0.439000966 | 0.660660836 | NA |
| Zrsr2 | 1585.658191 | 0.018486773 | 0.064562441 | 0.286339429 | 0.774618161 | 0.947238918 |
| Myo1c | 1011.679864 | 0.018489901 | 0.070095311 | 0.26378228 | 0.79194769 | 0.951728598 |
| Kcnv1 | 3273.894911 | 0.018493541 | 0.086724748 | 0.213244104 | 0.831136569 | 0.960162026 |
| Camsap3 | 1406.820521 | 0.018495321 | 0.052764016 | 0.350529058 | 0.725941687 | 0.933575664 |
| Pon2 | 2294.831359 | 0.018498469 | 0.060314008 | 0.306702706 | 0.75906967 | 0.942526199 |
| Slc35c1 | 780.4085517 | 0.018505326 | 0.066331902 | 0.278980778 | 0.780259576 | 0.948894514 |
| Ankef1 | 33.97826602 | 0.018515729 | 0.085178107 | 0.217376619 | 0.827914851 | 0.958723519 |
| Mtfr1 | 308.2393662 | 0.018518843 | 0.080086429 | 0.231235719 | 0.817131682 | 0.956302288 |
| Ccdc53 | 447.1501063 | 0.018520427 | 0.07454057 | 0.248461028 | 0.80377772 | 0.953895029 |
| Ddx42 | 4023.36249 | 0.018528843 | 0.050234274 | 0.36884863 | 0.712240555 | 0.930507332 |
| Pcbp2 | 8530.959251 | 0.018549731 | 0.042106136 | 0.440546979 | 0.659540995 | 0.915208181 |
| Tulp2 | 2.634622341 | 0.018550411 | 0.037434732 | 0.495540113 | 0.620218914 | NA |
| Tsx | 2.677119502 | 0.018574786 | 0.039833258 | 0.466313508 | 0.640991111 | NA |
| Zfp746 | 1114.524531 | 0.018591361 | 0.051621265 | 0.360149268 | 0.718735511 | 0.931758324 |
| Ubr4 | 8426.739225 | 0.018595022 | 0.079490478 | 0.233927667 | 0.815041134 | 0.956264994 |
| Sorbs3 | 1544.346509 | 0.018599718 | 0.078828366 | 0.235952094 | 0.813469844 | 0.956264994 |
| Hgd | 1.960486095 | 0.018602713 | 0.034123366 | 0.545160551 | 0.585643101 | NA |
| Gm10494 | 1.127933422 | 0.018609077 | 0.021609397 | 0.861156674 | 0.389151759 | NA |
| D630023F18Rik | 158.2330122 | 0.018615048 | 0.092830895 | 0.200526433 | 0.841068888 | 0.963094947 |
| Pik3r4 | 2273.824185 | 0.018622009 | 0.048023278 | 0.387770473 | 0.698185898 | 0.926409493 |
| Dusp23 | 155.4429452 | 0.018624404 | 0.094377636 | 0.19733917 | 0.843562128 | 0.964060034 |
| Slc35a2 | 1076.28585 | 0.018626033 | 0.066845485 | 0.278643096 | 0.780518736 | 0.948894514 |
| Usp22 | 12406.03501 | 0.018639728 | 0.044062768 | 0.423026717 | 0.672275768 | 0.919190306 |
| Wdr26 | 6211.28101 | 0.018639976 | 0.060912867 | 0.30601049 | 0.759596659 | 0.94272457 |
| Gm11127 | 55.21100843 | 0.018650041 | 0.094677081 | 0.196985802 | 0.843838649 | 0.964060034 |
| Rabep1 | 6829.230607 | 0.018657795 | 0.062438608 | 0.298818241 | 0.765078733 | 0.94510874 |
| Smim11 | 281.1019079 | 0.018663441 | 0.089497146 | 0.20853672 | 0.834809915 | 0.961348408 |
| Gm1110 | 2.910855909 | 0.018677339 | 0.033349354 | 0.56005101 | 0.575444644 | NA |
| Retnla | 0.543403425 | 0.018682949 | 0.016061777 | 1.16319309 | 0.24475117 | NA |
| 1810064F22Rik | 0.546021118 | 0.018728689 | 0.016081151 | 1.164636114 | 0.244166319 | NA |
| Strn3 | 2907.962242 | 0.018728782 | 0.07099748 | 0.263795026 | 0.791937868 | 0.951728598 |
| Crispld1 | 203.4479672 | 0.018745742 | 0.094301969 | 0.198784205 | 0.842431552 | 0.963252575 |
| Zbtb8os | 396.7528987 | 0.018747796 | 0.07345399 | 0.25523183 | 0.798544048 | 0.952978215 |
| Csmd3 | 1241.598349 | 0.018761209 | 0.067937821 | 0.276152637 | 0.782430833 | 0.949184473 |
| Patz1 | 1181.821587 | 0.018763384 | 0.058572974 | 0.320342 | 0.748709088 | 0.9410864 |
| Slc2a12 | 180.4482565 | 0.018773366 | 0.058494451 | 0.320942682 | NA | NA |
| Acss3 | 52.00767089 | 0.018776407 | 0.090356878 | 0.207802738 | 0.835382994 | 0.961504006 |
| Fam131b | 8051.693071 | 0.018779336 | 0.055890654 | 0.336001375 | 0.73686983 | 0.937206199 |
| 2310047M10Rik | 847.0577223 | 0.018782184 | 0.064713614 | 0.290235431 | 0.771636132 | 0.946528159 |
| Aldh1a2 | 302.0078489 | 0.018792632 | 0.074046676 | 0.253794404 | 0.799654397 | 0.953228631 |
| Chid1 | 2279.898182 | 0.01879705 | 0.070247881 | 0.267581737 | 0.789021291 | 0.951324462 |
| Ptdss2 | 1784.164498 | 0.018807233 | 0.059466485 | 0.316266098 | 0.751800542 | 0.941374879 |
| Prpf31 | 1344.733885 | 0.01880936 | 0.058242802 | 0.322947378 | 0.746735095 | 0.94097069 |
| 2310009A05Rik | 130.8533915 | 0.018817735 | 0.092377641 | 0.203704437 | 0.838584475 | 0.962110266 |
| Ptgis | 47.70772622 | 0.018828642 | 0.091337118 | 0.206144476 | 0.836678058 | 0.961729368 |
| Unc5c | 1563.304218 | 0.018839055 | 0.05494539 | 0.342868718 | 0.731697226 | 0.935424325 |
| AI597479 | 1429.046797 | 0.01884063 | 0.059578405 | 0.316232537 | 0.751826013 | 0.941374879 |
| Psma4 | 2923.900567 | 0.01884185 | 0.055903045 | 0.337045141 | 0.736082872 | 0.937120403 |
| Eprs | 4179.033859 | 0.01885362 | 0.056084678 | 0.336163469 | 0.736747599 | 0.937206199 |
| 2410066E13Rik | 1994.606962 | 0.01887771 | 0.065962727 | 0.286187534 | 0.774734491 | 0.947238918 |
| 4931428F04Rik | 1305.593642 | 0.018886076 | 0.067901776 | 0.278138177 | 0.780906291 | 0.948894514 |
| Tdrd1 | 34.96613852 | 0.018901681 | 0.075503615 | 0.250341401 | 0.802323342 | 0.95333733 |
| Gsk3a | 10343.82716 | 0.018915737 | 0.041432258 | 0.456546134 | 0.647997304 | 0.913155689 |
| Mapk1 | 28002.99232 | 0.018916551 | 0.071100474 | 0.266053797 | 0.790197778 | 0.951607109 |
| Grik2 | 1950.134638 | 0.018918887 | 0.06646301 | 0.284652874 | 0.775910098 | 0.947241001 |
| Tpt1 | 14978.53337 | 0.018944848 | 0.060489741 | 0.31319109 | 0.754135488 | 0.941923654 |
| Chst5 | 29.79830454 | 0.018952477 | 0.089158998 | 0.212569424 | 0.831662822 | 0.960389074 |
| Slc4a8 | 5422.833289 | 0.018962508 | 0.059274236 | 0.319911476 | 0.749035438 | 0.9410864 |
| 2410003L11Rik | 2.235867929 | 0.018966769 | 0.03434191 | 0.55229221 | 0.580748167 | NA |
| Odf3l1 | 2.937249825 | 0.018967814 | 0.036419504 | 0.520814729 | 0.602495842 | NA |
| Phospho2 | 923.0710993 | 0.018969119 | 0.082067717 | 0.231139836 | 0.817206169 | 0.956302288 |
| Bicd1 | 908.8114037 | 0.018971811 | 0.078277289 | 0.242366739 | 0.808496005 | 0.955716697 |
| Cpne8 | 1234.062279 | 0.018996527 | 0.060988363 | 0.31147789 | 0.755437349 | 0.942139348 |
| BC048609 | 1.850648943 | 0.019000658 | 0.026431168 | 0.718873161 | 0.472219075 | NA |
| Wdr47 | 6805.344373 | 0.01900355 | 0.059687177 | 0.318385807 | 0.750192303 | 0.9410864 |
| Slc33a1 | 1237.409157 | 0.019007407 | 0.071038144 | 0.267566218 | 0.789033237 | 0.951324462 |
| 1700010K23Rik | 1.446992607 | 0.019009717 | 0.022204148 | 0.856133616 | 0.391923876 | NA |
| Ccdc169 | 5.252289339 | 0.019012535 | 0.04770451 | 0.398547962 | 0.690226311 | NA |
| Fbln7 | 94.99374781 | 0.019019793 | 0.095089751 | 0.200019384 | 0.841465421 | 0.963133457 |
| Efemp2 | 377.4468579 | 0.019039305 | 0.091936399 | 0.207092127 | 0.835937911 | 0.961504006 |
| Plxna1 | 5302.096037 | 0.019046199 | 0.065357289 | 0.291416606 | 0.770732716 | 0.946250219 |
| Dnajb7 | 0.536670213 | 0.019047285 | 0.016216936 | 1.174530478 | 0.240182611 | NA |
| Narf | 2416.933334 | 0.019052454 | 0.058188262 | 0.327427782 | 0.743344356 | 0.939912916 |
| Asb16 | 34.05496244 | 0.019065775 | 0.08873314 | 0.21486645 | 0.829871443 | 0.959693701 |
| Rims1 | 7019.241446 | 0.019082418 | 0.058313918 | 0.327236073 | 0.743489339 | 0.939912916 |
| Tmcc1 | 3964.630832 | 0.019084189 | 0.068141387 | 0.280067513 | 0.779425708 | 0.948553773 |
| Ttc7 | 483.4525941 | 0.019094933 | 0.084304486 | 0.226499604 | 0.820812867 | 0.957604025 |
| Tmem2 | 625.38734 | 0.019099009 | 0.061053921 | 0.312821994 | 0.754415904 | 0.94196623 |
| Ap1s3 | 119.2128339 | 0.019101832 | 0.093647541 | 0.203975798 | 0.838372411 | 0.962077671 |
| Hadh | 886.5788698 | 0.019106158 | 0.064900588 | 0.294391134 | 0.768459041 | 0.94619074 |
| Snrpb2 | 744.4743191 | 0.019116387 | 0.065473885 | 0.291969647 | 0.770309831 | 0.94619074 |
| Bmp15 | 4.826487318 | 0.019133423 | 0.050158323 | 0.381460581 | 0.702861516 | NA |
| Tktl2 | 2.358548777 | 0.019138095 | 0.035147015 | 0.54451553 | 0.586086765 | NA |
| Cited2 | 959.0137624 | 0.019159621 | 0.075066896 | 0.255233955 | 0.798542407 | 0.952978215 |
| Dpep1 | 3.082210665 | 0.019168558 | 0.023987793 | 0.799096354 | NA | NA |
| Mta1 | 2643.631082 | 0.019174913 | 0.051014014 | 0.375875411 | 0.707009521 | 0.929707007 |
| Ndufa2 | 1690.770932 | 0.019179328 | 0.067480643 | 0.284219704 | 0.776242017 | 0.947241001 |
| Olfr446 | 0.543673105 | 0.019180825 | 0.016272792 | 1.178705262 | 0.23851556 | NA |
| Ntpcr | 606.3977133 | 0.019206986 | 0.067496048 | 0.284564598 | 0.775977737 | 0.947241001 |
| 1700096J18Rik | 0.43832756 | 0.019215673 | 0.016298341 | 1.178995635 | 0.238399914 | NA |
| Clec4d | 1.960996837 | 0.019232505 | 0.024230496 | 0.793731362 | 0.427351839 | NA |
| Trio | 7818.217123 | 0.019242748 | 0.063692768 | 0.30211826 | 0.762561913 | 0.944328589 |
| 2010204K13Rik | 462.0543593 | 0.019254476 | 0.072341942 | 0.266159239 | 0.790116574 | 0.951607109 |
| Ldlrad2 | 3.900073275 | 0.019264092 | 0.04363695 | 0.441462829 | 0.658877965 | NA |
| Ins2 | 1.539320337 | 0.019265456 | 0.027273036 | 0.70639208 | 0.479944345 | NA |
| Ccbl1 | 462.604712 | 0.019265533 | 0.079723005 | 0.241655875 | 0.809046823 | 0.955725085 |
| Cyth2 | 3184.611371 | 0.019268451 | 0.052087755 | 0.369922849 | 0.711439976 | 0.930507332 |
| 1700019L03Rik | 12.15822791 | 0.019269918 | 0.064425501 | 0.29910389 | 0.764860778 | NA |
| LOC100043315 | 0.548750089 | 0.019272087 | 0.016310961 | 1.181542093 | 0.237387442 | NA |
| Ift74 | 829.4137554 | 0.019276298 | 0.07041058 | 0.27376991 | 0.784261449 | 0.949699053 |
| Phc2 | 3359.676217 | 0.01927679 | 0.053090526 | 0.363092837 | 0.716535535 | 0.93125489 |
| Hadhb | 3142.315585 | 0.019311032 | 0.054827242 | 0.35221601 | 0.724676271 | 0.932965303 |
| Klc2 | 11693.53747 | 0.019311538 | 0.042147169 | 0.458192998 | 0.64681379 | 0.912750424 |
| Styxl1 | 6.073328644 | 0.019362844 | 0.051603114 | 0.375226265 | 0.707492199 | NA |
| Pea15a | 23765.23299 | 0.019370805 | 0.052286361 | 0.370475299 | 0.711028378 | 0.930507332 |
| Mynn | 630.5088272 | 0.019380676 | 0.066083097 | 0.293277356 | 0.76931016 | 0.94619074 |
| Krt80 | 165.6963024 | 0.019383222 | 0.0949729 | 0.204092136 | 0.838281499 | 0.962077671 |
| Gm4262 | 0.555591603 | 0.019386353 | 0.016358642 | 1.185083286 | 0.235984519 | NA |
| 4931402G19Rik | 0.587554335 | 0.01941968 | 0.01637094 | 1.186228737 | 0.23553198 | NA |
| Tnfsf13 | 18.44835723 | 0.019423992 | 0.077423626 | 0.250879385 | 0.801907363 | NA |
| Papd5 | 1664.056952 | 0.019425916 | 0.058098026 | 0.334364468 | 0.738104548 | 0.937539933 |
| Dennd6a | 1172.359513 | 0.019443018 | 0.060988631 | 0.318797411 | 0.749880141 | 0.9410864 |
| Ptrh1 | 97.3902771 | 0.01945025 | 0.094088506 | 0.206722913 | 0.836226262 | 0.961664 |
| Yipf3 | 1809.574892 | 0.019457827 | 0.05799201 | 0.335525995 | 0.737228339 | 0.937206199 |
| Ern2 | 15.78449249 | 0.019466268 | 0.057484679 | 0.338634018 | 0.734885453 | NA |
| Nt5c1a | 137.7849023 | 0.019473447 | 0.090439744 | 0.215319573 | 0.829518171 | 0.959526943 |
| Taf1 | 3574.728455 | 0.019499502 | 0.057079133 | 0.341622253 | 0.732635188 | 0.936076941 |
| Lmna | 1618.032975 | 0.019507894 | 0.06226139 | 0.313322491 | 0.754035665 | 0.941879961 |
| U2af2 | 5928.770422 | 0.019519048 | 0.040863154 | 0.477668661 | 0.632886053 | 0.906637358 |
| Mrpl48 | 1878.717643 | 0.019541175 | 0.058269855 | 0.335356501 | 0.737356177 | 0.937206199 |
| Chst12 | 837.0214946 | 0.019571232 | 0.073587632 | 0.265958175 | 0.790271421 | 0.951607109 |
| Dhrs7 | 919.8087509 | 0.019576824 | 0.061425408 | 0.318708903 | 0.749947263 | 0.9410864 |
| Sim2 | 5.442816465 | 0.019578294 | 0.050869408 | 0.384873635 | 0.70033104 | NA |
| Aco1 | 1454.117593 | 0.01959898 | 0.057146191 | 0.342962136 | 0.731626945 | 0.935424325 |
| Mphosph8 | 2216.805036 | 0.01959917 | 0.074307289 | 0.263758376 | 0.79196611 | 0.951728598 |
| Sorbs1 | 6792.458409 | 0.019604951 | 0.06149619 | 0.318799437 | 0.749878605 | 0.9410864 |
| Cngb3 | 0.855971059 | 0.019608973 | 0.016436001 | 1.193050113 | 0.232849747 | NA |
| Gm10637 | 0.568632945 | 0.01961709 | 0.016454259 | 1.192219605 | 0.233175148 | NA |
| Serpina12 | 0.763755743 | 0.019619424 | 0.020502119 | 0.95694616 | 0.338594431 | NA |
| Anxa5 | 2299.22525 | 0.019678686 | 0.063969965 | 0.307623838 | 0.758368579 | 0.942526199 |
| 1110059G10Rik | 410.793823 | 0.019680138 | 0.08152245 | 0.241407591 | 0.80923923 | 0.955725085 |
| Gm766 | 3.536026832 | 0.019683854 | 0.042451371 | 0.463680063 | 0.642876983 | NA |
| Crx | 1.113887639 | 0.019695619 | 0.022274415 | 0.884226129 | 0.376574156 | NA |
| Fam188a | 1277.61608 | 0.019698578 | 0.070765866 | 0.278362712 | 0.78073394 | 0.948894514 |
| Smap1 | 3815.804078 | 0.019709465 | 0.059351951 | 0.332077791 | 0.73983052 | 0.938583243 |
| Cactin | 1112.606658 | 0.019711076 | 0.063634311 | 0.309755478 | 0.756746911 | 0.942139348 |
| Mgam | 0.965269439 | 0.01974189 | 0.023426492 | 0.842716435 | 0.399387053 | NA |
| Farsa | 1324.830006 | 0.019747594 | 0.060999837 | 0.323731919 | 0.746141003 | 0.940755508 |
| 5330434G04Rik | 219.6216714 | 0.019751595 | 0.090618662 | 0.217963879 | 0.827457255 | 0.958657209 |
| Elovl1 | 695.5011703 | 0.019758057 | 0.079010385 | 0.250069118 | 0.802533897 | 0.95350952 |
| Ppp1r16b | 9051.981931 | 0.019760505 | 0.07339746 | 0.269226005 | 0.787755769 | 0.95096321 |
| Fzd7 | 338.1670163 | 0.01976669 | 0.077390902 | 0.255413612 | 0.798403659 | 0.952967259 |
| Mir1905 | 1.607791299 | 0.019812891 | 0.028138594 | 0.704118019 | 0.481359279 | NA |
| Pikfyve | 2315.787756 | 0.019815624 | 0.066096485 | 0.299798454 | 0.764330894 | 0.944872045 |
| Zkscan4 | 221.3423296 | 0.019818372 | 0.088244279 | 0.224585348 | 0.822301864 | 0.95778569 |
| Esrp1 | 4.255649014 | 0.019831569 | 0.044557843 | 0.445074698 | 0.656265773 | NA |
| Prss16 | 43.87131591 | 0.019832846 | 0.087229876 | 0.227362993 | 0.820141494 | 0.957571905 |
| Mis18a | 175.3863125 | 0.01985192 | 0.08792691 | 0.225777518 | 0.821374464 | 0.957699646 |
| Mir221 | 3.040170242 | 0.019884385 | 0.040132925 | 0.495463133 | 0.62027324 | NA |
| Gorab | 359.7611478 | 0.019902994 | 0.072699438 | 0.273770951 | 0.784260649 | 0.949699053 |
| AI507597 | 2.008287894 | 0.019908556 | 0.033074616 | 0.601928559 | 0.547221693 | NA |
| Selplg | 1532.777003 | 0.019909224 | 0.06603452 | 0.301497218 | 0.763035371 | 0.944431816 |
| Rap2c | 1047.083525 | 0.01994268 | 0.072932877 | 0.273438817 | 0.784515917 | 0.949699053 |
| Sac3d1 | 411.6348499 | 0.019959676 | 0.070797116 | 0.281927814 | 0.777998862 | 0.94784716 |
| Smtn | 378.185092 | 0.019962563 | 0.088050777 | 0.226716484 | 0.820644209 | 0.957571905 |
| H19 | 8.285261359 | 0.019997368 | 0.05172074 | 0.386641183 | 0.699021865 | NA |
| Rdm1 | 204.4538513 | 0.020003386 | 0.086549726 | 0.231120156 | 0.817221457 | 0.956302288 |
| Sqrdl | 233.8218677 | 0.020019984 | 0.09013594 | 0.222108787 | 0.824229198 | 0.957903916 |
| Wdr55 | 435.6905926 | 0.020021821 | 0.073696816 | 0.271678239 | 0.785869435 | 0.950484821 |
| Mib1 | 2866.751943 | 0.020029062 | 0.069323392 | 0.288922136 | 0.772640963 | 0.947121017 |
| Lrrc40 | 1039.431429 | 0.0200331 | 0.072498395 | 0.276324744 | 0.782298653 | 0.949184473 |
| Egln3 | 420.4351376 | 0.020047049 | 0.070361416 | 0.284915378 | 0.775708974 | 0.947241001 |
| Gmeb2 | 1103.799408 | 0.020047083 | 0.067693287 | 0.296145808 | 0.767118732 | 0.946118815 |
| Lhx9 | 67.62040579 | 0.020050962 | 0.059218058 | 0.338595394 | 0.734914553 | 0.936456337 |

| | | | | | | |
|---|---|---|---|---|---|---|
| Fam69a | 992.2106911 | -0.043268008 | 0.084590552 | -0.511499298 | 0.60900148 | 0.787503666 |
| Wbscr25 | 9.26874711 | 0.078853994 | 0.154212657 | 0.51133283 | 0.60911802 | 0.787588634 |
| Dio3 | 57.08990426 | -0.085556973 | 0.167393794 | -0.511111978 | 0.609272649 | 0.787624958 |
| Hc | 8.886618883 | 0.080904288 | 0.158296519 | 0.511093288 | 0.609285735 | 0.787624958 |
| Rpl22 | 3239.791195 | 0.027045474 | 0.052918807 | 0.511074893 | 0.609298615 | 0.787624958 |
| Prkacb | 21325.71257 | 0.034400718 | 0.067330424 | 0.510923834 | 0.609404391 | 0.787695974 |
| 4932411E22Rik | 106.238372 | 0.077424832 | 0.151626708 | 0.51062793 | 0.609611615 | 0.787891263 |
| Cnot8 | 2030.352198 | -0.032390805 | 0.063450388 | -0.510490267 | 0.609708032 | 0.787891263 |
| Cyp4f15 | 592.2127849 | -0.054744479 | 0.107235922 | -0.510505039 | 0.609697685 | 0.787891263 |
| Lias | 1053.786665 | 0.041999455 | 0.082367927 | 0.509900592 | 0.610121107 | 0.78826483 |
| Maml1 | 524.0835667 | 0.055819899 | 0.10945915 | 0.509961015 | 0.610078774 | 0.78826483 |
| Rlf | 1577.384865 | -0.0380831 | 0.074701671 | -0.509802514 | 0.610189824 | 0.78826483 |
| Slc39a9 | 2422.941793 | -0.023411018 | 0.045923126 | -0.509787109 | 0.610200618 | 0.78826483 |
| Agphd1 | 135.995157 | 0.061732892 | 0.121127049 | 0.509654059 | 0.610293844 | 0.788319535 |
| Itfg2 | 691.0574365 | 0.039473942 | 0.077470408 | 0.509535743 | 0.610376751 | 0.788360903 |
| Rbbp7 | 5257.589115 | 0.041533546 | 0.081525071 | 0.50945735 | 0.610431686 | 0.788366137 |
| Ddx54 | 1823.593484 | -0.029239776 | 0.057403987 | -0.509368387 | 0.610494031 | 0.788366656 |
| Igsf11 | 1502.429436 | -0.040889313 | 0.080283496 | -0.509311566 | 0.610533852 | 0.788366656 |
| Kdm1b | 349.8054666 | 0.041727701 | 0.081972736 | 0.509043651 | 0.610721628 | 0.788477702 |
| Rbck1 | 3305.827814 | -0.02832629 | 0.055644539 | -0.509057857 | 0.610711671 | 0.788477702 |
| Agmo | 259.1224963 | -0.047498044 | 0.093342715 | -0.50885646 | 0.610852841 | 0.788581396 |
| Ehd4 | 642.7232822 | 0.046853917 | 0.092126313 | 0.508583435 | 0.611044242 | 0.788697059 |
| Rsrc1 | 1504.216196 | 0.030216163 | 0.059410722 | 0.508597801 | 0.611034171 | 0.788697059 |
| 3830406C13Rik | 834.3781015 | -0.064507756 | 0.126908111 | -0.508302867 | 0.61124096 | 0.788819543 |
| Nudt12 | 241.6882444 | 0.046178455 | 0.090837229 | 0.508364859 | 0.611197492 | 0.788819543 |
| Igfbp7 | 662.2002832 | 0.046327845 | 0.091172309 | 0.508135038 | 0.611358645 | 0.788905708 |
| Ahr | 392.3705427 | -0.044437943 | 0.087510603 | -0.507800669 | 0.61159314 | 0.789142581 |
| Jtb | 872.9558435 | -0.040657499 | 0.080109294 | -0.507525369 | 0.611786241 | 0.789194573 |
| Mtmr10 | 894.9975315 | 0.045024736 | 0.088708566 | 0.50755793 | 0.611763401 | 0.789194573 |
| Slc46a3 | 266.3226753 | 0.049830425 | 0.098178111 | 0.507551273 | 0.61176807 | 0.789194573 |
| AV039307 | 18.02152348 | 0.084830314 | 0.167171774 | 0.507443998 | 0.611843321 | 0.789202498 |
| Gkap1 | 449.7771473 | 0.0559575 | 0.110308324 | 0.507282656 | 0.611956506 | 0.789217089 |
| Hs2st1 | 2226.048203 | 0.038530088 | 0.07594723 | 0.507327097 | 0.611925329 | 0.789217089 |
| Bicd2 | 3398.568428 | 0.031905894 | 0.062959104 | 0.506771729 | 0.612314995 | 0.789482277 |
| Pmm1 | 4135.479462 | -0.030846082 | 0.060862778 | -0.506813574 | 0.612285631 | 0.789482277 |
| Srrd | 514.4627484 | 0.039874869 | 0.078681282 | 0.506789766 | 0.612302338 | 0.789482277 |
| Eaf1 | 958.6380153 | -0.028425419 | 0.056107822 | -0.506621318 | 0.612420547 | 0.789486976 |
| Fosb | 880.3338224 | -0.084653084 | 0.167091056 | -0.506628457 | 0.612415538 | 0.789486976 |
| Cdh7 | 643.8437396 | -0.045129914 | 0.089102856 | -0.506492336 | 0.612511068 | 0.789511234 |
| Hmgxb4 | 665.9315276 | -0.045462537 | 0.089767205 | -0.506449294 | 0.612541277 | 0.789511234 |
| Tigd2 | 485.9670798 | 0.049546451 | 0.097848535 | 0.506358641 | 0.612604903 | 0.789527564 |
| Pnlip | 30.04926287 | 0.084538568 | 0.167023662 | 0.506147257 | 0.612753278 | 0.789587432 |
| Ttbk1 | 5301.237446 | -0.04602009 | 0.090913912 | -0.50619415 | 0.612720362 | 0.789587432 |
| 1700084C01Rik | 465.8140345 | -0.045264016 | 0.089577582 | -0.505305177 | 0.613344507 | 0.790241343 |
| Baiap3 | 1343.991877 | 0.079338609 | 0.157153156 | 0.504848973 | 0.613664916 | 0.790241343 |
| Cacna1a | 6106.130465 | -0.027066457 | 0.053610892 | -0.504868624 | 0.613651113 | 0.790241343 |
| Cdk6 | 23.68723309 | -0.083012259 | 0.164385073 | -0.504986603 | 0.613568246 | 0.790241343 |
| Gstp1 | 1583.387563 | -0.038249984 | 0.075765405 | -0.504847621 | 0.613665866 | 0.790241343 |
| Ppcdc | 432.9987021 | 0.045868965 | 0.090842175 | 0.504930282 | 0.613607804 | 0.790241343 |
| Prpf18 | 1951.620662 | -0.031370293 | 0.062115799 | -0.505029209 | 0.613538321 | 0.790241343 |
| Specc1l | 2616.77255 | 0.035188241 | 0.069701287 | 0.504843494 | 0.613668765 | 0.790241343 |
| Nlrp5-ps | 18.20773253 | 0.080738788 | 0.159991356 | 0.504644688 | 0.613808416 | 0.790355489 |
| Kif20b | 28.38733913 | -0.084707091 | 0.167918519 | -0.50445354 | 0.613942702 | 0.790462708 |
| 2310022A10Rik | 900.3087391 | 0.033423352 | 0.06627666 | 0.504300496 | 0.614050229 | 0.790490492 |
| Api5 | 3694.754164 | -0.033039719 | 0.065528346 | -0.504204983 | 0.614117339 | 0.790490492 |
| Itgb4 | 487.0040135 | 0.072639316 | 0.144053764 | 0.504251424 | 0.614084708 | 0.790490492 |
| 6820408C15Rik | 78.01011039 | 0.075231982 | 0.149408924 | 0.503530713 | 0.614591192 | 0.791034718 |
| Hpgd | 187.983353 | -0.066146275 | 0.131559073 | -0.50278763 | 0.615113592 | 0.791541695 |
| Sh2d4b | 12.79213699 | 0.083272381 | 0.165632978 | 0.502752423 | 0.615138347 | 0.791541695 |
| Trnp1 | 4578.937256 | -0.041877256 | 0.083290896 | -0.502783117 | 0.615116765 | 0.791541695 |
| Pkdrej | 85.1338863 | 0.066608109 | 0.132558861 | 0.502479493 | 0.615330274 | 0.791722908 |
| Dcaf4 | 774.2871877 | 0.037820794 | 0.075300369 | 0.502265726 | 0.615480615 | 0.791809747 |
| Pwp1 | 814.6656963 | -0.033405376 | 0.066513012 | -0.502238202 | 0.615499974 | 0.791809747 |
| AK010878 | 130.7434513 | -0.058204127 | 0.115984132 | -0.501828361 | 0.615788261 | 0.792073939 |
| Mlx | 1004.767174 | 0.035331393 | 0.070409186 | 0.501800898 | 0.615807582 | 0.792073939 |
| 4833417C18Rik | 10.52508099 | 0.079575372 | 0.158669616 | 0.501516132 | 0.616007927 | 0.79225276 |
| Nans | 489.1273568 | -0.04719701 | 0.094119579 | -0.501457937 | 0.616048874 | 0.79225276 |
| Matr3 | 13492.23547 | -0.035429334 | 0.070675888 | -0.501293091 | 0.616164867 | 0.792336165 |
| Mir343 | 8.169445386 | 0.073555497 | 0.146777519 | 0.501135992 | 0.616275418 | 0.792346803 |
| Pkd1 | 5634.166979 | 0.042835207 | 0.085476088 | 0.501136732 | 0.616274897 | 0.792346803 |
| Wdr53 | 302.3134645 | 0.043443621 | 0.086720082 | 0.500963792 | 0.616396606 | 0.792436857 |
| Serpinb1a | 424.8375972 | 0.074666584 | 0.149084195 | 0.50083501 | 0.616487244 | 0.792487626 |
| Zfp866 | 1114.712498 | -0.033918215 | 0.067750883 | -0.500631333 | 0.616630607 | 0.792606158 |
| Sav1 | 710.2840649 | 0.037241568 | 0.074409893 | 0.500492158 | 0.616728577 | 0.792666327 |
| Thap6 | 117.911951 | 0.06433808 | 0.128596462 | 0.500309877 | 0.6168569 | 0.792765495 |
| Chdh | 274.1768713 | -0.070860563 | 0.141660984 | -0.50021227 | 0.616925619 | 0.792788052 |
| Idh3a | 7739.965482 | -0.033371084 | 0.066741059 | -0.500008314 | 0.617069223 | 0.792841077 |
| Rims4 | 754.4587819 | -0.04480283 | 0.089597226 | -0.500047071 | 0.617041934 | 0.792841077 |
| 2810047C21Rik1 | 41.2318845 | 0.08392357 | 0.168057258 | 0.499374858 | 0.617515328 | 0.792926278 |
| Alpk1 | 190.8934208 | 0.068747193 | 0.137708937 | 0.499221003 | 0.6176237 | 0.792926278 |
| Cdkn1b | 812.4614429 | -0.046973047 | 0.094058655 | -0.499401645 | 0.617496461 | 0.792926278 |
| Gpr161 | 127.3023963 | 0.074031536 | 0.148223526 | 0.499458742 | 0.617456246 | 0.792926278 |
| Plxdc2 | 1212.886961 | -0.046419868 | 0.092937544 | -0.499473805 | 0.617445636 | 0.792926278 |
| Psmd3 | 4656.737295 | 0.028526985 | 0.057146834 | 0.499187501 | 0.617647299 | 0.792926278 |
| Rps6kc1 | 2260.823025 | 0.033324981 | 0.066754295 | 0.499218523 | 0.617625447 | 0.792926278 |
| Slc35e2 | 2068.476093 | -0.033272173 | 0.06658152 | -0.499720835 | 0.61727166 | 0.792926278 |
| Snora52 | 16.39225133 | 0.081809117 | 0.163840868 | 0.499320577 | 0.617553561 | 0.792926278 |
| Zfp296 | 67.67082881 | 0.078205341 | 0.156497935 | 0.499721233 | 0.61727138 | 0.792926278 |
| Sdpr | 392.3578933 | 0.054297244 | 0.108817405 | 0.498975724 | 0.617796486 | 0.793052092 |
| Dnajc6 | 13878.45464 | -0.033656692 | 0.067465244 | -0.498874539 | 0.617867772 | 0.793071598 |
| Gm5083 | 21.68296133 | -0.081342048 | 0.163072585 | -0.49880885 | 0.617914052 | 0.793071598 |
| Sgms1 | 1575.848696 | 0.03889691 | 0.078014549 | 0.498585341 | 0.618071534 | 0.793142318 |
| Trpc6 | 366.7489013 | -0.056691472 | 0.113696148 | -0.498622626 | 0.618045263 | 0.793142318 |
| Gpr157 | 139.8435857 | -0.062456867 | 0.125364511 | -0.498202135 | 0.618341578 | 0.793423139 |
| Gbf1 | 4199.01657 | 0.024798568 | 0.049801673 | 0.497946487 | 0.61852176 | 0.793588618 |
| BC037032 | 34.27856213 | -0.083671215 | 0.168068474 | -0.497840034 | 0.618596796 | 0.793592753 |
| Zbtb2 | 395.2091516 | 0.044160226 | 0.088711388 | 0.497796588 | 0.618627421 | 0.793592753 |
| Amz2 | 2034.68262 | -0.031826994 | 0.064039231 | -0.496992139 | 0.619194595 | 0.794188088 |
| C2cd4b | 27.38339722 | -0.082529569 | 0.166106688 | -0.496846753 | 0.619297123 | 0.794188088 |
| Mtbp | 82.86856437 | 0.075192918 | 0.151362149 | 0.496774909 | 0.619347791 | 0.794188088 |
| Rfc4 | 129.2673028 | -0.062956715 | 0.126696492 | -0.496909693 | 0.619252736 | 0.794188088 |
| Slc22a13b-ps | 17.59674693 | 0.081107373 | 0.163196266 | 0.496992821 | 0.619194114 | 0.794188088 |
| Psmd9 | 842.9677412 | 0.044340706 | 0.089304603 | 0.49651087 | 0.61953402 | 0.794361148 |
| Chrm2 | 316.0055412 | -0.071933242 | 0.144967127 | -0.496203814 | 0.61975062 | 0.794414831 |
| Fam86 | 145.1380965 | -0.064234813 | 0.129463706 | -0.496160781 | 0.619780979 | 0.794414831 |
| Hcfc2 | 871.4573886 | 0.038422176 | 0.07740906 | 0.496352446 | 0.61964577 | 0.794414831 |
| Lrrc8c | 640.6475225 | 0.055251395 | 0.111327082 | 0.496297888 | 0.619684256 | 0.794414831 |
| Wdr36 | 1102.116638 | -0.031199469 | 0.062913029 | -0.495914264 | 0.619954902 | 0.794572028 |
| Rtfdc1 | 2958.487745 | 0.032663688 | 0.065888868 | 0.495739103 | 0.620078495 | 0.794664698 |
| C1qbp | 1738.514956 | 0.04082244 | 0.082362205 | 0.4956453 | 0.620144686 | 0.794683795 |
| 6720489N17Rik | 82.92300074 | 0.078261731 | 0.158108563 | 0.494987302 | 0.620609084 | 0.795015325 |
| Cckar | 20.82064125 | -0.067407057 | 0.136145758 | -0.495109489 | 0.620522836 | 0.795015325 |
| Ptgs1 | 763.4948245 | 0.046127001 | 0.093184869 | 0.495005266 | 0.620596403 | 0.795015325 |
| Sec14l3 | 101.4660051 | -0.078185962 | 0.157978491 | -0.494915234 | 0.620659957 | 0.795015325 |
| Slc25a22 | 8664.2695 | -0.038362429 | 0.077511862 | -0.494923331 | 0.620654241 | 0.795015325 |
| Grik1 | 525.6646866 | 0.047450927 | 0.095965641 | 0.494457463 | 0.62098314 | 0.795363542 |
| Mettl6 | 511.026129 | 0.037411678 | 0.075708701 | 0.494152947 | 0.621198167 | 0.795573186 |
| N28178 | 11524.72701 | 0.034449617 | 0.06972632 | 0.494069048 | 0.621257416 | 0.795583305 |
| Gk5 | 73.95411687 | -0.073299729 | 0.148435096 | -0.493816697 | 0.62143564 | 0.795745769 |
| Ccndbp1 | 1787.864175 | -0.02925525 | 0.059273634 | -0.493562617 | 0.621615107 | 0.795909799 |
| 9530080O11Rik | 54.61233321 | 0.068693607 | 0.139371135 | 0.492882594 | 0.622095547 | 0.796459131 |
| Ina | 4755.291665 | 0.04501239 | 0.091355531 | 0.492716643 | 0.622212816 | 0.79654345 |
| Kbtbd4 | 1220.334743 | -0.043312808 | 0.087954439 | -0.492445963 | 0.622404114 | 0.796656699 |
| Mlycd | 890.1232525 | -0.037393334 | 0.07592338 | -0.492514083 | 0.622355969 | 0.796656699 |
| Wars | 3766.71202 | -0.026732175 | 0.054384224 | -0.491542819 | 0.623042575 | 0.797408034 |
| Ei24 | 4257.710656 | -0.030511106 | 0.062113674 | -0.491213999 | 0.623275098 | 0.797508 |
| Fastkd3 | 443.0948416 | -0.041714005 | 0.084913329 | -0.49125391 | 0.623246874 | 0.797508 |
| Trim47 | 130.614341 | -0.065780903 | 0.133901655 | -0.491262809 | 0.62324058 | 0.797508 |
| 1110020A21Rik | 23.16602162 | 0.082480142 | 0.168093298 | 0.490680729 | 0.623652278 | 0.797775617 |
| Baz2b | 1323.226655 | 0.043906446 | 0.08949046 | 0.490627114 | 0.623690206 | 0.797775617 |
| Ctsh | 492.7938271 | 0.052288663 | 0.106555469 | 0.490717775 | 0.623626073 | 0.797775617 |
| Rad51ap1 | 57.6017152 | 0.075242613 | 0.153349537 | 0.490660839 | 0.623666349 | 0.797775617 |
| Poc1b | 234.5608553 | 0.043416536 | 0.08853113 | 0.490409823 | 0.623843927 | 0.797906373 |
| Elavl1 | 2854.099808 | -0.025309829 | 0.051620689 | -0.490303976 | 0.623918814 | 0.797936286 |
| Pex11c | 12.99093749 | -0.080672044 | 0.164678016 | -0.489877434 | 0.624220633 | 0.79824887 |
| Zfp14 | 399.9540936 | 0.045222418 | 0.092325894 | 0.489812948 | 0.624266268 | 0.79824887 |
| Fam120aos | 479.6717005 | 0.039700411 | 0.081076687 | 0.48966494 | 0.624371015 | 0.798316926 |
| Rundc1 | 2024.472193 | -0.043011984 | 0.087971119 | -0.48893301 | 0.624889124 | 0.798913449 |
| Crtc1 | 9661.581225 | 0.03512708 | 0.071908554 | 0.48849654 | 0.625198176 | 0.799217394 |
| Irak3 | 61.01279577 | 0.075890718 | 0.155369998 | 0.488451562 | 0.625230027 | 0.799217394 |
| Kdm4c | 1158.809682 | 0.02814271 | 0.057645459 | 0.48820341 | 0.625405769 | 0.799244226 |
| Tmc3 | 18.55201597 | -0.07885378 | 0.161496857 | -0.4882682 | 0.625359883 | 0.799244226 |
| Unc5c | 1442.031136 | -0.041471312 | 0.084936754 | -0.488261093 | 0.625364917 | 0.799244226 |
| Zwint | 16037.34365 | -0.040401968 | 0.08277208 | -0.488111065 | 0.625471174 | 0.799261886 |
| Polr2b | 3018.961107 | 0.036225234 | 0.074301351 | 0.487544753 | 0.625872336 | 0.799642614 |
| Tmem259 | 3360.760967 | 0.034607501 | 0.070979378 | 0.487571205 | 0.625853596 | 0.799642614 |
| Ppp1r15b | 2559.183579 | 0.038022092 | 0.078014511 | 0.487372044 | 0.625994701 | 0.799667071 |
| Rab12 | 3158.092854 | 0.037385648 | 0.07670024 | 0.48742544 | 0.625956869 | 0.799667071 |
| Pbdc1 | 507.3006437 | 0.031533672 | 0.064727147 | 0.487178462 | 0.626131867 | 0.799776352 |
| Senp7 | 1172.903083 | 0.044169471 | 0.090700048 | 0.486983991 | 0.626269676 | 0.799820505 |
| Snrnp200 | 6163.730648 | 0.039245328 | 0.080587466 | 0.48699047 | 0.626265084 | 0.799820505 |
| Hcn1 | 2240.803533 | 0.046268868 | 0.095026946 | 0.486902606 | 0.626327352 | 0.799828236 |
| Rimbp3 | 117.7905104 | 0.061501661 | 0.126364253 | 0.48670142 | 0.626469939 | 0.799944391 |
| Gm11944 | 19.71670886 | 0.081734905 | 0.167990249 | 0.486545529 | 0.626580433 | 0.79996003 |
| Nubpl | 157.4451283 | 0.06135689 | 0.126127906 | 0.486465617 | 0.626637077 | 0.79996003 |
| Snord49a | 10.00875279 | 0.077216398 | 0.158718681 | 0.486498489 | 0.626613776 | 0.79996003 |
| Ccbl1 | 394.5870832 | 0.048737932 | 0.100379363 | 0.485537365 | 0.627295214 | 0.799960418 |
| Cdh26 | 9.193531413 | -0.075582973 | 0.155562262 | -0.485869592 | 0.627059629 | 0.799960418 |
| Ddx1 | 5637.070374 | 0.028103888 | 0.057870768 | 0.485631846 | 0.627228213 | 0.799960418 |
| Enpp3 | 19.97968512 | -0.079736776 | 0.164059783 | -0.486022685 | 0.626951082 | 0.799960418 |
| Gbp4 | 124.0646516 | 0.068543273 | 0.141106485 | 0.485755656 | 0.627140418 | 0.799960418 |
| Hist1h2bg | 8.629779784 | 0.075029339 | 0.154356684 | 0.486077681 | 0.626912091 | 0.799960418 |
| Mgat1 | 2180.038983 | -0.031998641 | 0.065868076 | -0.485798926 | 0.627109736 | 0.799960418 |
| Mstn | 12.4107045 | 0.078706883 | 0.162104085 | 0.485533 | 0.62729831 | 0.799960418 |
| Myl12a | 780.6169583 | 0.048713657 | 0.100163714 | 0.486340364 | 0.626725866 | 0.799960418 |
| Nol7 | 904.3571637 | -0.050142822 | 0.103276847 | -0.485518524 | 0.627308576 | 0.799960418 |
| Ocel1 | 172.3712685 | 0.055474294 | 0.114161472 | 0.485928335 | 0.627017978 | 0.799960418 |
| Olfm1 | 34448.53703 | -0.041748906 | 0.085977393 | -0.485580037 | 0.627264953 | 0.799960418 |
| Tle1 | 1397.26853 | -0.031392714 | 0.064567189 | -0.486202273 | 0.62682376 | 0.799960418 |
| Gtdc2 | 1144.758456 | -0.033081127 | 0.068174674 | -0.485240707 | 0.627505609 | 0.800135255 |
| Rnf113a1 | 129.737977 | -0.058705129 | 0.120996701 | -0.485179585 | 0.627548962 | 0.800135255 |
| A930006K02Rik | 18.86192921 | -0.081247997 | 0.167572541 | -0.484852689 | 0.627780844 | 0.800365046 |
| Clic5 | 503.8828523 | 0.050238567 | 0.103636537 | 0.484757296 | 0.627848518 | 0.800370404 |
| Syt7 | 13591.17518 | -0.031452501 | 0.064890504 | -0.484701136 | 0.627888361 | 0.800370404 |
| Oxnad1 | 585.6124477 | 0.03282735 | 0.067861354 | 0.483741454 | 0.62856937 | 0.801172575 |
| Atxn3 | 1219.881615 | 0.039749308 | 0.082326905 | 0.482822819 | 0.629221547 | 0.801937867 |
| Sh2d4a | 9.457385991 | -0.079018363 | 0.163706745 | -0.482682392 | 0.629321267 | 0.80199899 |
| Prrc2a | 20015.86181 | 0.025016776 | 0.051860087 | 0.482389787 | 0.629529075 | 0.802197836 |
| Chst7 | 47.86388514 | 0.07112757 | 0.14757028 | 0.481991157 | 0.629812228 | 0.802480427 |
| Faxc | 5539.462891 | -0.043375767 | 0.090011472 | -0.481891539 | 0.629882997 | 0.802480427 |
| Kctd5 | 866.1310624 | -0.035489372 | 0.07365097 | -0.481858852 | 0.629906219 | 0.802480427 |
| D330050G23Rik | 68.33213034 | 0.069044702 | 0.14334399 | 0.481671408 | 0.63003939 | 0.802584092 |
| Gga1 | 2641.466426 | 0.03803169 | 0.079045771 | 0.481135035 | 0.630420529 | 0.802712142 |
| Parp6 | 5636.092162 | -0.025909948 | 0.053835404 | -0.481280837 | 0.630316915 | 0.802712142 |
| Pdcd11 | 1096.934607 | -0.045792486 | 0.095163671 | -0.481197135 | 0.630376397 | 0.802712142 |
| Phlpp1 | 4528.613778 | -0.033292588 | 0.06921154 | -0.481026546 | 0.630497632 | 0.802712142 |
| Prim2 | 270.2410451 | 0.045044282 | 0.093643339 | 0.481019602 | 0.630502567 | 0.802712142 |
| Serhl | 167.5050927 | 0.062252648 | 0.129302878 | 0.481448278 | 0.630197931 | 0.802712142 |
| Tmem194b | 185.7876316 | 0.069936275 | 0.145315758 | 0.481271104 | 0.630323831 | 0.802712142 |
| Tagln2 | 439.2771569 | 0.05742314 | 0.119406052 | 0.480906446 | 0.630582992 | 0.802748572 |
| B4galt5 | 2576.209746 | -0.032869711 | 0.068362516 | -0.480814813 | 0.630648121 | 0.802765527 |
| Cep72 | 56.40899362 | 0.075539133 | 0.157148789 | 0.480685428 | 0.63074009 | 0.802816639 |
| 1700095B22Rik | 42.66254303 | 0.078672339 | 0.163738344 | 0.480475965 | 0.630888989 | 0.802873394 |
| E2f6 | 1306.476256 | -0.030428982 | 0.063358824 | -0.480264316 | 0.631039459 | 0.802873394 |
| Gpn1 | 913.2205586 | -0.028208092 | 0.058728321 | -0.480314966 | 0.631003448 | 0.802873394 |
| Lamtor3 | 1374.561926 | 0.037786124 | 0.078647099 | 0.480451596 | 0.630906314 | 0.802873394 |
| Ufsp2 | 1466.217874 | -0.03127833 | 0.065128146 | -0.480258251 | 0.631043771 | 0.802873394 |
| 1810013L24Rik | 1676.904736 | 0.032066051 | 0.066813671 | 0.479932487 | 0.6312754 | 0.802875181 |
| Bend6 | 5224.913692 | 0.033278877 | 0.069345259 | 0.479901264 | 0.631297602 | 0.802875181 |
| Klhl8 | 1348.671759 | 0.046262806 | 0.096402558 | 0.47989189 | 0.631304268 | 0.802875181 |
| Ptbp2 | 2706.003729 | 0.039697698 | 0.082702623 | 0.480005304 | 0.631223622 | 0.802875181 |
| Yy1 | 991.9881287 | 0.041597001 | 0.086643646 | 0.480092928 | 0.631161317 | 0.802875181 |
| Lyar | 296.0254457 | 0.040210221 | 0.083808623 | 0.479786208 | 0.631379421 | 0.802904855 |
| B3galtl | 823.5198714 | 0.031448769 | 0.065587694 | 0.47949191 | 0.631588722 | 0.802907422 |
| Bdkrb2 | 19.99913979 | 0.078976807 | 0.164686137 | 0.479559533 | 0.631540627 | 0.802907422 |
| Hoga1 | 27.29510821 | 0.080631299 | 0.168120509 | 0.479604182 | 0.631508872 | 0.802907422 |
| Tbccd1 | 533.0767839 | 0.039146652 | 0.08163636 | 0.479524707 | 0.631565396 | 0.802907422 |
| 6030407O03Rik | 10.30395131 | 0.072586031 | 0.151410387 | 0.479399285 | 0.631654602 | 0.802925294 |
| 3110007F17Rik | 53.34745158 | -0.069028704 | 0.144133387 | -0.478922377 | 0.631993851 | 0.803224734 |
| Pcdhb10 | 164.6923182 | -0.049382896 | 0.103110585 | -0.478931395 | 0.631987436 | 0.803224734 |
| Aspa | 540.3752395 | 0.06824639 | 0.142625499 | 0.478500624 | 0.632293931 | 0.803503065 |
| Ccser1 | 408.1849538 | 0.048068057 | 0.100462256 | 0.478468814 | 0.632316566 | 0.803503065 |
| Aard | 51.61656172 | -0.072927366 | 0.152457107 | -0.478346776 | 0.632403409 | 0.803547517 |
| 0610011F06Rik | 622.9309509 | 0.035313831 | 0.073843706 | 0.478223976 | 0.6324908 | 0.803592657 |
| Igsf1 | 310.3915469 | -0.072092913 | 0.150848543 | -0.477915874 | 0.632710083 | 0.803750179 |
| Sp3 | 1743.918816 | -0.037831906 | 0.079162145 | -0.477904003 | 0.632718532 | 0.803750179 |
| Acot1 | 191.2553657 | 0.049392266 | 0.103402015 | 0.477672184 | 0.632883545 | 0.80382799 |
| Zfp941 | 1268.396444 | 0.040564877 | 0.084920146 | 0.477682614 | 0.63287612 | 0.80382799 |
| 2700070H01Rik | 9.773139921 | 0.072692835 | 0.152207849 | 0.477589269 | 0.63294257 | 0.803837064 |
| Pigc | 432.2989833 | 0.035681176 | 0.074734576 | 0.477438656 | 0.633049793 | 0.803907343 |
| Add3 | 4566.887958 | -0.02369281 | 0.049658732 | -0.477112663 | 0.633281898 | 0.804136184 |
| Usp24 | 4120.736361 | -0.037172854 | 0.077954153 | -0.476855339 | 0.633465136 | 0.804302944 |
| Msi1 | 622.2950416 | 0.03817594 | 0.080075452 | 0.476749606 | 0.633540434 | 0.804332636 |
| Nyap1 | 3287.93037 | -0.029429713 | 0.061760871 | -0.476510652 | 0.633710621 | 0.804482782 |
| Fam168b | 15674.11167 | 0.026558372 | 0.055747212 | 0.476407183 | 0.633784318 | 0.804510423 |
| Ppp1r12a | 2709.059036 | -0.025807256 | 0.054189354 | -0.476242181 | 0.633901852 | 0.804593699 |
| Gm14015 | 11.86561769 | 0.075352651 | 0.158292407 | 0.476034531 | 0.634049778 | 0.804715535 |
| Kcnj16 | 1135.384627 | -0.049040157 | 0.103049449 | -0.475889559 | 0.634153061 | 0.804729607 |
| Pold3 | 1011.627632 | 0.028764296 | 0.060449941 | 0.47583662 | 0.634190778 | 0.804729607 |
| Wbp4 | 2158.69212 | 0.033633764 | 0.07068883 | 0.475800267 | 0.63421668 | 0.804729607 |
| 4930578C19Rik | 30.9467851 | -0.079363465 | 0.166828036 | -0.475720193 | 0.634273733 | 0.804736097 |
| Igsf9b | 686.5394148 | 0.055606091 | 0.11692159 | 0.475584454 | 0.634370453 | 0.804792909 |
| Asic1 | 3309.80094 | 0.032802755 | 0.068985314 | 0.47550345 | 0.634428175 | 0.804800241 |
| Mpv17l | 2447.938291 | 0.025568626 | 0.053785537 | 0.475381076 | 0.63451538 | 0.80484497 |
| Cluh | 4325.21593 | 0.02371903 | 0.049914087 | 0.475197117 | 0.634646482 | 0.804945366 |

| | | | | | | |
|---|---|---|---|---|---|---|
| A4galt | 98.42837766 | 0.020052643 | 0.09504738 | 0.210975234 | 0.832906597 | 0.960603511 |
| Klhl21 | 895.21471 | 0.020061575 | 0.070401204 | 0.284960672 | 0.775674272 | 0.947241001 |
| Lrrc3 | 723.2899482 | 0.020064927 | 0.079578938 | 0.252138664 | 0.800933886 | 0.953228631 |
| Gm13247 | 2.196852926 | 0.020067166 | 0.030902607 | 0.649368065 | 0.5161005 | NA |
| 1700001J03Rik | 3.185234464 | 0.020085637 | 0.042613474 | 0.471344733 | 0.637394576 | NA |
| Zfand6 | 1221.727536 | 0.020104932 | 0.063276204 | 0.317732908 | 0.750687547 | 0.9410864 |
| Scaf8 | 2806.283313 | 0.020107722 | 0.055261984 | 0.36386175 | 0.715961249 | 0.93125489 |
| Slc31a1 | 1016.882345 | 0.020108273 | 0.083308157 | 0.241372201 | 0.809266656 | 0.955725085 |
| Wdr77 | 2522.828476 | 0.020150701 | 0.050615416 | 0.398113904 | 0.690546226 | 0.923860706 |
| Mafg | 1079.825353 | 0.020152688 | 0.082383602 | 0.244620138 | 0.80675057 | 0.955587482 |
| 9430014N10Rik | 0.597814748 | 0.020160302 | 0.016674507 | 1.209049345 | 0.226643888 | NA |
| Spr | 779.7647022 | 0.020162421 | 0.071468695 | 0.282115429 | 0.777855003 | 0.94784716 |
| Chmp2a | 2623.674373 | 0.020190323 | 0.050299195 | 0.401404494 | 0.688122341 | 0.923860706 |
| Fmo4 | 2.521405816 | 0.020205947 | 0.033438939 | 0.604264008 | 0.545668131 | NA |
| Ube2i | 7606.254705 | 0.02021134 | 0.039472441 | 0.512036728 | 0.608625305 | 0.894374424 |
| Rab27a | 279.4633469 | 0.020216345 | 0.087720966 | 0.230461958 | 0.817732821 | 0.956663546 |
| DQ267102 | 11.08398629 | 0.020223075 | 0.067395396 | 0.300066125 | 0.764126718 | NA |
| Eif3a | 6822.743775 | 0.020229503 | 0.054451397 | 0.371514857 | 0.710254092 | 0.930478981 |
| Tapt1 | 2309.112017 | 0.020243679 | 0.057923006 | 0.349492901 | 0.7267193 | 0.9335794 |
| Gm5544 | 1.788290623 | 0.020244499 | 0.026601741 | 0.761021593 | 0.44664417 | NA |
| Ankrd33 | 1.131216723 | 0.020249091 | 0.021433641 | 0.944734062 | 0.344794663 | NA |
| Des | 77.99675585 | 0.020262161 | 0.09502112 | 0.2132385 | 0.83114094 | 0.960162026 |
| Tgfbr3 | 583.1386451 | 0.02027048 | 0.085686715 | 0.236565029 | 0.812994253 | 0.956264994 |
| Adcy2 | 7070.330304 | 0.020270716 | 0.050554731 | 0.400965765 | 0.688445329 | 0.923860706 |
| 4930488L21Rik | 10.01787041 | 0.020272715 | 0.063137017 | 0.321090796 | 0.748141585 | NA |
| Tmem258 | 733.7281752 | 0.020279255 | 0.068010084 | 0.298180124 | 0.765565691 | 0.945308333 |
| 1700028B04Rik | 2.068070408 | 0.020295285 | 0.030273231 | 0.670403663 | 0.5026005 | NA |
| 1700023E05Rik | 8.537211841 | 0.020296087 | 0.062655866 | 0.323929565 | 0.74599136 | NA |
| Tnnt3 | 2.988541184 | 0.020332155 | 0.035614001 | 0.570903403 | 0.568065124 | NA |
| Mettl25 | 204.7093029 | 0.020338549 | 0.087261142 | 0.233076818 | 0.815701756 | 0.956264994 |
| Syn2 | 23023.70974 | 0.020352404 | 0.061950918 | 0.32852466 | 0.742515002 | 0.939912916 |
| Dhx30 | 5114.721567 | 0.020357825 | 0.04981433 | 0.408674073 | 0.682778863 | 0.922882553 |
| Myrfl | 4.6581879 | 0.020372457 | 0.048079936 | 0.423720549 | 0.671769627 | NA |
| Steap4 | 20.18423418 | 0.020380678 | 0.065794423 | 0.309762993 | 0.756741195 | NA |
| Xkr8 | 283.5597063 | 0.020391385 | 0.086647611 | 0.23533695 | 0.813947219 | 0.956264994 |
| 2010109I03Rik | 12.0968058 | 0.02039468 | 0.062520602 | 0.326207348 | 0.744267482 | NA |
| Actr3b | 3699.222022 | 0.020412322 | 0.053054406 | 0.384743194 | 0.70042769 | 0.927053542 |
| Zfp712 | 256.0083457 | 0.020414682 | 0.092123245 | 0.22160185 | 0.824623842 | 0.957903916 |
| Hspa12b | 329.6672846 | 0.020416044 | 0.092074042 | 0.221735064 | 0.824520133 | 0.957903916 |
| Lrfn4 | 1434.932946 | 0.020416968 | 0.078803974 | 0.259085516 | 0.79556926 | 0.952636969 |
| Galnt6 | 835.4253515 | 0.020429738 | 0.092251542 | 0.22145687 | 0.824736716 | 0.957903916 |
| 3110045C21Rik | 22.98764411 | 0.02043452 | 0.085822861 | 0.238101131 | 0.811802657 | 0.956219756 |
| Abhd12 | 12355.33395 | 0.020439147 | 0.043634729 | 0.468414665 | 0.639488082 | 0.909767911 |
| Rtp2 | 4.62875023 | 0.020450335 | 0.041637916 | 0.491146929 | 0.623322532 | NA |
| Mir384 | 0.74312821 | 0.020453268 | 0.020406064 | 1.002313209 | 0.316192345 | NA |
| Sox6 | 1006.161787 | 0.020453694 | 0.07756711 | 0.263690287 | 0.792018581 | 0.951728598 |
| As3mt | 477.9433224 | 0.020465186 | 0.08841589 | 0.231465027 | 0.816953552 | 0.956302288 |
| Pwwp2b | 702.4520872 | 0.020466514 | 0.071116312 | 0.287789301 | 0.773508028 | 0.947121017 |
| Nqo1 | 182.6227743 | 0.020473585 | 0.09265367 | 0.220968955 | 0.825116606 | 0.957903916 |
| Mir491 | 2.72934465 | 0.020486623 | 0.040059048 | 0.511410642 | 0.609063545 | NA |
| Zfp943 | 514.265741 | 0.020505618 | 0.079720228 | 0.257219755 | 0.797009134 | 0.952885371 |
| Neil2 | 93.52987264 | 0.020510898 | 0.094895072 | 0.216142916 | 0.828876348 | 0.959237314 |
| C2cd2l | 8626.82246 | 0.020515415 | 0.066634485 | 0.307879846 | 0.758173762 | 0.942510233 |
| Hmgb1-rs17 | 1.496993781 | 0.020524115 | 0.028155668 | 0.728951443 | 0.466031365 | NA |
| Phyhip | 26751.42129 | 0.020531048 | 0.056733162 | 0.361887955 | 0.717435762 | 0.93125489 |
| Ankib1 | 2367.992256 | 0.020536793 | 0.063183833 | 0.325032405 | 0.745156547 | 0.940632024 |
| Tubg1 | 1513.832931 | 0.020541753 | 0.063410349 | 0.323949537 | 0.745976239 | 0.940755508 |
| Zcchc10 | 233.2471755 | 0.020562597 | 0.08666504 | 0.237265185 | 0.812451069 | 0.956219756 |
| Fmn2 | 3857.004185 | 0.020574642 | 0.053988263 | 0.381094723 | 0.703132963 | 0.927855161 |
| Nagk | 1247.100166 | 0.020585247 | 0.07198121 | 0.285980835 | 0.7748928 | 0.947238918 |
| Lrrn4cl | 63.6927127 | 0.020589283 | 0.094288593 | 0.218364512 | 0.827145114 | 0.958524924 |
| Sntn | 13.3729447 | 0.020592828 | 0.064248925 | 0.3205163 | 0.748576976 | NA |
| Zc3h12b | 474.8221754 | 0.020597299 | 0.073930321 | 0.278604211 | 0.780548581 | 0.948894514 |
| Dab2ip | 7119.213559 | 0.020602279 | 0.055066194 | 0.374136599 | 0.708302691 | 0.929959854 |
| Gm17745 | 1.317769838 | 0.020607821 | 0.025723828 | 0.801117961 | 0.423063359 | NA |
| Acan | 235.8600573 | 0.020613988 | 0.086285632 | 0.238904058 | 0.811179979 | 0.956135608 |
| Aldh18a1 | 978.8416549 | 0.02063145 | 0.058883041 | 0.350380177 | 0.726053401 | 0.933575664 |
| Rbp2 | 1.38813166 | 0.020648679 | 0.025077479 | 0.823395348 | 0.410283207 | NA |
| Cox6c | 5898.109783 | 0.020670713 | 0.046905908 | 0.440684642 | 0.659441317 | 0.915208181 |
| E330013P04Rik | 6.045918369 | 0.020674059 | 0.049198156 | 0.4202202 | 0.674324599 | NA |
| Gpr81 | 41.5017933 | 0.020692778 | 0.086817583 | 0.238347779 | 0.811611366 | 0.956161128 |
| Medag | 1180.488223 | 0.020715811 | 0.084653507 | 0.244712962 | 0.806678691 | 0.955587482 |
| Exosc7 | 595.0792705 | 0.020734887 | 0.064928894 | 0.319347611 | 0.749462933 | 0.9410864 |
| Btnl9 | 0.981826011 | 0.020760011 | 0.02347791 | 0.884235906 | 0.376568879 | NA |
| Rnf149 | 865.2568957 | 0.020768921 | 0.058288181 | 0.356314441 | 0.721605096 | 0.932513933 |
| Vrk2 | 71.16682616 | 0.020780743 | 0.094977812 | 0.218795764 | 0.826809146 | 0.958455003 |
| Clip1 | 4973.310359 | 0.020797423 | 0.058376616 | 0.356262913 | 0.72164368 | 0.932513933 |
| Gsg2 | 6.536854203 | 0.020810911 | 0.05240688 | 0.397102658 | 0.69129176 | NA |
| Nme8 | 0.893333255 | 0.020811006 | 0.0201828 | 1.031125796 | 0.302481832 | NA |
| Rtkn2 | 133.2368028 | 0.020820119 | 0.09213153 | 0.22598256 | 0.821214985 | 0.957611694 |
| Gm14379 | 14.30840983 | 0.020833738 | 0.062336049 | 0.334216531 | 0.73821617 | NA |
| Cox15 | 1214.126381 | 0.020840337 | 0.054304379 | 0.383768998 | 0.701149668 | 0.927502056 |
| Sema7a | 9010.790392 | 0.020846076 | 0.067791673 | 0.307502012 | 0.758461292 | 0.942526199 |
| Alox5 | 33.5229515 | 0.020882774 | 0.089179099 | 0.234166682 | 0.81485558 | 0.956264994 |
| 4931431C16Rik | 1.922618846 | 0.020906161 | 0.02623229 | 0.796962855 | 0.425472604 | NA |
| 2410018L13Rik | 108.1296907 | 0.020907814 | 0.092917742 | 0.225014225 | 0.821968208 | 0.95778569 |
| Pacs2 | 6982.382603 | 0.020932975 | 0.067241869 | 0.311308645 | 0.755565996 | 0.942139348 |
| Rpl7 | 8751.792771 | 0.020957047 | 0.053638122 | 0.390711797 | 0.696010278 | 0.925527709 |
| Mmd | 7810.58178 | 0.020969606 | 0.060707056 | 0.345422883 | 0.729776476 | 0.934125508 |
| Cant1 | 2188.746886 | 0.020969704 | 0.052161886 | 0.402011996 | 0.687675198 | 0.923860706 |
| Eed | 522.3326084 | 0.020970875 | 0.075240757 | 0.278716959 | 0.780462047 | 0.948894514 |
| Chrm1 | 9233.924903 | 0.020981207 | 0.066302033 | 0.316448919 | 0.751661791 | 0.941374879 |
| Snora33 | 7.139059484 | 0.020996349 | 0.057448712 | 0.365479885 | 0.714753219 | NA |
| 2610020C07Rik | 21.50471589 | 0.021002197 | 0.080813918 | 0.259883415 | 0.794953705 | NA |
| Zcchc4 | 228.3440087 | 0.021011353 | 0.084966889 | 0.247288715 | 0.804684792 | 0.954737506 |
| Cdk12 | 1163.485224 | 0.021012207 | 0.071117957 | 0.295455713 | 0.767645779 | 0.94619074 |
| Mavs | 405.1361502 | 0.02103632 | 0.090445996 | 0.232584316 | 0.816084207 | 0.956264994 |
| Ostn | 1.400948497 | 0.021040719 | 0.022028729 | 0.95514905 | 0.339502327 | NA |
| Dnajc7 | 6508.945282 | 0.021044932 | 0.048971228 | 0.429740737 | 0.667384247 | 0.916320387 |
| Gm12695 | 3.397062918 | 0.021048733 | 0.041365908 | 0.50884253 | 0.610862606 | NA |
| Ly6c2 | 1.642337728 | 0.021049717 | 0.025092598 | 0.83888153 | 0.401535793 | NA |
| Zfp68 | 1544.774309 | 0.021050658 | 0.076357794 | 0.275684472 | 0.782790423 | 0.949308206 |
| Hr | 2295.637585 | 0.021052056 | 0.073758617 | 0.285418259 | 0.775323721 | 0.947241001 |
| Mrpl53 | 662.4420921 | 0.021060717 | 0.065318569 | 0.322430782 | 0.747126368 | 0.94097069 |
| Gm20754 | 31.73011875 | 0.021063619 | 0.08791824 | 0.239581898 | 0.810654399 | 0.956073263 |
| Mark1 | 4567.746752 | 0.021072379 | 0.046967596 | 0.448657817 | 0.653678519 | 0.914683958 |
| Srd5a1 | 441.5359827 | 0.02108164 | 0.07420155 | 0.284113204 | 0.776323629 | 0.947241001 |
| Tbl3 | 1203.030503 | 0.021098419 | 0.062118764 | 0.339646469 | 0.734122779 | 0.936456337 |
| Pik3r1 | 6482.194144 | 0.021109859 | 0.056528083 | 0.373440214 | 0.708820834 | 0.930183885 |
| Pcsk2 | 19534.81616 | 0.021111273 | 0.058876074 | 0.358571337 | 0.719915792 | 0.93237311 |
| 9530048J24Rik | 0.658477857 | 0.021112121 | 0.017057385 | 1.237711468 | 0.215823065 | NA |
| Ndufa7 | 1762.67206 | 0.021118951 | 0.058705113 | 0.359746367 | 0.719036814 | 0.932065747 |
| Lbp | 180.1178385 | 0.021124114 | 0.043228588 | 0.488660744 | 0.6250819 | 0.901503145 |
| Rbm14 | 1301.126615 | 0.021126418 | 0.058561777 | 0.360754386 | 0.718283064 | 0.93125489 |
| Ralb | 711.0196377 | 0.021132588 | 0.072262551 | 0.292441767 | 0.769948878 | 0.94619074 |
| Tmem68 | 1179.90041 | 0.021135373 | 0.075538633 | 0.279795551 | 0.779634365 | 0.948728222 |
| Mat2b | 4653.343362 | 0.021144951 | 0.066014661 | 0.320306889 | 0.748735701 | 0.9410864 |
| Umps | 719.8507417 | 0.021148326 | 0.069730298 | 0.303287471 | 0.761670793 | 0.943656543 |
| Rps27 | 28.44165344 | 0.021152219 | 0.086885862 | 0.24344834 | 0.8076581 | 0.955716697 |
| Fam120c | 3417.49273 | 0.021172685 | 0.062017635 | 0.34139781 | 0.732804123 | 0.936094952 |
| Beta-s | 118.1808697 | 0.021174143 | 0.037955735 | 0.557864119 | 0.576937175 | 0.881236189 |
| Sept4 | 8225.011335 | 0.021182161 | 0.080220803 | 0.264048232 | 0.791742754 | 0.951728598 |
| Foxs1 | 13.39738723 | 0.021191098 | 0.073200847 | 0.289492528 | 0.772204497 | NA |
| Mbd5 | 2671.562558 | 0.021196773 | 0.058497849 | 0.362351316 | 0.717089515 | 0.93125489 |
| Ufd1l | 3117.908842 | 0.021213187 | 0.056208589 | 0.377401162 | 0.705875503 | 0.929184192 |
| Pwp1 | 876.6286458 | 0.02122464 | 0.058614638 | 0.36210477 | 0.717273739 | 0.93125489 |
| Tdh | 1.758036472 | 0.021237433 | 0.028575059 | 0.743215734 | 0.457351079 | NA |
| Fam217b | 1674.122277 | 0.021265556 | 0.061070029 | 0.348215909 | 0.727678041 | 0.9335794 |
| Fam159b | 18.45272737 | 0.021274493 | 0.082136101 | 0.259015134 | 0.795623564 | NA |
| Iltifb | 2.99550181 | 0.021279052 | 0.039549892 | 0.53803059 | 0.59055593 | NA |
| Chst1 | 11021.95561 | 0.021285857 | 0.066822933 | 0.3185412 | 0.750074448 | 0.9410864 |
| Naa60 | 4791.452585 | 0.021296823 | 0.039905328 | 0.533683696 | 0.593560391 | 0.886313552 |
| Nt5dc1 | 81.69760609 | 0.021307059 | 0.095060849 | 0.224141267 | 0.822647383 | 0.957821087 |
| Eif5 | 9610.98575 | 0.021315278 | 0.068327694 | 0.311956647 | 0.755073471 | 0.942123109 |
| Rilpl2 | 394.3579007 | 0.021321008 | 0.080922701 | 0.263473756 | 0.792185449 | 0.951728598 |
| 1700034F02Rik | 3.638767852 | 0.021327524 | 0.044977954 | 0.474177289 | 0.635373487 | NA |
| Nyap1 | 3618.903346 | 0.021333063 | 0.058884418 | 0.362287068 | 0.717137521 | 0.93125489 |
| Phpt1 | 1202.639008 | 0.021333521 | 0.068954079 | 0.309387372 | 0.757026875 | 0.942139348 |
| Rhno1 | 503.2603625 | 0.021338341 | 0.068964016 | 0.309412681 | 0.757007625 | 0.942139348 |
| Gm3336 | 1.205649186 | 0.021344625 | 0.027694425 | 0.770719212 | 0.440873382 | NA |
| Mis18bp1 | 8.049298573 | 0.021347361 | 0.059967654 | 0.355981257 | 0.721854602 | NA |
| Ube4b | 5507.336933 | 0.021352031 | 0.048386073 | 0.441284637 | 0.659006946 | 0.915208181 |
| Stk38l | 1516.03285 | 0.021356528 | 0.05063632 | 0.421763028 | 0.673197994 | 0.919353963 |
| Abcg5 | 1.070025063 | 0.021361044 | 0.022570586 | 0.946410671 | 0.343939165 | NA |
| Neil3 | 4.045797931 | 0.021381105 | 0.042949827 | 0.497815869 | 0.61861383 | NA |
| Bsnd | 0.623897432 | 0.021382981 | 0.017167026 | 1.24558446 | 0.21291699 | NA |
| Oat | 3335.263901 | 0.021385976 | 0.04976753 | 0.429717447 | 0.66740119 | 0.916320387 |
| Gm16845 | 27.63573501 | 0.021399647 | 0.082957208 | 0.257960068 | 0.796437725 | 0.952734447 |
| Cntn2 | 8266.706136 | 0.021400251 | 0.067234896 | 0.318290832 | 0.750264338 | 0.9410864 |
| Ankrd61 | 3.345195297 | 0.021401897 | 0.037034542 | 0.57789015 | 0.563338286 | NA |
| Fam104a | 1365.994267 | 0.021417515 | 0.060736777 | 0.35262844 | 0.724367013 | 0.932882346 |
| Grk6 | 3054.952951 | 0.021418821 | 0.054770554 | 0.391064524 | 0.695749543 | 0.925527709 |
| Tmcc2 | 8369.536368 | 0.02143501 | 0.056513098 | 0.379292777 | 0.704470463 | 0.928382774 |
| Smarcd1 | 8470.112969 | 0.021439213 | 0.052255699 | 0.410275114 | 0.681604146 | 0.922538401 |
| Tex22 | 1.628583663 | 0.021441351 | 0.028409663 | 0.754720365 | 0.450416785 | NA |
| B3galt6 | 465.5454521 | 0.021441446 | 0.078154065 | 0.274348437 | 0.783816865 | 0.949699053 |
| Cenpl | 64.85413096 | 0.021450717 | 0.094887658 | 0.226064355 | 0.821151368 | 0.957611694 |
| Ensa | 12617.45953 | 0.021452628 | 0.04103492 | 0.52278957 | 0.601120699 | 0.890297392 |
| Git2 | 1528.896096 | 0.021458315 | 0.054967253 | 0.390383625 | 0.696252895 | 0.92555371 |
| Gramd4 | 2432.291924 | 0.02146035 | 0.069072823 | 0.310691654 | 0.756035043 | 0.942139348 |
| Pdgfa | 833.2626315 | 0.021461337 | 0.070483761 | 0.304486258 | 0.76075746 | 0.943105713 |
| Slc17a7 | 54750.44893 | 0.021465954 | 0.063834251 | 0.336276426 | 0.736662425 | 0.937206199 |
| Zfp410 | 934.1518762 | 0.021476944 | 0.066502068 | 0.322951528 | 0.746731952 | 0.94097069 |
| Gtf2a2 | 544.0651869 | 0.021478089 | 0.073949259 | 0.290443594 | 0.771476897 | 0.946412752 |
| Pcdhb5 | 291.3369402 | 0.021478344 | 0.090471634 | 0.237404177 | 0.81234325 | 0.956219756 |
| Zfp512 | 2100.601185 | 0.021484786 | 0.054923425 | 0.391177092 | 0.69566634 | 0.925527709 |
| Wfikkn2 | 82.59592107 | 0.021499039 | 0.059073088 | 0.363939656 | 0.715903071 | 0.93125489 |
| Usmg5 | 3628.767415 | 0.021499499 | 0.074389985 | 0.289010661 | 0.772573219 | 0.947121017 |
| Tfap2a | 7.57410212 | 0.021503017 | 0.058216233 | 0.369364633 | 0.711855956 | NA |
| Gm10364 | 1.424602839 | 0.021514816 | 0.02530573 | 0.850195454 | 0.395216429 | NA |
| E130317F20Rik | 164.3759881 | 0.021533269 | 0.09145799 | 0.235444374 | 0.813863849 | 0.956264994 |
| Gm572 | 3.214112978 | 0.021561913 | 0.040544601 | 0.53180727 | 0.594859487 | NA |
| Polr2d | 477.3520165 | 0.021562836 | 0.075535062 | 0.285467907 | 0.775285688 | 0.947241001 |
| Stpg2 | 3.085868376 | 0.02157522 | 0.034505341 | 0.625271912 | 0.531792612 | NA |
| AI316807 | 873.0553844 | 0.021586786 | 0.067382056 | 0.320364015 | 0.748692401 | 0.9410864 |
| Spta1 | 0.959331758 | 0.021586889 | 0.019696337 | 1.095984953 | 0.27308536 | NA |
| Gm1332 | 1.557541471 | 0.021588034 | 0.030197969 | 0.71488366 | 0.474680934 | NA |
| Mgat4a | 2058.318255 | 0.021589886 | 0.056378053 | 0.382948424 | 0.701758006 | 0.92751465 |
| Glb1l2 | 28.66460502 | 0.021591544 | 0.066882453 | 0.322828224 | 0.746825337 | 0.94097069 |
| Ubxn7 | 1821.254951 | 0.021595024 | 0.066586376 | 0.324315956 | 0.745698841 | 0.940687916 |
| Irx6 | 8.569213613 | 0.021601755 | 0.058908384 | 0.366700864 | 0.713842162 | NA |
| Klk9 | 2.071874923 | 0.021605906 | 0.034868768 | 0.619634916 | 0.535498174 | NA |
| Papd7 | 2082.607239 | 0.021607972 | 0.062632289 | 0.344997313 | 0.73009639 | 0.934152665 |
| E2f2 | 32.73276693 | 0.021613687 | 0.085107164 | 0.253958498 | 0.799527621 | 0.953228631 |
| Gm7168 | 0.931597681 | 0.021623399 | 0.019909884 | 1.086063537 | 0.277450883 | NA |
| Mettl5 | 214.3073317 | 0.021657231 | 0.085738354 | 0.252596766 | 0.800579829 | 0.953228631 |
| Exo5 | 347.1928655 | 0.021665967 | 0.08231381 | 0.263211806 | 0.792387332 | 0.951728598 |
| Pdk2 | 5268.65198 | 0.021673972 | 0.043150894 | 0.502283255 | 0.615468287 | 0.896719391 |
| Gpr126 | 154.8421911 | 0.02168183 | 0.087981898 | 0.246435119 | 0.805345423 | 0.955131256 |
| Wnt5a | 312.8252163 | 0.021682655 | 0.081649773 | 0.265556832 | 0.790580535 | 0.951728598 |
| Agmo | 281.8501621 | 0.021684023 | 0.082948177 | 0.261416504 | 0.793771331 | 0.951728598 |
| Zfp324 | 489.0004982 | 0.021705723 | 0.069273526 | 0.313333591 | 0.754027232 | 0.941879961 |
| Ephx2 | 891.6867486 | 0.021713206 | 0.067620046 | 0.321106059 | 0.748130019 | 0.9410864 |
| Rad23b | 8418.823109 | 0.021752333 | 0.052864596 | 0.411472609 | 0.680726022 | 0.922355522 |
| Gsta4 | 1327.874971 | 0.021755089 | 0.056156598 | 0.387400411 | 0.6984598 | 0.926409493 |
| Macc1 | 4.16186299 | 0.021774526 | 0.043845156 | 0.496623286 | 0.619454729 | NA |
| Tbc1d9b | 12654.23887 | 0.021775168 | 0.059506928 | 0.365926607 | 0.714419842 | 0.931210733 |
| Slc25a22 | 9585.713366 | 0.021777451 | 0.071479479 | 0.304667177 | 0.76061965 | 0.943105713 |
| Ugt8a | 1973.313662 | 0.021780301 | 0.085162976 | 0.255748476 | 0.798145061 | 0.952967259 |
| Lmx1b | 1.240293532 | 0.021782687 | 0.025870009 | 0.842005385 | 0.399784939 | NA |
| Sord | 1036.389798 | 0.02180017 | 0.066541149 | 0.327619385 | 0.743199463 | 0.939912916 |
| Krt18 | 42.28911412 | 0.021806327 | 0.044150049 | 0.493913987 | 0.621366926 | 0.899589902 |
| Sptlc2 | 1073.555255 | 0.021815234 | 0.064481565 | 0.338317382 | 0.735124026 | 0.936456337 |
| Rab3gap2 | 4680.873836 | 0.021815345 | 0.053047771 | 0.411239622 | 0.680896838 | 0.922382492 |
| Ttc26 | 475.2209501 | 0.021817156 | 0.079362129 | 0.274906385 | 0.783388161 | 0.949478217 |
| Il12rb1 | 43.07853109 | 0.021831397 | 0.086797857 | 0.251520002 | 0.8014121 | 0.953228631 |
| Fabp7 | 625.7803171 | 0.021839424 | 0.076355521 | 0.286022857 | 0.774860615 | 0.947238918 |
| Ino80c | 1034.418129 | 0.021846983 | 0.061254421 | 0.356659694 | 0.721346583 | 0.932513933 |
| Rps7 | 7373.48833 | 0.021851754 | 0.05620986 | 0.388753038 | 0.697458844 | 0.9261158 |
| Cpa5 | 2.351345804 | 0.021856112 | 0.032654277 | 0.669318512 | 0.503292321 | NA |
| Eda | 312.6311316 | 0.021863037 | 0.086870625 | 0.251673529 | 0.801293419 | 0.953228631 |
| Samd8 | 3659.628401 | 0.021874208 | 0.069466191 | 0.314889988 | 0.752845183 | 0.94183464 |
| Prmt7 | 1603.129662 | 0.02188022 | 0.060116805 | 0.363961785 | 0.715886546 | 0.93125489 |
| Wnt16 | 14.92581312 | 0.02188438 | 0.072888541 | 0.300244452 | 0.7639907 | NA |
| Tmem199 | 1347.041587 | 0.021884523 | 0.054912705 | 0.39853296 | 0.690237367 | 0.923860706 |
| Slc26a9 | 1.289261018 | 0.021908425 | 0.021491627 | 1.019393481 | 0.3080162 | NA |
| Tsr2 | 3590.564462 | 0.021908532 | 0.056485002 | 0.387864591 | 0.698116243 | 0.926409493 |
| Tanc2 | 8520.550719 | 0.021908795 | 0.062818167 | 0.348765271 | 0.727265538 | 0.9335794 |
| Cdc42se2 | 6939.169026 | 0.021924711 | 0.050082817 | 0.437769119 | 0.661553659 | 0.915428982 |
| Pum2 | 8756.970725 | 0.021931182 | 0.067661296 | 0.324131869 | 0.745838201 | 0.94070033 |
| Unc50 | 1183.274345 | 0.021931794 | 0.061814635 | 0.354799385 | 0.722739889 | 0.93255988 |
| Schip1 | 154.7298963 | 0.02193575 | 0.094931618 | 0.231068961 | 0.817261228 | 0.956302288 |
| Mogs | 1111.601066 | 0.021937842 | 0.062853047 | 0.349033865 | 0.727063886 | 0.9335794 |
| Naa15 | 2328.903842 | 0.021947668 | 0.068222019 | 0.321709453 | 0.747672816 | 0.9410864 |
| Rasgrp1 | 24575.28858 | 0.021947896 | 0.073902358 | 0.296985057 | 0.766477918 | 0.946032648 |
| Wwc2 | 1871.585389 | 0.021952764 | 0.056128761 | 0.391114359 | 0.695712707 | 0.925527709 |
| Upk2 | 1.281566481 | 0.021954595 | 0.026367979 | 0.832623366 | 0.405057177 | NA |
| Il13ra2 | 4.470941776 | 0.021975331 | 0.038788261 | 0.566545913 | 0.571022731 | NA |
| Nmur1 | 3.599724091 | 0.021984464 | 0.042801781 | 0.513634324 | 0.607507677 | NA |
| Fmo5 | 224.9472015 | 0.021985125 | 0.086570209 | 0.253957165 | 0.799528651 | 0.953228631 |
| Cdk16 | 12558.35849 | 0.021986153 | 0.045479837 | 0.483426387 | 0.628793016 | 0.904346423 |
| Tor1aip2 | 2340.794684 | 0.021990358 | 0.059759306 | 0.367982153 | 0.712886542 | 0.930507332 |
| Fam63b | 3658.079968 | 0.021990915 | 0.064822236 | 0.339249562 | 0.734421736 | 0.936456337 |
| Akap5 | 4841.566334 | 0.021995861 | 0.079376867 | 0.277106691 | 0.781698184 | 0.948979619 |
| Srsf12 | 632.7183814 | 0.022003799 | 0.063163927 | 0.348360211 | 0.72756968 | 0.9335794 |

| | | | | | | |
|---|---|---|---|---|---|---|
| Crls1 | 805.5518575 | -0.030234166 | 0.06367124 | -0.474848086 | 0.634895256 | 0.805134487 |
| Tmem229b | 1359.901048 | 0.029941176 | 0.063055014 | 0.474842104 | 0.63489952 | 0.805134487 |
| Esr1 | 113.5684499 | 0.071690651 | 0.151035662 | 0.474660422 | 0.635029032 | 0.805232819 |
| Dnaja2 | 6513.125564 | -0.03016937 | 0.063595764 | -0.474392764 | 0.635219852 | 0.805306026 |
| Nipal2 | 255.5690212 | 0.055937573 | 0.117898888 | 0.474453782 | 0.635176349 | 0.805306026 |
| Pomt2 | 1117.46432 | 0.035557092 | 0.074957916 | 0.474360731 | 0.635242691 | 0.805306026 |
| Cd79b | 8.48564503 | -0.074215849 | 0.156505261 | -0.474206736 | 0.63535249 | 0.805379325 |
| A430035B10Rik | 75.17145507 | -0.071311549 | 0.150429448 | -0.474053116 | 0.635462031 | 0.805452283 |
| Ppip5k1 | 5988.750691 | 0.020960222 | 0.044238303 | 0.473802571 | 0.635640701 | 0.805612845 |
| Cr1l | 589.6359846 | 0.038225272 | 0.080722558 | 0.473538902 | 0.635828754 | 0.805715213 |
| Stam2 | 1073.711164 | 0.027878438 | 0.058881034 | 0.473470593 | 0.635877476 | 0.805715213 |
| Tbc1d20 | 1526.655238 | -0.043847879 | 0.092606286 | -0.473487072 | 0.635865722 | 0.805715213 |
| Crlf3 | 419.024549 | 0.040256449 | 0.085074166 | 0.473192405 | 0.636075916 | 0.805900748 |
| Maged1 | 19974.96192 | 0.031420306 | 0.066447588 | 0.472858483 | 0.636314147 | 0.806136663 |
| Asns | 3716.161975 | 0.029218937 | 0.061814672 | 0.472686113 | 0.636437136 | 0.806226554 |
| Clcn3 | 8209.459066 | -0.031903345 | 0.067523717 | -0.472476142 | 0.636586967 | 0.806350431 |
| Eif3a | 6309.261379 | 0.030162352 | 0.063869387 | 0.472250526 | 0.636747979 | 0.806488448 |
| Gfpt2 | 318.0836347 | 0.065642091 | 0.139058743 | 0.47204577 | 0.636894119 | 0.806541682 |
| Tmem14a | 972.6597829 | -0.043375238 | 0.091885708 | -0.472056422 | 0.636886516 | 0.806541682 |
| BC027231 | 205.2101524 | 0.054256436 | 0.114974191 | 0.471901002 | 0.636997453 | 0.806606615 |
| Cdkn2d | 96.0730874 | -0.060669831 | 0.128704038 | -0.471390266 | 0.637362066 | 0.806879358 |
| Coasy | 858.2951255 | 0.03304097 | 0.070094062 | 0.471380448 | 0.637369076 | 0.806879358 |
| Ppcs | 165.8454288 | -0.057812227 | 0.122619701 | -0.47147584 | 0.637300969 | 0.806879358 |
| Lrrc61 | 1055.723586 | 0.040841093 | 0.086700568 | 0.471059114 | 0.637598522 | 0.807097323 |
| Slc22a15 | 330.0559183 | 0.038261491 | 0.081235723 | 0.470993422 | 0.637645432 | 0.807097323 |
| Pcnxl2 | 4929.598415 | -0.036151506 | 0.076798576 | -0.470731466 | 0.637832511 | 0.807202232 |
| Pms1 | 136.2562566 | 0.052964061 | 0.112499873 | 0.47079218 | 0.63778915 | 0.807202232 |
| Ing5 | 728.3948221 | 0.032042498 | 0.068123889 | 0.47035626 | 0.638100509 | 0.807468732 |
| Thrb | 2558.303045 | -0.04054195 | 0.086219519 | -0.470217773 | 0.638199438 | 0.807468732 |
| Zfp558 | 121.4400737 | -0.06567017 | 0.139644133 | -0.470268015 | 0.638163546 | 0.807468732 |
| Ppp1r14a | 238.0493898 | -0.075451213 | 0.160559174 | -0.46992776 | 0.63840663 | 0.807664925 |
| 6430531B16Rik | 20.30496894 | -0.078290249 | 0.166786641 | -0.469403593 | 0.638781181 | 0.808051631 |
| Insc | 163.0756786 | -0.075888158 | 0.161686389 | -0.469354029 | 0.638816602 | 0.808051631 |
| E030019B06Rik | 63.2509931 | 0.069704359 | 0.148576651 | 0.469147466 | 0.638964233 | 0.808106426 |
| Zfand2a | 2717.425749 | 0.038891908 | 0.082892948 | 0.469182345 | 0.638939304 | 0.808106426 |
| Irs2 | 1557.312647 | 0.074600314 | 0.159069273 | 0.468980038 | 0.639083905 | 0.808191807 |
| Alg8 | 259.7344675 | 0.0578476 | 0.1233721 | 0.468887213 | 0.639150257 | 0.808209751 |
| 2010001E11Rik | 12.37412882 | 0.078074646 | 0.166582229 | 0.468685326 | 0.639294578 | 0.808326276 |
| Bcar3 | 581.860314 | -0.044159158 | 0.094300111 | -0.468283206 | 0.639582076 | 0.808434982 |
| Elmo3 | 65.58835888 | 0.067704816 | 0.144518226 | 0.468486352 | 0.639436829 | 0.808434982 |
| Exosc7 | 540.6850564 | -0.032380062 | 0.069147808 | -0.468273159 | 0.639589261 | 0.808434982 |
| Npm1 | 4408.65652 | 0.02708196 | 0.057823161 | 0.468358336 | 0.639528357 | 0.808434982 |
| Hist1h1e | 29.58430736 | -0.071986683 | 0.153831611 | -0.467957676 | 0.639814857 | 0.808654164 |
| Nsfl1c | 2866.65993 | 0.030182816 | 0.064509754 | 0.467879884 | 0.639870489 | 0.808658513 |
| Col9a1 | 21.84694913 | 0.077559735 | 0.165872226 | 0.467587232 | 0.640079798 | 0.808726883 |
| Lage3 | 322.1109161 | 0.043188736 | 0.092365469 | 0.467585302 | 0.640081178 | 0.808726883 |
| Mttp | 568.8631284 | 0.037590598 | 0.080378834 | 0.467667872 | 0.64002212 | 0.808726883 |
| Snora43 | 11.35438356 | 0.07516654 | 0.160794692 | 0.467469038 | 0.640164339 | 0.808766004 |
| Zfp828 | 1738.115065 | -0.027394807 | 0.058625849 | -0.467282053 | 0.640298095 | 0.808869032 |
| Pgm3 | 533.2770265 | 0.039935569 | 0.085521257 | 0.466966581 | 0.640523788 | 0.80903468 |
| Smarcb1 | 2130.257326 | -0.019520604 | 0.041804235 | -0.466952793 | 0.640533653 | 0.80903468 |
| Arg2 | 89.28678067 | 0.069168244 | 0.148211613 | 0.466685725 | 0.640724744 | 0.809078173 |
| Mfng | 136.8498088 | -0.052970887 | 0.113473793 | -0.466811642 | 0.640634646 | 0.809078173 |
| Txnrd1 | 2598.766296 | -0.025895551 | 0.055482052 | -0.466737438 | 0.640687741 | 0.809078173 |
| S100a10 | 748.0651074 | 0.047743427 | 0.102320977 | 0.466604486 | 0.640782877 | 0.80908564 |
| Orc3 | 1425.807233 | 0.032510318 | 0.069702206 | 0.466417347 | 0.640916797 | 0.809188791 |
| Kank4 | 269.9284852 | 0.061530062 | 0.132010896 | 0.466098357 | 0.6411451 | 0.809345134 |
| Rbbp8 | 180.146347 | 0.051116358 | 0.109655671 | 0.46615334 | 0.641105745 | 0.809345134 |
| Gm11762 | 41.57431306 | -0.072042719 | 0.154622865 | -0.465925389 | 0.641268907 | 0.809435475 |
| 1700008J07Rik | 64.48689863 | 0.069664507 | 0.149982351 | 0.464484699 | 0.642300519 | 0.809467217 |
| 5430416O09Rik | 35.48401167 | -0.075149949 | 0.161694505 | -0.464765014 | 0.642099745 | 0.809467217 |
| AI429214 | 106.2603508 | 0.063301095 | 0.136184494 | 0.464818666 | 0.642061319 | 0.809467217 |
| Ap1s3 | 113.0627882 | -0.05548598 | 0.119177306 | -0.465575051 | 0.641519705 | 0.809467217 |
| Atg4d | 883.3401051 | 0.041653789 | 0.089686055 | 0.464439968 | 0.64233256 | 0.809467217 |
| Cog6 | 1972.273183 | -0.027612334 | 0.059434837 | -0.464581632 | 0.642231088 | 0.809467217 |
| Ctbs | 303.0671159 | 0.043223427 | 0.093044433 | 0.464546086 | 0.642256549 | 0.809467217 |
| Exosc2 | 659.4288818 | 0.032920093 | 0.070725644 | 0.465461901 | 0.641600714 | 0.809467217 |
| Fgf12 | 6128.982329 | -0.037239721 | 0.08019612 | -0.464358138 | 0.642391177 | 0.809467217 |
| Kcnab1 | 4747.755296 | -0.044038515 | 0.094836166 | -0.464364147 | 0.642386873 | 0.809467217 |
| LOC106740 | 451.7221732 | -0.036432101 | 0.078422138 | -0.464563987 | 0.642243727 | 0.809467217 |
| Lrrc18 | 22.21560156 | -0.078098068 | 0.168101648 | -0.464588357 | 0.642226272 | 0.809467217 |
| Mmachc | 608.8961714 | -0.034772388 | 0.074882599 | -0.464358725 | 0.642390757 | 0.809467217 |
| P2ry2 | 28.73660034 | 0.077240212 | 0.166331618 | 0.4643748 | 0.642379241 | 0.809467217 |
| Pla2g15 | 1118.920937 | -0.029002057 | 0.062279892 | -0.465672882 | 0.641449666 | 0.809467217 |
| Ppm1g | 2569.78848 | 0.029259172 | 0.062871196 | 0.465382777 | 0.641657365 | 0.809467217 |
| Rbm24 | 646.4649537 | -0.036240709 | 0.077871031 | -0.465393978 | 0.641649346 | 0.809467217 |
| 3-Sep | 23623.84203 | 0.034826124 | 0.074864522 | 0.465188629 | 0.641796381 | 0.809467217 |
| Smn1 | 484.3892752 | 0.042986331 | 0.092375547 | 0.46534318 | 0.641685718 | 0.809467217 |
| Spock1 | 12375.56968 | 0.036230737 | 0.07789592 | 0.465117261 | 0.641847485 | 0.809467217 |
| Txndc11 | 1126.17895 | 0.037741768 | 0.081223483 | 0.464665714 | 0.642170865 | 0.809467217 |
| Idh3b | 7487.130642 | -0.017931151 | 0.038642692 | -0.464024371 | 0.642630285 | 0.809702662 |
| Litaf | 749.9752729 | 0.061288984 | 0.13210472 | 0.463942422 | 0.642688998 | 0.809704323 |
| Zfp874a | 506.9761365 | 0.039051574 | 0.084185255 | 0.463876652 | 0.642736122 | 0.809704323 |
| Ppp1r12c | 4117.033617 | 0.034214404 | 0.073779011 | 0.463741693 | 0.642832822 | 0.809760305 |
| Tdrd3 | 760.0894646 | 0.043109963 | 0.092996371 | 0.463566076 | 0.642958664 | 0.809852983 |
| Zfp280b | 939.3192414 | 0.032622734 | 0.070399926 | 0.463391597 | 0.643083701 | 0.809944632 |
| Kti12 | 353.5307854 | -0.04043999 | 0.087332835 | -0.463055957 | 0.643324259 | 0.810181749 |
| Kifc1 | 43.49081701 | -0.071098126 | 0.15363888 | -0.462761286 | 0.643535484 | 0.810250186 |
| Ogfod3 | 184.4510648 | 0.050951897 | 0.110083739 | 0.462846716 | 0.643474243 | 0.810250186 |
| Shisa7 | 7628.324715 | 0.032565087 | 0.070367398 | 0.462786568 | 0.64351736 | 0.810250186 |
| Tmem251 | 296.2046812 | 0.055662039 | 0.120338157 | 0.462546884 | 0.643689189 | 0.810377858 |
| Ascc1 | 527.2957859 | 0.034356631 | 0.074301184 | 0.462396822 | 0.643796778 | 0.810447455 |
| Kcnh5 | 839.7195182 | -0.041034811 | 0.088798372 | -0.462112201 | 0.644000861 | 0.810638504 |
| Atf6b | 2132.13656 | -0.038069855 | 0.082441046 | -0.461782779 | 0.644237101 | 0.810795483 |
| Cacul1 | 2645.685604 | -0.03651697 | 0.079126606 | -0.461500519 | 0.644439549 | 0.810795483 |
| Ccng2 | 1612.423112 | -0.045487203 | 0.098526465 | -0.461674977 | 0.644314418 | 0.810795483 |
| Stambpl1 | 546.95164 | -0.031224655 | 0.067650664 | -0.461557259 | 0.644398851 | 0.810795483 |
| Tprgl | 5487.78155 | 0.039783933 | 0.086202298 | 0.461518247 | 0.644426833 | 0.810795483 |
| Trim35 | 6438.632577 | -0.025194698 | 0.054558144 | -0.461795362 | 0.644228077 | 0.810795483 |
| Rbm41 | 352.0077338 | 0.038024619 | 0.082418553 | 0.46135995 | 0.64454038 | 0.8108565 |
| Arl1 | 3060.159073 | -0.029576671 | 0.064158992 | -0.460990274 | 0.644805583 | 0.810992589 |
| Mir453 | 8.11833049 | 0.071683246 | 0.155453174 | 0.461124367 | 0.644709381 | 0.810992589 |
| Prkrir | 3042.46139 | -0.031624395 | 0.068592867 | -0.46104495 | 0.644766356 | 0.810992589 |
| Rabgap1 | 4390.546154 | 0.031322135 | 0.067974765 | 0.460790625 | 0.644948828 | 0.811106911 |
| 2010003O02Rik | 48.97811894 | -0.070286749 | 0.15259041 | -0.46062363 | 0.645068655 | 0.811191765 |
| Clec2d | 152.480538 | -0.062926963 | 0.136849732 | -0.459825251 | 0.645641657 | 0.811715225 |
| Fbxo42 | 1831.312051 | 0.031972883 | 0.069519861 | 0.459910056 | 0.645580782 | 0.811715225 |
| Fhl2 | 1283.653666 | -0.040536104 | 0.088169563 | -0.459751677 | 0.645694472 | 0.811715225 |
| Scml2 | 30.08057921 | 0.073288376 | 0.159393222 | 0.459796062 | 0.64566261 | 0.811715225 |
| AA388235 | 141.9485314 | -0.059199505 | 0.128928199 | -0.459166463 | 0.646114631 | 0.812045758 |
| Hint3 | 355.0223802 | -0.045625599 | 0.099351294 | -0.459235078 | 0.646065363 | 0.812045758 |
| Zbtb3 | 148.6405204 | -0.05184315 | 0.112890686 | -0.459233192 | 0.646066718 | 0.812045758 |
| Hmgcll1 | 549.440076 | 0.036707509 | 0.07998431 | 0.458933864 | 0.64628166 | 0.812189799 |
| S100a4 | 51.76095752 | -0.074293386 | 0.161955619 | -0.458726822 | 0.646430351 | 0.812310775 |
| 1700049G17Rik | 235.2861814 | -0.043938461 | 0.095812596 | -0.458587526 | 0.646530396 | 0.812370607 |
| Dmp1 | 287.5943455 | -0.061533929 | 0.13425517 | -0.458335642 | 0.646711321 | 0.812466165 |
| Rasgrp3 | 818.928538 | -0.053778524 | 0.11732108 | -0.458387562 | 0.646674026 | 0.812466165 |
| Cdc25c | 16.86389724 | -0.075574705 | 0.165025781 | -0.457956957 | 0.646983365 | 0.812742035 |
| Ccdc105 | 12.42612336 | -0.074603093 | 0.162967989 | -0.457777589 | 0.647112239 | 0.812772132 |
| Tec | 93.23658821 | -0.060406851 | 0.131940251 | -0.457834894 | 0.647071065 | 0.812772132 |
| Ahdc1 | 2918.780824 | 0.045619895 | 0.099764807 | 0.457274426 | 0.647473809 | 0.81311793 |
| Cdkn2aipnl | 1485.392265 | 0.035777928 | 0.078258651 | 0.457175378 | 0.647544994 | 0.81311793 |
| Fam132a | 418.1455108 | -0.052599448 | 0.115052543 | -0.45717762 | 0.647543383 | 0.81311793 |
| Parg | 1424.952453 | -0.029365409 | 0.064262247 | -0.456962056 | 0.647698319 | 0.813244551 |
| D630039A03Rik | 10.10624168 | 0.073131712 | 0.160078831 | 0.456848116 | 0.647780219 | 0.813278284 |
| Eid2 | 1070.31375 | -0.028622927 | 0.06267827 | -0.456664276 | 0.647912372 | 0.813278284 |
| Ntpcr | 535.700454 | -0.039067674 | 0.085538126 | -0.456728198 | 0.64786642 | 0.813278284 |
| S1pr3 | 456.3812282 | 0.040779623 | 0.089305108 | 0.456632597 | 0.647935145 | 0.813278284 |
| Pigl | 235.2228964 | 0.049615178 | 0.108711873 | 0.456391532 | 0.648108454 | 0.813429922 |
| Thra | 21059.7299 | -0.045967965 | 0.10074598 | -0.456275921 | 0.648191576 | 0.813468353 |
| Gm5506 | 30.76186486 | 0.074593729 | 0.163577823 | 0.456013701 | 0.648380126 | 0.813573183 |
| Mcat | 528.371208 | 0.038626753 | 0.084693925 | 0.456074666 | 0.648336287 | 0.813573183 |
| Zc3h6 | 789.3637171 | 0.043587911 | 0.095615814 | 0.455865083 | 0.648487 | 0.813641394 |
| Zfp865 | 883.9341585 | 0.034379732 | 0.075437343 | 0.455738912 | 0.648577737 | 0.813689349 |
| Trim29 | 8.766087716 | -0.067707983 | 0.148618369 | -0.455582866 | 0.648689966 | 0.813764257 |
| Foxn3 | 544.0779441 | 0.046529371 | 0.102177047 | 0.455379875 | 0.648835971 | 0.81388152 |
| Rassf9 | 39.91020945 | 0.074496909 | 0.163673543 | 0.455155472 | 0.648997392 | 0.814018101 |
| Pmfbp1 | 18.42114727 | -0.076498483 | 0.168101281 | -0.455073768 | 0.649056169 | 0.814025926 |
| Tmem176b | 965.4574841 | 0.038017261 | 0.083564418 | 0.454945561 | 0.649148404 | 0.814075708 |
| Fgd2 | 205.1216026 | 0.069778304 | 0.153454823 | 0.454715615 | 0.649313845 | 0.814217281 |
| Nubp1 | 504.2843087 | -0.039714995 | 0.087378881 | -0.454514802 | 0.649458341 | 0.814332567 |
| Amer1 | 482.1731231 | -0.032362914 | 0.071256227 | -0.454176649 | 0.649701689 | 0.814571773 |
| Poldip2 | 3466.790439 | -0.025661248 | 0.056548882 | -0.45378878 | 0.64998086 | 0.81485585 |
| Lrrc8d | 1805.216571 | 0.032432912 | 0.071524254 | 0.453453339 | 0.650222336 | 0.815026686 |
| Rdh11 | 833.1999514 | -0.032153673 | 0.070906748 | -0.453464216 | 0.650214505 | 0.815026686 |
| 1700101I11Rik | 59.12270159 | 0.06277025 | 0.138721497 | 0.452491153 | 0.650915193 | 0.815240171 |
| Arhgef40 | 915.7118899 | 0.03356339 | 0.074153536 | 0.452620227 | 0.650822231 | 0.815240171 |
| Bcat2 | 390.3451884 | -0.042566175 | 0.094071781 | -0.452486122 | 0.650918817 | 0.815240171 |
| Cers3 | 9.894783385 | -0.069910092 | 0.154359736 | -0.452903679 | 0.650618102 | 0.815240171 |
| Eif2a | 1564.040355 | 0.032534938 | 0.071846286 | 0.452840918 | 0.650663297 | 0.815240171 |
| Ero1l | 2179.01182 | 0.032475295 | 0.071702753 | 0.452915592 | 0.650609523 | 0.815240171 |
| Gpatch3 | 204.7593501 | 0.051966695 | 0.11483736 | 0.452524294 | 0.650891324 | 0.815240171 |
| Irak4 | 67.12996675 | 0.062631244 | 0.138402896 | 0.452528426 | 0.650888348 | 0.815240171 |
| Lpar2 | 47.08296158 | -0.072310639 | 0.159615308 | -0.453030725 | 0.650526617 | 0.815240171 |
| Rmdn2 | 206.9554802 | 0.047544689 | 0.105021284 | 0.452714792 | 0.650754127 | 0.815240171 |
| Slc39a12 | 2032.137974 | 0.033747344 | 0.074623246 | 0.452236339 | 0.651098732 | 0.815376338 |
| Tpcn1 | 2014.925785 | -0.043742618 | 0.096735242 | -0.45218906 | 0.651132789 | 0.815376338 |
| Myl6b | 530.3553611 | 0.049067344 | 0.108626687 | 0.451706164 | 0.651480677 | 0.815691978 |
| Stx6 | 3167.10966 | -0.024999767 | 0.055347107 | -0.451690593 | 0.651491896 | 0.815691978 |
| Tshz2 | 1339.691874 | 0.063975908 | 0.141658725 | 0.451619964 | 0.651542785 | 0.815691978 |
| Glipr2 | 136.9618425 | 0.074793951 | 0.165707345 | 0.451361713 | 0.651728873 | 0.815859025 |
| Abr | 24718.65854 | 0.03105003 | 0.068817821 | 0.451191705 | 0.651851388 | 0.815946469 |
| Lrrc47 | 2403.8609 | 0.032174108 | 0.071361493 | 0.450860912 | 0.652089797 | 0.816178957 |
| 0610010F05Rik | 1557.816758 | 0.037724703 | 0.083720224 | 0.450604426 | 0.652274676 | 0.816214639 |
| Gm2382 | 107.8433377 | 0.056205258 | 0.124728617 | 0.450620385 | 0.652263172 | 0.816214639 |
| Trip11 | 1137.316174 | -0.039455817 | 0.087562432 | -0.450602114 | 0.652276343 | 0.816214639 |
| H2-Ke2 | 1238.780192 | 0.033041088 | 0.073340288 | 0.450517561 | 0.652337295 | 0.81622499 |
| Gm12359 | 13.38375729 | 0.071538028 | 0.158849406 | 0.450351243 | 0.652457195 | 0.816309091 |
| 4930519F09Rik | 26.74828577 | 0.075445713 | 0.167621402 | 0.450095942 | 0.652641263 | 0.816472811 |
| Pradc1 | 472.2996927 | 0.044539604 | 0.098971714 | 0.450023571 | 0.652693445 | 0.816472811 |
| Zfp408 | 477.5856246 | 0.043997839 | 0.097826035 | 0.449755927 | 0.65288644 | 0.8166483 |
| Pcdhgb1 | 272.4836591 | 0.0589293 | 0.131071675 | 0.449595998 | 0.653001774 | 0.81672663 |
| Sfrp4 | 50.09720636 | 0.074228866 | 0.165263391 | 0.449154925 | 0.653319901 | 0.817058564 |
| Tmem106b | 6553.444947 | 0.03241647 | 0.072221733 | 0.448846465 | 0.653542417 | 0.817270881 |
| Ntsr2 | 2971.02922 | 0.042239034 | 0.094154842 | 0.44861245 | 0.653711251 | 0.817350075 |
| Stx11 | 9.107791921 | 0.072160704 | 0.160849942 | 0.44862126 | 0.653704895 | 0.817350075 |
| Cnpy3 | 2088.929784 | -0.036423744 | 0.081249169 | -0.448296817 | 0.653938998 | 0.817485766 |
| Gm10653 | 27.69384023 | -0.072514605 | 0.161816506 | -0.448128601 | 0.654060387 | 0.817485766 |
| Hdgf | 5879.122691 | 0.021998027 | 0.049092169 | 0.448096459 | 0.654083583 | 0.817485766 |
| Lace1 | 668.8188396 | -0.033444079 | 0.074594206 | -0.44834687 | 0.653902879 | 0.817485766 |
| Zcrb1 | 1089.552523 | -0.039575854 | 0.088291846 | -0.44823906 | 0.653980676 | 0.817485766 |
| Gm16291 | 9.455070472 | 0.07212736 | 0.16101337 | 0.447958827 | 0.654182912 | 0.817543961 |
| Sema3e | 1080.066587 | -0.044398145 | 0.099236718 | -0.447396349 | 0.654588911 | 0.817985369 |
| Paqr7 | 2562.656074 | -0.032820397 | 0.073372063 | -0.447314621 | 0.654647912 | 0.817993125 |
| Svep1 | 76.95649176 | -0.072267353 | 0.161631741 | -0.447111147 | 0.654794811 | 0.818110701 |
| Cul5 | 2491.067696 | -0.027085456 | 0.060595852 | -0.446985323 | 0.654885657 | 0.818111807 |
| Tob1 | 1055.776651 | -0.041060821 | 0.091866129 | -0.446963658 | 0.6549013 | 0.818111807 |
| Ccdc74a | 536.1031166 | 0.041413375 | 0.0926869 | 0.446809364 | 0.655012709 | 0.818185015 |
| Scly | 513.5363888 | -0.048388588 | 0.108325395 | -0.44669662 | 0.655094123 | 0.818185942 |
| Tcta | 1073.94419 | 0.041282172 | 0.092423723 | 0.44666208 | 0.655119065 | 0.818185942 |
| Pigh | 384.060621 | -0.035081672 | 0.078600488 | -0.446328928 | 0.655359663 | 0.818420458 |
| Adam9 | 2494.625411 | 0.038199862 | 0.085605948 | 0.446229067 | 0.655431788 | 0.818444562 |
| Brca2 | 180.8978839 | 0.060614413 | 0.136032602 | 0.445587393 | 0.655895319 | 0.818891384 |
| Zfp758 | 197.0190867 | -0.069051582 | 0.154948764 | -0.445641385 | 0.655856311 | 0.818891384 |
| Gapdh | 48807.23821 | 0.033139709 | 0.07444133 | 0.445178889 | 0.656190482 | 0.819193886 |
| Fads6 | 364.153409 | 0.054338956 | 0.122147139 | 0.444864748 | 0.6564175 | 0.819411275 |
| 2900009J06Rik | 22.27651492 | -0.074648082 | 0.167850891 | -0.444728542 | 0.65651594 | 0.819468136 |
| Naa38 | 439.5377711 | -0.035635262 | 0.080149293 | -0.444611061 | 0.656600853 | 0.819508105 |
| Pigw | 134.5914783 | 0.049260062 | 0.110830745 | 0.444462066 | 0.656708549 | 0.819576501 |
| Add2 | 16689.42055 | 0.03121146 | 0.070298841 | 0.443982568 | 0.657055187 | 0.819629267 |
| Alms1 | 509.4697043 | -0.041780623 | 0.094121387 | -0.443901477 | 0.657113817 | 0.819629267 |
| Cyb561 | 3109.103695 | -0.028977994 | 0.065273367 | -0.443948198 | 0.657080037 | 0.819629267 |
| Phax | 1353.429561 | 0.038948649 | 0.08774123 | 0.443903616 | 0.65711227 | 0.819629267 |
| Poc1a | 326.96966 | -0.043654704 | 0.098335979 | -0.443934195 | 0.657090161 | 0.819629267 |
| Sin3a | 1747.839872 | 0.039176265 | 0.088171367 | 0.444319585 | 0.656811544 | 0.819629267 |
| Tjp2 | 1384.081889 | 0.029367771 | 0.066159816 | 0.443891365 | 0.657121128 | 0.819629267 |
| Siglech | 617.0011948 | -0.074283856 | 0.167396317 | -0.443760398 | 0.657215823 | 0.819681395 |
| 2310010M20Rik | 11.41739012 | -0.071824328 | 0.162022355 | -0.443298878 | 0.657549567 | 0.819772818 |
| Cdca4 | 252.2988388 | 0.041586973 | 0.093813695 | 0.443293203 | 0.657553671 | 0.819772818 |
| Chmp6 | 938.9167627 | 0.029873973 | 0.067374425 | 0.443402266 | 0.657474797 | 0.819772818 |
| Impa1 | 2415.723805 | 0.041544345 | 0.093658474 | 0.443572735 | 0.657351522 | 0.819772818 |
| Rps19bp1 | 474.0858653 | -0.035015874 | 0.078968702 | -0.443414584 | 0.657465889 | 0.819772818 |
| Dido1 | 3205.551805 | 0.02477542 | 0.055933145 | 0.442947013 | 0.657804061 | 0.819944081 |
| Ftsj1 | 776.2437647 | 0.030171656 | 0.068125449 | 0.442883774 | 0.657849804 | 0.819944081 |
| Nabp1 | 318.4965549 | 0.046069633 | 0.104016544 | 0.442906784 | 0.65783316 | 0.819944081 |
| Exoc2 | 2049.909998 | 0.034194599 | 0.07724005 | 0.442705552 | 0.657978726 | 0.820038802 |
| Rgs4 | 19846.32822 | -0.056317083 | 0.127232733 | -0.442630464 | 0.658033046 | 0.820040539 |
| Gria4 | 3805.706546 | 0.035597261 | 0.080464327 | 0.442398045 | 0.658201194 | 0.820133819 |
| Psd4 | 70.10342554 | 0.062411874 | 0.141081827 | 0.442380672 | 0.658213763 | 0.820133819 |
| Pir | 195.0371162 | -0.045399639 | 0.102651384 | -0.442270114 | 0.658293755 | 0.82016429 |
| Slc35a5 | 1722.931907 | -0.03219888 | 0.072827154 | -0.442127397 | 0.65839702 | 0.82016429 |
| Srp19 | 856.6904535 | -0.03617017 | 0.081802648 | -0.442163805 | 0.658370676 | 0.82016429 |
| Srp72 | 3866.597356 | -0.025177673 | 0.056958238 | -0.442037425 | 0.658462124 | 0.820179448 |
| 0610009B22Rik | 425.3823604 | -0.042064852 | 0.095201805 | -0.441849308 | 0.658598255 | 0.820224001 |
| Bpgm | 2144.097761 | -0.025306591 | 0.057347561 | -0.441284521 | 0.659007031 | 0.820224001 |
| Hspb7 | 16.85478504 | -0.074091056 | 0.167909247 | -0.441256555 | 0.659027275 | 0.820224001 |
| Ipp | 389.1948952 | -0.041386966 | 0.093719792 | -0.441603261 | 0.658776324 | 0.820224001 |
| Pdzd2 | 4433.401548 | -0.074207528 | 0.16812151 | -0.441392231 | 0.658929066 | 0.820224001 |
| Ssbp1 | 476.9496582 | 0.033570134 | 0.076008685 | 0.441661815 | 0.658733945 | 0.820224001 |
| Suv420h1 | 2105.269485 | -0.027383208 | 0.062040035 | -0.441379639 | 0.65893818 | 0.820224001 |
| Syp | 53687.97567 | -0.025026263 | 0.056687246 | -0.441479597 | 0.658865829 | 0.820224001 |
| Yipf3 | 1622.891288 | 0.025383006 | 0.057460957 | 0.441743529 | 0.658674807 | 0.820224001 |
| Zfp275 | 1303.542341 | -0.032993266 | 0.074683434 | -0.441774899 | 0.658652104 | 0.820224001 |
| Trank1 | 17780.32987 | 0.054604414 | 0.123827865 | 0.440970325 | 0.65923448 | 0.820415987 |
| 2210013O21Rik | 736.0575477 | -0.02988023 | 0.067844338 | -0.440423344 | 0.65963052 | 0.820537487 |

| | | | | | | |
|---|---|---|---|---|---|---|
| 1700028J19Rik | 45.18845501 | 0.022005786 | 0.090735712 | 0.242526182 | 0.808372472 | 0.955716697 |
| Acss2 | 2236.448779 | 0.022007482 | 0.057048406 | 0.385768574 | 0.699668072 | 0.926781946 |
| Mtmr1 | 1953.408088 | 0.022009199 | 0.071699126 | 0.306966072 | 0.758869197 | 0.942526199 |
| Zbtb7a | 5580.955499 | 0.022015541 | 0.063205363 | 0.34831761 | 0.72760167 | 0.9335794 |
| Gm1943 | 342.0960388 | 0.02202025 | 0.076603251 | 0.287458426 | 0.773761331 | 0.947121017 |
| Smad5 | 1663.912966 | 0.022045124 | 0.058904631 | 0.374251112 | 0.708217501 | 0.929959854 |
| Aqp7 | 1.536631908 | 0.022061047 | 0.028975234 | 0.76137599 | 0.446432524 | NA |
| Gm15348 | 0.666283812 | 0.022061902 | 0.017432894 | 1.265532984 | 0.205680332 | NA |
| L2hgdh | 879.184282 | 0.022079276 | 0.064327279 | 0.343233482 | 0.731422818 | 0.935350634 |
| Alg3 | 462.0286813 | 0.022104204 | 0.076790211 | 0.287851841 | 0.773460152 | 0.947121017 |
| Bmp6 | 358.350517 | 0.022114101 | 0.095101697 | 0.232531086 | 0.816125546 | 0.956264994 |
| Proser2 | 155.2120193 | 0.022116511 | 0.090370605 | 0.244731247 | 0.806664532 | 0.955587482 |
| Dtnbp1 | 1937.890772 | 0.022165269 | 0.048404519 | 0.457917353 | 0.647011819 | 0.912750424 |
| Slc22a23 | 4033.449442 | 0.022166087 | 0.055209575 | 0.4014899 | 0.688059473 | 0.923860706 |
| Armc10 | 898.1849763 | 0.022171213 | 0.065849637 | 0.336694534 | 0.736347184 | 0.93717515 |
| Lrp5 | 532.5823767 | 0.02218886 | 0.072565448 | 0.30577721 | 0.759774281 | 0.942724969 |
| U2surp | 2823.684805 | 0.022200827 | 0.059937897 | 0.370397169 | 0.711086583 | 0.930507332 |
| Nkd2 | 138.6003437 | 0.022207888 | 0.093504098 | 0.237507111 | 0.812263403 | 0.956219756 |
| Stpg1 | 68.19018347 | 0.022217492 | 0.094722402 | 0.234553724 | 0.814555131 | 0.956264994 |
| Kbtbd8 | 299.4242575 | 0.022241202 | 0.086128761 | 0.258232002 | 0.796227861 | 0.952734447 |
| Lsm7 | 707.4486536 | 0.022246909 | 0.084323374 | 0.263828495 | 0.791912076 | 0.951728598 |
| Prps2 | 1619.561725 | 0.02226279 | 0.069905264 | 0.318470866 | 0.750127791 | 0.9410864 |
| Gjb1 | 586.3641472 | 0.022295793 | 0.092263332 | 0.241653894 | 0.809048358 | 0.955725085 |
| Shox2 | 21.8812246 | 0.02229851 | 0.030982158 | 0.719721003 | 0.471696792 | NA |
| Cenpw | 64.5576932 | 0.022313166 | 0.09343041 | 0.238821234 | 0.811244204 | 0.956135608 |
| Tbrg3 | 1.057718031 | 0.022322322 | 0.024029566 | 0.928952332 | 0.35291379 | NA |
| Isx | 2.760722691 | 0.022349975 | 0.039740813 | 0.562393496 | 0.57384795 | NA |
| Anapc7 | 1562.940931 | 0.022351378 | 0.048296319 | 0.462796732 | 0.643510074 | 0.911258473 |
| Zfp944 | 347.5378677 | 0.022357449 | 0.089844013 | 0.248847394 | 0.803478829 | 0.953754272 |
| BC051226 | 109.1446425 | 0.022375507 | 0.094979939 | 0.235581399 | 0.813757508 | 0.956264994 |
| Emc8 | 2824.10095 | 0.022421439 | 0.049460937 | 0.453316097 | 0.650321143 | 0.914000665 |
| Usp34 | 6414.367507 | 0.022449512 | 0.071455405 | 0.314175145 | 0.753388019 | 0.94183464 |
| Secisbp2l | 6589.039596 | 0.022457902 | 0.064861717 | 0.346242786 | 0.729160262 | 0.933996867 |
| Smek2 | 1810.450033 | 0.022471044 | 0.081601671 | 0.275374804 | 0.783028299 | 0.949438153 |
| 4930404N11Rik | 3.326830391 | 0.022474632 | 0.036905857 | 0.608971961 | 0.542543024 | NA |
| Elmod1 | 10106.37323 | 0.022480311 | 0.068689416 | 0.327274749 | 0.743460089 | 0.939912916 |
| 4930593A02Rik | 2.444879053 | 0.022512884 | 0.033227568 | 0.677536322 | 0.498065733 | NA |
| Pcyox1l | 626.0822316 | 0.022521382 | 0.072720106 | 0.309699518 | 0.75678947 | 0.942139348 |
| Nfkbil1 | 287.0920814 | 0.022524846 | 0.078856073 | 0.285645034 | 0.775150008 | 0.947238918 |
| Gm2721 | 12.50821608 | 0.022536024 | 0.071172376 | 0.316640046 | 0.751516746 | NA |
| Prrt3 | 2076.388553 | 0.022547218 | 0.070954105 | 0.317771869 | 0.750657991 | 0.9410864 |
| Tesk1 | 4154.805602 | 0.022576909 | 0.057988643 | 0.389333279 | 0.697029621 | 0.925997259 |
| Dlgap5 | 7.278894875 | 0.02260857 | 0.05672639 | 0.398554704 | 0.690221343 | NA |
| Atp5j | 4228.157154 | 0.022609236 | 0.062195407 | 0.363519393 | 0.716216929 | 0.93125489 |
| Prr16 | 310.0110801 | 0.022615427 | 0.093757758 | 0.241211261 | 0.809391383 | 0.955725085 |
| Fam160a2 | 3795.860466 | 0.022624565 | 0.063380869 | 0.356962054 | 0.721120213 | 0.932513933 |
| Pklr | 4.148702938 | 0.022625359 | 0.044977737 | 0.503034616 | 0.614939935 | NA |
| Plxdc1 | 2028.460571 | 0.022649812 | 0.057340869 | 0.39500294 | 0.692840719 | 0.924216421 |
| Tmtc3 | 602.0030987 | 0.022652146 | 0.086308356 | 0.262455999 | 0.792969904 | 0.951728598 |
| Wfikkn1 | 6.71714547 | 0.022654796 | 0.055724431 | 0.406550505 | 0.684338156 | NA |
| Egfl7 | 995.6964821 | 0.022663628 | 0.079181319 | 0.286224432 | 0.774706232 | 0.947238918 |
| Kpnb1 | 9373.113242 | 0.022676518 | 0.056695977 | 0.399966967 | 0.689180847 | 0.923860706 |
| Zfp30 | 463.121794 | 0.022676893 | 0.072666845 | 0.31206657 | 0.754989932 | 0.942099798 |
| Prr18 | 1469.842065 | 0.022680692 | 0.082985821 | 0.27330804 | 0.784616435 | 0.949699053 |
| Otx2 | 72.66662828 | 0.022684811 | 0.040327703 | 0.56251186 | 0.573767325 | 0.879633264 |
| Sertad2 | 1442.249081 | 0.022695841 | 0.066489124 | 0.341346676 | 0.732842613 | 0.936094952 |
| Crygs | 0.530631763 | 0.022698452 | 0.017691822 | 1.282991185 | 0.199495164 | NA |
| Xab2 | 1675.25958 | 0.022701025 | 0.060764149 | 0.373592417 | 0.708707577 | 0.93015267 |
| Ercc8 | 429.8211814 | 0.022705932 | 0.073408008 | 0.309311376 | 0.757084678 | 0.942139348 |
| 4632415K11Rik | 409.7477116 | 0.022728675 | 0.070710947 | 0.321430778 | 0.747883962 | 0.9410864 |
| Ddx28 | 497.6684505 | 0.022730159 | 0.071067532 | 0.319838863 | 0.749090485 | 0.9410864 |
| Tmem230 | 988.032073 | 0.0227386 | 0.069034145 | 0.329381931 | 0.741867024 | 0.93956033 |
| Cast | 557.0704538 | 0.022740671 | 0.075623083 | 0.300710714 | 0.763635098 | 0.944771607 |
| Yrdc | 716.7426106 | 0.022760636 | 0.061408238 | 0.370644662 | 0.710902212 | 0.930507332 |
| Gabpb1 | 651.9286205 | 0.022780644 | 0.067225149 | 0.338870864 | 0.734707014 | 0.936456337 |
| Adcy5 | 12511.62248 | 0.02278219 | 0.074240345 | 0.306870752 | 0.758941752 | 0.942526199 |
| 2900093L17Rik | 3.222003427 | 0.022783919 | 0.03217361 | 0.708155484 | 0.478848707 | NA |
| Lbx2 | 1.972697308 | 0.022794599 | 0.030179853 | 0.755291924 | 0.450073844 | NA |
| Syt17 | 1317.18047 | 0.022794903 | 0.089662332 | 0.254230543 | 0.799317456 | 0.953196401 |
| Cnot6 | 1425.858772 | 0.022808261 | 0.075718989 | 0.30122247 | 0.763244857 | 0.944610615 |
| Tnc | 181.5854343 | 0.022808262 | 0.094731678 | 0.240767003 | 0.809735705 | 0.955876688 |
| Polr3e | 1564.639121 | 0.022816055 | 0.060040525 | 0.380010928 | 0.703937303 | 0.928127792 |
| Cyp4f39 | 0.745705075 | 0.02282669 | 0.017723662 | 1.287921777 | 0.197773194 | NA |
| Crtc3 | 787.9497277 | 0.022836673 | 0.066889336 | 0.341409768 | 0.732795122 | 0.936094952 |
| Col8a1 | 191.7011473 | 0.022856434 | 0.062197897 | 0.367479204 | 0.713261601 | 0.930507332 |
| Slc10a6 | 1.22971801 | 0.022873052 | 0.023106703 | 0.989888154 | 0.32222879 | NA |
| Rpl29 | 5317.366329 | 0.022873875 | 0.05256604 | 0.435145478 | 0.663456835 | 0.915428982 |
| Cdk8 | 920.2883986 | 0.022879126 | 0.065685001 | 0.348315831 | 0.727603006 | 0.9335794 |
| Zfp354c | 1898.580178 | 0.022893037 | 0.072897803 | 0.314042899 | 0.753488457 | 0.94183464 |
| Tmem237 | 840.2946033 | 0.022916663 | 0.072609113 | 0.315616903 | 0.752293306 | 0.941565791 |
| Prps1l3 | 593.845782 | 0.022920196 | 0.078812279 | 0.290820118 | 0.771188898 | 0.946250219 |
| Gpat2 | 18.46041544 | 0.022928917 | 0.073913758 | 0.310211764 | 0.756399925 | NA |
| D3Ertd751e | 659.4676532 | 0.022956113 | 0.076570242 | 0.299804629 | 0.764326184 | 0.944872045 |
| Irgc1 | 4.411005129 | 0.022956377 | 0.036739844 | 0.624835997 | 0.532078703 | NA |
| Cltc | 29028.76708 | 0.022958907 | 0.064851935 | 0.35402039 | 0.723323602 | 0.932617157 |
| Eif4ebp2 | 942.2518647 | 0.022967861 | 0.070621854 | 0.325223135 | 0.745012201 | 0.940632024 |
| Bik | 2.99264569 | 0.02298276 | 0.040166346 | 0.572189454 | 0.567193631 | NA |
| Vmo1 | 4.232400386 | 0.022988775 | 0.034590264 | 0.664602468 | 0.50630479 | NA |
| Itih5 | 2002.811949 | 0.023006702 | 0.081323282 | 0.28290425 | 0.777250236 | 0.947506677 |
| 2310035C23Rik | 2928.944225 | 0.023008079 | 0.065194681 | 0.352913445 | 0.724153331 | 0.932714927 |
| Fabp5 | 2735.544513 | 0.023012048 | 0.059183254 | 0.388827012 | 0.697404117 | 0.9261158 |
| Ccdc175 | 1.565342685 | 0.023012299 | 0.025942427 | 0.88705263 | 0.375050566 | NA |
| 2210416O15Rik | 2.824126492 | 0.023029475 | 0.039398146 | 0.584531945 | 0.55886248 | NA |
| Scgb3a1 | 2.59611355 | 0.023033629 | 0.039430957 | 0.584150896 | 0.559118795 | NA |
| Dedd | 1124.857376 | 0.023051792 | 0.053212355 | 0.433203751 | 0.664866754 | 0.915428982 |
| Mapre3 | 7756.615354 | 0.023056182 | 0.035379348 | 0.651684753 | 0.51460456 | 0.853195313 |
| 2810013P06Rik | 639.4029957 | 0.023064131 | 0.06277741 | 0.367395387 | 0.713324112 | 0.930507332 |
| Chmp7 | 4021.862568 | 0.02308186 | 0.045412824 | 0.508267455 | 0.61126579 | 0.894995149 |
| Tbcel | 2379.610695 | 0.023094935 | 0.057354593 | 0.40266931 | 0.687191514 | 0.923860706 |
| Slc1a7 | 4.297115518 | 0.023101447 | 0.048966021 | 0.471785265 | 0.63708007 | NA |
| Lamtor4 | 945.7629724 | 0.023107107 | 0.064903089 | 0.356024764 | 0.72182202 | 0.932513933 |
| 2310022A10Rik | 986.4513492 | 0.0231074 | 0.056365885 | 0.409953638 | 0.681839957 | 0.922544285 |
| Art4 | 3.398784008 | 0.023118987 | 0.040225731 | 0.5747313 | 0.565473036 | NA |
| Rwdd2b | 376.9818685 | 0.023120198 | 0.075536806 | 0.306078578 | 0.759544818 | 0.94272457 |
| Gm4349 | 25.20428387 | 0.023144844 | 0.081756411 | 0.283095162 | 0.77710389 | 0.947474493 |
| Pate2 | 8.388622204 | 0.023149862 | 0.064205341 | 0.360559752 | 0.718428582 | NA |
| Mcpt4 | 0.554788539 | 0.023153921 | 0.017865547 | 1.296009576 | 0.194972186 | NA |
| Daam2 | 4682.238942 | 0.023205309 | 0.075518487 | 0.307279844 | 0.758630376 | 0.942526199 |
| Sema6d | 3027.15409 | 0.023206266 | 0.077528625 | 0.299325128 | 0.764691984 | 0.944872045 |
| Fntb | 1331.650984 | 0.023212413 | 0.053002642 | 0.43794822 | 0.661423819 | 0.915428982 |
| Adap1 | 8298.297197 | 0.023231733 | 0.057282097 | 0.405567082 | 0.68506072 | 0.923835238 |
| Gch1 | 65.3091783 | 0.023253698 | 0.090984989 | 0.255577296 | 0.798277251 | 0.952967259 |
| Kcns1 | 919.9171472 | 0.023293136 | 0.079288563 | 0.293776741 | 0.76892851 | 0.94619074 |
| Ankrd40 | 5968.972367 | 0.023303639 | 0.050476687 | 0.46167133 | 0.644317034 | 0.912152041 |
| Snord57 | 5.688914333 | 0.023323226 | 0.053019859 | 0.439896038 | 0.660012406 | NA |
| Sf3a1 | 3634.542239 | 0.02333369 | 0.042098125 | 0.554269113 | 0.579394685 | 0.88248791 |
| Egfl6 | 117.1479493 | 0.023349293 | 0.093095579 | 0.250809904 | 0.801961084 | 0.95333733 |
| Armc9 | 1231.224553 | 0.023351182 | 0.052096708 | 0.448227584 | 0.653988957 | 0.914765873 |
| CK137956 | 94.87982515 | 0.02335236 | 0.093832717 | 0.248872253 | 0.803459599 | 0.953754272 |
| Slc7a8 | 3897.894192 | 0.023364539 | 0.056077707 | 0.416645769 | 0.676937527 | 0.921056062 |
| Msra | 1158.349555 | 0.023365644 | 0.05655811 | 0.413126326 | 0.679514062 | 0.921712697 |
| Sptb | 9619.602592 | 0.023375601 | 0.054680216 | 0.427496489 | 0.669017742 | 0.917484508 |
| Kbtbd11 | 6505.58947 | 0.023380403 | 0.053914911 | 0.433653751 | 0.664539896 | 0.915428982 |
| Ube2e3 | 2049.9747 | 0.023385998 | 0.064273263 | 0.363852663 | 0.715968035 | 0.93125489 |
| Ift57 | 1516.183053 | 0.023395722 | 0.061851162 | 0.378258406 | 0.705238641 | 0.928783753 |
| Mrps16 | 633.3032356 | 0.023397705 | 0.070365297 | 0.332517677 | 0.739498394 | 0.938418054 |
| Cyp1a1 | 4.659616567 | 0.0234061 | 0.048105559 | 0.486557061 | 0.626572259 | NA |
| Dusp9 | 43.9491792 | 0.023407297 | 0.092942666 | 0.251846633 | 0.801159611 | 0.953228631 |
| Add2 | 17912.42 | 0.023409474 | 0.055552613 | 0.421392848 | 0.673468241 | 0.919353963 |
| Asb12 | 0.989058214 | 0.023430978 | 0.02095495 | 1.11815956 | 0.263498853 | NA |
| Gpx2 | 2.931517556 | 0.023432331 | 0.039221468 | 0.597436351 | 0.55021609 | NA |
| Mrps17 | 1281.809489 | 0.023435358 | 0.051127161 | 0.45837393 | 0.646683818 | 0.912750424 |
| Ace | 1335.84005 | 0.023437112 | 0.073959813 | 0.316889822 | 0.751327205 | 0.9410864 |
| Epb4.9 | 17497.56027 | 0.023445491 | 0.058483978 | 0.400887415 | 0.688503016 | 0.923860706 |
| Tifa | 271.1922725 | 0.023457389 | 0.076682537 | 0.305902627 | 0.759678786 | 0.94272457 |
| Calm1 | 135982.678 | 0.023458414 | 0.063040015 | 0.372119421 | 0.709803937 | 0.930224827 |
| Pcgf1 | 415.2666676 | 0.023461503 | 0.073222087 | 0.320415655 | 0.74865326 | 0.9410864 |
| Rps24 | 8186.68895 | 0.023471248 | 0.073874219 | 0.317719069 | 0.750698045 | 0.9410864 |
| Hist1h4k | 10.32971507 | 0.023473689 | 0.064974981 | 0.361272729 | 0.717895578 | NA |
| Glyctk | 251.2299951 | 0.023474944 | 0.086890872 | 0.270165828 | 0.787032681 | 0.950722388 |
| E230016K23Rik | 3.566196814 | 0.023538358 | 0.043084847 | 0.546325672 | 0.584842092 | NA |
| Pop5 | 1371.083626 | 0.023553936 | 0.061800414 | 0.381129098 | 0.703107457 | 0.927855161 |
| Vmn1r206 | 3.215882515 | 0.02356743 | 0.036786954 | 0.64064641 | 0.52175244 | NA |
| Wdr74 | 915.4819834 | 0.023567839 | 0.061320205 | 0.384340519 | 0.70072608 | 0.92719503 |
| Rasl2-9 | 4.7721782 | 0.023574949 | 0.049948539 | 0.471984751 | 0.636937673 | NA |
| Ncor2 | 6690.949239 | 0.023578935 | 0.064165434 | 0.367470977 | 0.713267737 | 0.930507332 |
| Nop58 | 1090.33897 | 0.023607679 | 0.08022917 | 0.294253063 | 0.768564536 | 0.94619074 |
| Gpr182 | 68.95347525 | 0.02363312 | 0.092148358 | 0.256468161 | 0.797589363 | 0.952967259 |
| Slc41a3 | 237.5515732 | 0.023637933 | 0.093416378 | 0.253038426 | 0.80023852 | 0.953228631 |
| Raet1d | 0.584133578 | 0.023687208 | 0.018066562 | 1.311107696 | 0.189821381 | NA |
| Rasef | 4.299766666 | 0.023691842 | 0.042559055 | 0.556681581 | 0.577745004 | NA |
| Sars2 | 480.497672 | 0.023700462 | 0.080447266 | 0.294608675 | 0.768292834 | 0.94619074 |
| Uox | 26.57085437 | 0.023702703 | 0.084925308 | 0.279100582 | 0.780167637 | 0.948894514 |
| Gria3 | 10301.9165 | 0.023709418 | 0.064543596 | 0.367339586 | 0.71336573 | 0.930507332 |
| Ank2 | 20077.30876 | 0.023721301 | 0.057356893 | 0.413573672 | 0.679186358 | 0.921471438 |
| Rpl37 | 4554.883024 | 0.023721756 | 0.053245521 | 0.445516467 | 0.655946562 | 0.915208181 |
| Prnd | 7.235977917 | 0.023725526 | 0.058921325 | 0.402664498 | 0.687195054 | NA |
| Mzt1 | 1943.707297 | 0.023735609 | 0.059884797 | 0.396354506 | 0.691843523 | 0.923860706 |
| Zfp148 | 4427.75554 | 0.023746546 | 0.072192618 | 0.328933161 | 0.742206209 | 0.939873367 |
| Masp1 | 428.8092392 | 0.023746758 | 0.071331488 | 0.332907084 | 0.739204422 | 0.938362618 |
| Chia | 84.09555846 | 0.023749787 | 0.094589631 | 0.251082356 | 0.801750437 | 0.95333733 |
| Gm4788 | 3.384828324 | 0.023751532 | 0.040942184 | 0.580123719 | 0.56183119 | NA |
| Ramp1 | 2053.921042 | 0.023755638 | 0.052088416 | 0.456063733 | 0.648344148 | 0.91329006 |
| Ppp1r3b | 58.30968144 | 0.023756377 | 0.0896287 | 0.265053231 | 0.790968455 | 0.951728598 |
| 5730435O14Rik | 1.298720366 | 0.023769143 | 0.027309396 | 0.870365005 | 0.384100965 | NA |
| Mog | 2643.008173 | 0.023782822 | 0.085913559 | 0.276822691 | 0.781916257 | 0.948979619 |
| Trim40 | 4.971032975 | 0.023786152 | 0.049627703 | 0.479291817 | 0.631731042 | NA |
| Ces1g | 0.719463989 | 0.023800813 | 0.018095136 | 1.315315496 | 0.188403892 | NA |
| Cntnap5b | 598.0942971 | 0.02382705 | 0.092763122 | 0.256859077 | 0.797287562 | 0.952897915 |
| Dlgap1 | 21014.70375 | 0.023847536 | 0.059519792 | 0.400665653 | 0.688666302 | 0.923860706 |
| Pank3 | 2538.385381 | 0.023885694 | 0.06020122 | 0.396764292 | 0.691541285 | 0.923860706 |
| Nt5dc2 | 131.2597592 | 0.023903067 | 0.095098551 | 0.25135048 | 0.80154315 | 0.953268036 |
| Tbc1d5 | 2166.671719 | 0.023909471 | 0.058399571 | 0.409411755 | 0.682237515 | 0.922544285 |
| Mrm1 | 584.6644151 | 0.023918672 | 0.069671518 | 0.343306318 | 0.731368029 | 0.935350634 |
| Fam105b | 609.8190101 | 0.023942285 | 0.061719884 | 0.38791851 | 0.69807634 | 0.926409493 |
| Kcnk7 | 3.976956415 | 0.023963071 | 0.040999393 | 0.584473815 | 0.558901578 | NA |
| C8b | 3.119695849 | 0.023968605 | 0.042419168 | 0.565041851 | 0.572045298 | NA |
| Hist1h1e | 38.04432562 | 0.023970563 | 0.087154284 | 0.275035965 | 0.783288607 | 0.949478217 |
| Mir365-1 | 1.79566016 | 0.023982437 | 0.027152218 | 0.883258853 | 0.37709643 | NA |
| AY512931 | 5.63855703 | 0.023987368 | 0.042958567 | 0.558383801 | 0.576582332 | NA |
| Slc7a14 | 6587.15055 | 0.024008576 | 0.073166311 | 0.32813703 | 0.742808057 | 0.939912916 |
| Lrrc14 | 839.879937 | 0.024018273 | 0.069432199 | 0.345924137 | 0.729399729 | 0.933996867 |
| Taco1 | 465.3766403 | 0.024033516 | 0.066355369 | 0.362193989 | 0.717207071 | 0.93125489 |
| Ralgapa2 | 1230.757632 | 0.02403609 | 0.062844346 | 0.382470208 | 0.702112623 | 0.927762026 |
| 3110040N11Rik | 228.5261906 | 0.02404969 | 0.090338219 | 0.26621833 | 0.790071067 | 0.951607109 |
| Snrnp40 | 750.3586333 | 0.024052857 | 0.068808681 | 0.349561369 | 0.726667907 | 0.9335794 |
| Barhl1 | 4.660516633 | 0.024079839 | 0.035562798 | 0.677107566 | 0.498337709 | NA |
| 1700061G19Rik | 6.024746964 | 0.024085301 | 0.051717878 | 0.465705519 | 0.641426301 | NA |
| Palld | 98.49167647 | 0.024086298 | 0.095067713 | 0.253359392 | 0.799990505 | 0.953228631 |
| Slc35e2 | 2203.66353 | 0.024092597 | 0.072899091 | 0.330492427 | 0.741027914 | 0.939200025 |
| Ch25h | 15.55986639 | 0.024108772 | 0.073570074 | 0.327698083 | 0.743139953 | NA |
| Snph | 16326.58469 | 0.024113769 | 0.054184999 | 0.445026658 | 0.656300489 | 0.915208181 |
| Spata7 | 773.5487497 | 0.024123455 | 0.075960864 | 0.31757742 | 0.750805504 | 0.9410864 |
| Zfp637 | 993.9707072 | 0.024133837 | 0.062140887 | 0.388372911 | 0.697740088 | 0.926197242 |
| Sds | 6.973786394 | 0.024159127 | 0.04995144 | 0.483652267 | 0.628632674 | NA |
| Vps37c | 919.1336124 | 0.024167487 | 0.062422242 | 0.387161469 | 0.698636673 | 0.926409493 |
| Ribc1 | 13.05053646 | 0.024173352 | 0.069102748 | 0.349817524 | 0.726475646 | NA |
| Sh2b3 | 742.4839146 | 0.024207473 | 0.068265829 | 0.354606007 | 0.722884775 | 0.93255988 |
| Ccdc40 | 88.92466881 | 0.024208002 | 0.09445077 | 0.256302857 | 0.797716991 | 0.952967259 |
| Rpl28 | 4515.451775 | 0.024246335 | 0.06496218 | 0.373237712 | 0.70897153 | 0.930183885 |
| Amhr2 | 37.02119871 | 0.024255718 | 0.088470032 | 0.274168744 | 0.783954947 | 0.949699053 |
| Mir3091 | 2.273203209 | 0.02428218 | 0.03056087 | 0.794551329 | 0.42687454 | NA |
| Ocrl | 3778.350686 | 0.024295473 | 0.054836488 | 0.443053049 | 0.657727364 | 0.915208181 |
| Rab2b | 1865.848115 | 0.024297564 | 0.054887469 | 0.442679619 | 0.657997486 | 0.915208181 |
| Gnb2 | 10319.40907 | 0.024367338 | 0.064149075 | 0.379854864 | 0.704053153 | 0.928127792 |
| Crmp1 | 7886.429112 | 0.024367345 | 0.050966152 | 0.478108386 | 0.632573064 | 0.906637358 |
| Nubp2 | 899.798738 | 0.024370924 | 0.064772985 | 0.376251363 | 0.706730034 | 0.929476902 |
| Rit1 | 1070.317851 | 0.024371153 | 0.058155422 | 0.419069319 | 0.675165472 | 0.919907397 |
| Ttc19 | 3375.014419 | 0.024371592 | 0.069336924 | 0.351495146 | 0.725216914 | 0.933334512 |
| Dspp | 9.237751076 | 0.024386223 | 0.060736957 | 0.401505513 | 0.68804798 | NA |
| Gys1 | 896.1102894 | 0.024397904 | 0.078566276 | 0.31053914 | 0.756151 | 0.942139348 |
| Sept8 | 9478.225228 | 0.024399493 | 0.041855193 | 0.582950185 | 0.559926835 | 0.873713575 |
| Trim3 | 5302.547921 | 0.02440381 | 0.060890846 | 0.400779614 | 0.688582389 | 0.923860706 |
| Bax | 1044.168073 | 0.024409371 | 0.067969804 | 0.359120808 | 0.719504716 | 0.932251852 |
| Nfatc1 | 295.3470599 | 0.024411879 | 0.08329474 | 0.293078275 | 0.769462322 | 0.94619074 |
| Ms4a4d | 1.866797768 | 0.024418589 | 0.027747486 | 0.880028892 | 0.378843658 | NA |
| Usp33 | 6258.813554 | 0.024435562 | 0.059454373 | 0.410996876 | 0.681074826 | 0.922451017 |
| Grcc10 | 2489.403652 | 0.024454303 | 0.058792237 | 0.415944421 | 0.677450673 | 0.921064066 |
| Apc2 | 7163.901947 | 0.02446501 | 0.054401959 | 0.449708258 | 0.652920816 | 0.914683958 |
| Mapkap1 | 2053.297051 | 0.024472088 | 0.054765265 | 0.446854193 | 0.65498034 | 0.915208181 |
| Mgarp | 3.184530062 | 0.024476566 | 0.037182738 | 0.658277668 | 0.510359726 | NA |
| Atxn7l3 | 12622.98767 | 0.024479246 | 0.053910625 | 0.454070903 | 0.649777795 | 0.913891582 |
| Snhg8 | 339.3549297 | 0.024479435 | 0.074110589 | 0.330309547 | 0.74116608 | 0.939258605 |
| Tmem184a | 11.51737104 | 0.024485152 | 0.043846455 | 0.558429443 | 0.576551173 | NA |
| B930025P03Rik | 16.51731414 | 0.024507183 | 0.064442981 | 0.380292516 | 0.70372829 | NA |
| Dgkg | 3999.322969 | 0.02450874 | 0.066068929 | 0.370957123 | 0.710669468 | 0.930507332 |
| Mettl1 | 211.697519 | 0.024511591 | 0.08296701 | 0.295437808 | 0.767659455 | 0.94619074 |
| Cebpz | 1667.443777 | 0.024523579 | 0.068809785 | 0.356396688 | 0.72154351 | 0.932513933 |
| Abcb8 | 2049.627655 | 0.024534736 | 0.064839536 | 0.378391603 | 0.705139705 | 0.928783753 |
| Jup | 2165.810506 | 0.024535577 | 0.065212395 | 0.376240999 | 0.706737738 | 0.929476902 |
| Dctn1 | 18642.13113 | 0.024539474 | 0.047255328 | 0.519295394 | 0.603554765 | 0.891540503 |
| Mmp9 | 61.60077138 | 0.024550727 | 0.094482556 | 0.259844021 | 0.794984093 | 0.952171746 |
| Ssh2 | 1957.216504 | 0.02456054 | 0.074456088 | 0.32986611 | 0.741501132 | 0.939436288 |
| Klf1 | 7.439810736 | 0.024570712 | 0.056059248 | 0.438298991 | 0.661169558 | NA |
| Eepd1 | 864.7383722 | 0.024572937 | 0.068553536 | 0.358448861 | 0.720007432 | 0.932408661 |

| | | | | | | | | | | | | | |
|---|---|---|---|---|---|---|---|---|---|---|---|---|---|
| BC018507 | 5895.60377 | 0.048021587 | 0.109028701 | 0.440449044 | 0.659611911 | 0.820537487 | Sp2 | 749.7916449 | 0.024573246 | 0.063753767 | 0.385439906 | 0.699911521 | 0.926781946 |
| Bdh1 | 1710.099705 | -0.026931231 | 0.061153039 | -0.440390724 | 0.659654142 | 0.820537487 | Crb3 | 18.80099779 | 0.024579632 | 0.068288967 | 0.35993562 | 0.718895279 | NA |
| Car2 | 6400.215571 | -0.039021955 | 0.088575096 | -0.440552224 | 0.659537197 | 0.820537487 | Pygo1 | 471.8702692 | 0.024579828 | 0.086048457 | 0.285650997 | 0.77514544 | 0.947238918 |
| Casp7 | 99.61756201 | 0.062270044 | 0.141418851 | 0.440323506 | 0.659702819 | 0.820537487 | Zc3h7b | 10364.64474 | 0.024597788 | 0.05230013 | 0.47031983 | 0.638126532 | 0.908932006 |
| Gramd2 | 159.7254439 | 0.064004135 | 0.145232215 | 0.440702047 | 0.659428714 | 0.820537487 | B4galt3 | 1717.055847 | 0.024608659 | 0.055533792 | 0.44312945 | 0.657672105 | 0.915208181 |
| Sp6 | 23.35578097 | 0.073646472 | 0.167246017 | 0.440348136 | 0.659684982 | 0.820537487 | Klhl7 | 3108.543804 | 0.024610857 | 0.070619126 | 0.348501302 | 0.727463736 | 0.9335794 |
| Atg3 | 1498.330904 | -0.033298519 | 0.075649098 | -0.440170736 | 0.659813453 | 0.820543354 | Sri | 1928.940421 | 0.024614643 | 0.053714129 | 0.458252658 | 0.646770932 | 0.912750424 |
| Atp6ap1l | 351.2087245 | -0.048241864 | 0.109590405 | -0.440201525 | 0.659791155 | 0.820543354 | Opcml | 8410.674377 | 0.024622824 | 0.068080944 | 0.361669833 | 0.717598773 | 0.93125489 |
| Ccdc30 | 369.6223591 | 0.046228857 | 0.105058103 | 0.440031333 | 0.659914414 | 0.820603044 | Dynlrb1 | 4460.797246 | 0.024637277 | 0.059484719 | 0.414178249 | 0.67874357 | 0.921316909 |
| Memo1 | 635.4228854 | 0.048737795 | 0.110897672 | 0.439484379 | 0.660310598 | 0.820963923 | Ireb2 | 2613.282499 | 0.024658746 | 0.070361521 | 0.35045783 | 0.725995132 | 0.933575664 |
| Myl4 | 554.3424728 | -0.073618754 | 0.167484282 | -0.4395562 | 0.66025857 | 0.820963923 | Eif1 | 12720.74894 | 0.024667696 | 0.052346458 | 0.471239067 | 0.637470024 | 0.908932006 |
| Zfp644 | 2199.532098 | -0.031985698 | 0.072796012 | -0.43938805 | 0.660380384 | 0.820984808 | Gbf1 | 4642.931356 | 0.024669246 | 0.049449284 | 0.498879743 | 0.617864106 | 0.898138096 |
| Mrpl11 | 1262.73555 | -0.027274992 | 0.062085362 | -0.43931437 | 0.660433763 | 0.820985296 | Fkbp10 | 277.7961793 | 0.024716 | 0.081956861 | 0.301573293 | 0.76297737 | 0.944431816 |
| Kctd16 | 635.4272273 | -0.038013815 | 0.086599764 | -0.438959806 | 0.660690661 | 0.821172878 | Vamp8 | 272.1153111 | 0.024721664 | 0.094745758 | 0.260926347 | 0.794149306 | 0.951878276 |
| Ttc3 | 44297.92609 | -0.030094623 | 0.068550872 | -0.439011529 | 0.660653183 | 0.821172878 | Fam118a | 765.8215258 | 0.02472407 | 0.059528318 | 0.415332907 | 0.677898215 | 0.921155235 |
| 1700012D01Rik | 13.51378355 | -0.072424001 | 0.165068673 | -0.438750732 | 0.660842163 | 0.821229425 | Iqck | 64.35724011 | 0.024731881 | 0.09404322 | 0.262984202 | 0.792562756 | 0.951728598 |
| Gm5126 | 83.98629361 | 0.057808662 | 0.131757321 | 0.438751042 | 0.660841938 | 0.821229425 | E030019B13Rik | 19.86825102 | 0.024744717 | 0.075542876 | 0.327558583 | 0.743245442 | NA |
| Prkar1b | 36725.58424 | 0.033662713 | 0.076752218 | 0.438589451 | 0.660959042 | 0.821308797 | Klc3 | 78.06763474 | 0.024745184 | 0.09500532 | 0.26046103 | 0.794508172 | 0.951979493 |
| 4930451G09Rik | 121.2841846 | 0.060810001 | 0.13867221 | 0.438516128 | 0.661012181 | 0.821308961 | Cdk15 | 27.54721064 | 0.024774491 | 0.084014467 | 0.294883628 | 0.768082779 | 0.94619074 |
| Traf1 | 29.56528989 | -0.0736312 | 0.168014578 | -0.438242926 | 0.661210195 | 0.821489115 | Aimp1 | 1475.149809 | 0.024775412 | 0.058506408 | 0.423464923 | 0.671956086 | 0.919099256 |
| Fam89a | 24.49921398 | 0.073252606 | 0.167439637 | 0.437486651 | 0.661758455 | 0.822104354 | Lrrc56 | 176.6639672 | 0.024776718 | 0.091142048 | 0.271847281 | 0.785739449 | 0.950484821 |
| Capn12 | 29.29644343 | 0.073294317 | 0.167657976 | 0.437165704 | 0.66199118 | 0.822261612 | St18 | 459.1804387 | 0.024782835 | 0.09132094 | 0.271381731 | 0.786097452 | 0.950484821 |
| Zcchc24 | 3280.169977 | 0.035055057 | 0.080182189 | 0.437192561 | 0.661971704 | 0.822261612 | Cox7a2 | 4093.687619 | 0.024799278 | 0.05797718 | 0.427742056 | 0.668838928 | 0.91742722 |
| Zkscan7 | 118.9331238 | 0.069257742 | 0.158529021 | 0.436877371 | 0.662200284 | 0.822455407 | Ccdc148 | 693.7124892 | 0.024799893 | 0.073105242 | 0.339235492 | 0.734432334 | 0.936456337 |
| Kif5c | 40602.91736 | -0.028657895 | 0.065608428 | -0.436802046 | 0.662254915 | 0.822457331 | Tyrp1 | 4.059861114 | 0.024808276 | 0.048609379 | 0.510359863 | 0.609799371 | NA |
| Epn2 | 5804.92083 | -0.023044414 | 0.052782502 | -0.436591929 | 0.662407317 | 0.822580666 | Atg14 | 1032.366883 | 0.024816819 | 0.066143662 | 0.375195726 | 0.707514909 | 0.92990966 |
| Eprs | 3836.950151 | -0.021387173 | 0.048996843 | -0.436501049 | 0.662473238 | 0.822596598 | Dlg4 | 31099.60296 | 0.024818605 | 0.060146396 | 0.412636609 | 0.679872875 | 0.921823758 |
| Cebpz | 1534.359702 | -0.028665987 | 0.065683865 | -0.436423564 | 0.662529445 | 0.822600467 | Rab1b | 5925.596634 | 0.024835263 | 0.049834318 | 0.498356627 | 0.618232702 | 0.898314317 |
| Gm11127 | 44.66253358 | -0.072417545 | 0.166076139 | -0.436050269 | 0.662800258 | 0.822804839 | Erbb3 | 457.6064107 | 0.02483942 | 0.088202894 | 0.281616832 | 0.778237333 | 0.947901685 |
| Kbtbd3 | 245.4250349 | -0.05177782 | 0.1187321 | -0.436089479 | 0.66277181 | 0.822804839 | Gprc5a | 9.704609962 | 0.024851225 | 0.062553943 | 0.39727672 | 0.691163413 | NA |
| Islr2 | 2860.805204 | 0.053766859 | 0.123370914 | 0.435814711 | 0.66297117 | 0.822951074 | Mthfd1 | 1675.742466 | 0.02485978 | 0.046103413 | 0.539217785 | 0.58973659 | 0.885244777 |
| Ppp1r26 | 1305.724365 | 0.031386915 | 0.07205673 | 0.435586162 | 0.663137013 | 0.823090994 | Phb2 | 2321.231459 | 0.024860459 | 0.052128862 | 0.476903922 | 0.633430539 | 0.906932171 |
| Psmd7 | 3401.350144 | -0.030246107 | 0.069500078 | -0.435195295 | 0.663420677 | 0.823377121 | 2310044G17Rik | 4455.780238 | 0.024865277 | 0.062024391 | 0.400895144 | 0.688497326 | 0.923860706 |
| Rpusd4 | 302.7999289 | 0.033730686 | 0.077541 | 0.435004528 | 0.663559141 | 0.823483005 | Map2 | 24471.4282 | 0.024902888 | 0.061032543 | 0.408026385 | 0.683254304 | 0.922993772 |
| Fli1 | 161.2922819 | -0.048821016 | 0.11239652 | -0.434364123 | 0.664024046 | 0.823504463 | Pibf1 | 441.4693469 | 0.024905114 | 0.079393104 | 0.313693669 | 0.75375371 | 0.94183464 |
| Mob1b | 410.517406 | -0.039304769 | 0.090503745 | -0.434288871 | 0.664078685 | 0.823504463 | Supv3l1 | 903.2659824 | 0.02491661 | 0.063874488 | 0.390087037 | 0.696472188 | 0.92555371 |
| Naprt1 | 174.5882294 | 0.063097146 | 0.145206834 | 0.434532894 | 0.663901514 | 0.823504463 | Gpn3 | 547.8292056 | 0.02492134 | 0.06900412 | 0.361157276 | 0.717981878 | 0.93125489 |
| Nlrp1b | 26.08243885 | -0.071167018 | 0.163839653 | -0.434369926 | 0.664019833 | 0.823504463 | Gna11 | 5997.842192 | 0.024927459 | 0.055243596 | 0.451228032 | 0.651825208 | 0.914447303 |
| Peg10 | 1951.577894 | -0.073053613 | 0.168097894 | -0.434589701 | 0.663860272 | 0.823504463 | Ap1b1 | 9858.007086 | 0.024937838 | 0.049225015 | 0.506609044 | 0.612429161 | 0.895344807 |
| Serpinb6a | 1062.332578 | -0.027162265 | 0.062539976 | -0.43431844 | 0.664057215 | 0.823504463 | Zmat4 | 2609.941324 | 0.024957071 | 0.06529369 | 0.382227918 | 0.702292316 | 0.927855161 |
| Slc25a39 | 2064.625561 | -0.03090553 | 0.071079727 | -0.434800904 | 0.663706949 | 0.823504463 | Ssx2ip | 3658.834567 | 0.024991582 | 0.063915001 | 0.391012774 | 0.695787794 | 0.925527709 |
| Spred2 | 2410.178305 | -0.051288458 | 0.118108518 | -0.434248593 | 0.66410793 | 0.823504463 | Usp6nl | 854.3966154 | 0.02500975 | 0.072182113 | 0.346481264 | 0.728981062 | 0.933996867 |
| Ssr4 | 989.329928 | -0.031230269 | 0.071893319 | -0.434397376 | 0.663999903 | 0.823504463 | Mark2 | 5192.638462 | 0.025024655 | 0.059136407 | 0.423168339 | 0.672172445 | 0.919190306 |
| Tbc1d19 | 933.5075097 | 0.046031385 | 0.105898887 | 0.434672986 | 0.663799809 | 0.823504463 | Klrg1 | 7.076320156 | 0.025031011 | 0.058479006 | 0.428034143 | 0.668626263 | NA |
| Ccdc137 | 926.9431607 | 0.033287956 | 0.076698654 | 0.434009644 | 0.664281438 | 0.823653697 | Acot1 | 205.2443649 | 0.025048358 | 0.085515787 | 0.292909168 | 0.769591581 | 0.94619074 |
| Dicer1 | 2577.65294 | -0.030470405 | 0.070249101 | -0.433747965 | 0.664471472 | 0.823807252 | 4930529M08Rik | 40.80287016 | 0.025057057 | 0.09165824 | 0.27337485 | 0.784565084 | 0.949699053 |
| Faf1 | 1312.210893 | 0.029450667 | 0.067918228 | 0.433619477 | 0.664564789 | 0.823807252 | Il31ra | 7.699513683 | 0.025079736 | 0.048703495 | 0.514947368 | 0.606589799 | NA |
| Kcnj6 | 1147.719299 | -0.036721062 | 0.084684705 | -0.43362095 | 0.664563719 | 0.823807252 | St3gal5 | 6622.68357 | 0.025085751 | 0.055082477 | 0.455421631 | 0.648805936 | 0.913724196 |
| Ddx11 | 79.55756514 | -0.066983504 | 0.154544003 | -0.433426742 | 0.664704777 | 0.823829802 | Upk1a | 20.20476212 | 0.025093312 | 0.072908523 | 0.344175287 | 0.730714467 | NA |
| Gmpr2 | 702.5060629 | -0.039427114 | 0.090964103 | -0.433435969 | 0.664698074 | 0.823829802 | Msx3 | 1.765766832 | 0.025094908 | 0.028520217 | 0.879898908 | 0.378914076 | NA |
| Ripply2 | 55.23182112 | -0.067215765 | 0.155098457 | -0.433374817 | 0.664742493 | 0.823829802 | Ifltd1 | 20.98962547 | 0.025096019 | 0.077284857 | 0.324721043 | 0.745392207 | NA |
| 2900041M22Rik | 26.91761232 | -0.070781222 | 0.16347465 | -0.432979806 | 0.66502944 | 0.82397451 | 4930567H12Rik | 1.65398349 | 0.025106456 | 0.029726716 | 0.844575486 | 0.398347901 | NA |
| Elmsan1 | 1230.085004 | 0.048559769 | 0.112216694 | 0.432732128 | 0.665209386 | 0.82397451 | Pmpca | 4443.360325 | 0.025111198 | 0.038460437 | 0.652909837 | 0.513814407 | 0.853195313 |
| Ptms | 14660.56089 | 0.029352337 | 0.067835045 | 0.432701667 | 0.665231519 | 0.82397451 | Ptplb | 1555.183367 | 0.025114042 | 0.054147544 | 0.463807589 | 0.642785606 | 0.911258473 |
| Rhod | 17.59476613 | 0.072721638 | 0.168045086 | 0.432750756 | 0.665195852 | 0.82397451 | B3galtl | 916.7758885 | 0.025125126 | 0.060133983 | 0.417819081 | 0.676079399 | 0.920643418 |
| Spata5l1 | 64.7577502 | 0.060983039 | 0.140916036 | 0.432761523 | 0.665188029 | 0.82397451 | Cd163 | 58.30930589 | 0.025151551 | 0.08615253 | 0.291942107 | 0.770330888 | 0.94619074 |
| Tmod3 | 910.3499372 | -0.029455213 | 0.068007649 | -0.433116182 | 0.664930367 | 0.82397451 | Amot | 963.6202368 | 0.025178733 | 0.078962613 | 0.318869047 | 0.749825817 | 0.9410864 |
| Xkr7 | 116.7853846 | -0.062238588 | 0.143820361 | -0.432752272 | 0.665194751 | 0.82397451 | Astn1 | 13889.22965 | 0.02518289 | 0.061009196 | 0.412772026 | 0.679773648 | 0.921806574 |
| 2410006H16Rik | 385.6369425 | 0.047947586 | 0.110861425 | 0.432500178 | 0.665377923 | 0.824089971 | Zic2 | 436.5149283 | 0.025186328 | 0.094678514 | 0.266019464 | 0.790224219 | 0.951607109 |
| Caps2 | 8.66176849 | 0.06403012 | 0.148078978 | 0.432405198 | 0.665446941 | 0.824109576 | Ankrd34a | 2880.020847 | 0.025188063 | 0.05437196 | 0.463254654 | 0.643181845 | 0.911258473 |
| 2900002K06Rik | 9.300791402 | 0.069965871 | 0.162047258 | 0.431762139 | 0.665914299 | 0.824289729 | Plat | 1312.192342 | 0.025191736 | 0.069306111 | 0.363485065 | 0.716242567 | 0.93125489 |
| Dnttip2 | 943.6065321 | 0.039446226 | 0.091375719 | 0.431692645 | 0.665964814 | 0.824289729 | Zkscan6 | 577.6449231 | 0.025193054 | 0.068789397 | 0.366234552 | 0.714190062 | 0.93118435 |
| Nxt1 | 222.078285 | 0.045053516 | 0.10432217 | 0.431869045 | 0.665836594 | 0.824289729 | Gm15413 | 2.971967653 | 0.025199159 | 0.037741675 | 0.667674632 | 0.504341306 | NA |
| Slain2 | 938.3515146 | -0.030284584 | 0.070131568 | -0.431825279 | 0.665868405 | 0.824289729 | Hoxb4 | 1.65841631 | 0.025237632 | 0.028050018 | 0.899736755 | 0.368260358 | NA |
| Tex261 | 795.783763 | -0.029464522 | 0.068210743 | -0.431963063 | 0.665768259 | 0.824289729 | Pbx1 | 6943.50812 | 0.025256799 | 0.053862976 | 0.468908335 | 0.639135158 | 0.909490332 |
| Zfp607 | 38.94664459 | -0.072355068 | 0.167587331 | -0.431745453 | 0.665926428 | 0.824289729 | 1700001O22Rik | 177.907996 | 0.02525715 | 0.093913806 | 0.268939627 | 0.787976096 | 0.951071107 |
| Zscan2 | 266.9103225 | -0.044995635 | 0.104201536 | -0.431813549 | 0.665876931 | 0.824289729 | Eif2b1 | 961.0461242 | 0.025258817 | 0.053537525 | 0.471796508 | 0.637072044 | 0.908932006 |
| Adat1 | 270.2829925 | 0.04759798 | 0.11036029 | 0.431296258 | 0.666252971 | 0.824514659 | Med20 | 846.9367139 | 0.025269792 | 0.060644339 | 0.416688396 | 0.676906343 | 0.921056062 |
| BC005764 | 2301.52035 | 0.046697572 | 0.108266516 | 0.431320542 | 0.666235316 | 0.824514659 | Olah | 2.174097225 | 0.025274478 | 0.032854688 | 0.769280705 | 0.441726689 | NA |
| Sobp | 3303.739988 | 0.029496516 | 0.068409305 | 0.431176966 | 0.666339701 | 0.824556132 | Id4 | 1581.266745 | 0.02527619 | 0.069891995 | 0.361646427 | 0.717616266 | 0.93125489 |
| 2810468N07Rik | 896.4073398 | -0.042299047 | 0.098208688 | -0.430705756 | 0.666682333 | 0.824848364 | Neo1 | 5003.096134 | 0.025291051 | 0.054465674 | 0.46434845 | 0.642398117 | 0.911258473 |
| Ptk2 | 5572.36413 | -0.029974773 | 0.069590882 | -0.430728451 | 0.666665829 | 0.824848364 | Mpo | 4.007323397 | 0.025308525 | 0.036163846 | 0.699829473 | 0.484033806 | NA |
| Fam169a | 2410.100082 | 0.03360058 | 0.078069025 | 0.43039579 | 0.666907757 | 0.825012842 | Gen1 | 40.62602183 | 0.02530898 | 0.088827354 | 0.284923273 | 0.775702925 | 0.947241001 |
| Zfp429 | 61.47528034 | 0.061000359 | 0.141737189 | 0.430376528 | 0.666921766 | 0.825012842 | Ptpn5 | 11897.84777 | 0.025322391 | 0.062656608 | 0.404145568 | 0.686105678 | 0.923860706 |
| Ap1b1 | 9134.871198 | 0.032117106 | 0.074672945 | 0.430103641 | 0.667120251 | 0.825060833 | Gstm5 | 5391.326145 | 0.025346638 | 0.063610081 | 0.398468884 | 0.69028459 | 0.923860706 |
| Olfr464 | 15.24344999 | 0.063291619 | 0.147173769 | 0.430046872 | 0.667161546 | 0.825060833 | Opn5 | 1.046073401 | 0.025351705 | 0.021232669 | 1.193995229 | 0.232479834 | NA |
| Ptgr1 | 35.32123312 | -0.069072108 | 0.160621472 | -0.430030352 | 0.667173563 | 0.825060833 | 1700021N21Rik | 1.886347427 | 0.025371394 | 0.032114224 | 0.790036044 | 0.429506719 | NA |
| Zfp507 | 1225.348798 | -0.034412417 | 0.079983671 | -0.430243025 | 0.667018867 | 0.825060833 | Yaf2 | 2636.947887 | 0.025376616 | 0.066707152 | 0.380418222 | 0.70363499 | 0.928079892 |
| Meis1 | 245.0294757 | -0.067732913 | 0.157571093 | -0.429856214 | 0.667300238 | 0.825151627 | Srsf10 | 2117.541498 | 0.025389588 | 0.07103592 | 0.357419008 | 0.720778149 | 0.932513933 |
| Zfp938 | 551.7235508 | -0.035297306 | 0.082184742 | -0.429487342 | 0.667568604 | 0.8254176 | Ncbp1 | 2495.615462 | 0.025391879 | 0.061400907 | 0.413542408 | 0.679209259 | 0.921471438 |
| Sephs1 | 1023.780118 | -0.036346273 | 0.084646927 | -0.429386801 | 0.667641757 | 0.825442179 | Krt2 | 60.6929632 | 0.02539642 | 0.044827301 | 0.566539131 | 0.57102734 | 0.878246677 |
| Tmod1 | 2687.42237 | -0.025700202 | 0.059866038 | -0.429295186 | 0.66770842 | 0.825458729 | Pdgfb | 1267.80647 | 0.025448355 | 0.080029305 | 0.317987954 | 0.750494074 | 0.9410864 |
| 6430548M08Rik | 15780.44524 | -0.028368179 | 0.066103617 | -0.429147155 | 0.667816138 | 0.825526028 | Hes5 | 343.0352855 | 0.025450634 | 0.094624145 | 0.268965535 | 0.787956205 | 0.951071107 |
| H1f0 | 3398.65047 | -0.025889557 | 0.060416693 | -0.428516618 | 0.668275037 | 0.825961505 | Uchl5 | 1581.706537 | 0.025467301 | 0.074449099 | 0.342076684 | 0.732293183 | 0.935722272 |
| Igf2bp2 | 40.46602135 | -0.068767341 | 0.160470482 | -0.428535766 | 0.6682611 | 0.825961505 | Rbfox1 | 14778.77875 | 0.025469856 | 0.062224182 | 0.409324087 | 0.682301842 | 0.922544285 |
| D430020J02Rik | 112.7601341 | -0.05669348 | 0.132384123 | -0.428249842 | 0.668469232 | 0.826003851 | Txndc2 | 3.130159782 | 0.025479471 | 0.038756239 | 0.657428876 | 0.51090519 | NA |
| Dnajc1 | 951.9734186 | -0.034814096 | 0.081282953 | -0.428307465 | 0.668427285 | 0.826003851 | Slc17a8 | 257.5411261 | 0.025513124 | 0.087327852 | 0.292153347 | 0.77016938 | 0.94619074 |
| Polr1c | 670.9800609 | -0.038910356 | 0.090839729 | -0.428340735 | 0.668403066 | 0.826003851 | Stk30 | 308.8872255 | 0.025513619 | 0.074892495 | 0.340669901 | 0.7333521 | 0.936456337 |
| Otud6b | 2493.777066 | -0.028985555 | 0.067718497 | -0.428030105 | 0.668629203 | 0.826069761 | Ctdp1 | 766.9145408 | 0.025537262 | 0.060512881 | 0.422013659 | 0.673015048 | 0.919353963 |
| Tcf15 | 10.56466275 | -0.06840767 | 0.159793378 | -0.428100782 | 0.668577748 | 0.826069761 | Polr3k | 1380.428261 | 0.025550644 | 0.061458714 | 0.415736725 | 0.677602665 | 0.921093181 |
| Larp4 | 1542.208037 | 0.031993358 | 0.074784684 | 0.427806294 | 0.668792154 | 0.826205202 | Galnt14 | 928.073522 | 0.025552443 | 0.065020895 | 0.392988173 | 0.694328216 | 0.925473635 |
| Snta1 | 1110.505937 | 0.029239656 | 0.068378216 | 0.427616547 | 0.668930317 | 0.82631 | Tmem126a | 556.1418991 | 0.025555743 | 0.062845516 | 0.406643861 | 0.684269578 | 0.923655038 |
| Pih1d1 | 653.0168881 | -0.037998836 | 0.088914499 | -0.427363774 | 0.669114389 | 0.826471488 | Trim8 | 2799.340608 | 0.025566992 | 0.061686105 | 0.414469223 | 0.678530502 | 0.921316909 |
| Agtr1a | 26.77549627 | -0.071408016 | 0.167262577 | -0.426921654 | 0.669436393 | 0.826737404 | A530050N04Rik | 6.075994793 | 0.025581291 | 0.049807179 | 0.513606506 | 0.60752713 | NA |
| Egr1 | 11151.10528 | -0.071335836 | 0.167074078 | -0.426971301 | 0.669400231 | 0.826737404 | Tmf1 | 1602.96625 | 0.025585096 | 0.067503984 | 0.379016088 | 0.704675917 | 0.928530694 |
| Rab1b | 5317.849902 | -0.027667973 | 0.064827471 | -0.426793969 | 0.6695294 | 0.826786365 | Nt5c2 | 3069.938243 | 0.025588778 | 0.049538439 | 0.516543885 | 0.605474597 | 0.892362418 |
| Eaf2 | 37.33129048 | 0.06731804 | 0.157770894 | 0.42668225 | 0.669610781 | 0.826820962 | Scrt2 | 1029.459953 | 0.025600323 | 0.081190116 | 0.3153133 | 0.752523788 | 0.941691983 |
| Nsun3 | 181.6446103 | 0.05257777 | 0.12328435 | 0.426475619 | 0.66976131 | 0.82694093 | Aldh16a1 | 262.2093656 | 0.025614609 | 0.082943656 | 0.308819394 | 0.757458915 | 0.942139348 |
| Zkscan8 | 827.7428554 | 0.043505039 | 0.1020363 | 0.426368254 | 0.66983953 | 0.826971608 | Ncaph2 | 4746.006042 | 0.025649246 | 0.055398238 | 0.4629975 | 0.643366159 | 0.911258473 |
| Ermp1 | 2702.450553 | 0.027285228 | 0.064016163 | 0.42622405 | 0.669944594 | 0.827035419 | Cdadc1 | 2388.935397 | 0.025656409 | 0.054648102 | 0.469484002 | 0.638723718 | 0.909380945 |
| 5730559C18Rik | 90.1446376 | 0.058978895 | 0.138449147 | 0.425996813 | 0.670110168 | 0.827173912 | Pde1a | 9866.673962 | 0.025673106 | 0.071512737 | 0.359000459 | 0.719594746 | 0.932251852 |
| Eif4a3 | 1548.534764 | -0.026211035 | 0.061553188 | -0.425827425 | 0.670233601 | 0.827194473 | Syce2 | 28.1511615 | 0.025684254 | 0.086251309 | 0.297783935 | 0.765868077 | 0.945453142 |
| Mpzl3 | 53.23578763 | 0.060940781 | 0.143103906 | 0.425849876 | 0.670217241 | 0.827194473 | Dlx6 | 293.9240847 | 0.02568989 | 0.092753956 | 0.276968129 | 0.781804579 | 0.948979619 |
| Slc30a6 | 357.7358486 | -0.039407253 | 0.092561654 | -0.425740584 | 0.670296886 | 0.827206687 | Actg1 | 49361.89195 | 0.025691408 | 0.042858631 | 0.599445373 | 0.548875928 | 0.868267067 |
| Nog | 173.8712881 | -0.05419453 | 0.12740214 | -0.425381628 | 0.670558497 | 0.827463631 | Rab40b | 1007.969566 | 0.02569528 | 0.082983268 | 0.309644107 | 0.756831611 | 0.942139348 |
| Bves | 33.25624571 | -0.066537377 | 0.156512509 | -0.425124978 | 0.67074557 | 0.827527874 | Hmx3 | 1.451804699 | 0.025708222 | 0.018769909 | 1.36965091 | 0.170795899 | NA |
| Grk6 | 2871.794578 | -0.026419301 | 0.062149843 | -0.425090385 | 0.670770787 | 0.827527874 | 9430091E24Rik | 54.23804104 | 0.025709247 | 0.094167911 | 0.273014945 | 0.784841727 | 0.949834246 |
| Smad2 | 1640.372517 | -0.026269432 | 0.061779318 | -0.425214022 | 0.670680664 | 0.827527874 | Reep6 | 464.2920379 | 0.025715697 | 0.074307636 | 0.346070711 | 0.729289574 | 0.933996867 |
| Adamts18 | 91.11222444 | -0.064276276 | 0.151334397 | -0.424730118 | 0.671033425 | 0.827720077 | Ctnna1 | 3628.537208 | 0.025717495 | 0.045245111 | 0.568403831 | 0.56976079 | 0.87770886 |
| Rnf19b | 1702.067611 | -0.035175385 | 0.082807487 | -0.424785084 | 0.670993352 | 0.827720077 | Plk1s1 | 619.0908794 | 0.025761179 | 0.081804579 | 0.314911209 | 0.752829071 | 0.94183464 |
| Snai3 | 23.54896297 | 0.071232435 | 0.168073927 | 0.423816089 | 0.671699943 | 0.828476272 | Nsmce1 | 395.5379807 | 0.025770614 | 0.081632083 | 0.315692227 | 0.752236127 | 0.941565791 |
| Cd4 | 602.259641 | -0.052523266 | 0.12398252 | -0.42363444 | 0.671832434 | 0.828507779 | Etaa1 | 397.6356917 | 0.025773151 | 0.08705951 | 0.296040623 | 0.767199058 | 0.946118815 |
| Tnni2 | 8.924590337 | 0.068047852 | 0.16061271 | 0.42367663 | 0.671801661 | 0.828507779 | Cenpp | 23.99808206 | 0.025774469 | 0.082930287 | 0.310796807 | 0.755955097 | 0.942139348 |
| Epor | 306.2094584 | 0.033669758 | 0.079574083 | 0.423124677 | 0.672204299 | 0.828834427 | Epha6 | 810.9172101 | 0.025803431 | 0.089339551 | 0.288824274 | 0.772715856 | 0.947121017 |
| Trappc10 | 5278.031982 | 0.033617909 | 0.079445059 | 0.423159217 | 0.6721791 | 0.828834427 | Mir3061 | 41.01418997 | 0.025811351 | 0.089825506 | 0.287349906 | 0.773844413 | 0.947121017 |
| Adrb1 | 867.5492348 | 0.048751679 | 0.115258964 | 0.422975168 | 0.672313379 | 0.82890296 | Tmem106c | 1131.354572 | 0.025812351 | 0.06085497 | 0.424161766 | 0.671447842 | 0.91883668 |
| Itm2b | 19310.33097 | 0.021495044 | 0.050829089 | 0.422888638 | 0.672376513 | 0.828914839 | 4933417O13Rik | 12.21227934 | 0.025812689 | 0.065764605 | 0.392501236 | 0.694687898 | NA |
| Myzap | 30.33260973 | 0.068693778 | 0.162502756 | 0.422723773 | 0.672496809 | 0.82899718 | Gp49a | 5.613307118 | 0.025858697 | 0.034384855 | 0.752037395 | 0.452028571 | NA |
| Maf1 | 2356.112032 | 0.027621725 | 0.065392652 | 0.422397999 | 0.672734538 | 0.829205242 | Cstb | 618.3963443 | 0.025862141 | 0.07475407 | 0.345962987 | 0.729370531 | 0.933996867 |
| Pcdh1 | 4811.754715 | -0.039356691 | 0.093185939 | -0.422345806 | 0.672772628 | 0.829205242 | Tmem8b | 4454.966356 | 0.025881372 | 0.057510406 | 0.450029374 | 0.652689261 | 0.914683958 |
| Slc33a1 | 1139.42915 | -0.024068338 | 0.057123965 | -0.421335207 | 0.673510325 | 0.82998242 | Ptrh2 | 502.6035792 | 0.025891024 | 0.069098114 | 0.374699432 | 0.707884014 | 0.929959854 |
| Slc41a2 | 951.0694831 | -0.036709122 | 0.087114985 | -0.421386998 | 0.673472512 | 0.82998242 | Macrod2 | 1075.7308 | 0.025901146 | 0.07760845 | 0.333741321 | 0.738574766 | 0.937973164 |
| Hpse | 54.31243035 | 0.064145222 | 0.152470416 | 0.420706023 | 0.673969763 | 0.830416501 | Noa1 | 705.9277347 | 0.025908386 | 0.063469443 | 0.408202512 | 0.683125004 | 0.922993772 |
| Mphosph10 | 636.6394307 | 0.031681754 | 0.075302663 | 0.420725549 | 0.673955503 | 0.830416501 | Relt | 829.6282319 | 0.025909994 | 0.065980721 | 0.392690374 | 0.694548181 | 0.925527709 |
| Actrt3 | 29.59252788 | 0.070068244 | 0.166580949 | 0.420625791 | 0.674028358 | 0.830417276 | Eapp | 790.6372851 | 0.02591845 | 0.065778619 | 0.394025456 | 0.693562245 | 0.924707221 |
| Pex1 | 893.831509 | 0.02559368 | 0.060867046 | 0.420485008 | 0.674131179 | 0.830417276 | Onecut3 | 6.492253781 | 0.025919632 | 0.019148194 | 1.35363325 | NA | NA |
| Zfp119b | 82.76413208 | -0.064022862 | 0.152251017 | -0.420508599 | 0.674113949 | 0.830417276 | Cldn10 | 1149.881478 | 0.025925169 | 0.071093284 | 0.364664114 | 0.71536215 | 0.93125489 |
| Nlrx1 | 359.4824604 | 0.039993616 | 0.095134884 | 0.420388549 | 0.674201632 | 0.830438038 | Tmbim4 | 1090.074995 | 0.025930025 | 0.057204848 | 0.453283698 | 0.65034447 | 0.914000665 |
| Arhgap30 | 167.2045975 | 0.050245828 | 0.119626063 | 0.420024087 | 0.674467858 | 0.830567874 | Duoxa1 | 1.99090577 | 0.025936457 | 0.029311968 | 0.884841912 | 0.376241901 | NA |
| Eif3b | 3586.265112 | 0.028509209 | 0.067872563 | 0.420040264 | 0.67445604 | 0.830567874 | Faf1 | 1463.780132 | 0.025954648 | 0.056419809 | 0.460027233 | 0.645496673 | 0.912363974 |
| Trip12 | 8072.241072 | -0.03467379 | 0.082549776 | -0.420034944 | 0.674459927 | 0.830567874 | Taf6 | 1455.581009 | 0.025958162 | 0.064474979 | 0.402608304 | 0.6872364 | 0.923860706 |
| Galnt7 | 487.3457676 | -0.030760224 | 0.073274394 | -0.419795 | 0.674635218 | 0.830575929 | Thap3 | 402.6732951 | 0.025960175 | 0.07574551 | 0.342728892 | 0.731802425 | 0.935424325 |
| Gm14379 | 10.2125842 | 0.06381314 | 0.151991841 | 0.419845828 | 0.674598084 | 0.830575929 | Gm5868 | 2.147800347 | 0.025962826 | 0.025753468 | 1.008129296 | 0.313392395 | NA |
| Nup205 | 1307.013797 | 0.03330795 | 0.079326423 | 0.419884679 | 0.674569701 | 0.830575929 | Arc | 11827.65662 | 0.025967318 | 0.054756368 | 0.474233756 | 0.635333225 | 0.90754749 |
| Grb10 | 2062.083747 | -0.042115717 | 0.100461873 | -0.419220905 | 0.675054696 | 0.830976799 | Pmch | 252.5295557 | 0.025970199 | 0.021827577 | 1.189788469 | NA | NA |
| Phldb1 | 4435.029842 | 0.051977994 | 0.123992543 | 0.419202577 | 0.675068089 | 0.830976799 | 0610010B08Rik | 9.730001072 | 0.025991292 | 0.064879761 | 0.400607079 | 0.688709433 | NA |
| Ccdc129 | 63.36399785 | -0.069119003 | 0.164930974 | -0.419078366 | 0.675158861 | 0.831022513 | Nrsn1 | 17735.90897 | 0.025994977 | 0.060189239 | 0.431887454 | 0.665823213 | 0.915591677 |
| Znhit2 | 551.6008514 | -0.040613708 | 0.096932236 | -0.418990722 | 0.675222913 | 0.831035334 | Usp1 | 1054.99503 | 0.026003377 | 0.072454108 | 0.358894439 | 0.71967406 | 0.932251852 |
| 9130019O22Rik | 143.2438487 | -0.052760179 | 0.126005998 | -0.418711649 | 0.675426881 | 0.831055994 | Rasa1 | 1942.2559 | 0.026020772 | 0.070736599 | 0.367854436 | 0.712981776 | 0.930507332 |
| Fam149a | 1223.616837 | -0.03123285 | 0.074609551 | -0.418617315 | 0.675495833 | 0.831055994 | Diras2 | 15233.72614 | 0.026026952 | 0.058539272 | 0.444606689 | 0.656604013 | 0.915208181 |
| Pcdha2 | 102.6958712 | -0.057005871 | 0.136201416 | -0.418540958 | 0.675551647 | 0.831055994 | Gpr158 | 9090.89194 | 0.026036356 | 0.069567779 | 0.374258838 | 0.708211753 | 0.929959854 |
| Rpap2 | 674.5631901 | 0.031260253 | 0.074703838 | 0.41845578 | 0.675613911 | 0.831055994 | Gm16551 | 2.257490815 | 0.026038323 | 0.028134616 | 0.925490641 | 0.35471075 | NA |
| S100a13 | 427.7237441 | -0.039196569 | 0.093652105 | -0.418533773 | 0.675556899 | 0.831055994 | Gm10007 | 2.190538874 | 0.026046227 | 0.032723469 | 0.795949469 | 0.426061406 | NA |
| Sfxn5 | 6937.514704 | -0.050376441 | 0.120358517 | -0.418553193 | 0.675542703 | 0.831055994 | Neu1 | 2315.841425 | 0.026048762 | 0.062151668 | 0.41911606 | 0.675131314 | 0.919907397 |
| Zfp943 | 432.0137575 | 0.048637919 | 0.116232395 | 0.41845407 | 0.675615161 | 0.831055994 | Shf | 1201.896599 | 0.026049719 | 0.062580512 | 0.416259277 | 0.677220288 | 0.921064066 |
| Ppef1 | 24.01062762 | -0.069503469 | 0.166197912 | -0.418197004 | 0.675803086 | 0.831221165 | Lpo | 37.75473909 | 0.026076011 | 0.092178443 | 0.282886221 | 0.777264057 | 0.947506677 |
| Pdcd6ip | 3498.002413 | 0.022819034 | 0.054648357 | 0.417561205 | 0.676267967 | 0.831660915 | Med9 | 1976.026139 | 0.026097334 | 0.048415559 | 0.539027838 | 0.589867646 | 0.885258542 |
| Plxnb1 | 4677.240226 | -0.024892522 | 0.059611801 | -0.417577077 | 0.676256359 | 0.831660915 | Zer1 | 8544.117056 | 0.026098116 | 0.052214675 | 0.49982339 | 0.617199439 | 0.897621051 |
| Fam149b | 1069.503648 | -0.027126659 | 0.06498795 | -0.417410597 | 0.676378105 | 0.831730346 | Jam2 | 2125.005234 | 0.026114407 | 0.061323559 | 0.425846241 | 0.67021989 | 0.91809932 |
| Fbl | 988.5161814 | -0.030124828 | 0.072211959 | -0.417172283 | 0.676552397 | 0.831746635 | Rmi2 | 9.294668307 | 0.026130346 | 0.057289402 | 0.456111341 | 0.648309915 | NA |
| Gabrg2 | 6528.350616 | 0.038369908 | 0.091971985 | 0.417191262 | 0.676538516 | 0.831746635 | Pik3r5 | 190.249903 | 0.026150093 | 0.085917578 | 0.304362548 | 0.760851697 | 0.943105713 |
| Irgq | 6052.776393 | -0.031840709 | 0.076318482 | -0.417208359 | 0.676526012 | 0.831746635 | Psmd5 | 1809.073795 | 0.026166541 | 0.055654796 | 0.470157888 | 0.638242218 | 0.908932006 |
| Psmc5 | 5800.996883 | -0.033427916 | 0.080151177 | -0.417060819 | 0.676633923 | 0.831780863 | Cygb | 2590.194946 | 0.026171393 | 0.051964777 | 0.503637175 | 0.614516364 | 0.89611764 |
| 4930539E08Rik | 431.6877385 | -0.048340777 | 0.11593116 | -0.416978291 | 0.676694287 | 0.831789074 | 2900097C17Rik | 69371.02063 | 0.026180389 | 0.059304505 | 0.441457011 | 0.658882177 | 0.915208181 |
| Gm1078 | 56.91073823 | 0.065940065 | 0.158189416 | 0.41684246 | 0.676793643 | 0.831845209 | Tfpt | 674.7398583 | 0.026220742 | 0.071203069 | 0.368252973 | 0.712684614 | 0.930507332 |
| Neurl3 | 45.36199933 | 0.068437687 | 0.164385587 | 0.416324132 | 0.677172836 | 0.832192166 | Orc1 | 6.230791539 | 0.02623516 | 0.049055801 | 0.534802398 | 0.592786505 | NA |
| Slc6a7 | 3208.578681 | -0.038042387 | 0.091380003 | -0.416309761 | 0.677183351 | 0.832192166 | B230209E15Rik | 1129.751398 | 0.026236494 | 0.075218824 | 0.34880224 | 0.727237782 | 0.9335794 |
| Ctdspl2 | 830.0826495 | -0.035385283 | 0.08503937 | -0.416104716 | 0.677333379 | 0.832310523 | Tmem125 | 368.4495179 | 0.02623678 | 0.086007591 | 0.305051911 | 0.760326616 | 0.943070948 |
| Fam20a | 446.0180101 | -0.048977651 | 0.117757509 | -0.415919556 | 0.677468869 | 0.832362764 | Fos | 1219.200699 | 0.026306488 | 0.060083283 | 0.437833728 | 0.66150682 | 0.915428982 |
| Uqcrc1 | 8117.058384 | -0.028359583 | 0.068188502 | -0.415899786 | 0.677483336 | 0.832362764 | Ffar4 | 2.557899846 | 0.026316911 | 0.032399495 | 0.812262987 | 0.416640744 | NA |
| Calr4 | 15.88165734 | -0.069628797 | 0.167486492 | -0.415727836 | 0.67760917 | 0.832451354 | 4931417G12Rik | 3.2449091 | 0.026321776 | 0.036991845 | 0.711556178 | 0.476739651 | NA |
| M6pr | 3902.498128 | -0.020148604 | 0.048492246 | -0.415501562 | 0.677774772 | 0.832588783 | Mon1a | 819.6110487 | 0.02633482 | 0.071926457 | 0.366135367 | 0.714264068 | 0.931197258 |
| Gm1141 | 13.95942987 | 0.069234781 | 0.16685795 | 0.414932469 | 0.67819134 | 0.832842534 | Wapal | 1807.561045 | 0.026337199 | 0.068731687 | 0.383188607 | 0.701579925 | 0.92751465 |
| Naa10 | 758.6089307 | -0.02849518 | 0.06866557 | -0.41498497 | 0.678152906 | 0.832842534 | Rab11fip5 | 5190.927647 | 0.026352342 | 0.038810029 | 0.679008566 | 0.497132434 | 0.845038521 |
| Nkx6-2 | 278.3828384 | -0.053523058 | 0.128940283 | -0.415099584 | 0.678069004 | 0.832842534 | C330022C24Rik | 1.677133784 | 0.026358185 | 0.030583644 | 0.861839277 | 0.388775968 | NA |
| Pip4k2a | 3801.823976 | -0.029766304 | 0.07173889 | -0.414925625 | 0.67819635 | 0.832842534 | Fam186b | 46.56080866 | 0.026367771 | 0.09241167 | 0.285329449 | 0.775391753 | 0.947241001 |
| Gzmm | 52.18773383 | -0.062038578 | 0.149550732 | -0.414832993 | 0.678264165 | 0.832859802 | Chrna3 | 33.24641455 | 0.026368252 | 0.085064643 | 0.309978988 | 0.756576935 | 0.942139348 |
| Ptprk | 2786.966436 | 0.032125973 | 0.077489515 | 0.414584768 | 0.6784459 | 0.833016941 | Speer4b | 3.764123392 | 0.026380463 | 0.043419312 | 0.607574415 | 0.543469774 | NA |
| Ing1 | 1196.328541 | -0.024370669 | 0.058809273 | -0.414401815 | 0.67857986 | 0.8331154 | Rpl21 | 10665.76231 | 0.026389674 | 0.06578616 | 0.401143242 | 0.688314666 | 0.923860706 |
| Erich2 | 14.03996946 | 0.068119284 | 0.164536998 | 0.414005877 | 0.678869803 | 0.833141285 | Trim30e-ps1 | 3.819995155 | 0.026392912 | 0.038971309 | 0.677239568 | 0.498253967 | NA |
| Got1 | 13979.28938 | 0.025833826 | 0.062396673 | 0.414025704 | 0.678855283 | 0.833141285 | Kir3dl2 | 9.851000786 | 0.026411048 | 0.061745933 | 0.427737456 | 0.668842276 | NA |
| Pcm1 | 4244.235441 | -0.036043109 | 0.087054898 | -0.414027356 | 0.678854073 | 0.833141285 | Jhdm1d | 2549.879036 | 0.026411338 | 0.089541547 | 0.29496182 | 0.768023046 | 0.94619074 |
| Sgpp2 | 1175.770352 | 0.038099865 | 0.092011673 | 0.41407643 | 0.678818134 | 0.833141285 | Rnf187 | 11503.69444 | 0.026432014 | 0.040549558 | 0.651844707 | 0.514501358 | 0.853195313 |
| Thsd4 | 429.2092741 | -0.060368752 | 0.145808002 | -0.414029076 | 0.678852813 | 0.833141285 | Slc13a1 | 6.266668738 | 0.026464675 | 0.05437216 | 0.486732092 | 0.6264482 | NA |
| Polr3f | 971.9951725 | 0.03221143 | 0.077865489 | 0.413680448 | 0.679108148 | 0.833367784 | 7530420F21Rik | 14.24436944 | 0.026471014 | 0.074857956 | 0.353616573 | 0.723626252 | NA |
| Ypel5 | 3604.148929 | -0.032937368 | 0.079650648 | -0.413522913 | 0.679223539 | 0.833443376 | Ntn3 | 401.0350304 | 0.026493966 | 0.080754105 | 0.328081967 | 0.742849688 | 0.939912916 |
| Adamts15 | 250.6182692 | -0.038684573 | 0.093599321 | -0.413299719 | 0.679387035 | 0.833506033 | Parp1 | 3327.02438 | 0.026502761 | 0.058979145 | 0.449358171 | 0.6531733 | 0.914683958 |
| Dusp7 | 2576.178196 | -0.03368325 | 0.081642093 | -0.412572099 | 0.679920146 | 0.833506033 | Mgat1 | 2407.477624 | 0.026530352 | 0.059662571 | 0.444673293 | 0.656555872 | 0.915208181 |
| Ecm2 | 150.5324112 | -0.056678243 | 0.137287779 | -0.412842593 | 0.679721943 | 0.833506033 | Bhlhe22 | 997.3186921 | 0.026543294 | 0.082701413 | 0.320953331 | 0.748245758 | 0.9410864 |
| Esm1 | 11.91931579 | -0.068266928 | 0.165428 | -0.412668522 | 0.679849491 | 0.833506033 | Smim13 | 11035.80032 | 0.026553451 | 0.072693939 | 0.365277374 | 0.714904366 | 0.93125489 |
| Gabarap | 4629.714203 | -0.019388335 | 0.046992855 | -0.412580485 | 0.679914001 | 0.833506033 | Gm20744 | 0.684132458 | 0.026563716 | 0.019110759 | 1.389987503 | 0.164532672 | NA |
| Got2 | 11586.04955 | 0.031151311 | 0.075494702 | 0.412629104 | 0.679878375 | 0.833506033 | Wdr11 | 2466.112379 | 0.026569025 | 0.04882354 | 0.544184744 | 0.586314349 | 0.884588669 |
| Metap1 | 1573.517111 | 0.027838949 | 0.067454133 | 0.412709309 | 0.679819603 | 0.833506033 | Zfp955a | 457.8265945 | 0.026606287 | 0.080785262 | 0.329345804 | 0.741894327 | 0.93956033 |
| Rnf2 | 895.9670994 | 0.040274253 | 0.097537496 | 0.412910469 | 0.67967221 | 0.833506033 | Actb | 39092.0296 | 0.026608846 | 0.053022864 | 0.501837212 | 0.615782035 | 0.896906632 |
| Snora28 | 35.56134574 | 0.06649318 | 0.160855103 | 0.413373146 | 0.679333246 | 0.833506033 | Hcrt | 21.47296838 | 0.026611066 | 0.021465925 | 1.239688788 | NA | NA |
| Syf2 | 1426.741235 | 0.033282253 | 0.080560676 | 0.413132736 | 0.679509367 | 0.833506033 | Nek7 | 1830.046854 | 0.026644674 | 0.068150697 | 0.390967007 | 0.695821623 | 0.925527709 |
| Tmem248 | 2060.144527 | 0.03147079 | 0.076180287 | 0.413109371 | 0.679526484 | 0.833506033 | Grid2ip | 155.894082 | 0.026658086 | 0.095102521 | 0.280308931 | 0.779240499 | 0.948407834 |
| Unc5b | 1323.874436 | -0.068188771 | 0.165225327 | -0.412701684 | 0.679825191 | 0.833506033 | Adad2 | 3.406875155 | 0.026660396 | 0.042370591 | 0.629219341 | 0.529205469 | NA |
| Ube4a | 3101.553318 | -0.02872725 | 0.069661652 | -0.412382557 | 0.680059046 | 0.833610353 | Rspo1 | 645.2136961 | 0.026670765 | 0.078453425 | 0.33995667 | 0.733889159 | 0.936456337 |
| Gpatch11 | 1061.399188 | -0.025453797 | 0.061787654 | -0.411956038 | 0.680371646 | 0.833798326 | B3gnt6 | 14.01018673 | 0.026671624 | 0.07406805 | 0.360096204 | 0.718775191 | NA |
| Pcdh8 | 1208.425982 | 0.048354583 | 0.117368133 | 0.41199073 | 0.680346218 | 0.833798326 | Rgl1 | 5323.274968 | 0.026682335 | 0.061307364 | 0.435222352 | 0.66340104 | 0.915428982 |
| Phf19 | 40.98642829 | 0.063172588 | 0.153349001 | 0.411953049 | 0.680373837 | 0.833798326 | Ccdc135 | 119.0861456 | 0.02668373 | 0.052974574 | 0.503708253 | 0.614466408 | 0.89611764 |
| Gpr3 | 257.4173278 | -0.065311378 | 0.158602963 | -0.411791663 | 0.680492133 | 0.833877342 | Brf1 | 1495.667284 | 0.026684631 | 0.048572708 | 0.549374992 | 0.582748133 | 0.88406516 |
| Sfmbt2 | 109.2970527 | -0.061898236 | 0.150359849 | -0.411667323 | 0.680583279 | 0.833923079 | Stbd1 | 51.4973384 | 0.026687821 | 0.089731106 | 0.297419953 | 0.766145912 | 0.945703223 |
| Ankmy1 | 15.39265554 | 0.068861152 | 0.16738576 | 0.411391939 | 0.680785164 | 0.834104485 | Tnni1 | 128.838898 | 0.026701172 | 0.093066927 | 0.286902904 | 0.774186667 | 0.947238918 |
| Gprc5a | 13.00459215 | 0.060073292 | 0.146059266 | 0.411293951 | 0.680857005 | 0.834126546 | Uncx | 6.865153869 | 0.026720745 | 0.049643004 | 0.538258023 | 0.590398927 | NA |
| Cnot11 | 1171.390626 | 0.02685076 | 0.06531239 | 0.411112806 | 0.68098982 | 0.834127781 | Fbln5 | 279.381418 | 0.026732384 | 0.086081916 | 0.310545879 | 0.756145877 | 0.942139348 |
| Fam76a | 2070.424221 | 0.020548532 | 0.04997343 | 0.411189139 | 0.680933851 | 0.834127781 | Wfdc2 | 18.47947389 | 0.026742948 | 0.035228852 | 0.759120611 | NA | NA |
| Rpl21 | 9906.652497 | 0.028882592 | 0.070261587 | 0.411072302 | 0.681019519 | 0.834127781 | Etnk1 | 16398.34267 | 0.026752524 | 0.060761413 | 0.440288052 | 0.659728493 | 0.915208181 |
| 6030458C11Rik | 1663.573487 | 0.023457165 | 0.057079583 | 0.410955434 | 0.681105214 | 0.834166799 | Dlx2 | 228.8366357 | 0.026772556 | 0.088329452 | 0.303098851 | 0.76181453 | 0.943724873 |
| Pacs1 | 4444.263963 | -0.026494292 | 0.064501462 | -0.410754906 | 0.681252262 | 0.834215012 | Apobec2 | 4.194527813 | 0.02679462 | 0.048012483 | 0.558076127 | 0.576792402 | NA |
| Spats2 | 1031.01526 | -0.034294694 | 0.08348009 | -0.410812859 | 0.681209763 | 0.834215012 | Mag | 8433.319032 | 0.026795538 | 0.081302781 | 0.329577142 | 0.741719496 | 0.939502738 |
| Dennd2d | 15.34345509 | -0.066004395 | 0.160824054 | -0.410413699 | 0.681502499 | 0.834389547 | Mcc | 690.6135849 | 0.026796218 | 0.07198404 | 0.372252214 | 0.709705074 | 0.930224827 |
| Txnl4a | 4107.043261 | 0.029601396 | 0.07212004 | 0.410446192 | 0.681478668 | 0.834389547 | Stxbp6 | 2842.721118 | 0.02680464 | 0.062629812 | 0.427985315 | 0.668661812 | 0.917270893 |
| Aldh1l2 | 158.0304863 | -0.045691695 | 0.111426619 | -0.410060854 | 0.681761308 | 0.83457452 | Phf13 | 765.4796978 | 0.026832831 | 0.076358643 | 0.351405288 | 0.725284316 | 0.933334512 |
| Bphl | 788.699158 | -0.033124039 | 0.080775707 | -0.410074273 | 0.681751465 | 0.83457452 | Cpq | 539.7141006 | 0.026834746 | 0.07975934 | 0.336446446 | 0.73653423 | 0.937206199 |
| 2010107G23Rik | 795.435503 | -0.03071787 | 0.07504445 | -0.409329008 | 0.68229823 | 0.834770118 | Maml3 | 497.3678703 | 0.026838212 | 0.093376337 | 0.287419846 | 0.773790867 | 0.947121017 |
| A930012L18Rik | 79.56746871 | 0.066324821 | 0.161962891 | 0.40950628 | 0.682168159 | 0.834770118 | Ldlrad4 | 1421.229055 | 0.026839494 | 0.07371999 | 0.36407349 | 0.715803132 | 0.93125489 |
| Amn | 99.27541316 | -0.057702752 | 0.140947664 | -0.409391334 | 0.682252498 | 0.834770118 | Gm5087 | 2.76313607 | 0.026840366 | 0.035615263 | 0.75361975 | 0.451077582 | NA |
| Ano5 | 54.65094741 | -0.065587281 | 0.160172649 | -0.409478658 | 0.682188426 | 0.834770118 | Ccdc43 | 763.6764426 | 0.026859084 | 0.0677288 | 0.396568136 | 0.691685954 | 0.923860706 |
| Casp8ap2 | 573.727199 | 0.036309526 | 0.088690013 | 0.409398136 | 0.682247508 | 0.834770118 | Dhx32 | 1455.253134 | 0.026894399 | 0.058257369 | 0.46164802 | 0.644333753 | 0.912152041 |
| Ppp1r1a | 2731.781984 | -0.03014624 | 0.073638491 | -0.409381559 | 0.682259671 | 0.834770118 | Taf9 | 2574.138369 | 0.026899261 | 0.061540802 | 0.437096361 | 0.662041466 | 0.915428982 |
| Yipf4 | 1095.404243 | -0.038188722 | 0.09326966 | -0.409444203 | 0.682213706 | 0.834770118 | Gstt4 | 1.223288234 | 0.026930903 | 0.025307421 | 1.064150452 | 0.287260549 | NA |
| Pcdha10 | 100.6843495 | -0.059437697 | 0.145255475 | -0.409194197 | 0.682397153 | 0.834825226 | Herc1 | 9642.374489 | 0.026938647 | 0.068055469 | 0.395833686 | 0.692227724 | 0.923860706 |
| 2310009B15Rik | 128.9386165 | 0.046550827 | 0.113800209 | 0.409057485 | 0.682497476 | 0.83487957 | Sebox | 72.45696277 | 0.026938749 | 0.094818837 | 0.284107568 | 0.776327948 | 0.947241001 |
| Pld3 | 25469.9894 | -0.03587209 | 0.087709651 | -0.408986808 | 0.682549342 | 0.83487957 | Nova1 | 2529.555626 | 0.026946367 | 0.07649238 | 0.352275184 | 0.724631897 | 0.932965303 |
| C1rl | 20.98685579 | 0.064222249 | 0.15706913 | 0.408878873 | 0.682628554 | 0.834910548 | Gm15441 | 2.796897423 | 0.026952028 | 0.037620999 | 0.716409164 | 0.473738738 | NA |
| Amotl2 | 789.8739848 | 0.043092997 | 0.105482066 | 0.408533873 | 0.682881767 | 0.835099429 | Ephx4 | 2125.282538 | 0.026961586 | 0.072489543 | 0.371937589 | 0.709939318 | 0.930318316 |
| Cad | 912.7019598 | -0.042352184 | 0.103705673 | -0.408388306 | 0.682988618 | 0.835099429 | Smarca4 | 7084.989487 | 0.026962107 | 0.049174869 | 0.548290368 | 0.58349254 | 0.884313345 |
| Cmklr1 | 167.0508792 | 0.052072633 | 0.127482752 | 0.408468064 | 0.682930072 | 0.835099429 | Gid8 | 1858.276807 | 0.026963696 | 0.057181719 | 0.471543995 | 0.63725231 | 0.908932006 |
| Map1lc3b | 7861.185546 | 0.02381103 | 0.0583173 | 0.408301312 | 0.683052476 | 0.835099429 | Tab3 | 1142.917156 | 0.02696522 | 0.073392416 | 0.367411532 | 0.713312071 | 0.930507332 |
| Tex30 | 98.51765765 | 0.058672218 | 0.143676377 | 0.408363707 | 0.683006674 | 0.835099429 | Adipor1 | 3183.892681 | 0.0269673 | 0.048321117 | 0.558085187 | 0.576786215 | 0.881151498 |
| Manea | 1372.564735 | -0.037703714 | 0.092416128 | -0.407977647 | 0.683290086 | 0.835324017 | Otp | 15.53315368 | 0.026976496 | 0.031902944 | 0.845580164 | NA | NA |
| A630033H20Rik | 10.0229666 | -0.06385784 | 0.156590368 | -0.40780184 | 0.683419162 | 0.835349992 | Ica1 | 1917.557601 | 0.026978155 | 0.056712041 | 0.475704183 | 0.634285141 | 0.907220906 |
| Tsacc | 26.71589533 | 0.067829382 | 0.166321004 | 0.407822107 | 0.683404282 | 0.835349992 | Pde6b | 10.76250585 | 0.026982566 | 0.06209651 | 0.434526282 | 0.663906314 | NA |
| Gm11627 | 73.93308866 | -0.059754313 | 0.14670314 | -0.407314477 | 0.683777031 | 0.83565557 | Nme9 | 10.8260449 | 0.026986345 | 0.049904818 | 0.540756301 | 0.588675567 | NA |
| Kdm3a | 2216.805962 | 0.041904546 | 0.102873567 | 0.407340267 | 0.683758092 | 0.83565557 | Icam2 | 98.39521827 | 0.027000339 | 0.094855848 | 0.284646016 | 0.775915353 | 0.947241001 |
| Rassf8 | 329.7976563 | 0.0619367 | 0.152106005 | 0.407194311 | 0.683865279 | 0.835697503 | Sept6 | 3990.18888 | 0.027003368 | 0.05284299 | 0.511011362 | 0.609343101 | 0.89440021 |
| Yy2 | 38.90033311 | -0.061939187 | 0.152196998 | -0.406967206 | 0.684032075 | 0.835835407 | Arpc5 | 2892.799947 | 0.027007254 | 0.070091762 | 0.385312812 | 0.70000567 | 0.926781946 |
| 1700030C10Rik | 41.75334185 | 0.067525145 | 0.166045509 | 0.406666493 | 0.684252953 | 0.835958036 | Hddc2 | 337.2468707 | 0.027017805 | 0.079663513 | 0.339149047 | 0.734497451 | 0.936456337 |
| Cxcr2 | 20.95648698 | -0.06740143 | 0.165764232 | -0.406610216 | 0.684294292 | 0.835958036 | Trim27 | 989.2562488 | 0.027021402 | 0.059849014 | 0.451492845 | 0.651634381 | 0.914447303 |
| Sytl3 | 8.126615697 | 0.063615322 | 0.156409218 | 0.40672361 | 0.684210998 | 0.835958036 | Ranbp2 | 6111.813253 | 0.027029435 | 0.083858868 | 0.32232053 | 0.747209882 | 0.94097069 |
| Bckdk | 1456.264235 | 0.029875973 | 0.07351349 | 0.406401241 | 0.684447808 | 0.836079655 | Hsd17b4 | 2495.058912 | 0.027036468 | 0.044884952 | 0.602350376 | 0.546940935 | 0.867343234 |
| Nkap | 594.8974103 | -0.031904205 | 0.078518468 | -0.406327401 | 0.684502055 | 0.836080003 | 2210409D07Rik | 1.583313647 | 0.027043396 | 0.027668432 | 0.977409794 | 0.328366316 | NA |
| Klhl25 | 410.6960091 | -0.038728145 | 0.095345092 | -0.406189179 | 0.684603604 | 0.836138124 | Bsn | 20513.01519 | 0.027049133 | 0.077803198 | 0.347660946 | 0.72809483 | 0.93397258 |

| | | | | | | |
|---|---|---|---|---|---|---|
| Aga | 181.2810359 | 0.045689568 | 0.112523673 | 0.406044053 | 0.684710232 | 0.836202439 |
| Kbtbd8 | 267.8250472 | -0.048316193 | 0.119084397 | -0.405730675 | 0.684940501 | 0.836417727 |
| Lrrc10b | 1828.373756 | 0.044464635 | 0.109643123 | 0.405539664 | 0.68508087 | 0.836523208 |
| Pllp | 1571.60785 | 0.067917795 | 0.167506536 | 0.405463554 | 0.685136803 | 0.836525582 |
| Rangap1 | 10335.5674 | -0.025702348 | 0.063450772 | -0.405075424 | 0.685422071 | 0.836796779 |
| Xrn2 | 1863.653306 | -0.025624681 | 0.063280051 | -0.404940912 | 0.685520945 | 0.836796779 |
| Zfp354a | 300.3412002 | 0.040246773 | 0.099374148 | 0.405002446 | 0.685475713 | 0.836796779 |
| Mfsd9 | 227.6935512 | -0.046850541 | 0.115767722 | -0.404694331 | 0.68570221 | 0.836952107 |
| Ccdc53 | 414.3496882 | 0.035158573 | 0.086908667 | 0.404546224 | 0.685811094 | 0.837009587 |
| Dtna | 5221.727049 | -0.017857933 | 0.044158009 | -0.404409837 | 0.685911367 | 0.837009587 |
| Fam179a | 146.8151778 | 0.053414648 | 0.132071751 | 0.404436583 | 0.685891703 | 0.837009587 |
| Arc | 9540.124169 | 0.060139412 | 0.148748453 | 0.404302777 | 0.685990084 | 0.83703972 |
| Psme2 | 772.3363048 | -0.028931852 | 0.071612342 | -0.4040065 | 0.686207939 | 0.83723961 |
| Smarca5 | 2757.44182 | 0.032067671 | 0.079436033 | 0.403691747 | 0.686439409 | 0.837456079 |
| Rnf220 | 5440.385001 | 0.029801739 | 0.073854293 | 0.403520746 | 0.686565176 | 0.837543567 |
| Mgst3 | 2227.095869 | -0.039387021 | 0.097649408 | -0.403351354 | 0.686689768 | 0.837599793 |
| Plagl2 | 418.5569075 | 0.037298967 | 0.092481878 | 0.403311088 | 0.686719386 | 0.837599793 |
| 1700003M07Rik | 660.3946791 | -0.038614474 | 0.095838055 | -0.402913794 | 0.687011643 | 0.83763747 |
| Bcan | 9766.838403 | 0.031938297 | 0.079270706 | 0.402901638 | 0.687020586 | 0.83763747 |
| Fam192a | 1543.191143 | 0.020741125 | 0.051478878 | 0.402905544 | 0.687017712 | 0.83763747 |
| Neurl2 | 60.00124942 | 0.057442313 | 0.142481912 | 0.403155125 | 0.68683411 | 0.83763747 |
| Wdr5 | 1106.548806 | 0.025343042 | 0.06288924 | 0.402978986 | 0.686963683 | 0.83763747 |
| Ptpla | 193.4111578 | -0.043673695 | 0.108511975 | -0.402478115 | 0.687332191 | 0.837951449 |
| Vit | 82.39122005 | -0.055692689 | 0.138524453 | -0.402042298 | 0.687652898 | 0.838276475 |
| Abcf2 | 4194.718615 | 0.031491521 | 0.078369054 | 0.401836171 | 0.6878046 | 0.83832949 |
| Dnmt3b | 25.11731642 | 0.067067145 | 0.166871628 | 0.401908617 | 0.687751281 | 0.83832949 |
| Cdc42ep1 | 821.9142013 | 0.053383909 | 0.132941511 | 0.401559368 | 0.688008338 | 0.838511853 |
| Pofut1 | 934.8873864 | -0.029263613 | 0.072893089 | -0.401459367 | 0.688081948 | 0.838535606 |
| Gpx3 | 386.4456488 | -0.067225304 | 0.167503861 | -0.401335845 | 0.688172876 | 0.838580459 |
| 1600002K03Rik | 71.03084095 | 0.058013124 | 0.144772723 | 0.40071861 | 0.688627308 | 0.838994182 |
| Hexb | 4530.336024 | 0.054426439 | 0.135842691 | 0.400657839 | 0.688672055 | 0.838994182 |
| Sfpi1 | 107.2398593 | 0.055718714 | 0.139069387 | 0.400654054 | 0.688674843 | 0.838994182 |
| Agpat6 | 3763.62684 | -0.023944907 | 0.059827772 | -0.400230637 | 0.688986651 | 0.839242074 |
| Asf1a | 355.9358031 | -0.035580053 | 0.088884471 | -0.400295486 | 0.688938892 | 0.839242074 |
| Ormdl1 | 278.3572034 | -0.039656793 | 0.099104302 | -0.400152078 | 0.689044509 | 0.83924657 |
| Ccdc79 | 37.12780319 | -0.065955654 | 0.164864214 | -0.400060466 | 0.689111982 | 0.839262777 |
| Fcrl1 | 17.80957779 | 0.066442633 | 0.166127649 | 0.399949278 | 0.689193876 | 0.839296542 |
| Rtn4rl2 | 832.2749924 | -0.04348449 | 0.108745749 | -0.399873011 | 0.689250052 | 0.839298986 |
| Chaf1a | 208.4733668 | 0.042383003 | 0.106101926 | 0.399455553 | 0.689557567 | 0.839342364 |
| Dmc1 | 25.06697163 | -0.067073538 | 0.167966988 | -0.399325716 | 0.689653221 | 0.839342364 |
| Efha1 | 825.3441301 | 0.029182917 | 0.073072996 | 0.399366638 | 0.689623072 | 0.839342364 |
| Nfatc1 | 273.447019 | -0.039985787 | 0.100063617 | -0.399603657 | 0.689448463 | 0.839342364 |
| Parp1 | 3060.247051 | 0.028112605 | 0.070389922 | 0.399383951 | 0.689610318 | 0.839342364 |
| Phkg1 | 546.0504954 | 0.060236702 | 0.150858144 | 0.399293674 | 0.689676828 | 0.839342364 |
| Scaf8 | 2646.538638 | 0.0296285 | 0.074219712 | 0.399199874 | 0.689745936 | 0.839342364 |
| Tgfb3 | 609.7779621 | 0.05113336 | 0.12810551 | 0.399150356 | 0.68978242 | 0.839342364 |
| Tnfrsf22 | 37.57004468 | -0.066271884 | 0.166057775 | -0.399089314 | 0.689827395 | 0.839342364 |
| Zfand5 | 5774.781522 | -0.029129811 | 0.072978563 | -0.399155727 | 0.689778463 | 0.839342364 |
| Mfi2 | 33.32889935 | 0.066366362 | 0.166363551 | 0.398923688 | 0.689949434 | 0.839424934 |
| Gpr179 | 36.0180086 | 0.064177778 | 0.161075798 | 0.398432159 | 0.690311657 | 0.839697927 |
| Tmem74b | 126.2159995 | -0.056264341 | 0.141226258 | -0.398398585 | 0.690336401 | 0.839697927 |
| Zfp784 | 196.996887 | -0.03802579 | 0.095421127 | -0.398504938 | 0.690258019 | 0.839697927 |
| Olfr287 | 149.0922863 | -0.052759501 | 0.13248547 | -0.398228584 | 0.690461698 | 0.839784406 |
| Gm16062 | 34.96471668 | 0.06516609 | 0.163853342 | 0.397709863 | 0.690844066 | 0.840183513 |
| Snora65 | 38.85525724 | 0.066231292 | 0.166665587 | 0.397390324 | 0.69107965 | 0.840404057 |
| Fam222a | 391.3833895 | -0.049039226 | 0.123476484 | -0.39715438 | 0.691253621 | 0.840515881 |
| Mllt1 | 3411.091759 | -0.035317246 | 0.088933779 | -0.397118471 | 0.6912801 | 0.840515881 |
| Top2b | 4331.857518 | -0.024492177 | 0.061692551 | -0.397003792 | 0.691364664 | 0.840552739 |
| Mettl25 | 177.0620277 | -0.053379306 | 0.134525808 | -0.396796024 | 0.691517883 | 0.840673055 |
| Pianp | 6769.683666 | 0.023412932 | 0.05906214 | 0.396411856 | 0.691801221 | 0.840951524 |
| Lsm14a | 2436.089014 | 0.029168993 | 0.073624027 | 0.396188504 | 0.691965971 | 0.841034242 |
| Mapk8ip2 | 11704.49379 | 0.027230757 | 0.068734609 | 0.396172427 | 0.691977831 | 0.841034242 |
| Aldh3a2 | 2160.572348 | 0.017036943 | 0.043053913 | 0.395711849 | 0.692317613 | 0.841315224 |
| Hfe | 154.6631992 | -0.05146739 | 0.130041332 | -0.395777165 | 0.692269424 | 0.841315224 |
| 2010107G12Rik | 15.92267058 | 0.065523624 | 0.165653057 | 0.395547328 | 0.692439 | 0.841396743 |
| 1700003M02Rik | 56.26096886 | 0.062651637 | 0.158493806 | 0.395293914 | 0.692625991 | 0.841530804 |
| Mtag2 | 32.39563154 | 0.063263813 | 0.160179311 | 0.394956207 | 0.692875207 | 0.841530804 |
| Pfkp | 11400.6714 | -0.020219374 | 0.051161958 | -0.395203287 | 0.692692867 | 0.841530804 |
| Rab3ip | 1317.752991 | -0.034754612 | 0.087990749 | -0.394980297 | 0.692857429 | 0.841530804 |
| Snora7a | 24.12184307 | -0.066360765 | 0.167950999 | -0.395119798 | 0.692754478 | 0.841530804 |
| Zfp91 | 3705.715971 | -0.033603239 | 0.085062009 | -0.395044027 | 0.692810396 | 0.841530804 |
| Wdr12 | 1248.92305 | 0.026173169 | 0.066348901 | 0.39447781 | 0.693228307 | 0.841893666 |
| Aif1 | 206.2758759 | 0.064627624 | 0.1638966 | 0.394319493 | 0.693345173 | 0.841969599 |
| 2310067B10Rik | 3429.968763 | 0.02238125 | 0.056827085 | 0.393848289 | 0.69369305 | 0.842326028 |
| Anp32b | 1811.987244 | -0.030006641 | 0.076231737 | -0.393623995 | 0.693858663 | 0.842461102 |
| Zfp418 | 141.8325229 | -0.04409291 | 0.112084624 | -0.393389467 | 0.694031848 | 0.842605348 |
| 4933412E12Rik | 168.805319 | -0.048073042 | 0.122264864 | -0.393187707 | 0.694180848 | 0.842644671 |
| Chd2 | 2865.842598 | -0.043940338 | 0.11176937 | -0.393133985 | 0.694220524 | 0.842644671 |
| Mat2b | 4187.440749 | 0.028264945 | 0.07189817 | 0.393124685 | 0.694227393 | 0.842644671 |
| Ube2r2 | 3125.27766 | 0.020939221 | 0.053289438 | 0.39293379 | 0.694368384 | 0.842749784 |
| Alg6 | 291.4031744 | 0.044408702 | 0.113040349 | 0.392857084 | 0.69442504 | 0.842752531 |
| Abhd3 | 1836.469352 | 0.029619715 | 0.075442838 | 0.39261136 | 0.694606548 | 0.842774773 |
| Bmi1 | 1313.96627 | 0.027527885 | 0.070110763 | 0.392634218 | 0.694589663 | 0.842774773 |
| Tug1 | 3967.912928 | -0.027551446 | 0.070159533 | -0.392697115 | 0.694543202 | 0.842774773 |
| Nup210l | 20.13740858 | -0.065542483 | 0.167019857 | -0.3924233 | 0.694745473 | 0.842877328 |
| AA986860 | 146.5955933 | 0.0460316 | 0.117407751 | 0.392066109 | 0.695009368 | 0.843085508 |
| C8g | 31.25616881 | 0.065244137 | 0.166467579 | 0.391932997 | 0.695107722 | 0.843085508 |
| Grb7 | 90.33636937 | 0.047807846 | 0.121973216 | 0.391953637 | 0.695092471 | 0.843085508 |
| Zfp773 | 85.10809361 | 0.049244226 | 0.125656218 | 0.391896457 | 0.695134721 | 0.843085508 |
| 2610100L16Rik | 185.5937378 | -0.045279267 | 0.115594281 | -0.391708539 | 0.69527358 | 0.843187919 |
| Mc5r | 12.8930283 | 0.063131737 | 0.161278913 | 0.391444462 | 0.695468663 | 0.843303626 |
| Zic5 | 49.68080126 | -0.061336979 | 0.156698886 | -0.391432132 | 0.695477845 | 0.843303626 |
| Sst | 5341.811177 | 0.030222106 | 0.077237779 | 0.391286574 | 0.695585422 | 0.843368067 |
| Arl5b | 366.6798005 | -0.041079447 | 0.10500875 | -0.391200222 | 0.695649244 | 0.843379451 |
| N6amt1 | 718.4268342 | -0.029724888 | 0.07601353 | -0.391047338 | 0.695762246 | 0.843384465 |
| Spry2 | 1909.683635 | -0.047083299 | 0.120395533 | -0.391071809 | 0.695744157 | 0.843384465 |
| Tstd3 | 704.4772325 | 0.025840903 | 0.0661138 | 0.390854896 | 0.695904495 | 0.843490905 |
| Smagp | 42.09226627 | -0.059086622 | 0.151201547 | -0.39078054 | 0.69595946 | 0.843491543 |
| Srgap3 | 16963.41267 | -0.028421465 | 0.072766935 | -0.390582139 | 0.69610613 | 0.843603315 |
| 9030025P20Rik | 58.63900224 | 0.064237722 | 0.16452512 | 0.390443248 | 0.696208813 | 0.843661766 |
| Dtx4 | 2073.992369 | 0.031096735 | 0.079693813 | 0.390202624 | 0.696386721 | 0.84377249 |
| Gpr133 | 269.2582667 | 0.056524612 | 0.144870879 | 0.390172358 | 0.6964091 | 0.84377249 |
| Fam73a | 4280.187357 | 0.025325969 | 0.064972193 | 0.389797042 | 0.696686631 | 0.843827087 |
| Fuz | 439.7195122 | -0.056198743 | 0.14422155 | -0.389669528 | 0.696780932 | 0.843827087 |
| Hsd17b4 | 2302.102311 | 0.023866663 | 0.061232895 | 0.389768651 | 0.696707627 | 0.843827087 |
| Plxnb3 | 1186.867762 | 0.056628476 | 0.145244253 | 0.389884454 | 0.69662199 | 0.843827087 |
| Rab26 | 1197.178635 | -0.032308076 | 0.082909254 | -0.389679976 | 0.696773205 | 0.843827087 |
| Zfp81 | 341.8961846 | -0.04418349 | 0.113335699 | -0.389846191 | 0.696650286 | 0.843827087 |
| Arl6ip1 | 9707.698232 | -0.028794815 | 0.073926905 | -0.389503861 | 0.696903455 | 0.843843555 |
| Commd5 | 425.3007849 | -0.033547374 | 0.086119502 | -0.389544444 | 0.69687344 | 0.843843555 |
| Clcn6 | 2520.924009 | -0.020796594 | 0.053445316 | -0.389119112 | 0.697188036 | 0.844056214 |
| Kcnmb1 | 45.78864986 | -0.065392905 | 0.168047908 | -0.389132516 | 0.697178121 | 0.844056214 |
| Usf1 | 1790.081057 | 0.026916393 | 0.06922647 | 0.388816495 | 0.697411898 | 0.844261266 |
| Ltn1 | 2659.250855 | -0.023786881 | 0.061191174 | -0.388730582 | 0.697475457 | 0.844272245 |
| 9130017N09Rik | 9.187396398 | 0.060423061 | 0.15556757 | 0.388403966 | 0.69771711 | 0.844441138 |
| Ankrd46 | 7994.453846 | 0.019788812 | 0.050950634 | 0.388391877 | 0.697726055 | 0.844441138 |
| Fbxo33 | 1961.716611 | 0.035248455 | 0.090788653 | 0.388247359 | 0.697832989 | 0.844441138 |
| Rasgrf2 | 8122.328737 | 0.029229992 | 0.075274904 | 0.388309927 | 0.697786692 | 0.844441138 |
| Mon1a | 738.4380455 | -0.026846981 | 0.06919922 | -0.387966523 | 0.698040808 | 0.844626652 |
| Jak2 | 2195.093135 | -0.036377935 | 0.09390351 | -0.387396973 | 0.698462345 | 0.845070714 |
| Fabp3 | 2156.524791 | 0.029039182 | 0.075006478 | 0.387155648 | 0.698640982 | 0.845220846 |
| Gpr139 | 41.83853061 | 0.062438676 | 0.161548223 | 0.386501779 | 0.699125085 | 0.845410448 |
| Ispd | 249.1273609 | -0.050746475 | 0.131277369 | -0.3865592 | 0.699082568 | 0.845410448 |
| Mark3 | 2723.965011 | -0.029445165 | 0.076142225 | -0.386712699 | 0.698968914 | 0.845410448 |
| Rasip1 | 334.2003882 | 0.039342935 | 0.101731422 | 0.386733366 | 0.698953612 | 0.845410448 |
| Rsu1 | 759.3770822 | -0.031148325 | 0.080576545 | -0.386568146 | 0.699075944 | 0.845410448 |
| Taf9b | 1138.169589 | -0.028338511 | 0.073254365 | -0.386850811 | 0.698866658 | 0.845410448 |
| Bid | 395.5447106 | 0.037935109 | 0.098230812 | 0.386183397 | 0.699360849 | 0.84548184 |
| Them6 | 1651.046142 | -0.035819261 | 0.092765421 | -0.386127293 | 0.699402398 | 0.84548184 |
| Tor1aip2 | 2140.205571 | 0.022300524 | 0.057744035 | 0.386196149 | 0.699351405 | 0.84548184 |
| Zfp868 | 448.0792393 | -0.034222656 | 0.08861437 | -0.386197584 | 0.699350343 | 0.84548184 |
| Pcmt1 | 4767.629079 | -0.025948313 | 0.067250235 | -0.385847172 | 0.699609857 | 0.845666651 |
| Brs3 | 9.319675715 | 0.048610902 | 0.126043657 | 0.385667179 | 0.699743173 | 0.845695844 |
| Krtcap2 | 711.2375452 | 0.025658781 | 0.06652565 | 0.385697562 | 0.699720669 | 0.845695844 |
| Nme7 | 607.0840906 | -0.026452246 | 0.068604265 | -0.385577285 | 0.699809758 | 0.84571035 |
| B230217O12Rik | 371.3139376 | -0.036975898 | 0.095999573 | -0.385167321 | 0.700113453 | 0.846011375 |
| Bhlhb9 | 2573.006799 | -0.029201478 | 0.075872705 | -0.384874617 | 0.700330313 | 0.846207429 |
| Gdap2 | 1286.869663 | 0.037675469 | 0.097912069 | 0.384788811 | 0.70039389 | 0.846218257 |
| Nol11 | 1013.98569 | -0.029049945 | 0.075546685 | -0.384529713 | 0.700585878 | 0.846318228 |
| Ppp3cb | 9947.016993 | -0.033040943 | 0.085916135 | -0.384572033 | 0.700554518 | 0.846318228 |
| Rbm27 | 1572.999398 | -0.026981311 | 0.070246418 | -0.384095187 | 0.700907899 | 0.846575765 |
| Zfp593 | 279.453477 | -0.03957743 | 0.103040844 | -0.384094583 | 0.700908346 | 0.846575765 |
| Gm10790 | 58.58123094 | 0.062580504 | 0.163030761 | 0.383857031 | 0.701084415 | 0.84672242 |
| Csnk1g1 | 1527.726187 | -0.021563721 | 0.05620739 | -0.383645649 | 0.701241102 | 0.846734721 |
| Itih3 | 1230.272969 | 0.064399527 | 0.167825911 | 0.383728155 | 0.701179943 | 0.846734721 |
| Kif24 | 32.77392906 | 0.060248353 | 0.157051448 | 0.383621758 | 0.701258811 | 0.846734721 |
| Tpr | 4758.115382 | -0.030136161 | 0.078571991 | -0.383548392 | 0.701313197 | 0.846734721 |
| Ndufaf7 | 1213.227337 | 0.020498002 | 0.053494447 | 0.383179999 | 0.701586307 | 0.84699846 |
| Stoml2 | 1499.572987 | 0.026807907 | 0.069978922 | 0.383085457 | 0.701656402 | 0.847017086 |
| Ppil3 | 491.9346333 | -0.03359692 | 0.087727436 | -0.382969358 | 0.701742484 | 0.847055005 |
| Adap1 | 7550.623652 | -0.033917203 | 0.088592482 | -0.382845159 | 0.701834575 | 0.847100172 |
| Col4a5 | 190.3006083 | -0.053208792 | 0.139131099 | -0.38243637 | 0.702137718 | 0.847268053 |
| Rsl24d1 | 1280.848223 | -0.025595047 | 0.066913968 | -0.382506784 | 0.702085498 | 0.847268053 |
| Srek1ip1 | 488.5347867 | -0.033013907 | 0.086305935 | -0.382521864 | 0.702074314 | 0.847268053 |
| Epx | 11.18929245 | 0.061630194 | 0.161225498 | 0.38226084 | 0.702267898 | 0.847271714 |
| Fam53a | 611.20584 | 0.030691363 | 0.080314977 | 0.382137483 | 0.70235939 | 0.847271714 |
| Ocln | 342.3514371 | -0.03117342 | 0.081576481 | -0.382137352 | 0.702359487 | 0.847271714 |
| Wdr37 | 2972.896414 | 0.027708478 | 0.072505721 | 0.382155749 | 0.702345843 | 0.847271714 |
| Ppp2r2d | 2126.69928 | -0.032222859 | 0.084359731 | -0.381969674 | 0.702483859 | 0.847289811 |
| Smndc1 | 865.4516355 | -0.029340821 | 0.076806056 | -0.382011812 | 0.702452603 | 0.847289811 |
| Otof | 1550.687555 | 0.050984659 | 0.133526591 | 0.381831503 | 0.70258635 | 0.847347467 |
| 2010320M18Rik | 135.5676894 | -0.045816035 | 0.120049253 | -0.381643651 | 0.702725702 | 0.847383611 |
| Fbln5 | 280.9131924 | -0.054149807 | 0.141866258 | -0.381696165 | 0.702686745 | 0.847383611 |
| Arl8a | 7663.810906 | 0.020542989 | 0.053858527 | 0.381425007 | 0.702887908 | 0.847513249 |
| Klf6 | 2067.493996 | 0.026483878 | 0.069494044 | 0.381095657 | 0.70313227 | 0.847741918 |
| Uba2 | 3359.460742 | -0.028045681 | 0.073650905 | -0.380792083 | 0.703357534 | 0.847947529 |
| Aff2 | 504.5553694 | 0.038694218 | 0.101760258 | 0.380248826 | 0.703760719 | 0.848206358 |
| Gpr101 | 348.2286257 | 0.055498738 | 0.145907217 | 0.380370071 | 0.703670728 | 0.848206358 |
| Trap1 | 2511.036339 | 0.029011226 | 0.076303616 | 0.380207751 | 0.703791206 | 0.848206358 |
| Uhrf1bp1l | 4180.839776 | 0.038827242 | 0.102102742 | 0.380276194 | 0.703740405 | 0.848206358 |
| Ankrd34a | 2536.322731 | 0.026177021 | 0.068863187 | 0.380130837 | 0.703848296 | 0.848209186 |
| Adprhl2 | 640.9688262 | 0.025645905 | 0.067504638 | 0.379913231 | 0.704009826 | 0.84827189 |
| Gng8 | 37.21611361 | 0.063615578 | 0.167416468 | 0.379983996 | 0.703957295 | 0.84827189 |
| Grhl3 | 47.40409557 | 0.059924792 | 0.157863917 | 0.37959778 | 0.704244009 | 0.848488078 |
| Bcs1l | 666.1490812 | -0.045269534 | 0.11942293 | -0.379069031 | 0.704636603 | 0.848723455 |
| Dbil5 | 12.64777304 | 0.062051921 | 0.163640243 | 0.379197193 | 0.704541436 | 0.848723455 |
| Stk39 | 3028.607662 | 0.033032404 | 0.087139605 | 0.379074516 | 0.70463253 | 0.848723455 |
| Ube3c | 4442.656102 | -0.022501689 | 0.059365012 | -0.379039567 | 0.704658482 | 0.848723455 |
| Eda2r | 17.08784489 | 0.063115673 | 0.166611154 | 0.378820214 | 0.704821376 | 0.848818156 |
| Snapc1 | 999.5470086 | 0.026354631 | 0.069576545 | 0.378786147 | 0.704846675 | 0.848818156 |
| Gorasp1 | 1363.655214 | -0.022381743 | 0.059146764 | -0.378410272 | 0.705125839 | 0.849088346 |
| Cuzd1 | 8.582796885 | 0.058118933 | 0.15361726 | 0.378335958 | 0.705181037 | 0.849088824 |
| Csf1r | 4813.85558 | 0.038719839 | 0.102387454 | 0.378169759 | 0.705304489 | 0.849105497 |
| Rin1 | 3215.854483 | 0.032166781 | 0.085057851 | 0.37817533 | 0.705300351 | 0.849105497 |
| Pdap1 | 3467.31506 | 0.029755502 | 0.078712587 | 0.378027241 | 0.705410357 | 0.849166971 |
| Ift20 | 1240.024748 | 0.022663745 | 0.059992209 | 0.377778141 | 0.705595413 | 0.849191808 |
| Tox4 | 3198.239775 | 0.021593571 | 0.057159243 | 0.377779159 | 0.705594657 | 0.849191808 |
| Zfp618 | 79.27789691 | -0.059639818 | 0.157835378 | -0.377860897 | 0.705533932 | 0.849191808 |
| Egfl6 | 96.39370506 | -0.057913811 | 0.153334885 | -0.377694947 | 0.705657222 | 0.849200233 |
| Pcbp4 | 4421.508976 | 0.030369633 | 0.080471144 | 0.3773978 | 0.705878002 | 0.84939995 |
| 4930579K19Rik | 14.1102877 | 0.062157446 | 0.164778785 | 0.377217526 | 0.706011957 | 0.849429202 |
| Tnfsf13 | 13.30771191 | 0.060548647 | 0.160487252 | 0.377280101 | 0.705965458 | 0.849429202 |
| Lrrn1 | 2747.226897 | -0.030951237 | 0.082088794 | -0.377045823 | 0.706139551 | 0.849450793 |
| Rab11fip1 | 161.312349 | 0.050893968 | 0.134977548 | 0.377055066 | 0.706132683 | 0.849450793 |
| Dhx8 | 1862.569677 | 0.023418723 | 0.062224038 | 0.37636135 | 0.706648276 | 0.84980222 |
| Nhej1 | 58.00484063 | 0.057226202 | 0.152052743 | 0.37635758 | 0.706651078 | 0.84980222 |
| Pcgf3 | 1380.565886 | 0.037299055 | 0.09906375 | 0.376515675 | 0.706533564 | 0.84980222 |
| Zswim3 | 332.9282698 | -0.038112889 | 0.101257182 | -0.376396892 | 0.706621856 | 0.84980222 |
| Lrrn2 | 4845.566684 | 0.032390796 | 0.086101701 | 0.37619229 | 0.706773947 | 0.849884015 |
| Fmod | 1225.307613 | 0.063018581 | 0.167606159 | 0.375992035 | 0.706922818 | 0.849997062 |
| Otub2 | 1393.675464 | 0.03696995 | 0.098409536 | 0.375674463 | 0.707158925 | 0.850192579 |
| Rab2b | 1609.713451 | 0.026732494 | 0.071167901 | 0.375625716 | 0.70719517 | 0.850192579 |
| Trim26 | 788.4999411 | -0.026460061 | 0.07048361 | -0.375407287 | 0.707357586 | 0.850321859 |
| Raly | 2723.950879 | -0.02349248 | 0.062591658 | -0.375329255 | 0.707415611 | 0.850325639 |
| Tmem129 | 995.40102 | 0.02389205 | 0.063679026 | 0.37519496 | 0.707515478 | 0.850379708 |
| AW551984 | 527.8791313 | 0.061020023 | 0.162798798 | 0.374818635 | 0.707795354 | 0.850403021 |
| Anapc2 | 3495.689469 | -0.021861754 | 0.058329138 | -0.374799879 | 0.707809304 | 0.850403021 |
| Aqr | 1995.205657 | 0.02482254 | 0.066180105 | 0.375075555 | 0.707604277 | 0.850403021 |
| Atp2a2 | 39242.75966 | -0.022093619 | 0.058942614 | -0.374832697 | 0.707784895 | 0.850403021 |
| Herc4 | 1218.566154 | -0.035370802 | 0.094353272 | -0.374876267 | 0.70775249 | 0.850403021 |
| Cpn1 | 12.60344936 | 0.06219576 | 0.166043579 | 0.374574917 | 0.707976631 | 0.850538103 |
| Pum1 | 4424.690301 | -0.027524402 | 0.07351877 | -0.374386056 | 0.708117116 | 0.85064092 |
| Mapk3 | 7065.011126 | -0.02758795 | 0.073740793 | -0.374120597 | 0.708314595 | 0.850812182 |
| Cyyr1 | 129.675924 | -0.049720038 | 0.13298096 | -0.373888401 | 0.708487347 | 0.850953717 |
| 2310033P09Rik | 408.34126 | -0.032738151 | 0.087627654 | -0.373605249 | 0.708698029 | 0.850973901 |
| 2810403D21Rik | 8.541697037 | -0.056505798 | 0.151286654 | -0.373501541 | 0.708775198 | 0.850973901 |
| Btc | 15.5488052 | 0.062617156 | 0.167534388 | 0.373757032 | 0.70858509 | 0.850973901 |
| Dnm1l | 8928.123971 | -0.022806773 | 0.061062843 | -0.373496747 | 0.708778766 | 0.850973901 |
| Prrg1 | 117.0713054 | 0.05829068 | 0.15602729 | 0.37359285 | 0.708707254 | 0.850973901 |
| Hdac1 | 934.2778922 | -0.024674485 | 0.066080342 | -0.373401287 | 0.708849802 | 0.850993246 |
| Gal3st3 | 1754.830002 | 0.033857842 | 0.090719147 | 0.373216055 | 0.708987648 | 0.851092788 |
| 1700113A16Rik | 35.5575411 | 0.062712493 | 0.168119141 | 0.373024112 | 0.709130498 | 0.851132384 |
| Mterfd1 | 494.2881501 | 0.034486468 | 0.092435419 | 0.373087161 | 0.709083574 | 0.851132384 |
| Cxx1c | 3927.90585 | 0.023049154 | 0.061887956 | 0.372433598 | 0.709570043 | 0.851593977 |
| Ccdc59 | 558.5149376 | 0.030063618 | 0.080753151 | 0.372290345 | 0.709676686 | 0.85160178 |
| Tmem173 | 169.5640962 | 0.051440882 | 0.138178974 | 0.372277202 | 0.709686471 | 0.85160178 |
| Crybb3 | 21.8635798 | -0.062506166 | 0.167964804 | -0.372138473 | 0.709789753 | 0.85163609 |
| Phf3 | 3743.114935 | 0.0327928 | 0.088131097 | 0.372091137 | 0.709824995 | 0.85163609 |
| Rnf113a2 | 289.042663 | 0.036400475 | 0.097851425 | 0.371997391 | 0.709894792 | 0.851653883 |
| Cxxc5 | 2718.412233 | -0.033295937 | 0.089564763 | -0.37175264 | 0.710077028 | 0.851806555 |
| Ubxn4 | 2507.749842 | 0.02162758 | 0.058190217 | 0.37167038 | 0.710138281 | 0.851814084 |
| Mir9-2 | 52.88681209 | -0.06008887 | 0.161983999 | -0.370955591 | 0.710670609 | 0.852386624 |
| Kctd7 | 611.7363298 | -0.031145514 | 0.084009329 | -0.370738754 | 0.710832122 | 0.852448367 |
| Ubtd1 | 312.8229857 | 0.034847745 | 0.093991076 | 0.370755897 | 0.710819353 | 0.852448367 |
| Dfna5 | 389.4453581 | -0.032274116 | 0.087145709 | -0.370346593 | 0.711124262 | 0.852732707 |

| | | | | | | |
|---|---|---|---|---|---|---|
| Marf1 | 7072.053541 | 0.027070697 | 0.06996064 | 0.386941806 | 0.698799291 | 0.926409493 |
| Bcl9 | 2544.904686 | 0.02707555 | 0.060389696 | 0.448347189 | 0.653902649 | 0.914733232 |
| Mlkl | 10.4517768 | 0.027082982 | 0.063869256 | 0.424037846 | 0.671538213 | NA |
| Exosc6 | 259.940508 | 0.027094134 | 0.085090673 | 0.318414848 | 0.750170277 | 0.9410864 |
| Rcbtb1 | 2124.003821 | 0.02709952 | 0.063648573 | 0.425767911 | 0.670276971 | 0.91809932 |
| Adcyap1r1 | 10004.48219 | 0.027106256 | 0.064099108 | 0.422880396 | 0.672382526 | 0.919249749 |
| Plekhm3 | 2669.989955 | 0.027109125 | 0.06343947 | 0.42732269 | 0.669144308 | 0.917499549 |
| Acn9 | 177.3891692 | 0.027115143 | 0.087554305 | 0.30969514 | 0.756792799 | 0.942139348 |
| Asap2 | 2252.773331 | 0.027135992 | 0.074281789 | 0.365311502 | 0.714878893 | 0.93125489 |
| Serpine1 | 73.06677021 | 0.027152426 | 0.086987744 | 0.312140829 | 0.754933498 | 0.942099798 |
| Snord73a | 2.36658174 | 0.027186088 | 0.03095344 | 0.878289706 | 0.379786521 | NA |
| Pcdhgb8 | 30.14092795 | 0.027189298 | 0.086669429 | 0.313712669 | 0.753739278 | 0.94183464 |
| 2610528J11Rik | 1.604340883 | 0.027192576 | 0.030991607 | 0.877417411 | 0.38025996 | NA |
| Rpp38 | 202.2329828 | 0.027208125 | 0.089688587 | 0.303362177 | 0.761613867 | 0.943656543 |
| Megf6 | 324.6525616 | 0.027208632 | 0.094635355 | 0.287510219 | 0.773721678 | 0.947121017 |
| Naca | 6008.750296 | 0.02721581 | 0.057233665 | 0.475521009 | 0.634415662 | 0.907220906 |
| Pacrgl | 226.9400187 | 0.027236817 | 0.085151873 | 0.319861631 | 0.749073225 | 0.9410864 |
| Tbrg1 | 1967.670236 | 0.027238878 | 0.04973691 | 0.547659228 | 0.583925913 | 0.884313345 |
| Gm14207 | 2.709108448 | 0.027245098 | 0.035342421 | 0.770889429 | 0.440772473 | NA |
| Rpl18a | 7142.635575 | 0.027282053 | 0.054979289 | 0.496224186 | 0.619736249 | 0.899216177 |
| Vps37b | 1310.962151 | 0.027293097 | 0.067767068 | 0.402748671 | 0.687133125 | 0.923860706 |
| Epc1 | 1758.733766 | 0.027299764 | 0.048882308 | 0.558479434 | 0.576517044 | 0.880968503 |
| Lyve1 | 34.99872084 | 0.027317502 | 0.080242391 | 0.340437291 | 0.733527239 | 0.936456337 |
| Camsap1 | 5603.648446 | 0.027321858 | 0.053506736 | 0.510624632 | 0.609613924 | 0.89440021 |
| Ubox5 | 811.9144691 | 0.027325976 | 0.068530694 | 0.398740686 | 0.690084286 | 0.923860706 |
| Dpp8 | 9723.264526 | 0.027330192 | 0.057907469 | 0.471963171 | NA | NA |
| Dynll1 | 7325.422381 | 0.027337139 | 0.050004911 | 0.546689089 | 0.584592352 | 0.884313345 |
| Bhlhb9 | 2827.598845 | 0.027347828 | 0.070686157 | 0.386890862 | 0.698837007 | 0.926409493 |
| Taok3 | 2523.449173 | 0.027348435 | 0.050920886 | 0.53707696 | 0.591214455 | 0.8858275 |
| Tpd52l1 | 672.686288 | 0.027349068 | 0.068809846 | 0.397458645 | 0.691029277 | 0.923860706 |
| Jmy | 2634.920724 | 0.027355793 | 0.065730038 | 0.416184046 | 0.677275333 | 0.921064066 |
| Dpys | 14.05175839 | 0.027359848 | 0.073370555 | 0.372899559 | 0.7092232 | NA |
| Il1f9 | 55.55358002 | 0.027364535 | 0.093679271 | 0.292108755 | 0.770203473 | 0.94619074 |
| 4933403O08Rik | 4.029635093 | 0.027385753 | 0.034863019 | 0.785524427 | 0.432146154 | NA |
| Nr0b1 | 1.195404933 | 0.027412774 | 0.025456967 | 1.076827973 | 0.281557126 | NA |
| Dhx38 | 1673.409398 | 0.027463129 | 0.059333771 | 0.46285831 | 0.643465932 | 0.911258473 |
| Clcnka | 2.341993474 | 0.027466925 | 0.025190185 | 1.090382008 | 0.275544904 | NA |
| Sobp | 3600.309384 | 0.027481221 | 0.053578743 | 0.512912758 | 0.608012349 | 0.894004456 |
| Gm1653 | 21.2467651 | 0.027481362 | 0.079830404 | 0.344246817 | 0.730660677 | NA |
| Drd2 | 1353.429454 | 0.027557788 | 0.091116978 | 0.302444055 | 0.762313575 | 0.944182042 |
| Gulo | 3.444514516 | 0.027558312 | 0.043003824 | 0.640833986 | 0.521630549 | NA |
| 2310007L24Rik | 6.803695725 | 0.027566647 | 0.056175907 | 0.490720108 | 0.623624423 | NA |
| Slc35b1 | 1551.572583 | 0.027568265 | 0.05504377 | 0.500842606 | 0.616481898 | 0.897296158 |
| 4930431P03Rik | 1.853216132 | 0.027579792 | 0.029316647 | 0.94075534 | 0.346830252 | NA |
| Mtrf1l | 260.9656684 | 0.027590518 | 0.081684559 | 0.337769082 | 0.735537211 | 0.936736238 |
| Phgdh | 2934.791273 | 0.027597743 | 0.06651024 | 0.414939768 | 0.678185997 | 0.921316909 |
| Gm5878 | 1.080185468 | 0.027606514 | 0.026281385 | 1.050420826 | 0.293524675 | NA |
| Leng9 | 248.1971712 | 0.027622002 | 0.088960693 | 0.310496704 | 0.756183266 | 0.942139348 |
| Mbnl3 | 4.206337952 | 0.027622015 | 0.041162715 | 0.671044545 | 0.502192153 | NA |
| Cetn3 | 1335.523653 | 0.027645314 | 0.060065173 | 0.460255287 | 0.645332991 | 0.912363974 |
| Snx27 | 7077.109587 | 0.027645849 | 0.061133206 | 0.452223119 | 0.651108255 | 0.914343204 |
| Sh3rf2 | 537.5569452 | 0.027667515 | 0.094337246 | 0.293283045 | 0.769305813 | 0.94619074 |
| Ss18l1 | 2321.18701 | 0.027667975 | 0.049580853 | 0.558037497 | 0.57681878 | 0.881151498 |
| Avp | 265.6136494 | 0.027670559 | 0.026701423 | 1.036295318 | NA | NA |
| Gm648 | 1.803776798 | 0.027670623 | 0.029575591 | 0.935589838 | 0.349484409 | NA |
| Azgp1 | 0.949664049 | 0.027726544 | 0.022343884 | 1.240900797 | 0.214642398 | NA |
| Kifc3 | 2161.349083 | 0.027732873 | 0.065647717 | 0.422449922 | 0.672696646 | 0.919353963 |
| Otx2os1 | 1.819187122 | 0.027737557 | 0.025283976 | 1.09704098 | 0.272623478 | NA |
| Commd2 | 379.7697922 | 0.02776097 | 0.078266736 | 0.354696911 | 0.722816666 | 0.93255988 |
| Gjc1 | 174.3377636 | 0.027771182 | 0.091995278 | 0.301876168 | 0.762746464 | 0.944431816 |
| Satb2 | 2447.434984 | 0.027786779 | 0.0766271 | 0.362623397 | 0.71688623 | 0.93125489 |
| Fam189a2 | 539.7591073 | 0.027787972 | 0.080046043 | 0.347149853 | 0.728478742 | 0.933996867 |
| Nprl2 | 1259.52434 | 0.027790571 | 0.062610237 | 0.443866248 | 0.657139288 | 0.915208181 |
| Dnajc11 | 2756.87791 | 0.027812812 | 0.050875428 | 0.546684569 | 0.584595457 | 0.884313345 |
| Itpripl1 | 95.76589268 | 0.027816648 | 0.094030772 | 0.295824945 | 0.767363772 | 0.94619074 |
| Stk11 | 2285.012166 | 0.027827393 | 0.055582558 | 0.500649726 | 0.61661766 | 0.897296158 |
| Tmem109 | 1728.013501 | 0.027842411 | 0.060760856 | 0.458229401 | 0.646787639 | 0.912750424 |
| S1pr5 | 597.2507353 | 0.02785233 | 0.094069202 | 0.296083408 | 0.767166385 | 0.946118815 |
| Gabrb1 | 798.8713675 | 0.027871414 | 0.071359802 | 0.390575834 | 0.696110791 | 0.925527709 |
| Fchsd1 | 366.9271334 | 0.027878299 | 0.079449877 | 0.350891655 | 0.725669632 | 0.933575664 |
| Vps53 | 3132.143895 | 0.027887541 | 0.051870612 | 0.537636628 | 0.590827938 | 0.885693151 |
| Cldn1 | 89.18507606 | 0.027901617 | 0.06206131 | 0.449581503 | 0.653012228 | 0.914683958 |
| Vkorc1 | 410.1066254 | 0.027925036 | 0.077310302 | 0.361207183 | 0.717944573 | 0.93125489 |
| Vps4a | 4687.123892 | 0.027926583 | 0.0462474 | 0.603851971 | 0.545942063 | 0.866989658 |
| Slain2 | 1011.342311 | 0.02793055 | 0.064497056 | 0.43305154 | 0.664977327 | 0.915428982 |
| Dusp7 | 2864.873115 | 0.02793649 | 0.060720838 | 0.460080777 | 0.645458242 | 0.912363974 |
| Anxa11 | 2198.718769 | 0.027939427 | 0.076446038 | 0.365479073 | 0.714753826 | 0.93125489 |
| Vmn2r-ps54 | 1.634148256 | 0.027950197 | 0.029765743 | 0.93900551 | 0.347727915 | NA |
| Alas2 | 11.20152991 | 0.027964297 | 0.042892911 | 0.651956142 | 0.514429466 | NA |
| Lgals2 | 3.661322689 | 0.027970501 | 0.041651711 | 0.671533064 | 0.501881004 | NA |
| 3110009E18Rik | 34.12500378 | 0.027975983 | 0.085363464 | 0.327727835 | 0.743117456 | 0.939912916 |
| Ilf3 | 3784.795342 | 0.027976535 | 0.045958239 | 0.608738201 | 0.542697982 | 0.865148254 |
| 4933431E20Rik | 5657.442047 | 0.027978499 | 0.047065615 | 0.594457322 | 0.552206279 | 0.870265825 |
| Rpl35 | 3527.452421 | 0.027997342 | 0.064390238 | 0.434807244 | 0.663702346 | 0.915428982 |
| Ccdc28a | 646.4057603 | 0.028002913 | 0.07098357 | 0.394498515 | 0.693213023 | 0.924482501 |
| Zfp609 | 2600.118983 | 0.028005125 | 0.064113795 | 0.436803417 | 0.662253921 | 0.915428982 |
| Junb | 1824.061189 | 0.028017003 | 0.078859409 | 0.355277876 | 0.722381428 | 0.93255988 |
| Slc4a10 | 9665.719981 | 0.028024234 | 0.067596361 | 0.414581991 | 0.678447934 | 0.921316909 |
| Sparcl1 | 95552.39038 | 0.028033921 | 0.054832131 | 0.511268134 | 0.609163315 | 0.89440021 |
| Kansl1 | 4712.068806 | 0.028036269 | 0.044670206 | 0.627627924 | 0.530247712 | 0.857915133 |
| Rhoq | 1655.193187 | 0.028045596 | 0.068771236 | 0.40780997 | 0.683413193 | 0.923036732 |
| Suco | 1617.833245 | 0.028045751 | 0.068346558 | 0.410346212 | 0.681551997 | 0.922538401 |
| Rif1 | 1124.871273 | 0.028052821 | 0.081208304 | 0.345442765 | 0.729761532 | 0.934125508 |
| Zfp870 | 494.2907812 | 0.028054472 | 0.077088811 | 0.36392404 | 0.715914733 | 0.93125489 |
| Xpnpep2 | 13.02434224 | 0.028055392 | 0.071947865 | 0.389940579 | 0.696580487 | NA |
| Rpl23 | 6028.662333 | 0.028075558 | 0.069144769 | 0.406040235 | 0.684713037 | 0.923762012 |
| Ganab | 5678.249337 | 0.028086859 | 0.045098342 | 0.622791378 | 0.533421625 | 0.859128786 |
| Insm1 | 169.0599213 | 0.028089207 | 0.094079217 | 0.29856973 | 0.765268364 | 0.945182234 |
| Tpm3 | 6702.791957 | 0.02810621 | 0.039057431 | 0.719612366 | 0.471763696 | 0.833304495 |
| Rpl10a | 4451.543823 | 0.028125733 | 0.053182348 | 0.528854674 | 0.596906267 | 0.887922451 |
| Pde7b | 3295.230313 | 0.028162781 | 0.095094144 | 0.296156838 | 0.76711031 | 0.946118815 |
| Foxc2 | 27.5237018 | 0.02817131 | 0.077716535 | 0.362487987 | 0.716987399 | 0.93125489 |
| Ubn1 | 2992.444611 | 0.028177239 | 0.058339182 | 0.482989959 | 0.629102866 | 0.904428764 |
| Mapk3 | 7574.559889 | 0.028212507 | 0.053887057 | 0.523548849 | 0.600592366 | 0.890043329 |
| Tnr | 3379.038304 | 0.028229667 | 0.05703605 | 0.494944288 | 0.620639447 | 0.899412757 |
| Cul4b | 2178.667822 | 0.02822979 | 0.075840646 | 0.372225074 | 0.709725279 | 0.930224827 |
| Ankrd29 | 777.7050138 | 0.028236354 | 0.066746538 | 0.423038478 | 0.672267188 | 0.919190306 |
| Usp37 | 1158.450192 | 0.028236481 | 0.079398577 | 0.355629562 | 0.722118002 | 0.93255988 |
| Trappc2l | 1065.283706 | 0.028236699 | 0.057950399 | 0.487256332 | 0.62607669 | 0.902402691 |
| Pcf11 | 1365.102876 | 0.028242434 | 0.084132831 | 0.335688623 | 0.737105686 | 0.937206199 |
| Patl1 | 1169.233439 | 0.028256654 | 0.068817413 | 0.410603259 | 0.681363474 | 0.922538401 |
| Nhlh2 | 17.57108475 | 0.028279765 | 0.067360365 | 0.419827966 | 0.674611133 | NA |
| N4bp2l1 | 946.3497619 | 0.028297 | 0.066272697 | 0.426978247 | 0.669395172 | 0.91770119 |
| Rpn2 | 5699.977562 | 0.028297211 | 0.051737736 | 0.546935618 | 0.584422965 | 0.884313345 |
| Agbl1 | 11.5398979 | 0.028299056 | 0.067137207 | 0.421510766 | 0.673382151 | NA |
| Psmf1 | 1575.218612 | 0.02831211 | 0.067799264 | 0.41758728 | 0.676248898 | 0.920643418 |
| Gm10684 | 7.558916914 | 0.028313328 | 0.053831936 | 0.525957825 | 0.59891751 | NA |
| Ankrd63 | 2098.627005 | 0.028319891 | 0.077090434 | 0.367359343 | 0.713350994 | 0.930507332 |
| Slc30a5 | 1867.518602 | 0.028343824 | 0.062986048 | 0.450001627 | 0.652709267 | 0.914683958 |
| Myadml2 | 265.3926673 | 0.028344044 | 0.087891511 | 0.32248898 | 0.747082285 | 0.94097069 |
| Clec12b | 0.966868033 | 0.028361679 | 0.022552353 | 1.257592921 | 0.208539015 | NA |
| Gm16325 | 1.500677145 | 0.028400839 | 0.027649162 | 1.027186242 | 0.304332775 | NA |
| Ephb4 | 275.9549459 | 0.028404263 | 0.079607488 | 0.356803906 | 0.721238612 | 0.932513933 |
| 4933427D06Rik | 1.027095484 | 0.028405318 | 0.023094885 | 1.229939791 | 0.218719652 | NA |
| Hif1a | 2910.067772 | 0.028409314 | 0.061822709 | 0.459528773 | 0.645854494 | 0.912363974 |
| Zbtb3 | 168.3752935 | 0.028432892 | 0.0870531 | 0.3266155 | 0.743958718 | 0.939962107 |
| Papola | 4359.404446 | 0.028454279 | 0.075131093 | 0.378728403 | 0.704889559 | 0.928727958 |
| Ptp4a3 | 2307.548644 | 0.028456969 | 0.065209272 | 0.43639452 | 0.662550514 | 0.915428982 |
| Git1 | 17368.26165 | 0.028468728 | 0.063991349 | 0.444884015 | 0.656403575 | 0.915208181 |
| Iffo2 | 1273.298386 | 0.028472173 | 0.053911725 | 0.528125807 | 0.59741202 | 0.888165441 |
| Gpn1 | 1018.430583 | 0.028475994 | 0.059986078 | 0.47471005 | 0.634993654 | 0.907297112 |
| Gdi1 | 28373.95432 | 0.028486108 | 0.03784896 | 0.7526259 | 0.451674749 | 0.824558475 |
| Mcoln2 | 1.739256276 | 0.028515107 | 0.029925489 | 0.952870214 | 0.340655834 | NA |
| C1ql3 | 4464.274178 | 0.028553972 | 0.082727463 | 0.345157115 | 0.729976257 | 0.934152665 |
| Neto1 | 2810.44728 | 0.028556387 | 0.068467632 | 0.41707864 | 0.676620888 | 0.920884008 |
| Rplp2 | 3432.831938 | 0.028559286 | 0.054881754 | 0.520378529 | 0.602799772 | 0.891281205 |
| Cdk20 | 383.6576553 | 0.028566255 | 0.077641464 | 0.367925244 | 0.712928977 | 0.930507332 |
| Klhl28 | 552.3376202 | 0.028566914 | 0.074962335 | 0.381083572 | 0.703141237 | 0.927855161 |
| Pm20d2 | 440.1140099 | 0.028579191 | 0.070253058 | 0.406803519 | 0.684152302 | 0.923605608 |
| H13 | 2822.159125 | 0.028584154 | 0.054357219 | 0.525857549 | 0.598987186 | 0.889150487 |
| Tbc1d30 | 5785.382973 | 0.028608338 | 0.064630011 | 0.442647889 | 0.65802044 | 0.915208181 |
| Rbm4b | 420.2433066 | 0.028609502 | 0.076414687 | 0.374397946 | 0.708108271 | 0.929959854 |
| Ccdc89 | 25.82032089 | 0.028615686 | 0.085398192 | 0.335085385 | 0.737560677 | 0.937206199 |
| Slc34a2 | 2.042086693 | 0.028627916 | 0.033561842 | 0.852990014 | 0.393664842 | NA |
| Casq1 | 27.94500136 | 0.028630078 | 0.077636226 | 0.368772147 | 0.712297567 | 0.930507332 |
| Mfap3 | 1234.658304 | 0.028667792 | 0.081571543 | 0.351443538 | 0.725255625 | 0.933334512 |
| BC017612 | 66.83134447 | 0.028673505 | 0.095101137 | 0.301505391 | 0.76302914 | 0.944431816 |
| Polr2j | 796.5881912 | 0.028689577 | 0.064176798 | 0.447039704 | 0.654846393 | 0.915208181 |
| 1700009P17Rik | 72.46294828 | 0.028689744 | 0.084475639 | 0.33962151 | 0.734141577 | 0.936456337 |
| Klhl33 | 131.9150315 | 0.028691794 | 0.091508918 | 0.31354096 | 0.753869707 | 0.94183464 |
| Cps1 | 17.74640388 | 0.028694549 | 0.07101095 | 0.404086259 | 0.686149289 | NA |
| Dgkb | 6243.640685 | 0.028695177 | 0.086064266 | 0.333415691 | 0.738820521 | 0.938076299 |
| Hdac8 | 496.8528209 | 0.028701645 | 0.076737141 | 0.374025475 | 0.708385363 | 0.929959854 |
| 2610018G03Rik | 218.2087193 | 0.028747501 | 0.089581935 | 0.320907347 | 0.748280606 | 0.9410864 |
| 2410004I01Rik | 14.04308908 | 0.028757878 | 0.074424587 | 0.386402917 | 0.69919829 | NA |
| Foxj2 | 1922.068892 | 0.028762226 | 0.051153066 | 0.562277663 | 0.573926855 | 0.879633264 |
| Fgf14 | 1800.77957 | 0.028768866 | 0.07969464 | 0.360988727 | 0.718107874 | 0.93125489 |
| Slc4a1ap | 1207.091011 | 0.02876992 | 0.049789808 | 0.577827489 | 0.563380594 | 0.875108781 |
| 1700065J18Rik | 0.933090672 | 0.028782642 | 0.022803569 | 1.262199 | 0.206877188 | NA |
| Itm2c | 32996.07567 | 0.028792641 | 0.047625688 | 0.604561154 | 0.545470624 | 0.866714755 |
| Trappc12 | 3437.1841 | 0.028811559 | 0.042907794 | 0.671476109 | 0.501917274 | 0.8469854 |
| 9430016H08Rik | 405.5448671 | 0.028812418 | 0.075571387 | 0.381260938 | 0.703009636 | 0.927855161 |
| 4930465K10Rik | 1.907950431 | 0.028821705 | 0.032267187 | 0.893220244 | 0.371739239 | NA |
| Katnb1 | 2060.248964 | 0.028836201 | 0.062974096 | 0.457905761 | 0.647020148 | 0.912750424 |
| Pigc | 481.2161915 | 0.028836548 | 0.064104138 | 0.449839106 | 0.652826458 | 0.914683958 |
| Gpr88 | 8533.691 | 0.028845611 | 0.09296116 | 0.310297454 | 0.756334767 | 0.942139348 |
| Smim18 | 173.7006034 | 0.028856412 | 0.084225336 | 0.342609636 | 0.731892152 | 0.9354567 |
| Bzw2 | 1307.094933 | 0.028857613 | 0.058340554 | 0.494640711 | 0.62085376 | 0.899423727 |
| Tle3 | 1762.80627 | 0.028879938 | 0.067037746 | 0.430801145 | 0.666612967 | 0.916017403 |
| Kcnt2 | 974.5968447 | 0.028902445 | 0.083016112 | 0.348154642 | 0.72772405 | 0.9335794 |
| Dzip3 | 2684.340463 | 0.028903366 | 0.058785559 | 0.491674596 | 0.6229494 | 0.900360726 |
| Asb7 | 853.7700626 | 0.028908627 | 0.069213256 | 0.417674712 | 0.676184964 | 0.920643418 |
| D430036J16Rik | 104.4409099 | 0.028912882 | 0.093667716 | 0.308675001 | 0.757568762 | 0.942139348 |
| Gins2 | 28.36377864 | 0.028949071 | 0.088729173 | 0.326263275 | 0.744225172 | 0.940096833 |
| Cpox | 1471.049512 | 0.028963827 | 0.059591471 | 0.486039813 | 0.626938938 | 0.902926336 |
| Iapp | 1.033043189 | 0.028977438 | 0.019924447 | 1.454365994 | 0.145844863 | NA |
| Tas1r3 | 14.58962066 | 0.028979415 | 0.071065851 | 0.407782556 | 0.683433322 | NA |
| Ttc9c | 1433.001892 | 0.028985263 | 0.064155517 | 0.451796889 | 0.651415311 | 0.914343204 |
| Ints8 | 1112.859722 | 0.028998856 | 0.067025968 | 0.432651057 | 0.665268291 | 0.915428982 |
| B4galt5 | 2724.848681 | 0.029018241 | 0.05004049 | 0.579895224 | 0.561985276 | 0.874306322 |
| Sec14l1 | 11248.84812 | 0.02901831 | 0.047786016 | 0.607255264 | 0.543681522 | 0.86604998 |
| Nrl | 12.95202956 | 0.029051367 | 0.0737716 | 0.393801499 | 0.693727597 | NA |
| Gtsf1 | 2.826755307 | 0.029055897 | 0.035359602 | 0.821725776 | 0.411232985 | NA |
| Spry1 | 344.5534619 | 0.029057538 | 0.081804337 | 0.355207791 | 0.722433929 | 0.93255988 |
| Gorasp1 | 1512.549264 | 0.029060774 | 0.056924248 | 0.510516607 | 0.609689583 | 0.89440021 |
| Gpc1 | 3529.527674 | 0.029062568 | 0.065189906 | 0.445813933 | 0.655731656 | 0.915208181 |
| 1700021K19Rik | 3982.560209 | 0.029064504 | 0.056703692 | 0.51256566 | 0.608255179 | 0.894011407 |
| Vmn2r29 | 214.5813051 | 0.029079136 | 0.086651507 | 0.335587198 | 0.737182179 | 0.937206199 |
| Selenbp1 | 390.4675331 | 0.029080216 | 0.085872762 | 0.338643075 | 0.734878629 | 0.936456337 |
| Ptov1 | 5068.583792 | 0.029101398 | 0.058181968 | 0.500178996 | 0.616949047 | 0.897436596 |
| Ddx19b | 1637.38085 | 0.029113935 | 0.077364029 | 0.376323929 | 0.706676092 | 0.929476902 |
| Mir126 | 3.261818363 | 0.029115578 | 0.041687786 | 0.698419872 | 0.484914652 | NA |
| 6820408C15Rik | 92.84907989 | 0.029119919 | 0.094631636 | 0.307718642 | 0.758296433 | 0.942526199 |
| Rpgrip1 | 28.10911649 | 0.029120081 | 0.083069585 | 0.350550452 | 0.725925634 | 0.933575664 |
| Zfp839 | 554.1431413 | 0.029128014 | 0.066696651 | 0.436723791 | 0.662311673 | 0.915428982 |
| Gucy2c | 13.83646313 | 0.029130523 | 0.071245664 | 0.408874319 | 0.682631896 | NA |
| Pde4a | 5372.665172 | 0.029132047 | 0.05573273 | 0.522709839 | 0.601176191 | 0.890297392 |
| Fam114a1 | 259.8087014 | 0.029160506 | 0.08119656 | 0.359134742 | 0.719494292 | 0.932251852 |
| Zbtb41 | 861.0598178 | 0.029170028 | 0.079160824 | 0.368490713 | 0.712507369 | 0.930507332 |
| Tesk2 | 292.2783278 | 0.02918189 | 0.080143308 | 0.364121356 | 0.71576739 | 0.93125489 |
| Dcaf4 | 880.2003357 | 0.029182225 | 0.066259425 | 0.440423756 | 0.659630222 | 0.915208181 |
| Tada1 | 1252.211826 | 0.029189877 | 0.060492785 | 0.48253484 | 0.629426055 | 0.904624802 |
| Saal1 | 414.255196 | 0.029193474 | 0.068736691 | 0.424714574 | 0.671044758 | 0.918544681 |
| Ppp2r5b | 6227.352538 | 0.029202181 | 0.055548287 | 0.525708038 | 0.599091078 | 0.889150487 |
| Aldh6a1 | 2964.156377 | 0.029218553 | 0.060482452 | 0.483091408 | 0.629030835 | 0.904428764 |
| Mettl11b | 22.89154615 | 0.029222161 | 0.081643603 | 0.357923458 | 0.720400595 | 0.932513933 |
| Capn5 | 2665.903296 | 0.029239559 | 0.058956047 | 0.495955206 | 0.619926015 | 0.899216177 |
| Vps13b | 3019.33549 | 0.029241398 | 0.065348474 | 0.447468725 | 0.654536664 | 0.915091394 |
| Tmem229a | 2307.726268 | 0.029279958 | 0.064312638 | 0.455275343 | 0.648911163 | 0.913724196 |
| Fut8 | 2524.530269 | 0.029302315 | 0.067038614 | 0.437096078 | 0.662041671 | 0.915428982 |
| Gtf2f1 | 2556.536917 | 0.029312423 | 0.047528024 | 0.616739785 | 0.537406375 | 0.861996453 |
| Tril | 2251.460726 | 0.02931765 | 0.052125384 | 0.562444768 | 0.573813025 | 0.879633264 |
| 1810019J16Rik | 4.635627769 | 0.029321901 | 0.042629878 | 0.687825112 | 0.491562921 | NA |
| Gprin2 | 19.43680601 | 0.029325878 | 0.078292071 | 0.374570218 | 0.707980126 | NA |
| Epb4.1l1 | 26497.45911 | 0.029329088 | 0.043907053 | 0.667981235 | 0.504145571 | 0.847807553 |
| 9130221H12Rik | 304.159099 | 0.029334989 | 0.081063985 | 0.361874496 | 0.717445819 | 0.93125489 |
| Hmga1 | 15.5043637 | 0.029336961 | 0.078318908 | 0.374583375 | 0.707970339 | NA |
| Fxyd2 | 86.41771495 | 0.029356613 | 0.095056586 | 0.308833021 | 0.757448549 | 0.942139348 |
| Gpd2 | 5447.839173 | 0.029368623 | 0.046905557 | 0.626122475 | 0.531234612 | 0.858328942 |
| Cops4 | 2498.06974 | 0.029384935 | 0.054051463 | 0.543647368 | 0.586684157 | 0.884638284 |
| Itpr1 | 33699.14409 | 0.029393701 | 0.065852274 | 0.44635818 | 0.655338536 | 0.915208181 |
| Gm525 | 2.11527068 | 0.029415004 | 0.030727513 | 0.957285538 | 0.338423153 | NA |
| B230312C02Rik | 39.55334325 | 0.029431038 | 0.086935719 | 0.338537929 | 0.734957849 | 0.936456337 |
| Fastkd3 | 471.0320477 | 0.029435632 | 0.069767992 | 0.421907398 | 0.67309261 | 0.919353963 |
| Paip2 | 4320.151563 | 0.029436446 | 0.063923217 | 0.460496942 | 0.645159566 | 0.912363974 |
| Elmod3 | 945.246657 | 0.029448019 | 0.059581887 | 0.494244485 | 0.621133526 | 0.899589902 |
| Dars | 2006.621147 | 0.029450422 | 0.050753532 | 0.580263498 | 0.561736939 | 0.874107207 |
| Trim9 | 12978.39311 | 0.029478812 | 0.07421919 | 0.397185845 | 0.69123042 | 0.923860706 |
| Rasgrf2 | 8774.125166 | 0.029493548 | 0.074114743 | 0.397944417 | 0.690671158 | 0.923860706 |
| Dnase1l1 | 75.00606145 | 0.029496073 | 0.095030307 | 0.310385958 | 0.756267471 | 0.942139348 |
| Dda1 | 2175.793023 | 0.029507138 | 0.061999404 | 0.475926155 | 0.634126988 | 0.907220906 |
| Mpdz | 2847.237261 | 0.029513028 | 0.05109111 | 0.577654861 | 0.56349716 | 0.875108781 |
| Hk1 | 18313.52239 | 0.029525446 | 0.04799362 | 0.615195227 | 0.5384258 | 0.862082844 |
| Mtmr12 | 2358.657323 | 0.029532692 | 0.066361653 | 0.445026459 | 0.656300633 | 0.915208181 |
| Msl3l2 | 433.39366 | 0.029538378 | 0.075224695 | 0.392668624 | 0.694564248 | 0.925527709 |

| | | | | | | | | | | | | | |
|---|---|---|---|---|---|---|---|---|---|---|---|---|---|
| Mc3r | 19.25439407 | -0.056691494 | 0.153114044 | -0.370256658 | 0.711191265 | 0.852747056 | Ppp3ca | 36009.71848 | 0.029546103 | 0.072007139 | 0.410321851 | 0.681569865 | 0.922538401 |
| Pvrl1 | 1996.919697 | -0.03682486 | 0.099535343 | -0.369967681 | 0.711406572 | 0.852939211 | Rab8b | 2052.392587 | 0.029550875 | 0.073641976 | 0.401277596 | 0.688215757 | 0.923860706 |
| 2410018L13Rik | 86.02727569 | 0.059582139 | 0.161161469 | 0.369704618 | 0.711602591 | 0.853108213 | 1700001C19Rik | 3.612210519 | 0.029570778 | 0.04052432 | 0.729704466 | 0.465570851 | NA |
| 2700054A10Rik | 194.575206 | -0.038662399 | 0.104624809 | -0.369533756 | 0.711729917 | 0.853128836 | Ten1 | 626.456009 | 0.029576023 | 0.072800512 | 0.406261189 | 0.684550699 | 0.923762012 |
| Ascl1 | 136.3006672 | 0.04774349 | 0.129198338 | 0.369536408 | 0.711727941 | 0.853128836 | Fkbp8 | 11878.36172 | 0.029603316 | 0.0581408 | 0.509165951 | 0.610635907 | 0.894800514 |
| 1110008F13Rik | 1252.470317 | -0.025444399 | 0.068906263 | -0.369261053 | 0.711933152 | 0.853306433 | Dtd2 | 321.3887439 | 0.029606492 | 0.080697143 | 0.366884014 | 0.713705536 | 0.930719689 |
| Atg4b | 2507.266747 | -0.018497429 | 0.05019876 | -0.368483793 | 0.712512527 | 0.853612884 | Tdrd7 | 1713.364556 | 0.029613527 | 0.061383282 | 0.482436362 | 0.629495995 | 0.904635816 |
| D130017N08Rik | 449.1845871 | 0.026694792 | 0.072412376 | 0.368649583 | 0.712388933 | 0.853612884 | Rnf5 | 2138.50311 | 0.029614681 | 0.04927635 | 0.60099178 | 0.547845461 | 0.867640991 |
| Mettl11b | 22.23890637 | -0.061892815 | 0.167970376 | -0.368474589 | 0.71251939 | 0.853612884 | Serpina3h | 2.85656553 | 0.029626394 | 0.035562999 | 0.833067928 | 0.40480642 | NA |
| Pus7l | 105.9122602 | -0.046440173 | 0.126011959 | -0.368537823 | 0.712472248 | 0.853612884 | Abcg4 | 3172.312505 | 0.029627795 | 0.067085859 | 0.441639947 | 0.658749772 | 0.915208181 |
| Slc4a7 | 741.987174 | -0.037062304 | 0.100535721 | -0.368648111 | 0.71239003 | 0.853612884 | Tsen2 | 560.9298955 | 0.029634816 | 0.066665855 | 0.44452766 | 0.656661136 | 0.915208181 |
| Zfp523 | 3718.260338 | 0.026338448 | 0.071423184 | 0.368766088 | 0.712302084 | 0.853612884 | 1700001G17Rik | 8.904351366 | 0.029635553 | 0.061040878 | 0.485503392 | 0.627319307 | NA |
| Pigx | 295.3677739 | -0.031127037 | 0.084498136 | -0.36837543 | 0.712593315 | 0.853635443 | Amfr | 7522.171861 | 0.029636977 | 0.04238835 | 0.699177423 | 0.484441158 | 0.838713365 |
| Ntan1 | 1315.933607 | -0.028264974 | 0.076780372 | -0.368127599 | 0.712778092 | 0.853724781 | Fam134c | 3206.731703 | 0.029639044 | 0.055271597 | 0.536243664 | 0.59179016 | 0.886226713 |
| Pqlc3 | 34.36435913 | 0.059888839 | 0.162667131 | 0.368168042 | 0.712747938 | 0.853724781 | Pi4k2a | 3997.779369 | 0.029642991 | 0.044217795 | 0.670386004 | 0.502611755 | 0.84739426 |
| Cacng6 | 16.23484161 | -0.060784381 | 0.165403313 | -0.36749192 | 0.713252117 | 0.853949516 | LOC100504608 | 2.507285137 | 0.029651218 | 0.030677346 | 0.966550946 | 0.333768567 | NA |
| Cttnbp2nl | 1506.722623 | 0.033779742 | 0.091864595 | 0.367712302 | 0.713087767 | 0.853949516 | Fkbp1a | 19507.56803 | 0.029655644 | 0.046776064 | 0.633991877 | 0.526086123 | 0.855892499 |
| Ddx28 | 471.9630016 | -0.030581987 | 0.083194637 | -0.367595653 | 0.713174756 | 0.853949516 | 1700022P22Rik | 1.533755251 | 0.029668545 | 0.024999453 | 1.186767739 | 0.235319247 | NA |
| Pold2 | 434.3870496 | 0.027602677 | 0.075067935 | 0.367702621 | 0.713094986 | 0.853949516 | L3mbtl2 | 2120.13473 | 0.029675421 | 0.060331998 | 0.491868689 | 0.622812174 | 0.900285565 |
| Ppp1cc | 8068.296299 | -0.040447791 | 0.110082227 | -0.367432525 | 0.713296414 | 0.853949516 | Mtch2 | 3082.504962 | 0.029693582 | 0.054488992 | 0.544946433 | 0.58579036 | 0.884588669 |
| Tmem245 | 1256.132194 | 0.033636339 | 0.091509882 | 0.36757056 | 0.71319347 | 0.853949516 | Epn1 | 14982.43468 | 0.029715318 | 0.063016164 | 0.471550733 | 0.637247499 | 0.908932006 |
| Esyt3 | 96.54783528 | 0.057862833 | 0.157615956 | 0.36711279 | 0.713534888 | 0.854047893 | Mrps5 | 1319.754416 | 0.029724639 | 0.05620113 | 0.528897524 | 0.59687654 | 0.887922451 |
| Gpr176 | 612.3331152 | 0.027193203 | 0.074075607 | 0.367100639 | 0.713543951 | 0.854047893 | Rfwd3 | 1566.267095 | 0.02974526 | 0.056614192 | 0.5254029 | 0.599303138 | 0.889150487 |
| Gstp2 | 10.1928301 | -0.058140685 | 0.158350232 | -0.367165139 | 0.713495841 | 0.854047893 | Ipcef1 | 4819.100153 | 0.029757226 | 0.075343308 | 0.394955129 | 0.692876003 | 0.924216421 |
| Nmnat3 | 112.4271343 | -0.041862457 | 0.114080914 | -0.366954079 | 0.713653272 | 0.85411276 | Manbal | 1492.350986 | 0.029760427 | 0.060034832 | 0.495719327 | 0.62009245 | 0.899216177 |
| Chchd7 | 403.0671475 | 0.03437352 | 0.093812768 | 0.366405557 | 0.714062473 | 0.854404507 | Vps41 | 8954.391805 | 0.029785204 | 0.053530195 | 0.556418744 | 0.577924628 | 0.881703921 |
| Eif4e3 | 586.1100906 | -0.030614361 | 0.083538921 | -0.366468231 | 0.714015714 | 0.854404507 | Acad9 | 1818.043312 | 0.029801476 | 0.044643646 | 0.667541264 | 0.504426461 | 0.847807553 |
| Lynx1 | 18057.98361 | -0.033634925 | 0.091776444 | -0.36648756 | 0.714001294 | 0.854404507 | Bend7 | 111.066064 | 0.02980831 | 0.093961176 | 0.317240712 | 0.75106096 | 0.9410864 |
| Hyal2 | 318.0637515 | -0.029185294 | 0.079712516 | -0.366131887 | 0.714266665 | 0.854582834 | Flot1 | 4075.040378 | 0.029814973 | 0.060775062 | 0.49057906 | 0.6237242 | 0.900733873 |
| Pnpo | 2070.508823 | 0.026462011 | 0.072295345 | 0.366026479 | 0.714345318 | 0.854610945 | Tmem232 | 87.54835569 | 0.029817063 | 0.094929231 | 0.314097806 | 0.753446756 | 0.94183464 |
| Rmnd5b | 1848.063911 | 0.025085883 | 0.068581012 | 0.365784677 | 0.714525755 | 0.854760813 | Tmem168 | 631.8457754 | 0.029841304 | 0.071174608 | 0.419268959 | 0.67501958 | 0.919907397 |
| Trpc4 | 601.3768542 | -0.043220159 | 0.118307158 | -0.365321592 | 0.714871362 | 0.855108229 | Rprm | 2076.397932 | 0.02987114 | 0.06025266 | 0.495764677 | 0.620060449 | 0.899216177 |
| Dab2 | 449.0587846 | -0.046981303 | 0.128689547 | -0.365074741 | 0.715055616 | 0.855109657 | Esr2 | 22.54296179 | 0.029872778 | 0.074779014 | 0.39948077 | 0.68953899 | 0.923860706 |
| Dock8 | 268.6816601 | -0.034356433 | 0.094140032 | -0.364950302 | 0.715148505 | 0.855109657 | Mvd | 685.1178201 | 0.029876598 | 0.074876106 | 0.399013771 | 0.689883057 | 0.923860706 |
| Kcnn1 | 682.6949707 | -0.032875071 | 0.0900674 | -0.365005214 | 0.715107514 | 0.855109657 | Chac1 | 218.7731004 | 0.029891109 | 0.088041042 | 0.339513354 | 0.734223039 | 0.936456337 |
| Map3k7 | 2590.583368 | 0.021760282 | 0.059597422 | 0.365121192 | 0.715020942 | 0.855109657 | Fbxl21 | 224.6001 | 0.029898845 | 0.092086774 | 0.324681208 | 0.745422359 | 0.940632024 |
| Tnni3k | 22.62762996 | -0.061195781 | 0.167616339 | -0.365094366 | 0.715040966 | 0.855109657 | Mmd2 | 5951.510708 | 0.02990598 | 0.050849581 | 0.588126371 | 0.556447474 | 0.871665697 |
| Specc1 | 4106.055133 | 0.030698386 | 0.084155724 | 0.364780719 | 0.715275099 | 0.85515718 | Jak1 | 8130.569314 | 0.029910008 | 0.056656623 | 0.527917246 | 0.597556774 | 0.888198349 |
| Usp4 | 3907.474046 | 0.015776828 | 0.043253909 | 0.364749191 | 0.715298635 | 0.85515718 | Zfp459 | 120.211093 | 0.029911598 | 0.09508493 | 0.314577698 | 0.753082315 | 0.94183464 |
| Cdca8 | 42.98309132 | -0.059380725 | 0.162924094 | -0.364468653 | 0.715508078 | 0.85529714 | 9230110C19Rik | 336.3578769 | 0.029927759 | 0.081418855 | 0.367577746 | 0.713188111 | 0.930507332 |
| Mlf2 | 14957.98408 | -0.029087926 | 0.079814419 | -0.364444503 | 0.715526109 | 0.85529714 | Pou2f3 | 11.64641492 | 0.029929606 | 0.060234584 | 0.496884088 | 0.619270793 | NA |
| Cyb5rl | 102.4248412 | 0.050103877 | 0.137530884 | 0.364310007 | 0.715626528 | 0.855351186 | Cldn26 | 34.53931205 | 0.029940966 | 0.088838607 | 0.337026511 | 0.736096915 | 0.937120403 |
| Ctcf | 2628.789448 | 0.024850325 | 0.06826695 | 0.364016925 | 0.715845371 | 0.855546759 | Traf6 | 920.3990901 | 0.029986812 | 0.066740971 | 0.449301397 | 0.65321425 | 0.914683958 |
| Ctdsp1 | 1122.964765 | 0.022321903 | 0.061347454 | 0.363860303 | 0.715962329 | 0.855620542 | Fam213b | 2031.652552 | 0.029992122 | 0.066173856 | 0.453232189 | 0.650381556 | 0.914000665 |
| 2610301B20Rik | 1402.338923 | 0.026178152 | 0.072011924 | 0.363525248 | 0.716212556 | 0.855787564 | Tmem205 | 599.0355709 | 0.030025281 | 0.064258933 | 0.467254587 | 0.640317744 | 0.910092095 |
| Hat1 | 553.594975 | -0.03104853 | 0.085400261 | -0.36356482 | 0.716183001 | 0.855787564 | Snai3 | 30.63580156 | 0.030040702 | 0.081784048 | 0.367317373 | 0.713382297 | 0.930507332 |
| 2210016F16Rik | 306.7016654 | -0.043908192 | 0.120852222 | -0.363321348 | 0.716364848 | 0.855903528 | Cyhr1 | 5852.625988 | 0.030056667 | 0.047027861 | 0.639124684 | 0.522741823 | 0.854337225 |
| 3930402G23Rik | 13.26162826 | -0.05947501 | 0.163790112 | -0.363117219 | 0.716517322 | 0.855923913 | Chrna5 | 94.50771782 | 0.030057779 | 0.094776191 | 0.317144827 | 0.751133712 | 0.9410864 |
| Ccdc103 | 98.58619842 | 0.045043597 | 0.124077339 | 0.363028391 | 0.716583676 | 0.855923913 | Gan | 212.4499467 | 0.030065004 | 0.084942105 | 0.353947007 | 0.723378598 | 0.932617157 |
| Slc35e1 | 3929.044079 | 0.02288207 | 0.063014093 | 0.363126235 | 0.716510588 | 0.855923913 | Ttll11 | 1961.386907 | 0.030065024 | 0.05583505 | 0.538461488 | 0.590258486 | 0.885570623 |
| Slc48a1 | 5558.591535 | 0.025719875 | 0.070853126 | 0.363002685 | 0.716602878 | 0.855923913 | Hsf3 | 2.457946924 | 0.030077681 | 0.038160813 | 0.788182389 | 0.430590029 | NA |
| Paqr9 | 856.0129954 | -0.029183669 | 0.080413003 | -0.362922269 | 0.716662951 | 0.855929682 | Zfp110 | 751.2346559 | 0.030077902 | 0.070231577 | 0.428267496 | 0.66845638 | 0.917162311 |
| Cdh13 | 2552.047051 | -0.03431623 | 0.094670345 | -0.362481303 | 0.716992393 | 0.85599323 | Rdh16 | 6.818557127 | 0.030078355 | 0.056480605 | 0.53254307 | 0.59434992 | NA |
| Fibp | 1137.313197 | 0.027596319 | 0.076091975 | 0.362670555 | 0.716850998 | 0.85599323 | Sec14l2 | 2075.243697 | 0.030085277 | 0.061219485 | 0.491433027 | 0.623120209 | 0.900462014 |
| Gm11346 | 9.504323874 | 0.054617504 | 0.150651317 | 0.362542491 | 0.716946677 | 0.85599323 | Tslp | 4.162648845 | 0.030112507 | 0.046265925 | 0.650857126 | 0.51513872 | NA |
| Gm1604b | 15.63049129 | -0.059469929 | 0.164043063 | -0.362526324 | 0.716958756 | 0.85599323 | Stoml2 | 1631.313808 | 0.030115224 | 0.052575294 | 0.572801823 | 0.566778884 | 0.875959314 |
| Pcdhgb6 | 563.6521109 | -0.029672174 | 0.081841495 | -0.362556603 | 0.716936134 | 0.85599323 | Pqlc1 | 662.5895276 | 0.030117899 | 0.067477157 | 0.44634215 | 0.655350114 | 0.915208181 |
| Sumo3 | 5204.251865 | -0.030780368 | 0.084986419 | -0.362179837 | 0.717217646 | 0.856196179 | Map4k3 | 3338.460901 | 0.030130157 | 0.05919751 | 0.508976758 | 0.610768515 | 0.894800514 |
| Kcnh2 | 1589.47047 | -0.031252237 | 0.086323324 | -0.362037 | 0.717324381 | 0.856257624 | Rabl6 | 6754.425239 | 0.030158204 | 0.044858034 | 0.672303302 | 0.501390628 | 0.846601651 |
| Mthfs | 49.43999889 | -0.055574692 | 0.153605961 | -0.361800361 | 0.717501222 | 0.856364623 | Sephs1 | 1058.792201 | 0.030176781 | 0.062765894 | 0.480783103 | 0.630670661 | 0.905249211 |
| Rapgef4 | 24482.07475 | -0.031759053 | 0.087788178 | -0.361769133 | 0.71752456 | 0.856364623 | Ccser2 | 4922.800074 | 0.030177502 | 0.056405746 | 0.535007578 | 0.592644617 | 0.886226713 |
| Dnajc10 | 3333.191 | 0.02695325 | 0.074743854 | 0.360608238 | 0.71839233 | 0.857031081 | Fam117a | 211.8257745 | 0.03020353 | 0.090237785 | 0.334710458 | 0.737843512 | 0.937372297 |
| Rab3gap1 | 4366.750005 | 0.02710996 | 0.075184707 | 0.360578118 | 0.718414849 | 0.857031081 | Morn4 | 2450.819665 | 0.030208763 | 0.06208539 | 0.486567991 | 0.626564512 | 0.902822404 |
| Snip1 | 505.3550265 | 0.024290437 | 0.067362621 | 0.360592224 | 0.718404303 | 0.857031081 | Dnalc4 | 1844.547929 | 0.030211999 | 0.073893952 | 0.408856186 | 0.682645204 | 0.922857537 |
| Tada2b | 1455.415048 | -0.034171762 | 0.094763949 | -0.360598756 | 0.71839942 | 0.857031081 | Fam110c | 1.549691887 | 0.030214096 | 0.027400647 | 1.102678171 | 0.27016695 | NA |
| Tmem44 | 3010.115454 | -0.035398579 | 0.098101151 | -0.360837555 | 0.718220887 | 0.857031081 | 5730422E09Rik | 6.063063076 | 0.030220985 | 0.048171307 | 0.627364863 | 0.530420095 | NA |
| Xpo7 | 4815.894444 | 0.025228833 | 0.069957581 | 0.360630436 | 0.718375734 | 0.857031081 | Pcdhb14 | 347.5135326 | 0.030237564 | 0.084401543 | 0.358258432 | 0.720149923 | 0.932426935 |
| Opalin | 572.3601181 | -0.056439292 | 0.156618724 | -0.360361078 | 0.718577129 | 0.857158676 | Upk3bl | 4.960631809 | 0.030263684 | 0.052870719 | 0.572409151 | 0.567044818 | NA |
| Fam72a | 41.64150374 | -0.060425266 | 0.167869981 | -0.359952781 | 0.718882445 | 0.85745686 | Azi2 | 2415.718208 | 0.030270419 | 0.061303865 | 0.493776681 | 0.621463903 | 0.899589902 |
| Fbxo11 | 5194.978154 | -0.025223322 | 0.070158906 | -0.359517041 | 0.719208332 | 0.857779532 | Stag2 | 2683.478246 | 0.03027954 | 0.066369719 | 0.456225235 | 0.648228021 | 0.91329006 |
| Pbld1 | 19.82299317 | -0.059896621 | 0.166650898 | -0.359413729 | 0.719285606 | 0.857805663 | Egr2 | 417.2830477 | 0.030280459 | 0.051723423 | 0.585430303 | 0.558258418 | 0.872714355 |
| D930020B18Rik | 18.57144959 | -0.059397065 | 0.165334523 | -0.359253859 | 0.719405189 | 0.857816222 | Lsm1 | 489.6887922 | 0.030312959 | 0.070915594 | 0.427451243 | 0.66905069 | 0.917484508 |
| Nexn | 395.0169962 | -0.05487608 | 0.15273344 | -0.359293157 | 0.719375793 | 0.857816222 | Smarcb1 | 2339.505658 | 0.030313297 | 0.051604744 | 0.587412982 | 0.556926377 | 0.871851768 |
| Lrrc1 | 411.7669936 | -0.040987195 | 0.114212236 | -0.358868686 | 0.719693326 | 0.858093759 | Dnph1 | 89.57710358 | 0.030317656 | 0.093326913 | 0.32485437 | 0.745291293 | 0.940632024 |
| Tubb6 | 51.81741495 | -0.059736246 | 0.166538237 | -0.358693878 | 0.719824109 | 0.858134425 | 6720456H20Rik | 1179.963032 | 0.030340081 | 0.064466384 | 0.470634133 | 0.637902028 | 0.908932006 |
| Vil1 | 23.85968561 | 0.060204957 | 0.167888419 | 0.358601014 | 0.719893588 | 0.858134425 | Man2a1 | 1031.985349 | 0.030341663 | 0.076814079 | 0.395001318 | 0.692841915 | 0.924216421 |
| Zfp719 | 661.2410681 | -0.032598889 | 0.090888023 | -0.3586709 | 0.7198413 | 0.858134425 | Ndufaf6 | 190.9813776 | 0.030388394 | 0.087112965 | 0.348838939 | 0.727210228 | 0.9335794 |
| Spg11 | 1122.138438 | 0.025845463 | 0.072306798 | 0.357441675 | 0.720761182 | 0.859102528 | Npy5r | 155.4980991 | 0.030394662 | 0.089873497 | 0.338193827 | 0.735217127 | 0.936492809 |
| Slc25a26 | 562.0719877 | -0.030457347 | 0.085229937 | -0.357355037 | 0.720826033 | 0.859113735 | Agbl4 | 389.620156 | 0.030407673 | 0.073351137 | 0.414549444 | 0.678471764 | 0.921316909 |
| 1700040L02Rik | 70.02454988 | -0.051432668 | 0.144041018 | -0.357069596 | 0.721039705 | 0.859218836 | Krt25 | 2.46567565 | 0.030420017 | 0.031719157 | 0.959042402 | 0.337537384 | NA |
| Ngly1 | 871.0853123 | -0.028469248 | 0.079742443 | -0.357014993 | 0.721080581 | 0.859218836 | Tubb2b | 3947.512118 | 0.030448824 | 0.070741572 | 0.430423342 | 0.666887719 | 0.916077199 |
| Tead4 | 29.75796076 | -0.059477577 | 0.166542792 | -0.357130901 | 0.720993812 | 0.859218836 | Dnahc5 | 199.361247 | 0.030464337 | 0.093128474 | 0.32712162 | 0.7435759 | 0.939912916 |
| Mzf1 | 252.6883157 | -0.036252273 | 0.101633065 | -0.356697629 | 0.721318181 | 0.859435858 | Gm18409 | 0.801396501 | 0.030468423 | 0.020440004 | 1.490627071 | 0.136059433 | NA |
| Dmtf1 | 2396.924026 | -0.028111131 | 0.078840759 | -0.356555813 | 0.721424362 | 0.859467364 | Sec63 | 4151.258461 | 0.030469813 | 0.05758608 | 0.529117672 | 0.596723823 | 0.887922451 |
| Wiz | 1925.727432 | 0.024490863 | 0.068695349 | 0.356514139 | 0.721455566 | 0.859467364 | Setd8 | 1700.175687 | 0.030476148 | 0.058871341 | 0.517673751 | 0.604685915 | 0.891988593 |
| Trim24 | 1084.178235 | 0.0271531 | 0.07619455 | 0.356365381 | 0.721566952 | 0.85953397 | C1qtnf3 | 4.104191035 | 0.030504569 | 0.035685988 | 0.854805228 | 0.392658983 | NA |
| Bok | 970.617301 | -0.040649146 | 0.114149982 | -0.356102957 | 0.721763463 | 0.859604432 | Snx24 | 570.3194726 | 0.030516454 | 0.067643335 | 0.45113763 | 0.651890358 | 0.914447303 |
| Htr1f | 101.581009 | -0.043492956 | 0.122140484 | -0.356089601 | 0.721773464 | 0.859604432 | Aen | 1011.007635 | 0.030522389 | 0.058745946 | 0.519565869 | 0.603366192 | 0.891540503 |
| Taf9 | 2439.287394 | 0.028840178 | 0.080997147 | 0.356064125 | 0.721792543 | 0.859604432 | Btbd9 | 4136.766328 | 0.030528312 | 0.064426897 | 0.473844213 | 0.635611003 | 0.907676507 |
| Slc35e3 | 1352.309264 | -0.017858622 | 0.050192247 | -0.355804386 | 0.721987064 | 0.859770008 | Hmgn2 | 2752.527149 | 0.030528997 | 0.059637079 | 0.511913011 | 0.608711892 | 0.89440021 |
| Hhipl1 | 183.5973184 | 0.042515692 | 0.119680718 | 0.355242619 | 0.722407839 | 0.860204968 | Trim29 | 6.863682406 | 0.030529451 | 0.054860195 | 0.556495496 | 0.577872173 | NA |
| Dcaf6 | 3831.216018 | -0.026486771 | 0.074617059 | -0.354969383 | 0.722612528 | 0.86038258 | Krt28 | 4.287144234 | 0.030535663 | 0.045723565 | 0.667832083 | 0.504240785 | NA |
| Trim12c | 128.4595416 | 0.057152878 | 0.161044871 | 0.354887913 | 0.722673564 | 0.860389135 | Slc35f6 | 1597.487692 | 0.030541778 | 0.053135615 | 0.574789199 | 0.565433873 | 0.875364912 |
| Itga2b | 90.21095094 | 0.056716037 | 0.159876871 | 0.354748232 | 0.722778214 | 0.86044761 | Paqr7 | 2840.539603 | 0.030542959 | 0.046449069 | 0.657558046 | 0.51082216 | 0.85162432 |
| Ccdc111 | 151.9455901 | 0.052856415 | 0.149151515 | 0.35438068 | 0.723053611 | 0.860456111 | Apex2 | 139.1705835 | 0.030544475 | 0.093793501 | 0.325656622 | 0.744684167 | 0.940439377 |
| Col6a1 | 4536.20896 | -0.05641591 | 0.159214182 | -0.354339728 | 0.723084298 | 0.860456111 | Dis3l2 | 1205.049517 | 0.030545048 | 0.055469208 | 0.550666738 | 0.58186215 | 0.884020489 |
| Cyp2e1 | 20.29962298 | -0.052300098 | 0.147486292 | -0.354609889 | 0.722881867 | 0.860456111 | Zfp319 | 1013.948839 | 0.030546251 | 0.061370045 | 0.497738774 | 0.618668175 | 0.898575807 |
| Dlg3 | 8949.812473 | -0.02439447 | 0.068892181 | -0.354096346 | 0.723266681 | 0.860456111 | Pcdhb6 | 113.762484 | 0.030588636 | 0.095067982 | 0.321755389 | 0.747638013 | 0.9410864 |
| Eva1b | 48.05962262 | -0.057066099 | 0.161167594 | -0.354079241 | 0.723279499 | 0.860456111 | Shisa5 | 2460.162017 | 0.030591597 | 0.057452879 | 0.532464127 | 0.594404581 | 0.887209303 |
| Gsn | 2395.39108 | 0.033125283 | 0.093524271 | 0.354189158 | 0.723197128 | 0.860456111 | Gemin7 | 688.1867338 | 0.030595061 | 0.059800924 | 0.51161519 | 0.608920352 | 0.89440021 |
| Lrrc8b | 2659.863119 | 0.027352104 | 0.077170523 | 0.354437198 | 0.723011262 | 0.860456111 | Eef1a1 | 88995.03367 | 0.030595787 | 0.053173374 | 0.57539676 | 0.565022995 | 0.875364912 |
| Slc35g2 | 540.3725439 | -0.028387753 | 0.080175159 | -0.354071677 | 0.723285167 | 0.860456111 | Acsl1 | 2597.754848 | 0.03060114 | 0.051916168 | 0.589433714 | 0.555570366 | 0.871360989 |
| Tgfbi | 173.0605788 | 0.049507631 | 0.139675868 | 0.354446565 | 0.723004243 | 0.860456111 | Mrpl28 | 1316.683761 | 0.03060137 | 0.063538459 | 0.481619648 | 0.630076166 | 0.905068772 |
| Fsbp | 11.73931349 | -0.058832099 | 0.166317329 | -0.353734032 | 0.723538215 | 0.860691064 | Apeh | 789.4639851 | 0.030617484 | 0.072568311 | 0.421912586 | 0.673088823 | 0.919353963 |
| Tmem163 | 589.0797543 | 0.032103602 | 0.090790117 | 0.353602383 | 0.723636888 | 0.860742357 | Fcamr | 0.812623213 | 0.030621527 | 0.020490058 | 1.4944578 | 0.135056004 | NA |
| Sgsh | 311.9617191 | 0.045524263 | 0.128948074 | 0.353043369 | 0.724055928 | 0.861174679 | Srr | 4600.389344 | 0.030623513 | 0.051420258 | 0.59555347 | 0.551473568 | 0.869755927 |
| 1700008O03Rik | 20.55105543 | 0.059141992 | 0.167704402 | 0.352656171 | 0.724346221 | 0.861321592 | Tbcc | 485.2069964 | 0.030626071 | 0.081598993 | 0.375324135 | 0.707419419 | 0.92986829 |
| Abcg3 | 21.8414568 | 0.05898381 | 0.167225098 | 0.352721041 | 0.724297584 | 0.861321592 | Peli1 | 1413.263538 | 0.030643014 | 0.068220566 | 0.449175605 | 0.653304984 | 0.914683958 |
| Ncor2 | 6540.951016 | -0.031606109 | 0.089622101 | -0.352659772 | 0.724343522 | 0.861321592 | Rgs8 | 6027.649281 | 0.030644613 | 0.061829242 | 0.495633013 | 0.620153357 | 0.899216177 |
| Mllt4 | 4058.749112 | -0.024267406 | 0.068828883 | -0.352575911 | 0.7244064 | 0.861327047 | Tc2n | 8.119880446 | 0.030645448 | 0.036494274 | 0.839733036 | 0.401058087 | NA |
| Adamts4 | 766.4677925 | -0.042240392 | 0.120035031 | -0.351900541 | 0.724912853 | 0.861686848 | Cyp4f41-ps | 4.719036762 | 0.030659892 | 0.050105568 | 0.611905892 | 0.540600023 | NA |
| Ap4b1 | 621.9431798 | 0.024394737 | 0.069362802 | 0.351697684 | 0.725064998 | 0.861686848 | Aire | 5.872841891 | 0.030701788 | 0.049161988 | 0.624502585 | 0.532297573 | NA |
| Btf3l4 | 2774.364396 | 0.025263375 | 0.071855291 | 0.351586842 | 0.725148135 | 0.861686848 | Gab3 | 61.37936639 | 0.03072241 | 0.094863321 | 0.32385974 | 0.746044226 | 0.940755508 |
| Lgals1 | 349.2201738 | -0.033445984 | 0.095130743 | -0.35157913 | 0.725153919 | 0.861686848 | Il34 | 2156.430176 | 0.030731651 | 0.072195789 | 0.425670964 | 0.670347622 | 0.918109519 |
| Pgap2 | 322.7726639 | -0.026886116 | 0.076394656 | -0.351937126 | 0.724885416 | 0.861686848 | Cnot7 | 3385.306766 | 0.03073814 | 0.070913086 | 0.433462163 | 0.664679048 | 0.915428982 |
| Pou2f3 | 13.88675794 | 0.05353565 | 0.152246961 | 0.351636901 | 0.725110587 | 0.861686848 | Mapk8ip3 | 23485.13487 | 0.03076422 | 0.046398351 | 0.663045541 | 0.507301387 | 0.848958235 |
| Tspan18 | 146.0749015 | 0.055250208 | 0.157079863 | 0.351733232 | 0.725038336 | 0.861686848 | Tmed3 | 924.044474 | 0.030769669 | 0.064486606 | 0.477148211 | 0.633256586 | 0.906932171 |
| Tspyl1 | 7061.744047 | -0.020636963 | 0.058651287 | -0.35185865 | 0.724944271 | 0.861686848 | Exoc7 | 3151.697393 | 0.030778485 | 0.04752738 | 0.64759482 | 0.517247048 | 0.853297835 |
| Pccb | 1897.269827 | 0.021339117 | 0.060708664 | 0.351500353 | 0.725213008 | 0.861690977 | Agtr1a | 29.92458302 | 0.030804472 | 0.082580926 | 0.37302164 | 0.709132338 | 0.930183885 |
| Dnd1 | 34.02841686 | 0.058986871 | 0.1679657 | 0.35118403 | 0.725450291 | 0.861896471 | Foxp4 | 1636.062095 | 0.030824854 | 0.076956736 | 0.400547839 | 0.688753055 | 0.923860706 |
| Lhcgr | 17.49059571 | -0.05626589 | 0.160246217 | -0.351121485 | 0.72549721 | 0.861896471 | Duox1 | 3.647264892 | 0.03082956 | 0.045359817 | 0.679666754 | 0.496715492 | NA |
| Ddx4 | 97.52253123 | -0.054436128 | 0.155191773 | -0.350766842 | 0.725763275 | 0.862146452 | Synm | 2114.063102 | 0.03083496 | 0.046641434 | 0.661106609 | 0.508543947 | 0.850050333 |
| Arpc2 | 6530.824311 | 0.033108411 | 0.094490643 | 0.350388248 | 0.726047345 | 0.862162734 | Keap1 | 2927.65966 | 0.030844204 | 0.048067176 | 0.641689527 | 0.521074791 | 0.85372574 |
| Dnalc4 | 1614.750679 | 0.030024189 | 0.085690905 | 0.350377772 | 0.726055206 | 0.862162734 | Sh3rf3 | 1476.488946 | 0.030868823 | 0.079683201 | 0.387394367 | 0.698464273 | 0.926409493 |
| Fmnl3 | 660.8887709 | 0.031787232 | 0.090689632 | 0.350505694 | 0.725959218 | 0.862162734 | Dctn4 | 4263.605287 | 0.030892959 | 0.0509779 | 0.6060069 | 0.544510171 | 0.86624804 |
| Rbm11 | 220.7235669 | -0.041650217 | 0.118856439 | -0.350424573 | 0.726020087 | 0.862162734 | Polr3c | 795.4062611 | 0.030903236 | 0.057108468 | 0.541132283 | 0.588416408 | 0.885188411 |
| Zfp948 | 337.6644491 | 0.036261806 | 0.103428118 | 0.350599109 | 0.725889125 | 0.862162734 | Xpnpep1 | 1543.474683 | 0.03090785 | 0.051018955 | 0.605811117 | 0.544640187 | 0.86624804 |
| Sipa1 | 550.6994291 | 0.027270032 | 0.077933161 | 0.349915642 | 0.726402008 | 0.862508445 | 2210407C18Rik | 1.070974855 | 0.030927996 | 0.023792982 | 1.299878915 | 0.193642473 | NA |
| Gm10069 | 38.61119159 | -0.057489845 | 0.164386269 | -0.349724131 | 0.726545742 | 0.862613006 | Akap9 | 3850.443199 | 0.030936307 | 0.0806904 | 0.383395139 | 0.701426807 | 0.92751465 |
| Asb7 | 763.4842776 | 0.023269157 | 0.06658079 | 0.349487543 | 0.726723321 | 0.862625537 | Susd2 | 709.458078 | 0.030939004 | 0.08393919 | 0.368588307 | 0.712434612 | 0.930507332 |
| Fam45a | 1038.915575 | 0.026436935 | 0.075628437 | 0.349563416 | 0.726666371 | 0.862625537 | Pmf1 | 98.00850234 | 0.030945887 | 0.094743376 | 0.326628499 | 0.743948886 | 0.939962107 |
| Gstm4 | 343.9333416 | 0.030947479 | 0.088533548 | 0.349556524 | 0.726671544 | 0.862625537 | Nfasc | 12948.48597 | 0.030951232 | 0.058839193 | 0.526030877 | 0.598866754 | 0.889150487 |
| Chd4 | 8806.354994 | 0.031293596 | 0.089582499 | 0.349327119 | 0.726843741 | 0.862694072 | D830031N03Rik | 323.612467 | 0.030974549 | 0.085217698 | 0.36347554 | 0.716249682 | 0.93125489 |
| Scfd2 | 664.3973305 | 0.026703269 | 0.076456209 | 0.349262275 | 0.726892417 | 0.862694072 | Ccdc11 | 16.47753607 | 0.030983643 | 0.071894986 | 0.430956943 | 0.666499678 | NA |
| Adcy4 | 170.0421125 | 0.046029161 | 0.131996035 | 0.348716242 | 0.727302349 | 0.863081236 | 1700016K19Rik | 68.17025686 | 0.030991426 | 0.094698105 | 0.327265533 | 0.743467059 | 0.939912916 |
| Cmtm7 | 71.68286398 | -0.052886774 | 0.151772255 | -0.348461411 | 0.727493689 | 0.863081236 | Eif3b | 3850.415667 | 0.031033264 | 0.04281435 | 0.72483323 | 0.468554347 | 0.832446623 |
| Gm20754 | 33.03341295 | -0.058059063 | 0.166565507 | -0.348565944 | 0.727415199 | 0.863081236 | Taok1 | 5593.101119 | 0.031033456 | 0.071961027 | 0.431253659 | 0.666283942 | 0.915911852 |
| Mbtps2 | 1104.842195 | -0.02951621 | 0.084705508 | -0.348456794 | 0.727497156 | 0.863081236 | Rps26 | 2871.064861 | 0.031041517 | 0.069130244 | 0.449029467 | 0.6534104 | 0.914683958 |
| Rac1 | 11034.29477 | 0.01604279 | 0.046015458 | 0.348639143 | 0.727360237 | 0.863081236 | Fgf13 | 4360.349527 | 0.031051752 | 0.067091466 | 0.462827144 | 0.643488273 | 0.911258473 |
| Ctcfl | 19.05918266 | -0.058482329 | 0.168098706 | -0.347904696 | 0.727911758 | 0.863374782 | Tmem100 | 403.6363953 | 0.031061256 | 0.080591307 | 0.385416947 | 0.699928528 | 0.926781946 |
| Ifnlr1 | 10.28598187 | -0.049561526 | 0.142421309 | -0.34799235 | 0.727845929 | 0.863374782 | Galnt7 | 530.5643584 | 0.031061395 | 0.080099932 | 0.387783034 | 0.698176602 | 0.926409493 |
| Laptm4b | 3333.20333 | 0.019474058 | 0.055964278 | 0.347973012 | 0.727860451 | 0.863374782 | Rbp4 | 345.0147695 | 0.031086279 | 0.082222828 | 0.378073583 | 0.705375932 | 0.928783753 |
| Itch | 2810.052805 | 0.030512989 | 0.087741768 | 0.347758989 | 0.728021191 | 0.863398894 | Kif1b | 38937.04614 | 0.031104347 | 0.0616542 | 0.504496806 | 0.613912306 | 0.895890111 |
| Mlf1 | 30.96103818 | 0.058440718 | 0.16806386 | 0.347729238 | 0.728043536 | 0.863398894 | Glod5 | 6.494854489 | 0.031113233 | 0.054323007 | 0.57274505 | 0.56681733 | NA |
| Gm13152 | 60.84356989 | -0.055428356 | 0.159443518 | -0.347636311 | 0.728113333 | 0.863415581 | Jakmip1 | 3250.050842 | 0.031113411 | 0.043745156 | 0.711242443 | 0.476934012 | 0.835195112 |
| Mylk | 828.7129317 | -0.031073492 | 0.089412036 | -0.347531425 | 0.728192114 | 0.863442919 | C2cd4d | 3.727446145 | 0.031115958 | 0.040397744 | 0.77023999 | 0.441157547 | NA |
| 4632415L05Rik | 928.7117235 | -0.023128487 | 0.066569749 | -0.347432396 | 0.728266499 | 0.863465039 | Rpl36al | 1720.17208 | 0.031128724 | 0.072162024 | 0.431372657 | 0.666197428 | 0.915879599 |
| Lpo | 30.84883934 | -0.058317567 | 0.16801885 | -0.34708943 | 0.728524134 | 0.863638327 | Htatsf1 | 3244.153662 | 0.031138879 | 0.051625322 | 0.60317065 | 0.54639517 | 0.867128614 |
| Prrt3 | 1881.010398 | 0.02515609 | 0.072476868 | 0.347091297 | 0.728522732 | 0.863638327 | Slc35b2 | 915.3244772 | 0.031140004 | 0.059297121 | 0.525152044 | 0.599477499 | 0.889207613 |
| Ccdc96 | 72.82090446 | 0.046784519 | 0.134824622 | 0.347002784 | 0.728589227 | 0.863649414 | Rpl13a | 8693.027279 | 0.031140729 | 0.061877439 | 0.503264666 | 0.614778206 | 0.896163393 |
| Gnaz | 2385.102323 | -0.026481756 | 0.076351431 | -0.346840341 | 0.728711268 | 0.863727998 | Wdr41 | 1475.84898 | 0.031147751 | 0.053503015 | 0.582168141 | 0.56045343 | 0.873713575 |
| Tmem230 | 834.4622663 | 0.032450726 | 0.093626774 | 0.346596645 | 0.728894367 | 0.863878935 | Tnfrsf18 | 2.229469499 | 0.031152733 | 0.036629196 | 0.85048912 | 0.395053207 | NA |
| 4933417D19Rik | 14.66866737 | 0.057840978 | 0.167137561 | 0.346068099 | 0.729291537 | 0.864151352 | Adar | 9583.344352 | 0.031152989 | 0.050618768 | 0.61544344 | 0.538261911 | 0.862082844 |
| Scamp4 | 1725.596758 | 0.02987764 | 0.086321262 | 0.346121443 | 0.729251449 | 0.864151352 | 2510039O18Rik | 1040.104313 | 0.031177659 | 0.057804005 | 0.539368502 | 0.589632611 | 0.885188411 |
| Smim4 | 111.5682331 | -0.047319331 | 0.136717545 | -0.34611016 | 0.729259928 | 0.864151352 | Nbn | 775.8664975 | 0.031180527 | 0.075231138 | 0.414463049 | 0.678535023 | 0.921316909 |
| Rfx8 | 8.163543776 | -0.051202944 | 0.148051484 | -0.345845532 | 0.729458804 | 0.864253503 | Bcas2 | 1611.087504 | 0.031199804 | 0.049184224 | 0.634345753 | 0.525855203 | 0.855892499 |
| Ubr5 | 7064.848839 | -0.027188539 | 0.078623916 | -0.345804948 | 0.729489306 | 0.864253503 | 4932411E22Rik | 117.7389976 | 0.031201832 | 0.095000837 | 0.328437439 | 0.742580939 | 0.939912916 |
| Suz12 | 1498.348722 | -0.031044729 | 0.089796479 | -0.345723236 | 0.72955072 | 0.864260177 | Tubb1 | 1.798813337 | 0.031228877 | 0.029685014 | 1.052008154 | 0.292795809 | NA |
| Btbd10 | 3329.755175 | -0.037180141 | 0.107575007 | -0.34562062 | 0.729627847 | 0.864285464 | Ell2 | 1279.667192 | 0.031230903 | 0.078719512 | 0.396736494 | 0.691561786 | 0.923860706 |
| Lmtk2 | 9799.599148 | 0.025978649 | 0.075182621 | 0.345540612 | 0.729687985 | 0.864290622 | Ltb | 13.45619781 | 0.031256433 | 0.070548631 | 0.443048041 | 0.657730987 | NA |
| Apold1 | 269.2489309 | 0.050202694 | 0.145495579 | 0.345046182 | 0.730059652 | 0.864400447 | Shisa4 | 3874.67045 | 0.031257436 | 0.065413095 | 0.477846773 | 0.632759268 | 0.906637358 |
| Cbfa2t2 | 2673.953218 | -0.02421978 | 0.070169906 | -0.345159069 | 0.729974788 | 0.864400447 | D19Ertd737e | 470.6863504 | 0.031260987 | 0.072366671 | 0.431980455 | 0.665755618 | 0.915591677 |
| Mettl8 | 430.5957936 | 0.027807078 | 0.080587964 | 0.3450525 | 0.730054902 | 0.864400447 | Lurap1 | 552.3042711 | 0.031263328 | 0.072230908 | 0.432824793 | 0.66514206 | 0.915428982 |
| Negr1 | 9060.383222 | 0.025623179 | 0.074245733 | 0.345113155 | 0.730009304 | 0.864400447 | Rab25 | 2.848920711 | 0.031283722 | 0.03882352 | 0.805793033 | 0.420362196 | NA |
| Plekhg1 | 771.2916429 | -0.031657595 | 0.091739152 | -0.345082712 | 0.73003219 | 0.864400447 | Stard7 | 4933.793819 | 0.031301579 | 0.045590191 | 0.686585814 | 0.492343771 | 0.842340249 |
| Nostrin | 151.2481392 | 0.053659323 | 0.155638012 | 0.344770033 | 0.730267263 | 0.864580192 | Mir339 | 1.360556778 | 0.031306093 | 0.026142862 | 1.197500596 | 0.231111496 | NA |
| Slfn3 | 20.03007843 | -0.057864353 | 0.167887876 | -0.344660698 | 0.730349468 | 0.86461145 | Zfp365 | 23209.16665 | 0.031321817 | 0.062917889 | 0.497820527 | 0.618610547 | 0.898575807 |
| Brd3 | 2160.017833 | 0.023675528 | 0.068736247 | 0.344440223 | 0.730515243 | 0.864620115 | Ppp1r10 | 2212.752825 | 0.031322081 | 0.04727331 | 0.662574311 | 0.507603227 | 0.849139846 |
| Med7 | 455.5872962 | -0.029143654 | 0.084592937 | -0.344516394 | 0.730457969 | 0.864620115 | Ppp2r2c | 29407.02663 | 0.03132452 | 0.057266818 | 0.546992503 | 0.584383883 | 0.884313345 |
| Pcna | 1460.851695 | 0.027652107 | 0.08028407 | 0.344428315 | 0.730524198 | 0.864620115 | Mmp13 | 2.536290848 | 0.03132582 | 0.032368678 | 0.967781867 | 0.333153322 | NA |
| C2cd5 | 2221.632983 | 0.031169825 | 0.090636302 | 0.343900009 | 0.730921485 | 0.864877663 | Rnase4 | 632.3964794 | 0.031334865 | 0.071667664 | 0.437224596 | 0.661948474 | 0.915428982 |
| Cstf1 | 841.2285045 | 0.021111964 | 0.061383625 | 0.343934788 | 0.730895329 | 0.864877663 | Rps23 | 6104.436091 | 0.031337817 | 0.058772711 | 0.533203535 | 0.593892695 | 0.886549756 |
| Orai1 | 183.4613945 | 0.032373954 | 0.094129095 | 0.343931429 | 0.730897856 | 0.864877663 | Gck | 70.06369567 | 0.031341165 | 0.061851523 | 0.506716149 | 0.612353998 | 0.895344807 |
| Slc10a3 | 117.5178445 | 0.039476765 | 0.114810757 | 0.343842039 | 0.730965083 | 0.864877663 | Gucd1 | 612.0142677 | 0.031341299 | 0.068975781 | 0.454381212 | 0.649554473 | 0.913724196 |
| Btrc | 5301.500974 | 0.019835283 | 0.057756053 | 0.343432107 | 0.731273409 | 0.864988796 | Rpl27 | 2911.385572 | 0.031343219 | 0.048224025 | 0.649950299 | 0.515724326 | 0.853297835 |
| Coq9 | 2049.315332 | 0.02490255 | 0.072488804 | 0.343536502 | 0.731194885 | 0.864988796 | Ric8b | 2849.531979 | 0.031371523 | 0.064382698 | 0.487266366 | 0.626069579 | 0.902402691 |
| Dgkd | 4958.494059 | 0.022361845 | 0.065143259 | 0.343271828 | 0.731393973 | 0.864988796 | Avil | 12.63040505 | 0.031371549 | 0.067041288 | 0.467943717 | 0.639824839 | NA |
| Pja2 | 24020.60627 | -0.025777128 | 0.075076657 | -0.343344117 | 0.731339595 | 0.864988796 | Rexo2 | 1344.878489 | 0.031373548 | 0.060033823 | 0.522597868 | 0.601254125 | 0.890297392 |
| Trp53rk | 111.367297 | 0.042657287 | 0.124213717 | 0.343418486 | 0.731283655 | 0.864988796 | Znrf3 | 589.4041584 | 0.031378618 | 0.065233346 | 0.481021129 | 0.630501482 | 0.905249211 |
| Zfp354b | 99.15564212 | 0.047452333 | 0.138110591 | 0.343582147 | 0.731160553 | 0.864988796 | Zfp648 | 15.93789832 | 0.031380863 | 0.068530781 | 0.457909019 | 0.647017807 | NA |
| Ppp1r10 | 2030.734903 | -0.016909313 | 0.049270695 | -0.343192094 | 0.731453952 | 0.864993706 | Maob | 1186.620419 | 0.031400574 | 0.066755117 | 0.470384527 | 0.638080317 | 0.908932006 |
| Fam167a | 91.90246557 | -0.050243604 | 0.146504704 | -0.342948743 | 0.731637022 | 0.865144166 | Trim2 | 11434.13753 | 0.031408144 | 0.058317979 | 0.538567088 | 0.590185602 | 0.885570623 |
| Aldh3b1 | 181.9754703 | -0.040863138 | 0.119343212 | -0.342400183 | 0.732049751 | 0.865272295 | Vstm2l | 1982.502138 | 0.031427548 | 0.068291476 | 0.460197234 | 0.645374656 | 0.912363974 |
| Ccdc102a | 54.89448219 | 0.051252741 | 0.149750864 | 0.342253389 | 0.73216021 | 0.865272295 | Fhl5 | 3.349380727 | 0.031434051 | 0.04131624 | 0.76081587 | 0.446767054 | NA |
| Fam186b | 49.81298711 | 0.0565383 | 0.165075447 | 0.342499756 | 0.731974827 | 0.865272295 | Tnfaip1 | 2538.141721 | 0.031449613 | 0.045920707 | 0.684867772 | 0.493427365 | 0.843139148 |
| Fam199x | 365.0926378 | -0.037334102 | 0.109035833 | -0.342402131 | 0.732048285 | 0.865272295 | Aagab | 2475.645321 | 0.031463909 | 0.04923459 | 0.639061057 | 0.522783213 | 0.854337225 |
| Lmbrd2 | 1538.315373 | -0.032077711 | 0.093703573 | -0.342331783 | 0.73210122 | 0.865272295 | AW554918 | 669.0164213 | 0.031464744 | 0.064712587 | 0.486222935 | 0.626809112 | 0.902918401 |
| Mfap1a | 1101.280602 | -0.027407263 | 0.079998277 | -0.342598164 | 0.731900783 | 0.865272295 | Ttc30b | 457.5661767 | 0.031464825 | 0.088893341 | 0.353961556 | 0.723367694 | 0.932617157 |
| Msln | 50.56342227 | 0.056348289 | 0.16453837 | 0.342462912 | 0.73200255 | 0.865272295 | Bdkrb1 | 3.190715645 | 0.031466222 | 0.040417402 | 0.77853152 | 0.436255732 | NA |
| Rsph3a | 163.4196193 | 0.034250786 | 0.100086762 | 0.342210953 | 0.732192143 | 0.865272295 | Lta | 5.631046331 | 0.031468528 | 0.05147674 | 0.611315476 | 0.540990748 | NA |
| Gmds | 284.2111233 | -0.032797477 | 0.095863478 | -0.342126926 | 0.732255375 | 0.865281023 | Gal3st1 | 403.906242 | 0.031470314 | 0.078622336 | 0.400271931 | 0.68895624 | 0.923860706 |
| Med13l | 5431.808213 | 0.037757868 | 0.11040076 | 0.342007318 | 0.732345385 | 0.86532139 | Rbm27 | 1667.073722 | 0.031474388 | 0.068799167 | 0.457482102 | 0.647324564 | 0.912750424 |
| Arhgap26 | 4977.694183 | 0.033116562 | 0.097091582 | 0.341085819 | 0.733038977 | 0.865332757 | Dlg2 | 19247.23475 | 0.031488161 | 0.060103489 | 0.523899043 | 0.600348761 | 0.889841393 |
| Btaf1 | 1672.089714 | -0.030148177 | 0.088266098 | -0.341560094 | 0.732681974 | 0.865332757 | Grem1 | 34.47172527 | 0.031494134 | 0.078222606 | 0.402621902 | 0.687226395 | 0.923860706 |
| Btbd19 | 250.1686548 | 0.039139387 | 0.114783288 | 0.340985063 | 0.733114827 | 0.865332757 | Get4 | 1852.306889 | 0.03149576 | 0.058059462 | 0.5424742 | 0.587491876 | 0.88480114 |
| Ddhd1 | 2762.002079 | -0.020193397 | 0.059111933 | -0.341612875 | 0.732642247 | 0.865332757 | Msrb1 | 672.8602655 | 0.0315164 | 0.073206538 | 0.430513458 | 0.666822179 | 0.916077199 |
| Glod4 | 2752.528443 | -0.018482786 | 0.054107505 | -0.341593764 | 0.732656631 | 0.865332757 | 2610019F03Rik | 823.8171039 | 0.0315392 | 0.07157122 | 0.440668754 | 0.65945282 | 0.915208181 |
| Gm14634 | 29.08526529 | -0.056070133 | 0.164173317 | -0.341530124 | 0.732704531 | 0.865332757 | D330045A20Rik | 10.59280126 | 0.031553688 | 0.061892693 | 0.509812812 | 0.610182609 | NA |
| Mad2l1 | 244.957894 | 0.030570847 | 0.089508863 | 0.341539889 | 0.732697181 | 0.865332757 | St3gal4 | 897.0512347 | 0.031565373 | 0.072128696 | 0.43762573 | 0.661657616 | 0.915428982 |
| Map2k2 | 2449.132867 | 0.024575821 | 0.072061438 | 0.341039833 | 0.733073595 | 0.865332757 | Fam159a | 2.640132242 | 0.031569744 | 0.028990945 | 1.088951896 | 0.276175099 | NA |
| Pik3r1 | 6156.615693 | 0.030501356 | 0.089398911 | 0.341182639 | 0.732966092 | 0.865332757 | Ehbp1 | 2522.165322 | 0.031576683 | 0.058760398 | 0.53738034 | 0.591004921 | 0.8858275 |
| Qdpr | 5190.034086 | -0.052893157 | 0.155132031 | -0.340955744 | 0.733136899 | 0.865332757 | Ankrd13c | 2100.107927 | 0.031598479 | 0.072961246 | 0.433085786 | 0.664952449 | 0.915428982 |
| Snx6 | 1226.815068 | -0.02077683 | 0.060782896 | -0.341820338 | 0.732486103 | 0.865332757 | Cd276 | 491.8485862 | 0.031599316 | 0.072502782 | 0.435835907 | 0.66295579 | 0.915428982 |
| Spdl1 | 21.23886745 | 0.057278649 | 0.167881303 | 0.341185397 | 0.732964016 | 0.865332757 | Orc3 | 1644.42117 | 0.0316234 | 0.061059936 | 0.517907511 | 0.604522801 | 0.891930651 |
| Thbs4 | 231.5327989 | 0.055478779 | 0.162600348 | 0.341197172 | 0.732955152 | 0.865332757 | Lpcat2b | 1.246483544 | 0.031635866 | 0.027252719 | 1.160833365 | 0.245709671 | NA |
| Vmn2r57 | 22.07518669 | -0.055672752 | 0.16327707 | -0.340971036 | 0.733125386 | 0.865332757 | Add1 | 19769.02436 | 0.031637643 | 0.04380783 | 0.722191522 | 0.470176735 | 0.833245343 |
| A330076C08Rik | 14.55609804 | -0.055819023 | 0.163990714 | -0.340379171 | 0.733571003 | 0.865510426 | Gm101 | 14.92818003 | 0.031640206 | 0.061749971 | 0.512392248 | 0.608376515 | NA |
| Amica1 | 20.82008421 | 0.056868006 | 0.167008537 | 0.340509573 | 0.733472815 | 0.865510426 | Tmem97 | 624.066566 | 0.031655889 | 0.069550214 | 0.455151563 | 0.649000205 | 0.913724196 |
| Eml1 | 2424.6355 | 0.025614263 | 0.075276101 | 0.340270849 | 0.733652568 | 0.865510426 | Mospd3 | 527.8526164 | 0.031659284 | 0.070650745 | 0.448109704 | 0.654074025 | 0.914796773 |
| Inpp1 | 754.489519 | 0.023638965 | 0.069396811 | 0.340634742 | 0.733378572 | 0.865510426 | Tex30 | 105.3363976 | 0.031674357 | 0.094415558 | 0.33547815 | 0.737264425 | 0.937206199 |
| Isca1 | 6903.527287 | -0.02069802 | 0.060790348 | -0.340482011 | 0.733493568 | 0.865510426 | Tssk3 | 3.501972258 | 0.031684848 | 0.044042524 | 0.719414882 | 0.47188533 | NA |
| Psma8 | 8.264657935 | 0.051211141 | 0.150549059 | 0.340162476 | 0.733734175 | 0.865510426 | Clic6 | 517.7450935 | 0.031690441 | 0.045627846 | 0.694541688 | 0.487342566 | 0.839801621 |
| Spag9 | 17926.11447 | -0.022076264 | 0.064848335 | -0.340429154 | 0.733533367 | 0.865510426 | Gpr133 | 307.2523333 | 0.031698196 | 0.093764085 | 0.338063301 | 0.735315485 | 0.93653597 |
| Ttc30b | 404.298401 | -0.03701469 | 0.108814794 | -0.340162292 | 0.733734313 | 0.865510426 | Gosr2 | 4441.795297 | 0.031724289 | 0.042823542 | 0.740814228 | 0.458806087 | 0.828211411 |
| Atf2 | 4710.102518 | -0.021271527 | 0.06257795 | -0.33992049 | 0.733916406 | 0.865637095 | A630077J23Rik | 2.550528828 | 0.031736381 | 0.037780485 | 0.840020485 | 0.400896901 | NA |
| Selm | 1826.322815 | 0.02759373 | 0.081188758 | 0.339871321 | 0.733953435 | 0.865637095 | Cog6 | 2107.264933 | 0.031737144 | 0.051472649 | 0.616582684 | 0.537510019 | 0.861996453 |
| Edaradd | 12.02261461 | -0.053900075 | 0.158696585 | -0.339642313 | 0.734125909 | 0.865774611 | Trim23 | 2165.148236 | 0.031738147 | 0.076532989 | 0.414698908 | 0.678362332 | 0.921316909 |
| Faim | 520.295638 | -0.02384171 | 0.07028601 | -0.33920989 | 0.734451619 | 0.865899032 | Ifnlr1 | 12.89042552 | 0.031739994 | 0.06157362 | 0.515480403 | 0.60621736 | NA |
| Rep15 | 23.39507524 | 0.05672322 | 0.167223783 | 0.339205461 | 0.734454956 | 0.865899032 | Wipf3 | 8038.122761 | 0.031753573 | 0.073498975 | 0.432027423 | 0.665721482 | 0.915591677 |
| Sgca | 12.2212277 | 0.056546062 | 0.166689708 | 0.339229473 | 0.734436868 | 0.865899032 | Ece1 | 2822.164523 | 0.031758715 | 0.059250721 | 0.536005539 | 0.591954722 | 0.886226713 |
| Ube2f | 1073.730474 | 0.019287664 | 0.056824703 | 0.339423932 | 0.734290393 | 0.865899032 | Dctn3 | 2403.652134 | 0.031761342 | 0.047089698 | 0.674485995 | 0.500002387 | 0.846241311 |
| Klk14 | 28.65982534 | -0.055709342 | 0.164333895 | -0.339000923 | 0.734609035 | 0.86601479 | Rfx4 | 697.6099249 | 0.031768891 | 0.072198096 | 0.440023945 | 0.659919764 | 0.9153055 |
| C230004F18Rik | 165.7937705 | 0.036333128 | 0.10722639 | 0.338845017 | 0.734726486 | 0.866021467 | Zfp658 | 288.1045133 | 0.031806288 | 0.087472459 | 0.363614879 | 0.716145614 | 0.93125489 |
| Men1 | 2415.984577 | 0.023460083 | 0.069227089 | 0.338885881 | 0.734695701 | 0.866021467 | 0610031J06Rik | 1494.254466 | 0.031806535 | 0.059346682 | 0.53594461 | 0.591996833 | 0.886226713 |
| Ppp4r2 | 2692.462702 | -0.026562473 | 0.078472675 | -0.33849328 | 0.73499149 | 0.866267926 | Arid1b | 3797.719548 | 0.031819572 | 0.045266726 | 0.702935148 | 0.482096166 | 0.837477954 |
| Atp1b3 | 3771.220246 | 0.022618485 | 0.066894043 | 0.338124055 | 0.735269704 | 0.866398113 | Usp47 | 4964.281868 | 0.031823357 | 0.061797609 | 0.51496097 | 0.606580294 | 0.893085886 |
| Eif4ebp1 | 94.05779067 | -0.049550081 | 0.146542324 | -0.338128121 | 0.73526664 | 0.866398113 | Ssbp3 | 5175.466065 | 0.031834234 | 0.044474594 | 0.7157847 | 0.474124301 | 0.833994865 |
| Npas3 | 411.289914 | -0.040602758 | 0.120073518 | -0.338149152 | 0.735250792 | 0.866398113 | Eps8l1 | 178.6142132 | 0.031838452 | 0.041948812 | 0.758983407 | 0.447862482 | 0.822771016 |
| Gm14057 | 40.20866927 | 0.056448889 | 0.167048917 | 0.337918319 | 0.735424743 | 0.866514902 | Mdk | 399.6866949 | 0.031869974 | 0.076674062 | 0.415655214 | 0.677662318 | 0.921093181 |
| Cotl1 | 2145.843974 | 0.021474212 | 0.063619375 | 0.337542017 | 0.735708344 | 0.866717235 | Bhlha9 | 45.16825961 | 0.031873698 | 0.088313545 | 0.360915172 | 0.718162861 | 0.93125489 |
| Eftud1 | 596.7551554 | 0.034461089 | 0.102071868 | 0.337615938 | 0.73565263 | 0.866717235 | 2010005H15Rik | 1.140093182 | 0.03187787 | 0.023397187 | 1.362465882 | 0.173050904 | NA |
| A230072E10Rik | 24.84955612 | 0.056477327 | 0.167708804 | 0.33675827 | 0.736299133 | 0.867149496 | Zfp707 | 354.0981903 | 0.03188262 | 0.075500099 | 0.4222858 | 0.672816422 | 0.919353963 |
| Celf5 | 11000.68403 | -0.025039439 | 0.074341547 | -0.336816216 | 0.736255448 | 0.867149496 | Asph | 3724.573828 | 0.031889873 | 0.049146643 | 0.648871856 | 0.516421208 | 0.853297835 |
| Gli2 | 211.6036066 | -0.045321458 | 0.134562833 | -0.336805173 | 0.736263773 | 0.867149496 | Fbxw11 | 6398.42148 | 0.031892885 | 0.048058209 | 0.663630323 | 0.506926945 | 0.848958235 |
| Gosr2 | 4135.373615 | -0.018378971 | 0.054547849 | -0.336933008 | 0.736167402 | 0.867149496 | Ash1l | 7256.629629 | 0.031893759 | 0.074618464 | 0.427424484 | 0.669070177 | 0.917484508 |

| | | | | | | |
|---|---|---|---|---|---|---|
| Ckap2 | 24.35405877 | -0.056368467 | 0.16753916 | -0.336449502 | 0.736531926 | 0.867298501 |
| Cpne3 | 632.7993878 | 0.029129623 | 0.086581421 | 0.336441967 | 0.736537607 | 0.867298501 |
| Daam1 | 2839.69438 | -0.017914043 | 0.053271535 | -0.336277948 | 0.736661278 | 0.867312298 |
| Marveld2 | 38.81004968 | -0.052316338 | 0.155555897 | -0.336318577 | 0.736630642 | 0.867312298 |
| Rbks | 54.81880893 | -0.05290575 | 0.157362467 | -0.336203101 | 0.736717715 | 0.867312839 |
| Arid3c | 8.671034986 | 0.053323558 | 0.158839878 | 0.335706367 | 0.737092304 | 0.867687902 |
| 1810043G02Rik | 1065.549107 | 0.022216711 | 0.06623775 | 0.335408608 | 0.737316875 | 0.867845457 |
| Efemp2 | 333.4872041 | -0.036061168 | 0.107523191 | -0.335380376 | 0.737338169 | 0.867845457 |
| Thoc3 | 1056.491454 | -0.024121314 | 0.071945362 | -0.33527268 | 0.7374194 | 0.867875138 |
| D2Wsu81e | 1134.606289 | -0.023277702 | 0.069492757 | -0.334965872 | 0.737650831 | 0.867949724 |
| Gm16432 | 23.58158421 | 0.055214976 | 0.164820902 | 0.334999841 | 0.737625206 | 0.867949724 |
| Hist2h3c2 | 36.91056024 | -0.054266252 | 0.161935012 | -0.335111297 | 0.737541131 | 0.867949724 |
| Paqr4 | 2419.366756 | -0.030636098 | 0.091578893 | -0.334532307 | 0.737977917 | 0.868136804 |
| Rhot1 | 3174.185746 | -0.020014936 | 0.059805509 | -0.334667095 | 0.737876226 | 0.868136804 |
| Tctex1d2 | 258.5524557 | 0.032193875 | 0.096229449 | 0.334553248 | 0.737962117 | 0.868136804 |
| Arhgef16 | 27.12201044 | -0.055649893 | 0.1665622 | -0.334108776 | 0.738297478 | 0.86844679 |
| Hp1bp3 | 10147.57499 | 0.021844245 | 0.065413012 | 0.333943427 | 0.738422249 | 0.868527619 |
| Ap3m2 | 6943.140625 | 0.019128639 | 0.057308271 | 0.33378497 | 0.738541826 | 0.868561308 |
| Cbx4 | 1613.814629 | 0.022332648 | 0.066912916 | 0.333756902 | 0.738563007 | 0.868561308 |
| BC049352 | 30.91641004 | 0.054707204 | 0.16395963 | 0.33366265 | 0.738634137 | 0.868579032 |
| Pdgfb | 1146.216983 | -0.03379265 | 0.101341146 | -0.333454394 | 0.73879131 | 0.868697924 |
| Esrra | 1752.360015 | -0.021656955 | 0.064969626 | -0.333339697 | 0.738877877 | 0.868733786 |
| Nacad | 1944.206602 | 0.025837314 | 0.077565102 | 0.333104882 | 0.739055115 | 0.868810315 |
| Snrpd1 | 817.1731181 | -0.020198387 | 0.060629992 | -0.333141842 | 0.739027216 | 0.868810315 |
| 2410066E13Rik | 1786.320262 | -0.024874012 | 0.074900586 | -0.332093694 | 0.739818512 | 0.869641758 |
| Ndufs1 | 5778.796976 | 0.019044253 | 0.057383635 | 0.331876039 | 0.739982864 | 0.869768965 |
| 9330133O14Rik | 287.649951 | -0.029965612 | 0.090349306 | -0.331663996 | 0.74014299 | 0.869891185 |
| Arhgap11a | 157.2080622 | 0.041561997 | 0.1255517 | 0.331034919 | 0.740618109 | 0.870317557 |
| Srp54a | 117.988052 | 0.038475709 | 0.116211698 | 0.331082929 | 0.740581846 | 0.870317557 |
| Oma1 | 240.2759288 | 0.031388434 | 0.094889988 | 0.330787628 | 0.740804907 | 0.870471047 |
| Fndc1 | 230.1010035 | 0.050388316 | 0.152442333 | 0.330540181 | 0.740991837 | 0.87062467 |
| AI847159 | 11.61781671 | -0.052631922 | 0.159306009 | -0.330382526 | 0.741110943 | 0.870698586 |
| Kbtbd7 | 1264.054909 | -0.029155222 | 0.088309601 | -0.330147813 | 0.741288277 | 0.870751603 |
| Rpl7 | 8331.632546 | -0.022929385 | 0.06946201 | -0.33009965 | 0.741324667 | 0.870751603 |
| Scamp3 | 1538.93972 | 0.021852688 | 0.066188159 | 0.330160083 | 0.741279006 | 0.870751603 |
| Aplp1 | 47280.54444 | -0.020483442 | 0.062105495 | -0.329816902 | 0.741538315 | 0.870758226 |
| C030016D13Rik | 15.16833982 | -0.055151078 | 0.167265933 | -0.329720926 | 0.741610841 | 0.870758226 |
| Exog | 1092.216366 | 0.026919149 | 0.081628163 | 0.32977772 | 0.741567924 | 0.870758226 |
| Hsd17b11 | 1473.618405 | 0.019593955 | 0.059393064 | 0.329903079 | 0.741473197 | 0.870758226 |
| Wbscr27 | 323.5035875 | 0.036585998 | 0.110960705 | 0.329720312 | 0.741611304 | 0.870758226 |
| Slc35a1 | 593.5531464 | -0.031506146 | 0.095726714 | -0.329125954 | 0.742060488 | 0.871219611 |
| Hyi | 151.8503011 | -0.034566651 | 0.105106786 | -0.328871731 | 0.742252642 | 0.871313163 |
| Klhl12 | 1816.372597 | -0.017765545 | 0.0540135 | -0.328909341 | 0.742224213 | 0.871313163 |
| Loxl3 | 164.9779882 | 0.03919079 | 0.119288928 | 0.328536694 | 0.742505905 | 0.871467912 |
| Rad51c | 122.6735446 | -0.036638465 | 0.111531654 | -0.328502836 | 0.7425315 | 0.871467912 |
| Rangrf | 283.6757771 | -0.042722945 | 0.130064871 | -0.328474124 | 0.742553206 | 0.871467912 |
| Ak2 | 679.7522014 | 0.032243224 | 0.098449797 | 0.327509298 | 0.743282712 | 0.871861787 |
| H2-T10 | 50.77292073 | 0.052955554 | 0.161605304 | 0.327684504 | 0.743150221 | 0.871861787 |
| Prps2 | 1343.299138 | 0.036272727 | 0.110750177 | 0.327518454 | 0.743275788 | 0.871861787 |
| Ranbp1 | 1285.713858 | 0.022502492 | 0.068673698 | 0.327672637 | 0.743159194 | 0.871861787 |
| Rasa3 | 3611.415396 | 0.022564779 | 0.068885732 | 0.327568256 | 0.743238127 | 0.871861787 |
| Rps6kl1 | 1396.054718 | -0.025842233 | 0.078884464 | -0.327595976 | 0.743217165 | 0.871861787 |
| Snapc2 | 1012.315525 | 0.021451455 | 0.065449948 | 0.327753584 | 0.743097985 | 0.871861787 |
| Gna13 | 2860.771725 | 0.021852536 | 0.066753159 | 0.327363317 | 0.743393108 | 0.871894162 |
| Igsf8 | 6794.012464 | 0.025219452 | 0.077063522 | 0.327255383 | 0.743474735 | 0.871894162 |
| Ube2q2 | 2518.42581 | -0.018384086 | 0.056177571 | -0.32724957 | 0.743479132 | 0.871894162 |
| Cox6b2 | 56.03317897 | 0.049863796 | 0.152421585 | 0.327143928 | 0.743559028 | 0.871921863 |
| Btbd6 | 1870.13243 | 0.021724814 | 0.066480453 | 0.326784986 | 0.743830516 | 0.87217421 |
| Kcna2 | 13619.57716 | -0.024837525 | 0.076043288 | -0.326623497 | 0.74395267 | 0.872186452 |
| Vtcn1 | 9.615922797 | 0.050647125 | 0.155063256 | 0.326622347 | 0.74395354 | 0.872186452 |
| Adsl | 920.1517154 | -0.023062421 | 0.070659821 | -0.326386632 | 0.744131851 | 0.872202723 |
| Exosc1 | 457.1513466 | 0.026632061 | 0.081598137 | 0.326380753 | 0.744136297 | 0.872202723 |
| Ncoa5 | 2082.659723 | -0.022723156 | 0.069591899 | -0.326520135 | 0.744030858 | 0.872202723 |
| Cdh12 | 445.41938 | 0.036423907 | 0.111630011 | 0.32629135 | 0.744203932 | 0.872216016 |
| Atp2b2 | 30955.47886 | -0.023258484 | 0.07139818 | -0.325757388 | 0.744607921 | 0.872491502 |
| Gpr180 | 839.2537099 | -0.021200882 | 0.065074002 | -0.325796493 | 0.744578332 | 0.872491502 |
| Nck1 | 702.3564888 | 0.023186494 | 0.071145718 | 0.325901474 | 0.744498901 | 0.872491502 |
| Nrl | 11.43725739 | -0.054039002 | 0.16604577 | -0.325446422 | 0.744843226 | 0.872701221 |
| Nup160 | 558.8627217 | 0.027497558 | 0.084578224 | 0.325113916 | 0.745094857 | 0.872930035 |
| Klhl10 | 8.328962422 | -0.048219853 | 0.148382538 | -0.324969861 | 0.745203882 | 0.872982115 |
| Prr18 | 1194.081594 | -0.054096886 | 0.166499965 | -0.324906289 | 0.745251997 | 0.872982115 |
| Nacc1 | 5961.913637 | 0.024233346 | 0.074701595 | 0.324401993 | 0.745633711 | 0.873297205 |
| Tpd52l2 | 1772.891706 | -0.020957439 | 0.064596899 | -0.324434136 | 0.745609379 | 0.873297205 |
| Lrrc42 | 686.7737961 | 0.026227612 | 0.080871001 | 0.324314172 | 0.745700191 | 0.873309052 |
| Ap4s1 | 1102.478813 | -0.024746341 | 0.076364259 | -0.32405658 | 0.745895199 | 0.873471409 |
| Tlr9 | 58.65205663 | 0.051752438 | 0.159878434 | 0.323698678 | 0.746166172 | 0.873722693 |
| Lef1 | 216.4999994 | 0.041776453 | 0.129152525 | 0.32346602 | 0.746342337 | 0.873862932 |
| A630072M18Rik | 248.651983 | 0.02833527 | 0.087703893 | 0.323078812 | 0.746635556 | 0.873899395 |
| A830080D01Rik | 234.7024779 | -0.037205585 | 0.115269352 | -0.322770836 | 0.746868802 | 0.873899395 |
| Acsf2 | 692.3929229 | 0.026820628 | 0.083156693 | 0.322531195 | 0.747050309 | 0.873899395 |
| Akap13 | 1856.759863 | -0.033105212 | 0.102628709 | -0.322572627 | 0.747018927 | 0.873899395 |
| Fut10 | 425.7383717 | 0.026927149 | 0.083475278 | 0.322576336 | 0.747016118 | 0.873899395 |
| Gm13826 | 65.29880927 | -0.054234087 | 0.168048039 | -0.322729661 | 0.746899987 | 0.873899395 |
| Gm6313 | 27.61326038 | -0.053263189 | 0.164769917 | -0.323257967 | 0.746499884 | 0.873899395 |
| Grap2 | 12.91169784 | -0.05373861 | 0.16627204 | -0.323196912 | 0.746546119 | 0.873899395 |
| Inppl1 | 1648.337644 | 0.020402154 | 0.063175901 | 0.322942034 | 0.746739142 | 0.873899395 |
| Pmpca | 3970.37913 | 0.013781858 | 0.042723722 | 0.322580941 | 0.747012629 | 0.873899395 |
| Rims1 | 6072.698806 | 0.029864716 | 0.092372596 | 0.323307101 | 0.746462677 | 0.873899395 |
| Rnf130 | 1861.552785 | -0.018543139 | 0.057456197 | -0.322735235 | 0.746895765 | 0.873899395 |
| Pak2 | 2141.887296 | 0.020434673 | 0.06339492 | 0.322339289 | 0.747195672 | 0.874003453 |
| Ass1 | 1682.123934 | 0.02410641 | 0.07481603 | 0.322209159 | 0.747294246 | 0.874052771 |
| Aqp4 | 4582.598944 | -0.028565737 | 0.08870562 | -0.322028487 | 0.747431114 | 0.874111615 |
| P4ha2 | 417.7503252 | -0.033362311 | 0.103611654 | -0.321993803 | 0.747457389 | 0.874111615 |
| Rpia | 332.3970037 | 0.035781448 | 0.111169154 | 0.321864894 | 0.74755505 | 0.874159844 |
| Kdm5a | 2096.379063 | -0.024892692 | 0.077418943 | -0.32153231 | 0.747807031 | 0.87438851 |
| Dnm3 | 7535.458429 | -0.023963957 | 0.07454962 | -0.321449755 | 0.747869583 | 0.874395662 |
| Arhgap40 | 9.341655097 | -0.050629512 | 0.157560375 | -0.321334043 | 0.747957261 | 0.874432189 |
| B3galt5 | 2414.65666 | -0.022544355 | 0.070237643 | -0.32097255 | 0.748231193 | 0.874579799 |
| Gm684 | 51.86542157 | -0.053146855 | 0.165595437 | -0.320943959 | 0.74825286 | 0.874579799 |
| Srpk2 | 9243.14002 | 0.015771759 | 0.049134068 | 0.320994358 | 0.748214667 | 0.874579799 |
| Mrps22 | 542.6896349 | -0.030971993 | 0.096542651 | -0.320811499 | 0.748353245 | 0.874631152 |
| Pus1 | 469.9164399 | 0.026375179 | 0.082439448 | 0.319933963 | 0.749018391 | 0.875342506 |
| Epdr1 | 4923.01199 | -0.019700179 | 0.061590814 | -0.319855801 | 0.749077645 | 0.87534573 |
| Pon2 | 2221.097596 | 0.029466126 | 0.092186793 | 0.31963501 | 0.749245031 | 0.875475303 |
| C530005A16Rik | 249.0886477 | 0.045339465 | 0.141882302 | 0.319556875 | 0.74930427 | 0.875478498 |
| Gemin6 | 164.8311605 | 0.03698991 | 0.115797044 | 0.319437428 | 0.749394833 | 0.875518288 |
| Ttc39b | 3043.251009 | 0.023872737 | 0.074771037 | 0.319277859 | 0.749515821 | 0.875593616 |
| Fibin | 100.7398516 | 0.053105493 | 0.16643118 | 0.319083797 | 0.74966297 | 0.875699492 |
| B930025P03Rik | 12.59728235 | -0.051352892 | 0.161340584 | -0.318288747 | 0.750265919 | 0.876337743 |
| Loxl2 | 98.41123997 | 0.046401348 | 0.145990742 | 0.317837608 | 0.750608122 | 0.876671358 |
| Pou2f2 | 225.566791 | 0.038086189 | 0.119859099 | 0.317758014 | 0.750668501 | 0.876675794 |
| 3110082J24Rik | 30.79067243 | -0.053122968 | 0.167587477 | -0.316986504 | 0.751253843 | 0.876924335 |
| Fsd1l | 1063.504579 | 0.023534441 | 0.074240894 | 0.317001045 | 0.751242809 | 0.876924335 |
| Galnt9 | 2924.626317 | -0.044087415 | 0.139084771 | -0.316982335 | 0.751257006 | 0.876924335 |
| Mpst | 698.6365587 | -0.035718466 | 0.112692432 | -0.316955323 | 0.751277503 | 0.876924335 |
| Picalm | 5658.954887 | 0.020250563 | 0.063805089 | 0.317381636 | 0.75095404 | 0.876924335 |
| Snrnp40 | 692.9643211 | -0.026335313 | 0.083073489 | -0.317012245 | 0.751234311 | 0.876924335 |
| Tmem241 | 236.2524372 | 0.035792754 | 0.112853504 | 0.317161209 | 0.751121282 | 0.876924335 |
| Trmt112 | 693.3137594 | -0.021366669 | 0.067443889 | -0.316806602 | 0.751390354 | 0.876989993 |
| Atg9b | 194.579822 | 0.04119628 | 0.13024734 | 0.316292677 | 0.751780369 | 0.877114837 |
| Col5a2 | 201.5184001 | -0.046561305 | 0.147089705 | -0.316550399 | 0.751584778 | 0.877114837 |
| Kctd12b | 176.0821419 | -0.036642688 | 0.115845014 | -0.316307859 | 0.751768847 | 0.877114837 |
| Prmt3 | 829.8614498 | 0.021365489 | 0.067537175 | 0.316351528 | 0.751735704 | 0.877114837 |
| Tmem115 | 1095.15483 | -0.018385199 | 0.058118434 | -0.316340235 | 0.751744275 | 0.877114837 |
| Serbp1 | 5035.74888 | 0.024687245 | 0.078117149 | 0.3160285 | 0.751980876 | 0.877216661 |
| Zswim7 | 249.4677752 | 0.032268396 | 0.102083633 | 0.316097644 | 0.751928395 | 0.877216661 |
| 2900076A07Rik | 123.717829 | 0.040815082 | 0.129196393 | 0.31591503 | 0.752067003 | 0.877251084 |
| Acss1 | 1073.232393 | -0.040663143 | 0.128918297 | -0.315417935 | 0.752444351 | 0.87730433 |
| Cep120 | 2177.01557 | 0.025628132 | 0.081302573 | 0.315219204 | 0.752595226 | 0.87730433 |
| Gpr37 | 2419.471675 | 0.049652545 | 0.157463483 | 0.315327362 | 0.752513112 | 0.87730433 |
| Pitpnm3 | 5699.723898 | 0.027544156 | 0.087377929 | 0.315230128 | 0.752586932 | 0.87730433 |
| Ptov1 | 4545.89114 | -0.017704385 | 0.056117269 | -0.315489068 | 0.75239035 | 0.87730433 |
| Tmx1 | 1109.275501 | 0.028772074 | 0.091191597 | 0.315512338 | 0.752372685 | 0.87730433 |
| Vkorc1 | 346.606721 | -0.037874492 | 0.120019323 | -0.315569947 | 0.752328951 | 0.87730433 |
| Zfp605 | 555.0832218 | -0.023881988 | 0.075771669 | -0.315183608 | 0.752622251 | 0.87730433 |
| Zfp647 | 95.11270836 | -0.04328718 | 0.137196941 | -0.31551126 | 0.752373503 | 0.87730433 |
| Cdk12 | 1108.151734 | -0.026355387 | 0.083648909 | -0.315071496 | 0.75270737 | 0.877337546 |
| Ip6k1 | 11115.03151 | 0.022774788 | 0.072327915 | 0.314882408 | 0.752850939 | 0.877372881 |
| Phkb | 1570.296888 | 0.039519987 | 0.125486239 | 0.314934825 | 0.75281114 | 0.877372881 |
| 4930529M08Rik | 31.25026332 | 0.052703486 | 0.167437725 | 0.314764707 | 0.752940311 | 0.877408124 |
| Rpl23 | 5647.02169 | -0.020945931 | 0.066559799 | -0.314693423 | 0.752994438 | 0.877408124 |
| BC049715 | 16.82902151 | -0.052830637 | 0.168117865 | -0.314247606 | 0.753332988 | 0.877736601 |
| Bcl9l | 1782.408814 | 0.02101478 | 0.067030286 | 0.313511717 | 0.75389192 | 0.878261353 |
| Crtap | 197.589956 | 0.035452143 | 0.113083045 | 0.313505381 | 0.753896734 | 0.878261353 |
| Exo5 | 304.325051 | -0.035581763 | 0.113611967 | -0.31318675 | 0.754138784 | 0.878477282 |
| Dctn2 | 8988.493529 | 0.023062266 | 0.073728099 | 0.312801579 | 0.754431416 | 0.878686038 |
| Ttll1 | 1567.328126 | 0.020274857 | 0.064807801 | 0.312845938 | 0.754397712 | 0.878686038 |
| 0610040B10Rik | 39.07994039 | -0.048487587 | 0.155195302 | -0.312429479 | 0.75471415 | 0.878883205 |
| Abi1 | 4114.116412 | -0.024630263 | 0.078826962 | -0.312459874 | 0.754691053 | 0.878883205 |
| Mir707 | 10.6946565 | 0.049148616 | 0.157352536 | 0.312347146 | 0.754776714 | 0.878890005 |
| Man2a1 | 965.5910422 | -0.025500006 | 0.081795215 | -0.311754256 | 0.755227291 | 0.879348587 |
| Ttc7b | 8006.847535 | -0.021038767 | 0.067572737 | -0.311349938 | 0.755534607 | 0.879640307 |
| Abcd4 | 453.1293278 | 0.037973193 | 0.122207819 | 0.31072638 | 0.756008641 | 0.879680258 |
| Ccdc66 | 562.8360292 | 0.027725163 | 0.089130975 | 0.311060914 | 0.755754314 | 0.879680258 |
| Desi1 | 5137.662373 | -0.025033663 | 0.080535877 | -0.310838651 | 0.755923285 | 0.879680258 |
| Dlx1as | 574.1928372 | -0.027146056 | 0.087295312 | -0.31096809 | 0.75582488 | 0.879680258 |
| Ercc6 | 1268.420034 | -0.022121818 | 0.0712539 | -0.310464665 | 0.756207626 | 0.879680258 |
| Fbxo3 | 2782.695023 | -0.018039865 | 0.058079832 | -0.310604636 | 0.756101202 | 0.879680258 |
| Lrfn1 | 1371.266007 | 0.027613717 | 0.088935208 | 0.310492523 | 0.756186445 | 0.879680258 |
| Me3 | 1888.54766 | 0.022448348 | 0.072242867 | 0.310734456 | 0.756002501 | 0.879680258 |
| Nutf2 | 476.9424126 | -0.023966927 | 0.07721089 | -0.310408635 | 0.756250228 | 0.879680258 |
| Srpx | 19.20947324 | -0.052129024 | 0.167850824 | -0.310567577 | 0.756129379 | 0.879680258 |
| Trip13 | 21.2721915 | 0.052210029 | 0.167907146 | 0.310945844 | 0.755841792 | 0.879680258 |
| Xpa | 189.8406325 | 0.041275818 | 0.132667322 | 0.31112272 | 0.755707329 | 0.879680258 |
| Dnajb5 | 4907.142787 | -0.036929444 | 0.119020171 | -0.310278868 | 0.7563489 | 0.879728988 |
| Strip1 | 3124.122388 | -0.0220394 | 0.071064899 | -0.310130603 | 0.756461641 | 0.879794075 |
| Efcab14 | 2396.544081 | 0.019323912 | 0.062372759 | 0.309813331 | 0.756702914 | 0.879942581 |
| Meox1 | 21.57115546 | -0.052022422 | 0.167899381 | -0.309842843 | 0.756680469 | 0.879942581 |
| Tnfrsf21 | 2892.729433 | -0.01938768 | 0.062630311 | -0.309557468 | 0.756897504 | 0.880102809 |
| Ptgds | 25568.77765 | 0.049043388 | 0.15857036 | 0.309284713 | 0.757104959 | 0.88027797 |
| Prrg2 | 139.3613697 | 0.037818227 | 0.122436607 | 0.308880064 | 0.757412763 | 0.880569772 |
| 1700086O06Rik | 117.7000298 | -0.037700145 | 0.122097471 | -0.308770896 | 0.757495809 | 0.880600245 |
| Dhdh | 492.9445713 | -0.032732911 | 0.106069733 | -0.30859803 | 0.75762732 | 0.880620982 |
| Ints5 | 876.266885 | 0.027967105 | 0.090622015 | 0.308612705 | 0.757616155 | 0.880620982 |
| Pfdn2 | 1709.104182 | -0.026293557 | 0.085274572 | -0.308339943 | 0.757823675 | 0.880783139 |
| A430105I19Rik | 343.7695633 | 0.034634966 | 0.11240313 | 0.308131689 | 0.757982129 | 0.880865123 |
| Col4a3bp | 3236.718876 | 0.025935535 | 0.084179553 | 0.308097796 | 0.758007919 | 0.880865123 |
| Crebbp | 2235.255626 | 0.0329432 | 0.107050753 | 0.307734405 | 0.758284438 | 0.881093622 |
| Mybl2 | 39.80392499 | 0.048790022 | 0.158568795 | 0.307689932 | 0.758318282 | 0.881093622 |
| Crnkl1 | 721.7406018 | 0.023464738 | 0.076337332 | 0.307382217 | 0.758552462 | 0.881299629 |
| Eif3l | 2483.004582 | 0.024322238 | 0.079197021 | 0.307110513 | 0.758759256 | 0.881435884 |
| Nudcd2 | 373.4802144 | -0.029636904 | 0.096512425 | -0.307078638 | 0.758783517 | 0.881435884 |
| C2cd2l | 7997.75983 | -0.022200314 | 0.072372411 | -0.30675106 | 0.759032862 | 0.881587457 |
| Cklf | 46.78292553 | 0.047346221 | 0.154461724 | 0.306523971 | 0.759205731 | 0.881587457 |
| Gm4944 | 308.1623449 | -0.044267741 | 0.144427559 | -0.306504806 | 0.759220321 | 0.881587457 |
| Itpka | 4856.380867 | -0.032574978 | 0.106320755 | -0.306383999 | 0.75931229 | 0.881587457 |
| Mfsd6 | 4503.544478 | -0.015295309 | 0.049914136 | -0.30643242 | 0.759275427 | 0.881587457 |
| Tmem87b | 1557.173958 | 0.019801825 | 0.064610573 | 0.30647963 | 0.759239487 | 0.881587457 |
| Wdr55 | 389.7170651 | -0.024465578 | 0.079782875 | -0.306651999 | 0.75910827 | 0.881587457 |
| Gins2 | 25.34136488 | -0.050593779 | 0.165175155 | -0.306303806 | 0.759373342 | 0.881592279 |
| Akt1s1 | 1179.126679 | 0.025195572 | 0.082303722 | 0.306129196 | 0.759506279 | 0.881680549 |
| Gigyf2 | 2902.176205 | 0.021076429 | 0.068885586 | 0.305962827 | 0.759632949 | 0.881761531 |
| Ankrd49 | 338.8693129 | -0.023908916 | 0.078228826 | -0.305627948 | 0.759887938 | 0.881991439 |
| Cpsf2 | 2200.8284 | -0.021293947 | 0.069703115 | -0.305494905 | 0.75998925 | 0.882042954 |
| Fdx1l | 23.64457283 | -0.050463184 | 0.165248538 | -0.305377493 | 0.760078662 | 0.882080652 |
| Ly96 | 22.78309308 | 0.051133572 | 0.167641191 | 0.305017949 | 0.760352482 | 0.882332336 |
| Col13a1 | 55.00126042 | -0.046888151 | 0.153765654 | -0.304932539 | 0.760417533 | 0.88234174 |
| Fxyd7 | 1809.908539 | 0.038480482 | 0.126565237 | 0.304036742 | 0.761099899 | 0.882736869 |
| Lap3 | 1822.310906 | -0.028173721 | 0.092567956 | -0.304357167 | 0.760855796 | 0.882736869 |
| Pecr | 145.3310414 | 0.036576014 | 0.120282298 | 0.304084761 | 0.761063317 | 0.882736869 |
| Pole2 | 14.41585235 | 0.048699786 | 0.160088962 | 0.304204519 | 0.760972082 | 0.882736869 |
| Sdcbp | 6322.274343 | 0.016592555 | 0.054527766 | 0.304295524 | 0.760902755 | 0.882736869 |
| Zfml | 4458.901212 | -0.020098389 | 0.066096661 | -0.304075712 | 0.761070211 | 0.882736869 |
| Spata24 | 110.5729162 | -0.040149415 | 0.132096936 | -0.30393903 | 0.761174342 | 0.882757129 |
| Slco2b1 | 1052.869527 | -0.020693403 | 0.068125059 | -0.303756117 | 0.761313701 | 0.882852667 |
| Itpa | 1704.06511 | 0.020350795 | 0.06703579 | 0.303581039 | 0.761447099 | 0.882875204 |
| Wdr31 | 148.761649 | 0.040488688 | 0.133342828 | 0.303643534 | 0.761399481 | 0.882875204 |
| Cep170b | 12215.13351 | -0.026116892 | 0.086229387 | -0.302876934 | 0.76198365 | 0.883190125 |
| Rnf219 | 580.8321946 | 0.022988912 | 0.075886073 | 0.302939799 | 0.761935741 | 0.883190125 |
| Slc25a38 | 654.6583019 | -0.022640849 | 0.074721003 | -0.303005149 | 0.761885938 | 0.883190125 |
| Tcte1 | 107.0245045 | 0.041370885 | 0.136604928 | 0.302850604 | 0.762003717 | 0.883190125 |
| Tmem218 | 502.1212783 | -0.023738924 | 0.078374926 | -0.302889274 | 0.761974246 | 0.883190125 |
| Sh2d5 | 4824.921479 | -0.027381039 | 0.090445762 | -0.302734357 | 0.762092313 | 0.88322674 |
| Cwc25 | 678.065601 | -0.024772424 | 0.081877284 | -0.302555519 | 0.762228617 | 0.883300031 |
| Tbc1d10a | 871.9608594 | -0.024532393 | 0.081098341 | -0.302501789 | 0.76226957 | 0.883300031 |
| Elmod2 | 585.4417674 | 0.025536226 | 0.084441749 | 0.302412334 | 0.762337753 | 0.883312978 |
| Kcne4 | 64.94067904 | 0.049029375 | 0.162254771 | 0.302175246 | 0.762518473 | 0.883395148 |
| Ncoa7 | 3112.174392 | 0.022458226 | 0.074323222 | 0.302169702 | 0.762522699 | 0.883395148 |
| Gpr1 | 26.14264596 | 0.050334655 | 0.166821374 | 0.301727851 | 0.762859535 | 0.883467684 |
| Nop58 | 978.3830433 | 0.023809763 | 0.078925464 | 0.301674028 | 0.762900569 | 0.883467684 |
| Psmg2 | 507.0191698 | 0.026247864 | 0.087008205 | 0.301671132 | 0.762902777 | 0.883467684 |
| Taf5 | 236.4150467 | 0.033557008 | 0.111197317 | 0.301778941 | 0.762820585 | 0.883467684 |
| Tlr12 | 39.68963515 | 0.050643855 | 0.167895694 | 0.301638795 | 0.762927431 | 0.883467684 |
| Ube2j1 | 4046.004225 | 0.022708143 | 0.075264338 | 0.301711852 | 0.762871733 | 0.883467684 |
| Rnf24 | 2476.544661 | -0.024588548 | 0.081556228 | -0.301491973 | 0.76303937 | 0.883531276 |
| Abca17 | 32.03836886 | 0.050585711 | 0.168009383 | 0.301088603 | 0.763346932 | 0.883545658 |
| Bricd5 | 14.13602637 | -0.050140654 | 0.166388478 | -0.301346912 | 0.763149972 | 0.883545658 |
| Cct2 | 5035.270111 | -0.020173456 | 0.067043486 | -0.300901056 | 0.763489945 | 0.883545658 |
| Colec12 | 656.2896989 | 0.037987746 | 0.126319354 | 0.300727836 | 0.763622041 | 0.883545658 |
| Gm16023 | 12.20423533 | -0.050303071 | 0.167231448 | -0.300799113 | 0.763567684 | 0.883545658 |
| Il1a | 38.19105039 | 0.050353816 | 0.167343633 | 0.300900696 | 0.763490219 | 0.883545658 |
| Insr | 2606.667835 | -0.020184344 | 0.067083439 | -0.30088415 | 0.763502837 | 0.883545658 |
| Nudt16l1 | 1384.285795 | -0.019189135 | 0.063788804 | -0.300822926 | 0.763549525 | 0.883545658 |
| Slamf6 | 8.921558944 | -0.047126082 | 0.156455033 | -0.301211671 | 0.763253091 | 0.883545658 |
| Tmprss7 | 84.63888987 | 0.041436548 | 0.137647208 | 0.301034424 | 0.763388245 | 0.883545658 |

| | | | | | | |
|---|---|---|---|---|---|---|
| 1810026J23Rik | 3110.388477 | 0.031896767 | 0.049658203 | 0.642326244 | 0.520661378 | 0.853626092 |
| Pla2g16 | 1937.548566 | 0.031909803 | 0.059472964 | 0.536543016 | 0.591583315 | 0.88618534 |
| Pip5k1a | 2277.641769 | 0.031925754 | 0.06786797 | 0.470409743 | 0.638062305 | 0.908932006 |
| Map1b | 35301.17457 | 0.031934005 | 0.071225486 | 0.44835082 | 0.653900029 | 0.914733232 |
| Ncmap | 28.29647334 | 0.031935239 | 0.088123301 | 0.362392672 | 0.717058615 | 0.93125489 |
| Ckb | 49969.98867 | 0.031940839 | 0.050957857 | 0.62680891 | 0.530784502 | 0.858116381 |
| Cdh6 | 451.8518178 | 0.031945788 | 0.077374465 | 0.412872485 | 0.679700041 | 0.921792811 |
| Parva | 3776.591033 | 0.031954026 | 0.06219624 | 0.513761383 | 0.60741883 | 0.893686409 |
| 2010300C02Rik | 11894.21745 | 0.031956595 | 0.06314515 | 0.506081535 | 0.612799413 | 0.895428201 |
| Rfesd | 417.2865474 | 0.031964915 | 0.071362142 | 0.447925385 | 0.654207047 | 0.914894731 |
| Klhl12 | 1969.221069 | 0.031981071 | 0.053498913 | 0.597789166 | 0.549980621 | 0.869221902 |
| Jmjd8 | 1479.815022 | 0.031983927 | 0.06772866 | 0.472236237 | 0.636758177 | 0.908778667 |
| Cnn3 | 2731.482496 | 0.032002599 | 0.061701022 | 0.518672106 | 0.603989417 | 0.891718993 |
| Tom1l2 | 17058.29249 | 0.032003935 | 0.050296205 | 0.636309145 | 0.524574946 | 0.855487854 |
| Asb8 | 1880.713549 | 0.032008853 | 0.047738923 | 0.670498013 | 0.502540373 | 0.84739426 |
| Pdcd6ip | 3843.519286 | 0.032020846 | 0.057005032 | 0.561719647 | 0.574307046 | 0.879791496 |
| 1700101I11Rik | 66.6860692 | 0.03204477 | 0.094488458 | 0.339139514 | 0.734504633 | 0.936456337 |
| Gmpr | 1633.466118 | 0.03204971 | 0.063934628 | 0.501288754 | 0.616167919 | 0.89720439 |
| Mttp | 636.2254763 | 0.032051047 | 0.082923255 | 0.386514574 | 0.699115611 | 0.926567041 |
| Slc37a3 | 2494.096341 | 0.032061131 | 0.056298613 | 0.569483497 | 0.569028067 | 0.87745335 |
| Nacc2 | 6757.798505 | 0.032100882 | 0.047602662 | 0.67435056 | 0.500088467 | 0.846258745 |
| Rabl5 | 1422.308888 | 0.032125104 | 0.052893972 | 0.607349061 | 0.543619286 | 0.866046693 |
| Ranbp3 | 3290.02563 | 0.032128646 | 0.059029891 | 0.544277569 | 0.58625048 | 0.884588669 |
| Vps26b | 6106.798283 | 0.032130564 | 0.058812219 | 0.546324634 | 0.584842806 | 0.884313345 |
| Fgf23 | 17.94398926 | 0.032130968 | 0.065718659 | 0.488916975 | 0.624900478 | NA |
| Fam108b | 881.8077151 | 0.032134009 | 0.066360694 | 0.484232556 | 0.628220834 | 0.903965908 |
| Cyp4f16 | 150.9165522 | 0.032136438 | 0.092582415 | 0.347111684 | 0.728507416 | 0.933996867 |
| 6430573F11Rik | 351.9590568 | 0.032148386 | 0.082191895 | 0.391138147 | 0.695695125 | 0.925527709 |
| Coro2b | 9845.907753 | 0.032154869 | 0.040482929 | 0.794282189 | 0.427031171 | 0.810328132 |
| Lcmt1 | 2291.03139 | 0.032173902 | 0.0497848 | 0.646259536 | 0.518111288 | 0.853297835 |
| Adsl | 1018.072274 | 0.032176661 | 0.062367187 | 0.515922918 | 0.605908247 | 0.892639269 |
| Ttc36 | 2.108329158 | 0.032190502 | 0.03314844 | 0.97110158 | 0.331497697 | NA |
| Zfp516 | 793.040433 | 0.032211634 | 0.064592377 | 0.498690954 | 0.617997118 | 0.89815169 |
| Adrm1 | 2533.798306 | 0.032240891 | 0.058474559 | 0.551366122 | 0.58138272 | 0.884020489 |
| Rhbdl2 | 9.318544184 | 0.032248755 | 0.058042294 | 0.555607864 | 0.578478954 | NA |
| Ibtk | 1243.175737 | 0.032278585 | 0.067414082 | 0.478810712 | 0.632073295 | 0.906456381 |
| Ctu1 | 325.250607 | 0.032280524 | 0.078833896 | 0.409475182 | 0.682190976 | 0.922544285 |
| Hectd1 | 6625.172399 | 0.032286331 | 0.056627964 | 0.57014818 | 0.568577199 | 0.877419107 |
| Mrfap1 | 9980.236079 | 0.032291937 | 0.052133475 | 0.619408878 | 0.535647034 | 0.860926435 |
| Hacl1 | 240.2417037 | 0.032292161 | 0.084336885 | 0.382894879 | 0.701797709 | 0.92751465 |
| Gm9767 | 13.84558561 | 0.032323838 | 0.072424317 | 0.446311949 | 0.655371926 | NA |
| E130112N10Rik | 21.87284141 | 0.032336263 | 0.081833657 | 0.395146257 | 0.692734952 | NA |
| Slc4a2 | 1517.699465 | 0.032340113 | 0.093308309 | 0.346594144 | 0.728896246 | 0.933996867 |
| Anks4b | 8.388243669 | 0.032342792 | 0.05167188 | 0.625926355 | 0.531363247 | NA |
| 5730455P16Rik | 3618.322984 | 0.032343122 | 0.06089458 | 0.531133022 | 0.595326602 | 0.887381665 |
| Tekt2 | 41.883037 | 0.032360629 | 0.087956255 | 0.367917314 | 0.712934889 | 0.930507332 |
| Sept3 | 25264.01924 | 0.032363689 | 0.035800337 | 0.904005151 | 0.36599267 | 0.778210102 |
| Map3k7 | 2824.394722 | 0.032378069 | 0.054502597 | 0.594064715 | 0.55246883 | 0.870265825 |
| Clca1 | 9.099916493 | 0.032392674 | 0.059761286 | 0.542034421 | 0.587794794 | NA |
| Myo5b | 2000.474351 | 0.032416426 | 0.057563859 | 0.563138519 | 0.573340564 | 0.879352236 |
| Nav3 | 2945.86555 | 0.032429106 | 0.059516378 | 0.544877001 | 0.585838115 | 0.884588669 |
| Pla2g4d | 3.894030992 | 0.032439236 | 0.04538227 | 0.714799781 | 0.47473277 | NA |
| Gp9 | 9.686344736 | 0.032443333 | 0.062537887 | 0.518778845 | 0.603914972 | NA |
| Glt1d1 | 682.1278789 | 0.032445009 | 0.061935902 | 0.523848179 | 0.60038414 | 0.889841393 |
| Tmem117 | 263.5637299 | 0.032500272 | 0.087894957 | 0.369762644 | 0.711559351 | 0.930507332 |
| Dbi | 3334.027272 | 0.032512703 | 0.078549225 | 0.413915011 | 0.67893635 | 0.921359251 |
| Nicn1 | 4695.002435 | 0.032519948 | 0.046808681 | 0.69474182 | 0.487217115 | 0.839801621 |
| A730085K08Rik | 6.659131101 | 0.032526263 | 0.055370257 | 0.587432042 | 0.556913579 | NA |
| Sel1l | 9475.377267 | 0.032527372 | 0.055147668 | 0.589823158 | 0.555309215 | 0.871360989 |
| Ncbp2 | 2101.159569 | 0.032528834 | 0.055539296 | 0.585690422 | 0.558083571 | 0.872673743 |
| Pde6d | 1196.248058 | 0.032535607 | 0.063491318 | 0.512441828 | 0.608341823 | 0.894048292 |
| Zfp526 | 448.5382381 | 0.032537266 | 0.079850686 | 0.407476348 | 0.683658162 | 0.923110101 |
| Hapln2 | 1081.700083 | 0.032570879 | 0.090124132 | 0.361400195 | 0.717800302 | 0.93125489 |
| Rcn3 | 147.9949088 | 0.032581019 | 0.095058919 | 0.342745523 | 0.731789912 | 0.935424325 |
| Zfand5 | 6485.344159 | 0.032595186 | 0.0722581 | 0.451093876 | 0.651921891 | 0.914447303 |
| Aven | 341.6808024 | 0.032599964 | 0.081168803 | 0.401631693 | 0.687955102 | 0.923860706 |
| Bcl11a | 3219.854303 | 0.032600747 | 0.060352879 | 0.540168885 | 0.589080569 | 0.885188411 |
| H2-T22 | 1.762293184 | 0.032604282 | 0.031787578 | 1.025692532 | 0.305036535 | NA |
| Cyld | 8073.553005 | 0.032607692 | 0.076525357 | 0.426103103 | 0.670032719 | 0.918093366 |
| Esf1 | 1206.613117 | 0.032612234 | 0.071294866 | 0.457427523 | 0.647363785 | 0.912750424 |
| Supt4a | 1163.845714 | 0.03261377 | 0.057013559 | 0.572035333 | 0.567298038 | 0.876376826 |
| Arg1 | 3.22235964 | 0.032617184 | 0.040397342 | 0.807409159 | 0.419430792 | NA |
| Gprc5b | 8119.836315 | 0.032626726 | 0.06465177 | 0.504653254 | 0.613802399 | 0.895819717 |
| Slc24a1 | 9.098563725 | 0.03263224 | 0.057123723 | 0.571255482 | 0.567826474 | NA |
| Mast4 | 3053.898699 | 0.032632877 | 0.07659687 | 0.426034078 | 0.670083014 | 0.918093366 |
| Txnrd3 | 213.6602322 | 0.032635223 | 0.088319649 | 0.369512602 | 0.711745681 | 0.930507332 |
| AI661453 | 25.3059165 | 0.032643305 | 0.075452322 | 0.432634862 | 0.665280059 | 0.915428982 |
| Ttbk1 | 5535.607757 | 0.032738796 | 0.06653864 | 0.492026833 | 0.622700375 | 0.900224603 |
| Vapb | 7256.492685 | 0.032747135 | 0.041214843 | 0.794547112 | 0.426876993 | 0.810246336 |
| Hist3h2a | 986.5866838 | 0.032761512 | 0.070125622 | 0.467183194 | 0.640368816 | 0.910092095 |
| Mab21l2 | 12.94346964 | 0.032776368 | 0.05268707 | 0.622095104 | 0.533879334 | NA |
| Isy1 | 677.3220545 | 0.032792666 | 0.064847665 | 0.505687693 | 0.613075909 | 0.895428201 |
| Atg9a | 7701.874643 | 0.032811939 | 0.045565611 | 0.720103135 | 0.471461498 | 0.833245343 |
| Man1a2 | 6144.086133 | 0.032838423 | 0.067500397 | 0.486492297 | 0.626618166 | 0.902822404 |
| Dapk1 | 5507.924766 | 0.03284799 | 0.066016194 | 0.497574725 | 0.618783822 | 0.898575807 |
| Zc3h14 | 3640.903968 | 0.032861554 | 0.064952413 | 0.505932763 | 0.612903851 | 0.895428201 |
| Brd3 | 2337.284655 | 0.032884047 | 0.053171921 | 0.6184476 | 0.536280329 | 0.861250199 |
| Tmem19 | 1602.191049 | 0.032885891 | 0.051520187 | 0.638310788 | 0.523271391 | 0.854628731 |
| Crabp2 | 82.22443329 | 0.032910443 | 0.089228373 | 0.368833828 | 0.712251589 | 0.930507332 |
| Plxna4 | 4677.338901 | 0.032920032 | 0.071490052 | 0.460484098 | 0.645168783 | 0.912363974 |
| Padi6 | 17.00937294 | 0.032934919 | 0.073024051 | 0.451014674 | 0.651978973 | NA |
| Rnf167 | 2904.971766 | 0.032947815 | 0.05078423 | 0.648780444 | 0.5164803 | 0.853297835 |
| Lats2 | 1090.915133 | 0.032969854 | 0.077027847 | 0.428025126 | 0.668632828 | 0.917270893 |
| Atp6v0d2 | 2.790396319 | 0.032977469 | 0.024258659 | 1.359410206 | 0.174016639 | NA |
| Purb | 18992.73068 | 0.032983578 | 0.07462556 | 0.44198768 | 0.658498121 | 0.915208181 |
| Mapk8 | 4551.62203 | 0.03299887 | 0.06693303 | 0.493013234 | 0.622003237 | 0.900011855 |
| 1190002F15Rik | 1.531935788 | 0.033002522 | 0.026178881 | 1.260654404 | 0.207433388 | NA |
| Bhmt | 4.343513701 | 0.033004579 | 0.036840183 | 0.895885334 | 0.370314008 | NA |
| Akirin2 | 2119.745569 | 0.033018324 | 0.060104229 | 0.549351092 | 0.582764531 | 0.88406516 |
| Smim9 | 1.829118544 | 0.033019071 | 0.028586983 | 1.155038653 | 0.248074598 | NA |
| Lasp1 | 7423.273949 | 0.033060036 | 0.043158496 | 0.76601456 | 0.443667639 | 0.820717992 |
| Crebrf | 2604.462717 | 0.033065781 | 0.068767284 | 0.480835924 | 0.630633116 | 0.905249211 |
| Mmp14 | 411.7851467 | 0.033076152 | 0.094279959 | 0.35082909 | 0.725716572 | 0.933575664 |
| Mkl2 | 12853.78058 | 0.033078266 | 0.072702475 | 0.454981284 | 0.649122703 | 0.913724196 |
| Cyth3 | 1738.37911 | 0.033104073 | 0.049657789 | 0.666644116 | 0.504999483 | 0.848183273 |
| Syt6 | 1559.464452 | 0.033110661 | 0.084857375 | 0.390191904 | 0.696394648 | 0.92555371 |
| Arhgef9 | 20715.92114 | 0.033123602 | 0.060508207 | 0.54742329 | 0.584087958 | 0.884313345 |
| Myot | 1.669919046 | 0.033134325 | 0.025486936 | 1.300051337 | 0.193583375 | NA |
| Tbc1d22a | 1908.940176 | 0.033159444 | 0.058830926 | 0.563639681 | 0.572999375 | 0.879310647 |
| Impdh1 | 2636.700416 | 0.033165977 | 0.060809569 | 0.545407207 | 0.585473485 | 0.884512516 |
| Sfn | 67.91907469 | 0.033167123 | 0.094996458 | 0.349140628 | 0.726983737 | 0.9335794 |
| Cbll1 | 760.4269545 | 0.033178374 | 0.080560565 | 0.411843857 | 0.680453874 | 0.922298495 |
| Smad4 | 1649.336654 | 0.033178943 | 0.07576952 | 0.437893006 | 0.661463846 | 0.915428982 |
| 4930455F23Rik | 972.9030029 | 0.033185441 | 0.065609258 | 0.505804234 | 0.612994086 | 0.895428201 |
| Rassf10 | 32.84561343 | 0.033200356 | 0.089100808 | 0.372615661 | 0.709434515 | 0.930224827 |
| Spink13 | 1.549054206 | 0.033202734 | 0.025561366 | 1.298942122 | 0.193963793 | NA |
| C1ql2 | 148.9309008 | 0.033208363 | 0.072120913 | 0.460454001 | 0.645190382 | 0.912363974 |
| Cask | 3734.499423 | 0.033211505 | 0.051357806 | 0.646669062 | 0.51784615 | 0.853297835 |
| Abcc4 | 680.8562222 | 0.033222682 | 0.073539017 | 0.451769452 | 0.651435079 | 0.914343204 |
| Smo | 665.4025983 | 0.033240105 | 0.083436663 | 0.398387275 | 0.690344737 | 0.923860706 |
| Syn3 | 1987.677941 | 0.03324092 | 0.06716774 | 0.494894133 | 0.620674852 | 0.899412757 |
| Gstt1 | 354.3711448 | 0.033248909 | 0.083880239 | 0.396385484 | 0.691820673 | 0.923860706 |
| Serp1 | 1428.499196 | 0.033250065 | 0.071615789 | 0.464284007 | 0.642444281 | 0.911258473 |
| Chrnb3 | 91.35448139 | 0.033252489 | 0.095089814 | 0.349695593 | 0.726567162 | 0.9335794 |
| Neurog2 | 10.58013349 | 0.033283849 | 0.06460311 | 0.515205064 | 0.606409731 | NA |
| C87436 | 977.7366355 | 0.033293877 | 0.05546115 | 0.600309894 | 0.548299726 | 0.86798189 |
| Gid4 | 3702.575253 | 0.033300742 | 0.054929175 | 0.606248721 | 0.544349604 | 0.86624804 |
| Nr4a2 | 2329.717604 | 0.0333097 | 0.086449518 | 0.385308106 | 0.700009157 | 0.926781946 |
| Nucks1 | 4895.875906 | 0.033341998 | 0.063974171 | 0.52117906 | 0.60224204 | 0.89112297 |
| Megf10 | 1329.46255 | 0.033380235 | 0.074268776 | 0.449451804 | 0.653105768 | 0.914683958 |
| Aldh1b1 | 384.6247408 | 0.033394683 | 0.08710289 | 0.38339351 | 0.701428015 | 0.92751465 |
| Fnip2 | 947.1811742 | 0.033397821 | 0.059921849 | 0.557356322 | 0.577284001 | 0.881375635 |
| Pcdhga5 | 533.179367 | 0.033420933 | 0.082510985 | 0.405048288 | 0.685442017 | 0.923860706 |
| Elp4 | 544.531901 | 0.033444514 | 0.067468931 | 0.495702448 | 0.62010436 | 0.899216177 |
| Comp | 12.94748285 | 0.033455834 | 0.068560177 | 0.487977654 | 0.625565669 | NA |
| Filip1l | 159.3397364 | 0.033477987 | 0.094837662 | 0.353003085 | 0.724086128 | 0.932714927 |
| Foxd2 | 9.614899422 | 0.033481731 | 0.055543334 | 0.602803768 | 0.54663924 | NA |
| Sep15 | 4827.64494 | 0.033483535 | 0.055897108 | 0.599020884 | 0.549158958 | 0.868365902 |
| Igsf10 | 186.255584 | 0.033487941 | 0.094797661 | 0.35325704 | 0.723895749 | 0.932714927 |
| Chmp1a | 4010.33592 | 0.033499074 | 0.056108525 | 0.59704073 | 0.550480188 | 0.869565155 |
| Pitpnb | 3674.827026 | 0.033506257 | 0.051688007 | 0.648240449 | 0.516829444 | 0.853297835 |
| Lurap1l | 331.0264656 | 0.033525152 | 0.085799956 | 0.390736237 | 0.695992211 | 0.925527709 |
| Psap | 85413.27367 | 0.033528146 | 0.046937962 | 0.71430766 | 0.475036957 | 0.834699977 |
| Fam64a | 7.351570161 | 0.033528927 | 0.056307319 | 0.595463038 | 0.551533999 | NA |
| Ptprt | 4017.176099 | 0.033543036 | 0.070771716 | 0.473961044 | 0.635527687 | 0.907676507 |
| Atrn | 11077.82109 | 0.033560537 | 0.067326712 | 0.498472837 | 0.61815081 | 0.89828518 |
| Slx4 | 1330.91441 | 0.033580372 | 0.065758284 | 0.510663748 | 0.609586529 | 0.89440021 |
| Immp1l | 489.2569448 | 0.033592133 | 0.08274876 | 0.40595331 | 0.684776907 | 0.923762012 |
| Apopt1 | 833.1010329 | 0.03359523 | 0.064952813 | 0.517225177 | 0.604998979 | 0.892113813 |
| D2Wsu81e | 1270.826053 | 0.0335969 | 0.063703337 | 0.527396228 | 0.597918462 | 0.888528244 |
| Osbpl11 | 947.7527536 | 0.033602077 | 0.061687058 | 0.544718422 | 0.585947193 | 0.884588669 |
| Synj1 | 18192.74367 | 0.033624442 | 0.055923871 | 0.60125384 | 0.547670929 | 0.867553753 |
| Dapk3 | 1042.095434 | 0.033648882 | 0.064579721 | 0.521044094 | 0.602336055 | 0.89112297 |
| Bcl11b | 3719.918351 | 0.03364927 | 0.083243346 | 0.404227743 | 0.686045254 | 0.923860706 |
| Fam187a | 3.830482669 | 0.033660581 | 0.043839077 | 0.767821399 | 0.442593299 | NA |
| Denr | 2322.584796 | 0.033671079 | 0.064612428 | 0.521123882 | 0.602280475 | 0.89112297 |
| Arhgef17 | 11788.18575 | 0.033676865 | 0.057458224 | 0.58611045 | 0.557801294 | 0.872673743 |
| Tenm3 | 3557.641161 | 0.033678349 | 0.068765536 | 0.48975621 | 0.624306421 | 0.901010936 |
| Sdf4 | 12663.99184 | 0.033712644 | 0.043023082 | 0.783594353 | 0.433278171 | 0.813783127 |
| Iqcf5 | 2.986798258 | 0.033731908 | 0.03471307 | 0.971735085 | 0.331182358 | NA |
| D130017N08Rik | 511.5963103 | 0.033758303 | 0.070838244 | 0.476554767 | 0.6336792 | 0.906932171 |
| Nfs1 | 1520.615002 | 0.033796593 | 0.054567538 | 0.619353459 | 0.535683534 | 0.860926435 |
| Tmem37 | 50.01888128 | 0.033825709 | 0.094819799 | 0.356736774 | 0.721288873 | 0.932513933 |
| Ogfr | 1421.363754 | 0.033836567 | 0.05500314 | 0.615175199 | 0.538439025 | 0.862082844 |
| Ccdc117 | 806.0503039 | 0.033850704 | 0.077581103 | 0.436326663 | 0.66259974 | 0.915428982 |
| Surf1 | 2135.032187 | 0.033855612 | 0.055782027 | 0.606926889 | 0.543899433 | 0.86604998 |
| Zbbx | 16.07888367 | 0.033884796 | 0.072740549 | 0.465830912 | 0.641336537 | NA |
| Ddx10 | 796.650767 | 0.033886131 | 0.065551955 | 0.516935476 | 0.605201202 | 0.892231063 |
| Fnta | 1886.784149 | 0.033901238 | 0.057460067 | 0.589996494 | 0.555193 | 0.871360989 |
| Prickle2 | 9784.658351 | 0.033920864 | 0.066874311 | 0.507233096 | 0.611991276 | 0.895245244 |
| Dcun1d4 | 4603.67483 | 0.03392153 | 0.053503141 | 0.634010074 | 0.526074248 | 0.855892499 |
| Mef2d | 5409.4268 | 0.03394459 | 0.064460331 | 0.526596582 | 0.598473766 | 0.888717024 |
| Mtor | 5625.338735 | 0.033969318 | 0.048822961 | 0.695765215 | 0.486575877 | 0.839801621 |
| Acsbg1 | 6981.680987 | 0.033984615 | 0.06177502 | 0.550135223 | 0.582226629 | 0.884020489 |
| 1700088E04Rik | 143.1374331 | 0.034016959 | 0.094288938 | 0.360773596 | 0.718268703 | 0.93125489 |
| Brd7 | 1809.589395 | 0.034023709 | 0.052849195 | 0.643788608 | 0.519712522 | 0.853297835 |
| Supt6 | 8116.412477 | 0.034061864 | 0.047577321 | 0.715926488 | 0.474036742 | 0.833994865 |
| Rad51c | 134.6666012 | 0.034068421 | 0.089169225 | 0.382064786 | 0.70241331 | 0.927855161 |
| Eps15 | 8837.963978 | 0.034089941 | 0.065533557 | 0.520190612 | 0.602930728 | 0.891281205 |
| Rpgr | 733.3382894 | 0.034114117 | 0.077465191 | 0.440379956 | 0.65966194 | 0.915208181 |
| Lrrc18 | 24.4443545 | 0.034120087 | 0.075004964 | 0.454904391 | 0.649178024 | 0.913724196 |
| Fosl1 | 3.543819132 | 0.034128629 | 0.045672645 | 0.747244418 | 0.454916037 | NA |
| Lix1l | 851.3488499 | 0.034137531 | 0.060027561 | 0.568697622 | 0.569561362 | 0.877633217 |
| Mab21l1 | 47.28291418 | 0.034156745 | 0.089796292 | 0.380380352 | 0.703663097 | 0.928079892 |
| Mrpl45 | 1371.286693 | 0.034168496 | 0.055649719 | 0.613992254 | 0.539220446 | 0.862815409 |
| Dlgap4 | 11716.3133 | 0.034174706 | 0.053562795 | 0.638030662 | 0.523453721 | 0.854731187 |
| Gclc | 2728.399142 | 0.034184737 | 0.051850322 | 0.659296536 | 0.509705366 | 0.850457637 |
| 2210015D19Rik | 524.7517609 | 0.034201491 | 0.062807087 | 0.544548281 | 0.586064234 | 0.884588669 |
| Zfp111 | 481.1942115 | 0.034218292 | 0.075264704 | 0.454639296 | 0.649368759 | 0.913724196 |
| Cd209b | 14.13743795 | 0.034231317 | 0.06912884 | 0.495181415 | 0.620472068 | NA |
| Zbtb21 | 854.5866132 | 0.03423478 | 0.06374109 | 0.537091231 | 0.591204597 | 0.8858275 |
| Plbd1 | 6.918319111 | 0.034274427 | 0.045538679 | 0.752644294 | 0.451663693 | NA |
| 9230114K14Rik | 84.3271121 | 0.034276848 | 0.094960476 | 0.360959107 | 0.718130017 | 0.93125489 |
| Lancl1 | 8647.956352 | 0.034280806 | 0.051582238 | 0.664585476 | 0.506315662 | 0.848908639 |
| Ppp1r7 | 4320.688302 | 0.034282787 | 0.062075356 | 0.552276928 | 0.580758636 | 0.8837314 |
| Xkr4 | 782.0496379 | 0.034285631 | 0.083433881 | 0.410931755 | 0.681122577 | 0.922451017 |
| Amph | 10572.47176 | 0.034285954 | 0.042720018 | 0.802573491 | 0.422221296 | 0.808695226 |
| Pcdhb10 | 174.3146878 | 0.03429545 | 0.090191507 | 0.38025144 | 0.703758778 | 0.928079892 |
| Phf2 | 2613.648702 | 0.03429765 | 0.045426495 | 0.755014211 | 0.450240455 | 0.823365451 |
| Nrn1 | 6797.787259 | 0.034305528 | 0.058877417 | 0.582660199 | 0.560122072 | 0.873713575 |
| Axin1 | 1195.636397 | 0.034311553 | 0.058178914 | 0.589759253 | 0.555352064 | 0.871360989 |
| 2610002J02Rik | 736.6321611 | 0.034342403 | 0.07570666 | 0.453624594 | 0.650099049 | 0.914000665 |
| Ddx6 | 7483.665534 | 0.034347212 | 0.064605272 | 0.5316472 | 0.594970368 | 0.88732439 |
| Pla2g5 | 107.3205902 | 0.03437029 | 0.091938313 | 0.373840766 | 0.708522788 | 0.93005621 |
| Rab22a | 1220.573556 | 0.03437286 | 0.063722496 | 0.539414832 | 0.589600649 | 0.885188411 |
| 5430417L22Rik | 1538.067701 | 0.034373491 | 0.063575843 | 0.540669061 | 0.588735707 | 0.885188411 |
| Rbfox2 | 12535.67322 | 0.034379171 | 0.054492772 | 0.630894149 | 0.528109738 | 0.856965327 |
| Rtn2 | 3668.26897 | 0.034388021 | 0.051572792 | 0.666786101 | 0.504908772 | 0.848183273 |
| Pola1 | 183.4866778 | 0.034399002 | 0.08756802 | 0.392826082 | 0.694447939 | 0.925527709 |
| Cetn2 | 1159.612994 | 0.034406733 | 0.063769762 | 0.539546205 | 0.589510024 | 0.885188411 |
| C1qtnf2 | 29.32233302 | 0.034414503 | 0.084995602 | 0.404897451 | 0.685552892 | 0.923860706 |
| Plekhj1 | 725.218458 | 0.034418653 | 0.064679663 | 0.532140263 | 0.594628853 | 0.887269096 |
| Prkd1 | 497.8617416 | 0.034420086 | 0.074343929 | 0.462984484 | 0.643375489 | 0.911258473 |
| Rnf181 | 2668.684219 | 0.034421471 | 0.044180958 | 0.779101969 | 0.435919651 | 0.816020977 |
| Uba52 | 9305.035077 | 0.034422567 | 0.057470607 | 0.59895951 | 0.549199886 | 0.868365902 |
| Slc5a7 | 351.9755979 | 0.034427173 | 0.09485376 | 0.362950004 | 0.716642232 | 0.93125489 |
| Stx18 | 900.6309211 | 0.034433036 | 0.054860613 | 0.627645854 | 0.530235963 | 0.857915133 |
| Pnpla1 | 27.1947314 | 0.034443196 | 0.08595968 | 0.400690141 | 0.68864827 | 0.923860706 |
| Gm8979 | 1.305524027 | 0.034460502 | 0.027421503 | 1.256696344 | 0.208863613 | NA |
| Dctn5 | 3213.869024 | 0.034466484 | 0.045525078 | 0.757087871 | 0.44899722 | 0.823263966 |
| Mphosph6 | 384.8102637 | 0.034468587 | 0.077101748 | 0.44705325 | 0.654836612 | 0.915208181 |
| Map3k4 | 2406.184743 | 0.034476938 | 0.055458325 | 0.621672901 | 0.534156974 | 0.859551798 |
| Bsx | 3.740478274 | 0.034513251 | 0.025058813 | 1.377289953 | NA | NA |
| H1f0 | 3829.460374 | 0.034531159 | 0.057148838 | 0.604232049 | 0.545689376 | 0.866967503 |
| Pah | 5.588214659 | 0.034558588 | 0.05067429 | 0.68197478 | 0.495254898 | NA |
| Rtkn | 2633.854717 | 0.034566771 | 0.058168881 | 0.594248507 | 0.552345913 | 0.870265825 |
| Nek10 | 68.8962501 | 0.034569346 | 0.093654146 | 0.36911709 | 0.712040451 | 0.930507332 |
| Dexi | 1272.578744 | 0.034571435 | 0.061089558 | 0.565913985 | 0.571452254 | 0.878389918 |
| Pip4k2c | 7301.830824 | 0.034574094 | 0.061266832 | 0.56431992 | 0.572536427 | 0.879277718 |
| Fert2 | 655.3576907 | 0.034605821 | 0.07439807 | 0.465144068 | 0.641828289 | 0.911180388 |
| Ids | 18601.13721 | 0.034612413 | 0.053936527 | 0.641724916 | 0.521051808 | 0.85372574 |
| Dock5 | 640.9229967 | 0.034623664 | 0.089424224 | 0.387184397 | 0.6986197 | 0.926409493 |
| Ndufaf7 | 1377.741036 | 0.034643034 | 0.048467432 | 0.714769329 | 0.47475159 | 0.834491965 |
| Brk1 | 5586.905557 | 0.034647694 | 0.043724609 | 0.792407171 | 0.428123299 | 0.810486244 |
| Gdf7 | 7.559636392 | 0.034671873 | 0.058105643 | 0.596704059 | 0.550704982 | NA |

| | | | | | | |
|---|---|---|---|---|---|---|
| Exosc9 | 1175.019969 | 0.020184691 | 0.067187655 | 0.300422609 | 0.76385482 | 0.88359866 |
| Kdelc2 | 364.5407453 | -0.026587043 | 0.088482565 | -0.30047776 | 0.763812758 | 0.88359866 |
| Klf13 | 8368.027943 | -0.03257404 | 0.108405842 | -0.300482335 | 0.763809269 | 0.88359866 |
| Setmar | 248.7700971 | 0.027867366 | 0.092777207 | 0.300368665 | 0.763895963 | 0.88359866 |
| 4930511M06Rik | 8.185131572 | -0.044233494 | 0.147334258 | -0.300225452 | 0.764005192 | 0.883659036 |
| Ncaph2 | 4285.153713 | -0.016732277 | 0.055769654 | -0.300024762 | 0.764158268 | 0.883770113 |
| Prps1l3 | 550.3667741 | -0.025938107 | 0.086530518 | -0.299756751 | 0.764362706 | 0.883940571 |
| Krt80 | 162.7778352 | 0.049138404 | 0.163980649 | 0.299659774 | 0.764436685 | 0.883960146 |
| Ltbp4 | 2584.522188 | 0.038631867 | 0.129009047 | 0.299450837 | 0.764596078 | 0.884012509 |
| Olfr316 | 38.6680573 | -0.047331036 | 0.158046617 | -0.299475162 | 0.76457752 | 0.884012509 |
| Isy1 | 638.6514619 | -0.024436665 | 0.081674545 | -0.299195603 | 0.764790804 | 0.884105702 |
| Klf8 | 204.687072 | 0.031901723 | 0.10660281 | 0.299257807 | 0.764743345 | 0.884105702 |
| Asrgl1 | 5807.253655 | 0.024514627 | 0.082033179 | 0.298837947 | 0.765063696 | 0.884223251 |
| Idnk | 433.7869275 | 0.025092054 | 0.083935445 | 0.298944667 | 0.764982265 | 0.884223251 |
| Pepd | 848.7736388 | -0.021987174 | 0.073574126 | -0.298843831 | 0.765059206 | 0.884223251 |
| Hnrpll | 1770.995578 | -0.019131344 | 0.064078454 | -0.298561267 | 0.765274822 | 0.884342751 |
| Mfn2 | 9641.225098 | 0.015971496 | 0.053496377 | 0.298552851 | 0.765281245 | 0.884342751 |
| Psmc2 | 3680.664745 | 0.023923112 | 0.080198453 | 0.298298924 | 0.765475026 | 0.884500712 |
| Calm3 | 52351.66938 | -0.022042115 | 0.074055484 | -0.297643248 | 0.765975463 | 0.884503851 |
| Casp2 | 475.7705935 | 0.025897173 | 0.086994429 | 0.297687718 | 0.765941519 | 0.884503851 |
| Ddit3 | 844.3870488 | 0.024836372 | 0.083294828 | 0.298174244 | 0.765570179 | 0.884503851 |
| Dnahc17 | 14.61769339 | -0.050077466 | 0.168004751 | -0.298071726 | 0.76564842 | 0.884503851 |
| H2-Ke6 | 546.4619963 | -0.030992668 | 0.104167318 | -0.29752775 | 0.766063625 | 0.884503851 |
| Isoc2a | 587.927983 | 0.029004883 | 0.097419879 | 0.297730641 | 0.765908755 | 0.884503851 |
| Jakmip1 | 2990.335259 | 0.019182597 | 0.064415792 | 0.297793388 | 0.765860861 | 0.884503851 |
| Mtpap | 897.0318741 | 0.019774764 | 0.066475908 | 0.297472634 | 0.766105698 | 0.884503851 |
| Ptpn18 | 32.03810674 | -0.049025207 | 0.164733536 | -0.297603073 | 0.766006129 | 0.884503851 |
| Rec8 | 182.2562181 | -0.04447448 | 0.149476264 | -0.297535402 | 0.766057784 | 0.884503851 |
| Rps15a | 5554.302223 | -0.019466631 | 0.065388361 | -0.297707896 | 0.765926116 | 0.884503851 |
| Adck4 | 847.6899211 | 0.02234379 | 0.075154905 | 0.297303157 | 0.766235071 | 0.884521397 |
| Gm9776 | 120.3672513 | -0.03722929 | 0.125219056 | -0.29731329 | 0.766227336 | 0.884521397 |
| Dusp18 | 1576.996776 | 0.024465691 | 0.082354805 | 0.297076671 | 0.766407975 | 0.884589181 |
| Mamstr | 87.92722086 | 0.0412269 | 0.138753276 | 0.297123792 | 0.766372001 | 0.884589181 |
| 2810001G20Rik | 246.0069339 | -0.038476869 | 0.129664648 | -0.296741399 | 0.766663948 | 0.884818712 |
| Phf6 | 922.692861 | 0.027549908 | 0.092929726 | 0.296459588 | 0.766879125 | 0.884935217 |
| Wrnip1 | 1939.183233 | 0.019826427 | 0.066864348 | 0.296517164 | 0.766835161 | 0.884935217 |
| Slc16a13 | 180.7554453 | -0.030920351 | 0.104363351 | -0.29627595 | 0.767019351 | 0.884965212 |
| Sox5 | 1524.364923 | -0.024077987 | 0.081267305 | -0.296281355 | 0.767015224 | 0.884965212 |
| Tomm6 | 2195.278163 | -0.019270207 | 0.06507321 | -0.296131196 | 0.767129891 | 0.885026845 |
| Ntmt1 | 595.9173927 | -0.024984962 | 0.084413003 | -0.295984753 | 0.767241726 | 0.885089963 |
| Sgpp1 | 1187.96751 | -0.022171115 | 0.075022852 | -0.295524825 | 0.767592991 | 0.885429259 |
| Clvs2 | 1077.803794 | -0.033500185 | 0.113612723 | -0.294862974 | 0.768098557 | 0.885915696 |
| Ppp1r21 | 3008.228931 | 0.025294597 | 0.085795856 | 0.294823062 | 0.768129048 | 0.885915696 |
| Ppp2r1a | 22225.32532 | 0.021131643 | 0.071711206 | 0.294677005 | 0.768240631 | 0.88597844 |
| Ccdc34 | 239.4850173 | -0.034403027 | 0.116796471 | -0.294555363 | 0.768333565 | 0.886015592 |
| Pigyl | 452.9574557 | -0.02356638 | 0.0800257 | -0.294485143 | 0.768387215 | 0.886015592 |
| Cited2 | 915.6787917 | 0.032755534 | 0.111349712 | 0.294168102 | 0.768629454 | 0.886163015 |
| Stx1b | 9293.042055 | -0.01554056 | 0.052825551 | -0.294186429 | 0.76861545 | 0.886163015 |
| Dolpp1 | 637.1222478 | 0.02545981 | 0.086717444 | 0.293595029 | 0.769067374 | 0.886601927 |
| Mir3061 | 40.65219261 | 0.048060105 | 0.16374666 | 0.293502811 | 0.769137852 | 0.886617206 |
| Gm1976 | 851.3978434 | -0.02505219 | 0.085420464 | -0.29328089 | 0.769307459 | 0.886746747 |
| Bcat1 | 3867.723807 | 0.024863687 | 0.084845536 | 0.293046496 | 0.769486613 | 0.88688727 |
| Lrif1 | 376.2870324 | -0.029724326 | 0.101482565 | -0.292900815 | 0.769597966 | 0.886949634 |
| Cxx1a | 1045.486083 | -0.01721869 | 0.058824322 | -0.292713789 | 0.769740929 | 0.886958984 |
| Itga11 | 97.28021762 | 0.038419736 | 0.131228261 | 0.292770287 | 0.769697741 | 0.886958984 |
| Prss36 | 218.7301914 | 0.039431357 | 0.134760427 | 0.292603385 | 0.769825326 | 0.886958984 |
| Ttc39c | 873.6800332 | 0.040523694 | 0.138499621 | 0.292590652 | 0.769835059 | 0.886958984 |
| Vdac2 | 6375.710804 | -0.016742134 | 0.057283502 | -0.292267988 | 0.770081733 | 0.887177215 |
| Gapvd1 | 3619.632255 | -0.014566427 | 0.049892494 | -0.291956276 | 0.770320055 | 0.887384277 |
| Pced1b | 167.2704856 | 0.030531959 | 0.104603381 | 0.291883093 | 0.77037601 | 0.887384277 |
| Gm14403 | 279.9965452 | -0.031924569 | 0.109432249 | -0.291729079 | 0.770493774 | 0.88745395 |
| E330011O21Rik | 14.93124897 | 0.047961373 | 0.164511075 | 0.291538872 | 0.770639219 | 0.887532087 |
| Lysmd2 | 1116.330546 | 0.025968556 | 0.089111748 | 0.291415633 | 0.77073346 | 0.887532087 |
| Pvrl2 | 205.3598162 | -0.031569922 | 0.108331698 | -0.291419064 | 0.770730836 | 0.887532087 |
| Coprs | 609.4114605 | 0.024079612 | 0.082750322 | 0.290991162 | 0.771058079 | 0.887773937 |
| Pbx4 | 229.5874826 | -0.0316704 | 0.108826528 | -0.291017275 | 0.771038108 | 0.887773937 |
| 3110043O21Rik | 940.9841575 | -0.020751245 | 0.071376733 | -0.290728415 | 0.771259038 | 0.887901166 |
| Gm9958 | 44.37979541 | -0.046568666 | 0.160196674 | -0.290696835 | 0.771283193 | 0.887901166 |
| Dhx38 | 1559.743586 | 0.02271191 | 0.078226687 | 0.290334556 | 0.771560305 | 0.887979926 |
| Igfbp6 | 1164.660539 | 0.029998042 | 0.103302935 | 0.290389055 | 0.771518616 | 0.887979926 |
| Kif23 | 54.25964979 | -0.044564784 | 0.153532047 | -0.290263724 | 0.771614489 | 0.887979926 |
| Tor2a | 742.4029251 | 0.0212208 | 0.07310215 | 0.290289689 | 0.771594626 | 0.887979926 |
| Usp37 | 1062.485511 | -0.017575404 | 0.060571871 | -0.290157856 | 0.771695476 | 0.887979926 |
| Zfp281 | 1564.812668 | 0.021059528 | 0.072578314 | 0.290162819 | 0.771691679 | 0.887979926 |
| Hmgn5 | 336.9339212 | 0.029442042 | 0.101574125 | 0.289857697 | 0.771925106 | 0.888178197 |
| Setd2 | 3149.6809 | 0.021556882 | 0.074408833 | 0.289708647 | 0.772039141 | 0.888243443 |
| Farsb | 1985.79909 | 0.018547933 | 0.064057813 | 0.2895499 | 0.7721606 | 0.888308362 |
| Ybey | 245.789461 | 0.026698444 | 0.092227371 | 0.289485033 | 0.772210232 | 0.888308362 |
| Tmem184b | 3505.343241 | 0.01833948 | 0.063441345 | 0.289077723 | 0.772521901 | 0.888600915 |
| Snrpd3 | 1190.349532 | -0.021158553 | 0.073216494 | -0.28898615 | 0.772591976 | 0.888615551 |
| Fam32a | 2910.001655 | -0.015470884 | 0.053564258 | -0.288828494 | 0.772712626 | 0.888622387 |
| Tph1 | 14.6630428 | -0.045322825 | 0.156881923 | -0.288897692 | 0.77265967 | 0.888622387 |
| Akr1e1 | 895.3758632 | 0.024601442 | 0.08523442 | 0.288632718 | 0.772862456 | 0.888688505 |
| Trim33 | 2594.32056 | 0.023221288 | 0.080460873 | 0.288603478 | 0.772884833 | 0.888688505 |
| Becn1 | 4206.227103 | -0.013231618 | 0.0458968 | -0.288290638 | 0.773124273 | 0.888897854 |
| Ccs | 499.6885177 | 0.02225488 | 0.077234318 | 0.288147552 | 0.773233794 | 0.888918104 |
| Eif2s2 | 2471.138503 | 0.021482373 | 0.074693961 | 0.287605216 | 0.773648952 | 0.888918104 |
| Fhl4 | 32.35717745 | -0.047948763 | 0.166534851 | -0.287920294 | 0.773407752 | 0.888918104 |
| Galnt2 | 3082.777041 | 0.016542961 | 0.057503781 | 0.287684747 | 0.773588067 | 0.888918104 |
| Insrr | 22.66198715 | 0.048277034 | 0.16767866 | 0.287914004 | 0.773412567 | 0.888918104 |
| Kcmf1 | 2148.387075 | 0.022882912 | 0.079538872 | 0.2876947 | 0.773580447 | 0.888918104 |
| Mgme1 | 244.3784871 | 0.031820849 | 0.110577357 | 0.287770024 | 0.773522784 | 0.888918104 |
| P2rx3 | 101.2352895 | -0.035403594 | 0.123103073 | -0.287593095 | 0.773658231 | 0.888918104 |
| Pdzk1ip1 | 9.49998641 | -0.042734176 | 0.148541135 | -0.287692535 | 0.773582105 | 0.888918104 |
| Sec31b | 15.42932273 | -0.047676552 | 0.165960972 | -0.287275687 | 0.773901237 | 0.889131379 |
| 1200011I18Rik | 1886.887827 | -0.019454601 | 0.067907005 | -0.286488863 | 0.774503722 | 0.889236585 |
| Abt1 | 506.2398727 | -0.02212829 | 0.077133089 | -0.286884526 | 0.774200739 | 0.889236585 |
| Ccno | 67.12171089 | -0.045877911 | 0.159870634 | -0.28696897 | 0.77413608 | 0.889236585 |
| Cks1b | 58.90302479 | -0.048065384 | 0.167822204 | -0.286406584 | 0.774566732 | 0.889236585 |
| Dlc1 | 3070.282061 | 0.019930228 | 0.069556795 | 0.286531718 | 0.774470904 | 0.889236585 |
| Ifi47 | 44.3159028 | -0.047952778 | 0.167298164 | -0.286630628 | 0.77439516 | 0.889236585 |
| Pdlim1 | 319.4947746 | -0.036674986 | 0.128029121 | -0.286458157 | 0.774527237 | 0.889236585 |
| Snord96a | 51.7638111 | 0.044284565 | 0.154479015 | 0.286670424 | 0.774364686 | 0.889236585 |
| Tbc1d12 | 561.7835412 | 0.025605359 | 0.089401488 | 0.286408643 | 0.774565156 | 0.889236585 |
| Wwp1 | 2080.366097 | -0.021718094 | 0.075688605 | -0.286940075 | 0.774158205 | 0.889236585 |
| 6330409D20Rik | 59.57369671 | 0.043076654 | 0.150716293 | 0.285812856 | 0.775021461 | 0.889641698 |
| Arsk | 239.8758108 | -0.033071854 | 0.115734897 | -0.285755248 | 0.775065587 | 0.889641698 |
| Hace1 | 1923.585569 | 0.021595145 | 0.07560108 | 0.285645986 | 0.775149279 | 0.889641698 |
| Plxna1 | 5055.633147 | 0.026423678 | 0.092484258 | 0.285710001 | 0.775100244 | 0.889641698 |
| Ephb3 | 942.4561403 | -0.036428702 | 0.127567146 | -0.28556492 | 0.775211375 | 0.889647066 |
| Mir3064 | 37.45961149 | 0.047032251 | 0.164816995 | 0.285360444 | 0.77536801 | 0.889695026 |
| Unc50 | 1089.507123 | 0.020075875 | 0.070336492 | 0.285426157 | 0.77531767 | 0.889695026 |
| Grwd1 | 602.3074049 | 0.022191332 | 0.077802292 | 0.285227229 | 0.775470061 | 0.889746232 |
| Rad54l2 | 1465.062182 | -0.020851055 | 0.073150555 | -0.285043022 | 0.775611181 | 0.889842254 |
| Cdk5 | 4007.767527 | 0.015029167 | 0.052740157 | 0.284966284 | 0.775669973 | 0.889843815 |
| Zfp280d | 1450.307783 | -0.023831404 | 0.083675932 | -0.28480596 | 0.775792806 | 0.889918837 |
| Aqp9 | 63.190852 | -0.040193331 | 0.141258987 | -0.284536454 | 0.775999302 | 0.890089812 |
| 2310057M21Rik | 586.4641659 | -0.021851674 | 0.076819893 | -0.284453322 | 0.776063001 | 0.890092881 |
| Angptl4 | 94.05376116 | 0.047341926 | 0.166483379 | 0.284364279 | 0.776131231 | 0.890092881 |
| Bend5 | 283.9789911 | -0.036051316 | 0.126803714 | -0.284308047 | 0.77617432 | 0.890092881 |
| Smg8 | 698.7915834 | 0.022498997 | 0.079180679 | 0.284147562 | 0.776297299 | 0.890168025 |
| Chrm1 | 8391.945492 | 0.023766565 | 0.08377077 | 0.283709524 | 0.776632996 | 0.890235035 |
| Dazap2 | 5009.593901 | -0.013163688 | 0.046419507 | -0.283580957 | 0.776731532 | 0.890235035 |
| Gan | 204.9582929 | 0.034468743 | 0.121431073 | 0.283854389 | 0.776521972 | 0.890235035 |
| Gid8 | 1707.407579 | 0.017710763 | 0.062445575 | 0.283619177 | 0.776702239 | 0.890235035 |
| Myo5b | 1925.539543 | 0.028626117 | 0.100984092 | 0.283471545 | 0.776815391 | 0.890235035 |
| Sgk2 | 69.86783744 | 0.046668842 | 0.164515625 | 0.283674221 | 0.776660052 | 0.890235035 |
| Stx1a | 9710.462948 | 0.036632172 | 0.129204129 | 0.283521687 | 0.77677696 | 0.890235035 |
| Tnpo2 | 8398.545695 | 0.014249308 | 0.050260857 | 0.283507064 | 0.776788168 | 0.890235035 |
| 5430435G22Rik | 71.78652017 | -0.043416699 | 0.153416178 | -0.28299948 | 0.777177236 | 0.890235241 |
| Als2 | 1988.688954 | 0.021486671 | 0.075907617 | 0.283063443 | 0.777128204 | 0.890235241 |
| Calcr | 39.84509933 | -0.0304389 | 0.10744218 | -0.283304939 | 0.776943092 | 0.890235241 |
| Heca | 949.2197709 | -0.032996982 | 0.116619115 | -0.282946604 | 0.777217769 | 0.890235241 |
| Setd3 | 6793.818741 | 0.01839059 | 0.064988296 | 0.282983111 | 0.777189784 | 0.890235241 |
| Sipa1l3 | 2536.48869 | 0.02458359 | 0.086878206 | 0.282966132 | 0.777202799 | 0.890235241 |
| Sqrdl | 205.9951386 | -0.029694681 | 0.104840633 | -0.283236374 | 0.776995647 | 0.890235241 |
| Pln | 55.51473402 | 0.047517595 | 0.167984987 | 0.282868106 | 0.777277944 | 0.890238355 |
| Arap3 | 512.7888566 | -0.024704428 | 0.087381812 | -0.282718194 | 0.777392868 | 0.890304169 |
| Nptn | 20576.01339 | 0.019935845 | 0.070575451 | 0.282475637 | 0.777578825 | 0.890451316 |
| 2010012O05Rik | 861.5766989 | 0.022210538 | 0.078796253 | 0.281873026 | 0.778040873 | 0.890481986 |
| Asph | 3212.173557 | -0.023465136 | 0.0832443 | -0.28188279 | 0.778033386 | 0.890481986 |
| B230120H23Rik | 408.8499808 | 0.026644062 | 0.094488216 | 0.281982909 | 0.777956615 | 0.890481986 |
| D130020L05Rik | 86.87235258 | 0.035714752 | 0.126753187 | 0.281766107 | 0.778122861 | 0.890481986 |
| Gabre | 22.44163423 | 0.044420205 | 0.157649102 | 0.281766306 | 0.778122708 | 0.890481986 |
| Ipo5 | 6515.633211 | 0.020929028 | 0.074212876 | 0.282013437 | 0.777933207 | 0.890481986 |
| Lrriq1 | 89.36996211 | -0.046826666 | 0.165863823 | -0.28231995 | 0.77769819 | 0.890481986 |
| Parl | 770.8607167 | -0.021256311 | 0.075399011 | -0.281917643 | 0.778006661 | 0.890481986 |
| Tfg | 1696.969997 | -0.020800307 | 0.073715003 | -0.282171965 | 0.777811654 | 0.890481986 |
| Ccdc85a | 1705.205901 | -0.028826201 | 0.102409924 | -0.281478587 | 0.778343349 | 0.890668527 |
| Rnf17 | 24.30282666 | 0.047290321 | 0.168087401 | 0.281343639 | 0.778446842 | 0.890675607 |
| Shroom1 | 292.2662869 | -0.029841284 | 0.106075715 | -0.281320606 | 0.778464507 | 0.890675607 |
| Palld | 87.26360503 | 0.037311905 | 0.132854023 | 0.280848894 | 0.7788263 | 0.890957966 |
| Ppm1f | 3855.923264 | 0.019781586 | 0.070416823 | 0.280921304 | 0.77877076 | 0.890957966 |
| AI115009 | 80.29800702 | 0.042833063 | 0.152560279 | 0.280761569 | 0.778893282 | 0.890968808 |
| Galnt18 | 1276.170404 | -0.022741001 | 0.081033612 | -0.280636647 | 0.778989105 | 0.890983874 |
| Lama1 | 177.1675749 | -0.03881344 | 0.138363909 | -0.280517085 | 0.779080819 | 0.890983874 |
| Mill2 | 35.4384986 | -0.045517765 | 0.162436019 | -0.280219654 | 0.779308989 | 0.890983874 |
| Mrps30 | 706.2789369 | -0.023091752 | 0.082416072 | -0.280185058 | 0.77933553 | 0.890983874 |
| Nipsnap1 | 2558.592551 | -0.014970008 | 0.053436698 | -0.280144698 | 0.779366493 | 0.890983874 |
| Pigk | 2924.147807 | 0.013138905 | 0.04685452 | 0.280419166 | 0.779155935 | 0.890983874 |
| Rrs1 | 620.4528669 | 0.01839026 | 0.065644993 | 0.280147185 | 0.779364585 | 0.890983874 |
| Rtn1 | 37747.73286 | -0.019375575 | 0.069090361 | -0.280438174 | 0.779141353 | 0.890983874 |
| Hmgn3 | 1100.811943 | 0.023432164 | 0.083674445 | 0.280039675 | 0.779447066 | 0.891010243 |
| Adat2 | 190.8223108 | 0.026897194 | 0.096151824 | 0.279736703 | 0.779679517 | 0.89114959 |
| Lig4 | 504.5315982 | -0.02138826 | 0.076460137 | -0.279730865 | 0.779683997 | 0.89114959 |
| Cntfr | 2746.717561 | 0.013828274 | 0.049557888 | 0.279032754 | 0.780219689 | 0.891299632 |
| Dcaf17 | 561.4762993 | 0.027839182 | 0.099824824 | 0.278880349 | 0.78033665 | 0.891299632 |
| Fam132b | 25.52662574 | 0.046745028 | 0.167669639 | 0.278792443 | 0.780404114 | 0.891299632 |
| Gad1 | 19339.09233 | -0.03235108 | 0.115992703 | -0.278906164 | 0.780316838 | 0.891299632 |
| Isoc2b | 147.2186409 | 0.04069316 | 0.145821409 | 0.279061629 | 0.78019753 | 0.891299632 |
| Lrfn4 | 1316.480953 | -0.022984403 | 0.082355458 | -0.27908779 | 0.780177453 | 0.891299632 |
| Mrpl24 | 1097.081974 | 0.02296467 | 0.082287299 | 0.279079161 | 0.780184075 | 0.891299632 |
| Ptpn12 | 1249.98716 | 0.026177429 | 0.093772698 | 0.27915832 | 0.780123328 | 0.891299632 |
| Setd1b | 1984.839387 | -0.021854059 | 0.078314519 | -0.279055017 | 0.780202604 | 0.891299632 |
| Srcin1 | 13775.71067 | -0.020315218 | 0.072883577 | -0.278735194 | 0.780448051 | 0.891299632 |
| Vwc2l | 310.1385039 | -0.035906124 | 0.128705446 | -0.278979057 | 0.780260897 | 0.891299632 |
| B230209E15Rik | 987.9569785 | 0.029120052 | 0.104536168 | 0.278564376 | 0.780579155 | 0.891305125 |
| Mnat1 | 379.8134247 | -0.028124889 | 0.100985557 | -0.278504073 | 0.780625439 | 0.891305125 |
| Rd3 | 15.99540595 | -0.044133356 | 0.158456211 | -0.278520833 | 0.780612575 | 0.891305125 |
| 5033404E19Rik | 15.44656618 | -0.046719549 | 0.167870615 | -0.278306893 | 0.780776785 | 0.89141224 |
| Klhdc2 | 4523.563465 | 0.017986548 | 0.064718622 | 0.277919209 | 0.781074378 | 0.891554914 |
| Osbpl10 | 1225.832075 | 0.031478805 | 0.113214773 | 0.278045032 | 0.780977791 | 0.891554914 |
| Rcan1 | 2503.90635 | 0.02593748 | 0.093306385 | 0.277981836 | 0.781026302 | 0.891554914 |
| Gm13251 | 26.33329925 | -0.046710045 | 0.168121527 | -0.277835003 | 0.78113902 | 0.891563018 |
| Aldh7a1 | 3120.493645 | -0.026565878 | 0.095735453 | -0.277492585 | 0.781401899 | 0.891666005 |
| Nsun2 | 2938.212673 | 0.019430205 | 0.069992962 | 0.277602265 | 0.781317693 | 0.891666005 |
| Tdrd1 | 28.17556201 | -0.045070928 | 0.162389622 | -0.277548082 | 0.781359291 | 0.891666005 |
| Ndel1 | 3895.00482 | -0.023830187 | 0.085977988 | -0.277166143 | 0.781652536 | 0.891886323 |
| Cope | 2528.779574 | -0.017523485 | 0.063295624 | -0.276851441 | 0.78189418 | 0.892042002 |
| Slc4a10 | 8714.103571 | 0.019312344 | 0.069760325 | 0.276838497 | 0.781904119 | 0.892042002 |
| Thap11 | 1067.494443 | 0.020877111 | 0.075506101 | 0.276495682 | 0.782167376 | 0.89227664 |
| Atp7a | 422.9781785 | -0.030629624 | 0.11082083 | -0.276388691 | 0.782249542 | 0.892304675 |
| 2510039O18Rik | 931.4140059 | 0.020370448 | 0.07379391 | 0.276045114 | 0.782513416 | 0.892461248 |
| Tssk4 | 14.31179391 | 0.046035621 | 0.166804809 | 0.275984977 | 0.782559605 | 0.892461248 |
| Vasp | 509.0329317 | 0.02547592 | 0.092265938 | 0.276114035 | 0.782460482 | 0.892461248 |
| Pdpr | 803.6551546 | 0.022809692 | 0.082686189 | 0.275858541 | 0.782656718 | 0.892506307 |
| H2-M3 | 117.089485 | -0.034741341 | 0.125977348 | -0.275774507 | 0.782721265 | 0.892514224 |
| Fam109a | 411.6674098 | 0.024045564 | 0.087339582 | 0.275311183 | 0.783077173 | 0.892809779 |
| Zfp786 | 193.0017087 | -0.031823069 | 0.115599584 | -0.275287057 | 0.783095708 | 0.892809779 |
| Blvra | 537.7480437 | -0.023260277 | 0.084523891 | -0.275191743 | 0.78316893 | 0.892827563 |
| Mbd5 | 2679.117595 | -0.028307745 | 0.10290354 | -0.275090102 | 0.783247016 | 0.892850888 |
| Gm16894 | 34.25461438 | -0.045382331 | 0.165037607 | -0.274981754 | 0.783330257 | 0.892880085 |
| Ajuba | 82.96797721 | 0.037989508 | 0.138483462 | 0.274325232 | 0.783834695 | 0.893282702 |
| Gm16515 | 597.0403147 | -0.0199552 | 0.072730158 | -0.274373116 | 0.783797901 | 0.893282702 |
| Myo1g | 22.87287971 | 0.046037681 | 0.167838842 | 0.27429694 | 0.783856436 | 0.893282702 |
| Hs3st1 | 658.363842 | -0.026559199 | 0.096855955 | -0.274213387 | 0.783920641 | 0.893290168 |
| Dlgap1 | 19578.01017 | -0.017585033 | 0.06416161 | -0.274074062 | 0.784027706 | 0.893346469 |
| Thap7 | 590.8718038 | 0.021665473 | 0.079205091 | 0.273536368 | 0.78444094 | 0.893751594 |
| Eda | 276.4629197 | -0.030135514 | 0.110304657 | -0.273202558 | 0.784697514 | 0.893758522 |
| Ippk | 776.8478436 | 0.028593207 | 0.104658755 | 0.273204154 | 0.784696287 | 0.893758522 |
| Josd2 | 899.8417534 | 0.040030974 | 0.146515366 | 0.273220309 | 0.78468387 | 0.893758522 |
| Lrrc40 | 916.1603725 | 0.019681806 | 0.072054302 | 0.273152412 | 0.784736059 | 0.893758522 |
| Rpl30 | 8.556208069 | -0.043314411 | 0.158615427 | -0.273078172 | 0.784793125 | 0.893758522 |
| Trim14 | 15.2114696 | 0.045611721 | 0.166921285 | 0.273252875 | 0.784658838 | 0.893758522 |
| Dazap1 | 1235.727374 | 0.020877227 | 0.076491626 | 0.272934798 | 0.784903337 | 0.893818337 |
| Dbt | 780.5452209 | -0.018927367 | 0.069504889 | -0.272317051 | 0.785378247 | 0.894293419 |
| Entpd3 | 566.0440671 | -0.026299315 | 0.096688069 | -0.272001654 | 0.785620748 | 0.894503812 |
| Guca1b | 19.17304051 | 0.045494177 | 0.167627148 | 0.271401009 | 0.786082626 | 0.894963936 |
| 4632427E13Rik | 12.35297793 | -0.043359733 | 0.159995362 | -0.271006189 | 0.786386272 | 0.895243858 |
| Cpne1 | 1554.097881 | 0.017747336 | 0.065506244 | 0.27092587 | 0.786448047 | 0.895248405 |
| 1700123M08Rik | 67.99566086 | -0.0364712 | 0.134717271 | -0.270724011 | 0.786603308 | 0.895359364 |
| Pigv | 246.8083063 | 0.027342101 | 0.101042158 | 0.270600926 | 0.786697983 | 0.895391868 |
| Zfp846 | 392.1786724 | 0.029344886 | 0.108469182 | 0.270536626 | 0.786747443 | 0.895391868 |
| Ska2 | 272.2157243 | -0.025847542 | 0.095609453 | -0.270345049 | 0.786894811 | 0.895493808 |
| Fstl4 | 809.1844048 | 0.023367837 | 0.086517976 | 0.27009228 | 0.787089261 | 0.89564931 |
| Prom2 | 18.32913512 | 0.045055893 | 0.166904196 | 0.269950628 | 0.787198237 | 0.895707533 |
| Cd300a | 58.37782299 | 0.039027132 | 0.144692528 | 0.269724587 | 0.787372143 | 0.895804021 |
| Hecw1 | 2700.168424 | -0.024387752 | 0.090428795 | -0.269690113 | 0.787398668 | 0.895804021 |
| Ahi1 | 8857.041659 | 0.030570579 | 0.113464861 | 0.269427723 | 0.787600555 | 0.895932558 |
| Cntnap1 | 11866.67093 | -0.021149221 | 0.078558474 | -0.269216286 | 0.787763248 | 0.895932558 |
| Pi4k2b | 72.43232759 | 0.040240184 | 0.149416085 | 0.269316277 | 0.787686308 | 0.895932558 |
| Tnfrsf1b | 234.8457863 | 0.029775882 | 0.110534531 | 0.269380819 | 0.787636645 | 0.895932558 |
| Zfp39 | 476.6660527 | -0.026775409 | 0.099474888 | -0.269167521 | 0.787800772 | 0.895932558 |
| Znf41-ps | 42.23565308 | -0.044237989 | 0.164458385 | -0.268991997 | 0.787935841 | 0.896020398 |
| Ube2h | 5326.312223 | 0.013445897 | 0.050087268 | 0.268449398 | 0.78835342 | 0.896363682 |

| | | | | | | |
|---|---|---|---|---|---|---|
| Cpsf4l | 1.224820421 | 0.034672892 | 0.024556846 | 1.411944016 | 0.157966446 | NA |
| Neu4 | 388.5525933 | 0.034675599 | 0.078906278 | 0.439452973 | 0.66033335 | 0.915428982 |
| D10Jhu81e | 2408.169184 | 0.034687778 | 0.047873711 | 0.724568405 | 0.468716848 | 0.832446623 |
| Usp42 | 1091.588729 | 0.034716886 | 0.056326715 | 0.616348493 | 0.53766454 | 0.862037665 |
| Dnali1 | 60.54585601 | 0.034730395 | 0.087734136 | 0.395859541 | 0.69220865 | 0.923860706 |
| Rplp0 | 6413.227458 | 0.034747013 | 0.050848691 | 0.683341349 | 0.494391172 | 0.843503724 |
| Capns1 | 5410.627837 | 0.034769575 | 0.060594565 | 0.573806824 | 0.56609853 | 0.87575243 |
| Copa | 11806.98066 | 0.03477365 | 0.041828055 | 0.831347534 | 0.405777329 | 0.800733644 |
| C330018D20Rik | 375.0079947 | 0.03478033 | 0.074717616 | 0.465490359 | 0.641580339 | 0.911095592 |
| Gsn | 2603.920419 | 0.034780633 | 0.077452668 | 0.449056619 | 0.653390814 | 0.914683958 |
| Sdcbp | 6893.521911 | 0.034780685 | 0.05110572 | 0.680563459 | 0.496147755 | 0.844707254 |
| Ckm | 2.457932771 | 0.034789494 | 0.034875311 | 0.997539314 | 0.318502801 | NA |
| Cd80 | 17.99361608 | 0.034791262 | 0.078443923 | 0.443517619 | 0.657391378 | NA |
| Eif3m | 2165.208712 | 0.034800136 | 0.05639148 | 0.617116922 | 0.537157608 | 0.861996453 |
| Plcb1 | 12848.0359 | 0.034809305 | 0.072666153 | 0.479030517 | 0.631916919 | 0.906377074 |
| Echdc3 | 35.16945818 | 0.034811611 | 0.082872938 | 0.420060056 | 0.674441582 | 0.919867577 |
| Arfgef1 | 5610.477091 | 0.034824971 | 0.075831023 | 0.459244373 | 0.646058689 | 0.912548475 |
| 4933417D19Rik | 17.9403931 | 0.034830393 | 0.077016687 | 0.452244752 | 0.651092672 | NA |
| Copg1 | 15157.77424 | 0.034835083 | 0.04310709 | 0.808105655 | 0.419029762 | 0.806975048 |
| Mfap1a | 1219.592671 | 0.034835675 | 0.07339793 | 0.474613861 | 0.635062225 | 0.907297112 |
| Zfp831 | 1404.375448 | 0.034837087 | 0.087765027 | 0.396935871 | 0.691414751 | 0.923860706 |
| Tmem169 | 661.7897955 | 0.034837421 | 0.063396184 | 0.549519207 | 0.582649187 | 0.88406516 |
| Gm14295 | 1938.45606 | 0.034839062 | 0.086794935 | 0.401395108 | 0.688129251 | 0.923860706 |
| Elac1 | 1910.310858 | 0.034856266 | 0.06135876 | 0.568073183 | 0.569985276 | 0.8778444 |
| Pon1 | 5.488229233 | 0.034864427 | 0.028547616 | 1.221272816 | 0.221982742 | NA |
| Med19 | 1345.32423 | 0.034878253 | 0.057751488 | 0.603936875 | 0.545885611 | 0.866989658 |
| Agrp | 6.924577723 | 0.034906647 | 0.041735852 | 0.836370772 | NA | NA |
| Alad | 1146.885414 | 0.034938113 | 0.054141983 | 0.64530539 | 0.5187293 | 0.853297835 |
| Alox8 | 320.4272682 | 0.034964638 | 0.088269117 | 0.396114058 | 0.692020888 | 0.923860706 |
| Insm2 | 5.212682721 | 0.03498042 | 0.043277357 | 0.808284574 | 0.418926781 | NA |
| Tbc1d17 | 2175.508884 | 0.034982441 | 0.057251364 | 0.611032456 | 0.541178094 | 0.863862966 |
| Sec13 | 1716.225172 | 0.035025441 | 0.05917823 | 0.591863618 | 0.553941918 | 0.871028394 |
| Snrpb | 2195.882687 | 0.035035251 | 0.064515786 | 0.543049273 | 0.587095878 | 0.884663353 |
| Cdc123 | 2383.029745 | 0.035045453 | 0.046998626 | 0.745669735 | 0.455866946 | 0.826603827 |
| Fam174a | 862.0156519 | 0.035051012 | 0.070496754 | 0.497200368 | 0.619047762 | 0.898695177 |
| Ywhae | 25384.54213 | 0.035059741 | 0.046412706 | 0.755391017 | 0.450014401 | 0.823263966 |
| Ndufb9 | 4590.030676 | 0.035097067 | 0.052990918 | 0.662322303 | 0.507764686 | 0.849201121 |
| St8sia5 | 1957.099554 | 0.035101597 | 0.071714583 | 0.489462462 | 0.624514324 | 0.901133756 |
| Akr1b8 | 43.63937458 | 0.035115375 | 0.089565175 | 0.39206505 | 0.695010151 | 0.925527709 |
| Dpt | 4.74169612 | 0.035137042 | 0.048934768 | 0.718038401 | 0.472733609 | NA |
| Paip2b | 3715.459485 | 0.035150313 | 0.061141596 | 0.57490016 | 0.565358822 | 0.875364912 |
| Hiatl1 | 1583.161187 | 0.035163692 | 0.063872833 | 0.550526576 | 0.581958254 | 0.884020489 |
| Adora1 | 11969.92825 | 0.035175347 | 0.05785207 | 0.608022275 | 0.5431727 | 0.865558163 |
| Dmxl1 | 2477.941964 | 0.035181232 | 0.078832975 | 0.446275589 | 0.655398187 | 0.915208181 |
| Zdhhc20 | 1927.717238 | 0.035191365 | 0.062321396 | 0.564675496 | 0.572294504 | 0.879277718 |
| Atxn7l1 | 1047.350242 | 0.035247898 | 0.058439993 | 0.60314686 | 0.546410995 | 0.867128614 |
| Dcaf5 | 4420.56428 | 0.035258241 | 0.055445938 | 0.635903046 | 0.524839617 | 0.855551597 |
| Cul1 | 5708.404643 | 0.035261148 | 0.058254929 | 0.605290385 | 0.544986069 | 0.866418697 |
| Gucy1a2 | 2357.988973 | 0.035288297 | 0.086518651 | 0.40786925 | 0.68336967 | 0.923036732 |
| Sim1 | 21.02888898 | 0.035297961 | 0.032087711 | 1.100046082 | 0.271312044 | NA |
| Etfb | 969.2403766 | 0.035304436 | 0.062594778 | 0.564015664 | 0.572743472 | 0.879310647 |
| Mat1a | 1.880538289 | 0.035317475 | 0.032690082 | 1.080372798 | 0.279976204 | NA |
| Gm9958 | 51.76059818 | 0.035327359 | 0.094201528 | 0.375018962 | 0.707646365 | 0.929959854 |
| Nudc | 4642.277746 | 0.035352863 | 0.04516248 | 0.782792777 | 0.43374881 | 0.814351197 |
| Mlst8 | 1899.537259 | 0.035366575 | 0.062382659 | 0.566929581 | 0.570762025 | 0.878246677 |
| Sos2 | 1722.276081 | 0.035371367 | 0.069182476 | 0.511276389 | 0.609157535 | 0.89440021 |
| Kdelr1 | 2246.175163 | 0.035373486 | 0.050959937 | 0.694143053 | 0.487592501 | 0.839801621 |
| Lrrc7 | 2448.769855 | 0.035390214 | 0.075269307 | 0.470181214 | 0.638225555 | 0.908932006 |
| Ipo11 | 2539.709815 | 0.035394239 | 0.069451542 | 0.509624955 | 0.610314238 | 0.894800514 |
| Ttc17 | 1965.370352 | 0.035397609 | 0.0532301 | 0.664992347 | 0.506055388 | 0.848908639 |
| Pomgnt1 | 3328.139462 | 0.035403911 | 0.056618061 | 0.625311265 | 0.531766788 | 0.858328942 |
| Nme6 | 161.7417226 | 0.035410871 | 0.08901043 | 0.397828335 | 0.690756729 | 0.923860706 |
| Fmn1 | 1075.232518 | 0.0354182 | 0.080051883 | 0.442440563 | 0.658170432 | 0.915208181 |
| Ubash3b | 1389.855187 | 0.035428692 | 0.063905524 | 0.554391698 | 0.579310807 | 0.882452682 |
| BC025920 | 169.8906753 | 0.035439736 | 0.089040722 | 0.398017163 | 0.690617535 | 0.923860706 |
| Hells | 65.96714403 | 0.0354416 | 0.093194229 | 0.380298227 | 0.703724052 | 0.928079892 |
| Cir1 | 1291.543461 | 0.035464443 | 0.064492514 | 0.549900142 | 0.582387867 | 0.88406516 |
| Suds3 | 3755.615839 | 0.035472289 | 0.045683189 | 0.776484521 | 0.437462957 | 0.816715817 |
| Bbox1 | 75.72484481 | 0.035474969 | 0.092101126 | 0.385174108 | 0.700108425 | 0.92679987 |
| Chtop | 5182.952176 | 0.035482412 | 0.048349461 | 0.733873983 | 0.463025545 | 0.831014288 |
| Cacna2d1 | 8105.616857 | 0.035487932 | 0.075322169 | 0.471148574 | 0.63753464 | 0.908932006 |
| Ergic2 | 1887.519071 | 0.035494317 | 0.06570954 | 0.540169913 | 0.589079859 | 0.885188411 |
| Zfp281 | 1769.42323 | 0.035494815 | 0.073083358 | 0.485675748 | 0.627197081 | 0.903033871 |
| Ift46 | 1446.78044 | 0.035500181 | 0.060161977 | 0.59007671 | 0.555139222 | 0.871360989 |
| Cdkn2a | 9.82797659 | 0.03550637 | 0.049416921 | 0.718506321 | 0.472445152 | NA |
| 4930523C07Rik | 79.25576008 | 0.035506545 | 0.094541655 | 0.375565092 | 0.707240247 | 0.92986829 |
| Arhgap32 | 20119.87912 | 0.035555539 | 0.078684902 | 0.451872446 | 0.651360875 | 0.914343204 |
| Csrnp2 | 1371.523593 | 0.035556651 | 0.068030181 | 0.522659942 | 0.601210919 | 0.890297392 |
| Slc9a3r2 | 1504.747261 | 0.035565677 | 0.073944891 | 0.480975448 | 0.630533949 | 0.905249211 |
| Eif2b4 | 1193.599382 | 0.035579551 | 0.055128893 | 0.645388466 | 0.518675476 | 0.853297835 |
| Trip4 | 1733.957382 | 0.035581317 | 0.056954491 | 0.624732426 | 0.532146687 | 0.858328942 |
| Nr5a1 | 9.910560834 | 0.035594537 | 0.02610966 | 1.363270791 | NA | NA |
| Cog8 | 875.6406049 | 0.035626332 | 0.061916633 | 0.575391947 | 0.565026249 | 0.875364912 |
| Gm14446 | 162.5608299 | 0.035643462 | 0.091954262 | 0.387621641 | 0.698296051 | 0.926409493 |
| Foxa1 | 1.201777077 | 0.035665462 | 0.024516135 | 1.454775026 | 0.145731554 | NA |
| Col6a5 | 86.77813917 | 0.035669713 | 0.093422032 | 0.381812641 | 0.702600342 | 0.927855161 |
| Runx1t1 | 815.9607604 | 0.035688411 | 0.084448291 | 0.422606665 | 0.672582263 | 0.919353963 |
| Tekt1 | 76.18079098 | 0.03569563 | 0.095080645 | 0.375424776 | 0.707344582 | 0.92986829 |
| Hrh1 | 777.7074855 | 0.035748958 | 0.078102916 | 0.457716047 | 0.647156458 | 0.912750424 |
| Scnn1g | 6.224211241 | 0.035763881 | 0.051889762 | 0.689228076 | 0.490679751 | NA |
| Auts2 | 2886.007818 | 0.035769837 | 0.075983154 | 0.470760101 | 0.63781206 | 0.908932006 |
| Bpifa1 | 11.17320953 | 0.035779652 | 0.041704065 | 0.857941598 | 0.390924713 | NA |
| Syne1 | 18771.97789 | 0.035784319 | 0.061884611 | 0.578242615 | 0.563100331 | 0.87510382 |
| Lyplal1 | 162.8974637 | 0.035784398 | 0.090485602 | 0.395470624 | 0.692495597 | 0.924048415 |
| Fbn2 | 84.8601326 | 0.035792792 | 0.095064581 | 0.376510283 | 0.706537572 | 0.929476902 |
| Nucb2 | 636.1071033 | 0.035808965 | 0.077104466 | 0.464421418 | 0.642345848 | 0.911258473 |
| Cbx3 | 3589.586358 | 0.035818649 | 0.058234153 | 0.615079768 | 0.538502043 | 0.862082844 |
| Nr0b2 | 1.012840553 | 0.035822947 | 0.022118521 | 1.619590505 | 0.105320271 | NA |
| Clec2l | 1332.169927 | 0.035832077 | 0.059090719 | 0.606390946 | 0.544255179 | 0.86624804 |
| Satb1 | 5160.266937 | 0.035836654 | 0.070960217 | 0.505024574 | 0.613541577 | 0.895819717 |
| Zc3hav1l | 287.5552538 | 0.035852452 | 0.079796315 | 0.449299589 | 0.653215554 | 0.914683958 |
| Usp50 | 4.627395278 | 0.03585309 | 0.037820238 | 0.947986901 | 0.343136123 | NA |
| Sec22a | 595.3917864 | 0.035863629 | 0.066697716 | 0.537703998 | 0.590781419 | 0.885693151 |
| Ddx49 | 982.4630439 | 0.03587841 | 0.065562268 | 0.547241749 | 0.584212658 | 0.884313345 |
| Asgr1 | 30.54472829 | 0.035879056 | 0.079931851 | 0.448870584 | 0.653525017 | 0.914683958 |
| Dtwd1 | 245.0863596 | 0.035879652 | 0.080900735 | 0.443502172 | 0.657402549 | 0.915208181 |
| Bscl2 | 3235.318365 | 0.035899228 | 0.050532889 | 0.710413137 | 0.477447979 | 0.835289575 |
| 1200011I18Rik | 2053.083574 | 0.035905931 | 0.059832536 | 0.600107128 | 0.548434842 | 0.868006594 |
| 3110079O15Rik | 9.48931331 | 0.03591239 | 0.061536288 | 0.583596957 | 0.559491508 | NA |
| Hey2 | 165.4119558 | 0.035928697 | 0.090161223 | 0.398493898 | 0.690266155 | 0.923860706 |
| Dhrs2 | 2.342921304 | 0.035956013 | 0.036259636 | 0.991626404 | 0.321379804 | NA |
| Twist1 | 13.81463304 | 0.035966112 | 0.072789486 | 0.494111362 | 0.621227534 | NA |
| Kcne1l | 159.3121094 | 0.03597985 | 0.095096464 | 0.378351079 | 0.705169805 | 0.928783753 |
| Mon1b | 1924.403994 | 0.035988572 | 0.051616716 | 0.697227085 | 0.485660691 | 0.839446359 |
| Fth1 | 55905.41649 | 0.036000069 | 0.081663369 | 0.440834975 | 0.659332471 | 0.915208181 |
| Snx9 | 434.3776852 | 0.036004734 | 0.068177838 | 0.528100263 | 0.597429748 | 0.888165441 |
| Nop10 | 1051.846636 | 0.036006178 | 0.075365136 | 0.477756429 | 0.632823576 | 0.906637358 |
| Cisd1 | 2350.291021 | 0.036037259 | 0.046608947 | 0.773183276 | 0.439413922 | 0.817545617 |
| Snrpa1 | 702.8857397 | 0.036042296 | 0.060812783 | 0.592676312 | 0.553397797 | 0.870600462 |
| Atg4c | 1802.674312 | 0.036044928 | 0.061673309 | 0.584449393 | 0.558918004 | 0.873275564 |
| 9630028B13Rik | 14.62226931 | 0.036045652 | 0.064819688 | 0.556091102 | 0.578148578 | NA |
| Upf2 | 2669.223139 | 0.03605769 | 0.073668207 | 0.489460667 | 0.624515595 | 0.901133756 |
| Fat4 | 570.4319975 | 0.036059405 | 0.092130963 | 0.391392897 | 0.695506842 | 0.925527709 |
| Nebl | 1433.551425 | 0.036065117 | 0.062761681 | 0.574635935 | 0.565537544 | 0.875426083 |
| Lrp11 | 8240.367894 | 0.036098145 | 0.058545195 | 0.616585952 | 0.537507863 | 0.861996453 |
| Dym | 2518.918777 | 0.036118451 | 0.053179672 | 0.679177764 | 0.497025234 | 0.845038521 |
| Suox | 552.0955786 | 0.036126373 | 0.069449709 | 0.520180333 | 0.602937891 | 0.891281205 |
| Hmgn5 | 358.7031296 | 0.036128657 | 0.083223637 | 0.434115338 | 0.664204688 | 0.915428982 |
| Myh15 | 76.44030467 | 0.03613562 | 0.094814943 | 0.381117354 | 0.703116171 | 0.927855161 |
| Slc25a30 | 235.6215925 | 0.036146905 | 0.091226038 | 0.396234511 | 0.691932034 | 0.923860706 |
| Wars2 | 338.9216324 | 0.036147832 | 0.081658079 | 0.44267307 | 0.658002224 | 0.915208181 |
| Sox1 | 530.3994289 | 0.036149029 | 0.089119428 | 0.405624561 | 0.685018479 | 0.923835238 |
| F930015N05Rik | 76.12422523 | 0.036152117 | 0.094938001 | 0.380797116 | 0.703353799 | 0.927967056 |
| Churc1 | 676.9848993 | 0.036163686 | 0.066536591 | 0.543515764 | 0.58677474 | 0.884638284 |
| Dusp1 | 1507.804976 | 0.036165756 | 0.077111119 | 0.469008321 | 0.639063688 | 0.909490332 |
| Trnt1 | 1275.858953 | 0.036166806 | 0.071362485 | 0.506804177 | 0.612292226 | 0.895344807 |
| Prpf19 | 8673.016183 | 0.036167357 | 0.040410578 | 0.894997265 | 0.37078855 | 0.780761257 |
| Pak6 | 3992.494436 | 0.036188976 | 0.05908644 | 0.612475153 | 0.540223433 | 0.863544778 |
| Prx | 23.9803642 | 0.036191363 | 0.0841291 | 0.430188402 | 0.667058598 | 0.916077199 |
| Osbpl9 | 1901.94764 | 0.036192387 | 0.061863841 | 0.585032972 | 0.558525547 | 0.872850145 |
| Prlr | 276.2364173 | 0.036192463 | 0.046812989 | 0.773128651 | 0.439446247 | 0.817545617 |
| Lhx8 | 76.99132947 | 0.036195115 | 0.086195357 | 0.419919549 | 0.674544226 | 0.919867577 |
| Gsta2 | 2.645002416 | 0.036214514 | 0.036986507 | 0.979127707 | 0.327516883 | NA |
| 2900052N01Rik | 1712.447503 | 0.036221812 | 0.075063293 | 0.482550266 | 0.6294151 | 0.904624802 |
| 2810004N23Rik | 442.6856408 | 0.036269548 | 0.077663458 | 0.467009182 | 0.640493309 | 0.910174702 |
| Depdc7 | 58.01859656 | 0.036276572 | 0.094551686 | 0.383669226 | 0.701223624 | 0.92751465 |
| 2700089E24Rik | 3890.883384 | 0.036287799 | 0.067906064 | 0.534382305 | 0.593077059 | 0.886253111 |
| E030003E18Rik | 338.8016284 | 0.036299175 | 0.094860886 | 0.382656924 | 0.701974158 | 0.927663448 |
| Saa1 | 1.718321026 | 0.036325035 | 0.029429026 | 1.2343268 | 0.217081153 | NA |
| Pde9a | 672.8891056 | 0.036358053 | 0.066142886 | 0.549689548 | 0.582532327 | 0.88406516 |
| Scrn2 | 129.3806998 | 0.036367091 | 0.09280018 | 0.391885996 | 0.695142451 | 0.925527709 |
| Rbm42 | 1653.921983 | 0.036400142 | 0.059025321 | 0.616686891 | 0.537441269 | 0.861996453 |
| Nos1ap | 3601.636947 | 0.036413434 | 0.062515533 | 0.582470186 | 0.560250018 | 0.873713575 |
| 2410004B18Rik | 534.2046695 | 0.036413497 | 0.077338385 | 0.470833433 | 0.637759687 | 0.908932006 |
| Spn | 7.032922148 | 0.036417816 | 0.057789284 | 0.630182861 | 0.528574952 | NA |
| Ccdc136 | 3238.855686 | 0.036438294 | 0.043889279 | 0.830232233 | 0.406407494 | 0.800759397 |
| Wdr8 | 817.9945723 | 0.036448891 | 0.073003811 | 0.499273811 | 0.617586502 | 0.898001714 |
| Evl | 4792.552457 | 0.036461288 | 0.045927735 | 0.793883863 | 0.427263045 | 0.810334558 |
| Sept5 | 27566.69083 | 0.036466837 | 0.059723782 | 0.610591549 | 0.541470019 | 0.864139035 |
| Aim1 | 61.89689853 | 0.03647663 | 0.094601006 | 0.385583955 | 0.699804818 | 0.926781946 |
| Polr3h | 1551.535296 | 0.036485881 | 0.05970548 | 0.611097686 | 0.541134911 | 0.863862966 |
| Tmed7 | 3154.944175 | 0.036485908 | 0.070915871 | 0.514495664 | 0.606905491 | 0.893150688 |
| Epn3 | 107.8147269 | 0.036487645 | 0.056154034 | 0.649777808 | 0.515835756 | 0.853297835 |
| Nrg3 | 1863.906284 | 0.036503818 | 0.069615462 | 0.524363652 | 0.600025633 | 0.889748095 |
| Zfand3 | 3205.45094 | 0.036533015 | 0.04888363 | 0.747346617 | 0.45485436 | 0.826168452 |
| Amz1 | 726.7206732 | 0.036535054 | 0.086257821 | 0.423556418 | 0.671889345 | 0.919094512 |
| Magi1 | 4750.490204 | 0.036541914 | 0.044107745 | 0.828469314 | 0.407404767 | 0.801055189 |
| D630029K05Rik | 2.002195999 | 0.036551612 | 0.031179718 | 1.172288086 | 0.241081413 | NA |
| BC055111 | 5.303674951 | 0.036552988 | 0.045676519 | 0.800257754 | 0.423561474 | NA |
| Ecd | 1783.282302 | 0.036602023 | 0.050742918 | 0.721322793 | 0.470710936 | 0.833245343 |
| Zfp202 | 206.2649099 | 0.03660427 | 0.083407064 | 0.438862944 | 0.660760849 | 0.915428982 |
| Tnfrsf26 | 1.803338207 | 0.03661303 | 0.029333729 | 1.248154643 | 0.211974431 | NA |
| Elovl5 | 2882.541524 | 0.036622133 | 0.058278207 | 0.628401848 | 0.529740727 | 0.857915133 |
| Thoc5 | 1334.102972 | 0.036631679 | 0.052728097 | 0.694727875 | 0.487225855 | 0.839801621 |
| Coq10a | 1329.075156 | 0.036644159 | 0.051160439 | 0.716259669 | 0.473831025 | 0.833983942 |
| Drosha | 5441.449828 | 0.036646991 | 0.040507911 | 0.90468725 | 0.365631098 | 0.777735733 |
| Rell2 | 5067.287949 | 0.036649471 | 0.062329917 | 0.587991658 | 0.556537892 | 0.871666144 |
| Irgq | 6418.786199 | 0.03666716 | 0.055159651 | 0.664746046 | 0.506212937 | 0.848908639 |
| Lnpep | 1765.988565 | 0.036693149 | 0.086996117 | 0.421779155 | 0.673186222 | 0.919353963 |
| Snx14 | 2383.831237 | 0.036696305 | 0.06678409 | 0.549476755 | 0.582678312 | 0.88406516 |
| Slc16a12 | 141.9235147 | 0.036700074 | 0.095096831 | 0.385923203 | 0.699553545 | 0.926781946 |
| 8030423F21Rik | 2.702957672 | 0.036712257 | 0.038073183 | 0.964255001 | 0.334918095 | NA |
| Bcdin3d | 103.5321561 | 0.036724032 | 0.092373683 | 0.397559468 | 0.690954943 | 0.923860706 |
| Gpr55 | 5.296343738 | 0.036729036 | 0.05167457 | 0.710775837 | 0.477223156 | NA |
| Neurod6 | 2024.986081 | 0.036750126 | 0.091554278 | 0.401402606 | 0.688123731 | 0.923860706 |
| Atp2b1 | 27462.82502 | 0.036769172 | 0.079193588 | 0.464294808 | 0.642436543 | 0.911258473 |
| Cpne7 | 2449.14296 | 0.036806773 | 0.093218503 | 0.394844064 | 0.692957973 | 0.924240888 |
| Zfyve20 | 3069.757984 | 0.036820656 | 0.051145187 | 0.719924165 | 0.471571689 | 0.833245343 |
| 1500009C09Rik | 3577.375291 | 0.036829318 | 0.057816844 | 0.636999803 | 0.524124972 | 0.855442623 |
| Acaca | 4608.817875 | 0.036832032 | 0.055103745 | 0.668412511 | 0.503870312 | 0.847807553 |
| Tigd5 | 233.5608633 | 0.036836447 | 0.084094518 | 0.438036243 | 0.661360011 | 0.915428982 |
| Elovl2 | 701.7164468 | 0.036836498 | 0.073600587 | 0.50049191 | 0.616728751 | 0.897296158 |
| A430093F15Rik | 4.098456256 | 0.036858255 | 0.041150745 | 0.89568864 | 0.370419079 | NA |
| Adck2 | 941.0342725 | 0.036872143 | 0.066685502 | 0.552925933 | 0.580314129 | 0.883332604 |
| Hspb11 | 196.1545947 | 0.036873195 | 0.084891442 | 0.434357029 | 0.664029197 | 0.915428982 |
| E130309D14Rik | 2013.405399 | 0.03688187 | 0.055932614 | 0.659398292 | 0.509640039 | 0.850457637 |
| L3mbtl3 | 937.4724868 | 0.036896417 | 0.064642703 | 0.570774669 | 0.568152396 | 0.877136965 |
| Mtmr4 | 4922.569922 | 0.036913538 | 0.055664117 | 0.663147832 | 0.507235879 | 0.848958235 |
| Qpctl | 726.6513806 | 0.036916133 | 0.063328534 | 0.582930486 | 0.559940097 | 0.873713575 |
| Mapk8ip2 | 12672.88845 | 0.036921447 | 0.046120629 | 0.800540848 | 0.423397507 | 0.808999423 |
| Fam118b | 582.6463534 | 0.036936516 | 0.067874963 | 0.54418469 | 0.586314386 | 0.884588669 |
| 4932412D23Rik | 2.915764801 | 0.036945388 | 0.040757816 | 0.906461422 | 0.364691675 | NA |
| Ndufs6 | 1352.204156 | 0.036968278 | 0.062342647 | 0.592985369 | 0.553190944 | 0.870600462 |
| Gm8773 | 3.473818726 | 0.036981661 | 0.042376916 | 0.872684097 | 0.382835288 | NA |
| Sec22b | 2301.213228 | 0.036993649 | 0.056766932 | 0.651676035 | 0.514610186 | 0.853195313 |
| Ccz1 | 1297.994928 | 0.037007446 | 0.061889363 | 0.597961337 | 0.549865731 | 0.869134887 |
| Gm11529 | 2.860514319 | 0.037027156 | 0.039779461 | 0.930810904 | 0.351951388 | NA |
| Slc37a1 | 279.9556221 | 0.037042879 | 0.085042298 | 0.435581824 | 0.66314016 | 0.915428982 |
| Chmp4b | 6587.470511 | 0.037055126 | 0.041318574 | 0.896815226 | 0.369817522 | 0.780397655 |
| Acadsb | 2329.275907 | 0.03706257 | 0.0570715 | 0.649405913 | 0.516076043 | 0.853297835 |
| Fbxo15 | 6.795287828 | 0.03706822 | 0.052543071 | 0.70548255 | 0.480509987 | NA |
| Cdh20 | 964.4008414 | 0.037099212 | 0.072658113 | 0.510599722 | 0.609631371 | 0.89440021 |
| Pcdhgb5 | 223.2947254 | 0.037101748 | 0.08449148 | 0.439118219 | 0.660575878 | 0.915428982 |
| Phf23 | 1779.092683 | 0.037112072 | 0.051161979 | 0.725383818 | 0.468216598 | 0.832372329 |
| Prrg2 | 163.6529354 | 0.037152083 | 0.090779017 | 0.409258483 | 0.68234998 | 0.922544285 |
| Slc5a8 | 3.665361673 | 0.037157695 | 0.042819052 | 0.86778416 | 0.385512503 | NA |
| Oaz2 | 7289.869475 | 0.037159322 | 0.051538209 | 0.7210053 | 0.470906254 | 0.833245343 |
| Ppap2c | 130.2914855 | 0.037159768 | 0.094691112 | 0.392431418 | 0.694739476 | 0.925527709 |
| Ppp1r15a | 643.6401668 | 0.037186878 | 0.077833765 | 0.477773077 | 0.632811725 | 0.906637358 |
| Ccdc162 | 29.82213276 | 0.03719028 | 0.086769804 | 0.428608548 | 0.668208124 | 0.916994918 |
| St7 | 696.9975574 | 0.037199604 | 0.071880096 | 0.517523012 | 0.604791109 | 0.891988593 |
| Cyb5r3 | 3033.814293 | 0.037200109 | 0.062925274 | 0.591179145 | 0.554400395 | 0.871360989 |
| Dhx8 | 2027.217469 | 0.037205812 | 0.051017677 | 0.729272961 | 0.465834708 | 0.831904964 |
| Mme | 597.7960825 | 0.037227562 | 0.092521278 | 0.402367567 | 0.687413535 | 0.923860706 |
| Usp9x | 11182.577 | 0.037243026 | 0.072142154 | 0.516244991 | 0.605683311 | 0.892498451 |
| 5830473C10Rik | 3.20689873 | 0.037265069 | 0.041328895 | 0.901671066 | 0.367231628 | NA |
| Tmem170b | 3350.451087 | 0.037273897 | 0.065958412 | 0.565112096 | 0.571997521 | 0.879055861 |
| Lrrfip2 | 1462.78372 | 0.037305531 | 0.064045583 | 0.582484054 | 0.560240679 | 0.873713575 |
| Rnase6 | 13.18818965 | 0.037319865 | 0.063058897 | 0.591825528 | 0.553967427 | NA |
| Emc1 | 3991.314413 | 0.037328164 | 0.047179361 | 0.791196894 | 0.428829103 | 0.810705511 |
| Rab5b | 7717.139106 | 0.037347422 | 0.041636535 | 0.896986789 | 0.369725967 | 0.780397655 |
| Cacfd1 | 1535.679999 | 0.03735474 | 0.051932937 | 0.719288027 | 0.471963471 | 0.83342346 |
| Kars | 3008.559926 | 0.037355062 | 0.055067234 | 0.678353708 | 0.497547453 | 0.845236583 |
| Slc15a5 | 2.056374142 | 0.037361771 | 0.034842805 | 1.072295152 | 0.283587483 | NA |
| Apold1 | 234.8755958 | 0.037366024 | 0.074621989 | 0.500737438 | 0.616555921 | 0.897296158 |

| | | | | | | | | | | | | | | | |
|---|---|---|---|---|---|---|---|---|---|---|---|---|---|---|---|
| Znrd1 | 440.343332 | 0.022391346 | 0.083390047 | 0.268513413 | 0.788304152 | 0.896363682 | | | Lrrc8d | 2088.047168 | 0.037371266 | 0.066793875 | 0.559501398 | 0.575819577 | 0.880811222 |
| Magel2 | 143.6801864 | 0.03961851 | 0.147864511 | 0.267937926 | 0.7887471 | 0.896745492 | | | Selm | 1961.786054 | 0.037384269 | 0.063241509 | 0.591134983 | 0.554429982 | 0.871360989 |
| Creb3l4 | 11.23549125 | 0.043955628 | 0.164305323 | 0.267524065 | 0.789065688 | 0.896976066 | | | Rab4b | 2186.126537 | 0.037388289 | 0.055771009 | 0.670389325 | 0.502609638 | 0.84739426 |
| Rpl6 | 9126.420504 | 0.018233499 | 0.068142323 | 0.267579653 | 0.789022895 | 0.896976066 | | | Gbgt1 | 33.60790895 | 0.037388671 | 0.088697821 | 0.421528628 | 0.673369111 | 0.919353963 |
| Pds5a | 1835.559765 | 0.022757815 | 0.085103977 | 0.26741188 | 0.789152053 | 0.89700843 | | | Tmem161a | 1059.273809 | 0.037389176 | 0.063109047 | 0.592453498 | 0.553546952 | 0.870690094 |
| Trabd2b | 268.567508 | -0.04053713 | 0.151671312 | -0.267269591 | 0.789261598 | 0.897067136 | | | Gm7173 | 8.556949004 | 0.03739756 | 0.060414091 | 0.619020479 | 0.535902868 | NA |
| Nhlh2 | 15.12859802 | 0.042954127 | 0.160782349 | 0.267156978 | 0.789348299 | 0.897099872 | | | Trp63 | 27.38522602 | 0.037398657 | 0.085301656 | 0.438428262 | 0.661075863 | 0.915428982 |
| Dner | 6628.236437 | -0.024096925 | 0.090352611 | -0.266698715 | 0.789701143 | 0.897435053 | | | Rad51d | 1095.186318 | 0.037423551 | 0.060198238 | 0.621671868 | 0.534157654 | 0.859551798 |
| 5031414D18Rik | 72.77979042 | 0.037618641 | 0.141197717 | 0.266425278 | 0.7899117 | 0.897568135 | | | Cdc42bpa | 14426.72585 | 0.03743006 | 0.061382934 | 0.609779576 | 0.542007833 | 0.864560611 |
| Gcnt2 | 1771.547953 | -0.026611942 | 0.099952559 | -0.266245726 | 0.790049969 | 0.897568135 | | | Gabre | 34.58322732 | 0.037435734 | 0.041120897 | 0.910382221 | 0.36262097 | 0.776449324 |
| Plekhm3 | 2587.265647 | -0.022516267 | 0.084567366 | -0.266252437 | 0.790044801 | 0.897568135 | | | Srsf4 | 2239.573156 | 0.037439864 | 0.051497205 | 0.727027112 | 0.467209347 | 0.832211278 |
| Prmt7 | 1457.117018 | 0.01831571 | 0.06877224 | 0.266324176 | 0.789989555 | 0.897568135 | | | Inip | 805.6205584 | 0.03744274 | 0.069007547 | 0.542589057 | 0.587412775 | 0.884773743 |
| E330033B04Rik | 43.14432678 | 0.043079724 | 0.161927494 | 0.266043294 | 0.790205867 | 0.897679426 | | | Agphd1 | 164.819238 | 0.037456543 | 0.090178912 | 0.415358118 | 0.677879762 | 0.921155235 |
| Nt5dc2 | 142.2024771 | 0.041921697 | 0.15775013 | 0.265747463 | 0.790433707 | 0.897872424 | | | Bap1 | 6634.385386 | 0.03748244 | 0.044720754 | 0.838144198 | 0.401949722 | 0.799277436 |
| Rab21 | 1524.996577 | 0.018047305 | 0.068017761 | 0.265332237 | 0.790753533 | 0.898169873 | | | Clock | 3366.772998 | 0.037486315 | 0.073597064 | 0.509345252 | 0.610510244 | 0.894800514 |
| Cenpp | 20.92333691 | -0.044265274 | 0.16692753 | -0.265176595 | 0.790873424 | 0.898170716 | | | Igf2 | 3280.12683 | 0.037488557 | 0.069079677 | 0.542685755 | 0.587346184 | 0.884773743 |
| Chst1 | 10613.64237 | 0.026885978 | 0.101444672 | 0.265030953 | 0.790985617 | 0.898170716 | | | Akt1 | 7194.594152 | 0.037499157 | 0.046557572 | 0.805436258 | 0.420567976 | 0.8078776 |
| Efhb | 10.095097 | 0.042613942 | 0.160788966 | 0.265030261 | 0.79098615 | 0.898170716 | | | Fbxl17 | 4113.752539 | 0.037510041 | 0.058286302 | 0.643548133 | 0.519868493 | 0.853297835 |
| Lysmd4 | 999.2058148 | -0.02020682 | 0.076241405 | -0.265037358 | 0.790980683 | 0.898170716 | | | Nudt18 | 1459.118101 | 0.037513319 | 0.072091154 | 0.520359522 | 0.602813017 | 0.891281205 |
| A330076H08Rik | 80.31321737 | 0.041396467 | 0.156361451 | 0.264748544 | 0.791203178 | 0.898211972 | | | Nr2c2ap | 536.9398693 | 0.037517389 | 0.074161844 | 0.505885331 | 0.61293715 | 0.895428201 |
| Cry1 | 757.6069619 | -0.033695351 | 0.127297639 | -0.264697373 | 0.791242601 | 0.898211972 | | | Samd7 | 2.24010455 | 0.037520439 | 0.030229627 | 1.241181012 | 0.214538888 | NA |
| Ict1 | 929.3242243 | 0.016940804 | 0.064034156 | 0.264558873 | 0.791349306 | 0.898211972 | | | 5930412G12Rik | 97.831882 | 0.037527594 | 0.094676039 | 0.396379004 | 0.691825453 | 0.923860706 |
| Rnd2 | 2085.071817 | -0.020120338 | 0.076051455 | -0.264562173 | 0.791346763 | 0.898211972 | | | Tdrkh | 1128.905826 | 0.037528479 | 0.057254625 | 0.655466343 | 0.512167549 | 0.852888435 |
| Tceal8 | 915.333067 | -0.017550313 | 0.066287804 | -0.264759299 | 0.791194893 | 0.898211972 | | | Ankrd45 | 3385.170126 | 0.03756312 | 0.06888653 | 0.545289763 | 0.585554244 | 0.884542528 |
| Zfp759 | 264.2109664 | 0.02858673 | 0.108065461 | 0.264531607 | 0.791370313 | 0.898211972 | | | A230056P14Rik | 100.7843497 | 0.037573868 | 0.094764171 | 0.396498675 | 0.691737185 | 0.923860706 |
| Hmox2 | 2548.1047 | -0.016156485 | 0.061253954 | -0.263762326 | 0.791963066 | 0.898802261 | | | Timp1 | 10.32580293 | 0.037634766 | 0.066650827 | 0.564655648 | 0.572308007 | NA |
| Ptcd1 | 1070.810211 | 0.021854635 | 0.082874976 | 0.263706085 | 0.792006407 | 0.898802261 | | | Kctd5 | 928.9931265 | 0.037643771 | 0.055653508 | 0.676395301 | 0.4987897 | 0.846052342 |
| Cyth2 | 3027.412243 | -0.018298409 | 0.069461189 | -0.263433565 | 0.792216423 | 0.898898281 | | | Kcne4 | 64.65174636 | 0.037685396 | 0.093209191 | 0.404309875 | 0.685984865 | 0.923860706 |
| Maea | 3095.757541 | -0.015422299 | 0.058557437 | -0.263370448 | 0.792265066 | 0.898898281 | | | Slc18a1 | 5.995618548 | 0.037699422 | 0.055118264 | 0.683973309 | 0.49399202 | NA |
| Ssbp3 | 4795.561219 | 0.018747014 | 0.071142327 | 0.263514201 | 0.792154279 | 0.898898281 | | | Sult1c1 | 8.933841044 | 0.037704026 | 0.035922001 | 1.049608188 | 0.293898291 | NA |
| Lnp | 2270.636896 | 0.02603539 | 0.098917338 | 0.263203509 | 0.792393726 | 0.898978428 | | | Mapre1 | 3938.325166 | 0.03770807 | 0.057809911 | 0.652276917 | 0.514222548 | 0.853195313 |
| Vmp1 | 3654.976116 | -0.019472616 | 0.07410662 | -0.262764865 | 0.792731818 | 0.899296148 | | | Rcc2 | 4445.071524 | 0.037726251 | 0.053204613 | 0.709078579 | 0.478275715 | 0.835936459 |
| Polr3gl | 628.0053753 | 0.020719883 | 0.078887467 | 0.262651142 | 0.792819478 | 0.899329745 | | | Tmem147 | 1875.510172 | 0.037735012 | 0.058926113 | 0.640378436 | 0.521926599 | 0.854113377 |
| Xrn1 | 733.6349894 | -0.027293602 | 0.104026468 | -0.262371703 | 0.793034886 | 0.899508238 | | | Vps72 | 972.708412 | 0.037758542 | 0.058709036 | 0.643147031 | 0.5201287 | 0.853297835 |
| Cuedc1 | 1013.776411 | 0.023637883 | 0.09013445 | 0.262251367 | 0.793127654 | 0.899547608 | | | Rnase1 | 17.0044005 | 0.037767371 | 0.068351001 | 0.552550372 | 0.580571333 | NA |
| 1110034G24Rik | 241.3123629 | 0.027853759 | 0.106393815 | 0.261798668 | 0.793476666 | 0.899577456 | | | Slc20a2 | 2429.07958 | 0.037788237 | 0.056601944 | 0.667613769 | 0.504380166 | 0.847807553 |
| Adrb2 | 77.65653248 | 0.03476209 | 0.132812528 | 0.261738032 | 0.793523417 | 0.899577456 | | | Pygo2 | 2275.277368 | 0.037802389 | 0.068943472 | 0.54830992 | 0.583479117 | 0.884313345 |
| Dpp6 | 11489.48664 | -0.016651203 | 0.063566122 | -0.261950909 | 0.79335929 | 0.899577456 | | | Fam198a | 349.225277 | 0.037820748 | 0.077631358 | 0.487183908 | 0.626128008 | 0.902402691 |
| Eif3d | 2533.167786 | -0.01652823 | 0.06313186 | -0.261804895 | 0.793471865 | 0.899577456 | | | Kcnq5 | 4116.736189 | 0.037852458 | 0.072968684 | 0.518749354 | 0.60393554 | 0.891718993 |
| Poll | 388.8964172 | -0.026884591 | 0.102796009 | -0.261533412 | 0.793681187 | 0.899577456 | | | 4833418N02Rik | 28.90428122 | 0.037887652 | 0.087012959 | 0.435425394 | 0.663253682 | 0.915428982 |
| Tm7sf2 | 569.2492683 | 0.034528564 | 0.131990145 | 0.261599565 | 0.79363018 | 0.899577456 | | | Adal | 659.9342684 | 0.037924341 | 0.06930575 | 0.547203379 | 0.584239015 | 0.884313345 |
| Txndc9 | 1551.095204 | -0.01941339 | 0.074107996 | -0.2619608 | 0.793351664 | 0.899577456 | | | Fuom | 571.0488298 | 0.037924391 | 0.069821438 | 0.543162565 | 0.587017879 | 0.884663353 |
| Vamp2 | 48121.43284 | -0.011969191 | 0.045775287 | -0.261477138 | 0.793724577 | 0.899577456 | | | Tor1a | 997.8321903 | 0.037929621 | 0.055451 | 0.684020499 | 0.493962221 | 0.843404385 |
| Vwa7 | 41.05197614 | 0.043776892 | 0.167425043 | 0.261471588 | 0.793728857 | 0.899577456 | | | Ngly1 | 954.6577561 | 0.037935299 | 0.069787417 | 0.543583655 | 0.58672801 | 0.884638284 |
| Zcchc6 | 2626.625449 | 0.019331296 | 0.073934778 | 0.261464181 | 0.793734568 | 0.899577456 | | | Slc2a3 | 7196.587038 | 0.037937838 | 0.066003706 | 0.574783445 | 0.565437764 | 0.875364912 |
| Bcdin3d | 96.70941969 | -0.031976003 | 0.122481757 | -0.261067473 | 0.794040475 | 0.899794369 | | | Htr2b | 3.676363584 | 0.037941789 | 0.045406671 | 0.835599441 | 0.403380283 | NA |
| Extl3 | 8191.856638 | 0.016141388 | 0.061840303 | 0.261017281 | 0.794079181 | 0.899794369 | | | Arsj | 118.8941233 | 0.037947009 | 0.093454202 | 0.406049249 | 0.684706415 | 0.923762012 |
| Tmem88b | 2417.102433 | 0.042381573 | 0.162387697 | 0.26099005 | 0.794100181 | 0.899794369 | | | Efcab10 | 49.80047629 | 0.037963355 | 0.090076103 | 0.421458676 | 0.673420181 | 0.919353963 |
| Chodl | 44.57409464 | 0.042111588 | 0.16149468 | 0.260761454 | 0.794276472 | 0.899928311 | | | 2510049J12Rik | 2.521736267 | 0.037969207 | 0.035443573 | 1.071257895 | 0.284053486 | NA |
| 2210018M11Rik | 1964.574117 | 0.017925558 | 0.068764238 | 0.260681399 | 0.794338212 | 0.899932455 | | | Ndufv3 | 1733.566461 | 0.037977032 | 0.062804222 | 0.60468916 | 0.545385552 | 0.866680785 |
| Il1rapl2 | 143.6440427 | 0.034757001 | 0.133704315 | 0.259954222 | 0.794899086 | 0.900502042 | | | Anks1b | 12133.90948 | 0.037985947 | 0.060007393 | 0.633021118 | 0.526719852 | 0.85599981 |
| Ndufs3 | 3573.91356 | -0.016146162 | 0.06215408 | -0.259776379 | 0.795036272 | 0.900528223 | | | Rnf10 | 10109.82369 | 0.038001638 | 0.044789194 | 0.848455508 | 0.39618434 | 0.796565439 |
| Siglece | 38.91030855 | 0.041086077 | 0.15816111 | 0.259773572 | 0.795038438 | 0.900528223 | | | Ykt6 | 3987.132807 | 0.038003441 | 0.05145426 | 0.738586876 | 0.460157896 | 0.829022384 |
| Dek | 1760.98955 | 0.021167776 | 0.081592213 | 0.259433771 | 0.795300576 | 0.900638073 | | | Zfp37 | 793.7462444 | 0.038006191 | 0.07911838 | 0.480371198 | 0.630963471 | 0.905472959 |
| Iah1 | 302.8662738 | 0.022009542 | 0.08483482 | 0.259439957 | 0.795295804 | 0.900638073 | | | Tbl2 | 1182.389318 | 0.038037071 | 0.050326422 | 0.755807181 | 0.44976481 | 0.823263966 |
| Zdhhc9 | 2506.767678 | -0.018587327 | 0.071649052 | -0.259421809 | 0.795309805 | 0.900638073 | | | Fam212a | 60.47212867 | 0.038038 | 0.09416719 | 0.403941113 | 0.686256022 | 0.923860706 |
| Rbm14 | 1179.990966 | 0.01727389 | 0.066626544 | 0.259264381 | 0.79543126 | 0.900681926 | | | Sys1 | 985.8547656 | 0.03806577 | 0.064434138 | 0.590770226 | 0.554674387 | 0.871360989 |
| Zfp869 | 1151.939719 | -0.019567954 | 0.075487557 | -0.259220919 | 0.795464791 | 0.900681926 | | | Dennd1b | 505.6277922 | 0.03807056 | 0.077906254 | 0.488671423 | 0.625074338 | 0.901503145 |
| Cnot4 | 1335.05416 | 0.02079997 | 0.080330411 | 0.258930207 | 0.795689091 | 0.900804236 | | | Cyp2b10 | 2.206759211 | 0.038075077 | 0.030988384 | 1.228688702 | 0.219188544 | NA |
| Socs1 | 21.6289523 | -0.0433299 | 0.167332917 | -0.258944268 | 0.795678242 | 0.900804236 | | | Qrich1 | 3703.13536 | 0.038079097 | 0.045665937 | 0.833862165 | 0.40435866 | 0.800290023 |
| Pex6 | 2017.418013 | -0.030577602 | 0.118219747 | -0.258650546 | 0.79590488 | 0.90091781 | | | Klhdc2 | 4890.571383 | 0.038084509 | 0.05589337 | 0.681377931 | 0.495632383 | 0.844369474 |
| Psme3 | 4400.86656 | -0.01507503 | 0.058283627 | -0.258649477 | 0.795905704 | 0.90091781 | | | Slc7a4 | 2869.482937 | 0.038085648 | 0.076078746 | 0.500608255 | 0.616646852 | 0.897296158 |
| Ccdc86 | 314.61713 | 0.025821007 | 0.099876944 | 0.258528206 | 0.795999283 | 0.900934512 | | | Krt23 | 1.465321169 | 0.038095313 | 0.024884515 | 1.530884266 | 0.125797998 | NA |
| Gemin5 | 1196.74765 | 0.01592868 | 0.061624505 | 0.258479648 | 0.796036754 | 0.900934512 | | | Ano10 | 1059.778182 | 0.038099401 | 0.067926341 | 0.560892879 | 0.574870565 | 0.880304641 |
| Cenpm | 60.43224162 | -0.040662624 | 0.157559368 | -0.25807811 | 0.796346625 | 0.901219386 | | | Nlgn1 | 2887.615208 | 0.038127211 | 0.075103674 | 0.507661068 | 0.611691056 | 0.895166661 |
| Dhx15 | 3385.917432 | 0.022400925 | 0.086833567 | 0.257975406 | 0.796425888 | 0.90124326 | | | Tbcb | 2023.565176 | 0.038138058 | 0.054331542 | 0.701950595 | 0.482709975 | 0.837880119 |
| Nudcd3 | 6683.59845 | 0.014681675 | 0.056937977 | 0.257853812 | 0.796519732 | 0.901283629 | | | Adamts19 | 9.335933272 | 0.038150637 | 0.049770773 | 0.766526918 | 0.443362842 | NA |
| Arfgap2 | 2877.693048 | 0.011207503 | 0.043543953 | 0.257383682 | 0.796882599 | 0.901493186 | | | Pih1d1 | 729.8706901 | 0.038197663 | 0.068069011 | 0.561160843 | 0.574687894 | 0.880182448 |
| Arsj | 101.7584186 | 0.039324169 | 0.152862332 | 0.257252182 | 0.796984103 | 0.901493186 | | | Cyfip2 | 60715.76747 | 0.038211746 | 0.053621569 | 0.712618941 | 0.476081588 | 0.835195112 |
| BC017612 | 54.75053544 | -0.041468195 | 0.161206203 | -0.257236968 | 0.796995848 | 0.901493186 | | | Fbxl8 | 29.131312 | 0.03822193 | 0.086627646 | 0.441220924 | 0.659053066 | 0.915208181 |
| E430018J23Rik | 194.071015 | 0.029494142 | 0.114598624 | 0.257369077 | 0.796893872 | 0.901493186 | | | Pfdn5 | 3856.703368 | 0.038236124 | 0.057390511 | 0.6662447 | 0.505254706 | 0.848183273 |
| Polm | 276.0519778 | 0.027912317 | 0.108444273 | 0.257388575 | 0.796878822 | 0.901493186 | | | Ggt5 | 112.2145457 | 0.038241807 | 0.092310768 | 0.414272437 | 0.678674597 | 0.921316909 |
| Chmp7 | 3678.19903 | -0.011082504 | 0.043159143 | -0.256782309 | 0.797346828 | 0.901824348 | | | Hivep2 | 12754.31193 | 0.038250031 | 0.077741437 | 0.492016007 | 0.622708028 | 0.900224603 |
| Echdc2 | 338.4211601 | 0.036500378 | 0.142244957 | 0.256602264 | 0.797485827 | 0.901915723 | | | Vgll2 | 4.274707362 | 0.038268901 | 0.047853042 | 0.79971719 | 0.423874671 | NA |
| E230029C05Rik | 29.30619119 | 0.04291006 | 0.167325137 | 0.25644718 | 0.797605561 | 0.901985298 | | | Agps | 1835.596191 | 0.038276296 | 0.064185631 | 0.596337465 | 0.550949808 | 0.869642174 |
| 4930579G24Rik | 57.1945659 | -0.03936036 | 0.153729779 | -0.256036013 | 0.79792303 | 0.902278457 | | | Il12a | 96.52560462 | 0.038283633 | 0.094455653 | 0.40530801 | 0.68525112 | 0.923860706 |
| Aaed1 | 193.5681617 | -0.027446668 | 0.107355124 | -0.255662391 | 0.798211538 | 0.902312383 | | | Gak | 5662.56018 | 0.038285684 | 0.038455883 | 0.99557419 | 0.319457081 | 0.749427924 |
| Fbxw7 | 8382.801364 | -0.016151384 | 0.063141324 | -0.255797354 | 0.798107317 | 0.902312383 | | | Gm9962 | 70.00460941 | 0.038288446 | 0.095094485 | 0.402635814 | 0.687216159 | 0.923860706 |
| Myh4 | 22.2132619 | 0.042968763 | 0.168096195 | 0.255620079 | 0.798244213 | 0.902312383 | | | Cyp2j13 | 2.577506032 | 0.038299148 | 0.030642175 | 1.24988346 | 0.211342122 | NA |
| Ubap2 | 2014.848389 | 0.014360149 | 0.056147024 | 0.255759754 | 0.798136352 | 0.902312383 | | | 5330426P16Rik | 199.4362824 | 0.038317265 | 0.094862995 | 0.403922151 | 0.686269967 | 0.923860706 |
| Vps72 | 888.9115345 | 0.01636018 | 0.063981486 | 0.255701784 | 0.798181117 | 0.902312383 | | | Zfp91 | 3881.186594 | 0.038321013 | 0.065518915 | 0.584884733 | 0.558625224 | 0.872912006 |
| Spag7 | 1544.127887 | 0.013545848 | 0.053016735 | 0.255501365 | 0.79833589 | 0.90235018 | | | Ero1l | 2386.528937 | 0.038336916 | 0.055825042 | 0.686733302 | 0.492250807 | 0.842340249 |
| Angptl1 | 32.38231405 | 0.042304591 | 0.165661392 | 0.255367833 | 0.798439013 | 0.902400909 | | | Sgpp1 | 1269.901095 | 0.038345584 | 0.065921504 | 0.581685507 | 0.560778535 | 0.873713575 |
| Scamp5 | 19054.15493 | -0.011285818 | 0.044207569 | -0.255291529 | 0.798497942 | 0.902401686 | | | Tle4 | 2549.456615 | 0.038348192 | 0.070669264 | 0.542643143 | 0.587375528 | 0.884773743 |
| Ubc | 5276.473282 | -0.022274722 | 0.087279673 | -0.255210881 | 0.798560228 | 0.902406255 | | | 4933424G06Rik | 9.494021602 | 0.038348835 | 0.063518683 | 0.603741026 | 0.546015833 | NA |
| AU022252 | 561.5916079 | -0.021962725 | 0.086196332 | -0.254798832 | 0.798878477 | 0.902436799 | | | Rel | 63.67588292 | 0.038349865 | 0.094449895 | 0.406033958 | 0.68471765 | 0.923762012 |
| Dnajb13 | 12.90382355 | 0.041199476 | 0.161672786 | 0.254832472 | 0.798852494 | 0.902436799 | | | Rnf19b | 1793.699392 | 0.038354681 | 0.054839286 | 0.699401545 | 0.484301123 | 0.838713365 |
| Mospd2 | 347.0758614 | 0.020315138 | 0.079713638 | 0.254851479 | 0.798837813 | 0.902436799 | | | Bptf | 4632.690115 | 0.038362462 | 0.071590104 | 0.535862635 | 0.59205349 | 0.886226713 |
| Spast | 1963.878183 | 0.018216461 | 0.071433175 | 0.255014023 | 0.798712269 | 0.902436799 | | | Gcsh | 1659.720252 | 0.038363135 | 0.063948457 | 0.599907121 | 0.548568136 | 0.868028404 |
| Yif1a | 441.3166213 | -0.029719748 | 0.116556781 | -0.254980856 | 0.798737885 | 0.902436799 | | | Ric3 | 4145.216632 | 0.038367424 | 0.053305232 | 0.719768445 | 0.471667576 | 0.833304495 |
| BC068281 | 315.2953418 | 0.023221547 | 0.091283724 | 0.254388689 | 0.799195288 | 0.902499395 | | | Rimkla | 731.3550249 | 0.038409814 | 0.075529379 | 0.50854137 | 0.611073734 | 0.89489435 |
| Eva1a | 307.9052506 | -0.024733133 | 0.097153023 | -0.254579135 | 0.799048176 | 0.902499395 | | | Mras | 10814.17554 | 0.038425482 | 0.052797618 | 0.727788176 | 0.466743264 | 0.832095317 |
| Hdac8 | 440.9794588 | 0.020700658 | 0.08138649 | 0.254350056 | 0.799225132 | 0.902499395 | | | Bet3l | 3.839309565 | 0.038432953 | 0.047527637 | 0.808644307 | 0.418719771 | NA |
| Lrrc17 | 251.7539354 | -0.034157285 | 0.134212402 | -0.254501705 | 0.799107987 | 0.902499395 | | | Mthfs | 55.11688784 | 0.038439506 | 0.092715883 | 0.414594618 | 0.678438689 | 0.921316909 |
| Lyn | 415.821327 | -0.029451314 | 0.115754644 | -0.254428786 | 0.799164314 | 0.902499395 | | | Setd1b | 2073.4321 | 0.03845686 | 0.06592489 | 0.583343561 | 0.559662044 | 0.873713575 |
| Lrrc28 | 848.8341289 | 0.017974266 | 0.070693934 | 0.254254715 | 0.799298783 | 0.902516786 | | | Ddx59 | 326.562234 | 0.038463243 | 0.082764001 | 0.464733972 | 0.642121977 | 0.911258473 |
| Cog8 | 803.2138014 | 0.019063593 | 0.075071773 | 0.253938223 | 0.799543285 | 0.902575325 | | | Yy1 | 1059.946247 | 0.038464158 | 0.072741839 | 0.528776261 | 0.596960667 | 0.887922451 |
| Fam229b | 261.6089568 | -0.02994843 | 0.11801689 | -0.253763933 | 0.799677939 | 0.902575325 | | | Rbm43 | 732.7563893 | 0.038464558 | 0.068708906 | 0.559819106 | 0.57560283 | 0.880811222 |
| Klhl18 | 1525.119764 | -0.015791033 | 0.062159328 | -0.254041235 | 0.799463702 | 0.902575325 | | | Gtf3c5 | 857.5217751 | 0.038478734 | 0.068510032 | 0.561651092 | 0.574353763 | 0.879791496 |
| Nsun4 | 482.1980431 | -0.019628913 | 0.077382833 | -0.253659792 | 0.799758399 | 0.902575325 | | | Rab3ip | 1377.595051 | 0.038480087 | 0.059210307 | 0.649888324 | 0.515764361 | 0.853297835 |
| Nt5c1a | 130.7807405 | -0.029630497 | 0.116789331 | -0.253708937 | 0.799720429 | 0.902575325 | | | Fam63a | 1009.562378 | 0.038502054 | 0.059571348 | 0.646318325 | 0.518073223 | 0.853297835 |
| Tcf12 | 2004.053621 | -0.019747112 | 0.077773205 | -0.253906372 | 0.799567892 | 0.902575325 | | | Ufc1 | 2000.184659 | 0.038503748 | 0.045237018 | 0.851155756 | 0.394682836 | 0.795437883 |
| Twf2 | 1461.227018 | 0.021322782 | 0.084005724 | 0.253825346 | 0.799630491 | 0.902575325 | | | Aktip | 4485.472246 | 0.038508319 | 0.05815287 | 0.662191198 | 0.507848695 | 0.849201121 |
| AA465934 | 36.49250901 | 0.042069891 | 0.166010957 | 0.253416352 | 0.799946493 | 0.902665401 | | | Nom1 | 1217.881048 | 0.038529612 | 0.061942645 | 0.622020781 | 0.533928203 | 0.859470725 |
| BC051226 | 95.50094421 | 0.03192256 | 0.125974126 | 0.253405689 | 0.799954732 | 0.902665401 | | | Hist1h1c | 521.3730531 | 0.038539925 | 0.094810567 | 0.406493985 | 0.684379676 | 0.923655038 |
| Ccdc162 | 26.54197927 | -0.042429724 | 0.167569588 | -0.25320659 | 0.800108575 | 0.902703835 | | | 4933403G14Rik | 180.3540019 | 0.038556372 | 0.094399262 | 0.408439343 | 0.682951154 | 0.922927571 |
| Nrsn2 | 3560.574223 | -0.023647204 | 0.093500758 | -0.252909224 | 0.800338362 | 0.902703835 | | | Grin3b | 8.094519872 | 0.038563096 | 0.058457336 | 0.659679335 | 0.509459631 | NA |
| Phgdh | 2506.51496 | -0.025155296 | 0.099460788 | -0.252916717 | 0.800332571 | 0.902703835 | | | Artn | 16.38987981 | 0.03857233 | 0.067844083 | 0.568543755 | 0.569665804 | NA |
| Rassf2 | 3021.817833 | -0.01748886 | 0.069140323 | -0.252947334 | 0.800308911 | 0.902703835 | | | Coasy | 988.5220244 | 0.038575441 | 0.07642751 | 0.504732407 | 0.613746796 | 0.895819717 |
| Rb1 | 1193.018664 | -0.021210654 | 0.08377548 | -0.253184511 | 0.800125635 | 0.902703835 | | | Hnrnpul2 | 11648.1059 | 0.038575808 | 0.059123676 | 0.652459564 | 0.51410475 | 0.853195313 |
| Smyd1 | 91.45201895 | -0.042465103 | 0.167812016 | -0.253051622 | 0.800228322 | 0.902703835 | | | Fbxo28 | 2423.43761 | 0.03859144 | 0.061144853 | 0.631147805 | 0.527943886 | 0.856965327 |
| Pick1 | 1636.407717 | -0.01567441 | 0.062063189 | -0.252555659 | 0.800611598 | 0.902893264 | | | Rassf4 | 567.4108254 | 0.03859222 | 0.086855982 | 0.444324261 | 0.656808163 | 0.915208181 |
| Tor1a | 923.1722241 | 0.019891716 | 0.078766257 | 0.252541091 | 0.800622857 | 0.902893264 | | | Prdm13 | 3.048187347 | 0.0385994 | 0.027639073 | 1.396551942 | 0.162548351 | NA |
| Fam129c | 61.2611326 | 0.041966813 | 0.166312257 | 0.252337464 | 0.800780233 | 0.903005016 | | | Nlk | 5254.564151 | 0.038600272 | 0.075785348 | 0.50933687 | 0.610516118 | 0.894800514 |
| Dnajb12 | 1329.805461 | -0.020525529 | 0.081533691 | -0.25174292 | 0.80123978 | 0.903335626 | | | Trf | 16025.29359 | 0.038607446 | 0.087200955 | 0.44274109 | 0.657953018 | 0.915208181 |
| Gpc5 | 1535.771489 | 0.021982806 | 0.087311852 | 0.251773447 | 0.801216182 | 0.903335626 | | | Ppargc1b | 362.424251 | 0.038615213 | 0.083418336 | 0.462910377 | 0.643428609 | 0.911258473 |
| Impdh1 | 2298.052411 | -0.021363981 | 0.084868005 | -0.251731869 | 0.801248322 | 0.903335626 | | | Pak2 | 2368.191581 | 0.038638823 | 0.060363624 | 0.640101116 | 0.522106863 | 0.854113377 |
| Golim4 | 943.5162319 | -0.019794654 | 0.078742338 | -0.251385149 | 0.801516349 | 0.903572054 | | | Ranbp9 | 3938.882557 | 0.038647443 | 0.061041284 | 0.633136144 | 0.526644741 | 0.85599981 |
| Gins1 | 26.87842405 | 0.041720872 | 0.166053174 | 0.251250071 | 0.801620775 | 0.903624029 | | | Pex26 | 823.2083883 | 0.038649769 | 0.066363852 | 0.582391891 | 0.560302742 | 0.873713575 |
| A4gnt | 8.878031344 | -0.039058858 | 0.155601257 | -0.251018913 | 0.801799487 | 0.903669159 | | | Trp53bp1 | 4594.611014 | 0.03865846 | 0.06826724 | 0.566281279 | 0.571202585 | 0.878374234 |
| Cdh18 | 896.5594038 | -0.023279017 | 0.092755445 | -0.250971968 | 0.801835782 | 0.903669159 | | | Atp12a | 4.121024504 | 0.038664986 | 0.03966269 | 0.97484526 | 0.329637023 | NA |
| Snhg8 | 336.7867829 | -0.028925283 | 0.115199979 | -0.251087571 | 0.801746405 | 0.903669159 | | | Gm16796 | 8.423860254 | 0.038674692 | 0.062342258 | 0.620360779 | 0.535020288 | NA |
| Gm2a | 2334.528424 | -0.016786299 | 0.066972518 | -0.250644585 | 0.802088908 | 0.903888686 | | | Dpp9 | 3240.247971 | 0.03868399 | 0.053295369 | 0.725841479 | 0.467935956 | 0.832353544 |
| Nop9 | 415.5390289 | 0.022137514 | 0.088367587 | 0.250516215 | 0.802188166 | 0.903934796 | | | Pfkl | 7480.034835 | 0.038690497 | 0.076318534 | 0.506960688 | 0.612182402 | 0.895344807 |
| Cryl1 | 464.2802902 | 0.02511184 | 0.100321941 | 0.250312545 | 0.802345656 | 0.903980773 | | | Pde10a | 12955.69318 | 0.038695394 | 0.094957842 | 0.407500771 | 0.683640228 | 0.923110101 |
| Polr3d | 684.2946357 | 0.015451801 | 0.061714412 | 0.250375897 | 0.802296668 | 0.903980773 | | | Ndufv2 | 4270.231008 | 0.038707407 | 0.071687645 | 0.539945299 | 0.589234757 | 0.885188411 |
| 2210404O09Rik | 153.6185303 | 0.033999007 | 0.135953232 | 0.250078694 | 0.802526492 | 0.904053037 | | | Zfp719 | 751.0793206 | 0.03871284 | 0.070326124 | 0.55047595 | 0.581992968 | 0.884020489 |
| Arl6ip5 | 4772.529839 | -0.013447536 | 0.05375837 | -0.250147766 | 0.802473078 | 0.904053037 | | | Nrp1 | 1194.038201 | 0.038723821 | 0.083459588 | 0.463982889 | 0.642660005 | 0.911258473 |
| Ryk | 1144.263596 | -0.015655842 | 0.062634724 | -0.249954677 | 0.802622399 | 0.904095344 | | | AA467197 | 5.030761406 | 0.038744375 | 0.047348363 | 0.818283304 | 0.413195438 | NA |
| Crispld2 | 123.8667687 | 0.036515541 | 0.14614793 | 0.249853287 | 0.802700809 | 0.904117937 | | | 4931429I11Rik | 18.86230472 | 0.038790956 | 0.073449923 | 0.528127936 | 0.597410543 | NA |
| Plod2 | 164.0105132 | 0.034370283 | 0.137656364 | 0.249681758 | 0.802833467 | 0.904201623 | | | Sntb1 | 81.58741308 | 0.038818535 | 0.095102745 | 0.408174707 | 0.683145416 | 0.922993772 |
| Fra10ac1 | 415.5731097 | -0.029349042 | 0.117605542 | -0.249554926 | 0.80293156 | 0.904246372 | | | Pdxdc1 | 4410.610855 | 0.038828942 | 0.049163264 | 0.789795849 | 0.429647004 | 0.811442206 |
| Hddc3 | 354.1106827 | 0.02265633 | 0.090965157 | 0.249066024 | 0.80330971 | 0.90453926 | | | Mrpl37 | 1530.754725 | 0.038851195 | 0.051306888 | 0.757231562 | 0.448911144 | 0.823263966 |
| Pcca | 1025.357392 | -0.017338339 | 0.069622287 | -0.249034318 | 0.803334235 | 0.90453926 | | | Anapc15 | 403.2932147 | 0.038854446 | 0.075108584 | 0.517310326 | 0.604939547 | 0.892113813 |
| Ube2t | 36.89201971 | -0.041783996 | 0.167812432 | -0.248992256 | 0.803366772 | 0.90453926 | | | Kctd1 | 3554.850212 | 0.038867667 | 0.074988812 | 0.518312882 | 0.604239987 | 0.891718993 |
| Lrrc51 | 87.81519744 | -0.040922811 | 0.164552793 | -0.248691077 | 0.803599752 | 0.904735834 | | | Cc2d1a | 2305.969251 | 0.0388719 | 0.060411422 | 0.643452811 | 0.519930325 | 0.853297835 |
| Ift43 | 224.2838269 | -0.022297555 | 0.089701428 | -0.248575248 | 0.803689357 | 0.904770972 | | | Fam171a2 | 2852.710484 | 0.038883727 | 0.060022528 | 0.647818887 | 0.517102098 | 0.853297835 |
| Rtn3 | 47567.32987 | -0.013424615 | 0.054029055 | -0.248470293 | 0.803770552 | 0.904796639 | | | Il17re | 11.41325271 | 0.038895541 | 0.059856223 | 0.64981616 | 0.515810979 | NA |
| Lmbr1 | 1351.937033 | -0.015728532 | 0.063392639 | -0.24811291 | 0.804047047 | 0.904978122 | | | Ccdc85c | 548.6124088 | 0.038926291 | 0.079074952 | 0.492270815 | 0.622527909 | 0.900143381 |
| Snrpb | 2121.638785 | 0.024755684 | 0.099776681 | 0.248110918 | 0.804048588 | 0.904978122 | | | Xrcc4 | 307.0890215 | 0.038929444 | 0.078273314 | 0.497352692 | 0.61894036 | 0.898695177 |
| Cep89 | 496.3590553 | 0.023484967 | 0.094721008 | 0.247938311 | 0.804182137 | 0.905062688 | | | Aes | 15605.57331 | 0.038941884 | 0.046159603 | 0.843635592 | 0.398873069 | 0.797789838 |
| Dync2li1 | 615.5838307 | -0.020819446 | 0.084151278 | -0.247404989 | 0.804594814 | 0.90542198 | | | Chsy3 | 492.8939178 | 0.038948041 | 0.073597877 | 0.529200608 | 0.596666295 | 0.887922451 |
| Tsc22d2 | 1260.71676 | -0.021040171 | 0.085053852 | -0.247374697 | 0.804618255 | 0.90542198 | | | Fcrlb | 14.66400033 | 0.038954107 | 0.074981689 | 0.519514931 | 0.603401704 | NA |
| B230319C09Rik | 13.20432393 | -0.04096216 | 0.165836757 | -0.247002903 | 0.804905978 | 0.905484189 | | | Bcas1 | 4716.833287 | 0.03898823 | 0.089448693 | 0.435872549 | 0.662929203 | 0.915428982 |
| Ccdc126 | 333.5876204 | -0.025793365 | 0.10442104 | -0.247013105 | 0.804898082 | 0.905484189 | | | Dkk3 | 12643.19957 | 0.03899387 | 0.06038419 | 0.645762903 | 0.518432916 | 0.853297835 |
| Ppp1r3e | 258.2518698 | 0.023143751 | 0.093698943 | 0.247001192 | 0.804907301 | 0.905484189 | | | Amer3 | 1676.415074 | 0.039047771 | 0.064490064 | 0.605485079 | 0.544856736 | 0.866307897 |
| Zmat2 | 7168.934951 | -0.014607992 | 0.05914127 | -0.247001658 | 0.804906941 | 0.905484189 | | | Cnot2 | 2460.178025 | 0.039055301 | 0.047759867 | 0.817743078 | 0.413503907 | 0.80409073 |
| Bckdha | 1363.433234 | 0.017454409 | 0.070781905 | 0.246594219 | 0.805222279 | 0.905535571 | | | Chgb | 20595.60617 | 0.039078337 | 0.061406644 | 0.636386139 | 0.524524773 | 0.855487854 |
| Blnk | 399.808407 | 0.034324968 | 0.139213767 | 0.246563033 | 0.805246417 | 0.905535571 | | | Srf | 2467.390456 | 0.039086954 | 0.070185326 | 0.556910631 | 0.577588491 | 0.881493005 |
| Epn1 | 14098.95882 | 0.023637474 | 0.095896629 | 0.246489099 | 0.805303642 | 0.905535571 | | | Cog5 | 1060.625363 | 0.039105634 | 0.061536039 | 0.635491578 | 0.525107858 | 0.855700779 |
| Hcn2 | 4748.070076 | 0.022103975 | 0.089559807 | 0.246806861 | 0.8050577 | 0.905535571 | | | Synpo | 11437.95751 | 0.039112618 | 0.079019077 | 0.494976902 | 0.620616425 | 0.899412757 |
| Ier5 | 2676.015341 | -0.031114767 | 0.126087062 | -0.246772081 | 0.805084618 | 0.905535571 | | | Slc6a13 | 380.4688131 | 0.039113974 | 0.084450702 | 0.463157478 | 0.643251493 | 0.911258473 |
| Mylpf | 19.17409249 | 0.041350805 | 0.167647144 | 0.246653798 | 0.805176166 | 0.905535571 | | | Sssca1 | 534.7077253 | 0.039117452 | 0.067642388 | 0.578297919 | 0.563062999 | 0.87510382 |
| Hsf5 | 15.43381094 | 0.040253905 | 0.163366625 | 0.246402255 | 0.805370861 | 0.905543596 | | | 1600023N17Rik | 14.69097073 | 0.039118878 | 0.074249739 | 0.526855426 | 0.59829399 | NA |
| Wee1 | 978.6111253 | 0.020445547 | 0.08300102 | 0.246328863 | 0.805427669 | 0.905543596 | | | Shoc2 | 2545.503623 | 0.039119242 | 0.066352445 | 0.589567445 | 0.555480683 | 0.871360989 |
| Proca1 | 92.43109062 | 0.038367841 | 0.15582008 | 0.246231684 | 0.80550289 | 0.905562456 | | | Atxn1 | 3469.274185 | 0.039165486 | 0.071450931 | 0.548145212 | 0.583592198 | 0.884313345 |
| Pigq | 4166.684369 | -0.027546348 | 0.111912193 | -0.246142508 | 0.805571918 | 0.905574353 | | | BC125332 | 5.227761903 | 0.039185388 | 0.049712422 | 0.788241378 | 0.43055553 | NA |
| 5-Mar | 2731.516544 | -0.01481606 | 0.060214912 | -0.246053009 | 0.805641198 | 0.905583862 | | | Vps11 | 3655.071193 | 0.039194437 | 0.048890505 | 0.801677894 | 0.422739306 | 0.808946274 |
| Osbpl9 | 1747.035349 | 0.018888632 | 0.076789122 | 0.245980571 | 0.805697272 | 0.905583862 | | | Mcm5 | 262.3850434 | 0.039216214 | 0.085110919 | 0.460765955 | 0.64496653 | 0.912363974 |
| Fbxw8 | 1947.195259 | -0.015298292 | 0.062226343 | -0.245849118 | 0.805799033 | 0.905632542 | | | Rbm20 | 252.8014645 | 0.039217673 | 0.092236519 | 0.425185965 | 0.670701115 | 0.918333904 |
| Tlcd2 | 10.17774983 | 0.038793143 | 0.157872424 | 0.245724633 | 0.805895402 | 0.905675155 | | | Kpna3 | 3892.249974 | 0.039243614 | 0.068062468 | 0.576582303 | 0.564221657 | 0.875323716 |
| Enpp4 | 1379.939745 | -0.019523561 | 0.079573938 | -0.245351201 | 0.80618451 | 0.9057368 | | | Epha5 | 3231.079127 | 0.039244009 | 0.059895072 | 0.655212655 | 0.512330848 | 0.852934333 |
| Gnao1 | 28590.1635 | 0.015708462 | 0.063988955 | 0.245487082 | 0.806079309 | 0.9057368 | | | Sumo1 | 3084.116667 | 0.039244974 | 0.058108214 | 0.675377393 | 0.499436025 | 0.846241311 |
| Nop16 | 606.0944139 | -0.015899021 | 0.064797958 | -0.245363 | 0.806175375 | 0.9057368 | | | Lcor | 156.7969155 | 0.039251177 | 0.093462257 | 0.41996821 | 0.674508677 | 0.919867577 |
| Prkag2 | 3259.603873 | -0.013320769 | 0.054300156 | -0.245317322 | 0.80621074 | 0.9057368 | | | Fig4 | 1541.341694 | 0.039253329 | 0.061677475 | 0.63642892 | 0.524496896 | 0.855487854 |
| Trio | 7767.073632 | -0.0234605 | 0.095649293 | -0.245276247 | 0.806242542 | 0.9057368 | | | Siah1b | 138.2267667 | 0.039270546 | 0.088123865 | 0.445628959 | 0.655865288 | 0.915208181 |
| Csgalnact2 | 462.2501273 | -0.021322509 | 0.087118622 | -0.244752595 | 0.806648002 | 0.906126596 | | | Hmgcs2 | 438.1283012 | 0.039279682 | 0.094848487 | 0.414130826 | 0.678778298 | 0.921316909 |
| Sh2b1 | 3208.352808 | 0.01631594 | 0.066732215 | 0.244498697 | 0.806844611 | 0.906281746 | | | Cdkn2d | 103.2958927 | 0.03928205 | 0.093336794 | 0.420863495 | 0.673854763 | 0.919618028 |
| Lysmd3 | 247.8475825 | 0.030085765 | 0.12337949 | 0.243847381 | 0.807349023 | 0.906773168 | | | Zbtb20 | 92.19674938 | 0.039286518 | 0.0863528 | 0.45495361 | 0.649142613 | 0.913724196 |
| Maml2 | 429.1354292 | -0.025290163 | 0.103740421 | -0.243782644 | 0.807399163 | 0.906773168 | | | Peli3 | 627.8951374 | 0.039304426 | 0.066888057 | 0.587615005 | 0.556790737 | 0.871733374 |
| Nbeal1 | 594.7405356 | 0.026996131 | 0.110901803 | 0.243423731 | 0.807677162 | 0.90698013 | | | Tm4sf5 | 10.04134989 | 0.039305703 | 0.058444266 | 0.672533105 | 0.501244372 | NA |
| Vcp | 13642.10763 | 0.01182067 | 0.048566072 | 0.243393578 | 0.807700519 | 0.90698013 | | | G6pc3 | 1902.431932 | 0.039338561 | 0.073659809 | 0.534057329 | 0.593301871 | 0.886291956 |
| Mef2a | 3269.672315 | -0.018947969 | 0.077897149 | -0.243243426 | 0.807816828 | 0.907044998 | | | Trap1 | 2960.171148 | 0.039342889 | 0.046527222 | 0.845588597 | 0.397782289 | 0.797651232 |
| Asb16 | 29.41225114 | 0.040242204 | 0.165565354 | 0.24305933 | 0.807959437 | 0.907139385 | | | Braf | 5472.075405 | 0.039376882 | 0.070178159 | 0.561098816 | 0.574730175 | 0.880182448 |
| Ddx25 | 1680.872346 | 0.01705652 | 0.07025461 | 0.242781507 | 0.808174662 | 0.907315283 | | | Spink10 | 39.2490164 | 0.03940147 | 0.091247576 | 0.431808399 | 0.665880674 | 0.915591677 |
| D7Ertd715e | 141.6731363 | 0.034304345 | 0.141421018 | 0.242568933 | 0.80833935 | 0.907434422 | | | Tmem132a | 10450.60301 | 0.039407103 | 0.079652286 | 0.494739139 | 0.62078427 | 0.899412757 |
| 1810022K09Rik | 422.6895378 | 0.024593865 | 0.101480661 | 0.242350265 | 0.808508768 | 0.907558854 | | | Dpysl2 | 6304.401706 | 0.039420475 | 0.056612136 | 0.696325523 | 0.486224993 | 0.839801621 |
| C2cd2 | 632.7923645 | 0.019562623 | 0.080838367 | 0.241996764 | 0.808782671 | 0.907669035 | | | Arhgap40 | 12.03655567 | 0.039420714 | 0.062677228 | 0.628947957 | 0.529383128 | NA |
| Gm14326 | 732.6571993 | 0.025491944 | 0.105281356 | 0.242131606 | 0.808678189 | 0.907669035 | | | Unc5d | 2605.060235 | 0.039422171 | 0.082078094 | 0.480300764 | 0.631013546 | 0.905472959 |
| Rbmx2 | 206.6897772 | 0.022295421 | 0.092124119 | 0.242015022 | 0.808768524 | 0.907669035 | | | Dmp1 | 341.0126752 | 0.039423118 | 0.089432959 | 0.440811959 | 0.659349135 | 0.915208181 |
| P2rx7 | 460.5599437 | 0.027883889 | 0.115300254 | 0.241837189 | 0.808906321 | 0.907742054 | | | Chmp3 | 4818.785208 | 0.039446725 | 0.053573245 | 0.7363139 | 0.461539689 | 0.830270578 |
| Gm10416 | 10.56428286 | 0.039244343 | 0.162400777 | 0.241651203 | 0.809050443 | 0.907838034 | | | Oxa1l | 1818.086703 | 0.039457681 | 0.051830421 | 0.761284219 | 0.446487324 | 0.822210607 |
| Ccnj | 335.8546819 | -0.02132529 | 0.088418109 | -0.241186905 | 0.80941026 | 0.907978755 | | | Zfr | 13881.19244 | 0.039468336 | 0.058240927 | 0.677673557 | 0.497978696 | 0.845489269 |
| Rprm | 1794.654288 | 0.021907808 | 0.090823312 | 0.241213488 | 0.809389658 | 0.907978755 | | | 4921536K21Rik | 20.06948696 | 0.039473386 | 0.082257619 | 0.479875134 | 0.631316183 | NA |
| Skap2 | 435.8208284 | 0.021055146 | 0.087253543 | 0.241309926 | 0.809314918 | 0.907978755 | | | Fn1 | 2002.061496 | 0.039491187 | 0.077164756 | 0.51177751 | 0.608806732 | 0.89440021 |
| Socs2 | 447.5557506 | 0.024599947 | 0.101938138 | 0.241322309 | 0.809305321 | 0.907978755 | | | Yipf6 | 1798.166531 | 0.039508626 | 0.062849893 | 0.628618832 | 0.529598628 | 0.857915133 |
| 8430408G22Rik | 45.14930867 | 0.037977505 | 0.157594194 | 0.240982896 | 0.809568373 | 0.90802464 | | | Ttc27 | 805.9985363 | 0.039536034 | 0.070771354 | 0.558644587 | 0.576404304 | 0.880966011 |
| Rhbdf2 | 67.14118425 | 0.037041503 | 0.153681788 | 0.241027281 | 0.809533972 | 0.90802464 | | | Slc27a5 | 16.30839846 | 0.039539632 | 0.079314767 | 0.498515387 | 0.618120826 | NA |
| 9130011E15Rik | 820.5687599 | -0.022410709 | 0.093116655 | -0.240673478 | 0.809808196 | 0.908187085 | | | Ube2e2 | 2404.979575 | 0.039543041 | 0.049846017 | 0.793303928 | 0.427600769 | 0.810334558 |
| Slc41a3 | 202.0083861 | 0.030241958 | 0.125670531 | 0.240644787 | 0.809830435 | 0.908187085 | | | Cyp2j8 | 3.484359187 | 0.039553871 | 0.042656221 | 0.927270879 | 0.35378591 | NA |
| Dbx2 | 546.0121797 | 0.026670073 | 0.11091535 | 0.240454701 | 0.809978087 | 0.908286928 | | | Pebp1 | 15242.63197 | 0.039560461 | 0.037132559 | 1.065384719 | 0.286701869 | 0.722148665 |
| Abcc3 | 144.0955549 | -0.029896562 | 0.1244421 | -0.240244755 | 0.81014052 | 0.908361319 | | | Fam73a | 4677.19906 | 0.039562081 | 0.0571285 | 0.692510409 | 0.488616852 | 0.840356191 |
| Mgmt | 26.83399539 | -0.040187817 | 0.167359771 | -0.240128296 | 0.810230799 | 0.908361319 | | | Snrpd2 | 1394.798995 | 0.039566755 | 0.062313553 | 0.634962262 | 0.525453028 | 0.855879197 |
| Myt1 | 440.1319134 | -0.025879841 | 0.107913178 | -0.239820955 | 0.810469062 | 0.908361319 | | | Tnfaip8l1 | 281.9734565 | 0.039582235 | 0.085311227 | 0.463974515 | 0.642666005 | 0.911258473 |
| Selplg | 1250.746319 | -0.033790217 | 0.140719857 | -0.240124018 | 0.810234115 | 0.908361319 | | | St6galnac4 | 1223.657706 | 0.03963396 | 0.053816818 | 0.736460487 | 0.461450506 | 0.830270578 |
| Tmem240 | 728.5438775 | -0.023813098 | 0.099226077 | -0.239988308 | 0.810339321 | 0.908361319 | | | Rpl6 | 9806.062646 | 0.03965496 | 0.054688081 | 0.725111567 | 0.468383589 | 0.832446623 |
| Tubg1 | 1406.213298 | 0.020160084 | 0.084040653 | 0.239884908 | 0.810419481 | 0.908361319 | | | Poldip3 | 3470.759358 | 0.039667393 | 0.042237061 | 0.939160825 | 0.347648179 | 0.766038372 |
| Wdr75 | 1053.119343 | 0.0163205 | 0.0680691 | 0.239763713 | 0.81051344 | 0.908361319 | | | Cyb5r4 | 1799.251807 | 0.0396795 | 0.073337628 | 0.541052409 | 0.58847146 | 0.885188411 |
| Zfp667 | 688.2143782 | 0.025968258 | 0.108290284 | 0.23980229 | 0.810483532 | 0.908361319 | | | Ncoa5 | 2222.023306 | 0.039681658 | 0.055099655 | 0.720179785 | 0.471414309 | 0.833245343 |
| Gm7694 | 140.4494037 | -0.033977762 | 0.141938874 | -0.239383059 | 0.810808565 | 0.908618954 | | | H2-Q10 | 3.534138123 | 0.039682142 | 0.042124202 | 0.942027163 | 0.346178733 | NA |
| Rnf114 | 2365.282758 | -0.014063509 | 0.058765451 | -0.239315936 | 0.81086061 | 0.908618954 | | | Tmem135 | 2537.888349 | 0.039698681 | 0.056542086 | 0.702108532 | 0.482611482 | 0.837880119 |
| Zfyve9 | 2509.064972 | 0.019933558 | 0.083350735 | 0.239152759 | 0.810987133 | 0.908695013 | | | Cdh3 | 15.66940397 | 0.039705888 | 0.055026371 | 0.721579261 | 0.470553194 | NA |
| 5730405O15Rik | 36.13177291 | -0.038183137 | 0.160083456 | -0.238520196 | 0.811477653 | 0.909178882 | | | Fam168b | 17363.14694 | 0.039747737 | 0.050638853 | 0.784925704 | 0.43249713 | 0.813262565 |
| Ccz1 | 1146.622472 | -0.015432467 | 0.064839978 | -0.238008524 | 0.811874483 | 0.909426203 | | | 2310061I04Rik | 1706.823394 | 0.039750519 | 0.056948003 | 0.698014273 | 0.485168268 | 0.838982004 |
| Fndc3a | 2365.846673 | 0.021832856 | 0.091717968 | 0.238043391 | 0.81184744 | 0.909426203 | | | Fam172a | 1842.565285 | 0.03975105 | 0.064964322 | 0.611890481 | 0.54061022 | 0.863847445 |
| Kat5 | 2056.108397 | 0.012601636 | 0.052943257 | 0.238021542 | 0.811864386 | 0.909426203 | | | 9030612E09Rik | 49.51962606 | 0.039752639 | 0.094142816 | 0.422258867 | 0.672836078 | 0.919353963 |
| Fzd7 | 308.3830433 | 0.020487271 | 0.086165885 | 0.237765454 | 0.812063014 | 0.909490489 | | | Tox3 | 567.1624606 | 0.039784222 | 0.084833084 | 0.468970601 | 0.639090651 | 0.909490332 |
| Gm5535 | 13.29186154 | 0.038229895 | 0.160827472 | 0.237707493 | 0.812107972 | 0.909490489 | | | Cenpb | 5443.275758 | 0.039802818 | 0.061381893 | 0.648445581 | 0.516696798 | 0.853297835 |
| Zfp84 | 813.8602586 | -0.01678109 | 0.07058587 | -0.237740073 | 0.812082701 | 0.909490489 | | | Atp2b2 | 34053.61818 | 0.039806815 | 0.065095012 | 0.61151866 | 0.540856269 | 0.863847445 |
| Ccdc19 | 62.52335278 | -0.035124605 | 0.147920055 | -0.237456679 | 0.812302523 | 0.90957688 | | | Rhpn2 | 336.8227755 | 0.039808922 | 0.082413195 | 0.483040632 | 0.629066886 | 0.904428764 |
| Pon3 | 86.29999124 | 0.03987509 | 0.167873928 | 0.237529977 | 0.812245667 | 0.90957688 | | | Slc25a38 | 698.8869466 | 0.039823337 | 0.065509561 | 0.607901134 | 0.543253048 | 0.865558163 |
| Acadl | 1250.297944 | -0.016070873 | 0.067704327 | -0.23736847 | 0.812370948 | 0.909587364 | | | Eaf1 | 1064.410219 | 0.039837781 | 0.062510151 | 0.63730099 | 0.523928807 | 0.855218546 |
| Eme2 | 545.1727203 | 0.029926144 | 0.126152539 | 0.237221893 | 0.812484653 | 0.909587364 | | | Taf13 | 784.6500024 | 0.039849588 | 0.070928375 | 0.561828568 | 0.574232826 | 0.879791496 |
| Fam212a | 48.04433735 | 0.037541272 | 0.158349011 | 0.237079297 | 0.812595273 | 0.909587364 | | | Acox1 | 10123.43949 | 0.039853268 | 0.050767238 | 0.785019423 | 0.432442181 | 0.813262565 |
| Frg1 | 733.8853626 | 0.017680713 | 0.074581323 | 0.237066225 | 0.812605414 | 0.909587364 | | | Nkap | 651.0486679 | 0.039857896 | 0.073270227 | 0.543984879 | 0.586451878 | 0.884593012 |
| Glis1 | 18.42866847 | 0.039805058 | 0.167906716 | 0.237066502 | 0.812605199 | 0.909587364 | | | Lrba | 1736.386791 | 0.039866163 | 0.068071919 | 0.585647701 | 0.558112286 | 0.872673743 |
| Abcb8 | 1765.224281 | -0.022043598 | 0.093089013 | -0.236801285 | 0.812810954 | 0.909620291 | | | Spon2 | 2.607883662 | 0.039889285 | 0.0348561 | 1.144398985 | 0.252458209 | NA |
| Prpf31 | 1207.418397 | -0.01403338 | 0.059251443 | -0.236844528 | 0.812777406 | 0.909620291 | | | Fam13b | 3325.002583 | 0.039910626 | 0.074817357 | 0.533440732 | 0.593728528 | 0.886473491 |
| Zfand3 | 2970.793702 | -0.014162939 | 0.059782326 | -0.236908466 | 0.812727802 | 0.909620291 | | | Gabra4 | 5052.292896 | 0.039913126 | 0.084012242 | 0.475087018 | 0.634724951 | 0.907297112 |
| Gjd3 | 9.349774015 | 0.037346362 | 0.157828569 | 0.236626118 | 0.812946856 | 0.909640976 | | | Gnal | 14501.02437 | 0.039915316 | 0.073963335 | 0.53966355 | 0.589429082 | 0.885188411 |
| Gpr183 | 64.07246037 | -0.035687294 | 0.150782364 | -0.23668082 | 0.812904416 | 0.909640976 | | | Zdhhc14 | 1823.234888 | 0.039973368 | 0.077920384 | 0.513002704 | 0.607949429 | 0.894004456 |
| Rptor | 3441.702703 | -0.012604632 | 0.053286835 | -0.236543082 | 0.813011281 | 0.90964737 | | | Tada3 | 1346.111409 | 0.03998609 | 0.052450805 | 0.762354163 | 0.445848655 | 0.821773578 |
| Upp1 | 55.91329225 | -0.035120577 | 0.148742565 | -0.23611652 | 0.813342255 | 0.909951975 | | | Atp6v1c2 | 59.15687189 | 0.040002386 | 0.094236348 | 0.424489984 | 0.671208508 | 0.918682257 |
| Reep6 | 390.8513979 | -0.028720251 | 0.12169528 | -0.236001352 | 0.813431621 | 0.909960249 | | | Apbb1 | 17908.69019 | 0.040011792 | 0.05234453 | 0.764393002 | 0.444633078 | 0.821493054 |
| Rimkla | 671.3257058 | -0.023916551 | 0.101360378 | -0.235955617 | 0.81346711 | 0.909960249 | | | Pkig | 2033.509431 | 0.040016254 | 0.066468363 | 0.602034592 | 0.547151111 | 0.867487125 |
| Acaa1a | 1180.3101 | 0.031708261 | 0.134450838 | 0.235835351 | 0.813560436 | 0.909998945 | | | 3300002I08Rik | 24.08072593 | 0.040061825 | 0.08553408 | 0.468372661 | 0.639518115 | 0.909767911 |
| Dak | 407.2197754 | -0.027952226 | 0.118613702 | -0.235657644 | 0.813698339 | 0.910021804 | | | Mut | 1791.671159 | 0.040074097 | 0.067190388 | 0.596426036 | 0.550890652 | 0.869642174 |
| Etl4 | 4749.685033 | 0.020836305 | 0.088411378 | 0.23567447 | 0.813685282 | 0.910021804 | | | B4galt1 | 283.7322375 | 0.04007997 | 0.089759241 | 0.446527507 | 0.655216248 | 0.915208181 |
| Mtif2 | 927.2501011 | 0.014519696 | 0.061656082 | 0.235494947 | 0.8138246 | 0.910097319 | | | Rpl4 | 17930.86843 | 0.040097282 | 0.048176649 | 0.832297031 | 0.405241306 | 0.8005027 |
| BC065397 | 101.5208191 | -0.03940141 | 0.16745964 | -0.235288992 | 0.813984438 | 0.910144684 | | | Naa30 | 1064.978752 | 0.04011755 | 0.08202064 | 0.489115296 | 0.624760073 | 0.90130746 |
| Dph2 | 413.0690843 | 0.021890244 | 0.093012767 | 0.235346661 | 0.813939682 | 0.910144684 | | | A830082N09Rik | 1046.350898 | 0.040137589 | 0.077150437 | 0.520250963 | 0.602888669 | 0.891281205 |
| 2610020H08Rik | 26.82693275 | 0.039173815 | 0.166666383 | 0.235043289 | 0.814175134 | 0.910226534 | | | Spop | 8966.12308 | 0.040142297 | 0.043060358 | 0.932233247 | 0.351215997 | 0.769244462 |
| Timm50 | 771.7919499 | 0.016365104 | 0.069615794 | 0.235077458 | 0.814148614 | 0.910226534 | | | Il11 | 15.82387403 | 0.040153796 | 0.075195936 | 0.533988908 | 0.593349208 | NA |
| Stra13 | 247.5100693 | -0.026247974 | 0.111774783 | -0.234829123 | 0.814341363 | 0.910346686 | | | Pacs1 | 4929.724261 | 0.040154168 | 0.048231631 | 0.83252769 | 0.405111155 | 0.8005027 |
| Gfra4 | 1661.769917 | -0.021220805 | 0.090498283 | -0.234488487 | 0.81460577 | 0.910517271 | | | Gys2 | 2.767438859 | 0.040156196 | 0.038747426 | 1.036357781 | 0.300035259 | NA |

| | | | | | | |
|---|---|---|---|---|---|---|
| Pitpnm2 | 12354.86163 | -0.019189518 | 0.081864653 | -0.234405416 | 0.814670255 | 0.910517271 |
| Slc24a6 | 269.630771 | 0.030227373 | 0.128953061 | 0.234406014 | 0.81466979 | 0.910517271 |
| Fbxo45 | 2777.918009 | -0.013736659 | 0.058653458 | -0.234200321 | 0.814829466 | 0.910563847 |
| Vav3 | 301.1180343 | 0.027862823 | 0.118952268 | 0.234235324 | 0.814802293 | 0.910563847 |
| 8-Sep | 8313.202878 | 0.016290897 | 0.069754941 | 0.233544696 | 0.815338466 | 0.911066941 |
| Panx2 | 3515.223801 | -0.014902183 | 0.063829811 | -0.233467445 | 0.815398446 | 0.911068258 |
| 2010001M06Rik | 70.87226574 | -0.035731332 | 0.153551066 | -0.232699992 | 0.815994375 | 0.911208422 |
| Apex2 | 132.657123 | -0.030574111 | 0.131220613 | -0.232997778 | 0.815763131 | 0.911208422 |
| Bnip1 | 579.2204776 | 0.018705499 | 0.080353195 | 0.232790984 | 0.815923714 | 0.911208422 |
| Cbr1 | 1818.429086 | -0.014332999 | 0.061484219 | -0.233116704 | 0.815670784 | 0.911208422 |
| Fbln2 | 293.6164151 | 0.030466785 | 0.13081024 | 0.232908255 | 0.815832648 | 0.911208422 |
| Fbxo2 | 2319.33533 | 0.023006039 | 0.098835488 | 0.232771041 | 0.815939202 | 0.911208422 |
| Rc3h2 | 2340.179596 | 0.022122028 | 0.09500817 | 0.232843426 | 0.81588299 | 0.911208422 |
| Sh3glb2 | 5779.828244 | -0.015530388 | 0.06668869 | -0.232878884 | 0.815855456 | 0.911208422 |
| Jph3 | 10255.47169 | 0.017815535 | 0.076627597 | 0.23249502 | 0.816153555 | 0.911320494 |
| Qpctl | 655.664884 | 0.014854835 | 0.063923596 | 0.232384223 | 0.816239602 | 0.911350897 |
| Gm10432 | 109.6548149 | 0.030360571 | 0.131253851 | 0.231311849 | 0.817072542 | 0.912215158 |
| Klf4 | 283.8481897 | 0.035243143 | 0.152560268 | 0.231011281 | 0.817306038 | 0.912344363 |
| Nphs1as | 9.720336112 | -0.031418047 | 0.135980819 | -0.231047638 | 0.817277793 | 0.912344363 |
| Rp9 | 682.0548254 | -0.018334968 | 0.079452769 | -0.23076562 | 0.817496892 | 0.912425936 |
| Snapin | 1731.801023 | -0.017664978 | 0.076539436 | -0.230795766 | 0.81747347 | 0.912425936 |
| Ubxn1 | 1877.978809 | 0.018960181 | 0.082221736 | 0.230598155 | 0.817627001 | 0.912505421 |
| Ets2 | 3283.047716 | 0.025878867 | 0.11235705 | 0.230327043 | 0.817837648 | 0.912551433 |
| Fam185a | 620.8499388 | -0.020338325 | 0.088329985 | -0.230253912 | 0.817894471 | 0.912551433 |
| Psip1 | 5497.157328 | -0.014363534 | 0.062353425 | -0.230356781 | 0.817814542 | 0.912551433 |
| Zfand2b | 524.7684696 | -0.021338639 | 0.092709747 | -0.230166085 | 0.817962715 | 0.912551433 |
| Zfp953 | 48.12744804 | -0.036018179 | 0.156485086 | -0.230170045 | 0.817959638 | 0.912551433 |
| Mettl3 | 816.4399769 | 0.023155746 | 0.100841828 | 0.229624416 | 0.818383633 | 0.912870298 |
| Nme6 | 144.4761539 | -0.025111169 | 0.109383104 | -0.229570822 | 0.818425282 | 0.912870298 |
| Zfp120 | 515.5925515 | -0.021384604 | 0.09314998 | -0.229571749 | 0.818424561 | 0.912870298 |
| Gpr150 | 130.0591477 | 0.031305161 | 0.136411199 | 0.229491135 | 0.81848721 | 0.912873656 |
| Ykt6 | 3756.089337 | -0.015977197 | 0.069657963 | -0.229366418 | 0.818584135 | 0.912916043 |
| Txnrd2 | 468.1251824 | 0.017698934 | 0.077217039 | 0.229210215 | 0.818705534 | 0.912985716 |
| Ace2 | 73.33532919 | -0.032306672 | 0.141008883 | -0.229110903 | 0.81878272 | 0.913006079 |
| Casp6 | 92.26922438 | -0.031585424 | 0.13797199 | -0.228926351 | 0.818926159 | 0.913095376 |
| Kif7 | 97.70569441 | -0.034206736 | 0.149468236 | -0.228856224 | 0.818980666 | 0.913095376 |
| Adam17 | 682.3202812 | -0.021555372 | 0.094280579 | -0.228630031 | 0.819156481 | 0.913111003 |
| Smpd5 | 16.56457916 | -0.038388687 | 0.16792163 | -0.228610733 | 0.819171482 | 0.913111003 |
| Zbtb37 | 144.2714277 | -0.030481585 | 0.133333349 | -0.228611859 | 0.819170606 | 0.913111003 |
| A230001M10Rik | 132.6031211 | -0.037426568 | 0.164137176 | -0.228020058 | 0.819630648 | 0.913228629 |
| Aar2 | 1775.926419 | 0.015982571 | 0.070348458 | 0.227191488 | 0.820274846 | 0.913228629 |
| Acbd6 | 1269.956756 | 0.015438353 | 0.067998802 | 0.227038607 | 0.820393722 | 0.913228629 |
| BC030476 | 12.03320977 | 0.036225289 | 0.158928763 | 0.22793413 | 0.819697449 | 0.913228629 |
| Cgn | 207.4320808 | -0.030766322 | 0.134776445 | -0.228276703 | 0.819431135 | 0.913228629 |
| D11Wsu47e | 311.5929534 | 0.024077341 | 0.105895611 | 0.227368641 | 0.820137103 | 0.913228629 |
| Emr1 | 311.1617796 | 0.034716481 | 0.152914274 | 0.227032309 | 0.820398619 | 0.913228629 |
| Fancb | 93.61499575 | -0.029538151 | 0.129696617 | -0.227748047 | 0.819842118 | 0.913228629 |
| Gin1 | 202.448412 | 0.023495471 | 0.102990159 | 0.228133169 | 0.819542715 | 0.913228629 |
| Ift80 | 1001.891828 | -0.015087886 | 0.066478541 | -0.226958743 | 0.820455823 | 0.913228629 |
| Kbtbd13 | 9.75476809 | 0.034117553 | 0.15006488 | 0.227352015 | 0.82015003 | 0.913228629 |
| Lin7c | 2396.304023 | -0.017735803 | 0.07784205 | -0.227843477 | 0.819767926 | 0.913228629 |
| Psmd1 | 5930.389645 | -0.015180018 | 0.066836594 | -0.227121357 | 0.820329378 | 0.913228629 |
| Serpine1 | 73.92736112 | 0.037184821 | 0.163477217 | 0.227461793 | 0.820064676 | 0.913228629 |
| Slc29a1 | 865.8956978 | -0.022505962 | 0.099013553 | -0.227301836 | 0.820189045 | 0.913228629 |
| Snrpn | 52.58330523 | 0.03812647 | 0.167283391 | 0.227915454 | 0.819711968 | 0.913228629 |
| Ssh1 | 499.5685849 | 0.018447213 | 0.080983604 | 0.227789484 | 0.819809902 | 0.913228629 |
| Tmem131 | 3696.317512 | -0.012648549 | 0.055657091 | -0.227258539 | 0.82022271 | 0.913228629 |
| Zfp13 | 578.302945 | 0.020042739 | 0.088062531 | 0.227596676 | 0.819959804 | 0.913228629 |
| Zfp830 | 342.4723617 | -0.017693574 | 0.077660194 | -0.227833244 | 0.819775881 | 0.913228629 |
| Apex1 | 1189.181777 | -0.012294239 | 0.054218614 | -0.226753107 | 0.820615729 | 0.913341003 |
| Acot3 | 80.16038207 | -0.03229615 | 0.142685268 | -0.226345369 | 0.820932814 | 0.913628288 |
| 0610037L13Rik | 961.0195021 | 0.028150491 | 0.124414214 | 0.226264265 | 0.82099589 | 0.91363286 |
| Clstn3 | 18337.68636 | -0.012933957 | 0.057199697 | -0.226119325 | 0.821108615 | 0.91369268 |
| Gtf3c6 | 534.8002293 | -0.018870975 | 0.083519179 | -0.225947796 | 0.821242023 | 0.913775505 |
| Cnbd2 | 176.9133733 | 0.024963457 | 0.11054244 | 0.225826906 | 0.82133605 | 0.913814502 |
| Fam175b | 1221.178758 | -0.012544703 | 0.055570421 | -0.225744241 | 0.821400347 | 0.913820419 |
| Slc35a3 | 464.3128657 | -0.020108784 | 0.089131417 | -0.225608257 | 0.821506118 | 0.913872472 |
| AW011738 | 159.0116633 | -0.025185838 | 0.111723665 | -0.225429752 | 0.821644969 | 0.913961314 |
| Pskh1 | 442.4544604 | -0.01945868 | 0.086497008 | -0.224963623 | 0.822007574 | 0.914233389 |
| Zfp119a | 92.17451907 | 0.033852444 | 0.150430878 | 0.225036536 | 0.821950851 | 0.914233389 |
| Pkd2 | 1478.201243 | 0.017296769 | 0.077079106 | 0.224402825 | 0.822443872 | 0.914652983 |
| Cox7a1 | 36.86101703 | -0.037672863 | 0.168041956 | -0.224187246 | 0.822611607 | 0.914708213 |
| Cyp4f16 | 135.6896764 | -0.028713296 | 0.128060833 | -0.224216066 | 0.822589183 | 0.914708213 |
| Ezr | 2520.595334 | 0.02595503 | 0.115944696 | 0.223856985 | 0.822868587 | 0.91486265 |
| Ppapdc3 | 914.4584521 | 0.016735207 | 0.074754475 | 0.223868966 | 0.822859264 | 0.91486265 |
| Pkm | 50060.38061 | 0.015551355 | 0.069557631 | 0.223575109 | 0.823087933 | 0.915040858 |
| Gm10125 | 48.28660933 | -0.035837892 | 0.160397293 | -0.223432025 | 0.823199282 | 0.915098987 |
| Lrmp | 67.01253871 | -0.03374186 | 0.15113401 | -0.223257889 | 0.8233348 | 0.915183972 |
| Cd209b | 9.024613783 | 0.03069995 | 0.13761002 | 0.223093855 | 0.823462461 | 0.915205309 |
| Mog | 1972.154735 | 0.037406527 | 0.167681043 | 0.223081428 | 0.823472132 | 0.915205309 |
| Slc35b2 | 815.8139418 | -0.016605151 | 0.074482858 | -0.222939231 | 0.823582803 | 0.915262656 |
| A230072C01Rik | 299.285 | 0.027294379 | 0.122714084 | 0.22242255 | 0.823984959 | 0.915643904 |
| Ap2a2 | 10279.06416 | -0.015042637 | 0.067819235 | -0.221804881 | 0.82446578 | 0.915701468 |
| Apoa1bp | 1451.300584 | 0.014152412 | 0.06388748 | 0.2215209 | 0.824686865 | 0.915701468 |
| Calcrl | 194.14616 | -0.028580848 | 0.128728412 | -0.222024399 | 0.82429489 | 0.915701468 |
| Cetn3 | 1171.451407 | 0.015071566 | 0.067970755 | 0.221736047 | 0.824519367 | 0.915701468 |
| Dmrta1 | 9.124115221 | -0.032765252 | 0.14777958 | -0.22171705 | 0.824534157 | 0.915701468 |
| Eif2b5 | 2208.508654 | -0.017147889 | 0.077396606 | -0.22155867 | 0.824657459 | 0.915701468 |
| Elmo1 | 6678.457418 | -0.018879506 | 0.085186114 | -0.221626568 | 0.824604599 | 0.915701468 |
| Pex14 | 1775.004927 | 0.016930041 | 0.076253135 | 0.22202419 | 0.824295052 | 0.915701468 |
| Pnpla1 | 23.490394 | -0.036743955 | 0.16572521 | -0.221716145 | 0.824534861 | 0.915701468 |
| Slc31a1 | 884.4683802 | 0.019581862 | 0.088287573 | 0.221796352 | 0.82447242 | 0.915701468 |
| Zfp524 | 169.7678777 | -0.023540157 | 0.106180739 | -0.221698939 | 0.824548256 | 0.915701468 |
| Sos2 | 1631.220216 | -0.018247847 | 0.082407788 | -0.22143352 | 0.824754895 | 0.915711382 |
| 8430410A17Rik | 586.9472905 | -0.022126569 | 0.100051156 | -0.221152561 | 0.824973646 | 0.915793302 |
| Ech1 | 1305.745837 | 0.015069385 | 0.068199855 | 0.220959191 | 0.825124209 | 0.915793302 |
| Noc3l | 610.2383296 | 0.019216023 | 0.086960199 | 0.220974916 | 0.825111964 | 0.915793302 |
| Tmem192 | 554.5327578 | 0.019956735 | 0.090234149 | 0.221166106 | 0.824963099 | 0.915793302 |
| Zfp68 | 1415.394268 | -0.01553688 | 0.070306083 | -0.220989127 | 0.825100899 | 0.915793302 |
| Cacna1d | 2213.955075 | -0.019713415 | 0.089312035 | -0.220725181 | 0.825306423 | 0.915825236 |
| Dffa | 1658.795263 | 0.013582649 | 0.061546547 | 0.220689048 | 0.825334559 | 0.915825236 |
| Mastl | 43.73873704 | 0.033599777 | 0.152350442 | 0.220542697 | 0.825448523 | 0.915825236 |
| N4bp2l2 | 1212.088554 | -0.020065252 | 0.090951151 | -0.220615706 | 0.825391671 | 0.915825236 |
| Usp11 | 6887.169032 | 0.014261618 | 0.064633294 | 0.220654355 | 0.825361575 | 0.915825236 |
| Hhat | 26.66258117 | 0.036590543 | 0.16637129 | 0.219933036 | 0.825923307 | 0.91628639 |
| Fermt2 | 3400.889783 | -0.011483916 | 0.052293241 | -0.219606135 | 0.826177913 | 0.916503228 |
| Mphosph6 | 358.5510027 | -0.018627255 | 0.084953882 | -0.219263143 | 0.826445071 | 0.916733959 |
| Uros | 757.3493189 | -0.024888297 | 0.113645854 | -0.218998723 | 0.826651043 | 0.91689679 |
| A430005L14Rik | 456.9157829 | 0.022544573 | 0.103241431 | 0.218367496 | 0.827142789 | 0.916976744 |
| Anp32e | 4071.882198 | -0.011881593 | 0.054312982 | -0.21876157 | 0.826835784 | 0.916976744 |
| Arhgef9 | 18781.64348 | 0.012619225 | 0.057767821 | 0.218447317 | 0.827080602 | 0.916976744 |
| Arid3a | 279.5531646 | -0.018068853 | 0.082824181 | -0.21815915 | 0.827305112 | 0.916976744 |
| Clec10a | 29.32471976 | -0.036271345 | 0.165873006 | -0.218669365 | 0.826907615 | 0.916976744 |
| Fgfbp3 | 127.2092524 | -0.028315557 | 0.129568366 | -0.218537576 | 0.827010284 | 0.916976744 |
| Fus | 7776.590099 | 0.021780879 | 0.099845184 | 0.218146518 | 0.827314954 | 0.916976744 |
| Grtp1 | 90.68636162 | 0.029594553 | 0.13546735 | 0.218462622 | 0.827068678 | 0.916976744 |
| Kpna3 | 3486.819845 | -0.017076161 | 0.078247853 | -0.21823169 | 0.827248595 | 0.916976744 |
| Slc25a17 | 1080.913507 | -0.014701002 | 0.067318804 | -0.218378832 | 0.827133957 | 0.916976744 |
| H2afj | 594.9409748 | -0.0251402 | 0.115376466 | -0.217897126 | 0.827509267 | 0.917126508 |
| Ece1 | 2557.885601 | -0.011187121 | 0.051427615 | -0.217531392 | 0.827794245 | 0.917158757 |
| Fgfbp1 | 81.39850415 | 0.036497573 | 0.167899248 | 0.217377822 | 0.827913914 | 0.917158757 |
| Kif13a | 1513.942364 | 0.015240475 | 0.069983873 | 0.217771237 | 0.827607356 | 0.917158757 |
| Tchh | 250.5199436 | 0.022654972 | 0.104188056 | 0.217443082 | 0.82786306 | 0.917158757 |
| Tmem168 | 555.349853 | -0.017334725 | 0.079762956 | -0.217328016 | 0.827952725 | 0.917158757 |
| Tmem48 | 439.3963653 | 0.019272362 | 0.088631276 | 0.217444258 | 0.827862144 | 0.917158757 |
| Zfp746 | 1047.750062 | 0.015811235 | 0.072665163 | 0.217590304 | 0.82774834 | 0.917158757 |
| Zbtb26 | 319.0922871 | 0.020099973 | 0.092662154 | 0.216916748 | 0.828273226 | 0.917448195 |
| Ubash3b | 1253.519859 | 0.016701404 | 0.07708067 | 0.216674354 | 0.828462135 | 0.917591845 |
| Gfm1 | 1653.999973 | -0.014109718 | 0.065151547 | -0.216567663 | 0.828545288 | 0.917618349 |
| Zfp772 | 445.3973207 | -0.021030623 | 0.09741382 | -0.215889518 | 0.829073867 | 0.918138125 |
| 5930412G12Rik | 70.66179274 | 0.036231139 | 0.167991448 | 0.21567252 | 0.829243023 | 0.91825982 |
| 9030624J02Rik | 2154.927746 | -0.012903503 | 0.059884267 | -0.215474012 | 0.829397772 | 0.918365546 |
| Igf2as | 10.84695316 | -0.033791035 | 0.157088337 | -0.215108492 | 0.829682733 | 0.918615425 |
| Keap1 | 2720.706465 | 0.013782257 | 0.064141981 | 0.214871083 | 0.829867831 | 0.91875471 |
| Creb3l1 | 117.2056915 | 0.035009217 | 0.163265396 | 0.214431337 | 0.830210707 | 0.918979996 |
| Fcna | 10.64088507 | 0.029282414 | 0.136497986 | 0.214526344 | 0.830136625 | 0.918979996 |
| Ubr3 | 8552.910562 | -0.017065179 | 0.07960177 | -0.214381898 | 0.830249257 | 0.918979996 |
| Taf12 | 466.4672604 | -0.019114225 | 0.089223697 | -0.214228123 | 0.830369166 | 0.919047065 |
| Ghr | 352.1717997 | 0.022913285 | 0.107035289 | 0.214072249 | 0.830490716 | 0.91911594 |
| Ctr9 | 2314.529318 | -0.016072072 | 0.075189798 | -0.213753357 | 0.8307394 | 0.919325496 |
| Shprh | 1739.053546 | 0.019407841 | 0.090898692 | 0.213510672 | 0.830928666 | 0.919469273 |
| Dnahc7b | 165.7432123 | 0.035021406 | 0.164214317 | 0.213266462 | 0.831119131 | 0.919483028 |
| Oprd1 | 1006.069683 | -0.017656512 | 0.082779324 | -0.213296157 | 0.831095971 | 0.919483028 |
| Sacs | 299.8026036 | -0.027650661 | 0.129597284 | -0.213358334 | 0.831047477 | 0.919483028 |
| Bag5 | 1803.76061 | -0.020271723 | 0.095099351 | -0.213163635 | 0.831199331 | 0.9195061 |
| Atp6v1a | 20267.80659 | 0.015979285 | 0.074989406 | 0.213087229 | 0.831258925 | 0.919506375 |
| Uri1 | 1439.924126 | -0.019665783 | 0.092333171 | -0.212987191 | 0.831336953 | 0.919527039 |
| Cd3eap | 824.7792293 | 0.015131191 | 0.071068581 | 0.212909707 | 0.831397391 | 0.919528244 |
| 2310030G06Rik | 84.12084755 | 0.027802062 | 0.130817285 | 0.212525909 | 0.831696767 | 0.919597921 |
| 9430023L20Rik | 812.3570065 | -0.019856362 | 0.093497838 | -0.212372419 | 0.831816501 | 0.919597921 |
| Cml5 | 57.47623243 | -0.034976549 | 0.164406803 | -0.212743927 | 0.831526702 | 0.919597921 |
| Tbc1d13 | 2051.516885 | -0.01388035 | 0.06533844 | -0.21243773 | 0.831765553 | 0.919597921 |
| Wbscr16 | 437.0924488 | 0.017751642 | 0.083536779 | 0.212500916 | 0.831716263 | 0.919597921 |
| Zdhhc3 | 3233.997734 | 0.010844375 | 0.050996186 | 0.212650708 | 0.831599416 | 0.919597921 |
| Wbp1l | 1739.545726 | -0.014667344 | 0.069161712 | -0.212073175 | 0.832049947 | 0.919790374 |
| B3gnt9 | 128.2517424 | 0.026427958 | 0.124808178 | 0.211748607 | 0.832303165 | 0.919939024 |
| Exosc3 | 197.2737878 | -0.020182908 | 0.095296818 | -0.211789947 | 0.832270912 | 0.919939024 |
| Kif14 | 15.81632744 | -0.035101413 | 0.166075669 | -0.211357951 | 0.832607966 | 0.920210273 |
| 1810011O10Rik | 35.76285289 | 0.035498476 | 0.168100468 | 0.211174167 | 0.832751368 | 0.920237479 |
| Stk38l | 1409.986719 | 0.012957765 | 0.061354675 | 0.211194417 | 0.832735567 | 0.920237479 |
| Patz1 | 1092.522772 | -0.01448719 | 0.068720439 | -0.210813406 | 0.833032877 | 0.920482912 |
| Stat5a | 76.86898896 | 0.030553571 | 0.145030427 | 0.210670079 | 0.833144724 | 0.920488074 |
| Zbtb45 | 909.1845817 | -0.014657752 | 0.069581739 | -0.21065516 | 0.833156367 | 0.920488074 |
| Pcyt2 | 2424.828928 | 0.017239281 | 0.081895789 | 0.210502656 | 0.833275379 | 0.920520216 |
| Wdr26 | 5847.944787 | 0.016757262 | 0.07961995 | 0.21046562 | 0.833304282 | 0.920520216 |
| Eif2c4 | 663.3262626 | 0.017871374 | 0.085031281 | 0.210174108 | 0.833531787 | 0.920705889 |
| Dnph1 | 79.79991871 | -0.026495651 | 0.126241235 | -0.209881116 | 0.833760461 | 0.920864051 |
| Fhl3 | 34.28614244 | 0.034266301 | 0.163357842 | 0.209762203 | 0.833853274 | 0.920864051 |
| Smim11 | 251.8271336 | 0.025670403 | 0.122344002 | 0.209821506 | 0.833806988 | 0.920864051 |
| Prdm16 | 533.4657366 | -0.018464537 | 0.08817614 | -0.20940514 | 0.834131981 | 0.921106187 |
| Ccdc15 | 79.06884742 | 0.029632725 | 0.141722755 | 0.209089393 | 0.834378455 | 0.921120761 |
| Egln2 | 2669.206408 | 0.017156881 | 0.082087415 | 0.209007442 | 0.834442429 | 0.921120761 |
| Praf2 | 2754.444032 | -0.012605055 | 0.06028999 | -0.209073761 | 0.834390657 | 0.921120761 |
| Rnmtl1 | 350.6032545 | 0.017689499 | 0.084601629 | 0.20909171 | 0.834376646 | 0.921120761 |
| Whsc1 | 3905.255635 | -0.014087498 | 0.067305862 | -0.209305667 | 0.834209628 | 0.921120761 |
| Gm9866 | 938.384012 | 0.014150273 | 0.067736359 | 0.208902177 | 0.834524605 | 0.921145845 |
| Blzf1 | 819.136785 | 0.013063865 | 0.062564455 | 0.208806506 | 0.834599293 | 0.921162661 |
| Taf3 | 774.0498192 | 0.020160973 | 0.096642962 | 0.208612942 | 0.834750406 | 0.921258727 |
| Vrk1 | 609.9927974 | 0.018709646 | 0.089716144 | 0.208542695 | 0.834805249 | 0.921258727 |
| D3Ertd254e | 1196.647647 | 0.013372334 | 0.064330296 | 0.207869934 | 0.835330525 | 0.921732596 |
| Klf11 | 345.8746993 | -0.025068645 | 0.12061491 | -0.207840347 | 0.835353628 | 0.921732596 |
| Zhx1 | 3147.920426 | 0.011156568 | 0.053700954 | 0.20775363 | 0.83542134 | 0.921741668 |
| Ephx4 | 1872.042929 | 0.017384738 | 0.08389211 | 0.20722733 | 0.835832324 | 0.922129452 |
| Rfc5 | 333.0806462 | -0.019786364 | 0.095521352 | -0.207140746 | 0.835899942 | 0.922138391 |
| Zfp62 | 1244.622908 | 0.017404757 | 0.084112472 | 0.206922423 | 0.836070445 | 0.922260819 |
| P2ry14 | 93.23992548 | 0.027735606 | 0.134142432 | 0.206762364 | 0.83619545 | 0.922262177 |
| Pipox | 71.6765521 | -0.034739045 | 0.168071385 | -0.206692204 | 0.836250246 | 0.922262177 |
| Rab2a | 8426.314297 | -0.01244967 | 0.060195116 | -0.20682193 | 0.836148929 | 0.922262177 |
| Dpys | 11.14665628 | 0.033847309 | 0.16387341 | 0.206545458 | 0.83636486 | 0.922322928 |
| Kctd11 | 83.42781557 | -0.027842401 | 0.134898816 | -0.206394704 | 0.836482607 | 0.922387126 |
| Pop4 | 774.3721191 | -0.013433448 | 0.0651506 | -0.206190709 | 0.836641945 | 0.922497175 |
| Fam217a | 31.35323975 | 0.034145353 | 0.165813146 | 0.205926694 | 0.836848174 | 0.922658906 |
| Mogs | 1004.493087 | -0.012914657 | 0.062741777 | -0.205838244 | 0.836917266 | 0.922669427 |
| Cyb5d2 | 541.4193384 | -0.019348723 | 0.094058563 | -0.205709318 | 0.83701798 | 0.922697724 |
| Fezf2 | 845.3938196 | -0.02729931 | 0.132744576 | -0.205652919 | 0.837062038 | 0.922697724 |
| BC052688 | 25.97601984 | 0.034506101 | 0.168095561 | 0.205276693 | 0.837355952 | 0.922767037 |
| Gpr182 | 77.84557444 | 0.034045223 | 0.166275571 | 0.204751803 | 0.837766045 | 0.922767037 |
| Kdm3b | 2668.277346 | -0.011036058 | 0.053892585 | -0.204778786 | 0.837744962 | 0.922767037 |
| Lor | 122.8689931 | 0.0306024 | 0.149518053 | 0.204673612 | 0.837827138 | 0.922767037 |
| Nln | 1236.738217 | -0.014556002 | 0.07103154 | -0.204923078 | 0.837632224 | 0.922767037 |
| Nova1 | 2308.875951 | -0.014838123 | 0.072398483 | -0.204950739 | 0.837610613 | 0.922767037 |
| Prkcg | 11518.57231 | -0.017296681 | 0.084465682 | -0.204777614 | 0.837745878 | 0.922767037 |
| Rad17 | 536.5242237 | -0.013012783 | 0.063583169 | -0.20465767 | 0.837839594 | 0.922767037 |
| Rgs3 | 1143.772897 | -0.01932083 | 0.094237874 | -0.205021921 | 0.837554998 | 0.922767037 |
| Sftpc | 26.13468982 | -0.034442023 | 0.168091537 | -0.204900398 | 0.837649945 | 0.922767037 |
| Spty2d1 | 850.3143967 | 0.018193092 | 0.088763003 | 0.204962559 | 0.837601377 | 0.922767037 |
| Tiparp | 676.3074774 | 0.029035233 | 0.141764924 | 0.204812531 | 0.837718596 | 0.922767037 |
| Gpr107 | 2040.234263 | -0.010773115 | 0.052662226 | -0.204570069 | 0.837908042 | 0.922776829 |
| Cct8 | 6209.580887 | -0.011888253 | 0.058287215 | -0.203959869 | 0.83838486 | 0.922846414 |
| Fam13b | 3080.89765 | 0.016367692 | 0.080261258 | 0.203930176 | 0.838408063 | 0.922846414 |
| Pik3r5 | 168.5248051 | 0.020483429 | 0.100377886 | 0.204063167 | 0.838304137 | 0.922846414 |
| Prr15 | 29.20239371 | -0.033680989 | 0.165162709 | -0.203926113 | 0.838411238 | 0.922846414 |
| Rgmb | 1773.137858 | -0.014780339 | 0.072399842 | -0.204148769 | 0.838237244 | 0.922846414 |
| Stpg1 | 55.08842667 | 0.033551728 | 0.164566518 | 0.203879431 | 0.838447719 | 0.922846414 |
| Trub2 | 899.9603082 | -0.012421951 | 0.06091361 | -0.203927345 | 0.838410276 | 0.922846414 |
| Zhx3 | 1364.20454 | -0.024919567 | 0.122048912 | -0.20417689 | 0.83821527 | 0.922846414 |
| Hpgds | 243.5163283 | -0.030655793 | 0.150473297 | -0.203729124 | 0.838565182 | 0.922910138 |
| Nudt7 | 145.790949 | -0.024818126 | 0.12196453 | -0.203486426 | 0.838754854 | 0.922948524 |
| Rrp12 | 1550.121614 | -0.016532463 | 0.081255207 | -0.203463423 | 0.838772832 | 0.922948524 |
| Timp1 | 14.35168281 | 0.030730609 | 0.151043143 | 0.203455833 | 0.838778764 | 0.922948524 |
| Atoh7 | 12.08333944 | -0.032941739 | 0.162168381 | -0.203132933 | 0.839031132 | 0.923095104 |
| Dnajb6 | 5090.944604 | -0.009920257 | 0.048832859 | -0.203147176 | 0.839019999 | 0.923095104 |
| Ankrd34c | 318.3573853 | -0.022384585 | 0.110400978 | -0.202757121 | 0.839324874 | 0.923352712 |
| Amigo1 | 5297.982592 | -0.007814362 | 0.038659396 | -0.202133587 | 0.839812291 | 0.923775413 |
| Gm2897 | 9.349600119 | 0.032099766 | 0.158820852 | 0.202113044 | 0.839828351 | 0.923775413 |
| Camsap1 | 5447.76945 | -0.017366465 | 0.086078348 | -0.20175184 | 0.840110733 | 0.923777635 |
| Fggy | 407.6595333 | -0.028003081 | 0.138815249 | -0.201729141 | 0.840128479 | 0.923777635 |
| Gjc2 | 457.5489394 | -0.033103777 | 0.163944241 | -0.201920949 | 0.839978525 | 0.923777635 |
| Inpp5d | 752.5291971 | -0.017911914 | 0.088764424 | -0.201791593 | 0.840079654 | 0.923777635 |
| Ttc38 | 355.9054899 | 0.017619569 | 0.08729119 | 0.201848199 | 0.8400354 | 0.923777635 |
| Rab3il1 | 388.1476444 | -0.022510052 | 0.111777417 | -0.201382828 | 0.840399241 | 0.924009782 |
| Gipc2 | 48.14432263 | 0.029903241 | 0.148723883 | 0.201065493 | 0.840647364 | 0.924217005 |
| Dhx30 | 4709.26039 | 0.011286047 | 0.056246284 | 0.200654085 | 0.840969064 | 0.924293811 |
| Foxo4 | 785.9827068 | -0.017644205 | 0.087992941 | -0.200518419 | 0.841075154 | 0.924293811 |
| Gpr153 | 1197.687537 | -0.024332031 | 0.121209407 | -0.200743752 | 0.840898947 | 0.924293811 |

| | | | | | | |
|---|---|---|---|---|---|---|
| Tyro3 | 11049.08311 | 0.040164724 | 0.070354232 | 0.570892802 | 0.56807231 | 0.877106545 |
| Mcm2 | 213.3250939 | 0.04017676 | 0.093349125 | 0.430392468 | 0.666910173 | 0.916077199 |
| Psip1 | 6134.342547 | 0.040181013 | 0.059973949 | 0.669974438 | 0.502874086 | 0.847418674 |
| Tmem47 | 3358.238133 | 0.040182432 | 0.059627937 | 0.673886007 | 0.500383789 | 0.846268338 |
| Skp1a | 11697.12948 | 0.040198501 | 0.065400344 | 0.61465274 | 0.538784077 | 0.862393479 |
| Copb1 | 2748.780706 | 0.040200633 | 0.063447384 | 0.633605841 | 0.526338088 | 0.855977056 |
| Elavl1 | 3163.139415 | 0.040217836 | 0.055246115 | 0.727975818 | 0.46662839 | 0.832095317 |
| Pecam1 | 940.1205918 | 0.040227393 | 0.079230355 | 0.507727029 | 0.61164479 | 0.895166661 |
| Rabgap1 | 4751.349746 | 0.040234117 | 0.056284868 | 0.714830086 | 0.474714042 | 0.834491965 |
| Hist1h2be | 73.53797018 | 0.040258694 | 0.095038419 | 0.423604418 | 0.671854333 | 0.919094512 |
| Usp10 | 2397.539498 | 0.040276337 | 0.045536082 | 0.884492811 | 0.376430241 | 0.783947144 |
| Maml1 | 566.0718395 | 0.040297053 | 0.069538529 | 0.579492457 | 0.562256935 | 0.874635279 |
| Cd47 | 2415.201853 | 0.040308852 | 0.072124701 | 0.558877219 | 0.576245518 | 0.880966011 |
| Nudt9 | 1738.601295 | 0.040311279 | 0.056548205 | 0.712865769 | 0.475928823 | 0.835195112 |
| Ttc21a | 95.80847237 | 0.040313102 | 0.093160312 | 0.432728283 | 0.66521218 | 0.915428982 |
| Mrps11 | 607.1378782 | 0.040320727 | 0.06681116 | 0.603502867 | 0.546174209 | 0.867128614 |
| Rab15 | 12818.25421 | 0.040328444 | 0.062527445 | 0.644971877 | 0.518945411 | 0.853297835 |
| Dhcr7 | 1410.856342 | 0.040340107 | 0.072952377 | 0.552964952 | 0.58028741 | 0.883332604 |
| Nr4a1 | 4042.832383 | 0.040358992 | 0.069795742 | 0.578244323 | 0.563099178 | 0.87510382 |
| Ppp2r5c | 8689.531824 | 0.040375775 | 0.048060474 | 0.840103553 | 0.400850328 | 0.799169897 |
| 1300017J02Rik | 2.471300781 | 0.040393216 | 0.036901097 | 1.094634559 | 0.273676769 | NA |
| Tcf20 | 5061.403951 | 0.040400387 | 0.065942172 | 0.612663881 | 0.540098609 | 0.863544778 |
| Atp10a | 781.7540844 | 0.040408338 | 0.084796803 | 0.476531387 | 0.633695852 | 0.906932171 |
| Cd164l2 | 127.5729421 | 0.040427872 | 0.091352655 | 0.44254732 | 0.658093196 | 0.915208181 |
| Cdk5r1 | 13027.1673 | 0.040445446 | 0.061734647 | 0.655149872 | 0.512371265 | 0.852934333 |
| Uchl3 | 773.9448398 | 0.040461246 | 0.076816696 | 0.52672463 | 0.598384829 | 0.888717024 |
| Fam69a | 1070.903349 | 0.040463201 | 0.0799354 | 0.506198766 | 0.612717121 | 0.895428201 |
| Rp9 | 732.2698952 | 0.040466267 | 0.070608347 | 0.573108826 | 0.566571012 | 0.875959314 |
| Igf1r | 3143.198715 | 0.040466999 | 0.072334867 | 0.559439736 | 0.575861649 | 0.880811222 |
| Arhgef18 | 2528.739716 | 0.040482181 | 0.048396921 | 0.836461925 | 0.402895088 | 0.800009209 |
| Zbtb2 | 430.9781667 | 0.040491493 | 0.079443224 | 0.509690955 | 0.610267991 | 0.894800514 |
| Abi2 | 9618.214371 | 0.040549822 | 0.071861549 | 0.564277044 | 0.572565602 | 0.879277718 |
| Cmc1 | 346.3058325 | 0.040595504 | 0.076593118 | 0.530015035 | 0.596101507 | 0.88782636 |
| Stub1 | 4233.447462 | 0.040597821 | 0.044950563 | 0.903166023 | 0.366437787 | 0.778210102 |
| Btbd17 | 534.9062839 | 0.040657541 | 0.079275681 | 0.51286271 | 0.60804736 | 0.894004456 |
| Trmt112 | 762.4916843 | 0.040659358 | 0.069819414 | 0.58235032 | 0.560330737 | 0.873713575 |
| Ube2f | 1189.314176 | 0.040665566 | 0.058198663 | 0.698737122 | 0.48471633 | 0.838713365 |
| BC031361 | 259.2318633 | 0.040667173 | 0.082334998 | 0.493923288 | 0.621360357 | 0.899589902 |
| Klhdc8b | 1044.528724 | 0.040691219 | 0.076213182 | 0.533913132 | 0.593401636 | 0.886313552 |
| Gm16861 | 31.75664318 | 0.040694394 | 0.084107433 | 0.483838256 | 0.628500663 | 0.904099884 |
| Mep1a | 2.830565892 | 0.040697417 | 0.035489596 | 1.146742197 | 0.251488185 | NA |
| Mrpl10 | 2045.047854 | 0.040714385 | 0.054679911 | 0.744594939 | 0.456516629 | 0.82674985 |
| Psmb3 | 2002.353287 | 0.040736095 | 0.057522341 | 0.708178663 | 0.478834315 | 0.835936459 |
| Wt1 | 5.622697468 | 0.040779382 | 0.043018789 | 0.947943518 | 0.343158209 | NA |
| Stil | 32.77807842 | 0.040793501 | 0.09053666 | 0.450574396 | 0.652296324 | 0.914611624 |
| Fech | 2563.40109 | 0.040793603 | 0.047450228 | 0.859713515 | 0.389946983 | 0.792995922 |
| Gna12 | 2189.563876 | 0.040831929 | 0.067748907 | 0.602695015 | 0.546711599 | 0.867128614 |
| Cdkl1 | 586.1453496 | 0.040884736 | 0.074821624 | 0.54642941 | 0.584770798 | 0.884313345 |
| Htr5a | 1224.273156 | 0.040964679 | 0.063284326 | 0.647311618 | 0.517430284 | 0.853297835 |
| Ndufb6 | 1618.016413 | 0.040964807 | 0.059238855 | 0.69151923 | 0.4892393 | 0.840836703 |
| Nxf2 | 2.781649977 | 0.040968716 | 0.037990474 | 1.07839443 | 0.280857772 | NA |
| Zfp622 | 1188.747025 | 0.040973171 | 0.064894154 | 0.631384619 | 0.52778907 | 0.856965327 |
| AW146154 | 154.9836677 | 0.040974541 | 0.089515082 | 0.457738963 | 0.647139992 | 0.912750424 |
| Naf1 | 456.1947769 | 0.040981437 | 0.084572387 | 0.48457231 | 0.627979759 | 0.903708538 |
| Gpsm1 | 2152.435043 | 0.040985857 | 0.057659727 | 0.710822936 | 0.477193966 | 0.835247566 |
| Tuba4a | 24712.82242 | 0.040992911 | 0.052307237 | 0.783694816 | 0.433219205 | 0.813783127 |
| Hira | 2847.834971 | 0.040998725 | 0.050156965 | 0.81740841 | 0.413695071 | 0.804247137 |
| Ripply2 | 58.78617703 | 0.041004803 | 0.094493619 | 0.433942558 | 0.664330154 | 0.915428982 |
| Col10a1 | 16.75788107 | 0.041010318 | 0.075961361 | 0.539883928 | 0.589277082 | NA |
| Snx2 | 2214.221924 | 0.041017544 | 0.065330745 | 0.627844417 | 0.530105866 | 0.857915133 |
| Samd4b | 3655.579713 | 0.041025902 | 0.072065662 | 0.569285024 | 0.569162729 | 0.87745335 |
| Abcf3 | 3078.36581 | 0.041045209 | 0.044521381 | 0.921921276 | 0.356569641 | 0.773064269 |
| Ext2 | 1642.06644 | 0.04106843 | 0.049409135 | 0.831191037 | 0.405865717 | 0.800733644 |
| Lamp1 | 13801.0205 | 0.041070378 | 0.041271976 | 0.995115372 | 0.319680155 | 0.749632194 |
| Myl2 | 11.47400791 | 0.041087334 | 0.070802747 | 0.58030706 | 0.561707567 | NA |
| Ndufa6 | 2063.605394 | 0.04110073 | 0.064653625 | 0.635706506 | 0.524967735 | 0.855664421 |
| Grsf1 | 4335.633866 | 0.041110607 | 0.066446202 | 0.61870515 | 0.536110617 | 0.861072846 |
| Ube2d2a | 5972.446498 | 0.041113235 | 0.053281541 | 0.771622474 | 0.44033806 | 0.818470973 |
| Tspan1 | 4.926805836 | 0.041132059 | 0.047784147 | 0.860788809 | 0.38935437 | NA |
| Tusc1 | 74.93072418 | 0.041160684 | 0.094652593 | 0.434860605 | 0.663663611 | 0.915428982 |
| Eif3e | 2830.372619 | 0.041203037 | 0.062881887 | 0.655244922 | 0.512310076 | 0.852934333 |
| Mrps26 | 1301.807157 | 0.041203434 | 0.05963317 | 0.690948242 | 0.489598068 | 0.841119426 |
| Fam221b | 54.04660429 | 0.041236049 | 0.094702633 | 0.435426634 | 0.663252781 | 0.915428982 |
| Rprd1a | 5534.724745 | 0.041257325 | 0.062127576 | 0.664074278 | 0.506642772 | 0.848943697 |
| Fxr2 | 5451.267112 | 0.041291746 | 0.044816244 | 0.921356697 | 0.356864229 | 0.773088648 |
| Prss54 | 9.870856792 | 0.04129451 | 0.065107762 | 0.634248654 | 0.52591856 | NA |
| Car14 | 563.5479475 | 0.041313252 | 0.094670531 | 0.436389772 | 0.662553958 | 0.915428982 |
| Asb18 | 13.53016113 | 0.041329293 | 0.075540725 | 0.547112745 | 0.584301277 | NA |
| Tmed9 | 3910.371599 | 0.041347551 | 0.047849334 | 0.864119669 | 0.387522155 | 0.791583702 |
| 9530080O11Rik | 62.95875076 | 0.041409435 | 0.094926785 | 0.436224987 | 0.6626735 | 0.915428982 |
| Gdpd2 | 363.143587 | 0.041431818 | 0.091115945 | 0.454715342 | 0.649314042 | 0.913724196 |
| Ndufb7 | 1664.546242 | 0.041433003 | 0.056581854 | 0.732266626 | 0.464005845 | 0.83159896 |
| Sh3d21 | 56.77660645 | 0.041434006 | 0.094117173 | 0.440238526 | 0.659764359 | 0.915208181 |
| Apoa1bp | 1593.065319 | 0.041441266 | 0.052040179 | 0.796332125 | 0.425839018 | 0.809519334 |
| Arl2bp | 5161.581138 | 0.041446496 | 0.042714233 | 0.970320513 | 0.331886756 | 0.756958483 |
| Klhdc9 | 178.3624245 | 0.041497701 | 0.094836081 | 0.437572923 | 0.661695903 | 0.915428982 |
| Serpina3c | 1.816701476 | 0.041500322 | 0.028092581 | 1.477269825 | 0.139603323 | NA |
| Ncs1 | 15706.94283 | 0.041514433 | 0.052735622 | 0.787218029 | 0.431154245 | 0.812181606 |
| Zfp553 | 1013.253898 | 0.041523485 | 0.061509404 | 0.675075386 | 0.499627871 | 0.846241311 |
| Wdr7 | 10078.98971 | 0.041557515 | 0.056546524 | 0.734926088 | 0.462384509 | 0.83085074 |
| Arhgef10l | 2123.39725 | 0.041579759 | 0.064575805 | 0.643890675 | 0.519646329 | 0.853297835 |
| Xpo1 | 2931.18204 | 0.041615693 | 0.075546834 | 0.550859524 | 0.581729976 | 0.884020489 |
| Fabp4 | 4.619292328 | 0.04163797 | 0.046247627 | 0.900326635 | 0.367946451 | NA |
| Rnf122 | 150.2536487 | 0.041639871 | 0.093026304 | 0.447613943 | 0.654431839 | 0.915032909 |
| Senp6 | 3112.176232 | 0.041643162 | 0.068277811 | 0.609907691 | 0.541922957 | 0.864560611 |
| Nr1i3 | 4.819949339 | 0.041665551 | 0.047773654 | 0.872144931 | 0.383129317 | NA |
| Gm5127 | 4.841476578 | 0.04170756 | 0.047169166 | 0.884212365 | 0.376581585 | NA |
| Kcnb2 | 401.9151345 | 0.04171306 | 0.084995676 | 0.490766845 | 0.623591362 | 0.900694853 |
| E130114P18Rik | 129.3936959 | 0.041743948 | 0.093812055 | 0.44497424 | 0.65633837 | 0.915208181 |
| Elp6 | 345.0339147 | 0.041744231 | 0.088187813 | 0.473356008 | 0.63595921 | 0.907889736 |
| Gatsl3 | 134.6397733 | 0.041749849 | 0.093804208 | 0.44507437 | 0.65626601 | 0.915208181 |
| 1700020I14Rik | 2869.59612 | 0.041778036 | 0.080857618 | 0.516686449 | 0.605375057 | 0.892306249 |
| Eif2s2 | 2672.910255 | 0.041780579 | 0.062525556 | 0.668216037 | 0.5039957 | 0.847807553 |
| Gyk | 914.9462957 | 0.041802193 | 0.063633357 | 0.656922639 | 0.51123066 | 0.852011998 |
| Ttc21b | 568.6819447 | 0.041807614 | 0.076961689 | 0.5432263 | 0.586974001 | 0.884663353 |
| Fkbp9 | 1987.664237 | 0.041810284 | 0.058812363 | 0.710909779 | 0.477140146 | 0.835247566 |
| 6030458C11Rik | 1843.896651 | 0.041810889 | 0.056221611 | 0.74368003 | 0.457070074 | 0.827236468 |
| Vtcn1 | 9.503312626 | 0.041857753 | 0.062206799 | 0.672880672 | 0.501023209 | NA |
| E2f6 | 1429.334547 | 0.041862207 | 0.06777215 | 0.61769041 | 0.536779435 | 0.86167069 |
| Usp31 | 5864.895531 | 0.0418666 | 0.072703081 | 0.575857306 | 0.564711635 | 0.875364912 |
| Tmem202 | 70.58762089 | 0.04189167 | 0.095100991 | 0.44049667 | 0.659577424 | 0.915208181 |
| 1810034E14Rik | 8.11036596 | 0.041896863 | 0.052762344 | 0.794067514 | 0.427156129 | NA |
| Actg2 | 3.014386258 | 0.04189716 | 0.032850095 | 1.275404545 | 0.202166091 | NA |
| C130080G10Rik | 6.221132432 | 0.041910145 | 0.051090587 | 0.820310492 | 0.412039127 | NA |
| Dnajb6 | 5594.444838 | 0.041912674 | 0.054510654 | 0.76888959 | 0.441958858 | 0.819321835 |
| S100a10 | 782.2015871 | 0.041915447 | 0.065375299 | 0.641151138 | 0.521424492 | 0.854009716 |
| Hyal1 | 186.4544206 | 0.041935267 | 0.092785354 | 0.451959977 | 0.651297816 | 0.914343204 |
| Ap2s1 | 3990.713078 | 0.041956552 | 0.048559892 | 0.864016581 | 0.387578782 | 0.791583702 |
| Nap1l1 | 10159.15565 | 0.041990923 | 0.069744657 | 0.602066522 | 0.547129858 | 0.867487125 |
| 1600014C10Rik | 1392.087473 | 0.041993022 | 0.054617718 | 0.768853467 | 0.441980304 | 0.819321835 |
| Tsn | 3269.542276 | 0.042038796 | 0.048332454 | 0.869784023 | 0.384418444 | 0.78893223 |
| Rbm34 | 769.6456439 | 0.042044469 | 0.06711595 | 0.626445267 | 0.531022926 | 0.858310785 |
| Dnm1l | 9942.944337 | 0.042061402 | 0.057726068 | 0.728637918 | 0.466223178 | 0.832050282 |
| Vamp4 | 2584.076745 | 0.042088286 | 0.056682304 | 0.742529549 | 0.457766555 | 0.827642075 |
| Bmpr2 | 2291.268515 | 0.042088464 | 0.075322985 | 0.558773183 | 0.576316527 | 0.880966011 |
| Eng | 1161.075779 | 0.042095827 | 0.079202726 | 0.531494677 | 0.595076029 | 0.887381665 |
| Kcnmb1 | 58.44556278 | 0.0420969 | 0.090861963 | 0.463306079 | 0.643144989 | 0.911258473 |
| Aimp2 | 769.3677838 | 0.042101108 | 0.060009294 | 0.701576455 | 0.482943339 | 0.837940151 |
| Rala | 2296.559456 | 0.042114307 | 0.059872069 | 0.703404904 | 0.481803451 | 0.837383765 |
| Gm16907 | 9.282788243 | 0.042127644 | 0.058626784 | 0.718573341 | 0.472403844 | NA |
| Grap | 169.3459996 | 0.042157328 | 0.093738763 | 0.449732068 | 0.652903646 | 0.914683958 |
| Tex261 | 886.5577403 | 0.042176108 | 0.057379787 | 0.735034235 | 0.462318645 | 0.830835164 |
| Lrrc3b | 551.619615 | 0.042182741 | 0.074358148 | 0.567291438 | 0.570516193 | 0.878246677 |
| Isg20l2 | 607.9341382 | 0.042190042 | 0.07304154 | 0.577617095 | 0.563522663 | 0.875108781 |
| Btc | 18.96012651 | 0.04223326 | 0.079139231 | 0.533657701 | 0.593578379 | NA |
| Apc | 12874.77053 | 0.042251514 | 0.065748161 | 0.64262656 | 0.520466445 | 0.8535556 |
| Dtwd2 | 114.0674526 | 0.042314037 | 0.093045978 | 0.454764818 | 0.649278443 | 0.913724196 |
| Tacc2 | 3579.037098 | 0.042323688 | 0.046554708 | 0.909117246 | 0.36328824 | 0.776821522 |
| Esr1 | 133.6615975 | 0.0423239 | 0.073446617 | 0.576253914 | 0.564443568 | 0.875364912 |
| Dph5 | 340.4255467 | 0.042325085 | 0.073763892 | 0.573791366 | 0.566108992 | 0.87575243 |
| Actr8 | 1991.324311 | 0.042340856 | 0.045597464 | 0.928579184 | 0.353107214 | 0.770519168 |
| Lrrk2 | 3319.544875 | 0.042347303 | 0.087825342 | 0.482176344 | 0.62968068 | 0.904807492 |
| Zfp438 | 264.8196908 | 0.042360207 | 0.079848142 | 0.530509607 | 0.59575865 | 0.887497708 |
| Chst4 | 6.704431706 | 0.04239438 | 0.050781165 | 0.834844581 | 0.403805221 | NA |
| Lrrc73 | 1195.646956 | 0.042453943 | 0.065600081 | 0.647162967 | 0.517526476 | 0.853297835 |
| Mdga1 | 1152.557439 | 0.042456799 | 0.083964858 | 0.505649628 | 0.613102636 | 0.895428201 |
| Prmt8 | 4218.65397 | 0.042465414 | 0.054645608 | 0.777105701 | 0.437096411 | 0.816564673 |
| Naa20 | 1579.392049 | 0.042483726 | 0.060151871 | 0.706274387 | 0.480017519 | 0.836914993 |
| Tti1 | 1159.076545 | 0.04248663 | 0.052357288 | 0.811474992 | 0.417092949 | 0.806153084 |
| Pfdn1 | 1520.077688 | 0.042489231 | 0.054708741 | 0.776644289 | 0.437368664 | 0.816715817 |
| Fmo2 | 183.6569665 | 0.042490565 | 0.0764171 | 0.556034773 | 0.578187084 | 0.881850649 |
| Cep120 | 2315.040653 | 0.042499603 | 0.059739584 | 0.71141445 | 0.476827447 | 0.835195112 |
| Fgf16 | 19.04434867 | 0.042510777 | 0.072182968 | 0.588930851 | 0.555907661 | NA |
| Cdh2 | 5159.925315 | 0.042522604 | 0.056854286 | 0.747922567 | 0.454506865 | 0.826168452 |
| Myo1d | 1104.629728 | 0.042525892 | 0.079665514 | 0.533805523 | 0.593476092 | 0.886313552 |
| Acad8 | 989.6005796 | 0.042540562 | 0.076554796 | 0.555687745 | 0.578424336 | 0.881901419 |
| Cdc16 | 1988.251122 | 0.042551678 | 0.053051339 | 0.802084892 | 0.422503854 | 0.808695226 |
| Ccdc124 | 2165.5407 | 0.042558831 | 0.053273161 | 0.798879396 | 0.424360347 | 0.809162684 |
| Aqr | 2160.438812 | 0.042567584 | 0.056963474 | 0.74727859 | 0.454895413 | 0.826168452 |
| Hspb8 | 590.054229 | 0.04259306 | 0.087336434 | 0.487689483 | 0.625769802 | 0.902316076 |
| Tpmt | 402.9915102 | 0.042599293 | 0.075175931 | 0.566661325 | 0.570944302 | 0.878246677 |
| AK129341 | 779.6873915 | 0.042600888 | 0.085340294 | 0.499188434 | 0.617646642 | 0.898001714 |
| Riok2 | 935.5331215 | 0.042604983 | 0.06759892 | 0.630261302 | 0.528523637 | 0.857228044 |
| Tomm70a | 8423.242691 | 0.042626572 | 0.078028297 | 0.546296326 | 0.584862261 | 0.884313345 |
| Rtn4r | 3476.872765 | 0.042647371 | 0.069329329 | 0.615141839 | 0.538461054 | 0.862082844 |
| Slc25a51 | 3829.582885 | 0.042669836 | 0.050413292 | 0.846400492 | 0.397329364 | 0.797340699 |
| Cnot11 | 1286.641683 | 0.042676833 | 0.063522074 | 0.671842568 | 0.501683926 | 0.846689985 |
| Evi5l | 1581.017097 | 0.042687144 | 0.048799716 | 0.874741635 | 0.3817145 | 0.786663048 |
| C630020P19Rik | 161.2830284 | 0.042687857 | 0.092076713 | 0.463611872 | 0.642925847 | 0.911258473 |
| Wdr70 | 512.7918887 | 0.042688022 | 0.067257384 | 0.634696431 | 0.525626422 | 0.855892499 |
| Slc22a13 | 1.715931041 | 0.042701597 | 0.028704312 | 1.487637019 | 0.136846645 | NA |
| Ctbs | 361.2927296 | 0.042716188 | 0.076855672 | 0.555797466 | 0.578349318 | 0.881901419 |
| Zfp772 | 508.9897088 | 0.042721467 | 0.082755735 | 0.516235729 | 0.605689779 | 0.892498451 |
| Vamp2 | 52658.47258 | 0.042746114 | 0.039497963 | 1.082235908 | 0.279147717 | 0.718571634 |
| Gm2a | 2675.517015 | 0.042756108 | 0.058762295 | 0.727611276 | 0.466851576 | 0.832107746 |
| Kif16b | 793.6921808 | 0.042776267 | 0.066766313 | 0.64068638 | 0.521726465 | 0.854113377 |
| Lyrm1 | 266.1242574 | 0.042776877 | 0.079441331 | 0.538471303 | 0.590251712 | 0.885570623 |
| Elmod2 | 662.7499074 | 0.042801889 | 0.074530544 | 0.574286547 | 0.565773911 | 0.875605352 |
| Tnn | 18.93388326 | 0.042813105 | 0.079792148 | 0.53655787 | 0.591573052 | NA |
| Taf7l | 2.813960406 | 0.042845011 | 0.033359036 | 1.284359962 | 0.199016035 | NA |
| Senp5 | 3310.27789 | 0.042851636 | 0.065211215 | 0.657120647 | 0.511103344 | 0.851995315 |
| 2410057H14Rik | 4.173405219 | 0.042866337 | 0.038902565 | 1.101889744 | 0.270509608 | NA |
| Ddb1 | 11658.2777 | 0.042868131 | 0.039525187 | 1.084577552 | 0.278108802 | 0.718319852 |
| Fam196b | 141.2264603 | 0.04288258 | 0.075248069 | 0.569882801 | 0.568757191 | 0.877419107 |
| Gpkow | 1792.081372 | 0.042886789 | 0.050283148 | 0.852905821 | 0.393711534 | 0.794754188 |
| D6Ertd474e | 20.77966601 | 0.042898075 | 0.076306518 | 0.562181003 | 0.573992703 | NA |
| Dnajc30 | 1293.297223 | 0.042900034 | 0.057199265 | 0.750010232 | 0.453248542 | 0.82562629 |
| Ppp2r4 | 7480.345278 | 0.042904631 | 0.049381782 | 0.868835217 | 0.384937268 | 0.789016184 |
| Orai1 | 205.9631901 | 0.042931808 | 0.087191138 | 0.492387288 | 0.622445584 | 0.900114632 |
| Myo1h | 83.67972504 | 0.042935912 | 0.093854369 | 0.457473767 | 0.647330554 | 0.912750424 |
| R3hdm2 | 12523.14988 | 0.042995501 | 0.057881427 | 0.742820324 | 0.457590469 | 0.827642075 |
| C130050O18Rik | 7.584762748 | 0.04300196 | 0.057171212 | 0.752161074 | 0.451954199 | NA |
| Setx | 3426.636317 | 0.04300648 | 0.062503953 | 0.688060159 | 0.491414898 | 0.841892493 |
| Clpb | 2254.761775 | 0.043019695 | 0.045283049 | 0.950017636 | 0.342103292 | 0.763752913 |
| Fam46b | 3.493317917 | 0.043028374 | 0.040967118 | 1.050314884 | 0.293573364 | NA |
| Snx13 | 3805.381011 | 0.043033042 | 0.065344366 | 0.658557808 | 0.510179764 | 0.850846298 |
| Serinc2 | 250.6216452 | 0.043055141 | 0.093658294 | 0.459704515 | 0.645728329 | 0.912363974 |
| Cryba4 | 20.49017498 | 0.043057359 | 0.0768289 | 0.560431797 | 0.575184948 | NA |
| Zfp706 | 8317.970152 | 0.043060088 | 0.056658489 | 0.759993574 | 0.447258426 | 0.822458097 |
| Tdrd3 | 828.9038367 | 0.043060536 | 0.075345874 | 0.571504898 | 0.567657441 | 0.876652349 |
| Calcr | 71.67556096 | 0.043065892 | 0.036010931 | 1.195911656 | 0.231731036 | 0.683754831 |
| Fam219a | 7299.216877 | 0.043066629 | 0.044808562 | 0.961124988 | 0.336489327 | 0.75970798 |
| Ptprm | 2015.425282 | 0.04306773 | 0.069895375 | 0.616174251 | 0.537779521 | 0.862037665 |
| Eml2 | 5820.892064 | 0.043079043 | 0.062883987 | 0.685055841 | 0.493308684 | 0.843035523 |
| Ubxn10 | 94.48964215 | 0.043088377 | 0.087386926 | 0.493075779 | 0.621959044 | 0.900011855 |
| Srpx | 23.12686749 | 0.043120685 | 0.085178346 | 0.506239992 | 0.612688183 | 0.895428201 |
| Wipi2 | 5439.478921 | 0.043125196 | 0.038672085 | 1.11515052 | 0.264785926 | 0.707370515 |
| Grk5 | 572.4293992 | 0.043148919 | 0.062632739 | 0.688919566 | 0.490873886 | 0.841866734 |
| Pip4k2b | 8379.547198 | 0.043183773 | 0.048813623 | 0.884666419 | 0.376336573 | 0.783947144 |
| Vax1 | 16.70707024 | 0.043202241 | 0.064095322 | 0.674031104 | 0.500291539 | NA |
| Saysd1 | 201.6698073 | 0.0432164 | 0.083505539 | 0.517527343 | 0.604788087 | 0.891988593 |
| Btd | 381.7954214 | 0.043221223 | 0.081417802 | 0.530857156 | 0.595517769 | 0.887497708 |
| Grpr | 29.1294315 | 0.043222298 | 0.080625622 | 0.536086389 | 0.591898846 | 0.886226713 |
| Parg | 1568.447523 | 0.043239029 | 0.067020059 | 0.645165491 | 0.518819947 | 0.853297835 |
| Kntc1 | 17.95387604 | 0.043265599 | 0.078343181 | 0.552257376 | 0.580772029 | NA |
| Rnft2 | 4777.991895 | 0.043294372 | 0.050020304 | 0.865535953 | 0.386744691 | 0.790912767 |
| Tomm40l | 2714.700796 | 0.043294748 | 0.05838012 | 0.741600869 | 0.458329197 | 0.827967904 |
| BC030500 | 278.3131111 | 0.043304557 | 0.082737386 | 0.523397688 | 0.600697533 | 0.890108371 |
| Ccdc86 | 364.2331175 | 0.043317158 | 0.081538137 | 0.531250278 | 0.595245356 | 0.887381665 |
| Zfp777 | 1290.086799 | 0.043349 | 0.061486073 | 0.705021447 | 0.48079689 | 0.836914993 |
| Zfp119b | 91.30390689 | 0.04337774 | 0.094838916 | 0.457383335 | 0.647395541 | 0.912750424 |
| St6galnac2 | 202.76118 | 0.04338701 | 0.093236114 | 0.46534554 | 0.641684027 | 0.911095592 |
| Zhx1 | 3481.895013 | 0.043394737 | 0.059192521 | 0.733111824 | 0.463490229 | 0.831397761 |
| Pou3f4 | 130.4732084 | 0.043395422 | 0.094390391 | 0.459744063 | 0.645699938 | 0.912363974 |
| Kif5a | 154107.5859 | 0.04342651 | 0.058983088 | 0.736253587 | 0.461576386 | 0.830270578 |
| Fndc3a | 2498.566194 | 0.043442053 | 0.069178789 | 0.627967821 | 0.53002502 | 0.857915133 |
| Lrig1 | 1781.006751 | 0.043457089 | 0.080334166 | 0.540954009 | 0.588539283 | 0.885188411 |
| Nqo2 | 415.8770042 | 0.043475672 | 0.081708207 | 0.532084512 | 0.594667464 | 0.887269096 |
| Trim14 | 20.6128322 | 0.043484537 | 0.078704818 | 0.552501583 | 0.58060475 | NA |
| Dctn2 | 9914.24023 | 0.043497741 | 0.053951854 | 0.806232548 | 0.420108775 | 0.807657282 |
| Commd3 | 1522.911871 | 0.043510605 | 0.053461889 | 0.813862084 | 0.415723964 | 0.805633085 |
| Bcl10 | 493.3104382 | 0.043513512 | 0.076367281 | 0.569792599 | 0.568818376 | 0.877419107 |
| Chchd1 | 624.0586705 | 0.043515579 | 0.070027531 | 0.621406722 | 0.53433205 | 0.859648542 |
| Rad23a | 2264.909881 | 0.043521824 | 0.048010856 | 0.906499654 | 0.364671448 | 0.777402033 |
| Fitm2 | 645.3870066 | 0.043523751 | 0.058796601 | 0.740242645 | 0.459152776 | 0.828381305 |

| | | | | | | | | | | | | | |
|---|---|---|---|---|---|---|---|---|---|---|---|---|---|
| Gstt2 | 74.08820043 | -0.02868563 | 0.142884172 | -0.200761423 | 0.840885129 | 0.924293811 | Fam175b | 1335.367393 | 0.043532769 | 0.059460216 | 0.732132705 | 0.464087573 | 0.83159896 |
| Kcnrg | 18.47035517 | -0.033143681 | 0.165250239 | -0.200566618 | 0.841037463 | 0.924293811 | Qsox2 | 902.656472 | 0.043554023 | 0.064558016 | 0.674649342 | 0.499898576 | 0.846241311 |
| Tram1 | 866.7421984 | 0.01216284 | 0.060559817 | 0.20084011 | 0.840823599 | 0.924293811 | Ccdc127 | 3596.312014 | 0.043562874 | 0.054191719 | 0.803865889 | 0.421474432 | 0.808555694 |
| Fkbpl | 252.9610019 | 0.026551754 | 0.132477973 | 0.200423916 | 0.841149057 | 0.924309467 | Nmd3 | 790.7195706 | 0.043564702 | 0.067384733 | 0.646507014 | 0.517951056 | 0.853297835 |
| A330035P11Rik | 23.35424737 | 0.032955813 | 0.164923502 | 0.199824841 | 0.841617573 | 0.924555649 | Tcf23 | 22.82379261 | 0.043567769 | 0.076946431 | 0.5662091 | 0.571251645 | 0.878374234 |
| BC020402 | 133.750866 | 0.025546446 | 0.127708838 | 0.200036631 | 0.841451933 | 0.924555649 | Wdr45b | 2808.480853 | 0.043594095 | 0.055408688 | 0.786773636 | 0.431414389 | 0.812181606 |
| Camk1g | 1627.500206 | -0.027427099 | 0.137303035 | -0.199755956 | 0.841671449 | 0.924555649 | Tnks2 | 6593.270637 | 0.043608386 | 0.071331035 | 0.611352205 | 0.540966437 | 0.863847445 |
| Ddx19b | 1465.477765 | -0.013979938 | 0.069910806 | -0.199968198 | 0.841505453 | 0.924555649 | Mn1 | 1883.530982 | 0.043622497 | 0.075625951 | 0.576819152 | 0.56406163 | 0.875251366 |
| Zfp398 | 526.993776 | 0.015029909 | 0.07520725 | 0.199846541 | 0.841600601 | 0.924555649 | Clip2 | 7633.273073 | 0.043623084 | 0.052227013 | 0.835259032 | 0.403571879 | 0.800224169 |
| Nt5dc1 | 68.25857394 | -0.031338288 | 0.15710204 | -0.199477284 | 0.841889411 | 0.924729515 | Trp53i11 | 4444.985621 | 0.043636727 | 0.080891865 | 0.539445186 | 0.589579709 | 0.885188411 |
| Snrpa1 | 659.1078367 | -0.015461882 | 0.07754719 | -0.199386747 | 0.841960227 | 0.924741742 | Btbd3 | 7228.764526 | 0.043642366 | 0.067767695 | 0.643999563 | 0.519575718 | 0.853297835 |
| Rasal1 | 3560.212332 | -0.020937242 | 0.105057815 | -0.199292568 | 0.842033892 | 0.924757097 | Cd151 | 1006.814956 | 0.043646574 | 0.068322447 | 0.638832124 | 0.522932148 | 0.854337225 |
| Aqp6 | 12.38268249 | -0.031418642 | 0.15795969 | -0.198902912 | 0.842338692 | 0.924950886 | Gfod1 | 6975.219711 | 0.043653239 | 0.086592535 | 0.50412243 | 0.614175346 | 0.896093921 |
| Fam194a | 74.8162093 | 0.032338688 | 0.162638335 | 0.19883804 | 0.842389438 | 0.924950886 | Rab10 | 5913.93494 | 0.043675911 | 0.05824169 | 0.749908033 | 0.453310096 | 0.82562629 |
| Tesk1 | 3904.763772 | -0.014274921 | 0.07177418 | -0.198886573 | 0.842351473 | 0.924950886 | Setd3 | 7327.255034 | 0.043691153 | 0.044976139 | 0.971429593 | 0.331334398 | 0.75639847 |
| Dnaja3 | 2653.465148 | 0.009887132 | 0.049745159 | 0.198755673 | 0.842453872 | 0.924956086 | Calr4 | 18.68087361 | 0.043703375 | 0.074360306 | 0.587724516 | 0.556717217 | NA |
| Adrbk1 | 10165.95731 | 0.017915739 | 0.090288307 | 0.198428122 | 0.842710116 | 0.925040775 | Irs1 | 795.9643901 | 0.043722062 | 0.074010775 | 0.590752659 | 0.554686159 | 0.871360989 |
| Tatdn1 | 255.6895049 | 0.022954524 | 0.115678283 | 0.198434173 | 0.842705383 | 0.925040775 | Pkdrej | 99.70849658 | 0.043723199 | 0.095089871 | 0.459809217 | 0.645653167 | 0.912363974 |
| Zfp609 | 2542.565259 | 0.019934032 | 0.10038408 | 0.198577628 | 0.842593156 | 0.925040775 | Pawr | 70.889627 | 0.04372397 | 0.095018761 | 0.460161444 | 0.645400344 | 0.912363974 |
| Trnau1ap | 460.4308242 | -0.017390173 | 0.087773491 | -0.198125573 | 0.842946817 | 0.925235051 | Mlph | 132.4334585 | 0.043755643 | 0.094906166 | 0.461041099 | 0.644769119 | 0.912353932 |
| Hnrnpul1 | 3451.958726 | 0.011843657 | 0.059803334 | 0.198043416 | 0.843011095 | 0.925240059 | Phf20 | 4099.1054 | 0.043768821 | 0.056959478 | 0.768420333 | 0.442237505 | 0.819589435 |
| Fitm2 | 572.6114104 | 0.012373242 | 0.062550445 | 0.197812211 | 0.843191992 | 0.925307509 | Pik3cb | 1659.310372 | 0.043794506 | 0.066940241 | 0.654232865 | 0.512961791 | 0.85302356 |
| Spata21 | 24.57871801 | 0.033239722 | 0.168011559 | 0.197841876 | 0.843168782 | 0.925307509 | Grxcr2 | 11.53202877 | 0.043798539 | 0.047058954 | 0.930716371 | 0.352000299 | NA |
| Dtd2 | 284.7222374 | -0.017055597 | 0.086361161 | -0.197491524 | 0.843442914 | 0.925451775 | Thumpd3 | 913.4852124 | 0.043810535 | 0.059107295 | 0.741203518 | 0.458570051 | 0.828043599 |
| Rfwd3 | 1455.186398 | -0.01192125 | 0.060344403 | -0.197553526 | 0.843394399 | 0.925451775 | Marc2 | 1882.385599 | 0.043826135 | 0.06685017 | 0.655587485 | 0.51208958 | 0.852888435 |
| Necab3 | 2625.21566 | -0.017624998 | 0.089444064 | -0.197050507 | 0.843788014 | 0.92558548 | Sdr39u1 | 2173.377465 | 0.043826521 | 0.061588418 | 0.711603283 | 0.476710474 | 0.835195112 |
| Ttn | 17.95583686 | -0.032644632 | 0.165549319 | -0.197189768 | 0.843679038 | 0.92558548 | Foxh1 | 1.867489414 | 0.043831542 | 0.031804184 | 1.378169047 | 0.168151101 | NA |
| Vcl | 1709.549346 | 0.01746234 | 0.08862763 | 0.19703043 | 0.843803725 | 0.92558548 | Rprml | 1158.558784 | 0.043841278 | 0.075511299 | 0.580592295 | 0.561515266 | 0.873934985 |
| Xpnpep1 | 1422.339839 | 0.011844306 | 0.060079331 | 0.197144439 | 0.843714509 | 0.92558548 | Eif6 | 1384.007953 | 0.04385806 | 0.064523667 | 0.67972051 | 0.496681447 | 0.8449088 |
| 4930455F23Rik | 835.7737628 | 0.015665281 | 0.079627398 | 0.196732303 | 0.84403703 | 0.925644801 | Fbp1 | 10.72618356 | 0.043871904 | 0.064749195 | 0.677566787 | 0.49804641 | NA |
| Bend3 | 274.8563567 | -0.019224806 | 0.097644908 | -0.196884875 | 0.84391763 | 0.925644801 | 9830001H06Rik | 2546.660144 | 0.043893757 | 0.079917713 | 0.549236899 | 0.582842885 | 0.88406516 |
| Dennd1b | 470.9442163 | -0.018249141 | 0.092758931 | -0.196737295 | 0.844033124 | 0.925644801 | Sp3 | 1855.246975 | 0.043925921 | 0.070251485 | 0.625266798 | 0.531795967 | 0.858328942 |
| Snx25 | 2484.099202 | 0.018655494 | 0.095098724 | 0.196169757 | 0.844477299 | 0.92606209 | Gmeb1 | 960.2058012 | 0.043947359 | 0.056789075 | 0.773869954 | 0.439007699 | 0.817545617 |
| Sap30bp | 1530.187915 | 0.015403719 | 0.078593656 | 0.195991888 | 0.844616516 | 0.926149207 | Tmem52 | 20.46076827 | 0.043961173 | 0.078833322 | 0.557647098 | 0.577085388 | NA |
| Adamts7 | 14.25209264 | -0.031825256 | 0.163225321 | -0.194977445 | 0.845410605 | 0.926167851 | Ube2m | 4519.868387 | 0.043975999 | 0.052101672 | 0.844041979 | 0.398645948 | 0.797789838 |
| Arsb | 5349.719046 | -0.011769654 | 0.060233767 | -0.195399607 | 0.845080124 | 0.926167851 | Cap1 | 11124.4634 | 0.04397837 | 0.051412686 | 0.855399177 | 0.392330199 | 0.794184423 |
| Asb6 | 1079.897126 | 0.015217288 | 0.077798323 | 0.195599175 | 0.844923906 | 0.926167851 | Rab20 | 16.76572115 | 0.043996295 | 0.066262564 | 0.6639691 | 0.506710088 | NA |
| Cox11 | 507.0562127 | 0.015523617 | 0.079528095 | 0.195196642 | 0.845239008 | 0.926167851 | Rpl19 | 10130.44555 | 0.044013236 | 0.053437912 | 0.823633163 | 0.410148027 | 0.80184319 |
| Dnajb14 | 754.8567296 | -0.016525466 | 0.084711714 | -0.19507887 | 0.845331204 | 0.926167851 | Sema5a | 1891.247813 | 0.044013751 | 0.065409885 | 0.672891426 | 0.501016367 | 0.846601651 |
| Dok2 | 22.33949463 | 0.032811367 | 0.168004801 | 0.195300175 | 0.84515796 | 0.926167851 | Ifngr2 | 1842.751798 | 0.04403189 | 0.050700741 | 0.868466404 | 0.385139056 | 0.789016184 |
| Grem1 | 40.53111566 | -0.030539214 | 0.156584509 | -0.19503343 | 0.845366777 | 0.926167851 | Pex19 | 3583.470828 | 0.044045704 | 0.041808739 | 1.05350473 | 0.292109728 | 0.727847831 |
| Ints4 | 1309.395563 | 0.012400662 | 0.063382579 | 0.195647797 | 0.844885847 | 0.926167851 | Enc1 | 39062.24961 | 0.044055262 | 0.077824792 | 0.566082618 | 0.571337619 | 0.878389918 |
| Mdh1b | 47.13552877 | -0.032626121 | 0.167037662 | -0.195321941 | 0.845140921 | 0.926167851 | Larp1b | 559.2875416 | 0.044078768 | 0.075930208 | 0.580516892 | 0.561566099 | 0.873934985 |
| Pex3 | 979.7850956 | -0.013991136 | 0.071603708 | -0.195396804 | 0.845082318 | 0.926167851 | Cnot4 | 1447.677367 | 0.044089534 | 0.071650451 | 0.615342037 | 0.538328862 | 0.862082844 |
| Samd14 | 1743.403481 | 0.024022417 | 0.122813925 | 0.195600108 | 0.844923175 | 0.926167851 | Adora2b | 146.2352356 | 0.044089802 | 0.089886368 | 0.490505996 | 0.623775888 | 0.900733873 |
| Soat1 | 814.0589817 | -0.013771564 | 0.070549304 | -0.195204819 | 0.845232607 | 0.926167851 | Hipk1 | 6506.15531 | 0.044094714 | 0.058270301 | 0.756727067 | 0.449213395 | 0.823263966 |
| Taco1 | 418.6189808 | -0.015836213 | 0.080852296 | -0.195865962 | 0.84471508 | 0.926167851 | Hapln3 | 23.67634687 | 0.04410077 | 0.077486056 | 0.569144594 | 0.569258018 | 0.87745335 |
| A830019L24Rik | 13.27229272 | -0.032544112 | 0.166993928 | -0.194882011 | 0.845485317 | 0.926174459 | Scarb2 | 3376.609283 | 0.044103661 | 0.056403686 | 0.7819287 | 0.434256477 | 0.814883037 |
| Cnpy4 | 356.6338269 | -0.015393897 | 0.079017205 | -0.19481703 | 0.84553619 | 0.926174459 | Hrct1 | 27.93839499 | 0.044106288 | 0.086917774 | 0.507448431 | 0.611840212 | 0.895245244 |
| Erf | 2011.426768 | -0.01359097 | 0.069828412 | -0.194633815 | 0.845679628 | 0.926203011 | Ccdc36 | 29.77106284 | 0.044108525 | 0.089302635 | 0.493921879 | 0.621361352 | 0.899589902 |
| Thap4 | 757.8091901 | -0.015652824 | 0.080423067 | -0.194631026 | 0.845681812 | 0.926203011 | Serpine2 | 7949.444971 | 0.044110356 | 0.052307367 | 0.843291457 | 0.399065459 | 0.797801385 |
| Rdh10 | 309.1945007 | 0.019176912 | 0.098623471 | 0.194445726 | 0.845826889 | 0.926296425 | Gpatch4 | 739.5136368 | 0.044118555 | 0.058634075 | 0.752438831 | 0.451787202 | 0.824558475 |
| E330009J07Rik | 674.6469564 | 0.02037472 | 0.105092441 | 0.193874269 | 0.846274332 | 0.926589958 | Gstm1 | 8579.775442 | 0.044124874 | 0.074456257 | 0.592628153 | 0.553430034 | 0.870600462 |
| Kcnq3 | 1884.500857 | -0.016938677 | 0.087312018 | -0.194001664 | 0.846174579 | 0.926589958 | Cntnap3 | 499.5988788 | 0.044141673 | 0.076398642 | 0.577780859 | 0.563412079 | 0.875108781 |
| Srsf4 | 2040.587141 | 0.011379428 | 0.05868343 | 0.193912119 | 0.846244694 | 0.926589958 | Rpp25 | 522.0898787 | 0.044150684 | 0.071772474 | 0.615147856 | 0.53845708 | 0.862082844 |
| A330032B11Rik | 8.622986397 | 0.030440849 | 0.157459087 | 0.19332545 | 0.846704096 | 0.926969536 | Gm14204 | 424.8587609 | 0.044164815 | 0.084082398 | 0.52525637 | 0.599404983 | 0.889190848 |
| Adam23 | 6781.288896 | 0.011789072 | 0.060995177 | 0.193278755 | 0.846740664 | 0.926969536 | Slc46a3 | 309.3021528 | 0.044176699 | 0.083030564 | 0.532053465 | 0.594688966 | 0.887269096 |
| Tmed1 | 367.7612492 | -0.02036904 | 0.105567217 | -0.192948535 | 0.846999274 | 0.927187138 | Adrbk1 | 10986.45263 | 0.044194619 | 0.059331014 | 0.744882244 | 0.45634291 | 0.82674985 |
| Dhx32 | 1385.512775 | 0.017535956 | 0.091071875 | 0.192550727 | 0.847310837 | 0.927331646 | Rpe | 681.6518674 | 0.044196813 | 0.079980029 | 0.552598109 | 0.580538637 | 0.883489182 |
| Ncoa1 | 6428.528386 | -0.017226121 | 0.089445292 | -0.192588343 | 0.847281375 | 0.927331646 | Dhx40 | 1249.71941 | 0.044218447 | 0.064627963 | 0.684199919 | 0.493848933 | 0.843404385 |
| Nfil3 | 337.4089507 | 0.025167509 | 0.130664363 | 0.192611884 | 0.847262937 | 0.927331646 | Ptpn3 | 3381.431787 | 0.044229271 | 0.060670701 | 0.729005447 | 0.465998331 | 0.832050282 |
| Pxmp4 | 388.5847757 | -0.022598696 | 0.117422071 | -0.192456972 | 0.847384269 | 0.927346508 | Polr1d | 1423.71042 | 0.044232779 | 0.06122566 | 0.722454905 | 0.470014841 | 0.833245343 |
| Kpna4 | 2773.296383 | -0.014550523 | 0.075911714 | -0.191676906 | 0.847995297 | 0.927884119 | Ankrd17 | 6670.933469 | 0.044236631 | 0.067030748 | 0.659945362 | 0.509288893 | 0.850336834 |
| Zfp212 | 595.0481712 | 0.015553825 | 0.08112725 | 0.191721339 | 0.84796049 | 0.927884119 | Rnf180 | 417.7981334 | 0.044251436 | 0.076090419 | 0.581563838 | 0.560860507 | 0.873713575 |
| B230209K01Rik | 63.1950679 | 0.028309981 | 0.147807756 | 0.191532447 | 0.848108462 | 0.927924127 | Clec4n | 4.67078998 | 0.044253722 | 0.047932956 | 0.923242089 | 0.35588106 | NA |
| Gpr83 | 1037.447373 | -0.019242718 | 0.100565316 | -0.191345471 | 0.848254938 | 0.927924127 | Srebf2 | 9405.293803 | 0.044255014 | 0.04939091 | 0.896015354 | 0.370244563 | 0.780463289 |
| Npas4 | 1200.606961 | -0.03012002 | 0.157429025 | -0.191324437 | 0.848271417 | 0.927924127 | Mup5 | 10.4859403 | 0.044275738 | 0.06211923 | 0.712754134 | 0.475997912 | NA |
| Zfp386 | 573.832646 | -0.017657632 | 0.092260678 | -0.191388496 | 0.848221233 | 0.927924127 | Zfp12 | 1588.944215 | 0.044279727 | 0.053593953 | 0.826207532 | 0.408686377 | 0.801055189 |
| Gtf2b | 1081.669442 | -0.018729953 | 0.097991018 | -0.191139485 | 0.848416313 | 0.92801711 | Rhoa | 6160.156382 | 0.044290492 | 0.048114045 | 0.920531461 | 0.3572951 | 0.773597947 |
| Gss | 443.3035945 | 0.013777695 | 0.072338027 | 0.190462688 | 0.848946579 | 0.928531575 | Tbl1x | 1628.922442 | 0.044319998 | 0.057572706 | 0.769809193 | 0.441413086 | 0.819001978 |
| Apoa2 | 20.9844396 | -0.031848781 | 0.167511403 | -0.190129032 | 0.849208021 | 0.928542254 | Ankrd46 | 8837.500308 | 0.044324842 | 0.044941212 | 0.986284959 | 0.323993299 | 0.752925264 |
| Ing2 | 979.5376707 | -0.014571501 | 0.076627813 | -0.190159428 | 0.849184203 | 0.928542254 | Sdhc | 3109.837334 | 0.044340112 | 0.050540604 | 0.877316614 | 0.380314691 | 0.786275719 |
| Rrp9 | 619.674667 | 0.017565374 | 0.092270698 | 0.190367846 | 0.849020892 | 0.928542254 | Dkk2 | 20.60244448 | 0.044347668 | 0.081198169 | 0.54616587 | 0.584951924 | NA |
| Serinc2 | 225.1979403 | 0.030234624 | 0.159048416 | 0.190096981 | 0.849233136 | 0.928542254 | Ocm | 24.283821 | 0.044368584 | 0.083045045 | 0.534271301 | 0.593153845 | 0.886253111 |
| Suv39h1 | 1187.937113 | 0.01579404 | 0.083096864 | 0.190067817 | 0.849255989 | 0.928542254 | Akirin1 | 2148.36703 | 0.044390296 | 0.061496656 | 0.721832689 | 0.470397349 | 0.833245343 |
| Ido2 | 76.15362988 | 0.028316441 | 0.149081247 | 0.189939659 | 0.849356415 | 0.928586529 | B230323A14Rik | 3.064303286 | 0.044390579 | 0.036454969 | 1.217682524 | 0.223344641 | NA |
| Armc10 | 818.6670109 | 0.012936698 | 0.068212668 | 0.18965243 | 0.8495815 | 0.928767075 | Isoc2a | 705.626877 | 0.044398919 | 0.068245376 | 0.650577685 | 0.515319139 | 0.853297835 |
| Hs6st1 | 2689.588529 | 0.016665209 | 0.087931425 | 0.189525065 | 0.849681312 | 0.928810656 | Pls1 | 608.9812672 | 0.044409413 | 0.078676417 | 0.564456479 | 0.57244351 | 0.879277718 |
| Mios | 907.7136053 | 0.013856363 | 0.073218165 | 0.189247617 | 0.849898749 | 0.928982802 | Coa4 | 80.24238244 | 0.044411909 | 0.0950481 | 0.467257195 | 0.640315878 | 0.910092095 |
| Cenph | 19.03299071 | 0.031760951 | 0.168098096 | 0.188942957 | 0.850137525 | 0.929047173 | Golph3l | 919.1420795 | 0.044424258 | 0.062243444 | 0.713717858 | 0.475401662 | 0.834927401 |
| Isg20l2 | 552.0488136 | -0.014355722 | 0.075967855 | -0.188971006 | 0.850115541 | 0.929047173 | Yes1 | 424.6763001 | 0.044468119 | 0.075375055 | 0.589958028 | 0.555218789 | 0.871360989 |
| Myh14 | 1477.210071 | 0.024081433 | 0.127399218 | 0.189023395 | 0.850074481 | 0.929047173 | Sybu | 3415.872903 | 0.044476223 | 0.052957897 | 0.839841195 | 0.400997432 | 0.799277436 |
| 4833412C05Rik | 27.76268487 | -0.031603568 | 0.167742076 | -0.188405731 | 0.850558608 | 0.929441785 | Mrps6 | 614.6598125 | 0.044488887 | 0.062661873 | 0.709983353 | 0.477714459 | 0.835457925 |
| Zfp53 | 168.2957928 | -0.019424675 | 0.103144467 | -0.188324938 | 0.850621938 | 0.929445437 | Carm1 | 3486.052182 | 0.044506472 | 0.044928272 | 0.990611704 | 0.32187522 | 0.751214565 |
| Gbe1 | 453.2431497 | -0.017221079 | 0.09166574 | -0.187868216 | 0.850979959 | 0.929771065 | Marcks | 3502.891669 | 0.044524195 | 0.072073767 | 0.617758683 | 0.536734423 | 0.86167069 |
| Prkag1 | 1877.766315 | 0.016572308 | 0.088275992 | 0.187732895 | 0.851086042 | 0.929821403 | Scnn1b | 1.824694551 | 0.044532629 | 0.028566859 | 1.55889134 | 0.1190221 | NA |
| Ccdc85b | 1943.503213 | -0.018104509 | 0.096518048 | -0.187576409 | 0.851208721 | 0.929876317 | Myo5a | 24015.03842 | 0.044544645 | 0.063750035 | 0.698739143 | 0.484715067 | 0.838713365 |
| Uxt | 170.7777474 | -0.019350028 | 0.103191524 | -0.187515672 | 0.851256337 | 0.929876317 | Slc5a6 | 1284.201278 | 0.044572785 | 0.059895536 | 0.744175414 | 0.45677036 | 0.827003147 |
| Nisch | 17129.72234 | -0.010048584 | 0.053639938 | -0.187333999 | 0.851398767 | 0.929966338 | Smim7 | 3574.660152 | 0.044594646 | 0.054047996 | 0.82509342 | 0.409318555 | 0.801055189 |
| Arfip1 | 623.7055757 | 0.014701619 | 0.07856456 | 0.187127874 | 0.851560373 | 0.930000411 | St3gal2 | 5290.971887 | 0.044596666 | 0.044280525 | 1.007139489 | 0.313867747 | 0.745923449 |
| Ccdc115 | 589.8750615 | 0.015187064 | 0.0811403 | 0.187170413 | 0.851527022 | 0.930000411 | Tmem132b | 7390.676506 | 0.044610709 | 0.05163601 | 0.863945716 | 0.387617711 | 0.791583702 |
| Cog4 | 2012.736954 | 0.010996807 | 0.058786166 | 0.187064538 | 0.851610032 | 0.930000411 | Ginm1 | 456.5552701 | 0.044631148 | 0.070984174 | 0.628747877 | 0.529514129 | 0.857915133 |
| Larp4b | 2626.189986 | -0.011652162 | 0.062320862 | -0.186970484 | 0.851683775 | 0.930015393 | Eno4 | 141.1017829 | 0.044635462 | 0.093290095 | 0.478458751 | 0.632323726 | 0.906498456 |
| Galm | 100.0177695 | 0.025618158 | 0.137184164 | 0.186742825 | 0.851862277 | 0.930144758 | Utp6 | 1859.445799 | 0.044645281 | 0.063372013 | 0.704495229 | 0.48112442 | 0.8371092 |
| Zscan12 | 271.5163544 | 0.015552226 | 0.083318167 | 0.18666068 | 0.851926687 | 0.930149537 | Ccndbp1 | 1928.43091 | 0.044695842 | 0.051129277 | 0.874173183 | 0.382023948 | 0.786663048 |
| Il11ra1 | 572.0125643 | -0.019851818 | 0.106421323 | -0.18653985 | 0.852021432 | 0.930187434 | Tmem129 | 1151.329135 | 0.044706783 | 0.056120231 | 0.796625066 | 0.425668816 | 0.809519334 |
| Chst5 | 26.48265673 | 0.03130002 | 0.168074849 | 0.186226675 | 0.852267006 | 0.930279 | Ddo | 303.6005966 | 0.04472642 | 0.087497436 | 0.511174067 | 0.609229176 | 0.89440021 |
| Nell2 | 21824.50246 | 0.015467306 | 0.083064499 | 0.186208383 | 0.85228135 | 0.930279 | Emc9 | 565.8434124 | 0.044728822 | 0.071901113 | 0.622088036 | 0.533883982 | 0.859470725 |
| Psen2 | 521.0037318 | 0.013517266 | 0.072615627 | 0.18614817 | 0.852328567 | 0.930279 | Ndrg4 | 81956.22306 | 0.044738088 | 0.058533506 | 0.764315875 | 0.444679027 | 0.821493054 |
| Zfp422 | 776.668348 | -0.013068961 | 0.070215431 | -0.186126619 | 0.852345468 | 0.930279 | Slc39a12 | 2374.079635 | 0.044748958 | 0.063189528 | 0.708170472 | 0.478839401 | 0.835936459 |
| Rpl3l | 12.82023564 | -0.03095389 | 0.166417576 | -0.18600133 | 0.852443718 | 0.9303207 | Uqcrb | 2312.6422 | 0.044752172 | 0.072420087 | 0.617952477 | 0.536606666 | 0.861583776 |
| Mccc2 | 625.639699 | 0.016437901 | 0.088455551 | 0.185832332 | 0.852576249 | 0.930345794 | Trappc10 | 5613.073022 | 0.044756801 | 0.043620459 | 1.026050657 | 0.304867707 | 0.737453291 |
| Trmt10a | 214.4803079 | -0.022996337 | 0.123756736 | -0.185818874 | 0.852586803 | 0.930345794 | Chmp2b | 1570.988367 | 0.04476616 | 0.064736592 | 0.691512465 | 0.48924355 | 0.840836703 |
| AA543186 | 8.427306637 | -0.027843815 | 0.150265941 | -0.185296913 | 0.852996159 | 0.930648992 | Ttc30a2 | 1.986946704 | 0.0447706 | 0.030579427 | 1.464075827 | 0.143173231 | NA |
| Abra | 8.25901321 | -0.029367637 | 0.159033834 | -0.184662825 | 0.853493506 | 0.930648992 | Insig1 | 3059.414677 | 0.044793605 | 0.085975503 | 0.521004277 | 0.602363792 | 0.89112297 |
| Cpd | 4869.484081 | -0.010740822 | 0.058085609 | -0.184913647 | 0.853296767 | 0.930648992 | Cmya5 | 414.7496218 | 0.044801591 | 0.08140449 | 0.550357737 | 0.58207403 | 0.884020489 |
| Elovl7 | 726.9286716 | 0.025928214 | 0.140450701 | 0.184607224 | 0.853537119 | 0.930648992 | Nbas | 2370.687572 | 0.044837927 | 0.057668295 | 0.777514348 | 0.436855372 | 0.816490111 |
| Fam118a | 699.0848997 | -0.013104741 | 0.070783137 | -0.185139304 | 0.853119774 | 0.930648992 | 1700003M02Rik | 70.19258166 | 0.044845705 | 0.094498112 | 0.474567205 | 0.635095486 | 0.907297112 |
| Foxr2 | 21.14520136 | -0.030960789 | 0.167544743 | -0.184791166 | 0.853392837 | 0.930648992 | Atxn3 | 1324.99662 | 0.044855561 | 0.068133846 | 0.658344771 | 0.510316616 | 0.850976829 |
| Lsm7 | 651.2409455 | -0.014629177 | 0.079142634 | -0.184845713 | 0.853350052 | 0.930648992 | Fam108a | 3532.431138 | 0.044865958 | 0.065156759 | 0.68858486 | 0.491084551 | 0.841866734 |
| Rgma | 2021.873058 | 0.01012398 | 0.054728917 | 0.184984111 | 0.853241498 | 0.930648992 | Rc3h1 | 1765.830521 | 0.044884631 | 0.073623169 | 0.609653615 | 0.542091288 | 0.864560611 |
| Shcbp1 | 26.20193717 | -0.030309907 | 0.164240729 | -0.184545621 | 0.853585441 | 0.930648992 | Nt5c3l | 1211.298709 | 0.04491428 | 0.064523568 | 0.696091081 | 0.486371792 | 0.839801621 |
| Smarcad1 | 1025.307903 | 0.015363266 | 0.083023644 | 0.185046876 | 0.853192269 | 0.930648992 | Tas1r1 | 6.482190712 | 0.044919166 | 0.051713112 | 0.868622373 | 0.385053713 | NA |
| Strn3 | 2711.490628 | 0.013184776 | 0.071372099 | 0.184732916 | 0.853438527 | 0.930648992 | C1qbp | 1888.3787 | 0.044932364 | 0.067626241 | 0.664422034 | 0.506420234 | 0.848908639 |
| Vwa9 | 807.2230178 | 0.013534621 | 0.073335291 | 0.184558084 | 0.853575665 | 0.930648992 | Laptm4a | 6802.187437 | 0.044948954 | 0.055022257 | 0.816923134 | 0.413972356 | 0.804285711 |
| Foxf1 | 33.81081508 | 0.029991954 | 0.162748812 | 0.184283705 | 0.853790896 | 0.930807497 | Hsp90ab1 | 67936.94723 | 0.044968559 | 0.033928462 | 1.325393351 | 0.185040721 | 0.637780733 |
| Gm15545 | 14.60392867 | -0.030572763 | 0.165970727 | -0.184205753 | 0.853852047 | 0.930808669 | Mblac2 | 908.3606643 | 0.0449749 | 0.062785577 | 0.71632534 | 0.473790484 | 0.833983942 |
| Adh5 | 1683.810814 | 0.009814553 | 0.05340946 | 0.183760575 | 0.854201286 | 0.930861916 | 4932441J04Rik | 29.20089624 | 0.044979802 | 0.08987124 | 0.500491617 | 0.616728958 | 0.897296158 |
| Coil | 784.7968806 | 0.011216755 | 0.061039823 | 0.183761266 | 0.854200744 | 0.930861916 | Kcnma1 | 4212.108782 | 0.044995226 | 0.066606132 | 0.675541795 | 0.499331607 | 0.846241311 |
| Fstl3 | 69.54222957 | 0.024693306 | 0.134366951 | 0.183775145 | 0.854189856 | 0.930861916 | Otub1 | 7709.163798 | 0.045001995 | 0.045216382 | 0.995258641 | 0.319610488 | 0.749632194 |
| Katnal1 | 3971.691127 | 0.008739849 | 0.047555446 | 0.183782303 | 0.85418424 | 0.930861916 | Fbxl4 | 798.6534185 | 0.045014602 | 0.06355876 | 0.708235991 | 0.478798719 | 0.835936459 |
| Xrcc2 | 261.2427055 | 0.017718479 | 0.096356492 | 0.18388464 | 0.854103955 | 0.930861916 | Acbd5 | 2832.858341 | 0.045023037 | 0.067564818 | 0.666368063 | 0.505175871 | 0.848183273 |
| Fam60a | 88.72269921 | -0.025180752 | 0.137140303 | -0.183613072 | 0.854317007 | 0.930922548 | Txndc11 | 1214.490965 | 0.045031285 | 0.058033178 | 0.775957591 | 0.437774027 | 0.816984132 |
| Cant1 | 1932.726305 | 0.010213039 | 0.05572207 | 0.18328534 | 0.854574136 | 0.931137248 | C1qtnf6 | 80.07360459 | 0.045032983 | 0.095102193 | 0.473522025 | 0.635840791 | 0.907889736 |
| Gimap8 | 60.41812464 | 0.026956229 | 0.14727823 | 0.183029283 | 0.854775043 | 0.931269015 | Hdgf | 6494.240487 | 0.04503816 | 0.038614777 | 1.166345203 | 0.243474904 | 0.692772033 |
| Tmem160 | 593.9720746 | -0.016131478 | 0.088160752 | -0.182978002 | 0.854815279 | 0.931269015 | Zc4h2 | 723.2434763 | 0.045042576 | 0.078359237 | 0.574821521 | 0.56541201 | 0.875364912 |
| Ehmt2 | 7905.799161 | 0.011900013 | 0.065116166 | 0.182750515 | 0.854993778 | 0.931332508 | Osbpl3 | 1679.310054 | 0.045056165 | 0.082837032 | 0.543913321 | 0.586501122 | 0.884593012 |
| Fam151b | 177.2680439 | -0.021716939 | 0.118828978 | -0.182757939 | 0.854987953 | 0.931332508 | Papss2 | 839.2769651 | 0.045064115 | 0.074494934 | 0.604928582 | 0.545226451 | 0.866611164 |
| Midn | 2183.873371 | -0.024589066 | 0.134666581 | -0.182592189 | 0.855118014 | 0.931402355 | Mul1 | 2047.812084 | 0.04509127 | 0.047528976 | 0.948711166 | 0.342767533 | 0.763762832 |
| Mmp28 | 111.1680894 | -0.021838346 | 0.119806537 | -0.182280091 | 0.855362922 | 0.931603621 | Dcstamp | 4.176112139 | 0.045117702 | 0.039746609 | 1.135133368 | 0.256319442 | NA |
| Naif1 | 96.22778709 | 0.024516392 | 0.13462835 | 0.182104227 | 0.855500931 | 0.931642409 | Ip6k1 | 11946.93314 | 0.045145969 | 0.048866054 | 0.923871786 | 0.355553074 | 0.772370994 |
| Senp2 | 4274.369935 | -0.010447225 | 0.057376653 | -0.182081465 | 0.855518794 | 0.931642409 | Tnpo3 | 4183.72099 | 0.045159374 | 0.049689227 | 0.908836319 | 0.363436532 | 0.776821522 |
| 4930481A15Rik | 64.08803779 | -0.026740508 | 0.147335511 | -0.181493981 | 0.855979857 | 0.931766735 | Rarres2 | 213.6741156 | 0.045163192 | 0.095056774 | 0.47511808 | 0.634702812 | 0.907297112 |
| Fam189a1 | 3209.389555 | 0.012739998 | 0.070103856 | 0.181730339 | 0.855794355 | 0.931766735 | Cacna2d3 | 5442.984278 | 0.045164289 | 0.068544246 | 0.658907078 | 0.50995544 | 0.850575622 |
| Fgfr2 | 2224.543239 | -0.015813242 | 0.087136711 | -0.181476235 | 0.855993785 | 0.931766735 | Ksr2 | 965.5400077 | 0.04520676 | 0.072318396 | 0.62510733 | 0.531900617 | 0.858328942 |
| Gjc3 | 1671.650198 | 0.023061021 | 0.127021183 | 0.181552559 | 0.855933883 | 0.931766735 | Grm3 | 3024.754287 | 0.045215607 | 0.064299751 | 0.70320035 | 0.481930901 | 0.837383765 |
| Pfkfb3 | 1926.590514 | 0.022420726 | 0.123319732 | 0.181809723 | 0.855732054 | 0.931766735 | Thrap3 | 5817.448631 | 0.045217327 | 0.0426607 | 1.059929343 | 0.289176745 | 0.724444298 |
| Snx2 | 2017.458977 | -0.012094036 | 0.066592035 | -0.181613851 | 0.855885778 | 0.931766735 | Csnk2a2 | 1472.416881 | 0.045217548 | 0.051969803 | 0.87007349 | 0.384260244 | 0.788718992 |
| Aimp1 | 1379.377835 | -0.012430511 | 0.06875195 | -0.180802305 | 0.856522753 | 0.932277031 | Tsga14 | 334.2112915 | 0.045219733 | 0.078448304 | 0.576427161 | 0.56432649 | 0.875323716 |
| H2-DMa | 504.0634724 | -0.024259737 | 0.134252294 | -0.180702591 | 0.856601025 | 0.932280843 | Vrtn | 4.278817104 | 0.045236145 | 0.048817359 | 0.926640555 | 0.354113191 | NA |
| Slc35f5 | 675.4444576 | 0.012568278 | 0.069589548 | 0.180605835 | 0.856676976 | 0.932280843 | Ehd3 | 5963.304865 | 0.045242612 | 0.057598927 | 0.785476644 | 0.432174159 | 0.813254665 |
| Tlr3 | 584.2735962 | 0.015604809 | 0.086420736 | 0.180567884 | 0.856706766 | 0.932280843 | 5033406O09Rik | 7.494689832 | 0.045245195 | 0.055251462 | 0.818895885 | 0.412845818 | NA |
| Ly6c1 | 1011.477383 | 0.020797984 | 0.115327064 | 0.180339144 | 0.856886327 | 0.932407044 | Fkrp | 2446.954951 | 0.045249799 | 0.057034553 | 0.793375184 | 0.427559264 | 0.810334558 |
| Naa60 | 4317.511343 | -0.006978166 | 0.038710205 | -0.18026683 | 0.856943095 | 0.932407044 | Pcnp | 5536.58297 | 0.045280704 | 0.053466523 | 0.846898248 | 0.397051839 | 0.797284168 |
| Ifrd1 | 1410.914111 | 0.020552304 | 0.114104536 | 0.180118206 | 0.85705977 | 0.932468512 | Pcbd2 | 174.9820814 | 0.045281632 | 0.093557947 | 0.483995569 | 0.628389014 | 0.904028802 |
| Jmjd8 | 1302.631359 | -0.011971617 | 0.066574244 | -0.179823544 | 0.857291099 | 0.93252375 | B3gnt5 | 28.36080055 | 0.045289136 | 0.084643684 | 0.535056298 | 0.592610928 | 0.886226713 |
| Larp1 | 6294.587717 | -0.012919946 | 0.071840312 | -0.179842572 | 0.857276161 | 0.93252375 | Cuedc2 | 3873.381626 | 0.045318258 | 0.044967338 | 1.007803891 | 0.313548618 | 0.745844951 |
| Trp63 | 22.58944417 | -0.029807007 | 0.16573154 | -0.179851143 | 0.857269432 | 0.93252375 | Zfp143 | 490.4812056 | 0.045322409 | 0.078722568 | 0.575723202 | 0.56480229 | 0.875364912 |
| Hnrnpr | 2528.107511 | -0.010524565 | 0.058660523 | -0.179414783 | 0.857612024 | 0.932741872 | Gas7 | 26669.47141 | 0.045355054 | 0.053424033 | 0.848963495 | 0.395901606 | 0.796351698 |
| Pgr | 974.4240757 | 0.019250236 | 0.107290461 | 0.179421691 | 0.857606601 | 0.932741872 | Arl8b | 9487.114692 | 0.045362921 | 0.069400097 | 0.653643484 | 0.513341522 | 0.853195313 |
| Nek3 | 57.00170093 | 0.026706281 | 0.148989758 | 0.179249107 | 0.857742106 | 0.932778042 | Rab39b | 1724.127224 | 0.045364682 | 0.076996548 | 0.589178128 | 0.555741788 | 0.87140103 |
| Zzz3 | 2356.201211 | 0.010039344 | 0.05601716 | 0.179219076 | 0.857765686 | 0.932778042 | Zfp275 | 1480.222797 | 0.045369887 | 0.07270124 | 0.624059333 | 0.53258862 | 0.858328942 |
| Ski | 6830.00348 | -0.009068559 | 0.050689819 | -0.178902971 | 0.85801389 | 0.932982469 | E030013I19Rik | 51.28353327 | 0.04537052 | 0.093441543 | 0.485549771 | 0.627286417 | 0.903068631 |
| B230118H07Rik | 732.6164316 | 0.01576599 | 0.08820319 | 0.178746259 | 0.858136944 | 0.932987515 | Rdh11 | 903.3951386 | 0.045381643 | 0.065518921 | 0.692649423 | 0.488529587 | 0.840318276 |
| Rabac1 | 2860.612652 | 0.013428402 | 0.075191098 | 0.178590322 | 0.858259394 | 0.932987515 | Tmem101 | 698.9130905 | 0.045407958 | 0.062796249 | 0.723099845 | 0.469618544 | 0.833245343 |
| Sertad3 | 82.50069842 | -0.025859823 | 0.144798329 | -0.178592 | 0.858258077 | 0.932987515 | Adam22 | 10178.21649 | 0.045415683 | 0.060073198 | 0.756005744 | 0.449645752 | 0.823263966 |
| Znrd1as | 104.2594589 | 0.025216151 | 0.14111337 | 0.17869427 | 0.858177769 | 0.932987515 | Lrp12 | 2548.397071 | 0.045416377 | 0.058153468 | 0.780974535 | 0.434817472 | 0.815326688 |
| Dip2b | 4044.149763 | -0.008806491 | 0.049362686 | -0.178403809 | 0.858405858 | 0.933081266 | Gm4477 | 2.814227903 | 0.045446152 | 0.039516928 | 1.150042642 | 0.250126309 | NA |
| Get4 | 1755.022801 | 0.01446647 | 0.081176228 | 0.178210665 | 0.858557534 | 0.933180669 | Hp1bp3 | 10974.56032 | 0.045448944 | 0.046767697 | 0.971802052 | 0.331149035 | 0.756212998 |
| Fzd1 | 420.3180396 | -0.021516984 | 0.120814396 | -0.178099504 | 0.858644831 | 0.933210088 | Snrnp27 | 1097.872609 | 0.045465263 | 0.06692373 | 0.679359363 | 0.496910192 | 0.845038521 |
| Syap1 | 868.6686939 | 0.011848808 | 0.066631075 | 0.17782706 | 0.858858795 | 0.93337716 | Slc23a1 | 9.653164508 | 0.045469828 | 0.06756512 | 0.672977826 | 0.500961398 | NA |
| Eml6 | 1171.415609 | 0.015660125 | 0.088202961 | 0.177546474 | 0.859079164 | 0.933420236 | Adamtsl3 | 50.51023068 | 0.045476035 | 0.089647052 | 0.507278643 | 0.611959321 | 0.895245244 |
| Golt1b | 1347.891973 | -0.0118331 | 0.066647134 | -0.177548528 | 0.85907755 | 0.933420236 | Mex3c | 1629.261196 | 0.045493935 | 0.077637439 | 0.585979333 | 0.557889403 | 0.872673743 |
| Kdm4a | 2273.818255 | -0.014287363 | 0.080425704 | -0.17764672 | 0.859000431 | 0.933420236 | Arhgef7 | 5862.864316 | 0.045494322 | 0.056000584 | 0.812390135 | 0.416567805 | 0.806153084 |
| Rhot2 | 3389.89882 | -0.016421563 | 0.092633898 | -0.177273796 | 0.859293332 | 0.933587469 | Ccdc105 | 13.01339065 | 0.045497599 | 0.070564235 | 0.644768543 | 0.51907719 | NA |
| Ube2i | 7013.49087 | -0.007844319 | 0.044317392 | -0.177003172 | 0.859505898 | 0.933752937 | Fam76a | 2283.434185 | 0.045530338 | 0.047417448 | 0.960202187 | 0.336953467 | 0.760113439 |
| Atad2b | 704.6344387 | -0.017217593 | 0.097426468 | -0.176723976 | 0.859725207 | 0.933829543 | Hip1 | 3005.812957 | 0.045544811 | 0.058880211 | 0.773516445 | 0.4392168 | 0.817545617 |
| Lsm11 | 1303.944447 | -0.019415971 | 0.109811988 | -0.176811036 | 0.85965682 | 0.933829543 | Vps36 | 547.2749956 | 0.045547386 | 0.069875514 | 0.651836153 | 0.514506876 | 0.853195313 |
| Sgta | 4380.261946 | 0.011842485 | 0.067026652 | 0.176683217 | 0.859757225 | 0.933829543 | Tmem48 | 488.6741707 | 0.045565459 | 0.078474521 | 0.580640164 | 0.561482997 | 0.873934985 |
| E130012A19Rik | 2263.404472 | -0.022373602 | 0.126776155 | -0.176481152 | 0.859915955 | 0.933871017 | Apoc3 | 13.81388057 | 0.045569123 | 0.070139214 | 0.649695381 | 0.515889008 | NA |
| Mal | 8265.278186 | -0.017999683 | 0.101958451 | -0.176539392 | 0.859870204 | 0.933871017 | Arl6 | 801.9515331 | 0.045581614 | 0.066201168 | 0.688531872 | 0.491117906 | 0.841866734 |
| Acy1 | 402.5595259 | -0.018339215 | 0.104278165 | -0.175868217 | 0.860397475 | 0.93408313 | Cpsf3l | 1046.438122 | 0.045582705 | 0.062289425 | 0.731788813 | 0.464297478 | 0.83159896 |
| Atp6v0e2 | 14821.55807 | 0.01059722 | 0.06026324 | 0.175848834 | 0.860412704 | 0.93408313 | Dlx4 | 14.29464018 | 0.045584425 | 0.065122306 | 0.699981745 | 0.483938705 | NA |
| B4galt3 | 1553.322516 | 0.01094893 | 0.062259175 | 0.175860507 | 0.860403533 | 0.93408313 | Tmbim6 | 13537.32555 | 0.04559131 | 0.039217564 | 1.162522727 | 0.245023199 | 0.694614946 |
| Fa2h | 1266.646751 | 0.024079557 | 0.136802506 | 0.176016928 | 0.860280643 | 0.93408313 | Lin52 | 1128.827138 | 0.045637196 | 0.061343935 | 0.743956123 | 0.45690302 | 0.827140238 |
| Shroom4 | 180.2670878 | -0.02326056 | 0.132198268 | -0.175952079 | 0.86033159 | 0.93408313 | Hs3st2 | 1577.557515 | 0.045659288 | 0.069830332 | 0.653860393 | 0.513201753 | 0.853143588 |
| Ehmt1 | 1302.023168 | 0.010291587 | 0.058621763 | 0.175559148 | 0.860640299 | 0.934210484 | Ept1 | 3398.314274 | 0.045686068 | 0.064752707 | 0.705546843 | 0.480469991 | 0.836914993 |
| Gm4925 | 12.29861451 | -0.029195699 | 0.166313636 | -0.175546033 | 0.860650603 | 0.934210484 | Psmg2 | 589.135085 | 0.045691501 | 0.063545475 | 0.719036265 | 0.472118576 | 0.833441697 |
| Golph3l | 823.0960671 | -0.010964919 | 0.062550361 | -0.17529746 | 0.860845908 | 0.934292638 | Rpp14 | 1341.261577 | 0.045708498 | 0.064203157 | 0.711935363 | 0.476504802 | 0.835195112 |
| Pcdha11 | 122.1904716 | 0.026130243 | 0.149178591 | 0.175160814 | 0.860953274 | 0.934292638 | Topbp1 | 1012.20112 | 0.045714583 | 0.071235472 | 0.641739038 | 0.521042638 | 0.85372574 |
| Rad23b | 7500.137476 | 0.010797137 | 0.061625362 | 0.175206063 | 0.86091772 | 0.934292638 | Gpr150 | 154.9789695 | 0.045722466 | 0.089554984 | 0.510551892 | 0.60966487 | 0.89440021 |
| Rrm2b | 1442.642542 | -0.014218375 | 0.08118165 | -0.175142723 | 0.860967489 | 0.934292638 | Insl6 | 11.67445781 | 0.045737944 | 0.062512642 | 0.731659108 | 0.464376661 | NA |
| Gtpbp6 | 825.2574961 | 0.016903531 | 0.096694238 | 0.174814258 | 0.861225584 | 0.934441821 | Mre11a | 402.6208153 | 0.045745819 | 0.075051494 | 0.609525763 | 0.542176002 | 0.86460079 |
| Slc35a2 | 930.2609508 | 0.015007002 | 0.085820179 | 0.174865652 | 0.8611852 | 0.934441821 | Adamts8 | 153.9663283 | 0.045752421 | 0.09028175 | 0.506773752 | 0.612313575 | 0.895344807 |
| Ahcy | 1374.559168 | -0.016828835 | 0.096468134 | -0.174449684 | 0.86151207 | 0.934625736 | Gprin1 | 5979.255856 | 0.045764093 | 0.064840522 | 0.705794643 | 0.480315854 | 0.836914993 |
| Col22a1 | 107.1362851 | 0.023478159 | 0.134587726 | 0.174445024 | 0.861515733 | 0.934625736 | Angptl7 | 8.152985034 | 0.045766873 | 0.059600794 | 0.76789032 | 0.442552349 | NA |
| Sigirr | 56.1650546 | -0.025129764 | 0.144159729 | -0.174318892 | 0.861614853 | 0.934667824 | Fut4 | 13.954792 | 0.045790123 | 0.061513627 | 0.744389906 | 0.456640624 | NA |
| Slc6a17 | 26027.30972 | -0.013439028 | 0.077156152 | -0.174179608 | 0.861724311 | 0.934721119 | Tlk1 | 5023.081272 | 0.045802674 | 0.068904294 | 0.664728873 | 0.506223923 | 0.848908639 |
| Arl2 | 1502.057455 | -0.013260248 | 0.076176907 | -0.174071762 | 0.861809065 | 0.934747613 | Rpl10 | 8775.641122 | 0.04582801 | 0.043210852 | 1.060567141 | 0.288886662 | 0.724444298 |
| Fam117b | 3117.335118 | 0.013505229 | 0.077675446 | 0.173867408 | 0.861969667 | 0.93479093 | Mis12 | 1069.869418 | 0.045841097 | 0.069026296 | 0.664110637 | 0.506619503 | 0.848943697 |
| Mir3082 | 8.646482355 | 0.023990345 | 0.137928954 | 0.173932622 | 0.861918414 | 0.93479093 | Sgip1 | 5922.293603 | 0.045849454 | 0.051939432 | 0.882748459 | 0.377372194 | 0.783997209 |
| Gm4787 | 15.75465411 | 0.028962348 | 0.1667884 | 0.173647257 | 0.862142691 | 0.934840346 | Fads3 | 1467.440238 | 0.045852918 | 0.067984795 | 0.674458428 | 0.500019907 | 0.846241311 |
| U2af1l4 | 872.8100903 | -0.015304611 | 0.08810341 | -0.173711907 | 0.862091879 | 0.934840346 | Pom121 | 3919.997651 | 0.045854256 | 0.050304784 | 0.91152874 | 0.36201685 | 0.776449324 |
| Vim | 1536.529387 | -0.016590545 | 0.09557915 | -0.173579123 | 0.86219624 | 0.934840346 | Cd59b | 23.37160762 | 0.045916258 | 0.082075605 | 0.559438554 | 0.575862455 | 0.880811222 |
| Epas1 | 6564.44705 | -0.010289574 | 0.059322564 | -0.173451261 | 0.862296735 | 0.934842752 | Hsd17b13 | 9.089481088 | 0.045956012 | 0.051852734 | 0.886279436 | 0.375466967 | NA |
| Irs3 | 9.760925453 | 0.025720749 | 0.148312411 | 0.173422677 | 0.862319131 | 0.934842752 | 4930426D05Rik | 37.79481697 | 0.046005186 | 0.089205915 | 0.515719009 | 0.606050676 | 0.892684114 |
| F930015N05Rik | 66.58481139 | -0.024917239 | 0.143817773 | -0.17325563 | 0.862450499 | 0.934919753 | Mpp5 | 1646.886733 | 0.046023545 | 0.061905341 | 0.743450318 | 0.45720909 | 0.827384981 |
| Trim25 | 349.6181088 | 0.021487266 | 0.124100372 | 0.173144248 | 0.862538045 | 0.934949242 | Hcfc2 | 969.1486022 | 0.04603306 | 0.063040144 | 0.730218191 | 0.465256825 | 0.831682622 |
| Dgkz | 20930.5799 | -0.016876264 | 0.097625907 | -0.172866651 | 0.862756245 | 0.93512034 | Gatad2b | 1173.615804 | 0.046034562 | 0.073262737 | 0.628348922 | 0.52977539 | 0.857915133 |
| 2310044G17Rik | 4038.125675 | -0.008549817 | 0.049585618 | -0.172425332 | 0.863103158 | 0.935401714 | Ift80 | 1121.762585 | 0.046040179 | 0.058369062 | 0.788777097 | 0.430242298 | 0.811860174 |
| Arl6ip6 | 235.7356215 | 0.015949672 | 0.092524729 | 0.172382806 | 0.863136589 | 0.935401714 | Lamp2 | 2677.796491 | 0.046042601 | 0.066073748 | 0.696836524 | 0.485905106 | 0.839716418 |
| Igf2 | 3141.66255 | 0.027784624 | 0.161361082 | 0.172189126 | 0.863288847 | 0.935501286 | Itgbl1 | 58.09859479 | 0.046058628 | 0.091684768 | 0.502358557 | 0.615415325 | 0.896719391 |
| Psmd6 | 2014.471549 | -0.01100942 | 0.064002729 | -0.17201486 | 0.863425847 | 0.935584311 | C030037D09Rik | 172.7147809 | 0.046061999 | 0.091141201 | 0.505391619 | 0.613283805 | 0.895602749 |
| Nphs1 | 81.4127743 | 0.026257464 | 0.152759697 | 0.17188738 | 0.863526068 | 0.935627475 | Slc5a11 | 25.12181864 | 0.04606376 | 0.078623709 | 0.585876206 | 0.557958708 | 0.872673743 |
| Asf1b | 28.25143863 | -0.028591719 | 0.1664508 | -0.171772794 | 0.863616155 | 0.935659653 | Odf3l2 | 10.26046263 | 0.046068585 | 0.066825327 | 0.689388107 | 0.490579065 | NA |
| Hnrnpa3 | 6121.153902 | -0.017976562 | 0.104766768 | -0.171586492 | 0.863762627 | 0.935752912 | Wrnip1 | 2107.647197 | 0.046069782 | 0.052176246 | 0.882964677 | 0.377255357 | 0.783997209 |
| Atmin | 4509.069956 | 0.014056039 | 0.082135603 | 0.171132108 | 0.864119889 | 0.935781262 | Kcnq4 | 280.9423353 | 0.046093231 | 0.086947976 | 0.530124249 | 0.596025787 | 0.887804613 |
| Dcdc2c | 124.5243508 | -0.021553907 | 0.125908439 | -0.171187154 | 0.864076608 | 0.935781262 | Fam84b | 389.9520169 | 0.046097931 | 0.09382792 | 0.491302915 | 0.623212218 | 0.900470921 |
| Gm4349 | 20.24079729 | 0.028633696 | 0.167205481 | 0.171248548 | 0.864028336 | 0.935781262 | Thrsp | 618.7899953 | 0.046116852 | 0.074324132 | 0.620482893 | 0.534939913 | 0.860334055 |
| Psd2 | 2463.7829 | -0.015767126 | 0.092163331 | -0.171078085 | 0.864162368 | 0.935781262 | P2rx2 | 4.341453551 | 0.04611963 | 0.047839614 | 0.96404686 | 0.335022433 | NA |
| Tcp1 | 7167.527191 | -0.010465955 | 0.061198863 | -0.171015509 | 0.864211571 | 0.935781262 | Matn3 | 2.811696213 | 0.046127791 | 0.040475731 | 1.139640711 | 0.254436017 | NA |
| Wdr65 | 45.16773207 | 0.028799916 | 0.168086999 | 0.171339343 | 0.863956946 | 0.935781262 | Slc25a32 | 325.0354166 | 0.046158721 | 0.085846003 | 0.537692134 | 0.590789611 | 0.885693151 |
| Zfp526 | 403.2339095 | 0.015865082 | 0.092731292 | 0.171086605 | 0.864155668 | 0.935781262 | Fundc2 | 1653.157992 | 0.046168962 | 0.062188132 | 0.742407929 | 0.457840217 | 0.827642075 |
| Rce1 | 521.4163589 | 0.015598805 | 0.091309845 | 0.170833769 | 0.864354476 | 0.935870597 | Car12 | 1000.483699 | 0.046173215 | 0.085407625 | 0.540621689 | 0.588768365 | 0.885188411 |
| Abracl | 198.5400436 | 0.022395558 | 0.13117762 | 0.170726974 | 0.864438452 | 0.93589612 | Ube2h | 5839.538812 | 0.046174462 | 0.048652914 | 0.949058514 | 0.342590853 | 0.763762832 |
| Clip4 | 2506.099618 | -0.018294823 | 0.10729707 | -0.17050627 | 0.864612004 | 0.936018613 | Prok2 | 7.155140366 | 0.046230248 | 0.05643805 | 0.819132628 | 0.412710748 | NA |
| Hk1 | 16890.39822 | -0.012041658 | 0.070721709 | -0.170268199 | 0.864799219 | 0.936113441 | Dnajb14 | 804.8074676 | 0.046236216 | 0.064223647 | 0.71992511 | 0.471571106 | 0.833245343 |
| Tab3 | 1069.814722 | -0.014374682 | 0.084437138 | -0.170241223 | 0.864820433 | 0.936113441 | Mrpl3 | 1898.789156 | 0.046244037 | 0.064053211 | 0.721962826 | 0.470317333 | 0.833245343 |
| 1810062O18Rik | 34.12228254 | -0.027184373 | 0.160096206 | -0.16980023 | 0.865167246 | 0.936273888 | 9530052E02Rik | 13.79833074 | 0.046247707 | 0.066115938 | 0.699494079 | 0.484243312 | NA |
| 2610524H06Rik | 368.6887947 | -0.013268687 | 0.078428042 | -0.169182945 | 0.865652745 | 0.936273888 | Lamtor1 | 2409.271538 | 0.046248909 | 0.051965009 | 0.89000098 | 0.37346536 | 0.782348609 |
| Ampd2 | 2899.248292 | -0.017107751 | 0.100912845 | -0.169529966 | 0.865379804 | 0.936273888 | Serpinb6a | 1191.053835 | 0.046261313 | 0.057218356 | 0.808504752 | 0.418800072 | 0.806975048 |
| Cdkal1 | 421.2800841 | -0.014151287 | 0.083496878 | -0.169482832 | 0.865416875 | 0.936273888 | Dbt | 869.4177858 | 0.046279244 | 0.058942454 | 0.78515977 | 0.432359899 | 0.813262565 |
| 9-Mar | 615.0578912 | -0.011291036 | 0.06667253 | -0.169350644 | 0.865520843 | 0.936273888 | Gpr4 | 146.6471352 | 0.046281621 | 0.089918301 | 0.514707464 | 0.606757457 | 0.893150688 |
| Mdp1 | 1424.19307 | -0.010334746 | 0.061128224 | -0.169066688 | 0.865744187 | 0.936273888 | Gdap1 | 5040.736906 | 0.046306435 | 0.065876467 | 0.702928328 | 0.482100416 | 0.837477954 |
| Mtrf1l | 239.6337421 | -0.015979038 | 0.094392104 | -0.169283632 | 0.86557355 | 0.936273888 | Gm2518 | 12.24174935 | 0.046312222 | 0.069270782 | 0.668567914 | 0.503771146 | NA |
| Ppp3r1 | 28685.26238 | -0.013233251 | 0.078046085 | -0.169556882 | 0.865358635 | 0.936273888 | Thop1 | 1659.378584 | 0.046316006 | 0.064426326 | 0.718898758 | 0.472203302 | 0.833441697 |
| Set | 5879.256784 | -0.009164455 | 0.053941831 | -0.169895136 | 0.865092606 | 0.936273888 | Cltb | 5343.39248 | 0.046319589 | 0.043188589 | 1.072495994 | 0.283497312 | 0.721034821 |
| Tlr6 | 21.81002346 | 0.028158953 | 0.166096093 | 0.16953411 | 0.865376545 | 0.936273888 | Bcl7b | 1801.100569 | 0.046319824 | 0.053979332 | 0.858102962 | 0.390835612 | 0.794184423 |
| Toe1 | 438.9140672 | 0.013977515 | 0.082628399 | 0.169161146 | 0.865669891 | 0.936273888 | Sgtb | 6196.746426 | 0.046327633 | 0.063795885 | 0.726185274 | 0.467725198 | 0.832353544 |
| Ttyh1 | 19043.01706 | -0.007815355 | 0.046045622 | -0.169730687 | 0.865221939 | 0.936273888 | Scly | 606.9125546 | 0.046340396 | 0.065934401 | 0.702825756 | 0.482164344 | 0.837488851 |
| Zmym2 | 3695.228136 | -0.011353688 | 0.067160159 | -0.169053923 | 0.865754227 | 0.936273888 | Atf2 | 5146.090263 | 0.046344308 | 0.067460784 | 0.686981462 | 0.492094411 | 0.842340249 |
| 1110054M08Rik | 111.5220471 | 0.022146323 | 0.131121525 | 0.16889922 | 0.865875912 | 0.93634013 | Cebpd | 72.12625372 | 0.046369391 | 0.093088578 | 0.498121169 | 0.618398641 | 0.898465559 |
| Fam53b | 1897.639704 | 0.019248602 | 0.114020447 | 0.16881711 | 0.865940499 | 0.936344623 | Ppp3r1 | 31725.43799 | 0.046376642 | 0.079073567 | 0.586499932 | 0.557539607 | 0.872623676 |
| 4930556M19Rik | 9.227431491 | 0.026757754 | 0.159243386 | 0.168030548 | 0.866559247 | 0.936411385 | Kcnab3 | 4350.224734 | 0.046391626 | 0.074348832 | 0.623972495 | 0.532645649 | 0.858328942 |
| Abhd6 | 2074.904578 | -0.014446968 | 0.085792103 | -0.168395073 | 0.866272484 | 0.936411385 | Rpl3 | 18791.30444 | 0.046396371 | 0.047685586 | 0.972964257 | 0.330571067 | 0.756141871 |

| | | | | | | | | | | | | | |
|---|---|---|---|---|---|---|---|---|---|---|---|---|---|
| Diap1 | 2418.892902 | 0.019786781 | 0.117799254 | 0.167970341 | 0.866606612 | 0.936411385 | Irf2bp2 | 1449.896811 | 0.046407226 | 0.076972156 | 0.602909272 | 0.546569048 | 0.867128614 |
| Gorasp2 | 3845.741731 | 0.011352127 | 0.067424972 | 0.168366801 | 0.866294724 | 0.936411385 | Bcl7c | 316.0941806 | 0.046415132 | 0.077960759 | 0.595365321 | 0.551599301 | 0.869755927 |
| Ipo13 | 4119.97244 | 0.010286325 | 0.061215714 | 0.168034058 | 0.866556485 | 0.936411385 | Emid1 | 135.0211432 | 0.046424563 | 0.094555665 | 0.490976015 | 0.623443412 | 0.900570704 |
| Olfr78 | 44.40332962 | -0.026908616 | 0.159705959 | -0.168488489 | 0.866198999 | 0.936411385 | Fkbp3 | 3872.051503 | 0.04642505 | 0.068393932 | 0.67878901 | 0.497271558 | 0.845038521 |
| Riiad1 | 48.29344377 | 0.026391183 | 0.156876712 | 0.168228811 | 0.866403276 | 0.936411385 | Fbxo40 | 8.094448538 | 0.046433742 | 0.052885263 | 0.878009112 | 0.379938774 | NA |
| Sertad2 | 1327.933488 | -0.013266656 | 0.078713369 | -0.168543871 | 0.866155434 | 0.936411385 | Six6 | 5.085214279 | 0.046434107 | 0.030518879 | 1.52148795 | NA | NA |
| Slc25a15 | 496.9165571 | -0.012732928 | 0.075719383 | -0.168159422 | 0.866457863 | 0.936411385 | Fam108c | 976.5965541 | 0.046483277 | 0.066166004 | 0.702525066 | 0.482351775 | 0.837629227 |
| Tgfb1 | 225.5138237 | -0.01894941 | 0.112569822 | -0.168334728 | 0.866319954 | 0.936411385 | Chn1 | 46967.97579 | 0.046539425 | 0.060026207 | 0.775318428 | 0.438151523 | 0.816984132 |
| Zbtb25 | 353.6152405 | 0.015267709 | 0.091038583 | 0.167705914 | 0.866814643 | 0.93656606 | Psma3 | 3206.687555 | 0.046564389 | 0.055504393 | 0.838931593 | 0.401507698 | 0.799277436 |
| Zfp874b | 447.417298 | 0.013014108 | 0.077633725 | 0.167634725 | 0.866870651 | 0.93656606 | Atp8a1 | 11217.54984 | 0.046578243 | 0.060718045 | 0.767123566 | 0.443008052 | 0.820159899 |
| Ccdc28a | 585.1135855 | 0.011614931 | 0.069369334 | 0.1674361 | 0.867026923 | 0.936619688 | Pdk4 | 538.2805523 | 0.046609935 | 0.084666071 | 0.55051492 | 0.581966246 | 0.884020489 |
| Wdr34 | 1019.914335 | -0.01243662 | 0.074284857 | -0.167417967 | 0.867041189 | 0.936619688 | Neurog1 | 5.445158285 | 0.046610454 | 0.044609388 | 1.044857494 | 0.296088838 | NA |
| Morc4 | 155.3713355 | -0.022746238 | 0.135991738 | -0.167261911 | 0.867163973 | 0.936621721 | Sra1 | 1002.011613 | 0.046612607 | 0.054677717 | 0.852497319 | 0.393938126 | 0.794754188 |
| Ric8 | 2169.884845 | 0.010002849 | 0.059792811 | 0.167291832 | 0.867140431 | 0.936621721 | Cbx1 | 726.8946924 | 0.046627696 | 0.066797674 | 0.698043707 | 0.485149861 | 0.838982004 |
| Dpy19l4 | 635.531608 | -0.015182925 | 0.091012606 | -0.166822222 | 0.867509933 | 0.936799477 | Il22ra1 | 4.127623594 | 0.046634623 | 0.046536714 | 1.002103894 | 0.316293418 | NA |
| Dusp28 | 674.7630908 | 0.016658425 | 0.099802143 | 0.166914498 | 0.867437325 | 0.936799477 | Fmnl2 | 2158.147751 | 0.046669464 | 0.056382148 | 0.827734777 | 0.407820719 | 0.801055189 |
| Zdhhc5 | 2646.557679 | -0.013466238 | 0.080710513 | -0.16684614 | 0.867491113 | 0.936799477 | Uba1 | 15890.43162 | 0.046673632 | 0.037133168 | 1.256925668 | 0.208780553 | 0.66358495 |
| Hexa | 2136.531987 | 0.017103872 | 0.102663372 | 0.166601507 | 0.867683608 | 0.936921724 | Lmo3 | 3682.135072 | 0.046688593 | 0.075872771 | 0.615353726 | 0.538321144 | 0.862082844 |
| 9130227L01Rik | 9.932814981 | 0.026764769 | 0.160754641 | 0.166494535 | 0.867767783 | 0.936947319 | Fam178b | 7.044463249 | 0.046700274 | 0.047194266 | 0.989532805 | 0.322402528 | NA |
| Pigp | 651.4541943 | -0.014333709 | 0.086135268 | -0.16640929 | 0.867834864 | 0.936954455 | Poc5 | 1381.643238 | 0.046718394 | 0.057510027 | 0.812352145 | 0.416589597 | 0.806153084 |
| Zfp444 | 1031.943148 | -0.012546151 | 0.075519971 | -0.166130247 | 0.868054451 | 0.937126231 | Sumo3 | 5379.947736 | 0.046747513 | 0.049363486 | 0.947005924 | 0.34363576 | 0.763762832 |
| Gnpnat1 | 320.5264742 | -0.015922907 | 0.095989505 | -0.165881748 | 0.868250011 | 0.93715637 | Pcna | 1566.822678 | 0.046754134 | 0.055241547 | 0.846358154 | 0.397352975 | 0.797340699 |
| Mad2l1bp | 302.0338083 | 0.016423135 | 0.099015554 | 0.165864196 | 0.868263825 | 0.93715637 | Eci2 | 1204.114626 | 0.0467819 | 0.070182115 | 0.666578653 | 0.505041308 | 0.848183273 |
| Zfp219 | 1163.620912 | 0.014398027 | 0.086752279 | 0.165967127 | 0.86818282 | 0.93715637 | Naa11 | 2.131511852 | 0.046836663 | 0.034419832 | 1.360746417 | 0.173593841 | NA |
| Cd79a | 12.35307453 | 0.026221886 | 0.158731072 | 0.165196929 | 0.868788982 | 0.937641239 | Rad21 | 5399.474624 | 0.046841316 | 0.057086581 | 0.820531121 | 0.411913396 | 0.802795661 |
| Fam167b | 11.85442749 | -0.025027567 | 0.151578676 | -0.16511272 | 0.868855261 | 0.937641239 | Arpc2 | 6863.342616 | 0.046872135 | 0.057981381 | 0.808399773 | 0.418860483 | 0.806975048 |
| Leng9 | 221.9550056 | -0.015918412 | 0.09648146 | -0.164989338 | 0.868952374 | 0.937641239 | Pik3c3 | 2522.269866 | 0.046889421 | 0.058784029 | 0.797655796 | 0.425070262 | 0.809421961 |
| Polg2 | 67.75887767 | -0.026285431 | 0.159393578 | -0.164908971 | 0.869015631 | 0.937641239 | Lrrc4 | 5048.97064 | 0.046896084 | 0.053422996 | 0.877825788 | 0.380038268 | 0.786275719 |
| Ywhab | 24096.32406 | -0.007528205 | 0.045637418 | -0.164956856 | 0.868977941 | 0.937641239 | Atp5g2 | 3372.638699 | 0.046899606 | 0.055863705 | 0.839536258 | 0.401168452 | 0.799277436 |
| Col6a4 | 215.4654053 | 0.027025995 | 0.164060037 | 0.164732349 | 0.869154654 | 0.937725939 | Mrps9 | 957.0270195 | 0.046904482 | 0.060284326 | 0.77805435 | 0.436536973 | 0.816420249 |
| Ing3 | 562.1509928 | 0.014055574 | 0.085463727 | 0.164462453 | 0.869367102 | 0.937824541 | 4930452B06Rik | 566.8694402 | 0.046905649 | 0.075751604 | 0.619203376 | 0.535782388 | 0.860926435 |
| Snapc5 | 580.3158523 | 0.01448811 | 0.088092816 | 0.164464143 | 0.869365772 | 0.937824541 | Leprotl1 | 3127.497582 | 0.046915679 | 0.056100315 | 0.836281919 | 0.402996323 | 0.800009209 |
| Etv4 | 310.5063925 | 0.022191709 | 0.135089388 | 0.164274258 | 0.869515246 | 0.937919049 | Tiam1 | 5361.423897 | 0.046921982 | 0.067092129 | 0.699366417 | 0.48432307 | 0.838713365 |
| Ndufs2 | 6986.995156 | -0.0102067 | 0.062214235 | -0.164057305 | 0.869686032 | 0.938037966 | Fis1 | 3826.085012 | 0.046927493 | 0.049573619 | 0.946622299 | 0.343831277 | 0.763762832 |
| Itgb7 | 18.59006237 | 0.027544647 | 0.167982683 | 0.163973132 | 0.869752295 | 0.938044136 | Zcchc3 | 381.3339381 | 0.046943307 | 0.070657215 | 0.664380941 | 0.506446528 | 0.848908639 |
| 1700011J10Rik | 278.0213912 | 0.014985983 | 0.091496951 | 0.163786693 | 0.869899068 | 0.93813713 | F630042J09Rik | 31.03482154 | 0.046956135 | 0.090579571 | 0.51839653 | 0.604181635 | 0.891718993 |
| Hspa12a | 9771.55351 | -0.009691682 | 0.059310609 | -0.163405532 | 0.870199147 | 0.938395434 | Ipo8 | 2372.290738 | 0.046972515 | 0.065384125 | 0.718408548 | 0.472505417 | 0.833671043 |
| Ssr3 | 4804.285719 | -0.012905814 | 0.079089733 | -0.163179383 | 0.870377199 | 0.938522118 | Plk2 | 9175.85058 | 0.046980444 | 0.067158572 | 0.699545008 | 0.484211496 | 0.838713365 |
| Ly6g6f | 13.0132176 | 0.024712933 | 0.151615148 | 0.162997781 | 0.870520182 | 0.938610974 | Mex3a | 176.6476821 | 0.04698804 | 0.089625683 | 0.524269806 | 0.600090895 | 0.889754041 |
| Pdcd5 | 932.4693285 | 0.015109821 | 0.092783149 | 0.162850918 | 0.870635816 | 0.938670332 | Atg3 | 1578.929811 | 0.046994588 | 0.056590135 | 0.830437821 | 0.406291289 | 0.800759397 |
| 4931403E22Rik | 27.13302725 | 0.024627988 | 0.151418141 | 0.162648859 | 0.870794915 | 0.938711223 | Alx3 | 25.26391443 | 0.04699727 | 0.072334575 | 0.649720689 | 0.515872658 | 0.853297835 |
| Sde2 | 858.7217018 | -0.013972244 | 0.085883631 | -0.162688096 | 0.87076402 | 0.938711223 | Dpep2 | 2.742759324 | 0.047008329 | 0.037885081 | 1.240813736 | 0.214674566 | NA |
| Dbn1 | 9118.412048 | 0.010527404 | 0.064790162 | 0.162484608 | 0.870924248 | 0.938785328 | Rph3a | 21853.62779 | 0.047014144 | 0.056545511 | 0.831439014 | 0.405725667 | 0.800733644 |
| Fam175a | 146.504264 | 0.02111675 | 0.130151134 | 0.162247914 | 0.87111063 | 0.938790297 | Gspt1 | 5034.92348 | 0.047014748 | 0.066997178 | 0.701742212 | 0.482839943 | 0.837880119 |
| Fbrsl1 | 1661.515585 | -0.012921422 | 0.079570505 | -0.162389588 | 0.87099907 | 0.938790297 | Itgb4 | 629.0901408 | 0.047014795 | 0.088525864 | 0.531085416 | 0.59535959 | 0.887381665 |
| Zfp273 | 195.1523405 | -0.019073015 | 0.117525479 | -0.16228834 | 0.871078797 | 0.938790297 | Arpc5l | 2522.980218 | 0.047036114 | 0.05883087 | 0.799514173 | 0.423992332 | 0.808999423 |
| Frmd4b | 892.7113225 | 0.012038995 | 0.074399377 | 0.161815803 | 0.871450908 | 0.939091694 | Ispd | 298.1157838 | 0.047044891 | 0.081494294 | 0.577278351 | 0.563751436 | 0.875120313 |
| Zfp72 | 138.4753732 | 0.020322321 | 0.126156442 | 0.161088258 | 0.872023889 | 0.939643795 | Myoz2 | 4.160797024 | 0.047051644 | 0.047911451 | 0.982054262 | 0.326073117 | NA |
| Nsmce4a | 1049.429617 | 0.014735959 | 0.091580862 | 0.160906534 | 0.872167016 | 0.939732666 | Chmp5 | 2487.650144 | 0.047056908 | 0.051465049 | 0.914346897 | 0.360534595 | 0.776337125 |
| Cenpn | 35.67927986 | -0.02696999 | 0.167864943 | -0.160664814 | 0.872357404 | 0.939807093 | Cav3 | 4.642390565 | 0.047061413 | 0.045782935 | 1.027924773 | 0.303985215 | NA |
| Tpst2 | 750.4762317 | 0.01416992 | 0.088188898 | 0.16067691 | 0.872347876 | 0.939807093 | Xrcc3 | 366.9843771 | 0.047065633 | 0.091280502 | 0.515615404 | 0.606123049 | 0.892684114 |
| Coq7 | 646.4501694 | 0.014124141 | 0.088114113 | 0.160293749 | 0.872649683 | 0.940056608 | Kif3a | 8945.720867 | 0.047068231 | 0.050786524 | 0.926785838 | 0.354037739 | 0.771423785 |
| Mir682 | 56.23566017 | 0.026865879 | 0.167955249 | 0.159958556 | 0.872913721 | 0.940275667 | Cyb5d1 | 454.687513 | 0.047092456 | 0.065926716 | 0.71431521 | 0.475032289 | 0.834699977 |
| D830015G02Rik | 14.16509282 | 0.025456755 | 0.159235324 | 0.159868765 | 0.872984454 | 0.940286488 | Prkcz | 12219.44863 | 0.04710345 | 0.046267059 | 1.018077469 | 0.308641142 | 0.740922364 |
| Chmp4b | 6094.359911 | -0.009062981 | 0.056846709 | -0.159428426 | 0.873331345 | 0.940594735 | Mpnd | 2481.101488 | 0.047113091 | 0.06682106 | 0.705063507 | 0.480770716 | 0.836914993 |
| Dtymk | 535.7868936 | -0.017931661 | 0.112630447 | -0.159207939 | 0.873505049 | 0.94068029 | Ppfia2 | 3852.872739 | 0.04711314 | 0.054713536 | 0.86108745 | 0.389189881 | 0.79256585 |
| Phc2 | 3118.060063 | 0.010196562 | 0.064059432 | 0.159173468 | 0.873532207 | 0.94068029 | Ywhab | 27181.18443 | 0.047130186 | 0.051663855 | 0.91224679 | 0.361638818 | 0.776449324 |
| Paip1 | 2364.819852 | 0.009681235 | 0.060879272 | 0.159023494 | 0.873650364 | 0.940742146 | Il4i1 | 1.064684283 | 0.047135859 | 0.025301501 | 1.862966907 | 0.062466922 | NA |
| A430033K04Rik | 669.2723637 | -0.011347727 | 0.071457435 | -0.158803999 | 0.873823298 | 0.940752848 | Rnf214 | 2692.522217 | 0.0471379 | 0.056524566 | 0.833936519 | 0.404316757 | 0.800290023 |
| Ndufa4l2 | 80.71192522 | 0.02253701 | 0.142019313 | 0.158689757 | 0.873913308 | 0.940752848 | Acp1 | 2252.984393 | 0.047167886 | 0.053890296 | 0.875257498 | 0.381433814 | 0.786663048 |
| Nupr1l | 12.20577834 | -0.025195334 | 0.15883527 | -0.158625562 | 0.873963888 | 0.940752848 | 9930021J03Rik | 2943.656426 | 0.047197054 | 0.076489729 | 0.617037798 | 0.537209795 | 0.861996453 |
| Rab10 | 5226.402416 | -0.010594458 | 0.066759868 | -0.158695005 | 0.873909174 | 0.940752848 | Nkx6-2 | 320.7178134 | 0.047222037 | 0.094288383 | 0.500825605 | 0.616493864 | 0.897296158 |
| Tdo2 | 12.27823485 | -0.025298892 | 0.159335913 | -0.158777085 | 0.873844503 | 0.940752848 | Ap3m2 | 7879.773524 | 0.047232872 | 0.04411614 | 1.070648327 | 0.284327585 | 0.721548306 |
| Tmem125 | 273.8593757 | 0.026370475 | 0.166491276 | 0.158389529 | 0.874149864 | 0.940887669 | Gtf3c4 | 2317.169582 | 0.047234049 | 0.058453987 | 0.808055219 | 0.419058795 | 0.806975048 |
| 1700048O20Rik | 309.413122 | 0.021783935 | 0.137701808 | 0.158196437 | 0.874302012 | 0.940905765 | Rabac1 | 3178.400989 | 0.047247027 | 0.053378858 | 0.885126223 | 0.376088558 | 0.783947144 |
| Arhgap25 | 417.2620577 | -0.022531239 | 0.142544686 | -0.158064392 | 0.874406059 | 0.940905765 | Bet1l | 1074.042338 | 0.047291196 | 0.057638944 | 0.820472979 | 0.411946527 | 0.802795661 |
| Rqcd1 | 1239.515105 | -0.010165519 | 0.064277695 | -0.158150019 | 0.874338587 | 0.940905765 | Mgat5 | 1827.879643 | 0.047319975 | 0.075738743 | 0.624778991 | 0.532116121 | 0.858328942 |
| Ttc5 | 1312.680461 | 0.013246464 | 0.083806597 | 0.158059917 | 0.874409585 | 0.940905765 | Tbpl1 | 2501.372182 | 0.047320834 | 0.044991619 | 1.051769973 | 0.292905099 | 0.728526371 |
| Noa1 | 666.1895431 | -0.012939446 | 0.082039366 | -0.157722401 | 0.874675548 | 0.941126593 | Slc16a9 | 138.0628451 | 0.047325989 | 0.089760291 | 0.527248609 | 0.598020957 | 0.88858965 |
| Zfp939 | 236.7884263 | 0.016306482 | 0.103530082 | 0.157504769 | 0.874847049 | 0.941245759 | Abca8a | 612.5531924 | 0.047330608 | 0.082650864 | 0.572657147 | 0.566876858 | 0.875959314 |
| Fgd5 | 384.7542704 | 0.01753349 | 0.111466471 | 0.157298332 | 0.875009735 | 0.941355425 | Rpl7a | 8086.812699 | 0.047334574 | 0.04721673 | 1.0024958 | 0.316104194 | 0.745993052 |
| Aste1 | 202.5166763 | -0.016480473 | 0.105029099 | -0.156913397 | 0.875313102 | 0.941551041 | Timmdc1 | 687.6647289 | 0.047343779 | 0.065311933 | 0.724887113 | 0.468521287 | 0.832446623 |
| Kctd15 | 416.5116248 | -0.016300842 | 0.103845649 | -0.156971839 | 0.875267043 | 0.941551041 | Usp15 | 2710.041592 | 0.047379204 | 0.068141152 | 0.695309697 | 0.486861239 | 0.839801621 |
| Il6st | 3669.446005 | 0.010612232 | 0.067673785 | 0.156814514 | 0.875391034 | 0.941569502 | Eif2ak3 | 1498.699783 | 0.047388162 | 0.056946298 | 0.832155265 | 0.405321311 | 0.8005027 |
| 4930426L09Rik | 23.66841967 | -0.026295945 | 0.167789507 | -0.156719839 | 0.875465651 | 0.941584395 | Klhl30 | 7.677939076 | 0.047406261 | 0.053809813 | 0.880996576 | 0.378319673 | NA |
| Gna15 | 76.39278047 | -0.023150763 | 0.148036728 | -0.156385268 | 0.875729349 | 0.941802633 | Trim35 | 7009.517827 | 0.04740717 | 0.052827588 | 0.897394163 | 0.369508626 | 0.780397655 |
| Fbln7 | 79.98409254 | -0.022624109 | 0.14507332 | -0.155949484 | 0.87607284 | 0.942041265 | Eef1d | 2395.418703 | 0.04741809 | 0.053907524 | 0.879619136 | 0.379065669 | 0.785380516 |
| Hnrnpf | 3236.496776 | -0.010771316 | 0.069043957 | -0.156006645 | 0.876027784 | 0.942041265 | Slc19a2 | 678.0016153 | 0.047426866 | 0.061330165 | 0.773304077 | 0.439342444 | 0.817545617 |
| Bag4 | 2823.315215 | -0.011408582 | 0.073296242 | -0.155650298 | 0.876308676 | 0.94205372 | G6pdx | 1680.788967 | 0.047433191 | 0.059629314 | 0.795467662 | 0.426341515 | 0.809717746 |
| Kcnk5 | 36.15723668 | -0.026138712 | 0.167944226 | -0.155639241 | 0.876317392 | 0.94205372 | Gm10789 | 9.734770095 | 0.047439897 | 0.061637216 | 0.7696632 | 0.441499705 | NA |
| Med4 | 811.3447634 | 0.014993058 | 0.096411585 | 0.155510954 | 0.876418519 | 0.94205372 | Ppp1cc | 8080.254641 | 0.04744481 | 0.069778199 | 0.679937442 | 0.496544072 | 0.844829674 |
| Por | 2798.563722 | 0.012610459 | 0.080944767 | 0.155790916 | 0.876197831 | 0.94205372 | Katnal1 | 4484.73615 | 0.04745993 | 0.053701557 | 0.883771953 | 0.37681933 | 0.783969863 |
| Tpm3 | 6104.833985 | -0.008905319 | 0.057279245 | -0.155471995 | 0.87644923 | 0.94205372 | Zfp946 | 550.3286806 | 0.047467654 | 0.082230671 | 0.577249984 | 0.563770596 | 0.875120313 |
| Zfp322a | 1271.928996 | -0.012624075 | 0.081188878 | -0.155490199 | 0.87643488 | 0.94205372 | Klhdc3 | 7473.747242 | 0.047469574 | 0.048128875 | 0.986301343 | 0.323985261 | 0.752925264 |
| 2310047M10Rik | 785.9265885 | 0.011704978 | 0.075416122 | 0.155205254 | 0.876659506 | 0.942136042 | Pigr | 4.08118176 | 0.047477549 | 0.04476575 | 1.060577548 | 0.288881931 | NA |
| Prob1 | 239.5822716 | -0.014272987 | 0.091998654 | -0.155143437 | 0.876708239 | 0.942136042 | Gja1 | 7234.807969 | 0.047481603 | 0.069637724 | 0.681837377 | 0.495341787 | 0.844369474 |
| Trp53i13 | 115.2111705 | -0.023514704 | 0.151426869 | -0.155287529 | 0.876594647 | 0.942136042 | Slc3a2 | 4633.022832 | 0.047485209 | 0.043546389 | 1.090451118 | 0.275514475 | 0.717137213 |
| Thumpd2 | 143.2234464 | -0.018081733 | 0.116691127 | -0.154953797 | 0.876857742 | 0.942231351 | Ces1d | 7.94638497 | 0.047489545 | 0.050607375 | 0.938391799 | 0.348043099 | NA |
| 1600014C10Rik | 1285.164816 | 0.010307636 | 0.066681858 | 0.154579324 | 0.877152971 | 0.942287189 | Fam211a | 694.4062806 | 0.047493718 | 0.065963371 | 0.720001384 | 0.471524144 | 0.833245343 |
| E130308A19Rik | 366.9387067 | 0.019118436 | 0.123641097 | 0.154628492 | 0.877114207 | 0.942287189 | Ndufa5 | 2222.605013 | 0.047502414 | 0.069307156 | 0.685389753 | 0.493098008 | 0.842873814 |
| Sash3 | 90.12942851 | 0.023456062 | 0.15172531 | 0.154595579 | 0.877140156 | 0.942287189 | Prob1 | 269.0963437 | 0.047509289 | 0.091657151 | 0.518336963 | 0.604223188 | 0.891718993 |
| Taf7 | 258.6372587 | -0.015061469 | 0.097316214 | -0.154768343 | 0.877003949 | 0.942287189 | Tatdn2 | 1749.025391 | 0.047509751 | 0.053403476 | 0.889637811 | 0.373660396 | 0.782348609 |
| Tbc1d9b | 11676.7819 | 0.011772268 | 0.07619679 | 0.154498221 | 0.877216914 | 0.942290548 | Pard6b | 225.482892 | 0.047526717 | 0.084910379 | 0.559727998 | 0.575664982 | 0.880811222 |
| Camsap2 | 7914.872972 | -0.013393235 | 0.086905114 | -0.154113316 | 0.877520389 | 0.942518136 | Sh3gl1 | 1345.788856 | 0.047529874 | 0.07209302 | 0.659285381 | 0.509712528 | 0.850457637 |
| Gm16197 | 18.97442506 | 0.02583713 | 0.167859801 | 0.153920893 | 0.87767211 | 0.942518136 | Thnsl1 | 911.4655345 | 0.047546735 | 0.062259018 | 0.76369234 | 0.445050607 | 0.821493054 |
| Osr1 | 73.55707676 | 0.023458631 | 0.152362272 | 0.153966141 | 0.877636433 | 0.942518136 | Peo1 | 700.8135561 | 0.047576752 | 0.06472479 | 0.735062283 | 0.462301563 | 0.830835164 |
| Tcf4 | 9297.539176 | 0.008420636 | 0.054702622 | 0.153934774 | 0.877661166 | 0.942518136 | 1700007G11Rik | 21.93590021 | 0.04758097 | 0.074097629 | 0.6421389 | 0.520783001 | NA |
| Slc26a5 | 10.66626192 | -0.024122216 | 0.157041592 | -0.153603998 | 0.877921984 | 0.942721133 | Idh1 | 1534.436858 | 0.04758741 | 0.058883215 | 0.808165958 | 0.418995052 | 0.806975048 |
| Rnf185 | 1447.286638 | 0.011677565 | 0.076160135 | 0.153329101 | 0.878138752 | 0.942888554 | Map4 | 19264.97133 | 0.04762083 | 0.050277108 | 0.947167247 | 0.343553562 | 0.763762832 |
| Cers6 | 1315.767097 | -0.013630231 | 0.089339762 | -0.152566233 | 0.878740356 | 0.943469134 | Ralgds | 4700.522137 | 0.047629197 | 0.063717145 | 0.747509908 | 0.454755824 | 0.826168452 |
| Cyth4 | 487.6159804 | 0.01941896 | 0.127650407 | 0.152126109 | 0.879087472 | 0.943698723 | Trim10 | 2.150013696 | 0.047652282 | 0.034141281 | 1.39573796 | 0.162793419 | NA |
| Pde2a | 22668.44252 | -0.010128869 | 0.066609506 | -0.15206342 | 0.879136915 | 0.943698723 | Fam126b | 3396.38772 | 0.047657145 | 0.084621228 | 0.563181909 | 0.573311021 | 0.879352236 |
| Rpl14 | 6660.641903 | 0.010923351 | 0.07179587 | 0.152144564 | 0.879072917 | 0.943698723 | Ppih | 367.6000354 | 0.047673708 | 0.072832392 | 0.654567387 | 0.512746327 | 0.852934798 |
| Gm10334 | 13.37430389 | -0.023400189 | 0.1542345 | -0.151718255 | 0.879409159 | 0.943925563 | Krt75 | 1.20445185 | 0.047683262 | 0.025431483 | 1.874969783 | 0.060796881 | NA |
| Fzd6 | 395.1158759 | -0.017723871 | 0.117036379 | -0.151438987 | 0.879629437 | 0.943965819 | Ndufaf2 | 460.8916777 | 0.047685366 | 0.073710035 | 0.646931801 | 0.517676083 | 0.853297835 |
| Hltf | 898.5618034 | -0.017884193 | 0.11806619 | -0.151475989 | 0.87960025 | 0.943965819 | Mllt1 | 3605.110729 | 0.047691004 | 0.068124414 | 0.700057451 | 0.483891427 | 0.838713365 |
| Tmem219 | 330.7484836 | -0.013630865 | 0.089947909 | -0.151541768 | 0.879548365 | 0.943965819 | Dnajc18 | 5229.414324 | 0.047707017 | 0.049308452 | 0.967522105 | 0.333283096 | 0.757730862 |
| Gm15612 | 14.98544943 | -0.022533292 | 0.149110892 | -0.151117682 | 0.879882884 | 0.944172409 | Sync | 56.60589655 | 0.047709302 | 0.093373475 | 0.510951341 | 0.60938513 | 0.89440021 |
| Arap1 | 715.4586533 | 0.010510447 | 0.069615465 | 0.150978618 | 0.879992583 | 0.944181141 | Eif4enif1 | 2523.070068 | 0.047710278 | 0.048724307 | 0.979188443 | 0.327486878 | 0.754492369 |
| Cnn3 | 2522.322828 | 0.010105897 | 0.066947367 | 0.150952863 | 0.880012899 | 0.944181141 | Scn5a | 470.6619929 | 0.047730034 | 0.094644238 | 0.504309978 | 0.614043567 | 0.895991647 |
| Mis12 | 984.6210667 | 0.012404985 | 0.082530873 | 0.150307209 | 0.880522246 | 0.944662213 | Ier5 | 3092.861158 | 0.047738742 | 0.094814346 | 0.503497035 | 0.614614864 | 0.89611764 |
| Cbln3 | 36.0143865 | 0.024832732 | 0.16553768 | 0.150012569 | 0.880754699 | 0.94471823 | H2-Ke2 | 1354.50902 | 0.047743719 | 0.050952344 | 0.937026937 | 0.348744702 | 0.767555385 |
| Cytl1 | 25.07304251 | -0.025136332 | 0.167513636 | -0.150055441 | 0.880720875 | 0.94471823 | Pou3f1 | 759.3548512 | 0.047769191 | 0.088566343 | 0.539360542 | 0.589638102 | 0.885188411 |
| Suox | 529.9425783 | 0.015083354 | 0.100549546 | 0.150009172 | 0.880757379 | 0.94471823 | Med13 | 2694.629995 | 0.047788413 | 0.078382988 | 0.609678377 | 0.542074881 | 0.864560611 |
| Gm20597 | 12.22818238 | 0.024518703 | 0.163584025 | 0.14988446 | 0.880855773 | 0.944758365 | Rgs12 | 2815.904419 | 0.047799998 | 0.07492864 | 0.637940282 | 0.523512555 | 0.854731187 |
| Prr22 | 34.40837143 | 0.025001901 | 0.167027748 | 0.14968711 | 0.881011479 | 0.94485996 | Zdhhc22 | 678.7872955 | 0.047807844 | 0.063988549 | 0.747131245 | 0.454984342 | 0.826168452 |
| Fam188a | 1158.275097 | 0.008587126 | 0.057468507 | 0.149423169 | 0.881219731 | 0.944968814 | Cnp | 13254.7067 | 0.047809345 | 0.074312078 | 0.643359009 | 0.519991175 | 0.853297835 |
| Olfm2 | 3697.667851 | -0.01704423 | 0.114140632 | -0.149326577 | 0.881295945 | 0.944968814 | Stt3a | 1460.620194 | 0.047836321 | 0.060431484 | 0.791579448 | 0.428605934 | 0.810486244 |
| Rpl36a | 2433.186553 | 0.011617202 | 0.077772784 | 0.149373612 | 0.881258833 | 0.944968814 | Slc1a4 | 2977.968852 | 0.047846944 | 0.061536479 | 0.777537881 | 0.436841494 | 0.816490111 |
| Mrm1 | 519.0977907 | -0.014576433 | 0.097724018 | -0.149159164 | 0.881428042 | 0.944987803 | Vamp3 | 1049.342801 | 0.047866272 | 0.062833179 | 0.761799308 | 0.446179794 | 0.822210607 |
| Nsun6 | 269.5786637 | 0.014842172 | 0.099563619 | 0.149072246 | 0.881496625 | 0.944987803 | Zfhx4 | 547.5503279 | 0.04787304 | 0.094270613 | 0.507825703 | 0.611575582 | 0.895166661 |
| Vezf1 | 1580.460218 | -0.01176952 | 0.078915033 | -0.149141668 | 0.881441847 | 0.944987803 | C230081A13Rik | 1816.429615 | 0.047873083 | 0.068051196 | 0.703486293 | 0.481752746 | 0.837383765 |
| 1700019L03Rik | 8.502020403 | 0.021972329 | 0.147567974 | 0.148896323 | 0.881635443 | 0.945021542 | Lrrc8b | 3012.355475 | 0.047904214 | 0.060030479 | 0.797998209 | 0.424871529 | 0.809188912 |
| Myh1 | 24.65241061 | 0.025017723 | 0.168042031 | 0.148877769 | 0.881650084 | 0.945021542 | Egfem1 | 200.1171509 | 0.047905072 | 0.089689312 | 0.534122413 | 0.593256844 | 0.886291956 |
| Snd1 | 3064.39746 | -0.007636581 | 0.051324973 | -0.148788796 | 0.881720292 | 0.94503142 | Il1rapl1 | 311.0167253 | 0.047910196 | 0.092263036 | 0.519278334 | 0.60356666 | 0.891540503 |
| Trib1 | 859.966982 | -0.024983689 | 0.168014526 | -0.148699577 | 0.881790695 | 0.945041504 | Tmem56 | 2851.762777 | 0.047911316 | 0.082388211 | 0.581531206 | 0.560882493 | 0.873713575 |
| Lsm12 | 533.4127831 | -0.011556167 | 0.077882013 | -0.148380433 | 0.882042541 | 0.945246032 | BC100451 | 6.913574096 | 0.047916715 | 0.055378847 | 0.865253025 | 0.386899928 | NA |
| 4833422C13Rik | 133.6161986 | -0.018892619 | 0.127548957 | -0.148120533 | 0.882247645 | 0.945269692 | Kif26b | 315.9457154 | 0.047917843 | 0.086882207 | 0.551526535 | 0.581272782 | 0.883958169 |
| Fez2 | 1431.770481 | -0.010151042 | 0.068491529 | -0.148208726 | 0.882178046 | 0.945269692 | Fen1 | 586.1066246 | 0.047918434 | 0.067401485 | 0.710940335 | 0.47712121 | 0.835247566 |
| Mrps15 | 821.8131202 | 0.013524767 | 0.091270395 | 0.148183508 | 0.882197947 | 0.945269692 | Aox4 | 41.60249158 | 0.047922249 | 0.091692837 | 0.522638957 | 0.601225525 | 0.890297392 |
| Asic2 | 2476.611449 | -0.009294476 | 0.062869443 | -0.147837733 | 0.88247083 | 0.945443441 | Padi1 | 3.123640962 | 0.047950661 | 0.037672567 | 1.272827017 | 0.203079432 | NA |
| Abcb9 | 2065.501332 | -0.014722898 | 0.100129344 | -0.147038795 | 0.883101399 | 0.945450326 | B3gnt4 | 23.48582515 | 0.047951188 | 0.07557752 | 0.634463634 | 0.525778292 | 0.855892499 |
| B630005N14Rik | 710.6606425 | -0.014294388 | 0.097134512 | -0.147160749 | 0.883005141 | 0.945450326 | 0610007P14Rik | 1201.098142 | 0.047952886 | 0.053574478 | 0.89506959 | 0.370749889 | 0.780761257 |
| Bet1l | 955.4434053 | 0.012144761 | 0.08278807 | 0.146696992 | 0.883371193 | 0.945450326 | Sv2b | 24553.25358 | 0.047955912 | 0.07492399 | 0.640060848 | 0.522133041 | 0.854113377 |
| C2cd3 | 1778.371188 | -0.010931797 | 0.074472784 | -0.146789156 | 0.883298444 | 0.945450326 | Exosc9 | 1289.148943 | 0.047963652 | 0.0641509 | 0.747669202 | 0.454659712 | 0.826168452 |
| C2cd4c | 773.9773914 | -0.018946458 | 0.128810568 | -0.147087761 | 0.88306275 | 0.945450326 | Scyl1 | 3104.49187 | 0.047983032 | 0.052725923 | 0.910046323 | 0.362798081 | 0.776599605 |
| Chmp2a | 2398.925999 | 0.008172516 | 0.055610913 | 0.146958858 | 0.883164494 | 0.945450326 | Rnf4 | 3804.186659 | 0.04798718 | 0.053637106 | 0.894663842 | 0.370966813 | 0.780761257 |
| Cth | 118.8401007 | 0.019616487 | 0.13358341 | 0.14684823 | 0.883251815 | 0.945450326 | Ankrd54 | 527.228082 | 0.048002737 | 0.076563947 | 0.626962676 | 0.530683701 | 0.858116381 |
| Eya1 | 428.3649427 | 0.013572443 | 0.092537335 | 0.146669919 | 0.883392563 | 0.945450326 | Bsg | 14125.52614 | 0.048007508 | 0.061124178 | 0.785409465 | 0.432213533 | 0.813254665 |
| Ifnar2 | 617.4991832 | -0.0128693 | 0.087360973 | -0.147311774 | 0.882885939 | 0.945450326 | BC029214 | 1075.448506 | 0.048007994 | 0.074427514 | 0.645030197 | 0.518907617 | 0.853297835 |
| Rab5a | 3672.750555 | -0.009563287 | 0.065106855 | -0.146886022 | 0.883221985 | 0.945450326 | Rnf115 | 2116.303904 | 0.048011676 | 0.060708539 | 0.790855412 | 0.42902837 | 0.810833477 |
| Rad21l | 17.42333812 | -0.024784342 | 0.167786821 | -0.14771328 | 0.882569051 | 0.945450326 | Wdr33 | 2555.275219 | 0.048026529 | 0.049280933 | 0.974545845 | 0.329785588 | 0.756141871 |
| Rad51ap2 | 31.03090573 | 0.024612464 | 0.167346568 | 0.147074804 | 0.883072977 | 0.945450326 | Zdhhc7 | 1422.281772 | 0.048068062 | 0.061267934 | 0.784554969 | 0.432714541 | 0.813460776 |
| Rhbdd2 | 3201.831224 | -0.008843938 | 0.060149514 | -0.147032583 | 0.883106302 | 0.945450326 | Fam171a1 | 4095.646063 | 0.048073657 | 0.063777579 | 0.753770491 | 0.450987046 | 0.823992799 |
| Ubqln2 | 6647.014602 | 0.008731367 | 0.059348605 | 0.147120011 | 0.883037295 | 0.945450326 | Prss8 | 4.439235943 | 0.048074266 | 0.048129732 | 0.998847574 | 0.317868536 | NA |
| Zfp612 | 2380.739916 | 0.010984234 | 0.074793392 | 0.146861026 | 0.883241715 | 0.945450326 | Trim44 | 16237.15466 | 0.048081743 | 0.054877792 | 0.876160305 | NA | NA |
| Actr5 | 243.2720842 | -0.012804967 | 0.08759251 | -0.146187924 | 0.883773039 | 0.945726878 | Rrm2b | 1630.948967 | 0.048098124 | 0.069243351 | 0.694624447 | 0.487290687 | 0.839801621 |
| Psmd13 | 2530.165012 | -0.011518748 | 0.07876229 | -0.146246998 | 0.883726405 | 0.945726878 | Lrrc28 | 941.815737 | 0.048100784 | 0.064338624 | 0.747619096 | 0.454689943 | 0.826168452 |
| Cul9 | 3362.681816 | 0.009571489 | 0.06552039 | 0.146084127 | 0.883854977 | 0.945749241 | Nat14 | 2264.713393 | 0.048105277 | 0.058179555 | 0.826841609 | 0.408326843 | 0.801055189 |
| Pxdc1 | 140.863038 | -0.018766531 | 0.128782811 | -0.145722325 | 0.884140597 | 0.945858897 | Ubl5 | 2733.87901 | 0.048114096 | 0.062452836 | 0.770406914 | 0.441058553 | 0.818962636 |
| Rbms2 | 608.1929993 | -0.012202837 | 0.083673485 | -0.145838753 | 0.884048683 | 0.945858897 | Creb3 | 1609.118852 | 0.048132337 | 0.046915126 | 1.025944949 | 0.304917534 | 0.737453291 |
| Slc52a2 | 493.9896092 | 0.012311651 | 0.084456003 | 0.145775914 | 0.88409829 | 0.945858897 | Dnalc1 | 5765.663337 | 0.048134792 | 0.062659704 | 0.768193734 | 0.442372096 | 0.819629727 |
| Lsm10 | 207.2721434 | 0.014712808 | 0.10115299 | 0.145451047 | 0.884354763 | 0.945957384 | Olfr658 | 4.357951792 | 0.048140379 | 0.047067655 | 1.022791115 | 0.30640662 | NA |
| Tpk1 | 85.35820468 | -0.018330993 | 0.12597976 | -0.145507443 | 0.884310239 | 0.945957384 | Dazap2 | 5582.617312 | 0.048143788 | 0.049910716 | 0.964598225 | 0.334746089 | 0.758238352 |
| Dqx1 | 29.89861834 | -0.023846921 | 0.165143384 | -0.14440131 | 0.885183582 | 0.946647876 | Atp5h | 6144.417114 | 0.048156236 | 0.057512969 | 0.837310909 | 0.402417829 | 0.79955155 |
| Gal3st4 | 161.6075657 | 0.021447541 | 0.148501714 | 0.144426217 | 0.885163915 | 0.946647876 | Lmbrd2 | 1637.994182 | 0.048193351 | 0.077490189 | 0.62192842 | 0.533988936 | 0.859470725 |
| Urm1 | 1298.505535 | -0.009146832 | 0.063287642 | -0.14452794 | 0.885083595 | 0.946647876 | Hyal2 | 353.5065767 | 0.048209658 | 0.074394351 | 0.64802847 | 0.516966536 | 0.853297835 |
| Ccl27a | 1563.130936 | 0.019039047 | 0.132012267 | 0.144221802 | 0.885325326 | 0.946712918 | Slc44a1 | 3778.883667 | 0.048220238 | 0.07587695 | 0.635505748 | 0.525098619 | 0.855700779 |
| F12 | 23.74888089 | 0.023963177 | 0.16623149 | 0.14415546 | 0.885377711 | 0.946712918 | Pex13 | 1241.952532 | 0.048220833 | 0.066878494 | 0.721021518 | 0.470896276 | 0.833245343 |
| Mafg | 973.5102133 | 0.01337885 | 0.092849266 | 0.144092146 | 0.885427707 | 0.946712918 | Slc48a1 | 6386.780429 | 0.048222053 | 0.049997004 | 0.964498851 | 0.334795884 | 0.758238352 |
| Anxa8 | 17.78290713 | 0.024111675 | 0.168038983 | 0.143488583 | 0.885904328 | 0.947157166 | Pcdhga4 | 595.6335306 | 0.048224892 | 0.075257375 | 0.640799554 | 0.521652923 | 0.854113377 |
| Cryzl1 | 1627.212727 | -0.009463941 | 0.066029748 | -0.143328444 | 0.886030792 | 0.947171527 | Mycl1 | 519.5995909 | 0.048228789 | 0.079587641 | 0.605983393 | 0.544525781 | 0.86624804 |
| Npepl1 | 452.0506384 | -0.017050521 | 0.118970881 | -0.143316756 | 0.886040023 | 0.947171527 | 4930579G24Rik | 63.23709868 | 0.048229753 | 0.094947082 | 0.507964556 | 0.6114782 | 0.895166661 |
| Acot13 | 1414.828958 | -0.012606161 | 0.088094181 | -0.143098683 | 0.886212245 | 0.947254819 | Clip3 | 24785.60995 | 0.048233037 | 0.042261774 | 1.14129229 | 0.25374831 | 0.701619624 |
| Tex264 | 2099.229709 | 0.009037454 | 0.063171032 | 0.14306327 | 0.886240213 | 0.947254819 | Sigmar1 | 1383.255958 | 0.04823406 | 0.055222955 | 0.873442206 | 0.382422094 | 0.786954445 |
| Vezt | 1829.219662 | -0.007576155 | 0.053084241 | -0.142719478 | 0.886511733 | 0.947479671 | Tlk2 | 2443.836372 | 0.048237662 | 0.068792351 | 0.701206767 | 0.483173988 | 0.838104566 |
| Nell1 | 1934.637275 | 0.010922752 | 0.076578643 | 0.14263444 | 0.886578896 | 0.947486096 | Nme2 | 2265.878812 | 0.048241698 | 0.056069085 | 0.860397455 | 0.389569988 | 0.792715562 |
| Flywch2 | 228.3754321 | 0.014599729 | 0.10241519 | 0.142554328 | 0.88664217 | 0.947488363 | Klhl3 | 769.4199014 | 0.048254333 | 0.091337352 | 0.528308868 | 0.597284977 | 0.888165441 |
| Fzd8 | 118.166857 | -0.019708393 | 0.138558891 | -0.142238385 | 0.886891713 | 0.947689669 | Exosc1 | 504.1819983 | 0.048259266 | 0.065156584 | 0.740665994 | 0.458895983 | 0.828270757 |
| Axl | 2324.856205 | -0.008711061 | 0.061300293 | -0.142104722 | 0.886997289 | 0.947715232 | Mettl14 | 855.3611547 | 0.048294064 | 0.063206606 | 0.764066714 | 0.444827487 | 0.821493054 |
| Slc13a4 | 581.6961566 | 0.019147704 | 0.13486599 | 0.141975782 | 0.887099135 | 0.947715232 | Evpl | 245.8334212 | 0.048313114 | 0.088560325 | 0.545539033 | 0.585382842 | 0.884512516 |
| Zfp960 | 42.15163643 | -0.022600511 | 0.159156158 | -0.142002114 | 0.887078336 | 0.947715232 | Mxd4 | 1095.986639 | 0.048332764 | 0.069603898 | 0.694397372 | 0.487433042 | 0.839801621 |
| Camk2a | 133864.2327 | 0.012878527 | 0.090792969 | 0.141844982 | 0.887202453 | 0.947760259 | Emc7 | 2508.717385 | 0.048336026 | 0.056832382 | 0.850501506 | 0.395046323 | 0.795790167 |
| Mplkip | 213.8935559 | 0.015674491 | 0.110569771 | 0.141761088 | 0.887268721 | 0.947765705 | Hhip | 436.432172 | 0.04834592 | 0.086788488 | 0.557054527 | 0.577490175 | 0.881435566 |
| Tomm20 | 15656.25893 | -0.011693676 | 0.082545959 | -0.141662616 | 0.887346505 | 0.947783452 | Sgms2 | 168.7468205 | 0.048346922 | 0.09491126 | 0.509390794 | 0.610478328 | 0.894800514 |
| Slc25a25 | 2660.084245 | -0.013145657 | 0.092941241 | -0.141440518 | 0.887521947 | 0.947905498 | Rragc | 3379.268249 | 0.048378786 | 0.058043538 | 0.833491328 | 0.404567687 | 0.800290023 |
| Dcaf10 | 946.929872 | 0.012733841 | 0.090127444 | 0.14128705 | 0.88764318 | 0.947969633 | Btn1a1 | 3.336976282 | 0.048394921 | 0.03746538 | 1.291723772 | 0.196452822 | NA |
| Cmip | 13605.92313 | 0.007802127 | 0.05534535 | 0.140971672 | 0.887892322 | 0.948067112 | Lnp | 2714.706288 | 0.048399682 | 0.077441969 | 0.624980009 | 0.531984179 | 0.858328942 |
| Spata17 | 14.13120632 | -0.023027916 | 0.163389077 | -0.140939137 | 0.887918024 | 0.948067112 | Nell2 | 23909.52563 | 0.048407663 | 0.063723612 | 0.759650339 | 0.447463621 | 0.822458097 |
| Syncrip | 2513.80012 | -0.013573633 | 0.096267255 | -0.140999483 | 0.887870351 | 0.948067112 | Exoc8 | 1571.457524 | 0.048415364 | 0.056685075 | 0.854111312 | 0.393043317 | 0.794620351 |
| Rmnd5a | 6258.636161 | -0.010292076 | 0.073143968 | -0.140709836 | 0.888099175 | 0.948129857 | Nptn | 23427.94185 | 0.048418855 | 0.065903284 | 0.734695634 | 0.46252488 | 0.83094391 |
| Zfp157 | 963.8884807 | 0.01145286 | 0.081391872 | 0.140712577 | 0.888097009 | 0.948129857 | Chad | 5.366179878 | 0.048433682 | 0.049715815 | 0.974210756 | 0.329951905 | NA |
| Plaa | 1717.741828 | 0.013763256 | 0.097941519 | 0.140525249 | 0.888245005 | 0.948220209 | Mcl1 | 2770.663348 | 0.048436286 | 0.066478917 | 0.728596193 | 0.466248708 | 0.832050282 |
| Armcx4 | 3263.543913 | -0.012037519 | 0.085748317 | -0.140381984 | 0.888358192 | 0.948275702 | Gm3219 | 23.59290988 | 0.048443098 | 0.08109547 | 0.597358861 | 0.550267814 | 0.869486595 |
| Phospho2 | 904.4462402 | 0.015476922 | 0.110323327 | 0.140286937 | 0.888433285 | 0.948290529 | Scrg1 | 800.3154471 | 0.048446353 | 0.085973322 | 0.563504486 | 0.573091406 | 0.879310647 |
| Ddb1 | 10855.09087 | -0.008820483 | 0.062921797 | -0.140181678 | 0.888516448 | 0.948313966 | Gpr50 | 27.82497676 | 0.04845855 | 0.040462001 | 1.197631075 | NA | NA |
| Cd53 | 252.0220639 | 0.017764764 | 0.126978142 | 0.139904107 | 0.888735756 | 0.948414813 | Ythdf1 | 2522.534626 | 0.048463218 | 0.053769245 | 0.901318563 | 0.367418967 | 0.778801679 |
| Tmem82 | 23.57419305 | 0.023456886 | 0.167753286 | 0.139829669 | 0.888794571 | 0.948414813 | Phf6 | 1002.65708 | 0.048479372 | 0.078995968 | 0.613694251 | 0.539417388 | 0.862853979 |
| Trim36 | 268.6231078 | 0.019991874 | 0.142864123 | 0.139936282 | 0.888710334 | 0.948414813 | Fbxo2 | 2461.630796 | 0.048493129 | 0.062224153 | 0.779329675 | 0.435785538 | 0.815960393 |
| N4bp2 | 260.751339 | 0.021026107 | 0.150590677 | 0.139624229 | 0.888956896 | 0.948457385 | Emp1 | 95.23179129 | 0.048508169 | 0.095037557 | 0.510410522 | 0.609763887 | 0.894418885 |
| Tead1 | 808.9096693 | -0.018800591 | 0.134618837 | -0.139657951 | 0.888930251 | 0.948457385 | Gstm2 | 90.11721129 | 0.048518403 | 0.090199898 | 0.537898649 | 0.590647021 | 0.885693151 |
| Amz1 | 624.5352566 | -0.015178556 | 0.108921084 | -0.139353697 | 0.88917066 | 0.948501164 | Rnf165 | 2977.436261 | 0.048520422 | 0.090387663 | 0.536803588 | 0.591403293 | 0.886007038 |
| Pigm | 545.2732085 | -0.012875428 | 0.092403041 | -0.139339877 | 0.88918158 | 0.948501164 | Gpr64 | 94.60542156 | 0.048522894 | 0.094013933 | 0.516124492 | 0.605767463 | 0.892522382 |
| Usp47 | 4655.399538 | -0.009196692 | 0.065945957 | -0.139458009 | 0.889088236 | 0.948501164 | Txndc9 | 1655.2583 | 0.04855007 | 0.053967576 | 0.899615534 | 0.368324888 | 0.779243711 |
| Prdm4 | 1471.746885 | 0.006626593 | 0.047662126 | 0.139032671 | 0.889424332 | 0.948694796 | Klf4 | 304.0337988 | 0.048552703 | 0.082805375 | 0.58634724 | 0.557642192 | 0.872673743 |
| Gtf2h2 | 574.5250723 | 0.012275773 | 0.088458488 | 0.138774393 | 0.889628429 | 0.948727446 | Fndc7 | 8.500584234 | 0.048554075 | 0.059187931 | 0.820337424 | 0.412023778 | NA |
| Sdhaf1 | 373.4587599 | -0.011329334 | 0.081646103 | -0.138761475 | 0.889638638 | 0.948727446 | Uqcr10 | 2198.325799 | 0.048560297 | 0.057916855 | 0.838448447 | 0.40177889 | 0.799277436 |
| Tpx2 | 138.4942745 | -0.016017427 | 0.115400112 | -0.138799058 | 0.889608938 | 0.948727446 | Ubxn6 | 2782.362895 | 0.048565449 | 0.052410277 | 0.926639815 | 0.354113575 | 0.771423785 |
| Hydin | 79.45290013 | -0.022211309 | 0.160529204 | -0.138363042 | 0.889953504 | 0.948997908 | Fancl | 196.0795827 | 0.048573784 | 0.086810736 | 0.559536598 | 0.575795561 | 0.880811222 |
| Oxgr1 | 13.71520652 | -0.022313043 | 0.161497284 | -0.138163581 | 0.890111137 | 0.949100678 | Mpi | 2125.942331 | 0.048585385 | 0.05930101 | 0.81930113 | 0.412614628 | 0.803544722 |
| Stk25 | 8821.653956 | -0.008145688 | 0.059005848 | -0.138048832 | 0.890201825 | 0.949132058 | Itgb7 | 21.00765731 | 0.048590995 | 0.074973507 | 0.648108867 | 0.51691454 | NA |
| 4930513N10Rik | 26.34595695 | 0.022874549 | 0.165944722 | 0.137844389 | 0.890363401 | 0.949197578 | Kctd7 | 683.2730962 | 0.048612999 | 0.074326189 | 0.654049401 | 0.513079979 | 0.853038774 |
| Wwp2 | 1724.72234 | 0.010201599 | 0.0740233 | 0.137816049 | 0.8903858 | 0.949197578 | Limd1 | 564.306984 | 0.048613462 | 0.066590384 | 0.730037262 | 0.465367408 | 0.831682622 |
| S100a11 | 152.3316644 | -0.015323827 | 0.111355116 | -0.137612242 | 0.89054688 | 0.949238674 | 2310011J03Rik | 1111.564964 | 0.048622906 | 0.058058982 | 0.837474319 | 0.402326006 | 0.79955155 |
| Vwc2 | 211.4320399 | -0.014005566 | 0.101764941 | -0.137626631 | 0.890535507 | 0.949238674 | Gcnt2 | 1899.370583 | 0.048625031 | 0.082658287 | 0.588265651 | 0.556353998 | 0.871613262 |
| Fam168a | 14558.70205 | 0.006752528 | 0.049134441 | 0.137429639 | 0.890691205 | 0.949261902 | Spag9 | 19197.79926 | 0.048635049 | 0.063431368 | 0.766734974 | 0.443239105 | 0.820398986 |
| Rhof | 1704.272908 | 0.01099219 | 0.079964174 | 0.13746394 | 0.890664094 | 0.949261902 | Cry2 | 7734.337601 | 0.048642877 | 0.060622766 | 0.802386313 | 0.422329528 | 0.808695226 |
| Fancd2 | 192.1573724 | -0.015553346 | 0.11344008 | -0.137106272 | 0.890946795 | 0.949468989 | E430025E21Rik | 2347.891125 | 0.048654183 | 0.057076413 | 0.852439399 | 0.39397026 | 0.794754188 |
| Cmtm5 | 1197.720742 | -0.022679606 | 0.166193847 | -0.136464778 | 0.891453867 | 0.949944032 | Klrg2 | 9.928711501 | 0.048659079 | 0.057940834 | 0.839806334 | 0.401016982 | NA |
| Crtc3 | 736.0602214 | -0.010951644 | 0.080322746 | -0.136345495 | 0.89154816 | 0.949979176 | Vopp1 | 8373.689951 | 0.048712607 | 0.094864405 | 0.51349721 | 0.607603562 | 0.893867666 |
| Slc35b1 | 1413.961795 | 0.007856621 | 0.05766349 | 0.136249495 | 0.891624049 | 0.949994706 | Hmmr | 38.39553436 | 0.048731835 | 0.089297976 | 0.545721609 | 0.585257316 | 0.884512516 |
| Nhlh1 | 19.30096348 | -0.022767078 | 0.167314743 | -0.136073355 | 0.891763291 | 0.950074875 | BC089491 | 32.0808767 | 0.048754266 | 0.081851615 | 0.595642078 | 0.55141436 | 0.869755927 |
| Zfp319 | 906.8293593 | -0.008973745 | 0.065983816 | -0.135999179 | 0.89182193 | 0.950074875 | Hgs | 3591.652971 | 0.048798296 | 0.05228151 | 0.933375789 | 0.350625976 | 0.768667813 |
| Cops2 | 2772.235842 | 0.01093053 | 0.080464827 | 0.135842334 | 0.891945923 | 0.950141639 | Tppp | 29335.93543 | 0.048800577 | 0.047430265 | 1.028891106 | 0.303530848 | 0.736936169 |
| Akr1c13 | 9.949949216 | 0.020615761 | 0.152154401 | 0.135492374 | 0.892222593 | 0.950199203 | Pcdhb8 | 156.514974 | 0.048802194 | 0.090377875 | 0.539979429 | 0.589211219 | 0.885188411 |
| BC033916 | 15.55111826 | 0.021943659 | 0.162932534 | 0.134679421 | 0.892865344 | 0.950199203 | Chrna6 | 29.03269317 | 0.048802236 | 0.06307884 | 0.773670474 | NA | NA |
| Cd80 | 16.39149383 | 0.022685861 | 0.168116002 | 0.13494171 | 0.892657961 | 0.950199203 | Vps33a | 6071.359379 | 0.048809915 | 0.048240406 | 1.01180564 | 0.311630993 | 0.74424258 |
| Cstf3 | 924.6161184 | -0.00973183 | 0.072041229 | -0.135086947 | 0.89254313 | 0.950199203 | 8430408G22Rik | 65.36938146 | 0.048817138 | 0.074387338 | 0.656256012 | 0.511659415 | 0.852417908 |
| Ddx56 | 1432.753155 | 0.007971567 | 0.059456586 | 0.134073749 | 0.893344257 | 0.950199203 | Gm2762 | 20.46052548 | 0.048817312 | 0.078244362 | 0.623908363 | 0.532687767 | NA |
| Ebf3 | 41.58639907 | 0.022091859 | 0.164392424 | 0.134384899 | 0.893098222 | 0.950199203 | Oxt | 130.0454043 | 0.048821379 | 0.029397877 | 1.660711062 | 0.096771491 | 0.516802454 |
| Eif4e2 | 2134.786825 | -0.007124623 | 0.052601597 | -0.135444987 | 0.892260057 | 0.950199203 | Dcun1d2 | 1903.148625 | 0.048847494 | 0.05477243 | 0.891826296 | 0.372486045 | 0.781446026 |
| Fgd4 | 545.1506482 | -0.014466169 | 0.107424257 | -0.134663893 | 0.892877622 | 0.950199203 | Zfp617 | 1363.988524 | 0.048849134 | 0.06864091 | 0.711662095 | 0.476674045 | 0.835195112 |
| Galnt14 | 799.7241445 | 0.012044727 | 0.089740688 | 0.134217013 | 0.893230973 | 0.950199203 | Rspry1 | 1018.228248 | 0.048849283 | 0.060200338 | 0.811445333 | 0.417109974 | 0.806153084 |
| Gnai2 | 6623.334358 | -0.006613817 | 0.049255298 | -0.134276248 | 0.893184135 | 0.950199203 | Slc43a2 | 3460.132762 | 0.048875464 | 0.061337836 | 0.796824067 | 0.425553216 | 0.809519334 |
| Gp5 | 26.03207002 | 0.02215906 | 0.165056146 | 0.13425165 | 0.893203585 | 0.950199203 | Lmbr1 | 1499.319357 | 0.048890502 | 0.068874844 | 0.709845565 | 0.47779991 | 0.835457925 |
| Grik5 | 8809.341348 | 0.009723477 | 0.071796825 | 0.135430456 | 0.892271546 | 0.950199203 | Adra1b | 829.6518513 | 0.048901092 | 0.077718617 | 0.629206931 | 0.529213592 | 0.857868166 |
| Lhx6 | 1025.496852 | -0.01556513 | 0.115676482 | -0.134557431 | 0.8929618 | 0.950199203 | Lrrc14b | 9.877003295 | 0.048906665 | 0.058985798 | 0.829126107 | 0.407033052 | NA |
| Mnd1 | 12.08354297 | -0.022266144 | 0.165249734 | -0.134742387 | 0.892815559 | 0.950199203 | Slc26a2 | 188.7104057 | 0.04890959 | 0.086750243 | 0.563797725 | 0.5728918 | 0.879310647 |
| Mrps2 | 1162.576365 | -0.00810478 | 0.059975345 | -0.135135187 | 0.89250499 | 0.950199203 | Spata22 | 2.45023949 | 0.048909944 | 0.036091725 | 1.355156747 | 0.175367622 | NA |
| Nnat | 4762.766555 | -0.0168729 | 0.12507643 | -0.13490072 | 0.892690371 | 0.950199203 | Nrxn1 | 12651.71961 | 0.048912065 | 0.066984667 | 0.730197926 | 0.46526921 | 0.831682622 |
| Oasl1 | 10.02168297 | 0.021017553 | 0.155478588 | 0.135179725 | 0.892469777 | 0.950199203 | Kif3b | 4378.858949 | 0.048933737 | 0.055084204 | 0.888344265 | 0.374355596 | 0.78293624 |
| Plce1 | 683.2283187 | 0.022339489 | 0.165414542 | 0.135051545 | 0.892571121 | 0.950199203 | Lpcat3 | 677.1899507 | 0.048934587 | 0.080109493 | 0.610846297 | 0.54130134 | 0.863964758 |
| Ptpre | 1341.6084 | 0.006575796 | 0.048942296 | 0.134358149 | 0.893119374 | 0.950199203 | Cks1b | 50.97863319 | 0.048936454 | 0.090673348 | 0.539700532 | 0.589403573 | 0.885188411 |
| R3hdm2 | 11071.28847 | 0.008816192 | 0.065296758 | 0.135017304 | 0.892598193 | 0.950199203 | Ppic | 52.69714413 | 0.048938541 | 0.09380209 | 0.521721223 | 0.601864445 | 0.890837878 |
| Shank1 | 15649.75751 | -0.013244302 | 0.098822735 | -0.1340208 | 0.893386127 | 0.950199203 | Dao | 6.392253965 | 0.048947031 | 0.051882196 | 0.943426361 | 0.345462865 | NA |

| | | | | | | | | | | | | | | | |
|---|---|---|---|---|---|---|---|---|---|---|---|---|---|---|---|
| Srsf9 | 1112.756146 | -0.011188332 | 0.083290734 | -0.134328645 | 0.893142703 | 0.950199203 | | | Clec4g | 4.159828353 | 0.048953986 | 0.043077175 | 1.136425179 | 0.255778663 | NA |
| Stx12 | 4461.650201 | 0.007150024 | 0.053362367 | 0.133990002 | 0.893410481 | 0.950199203 | | | Svop | 8344.723636 | 0.048962822 | 0.063492594 | 0.771158002 | 0.440613284 | 0.818563572 |
| Zfp738 | 559.3449046 | 0.014268906 | 0.106591608 | 0.133865193 | 0.893509175 | 0.950238942 | | | Sult1c2 | 8.112205283 | 0.048965101 | 0.039291204 | 1.246210237 | 0.212687222 | NA |
| Casp12 | 38.08740145 | 0.02235555 | 0.167277637 | 0.133643388 | 0.893684573 | 0.950305245 | | | Emg1 | 882.1332981 | 0.048976577 | 0.062825846 | 0.779560965 | 0.43564934 | 0.815960393 |
| Fam26e | 221.5847582 | -0.014632546 | 0.109563035 | -0.133553673 | 0.89375552 | 0.950305245 | | | Tmed5 | 756.2455296 | 0.048994233 | 0.079818362 | 0.613821585 | 0.539333232 | 0.862815409 |
| Gpt2 | 2410.923909 | 0.009898594 | 0.074112196 | 0.133562275 | 0.893748717 | 0.950305245 | | | Ppp1ca | 8337.210691 | 0.048995375 | 0.051473309 | 0.951859823 | 0.341168078 | 0.762819666 |
| Ppp5c | 4285.859346 | -0.007256878 | 0.054396744 | -0.133406477 | 0.893871923 | 0.950328085 | | | Ddah2 | 283.3190953 | 0.049009057 | 0.085637614 | 0.572284236 | 0.567129428 | 0.876209537 |
| Slc25a40 | 348.0477285 | -0.017495065 | 0.131328292 | -0.133216272 | 0.894022342 | 0.950328085 | | | Slc16a14 | 238.2213052 | 0.049017754 | 0.093038621 | 0.526853834 | 0.598295095 | 0.888717024 |
| Usp51 | 49.87343534 | -0.022266212 | 0.167125046 | -0.13323085 | 0.894010813 | 0.950328085 | | | D10Wsu52e | 4741.037952 | 0.04903243 | 0.045413626 | 1.079685427 | 0.280282286 | 0.71983389 |
| Zfp513 | 828.8889855 | 0.010045251 | 0.075362625 | 0.13329221 | 0.893962288 | 0.950328085 | | | Golga3 | 2417.536845 | 0.049033409 | 0.05331169 | 0.919749676 | 0.357703587 | 0.773905615 |
| Car15 | 186.9197194 | -0.01556325 | 0.117063314 | -0.132947287 | 0.894235069 | 0.950489 | | | Prdx1 | 5178.568046 | 0.049038405 | 0.05535851 | 0.885833185 | 0.375707425 | 0.783947144 |
| Mtf1 | 1388.641237 | 0.011145645 | 0.083939875 | 0.132781287 | 0.894366354 | 0.950563334 | | | Wasf3 | 2974.261158 | 0.049046262 | 0.050855165 | 0.964430301 | 0.334830237 | 0.758238352 |
| Il17rb | 53.02902458 | -0.019307071 | 0.145631729 | -0.132574621 | 0.894529805 | 0.950606639 | | | Mir1894 | 3.493832003 | 0.049049938 | 0.044241974 | 1.108674265 | 0.267570728 | NA |
| Tmem41b | 1435.130404 | -0.008409024 | 0.063423796 | -0.132584685 | 0.894521845 | 0.950606639 | | | Rab5a | 4012.876439 | 0.049056957 | 0.073263317 | 0.669597811 | 0.50311421 | 0.847567618 |
| A330041J22Rik | 39.1580887 | -0.021289209 | 0.160684069 | -0.132491099 | 0.894595863 | 0.950611638 | | | Enpp4 | 1538.339634 | 0.049057844 | 0.073884612 | 0.663979175 | 0.50670364 | 0.84894776 |
| Rb1cc1 | 3929.369159 | 0.008998143 | 0.067978003 | 0.132368448 | 0.89469287 | 0.950638867 | | | Mbtps2 | 1247.552409 | 0.049065925 | 0.075130086 | 0.653079589 | 0.51370497 | 0.853195313 |
| Thsd7b | 263.5779133 | -0.01626495 | 0.122936609 | -0.132303553 | 0.894744197 | 0.950638867 | | | Acot9 | 715.5236344 | 0.04906975 | 0.068302272 | 0.71842047 | 0.472498069 | 0.833671043 |
| Iws1 | 1395.55833 | 0.010003335 | 0.075756688 | 0.132045569 | 0.894948249 | 0.950790466 | | | Tmem119 | 1052.632164 | 0.049089966 | 0.075016067 | 0.654392698 | 0.512858838 | 0.852963989 |
| Babam1 | 2320.239589 | 0.007418353 | 0.056253759 | 0.131873011 | 0.895084736 | 0.950796033 | | | Cacna2d2 | 2902.459956 | 0.049113332 | 0.082672182 | 0.59407325 | 0.552463121 | 0.870265825 |
| Hspa1b | 200.294678 | -0.019842128 | 0.150702695 | -0.131664053 | 0.89525002 | 0.950796033 | | | Paqr3 | 361.9867078 | 0.049119599 | 0.07580047 | 0.648011803 | 0.516977316 | 0.853297835 |
| Lox | 24.04041747 | -0.02182607 | 0.166016766 | -0.131469069 | 0.895404254 | 0.950796033 | | | Sh2b1 | 3527.037226 | 0.049120857 | 0.061550789 | 0.798054056 | 0.424839122 | 0.809188912 |
| Malat1 | 9205.227744 | -0.018504491 | 0.140409697 | -0.131789269 | 0.895150975 | 0.950796033 | | | Dctpp1 | 264.955618 | 0.049173592 | 0.079060103 | 0.621977336 | 0.533956771 | 0.859470725 |
| Nupl1 | 1339.042615 | -0.011075942 | 0.083939253 | -0.131951873 | 0.895022358 | 0.950796033 | | | Senp8 | 284.4279286 | 0.049199262 | 0.083485833 | 0.589312706 | 0.555651523 | 0.871360989 |
| Pkib | 295.324103 | 0.013351484 | 0.101595316 | 0.131418301 | 0.895444412 | 0.950796033 | | | Styk1 | 13.01413797 | 0.049211791 | 0.061190532 | 0.804238653 | 0.42125916 | NA |
| Wfs1 | 6854.469288 | 0.017498895 | 0.133090224 | 0.131481446 | 0.895394463 | 0.950796033 | | | Abhd8 | 8562.653295 | 0.049251326 | 0.064940091 | 0.758411718 | 0.448204545 | 0.823087132 |
| Zbtb21 | 749.7603584 | -0.009431491 | 0.071725267 | -0.131494676 | 0.895383998 | 0.950796033 | | | Zdhhc6 | 1091.707145 | 0.049260973 | 0.059543481 | 0.827310937 | 0.408060846 | 0.801055189 |
| Whsc1l1 | 3505.082883 | 0.007970818 | 0.060736476 | 0.131236094 | 0.895588544 | 0.95088391 | | | Npepps | 7205.206329 | 0.049266079 | 0.051167147 | 0.962845937 | 0.335624844 | 0.759126439 |
| Efna4 | 8.988549763 | -0.020414968 | 0.155885272 | -0.130961491 | 0.895805771 | 0.950984215 | | | Dmrt2 | 35.66620313 | 0.049266263 | 0.092053066 | 0.535194154 | 0.592515608 | 0.886226713 |
| Rpp38 | 178.2689245 | -0.015094259 | 0.115251105 | -0.130968451 | 0.895800265 | 0.950984215 | | | 9530026P05Rik | 12.89476998 | 0.049269322 | 0.063692081 | 0.773554911 | 0.439194044 | NA |
| Efcab11 | 9.084649627 | 0.020969714 | 0.160543606 | 0.130616935 | 0.896078346 | 0.951208405 | | | Gab2 | 1188.732872 | 0.049269561 | 0.05966768 | 0.825732809 | 0.408955677 | 0.801055189 |
| Peo1 | 621.5958862 | -0.009333946 | 0.071635705 | -0.130297392 | 0.896331143 | 0.951411573 | | | Prrc2b | 26688.02249 | 0.049288871 | 0.055807194 | 0.883199235 | 0.377128635 | 0.783997209 |
| Inip | 745.0784196 | 0.0089172 | 0.068534638 | 0.130112305 | 0.896477575 | 0.951501818 | | | Ppa2 | 1298.709846 | 0.049290058 | 0.065120647 | 0.756903694 | 0.449107562 | 0.823263966 |
| E130215H24Rik | 14.35097443 | -0.021325141 | 0.164313867 | -0.129782965 | 0.896738141 | 0.95158282 | | | Hat1 | 605.623402 | 0.049292391 | 0.076383882 | 0.645324502 | 0.518716918 | 0.853297835 |
| Mdh2 | 9992.289569 | 0.008592264 | 0.066199651 | 0.129793195 | 0.896730047 | 0.95158282 | | | Mrpl2 | 921.5959112 | 0.049298662 | 0.067418829 | 0.731229874 | 0.464638756 | 0.83159896 |
| Shisa4 | 3170.291312 | -0.017621961 | 0.135666764 | -0.129891512 | 0.89665226 | 0.95158282 | | | Tef | 19815.82884 | 0.049302338 | 0.061949401 | 0.795848502 | 0.426120097 | 0.809519334 |
| Pdik1l | 543.787077 | -0.009762427 | 0.075267292 | -0.129703449 | 0.896801054 | 0.951584408 | | | Mrpl32 | 442.5518272 | 0.049330231 | 0.073815196 | 0.668293705 | 0.503946131 | 0.847807553 |
| Cstb | 599.1638882 | 0.01352667 | 0.104529639 | 0.129405118 | 0.897037098 | 0.951704519 | | | Yae1d1 | 1096.020737 | 0.049340712 | 0.057281001 | 0.861380069 | 0.389028749 | 0.792459909 |
| Mettl1 | 183.5823193 | 0.013770763 | 0.106390783 | 0.12943568 | 0.897012916 | 0.951704519 | | | Speer4d | 1.221072351 | 0.049375022 | 0.025867673 | 1.908753941 | 0.056293841 | NA |
| Nfkbie | 96.30878828 | 0.016742832 | 0.129463243 | 0.129324982 | 0.897100505 | 0.951706622 | | | Myrf | 3175.776699 | 0.049381425 | 0.084364433 | 0.585334639 | 0.558322728 | 0.872720967 |
| 2900079G21Rik | 208.6325909 | 0.013409575 | 0.103761734 | 0.1292343 | 0.897172256 | 0.951717577 | | | Ntf3 | 50.06571224 | 0.04938382 | 0.08567025 | 0.576440712 | 0.564317333 | 0.875323716 |
| Pirt | 8.396537137 | -0.019791813 | 0.153287778 | -0.129115399 | 0.897266337 | 0.951752217 | | | Tmem29 | 519.2715152 | 0.049408171 | 0.070040032 | 0.705427583 | 0.480544184 | 0.836914993 |
| Trh | 268.6055651 | 0.016998232 | 0.131817065 | 0.128953198 | 0.897394682 | 0.951823193 | | | Gm3558 | 35.07799301 | 0.049418751 | 0.083370732 | 0.592758987 | 0.553342459 | 0.870600462 |
| Klhl15 | 209.882254 | 0.015603453 | 0.121099064 | 0.128848663 | 0.897477399 | 0.951845767 | | | Tubb4a | 33893.0329 | 0.049432374 | 0.0547887 | 0.902236656 | 0.366931166 | 0.778335591 |
| Serpinb9 | 363.2894913 | -0.016158566 | 0.125580035 | -0.128671456 | 0.897617622 | 0.951916874 | | | Rpl15 | 7214.929103 | 0.049443162 | 0.037158007 | 1.330619339 | 0.183314296 | 0.63672274 |
| Tspyl4 | 21133.37498 | -0.006284679 | 0.048866692 | -0.128608652 | 0.89766732 | 0.951916874 | | | Napsa | 2.854951619 | 0.049456754 | 0.040397396 | 1.22425599 | 0.220855674 | NA |
| Hif1a | 2660.89268 | 0.009629944 | 0.075049869 | 0.128313941 | 0.897900533 | 0.952099019 | | | Aip | 2679.825726 | 0.049459693 | 0.042615027 | 1.160616261 | 0.245797989 | 0.695249198 |
| 4632428N05Rik | 1314.845776 | -0.010116115 | 0.079104561 | -0.127882834 | 0.898241696 | 0.952125419 | | | Tmem9b | 3772.70643 | 0.04947074 | 0.062361186 | 0.793293767 | 0.427606688 | 0.810334558 |
| Asap3 | 238.0782798 | -0.017095858 | 0.133630779 | -0.127933538 | 0.898201569 | 0.952125419 | | | Pcyt1a | 2007.202353 | 0.049471191 | 0.062152504 | 0.795964579 | 0.426052623 | 0.809519334 |
| Hook3 | 2926.548077 | 0.009693743 | 0.075663759 | 0.128116058 | 0.898057128 | 0.952125419 | | | Dnajc16 | 3140.019683 | 0.049473851 | 0.052268804 | 0.946527317 | 0.343879696 | 0.763762832 |
| Ly6a | 840.7973301 | -0.011866088 | 0.092693419 | -0.128014353 | 0.898137614 | 0.952125419 | | | Slc35e3 | 1507.801721 | 0.049502515 | 0.05358304 | 0.923846706 | 0.355566134 | 0.772370994 |
| Prkaca | 7682.850179 | 0.006129347 | 0.047874688 | 0.128028971 | 0.898126046 | 0.952125419 | | | Slc30a9 | 6540.362179 | 0.049511684 | 0.047746551 | 1.036968811 | 0.299750393 | 0.733704078 |
| Rgs17 | 3085.962909 | 0.012160141 | 0.095137437 | 0.127816568 | 0.898294138 | 0.952125419 | | | Dap3 | 2718.75506 | 0.049523255 | 0.05179562 | 0.956128241 | 0.339007448 | 0.761116259 |
| Erlin1 | 1135.659156 | -0.009407409 | 0.073695378 | -0.127652632 | 0.898423878 | 0.952132665 | | | Map3k14 | 272.2316413 | 0.049532466 | 0.090015267 | 0.550267386 | 0.58213599 | 0.884020489 |
| Mettl15 | 128.0784093 | 0.016147776 | 0.12645803 | 0.127692769 | 0.898392113 | 0.952132665 | | | Hmgb1 | 7109.140258 | 0.04953871 | 0.062056291 | 0.79828667 | 0.424704152 | 0.809188912 |
| Acpl2 | 575.0811664 | -0.010419503 | 0.081676942 | -0.127569695 | 0.898489515 | 0.9521371 | | | Ssr3 | 5123.979975 | 0.049554038 | 0.065674059 | 0.754545074 | 0.450521991 | 0.823490932 |
| Ckb | 46198.40988 | -0.007916275 | 0.062138119 | -0.127398038 | 0.898625369 | 0.952215939 | | | Slit2 | 1351.024103 | 0.04956418 | 0.074171439 | 0.668238089 | 0.503981625 | 0.847807553 |
| Bin2 | 244.5924536 | 0.019614738 | 0.154251452 | 0.1271608 | 0.898813131 | 0.952349767 | | | Xrn2 | 2046.313461 | 0.049565215 | 0.059723484 | 0.829911651 | 0.406588737 | 0.800917161 |
| Ngrn | 1082.374322 | -0.008031186 | 0.063287437 | -0.126900164 | 0.899019417 | 0.952438075 | | | Rnf152 | 939.9366662 | 0.049581375 | 0.074769871 | 0.663119704 | 0.507253892 | 0.848958235 |
| Rnls | 22.99296527 | -0.021316199 | 0.167880893 | -0.126972154 | 0.898962438 | 0.952438075 | | | H2afz | 3022.699343 | 0.04960472 | 0.049899355 | 0.994095416 | 0.320176417 | 0.750154748 |
| Derl2 | 798.0939639 | -0.011523458 | 0.091171327 | -0.126393445 | 0.899420491 | 0.952724376 | | | Fiz1 | 898.0147067 | 0.049608424 | 0.055137798 | 0.899717186 | 0.368270775 | 0.779243711 |
| Letm1 | 3202.125936 | 0.008313104 | 0.065749121 | 0.126436726 | 0.899386232 | 0.952724376 | | | Myocd | 10.17793859 | 0.049609451 | 0.067753315 | 0.732206994 | 0.464042236 | NA |
| Mfap4 | 62.80055167 | -0.019970421 | 0.158086786 | -0.126325679 | 0.89947413 | 0.952724376 | | | Ndrg2 | 33606.63464 | 0.04961117 | 0.068587935 | 0.72332211 | 0.469482012 | 0.833245343 |
| Tmem143 | 703.0138949 | 0.014471242 | 0.114707176 | 0.126158123 | 0.89960676 | 0.952799722 | | | Zswim7 | 271.9425458 | 0.049628604 | 0.079351005 | 0.625431328 | 0.531688006 | 0.858328942 |
| B3gnt6 | 13.87083927 | -0.020991617 | 0.166780011 | -0.125864108 | 0.899839494 | 0.952981075 | | | Gdi2 | 9130.207938 | 0.049628818 | 0.056237335 | 0.882488801 | 0.377512533 | 0.784176613 |
| Rbm20 | 242.306444 | -0.018727476 | 0.148917871 | -0.125757072 | 0.899924223 | 0.952991929 | | | Ap3s2 | 4330.097968 | 0.049641753 | 0.043410048 | 1.143554425 | 0.252808472 | 0.700378199 |
| Zfp71-rs1 | 494.0088994 | -0.010904564 | 0.086753633 | -0.125695761 | 0.899972758 | 0.952991929 | | | Snrpf | 479.975422 | 0.049660857 | 0.073325844 | 0.677262664 | 0.498239315 | 0.845625663 |
| Wnt10a | 304.5620225 | 0.017328798 | 0.138100367 | 0.125479738 | 0.900143765 | 0.953107872 | | | D3Bwg0562e | 8979.751846 | 0.049692959 | 0.072955353 | 0.681142061 | 0.495781605 | 0.844379526 |
| Nudcd1 | 538.4244488 | -0.010131925 | 0.080885541 | -0.125262493 | 0.900315745 | 0.953224829 | | | Nrbf2 | 813.0591272 | 0.04969911 | 0.061174719 | 0.81241256 | 0.416554941 | 0.806153084 |
| Musk | 78.52697804 | 0.019019062 | 0.152092259 | 0.125049506 | 0.900484358 | 0.953338206 | | | Igfbp2 | 1285.98226 | 0.049701797 | 0.085286619 | 0.582761958 | 0.560053558 | 0.873713575 |
| Acbd3 | 1366.151762 | 0.008282871 | 0.066344157 | 0.124847027 | 0.900644656 | 0.953358248 | | | Zbtb18 | 11090.63823 | 0.049702474 | 0.071720027 | 0.693006905 | 0.488305218 | 0.840317688 |
| Calhm2 | 158.3646228 | 0.017919945 | 0.144086167 | 0.124369642 | 0.901022609 | 0.953358248 | | | 2310069G16Rik | 93.24951268 | 0.049712501 | 0.095012883 | 0.523218524 | 0.600822192 | 0.890202281 |
| Fam49b | 5641.237353 | -0.008534813 | 0.06849423 | -0.1246063 | 0.90083524 | 0.953358248 | | | Wscd1 | 3358.000689 | 0.049767481 | 0.06898089 | 0.721467655 | 0.470621834 | 0.833245343 |
| Ikbkg | 1791.045074 | -0.007376092 | 0.05938506 | -0.124207869 | 0.901150692 | 0.953358248 | | | Wdr1 | 6441.028026 | 0.04977081 | 0.051903347 | 0.958913317 | 0.337602415 | 0.760312044 |
| Nfe2l3 | 145.5723182 | -0.017942915 | 0.144210292 | -0.124421875 | 0.900981254 | 0.953358248 | | | Snap91 | 13767.80365 | 0.049771338 | 0.056681888 | 0.878081863 | 0.379899295 | 0.786275719 |
| Pcdhga3 | 587.4198588 | -0.012075166 | 0.097297211 | -0.124105984 | 0.90123136 | 0.953358248 | | | Copg2 | 3890.38889 | 0.049781366 | 0.058867547 | 0.845650417 | 0.397747791 | 0.797651232 |
| Rrbp1 | 2195.598937 | 0.005487427 | 0.043982308 | 0.124764407 | 0.900710066 | 0.953358248 | | | 2010001M06Rik | 77.64222168 | 0.049814547 | 0.094792898 | 0.525509271 | 0.59922921 | 0.889150487 |
| Ruvbl1 | 1283.71855 | 0.008354773 | 0.067184135 | 0.124356337 | 0.901033143 | 0.953358248 | | | C2cd2 | 722.4431312 | 0.049835405 | 0.072575299 | 0.686671714 | 0.492289625 | 0.842340249 |
| Ruvbl2 | 836.4055891 | 0.008114127 | 0.065217884 | 0.12441568 | 0.900986159 | 0.953358248 | | | Ppil3 | 557.8932343 | 0.049855515 | 0.070775051 | 0.704422175 | 0.481169901 | 0.8371092 |
| Skida1 | 289.4352532 | -0.014756732 | 0.118916736 | -0.124092978 | 0.901241658 | 0.953358248 | | | Tigd3 | 133.7414706 | 0.049860103 | 0.09198326 | 0.542056272 | 0.587779741 | 0.885142911 |
| Spry4 | 640.8021378 | 0.020415338 | 0.164436582 | 0.124153264 | 0.901193926 | 0.953358248 | | | Msi2 | 4026.501098 | 0.049878323 | 0.055841242 | 0.893216573 | 0.371741204 | 0.781124365 |
| Top1 | 3006.688218 | 0.008854064 | 0.071071455 | 0.124579744 | 0.900856265 | 0.953358248 | | | Jak2 | 2287.642963 | 0.049886564 | 0.078950812 | 0.631868917 | 0.527472536 | 0.856575299 |
| Gm20257 | 31.09443483 | -0.020544662 | 0.1657685 | -0.123935865 | 0.901366056 | 0.953424745 | | | Hdlbp | 9996.648839 | 0.049889553 | 0.042573984 | 1.17183192 | 0.241264545 | 0.692393571 |
| Dtx1 | 2745.514973 | 0.006678579 | 0.053948539 | 0.123795365 | 0.901477302 | 0.95343761 | | | Clec16a | 4805.594631 | 0.049896299 | 0.050363351 | 0.990726348 | 0.321819222 | 0.751204566 |
| Mafa | 14.5686718 | -0.020696634 | 0.167225167 | -0.12376507 | 0.90150129 | 0.95343761 | | | Camkk1 | 7183.229903 | 0.049912613 | 0.077452807 | 0.644426137 | 0.51929914 | 0.853297835 |
| Snap47 | 13389.23683 | 0.007507171 | 0.060813325 | 0.123446148 | 0.901753816 | 0.953639589 | | | Pus1 | 525.1064458 | 0.049922184 | 0.070904717 | 0.704074226 | 0.48138655 | 0.837138876 |
| Mpg | 426.9234781 | -0.009584608 | 0.077731163 | -0.123304577 | 0.901865917 | 0.953693046 | | | Pgrmc2 | 3387.956748 | 0.049922819 | 0.050061129 | 0.997237191 | 0.318649393 | 0.74851037 |
| 1700066M21Rik | 779.9621018 | -0.011243137 | 0.091277492 | -0.123175354 | 0.901968241 | 0.953736158 | | | Zscan12 | 307.6519267 | 0.049938495 | 0.074076445 | 0.674148107 | 0.500217157 | 0.846268338 |
| Btbd7 | 651.358974 | -0.009751146 | 0.079267978 | -0.123014947 | 0.902095262 | 0.953805376 | | | Gpx8 | 117.2531456 | 0.049956622 | 0.090509582 | 0.551948439 | 0.58098368 | 0.883796098 |
| Pfn1 | 2882.110852 | -0.00848593 | 0.069228154 | -0.12257918 | 0.902440342 | 0.953858521 | | | Rufy3 | 10205.35894 | 0.049973534 | 0.050828094 | 0.983187243 | 0.325515292 | 0.75293408 |
| Slc22a17 | 22120.49638 | -0.007807489 | 0.063586446 | -0.122785433 | 0.90227701 | 0.953858521 | | | Ythdf3 | 2216.958067 | 0.049978559 | 0.074183397 | 0.673716241 | 0.500491735 | 0.846307849 |
| Spata2l | 2011.458395 | -0.011538775 | 0.094028553 | -0.122715645 | 0.902332275 | 0.953858521 | | | Tead1 | 768.502256 | 0.049997161 | 0.07886531 | 0.633956319 | 0.52610933 | 0.855892499 |
| Trdmt1 | 157.4064013 | 0.015349987 | 0.125011144 | 0.122788951 | 0.902274224 | 0.953858521 | | | Clcn3 | 8891.236638 | 0.050014989 | 0.058048773 | 0.861602864 | 0.388906094 | 0.79235741 |
| Usp45 | 1768.588964 | -0.009774961 | 0.079754735 | -0.122562765 | 0.902453341 | 0.953858521 | | | Tnfrsf8 | 55.29894472 | 0.05001883 | 0.092489917 | 0.540803051 | 0.588643339 | 0.885188411 |
| Rpn2 | 5011.598066 | 0.006718546 | 0.054891845 | 0.122396073 | 0.902585349 | 0.953932974 | | | Bcmo1 | 4.339483503 | 0.050020941 | 0.037650045 | 1.328575887 | 0.183987932 | NA |
| Amer2 | 845.8173871 | -0.01184249 | 0.096838642 | -0.122290953 | 0.902668596 | 0.953955885 | | | Tctex1d2 | 290.3038354 | 0.050030401 | 0.08091926 | 0.61827556 | 0.536393709 | 0.861337058 |
| 4931429I11Rik | 14.8141693 | -0.020111119 | 0.16480017 | -0.122033366 | 0.902872594 | 0.9539997 | | | Ly6d | 16.91919721 | 0.050032565 | 0.067138259 | 0.7452169 | 0.456140608 | NA |
| Mapk12 | 228.8538536 | -0.014439364 | 0.118350244 | -0.122005361 | 0.902894772 | 0.9539997 | | | Olfr1344 | 27.38216268 | 0.05003474 | 0.085981291 | 0.581925894 | 0.560616598 | 0.873713575 |
| Tfpi | 165.9017495 | 0.014867838 | 0.121730097 | 0.122137729 | 0.902789942 | 0.9539997 | | | Prpf6 | 2918.388802 | 0.050034962 | 0.041751842 | 1.198389342 | 0.23076548 | 0.683402059 |
| Eif2c1 | 4114.239159 | -0.008885916 | 0.072887048 | -0.121913507 | 0.902967518 | 0.954011506 | | | Mrpl55 | 720.534564 | 0.050049355 | 0.065545134 | 0.76358613 | 0.445113917 | 0.821493054 |
| Dgke | 2985.479348 | -0.008015733 | 0.065832748 | -0.121759056 | 0.903089841 | 0.954075685 | | | 1700007F19Rik | 9.122245547 | 0.050092497 | 0.059418242 | 0.843049125 | 0.39920097 | NA |
| Chic1 | 1330.480205 | 0.008255748 | 0.067873933 | 0.121633553 | 0.903189239 | 0.954113462 | | | Anapc16 | 1611.637967 | 0.05009406 | 0.04897161 | 1.022920431 | 0.306345469 | 0.738584943 |
| Kbtbd2 | 2139.009392 | -0.009301334 | 0.076517409 | -0.121558403 | 0.903248758 | 0.954113462 | | | Ica1l | 1608.044001 | 0.050116412 | 0.066121067 | 0.757949235 | 0.448481374 | 0.823263966 |
| Ppargc1a | 2193.101935 | -0.014012445 | 0.11543082 | -0.121392577 | 0.903380096 | 0.954122826 | | | Gng5 | 434.9680834 | 0.050123858 | 0.075267275 | 0.665944901 | 0.505446319 | 0.848406891 |
| Rgs12 | 2446.854962 | -0.010629988 | 0.087567657 | -0.12139171 | 0.903380783 | 0.954122826 | | | Snx19 | 4306.221232 | 0.050168582 | 0.054455444 | 0.921277617 | 0.356905504 | 0.773088648 |
| Pik3r6 | 172.9471594 | -0.014130104 | 0.116508989 | -0.121279092 | 0.90346998 | 0.954123446 | | | Rsbn1 | 1095.541202 | 0.05017455 | 0.074854459 | 0.670294743 | 0.502669918 | 0.84739426 |
| Trim59 | 580.067647 | 0.016578397 | 0.136745438 | 0.12123547 | 0.90350453 | 0.954123446 | | | Entpd7 | 510.8477122 | 0.050176161 | 0.072442147 | 0.69263768 | 0.488536957 | 0.840318276 |
| B430212C06Rik | 9.053560191 | 0.018923953 | 0.156286665 | 0.121084888 | 0.903623799 | 0.954184362 | | | Shb | 464.1924432 | 0.050191093 | 0.080675598 | 0.622134754 | 0.533853264 | 0.859470725 |
| Cyp27b1 | 12.6113855 | -0.019311519 | 0.159953851 | -0.12073182 | 0.903903455 | 0.954249731 | | | Nckap1 | 23486.7415 | 0.050213573 | 0.068641874 | 0.731529754 | 0.464455637 | 0.83159896 |
| Gm3414 | 68.06144855 | 0.018933623 | 0.156676822 | 0.120845081 | 0.903813743 | 0.954249731 | | | Piezo2 | 39.419641 | 0.050214579 | 0.088198944 | 0.569333107 | 0.569130103 | 0.87745335 |
| Pkn2 | 1091.404654 | 0.008720749 | 0.072177055 | 0.120824399 | 0.903830124 | 0.954249731 | | | Acsm5 | 8.175990469 | 0.050223096 | 0.057783852 | 0.869154512 | 0.384762624 | NA |
| Pltp | 2624.837207 | -0.019890289 | 0.164903216 | -0.120617956 | 0.903993646 | 0.954249731 | | | Slc39a7 | 3757.58495 | 0.050225799 | 0.053308368 | 0.94217476 | 0.346103174 | 0.76510344 |
| Pycard | 81.33847453 | 0.019726985 | 0.163505325 | 0.120650414 | 0.903967936 | 0.954249731 | | | Myo1e | 941.8448037 | 0.050229138 | 0.066304116 | 0.757556862 | 0.448716313 | 0.823263966 |
| Cyb5d1 | 411.6726485 | 0.009079058 | 0.075511015 | 0.120234889 | 0.904297081 | 0.954439985 | | | Hsd17b7 | 950.5711047 | 0.050231277 | 0.0724374 | 0.693443957 | 0.488030985 | 0.839945271 |
| Sh3bgrl3 | 3408.3619 | -0.009485502 | 0.078872594 | -0.120263601 | 0.904274337 | 0.954439985 | | | Cope | 2789.474421 | 0.050233301 | 0.062634969 | 0.802000893 | 0.422552442 | 0.808695226 |
| Tmem87a | 1179.998807 | 0.007502279 | 0.062471502 | 0.120091227 | 0.904410882 | 0.954495075 | | | Zfp551 | 232.9542769 | 0.05023398 | 0.084343763 | 0.595586183 | 0.551451709 | 0.869755927 |
| Cyp4f17 | 74.3098625 | -0.017314129 | 0.144480974 | -0.119836742 | 0.904612475 | 0.95457779 | | | Exoc2 | 2398.884347 | 0.050250132 | 0.058668056 | 0.85651605 | 0.391712399 | 0.794184423 |
| Kdelr3 | 20.8035302 | -0.020146817 | 0.168056616 | -0.119881127 | 0.904577315 | 0.95457779 | | | Megf11 | 829.0870972 | 0.050252382 | 0.085247703 | 0.589486638 | 0.555534873 | 0.871360989 |
| Npy1r | 1008.81523 | 0.008117226 | 0.067971882 | 0.119420351 | 0.904942339 | 0.954860842 | | | Cdc42ep4 | 1389.80762 | 0.050256487 | 0.067447801 | 0.745116765 | 0.456201135 | 0.82674985 |
| Pole3 | 383.5262326 | -0.013832844 | 0.116116483 | -0.11912903 | 0.905173132 | 0.955010028 | | | Rnf13 | 4476.977457 | 0.050260658 | 0.066763532 | 0.752816043 | 0.451560465 | 0.824520467 |
| Zfp839 | 495.6389862 | 0.010819577 | 0.090854938 | 0.11908628 | 0.905207001 | 0.955010028 | | | Pcsk6 | 957.9676874 | 0.050263985 | 0.079695382 | 0.630701352 | 0.528235815 | 0.856965327 |
| Aox4 | 36.63004758 | -0.01961695 | 0.164928067 | -0.118942457 | 0.905320945 | 0.955065208 | | | Erh | 1616.933183 | 0.050315752 | 0.049031858 | 1.026184899 | 0.304804438 | 0.737453291 |
| C5ar2 | 46.08286919 | -0.018673785 | 0.157290577 | -0.118721574 | 0.905495944 | 0.955067686 | | | Dcn | 537.0362102 | 0.0503436 | 0.084662395 | 0.594639445 | 0.552084507 | 0.870265825 |
| Polr3h | 1344.222246 | -0.010183221 | 0.085699565 | -0.118824653 | 0.905414277 | 0.955067686 | | | Mfsd12 | 655.1790922 | 0.050344442 | 0.070507455 | 0.714030058 | 0.475208593 | 0.834790183 |
| Tmem151a | 8629.333015 | -0.010133825 | 0.085369047 | -0.118706083 | 0.905508217 | 0.955067686 | | | Trp53rk | 128.4257079 | 0.050350394 | 0.094688935 | 0.531745279 | 0.594902427 | 0.887314166 |
| Agpat3 | 8408.979351 | 0.006970984 | 0.059136288 | 0.117879964 | 0.906162769 | 0.955432864 | | | Trappc4 | 1632.86807 | 0.050387845 | 0.046214463 | 1.090304675 | 0.275578957 | 0.717137213 |
| Fndc4 | 2970.414117 | -0.00816436 | 0.069257962 | -0.11788334 | 0.906160094 | 0.955432864 | | | Mtmr10 | 974.8946923 | 0.050393524 | 0.071733421 | 0.702511086 | 0.48236049 | 0.837629227 |
| Hexim2 | 321.535283 | 0.013202657 | 0.111717412 | 0.118179044 | 0.905925794 | 0.955432864 | | | Arhgap1 | 3609.576663 | 0.050397322 | 0.048120228 | 1.047320934 | 0.294951584 | 0.73000898 |
| Nrep | 2158.152474 | -0.019463996 | 0.165113836 | -0.117882283 | 0.906160931 | 0.955432864 | | | Ripk4 | 32.07509919 | 0.050409216 | 0.088177508 | 0.571678847 | 0.567539568 | 0.876563504 |
| Tmem161a | 942.0641698 | -0.008868627 | 0.075117872 | -0.118062814 | 0.906017888 | 0.955432864 | | | AK010878 | 140.4038519 | 0.050438883 | 0.088920496 | 0.567235735 | 0.570554032 | 0.878246677 |
| Pdcd2l | 392.6584843 | 0.008908807 | 0.07567425 | 0.11772574 | 0.906284971 | 0.955461008 | | | Snapc5 | 674.8120294 | 0.050451518 | 0.074750911 | 0.674928464 | 0.499721215 | 0.846241311 |
| Zbtb22 | 1396.52152 | 0.009818406 | 0.083425554 | 0.117690626 | 0.906312794 | 0.955461008 | | | Bri3 | 985.2649874 | 0.050451751 | 0.061269412 | 0.823441072 | 0.410257215 | 0.80184319 |
| Gemin8 | 343.2293741 | 0.010165546 | 0.086915963 | 0.116958334 | 0.906893072 | 0.956007705 | | | Pgr15l | 26.80586984 | 0.050457193 | 0.038483038 | 1.311154108 | NA | NA |
| Puf60 | 4792.285115 | -0.006761095 | 0.057893414 | -0.116785217 | 0.907030259 | 0.956029949 | | | Zfp474 | 18.97248162 | 0.050480626 | 0.069627891 | 0.725005815 | 0.468448463 | NA |
| Seh1l | 1486.889284 | 0.007214505 | 0.061780732 | 0.11677598 | 0.907037579 | 0.956029949 | | | Mmel1 | 40.06437495 | 0.050501803 | 0.092047767 | 0.548647779 | 0.58324719 | 0.884313345 |
| Trpm2 | 1559.0131 | 0.012006891 | 0.102942207 | 0.1166372 | 0.907147558 | 0.956080828 | | | Rgcc | 374.2452108 | 0.050522186 | 0.084736757 | 0.596225156 | 0.551024822 | 0.869642174 |
| Tmtc3 | 547.1921766 | -0.010911905 | 0.093634459 | -0.116537278 | 0.907226744 | 0.95609925 | | | Gstk1 | 349.1316588 | 0.050538838 | 0.089094004 | 0.567252965 | 0.570542328 | 0.878246677 |
| Adh1 | 14.13129829 | -0.018381701 | 0.157848268 | -0.116451714 | 0.907294552 | 0.956105679 | | | Abca2 | 14392.90496 | 0.050548758 | 0.059067372 | 0.855781399 | 0.392118706 | 0.794184423 |
| Lonrf1 | 1321.721782 | -0.008894707 | 0.07647629 | -0.116306728 | 0.907409454 | 0.95616173 | | | Gspt2 | 1328.582331 | 0.050549861 | 0.060370953 | 0.837320905 | 0.402412212 | 0.79955155 |
| Mrpl16 | 843.8670443 | -0.008997183 | 0.077523798 | -0.116057044 | 0.907607332 | 0.956240175 | | | Ednra | 178.5656625 | 0.05056185 | 0.089251615 | 0.566509073 | 0.571047767 | 0.878246677 |
| Paqr8 | 5627.920049 | -0.00913367 | 0.078660804 | -0.116114629 | 0.907561695 | 0.956240175 | | | Tox4 | 3499.018986 | 0.050575181 | 0.050507701 | 1.001336023 | 0.316664383 | 0.746005 |
| Slc6a12 | 47.73186695 | 0.019414982 | 0.167685241 | 0.115782292 | 0.907825085 | 0.95640456 | | | Agpat3 | 9118.494202 | 0.050592466 | 0.040895464 | 1.237116833 | 0.216043711 | 0.670049713 |
| Ahcyl2 | 7230.650512 | 0.007863996 | 0.06829816 | 0.115142138 | 0.908332461 | 0.956463738 | | | Ebag9 | 667.1245661 | 0.050602721 | 0.07754734 | 0.652539745 | 0.514053042 | 0.853195313 |
| Arhgef10l | 2156.661495 | 0.014215295 | 0.123328713 | 0.115263469 | 0.908236293 | 0.956463738 | | | Mipep | 1225.524303 | 0.050605043 | 0.051731519 | 0.978224571 | 0.327963266 | 0.754778534 |
| CK137956 | 81.90248776 | 0.015827107 | 0.138573029 | 0.114214913 | 0.90906743 | 0.956463738 | | | Smap2 | 8620.253198 | 0.050633256 | 0.047213837 | 1.072424085 | 0.283529595 | 0.721034821 |
| Cat | 1780.542594 | -0.01294155 | 0.113410297 | -0.114112656 | 0.90914849 | 0.956463738 | | | Snca | 8558.941242 | 0.050646969 | 0.058307327 | 0.868620997 | 0.385054466 | 0.789016184 |
| Coq4 | 613.7277962 | -0.011694159 | 0.101769923 | -0.114907814 | 0.908518192 | 0.956463738 | | | Terf2 | 1752.22177 | 0.050649536 | 0.052065367 | 0.972806667 | 0.330649399 | 0.756141871 |
| Dcxr | 150.4381454 | 0.014865078 | 0.129187926 | 0.115065536 | 0.908393177 | 0.956463738 | | | Tmem151a | 9943.82236 | 0.050674756 | 0.053283146 | 0.951046623 | 0.341580709 | 0.76303628 |
| Gnb1l | 158.6264512 | 0.012256131 | 0.106536814 | 0.115041272 | 0.908412409 | 0.956463738 | | | Ccdc90b | 434.0238802 | 0.050681802 | 0.0731691 | 0.69266674 | 0.488518717 | 0.840318276 |
| Homez | 704.0498184 | 0.007922229 | 0.06862874 | 0.115436026 | 0.908099525 | 0.956463738 | | | Zfp775 | 338.0714683 | 0.050682961 | 0.085493306 | 0.59282959 | 0.553295203 | 0.870600462 |
| Hsp90ab1 | 62236.55288 | -0.006212361 | 0.053911814 | -0.115231909 | 0.908261307 | 0.956463738 | | | Bai3 | 4759.591146 | 0.050708472 | 0.056148741 | 0.903109694 | 0.36646768 | 0.778210102 |
| Kcp | 63.51843393 | -0.018948426 | 0.164377715 | -0.115273695 | 0.908228188 | 0.956463738 | | | Senp7 | 1268.732417 | 0.050718988 | 0.059202878 | 0.856698014 | 0.391611801 | 0.794184423 |
| Pnck | 2488.505235 | 0.01004438 | 0.087365191 | 0.11497005 | 0.908468861 | 0.956463738 | | | Ppp2r2d | 2211.522503 | 0.050723154 | 0.063171553 | 0.802942955 | 0.422007707 | 0.808695226 |
| Pno1 | 762.4529719 | -0.008465338 | 0.073423137 | -0.11529524 | 0.908211111 | 0.956463738 | | | Wdr92 | 592.640039 | 0.050756414 | 0.069320354 | 0.732200741 | 0.464046052 | 0.83159896 |
| Slc16a14 | 209.0704238 | 0.013808256 | 0.120244563 | 0.114834762 | 0.908576095 | 0.956463738 | | | Atp1a3 | 106168.954 | 0.050785875 | 0.047186179 | 1.076287095 | 0.281798878 | 0.720163706 |
| Slc39a7 | 3402.003784 | -0.005738184 | 0.050301449 | -0.114075917 | 0.909177613 | 0.956463738 | | | Atg4d | 986.9685738 | 0.050793341 | 0.074098144 | 0.685487355 | 0.493036437 | 0.842867752 |
| Sort1 | 27082.2747 | 0.008538676 | 0.074808586 | 0.114140317 | 0.909126563 | 0.956463738 | | | Zmiz1 | 6664.247321 | 0.050819645 | 0.061581104 | 0.82524738 | 0.409231159 | 0.801055189 |
| St3gal5 | 6035.551493 | -0.007349071 | 0.064309106 | -0.114277306 | 0.909017972 | 0.956463738 | | | Rnf121 | 924.7430275 | 0.050825765 | 0.054237485 | 0.937096629 | 0.348708855 | 0.767555385 |
| Tmem204 | 307.6590033 | -0.010375588 | 0.090561681 | -0.1145693 | 0.908786514 | 0.956463738 | | | Hao1 | 3.93938632 | 0.050835104 | 0.043906789 | 1.157795975 | 0.246947306 | NA |
| Uck1 | 1485.092848 | -0.007293255 | 0.063923156 | -0.114094106 | 0.909163194 | 0.956463738 | | | Wdr18 | 2451.001299 | 0.050851632 | 0.054172601 | 0.938696512 | 0.347886585 | 0.766247237 |
| Zbtb1 | 460.9006936 | -0.011269535 | 0.098670311 | -0.114214045 | 0.909068119 | 0.956463738 | | | Yy2 | 42.8618466 | 0.050855041 | 0.093096853 | 0.546259511 | 0.584887564 | 0.884313345 |
| Zfhx2 | 2551.685006 | -0.008441085 | 0.073724874 | -0.114494394 | 0.90884589 | 0.956463738 | | | Lman2l | 1538.335193 | 0.050904957 | 0.07153416 | 0.711617452 | 0.476701697 | 0.835195112 |
| Zfp959 | 155.2584251 | 0.015049763 | 0.131141944 | 0.114759343 | 0.908635875 | 0.956463738 | | | 6030419C18Rik | 6202.466997 | 0.050905092 | 0.06384142 | 0.797367794 | 0.425237458 | 0.809519334 |
| Cdkl1 | 507.6461099 | 0.00977873 | 0.085785355 | 0.113990665 | 0.909245193 | 0.956469891 | | | Zfp644 | 2359.999673 | 0.050908375 | 0.071049371 | 0.716521128 | 0.473669626 | 0.833983942 |
| Aldh18a1 | 903.8181963 | -0.007498777 | 0.06594477 | -0.113712988 | 0.909465317 | 0.956511805 | | | Rock2 | 10632.62065 | 0.050926358 | 0.080065152 | 0.636061458 | 0.524736366 | 0.855487854 |
| Fam134a | 8263.703757 | 0.006266921 | 0.05509244 | 0.11375283 | 0.909433732 | 0.956511805 | | | Alg14 | 298.5990896 | 0.050945565 | 0.07645973 | 0.666305844 | 0.505215631 | 0.848183273 |
| Mcm3 | 133.9981209 | -0.015427611 | 0.135678904 | -0.113706776 | 0.909470241 | 0.956511805 | | | Arfgef2 | 3991.424462 | 0.050945571 | 0.05531046 | 0.921083845 | 0.357006654 | 0.773088648 |
| Armc4 | 26.7379031 | 0.018851512 | 0.165925659 | 0.113614207 | 0.909543625 | 0.956524056 | | | Mypop | 1573.842797 | 0.050970457 | 0.049148451 | 1.037071489 | 0.299702541 | 0.733704078 |
| Bean1 | 566.6786218 | -0.010087987 | 0.089104443 | -0.113215312 | 0.909859857 | 0.956579732 | | | C530044C16Rik | 7.76552431 | 0.050984082 | 0.051802263 | 0.984205698 | 0.325014385 | NA |
| Dedd | 1070.650126 | 0.00932881 | 0.082376704 | 0.113245731 | 0.909835741 | 0.956579732 | | | Apba2 | 8985.215446 | 0.051001321 | 0.060786366 | 0.839025672 | 0.401454904 | 0.799277436 |
| Prr16 | 289.7261963 | 0.013257001 | 0.117146865 | 0.113165647 | 0.90989923 | 0.956579732 | | | Mnat1 | 446.239477 | 0.051003926 | 0.078838468 | 0.646942121 | 0.517669404 | 0.853297835 |
| Wdr82 | 6249.904103 | 0.005107405 | 0.045101436 | 0.113242628 | 0.909838201 | 0.956579732 | | | Krt26 | 1.845863774 | 0.051004945 | 0.028137004 | 1.812735428 | 0.069872637 | NA |
| Zfp433 | 41.25529608 | 0.018182206 | 0.160679754 | 0.113158042 | 0.909905259 | 0.956579732 | | | Trove2 | 1824.928792 | 0.051013726 | 0.07650855 | 0.666771565 | 0.504918058 | 0.848183273 |
| Klhdc10 | 4548.620363 | -0.00617608 | 0.054624484 | -0.113064313 | 0.909979568 | 0.956591808 | | | Rai2 | 432.3785076 | 0.051040191 | 0.08529758 | 0.598377952 | 0.549587774 | 0.868884633 |
| Taf4b | 227.0967711 | 0.010774342 | 0.09535845 | 0.112987802 | 0.910040226 | 0.956591808 | | | Elk4 | 383.8420086 | 0.051067204 | 0.082337124 | 0.620220893 | 0.535112368 | 0.860420883 |
| Dkkl1 | 639.0562889 | -0.014680317 | 0.130055341 | -0.112877467 | 0.910127701 | 0.956618859 | | | Plekhf2 | 306.6307872 | 0.051072625 | 0.081413604 | 0.627322983 | 0.530447541 | 0.857953566 |
| C130060C02Rik | 10.85089783 | -0.018507212 | 0.164266371 | -0.112665862 | 0.910295467 | 0.956675494 | | | Tm9sf4 | 4630.078282 | 0.051080997 | 0.048224666 | 1.059229676 | 0.289495193 | 0.724785794 |
| Gipr | 56.31207524 | 0.018313971 | 0.162568682 | 0.112653745 | 0.910305074 | 0.956675494 | | | Cldn3 | 6.534962883 | 0.051082785 | 0.048784934 | 1.047101651 | 0.295052698 | NA |
| 2610044O15Rik8 | 274.3519795 | -0.011255253 | 0.100356178 | -0.112153065 | 0.910702043 | 0.956833082 | | | 2410002F23Rik | 2425.453917 | 0.051088683 | 0.054270786 | 0.941366191 | 0.346517233 | 0.765286747 |
| C330027C09Rik | 110.9948917 | -0.016454939 | 0.146508223 | -0.112314101 | 0.910574362 | 0.956833082 | | | G0s2 | 105.9997458 | 0.051091796 | 0.093884766 | 0.544196875 | 0.586306002 | 0.884588669 |
| Lipo1 | 172.7298145 | 0.015415541 | 0.137427483 | 0.112172185 | 0.910686883 | 0.956833082 | | | Pafah1b1 | 26402.36539 | 0.051092706 | 0.064268814 | 0.794984414 | 0.426622568 | 0.809869241 |
| Tra2a | 1697.730144 | 0.013100424 | 0.116661043 | 0.112294765 | 0.910589693 | 0.956833082 | | | Psma6 | 2917.066034 | 0.051099726 | 0.044215964 | 1.155684984 | 0.247810033 | 0.696944196 |
| Tmem109 | 1501.349548 | -0.010448655 | 0.093274268 | -0.112020768 | 0.910806939 | 0.956878405 | | | BC039966 | 16.51023395 | 0.05111467 | 0.078996922 | 0.64704635 | 0.517601946 | NA |
| Hnrnpl | 5373.954906 | 0.008185926 | 0.073251516 | 0.111750938 | 0.91102089 | 0.957038285 | | | Glud1 | 10774.20738 | 0.051138422 | 0.050353818 | 1.015581818 | 0.309828567 | 0.742545745 |
| 0610043K17Rik | 13.43557615 | 0.018251914 | 0.163727531 | 0.111477364 | 0.911237814 | 0.957071495 | | | Oard1 | 454.3637353 | 0.05116237 | 0.066827976 | 0.765583111 | 0.443924397 | 0.820935174 |
| Cenpl | 56.04421654 | -0.016430989 | 0.147354997 | -0.111506157 | 0.911214984 | 0.957071495 | | | Topors | 1030.917881 | 0.051197034 | 0.068618558 | 0.746110603 | 0.455600605 | 0.826603827 |
| Rab13 | 158.6167318 | 0.011962421 | 0.107235387 | 0.111552927 | 0.911177898 | 0.957071495 | | | Prss56 | 8.186573763 | 0.051198289 | 0.062239774 | 0.82259761 | 0.410736858 | NA |
| Alyref2 | 81.42775452 | -0.016331463 | 0.147017587 | -0.1110851 | 0.911548864 | 0.957333295 | | | Myl12a | 827.6842992 | 0.051198654 | 0.073141904 | 0.699990711 | 0.483933105 | 0.838713365 |
| Zfp317 | 1045.148324 | 0.009010826 | 0.081262943 | 0.110884809 | 0.911707692 | 0.957435203 | | | Pcnt | 1609.595764 | 0.051231304 | 0.054429871 | 0.941235086 | 0.3465844 | 0.765286747 |
| Cxcl5 | 9.024761255 | 0.017767527 | 0.160579034 | 0.110646617 | 0.91189658 | 0.957533205 | | | Rnf185 | 1554.603821 | 0.051232747 | 0.054562894 | 0.938966803 | 0.347747789 | 0.766057685 |
| Gtf3c5 | 776.9756172 | 0.008577469 | 0.077546072 | 0.110611264 | 0.911924615 | 0.957533205 | | | Lrrc4c | 4815.938085 | 0.051235712 | 0.068119127 | 0.752148691 | 0.451961645 | 0.824608086 |
| Nsa2 | 1669.867427 | 0.007514802 | 0.068217907 | 0.110158787 | 0.912283446 | 0.957845071 | | | Gstp1 | 1778.936799 | 0.051267808 | 0.063471797 | 0.8077258 | 0.419248448 | 0.807114502 |
| Gltscr1 | 828.6145941 | 0.008766322 | 0.079713801 | 0.109972453 | 0.912431222 | 0.957935313 | | | Dmwd | 5259.762114 | 0.05126978 | 0.059971985 | 0.854895503 | 0.392609001 | 0.794184423 |
| 1200014J11Rik | 851.4867616 | -0.010445507 | 0.095216536 | -0.109702656 | 0.912645194 | 0.958095036 | | | Bambi | 147.8322686 | 0.051270186 | 0.093742994 | 0.546922861 | 0.584431729 | 0.884313345 |
| Ern2 | 13.93636741 | -0.018139584 | 0.166426952 | -0.108994268 | 0.913207036 | 0.958554964 | | | Cdc42ep3 | 512.6752049 | 0.051273792 | 0.074235334 | 0.690692539 | 0.489758779 | 0.841119426 |
| Nr2c2ap | 490.9915276 | -0.008931279 | 0.081923693 | -0.109019127 | 0.913187319 | 0.958554964 | | | Fam65b | 2826.288801 | 0.051285071 | 0.073747134 | 0.695417816 | 0.486793498 | 0.839801621 |
| Lca5 | 371.168883 | 0.013096488 | 0.120559642 | 0.108630783 | 0.913495344 | 0.958741946 | | | Atg7 | 1149.829656 | 0.051285775 | 0.058477367 | 0.877019224 | 0.380476197 | 0.786275719 |
| Slc18b1 | 793.2573755 | -0.008703264 | 0.080130475 | -0.108613655 | 0.91350893 | 0.958741946 | | | Dpy19l3 | 2743.488024 | 0.051286768 | 0.071148666 | 0.7208395 | 0.47100827 | 0.833245343 |
| B930003M22Rik | 25.38807303 | -0.01800604 | 0.16661423 | -0.10807024 | 0.913939975 | 0.958938792 | | | Ano7 | 16.69086913 | 0.051310865 | 0.071850866 | 0.714130092 | 0.47514674 | NA |
| Dennd4c | 952.7512021 | -0.009799404 | 0.090680559 | -0.1080651 | 0.913944052 | 0.958938792 | | | Tmem106b | 7221.466313 | 0.051312936 | 0.065220102 | 0.786765658 | 0.43141906 | 0.812181606 |
| Nr3c2 | 1286.47972 | 0.009286106 | 0.085758376 | 0.108282207 | 0.913771836 | 0.958938792 | | | Phactr4 | 593.0931151 | 0.051318998 | 0.078845479 | 0.650880673 | 0.515123519 | 0.853297835 |
| Tmem69 | 467.3109662 | 0.009899799 | 0.091573342 | 0.108107866 | 0.913910128 | 0.958938792 | | | Dusp19 | 379.4090556 | 0.051336573 | 0.081576709 | 0.629304288 | 0.529149865 | 0.857868166 |
| Ckap5 | 7591.947189 | -0.006649455 | 0.061834647 | -0.107536074 | 0.914363708 | 0.959068549 | | | Polr3d | 770.755334 | 0.051353308 | 0.058657738 | 0.875473716 | 0.381316205 | 0.786663048 |
| Fastkd2 | 695.0841194 | -0.009306513 | 0.086745735 | -0.10728496 | 0.914562916 | 0.959068549 | | | Cog4 | 2217.900071 | 0.05137068 | 0.047133488 | 1.089897714 | 0.275758204 | 0.717346645 |
| Fkbp14 | 397.6364237 | -0.009241841 | 0.085970513 | -0.10750013 | 0.914392222 | 0.959068549 | | | Pla2g12a | 559.9562487 | 0.051420217 | 0.071227055 | 0.721919739 | 0.470343825 | 0.833245343 |
| Gstm2 | 92.35188109 | -0.016121525 | 0.149929805 | -0.107527155 | 0.914370784 | 0.959068549 | | | Chm | 2007.146936 | 0.051425543 | 0.082548795 | 0.622971455 | 0.53330328 | 0.859128696 |
| Kdelr2 | 745.2885157 | -0.006662552 | 0.062007873 | -0.107446865 | 0.914434477 | 0.959068549 | | | Tmed8 | 2974.9104 | 0.051435513 | 0.058085084 | 0.885520167 | 0.375876148 | 0.783947144 |
| Rpl28 | 4181.10371 | -0.007003841 | 0.065238825 | -0.107356943 | 0.914505811 | 0.959068549 | | | Sars | 7387.941725 | 0.051440477 | 0.044554672 | 1.15454732 | 0.24827585 | 0.697708453 |
| Stk24 | 1889.851866 | -0.008784906 | 0.08165155 | -0.107590191 | 0.914320778 | 0.959068549 | | | Brcc3 | 1016.768983 | 0.051462067 | 0.056708025 | 0.907491785 | 0.364146792 | 0.777191869 |
| Taf15 | 1555.251379 | 0.013656929 | 0.127287823 | 0.107291715 | 0.914557557 | 0.959068549 | | | Igsf8 | 7891.669454 | 0.051472185 | 0.05758644 | 0.893824734 | 0.371415673 | 0.780761257 |
| Ncoa4 | 3930.470419 | 0.005568666 | 0.051950364 | 0.107192059 | 0.914636615 | 0.959080923 | | | Dlx6as1 | 619.8218595 | 0.051501193 | 0.079962526 | 0.644066618 | 0.519532236 | 0.853297835 |
| Arhgap27 | 705.1589356 | 0.009341777 | 0.087282287 | 0.107029465 | 0.914765604 | 0.959140337 | | | Tmcc3 | 2446.992838 | 0.051508963 | 0.077992776 | 0.660432481 | 0.508976334 | 0.850095619 |
| Cnp | 10698.14211 | 0.013912745 | 0.130275889 | 0.106794471 | 0.914952034 | 0.959140337 | | | Peg3 | 11309.99947 | 0.051521156 | 0.082911352 | 0.621400513 | 0.534336134 | 0.859648542 |
| Cntnap5a | 942.3713868 | -0.012237886 | 0.114629235 | -0.106760602 | 0.914978904 | 0.959140337 | | | Slc30a2 | 3.227866408 | 0.051521863 | 0.042887364 | 1.201329688 | 0.229623338 | NA |
| Lrch2 | 612.1794191 | 0.010787176 | 0.100897411 | 0.106912316 | 0.914858542 | 0.959140337 | | | Zic1 | 434.1994382 | 0.051523067 | 0.085155575 | 0.605046317 | 0.545148222 | 0.866581645 |
| Slk | 4355.104365 | -0.009469322 | 0.088721815 | -0.106730486 | 0.915002796 | 0.959140337 | | | Gpr137 | 3289.924922 | 0.051528906 | 0.06037088 | 0.85353908 | 0.393360425 | 0.794754188 |
| Atp6v1h | 6305.724868 | 0.006648692 | 0.062664232 | 0.106100279 | 0.91550279 | 0.959270321 | | | A930004D18Rik | 119.3633853 | 0.051535967 | 0.093491077 | 0.551239423 | 0.581469558 | 0.884020489 |
| Cnih4 | 1379.421505 | 0.007194052 | 0.067867299 | 0.106001743 | 0.915580969 | 0.959270321 | | | Egr1 | 12389.93262 | 0.05153618 | 0.07852594 | 0.656294973 | 0.511634351 | 0.852417908 |
| Gm14420 | 1019.792743 | -0.008980748 | 0.084423044 | -0.106377918 | 0.915282513 | 0.959270321 | | | Atp5j2 | 3134.284665 | 0.051544177 | 0.051924047 | 0.992684128 | 0.320863912 | 0.750352874 |
| Ndufa10 | 4163.439181 | -0.005384895 | 0.050732325 | -0.106143272 | 0.915468679 | 0.959270321 | | | Rundc3b | 1692.040264 | 0.051591753 | 0.064313467 | 0.802192056 | 0.422441871 | 0.808695226 |
| Nfatc4 | 74.22616369 | -0.017285484 | 0.163147702 | -0.105949906 | 0.915622098 | 0.959270321 | | | Chrne | 3.430925882 | 0.051612441 | 0.034979828 | 1.475491553 | 0.140080438 | NA |
| Tctex1d1 | 8.886442323 | 0.016591308 | 0.156567577 | 0.105968992 | 0.915606955 | 0.959270321 | | | Pak1 | 24175.81546 | 0.051628108 | 0.057957371 | 0.89079451 | 0.373039421 | 0.782226671 |
| Thap2 | 453.9801629 | 0.009895664 | 0.093111503 | 0.106277569 | 0.915362128 | 0.959270321 | | | Col9a2 | 265.0029341 | 0.051632329 | 0.093614747 | 0.551540544 | 0.581263182 | 0.883958169 |
| Yod1 | 357.2769674 | 0.009250261 | 0.08689027 | 0.106459114 | 0.915218093 | 0.959270321 | | | Armcx1 | 2873.811515 | 0.051662631 | 0.053602475 | 0.963810566 | 0.335140909 | 0.75853176 |
| Axin1 | 1125.106509 | 0.008208521 | 0.078036638 | 0.105188043 | 0.916226598 | 0.959773843 | | | Apaf1 | 740.1200539 | 0.051669685 | 0.092219236 | 0.560291841 | 0.575280391 | 0.880746518 |
| Ppp1r16b | 8477.186128 | 0.009375605 | 0.089082916 | 0.105245822 | 0.916180752 | 0.959773843 | | | Mad1l1 | 629.725253 | 0.051673821 | 0.063771129 | 0.810301171 | 0.417767102 | 0.80621897 |
| Ap1m1 | 2427.713154 | 0.007119125 | 0.067756201 | 0.105069719 | 0.916320487 | 0.959782428 | | | Hpcal4 | 50641.00071 | 0.051706187 | 0.063479232 | 0.814537058 | 0.415337352 | 0.805633085 |
| Fgf2 | 21.90388538 | -0.017633163 | 0.168080348 | -0.104909128 | 0.916447916 | 0.959782428 | | | Zxdc | 1253.162486 | 0.051713007 | 0.055802115 | 0.926721284 | 0.354071264 | 0.771423785 |
| Rras | 295.6393632 | 0.010860285 | 0.103564 | 0.104865449 | 0.916482576 | 0.959782428 | | | H2afy | 2482.03584 | 0.051720327 | 0.05993331 | 0.862964638 | 0.38815691 | 0.791889115 |
| Tmem216 | 189.8635601 | 0.012106077 | 0.115398251 | 0.104906934 | 0.916449657 | 0.959782428 | | | Rint1 | 732.0645556 | 0.051726698 | 0.058493522 | 0.884314982 | 0.376526203 | 0.783953619 |
| Pcdhga9 | 438.3674225 | 0.010783577 | 0.102963523 | 0.104732015 | 0.916588458 | 0.959828438 | | | Gart | 1523.357011 | 0.051744385 | 0.050570761 | 1.023207561 | 0.306209719 | 0.738584943 |
| Cxadr | 941.1407024 | 0.009268007 | 0.08856428 | 0.104647239 | 0.916655729 | 0.959834011 | | | Stam | 2398.54688 | 0.05174622 | 0.060696352 | 0.852542505 | 0.393913058 | 0.794754188 |
| Gdpgp1 | 341.3943283 | 0.009358514 | 0.089682532 | 0.10435158 | 0.916890346 | 0.960014801 | | | Pianp | 7731.401596 | 0.051748722 | 0.047212141 | 1.096089287 | 0.273039702 | 0.714368645 |
| Atic | 1135.888063 | 0.006858914 | 0.06590389 | 0.104074497 | 0.91711023 | 0.96018014 | | | Ddx56 | 1579.473256 | 0.05174973 | 0.053612143 | 0.96526136 | 0.33441392 | 0.758238352 |
| Zfp27 | 727.8884507 | 0.008922646 | 0.085836934 | 0.103948794 | 0.917209985 | 0.960219696 | | | Mbtps1 | 4206.309889 | 0.051763612 | 0.044131996 | 1.172927047 | 0.240825062 | 0.691863635 |
| Espl1 | 51.19082012 | -0.017126746 | 0.165379119 | -0.103560511 | 0.917518127 | 0.960447004 | | | Eef2 | 45690.34342 | 0.051775117 | 0.053665792 | 0.964769465 | 0.334660293 | 0.758238352 |

| | | | | | | |
|---|---|---|---|---|---|---|
| Rprd2 | 2219.766922 | -0.006812227 | 0.065806551 | -0.103518979 | 0.917551087 | 0.960447004 |
| Tmem132b | 7073.334779 | -0.009188575 | 0.088966727 | -0.103281028 | 0.917739932 | 0.960579782 |
| Gdi2 | 8218.755862 | 0.005240927 | 0.050884374 | 0.102996789 | 0.917965519 | 0.960750997 |
| Osgin2 | 605.3389432 | -0.008976945 | 0.08777697 | -0.102269935 | 0.918542418 | 0.96128985 |
| Ccdc104 | 2972.470668 | -0.005938486 | 0.058213307 | -0.102012514 | 0.918746742 | 0.961396449 |
| Spopl | 335.5293538 | 0.008211547 | 0.080516996 | 0.101985258 | 0.918768376 | 0.961396449 |
| Mpc1 | 2852.030273 | -0.004863468 | 0.047799363 | -0.101747552 | 0.918957057 | 0.961504568 |
| Nrn1 | 5992.384935 | 0.009717182 | 0.09554871 | 0.101698725 | 0.918995815 | 0.961504568 |
| Trappc1 | 1036.212537 | -0.007871819 | 0.077476097 | -0.101603197 | 0.919071642 | 0.961518975 |
| Clint1 | 2080.405397 | 0.007505495 | 0.074186746 | 0.101170298 | 0.919415274 | 0.961813535 |
| Mknk2 | 1738.556951 | 0.009457847 | 0.093587666 | 0.101058688 | 0.919503872 | 0.961841277 |
| Dync1li1 | 2535.881072 | 0.006003032 | 0.059500814 | 0.100889922 | 0.919637843 | 0.961851542 |
| Prss50 | 20.46506222 | 0.016958587 | 0.168081258 | 0.100895167 | 0.919633679 | 0.961851542 |
| Dscr3 | 1303.525707 | -0.005959997 | 0.059183547 | -0.100703618 | 0.919785739 | 0.961876366 |
| Rnf10 | 9304.627882 | 0.006490959 | 0.064417743 | 0.100763529 | 0.919738179 | 0.961876366 |
| Pibf1 | 383.5208723 | -0.010104772 | 0.10067556 | -0.100369668 | 0.92005085 | 0.962082111 |
| Zfp560 | 265.8422889 | -0.010844311 | 0.108119445 | -0.100299356 | 0.920106669 | 0.962082111 |
| Tspo | 101.1323323 | -0.012317522 | 0.123032321 | -0.100116144 | 0.920252119 | 0.962169265 |
| Gt(ROSA)26Sor | 100.8166139 | 0.014085894 | 0.141035047 | 0.099875133 | 0.920443459 | 0.962304383 |
| Rhobtb3 | 921.3101975 | 0.011464106 | 0.115027427 | 0.099664107 | 0.920610997 | 0.962411754 |
| Sphk1 | 148.0795913 | 0.01553883 | 0.156029109 | 0.099589299 | 0.92067039 | 0.962411754 |
| 3110082I17Rik | 277.8307144 | -0.009020973 | 0.090904605 | -0.099235602 | 0.920951208 | 0.962640357 |
| Gm5077 | 9.747069564 | 0.015290097 | 0.154367877 | 0.099049733 | 0.921098783 | 0.962729664 |
| Atg2b | 2937.258318 | -0.009752592 | 0.098692747 | -0.098817713 | 0.921283004 | 0.962792315 |
| Rad21 | 4971.8104 | 0.006053248 | 0.061247519 | 0.098832538 | 0.921271234 | 0.962792315 |
| Dsc3 | 20.36601679 | -0.016509183 | 0.167599826 | -0.098503581 | 0.921532429 | 0.962988025 |
| 4930519G04Rik | 9.315846225 | -0.015921966 | 0.162048462 | -0.098254345 | 0.92173033 | 0.963129871 |
| Ankrd11 | 8780.980855 | -0.007471755 | 0.07650391 | -0.097665012 | 0.922198299 | 0.963164169 |
| Arsi | 19.21450399 | -0.016419574 | 0.167564068 | -0.097989825 | 0.921940373 | 0.963164169 |
| Dtnbp1 | 1879.735553 | 0.009495668 | 0.097038617 | 0.09785453 | 0.922047806 | 0.963164169 |
| Rad54l | 22.45863722 | 0.016456679 | 0.168055593 | 0.097924019 | 0.921992628 | 0.963164169 |
| Scrn2 | 107.1911142 | -0.013495625 | 0.137897583 | -0.097867014 | 0.922037893 | 0.963164169 |
| Slc25a47 | 72.36580198 | 0.012846491 | 0.131500273 | 0.097691743 | 0.922177072 | 0.963164169 |
| Zfp143 | 458.6307551 | -0.008637859 | 0.088418566 | -0.09769282 | 0.922176217 | 0.963164169 |
| Alkbh6 | 922.1739909 | 0.007941385 | 0.081506005 | 0.097433125 | 0.92238244 | 0.963291556 |
| Alkbh4 | 209.1106547 | -0.010273647 | 0.105537529 | -0.097345913 | 0.922451696 | 0.963298954 |
| Ankrd52 | 3504.184953 | -0.005982595 | 0.06152692 | -0.097235404 | 0.922539453 | 0.96332567 |
| Angpt2 | 70.5707085 | 0.015714696 | 0.161792786 | 0.097128533 | 0.922624322 | 0.963349366 |
| Chek2 | 50.25845472 | -0.014473256 | 0.149273114 | -0.096958223 | 0.922759571 | 0.96342566 |
| Zfp109 | 308.1359012 | 0.008482579 | 0.087601469 | 0.09683147 | 0.922860232 | 0.963465833 |
| Rpl18a | 6791.102507 | -0.007235561 | 0.074803647 | -0.096727377 | 0.922942898 | 0.963487216 |
| Ino80e | 909.081151 | -0.006361709 | 0.065851218 | -0.096607311 | 0.92303825 | 0.963521839 |
| Avp | 98.95776403 | 0.010308915 | 0.107210402 | 0.09615592 | 0.923396739 | 0.963701272 |
| BC016423 | 1398.750865 | -0.007401878 | 0.076917132 | -0.096231855 | 0.923336431 | 0.963701272 |
| Det1 | 134.4439488 | -0.01234593 | 0.128268653 | -0.096250559 | 0.923321577 | 0.963701272 |
| Tmem202 | 67.22361334 | 0.013716598 | 0.142861517 | 0.096013249 | 0.92351005 | 0.963754611 |
| Hpdl | 38.20390556 | -0.01552926 | 0.161988137 | -0.095866648 | 0.923626483 | 0.963811202 |
| Cenpe | 63.03364068 | -0.014939392 | 0.156007101 | -0.095760974 | 0.923710413 | 0.963833871 |
| Atl1 | 4497.646216 | 0.007135811 | 0.074894402 | 0.095278296 | 0.924093782 | 0.964025796 |
| Egf | 53.65474773 | -0.015608856 | 0.16393095 | -0.095216044 | 0.924143227 | 0.964025796 |
| Fsd1 | 1439.695119 | 0.006182816 | 0.064798681 | 0.095415774 | 0.923984587 | 0.964025796 |
| Gtf3c1 | 6529.694508 | 0.007323265 | 0.076850026 | 0.095292947 | 0.924082145 | 0.964025796 |
| Akap12 | 1310.823654 | 0.011906066 | 0.125395172 | 0.094948362 | 0.924355843 | 0.964182673 |
| Sec16a | 3243.065085 | 0.007692294 | 0.081173332 | 0.094763807 | 0.924502435 | 0.964270665 |
| Stx8 | 680.4846079 | 0.005976103 | 0.063116102 | 0.094684279 | 0.924565605 | 0.96427164 |
| Chn1 | 43142.48266 | -0.007424474 | 0.078715853 | -0.094319935 | 0.924855015 | 0.964378734 |
| Fuca2 | 1472.079957 | -0.006831982 | 0.072358263 | -0.094418826 | 0.924776462 | 0.964378734 |
| Gtpbp8 | 343.9583588 | -0.009479067 | 0.100424836 | -0.094389668 | 0.924799623 | 0.964378734 |
| Coro2a | 1557.258025 | -0.008966294 | 0.095310079 | -0.094074985 | 0.925049592 | 0.964386918 |
| Ggta1 | 207.905302 | -0.011497537 | 0.122152207 | -0.094124678 | 0.925010117 | 0.964386918 |
| Rin3 | 145.4148716 | 0.01389528 | 0.147647147 | 0.094111402 | 0.925020663 | 0.964386918 |
| 1810010H24Rik | 93.82252502 | 0.012389605 | 0.13213979 | 0.093761346 | 0.925298738 | 0.964581756 |
| Akt1 | 6646.963061 | 0.006313688 | 0.06760388 | 0.093392397 | 0.92559183 | 0.964692796 |
| Gdf9 | 27.08459714 | 0.015665872 | 0.16788384 | 0.093313755 | 0.925654305 | 0.964692796 |
| Poldip3 | 3166.60469 | -0.004885659 | 0.052246727 | -0.093511291 | 0.92549738 | 0.964692796 |
| Tceanc2 | 729.536223 | -0.005796336 | 0.062102608 | -0.09333482 | 0.92563757 | 0.964692796 |
| 1110008P14Rik | 2727.88039 | -0.011954374 | 0.12850458 | -0.093026834 | 0.925882243 | 0.964696902 |
| 3110001I22Rik | 56.60339567 | 0.014207755 | 0.153037229 | 0.092838556 | 0.92603182 | 0.964696902 |
| Gid4 | 3346.752378 | -0.004587189 | 0.049339919 | -0.092971157 | 0.925926475 | 0.964696902 |
| Heatr1 | 1104.786695 | -0.007506866 | 0.080692305 | -0.093030752 | 0.92587913 | 0.964696902 |
| Map9 | 5159.779301 | -0.006459223 | 0.069366609 | -0.093117179 | 0.92581047 | 0.964696902 |
| Tmem8c | 12.46530337 | -0.014719441 | 0.158524403 | -0.092852842 | 0.92602047 | 0.964696902 |
| Ggps1 | 1556.760242 | 0.00655224 | 0.070637933 | 0.092758089 | 0.926095747 | 0.964698636 |
| 3110040N11Rik | 200.2479906 | -0.010311439 | 0.111887495 | -0.092158995 | 0.926571716 | 0.964884184 |
| Atp6v1g2 | 18419.25597 | 0.00563271 | 0.061011689 | 0.092321818 | 0.926442354 | 0.964884184 |
| Fbxw11 | 5840.817051 | 0.005025535 | 0.054393946 | 0.092391446 | 0.926387036 | 0.964884184 |
| Gnptab | 3552.563981 | -0.005160593 | 0.05597666 | -0.092191876 | 0.926545593 | 0.964884184 |
| Snhg10 | 48.81038557 | -0.014813644 | 0.160769776 | -0.092141968 | 0.926585245 | 0.964884184 |
| Bdp1 | 1652.553161 | -0.008134477 | 0.088435951 | -0.091981567 | 0.926712685 | 0.964952039 |
| Iqsec2 | 8981.57022 | -0.008630984 | 0.094373746 | -0.091455347 | 0.927130785 | 0.965037944 |
| Mrpl4 | 1368.996757 | 0.006533102 | 0.071311346 | 0.091613784 | 0.927004899 | 0.965037944 |
| Nup50 | 1760.151316 | 0.005786979 | 0.063208041 | 0.091554471 | 0.927052026 | 0.965037944 |
| Pcsk1n | 7834.327333 | 0.014376645 | 0.1572758 | 0.091410405 | 0.927166495 | 0.965037944 |
| Prcc | 1597.469384 | -0.007103005 | 0.077514137 | -0.091634958 | 0.926988075 | 0.965037944 |
| Xlr3a | 19.19988855 | 0.013191312 | 0.144313413 | 0.091407386 | 0.927168893 | 0.965037944 |
| Haus8 | 237.5863527 | -0.009455107 | 0.103644752 | -0.091226103 | 0.927312934 | 0.965058208 |
| Wsb2 | 12312.67374 | -0.008687331 | 0.095203949 | -0.091249691 | 0.927294192 | 0.965058208 |
| Mipol1 | 93.46203294 | 0.012756516 | 0.140047371 | 0.091087148 | 0.927423344 | 0.965108288 |
| D19Bwg1357e | 1163.727505 | 0.006697117 | 0.073743056 | 0.090816912 | 0.927638072 | 0.965162524 |
| Fancc | 326.8671602 | -0.009188663 | 0.101211927 | -0.09078637 | 0.927662341 | 0.965162524 |
| Syt12 | 1819.121522 | -0.007615453 | 0.083766944 | -0.090912385 | 0.927562209 | 0.965162524 |
| 1190007I07Rik | 143.3317598 | 0.011740821 | 0.130194697 | 0.090178949 | 0.928145012 | 0.96528002 |
| Aamp | 4706.653545 | -0.007936662 | 0.087951122 | -0.090239464 | 0.928096924 | 0.96528002 |
| Cbr4 | 341.7830142 | -0.00829273 | 0.091743701 | -0.090390189 | 0.927977153 | 0.96528002 |
| Chst14 | 71.98985693 | 0.01309629 | 0.145058638 | 0.090282728 | 0.928062545 | 0.96528002 |
| Fam213b | 1812.453392 | 0.007492157 | 0.082922882 | 0.090350898 | 0.928008375 | 0.96528002 |
| Vrk2 | 61.55109073 | -0.012921861 | 0.143299449 | -0.090173839 | 0.928149073 | 0.96528002 |
| Ctsk | 133.4660094 | -0.011386127 | 0.126539442 | -0.08998085 | 0.928302433 | 0.965309926 |
| Unc119 | 794.7407132 | 0.006478043 | 0.071961311 | 0.090021195 | 0.928270372 | 0.965309926 |
| 4930539J05Rik | 20.49971388 | 0.015093239 | 0.167997421 | 0.089842088 | 0.928412702 | 0.965352267 |
| Ptchd2 | 1779.008469 | 0.006707008 | 0.074710907 | 0.089772803 | 0.928467761 | 0.965352267 |
| Fam155a | 3259.964307 | -0.004650443 | 0.051887305 | -0.089625842 | 0.928584548 | 0.96540891 |
| Nos1ap | 3410.102435 | 0.0093835 | 0.104803737 | 0.089534021 | 0.928657517 | 0.965419992 |
| Aftph | 3434.620288 | -0.005519066 | 0.061823566 | -0.089271233 | 0.928866356 | 0.965507535 |
| Sox10 | 1522.089714 | 0.0114482 | 0.12813357 | 0.089345827 | 0.928807075 | 0.965507535 |
| 9130221H12Rik | 277.362702 | 0.008514152 | 0.095823147 | 0.088852765 | 0.929198923 | 0.965521426 |
| F730043M19Rik | 274.6463471 | 0.01003784 | 0.113015163 | 0.088818523 | 0.929226137 | 0.965521426 |
| Mcrs1 | 2498.203394 | -0.005879474 | 0.066222259 | -0.088783949 | 0.929253614 | 0.965521426 |
| Pgm2 | 2626.346154 | -0.007514407 | 0.084429805 | -0.089001825 | 0.92908046 | 0.965521426 |
| Synpo | 10628.55799 | -0.009221126 | 0.103665857 | -0.08895046 | 0.929121282 | 0.965521426 |
| Wipi2 | 5049.482316 | 0.005763065 | 0.064846973 | 0.088871765 | 0.929183823 | 0.965521426 |
| Zfp809 | 377.5169647 | 0.008850475 | 0.100041454 | 0.08846808 | 0.929504653 | 0.965717501 |
| Henmt1 | 50.54880876 | 0.013676025 | 0.154734011 | 0.08838409 | 0.929571406 | 0.965722098 |
| 9330179D12Rik | 79.86706808 | -0.013741492 | 0.155928615 | -0.088126813 | 0.929775885 | 0.965740267 |
| Mypop | 1387.588612 | 0.005771083 | 0.065476182 | 0.088140183 | 0.929765259 | 0.965740267 |
| Ythdf2 | 1410.257702 | -0.006109565 | 0.069315558 | -0.088141326 | 0.929764351 | 0.965740267 |
| Gpr156 | 47.35591155 | -0.013033328 | 0.148409143 | -0.087820252 | 0.930019541 | 0.965928594 |
| Kcng3 | 65.71152108 | 0.013937824 | 0.159380392 | 0.087450055 | 0.930313784 | 0.965975189 |
| Mir132 | 12.86860148 | 0.012978124 | 0.148302017 | 0.087511447 | 0.930264987 | 0.965975189 |
| Onecut2 | 208.9002343 | 0.011188251 | 0.127801619 | 0.087543893 | 0.930239197 | 0.965975189 |
| Vgll4 | 419.589731 | -0.006364854 | 0.072625522 | -0.087639362 | 0.930163316 | 0.965975189 |
| Bcl6 | 1234.413767 | -0.011561643 | 0.132451699 | -0.0872895 | 0.9304414 | 0.965986374 |
| Spryd3 | 6716.414232 | 0.006870055 | 0.078713152 | 0.087279627 | 0.930449248 | 0.965986374 |
| Actr1a | 8736.460976 | 0.004481038 | 0.051428965 | 0.087130625 | 0.930567683 | 0.966044602 |
| Adnp | 5772.486053 | -0.004839853 | 0.055780969 | -0.086765304 | 0.930858067 | 0.966194882 |
| Lrp5 | 510.8987494 | -0.008287535 | 0.095574138 | -0.086713152 | 0.930899522 | 0.966194882 |
| Pik3cd | 1209.048638 | 0.006330809 | 0.072911159 | 0.086829084 | 0.930807369 | 0.966194882 |
| Lrriq3 | 16.26361541 | 0.01407704 | 0.163359225 | 0.086172298 | 0.931329452 | 0.966576363 |
| D130040H23Rik | 98.53539653 | -0.012356729 | 0.144169568 | -0.085709691 | 0.931697199 | 0.966893262 |
| Eri3 | 4486.789116 | 0.005750613 | 0.067439399 | 0.085270818 | 0.932046091 | 0.967125065 |
| Kcnu1 | 127.0979844 | -0.012084503 | 0.141793856 | -0.085225855 | 0.932081836 | 0.967125065 |
| Pafah2 | 310.1529266 | -0.008299969 | 0.09742529 | -0.085193168 | 0.932107822 | 0.967125065 |
| Vps26b | 5657.925355 | -0.004826864 | 0.056807239 | -0.084969177 | 0.932285896 | 0.967245056 |
| Prima1 | 173.4761502 | 0.009171923 | 0.108782303 | 0.084314478 | 0.932806401 | 0.96772028 |
| 1600012H06Rik | 527.922023 | -0.00771546 | 0.09180318 | -0.084043492 | 0.933021852 | 0.967755144 |
| Ctnna3 | 70.19221622 | -0.013040146 | 0.155172356 | -0.084036531 | 0.933027387 | 0.967755144 |
| Eif4g1 | 11861.20377 | 0.004938552 | 0.058677818 | 0.084163872 | 0.932926142 | 0.967755144 |
| 4932441J04Rik | 23.10635807 | 0.013840736 | 0.167442492 | 0.082659638 | 0.934122179 | 0.967854022 |
| BC004004 | 2056.246026 | -0.005113492 | 0.061846985 | -0.082679723 | 0.934106208 | 0.967854022 |
| Cyb5r1 | 738.7952058 | 0.008211656 | 0.098840666 | 0.083079727 | 0.933788145 | 0.967854022 |
| Glt28d2 | 13.25781499 | -0.013801469 | 0.164878393 | -0.083706962 | 0.933289422 | 0.967854022 |
| Mrgprf | 44.801966 | -0.013951182 | 0.167947905 | -0.083068507 | 0.933797067 | 0.967854022 |
| Pank2 | 2071.923451 | 0.006394251 | 0.077202377 | 0.08282453 | 0.933991064 | 0.967854022 |
| Papss1 | 2600.48444 | -0.004154017 | 0.049698383 | -0.083584556 | 0.933386747 | 0.967854022 |
| Phip | 1864.166766 | -0.007299971 | 0.08769089 | -0.083246627 | 0.933655438 | 0.967854022 |
| Sgol2 | 41.79432965 | -0.013154784 | 0.158906707 | -0.082783062 | 0.934024038 | 0.967854022 |
| Slc4a1ap | 1124.905011 | 0.00564399 | 0.067781506 | 0.083267411 | 0.933638912 | 0.967854022 |
| Smpd3 | 4604.49502 | 0.007164874 | 0.085597611 | 0.083704133 | 0.933291671 | 0.967854022 |
| Stk4 | 2041.696668 | 0.006361011 | 0.076279282 | 0.083391066 | 0.933540592 | 0.967854022 |
| Tinagl1 | 328.052975 | -0.01167691 | 0.140865486 | -0.082894045 | 0.93393579 | 0.967854022 |
| Ttc23 | 137.6945784 | -0.010284783 | 0.123023238 | -0.083600327 | 0.933374207 | 0.967854022 |
| Urgcp | 3087.921149 | -0.00587621 | 0.070483796 | -0.083369661 | 0.933557611 | 0.967854022 |
| Vmn2r-ps129 | 13.95260298 | -0.013731065 | 0.165510829 | -0.08296173 | 0.933881969 | 0.967854022 |
| Dcaf5 | 4175.404969 | -0.006379166 | 0.077260758 | -0.0825667 | 0.93419608 | 0.967865869 |
| Chp1 | 6534.915863 | -0.005715103 | 0.069487122 | -0.082246936 | 0.93445035 | 0.968064571 |
| 7530420F21Rik | 16.91309614 | -0.013321429 | 0.162129686 | -0.082165269 | 0.934515291 | 0.968067121 |
| 1700045I19Rik | 18.92302234 | -0.01375968 | 0.168048353 | -0.081879291 | 0.934742703 | 0.968108518 |
| Ilk | 2682.990721 | -0.005732029 | 0.069883527 | -0.082022611 | 0.934628733 | 0.968108518 |
| Wdtc1 | 3136.325127 | -0.007285369 | 0.088903965 | -0.081946498 | 0.934689259 | 0.968108518 |
| Sdsl | 71.29409587 | 0.0133257 | 0.163461476 | 0.081521962 | 0.93502686 | 0.96833809 |
| Nmrk1 | 179.478278 | 0.011208014 | 0.137754703 | 0.081362119 | 0.935153974 | 0.968405004 |
| Uspl1 | 1583.709729 | 0.004329826 | 0.053481699 | 0.080959023 | 0.93547454 | 0.968672227 |
| 1700030K09Rik | 326.6655629 | 0.00744176 | 0.092068422 | 0.080828587 | 0.935578273 | 0.9687149 |
| 8430427H17Rik | 2717.094938 | 0.004995738 | 0.062041233 | 0.080522874 | 0.935821404 | 0.968837153 |
| Clvs1 | 2400.068038 | -0.006471659 | 0.080292351 | -0.080601186 | 0.935759123 | 0.968837153 |
| 4930447C04Rik | 75.62695959 | -0.013416989 | 0.167294313 | -0.080199909 | 0.936078262 | 0.969038324 |
| 1110017D15Rik | 61.5582291 | -0.011622887 | 0.145329946 | -0.079975857 | 0.936256458 | 0.969093297 |
| Pi15 | 9.523208202 | -0.012417621 | 0.155265038 | -0.07997693 | 0.936255605 | 0.969093297 |
| Rhoh | 30.89465425 | 0.01319374 | 0.165262233 | 0.079835181 | 0.936368343 | 0.969144363 |
| Adcy10 | 8.387214761 | -0.011917851 | 0.149673539 | -0.07962564 | 0.936535002 | 0.969252109 |
| Vps52 | 3074.097243 | -0.005881734 | 0.074192647 | -0.079276512 | 0.936812688 | 0.969474739 |
| Scarf2 | 100.4106626 | 0.010525248 | 0.133099913 | 0.079077796 | 0.936970744 | 0.969573546 |
| Eif3e | 2656.774389 | -0.006027796 | 0.076577872 | -0.078714592 | 0.937259638 | 0.969742961 |
| H6pd | 416.8723981 | -0.008288047 | 0.10528719 | -0.078718473 | 0.937256552 | 0.969742961 |
| Zfp184 | 114.8890868 | 0.009987746 | 0.127216188 | 0.078510024 | 0.937422357 | 0.969846554 |
| Tmsb15b1 | 10.78356615 | -0.012612815 | 0.161053046 | -0.078314666 | 0.937577751 | 0.969942557 |
| Gtf3a | 661.6806021 | 0.005399073 | 0.069260842 | 0.077952742 | 0.937865645 | 0.970068747 |
| Oxsm | 419.4854133 | -0.007842087 | 0.100549571 | -0.077992247 | 0.937834221 | 0.970068747 |
| Tmem136 | 871.5745593 | -0.007008244 | 0.089935536 | -0.077925194 | 0.937887559 | 0.970068747 |
| 1700019G17Rik | 486.5187861 | -0.010241584 | 0.13207937 | -0.077541132 | 0.938193071 | 0.970115459 |
| Arl5c | 73.47917961 | -0.012190556 | 0.157335275 | -0.077481393 | 0.938240593 | 0.970115459 |
| Fmnl2 | 1955.363534 | 0.00469223 | 0.060745927 | 0.077243536 | 0.938429808 | 0.970115459 |
| Klhl24 | 2822.895878 | 0.006151223 | 0.079320336 | 0.077549135 | 0.938186705 | 0.970115459 |
| Prkab1 | 546.0617654 | -0.006280806 | 0.081316775 | -0.077238745 | 0.93843362 | 0.970115459 |
| Stil | 31.08527343 | -0.012647669 | 0.163289113 | -0.077455678 | 0.938261049 | 0.970115459 |
| Tnfrsf13c | 12.39775922 | 0.012678673 | 0.163185513 | 0.077694846 | 0.938070794 | 0.970115459 |
| Zfp579 | 404.6656401 | 0.010306945 | 0.133172238 | 0.0773956 | 0.938308841 | 0.970115459 |
| Olfml1 | 504.5213273 | 0.008217022 | 0.106817137 | 0.076926067 | 0.93868236 | 0.970307858 |
| Fam198a | 297.4867065 | 0.008182682 | 0.106835247 | 0.076591595 | 0.938948445 | 0.97051816 |
| Tprkb | 2298.747064 | 0.006542309 | 0.085789411 | 0.076260094 | 0.939212174 | 0.970725997 |
| Mtfp1 | 2056.671375 | -0.006346638 | 0.083352079 | -0.076142533 | 0.939305702 | 0.970757908 |
| Rap1gap | 10353.71211 | 0.004801876 | 0.063256824 | 0.075910792 | 0.939490071 | 0.970883689 |
| Med9 | 1749.001415 | 0.004111811 | 0.054289312 | 0.075738858 | 0.939626861 | 0.97096029 |
| Gtf2i | 9835.058737 | -0.003246493 | 0.042922409 | -0.075636316 | 0.939708443 | 0.970979835 |
| Hmgcl | 758.9093466 | -0.005756765 | 0.076278603 | -0.075470244 | 0.939840572 | 0.971043809 |
| Ppih | 342.7335673 | -0.00652199 | 0.086516524 | -0.075384329 | 0.939908928 | 0.971043809 |
| Tm2d3 | 959.3000823 | 0.004836901 | 0.064216151 | 0.075322181 | 0.939958374 | 0.971043809 |
| Gcc1 | 896.1615592 | -0.004837321 | 0.064612797 | -0.074866293 | 0.940321095 | 0.97135376 |
| Lrpprc | 2867.53332 | -0.005056311 | 0.067643248 | -0.074749675 | 0.940413883 | 0.971384847 |
| Fam154b | 56.91552099 | 0.011806756 | 0.158451531 | 0.074513357 | 0.940601913 | 0.971514302 |
| Necab2 | 3048.566826 | -0.008009581 | 0.107909799 | -0.074224784 | 0.940831526 | 0.971686686 |
| Gimap4 | 12.28994306 | -0.012333638 | 0.166836841 | -0.073926348 | 0.94106899 | 0.971737617 |
| Psmc3 | 4912.858166 | -0.005200973 | 0.070219034 | -0.074067852 | 0.940956395 | 0.971737617 |
| Tmem88 | 95.43750643 | -0.011243905 | 0.151957365 | -0.073993816 | 0.941015306 | 0.971737617 |
| Arhgap1 | 3366.549017 | -0.005084057 | 0.069304024 | -0.073358755 | 0.941520638 | 0.972074416 |
| Pde12 | 466.9658067 | 0.005070808 | 0.06908819 | 0.073396158 | 0.941490875 | 0.972074416 |
| Tnfaip2 | 43.05045635 | -0.011571267 | 0.157972542 | -0.073248594 | 0.941608297 | 0.972100144 |
| Mak16 | 1080.603763 | -0.005316367 | 0.07277219 | -0.073054929 | 0.941762408 | 0.972175724 |
| Naca | 5629.003993 | -0.004692384 | 0.064280201 | -0.072998894 | 0.941806997 | 0.972175724 |
| Nin | 2260.93216 | -0.005293064 | 0.07265865 | -0.072848372 | 0.941926778 | 0.972234595 |
| 4921524J17Rik | 260.9266079 | 0.010021695 | 0.13782981 | 0.072710651 | 0.942036373 | 0.972282944 |
| Nat2 | 25.73834198 | 0.012070128 | 0.166313633 | 0.072574498 | 0.942144721 | 0.972330001 |
| Ndufv1 | 5770.351763 | -0.005912573 | 0.081716599 | -0.072354614 | 0.942319703 | 0.972445816 |
| Cyp4f14 | 320.5134533 | 0.00866893 | 0.120258617 | 0.072085733 | 0.94253368 | 0.972601855 |
| Hagh | 2275.049144 | 0.006813964 | 0.094629649 | 0.072006648 | 0.942596617 | 0.972602024 |
| Gm10638 | 24.4786567 | 0.012088469 | 0.168104581 | 0.071910406 | 0.942673208 | 0.972616281 |
| Celf1 | 9904.330412 | 0.004293896 | 0.059792174 | 0.071813682 | 0.942750184 | 0.972629446 |
| Zbtb40 | 650.5554704 | -0.008061655 | 0.112378522 | -0.071736614 | 0.942811517 | 0.972629446 |
| 4932416H05Rik | 257.4662096 | -0.00660409 | 0.092408737 | -0.071466079 | 0.943026821 | 0.972722027 |
| Muc2 | 8.862724349 | -0.010592989 | 0.148168998 | -0.071492612 | 0.943005704 | 0.972722027 |
| 2810474O19Rik | 493.7560424 | 0.006040051 | 0.085117943 | 0.070960963 | 0.943428824 | 0.972853049 |
| Fam220a | 1357.815134 | -0.006842247 | 0.096272255 | -0.071071852 | 0.94334057 | 0.972853049 |
| Gm2721 | 10.79135489 | 0.011624016 | 0.163297111 | 0.071183229 | 0.943251929 | 0.972853049 |
| Rras2 | 402.9845218 | 0.008138882 | 0.11474421 | 0.070930652 | 0.943452948 | 0.972853049 |
| Sap30l | 739.2251498 | 0.007201341 | 0.101553197 | 0.070912006 | 0.943467788 | 0.972853049 |
| Ino80b | 314.043753 | 0.005961579 | 0.084357903 | 0.070670066 | 0.943660345 | 0.97298685 |
| Eif2b1 | 892.1375464 | 0.004909488 | 0.069784713 | 0.07035191 | 0.943913567 | 0.973183178 |
| Acot12 | 12.38480903 | -0.011311877 | 0.16104794 | -0.070239192 | 0.94400328 | 0.973210914 |
| Trim13 | 263.072493 | 0.007534811 | 0.107508434 | 0.070085768 | 0.944125394 | 0.973272046 |
| Sox7 | 86.25078052 | -0.009433982 | 0.135100449 | -0.069829391 | 0.944329453 | 0.97341764 |
| Gpr160 | 37.12839143 | -0.010870989 | 0.155902932 | -0.069729218 | 0.944409186 | 0.973435067 |
| Marveld1 | 434.4174166 | -0.006443494 | 0.092539691 | -0.069629519 | 0.944488541 | 0.973452102 |
| 2700038G22Rik | 30.97042964 | -0.01168773 | 0.168108256 | -0.069525021 | 0.944571717 | 0.973473073 |
| Hrc | 13.38579228 | -0.011367328 | 0.16463901 | -0.069043954 | 0.944954633 | 0.973738167 |
| Sik1 | 768.448703 | 0.011449437 | 0.165700345 | 0.069097243 | 0.944912215 | 0.973738167 |

| | | | | | | |
|---|---|---|---|---|---|---|
| Gabrr2 | 39.86393692 | 0.051780018 | 0.085919948 | 0.602654204 | 0.546738753 | 0.867128614 |
| Enox1 | 1157.572031 | 0.051799468 | 0.062676743 | 0.826454368 | 0.408546393 | 0.801055189 |
| Casc4 | 4107.250857 | 0.05183137 | 0.061874326 | 0.837687826 | 0.402206053 | 0.79955155 |
| 9430020K01Rik | 7191.252919 | 0.05186304 | 0.072448808 | 0.715857736 | 0.474079198 | 0.833994865 |
| Foxo1 | 1562.688142 | 0.051879721 | 0.06218715 | 0.834251452 | 0.404139303 | 0.800290023 |
| Ghdc | 309.0362297 | 0.05189567 | 0.081212416 | 0.639011523 | 0.522815436 | 0.854337225 |
| Mterfd2 | 490.7996349 | 0.051897905 | 0.069923849 | 0.742206074 | 0.457962489 | 0.827642075 |
| Cops6 | 3369.72653 | 0.051912023 | 0.047941795 | 1.082813503 | 0.278891211 | 0.71849589 |
| Psma8 | 12.68257091 | 0.051926432 | 0.069089088 | 0.751586592 | 0.452299709 | NA |
| Aff4 | 9748.687109 | 0.051963586 | 0.068535525 | 0.758199286 | 0.448331688 | 0.823216547 |
| Ccdc71 | 1207.031178 | 0.051993925 | 0.053399957 | 0.973669783 | 0.330220525 | 0.756141871 |
| Eva1c | 67.47001664 | 0.052030573 | 0.09505339 | 0.547382609 | 0.5841159 | 0.884313345 |
| Glg1 | 5700.177223 | 0.052042166 | 0.06381714 | 0.815488848 | 0.414792545 | 0.805087722 |
| Cnbp | 8508.111433 | 0.052047473 | 0.046170817 | 1.12728075 | 0.259623799 | 0.703893404 |
| Trpm3 | 1983.67992 | 0.052056581 | 0.094696873 | 0.549718058 | 0.582512769 | 0.88406516 |
| Rhot1 | 3456.474528 | 0.052058004 | 0.059353785 | 0.877079774 | 0.38044331 | 0.786275719 |
| Mylk3 | 190.0963092 | 0.052059504 | 0.09316151 | 0.558809143 | 0.576291982 | 0.880966011 |
| Max | 2073.222817 | 0.052061384 | 0.049857995 | 1.044193298 | 0.296395967 | 0.731596911 |
| Olig2 | 1007.657254 | 0.052082436 | 0.081991596 | 0.635216762 | 0.525287053 | 0.855800777 |
| Atp5g3 | 7057.107928 | 0.052087328 | 0.047392183 | 1.099070028 | 0.271737522 | 0.713790099 |
| Gm5 | 7.162955226 | 0.052152894 | 0.057249605 | 0.910973859 | 0.362309147 | NA |
| Capza1 | 2008.85684 | 0.052169985 | 0.065163462 | 0.800601797 | 0.42336221 | 0.808999423 |
| Lhfpl4 | 6876.209318 | 0.052171946 | 0.051437119 | 1.014285936 | 0.310446333 | 0.743290495 |
| Ube2w | 2093.497837 | 0.052185945 | 0.071697316 | 0.727864703 | 0.466696412 | 0.832095317 |
| D3Ertd254e | 1336.325613 | 0.052198631 | 0.074457289 | 0.701054688 | 0.483268886 | 0.838104566 |
| Prkce | 18609.37405 | 0.052207058 | 0.048636663 | 1.073409531 | 0.28308741 | 0.720920007 |
| Tom1 | 1860.478475 | 0.052218238 | 0.069911516 | 0.746918976 | 0.455112471 | 0.826267186 |
| Ttc7b | 8638.808872 | 0.052225472 | 0.048891861 | 1.068183354 | 0.285437812 | 0.722148665 |
| Ggps1 | 1710.688767 | 0.052237007 | 0.064758047 | 0.806648896 | 0.419868795 | 0.807657282 |
| Cops8 | 3852.302818 | 0.052241476 | 0.044057749 | 1.185749986 | 0.235721048 | 0.686877421 |
| Rnaseh2a | 707.274422 | 0.052262759 | 0.059439 | 0.87926712 | 0.379256459 | 0.785552449 |
| Eif4e2 | 2345.62571 | 0.052296275 | 0.046470337 | 1.125368966 | 0.260432716 | 0.704058139 |
| Osbpl1a | 14364.4813 | 0.052320912 | 0.06789016 | 0.770670031 | 0.44090254 | 0.818786793 |
| Atp5s | 511.292725 | 0.052334484 | 0.06888083 | 0.759783014 | 0.447384298 | 0.822458097 |
| Zik1 | 758.14953 | 0.052339743 | 0.078626326 | 0.665677081 | 0.505617526 | 0.84841357 |
| Toe1 | 484.9837271 | 0.052360089 | 0.065246661 | 0.80249453 | 0.422266951 | 0.808695226 |
| Ilf2 | 3425.057396 | 0.052362473 | 0.050777425 | 1.031215599 | 0.302439727 | 0.735974513 |
| Cfl1 | 27611.35003 | 0.052380182 | 0.036864677 | 1.420877278 | 0.15535244 | 0.604578396 |
| Gm5535 | 19.99808624 | 0.05239694 | 0.074522727 | 0.703100141 | 0.481993344 | NA |
| Zfp748 | 487.7329107 | 0.052401934 | 0.079280069 | 0.660972354 | 0.508630043 | 0.850050333 |
| Snupn | 488.7591427 | 0.052408468 | 0.076947894 | 0.681090349 | 0.495814323 | 0.844379526 |
| Cdk18 | 1753.158895 | 0.052412614 | 0.076958389 | 0.681051344 | 0.495839003 | 0.844379526 |
| 9030617O03Rik | 368.5386635 | 0.052423783 | 0.085405707 | 0.613820607 | 0.539333878 | 0.862815409 |
| Tbc1d4 | 700.836372 | 0.05245282 | 0.076250271 | 0.687903391 | 0.491513621 | 0.841892493 |
| Nsun4 | 540.4878928 | 0.052478836 | 0.070960529 | 0.739549677 | 0.459573286 | 0.828566657 |
| Ntm | 5651.835564 | 0.052483697 | 0.048770741 | 1.076130805 | 0.281868761 | 0.720215588 |
| 1200014J11Rik | 897.2719695 | 0.052493858 | 0.071187767 | 0.737399972 | 0.460879153 | 0.829479689 |
| Qdpr | 6348.612752 | 0.052498831 | 0.065916745 | 0.796441496 | 0.425775467 | 0.809519334 |
| Bdh2 | 57.46826962 | 0.052525261 | 0.092143928 | 0.570034973 | 0.568653978 | 0.877419107 |
| Jkamp | 1681.531817 | 0.052527873 | 0.05108902 | 1.028163631 | 0.303872863 | 0.737396794 |
| Pccb | 2104.179748 | 0.052528501 | 0.054745475 | 0.959503976 | 0.337304917 | 0.760113439 |
| Ddx25 | 1819.778613 | 0.052531321 | 0.05782872 | 0.908395017 | 0.363669558 | 0.776984985 |
| Vwa5b1 | 219.5862542 | 0.052531608 | 0.094459214 | 0.556130057 | 0.578121949 | 0.881843897 |
| Lysmd1 | 883.2985224 | 0.052547752 | 0.055644915 | 0.944340584 | 0.344995633 | 0.764633288 |
| Cspg4 | 857.3446284 | 0.052550285 | 0.081585858 | 0.644110225 | 0.519503961 | 0.853297835 |
| Ppp1r13b | 3925.401745 | 0.052551944 | 0.051233083 | 1.02574237 | 0.305013037 | 0.737453291 |
| Sall2 | 3175.644506 | 0.052571679 | 0.057294475 | 0.917569787 | 0.358844141 | 0.774757728 |
| Ncoa7 | 3389.027683 | 0.052574478 | 0.067345668 | 0.78066606 | 0.434998928 | 0.815326688 |
| Arl16 | 820.1881203 | 0.052588628 | 0.062298013 | 0.844146152 | 0.398587741 | 0.797789838 |
| Atl2 | 2476.163667 | 0.052590953 | 0.063661334 | 0.826105107 | 0.408744471 | 0.801055189 |
| Nrap | 59.02367751 | 0.052593774 | 0.093564335 | 0.562113485 | 0.574038701 | 0.879633264 |
| Fam49a | 10779.65902 | 0.05259517 | 0.076772129 | 0.685081557 | 0.493292457 | 0.843035523 |
| Gria1 | 17289.37909 | 0.05260368 | 0.052238768 | 1.006985475 | 0.313941754 | 0.745923449 |
| Golga7 | 3026.161313 | 0.052648657 | 0.055492249 | 0.948756964 | 0.342744235 | 0.763762832 |
| Atp13a3 | 2927.098222 | 0.052657175 | 0.082433082 | 0.638786928 | 0.522961553 | 0.854337225 |
| Tbc1d2b | 1090.465474 | 0.052662392 | 0.055087787 | 0.955972174 | 0.339086293 | 0.761175629 |
| Ube2q1 | 4232.671677 | 0.052664701 | 0.050908628 | 1.034494599 | 0.300905001 | 0.734983359 |
| Heatr3 | 996.0667476 | 0.052666445 | 0.072151271 | 0.729944791 | 0.465423932 | 0.831682622 |
| Zbtb5 | 508.0232383 | 0.052668732 | 0.067201864 | 0.783739152 | 0.433193184 | 0.813783127 |
| Sec23ip | 1910.857062 | 0.052676412 | 0.06123537 | 0.860228525 | 0.389663084 | 0.792715562 |
| Cdk7 | 2677.038501 | 0.052680691 | 0.065832956 | 0.800217619 | 0.423584723 | 0.808999423 |
| Usf1 | 1960.666402 | 0.052692145 | 0.059427444 | 0.886663493 | 0.375260099 | 0.783489748 |
| Dhx15 | 3629.886969 | 0.052705562 | 0.068152102 | 0.773351972 | 0.439314106 | 0.817545617 |
| Smc1a | 4187.087663 | 0.052717881 | 0.056680304 | 0.930091725 | 0.352323595 | 0.770257962 |
| Sil1 | 782.3109718 | 0.052763129 | 0.073548903 | 0.717388383 | 0.473134486 | 0.833983942 |
| Arl8a | 8534.214947 | 0.052771965 | 0.042029946 | 1.255580137 | 0.209268237 | 0.664196937 |
| Csgalnact2 | 496.0452852 | 0.052821 | 0.071747428 | 0.736207584 | 0.461604378 | 0.830270578 |
| Itpa | 1876.580369 | 0.0528637 | 0.056220514 | 0.940291995 | 0.347067805 | 0.765953651 |
| Ppat | 715.0373372 | 0.052906829 | 0.06926094 | 0.763876855 | 0.444940631 | 0.821493054 |
| Cyp19a1 | 13.20762524 | 0.052917346 | 0.064472186 | 0.820777907 | 0.411772785 | NA |
| Bmpr1a | 2471.165121 | 0.052917569 | 0.064054154 | 0.826137978 | 0.408725826 | 0.801055189 |
| AI115009 | 102.1412007 | 0.052923993 | 0.093719326 | 0.564707364 | 0.572272825 | 0.879277718 |
| Capn10 | 956.1157049 | 0.052943919 | 0.077645147 | 0.681870296 | 0.49532097 | 0.844369474 |
| Vps13a | 2475.684744 | 0.052946622 | 0.075288012 | 0.703254349 | 0.481897255 | 0.837383765 |
| Rps9 | 5663.688088 | 0.052982256 | 0.052663094 | 1.006060465 | 0.314386485 | 0.745993052 |
| Afg3l2 | 3627.167781 | 0.053021028 | 0.03908894 | 1.356420219 | 0.174965504 | 0.628026741 |
| Sdf2 | 1634.040642 | 0.053033532 | 0.05813755 | 0.912207892 | 0.36165929 | 0.776449324 |
| Hhat | 30.44325075 | 0.053036393 | 0.089750241 | 0.590933152 | 0.554565212 | 0.871360989 |
| Cntnap2 | 4295.383354 | 0.053042983 | 0.062789007 | 0.844781359 | 0.398232924 | 0.797789838 |
| Ccdc132 | 3086.77643 | 0.053066333 | 0.072617922 | 0.730760837 | 0.464925248 | 0.831682622 |
| Lyrm4 | 738.8070238 | 0.053074355 | 0.07515113 | 0.706234951 | 0.480042038 | 0.836914993 |
| Tfam | 1321.716206 | 0.053089698 | 0.06050441 | 0.877451699 | 0.380241343 | 0.786275719 |
| Dgat2 | 2718.270851 | 0.053121209 | 0.064246256 | 0.826837425 | 0.408329215 | 0.801055189 |
| Desi1 | 5944.351073 | 0.053136729 | 0.055630847 | 0.955166639 | 0.339493434 | 0.761618782 |
| Asic1 | 3833.652809 | 0.053148931 | 0.062994392 | 0.84370892 | 0.398832082 | 0.797789838 |
| Aldh3a2 | 2429.358575 | 0.053149482 | 0.047212799 | 1.125743084 | 0.260274282 | 0.704058139 |
| A330050F15Rik | 292.3073731 | 0.053160974 | 0.08496333 | 0.625693153 | 0.531516225 | 0.858328942 |
| Nmt1 | 4675.311893 | 0.053164174 | 0.053846969 | 0.987319705 | 0.323485934 | 0.752925264 |
| Dennd5b | 3603.414314 | 0.053167429 | 0.077531623 | 0.685751531 | 0.492869805 | 0.842781259 |
| Rnf24 | 2614.386453 | 0.053169267 | 0.060725219 | 0.875571443 | 0.381263056 | 0.786663048 |
| Brs3 | 13.73253843 | 0.053172463 | 0.051206817 | 1.038386425 | 0.299090187 | NA |
| Cggbp1 | 2268.672101 | 0.053184917 | 0.062101901 | 0.856413674 | 0.391769004 | 0.794184423 |
| Abce1 | 3781.612312 | 0.053201861 | 0.054589947 | 0.974572494 | 0.329772363 | 0.756141871 |
| Nadkd1 | 1394.695373 | 0.053211899 | 0.052970863 | 1.004550348 | 0.315113416 | 0.745993052 |
| Ppp2r3c | 1141.669163 | 0.053221722 | 0.070046143 | 0.759809467 | 0.447368483 | 0.822458097 |
| Rgmb | 1890.494421 | 0.053231087 | 0.054253073 | 0.981162604 | 0.326512562 | 0.753373722 |
| Rnmt | 2775.54388 | 0.053247369 | 0.056901886 | 0.935775128 | 0.34938898 | 0.768268271 |
| Wif1 | 29.11675392 | 0.053255099 | 0.082127488 | 0.648444273 | 0.516697643 | 0.853297835 |
| Rab5c | 5730.428263 | 0.053260095 | 0.038176446 | 1.39510354 | 0.162984619 | 0.614351408 |
| Kdm5c | 3416.711453 | 0.053263366 | 0.056319639 | 0.945733446 | 0.344284559 | 0.76410657 |
| Gpr20 | 7.753135342 | 0.053275714 | 0.057046404 | 0.933901352 | 0.350354781 | NA |
| Nacc1 | 6470.869709 | 0.053281436 | 0.048274201 | 1.103724865 | 0.269712506 | 0.71207631 |
| Cnpy1 | 3.608850288 | 0.053283237 | 0.038891373 | 1.370052879 | 0.170670395 | NA |
| Stk32c | 7229.015864 | 0.053289616 | 0.068438414 | 0.778650654 | 0.436185532 | 0.816078186 |
| Srpk2 | 10303.57613 | 0.053296462 | 0.056270153 | 0.947153313 | 0.343560661 | 0.763762832 |
| Pde4b | 5750.642139 | 0.053316271 | 0.063816835 | 0.835457769 | 0.403460015 | 0.800224169 |
| Tbc1d25 | 606.2077326 | 0.05331918 | 0.063853426 | 0.835024576 | 0.403703871 | 0.800224169 |
| Btrc | 5824.703144 | 0.053330774 | 0.049566794 | 1.075937522 | 0.2819552 | 0.720294902 |
| Vipas39 | 2416.619174 | 0.053336363 | 0.053809487 | 0.991207425 | 0.321584306 | 0.751018267 |
| Edc3 | 1305.808459 | 0.053337154 | 0.058043059 | 0.918923896 | 0.35813538 | 0.774147996 |
| Cd3eap | 901.7751579 | 0.053339226 | 0.063595273 | 0.838729419 | 0.401621166 | 0.799277436 |
| Sfxn1 | 2422.935702 | 0.053342661 | 0.054174602 | 0.98464335 | 0.324799288 | 0.75293408 |
| Lamtor3 | 1485.262467 | 0.053348803 | 0.059873391 | 0.891026915 | 0.372914731 | 0.782226671 |
| Mkrn3 | 22.78616889 | 0.053353913 | 0.079922287 | 0.667572405 | 0.504406577 | 0.847807553 |
| Leprel4 | 386.6313519 | 0.053357283 | 0.0890427 | 0.599232534 | 0.549017831 | 0.868267067 |
| Angptl2 | 241.7269622 | 0.053401192 | 0.077784385 | 0.686528429 | 0.492379943 | 0.842340249 |
| Polr2b | 3279.652756 | 0.053415582 | 0.056085063 | 0.952402991 | 0.340892643 | 0.76263292 |
| Gm6277 | 165.1035631 | 0.053427202 | 0.091930877 | 0.581167109 | 0.561127833 | 0.87387211 |
| Trak2 | 9143.460176 | 0.053428263 | 0.053260481 | 1.003150218 | 0.315788388 | 0.745993052 |
| Adnp | 6370.437369 | 0.053436574 | 0.052002491 | 1.027577205 | 0.304148751 | 0.737453291 |
| Cap2 | 10577.22333 | 0.053439928 | 0.057076726 | 0.9362823 | 0.349127859 | 0.768165901 |
| Grik1 | 616.9820497 | 0.053441965 | 0.084730028 | 0.630732296 | 0.528215578 | 0.856965327 |
| Dhfr | 242.5307081 | 0.053443609 | 0.084316737 | 0.63384342 | 0.526183013 | 0.855916512 |
| Map3k10 | 6897.374397 | 0.053450101 | 0.066011891 | 0.809704132 | 0.418110243 | 0.80646254 |
| Mrpl30 | 1430.491034 | 0.053455515 | 0.060094422 | 0.889525406 | 0.373720775 | 0.782348609 |
| Itga3 | 980.4745997 | 0.053462453 | 0.068255979 | 0.783264033 | 0.43347208 | 0.813936845 |
| Pcdhb13 | 301.4471925 | 0.053464109 | 0.083113696 | 0.643264724 | 0.520052342 | 0.853297835 |
| Klhdc1 | 528.0980603 | 0.05347114 | 0.07460007 | 0.716770637 | 0.473515632 | 0.833983942 |
| Xpo7 | 5228.282512 | 0.053475459 | 0.058824234 | 0.909071919 | 0.363312164 | 0.776821522 |
| Dnajc25 | 104.9663613 | 0.053482033 | 0.094880971 | 0.56367502 | 0.572975321 | 0.879310647 |
| Msl2 | 2151.242394 | 0.053492168 | 0.057781804 | 0.925761466 | 0.354569957 | 0.77200441 |
| Ube2l3 | 7198.042473 | 0.053504366 | 0.045408813 | 1.178281543 | 0.238684383 | 0.689537017 |
| Fam183b | 77.89235173 | 0.053507509 | 0.090879424 | 0.588774738 | 0.556012394 | 0.871546692 |
| Nkx2-1 | 63.39790507 | 0.053510305 | 0.088792643 | 0.60264346 | 0.546745902 | 0.867128614 |
| Sox15 | 2.792802749 | 0.053512231 | 0.038829415 | 1.378136405 | 0.168161178 | NA |
| Cntn6 | 889.7674486 | 0.053515085 | 0.070498379 | 0.759096669 | 0.447794732 | 0.822750604 |
| Ccs | 563.7553511 | 0.053521632 | 0.073744624 | 0.725769949 | 0.467979812 | 0.832353544 |
| Mrpl27 | 1210.001676 | 0.053531315 | 0.064061362 | 0.835625624 | 0.403365549 | 0.800224169 |
| Atp6v1f | 3287.146837 | 0.053532671 | 0.049935267 | 1.072041359 | 0.283701456 | 0.721155194 |
| Pcdhga9 | 500.020102 | 0.053536898 | 0.082980127 | 0.645177347 | 0.518812264 | 0.853297835 |
| BC017158 | 998.2834206 | 0.053601427 | 0.073771871 | 0.726583545 | 0.467481112 | 0.832353544 |
| Ttc39b | 3331.717801 | 0.05361172 | 0.068787755 | 0.779378835 | 0.435756588 | 0.815960393 |
| Tomm20 | 16551.82095 | 0.053611774 | 0.063864287 | 0.839464065 | 0.401208946 | 0.799277436 |
| Cplx4 | 2.211179868 | 0.053626705 | 0.036165164 | 1.482827625 | 0.138120214 | NA |
| Sh3gl3 | 1189.490956 | 0.053637739 | 0.06724931 | 0.797595371 | 0.425105338 | 0.809421961 |
| Ube2s | 2058.72085 | 0.053644028 | 0.052396289 | 1.023813495 | 0.305923375 | 0.738179891 |
| Clint1 | 2227.856707 | 0.053646391 | 0.049035386 | 1.094034244 | 0.27393996 | 0.716079153 |
| Sumo2 | 4830.276763 | 0.05365731 | 0.060328505 | 0.889418869 | 0.373778008 | 0.782348609 |
| Scamp5 | 21340.20422 | 0.053665051 | 0.032336085 | 1.659602593 | 0.096994426 | 0.517612431 |
| Picalm | 6284.066073 | 0.053670773 | 0.059414989 | 0.90332042 | 0.366355862 | 0.778210102 |
| Cox7a2l | 5273.18288 | 0.053716005 | 0.047196907 | 1.138125532 | 0.255068068 | 0.701727038 |
| Ncoa4 | 4332.564655 | 0.053741804 | 0.051281006 | 1.047986547 | 0.294644805 | 0.729630146 |
| Selrc1 | 509.8833535 | 0.053775889 | 0.07536039 | 0.713582943 | 0.475485108 | 0.834973002 |
| Hs2st1 | 2451.787046 | 0.053784131 | 0.06138611 | 0.876161244 | 0.380942382 | 0.786576978 |
| Acsl6 | 7157.849071 | 0.053793477 | 0.047501012 | 1.132470141 | 0.257436831 | 0.703271881 |
| Wdfy3 | 8573.562693 | 0.053805755 | 0.073433988 | 0.732709151 | 0.463735841 | 0.83159896 |
| Pard3b | 178.5138623 | 0.053811071 | 0.089714403 | 0.599804151 | 0.548636766 | 0.868042443 |
| 1110008P14Rik | 3264.994854 | 0.053812449 | 0.066410441 | 0.810301024 | 0.417767187 | 0.80621897 |
| 4931440J10Rik | 4.31903238 | 0.053814228 | 0.046641175 | 1.153792271 | 0.248585344 | NA |
| Ogg1 | 226.6285956 | 0.053820896 | 0.087436583 | 0.615542072 | 0.538196794 | 0.862082844 |
| 4732415M23Rik | 26.35568566 | 0.053827747 | 0.086123696 | 0.625005074 | 0.531967728 | 0.858328942 |
| Stox2 | 5930.490023 | 0.053842686 | 0.067344101 | 0.799515994 | 0.423991276 | 0.808999423 |
| Nkain1 | 2879.051524 | 0.053843342 | 0.059604296 | 0.903346655 | 0.366341942 | 0.778210102 |
| Grin2a | 1618.02739 | 0.053845416 | 0.08763419 | 0.614433893 | 0.538928645 | 0.862452076 |
| Sspn | 578.0131677 | 0.053850235 | 0.079203324 | 0.679898678 | 0.496568619 | 0.844829674 |
| Serinc3 | 13280.33102 | 0.053854422 | 0.057791233 | 0.931878748 | 0.351399192 | 0.769244462 |
| Tmem184c | 2909.542566 | 0.053860691 | 0.065732433 | 0.819392929 | 0.412562268 | 0.803544722 |
| Stmn2 | 14804.05942 | 0.053863867 | 0.048474371 | 1.111182397 | 0.266489847 | 0.709239926 |
| 1500016L03Rik | 3.555802094 | 0.053873473 | 0.037228762 | 1.447092768 | 0.147870943 | NA |
| Folh1 | 314.6547868 | 0.053874864 | 0.087052859 | 0.618875289 | 0.535998519 | 0.860988 |
| Adam5 | 10.4070559 | 0.053888306 | 0.069458917 | 0.775829926 | 0.437849412 | NA |
| Surf6 | 741.6188692 | 0.053893318 | 0.063484878 | 0.848915836 | 0.395928126 | 0.796351698 |
| Ppp2r5e | 3900.520329 | 0.053896455 | 0.063300833 | 0.851433578 | 0.394528546 | 0.795437883 |
| 2700081L22Rik | 7.690517445 | 0.053905729 | 0.058861962 | 0.915799055 | 0.359772299 | NA |
| Ppp3cb | 11080.48906 | 0.053925021 | 0.074904445 | 0.719917499 | 0.471575793 | 0.833245343 |
| Rbm18 | 1782.862473 | 0.053925261 | 0.070652716 | 0.763243987 | 0.4453179 | 0.821629343 |
| Pcdhga3 | 635.4462423 | 0.05393449 | 0.078520403 | 0.686885045 | 0.492155172 | 0.842340249 |
| Tsen34 | 1780.339706 | 0.053938242 | 0.047609805 | 1.132922978 | 0.2572466 | 0.703199912 |
| Gabbr1 | 33901.78661 | 0.05397438 | 0.043297902 | 1.24658188 | 0.212550849 | 0.666270994 |
| Snrpa | 1736.399821 | 0.053976629 | 0.060857112 | 0.886940354 | 0.375111014 | 0.783489748 |
| Dynlrb2 | 48.78579972 | 0.05399179 | 0.085132633 | 0.634207919 | 0.52594514 | 0.855892499 |
| Zfp563 | 595.3001898 | 0.053992926 | 0.077564819 | 0.696100712 | 0.486365761 | 0.839801621 |
| Dnajc24 | 180.7641425 | 0.053999043 | 0.093416065 | 0.578048786 | 0.563231182 | 0.875108781 |
| Lysmd3 | 275.1516301 | 0.054009103 | 0.086712026 | 0.62285597 | 0.533379174 | 0.859128786 |
| Scn2a1 | 13451.62779 | 0.054022946 | 0.071006667 | 0.760815128 | 0.446767498 | 0.822210607 |
| Nmur2 | 3.957705342 | 0.054064988 | 0.031112128 | 1.737746379 | 0.082255513 | NA |
| Homer2 | 2656.123097 | 0.05410962 | 0.075017114 | 0.72129701 | 0.470726796 | 0.833245343 |
| Smad3 | 4334.874365 | 0.054115981 | 0.084579423 | 0.639824423 | 0.522286753 | 0.854113377 |
| Lamtor2 | 1010.401644 | 0.054122296 | 0.0662892 | 0.816457214 | 0.414238685 | 0.804541293 |
| Drd1a | 2137.950667 | 0.054123301 | 0.094497366 | 0.572749307 | 0.566814447 | 0.875959314 |
| Scara5 | 28.79602406 | 0.05415076 | 0.06779818 | 0.798705209 | 0.424461366 | 0.809162684 |
| Hs3st3a1 | 82.98658041 | 0.054153224 | 0.094211383 | 0.574805533 | 0.565422824 | 0.875364912 |
| Fam222b | 1724.690129 | 0.054158113 | 0.049054979 | 1.104028868 | 0.269580615 | 0.711972822 |
| Clstn1 | 53421.70596 | 0.054164424 | 0.050772865 | 1.066798643 | 0.286062772 | 0.722148665 |
| Ncoa2 | 4926.514833 | 0.054165053 | 0.05971327 | 0.907085687 | 0.364361486 | 0.777237064 |
| Vta1 | 1445.067913 | 0.054189353 | 0.050209249 | 1.079270335 | 0.280467233 | 0.720080692 |
| Lcmt2 | 280.5618335 | 0.054226037 | 0.092016093 | 0.589310364 | 0.555653094 | 0.871360989 |
| Grina | 16946.74044 | 0.054228243 | 0.050030389 | 1.083906098 | 0.278406436 | 0.71849589 |
| Ush1g | 33.00694938 | 0.054233289 | 0.082905899 | 0.654154761 | 0.513012104 | 0.85302356 |
| Rnd3 | 598.3951675 | 0.054237302 | 0.090379355 | 0.600107205 | 0.548434791 | 0.868006594 |
| Gyg | 1172.456874 | 0.054237348 | 0.064707126 | 0.838197446 | 0.401919821 | 0.799277436 |
| Exosc2 | 727.9633689 | 0.054247884 | 0.065186584 | 0.832194008 | 0.405299445 | 0.8005027 |
| BC003965 | 1048.920872 | 0.054253721 | 0.059473569 | 0.912232486 | 0.361646346 | 0.776449324 |
| Cnst | 2455.102067 | 0.054266113 | 0.062418761 | 0.869387852 | 0.384635025 | 0.789016184 |
| Mpp1 | 1713.405663 | 0.054299006 | 0.05868972 | 0.925187682 | 0.35486829 | 0.772141519 |
| Crtac1 | 2643.917884 | 0.054304528 | 0.070070533 | 0.774998068 | 0.438340801 | 0.817094139 |
| Pigz | 473.9264018 | 0.054313718 | 0.091170848 | 0.595735581 | 0.551351885 | 0.869755927 |
| Plxna2 | 6935.368802 | 0.054314731 | 0.061496549 | 0.883215908 | 0.377119628 | 0.783997209 |
| Ptger2 | 21.50792901 | 0.054315399 | 0.080208402 | 0.677178428 | 0.498292753 | NA |
| Zfyve1 | 2203.541697 | 0.054320143 | 0.052916709 | 1.026521566 | 0.304645803 | 0.737453291 |
| Ndufb4 | 1792.126137 | 0.054331329 | 0.066451507 | 0.817608694 | 0.413580662 | 0.804132338 |
| Cyp20a1 | 377.3691754 | 0.054341422 | 0.07139565 | 0.761130711 | 0.446578999 | 0.822210607 |
| Pphln1 | 1632.405479 | 0.05434829 | 0.066396464 | 0.818541932 | 0.413047809 | 0.803849473 |
| Glt8d2 | 658.066425 | 0.05435973 | 0.072486973 | 0.749924119 | 0.453300407 | 0.82562629 |
| Efnb2 | 2446.109008 | 0.05438831 | 0.072961421 | 0.745439285 | 0.456006202 | 0.826649955 |
| Dnajc14 | 2753.266975 | 0.054412365 | 0.051842413 | 1.049572372 | 0.293914765 | 0.729288721 |
| Pcdha6 | 123.6958951 | 0.054414144 | 0.094933924 | 0.573179129 | 0.566523414 | 0.875959314 |
| Creld1 | 3008.440181 | 0.054418312 | 0.060221428 | 0.903637036 | 0.366187896 | 0.778210102 |
| Shisa3 | 37.41123878 | 0.054419666 | 0.090713201 | 0.599909007 | 0.548566879 | 0.868028404 |
| Uqcrh | 4477.943128 | 0.054432974 | 0.062390087 | 0.872461894 | 0.382956447 | 0.787600928 |
| Vps26a | 2209.289948 | 0.05445087 | 0.059578384 | 0.913936668 | 0.360750125 | 0.776337125 |
| Adra2b | 47.8449036 | 0.054464093 | 0.091648228 | 0.594273286 | 0.552329343 | 0.870265825 |
| Lrsam1 | 1770.587972 | 0.054474784 | 0.06291255 | 0.865881037 | 0.386555403 | 0.790748315 |
| Rpsa | 13392.24864 | 0.054487733 | 0.052452026 | 1.038810835 | 0.298892721 | 0.733276458 |
| Lemd3 | 1495.260578 | 0.054490053 | 0.07035213 | 0.774533101 | 0.438615602 | 0.817139233 |

| | | | | | | | | | | | | | |
|---|---|---|---|---|---|---|---|---|---|---|---|---|---|
| Fam122b | 212.0208943 | -0.007688263 | 0.112300965 | -0.068461238 | 0.945418475 | 0.97415135 | Tln2 | 5780.046072 | 0.05449145 | 0.061994614 | 0.878970715 | NA | NA |
| 9430083A17Rik | 102.4786737 | -0.009589845 | 0.141621933 | -0.067714404 | 0.946012983 | 0.974313896 | Gm4944 | 364.9740824 | 0.054498101 | 0.091789539 | 0.593728885 | 0.55269346 | 0.870289995 |
| Abhd14a | 847.5860223 | -0.006352051 | 0.093736824 | -0.067764734 | 0.945972917 | 0.974313896 | Dtx1 | 3033.08434 | 0.054500758 | 0.05309902 | 1.026398561 | 0.304703756 | 0.737453291 |
| BC055402 | 10.74711078 | -0.011053039 | 0.162740628 | -0.06791813 | 0.945850806 | 0.974313896 | Scn2b | 15756.20764 | 0.054518269 | 0.055465272 | 0.982926187 | 0.325643768 | 0.75293408 |
| Harbi1 | 213.1063559 | 0.006355709 | 0.09386649 | 0.0677101 | 0.946016409 | 0.974313896 | Ppp1r12b | 2043.516944 | 0.054531551 | 0.067242821 | 0.810964657 | 0.417385967 | 0.806153084 |
| Sec22b | 2119.657405 | -0.004162503 | 0.06115745 | -0.06806207 | 0.945736224 | 0.974313896 | Tctn2 | 965.4424457 | 0.054568758 | 0.060085759 | 0.908181217 | 0.363782486 | 0.777103587 |
| Sema3g | 301.0340115 | -0.008717249 | 0.128605984 | -0.067782607 | 0.94595869 | 0.974313896 | Nol12 | 660.6952756 | 0.05459495 | 0.066018366 | 0.82696609 | 0.408256282 | 0.801055189 |
| Zfp954 | 430.6654951 | -0.005826521 | 0.085911312 | -0.067820189 | 0.945928773 | 0.974313896 | Eif3l | 2655.700377 | 0.054643283 | 0.043653868 | 1.251739778 | 0.2106647 | 0.6650979 |
| Lin52 | 1022.319749 | -0.004289962 | 0.063595218 | -0.067457301 | 0.946217654 | 0.974411039 | Mrpl35 | 1132.95293 | 0.054649455 | 0.058592338 | 0.932706498 | 0.350971529 | 0.76907715 |
| Tekt2 | 34.85591462 | 0.011294409 | 0.16768575 | 0.067354614 | 0.9462994 | 0.974411039 | Cabp5 | 8.288640541 | 0.054672701 | 0.057294347 | 0.95424249 | 0.339960912 | NA |
| Zfp654 | 567.9641119 | -0.006802611 | 0.100914843 | -0.067409417 | 0.946255772 | 0.974411039 | Gm5643 | 6.414334197 | 0.054684263 | 0.050195531 | 1.089424927 | 0.275966544 | NA |
| Cactin | 989.4322206 | 0.005168289 | 0.07716528 | 0.066976871 | 0.946600116 | 0.974464907 | Dgki | 1512.431845 | 0.054699602 | 0.079740594 | 0.685969332 | 0.492732448 | 0.842744794 |
| Cd82 | 680.0478734 | -0.008735052 | 0.130093636 | -0.067144343 | 0.946466793 | 0.974464907 | Cln6 | 563.1927362 | 0.054723666 | 0.074817066 | 0.731432938 | 0.464514753 | 0.83159896 |
| Cul4b | 1963.434461 | -0.005430359 | 0.081082938 | -0.06697289 | 0.946603286 | 0.974464907 | Ccl24 | 4.457229419 | 0.054746025 | 0.046837186 | 1.168858109 | 0.242460804 | NA |
| S1pr2 | 112.6307831 | -0.007923385 | 0.118097703 | -0.067091783 | 0.946508636 | 0.974464907 | Cdc14b | 497.8220295 | 0.054752078 | 0.078777798 | 0.695019146 | 0.487043303 | 0.839801621 |
| Bloc1s2a | 844.5482714 | -0.004421678 | 0.06620942 | -0.066783223 | 0.94675428 | 0.974490853 | Fbxo31 | 6093.084723 | 0.054790227 | 0.057195347 | 0.957949038 | 0.338088457 | 0.760581219 |
| Epb4.1l4a | 130.8769088 | -0.010586295 | 0.158419553 | -0.066824422 | 0.946721481 | 0.974490853 | Paqr4 | 2819.103881 | 0.05479151 | 0.05800008 | 0.9446799 | 0.344822322 | 0.764561711 |
| Cacnb3 | 11473.69893 | -0.004923024 | 0.074158438 | -0.066385213 | 0.947071143 | 0.974575798 | Nipsnap1 | 2808.491747 | 0.054799945 | 0.048587514 | 1.127860628 | 0.259378784 | 0.703893404 |
| Fzr1 | 2485.594993 | -0.004620119 | 0.069618341 | -0.066363527 | 0.947088407 | 0.974575798 | Ephb3 | 1082.27588 | 0.054823757 | 0.081798849 | 0.670226501 | 0.502713412 | 0.84739426 |
| Mtrf1 | 265.3600913 | 0.005414439 | 0.081328258 | 0.066575121 | 0.946919952 | 0.974575798 | Gm14492 | 3.577989769 | 0.054836955 | 0.040225178 | 1.363249533 | 0.172803881 | NA |
| Trappc6a | 186.033769 | 0.007531103 | 0.113317436 | 0.066460231 | 0.947011419 | 0.974575798 | Krt1 | 111.3443878 | 0.054882212 | 0.095079144 | 0.577226617 | 0.563786379 | 0.875120313 |
| 1500015L24Rik | 19.24326038 | 0.010994645 | 0.166729753 | 0.065942911 | 0.947423276 | 0.974855641 | Plaa | 1796.127145 | 0.054891234 | 0.070582213 | 0.777692154 | 0.436750519 | 0.816490111 |
| Rbfox2 | 11969.74879 | 0.005512853 | 0.0837307 | 0.065840287 | 0.947504981 | 0.97487497 | Eif4g1 | 12867.15742 | 0.054893764 | 0.038059578 | 1.442311446 | 0.149214533 | 0.597429754 |
| Anapc1 | 3795.769805 | -0.003851175 | 0.058664727 | -0.065647204 | 0.947658707 | 0.974968394 | Fip1l1 | 2293.245223 | 0.054897769 | 0.060471413 | 0.907830091 | 0.363967997 | 0.777191869 |
| Bfar | 1396.121496 | 0.003918516 | 0.059796093 | 0.065531312 | 0.947750976 | 0.974998581 | Fam120b | 3615.372369 | 0.054898503 | 0.055492386 | 0.989297933 | 0.322517395 | 0.752229878 |
| Zfp148 | 4125.639607 | 0.004838061 | 0.073999896 | 0.065379299 | 0.947872006 | 0.975058349 | 2900056M20Rik | 885.1726572 | 0.054930133 | 0.076800784 | 0.7152288 | 0.474467676 | 0.834396787 |
| A130010J15Rik | 218.2191523 | 0.006219038 | 0.09538217 | 0.065201265 | 0.948013754 | 0.975127361 | Mmadhc | 1460.132399 | 0.054954432 | 0.058303892 | 0.942551692 | 0.345910259 | 0.765097832 |
| Ccdc58 | 190.2807117 | 0.007841222 | 0.120447414 | 0.065100788 | 0.948093753 | 0.975127361 | Ptp4a2 | 4967.961582 | 0.054956778 | 0.061785287 | 0.889480013 | 0.37374516 | 0.782348609 |
| Ppp2r5d | 4658.48742 | -0.005506074 | 0.084633436 | -0.065057899 | 0.948127901 | 0.975127361 | Mtss1 | 1398.216377 | 0.054958875 | 0.076252142 | 0.720751879 | 0.471062187 | 0.833245343 |
| Fam98b | 1571.510892 | 0.003411073 | 0.052649147 | 0.064788757 | 0.948342193 | 0.975218288 | 9130401M01Rik | 909.9953759 | 0.054973953 | 0.058206733 | 0.944460389 | 0.344934435 | 0.764614314 |
| Plp1 | 49858.16529 | 0.010048777 | 0.155038674 | 0.06481465 | 0.948321577 | 0.975218288 | Taf7 | 293.2422514 | 0.054974512 | 0.086926587 | 0.632424596 | 0.527109466 | 0.856196078 |
| Acad9 | 1584.367166 | 0.004156651 | 0.064318371 | 0.064626193 | 0.94847163 | 0.975243916 | Nabp2 | 1730.64675 | 0.054984995 | 0.058420725 | 0.941189871 | 0.346607566 | 0.765286747 |
| Snord99 | 14.02318418 | 0.010449128 | 0.16175283 | 0.06459935 | 0.948493002 | 0.975243916 | Fbxl18 | 662.5350711 | 0.05500481 | 0.074063121 | 0.742674754 | 0.457678618 | 0.827642075 |
| Xrcc3 | 296.7279934 | -0.008774742 | 0.136019014 | -0.064511141 | 0.948563236 | 0.975251412 | Cmip | 15078.90582 | 0.055029525 | 0.049685144 | 1.107564968 | 0.26804974 | 0.71037507 |
| Angptl2 | 173.1158367 | 0.010616632 | 0.165221774 | 0.064256859 | 0.948765704 | 0.97538782 | Glrx3 | 329.5228097 | 0.055041902 | 0.085704652 | 0.642227712 | 0.520725342 | 0.853634615 |
| Ier5l | 317.6297975 | -0.009154008 | 0.14261604 | -0.064186387 | 0.948821817 | 0.97538782 | Atp5d | 5763.485063 | 0.05504871 | 0.057570377 | 0.956198524 | 0.338971945 | 0.761116259 |
| Hiatl1 | 1436.60328 | -0.003559058 | 0.055823757 | -0.063755258 | 0.949165105 | 0.975508052 | Ppm1h | 4513.184505 | 0.055067143 | 0.066267489 | 0.830982793 | 0.40598335 | 0.800733644 |
| Lsm2 | 196.8088213 | 0.007301621 | 0.114583372 | 0.06372322 | 0.949190616 | 0.975508052 | Usp54 | 4136.89888 | 0.055095439 | 0.079244367 | 0.695260002 | 0.486892376 | 0.839801621 |
| Nrd1 | 6064.886265 | -0.003712292 | 0.058182628 | -0.063804126 | 0.949126193 | 0.975508052 | Psmd8 | 3455.655223 | 0.055112414 | 0.043098323 | 1.278760081 | 0.200981555 | 0.654716629 |
| Pdgfd | 83.22977343 | 0.008639413 | 0.135285865 | 0.063860425 | 0.949081364 | 0.975508052 | Psd3 | 22118.62539 | 0.05511411 | 0.069485362 | 0.793175834 | 0.427675385 | 0.810334558 |
| Vwce | 14.9139325 | 0.010522902 | 0.166318202 | 0.063269696 | 0.949551746 | 0.975814469 | Smim12 | 1259.862754 | 0.055129935 | 0.060935174 | 0.904730906 | 0.365607964 | 0.777735733 |
| Amigo2 | 436.8699894 | -0.009431617 | 0.150147576 | -0.062815648 | 0.949913306 | 0.975997623 | Mbnl2 | 8145.314078 | 0.055146853 | 0.069544265 | 0.792974842 | 0.42779248 | 0.810334558 |
| Bmf | 171.1240867 | -0.008605235 | 0.137007358 | -0.062808564 | 0.949918946 | 0.975997623 | Eea1 | 2019.918117 | 0.055171526 | 0.060239987 | 0.915862182 | 0.359739184 | 0.776027411 |
| E030011O05Rik | 18.15322911 | -0.010538072 | 0.167702491 | -0.062837899 | 0.949895587 | 0.975997623 | Rps6ka5 | 1014.088615 | 0.055174587 | 0.080149127 | 0.688399103 | 0.491201488 | 0.841892493 |
| 8030462N17Rik | 1074.09818 | -0.005646903 | 0.090393257 | -0.062470404 | 0.95018823 | 0.976122789 | Apitd1 | 74.17766495 | 0.055214664 | 0.095088754 | 0.580664503 | 0.56146659 | 0.873934985 |
| Meaf6 | 1341.154369 | -0.003970438 | 0.063610212 | -0.062418242 | 0.950229768 | 0.976122789 | P2rx5 | 39.36389795 | 0.05521955 | 0.085380297 | 0.646748152 | 0.517794953 | 0.853297835 |
| Trim23 | 1992.977334 | -0.005689204 | 0.091071525 | -0.062469621 | 0.950188854 | 0.976122789 | Asphd1 | 610.1073789 | 0.05524459 | 0.072939056 | 0.757407518 | 0.448805753 | 0.823263966 |
| Fhl1 | 3417.754905 | -0.006312011 | 0.101329014 | -0.06229224 | 0.950330108 | 0.976161144 | St3gal3 | 2211.205826 | 0.055252187 | 0.052393329 | 1.054565311 | 0.291624176 | 0.727847831 |
| Ctu1 | 286.7803078 | 0.006027457 | 0.096956558 | 0.062166574 | 0.950430181 | 0.976199219 | Cstf2t | 3550.870806 | 0.055258241 | 0.057438152 | 0.962047683 | 0.336025652 | 0.759375714 |
| Foxj3 | 3362.595721 | -0.003815194 | 0.062118628 | -0.061417871 | 0.95102642 | 0.976746875 | Col25a1 | 2176.426035 | 0.055263428 | 0.084434667 | 0.654511115 | 0.512782568 | 0.852934798 |
| Cubn | 36.03025949 | -0.010266064 | 0.167606967 | -0.061250816 | 0.95115946 | 0.976761844 | Phka1 | 1276.536408 | 0.055278979 | 0.073814217 | 0.748893385 | 0.453921468 | 0.825985606 |
| Hacl1 | 200.0825677 | 0.007474242 | 0.122203836 | 0.061162093 | 0.951230119 | 0.976761844 | Wsb2 | 12913.47007 | 0.055279275 | 0.055439013 | 0.997118665 | 0.318706914 | 0.74851037 |
| Rab42 | 30.2532644 | 0.009790224 | 0.159938354 | 0.061212484 | 0.951189987 | 0.976761844 | Slc6a3 | 67.01662748 | 0.055304608 | 0.029602896 | 1.86821614 | NA | NA |
| Dynlt1a | 41.20297215 | -0.009665877 | 0.158450782 | -0.061002395 | 0.951357302 | 0.976827702 | Mrpl38 | 1438.179278 | 0.055309453 | 0.066563546 | 0.830927083 | 0.406014823 | 0.800733644 |
| Nek11 | 66.01732707 | 0.008994637 | 0.147751071 | 0.060876969 | 0.951457191 | 0.976847879 | Smyd1 | 119.4640895 | 0.055311975 | 0.09507629 | 0.581764131 | 0.560725567 | 0.873713575 |
| Yrdc | 678.0361482 | 0.004914833 | 0.080810287 | 0.060819393 | 0.951503045 | 0.976847879 | Naa16 | 213.272083 | 0.055317443 | 0.087881042 | 0.629458198 | 0.529049127 | 0.857868166 |
| Rnf34 | 2139.328822 | 0.005572195 | 0.091772486 | 0.060717492 | 0.9515842 | 0.976866468 | Slc26a3 | 6.939871194 | 0.055338 | 0.058427138 | 0.947128383 | 0.343573363 | NA |
| Lipa | 888.3373778 | -0.003984588 | 0.065838185 | -0.060520922 | 0.951740753 | 0.976913781 | Apol9b | 6.63364543 | 0.055340578 | 0.053637146 | 1.031758437 | 0.302185294 | NA |
| Sfr1 | 2837.817735 | 0.004144172 | 0.068497258 | 0.060501287 | 0.951756391 | 0.976913781 | 2810025M15Rik | 453.7925597 | 0.055348316 | 0.075396229 | 0.734099265 | 0.462888241 | 0.831014288 |
| Btbd16 | 30.44447211 | -0.009825854 | 0.163092197 | -0.060247236 | 0.951958725 | 0.976992017 | 2810047C21Rik1 | 57.0214405 | 0.055391141 | 0.081437011 | 0.680171585 | 0.496395822 | 0.844826506 |
| Npas2 | 2211.395207 | -0.006027003 | 0.100032487 | -0.06025046 | 0.951956158 | 0.976992017 | 2900008C10Rik | 174.412632 | 0.055395737 | 0.087295262 | 0.63457897 | 0.525703047 | 0.855892499 |
| Naa30 | 941.3673321 | 0.006539833 | 0.108916858 | 0.06004427 | 0.952120376 | 0.977093198 | Klhdc5 | 1461.714013 | 0.055402546 | 0.070141983 | 0.789862837 | 0.429607877 | 0.811442206 |
| A630066F11Rik | 25.74618602 | -0.009898697 | 0.16639836 | -0.059487951 | 0.952563463 | 0.977224279 | H2afv | 991.4426575 | 0.055408777 | 0.05552828 | 0.997847883 | 0.318353127 | 0.74851037 |
| Ankef1 | 27.39198519 | 0.009975288 | 0.167203563 | 0.059659544 | 0.952426794 | 0.977224279 | Ik | 4905.7423 | 0.055419479 | 0.060669551 | 0.913464462 | 0.360998316 | 0.776449324 |
| Nubp2 | 794.0849247 | 0.004194273 | 0.070367914 | 0.059604908 | 0.952470309 | 0.977224279 | 1110054M08Rik | 134.4103889 | 0.055423919 | 0.094612304 | 0.585800333 | 0.5580097 | 0.872673743 |
| Pgrmc2 | 3100.645419 | 0.003647188 | 0.061154992 | 0.059638435 | 0.952443606 | 0.977224279 | Pdpk1 | 7617.298576 | 0.055428265 | 0.065757867 | 0.842914584 | 0.399276216 | 0.798004076 |
| R74862 | 163.6936304 | -0.006186852 | 0.103888555 | -0.059552776 | 0.952511831 | 0.977224279 | Rnf217 | 434.1666649 | 0.055436959 | 0.083415311 | 0.664589734 | 0.506312937 | 0.848908639 |
| 2700046G09Rik | 96.18597442 | 0.00823954 | 0.139166834 | 0.059206207 | 0.952787866 | 0.977246133 | Dnaja3 | 2923.744921 | 0.055441162 | 0.046320966 | 1.196891314 | 0.23134892 | 0.683460324 |
| 4933411K20Rik | 1254.382468 | -0.004104786 | 0.070567674 | -0.058168085 | 0.953614742 | 0.977246133 | Cnnm4 | 629.9917767 | 0.055461809 | 0.072223303 | 0.767921245 | 0.442533975 | 0.819825088 |
| Abcc1 | 1059.609969 | 0.004113347 | 0.070681532 | 0.058195494 | 0.95359291 | 0.977246133 | Cbx5 | 9224.183557 | 0.055467769 | 0.057422771 | 0.965954238 | 0.33406708 | 0.758212399 |
| Apbb1 | 16349.25817 | 0.004680395 | 0.079391335 | 0.058953472 | 0.952989168 | 0.977246133 | Zfand2a | 2944.117406 | 0.055504438 | 0.053542934 | 1.036634213 | 0.299906362 | 0.733704078 |
| Cdk13 | 2154.752081 | -0.004412785 | 0.074680995 | -0.059088466 | 0.952881645 | 0.977246133 | Ralgps2 | 1816.028591 | 0.055521962 | 0.064425032 | 0.861807288 | 0.388793574 | 0.79235741 |
| Cep57l1 | 155.8028308 | -0.006931669 | 0.118580506 | -0.058455382 | 0.953385902 | 0.977246133 | Tspan8 | 11.48362848 | 0.05553769 | 0.062145853 | 0.893666885 | 0.371500148 | NA |
| Cops7b | 805.7245426 | 0.00462374 | 0.078701722 | 0.058750179 | 0.953151091 | 0.977246133 | Gpr107 | 2260.452708 | 0.055539765 | 0.055445085 | 1.001707648 | 0.316484812 | 0.745993052 |
| Gemin7 | 632.4105373 | -0.003999334 | 0.067970227 | -0.058839501 | 0.953079945 | 0.977246133 | Pfkp | 12577.65326 | 0.055545634 | 0.042513123 | 1.306552661 | 0.191364683 | 0.646682644 |
| Hsd3b7 | 257.0368928 | 0.006980037 | 0.119659715 | 0.058332388 | 0.95348387 | 0.977246133 | Mrps14 | 958.6216779 | 0.055551084 | 0.062840001 | 0.884008316 | 0.376691724 | 0.783969863 |
| Hsp90aa1 | 43613.85765 | 0.003880922 | 0.06598576 | 0.058814534 | 0.953099832 | 0.977246133 | Atp2a2 | 43170.81825 | 0.055574908 | 0.058282506 | 0.95354356 | 0.340314737 | 0.76228402 |
| Kcng2 | 168.8928808 | -0.007982291 | 0.137246599 | -0.058160209 | 0.953621015 | 0.977246133 | Frs3 | 713.8735248 | 0.05558176 | 0.081895078 | 0.678694755 | 0.49733129 | 0.845038521 |
| Nme5 | 270.0574552 | -0.005900529 | 0.100452812 | -0.058739308 | 0.95315975 | 0.977246133 | Fam216a | 1516.370768 | 0.055582976 | 0.060814687 | 0.913972901 | 0.360731085 | 0.776337125 |
| Sarnp | 1077.007126 | 0.004310534 | 0.073686096 | 0.058498605 | 0.953351473 | 0.977246133 | Syf2 | 1530.957202 | 0.055582989 | 0.055493011 | 1.001621426 | 0.316526469 | 0.745993052 |
| Sco1 | 378.8551667 | -0.005293503 | 0.090919272 | -0.058222013 | 0.953571787 | 0.977246133 | Dync2li1 | 702.0541146 | 0.055591167 | 0.064832846 | 0.85745375 | 0.391194165 | 0.794184423 |
| Scyl1 | 2810.376304 | 0.003201255 | 0.054071517 | 0.059204089 | 0.952789553 | 0.977246133 | Gpr155 | 3323.475644 | 0.055592422 | 0.083019037 | 0.669634631 | 0.503090732 | 0.847567618 |
| Slc26a6 | 268.9945167 | 0.005626126 | 0.095199947 | 0.059097992 | 0.952874058 | 0.977246133 | Habp4 | 9677.312808 | 0.055596191 | 0.041096919 | 1.352806774 | 0.176117368 | 0.628879275 |
| Tmbim4 | 968.1389399 | -0.003457344 | 0.059491382 | -0.058115035 | 0.953656998 | 0.977246133 | Psmd3 | 5201.791336 | 0.055618563 | 0.052198389 | 1.065522587 | 0.28663951 | 0.722148665 |
| Rabep1 | 6210.979818 | -0.003077299 | 0.053162054 | -0.057885251 | 0.953840031 | 0.977369052 | Ppp1r26 | 1500.51084 | 0.055651369 | 0.064782709 | 0.859046656 | 0.390314776 | 0.793521685 |
| Pex16 | 626.9319964 | -0.004925009 | 0.08523702 | -0.057780159 | 0.953923743 | 0.977390191 | Corin | 10.86330466 | 0.055658772 | 0.058428795 | 0.952591474 | 0.340797099 | NA |
| Dach2 | 21.94758897 | 0.009619436 | 0.166821333 | 0.057663107 | 0.954016981 | 0.977421087 | Syt1 | 72163.96653 | 0.055660762 | 0.059253209 | 0.939371266 | 0.34754016 | 0.766038372 |
| Stag2 | 2476.029157 | 0.004233482 | 0.073795191 | 0.057367998 | 0.954252055 | 0.977597285 | Snapc1 | 1113.25254 | 0.055677082 | 0.067344197 | 0.82675397 | 0.408376525 | 0.801055189 |
| Pcbp1 | 4028.49714 | 0.003697711 | 0.064646813 | 0.057198654 | 0.95438695 | 0.977639006 | Lonrf3 | 1057.417065 | 0.055689229 | 0.082639262 | 0.673883427 | 0.50038543 | 0.846268338 |
| Slc7a14 | 6549.082324 | 0.006396928 | 0.111915705 | 0.057158451 | 0.954418975 | 0.977639006 | Grpel1 | 1636.217454 | 0.055703993 | 0.052027269 | 1.07066917 | 0.28431821 | 0.721548306 |
| Agbl1 | 12.74074008 | 0.009210792 | 0.162521243 | 0.056674387 | 0.954804578 | 0.977713021 | Arf5 | 3499.716555 | 0.055704243 | 0.058344844 | 0.954741482 | 0.339708447 | 0.761679979 |
| Gpr25 | 251.9183422 | 0.005959122 | 0.105216433 | 0.056636803 | 0.954834517 | 0.977713021 | Alox12e | 5.988396507 | 0.055717579 | 0.049113259 | 1.134471212 | 0.256596943 | NA |
| Mettl2 | 541.840372 | 0.004291354 | 0.07569776 | 0.056690632 | 0.954791637 | 0.977713021 | Ahcyl2 | 7881.915158 | 0.055718934 | 0.052253209 | 1.0663256 | 0.286276481 | 0.722148665 |
| Olig2 | 877.0502211 | 0.004965607 | 0.087174494 | 0.056961699 | 0.954575705 | 0.977713021 | Kif26a | 336.0786843 | 0.055721097 | 0.081010978 | 0.68782156 | 0.491565157 | 0.841892493 |
| Pacsin1 | 20078.34289 | -0.003778743 | 0.066477968 | -0.056842033 | 0.954671031 | 0.977713021 | Gabrb3 | 12095.47701 | 0.055722769 | 0.063072014 | 0.883478502 | 0.376977793 | 0.783997209 |
| Psen1 | 2561.575049 | -0.003260898 | 0.057628242 | -0.05658507 | 0.954875728 | 0.977713021 | Fam122a | 394.1339247 | 0.055734279 | 0.089301473 | 0.624113772 | 0.53255287 | 0.858328942 |
| Snrpa | 1522.537082 | 0.00379388 | 0.067132597 | 0.056513236 | 0.954932952 | 0.977713021 | Zdhhc2 | 1286.90106 | 0.055753519 | 0.076250971 | 0.731184378 | 0.464666541 | 0.83159896 |
| Far1 | 2281.220372 | 0.003728609 | 0.066288439 | 0.056248261 | 0.955144035 | 0.977864522 | D17Wsu92e | 13171.73342 | 0.055768152 | 0.051401183 | 1.084958525 | 0.277940025 | 0.718263085 |
| Nap1l5 | 9994.425805 | -0.005011961 | 0.089596017 | -0.055939553 | 0.955389962 | 0.977922446 | Idnk | 483.8567001 | 0.055780352 | 0.071567041 | 0.779413982 | 0.43573589 | 0.815960393 |
| Rpp30 | 284.6175693 | -0.005691787 | 0.101722529 | -0.055954046 | 0.955378416 | 0.977922446 | Smim1 | 490.1394891 | 0.055791856 | 0.083250468 | 0.670168684 | 0.502750265 | 0.84739426 |
| Siah1a | 806.2354318 | 0.003639596 | 0.064997774 | 0.055995699 | 0.955345234 | 0.977922446 | Ccni | 8370.714697 | 0.055831392 | 0.056210632 | 0.993253235 | 0.320586561 | 0.750352874 |
| Mgat4b | 2072.041188 | 0.004750535 | 0.086566237 | 0.054877452 | 0.956236096 | 0.978659229 | Ccdc97 | 2432.423734 | 0.055841232 | 0.053106819 | 1.051488931 | 0.293034091 | 0.728526371 |
| Sox17 | 169.1801436 | 0.008991289 | 0.163767533 | 0.054902757 | 0.956215935 | 0.978659229 | Irak1bp1 | 463.6792055 | 0.0558601 | 0.074155205 | 0.753286307 | 0.451277885 | 0.824250291 |
| Diap3 | 42.3746058 | 0.009159237 | 0.167149154 | 0.054796789 | 0.956300359 | 0.978660354 | Nxpe2 | 9.965410442 | 0.055891498 | 0.061722779 | 0.905524646 | 0.365187509 | NA |
| Atp6ap1 | 15633.25101 | -0.002813573 | 0.051464297 | -0.054670388 | 0.956401061 | 0.978698768 | Prpf18 | 2104.458144 | 0.055927622 | 0.062487252 | 0.895024506 | 0.370773988 | 0.780761257 |
| Zdhhc15 | 430.8799153 | -0.005830956 | 0.107325475 | -0.054329656 | 0.956672522 | 0.978911905 | Dr1 | 1070.264524 | 0.055935294 | 0.07162193 | 0.780979992 | 0.434814263 | 0.815326688 |
| Rgag4 | 1276.380522 | 0.005445396 | 0.100820139 | 0.054010999 | 0.956926401 | 0.979107024 | Cacng3 | 4551.092932 | 0.055950267 | 0.059579454 | 0.939086596 | 0.347686286 | 0.766038372 |
| Nr4a1 | 3817.794586 | 0.008555188 | 0.158634597 | 0.053930151 | 0.956990815 | 0.979108273 | Sncb | 16574.42666 | 0.055950559 | 0.03864442 | 1.447830207 | 0.147664542 | 0.594835037 |
| Tmed9 | 3613.415968 | 0.003549217 | 0.065950874 | 0.053816074 | 0.957081703 | 0.979136606 | Rap1gap | 11284.05259 | 0.055954546 | 0.064199262 | 0.871576163 | 0.383439639 | 0.787702475 |
| Mterfd2 | 439.9342985 | -0.004534303 | 0.085155318 | -0.05324744 | 0.957534757 | 0.979524959 | Bnip3l | 5460.576874 | 0.055955415 | 0.061909427 | 0.903827057 | 0.366087112 | 0.778210102 |
| Tgs1 | 1329.798506 | -0.003488217 | 0.0655915 | -0.053180933 | 0.957587747 | 0.979524959 | Zfp385b | 1546.082354 | 0.05599733 | 0.068389791 | 0.818796632 | 0.412902453 | 0.803781966 |
| Gm5468 | 120.8048269 | -0.00718032 | 0.136221637 | -0.052710569 | 0.957962518 | 0.979843626 | Sptlc3 | 6.629747119 | 0.05601634 | 0.041102698 | 1.362838492 | 0.172933417 | NA |
| Lama2 | 679.8431429 | -0.00447044 | 0.085047467 | -0.052564062 | 0.958079252 | 0.979898339 | Gipc2 | 56.72068496 | 0.056026493 | 0.094759634 | 0.591248521 | 0.554353917 | 0.871360989 |
| Cpeb4 | 2154.009699 | 0.005145618 | 0.098334173 | 0.052327874 | 0.958267444 | 0.97996401 | Galm | 111.5457009 | 0.056035158 | 0.093981617 | 0.596235306 | 0.551018042 | 0.869642174 |
| Polr3b | 741.6605126 | 0.003302725 | 0.063119784 | 0.052324719 | 0.958269957 | 0.97996401 | Zfp330 | 841.5815714 | 0.056037506 | 0.065041112 | 0.861570548 | 0.388923884 | 0.79235741 |
| Arfrp1 | 1179.621703 | 0.00513677 | 0.098600303 | 0.052096894 | 0.958451487 | 0.980084962 | 9330159F19Rik | 3337.18 | 0.056047183 | 0.074387362 | 0.753450329 | 0.451179349 | 0.824173947 |
| Slc6a9 | 2037.336911 | -0.006062075 | 0.116584707 | -0.051997176 | 0.958530944 | 0.980101527 | BC048546 | 337.2759692 | 0.056052295 | 0.089256322 | 0.627992439 | 0.530008893 | 0.857915133 |
| Ropn1l | 15.23705801 | -0.008632749 | 0.16648075 | -0.05185434 | 0.958644757 | 0.980153218 | Kdelr2 | 848.2293423 | 0.056066507 | 0.056886574 | 0.98558417 | 0.324337209 | 0.75293408 |
| Cdadc1 | 2148.249051 | 0.002758667 | 0.05370501 | 0.051367028 | 0.959033058 | 0.980168304 | Fam89b | 807.1961456 | 0.056070378 | 0.066081387 | 0.848504869 | 0.396156862 | 0.796565439 |
| Chd1l | 367.5512368 | 0.004185183 | 0.081331214 | 0.051458509 | 0.958960163 | 0.980168304 | Akt3 | 7124.061253 | 0.056072493 | 0.066879525 | 0.838410462 | 0.401800216 | 0.799277436 |
| Efcab5 | 149.4599485 | -0.005906504 | 0.114962165 | -0.05137781 | 0.959024466 | 0.980168304 | Dlx1 | 871.9934522 | 0.056074054 | 0.082039914 | 0.683497233 | 0.494292698 | 0.843503724 |
| Efhd2 | 9789.435918 | 0.005075965 | 0.098985112 | 0.051280081 | 0.95910234 | 0.980168304 | Eif2a | 1716.966961 | 0.056090508 | 0.063537246 | 0.882797276 | 0.377345813 | 0.783997209 |
| Ggcx | 588.9321664 | -0.006473137 | 0.126128802 | -0.051321637 | 0.959069227 | 0.980168304 | Lmln | 543.5146396 | 0.056098959 | 0.068330762 | 0.820991259 | 0.411651247 | 0.802795661 |
| Pctp | 77.35417305 | -0.008259466 | 0.159696003 | -0.051719926 | 0.958751859 | 0.980168304 | Pex5l | 8945.396087 | 0.056107396 | 0.055901718 | 1.003679279 | 0.315533228 | 0.745993052 |
| Pdgfa | 738.0017226 | -0.004123908 | 0.080343637 | -0.051328373 | 0.959063859 | 0.980168304 | Taf5l | 877.2848025 | 0.056110909 | 0.063274847 | 0.886780634 | 0.375197016 | 0.783489748 |
| Clec14a | 166.7864288 | -0.006292612 | 0.122919662 | -0.051192882 | 0.959171824 | 0.980174663 | Rbfa | 966.5041841 | 0.056113016 | 0.059965281 | 0.935758419 | 0.349397585 | 0.768268271 |
| Fam104a | 1285.607952 | 0.003851724 | 0.075381455 | 0.051096434 | 0.959248677 | 0.980188551 | Ppp1r15b | 2765.588894 | 0.056114554 | 0.070493213 | 0.796027747 | 0.426015908 | 0.809519334 |
| Nkrf | 1292.220283 | -0.003720215 | 0.073125956 | -0.050874069 | 0.959425869 | 0.98030496 | Dlg5 | 1841.303824 | 0.056135838 | 0.073756752 | 0.761094224 | 0.44660079 | 0.822210607 |
| 2310039L15Rik | 17.82082552 | -0.008428966 | 0.168096725 | -0.050143544 | 0.960008 | 0.98049586 | Zfp654 | 624.5531815 | 0.056145726 | 0.08648648 | 0.649184999 | 0.516218806 | 0.853297835 |
| Btbd2 | 2909.317488 | 0.003253568 | 0.064522285 | 0.050425497 | 0.959783319 | 0.98049586 | Nab1 | 1137.920649 | 0.056159226 | 0.071312289 | 0.787511205 | 0.430982672 | 0.812181606 |
| Cideb | 149.1887687 | 0.008304306 | 0.165237388 | 0.050256824 | 0.95991773 | 0.98049586 | Mrgprf | 40.48612488 | 0.056166752 | 0.088737687 | 0.632952625 | 0.526764581 | 0.85599981 |
| Dusp27 | 27.1016467 | 0.008228596 | 0.163678439 | 0.050272938 | 0.959904889 | 0.98049586 | 2310022B05Rik | 3287.896357 | 0.056171364 | 0.079624873 | 0.705449962 | 0.480530261 | 0.836914993 |
| Dut | 693.1935947 | 0.003481451 | 0.069733771 | 0.049924889 | 0.960182244 | 0.98049586 | Esco1 | 876.4277949 | 0.056180477 | 0.080898986 | 0.69445217 | 0.487398686 | 0.839801621 |
| Man2a2 | 11034.22239 | 0.002270259 | 0.045385774 | 0.050021379 | 0.960105351 | 0.98049586 | Srxn1 | 4972.62886 | 0.056201786 | 0.052124369 | 1.078224765 | 0.280933462 | 0.720080692 |
| Ror2 | 36.42642384 | -0.00834492 | 0.165972723 | -0.050278866 | 0.959900165 | 0.98049586 | Zfp941 | 1454.252698 | 0.056207208 | 0.069741224 | 0.805939513 | 0.420277727 | 0.807657282 |
| Slc25a46 | 3359.500275 | 0.003826131 | 0.076156218 | 0.050240564 | 0.959930687 | 0.98049586 | Zfp612 | 2738.078057 | 0.056208031 | 0.069195508 | 0.812307511 | 0.416615202 | 0.806153084 |
| Tsen15 | 546.9296642 | -0.003690376 | 0.073805512 | -0.050001356 | 0.960121308 | 0.98049586 | March5 | 3008.401717 | 0.056215187 | 0.060470174 | 0.929634946 | 0.352560127 | 0.770466136 |
| Axin2 | 1081.939312 | 0.003516955 | 0.071109375 | 0.049458383 | 0.960554002 | 0.980746208 | Rfwd2 | 3150.224455 | 0.056239257 | 0.072569976 | 0.774965897 | 0.438359812 | 0.817094139 |
| Ptafr | 36.16946345 | 0.008115902 | 0.164026793 | 0.049479122 | 0.960537475 | 0.980746208 | Map9 | 5566.631945 | 0.056241761 | 0.065590377 | 0.857469702 | 0.391185352 | 0.794184423 |
| Suclg2 | 1009.37556 | -0.00297963 | 0.060735037 | -0.049059488 | 0.960871888 | 0.98100613 | Prr14l | 3091.086824 | 0.056265208 | 0.076024308 | 0.740094973 | 0.459242369 | 0.828381305 |
| Fastkd1 | 479.97532 | -0.004935354 | 0.100919764 | -0.048903741 | 0.960996008 | 0.981068205 | 4933411K20Rik | 1410.738907 | 0.056268612 | 0.068759468 | 0.818339843 | 0.413163161 | 0.803854924 |
| Fut11 | 719.7919859 | 0.003251783 | 0.066649446 | 0.048789354 | 0.961087166 | 0.981096624 | Casd1 | 2650.557358 | 0.056280177 | 0.065745292 | 0.856033572 | 0.39197921 | 0.794184423 |
| Pde11a | 99.96411309 | 0.007482275 | 0.154423856 | 0.048452846 | 0.961355344 | 0.981305731 | Reep1 | 5729.254141 | 0.056289474 | 0.054188654 | 1.038768637 | 0.298912351 | 0.733276458 |
| Samsn1 | 63.37936516 | 0.007661068 | 0.159379578 | 0.048068067 | 0.961661996 | 0.981554082 | Ubxn2b | 1710.044016 | 0.056291074 | 0.074419476 | 0.756402456 | 0.449407936 | 0.823263966 |
| Comp | 10.00674596 | 0.007257308 | 0.15378689 | 0.047190682 | 0.962361254 | 0.98200904 | D030056L22Rik | 791.7127024 | 0.056296135 | 0.069742102 | 0.807204444 | 0.419548706 | 0.807483875 |
| Rrm1 | 777.3881959 | 0.00413322 | 0.087563626 | 0.047202479 | 0.962351852 | 0.98200904 | Meaf6 | 1468.924919 | 0.056301882 | 0.058770503 | 0.957995587 | 0.338064984 | 0.760581219 |
| Rs1 | 8.459530939 | -0.007195348 | 0.152344108 | -0.04723089 | 0.962329209 | 0.98200904 | Tspan3 | 11131.18746 | 0.056314639 | 0.039514669 | 1.425157809 | 0.154111589 | 0.60340295 |
| Stom | 570.183771 | 0.004323496 | 0.091157302 | 0.047428964 | 0.962171345 | 0.98200904 | Tomm6 | 2467.727167 | 0.056329492 | 0.05371448 | 1.048683559 | 0.294323783 | 0.729576944 |
| 1110008J03Rik | 487.1465197 | -0.004010393 | 0.087640028 | -0.045759837 | 0.963501671 | 0.982137951 | 8030462N17Rik | 1127.476326 | 0.056337703 | 0.076959686 | 0.732041749 | 0.464143086 | 0.83159896 |
| AF529169 | 158.6245214 | -0.007075104 | 0.155685073 | -0.045444971 | 0.963752637 | 0.982137951 | Snrpc | 637.6955396 | 0.056396607 | 0.062558669 | 0.901499467 | 0.367322817 | 0.778711506 |
| Arl4d | 866.6304843 | 0.006698774 | 0.143216735 | 0.04677368 | 0.962693606 | 0.982137951 | Spp1 | 503.9173172 | 0.05639672 | 0.089764058 | 0.628277294 | 0.529822303 | 0.857915133 |
| Cbx8 | 255.483013 | -0.00443187 | 0.096445481 | -0.045952078 | 0.963348446 | 0.982137951 | Lck | 59.2696806 | 0.056407346 | 0.094282251 | 0.598281704 | 0.549651983 | 0.868891577 |
| Chd8 | 4609.269977 | 0.002865937 | 0.06215293 | 0.04611105 | 0.963221739 | 0.982137951 | Inpp5a | 2738.169681 | 0.056418075 | 0.055604607 | 1.014629515 | 0.310282465 | 0.743039093 |
| Disp2 | 15021.49942 | 0.002665886 | 0.057100767 | 0.046687391 | 0.96276238 | 0.982137951 | Scnn1a | 275.2606305 | 0.056419403 | 0.080401085 | 0.701724392 | 0.482851059 | 0.837880119 |
| Fbxo44 | 3575.105884 | -0.0028674 | 0.063232312 | -0.045347073 | 0.963830667 | 0.982137951 | Fut2 | 8.902867462 | 0.056420039 | 0.061228796 | 0.921462482 | 0.35680902 | NA |
| Frmd8 | 889.5579279 | 0.005251624 | 0.116312405 | 0.045151019 | 0.963986936 | 0.982137951 | Mrpl13 | 907.8816668 | 0.056442611 | 0.070232668 | 0.803651811 | 0.421598092 | 0.808573261 |
| Fsip1 | 12.82421761 | 0.007634898 | 0.166329954 | 0.045902124 | 0.963388261 | 0.982137951 | Sephs2 | 1207.621649 | 0.056457887 | 0.060556195 | 0.932322232 | 0.351170021 | 0.769244462 |
| Gse1 | 2850.179647 | -0.003778298 | 0.082246097 | -0.045938933 | 0.963358923 | 0.982137951 | Timm13 | 1534.012172 | 0.05646524 | 0.066894807 | 0.844090024 | 0.398619102 | 0.797789838 |
| Inhbe | 8.131688114 | -0.006905114 | 0.150797791 | -0.04579055 | 0.963477191 | 0.982137951 | Syt13 | 11629.25394 | 0.056514151 | 0.056406146 | 1.001914787 | 0.31638475 | 0.745993052 |
| Kctd4 | 690.0822562 | -0.003944364 | 0.085006513 | -0.04640073 | 0.962990855 | 0.982137951 | Rwdd1 | 1412.193811 | 0.056514611 | 0.064020282 | 0.882761049 | 0.37736539 | 0.783997209 |
| Lysmd1 | 827.4086473 | -0.003479754 | 0.076838085 | -0.045286839 | 0.963878678 | 0.982137951 | Ctxn3 | 73.23637755 | 0.056525496 | 0.091756904 | 0.616035347 | 0.537871191 | 0.862082844 |
| Mamdc2 | 49.91430138 | 0.006741999 | 0.144846045 | 0.046545968 | 0.962875096 | 0.982137951 | Coq3 | 654.689873 | 0.05653823 | 0.066249239 | 0.853417049 | 0.39342807 | 0.794754188 |
| Mto1 | 815.1214825 | 0.003548518 | 0.077095049 | 0.046027834 | 0.963288065 | 0.982137951 | Arhgdig | 1809.750878 | 0.05656606 | 0.060495591 | 0.935044336 | 0.349765451 | 0.768521092 |
| Obfc1 | 426.6985832 | 0.003915796 | 0.084021844 | 0.046604495 | 0.96282845 | 0.982137951 | Sesn3 | 1860.815858 | 0.05656784 | 0.074279752 | 0.761551275 | 0.446327865 | 0.822210607 |
| Pde1a | 8720.519525 | -0.004357806 | 0.095476325 | -0.045642787 | 0.963594965 | 0.982137951 | Drg1 | 2757.967359 | 0.05657654 | 0.04846066 | 1.167473566 | 0.243019177 | 0.692772033 |
| Psmb10 | 1049.272001 | 0.00406844 | 0.089974962 | 0.045217467 | 0.963933972 | 0.982137951 | Sh3bp5l | 2155.749485 | 0.056586097 | 0.065604556 | 0.862533051 | 0.388394253 | 0.792167218 |
| Rnf152 | 809.4292148 | -0.004721924 | 0.104644556 | -0.045123457 | 0.964008904 | 0.982137951 | Sidt1 | 2841.12004 | 0.056598131 | 0.065389888 | 0.865548676 | 0.386737712 | 0.790912767 |
| Six3os1 | 188.1669923 | -0.007422247 | 0.162999395 | -0.045535426 | 0.963680538 | 0.982137951 | Tacstd2 | 7.87253979 | 0.056609017 | 0.058529019 | 0.967195714 | 0.333446205 | NA |
| Spen | 3396.743117 | 0.004501472 | 0.098446264 | 0.045725169 | 0.963529303 | 0.982137951 | Prrg4 | 37.51370285 | 0.056615189 | 0.090742993 | 0.623907011 | 0.532688656 | 0.858328942 |
| Ttc1 | 1397.041825 | -0.003385809 | 0.073702243 | -0.045939029 | 0.963358846 | 0.982137951 | Vps28 | 3066.578329 | 0.056646548 | 0.058497869 | 0.968352336 | 0.332868435 | 0.757641999 |
| Ube3b | 6260.927891 | 0.003159855 | 0.068567306 | 0.046083991 | 0.963243306 | 0.982137951 | Prpsap2 | 1301.212155 | 0.056670648 | 0.058521524 | 0.968372724 | 0.332858256 | 0.757641999 |
| Zmym5 | 1384.54329 | 0.003219723 | 0.070199185 | 0.045865531 | 0.963417427 | 0.982137951 | Kti12 | 397.710618 | 0.056695435 | 0.074874607 | 0.757205107 | 0.448926991 | 0.823263966 |
| Zfp493 | 133.1399687 | 0.005725169 | 0.128558279 | 0.044533645 | 0.964479034 | 0.982552314 | Sorl1 | 8356.353472 | 0.056718165 | 0.063081309 | 0.899127906 | 0.368584536 | 0.779452463 |
| Snx21 | 919.2078478 | -0.003390138 | 0.076275026 | -0.044446244 | 0.964548701 | 0.982558683 | Atp6v1e1 | 13924.93199 | 0.056725623 | 0.046104667 | 1.230366198 | 0.218560005 | 0.671604514 |
| Tmem206 | 387.6941321 | 0.003618075 | 0.081786304 | 0.044238154 | 0.96471457 | 0.982663043 | Frmpd4 | 3872.182689 | 0.056727611 | 0.071506975 | 0.793315775 | 0.427593868 | 0.810334558 |
| Gm5124 | 697.063281 | -0.003199907 | 0.072756585 | -0.043980989 | 0.964919557 | 0.98274263 | Ccnc | 816.7597675 | 0.056758506 | 0.068977588 | 0.822854317 | 0.410590844 | 0.802184613 |
| Pigs | 2204.547156 | 0.003321502 | 0.075456974 | 0.04401849 | 0.964889665 | 0.98274263 | Acot2 | 207.4409855 | 0.056761659 | 0.080682229 | 0.703521207 | 0.481730996 | 0.837383765 |
| 2210015D19Rik | 475.4594587 | -0.003065715 | 0.070105717 | -0.043729884 | 0.965119718 | 0.982881879 | Wrb | 2166.33031 | 0.056773467 | 0.055667188 | 1.019873086 | 0.307788655 | 0.740098991 |
| Capsl | 39.03536385 | 0.006912649 | 0.159480618 | 0.04334476 | 0.965426712 | 0.983126192 | Abcb10 | 1258.086358 | 0.056800684 | 0.053936572 | 1.053101491 | 0.29229448 | 0.728154624 |
| Evpl | 216.1698738 | 0.004426387 | 0.102297543 | 0.043269731 | 0.96548652 | 0.983126192 | Fahd1 | 485.3630308 | 0.056814968 | 0.071675363 | 0.792670814 | 0.427969639 | 0.810464138 |
| Sv2a | 19617.77277 | 0.001596253 | 0.036984334 | 0.043160239 | 0.9655738 | 0.983128201 | Cbx4 | 1767.639275 | 0.056816982 | 0.057026569 | 0.996324732 | 0.319092391 | 0.748931341 |
| Wdr47 | 6127.093932 | -0.002318444 | 0.05378215 | -0.043108056 | 0.965615398 | 0.983128201 | Tmco3 | 1708.603178 | 0.056842113 | 0.047130284 | 1.206063466 | 0.22779302 | 0.680415543 |
| Senp6 | 2874.706153 | 0.002979048 | 0.069511364 | 0.04285699 | 0.965815534 | 0.983267355 | Slc36a4 | 1159.198057 | 0.056845801 | 0.065006729 | 0.874460258 | 0.381867654 | 0.786663048 |
| Dcun1d2 | 1702.580519 | -0.002520898 | 0.059217931 | -0.042569835 | 0.966044442 | 0.983435781 | Zfp583 | 215.8113838 | 0.056855592 | 0.084740114 | 0.670940705 | 0.502258304 | 0.84736401 |
| Ncoa6 | 3536.951371 | 0.003325574 | 0.078669356 | 0.042272803 | 0.966281226 | 0.983612201 | Ttyh2 | 1928.785964 | 0.056872674 | 0.084255657 | 0.675001252 | 0.49967497 | 0.846241311 |
| Tspan2 | 3678.412705 | 0.004323105 | 0.102722713 | 0.042085195 | 0.966430782 | 0.983635193 | Thnsl2 | 194.162328 | 0.056876721 | 0.090844619 | 0.62608795 | 0.531257256 | 0.858328942 |
| Zbbx | 11.49861351 | -0.006503412 | 0.154510209 | -0.042090503 | 0.966426551 | 0.983635193 | Pbrm1 | 2244.85273 | 0.056884944 | 0.069341618 | 0.820357905 | 0.412012105 | 0.802795661 |
| Mars2 | 337.7014098 | 0.003673713 | 0.08783402 | 0.041825622 | 0.96663771 | 0.983711068 | Ncrna00086 | 950.6696736 | 0.056893737 | 0.056533026 | 1.006380522 | 0.314232559 | 0.745993052 |
| Rpp40 | 119.6578433 | 0.00526221 | 0.125917736 | 0.041790857 | 0.966665423 | 0.983711068 | Gm4847 | 4.393942755 | 0.056908691 | 0.029501637 | 1.929001127 | NA | NA |
| Znrf1 | 6104.394776 | 0.003409104 | 0.081649794 | 0.041752753 | 0.9666958 | 0.983711068 | Atg2b | 3011.223506 | 0.056914546 | 0.060341415 | 0.943208669 | 0.34557418 | 0.765097832 |
| Wac | 4679.775774 | -0.003112016 | 0.074718909 | -0.041649651 | 0.966777992 | 0.983730098 | Ppil1 | 565.2194004 | 0.056927473 | 0.07011465 | 0.811919797 | 0.416837654 | 0.806153084 |
| Gstm7 | 593.5406854 | 0.00455294 | 0.110112368 | 0.041348125 | 0.967018368 | 0.983852768 | 4833439L19Rik | 5964.124729 | 0.056942915 | 0.041199616 | 1.382122475 | 0.166934099 | 0.617356019 |
| Zfp36l2 | 996.9950571 | 0.0057473 | 0.139028106 | 0.041339121 | 0.967025545 | 0.983852768 | Entpd5 | 1925.334834 | 0.05694321 | 0.051604255 | 1.103459598 | 0.269827627 | 0.71216373 |
| Adamts6 | 146.8056434 | 0.005662724 | 0.138581031 | 0.040862185 | 0.967405765 | 0.984110362 | Phax | 1450.365537 | 0.056948119 | 0.062380087 | 0.912921443 | 0.361283859 | 0.776449324 |
| Slc24a4 | 1145.890085 | 0.004733738 | 0.115628632 | 0.040939151 | 0.967344406 | 0.984110362 | Cs | 13075.76874 | 0.056965273 | 0.038844053 | 1.466512075 | 0.142508834 | 0.588697979 |
| Gm13399 | 40.15132026 | -0.006669718 | 0.164267058 | -0.040602892 | 0.967612478 | 0.984256023 | Spcs2 | 2927.906277 | 0.056973397 | 0.053106669 | 1.072810586 | 0.28335611 | 0.721034821 |
| Acyp1 | 341.6497221 | -0.003122839 | 0.077612893 | -0.040236084 | 0.96790491 | 0.984424227 | Fbxo44 | 3909.458502 | 0.056986596 | 0.052360259 | 1.088355874 | 0.276438033 | 0.717475105 |
| Bre | 1266.240981 | -0.002829513 | 0.07023688 | -0.040285291 | 0.96786568 | 0.984424227 | Camk2n2 | 3595.91626 | 0.057010047 | 0.066253444 | 0.860484283 | 0.389522144 | 0.792715562 |
| Ap1s1 | 8167.578864 | -0.003501231 | 0.087715098 | -0.039915946 | 0.968160138 | 0.984489938 | 9330175M20Rik | 12.09548082 | 0.057033723 | 0.066097368 | 0.862874343 | 0.388206559 | NA |
| Cldn11 | 3438.816519 | 0.004714588 | 0.118098054 | 0.039920965 | 0.968156137 | 0.984489938 | Dcxr | 172.772978 | 0.057052683 | 0.088394311 | 0.645433884 | 0.518646051 | 0.853297835 |
| Sox2ot | 1187.128559 | -0.006427447 | 0.160837404 | -0.039962387 | 0.968123114 | 0.984489938 | Xpot | 4214.112191 | 0.057056733 | 0.055511623 | 1.027834005 | 0.304027918 | 0.737453291 |
| Trove2 | 1663.904543 | 0.003225374 | 0.08121899 | 0.039712065 | 0.968322683 | 0.984590605 | Ppp4c | 1127.648505 | 0.057081688 | 0.059028733 | 0.967015298 | 0.333536386 | 0.757852268 |
| Edem3 | 2190.712256 | -0.003613864 | 0.091825824 | -0.039355644 | 0.968606843 | 0.984750291 | Dlst | 5427.784385 | 0.057083961 | 0.038881352 | 1.46815782 | 0.142061359 | 0.587991041 |
| Sec11c | 1295.041111 | -0.004041508 | 0.102511083 | -0.039425079 | 0.968551485 | 0.984750291 | Efna5 | 1481.917923 | 0.05708893 | 0.076197865 | 0.749219542 | 0.453724892 | 0.825985606 |
| Zfyve19 | 553.0646934 | 0.003093714 | 0.078945153 | 0.039188145 | 0.968740385 | 0.984821437 | Ank | 9859.875926 | 0.057097323 | 0.047893548 | 1.192171497 | 0.233194007 | 0.685064073 |
| Cnot1 | 5610.769466 | 0.00343527 | 0.088684359 | 0.03873592 | 0.969100935 | 0.985123336 | E330034G19Rik | 6.454303816 | 0.057107534 | 0.05836443 | 0.978464698 | 0.327844543 | NA |
| Actl6a | 433.003353 | -0.003766829 | 0.097850611 | -0.038495718 | 0.969292446 | 0.985253372 | Mtif3 | 668.8937555 | 0.057110174 | 0.068304 | 0.83611757 | 0.403088765 | 0.800009209 |
| Coro1a | 3037.449818 | -0.003785358 | 0.099312325 | -0.038115697 | 0.969595436 | 0.98536742 | Zfp964 | 81.87597017 | 0.057131486 | 0.09479105 | 0.602709707 | 0.546701823 | 0.867128614 |
| Mab21l2 | 8.138141471 | -0.004338139 | 0.113370529 | -0.038265139 | 0.969476285 | 0.98536742 | Arf4 | 3287.815003 | 0.05713696 | 0.062293233 | 0.917225793 | 0.359024333 | 0.774860309 |
| Sec14l1 | 10262.44675 | 0.001754777 | 0.045963057 | 0.038177987 | 0.969545772 | 0.98536742 | Galk1 | 370.2363397 | 0.057150229 | 0.084473823 | 0.676543657 | 0.498695537 | 0.846052342 |
| Mapk14 | 3956.92498 | 0.001854529 | 0.049049087 | 0.037809646 | 0.969839454 | 0.985550764 | Dkc1 | 1071.570197 | 0.05716829 | 0.063782453 | 0.896301217 | 0.370091909 | 0.780397655 |
| Rcsd1 | 247.4616877 | 0.004167505 | 0.110750411 | 0.037629702 | 0.969982926 | 0.985631916 | AU022252 | 592.4238723 | 0.057181773 | 0.066976014 | 0.85376494 | 0.393235244 | 0.794754188 |
| Vprbp | 1521.806675 | 0.003208061 | 0.085907482 | 0.037343203 | 0.970211358 | 0.985799382 | 1700021F05Rik | 527.7904998 | 0.057187505 | 0.079704852 | 0.717490891 | 0.473071255 | 0.833983942 |
| Agpat4 | 1777.268959 | 0.002914364 | 0.079260148 | 0.036769596 | 0.970668716 | 0.986190316 | Mpp2 | 7998.30909 | 0.057187613 | 0.057104995 | 1.001446786 | 0.316610855 | 0.745999882 |
| Armc7 | 385.3887491 | -0.003305575 | 0.090067699 | -0.036701003 | 0.970723409 | 0.986190316 | Slc1a6 | 389.7458701 | 0.057193039 | 0.081022356 | 0.705892073 | 0.480255257 | 0.836914993 |
| Hccs | 747.401672 | -0.002930734 | 0.080197339 | -0.036544033 | 0.970848569 | 0.986252801 | Bckdhb | 790.4720255 | 0.057202468 | 0.060224521 | 0.94982022 | 0.34220361 | 0.763762832 |
| Slc35d2 | 25.41426128 | -0.006025861 | 0.168005418 | -0.035867065 | 0.971388357 | 0.986736459 | Klf12 | 1574.795042 | 0.057218084 | 0.068073207 | 0.840537517 | 0.400607076 | 0.799169897 |
| Clec5a | 102.0897927 | 0.005342752 | 0.149441873 | 0.035751369 | 0.97148061 | 0.986765476 | Rpusd1 | 2529.715019 | 0.057223631 | 0.050931112 | 1.123549603 | 0.261204146 | 0.704173028 |
| 10-Sep | 178.4881779 | -0.004935324 | 0.138971798 | -0.035513132 | 0.971670575 | 0.986893732 | Fh1 | 2814.765895 | 0.057226028 | 0.045339648 | 1.262163037 | 0.206890126 | 0.661374554 |
| Vat1 | 1880.065325 | -0.003754861 | 0.10597752 | -0.035430733 | 0.971736279 | 0.986895772 | Foxb1 | 4.825874593 | 0.057232918 | 0.043967225 | 1.301717781 | 0.193012879 | NA |
| 2010300C02Rik | 11787.69074 | 0.003723328 | 0.106166639 | 0.035070599 | 0.972023446 | 0.98706365 | Sdhaf1 | 416.0814894 | 0.057245159 | 0.084328664 | 0.678833935 | 0.497243089 | 0.845038521 |
| Adnp2 | 1096.30649 | -0.002758645 | 0.078892178 | -0.034967286 | 0.972105827 | 0.98706365 | Fam227b | 2.743729671 | 0.05725251 | 0.034945778 | 1.638324092 | 0.101354104 | NA |
| Cryz | 594.1064462 | -0.003477279 | 0.099624496 | -0.03490386 | 0.972156403 | 0.98706365 | Ptgfr | 38.25095048 | 0.057261516 | 0.087350127 | 0.655540162 | 0.512120037 | 0.852888435 |
| Sema4c | 657.1191632 | 0.003085459 | 0.088259259 | 0.03495904 | 0.972112402 | 0.98706365 | Ncl | 11344.7469 | 0.057262919 | 0.055093265 | 1.039381472 | 0.298627357 | 0.732994423 |
| Xpnpep2 | 10.9238657 | 0.005762924 | 0.165646731 | 0.03479045 | 0.972246836 | 0.987090785 | Camsap2 | 8275.832725 | 0.057265379 | 0.064726581 | 0.884727393 | 0.376303678 | 0.783947144 |
| Atp6v0d1 | 8909.064425 | 0.002169196 | 0.062519447 | 0.034696334 | 0.972321884 | 0.987102298 | Rem1 | 16.10280817 | 0.057284541 | 0.073851309 | 0.77567401 | 0.437941489 | NA |
| Isg20 | 17.28502391 | -0.005781609 | 0.167831847 | -0.034448821 | 0.972519253 | 0.987237981 | Tex15 | 41.8687796 | 0.057295512 | 0.092785222 | 0.617506868 | 0.536900453 | 0.861769721 |
| Dapk1 | 5580.837138 | 0.003849414 | 0.115989933 | 0.033187479 | 0.973525083 | 0.987751553 | Tceal6 | 2325.855021 | 0.057305091 | 0.059695357 | 0.959958937 | 0.337075882 | 0.760113439 |
| Gm16998 | 19.32956427 | -0.005611498 | 0.167908013 | -0.033420074 | 0.973339602 | 0.987751553 | Cacng2 | 2206.437561 | 0.057307037 | 0.055273392 | 1.036792478 | 0.299832582 | 0.733704078 |
| Hhex | 73.58037285 | 0.004801122 | 0.144418124 | 0.033244597 | 0.973479534 | 0.987751553 | Gm17801 | 4.682773056 | 0.057314924 | 0.04942557 | 1.159620902 | 0.246203187 | NA |
| Map4 | 17264.18494 | -0.001629119 | 0.048301905 | -0.033727837 | 0.973094181 | 0.987751553 | Zscan22 | 434.4183922 | 0.057332497 | 0.067969905 | 0.843498257 | 0.39894984 | 0.797789838 |
| Padi2 | 1366.575876 | 0.004144529 | 0.124929946 | 0.033174827 | 0.973535172 | 0.987751553 | Rps6 | 8791.384766 | 0.057356058 | 0.064875779 | 0.884090467 | 0.376647379 | 0.783969863 |
| Rsl1d1 | 1767.970769 | -0.002245538 | 0.06678787 | -0.033621946 | 0.973178622 | 0.987751553 | 1110004E09Rik | 892.4639649 | 0.057361637 | 0.059782488 | 0.959505673 | 0.337304063 | 0.760113439 |
| Rwdd4a | 933.008118 | 0.002073982 | 0.06228831 | 0.033296493 | 0.973438151 | 0.987751553 | Gm19461 | 34.73284799 | 0.057371435 | 0.072094544 | 0.795780548 | 0.4261596 | 0.809519334 |
| Swap70 | 549.3918771 | -0.003226478 | 0.096667396 | -0.033377104 | 0.973373868 | 0.987751553 | Gal | 47.98962653 | 0.05738741 | 0.034086602 | 1.68357674 | NA | NA |

| | | | | | | |
|---|---|---|---|---|---|---|
| Ttl | 3719.099822 | -0.002314237 | 0.070327071 | -0.032906774 | 0.973748931 | 0.987903741 |
| Acot6 | 129.0098936 | -0.004092168 | 0.125852082 | -0.032515693 | 0.974060801 | 0.988155441 |
| Lrp2bp | 17.00275331 | -0.005298453 | 0.165548536 | -0.032005437 | 0.974467715 | 0.988503521 |
| Sil1 | 693.5147739 | -0.002260879 | 0.070864069 | -0.031904441 | 0.974548257 | 0.988520503 |
| Abcf1 | 3444.169422 | 0.001334993 | 0.042315921 | 0.031548246 | 0.974832317 | 0.98858135 |
| Aqp1 | 91.21261106 | 0.003280536 | 0.103718952 | 0.031629091 | 0.974767844 | 0.98858135 |
| Ccdc50 | 2262.571162 | -0.00262221 | 0.083220457 | -0.031509195 | 0.97486346 | 0.98858135 |
| Nsf | 38819.19215 | -0.001928306 | 0.060856526 | -0.031686106 | 0.974722375 | 0.98858135 |
| Cd1d1 | 32.54327098 | -0.005093142 | 0.163182207 | -0.031211382 | 0.975100963 | 0.988757481 |
| Mrpl37 | 1369.966361 | 0.001892326 | 0.061511563 | 0.030763753 | 0.975457948 | 0.988883873 |
| Prickle1 | 3462.182822 | 0.002618281 | 0.085087885 | 0.030771495 | 0.975451773 | 0.988883873 |
| Tango6 | 248.1212418 | 0.003314154 | 0.107951024 | 0.030700537 | 0.975508363 | 0.988883873 |
| Usp15 | 2450.505048 | 0.002000606 | 0.064661848 | 0.03093951 | 0.97531778 | 0.988883873 |
| Zfp128 | 357.9806866 | -0.00254551 | 0.083037526 | -0.03065494 | 0.975544727 | 0.988883873 |
| Celf3 | 5659.13181 | 0.00269921 | 0.088477276 | 0.030507382 | 0.975662406 | 0.988938462 |
| Mrps26 | 1189.58237 | -0.001858848 | 0.06167794 | -0.030137965 | 0.975957023 | 0.989172376 |
| Fadd | 111.7609819 | 0.004711326 | 0.158478851 | 0.029728423 | 0.976283644 | 0.989438695 |
| Kif21a | 8666.123326 | 0.001674923 | 0.057035947 | 0.029366102 | 0.976572608 | 0.989551211 |
| Mgst1 | 211.8973778 | -0.00373632 | 0.126859479 | -0.029452431 | 0.976503757 | 0.989551211 |
| Paip2b | 3515.682181 | 0.002360915 | 0.080442824 | 0.02934898 | 0.976586264 | 0.989551211 |
| Lyve1 | 47.28689215 | -0.004872152 | 0.166685596 | -0.029229591 | 0.976681481 | 0.989582976 |
| Coq5 | 1367.132236 | 0.001647332 | 0.056608635 | 0.029100364 | 0.976784545 | 0.98960427 |
| Kctd10 | 2514.235073 | 0.001762763 | 0.060694797 | 0.029043073 | 0.976830238 | 0.98960427 |
| Aass | 199.9081355 | 0.003573214 | 0.124632008 | 0.028670113 | 0.977127693 | 0.989687009 |
| Kcnd1 | 267.568035 | 0.003421789 | 0.119139533 | 0.02872085 | 0.977087227 | 0.989687009 |
| Rnaseh2a | 653.4693222 | 0.002201751 | 0.076929657 | 0.028620313 | 0.977167411 | 0.989687009 |
| Tmem26 | 13.05834903 | -0.004356366 | 0.15095382 | -0.028858932 | 0.976977099 | 0.989687009 |
| Bco2 | 154.7218001 | -0.003870107 | 0.13595729 | -0.028465606 | 0.9772908 | 0.989703857 |
| Metap2 | 2494.62294 | -0.001629565 | 0.057337225 | -0.028420721 | 0.977326598 | 0.989703857 |
| Pcdh7 | 4754.351195 | 0.002609328 | 0.092009986 | 0.028359187 | 0.977375675 | 0.989703857 |
| Commd10 | 749.7467277 | 0.002324648 | 0.083314259 | 0.027902162 | 0.977740184 | 0.989854996 |
| Rab15 | 12222.68083 | -0.002641388 | 0.094788243 | -0.027866202 | 0.977768865 | 0.989854996 |
| Rfc2 | 1038.923328 | 0.002585927 | 0.092567043 | 0.027935716 | 0.977713422 | 0.989854996 |
| Vps37b | 1198.68308 | -0.002144972 | 0.077014194 | -0.027851643 | 0.977780477 | 0.989854996 |
| 5330413P13Rik | 90.78374376 | -0.00372425 | 0.135079343 | -0.027570836 | 0.978004443 | 0.989914466 |
| Arhgef4 | 8839.262913 | 0.002123502 | 0.077112625 | 0.027537674 | 0.978030892 | 0.989914466 |
| Nt5c | 854.1199201 | 0.002257567 | 0.081977984 | 0.027538704 | 0.97803007 | 0.989914466 |
| Nup155 | 886.5601017 | -0.00211878 | 0.078034319 | -0.027151899 | 0.97833858 | 0.990096536 |
| Rag1 | 10.12924035 | -0.004139332 | 0.152280533 | -0.027182278 | 0.978314351 | 0.990096536 |
| 1600023N17Rik | 13.30493131 | -0.004499118 | 0.167417626 | -0.026873619 | 0.978560535 | 0.990191805 |
| Rasd2 | 8500.339938 | -0.002628553 | 0.097619176 | -0.02692661 | 0.97851827 | 0.990191805 |
| Rbmxl1 | 1487.789283 | -0.002066028 | 0.077445511 | -0.026677186 | 0.97871721 | 0.990285669 |
| Plscr4 | 132.2827043 | 0.003571602 | 0.134521333 | 0.026550447 | 0.978818297 | 0.990323279 |
| Aldh9a1 | 1303.543604 | -0.002770205 | 0.106124073 | -0.026103459 | 0.979174818 | 0.990326427 |
| Cab39 | 7065.999796 | -0.001858733 | 0.07042627 | -0.026392606 | 0.978944192 | 0.990326427 |
| Il27ra | 15.85672355 | -0.004121346 | 0.158771101 | -0.025957787 | 0.979291008 | 0.990326427 |
| Nadkd1 | 1323.673688 | -0.00229767 | 0.088947468 | -0.025831768 | 0.979391523 | 0.990326427 |
| Neurl4 | 5312.826621 | -0.001665767 | 0.063877725 | -0.026077429 | 0.97919558 | 0.990326427 |
| Rarres1 | 21.18027367 | 0.004405938 | 0.16772645 | 0.026268594 | 0.979043105 | 0.990326427 |
| Rnase6 | 12.00093778 | -0.004047316 | 0.156718844 | -0.025825329 | 0.979396659 | 0.990326427 |
| Tmem236 | 9.708465889 | -0.0039695 | 0.151812044 | -0.026147466 | 0.979139718 | 0.990326427 |
| Uhrf2 | 1747.800848 | 0.001914713 | 0.073902783 | 0.025908531 | 0.979330296 | 0.990326427 |
| Myh13 | 23.58856819 | 0.004298491 | 0.168086438 | 0.025573096 | 0.979597845 | 0.990465219 |
| Kcnk12 | 195.6128008 | 0.002987966 | 0.120127797 | 0.02487323 | 0.98015608 | 0.990964979 |
| Dennd2c | 38.62710557 | -0.004107979 | 0.165768984 | -0.024781349 | 0.980229368 | 0.990974411 |
| Fcrls | 1049.299538 | 0.00386032 | 0.156356063 | 0.024689287 | 0.9803028 | 0.990983988 |
| Atp2a3 | 434.5558618 | 0.002851754 | 0.117091652 | 0.024354886 | 0.980569533 | 0.991124297 |
| Smarcd1 | 7756.527655 | -0.001173761 | 0.048118719 | -0.024393033 | 0.980539106 | 0.991124297 |
| Tmem120a | 473.5902356 | -0.002058137 | 0.085616241 | -0.024039093 | 0.980821426 | 0.991314232 |
| Gm16386 | 25.04830294 | 0.003967832 | 0.166506675 | 0.02382987 | 0.980988314 | 0.991418234 |
| Kirrel2 | 352.2542595 | -0.003447814 | 0.145494654 | -0.023697184 | 0.981094153 | 0.991460527 |
| Piezo1 | 341.5240115 | 0.002136649 | 0.09128698 | 0.023405847 | 0.981326541 | 0.991566025 |
| Slc43a1 | 8.342027049 | -0.003471896 | 0.148112058 | -0.02344101 | 0.981298492 | 0.991566025 |
| Pemt | 62.88058871 | 0.00323902 | 0.138915275 | 0.023316512 | 0.981397801 | 0.991573363 |
| Wbp5 | 1523.56789 | -0.002090061 | 0.09028048 | -0.023150747 | 0.981530026 | 0.991642294 |
| Ccdc6 | 2516.882976 | -0.0025616 | 0.112144761 | -0.022841908 | 0.981776379 | 0.991657986 |
| Ddx51 | 802.8650365 | -0.002052875 | 0.094147373 | -0.021804905 | 0.982603581 | 0.991657986 |
| E130307A14Rik | 547.485058 | -0.001841821 | 0.083705085 | -0.02200369 | 0.982445012 | 0.991657986 |
| Egfem1 | 187.2674802 | -0.0026555 | 0.116763893 | -0.022742475 | 0.981855695 | 0.991657986 |
| Fam221b | 45.74367774 | -0.00339704 | 0.156416615 | -0.0217179 | 0.982672985 | 0.991657986 |
| Hdac4 | 1274.619603 | 0.00222812 | 0.098441051 | 0.022634052 | 0.981942181 | 0.991657986 |
| Mir212 | 13.63421025 | 0.00357123 | 0.160901731 | 0.022195097 | 0.982292329 | 0.991657986 |
| Mmp19 | 27.56480974 | 0.003743919 | 0.167245996 | 0.022385704 | 0.982140284 | 0.991657986 |
| Ntng2 | 642.6352997 | 0.002780037 | 0.127449164 | 0.021812908 | 0.982597197 | 0.991657986 |
| Plcb2 | 170.3644098 | 0.003024491 | 0.134237082 | 0.022530964 | 0.982024413 | 0.991657986 |
| Pogz | 2689.165033 | -0.001430227 | 0.064274557 | -0.022251843 | 0.982247063 | 0.991657986 |
| Slc9a2 | 238.8313981 | 0.002851884 | 0.125619273 | 0.022702602 | 0.981887501 | 0.991657986 |
| Tbx1 | 44.81016938 | -0.003622845 | 0.158988405 | -0.022786848 | 0.981820299 | 0.991657986 |
| Tmem178 | 2281.969434 | 0.002656471 | 0.120365645 | 0.022070013 | 0.982392107 | 0.991657986 |
| Tmem41a | 507.4769731 | 0.001802434 | 0.083111142 | 0.021687034 | 0.982697607 | 0.991657986 |
| Tnpo1 | 1923.527351 | 0.002175728 | 0.095662554 | 0.022743775 | 0.981854658 | 0.991657986 |
| Zdhhc18 | 1163.638478 | 0.001246604 | 0.056438259 | 0.022087926 | 0.982377818 | 0.991657986 |
| Zmynd19 | 661.0664706 | 0.001650568 | 0.074941837 | 0.022024649 | 0.982428293 | 0.991657986 |
| Rgn | 10.15687085 | 0.003405989 | 0.158195522 | 0.021530252 | 0.982822672 | 0.991719601 |
| C030034I22Rik | 82.23014544 | -0.003228001 | 0.150816747 | -0.021403462 | 0.982923812 | 0.991757068 |
| Epha4 | 7563.200335 | -0.001738958 | 0.08179665 | -0.021259522 | 0.983038634 | 0.991808334 |
| Magix | 19.19766925 | 0.003203462 | 0.152765067 | 0.020969857 | 0.983269701 | 0.991976869 |
| Kcnk1 | 3415.837048 | -0.001354949 | 0.065733732 | -0.020612697 | 0.983554612 | 0.992174822 |
| Pdzk1 | 10.90663115 | 0.003238757 | 0.157501546 | 0.020563335 | 0.983593989 | 0.992174822 |
| Usp3 | 620.5620409 | 0.00204291 | 0.099945057 | 0.020440331 | 0.983692111 | 0.992209203 |
| Tm9sf4 | 4023.717384 | -0.001223899 | 0.060619961 | -0.020189709 | 0.983892037 | 0.992281665 |
| Zfp672 | 1071.752725 | -0.001572108 | 0.07758748 | -0.020262397 | 0.983834053 | 0.992281665 |
| Nme4 | 61.49123493 | 0.003113063 | 0.155693431 | 0.019994825 | 0.984047501 | 0.992373859 |
| Ola1 | 3219.105232 | 0.001461761 | 0.07422509 | 0.019693629 | 0.984287773 | 0.992487471 |
| Tmem28 | 278.9204815 | -0.002536698 | 0.128812145 | -0.019693004 | 0.984288271 | 0.992487471 |
| Bai2 | 17780.92237 | 0.001519213 | 0.078021277 | 0.019471773 | 0.984464755 | 0.992536238 |
| Pacs2 | 5988.197758 | -0.001769458 | 0.090853946 | -0.019475854 | 0.984461499 | 0.992536238 |
| Fbxo41 | 6372.723859 | 0.001401646 | 0.072975559 | 0.019207056 | 0.984675929 | 0.99268455 |
| Pcdhgb5 | 183.0302292 | 0.002506155 | 0.131344967 | 0.019080711 | 0.984776719 | 0.992699952 |
| Sgcb | 2422.764933 | 0.000868053 | 0.045652275 | 0.019014452 | 0.984829576 | 0.992699952 |
| Tmem98 | 347.7060939 | 0.003177439 | 0.167701803 | 0.018946961 | 0.984883417 | 0.992699952 |
| Spata3 | 11.59395914 | -0.003014432 | 0.16026466 | -0.018809086 | 0.984993405 | 0.992746232 |
| Arfgef2 | 3861.951108 | -0.001701558 | 0.092112444 | -0.01847262 | 0.98526182 | 0.992823012 |
| Scp2 | 3809.812125 | 0.000836352 | 0.044963602 | 0.018600644 | 0.985159689 | 0.992823012 |
| Tceanc | 71.20026113 | 0.002751489 | 0.148445861 | 0.018535303 | 0.985211815 | 0.992823012 |
| 1110004F10Rik | 2800.142859 | -0.001281422 | 0.07079693 | -0.018099968 | 0.985559104 | 0.993057993 |
| Golt1a | 33.23396233 | -0.003003961 | 0.166963962 | -0.017991675 | 0.985645495 | 0.993080459 |
| Zfp668 | 480.4279658 | 0.001305532 | 0.073554825 | 0.017749104 | 0.985839007 | 0.993210844 |
| Adora2b | 116.3628664 | -0.002579305 | 0.147382411 | -0.017500768 | 0.98603712 | 0.993345848 |
| Atxn1l | 2460.056662 | -0.00152341 | 0.088108897 | -0.017290077 | 0.986205201 | 0.993385996 |
| Kcnq2 | 7046.441661 | 0.001090031 | 0.063000029 | 0.017302079 | 0.986195627 | 0.993385996 |
| AU040320 | 3338.256367 | 0.000963815 | 0.059182281 | 0.016285525 | 0.987006605 | 0.993681113 |
| Erbb2 | 136.7994706 | 0.002106182 | 0.127113835 | 0.016569259 | 0.986780249 | 0.993681113 |
| Frat2 | 193.9293929 | 0.001966083 | 0.119653138 | 0.016431524 | 0.986890131 | 0.993681113 |
| Lztr1 | 6350.258598 | 0.001455694 | 0.087417554 | 0.016652188 | 0.98671409 | 0.993681113 |
| Mir5133 | 10.68092198 | 0.002350102 | 0.14507416 | 0.016199312 | 0.987075385 | 0.993681113 |
| Olfml3 | 870.0806096 | 0.002692105 | 0.162431527 | 0.016573781 | 0.986776641 | 0.993681113 |
| Osbpl3 | 1459.937824 | -0.00141628 | 0.086745438 | -0.016326849 | 0.986973638 | 0.993681113 |
| Rcor2 | 472.2407876 | -0.001624162 | 0.099837223 | -0.016268097 | 0.987020509 | 0.993681113 |
| Tsg101 | 2392.474888 | -0.001490636 | 0.091301186 | -0.016326583 | 0.98697385 | 0.993681113 |
| Git1 | 16622.79819 | 0.001599394 | 0.099729211 | 0.016037368 | 0.98720458 | 0.993746606 |
| Bckdhb | 700.0788937 | -0.001075777 | 0.06806251 | -0.015805714 | 0.98738939 | 0.993868071 |
| Tgfbr3 | 566.5736921 | 0.001678441 | 0.107970966 | 0.015545301 | 0.987597144 | 0.994012612 |
| Ncbp2 | 1875.304417 | -0.000829732 | 0.053664783 | -0.015461393 | 0.987664085 | 0.994015416 |
| Fnta | 1717.65578 | 0.00090605 | 0.059794542 | 0.015152724 | 0.987910338 | 0.994198673 |
| Gen1 | 37.2093745 | 0.002336947 | 0.156403349 | 0.014941798 | 0.988078614 | 0.994243474 |
| Mus81 | 699.6740013 | -0.001272845 | 0.086036196 | -0.014794299 | 0.988196288 | 0.994243474 |
| Ptcd3 | 1808.491483 | -0.001434729 | 0.097103948 | -0.014775189 | 0.988211534 | 0.994243474 |
| Ube2d1 | 1539.912066 | 0.001080004 | 0.072437366 | 0.01490949 | 0.988104389 | 0.994243474 |
| Ankrd40 | 5652.590667 | 0.001049128 | 0.072294241 | 0.014511911 | 0.988421577 | 0.994390229 |
| Hcrtr1 | 93.29354779 | -0.002152407 | 0.149276866 | -0.01441889 | 0.988495789 | 0.994400322 |
| Gm14325 | 796.1449138 | 0.001454586 | 0.101456665 | 0.014337022 | 0.988561103 | 0.994401463 |
| Slc12a5 | 27128.00226 | 0.000767148 | 0.054094762 | 0.014181557 | 0.988685134 | 0.994461664 |
| Akap2 | 3477.887049 | -0.000885137 | 0.064873061 | -0.013644136 | 0.989113892 | 0.994505822 |
| Irx1 | 11.85203082 | 0.002281948 | 0.163506049 | 0.013956351 | 0.988864805 | 0.994505822 |
| Kif3c | 11925.10338 | 0.000961997 | 0.070952256 | 0.013558367 | 0.98918232 | 0.994505822 |
| Prpf8 | 10564.95747 | -0.001020504 | 0.075688785 | -0.0134829 | 0.989242528 | 0.994505822 |
| Ptger2 | 17.25324598 | 0.002280266 | 0.1673403 | 0.013626522 | 0.989127945 | 0.994505822 |
| Rbl1 | 89.18921211 | 0.001790145 | 0.132524714 | 0.013508011 | 0.989222494 | 0.994505822 |
| Rbpms2 | 95.1584948 | -0.00175344 | 0.127388021 | -0.013764562 | 0.989017816 | 0.994505822 |
| Rnf41 | 3283.745133 | 0.000947558 | 0.069936905 | 0.013548749 | 0.989189993 | 0.994505822 |
| Cdc27 | 2473.86381 | -0.000936024 | 0.069933918 | -0.01338441 | 0.989321105 | 0.994520288 |
| Pabpc5 | 54.1662827 | 0.002040712 | 0.156791984 | 0.013015409 | 0.989615499 | 0.99475169 |
| Myh15 | 60.90675207 | 0.002057085 | 0.164094185 | 0.012536002 | 0.989997979 | 0.995071599 |
| 4632434I11Rik | 25.98516669 | -0.002020526 | 0.167527571 | -0.012060855 | 0.990377063 | 0.995108658 |
| Auh | 1526.627737 | 0.0007585 | 0.06385699 | 0.011878097 | 0.990522873 | 0.995108658 |
| Ccdc87 | 131.6507694 | -0.001811512 | 0.152924596 | -0.011845783 | 0.990548654 | 0.995108658 |
| Lgi4 | 1434.531332 | -0.001551298 | 0.128349622 | -0.012086499 | 0.990356604 | 0.995108658 |
| Lrrc6 | 247.0874761 | -0.001407613 | 0.118687475 | -0.011859829 | 0.990537447 | 0.995108658 |
| Lyst | 3153.545997 | -0.001600931 | 0.132486392 | -0.012083741 | 0.990358804 | 0.995108658 |
| Pla2g2d | 16.24653744 | -0.001999872 | 0.163386675 | -0.012240114 | 0.990234046 | 0.995108658 |
| Terf2ip | 2328.154726 | 0.000799079 | 0.067053076 | 0.011917116 | 0.990491742 | 0.995108658 |
| Hes1 | 248.3937809 | 0.001428428 | 0.122410824 | 0.011669133 | 0.99068959 | 0.9951212 |
| Rap1gap2 | 12501.18537 | -0.001025869 | 0.087634889 | -0.011706173 | 0.990660039 | 0.9951212 |
| Acad12 | 79.61071545 | -0.001665999 | 0.145267747 | -0.011468472 | 0.990849684 | 0.99521749 |
| Akna | 261.9427128 | -0.001149356 | 0.102197599 | -0.011246406 | 0.991026855 | 0.995252438 |
| Dpp8 | 8896.008392 | -0.0005846 | 0.052240346 | -0.011190587 | 0.99107139 | 0.995252438 |
| Gpr34 | 366.3839922 | 0.001867766 | 0.16811907 | 0.01110978 | 0.99113586 | 0.995252438 |
| Phc3 | 2744.1408 | -0.000797074 | 0.072185538 | -0.011042011 | 0.991189929 | 0.995252438 |
| Snrpb2 | 652.4126242 | 0.000950131 | 0.086200805 | 0.011022304 | 0.991205652 | 0.995252438 |
| Mterf | 59.80421933 | 0.001637926 | 0.151958411 | 0.010778777 | 0.991399946 | 0.995318523 |
| Stbd1 | 46.14457439 | 0.001736364 | 0.16040784 | 0.010824684 | 0.991363321 | 0.995318523 |
| Avil | 10.92911177 | -0.001614692 | 0.15745005 | -0.010255268 | 0.991817623 | 0.995593735 |
| Mkrn2 | 1157.719147 | 0.000732318 | 0.072349977 | 0.010121885 | 0.991924042 | 0.995593735 |
| Papola | 4072.027534 | -0.000830241 | 0.082096145 | -0.010113037 | 0.991931101 | 0.995593735 |
| 2-Sep | 2153.618269 | -0.000864322 | 0.084675078 | -0.010207519 | 0.99185572 | 0.995593735 |
| Gna11 | 5692.958845 | 0.000823935 | 0.083820209 | 0.009829794 | 0.992157085 | 0.995756048 |
| B3gnt7 | 40.69134208 | 0.001586532 | 0.168070111 | 0.009439702 | 0.992468319 | 0.995905917 |
| Fam101a | 17.36029988 | -0.001582826 | 0.168095498 | -0.00941623 | 0.992487047 | 0.995905917 |
| Gm11837 | 25.83735658 | 0.001558223 | 0.165751743 | 0.009400944 | 0.992499242 | 0.995905917 |
| Mxra8 | 218.7329288 | -0.001058822 | 0.113689213 | -0.009313306 | 0.992569164 | 0.995911581 |
| Mycl1 | 526.5835398 | -0.001164943 | 0.128205654 | -0.009086522 | 0.992750105 | 0.995992825 |
| Omd | 35.65791024 | 0.001509393 | 0.166771188 | 0.00905068 | 0.9927787 | 0.995992825 |
| Ppp1r17 | 18.66143946 | -0.001434873 | 0.16153546 | -0.008882714 | 0.992912713 | 0.996062776 |
| Caml | 843.456301 | 0.000472014 | 0.058882645 | 0.008016185 | 0.993604079 | 0.996562765 |
| Pde5a | 514.8273888 | -0.000758537 | 0.092836744 | -0.008170651 | 0.993480836 | 0.996562765 |
| Zfp942 | 457.6363565 | 0.001074722 | 0.132902675 | 0.008086537 | 0.993547947 | 0.996562765 |
| Golph3 | 2958.782088 | -0.00066526 | 0.085508761 | -0.007780025 | 0.993792501 | 0.996622719 |
| Slc35a4 | 2041.410042 | -0.000466242 | 0.059812484 | -0.007795061 | 0.993780504 | 0.996622719 |
| Acp1 | 1994.917265 | 0.000388099 | 0.05408669 | 0.007175502 | 0.994274827 | 0.996807126 |
| Bmx | 61.26489296 | -0.00116046 | 0.157502282 | -0.007367892 | 0.994121326 | 0.996807126 |
| Fam43b | 849.2272051 | -0.000613415 | 0.089245032 | -0.006873381 | 0.994515879 | 0.996807126 |
| Gm5918 | 130.1198713 | -0.00087513 | 0.129396702 | -0.006763156 | 0.994603824 | 0.996807126 |
| Jag1 | 352.4823198 | 0.000649683 | 0.095392869 | 0.006810608 | 0.994565963 | 0.996807126 |
| Mfsd3 | 175.8922193 | -0.000745592 | 0.108627142 | -0.006863774 | 0.994523544 | 0.996807126 |
| Plrg1 | 1346.263033 | -0.000587879 | 0.0858402 | -0.006848532 | 0.994535705 | 0.996807126 |
| Ppapdc1b | 598.3588504 | 0.000737008 | 0.099627551 | 0.00739763 | 0.994097599 | 0.996807126 |
| Samd9l | 278.0205214 | 0.00095588 | 0.133756259 | 0.007146429 | 0.994298023 | 0.996807126 |
| Timm9 | 828.7492188 | 0.000530912 | 0.078732755 | 0.006743214 | 0.994619735 | 0.996807126 |
| E2f1 | 613.2955741 | -0.000684424 | 0.111908831 | -0.006115907 | 0.995120243 | 0.99724423 |
| Nptxr | 128.1616818 | -0.00063963 | 0.11451604 | -0.005585502 | 0.995543437 | 0.997539288 |
| Pcdhac1 | 135.5874904 | 0.000832668 | 0.148084182 | 0.005622935 | 0.995513571 | 0.997539288 |
| Tmem56 | 2489.703399 | -0.000477591 | 0.098601722 | -0.004843642 | 0.996135348 | 0.99806784 |
| Psmb3 | 1875.521334 | 0.000353337 | 0.076583755 | 0.004613735 | 0.996318785 | 0.998122543 |
| Rps19-ps3 | 46.27803518 | -0.000694092 | 0.148410179 | -0.004676849 | 0.996268428 | 0.998122543 |
| Gstk1 | 299.8220455 | 0.000451616 | 0.099756995 | 0.004527161 | 0.99638786 | 0.998127207 |
| Hkdc1 | 547.9840648 | -0.000668757 | 0.15261185 | -0.004382081 | 0.996503617 | 0.998178629 |
| Ninj2 | 69.13077651 | -0.000681194 | 0.165557552 | -0.004114547 | 0.996717076 | 0.998327906 |
| Acaa1b | 12.80983927 | 0.000559127 | 0.15975484 | 0.003499907 | 0.997207484 | 0.998496366 |
| Adamts13 | 33.87468898 | -0.000591131 | 0.162275631 | -0.003642758 | 0.997093506 | 0.998496366 |
| Eif3k | 2610.413977 | 0.000187263 | 0.051557469 | 0.00363213 | 0.997101986 | 0.998496366 |
| Nr4a3 | 376.2466614 | -0.000576655 | 0.163903347 | -0.003518261 | 0.99719284 | 0.998496366 |
| Sf3b3 | 3715.511069 | 0.000158634 | 0.044406627 | 0.003572294 | 0.997149728 | 0.998496366 |
| Tbl1x | 1552.544523 | -0.000271419 | 0.086524969 | -0.003136891 | 0.997497127 | 0.998678719 |
| Tysnd1 | 512.4210779 | 0.000347951 | 0.111878406 | 0.003110086 | 0.997518514 | 0.998678719 |
| Isca2 | 801.6239039 | 0.000213654 | 0.070766964 | 0.003019118 | 0.997591096 | 0.998686854 |
| Srp54c | 562.3280563 | -0.000228141 | 0.079550465 | -0.002867874 | 0.997711771 | 0.998743131 |
| Sim1 | 10.55397087 | 0.000308483 | 0.110836533 | 0.002783224 | 0.997779311 | 0.998746214 |
| AF251705 | 117.7600714 | 0.00035095 | 0.139450625 | 0.002516661 | 0.997991997 | 0.998765534 |
| Rsg1 | 36.59120384 | -0.000413558 | 0.157190115 | -0.002630941 | 0.997900815 | 0.998765534 |
| Uso1 | 3796.483903 | -0.000201437 | 0.07785354 | -0.002587378 | 0.997935573 | 0.998765534 |
| 2810408I11Rik | 12.93392093 | -0.000273083 | 0.166876541 | -0.001636437 | 0.998694313 | 0.999065354 |
| 6530402F18Rik | 617.9998662 | -0.000170737 | 0.103713633 | -0.00164623 | 0.998686499 | 0.999065354 |
| Arhgap36 | 156.6084681 | -0.000217707 | 0.140817256 | -0.001546024 | 0.998766452 | 0.999065354 |
| Clcf1 | 15.12012217 | 0.000262736 | 0.168121392 | 0.001562776 | 0.998753085 | 0.999065354 |
| Hist1h4h | 8.883851178 | -0.000285968 | 0.151345079 | -0.001889511 | 0.998492389 | 0.999065354 |
| Magi2 | 4732.345226 | 0.000100652 | 0.073438427 | 0.001370561 | 0.998906451 | 0.999065354 |
| Mgat5b | 4226.267606 | -0.000137794 | 0.09467706 | -0.001455412 | 0.99883875 | 0.999065354 |
| Plekhg6 | 10.90760365 | -0.000225978 | 0.157194825 | -0.001437567 | 0.998852988 | 0.999065354 |
| Smcr8 | 2495.934167 | -0.00016619 | 0.081525538 | -0.002038508 | 0.998373507 | 0.999065354 |
| Zfp511 | 451.450827 | 0.000104494 | 0.083447117 | 0.00125222 | 0.999000874 | 0.999065354 |
| Zfp677 | 63.29793703 | -0.000204645 | 0.156325169 | -0.0013091 | 0.99895549 | 0.999065354 |
| Sumf1 | 916.2325874 | 3.95E-05 | 0.088585964 | 0.000445386 | 0.999644634 | 0.999644634 |
| 0610005C13Rik | 0.787035502 | -0.008368886 | 0.067921826 | -0.123213505 | 0.901938032 | NA |
| 0610008F07Rik | 0 | NA | NA | NA | NA | NA |
| 0610009B14Rik | 0.400957162 | -0.004374601 | 0.047713229 | -0.091685289 | 0.926948086 | NA |
| 0610012H03Rik | 0 | NA | NA | NA | NA | NA |
| 0610031O16Rik | 0 | NA | NA | NA | NA | NA |
| 0610038L08Rik | 0.159967308 | 0.0240345 | 0.030870969 | 0.778547006 | 0.436246606 | NA |
| 0610039K10Rik | 1.397022734 | 0.013049902 | 0.067191501 | 0.194219532 | 0.84600399 | NA |
| 1100001G20Rik | 0.503646335 | -0.052098212 | 0.044076498 | -1.181995275 | 0.237207576 | NA |
| 1110025L11Rik | 0 | NA | NA | NA | NA | NA |
| 1110028F11Rik | 0.2777807 | 0.011208478 | 0.038523646 | 0.290950592 | 0.771089108 | NA |
| 1110028F18Rik | 0 | NA | NA | NA | NA | NA |
| 1110036E04Rik | 0.099009884 | 0.018205726 | 0.027896865 | 0.652608315 | 0.514008823 | NA |

| | | | | | | |
|---|---|---|---|---|---|---|
| Rmnd5a | 6754.819649 | 0.057398855 | 0.072416107 | 0.792625522 | 0.427996035 | 0.810464138 |
| Aprt | 728.6856793 | 0.057416559 | 0.070679893 | 0.81234643 | 0.416592876 | 0.806153084 |
| Ccdc67 | 16.39275236 | 0.057428719 | 0.065612001 | 0.875277666 | 0.381422843 | NA |
| 4930506M07Rik | 1390.457924 | 0.057435473 | 0.073205961 | 0.784573716 | 0.432703546 | 0.813460776 |
| Dst | 14511.95524 | 0.057450139 | 0.081117165 | 0.708236522 | 0.478798389 | 0.835936459 |
| Cpeb3 | 2875.080441 | 0.057470406 | 0.075846639 | 0.757718552 | 0.44861949 | 0.823263966 |
| Fam35a | 239.4821845 | 0.057476511 | 0.089787509 | 0.640139272 | 0.522082059 | 0.854113377 |
| Map6 | 5643.015192 | 0.057481101 | 0.040678879 | 1.413045333 | 0.157642395 | 0.608141173 |
| Dbndd1 | 613.2349542 | 0.057508446 | 0.069521072 | 0.827208849 | 0.408118697 | 0.801055189 |
| Ptgds | 21239.52221 | 0.057520432 | 0.090529004 | 0.635381252 | 0.525179792 | 0.855722004 |
| Anapc2 | 3906.988553 | 0.057531463 | 0.052254987 | 1.100975563 | 0.270907293 | 0.712640228 |
| Tut1 | 1004.334645 | 0.05754772 | 0.057481985 | 1.001143582 | 0.316757398 | 0.746103123 |
| Mta3 | 2303.530504 | 0.057563726 | 0.062094084 | 0.927040425 | 0.353905546 | 0.771317624 |
| Alk | 328.6680858 | 0.057582737 | 0.087130896 | 0.660876219 | 0.508691698 | 0.850050333 |
| Rps3 | 8622.623395 | 0.057598059 | 0.04708769 | 1.223208416 | 0.221250988 | 0.672757795 |
| 2310057M21Rik | 631.1492045 | 0.057609352 | 0.067973659 | 0.84752466 | 0.396702748 | 0.796916625 |
| Fam49b | 6191.594442 | 0.057617991 | 0.071038594 | 0.811080123 | 0.41731966 | 0.806153084 |
| Gsk3b | 8614.97969 | 0.057632525 | 0.060206336 | 0.95725016 | 0.338441005 | 0.760991092 |
| Fktn | 1420.590095 | 0.057646888 | 0.079967644 | 0.720877661 | 0.470984788 | 0.833245343 |
| Pcdhac1 | 170.611437 | 0.057657724 | 0.089117241 | 0.64698731 | 0.517640156 | 0.853297835 |
| Usp4 | 4348.846473 | 0.057670057 | 0.04183007 | 1.378674642 | 0.167995091 | 0.617953076 |
| Def8 | 2557.571065 | 0.057682169 | 0.047090632 | 1.224918134 | 0.220606067 | 0.672757795 |
| Vwa2 | 1.923373596 | 0.057692003 | 0.031861434 | 1.810715854 | 0.070184847 | NA |
| Arhgef4 | 9928.076527 | 0.057712827 | 0.055517526 | 1.039542484 | 0.29855251 | 0.732994423 |
| Phf17 | 1699.710454 | 0.057723861 | 0.058063816 | 0.994145154 | 0.320152205 | 0.750154748 |
| Asb4 | 58.97699677 | 0.057732217 | 0.040042318 | 1.441780086 | NA | NA |
| Gucy1b3 | 7564.613198 | 0.057753627 | 0.070582213 | 0.818246187 | 0.413216627 | 0.803854924 |
| Thap7 | 654.3745925 | 0.057756836 | 0.075921574 | 0.760743401 | 0.446810347 | 0.822210607 |
| Hap1 | 9885.436271 | 0.057764397 | 0.094830965 | 0.609130122 | 0.542438193 | 0.864833084 |
| Spata6 | 169.6138203 | 0.057773893 | 0.094379853 | 0.612142218 | 0.540443667 | 0.863734703 |
| Twf2 | 1636.708815 | 0.057777561 | 0.058460489 | 0.988318131 | 0.322996868 | 0.752792382 |
| 2610028E06Rik | 13.97994078 | 0.05777913 | 0.071754034 | 0.805238775 | 0.420681905 | NA |
| Ogfrl1 | 4750.133711 | 0.057791795 | 0.05939311 | 0.973038711 | 0.330534063 | 0.756141871 |
| Dffa | 1828.956757 | 0.057794736 | 0.052755325 | 1.095524204 | 0.273287048 | 0.714887026 |
| Ppdpf | 1267.634505 | 0.057819098 | 0.063752921 | 0.906924688 | 0.364446625 | 0.777237064 |
| Ube2q2 | 2864.024775 | 0.057819162 | 0.058766301 | 0.983882952 | 0.325173067 | 0.75293408 |
| Rac1 | 12289.47755 | 0.057830255 | 0.040619938 | 1.423691365 | 0.154535834 | 0.604361735 |
| Igfbp4 | 3271.215952 | 0.057855683 | 0.078662234 | 0.73549505 | 0.462038054 | 0.830604784 |
| Myc | 140.1656095 | 0.05785674 | 0.09265005 | 0.624465287 | 0.53232206 | 0.858328942 |
| Zfp87 | 657.5178677 | 0.057870525 | 0.067015471 | 0.863539782 | 0.387840756 | 0.791889115 |
| 2810408M09Rik | 503.2475503 | 0.057873246 | 0.071164777 | 0.813228798 | 0.416086891 | 0.805984555 |
| Lrtm2 | 3971.714544 | 0.057894257 | 0.055697376 | 1.039443179 | 0.298598672 | 0.732994423 |
| Ccny | 5140.962596 | 0.057895364 | 0.068023206 | 0.851111958 | 0.394707164 | 0.795437883 |
| Myt1l | 11277.89934 | 0.057925897 | 0.063047988 | 0.918758852 | 0.358221719 | 0.774201531 |
| Oprl1 | 2390.430432 | 0.0579615 | 0.060971118 | 0.95063863 | 0.341787851 | 0.763229874 |
| Pdap1 | 3759.029649 | 0.05799687 | 0.054694436 | 1.060379701 | 0.288971893 | 0.724444298 |
| Pcmt1 | 5409.20742 | 0.058003324 | 0.057678854 | 1.005625459 | 0.314595772 | 0.745993052 |
| Wdr5b | 187.8663093 | 0.058004858 | 0.085714162 | 0.676724325 | 0.49858088 | 0.846052342 |
| Ppp1cb | 8133.580152 | 0.058009551 | 0.076855279 | 0.754789413 | 0.450375347 | 0.823365451 |
| Dad1 | 2049.219719 | 0.058026593 | 0.048981283 | 1.184668696 | 0.236148464 | 0.687039131 |
| Pnrc2 | 1271.789725 | 0.058027044 | 0.068937159 | 0.84173826 | 0.399934478 | 0.798662339 |
| BC005624 | 2975.489165 | 0.058030661 | 0.049862949 | 1.163803245 | 0.244503757 | 0.694361988 |
| Rbx1 | 3532.351187 | 0.058038091 | 0.041593222 | 1.39537378 | 0.162903154 | 0.614351408 |
| Myom2 | 7.308155604 | 0.058056146 | 0.058178974 | 0.997888797 | 0.318333285 | NA |
| Gm14023 | 9.967836739 | 0.058067441 | 0.066496197 | 0.873244542 | 0.382529802 | NA |
| Gareml | 636.2626547 | 0.05807869 | 0.076435696 | 0.759837258 | 0.447351869 | 0.822458097 |
| Taf5 | 259.3027581 | 0.058080001 | 0.091124241 | 0.637371571 | 0.523882842 | 0.855218546 |
| Lsm6 | 1284.17746 | 0.058109878 | 0.062528723 | 0.929330956 | 0.352717597 | 0.770519168 |
| Tmem242 | 1299.104209 | 0.058120457 | 0.053509046 | 1.086180026 | 0.277399352 | 0.718015501 |
| Kcnh5 | 906.9526816 | 0.058133602 | 0.083736656 | 0.694243181 | 0.487529717 | 0.839801621 |
| Cep19 | 2411.332976 | 0.058149924 | 0.062401385 | 0.931869124 | 0.351404166 | 0.769244462 |
| Mmp16 | 862.3429657 | 0.058156869 | 0.07086058 | 0.820722455 | 0.411804377 | 0.802795661 |
| Uso1 | 4057.965747 | 0.058171661 | 0.05892873 | 0.987152806 | 0.323567734 | 0.752925264 |
| Slc39a3 | 3381.566683 | 0.058182701 | 0.058017405 | 1.002849079 | 0.315933685 | 0.745993052 |
| 2410015M20Rik | 1267.844495 | 0.05818645 | 0.064263886 | 0.905429992 | 0.365237633 | 0.777735733 |
| Ccnh | 1317.370645 | 0.05819096 | 0.071215253 | 0.817113719 | 0.413863444 | 0.804285711 |
| Ndufa8 | 3069.848201 | 0.058214269 | 0.046683787 | 1.246991139 | 0.212400747 | 0.666270994 |
| Homer1 | 6575.036986 | 0.058216802 | 0.089317742 | 0.651794379 | 0.514533828 | 0.853195313 |
| Atg12 | 2611.715565 | 0.058222841 | 0.063136823 | 0.92216932 | 0.356440263 | 0.773064269 |
| Aars | 7940.137466 | 0.058236826 | 0.043107708 | 1.35096086 | 0.176707971 | 0.629083076 |
| Zfp804b | 57.20184686 | 0.058240558 | 0.090649181 | 0.642483004 | 0.520559621 | 0.8535556 |
| Tpd52l2 | 1911.520936 | 0.058256131 | 0.048796968 | 1.193847365 | 0.23253768 | 0.684327865 |
| Nme7 | 667.6579572 | 0.058270003 | 0.0644022 | 0.904782811 | 0.36558046 | 0.777735733 |
| Slc25a29 | 474.9895771 | 0.058276818 | 0.072750253 | 0.80105314 | 0.423100088 | 0.808999423 |
| Dcbld1 | 392.7101598 | 0.058280613 | 0.074697463 | 0.780222123 | 0.435260142 | 0.815502235 |
| Eif4a2 | 24646.79695 | 0.058307401 | 0.059512963 | 0.979742865 | 0.327213061 | 0.754100734 |
| Zfp13 | 639.4795662 | 0.058316355 | 0.070505503 | 0.827117778 | 0.408170309 | 0.801055189 |
| Erp29 | 2474.579857 | 0.058320386 | 0.059995392 | 0.972081096 | 0.331010206 | 0.756212998 |
| Fgf22 | 37.18324962 | 0.058324324 | 0.085494741 | 0.682197796 | 0.495113888 | 0.844332512 |
| Plin3 | 704.4615415 | 0.058330895 | 0.077894262 | 0.748847128 | 0.453949351 | 0.825985606 |
| Ephx1 | 1207.60371 | 0.058336277 | 0.070470061 | 0.82781648 | 0.40777444 | 0.801055189 |
| Calb1 | 5032.35623 | 0.058345284 | 0.081146177 | 0.719014584 | 0.472131934 | 0.833441697 |
| Prep | 989.3502471 | 0.058350023 | 0.058227759 | 1.002099741 | 0.316295423 | 0.745993052 |
| G630093K05Rik | 9.26186484 | 0.058351986 | 0.059490101 | 0.980868831 | 0.32665743 | NA |
| Mien1 | 1744.956767 | 0.058355401 | 0.052939648 | 1.102300528 | 0.27033104 | 0.71216373 |
| Rpl11 | 6379.020324 | 0.058377504 | 0.06019906 | 0.969741126 | 0.332175546 | 0.757166892 |
| Sipa1l2 | 2724.398469 | 0.058386734 | 0.066060071 | 0.883843043 | 0.376780947 | 0.783969863 |
| Kat2b | 1183.077278 | 0.058391467 | 0.072696364 | 0.80322403 | 0.421845259 | 0.808695226 |
| Tmem50a | 2992.071349 | 0.058409006 | 0.044949476 | 1.29943685 | 0.193794052 | 0.649958942 |
| Slc19a3 | 156.7515836 | 0.058410321 | 0.094329884 | 0.619213322 | 0.535775837 | 0.860926435 |
| Mlec | 7117.103988 | 0.05841764 | 0.056774739 | 1.028937177 | 0.303509197 | 0.736936169 |
| Smim14 | 4212.661797 | 0.058425312 | 0.04936295 | 1.183586307 | 0.236576863 | 0.687308962 |
| Steap3 | 223.414919 | 0.05845278 | 0.090884473 | 0.643154743 | 0.520123696 | 0.853297835 |
| Ppp3r2 | 5.322600447 | 0.058454225 | 0.052093884 | 1.122093808 | 0.261822557 | NA |
| Ppm1b | 4285.478556 | 0.058465 | 0.058340925 | 1.002126731 | 0.316282389 | 0.745993052 |
| Pex5 | 2346.030101 | 0.058486063 | 0.052402581 | 1.116091256 | 0.264383074 | 0.707261241 |
| Terf2ip | 2556.706724 | 0.058497617 | 0.054431006 | 1.074711312 | 0.282503997 | 0.720824357 |
| Scn9a | 127.1963583 | 0.058498307 | 0.06178479 | 0.94680756 | 0.343736849 | 0.763762832 |
| Lrrc47 | 2637.521065 | 0.058501772 | 0.053223501 | 1.099171831 | 0.271693123 | 0.713790099 |
| Eif3h | 4317.106618 | 0.058522504 | 0.055130605 | 1.061524779 | 0.288451478 | 0.724444298 |
| Btf3l4 | 3095.723011 | 0.058535401 | 0.060599404 | 0.965940205 | 0.334074102 | 0.758212399 |
| F2rl2 | 88.43269118 | 0.05853762 | 0.094564367 | 0.619024083 | 0.535900494 | 0.860988 |
| Spc24 | 67.91946935 | 0.058584437 | 0.095034562 | 0.616454014 | 0.537594914 | 0.862027124 |
| Tmem43 | 1770.52713 | 0.058595205 | 0.054389236 | 1.077330895 | 0.281332465 | 0.720080692 |
| 4932438H23Rik | 51.30655632 | 0.058607026 | 0.088724715 | 0.660549044 | 0.508901556 | 0.850080958 |
| Cacng7 | 3114.178893 | 0.058657581 | 0.056292135 | 1.042020895 | 0.297401992 | 0.732415975 |
| 1110018G07Rik | 8007.702185 | 0.058661928 | 0.055351589 | 1.059805659 | 0.289233022 | 0.724444298 |
| Pex7 | 845.7996189 | 0.05867604 | 0.061333629 | 0.956669959 | 0.338733865 | 0.761006691 |
| Serpinb6b | 54.99398414 | 0.058676519 | 0.093505654 | 0.627518403 | 0.530319477 | 0.857915133 |
| Cby1 | 1215.407912 | 0.058687737 | 0.060619857 | 0.968127286 | 0.332980804 | 0.75765651 |
| Olfm2 | 4335.832509 | 0.058698079 | 0.070134273 | 0.836938584 | 0.402627091 | 0.799748547 |
| Wdtc1 | 3367.580455 | 0.058699192 | 0.056852043 | 1.032490463 | 0.301842413 | 0.735974432 |
| Tnip1 | 1238.210685 | 0.058718802 | 0.059598481 | 0.985239911 | 0.324506241 | 0.75293408 |
| Frat2 | 215.7218289 | 0.058748008 | 0.087403284 | 0.672148749 | 0.501489005 | 0.846601651 |
| Rab6a | 21026.24767 | 0.058809835 | 0.062946115 | 0.934288558 | 0.350155065 | 0.768667813 |
| Lrg1 | 46.51723309 | 0.058818942 | 0.093024651 | 0.632294135 | 0.527194695 | 0.856219828 |
| Dhx36 | 3413.794205 | 0.058822234 | 0.062320712 | 0.943863306 | 0.345239505 | 0.764823708 |
| Fbxo11 | 5615.694671 | 0.058825857 | 0.066834524 | 0.880171697 | 0.378766303 | 0.785380516 |
| Kctd10 | 2756.050876 | 0.058835035 | 0.048157118 | 1.221730815 | 0.221809439 | 0.672987912 |
| Poldip2 | 3786.933966 | 0.058838632 | 0.04957699 | 1.186813304 | 0.23530127 | 0.686276191 |
| Endog | 280.7444189 | 0.058874524 | 0.08049315 | 0.731422787 | 0.464520951 | 0.83159896 |
| Ubac1 | 1611.599458 | 0.058881549 | 0.0497875 | 1.182657269 | 0.236945006 | 0.687684946 |
| Adk | 2149.417007 | 0.0588823 | 0.054252464 | 1.085338732 | 0.277771658 | 0.718083484 |
| Zfp597 | 939.1581275 | 0.058937391 | 0.079731867 | 0.739194924 | 0.459788643 | 0.828852051 |
| Fbxo3 | 3029.43806 | 0.05899242 | 0.058748073 | 1.00415923 | 0.31530187 | 0.745993052 |
| Sap18 | 1249.697788 | 0.058999638 | 0.062944627 | 0.937326037 | 0.348590874 | 0.767449383 |
| Hipk3 | 4041.295141 | 0.059004096 | 0.05938928 | 0.99351425 | 0.320459409 | 0.750321306 |
| Calm3 | 59475.43716 | 0.059014232 | 0.046433912 | 1.270929579 | 0.203753702 | 0.657622144 |
| Pdia4 | 1715.794411 | 0.059046112 | 0.064963285 | 0.908915119 | 0.363394933 | 0.776821522 |
| Myl6b | 650.6952316 | 0.059059137 | 0.062690817 | 0.942069987 | 0.346156809 | 0.765105571 |
| Fam19a5 | 4645.213229 | 0.059063555 | 0.048128116 | 1.227215205 | 0.219741716 | 0.672421385 |
| Nxt2 | 1089.310603 | 0.059063688 | 0.061977378 | 0.952987856 | 0.340596224 | 0.762546123 |
| Etv6 | 945.4848535 | 0.059073249 | 0.074289182 | 0.795179699 | 0.426508979 | 0.809764424 |
| Nploc4 | 3402.602247 | 0.059075888 | 0.042428063 | 1.39237769 | 0.163808057 | 0.614351408 |
| Ralgapb | 7376.947074 | 0.059108765 | 0.059276482 | 0.997170597 | 0.31868171 | 0.74851037 |
| Gm20300 | 1708.167474 | 0.059115401 | 0.072745045 | 0.812638181 | 0.416425534 | 0.806153084 |
| Vti1b | 4881.474513 | 0.059123891 | 0.049772594 | 1.187880441 | 0.234880516 | 0.686217695 |
| Icmt | 3910.656538 | 0.059125892 | 0.044304723 | 1.334527964 | 0.182030889 | 0.635245711 |
| Rnf17 | 33.31415361 | 0.059126125 | 0.081819969 | 0.722636854 | 0.46990302 | 0.833245343 |
| Rnls | 30.14557899 | 0.059129574 | 0.085654542 | 0.690326197 | 0.489989077 | 0.841321671 |
| AI837181 | 2137.540521 | 0.05914596 | 0.052456152 | 1.127531422 | 0.259517863 | 0.703893404 |
| Ndnl2 | 662.5974457 | 0.059153089 | 0.06294612 | 0.939741632 | 0.347350105 | 0.766038372 |
| C1d | 922.9927176 | 0.059153201 | 0.069930531 | 0.845885203 | 0.397616788 | 0.797649698 |
| Plod1 | 910.0275586 | 0.059155782 | 0.074567223 | 0.793321513 | 0.427590526 | 0.810334558 |
| Sulf1 | 1106.627434 | 0.059167581 | 0.093010735 | 0.636137118 | 0.524687055 | 0.855487854 |
| Vmn2r85 | 9.407297401 | 0.059184546 | 0.057486122 | 1.029544948 | 0.30322367 | NA |
| Ttll1 | 1749.389657 | 0.059189274 | 0.057465174 | 1.030002521 | 0.303008822 | 0.73653048 |
| Npm1 | 4909.19626 | 0.059203382 | 0.05961199 | 0.993145547 | 0.32063903 | 0.750352874 |
| Tox | 667.1458423 | 0.059227542 | 0.082160646 | 0.720874835 | 0.470986527 | 0.833245343 |
| Zbtb22 | 1521.383231 | 0.059229532 | 0.074870866 | 0.791089172 | 0.428891957 | 0.810705511 |
| Fam210a | 1871.731944 | 0.059233695 | 0.063448091 | 0.93357725 | 0.350522005 | 0.768667813 |
| Cox6b1 | 6048.30261 | 0.059236074 | 0.05769778 | 1.02666123 | 0.304580011 | 0.737453291 |
| Plekhd1 | 193.7042958 | 0.059261351 | 0.093929175 | 0.630915272 | 0.528095925 | 0.856965327 |
| Zbtb38 | 1510.055168 | 0.059277272 | 0.068914112 | 0.86016159 | 0.389699974 | 0.792715562 |
| Smim15 | 1152.427635 | 0.059308038 | 0.06703815 | 0.884690851 | 0.376323391 | 0.783947144 |
| Napb | 24865.46334 | 0.059314715 | 0.060828513 | 0.975113672 | 0.329503878 | 0.756141871 |
| Acat2 | 1274.60324 | 0.059333386 | 0.060417626 | 0.982054244 | 0.326073125 | 0.753049596 |
| Mtrf1 | 296.6505976 | 0.059334654 | 0.074945984 | 0.791698917 | 0.428536254 | 0.810486244 |
| Spata5 | 521.82967 | 0.059341208 | 0.077712886 | 0.76359547 | 0.445108349 | 0.821493054 |
| Ddx1 | 6232.239062 | 0.059344198 | 0.055548252 | 1.06833601 | 0.28536897 | 0.722148665 |
| Dnajc3 | 2560.571912 | 0.059345308 | 0.055866985 | 1.062260811 | 0.2881173 | 0.724103151 |
| Rnaseh2b | 296.9996688 | 0.059358722 | 0.084899713 | 0.699162813 | 0.484450288 | 0.838713365 |
| Hdac4 | 1353.182725 | 0.059361006 | 0.07516623 | 0.789729723 | 0.429685629 | 0.811442206 |
| Tiam2 | 2792.513929 | 0.059408979 | 0.077146947 | 0.770075564 | 0.44125507 | 0.818962636 |
| Comt | 3679.304222 | 0.059419642 | 0.046759224 | 1.270757659 | 0.203814876 | 0.657622144 |
| Snx3 | 5302.910604 | 0.059430981 | 0.051948893 | 1.144027856 | 0.252612085 | 0.700378199 |
| Zc3h10 | 629.1851297 | 0.059433147 | 0.065541694 | 0.906799076 | 0.364513058 | 0.777237064 |
| Pink1 | 16539.76894 | 0.059457641 | 0.041509804 | 1.432375856 | 0.152036284 | 0.600023818 |
| Dcaf6 | 4088.089962 | 0.059463168 | 0.058005982 | 1.025121298 | 0.305305957 | 0.737793102 |
| Asb11 | 167.177861 | 0.05948811 | 0.085450222 | 0.696172675 | 0.486320698 | 0.839801621 |
| Gnaz | 2626.301107 | 0.059513608 | 0.055501023 | 1.0722975 | 0.283586429 | 0.72105312 |
| Dnaja2 | 7142.598067 | 0.059517408 | 0.061926661 | 0.961095059 | 0.336504374 | 0.75970798 |
| Xiap | 3706.010268 | 0.059520241 | 0.066719153 | 0.892101264 | 0.372338658 | 0.781446026 |
| Uqcrfs1 | 3442.393437 | 0.059527438 | 0.048835511 | 1.218937559 | 0.222867893 | 0.67403269 |
| Apip | 356.9861328 | 0.059550552 | 0.082225165 | 0.724237626 | 0.468919862 | 0.832603584 |
| Trappc3 | 2255.91596 | 0.059603691 | 0.05078639 | 1.173615418 | 0.240549103 | 0.691787026 |
| Rnf11 | 6596.111337 | 0.059620623 | 0.065478592 | 0.910536116 | 0.362539844 | 0.776449324 |
| Rrn3 | 2879.715981 | 0.05962373 | 0.05477705 | 1.088480129 | 0.276383204 | 0.717475105 |
| Mapk10 | 20050.86103 | 0.059640019 | 0.046632625 | 1.27893333 | 0.200920534 | 0.654716629 |
| Rdx | 3146.41282 | 0.059645359 | 0.058229534 | 1.024314542 | 0.305686731 | 0.738091018 |
| Wbp1l | 1878.191305 | 0.059659229 | 0.052745625 | 1.131074454 | 0.258023755 | 0.703616072 |
| Epdr1 | 5575.033613 | 0.059667191 | 0.052712668 | 1.131932661 | 0.257662746 | 0.703403317 |
| Agk | 1736.406618 | 0.059684111 | 0.053199777 | 1.121886492 | 0.261910705 | 0.704317239 |
| Eef1g | 11097.45697 | 0.059686884 | 0.056872512 | 1.049485631 | 0.293954665 | 0.729288721 |
| Creb3l1 | 110.1748762 | 0.05969399 | 0.095086295 | 0.627787524 | 0.53014314 | 0.857915133 |
| Bmp2 | 124.8062992 | 0.059714129 | 0.091511888 | 0.652528649 | 0.514060198 | 0.853195313 |
| Trappc8 | 2843.956723 | 0.059728176 | 0.066693386 | 0.895563711 | 0.370485824 | 0.780632011 |
| Smad7 | 808.5485568 | 0.05973465 | 0.077479414 | 0.770974474 | 0.440722062 | 0.818660955 |
| Ciao1 | 2182.309198 | 0.059743927 | 0.056711041 | 1.053479639 | 0.292121221 | 0.727847831 |
| Glyr1 | 10006.71581 | 0.059747606 | 0.042992252 | 1.389729632 | 0.164610993 | 0.614688762 |
| Atp5sl | 1701.822389 | 0.059752565 | 0.054866864 | 1.089046487 | 0.276133386 | 0.717475105 |
| Slc25a48 | 6.135086484 | 0.059756556 | 0.049359377 | 1.210642432 | 0.226032475 | NA |
| Cadps | 12491.5789 | 0.059762567 | 0.054040826 | 1.105878122 | 0.268779276 | 0.710903001 |
| AI467606 | 75.53129156 | 0.059772006 | 0.093963969 | 0.636116228 | 0.524700669 | 0.855487854 |
| Eif4a1 | 10970.88632 | 0.059783351 | 0.045121253 | 1.324948806 | 0.185188132 | 0.637780733 |
| Llph | 997.4243182 | 0.059786258 | 0.068344737 | 0.874774855 | 0.381696421 | 0.786663048 |
| Myh1 | 29.11644539 | 0.059800306 | 0.090113275 | 0.663612611 | 0.506938284 | 0.848958235 |
| Slc25a1 | 1350.115354 | 0.059801802 | 0.067788088 | 0.882187484 | 0.377675429 | 0.784178571 |
| Stx12 | 4962.126657 | 0.059820843 | 0.060273049 | 0.992497386 | 0.320954954 | 0.750352874 |
| Meis1 | 266.601734 | 0.059829336 | 0.092672824 | 0.645597419 | 0.51854011 | 0.853297835 |
| Snap29 | 1293.699706 | 0.059835645 | 0.067575886 | 0.885458546 | 0.375909368 | 0.783947144 |
| Lyzl4 | 21.02682536 | 0.059877229 | 0.077661834 | 0.770999425 | 0.440707272 | NA |
| Gpx1 | 1942.661347 | 0.059881847 | 0.053971842 | 1.109501651 | 0.267213833 | 0.709971477 |
| Eif2b5 | 2359.64404 | 0.059896429 | 0.047263865 | 1.267277436 | 0.205056106 | 0.658534108 |
| Asb1 | 2696.664444 | 0.059902127 | 0.060845913 | 0.984488917 | 0.324875178 | 0.75293408 |
| P4ha2 | 450.0449871 | 0.05991677 | 0.084536357 | 0.708769249 | 0.478467684 | 0.835936459 |
| Tbc1d9 | 5240.074349 | 0.059924678 | 0.048912749 | 1.225134119 | 0.220524692 | 0.672757795 |
| Slc39a14 | 1658.432656 | 0.059925255 | 0.057621575 | 1.039979455 | 0.298349446 | 0.732994423 |
| Zfp592 | 1673.355306 | 0.059940421 | 0.06075682 | 0.986562843 | 0.323856993 | 0.752925264 |
| Tenm4 | 3283.136285 | 0.059942384 | 0.07320571 | 0.818821163 | 0.412888455 | 0.803781966 |
| Sv2a | 21942.79796 | 0.059942875 | 0.036525461 | 1.641125748 | 0.100771315 | 0.521542547 |
| Mrps34 | 994.6701151 | 0.059975014 | 0.059195428 | 1.013169702 | 0.31097911 | 0.743730778 |
| Kcnn2 | 1195.943831 | 0.059981008 | 0.079402113 | 0.755408205 | 0.450004091 | 0.823263966 |
| Mff | 6757.570028 | 0.059990959 | 0.051318144 | 1.169000957 | 0.242403246 | 0.692772033 |
| Ube3a | 1981.932883 | 0.059991553 | 0.07313666 | 0.820266514 | 0.412064192 | 0.802795661 |
| Aasdhppt | 749.0952049 | 0.060028023 | 0.073524065 | 0.816440483 | 0.414248251 | 0.804541293 |
| Rqcd1 | 1361.020674 | 0.060039065 | 0.059722005 | 1.005308924 | 0.314748119 | 0.745993052 |
| Sh3bgrl2 | 1297.067861 | 0.060048372 | 0.068873544 | 0.871864112 | 0.383282514 | 0.787602461 |
| 4930432K21Rik | 135.5021809 | 0.060053582 | 0.091982604 | 0.652879779 | 0.513833786 | 0.853195313 |
| Cox10 | 1120.894586 | 0.060053935 | 0.060513303 | 0.992408807 | 0.320998145 | 0.750352874 |
| Tmem41b | 1576.260596 | 0.060089498 | 0.062316997 | 0.964255346 | 0.334917922 | 0.758263364 |
| Mrpl54 | 488.5930813 | 0.060094994 | 0.084393205 | 0.712083324 | 0.47641318 | 0.835195112 |
| Mrps35 | 568.8587691 | 0.060096884 | 0.072372302 | 0.830385143 | 0.406321063 | 0.800759397 |
| Eci1 | 464.9017671 | 0.060097709 | 0.082907584 | 0.724875893 | 0.468528171 | 0.832446623 |
| Ogfod1 | 6590.582051 | 0.060110051 | 0.056626881 | 1.061510897 | 0.288457783 | 0.724444298 |
| Draxin | 26.49340856 | 0.060140626 | 0.087314454 | 0.688782018 | 0.490960453 | 0.841866734 |
| Spa17 | 75.88814891 | 0.060143428 | 0.095017284 | 0.632973553 | 0.526750914 | 0.85599981 |
| Mkln1 | 2524.590958 | 0.060146559 | 0.077765116 | 0.773438807 | NA | NA |
| Got2 | 12495.49033 | 0.060152549 | 0.046639432 | 1.289735886 | 0.197142375 | 0.653065027 |
| Usp30 | 1787.147395 | 0.060186846 | 0.049053398 | 1.226965895 | 0.219835409 | 0.672421385 |
| Sfrp2 | 60.84432214 | 0.060187735 | 0.094305998 | 0.63821746 | 0.523332133 | 0.854628731 |
| C1qtnf9 | 16.28389737 | 0.06021252 | 0.077083528 | 0.781133424 | 0.434724025 | NA |
| Syde2 | 145.6925192 | 0.060223529 | 0.090803152 | 0.663231703 | 0.50718217 | 0.848958235 |
| Polr2g | 2037.095245 | 0.060254766 | 0.054415434 | 1.107310212 | 0.268159831 | 0.71037507 |
| Aldh3b2 | 68.08524532 | 0.060299536 | 0.092101045 | 0.654710658 | 0.512654062 | 0.852934798 |

| | | | | | | | | | | | | | | | |
|---|---|---|---|---|---|---|---|---|---|---|---|---|---|---|---|
| 1190002F15Rik | 2.065659253 | -0.174155995 | 0.097164863 | -1.792376267 | 0.073072714 | NA | | | Acy3 | 104.3460704 | 0.060301281 | 0.094024842 | 0.641333499 | 0.521306029 | 0.853911973 |
| 1190003K10Rik | 0.17833704 | -0.002390466 | 0.031282281 | -0.076415987 | 0.939088151 | NA | | | Npr3 | 491.947377 | 0.060303715 | 0.091220357 | 0.661077387 | 0.508562686 | 0.850050333 |
| 1300002K09Rik | 4.318427049 | 0.218488128 | 0.129086896 | 1.692566282 | 0.090538054 | NA | | | Ctdnep1 | 2785.643481 | 0.060307667 | 0.048258822 | 1.249671356 | 0.211419625 | 0.666270994 |
| 1300015D01Rik | 0.640586365 | -0.082316043 | 0.054836985 | -1.501104465 | 0.133328544 | NA | | | Alpk2 | 19.38808774 | 0.060308465 | 0.081858256 | 0.736742613 | 0.461278887 | NA |
| 1300017J02Rik | 3.063965952 | -0.141407668 | 0.118448093 | -1.193836595 | 0.232541894 | NA | | | Tmem198 | 2333.325568 | 0.060319605 | 0.079804455 | 0.755842572 | 0.449743589 | 0.823263966 |
| 1300018J18Rik | 838.4683718 | 0.221761906 | 0.092208535 | 2.405004119 | NA | NA | | | Smcr8 | 2637.69212 | 0.060341891 | 0.055609452 | 1.085101341 | 0.277876773 | 0.718227404 |
| 1500002O10Rik | 5.66066331 | -0.019385632 | 0.142637895 | -0.135908007 | 0.891894005 | NA | | | Grb14 | 832.5957558 | 0.060348218 | 0.070563519 | 0.855232553 | 0.392422418 | 0.794184423 |
| 1500016L03Rik | 2.023832634 | -0.027536726 | 0.096273574 | -0.286025798 | 0.774858363 | NA | | | Ctnnb1 | 19029.40035 | 0.060361199 | 0.045533381 | 1.325647216 | 0.184956579 | 0.637780733 |
| 1600002D24Rik | 0.496539307 | -0.049785487 | 0.052778288 | -0.943294857 | 0.345530106 | NA | | | Cryl1 | 545.0190727 | 0.060367116 | 0.086024874 | 0.701740246 | 0.482841169 | 0.837880119 |
| 1600010M07Rik | 1.65989692 | 0.123419835 | 0.094419236 | 1.307147157 | 0.191162738 | NA | | | Iqsec1 | 15142.34491 | 0.060382194 | 0.037039419 | 1.630214383 | 0.103056195 | 0.522975271 |
| 1600014C23Rik | 2.580700065 | 0.038362432 | 0.11349426 | 0.338012092 | 0.735354076 | NA | | | Sec61g | 767.8473452 | 0.060399898 | 0.079774173 | 0.757135994 | 0.448968392 | 0.823263966 |
| 1600014K23Rik | 0.347667565 | 0.012696751 | 0.035421431 | 0.358448274 | 0.720007871 | NA | | | Kcmf1 | 2307.880175 | 0.060403347 | 0.052656061 | 1.14713 | 0.251327897 | 0.699465245 |
| 1600015I10Rik | 0.107716603 | -0.020477583 | 0.027954224 | -0.732539849 | 0.46383913 | NA | | | Sppl3 | 3528.074657 | 0.06041216 | 0.053593101 | 1.12723763 | 0.259642024 | 0.703893404 |
| 1600016N20Rik | 5.956205954 | 0.142192876 | 0.143802093 | 0.988809503 | 0.322756354 | NA | | | Zfp85-rs1 | 152.3697686 | 0.060423333 | 0.092068697 | 0.656285307 | 0.51164057 | 0.852417908 |
| 1600019K03Rik | 0 | NA | NA | NA | NA | NA | | | 9530053A07Rik | 7.95183268 | 0.060433303 | 0.051743254 | 1.16794555 | 0.242828729 | NA |
| 1600025M17Rik | 0 | NA | NA | NA | NA | NA | | | Tmem57 | 2812.267771 | 0.060436189 | 0.054985804 | 1.099123503 | 0.2717142 | 0.713790099 |
| 1600029D21Rik | 5.514682936 | 0.320197518 | 0.141155291 | 2.268406061 | 0.023304469 | NA | | | Rtn4ip1 | 604.0147871 | 0.060448315 | 0.069904698 | 0.864724651 | 0.387189936 | 0.791583702 |
| 1600029I14Rik | 2.401394009 | 0.039527292 | 0.10482021 | 0.377096101 | 0.706102189 | NA | | | Cdv3 | 4001.5089 | 0.060466587 | 0.054332182 | 1.112905539 | 0.265749001 | 0.708046586 |
| 1700001C19Rik | 3.199909664 | -0.063890173 | 0.124721809 | -0.512261439 | 0.608468049 | NA | | | Mfn2 | 10744.93433 | 0.060496952 | 0.046873921 | 1.290631349 | 0.19683154 | 0.652874709 |
| 1700001D01Rik | 0 | NA | NA | NA | NA | NA | | | Ap5b1 | 150.934432 | 0.060497193 | 0.094692389 | 0.638881257 | 0.522900182 | 0.854337225 |
| 1700001F09Rik | 0.192142086 | -0.026384592 | 0.031527717 | -0.836869715 | 0.402665805 | NA | | | Slc24a2 | 29718.16885 | 0.060503916 | 0.077479635 | 0.78090089 | 0.434860789 | 0.815326688 |
| 1700001G11Rik | 6.790179138 | 0.042434055 | 0.150812379 | 0.281369843 | 0.778426746 | NA | | | Ptbp2 | 2955.577599 | 0.060505937 | 0.063418306 | 0.954076842 | 0.340044747 | 0.761914237 |
| 1700001G17Rik | 7.440824058 | 0.098663269 | 0.152781807 | 0.64577891 | 0.518422548 | NA | | | Zfp106 | 12919.22506 | 0.060516646 | 0.059043877 | 1.024943637 | 0.305389783 | 0.737872918 |
| 1700001J03Rik | 2.254932462 | -0.035214803 | 0.100425809 | -0.350654906 | 0.725847259 | NA | | | Rptor | 3791.592767 | 0.060520822 | 0.047481037 | 1.274631435 | 0.202439726 | 0.65606599 |
| 1700001J11Rik | 2.485277953 | -0.006079885 | 0.106896884 | -0.056876165 | 0.954643841 | NA | | | Dynlt3 | 3879.281735 | 0.060521649 | 0.063053667 | 0.959843446 | 0.337134013 | 0.760113439 |
| 1700001K19Rik | 0.352713512 | 0.01988513 | 0.041853977 | 0.475107309 | 0.634710489 | NA | | | Stac2 | 5334.020699 | 0.060543806 | 0.066011284 | 0.917173592 | 0.359051682 | 0.774860309 |
| 1700001K23Rik | 0.107716603 | -0.020477583 | 0.027954224 | -0.732539849 | 0.46383913 | NA | | | Yme1l1 | 2871.005648 | 0.060576948 | 0.067848304 | 0.892829217 | 0.371948638 | 0.781368063 |
| 1700001P01Rik | 5.989583001 | 0.256745114 | 0.145391429 | 1.765888917 | 0.077414487 | NA | | | Rnf19a | 1603.205691 | 0.060614681 | 0.069095448 | 0.87726011 | 0.380345373 | 0.786275719 |
| 1700003C15Rik | 0 | NA | NA | NA | NA | NA | | | Sh3bgrl | 1643.953609 | 0.060619035 | 0.078258759 | 0.77459745 | 0.438577564 | 0.817139233 |
| 1700003F12Rik | 5.668743844 | 0.190346875 | 0.146811413 | 1.296540045 | 0.194789493 | NA | | | Mogat1 | 5.053378446 | 0.060638758 | 0.044979376 | 1.348145826 | 0.177611485 | NA |
| 1700003F17Rik | 0 | NA | NA | NA | NA | NA | | | Gstt2 | 88.25032805 | 0.060650327 | 0.094797659 | 0.63978718 | 0.522310968 | 0.854113377 |
| 1700003G13Rik | 0 | NA | NA | NA | NA | NA | | | Ranbp6 | 1952.751867 | 0.060652578 | 0.067136688 | 0.903419276 | 0.366303414 | 0.778210102 |
| 1700003G18Rik | 0.094392817 | 0.017895942 | 0.027818137 | 0.643319236 | 0.520016977 | NA | | | Ak7 | 118.0907536 | 0.060664087 | 0.087965312 | 0.689636468 | 0.490422828 | 0.841767931 |
| 1700003H04Rik | 0 | NA | NA | NA | NA | NA | | | Tmem134 | 781.9775544 | 0.060677767 | 0.069928567 | 0.867710717 | 0.385552718 | 0.789252928 |
| 1700003L19Rik | 1.181434631 | 0.01240029 | 0.079220911 | 0.156527991 | 0.875616858 | NA | | | Sort1 | 29328.97443 | 0.060678297 | 0.05656987 | 1.072625716 | 0.283439082 | 0.721034821 |
| 1700003P14Rik | 0 | NA | NA | NA | NA | NA | | | Timp2 | 17050.32347 | 0.060687673 | 0.079595742 | 0.762448742 | 0.445792225 | 0.821773578 |
| 1700006A11Rik | 0 | NA | NA | NA | NA | NA | | | Ppp1r11 | 2192.794445 | 0.060707552 | 0.05591665 | 1.085679353 | 0.277620879 | 0.71807708 |
| 1700006E09Rik | 0 | NA | NA | NA | NA | NA | | | Ccdc47 | 4784.637751 | 0.060710251 | 0.049088659 | 1.236746986 | 0.216181029 | 0.670049713 |
| 1700006F04Rik | 0.298391179 | 0.008861245 | 0.039220874 | 0.225931861 | 0.821254417 | NA | | | Dio3os | 8.385694236 | 0.060715971 | 0.056088668 | 1.08249978 | 0.279030513 | NA |
| 1700006H21Rik | 0 | NA | NA | NA | NA | NA | | | Dlg1 | 8160.710735 | 0.060725095 | 0.067039557 | 0.905809917 | 0.365036471 | 0.777634019 |
| 1700007B14Rik | 5.000454671 | 0.271909199 | 0.120059613 | 2.264784899 | 0.023525883 | NA | | | Kif5c | 45120.6547 | 0.060744292 | 0.052503474 | 1.156957571 | 0.247289696 | 0.696744638 |
| 1700007J10Rik | 4.664008559 | 0.177386577 | 0.132557753 | 1.338183347 | 0.180836676 | NA | | | 4732456N10Rik | 6.092106704 | 0.060760431 | 0.036773575 | 1.652285137 | 0.098476438 | NA |
| 1700007K09Rik | 5.328206121 | 0.020225251 | 0.133870142 | 0.151081122 | 0.879911724 | NA | | | Rhobtb3 | 1108.576675 | 0.060794008 | 0.065448966 | 0.928876515 | 0.352953085 | 0.770519168 |
| 1700008A04Rik | 0 | NA | NA | NA | NA | NA | | | Tlr5 | 18.58821657 | 0.060800915 | 0.080672454 | 0.753676284 | 0.451043626 | NA |
| 1700008F21Rik | 0 | NA | NA | NA | NA | NA | | | Ppp1r9a | 17055.89033 | 0.060801238 | 0.072807808 | 0.835092277 | 0.403665755 | 0.800224169 |
| 1700008I05Rik | 0.201994201 | 0.000227241 | 0.031727614 | 0.00716224 | 0.994285408 | NA | | | Slc52a3 | 99.73116728 | 0.060816068 | 0.094353225 | 0.644557378 | 0.519214063 | 0.853297835 |
| 1700008K24Rik | 0 | NA | NA | NA | NA | NA | | | Gnai3 | 1273.06518 | 0.060840243 | 0.06931864 | 0.877689501 | 0.380112244 | 0.786275719 |
| 1700008P02Rik | 0 | NA | NA | NA | NA | NA | | | Ndufs3 | 3870.133386 | 0.060846057 | 0.042395294 | 1.435207817 | 0.151227887 | 0.598846773 |
| 1700009C05Rik | 1.413023452 | 0.042920249 | 0.080989815 | 0.529946255 | 0.596149195 | NA | | | Ppapdc1a | 182.6902633 | 0.060858893 | 0.090790372 | 0.670323196 | 0.502651783 | 0.84739426 |
| 1700009J07Rik | 1.022906055 | -0.048349521 | 0.077944357 | -0.620308166 | 0.53505492 | NA | | | Penk | 12888.26142 | 0.060865754 | 0.082761841 | 0.735432576 | 0.462076088 | 0.830604784 |
| 1700009N14Rik | 0 | NA | NA | NA | NA | NA | | | Tchh | 272.0762785 | 0.060881704 | 0.083647021 | 0.72784068 | 0.466711119 | 0.832095317 |
| 1700010B08Rik | 0 | NA | NA | NA | NA | NA | | | Tbca | 1126.067798 | 0.060907343 | 0.083873653 | 0.726179689 | 0.467728622 | 0.832353544 |
| 1700010D01Rik | 0 | NA | NA | NA | NA | NA | | | Trmt12 | 377.8811517 | 0.060911675 | 0.073024146 | 0.83413061 | 0.404207388 | 0.800290023 |
| 1700010I02Rik | 0 | NA | NA | NA | NA | NA | | | Rbm8a | 1803.305449 | 0.060923133 | 0.058462284 | 1.042092927 | 0.297368597 | 0.732415975 |
| 1700010J16Rik | 0 | NA | NA | NA | NA | NA | | | Gabra6 | 12.95759981 | 0.060932661 | 0.042937847 | 1.419089824 | 0.15587283 | NA |
| 1700010K23Rik | 0.843194396 | 0.001915388 | 0.057336671 | 0.033405992 | 0.973350831 | NA | | | Pa2g4 | 2935.840477 | 0.060940918 | 0.044771479 | 1.361154886 | 0.173464748 | 0.625247872 |
| 1700011A15Rik | 0.28821313 | -0.029561117 | 0.033427731 | -0.884329147 | 0.376518559 | NA | | | 1700026D08Rik | 40.77998813 | 0.060941156 | 0.086894887 | 0.70132039 | 0.483103092 | 0.838104566 |
| 1700011B04Rik | 0 | NA | NA | NA | NA | NA | | | Stard3nl | 1314.109297 | 0.060943503 | 0.053164432 | 1.146320965 | 0.251662371 | 0.699465245 |
| 1700011E24Rik | 1.014018918 | -0.01008284 | 0.071659643 | -0.140704577 | 0.888103329 | NA | | | Pou6f2 | 44.83223655 | 0.060960838 | 0.086198694 | 0.707213014 | 0.479434112 | 0.836346813 |
| 1700011H14Rik | 0.181709657 | 0.024794574 | 0.031325423 | 0.791516008 | 0.428642938 | NA | | | Ier3ip1 | 1098.531755 | 0.06098696 | 0.062731774 | 0.972186122 | 0.330957963 | 0.756212998 |
| 1700011I03Rik | 3.089749683 | -0.036099319 | 0.115858701 | -0.311580556 | 0.755359313 | NA | | | Jazf1 | 1207.873628 | 0.060998932 | 0.054397947 | 1.121346212 | 0.262140522 | 0.704505684 |
| 1700011L22Rik | 0.287632815 | 0.04259931 | 0.040911109 | 1.041265103 | 0.297752528 | NA | | | Fkbp11 | 19.21931376 | 0.061002124 | 0.070172074 | 0.86932195 | 0.38467106 | NA |
| 1700011M02Rik | 0 | NA | NA | NA | NA | NA | | | Eif4g2 | 26798.44199 | 0.061009435 | 0.055801768 | 1.093324412 | 0.274251388 | 0.716185716 |
| 1700012A03Rik | 0 | NA | NA | NA | NA | NA | | | Hrh3 | 4619.972979 | 0.061023871 | 0.075216465 | 0.811310013 | 0.417187661 | 0.806153084 |
| 1700012B07Rik | 0 | NA | NA | NA | NA | NA | | | Ffar2 | 2.783664535 | 0.061026479 | 0.037932509 | 1.608817363 | 0.107656283 | NA |
| 1700012I11Rik | 0 | NA | NA | NA | NA | NA | | | A430033K04Rik | 739.1486408 | 0.061030999 | 0.067834302 | 0.899707036 | 0.368276178 | 0.779243711 |
| 1700012L04Rik | 0.333037401 | -0.016093237 | 0.040545167 | -0.396921218 | 0.691425557 | NA | | | Strip1 | 3434.407204 | 0.061044497 | 0.054244111 | 1.125366336 | 0.26043383 | 0.704058139 |
| 1700012P22Rik | 0.292249882 | 0.042980062 | 0.04108841 | 1.046038582 | 0.29554322 | NA | | | Prodh | 1220.572305 | 0.061049257 | 0.07860829 | 0.776626187 | 0.437379347 | 0.816715817 |
| 1700013D24Rik | 0.188785634 | 0.025025111 | 0.031461603 | 0.79541754 | 0.42637066 | NA | | | Wbp2 | 18407.93541 | 0.061052623 | 0.040802874 | 1.496282423 | 0.134580074 | 0.577862888 |
| 1700013G24Rik | 0 | NA | NA | NA | NA | NA | | | Dfna5 | 443.6536987 | 0.06109539 | 0.072375897 | 0.844140008 | 0.398591173 | 0.797789838 |
| 1700013H16Rik | 0.653760176 | -0.030006487 | 0.054578099 | -0.549789885 | 0.582463497 | NA | | | Pomp | 3108.002956 | 0.061101164 | 0.046995065 | 1.3001613 | 0.193545692 | 0.649744561 |
| 1700015E13Rik | 0.094392817 | 0.017895942 | 0.027818137 | 0.643319236 | 0.520016977 | NA | | | Dnajc21 | 1218.6789 | 0.061109443 | 0.078213943 | 0.781311372 | 0.434619383 | 0.815326688 |
| 1700015F17Rik | 2.761763786 | 0.067473314 | 0.109707585 | 0.615028702 | 0.538535766 | NA | | | Vac14 | 1293.269055 | 0.061112679 | 0.062814567 | 0.972906158 | 0.330599944 | 0.756141871 |
| 1700015G11Rik | 0 | NA | NA | NA | NA | NA | | | Hbegf | 546.505629 | 0.061128626 | 0.075465874 | 0.810016809 | 0.417930515 | 0.80622293 |
| 1700016C15Rik | 0 | NA | NA | NA | NA | NA | | | Pcdh18 | 296.6320954 | 0.061139011 | 0.09040424 | 0.676284772 | 0.498859859 | 0.846052342 |
| 1700016D06Rik | 0 | NA | NA | NA | NA | NA | | | Tiprl | 1613.516915 | 0.061165413 | 0.068049461 | 0.898837588 | 0.368739176 | 0.779573594 |
| 1700016G22Rik | 0.499641834 | 0.064869181 | 0.050070714 | 1.295551338 | 0.195130103 | NA | | | Frmpd1 | 650.2935885 | 0.061173319 | 0.08473603 | 0.721928083 | 0.470338694 | 0.833245343 |
| 1700016H13Rik | 0.410728284 | -0.005925811 | 0.04256373 | -0.139222079 | 0.889274662 | NA | | | Rbks | 56.17545918 | 0.061181684 | 0.094645603 | 0.646429228 | 0.518001417 | 0.853297835 |
| 1700016L04Rik | 1.09910115 | 0.022207625 | 0.066299534 | 0.334958993 | 0.73765602 | NA | | | Sos1 | 2752.380641 | 0.061195705 | 0.072057959 | 0.849256707 | 0.395738465 | 0.796301004 |
| 1700016L21Rik | 0 | NA | NA | NA | NA | NA | | | Rhov | 336.224113 | 0.061197211 | 0.082280712 | 0.743761318 | 0.457020886 | 0.827236468 |
| 1700016P04Rik | 0 | NA | NA | NA | NA | NA | | | Cox4i1 | 13085.91229 | 0.061209699 | 0.043701436 | 1.400633578 | 0.161323674 | 0.612970886 |
| 1700017D01Rik | 0 | NA | NA | NA | NA | NA | | | Pcgf3 | 1464.532127 | 0.061219702 | 0.07493085 | 0.817015992 | 0.413919289 | 0.804285711 |
| 1700017G19Rik | 0.402750607 | -0.037318681 | 0.048991399 | -0.76173946 | 0.446215519 | NA | | | Pdik1l | 608.5190103 | 0.061226706 | 0.0764731 | 0.800630629 | 0.423345514 | 0.808999423 |
| 1700017J07Rik | 0.239950962 | 0.027417169 | 0.032831728 | 0.835081494 | 0.403671825 | NA | | | Dhcr24 | 5718.990761 | 0.061234404 | 0.061253351 | 0.99969067 | 0.317460229 | 0.747273907 |
| 1700017N19Rik | 0 | NA | NA | NA | NA | NA | | | Amer2 | 891.4527147 | 0.06127103 | 0.074673212 | 0.82052223 | 0.411918462 | 0.802795661 |
| 1700018A04Rik | 1.349866079 | 0.002120246 | 0.087195859 | 0.024315901 | 0.98060063 | NA | | | Lao1 | 8.050104979 | 0.06128545 | 0.048097188 | 1.274200285 | 0.202592444 | NA |
| 1700018B08Rik | 0.094392817 | 0.017895942 | 0.027818137 | 0.643319236 | 0.520016977 | NA | | | Sfmbt2 | 123.4476883 | 0.061373057 | 0.094994634 | 0.646068676 | 0.518234881 | 0.853297835 |
| 1700018B24Rik | 0 | NA | NA | NA | NA | NA | | | Dcun1d1 | 1792.300614 | 0.061387305 | 0.073202974 | 0.838590315 | 0.401699247 | 0.799277436 |
| 1700018C11Rik | 0 | NA | NA | NA | NA | NA | | | Tmed4 | 2554.19138 | 0.061387513 | 0.046002226 | 1.334446567 | 0.182057547 | 0.635245711 |
| 1700018F24Rik | 0 | NA | NA | NA | NA | NA | | | Fbxo9 | 3219.305294 | 0.061403123 | 0.047956752 | 1.280385354 | 0.200409642 | 0.654475062 |
| 1700018G05Rik | 0 | NA | NA | NA | NA | NA | | | Txnl1 | 2604.432853 | 0.061433602 | 0.05899577 | 1.041322145 | 0.297726062 | 0.73263913 |
| 1700019A02Rik | 1.732172825 | 0.01584073 | 0.093920608 | 0.168660855 | 0.866063411 | NA | | | Pcdhb12 | 326.9293572 | 0.061437293 | 0.08098756 | 0.758601612 | 0.448090908 | 0.822982492 |
| 1700019B03Rik | 5.325715112 | 0.11904574 | 0.130617208 | 0.91140931 | 0.36207975 | NA | | | Aldh2 | 2824.1754 | 0.061438321 | 0.066928201 | 0.917973591 | 0.358632692 | 0.774627234 |
| 1700019B21Rik | 0 | NA | NA | NA | NA | NA | | | Tcaim | 744.5200474 | 0.061440328 | 0.074786899 | 0.821538655 | 0.411339515 | 0.802795661 |
| 1700019E08Rik | 0 | NA | NA | NA | NA | NA | | | Msh3 | 1010.487693 | 0.061444332 | 0.063372923 | 0.969567575 | 0.332262082 | 0.757166892 |
| 1700019G24Rik | 2.199194789 | 0.093608401 | 0.091510596 | 1.022924175 | 0.306343698 | NA | | | Lyg2 | 4.663625196 | 0.061451145 | 0.047218918 | 1.301409435 | 0.193118346 | NA |
| 1700019M22Rik | 0 | NA | NA | NA | NA | NA | | | Dsc3 | 25.10332378 | 0.061458569 | 0.083717743 | 0.734116411 | 0.462877792 | 0.831014288 |
| 1700019N19Rik | 6.154910495 | -0.063797621 | 0.146511642 | -0.43544404 | 0.66324015 | NA | | | Peli2 | 1058.558474 | 0.061467201 | 0.063247332 | 0.971854456 | 0.33112296 | 0.756212998 |
| 1700019O17Rik | 0 | NA | NA | NA | NA | NA | | | Zmat5 | 516.2055326 | 0.061483465 | 0.072201142 | 0.851558057 | 0.394459427 | 0.795437883 |
| 1700020A23Rik | 0.094392817 | 0.017895942 | 0.027818137 | 0.643319236 | 0.520016977 | NA | | | Robo1 | 1626.330009 | 0.06149066 | 0.06904225 | 0.890623649 | 0.373131108 | 0.782226671 |
| 1700020G17Rik | 0.204276544 | 2.70E-05 | 0.031767541 | 0.000850493 | 0.999321405 | NA | | | Tcp1 | 7801.027021 | 0.061496017 | 0.055384338 | 1.110350314 | 0.2668481 | 0.709441589 |
| 1700020L24Rik | 6.022953304 | 0.19663498 | 0.139041814 | 1.414214722 | 0.157298867 | NA | | | Gemin4 | 494.8568323 | 0.061520496 | 0.074467397 | 0.82614001 | 0.408724674 | 0.801055189 |
| 1700020M21Rik | 0 | NA | NA | NA | NA | NA | | | Tmx1 | 1197.548909 | 0.061521481 | 0.063302942 | 0.971858159 | 0.331121118 | 0.756212998 |
| 1700020N01Rik | 0.620554997 | 0.021634778 | 0.053065255 | 0.40770138 | 0.683492924 | NA | | | Fam43a | 578.6182751 | 0.061531544 | 0.08686488 | 0.70835928 | 0.478722172 | 0.835936459 |
| 1700020N15Rik | 0 | NA | NA | NA | NA | NA | | | Mtrr | 533.70605 | 0.061547524 | 0.081797237 | 0.75244014 | 0.451786415 | 0.824558475 |
| 1700020N18Rik | 1.495483865 | 0.062587367 | 0.087569837 | 0.714713756 | 0.474785936 | NA | | | Med21 | 531.5227941 | 0.061562164 | 0.079800659 | 0.771449325 | 0.440440648 | 0.818556916 |
| 1700021F07Rik | 1.755178943 | -0.177768483 | 0.087435428 | -2.033140181 | 0.042038362 | NA | | | Dusp3 | 9168.745663 | 0.061587579 | 0.054492332 | 1.130206344 | 0.258389287 | 0.703616072 |
| 1700021N21Rik | 2.167429004 | -0.162674127 | 0.106199892 | -1.531772994 | 0.125578462 | NA | | | Stim1 | 4242.418292 | 0.061595875 | 0.04773217 | 1.290447819 | 0.196895218 | 0.652874709 |
| 1700022A21Rik | 0 | NA | NA | NA | NA | NA | | | Rbms2 | 648.6625673 | 0.061620036 | 0.062760795 | 0.981823714 | 0.326186704 | 0.753049596 |
| 1700022A22Rik | 0 | NA | NA | NA | NA | NA | | | Zfp532 | 2445.603619 | 0.061620553 | 0.05436768 | 1.133404132 | 0.25704458 | 0.703199912 |
| 1700022E09Rik | 0 | NA | NA | NA | NA | NA | | | Gbas | 2806.405437 | 0.061626231 | 0.05192618 | 1.186804634 | 0.235304691 | 0.686276191 |
| 1700022H16Rik | 0.119397641 | -0.020477583 | 0.027954224 | -0.732539845 | 0.463839132 | NA | | | Fev | 1.697717137 | 0.061628796 | 0.03091065 | 1.993772266 | 0.046176948 | NA |
| 1700022I11Rik | 0 | NA | NA | NA | NA | NA | | | Vps45 | 1589.686145 | 0.061631834 | 0.064706061 | 0.952489349 | 0.340848865 | 0.76263292 |
| 1700022P22Rik | 0.294090812 | 0.009484222 | 0.033616198 | 0.282132504 | 0.77784191 | NA | | | Gde1 | 7272.4787 | 0.061639758 | 0.050451416 | 1.221764674 | 0.221796631 | 0.672987912 |
| 1700023A16Rik | 2.179656538 | 0.082062822 | 0.086572956 | 0.947903666 | 0.343178498 | NA | | | Bcam | 825.1390905 | 0.061642872 | 0.084757184 | 0.727287874 | 0.467049625 | 0.832211278 |
| 1700023C21Rik | 0 | NA | NA | NA | NA | NA | | | Zfp449 | 236.934497 | 0.061644256 | 0.082511022 | 0.747103293 | 0.455001213 | 0.826168452 |
| 1700023E05Rik | 7.707835974 | -0.111299066 | 0.158723178 | -0.701214954 | 0.483168879 | NA | | | Ube4a | 3319.742511 | 0.061675044 | 0.05766619 | 1.06951826 | 0.284836206 | 0.721641599 |
| 1700023F02Rik | 0.978055737 | 0.13187226 | 0.070074519 | 1.881886059 | 0.059851488 | NA | | | Acadl | 1427.329663 | 0.061675916 | 0.067514878 | 0.913515923 | 0.360971263 | 0.776449324 |
| 1700023L04Rik | 4.933102782 | 0.176106676 | 0.142383379 | 1.23684855 | 0.216143314 | NA | | | Asb3 | 574.6769187 | 0.061677611 | 0.07030278 | 0.877313962 | 0.380316131 | 0.786275719 |
| 1700024B18Rik | 0 | NA | NA | NA | NA | NA | | | C77080 | 1226.815923 | 0.061679019 | 0.075722452 | 0.814540698 | 0.415335267 | 0.805633085 |
| 1700024G13Rik | 5.754891192 | -0.009231793 | 0.13331388 | -0.069248554 | 0.944791775 | NA | | | Glp1r | 24.55941584 | 0.061683888 | 0.054694219 | 1.127795386 | NA | NA |
| 1700024P03Rik | 0 | NA | NA | NA | NA | NA | | | Rwdd4a | 1048.936827 | 0.061685927 | 0.0694893 | 0.887703964 | 0.374700013 | 0.783149984 |
| 1700024P04Rik | 0 | NA | NA | NA | NA | NA | | | Ndufb5 | 3366.529859 | 0.061689798 | 0.055957146 | 1.102447175 | 0.270267312 | 0.71216373 |
| 1700025B11Rik | 0 | NA | NA | NA | NA | NA | | | Snx4 | 1563.230313 | 0.061690483 | 0.060117048 | 1.026172853 | 0.304810115 | 0.737453291 |
| 1700025C18Rik | 0 | NA | NA | NA | NA | NA | | | Prune2 | 4721.558093 | 0.061700444 | 0.067582402 | 0.912966134 | 0.361260353 | 0.776449324 |
| 1700025E21Rik | 0.785077497 | 0.025108046 | 0.067164331 | 0.373830062 | 0.708530752 | NA | | | Gadd45gip1 | 989.685134 | 0.061704557 | 0.05496524 | 1.12261052 | 0.261602945 | 0.704173028 |
| 1700025F22Rik | 0 | NA | NA | NA | NA | NA | | | Myod1 | 7.074066916 | 0.061711946 | 0.056025788 | 1.1014918 | 0.270682671 | NA |
| 1700025F24Rik | 0.425134012 | -0.003256152 | 0.042864108 | -0.075964537 | 0.939447312 | NA | | | Tuba1b | 10246.85473 | 0.061737777 | 0.044594159 | 1.384436403 | 0.166224869 | 0.616594087 |
| 1700025K24Rik | 0 | NA | NA | NA | NA | NA | | | Iscu | 2207.896892 | 0.06175409 | 0.071433401 | 0.864498812 | 0.387313933 | 0.791583702 |
| 1700025M24Rik | 0.116696964 | -0.020477583 | 0.027954224 | -0.732539847 | 0.463839131 | NA | | | Stk32a | 332.320069 | 0.061783473 | 0.089462196 | 0.690609837 | 0.489810764 | 0.841119426 |
| 1700025N23Rik | 0.198019769 | 0.025308392 | 0.031628095 | 0.80018706 | 0.423602426 | NA | | | Kpna6 | 5832.198914 | 0.061784047 | 0.04500337 | 1.372876 | 0.169790899 | 0.62000568 |
| 1700026D11Rik | 0.199381295 | -0.004055078 | 0.031639813 | -0.128163771 | 0.89801937 | NA | | | Rgs2 | 2590.691165 | 0.061787181 | 0.081346852 | 0.759552216 | 0.447522291 | 0.822458097 |
| 1700026F02Rik | 0 | NA | NA | NA | NA | NA | | | Fbxo21 | 6541.096973 | 0.061798779 | 0.04048973 | 1.526282819 | 0.126939446 | 0.563718247 |
| 1700026L06Rik | 4.384532908 | 0.260504994 | 0.122357671 | 2.12904505 | 0.033250533 | NA | | | Tomm7 | 1414.1116 | 0.061804561 | 0.050746226 | 1.217914422 | 0.223256495 | 0.674214152 |
| 1700027A15Rik | 0.079983654 | 0.016740032 | 0.027674178 | 0.604897175 | 0.54524732 | NA | | | Idh3b | 8393.420495 | 0.061822823 | 0.042195989 | 1.465135057 | 0.142884074 | 0.589284353 |
| 1700027F09Rik | 0 | NA | NA | NA | NA | NA | | | 4930515G01Rik | 9.873490607 | 0.061852038 | 0.065490935 | 0.944436636 | 0.344946567 | NA |
| 1700027H10Rik | 0.690975005 | 0.005249197 | 0.047660626 | 0.110136963 | 0.912300754 | NA | | | Gcc2 | 3650.888607 | 0.061856361 | 0.065209452 | 0.948579677 | 0.342834431 | 0.763762832 |
| 1700027I24Rik | 0.222620122 | -0.001447027 | 0.032059667 | -0.04513545 | 0.963999345 | NA | | | Il17rd | 233.8025272 | 0.061860505 | 0.087426248 | 0.707573604 | 0.47921009 | 0.836180652 |
| 1700027J07Rik | 0.510163613 | 0.009578335 | 0.050846372 | 0.188377952 | 0.850580382 | NA | | | Aifm1 | 959.9196346 | 0.061876099 | 0.056874865 | 1.08793399 | 0.276624249 | 0.717475105 |
| 1700028B04Rik | 1.206882925 | 0.053053473 | 0.077520621 | 0.684378841 | 0.493735974 | NA | | | Exosc10 | 1855.397905 | 0.061893391 | 0.051280959 | 1.206946819 | 0.227452625 | 0.680203999 |
| 1700028D13Rik | 0 | NA | NA | NA | NA | NA | | | C1galt1 | 326.160726 | 0.061894486 | 0.088551888 | 0.698962916 | 0.484575207 | 0.838713365 |
| 1700028E10Rik | 7.104082621 | 0.015533361 | 0.156379048 | 0.099331474 | 0.92087509 | NA | | | Manea | 1578.385944 | 0.061927835 | 0.065320248 | 0.948064912 | 0.343096409 | 0.763762832 |
| 1700028I16Rik | 0.544515322 | -0.071444103 | 0.051231676 | -1.394529886 | 0.163157651 | NA | | | Bcl6 | 1205.100555 | 0.061932081 | 0.083563491 | 0.741138024 | 0.458609756 | 0.828043599 |
| 1700028M03Rik | 0.116696964 | -0.020477583 | 0.027954224 | -0.732539847 | 0.463839131 | NA | | | Zfp296 | 82.59661152 | 0.061949483 | 0.095068047 | 0.651633062 | 0.514637914 | 0.853195313 |
| 1700028P15Rik | 0 | NA | NA | NA | NA | NA | | | Rab12 | 3414.613538 | 0.06196007 | 0.052612526 | 1.17766764 | 0.238929132 | 0.689537017 |
| 1700029B22Rik | 0 | NA | NA | NA | NA | NA | | | S100pbp | 719.0841065 | 0.061985822 | 0.075148722 | 0.824842005 | 0.409461296 | 0.801055189 |
| 1700029F12Rik | 0 | NA | NA | NA | NA | NA | | | Rims4 | 843.1632335 | 0.061986609 | 0.071718811 | 0.864300569 | 0.387422797 | 0.791583702 |
| 1700029H14Rik | 0.571399219 | 0.021750442 | 0.053110482 | 0.409532001 | 0.682149287 | NA | | | Baalc | 9988.794845 | 0.062008641 | 0.048384855 | 1.281571283 | 0.199993079 | 0.654380963 |
| 1700029J03Rik | 0 | NA | NA | NA | NA | NA | | | Cse1l | 3062.585649 | 0.062022533 | 0.073755374 | 0.840922226 | 0.400391506 | 0.799132594 |
| 1700029M20Rik | 1.763654503 | -0.237893836 | 0.094151439 | -2.526714805 | 0.011513496 | NA | | | Cnksr3 | 215.7862117 | 0.062025619 | 0.092084642 | 0.673571815 | 0.500583577 | 0.846307849 |
| 1700029N11Rik | 0 | NA | NA | NA | NA | NA | | | Cntn4 | 2288.37415 | 0.062045806 | 0.077514967 | 0.800436474 | 0.423457956 | 0.808999423 |
| 1700029P11Rik | 0 | NA | NA | NA | NA | NA | | | Hrk | 2473.255116 | 0.062046775 | 0.080844479 | 0.767483148 | 0.442794309 | 0.819995323 |
| 1700030A11Rik | 1.17684774 | 0.072940977 | 0.07688226 | 0.948736124 | 0.342754837 | NA | | | AA986860 | 173.6189217 | 0.062056566 | 0.09200866 | 0.674464402 | 0.50001611 | 0.846241311 |
| 1700030F04Rik | 0.880480036 | -0.025404946 | 0.074833042 | -0.339488353 | 0.734241869 | NA | | | Pcdhga7 | 844.6752429 | 0.06206926 | 0.070981643 | 0.874441017 | 0.381878129 | 0.786663048 |
| 1700030F18Rik | 0.276690427 | -0.012033946 | 0.033317809 | -0.361186582 | 0.717959972 | NA | | | Taf3 | 819.8214554 | 0.062073693 | 0.077995445 | 0.79586306 | 0.426111634 | 0.809519334 |
| 1700030M09Rik | 0.098353386 | -0.020212831 | 0.02786466 | -0.725393061 | 0.468210929 | NA | | | Taf11 | 1237.001955 | 0.062091564 | 0.061961722 | 1.002095523 | 0.31629746 | 0.745993052 |
| 1700030N03Rik | 0 | NA | NA | NA | NA | NA | | | Zim1 | 166.340216 | 0.062121795 | 0.094004488 | 0.660838602 | 0.508715824 | 0.850050333 |
| 1700030O20Rik | 0 | NA | NA | NA | NA | NA | | | Nab2 | 1946.717768 | 0.062177496 | 0.090001804 | 0.690847213 | 0.489661562 | 0.841119426 |
| 1700031A10Rik | 0 | NA | NA | NA | NA | NA | | | Eri3 | 4970.993266 | 0.062199112 | 0.048528235 | 1.281709757 | 0.199944481 | 0.654380963 |
| 1700031F05Rik | 0 | NA | NA | NA | NA | NA | | | Gm13749 | 16.83532029 | 0.062213275 | 0.074412496 | 0.836059504 | 0.403121429 | NA |
| 1700031M16Rik | 0.395349417 | 0.027211441 | 0.048374467 | 0.562516604 | 0.573764094 | NA | | | Brms1l | 2701.797083 | 0.062221354 | 0.059340382 | 1.048549934 | 0.294385309 | 0.729576944 |
| 1700031P21Rik | 0 | NA | NA | NA | NA | NA | | | Wdr36 | 1247.389238 | 0.062238152 | 0.06225062 | 0.999799709 | 0.317407447 | 0.747270757 |
| 1700034E13Rik | 0.119397641 | -0.020477583 | 0.027954224 | -0.732539845 | 0.463839132 | NA | | | Nupl1 | 1425.800427 | 0.062248583 | 0.068601308 | 0.907396443 | 0.36419719 | 0.777191869 |
| 1700034F02Rik | 3.160626376 | -0.105399317 | 0.124784832 | -0.844648464 | 0.398307142 | NA | | | Pcnx | 5937.974144 | 0.062250604 | 0.061126735 | 1.018385881 | 0.308494609 | 0.740806186 |
| 1700034G24Rik | 0.188785634 | 0.025025111 | 0.031461603 | 0.79541754 | 0.42637066 | NA | | | Cand1 | 6742.394693 | 0.062294451 | 0.058397387 | 1.066733525 | 0.286092184 | 0.722148665 |
| 1700034I23Rik | 0.46990427 | 0.035927024 | 0.051562175 | 0.696770909 | 0.485946174 | NA | | | Aldh5a1 | 5273.408651 | 0.062298116 | 0.061376005 | 1.015023967 | 0.310094403 | 0.743030104 |
| 1700034J05Rik | 0.567794913 | -0.037435068 | 0.058745955 | -0.637236527 | 0.523970788 | NA | | | Rpa1 | 2097.611207 | 0.062301601 | 0.043667341 | 1.426732181 | 0.153657106 | 0.60340295 |
| 1700034K08Rik | 0 | NA | NA | NA | NA | NA | | | A330040F15Rik | 10.12829481 | 0.062304942 | 0.062078145 | 1.00365341 | 0.315545701 | NA |
| 1700034O15Rik | 0 | NA | NA | NA | NA | NA | | | Usp46 | 4549.777705 | 0.062307357 | 0.058474981 | 1.065538733 | 0.286632208 | 0.722148665 |
| 1700034P13Rik | 2.790014232 | 0.094678183 | 0.115412086 | 0.820348946 | 0.412017211 | NA | | | Dmrtc1a | 196.1539943 | 0.062313652 | 0.093611188 | 0.66566458 | 0.505625518 | 0.84841357 |
| 1700036G14Rik | 0.08731684 | 0.017365056 | 0.027733412 | 0.626142072 | 0.531221759 | NA | | | Dcaf7 | 12022.97118 | 0.062330546 | 0.04990814 | 1.248905404 | 0.211699675 | 0.666270994 |
| 1700039E15Rik | 1.781638371 | 0.01592076 | 0.072687075 | 0.219031518 | 0.826625496 | NA | | | Pla1a | 63.81061276 | 0.062402986 | 0.095100551 | 0.65617902 | 0.511708946 | 0.852417908 |
| 1700039E22Rik | 0 | NA | NA | NA | NA | NA | | | Mesdc1 | 1161.295749 | 0.062417522 | 0.073405874 | 0.850306913 | 0.395154474 | 0.795857764 |
| 1700041C23Rik | 0 | NA | NA | NA | NA | NA | | | Zfp791 | 189.6368672 | 0.062418394 | 0.085682526 | 0.72848452 | 0.466317042 | 0.832050282 |
| 1700041M19Rik | 0.098353386 | -0.020212831 | 0.02786466 | -0.725393061 | 0.468210929 | NA | | | Cdc42 | 11081.24545 | 0.062418464 | 0.05513435 | 1.132115709 | 0.257585791 | 0.703325414 |
| 1700042B14Rik | 0.096071043 | -0.020098795 | 0.027825948 | -0.722304036 | 0.470107572 | NA | | | Pmvk | 1225.543727 | 0.062431875 | 0.072536753 | 0.860692988 | 0.389407156 | 0.792715562 |
| 1700042G07Rik | 0 | NA | NA | NA | NA | NA | | | Nxph2 | 72.06861153 | 0.062452044 | 0.090362381 | 0.691128802 | 0.489484602 | 0.841119426 |
| 1700042G15Rik | 0 | NA | NA | NA | NA | NA | | | Iqcg | 179.2352012 | 0.062458043 | 0.095065037 | 0.657003298 | 0.511178796 | 0.852011998 |
| 1700042O10Rik | 0 | NA | NA | NA | NA | NA | | | Dnajb9 | 1408.333355 | 0.062483628 | 0.075746211 | 0.824907643 | 0.409424027 | 0.801055189 |
| 1700044C05Rik | 0 | NA | NA | NA | NA | NA | | | Vstm2b | 1029.586552 | 0.062506996 | 0.059693317 | 1.047135578 | 0.295037053 | 0.73000898 |
| 1700044K03Rik | 0 | NA | NA | NA | NA | NA | | | Ltv1 | 639.4773597 | 0.062512912 | 0.065594099 | 0.953026475 | 0.340576657 | 0.762546123 |
| 1700045H11Rik | 0.079983654 | 0.016740032 | 0.027674178 | 0.604897175 | 0.54524732 | NA | | | Wdr37 | 3243.682756 | 0.062514752 | 0.058946065 | 1.060541568 | 0.28889829 | 0.724444298 |
| 1700046C09Rik | 0 | NA | NA | NA | NA | NA | | | Rgs19 | 610.7127286 | 0.062547062 | 0.066899696 | 0.934937908 | 0.349820299 | 0.768525358 |
| 1700047E10Rik | 0.30236719 | -0.01694275 | 0.039582453 | -0.428036895 | 0.668624259 | NA | | | Dgcr2 | 6337.521283 | 0.062560748 | 0.055299986 | 1.131297713 | 0.257929806 | 0.703616072 |
| 1700047G03Rik | 0 | NA | NA | NA | NA | NA | | | Napepld | 872.409783 | 0.062580685 | 0.064792469 | 0.965863561 | 0.334112458 | 0.758212399 |
| 1700047I17Rik2 | 0 | NA | NA | NA | NA | NA | | | Zbtb46 | 257.56855 | 0.062583809 | 0.086290215 | 0.725271211 | 0.468285664 | 0.832393239 |
| 1700047L14Rik | 0 | NA | NA | NA | NA | NA | | | Zdhhc3 | 3617.501925 | 0.062591634 | 0.047597535 | 1.315018415 | 0.188503714 | 0.64225749 |
| 1700048M11Rik | 0 | NA | NA | NA | NA | NA | | | Myl6 | 8971.997501 | 0.06261412 | 0.057092343 | 1.096716593 | 0.272765301 | 0.714142044 |
| 1700049E15Rik | 0 | NA | NA | NA | NA | NA | | | Ppp4r2 | 2868.505732 | 0.062614241 | 0.077997533 | 0.802772072 | 0.422106487 | 0.808695226 |
| 1700049E22Rik | 0 | NA | NA | NA | NA | NA | | | St3gal1 | 1177.778037 | 0.062626811 | 0.068432405 | 0.9151631 | 0.360106013 | 0.776328279 |
| 1700049L16Rik | 0 | NA | NA | NA | NA | NA | | | Erc2 | 7315.038933 | 0.062630205 | 0.063675228 | 0.98358823 | 0.325318014 | 0.75293408 |
| 1700051A21Rik | 1.76604708 | 0.096531081 | 0.095244804 | 1.013504957 | 0.31081903 | NA | | | Dach1 | 560.4255589 | 0.062659112 | 0.093741567 | 0.668423982 | 0.503862991 | 0.847807553 |
| 1700052I22Rik | 0 | NA | NA | NA | NA | NA | | | Gal3st3 | 1972.708896 | 0.062676331 | 0.071605742 | 0.875297556 | 0.381412024 | 0.786663048 |
| 1700054A03Rik | 0.119397641 | -0.020477583 | 0.027954224 | -0.732539845 | 0.463839132 | NA | | | Gm694 | 72.25480243 | 0.062684065 | 0.08523079 | 0.735462669 | 0.462057767 | 0.830604784 |
| 1700054K19Rik | 0 | NA | NA | NA | NA | NA | | | Slc35e1 | 4289.669986 | 0.062691291 | 0.04443359 | 1.410898639 | 0.158274504 | 0.609156864 |
| 1700054M17Rik | 0.651627236 | -0.059904957 | 0.056388213 | -1.062366654 | 0.288069266 | NA | | | Prss35 | 107.196387 | 0.062716774 | 0.094862646 | 0.661132461 | 0.50852737 | 0.850050333 |
| 1700054O13Rik | 0 | NA | NA | NA | NA | NA | | | Rabgef1 | 1621.840128 | 0.062751289 | 0.055210178 | 1.136589149 | 0.255710078 | 0.701952734 |
| 1700055C04Rik | 0 | NA | NA | NA | NA | NA | | | Eif2b2 | 1169.748277 | 0.06275281 | 0.053779073 | 1.166862997 | 0.243265701 | 0.692772033 |
| 1700055N04Rik | 2.253040213 | -0.154269086 | 0.094692377 | -1.629160568 | 0.103279032 | NA | | | Haus4 | 80.47024945 | 0.062802556 | 0.095059206 | 0.660667791 | 0.508825384 | 0.850051469 |
| 1700056E22Rik | 2.635905444 | 0.037144121 | 0.112075612 | 0.331420196 | 0.740327112 | NA | | | Smcr7l | 1296.828766 | 0.062831618 | 0.062563564 | 1.004284509 | 0.315241498 | 0.745993052 |
| 1700057G04Rik | 0 | NA | NA | NA | NA | NA | | | 2410018M08Rik | 246.9079346 | 0.062879766 | 0.087247064 | 0.720709247 | 0.471088422 | 0.833245343 |
| 1700057H15Rik | 0.098353386 | -0.020212831 | 0.02786466 | -0.725393061 | 0.468210929 | NA | | | Srsf3 | 4027.577882 | 0.06289126 | 0.054119584 | 1.162079521 | 0.245203165 | 0.694718388 |
| 1700058G18Rik | 5.410811809 | 0.122361825 | 0.125293175 | 0.976604074 | 0.328765202 | NA | | | Nudcd3 | 7560.839367 | 0.062905122 | 0.041258089 | 1.524673661 | 0.127340514 | 0.564894808 |
| 1700060C16Rik | 0 | NA | NA | NA | NA | NA | | | Ywhah | 36175.1318 | 0.062956499 | 0.055487001 | 1.134617072 | 0.256535797 | 0.702836475 |
| 1700060C20Rik | 0.197363271 | -0.000561558 | 0.031650166 | -0.017742642 | 0.985844163 | NA | | | Cdh12 | 493.1495629 | 0.062956693 | 0.090690883 | 0.694189875 | 0.487563141 | 0.839801621 |
| 1700061F12Rik | 0 | NA | NA | NA | NA | NA | | | Als2 | 2161.274413 | 0.062966107 | 0.061966687 | 1.016128348 | 0.309568271 | 0.742289322 |
| 1700061G19Rik | 5.80828273 | -0.169261873 | 0.140020175 | -1.208839177 | 0.226724637 | NA | | | Ndrg3 | 25733.7713 | 0.062997345 | 0.039594521 | 1.59106219 | 0.111595581 | 0.538992394 |
| 1700061I17Rik | 0.285336609 | 0.031742875 | 0.03528833 | 0.899528977 | 0.368370968 | NA | | | Bst1 | 5.469989599 | 0.063012093 | 0.046710665 | 1.348987289 | 0.177341049 | NA |
| 1700063A18Rik | 0 | NA | NA | NA | NA | NA | | | Ddx47 | 1771.279005 | 0.063015847 | 0.051409112 | 1.225771947 | 0.220284507 | 0.672757795 |
| 1700063O14Rik | 1.386926346 | -0.16451203 | 0.088094712 | -1.867445009 | 0.061839469 | NA | | | Gm16897 | 14.79302018 | 0.06301833 | 0.074335512 | 0.847755379 | 0.396574218 | NA |
| 1700064J06Rik | 0 | NA | NA | NA | NA | NA | | | Pmp22 | 1082.570996 | 0.063033981 | 0.082553014 | 0.763557598 | 0.445130926 | 0.821493054 |
| 1700064M15Rik | 1.360944942 | 0.053265395 | 0.088665645 | 0.600744463 | 0.5480102 | NA | | | Cwc25 | 730.0717149 | 0.063046467 | 0.063867614 | 0.987142984 | 0.323572548 | 0.752925264 |
| 1700065D16Rik | 1.437427219 | 0.011840258 | 0.087890685 | 0.134715731 | 0.892836635 | NA | | | Nr4a3 | 393.1229689 | 0.063048523 | 0.069647906 | 0.905246499 | 0.365334813 | 0.777735733 |
| 1700065I16Rik | 0 | NA | NA | NA | NA | NA | | | A730056A06Rik | 140.7574483 | 0.063058142 | 0.092407111 | 0.682394907 | 0.494989275 | 0.844219116 |
| 1700065I17Rik | 0 | NA | NA | NA | NA | NA | | | Ubl7 | 3060.986888 | 0.063078882 | 0.049361156 | 1.277905285 | 0.201282824 | 0.654716629 |
| 1700065J11Rik | 0.08731684 | 0.017365056 | 0.027733412 | 0.626142072 | 0.531221759 | NA | | | Sox5 | 1692.413849 | 0.063080451 | 0.067583041 | 0.93337692 | 0.350625393 | 0.768667813 |
| 1700065J18Rik | 0.282093074 | 0.009461846 | 0.038656194 | 0.244769212 | 0.806635134 | NA | | | Uri1 | 1518.548269 | 0.063092996 | 0.06824429 | 0.924516845 | 0.355217285 | 0.772206867 |
| 1700065O20Rik | 0 | NA | NA | NA | NA | NA | | | Gltscr2 | 2265.139876 | 0.063094042 | 0.05776563 | 1.092241913 | 0.274726785 | 0.716459834 |
| 1700066B17Rik | 0.199381295 | -0.004055078 | 0.031639813 | -0.128163771 | 0.89801937 | NA | | | Ptprj | 7011.397171 | 0.063100501 | 0.071725731 | 0.879747064 | 0.378996347 | 0.785380516 |
| 1700066B19Rik | 1.988627642 | -0.036136824 | 0.088810508 | -0.406898072 | 0.684082852 | NA | | | Ptcra | 9.517173675 | 0.063105116 | 0.062823389 | 1.004484424 | 0.315145175 | NA |
| 1700066J24Rik | 4.91723478 | 0.019788423 | 0.106371591 | 0.186031088 | 0.852420382 | NA | | | Socs7 | 6749.84427 | 0.063108402 | 0.069238565 | 0.911463168 | 0.362051384 | 0.776449324 |
| 1700066N21Rik | 0 | NA | NA | NA | NA | NA | | | Nthl1 | 93.14107384 | 0.063112723 | 0.09447676 | 0.668023785 | 0.50411841 | 0.847807553 |
| 1700066O22Rik | 0 | NA | NA | NA | NA | NA | | | Fbxw8 | 2203.642126 | 0.063116476 | 0.05489349 | 1.149798919 | 0.250226701 | 0.698461386 |
| 1700067G17Rik | 0 | NA | NA | NA | NA | NA | | | A730098P11Rik | 8.658001106 | 0.063127762 | 0.062387541 | 1.011864875 | 0.311602666 | NA |
| 1700067K01Rik | 1.22238232 | -0.038266874 | 0.075601477 | -0.506165695 | 0.612740335 | NA | | | Slc4a4 | 7230.561922 | 0.06313794 | 0.068284654 | 0.924628536 | 0.355159164 | 0.772206867 |
| 1700067P10Rik | 1.352373962 | 0.01319769 | 0.082785853 | 0.159419633 | 0.873338272 | NA | | | Ssu72 | 2100.945725 | 0.0631747 | 0.049967294 | 1.264321017 | 0.206114829 | 0.660203749 |
| 1700069L16Rik | 0 | NA | NA | NA | NA | NA | | | Rtn4 | 28672.34743 | 0.06318593 | 0.064927197 | 0.973181242 | 0.330463232 | 0.756141871 |
| 1700069P05Rik | 0 | NA | NA | NA | NA | NA | | | Cldn25 | 2475.4394 | 0.063191205 | 0.049115957 | 1.28657181 | 0.198243575 | 0.653581647 |
| 1700071K01Rik | 0 | NA | NA | NA | NA | NA | | | Areg | 3.383278858 | 0.06319761 | 0.041909279 | 1.507962239 | 0.131564192 | NA |
| 1700071M16Rik | 1.025937788 | -0.019215963 | 0.066067457 | -0.290853686 | 0.771163224 | NA | | | Fbxo42 | 2005.240222 | 0.063199962 | 0.051054693 | 1.237887406 | 0.215757813 | 0.669822181 |
| 1700072B07Rik | 0 | NA | NA | NA | NA | NA | | | Nudt16l1 | 1535.975348 | 0.063209026 | 0.060972901 | 1.036640069 | 0.299903632 | 0.733704078 |
| 1700072O05Rik | 0 | NA | NA | NA | NA | NA | | | Fam187b | 3.085768913 | 0.06323424 | 0.037582946 | 1.682524831 | 0.092467114 | NA |
| 1700073E17Rik | 0.079983654 | 0.016740032 | 0.027674178 | 0.604897175 | 0.54524732 | NA | | | Ndufa1 | 1428.3963 | 0.06323626 | 0.064269569 | 0.983922272 | 0.325153732 | 0.75293408 |
| 1700074P13Rik | 0 | NA | NA | NA | NA | NA | | | Tcea3 | 16.16589703 | 0.063237909 | 0.072027409 | 0.877970061 | 0.379959966 | NA |
| 1700080E11Rik | 0 | NA | NA | NA | NA | NA | | | 8430419L09Rik | 14249.83385 | 0.063239274 | 0.048266886 | 1.310200022 | 0.190128178 | 0.644966498 |

| | | | | | | | | | | | | | | | |
|---|---|---|---|---|---|---|---|---|---|---|---|---|---|---|---|
| 1700080N15Rik | 0 | NA | NA | NA | NA | NA | | | Slc25a20 | 1446.197791 | 0.063239612 | 0.053379348 | 1.184720577 | 0.236127944 | 0.687039131 |
| 1700080O16Rik | 0.099009884 | 0.018205726 | 0.027896865 | 0.652608315 | 0.514008823 | NA | | | Abcf1 | 3821.328531 | 0.063266666 | 0.039489356 | 1.602119479 | 0.109129191 | 0.535010276 |
| 1700081H04Rik | 0 | NA | NA | NA | NA | NA | | | Poglut1 | 2210.157318 | 0.063293908 | 0.057703357 | 1.096884318 | 0.272691965 | 0.714142044 |
| 1700081H22Rik | 0 | NA | NA | NA | NA | NA | | | Sf3b2 | 10948.94095 | 0.063323707 | 0.043358875 | 1.460455497 | 0.14416493 | 0.589967032 |
| 1700084F23Rik | 0 | NA | NA | NA | NA | NA | | | Camta1 | 11932.51425 | 0.06336493 | 0.060257948 | 1.051561373 | 0.293000838 | 0.728526371 |
| 1700084J12Rik | 0.098353386 | -0.020212831 | 0.02786466 | -0.725393061 | 0.468210929 | NA | | | Cd82 | 842.8715933 | 0.063396121 | 0.087016558 | 0.728552389 | 0.466275511 | 0.832050282 |
| 1700085B03Rik | 0 | NA | NA | NA | NA | NA | | | Itsn1 | 7098.836112 | 0.063399474 | 0.062724871 | 1.010754951 | 0.312133729 | 0.744800424 |
| 1700085C21Rik | 0.394855623 | -0.002402222 | 0.042167018 | -0.056969207 | 0.954569725 | NA | | | Gfm1 | 1797.379044 | 0.063400241 | 0.060223132 | 1.052755616 | 0.292453012 | 0.728310617 |
| 1700091E21Rik | 1.870113224 | 0.15100164 | 0.099934872 | 1.511000488 | 0.130788327 | NA | | | Lipt1 | 167.2489064 | 0.063414675 | 0.089900533 | 0.705387084 | 0.48056938 | 0.836914993 |
| 1700091H14Rik | 7.58037232 | 0.00378325 | 0.151503259 | 0.024971409 | 0.980077769 | NA | | | Rab6b | 48049.33502 | 0.063437969 | 0.045808683 | 1.384845963 | 0.166099573 | 0.61635484 |
| 1700092C02Rik | 0.08731684 | 0.017365056 | 0.027733412 | 0.626142072 | 0.531221759 | NA | | | Crem | 505.3920905 | 0.063440196 | 0.083824278 | 0.756823643 | 0.449155526 | 0.823263966 |
| 1700092C10Rik | 0 | NA | NA | NA | NA | NA | | | 1700007K13Rik | 77.13125273 | 0.063450203 | 0.095053959 | 0.667517725 | 0.504441492 | 0.847807553 |
| 1700092E19Rik | 0 | NA | NA | NA | NA | NA | | | Nhp2 | 854.9009451 | 0.063502198 | 0.062103068 | 1.022529156 | 0.30653052 | 0.738806192 |
| 1700092K14Rik | 2.149771734 | 0.043087493 | 0.091281036 | 0.472031155 | 0.636904551 | NA | | | Hdgfrp3 | 4927.369758 | 0.063539801 | 0.069232357 | 0.917776069 | 0.358736114 | 0.774639629 |
| 1700092M07Rik | 1.382630839 | 0.038598511 | 0.077139913 | 0.500370169 | 0.616814454 | NA | | | Lrpap1 | 7849.252989 | 0.063547226 | 0.047329967 | 1.3426425 | 0.179387762 | 0.633240978 |
| 1700093K21Rik | 5.207369283 | 0.198452885 | 0.126825587 | 1.564770083 | 0.117636829 | NA | | | Csnk2b | 3037.11541 | 0.063548039 | 0.051954966 | 1.223136961 | 0.22127797 | 0.672757795 |
| 1700094J05Rik | 1.235265611 | 0.066574415 | 0.071141067 | 0.935808499 | 0.349371795 | NA | | | Serbp1 | 5437.705046 | 0.063577205 | 0.056256725 | 1.130126307 | 0.258423006 | 0.703616072 |
| 1700094M24Rik | 0 | NA | NA | NA | NA | NA | | | Pcyox1 | 6091.937461 | 0.063583704 | 0.050024172 | 1.271059591 | 0.20370745 | 0.657622144 |
| 1700095A21Rik | 0 | NA | NA | NA | NA | NA | | | Ttpal | 3629.86097 | 0.063587292 | 0.060429265 | 1.052259904 | 0.292680322 | 0.728496032 |
| 1700095B10Rik | 0.352791569 | -0.038810225 | 0.038158703 | -1.017074017 | 0.30911822 | NA | | | Nsg2 | 27605.96569 | 0.063599337 | 0.055636869 | 1.143114946 | 0.252990869 | 0.700378199 |
| 1700096J18Rik | 0.096071043 | -0.020098795 | 0.027825948 | -0.722304036 | 0.470107572 | NA | | | Capzb | 8003.255689 | 0.063599636 | 0.044706706 | 1.422597229 | 0.154852948 | 0.604489543 |
| 1700097N02Rik | 0.311993147 | -0.014421208 | 0.039860783 | -0.361789373 | 0.717509434 | NA | | | Mpv17l | 2804.856987 | 0.063610375 | 0.057078699 | 1.114432805 | 0.265093556 | 0.707370515 |
| 1700100L14Rik | 0 | NA | NA | NA | NA | NA | | | LOC100861615 | 6.141076021 | 0.063642805 | 0.056427364 | 1.127871302 | 0.259374275 | NA |
| 1700101E01Rik | 6.980296523 | 0.342701974 | 0.153761258 | 2.228792724 | 0.025827699 | NA | | | Ankrd6 | 1781.811736 | 0.063676894 | 0.073769261 | 0.863190066 | 0.388032974 | 0.791889115 |
| 1700101O22Rik | 0 | NA | NA | NA | NA | NA | | | A930017M01Rik | 584.331785 | 0.063693565 | 0.076425375 | 0.833408603 | 0.404614325 | 0.800290023 |
| 1700102H20Rik | 5.830630321 | -0.068945968 | 0.136910925 | -0.503582663 | 0.614554678 | NA | | | Mpped1 | 8852.494075 | 0.063704878 | 0.058277713 | 1.093125907 | 0.274338523 | 0.716185716 |
| 1700104A03Rik | 0.105923158 | 0.018524693 | 0.027977518 | 0.662127842 | 0.507889294 | NA | | | Epc2 | 2140.43699 | 0.063710459 | 0.059970857 | 1.062356992 | 0.28807365 | 0.724103151 |
| 1700105P06Rik | 3.909002408 | 0.146085372 | 0.133234882 | 1.096449896 | 0.272881938 | NA | | | Stambp | 1484.918157 | 0.063743784 | 0.05377054 | 1.185477846 | 0.235828569 | 0.687039131 |
| 1700106J16Rik | 0 | NA | NA | NA | NA | NA | | | Ecsit | 1564.704508 | 0.063750227 | 0.054727356 | 1.164869489 | 0.244071826 | 0.69367782 |
| 1700108F19Rik | 0.495097952 | -0.018652806 | 0.05081183 | -0.367095744 | 0.713547602 | NA | | | Rtl1 | 34.31035068 | 0.063775695 | 0.07163156 | 0.890329567 | 0.37328895 | 0.782226671 |
| 1700108J01Rik | 0.119397641 | -0.020477583 | 0.027954224 | -0.732539845 | 0.463839132 | NA | | | Pnma1 | 171.4126621 | 0.063803043 | 0.091682726 | 0.695911273 | 0.486484397 | 0.839801621 |
| 1700109F18Rik | 0.323149808 | -0.030167563 | 0.033779321 | -0.893077861 | 0.371815478 | NA | | | Rnf216 | 2433.823386 | 0.063807255 | 0.053275493 | 1.197684922 | 0.231039702 | 0.683402059 |
| 1700109G14Rik | 1.172532083 | -0.100619872 | 0.080734232 | -1.246309893 | 0.212650647 | NA | | | Nr2c2 | 7033.064735 | 0.063843631 | 0.059587448 | 1.0714275 | 0.283977253 | 0.721415456 |
| 1700109G15Rik | 0 | NA | NA | NA | NA | NA | | | Snx6 | 1403.770931 | 0.06385707 | 0.060808316 | 1.050137125 | 0.293655072 | 0.729168132 |
| 1700109I08Rik | 0 | NA | NA | NA | NA | NA | | | Pgam5 | 1989.944492 | 0.063874998 | 0.054891937 | 1.163649913 | 0.244565915 | 0.694402753 |
| 1700110I01Rik | 0 | NA | NA | NA | NA | NA | | | Amn1 | 324.1682098 | 0.063875218 | 0.078370156 | 0.815045186 | 0.415046446 | 0.805365305 |
| 1700110K17Rik | 1.860880301 | 0.07121156 | 0.08778855 | 0.811171393 | 0.417267251 | NA | | | Inpp4a | 7425.757087 | 0.063881615 | 0.061542819 | 1.038002755 | 0.299268772 | 0.733704078 |
| 1700110M21Rik | 0.099009884 | 0.018205726 | 0.027896865 | 0.652608315 | 0.514008823 | NA | | | Cul2 | 2352.356581 | 0.063886992 | 0.063157984 | 1.011542621 | 0.311756792 | 0.74424258 |
| 1700111N16Rik | 0.188785634 | 0.025025111 | 0.031461603 | 0.79541754 | 0.42637066 | NA | | | Zc2hc1a | 2173.234254 | 0.063888759 | 0.066244448 | 0.964439455 | 0.334825649 | 0.758238352 |
| 1700112H15Rik | 0 | NA | NA | NA | NA | NA | | | Mxi1 | 3532.067226 | 0.063912739 | 0.059594858 | 1.072453912 | 0.283516204 | 0.721034821 |
| 1700112J05Rik | 0 | NA | NA | NA | NA | NA | | | Psmc4 | 3494.689964 | 0.063951421 | 0.045861187 | 1.394456277 | 0.163179864 | 0.614351408 |
| 1700113H08Rik | 0.66421776 | -0.022294278 | 0.060705401 | -0.367253607 | 0.713429856 | NA | | | Creb1 | 2236.596178 | 0.063962416 | 0.076279862 | 0.838522966 | 0.401737055 | 0.799277436 |
| 1700119H24Rik | 0.496818212 | 0.030597582 | 0.045890909 | 0.666746028 | 0.504934373 | NA | | | Sntg1 | 981.6187438 | 0.06396341 | 0.069841531 | 0.915836303 | 0.35975276 | 0.776027411 |
| 1700120E14Rik | 0 | NA | NA | NA | NA | NA | | | Btbd1 | 5306.902157 | 0.063990755 | 0.055545568 | 1.152040695 | 0.249304351 | 0.697864808 |
| 1700120G07Rik | 0.247284148 | 0.028796741 | 0.033643376 | 0.855940884 | 0.392030479 | NA | | | Pitpnc1 | 3875.589455 | 0.063991781 | 0.06624498 | 0.965986874 | 0.334050749 | 0.758212399 |
| 1700120K04Rik | 1.267252396 | 0.099744843 | 0.074046513 | 1.347056583 | 0.17796201 | NA | | | Foxred2 | 1028.714544 | 0.064004389 | 0.07273047 | 0.880021666 | 0.378847573 | 0.785380516 |
| 1700121L16Rik | 0 | NA | NA | NA | NA | NA | | | Pole3 | 391.5684666 | 0.064007443 | 0.075077193 | 0.852555092 | 0.393906075 | 0.794754188 |
| 1700121N20Rik | 1.017859841 | -0.064033937 | 0.073737633 | -0.868402388 | 0.385174088 | NA | | | Zfp385a | 3383.174848 | 0.06401033 | 0.054777931 | 1.168542306 | 0.242588085 | 0.692772033 |
| 1700122O11Rik | 0 | NA | NA | NA | NA | NA | | | Tmem50b | 3839.535159 | 0.064029567 | 0.053162514 | 1.204411955 | 0.228430394 | 0.681631039 |
| 1700123I01Rik | 2.289901513 | 0.025548849 | 0.106337215 | 0.240262538 | 0.810126735 | NA | | | Yipf2 | 870.0851972 | 0.064031327 | 0.065279378 | 0.980881393 | 0.326651234 | 0.753403593 |
| 1700123K08Rik | 0.569095902 | 0.022553468 | 0.05830359 | 0.386828113 | 0.698883463 | NA | | | Hspa12a | 11063.82947 | 0.064036016 | 0.05515284 | 1.161064717 | 0.245615582 | 0.69504627 |
| 1700123L14Rik | 1.304710972 | 0.030808095 | 0.071729445 | 0.429504154 | 0.667556372 | NA | | | Fam3c | 2579.360557 | 0.064037401 | 0.061721903 | 1.037515017 | 0.299495901 | 0.733704078 |
| 1700123O12Rik | 0 | NA | NA | NA | NA | NA | | | Snx17 | 3266.924324 | 0.06404338 | 0.0432825 | 1.479659921 | 0.138964027 | 0.583135914 |
| 1700123O21Rik | 0.199381295 | -0.004055078 | 0.031639813 | -0.128163771 | 0.89801937 | NA | | | Cacnb1 | 4443.635518 | 0.0640705 | 0.066959039 | 0.956861098 | 0.338637369 | 0.760991092 |
| 1700125G02Rik | 0.38258743 | 0.019046329 | 0.042646787 | 0.446606435 | 0.655159248 | NA | | | Acot13 | 1503.917596 | 0.064071546 | 0.063020215 | 1.016682446 | 0.309304519 | 0.742145027 |
| 1700125G22Rik | 0.391976801 | -0.003620343 | 0.047492484 | -0.076229803 | 0.939236272 | NA | | | Cox16 | 304.7187999 | 0.064071856 | 0.082361601 | 0.777933591 | 0.436608164 | 0.816448274 |
| 1700125H03Rik | 1.302823544 | -0.056690712 | 0.074863899 | -0.757250325 | 0.448899905 | NA | | | Erlin2 | 2559.467494 | 0.064078234 | 0.054232598 | 1.181544609 | 0.237386443 | 0.688415946 |
| 1700125H20Rik | 2.701148872 | 0.116217923 | 0.115881161 | 1.002906098 | 0.31590617 | NA | | | Stt3b | 4295.42964 | 0.064083953 | 0.062412689 | 1.026777622 | 0.304525189 | 0.737453291 |
| 1700126H18Rik | 0 | NA | NA | NA | NA | NA | | | Zfp507 | 1314.407529 | 0.064106423 | 0.055574595 | 1.153520298 | 0.248696891 | 0.697784358 |
| 1700128A07Rik | 0 | NA | NA | NA | NA | NA | | | Pgap2 | 362.6601981 | 0.064110288 | 0.071031925 | 0.902555965 | 0.366761606 | 0.778316887 |
| 1700128F08Rik | 0 | NA | NA | NA | NA | NA | | | Abcd3 | 3023.997832 | 0.064138684 | 0.054364229 | 1.179795703 | 0.238081479 | 0.688781431 |
| 1700129C05Rik | 0 | NA | NA | NA | NA | NA | | | Hddc3 | 409.9594481 | 0.064149183 | 0.080567476 | 0.796216866 | 0.425905996 | 0.809519334 |
| 1810006J02Rik | 0 | NA | NA | NA | NA | NA | | | Rsph9 | 357.5396842 | 0.064152957 | 0.080138962 | 0.800521429 | 0.423408753 | 0.808999423 |
| 1810007C17Rik | 0 | NA | NA | NA | NA | NA | | | Fgl1 | 9.77194487 | 0.064170587 | 0.063959014 | 1.003307948 | 0.315712303 | NA |
| 1810007D17Rik | 1.365135949 | 0.016537606 | 0.084668701 | 0.195321361 | 0.845141375 | NA | | | Ndufa13 | 3693.434647 | 0.06418445 | 0.059781071 | 1.07365842 | 0.282975804 | 0.720920007 |
| 1810007I06Rik | 0 | NA | NA | NA | NA | NA | | | Clmn | 4733.764666 | 0.064195151 | 0.075402433 | 0.8513671 | 0.394565462 | 0.795437883 |
| 1810008I18Rik | 0 | NA | NA | NA | NA | NA | | | Dohh | 1820.862141 | 0.064195854 | 0.074466204 | 0.862080393 | 0.38864328 | 0.79235741 |
| 1810009J06Rik | 0 | NA | NA | NA | NA | NA | | | Copz2 | 158.5035938 | 0.064207921 | 0.092694394 | 0.69268397 | 0.488507901 | 0.840318276 |
| 1810010D01Rik | 4.821765732 | 0.036112285 | 0.121594504 | 0.296989449 | 0.766474564 | NA | | | Cdk14 | 8884.548188 | 0.064209449 | 0.051862882 | 1.238061724 | 0.215693175 | 0.669812326 |
| 1810012K16Rik | 5.815315053 | 0.209559767 | 0.143749318 | 1.45781399 | 0.144891821 | NA | | | Rb1cc1 | 4343.638024 | 0.064212156 | 0.075468354 | 0.850848764 | 0.394853369 | 0.795622145 |
| 1810013A23Rik | 0 | NA | NA | NA | NA | NA | | | Fam122b | 243.288841 | 0.064224574 | 0.081906152 | 0.784123925 | 0.432967398 | 0.813783127 |
| 1810018F18Rik | 0 | NA | NA | NA | NA | NA | | | Zfp933 | 585.3369672 | 0.064237457 | 0.064764357 | 0.991864352 | 0.321263701 | 0.750510858 |
| 1810019D21Rik | 2.62439483 | 0.164416764 | 0.105783488 | 1.554276254 | 0.120118535 | NA | | | Gnptab | 3873.801898 | 0.064250364 | 0.048676548 | 1.319944952 | 0.186853398 | 0.641054259 |
| 1810019J16Rik | 4.780621242 | -0.028542864 | 0.133833793 | -0.213270977 | 0.83111561 | NA | | | Zscan21 | 711.8900308 | 0.064278435 | 0.06546413 | 0.981887869 | 0.326155093 | 0.753049596 |
| 1810034E14Rik | 6.08985207 | -0.097671016 | 0.140143226 | -0.696937117 | 0.485842148 | NA | | | Btbd2 | 3229.529778 | 0.064297315 | 0.061341893 | 1.048179502 | 0.294555913 | 0.72962173 |
| 1810035I16Rik | 0 | NA | NA | NA | NA | NA | | | A730090H04Rik | 17.38644728 | 0.064303716 | 0.079626722 | 0.807564534 | 0.41934131 | NA |
| 1810046K07Rik | 0.294043327 | 0.010508917 | 0.039092051 | 0.2688249 | 0.788064431 | NA | | | Nucb1 | 3357.893159 | 0.064310435 | 0.056416888 | 1.139914613 | 0.254321876 | 0.701619624 |
| 1810053B23Rik | 0 | NA | NA | NA | NA | NA | | | Thrb | 2834.719377 | 0.064318798 | 0.065773307 | 0.977886035 | 0.328130693 | 0.754778534 |
| 1810062G17Rik | 3.223892963 | 0.062676836 | 0.125600841 | 0.499016055 | 0.617768073 | NA | | | Csk | 1182.876189 | 0.064324967 | 0.054834313 | 1.173078747 | 0.240764228 | 0.691863635 |
| 1810064F22Rik | 0.284856677 | 0.008904946 | 0.033416532 | 0.266483248 | 0.789867059 | NA | | | 4933426M11Rik | 2473.335655 | 0.064361042 | 0.05390041 | 1.19407332 | 0.232449289 | 0.684327865 |
| 1810065E05Rik | 0.308183275 | 0.007047773 | 0.033991235 | 0.207340891 | 0.835743642 | NA | | | Suv39h1 | 1288.646807 | 0.064365515 | 0.066905966 | 0.962029531 | 0.33603477 | 0.759375714 |
| 2010003K11Rik | 0.308183275 | 0.007047773 | 0.033991235 | 0.207340891 | 0.835743642 | NA | | | Faf2 | 2483.602393 | 0.064371165 | 0.047164854 | 1.36481215 | 0.172312099 | 0.623258355 |
| 2010005H15Rik | 1.375106598 | -0.141443519 | 0.075358259 | -1.876947794 | 0.060525249 | NA | | | Ndufs2 | 7542.694531 | 0.064374118 | 0.040817086 | 1.577136518 | 0.114764116 | 0.543130934 |
| 2010009K17Rik | 0 | NA | NA | NA | NA | NA | | | A4gnt | 9.347932523 | 0.064386711 | 0.063767257 | 1.009714293 | 0.312632193 | NA |
| 2010010A06Rik | 0 | NA | NA | NA | NA | NA | | | Eif3c | 8001.520838 | 0.064387959 | 0.048305557 | 1.332930674 | 0.182554555 | 0.635424885 |
| 2010016I18Rik | 0.408673203 | -0.002822473 | 0.042495157 | -0.066418693 | 0.947044488 | NA | | | Mrpl14 | 366.3344351 | 0.064389679 | 0.080090993 | 0.803956552 | 0.421422068 | 0.808555694 |
| 2010106C02Rik | 0.079983654 | 0.016740032 | 0.027674178 | 0.604897175 | 0.54524732 | NA | | | Galnt1 | 2503.410677 | 0.064396828 | 0.058695027 | 1.097142835 | 0.272578958 | 0.714142044 |
| 2010106E10Rik | 0.398790038 | -0.006260381 | 0.047666372 | -0.131337473 | 0.895508349 | NA | | | Mamld1 | 1354.721044 | 0.064401376 | 0.074337269 | 0.866340361 | 0.386303539 | 0.790344395 |
| 2010109A12Rik | 1.249209019 | -0.013485133 | 0.089181369 | -0.151210208 | 0.879809898 | NA | | | Sae1 | 3776.683371 | 0.06440322 | 0.04586012 | 1.404340404 | 0.160217508 | 0.612291377 |
| 2010308F09Rik | 6.56425722 | 0.305770883 | 0.151325555 | 2.020616298 | 0.0433195 | NA | | | Snw1 | 3377.999625 | 0.064419573 | 0.059776248 | 1.077678435 | 0.281177286 | 0.720080692 |
| 2010310C07Rik | 0 | NA | NA | NA | NA | NA | | | Glrx | 1422.542194 | 0.064431552 | 0.060475904 | 1.06540866 | 0.28669104 | 0.722148665 |
| 2200002D01Rik | 2.751391874 | -0.1236684 | 0.111516433 | -1.10897019 | 0.267443042 | NA | | | Ano4 | 954.4957674 | 0.064440993 | 0.060617056 | 1.063083507 | 0.287744083 | 0.723769792 |
| 2200002J24Rik | 0 | NA | NA | NA | NA | NA | | | Gm1976 | 963.0075212 | 0.064445416 | 0.07488903 | 0.860545471 | 0.38948843 | 0.792715562 |
| 2210010C04Rik | 0.430100701 | -0.046155006 | 0.041541744 | -1.111051218 | 0.266546304 | NA | | | Rsrc1 | 1739.978208 | 0.064468734 | 0.054033112 | 1.193133822 | 0.232816968 | 0.684628272 |
| 2210019I11Rik | 0 | NA | NA | NA | NA | NA | | | Edem3 | 2318.533988 | 0.064474788 | 0.070124055 | 0.919438963 | 0.357866018 | 0.77402647 |
| 2210407C18Rik | 1.119479332 | -0.053604557 | 0.078463285 | -0.683180128 | 0.494493028 | NA | | | Rtn3 | 54052.64243 | 0.064493014 | 0.054320894 | 1.187259797 | 0.235125161 | 0.686276191 |
| 2210409D07Rik | 1.094514179 | -0.05351922 | 0.067126426 | -0.79728987 | 0.425282702 | NA | | | Adi1 | 1210.517759 | 0.064494424 | 0.053417933 | 1.207355303 | 0.227295341 | 0.680203999 |
| 2210409E12Rik | 0.188785634 | 0.025025111 | 0.031461603 | 0.79541754 | 0.42637066 | NA | | | Ldlr | 1116.443046 | 0.06453303 | 0.094709682 | 0.681377331 | 0.495632763 | 0.844369474 |
| 2210414B05Rik | 0 | NA | NA | NA | NA | NA | | | Mir703 | 11.36090892 | 0.064582386 | 0.058032452 | 1.112866756 | 0.26576566 | NA |
| 2210416O15Rik | 1.648555063 | 0.089082826 | 0.096799845 | 0.920278602 | 0.357427188 | NA | | | Rpp25l | 396.6345551 | 0.064613466 | 0.078475752 | 0.823355804 | 0.410305688 | 0.80184319 |
| 2210417A02Rik | 0.159967308 | 0.0240345 | 0.030870969 | 0.778547006 | 0.436246606 | NA | | | Lrpprc | 3268.486324 | 0.064620628 | 0.05565261 | 1.161142818 | 0.245583824 | 0.69504627 |
| 2210420H20Rik | 0.098353386 | -0.020212831 | 0.02786466 | -0.725393061 | 0.468210929 | NA | | | Gpr180 | 937.7739912 | 0.064653129 | 0.060059071 | 1.076492324 | 0.281707132 | 0.720163706 |
| 2300002M23Rik | 0.824582689 | -0.009713019 | 0.062308342 | -0.155886342 | 0.876122611 | NA | | | Prim2 | 299.9606647 | 0.064672389 | 0.088858995 | 0.727809142 | 0.466730427 | 0.832095317 |
| 2300003K06Rik | 0 | NA | NA | NA | NA | NA | | | Slbp | 847.7891587 | 0.064679449 | 0.064427894 | 1.003904436 | 0.315424678 | 0.745993052 |
| 2300005B03Rik | 0.768098354 | -0.017953171 | 0.049913376 | -0.359686576 | 0.719081532 | NA | | | Zbtb11 | 2332.297816 | 0.064697312 | 0.080967493 | 0.799052925 | 0.424259724 | 0.809162684 |
| 2310001H17Rik | 5.601893641 | 0.034433326 | 0.127895942 | 0.26922923 | 0.787753288 | NA | | | Wrap53 | 468.8541712 | 0.064699174 | 0.081580186 | 0.793074616 | 0.427734351 | 0.810334558 |
| 2310001K24Rik | 0.107716603 | -0.020477583 | 0.027954224 | -0.732539849 | 0.46383913 | NA | | | Tnfrsf12a | 148.4324896 | 0.064713178 | 0.093506865 | 0.692068736 | 0.488894164 | 0.840634071 |
| 2310002D06Rik | 0.20671448 | -0.003314027 | 0.03178208 | -0.10427343 | 0.916952363 | NA | | | Slc35c2 | 2070.874074 | 0.064721 | 0.058715764 | 1.102276379 | 0.270341535 | 0.71216373 |
| 2310002F09Rik | 6.146738874 | 0.343511533 | 0.134253687 | 2.558674865 | 0.010507196 | NA | | | Gpc5 | 1630.89767 | 0.064742692 | 0.056171487 | 1.15258996 | 0.249078726 | 0.697784358 |
| 2310002J15Rik | 0.867184421 | 0.09414792 | 0.068136439 | 1.381755799 | 0.167046695 | NA | | | A230001M10Rik | 173.113857 | 0.06475362 | 0.095039145 | 0.681336305 | 0.495658716 | 0.844369474 |
| 2310002L09Rik | 0.105923158 | 0.018524693 | 0.027977518 | 0.662127842 | 0.507889294 | NA | | | Pgm3 | 598.7955767 | 0.064782373 | 0.072575495 | 0.892620481 | 0.372060447 | 0.781371686 |
| 2310005A03Rik | 0.094392817 | 0.017895942 | 0.027818137 | 0.643319236 | 0.520016977 | NA | | | Tmem183a | 2221.324488 | 0.06482197 | 0.058091365 | 1.1158624 | 0.264481039 | 0.707370515 |
| 2310005E17Rik | 0 | NA | NA | NA | NA | NA | | | Slc35a4 | 2298.56425 | 0.0648294 | 0.050260976 | 1.289855591 | 0.197100802 | 0.653065027 |
| 2310005G13Rik | 1.542146306 | 0.042012054 | 0.090068075 | 0.466447787 | 0.640895013 | NA | | | Aplp2 | 26177.45044 | 0.064845083 | 0.045049532 | 1.439417446 | 0.150032284 | 0.598234686 |
| 2310007B03Rik | 1.318607928 | 0.094192747 | 0.080628132 | 1.168236742 | 0.242711284 | NA | | | 2810429I04Rik | 28.44363196 | 0.064865032 | 0.085251203 | 0.760869404 | 0.446735075 | 0.822210607 |
| 2310007L24Rik | 3.804890523 | 0.232236128 | 0.131254596 | 1.769356159 | 0.076834457 | NA | | | Pmepa1 | 2491.679021 | 0.064866868 | 0.069854352 | 0.928601669 | 0.353095556 | 0.770519168 |
| 2310008N11Rik | 0.17463368 | 0.024560283 | 0.031186242 | 0.787535833 | 0.430968261 | NA | | | Ociad2 | 3690.307749 | 0.064871268 | 0.060927778 | 1.064724002 | 0.287000845 | 0.722400671 |
| 2310014F07Rik | 1.484926722 | 0.099947604 | 0.090094341 | 1.109366068 | 0.267272294 | NA | | | Pdzd8 | 3081.970805 | 0.064875878 | 0.067952078 | 0.95472986 | 0.339714326 | 0.761679979 |
| 2310015B20Rik | 2.653928998 | 0.017448554 | 0.106482186 | 0.163863598 | 0.869838524 | NA | | | Ak8 | 93.4145262 | 0.064880385 | 0.094535522 | 0.68630695 | 0.492519567 | 0.842479891 |
| 2310015D24Rik | 5.901268509 | 0.262936099 | 0.142809491 | 1.841166836 | 0.065597114 | NA | | | Trappc9 | 3660.918189 | 0.064894961 | 0.050652993 | 1.281167335 | 0.200134897 | 0.654470849 |
| 2310016D03Rik | 0 | NA | NA | NA | NA | NA | | | Cnot6l | 675.0700039 | 0.064903481 | 0.085766837 | 0.756743321 | 0.449203656 | 0.823263966 |
| 2310020H05Rik | 0 | NA | NA | NA | NA | NA | | | Nop14 | 1040.10871 | 0.064927588 | 0.066762833 | 0.97251098 | 0.330796405 | 0.756212998 |
| 2310030A07Rik | 0.096071043 | -0.020098795 | 0.027825948 | -0.722304036 | 0.470107572 | NA | | | Cdpf1 | 322.4609421 | 0.064942521 | 0.084341772 | 0.769992371 | 0.441304418 | 0.818962636 |
| 2310034C09Rik | 0 | NA | NA | NA | NA | NA | | | Fndc4 | 3402.041871 | 0.064955196 | 0.056285846 | 1.154023639 | 0.248490478 | 0.697784358 |
| 2310034O05Rik | 4.304087238 | 0.013054971 | 0.128955331 | 0.101236377 | 0.91936282 | NA | | | Commd5 | 471.4291037 | 0.064977974 | 0.07538763 | 0.861918254 | 0.388732503 | 0.79235741 |
| 2310043L19Rik | 1.683619574 | -0.027326437 | 0.085849419 | -0.318306603 | 0.750252376 | NA | | | Susd4 | 4078.104016 | 0.06498153 | 0.049792968 | 1.305034269 | 0.191881179 | 0.646997681 |
| 2310043O21Rik | 0.094392817 | 0.017895942 | 0.027818137 | 0.643319236 | 0.520016977 | NA | | | Smarce1 | 2149.81454 | 0.0649831 | 0.049776777 | 1.305490311 | 0.191725944 | 0.646997681 |
| 2310050C09Rik | 0 | NA | NA | NA | NA | NA | | | Trim32 | 8998.297681 | 0.064998787 | 0.05390725 | 1.205752232 | 0.227913039 | 0.680415543 |
| 2310057J18Rik | 0 | NA | NA | NA | NA | NA | | | Pold2 | 499.4649789 | 0.065007201 | 0.072812798 | 0.892799109 | 0.371964764 | 0.781368063 |
| 2310057N15Rik | 0 | NA | NA | NA | NA | NA | | | Wipf2 | 4780.804919 | 0.065020042 | 0.066455605 | 0.978398165 | 0.327877435 | 0.754778534 |
| 2310061N02Rik | 0 | NA | NA | NA | NA | NA | | | Zfp334 | 986.7695268 | 0.065035399 | 0.0600154 | 1.083645173 | 0.278522154 | 0.71849589 |
| 2310065F04Rik | 3.373993267 | 0.025467206 | 0.12625687 | 0.201709471 | 0.840143858 | NA | | | Hnrnpk | 13442.92726 | 0.065036209 | 0.052666497 | 1.234868689 | 0.216879378 | 0.670185967 |
| 2310068J16Rik | 6.350782555 | -0.130231293 | 0.150385883 | -0.865980837 | 0.386500671 | NA | | | Wibg | 603.8396559 | 0.06508218 | 0.068729773 | 0.946928494 | 0.343675217 | 0.763762832 |
| 2310069B03Rik | 0.222620122 | -0.001447027 | 0.032059667 | -0.04513545 | 0.963999345 | NA | | | Fam72a | 54.50845018 | 0.065082747 | 0.093138859 | 0.698771144 | 0.484695065 | 0.838713365 |
| 2310079G19Rik | 0 | NA | NA | NA | NA | NA | | | 1110057K04Rik | 1771.079259 | 0.065102735 | 0.052577979 | 1.238212968 | 0.215637104 | 0.669812326 |
| 2310081J21Rik | 0.079983654 | 0.016740032 | 0.027674178 | 0.604897175 | 0.54524732 | NA | | | Cdkl5 | 2591.540022 | 0.065120724 | 0.076169041 | 0.85495004 | 0.392578806 | 0.794184423 |
| 2410003L11Rik | 1.778094025 | 0.009087409 | 0.098055755 | 0.092675939 | 0.926161013 | NA | | | Rap1gds1 | 18058.04604 | 0.065139447 | 0.062414917 | 1.043651896 | 0.296646473 | 0.73196648 |
| 2410007B07Rik | 0.494509999 | 0.03737204 | 0.052072961 | 0.717686096 | 0.472950858 | NA | | | Lingo2 | 943.0089269 | 0.065169829 | 0.083710198 | 0.778517206 | 0.436264167 | 0.816120192 |
| 2410012E07Rik | 0 | NA | NA | NA | NA | NA | | | Skiv2l2 | 2990.258478 | 0.065182514 | 0.057708478 | 1.12951366 | 0.258681211 | 0.703616072 |
| 2410012M07Rik | 0.08731684 | 0.017365056 | 0.027733412 | 0.626142072 | 0.531221759 | NA | | | Uap1l1 | 602.9527432 | 0.065184037 | 0.074086628 | 0.879835383 | 0.378948494 | 0.785380516 |
| 2410017I17Rik | 0 | NA | NA | NA | NA | NA | | | Ppp6c | 1330.257402 | 0.065191426 | 0.069957436 | 0.931872712 | 0.351402311 | 0.769244462 |
| 2410021H03Rik | 0.159967308 | 0.0240345 | 0.030870969 | 0.778547006 | 0.436246606 | NA | | | Sh2d4a | 10.66255757 | 0.065194843 | 0.064445946 | 1.011620544 | 0.311719519 | NA |
| 2410057H14Rik | 3.283049985 | -0.087052309 | 0.12333788 | -0.705803515 | 0.480310336 | NA | | | Arhgap44 | 6756.842509 | 0.065229868 | 0.045620041 | 1.429851163 | 0.152759741 | 0.601351762 |
| 2410076I21Rik | 4.680900391 | -0.06182659 | 0.121138272 | -0.510380318 | 0.609785043 | NA | | | Gabarap | 5130.223609 | 0.065243867 | 0.056693396 | 1.150819532 | 0.249806488 | 0.698227639 |
| 2410088K16Rik | 0 | NA | NA | NA | NA | NA | | | Tssc1 | 1624.957373 | 0.065246489 | 0.067769829 | 0.962766026 | 0.335664954 | 0.759126439 |
| 2410114N07Rik | 0 | NA | NA | NA | NA | NA | | | Nt5e | 361.2258381 | 0.065247524 | 0.091283016 | 0.714782734 | 0.474743306 | 0.834491965 |
| 2410124H12Rik | 1.206912129 | -0.098297715 | 0.071099962 | -1.382528372 | 0.166809525 | NA | | | Tbc1d14 | 4529.808611 | 0.065274977 | 0.052711266 | 1.238349635 | 0.215586447 | 0.669812326 |
| 2410137F16Rik | 1.577581099 | 0.043788493 | 0.086383756 | 0.506906564 | 0.61222038 | NA | | | Tmem179 | 5593.783962 | 0.06530085 | 0.06202906 | 1.052746098 | 0.292457375 | 0.728310617 |
| 2410141K09Rik | 0 | NA | NA | NA | NA | NA | | | Minpp1 | 2565.450014 | 0.065311947 | 0.052047566 | 1.254851144 | 0.209532802 | 0.664226905 |
| 2510003D18Rik | 0.639497534 | 0.043360211 | 0.050060413 | 0.866157682 | 0.386403697 | NA | | | Agpat6 | 4066.440426 | 0.065319571 | 0.039097502 | 1.670683978 | 0.094784115 | 0.512787358 |
| 2510049J12Rik | 1.751491653 | 0.009023362 | 0.079900997 | 0.112931778 | 0.910084642 | NA | | | Chic2 | 201.6589089 | 0.065329 | 0.084945669 | 0.76906805 | 0.441852914 | 0.819294796 |
| 2600006L11Rik | 0 | NA | NA | NA | NA | NA | | | Zfp292 | 1714.158627 | 0.065335835 | 0.073483932 | 0.889117301 | 0.373940042 | 0.782462265 |
| 2610016A17Rik | 0.660333954 | -0.101834242 | 0.060586068 | -1.680819444 | 0.092797991 | NA | | | Atp1b2 | 24295.2928 | 0.065336984 | 0.068909708 | 0.948153546 | 0.343051292 | 0.763762832 |
| 2610027K06Rik | 2.452697712 | 0.02012729 | 0.114137613 | 0.176342303 | 0.86002503 | NA | | | Dnahc6 | 121.4181679 | 0.065344112 | 0.090522782 | 0.721852671 | 0.470385062 | 0.833245343 |
| 2610028H24Rik | 2.231059021 | 0.134326748 | 0.102585819 | 1.309408549 | 0.190395997 | NA | | | Zmynd19 | 727.4106122 | 0.065363114 | 0.062007938 | 1.054108819 | 0.291833099 | 0.727847831 |
| 2610035F20Rik | 6.484895822 | -0.102606829 | 0.148692803 | -0.690059149 | 0.490156992 | NA | | | Ip6k2 | 2616.516774 | 0.065366183 | 0.058204702 | 1.123039551 | 0.261420697 | 0.704173028 |
| 2610037D02Rik | 2.54646873 | 0.128129051 | 0.112391189 | 1.140027538 | 0.254274829 | NA | | | Ddx3x | 12272.80136 | 0.065367824 | 0.063275493 | 1.033067007 | 0.301572542 | 0.73587145 |
| 2610206C17Rik | 1.068033753 | 0.068003709 | 0.080072483 | 0.84927689 | 0.395727237 | NA | | | Zfp938 | 615.7400012 | 0.065367997 | 0.07852872 | 0.832408796 | 0.405178239 | 0.8005027 |
| 2610307P16Rik | 7.718447562 | -0.385415747 | 0.147331387 | -2.615978546 | 0.008897215 | NA | | | Tmem255b | 53.58792924 | 0.065372881 | 0.091884422 | 0.711468602 | 0.476793901 | 0.835195112 |
| 2610528J11Rik | 1.22007994 | -0.050163986 | 0.086589842 | -0.57932876 | 0.562367363 | NA | | | Ptpn1 | 2173.654056 | 0.065382073 | 0.078865062 | 0.829037231 | 0.40708334 | 0.800975798 |
| 2700086A05Rik | 0 | NA | NA | NA | NA | NA | | | Fdx1 | 215.459652 | 0.06538658 | 0.085244536 | 0.767047168 | 0.443053472 | 0.820159899 |
| 2700089I24Rik | 0.107716603 | -0.020477583 | 0.027954224 | -0.732539849 | 0.46383913 | NA | | | Nek2 | 69.47983999 | 0.065403085 | 0.094764593 | 0.690163735 | 0.490091226 | 0.841397607 |
| 2810007J24Rik | 0 | NA | NA | NA | NA | NA | | | Erc1 | 2701.031583 | 0.065425523 | 0.060394291 | 1.083306419 | 0.278672437 | 0.71849589 |
| 2810404M03Rik | 4.904431192 | 0.047550992 | 0.130091949 | 0.365518331 | 0.714724526 | NA | | | Cbl | 1362.001873 | 0.065430623 | 0.085684369 | 0.76362379 | 0.445091468 | 0.821493054 |
| 2810405F15Rik | 2.621914376 | -0.036617563 | 0.107992482 | -0.339075109 | 0.734553149 | NA | | | Ntmt1 | 661.0942672 | 0.065445015 | 0.076582241 | 0.854571684 | 0.392788309 | 0.794436505 |
| 2810442I21Rik | 1.913210347 | -0.036894559 | 0.089449939 | -0.412460413 | 0.680001991 | NA | | | Anxa8 | 23.55428055 | 0.065447804 | 0.082889179 | 0.789582002 | 0.429771923 | 0.811499668 |
| 2810442N19Rik | 0.096071043 | -0.020098795 | 0.027825948 | -0.722304036 | 0.470107572 | NA | | | Magel2 | 203.5709119 | 0.065458445 | 0.058543387 | 1.118118515 | 0.26351638 | 0.706200557 |
| 2810471M01Rik | 0 | NA | NA | NA | NA | NA | | | A030009H04Rik | 4598.088121 | 0.065459159 | 0.060138561 | 1.088472312 | 0.276386653 | 0.717475105 |
| 2900057B20Rik | 0 | NA | NA | NA | NA | NA | | | Luzp2 | 6396.981403 | 0.065468913 | 0.056199905 | 1.164929233 | 0.244047639 | 0.69367782 |
| 2900060B14Rik | 0.096071043 | -0.020098795 | 0.027825948 | -0.722304036 | 0.470107572 | NA | | | D430041D05Rik | 9268.905441 | 0.065541189 | 0.071297662 | 0.919261412 | 0.357958856 | 0.774087808 |
| 2900092C05Rik | 0.105923158 | 0.018524693 | 0.027977518 | 0.662127842 | 0.507889294 | NA | | | Igdcc3 | 68.32232789 | 0.065552172 | 0.092346509 | 0.709850035 | 0.477797138 | 0.835457925 |
| 2900093L17Rik | 1.048127784 | 0.10735569 | 0.071642996 | 1.498481295 | 0.134008249 | NA | | | Ipo7 | 4807.592728 | 0.06555955 | 0.07268959 | 0.901911127 | 0.36710408 | 0.778490943 |
| 3100003L05Rik | 1.071449219 | -0.042018488 | 0.076873758 | -0.546590785 | 0.584659902 | NA | | | Fam195b | 2091.122114 | 0.065589956 | 0.064933281 | 1.010113082 | 0.312441115 | 0.744874388 |
| 3110009F21Rik | 0.902191072 | 0.013251232 | 0.063848263 | 0.207542558 | 0.835586159 | NA | | | B230219D22Rik | 4365.181058 | 0.065601654 | 0.067252223 | 0.975457026 | 0.329333608 | 0.756141871 |
| 3110015C05Rik | 1.176732261 | 0.066978284 | 0.07561092 | 0.885828185 | 0.375710119 | NA | | | Ift43 | 254.7814741 | 0.065601803 | 0.079216584 | 0.828132186 | 0.407595644 | 0.801055189 |
| 3110021A11Rik | 5.658365749 | 0.139867743 | 0.140350995 | 0.99655683 | 0.318979669 | NA | | | Zmat3 | 4692.980104 | 0.065607623 | 0.067564491 | 0.971037038 | 0.331529835 | 0.756606816 |
| 3110079O15Rik | 6.442475272 | -0.011307627 | 0.141434464 | -0.07994959 | 0.936277349 | NA | | | Slc25a19 | 1616.561762 | 0.06561445 | 0.060041598 | 1.092816508 | 0.274474372 | 0.716185716 |
| 3110099E03Rik | 7.584837601 | -0.210768864 | 0.133039148 | -1.584261978 | 0.113134114 | NA | | | Aig1 | 918.7739701 | 0.065631036 | 0.060746128 | 1.080415143 | 0.279957356 | 0.719332628 |
| 3200001D21Rik | 0.211846316 | 0.025707621 | 0.031860375 | 0.806883811 | 0.419733426 | NA | | | Snx31 | 8.724798734 | 0.065633318 | 0.062596296 | 1.04851761 | 0.294400193 | NA |
| 3300005D01Rik | 0.313786592 | -0.036039695 | 0.036798757 | -0.979372609 | 0.327395906 | NA | | | Spen | 3518.302242 | 0.065646741 | 0.070027549 | 0.937441652 | 0.348531424 | 0.767434814 |
| 3830403N18Rik | 2.284420513 | -0.010745656 | 0.097777774 | -0.109898762 | 0.912489664 | NA | | | Oaf | 429.4222265 | 0.065657027 | 0.078139675 | 0.840252116 | 0.400767043 | 0.799169897 |
| 3830417A13Rik | 0 | NA | NA | NA | NA | NA | | | Pycr2 | 749.0113552 | 0.065672622 | 0.072067937 | 0.911259913 | 0.362158443 | 0.776449324 |
| 4732416N19Rik | 7.177832134 | 0.12671669 | 0.146110089 | 0.867268586 | 0.385794865 | NA | | | Hgsnat | 2559.376835 | 0.065677957 | 0.051543566 | 1.27422222 | 0.202584673 | 0.65606599 |
| 4732456N10Rik | 4.55523015 | 0.089755881 | 0.10153235 | 0.884012637 | 0.376689391 | NA | | | Tbc1d8b | 246.1292695 | 0.065680573 | 0.090645845 | 0.72458449 | 0.468706977 | 0.832446623 |
| 4732490B19Rik | 0.539953942 | 0.069964435 | 0.051895658 | 1.348175111 | 0.177602068 | NA | | | 6430548M08Rik | 17036.46239 | 0.065686743 | 0.052152817 | 1.259505177 | 0.207847922 | 0.662965736 |
| 4833403I15Rik | 3.239762966 | 0.126166008 | 0.101478368 | 1.243279829 | 0.213764734 | NA | | | Chrm4 | 2155.539563 | 0.065696597 | 0.074520726 | 0.88158826 | 0.377999506 | 0.784627154 |
| 4833419F23Rik | 5.458823561 | 0.033687057 | 0.141094359 | 0.238755518 | 0.811295164 | NA | | | Ptprf | 3917.438714 | 0.06569772 | 0.070252313 | 0.935168072 | 0.34970169 | 0.768497238 |
| 4833423E24Rik | 5.246093872 | -0.129244546 | 0.134098976 | -0.963799647 | 0.335146384 | NA | | | Gins4 | 540.988037 | 0.065699128 | 0.077017306 | 0.853043714 | 0.393635063 | 0.794754188 |
| 4833427F10Rik | 0.285336609 | 0.031742875 | 0.03528833 | 0.899528977 | 0.368370968 | NA | | | Kif21a | 9545.683983 | 0.065734853 | 0.048040308 | 1.36832704 | 0.17120973 | 0.621282174 |
| 4833427G06Rik | 5.338523982 | 0.267809996 | 0.144546026 | 1.852766231 | 0.063915872 | NA | | | Dbc1 | 10570.58606 | 0.065740069 | 0.05705084 | 1.15230677 | 0.249195036 | 0.697861402 |
| 4833428L15Rik | 0.969874391 | -0.007538203 | 0.071551649 | -0.105353313 | 0.916095461 | NA | | | Tmem70 | 1165.906088 | 0.065756079 | 0.068736507 | 0.956639809 | 0.338749089 | 0.761006691 |
| 4921501E09Rik | 0.279364949 | 0.005060019 | 0.033327057 | 0.151829152 | 0.879321689 | NA | | | Oxgr1 | 13.54600362 | 0.06575966 | 0.066406557 | 0.990258543 | 0.322047764 | NA |
| 4921504E06Rik | 0.502404983 | 0.007339048 | 0.055675222 | 0.13181893 | 0.895127513 | NA | | | Slitrk6 | 64.60520279 | 0.065829578 | 0.077795779 | 0.846184452 | 0.397449854 | 0.797424942 |
| 4921506M07Rik | 0 | NA | NA | NA | NA | NA | | | Pigu | 933.3138883 | 0.065839043 | 0.056869526 | 1.157720976 | 0.246977921 | 0.696511265 |
| 4921507L20Rik | 0.345637534 | 0.019330126 | 0.041637051 | 0.464253003 | 0.642466491 | NA | | | Matn4 | 290.632908 | 0.065849028 | 0.086560933 | 0.760724567 | 0.446821598 | 0.822210607 |
| 4921508D12Rik | 0.740533532 | 0.01739619 | 0.062889035 | 0.276617211 | 0.782074048 | NA | | | Prdm11 | 122.5796996 | 0.065869186 | 0.090022669 | 0.731695549 | 0.464354414 | 0.83159896 |
| 4921509C19Rik | 0.238795281 | -0.027601606 | 0.032267774 | -0.855392297 | 0.392334007 | NA | | | Cacna1i | 5366.095026 | 0.065881892 | 0.061258786 | 1.075468463 | 0.282165045 | 0.720537371 |
| 4921509O07Rik | 1.096585181 | -0.028447705 | 0.083638729 | -0.340125984 | 0.733761655 | NA | | | Tspan9 | 2185.312416 | 0.065904643 | 0.048033549 | 1.372054413 | 0.1700465 | 0.620316915 |
| 4921511C10Rik | 0.496152747 | 0.00873411 | 0.050596247 | 0.172623666 | 0.862947248 | NA | | | 1810009A15Rik | 372.5924978 | 0.065909833 | 0.079092797 | 0.833322821 | 0.404662689 | 0.800290023 |
| 4921511C20Rik | 0 | NA | NA | NA | NA | NA | | | Map1lc3b | 8770.993507 | 0.065966749 | 0.043629799 | 1.511965462 | 0.130542649 | 0.569619261 |
| 4921511H03Rik | 0 | NA | NA | NA | NA | NA | | | 9930014A18Rik | 92.62978358 | 0.065985781 | 0.094787201 | 0.696146525 | 0.486337073 | 0.839801621 |
| 4921511I17Rik | 0.283178451 | 0.028390488 | 0.033368195 | 0.850824814 | 0.394866675 | NA | | | Api5 | 4076.523725 | 0.06598843 | 0.060022561 | 1.099393787 | 0.27159634 | 0.713790099 |
| 4921511M17Rik | 0 | NA | NA | NA | NA | NA | | | Cnnm2 | 1253.078455 | 0.06599525 | 0.054943191 | 1.2011543 | 0.229691353 | 0.682724729 |
| 4921513I03Rik | 0 | NA | NA | NA | NA | NA | | | Myg1 | 930.0516972 | 0.065999662 | 0.061195716 | 1.078501345 | 0.280810082 | 0.720080692 |
| 4921515E04Rik | 0.302603776 | 0.00963845 | 0.039370813 | 0.244812063 | 0.806601953 | NA | | | Ddx24 | 8997.721972 | 0.066034 | 0.046130531 | 1.43145978 | 0.152298486 | 0.600512434 |
| 4921517D22Rik | 0.238795281 | -0.027601606 | 0.032267774 | -0.855392297 | 0.392334007 | NA | | | Aarsd1 | 1900.21336 | 0.066041838 | 0.0602034 | 1.096978542 | 0.272650772 | 0.714142044 |
| 4921524L21Rik | 0 | NA | NA | NA | NA | NA | | | Tmc1 | 14.99237296 | 0.066050167 | 0.072076444 | 0.916390473 | 0.35946213 | NA |
| 4921525B02Rik | 0.470066974 | 0.035966715 | 0.05156661 | 0.697480692 | 0.485502018 | NA | | | Polr2e | 1712.271554 | 0.066054014 | 0.056783673 | 1.163257159 | 0.244725182 | 0.694476029 |
| 4921530D09Rik | 0.107716603 | -0.020477583 | 0.027954224 | -0.732539849 | 0.46383913 | NA | | | 6330403A02Rik | 14847.70288 | 0.066066198 | 0.074078798 | 0.891836797 | 0.372480416 | 0.781446026 |
| 4921530L18Rik | 7.312735488 | 0.072810329 | 0.155593381 | 0.467952615 | 0.639818476 | NA | | | Gnb1 | 26820.25464 | 0.066095788 | 0.040790926 | 1.620355168 | 0.105156004 | 0.526359793 |
| 4921530L21Rik | 1.817384064 | 0.014184023 | 0.080545876 | 0.176098685 | 0.860216414 | NA | | | 9030624J02Rik | 2454.331406 | 0.066096803 | 0.055200843 | 1.197387563 | 0.231155529 | 0.683402059 |
| 4921531P14Rik | 0 | NA | NA | NA | NA | NA | | | Susd5 | 321.2174868 | 0.066099464 | 0.08276207 | 0.798668566 | 0.424482618 | 0.809162684 |
| 4921533I20Rik | 0 | NA | NA | NA | NA | NA | | | Slc35a5 | 1830.40936 | 0.066101323 | 0.06133329 | 1.077739734 | 0.281149921 | 0.720080692 |
| 4921539E11Rik | 0 | NA | NA | NA | NA | NA | | | Mknk2 | 1884.511799 | 0.066106707 | 0.071088498 | 0.929921285 | 0.352411842 | 0.770257962 |
| 4922502D21Rik | 0 | NA | NA | NA | NA | NA | | | Zfp444 | 1112.267742 | 0.066119595 | 0.057196321 | 1.156011325 | 0.247676525 | 0.696944196 |
| 4922502H24Rik | 0 | NA | NA | NA | NA | NA | | | Ttc3 | 47349.97241 | 0.066129742 | 0.055540508 | 1.190657853 | 0.23378793 | 0.685064073 |
| 4922502N22Rik | 0.430253841 | 0.022125686 | 0.043852981 | 0.504542347 | 0.613880312 | NA | | | Gm5124 | 809.7497056 | 0.066134389 | 0.071327569 | 0.927192522 | 0.353826585 | 0.77126125 |
| 4922505E12Rik | 0.555364211 | 0.060011321 | 0.059945413 | 1.001099473 | 0.31677872 | NA | | | Wdr82 | 6918.831816 | 0.066141245 | 0.045391007 | 1.45714424 | 0.145076569 | 0.590634035 |
| 4930401C15Rik | 0 | NA | NA | NA | NA | NA | | | Zfp770 | 1035.512961 | 0.066152384 | 0.082027434 | 0.806466579 | 0.419973871 | 0.807657282 |
| 4930401O10Rik | 0 | NA | NA | NA | NA | NA | | | Tcf25 | 29243.05939 | 0.066152609 | 0.037563159 | 1.761103477 | 0.078220891 | 0.475440315 |
| 4930401O12Rik | 0 | NA | NA | NA | NA | NA | | | Myo16 | 1426.063034 | 0.066162228 | 0.065361196 | 1.012255474 | 0.311415918 | 0.744017702 |
| 4930402F06Rik | 0.56395571 | 0.017428757 | 0.047045094 | 0.370469173 | 0.711032941 | NA | | | Lamtor5 | 1213.304515 | 0.066186164 | 0.058414399 | 1.133045365 | 0.257195203 | 0.703199912 |
| 4930402F11Rik | 0 | NA | NA | NA | NA | NA | | | Arl2 | 1753.651004 | 0.066197813 | 0.056598341 | 1.169606954 | 0.242159177 | 0.692772033 |
| 4930402K13Rik | 0 | NA | NA | NA | NA | NA | | | Elof1 | 1132.889576 | 0.066216002 | 0.059843355 | 1.106488794 | 0.268515012 | 0.710723225 |
| 4930404A05Rik | 0 | NA | NA | NA | NA | NA | | | Srp54a | 136.7049572 | 0.066229526 | 0.089843114 | 0.737168637 | 0.461019804 | 0.829630038 |
| 4930404A10Rik | 0.192746203 | -0.000975302 | 0.031567365 | -0.030895906 | 0.975352555 | NA | | | Nrip2 | 744.9196021 | 0.066255635 | 0.089835627 | 0.7375207 | 0.460805759 | 0.829450367 |
| 4930404H11Rik | 2.137369726 | 0.104910205 | 0.10463188 | 1.002660038 | 0.316024917 | NA | | | Thg1l | 211.3284299 | 0.066300901 | 0.082904094 | 0.799730116 | 0.42386718 | 0.808999423 |
| 4930404I05Rik | 0.761449927 | -0.003542071 | 0.055175041 | -0.064196988 | 0.948813375 | NA | | | Srpx2 | 79.37463046 | 0.066303205 | 0.094903893 | 0.698635254 | 0.484780006 | 0.838722531 |
| 4930404N11Rik | 4.061198555 | -0.06188911 | 0.102513511 | -0.60371662 | 0.546032062 | NA | | | Gstm4 | 403.889741 | 0.066325898 | 0.077980674 | 0.850542765 | 0.395023395 | 0.795790167 |
| 4930405A10Rik | 0 | NA | NA | NA | NA | NA | | | Tbc1d12 | 623.5509863 | 0.066347074 | 0.081313298 | 0.81594371 | 0.414532331 | 0.804797733 |
| 4930405D11Rik | 0.107716603 | -0.020477583 | 0.027954224 | -0.732539849 | 0.46383913 | NA | | | Atp6v0a4 | 13.88961335 | 0.066359221 | 0.068482442 | 0.968996119 | 0.332547125 | NA |
| 4930405J17Rik | 0 | NA | NA | NA | NA | NA | | | Mcm10 | 24.72572504 | 0.066381083 | 0.086076394 | 0.771188005 | 0.440595503 | 0.818563572 |
| 4930405P13Rik | 0.20210942 | -0.001755803 | 0.031727492 | -0.055340098 | 0.955867518 | NA | | | 2610035D17Rik | 607.6685015 | 0.066415784 | 0.073820147 | 0.899697255 | 0.368281385 | 0.779243711 |
| 4930406D18Rik | 0 | NA | NA | NA | NA | NA | | | Zmpste24 | 1353.600645 | 0.066423647 | 0.052407729 | 1.267439909 | 0.204998037 | 0.658514261 |
| 4930407I10Rik | 0.094392817 | 0.017895942 | 0.027818137 | 0.643319236 | 0.520016977 | NA | | | Larp1 | 6702.731412 | 0.066434186 | 0.042835937 | 1.550898376 | 0.12092604 | 0.555002664 |

| | | | | | | | | | | | | | | | |
|---|---|---|---|---|---|---|---|---|---|---|---|---|---|---|---|
| 4930408F14Rik | 0 | NA | NA | NA | NA | NA | | | Paip1 | 2662.877352 | 0.066446072 | 0.072296365 | 0.919079012 | 0.358054246 | 0.774087808 |
| 4930412B13Rik | 0 | NA | NA | NA | NA | NA | | | Stx8 | 763.2681556 | 0.066465384 | 0.061713777 | 1.076994271 | 0.281482825 | 0.720080692 |
| 4930412O13Rik | 0.66557215 | 0.04559458 | 0.049811598 | 0.915340648 | 0.360012826 | NA | | | Rnf7 | 2227.328874 | 0.066488499 | 0.060209274 | 1.104289994 | 0.269467362 | 0.711972822 |
| 4930413E15Rik | 0 | NA | NA | NA | NA | NA | | | 2610306M01Rik | 176.9390866 | 0.066491859 | 0.087131056 | 0.76312468 | 0.445389042 | 0.821656239 |
| 4930413F20Rik | 3.863764201 | 0.021671767 | 0.125973555 | 0.172034259 | 0.863410596 | NA | | | Rrp36 | 500.706703 | 0.066501711 | 0.067292176 | 0.988253241 | 0.323028639 | 0.752792382 |
| 4930413G21Rik | 3.735394276 | -0.053166696 | 0.126238523 | -0.421160631 | 0.67363779 | NA | | | Ppp1r32 | 54.73790139 | 0.066550593 | 0.090107013 | 0.738572839 | 0.460166422 | 0.829022384 |
| 4930413M19Rik | 0 | NA | NA | NA | NA | NA | | | Flot2 | 5091.606352 | 0.066577955 | 0.052409048 | 1.270352311 | 0.203959162 | 0.65778952 |
| 4930414N06Rik | 0.574278041 | 0.012588346 | 0.03989752 | 0.315517008 | 0.75236914 | NA | | | Praf2 | 3131.873801 | 0.066583734 | 0.068289959 | 0.975014991 | 0.329552824 | 0.756141871 |
| 4930415F15Rik | 6.800379707 | 0.161059409 | 0.128369336 | 1.254656401 | 0.209603519 | NA | | | Ptpn14 | 755.9632099 | 0.06661518 | 0.094509066 | 0.704854919 | 0.480900527 | 0.836995094 |
| 4930415L06Rik | 0 | NA | NA | NA | NA | NA | | | Psmc2 | 3938.541794 | 0.066643344 | 0.045311797 | 1.47077249 | 0.141352655 | 0.586989325 |
| 4930415O20Rik | 1.339143676 | -0.02737891 | 0.081591324 | -0.335561533 | 0.737201536 | NA | | | Xrcc2 | 293.566991 | 0.066650973 | 0.083322116 | 0.799919359 | 0.42375752 | 0.808999423 |
| 4930417O13Rik | 4.964691473 | -0.356244184 | 0.141475704 | -2.518059094 | 0.011800352 | NA | | | Rassf8 | 330.3216505 | 0.066675613 | 0.08725283 | 0.764165619 | 0.444768552 | 0.821493054 |
| 4930417O22Rik | 1.440912637 | 0.05008834 | 0.082749727 | 0.605299152 | 0.544980245 | NA | | | Fkbp14 | 440.07528 | 0.066690127 | 0.077990701 | 0.855103564 | 0.392493817 | 0.794184423 |
| 4930419G24Rik | 1.09495344 | 0.103030006 | 0.078097338 | 1.319251196 | 0.187085148 | NA | | | Psma5 | 2362.589153 | 0.066697626 | 0.050683681 | 1.315958609 | 0.188187935 | 0.642130211 |
| 4930423M02Rik | 0 | NA | NA | NA | NA | NA | | | Clptm1 | 7576.310765 | 0.066722693 | 0.045876447 | 1.454399757 | 0.145835507 | 0.592514474 |
| 4930425K10Rik | 0.17463368 | 0.024560283 | 0.031186242 | 0.787535833 | 0.430968261 | NA | | | Lyrm7 | 471.4010191 | 0.066742892 | 0.074171903 | 0.899840633 | 0.368205067 | 0.779243711 |
| 4930425O10Rik | 0.593519161 | -0.035580971 | 0.059082103 | -0.60222925 | 0.547021548 | NA | | | Ywhaz | 47863.67434 | 0.066754892 | 0.056719421 | 1.176931836 | 0.239222714 | 0.689747746 |
| 4930428D18Rik | 0 | NA | NA | NA | NA | NA | | | Cep170b | 13060.53013 | 0.066783253 | 0.059660142 | 1.119394812 | 0.262971743 | 0.70568202 |
| 4930428E07Rik | 0.116696964 | -0.020477583 | 0.027954224 | -0.732539847 | 0.463839131 | NA | | | Sdc2 | 2040.638072 | 0.066792613 | 0.070094187 | 0.952898043 | 0.340641732 | 0.762546123 |
| 4930428G15Rik | 0.712869317 | -0.061164802 | 0.067168122 | -0.91062248 | 0.362494321 | NA | | | Armc6 | 946.9346066 | 0.066795745 | 0.066609883 | 1.002790309 | 0.315962045 | 0.745993052 |
| 4930428O21Rik | 0 | NA | NA | NA | NA | NA | | | D5Ertd579e | 6869.052419 | 0.066797928 | 0.072310991 | 0.923758985 | 0.355611814 | 0.772370994 |
| 4930429D17Rik | 0 | NA | NA | NA | NA | NA | | | Apln | 1177.523283 | 0.066802182 | 0.08434411 | 0.792019525 | 0.428349292 | 0.810486244 |
| 4930429F11Rik | 0 | NA | NA | NA | NA | NA | | | Pias2 | 2562.443234 | 0.066837043 | 0.061278502 | 1.090709475 | 0.27540074 | 0.717061772 |
| 4930429F24Rik | 1.878795999 | -0.006604545 | 0.109082731 | -0.060546206 | 0.951720616 | NA | | | Rsl1d1 | 1937.993221 | 0.066874421 | 0.061355241 | 1.089954492 | 0.275733191 | 0.717346645 |
| 4930430A15Rik | 0 | NA | NA | NA | NA | NA | | | Elavl3 | 10388.289 | 0.066888209 | 0.050310977 | 1.329495331 | 0.183684605 | 0.637355821 |
| 4930430D24Rik | 0 | NA | NA | NA | NA | NA | | | Lefty2 | 17.65535745 | 0.066936662 | 0.073624229 | 0.909166219 | 0.363262393 | NA |
| 4930430F21Rik | 0.401377075 | -0.004225889 | 0.047720475 | -0.08855505 | 0.929435533 | NA | | | Irf2bp1 | 1667.780787 | 0.066949941 | 0.048552981 | 1.378904861 | 0.167924089 | 0.617953076 |
| 4930430J02Rik | 0 | NA | NA | NA | NA | NA | | | Hnrnph2 | 5113.969162 | 0.066954524 | 0.048093977 | 1.39216027 | 0.163873871 | 0.614351408 |
| 4930431F12Rik | 2.357898175 | 0.060875398 | 0.098157387 | 0.620181528 | 0.535138281 | NA | | | Itgb1bp1 | 826.6367653 | 0.066968857 | 0.058430498 | 1.146128466 | 0.251742001 | 0.699465245 |
| 4930431P03Rik | 1.064313701 | -0.021011496 | 0.076941904 | -0.273082613 | 0.784789711 | NA | | | Tpbg | 581.4412408 | 0.066994439 | 0.083383245 | 0.803452055 | 0.421713498 | 0.808687824 |
| 4930432J09Rik | 0.649580461 | 0.072845422 | 0.063527461 | 1.146676111 | 0.251515507 | NA | | | Nudt10 | 500.7109192 | 0.066995588 | 0.081193658 | 0.825133267 | 0.409295935 | 0.801055189 |
| 4930432K09Rik | 0 | NA | NA | NA | NA | NA | | | Nog | 205.7963875 | 0.067070754 | 0.08879907 | 0.755309203 | 0.450063478 | 0.823263966 |
| 4930432M17Rik | 0 | NA | NA | NA | NA | NA | | | Prdx4 | 444.2162913 | 0.067107078 | 0.075468472 | 0.889206795 | 0.373891952 | 0.782462265 |
| 4930433B08Rik | 0 | NA | NA | NA | NA | NA | | | C78339 | 972.2169703 | 0.067114992 | 0.071206672 | 0.942537975 | 0.345917278 | 0.765097832 |
| 4930433I11Rik | 0.185906812 | 0.024895454 | 0.031385107 | 0.793225068 | 0.427646705 | NA | | | Nsun2 | 3189.606666 | 0.067126572 | 0.043570826 | 1.540631144 | 0.123406584 | 0.558269708 |
| 4930433N12Rik | 0.094392817 | 0.017895942 | 0.027818137 | 0.643319236 | 0.520016977 | NA | | | Nedd4 | 23601.30461 | 0.06721919 | 0.053357618 | 1.259786186 | 0.207746504 | 0.662799412 |
| 4930434J06Rik | 0.17833704 | -0.002390466 | 0.031282281 | -0.076415987 | 0.939088151 | NA | | | Pex16 | 739.1143716 | 0.067219447 | 0.070826934 | 0.949066164 | 0.342586962 | 0.763762832 |
| 4930435E12Rik | 0 | NA | NA | NA | NA | NA | | | Fhad1 | 281.7001767 | 0.067232943 | 0.091679819 | 0.733345073 | 0.463347991 | 0.831245266 |
| 4930438E09Rik | 0 | NA | NA | NA | NA | NA | | | Kcna2 | 14557.04114 | 0.067293174 | 0.068258924 | 0.985851666 | 0.324205908 | 0.752925264 |
| 4930440C22Rik | 0 | NA | NA | NA | NA | NA | | | Gpx4 | 8268.645602 | 0.067309629 | 0.050901251 | 1.322357055 | 0.186049282 | 0.639960187 |
| 4930441J16Rik | 0 | NA | NA | NA | NA | NA | | | Vcp | 15351.01958 | 0.067343033 | 0.043067819 | 1.563650868 | 0.117899582 | 0.548913246 |
| 4930442J19Rik | 0 | NA | NA | NA | NA | NA | | | Hyal3 | 28.9094863 | 0.067356816 | 0.088037678 | 0.765090788 | 0.444217485 | 0.821372605 |
| 4930442L01Rik | 0 | NA | NA | NA | NA | NA | | | Hnrnpf | 3467.150263 | 0.067356985 | 0.051763781 | 1.301237741 | 0.193177091 | 0.649744561 |
| 4930443O20Rik | 0.185670226 | -0.001645086 | 0.031432796 | -0.052336624 | 0.958260472 | NA | | | Copz1 | 3746.707802 | 0.06738345 | 0.042653813 | 1.579775532 | 0.114158281 | 0.542980743 |
| 4930444F02Rik | 0 | NA | NA | NA | NA | NA | | | Ptplad1 | 7755.320922 | 0.06739126 | 0.051892413 | 1.298672691 | 0.19405628 | 0.65013848 |
| 4930444G20Rik | 0.380940254 | 0.025976956 | 0.047990576 | 0.54129287 | 0.588305734 | NA | | | Nlrp10 | 10.63405449 | 0.067405204 | 0.062113738 | 1.08518995 | 0.277837534 | NA |
| 4930444M15Rik | 2.513581065 | 0.181080302 | 0.110869587 | 1.633272988 | 0.102411595 | NA | | | Slc27a4 | 4262.290948 | 0.067412642 | 0.053173088 | 1.267796243 | 0.204870724 | 0.658461308 |
| 4930447A16Rik | 0 | NA | NA | NA | NA | NA | | | 2410016O06Rik | 529.9190121 | 0.067418862 | 0.06959215 | 0.968771078 | 0.33265942 | 0.757637305 |
| 4930447J18Rik | 0 | NA | NA | NA | NA | NA | | | Mrps7 | 1489.562191 | 0.067440714 | 0.057513892 | 1.172598687 | 0.240956776 | 0.691863635 |
| 4930447K03Rik | 7.44066142 | 0.025083196 | 0.155849513 | 0.160944977 | 0.872136737 | NA | | | Fam115a | 9072.443785 | 0.067449541 | 0.043681172 | 1.54413304 | 0.122556106 | 0.557198747 |
| 4930448C13Rik | 0 | NA | NA | NA | NA | NA | | | Stxbp1 | 46683.02007 | 0.06745241 | 0.040840218 | 1.651617305 | 0.09861259 | 0.520050241 |
| 4930448F12Rik | 0.08731684 | 0.017365056 | 0.027733412 | 0.626142072 | 0.531221759 | NA | | | Adam19 | 1067.919936 | 0.067455357 | 0.074162422 | 0.909562488 | 0.363053289 | 0.776821522 |
| 4930448H16Rik | 0 | NA | NA | NA | NA | NA | | | Apoa2 | 25.11865789 | 0.067472187 | 0.084675118 | 0.79683606 | 0.42554625 | 0.809519334 |
| 4930448I06Rik | 0 | NA | NA | NA | NA | NA | | | Zfp704 | 3341.806908 | 0.06747345 | 0.07263533 | 0.928934317 | 0.352923127 | 0.770519168 |
| 4930448I18Rik | 0 | NA | NA | NA | NA | NA | | | Arih1 | 4352.880523 | 0.067501299 | 0.07269425 | 0.928564483 | 0.353114835 | 0.770519168 |
| 4930448K20Rik | 0 | NA | NA | NA | NA | NA | | | Msantd3 | 366.0076895 | 0.067525483 | 0.077002522 | 0.87692561 | 0.380527045 | 0.786275719 |
| 4930449E01Rik | 0.08731684 | 0.017365056 | 0.027733412 | 0.626142072 | 0.531221759 | NA | | | Fam134a | 9301.710451 | 0.067545534 | 0.040795689 | 1.655702755 | 0.097782029 | 0.518456434 |
| 4930449E18Rik | 0 | NA | NA | NA | NA | NA | | | Nolc1 | 3019.987935 | 0.067550974 | 0.045408814 | 1.487618107 | 0.136851635 | 0.581089406 |
| 4930449I24Rik | 0 | NA | NA | NA | NA | NA | | | Ush2a | 5.553128838 | 0.067570226 | 0.04704939 | 1.43615519 | 0.150958186 | NA |
| 4930451I11Rik | 2.411114941 | -0.043380949 | 0.11076614 | -0.391644498 | 0.695320905 | NA | | | Atad1 | 4481.84066 | 0.067576852 | 0.059802352 | 1.130003247 | 0.258474856 | 0.703616072 |
| 4930452A19Rik | 0.430253841 | 0.022125686 | 0.043852981 | 0.504542347 | 0.613880312 | NA | | | Zswim1 | 1193.107524 | 0.067598492 | 0.056832444 | 1.189434889 | 0.234268578 | 0.68557298 |
| 4930452N14Rik | 1.232026643 | -0.00778314 | 0.066195024 | -0.117578928 | 0.906401302 | NA | | | Pik3c2a | 1107.504017 | 0.067615335 | 0.068500348 | 0.98708017 | 0.323603338 | 0.752925264 |
| 4930453H23Rik | 0.079983654 | 0.016740032 | 0.027674178 | 0.604897175 | 0.54524732 | NA | | | Lrif1 | 420.3430366 | 0.06763452 | 0.084430167 | 0.801070547 | 0.423090806 | 0.808999423 |
| 4930453L07Rik | 0.212768007 | -0.02694223 | 0.031868764 | -0.845411806 | 0.397880955 | NA | | | Strn | 2837.023923 | 0.06763509 | 0.08218834 | 0.822928054 | 0.410548908 | 0.802184613 |
| 4930455B14Rik | 0.832005321 | -0.09663103 | 0.066726197 | -1.4481723 | 0.147568869 | NA | | | Anp32e | 4576.2442 | 0.067655125 | 0.052743741 | 1.282713817 | 0.199592357 | 0.654021787 |
| 4930455C13Rik | 0.543893889 | 0.070201746 | 0.051980176 | 1.350548462 | 0.17684012 | NA | | | Arpc1a | 6032.413455 | 0.067702848 | 0.040189415 | 1.68459402 | 0.092066926 | 0.505579538 |
| 4930455D15Rik | 0.119397641 | -0.020477583 | 0.027954224 | -0.732539845 | 0.463839132 | NA | | | Atpif1 | 3983.457429 | 0.06771174 | 0.054507833 | 1.242238712 | 0.214148501 | 0.668253731 |
| 4930455F16Rik | 0.410740291 | -0.005062362 | 0.047943713 | -0.105589704 | 0.915907894 | NA | | | Dpy30 | 598.2482697 | 0.067729891 | 0.07776784 | 0.870924156 | 0.383795566 | 0.78787654 |
| 4930455H04Rik | 0.861910856 | 0.04703845 | 0.058004879 | 0.810939544 | 0.41740039 | NA | | | Arl10 | 405.0621861 | 0.067741656 | 0.086471331 | 0.783400168 | 0.433392158 | 0.813891982 |
| 4930455J16Rik | 0 | NA | NA | NA | NA | NA | | | C1qtnf1 | 410.6121988 | 0.067756294 | 0.073817535 | 0.917888876 | 0.358677046 | 0.774627234 |
| 4930456L15Rik | 0 | NA | NA | NA | NA | NA | | | Igsf3 | 793.4124178 | 0.067767397 | 0.063324048 | 1.070168441 | 0.284543497 | 0.721641599 |
| 4930459L07Rik | 0 | NA | NA | NA | NA | NA | | | Coq5 | 1514.554985 | 0.067774574 | 0.054162488 | 1.251319441 | 0.210817955 | 0.665414392 |
| 4930461G14Rik | 0.462752832 | 0.005884822 | 0.049969805 | 0.117767555 | 0.906251838 | NA | | | Pcyt1b | 2767.012006 | 0.067779086 | 0.052524857 | 1.29041922 | 0.196905142 | 0.652874709 |
| 4930465A12Rik | 0 | NA | NA | NA | NA | NA | | | Itih2 | 149.4433111 | 0.067782269 | 0.092264316 | 0.734653137 | 0.462550768 | 0.83094391 |
| 4930465K10Rik | 1.060764143 | -0.020894344 | 0.066152052 | -0.315853302 | 0.752113858 | NA | | | Uqcr11 | 2486.795524 | 0.067794891 | 0.0718251 | 0.943888568 | 0.345226594 | 0.764823708 |
| 4930465M20Rik | 0.386222787 | 0.053439715 | 0.045634083 | 1.171048305 | 0.241579361 | NA | | | Fzd9 | 179.3480099 | 0.067812621 | 0.092804183 | 0.730706511 | 0.464958438 | 0.831682622 |
| 4930467D21Rik | 2.016869231 | 0.118619644 | 0.094075112 | 1.260903567 | 0.207343593 | NA | | | Rangap1 | 11689.09376 | 0.067846031 | 0.046970896 | 1.444427032 | 0.148618891 | 0.596199396 |
| 4930467E23Rik | 0 | NA | NA | NA | NA | NA | | | Wwc1 | 3544.21486 | 0.067890673 | 0.05565777 | 1.219787861 | 0.222545304 | 0.67374894 |
| 4930467K11Rik | 2.686659219 | -0.053715893 | 0.107438707 | -0.499967794 | 0.617097755 | NA | | | Lrrc49 | 3466.944977 | 0.067900383 | 0.055677706 | 1.219525503 | 0.222644802 | 0.67374894 |
| 4930468A15Rik | 0.38092772 | -0.000766687 | 0.035977 | -0.021310467 | 0.982997994 | NA | | | Nudt17 | 284.8727331 | 0.067918543 | 0.087472945 | 0.776451995 | 0.437482154 | 0.816715817 |
| 4930469G21Rik | 0.258320694 | 0.006870708 | 0.033097595 | 0.207589347 | 0.835549623 | NA | | | Kndc1 | 6326.594644 | 0.067947913 | 0.067586492 | 1.005347532 | 0.314729535 | 0.745993052 |
| 4930470P17Rik | 0 | NA | NA | NA | NA | NA | | | Adamtsl1 | 352.0561341 | 0.06795086 | 0.087871676 | 0.773296507 | 0.439346923 | 0.817545617 |
| 4930471C04Rik | 6.068235747 | 0.148690851 | 0.13758957 | 1.080684028 | 0.27983769 | NA | | | H2afy2 | 683.0636943 | 0.06796715 | 0.065301108 | 1.04082691 | 0.297955888 | 0.732916829 |
| 4930471G03Rik | 0.079983654 | 0.016740032 | 0.027674178 | 0.604897175 | 0.54524732 | NA | | | Hiat1 | 2235.002827 | 0.067998806 | 0.065163434 | 1.043511697 | 0.296711366 | 0.73200226 |
| 4930471I20Rik | 0.815331231 | 0.023730175 | 0.06737475 | 0.352211695 | 0.724679507 | NA | | | Cttnbp2nl | 1602.565155 | 0.068082399 | 0.053257388 | 1.278365326 | 0.201120644 | 0.654716629 |
| 4930471M09Rik | 0.820973202 | -0.049337952 | 0.072961054 | -0.676223119 | 0.498898996 | NA | | | Lysmd2 | 1189.765707 | 0.068094712 | 0.055890752 | 1.21835383 | 0.223089542 | 0.674141604 |
| 4930473O22Rik | 0.265890466 | 0.030259369 | 0.03447172 | 0.877802699 | 0.3800508 | NA | | | Cebpa | 385.6366623 | 0.068095924 | 0.085166844 | 0.799559085 | 0.423966301 | 0.808999423 |
| 4930474G06Rik | 6.829476927 | -0.008029662 | 0.144632649 | -0.05551763 | 0.955726085 | NA | | | Slc9a6 | 6194.681242 | 0.068099422 | 0.062864112 | 1.083279793 | 0.278684251 | 0.71849589 |
| 4930474H20Rik | 0 | NA | NA | NA | NA | NA | | | Kat7 | 4346.627133 | 0.068108902 | 0.046070011 | 1.478378246 | 0.139306565 | 0.583665174 |
| 4930474M22Rik | 0 | NA | NA | NA | NA | NA | | | Map2k1 | 12014.39273 | 0.068114796 | 0.058977021 | 1.154937892 | 0.248115861 | 0.697476023 |
| 4930474N05Rik | 0 | NA | NA | NA | NA | NA | | | Atp10b | 110.6800002 | 0.068121031 | 0.09506608 | 0.716565064 | 0.473642508 | 0.833983942 |
| 4930474N09Rik | 3.418441142 | -0.06720669 | 0.121752223 | -0.551995593 | 0.580951372 | NA | | | Mphosph10 | 724.6335733 | 0.068145917 | 0.082520287 | 0.825808048 | 0.408912988 | 0.801055189 |
| 4930478L05Rik | 0 | NA | NA | NA | NA | NA | | | Avpr1a | 13.19834505 | 0.068148503 | 0.053423323 | 1.275632045 | NA | NA |
| 4930478P22Rik | 0 | NA | NA | NA | NA | NA | | | Csde1 | 9247.926197 | 0.068156302 | 0.057994009 | 1.175230034 | 0.239902698 | 0.690751361 |
| 4930479M11Rik | 0.08731684 | 0.017365056 | 0.027733412 | 0.626142072 | 0.531221759 | NA | | | Cpa6 | 9.879624675 | 0.068161363 | 0.061984076 | 1.099659252 | 0.271480615 | NA |
| 4930480E11Rik | 0 | NA | NA | NA | NA | NA | | | Sdhaf2 | 1601.830204 | 0.068180324 | 0.055228548 | 1.234512325 | 0.217012057 | 0.670185967 |
| 4930480G23Rik | 0.094392817 | 0.017895942 | 0.027818137 | 0.643319236 | 0.520016977 | NA | | | Mlf2 | 15778.97725 | 0.068190061 | 0.046032862 | 1.481334381 | 0.138517491 | 0.583109099 |
| 4930480K15Rik | 0 | NA | NA | NA | NA | NA | | | Rab11b | 10097.97235 | 0.068202115 | 0.037089506 | 1.838852059 | 0.065936949 | 0.446279967 |
| 4930480M12Rik | 0 | NA | NA | NA | NA | NA | | | Alg10b | 1247.728985 | 0.06820329 | 0.076035815 | 0.896989001 | 0.369724786 | 0.780397655 |
| 4930483J18Rik | 1.04385259 | 0.004672836 | 0.066845675 | 0.069904832 | 0.944269407 | NA | | | Ctnnbl1 | 1130.314371 | 0.06820837 | 0.05663938 | 1.204256995 | 0.228490263 | 0.681669657 |
| 4930483K19Rik | 0.239950962 | 0.027417169 | 0.032831728 | 0.835081494 | 0.403671825 | NA | | | Wdr12 | 1413.923785 | 0.068209953 | 0.056347936 | 1.210513795 | 0.2260818 | 0.678525668 |
| 4930483O08Rik | 1.313476444 | 0.109374984 | 0.086680888 | 1.261811991 | 0.207016446 | NA | | | Trappc6b | 2322.942246 | 0.068210017 | 0.056402393 | 1.209346162 | 0.226529883 | 0.679565252 |
| 4930486F22Rik | 0 | NA | NA | NA | NA | NA | | | Lipa | 1010.983124 | 0.06821836 | 0.057302944 | 1.19048613 | 0.233855378 | 0.685064073 |
| 4930486I03Rik | 2.188966278 | -0.12387563 | 0.109669576 | -1.12953505 | 0.258672193 | NA | | | Sdk2 | 554.5047146 | 0.068226591 | 0.087941321 | 0.775819494 | 0.437855572 | 0.816984132 |
| 4930486L24Rik | 6.815623883 | 0.105972792 | 0.134994638 | 0.785014825 | 0.432444876 | NA | | | Maats1 | 69.76084729 | 0.068235079 | 0.094845002 | 0.719437793 | 0.471871218 | 0.833361862 |
| 4930487D11Rik | 0.096071043 | -0.020098795 | 0.027825948 | -0.722304036 | 0.470107572 | NA | | | Zfp35 | 975.9423863 | 0.06824982 | 0.077298492 | 0.882938562 | 0.377269467 | 0.783997209 |
| 4930487H11Rik | 0.187700257 | -0.003168524 | 0.031449734 | -0.100748845 | 0.919749836 | NA | | | Tfb2m | 969.69367 | 0.068252397 | 0.06344105 | 1.075839653 | 0.281998975 | 0.720294902 |
| 4930488B22Rik | 0 | NA | NA | NA | NA | NA | | | Ctnnbip1 | 1387.161585 | 0.068266346 | 0.062864476 | 1.085928808 | 0.27751049 | 0.71807708 |
| 4930488L21Rik | 8.031833972 | 0.012865558 | 0.153101826 | 0.084032689 | 0.933030442 | NA | | | Ube2e1 | 2125.513654 | 0.068282293 | 0.049822293 | 1.37051685 | 0.170525619 | 0.620660272 |
| 4930500F04Rik | 0 | NA | NA | NA | NA | NA | | | Ttc33 | 1492.726429 | 0.068283608 | 0.06086632 | 1.121861958 | 0.261921138 | 0.704317239 |
| 4930500J02Rik | 7.694952451 | 0.155917232 | 0.145144368 | 1.074221712 | 0.282723323 | NA | | | Il25 | 3.995688887 | 0.068292095 | 0.038079868 | 1.793391053 | 0.072910417 | NA |
| 4930500L23Rik | 0.40780145 | -0.036172458 | 0.049100092 | -0.736708546 | 0.461299609 | NA | | | Ppp1r8 | 1361.276789 | 0.068323339 | 0.057041131 | 1.197790754 | 0.230998488 | 0.683402059 |
| 4930502A04Rik | 0 | NA | NA | NA | NA | NA | | | Ddost | 3644.25303 | 0.068333229 | 0.053307381 | 1.281871806 | 0.19988762 | 0.654380963 |
| 4930502E09Rik | 0.099009884 | 0.018205726 | 0.027896865 | 0.652608315 | 0.514008823 | NA | | | Sesn1 | 2731.704333 | 0.068334262 | 0.058442074 | 1.16926485 | 0.24229694 | 0.692772033 |
| 4930503B20Rik | 2.495604498 | 0.020978341 | 0.111466346 | 0.18820336 | 0.850717239 | NA | | | Ly6h | 4944.000726 | 0.068391 | 0.074168249 | 0.922106165 | 0.356473201 | 0.773064269 |
| 4930503E14Rik | 1.459175472 | 0.008307894 | 0.078779715 | 0.10545728 | 0.916012967 | NA | | | Ap1g1 | 5333.316232 | 0.068391521 | 0.058847022 | 1.162191706 | 0.245157603 | 0.694718388 |
| 4930503E24Rik | 0 | NA | NA | NA | NA | NA | | | Mtpn | 13536.04364 | 0.068448007 | 0.073746983 | 0.928146534 | 0.353331564 | 0.770624606 |
| 4930503H13Rik | 0.176054697 | -0.002190495 | 0.031239146 | -0.070120197 | 0.944097991 | NA | | | Matn2 | 1162.965927 | 0.068467547 | 0.069395164 | 0.986632832 | 0.323822669 | 0.752925264 |
| 4930503O07Rik | 0 | NA | NA | NA | NA | NA | | | Zfp9 | 1145.238766 | 0.068469289 | 0.057151558 | 1.198030141 | 0.230905283 | 0.683402059 |
| 4930504O13Rik | 0 | NA | NA | NA | NA | NA | | | A330076C08Rik | 18.65219595 | 0.068503273 | 0.072605934 | 0.943494128 | 0.345428217 | NA |
| 4930505A04Rik | 6.08102324 | -0.060985046 | 0.15135395 | -0.402929993 | 0.686999726 | NA | | | Chaf1b | 17.65788618 | 0.068515533 | 0.078974839 | 0.867561538 | 0.385634409 | NA |
| 4930505G20Rik | 0 | NA | NA | NA | NA | NA | | | Siah3 | 83.9924798 | 0.068517009 | 0.094947262 | 0.721632273 | 0.470520592 | 0.833245343 |
| 4930509E16Rik | 1.005958322 | 0.045463035 | 0.072765671 | 0.624786867 | 0.532110951 | NA | | | Eftud2 | 4504.08206 | 0.068529637 | 0.053815599 | 1.273415851 | 0.202870516 | 0.656606302 |
| 4930509J09Rik | 0 | NA | NA | NA | NA | NA | | | Ubac2 | 1307.471024 | 0.068553562 | 0.053424543 | 1.283184824 | 0.199427332 | 0.65377551 |
| 4930509K18Rik | 0 | NA | NA | NA | NA | NA | | | N4bp1 | 3181.350204 | 0.068554552 | 0.063548857 | 1.078769229 | 0.280690615 | 0.720080692 |
| 4930511A02Rik | 2.134322429 | 0.023974154 | 0.087649941 | 0.273521619 | 0.784452276 | NA | | | Cacnb2 | 2206.629121 | 0.068566007 | 0.057738373 | 1.187529261 | 0.235018922 | 0.686276191 |
| 4930511E03Rik | 0 | NA | NA | NA | NA | NA | | | Cops5 | 2455.900228 | 0.068573093 | 0.048767626 | 1.40611916 | 0.159688744 | 0.611570723 |
| 4930511M11Rik | 0 | NA | NA | NA | NA | NA | | | Txndc5 | 2663.715615 | 0.068575108 | 0.059888621 | 1.145044038 | 0.252190915 | 0.69992334 |
| 4930512B01Rik | 1.54174701 | 0.130241365 | 0.088845497 | 1.465930961 | 0.142667096 | NA | | | Tug1 | 4490.365497 | 0.068588721 | 0.058658433 | 1.169290033 | 0.242286797 | 0.692772033 |
| 4930513D17Rik | 0.929186171 | -0.000153963 | 0.058545282 | -0.002629812 | 0.997901716 | NA | | | Idi1 | 1713.710739 | 0.068602302 | 0.073919948 | 0.928062093 | 0.353375361 | 0.770624606 |
| 4930513O06Rik | 0 | NA | NA | NA | NA | NA | | | Ntn1 | 527.119091 | 0.068618913 | 0.084271583 | 0.814259221 | 0.415496466 | 0.805633085 |
| 4930515B02Rik | 0 | NA | NA | NA | NA | NA | | | Agtrap | 522.3748529 | 0.068621382 | 0.088606707 | 0.774449075 | 0.438665272 | 0.817139233 |
| 4930515G01Rik | 7.402560496 | -0.000999662 | 0.149758177 | -0.006675172 | 0.994674023 | NA | | | Stk35 | 1688.099703 | 0.068627429 | 0.065539876 | 1.047109528 | 0.295049065 | 0.73000898 |
| 4930515G16Rik | 0 | NA | NA | NA | NA | NA | | | Psmc3 | 5329.132943 | 0.068629547 | 0.051531569 | 1.331796189 | 0.182927171 | 0.636150366 |
| 4930515L03Rik | 1.09530653 | 0.005333132 | 0.072668172 | 0.073390207 | 0.94149561 | NA | | | Tcp11l1 | 478.8138155 | 0.068634877 | 0.081355671 | 0.843639731 | 0.398870756 | 0.797789838 |
| 4930515L19Rik | 0 | NA | NA | NA | NA | NA | | | Ccdc126 | 365.5588396 | 0.068659147 | 0.08716563 | 0.787686003 | 0.430880395 | 0.812181606 |
| 4930517E11Rik | 0 | NA | NA | NA | NA | NA | | | Pgam1 | 22164.98758 | 0.068666825 | 0.031413486 | 2.185902762 | 0.02882272 | 0.316461703 |
| 4930518P08Rik | 0 | NA | NA | NA | NA | NA | | | Sdf2l1 | 298.2674253 | 0.068702504 | 0.090106371 | 0.762460005 | 0.445785505 | 0.821773578 |
| 4930519D14Rik | 0.094392817 | 0.017895942 | 0.027818137 | 0.643319236 | 0.520016977 | NA | | | Ctdspl2 | 886.5322965 | 0.068712848 | 0.082467126 | 0.833215024 | 0.404723471 | 0.800301271 |
| 4930519F16Rik | 1.681417607 | 0.226274572 | 0.095808368 | 2.361741218 | 0.018189334 | NA | | | Slitrk5 | 3009.392674 | 0.068737406 | 0.056047385 | 1.22641592 | 0.220042198 | 0.672757795 |
| 4930519F24Rik | 0.119397641 | -0.020477583 | 0.027954224 | -0.732539845 | 0.463839132 | NA | | | Tenm2 | 2612.347159 | 0.068737478 | 0.077071309 | 0.891868572 | 0.372463382 | 0.781446026 |
| 4930519H02Rik | 0.107716603 | -0.020477583 | 0.027954224 | -0.732539849 | 0.46383913 | NA | | | Mkrn1 | 5056.16282 | 0.068745197 | 0.055168967 | 1.246084524 | 0.212733366 | 0.666270994 |
| 4930520P13Rik | 0 | NA | NA | NA | NA | NA | | | Esrp2 | 3.655329606 | 0.068749042 | 0.042030064 | 1.635711109 | 0.101900063 | NA |
| 4930521E06Rik | 0 | NA | NA | NA | NA | NA | | | Rassf5 | 1359.212837 | 0.06875831 | 0.061576759 | 1.116627614 | 0.26415358 | 0.707085906 |
| 4930522H14Rik | 0 | NA | NA | NA | NA | NA | | | Wfdc1 | 210.6951663 | 0.068759648 | 0.081075123 | 0.848097974 | 0.39638341 | 0.796715845 |
| 4930522O17Rik | 0.119397641 | -0.020477583 | 0.027954224 | -0.732539845 | 0.463839132 | NA | | | Flt4 | 199.4383297 | 0.068799874 | 0.092839622 | 0.741061553 | 0.45865612 | 0.828043599 |
| 4930523O13Rik | 0 | NA | NA | NA | NA | NA | | | Hcfc1r1 | 2489.989044 | 0.068827984 | 0.067367768 | 1.021675297 | 0.306934607 | 0.739163119 |
| 4930524B15Rik | 6.554640627 | 0.05191124 | 0.144854754 | 0.358367525 | 0.720068291 | NA | | | Gabrg3 | 233.9012017 | 0.068857803 | 0.09440856 | 0.729359746 | 0.465781634 | 0.831904964 |
| 4930524C18Rik | 0 | NA | NA | NA | NA | NA | | | Fbxo47 | 32.37521958 | 0.068870252 | 0.083519497 | 0.824600903 | 0.409598209 | 0.801196796 |
| 4930524N10Rik | 0 | NA | NA | NA | NA | NA | | | Ndufs4 | 2193.709452 | 0.068894079 | 0.056275799 | 1.224222144 | 0.220868438 | 0.672757795 |
| 4930524O05Rik | 0 | NA | NA | NA | NA | NA | | | Csnk1e | 4529.457715 | 0.068910097 | 0.046521749 | 1.481244751 | 0.138541365 | 0.583109099 |
| 4930524O08Rik | 0 | NA | NA | NA | NA | NA | | | Rnf150 | 3982.987324 | 0.068929348 | 0.079143058 | 0.870946233 | 0.383783511 | 0.78787654 |
| 4930525D18Rik | 0.882749213 | 0.107936301 | 0.072744355 | 1.483775627 | 0.137868455 | NA | | | Anxa7 | 4177.441217 | 0.068935095 | 0.04809485 | 1.433315526 | 0.151767687 | 0.59939571 |
| 4930525M21Rik | 0 | NA | NA | NA | NA | NA | | | Ermn | 2834.863886 | 0.068938843 | 0.089518106 | 0.770110611 | 0.441234282 | 0.818962636 |
| 4930526D03Rik | 0 | NA | NA | NA | NA | NA | | | I7Rn6 | 937.9543132 | 0.068948576 | 0.06431848 | 1.071987011 | 0.283725866 | 0.721155194 |
| 4930526L06Rik | 0.978497497 | 0.043411735 | 0.06751127 | 0.643029462 | 0.520204983 | NA | | | Rdh12 | 89.66463097 | 0.06896914 | 0.093984178 | 0.733837776 | 0.463047614 | 0.831014288 |
| 4930527F14Rik | 0.107716603 | -0.020477583 | 0.027954224 | -0.732539849 | 0.46383913 | NA | | | Tfip11 | 2396.274955 | 0.069003317 | 0.061110016 | 1.129165421 | 0.258828058 | 0.703616072 |
| 4930527G23Rik | 0.098353386 | -0.020212831 | 0.02786466 | -0.725393061 | 0.468210929 | NA | | | Dnajb4 | 2817.04606 | 0.069021767 | 0.076221835 | 0.905537977 | 0.36518045 | 0.777735733 |
| 4930528D03Rik | 1.522758917 | 0.143613366 | 0.093881166 | 1.529735638 | 0.126082178 | NA | | | Cml5 | 73.57518549 | 0.069058867 | 0.094064984 | 0.734161267 | 0.462850457 | 0.831014288 |
| 4930528P14Rik | 0.618726074 | 0.039228334 | 0.048427554 | 0.810041617 | 0.417916257 | NA | | | Srp9 | 2282.325396 | 0.069066891 | 0.047815608 | 1.444442398 | 0.148614572 | 0.596199396 |
| 4930529C04Rik | 0 | NA | NA | NA | NA | NA | | | Ostm1 | 2451.759829 | 0.069068062 | 0.059983721 | 1.151446784 | 0.249548476 | 0.698043744 |
| 4930529F24Rik | 0 | NA | NA | NA | NA | NA | | | Pvr | 319.5300256 | 0.069075572 | 0.081581389 | 0.846707478 | 0.397158189 | 0.797284168 |
| 4930529K09Rik | 0.079983654 | 0.016740032 | 0.027674178 | 0.604897175 | 0.54524732 | NA | | | Gcc1 | 1001.496428 | 0.069079262 | 0.063597597 | 1.086192955 | 0.277393634 | 0.718015501 |
| 4930529L06Rik | 0 | NA | NA | NA | NA | NA | | | Ppp2r2a | 4567.930386 | 0.069085156 | 0.070235784 | 0.983617649 | 0.325303544 | 0.75293408 |
| 4930533B01Rik | 0 | NA | NA | NA | NA | NA | | | Ube2ql1 | 7668.109934 | 0.069089271 | 0.056643159 | 1.219728412 | 0.222567847 | 0.67374894 |
| 4930533P14Rik | 0.323149808 | -0.030167563 | 0.033779321 | -0.893077861 | 0.371815478 | NA | | | LOC106740 | 505.4102921 | 0.069099969 | 0.070354953 | 0.982162104 | 0.326019994 | 0.753049596 |
| 4930539C22Rik | 0 | NA | NA | NA | NA | NA | | | Zfp647 | 99.28900146 | 0.069110779 | 0.094334272 | 0.732615807 | 0.463792787 | 0.83159896 |
| 4930539M17Rik | 5.329181811 | 0.014030299 | 0.140677651 | 0.099733673 | 0.920555766 | NA | | | Gpr56 | 2823.990853 | 0.069118362 | 0.082242212 | 0.840424397 | 0.400670475 | 0.799169897 |
| 4930539N22Rik | 0 | NA | NA | NA | NA | NA | | | Dpyd | 258.7254624 | 0.069120294 | 0.084668019 | 0.816368386 | 0.414289472 | 0.804541293 |
| 4930540M03Rik | 0 | NA | NA | NA | NA | NA | | | Gatad1 | 2443.114777 | 0.069188814 | 0.056535443 | 1.223813083 | 0.221022748 | 0.672757795 |
| 4930542C21Rik | 0 | NA | NA | NA | NA | NA | | | Fgf17 | 24.35879781 | 0.069229805 | 0.082707254 | 0.837046355 | 0.402566512 | 0.799737577 |
| 4930542D17Rik | 0.079983654 | 0.016740032 | 0.027674178 | 0.604897175 | 0.54524732 | NA | | | Psmb7 | 4476.414775 | 0.069233147 | 0.051241691 | 1.35110974 | 0.176660282 | 0.629083076 |
| 4930542H20Rik | 0 | NA | NA | NA | NA | NA | | | Ptdss1 | 3262.927915 | 0.069261178 | 0.043557839 | 1.590096748 | 0.111812999 | 0.538992394 |
| 4930543E12Rik | 0.488510588 | 0.029306206 | 0.046230877 | 0.633909798 | 0.526139691 | NA | | | Twsg1 | 1285.260338 | 0.069266027 | 0.075953838 | 0.911948999 | 0.361795566 | 0.776449324 |
| 4930544D05Rik | 0.331243956 | 0.008596723 | 0.034710864 | 0.247666645 | 0.804392342 | NA | | | Stat5b | 1599.268525 | 0.069300258 | 0.062506866 | 1.108682337 | 0.267567244 | 0.710152314 |
| 4930544G11Rik | 0 | NA | NA | NA | NA | NA | | | C230052I12Rik | 190.1832651 | 0.069355516 | 0.083162058 | 0.833980266 | 0.404292104 | 0.800290023 |
| 4930544M13Rik | 0 | NA | NA | NA | NA | NA | | | Trpc4ap | 7430.420812 | 0.069378811 | 0.036264256 | 1.913145875 | 0.05572938 | 0.413057568 |
| 4930545E07Rik | 0 | NA | NA | NA | NA | NA | | | Slc38a10 | 3431.298764 | 0.069393538 | 0.053068442 | 1.307623413 | 0.191001072 | 0.646547945 |
| 4930545H06Rik | 0.813882095 | 0.031557003 | 0.065495559 | 0.481818976 | 0.629934548 | NA | | | Kif3c | 12909.76968 | 0.069409121 | 0.038636215 | 1.796478279 | 0.072418484 | 0.462192839 |
| 4930545L23Rik | 0.425621188 | 0.027403281 | 0.044342756 | 0.617987764 | 0.536583404 | NA | | | Mboat7 | 5438.495463 | 0.069422782 | 0.05372357 | 1.292222054 | 0.196280256 | 0.651844308 |
| 4930546C10Rik | 2.10939122 | -0.057982471 | 0.099752708 | -0.581262119 | 0.561063807 | NA | | | Mlycd | 962.1934721 | 0.069426342 | 0.067383175 | 1.03032162 | 0.302859053 | 0.736289641 |
| 4930546K05Rik | 0.565738727 | -0.007881918 | 0.057869715 | -0.13620108 | 0.891662321 | NA | | | Usp53 | 839.3599631 | 0.069445108 | 0.07297557 | 0.951621313 | 0.341289069 | 0.762819666 |
| 4930547C10Rik | 0 | NA | NA | NA | NA | NA | | | Esrra | 1930.98202 | 0.069467428 | 0.073184174 | 0.949213797 | 0.342511886 | 0.763762832 |
| 4930547E08Rik | 0 | NA | NA | NA | NA | NA | | | Slc35b4 | 3406.90302 | 0.069494878 | 0.047122646 | 1.47476606 | 0.140275449 | 0.585529073 |
| 4930547E14Rik | 0 | NA | NA | NA | NA | NA | | | Spire1 | 8944.91141 | 0.0695101 | 0.055059819 | 1.26244693 | 0.206788011 | 0.661193627 |
| 4930548G14Rik | 0 | NA | NA | NA | NA | NA | | | Pin1 | 2659.625089 | 0.069516713 | 0.050444163 | 1.378092302 | 0.168174792 | 0.618133132 |
| 4930548H24Rik | 0.380940254 | 0.025976956 | 0.047990576 | 0.54129287 | 0.588305734 | NA | | | Gars | 5594.50856 | 0.069529143 | 0.053186957 | 1.307259292 | 0.191124665 | 0.646547945 |
| 4930548J01Rik | 0 | NA | NA | NA | NA | NA | | | Pmfbp1 | 23.20620155 | 0.069579691 | 0.070745109 | 0.983526523 | 0.325348368 | 0.75293408 |
| 4930548K13Rik | 0 | NA | NA | NA | NA | NA | | | Bend6 | 6011.222302 | 0.069581071 | 0.062667255 | 1.110325807 | 0.266858656 | 0.709441589 |
| 4930549C01Rik | 0 | NA | NA | NA | NA | NA | | | Vps29 | 2383.36857 | 0.069581861 | 0.056425745 | 1.233158049 | 0.217516804 | 0.670311138 |
| 4930549G23Rik | 1.00591031 | 0.019068594 | 0.071772183 | 0.265682233 | 0.790483948 | NA | | | Disp2 | 16945.79879 | 0.06959685 | 0.066586214 | 1.045214117 | 0.29592402 | 0.730928808 |
| 4930550L24Rik | 0.159967308 | 0.0240345 | 0.030870969 | 0.778547006 | 0.436246606 | NA | | | Gm10778 | 14.8010387 | 0.069627164 | 0.070075375 | 0.993603878 | 0.320415755 | NA |
| 4930552N02Rik | 0 | NA | NA | NA | NA | NA | | | 4922501L14Rik | 267.713592 | 0.069630084 | 0.085605942 | 0.813379095 | 0.416000741 | 0.805925149 |
| 4930552P12Rik | 0.931403597 | -0.00929293 | 0.064794173 | -0.143422307 | 0.885956667 | NA | | | Speer7-ps1 | 5.297765712 | 0.069630477 | 0.05101513 | 1.364898551 | 0.172284938 | NA |
| 4930553E22Rik | 0.08731684 | 0.017365056 | 0.027733412 | 0.626142072 | 0.531221759 | NA | | | Atg4b | 2778.554339 | 0.069644533 | 0.047448484 | 1.467792569 | 0.142160577 | 0.587991041 |
| 4930554C24Rik | 0 | NA | NA | NA | NA | NA | | | Bop1 | 2068.641063 | 0.06965541 | 0.06941798 | 1.003420295 | 0.315658116 | 0.745993052 |
| 4930555B11Rik | 0 | NA | NA | NA | NA | NA | | | Lmf1 | 619.919614 | 0.069662799 | 0.068379678 | 1.018764649 | 0.308314712 | 0.74062833 |
| 4930555G01Rik | 5.980794058 | -0.133563684 | 0.15019306 | -0.889279997 | 0.373852619 | NA | | | Dctn6 | 2037.150579 | 0.069672487 | 0.052949009 | 1.315841203 | 0.188227347 | 0.642130211 |
| 4930556C24Rik | 0 | NA | NA | NA | NA | NA | | | D6Wsu163e | 1397.929567 | 0.069672919 | 0.05448686 | 1.278710485 | 0.200999026 | 0.654716629 |
| 4930556G01Rik | 0 | NA | NA | NA | NA | NA | | | Fzr1 | 2770.037971 | 0.069692943 | 0.058554601 | 1.190221467 | 0.233959358 | 0.685180168 |
| 4930556J02Rik | 0 | NA | NA | NA | NA | NA | | | Fam212b | 6679.721286 | 0.069707707 | 0.069573043 | 1.001935568 | 0.316374713 | 0.745993052 |
| 4930556N09Rik | 0.32048461 | -0.048989879 | 0.042627894 | -1.149244659 | 0.25045511 | NA | | | Ropn1 | 3.393929158 | 0.06970965 | 0.043216346 | 1.613038976 | 0.10673604 | NA |
| 4930557A04Rik | 0 | NA | NA | NA | NA | NA | | | Nmbr | 43.07922016 | 0.069721887 | 0.089665971 | 0.777573547 | 0.436820461 | 0.816490111 |
| 4930557J02Rik | 0 | NA | NA | NA | NA | NA | | | Epas1 | 7309.736955 | 0.069731754 | 0.06537927 | 1.066572842 | 0.286164769 | 0.722148665 |
| 4930558C23Rik | 0 | NA | NA | NA | NA | NA | | | Serp2 | 1897.907762 | 0.06974348 | 0.063132777 | 1.104711109 | 0.269284788 | 0.711922711 |
| 4930558G05Rik | 0 | NA | NA | NA | NA | NA | | | Rapgef2 | 6139.684569 | 0.06975494 | 0.06150342 | 1.134163599 | 0.25672593 | 0.703000737 |
| 4930558J18Rik | 1.445046221 | 0.012865077 | 0.080275315 | 0.160261929 | 0.872674748 | NA | | | Zfp367 | 918.4298852 | 0.069760404 | 0.074699454 | 0.933881035 | 0.350365263 | 0.768667813 |
| 4930558K02Rik | 0.6359905 | 0.05770125 | 0.047097028 | 1.225156918 | 0.220516103 | NA | | | Vil1 | 27.57050787 | 0.069766129 | 0.084085342 | 0.829706194 | 0.406704919 | 0.800917161 |
| 4930562C15Rik | 0.17833704 | -0.002390466 | 0.031282281 | -0.076415987 | 0.939088151 | NA | | | Pcdha11 | 151.4618078 | 0.069778228 | 0.090234166 | 0.773301641 | 0.439343885 | 0.817545617 |
| 4930562F07Rik | 0.795601132 | -0.041375411 | 0.068722503 | -0.602064955 | 0.547130901 | NA | | | Tmem104 | 897.804177 | 0.069788922 | 0.056656359 | 1.231793269 | 0.218026318 | 0.670929686 |
| 4930563D23Rik | 0.08731684 | 0.017365056 | 0.027733412 | 0.626142072 | 0.531221759 | NA | | | 2610034B18Rik | 787.1820655 | 0.069792612 | 0.070623979 | 0.988228263 | 0.32304087 | 0.752792382 |
| 4930563E18Rik | 1.896076537 | 0.04518369 | 0.096495718 | 0.468245545 | 0.639609006 | NA | | | Cyp4f15 | 612.5017202 | 0.069823616 | 0.085795712 | 0.813835733 | 0.415739062 | 0.805633085 |
| 4930563F08Rik | 0 | NA | NA | NA | NA | NA | | | Exog | 1188.821768 | 0.069834143 | 0.062887672 | 1.110458382 | 0.266801552 | 0.709441589 |
| 4930563M20Rik | 0 | NA | NA | NA | NA | NA | | | Slc5a5 | 991.2797174 | 0.069836589 | 0.06464279 | 1.080346145 | 0.279988068 | 0.719332628 |
| 4930564B18Rik | 0.105923158 | 0.018524693 | 0.027977518 | 0.662127842 | 0.507889294 | NA | | | Rhog | 1190.624217 | 0.069842581 | 0.076663316 | 0.911030009 | 0.362279563 | 0.776449324 |
| 4930564C03Rik | 0 | NA | NA | NA | NA | NA | | | Coch | 1916.920801 | 0.069876183 | 0.083243743 | 0.839416643 | 0.401235548 | 0.799277436 |
| 4930564D02Rik | 0 | NA | NA | NA | NA | NA | | | Prdx3 | 2343.64249 | 0.069884635 | 0.060283443 | 1.159267485 | 0.246347171 | 0.696058939 |
| 4930564G21Rik | 0 | NA | NA | NA | NA | NA | | | Calm2 | 53347.05689 | 0.069893612 | 0.061722161 | 1.132390868 | 0.257470143 | 0.703271881 |
| 4930564K09Rik | 0 | NA | NA | NA | NA | NA | | | Crk | 5978.367736 | 0.069895266 | 0.069032097 | 1.012503868 | 0.311297198 | 0.743856408 |
| 4930565D16Rik | 0 | NA | NA | NA | NA | NA | | | Mvk | 709.1090954 | 0.069903715 | 0.079455184 | 0.879787963 | 0.378974187 | 0.785380516 |
| 4930565N06Rik | 0 | NA | NA | NA | NA | NA | | | Lrrc61 | 1260.197272 | 0.069920115 | 0.071595813 | 0.976595027 | 0.328769682 | 0.755788917 |
| 4930567H12Rik | 0.688757862 | 0.006342159 | 0.055170474 | 0.114955679 | 0.908480252 | NA | | | Rmrp | 4.131041172 | 0.069953242 | 0.043600385 | 1.604418003 | 0.108621945 | NA |
| 4930567H17Rik | 0 | NA | NA | NA | NA | NA | | | Vapa | 4465.055123 | 0.069955944 | 0.064433161 | 1.085713354 | 0.277605831 | 0.71807708 |
| 4930567J20Rik | 0 | NA | NA | NA | NA | NA | | | Rcan1 | 2685.903935 | 0.06995945 | 0.064575741 | 1.083370466 | 0.27864402 | 0.71849589 |
| 4930567K20Rik | 0.096071043 | -0.020098795 | 0.027825948 | -0.722304036 | 0.470107572 | NA | | | Ndufs1 | 6494.930839 | 0.069961937 | 0.051517312 | 1.358027712 | 0.174454891 | 0.627260333 |
| 4930568D16Rik | 0 | NA | NA | NA | NA | NA | | | Nrxn3 | 8655.758785 | 0.069979934 | 0.059374948 | 1.178610461 | 0.238553324 | 0.689459926 |
| 4930568E12Rik | 0 | NA | NA | NA | NA | NA | | | Ly6g6d | 10.93927542 | 0.069986288 | 0.062764443 | 1.115062682 | 0.264823562 | NA |
| 4930568G15Rik | 1.328994491 | 0.001593344 | 0.086013489 | 0.018524352 | 0.985220551 | NA | | | Cnga3 | 6.836651156 | 0.070017443 | 0.040670994 | 1.721557183 | 0.085149765 | NA |
| 4930568K20Rik | 0 | NA | NA | NA | NA | NA | | | Psenen | 1446.834647 | 0.070023349 | 0.054396396 | 1.287279208 | 0.197996988 | 0.653535282 |
| 4930571K23Rik | 0 | NA | NA | NA | NA | NA | | | Pcbp3 | 2295.292946 | 0.070039119 | 0.059464275 | 1.177835243 | 0.238862296 | 0.689537017 |
| 4930572K03Rik | 0.238795281 | -0.027601606 | 0.032267774 | -0.855392297 | 0.392334007 | NA | | | Ano3 | 6424.921394 | 0.070053564 | 0.087076339 | 0.804507454 | 0.421103966 | 0.808470525 |
| 4930572O03Rik | 0.705750341 | 0.031598865 | 0.054521254 | 0.579569665 | 0.562204855 | NA | | | Tmem218 | 569.4828748 | 0.070062897 | 0.071163558 | 0.984533369 | 0.324853333 | 0.75293408 |
| 4930572O13Rik | 0.380940254 | 0.025976956 | 0.047990576 | 0.54129287 | 0.588305734 | NA | | | Spin1 | 9660.627803 | 0.070078281 | 0.062214385 | 1.126399957 | 0.259996265 | 0.704052663 |
| 4930573O16Rik | 0 | NA | NA | NA | NA | NA | | | Nrip3 | 5387.897395 | 0.0700821 | 0.069107994 | 1.01409542 | 0.310537223 | 0.743385581 |
| 4930578E11Rik | 0.176054697 | -0.002190495 | 0.031239146 | -0.070120197 | 0.944097991 | NA | | | Fam131a | 11379.59752 | 0.070106379 | 0.071289657 | 0.983401837 | 0.325409707 | 0.75293408 |

| | | | | | | |
|---|---|---|---|---|---|---|
| 4930578I06Rik | 0 | NA | NA | NA | NA | NA |
| 4930578N18Rik | 0 | NA | NA | NA | NA | NA |
| 4930579F01Rik | 0 | NA | NA | NA | NA | NA |
| 4930579G18Rik | 3.406179514 | 0.084716588 | 0.115538925 | 0.733229847 | 0.463418254 | NA |
| 4930583K01Rik | 0.974763109 | 0.077267739 | 0.069555094 | 1.110885411 | 0.266617676 | NA |
| 4930583P06Rik | 0 | NA | NA | NA | NA | NA |
| 4930584F24Rik | 0 | NA | NA | NA | NA | NA |
| 4930590L20Rik | 0 | NA | NA | NA | NA | NA |
| 4930591A17Rik | 0 | NA | NA | NA | NA | NA |
| 4930592A05Rik | 0.68657355 | 0.032306041 | 0.060686154 | 0.532346161 | 0.594486266 | NA |
| 4930592I03Rik | 0 | NA | NA | NA | NA | NA |
| 4930593A02Rik | 1.28787035 | 0.023814336 | 0.075742054 | 0.314413659 | 0.753206884 | NA |
| 4930593C16Rik | 0.49242343 | -0.024792866 | 0.039435663 | -0.6286915 | 0.529551044 | NA |
| 4930594C11Rik | 0.751088818 | -0.011157514 | 0.061845162 | -0.180410466 | 0.856830339 | NA |
| 4930595M18Rik | 0 | NA | NA | NA | NA | NA |
| 4930596D02Rik | 0 | NA | NA | NA | NA | NA |
| 4930596M17Rik | 0.605945404 | -0.004798112 | 0.052429254 | -0.091515938 | 0.927082643 | NA |
| 4930597G03Rik | 0 | NA | NA | NA | NA | NA |
| 4930598F16Rik | 0 | NA | NA | NA | NA | NA |
| 4930599N23Rik | 5.22008009 | 0.087699406 | 0.134255255 | 0.653228851 | 0.513608753 | NA |
| 4931402G19Rik | 0.997312756 | -0.104805702 | 0.06156002 | -1.702496218 | 0.088662387 | NA |
| 4931403G20Rik | 0.19442443 | -0.026451352 | 0.031568725 | -0.837897386 | 0.402088337 | NA |
| 4931406B18Rik | 0 | NA | NA | NA | NA | NA |
| 4931406H21Rik | 0.789628455 | 0.016746707 | 0.050267784 | 0.333149899 | 0.739021135 | NA |
| 4931408C20Rik | 0.096071043 | -0.020098795 | 0.027825948 | -0.722304036 | 0.470107572 | NA |
| 4931409K22Rik | 5.124992282 | 0.145853595 | 0.141286002 | 1.032328699 | 0.301918161 | NA |
| 4931417E11Rik | 0 | NA | NA | NA | NA | NA |
| 4931417G12Rik | 2.927860015 | -0.140760812 | 0.117456152 | -1.198411569 | 0.230756831 | NA |
| 4931419H13Rik | 0.430483556 | -0.061355922 | 0.047577744 | -1.289592921 | 0.197192035 | NA |
| 4931420L22Rik | 0 | NA | NA | NA | NA | NA |
| 4931423N10Rik | 7.255305665 | 0.135750534 | 0.130148345 | 1.043044647 | 0.296927615 | NA |
| 4931428L18Rik | 0.466626353 | 0.089501577 | 0.058821838 | 1.521570567 | 0.128116717 | NA |
| 4931429L15Rik | 3.548963464 | 0.10799846 | 0.122005073 | 0.885196474 | 0.376050673 | NA |
| 4931429P17Rik | 0.185670226 | -0.001645086 | 0.031432796 | -0.052336624 | 0.958260472 | NA |
| 4931430N09Rik | 0.287795519 | 0.031925978 | 0.035387605 | 0.902179657 | 0.366961439 | NA |
| 4931431B13Rik | 0.094392817 | 0.017895942 | 0.027818137 | 0.643319236 | 0.520016977 | NA |
| 4931431C16Rik | 0.900243631 | 0.028123536 | 0.058318278 | 0.482242223 | 0.629633886 | NA |
| 4931431F19Rik | 0 | NA | NA | NA | NA | NA |
| 4931440J10Rik | 5.291001609 | -0.206325516 | 0.135621765 | -1.521330421 | 0.12817694 | NA |
| 4931440L10Rik | 0.456340463 | 0.030209904 | 0.045608803 | 0.662370032 | 0.507734105 | NA |
| 4931440P22Rik | 7.476089336 | 0.288538908 | 0.157216927 | 1.835291613 | 0.066462493 | NA |
| 4932411N23Rik | 0.096071043 | -0.020098795 | 0.027825948 | -0.722304036 | 0.470107572 | NA |
| 4932412D23Rik | 3.061015359 | -0.208674211 | 0.121981904 | -1.710698097 | 0.087136859 | NA |
| 4932413F04Rik | 0 | NA | NA | NA | NA | NA |
| 4932414J04Rik | 0 | NA | NA | NA | NA | NA |
| 4932414N04Rik | 0 | NA | NA | NA | NA | NA |
| 4932415M13Rik | 0.450736154 | 0.045589387 | 0.042063495 | 1.083823084 | 0.278443248 | NA |
| 4932416K20Rik | 0.107716603 | -0.020477583 | 0.027954224 | -0.732539849 | 0.46383913 | NA |
| 4932429P05Rik | 0 | NA | NA | NA | NA | NA |
| 4932435O22Rik | 5.460629182 | -0.040521896 | 0.148776287 | -0.272367973 | 0.785339096 | NA |
| 4932702P03Rik | 1.398967995 | -0.087155862 | 0.079085499 | -1.10204605 | 0.270441652 | NA |
| 4933400A11Rik | 0 | NA | NA | NA | NA | NA |
| 4933400B14Rik | 0.197363271 | -0.000561558 | 0.031650166 | -0.017742642 | 0.985844163 | NA |
| 4933400C23Rik | 0.62365482 | 0.069517633 | 0.062256133 | 1.116639103 | 0.264148665 | NA |
| 4933400F21Rik | 0.096071043 | -0.020098795 | 0.027825948 | -0.722304036 | 0.470107572 | NA |
| 4933400L20Rik | 0.295060159 | -0.029687054 | 0.033501022 | -0.886153701 | 0.375534709 | NA |
| 4933401B06Rik | 0 | NA | NA | NA | NA | NA |
| 4933401D09Rik | 0 | NA | NA | NA | NA | NA |
| 4933401H06Rik | 0.174376471 | 0.024538273 | 0.031173126 | 0.787161136 | 0.431187544 | NA |
| 4933401P06Rik | 0 | NA | NA | NA | NA | NA |
| 4933402C06Rik | 0.947736072 | -0.037611715 | 0.058791758 | -0.639744696 | 0.522338592 | NA |
| 4933402D24Rik | 7.163047344 | 0.223585585 | 0.155171886 | 1.440889781 | 0.149615824 | NA |
| 4933402E13Rik | 0.096071043 | -0.020098795 | 0.027825948 | -0.722304036 | 0.470107572 | NA |
| 4933402J07Rik | 1.042779188 | 0.116034349 | 0.074231793 | 1.563135483 | 0.118020731 | NA |
| 4933402J10Rik | 0 | NA | NA | NA | NA | NA |
| 4933402J15Rik | 1.208531101 | -0.036599839 | 0.079628192 | -0.45963418 | 0.645778821 | NA |
| 4933402N03Rik | 0 | NA | NA | NA | NA | NA |
| 4933402N22Rik | 0 | NA | NA | NA | NA | NA |
| 4933402P03Rik | 0 | NA | NA | NA | NA | NA |
| 4933403O08Rik | 2.848565718 | -0.00935093 | 0.116946144 | -0.079959287 | 0.936269636 | NA |
| 4933404G15Rik | 0 | NA | NA | NA | NA | NA |
| 4933404K08Rik | 0.222620122 | -0.001447027 | 0.032059667 | -0.04513545 | 0.963999345 | NA |
| 4933405D12Rik | 1.532017869 | 0.163082275 | 0.090868546 | 1.794705448 | 0.072700642 | NA |
| 4933405E24Rik | 0 | NA | NA | NA | NA | NA |
| 4933405L10Rik | 7.40056411 | 0.127715269 | 0.153739476 | 0.830725274 | 0.406128845 | NA |
| 4933405O20Rik | 0.212768007 | -0.02694223 | 0.031868764 | -0.845411806 | 0.397880955 | NA |
| 4933406D12Rik | 0 | NA | NA | NA | NA | NA |
| 4933406F09Rik | 0 | NA | NA | NA | NA | NA |
| 4933406G16Rik | 0 | NA | NA | NA | NA | NA |
| 4933406J08Rik | 0 | NA | NA | NA | NA | NA |
| 4933406J10Rik | 0.119397641 | -0.020477583 | 0.027954224 | -0.732539845 | 0.463839132 | NA |
| 4933406K04Rik | 0 | NA | NA | NA | NA | NA |
| 4933406M09Rik | 0.105923158 | 0.018524693 | 0.027977518 | 0.662127842 | 0.507889294 | NA |
| 4933407E24Rik | 0.896399808 | -0.000816113 | 0.069363515 | -0.011765745 | 0.99061251 | NA |
| 4933407G14Rik | 0.096071043 | -0.020098795 | 0.027825948 | -0.722304036 | 0.470107572 | NA |
| 4933407I05Rik | 2.918431594 | -0.065131308 | 0.103673378 | -0.628235609 | 0.529849606 | NA |
| 4933408J17Rik | 1.306557241 | -0.07162614 | 0.091615212 | -0.781814924 | 0.434323349 | NA |
| 4933408N05Rik | 0.496037528 | 0.012714628 | 0.055562508 | 0.228834661 | 0.818997426 | NA |
| 4933409G03Rik | 0 | NA | NA | NA | NA | NA |
| 4933409K07Rik | 1.09986382 | 0.036652755 | 0.083827004 | 0.437242813 | 0.661935264 | NA |
| 4933411E08Rik | 1.335811433 | -0.057819597 | 0.082562911 | -0.700309579 | 0.483733991 | NA |
| 4933411G06Rik | 0 | NA | NA | NA | NA | NA |
| 4933411G11Rik | 0.099009884 | 0.018205726 | 0.027896865 | 0.652608315 | 0.514008823 | NA |
| 4933411K16Rik | 1.808784137 | -0.094933916 | 0.092210675 | -1.029532823 | 0.303229364 | NA |
| 4933412E24Rik | 0 | NA | NA | NA | NA | NA |
| 4933413J09Rik | 0 | NA | NA | NA | NA | NA |
| 4933413L06Rik | 0 | NA | NA | NA | NA | NA |
| 4933415F23Rik | 0.489904938 | 0.03017109 | 0.045748741 | 0.659495523 | 0.50957762 | NA |
| 4933416C03Rik | 0 | NA | NA | NA | NA | NA |
| 4933416E03Rik | 0 | NA | NA | NA | NA | NA |
| 4933416I08Rik | 0.723371454 | 0.006844428 | 0.061016294 | 0.112173772 | 0.910685625 | NA |
| 4933416M06Rik | 0 | NA | NA | NA | NA | NA |
| 4933416M07Rik | 0.094392817 | 0.017895942 | 0.027818137 | 0.643319236 | 0.520016977 | NA |
| 4933417A18Rik | 0 | NA | NA | NA | NA | NA |
| 4933417E11Rik | 0.408720688 | -0.001528354 | 0.047884645 | -0.03191741 | 0.974537915 | NA |
| 4933417G07Rik | 0 | NA | NA | NA | NA | NA |
| 4933421I07Rik | 0 | NA | NA | NA | NA | NA |
| 4933422A05Rik | 0.524867658 | 0.066447841 | 0.050604724 | 1.313075849 | 0.189157391 | NA |
| 4933422H20Rik | 2.50170239 | 0.023268841 | 0.107765362 | 0.215921334 | 0.829049067 | NA |
| 4933424G05Rik | 0.832944429 | -0.043859678 | 0.073013575 | -0.600705798 | 0.548035956 | NA |
| 4933424G06Rik | 7.064976964 | 0.003371442 | 0.153226087 | 0.022003052 | 0.982445521 | NA |
| 4933425B07Rik | 0 | NA | NA | NA | NA | NA |
| 4933425H06Rik | 0.079983654 | 0.016740032 | 0.027674178 | 0.604897175 | 0.54524732 | NA |
| 4933425L06Rik | 0.531154764 | -0.070672053 | 0.050962944 | -1.38673411 | 0.165522855 | NA |
| 4933426D04Rik | 0.116696964 | -0.020477583 | 0.027954224 | -0.732539847 | 0.463839131 | NA |
| 4933427D06Rik | 2.117011045 | -0.056698448 | 0.085191871 | -0.665538244 | 0.505706291 | NA |
| 4933427E11Rik | 2.800614345 | 0.195179288 | 0.104946691 | 1.859794581 | 0.062914594 | NA |
| 4933427E13Rik | 0.55073328 | 0.049633073 | 0.054814118 | 0.905479733 | 0.365211292 | NA |
| 4933428C19Rik | 0.195080928 | -0.000361156 | 0.031609476 | -0.011425558 | 0.990883922 | NA |
| 4933429K18Rik | 0 | NA | NA | NA | NA | NA |
| 4933429O19Rik | 1.152370116 | 0.065394523 | 0.061053146 | 1.071108167 | 0.284120797 | NA |
| 4933430H16Rik | 0.105923158 | 0.018524693 | 0.027977518 | 0.662127842 | 0.507889294 | NA |
| 4933430M04Rik | 1.602568339 | 0.04362618 | 0.087617457 | 0.497916531 | 0.618542875 | NA |
| 4933430N04Rik | 0 | NA | NA | NA | NA | NA |
| 4933431G14Rik | 0 | NA | NA | NA | NA | NA |
| 4933432G23Rik | 0 | NA | NA | NA | NA | NA |
| 4933432I03Rik | 0.19046386 | -0.000774927 | 0.031526133 | -0.024580468 | 0.980389599 | NA |
| 4933433C11Rik | 1.372710149 | 0.073171619 | 0.085220248 | 0.85861777 | 0.390551431 | NA |
| 4933433F19Rik | 0 | NA | NA | NA | NA | NA |
| 4933433G15Rik | 5.462890164 | 0.181224023 | 0.127936922 | 1.416510729 | 0.156626025 | NA |
| 4933433G19Rik | 3.338064257 | 0.013275869 | 0.109267928 | 0.121498311 | 0.903296353 | NA |
| 4933433H22Rik | 0.421503194 | -0.044552504 | 0.040840123 | -1.090900344 | 0.275316736 | NA |
| 4933434I20Rik | 0 | NA | NA | NA | NA | NA |
| 4933436E23Rik | 0 | NA | NA | NA | NA | NA |
| 4933436H12Rik | 0.079983654 | 0.016740032 | 0.027674178 | 0.604897175 | 0.54524732 | NA |
| 4933436I01Rik | 0 | NA | NA | NA | NA | NA |
| 4933438B17Rik | 0.566356902 | 0.035306671 | 0.036986662 | 0.954578446 | 0.339790921 | NA |
| 4933438K21Rik | 0.294043327 | 0.010508917 | 0.039092051 | 0.2688249 | 0.788064431 | NA |
| 4933439K11Rik | 4.788981835 | 0.192743155 | 0.141262722 | 1.364430423 | 0.172432138 | NA |
| 4933440J02Rik | 0 | NA | NA | NA | NA | NA |
| 4933440M02Rik | 0.617197563 | -0.009986935 | 0.063001754 | -0.158518368 | 0.874048348 | NA |
| 5031410I06Rik | 1.406075504 | 0.071744676 | 0.086475795 | 0.82965038 | 0.406736484 | NA |
| 5031425F14Rik | 3.624529577 | -0.156543277 | 0.119421821 | -1.310843152 | 0.18991076 | NA |
| 5031426D15Rik | 7.783855974 | -0.041969229 | 0.14812144 | -0.283343376 | 0.77691363 | NA |
| 5031434C07Rik | 0.529876295 | -0.026705656 | 0.04018514 | -0.664565453 | 0.506328472 | NA |
| 5033403H07Rik | 0 | NA | NA | NA | NA | NA |
| 5033406O09Rik | 6.163381864 | -0.067236878 | 0.148473129 | -0.452855535 | 0.650652771 | NA |
| 5330411J11Rik | 3.284653191 | -0.067891171 | 0.124746327 | -0.544233826 | 0.586280577 | NA |
| 5330439B14Rik | 0 | NA | NA | NA | NA | NA |
| 5430401F13Rik | 1.104555619 | 0.076855026 | 0.069738205 | 1.102050531 | 0.270439704 | NA |
| 5430402E10Rik | 1.709813011 | 0.036245416 | 0.04818324 | 0.75224115 | NA | NA |
| 5430403N17Rik | 0.990847688 | -0.007537041 | 0.064922126 | -0.11609356 | 0.907578393 | NA |
| 5430419D17Rik | 0.402033573 | 0.020557702 | 0.043166232 | 0.476245001 | 0.633899843 | NA |
| 5430421F17Rik | 4.448672424 | 0.010685712 | 0.125815844 | 0.084931372 | 0.932315951 | NA |
| 5430421N21Rik | 0.331747315 | -0.03734372 | 0.03744515 | -0.99729124 | 0.318623164 | NA |
| 5430425J12Rik | 0 | NA | NA | NA | NA | NA |
| 5430427M07Rik | 0.233393928 | -0.027475306 | 0.032191697 | -0.85349047 | 0.39338737 | NA |
| 5430428K19Rik | 0 | NA | NA | NA | NA | NA |
| 5430434I15Rik | 0 | NA | NA | NA | NA | NA |
| 5430437J10Rik | 0.923168337 | -0.053697833 | 0.07081308 | -0.758303882 | 0.448269084 | NA |
| 5430440P10Rik | 2.025330369 | -0.039732542 | 0.096551468 | -0.411516702 | 0.680693697 | NA |
| 5530400C23Rik | 0 | NA | NA | NA | NA | NA |
| 5530401A14Rik | 1.195750858 | -0.076546543 | 0.064444899 | -1.1877828 | 0.234918993 | NA |
| 5730403I07Rik | 2.921794355 | -0.096465799 | 0.092218981 | -1.046051453 | 0.295537277 | NA |
| 5730412P04Rik | 1.225651752 | -0.000777335 | 0.074383427 | -0.010450375 | 0.991661959 | NA |
| 5730416F02Rik | 0 | NA | NA | NA | NA | NA |
| 5730420D15Rik | 2.378564255 | 0.06379947 | 0.106297844 | 0.60019533 | 0.548376066 | NA |
| 5730422E09Rik | 5.36260156 | -0.101837742 | 0.131950107 | -0.771789769 | 0.440238952 | NA |
| 5730435O14Rik | 1.274846761 | -0.063631618 | 0.08842494 | -0.71961166 | 0.471764131 | NA |
| 5730457N03Rik | 0 | NA | NA | NA | NA | NA |
| 5730460C07Rik | 0 | NA | NA | NA | NA | NA |
| 5730488B01Rik | 0 | NA | NA | NA | NA | NA |
| 5830403M04Rik | 1.966094221 | 0.013819393 | 0.098611424 | 0.140139876 | 0.888549475 | NA |
| 5830411N06Rik | 1.463700863 | 0.137456779 | 0.093695528 | 1.467058059 | 0.142360262 | NA |
| 5830416I19Rik | 4.743051969 | 0.209804681 | 0.137766789 | 1.522897369 | 0.127784384 | NA |
| 5830428M24Rik | 0 | NA | NA | NA | NA | NA |
| 5830432E09Rik | 3.127986928 | 0.075015536 | 0.116468975 | 0.644081707 | 0.519522452 | NA |
| 5830473C10Rik | 2.529883987 | -0.019438374 | 0.111860035 | -0.173774073 | 0.862043021 | NA |
| 6030408B16Rik | 6.325265018 | 0.100117471 | 0.144425714 | 0.693210841 | 0.488177246 | NA |
| 6030440G07Rik | 0 | NA | NA | NA | NA | NA |
| 6030466F02Rik | 1.43368476 | 0.060713864 | 0.090463583 | 0.671141489 | 0.502130399 | NA |
| 6030468B19Rik | 2.58199959 | 0.13739291 | 0.106765721 | 1.286863507 | 0.198141867 | NA |
| 6030498E09Rik | 0.558783501 | -0.006176017 | 0.052494503 | -0.117650732 | 0.906344406 | NA |
| 6330407A03Rik | 0.107716603 | -0.020477583 | 0.027954224 | -0.732539849 | 0.46383913 | NA |
| 6330410L21Rik | 6.399064506 | 0.07381682 | 0.137644779 | 0.536284921 | 0.59176165 | NA |
| 6330415B21Rik | 0 | NA | NA | NA | NA | NA |
| 6430562O15Rik | 1.492400801 | 0.032328451 | 0.083519906 | 0.3870748 | 0.698700833 | NA |
| 6430706D22Rik | 0 | NA | NA | NA | NA | NA |
| 7420426K07Rik | 0 | NA | NA | NA | NA | NA |
| 7420461P10Rik | 0.096071043 | -0.020098795 | 0.027825948 | -0.722304036 | 0.470107572 | NA |
| 7420700N18Rik | 0 | NA | NA | NA | NA | NA |
| 7420701I03Rik | 0 | NA | NA | NA | NA | NA |
| 7530416G11Rik | 0.186326724 | 0.024939385 | 0.031411053 | 0.793968451 | 0.427213798 | NA |
| 7630403G23Rik | 6.137286725 | 0.0464188 | 0.104719735 | 0.443266975 | 0.65757264 | NA |
| 8030411F24Rik | 0 | NA | NA | NA | NA | NA |
| 8030423F21Rik | 2.141433242 | 0.034766227 | 0.097894192 | 0.355140854 | 0.722484072 | NA |
| 8030423J24Rik | 1.331013364 | 0.035204865 | 0.084182414 | 0.418197386 | 0.675802807 | NA |
| 8030442B05Rik | 2.133971828 | 0.051578361 | 0.100061993 | 0.515464057 | 0.60622878 | NA |
| 8030443G20Rik | 0.663524823 | -0.038052693 | 0.059997452 | -0.634238477 | 0.5259252 | NA |
| 8430403D17Rik | 0.098353386 | -0.020212831 | 0.02786466 | -0.725393061 | 0.468210929 | NA |
| 8430422H06Rik | 0 | NA | NA | NA | NA | NA |
| 8430423G03Rik | 0 | NA | NA | NA | NA | NA |
| 8430431K14Rik | 0 | NA | NA | NA | NA | NA |
| 8430436N08Rik | 0.302785524 | -0.016957309 | 0.039595455 | -0.428264025 | 0.668458907 | NA |
| 8430437L04Rik | 0.098353386 | -0.020212831 | 0.02786466 | -0.725393061 | 0.468210929 | NA |
| 9030204H09Rik | 0.362076728 | 0.018991228 | 0.042095091 | 0.451150657 | 0.65188097 | NA |
| 9030404E10Rik | 0.185906812 | 0.024895454 | 0.031385107 | 0.793225068 | 0.427646705 | NA |
| 9030619P08Rik | 0.309710804 | -0.014226733 | 0.039782345 | -0.357614233 | 0.720632025 | NA |
| 9130008F23Rik | 4.619969587 | -0.064091833 | 0.121275204 | -0.528482583 | 0.597164432 | NA |
| 9130015A21Rik | 1.12994131 | 0.036923979 | 0.07408221 | 0.498418966 | 0.618188772 | NA |
| 9130015L21Rik | 0.283178451 | 0.028390488 | 0.033368195 | 0.850824814 | 0.394866675 | NA |
| 9130019P16Rik | 7.620208972 | 0.006877866 | 0.155372474 | 0.044266952 | 0.964691614 | NA |
| 9130204L05Rik | 0 | NA | NA | NA | NA | NA |
| 9130206I24Rik | 5.39408971 | -0.015742662 | 0.133937251 | -0.117537595 | 0.906434054 | NA |
| 9130209A04Rik | 0 | NA | NA | NA | NA | NA |
| 9130221F21Rik | 0 | NA | NA | NA | NA | NA |
| 9130230L23Rik | 0 | NA | NA | NA | NA | NA |
| 9130409I23Rik | 0.192142086 | -0.026384592 | 0.031527717 | -0.836869715 | 0.402665805 | NA |
| 9230009I02Rik | 6.460502141 | 0.126379548 | 0.1389964 | 0.909228929 | 0.363229297 | NA |
| 9230102K24Rik | 1.840445218 | -0.054323261 | 0.094912351 | -0.572351865 | 0.567083619 | NA |
| 9230102O04Rik | 0.159967308 | 0.0240345 | 0.030870969 | 0.778547006 | 0.436246606 | NA |
| 9230104L09Rik | 0.833822031 | 0.048595634 | 0.059165953 | 0.821344557 | 0.411450034 | NA |
| 9230105E05Rik | 0 | NA | NA | NA | NA | NA |
| 9230110F15Rik | 0.094392817 | 0.017895942 | 0.027818137 | 0.643319236 | 0.520016977 | NA |
| 9230112D13Rik | 0 | NA | NA | NA | NA | NA |
| 9230115E21Rik | 0.389426822 | -0.00542245 | 0.0474348 | -0.114313744 | 0.908989088 | NA |
| 9330111N05Rik | 6.798952139 | -0.025466348 | 0.155174401 | -0.164114364 | 0.869641115 | NA |
| 9330162B11Rik | 0.289311041 | 0.012154199 | 0.038946437 | 0.312074719 | 0.754983739 | NA |
| 9330175E14Rik | 1.46230452 | 0.156422717 | 0.081557834 | 1.917936127 | 0.055119111 | NA |
| 9330178D15Rik | 0 | NA | NA | NA | NA | NA |
| 9330182O14Rik | 0 | NA | NA | NA | NA | NA |
| 9430007A20Rik | 0 | NA | NA | NA | NA | NA |
| 9430014N10Rik | 0.423821016 | -0.060879619 | 0.047397786 | -1.284440147 | 0.198987993 | NA |
| 9430018G01Rik | 1.1451679 | 0.008926323 | 0.072644483 | 0.122876814 | 0.902204646 | NA |
| 9430019J16Rik | 0 | NA | NA | NA | NA | NA |

| | | | | | | |
|---|---|---|---|---|---|---|
| Nt5c | 930.9865909 | 0.070127695 | 0.066561434 | 1.05357849 | 0.292075941 | 0.727847831 |
| Ccdc115 | 650.314827 | 0.070131823 | 0.067326062 | 1.041674221 | 0.297562745 | 0.732415975 |
| Syncrip | 2643.303813 | 0.070136804 | 0.07860143 | 0.892309523 | 0.372227051 | 0.781446026 |
| Adam18 | 23.43330585 | 0.070144023 | 0.084010225 | 0.834946263 | 0.403747965 | 0.800224169 |
| Wnt5b | 75.13131147 | 0.070144337 | 0.095062537 | 0.737875716 | 0.460589977 | 0.829267477 |
| Ciapin1 | 2439.991056 | 0.070147531 | 0.055247136 | 1.269704402 | 0.204189944 | 0.65778952 |
| Camkmt | 295.8258925 | 0.07017215 | 0.081682359 | 0.859085745 | 0.390293211 | 0.793521685 |
| Fndc5 | 3823.986315 | 0.070182142 | 0.061830385 | 1.135075294 | 0.256343772 | 0.702442866 |
| Snx21 | 1002.04595 | 0.070195126 | 0.065757151 | 1.067490378 | 0.285750456 | 0.722148665 |
| BC027072 | 16.12087067 | 0.070197149 | 0.075423624 | 0.930705065 | 0.352006149 | NA |
| Gm20748 | 27.91853773 | 0.07020105 | 0.085151717 | 0.824423183 | 0.409699146 | 0.801196796 |
| Pfkm | 19955.37311 | 0.070217719 | 0.050230899 | 1.397898915 | 0.162143425 | 0.613957643 |
| Mrpl47 | 433.8482711 | 0.070231983 | 0.077121156 | 0.910670777 | 0.362468866 | 0.776449324 |
| Rcn2 | 4249.444776 | 0.070273155 | 0.050096845 | 1.402746104 | 0.160692564 | 0.612387888 |
| Ttc1 | 1514.306948 | 0.070301257 | 0.057935585 | 1.213438289 | 0.224962295 | 0.67664265 |
| Glo1 | 3551.456179 | 0.070302817 | 0.046728427 | 1.504497837 | 0.13245323 | 0.574161628 |
| Cisd2 | 1588.63032 | 0.070303985 | 0.059023722 | 1.191114045 | 0.233608817 | 0.685064073 |
| Pik3ca | 3428.74435 | 0.070309159 | 0.069398846 | 1.013117126 | 0.311004219 | 0.743730778 |
| Prss12 | 770.3389609 | 0.070349258 | 0.075502357 | 0.931749158 | 0.351466175 | 0.769244462 |
| Ipo5 | 7103.601652 | 0.070358756 | 0.058445321 | 1.203838987 | 0.228651818 | 0.68173159 |
| Pcdhgb1 | 317.7048591 | 0.070386228 | 0.094373821 | 0.745823654 | 0.455773949 | 0.826603827 |
| 0610011F06Rik | 724.1337056 | 0.070412082 | 0.067644424 | 1.040914801 | 0.297915092 | 0.732916829 |
| Ebf2 | 36.50106641 | 0.070423551 | 0.084595202 | 0.832476902 | 0.40513981 | 0.8005027 |
| Cyb5 | 1680.104763 | 0.070465718 | 0.053383261 | 1.319996494 | 0.186836189 | 0.641054259 |
| Emc2 | 2158.64419 | 0.070472622 | 0.054389112 | 1.295711951 | 0.195074743 | 0.65093872 |
| Hecw1 | 2968.508811 | 0.070474215 | 0.075495113 | 0.933493731 | 0.350565106 | 0.768667813 |
| Pds5b | 4477.816906 | 0.070477125 | 0.065112168 | 1.082395608 | 0.279076779 | 0.718516358 |
| Cd81 | 10156.31008 | 0.070488608 | 0.064265955 | 1.096826584 | 0.272717207 | 0.714142044 |
| Srp14 | 2308.149641 | 0.070506084 | 0.061177139 | 1.152490704 | 0.249119488 | 0.697784358 |
| Arhgdia | 12099.5019 | 0.070514446 | 0.056262868 | 1.253303439 | 0.210095297 | 0.664608511 |
| Depdc5 | 2900.832751 | 0.07051627 | 0.052554495 | 1.341774287 | 0.179669192 | 0.633459544 |
| Fndc3b | 842.1388725 | 0.070601371 | 0.063912884 | 1.104650049 | 0.269311255 | 0.711922711 |
| Smarca2 | 12135.33654 | 0.070606209 | 0.05975219 | 1.181650551 | 0.237344387 | 0.688415946 |
| Cyp2j9 | 1236.884403 | 0.070606323 | 0.085468862 | 0.826105811 | 0.408744072 | 0.801055189 |
| Kcnip3 | 6584.170717 | 0.07061133 | 0.059576104 | 1.185229062 | 0.235926892 | 0.687039131 |
| Sar1a | 5713.453125 | 0.070664856 | 0.045899972 | 1.539540275 | 0.123672453 | 0.558428989 |
| Slc39a11 | 682.2385319 | 0.070671377 | 0.070017837 | 1.00933391 | 0.312814523 | 0.745030949 |
| Abcf2 | 4539.1199 | 0.070689281 | 0.038140212 | 1.853405569 | 0.06382425 | 0.441368101 |
| Ubqln2 | 7488.477001 | 0.070701974 | 0.04297773 | 1.645083952 | 0.0999525 | 0.521115714 |
| Ldoc1 | 55.82609096 | 0.070721418 | 0.093106413 | 0.759576231 | 0.447507931 | 0.822458097 |
| B230216N24Rik | 133.1087361 | 0.070723458 | 0.091213873 | 0.775358571 | 0.438127809 | 0.816984132 |
| Psen2 | 584.871434 | 0.070725584 | 0.069576391 | 1.016516999 | 0.309383257 | 0.742145027 |
| Zmynd8 | 6001.194261 | 0.070765677 | 0.057314776 | 1.23468469 | 0.216947876 | 0.670185967 |
| Nanp | 584.5423537 | 0.070791336 | 0.064563057 | 1.096468148 | 0.272873955 | 0.714192265 |
| Lypla2 | 2763.008458 | 0.070791404 | 0.066573287 | 1.063360511 | 0.287618493 | 0.723579189 |
| Dstn | 6049.520356 | 0.070811051 | 0.056053661 | 1.263272533 | 0.206491254 | 0.660681047 |
| Slc28a3 | 7.164824504 | 0.070812469 | 0.045863928 | 1.543968707 | 0.122595914 | NA |
| Snx8 | 586.4766663 | 0.070828217 | 0.08509565 | 0.83233652 | 0.405219022 | 0.8005027 |
| D430020J02Rik | 131.8205057 | 0.070846892 | 0.094373535 | 0.7507072 | 0.452828889 | 0.82547056 |
| Gm5820 | 38.52581085 | 0.070863853 | 0.087031144 | 0.814235561 | 0.415510017 | 0.805633085 |
| Fam169b | 23.75927967 | 0.070867239 | 0.076283607 | 0.928996958 | 0.352890663 | 0.770519168 |
| Sertad4 | 835.1048019 | 0.070881799 | 0.077869626 | 0.910262477 | 0.362684102 | 0.776470047 |
| Pld3 | 28867.81376 | 0.070884537 | 0.057324749 | 1.236543342 | 0.216256666 | 0.670049713 |
| Agpat1 | 5249.82963 | 0.070902443 | 0.053280873 | 1.330729742 | 0.183277953 | 0.63672274 |
| Tmie | 229.4721811 | 0.070915272 | 0.094988137 | 0.746569772 | 0.455323301 | 0.826546647 |
| Gdnf | 46.41318351 | 0.070937896 | 0.092939129 | 0.763272654 | 0.445300807 | 0.821629343 |
| Rsph4a | 227.3965194 | 0.070938646 | 0.089185101 | 0.795409155 | 0.426375536 | 0.809717746 |
| Nrip1 | 1017.411928 | 0.070939735 | 0.072537761 | 0.977969734 | 0.328089293 | 0.754778534 |
| Pdgfd | 90.63040677 | 0.070943548 | 0.095098031 | 0.746004386 | 0.455664766 | 0.826603827 |
| Wwp2 | 1891.50686 | 0.07097223 | 0.055468437 | 1.279506582 | 0.200718723 | 0.654716629 |
| Rcsd1 | 264.3387952 | 0.07097431 | 0.082498756 | 0.860307639 | 0.389619483 | 0.792715562 |
| Eri1 | 610.877321 | 0.070996066 | 0.085963185 | 0.825889201 | 0.408866947 | 0.801055189 |
| Cyp2b23 | 2.310920538 | 0.07099733 | 0.032549341 | 2.181221712 | 0.029167022 | NA |
| Gpm6a | 38343.45946 | 0.071009367 | 0.072861115 | 0.974585241 | 0.329766037 | 0.756141871 |
| Eif3g | 2183.800845 | 0.071049243 | 0.047803036 | 1.486291425 | 0.137202054 | 0.581727097 |
| Shc3 | 1132.804364 | 0.07105158 | 0.074123906 | 0.958551495 | 0.337784738 | 0.760581219 |
| Zfp934 | 73.70059542 | 0.071074886 | 0.093998329 | 0.756129251 | 0.449571706 | 0.823263966 |
| Atp6v0a1 | 25402.80009 | 0.071092293 | 0.043712398 | 1.626364525 | 0.103872133 | 0.523645898 |
| Eif1ax | 1825.554996 | 0.071093588 | 0.07194471 | 0.98816977 | 0.32306951 | 0.752792382 |
| Arf1 | 16005.64003 | 0.071095789 | 0.038129191 | 1.864602623 | 0.062237125 | 0.436873272 |
| Sec22c | 2309.318821 | 0.071120273 | 0.061647937 | 1.153652125 | 0.248642819 | 0.697784358 |
| Gtf2f2 | 440.5862172 | 0.071150645 | 0.072956057 | 0.975253433 | 0.329434563 | 0.756141871 |
| Tomm22 | 2136.873924 | 0.071150978 | 0.055278806 | 1.287129443 | 0.198049175 | 0.653535282 |
| Lrrc59 | 4408.116712 | 0.071152993 | 0.048749359 | 1.459567773 | 0.144408902 | 0.589967032 |
| Pde11a | 96.82414906 | 0.071176966 | 0.095069413 | 0.748684191 | 0.454047574 | 0.825985606 |
| Gss | 511.7818451 | 0.071177635 | 0.066301429 | 1.073546011 | 0.283026206 | 0.720920007 |
| Cacnb4 | 9153.097186 | 0.071188051 | 0.079237911 | 0.898408981 | 0.36896755 | 0.779694884 |
| Shd | 847.1482601 | 0.071216415 | 0.074122236 | 0.960796908 | 0.336654294 | 0.75970798 |
| Dsg1c | 10.17379232 | 0.07122315 | 0.06248093 | 1.139918221 | 0.254320373 | NA |
| A630007B06Rik | 1137.152647 | 0.071285538 | 0.079765962 | 0.893683672 | 0.371491164 | 0.780761257 |
| AI597468 | 1175.883627 | 0.071313414 | 0.077236445 | 0.923313005 | 0.355844113 | 0.772643677 |
| Zfp426 | 1130.635745 | 0.071320162 | 0.075774657 | 0.941213913 | 0.346595248 | 0.765286747 |
| Ak1 | 3955.673936 | 0.071323177 | 0.058104621 | 1.227495778 | 0.219636308 | 0.672421385 |
| Pdcd6 | 2014.775614 | 0.071328849 | 0.053670767 | 1.32900746 | 0.183845509 | 0.637355821 |
| Aard | 58.11741459 | 0.0713942 | 0.092359621 | 0.773002301 | 0.439521019 | 0.817580013 |
| Pak1ip1 | 1562.194914 | 0.071396275 | 0.055934946 | 1.27641628 | 0.201808404 | 0.655168687 |
| Nono | 5363.146507 | 0.07140361 | 0.047603683 | 1.499959767 | 0.133624825 | 0.57588358 |
| Sc5d | 1801.222618 | 0.071415231 | 0.065106245 | 1.096902933 | 0.272683826 | 0.714142044 |
| 1500011B03Rik | 3262.675584 | 0.071424025 | 0.051097055 | 1.397810983 | 0.162169835 | 0.613957643 |
| Usp14 | 6045.750312 | 0.071445262 | 0.068505594 | 1.042911348 | 0.296989354 | 0.732415975 |
| Figf | 91.05785371 | 0.071504276 | 0.095056551 | 0.752228811 | 0.45191347 | 0.824608086 |
| Aldoa | 63778.62338 | 0.071516966 | 0.045330627 | 1.577674325 | 0.114640448 | 0.543130934 |
| Ubtd2 | 1557.637216 | 0.071524709 | 0.074959809 | 0.954174101 | 0.339995522 | 0.761914237 |
| Dner | 7463.302369 | 0.071537824 | 0.063728201 | 1.122545786 | 0.261630452 | 0.704173028 |
| Pgs1 | 3158.641893 | 0.071540708 | 0.053623252 | 1.334135961 | 0.182159302 | 0.635245711 |
| Ercc3 | 1365.771599 | 0.071578131 | 0.057378064 | 1.247482517 | 0.212220627 | 0.666270994 |
| Adprh | 3415.308842 | 0.071579129 | 0.047853518 | 1.495796606 | 0.134706666 | 0.577947814 |
| Zfp423 | 806.5306003 | 0.07161542 | 0.085004096 | 0.842493753 | 0.399511635 | 0.798363738 |
| Fam192a | 1743.137961 | 0.071635606 | 0.048745071 | 1.469596919 | 0.141670955 | 0.587317993 |
| Eif2ak1 | 3572.986522 | 0.071643152 | 0.052947914 | 1.353087344 | 0.176027728 | 0.62871374 |
| Araf | 5262.505894 | 0.071651355 | 0.046877145 | 1.528492297 | 0.126390355 | 0.563001009 |
| Nsmce4a | 1113.572872 | 0.071702084 | 0.065047656 | 1.102300811 | 0.270330917 | 0.71216373 |
| Zfp280b | 1047.607389 | 0.071711043 | 0.064247422 | 1.116169969 | 0.264349387 | 0.707261241 |
| Chchd6 | 1417.396167 | 0.071732126 | 0.056005739 | 1.280799576 | 0.200264073 | 0.654475062 |
| 1110015O18Rik | 11.89367556 | 0.071733923 | 0.057230806 | 1.253414514 | 0.210054892 | NA |
| Lipe | 424.3891044 | 0.07173417 | 0.075374646 | 0.95170158 | 0.341248348 | 0.762819666 |
| Ube2k | 4452.124036 | 0.071737885 | 0.061396074 | 1.168444177 | 0.242627644 | 0.692772033 |
| Hspa14 | 1073.417694 | 0.071740478 | 0.06202576 | 1.156623938 | 0.247426038 | 0.696744638 |
| Mmgt1 | 1676.177443 | 0.071744908 | 0.063535565 | 1.129208617 | 0.25880984 | 0.703616072 |
| Stk3 | 619.0792355 | 0.071754064 | 0.06827813 | 1.050908451 | 0.293300638 | 0.728786362 |
| Mrpl12 | 1621.573403 | 0.071766789 | 0.051543024 | 1.392366668 | 0.163811393 | 0.614351408 |
| Lamb3 | 90.20870048 | 0.071768501 | 0.095021743 | 0.75528504 | 0.450077973 | 0.823263966 |
| Sbspon | 15.19441133 | 0.071778908 | 0.076372589 | 0.939851695 | 0.347293638 | NA |
| Sec11a | 686.2205541 | 0.071806993 | 0.065369411 | 1.098480046 | 0.271994926 | 0.714078854 |
| Med18 | 114.5724284 | 0.071826079 | 0.091827741 | 0.782182786 | 0.434107159 | 0.814708087 |
| Pcdhac2 | 1555.088788 | 0.071838495 | 0.065277644 | 1.10050686 | 0.271111342 | 0.712918783 |
| Itga9 | 506.9769934 | 0.071857597 | 0.085168748 | 0.843708503 | 0.398832315 | 0.797789838 |
| 4930519F09Rik | 28.68851355 | 0.071867477 | 0.087116021 | 0.824962804 | 0.409392708 | 0.801055189 |
| Hspbp1 | 2474.504057 | 0.071920623 | 0.058461212 | 1.230228046 | 0.21861172 | 0.671604514 |
| Vpreb1 | 3.016205768 | 0.071921276 | 0.040550188 | 1.773636082 | 0.076123365 | NA |
| Hdc | 125.4646549 | 0.07194132 | 0.063804271 | 1.127531414 | 0.259517867 | 0.703893404 |
| Wnk3 | 1698.643716 | 0.071948832 | 0.073563709 | 0.978047921 | 0.328050624 | 0.754778534 |
| Rpl18 | 3226.862672 | 0.071962382 | 0.059081964 | 1.218009295 | 0.22322044 | 0.674214152 |
| Tacr1 | 504.2905649 | 0.071969028 | 0.091671826 | 0.78507248 | 0.432411074 | 0.813262565 |
| Pi16 | 38.9080401 | 0.071998401 | 0.072979245 | 0.986559965 | 0.323858405 | 0.752925264 |
| Lypla1 | 797.6182795 | 0.072016558 | 0.075369932 | 0.955507795 | 0.339320966 | 0.76146707 |
| Pdhb | 4512.572357 | 0.072022092 | 0.047958259 | 1.50176619 | 0.133157503 | 0.575193212 |
| Nsf | 42381.32647 | 0.072031458 | 0.038351712 | 1.878181031 | 0.060356404 | 0.430493689 |
| Ogfod3 | 210.80085 | 0.072041413 | 0.090943308 | 0.79215738 | 0.428268916 | 0.810486244 |
| Gm1661 | 19.82913182 | 0.072043302 | 0.077996017 | 0.923679244 | 0.355653342 | NA |
| Ccdc60 | 24.45548615 | 0.072046244 | 0.077882029 | 0.925068914 | 0.354930062 | 0.772160263 |
| Fbxo24 | 8.470166489 | 0.072053208 | 0.058914868 | 1.223005504 | 0.221327617 | NA |
| Gm11992 | 58.02173373 | 0.072068262 | 0.093658763 | 0.76947698 | 0.441610205 | 0.819100918 |
| Casp8ap2 | 638.3047575 | 0.072072936 | 0.086449809 | 0.833696881 | 0.404451816 | 0.800290023 |
| Hrasls | 1302.88489 | 0.072085958 | 0.0677856 | 1.063440584 | 0.287582197 | 0.723579189 |
| Armc1 | 3211.537618 | 0.072087862 | 0.053754109 | 1.341067012 | 0.179898697 | 0.633658697 |
| Kcnq3 | 2021.236095 | 0.072112303 | 0.073199226 | 0.985151166 | 0.324549824 | 0.75293408 |
| Ugdh | 959.7120464 | 0.072116983 | 0.056697782 | 1.271954224 | 0.203389384 | 0.657417489 |
| Tph2 | 40.52028626 | 0.072137935 | 0.089491556 | 0.806086497 | 0.420192977 | 0.807657282 |
| Rtfdc1 | 3305.120197 | 0.072142694 | 0.047636972 | 1.514426522 | 0.129917697 | 0.56891406 |
| Vav3 | 319.3728672 | 0.072173549 | 0.092266957 | 0.782225311 | 0.434082172 | 0.814708087 |
| Eef1a2 | 40259.61988 | 0.072173664 | 0.050698264 | 1.423592409 | 0.154564494 | 0.604361735 |
| Ccdc55 | 715.9519455 | 0.072173904 | 0.066346553 | 1.087832004 | 0.276669278 | 0.717475105 |
| H2afx | 830.2712995 | 0.07219821 | 0.06668055 | 1.082747665 | 0.278920441 | 0.71849589 |
| Eps8l2 | 243.3265256 | 0.072199583 | 0.086942199 | 0.830431987 | 0.406294587 | 0.800759397 |
| Galnt2 | 3487.605994 | 0.072208066 | 0.054553452 | 1.323620461 | 0.185629126 | 0.638817505 |
| Vprbp | 1622.949327 | 0.072227344 | 0.066084533 | 1.092953836 | 0.274414069 | 0.716185716 |
| Pnpla8 | 4483.503475 | 0.072229493 | 0.06698433 | 1.078304334 | 0.280897963 | 0.720080692 |
| Gadd45g | 334.4906166 | 0.072246624 | 0.086649653 | 0.833778576 | 0.40440577 | 0.800290023 |
| Lonrf1 | 1468.410044 | 0.072308291 | 0.076340231 | 0.9471846 | 0.343544721 | 0.763762832 |
| Ppp2r2b | 5452.214812 | 0.072310493 | 0.04483386 | 1.612854501 | 0.106776122 | 0.530447999 |
| Eif4e | 3911.510224 | 0.072344089 | 0.055406446 | 1.305698053 | 0.19165526 | 0.646997681 |
| Igsf1 | 408.7876096 | 0.072351781 | 0.0614293 | 1.177805724 | 0.238874066 | 0.689537017 |
| Gdf9 | 37.00989605 | 0.072362565 | 0.092309019 | 0.783916527 | 0.433089092 | 0.813783127 |
| Rab11a | 3334.297483 | 0.072367762 | 0.064785816 | 1.117030959 | 0.263981088 | 0.706954491 |
| Phex | 27.54317274 | 0.072370161 | 0.080157228 | 0.902852582 | 0.36660414 | 0.778210102 |
| Pdha1 | 9971.562582 | 0.072408403 | 0.048136704 | 1.504224364 | 0.132523606 | 0.574295318 |
| D630039A03Rik | 12.6205393 | 0.072431854 | 0.064370284 | 1.125237445 | 0.26048843 | NA |
| Eno2 | 46505.65641 | 0.07244307 | 0.042917634 | 1.68795582 | 0.09141971 | 0.504162 |
| Prmt5 | 2413.694046 | 0.072532893 | 0.044217348 | 1.640371846 | 0.100927876 | 0.521542547 |
| Vps18 | 2504.571297 | 0.072538717 | 0.053738121 | 1.349855845 | 0.177062227 | 0.629241039 |
| Nme3 | 542.7119345 | 0.072550621 | 0.06711955 | 1.08091638 | 0.279734312 | 0.718934998 |
| Brix1 | 775.9668915 | 0.072553817 | 0.064213978 | 1.129875752 | 0.258528583 | 0.703616072 |
| Sytl4 | 63.80699916 | 0.072592 | 0.044232776 | 1.64113598 | NA | NA |
| Zfp788 | 1372.347446 | 0.072598457 | 0.064366782 | 1.127887006 | 0.259367642 | 0.703893404 |
| Ube2b | 5183.109642 | 0.072617363 | 0.058171638 | 1.24832934 | 0.211910474 | 0.666270994 |
| Nudt21 | 884.7805253 | 0.072702547 | 0.064872903 | 1.120692058 | 0.262418964 | 0.704862772 |
| Osbpl10 | 1283.052683 | 0.072706239 | 0.061822529 | 1.176047638 | 0.23957584 | 0.690203878 |
| Pthlh | 106.2116703 | 0.07270871 | 0.095094003 | 0.764598265 | 0.444510803 | 0.821493054 |
| 4632415L05Rik | 1007.194435 | 0.072714987 | 0.060598054 | 1.199955806 | 0.230156505 | 0.683133099 |
| Psma1 | 2696.475169 | 0.072721993 | 0.061069919 | 1.190798918 | 0.233732534 | 0.685064073 |
| Ndst4 | 275.1989516 | 0.072732271 | 0.094357675 | 0.770814576 | 0.440816846 | 0.818732324 |
| Gpx6 | 39.78474463 | 0.072752806 | 0.081059957 | 0.897518436 | 0.36944234 | 0.780397655 |
| D10Bwg1379e | 2606.834587 | 0.072786657 | 0.072048145 | 1.010250265 | 0.312375402 | 0.74483997 |
| Crybg3 | 318.6719045 | 0.072810237 | 0.085097034 | 0.855614282 | 0.392211167 | 0.794184423 |
| Gm2694 | 45.60092276 | 0.07283557 | 0.092626857 | 0.786333174 | 0.431672322 | 0.812553085 |
| Rab31 | 3938.760262 | 0.072880107 | 0.061390027 | 1.187165246 | 0.235162447 | 0.686276191 |
| 1810055G02Rik | 824.4212943 | 0.072898561 | 0.066356832 | 1.098584097 | 0.271949517 | 0.714078854 |
| Smyd2 | 2263.896485 | 0.072907661 | 0.050142883 | 1.453998168 | 0.145946814 | 0.592514474 |
| Abhd13 | 1780.513079 | 0.07292907 | 0.074572778 | 0.977958346 | 0.328094926 | 0.754778534 |
| Ptpra | 7503.281894 | 0.072943167 | 0.043395292 | 1.680900451 | 0.092782252 | 0.506868248 |
| Pithd1 | 2922.276534 | 0.07296127 | 0.049363792 | 1.478032108 | 0.139399184 | 0.583884819 |
| Mapk4 | 4369.782043 | 0.072967449 | 0.06580204 | 1.108893407 | 0.267476169 | 0.710152314 |
| Slc12a8 | 59.07772276 | 0.072970102 | 0.094608486 | 0.771284959 | 0.440538046 | 0.818563572 |
| Tars | 1271.444276 | 0.073007815 | 0.054609133 | 1.336915851 | 0.181250106 | 0.634469716 |
| Pkia | 6587.188655 | 0.073008175 | 0.07502327 | 0.973140391 | 0.330483532 | 0.756141871 |
| Efha2 | 3504.806222 | 0.073013551 | 0.070042332 | 1.042420332 | 0.297216844 | 0.732415975 |
| Mllt11 | 10874.70604 | 0.073035187 | 0.052474898 | 1.391811897 | 0.163979366 | 0.614351408 |
| Ergic3 | 3341.878652 | 0.073062832 | 0.065493754 | 1.115569462 | 0.26460647 | 0.707370515 |
| Twistnb | 361.8585693 | 0.073122856 | 0.078717925 | 0.928922551 | 0.352929225 | 0.770519168 |
| Tpgs2 | 2529.103205 | 0.073136257 | 0.058678611 | 1.246386979 | 0.212622359 | 0.666270994 |
| Bcar1 | 2138.492097 | 0.07315399 | 0.061273065 | 1.193901277 | 0.232516588 | 0.684327865 |
| Mtmr6 | 6627.085125 | 0.073165539 | 0.06511222 | 1.123683678 | 0.261147243 | 0.704173028 |
| Lynx1 | 20758.29343 | 0.073173394 | 0.058114526 | 1.259123979 | 0.207985554 | 0.66299025 |
| Lypd6b | 701.3095382 | 0.073177226 | 0.064956444 | 1.126558373 | 0.259929247 | 0.704052663 |
| Kctd13 | 4852.530545 | 0.073177503 | 0.041355312 | 1.769482569 | 0.076813377 | 0.472560915 |
| Aurkaip1 | 1699.031199 | 0.073187052 | 0.051790512 | 1.413136295 | 0.157615653 | 0.608141173 |
| Cox6b2 | 73.49383735 | 0.073224244 | 0.091616552 | 0.799246889 | 0.424147268 | 0.809082497 |
| Alkbh8 | 632.4334323 | 0.07322619 | 0.079819488 | 0.917397401 | 0.358934434 | 0.774837508 |
| Spryd3 | 7378.919815 | 0.073243048 | 0.057955452 | 1.263781835 | 0.206308343 | 0.660297797 |
| Glod4 | 3157.613902 | 0.073267232 | 0.054692243 | 1.33962749 | 0.180366482 | 0.634469716 |
| Pbx3 | 586.9501735 | 0.073280409 | 0.088772349 | 0.825486878 | 0.409095229 | 0.801055189 |
| Ndufaf5 | 1041.485861 | 0.073298859 | 0.058885522 | 1.244768767 | 0.21321676 | 0.666732456 |
| Gm17751 | 7.00940474 | 0.073318945 | 0.053401815 | 1.372967287 | 0.169762517 | NA |
| Dph3 | 1272.873016 | 0.07334337 | 0.060005929 | 1.222268717 | 0.221606025 | 0.67287816 |
| Prrg3 | 2507.966827 | 0.073350176 | 0.063677673 | 1.151897864 | 0.249363046 | 0.697864808 |
| Kcnd3 | 4289.439797 | 0.073363712 | 0.076014976 | 0.965121818 | 0.3344838 | 0.758238352 |
| Wac | 5085.862409 | 0.073368074 | 0.0646131 | 1.135498435 | 0.256166537 | 0.702218644 |
| March6 | 12220.5601 | 0.073377241 | 0.06174924 | 1.188310037 | 0.234711285 | 0.685908078 |
| Fstl1 | 1630.596736 | 0.073406208 | 0.070851357 | 1.036059306 | 0.300174475 | 0.733939477 |
| Sec23b | 1864.130103 | 0.073412925 | 0.047208012 | 1.555094614 | 0.119923537 | 0.553708672 |
| Slc35e4 | 1044.884672 | 0.073428131 | 0.073977666 | 0.992571615 | 0.320918763 | 0.750352874 |
| Kcnh8 | 24.73074827 | 0.073437954 | 0.075610505 | 0.97126654 | 0.331415566 | 0.756464884 |
| Tnni3k | 29.04161803 | 0.073438454 | 0.087940041 | 0.835096873 | 0.403663167 | 0.800224169 |
| Amica1 | 21.70011885 | 0.073438617 | 0.078598749 | 0.934348417 | 0.350124197 | NA |
| Lpgat1 | 14608.22065 | 0.0734439 | 0.064768769 | 1.133940034 | 0.256819703 | 0.703000737 |
| Snap25 | 116859.6071 | 0.073446169 | 0.062382461 | 1.177352878 | 0.23905469 | 0.689537017 |
| Fuca1 | 1187.416877 | 0.073473532 | 0.063187685 | 1.162782466 | 0.244917773 | 0.694568584 |
| Glipr2 | 127.1244699 | 0.073502855 | 0.094440966 | 0.778294187 | 0.436395602 | 0.816260946 |
| Slc39a10 | 10931.11749 | 0.0735039 | 0.07919505 | 0.928137563 | 0.353336217 | 0.770624606 |
| Mapk9 | 12157.16594 | 0.073513206 | 0.053231988 | 1.380996816 | 0.16727994 | 0.617511666 |
| Dnajb3 | 28.74850084 | 0.073533295 | 0.085069677 | 0.864389031 | 0.387374217 | 0.791583702 |
| Npffr1 | 21.71576102 | 0.07353923 | 0.074196885 | 0.991136349 | 0.321619006 | NA |
| P2rx3 | 118.1726567 | 0.073570892 | 0.09300209 | 0.791067077 | 0.42890485 | 0.810705511 |
| Csrp2bp | 1538.387215 | 0.0735986 | 0.04910805 | 1.498707432 | 0.133949548 | 0.576348084 |
| Clip4 | 2567.078921 | 0.073603088 | 0.057672629 | 1.27622218 | 0.20187699 | 0.655168687 |
| Eif2s3x | 8043.224867 | 0.073604096 | 0.053070595 | 1.386909206 | 0.16546945 | 0.615678595 |
| Gtf2e2 | 568.4405945 | 0.073610225 | 0.077083422 | 0.95494236 | 0.339606847 | 0.761679979 |
| Mrps2 | 1328.675318 | 0.073612765 | 0.056793076 | 1.296157391 | 0.194921266 | 0.650861767 |
| Fam210b | 1147.989043 | 0.073658678 | 0.076539897 | 0.962356636 | 0.335870488 | 0.759354819 |
| Syp | 60443.48884 | 0.073674903 | 0.045264095 | 1.62766765 | 0.103595376 | 0.523462117 |
| Cyc1 | 5281.680731 | 0.073684974 | 0.05250939 | 1.40327234 | 0.160535643 | 0.612387888 |
| Psmd9 | 912.3056086 | 0.073691284 | 0.06099063 | 1.208239441 | 0.226955175 | 0.67988263 |
| Gprasp2 | 6018.148391 | 0.073692662 | 0.076204757 | 0.967034932 | 0.333526572 | 0.757852268 |
| Ssbp2 | 3899.289479 | 0.073702627 | 0.062948848 | 1.170833618 | 0.241665662 | 0.692393571 |

| | | | | | | |
|---|---|---|---|---|---|---|
| 9430060I03Rik | 7.122908069 | -0.012088913 | 0.143328609 | -0.084344032 | 0.932782905 | NA |
| 9430069I07Rik | 0 | NA | NA | NA | NA | NA |
| 9430076C15Rik | 1.2997562 | -0.001626253 | 0.081439436 | -0.01996886 | 0.984068214 | NA |
| 9430076G02Rik | 1.27944664 | -0.029819558 | 0.086373056 | -0.345241433 | 0.729912872 | NA |
| 9530002B09Rik | 0 | NA | NA | NA | NA | NA |
| 9530003J23Rik | 0.211846316 | 0.025707621 | 0.031860375 | 0.806883811 | 0.419733426 | NA |
| 9530026F06Rik | 0.203787646 | -0.02671144 | 0.031728022 | -0.841887971 | 0.399850664 | NA |
| 9530027J09Rik | 2.461712552 | 0.106928818 | 0.10883103 | 0.982521413 | 0.325843039 | NA |
| 9530048J24Rik | 0.521607058 | -0.053101502 | 0.044487442 | -1.193629028 | 0.232623114 | NA |
| 9530053A07Rik | 5.088403885 | 0.305295635 | 0.117933637 | 2.588707018 | 0.009633702 | NA |
| 9530059O14Rik | 3.90566549 | 0.047944508 | 0.131037513 | 0.365883834 | 0.71445176 | NA |
| 9630028H03Rik | 0.492226933 | -0.036718726 | 0.046334526 | -0.792470094 | 0.428086623 | NA |
| 9830107B12Rik | 0 | NA | NA | NA | NA | NA |
| 9830132P13Rik | 0.105923158 | 0.018524693 | 0.027977518 | 0.662127842 | 0.507889294 | NA |
| 9930111J21Rik2 | 0.254617334 | 0.029389883 | 0.033981526 | 0.864878265 | 0.387105608 | NA |
| A1bg | 0 | NA | NA | NA | NA | NA |
| A1cf | 0.08731684 | 0.017365056 | 0.027733412 | 0.626142072 | 0.531221759 | NA |
| A230020J21Rik | 0.187700257 | -0.003168524 | 0.031449734 | -0.100748845 | 0.919749836 | NA |
| A230028O05Rik | 7.252867673 | -0.008809902 | 0.133754343 | -0.065866289 | 0.94748428 | NA |
| A230108P19Rik | 0.346294032 | 0.050460286 | 0.044390535 | 1.136735257 | 0.255648976 | NA |
| A330093E20Rik | 6.749109249 | 0.016053548 | 0.142900097 | 0.11234106 | 0.910552987 | NA |
| A3galt2 | 7.237645065 | 0.021751122 | 0.145357671 | 0.149638626 | 0.881049732 | NA |
| A430071A18Rik | 0.483152594 | -0.021244376 | 0.05063691 | -0.419543292 | 0.674819123 | NA |
| A430088P11Rik | 0.174376471 | 0.024538273 | 0.031173126 | 0.787161136 | 0.431187544 | NA |
| A430089I19Rik | 0 | NA | NA | NA | NA | NA |
| A430090L17Rik | 0 | NA | NA | NA | NA | NA |
| A430092G05Rik | 0.379842343 | -0.027920334 | 0.043022777 | -0.648966337 | 0.516360135 | NA |
| A430093F15Rik | 2.465706733 | 0.010129674 | 0.111331717 | 0.090986418 | 0.927503383 | NA |
| A530006G24Rik | 0 | NA | NA | NA | NA | NA |
| A530013C23Rik | 2.442407716 | 0.087283208 | 0.097467933 | 0.895506915 | 0.370516171 | NA |
| A530016L24Rik | 0.520737658 | -0.0194622 | 0.051269202 | -0.379608015 | 0.70423641 | NA |
| A530032D15Rik | 0.506479192 | -0.018468102 | 0.045386828 | -0.406904433 | 0.68407818 | NA |
| A530046M15Rik | 0 | NA | NA | NA | NA | NA |
| A530050N04Rik | 5.660482064 | 0.045064192 | 0.12986375 | 0.347011328 | 0.728582809 | NA |
| A530053G22Rik | 1.112233245 | 0.118486707 | 0.073791026 | 1.605706178 | 0.108338483 | NA |
| A530064D06Rik | 1.664664629 | 0.131011716 | 0.079362674 | 1.650797653 | 0.098779899 | NA |
| A530065N20Rik | 0.198019769 | 0.025308392 | 0.031628095 | 0.80018706 | 0.423602426 | NA |
| A530088E08Rik | 1.489254681 | 0.099355415 | 0.093643733 | 1.060993744 | 0.288692744 | NA |
| A530099J19Rik | 0.204013804 | -0.003115899 | 0.031740777 | -0.098167077 | 0.921799625 | NA |
| A630010A05Rik | 0 | NA | NA | NA | NA | NA |
| A630012P03Rik | 0.096071043 | -0.020098795 | 0.027825948 | -0.722304036 | 0.470107572 | NA |
| A630020A06 | 1.158197331 | -0.063360987 | 0.07979942 | -0.794003111 | 0.42719362 | NA |
| A630023A22Rik | 0 | NA | NA | NA | NA | NA |
| A630073D07Rik | 0 | NA | NA | NA | NA | NA |
| A630075F10Rik | 0.079983654 | 0.016740032 | 0.027674178 | 0.604897175 | 0.54524732 | NA |
| A630076J17Rik | 0.501077839 | -0.018122177 | 0.045283609 | -0.400192852 | 0.689014479 | NA |
| A630077J23Rik | 3.174756188 | -0.231080716 | 0.12488884 | -1.850291157 | 0.064271597 | NA |
| A630095E13Rik | 0.192142086 | -0.026384592 | 0.031527717 | -0.836869715 | 0.402665805 | NA |
| A630095N17Rik | 1.318866669 | -0.014521743 | 0.072030855 | -0.201604475 | 0.840225947 | NA |
| A730008H23Rik | 0 | NA | NA | NA | NA | NA |
| A730018C14Rik | 8.063125415 | -0.195203337 | 0.14620121 | -1.335169092 | 0.18182101 | NA |
| A730082K24Rik | 0 | NA | NA | NA | NA | NA |
| A730085A09Rik | 3.397070152 | 0.259675453 | 0.108332692 | 2.397018369 | 0.016529095 | NA |
| A730090N16Rik | 1.353239558 | 0.100290753 | 0.081106574 | 1.236530504 | 0.216261434 | NA |
| A730098P11Rik | 7.212616839 | -0.145312751 | 0.154259555 | -0.942001623 | 0.346191809 | NA |
| A930001A20Rik | 4.837739006 | 0.012436999 | 0.141469758 | 0.087912777 | 0.929946002 | NA |
| A930001C03Rik | 2.223242678 | 0.121634856 | 0.098371412 | 1.236485818 | 0.216278034 | NA |
| A930006I01Rik | 0.079983654 | 0.016740032 | 0.027674178 | 0.604897175 | 0.54524732 | NA |
| A930018P22Rik | 1.843354373 | 0.069817574 | 0.092858753 | 0.751868533 | 0.452130123 | NA |
| A930019D19Rik | 0.536358603 | 0.018999954 | 0.051864556 | 0.366337928 | 0.714112931 | NA |
| A930041C12Rik | 1.176855315 | 0.004953334 | 0.066942803 | 0.073993531 | 0.941015533 | NA |
| AA467197 | 3.381262845 | 0.031271476 | 0.118097646 | 0.26479339 | 0.791168629 | NA |
| AA536875 | 0.482409049 | 0.03844785 | 0.05183958 | 0.741669784 | 0.458287431 | NA |
| AA545190 | 0 | NA | NA | NA | NA | NA |
| AA619741 | 0.159967308 | 0.0240345 | 0.030870969 | 0.778547006 | 0.436246606 | NA |
| AA792892 | 0 | NA | NA | NA | NA | NA |
| AB099516 | 0.195033442 | -0.002424765 | 0.031596358 | -0.076741901 | 0.93882887 | NA |
| AF357355 | 3.574150081 | 0.111350073 | 0.106999133 | 1.040663321 | 0.298031832 | NA |
| AF357399 | 6.262553216 | 0.142496222 | 0.145564736 | 0.97891994 | 0.327619539 | NA |
| AF357425 | 4.230150771 | 0.244130117 | 0.130024704 | 1.877567183 | 0.060440399 | NA |
| AF357426 | 5.609738731 | 0.096073778 | 0.137180076 | 0.7003479 | 0.483710064 | NA |
| AF366264 | 0 | NA | NA | NA | NA | NA |
| AI314831 | 0.099009884 | 0.018205726 | 0.027896865 | 0.652608315 | 0.514008823 | NA |
| AI427809 | 5.07471216 | -0.044295103 | 0.136950375 | -0.32343908 | 0.746362737 | NA |
| AI463170 | 0.48794879 | 0.064131883 | 0.049799661 | 1.287797577 | 0.197816436 | NA |
| AI506816 | 2.116057961 | -0.06180638 | 0.092132999 | -0.670838684 | 0.502323301 | NA |
| AI507597 | 2.297210766 | -0.060771891 | 0.104072412 | -0.583938528 | 0.559261671 | NA |
| AI646519 | 0 | NA | NA | NA | NA | NA |
| AI662270 | 4.790869034 | -0.039136686 | 0.117567014 | -0.33288832 | 0.739218587 | NA |
| AI747448 | 0.263371537 | 0.009921338 | 0.037961873 | 0.261350055 | 0.793822569 | NA |
| AU015228 | 0.297029653 | 0.02866449 | 0.033517577 | 0.855207712 | 0.392436167 | NA |
| AU015836 | 0.196680618 | -0.003857632 | 0.031597546 | -0.122086447 | 0.902830556 | NA |
| AU016765 | 0 | NA | NA | NA | NA | NA |
| AU018091 | 0 | NA | NA | NA | NA | NA |
| AU018829 | 0 | NA | NA | NA | NA | NA |
| AU019990 | 0.200315975 | 0.025370485 | 0.031664208 | 0.801235428 | 0.422995365 | NA |
| AU021063 | 1.577856553 | 0.091196744 | 0.083203315 | 1.096071045 | 0.273047685 | NA |
| AU022751 | 0.722087843 | -0.018583788 | 0.054979351 | -0.33801396 | 0.735352668 | NA |
| AU022793 | 5.942553399 | 0.331664192 | 0.139439668 | 2.378549788 | 0.01738089 | NA |
| AU023871 | 0.824694589 | 0.002268418 | 0.057226179 | 0.039639519 | 0.96838052 | NA |
| AV051173 | 4.482025111 | -0.090561204 | 0.134030697 | -0.675675093 | 0.499246953 | NA |
| AV320801 | 0 | NA | NA | NA | NA | NA |
| AY074887 | 1.746146875 | 0.066616651 | 0.093763116 | 0.710478217 | 0.477407634 | NA |
| AY358078 | 0.600695175 | -0.01924069 | 0.047747217 | -0.40296987 | 0.68697039 | NA |
| AY512915 | 0.381360167 | 0.020495541 | 0.042644778 | 0.480610819 | 0.630793125 | NA |
| AY512931 | 3.937876758 | 0.045757036 | 0.124981246 | 0.366111214 | 0.71428209 | NA |
| AY761184 | 0 | NA | NA | NA | NA | NA |
| AY761185 | 0 | NA | NA | NA | NA | NA |
| Aadac | 0 | NA | NA | NA | NA | NA |
| Aadacl2 | 0 | NA | NA | NA | NA | NA |
| Aadacl3 | 0.08731684 | 0.017365056 | 0.027733412 | 0.626142072 | 0.531221759 | NA |
| Aadat | 6.836992621 | -0.119528564 | 0.150494808 | -0.794237129 | 0.427057397 | NA |
| Abca12 | 6.783017849 | 0.064861459 | 0.135527712 | 0.478584473 | 0.632234266 | NA |
| Abca14 | 1.030459691 | -0.007957983 | 0.064045404 | -0.124255332 | 0.901113113 | NA |
| Abca15 | 1.021931072 | 0.119306829 | 0.075489238 | 1.580448199 | 0.114004261 | NA |
| Abca16 | 0.08731684 | 0.017365056 | 0.027733412 | 0.626142072 | 0.531221759 | NA |
| Abcb11 | 0.781051818 | -0.037914647 | 0.067597685 | -0.56088677 | 0.57487473 | NA |
| Abcb5 | 0.192142086 | -0.026384592 | 0.031527717 | -0.836869715 | 0.402665805 | NA |
| Abcg5 | 1.062502086 | -0.033374209 | 0.06048551 | -0.551771973 | 0.581104592 | NA |
| Abcg8 | 0.215468684 | -0.027008432 | 0.031909031 | -0.846419662 | 0.397318674 | NA |
| Abhd12b | 6.56843143 | 0.215727634 | 0.116561549 | 1.850761555 | 0.064203864 | NA |
| Abhd16b | 0.098353386 | -0.020212831 | 0.02786466 | -0.725393061 | 0.468210929 | NA |
| Abo | 0 | NA | NA | NA | NA | NA |
| Abp1 | 1.087822263 | 0.131401943 | 0.077071413 | 1.704937519 | 0.088206078 | NA |
| Acap1 | 3.34557331 | -0.091379074 | 0.111964918 | -0.816140234 | 0.414419935 | NA |
| Acbd7 | 7.53369653 | 0.006552191 | 0.146003894 | 0.044876826 | 0.964205488 | NA |
| Ace3 | 0 | NA | NA | NA | NA | NA |
| Acer1 | 0.609905974 | -0.029829418 | 0.054035901 | -0.552029618 | 0.580928061 | NA |
| Acmsd | 0.87601496 | 0.003546103 | 0.064153208 | 0.055275543 | 0.955918946 | NA |
| Acnat1 | 4.202123611 | 0.108514166 | 0.124992797 | 0.868163353 | 0.385304913 | NA |
| Acnat2 | 0 | NA | NA | NA | NA | NA |
| Acot10 | 0.08731684 | 0.017365056 | 0.027733412 | 0.626142072 | 0.531221759 | NA |
| Acox2 | 5.121583048 | 0.190673171 | 0.12464736 | 1.529700845 | 0.126090794 | NA |
| Acoxl | 0.876552974 | 0.032384074 | 0.064824785 | 0.499563152 | 0.617382709 | NA |
| Acpt | 0 | NA | NA | NA | NA | NA |
| Acrv1 | 0 | NA | NA | NA | NA | NA |
| Acsbg2 | 1.704303594 | 0.033296316 | 0.090317104 | 0.368660143 | 0.71238106 | NA |
| Acsm1 | 0 | NA | NA | NA | NA | NA |
| Acsm2 | 0.341110531 | -0.038008058 | 0.037770309 | -1.006294602 | 0.314273876 | NA |
| Acsm3 | 4.98918845 | 0.251118734 | 0.130025547 | 1.931303044 | 0.053445587 | NA |
| Acsm4 | 1.474061535 | 0.040944629 | 0.079840068 | 0.512833097 | 0.608068076 | NA |
| Actbl2 | 0.571288896 | 0.022514058 | 0.053122995 | 0.423810033 | 0.671704361 | NA |
| Actc1 | 0 | NA | NA | NA | NA | NA |
| Actg2 | 3.660722762 | -0.190788708 | 0.123651213 | -1.542958637 | 0.122840812 | NA |
| Actl10 | 0.197363271 | -0.000561558 | 0.031650166 | -0.017742642 | 0.985844163 | NA |
| Actl11 | 0 | NA | NA | NA | NA | NA |
| Actl7a | 0 | NA | NA | NA | NA | NA |
| Actl7b | 0.910275419 | 0.002696959 | 0.070454943 | 0.038279207 | 0.969465069 | NA |
| Actl9 | 5.064570303 | -0.018445694 | 0.121375821 | -0.151971732 | 0.879209231 | NA |
| Actr8 | 1805.598313 | 0.1420983 | 0.080864198 | 1.757246147 | NA | NA |
| Actrt1 | 0 | NA | NA | NA | NA | NA |
| Actrt2 | 0 | NA | NA | NA | NA | NA |
| Ada | 20.83374261 | 0.2270041 | 0.157922884 | 1.437436386 | NA | NA |
| Adad1 | 0.215050351 | -0.027006021 | 0.031907566 | -0.846382992 | 0.397339123 | NA |
| Adad2 | 3.510446752 | -0.083303528 | 0.112342153 | -0.741516216 | 0.458380504 | NA |
| Adam1b | 0.193239998 | 0.025145508 | 0.031532422 | 0.797449311 | 0.42519013 | NA |
| Adam2 | 0.746849799 | -0.00504642 | 0.061759293 | -0.081711098 | 0.934876453 | NA |
| Adam20 | 0 | NA | NA | NA | NA | NA |
| Adam24 | 0 | NA | NA | NA | NA | NA |
| Adam25 | 0 | NA | NA | NA | NA | NA |
| Adam26a | 0 | NA | NA | NA | NA | NA |
| Adam26b | 0 | NA | NA | NA | NA | NA |
| Adam28 | 1.418099254 | 0.061945414 | 0.090504709 | 0.684444096 | 0.493694779 | NA |
| Adam29 | 0 | NA | NA | NA | NA | NA |
| Adam3 | 3.516109624 | 0.16018811 | 0.115274822 | 1.389619243 | 0.164644529 | NA |
| Adam30 | 0 | NA | NA | NA | NA | NA |
| Adam34 | 0 | NA | NA | NA | NA | NA |
| Adam39 | 0 | NA | NA | NA | NA | NA |
| Adam4 | 7.268415126 | -0.239545171 | 0.157467127 | -1.521239231 | 0.128199814 | NA |
| Adam5 | 7.351368263 | 0.003698338 | 0.156765848 | 0.023591476 | 0.981178471 | NA |
| Adam6a | 0 | NA | NA | NA | NA | NA |
| Adam6b | 0 | NA | NA | NA | NA | NA |
| Adam7 | 0 | NA | NA | NA | NA | NA |
| Adamdec1 | 0.098353386 | -0.020212831 | 0.02786466 | -0.725393061 | 0.468210929 | NA |
| Adamts12 | 8.020384222 | -0.177224349 | 0.150919601 | -1.174296437 | 0.24027631 | NA |
| Adh4 | 0 | NA | NA | NA | NA | NA |
| Adh6-ps1 | 0 | NA | NA | NA | NA | NA |
| Adh6a | 0 | NA | NA | NA | NA | NA |
| Adh7 | 3.457528501 | -0.027422988 | 0.106834914 | -0.256685635 | 0.797421461 | NA |
| Adig | 0.639411646 | 0.032045962 | 0.060168178 | 0.532606496 | 0.594306005 | NA |
| Adipoq | 1.743847353 | 0.074828092 | 0.098163715 | 0.762278525 | 0.445893788 | NA |
| Adipor2 | 3159.946284 | 0.27510772 | 0.160253584 | 1.716702445 | NA | NA |
| Adm2 | 0 | NA | NA | NA | NA | NA |
| Adprhl1 | 4.519450566 | 0.05331919 | 0.137129957 | 0.388822334 | 0.697407578 | NA |
| Afm | 0 | NA | NA | NA | NA | NA |
| Afp | 4.10379429 | -0.083586349 | 0.117267397 | -0.712784212 | 0.475979297 | NA |
| Agr2 | 1.226054931 | -0.039562708 | 0.075085596 | -0.526901426 | 0.598262043 | NA |
| Agr3 | 0.977524795 | -0.093181919 | 0.064611519 | -1.442187401 | 0.149249514 | NA |
| Agrp | 4.094995815 | 0.004568953 | 0.130892431 | 0.03490617 | 0.972154561 | NA |
| Agt | 964.1615949 | 0.346699071 | 0.142928413 | 2.425683343 | NA | NA |
| Agtr1b | 4.917559474 | -0.162501733 | 0.141827691 | -1.145768723 | 0.25189086 | NA |
| Agxt | 0 | NA | NA | NA | NA | NA |
| Agxt2 | 0.299163155 | 0.032734233 | 0.035823547 | 0.913763036 | 0.360841373 | NA |
| Ahsg | 0.20671448 | -0.003314027 | 0.03178208 | -0.10427343 | 0.916952363 | NA |
| Aicda | 0.19442443 | -0.026451352 | 0.031568725 | -0.837897386 | 0.402088337 | NA |
| Aipl1 | 0.958438028 | -0.024468712 | 0.063760441 | -0.383760081 | 0.701156278 | NA |
| Aire | 6.525529669 | 0.251428125 | 0.119637062 | 2.101590597 | 0.035589154 | NA |
| Akap14 | 6.612560473 | 0.229692455 | 0.150132845 | 1.529928086 | 0.126034531 | NA |
| Akap4 | 0 | NA | NA | NA | NA | NA |
| Aknad1 | 0.915478062 | -0.025627691 | 0.074998556 | -0.341709124 | 0.732569805 | NA |
| Akp3 | 0.295060159 | -0.029687054 | 0.033501022 | -0.886153701 | 0.375534709 | NA |
| Akr1b3 | 0 | NA | NA | NA | NA | NA |
| Akr1b7 | 1.01527448 | 0.083273158 | 0.071468685 | 1.165169853 | 0.243950246 | NA |
| Akr1c12 | 1.073983606 | 0.05470851 | 0.060038048 | 0.911230649 | 0.362173858 | NA |
| Akr1c19 | 0.305079873 | -0.015054679 | 0.039642215 | -0.379763828 | 0.704120735 | NA |
| Akr1c20 | 0 | NA | NA | NA | NA | NA |
| Akr1c21 | 0.327267802 | 0.035667551 | 0.037337881 | 0.955264473 | 0.339443969 | NA |
| Akr1c6 | 0 | NA | NA | NA | NA | NA |
| Akr1cl | 0.206069989 | -0.026776453 | 0.031767726 | -0.842882251 | 0.399294301 | NA |
| Akr1d1 | 0.08731684 | 0.017365056 | 0.027733412 | 0.626142072 | 0.531221759 | NA |
| Alas2 | 24.17344701 | -0.175998604 | 0.091829495 | -1.916580334 | NA | NA |
| Alb | 0.702296617 | 0.005938661 | 0.054546958 | 0.10887246 | 0.913303651 | NA |
| Aldh8a1 | 1.984465515 | 0.031198082 | 0.097224269 | 0.320887796 | 0.748295423 | NA |
| Aldoart1 | 0.239950962 | 0.027417169 | 0.032831728 | 0.835081494 | 0.403671825 | NA |
| Aldoart2 | 0.977385348 | 0.0095598 | 0.058332584 | 0.163884395 | 0.869822152 | NA |
| Alms1-ps2 | 3.595120074 | 0.020417395 | 0.128248318 | 0.159202046 | 0.873509692 | NA |
| Alox12e | 4.988882264 | 0.033035474 | 0.128125116 | 0.257837613 | 0.796532235 | NA |
| Alox15 | 2.286331333 | 0.01030815 | 0.102305164 | 0.100758844 | 0.919741898 | NA |
| Alpi | 0.094392817 | 0.017895942 | 0.027818137 | 0.643319236 | 0.520016977 | NA |
| Alppl2 | 0 | NA | NA | NA | NA | NA |
| Als2cr11 | 0.488462861 | -0.011555605 | 0.044402921 | -0.26024426 | 0.794675365 | NA |
| Als2cr12 | 0.810586661 | 0.022431908 | 0.063397743 | 0.353828177 | 0.723467655 | NA |
| Alx1 | 4.136739917 | 0.074635799 | 0.129062468 | 0.578292049 | 0.563066962 | NA |
| Ambn | 0.256038351 | 0.007085343 | 0.03307033 | 0.214250763 | 0.830351512 | NA |
| Ambp | 0 | NA | NA | NA | NA | NA |
| Amd1 | 0 | NA | NA | NA | NA | NA |
| Amd2 | 0 | NA | NA | NA | NA | NA |
| Amelx | 0.376356809 | 0.021703132 | 0.042541541 | 0.510163288 | 0.60993707 | NA |
| Ampd1 | 0.116696964 | -0.020477583 | 0.027954224 | -0.732539847 | 0.463839131 | NA |
| Amtn | 0 | NA | NA | NA | NA | NA |
| Amy2a2 | 0 | NA | NA | NA | NA | NA |
| Amy2a3 | 0 | NA | NA | NA | NA | NA |
| Amy2a4 | 0 | NA | NA | NA | NA | NA |
| Amy2a5 | 0 | NA | NA | NA | NA | NA |
| Amy2b | 0.079983654 | 0.016740032 | 0.027674178 | 0.604897175 | 0.54524732 | NA |
| Ang2 | 0 | NA | NA | NA | NA | NA |
| Ang3 | 0 | NA | NA | NA | NA | NA |
| Ang4 | 0.098353386 | -0.020212831 | 0.02786466 | -0.725393061 | 0.468210929 | NA |
| Ang5 | 0 | NA | NA | NA | NA | NA |
| Ang6 | 0 | NA | NA | NA | NA | NA |
| Angpt4 | 1.123832365 | -0.079502269 | 0.075748052 | -1.049561891 | 0.293919586 | NA |
| Angptl3 | 6.915324161 | -0.035852231 | 0.154708243 | -0.231740923 | 0.816739244 | NA |
| Ankar | 0.599827183 | 0.017101474 | 0.046253359 | 0.369734744 | 0.711580142 | NA |
| Ankk1 | 0.472380367 | 0.010114997 | 0.055121332 | 0.183504213 | 0.854402413 | NA |
| Ankrd1 | 1.798583869 | 0.034259767 | 0.089266409 | 0.383792377 | 0.701132339 | NA |

| | | | | | | |
|---|---|---|---|---|---|---|
| Pdhx | 1779.536542 | 0.073704214 | 0.05283234 | 1.395058661 | 0.162998151 | 0.614351408 |
| Actr6 | 501.0616266 | 0.073724827 | 0.07775541 | 0.948163305 | 0.343046325 | 0.763762832 |
| Slc18a2 | 55.4905087 | 0.07372713 | 0.046646256 | 1.580558357 | 0.113979054 | 0.54248353 |
| Ube2v1 | 8981.914963 | 0.073730949 | 0.042799139 | 1.722720367 | 0.084939105 | 0.488499377 |
| Plekha3 | 1598.702769 | 0.073763511 | 0.057892288 | 1.274150898 | 0.202609943 | 0.65606599 |
| Zyg11b | 11260.90386 | 0.073818361 | 0.069705386 | 1.059005127 | 0.289597445 | 0.72491685 |
| Emc4 | 3446.759207 | 0.073819726 | 0.042675916 | 1.729774821 | 0.083670515 | 0.486580734 |
| Akap6 | 10170.44936 | 0.07382445 | 0.068897832 | 1.071506133 | 0.283941915 | 0.721415456 |
| Ntrk1 | 102.6440613 | 0.073824776 | 0.09453896 | 0.780892617 | 0.434865655 | 0.815326688 |
| Magi2 | 5193.750052 | 0.073827206 | 0.054925713 | 1.344128312 | 0.178906898 | 0.632619671 |
| Rtn1 | 41995.43105 | 0.073827287 | 0.054429803 | 1.356376169 | 0.174979512 | 0.628026741 |
| Camk2d | 5114.463441 | 0.073842603 | 0.077160919 | 0.956994854 | 0.338569852 | 0.760991092 |
| Reps2 | 8885.740192 | 0.073883745 | 0.070229859 | 1.052027534 | 0.292786917 | 0.728526371 |
| Npnt | 698.5263599 | 0.073893637 | 0.082092242 | 0.900129347 | 0.36805142 | 0.779243711 |
| Tmem255a | 1274.473093 | 0.073898796 | 0.091768693 | 0.805272408 | 0.4206625 | 0.807944816 |
| Sertm1 | 1336.616456 | 0.073900319 | 0.078067384 | 0.9466222 | 0.343831327 | 0.763762832 |
| Agpat4 | 2055.838811 | 0.073915097 | 0.057895331 | 1.276702217 | 0.201707398 | 0.655168687 |
| Tceanc2 | 811.2170427 | 0.073942204 | 0.062879183 | 1.175940913 | 0.239618488 | 0.690203878 |
| Kcnb1 | 13815.29858 | 0.073949583 | 0.069533761 | 1.063506164 | 0.287552471 | 0.723579189 |
| Sf3b5 | 821.3050851 | 0.074016552 | 0.056913404 | 1.300511782 | 0.193425621 | 0.649744561 |
| Pum1 | 4730.226026 | 0.074035192 | 0.055394609 | 1.336505364 | 0.181384148 | 0.634469716 |
| Ing3 | 605.6728061 | 0.074041592 | 0.064237518 | 1.152622251 | 0.249065466 | 0.697784358 |
| Vti1a | 2991.907455 | 0.074044733 | 0.052722641 | 1.404420033 | 0.160193809 | 0.612291377 |
| Tax1bp1 | 6667.016245 | 0.074053332 | 0.061360047 | 1.206865638 | 0.227483893 | 0.680203999 |
| Bag2 | 339.2077925 | 0.074067963 | 0.076105961 | 0.973221563 | 0.330443196 | 0.756141871 |
| Atp6v1g2 | 20292.21403 | 0.074073297 | 0.048989596 | 1.512020983 | 0.130528525 | 0.569619261 |
| Hspa4l | 5388.261287 | 0.074077704 | 0.065776706 | 1.126199659 | 0.260081018 | 0.704052663 |
| Pvrl2 | 231.174141 | 0.074086723 | 0.091374532 | 0.810802761 | 0.417478948 | 0.806153084 |
| Rrad | 83.36854814 | 0.074099995 | 0.093063731 | 0.796228502 | 0.425899234 | 0.809519334 |
| AI854703 | 1416.499807 | 0.074108968 | 0.059367044 | 1.248318312 | 0.211914511 | 0.666270994 |
| Atf6 | 4921.29566 | 0.074123133 | 0.068862706 | 1.076390077 | 0.281752838 | 0.720163706 |
| C330006A16Rik | 3211.590967 | 0.074126883 | 0.051449743 | 1.440762932 | 0.149651669 | 0.598098567 |
| Adam23 | 7538.113085 | 0.074134709 | 0.056816684 | 1.304805275 | 0.191959162 | 0.647018138 |
| 9330111N05Rik | 9.001082341 | 0.074143146 | 0.063270605 | 1.171841901 | 0.241260537 | NA |
| Dhps | 1139.219003 | 0.074145395 | 0.056849972 | 1.304229235 | 0.192155435 | 0.647321393 |
| Acsl5 | 3928.803513 | 0.07418418 | 0.065232962 | 1.137219249 | 0.255446642 | 0.701952734 |
| Tbc1d13 | 2229.255033 | 0.074206774 | 0.051468527 | 1.441789334 | 0.149361813 | 0.59769413 |
| Hepacam | 4321.383488 | 0.074209086 | 0.071603632 | 1.036387175 | 0.300021551 | 0.733812604 |
| Reep5 | 18085.46623 | 0.074209177 | 0.049876712 | 1.487852224 | 0.136789869 | 0.580996972 |
| L3hypdh | 54.47056413 | 0.074210392 | 0.094322074 | 0.786776515 | 0.431412704 | 0.812181606 |
| Mfsd6 | 5086.404996 | 0.074216063 | 0.053736722 | 1.381105133 | 0.167246638 | 0.617511666 |
| Prr13 | 1389.21344 | 0.074217607 | 0.066981043 | 1.108038985 | 0.26784498 | 0.710243917 |
| Nkiras1 | 3453.731083 | 0.074222816 | 0.04964967 | 1.49493069 | 0.134932532 | 0.578521238 |
| Spag16 | 38.08820618 | 0.074272478 | 0.078037676 | 0.951751531 | 0.341223008 | 0.762819666 |
| Gipc1 | 2620.622055 | 0.074280674 | 0.073464179 | 1.011114191 | 0.311961779 | 0.744587053 |
| Ptges3 | 3277.057921 | 0.074300802 | 0.046368075 | 1.602412895 | 0.109064335 | 0.535010276 |
| Lgals3 | 42.42317168 | 0.07435192 | 0.075790977 | 0.981012821 | 0.326586419 | 0.753373722 |
| Zfp146 | 1425.527078 | 0.074376123 | 0.059273741 | 1.25479042 | 0.209554851 | 0.664226905 |
| Nol10 | 975.9402986 | 0.074404687 | 0.064250846 | 1.158034346 | 0.24685002 | 0.696511265 |
| Rabl3 | 347.8712514 | 0.074442106 | 0.078172199 | 0.952283644 | 0.340953151 | 0.76263292 |
| Set | 6580.399084 | 0.074451701 | 0.053975055 | 1.379372403 | 0.167779964 | 0.617953076 |
| Mgat2 | 947.6314625 | 0.074454834 | 0.065414797 | 1.138195583 | 0.255038823 | 0.701727038 |
| Hspa9 | 10110.92277 | 0.074492542 | 0.053788329 | 1.384920184 | 0.166076874 | 0.61635484 |
| Adck4 | 979.8631154 | 0.074533886 | 0.078114082 | 0.95416708 | 0.339999075 | 0.761914237 |
| Srd5a3 | 737.2633937 | 0.074543613 | 0.065033112 | 1.146240915 | 0.251695483 | 0.699465245 |
| Psmd4 | 3415.509955 | 0.074551827 | 0.047652878 | 1.564476889 | 0.117705616 | 0.548537627 |
| Prcc | 1725.958996 | 0.074588141 | 0.054569222 | 1.366853667 | 0.171671177 | 0.622234025 |
| Fbxo36 | 106.9655308 | 0.074591614 | 0.094414119 | 0.790047239 | 0.429500181 | 0.811442206 |
| Atg5 | 865.2977639 | 0.074596545 | 0.070465718 | 1.058621789 | 0.289772059 | 0.725139224 |
| Klhl14 | 119.7464575 | 0.074606774 | 0.090084409 | 0.828187419 | 0.407564368 | 0.801055189 |
| Mrpl41 | 1357.814261 | 0.074627515 | 0.055966863 | 1.33342324 | 0.18239295 | 0.635245711 |
| Myof | 227.6737549 | 0.074628011 | 0.093769048 | 0.795870407 | 0.426107363 | 0.809519334 |
| Dph2 | 480.4501538 | 0.074640244 | 0.081028679 | 0.921158346 | 0.356967762 | 0.773088648 |
| 9430023L20Rik | 882.1102561 | 0.074666394 | 0.072922431 | 1.023915316 | 0.305875275 | 0.738179891 |
| B3gat2 | 565.0558174 | 0.074678886 | 0.072982207 | 1.023247837 | 0.30619068 | 0.738584943 |
| Dnajc5 | 16585.20147 | 0.074681649 | 0.048320578 | 1.545545424 | 0.12221439 | 0.556819993 |
| Rhbdd2 | 3657.143109 | 0.074683654 | 0.057720731 | 1.293879207 | 0.195707148 | 0.651423791 |
| Slc16a13 | 200.9155014 | 0.074700365 | 0.090226418 | 0.82792121 | 0.407715122 | 0.801055189 |
| Lmbrd1 | 8667.37909 | 0.074710423 | 0.058364158 | 1.280073691 | 0.20051922 | 0.654475062 |
| Pcdhga6 | 945.7823497 | 0.074743425 | 0.075069311 | 0.995658869 | 0.319415921 | 0.749427924 |
| Tm9sf3 | 6955.598121 | 0.074753765 | 0.059696032 | 1.252240104 | 0.210482387 | 0.664955883 |
| Irs4 | 205.7260439 | 0.074760602 | 0.045294275 | 1.650553015 | NA | NA |
| 4930524L23Rik | 17.06832118 | 0.074790309 | 0.076187767 | 0.981657714 | 0.326268504 | NA |
| G530011O06Rik | 168.3457632 | 0.074791715 | 0.062424915 | 1.198106812 | 0.230875437 | 0.683402059 |
| Smarca5 | 3038.980075 | 0.074801592 | 0.077607917 | 0.963839709 | 0.335126295 | 0.75853176 |
| Sik3 | 6774.248664 | 0.074817033 | 0.047591936 | 1.572052739 | 0.115938321 | 0.54588186 |
| Pcif1 | 1932.132689 | 0.074817088 | 0.054121833 | 1.38238273 | 0.166854216 | 0.617356019 |
| Gm16701 | 92.31056591 | 0.07482904 | 0.095058634 | 0.787188251 | 0.431171674 | 0.812181606 |
| BC022687 | 533.1758843 | 0.074848374 | 0.079953556 | 0.936148155 | 0.349196912 | 0.768201476 |
| Rnf130 | 2071.30084 | 0.074859988 | 0.058722537 | 1.274808482 | 0.202377038 | 0.656043037 |
| Gtpbp4 | 1421.970142 | 0.074884287 | 0.061677357 | 1.214129315 | 0.224698347 | 0.676420098 |
| Fbxw2 | 3047.023294 | 0.074902347 | 0.051760598 | 1.447092013 | 0.147871155 | 0.594842535 |
| Josd1 | 4249.719417 | 0.074926327 | 0.054542755 | 1.373717305 | 0.169529462 | 0.619945581 |
| Rbm12 | 832.1186188 | 0.074930791 | 0.076167164 | 0.983767647 | 0.32522977 | 0.75293408 |
| Cyb5b | 8032.945354 | 0.074932075 | 0.040908197 | 1.831712977 | 0.066994194 | 0.447688159 |
| Mapt | 24603.75781 | 0.074948731 | 0.046598231 | 1.608402925 | 0.107746961 | 0.533015565 |
| Rnf2 | 957.4969753 | 0.074975564 | 0.069545662 | 1.078076788 | 0.28099949 | 0.720080692 |
| Sf3b3 | 4155.579579 | 0.074976221 | 0.045251405 | 1.656881608 | 0.097543414 | 0.518456434 |
| Gpr6 | 321.6066478 | 0.074995284 | 0.092462355 | 0.811089913 | 0.417314038 | 0.806153084 |
| Smtnl2 | 49.70303081 | 0.075025268 | 0.090900539 | 0.825355591 | 0.409169739 | 0.801055189 |
| Hspe1 | 1547.537039 | 0.075038325 | 0.055787222 | 1.345080867 | 0.178599121 | 0.632097108 |
| Gm5141 | 122.5258018 | 0.075052929 | 0.094932746 | 0.790590514 | 0.429182984 | 0.811020168 |
| Ubap1 | 2096.181693 | 0.075070534 | 0.049455468 | 1.517942052 | 0.129029008 | 0.567618221 |
| Pam | 9043.512383 | 0.075085981 | 0.052226795 | 1.437690776 | 0.150521811 | 0.598511432 |
| Synpo2l | 9.88536195 | 0.075087552 | 0.063577026 | 1.181048514 | 0.237583451 | NA |
| Csnk1g2 | 5800.137817 | 0.075092028 | 0.060895883 | 1.233121579 | 0.217530408 | 0.670311138 |
| Gramd1b | 4160.001306 | 0.075097311 | 0.069960675 | 1.073421765 | 0.283081923 | 0.720920007 |
| Aggf1 | 2812.029544 | 0.075118615 | 0.056558482 | 1.328158245 | 0.184125835 | 0.637571366 |
| Ebp | 624.9217604 | 0.075118994 | 0.060900346 | 1.233474014 | 0.217398966 | 0.670311138 |
| Cfl2 | 2387.032931 | 0.075123691 | 0.056282392 | 1.334763651 | 0.181953714 | 0.635245711 |
| Spag7 | 1756.931158 | 0.075140069 | 0.057959949 | 1.296413661 | 0.194833008 | 0.65082254 |
| Esyt3 | 128.6670753 | 0.075140955 | 0.068912078 | 1.090388758 | 0.275541932 | 0.717137213 |
| Sncg | 118.5122658 | 0.075141293 | 0.068986634 | 1.089215247 | 0.276058977 | 0.717475105 |
| Kcnn3 | 970.8322394 | 0.075142668 | 0.079930884 | 0.940095536 | 0.347168558 | 0.765974975 |
| Elfn1 | 1417.850479 | 0.075163158 | 0.062731278 | 1.198176732 | 0.230848222 | 0.683402059 |
| Rpf1 | 387.3146451 | 0.075170884 | 0.082349692 | 0.912825313 | 0.361334423 | 0.776449324 |
| Bcl2l2 | 6863.580407 | 0.075172425 | 0.04747029 | 1.583567862 | 0.113292092 | 0.540327972 |
| Ptch1 | 1692.131562 | 0.075179098 | 0.075048699 | 1.001737524 | 0.316470379 | 0.745993052 |
| Mmp12 | 13.41825555 | 0.075191745 | 0.040062212 | 1.876874548 | 0.06053529 | NA |
| Syt11 | 39148.85559 | 0.075213825 | 0.040403776 | 1.861554323 | 0.062665937 | 0.437847714 |
| Dnajc10 | 3684.866483 | 0.075271483 | 0.061512325 | 1.223681318 | 0.22107247 | 0.672757795 |
| Cacul1 | 2839.608649 | 0.075282864 | 0.058467901 | 1.287593086 | 0.197887647 | 0.653535282 |
| Akr1a1 | 11959.34314 | 0.075302082 | 0.046259709 | 1.627811424 | 0.103564878 | 0.523462117 |
| Plin5 | 29.82022386 | 0.075332977 | 0.077018098 | 0.978120448 | 0.328014756 | 0.754778534 |
| Upp2 | 326.4756794 | 0.075342054 | 0.085575511 | 0.880416055 | 0.378633962 | 0.785380516 |
| Tmem64 | 1994.492545 | 0.075346584 | 0.06707062 | 1.123391805 | 0.261271129 | 0.704173028 |
| Tpst1 | 1361.203929 | 0.075349164 | 0.056918278 | 1.323813131 | 0.185565114 | 0.638748542 |
| Zfp422 | 870.8523047 | 0.075389563 | 0.062188166 | 1.212281506 | 0.225404641 | 0.677510814 |
| Klhl18 | 1667.391587 | 0.075405691 | 0.047782979 | 1.578086842 | 0.114545661 | 0.543130934 |
| Sms | 4895.616278 | 0.075418552 | 0.066970779 | 1.126141174 | 0.260105768 | 0.704052663 |
| Kcnh7 | 1246.991139 | 0.075427846 | 0.089653901 | 0.841322514 | 0.400167281 | 0.798904608 |
| Laptm4b | 3837.876889 | 0.075441289 | 0.047359542 | 1.592948016 | 0.111171853 | 0.538114742 |
| Chtf8 | 3517.679086 | 0.07545129 | 0.054276245 | 1.390134654 | 0.164487992 | 0.614688762 |
| Copb2 | 4402.403848 | 0.075457235 | 0.048771942 | 1.547144351 | 0.121828441 | 0.556153344 |
| Rbbp4 | 5103.57892 | 0.075466251 | 0.05119521 | 1.474088132 | 0.140457863 | 0.585785508 |
| Atp6v1b2 | 33561.13915 | 0.075473935 | 0.046942508 | 1.607795106 | 0.107880061 | 0.533015565 |
| Lgi1 | 6964.864269 | 0.075482552 | 0.05070532 | 1.488651533 | 0.136579154 | 0.580611294 |
| Ckap4 | 898.1424583 | 0.075487977 | 0.059614815 | 1.266262052 | 0.205419278 | 0.659018859 |
| Stau2 | 4515.831692 | 0.075489422 | 0.062609939 | 1.20570989 | 0.22792937 | 0.680415543 |
| Slc46a1 | 389.7501173 | 0.075541299 | 0.080310828 | 0.940611624 | 0.346903923 | 0.76579125 |
| Fscn1 | 8062.839807 | 0.075584015 | 0.061749257 | 1.224047371 | 0.220934358 | 0.672757795 |
| 9530077C05Rik | 268.9377397 | 0.075594223 | 0.07911198 | 0.955534457 | 0.339307489 | 0.76146707 |
| Tecr | 12862.80772 | 0.075594458 | 0.050891984 | 1.48539027 | 0.137440472 | 0.582012447 |
| Rnf208 | 4277.136376 | 0.075595219 | 0.058667258 | 1.28854187 | 0.197557404 | 0.653498553 |
| Ptx4 | 6.286111793 | 0.075619818 | 0.054628855 | 1.384246789 | 0.166282901 | NA |
| Aldoc | 34640.54346 | 0.07562023 | 0.0665023 | 1.13710698 | 0.255493566 | 0.701952734 |
| Myh2 | 57.6902851 | 0.075630046 | 0.092471843 | 0.817871079 | 0.413430806 | 0.804056218 |
| Pdcd10 | 1216.994334 | 0.075643225 | 0.082121717 | 0.921111103 | 0.356992424 | 0.773088648 |
| Bdh1 | 1944.60528 | 0.075650963 | 0.061540231 | 1.229292801 | 0.218962045 | 0.671730236 |
| Pcdha2 | 109.5392377 | 0.075655345 | 0.094317191 | 0.802137384 | 0.422473492 | 0.808695226 |
| Tstd3 | 791.1208429 | 0.075667681 | 0.06516185 | 1.161226718 | 0.245549712 | 0.69504627 |
| Wbp11 | 3941.03823 | 0.075696923 | 0.047324763 | 1.599520375 | 0.109705025 | 0.535311472 |
| Gabrq | 76.52522193 | 0.075697621 | 0.047021841 | 1.609839586 | 0.107432881 | 0.532254443 |
| Larp4 | 1703.300649 | 0.075701883 | 0.071545603 | 1.05809274 | 0.290013164 | 0.725277354 |
| Ctr9 | 2483.776996 | 0.075702739 | 0.059348868 | 1.275554888 | 0.20211291 | 0.655443359 |
| Dcakd | 1062.5341 | 0.075724673 | 0.075114294 | 1.008126 | 0.313393977 | 0.745844951 |
| Crbn | 3272.868249 | 0.075731774 | 0.063441925 | 1.193718099 | 0.232588258 | 0.684338068 |
| Nefh | 3638.723414 | 0.075734482 | 0.087236564 | 0.868150676 | 0.385311852 | 0.789016184 |
| Entpd6 | 2256.551258 | 0.075749151 | 0.058366562 | 1.2978176 | 0.19435002 | 0.650198399 |
| Tmem60 | 541.4205692 | 0.07577328 | 0.078929442 | 0.960012874 | 0.337048735 | 0.760113439 |
| Dok7 | 491.2037876 | 0.075809611 | 0.08494125 | 0.892494644 | 0.372127862 | 0.781371686 |
| Psmd6 | 2218.497418 | 0.075820631 | 0.057795843 | 1.311869983 | 0.189564007 | 0.644118541 |
| Stac | 40.35693625 | 0.075830473 | 0.078701386 | 0.963521441 | 0.335285909 | 0.758741723 |
| Gnai1 | 5740.643234 | 0.075850801 | 0.054078551 | 1.402604175 | 0.160734906 | 0.612387888 |
| Fam179b | 2708.567775 | 0.075873964 | 0.076469294 | 0.992214784 | 0.321092762 | 0.750352874 |
| Stam2 | 1229.189055 | 0.075875767 | 0.064786145 | 1.171172746 | 0.241529348 | 0.692393571 |
| 2700054A10Rik | 206.1290538 | 0.075914487 | 0.081216012 | 0.934723152 | 0.349930992 | 0.768536074 |
| Psmd13 | 2707.256983 | 0.075920253 | 0.047776379 | 1.589075069 | 0.112043446 | 0.538992394 |
| Csrp1 | 11311.99089 | 0.075923985 | 0.074684414 | 1.016597456 | 0.309344965 | 0.742145027 |
| Ccdc94 | 254.9011773 | 0.075928221 | 0.087467729 | 0.868071251 | 0.385355328 | 0.789016184 |
| Lsm14a | 2668.717621 | 0.075933258 | 0.060687687 | 1.251213583 | 0.210856564 | 0.665414392 |
| Mrps36 | 668.5259063 | 0.075936419 | 0.075756992 | 1.00236845 | 0.316165674 | 0.745993052 |
| Snx25 | 2636.958146 | 0.07595438 | 0.064439974 | 1.178684197 | 0.238523951 | 0.689459926 |
| Fbxo45 | 3096.504317 | 0.075954972 | 0.062130119 | 1.222514501 | 0.221513124 | 0.672757795 |
| Enpp5 | 10572.3316 | 0.07600891 | 0.057280939 | 1.326949433 | 0.184525411 | 0.637780733 |
| Eif1b | 3107.627447 | 0.076039273 | 0.064813078 | 1.17320878 | 0.240712092 | 0.691863635 |
| Pcbp1 | 4410.849631 | 0.076078745 | 0.040377239 | 1.884198766 | 0.059538094 | 0.428234899 |
| Fam163a | 361.6763684 | 0.076084949 | 0.086782789 | 0.876728552 | 0.380634095 | 0.786275719 |
| Hspa4 | 11379.9623 | 0.076124943 | 0.058633711 | 1.298313586 | 0.1941796 | 0.65013848 |
| Mrpl50 | 1335.729573 | 0.076132526 | 0.05880259 | 1.294713813 | 0.195418973 | 0.651003086 |
| Cdsn | 111.2833396 | 0.076133756 | 0.092633958 | 0.821877395 | 0.41114668 | 0.802622856 |
| Olfr1372-ps1 | 9.365635562 | 0.076149352 | 0.062161924 | 1.225016003 | 0.220569191 | NA |
| Nars | 8630.069556 | 0.076149374 | 0.039997024 | 1.903876001 | 0.056926335 | 0.418112798 |
| Gbe1 | 504.3321159 | 0.076165012 | 0.073206416 | 1.040414437 | 0.298147397 | 0.732930969 |
| Fat2 | 22.35304398 | 0.076205833 | 0.052459277 | 1.452666468 | 0.146316381 | 0.593022254 |
| Fads2 | 3884.422893 | 0.076220943 | 0.070284536 | 1.084462481 | 0.278159794 | 0.718319852 |
| Plcb4 | 2518.398446 | 0.076227929 | 0.067543667 | 1.128572556 | 0.259078193 | 0.703828284 |
| Exoc4 | 3026.058104 | 0.076275142 | 0.053025151 | 1.438471018 | 0.150300455 | 0.598511432 |
| Sgk3 | 545.6037489 | 0.076284584 | 0.089492518 | 0.852412982 | 0.393984917 | 0.794754188 |
| Rmdn3 | 1903.948619 | 0.076290316 | 0.049221939 | 1.549925031 | 0.121159511 | 0.555321126 |
| Nipa1 | 2195.766785 | 0.076307815 | 0.061211733 | 1.246620725 | 0.212536599 | 0.666270994 |
| Camk4 | 11348.35482 | 0.076362095 | 0.074244152 | 1.028526744 | 0.303702117 | 0.73722877 |
| Rab18 | 3678.106911 | 0.076369789 | 0.067578516 | 1.130089762 | 0.258438403 | 0.703616072 |
| Casr | 42.59387738 | 0.076370422 | 0.079708893 | 0.958116707 | 0.338003912 | 0.760581219 |
| Atp6v0b | 7148.792276 | 0.076386818 | 0.060708024 | 1.258265592 | 0.20829572 | 0.663292927 |
| Cbwd1 | 441.3245274 | 0.076395555 | 0.077926803 | 0.980350173 | 0.326913296 | 0.753888322 |
| Gm826 | 3.661728426 | 0.076395611 | 0.042095465 | 1.814818094 | 0.069551869 | NA |
| Dlk1 | 647.2709613 | 0.076415898 | 0.054944074 | 1.390794179 | 0.164287849 | 0.614588023 |
| 4930579K19Rik | 15.76751413 | 0.076424148 | 0.074401627 | 1.027183833 | 0.304333909 | NA |
| Tango2 | 2667.494885 | 0.076425288 | 0.0607013 | 1.259038739 | 0.20801634 | 0.66299025 |
| Cilp | 5.825388647 | 0.076427783 | 0.044317341 | 1.724557049 | 0.08460733 | NA |
| Cetn4 | 241.5900601 | 0.0764335 | 0.086860648 | 0.879955446 | 0.378883446 | 0.785380516 |
| Pfdn4 | 472.5566547 | 0.07644192 | 0.077434677 | 0.987179424 | 0.323554687 | 0.752925264 |
| P4hb | 4890.533973 | 0.076455719 | 0.046845252 | 1.632091093 | 0.102660298 | 0.522975271 |
| Rgs7bp | 13059.16152 | 0.076466586 | 0.082187843 | 0.930388045 | 0.352170206 | 0.770193378 |
| Cyp4f17 | 83.40454236 | 0.076500315 | 0.095100064 | 0.804419183 | 0.421154926 | 0.808470525 |
| Pcgf5 | 339.9232043 | 0.07650744 | 0.078940303 | 0.969180982 | 0.332454898 | 0.757290239 |
| Zbtb10 | 326.7619706 | 0.076532556 | 0.088153484 | 0.868173926 | 0.385299126 | 0.789016184 |
| Phactr3 | 5582.731063 | 0.076565821 | 0.050479669 | 1.516765499 | 0.129325901 | 0.568274875 |
| Mea1 | 2177.96862 | 0.076568932 | 0.059995055 | 1.276254044 | 0.20186573 | 0.655168687 |
| Chd5 | 12701.51623 | 0.076570151 | 0.057969569 | 1.320868033 | 0.18654537 | 0.640826269 |
| Slc25a14 | 2019.543258 | 0.076603627 | 0.061598067 | 1.243604387 | 0.213645201 | 0.667262579 |
| Mrpl18 | 1067.727661 | 0.076612351 | 0.053826791 | 1.42331263 | 0.154645547 | 0.604402956 |
| Rgag4 | 1451.827447 | 0.076622213 | 0.074638766 | 1.026573948 | 0.304621126 | 0.737453291 |
| Ube2z | 4514.000159 | 0.076624045 | 0.040148011 | 1.908539008 | 0.056321586 | 0.416282454 |
| Dlx1as | 618.7221959 | 0.076630845 | 0.079911537 | 0.95894595 | 0.337585975 | 0.760312044 |
| Matr3 | 14816.56003 | 0.076670619 | 0.070294305 | 1.090708839 | 0.27540102 | 0.717061772 |
| Txndc15 | 2139.404086 | 0.076679576 | 0.053387819 | 1.436274756 | 0.150924174 | 0.598846773 |
| Usp11 | 7947.496259 | 0.076680437 | 0.058973993 | 1.300241567 | 0.193518188 | 0.649744561 |
| Cox7c | 3208.653348 | 0.076681785 | 0.051572505 | 1.486873386 | 0.137048255 | 0.581414412 |
| M6pr | 4314.786775 | 0.076682512 | 0.047450798 | 1.616042627 | 0.106085097 | 0.52850889 |
| Ak5 | 8514.663961 | 0.076687076 | 0.060401106 | 1.26963033 | 0.204216341 | 0.65778952 |
| Tmem28 | 289.7583628 | 0.07670975 | 0.086235375 | 0.889539238 | 0.373713345 | 0.782348609 |
| 1500017E21Rik | 12.72435476 | 0.076713055 | 0.067532914 | 1.13593581 | 0.25598343 | NA |
| AI504432 | 1077.13128 | 0.076725719 | 0.083248335 | 0.9216487 | 0.356711847 | 0.773088648 |
| Sap30bp | 1655.346027 | 0.076731354 | 0.04968409 | 1.544384812 | 0.122495137 | 0.557095919 |
| Asic2 | 2824.856602 | 0.076731428 | 0.047044991 | 1.631022274 | 0.10288562 | 0.522975271 |
| Fam189a1 | 3618.714526 | 0.07674686 | 0.046453166 | 1.652134106 | 0.098507216 | 0.520050241 |
| Fam155a | 3678.797636 | 0.076766892 | 0.046273736 | 1.658973307 | 0.097121171 | 0.517908272 |
| Trak1 | 8494.42346 | 0.076783015 | 0.065837705 | 1.16624683 | 0.243514664 | 0.692772033 |
| Otud6b | 2715.267217 | 0.076789261 | 0.063616017 | 1.207074331 | 0.227403519 | 0.680203999 |
| Rab21 | 1693.249215 | 0.076802282 | 0.061516071 | 1.248491342 | 0.211851177 | 0.666270994 |
| Srp19 | 917.26169 | 0.076819395 | 0.061235722 | 1.254486637 | 0.20966518 | 0.664226905 |
| Elfn2 | 4658.009703 | 0.076833606 | 0.075958344 | 1.011522915 | 0.311766219 | 0.74424258 |
| Paics | 4063.189962 | 0.076834444 | 0.055126049 | 1.393795602 | 0.163379337 | 0.614351408 |
| Rbbp7 | 5730.861776 | 0.076848445 | 0.060606011 | 1.268000383 | 0.204797814 | 0.658461308 |
| Prtn3 | 39.77644207 | 0.076872671 | 0.089088472 | 0.862880112 | 0.388203386 | 0.791889115 |
| Cnpy3 | 2429.083737 | 0.076873308 | 0.055853367 | 1.376341533 | 0.16871592 | 0.619205089 |
| Sec23a | 4500.418186 | 0.076886227 | 0.06309613 | 1.218556943 | 0.2230124 | 0.674048714 |
| Zfp655 | 1626.959197 | 0.076899651 | 0.07220559 | 1.065009655 | 0.286871561 | 0.722256881 |
| Csnk1g3 | 2512.461737 | 0.076909988 | 0.063245292 | 1.216058713 | 0.223962557 | 0.675094439 |
| B230217O12Rik | 405.4880799 | 0.07693137 | 0.074553502 | 1.031894792 | 0.302121406 | 0.735974432 |
| Ttc39d | 3.452478373 | 0.076952445 | 0.039631305 | 1.941708572 | 0.05217239 | NA |
| Grin2b | 1839.507671 | 0.077003993 | 0.092477554 | 0.832677665 | 0.405026544 | 0.8005027 |
| Unc80 | 12761.92811 | 0.077034424 | 0.071106083 | 1.083373183 | 0.278642814 | 0.71849589 |

| | | | | | | | | | | | | | |
|---|---|---|---|---|---|---|---|---|---|---|---|---|---|
| Ankrd22 | 1.287552525 | 0.078051246 | 0.076533001 | 1.019837784 | 0.3078054 | NA | Cngb1 | 23.85274932 | 0.077038794 | 0.053501313 | 1.439942105 | NA | NA |
| Ankrd33 | 0.563199176 | -0.006483013 | 0.052564167 | -0.123335213 | 0.901841657 | NA | Rhob | 9163.563858 | 0.077047041 | 0.05798743 | 1.328685211 | 0.183951846 | 0.637355821 |
| Ankrd36 | 0.116696964 | -0.020477583 | 0.027954224 | -0.732539847 | 0.463839131 | NA | Gm7271 | 3.470884289 | 0.077060258 | 0.042759967 | 1.802158974 | 0.071520397 | NA |
| Ankrd53 | 0 | NA | NA | NA | NA | NA | Cmtm4 | 8458.842364 | 0.077061413 | 0.064044791 | 1.203242474 | 0.228882503 | 0.681764163 |
| Ankrd60 | 0.096071043 | -0.020098795 | 0.027825948 | -0.722304036 | 0.470107572 | NA | Acbd3 | 1498.978765 | 0.077092753 | 0.055916893 | 1.378702368 | 0.167986539 | 0.617953076 |
| Ankrd61 | 2.998291199 | -0.07531032 | 0.118007854 | -0.638180576 | 0.52335614 | NA | Sspo | 136.8729808 | 0.077097351 | 0.095081612 | 0.810854478 | 0.417449245 | 0.806153084 |
| Ankrd7 | 2.336743905 | -0.076060354 | 0.113097251 | -0.672521688 | 0.501251638 | NA | Slc29a4 | 530.6355972 | 0.07711933 | 0.088933021 | 0.867161934 | 0.38585329 | 0.78964566 |
| Anks4b | 7.515308668 | 0.121738445 | 0.144283321 | 0.843745797 | 0.39881147 | NA | Nfe2l1 | 17287.5355 | 0.077119915 | 0.035146255 | 2.194256967 | 0.028216941 | 0.312646292 |
| Ano9 | 0.54939413 | 0.01643428 | 0.052034848 | 0.315832191 | 0.752129883 | NA | Efhd2 | 10865.1192 | 0.077142133 | 0.05138445 | 1.501273874 | 0.133284739 | 0.575362296 |
| Antxrl | 0 | NA | NA | NA | NA | NA | Ttpa | 109.7038733 | 0.077164881 | 0.092990163 | 0.829817686 | 0.40664187 | 0.800917161 |
| Anxa10 | 2.311589427 | -0.178605716 | 0.108068424 | -1.65270955 | 0.09838999 | NA | Zfpm2 | 642.8478396 | 0.077174157 | 0.066728149 | 1.156545752 | 0.247457997 | 0.696744638 |
| Anxa13 | 1.378542077 | -0.040044204 | 0.077213795 | -0.518614634 | 0.604029502 | NA | Slc45a1 | 2395.776975 | 0.077174891 | 0.051596169 | 1.495748472 | 0.134719214 | 0.577947814 |
| Aox3l1 | 0.48472991 | 0.029813289 | 0.045640658 | 0.653217767 | 0.513615898 | NA | Hmgcll1 | 621.4183225 | 0.077201249 | 0.070626862 | 1.093086213 | 0.274355949 | 0.716185716 |
| Apcs | 0 | NA | NA | NA | NA | NA | Eltd1 | 937.0786802 | 0.077211251 | 0.088215384 | 0.875258342 | 0.381433355 | 0.786663048 |
| Apoa4 | 0.215050351 | -0.027006021 | 0.031907566 | -0.846382992 | 0.397339123 | NA | Nanos1 | 1200.633553 | 0.077256668 | 0.072865725 | 1.060260741 | 0.289025994 | 0.724444298 |
| Apoa5 | 0.107716603 | -0.020477583 | 0.027954224 | -0.732539849 | 0.46383913 | NA | Elavl2 | 2761.412839 | 0.077283602 | 0.060995289 | 1.267042146 | 0.20514022 | 0.658534108 |
| Apob | 1.711512134 | -0.016972912 | 0.088064362 | -0.192733039 | 0.847168047 | NA | Ogdhl | 5555.216912 | 0.07730408 | 0.060166181 | 1.284842734 | 0.198847246 | 0.65377551 |
| Apobec2 | 2.468537556 | 0.048974147 | 0.110988758 | 0.441253219 | 0.659029689 | NA | Lancl2 | 4268.142315 | 0.077320111 | 0.068860433 | 1.122852533 | 0.26150013 | 0.704173028 |
| Apobec4 | 0.493412615 | 0.00310454 | 0.044827045 | 0.069255952 | 0.944785887 | NA | Cxx1b | 2037.078947 | 0.07732998 | 0.06318805 | 1.223807025 | 0.221025034 | 0.672757795 |
| Apoc2 | 0.912251645 | 0.081212828 | 0.06270407 | 1.295176341 | 0.195259404 | NA | Ghitm | 18209.57201 | 0.077355438 | 0.060258554 | 1.28372543 | 0.199238043 | 0.65377551 |
| Apoc4 | 0.405897065 | -0.027138935 | 0.04364914 | -0.621751886 | 0.534105028 | NA | Sgce | 1011.990903 | 0.077358549 | 0.060531864 | 1.277980625 | 0.201256258 | 0.654716629 |
| Apod | 10116.30964 | 0.302350263 | 0.144000193 | 2.099651796 | NA | NA | Xk | 947.6192772 | 0.077374153 | 0.072589887 | 1.065908152 | 0.286465164 | 0.722148665 |
| Apof | 3.089470404 | -0.107454105 | 0.121800419 | -0.882214582 | 0.377660778 | NA | Klhl15 | 234.4710552 | 0.07737946 | 0.092305356 | 0.838298703 | 0.401862964 | 0.799277436 |
| Apol10a | 0.507828167 | -0.069254262 | 0.050465732 | -1.372302744 | 0.169969212 | NA | Prkag1 | 1995.653115 | 0.077399605 | 0.054788216 | 1.412705337 | 0.157742382 | 0.608141173 |
| Apol11a | 0 | NA | NA | NA | NA | NA | Tex38 | 5.739947728 | 0.077407281 | 0.053701485 | 1.441436506 | 0.149461403 | NA |
| Apol11b | 1.366416063 | -0.020011337 | 0.07350953 | -0.272227785 | 0.785446879 | NA | Mob4 | 1850.649568 | 0.077426693 | 0.060372012 | 1.282493165 | 0.199669701 | 0.654127668 |
| Apol6 | 5.585729203 | 0.268061819 | 0.136635685 | 1.961872689 | 0.049777308 | NA | Atp6v1g1 | 2705.23013 | 0.077427213 | 0.048829521 | 1.585663976 | 0.112815555 | 0.540309513 |
| Apol7b | 4.031407462 | 0.29226632 | 0.12256045 | 2.384670752 | 0.017094413 | NA | Mllt3 | 1382.972384 | 0.077446537 | 0.067897494 | 1.140639099 | 0.25402014 | 0.701619624 |
| Apol7c | 0 | NA | NA | NA | NA | NA | Idh3a | 8301.072174 | 0.077477734 | 0.046093069 | 1.680897732 | 0.092782781 | 0.506868248 |
| Apol7d | 0.781403623 | -0.009451213 | 0.067897486 | -0.139198272 | 0.889293474 | NA | Pnck | 2894.289033 | 0.077532172 | 0.079993163 | 0.969234976 | 0.332427964 | 0.757290239 |
| Apol7e | 0 | NA | NA | NA | NA | NA | Cerk | 3846.688005 | 0.077538546 | 0.05304474 | 1.461757492 | 0.143807676 | 0.589967032 |
| Apol9a | 3.435832313 | 0.074800035 | 0.131540197 | 0.568647736 | 0.569595223 | NA | Cdip1 | 11040.3728 | 0.077542528 | 0.035513803 | 2.18344762 | 0.029002862 | 0.317240647 |
| Apol9b | 5.046188569 | -0.118540898 | 0.135451484 | -0.875153922 | 0.381490161 | NA | Nat8l | 22445.44556 | 0.077549519 | 0.044844521 | 1.729297507 | 0.083755863 | 0.486580734 |
| Apom | 1.537810199 | -0.145311625 | 0.097659617 | -1.487939734 | 0.136766788 | NA | Kcnd2 | 4327.19521 | 0.077558694 | 0.071077839 | 1.091179687 | 0.275193824 | 0.716917118 |
| Apon | 0.668229972 | 0.034229313 | 0.060509279 | 0.565687011 | 0.571606567 | NA | Eif2b3 | 578.4021489 | 0.07760264 | 0.064787111 | 1.197809844 | 0.230991054 | 0.683402059 |
| Apoo-ps | 0 | NA | NA | NA | NA | NA | Klf7 | 368.6588933 | 0.077604453 | 0.075232059 | 1.031534352 | 0.302290308 | 0.735974513 |
| Aqp12 | 0 | NA | NA | NA | NA | NA | Pdk3 | 1237.852272 | 0.077618527 | 0.066110927 | 1.174065033 | 0.240368978 | 0.6914669 |
| Aqp2 | 1.788037816 | -0.082425166 | 0.10200462 | -0.808053268 | 0.419059918 | NA | Mycbp | 167.0846487 | 0.077633771 | 0.093123565 | 0.833664075 | 0.404470308 | 0.800290023 |
| Aqp3 | 1.584931511 | -0.02089577 | 0.09401094 | -0.22226956 | 0.824104047 | NA | Cript | 1485.38317 | 0.077658983 | 0.06425285 | 1.208646511 | 0.226798679 | 0.679834424 |
| Aqp5 | 7.883463462 | -0.118305929 | 0.157776108 | -0.74983425 | 0.453354538 | NA | Sparc | 12201.39713 | 0.077664818 | 0.08906601 | 0.871991654 | 0.38321293 | 0.787602461 |
| Aqp7 | 0.967893051 | 0.018368493 | 0.072823062 | 0.252234557 | 0.800859769 | NA | Ndfip1 | 11585.41847 | 0.077674207 | 0.059234636 | 1.311297108 | 0.189757405 | 0.644323531 |
| Aqp8 | 0.312649645 | 0.012040436 | 0.039723828 | 0.303103628 | 0.761810889 | NA | Smndc1 | 970.5266762 | 0.077682372 | 0.064702515 | 1.200608226 | 0.229903208 | 0.683079159 |
| Areg | 5.148644673 | -0.121385393 | 0.115016147 | -1.055376966 | 0.291252953 | NA | Mfhas1 | 2443.776761 | 0.077684266 | 0.060298696 | 1.288324148 | 0.197633151 | 0.653498553 |
| Arg1 | 1.344317085 | 0.139473702 | 0.085254256 | 1.635973488 | 0.101845136 | NA | Ergic1 | 6427.809363 | 0.077716962 | 0.04185326 | 1.856891482 | 0.063326595 | 0.440607157 |
| Arhgap8 | 5.170951952 | 0.051176054 | 0.139942827 | 0.365692586 | 0.714594479 | NA | Gm5148 | 99.90802547 | 0.07771962 | 0.094475433 | 0.822643695 | 0.410710643 | 0.802310759 |
| Arhgef11 | 4184.845402 | 0.070485036 | 0.09767587 | 0.721621794 | NA | NA | Gdap1l1 | 2891.989408 | 0.077731134 | 0.055034868 | 1.412397945 | 0.157832821 | 0.608299168 |
| Arhgef38 | 0.336022005 | 0.013628574 | 0.035131903 | 0.387925872 | 0.698070891 | NA | Filip1 | 729.1666579 | 0.077735394 | 0.076734577 | 1.013042579 | 0.311039823 | 0.743730778 |
| Arid5b | 941.6196678 | 0.286301178 | 0.133880563 | 2.138482028 | NA | NA | Ccnd1 | 2590.730164 | 0.077744648 | 0.079007248 | 0.984019194 | 0.325106076 | 0.75293408 |
| Arl13a | 0.107716603 | -0.020477583 | 0.027954224 | -0.732539849 | 0.46383913 | NA | Syngr3 | 6599.134168 | 0.077764134 | 0.05468375 | 1.422070263 | 0.155005855 | 0.604489543 |
| Arl14 | 0 | NA | NA | NA | NA | NA | Emc3 | 3108.815265 | 0.077771603 | 0.052013994 | 1.495205357 | 0.134860856 | 0.578384645 |
| Arl14epl | 0.211089781 | -0.002448206 | 0.031868916 | -0.076821119 | 0.93876585 | NA | Ccdc25 | 1917.416365 | 0.077793791 | 0.060775933 | 1.280009816 | 0.200541684 | 0.654475062 |
| Arl9 | 3.583664441 | -0.020384064 | 0.129789218 | -0.157055144 | 0.875201389 | NA | Rprd1b | 1381.262039 | 0.077808486 | 0.061295588 | 1.269397827 | 0.204299211 | 0.65778952 |
| Armc12 | 0.664808826 | 0.07862935 | 0.054816974 | 1.434397841 | 0.151458764 | NA | Ing1 | 1326.788792 | 0.077843696 | 0.059067121 | 1.317885385 | 0.187542019 | 0.641776649 |
| Arrdc5 | 0.098353386 | -0.020212831 | 0.02786466 | -0.725393061 | 0.468210929 | NA | Thap11 | 1181.388088 | 0.077859635 | 0.062847187 | 1.23887222 | 0.215392823 | 0.669690956 |
| Art1 | 1.104411432 | 0.039149796 | 0.074125616 | 0.528154746 | 0.597391936 | NA | Mxd3 | 29.13045532 | 0.077862733 | 0.081038059 | 0.960816856 | 0.336644262 | 0.75970798 |
| Art2a-ps | 0 | NA | NA | NA | NA | NA | Ube2j1 | 4426.341397 | 0.077871671 | 0.053465877 | 1.456474219 | 0.145261572 | 0.591055907 |
| Art2b | 0 | NA | NA | NA | NA | NA | Zfp931 | 142.9720142 | 0.077919733 | 0.091951727 | 0.847398259 | 0.396773174 | 0.796947889 |
| Art4 | 3.142597282 | -0.072302224 | 0.091704501 | -0.788426123 | 0.430447496 | NA | Cystm1 | 639.176107 | 0.077935585 | 0.068416437 | 1.139135398 | 0.254646685 | 0.701619624 |
| Art5 | 0.685174529 | -0.047276997 | 0.061327002 | -0.77090018 | 0.440766101 | NA | Frrs1l | 8881.919733 | 0.077936994 | 0.078746401 | 0.989721348 | 0.322310337 | 0.751988429 |
| Asb12 | 0.634307402 | -0.006355127 | 0.053469976 | -0.118854125 | 0.905390927 | NA | Cftr | 60.39311418 | 0.077937186 | 0.095032509 | 0.820110789 | 0.412152952 | 0.802860907 |
| Asb14 | 3.916359886 | 0.143258216 | 0.136141293 | 1.052276007 | 0.292672936 | NA | Golph3 | 3170.186263 | 0.077947303 | 0.068541221 | 1.13723249 | 0.255441108 | 0.701952734 |
| Asb15 | 2.114224913 | 0.009215635 | 0.096415217 | 0.095582784 | 0.923851939 | NA | Rnf157 | 17133.33016 | 0.077961849 | 0.046380734 | 1.680910198 | 0.092780359 | 0.506868248 |
| Asb17 | 0.099009884 | 0.018205726 | 0.027896865 | 0.652608315 | 0.514008823 | NA | Hes1 | 268.4224395 | 0.077964452 | 0.089797751 | 0.868222768 | 0.385272393 | 0.789016184 |
| Asb9 | 0.498228802 | 0.006953757 | 0.044429446 | 0.156512357 | 0.875629179 | NA | Car10 | 6162.508541 | 0.077979196 | 0.057391462 | 1.358724681 | 0.174233848 | 0.627072384 |
| Ascl2 | 7.976169801 | 0.042936466 | 0.148797997 | 0.288555405 | 0.772921626 | NA | Tarsl2 | 1988.905996 | 0.077988129 | 0.069801785 | 1.117279864 | NA | NA |
| Ascl3 | 3.163888827 | 0.043878647 | 0.11099558 | 0.395318869 | 0.692607576 | NA | Cish | 352.4925579 | 0.07799641 | 0.072675193 | 1.073219158 | 0.283172797 | 0.720995619 |
| Ascl4 | 5.229341579 | 0.086470426 | 0.133779624 | 0.6463647 | 0.518043196 | NA | AA387883 | 82.15072352 | 0.078004798 | 0.094954727 | 0.821494625 | 0.411364584 | 0.802795661 |
| Asgr2 | 3.05986852 | 0.106795161 | 0.108520943 | 0.984097239 | 0.325067704 | NA | Sept9 | 3420.557151 | 0.07801198 | 0.05913266 | 1.319270588 | 0.187078667 | 0.641526137 |
| Asic5 | 0.20210942 | -0.001755803 | 0.031727492 | -0.055340098 | 0.955867518 | NA | Men1 | 2639.718043 | 0.078014933 | 0.042197223 | 1.848816772 | 0.064484275 | 0.443022969 |
| Astl | 0.183387883 | -0.001444816 | 0.031390672 | -0.046026918 | 0.963288796 | NA | Fam149a | 1401.334263 | 0.078070878 | 0.058748949 | 1.328889791 | 0.183884332 | 0.637355821 |
| Asz1 | 0.105923158 | 0.018524693 | 0.027977518 | 0.662127842 | 0.507889294 | NA | Ppp2r1a | 24334.62988 | 0.078084687 | 0.043403759 | 1.799030533 | 0.072013851 | 0.460419542 |
| Atf7ip2 | 0.776419309 | -0.033338817 | 0.062710421 | -0.531631219 | 0.594981438 | NA | Pdia3 | 8246.25866 | 0.078106066 | 0.065582896 | 1.190951776 | 0.233672517 | 0.685064073 |
| Atoh1 | 0 | NA | NA | NA | NA | NA | Fam129b | 1678.739922 | 0.078109589 | 0.075097095 | 1.040114649 | 0.298286638 | 0.732994423 |
| Atp12a | 4.689740524 | -0.232568477 | 0.128532634 | -1.809411894 | 0.070387037 | NA | Pelo | 480.9243971 | 0.078113275 | 0.067861944 | 1.151061552 | 0.249706914 | 0.698181362 |
| Atp1a4 | 3.813822435 | 0.092473371 | 0.132323449 | 0.698843416 | 0.484649892 | NA | Rnf41 | 3563.521923 | 0.078126392 | 0.04592833 | 1.701050162 | 0.088933568 | 0.49820432 |
| Atp1b4 | 0 | NA | NA | NA | NA | NA | Spock1 | 13826.20038 | 0.078146239 | 0.040015107 | 1.952918417 | 0.050829263 | 0.403266245 |
| Atp4b | 0 | NA | NA | NA | NA | NA | Canx | 14191.647 | 0.07815315 | 0.055777551 | 1.401157789 | 0.161166894 | 0.612724387 |
| Atp6v0c | 2.314865729 | -0.012024494 | 0.114414417 | -0.10509597 | 0.916299657 | NA | Otud1 | 657.7845662 | 0.078157502 | 0.081808685 | 0.95536925 | 0.339390999 | 0.761506588 |
| Atp6v0c-ps2 | 0 | NA | NA | NA | NA | NA | Naa38 | 474.835151 | 0.078176845 | 0.067377837 | 1.160275371 | 0.245936709 | 0.695335425 |
| Atp6v0d2 | 0.451040743 | 0.02612425 | 0.044851076 | 0.582466505 | 0.560252497 | NA | Xpr1 | 6590.654136 | 0.078180862 | 0.053381876 | 1.464558141 | 0.14304151 | 0.589285584 |
| Atp6v1b1 | 0.652296579 | 0.083224305 | 0.067807775 | 1.227356387 | 0.219688671 | NA | Pes1 | 1902.388603 | 0.078200785 | 0.05386396 | 1.451820198 | 0.146551606 | 0.59347885 |
| Atp6v1e2 | 1.495931226 | 0.071170774 | 0.091244895 | 0.779997327 | 0.435392449 | NA | Rabgap1l | 4336.257362 | 0.078213922 | 0.075044059 | 1.042240033 | 0.297300407 | 0.732415975 |
| Atp6v1g3 | 0.317012939 | 0.010888466 | 0.039852335 | 0.273220278 | 0.784683893 | NA | Pdxp | 6369.921958 | 0.078232793 | 0.052578809 | 1.487914889 | 0.136773341 | 0.580996972 |
| Atp8b3 | 6.755804481 | -0.138494199 | 0.141631099 | -0.977851615 | 0.328147719 | NA | Pld5 | 552.8031989 | 0.078248718 | 0.07950188 | 0.984237335 | 0.324998833 | 0.75293408 |
| Atp8b4 | 3.429688571 | -0.093214292 | 0.117465375 | -0.79354697 | 0.427459215 | NA | Cds2 | 31122.44181 | 0.078270039 | 0.051447613 | 1.521354133 | 0.128170993 | 0.566072313 |
| Aurkc | 4.002984267 | 0.171843579 | 0.1313428 | 1.308359338 | 0.190751458 | NA | Usp32 | 7933.86317 | 0.078285908 | 0.070631004 | 1.108378801 | 0.267698256 | 0.710152314 |
| Avpr1b | 1.493018595 | 0.124975862 | 0.073750772 | 1.694570211 | 0.090156982 | NA | Flrt1 | 1402.872398 | 0.078326271 | 0.068528721 | 1.142969983 | 0.253051054 | 0.700378199 |
| Avpr2 | 0 | NA | NA | NA | NA | NA | Drap1 | 3784.359968 | 0.078352509 | 0.055577879 | 1.409778677 | 0.158605045 | 0.609496529 |
| Awat1 | 0.099009884 | 0.018205726 | 0.027896865 | 0.652608315 | 0.514008823 | NA | Grb2 | 6910.897565 | 0.078358132 | 0.046122013 | 1.698931324 | 0.089332124 | 0.499459783 |
| Aym1 | 0.299858689 | -0.034980473 | 0.036264327 | -0.964597324 | 0.33474654 | NA | Islr2 | 3048.861764 | 0.078389167 | 0.084639614 | 0.926152227 | 0.354366876 | 0.771744113 |
| Azgp1 | 0.695983529 | -0.04154153 | 0.056086079 | -0.740674517 | 0.458890814 | NA | Siah2 | 662.4420464 | 0.07840571 | 0.075996734 | 1.031698425 | 0.302213416 | 0.735974513 |
| B020004C17Rik | 0 | NA | NA | NA | NA | NA | Rassf6 | 34.57196571 | 0.078442245 | 0.089056499 | 0.880814385 | 0.378418293 | 0.785272017 |
| B020004J07Rik | 7.203025663 | -0.048641878 | 0.151001513 | -0.322128417 | 0.747355411 | NA | Specc1 | 4427.686361 | 0.078445435 | 0.060329519 | 1.300282777 | 0.193504069 | 0.649744561 |
| B020014A21Rik | 0 | NA | NA | NA | NA | NA | Ttll5 | 1615.190491 | 0.078445724 | 0.063431145 | 1.23670673 | 0.216195979 | 0.670049713 |
| B020018J22Rik | 0.999349742 | 0.048801405 | 0.07392027 | 0.660189745 | 0.509132072 | NA | Large | 7447.765403 | 0.078445812 | 0.058580431 | 1.339112913 | 0.180533917 | 0.634469716 |
| B130006D01Rik | 0.419064155 | -0.029347008 | 0.044028886 | -0.666539874 | 0.505066086 | NA | Ccdc149 | 2581.669464 | 0.078523838 | 0.054622999 | 1.437559987 | 0.150558941 | 0.598511432 |
| B230119M05Rik | 0.818183462 | -0.07350399 | 0.07059028 | -1.041276359 | 0.297747306 | NA | Mettl22 | 614.4825177 | 0.078530236 | 0.06916611 | 1.135386045 | 0.256213604 | 0.702218644 |
| B230208H11Rik | 4.25069826 | 0.178552036 | 0.136488432 | 1.308184389 | 0.190810776 | NA | N4bp3 | 524.6189784 | 0.078538671 | 0.087081642 | 0.901896991 | 0.36711159 | 0.778490943 |
| B230214G05Rik | 0.099009884 | 0.018205726 | 0.027896865 | 0.652608315 | 0.514008823 | NA | Med4 | 864.6842326 | 0.078551633 | 0.069832032 | 1.124865361 | 0.260646092 | 0.704058139 |
| B230323A14Rik | 1.848492479 | -0.086778593 | 0.095571815 | -0.907993563 | 0.363881622 | NA | Fem1a | 2788.302274 | 0.078566318 | 0.05717932 | 1.374033802 | 0.169431189 | 0.619945581 |
| B430010I23Rik | 1.388852301 | -0.075926941 | 0.076505134 | -0.992442438 | 0.320981746 | NA | Fezf1 | 31.45713106 | 0.078624008 | 0.042494723 | 1.850206408 | NA | NA |
| B4galnt2 | 4.739173048 | 0.147341103 | 0.138913892 | 1.060664996 | 0.288842173 | NA | 1700045I19Rik | 22.50710758 | 0.078626678 | 0.074865577 | 1.05023806 | 0.293608675 | 0.729168132 |
| B930018H19Rik | 7.905492961 | 0.171055224 | 0.150025173 | 1.140176817 | 0.254212644 | NA | Ptrhd1 | 499.59164 | 0.078636746 | 0.06820622 | 1.152926318 | 0.248940629 | 0.697784358 |
| B930092H01Rik | 0.094392817 | 0.017895942 | 0.027818137 | 0.643319236 | 0.520016977 | NA | Aga | 212.3878963 | 0.07864745 | 0.08957776 | 0.877979652 | 0.379954761 | 0.786275719 |
| BB014433 | 0 | NA | NA | NA | NA | NA | Hsph1 | 13809.52882 | 0.078716857 | 0.062577921 | 1.257901437 | 0.208427404 | 0.663405095 |
| BB019430 | 0.21363976 | -0.000753674 | 0.031922632 | -0.02360938 | 0.98116419 | NA | Rusc2 | 6804.061672 | 0.0787189 | 0.05307197 | 1.483248108 | 0.138008504 | 0.582865025 |
| BB123696 | 0.639521788 | 0.057858035 | 0.047157191 | 1.226918608 | 0.219853184 | NA | Nrbp1 | 3402.692718 | 0.078725068 | 0.045278309 | 1.738692751 | 0.082088822 | 0.482582908 |
| BB283400 | 0 | NA | NA | NA | NA | NA | Immt | 5781.222406 | 0.078782522 | 0.053866717 | 1.462545466 | 0.143591795 | 0.589907432 |
| BB287469 | 0 | NA | NA | NA | NA | NA | Fam169a | 2637.963895 | 0.078785933 | 0.063101834 | 1.248552194 | 0.211828907 | 0.666270994 |
| BB557941 | 7.23918054 | 0.27121741 | 0.15688831 | 1.728729245 | 0.083857565 | NA | Fads1 | 10507.90044 | 0.078804379 | 0.062665293 | 1.25754425 | 0.208556626 | 0.663405095 |
| BC016579 | 6.725059 | 0.237332762 | 0.145391178 | 1.632373889 | 0.102600746 | NA | Mt2 | 2668.855975 | 0.078810492 | 0.095038104 | 0.829251523 | 0.406962096 | 0.800975798 |
| BC021614 | 0.096071043 | -0.020098795 | 0.027825948 | -0.722304036 | 0.470107572 | NA | Pknox2 | 3354.179219 | 0.078821502 | 0.04963236 | 1.588107077 | 0.112262128 | 0.538992394 |
| BC021767 | 3.096620928 | 0.227057757 | 0.119923008 | 1.893362757 | 0.058309645 | NA | Actr10 | 5211.666836 | 0.078823453 | 0.05728658 | 1.375949725 | 0.168837199 | 0.619267834 |
| BC024386 | 0 | NA | NA | NA | NA | NA | Tmem59l | 10839.8556 | 0.078862174 | 0.046395642 | 1.699775456 | 0.08917317 | 0.499163571 |
| BC024582 | 2.943682564 | 0.085967715 | 0.117368464 | 0.732460082 | 0.463887799 | NA | Timm8a1 | 497.3095096 | 0.078867898 | 0.075802628 | 1.040437513 | 0.298136681 | 0.732930969 |
| BC030870 | 0 | NA | NA | NA | NA | NA | Rftn1 | 411.651816 | 0.078875644 | 0.086748454 | 0.909245529 | 0.363220536 | 0.776821522 |
| BC039771 | 6.94654834 | -0.064269477 | 0.142735937 | -0.450268362 | 0.652516949 | NA | Ctbp1 | 7754.502305 | 0.078892703 | 0.045486253 | 1.734429579 | 0.082841891 | 0.484423387 |
| BC048502 | 0 | NA | NA | NA | NA | NA | Dixdc1 | 4073.25443 | 0.078923624 | 0.064485883 | 1.223889963 | 0.220993741 | 0.672757795 |
| BC048507 | 0.186326724 | 0.024939385 | 0.031411053 | 0.793968451 | 0.427213798 | NA | Ndst1 | 4721.258535 | 0.078936338 | 0.041203309 | 1.915776676 | 0.055393528 | 0.413057568 |
| BC048562 | 0.119397641 | -0.020477583 | 0.027954224 | -0.732539845 | 0.463839132 | NA | Gad2 | 10636.33808 | 0.078941999 | 0.077390706 | 1.020044951 | 0.307707143 | 0.740098991 |
| BC048602 | 0 | NA | NA | NA | NA | NA | Rab4a | 1557.677251 | 0.078950438 | 0.061419562 | 1.285428226 | 0.198642683 | 0.65377551 |
| BC048609 | 0.609009071 | 0.027901577 | 0.054375796 | 0.513124931 | 0.607863933 | NA | Tdgf1 | 21.73456736 | 0.078963855 | 0.077043451 | 1.024926246 | 0.305397989 | NA |
| BC048644 | 3.888042553 | -0.052750898 | 0.125030752 | -0.42190339 | 0.673095536 | NA | Fam115c | 33.68736915 | 0.078977961 | 0.085449535 | 0.924264371 | 0.355348688 | 0.772370994 |
| BC048671 | 0.119397641 | -0.020477583 | 0.027954224 | -0.732539845 | 0.463839132 | NA | Klhl9 | 3830.805735 | 0.078984727 | 0.063930014 | 1.235487406 | 0.216649161 | 0.670049713 |
| BC048679 | 0.096071043 | -0.020098795 | 0.027825948 | -0.722304036 | 0.470107572 | NA | Slu7 | 2860.774012 | 0.07900343 | 0.057606746 | 1.371426708 | 0.170241976 | 0.620316915 |
| BC049635 | 2.342311452 | 0.138256711 | 0.104499118 | 1.3230419 | 0.185821444 | NA | C030030A07Rik | 311.091108 | 0.079022271 | 0.088124532 | 0.896711384 | 0.369872944 | 0.780397655 |
| BC049730 | 0 | NA | NA | NA | NA | NA | Scoc | 4350.104393 | 0.079028979 | 0.069107116 | 1.143572237 | 0.252801081 | 0.700378199 |
| BC049762 | 3.134227431 | 0.205648659 | 0.113506461 | 1.811779322 | 0.070020301 | NA | Acvrl1 | 858.0658386 | 0.079030928 | 0.081471747 | 0.97004091 | 0.3320261 | 0.757144605 |
| BC051019 | 6.67735047 | 0.156031214 | 0.127020367 | 1.228395233 | 0.219298637 | NA | Tceal7 | 24.2213352 | 0.079038219 | 0.080174039 | 0.98583308 | 0.32421503 | 0.752925264 |
| BC051070 | 0.579950701 | 0.023389586 | 0.058461528 | 0.400085095 | 0.689093842 | NA | Slc25a11 | 4125.392369 | 0.079043237 | 0.051266275 | 1.541817438 | 0.123117963 | 0.557484892 |
| BC051537 | 2.482475864 | 0.067703871 | 0.086602696 | 0.781775561 | 0.434346486 | NA | 1700066M21Rik | 893.9267816 | 0.079043243 | 0.081142404 | 0.974129924 | 0.329992033 | 0.756141871 |
| BC051628 | 3.40388302 | -0.099261099 | 0.127053494 | -0.78125438 | 0.434652896 | NA | Trim26 | 869.5804525 | 0.079093175 | 0.069366581 | 1.140220173 | 0.254194585 | 0.701619624 |
| BC051665 | 0.461343821 | 0.036894045 | 0.051391044 | 0.717908062 | 0.472813976 | NA | Zbtb14 | 1212.036255 | 0.079145165 | 0.060968898 | 1.298123583 | 0.194244872 | 0.65013848 |
| BC053393 | 1.735623088 | 0.023509006 | 0.087789743 | 0.26778762 | 0.7888628 | NA | Pcdha3 | 104.9119319 | 0.079145319 | 0.093158037 | 0.849581229 | 0.395557952 | 0.796239418 |
| BC055111 | 3.288159444 | 0.158698338 | 0.119825494 | 1.324412139 | 0.185366205 | NA | Rilpl1 | 2326.249729 | 0.079152995 | 0.082300761 | 0.961752894 | 0.336173746 | 0.759567558 |
| BC061212 | 0 | NA | NA | NA | NA | NA | Chrnb2 | 3214.461829 | 0.079161749 | 0.048098944 | 1.645810549 | 0.09980277 | 0.521115714 |
| BC061237 | 0.098353386 | -0.020212831 | 0.02786466 | -0.725393061 | 0.468210929 | NA | Ccdc12 | 954.6078983 | 0.079180159 | 0.062921366 | 1.258398615 | 0.208247633 | 0.663292927 |
| BC080695 | 0 | NA | NA | NA | NA | NA | Morc2b | 58.45681588 | 0.079195179 | 0.094680592 | 0.836445751 | 0.402904184 | 0.800009209 |
| BC089597 | 0.288817247 | -0.014517254 | 0.039073354 | -0.37153846 | 0.710236516 | NA | 4930444A02Rik | 1576.567122 | 0.079198052 | 0.064381669 | 1.23013357 | 0.218647091 | 0.671604514 |
| BC090627 | 0 | NA | NA | NA | NA | NA | Scn10a | 31.61648218 | 0.07921164 | 0.086611191 | 0.914565886 | 0.360419574 | 0.776337125 |
| BC094916 | 0 | NA | NA | NA | NA | NA | Gm6682 | 30.22478552 | 0.079229642 | 0.090915042 | 0.871469019 | 0.383498115 | 0.787711201 |
| BC096441 | 0 | NA | NA | NA | NA | NA | Kif13a | 1721.15488 | 0.079232976 | 0.072994628 | 1.085463107 | 0.277716595 | 0.718083484 |
| BC100451 | 5.237384986 | -0.033081727 | 0.145372682 | -0.227564947 | 0.819984473 | NA | Gosr1 | 2347.842871 | 0.079242787 | 0.048233913 | 1.642885302 | 0.100406667 | 0.5211685 |
| BC100530 | 0 | NA | NA | NA | NA | NA | Mvb12b | 6136.781309 | 0.079243393 | 0.045596593 | 1.737923572 | 0.082224282 | 0.482972064 |
| BC107364 | 1.651366412 | -0.041334664 | 0.086431294 | -0.478237244 | 0.632481357 | NA | Commd6 | 665.6802608 | 0.079260466 | 0.070327047 | 1.127026788 | 0.259731155 | 0.703893404 |
| BC117090 | 0 | NA | NA | NA | NA | NA | Nlgn3 | 3094.930638 | 0.07926611 | 0.058325115 | 1.35903908 | 0.174134205 | 0.626964546 |
| BC125332 | 2.627048511 | 0.091651976 | 0.112830304 | 0.812299298 | 0.416619914 | NA | Dbr1 | 440.7555685 | 0.079302861 | 0.069394888 | 1.14277669 | 0.253131319 | 0.700378199 |
| Baat | 5.476431584 | 0.084660008 | 0.138559451 | 0.611001325 | 0.541198703 | NA | A530058N18Rik | 85.40079712 | 0.079306335 | 0.094844016 | 0.836176474 | 0.403055631 | 0.800009209 |
| Bag3 | 372.7242188 | 0.393999397 | 0.12888778 | 3.056918177 | NA | NA | Tubg2 | 2184.491002 | 0.079307408 | 0.055956922 | 1.417293961 | 0.156397 | 0.60719411 |
| Banf2 | 0 | NA | NA | NA | NA | NA | Lsmd1 | 726.4412722 | 0.079376165 | 0.064268467 | 1.235071713 | 0.216803816 | 0.670185967 |
| Barhl1 | 2.010134074 | 0.116289232 | 0.087097976 | 1.335154244 | 0.181825869 | NA | Uchl1 | 19027.9955 | 0.079381434 | 0.045747337 | 1.7352143 | 0.082702855 | 0.484021622 |
| Barx1 | 0 | NA | NA | NA | NA | NA | Blcap | 3738.659455 | 0.079411546 | 0.057961198 | 1.37008117 | 0.170661565 | 0.620885919 |
| Batf | 5.419938029 | -0.01380056 | 0.125152616 | -0.110269848 | 0.912195369 | NA | Kif2c | 7.480792864 | 0.079476813 | 0.055501624 | 1.431972753 | 0.152151619 | NA |
| Bc1 | 0 | NA | NA | NA | NA | NA | Mapkapk2 | 1431.093553 | 0.079479414 | 0.055297568 | 1.437303983 | 0.150631638 | 0.598511432 |
| Bcl2a1a | 6.646647669 | 0.301929753 | 0.147171219 | 2.051554339 | 0.040212994 | NA | Baiap3 | 1923.307258 | 0.079483187 | 0.07493149 | 1.060744773 | 0.288805906 | 0.724444298 |
| Bcl2a1c | 0 | NA | NA | NA | NA | NA | Hint2 | 416.6419243 | 0.079492325 | 0.078064781 | 1.018286657 | 0.308541747 | 0.740806186 |
| Bcl2a1d | 1.446188375 | 0.02523781 | 0.079787286 | 0.316313679 | 0.751764429 | NA | Mtmr9 | 2889.962594 | 0.07949636 | 0.056182815 | 1.414958656 | 0.157080618 | 0.607833353 |
| Bcl2l10 | 1.091020553 | 0.03623669 | 0.078890562 | 0.459328585 | 0.645998224 | NA | C920025E04Rik | 53.02889323 | 0.079523684 | 0.088685213 | 0.896696087 | 0.369881109 | 0.780397655 |
| Bcl2l14 | 0.167300494 | 0.024299763 | 0.031030539 | 0.783091891 | 0.433573153 | NA | 1600012H06Rik | 606.1495293 | 0.079553244 | 0.071762871 | 1.108557156 | 0.267621269 | 0.710152314 |
| Bcmo1 | 2.427182447 | 0.127250925 | 0.090455439 | 1.40678025 | 0.159492561 | NA | Bcap29 | 1485.330117 | 0.079555237 | 0.077066517 | 1.032293143 | 0.301934812 | 0.735974432 |
| Bdkrb1 | 2.029057989 | -0.014098727 | 0.10124509 | -0.139253442 | 0.889249879 | NA | Ucp2 | 1064.290598 | 0.079573595 | 0.094187404 | 0.844843281 | 0.398198346 | 0.797789838 |
| Best2 | 2.085187786 | 0.009796423 | 0.099723293 | 0.098236055 | 0.921744853 | NA | Cdkn3 | 11.08123189 | 0.07957634 | 0.06924484 | 1.149202441 | 0.250472515 | NA |
| Best3 | 7.278845561 | 0.256466239 | 0.1341041 | 1.912441446 | 0.055819596 | NA | Insig2 | 1804.509986 | 0.079632578 | 0.063969769 | 1.244847037 | 0.213187982 | 0.666732456 |
| Bet3l | 5.266578649 | -0.028364686 | 0.134085205 | -0.211542249 | 0.832464168 | NA | Arhgap36 | 219.7885606 | 0.079641204 | 0.05449271 | 1.461501997 | 0.143877728 | 0.589967032 |
| Beta-s | 296.0418766 | -0.188675608 | 0.093899957 | -2.009325826 | NA | NA | Ankfn1 | 103.794273 | 0.079687882 | 0.094758929 | 0.840953809 | 0.400373812 | 0.799132594 |
| Bex6 | 0 | NA | NA | NA | NA | NA | Plp1 | 64986.24614 | 0.079694309 | 0.077719935 | 1.025403705 | 0.305172741 | 0.737593883 |
| Bglap | 1.768971194 | -0.020394106 | 0.084638305 | -0.240955985 | 0.809589231 | NA | Ubfd1 | 2776.912894 | 0.079696148 | 0.059182283 | 1.34662172 | 0.178102095 | 0.631310648 |
| Bglap2 | 1.256570555 | 0.021432703 | 0.069188086 | 0.309774477 | 0.756732462 | NA | Asna1 | 3497.021696 | 0.079754921 | 0.051977935 | 1.534399561 | 0.124931386 | 0.560889589 |
| Bglap3 | 7.422566327 | 0.535619198 | 0.147779456 | 3.624449656 | 0.000289577 | NA | Hras1 | 4073.663734 | 0.079757435 | 0.044329894 | 1.799179489 | 0.071990293 | 0.460419542 |
| Bhlha15 | 4.158081394 | 0.118235459 | 0.129962801 | 0.90976386 | 0.362947058 | NA | Chst11 | 3124.08881 | 0.079772641 | 0.058988476 | 1.352342806 | 0.176265677 | 0.629083076 |
| Bhlhe23 | 2.453154771 | 0.019244711 | 0.097268501 | 0.197851415 | 0.843161318 | NA | Sla | 467.2570232 | 0.079799329 | 0.078632752 | 1.014835766 | 0.310184122 | 0.743030104 |
| Bhmt | 2.953675465 | 0.089629847 | 0.093638558 | 0.95718952 | 0.338471606 | NA | Krt222 | 1759.015995 | 0.079820869 | 0.065625658 | 1.216305819 | 0.223868446 | 0.675094439 |
| Bik | 2.824761723 | -0.008820612 | 0.11210762 | -0.078679862 | 0.937287264 | NA | Sez6l | 10063.62599 | 0.079843045 | 0.062003885 | 1.28771036 | 0.197846806 | 0.653535282 |
| Birc7 | 3.402648078 | 0.145769176 | 0.115999903 | 1.256631883 | 0.208886964 | NA | Ralbp1 | 3531.024216 | 0.079853283 | 0.050645596 | 1.576707333 | 0.114862882 | 0.543130934 |
| Blk | 0.421776838 | -0.034999817 | 0.049427494 | -0.708104206 | 0.478880548 | NA | Scaper | 1705.806083 | 0.079853966 | 0.060662911 | 1.316355659 | 0.188054698 | 0.641854955 |
| Bmp10 | 0 | NA | NA | NA | NA | NA | Sub1 | 7119.340532 | 0.079880646 | 0.063747957 | 1.253069899 | 0.21018027 | 0.664608511 |
| Bmp15 | 7.566656856 | -0.346971736 | 0.152251678 | -2.278935381 | 0.022670907 | NA | Pcdh10 | 7219.670581 | 0.079891866 | 0.054693119 | 1.460729758 | 0.144089619 | 0.589967032 |
| Bmp8a | 5.134289876 | 0.071577255 | 0.141882678 | 0.504481985 | 0.613922718 | NA | Kcnh2 | 1783.783108 | 0.079896925 | 0.075515962 | 1.058013736 | 0.29004918 | 0.725277354 |
| Bmp8b | 7.181411658 | -0.076495296 | 0.15240289 | -0.501928117 | 0.615718086 | NA | Anxa10 | 3.186994858 | 0.079898727 | 0.041622446 | 1.919606762 | 0.054907592 | NA |
| Bnc1 | 0.193239998 | 0.025145508 | 0.031532422 | 0.797449311 | 0.42519013 | NA | Pop7 | 467.7312279 | 0.079903872 | 0.064910448 | 1.230986301 | 0.218327988 | 0.671344538 |
| Boll | 0 | NA | NA | NA | NA | NA | Pgap3 | 446.2472421 | 0.079919211 | 0.071942056 | 1.11088306 | 0.266618688 | 0.709441589 |
| Bpi | 0.482559746 | 0.027836625 | 0.0455364 | 0.611304907 | 0.540997743 | NA | Rph3al | 36.11499381 | 0.079960023 | 0.086299869 | 0.926537014 | 0.354166971 | 0.771424414 |
| Bpifa2 | 0.119397641 | -0.020477583 | 0.027954224 | -0.732539845 | 0.463839132 | NA | Serpinb8 | 474.0693128 | 0.079981169 | 0.09003511 | 0.888333105 | 0.374361597 | 0.78293624 |
| Bpifa3 | 0 | NA | NA | NA | NA | NA | Kdm6b | 1297.977244 | 0.079982203 | 0.090427267 | 0.884492091 | 0.37643063 | 0.783947144 |
| Bpifa5 | 0.360960404 | 0.037959713 | 0.038487497 | 0.986286873 | 0.32399236 | NA | Stk4 | 2216.817596 | 0.080001497 | 0.064561018 | 1.23916102 | 0.215285873 | 0.669593089 |
| Bpifa6 | 0.105923158 | 0.018524693 | 0.027977518 | 0.662127842 | 0.507889294 | NA | Zap70 | 53.7361053 | 0.0800069 | 0.090523984 | 0.883819917 | 0.376793433 | 0.783969863 |
| Bpifb1 | 7.364245161 | -0.20212543 | 0.150245184 | -1.345303886 | 0.178527119 | NA | Mkrn2 | 1275.752305 | 0.080024724 | 0.061250602 | 1.306513256 | 0.191378074 | 0.646682644 |
| Bpifb2 | 0.288817247 | -0.014517254 | 0.039073354 | -0.37153846 | 0.710236516 | NA | Ptpru | 1997.330321 | 0.080047503 | 0.080552767 | 0.993727548 | 0.320355527 | 0.750259458 |
| Bpifb3 | 2.614841149 | -0.051131894 | 0.089706022 | -0.569994002 | 0.568681766 | NA | Hsdl1 | 2227.238256 | 0.080048906 | 0.056132289 | 1.426075921 | 0.153846428 | 0.60340295 |
| Bpifb4 | 4.235457987 | 0.272341675 | 0.126416023 | 2.154328774 | 0.031214398 | NA | Trappc2 | 984.1130138 | 0.08012563 | 0.070069775 | 1.143512028 | 0.252826064 | 0.700378199 |
| Bpifb5 | 1.33467163 | 0.081170829 | 0.078213963 | 1.037804842 | 0.299360922 | NA | Hdac2 | 2977.59833 | 0.080127973 | 0.058419396 | 1.371598799 | 0.170188368 | 0.620316915 |
| Bpifb6 | 1.204479727 | 0.022259603 | 0.079788357 | 0.278983096 | 0.780257797 | NA | 2610034M16Rik | 148.4834789 | 0.080171908 | 0.094593709 | 0.847539535 | 0.39669446 | 0.796916625 |
| Bpifb9a | 2.943751015 | 0.132924612 | 0.075548193 | 1.759467792 | 0.078498086 | NA | Pigs | 2605.706601 | 0.080172192 | 0.058027529 | 1.381623404 | 0.167087364 | 0.617356019 |
| Bpifb9b | 7.716286422 | 0.193258552 | 0.094622319 | 2.042420371 | 0.041109851 | NA | Spryd7 | 1374.202366 | 0.080183174 | 0.066415538 | 1.207295414 | 0.227318396 | 0.680203999 |
| Brat1 | 0.953203133 | -0.012833519 | 0.065301315 | -0.196527732 | 0.844197129 | NA | Crh | 148.5363309 | 0.080188668 | 0.091802706 | 0.873489156 | 0.382396514 | 0.786954445 |
| Bsnd | 0.292777816 | -0.03443077 | 0.035983808 | -0.956840623 | 0.338647705 | NA | Psmc6 | 2624.237006 | 0.08019504 | 0.069221651 | 1.158525378 | 0.246649701 | 0.696511265 |
| Bsph1 | 1.117230264 | 0.051571758 | 0.06768252 | 0.761965688 | 0.446080483 | NA | Scfd1 | 1274.900107 | 0.080199911 | 0.063568164 | 1.261636422 | 0.207079644 | 0.661689157 |
| Bsph2 | 0 | NA | NA | NA | NA | NA | Dlat | 3900.732607 | 0.080205752 | 0.05159419 | 1.554550081 | 0.12005326 | 0.553708672 |
| Bst1 | 6.492475 | 0.055254644 | 0.115978935 | 0.47641965 | 0.633775438 | NA | Katnal2 | 153.5936616 | 0.080216533 | 0.093805683 | 0.855135106 | 0.392476357 | 0.794184423 |
| Bsx | 0.675759004 | 0.004073513 | 0.060574347 | 0.06724815 | 0.946384154 | NA | Gm20187 | 12.6754678 | 0.080235629 | 0.073597018 | 1.090202171 | 0.275624097 | NA |
| Btg3 | 0 | NA | NA | NA | NA | NA | Polr2f | 783.1488773 | 0.080278252 | 0.058813129 | 1.36497161 | 0.172261973 | 0.623232235 |
| Btg4 | 0.297734681 | -0.017963657 | 0.03944387 | -0.455423286 | 0.648804746 | NA | Clybl | 660.442379 | 0.080330986 | 0.075268018 | 1.067265861 | 0.285851799 | 0.722148665 |
| Btn1a1 | 1.634812729 | -0.045320037 | 0.090787953 | -0.499185585 | 0.617648649 | NA | Cinp | 1603.433467 | 0.080362001 | 0.064734222 | 1.241414483 | 0.214452672 | 0.668484873 |
| Btnl1 | 0.119397641 | -0.020477583 | 0.027954224 | -0.732539845 | 0.463839132 | NA | Myadm | 2770.564171 | 0.080382501 | 0.055041901 | 1.460387425 | 0.144183627 | 0.589967032 |
| Btnl10 | 0.314060235 | -0.015766464 | 0.039947878 | -0.394675873 | 0.693082111 | NA | 2510003E04Rik | 7058.514774 | 0.080387111 | 0.04993463 | 1.609846912 | 0.107431281 | 0.532254443 |
| Btnl2 | 0.08731684 | 0.017365056 | 0.027733412 | 0.626142072 | 0.531221759 | NA | Adarb1 | 11079.32106 | 0.080395419 | 0.055584626 | 1.446360727 | 0.148076051 | 0.595162562 |
| Btnl4 | 0 | NA | NA | NA | NA | NA | Pank1 | 665.8333221 | 0.08040755 | 0.064823241 | 1.240412356 | 0.214822913 | 0.668817995 |
| Btnl5 | 0.462360653 | -0.013923057 | 0.043980692 | -0.316572026 | 0.751568365 | NA | 2810407C02Rik | 6146.307487 | 0.080423607 | 0.05867292 | 1.370710831 | 0.170465117 | 0.620595561 |
| Btnl6 | 0.198019769 | 0.025308392 | 0.031628095 | 0.80018706 | 0.423602426 | NA | Tamm41 | 283.0233369 | 0.080464621 | 0.077648859 | 1.036262755 | 0.300079577 | 0.733830965 |
| Btnl9 | 0.762823459 | -0.031698507 | 0.061350021 | -0.516682909 | 0.605377529 | NA | Ptcd1 | 1172.676065 | 0.080499426 | 0.063685658 | 1.264011844 | 0.206225776 | 0.660267934 |
| C030007H22Rik | 2.060017668 | -0.091560523 | 0.094327218 | -0.970669174 | 0.331713047 | NA | Pex2 | 878.8786157 | 0.080533128 | 0.058373473 | 1.37961858 | 0.167704114 | 0.617953076 |
| C030034L19Rik | 3.921997651 | 0.236487143 | 0.118383829 | 1.997630463 | 0.045756738 | NA | Impad1 | 3350.812867 | 0.080535163 | 0.070891681 | 1.136031224 | 0.255943497 | 0.701952734 |
| C130021I20Rik | 5.811273544 | -0.191888635 | 0.147708656 | -1.299102166 | 0.19390887 | NA | St8sia3 | 7756.41902 | 0.080540206 | 0.066767792 | 1.206273321 | 0.227712121 | 0.680415543 |
| C130026I21Rik | 1.924063848 | 0.102343837 | 0.102984531 | 0.99377873 | 0.320330604 | NA | Fam213a | 4319.33581 | 0.080570112 | 0.058852156 | 1.369025654 | 0.170991256 | 0.621020522 |
| C130050O18Rik | 6.790989308 | -0.127379167 | 0.14490064 | -0.879079397 | 0.379358228 | NA | Hsbp1 | 7209.135297 | 0.080573265 | 0.0429427 | 1.876297113 | 0.060614493 | 0.431188114 |
| C130079G13Rik | 0 | NA | NA | NA | NA | NA | Rbm15b | 1568.469615 | 0.080577478 | 0.062303068 | 1.293314776 | 0.195902212 | 0.651482309 |
| C130080G10Rik | 4.632132215 | 0.016732535 | 0.137320041 | 0.121850641 | 0.903017307 | NA | Slc25a47 | 85.76886298 | 0.080579059 | 0.094446263 | 0.853173609 | 0.393563036 | 0.794754188 |
| C130083M11Rik | 1.351018946 | -0.009805109 | 0.088731714 | -0.110502868 | 0.912010575 | NA | Helz | 1077.913573 | 0.080590346 | 0.081283141 | 0.99147676 | 0.321452835 | 0.750831947 |
| C1ql4 | 0.497028206 | -0.021472216 | 0.055798694 | -0.384815747 | 0.700373932 | NA | Rbp1 | 300.3835604 | 0.080606036 | 0.09353727 | 0.861753139 | 0.388823377 | 0.79235741 |
| C1qtnf3 | 2.839363405 | 0.018647936 | 0.10561413 | 0.176566674 | 0.859848773 | NA | Fbxo33 | 2126.161525 | 0.080647174 | 0.075398636 | 1.069610514 | 0.284794661 | 0.721641599 |
| C1qtnf5 | 0 | NA | NA | NA | NA | NA | Efhd1 | 1008.073996 | 0.080698511 | 0.085201728 | 0.947146409 | 0.343564179 | 0.763762832 |
| C1rb | 0.269026497 | 0.030520093 | 0.03461589 | 0.881678683 | 0.377950592 | NA | Psmb1 | 4142.98066 | 0.08070255 | 0.048913954 | 1.649888092 | 0.098965827 | 0.520271327 |
| C230024C17Rik | 1.292691633 | 0.079787348 | 0.077758798 | 1.026087726 | 0.304850235 | NA | Tmem132e | 914.7755479 | 0.080718955 | 0.089709684 | 0.899779729 | 0.368237484 | 0.779243711 |
| C230079O03Rik | 0 | NA | NA | NA | NA | NA | C130021I20Rik | 7.322352946 | 0.080744317 | 0.052588318 | 1.535404069 | 0.124684605 | NA |
| C2cd4a | 2.754249344 | -0.038932739 | 0.112024442 | -0.347537893 | 0.728187256 | NA | Bag4 | 3059.825128 | 0.080748043 | 0.067800422 | 1.190966676 | 0.233666667 | 0.685064073 |
| C2cd4d | 3.12534758 | -0.095787147 | 0.118604197 | -0.807620217 | 0.419309245 | NA | Arl1 | 3332.411309 | 0.080755777 | 0.063293062 | 1.275902525 | 0.201989979 | 0.655373337 |
| C330008G21Rik | 0 | NA | NA | NA | NA | NA | A730046J19Rik | 29.30870069 | 0.080820539 | 0.07475783 | 1.081097986 | 0.279653529 | 0.718934998 |
| C330011F03Rik | 0 | NA | NA | NA | NA | NA | Tmem128 | 731.7569816 | 0.080823153 | 0.066915171 | 1.207844968 | 0.227106901 | 0.680196875 |
| C330013F16Rik | 0.300084847 | -0.016747215 | 0.039501486 | -0.423964169 | 0.671591945 | NA | Mpped2 | 1288.648609 | 0.080849858 | 0.064422519 | 1.254993737 | 0.209481034 | 0.664226905 |
| C330021F23Rik | 2.771310574 | 0.021981985 | 0.112445875 | 0.195489471 | 0.845009779 | NA | Nkain3 | 28.97645787 | 0.080874129 | 0.088889689 | 0.909825755 | 0.36291441 | 0.776734156 |
| C330022C24Rik | 1.325156643 | 0.001301153 | 0.081950265 | 0.015877345 | 0.987332244 | NA | Zbtb4 | 8131.29642 | 0.080887514 | 0.046340066 | 1.745520031 | 0.080894389 | 0.479817024 |
| C330024C12Rik | 0.094392817 | 0.017895942 | 0.027818137 | 0.643319236 | 0.520016977 | NA | Ankrd37 | 143.7170454 | 0.080898578 | 0.092370718 | 0.875803284 | 0.381136984 | 0.786663048 |
| C330024D21Rik | 2.92332916 | 0.087083422 | 0.12020982 | 0.724428521 | 0.468802695 | NA | Zfp180 | 2806.235038 | 0.080905126 | 0.073166561 | 1.105766422 | 0.268827632 | 0.710903001 |
| C330046G13Rik | 0.119397641 | -0.020477583 | 0.027954224 | -0.732539845 | 0.463839132 | NA | Enkur | 103.0592239 | 0.080915341 | 0.094725717 | 0.854206693 | 0.392990476 | 0.794620351 |
| C430002E04Rik | 0 | NA | NA | NA | NA | NA | Atp8b3 | 5.832107403 | 0.080920739 | 0.054003473 | 1.498435837 | 0.134020051 | NA |

| | | | | | | |
|---|---|---|---|---|---|---|
| C430049B03Rik | 6.112321871 | -0.081808695 | 0.149951652 | -0.545567146 | 0.585363513 | NA |
| C4bp | 0.765844708 | 0.066775506 | 0.059510282 | 1.122083506 | 0.261826936 | NA |
| C4bp-ps1 | 4.112967842 | 0.187839373 | 0.131127314 | 1.432496158 | 0.152001877 | NA |
| C530044C16Rik | 6.273837178 | -0.074356482 | 0.133433124 | -0.557256545 | 0.577352161 | NA |
| C6 | 0.652231925 | -0.101419571 | 0.060468106 | -1.677240726 | 0.093495418 | NA |
| C630028M04Rik | 0 | NA | NA | NA | NA | NA |
| C730027H18Rik | 0.096071043 | -0.020098795 | 0.027825948 | -0.722304036 | 0.470107572 | NA |
| C730036E19Rik | 2.532110895 | -0.136872893 | 0.11569686 | -1.183030317 | 0.236797132 | NA |
| C86187 | 1.036144045 | 0.047346346 | 0.07129148 | 0.664123476 | 0.506611286 | NA |
| C86695 | 0.410466648 | -0.020907232 | 0.037449064 | -0.55828449 | 0.576650134 | NA |
| C87198 | 0.119397641 | -0.020477583 | 0.027954224 | -0.732539845 | 0.463839132 | NA |
| C87414 | 0 | NA | NA | NA | NA | NA |
| C87499 | 0.105923158 | 0.018524693 | 0.027977518 | 0.662127842 | 0.507889294 | NA |
| C87977 | 0 | NA | NA | NA | NA | NA |
| C8a | 1.560028588 | 0.060622267 | 0.078764365 | 0.769666169 | 0.441497943 | NA |
| C8b | 3.447200476 | 0.113633108 | 0.103216963 | 1.10091505 | 0.270933631 | NA |
| C9 | 0 | NA | NA | NA | NA | NA |
| C920009B18Rik | 1.47656618 | -0.00521879 | 0.078807297 | -0.06622217 | 0.947200946 | NA |
| Cabp2 | 0.08731684 | 0.017365056 | 0.027733412 | 0.626142072 | 0.531221759 | NA |
| Cabp4 | 1.249089198 | 0.115590482 | 0.08163889 | 1.415875226 | 0.15681204 | NA |
| Cabp5 | 6.461264216 | -0.095840601 | 0.144840321 | -0.661698343 | 0.508164567 | NA |
| Cabs1 | 0.797272051 | 0.015791619 | 0.056720173 | 0.27841275 | 0.780695533 | NA |
| Cacna1f | 7.407490114 | 0.167424236 | 0.137324519 | 1.219186769 | 0.222773313 | NA |
| Calcb | 3.460229114 | -0.11511207 | 0.101596372 | -1.133033276 | 0.257200279 | NA |
| Calcoco2 | 1.659453276 | 0.080678432 | 0.092747754 | 0.869869383 | 0.384371789 | NA |
| Calhm1 | 0.098353386 | -0.020212831 | 0.02786466 | -0.725393061 | 0.468210929 | NA |
| Calm4 | 0 | NA | NA | NA | NA | NA |
| Calm5 | 0 | NA | NA | NA | NA | NA |
| Calml3 | 0.181709657 | 0.024794574 | 0.031325423 | 0.791516008 | 0.428642938 | NA |
| Calml4 | 24.27553477 | 0.414091445 | 0.148607148 | 2.78648404 | NA | NA |
| Camp | 4.92462669 | -0.038932099 | 0.130681482 | -0.297915958 | 0.765767308 | NA |
| Capn13 | 0.577221778 | 0.048337644 | 0.054475647 | 0.887325731 | 0.374903556 | NA |
| Capn8 | 1.130778681 | 0.149419225 | 0.081821028 | 1.826171453 | 0.06782444 | NA |
| Capn9 | 7.137506979 | 0.094543834 | 0.150215766 | 0.629386891 | 0.529095798 | NA |
| Capns2 | 5.140823299 | -0.151525387 | 0.133278981 | -1.136903852 | 0.255578482 | NA |
| Capza3 | 0 | NA | NA | NA | NA | NA |
| Car1 | 4.365296727 | 0.287287806 | 0.138705328 | 2.07120959 | 0.038339214 | NA |
| Car6 | 0.686664619 | 0.040716927 | 0.066216252 | 0.614908356 | 0.538615244 | NA |
| Casc5 | 6.936931086 | -0.305600687 | 0.147197613 | -2.076125285 | 0.037882365 | NA |
| Casp14 | 0.099009884 | 0.018205726 | 0.027896865 | 0.652608315 | 0.514008823 | NA |
| Catsper1 | 0.283178451 | 0.028390488 | 0.033368195 | 0.850824814 | 0.394866675 | NA |
| Catsper4 | 0.711136086 | 0.059233718 | 0.058127284 | 1.019034662 | 0.308186511 | NA |
| Catsperb | 0 | NA | NA | NA | NA | NA |
| Catsperg2 | 2.428626099 | 0.037121406 | 0.102317898 | 0.362804624 | 0.716750837 | NA |
| Cav3 | 3.428642594 | 0.017420055 | 0.115867918 | 0.150344072 | 0.880493164 | NA |
| Cblc | 0.448020035 | 0.075829036 | 0.05417878 | 1.399607666 | 0.161630837 | NA |
| Ccdc144b | 0.098353386 | -0.020212831 | 0.02786466 | -0.725393061 | 0.468210929 | NA |
| Ccdc147 | 6.580716721 | 0.0815182 | 0.152400081 | 0.534896042 | 0.592721745 | NA |
| Ccdc150 | 4.960270646 | 0.201929436 | 0.139347613 | 1.449105807 | 0.147308036 | NA |
| Ccdc152 | 0.332165648 | -0.049934083 | 0.043026356 | -1.160546414 | 0.245826408 | NA |
| Ccdc154 | 0.937815091 | 0.042109344 | 0.066319649 | 0.634945222 | 0.525464142 | NA |
| Ccdc169 | 6.182358922 | -0.138730546 | 0.136232021 | -1.018340219 | 0.308516301 | NA |
| Ccdc172 | 1.914837824 | 0.003450926 | 0.094072603 | 0.036683645 | 0.970737249 | NA |
| Ccdc175 | 1.012519264 | -0.002321965 | 0.06532011 | -0.035547483 | 0.971643185 | NA |
| Ccdc178 | 0.432801378 | -0.061508142 | 0.047634613 | -1.291248906 | 0.196617382 | NA |
| Ccdc27 | 2.025826332 | 0.009484835 | 0.082766829 | 0.114597055 | 0.908764514 | NA |
| Ccdc38 | 3.560846977 | -0.036810944 | 0.127471875 | -0.288776988 | 0.772752043 | NA |
| Ccdc42b | 6.692989405 | 0.266806993 | 0.144128746 | 1.851171264 | 0.064144918 | NA |
| Ccdc54 | 0.19442443 | -0.026451352 | 0.031568725 | -0.837897386 | 0.402088337 | NA |
| Ccdc64b | 0.197363271 | -0.000561558 | 0.031650166 | -0.017742642 | 0.985844163 | NA |
| Ccdc68 | 3.776624343 | -0.052750183 | 0.117658022 | -0.448334772 | 0.653911609 | NA |
| Ccdc7 | 0 | NA | NA | NA | NA | NA |
| Ccdc70 | 0 | NA | NA | NA | NA | NA |
| Ccdc73 | 1.245860835 | -0.055379176 | 0.0814382 | -0.680014731 | 0.496495133 | NA |
| Ccdc83 | 0.874461425 | 0.034875448 | 0.068401208 | 0.509865971 | 0.610145364 | NA |
| Ccer1 | 0.198019769 | 0.025308392 | 0.031628095 | 0.80018706 | 0.423602426 | NA |
| Ccin | 1.246725701 | 0.063284555 | 0.059794406 | 1.058369155 | NA | NA |
| Ccl1 | 0 | NA | NA | NA | NA | NA |
| Ccl11 | 2.177316376 | 0.111290404 | 0.095260493 | 1.168274494 | 0.24269606 | NA |
| Ccl19 | 0.545128971 | 0.052193958 | 0.044892258 | 1.162649418 | 0.244971772 | NA |
| Ccl2 | 7.657881003 | 0.100684772 | 0.142001661 | 0.709039393 | 0.478300032 | NA |
| Ccl20 | 0.331747315 | -0.03734372 | 0.03744515 | -0.99729124 | 0.318623164 | NA |
| Ccl21a | 0 | NA | NA | NA | NA | NA |
| Ccl21b | 0 | NA | NA | NA | NA | NA |
| Ccl21c | 0 | NA | NA | NA | NA | NA |
| Ccl22 | 7.745376831 | 0.233001805 | 0.14669812 | 1.588308044 | 0.112216699 | NA |
| Ccl24 | 5.011820588 | -0.017888037 | 0.129120674 | -0.138537356 | 0.889815748 | NA |
| Ccl26 | 0 | NA | NA | NA | NA | NA |
| Ccl27b | 0 | NA | NA | NA | NA | NA |
| Ccl5 | 3.853689555 | 0.197422328 | 0.11401529 | 1.731542575 | 0.083355038 | NA |
| Ccl7 | 3.246695258 | -0.094285733 | 0.117179106 | -0.804629228 | 0.421033671 | NA |
| Ccl8 | 3.889292092 | 0.087408529 | 0.112510266 | 0.776893814 | 0.437221421 | NA |
| Ccnb1ip1 | 0.323674185 | -0.015335151 | 0.04024759 | -0.381020351 | 0.703188147 | NA |
| Ccnb3 | 0.159967308 | 0.0240345 | 0.030870969 | 0.778547006 | 0.436246606 | NA |
| Ccr10 | 4.373914362 | 0.133965387 | 0.127212233 | 1.053085725 | 0.292301705 | NA |
| Ccr1l1 | 1.053006083 | 0.006999787 | 0.0772405 | 0.090623268 | 0.927791942 | NA |
| Ccr3 | 1.577672267 | -0.049590578 | 0.095854562 | -0.517352299 | 0.604910252 | NA |
| Ccr4 | 0.159967308 | 0.0240345 | 0.030870969 | 0.778547006 | 0.436246606 | NA |
| Ccr6 | 7.210458705 | 0.070182679 | 0.153807148 | 0.456303104 | 0.648172032 | NA |
| Ccr7 | 3.227672923 | -0.017707268 | 0.125379273 | -0.141229627 | 0.887688542 | NA |
| Ccr8 | 0 | NA | NA | NA | NA | NA |
| Ccrl1 | 4.164982232 | 0.168830964 | 0.106907222 | 1.579228797 | 0.114283587 | NA |
| Cct6b | 5.668538221 | 0.041738134 | 0.133284779 | 0.313150038 | 0.754166675 | NA |
| Cct8l1 | 2.531438489 | 0.135460055 | 0.100688233 | 1.345341474 | 0.178514986 | NA |
| Cd160 | 0.079983654 | 0.016740032 | 0.027674178 | 0.604897175 | 0.54524732 | NA |
| Cd163l1 | 3.295479378 | 0.039348761 | 0.123032794 | 0.319823357 | 0.74910224 | NA |
| Cd177 | 3.22070232 | -0.182423077 | 0.106474589 | -1.71330154 | 0.086657079 | NA |
| Cd19 | 1.431574452 | -0.044563126 | 0.090335411 | -0.493307389 | 0.621795408 | NA |
| Cd1d2 | 0 | NA | NA | NA | NA | NA |
| Cd2 | 2.21153379 | -0.207226095 | 0.10275758 | -2.016650212 | 0.043732029 | NA |
| Cd200r2 | 1.931964708 | -0.008143172 | 0.08424728 | -0.096657978 | 0.922998012 | NA |
| Cd200r3 | 0 | NA | NA | NA | NA | NA |
| Cd200r4 | 6.705642094 | 0.412995008 | 0.140600688 | 2.937361216 | 0.003310183 | NA |
| Cd207 | 0.185670226 | -0.001645086 | 0.031432796 | -0.052336624 | 0.958260472 | NA |
| Cd209d | 2.214390628 | -0.033180086 | 0.100762333 | -0.329290567 | 0.741936073 | NA |
| Cd209e | 0.275064582 | 0.010932281 | 0.038417727 | 0.284563456 | 0.775978612 | NA |
| Cd209g | 5.493189024 | -0.037289074 | 0.130204764 | -0.286387935 | 0.774581015 | NA |
| Cd226 | 0.976978344 | 0.024011131 | 0.080907063 | 0.296774229 | 0.766638882 | NA |
| Cd244 | 2.627008601 | 0.11305792 | 0.09932635 | 1.138246996 | 0.25501736 | NA |
| Cd247 | 3.053224201 | -0.165473618 | 0.118748421 | -1.393480581 | 0.163474513 | NA |
| Cd27 | 0.488467756 | -0.020554715 | 0.055630496 | -0.369486469 | 0.711765157 | NA |
| Cd300c | 0.098353386 | -0.020212831 | 0.02786466 | -0.725393061 | 0.468210929 | NA |
| Cd300e | 4.939095036 | 0.126572946 | 0.134377331 | 0.941921865 | 0.346232645 | NA |
| Cd300lb | 2.653118307 | 0.042060645 | 0.113232116 | 0.37145508 | 0.710298607 | NA |
| Cd300ld | 3.963134501 | -0.112237914 | 0.124950874 | -0.898256334 | 0.369048907 | NA |
| Cd300lf | 6.429006114 | 0.235143999 | 0.104052941 | 2.259849607 | NA | NA |
| Cd300lg | 5.582805756 | 0.094471637 | 0.150070218 | 0.62951622 | 0.529011153 | NA |
| Cd300lh | 0.116696964 | -0.020477583 | 0.027954224 | -0.732539847 | 0.463839131 | NA |
| Cd3d | 0.879765535 | 0.027449169 | 0.061992718 | 0.442780535 | 0.657924484 | NA |
| Cd3e | 1.473692292 | 0.064687221 | 0.076260439 | 0.848240873 | 0.396303839 | NA |
| Cd3g | 0.588168521 | -0.033702544 | 0.053860702 | -0.625735339 | 0.53148855 | NA |
| Cd40lg | 0.198019769 | 0.025308392 | 0.031628095 | 0.80018706 | 0.423602426 | NA |
| Cd5 | 5.295899881 | 0.160072967 | 0.128744657 | 1.243336774 | 0.213743758 | NA |
| Cd5l | 0.48156345 | -0.012077111 | 0.044778632 | -0.269707011 | 0.787385666 | NA |
| Cd69 | 0.810921301 | 0.051807665 | 0.063085565 | 0.821228515 | 0.411516117 | NA |
| Cd70 | 1.912147608 | 0.005744978 | 0.094550521 | 0.060760932 | 0.951549604 | NA |
| Cd8a | 1.366151416 | -0.005040101 | 0.076024171 | -0.066296029 | 0.947142145 | NA |
| Cd8b1 | 0.884508339 | -0.04816716 | 0.06562523 | -0.733973201 | 0.462965071 | NA |
| Cd96 | 1.779697318 | -0.024466514 | 0.09673155 | -0.252932101 | 0.800320683 | NA |
| Cdcp2 | 0.098353386 | -0.020212831 | 0.02786466 | -0.725393061 | 0.468210929 | NA |
| Cdh16 | 0 | NA | NA | NA | NA | NA |
| Cdh17 | 2.472933667 | 0.014869444 | 0.112928582 | 0.131671217 | 0.895244353 | NA |
| Cdhr5 | 0.846679412 | 0.067107966 | 0.071303602 | 0.941158145 | 0.346623822 | NA |
| Cdkn1a | 1019.537859 | 0.11224457 | 0.152161752 | 0.737666122 | NA | NA |
| Cdkn2a | 4.153287324 | 0.340939665 | 0.125446185 | 2.717816134 | 0.006571434 | NA |
| Cdr1 | 1.191371392 | 0.085765618 | 0.070050215 | 1.224344827 | 0.220822173 | NA |
| Cdrt4 | 1.650567937 | -0.033895378 | 0.092427697 | -0.366723165 | 0.713825525 | NA |
| Cdx1 | 0 | NA | NA | NA | NA | NA |
| Cdx2 | 0 | NA | NA | NA | NA | NA |
| Cdx4 | 0 | NA | NA | NA | NA | NA |
| Ceacam-ps1 | 1.014480811 | 0.116186393 | 0.073507767 | 1.580600225 | 0.113969474 | NA |
| Ceacam10 | 4.137747377 | 0.061850514 | 0.123778286 | 0.499687921 | 0.617294839 | NA |
| Ceacam11 | 0 | NA | NA | NA | NA | NA |
| Ceacam12 | 0 | NA | NA | NA | NA | NA |
| Ceacam13 | 0 | NA | NA | NA | NA | NA |
| Ceacam14 | 0 | NA | NA | NA | NA | NA |
| Ceacam15 | 1.645313989 | -0.050038391 | 0.097491763 | -0.513257626 | 0.607771121 | NA |
| Ceacam16 | 0.730358311 | 0.059975352 | 0.057700588 | 1.039423589 | 0.298607778 | NA |
| Ceacam18 | 0.412387467 | -0.005395598 | 0.042599376 | -0.126659095 | 0.899210223 | NA |
| Ceacam19 | 0 | NA | NA | NA | NA | NA |
| Ceacam20 | 0 | NA | NA | NA | NA | NA |
| Ceacam3 | 1.08876445 | 0.018664533 | 0.066257674 | 0.281696177 | 0.778176486 | NA |
| Ceacam5 | 0 | NA | NA | NA | NA | NA |
| Cebpe | 0.307917359 | 0.010394288 | 0.03404619 | 0.305299606 | 0.760137976 | NA |
| Cel | 0.393805724 | -0.036573061 | 0.048755423 | -0.750133191 | 0.453174491 | NA |
| Cela2a | 0.258977192 | 0.029727007 | 0.03417262 | 0.869907163 | 0.38435114 | NA |
| Cela3b | 0 | NA | NA | NA | NA | NA |
| Cep55 | 8.057431714 | -0.029397394 | 0.156868106 | -0.187401985 | 0.851345466 | NA |
| Cer1 | 4.428946437 | -0.038589098 | 0.132362777 | -0.291540408 | 0.770638044 | NA |
| Cers1 | 0 | NA | NA | NA | NA | NA |
| Ces1a | 2.303102152 | -0.246464794 | 0.102371746 | -2.407547029 | 0.016060094 | NA |
| Ces1b | 0 | NA | NA | NA | NA | NA |
| Ces1c | 0 | NA | NA | NA | NA | NA |
| Ces1d | 4.568624031 | 0.036060239 | 0.140350017 | 0.256930775 | 0.797232213 | NA |
| Ces1e | 0 | NA | NA | NA | NA | NA |
| Ces1f | 3.2535107 | -0.019126911 | 0.109824549 | -0.174158798 | 0.861740665 | NA |
| Ces1g | 0.213790458 | -0.002646789 | 0.031909369 | -0.082947074 | 0.933893623 | NA |
| Ces2a | 0 | NA | NA | NA | NA | NA |
| Ces2b | 0.838780774 | 0.045559078 | 0.062686379 | 0.726777954 | 0.46736199 | NA |
| Ces2c | 0 | NA | NA | NA | NA | NA |
| Ces2d-ps | 0 | NA | NA | NA | NA | NA |
| Ces2e | 0.763784651 | -0.00317115 | 0.056229574 | -0.056396483 | 0.955025958 | NA |
| Ces2f | 0 | NA | NA | NA | NA | NA |
| Ces2h | 0 | NA | NA | NA | NA | NA |
| Ces3a | 0 | NA | NA | NA | NA | NA |
| Ces3b | 0 | NA | NA | NA | NA | NA |
| Ces4a | 0 | NA | NA | NA | NA | NA |
| Cetn1 | 0 | NA | NA | NA | NA | NA |
| Cfc1 | 0.203787646 | -0.02671144 | 0.031728022 | -0.841887971 | 0.399850664 | NA |
| Cfd | 0 | NA | NA | NA | NA | NA |
| Cfhr1 | 0 | NA | NA | NA | NA | NA |
| Cfhr2 | 0.572845755 | -0.029822974 | 0.053532279 | -0.557102627 | 0.577457312 | NA |
| Cfi | 0.871541253 | 0.00020978 | 0.064142277 | 0.003270542 | 0.99739049 | NA |
| Cga | 4.56743289 | 0.289719723 | 0.105259322 | 2.752437679 | 0.00591534 | NA |
| Chad | 3.328561322 | -0.006873845 | 0.126534448 | -0.054323906 | 0.956677103 | NA |
| Chi3l3 | 7.451930034 | -0.057053337 | 0.137564507 | -0.41473879 | 0.678333133 | NA |
| Chi3l4 | 0.098353386 | -0.020212831 | 0.02786466 | -0.725393061 | 0.468210929 | NA |
| Chit1 | 5.854022429 | 0.04452478 | 0.147620979 | 0.30161553 | 0.762945168 | NA |
| Chmp4c | 3.675516495 | 0.040397073 | 0.114601444 | 0.352500558 | 0.7244629 | NA |
| Chrdl2 | 5.33060177 | 0.084383174 | 0.143046078 | 0.589902044 | 0.555256323 | NA |
| Chrna9 | 0.217751027 | -0.027071875 | 0.031947577 | -0.847384294 | 0.396780956 | NA |
| Chrnd | 0 | NA | NA | NA | NA | NA |
| Chrne | 1.690479618 | -0.035285547 | 0.098151323 | -0.359501494 | 0.719219961 | NA |
| Chrng | 0.096071043 | -0.020098795 | 0.027825948 | -0.722304036 | 0.470107572 | NA |
| Chst13 | 1.103949313 | -0.080511457 | 0.075243076 | -1.070018145 | 0.284611141 | NA |
| Chst4 | 4.187077356 | -0.01092005 | 0.114141695 | -0.095670996 | 0.923781877 | NA |
| Cib3 | 0.470878708 | 0.028927796 | 0.04534639 | 0.637929415 | 0.523519629 | NA |
| Cib4 | 0 | NA | NA | NA | NA | NA |
| Cidec | 0.331243956 | 0.008596723 | 0.034710864 | 0.247666645 | 0.804392342 | NA |
| Ciita | 9.936993295 | 0.141291351 | 0.109518408 | 1.290115088 | NA | NA |
| Cilp | 5.520717217 | -0.435128263 | 0.144845084 | -3.004094107 | 0.002663729 | NA |
| Ckm | 0.969645158 | 0.020592352 | 0.076153894 | 0.270404457 | 0.786849112 | NA |
| Ckmt2 | 1.03630432 | 0.003079144 | 0.064486102 | 0.047748952 | 0.961916321 | NA |
| Clca2 | 2.012024816 | 0.049548493 | 0.09627614 | 0.514649766 | 0.606797782 | NA |
| Clca3 | 0.7238653 | 0.106446607 | 0.063481683 | 1.676808206 | 0.093579993 | NA |
| Clca4 | 0.192142086 | -0.026384592 | 0.031527717 | -0.836869715 | 0.402665805 | NA |
| Clca5 | 0.352713512 | 0.01988513 | 0.041853977 | 0.475107309 | 0.634710489 | NA |
| Clca6 | 0.105923158 | 0.018524693 | 0.027977518 | 0.662127842 | 0.507889294 | NA |
| Clcnka | 6.973684801 | 0.073460396 | 0.063858896 | 1.150354928 | NA | NA |
| Clcnkb | 1.171397964 | 0.084597537 | 0.085157829 | 0.993420544 | 0.320505054 | NA |
| Cldn13 | 0.236094605 | -0.028257939 | 0.032632891 | -0.865934257 | 0.386526215 | NA |
| Cldn15 | 5.074131718 | -0.051908619 | 0.133040948 | -0.390170243 | 0.696410664 | NA |
| Cldn16 | 0.105923158 | 0.018524693 | 0.027977518 | 0.662127842 | 0.507889294 | NA |
| Cldn17 | 0 | NA | NA | NA | NA | NA |
| Cldn18 | 0.195033442 | -0.002424765 | 0.031596358 | -0.076741901 | 0.93882887 | NA |
| Cldn2 | 30.84619758 | 0.280923164 | 0.117259105 | 2.395747128 | NA | NA |
| Cldn20 | 7.192744513 | 0.061794147 | 0.156725465 | 0.39428275 | 0.693372298 | NA |
| Cldn24 | 0.119397641 | -0.020477583 | 0.027954224 | -0.732539845 | 0.463839132 | NA |
| Cldn3 | 3.452382754 | 0.224953466 | 0.121639461 | 1.849346131 | 0.064407849 | NA |
| Cldn4 | 0.079983654 | 0.016740032 | 0.027674178 | 0.604897175 | 0.54524732 | NA |
| Cldn6 | 0.79738299 | 0.056420574 | 0.069491893 | 0.811901524 | 0.41684814 | NA |
| Cldn7 | 0.358192922 | -0.030696978 | 0.034083535 | -0.900639514 | 0.367780018 | NA |
| Cldn8 | 0 | NA | NA | NA | NA | NA |
| Cldn9 | 3.68848754 | 0.142388042 | 0.12567826 | 1.132956819 | 0.257232387 | NA |
| Cldnd2 | 0.68889895 | 0.008020532 | 0.060682319 | 0.132172472 | 0.894847874 | NA |
| Clec12b | 1.117520908 | -0.112944557 | 0.069043422 | -1.635848203 | 0.101871361 | NA |
| Clec1b | 2.16508377 | -0.053897296 | 0.107256072 | -0.502510441 | 0.61530851 | NA |
| Clec2e | 0.724560782 | -0.05162733 | 0.062704856 | -0.823338625 | 0.410315455 | NA |
| Clec2g | 0.500981137 | -0.05018208 | 0.04329235 | -1.159144275 | 0.246397381 | NA |
| Clec2h | 0.236094605 | -0.028257939 | 0.032632891 | -0.865934257 | 0.386526215 | NA |
| Clec2i | 1.473017754 | -0.004436983 | 0.091300537 | -0.048597552 | 0.96124002 | NA |
| Clec3a | 1.276319948 | 0.026881504 | 0.082439033 | 0.326077382 | 0.744365809 | NA |
| Clec4a4 | 0.079983654 | 0.016740032 | 0.027674178 | 0.604897175 | 0.54524732 | NA |
| Clec4b1 | 0.298180463 | -0.015304498 | 0.039412854 | -0.388312359 | 0.697784893 | NA |
| Clec4b2 | 0 | NA | NA | NA | NA | NA |

| | | | | | | |
|---|---|---|---|---|---|---|
| Smad9 | 818.0579377 | 0.080954761 | 0.069109018 | 1.171406613 | 0.241435375 | 0.692393571 |
| Ing2 | 1118.753983 | 0.080978536 | 0.07344398 | 1.102589166 | 0.270205617 | 0.71216373 |
| Atl1 | 4990.246012 | 0.080994324 | 0.063378066 | 1.277955112 | 0.201265254 | 0.654716629 |
| Med14 | 3767.419252 | 0.081000449 | 0.064671858 | 1.252483719 | 0.210393657 | 0.664820157 |
| Pcp2 | 11.80046778 | 0.081016499 | 0.048641896 | 1.66557033 | 0.095799025 | NA |
| Tkt | 6101.803659 | 0.081058906 | 0.061405268 | 1.32006435 | 0.186813534 | 0.641054259 |
| Ndel1 | 4114.894361 | 0.081087649 | 0.058600191 | 1.383743759 | 0.166436929 | 0.616902656 |
| Dennd2d | 20.14053706 | 0.081089285 | 0.082087447 | 0.987840261 | 0.323230887 | NA |
| Cdc34 | 1278.906297 | 0.081090984 | 0.062904597 | 1.28911062 | 0.197359632 | 0.653337743 |
| 2210016L21Rik | 4256.002858 | 0.081101928 | 0.05059469 | 1.602973104 | 0.108940592 | 0.534621387 |
| Syngr1 | 29505.42441 | 0.081106951 | 0.047666988 | 1.701532972 | 0.088842952 | 0.49789071 |
| Pim2 | 1837.573036 | 0.081128026 | 0.049923171 | 1.625057556 | 0.104150296 | 0.523645898 |
| Trmt11 | 270.6850803 | 0.081141404 | 0.083279264 | 0.974329024 | 0.329893198 | 0.756141871 |
| Cab39 | 7711.489134 | 0.081145726 | 0.056579195 | 1.434197253 | 0.151515981 | 0.598905575 |
| Stmn3 | 13586.52853 | 0.081159954 | 0.039183309 | 2.071288913 | 0.038331805 | 0.354869883 |
| Aff2 | 564.5936772 | 0.081161109 | 0.085490032 | 0.949363422 | 0.342435807 | 0.763762832 |
| Lbh | 2123.78035 | 0.081161668 | 0.065326744 | 1.242395731 | 0.21409059 | 0.668216569 |
| Yipf1 | 1188.421963 | 0.081163624 | 0.056156274 | 1.445317122 | 0.148368831 | 0.595689782 |
| Tspyl5 | 2293.416301 | 0.081176831 | 0.053404124 | 1.520047977 | 0.128498918 | 0.566659164 |
| Dync1li1 | 2857.555847 | 0.081179078 | 0.053585801 | 1.51493634 | 0.129788527 | 0.568885619 |
| Slc38a2 | 5628.950296 | 0.081197944 | 0.075541703 | 1.074875737 | 0.282430366 | 0.72081452 |
| Rab14 | 9290.168509 | 0.081213566 | 0.059565536 | 1.363432149 | 0.172746355 | 0.624056899 |
| Nt5dc3 | 7321.82067 | 0.081226463 | 0.052107521 | 1.558824169 | 0.119038002 | 0.551385848 |
| Yeats4 | 1081.547413 | 0.081275214 | 0.068743157 | 1.182302615 | 0.237085648 | 0.687955679 |
| Gm16702 | 184.1232918 | 0.08128175 | 0.094253006 | 0.862378322 | 0.388479366 | 0.792229575 |
| Bpnt1 | 2012.712104 | 0.081304535 | 0.045815267 | 1.774616615 | 0.075961212 | 0.471947205 |
| Lrrc34 | 7.556988423 | 0.081320163 | 0.052316007 | 1.554403092 | 0.120088296 | NA |
| Kbtbd2 | 2311.899907 | 0.08134854 | 0.074657047 | 1.089629761 | 0.275876268 | 0.717396826 |
| Cenpk | 24.15009258 | 0.081356171 | 0.08490358 | 0.958218388 | 0.337952647 | 0.760581219 |
| Pfkfb3 | 2022.751178 | 0.081371353 | 0.091047674 | 0.893722479 | 0.371470395 | 0.780761257 |
| B130024G19Rik | 16.98105827 | 0.081386712 | 0.068976547 | 1.179918632 | 0.238032579 | NA |
| Atp6v0e2 | 16826.7016 | 0.081390298 | 0.043336713 | 1.878091173 | 0.060368694 | 0.430493689 |
| Tspan33 | 985.456722 | 0.081396414 | 0.070315256 | 1.157592516 | 0.247030365 | 0.696511265 |
| Elp3 | 3903.957752 | 0.081427612 | 0.048598987 | 1.675500168 | 0.093836138 | 0.509556539 |
| Lamp5 | 8675.516963 | 0.081431712 | 0.074917631 | 1.086949906 | 0.277058949 | 0.717815689 |
| Gpr44 | 4.393186002 | 0.081476998 | 0.049535959 | 1.644805093 | 0.100010012 | NA |
| Rpl7l1 | 1589.410929 | 0.081482389 | 0.051451862 | 1.583662588 | 0.113270523 | 0.540327972 |
| Theg | 15.05706019 | 0.081507776 | 0.072246471 | 1.12819042 | 0.259239509 | NA |
| Dzank1 | 14206.39871 | 0.081513717 | 0.063039938 | 1.293048818 | 0.195994175 | 0.651490021 |
| Slc30a4 | 3427.624056 | 0.081520487 | 0.08383171 | 0.972430213 | 0.330836568 | 0.756212998 |
| Tmem33 | 3971.856095 | 0.081543993 | 0.064996582 | 1.254588945 | 0.209628019 | 0.664226905 |
| Cbr1 | 2009.427586 | 0.081559779 | 0.053042223 | 1.537638793 | 0.124136957 | 0.559205951 |
| Cdipt | 5713.994163 | 0.081561553 | 0.057315349 | 1.4230316 | 0.154726995 | 0.604402956 |
| Fam20b | 5170.843302 | 0.081569017 | 0.048095377 | 1.695984578 | 0.089888801 | 0.500949827 |
| Rnf168 | 1513.098342 | 0.081612785 | 0.065626279 | 1.24359914 | 0.213647133 | 0.667262579 |
| Rps6kc1 | 2528.111936 | 0.081653278 | 0.054801568 | 1.489980701 | 0.136229311 | 0.579855848 |
| Zfp300 | 121.0078867 | 0.08166271 | 0.094969679 | 0.859881928 | 0.389854131 | 0.792918106 |
| Arl5a | 4048.754827 | 0.081663789 | 0.05717118 | 1.428408313 | NA | NA |
| Trhr | 199.8131296 | 0.081683945 | 0.073635818 | 1.10929636 | 0.267302355 | 0.709971477 |
| Wnt10a | 339.8964078 | 0.081708046 | 0.087482134 | 0.933996952 | 0.350305465 | 0.768667813 |
| Rab8a | 1423.917367 | 0.081774878 | 0.051637392 | 1.583636876 | 0.113276377 | 0.540327972 |
| Acpl2 | 636.7795203 | 0.081787645 | 0.067054974 | 1.219710326 | 0.222574706 | 0.67374894 |
| Rps15a-ps4 | 14.50369155 | 0.081793629 | 0.074215856 | 1.102104498 | 0.270416244 | NA |
| Pde12 | 532.1722369 | 0.081839728 | 0.071055731 | 1.151768154 | 0.249416357 | 0.697864808 |
| Uhmk1 | 2383.265487 | 0.081849929 | 0.068683809 | 1.191691755 | 0.233382132 | 0.685064073 |
| Mb21d2 | 1917.30955 | 0.081866042 | 0.050608076 | 1.617647778 | 0.105738526 | 0.527979239 |
| Tpm4 | 1270.514664 | 0.081888894 | 0.071811467 | 1.14033172 | 0.254148128 | 0.701619624 |
| Gkap1 | 518.2502469 | 0.081957224 | 0.083095034 | 0.986307127 | 0.323982424 | 0.752925264 |
| Tex2 | 5126.287909 | 0.081995182 | 0.049293411 | 1.663410595 | 0.096230274 | 0.516109386 |
| Efcc1 | 270.7722985 | 0.082023099 | 0.083696026 | 0.980011877 | 0.327080256 | 0.754033931 |
| Aaed1 | 214.405573 | 0.082041967 | 0.085484816 | 0.959725616 | 0.337193328 | 0.760113439 |
| Zcchc17 | 2162.619925 | 0.082045297 | 0.054180976 | 1.514282381 | 0.129954235 | 0.56891406 |
| Smad1 | 854.750301 | 0.082065562 | 0.06513959 | 1.259841546 | 0.207726529 | 0.662799412 |
| Adam32 | 13.84849078 | 0.082076164 | 0.071734579 | 1.144164574 | 0.252555392 | NA |
| Itih3 | 1389.227926 | 0.082109937 | 0.075733785 | 1.084191648 | NA | NA |
| Cnrip1 | 2729.794564 | 0.082130881 | 0.057166006 | 1.436708384 | 0.150800872 | 0.598846773 |
| Ddx54 | 2012.712132 | 0.082145784 | 0.050551899 | 1.624979188 | 0.104166994 | 0.523645898 |
| Ttyh3 | 11442.48905 | 0.082150525 | 0.057819091 | 1.420820076 | 0.155369073 | 0.604578396 |
| Man1a | 749.4448074 | 0.08215378 | 0.073906728 | 1.111587294 | 0.266315638 | 0.709036099 |
| Chordc1 | 1628.71302 | 0.082176648 | 0.070780348 | 1.161009376 | 0.245638086 | 0.69504627 |
| Ctxn2 | 275.2576196 | 0.082255497 | 0.084996526 | 0.967751274 | 0.333168604 | 0.757730862 |
| Gabrg1 | 996.1057436 | 0.082271228 | 0.08020452 | 1.025767972 | 0.305000966 | 0.737453291 |
| Nudt12 | 282.852855 | 0.082286416 | 0.078026812 | 1.05459154 | 0.291612174 | 0.727847831 |
| Wars | 4304.605108 | 0.082295084 | 0.044608374 | 1.844834894 | 0.06506156 | 0.444536321 |
| Setd7 | 9696.638615 | 0.082302898 | 0.06364355 | 1.293185221 | 0.195947006 | 0.651482309 |
| Fgfr3 | 5085.562193 | 0.082304346 | 0.053761132 | 1.530926598 | 0.125787535 | 0.562787717 |
| Arcn1 | 5628.987344 | 0.082329965 | 0.056768906 | 1.450265121 | 0.146984601 | 0.594031723 |
| Crkl | 2231.219008 | 0.082364591 | 0.068791707 | 1.197304074 | 0.231188057 | 0.683402059 |
| Scp2 | 4260.485143 | 0.082393238 | 0.042216695 | 1.951674297 | 0.050976886 | 0.403266245 |
| Wdr35 | 2113.092058 | 0.082394169 | 0.058832602 | 1.400484873 | 0.16136817 | 0.612979613 |
| Plekhb1 | 11293.95654 | 0.082400913 | 0.08002323 | 1.029712413 | 0.303145027 | 0.73655042 |
| Erbb4 | 1075.581145 | 0.082449072 | 0.086501815 | 0.953148459 | 0.340514856 | 0.762546123 |
| Dpp10 | 7596.747983 | 0.082456233 | 0.072380974 | 1.139197626 | 0.254620735 | 0.701619624 |
| Pgm2l1 | 14064.81734 | 0.082461172 | 0.076147967 | 1.08290707 | 0.278849674 | 0.71849589 |
| Ndufb11 | 2548.119861 | 0.082487948 | 0.05048774 | 1.633821365 | 0.102296365 | 0.522856086 |
| Dcaf12 | 2113.902775 | 0.082490955 | 0.056805333 | 1.452169202 | 0.146454564 | 0.593251254 |
| Prkg2 | 570.205071 | 0.08249765 | 0.085544618 | 0.964381535 | 0.334854676 | 0.758238352 |
| Asf1b | 26.25214772 | 0.082501314 | 0.082331192 | 1.002066311 | 0.316311567 | 0.745993052 |
| Pdss2 | 390.3417781 | 0.082506141 | 0.076016797 | 1.085367234 | 0.277759038 | 0.718083484 |
| Rrp1 | 12144.12895 | 0.082511083 | 0.042053469 | 1.962051775 | 0.049756457 | 0.399757904 |
| Pex11b | 1687.408521 | 0.082512106 | 0.049548158 | 1.665291079 | 0.095854697 | 0.515046226 |
| Rgma | 2267.380066 | 0.0825893 | 0.060341673 | 1.368694235 | 0.171094873 | 0.621020522 |
| B3gat3 | 4202.738223 | 0.082620147 | 0.065359342 | 1.264090859 | 0.206197417 | 0.660267934 |
| Aftph | 3788.783783 | 0.08265655 | 0.070606536 | 1.170664279 | 0.241733749 | 0.692421615 |
| Zbtb26 | 371.8672875 | 0.082660693 | 0.089870286 | 0.919777789 | 0.357688893 | 0.773905615 |
| 1700086L19Rik | 843.2468835 | 0.08266624 | 0.084953779 | 0.97307314 | 0.330516953 | 0.756141871 |
| Dpp3 | 1559.908959 | 0.082699387 | 0.059650173 | 1.386406484 | 0.165622818 | 0.616091434 |
| Psma7 | 2358.18614 | 0.082724753 | 0.050194774 | 1.648075018 | 0.099337278 | 0.520411863 |
| Zfp354b | 124.1056714 | 0.082739464 | 0.091503211 | 0.904224707 | 0.365876262 | 0.778143276 |
| AI413582 | 907.536474 | 0.082757488 | 0.072164834 | 1.146784155 | 0.25147084 | 0.699465245 |
| Gabrg2 | 7909.983652 | 0.082761665 | 0.066261704 | 1.249012038 | 0.211660671 | 0.666270994 |
| Gm14015 | 16.0461305 | 0.082768625 | 0.070772813 | 1.169497463 | 0.242203263 | NA |
| Prpf38a | 686.1645718 | 0.082804126 | 0.070211859 | 1.179346722 | 0.238260141 | 0.689023852 |
| Tmem222 | 2831.533113 | 0.08280468 | 0.057313144 | 1.444776423 | 0.148520695 | 0.59613474 |
| Popdc3 | 31.52475347 | 0.082811817 | 0.087632217 | 0.944992834 | 0.344662535 | 0.764478237 |
| Csnk1d | 6775.670902 | 0.082872703 | 0.050545683 | 1.639560473 | 0.101096587 | 0.521924472 |
| Galnt18 | 1412.165473 | 0.082877949 | 0.072006633 | 1.15097659 | 0.249741867 | 0.698181362 |
| Adra1a | 404.5498637 | 0.082883198 | 0.079570951 | 1.041626329 | 0.297584957 | 0.732415975 |
| Grlf1 | 11459.87932 | 0.082884725 | 0.053416047 | 1.551682115 | 0.120738304 | 0.554612895 |
| Gpr139 | 52.35536328 | 0.082895101 | 0.088559824 | 0.936035069 | 0.349255132 | 0.768213211 |
| 1700013G23Rik | 17.76692386 | 0.082906538 | 0.07785475 | 1.064887347 | 0.286926911 | NA |
| Tmem189 | 1653.73224 | 0.082951305 | 0.056036023 | 1.480321065 | 0.138787584 | 0.583135914 |
| Fabp3 | 2402.067766 | 0.082953629 | 0.053598929 | 1.547673251 | 0.121700985 | 0.555997647 |
| Dip2c | 4275.984735 | 0.082961294 | 0.06119646 | 1.355655105 | 0.17520893 | 0.628026741 |
| Mettl6 | 585.084674 | 0.082997082 | 0.070410952 | 1.178752444 | 0.238496766 | 0.689459926 |
| Faxc | 5801.365171 | 0.083004784 | 0.071055425 | 1.168169552 | 0.24273838 | 0.692772033 |
| Pten | 8594.251715 | 0.083023492 | 0.073673584 | 1.126909921 | 0.259780569 | 0.703893404 |
| Tmem150c | 1587.241756 | 0.083044512 | 0.065285597 | 1.272018874 | 0.203366413 | 0.657417489 |
| 1700007P06Rik | 19.30882388 | 0.083059053 | 0.079763995 | 1.0413101 | 0.297731651 | NA |
| Fgf18 | 53.70592475 | 0.083091552 | 0.093546754 | 0.888235542 | 0.374414064 | 0.78293624 |
| D8Ertd82e | 1726.467339 | 0.083095471 | 0.072946324 | 1.139131706 | 0.254648225 | 0.701619624 |
| Nop2 | 1030.056657 | 0.083147082 | 0.062920328 | 1.321466128 | 0.186345989 | 0.640677359 |
| Ift52 | 1321.571298 | 0.08314801 | 0.049719723 | 1.672334532 | 0.094458374 | 0.511705397 |
| Srm | 1750.316817 | 0.083148312 | 0.06185605 | 1.344222797 | 0.178876352 | 0.632619671 |
| Ap2m1 | 22677.83199 | 0.083154696 | 0.045818866 | 1.814857155 | 0.069545864 | 0.455055507 |
| Ift27 | 668.0443243 | 0.083206332 | 0.0687089 | 1.210997878 | 0.225896218 | 0.678248958 |
| Tnfaip8 | 233.9844604 | 0.083211654 | 0.092651536 | 0.898114131 | 0.369124706 | 0.779913525 |
| Gpr125 | 883.6500047 | 0.083229252 | 0.067980734 | 1.224306456 | 0.220836642 | 0.672757795 |
| March4 | 1122.249799 | 0.083229522 | 0.057027299 | 1.459468071 | 0.144436322 | 0.589967032 |
| Snx29 | 329.1881835 | 0.083239904 | 0.07326694 | 1.136118189 | 0.255907104 | 0.701952734 |
| Gm10406 | 66.60979705 | 0.083240887 | 0.0945214 | 0.880656511 | 0.378503762 | 0.785337186 |
| Calr | 13796.90374 | 0.083288761 | 0.055454198 | 1.501937883 | 0.133113152 | 0.575193212 |
| Galnt13 | 1293.149979 | 0.083313959 | 0.088660658 | 0.939694793 | 0.347374137 | 0.766038372 |
| Ate1 | 3285.588373 | 0.083346879 | 0.054748045 | 1.522371787 | 0.12791595 | 0.565771339 |
| Naa50 | 2302.023933 | 0.083359587 | 0.076233061 | 1.093483405 | 0.274181611 | 0.716185716 |
| Atrnl1 | 6466.36453 | 0.083398726 | 0.072036501 | 1.15772871 | 0.246974764 | 0.696511265 |
| Pex11c | 17.59566597 | 0.083424841 | 0.072837259 | 1.145359413 | 0.252060304 | NA |
| Fam43b | 964.9042267 | 0.083444858 | 0.069432102 | 1.201819564 | 0.229433444 | 0.682503526 |
| Fam207a | 1423.328824 | 0.083461095 | 0.05769411 | 1.446613777 | 0.148005125 | 0.595114347 |
| 1110001A16Rik | 392.4718473 | 0.083471188 | 0.074408126 | 1.121802043 | 0.261946617 | 0.704317239 |
| Arl6ip5 | 5402.766538 | 0.083476793 | 0.049437049 | 1.688547238 | 0.091306229 | 0.503727415 |
| Angptl1 | 35.88989857 | 0.083493021 | 0.087547723 | 0.953685811 | 0.340242705 | 0.762240211 |
| Nts | 153.6098757 | 0.08349895 | 0.075067738 | 1.112314707 | 0.266002863 | 0.708592991 |
| Gmfb | 6847.586738 | 0.083517728 | 0.075829388 | 1.101389974 | 0.270726967 | 0.712294861 |
| Kcnj14 | 94.14024467 | 0.083524027 | 0.095099093 | 0.878284156 | 0.379789532 | 0.786275719 |
| Lcn2 | 11.95296539 | 0.083529824 | 0.054922835 | 1.520857827 | 0.128295519 | NA |
| Fbll1 | 855.920131 | 0.083568552 | 0.070941552 | 1.177991591 | 0.238799959 | 0.689537017 |
| Swsap1 | 542.8868675 | 0.083576329 | 0.064514002 | 1.295475805 | 0.195156142 | 0.65093872 |
| Srbd1 | 446.9008292 | 0.08358113 | 0.073246637 | 1.14109171 | 0.253831761 | 0.701619624 |
| Mtmr7 | 3616.279528 | 0.083627476 | 0.059394036 | 1.408011333 | 0.159127714 | 0.610858662 |
| Prkch | 366.601411 | 0.083646678 | 0.083037361 | 1.007337859 | 0.313772442 | 0.745844951 |
| Rufy4 | 3.038122281 | 0.083663887 | 0.038536857 | 2.171009594 | 0.029930447 | NA |
| Emx2 | 311.0382275 | 0.083716484 | 0.083586168 | 1.001559065 | 0.3165566 | 0.745993052 |
| Slc45a4 | 2108.80798 | 0.083732339 | 0.059563476 | 1.4057665 | 0.159793473 | 0.611797571 |
| Ebf3 | 58.43011339 | 0.083735138 | 0.075429089 | 1.110117323 | 0.266948474 | 0.709550508 |
| Kcnip1 | 1203.638963 | 0.083775591 | 0.085501495 | 0.979814343 | 0.327177771 | 0.754100734 |
| Timm8b | 1826.243628 | 0.083778807 | 0.060994867 | 1.373538632 | 0.16958496 | 0.619945581 |
| Fcgbp | 20.98965864 | 0.083779879 | 0.073454303 | 1.140571414 | 0.254048319 | NA |
| Gabarapl1 | 26218.28507 | 0.083802663 | 0.034120248 | 2.45609769 | 0.014045497 | 0.227385586 |
| Ctcf | 2899.360203 | 0.083806938 | 0.060233549 | 1.391366429 | 0.164114339 | 0.614351408 |
| Ghrh | 7.211534777 | 0.083829007 | 0.043986784 | 1.905777143 | 0.056679126 | NA |
| Pcdhga11 | 944.8951425 | 0.083844466 | 0.05338181 | 1.570656106 | 0.11626255 | 0.546307021 |
| Fpgt | 982.5347785 | 0.083853162 | 0.06580355 | 1.274295427 | 0.202558736 | 0.65606599 |
| Mthfd2 | 240.8989066 | 0.083891486 | 0.08558493 | 0.980213292 | 0.326980845 | 0.753924405 |
| Lyst | 3053.330445 | 0.083941358 | 0.073763116 | 1.137985512 | 0.255126532 | 0.701754971 |
| Calhm2 | 196.0583231 | 0.083970436 | 0.093019293 | 0.902720646 | 0.366674176 | 0.778245044 |
| Rnf113a1 | 142.5391582 | 0.083971162 | 0.093294342 | 0.900067042 | 0.368084574 | 0.779243711 |
| Lrrn3 | 2639.393133 | 0.083999506 | 0.062065223 | 1.353406973 | 0.175925651 | 0.628658305 |
| Gsto1 | 952.1495535 | 0.084019849 | 0.06705928 | 1.25291904 | 0.210235172 | 0.664608511 |
| Tmppe | 1038.824429 | 0.084049809 | 0.057647692 | 1.457990884 | 0.144843056 | 0.590269908 |
| Vps35 | 7451.349308 | 0.08405319 | 0.073336507 | 1.146130253 | 0.251741261 | 0.699465245 |
| Dpp6 | 12714.81085 | 0.084070385 | 0.061190083 | 1.373921737 | 0.169465981 | 0.619945581 |
| Trib1 | 979.817492 | 0.084113688 | 0.085607808 | 0.982546927 | 0.325830476 | 0.753033837 |
| Fkbp2 | 2292.30727 | 0.084153044 | 0.066841795 | 1.258988382 | 0.208034528 | 0.66299025 |
| Actr1a | 9762.794698 | 0.084155732 | 0.041464308 | 2.029594507 | 0.042397774 | 0.373028504 |
| Med27 | 775.3672512 | 0.084160625 | 0.072178957 | 1.165999462 | 0.243614663 | 0.692824881 |
| Pfdn2 | 1844.946409 | 0.084167683 | 0.069603469 | 1.209245522 | 0.226568533 | 0.679565252 |
| Pigt | 3689.18136 | 0.084177195 | 0.049013264 | 1.717437033 | 0.08589935 | 0.491218621 |
| Cct4 | 4022.609911 | 0.084179455 | 0.057993927 | 1.451521882 | 0.146634594 | 0.593534375 |
| Sucla2 | 5968.439682 | 0.084203047 | 0.062773655 | 1.341375563 | 0.179798548 | 0.633459544 |
| Grm1 | 2015.654262 | 0.084211469 | 0.066967826 | 1.257491456 | 0.208575731 | 0.663405095 |
| Usp39 | 1312.175654 | 0.084238308 | 0.053478829 | 1.575171148 | 0.115216945 | 0.543478516 |
| Hectd2 | 667.443337 | 0.084249243 | 0.08754287 | 0.962376985 | 0.33586027 | 0.759354819 |
| Far2 | 569.9657833 | 0.084255509 | 0.075252053 | 1.119643993 | 0.2628655 | 0.705670719 |
| Tmed2 | 4128.813841 | 0.084291767 | 0.058208224 | 1.44810752 | 0.147586983 | 0.594737171 |
| Kremen2 | 41.82555899 | 0.084318065 | 0.0930714 | 0.905950324 | 0.364962146 | 0.777634019 |
| Pcx | 3872.697909 | 0.084321554 | 0.05405837 | 1.559824197 | 0.118801432 | 0.550992847 |
| Nr1d2 | 5403.613271 | 0.084340317 | 0.08155912 | 1.034100371 | 0.301089243 | 0.735062578 |
| Uap1 | 2151.64394 | 0.084348091 | 0.065504137 | 1.287675785 | 0.197858846 | 0.653535282 |
| Pcdhgb6 | 652.2091989 | 0.084384972 | 0.072196992 | 1.168815627 | 0.242477923 | 0.692772033 |
| Usp5 | 9597.998221 | 0.0843867 | 0.047490515 | 1.77691691 | 0.075581912 | 0.470597021 |
| Ptges2 | 1220.606399 | 0.084403559 | 0.060473923 | 1.395701724 | 0.162804335 | 0.614351408 |
| Chga | 6516.638227 | 0.084418218 | 0.063706479 | 1.325111974 | 0.185134015 | 0.637780733 |
| Grwd1 | 688.8024045 | 0.084425734 | 0.062858929 | 1.343098506 | 0.179240079 | 0.633029218 |
| Qprt | 15.72880647 | 0.084457695 | 0.070843279 | 1.19217653 | 0.233192034 | NA |
| Birc7 | 8.494131837 | 0.084461153 | 0.06314048 | 1.337670427 | 0.181003895 | NA |
| Heph | 360.6813782 | 0.084467911 | 0.088322147 | 0.956361615 | 0.33888957 | 0.761116259 |
| Immp2l | 102.2109603 | 0.084468182 | 0.092888968 | 0.909345687 | 0.363167681 | 0.776821522 |
| Bace1 | 4372.096654 | 0.084480107 | 0.053344413 | 1.583673023 | 0.113268147 | 0.540327972 |
| Fam150b | 39.22457132 | 0.084541312 | 0.089414601 | 0.945497848 | 0.344404769 | 0.764139862 |
| Aar2 | 1954.484811 | 0.084592423 | 0.057541949 | 1.470100045 | 0.14153466 | 0.587141261 |
| B630005N14Rik | 791.0056192 | 0.0845982 | 0.079819722 | 1.059865881 | 0.28920562 | 0.724444298 |
| Ptma | 8748.120686 | 0.084599669 | 0.044169819 | 1.915327523 | 0.055450748 | 0.413057568 |
| 5730577I03Rik | 67.81872095 | 0.084623378 | 0.095023325 | 0.890553744 | 0.373168624 | 0.782226671 |
| Gm16677 | 22.92660709 | 0.084655918 | 0.07746209 | 1.092869003 | 0.27445132 | 0.716185716 |
| Atp6v1c1 | 10630.10081 | 0.084702522 | 0.050670502 | 1.671633759 | 0.094596564 | 0.511963369 |
| Sept7 | 14513.41068 | 0.084710402 | 0.073616719 | 1.150695161 | 0.249857668 | 0.698236338 |
| Tac2 | 354.1943015 | 0.084710586 | 0.094014931 | 0.901033323 | 0.367570602 | 0.778899851 |
| Trpc5 | 382.4622469 | 0.084720099 | 0.08738542 | 0.969499253 | 0.332296153 | 0.757166892 |
| Rybp | 2625.415409 | 0.084731529 | 0.07577101 | 1.118257879 | 0.263456871 | 0.706200557 |
| Pnma2 | 3109.840353 | 0.08473917 | 0.058056814 | 1.459590437 | 0.144402669 | 0.589967032 |
| Pqbp1 | 2239.848351 | 0.084746694 | 0.053456918 | 1.585326963 | 0.112892066 | 0.540327972 |
| Hist2h3c1 | 69.52019692 | 0.084785599 | 0.094148758 | 0.900549313 | 0.367827995 | 0.77910024 |
| Mcfd2 | 2670.903835 | 0.084810468 | 0.049153541 | 1.725419276 | 0.08445194 | 0.488499377 |
| A430005L14Rik | 498.6868919 | 0.084821546 | 0.079317352 | 1.069394583 | 0.284891907 | 0.721641599 |
| Tspan31 | 2375.177829 | 0.084859275 | 0.05556186 | 1.527293634 | 0.126688012 | 0.563635546 |
| Fcna | 8.27766832 | 0.084924289 | 0.061488887 | 1.381132308 | 0.167238283 | NA |
| Esd | 2219.540545 | 0.084933481 | 0.057647859 | 1.473315434 | 0.140666001 | 0.58648517 |
| Sema3c | 1083.476865 | 0.084991606 | 0.068788629 | 1.23554732 | 0.216626877 | 0.670049713 |
| BC029722 | 2401.273901 | 0.085003264 | 0.055582544 | 1.52931584 | 0.126186165 | 0.562787717 |
| Tceb2 | 3249.967871 | 0.085015272 | 0.05057343 | 1.681026435 | 0.09275778 | 0.506868248 |
| Syne3 | 105.8273005 | 0.085072428 | 0.095096426 | 0.894591223 | 0.371005645 | 0.780761257 |
| Tmem127 | 6313.310676 | 0.085145459 | 0.042329134 | 2.01150958 | 0.044271659 | 0.380589751 |
| Maged2 | 1815.205831 | 0.085157098 | 0.067169053 | 1.267802568 | 0.204868465 | 0.658461308 |
| Nipa2 | 1219.979795 | 0.085184386 | 0.080350737 | 1.060156883 | 0.289073233 | 0.724444298 |
| Bbs9 | 760.1745907 | 0.085185122 | 0.066321438 | 1.28442815 | 0.198992189 | 0.65377551 |
| Otud7b | 2952.429699 | 0.085199063 | 0.072954413 | 1.16783974 | 0.242871415 | 0.692772033 |
| Tob2 | 1394.767803 | 0.085204954 | 0.068251697 | 1.248393198 | 0.211887099 | 0.666270994 |
| Ubald1 | 3725.507584 | 0.085277444 | 0.049595449 | 1.719461052 | 0.085530454 | 0.490104683 |
| Pef1 | 2749.388397 | 0.085303351 | 0.05626263 | 1.516163578 | 0.129477995 | 0.568388444 |
| Cttn | 4965.778795 | 0.085341816 | 0.050405501 | 1.69310522 | 0.090435441 | 0.502238744 |
| Wasl | 4956.996373 | 0.085407585 | 0.063456464 | 1.345924109 | 0.178326993 | 0.631799487 |
| Ripply3 | 11.10919622 | 0.08541851 | 0.067732581 | 1.261114067 | 0.207267753 | NA |
| Pvt1 | 369.3005322 | 0.085420709 | 0.075150873 | 1.136656246 | 0.255682017 | 0.701952734 |
| Ndufv1 | 6185.171454 | 0.085477536 | 0.053831619 | 1.58786858 | 0.11231606 | 0.538992394 |

| | | | | | | | | | | | | | | |
|---|---|---|---|---|---|---|---|---|---|---|---|---|---|---|
| Clec4d | 1.084798614 | 0.060484599 | 0.072840762 | 0.830367456 | 0.40633106 | NA | | | Rnf223 | 36.36356256 | 0.085503618 | 0.091268872 | 0.936832199 | 0.348844879 | 0.767659554 |
| Clec4e | 0 | NA | NA | NA | NA | NA | | | Mgst3 | 2307.03499 | 0.085506423 | 0.053997688 | 1.583520078 | 0.113302974 | 0.540327972 |
| Clec4f | 0.094392817 | 0.017895942 | 0.027818137 | 0.643319236 | 0.520016977 | NA | | | Mpp6 | 1409.470282 | 0.085540347 | 0.071166896 | 1.201968218 | 0.229375842 | 0.682503526 |
| Clec4g | 3.369466128 | -0.135668073 | 0.107077407 | -1.267009323 | 0.205151956 | NA | | | Rcan3 | 2166.267965 | 0.085561998 | 0.071096832 | 1.203457255 | 0.228799423 | 0.681764163 |
| Clec4n | 6.992929713 | -0.475495151 | 0.148866014 | -3.194114892 | 0.001402603 | NA | | | Apba1 | 9401.40803 | 0.08556441 | 0.052216966 | 1.638632368 | 0.101289847 | 0.52249159 |
| Clec9a | 1.090837823 | -0.027094434 | 0.075375517 | -0.35945934 | 0.71925149 | NA | | | Ndufa12 | 2541.607643 | 0.085601463 | 0.061258845 | 1.397373121 | 0.162301398 | 0.613957643 |
| Clhc1 | 5.619247619 | 0.084229664 | 0.141310262 | 0.596061905 | 0.551133872 | NA | | | Tdrd12 | 10.59576165 | 0.085604345 | 0.06872905 | 1.245533667 | 0.212935648 | NA |
| Clic3 | 6.26346902 | 0.164607011 | 0.14159867 | 1.162489806 | 0.245036563 | NA | | | Lphn3 | 3854.946134 | 0.085604401 | 0.064042283 | 1.336685668 | 0.181325262 | 0.634469716 |
| Clnk | 2.97680312 | 0.080992202 | 0.116628595 | 0.694445493 | 0.487402872 | NA | | | Snx16 | 991.1614126 | 0.085605747 | 0.071860942 | 1.191269475 | 0.233547813 | 0.685064073 |
| Clps | 0.192746203 | -0.000975302 | 0.031567365 | -0.030895906 | 0.975352555 | NA | | | Plrg1 | 1435.288892 | 0.085652387 | 0.05995532 | 1.428603627 | 0.153118191 | 0.601784319 |
| Clpsl2 | 0 | NA | NA | NA | NA | NA | | | Pcdhb20 | 449.6009236 | 0.085665844 | 0.074591739 | 1.14846289 | 0.250777524 | 0.699068333 |
| Clrn2 | 7.996239079 | 0.25158323 | 0.156610172 | 1.606429689 | 0.108179532 | NA | | | Gpr85 | 1424.532415 | 0.085748882 | 0.066737757 | 1.284863114 | 0.198840122 | 0.65377551 |
| Clrn3 | 0.116696964 | -0.020477583 | 0.027954224 | -0.732539847 | 0.463839131 | NA | | | Mpc1 | 3181.559217 | 0.085751087 | 0.051793361 | 1.655638583 | 0.097795032 | 0.518456434 |
| Cma1 | 0.609727828 | -0.037051079 | 0.059330555 | -0.624485625 | 0.532308708 | NA | | | Mto1 | 891.9436817 | 0.085810243 | 0.057548842 | 1.491085485 | 0.135939053 | 0.579435507 |
| Cma2 | 0 | NA | NA | NA | NA | NA | | | C530008M17Rik | 1973.451997 | 0.085890309 | 0.075337297 | 1.140076869 | 0.254254278 | 0.701619624 |
| Cmah | 6.407250863 | 0.081367871 | 0.142703912 | 0.570186691 | 0.568551082 | NA | | | Fam46a | 311.9802498 | 0.085890999 | 0.093893477 | 0.914770678 | 0.360312031 | 0.776337125 |
| Cmtm1 | 0.392633299 | 0.026946 | 0.048301387 | 0.557872169 | 0.576931677 | NA | | | Ptcd2 | 1142.282667 | 0.085942237 | 0.063583498 | 1.351643739 | 0.176489311 | 0.629083076 |
| Cmtm2a | 0.096071043 | -0.020098795 | 0.027825948 | -0.722304036 | 0.470107572 | NA | | | Hsp90b1 | 15752.54307 | 0.085946784 | 0.060471697 | 1.421272896 | 0.15523744 | 0.604578396 |
| Cmtm2b | 0 | NA | NA | NA | NA | NA | | | Tm2d3 | 1074.154362 | 0.0859613 | 0.066061416 | 1.301233091 | 0.193178682 | 0.649744561 |
| Cnfn | 0.612124479 | 0.05749822 | 0.056841187 | 1.011559104 | 0.311748908 | NA | | | 4930513N10Rik | 29.95671132 | 0.086011935 | 0.088295113 | 0.974141504 | 0.329986284 | 0.756141871 |
| Cnga1 | 0 | NA | NA | NA | NA | NA | | | Stmn1 | 24211.67332 | 0.086015814 | 0.04618757 | 1.862315196 | 0.062558675 | 0.437519166 |
| Cnga3 | 7.394195149 | -0.432416454 | 0.135990071 | -3.179764904 | 0.001473946 | NA | | | Spty2d1 | 922.5526142 | 0.086038695 | 0.066633079 | 1.291230975 | 0.196623597 | 0.652536983 |
| Cngb3 | 0.233393928 | -0.027475306 | 0.032191697 | -0.85349047 | 0.39338737 | NA | | | Slc5a3 | 1060.568687 | 0.086067173 | 0.073688661 | 1.167983947 | 0.24281324 | 0.692772033 |
| Cnpy1 | 1.353565994 | 0.029979155 | 0.07482703 | 0.400646063 | 0.688680727 | NA | | | Ndufaf1 | 751.5937688 | 0.086092314 | 0.060399458 | 1.425382225 | 0.154046743 | 0.60340295 |
| Cntf | 3.262999941 | 0.107163825 | 0.098631722 | 1.086504656 | 0.277255782 | NA | | | Tmem251 | 316.279217 | 0.086094254 | 0.082129247 | 1.048277651 | 0.294510703 | 0.72962173 |
| Col8a2 | 118.1302481 | 0.154562689 | 0.161210953 | 0.95876047 | NA | NA | | | 2810001G20Rik | 247.3767091 | 0.086152591 | 0.080546332 | 1.069602911 | 0.284798084 | 0.721641599 |
| Colec10 | 0.094392817 | 0.017895942 | 0.027818137 | 0.643319236 | 0.520016977 | NA | | | Samd4 | 1411.282434 | 0.086158614 | 0.074386566 | 1.158255031 | 0.246759977 | 0.696511265 |
| Cox7b2 | 5.399662925 | -0.04210873 | 0.145802866 | -0.288805913 | 0.772729907 | NA | | | Drg2 | 2262.261811 | 0.086174513 | 0.053175769 | 1.620559802 | 0.105112079 | 0.526359793 |
| Cox8c | 0 | NA | NA | NA | NA | NA | | | Cdca3 | 22.69534928 | 0.086207175 | 0.083536598 | 1.031968947 | 0.302086665 | 0.735974432 |
| Cp | 859.6799956 | -0.035654034 | 0.143638879 | -0.248219943 | NA | NA | | | Sec24d | 690.3225815 | 0.086214668 | 0.074340306 | 1.159729795 | 0.246158835 | 0.695662109 |
| Cpa1 | 0 | NA | NA | NA | NA | NA | | | Slc6a1 | 20951.09253 | 0.086215384 | 0.03951887 | 2.181625738 | 0.029137166 | 0.317415456 |
| Cpa3 | 1.006239153 | 0.013014933 | 0.062058454 | 0.209720547 | 0.833885788 | NA | | | Ptp4a1 | 421.5572657 | 0.086242986 | 0.079261892 | 1.088076303 | 0.276561424 | 0.717475105 |
| Cpa5 | 1.71620555 | 0.041323327 | 0.096663668 | 0.427495957 | 0.669018129 | NA | | | Uvrag | 1742.796682 | 0.086262086 | 0.059814119 | 1.442169305 | 0.149254618 | 0.597429754 |
| Cpb1 | 5.316919271 | -0.100469728 | 0.146862974 | -0.684105226 | 0.493908722 | NA | | | Yars | 4185.921649 | 0.086268485 | 0.046565659 | 1.852620284 | 0.063936803 | 0.441368101 |
| Cpb2 | 0 | NA | NA | NA | NA | NA | | | Tmem8c | 18.75197626 | 0.08628089 | 0.072668 | 1.187329909 | 0.235097515 | NA |
| Cphx1 | 0 | NA | NA | NA | NA | NA | | | Rom1 | 193.8409752 | 0.086336158 | 0.093381746 | 0.924550697 | 0.355199669 | 0.772206867 |
| Cphx2 | 0 | NA | NA | NA | NA | NA | | | Pappa2 | 172.7907333 | 0.086336792 | 0.091685571 | 0.941661711 | 0.346365864 | 0.765286747 |
| Cplx4 | 2.388063154 | 0.001138628 | 0.084311275 | 0.013505048 | 0.989224859 | NA | | | Sez6l2 | 19002.18718 | 0.086362575 | 0.048177975 | 1.792573767 | 0.073041104 | 0.463924389 |
| Cpn2 | 0 | NA | NA | NA | NA | NA | | | Nsfl1c | 3215.548756 | 0.086380576 | 0.056401133 | 1.531539721 | 0.125636057 | 0.562573788 |
| Cpsf4l | 0.295716657 | -0.010253415 | 0.03372058 | -0.304069933 | 0.761074613 | NA | | | F8a | 494.3545362 | 0.086402507 | 0.08066558 | 1.071119895 | 0.284115524 | 0.721514352 |
| Cpt1b | 2.04416705 | 0.107359693 | 0.101147295 | 1.061419315 | 0.288499383 | NA | | | Bod1 | 2741.268763 | 0.086451575 | 0.046135435 | 1.873864973 | 0.060949038 | 0.432354503 |
| Cpvl | 0.350688377 | 0.02015342 | 0.04180392 | 0.482094028 | 0.629739152 | NA | | | Cmss1 | 339.4917291 | 0.086475793 | 0.073856269 | 1.170865993 | 0.241652646 | 0.692393571 |
| Cpxcr1 | 0 | NA | NA | NA | NA | NA | | | Zfp521 | 539.2766053 | 0.086477894 | 0.085711051 | 1.008946837 | 0.313000131 | 0.745350804 |
| Cpz | 7.517777699 | 0.086076124 | 0.150796912 | 0.57080827 | 0.568129616 | NA | | | Scn1b | 17860.08847 | 0.086481376 | 0.054028691 | 1.60065651 | 0.109453019 | 0.535243338 |
| Crct1 | 0.698757213 | 0.06007153 | 0.05689194 | 1.055888217 | 0.291019288 | NA | | | Six4 | 90.61461893 | 0.086515264 | 0.093781601 | 0.922518526 | 0.356258172 | 0.772965825 |
| Crisp1 | 0 | NA | NA | NA | NA | NA | | | Tmem167b | 2337.092422 | 0.086529066 | 0.053395948 | 1.620517468 | 0.105121165 | 0.526359793 |
| Crisp2 | 0 | NA | NA | NA | NA | NA | | | Pls3 | 7486.020904 | 0.086531376 | 0.065027401 | 1.330690993 | 0.183290708 | 0.63672274 |
| Crisp3 | 0 | NA | NA | NA | NA | NA | | | Abcc9 | 481.7000829 | 0.086531829 | 0.082752142 | 1.045674797 | 0.295711203 | 0.730775932 |
| Crisp4 | 0 | NA | NA | NA | NA | NA | | | Txndc16 | 3279.849748 | 0.086544993 | 0.065246201 | 1.326437281 | 0.184694897 | 0.637780733 |
| Crnn | 0 | NA | NA | NA | NA | NA | | | 9130011E15Rik | 867.8905073 | 0.086560584 | 0.059648267 | 1.451183543 | 0.146728758 | 0.593534375 |
| Crp | 0 | NA | NA | NA | NA | NA | | | Tmem14a | 1139.379778 | 0.086599452 | 0.063971836 | 1.353712143 | 0.175828232 | 0.628658305 |
| Crtam | 2.957490911 | 0.055928676 | 0.120427015 | 0.464419677 | 0.642347095 | NA | | | Napg | 6789.598311 | 0.086627488 | 0.052896587 | 1.637676307 | 0.101489236 | 0.522606076 |
| Crx | 0.508673765 | -0.022465145 | 0.056012253 | -0.401075549 | 0.688364502 | NA | | | Mctp1 | 2037.603057 | 0.086636578 | 0.084019796 | 1.031144823 | 0.302472911 | 0.735974513 |
| Crxos1 | 3.825199017 | 0.261274637 | 0.129334325 | 2.020149226 | 0.043367911 | NA | | | Vdac3 | 5628.43881 | 0.086640238 | 0.053292384 | 1.625752726 | 0.104002269 | 0.523645898 |
| Cryaa | 3.527884846 | 0.054035705 | 0.114948909 | 0.470084542 | 0.638294618 | NA | | | Avpi1 | 697.2754858 | 0.086653934 | 0.072125055 | 1.201440108 | 0.229580526 | 0.682539238 |
| Cryba1 | 1.263743685 | -9.22E-05 | 0.086134906 | -0.001070538 | 0.999145834 | NA | | | Gamt | 340.1396624 | 0.086694098 | 0.07773832 | 1.115204166 | 0.264762941 | 0.707370515 |
| Cryba2 | 6.796622269 | 0.184559996 | 0.13483829 | 1.368750641 | 0.171077235 | NA | | | Mt1 | 5018.895583 | 0.086695853 | 0.084665186 | 1.023984668 | 0.305842517 | 0.738179891 |
| Crybb2 | 0.098353386 | -0.020212831 | 0.02786466 | -0.725393061 | 0.468210929 | NA | | | Cytl1 | 31.06667745 | 0.086721331 | 0.07502898 | 1.155837789 | 0.247747513 | 0.696944196 |
| Crygb | 0 | NA | NA | NA | NA | NA | | | Mthfd2l | 149.0474252 | 0.086728611 | 0.094854961 | 0.914328681 | 0.360544164 | 0.776337125 |
| Crygc | 0 | NA | NA | NA | NA | NA | | | Ncapg2 | 225.4243477 | 0.086729848 | 0.086506583 | 1.002580909 | 0.316063111 | 0.745993052 |
| Crygd | 0 | NA | NA | NA | NA | NA | | | Clpx | 1250.72679 | 0.086731308 | 0.070532112 | 1.229671214 | 0.21882025 | 0.671604514 |
| Cryge | 0.079983654 | 0.016740032 | 0.027674178 | 0.604897175 | 0.54524732 | NA | | | Ajap1 | 2019.212675 | 0.086758164 | 0.067572355 | 1.28392986 | 0.199166498 | 0.65377551 |
| Crygf | 0 | NA | NA | NA | NA | NA | | | Eif3i | 2662.762687 | 0.086781569 | 0.049032178 | 1.769890158 | 0.076745441 | 0.472560915 |
| Crygs | 0.495946458 | 0.007147691 | 0.044404124 | 0.160969084 | 0.87211775 | NA | | | Efna3 | 1430.919074 | 0.086796564 | 0.061768397 | 1.405193742 | 0.159963675 | 0.612126538 |
| Csf2 | 0 | NA | NA | NA | NA | NA | | | Ssr2 | 2046.677488 | 0.086798766 | 0.051534674 | 1.684278937 | 0.092127774 | 0.505579538 |
| Csf3 | 0.295416859 | -0.013215976 | 0.033806277 | -0.390932592 | 0.695847062 | NA | | | Maea | 3413.970505 | 0.086804696 | 0.043719138 | 1.98550796 | 0.047087985 | 0.391304391 |
| Csl | 0 | NA | NA | NA | NA | NA | | | Ppargc1a | 2273.64967 | 0.086831102 | 0.067929146 | 1.278259889 | 0.201157806 | 0.654716629 |
| Csn1s1 | 0.276102474 | 0.031058433 | 0.034913626 | 0.889579115 | 0.373691925 | NA | | | Yipf4 | 1230.732479 | 0.08683229 | 0.074160145 | 1.170875395 | 0.241648867 | 0.692393571 |
| Csn1s2a | 0 | NA | NA | NA | NA | NA | | | Rfk | 2251.865167 | 0.086843077 | 0.057323232 | 1.514971741 | 0.129779561 | 0.568885619 |
| Csn1s2b | 0 | NA | NA | NA | NA | NA | | | Stx1b | 10289.50076 | 0.086858141 | 0.036425383 | 2.384549863 | 0.01710003 | 0.249184268 |
| Csn2 | 0.079983654 | 0.016740032 | 0.027674178 | 0.604897175 | 0.54524732 | NA | | | Kcnk1 | 3794.988084 | 0.086872322 | 0.06033726 | 1.439779038 | 0.149929924 | 0.598234686 |
| Csn3 | 0 | NA | NA | NA | NA | NA | | | Arhgap26 | 5313.781019 | 0.086905199 | 0.060976322 | 1.425228602 | 0.154091131 | 0.60340295 |
| Csnka2ip | 0.48794879 | 0.064131883 | 0.049799661 | 1.287797577 | 0.197816436 | NA | | | Six3 | 469.0312369 | 0.086957279 | 0.090481709 | 0.961048152 | 0.336527957 | 0.75970798 |
| Cspg5 | 10656.73371 | -0.232533607 | 0.113845531 | -2.042536101 | NA | NA | | | Krr1 | 1824.523853 | 0.086967169 | 0.062369939 | 1.394376366 | 0.163203981 | 0.614351408 |
| Csrp2bp | 1388.527159 | 0.203391239 | 0.102418302 | 1.985887628 | NA | NA | | | Setbp1 | 1661.148975 | 0.086970307 | 0.060556298 | 1.436189289 | 0.150948486 | 0.598846773 |
| Csrp3 | 0.321356363 | -0.01100297 | 0.034551013 | -0.318455781 | 0.750139232 | NA | | | Sema3e | 1228.362807 | 0.086971745 | 0.079988401 | 1.08730445 | 0.276902283 | 0.7177927 |
| Cst10 | 0 | NA | NA | NA | NA | NA | | | Scrn3 | 1336.159214 | 0.086971755 | 0.066554586 | 1.306773291 | 0.191289719 | 0.646682644 |
| Cst11 | 0.691297128 | 0.034620722 | 0.06074969 | 0.569891329 | 0.568751406 | NA | | | Thumpd1 | 2285.854121 | 0.086981982 | 0.056153221 | 1.549011446 | 0.121378969 | 0.555673148 |
| Cst12 | 0.800245648 | -0.038051497 | 0.068702165 | -0.553861688 | 0.579673506 | NA | | | Kcnab1 | 5216.535938 | 0.08698588 | 0.081888951 | 1.062241955 | 0.288125858 | 0.724103151 |
| Cst13 | 0 | NA | NA | NA | NA | NA | | | Letm1 | 3511.298388 | 0.086988186 | 0.043353368 | 2.006492014 | 0.04480378 | 0.381265206 |
| Cst8 | 0 | NA | NA | NA | NA | NA | | | Spon1 | 3256.201236 | 0.086990371 | 0.06249781 | 1.391894718 | 0.163954282 | 0.614351408 |
| Cst9 | 0 | NA | NA | NA | NA | NA | | | Mfsd5 | 1393.953558 | 0.087000683 | 0.051720788 | 1.682122153 | 0.092545155 | 0.506868248 |
| Csta | 0 | NA | NA | NA | NA | NA | | | Fbxo46 | 473.968943 | 0.087002744 | 0.071129925 | 1.223152485 | 0.221272108 | 0.672757795 |
| Cstl1 | 5.864419947 | 0.061379307 | 0.14608311 | 0.420167034 | 0.674363436 | NA | | | Crnkl1 | 795.1444645 | 0.087024407 | 0.070069358 | 1.241975238 | 0.214245698 | 0.668413448 |
| Ctag2 | 0 | NA | NA | NA | NA | NA | | | Mrpl20 | 1388.2695 | 0.087026465 | 0.058251232 | 1.493984974 | 0.135179547 | 0.578555714 |
| Ctf2 | 0 | NA | NA | NA | NA | NA | | | Cdyl2 | 1095.354892 | 0.087066997 | 0.094652659 | 0.91985791 | 0.357647016 | 0.773905615 |
| Ctla2a | 199.18628 | 0.282252577 | 0.162631217 | 1.735537511 | NA | NA | | | Prkar1b | 41686.5328 | 0.087089929 | 0.053957964 | 1.614032876 | 0.106520295 | 0.53002185 |
| Ctla4 | 2.082501219 | 0.057465537 | 0.099780127 | 0.575921667 | 0.564668129 | NA | | | Hnrnpab | 4230.260515 | 0.087150503 | 0.05409738 | 1.610993029 | 0.107181242 | 0.532011007 |
| Ctrb1 | 0.167300494 | 0.024299763 | 0.031030539 | 0.783091891 | 0.433573153 | NA | | | Rad21l | 24.28709945 | 0.08715581 | 0.078616829 | 1.108615181 | 0.267596226 | 0.710152314 |
| Ctrc | 0 | NA | NA | NA | NA | NA | | | Gprasp1 | 27616.76366 | 0.087196375 | 0.062068907 | 1.404831802 | 0.1600713 | 0.612215825 |
| Cts3 | 0 | NA | NA | NA | NA | NA | | | Map2k5 | 1353.735923 | 0.087234525 | 0.052917682 | 1.648494812 | 0.099251174 | 0.520390402 |
| Cts6 | 0 | NA | NA | NA | NA | NA | | | Mrpl1 | 563.4331806 | 0.087236606 | 0.064191358 | 1.359008571 | 0.174143873 | 0.626964546 |
| Cts7 | 0 | NA | NA | NA | NA | NA | | | Gabra3 | 7375.196708 | 0.08724251 | 0.065271726 | 1.336604919 | 0.181351633 | 0.634469716 |
| Cts8 | 0 | NA | NA | NA | NA | NA | | | Pcdhb21 | 86.36691118 | 0.087247619 | 0.094621917 | 0.922065645 | 0.356494336 | 0.773064269 |
| Cts8-ps | 0 | NA | NA | NA | NA | NA | | | 1110004F10Rik | 3058.448699 | 0.087247781 | 0.053504643 | 1.630658115 | 0.102962479 | 0.522975271 |
| Ctse | 3.374962363 | 0.129652481 | 0.101517341 | 1.277146145 | 0.201550656 | NA | | | Nt5c3 | 1731.78508 | 0.087255592 | 0.068533098 | 1.273189072 | 0.202950958 | 0.656606302 |
| Ctsg | 0.286698134 | 0.007861574 | 0.038801625 | 0.202609399 | 0.839440343 | NA | | | Hsd17b11 | 1663.063004 | 0.087285048 | 0.060585988 | 1.440680461 | 0.149674977 | 0.598098567 |
| Ctsj | 0 | NA | NA | NA | NA | NA | | | Gps1 | 5265.501852 | 0.087295075 | 0.04347302 | 2.008028766 | 0.044640235 | 0.380589751 |
| Ctsll3 | 0 | NA | NA | NA | NA | NA | | | Dusp26 | 3137.88749 | 0.087315192 | 0.067130656 | 1.300675392 | 0.193369589 | 0.649744561 |
| Ctsm | 0 | NA | NA | NA | NA | NA | | | Sult1a1 | 361.0962343 | 0.087354408 | 0.08187541 | 1.066918729 | 0.286008537 | 0.722148665 |
| Ctsq | 0 | NA | NA | NA | NA | NA | | | Chpf | 3308.311216 | 0.087355525 | 0.062647823 | 1.39439043 | 0.163199736 | 0.614351408 |
| Ctsr | 0 | NA | NA | NA | NA | NA | | | Rtf1 | 4677.623887 | 0.087362763 | 0.055115171 | 1.585094666 | 0.112944827 | 0.540327972 |
| Ctsw | 4.625655994 | -0.057633096 | 0.132025276 | -0.436530771 | 0.662451679 | NA | | | Mdh2 | 10944.66689 | 0.087376902 | 0.045145197 | 1.935463957 | 0.052933388 | 0.406864177 |
| Cxcl1 | 3.061675514 | 0.281468208 | 0.11865801 | 2.372096143 | 0.017687488 | NA | | | Slc41a1 | 3421.622285 | 0.087384065 | 0.061494506 | 1.421006047 | 0.155315002 | 0.604578396 |
| Cxcl11 | 0.946603521 | -0.013609344 | 0.077083244 | -0.176553859 | 0.85985884 | NA | | | 2700060E02Rik | 2032.891568 | 0.087404089 | 0.05780729 | 1.511990771 | 0.13053621 | 0.569619261 |
| Cxcl13 | 2.081694735 | -0.132821048 | 0.089437611 | -1.48506928 | 0.137525473 | NA | | | Prss23 | 716.5379386 | 0.087415992 | 0.065528292 | 1.334019079 | 0.182197604 | 0.635245711 |
| Cxcl17 | 0 | NA | NA | NA | NA | NA | | | Apoh | 15.83451376 | 0.08743866 | 0.076604287 | 1.141432987 | 0.253689784 | NA |
| Cxcl2 | 0.379249494 | 0.016397727 | 0.036387388 | 0.450643137 | 0.652246771 | NA | | | Gabrb2 | 13276.46822 | 0.087445029 | 0.075600322 | 1.156675364 | 0.247405019 | 0.696744638 |
| Cxcl3 | 0.096071043 | -0.020098795 | 0.027825948 | -0.722304036 | 0.470107572 | NA | | | Spata2l | 2172.480329 | 0.087458006 | 0.072237259 | 1.210704928 | 0.226008513 | 0.67844589 |
| Cxcl9 | 11.7388471 | 0.0620493 | 0.049980769 | 1.241463479 | NA | NA | | | Cirh1a | 1279.991974 | 0.087461908 | 0.060597714 | 1.443320259 | 0.148930275 | 0.596789291 |
| Cxcr1 | 0 | NA | NA | NA | NA | NA | | | Zfp935 | 269.2457117 | 0.08746786 | 0.093504594 | 0.935439174 | 0.349562016 | 0.768306529 |
| Cxcr3 | 2.801578802 | 0.029349885 | 0.103400683 | 0.283846142 | 0.776528292 | NA | | | Ywhaq | 13941.4118 | 0.087516376 | 0.054754411 | 1.598343862 | 0.109966471 | 0.535492107 |
| Cxcr5 | 0 | NA | NA | NA | NA | NA | | | Dld | 4292.888471 | 0.087520908 | 0.057200368 | 1.530075961 | 0.125997928 | 0.562787717 |
| Cxcr6 | 1.83817932 | -0.007441211 | 0.084213542 | -0.088361215 | 0.929589587 | NA | | | Nol11 | 1113.205864 | 0.087592723 | 0.06494562 | 1.348708704 | 0.177430549 | 0.629852432 |
| Cyct | 0.107716603 | -0.020477583 | 0.027954224 | -0.732539849 | 0.46383913 | NA | | | Taf10 | 839.0475278 | 0.087598725 | 0.071697378 | 1.221784227 | 0.221789235 | 0.672987912 |
| Cylc1 | 0 | NA | NA | NA | NA | NA | | | Synrg | 4257.245829 | 0.087611614 | 0.045966727 | 1.90597895 | 0.056652937 | 0.417225452 |
| Cylc2 | 0 | NA | NA | NA | NA | NA | | | 2810403D21Rik | 9.585207386 | 0.087633712 | 0.06250364 | 1.402057749 | 0.160898003 | NA |
| Cym | 1.731090764 | -0.014411747 | 0.092176278 | -0.156349844 | 0.87575727 | NA | | | Pcdhb22 | 208.1356312 | 0.087643176 | 0.0813599 | 1.077228173 | 0.281378342 | 0.720080692 |
| Cyp11b1 | 0.491454083 | 0.009574229 | 0.044346165 | 0.215897555 | 0.829067603 | NA | | | Cplx1 | 33806.59349 | 0.087651385 | 0.055975193 | 1.565896972 | 0.117372739 | 0.548468184 |
| Cyp11b2 | 0.211846316 | 0.025707621 | 0.031860375 | 0.806883811 | 0.419733426 | NA | | | Adh5 | 1949.050324 | 0.087672417 | 0.050110678 | 1.749575575 | 0.080191577 | 0.47917024 |
| Cyp17a1 | 0.289473745 | 0.012204471 | 0.038954522 | 0.313300481 | 0.754052385 | NA | | | Mtch1 | 14983.17587 | 0.087685514 | 0.051330997 | 1.708237106 | 0.087592356 | 0.495597906 |
| Cyp1a1 | 3.824364194 | 0.007771535 | 0.133483278 | 0.058221041 | 0.95357256 | NA | | | Calu | 3851.00062 | 0.087691342 | 0.049542089 | 1.770037219 | 0.076720941 | 0.472560915 |
| Cyp1a2 | 0.196680618 | -0.003857632 | 0.031597546 | -0.122086447 | 0.902830556 | NA | | | Amigo3 | 52.96610658 | 0.087699909 | 0.094141897 | 0.931571512 | 0.351558011 | 0.769244462 |
| Cyp21a1 | 1.666562999 | 0.129478863 | 0.086451318 | 1.497708378 | 0.134209034 | NA | | | Mgat3 | 8853.65715 | 0.087715572 | 0.053595386 | 1.636625451 | 0.101708754 | 0.522606076 |
| Cyp24a1 | 1.70862315 | 0.086528132 | 0.092714498 | 0.933275096 | 0.35067795 | NA | | | Pfn2 | 16175.77163 | 0.087716759 | 0.053443104 | 1.641311078 | 0.100732858 | 0.521542547 |
| Cyp26a1 | 4.792384107 | 0.049591066 | 0.128326215 | 0.386445325 | 0.699166888 | NA | | | Gapdh | 53989.88843 | 0.087751366 | 0.051177279 | 1.714654772 | 0.08640854 | 0.492030755 |
| Cyp26c1 | 0.167300494 | 0.024299763 | 0.031030539 | 0.783091891 | 0.433573153 | NA | | | Cd38 | 118.2301469 | 0.087782615 | 0.092974821 | 0.94415471 | 0.345090595 | 0.764727076 |
| Cyp2a12 | 0 | NA | NA | NA | NA | NA | | | Cml3 | 202.9223107 | 0.087783695 | 0.093224135 | 0.941641297 | 0.346376319 | 0.765286747 |
| Cyp2a22 | 0 | NA | NA | NA | NA | NA | | | Ociad1 | 11536.28788 | 0.087792089 | 0.048178322 | 1.822232196 | 0.068419761 | 0.451346709 |
| Cyp2a4 | 0.198019769 | 0.025308392 | 0.031628095 | 0.80018706 | 0.423602426 | NA | | | Mfge8 | 6569.319254 | 0.087803815 | 0.066677052 | 1.316852076 | 0.187888213 | 0.641781416 |
| Cyp2ab1 | 0.556822015 | 0.070925085 | 0.052229705 | 1.357945369 | 0.17448102 | NA | | | Ethe1 | 439.7883704 | 0.087805048 | 0.067148011 | 1.307634382 | 0.190997349 | 0.646547945 |
| Cyp2b10 | 1.93622122 | -0.10604424 | 0.079189005 | -1.339128328 | 0.1805289 | NA | | | G3bp1 | 2436.892835 | 0.08780554 | 0.057000874 | 1.540424452 | 0.123456925 | 0.558323565 |
| Cyp2b13 | 0 | NA | NA | NA | NA | NA | | | Mrps18a | 1397.951415 | 0.087811101 | 0.06197653 | 1.416844424 | 0.156528418 | 0.60719411 |
| Cyp2b19 | 0 | NA | NA | NA | NA | NA | | | Igf1 | 334.7252548 | 0.087831805 | 0.086651849 | 1.013617204 | 0.310765445 | 0.743686797 |
| Cyp2b23 | 0.389457872 | -0.034711543 | 0.048622122 | -0.713904319 | 0.475286347 | NA | | | St13 | 4256.461169 | 0.087846578 | 0.050226602 | 1.749004991 | 0.080290157 | 0.479231997 |
| Cyp2b9 | 0 | NA | NA | NA | NA | NA | | | Sdhb | 4338.001759 | 0.087863415 | 0.048914449 | 1.796267085 | 0.07245205 | 0.462203987 |
| Cyp2c29 | 0 | NA | NA | NA | NA | NA | | | Bub3 | 2552.057001 | 0.087867293 | 0.051487377 | 1.706579323 | 0.087900271 | 0.495973914 |
| Cyp2c37 | 0 | NA | NA | NA | NA | NA | | | Ttc28 | 818.2303 | 0.087875361 | 0.07211124 | 1.218608384 | 0.222992865 | 0.674048714 |
| Cyp2c38 | 0 | NA | NA | NA | NA | NA | | | Oxr1 | 11650.02434 | 0.087881657 | 0.066631379 | 1.318922996 | 0.187194857 | 0.641692692 |
| Cyp2c39 | 0 | NA | NA | NA | NA | NA | | | Il33 | 3007.669493 | 0.08790318 | 0.088462748 | 0.993674533 | 0.320381345 | 0.750259458 |
| Cyp2c40 | 0 | NA | NA | NA | NA | NA | | | Ppp2r5d | 4937.991577 | 0.087912739 | 0.038614657 | 2.276667642 | 0.02280608 | 0.2841176 |
| Cyp2c50 | 0 | NA | NA | NA | NA | NA | | | Mpp7 | 230.502208 | 0.087944391 | 0.087861917 | 1.000938672 | 0.316856459 | 0.746215455 |
| Cyp2c53-ps | 0 | NA | NA | NA | NA | NA | | | Ubxn2a | 2180.107869 | 0.087947339 | 0.073395631 | 1.198263958 | 0.230814273 | 0.683402059 |
| Cyp2c54 | 0 | NA | NA | NA | NA | NA | | | Znhit1 | 1114.694852 | 0.087958747 | 0.063644339 | 1.382035681 | 0.166960746 | 0.617356019 |
| Cyp2c55 | 0.755177424 | 0.04798083 | 0.063384139 | 0.756984806 | 0.449058965 | NA | | | Trmt61b | 308.3098543 | 0.087959432 | 0.075738431 | 1.161357989 | 0.245496346 | 0.69504627 |
| Cyp2c65 | 0.404910094 | -0.043499547 | 0.040365123 | -1.077651771 | 0.281189189 | NA | | | Acot3 | 95.90063736 | 0.087963954 | 0.094725428 | 0.92862028 | 0.353085908 | 0.770519168 |
| Cyp2c66 | 0 | NA | NA | NA | NA | NA | | | Atp5g1 | 4529.742687 | 0.087980616 | 0.05757964 | 1.527981348 | 0.126517169 | 0.563050456 |
| Cyp2c67 | 0 | NA | NA | NA | NA | NA | | | Cdh13 | 2807.203556 | 0.087982313 | 0.078970108 | 1.114121736 | 0.265226965 | 0.707434245 |
| Cyp2c68 | 0 | NA | NA | NA | NA | NA | | | Abi1 | 4462.462995 | 0.088026376 | 0.064664226 | 1.361283993 | 0.17342396 | 0.625247872 |
| Cyp2c69 | 0 | NA | NA | NA | NA | NA | | | Eif4h | 14478.46463 | 0.088027409 | 0.039819342 | 2.210669626 | 0.027058723 | 0.306595176 |
| Cyp2c70 | 0.116696964 | -0.020477583 | 0.027954224 | -0.732539847 | 0.463839131 | NA | | | Nipal3 | 6381.032078 | 0.088034399 | 0.045863942 | 1.919468636 | 0.054925052 | 0.412110852 |
| Cyp2d10 | 0 | NA | NA | NA | NA | NA | | | Tmx3 | 2317.274502 | 0.088045114 | 0.078371865 | 1.123427579 | 0.261255942 | 0.704173028 |
| Cyp2d11 | 0 | NA | NA | NA | NA | NA | | | Extl3 | 9215.509741 | 0.088057565 | 0.049130631 | 1.792314966 | 0.073082527 | 0.463984612 |
| Cyp2d12 | 0 | NA | NA | NA | NA | NA | | | Myo18b | 96.79263086 | 0.08805779 | 0.094329044 | 0.933517247 | 0.35055297 | 0.768667813 |
| Cyp2d13 | 0 | NA | NA | NA | NA | NA | | | Coa3 | 2016.328721 | 0.088075661 | 0.061987314 | 1.420865907 | 0.155355746 | 0.604578396 |
| Cyp2d26 | 1.875993474 | 0.071241832 | 0.093101132 | 0.765209084 | 0.444147052 | NA | | | 1810037I17Rik | 1371.377611 | 0.088226951 | 0.053475848 | 1.649846673 | 0.0989743 | 0.520271327 |
| Cyp2d34 | 0 | NA | NA | NA | NA | NA | | | Dnajc6 | 15129.80773 | 0.08823239 | 0.052030291 | 1.695788883 | 0.089925869 | 0.500949827 |
| Cyp2d37-ps | 0 | NA | NA | NA | NA | NA | | | Socs2 | 488.6314499 | 0.088234564 | 0.074506167 | 1.184258544 | 0.236310734 | 0.687217161 |
| Cyp2d40 | 0 | NA | NA | NA | NA | NA | | | Mlf1 | 40.1905445 | 0.088243868 | 0.084815715 | 1.040418842 | 0.298145352 | 0.732930969 |
| Cyp2d9 | 0 | NA | NA | NA | NA | NA | | | Bach2 | 470.4892537 | 0.088261328 | 0.088939581 | 0.992374006 | 0.321015114 | 0.750352874 |
| Cyp2g1 | 3.774254895 | -0.005822804 | 0.074314066 | -0.078353992 | 0.93754647 | NA | | | Exoc3 | 5163.121147 | 0.088268694 | 0.051681646 | 1.707931178 | 0.087649113 | 0.495597906 |
| Cyp2j11 | 0 | NA | NA | NA | NA | NA | | | Caly | 5312.493231 | 0.088271331 | 0.061777848 | 1.428850859 | 0.153047104 | 0.601704476 |
| Cyp2j13 | 2.94176132 | -0.210737144 | 0.115223812 | -1.828937432 | 0.067408982 | NA | | | Plekhg1 | 890.4909819 | 0.088275084 | 0.093786506 | 0.941234384 | 0.34658476 | 0.765286747 |
| Cyp2j5 | 0 | NA | NA | NA | NA | NA | | | Panx2 | 4005.646339 | 0.088278234 | 0.048475445 | 1.821091768 | 0.068592908 | 0.452078306 |
| Cyp2j8 | 3.210243929 | -0.128843303 | 0.115225143 | -1.118187402 | 0.263486964 | NA | | | 4930441O14Rik | 161.7037849 | 0.088280588 | 0.085495215 | 1.03257929 | 0.301800824 | 0.735974432 |
| Cyp2t4 | 0.094392817 | 0.017895942 | 0.027818137 | 0.643319236 | 0.520016977 | NA | | | Eif5a | 10577.08237 | 0.088329197 | 0.04030056 | 2.191761052 | 0.028396765 | 0.313443317 |
| Cyp2w1 | 0 | NA | NA | NA | NA | NA | | | Tmem196 | 447.4272723 | 0.088353774 | 0.072771075 | 1.214133141 | 0.224696887 | 0.676420098 |
| Cyp3a11 | 0 | NA | NA | NA | NA | NA | | | Pitpna | 15067.37498 | 0.08835814 | 0.047611265 | 1.855824266 | 0.063478611 | 0.440995266 |
| Cyp3a16 | 0 | NA | NA | NA | NA | NA | | | Nup35 | 607.2336138 | 0.088359085 | 0.067272447 | 1.313451336 | 0.189030909 | 0.643290796 |
| Cyp3a25 | 0 | NA | NA | NA | NA | NA | | | 4930453N24Rik | 575.0473276 | 0.088396443 | 0.079313249 | 1.114523032 | 0.265054869 | 0.707370515 |
| Cyp3a41a | 0 | NA | NA | NA | NA | NA | | | Poll | 463.4766388 | 0.088447697 | 0.07841595 | 1.127929934 | 0.259349511 | 0.703893404 |
| Cyp3a41b | 0 | NA | NA | NA | NA | NA | | | Eln | 317.1344407 | 0.088451034 | 0.091348184 | 0.968284539 | 0.332902284 | 0.757641999 |
| Cyp3a44 | 0 | NA | NA | NA | NA | NA | | | D10Wsu102e | 1130.866456 | 0.088456875 | 0.066332365 | 1.333540198 | 0.182354593 | 0.635245711 |
| Cyp3a57 | 1.037445604 | 0.04466278 | 0.065037715 | 0.686721228 | 0.492258417 | NA | | | Mecp2 | 7058.127029 | 0.088467667 | 0.067133875 | 1.317779824 | 0.187577364 | 0.641776649 |
| Cyp3a59 | 0 | NA | NA | NA | NA | NA | | | Csnk1a1 | 7095.191343 | 0.088474528 | 0.065445772 | 1.351875376 | 0.176415186 | 0.629083076 |
| Cyp4a10 | 0 | NA | NA | NA | NA | NA | | | Ndfip2 | 4230.045981 | 0.0885011 | 0.075311711 | 1.175130642 | 0.239942453 | 0.690751361 |
| Cyp4a12a | 0 | NA | NA | NA | NA | NA | | | Cd300lg | 9.523451092 | 0.088566592 | 0.058354239 | 1.517740511 | 0.129079827 | NA |
| Cyp4a12b | 3.14207922 | -0.096682315 | 0.116253188 | -0.831653017 | 0.405604827 | NA | | | Ghsr | 15.88990149 | 0.088572176 | 0.044265631 | 2.000924269 | NA | NA |
| Cyp4a14 | 0 | NA | NA | NA | NA | NA | | | Tceal3 | 3202.221898 | 0.088574254 | 0.070635506 | 1.253962198 | 0.209855746 | 0.664421221 |
| Cyp4a29-ps | 0 | NA | NA | NA | NA | NA | | | Dhdds | 3227.124544 | 0.088579737 | 0.053635079 | 1.651526185 | 0.098631178 | 0.520050241 |
| Cyp4a30b-ps | 0 | NA | NA | NA | NA | NA | | | 1810041L15Rik | 2953.868386 | 0.088583538 | 0.073947461 | 1.197925349 | 0.23094608 | 0.683402059 |
| Cyp4a31 | 0 | NA | NA | NA | NA | NA | | | Unc5a | 5471.624618 | 0.088585107 | 0.077921131 | 1.136856021 | 0.25559848 | 0.701952734 |
| Cyp4a32 | 0 | NA | NA | NA | NA | NA | | | Prdm2 | 3739.80308 | 0.08861503 | 0.069338918 | 1.277998451 | 0.201249972 | 0.654716629 |
| Cyp4f18 | 3.549067979 | 0.149919138 | 0.092241097 | 1.625296564 | 0.104099383 | NA | | | Dok5 | 799.5301928 | 0.088652164 | 0.069841112 | 1.269340671 | 0.204319587 | 0.65778952 |
| Cyp4f39 | 0.765385359 | -0.115948354 | 0.064415651 | -1.800002827 | 0.071860192 | NA | | | Timm22 | 1028.551786 | 0.088682235 | 0.054692002 | 1.621484539 | 0.104913763 | 0.525690695 |
| Cyp4f40 | 0 | NA | NA | NA | NA | NA | | | Limk1 | 2738.60956 | 0.088697593 | 0.052066392 | 1.703547895 | 0.088465583 | 0.497310781 |
| Cyp4f41-ps | 3.218777658 | 0.102065351 | 0.119120128 | 0.856827075 | 0.39154046 | NA | | | Ap2b1 | 19084.76379 | 0.088703351 | 0.044155168 | 2.008900797 | 0.044547655 | 0.380589751 |
| Cyp7a1 | 0.206069989 | -0.026776453 | 0.031767726 | -0.842882251 | 0.399294301 | NA | | | Zfp942 | 547.3519317 | 0.088721611 | 0.086436454 | 1.026437418 | 0.304685448 | 0.737453291 |
| Cyp8b1 | 0 | NA | NA | NA | NA | NA | | | Gabarapl2 | 5004.294183 | 0.088734108 | 0.041547767 | 2.135713038 | 0.032702811 | 0.330348423 |
| Cypt1 | 0 | NA | NA | NA | NA | NA | | | Ppm1a | 6948.59036 | 0.088739017 | 0.049647908 | 1.78736671 | 0.073878238 | 0.466430192 |
| Cypt10 | 0 | NA | NA | NA | NA | NA | | | Puf60 | 5330.702898 | 0.088787677 | 0.054550399 | 1.627626539 | 0.103604098 | 0.523462117 |
| Cypt12 | 0.258977192 | 0.029727007 | 0.03417262 | 0.869907163 | 0.38435114 | NA | | | Atp2b4 | 14382.58423 | 0.088803017 | 0.071936242 | 1.234468393 | 0.217028417 | 0.670185967 |
| Cypt14 | 0 | NA | NA | NA | NA | NA | | | Gpr146 | 1111.181766 | 0.088805432 | 0.063980133 | 1.388015738 | 0.165132253 | 0.615077498 |
| Cypt15 | 0 | NA | NA | NA | NA | NA | | | Dcps | 492.3686965 | 0.088834622 | 0.067359095 | 1.318821483 | 0.1872288 | 0.641692692 |
| Cypt2 | 0 | NA | NA | NA | NA | NA | | | Rnf14 | 15670.97569 | 0.08885741 | 0.050129708 | 1.772549915 | 0.076303317 | 0.472435997 |
| Cypt3 | 0 | NA | NA | NA | NA | NA | | | Tmx2 | 9177.878122 | 0.088870068 | 0.04455064 | 1.994810153 | 0.04606359 | 0.387222051 |
| Cypt4 | 0 | NA | NA | NA | NA | NA | | | Cited4 | 233.5489674 | 0.088873618 | 0.09229641 | 0.96291522 | 0.335590071 | 0.759126439 |
| Cypt7 | 0 | NA | NA | NA | NA | NA | | | Gdf10 | 382.3402594 | 0.088876768 | 0.087053319 | 1.020946347 | 0.307279859 | 0.739487613 |
| Cypt8 | 0 | NA | NA | NA | NA | NA | | | Neurod4 | 20.20722562 | 0.08889342 | 0.079446283 | 1.11891225 | 0.263177576 | NA |
| Cypt9 | 0 | NA | NA | NA | NA | NA | | | Acer2 | 396.7680386 | 0.088893877 | 0.075113434 | 1.183461754 | 0.236626196 | 0.687308962 |
| Cytip | 7.628773967 | 0.111026357 | 0.147352558 | 0.753474241 | 0.451164985 | NA | | | Dclk1 | 41359.77701 | 0.0889448 | 0.06856943 | 1.297149474 | 0.194579761 | 0.650659672 |
| D030018L15Rik | 0.401795408 | -0.00423014 | 0.047730791 | -0.088624972 | 0.929379961 | NA | | | Egln2 | 2929.720113 | 0.088977861 | 0.063348136 | 1.404585288 | 0.160144634 | 0.612291377 |
| D030024E09Rik | 0 | NA | NA | NA | NA | NA | | | Dcc | 987.4288771 | 0.088989828 | 0.071599348 | 1.242886015 | 0.21390984 | 0.667795903 |
| D030025E07Rik | 0.46655017 | -0.036459544 | 0.045831149 | -0.795518874 | 0.426311736 | NA | | | Wdr65 | 63.5095083 | 0.089017178 | 0.092754617 | 0.959706168 | 0.337203118 | 0.760113439 |
| D030025P21Rik | 0.412413621 | -0.026778838 | 0.043795168 | -0.611456443 | 0.540897446 | NA | | | Armcx2 | 4223.581534 | 0.089033124 | 0.049207607 | 1.809336581 | 0.070398729 | 0.456017009 |
| D030040B21Rik | 0.462334498 | 0.005887023 | 0.049959785 | 0.117835246 | 0.906198202 | NA | | | Birc5 | 34.10042573 | 0.089040362 | 0.088731241 | 1.003483794 | 0.315627492 | 0.745993052 |
| D030045P18Rik | 1.457485358 | -0.095836437 | 0.083616539 | -1.146142118 | 0.251736353 | NA | | | Hebp1 | 800.0581332 | 0.089041158 | 0.074054086 | 1.202380079 | 0.229216304 | 0.682442899 |
| D030047H15Rik | 5.035610353 | 0.160455316 | 0.132056806 | 1.21504768 | 0.224347907 | NA | | | Cfdp1 | 2870.268979 | 0.089049487 | 0.061937046 | 1.437741918 | 0.150507295 | 0.598511432 |
| D130009I18Rik | 0 | NA | NA | NA | NA | NA | | | Ildr2 | 7701.035073 | 0.089057649 | 0.061061885 | 1.458481823 | 0.144707782 | 0.590269908 |
| D130058E03 | 0.098353386 | -0.020212831 | 0.02786466 | -0.725393061 | 0.468210929 | NA | | | Tmem151b | 6275.11787 | 0.089062025 | 0.07676114 | 1.160248853 | 0.245947502 | 0.695335425 |
| D16Ertd519e | 0.317012939 | 0.010888466 | 0.039852335 | 0.273220278 | 0.784683893 | NA | | | Lgalsl | 3164.649639 | 0.089074301 | 0.063583407 | 1.400904806 | 0.161242541 | 0.612822908 |
| D17Ertd648e | 0.266310378 | 0.0407635 | 0.040043874 | 1.017970943 | 0.308691765 | NA | | | Cdr2l | 1234.485715 | 0.089091089 | 0.062734959 | 1.420118709 | 0.155573124 | 0.605210285 |
| D1Pas1 | 0 | NA | NA | NA | NA | NA | | | Psme3 | 4834.970964 | 0.089110208 | 0.049895982 | 1.785919518 | 0.07411229 | 0.467052116 |
| D230030E09Rik | 0.08731684 | 0.017365056 | 0.027733412 | 0.626142072 | 0.531221759 | NA | | | Vbp1 | 1499.768177 | 0.089131291 | 0.066697386 | 1.336353577 | 0.181433732 | 0.634469716 |
| D330022K07Rik | 2.055724125 | 0.162640569 | 0.098066497 | 1.6584723 | 0.097222174 | NA | | | Gorasp2 | 4209.114924 | 0.089185271 | 0.049524308 | 1.800838282 | 0.071728373 | 0.460037575 |
| D530049I02Rik | 0 | NA | NA | NA | NA | NA | | | Cops7a | 5543.562455 | 0.089238751 | 0.04432242 | 2.013399776 | 0.044072589 | 0.380217243 |
| D5Ertd577e | 0 | NA | NA | NA | NA | NA | | | Serinc1 | 35054.02552 | 0.089247255 | 0.065900111 | 1.354280802 | 0.175646809 | 0.628459435 |
| D5Ertd605e | 2.405648674 | -0.064213337 | 0.106402337 | -0.60349555 | 0.546179075 | NA | | | Higd1a | 2821.966518 | 0.089263191 | 0.057493064 | 1.55259062 | 0.120520968 | 0.554583368 |
| D630010B17Rik | 2.410794075 | -0.077905693 | 0.106244561 | -0.733267585 | 0.463395241 | NA | | | Tsc22d4 | 2688.805799 | 0.089287803 | 0.069904475 | 1.27728308 | 0.201502325 | 0.655108051 |
| D630013N20Rik | 3.010989214 | 0.126214459 | 0.110023053 | 1.147163757 | 0.251313947 | NA | | | Sqstm1 | 18338.90914 | 0.089291494 | 0.039311027 | 2.271410859 | 0.023122118 | 0.286019362 |
| D630029K05Rik | 1.675520834 | -0.144985409 | 0.094226611 | -1.538688569 | 0.123880344 | NA | | | Tmem229b | 1572.212926 | 0.08929369 | 0.059755817 | 1.494309597 | 0.135094718 | 0.578555714 |
| D630032N06Rik | 6.094611392 | 0.209464017 | 0.150812663 | 1.388902047 | 0.164862538 | NA | | | B3gnt1 | 4438.706786 | 0.089345436 | 0.043827356 | 2.038576899 | 0.041492273 | 0.368048911 |
| D630033O11Rik | 4.324882926 | 0.083293511 | 0.13742743 | 0.606090869 | 0.544454414 | NA | | | Car8 | 533.890716 | 0.089348215 | 0.090026209 | 0.992468922 | 0.320968832 | 0.750352874 |
| D630045M09Rik | 0.383033497 | -0.001876918 | 0.041875907 | -0.044820955 | 0.964250022 | NA | | | Cct3 | 4571.027001 | 0.08937252 | 0.0411821 | 2.170178774 | 0.029993305 | 0.318947844 |
| D6Ertd527e | 2.420643215 | -0.058708771 | 0.110519336 | -0.531208139 | 0.595274554 | NA | | | Pim3 | 1536.526028 | 0.089388175 | 0.071114876 | 1.256954662 | 0.208770053 | 0.66358495 |
| D730001G18Rik | 0.08731684 | 0.017365056 | 0.027733412 | 0.626142072 | 0.531221759 | NA | | | Ctnna2 | 6788.090262 | 0.089390171 | 0.049390519 | 1.809864977 | 0.070316728 | 0.456017009 |
| D730005E14Rik | 4.188776461 | 0.22374399 | 0.126599061 | 1.767343206 | 0.077170768 | NA | | | Yif1b | 969.1401045 | 0.089393009 | 0.074695416 | 1.196767008 | 0.231397381 | 0.683464489 |
| D730045A05Rik | 0.566001468 | -0.004543743 | 0.057856841 | -0.078534241 | 0.937403094 | NA | | | Mapk1ip1l | 1877.698815 | 0.089403665 | 0.055961598 | 1.597589568 | 0.11013435 | 0.535496439 |
| D730048I06Rik | 0 | NA | NA | NA | NA | NA | | | Syt16 | 2937.660993 | 0.089408118 | 0.05532687 | 1.615998112 | 0.106094721 | 0.52850889 |
| D730050B12Rik | 0 | NA | NA | NA | NA | NA | | | Snrpd3 | 1291.323822 | 0.089462072 | 0.05708501 | 1.567172751 | 0.117074318 | 0.548233894 |
| D7Ertd143e | 0 | NA | NA | NA | NA | NA | | | Scd2 | 76254.07008 | 0.089468835 | 0.064568428 | 1.385643686 | 0.165855732 | 0.616168889 |
| D830005E20Rik | 0 | NA | NA | NA | NA | NA | | | Ssr1 | 6041.725923 | 0.089503092 | 0.049478146 | 1.8089419 | 0.07046003 | 0.456017009 |
| D830013O20Rik | 0.341934174 | 0.0366924 | 0.037856746 | 0.969243346 | 0.332423789 | NA | | | Pkm | 55423.20069 | 0.089523306 | 0.050808591 | 1.761971823 | 0.078074058 | 0.475440315 |
| D830031N03Rik | 392.919938 | -0.083108005 | 0.157079395 | -0.52908279 | NA | NA | | | 1110008J03Rik | 538.0184866 | 0.08953554 | 0.072517901 | 1.234668118 | 0.216954046 | 0.670185967 |
| D830046C22Rik | 2.078483941 | 0.00507376 | 0.091802309 | 0.055268325 | 0.955924697 | NA | | | Nup62 | 1183.074619 | 0.0895744 | 0.058572287 | 1.52929662 | 0.126190927 | 0.562787717 |

| | | | | | | |
|---|---|---|---|---|---|---|
| D930007P13Rik | 0.079983654 | 0.016740032 | 0.027674178 | 0.604897175 | 0.54524732 | NA |
| D930015M05Rik | 4.259944431 | 0.137513506 | 0.130806399 | 1.051275065 | 0.293132276 | NA |
| D930028M14Rik | 5.369676831 | -0.050589606 | 0.134747415 | -0.375440274 | 0.707333057 | NA |
| D930032P07Rik | 0.397753127 | 0.04024206 | 0.039595247 | 1.016335617 | 0.309469593 | NA |
| DQ267100 | 7.292888916 | 0.171924681 | 0.142912322 | 1.203008098 | 0.228973187 | NA |
| DQ267101 | 0.619937596 | 0.079772058 | 0.055246832 | 1.443920948 | 0.148761213 | NA |
| DQ267102 | 7.417180309 | 0.118441321 | 0.155041872 | 0.763931187 | 0.444908251 | NA |
| DXBay18 | 0 | NA | NA | NA | NA | NA |
| Dab1 | 4052.864631 | -0.051749828 | 0.111027227 | -0.466100337 | NA | NA |
| Daf2 | 0.188785634 | 0.025025111 | 0.031461603 | 0.79541754 | 0.42637066 | NA |
| Dao | 3.940688519 | -0.018999632 | 0.131076522 | -0.144950685 | 0.884749808 | NA |
| Dbx1 | 0 | NA | NA | NA | NA | NA |
| Dcaf8 | 6303.052294 | 0.145013089 | 0.073143629 | 1.982579889 | NA | NA |
| Dcbld2 | 1685.706633 | 0.162802744 | 0.113311397 | 1.436772894 | NA | NA |
| Dclre1b | 341.6925532 | 0.17318999 | 0.128943557 | 1.343145746 | NA | NA |
| Dcpp1 | 0 | NA | NA | NA | NA | NA |
| Dcpp2 | 0 | NA | NA | NA | NA | NA |
| Dcpp3 | 0 | NA | NA | NA | NA | NA |
| Dcst1 | 1.016765751 | 0.015991548 | 0.071806909 | 0.222702077 | 0.823767385 | NA |
| Dcstamp | 2.851231242 | 0.075215276 | 0.092361064 | 0.81436129 | 0.415438008 | NA |
| Dct | 24.09630804 | -0.344028829 | 0.157984581 | -2.177610165 | NA | NA |
| Ddi1 | 0 | NA | NA | NA | NA | NA |
| Ddx43 | 5.904419629 | -0.033536154 | 0.145966234 | -0.229752819 | 0.818283849 | NA |
| Dear1 | 0.954295622 | 0.01559776 | 0.066390676 | 0.234939012 | 0.81425607 | NA |
| Defa-ps1 | 0 | NA | NA | NA | NA | NA |
| Defa-ps12 | 0 | NA | NA | NA | NA | NA |
| Defa-ps13 | 0 | NA | NA | NA | NA | NA |
| Defa-rs1 | 0.116696964 | -0.020477583 | 0.027954224 | -0.732539847 | 0.463839131 | NA |
| Defa-rs7 | 0 | NA | NA | NA | NA | NA |
| Defa17 | 0 | NA | NA | NA | NA | NA |
| Defa20 | 0 | NA | NA | NA | NA | NA |
| Defa21 | 0 | NA | NA | NA | NA | NA |
| Defa22 | 0 | NA | NA | NA | NA | NA |
| Defa23 | 0 | NA | NA | NA | NA | NA |
| Defa24 | 0 | NA | NA | NA | NA | NA |
| Defa25 | 0 | NA | NA | NA | NA | NA |
| Defa26 | 0 | NA | NA | NA | NA | NA |
| Defa3 | 0 | NA | NA | NA | NA | NA |
| Defa4 | 0 | NA | NA | NA | NA | NA |
| Defa5 | 0 | NA | NA | NA | NA | NA |
| Defa6 | 0 | NA | NA | NA | NA | NA |
| Defb10 | 0 | NA | NA | NA | NA | NA |
| Defb11 | 0.804457876 | 0.06189952 | 0.048640448 | 1.272593545 | 0.20316231 | NA |
| Defb12 | 0 | NA | NA | NA | NA | NA |
| Defb13 | 0 | NA | NA | NA | NA | NA |
| Defb14 | 0.204933042 | 0.025509582 | 0.031745334 | 0.803569494 | 0.421645647 | NA |
| Defb15 | 0 | NA | NA | NA | NA | NA |
| Defb18 | 0.098353386 | -0.020212831 | 0.02786466 | -0.725393061 | 0.468210929 | NA |
| Defb19 | 0 | NA | NA | NA | NA | NA |
| Defb2 | 0 | NA | NA | NA | NA | NA |
| Defb20 | 0.407030455 | 0.026116724 | 0.048660926 | 0.536708323 | 0.591469106 | NA |
| Defb21 | 0 | NA | NA | NA | NA | NA |
| Defb22 | 0 | NA | NA | NA | NA | NA |
| Defb23 | 0.094392817 | 0.017895942 | 0.027818137 | 0.643319236 | 0.520016977 | NA |
| Defb25 | 0.644647693 | -0.028208895 | 0.054433229 | -0.518229311 | 0.604298287 | NA |
| Defb26 | 0 | NA | NA | NA | NA | NA |
| Defb28 | 0.419482488 | -0.03706176 | 0.049390016 | -0.75038972 | 0.453020021 | NA |
| Defb29 | 0 | NA | NA | NA | NA | NA |
| Defb3 | 0 | NA | NA | NA | NA | NA |
| Defb30 | 0.498045124 | 0.003767638 | 0.044410484 | 0.084836674 | 0.932391237 | NA |
| Defb34 | 0 | NA | NA | NA | NA | NA |
| Defb35 | 0 | NA | NA | NA | NA | NA |
| Defb36 | 0 | NA | NA | NA | NA | NA |
| Defb37 | 0 | NA | NA | NA | NA | NA |
| Defb38 | 0 | NA | NA | NA | NA | NA |
| Defb39 | 0 | NA | NA | NA | NA | NA |
| Defb4 | 0 | NA | NA | NA | NA | NA |
| Defb40 | 0 | NA | NA | NA | NA | NA |
| Defb41 | 0.08731684 | 0.017365056 | 0.027733412 | 0.626142072 | 0.531221759 | NA |
| Defb42 | 0.556822015 | 0.070925085 | 0.052229705 | 1.357945369 | 0.17448102 | NA |
| Defb43 | 0 | NA | NA | NA | NA | NA |
| Defb44-ps | 0 | NA | NA | NA | NA | NA |
| Defb45 | 0 | NA | NA | NA | NA | NA |
| Defb46 | 0 | NA | NA | NA | NA | NA |
| Defb47 | 0.688621028 | 0.009994152 | 0.060688229 | 0.164680234 | 0.869195675 | NA |
| Defb48 | 0 | NA | NA | NA | NA | NA |
| Defb5 | 0 | NA | NA | NA | NA | NA |
| Defb50 | 0 | NA | NA | NA | NA | NA |
| Defb6 | 0.509507115 | -0.017094169 | 0.051034812 | -0.334951159 | 0.73766193 | NA |
| Defb7 | 0 | NA | NA | NA | NA | NA |
| Defb8 | 0 | NA | NA | NA | NA | NA |
| Defb9 | 0 | NA | NA | NA | NA | NA |
| Depdc1a | 1.971358144 | -0.047190598 | 0.092630525 | -0.509449751 | 0.610437011 | NA |
| Dfnb59 | 2.167006197 | -0.005730924 | 0.109633413 | -0.052273516 | 0.958310755 | NA |
| Dhh | 5.817106699 | 0.118128211 | 0.138605369 | 0.85226288 | 0.394068203 | NA |
| Dhrs2 | 1.268245534 | 0.002510205 | 0.091751101 | 0.027358849 | 0.97817352 | NA |
| Dhrs7c | 0.21363976 | -0.000753674 | 0.031922632 | -0.02360938 | 0.98116419 | NA |
| Dhrs9 | 2.54759367 | 0.067456978 | 0.112489165 | 0.599675332 | 0.548722631 | NA |
| Dio1 | 0.196706773 | -0.026517754 | 0.031609464 | -0.838918169 | 0.401515231 | NA |
| Dkk1 | 0.392799438 | 0.019887619 | 0.042925294 | 0.463307699 | 0.643143828 | NA |
| Dkk4 | 0 | NA | NA | NA | NA | NA |
| Dlx3 | 3.760782338 | -0.211117866 | 0.118229402 | -1.785662982 | 0.074153843 | NA |
| Dlx6as2 | 0.997141522 | -0.047699308 | 0.081438456 | -0.585709869 | 0.558070501 | NA |
| Dmbt1 | 0.278003423 | 0.03119216 | 0.03498768 | 0.891518392 | 0.372651129 | NA |
| Dmrt1 | 0 | NA | NA | NA | NA | NA |
| Dmrt3 | 3.573628683 | 0.294315984 | 0.113503582 | 2.593010534 | 0.009513986 | NA |
| Dmrtc1b | 0 | NA | NA | NA | NA | NA |
| Dmrtc1c1 | 0 | NA | NA | NA | NA | NA |
| Dmrtc1c2 | 0 | NA | NA | NA | NA | NA |
| Dmrtc2 | 0.484073412 | 0.009324742 | 0.050383912 | 0.185073807 | 0.853171145 | NA |
| Dnajb7 | 0.19442443 | -0.026451352 | 0.031568725 | -0.837897386 | 0.402088337 | NA |
| Dnajb8 | 0 | NA | NA | NA | NA | NA |
| Dnajc22 | 0.977141333 | -0.006109669 | 0.063937801 | -0.095556438 | 0.923872864 | NA |
| Dnajc5b | 1.34728483 | -0.129015236 | 0.080200045 | -1.608667881 | 0.107688983 | NA |
| Dnajc5g | 0.193239998 | 0.025145508 | 0.031532422 | 0.797449311 | 0.42519013 | NA |
| Dnase1 | 0.566503969 | 0.017827893 | 0.052996707 | 0.336396242 | 0.736572083 | NA |
| Dnase1l3 | 6.793814045 | 0.252224576 | 0.117630774 | 2.144205689 | 0.032016409 | NA |
| Dnase2b | 1.242544597 | 0.02098239 | 0.072567323 | 0.289143781 | 0.772471351 | NA |
| Dnm3os | 1.628968287 | 0.006649644 | 0.090978817 | 0.073090022 | 0.941734482 | NA |
| Dnmt3l | 3.539878472 | -0.008370061 | 0.128629357 | -0.065071158 | 0.948117344 | NA |
| Dntt | 0.312649645 | 0.012040436 | 0.039723828 | 0.303103628 | 0.761810889 | NA |
| Doxl2 | 0 | NA | NA | NA | NA | NA |
| Dpcr1 | 0.364243852 | 0.020744286 | 0.042194559 | 0.491634149 | 0.622977998 | NA |
| Dpep1 | 0.611217879 | 0.037217074 | 0.037942619 | 0.980877835 | NA | NA |
| Dpep2 | 2.906442131 | 0.012397415 | 0.100063396 | 0.123895602 | 0.901397935 | NA |
| Dpep3 | 0 | NA | NA | NA | NA | NA |
| Dppa1 | 1.738467432 | -0.069286381 | 0.097356143 | -0.711679603 | 0.476663201 | NA |
| Dppa2 | 0.871485526 | 0.006170912 | 0.065128794 | 0.09474937 | 0.924513903 | NA |
| Dppa3 | 0 | NA | NA | NA | NA | NA |
| Dppa4 | 0 | NA | NA | NA | NA | NA |
| Dppa5a | 3.237291537 | 0.04849488 | 0.110106154 | 0.440437504 | 0.659620267 | NA |
| Dpt | 4.263485727 | -0.116786472 | 0.130039205 | -0.89808664 | 0.369139361 | NA |
| Dpy19l2 | 7.935042318 | 0.072922031 | 0.153326317 | 0.475600223 | 0.634359216 | NA |
| Drap1 | 3856.029312 | -0.055052089 | 0.116475557 | -0.472649291 | NA | NA |
| Dsc1 | 2.589115619 | -0.063544332 | 0.107979268 | -0.588486412 | 0.556205852 | NA |
| Dscc1 | 3.842157774 | 0.129346656 | 0.123408192 | 1.048120503 | 0.294583091 | NA |
| Dsg1a | 1.78476557 | -0.039501569 | 0.082623023 | -0.478093976 | 0.632583319 | NA |
| Dsg1b | 1.318872376 | 0.046211558 | 0.068456056 | 0.675054338 | 0.499641243 | NA |
| Dsg3 | 1.289828268 | -0.073754435 | 0.078075202 | -0.944658915 | 0.344833038 | NA |
| Dsg4 | 0 | NA | NA | NA | NA | NA |
| Duox1 | 2.139866676 | 0.06955864 | 0.111811441 | 0.622106643 | 0.533871747 | NA |
| Duox2 | 2.520612473 | -0.124688883 | 0.100292205 | -1.243255967 | 0.213773525 | NA |
| Duoxa1 | 1.486072896 | -0.093221287 | 0.08844968 | -1.053947138 | 0.29190712 | NA |
| Duoxa2 | 0.466789057 | 0.062805705 | 0.049307408 | 1.273757982 | 0.202749201 | NA |
| Dupd1 | 2.002378788 | -0.028313667 | 0.110639196 | -0.255909914 | 0.798020399 | NA |
| Dusp13 | 7.723204011 | 0.200993409 | 0.150781543 | 1.333010691 | 0.182528295 | NA |
| Dusp21 | 0 | NA | NA | NA | NA | NA |
| Dux | 1.104854567 | 0.068644114 | 0.08014215 | 0.856529481 | 0.391704973 | NA |
| Duxbl1 | 0 | NA | NA | NA | NA | NA |
| Duxbl2 | 0 | NA | NA | NA | NA | NA |
| Duxbl3 | 0 | NA | NA | NA | NA | NA |
| Dydc1 | 1.163652955 | 0.016304391 | 0.079213808 | 0.205827638 | 0.836925551 | NA |
| Dynap | 0 | NA | NA | NA | NA | NA |
| Dynlt1c | 0 | NA | NA | NA | NA | NA |
| Dynlt1f | 0 | NA | NA | NA | NA | NA |
| Dyrk4 | 8.052629629 | 0.081786825 | 0.14085509 | 0.580645156 | 0.561479632 | NA |
| Dytn | 2.675503349 | 0.146201412 | 0.116647499 | 1.253360879 | 0.210074402 | NA |
| E030002O03Rik | 5.963835012 | -0.376582793 | 0.14013703 | -2.687246855 | 0.007204369 | NA |
| E030018B13Rik | 0 | NA | NA | NA | NA | NA |
| E030024N20Rik | 1.254731069 | 0.00474194 | 0.081515214 | 0.058172459 | 0.953611258 | NA |
| E030025P04Rik | 0.91606928 | -0.07312839 | 0.066468253 | -1.100200264 | 0.271244876 | NA |
| E030044B06Rik | 0.527043497 | -0.046865061 | 0.053304187 | -0.879200371 | 0.379292643 | NA |
| E130018N17Rik | 1.949237493 | 0.041858869 | 0.102735546 | 0.407442908 | 0.683682718 | NA |
| E130201H02Rik | 4.146769221 | -0.170755899 | 0.130373563 | -1.309743281 | 0.190282697 | NA |
| E130218I03Rik | 0.865351476 | 0.022878361 | 0.056461578 | 0.405202304 | 0.685328812 | NA |
| E130304I02Rik | 2.753772459 | 0.080587187 | 0.115470125 | 0.697905081 | 0.485236557 | NA |
| E130310I04Rik | 2.721459421 | 0.06797549 | 0.106170778 | 0.640246701 | 0.522012225 | NA |
| E230016K23Rik | 3.351906977 | 0.021074134 | 0.123095519 | 0.171201472 | 0.86406535 | NA |
| E230019M04Rik | 0.976884471 | 0.068046414 | 0.068004534 | 1.00061584 | 0.317012569 | NA |
| E330012B07Rik | 0.583727788 | 0.046764408 | 0.054560797 | 0.857106391 | 0.391386089 | NA |
| E330013P04Rik | 4.000472248 | 0.030423815 | 0.115649079 | 0.263070104 | 0.792496546 | NA |
| E330014E10Rik | 0 | NA | NA | NA | NA | NA |
| E330017A01Rik | 0 | NA | NA | NA | NA | NA |
| E330017L17Rik | 2.113500612 | 0.087607488 | 0.101099921 | 0.866543582 | 0.386192137 | NA |
| E330021D16Rik | 0 | NA | NA | NA | NA | NA |
| E330023G01Rik | 0.825193745 | 0.019755774 | 0.050840231 | 0.388585451 | 0.697582831 | NA |
| E330034G19Rik | 7.070014016 | -0.077931341 | 0.150930311 | -0.516339898 | 0.605617035 | NA |
| E430016F16Rik | 7.844773586 | -0.092996683 | 0.149346694 | -0.622689931 | 0.533488301 | NA |
| EU599041 | 0 | NA | NA | NA | NA | NA |
| Ear1 | 0.08731684 | 0.017365056 | 0.027733412 | 0.626142072 | 0.531221759 | NA |
| Ear10 | 0 | NA | NA | NA | NA | NA |
| Ear11 | 0.601913201 | 0.016398223 | 0.047539851 | 0.344936356 | 0.730142218 | NA |
| Ear12 | 0 | NA | NA | NA | NA | NA |
| Ear14 | 0 | NA | NA | NA | NA | NA |
| Ear2 | 0.19442443 | -0.026451352 | 0.031568725 | -0.837897386 | 0.402088337 | NA |
| Ear3 | 0 | NA | NA | NA | NA | NA |
| Ear4 | 0 | NA | NA | NA | NA | NA |
| Ear5 | 0 | NA | NA | NA | NA | NA |
| Ear6 | 0 | NA | NA | NA | NA | NA |
| Ear7 | 0 | NA | NA | NA | NA | NA |
| Eci3 | 2.696487227 | 0.082874488 | 0.113218419 | 0.731987681 | 0.464176087 | NA |
| Ect2l | 7.825503896 | 0.182843732 | 0.152792908 | 1.196676826 | 0.231432543 | NA |
| Eddm3b | 0 | NA | NA | NA | NA | NA |
| Edn2 | 1.77512903 | 0.006815807 | 0.101457163 | 0.067179155 | 0.946439079 | NA |
| Efcab8 | 0.378390275 | 0.024206488 | 0.047912756 | 0.50522012 | 0.61340424 | NA |
| Efcab9 | 7.840917801 | -0.055421878 | 0.158129876 | -0.350483285 | 0.725976032 | NA |
| Egfbp2 | 0 | NA | NA | NA | NA | NA |
| Ehf | 5.217180314 | 0.307076261 | 0.116888929 | 2.627077364 | 0.008612175 | NA |
| Eid3 | 3.601856976 | 0.083050739 | 0.130042195 | 0.638644548 | 0.523054194 | NA |
| Eif3j1 | 0 | NA | NA | NA | NA | NA |
| Eif3j2 | 0.119397641 | -0.020477583 | 0.027954224 | -0.732539845 | 0.463839132 | NA |
| Eif4e1b | 0.390895054 | 0.015442627 | 0.036628692 | 0.421599181 | 0.673317604 | NA |
| Elane | 1.329531748 | -0.068571592 | 0.08965843 | -0.764809201 | 0.444385168 | NA |
| Elf3 | 5.286250829 | -0.126652234 | 0.133251651 | -0.950474034 | 0.341871442 | NA |
| Ell3 | 3.148015308 | -0.034543619 | 0.123897159 | -0.278808803 | 0.780391558 | NA |
| Elovl3 | 0.766232536 | -0.054016814 | 0.063110901 | -0.855903078 | 0.392051392 | NA |
| Elovl4 | 1842.92971 | 0.113001034 | 0.121353031 | 0.931176039 | NA | NA |
| Eme1 | 4.324249447 | -0.274760079 | 0.13995426 | -1.963213398 | 0.049621383 | NA |
| Emilin3 | 5.724997593 | 0.106658926 | 0.135393543 | 0.787769665 | 0.430831448 | NA |
| Emr4 | 1.817799692 | -0.086568146 | 0.101525858 | -0.852670915 | 0.393841825 | NA |
| En1 | 1.78813753 | 0.088671736 | 0.086840617 | 1.021085979 | 0.307213706 | NA |
| Enam | 0 | NA | NA | NA | NA | NA |
| Eno1 | 2.611817104 | 0.013854512 | 0.098205679 | 0.141076485 | 0.88780952 | NA |
| Enpp2 | 12168.30993 | 0.298143001 | 0.166153531 | 1.794382581 | NA | NA |
| Enpp7 | 0 | NA | NA | NA | NA | NA |
| Enthd1 | 0.955421901 | 0.066721921 | 0.069484429 | 0.960242778 | 0.336933042 | NA |
| Entpd8 | 0.276690427 | -0.012033946 | 0.033317809 | -0.361186582 | 0.717959972 | NA |
| Ep300 | 3273.437727 | -0.059880036 | 0.09823326 | -0.609569872 | NA | NA |
| Epb4.2 | 0.494244083 | 0.041448004 | 0.052104748 | 0.795474608 | 0.426337476 | NA |
| Epcam | 6.107276653 | -0.068676491 | 0.146376 | -0.469178627 | 0.638941961 | NA |
| Epgn | 2.711438282 | 0.025508325 | 0.10108538 | 0.252344351 | 0.80077491 | NA |
| Epo | 0.59261035 | -0.06016431 | 0.055570839 | -1.082659739 | 0.278959481 | NA |
| Eppin | 0 | NA | NA | NA | NA | NA |
| Eppk1 | 6.623047455 | 0.15723547 | 0.151422488 | 1.038389157 | 0.299088916 | NA |
| Eps8l3 | 5.852289433 | 0.163762554 | 0.117493252 | 1.393803913 | 0.163376826 | NA |
| Epyc | 1.96563487 | 0.131724268 | 0.100098432 | 1.315947373 | 0.188191707 | NA |
| Eras | 0 | NA | NA | NA | NA | NA |
| Ereg | 0.198019769 | 0.025308392 | 0.031628095 | 0.80018706 | 0.423602426 | NA |
| Erp27 | 3.119599062 | 0.148250721 | 0.119277426 | 1.242906769 | 0.213902191 | NA |
| Esco2 | 6.581219312 | 0.002982056 | 0.151701691 | 0.019657365 | 0.984316702 | NA |
| Esrp1 | 3.052494327 | -0.035666884 | 0.108028346 | -0.330162273 | 0.741277352 | NA |
| Esrp2 | 2.340204657 | 0.01422642 | 0.0984252 | 0.144540421 | 0.885073741 | NA |
| Esx1 | 0 | NA | NA | NA | NA | NA |
| Etd | 0 | NA | NA | NA | NA | NA |
| Etv2 | 0.096071043 | -0.020098795 | 0.027825948 | -0.722304036 | 0.470107572 | NA |
| Evx1 | 0.105923158 | 0.018524693 | 0.027977518 | 0.662127842 | 0.507889294 | NA |
| Evx2 | 0 | NA | NA | NA | NA | NA |
| F10 | 0.608590881 | 0.025239369 | 0.042882565 | 0.588560172 | 0.556157176 | NA |
| F11 | 1.014523425 | -0.008946388 | 0.077241892 | -0.115823006 | 0.907792817 | NA |
| F13b | 0 | NA | NA | NA | NA | NA |
| F2 | 2.575664693 | 0.015768548 | 0.099068314 | 0.159168429 | 0.873536176 | NA |
| F2rl3 | 4.338555771 | 0.074491482 | 0.116644866 | 0.638617748 | 0.523071632 | NA |
| F5 | 90.66509134 | 0.306242572 | 0.130779031 | 2.34167948 | NA | NA |
| F630028O10Rik | 3.168600947 | -0.113463967 | 0.101133298 | -1.121924925 | 0.261894362 | NA |
| F630111L10Rik | 0 | NA | NA | NA | NA | NA |

| | | | | | | |
|---|---|---|---|---|---|---|
| Spg21 | 1343.345702 | 0.08959688 | 0.056402254 | 1.588533668 | 0.112165714 | 0.538992394 |
| Ptprg | 3065.153055 | 0.089599365 | 0.050652426 | 1.768905693 | 0.076909614 | 0.472576677 |
| Dnm3 | 8411.969349 | 0.089616454 | 0.066986428 | 1.337830015 | 0.180951855 | 0.634469716 |
| Trp53bp2 | 1457.055896 | 0.08963734 | 0.065096443 | 1.376992909 | 0.16851444 | 0.618923074 |
| Kcnc4 | 3245.25378 | 0.089658822 | 0.056347825 | 1.591167384 | 0.111571911 | 0.538992394 |
| A530054K11Rik | 474.9885696 | 0.089717141 | 0.083566956 | 1.073595898 | 0.283003837 | 0.720920007 |
| Gda | 11549.53277 | 0.089735863 | 0.074341658 | 1.207073736 | 0.227403749 | 0.680203999 |
| Bad | 772.3944833 | 0.089804136 | 0.067102948 | 1.338303878 | 0.180797398 | 0.634469716 |
| Srp68 | 3038.460287 | 0.08981335 | 0.049006912 | 1.83266698 | 0.066852111 | 0.447688159 |
| Imp4 | 777.9491588 | 0.089820086 | 0.06927904 | 1.296497263 | 0.194804223 | 0.65082254 |
| A830010M20Rik | 8095.231638 | 0.089836025 | 0.075057473 | 1.196896487 | 0.231346904 | 0.683460324 |
| Abl2 | 3482.953622 | 0.08984234 | 0.065184158 | 1.378284882 | 0.168115349 | 0.6180824 |
| Tmem126b | 566.2873938 | 0.089845331 | 0.065893438 | 1.363494371 | 0.172726757 | 0.624056899 |
| Tubb5 | 17841.21834 | 0.089898136 | 0.050101506 | 1.794320026 | 0.072762104 | 0.46336796 |
| Mc3r | 26.64443498 | 0.089923041 | 0.067700102 | 1.328255616 | 0.184093677 | 0.637571366 |
| Dirc2 | 3140.053643 | 0.090056514 | 0.050599852 | 1.779778209 | 0.075112265 | 0.468962081 |
| Sox11 | 470.8615444 | 0.090074082 | 0.074996936 | 1.201036821 | 0.229736918 | 0.682724729 |
| Psmd14 | 2781.675319 | 0.090086328 | 0.06005286 | 1.500117186 | 0.13358405 | 0.57588358 |
| Slc24a3 | 5470.550443 | 0.090110445 | 0.060900614 | 1.479631158 | 0.138971707 | 0.583135914 |
| Adcy9 | 3015.763352 | 0.090116095 | 0.071555418 | 1.259388845 | 0.207889917 | 0.662965736 |
| Frmpd3 | 169.1838599 | 0.090129418 | 0.090834361 | 0.992239253 | 0.321080829 | 0.750352874 |
| Ywhag | 50321.20208 | 0.090159837 | 0.038545187 | 2.339068611 | 0.019331883 | 0.263564471 |
| Smu1 | 2624.158638 | 0.090197417 | 0.055124811 | 1.636239929 | 0.101789383 | 0.522606076 |
| Gfpt1 | 3360.481044 | 0.090198433 | 0.061130584 | 1.475504206 | 0.140077038 | 0.585376897 |
| Synj2 | 4495.854977 | 0.090238795 | 0.078640938 | 1.14747862 | 0.251183864 | 0.699465245 |
| Ly6c1 | 1099.017821 | 0.090242592 | 0.08289351 | 1.088656898 | 0.276305216 | 0.717475105 |
| Tcea1 | 3289.484507 | 0.090267611 | 0.059520682 | 1.516575551 | 0.129373882 | 0.568274875 |
| Ptgfrn | 2200.037166 | 0.090322306 | 0.065176207 | 1.385817162 | 0.165802741 | 0.616168889 |
| Efnb3 | 5471.412496 | 0.090337745 | 0.073472654 | 1.229542417 | 0.218868505 | 0.671604514 |
| Fgf1 | 2645.397517 | 0.090339193 | 0.081825359 | 1.104048844 | 0.26957195 | 0.711972822 |
| Lrfn5 | 2135.818812 | 0.090355417 | 0.060959436 | 1.482222006 | 0.138281233 | 0.582904025 |
| Apod | 8109.46498 | 0.090362561 | 0.094870087 | 0.952487391 | 0.340849858 | 0.76263292 |
| Enah | 3098.111227 | 0.090363928 | 0.067088274 | 1.346940712 | 0.177999328 | 0.631188775 |
| Zcchc18 | 12950.05034 | 0.090366742 | 0.041737095 | 2.165142132 | 0.030376802 | 0.32021294 |
| Ermp1 | 3175.548393 | 0.090424439 | 0.057747182 | 1.565867566 | 0.117379624 | 0.548468184 |
| Socs5 | 5398.655832 | 0.090427255 | 0.061489322 | 1.470617211 | 0.141394667 | 0.586995979 |
| Atp5b | 45294.39078 | 0.090454423 | 0.038202575 | 2.367757236 | 0.017896274 | 0.255043174 |
| Kbtbd7 | 1390.901919 | 0.090459966 | 0.075709517 | 1.194829513 | 0.232153647 | 0.684268763 |
| Amacr | 367.7320397 | 0.090473022 | 0.08180601 | 1.105945917 | 0.268749929 | 0.710903001 |
| A330048O09Rik | 40.84209247 | 0.090510178 | 0.087894285 | 1.029761812 | 0.303121831 | 0.73655042 |
| Alyref2 | 88.93795727 | 0.090512745 | 0.094648181 | 0.956307282 | 0.338917012 | 0.761116259 |
| Alcam | 6409.684531 | 0.090524892 | 0.064053432 | 1.41327152 | 0.157575905 | 0.608141173 |
| Gria4 | 4579.412275 | 0.090550242 | 0.0557099 | 1.625388702 | 0.104079762 | 0.523645898 |
| Gopc | 2556.728171 | 0.09056553 | 0.065216235 | 1.388696087 | 0.164925184 | 0.615071433 |
| Asb13 | 2384.192693 | 0.090566073 | 0.054833947 | 1.651642427 | 0.098607466 | 0.520050241 |
| Atp6ap1 | 17447.159 | 0.090603555 | 0.040207126 | 2.253420315 | 0.02423266 | 0.291915574 |
| Them6 | 1756.445251 | 0.090653701 | 0.057875886 | 1.566346671 | 0.11726748 | 0.548468184 |
| Smg8 | 774.6402723 | 0.09065621 | 0.066081008 | 1.37189508 | 0.170096102 | 0.620316915 |
| Ubxn1 | 2033.47382 | 0.09066502 | 0.051239649 | 1.769430932 | 0.076821987 | 0.472560915 |
| Nhs | 229.3773049 | 0.090692633 | 0.093723177 | 0.967664945 | 0.333211731 | 0.757730862 |
| Magoh | 453.656887 | 0.090694691 | 0.072235468 | 1.255542377 | 0.209281934 | 0.664196937 |
| Myl1 | 10.27370433 | 0.090734412 | 0.062670811 | 1.447793801 | 0.147674727 | NA |
| Glul | 69300.96669 | 0.090747992 | 0.062057254 | 1.462326898 | 0.143651651 | 0.589907432 |
| Gnas | 51957.59659 | 0.0907793 | 0.05157226 | 1.760235058 | 0.07836796 | 0.475744405 |
| 1700025G04Rik | 4653.057972 | 0.090811634 | 0.051256153 | 1.771721608 | 0.076440781 | 0.472560915 |
| Nomo1 | 6213.586004 | 0.090821226 | 0.050656753 | 1.792875009 | 0.072992912 | 0.463821104 |
| Rcn1 | 1099.054125 | 0.090857593 | 0.081370908 | 1.116585719 | 0.2641715 | 0.707085906 |
| Ndufb10 | 3378.546648 | 0.090870523 | 0.048611971 | 1.869303388 | 0.061580617 | 0.435077842 |
| Kifap3 | 13367.37025 | 0.090879675 | 0.063496184 | 1.43126199 | 0.152355143 | 0.600557969 |
| Eid1 | 4955.580054 | 0.090892436 | 0.057486971 | 1.581096283 | 0.113856025 | 0.542263927 |
| Auh | 1685.521634 | 0.090913404 | 0.060100323 | 1.512694095 | 0.13035738 | 0.569619261 |
| Slc6a9 | 2515.510375 | 0.090924815 | 0.076995231 | 1.1809149 | 0.237636532 | 0.688529917 |
| Ccdc92 | 3704.111099 | 0.090930694 | 0.042461961 | 2.14146243 | 0.032236765 | 0.327711599 |
| Imp3 | 742.6083373 | 0.090975578 | 0.069185029 | 1.314960473 | 0.188523187 | 0.64225749 |
| Abat | 16573.93197 | 0.090992166 | 0.067983513 | 1.338444614 | 0.180751544 | 0.634469716 |
| Cx3cl1 | 32735.73569 | 0.09099946 | 0.059182636 | 1.53760403 | 0.124145462 | 0.559205951 |
| Park7 | 3140.114478 | 0.091059692 | 0.04957736 | 1.836719255 | 0.066251351 | 0.446279967 |
| Gm13152 | 77.82015888 | 0.091075308 | 0.095029273 | 0.958392134 | 0.33786506 | 0.760581219 |
| Sdha | 18069.20178 | 0.091079344 | 0.037425772 | 2.433599635 | 0.01494952 | 0.232501856 |
| Tmem59 | 7226.004225 | 0.091124916 | 0.054251058 | 1.679689189 | 0.093017804 | 0.507388894 |
| Med7 | 506.0736496 | 0.091126216 | 0.079951431 | 1.139769661 | 0.254382276 | 0.701619624 |
| Arl4c | 2454.617196 | 0.091144098 | 0.057152779 | 1.594744837 | 0.110769307 | 0.537406303 |
| Zfp763 | 779.0492561 | 0.091177355 | 0.082997151 | 1.09856006 | 0.271960007 | 0.714078854 |
| Dyrk2 | 684.1796817 | 0.091221031 | 0.077644268 | 1.174858546 | 0.240051313 | 0.690902589 |
| Lsm11 | 1448.385807 | 0.091256448 | 0.075404098 | 1.210231952 | 0.226189901 | 0.678709873 |
| Ndufa11 | 2132.943261 | 0.091288269 | 0.058810922 | 1.552233262 | 0.12060642 | 0.554583368 |
| Ogdh | 14131.17915 | 0.091303635 | 0.04887328 | 1.868170789 | 0.061738269 | 0.435133477 |
| Ak4 | 2389.606713 | 0.091311198 | 0.082864534 | 1.101933404 | 0.270490625 | 0.712294861 |
| Sgta | 4902.574164 | 0.091331294 | 0.057289306 | 1.594211905 | 0.110888581 | 0.537434808 |
| BC033916 | 17.06183678 | 0.091350289 | 0.071926933 | 1.27004288 | 0.204069356 | NA |
| Vat1 | 2089.559691 | 0.091369409 | 0.094714853 | 0.964678779 | 0.334705727 | 0.758238352 |
| Irf2bpl | 2773.590539 | 0.091379817 | 0.072309455 | 1.263732613 | 0.206326015 | 0.660297797 |
| Adam9 | 2716.25473 | 0.091384839 | 0.054238812 | 1.68486061 | 0.092015468 | 0.505579538 |
| Prepl | 21130.94236 | 0.091402988 | 0.057922049 | 1.578034447 | 0.114557696 | 0.543130934 |
| Alkbh4 | 236.4742494 | 0.091408035 | 0.07923145 | 1.15368372 | 0.248629861 | 0.697784358 |
| Nkiras2 | 369.8900942 | 0.091409446 | 0.073265527 | 1.247646049 | 0.212160707 | 0.666270994 |
| Frem3 | 111.1131637 | 0.091430144 | 0.063242079 | 1.445716927 | 0.148256615 | 0.595568469 |
| Pcdh19 | 1549.851102 | 0.091466342 | 0.079757077 | 1.146811611 | 0.25145949 | 0.699465245 |
| Apex1 | 1336.546402 | 0.091470592 | 0.052822359 | 1.731664287 | 0.083333353 | 0.48555968 |
| Polr2h | 571.0716747 | 0.091511376 | 0.066040226 | 1.385691434 | 0.165841145 | 0.616168889 |
| Hs6st3 | 542.3178099 | 0.091556391 | 0.078474808 | 1.166697855 | 0.243332409 | 0.692772033 |
| Rer1 | 2362.472884 | 0.091563768 | 0.052715349 | 1.736947023 | 0.082396522 | 0.483019453 |
| Prickle1 | 3910.302872 | 0.091573792 | 0.067180343 | 1.363103965 | 0.172849747 | 0.624135563 |
| Gcat | 394.0764756 | 0.091594951 | 0.083710255 | 1.094190328 | 0.273871512 | 0.716079153 |
| Cryab | 3885.595258 | 0.091606745 | 0.076500039 | 1.197473179 | 0.231122176 | 0.683402059 |
| Rrs1 | 711.4447262 | 0.091607053 | 0.067472897 | 1.357686676 | 0.174563126 | 0.627494178 |
| Tm2d2 | 1934.821684 | 0.091640404 | 0.047407973 | 1.933016716 | 0.053234139 | 0.407317206 |
| Midn | 2442.124344 | 0.091687561 | 0.094139502 | 0.973954168 | 0.330079296 | 0.756141871 |
| Mmp17 | 9912.905129 | 0.091708249 | 0.065382643 | 1.40263907 | 0.160724495 | 0.612387888 |
| 2700046A07Rik | 29.63482662 | 0.09171685 | 0.086695279 | 1.057922076 | 0.29009097 | 0.725277354 |
| Trim54 | 20.64769469 | 0.091769008 | 0.079591331 | 1.153002558 | 0.248909335 | NA |
| Eif1a | 1622.631862 | 0.091802856 | 0.063771861 | 1.439551148 | 0.149994429 | 0.598234686 |
| St8sia1 | 2494.507917 | 0.091812463 | 0.065024979 | 1.411956835 | 0.157962671 | 0.608476733 |
| Ahdc1 | 3159.577334 | 0.091821083 | 0.070974495 | 1.293719432 | 0.195762351 | 0.651423791 |
| Caprin1 | 8944.849851 | 0.09183071 | 0.05625634 | 1.632361968 | 0.102603256 | 0.522975271 |
| Rnasek | 4038.60441 | 0.091843783 | 0.048515792 | 1.893069833 | 0.058348583 | 0.423355579 |
| Uba3 | 2869.617247 | 0.091851298 | 0.061655468 | 1.489751042 | 0.136289708 | 0.579855848 |
| Acyp1 | 391.5395527 | 0.091862449 | 0.073374789 | 1.251962014 | 0.210583706 | 0.6650979 |
| Mapre2 | 15257.26421 | 0.091884824 | 0.050599495 | 1.81592372 | 0.069382074 | 0.454985124 |
| Thumpd2 | 156.6460917 | 0.091895952 | 0.088580361 | 1.037430321 | 0.299535354 | 0.733704078 |
| Bcas3 | 2919.233344 | 0.091901474 | 0.050073889 | 1.835317277 | 0.066458693 | 0.44672789 |
| Slc25a39 | 2361.069664 | 0.091912316 | 0.066489353 | 1.382361418 | 0.166860756 | 0.617356019 |
| Zfyve9 | 2725.618332 | 0.091926287 | 0.067060956 | 1.370787013 | 0.170441361 | 0.620595561 |
| Gm10509 | 173.4146698 | 0.091930936 | 0.092180813 | 0.997289281 | 0.318624115 | 0.74851037 |
| Tufm | 2433.355961 | 0.091941491 | 0.055162334 | 1.666744034 | 0.095565313 | 0.514803116 |
| Pmm1 | 4604.255351 | 0.091956385 | 0.054243602 | 1.695248505 | 0.090028289 | 0.500949827 |
| Prmt1 | 2351.33227 | 0.091960395 | 0.045017639 | 2.042763635 | 0.041075842 | 0.366278503 |
| Gas6 | 4190.759783 | 0.091963327 | 0.07146969 | 1.28674584 | 0.19818289 | 0.65354259 |
| Utp14b | 316.6823049 | 0.091979544 | 0.095099374 | 0.967193999 | 0.333447061 | 0.757852268 |
| BC052040 | 327.0796409 | 0.091984685 | 0.077719218 | 1.183551341 | 0.236590712 | 0.687308962 |
| Urod | 1494.053199 | 0.092013125 | 0.054382538 | 1.69196083 | 0.090653442 | 0.502789698 |
| 2900011O08Rik | 12361.1454 | 0.09211623 | 0.050228156 | 1.833956054 | 0.066660518 | 0.447318173 |
| Egr3 | 2608.748014 | 0.092127085 | 0.089496548 | 1.029392613 | 0.303295219 | 0.736610325 |
| 4930486L24Rik | 8.588095533 | 0.09216427 | 0.059721104 | 1.543244562 | 0.122771449 | NA |
| Rab3b | 2577.998487 | 0.092164506 | 0.05584607 | 1.650331091 | 0.098875237 | 0.520271327 |
| Dnahc8 | 21.46121323 | 0.092169282 | 0.074376486 | 1.239226085 | 0.215261783 | NA |
| Dnajc9 | 1023.235866 | 0.092170899 | 0.074649538 | 1.234714935 | 0.216936616 | 0.670185967 |
| Pgk1 | 18657.599 | 0.092182015 | 0.044556618 | 2.068873697 | 0.03855794 | 0.355824003 |
| Zfp619 | 221.0969102 | 0.092187181 | 0.085862147 | 1.073664988 | 0.282972859 | 0.720920007 |
| Pgap1 | 1532.732071 | 0.092209792 | 0.078729212 | 1.171227159 | 0.241507481 | 0.692393571 |
| Manf | 1513.108 | 0.092246438 | 0.064440248 | 1.43150346 | 0.152285976 | 0.600512434 |
| Klhl11 | 1080.696485 | 0.092261311 | 0.074767152 | 1.233981887 | 0.217209654 | 0.670311138 |
| 1700017B05Rik | 657.650475 | 0.092316474 | 0.073976799 | 1.247911172 | 0.212063589 | 0.666270994 |
| Degs1 | 3403.016293 | 0.092321297 | 0.049493832 | 1.865309124 | 0.062138087 | 0.436531911 |
| Tomm40 | 2343.95292 | 0.092324537 | 0.055087159 | 1.675972008 | 0.093743676 | 0.509378873 |
| Gm3604 | 48.09479956 | 0.092384909 | 0.090369992 | 1.022296306 | 0.306640681 | 0.738806192 |
| Clstn3 | 20819.20551 | 0.092410544 | 0.037422889 | 2.469358903 | 0.013535539 | 0.223175067 |
| Gdap2 | 1380.232387 | 0.092422258 | 0.060689437 | 1.522872217 | 0.127790678 | 0.565596401 |
| Ppp1r17 | 28.25592435 | 0.092423635 | 0.060664957 | 1.523509437 | 0.127631302 | 0.565235453 |
| Nif3l1 | 807.8588399 | 0.092463457 | 0.067649306 | 1.366805699 | 0.171686216 | 0.622234025 |
| Tsg101 | 2520.577723 | 0.092482163 | 0.04816291 | 1.920194675 | 0.054833314 | 0.412110852 |
| Gpx3 | 477.309003 | 0.092523715 | 0.065939989 | 1.403150291 | NA | NA |
| Fam110b | 1262.629062 | 0.092545112 | 0.062085633 | 1.490604295 | 0.136065416 | 0.579784756 |
| Gabra2 | 3710.165519 | 0.092547857 | 0.095051318 | 0.973661997 | 0.330224392 | 0.756141871 |
| Pnpo | 2287.37089 | 0.092574636 | 0.059828095 | 1.54734388 | 0.121780346 | 0.556108553 |
| Aco2 | 23504.93539 | 0.09261684 | 0.047029981 | 1.969314852 | 0.048916947 | 0.39652208 |
| Rnf6 | 2783.786354 | 0.092633752 | 0.059521169 | 1.556316085 | 0.119632948 | 0.552907479 |
| Nudt4 | 6825.888024 | 0.092639313 | 0.066988417 | 1.382915392 | 0.16669081 | 0.617356019 |
| Rpl8 | 8358.142175 | 0.092647835 | 0.058515583 | 1.583301903 | 0.113352669 | 0.540387554 |
| Dcaf12l1 | 1647.653784 | 0.092685609 | 0.069024932 | 1.342784507 | 0.179341762 | 0.633232481 |
| Sdcbp2 | 34.95598718 | 0.092718129 | 0.091936312 | 1.008503902 | 0.313212616 | 0.745612333 |
| Cep76 | 329.5109579 | 0.092787914 | 0.082717918 | 1.121738999 | 0.26197343 | 0.704317239 |
| Plekhb2 | 12739.84243 | 0.092802959 | 0.043921649 | 2.112920642 | 0.034607561 | 0.33922827 |
| Faim2 | 29969.82156 | 0.092849094 | 0.051786233 | 1.792930075 | 0.072984106 | 0.463821104 |
| Tgm4 | 15.52487044 | 0.092853931 | 0.077160972 | 1.203379494 | 0.228829499 | NA |
| Cops2 | 3156.898681 | 0.092859797 | 0.066668824 | 1.392851898 | 0.163664581 | 0.614351408 |
| Ppp5c | 4830.762121 | 0.092919356 | 0.052701902 | 1.763111993 | 0.077881603 | 0.475337606 |
| Gzmm | 62.95828639 | 0.092973951 | 0.094130528 | 0.987713054 | 0.323293199 | 0.752925264 |
| Kcnj16 | 1347.101363 | 0.092992352 | 0.069237265 | 1.343096851 | 0.179240615 | 0.633029218 |
| Pafah1b2 | 8430.692115 | 0.093011559 | 0.057002001 | 1.631724457 | 0.102737546 | 0.522975271 |
| Ppm1d | 602.6116995 | 0.093100477 | 0.068864655 | 1.351934139 | 0.176396385 | 0.629083076 |
| Gabra5 | 2072.204624 | 0.093127129 | 0.073657375 | 1.264328634 | 0.206112096 | 0.660203749 |
| Slc25a46 | 3807.521163 | 0.093132563 | 0.065486276 | 1.422169172 | 0.154977146 | 0.604489543 |
| Fbxo25 | 2539.706752 | 0.09314274 | 0.054088598 | 1.722040197 | 0.085062237 | 0.488771383 |
| Per2 | 2341.741477 | 0.093145756 | 0.084145803 | 1.106956652 | 0.26831267 | 0.710391551 |
| Ccdc113 | 120.2000192 | 0.09314887 | 0.094256478 | 0.988248999 | 0.323030717 | 0.752792382 |
| Trpc3 | 811.4107849 | 0.093154926 | 0.075373388 | 1.235912681 | 0.216491023 | 0.670049713 |
| Bcor | 1492.728439 | 0.093204036 | 0.085247236 | 1.093337921 | 0.274245459 | 0.716185716 |
| Agpat5 | 1638.658267 | 0.093241591 | 0.056656837 | 1.64572531 | 0.099820326 | 0.521115714 |
| Ncald | 22907.36797 | 0.093266255 | 0.043559406 | 2.141127798 | 0.032263734 | 0.327711599 |
| Klf14 | 29.49413904 | 0.093271583 | 0.087068581 | 1.071242705 | 0.284060314 | 0.721500285 |
| Enox2 | 417.7508755 | 0.093292498 | 0.076194879 | 1.224393282 | 0.220803902 | 0.672757795 |
| Clec12a | 29.47095516 | 0.093297128 | 0.084881991 | 1.099139259 | 0.271707328 | 0.713790099 |
| Nptx2 | 2494.699731 | 0.093333394 | 0.090411043 | 1.032322947 | 0.301920855 | 0.735974432 |
| Nr2f2 | 1013.33636 | 0.093361038 | 0.094229241 | 0.990786266 | 0.321789957 | 0.751204566 |
| Asns | 4199.982374 | 0.093377949 | 0.051524768 | 1.812292474 | 0.069941017 | 0.455201066 |
| Atxn10 | 10556.49163 | 0.093423463 | 0.051032379 | 1.830670354 | 0.067149761 | 0.447688159 |
| Ndufa9 | 3857.643713 | 0.093428253 | 0.044403983 | 2.104051188 | 0.035373982 | 0.342065168 |
| AW551984 | 819.1546187 | 0.0934727 | 0.060027188 | 1.557172731 | 0.119429481 | 0.55267048 |
| Podxl2 | 5774.139812 | 0.093477955 | 0.061782835 | 1.513008513 | 0.130277496 | 0.569619261 |
| Hmox2 | 2795.38223 | 0.093488979 | 0.049892536 | 1.873806907 | 0.060957044 | 0.432354503 |
| Rpn1 | 5446.249637 | 0.093495728 | 0.055351663 | 1.689122294 | 0.091195996 | 0.503727415 |
| Mrps21 | 985.4771378 | 0.0935054 | 0.063904642 | 1.463201999 | 0.143412114 | 0.589907432 |
| Wee1 | 1090.670465 | 0.0935065 | 0.078475959 | 1.191530527 | 0.23344538 | 0.685064073 |
| Tmem66 | 9931.541202 | 0.093539759 | 0.048816164 | 1.916163644 | 0.055344269 | 0.413057568 |
| Gstm7 | 734.2944795 | 0.093548367 | 0.088749005 | 1.054077919 | 0.291847245 | 0.727847831 |
| Htr5b | 46.64434564 | 0.093558475 | 0.087481295 | 1.069468333 | 0.284858691 | 0.721641599 |
| Scn11a | 20.10733417 | 0.093561803 | 0.079409973 | 1.178212245 | 0.238712002 | NA |
| Tmem159 | 419.5400146 | 0.093588312 | 0.080245861 | 1.16626965 | 0.24350544 | 0.692772033 |
| Zfp947 | 69.90886284 | 0.093592947 | 0.094159933 | 0.993978476 | 0.320233347 | 0.750154748 |
| Scfd2 | 786.918445 | 0.093628544 | 0.063174277 | 1.482067518 | 0.138322331 | 0.582904025 |
| E2f3 | 1396.285099 | 0.093634688 | 0.05910368 | 1.584244625 | 0.113138062 | 0.540327972 |
| Atp1b1 | 59295.23158 | 0.093651573 | 0.044068888 | 2.125117689 | 0.033576794 | 0.335456507 |
| Clptm1l | 4368.264133 | 0.093653178 | 0.044729364 | 2.093773991 | 0.036280118 | 0.345124424 |
| Serpinb1b | 69.18320478 | 0.093660204 | 0.08829966 | 1.060708546 | 0.288822375 | 0.724444298 |
| Atp6v1a | 22768.36613 | 0.093662084 | 0.062161628 | 1.506750838 | 0.131874535 | 0.572833273 |
| Gnl3l | 9949.833549 | 0.093709365 | 0.05360905 | 1.748013895 | 0.080461621 | 0.479417114 |
| Tasp1 | 765.7294994 | 0.093788339 | 0.07122595 | 1.316772038 | 0.187915048 | 0.641781416 |
| Spns2 | 2586.070195 | 0.093794787 | 0.077277367 | 1.213742016 | 0.224846254 | 0.676494758 |
| Ldhb | 20367.79021 | 0.093805328 | 0.046659386 | 2.010427817 | 0.044385929 | 0.380589751 |
| Gnb4 | 1402.306681 | 0.093822117 | 0.062265134 | 1.506816261 | 0.13185776 | 0.572833273 |
| Zfp874b | 516.1229943 | 0.093866387 | 0.082803646 | 1.133602099 | 0.256961492 | 0.703199912 |
| Taldo1 | 2899.631857 | 0.09389689 | 0.045663557 | 2.056276287 | 0.039755893 | 0.36080087 |
| 2900026A02Rik | 3081.60413 | 0.09397971 | 0.063216659 | 1.486628869 | 0.137112859 | 0.581518653 |
| Chn2 | 2140.899479 | 0.093980232 | 0.079192798 | 1.186727006 | 0.235335319 | 0.686276191 |
| Tpi1 | 24122.11141 | 0.093986901 | 0.046727679 | 2.011375331 | 0.044285827 | 0.380589751 |
| Mdga2 | 1534.84931 | 0.093998012 | 0.06387714 | 1.471543848 | 0.141144098 | 0.586989325 |
| Hnrnpc | 7184.531359 | 0.094031239 | 0.058771651 | 1.599942106 | 0.109611427 | 0.535311472 |
| Rnd1 | 952.8233762 | 0.094050848 | 0.073259702 | 1.283800581 | 0.19921174 | 0.65377551 |
| Lrrc32 | 279.8456611 | 0.094123179 | 0.083329333 | 1.129532361 | 0.258673326 | 0.703616072 |
| A730017C20Rik | 1673.97398 | 0.094130943 | 0.09506434 | 0.990181415 | NA | NA |
| Caln1 | 2814.775874 | 0.09415117 | 0.079986364 | 1.177090265 | 0.23915948 | 0.689702324 |
| BC031181 | 3192.022316 | 0.094177172 | 0.044579775 | 2.112553788 | 0.034638978 | 0.33922827 |
| Sox8 | 1947.678855 | 0.094200513 | 0.081831998 | 1.151145208 | 0.249672502 | 0.698181362 |
| Extl2 | 3686.921849 | 0.094204515 | 0.057459106 | 1.639505409 | 0.101108045 | 0.521924472 |
| Necab1 | 4790.977774 | 0.094207094 | 0.08163191 | 1.154047417 | 0.248480729 | 0.697784358 |
| Gdpd1 | 1676.744932 | 0.094211351 | 0.05773438 | 1.631806733 | 0.102720206 | 0.522975271 |
| Car13 | 43.47522351 | 0.094212491 | 0.081919642 | 1.150059841 | 0.250119225 | 0.698461386 |
| Hagh | 2432.764688 | 0.094273056 | 0.065170587 | 1.446558337 | 0.148020662 | 0.595114347 |
| Fam53b | 2144.404432 | 0.094294905 | 0.082902057 | 1.13742539 | 0.255360498 | 0.701952734 |
| Ormdl3 | 2378.949166 | 0.094318361 | 0.048153868 | 1.958687131 | 0.050149438 | 0.401582548 |
| Rnf128 | 465.328836 | 0.094323212 | 0.076284414 | 1.23646767 | 0.216284776 | 0.670049713 |
| Taf9b | 1296.709213 | 0.094356937 | 0.069864253 | 1.350575342 | 0.176831504 | 0.629241039 |
| Mrgpre | 581.8105472 | 0.094422954 | 0.087335509 | 1.081155934 | 0.279629535 | 0.718934998 |
| Ppid | 2502.175388 | 0.094449458 | 0.055910359 | 1.689301603 | 0.091161646 | 0.503727415 |
| Tcap | 167.7699443 | 0.094451253 | 0.094767885 | 0.99665887 | 0.31893012 | 0.748792456 |
| Pex14 | 1956.033731 | 0.094578912 | 0.048791734 | 1.898727572 | 0.057600304 | 0.419820382 |
| Metap2 | 2819.049468 | 0.09458291 | 0.051033344 | 1.853355151 | 0.063831471 | 0.441368101 |
| Gng2 | 8707.815231 | 0.09466098 | 0.046370737 | 2.04139476 | 0.041211605 | 0.367038487 |
| N6amt2 | 666.0738179 | 0.09469754 | 0.061668275 | 1.535595753 | 0.124637562 | 0.560776598 |
| Tm9sf2 | 5086.132671 | 0.094725752 | 0.069507826 | 1.362807007 | 0.172943342 | 0.62429796 |
| Stk39 | 3283.030164 | 0.094745431 | 0.061420015 | 1.542582355 | 0.122932142 | 0.557484892 |
| Kcnj6 | 1248.893749 | 0.094776182 | 0.079167038 | 1.197167212 | 0.231241387 | 0.683420627 |
| Dnajb11 | 2030.517939 | 0.094793836 | 0.06144135 | 1.54283453 | 0.122870929 | 0.557484892 |
| Vstm2a | 3151.2393 | 0.094872917 | 0.066822277 | 1.419779771 | 0.155671806 | 0.605432036 |

| | | | | | | | | | | | | | |
|---|---|---|---|---|---|---|---|---|---|---|---|---|---|
| F630206G17Rik | 0 | NA | NA | NA | NA | NA | Nudt3 | 9080.612077 | 0.094876624 | 0.043361424 | 2.188042193 | 0.028666529 | 0.315230284 |
| F7 | 3.810119778 | 0.055875058 | 0.107332469 | 0.520579268 | 0.602659895 | NA | Coro1c | 4056.083877 | 0.094890855 | 0.044931471 | 2.111901803 | 0.034694873 | 0.33922827 |
| F730035M05Rik | 0.862304613 | 0.072123711 | 0.060987028 | 1.18260741 | 0.236964775 | NA | Dcun1d3 | 843.8940995 | 0.094910085 | 0.06907542 | 1.37400664 | 0.169439621 | 0.619945581 |
| F830002L21Rik | 0 | NA | NA | NA | NA | NA | Igfbp3 | 460.7723442 | 0.094942596 | 0.085278389 | 1.113325398 | 0.265568702 | 0.707696014 |
| F830045P16Rik | 3.177521128 | -0.054521823 | 0.100498841 | -0.542511952 | 0.587465876 | NA | Fam131c | 110.7094655 | 0.094944357 | 0.095097477 | 0.998389863 | 0.318090347 | 0.748272127 |
| Fabp1 | 0 | NA | NA | NA | NA | NA | Mcm7 | 512.0253477 | 0.094972152 | 0.068257345 | 1.39138361 | 0.164109132 | 0.614351408 |
| Fabp12 | 0 | NA | NA | NA | NA | NA | 2610002M06Rik | 2624.318357 | 0.095000244 | 0.071002667 | 1.33798134 | 0.18090252 | 0.634469716 |
| Fabp2 | 0 | NA | NA | NA | NA | NA | Dio2 | 3249.051958 | 0.095001628 | 0.094825654 | 1.00185576 | 0.316413262 | 0.745993052 |
| Fabp4 | 2.614357751 | 0.013335283 | 0.118686773 | 0.112356947 | 0.910540391 | NA | Cct6a | 5553.718965 | 0.09500521 | 0.05215642 | 1.821543903 | 0.068524219 | 0.451830599 |
| Fabp6 | 0.238795281 | -0.027601606 | 0.032267774 | -0.855392297 | 0.392334007 | NA | Atp5l | 3807.233768 | 0.095012361 | 0.056394023 | 1.684794878 | 0.092028153 | 0.505579538 |
| Fabp9 | 0 | NA | NA | NA | NA | NA | Zcchc24 | 3614.830739 | 0.095018846 | 0.06362242 | 1.493480545 | 0.135311443 | 0.57866043 |
| Faim3 | 2.164581991 | 0.008983142 | 0.104935757 | 0.085606108 | 0.931779543 | NA | Gm5595 | 159.1372985 | 0.095019229 | 0.09473094 | 1.003043248 | 0.315839995 | 0.745993052 |
| Fam110c | 3.070669514 | -0.266438682 | 0.110310311 | -2.415356089 | 0.015719833 | NA | Cdkal1 | 483.8980883 | 0.095026717 | 0.078159733 | 1.215801452 | 0.224060565 | 0.675148076 |
| Fam115e | 0.116696964 | -0.020477583 | 0.027954224 | -0.732539847 | 0.463839131 | NA | Tet3 | 2652.316234 | 0.095037311 | 0.08263366 | 1.150104092 | 0.250101001 | 0.698461386 |
| Fam122c | 0 | NA | NA | NA | NA | NA | Odc1 | 2765.060226 | 0.095043241 | 0.054083322 | 1.757348414 | 0.07885844 | 0.476674706 |
| Fam124b | 1.211753037 | -0.015175223 | 0.08484861 | -0.178850585 | 0.858055024 | NA | Ext1 | 1738.094518 | 0.095052857 | 0.065683244 | 1.447140123 | 0.147857683 | 0.594842535 |
| Fam150a | 1.362713392 | -0.078483362 | 0.086579937 | -0.906484396 | 0.364679521 | NA | 4833424O15Rik | 1509.422047 | 0.095070498 | 0.076911381 | 1.23610442 | 0.216419752 | 0.670049713 |
| Fam151a | 3.3190959 | 0.04965642 | 0.112951216 | 0.439627135 | 0.660207184 | NA | Clic5 | 580.7018636 | 0.095083468 | 0.088786782 | 1.070919183 | 0.28420577 | 0.721548306 |
| Fam154a | 1.591349109 | -0.010914459 | 0.086180799 | -0.126646068 | 0.899220533 | NA | Vdac1 | 18000.96992 | 0.095100442 | 0.040712544 | 2.33590026 | 0.01949644 | 0.263578327 |
| Fam159a | 1.405242769 | -0.067305686 | 0.08552105 | -0.787007247 | 0.431277624 | NA | Acsl4 | 3122.823219 | 0.095104061 | 0.079757498 | 1.192415307 | 0.233098441 | 0.685064073 |
| Fam162b | 0 | NA | NA | NA | NA | NA | Sdc3 | 10453.98971 | 0.095114079 | 0.059809399 | 1.590286475 | 0.111770246 | 0.538992394 |
| Fam166a | 2.776405486 | 0.138982001 | 0.109737877 | 1.266490704 | 0.205337455 | NA | Pip4k2a | 4329.728961 | 0.095181881 | 0.067984799 | 1.400046525 | 0.161499387 | 0.613176432 |
| Fam170a | 0.08731684 | 0.017365056 | 0.027733412 | 0.626142072 | 0.531221759 | NA | Meis2 | 5451.105888 | 0.095184921 | 0.072179295 | 1.318728894 | 0.187259763 | 0.641692692 |
| Fam170b | 0.38968403 | -0.004310943 | 0.042055996 | -0.102504825 | 0.918355984 | NA | Trim37 | 18607.40739 | 0.095198404 | 0.062448114 | 1.524440021 | 0.127398829 | 0.564894808 |
| Fam177a | 0 | NA | NA | NA | NA | NA | Ccdc153 | 105.6966644 | 0.095251622 | 0.093437614 | 1.019414108 | 0.308006411 | 0.740254985 |
| Fam178b | 3.829658447 | -0.006411603 | 0.120401891 | -0.053251678 | 0.957531381 | NA | Arl3 | 2890.875837 | 0.09525522 | 0.051834737 | 1.837671532 | 0.066110821 | 0.446279967 |
| Fam187a | 2.566500152 | -0.003690655 | 0.106798931 | -0.034557039 | 0.972432959 | NA | Slc9a3r1 | 1633.505924 | 0.095263181 | 0.06241037 | 1.526399886 | 0.126910306 | 0.563718247 |
| Fam187b | 2.035303952 | -0.085013408 | 0.105238561 | -0.807816139 | 0.419196433 | NA | Heatr5b | 2357.337869 | 0.095269698 | 0.061896152 | 1.539186122 | 0.123758865 | 0.558645515 |
| Fam209 | 1.449700838 | -0.062223199 | 0.083969511 | -0.741021334 | 0.458680505 | NA | Heg1 | 913.3161586 | 0.095331428 | 0.070319406 | 1.355691594 | 0.175197315 | 0.628026741 |
| Fam213a | 3923.705842 | 0.107790065 | 0.084980665 | 1.268406934 | NA | NA | Tmprss6 | 24.85148337 | 0.095350129 | 0.080492156 | 1.184589074 | 0.236179959 | 0.687039131 |
| Fam227b | 1.101352946 | -0.029980011 | 0.083655862 | -0.3583731 | 0.72006412 | NA | Arhgef10 | 2361.119044 | 0.095391337 | 0.07143084 | 1.335436309 | 0.181733588 | 0.635192997 |
| Fam229a | 2.800884478 | 0.091358648 | 0.11708871 | 0.780251553 | 0.435242823 | NA | Ndrg1 | 5432.108449 | 0.095394795 | 0.074731202 | 1.276505559 | 0.201776863 | 0.655168687 |
| Fam24a | 0 | NA | NA | NA | NA | NA | Ppme1 | 10400.41949 | 0.095414488 | 0.050241849 | 1.899103839 | 0.057550824 | 0.419820382 |
| Fam25c | 0.333037401 | -0.016093237 | 0.040545167 | -0.396921218 | 0.691425557 | NA | Ttc30a1 | 148.3520825 | 0.095426112 | 0.09135324 | 1.04458377 | 0.296215384 | 0.731524085 |
| Fam26d | 0.665247938 | 0.039121182 | 0.061388665 | 0.63727045 | 0.523948696 | NA | Tspyl1 | 7929.130684 | 0.095447589 | 0.060868255 | 1.568101285 | 0.116857497 | 0.547395033 |
| Fam3b | 0 | NA | NA | NA | NA | NA | Lhfp | 919.6957953 | 0.09545261 | 0.061799723 | 1.544547542 | 0.122455743 | 0.557091176 |
| Fam46b | 3.030819446 | -0.069823169 | 0.110165979 | -0.633799742 | 0.526211521 | NA | Krt73 | 56.13520293 | 0.095467276 | 0.094560901 | 1.009585092 | 0.312694115 | 0.745030949 |
| Fam46d | 0.670842878 | 0.03854673 | 0.060574172 | 0.636355866 | 0.5245445 | NA | Ebpl | 320.9371989 | 0.095475642 | 0.086340333 | 1.10580581 | 0.26881058 | 0.710903001 |
| Fam47c | 0 | NA | NA | NA | NA | NA | Bpgm | 2395.308967 | 0.095476827 | 0.05761985 | 1.65701277 | 0.097516893 | 0.518456434 |
| Fam47e | 7.995450745 | 0.077393978 | 0.155229753 | 0.498576958 | 0.618077441 | NA | Gm9895 | 13.59746244 | 0.095481782 | 0.070554052 | 1.353313942 | 0.175955357 | NA |
| Fam50b | 0 | NA | NA | NA | NA | NA | Elp5 | 2294.160593 | 0.09551502 | 0.044694528 | 2.137062982 | 0.032592869 | 0.329798129 |
| Fam71a | 0 | NA | NA | NA | NA | NA | Mertk | 2079.525796 | 0.095551288 | 0.067952458 | 1.406149105 | 0.159679853 | 0.611570723 |
| Fam71b | 0.08731684 | 0.017365056 | 0.027733412 | 0.626142072 | 0.531221759 | NA | Gpr83 | 1189.988427 | 0.095553561 | 0.065137441 | 1.466952956 | 0.142388853 | 0.588600023 |
| Fam71d | 4.862649671 | 0.064585519 | 0.140295324 | 0.460354042 | 0.645262117 | NA | Gm9079 | 69.45632179 | 0.095574809 | 0.095068069 | 1.005330287 | 0.314737836 | 0.745993052 |
| Fam71e2 | 0.099009884 | 0.018205726 | 0.027896865 | 0.652608315 | 0.514008823 | NA | Txnl4b | 732.4136241 | 0.095625112 | 0.065372174 | 1.462780041 | 0.143527576 | 0.589907432 |
| Fam71f1 | 0 | NA | NA | NA | NA | NA | Bcl2l13 | 2701.947822 | 0.095633819 | 0.058477949 | 1.635382581 | 0.101968872 | 0.522723164 |
| Fam83a | 0.426633926 | -0.004170814 | 0.042905384 | -0.097209576 | 0.922559963 | NA | Map7d2 | 7576.269299 | 0.09566694 | 0.061204868 | 1.563060954 | 0.118038258 | 0.549241002 |
| Fam83b | 3.412064704 | -0.092500129 | 0.119400519 | -0.774704576 | 0.438514247 | NA | Abracl | 217.3400706 | 0.095738708 | 0.089845524 | 1.065592407 | 0.286607933 | 0.722148665 |
| Fam83c | 0.238795281 | -0.027601606 | 0.032267774 | -0.855392297 | 0.392334007 | NA | Etfdh | 1224.487096 | 0.09574782 | 0.055480828 | 1.725782106 | 0.08438662 | 0.488499377 |
| Fam83e | 3.100974519 | -0.146460362 | 0.126535299 | -1.15746644 | 0.247081843 | NA | Paqr9 | 943.4207947 | 0.095756964 | 0.070620103 | 1.355944848 | 0.175116717 | 0.628026741 |
| Fam83g | 7.944149917 | -0.110072069 | 0.148975254 | -0.738861431 | 0.459991145 | NA | Clpp | 1333.019528 | 0.095760094 | 0.061749867 | 1.550774096 | 0.120955831 | 0.555002664 |
| Fasl | 3.499930511 | 0.167464601 | 0.12045149 | 1.390307425 | 0.164435544 | NA | Lin7a | 3041.350817 | 0.095776907 | 0.061839039 | 1.548809747 | 0.121427462 | 0.555720009 |
| Fate1 | 3.933553786 | 0.091768208 | 0.121420309 | 0.755789615 | 0.449775344 | NA | Dleu7 | 155.6975325 | 0.095803974 | 0.087670023 | 1.092779164 | 0.274490772 | 0.716185716 |
| Fbp1 | 6.741081777 | 0.104716587 | 0.141920064 | 0.737856119 | 0.460601887 | NA | Khdrbs1 | 4457.945494 | 0.095809718 | 0.048309658 | 1.983241494 | 0.047340461 | 0.391951501 |
| Fbp2 | 0.678304087 | 0.007077915 | 0.060597108 | 0.116802858 | 0.907016279 | NA | Chrm5 | 37.81740929 | 0.095853243 | 0.076414669 | 1.254382754 | 0.209702918 | 0.664226905 |
| Fbxo15 | 4.952400966 | -0.052408258 | 0.130119018 | -0.402771698 | 0.687116184 | NA | Sycp3 | 10.03624008 | 0.095853456 | 0.058949677 | 1.62602172 | 0.103945035 | NA |
| Fbxo24 | 6.798809193 | -0.16018688 | 0.15314045 | -1.046012856 | 0.295555097 | NA | Galr1 | 22.65099282 | 0.095875448 | 0.068427108 | 1.401132556 | 0.161174438 | 0.612724387 |
| Fbxo39 | 0 | NA | NA | NA | NA | NA | Peg10 | 2341.023213 | 0.095878879 | 0.080540545 | 1.190442403 | 0.233872555 | 0.685064073 |
| Fbxo40 | 6.1891596 | 0.056783087 | 0.136295278 | 0.416618152 | 0.67695773 | NA | Has3 | 104.9971422 | 0.095885673 | 0.093731111 | 1.022986624 | 0.30631417 | 0.738584943 |
| Fbxo43 | 4.772613506 | 0.165549102 | 0.135142861 | 1.224993319 | 0.220577738 | NA | Smad6 | 62.47383324 | 0.095893344 | 0.094122651 | 1.01881262 | 0.308291933 | 0.74062833 |
| Fbxw13 | 1.218222965 | -0.060677382 | 0.06307571 | -0.961976984 | 0.336061165 | NA | Rpf2 | 582.2218958 | 0.095902791 | 0.066686884 | 1.438105743 | 0.150404053 | 0.598511432 |
| Fbxw14 | 0 | NA | NA | NA | NA | NA | Unc5cl | 8.797076338 | 0.095903334 | 0.060070847 | 1.596503772 | 0.110376364 | NA |
| Fbxw15 | 0.869516844 | 0.009508591 | 0.062816444 | 0.151371049 | 0.879683026 | NA | Gm561 | 325.8544624 | 0.095960179 | 0.075682364 | 1.267933158 | 0.204821822 | 0.658461308 |
| Fbxw16 | 0.094392817 | 0.017895942 | 0.027818137 | 0.643319236 | 0.520016977 | NA | Pcnxl4 | 1887.423036 | 0.096027123 | 0.060466507 | 1.588104351 | 0.112262745 | 0.538992394 |
| Fbxw18 | 0.204276544 | 2.70E-05 | 0.031767541 | 0.000850493 | 0.999321405 | NA | Scd3 | 217.6522718 | 0.096032099 | 0.083618555 | 1.148454419 | 0.250781019 | 0.699068333 |
| Fbxw19 | 0 | NA | NA | NA | NA | NA | Fam171b | 9992.470243 | 0.096038649 | 0.064178307 | 1.496434753 | 0.134540399 | 0.577862888 |
| Fbxw20 | 0 | NA | NA | NA | NA | NA | Ppp1r2 | 5178.062344 | 0.096070434 | 0.069675464 | 1.378827332 | 0.167947998 | 0.617953076 |
| Fbxw21 | 0 | NA | NA | NA | NA | NA | Vps25 | 1606.358034 | 0.09612357 | 0.055589049 | 1.729181775 | 0.083776567 | 0.486580734 |
| Fbxw22 | 0 | NA | NA | NA | NA | NA | Tead2 | 78.14707801 | 0.096149489 | 0.094934677 | 1.012796288 | 0.311157474 | 0.743823372 |
| Fbxw24 | 0.592206887 | -0.00501336 | 0.052942776 | -0.094693948 | 0.924557925 | NA | Fam117b | 3423.313327 | 0.096173321 | 0.050901557 | 1.88939842 | 0.058838462 | 0.425190581 |
| Fbxw26 | 0 | NA | NA | NA | NA | NA | Scd1 | 8228.779937 | 0.096191448 | 0.082174726 | 1.170572179 | 0.241770786 | 0.692421615 |
| Fbxw28 | 0 | NA | NA | NA | NA | NA | B3galnt1 | 1597.044895 | 0.096209558 | 0.054741187 | 1.75753512 | 0.078826641 | 0.476674706 |
| Fcamr | 0.772426891 | 0.028456539 | 0.048674378 | 0.584630763 | 0.558796018 | NA | Ralyl | 2428.283489 | 0.096234824 | 0.064182585 | 1.499391531 | 0.133772089 | 0.576096463 |
| Fcer1a | 0.314431083 | -0.017921647 | 0.039992366 | -0.448126696 | 0.654061762 | NA | Eps8 | 1272.72727 | 0.096238763 | 0.064447811 | 1.493282104 | 0.135363358 | 0.57866043 |
| Fcer2a | 0.722610492 | -0.050589987 | 0.061769794 | -0.819008519 | 0.412781553 | NA | Dusp5 | 197.3401857 | 0.096271242 | 0.073362399 | 1.312269554 | 0.189429201 | 0.644077615 |
| Fcnb | 0.373417968 | -0.003176841 | 0.047020903 | -0.067562317 | 0.946134054 | NA | Exosc3 | 226.2091564 | 0.096279696 | 0.086462209 | 1.113546576 | 0.265473756 | 0.707651445 |
| Fcrl5 | 0.203787646 | -0.02671144 | 0.031728022 | -0.841887971 | 0.399850664 | NA | Gdpgp1 | 396.8884283 | 0.09629996 | 0.08178112 | 1.177532898 | 0.238982875 | 0.689537017 |
| Fcrl6 | 1.1676572 | 0.012223131 | 0.075074963 | 0.162812343 | 0.870666189 | NA | Pja2 | 26391.161 | 0.096301293 | 0.066232543 | 1.453987551 | 0.145949758 | 0.592514474 |
| Fcrla | 0.440108409 | -0.030798726 | 0.044545461 | -0.691399874 | 0.489314283 | NA | Cdc5l | 2626.899919 | 0.096309011 | 0.062792812 | 1.53375851 | 0.125089074 | 0.561335769 |
| Fer1l4 | 0.421660619 | 0.058159795 | 0.047506763 | 1.224242436 | 0.220860785 | NA | Bre | 1475.790629 | 0.096311259 | 0.055097524 | 1.74801428 | 0.080461555 | 0.479417114 |
| Ferd3l | 0.709143477 | 0.005126568 | 0.054605503 | 0.09388372 | 0.925201526 | NA | Atp5c1 | 7568.670092 | 0.096336397 | 0.052720727 | 1.827296444 | 0.06765521 | 0.449158855 |
| Fetub | 2.839570631 | 0.077230581 | 0.10495259 | 0.735861596 | 0.461814931 | NA | Tsnax | 9032.506058 | 0.096393164 | 0.05353008 | 1.800728941 | 0.071745614 | 0.460037575 |
| Fev | 0.975805528 | 0.028297693 | 0.057303592 | 0.493820588 | 0.621432892 | NA | Sgcg | 5.675724316 | 0.096414682 | 0.053240794 | 1.810917419 | 0.070153636 | NA |
| Ffar1 | 0 | NA | NA | NA | NA | NA | Galnt11 | 1623.099852 | 0.096420457 | 0.060583694 | 1.591524889 | 0.111491499 | 0.538992394 |
| Ffar2 | 1.906626708 | -0.141934789 | 0.096254684 | -1.474575388 | 0.140326736 | NA | Atp2b3 | 4817.397347 | 0.096426831 | 0.055613076 | 1.733887752 | 0.082938001 | 0.484423387 |
| Ffar3 | 3.190905903 | -0.133988432 | 0.107408912 | -1.247461028 | 0.212228501 | NA | Aatf | 545.0507167 | 0.096485473 | 0.070572018 | 1.367191651 | 0.171565242 | 0.622106016 |
| Ffar4 | 1.74008905 | 0.008127778 | 0.099620747 | 0.081587199 | 0.934974981 | NA | Chchd4 | 1237.459649 | 0.096513164 | 0.051578869 | 1.87117641 | 0.061320633 | 0.434085536 |
| Fga | 0.61964287 | -0.002240474 | 0.053268353 | -0.042060124 | 0.966450769 | NA | Sfrp5 | 101.0748739 | 0.096578383 | 0.069326103 | 1.393102728 | 0.163588728 | 0.614351408 |
| Fgb | 0.200315975 | 0.025370485 | 0.031664208 | 0.801235428 | 0.422995365 | NA | Ppapdc3 | 1032.819432 | 0.096596715 | 0.061416983 | 1.572801354 | 0.115764822 | 0.545619664 |
| Fgf15 | 1.659710916 | -0.101587859 | 0.087624442 | -1.159355271 | 0.246311401 | NA | Pxdc1 | 159.0585758 | 0.096597216 | 0.090510933 | 1.067243626 | 0.285861837 | 0.722148665 |
| Fgf20 | 0.203787646 | -0.02671144 | 0.031728022 | -0.841887971 | 0.399850664 | NA | Gm14634 | 33.46517389 | 0.096646243 | 0.089986057 | 1.074013524 | 0.28281662 | 0.720920007 |
| Fgf21 | 0.079983654 | 0.016740032 | 0.027674178 | 0.604897175 | 0.54524732 | NA | Snrnp35 | 362.8354171 | 0.096651209 | 0.075427795 | 1.281373917 | 0.200062361 | 0.654380963 |
| Fgf4 | 0 | NA | NA | NA | NA | NA | Eftud1 | 643.6036202 | 0.096669218 | 0.070398821 | 1.373165288 | 0.169700968 | 0.619945581 |
| Fgf6 | 0 | NA | NA | NA | NA | NA | Nxph1 | 1436.010892 | 0.096706804 | 0.064388695 | 1.501922111 | 0.133117225 | 0.575193212 |
| Fgf8 | 0.390839854 | 0.053765426 | 0.045767491 | 1.174751441 | 0.240094173 | NA | Prkaca | 8825.434052 | 0.096711108 | 0.048899648 | 1.977746501 | 0.047957312 | 0.393352856 |
| Fgg | 0.098353386 | -0.020212831 | 0.02786466 | -0.725393061 | 0.468210929 | NA | Armcx4 | 3517.916265 | 0.096712818 | 0.059411671 | 1.627842074 | 0.103558377 | 0.523462117 |
| Fgl1 | 7.191908138 | -0.035068286 | 0.157767221 | -0.222278658 | 0.824096965 | NA | Opa3 | 1879.266386 | 0.096716915 | 0.052191489 | 1.853116614 | 0.063865646 | 0.441368101 |
| Fhl5 | 5.232462433 | -0.306394061 | 0.143279146 | -2.138441427 | 0.032480933 | NA | Sema4b | 715.5615369 | 0.096728001 | 0.075793255 | 1.276208567 | 0.201881801 | 0.655168687 |
| Figla | 0.841377836 | 0.042306321 | 0.058185242 | 0.727097102 | 0.467166474 | NA | Phf8 | 1409.554889 | 0.096733567 | 0.066581008 | 1.452870274 | 0.146259776 | 0.592958276 |
| Fitm1 | 0.533252617 | -0.072906618 | 0.051718649 | -1.409677544 | 0.158634918 | NA | Gphn | 3130.321217 | 0.096735731 | 0.04587973 | 2.108463405 | 0.034990925 | 0.34025917 |
| Fkbp5 | 2046.344257 | 0.210444595 | 0.157993516 | 1.331982481 | NA | NA | Lrrn1 | 2959.855618 | 0.096785072 | 0.059330158 | 1.631296382 | 0.102827796 | 0.522975271 |
| Fkbp6 | 1.41373376 | 0.013835613 | 0.076512541 | 0.180828044 | 0.85650255 | NA | Avl9 | 2741.159983 | 0.096786863 | 0.064810549 | 1.493381304 | 0.135337404 | 0.57866043 |
| Flg2 | 0 | NA | NA | NA | NA | NA | Rab2a | 9564.63237 | 0.096858031 | 0.065218615 | 1.485128619 | 0.137509756 | 0.582012447 |
| Fmo3 | 0.116696964 | -0.020477583 | 0.027954224 | -0.732539847 | 0.463839131 | NA | Rab3a | 21820.18913 | 0.09688102 | 0.038258649 | 2.532264559 | 0.011332845 | 0.202439555 |
| Fmo4 | 3.176147293 | 0.010691655 | 0.107144884 | 0.099786893 | 0.920513514 | NA | 9530068E07Rik | 3415.271996 | 0.096896527 | 0.042507856 | 2.279496934 | 0.022637542 | 0.28302679 |
| Fmo6 | 0.601929635 | 0.059495052 | 0.0606896 | 0.980317088 | 0.326929622 | NA | Chchd3 | 1680.613068 | 0.096929144 | 0.056865778 | 1.704525071 | 0.088283036 | 0.496728133 |
| Fmo9 | 0 | NA | NA | NA | NA | NA | Ino80e | 1020.68219 | 0.096934344 | 0.066896541 | 1.449018775 | 0.14733234 | 0.59448599 |
| Fmr1nb | 0.976635247 | -0.01263563 | 0.077170442 | -0.163736661 | 0.869938456 | NA | Sh3bgr | 37.20414397 | 0.096982907 | 0.085374979 | 1.135964047 | 0.255971612 | 0.701952734 |
| Fnd3c2 | 0 | NA | NA | NA | NA | NA | Lrrn2 | 5347.457364 | 0.096987774 | 0.065685568 | 1.476546165 | 0.139797332 | 0.585046396 |
| Fndc3c1 | 0.501157356 | 0.032769269 | 0.046039771 | 0.711760042 | 0.47661338 | NA | Dnajc27 | 7208.782797 | 0.097013763 | 0.047237822 | 2.053730645 | 0.040001769 | 0.361359262 |
| Fndc7 | 7.809630633 | -0.118504735 | 0.149768535 | -0.79125255 | 0.428796631 | NA | Lpar1 | 1154.151294 | 0.097032861 | 0.069144607 | 1.403332303 | 0.16051777 | 0.612387888 |
| Fndc8 | 0.287795519 | 0.031925978 | 0.035387605 | 0.902179657 | 0.366961439 | NA | Elmsan1 | 1288.056934 | 0.097057012 | 0.060526286 | 1.603551407 | 0.108812969 | 0.534621387 |
| Folr1 | 46.57336198 | 0.422003245 | 0.12823039 | 3.29097686 | NA | NA | Dll3 | 43.08684187 | 0.097082981 | 0.093465833 | 1.038700213 | 0.298944182 | 0.733276458 |
| Folr4 | 0.30279753 | -0.014859216 | 0.039561964 | -0.375593495 | 0.707219128 | NA | Shfm1 | 1147.914637 | 0.097084629 | 0.062083796 | 1.563767605 | 0.117872155 | 0.548913246 |
| Fosl1 | 2.366274081 | 0.256716804 | 0.098691656 | 2.601200693 | 0.009289809 | NA | Tnxb | 175.217121 | 0.097102319 | 0.095076307 | 1.021309329 | 0.307107908 | 0.739196267 |
| Foxa1 | 0.268769288 | 0.03049543 | 0.034602455 | 0.881308274 | 0.378150989 | NA | Rab11fip4 | 6477.947085 | 0.097114459 | 0.059482072 | 1.632667726 | 0.102538899 | 0.522975271 |
| Foxa2 | 0 | NA | NA | NA | NA | NA | Tspan6 | 495.6796091 | 0.097120677 | 0.079823175 | 1.216697742 | 0.223719239 | 0.675094439 |
| Foxa3 | 0 | NA | NA | NA | NA | NA | Slc27a3 | 167.297147 | 0.097162939 | 0.092895651 | 1.045936352 | 0.29559042 | 0.730624344 |
| Foxb1 | 2.808150522 | 0.01249967 | 0.113219953 | 0.11040165 | 0.912090845 | NA | 0610009B22Rik | 459.2832703 | 0.097172139 | 0.067029442 | 1.449693377 | 0.147144043 | 0.594031723 |
| Foxb2 | 0.770230356 | -0.005970955 | 0.061934977 | -0.096406831 | 0.923197467 | NA | Nup50 | 1959.262861 | 0.097183169 | 0.061534937 | 1.579316941 | 0.114263378 | 0.543130934 |
| Foxd3 | 3.520537522 | -0.178102128 | 0.080494249 | -2.212606862 | 0.026924762 | NA | Prelid1 | 3961.159083 | 0.097197526 | 0.046762478 | 2.078536692 | 0.037659955 | 0.351347789 |
| Foxd4 | 0.079983654 | 0.016740032 | 0.027674178 | 0.604897175 | 0.54524732 | NA | Nipal2 | 310.9845804 | 0.097218388 | 0.087181814 | 1.11512234 | 0.264797999 | 0.707370515 |
| Foxe1 | 0 | NA | NA | NA | NA | NA | Cgrrf1 | 671.7941936 | 0.097233427 | 0.065210579 | 1.4910683 | 0.135943564 | 0.579435507 |
| Foxe3 | 0 | NA | NA | NA | NA | NA | Atp5f1 | 7781.49038 | 0.097245486 | 0.053751902 | 1.809154316 | 0.070427033 | 0.456017009 |
| Foxh1 | 0.626277961 | -0.006339789 | 0.058658962 | -0.108078782 | 0.913933199 | NA | Rab1 | 6910.695407 | 0.097256125 | 0.047048339 | 2.067153196 | 0.038719721 | 0.356201815 |
| Foxi1 | 2.438620476 | 0.109743235 | 0.113148543 | 0.96990409 | 0.332094301 | NA | Mid2 | 1281.701213 | 0.097272714 | 0.06960844 | 1.397427007 | 0.162285203 | 0.613957643 |
| Foxi2 | 0.185670226 | -0.001645086 | 0.031432796 | -0.052336624 | 0.958260472 | NA | Ssna1 | 898.1755689 | 0.097295942 | 0.060285413 | 1.6139218 | 0.10654439 | 0.53002185 |
| Foxi3 | 0 | NA | NA | NA | NA | NA | Lipg | 293.5611412 | 0.097314536 | 0.08958327 | 1.086302566 | 0.277345152 | 0.718015501 |
| Foxl1 | 1.495943527 | -0.128616887 | 0.092341476 | -1.392839847 | 0.163668226 | NA | Irak2 | 841.2951888 | 0.097321969 | 0.069874535 | 1.392810263 | 0.163677174 | 0.614351408 |
| Foxn1 | 0.700186587 | 0.005332081 | 0.060807798 | 0.087687447 | 0.930125097 | NA | Tmem26 | 21.56230327 | 0.097323038 | 0.060125833 | 1.618655954 | 0.105521309 | NA |
| Foxn4 | 0.08731684 | 0.017365056 | 0.027733412 | 0.626142072 | 0.531221759 | NA | Angptl4 | 94.08156506 | 0.097324146 | 0.091879697 | 1.059256287 | 0.289483077 | 0.724785794 |
| Foxp3 | 3.104468821 | 0.137387151 | 0.116806182 | 1.176197596 | 0.239515925 | NA | Slc39a9 | 2762.851126 | 0.097335699 | 0.04360394 | 2.232268445 | 0.025597226 | 0.300343543 |
| Foxr1 | 0.528836942 | -0.072640212 | 0.051627513 | -1.407005851 | 0.159425654 | NA | Tmem178b | 2051.458862 | 0.097354273 | 0.079818592 | 1.219694186 | 0.222580826 | 0.67374894 |
| Fpr-rs3 | 0 | NA | NA | NA | NA | NA | A230056J06Rik | 39.20412072 | 0.097357586 | 0.092777233 | 1.049369368 | 0.294008151 | 0.729296858 |
| Fpr-rs4 | 0 | NA | NA | NA | NA | NA | Serf1 | 226.3717939 | 0.097388772 | 0.089601224 | 1.086913415 | 0.277075077 | 0.717815689 |
| Fpr-rs6 | 0 | NA | NA | NA | NA | NA | Tulp4 | 8135.880014 | 0.097395795 | 0.060134731 | 1.619626357 | 0.105312564 | 0.526780409 |
| Fpr1 | 1.538364536 | 0.048392463 | 0.076911902 | 0.629193423 | 0.529222434 | NA | Foxr2 | 26.4338135 | 0.097420891 | 0.081898567 | 1.189531095 | 0.234230742 | 0.68557298 |
| Fpr2 | 6.236296654 | 0.227658796 | 0.135529912 | 1.679767908 | 0.093002481 | NA | Trp53inp1 | 486.8551964 | 0.097423064 | 0.078133275 | 1.246883153 | 0.212440345 | 0.666270994 |
| Fpr3 | 0 | NA | NA | NA | NA | NA | Mospd1 | 383.6920243 | 0.09744821 | 0.077534885 | 1.256830531 | 0.208815008 | 0.66358495 |
| Frmpd1 | 622.6516966 | 0.204622233 | 0.136374933 | 1.50043874 | NA | NA | Arxes2 | 629.3982382 | 0.097499071 | 0.068185168 | 1.429916113 | 0.152741097 | 0.601351762 |
| Fscb | 0 | NA | NA | NA | NA | NA | Hprt | 4394.174498 | 0.097500894 | 0.063965335 | 1.524277074 | 0.127439512 | 0.564902761 |
| Fscn2 | 6.00751901 | 0.036516432 | 0.135683288 | 0.269129916 | 0.78782971 | NA | Jagn1 | 895.4174378 | 0.097513881 | 0.061447397 | 1.586948933 | 0.112524211 | 0.5393691 |
| Fscn3 | 0.498928155 | 0.04751468 | 0.042883458 | 1.107995533 | 0.267863745 | NA | Mall | 34.37002237 | 0.097523898 | 0.082648194 | 1.179988257 | 0.238004886 | 0.688758314 |
| Fshb | 0.192142086 | -0.026384592 | 0.031527717 | -0.836869715 | 0.402665805 | NA | Aff1 | 1100.134742 | 0.097531604 | 0.079590897 | 1.22541155 | 0.220420197 | 0.672757795 |
| Fshr | 0.431541485 | -0.028919407 | 0.044298965 | -0.652823532 | 0.513870051 | NA | Fkbpl | 260.5702435 | 0.097540958 | 0.084396876 | 1.155741331 | 0.247786977 | 0.696944196 |
| Ftcd | 3.16605085 | -0.077390674 | 0.122079584 | -0.633936248 | 0.526122428 | NA | Itch | 3050.060612 | 0.097549024 | 0.070074018 | 1.392085496 | 0.16389651 | 0.614351408 |
| Fthl17 | 0 | NA | NA | NA | NA | NA | Zdhhc9 | 2916.866403 | 0.097569811 | 0.055728578 | 1.750803878 | 0.079979698 | 0.478751136 |
| Ftmt | 0 | NA | NA | NA | NA | NA | Hsp90aa1 | 50276.28609 | 0.097588041 | 0.062147732 | 1.5702591 | 0.116354845 | 0.546450205 |
| Fut1 | 1.049546946 | 0.098197456 | 0.077943846 | 1.259848741 | 0.207723933 | NA | Rraga | 3550.711971 | 0.097695608 | 0.044868448 | 2.177378818 | 0.029452315 | 0.317771905 |
| Fut2 | 4.395458818 | 0.292207881 | 0.128666096 | 2.271055773 | 0.023143602 | NA | 1110051M20Rik | 3264.514111 | 0.097717946 | 0.057848253 | 1.689211696 | 0.091178868 | 0.503727415 |
| Fut4-ps1 | 0.820184021 | -0.034797393 | 0.063024181 | -0.552127644 | 0.580860904 | NA | Klhdc10 | 5028.887036 | 0.097747645 | 0.049302459 | 1.982611967 | 0.04741079 | 0.391951501 |
| Fxyd3 | 3.789960698 | 0.00572286 | 0.129084969 | 0.044334053 | 0.964638128 | NA | Plcl1 | 1943.281384 | 0.097751214 | 0.069081304 | 1.415016913 | 0.157063537 | 0.607833353 |
| Fxyd4 | 0.211089781 | -0.002448206 | 0.031868916 | -0.076821119 | 0.93876585 | NA | Txnrd1 | 2888.306394 | 0.097758633 | 0.060256852 | 1.622365435 | 0.104725125 | 0.525107757 |
| G630055G22Rik | 0.19046386 | -0.000774927 | 0.031526133 | -0.024580468 | 0.980389599 | NA | Rnd2 | 2243.273133 | 0.097771875 | 0.056724683 | 1.723621343 | 0.084776223 | 0.488499377 |
| G630071F17Rik | 0 | NA | NA | NA | NA | NA | Sik1 | 806.0338662 | 0.097794031 | 0.083456906 | 1.171790747 | 0.241281079 | 0.692393571 |
| G630090E17Rik | 1.842049084 | -0.193652633 | 0.103772397 | -1.866128554 | 0.062023382 | NA | Tmem185b | 570.3799743 | 0.097806304 | 0.074106094 | 1.319814591 | 0.186896929 | 0.641054259 |
| G630093K05Rik | 7.183520563 | -0.128233229 | 0.148141392 | -0.865613768 | 0.386702003 | NA | Dgkk | 177.885269 | 0.097809768 | 0.05735883 | 1.705225999 | 0.088152283 | 0.496559782 |
| G6b | 0 | NA | NA | NA | NA | NA | B4galt6 | 6811.098659 | 0.09783412 | 0.060009824 | 1.630301726 | 0.103037743 | 0.522975271 |
| G6pc | 0.843897386 | 0.005955343 | 0.070126262 | 0.084923151 | 0.932322486 | NA | Zfp808 | 51.7455227 | 0.09784008 | 0.093035822 | 1.051638802 | 0.292965298 | 0.728526371 |
| G6pc2 | 0.08731684 | 0.017365056 | 0.027733412 | 0.626142072 | 0.531221759 | NA | Ubqln1 | 7937.754046 | 0.097881486 | 0.045059231 | 2.172284898 | 0.02983418 | 0.318437312 |
| G6pd2 | 0 | NA | NA | NA | NA | NA | 3110035E14Rik | 14883.03804 | 0.097910667 | 0.078351175 | 1.249638796 | 0.211431524 | 0.666270994 |
| G730013B05Rik | 6.028322006 | 0.036204347 | 0.150148774 | 0.241123161 | 0.809459662 | NA | Soga3 | 2511.269222 | 0.097929925 | 0.062087053 | 1.577300268 | 0.114726451 | 0.543130934 |
| Gabra6 | 6.479264846 | 0.055393262 | 0.120871636 | 0.458281726 | 0.646750051 | NA | Banf1 | 1306.28899 | 0.098029484 | 0.053630791 | 1.827858247 | 0.067570829 | 0.448803776 |
| Gabrp | 0.185670226 | -0.001645086 | 0.031432796 | -0.052336624 | 0.958260472 | NA | Nek3 | 70.51091647 | 0.098030971 | 0.095100282 | 1.030816828 | 0.302626725 | 0.735974513 |
| Gabrr1 | 6.273610202 | 0.381262814 | 0.1369402 | 2.784155518 | 0.00536673 | NA | Skap1 | 60.81650945 | 0.09804389 | 0.095082023 | 1.031150644 | 0.302470182 | 0.735974513 |
| Gabrr2 | 39.57276487 | 0.578255091 | 0.155960107 | 3.70771156 | NA | NA | Farsb | 2339.58868 | 0.098076233 | 0.048809278 | 2.009376827 | 0.044497186 | 0.380589751 |
| Gabrr3 | 2.518757579 | -0.016978466 | 0.114334698 | -0.148497932 | 0.881949818 | NA | Ccdc170 | 20.28776809 | 0.098089297 | 0.071468722 | 1.372478671 | 0.169914474 | NA |
| Gal3st2 | 1.582316677 | 0.152511966 | 0.085650141 | 1.780638826 | 0.074971473 | NA | Btbd11 | 1433.131587 | 0.098099299 | 0.082478119 | 1.18939787 | 0.234283138 | 0.68557298 |
| Galnt16 | 6936.344997 | 0.097446571 | 0.165043944 | 0.590428033 | NA | NA | Isca1 | 7548.434374 | 0.098123313 | 0.051513792 | 1.90479693 | 0.056806473 | 0.417808014 |
| Galnt5 | 1.004129514 | 0.045353969 | 0.071174536 | 0.63722184 | 0.523980354 | NA | Chic1 | 1526.332113 | 0.098137818 | 0.070737043 | 1.387361038 | 0.165331699 | 0.615639135 |
| Galntl5 | 0 | NA | NA | NA | NA | NA | Ttll7 | 10352.65285 | 0.09814798 | 0.063642161 | 1.542184906 | 0.123028667 | 0.557484892 |
| Galp | 0.192142086 | -0.026384592 | 0.031527717 | -0.836869715 | 0.402665805 | NA | Slc7a5 | 3439.387918 | 0.098171698 | 0.058769374 | 1.670456738 | 0.094829032 | 0.512839359 |
| Galr2 | 0.573673924 | -0.001525252 | 0.046178452 | -0.033029522 | 0.973651045 | NA | Polr2m | 7275.627501 | 0.098214049 | 0.049701534 | 1.976076813 | 0.048146078 | 0.393789375 |
| Galr3 | 3.99998227 | -0.081628314 | 0.127764679 | -0.638895783 | 0.522890732 | NA | Grhl3 | 52.8985134 | 0.098251715 | 0.094314587 | 1.041744626 | 0.297530093 | 0.732415975 |
| Gapdhs | 0.105923158 | 0.018524693 | 0.027977518 | 0.662127842 | 0.507889294 | NA | Rcl1 | 626.2873029 | 0.098264927 | 0.072090039 | 1.363086064 | 0.172855388 | 0.624135563 |
| Gapt | 0.183387883 | -0.001444816 | 0.031390672 | -0.046026918 | 0.963288796 | NA | Atxn7l3b | 8682.103927 | 0.098265001 | 0.054193885 | 1.813211983 | 0.069799132 | 0.455201066 |
| Gast | 3.825093049 | -0.166486861 | 0.119860133 | -1.389009481 | 0.164829867 | NA | Adcy1 | 39493.82823 | 0.098265881 | 0.070309757 | 1.397613722 | 0.162229096 | 0.613957643 |
| Gata1 | 0.30883905 | -0.035660532 | 0.036608049 | -0.97411725 | 0.329998325 | NA | Nol4 | 3975.930586 | 0.098266035 | 0.071194209 | 1.380253209 | 0.167508696 | 0.617953076 |
| Gata3 | 3.030947112 | 0.230762804 | 0.092189626 | 2.503132016 | 0.012309962 | NA | Tusc3 | 6926.764975 | 0.098292523 | 0.052729786 | 1.864079683 | 0.062310515 | 0.436873272 |
| Gata4 | 0.094392817 | 0.017895942 | 0.027818137 | 0.643319236 | 0.520016977 | NA | Cxx1a | 1181.439099 | 0.098299017 | 0.064716691 | 1.51891289 | 0.128784424 | 0.567401439 |
| Gata5 | 0 | NA | NA | NA | NA | NA | Pip5k1b | 842.7611792 | 0.098301834 | 0.060003286 | 1.638274196 | 0.101364508 | 0.522606076 |
| Gata6 | 2.593282027 | 0.063245836 | 0.115025439 | 0.549842158 | 0.58242764 | NA | Tmem38b | 374.2449343 | 0.098309325 | 0.079175778 | 1.241659099 | 0.214362367 | 0.668471732 |
| Gatsl2 | 4780.810923 | 0.006269933 | 0.122850168 | 0.05103724 | NA | NA | Ache | 2560.752976 | 0.098312402 | 0.077551172 | 1.26771007 | 0.204901507 | 0.658461308 |
| Gbp1 | 0 | NA | NA | NA | NA | NA | Tmx4 | 11598.73543 | 0.098511191 | 0.06252083 | 1.575653917 | 0.115105583 | 0.543394118 |
| Gbp10 | 13.05975424 | 0.090124964 | 0.061635827 | 1.462217151 | NA | NA | Dlg3 | 9685.949002 | 0.098544172 | 0.045195556 | 2.180395175 | 0.029228181 | 0.317415456 |
| Gbp6 | 126.1925848 | 0.32540605 | 0.138247608 | 2.353791538 | NA | NA | Mrpl49 | 1418.23281 | 0.098548117 | 0.057718416 | 1.707394694 | 0.087748716 | 0.495775125 |
| Gbp8 | 9.717919696 | 0.15023243 | 0.102796668 | 1.461452329 | NA | NA | Tmeff2 | 2822.970069 | 0.098568238 | 0.047207125 | 2.087994942 | 0.036798291 | 0.347730275 |
| Gc | 1.827451186 | -0.108507875 | 0.088999008 | -1.219203195 | 0.22276708 | NA | Robo2 | 4895.781865 | 0.09857299 | 0.070728834 | 1.393674755 | 0.163415843 | 0.614351408 |
| Gcg | 0.707079824 | 0.006315794 | 0.06085822 | 0.103778824 | 0.917344871 | NA | Aldh1a3 | 39.09578032 | 0.098576536 | 0.092288597 | 1.068133437 | 0.285460325 | 0.722148665 |
| Gcgr | 0.593519161 | -0.035580971 | 0.059082103 | -0.60222925 | 0.547021548 | NA | Hcn4 | 175.8588439 | 0.098614712 | 0.094469966 | 1.043873684 | 0.296543835 | 0.731837538 |
| Gckr | 3.499604695 | 0.086851506 | 0.123832684 | 0.701361734 | 0.483077296 | NA | Cadm4 | 9944.230764 | 0.098633257 | 0.050545051 | 1.95139299 | 0.051010315 | 0.403266245 |
| Gcm1 | 0.159967308 | 0.0240345 | 0.030870969 | 0.778547006 | 0.436246606 | NA | Rslcan18 | 39.63508228 | 0.09863408 | 0.092564577 | 1.065570466 | 0.286617856 | 0.722148665 |
| Gcm2 | 0.08731684 | 0.017365056 | 0.027733412 | 0.626142072 | 0.531221759 | NA | Glb1 | 1138.176301 | 0.098649196 | 0.074231257 | 1.328944162 | 0.183866392 | 0.637355821 |
| Gcnt3 | 0 | NA | NA | NA | NA | NA | Rasgrf1 | 15912.5894 | 0.09866909 | 0.061073392 | 1.615582291 | 0.106184655 | 0.528775558 |
| Gcnt7 | 0.327786745 | -0.013119639 | 0.034805201 | -0.376944789 | 0.706214636 | NA | Dync1i1 | 5419.843574 | 0.09870996 | 0.04373901 | 2.256794545 | 0.024020916 | 0.29172307 |
| Gcsam | 0.195033442 | -0.002424765 | 0.031596358 | -0.076741901 | 0.93882887 | NA | Gnb5 | 5789.706147 | 0.098766184 | 0.063205535 | 1.562619224 | 0.118142184 | 0.549241002 |
| Gdf1 | 0.098353386 | -0.020212831 | 0.02786466 | -0.725393061 | 0.468210929 | NA | Kcna1 | 10999.59489 | 0.098780072 | 0.068664093 | 1.43859867 | 0.150264264 | 0.598511432 |
| Gdf15 | 1.598617737 | -0.019268878 | 0.101151324 | -0.190495563 | 0.84892082 | NA | Slc41a2 | 1050.023712 | 0.098857044 | 0.074853444 | 1.32067462 | 0.18660988 | 0.640826269 |
| Gdf2 | 0.098353386 | -0.020212831 | 0.02786466 | -0.725393061 | 0.468210929 | NA | Lingo1 | 14657.53753 | 0.098874566 | 0.071843471 | 1.376249846 | 0.168744295 | 0.619205089 |
| Gdf3 | 0.297029653 | 0.02866449 | 0.033517577 | 0.855207712 | 0.392436167 | NA | Pkp4 | 10054.8987 | 0.098963834 | 0.049871606 | 1.98437232 | 0.047214349 | 0.391827402 |
| Gdf6 | 3.613165193 | -0.133304603 | 0.102818013 | -1.2965102 | 0.194799768 | NA | Slc25a44 | 5141.893077 | 0.098974726 | 0.047951926 | 2.064040674 | 0.03901386 | 0.358227137 |
| Gdf7 | 5.801625888 | -0.027578161 | 0.144280865 | -0.191142191 | 0.848414193 | NA | Tmem41a | 567.7626192 | 0.099021668 | 0.068707979 | 1.441196044 | 0.149529306 | 0.598034883 |
| Gem | 89.20727141 | 0.181509619 | 0.152133927 | 1.193091 | NA | NA | H3f3b | 5847.095526 | 0.09905534 | 0.057750256 | 1.715236372 | 0.086301899 | 0.491660395 |
| Gfi1 | 0.615267569 | -0.003068902 | 0.053224915 | -0.057659123 | 0.954020155 | NA | Nedd8 | 2864.381167 | 0.09916139 | 0.050821658 | 1.951164016 | 0.051037538 | 0.403266245 |
| Gfi1b | 0.107716603 | -0.020477583 | 0.027954224 | -0.732539849 | 0.46383913 | NA | Scn1a | 7794.940686 | 0.099165631 | 0.070482155 | 1.406960823 | 0.159439006 | 0.61140734 |
| Gfra3 | 0.513772093 | -0.015418338 | 0.051085511 | -0.301814299 | 0.76279363 | NA | Bend4 | 456.6978383 | 0.099191 | 0.077826044 | 1.274521924 | 0.202478508 | 0.65606599 |
| Gfral | 0.099009884 | 0.018205726 | 0.027896865 | 0.652608315 | 0.514008823 | NA | Sod2 | 5251.63261 | 0.099196583 | 0.057658744 | 1.72040833 | 0.085358244 | 0.489697414 |
| Ggt6 | 3.368657239 | -0.096339354 | 0.116431997 | -0.82743023 | 0.407993252 | NA | Ank1 | 3575.952339 | 0.099202173 | 0.063235368 | 1.568776709 | 0.116699977 | 0.546833506 |
| Ghrh | 3.987779931 | -0.145342207 | 0.12010446 | -1.210131639 | 0.226228384 | NA | Pex10 | 399.3846228 | 0.099314628 | 0.076348393 | 1.30080837 | 0.193324056 | 0.649744561 |
| Ghrhr | 1.075850375 | 0.055512784 | 0.058551877 | 0.948095722 | 0.343080726 | NA | Rab7 | 5146.013305 | 0.099326905 | 0.054690927 | 1.816149592 | 0.069347428 | 0.454985124 |
| Ghrl | 2.238825209 | 0.169029556 | 0.097527166 | 1.733153566 | 0.083068378 | NA | Trh | 400.3082846 | 0.099328613 | 0.050807142 | 1.955012823 | 0.050581557 | 0.403264376 |
| Ghsr | 6.587346289 | 0.075959234 | 0.132407323 | 0.573678494 | 0.566185384 | NA | Zfp72 | 169.8288826 | 0.099402748 | 0.094626948 | 1.050469773 | 0.293502181 | 0.729162423 |
| Gif | 0 | NA | NA | NA | NA | NA | Zfp709 | 256.4531521 | 0.099433727 | 0.087463649 | 1.136857744 | 0.25559776 | 0.701952734 |
| Gimap3 | 3.510398947 | -0.113363313 | 0.105957268 | -1.069896525 | 0.284665888 | NA | Klhl32 | 396.8660416 | 0.099464655 | 0.086873408 | 1.144937878 | 0.252234892 | 0.69992334 |
| Gimap6 | 136.4938202 | 0.192641666 | 0.161529907 | 1.1926068 | NA | NA | Timm10 | 588.8626174 | 0.099490157 | 0.076479676 | 1.300870534 | 0.193302774 | 0.649744561 |
| Gimap7 | 0.295716657 | -0.010253415 | 0.03372058 | -0.304069933 | 0.761074613 | NA | Hspd1 | 7221.753944 | 0.099542681 | 0.053072493 | 1.875598367 | 0.060710451 | 0.431659328 |
| Gip | 0 | NA | NA | NA | NA | NA | Pp2d1 | 6.503277534 | 0.099546201 | 0.056456035 | 1.763251741 | 0.077858041 | NA |
| Gja10 | 0.517954385 | 0.0503077 | 0.044100896 | 1.140741003 | 0.253977718 | NA | Cops3 | 2454.449595 | 0.09957971 | 0.050478313 | 1.972722615 | 0.048527174 | 0.395027926 |
| Gja3 | 4.961887536 | -0.029290092 | 0.129110669 | -0.22686035 | 0.820532334 | NA | Hopx | 831.9402275 | 0.09958674 | 0.087235056 | 1.141590824 | 0.25362414 | 0.701608124 |
| Gja6 | 5.048565387 | 0.244324266 | 0.127284965 | 1.919506094 | 0.054920316 | NA | Tmem248 | 2261.480952 | 0.099603532 | 0.052621673 | 1.892823351 | 0.058381365 | 0.423355579 |
| Gja8 | 0.08731684 | 0.017365056 | 0.027733412 | 0.626142072 | 0.531221759 | NA | Sh3bp5 | 2206.68011 | 0.099631987 | 0.061341944 | 1.624206548 | 0.104331734 | 0.524144076 |
| Gjb3 | 3.015169002 | -0.044134795 | 0.111096007 | -0.397267158 | 0.691170463 | NA | Tmem17 | 191.8316393 | 0.099636178 | 0.085393739 | 1.166785523 | 0.243296995 | 0.692772033 |
| Gjb4 | 0.079983654 | 0.016740032 | 0.027674178 | 0.604897175 | 0.54524732 | NA | Uprt | 146.9649442 | 0.099641545 | 0.091187912 | 1.09270563 | 0.274523067 | 0.716185716 |
| Gjb5 | 0.400694422 | -0.006405997 | 0.042335824 | -0.151313856 | 0.879728139 | NA | Asb5 | 19.10611198 | 0.099665697 | 0.07799077 | 1.277916559 | 0.201278849 | NA |
| Gjd4 | 0.721947876 | -0.045751712 | 0.056502661 | -0.809726681 | 0.41809728 | NA | Kcna6 | 5513.77789 | 0.099680216 | 0.06034626 | 1.651804359 | 0.09857444 | 0.520050241 |
| Gje1 | 0.982925089 | 0.044350041 | 0.071094565 | 0.623817603 | 0.532747378 | NA | Ngf | 79.01724799 | 0.099718775 | 0.095078483 | 1.048804865 | 0.294267938 | 0.729567515 |
| Gk2 | 0.08731684 | 0.017365056 | 0.027733412 | 0.626142072 | 0.531221759 | NA | Msantd1 | 84.47720805 | 0.099752272 | 0.095091019 | 1.049018854 | 0.294169441 | 0.729447815 |
| Gkn1 | 2.904403723 | -0.02022624 | 0.119027508 | -0.169929123 | 0.865065877 | NA | Mrap | 11.84095858 | 0.099814719 | 0.068854661 | 1.449643599 | 0.147157931 | NA |
| Gkn2 | 0.119397641 | -0.020477583 | 0.027954224 | -0.732539845 | 0.463839132 | NA | Usp51 | 46.00038944 | 0.099819964 | 0.091763421 | 1.087796886 | 0.276684784 | 0.717475105 |
| Glb1l3 | 1.079110674 | 0.030086877 | 0.074340139 | 0.404719133 | 0.685683976 | NA | Dnaja1 | 11565.0829 | 0.099842215 | 0.065252385 | 1.530092967 | 0.125993719 | 0.562787717 |
| Glipr1l1 | 0.218407525 | -0.002234018 | 0.031987735 | -0.069839845 | 0.944321133 | NA | Atp6v0d1 | 9892.060536 | 0.099881814 | 0.042634335 | 2.342755274 | 0.019141934 | 0.263564471 |
| Glipr1l2 | 0 | NA | NA | NA | NA | NA | Ubiad1 | 443.7472002 | 0.099882217 | 0.074606948 | 1.338779033 | 0.180642619 | 0.634469716 |
| Glod5 | 3.668959318 | 0.230269405 | 0.12459466 | 1.84814827 | 0.064580896 | NA | Iars2 | 2397.818177 | 0.099907654 | 0.053178691 | 1.878715921 | 0.060283293 | 0.43030718 |
| Glra4 | 6.164400178 | -0.030199606 | 0.126484026 | -0.238762215 | 0.811289971 | NA | Strap | 4315.975215 | 0.099916434 | 0.054399627 | 1.836711723 | 0.066252464 | 0.446279967 |
| Glrp1 | 4.989901877 | 0.157099965 | 0.132768472 | 1.183262583 | 0.236705097 | NA | Gadd45a | 238.1080149 | 0.099917279 | 0.094253356 | 1.060092537 | 0.289102503 | 0.724444298 |

| | | | | | | | | | | | | | | | |
|---|---|---|---|---|---|---|---|---|---|---|---|---|---|---|---|
| Glt6d1 | 0 | NA | NA | NA | NA | NA | | | App | 72861.48143 | 0.099932758 | 0.040815393 | 2.448408568 | 0.014348884 | 0.229550543 |
| Gltpd2 | 0.119397641 | -0.020477583 | 0.027954224 | -0.732539845 | 0.463839132 | NA | | | Fam83h | 152.6974081 | 0.09993902 | 0.094601598 | 1.056419996 | 0.290776374 | 0.726615794 |
| Glyat | 0.477174001 | 0.011008699 | 0.05521377 | 0.199383206 | 0.841962996 | NA | | | Txn2 | 4295.450081 | 0.100007885 | 0.051357514 | 1.947288299 | 0.051500181 | 0.404373852 |
| Glyatl3 | 0.178993538 | 0.024685672 | 0.031260828 | 0.789667872 | 0.429721759 | NA | | | Azin1 | 6655.628201 | 0.100058376 | 0.072240654 | 1.385070182 | 0.166031009 | 0.61635484 |
| Glycam1 | 2.218913329 | 0.217720313 | 0.097647593 | 2.229653655 | 0.025770445 | NA | | | Edf1 | 2775.077818 | 0.100089652 | 0.055235003 | 1.81206929 | 0.069975491 | 0.455201066 |
| Gm10007 | 1.665083702 | -0.004579291 | 0.09434744 | -0.048536461 | 0.961288707 | NA | | | 2010320M18Rik | 148.4111201 | 0.100124798 | 0.092947416 | 1.077219818 | 0.281382074 | 0.720080692 |
| Gm10012 | 0.814547248 | -0.010378377 | 0.06227327 | -0.166658623 | 0.867638664 | NA | | | Adamts2 | 300.1633795 | 0.100126121 | 0.087339324 | 1.146403666 | 0.251628167 | 0.699465245 |
| Gm10024 | 0 | NA | NA | NA | NA | NA | | | Ap1s2 | 378.9447483 | 0.10013101 | 0.087482219 | 1.144586994 | 0.252380282 | 0.700033603 |
| Gm10046 | 2.368722812 | 0.040270498 | 0.110488348 | 0.364477329 | 0.7155016 | NA | | | Arf6 | 1448.021783 | 0.100139159 | 0.059283791 | 1.689149043 | 0.091190871 | 0.503727415 |
| Gm10052 | 0 | NA | NA | NA | NA | NA | | | Sdhd | 3060.881682 | 0.100173232 | 0.044788905 | 2.23656353 | 0.025314881 | 0.298582503 |
| Gm10057 | 0 | NA | NA | NA | NA | NA | | | Syt5 | 6086.151249 | 0.100197276 | 0.046019853 | 2.177261953 | 0.029461028 | 0.317771905 |
| Gm10058 | 0 | NA | NA | NA | NA | NA | | | Cd36 | 73.97923723 | 0.100198004 | 0.080230727 | 1.248873194 | 0.211711457 | 0.666270994 |
| Gm10081 | 0 | NA | NA | NA | NA | NA | | | Psmd11 | 2300.146664 | 0.100204232 | 0.053252594 | 1.881677949 | 0.059879755 | 0.428722425 |
| Gm10094 | 0 | NA | NA | NA | NA | NA | | | Klhl8 | 1453.565078 | 0.100227799 | 0.057034721 | 1.757311991 | 0.078864645 | 0.476674706 |
| Gm10096 | 0 | NA | NA | NA | NA | NA | | | Pmpcb | 1765.269315 | 0.100240322 | 0.057755125 | 1.735609119 | 0.082632973 | 0.484002515 |
| Gm10100 | 0 | NA | NA | NA | NA | NA | | | Ccng1 | 3587.237864 | 0.10024815 | 0.06773711 | 1.479959064 | 0.138884172 | 0.583135914 |
| Gm10104 | 0 | NA | NA | NA | NA | NA | | | Hnmt | 690.5537955 | 0.100252917 | 0.072676948 | 1.37943212 | 0.167761562 | 0.617953076 |
| Gm10142 | 0 | NA | NA | NA | NA | NA | | | Arf2 | 2080.706031 | 0.100254551 | 0.059936548 | 1.672678099 | 0.094390683 | 0.511705397 |
| Gm10147 | 0 | NA | NA | NA | NA | NA | | | Xbp1 | 3725.960326 | 0.100286469 | 0.075883797 | 1.321579475 | 0.186308222 | 0.640677359 |
| Gm10190 | 5.185794867 | -0.105169283 | 0.13768921 | -0.763816446 | 0.444976635 | NA | | | Carkd | 1344.265715 | 0.100290879 | 0.060382754 | 1.660919259 | 0.096729664 | 0.516802454 |
| Gm10220 | 2.282862464 | 0.093317574 | 0.110217915 | 0.846664303 | 0.397182261 | NA | | | Flrt3 | 1151.47031 | 0.100298945 | 0.080804689 | 1.241251543 | 0.214512839 | 0.668528965 |
| Gm10228 | 0 | NA | NA | NA | NA | NA | | | Efr3a | 6897.875722 | 0.100298958 | 0.068552252 | 1.463102305 | 0.143439387 | 0.589907432 |
| Gm10229 | 0 | NA | NA | NA | NA | NA | | | Cct8 | 6968.949747 | 0.100301653 | 0.045448973 | 2.206906918 | 0.027320563 | 0.308173879 |
| Gm10230 | 0 | NA | NA | NA | NA | NA | | | Phyh | 3090.140028 | 0.100327226 | 0.073433878 | 1.366225352 | 0.171868243 | 0.622583066 |
| Gm10248 | 0.414849978 | -0.030430887 | 0.043968299 | -0.692109732 | 0.488868421 | NA | | | Cmpk1 | 1876.177159 | 0.100385955 | 0.068742421 | 1.460320328 | 0.144202058 | 0.589967032 |
| Gm10256 | 0 | NA | NA | NA | NA | NA | | | Atp8a2 | 1318.699026 | 0.100394954 | 0.059741108 | 1.680500385 | 0.092859999 | 0.507099381 |
| Gm10267 | 0 | NA | NA | NA | NA | NA | | | Arid5a | 453.440432 | 0.100409561 | 0.086215197 | 1.164638776 | 0.244165241 | 0.693692714 |
| Gm10272 | 0 | NA | NA | NA | NA | NA | | | Pcdh7 | 5410.637317 | 0.100493309 | 0.071633663 | 1.402878275 | 0.16065314 | 0.612387888 |
| Gm10280 | 0.099009884 | 0.018205726 | 0.027896865 | 0.652608315 | 0.514008823 | NA | | | Rad18 | 188.7378379 | 0.100500769 | 0.089487891 | 1.123065564 | 0.26140965 | 0.704173028 |
| Gm10318 | 0 | NA | NA | NA | NA | NA | | | Asic4 | 1061.735998 | 0.100562422 | 0.075622345 | 1.329797722 | 0.183584927 | 0.637217359 |
| Gm10319 | 0 | NA | NA | NA | NA | NA | | | Dnajb12 | 1423.930332 | 0.100568816 | 0.055182335 | 1.822482063 | 0.068381873 | 0.451301721 |
| Gm10324 | 0.08731684 | 0.017365056 | 0.027733412 | 0.626142072 | 0.531221759 | NA | | | Zdhhc5 | 2866.36717 | 0.100598338 | 0.060871241 | 1.652641498 | 0.098403847 | 0.520050241 |
| Gm10352 | 0 | NA | NA | NA | NA | NA | | | Nhp2l1 | 4171.408381 | 0.100621218 | 0.054647471 | 1.8412786 | 0.065580742 | 0.4455608 |
| Gm10364 | 1.021338241 | -0.011381811 | 0.07725403 | -0.147329678 | 0.882871809 | NA | | | Sbds | 2490.967058 | 0.10062428 | 0.053842619 | 1.868859325 | 0.061642389 | 0.435129208 |
| Gm10373 | 0.806298578 | 0.016681995 | 0.049256587 | 0.338675417 | 0.734854262 | NA | | | Cmtm8 | 39.76824599 | 0.100651014 | 0.086596253 | 1.162302194 | 0.245112736 | 0.694718388 |
| Gm10375 | 0 | NA | NA | NA | NA | NA | | | Pnkd | 8518.101533 | 0.100669598 | 0.044708331 | 2.25169665 | 0.024341449 | 0.292457679 |
| Gm10377 | 0 | NA | NA | NA | NA | NA | | | Samd14 | 1809.334686 | 0.100674844 | 0.074471222 | 1.351862387 | 0.176419342 | 0.629083076 |
| Gm10389 | 2.031687473 | 0.01096503 | 0.109148051 | 0.100460152 | 0.919979017 | NA | | | Zfp213 | 328.7713105 | 0.100689866 | 0.075269741 | 1.337720378 | 0.180987606 | 0.634469716 |
| Gm10390 | 0.844386285 | 0.031879823 | 0.064648562 | 0.493125017 | 0.621924255 | NA | | | Ntrk2 | 25271.18833 | 0.100735871 | 0.058150611 | 1.732326948 | 0.083215367 | 0.485168543 |
| Gm10400 | 0.096071043 | -0.020098795 | 0.027825948 | -0.722304036 | 0.470107572 | NA | | | Zc3hc1 | 952.1529602 | 0.100747739 | 0.061339923 | 1.642449712 | 0.10049684 | 0.521177114 |
| Gm10408 | 0.440944058 | 0.059506558 | 0.048029448 | 1.238959851 | 0.215360367 | NA | | | Pcdhga8 | 582.2285774 | 0.100806413 | 0.089086061 | 1.13156212 | 0.257818573 | 0.703616072 |
| Gm10409 | 0 | NA | NA | NA | NA | NA | | | Tagln3 | 9297.751364 | 0.100819554 | 0.05241151 | 1.923614771 | 0.054402888 | 0.410949742 |
| Gm10413 | 0 | NA | NA | NA | NA | NA | | | Entpd3 | 626.8333617 | 0.100864964 | 0.075700624 | 1.332419189 | 0.18272248 | 0.635742932 |
| Gm10415 | 0.273386355 | 0.041384988 | 0.040339907 | 1.025906866 | 0.304935486 | NA | | | 4933430I17Rik | 15.56437843 | 0.100868527 | 0.073781715 | 1.367120929 | 0.171587404 | NA |
| Gm10436 | 0 | NA | NA | NA | NA | NA | | | Mrrf | 714.6948683 | 0.100923847 | 0.06157791 | 1.638961879 | 0.101221199 | 0.522322962 |
| Gm10439 | 0 | NA | NA | NA | NA | NA | | | Syt2 | 2133.005195 | 0.100936533 | 0.083828113 | 1.204089287 | 0.22855507 | 0.681722988 |
| Gm10440 | 2.269802738 | 0.113465519 | 0.106587171 | 1.064532609 | 0.28708749 | NA | | | Appbp2 | 1861.088909 | 0.10095096 | 0.072238803 | 1.397461684 | 0.162274781 | 0.613957643 |
| Gm10445 | 0 | NA | NA | NA | NA | NA | | | Prdx2 | 7445.240719 | 0.101029592 | 0.050134945 | 2.015153135 | 0.043888606 | 0.379253953 |
| Gm1045 | 0.678127538 | -0.001548773 | 0.04101571 | -0.037760472 | 0.969878661 | NA | | | Ftsjd1 | 264.299445 | 0.101044394 | 0.0872842 | 1.157648162 | 0.247007646 | 0.696511265 |
| Gm10466 | 0 | NA | NA | NA | NA | NA | | | Srpr | 3279.507345 | 0.101051311 | 0.049484213 | 2.042091925 | 0.041142414 | 0.366647057 |
| Gm10471 | 3.549274368 | 0.013462761 | 0.114199705 | 0.117887877 | 0.906156499 | NA | | | Spock2 | 32981.30078 | 0.101062821 | 0.05144195 | 1.964599329 | 0.04946063 | 0.39893239 |
| Gm10474 | 3.710584375 | 0.001827684 | 0.119330919 | 0.015316094 | 0.987780003 | NA | | | Npsr1 | 195.7970387 | 0.10109771 | 0.087959363 | 1.149368372 | 0.250404116 | 0.698688088 |
| Gm10485 | 0.301004085 | 0.013136417 | 0.039354285 | 0.333798913 | 0.738531304 | NA | | | Fto | 9756.982879 | 0.101122789 | 0.044040089 | 2.296153155 | 0.021667126 | 0.276408712 |
| Gm10486 | 0 | NA | NA | NA | NA | NA | | | Fem1b | 6906.590098 | 0.101133314 | 0.068966171 | 1.466419154 | 0.142534131 | 0.588697979 |
| Gm10487 | 0 | NA | NA | NA | NA | NA | | | Hibadh | 2083.484178 | 0.101140734 | 0.058331812 | 1.733886373 | 0.082938246 | 0.484423387 |
| Gm10488 | 0 | NA | NA | NA | NA | NA | | | Tmem115 | 1239.319519 | 0.101157049 | 0.054539599 | 1.854745031 | 0.063632645 | 0.440996088 |
| Gm10494 | 0.613100444 | -0.006928698 | 0.062938259 | -0.110087217 | 0.912340206 | NA | | | Tmem30a | 12720.46596 | 0.101175547 | 0.078141632 | 1.294771357 | 0.195399116 | 0.651003086 |
| Gm10510 | 0.256038351 | 0.007085343 | 0.03307033 | 0.214250763 | 0.830351512 | NA | | | Slc25a16 | 1719.159798 | 0.101199367 | 0.064081759 | 1.579222671 | 0.114284992 | 0.543130934 |
| Gm10512 | 0.107716603 | -0.020477583 | 0.027954224 | -0.732539849 | 0.46383913 | NA | | | Spint2 | 707.3367213 | 0.101233272 | 0.092568068 | 1.093608993 | 0.274126503 | 0.716185716 |
| Gm10536 | 1.848793241 | 0.060591749 | 0.106909603 | 0.566756847 | 0.570879392 | NA | | | 1700123O20Rik | 578.1495442 | 0.101263347 | 0.069772071 | 1.451345019 | 0.146683811 | 0.593534375 |
| Gm10538 | 0 | NA | NA | NA | NA | NA | | | Etl4 | 5421.192464 | 0.101299105 | 0.047181495 | 2.147009233 | 0.031792547 | 0.326301135 |
| Gm10549 | 0.945542035 | 0.081901904 | 0.07456633 | 1.098376493 | 0.272040123 | NA | | | Timm23 | 2311.203929 | 0.101304544 | 0.050978763 | 1.987191109 | 0.046901223 | 0.39102736 |
| Gm10556 | 0.615078183 | -0.001857088 | 0.053211811 | -0.034899925 | 0.97215954 | NA | | | Snx1 | 2966.863467 | 0.101320602 | 0.043405528 | 2.33427878 | 0.019581129 | 0.26397034 |
| Gm10578 | 0 | NA | NA | NA | NA | NA | | | B020004J07Rik | 9.358360774 | 0.10133877 | 0.065032411 | 1.558280984 | 0.119166655 | NA |
| Gm10584 | 5.185747891 | -0.037390519 | 0.135246782 | -0.276461433 | 0.782193678 | NA | | | Zfp708 | 142.9321592 | 0.101355901 | 0.09186573 | 1.103304809 | 0.269894819 | 0.71216373 |
| Gm10591 | 0 | NA | NA | NA | NA | NA | | | Nnat | 5669.7866 | 0.101389063 | 0.094124471 | 1.077180698 | 0.281399547 | 0.720080692 |
| Gm10619 | 0 | NA | NA | NA | NA | NA | | | Trhde | 1377.912931 | 0.101418734 | 0.068427134 | 1.482142075 | 0.138302495 | 0.582904025 |
| Gm10636 | 2.154172627 | 0.072376572 | 0.110402682 | 0.655568967 | 0.512101498 | NA | | | Pigf | 238.4987365 | 0.101438307 | 0.088649839 | 1.144258214 | 0.252516567 | 0.700277903 |
| Gm10637 | 0.482637559 | -0.049175986 | 0.042867026 | -1.14717514 | 0.251309244 | NA | | | Ankrd27 | 1839.694607 | 0.101439393 | 0.057582264 | 1.761643015 | 0.078129631 | 0.475440315 |
| Gm10639 | 0.479713553 | 0.008997466 | 0.050300534 | 0.178874167 | 0.858036507 | NA | | | Sult4a1 | 18853.22657 | 0.101490628 | 0.044471449 | 2.282152487 | 0.022480339 | 0.281750998 |
| Gm10640 | 6.972240599 | -0.029134361 | 0.151803251 | -0.191921853 | 0.84780342 | NA | | | 1700008O03Rik | 25.79014218 | 0.101567999 | 0.084236461 | 1.205748649 | 0.227914421 | 0.680415543 |
| Gm10649 | 6.142534515 | -0.083052187 | 0.142983268 | -0.580852493 | 0.561339873 | NA | | | Rerg | 560.8866757 | 0.101583678 | 0.08182001 | 1.241550548 | 0.214402437 | 0.668471732 |
| Gm10662 | 0 | NA | NA | NA | NA | NA | | | Gnl1 | 8222.838656 | 0.101653183 | 0.048689524 | 2.087783449 | 0.036817373 | 0.347730275 |
| Gm10664 | 0.394136287 | -0.017186747 | 0.03688538 | -0.465950104 | 0.641251216 | NA | | | Mrps23 | 1241.200586 | 0.10167709 | 0.049814802 | 2.041101966 | 0.041240693 | 0.367072492 |
| Gm10665 | 0 | NA | NA | NA | NA | NA | | | Khdrbs3 | 5337.640607 | 0.10168123 | 0.037217164 | 2.732105844 | 0.006293092 | 0.152102254 |
| Gm10666 | 0 | NA | NA | NA | NA | NA | | | Spata2 | 3374.979222 | 0.10170534 | 0.057558669 | 1.766985602 | 0.07723064 | 0.4735552 |
| Gm10670 | 0 | NA | NA | NA | NA | NA | | | Ccdc146 | 37.03253974 | 0.101714554 | 0.084757828 | 1.200060888 | 0.230115694 | 0.683133099 |
| Gm10677 | 6.693545595 | -0.04670418 | 0.125735101 | -0.371449016 | 0.710303123 | NA | | | Paqr6 | 283.0685876 | 0.101741725 | 0.092163356 | 1.103928179 | 0.269624294 | 0.711972822 |
| Gm10681 | 0 | NA | NA | NA | NA | NA | | | Fam199x | 412.9827982 | 0.101745361 | 0.084371363 | 1.205922925 | 0.22784721 | 0.680415543 |
| Gm10684 | 3.96246455 | 0.214059037 | 0.12896362 | 1.659840473 | 0.096946549 | NA | | | Psmd2 | 9939.602825 | 0.101879367 | 0.043339107 | 2.350749094 | 0.018735663 | 0.261184487 |
| Gm10696 | 0.107716603 | -0.020477583 | 0.027954224 | -0.732539849 | 0.46383913 | NA | | | Hars | 3971.040918 | 0.101888347 | 0.045642752 | 2.232300696 | 0.025595096 | 0.300343543 |
| Gm10697 | 0 | NA | NA | NA | NA | NA | | | Nop16 | 684.8504176 | 0.101976385 | 0.062503688 | 1.631525887 | 0.102779402 | 0.522975271 |
| Gm10714 | 5.966882616 | 0.222710908 | 0.141662646 | 1.572121616 | 0.115922349 | NA | | | Mki67ip | 945.1054064 | 0.101978913 | 0.062530984 | 1.630854123 | 0.102921104 | 0.522975271 |
| Gm10731 | 1.168665658 | 0.004111664 | 0.072666983 | 0.056582281 | 0.954877949 | NA | | | Fam107b | 335.1305349 | 0.102007013 | 0.075826329 | 1.345271681 | 0.178537515 | 0.632082854 |
| Gm10745 | 0 | NA | NA | NA | NA | NA | | | Hyou1 | 5364.034184 | 0.102059902 | 0.061034884 | 1.672156883 | 0.09449339 | 0.511705397 |
| Gm10768 | 1.119424133 | -0.021548962 | 0.077527442 | -0.277952703 | 0.781048666 | NA | | | Nlrp5-ps | 25.13263652 | 0.102071353 | 0.078464672 | 1.300857448 | 0.193307254 | 0.649744561 |
| Gm10782 | 0 | NA | NA | NA | NA | NA | | | Ftsj2 | 289.3321847 | 0.102081731 | 0.078853006 | 1.294582607 | 0.195464256 | 0.651003086 |
| Gm10787 | 0.116696964 | -0.020477583 | 0.027954224 | -0.732539847 | 0.463839131 | NA | | | Exoc5 | 2021.174596 | 0.102120151 | 0.073535967 | 1.388710247 | 0.164920877 | 0.615071433 |
| Gm10789 | 6.078012854 | 0.015137487 | 0.129637506 | 0.116767803 | 0.907044058 | NA | | | Amigo2 | 503.1895601 | 0.102127363 | 0.082764832 | 1.233946362 | 0.217222892 | 0.670311138 |
| Gm10791 | 7.070179103 | 0.046711269 | 0.141089308 | 0.331075893 | 0.740587161 | NA | | | Ube2n | 7704.799102 | 0.1021284 | 0.051583709 | 1.979857617 | 0.04771953 | 0.392678115 |
| Gm10804 | 0 | NA | NA | NA | NA | NA | | | Traf3 | 2535.95311 | 0.102169509 | 0.074307306 | 1.374959132 | 0.169144115 | 0.619512712 |
| Gm10818 | 2.069276574 | 0.069858037 | 0.094961905 | 0.735642754 | 0.461948137 | NA | | | Emc6 | 974.5942158 | 0.102170604 | 0.074362049 | 1.373961663 | 0.169453585 | 0.619945581 |
| Gm1082 | 2.166970131 | 0.011149646 | 0.097126817 | 0.114794721 | 0.908607833 | NA | | | Pdp1 | 6703.975173 | 0.102187471 | 0.095073083 | 1.07483073 | 0.28245052 | 0.72081452 |
| Gm10823 | 0 | NA | NA | NA | NA | NA | | | Fars2 | 596.5474434 | 0.10221101 | 0.072124902 | 1.417138973 | 0.1564423 | 0.60719411 |
| Gm10825 | 5.723088238 | -0.040102144 | 0.128624986 | -0.311775695 | 0.755210997 | NA | | | Umodl1 | 16.17612813 | 0.102219662 | 0.070823355 | 1.443304422 | 0.148934735 | NA |
| Gm10857 | 0.607229355 | 0.041512175 | 0.049138713 | 0.844795719 | 0.398224905 | NA | | | Clns1a | 2504.730721 | 0.102223543 | 0.047540666 | 2.150233743 | 0.03153673 | 0.32443523 |
| Gm10872 | 0.506202321 | -0.046639326 | 0.052906751 | -0.881538272 | 0.378026549 | NA | | | Psmb4 | 3719.480736 | 0.102231233 | 0.044261135 | 2.309729118 | 0.020903156 | 0.272322188 |
| Gm10921 | 0 | NA | NA | NA | NA | NA | | | Ppp2cb | 4528.981371 | 0.102251011 | 0.053145141 | 1.923995487 | 0.054355149 | 0.410949742 |
| Gm10922 | 0 | NA | NA | NA | NA | NA | | | Nxpe3 | 1212.16638 | 0.102268397 | 0.056323171 | 1.815742904 | 0.069409819 | 0.454985124 |
| Gm1110 | 1.351077591 | 0.096155621 | 0.083031192 | 1.158066254 | 0.246837 | NA | | | Lsamp | 2368.805213 | 0.102348574 | 0.055419245 | 1.846805649 | 0.064775313 | 0.443414795 |
| Gm11128 | 6.173766686 | -0.039183861 | 0.133097611 | -0.294399433 | 0.7684527 | NA | | | Nxph4 | 238.3032636 | 0.102390866 | 0.094680994 | 1.081429988 | 0.279505888 | 0.718934998 |
| Gm11149 | 4.404330934 | 0.067551017 | 0.101183254 | 0.667610641 | 0.504382164 | NA | | | Gpr101 | 462.2642137 | 0.102395063 | 0.066259634 | 1.545361138 | 0.122258935 | 0.556819993 |
| Gm11166 | 1.589607904 | 0.014514141 | 0.097236105 | 0.14926699 | 0.881342961 | NA | | | Lonrf2 | 7786.819566 | 0.102396653 | 0.063078333 | 1.623325287 | 0.104519887 | 0.524427522 |
| Gm11186 | 0 | NA | NA | NA | NA | NA | | | Dbndd2 | 5936.280301 | 0.102400906 | 0.062139858 | 1.64791022 | 0.099371096 | 0.520411863 |
| Gm11194 | 0 | NA | NA | NA | NA | NA | | | Kcnn1 | 779.9608141 | 0.102435868 | 0.075937362 | 1.348952166 | 0.177352331 | 0.629728662 |
| Gm11201 | 2.19009043 | -0.047749419 | 0.091201009 | -0.523562398 | 0.600582941 | NA | | | Ptpn9 | 2091.81978 | 0.102508741 | 0.071189956 | 1.439932643 | 0.149886457 | 0.598234686 |
| Gm11202 | 0.29182997 | 0.032201186 | 0.035537591 | 0.906116166 | 0.36487437 | NA | | | Sik2 | 1375.924179 | 0.102535973 | 0.091245074 | 1.123742559 | 0.261122256 | 0.704173028 |
| Gm11213 | 0 | NA | NA | NA | NA | NA | | | Enho | 1422.275599 | 0.102546919 | 0.066488236 | 1.542331784 | 0.122992989 | 0.557484892 |
| Gm1123 | 2.051606762 | 0.064583559 | 0.094182112 | 0.685730627 | 0.49288299 | NA | | | Grhpr | 1031.2325 | 0.102554263 | 0.07072739 | 1.449993598 | 0.147060304 | 0.594031723 |
| Gm11240 | 0.215468684 | -0.027008432 | 0.031909031 | -0.846419662 | 0.397318674 | NA | | | Ubc | 5673.517106 | 0.102639231 | 0.067097705 | 1.52969808 | 0.126091479 | 0.562787717 |
| Gm11351 | 0.198019769 | 0.025308392 | 0.031628095 | 0.80018706 | 0.423602426 | NA | | | Bcap31 | 2315.805919 | 0.1026466 | 0.061995737 | 1.655704169 | 0.097781743 | 0.518456434 |
| Gm1140 | 0 | NA | NA | NA | NA | NA | | | A830018L16Rik | 2979.518549 | 0.10264729 | 0.072219481 | 1.421324116 | 0.155222556 | 0.604578396 |
| Gm11413 | 7.376766201 | 0.069026574 | 0.148567712 | 0.46461356 | 0.64220822 | NA | | | Atp5o | 5567.383678 | 0.102648044 | 0.062687616 | 1.637453314 | 0.101535786 | 0.522606076 |
| Gm11426 | 0.08731684 | 0.017365056 | 0.027733412 | 0.626142072 | 0.531221759 | NA | | | Gm11549 | 2066.04018 | 0.102704971 | 0.074647681 | 1.375862852 | 0.168864098 | 0.619267834 |
| Gm11435 | 0.098353386 | -0.020212831 | 0.02786466 | -0.725393061 | 0.468210929 | NA | | | Aamp | 5143.013101 | 0.102720906 | 0.069788702 | 1.471884466 | 0.141052079 | 0.586989325 |
| Gm11437 | 0.293434314 | -0.014075032 | 0.039227749 | -0.358802961 | 0.719742498 | NA | | | Slc25a33 | 1167.586742 | 0.102732093 | 0.065198158 | 1.575690114 | 0.115097237 | 0.543394118 |
| Gm11468 | 2.177079743 | -0.213911707 | 0.103844481 | -2.059923699 | 0.039405836 | NA | | | Sulf2 | 4225.877953 | 0.102750526 | 0.067467932 | 1.522953546 | 0.127770328 | 0.565596401 |
| Gm11487 | 0 | NA | NA | NA | NA | NA | | | Ran | 7704.165734 | 0.102814795 | 0.048523198 | 2.118879192 | 0.034100678 | 0.336970372 |
| Gm11529 | 2.633445258 | -0.176207923 | 0.113559765 | -1.551675658 | 0.12073985 | NA | | | Ngfr | 202.8552945 | 0.102862154 | 0.08089483 | 1.271554112 | 0.20353159 | 0.657622144 |
| Gm11541 | 0 | NA | NA | NA | NA | NA | | | Kpna2 | 1313.979789 | 0.102894895 | 0.062621565 | 1.643122377 | 0.100357616 | 0.5211685 |
| Gm11544 | 0 | NA | NA | NA | NA | NA | | | Mrpl43 | 1130.444247 | 0.102938064 | 0.058567401 | 1.757599988 | 0.078815595 | 0.476674706 |
| Gm11545 | 6.538721738 | 0.519609241 | 0.148984531 | 3.48767243 | 0.000487245 | NA | | | Resp18 | 2680.184614 | 0.102960777 | 0.094653075 | 1.087770014 | 0.27669665 | 0.717475105 |
| Gm11548 | 0 | NA | NA | NA | NA | NA | | | Slc25a17 | 1233.66415 | 0.103040743 | 0.060174728 | 1.712359094 | 0.086830513 | 0.493466364 |
| Gm11554 | 0 | NA | NA | NA | NA | NA | | | Edil3 | 5534.306273 | 0.103050306 | 0.058756214 | 1.75386226 | 0.079454112 | 0.477013957 |
| Gm11559 | 0 | NA | NA | NA | NA | NA | | | Ube2g2 | 1887.431077 | 0.103053474 | 0.061303202 | 1.681045539 | 0.09275407 | 0.506868248 |
| Gm11562 | 0 | NA | NA | NA | NA | NA | | | Armcx3 | 4059.873981 | 0.103067797 | 0.057346418 | 1.797283967 | 0.07229055 | 0.461579135 |
| Gm11563 | 0 | NA | NA | NA | NA | NA | | | Ola1 | 3623.287017 | 0.103105546 | 0.063913836 | 1.613196011 | 0.10670193 | 0.530442242 |
| Gm11564 | 0 | NA | NA | NA | NA | NA | | | Ctnna3 | 78.39929691 | 0.103130013 | 0.093989747 | 1.097247482 | 0.272533222 | 0.714142044 |
| Gm11565 | 0 | NA | NA | NA | NA | NA | | | Kdm4a | 2450.191678 | 0.103144197 | 0.056063154 | 1.839785851 | 0.065799683 | 0.445805132 |
| Gm11567 | 0 | NA | NA | NA | NA | NA | | | Nsg1 | 12026.7152 | 0.10314885 | 0.046800211 | 2.204025304 | 0.027522565 | 0.308173879 |
| Gm11568 | 0 | NA | NA | NA | NA | NA | | | Mrto4 | 750.287953 | 0.103198162 | 0.062528605 | 1.650415227 | 0.098858039 | 0.520271327 |
| Gm11569 | 0 | NA | NA | NA | NA | NA | | | Prorsd1 | 66.8005408 | 0.103202742 | 0.091945382 | 1.122435291 | 0.261677406 | 0.704173028 |
| Gm11570 | 0 | NA | NA | NA | NA | NA | | | Slc39a6 | 2041.152423 | 0.103219453 | 0.060942448 | 1.693720165 | 0.090318471 | 0.50209189 |
| Gm11595 | 0 | NA | NA | NA | NA | NA | | | Ndufs7 | 2402.143527 | 0.103279062 | 0.064800698 | 1.59379551 | 0.110981844 | 0.537553271 |
| Gm11596 | 0.098353386 | -0.020212831 | 0.02786466 | -0.725393061 | 0.468210929 | NA | | | Cntfr | 3163.08383 | 0.103310282 | 0.04965591 | 2.080523391 | 0.037477552 | 0.350065015 |
| Gm11648 | 4.544781541 | 0.068948765 | 0.138959469 | 0.496178959 | 0.619768155 | NA | | | Cgref1 | 730.8393697 | 0.103368743 | 0.07372874 | 1.402014227 | 0.160910998 | 0.612415571 |
| Gm11710 | 0 | NA | NA | NA | NA | NA | | | Nfil3 | 359.0604024 | 0.103402345 | 0.087148178 | 1.186511837 | 0.23542023 | 0.686276191 |
| Gm11711 | 0 | NA | NA | NA | NA | NA | | | Stab2 | 28.11291031 | 0.10340603 | 0.087560304 | 1.180969291 | 0.237614923 | 0.688529917 |
| Gm11747 | 0.2777807 | 0.011208478 | 0.038523646 | 0.290950592 | 0.771089108 | NA | | | Dcp2 | 3672.514735 | 0.103409099 | 0.066931344 | 1.545002578 | 0.12234564 | 0.556819993 |
| Gm11756 | 0 | NA | NA | NA | NA | NA | | | Cldn12 | 1969.163834 | 0.103435848 | 0.065308587 | 1.583801664 | 0.113238861 | 0.540327972 |
| Gm11757 | 0 | NA | NA | NA | NA | NA | | | Mif | 3916.093695 | 0.103459276 | 0.057379366 | 1.803074574 | 0.071376504 | 0.459174089 |
| Gm11758 | 0 | NA | NA | NA | NA | NA | | | Itpk1 | 2774.904524 | 0.103525073 | 0.061425798 | 1.685367983 | 0.091917597 | 0.505527387 |
| Gm11937 | 0 | NA | NA | NA | NA | NA | | | Nov | 3392.544286 | 0.103529678 | 0.089134526 | 1.161499176 | 0.245438958 | 0.69504627 |
| Gm11938 | 0 | NA | NA | NA | NA | NA | | | Ctif | 5065.395647 | 0.103529874 | 0.048357291 | 2.140936183 | 0.032279185 | 0.327711599 |
| Gm11961 | 1.284420953 | -0.023135237 | 0.082292133 | -0.281135459 | 0.778606505 | NA | | | Ugcg | 3144.18393 | 0.10355382 | 0.064057429 | 1.616577821 | 0.105969443 | 0.52850889 |
| Gm11978 | 2.445186259 | 0.100952169 | 0.097897036 | 1.031207611 | 0.302443472 | NA | | | Hypk | 1083.625834 | 0.103581019 | 0.071909456 | 1.440436685 | 0.149743891 | 0.598098567 |
| Gm11981 | 0 | NA | NA | NA | NA | NA | | | Zfp59 | 355.6464229 | 0.103640602 | 0.073887611 | 1.402679022 | 0.160712576 | 0.612387888 |
| Gm11985 | 0 | NA | NA | NA | NA | NA | | | Fzd3 | 5143.528497 | 0.103658139 | 0.077085854 | 1.344710257 | 0.178718821 | 0.632262443 |
| Gm12 | 0 | NA | NA | NA | NA | NA | | | Grm5 | 6530.84366 | 0.103658927 | 0.076277696 | 1.358967719 | 0.174156818 | 0.626964546 |
| Gm12060 | 3.907643635 | 0.041831464 | 0.122959699 | 0.340204669 | 0.733702402 | NA | | | Cds1 | 5172.605435 | 0.103682157 | 0.058115554 | 1.784068974 | 0.074412458 | 0.46772625 |
| Gm12130 | 0 | NA | NA | NA | NA | NA | | | Srprb | 1711.861855 | 0.103708587 | 0.053813943 | 1.927169456 | 0.053958515 | 0.409506392 |
| Gm12159 | 0 | NA | NA | NA | NA | NA | | | Tex264 | 2412.686386 | 0.103718725 | 0.056042488 | 1.850715919 | 0.064210433 | 0.442047746 |
| Gm12169 | 0.107716603 | -0.020477583 | 0.027954224 | -0.732539849 | 0.46383913 | NA | | | Rora | 485.8677261 | 0.103725018 | 0.081696933 | 1.269631727 | 0.204215842 | 0.65778952 |
| Gm12171 | 0.20671448 | -0.003314027 | 0.03178208 | -0.10427343 | 0.916952363 | NA | | | Gpd1l | 6436.086605 | 0.1037361 | 0.043901475 | 2.362929716 | 0.018131108 | 0.256569934 |
| Gm12185 | 8.060386651 | 0.00437008 | 0.133305673 | 0.032782402 | 0.973848112 | NA | | | Slmo2 | 1601.198099 | 0.103762206 | 0.060389963 | 1.718202836 | 0.085759624 | 0.490643681 |
| Gm12191 | 0 | NA | NA | NA | NA | NA | | | Scamp1 | 10363.30544 | 0.103788864 | 0.060655967 | 1.711107223 | 0.087061321 | 0.494198026 |
| Gm12228 | 0 | NA | NA | NA | NA | NA | | | Siva1 | 190.2692243 | 0.103817192 | 0.093157697 | 1.114424203 | 0.265097245 | 0.707370515 |
| Gm12238 | 0.462717353 | 0.004293417 | 0.044257148 | 0.097010698 | 0.922717899 | NA | | | Tbc1d22b | 2224.341879 | 0.103846263 | 0.062065622 | 1.67316882 | 0.094294067 | 0.511469609 |
| Gm12250 | 22.08773812 | 0.203168811 | 0.139805838 | 1.453221225 | NA | NA | | | D15Ertd621e | 3359.223735 | 0.103847244 | 0.06461495 | 1.607170547 | 0.108016961 | 0.533425002 |
| Gm12253 | 0 | NA | NA | NA | NA | NA | | | Urgcp | 3426.48322 | 0.103902296 | 0.056358846 | 1.843584515 | 0.065243714 | 0.444691459 |
| Gm12295 | 1.534685539 | 0.015731773 | 0.084450203 | 0.186284614 | 0.852221573 | NA | | | Ubl3 | 6609.28289 | 0.103920246 | 0.054309444 | 1.91348389 | 0.055686134 | 0.413057568 |
| Gm12298 | 0 | NA | NA | NA | NA | NA | | | Gm17455 | 13.17172147 | 0.103929399 | 0.063904196 | 1.626331368 | 0.103879182 | NA |
| Gm12360 | 4.13031607 | 0.095576326 | 0.09404658 | 1.016265824 | 0.309502819 | NA | | | Xylt1 | 860.2847853 | 0.103934875 | 0.066981787 | 1.551688595 | 0.120736753 | 0.554612895 |
| Gm12409 | 0.368800901 | -0.003607529 | 0.046901082 | -0.07691782 | 0.938688921 | NA | | | Trappc13 | 1961.087758 | 0.103940683 | 0.058237691 | 1.784766537 | 0.074299193 | 0.467418832 |
| Gm12504 | 3.0697094 | -0.033102735 | 0.107900842 | -0.30678848 | 0.759004377 | NA | | | Lrfn2 | 1095.753781 | 0.103959865 | 0.088106434 | 1.179934988 | 0.238026073 | 0.688758314 |
| Gm12505 | 0 | NA | NA | NA | NA | NA | | | Glra3 | 204.1073634 | 0.103968416 | 0.092982114 | 1.118155008 | 0.263500796 | 0.706200557 |
| Gm12530 | 5.056708818 | 0.044885282 | 0.144270955 | 0.311117939 | 0.755710964 | NA | | | Mcts2 | 221.0839935 | 0.103986416 | 0.080348097 | 1.294198868 | 0.195596738 | 0.651212059 |
| Gm12597 | 0 | NA | NA | NA | NA | NA | | | Thbs4 | 340.5523734 | 0.103991061 | 0.09495903 | 1.095115028 | 0.273466246 | 0.715226987 |
| Gm12603 | 0.105923158 | 0.018524693 | 0.027977518 | 0.662127842 | 0.507889294 | NA | | | Isl1 | 104.568127 | 0.104110596 | 0.089268148 | 1.166268124 | 0.243506057 | 0.692772033 |
| Gm12633 | 0 | NA | NA | NA | NA | NA | | | Txnl4a | 4564.104754 | 0.104134094 | 0.046582311 | 2.235485786 | 0.025385474 | 0.298582503 |
| Gm12657 | 2.130322593 | 0.099612284 | 0.090831579 | 1.096670184 | 0.272785595 | NA | | | H2-T10 | 57.64705417 | 0.104144481 | 0.094920093 | 1.097180567 | 0.272562466 | 0.714142044 |
| Gm12669 | 0 | NA | NA | NA | NA | NA | | | Arhgap6 | 581.7381422 | 0.104153787 | 0.091279466 | 1.141042909 | 0.253852067 | 0.701619624 |
| Gm12695 | 2.027333788 | 0.006431673 | 0.090511717 | 0.07105901 | 0.943350791 | NA | | | Gstm6 | 273.1035258 | 0.104171823 | 0.092231048 | 1.129465887 | 0.258701352 | 0.703616072 |
| Gm12709 | 0.649215748 | 0.028699483 | 0.054876546 | 0.522982684 | 0.600986303 | NA | | | Cox8b | 21.55887925 | 0.10417812 | 0.059218938 | 1.759202787 | 0.078543071 | NA |
| Gm12718 | 0.107716603 | -0.020477583 | 0.027954224 | -0.732539849 | 0.46383913 | NA | | | Acta1 | 328.3779266 | 0.104224747 | 0.080044185 | 1.302090179 | 0.19288556 | 0.649602848 |
| Gm12789 | 0.119397641 | -0.020477583 | 0.027954224 | -0.732539845 | 0.463839132 | NA | | | Neto2 | 2539.318839 | 0.104226398 | 0.086615718 | 1.203319677 | 0.228852637 | 0.681764163 |
| Gm12794 | 0.198019769 | 0.025308392 | 0.031628095 | 0.80018706 | 0.423602426 | NA | | | Mesdc2 | 2260.582711 | 0.104267312 | 0.061638018 | 1.691607161 | 0.0907209 | 0.502789698 |
| Gm128 | 4.202001019 | 0.079275767 | 0.133426887 | 0.594151365 | 0.552410878 | NA | | | Glrx5 | 965.3516248 | 0.104285603 | 0.063521132 | 1.641746604 | 0.100642529 | 0.521542547 |
| Gm12830 | 1.402624803 | -0.010824009 | 0.069768248 | -0.155142337 | 0.876709106 | NA | | | Mrps31 | 638.6470024 | 0.104289272 | 0.072217679 | 1.444096154 | 0.14871193 | 0.596407922 |
| Gm12839 | 0 | NA | NA | NA | NA | NA | | | Ube2d3 | 4695.295466 | 0.104300823 | 0.067941187 | 1.535163383 | 0.1247437 | 0.560776598 |
| Gm12886 | 0 | NA | NA | NA | NA | NA | | | Atp6ap2 | 9301.666133 | 0.104304283 | 0.063405162 | 1.645044022 | 0.099960733 | 0.521115714 |
| Gm12887 | 0 | NA | NA | NA | NA | NA | | | AI846148 | 884.4784894 | 0.104315701 | 0.060007929 | 1.738365297 | 0.082146467 | 0.482710189 |
| Gm12888 | 0 | NA | NA | NA | NA | NA | | | Ndufa10 | 4725.745646 | 0.104323413 | 0.047012222 | 2.219070029 | 0.026481961 | 0.303950052 |
| Gm13011 | 0 | NA | NA | NA | NA | NA | | | Cdc37l1 | 2757.542099 | 0.104324454 | 0.067472172 | 1.546184918 | 0.122059915 | 0.55651046 |
| Gm13023 | 0 | NA | NA | NA | NA | NA | | | AI480653 | 1258.920962 | 0.104397135 | 0.057835695 | 1.80506406 | 0.071064658 | 0.458300422 |
| Gm13032 | 0.316760911 | -0.015973748 | 0.040037625 | -0.398968415 | 0.689916477 | NA | | | Lpar2 | 55.59011525 | 0.104499277 | 0.094320517 | 1.107916714 | 0.267897786 | 0.710254288 |
| Gm13034 | 0.261950519 | 0.027936351 | 0.033119325 | 0.843506059 | 0.398945478 | NA | | | Fa2h | 1629.162906 | 0.104515127 | 0.085326742 | 1.224881257 | 0.220619963 | 0.672757795 |
| Gm13037 | 0.192746203 | -0.000975302 | 0.031567365 | -0.030895906 | 0.975352555 | NA | | | Pth2r | 56.227155 | 0.104541214 | 0.088120723 | 1.186340852 | 0.23548772 | 0.686335196 |
| Gm13040 | 0 | NA | NA | NA | NA | NA | | | Srp72 | 4345.937837 | 0.104583889 | 0.058532176 | 1.786776045 | 0.073973693 | 0.466583525 |
| Gm13043 | 0 | NA | NA | NA | NA | NA | | | Map2k4 | 9630.538481 | 0.10467754 | 0.052882503 | 1.979436197 | 0.047766916 | 0.392678115 |
| Gm13051 | 3.005711473 | 0.048107853 | 0.097982051 | 0.490986386 | 0.623436076 | NA | | | Mrps22 | 574.5290449 | 0.104754065 | 0.069775517 | 1.50130116 | 0.133277685 | 0.575362296 |
| Gm13057 | 0 | NA | NA | NA | NA | NA | | | Ifrd1 | 1513.312934 | 0.10476776 | 0.081093893 | 1.291931563 | 0.196380846 | 0.652029296 |
| Gm13078 | 0 | NA | NA | NA | NA | NA | | | Cog7 | 2543.80885 | 0.104788526 | 0.044188065 | 2.371421498 | 0.01771981 | 0.254345817 |
| Gm13083 | 0 | NA | NA | NA | NA | NA | | | Ctps | 1852.536497 | 0.104828241 | 0.061363704 | 1.708310177 | 0.087578804 | 0.495597906 |
| Gm13084 | 0 | NA | NA | NA | NA | NA | | | Rundc1 | 2217.399086 | 0.104853761 | 0.079835008 | 1.313380727 | 0.189054689 | 0.643290796 |
| Gm13088 | 0 | NA | NA | NA | NA | NA | | | Lrrc33 | 315.7996311 | 0.104865205 | 0.094706048 | 1.107270414 | 0.268177032 | 0.71037507 |
| Gm13102 | 0 | NA | NA | NA | NA | NA | | | Trmt10b | 466.451101 | 0.104879334 | 0.072025022 | 1.456151354 | 0.145350784 | 0.591253286 |
| Gm13103 | 0 | NA | NA | NA | NA | NA | | | Dennd2a | 1162.726807 | 0.104936217 | 0.061155978 | 1.71587831 | 0.086184317 | 0.491660395 |
| Gm13109 | 0 | NA | NA | NA | NA | NA | | | Cdc37 | 4498.253983 | 0.104944537 | 0.04267151 | 2.459358428 | 0.013918558 | 0.226406466 |
| Gm13119 | 0 | NA | NA | NA | NA | NA | | | Kpna1 | 5852.434835 | 0.105041622 | 0.072539378 | 1.448063456 | 0.147599305 | 0.594737171 |
| Gm13124 | 0 | NA | NA | NA | NA | NA | | | Hcrtr1 | 115.5685759 | 0.105049688 | 0.094905127 | 1.106891598 | 0.268340799 | 0.710391551 |
| Gm13125 | 0 | NA | NA | NA | NA | NA | | | Pnoc | 343.0954923 | 0.105069642 | 0.093908043 | 1.118856697 | 0.263201279 | 0.705919827 |
| Gm13128 | 0 | NA | NA | NA | NA | NA | | | Lrrc8c | 690.8206575 | 0.105140112 | 0.065788643 | 1.598149893 | 0.110009622 | 0.535496439 |
| Gm13139 | 4.772703669 | 0.22596694 | 0.138142542 | 1.635752005 | 0.101891501 | NA | | | Ephb2 | 878.6866315 | 0.105160077 | 0.069786678 | 1.506878958 | 0.131841686 | 0.572833273 |
| Gm13154 | 0.853962284 | -0.033059725 | 0.056950476 | -0.58049954 | 0.561577797 | NA | | | Jsrp1 | 116.6918845 | 0.105161448 | 0.092823662 | 1.132916384 | 0.257249369 | 0.703199912 |
| Gm13177 | 0 | NA | NA | NA | NA | NA | | | Cdc25a | 300.1301301 | 0.105194787 | 0.085371121 | 1.232205754 | 0.217872234 | 0.670929686 |
| Gm13178 | 0.098353386 | -0.020212831 | 0.02786466 | -0.725393061 | 0.468210929 | NA | | | G3bp2 | 12083.91797 | 0.105259685 | 0.05908024 | 1.781639437 | 0.074808049 | 0.468388672 |
| Gm13212 | 3.161633673 | 0.139904147 | 0.096489302 | 1.449944645 | 0.147073956 | NA | | | Top1 | 3385.101512 | 0.10531025 | 0.073270317 | 1.437283956 | 0.150637326 | 0.598511432 |
| Gm1322 | 0 | NA | NA | NA | NA | NA | | | Tbccd1 | 594.5363998 | 0.105321424 | 0.062689326 | 1.680053529 | 0.092946901 | 0.507192595 |
| Gm13242 | 0.079983654 | 0.016740032 | 0.027674178 | 0.604897175 | 0.54524732 | NA | | | Nudt5 | 483.986821 | 0.105362368 | 0.072730665 | 1.448664981 | 0.147431165 | 0.594719549 |
| Gm13247 | 2.081103138 | -0.06677113 | 0.092140514 | -0.72466635 | 0.468656743 | NA | | | Pigk | 3383.311846 | 0.105405528 | 0.054004471 | 1.951792611 | 0.050962832 | 0.403266245 |
| Gm13271 | 0 | NA | NA | NA | NA | NA | | | Zdhhc15 | 523.7139974 | 0.105423677 | 0.078638204 | 1.34061655 | 0.180044981 | 0.634020218 |
| Gm13272 | 0 | NA | NA | NA | NA | NA | | | Cib2 | 693.6805191 | 0.105423689 | 0.068639788 | 1.535897644 | 0.124563486 | 0.560776598 |
| Gm13275 | 0 | NA | NA | NA | NA | NA | | | Phactr2 | 1636.561417 | 0.105443104 | 0.069946466 | 1.507482944 | 0.131686912 | 0.572734933 |
| Gm13276 | 0 | NA | NA | NA | NA | NA | | | Jrk | 469.5244119 | 0.105465455 | 0.085043976 | 1.240128467 | 0.214927881 | 0.668817995 |
| Gm13277 | 0 | NA | NA | NA | NA | NA | | | Polr2c | 1819.836529 | 0.105496597 | 0.059857496 | 1.762462589 | 0.077991172 | 0.475337606 |
| Gm13278 | 0 | NA | NA | NA | NA | NA | | | Rpap3 | 1050.366857 | 0.105612274 | 0.062979846 | 1.67692175 | 0.093557784 | 0.508805831 |
| Gm13279 | 0 | NA | NA | NA | NA | NA | | | Zfp830 | 383.6474421 | 0.105633668 | 0.074105092 | 1.42545762 | 0.154024962 | 0.60340295 |
| Gm13280 | 0 | NA | NA | NA | NA | NA | | | Zfp711 | 336.8441745 | 0.105648226 | 0.082533096 | 1.280071028 | 0.200520157 | 0.654475062 |
| Gm13283 | 0 | NA | NA | NA | NA | NA | | | Stx4a | 1821.855474 | 0.105658285 | 0.063395217 | 1.6666602 | 0.095581991 | 0.514803116 |
| Gm13285 | 0 | NA | NA | NA | NA | NA | | | Rab3gap1 | 4804.65865 | 0.105674197 | 0.052121875 | 2.027444264 | 0.042616999 | 0.37389754 |
| Gm13286 | 0 | NA | NA | NA | NA | NA | | | Ppp2ca | 14304.24727 | 0.10567995 | 0.055870011 | 1.891532634 | NA | NA |
| Gm13287 | 0 | NA | NA | NA | NA | NA | | | Tcf7l2 | 240.0164466 | 0.105698559 | 0.084054251 | 1.257504026 | 0.208571182 | 0.663405095 |
| Gm13288 | 0 | NA | NA | NA | NA | NA | | | Dusp6 | 2377.454242 | 0.105744549 | 0.093508948 | 1.130849527 | 0.25811843 | 0.703616072 |
| Gm13289 | 0 | NA | NA | NA | NA | NA | | | 2610318N02Rik | 34.96250787 | 0.105750638 | 0.090476797 | 1.168815003 | 0.242478174 | 0.692772033 |
| Gm13290 | 0 | NA | NA | NA | NA | NA | | | Prkar2b | 2690.899252 | 0.105763771 | 0.074787359 | 1.414193144 | 0.157305201 | 0.607878516 |
| Gm13298 | 0.482734261 | -0.021234089 | 0.050628366 | -0.4194109 | 0.67491586 | NA | | | Sfr1 | 3131.023416 | 0.105782515 | 0.05467863 | 1.934622629 | 0.053036621 | 0.40708779 |
| Gm13304 | 0 | NA | NA | NA | NA | NA | | | Negr1 | 10429.72397 | 0.105784807 | 0.073382325 | 1.441557039 | 0.149427375 | 0.597791807 |
| Gm13305 | 0 | NA | NA | NA | NA | NA | | | Fam134b | 1545.574591 | 0.105792978 | 0.060105396 | 1.760124471 | 0.078386704 | 0.475744405 |
| Gm13306 | 0 | NA | NA | NA | NA | NA | | | Psmb5 | 1846.434549 | 0.105795992 | 0.060672022 | 1.743736045 | 0.081205127 | 0.479898997 |
| Gm13308 | 0 | NA | NA | NA | NA | NA | | | Tgoln1 | 4185.144269 | 0.105796357 | 0.055406831 | 1.909446105 | 0.056204568 | 0.415696311 |
| Gm13315 | 0.585367496 | 0.027251914 | 0.062924522 | 0.433088935 | 0.664950161 | NA | | | Ahi1 | 10451.49133 | 0.105836512 | 0.094864834 | 1.115655908 | 0.264569451 | 0.707370515 |
| Gm1332 | 0.496526426 | 0.041177769 | 0.052144109 | 0.78969169 | 0.429707846 | NA | | | Garem | 1414.446302 | 0.105839901 | 0.069946482 | 1.513155468 | 0.130240173 | 0.569619261 |
| Gm13363 | 0 | NA | NA | NA | NA | NA | | | Nutf2 | 543.9446335 | 0.105847491 | 0.076556627 | 1.382603908 | 0.16678635 | 0.617356019 |
| Gm13490 | 0.37445586 | 0.021594193 | 0.042489861 | 0.508219911 | 0.611299129 | NA | | | Faim | 598.2527767 | 0.105901512 | 0.065154325 | 1.625394972 | 0.104078427 | 0.523645898 |

| | | | | | | | | | | | | | | | |
|---|---|---|---|---|---|---|---|---|---|---|---|---|---|---|---|
| Gm13497 | 0.899153043 | 0.057189445 | 0.063719345 | 0.897520924 | 0.369441013 | NA | | | Chp1 | 7148.898005 | 0.106032622 | 0.043015615 | 2.464979823 | 0.013702097 | 0.224208584 |
| Gm13498 | 0.276664272 | 0.005276976 | 0.033299614 | 0.158469592 | 0.87408678 | NA | | | Gbx2 | 8.666318005 | 0.106056395 | 0.062039024 | 1.709511018 | 0.087356333 | NA |
| Gm13539 | 0 | NA | NA | NA | NA | NA | | | Ppm1g | 2862.781001 | 0.106067153 | 0.045710362 | 2.32041812 | 0.02031827 | 0.269537155 |
| Gm13544 | 3.018081471 | -0.154871573 | 0.111627587 | -1.387395155 | 0.165321302 | NA | | | Morf4l1 | 12585.97659 | 0.106158923 | 0.057938185 | 1.832279067 | 0.066909854 | 0.447688159 |
| Gm13546 | 0.744732266 | 0.0208816 | 0.067843496 | 0.307790742 | 0.758241567 | NA | | | Tmem108 | 913.8428323 | 0.106290938 | 0.071559584 | 1.485348741 | 0.137451467 | 0.582012447 |
| Gm13547 | 3.314472048 | 0.200455762 | 0.113211871 | 1.770624943 | 0.076623091 | NA | | | Sec61b | 420.5226203 | 0.106381167 | 0.079975734 | 1.330168049 | 0.183462909 | 0.636946035 |
| Gm13580 | 0.08731684 | 0.017365056 | 0.027733412 | 0.626142072 | 0.531221759 | NA | | | Klf10 | 1245.697147 | 0.106384983 | 0.085778471 | 1.240229417 | 0.214890551 | 0.668817995 |
| Gm13582 | 0.079983654 | 0.016740032 | 0.027674178 | 0.604897175 | 0.54524732 | NA | | | Tuba1a | 26588.13307 | 0.106437887 | 0.049425447 | 2.153503757 | 0.031279109 | 0.32443523 |
| Gm136 | 7.486667715 | -0.332955286 | 0.140212455 | -2.374648435 | 0.017565672 | NA | | | Mal2 | 6535.906745 | 0.106459906 | 0.070047716 | 1.519819812 | 0.128556268 | 0.566740017 |
| Gm13710 | 4.935292295 | 0.197781296 | 0.13092614 | 1.510632606 | 0.130882082 | NA | | | 3110070M22Rik | 35.41209493 | 0.106466091 | 0.081490371 | 1.306486764 | 0.191387077 | 0.646682644 |
| Gm13718 | 0 | NA | NA | NA | NA | NA | | | Stk32b | 180.6844995 | 0.10649767 | 0.094858942 | 1.122695107 | 0.261567007 | 0.704173028 |
| Gm13752 | 0 | NA | NA | NA | NA | NA | | | Ormdl1 | 301.3183507 | 0.106516508 | 0.080304384 | 1.326409627 | 0.184704052 | 0.637780733 |
| Gm13769 | 0 | NA | NA | NA | NA | NA | | | Nmnat2 | 8385.882319 | 0.106531918 | 0.04360558 | 2.443079944 | 0.014562511 | 0.23143877 |
| Gm13807 | 0 | NA | NA | NA | NA | NA | | | Atp9a | 18025.22499 | 0.106553575 | 0.053168175 | 2.004085642 | 0.045060887 | 0.383004357 |
| Gm13871 | 0 | NA | NA | NA | NA | NA | | | Zfp184 | 125.6997692 | 0.106612647 | 0.092974812 | 1.146683117 | 0.251512611 | 0.699465245 |
| Gm13939 | 0 | NA | NA | NA | NA | NA | | | Loxhd1 | 10.15196697 | 0.106617135 | 0.068304361 | 1.560912556 | 0.118544384 | NA |
| Gm13944 | 0.17463368 | 0.024560283 | 0.031186242 | 0.787535833 | 0.430968261 | NA | | | Fam19a4 | 21.40448009 | 0.106627185 | 0.077855466 | 1.369552963 | 0.170826491 | NA |
| Gm14005 | 6.259972334 | -0.189276401 | 0.148707537 | -1.272809737 | 0.203085565 | NA | | | Gm5113 | 922.7197448 | 0.1066581 | 0.06658729 | 1.601778667 | 0.109204562 | 0.535010276 |
| Gm14023 | 6.950434306 | -0.065195302 | 0.14628259 | -0.445680525 | 0.655828034 | NA | | | Spry4 | 730.1003383 | 0.106710691 | 0.088288994 | 1.208652252 | 0.226796473 | 0.679834424 |
| Gm14047 | 2.932336519 | -0.067815572 | 0.119467639 | -0.567648046 | 0.570273976 | NA | | | Tmod2 | 22410.72088 | 0.106719066 | 0.058379379 | 1.828026731 | 0.06754554 | 0.448803776 |
| Gm14085 | 0 | NA | NA | NA | NA | NA | | | Ndufb8 | 3769.9787 | 0.106734567 | 0.052311472 | 2.040366323 | 0.041313854 | 0.367273584 |
| Gm14092 | 1.280276179 | -0.017309126 | 0.073661637 | -0.23498155 | 0.814223053 | NA | | | 2210013O21Rik | 811.3279508 | 0.106740226 | 0.065378398 | 1.632652807 | 0.102542038 | 0.522975271 |
| Gm14124 | 4.114478889 | 0.012894769 | 0.104654401 | 0.123212866 | 0.901938538 | NA | | | Prps1 | 3155.805415 | 0.106823562 | 0.052992154 | 2.015837327 | 0.043816989 | 0.378860465 |
| Gm14137 | 5.393404973 | -0.045281017 | 0.114638313 | -0.394990257 | 0.692850079 | NA | | | Slc35d3 | 258.2641998 | 0.106844248 | 0.091469941 | 1.168080427 | 0.242774325 | 0.692772033 |
| Gm14139 | 0.491740777 | -0.038091983 | 0.046390976 | -0.821107608 | 0.411584977 | NA | | | Rpusd2 | 342.5415737 | 0.106847259 | 0.074486587 | 1.434449654 | 0.151443987 | 0.598905575 |
| Gm14151 | 0.186326724 | 0.024939385 | 0.031411053 | 0.793968451 | 0.427213798 | NA | | | St7l | 869.7450796 | 0.106897175 | 0.071660877 | 1.491709004 | 0.135775448 | 0.579435507 |
| Gm14164 | 1.100526109 | 0.060038741 | 0.072132852 | 0.832335606 | 0.405219538 | NA | | | Itfg2 | 806.2369996 | 0.106902818 | 0.072708664 | 1.470289961 | 0.141483239 | 0.587141261 |
| Gm14207 | 1.481295904 | 0.039395746 | 0.088746968 | 0.443910889 | 0.657107012 | NA | | | Zkscan1 | 2664.512498 | 0.106914836 | 0.066271937 | 1.613274643 | 0.106684853 | 0.530442242 |
| Gm14288 | 0 | NA | NA | NA | NA | NA | | | Nkain2 | 2263.081525 | 0.106936186 | 0.06479087 | 1.65048233 | 0.098844324 | 0.520271327 |
| Gm14305 | 0 | NA | NA | NA | NA | NA | | | Ccdc32 | 1671.534796 | 0.106937067 | 0.052517955 | 2.036200116 | 0.041730265 | 0.36916798 |
| Gm14306 | 0 | NA | NA | NA | NA | NA | | | Pde4d | 3241.869874 | 0.106948234 | 0.074230225 | 1.440763971 | 0.149651375 | 0.598098567 |
| Gm14308 | 1.402071703 | 0.146343382 | 0.078928163 | 1.854133891 | 0.063720007 | NA | | | Clcn6 | 2810.800385 | 0.106949604 | 0.049177697 | 2.174758262 | 0.029648236 | 0.317771905 |
| Gm14318 | 0 | NA | NA | NA | NA | NA | | | Sult2b1 | 302.7605581 | 0.106962003 | 0.090538528 | 1.181397637 | 0.237444796 | 0.688447727 |
| Gm14327 | 4.815060157 | 0.072283916 | 0.136988002 | 0.527666036 | 0.59773115 | NA | | | Nrarp | 632.5462193 | 0.106990176 | 0.075513702 | 1.416831286 | 0.15653226 | 0.60719411 |
| Gm14345 | 0 | NA | NA | NA | NA | NA | | | Peg13 | 18894.70533 | 0.107051105 | 0.044096252 | 2.427668994 | 0.015196205 | 0.233769172 |
| Gm14346 | 0 | NA | NA | NA | NA | NA | | | Mtap | 448.4432552 | 0.107062861 | 0.068855858 | 1.554883834 | 0.119973738 | 0.553708672 |
| Gm14347 | 0 | NA | NA | NA | NA | NA | | | Lhx1 | 8.55942623 | 0.107078106 | 0.04774115 | 2.242889132 | 0.024903965 | NA |
| Gm14351 | 0 | NA | NA | NA | NA | NA | | | BC028528 | 59.29283911 | 0.107090031 | 0.093127115 | 1.149933952 | 0.250171075 | 0.698461386 |
| Gm14354 | 0 | NA | NA | NA | NA | NA | | | Fyttd1 | 2745.040308 | 0.107130298 | 0.069012873 | 1.552323405 | 0.12058486 | 0.554583368 |
| Gm14374 | 0 | NA | NA | NA | NA | NA | | | Tollip | 7439.439908 | 0.107136865 | 0.040008382 | 2.677860506 | 0.007409407 | 0.163894025 |
| Gm14391 | 0 | NA | NA | NA | NA | NA | | | Furin | 1583.895526 | 0.10713985 | 0.064239676 | 1.667814288 | 0.095352599 | 0.51433043 |
| Gm14430 | 0 | NA | NA | NA | NA | NA | | | Ankrd50 | 1681.66127 | 0.107148949 | 0.071433352 | 1.499984903 | 0.133618313 | 0.57588358 |
| Gm14431 | 0 | NA | NA | NA | NA | NA | | | Ado | 2342.422587 | 0.107158915 | 0.068642672 | 1.561112218 | 0.118497276 | 0.550284983 |
| Gm14434 | 0 | NA | NA | NA | NA | NA | | | Dio3 | 74.36877849 | 0.107178413 | 0.062580752 | 1.712641822 | 0.086778454 | 0.493466364 |
| Gm14436 | 0 | NA | NA | NA | NA | NA | | | Nfkbie | 112.017 | 0.107198343 | 0.093522969 | 1.146224764 | 0.251702164 | 0.699465245 |
| Gm14440 | 0 | NA | NA | NA | NA | NA | | | Atcay | 11732.74833 | 0.107221738 | 0.040938309 | 2.619105191 | 0.008816076 | 0.179358986 |
| Gm14458 | 0 | NA | NA | NA | NA | NA | | | Wnt7a | 424.0786088 | 0.107282227 | 0.080213313 | 1.337461613 | 0.181072004 | 0.634469716 |
| Gm14459 | 0 | NA | NA | NA | NA | NA | | | Bag5 | 1904.450722 | 0.107299453 | 0.06116913 | 1.754143862 | 0.07940586 | 0.477013957 |
| Gm14461 | 2.338466806 | -0.020661496 | 0.099826237 | -0.206974609 | 0.836029688 | NA | | | Dscam | 3166.689704 | 0.107331758 | 0.055972194 | 1.917590679 | 0.055162933 | 0.412614194 |
| Gm14474 | 0 | NA | NA | NA | NA | NA | | | AI464131 | 1668.549762 | 0.107355554 | 0.072494567 | 1.480877247 | 0.138639287 | 0.583109099 |
| Gm14475 | 0 | NA | NA | NA | NA | NA | | | Rad17 | 610.5109504 | 0.107393206 | 0.065647012 | 1.635919169 | 0.101856506 | 0.522606076 |
| Gm14476 | 0 | NA | NA | NA | NA | NA | | | 1700037H04Rik | 2126.14505 | 0.10741971 | 0.059419716 | 1.807812582 | 0.070635675 | 0.456632761 |
| Gm14477 | 0 | NA | NA | NA | NA | NA | | | 1700003E16Rik | 998.5377028 | 0.107428987 | 0.072348953 | 1.484872721 | 0.137577543 | 0.582129739 |
| Gm14478 | 0 | NA | NA | NA | NA | NA | | | Cacng5 | 648.5670185 | 0.107505849 | 0.085803656 | 1.252928542 | 0.210231713 | 0.664608511 |
| Gm14479 | 0 | NA | NA | NA | NA | NA | | | Alg11 | 1517.288033 | 0.107532392 | 0.07181274 | 1.497399921 | 0.134289229 | 0.57712584 |
| Gm14482 | 0 | NA | NA | NA | NA | NA | | | Snx10 | 3697.980186 | 0.107544105 | 0.061209537 | 1.756982807 | 0.07892074 | 0.476674706 |
| Gm14483 | 0.098353386 | -0.020212831 | 0.02786466 | -0.725393061 | 0.468210929 | NA | | | Ahnak | 1507.113792 | 0.107558645 | 0.092168779 | 1.16697483 | 0.243220534 | 0.692772033 |
| Gm14484 | 0 | NA | NA | NA | NA | NA | | | Cul3 | 4967.403187 | 0.107574959 | 0.071633933 | 1.501731847 | 0.133166376 | 0.575193212 |
| Gm14492 | 3.401484765 | -0.077609153 | 0.117692894 | -0.659420886 | 0.509625534 | NA | | | Wdr61 | 1339.475806 | 0.107578111 | 0.054341462 | 1.979669048 | 0.047740729 | 0.392678115 |
| Gm14496 | 0 | NA | NA | NA | NA | NA | | | Nrp2 | 1511.405105 | 0.107625121 | 0.085787946 | 1.254548295 | 0.209642783 | 0.664226905 |
| Gm14501 | 0 | NA | NA | NA | NA | NA | | | Kctd20 | 1926.167099 | 0.107647421 | 0.058262693 | 1.84762178 | 0.064657077 | 0.443022969 |
| Gm14511 | 0 | NA | NA | NA | NA | NA | | | D8Ertd738e | 1018.093266 | 0.107662778 | 0.061267445 | 1.757259148 | 0.078873648 | 0.476674706 |
| Gm14525 | 0 | NA | NA | NA | NA | NA | | | Zfp651 | 4113.652274 | 0.107738988 | 0.062458416 | 1.724971517 | 0.084532606 | 0.488499377 |
| Gm14548 | 0 | NA | NA | NA | NA | NA | | | Rn45s | 16791.75629 | 0.10774021 | 0.058198413 | 1.851256846 | 0.064132611 | 0.441721342 |
| Gm14625 | 0 | NA | NA | NA | NA | NA | | | Psmc5 | 6209.356671 | 0.10774871 | 0.045069869 | 2.390703851 | 0.016816111 | 0.246489231 |
| Gm14632 | 0 | NA | NA | NA | NA | NA | | | Gsr | 1006.611931 | 0.107749123 | 0.055270212 | 1.949497186 | 0.051236079 | 0.403266245 |
| Gm14635 | 0 | NA | NA | NA | NA | NA | | | Kctd3 | 3654.181805 | 0.107797545 | 0.052237847 | 2.063590885 | 0.039056523 | 0.358392323 |
| Gm14685 | 0 | NA | NA | NA | NA | NA | | | Kcnj8 | 156.4024681 | 0.107800248 | 0.094835759 | 1.136704638 | 0.25566178 | 0.701952734 |
| Gm14692 | 0 | NA | NA | NA | NA | NA | | | Rnf139 | 1283.466406 | 0.107824567 | 0.073633506 | 1.464341077 | 0.143100779 | 0.589362607 |
| Gm14718 | 0 | NA | NA | NA | NA | NA | | | Eif3d | 2836.794225 | 0.107840287 | 0.052448905 | 2.056101782 | 0.039772706 | 0.36080087 |
| Gm14725 | 0 | NA | NA | NA | NA | NA | | | Ptprn2 | 18312.04008 | 0.107926189 | 0.051643524 | 2.08983005 | 0.036633069 | 0.346440078 |
| Gm14743 | 2.913104147 | 0.132434134 | 0.069315834 | 1.910589925 | NA | NA | | | Eif2c2 | 2829.732244 | 0.107958387 | 0.077325986 | 1.396146269 | 0.162670453 | 0.614351408 |
| Gm14744 | 1.229087554 | 0.083549556 | 0.056066624 | 1.490183475 | 0.136176 | NA | | | Nceh1 | 6254.402864 | 0.108003228 | 0.063700947 | 1.695472875 | 0.089985752 | 0.500949827 |
| Gm14781 | 0 | NA | NA | NA | NA | NA | | | Snrk | 4118.879577 | 0.108021429 | 0.052827292 | 2.044803453 | 0.040874239 | 0.365377968 |
| Gm14812 | 0.079983654 | 0.016740032 | 0.027674178 | 0.604897175 | 0.54524732 | NA | | | Elovl6 | 2653.400992 | 0.108064994 | 0.076783747 | 1.407394121 | 0.159310553 | 0.611076076 |
| Gm14819 | 0 | NA | NA | NA | NA | NA | | | Ubl4 | 3346.348793 | 0.10806557 | 0.050616334 | 2.13499401 | 0.032761499 | 0.330481625 |
| Gm14850 | 0 | NA | NA | NA | NA | NA | | | Smoc1 | 800.0629246 | 0.108149375 | 0.093902551 | 1.151719237 | 0.249436465 | 0.697864808 |
| Gm14851 | 0 | NA | NA | NA | NA | NA | | | Zfp790 | 812.5874415 | 0.108153813 | 0.079007942 | 1.368897997 | 0.171031162 | 0.621020522 |
| Gm14858 | 0.386222787 | 0.053439715 | 0.045634083 | 1.171048305 | 0.241579361 | NA | | | Nppc | 100.7698103 | 0.108181708 | 0.093450993 | 1.157630379 | 0.247014907 | 0.696511265 |
| Gm14873 | 1.789798875 | -0.027703176 | 0.081284736 | -0.340816457 | 0.73324176 | NA | | | Syt10 | 294.1553375 | 0.108196649 | 0.090820995 | 1.191317593 | 0.23352893 | 0.685064073 |
| Gm14920 | 0 | NA | NA | NA | NA | NA | | | Uqcrc1 | 8886.647328 | 0.108212086 | 0.051025396 | 2.120749541 | 0.033942885 | 0.336326295 |
| Gm15008 | 0.310199702 | 0.010205801 | 0.034113052 | 0.299175856 | 0.76480587 | NA | | | Ak3 | 3584.321445 | 0.108218288 | 0.062759737 | 1.72432666 | 0.084648889 | 0.488499377 |
| Gm15023 | 0 | NA | NA | NA | NA | NA | | | Cbfb | 675.8876827 | 0.108244175 | 0.07757123 | 1.395416511 | 0.162890275 | 0.614351408 |
| Gm15056 | 0 | NA | NA | NA | NA | NA | | | Arxes1 | 477.7195351 | 0.10832122 | 0.081271544 | 1.332830835 | 0.182587324 | 0.635424885 |
| Gm15070 | 0.575738104 | 0.018512958 | 0.053137856 | 0.348394908 | 0.727543626 | NA | | | Ssb | 3826.952959 | 0.108361527 | 0.062052503 | 1.746287781 | 0.080760958 | 0.479417114 |
| Gm15080 | 0 | NA | NA | NA | NA | NA | | | Psmd1 | 6504.291334 | 0.108402944 | 0.04715326 | 2.298949099 | 0.02150783 | 0.275709409 |
| Gm15085 | 0 | NA | NA | NA | NA | NA | | | Yars2 | 547.6729949 | 0.108412222 | 0.076778069 | 1.412020687 | 0.15794387 | 0.608476733 |
| Gm15091 | 0 | NA | NA | NA | NA | NA | | | Kcnrg | 20.48787797 | 0.108423082 | 0.080441899 | 1.347843389 | 0.177708759 | NA |
| Gm15093 | 0 | NA | NA | NA | NA | NA | | | Cnpy2 | 1005.149383 | 0.108434329 | 0.068616362 | 1.58029843 | 0.114038539 | 0.54258887 |
| Gm15097 | 0 | NA | NA | NA | NA | NA | | | Bzw1 | 6708.941635 | 0.108437263 | 0.066962814 | 1.61936537 | 0.105368673 | 0.526879637 |
| Gm15104 | 4.111783717 | 0.20070567 | 0.12065897 | 1.663412759 | 0.096229841 | NA | | | Rac3 | 640.4748772 | 0.108455027 | 0.082751027 | 1.310618511 | 0.189986682 | 0.644800595 |
| Gm15107 | 0 | NA | NA | NA | NA | NA | | | Mchr1 | 906.7964038 | 0.108480088 | 0.063080509 | 1.71970851 | 0.08548544 | 0.490040057 |
| Gm15114 | 0 | NA | NA | NA | NA | NA | | | Nap1l4 | 4782.192198 | 0.108525945 | 0.049441992 | 2.195015612 | 0.028162478 | 0.312281033 |
| Gm15127 | 0 | NA | NA | NA | NA | NA | | | 0610009D07Rik | 1146.335862 | 0.108551464 | 0.056022341 | 1.937645976 | 0.052666429 | 0.406864177 |
| Gm15133 | 1.055623674 | 0.111406771 | 0.072854512 | 1.529167761 | 0.126222861 | NA | | | Ehd1 | 1595.309918 | 0.108559966 | 0.050766885 | 2.138401148 | 0.032484199 | 0.329285052 |
| Gm15140 | 0 | NA | NA | NA | NA | NA | | | Cdh22 | 1770.680554 | 0.108560191 | 0.074376852 | 1.459596474 | 0.144401009 | 0.589967032 |
| Gm15179 | 0.582829523 | 0.023716638 | 0.058499643 | 0.405415095 | 0.685172417 | NA | | | Ptges3l | 91.91072577 | 0.10860328 | 0.094839716 | 1.145124479 | 0.252157596 | 0.69992334 |
| Gm15217 | 0 | NA | NA | NA | NA | NA | | | Adm | 75.39743338 | 0.108607159 | 0.094988463 | 1.143372098 | 0.252884132 | 0.700378199 |
| Gm1527 | 0 | NA | NA | NA | NA | NA | | | Mpv17l2 | 435.6982943 | 0.108664116 | 0.077345655 | 1.404915592 | 0.16004638 | 0.612215825 |
| Gm15284 | 0 | NA | NA | NA | NA | NA | | | Shisa9 | 2812.682748 | 0.10869176 | 0.065142054 | 1.668534441 | 0.095209681 | 0.514323475 |
| Gm15292 | 0 | NA | NA | NA | NA | NA | | | Cnih | 2298.233143 | 0.108695794 | 0.063965164 | 1.699296729 | 0.089263289 | 0.499459783 |
| Gm15293 | 0 | NA | NA | NA | NA | NA | | | Flywch1 | 9607.769639 | 0.108799709 | 0.040473404 | 2.688177855 | 0.007184312 | 0.161546928 |
| Gm15299 | 0 | NA | NA | NA | NA | NA | | | Psmb2 | 1974.985633 | 0.108862675 | 0.04738189 | 2.297558727 | 0.021586917 | 0.276274499 |
| Gm15308 | 0 | NA | NA | NA | NA | NA | | | Aamdc | 649.9617597 | 0.109011685 | 0.061613476 | 1.769283146 | 0.076846635 | 0.472560915 |
| Gm15315 | 0 | NA | NA | NA | NA | NA | | | Fam5b | 5197.762044 | 0.109028636 | 0.061484082 | 1.773282334 | 0.076181935 | 0.472435997 |
| Gm15319 | 0 | NA | NA | NA | NA | NA | | | Esrrb | 145.9412624 | 0.109055377 | 0.090619711 | 1.203439915 | 0.22880613 | 0.681764163 |
| Gm15348 | 0 | NA | NA | NA | NA | NA | | | Gatsl2 | 4772.356991 | 0.109120397 | 0.05958046 | 1.831479588 | 0.067028992 | 0.447688159 |
| Gm15350 | 0.537538576 | 0.051328891 | 0.044528031 | 1.152732117 | 0.249020355 | NA | | | Leo1 | 933.673296 | 0.109130001 | 0.060782391 | 1.795421315 | 0.072586599 | 0.462655961 |
| Gm15386 | 0 | NA | NA | NA | NA | NA | | | Kctd21 | 890.6862418 | 0.109133378 | 0.058560113 | 1.863612822 | 0.062376096 | 0.436873272 |
| Gm15401 | 0.927537728 | 0.084647147 | 0.074477453 | 1.136547297 | 0.255727583 | NA | | | Ar | 349.2862274 | 0.109156591 | 0.087791859 | 1.24335664 | 0.213736441 | 0.667398009 |
| Gm15408 | 7.199789303 | -0.054936747 | 0.14384481 | -0.381916781 | 0.702523092 | NA | | | Sptssb | 111.0992747 | 0.109187698 | 0.094081986 | 1.160559023 | 0.245821278 | 0.695249198 |
| Gm15412 | 0 | NA | NA | NA | NA | NA | | | E130309F12Rik | 685.5043686 | 0.109203871 | 0.077355152 | 1.411720712 | 0.158032212 | 0.608583223 |
| Gm15413 | 2.044128532 | 0.119959044 | 0.106424412 | 1.127176011 | 0.259668071 | NA | | | Nmral1 | 443.9773331 | 0.109271812 | 0.079778837 | 1.369684189 | 0.170785506 | 0.620885919 |
| Gm15417 | 73.02544883 | 0.176601473 | 0.16747169 | 1.054515379 | NA | NA | | | Rnf26 | 1562.797909 | 0.109275464 | 0.062615818 | 1.745173462 | 0.080954679 | 0.479898997 |
| Gm15441 | 2.092731098 | 0.10037557 | 0.098003676 | 1.024202087 | 0.305739833 | NA | | | Acat1 | 3558.343485 | 0.109353305 | 0.053168491 | 2.056731401 | 0.03971207 | 0.36080087 |
| Gm15455 | 2.552545577 | 0.143903929 | 0.108635775 | 1.324645856 | 0.185288639 | NA | | | Cmas | 3313.379544 | 0.109412609 | 0.048189796 | 2.270451786 | 0.023180185 | 0.286020752 |
| Gm15471 | 2.881483846 | 0.074193832 | 0.092582574 | 0.801380094 | 0.422911636 | NA | | | Rhof | 1902.266352 | 0.109432788 | 0.063002001 | 1.73697321 | 0.0823919 | 0.483019453 |
| Gm1553 | 0.079983654 | 0.016740032 | 0.027674178 | 0.604897175 | 0.54524732 | NA | | | Prosc | 2950.538372 | 0.109485619 | 0.057656794 | 1.898919648 | 0.057575042 | 0.419820382 |
| Gm156 | 0 | NA | NA | NA | NA | NA | | | Ptpdc1 | 1359.416188 | 0.10955618 | 0.060542094 | 1.80958689 | 0.070359875 | 0.456017009 |
| Gm1564 | 0.211846316 | 0.025707621 | 0.031860375 | 0.806883811 | 0.419733426 | NA | | | Cenpq | 132.8411834 | 0.109616153 | 0.093923141 | 1.16708356 | 0.243176626 | 0.692772033 |
| Gm15698 | 0.098353386 | -0.020212831 | 0.02786466 | -0.725393061 | 0.468210929 | NA | | | Timp4 | 1002.295067 | 0.109617012 | 0.076377847 | 1.435193794 | 0.151231882 | 0.598846773 |
| Gm15713 | 0 | NA | NA | NA | NA | NA | | | Clcn4-2 | 9510.150246 | 0.109621432 | 0.059638831 | 1.838088214 | 0.066049408 | 0.446279967 |
| Gm15772 | 0 | NA | NA | NA | NA | NA | | | Pdzrn4 | 298.1133761 | 0.109622813 | 0.089667753 | 1.222544447 | 0.221501807 | 0.672757795 |
| Gm15787 | 6.597693041 | 0.006260648 | 0.151751078 | 0.04125604 | 0.967091778 | NA | | | Sod1 | 6047.24217 | 0.109624542 | 0.058810066 | 1.864043846 | 0.062315547 | 0.436873272 |
| Gm15850 | 1.979993066 | -0.122100993 | 0.091664009 | -1.332049459 | 0.182843937 | NA | | | Abhd6 | 2259.205042 | 0.10969456 | 0.061837626 | 1.773912876 | 0.076077563 | 0.472435997 |
| Gm1587 | 0.309826022 | -0.01193339 | 0.034225532 | -0.348669245 | 0.727337636 | NA | | | Spin4 | 25.24535738 | 0.109704127 | 0.082570444 | 1.328612537 | 0.183975834 | 0.637355821 |
| Gm15880 | 3.002832699 | -0.056348994 | 0.112379008 | -0.501419215 | 0.616076119 | NA | | | Emc10 | 7147.984889 | 0.10972018 | 0.056697068 | 1.935200246 | 0.052965728 | 0.406864177 |
| Gm15881 | 0 | NA | NA | NA | NA | NA | | | Got1 | 15572.72228 | 0.109737768 | 0.039572161 | 2.773105258 | 0.005552416 | 0.141004182 |
| Gm15915 | 0.587717662 | 0.024545414 | 0.058572002 | 0.419063942 | 0.675169402 | NA | | | Tsc22d1 | 16691.0519 | 0.109758372 | 0.049453006 | 2.219447928 | 0.026456266 | 0.303950052 |
| Gm15941 | 0.901370294 | -0.023544105 | 0.064578436 | -0.364581526 | 0.715423807 | NA | | | 2010107G23Rik | 919.6297053 | 0.109782937 | 0.063000881 | 1.74256193 | 0.081410165 | 0.480521762 |
| Gm15987 | 0.659787236 | 0.016087308 | 0.046903158 | 0.342989874 | 0.731606078 | NA | | | Scn3b | 5991.110544 | 0.109790127 | 0.069462601 | 1.580564574 | 0.113977631 | 0.54248353 |
| Gm16063 | 0 | NA | NA | NA | NA | NA | | | Bmper | 413.1133067 | 0.109836516 | 0.086513355 | 1.269590296 | 0.204230608 | 0.65778952 |
| Gm16065 | 0.651198668 | 0.010138487 | 0.065005828 | 0.155962742 | 0.876062389 | NA | | | Plcxd2 | 5910.421711 | 0.10984289 | 0.067545457 | 1.626206934 | 0.103905642 | 0.523645898 |
| Gm16130 | 0.079983654 | 0.016740032 | 0.027674178 | 0.604897175 | 0.54524732 | NA | | | Dnajb5 | 5490.543952 | 0.109851248 | 0.094908445 | 1.157444393 | 0.247090846 | 0.696534373 |
| Gm16157 | 0.652614779 | -0.082917553 | 0.055030012 | -1.506769682 | 0.131869703 | NA | | | Zc3h15 | 4085.223746 | 0.10985888 | 0.056303524 | 1.95119011 | 0.051034435 | 0.403266245 |
| Gm16294 | 5.002996032 | 0.039285375 | 0.137260593 | 0.286210149 | 0.774717171 | NA | | | Wdr54 | 570.4358012 | 0.109883191 | 0.070693061 | 1.554370253 | 0.120096125 | 0.553708672 |
| Gm1631 | 0.29109959 | -0.010630118 | 0.033578644 | -0.316573778 | 0.751567036 | NA | | | Steap2 | 699.9225502 | 0.109886792 | 0.081515907 | 1.348041087 | 0.177645168 | 0.630460228 |
| Gm16325 | 1.21199048 | -0.080292872 | 0.085386543 | -0.940345746 | 0.347040242 | NA | | | Ppia | 31521.712 | 0.109906972 | 0.049654262 | 2.213444875 | 0.02686699 | 0.306064716 |
| Gm16336 | 0 | NA | NA | NA | NA | NA | | | Ppa1 | 2733.588404 | 0.109937422 | 0.056500318 | 1.945784139 | 0.051680675 | 0.404936564 |
| Gm16367 | 0 | NA | NA | NA | NA | NA | | | Adssl1 | 950.4999389 | 0.109958822 | 0.076886888 | 1.430137514 | 0.152677555 | 0.601351762 |
| Gm16381 | 0 | NA | NA | NA | NA | NA | | | Rabif | 1304.75897 | 0.109983182 | 0.060018524 | 1.832487298 | 0.066878853 | 0.447688159 |
| Gm16390 | 0 | NA | NA | NA | NA | NA | | | Sf3a3 | 1242.746955 | 0.1099921 | 0.056766686 | 1.937617079 | 0.052669957 | 0.406864177 |
| Gm16404 | 0 | NA | NA | NA | NA | NA | | | Bsdc1 | 2859.334589 | 0.110018502 | 0.044186396 | 2.489872715 | 0.012778885 | 0.215303986 |
| Gm16405 | 0 | NA | NA | NA | NA | NA | | | Gimap5 | 20.05676061 | 0.11002086 | 0.072151908 | 1.524850326 | 0.127296434 | NA |
| Gm16430 | 0 | NA | NA | NA | NA | NA | | | Chst10 | 3375.405718 | 0.110029238 | 0.048450451 | 2.270964183 | 0.023149146 | 0.286019362 |
| Gm16442 | 0 | NA | NA | NA | NA | NA | | | 4931414P19Rik | 166.2483499 | 0.110083868 | 0.088389412 | 1.245441795 | 0.212969398 | 0.666579073 |
| Gm16445 | 0 | NA | NA | NA | NA | NA | | | Zfp536 | 610.5845141 | 0.110110513 | 0.079330509 | 1.387997065 | 0.165137939 | 0.615077498 |
| Gm16451 | 0 | NA | NA | NA | NA | NA | | | Mdh1 | 29272.27832 | 0.110194178 | 0.043341297 | 2.542475302 | 0.011007038 | 0.199361888 |
| Gm1647 | 0.840912053 | 0.002766328 | 0.063990717 | 0.043230145 | 0.965518076 | NA | | | Arhgap31 | 1721.422996 | 0.110217915 | 0.07862364 | 1.401841927 | 0.160962455 | 0.612415571 |
| Gm16497 | 0.227114243 | -0.027513412 | 0.032207244 | -0.854261616 | 0.39296005 | NA | | | Slc25a12 | 6993.898921 | 0.110237407 | 0.043862093 | 2.513272854 | 0.011961678 | 0.208339729 |
| Gm16501 | 0 | NA | NA | NA | NA | NA | | | Mrpl42 | 1150.204179 | 0.110244764 | 0.072068631 | 1.52971913 | 0.126086266 | 0.562787717 |
| Gm16523 | 0.404218839 | -0.002314653 | 0.036541701 | -0.063342793 | 0.94949354 | NA | | | Pnmal2 | 16599.49626 | 0.110322232 | 0.065296169 | 1.689566675 | 0.091110886 | 0.503727415 |
| Gm16548 | 1.429623161 | 0.050192426 | 0.084987252 | 0.590587707 | 0.554796704 | NA | | | Akap10 | 1338.924471 | 0.110420222 | 0.076481018 | 1.443759829 | 0.148806546 | 0.596622655 |
| Gm16551 | 0.946752622 | -0.006409306 | 0.071468774 | -0.089679802 | 0.928541667 | NA | | | Gad1 | 20625.89624 | 0.110438486 | 0.061576073 | 1.793529219 | 0.072888343 | 0.463573784 |
| Gm16617 | 0.401340017 | -0.003486686 | 0.042331768 | -0.082365706 | 0.934355906 | NA | | | Tmco1 | 1203.332133 | 0.110451931 | 0.065056944 | 1.697773116 | 0.089550592 | 0.500119914 |
| Gm16796 | 7.151077362 | 0.113291412 | 0.154698788 | 0.732335482 | 0.463963827 | NA | | | Ccdc91 | 1608.608868 | 0.110472947 | 0.057750824 | 1.912924163 | 0.055757762 | 0.413057568 |
| Gm16833 | 0.423692631 | 0.031061933 | 0.034784099 | 0.892992313 | 0.371861289 | NA | | | Crybb1 | 120.631174 | 0.110495191 | 0.094584868 | 1.168212135 | 0.242721207 | 0.692772033 |
| Gm16853 | 6.300706512 | 0.243538079 | 0.148707516 | 1.637698518 | 0.1014846 | NA | | | Abcb7 | 1456.020763 | 0.110499131 | 0.067799307 | 1.629797341 | 0.103144336 | 0.522975271 |
| Gm16863 | 0.539474011 | 0.04933016 | 0.054620124 | 0.903149904 | 0.366446341 | NA | | | Scg5 | 7806.697272 | 0.11050813 | 0.060432788 | 1.828612141 | 0.067457733 | 0.448803776 |
| Gm16880 | 1.013259476 | -0.016392334 | 0.071729171 | -0.228530927 | 0.819233516 | NA | | | Ngfrap1 | 5366.659253 | 0.110511844 | 0.060076928 | 1.839505585 | 0.065840857 | 0.445876127 |
| Gm16982 | 1.937057278 | 0.072024504 | 0.085701883 | 0.840407481 | 0.400679956 | NA | | | Ppm1l | 3064.522481 | 0.110580073 | 0.063836873 | 1.732228848 | 0.083232825 | 0.485168543 |
| Gm17019 | 0.813807001 | 0.049577646 | 0.06258422 | 0.792174864 | 0.428258723 | NA | | | Skil | 3017.793096 | 0.110603232 | 0.081683089 | 1.354053004 | 0.175719468 | 0.628540013 |
| Gm1715 | 3.988540077 | 0.003802252 | 0.116258024 | 0.032705289 | 0.973909606 | NA | | | Dlc1 | 3515.895399 | 0.110618581 | 0.068631149 | 1.611783896 | 0.107008974 | 0.531423026 |
| Gm1720 | 0.159967308 | 0.0240345 | 0.030870969 | 0.778547006 | 0.436246606 | NA | | | Nup62cl | 9.359482255 | 0.110626574 | 0.062902489 | 1.758699485 | 0.078628565 | NA |
| Gm17252 | 0 | NA | NA | NA | NA | NA | | | Lzts2 | 1716.032477 | 0.110651307 | 0.068307473 | 1.619900474 | 0.105253658 | 0.526667116 |
| Gm17359 | 0.19442443 | -0.026451352 | 0.031568725 | -0.837897386 | 0.402088337 | NA | | | Pcdhga1 | 289.0068331 | 0.110668794 | 0.092905935 | 1.191191872 | 0.23357827 | 0.685064073 |
| Gm17365 | 0 | NA | NA | NA | NA | NA | | | Glra4 | 6.424763043 | 0.110699917 | 0.058065579 | 1.906463682 | 0.056590073 | NA |
| Gm17384 | 5.411198423 | 0.176646725 | 0.142620572 | 1.238578157 | 0.215501762 | NA | | | Prkar1a | 28417.63793 | 0.110723877 | 0.047557012 | 2.328234501 | 0.019899655 | 0.26641695 |
| Gm17495 | 0.116696964 | -0.020477583 | 0.027954224 | -0.732539847 | 0.463839131 | NA | | | Cisd3 | 677.1743138 | 0.110755361 | 0.061154901 | 1.811062713 | 0.070131145 | 0.456009403 |
| Gm17660 | 6.825038994 | 0.286873783 | 0.151756259 | 1.890358815 | 0.058709988 | NA | | | Gadd45b | 426.8603743 | 0.1107899 | 0.093836711 | 1.180666914 | 0.23773507 | 0.688602119 |
| Gm17677 | 0.186326724 | 0.024939385 | 0.031411053 | 0.793968451 | 0.427213798 | NA | | | Per1 | 5332.737938 | 0.110837858 | 0.091369324 | 1.213075166 | NA | NA |
| Gm17689 | 0 | NA | NA | NA | NA | NA | | | Nip7 | 1008.380848 | 0.110848122 | 0.0625813 | 1.77126587 | 0.0765165 | 0.472560915 |
| Gm17727 | 0 | NA | NA | NA | NA | NA | | | Lrrn4 | 25.47533856 | 0.110884931 | 0.08354324 | 1.327275926 | 0.184417424 | 0.637780733 |
| Gm17745 | 1.856371848 | -0.137630911 | 0.089636171 | -1.535439431 | 0.124675924 | NA | | | Rnf220 | 6073.003461 | 0.110911411 | 0.044625594 | 2.485376665 | 0.012941442 | 0.216344675 |
| Gm17746 | 0.914430724 | 0.026377469 | 0.058012971 | 0.454682264 | 0.649337842 | NA | | | Tmem55a | 1522.057584 | 0.110922333 | 0.06184624 | 1.793517819 | 0.072890164 | 0.463573784 |
| Gm17751 | 7.248427673 | -0.336861661 | 0.152608484 | -2.207358666 | 0.027289012 | NA | | | Dpcd | 1510.483475 | 0.110960488 | 0.05473822 | 2.027111744 | 0.042650986 | 0.37389754 |
| Gm17757 | 0 | NA | NA | NA | NA | NA | | | Alg2 | 6092.892262 | 0.110989966 | 0.056917102 | 1.950028406 | 0.051172733 | 0.403266245 |
| Gm17801 | 3.177281092 | -0.082888088 | 0.124350382 | -0.666568823 | 0.505047589 | NA | | | Gfod2 | 764.6225987 | 0.111020938 | 0.060986685 | 1.820412728 | 0.068696176 | 0.452553581 |
| Gm17830 | 0.426633926 | -0.004170814 | 0.042905384 | -0.097209576 | 0.922559963 | NA | | | Pim1 | 123.7989194 | 0.111023488 | 0.095081215 | 1.167670055 | 0.24293988 | 0.692772033 |
| Gm18409 | 1.082787731 | -0.073424906 | 0.057334288 | -1.280645646 | 0.20031816 | NA | | | Gpr19 | 505.7297864 | 0.111023726 | 0.075432684 | 1.471825205 | 0.141068085 | 0.586989325 |
| Gm18853 | 0 | NA | NA | NA | NA | NA | | | Grik3 | 5968.400168 | 0.111040974 | 0.066159798 | 1.678375345 | 0.093273848 | 0.50821302 |
| Gm19276 | 0.275017096 | 0.008844542 | 0.038389071 | 0.2303922 | 0.817787022 | NA | | | Swi5 | 3509.847136 | 0.11106457 | 0.048679442 | 2.281549779 | 0.022515935 | 0.28195385 |
| Gm19277 | 0 | NA | NA | NA | NA | NA | | | Thoc3 | 1164.185172 | 0.111101843 | 0.061718029 | 1.800152158 | 0.071836616 | 0.460196881 |
| Gm19299 | 0.176054697 | -0.002190495 | 0.031239146 | -0.070120197 | 0.944097991 | NA | | | Cct7 | 7176.798944 | 0.111114479 | 0.041113497 | 2.702627736 | 0.006879375 | 0.15963226 |
| Gm19303 | 0.282397767 | 0.011595411 | 0.038695572 | 0.299657307 | 0.764438567 | NA | | | Gpbp1 | 3672.725782 | 0.111157803 | 0.069681356 | 1.595230188 | 0.11066077 | 0.537406303 |
| Gm19395 | 0 | NA | NA | NA | NA | NA | | | Ccser1 | 453.684784 | 0.111184436 | 0.074564829 | 1.49111099 | 0.135932357 | 0.579435507 |
| Gm19402 | 0.098353386 | -0.020212831 | 0.02786466 | -0.725393061 | 0.468210929 | NA | | | Cd302 | 504.167 | 0.111197785 | 0.075692348 | 1.469075638 | 0.141812275 | 0.587554222 |
| Gm19424 | 0.360303906 | 0.01488513 | 0.035847896 | 0.415230234 | 0.677973368 | NA | | | Arl6ip1 | 10622.56379 | 0.11122708 | 0.060757012 | 1.830687121 | 0.067147256 | 0.447688159 |
| Gm19434 | 1.594634258 | -0.015377979 | 0.093891344 | -0.163784838 | 0.869900528 | NA | | | Nfkbib | 793.8737636 | 0.111250625 | 0.071603241 | 1.553709346 | 0.120253763 | 0.554014004 |
| Gm19466 | 0.338960846 | 0.036479516 | 0.037750094 | 0.966342396 | 0.333872878 | NA | | | Tenm1 | 1003.244866 | 0.111278087 | 0.080433968 | 1.383471308 | 0.166520399 | 0.616902656 |
| Gm19510 | 0.187700257 | -0.003168524 | 0.031449734 | -0.100748845 | 0.919749836 | NA | | | Oxct1 | 12418.4517 | 0.111358768 | 0.051487723 | 2.162821782 | 0.03055489 | 0.320602541 |
| Gm19557 | 2.938296458 | 0.070952002 | 0.1066624 | 0.665201628 | 0.505921539 | NA | | | Lrrtm4 | 1101.401546 | 0.111360928 | 0.065876156 | 1.690458795 | 0.090940215 | 0.503727415 |
| Gm19583 | 0.21363976 | -0.000753674 | 0.031922632 | -0.02360938 | 0.98116419 | NA | | | Mmp24 | 2053.985057 | 0.111389536 | 0.071558204 | 1.556628449 | 0.119558725 | 0.55282356 |
| Gm19589 | 0.107716603 | -0.020477583 | 0.027954224 | -0.732539849 | 0.46383913 | NA | | | Sap30 | 139.2768409 | 0.111433935 | 0.091601592 | 1.216506531 | 0.223792025 | 0.675094439 |
| Gm19619 | 8.065104231 | 0.164940638 | 0.156455409 | 1.054234169 | 0.29177572 | NA | | | Sgsh | 402.8846555 | 0.111459505 | 0.087759226 | 1.270060252 | 0.204063169 | 0.65778952 |
| Gm1965 | 0.188785634 | 0.025025111 | 0.031461603 | 0.79541754 | 0.42637066 | NA | | | D17H6S53E | 607.735037 | 0.111488352 | 0.064771431 | 1.721258131 | 0.085203993 | 0.489390873 |
| Gm19668 | 0 | NA | NA | NA | NA | NA | | | Agpat9 | 247.2578 | 0.111542782 | 0.091477971 | 1.219340369 | 0.222715033 | 0.67374894 |
| Gm1968 | 0.105923158 | 0.018524693 | 0.027977518 | 0.662127842 | 0.507889294 | NA | | | C330007P06Rik | 1993.078756 | 0.11156069 | 0.062929781 | 1.772780517 | 0.076265083 | 0.472435997 |
| Gm19689 | 2.262677967 | 0.052066007 | 0.10442818 | 0.498581957 | 0.618073919 | NA | | | Rnf34 | 2285.808367 | 0.111591535 | 0.053388782 | 2.090168213 | 0.036602692 | 0.346440078 |
| Gm19710 | 2.050198435 | -0.037167857 | 0.104183566 | -0.356753102 | 0.721276611 | NA | | | Myh7 | 704.7068128 | 0.111640906 | 0.086115405 | 1.296423948 | 0.194829466 | 0.65082254 |
| Gm19782 | 0.305079873 | -0.015054679 | 0.039642215 | -0.379763828 | 0.704120735 | NA | | | Unc13c | 2723.02985 | 0.111718814 | 0.049498368 | 2.176007721 | 0.029591791 | 0.692038786 |
| Gm19784 | 0 | NA | NA | NA | NA | NA | | | Ylpf5 | 1466.195301 | 0.111758526 | 0.064618634 | 1.741157331 | 0.081656004 | 0.481581452 |
| Gm1987 | 0 | NA | NA | NA | NA | NA | | | Bud31 | 1189.150247 | 0.11179084 | 0.063405698 | 1.763104016 | 0.077882948 | 0.475337606 |
| Gm19897 | 2.557211408 | 0.060659439 | 0.097579969 | 0.621638224 | 0.534179781 | NA | | | Cradd | 191.3212028 | 0.111812922 | 0.093310212 | 1.198292445 | 0.230803187 | 0.683402059 |
| Gm1993 | 0 | NA | NA | NA | NA | NA | | | Ahsa1 | 4364.075358 | 0.111873323 | 0.04679463 | 2.389970254 | 0.016849737 | 0.24653656 |
| Gm1995 | 0 | NA | NA | NA | NA | NA | | | Mrps25 | 1349.032936 | 0.11187819 | 0.057672238 | 1.93989981 | 0.052392232 | 0.406355487 |
| Gm19990 | 1.264991571 | -0.0324123 | 0.085073853 | -0.380990153 | 0.703210555 | NA | | | Nans | 522.342846 | 0.111879916 | 0.070951112 | 1.576859226 | 0.11482792 | 0.541130934 |
| Gm2002 | 0 | NA | NA | NA | NA | NA | | | Dnajc28 | 578.8346001 | 0.111918139 | 0.073667826 | 1.519226838 | 0.128705408 | 0.567225351 |
| Gm20098 | 0.215433205 | -0.027029776 | 0.031922004 | -0.84674434 | 0.397137638 | NA | | | Fam92a | 1493.784776 | 0.111921764 | 0.061381988 | 1.823364926 | 0.068248139 | 0.451155091 |
| Gm20110 | 0.294090812 | -0.00484422 | 0.03361619 | 0.282132504 | 0.77784191 | NA | | | Pygm | 1787.068689 | 0.111937023 | 0.068768326 | 1.627740981 | 0.10357982 | 0.523462117 |
| Gm2012 | 0 | NA | NA | NA | NA | NA | | | Ppap2b | 846.6607414 | 0.111965051 | 0.075886128 | 1.475435018 | 0.140095627 | 0.585376897 |
| Gm20125 | 1.627670774 | 0.215002718 | 0.093678218 | 2.295119629 | 0.021726269 | NA | | | Orc5 | 969.218877 | 0.112025446 | 0.056833206 | 1.971161327 | 0.048705426 | 0.395027926 |
| Gm20139 | 1.140243808 | -0.052958208 | 0.079629872 | -0.66505455 | 0.506015603 | NA | | | Vdac3 | 7090.679269 | 0.112034037 | 0.048738671 | 2.298662823 | 0.021523783 | 0.275709409 |
| Gm2016 | 0 | NA | NA | NA | NA | NA | | | Ythdf2 | 1556.631747 | 0.112149797 | 0.063606578 | 1.763179221 | 0.077870267 | 0.475337606 |
| Gm20172 | 0 | NA | NA | NA | NA | NA | | | Alg8 | 840.4244159 | 0.112153875 | 0.061632011 | 1.819734147 | 0.068799501 | 0.452823537 |
| Gm20199 | 0.176054697 | -0.002190495 | 0.031239146 | -0.070120197 | 0.944097991 | NA | | | Deptor | 452.0526655 | 0.11221544 | 0.0857915 | 1.30800185 | 0.190872682 | 0.646448258 |
| Gm2022 | 0 | NA | NA | NA | NA | NA | | | Cmbl | 399.0173737 | 0.112224399 | 0.085230291 | 1.316716951 | 0.187932614 | 0.641781416 |
| Gm20268 | 0 | NA | NA | NA | NA | NA | | | Tmem178 | 2505.288308 | 0.112292692 | 0.07343359 | 1.529137388 | 0.126221466 | 0.562787717 |
| Gm2027 | 5.893859806 | -0.051956358 | 0.138081687 | -0.376272616 | 0.706714235 | NA | | | Cenpn | 37.07319102 | 0.112318629 | 0.084317623 | 1.332089612 | 0.182830744 | 0.635967285 |
| Gm2030 | 0 | NA | NA | NA | NA | NA | | | Gpr98 | 79.6564586 | 0.112336858 | 0.094770486 | 1.185356985 | 0.235837631 | 0.687039131 |
| Gm20324 | 7.74618319 | 0.049083581 | 0.157553918 | 0.311535132 | 0.755393839 | NA | | | Zbtb7b | 703.2548198 | 0.112370822 | 0.070083539 | 1.603383971 | 0.108849908 | 0.534621387 |
| Gm20356 | 0 | NA | NA | NA | NA | NA | | | Atp6v1d | 9754.728429 | 0.112453389 | 0.058692252 | 1.915983531 | 0.055367192 | 0.413057568 |
| Gm20362 | 1.746981493 | -0.005619916 | 0.100535985 | -0.055899543 | 0.955421835 | NA | | | Actrt3 | 34.85827009 | 0.112533282 | 0.091806718 | 1.225763046 | 0.220287857 | 0.672757795 |
| Gm20408 | 0 | NA | NA | NA | NA | NA | | | Haus8 | 288.3939228 | 0.112536772 | 0.082078641 | 1.37108474 | 0.170348541 | 0.620482174 |
| Gm20410 | 0 | NA | NA | NA | NA | NA | | | Luzp1 | 5599.610212 | 0.112587915 | 0.075450468 | 1.492209635 | 0.135644197 | 0.579179191 |
| Gm2042 | 0 | NA | NA | NA | NA | NA | | | Ect2l | 14.96070415 | 0.112604494 | 0.068384807 | 1.646630267 | 0.099634065 | NA |
| Gm20516 | 0 | NA | NA | NA | NA | NA | | | Ccnd3 | 942.0067076 | 0.112642648 | 0.067758363 | 1.712457467 | 0.086812337 | 0.493466364 |
| Gm20554 | 1.055791187 | 0.040993609 | 0.080037269 | 0.51218151 | 0.608523982 | NA | | | Egln1 | 3971.342359 | 0.112732677 | 0.068613208 | 1.643017147 | 0.100379386 | 0.5211685 |
| Gm20556 | 0.096071043 | -0.002009795 | 0.027825948 | -0.722340436 | 0.470107572 | NA | | | Mrpl15 | 983.0350172 | 0.112740033 | 0.066506806 | 1.695165346 | 0.090044059 | 0.500949827 |
| Gm20579 | 0 | NA | NA | NA | NA | NA | | | Mettl18 | 165.0242488 | 0.112808922 | 0.087837522 | 1.28429081 | 0.199040221 | 0.65377551 |
| Gm20580 | 0 | NA | NA | NA | NA | NA | | | Gm129 | 856.2818815 | 0.112817373 | 0.088620244 | 1.273042909 | 0.203008216 | 0.656606302 |
| Gm20581 | 0 | NA | NA | NA | NA | NA | | | Ttc39c | 882.0724182 | 0.112842558 | 0.072583391 | 1.554660873 | 0.120026858 | 0.553708672 |
| Gm20594 | 1.362017742 | -0.00396715 | 0.073343854 | -0.049948753 | 0.964148157 | NA | | | Rheb | 3351.622749 | 0.112889599 | 0.069993486 | 1.612858639 | 0.106775223 | 0.530447999 |
| Gm20604 | 0 | NA | NA | NA | NA | NA | | | Cept1 | 1330.661152 | 0.11291086 | 0.065549382 | 1.722673194 | 0.084947641 | 0.488499377 |
| Gm20611 | 0.281741269 | -0.015233555 | 0.038833149 | -0.392282259 | 0.69484967 | NA | | | Pcdhb18 | 168.4858115 | 0.112917372 | 0.091291798 | 1.236884081 | 0.216130121 | 0.670049713 |

| | | | | | | | | | | | | | | |
|---|---|---|---|---|---|---|---|---|---|---|---|---|---|---|
| Gm20735 | 1.169054187 | -0.010860347 | 0.079141224 | -0.137227436 | 0.890851025 | NA | | | Psen1 | 2859.812873 | 0.112926339 | 0.05077988 | 2.223840225 | 0.026159196 | 0.302373406 |
| Gm20740 | 0 | NA | NA | NA | NA | NA | | | Mtf1 | 1532.249891 | 0.113078386 | 0.074195757 | 1.524054628 | 0.127495066 | 0.564976608 |
| Gm20741 | 0 | NA | NA | NA | NA | NA | | | Fosb | 840.5505186 | 0.113091681 | 0.091801595 | 1.231914121 | 0.217981166 | 0.670929686 |
| Gm20744 | 0.224413567 | -0.027287719 | 0.032077135 | -0.850690659 | 0.394941214 | NA | | | Rnf103 | 1770.206256 | 0.113124629 | 0.065814253 | 1.718846968 | 0.085642241 | 0.490358371 |
| Gm20745 | 0.159967308 | 0.0240345 | 0.030870969 | 0.778547006 | 0.436246606 | NA | | | Sacm1l | 3507.399694 | 0.113169353 | 0.066816729 | 1.693727835 | 0.090317013 | 0.50209189 |
| Gm20750 | 0.296373155 | 0.009272162 | 0.033639931 | 0.275629621 | 0.782832556 | NA | | | Blmh | 2373.356012 | 0.113178974 | 0.049412314 | 2.290501387 | 0.021992268 | 0.278517602 |
| Gm20752 | 4.973363912 | 0.084306828 | 0.122115078 | 0.690388352 | 0.48995 | NA | | | Rab9b | 1241.718515 | 0.113218103 | 0.075017128 | 1.509230037 | 0.131240007 | 0.571460533 |
| Gm20755 | 0.619937596 | 0.079772058 | 0.055246832 | 1.443920948 | 0.148761213 | NA | | | Snx22 | 301.5839025 | 0.11324002 | 0.092758735 | 1.220801687 | 0.222161115 | 0.673437469 |
| Gm20756 | 0.403835985 | -0.003083065 | 0.042384291 | -0.072740739 | 0.94201243 | NA | | | Mrpl17 | 1032.48986 | 0.113248284 | 0.055362067 | 2.045593495 | 0.040796382 | 0.365377968 |
| Gm20757 | 0.32854328 | 0.00879128 | 0.034635881 | 0.253820007 | 0.799634616 | NA | | | Vcpip1 | 2477.210919 | 0.11332003 | 0.071841892 | 1.577353094 | 0.114714302 | 0.543130934 |
| Gm20758 | 0 | NA | NA | NA | NA | NA | | | Chsy1 | 591.3559668 | 0.113352125 | 0.069269576 | 1.636391204 | 0.101757739 | 0.522606076 |
| Gm20759 | 0 | NA | NA | NA | NA | NA | | | Cckar | 33.0881941 | 0.113360971 | 0.059447157 | 1.906919966 | 0.056530952 | 0.416806094 |
| Gm20765 | 0 | NA | NA | NA | NA | NA | | | Nsdhl | 1044.978714 | 0.113383854 | 0.05641821 | 2.009703159 | 0.044462615 | 0.380589751 |
| Gm2083 | 0 | NA | NA | NA | NA | NA | | | Spsb1 | 903.7217685 | 0.113407035 | 0.080225383 | 1.413605392 | 0.157477797 | 0.608141173 |
| Gm20878 | 0 | NA | NA | NA | NA | NA | | | Mrpl21 | 595.0490793 | 0.113432268 | 0.068534404 | 1.655114239 | 0.097901327 | 0.518695741 |
| Gm21002 | 0 | NA | NA | NA | NA | NA | | | Tprn | 879.2536157 | 0.113440478 | 0.074764669 | 1.517300604 | 0.129190806 | 0.568157931 |
| Gm21057 | 1.653400267 | -0.008468466 | 0.100130603 | -0.084574206 | 0.932599906 | NA | | | Adc | 1198.999713 | 0.113636341 | 0.063645474 | 1.785458321 | 0.074187006 | 0.467117664 |
| Gm2109 | 2.980264325 | 0.06341751 | 0.11607839 | 0.546333468 | 0.584836734 | NA | | | Aph1b | 1189.514552 | 0.113646266 | 0.062272486 | 1.824983592 | 0.068003505 | 0.450415406 |
| Gm21119 | 0 | NA | NA | NA | NA | NA | | | Xylt2 | 1102.917814 | 0.113659079 | 0.058380376 | 1.946871303 | 0.051550166 | 0.404394022 |
| Gm21276 | 0 | NA | NA | NA | NA | NA | | | Stard4 | 931.0726841 | 0.113732141 | 0.063992413 | 1.777275408 | 0.075522938 | 0.470431472 |
| Gm21293 | 0 | NA | NA | NA | NA | NA | | | Slc6a15 | 2757.264872 | 0.113761989 | 0.055389646 | 2.053849377 | 0.039990272 | 0.361359262 |
| Gm21304 | 0 | NA | NA | NA | NA | NA | | | Stx7 | 3956.888454 | 0.113819024 | 0.054276233 | 2.09703248 | 0.0359907 | 0.344080613 |
| Gm21312 | 0 | NA | NA | NA | NA | NA | | | BC004004 | 2301.903003 | 0.113846595 | 0.052386805 | 2.173192194 | 0.029765855 | 0.318159537 |
| Gm21319 | 0 | NA | NA | NA | NA | NA | | | Smyd3 | 1073.198134 | 0.113858666 | 0.061277759 | 1.858074916 | 0.063158376 | 0.440342223 |
| Gm21498 | 0 | NA | NA | NA | NA | NA | | | Ptpn11 | 9081.521826 | 0.113885046 | 0.043468871 | 2.619921857 | 0.008794992 | 0.179191559 |
| Gm21541 | 0 | NA | NA | NA | NA | NA | | | Mob3a | 516.2944968 | 0.113909758 | 0.086801337 | 1.312304183 | 0.189417521 | 0.644077615 |
| Gm21586 | 0 | NA | NA | NA | NA | NA | | | Zfp593 | 335.9758142 | 0.113961029 | 0.076242253 | 1.494722745 | 0.134986816 | 0.578547014 |
| Gm21637 | 0 | NA | NA | NA | NA | NA | | | Chodl | 56.31277321 | 0.113980881 | 0.088347162 | 1.29014762 | 0.196999409 | 0.653038204 |
| Gm21677 | 0 | NA | NA | NA | NA | NA | | | Prkacb | 24312.29171 | 0.114110539 | 0.063204793 | 1.805409576 | 0.071010614 | 0.458300422 |
| Gm21693 | 0 | NA | NA | NA | NA | NA | | | Adamts13 | 44.6965783 | 0.114213628 | 0.092414063 | 1.235890126 | 0.216499408 | 0.670049713 |
| Gm21704 | 0 | NA | NA | NA | NA | NA | | | Tmem9 | 2111.890346 | 0.11424113 | 0.070998263 | 1.60906936 | 0.107601176 | 0.532801865 |
| Gm21708 | 0 | NA | NA | NA | NA | NA | | | P4ha1 | 2147.692088 | 0.11424387 | 0.088219365 | 1.29499765 | 0.19532104 | 0.651003086 |
| Gm2176 | 0 | NA | NA | NA | NA | NA | | | Ston1 | 176.6388677 | 0.114265738 | 0.085663645 | 1.333888346 | 0.182240451 | 0.635245711 |
| Gm21945 | 0 | NA | NA | NA | NA | NA | | | Sugt1 | 2221.110583 | 0.114284732 | 0.071191461 | 1.605315176 | 0.108424461 | 0.534070438 |
| Gm21946 | 0 | NA | NA | NA | NA | NA | | | Hdhd2 | 3031.820926 | 0.11429999 | 0.049596912 | 2.304578755 | 0.021190177 | 0.274094844 |
| Gm21947 | 0 | NA | NA | NA | NA | NA | | | Ninj1 | 407.287414 | 0.114314142 | 0.081948634 | 1.394948716 | 0.163031305 | 0.614351408 |
| Gm21949 | 0 | NA | NA | NA | NA | NA | | | Mbp | 75747.34624 | 0.114314726 | 0.085543366 | 1.33633655 | 0.181439295 | 0.634469716 |
| Gm21950 | 0 | NA | NA | NA | NA | NA | | | Gpr137c | 839.1832896 | 0.114316951 | 0.070161397 | 1.629342568 | 0.10324052 | 0.522975271 |
| Gm21951 | 0 | NA | NA | NA | NA | NA | | | Nfu1 | 1223.426483 | 0.114321991 | 0.06385293 | 1.790395388 | 0.073390371 | 0.464720371 |
| Gm2381 | 1.105618087 | 0.098496357 | 0.077202422 | 1.275819518 | 0.202019326 | NA | | | Itfg1 | 11150.74297 | 0.114426572 | 0.066668913 | 1.716340762 | 0.086099692 | 0.491621118 |
| Gm2447 | 0.505546546 | 0.009202456 | 0.05076589 | 0.181272426 | 0.856153749 | NA | | | Cpsf2 | 2419.025691 | 0.114502623 | 0.060180939 | 1.902639371 | 0.057087618 | 0.418604109 |
| Gm2506 | 0 | NA | NA | NA | NA | NA | | | Exoc6b | 5186.665773 | 0.114649084 | 0.049954963 | 2.29504894 | 0.02173032 | 0.276647349 |
| Gm2516 | 0 | NA | NA | NA | NA | NA | | | Sar1b | 1303.027518 | 0.114793439 | 0.067175799 | 1.708851105 | 0.087478534 | 0.495597906 |
| Gm2663 | 0 | NA | NA | NA | NA | NA | | | Smim3 | 335.3800915 | 0.114812241 | 0.093287211 | 1.23073934 | 0.218420369 | 0.671344538 |
| Gm2696 | 0 | NA | NA | NA | NA | NA | | | Prdx5 | 5559.963931 | 0.114830698 | 0.055055967 | 2.085708514 | 0.037005035 | 0.348173348 |
| Gm2799 | 0 | NA | NA | NA | NA | NA | | | Diras1 | 5451.239281 | 0.114872362 | 0.051372735 | 2.236056962 | 0.02534804 | 0.298582503 |
| Gm2825 | 0 | NA | NA | NA | NA | NA | | | Fam69b | 1946.793305 | 0.114911977 | 0.062968189 | 1.824921117 | 0.068012934 | 0.450415406 |
| Gm2837 | 0 | NA | NA | NA | NA | NA | | | Rangrf | 349.1368184 | 0.11492405 | 0.079661102 | 1.442662056 | 0.149115693 | 0.597202801 |
| Gm2848 | 0.099009884 | 0.018205726 | 0.027896865 | 0.652608315 | 0.514008823 | NA | | | Trp53inp2 | 8184.98424 | 0.114930656 | 0.061825804 | 1.858943168 | 0.063035194 | 0.440004437 |
| Gm2863 | 0 | NA | NA | NA | NA | NA | | | Klhl25 | 449.2875211 | 0.114939995 | 0.07236601 | 1.588314669 | 0.112215202 | 0.538992394 |
| Gm2913 | 0 | NA | NA | NA | NA | NA | | | Plk3 | 638.339206 | 0.114970637 | 0.093914431 | 1.224206288 | 0.220874418 | 0.672757795 |
| Gm2933 | 0 | NA | NA | NA | NA | NA | | | Etf1 | 3511.59025 | 0.114983973 | 0.056597039 | 2.031625239 | 0.04219161 | 0.371439588 |
| Gm3020 | 0 | NA | NA | NA | NA | NA | | | Tmem11 | 786.1756962 | 0.115017819 | 0.059028173 | 1.948524104 | 0.051352283 | 0.403430649 |
| Gm3139 | 0 | NA | NA | NA | NA | NA | | | Lhx5 | 11.93645414 | 0.115086542 | 0.056891635 | 2.02290797 | 0.043082637 | NA |
| Gm3143 | 0.079983654 | 0.016740032 | 0.027674178 | 0.604897175 | 0.54524732 | NA | | | Vsnl1 | 23661.49269 | 0.115139901 | 0.064944779 | 1.772889263 | 0.076247058 | 0.472435997 |
| Gm3238 | 0 | NA | NA | NA | NA | NA | | | 4930555G01Rik | 9.366760938 | 0.115141832 | 0.06513041 | 1.767865929 | 0.07708332 | NA |
| Gm3258 | 0.295060159 | -0.029687054 | 0.033501022 | -0.886153701 | 0.375534709 | NA | | | Lrrc24 | 1143.920524 | 0.115166083 | 0.071609907 | 1.608242317 | 0.107782119 | 0.533015565 |
| Gm3259 | 0 | NA | NA | NA | NA | NA | | | Fdps | 2348.381945 | 0.115171196 | 0.050432869 | 2.283653478 | 0.022391904 | 0.281750998 |
| Gm3264 | 1.613866166 | 0.026648896 | 0.096497914 | 0.276160333 | 0.782424922 | NA | | | Surf4 | 3165.405312 | 0.115184931 | 0.043343621 | 2.657482855 | 0.007872659 | 0.169996349 |
| Gm3285 | 0 | NA | NA | NA | NA | NA | | | Tmed10 | 6588.343332 | 0.115191705 | 0.051815213 | 2.223125176 | 0.02620736 | 0.302373406 |
| Gm3286 | 0 | NA | NA | NA | NA | NA | | | Zhx2 | 684.0154422 | 0.115276993 | 0.073610351 | 1.566043255 | 0.117338491 | 0.548468184 |
| Gm3317 | 0 | NA | NA | NA | NA | NA | | | Crhbp | 766.7828428 | 0.115315599 | 0.070547343 | 1.634584568 | 0.102136165 | 0.522723164 |
| Gm3336 | 0.589328065 | -0.006539609 | 0.058189931 | -0.112383855 | 0.910519056 | NA | | | Hdac9 | 1347.020357 | 0.115356845 | 0.065115114 | 1.771583254 | 0.076463761 | 0.472560915 |
| Gm3376 | 0 | NA | NA | NA | NA | NA | | | Slc7a3 | 409.7692653 | 0.115380748 | 0.091581394 | 1.259871053 | 0.207715883 | 0.662799412 |
| Gm3383 | 0 | NA | NA | NA | NA | NA | | | Myl12b | 7901.317927 | 0.115397733 | 0.050517364 | 2.28431817 | 0.022352839 | 0.281750998 |
| Gm3402 | 0 | NA | NA | NA | NA | NA | | | Mrpl22 | 444.2835168 | 0.115485212 | 0.074081487 | 1.558894362 | 0.119021385 | 0.551385848 |
| Gm3404 | 0 | NA | NA | NA | NA | NA | | | Aldh1a1 | 2479.789611 | 0.115486983 | 0.088819617 | 1.300241851 | 0.193518091 | 0.649744561 |
| Gm3409 | 0 | NA | NA | NA | NA | NA | | | Zfp828 | 1932.18921 | 0.115501068 | 0.046140206 | 2.503262971 | 0.012305407 | 0.212038374 |
| Gm3415 | 0 | NA | NA | NA | NA | NA | | | Ndufb3 | 1683.00762 | 0.115513527 | 0.054702497 | 2.111668278 | 0.034714912 | 0.33922827 |
| Gm3417 | 0 | NA | NA | NA | NA | NA | | | Chl1 | 5342.755183 | 0.115561632 | 0.062392589 | 1.852169197 | 0.06400153 | 0.441368101 |
| Gm3428 | 0 | NA | NA | NA | NA | NA | | | Eif2s1 | 1565.471226 | 0.115601921 | 0.071196742 | 1.623696774 | 0.10444054 | 0.524404868 |
| Gm3434 | 0.116696964 | -0.020477583 | 0.027954224 | -0.732539847 | 0.463839131 | NA | | | Scrn1 | 5482.763182 | 0.115685405 | 0.061234875 | 1.889207831 | 0.058863986 | 0.425190581 |
| Gm3448 | 0 | NA | NA | NA | NA | NA | | | Zfp689 | 292.5122166 | 0.115726889 | 0.07846537 | 1.474878521 | 0.140245206 | 0.585529073 |
| Gm3494 | 0 | NA | NA | NA | NA | NA | | | Snap47 | 15247.82908 | 0.115772719 | 0.04464793 | 2.593014276 | 0.009513882 | 0.186251552 |
| Gm3500 | 0 | NA | NA | NA | NA | NA | | | Syt4 | 8476.126474 | 0.115836159 | 0.070499243 | 1.643083732 | 0.10036561 | 0.5211685 |
| Gm362 | 0 | NA | NA | NA | NA | NA | | | Epb4.1l3 | 8982.149257 | 0.115838306 | 0.057012647 | 2.031800185 | 0.042173888 | 0.371439588 |
| Gm364 | 0.094392817 | 0.017895942 | 0.027818137 | 0.643319236 | 0.520016977 | NA | | | Coq10b | 876.6332329 | 0.115850154 | 0.085510938 | 1.354799237 | 0.17548153 | 0.628307789 |
| Gm3646 | 0 | NA | NA | NA | NA | NA | | | Dnmt3b | 34.18673748 | 0.115914328 | 0.083751386 | 1.384028769 | 0.166349646 | 0.616741949 |
| Gm3696 | 1.101142027 | 0.008536422 | 0.072641135 | 0.117514982 | 0.906451972 | NA | | | Slc25a5 | 7927.026801 | 0.115919011 | 0.054970438 | 2.108751817 | 0.03496601 | 0.34025917 |
| Gm3701 | 0 | NA | NA | NA | NA | NA | | | Impa1 | 2621.57463 | 0.115925064 | 0.063190963 | 1.834519656 | 0.066576893 | 0.447108619 |
| Gm3706 | 0 | NA | NA | NA | NA | NA | | | Slc18a3 | 92.61240397 | 0.115948566 | 0.094964259 | 1.220970587 | 0.222097157 | 0.673384115 |
| Gm3716 | 4.910453018 | -0.097938929 | 0.120135335 | -0.815238321 | 0.414935907 | NA | | | Pfn4 | 219.8162655 | 0.116018748 | 0.086860766 | 1.335686442 | 0.181651783 | 0.635059878 |
| Gm3750 | 0 | NA | NA | NA | NA | NA | | | Clvs1 | 2726.460098 | 0.116127483 | 0.071049471 | 1.634459503 | 0.102162403 | 0.522723164 |
| Gm3763 | 0 | NA | NA | NA | NA | NA | | | Uxs1 | 1528.570298 | 0.116207464 | 0.05569943 | 2.086331273 | 0.036948626 | 0.348173348 |
| Gm3776 | 0.413840517 | 0.018406376 | 0.037259486 | 0.494005092 | 0.621302583 | NA | | | Prkcq | 571.2505346 | 0.116222184 | 0.086915347 | 1.337188287 | 0.181161184 | 0.634469716 |
| Gm382 | 0 | NA | NA | NA | NA | NA | | | Tmem200a | 600.9512971 | 0.11623524 | 0.073932631 | 1.57217777 | 0.115909329 | 0.54588186 |
| Gm3833 | 0 | NA | NA | NA | NA | NA | | | Agap1 | 6162.315011 | 0.116238259 | 0.057645931 | 2.016417423 | 0.043756345 | 0.37878705 |
| Gm3925 | 0 | NA | NA | NA | NA | NA | | | 1500009L16Rik | 413.8167108 | 0.116309947 | 0.076481245 | 1.520764291 | 0.128318999 | 0.566288196 |
| Gm3985 | 0.197363271 | -0.000561558 | 0.031650166 | -0.017742642 | 0.985844163 | NA | | | Dnttip2 | 1033.816921 | 0.116365349 | 0.070592862 | 1.648401063 | 0.099270398 | 0.520390402 |
| Gm4013 | 7.753198208 | -0.02975479 | 0.146804805 | -0.202682671 | 0.839383069 | NA | | | Psmd7 | 3695.8742 | 0.116378122 | 0.039169994 | 2.971103897 | 0.002967314 | 0.09427942 |
| Gm4027 | 0 | NA | NA | NA | NA | NA | | | Tmem252 | 38.17854281 | 0.116419799 | 0.086877925 | 1.340038904 | 0.180232697 | 0.63422 |
| Gm4064 | 0 | NA | NA | NA | NA | NA | | | Ism1 | 250.2263968 | 0.116536422 | 0.085557467 | 1.362083587 | 0.173171507 | 0.624655902 |
| Gm4070 | 0 | NA | NA | NA | NA | NA | | | Nefm | 10432.9716 | 0.11656484 | 0.071352529 | 1.63364693 | 0.102333008 | 0.522859399 |
| Gm41 | 0 | NA | NA | NA | NA | NA | | | Hdhd3 | 141.8154252 | 0.11658156 | 0.091565322 | 1.27320647 | 0.202944785 | 0.656606302 |
| Gm4133 | 0 | NA | NA | NA | NA | NA | | | Elmo1 | 7226.94836 | 0.116692207 | 0.056627428 | 2.060701158 | 0.039331559 | 0.35977974 |
| Gm4141 | 0 | NA | NA | NA | NA | NA | | | Trim45 | 443.6801179 | 0.116716375 | 0.076016125 | 1.535416001 | 0.124681676 | 0.560776598 |
| Gm4172 | 0 | NA | NA | NA | NA | NA | | | Fbxo7 | 1136.231077 | 0.116826783 | 0.049993759 | 2.336827328 | 0.019448164 | 0.263564471 |
| Gm4175 | 0 | NA | NA | NA | NA | NA | | | Uqcrc2 | 8409.644389 | 0.116935102 | 0.048398244 | 2.416102163 | 0.015687659 | 0.237621413 |
| Gm4177 | 0 | NA | NA | NA | NA | NA | | | Atp5a1 | 35358.49742 | 0.116965915 | 0.040652871 | 2.877187094 | 0.004012376 | 0.113864147 |
| Gm4179 | 0 | NA | NA | NA | NA | NA | | | Tspan11 | 359.1697115 | 0.116972321 | 0.073340797 | 1.594914764 | 0.110731298 | 0.537406303 |
| Gm4187 | 0 | NA | NA | NA | NA | NA | | | Cd99l2 | 4666.777429 | 0.116985412 | 0.063398763 | 1.845231762 | 0.065003832 | 0.4443509 |
| Gm4201 | 0 | NA | NA | NA | NA | NA | | | E330009J07Rik | 740.4031864 | 0.11706226 | 0.085757286 | 1.365041558 | 0.172239989 | 0.623232235 |
| Gm4214 | 0 | NA | NA | NA | NA | NA | | | Zfp74 | 760.0134155 | 0.11706622 | 0.061721067 | 1.89669792 | 0.057867818 | 0.420925353 |
| Gm4216 | 0 | NA | NA | NA | NA | NA | | | Acot7 | 9027.936893 | 0.117086193 | 0.058922609 | 1.987118288 | 0.04690929 | 0.39102736 |
| Gm4220 | 0 | NA | NA | NA | NA | NA | | | Ppard | 1146.326812 | 0.117147026 | 0.064795065 | 1.807962175 | 0.070612388 | 0.456632761 |
| Gm4224 | 0 | NA | NA | NA | NA | NA | | | Plxnc1 | 3006.303328 | 0.117158885 | 0.076949307 | 1.522546338 | 0.127872244 | 0.565771339 |
| Gm4251 | 0 | NA | NA | NA | NA | NA | | | Kcnip4 | 2166.222137 | 0.117200786 | 0.068034188 | 1.72267487 | 0.084947337 | 0.488499377 |
| Gm4262 | 0.487806363 | -0.025218575 | 0.039384805 | -0.640312313 | 0.521969578 | NA | | | Pcdhgb7 | 363.3764578 | 0.117204923 | 0.089642967 | 1.307463665 | 0.191055287 | 0.646547945 |
| Gm4265 | 0.371083244 | -0.002928723 | 0.041579259 | -0.070437114 | 0.943845752 | NA | | | Astn2 | 2207.425747 | 0.117233846 | 0.058557798 | 2.002019363 | 0.045282649 | 0.383869599 |
| Gm4278 | 0 | NA | NA | NA | NA | NA | | | Lzts1 | 1702.330054 | 0.11724301 | 0.083279689 | 1.407822387 | 0.15918367 | 0.610912017 |
| Gm428 | 0 | NA | NA | NA | NA | NA | | | Fam78a | 405.6506848 | 0.117281511 | 0.085032759 | 1.379250921 | 0.167817403 | 0.617953076 |
| Gm4297 | 0 | NA | NA | NA | NA | NA | | | Ascc1 | 611.910022 | 0.117419912 | 0.071740403 | 1.636733387 | 0.10168619 | 0.522606076 |
| Gm4301 | 0 | NA | NA | NA | NA | NA | | | Sema3a | 1086.555683 | 0.117424685 | 0.07185719 | 1.634139665 | 0.102229528 | 0.522856086 |
| Gm4302 | 0 | NA | NA | NA | NA | NA | | | Nap1l3 | 1884.967831 | 0.117440338 | 0.071276063 | 1.647682714 | 0.099417797 | 0.520411863 |
| Gm4303 | 0 | NA | NA | NA | NA | NA | | | Cnot8 | 2239.411061 | 0.117501749 | 0.05210196 | 2.255227051 | 0.024119081 | 0.29178139 |
| Gm4305 | 0 | NA | NA | NA | NA | NA | | | Opn3 | 180.122375 | 0.117513636 | 0.089732869 | 1.309594094 | 0.190333188 | 0.645373455 |
| Gm4307 | 0 | NA | NA | NA | NA | NA | | | Gjd2 | 255.6622307 | 0.117560327 | 0.089472357 | 1.313929031 | 0.188870088 | 0.643114603 |
| Gm4312 | 0 | NA | NA | NA | NA | NA | | | Slc36a1 | 4282.048322 | 0.117570206 | 0.069280984 | 1.697005424 | 0.089695635 | 0.500737431 |
| Gm4340 | 0 | NA | NA | NA | NA | NA | | | Tra2b | 2162.143002 | 0.117591032 | 0.048727735 | 2.413225879 | 0.015812017 | 0.239006616 |
| Gm4345 | 0 | NA | NA | NA | NA | NA | | | Supt7l | 765.0733134 | 0.117604666 | 0.065758982 | 1.788419802 | 0.073708304 | 0.465836525 |
| Gm436 | 0 | NA | NA | NA | NA | NA | | | Klhl1 | 236.4634064 | 0.117609518 | 0.094028747 | 1.250782571 | 0.211013815 | 0.665621429 |
| Gm438 | 0 | NA | NA | NA | NA | NA | | | Slc45a3 | 57.05666145 | 0.117632848 | 0.094307144 | 1.247337615 | 0.212273731 | 0.666270994 |
| Gm44 | 0 | NA | NA | NA | NA | NA | | | Nostrin | 165.2980796 | 0.117795388 | 0.094714151 | 1.243693652 | 0.213612333 | 0.667262579 |
| Gm4461 | 2.006666071 | 0.020792586 | 0.103606388 | 0.200688264 | 0.840942337 | NA | | | Zbtb42 | 48.93665689 | 0.117879185 | 0.094455098 | 1.247991772 | 0.21203407 | 0.666270994 |
| Gm4477 | 1.064432019 | 0.028665342 | 0.074319474 | 0.385704324 | 0.69971566 | NA | | | Gpatch2l | 1684.598212 | 0.117883938 | 0.052249184 | 2.256187175 | 0.024058911 | 0.29172307 |
| Gm4489 | 0.479444338 | 0.008662674 | 0.055255961 | 0.156773568 | 0.875423305 | NA | | | Fam5c | 1664.899201 | 0.117889044 | 0.06532026 | 1.804785285 | 0.071108288 | 0.458300422 |
| Gm4498 | 0 | NA | NA | NA | NA | NA | | | Epm2aip1 | 5918.878465 | 0.117895289 | 0.055066774 | 2.14095144 | 0.032277954 | 0.327711599 |
| Gm4541 | 0 | NA | NA | NA | NA | NA | | | Suclg1 | 3490.521047 | 0.117930465 | 0.046288441 | 2.547730314 | 0.010842626 | 0.198744008 |
| Gm4559 | 0.094392817 | 0.017895942 | 0.027818137 | 0.643319236 | 0.520016977 | NA | | | Tspyl3 | 1588.911426 | 0.117947856 | 0.068185496 | 1.729820591 | 0.083664479 | 0.486580734 |
| Gm4566 | 0.38134816 | 0.013258311 | 0.036322589 | 0.365015595 | 0.715099765 | NA | | | Adra2c | 758.8377062 | 0.118057346 | 0.073858026 | 1.598436253 | 0.109945922 | 0.535492107 |
| Gm4567 | 0 | NA | NA | NA | NA | NA | | | Fgf2 | 24.95742788 | 0.118072268 | 0.084454556 | 1.398056817 | 0.162096006 | 0.613957643 |
| Gm4598 | 3.777051371 | -0.147144992 | 0.120949555 | -1.216581515 | 0.223763479 | NA | | | Nus1 | 3916.543561 | 0.118092951 | 0.073404417 | 1.60879898 | 0.107660304 | 0.532835971 |
| Gm4710 | 0.283178451 | 0.028390488 | 0.033368195 | 0.850824814 | 0.394866675 | NA | | | Spdya | 43.11098216 | 0.118094326 | 0.093316639 | 1.26552271 | 0.205684013 | 0.659550987 |
| Gm4719 | 0 | NA | NA | NA | NA | NA | | | Inpp5f | 4599.175924 | 0.11812942 | 0.068192732 | 1.732287529 | 0.083222381 | 0.485168543 |
| Gm4724 | 0 | NA | NA | NA | NA | NA | | | Derl1 | 2920.692473 | 0.118152283 | 0.053105035 | 2.224879109 | 0.026089355 | 0.302212096 |
| Gm4736 | 0 | NA | NA | NA | NA | NA | | | Csrp2 | 130.316805 | 0.118159372 | 0.094938989 | 1.244582151 | 0.213285385 | 0.666732456 |
| Gm4745 | 0 | NA | NA | NA | NA | NA | | | Plcz1 | 62.93226129 | 0.118248748 | 0.093767887 | 1.261079364 | 0.207280255 | 0.661893379 |
| Gm4759 | 2.439005598 | 0.006430841 | 0.091774583 | 0.07007213 | 0.944136249 | NA | | | Txn1 | 1573.28178 | 0.118309258 | 0.050932165 | 2.322879032 | 0.020185652 | 0.26886923 |
| Gm4763 | 0.227114243 | -0.027513412 | 0.032207244 | -0.854261616 | 0.39296005 | NA | | | Foxp2 | 926.6374136 | 0.118335482 | 0.090192178 | 1.312037084 | 0.189507623 | 0.644077615 |
| Gm4776 | 0.098353386 | -0.020212831 | 0.02786466 | -0.725393061 | 0.468210929 | NA | | | Eef1e1 | 602.3583082 | 0.118342848 | 0.07246298 | 1.633149071 | 0.102437648 | 0.522975271 |
| Gm4788 | 3.340403103 | -0.141652066 | 0.126845024 | -1.116733335 | 0.26410836 | NA | | | Serpini1 | 10679.45 | 0.118362616 | 0.076947309 | 1.538229432 | 0.123992527 | 0.559205951 |
| Gm4791 | 0.285336609 | 0.031742875 | 0.03528833 | 0.899528977 | 0.368370968 | NA | | | Ezr | 2661.976918 | 0.118367562 | 0.072547767 | 1.631581043 | 0.102767774 | 0.522975271 |
| Gm4792 | 0.309710804 | -0.014226733 | 0.039782345 | -0.357614233 | 0.720632025 | NA | | | Mettl7a1 | 1222.799213 | 0.118394175 | 0.085141508 | 1.390557637 | 0.16435961 | 0.614641427 |
| Gm4794 | 0 | NA | NA | NA | NA | NA | | | Gmppb | 419.5956503 | 0.118425554 | 0.077504698 | 1.527979049 | 0.12651774 | 0.563050456 |
| Gm4814 | 3.46518233 | 0.066414662 | 0.130114548 | 0.510432251 | 0.609748667 | NA | | | Maged1 | 22501.86593 | 0.118479978 | 0.060062933 | 1.972597278 | 0.048541463 | 0.395027926 |
| Gm4827 | 2.326590804 | -0.057497865 | 0.096373257 | -0.596616389 | 0.550763527 | NA | | | Fam26e | 243.9636622 | 0.11848865 | 0.086363625 | 1.371974019 | 0.170071526 | 0.620316915 |
| Gm4832 | 0.296373155 | 0.009272162 | 0.033639931 | 0.275629621 | 0.782832556 | NA | | | Gskip | 1316.288008 | 0.118489287 | 0.06604944 | 1.793948395 | 0.072821406 | 0.463542395 |
| Gm4836 | 0 | NA | NA | NA | NA | NA | | | Tmem164 | 1451.758121 | 0.118489308 | 0.062588195 | 1.89315747 | 0.058336931 | 0.423355579 |
| Gm4841 | 9.063191654 | 0.269685464 | 0.111962688 | 2.408708371 | NA | NA | | | Fam84a | 2780.703902 | 0.11854753 | 0.070414606 | 1.683564495 | 0.092265865 | 0.505947134 |
| Gm4846 | 0 | NA | NA | NA | NA | NA | | | Sf3b4 | 1873.634876 | 0.118552846 | 0.054089596 | 2.19178648 | 0.028394928 | 0.313443317 |
| Gm4847 | 0.927575411 | 0.00625276 | 0.069430007 | 0.090058465 | 0.928240755 | NA | | | Itpkc | 408.807499 | 0.118598767 | 0.079965786 | 1.483118887 | 0.138042827 | 0.582865025 |
| Gm4850 | 0 | NA | NA | NA | NA | NA | | | Hes6 | 379.405116 | 0.11868345 | 0.08481287 | 1.399356598 | 0.161706075 | 0.613781669 |
| Gm4858 | 0 | NA | NA | NA | NA | NA | | | Hipk2 | 1418.640082 | 0.118762485 | 0.083749069 | 1.418075287 | 0.156168785 | 0.606877415 |
| Gm4861 | 1.64668743 | 0.047849013 | 0.068404387 | 0.699502111 | 0.484238294 | NA | | | Lonp2 | 3198.504055 | 0.1187696 | 0.060052042 | 1.977777877 | 0.04795377 | 0.393352856 |
| Gm4871 | 0 | NA | NA | NA | NA | NA | | | Hspa8 | 89369.25306 | 0.118776391 | 0.050152073 | 2.368324658 | 0.017868848 | 0.255043174 |
| Gm4872 | 0 | NA | NA | NA | NA | NA | | | Gm16039 | 1228.728655 | 0.118779939 | 0.057506152 | 2.065517073 | 0.038874102 | 0.357169642 |
| Gm4884 | 0 | NA | NA | NA | NA | NA | | | Crebl2 | 1521.161969 | 0.118802798 | 0.051228398 | 2.319080871 | 0.020390652 | 0.269758295 |
| Gm4894 | 0.409448625 | -0.030061717 | 0.043829456 | -0.685879317 | 0.492789214 | NA | | | Bend5 | 291.137294 | 0.118828489 | 0.083355998 | 1.425554156 | 0.153997077 | 0.60340295 |
| Gm4906 | 0 | NA | NA | NA | NA | NA | | | Uba2 | 3645.001772 | 0.118851972 | 0.05217105 | 2.278121147 | 0.022719361 | 0.283293487 |
| Gm4907 | 0 | NA | NA | NA | NA | NA | | | Net1 | 1994.40915 | 0.118954355 | 0.075739321 | 1.570575935 | 0.116281183 | 0.546307021 |
| Gm4922 | 0 | NA | NA | NA | NA | NA | | | Gpr149 | 127.3449685 | 0.119066421 | 0.094351979 | 1.261938777 | 0.206970817 | 0.661486929 |
| Gm4937 | 0 | NA | NA | NA | NA | NA | | | Psmc1 | 5905.22531 | 0.119073321 | 0.041984366 | 2.836134752 | 0.004566318 | 0.125625622 |
| Gm4952 | 5.288185368 | 0.333357223 | 0.146925878 | 2.268880256 | 0.023275609 | NA | | | Vgll3 | 84.78048203 | 0.119170506 | 0.081667676 | 1.45921265 | 0.144506588 | 0.589967032 |
| Gm4956 | 0.188785634 | 0.025025111 | 0.031461603 | 0.79541754 | 0.42637066 | NA | | | Palm2 | 2859.497511 | 0.119216027 | 0.072634434 | 1.641315578 | 0.100731924 | 0.521542547 |
| Gm4961 | 5.368778638 | 0.018221312 | 0.147291842 | 0.123708901 | 0.901545764 | NA | | | Bnip3 | 4047.164247 | 0.119252869 | 0.064247008 | 1.856162216 | 0.06343044 | 0.440995266 |
| Gm4971 | 0.215050351 | -0.027006021 | 0.031907566 | -0.846382992 | 0.397339123 | NA | | | Cadm1 | 6397.831807 | 0.119254931 | 0.0554023 | 2.15252671 | 0.031355894 | 0.32443523 |
| Gm4975 | 0.561778158 | 0.022427986 | 0.058194018 | 0.385400203 | 0.699940932 | NA | | | Fkbp4 | 4651.812433 | 0.119294613 | 0.050497354 | 2.36239334 | 0.018157366 | 0.256569934 |
| Gm4981 | 0.844193991 | 0.068484887 | 0.07127605 | 0.960840099 | 0.336632573 | NA | | | Trpc4 | 617.3699563 | 0.119461928 | 0.072992559 | 1.636631603 | 0.101707468 | 0.522606076 |
| Gm4984 | 0.283178451 | 0.028390488 | 0.033368195 | 0.850824814 | 0.394866675 | NA | | | Cct5 | 5809.621035 | 0.119480639 | 0.046740062 | 2.556278986 | 0.010579826 | 0.196611595 |
| Gm5 | 5.809474054 | 0.011902801 | 0.145173819 | 0.081989999 | 0.934654667 | NA | | | B3galt2 | 1512.850995 | 0.119483205 | 0.078533793 | 1.521424098 | 0.128153445 | 0.566072313 |
| Gm5039 | 0 | NA | NA | NA | NA | NA | | | Kitl | 1895.328179 | 0.119601689 | 0.086120586 | 1.388770034 | 0.16490269 | 0.615071433 |
| Gm5065 | 0.105923158 | 0.018524693 | 0.027977518 | 0.662127842 | 0.507889294 | NA | | | Micu1 | 3702.047633 | 0.119606054 | 0.043881283 | 2.725673587 | 0.006417042 | 0.154049193 |
| Gm5071 | 0 | NA | NA | NA | NA | NA | | | Necap1 | 8477.885494 | 0.119648888 | 0.055324205 | 2.16268609 | 0.030565332 | 0.320602541 |
| Gm5072 | 0 | NA | NA | NA | NA | NA | | | Grin3a | 2264.129697 | 0.119649079 | 0.081486579 | 1.46832866 | 0.14201497 | 0.587991041 |
| Gm5082 | 0.096071043 | -0.020098795 | 0.027825948 | -0.722304036 | 0.470107572 | NA | | | Prmt2 | 2018.358488 | 0.119687109 | 0.07404295 | 1.616455162 | 0.10599594 | 0.52850889 |
| Gm5084 | 2.61041684 | 0.186756539 | 0.111605932 | 1.673356747 | 0.094257087 | NA | | | Mrpl39 | 936.164238 | 0.11974465 | 0.069392717 | 1.725608328 | 0.0844179 | 0.488499377 |
| Gm5087 | 1.871052239 | 0.05152163 | 0.100714692 | 0.511560222 | 0.608958831 | NA | | | Hpdl | 44.09153172 | 0.119781103 | 0.091370973 | 1.310931669 | 0.18988085 | 0.644592015 |
| Gm5088 | 0.185906812 | 0.024895454 | 0.031385107 | 0.793225068 | 0.427646705 | NA | | | Alox12b | 298.4545643 | 0.119784496 | 0.090973138 | 1.316701816 | 0.187938595 | 0.641781416 |
| Gm5091 | 0 | NA | NA | NA | NA | NA | | | Pard3 | 993.9469024 | 0.119793072 | 0.061924358 | 1.934506471 | 0.053050887 | 0.40708779 |
| Gm5095 | 0.116696964 | -0.020477583 | 0.027954224 | -0.732539847 | 0.463839131 | NA | | | D17Wsu104e | 1010.517933 | 0.119811651 | 0.072324237 | 1.65659059 | 0.097602276 | 0.518456434 |
| Gm5105 | 7.748551166 | -0.000784878 | 0.133317147 | -0.005887296 | 0.995302644 | NA | | | Abcg1 | 3870.052705 | 0.119836036 | 0.046676278 | 2.56738631 | 0.010246838 | 0.193809343 |
| Gm5108 | 0.183387883 | -0.001444816 | 0.031390672 | -0.046026918 | 0.963288796 | NA | | | Ldoc1l | 3538.025691 | 0.119838581 | 0.05847832 | 2.049282215 | 0.040434527 | 0.363685409 |
| Gm5114 | 0 | NA | NA | NA | NA | NA | | | Tmem204 | 363.0165172 | 0.119848901 | 0.081638212 | 1.468049065 | 0.142090896 | 0.587991041 |
| Gm5122 | 0.913584581 | 0.006419397 | 0.078297819 | 0.081986924 | 0.934657112 | NA | | | Mettl16 | 1006.342627 | 0.119916864 | 0.072800378 | 1.647201118 | 0.099516714 | 0.520674 |
| Gm5127 | 5.072092974 | -0.017973835 | 0.13035956 | -0.137878914 | 0.890336116 | NA | | | Tfg | 1871.175527 | 0.11992129 | 0.054528243 | 2.19925095 | 0.027860083 | 0.310587544 |
| Gm5128 | 0 | NA | NA | NA | NA | NA | | | Sec62 | 6059.649482 | 0.120024725 | 0.056408963 | 2.127759807 | 0.033357003 | 0.334860976 |
| Gm5129 | 0.266310378 | 0.0407635 | 0.040043874 | 1.017970943 | 0.308691765 | NA | | | Pnmal1 | 3494.502714 | 0.120112201 | 0.056420535 | 2.128873851 | 0.033264699 | 0.334628126 |
| Gm5132 | 0.215706849 | -0.002035226 | 0.0319478 | -0.063704747 | 0.949205325 | NA | | | Nhsl2 | 2838.33764 | 0.120125619 | 0.077277522 | 1.554470372 | 0.120072259 | 0.553708672 |
| Gm5134 | 2.979985258 | 0.011748176 | 0.101498697 | 0.11574706 | 0.907853008 | NA | | | Psma2 | 2835.465863 | 0.120125759 | 0.048972567 | 2.45291939 | 0.014170209 | 0.228200061 |
| Gm5136 | 0.096071043 | -0.020098795 | 0.027825948 | -0.722304036 | 0.470107572 | NA | | | Gltpd1 | 929.1307266 | 0.120247631 | 0.063888035 | 1.882161982 | 0.059814026 | 0.428642594 |
| Gm5142 | 0 | NA | NA | NA | NA | NA | | | Gne | 1724.886544 | 0.120249072 | 0.052436069 | 2.2932511 | 0.021833553 | 0.277716141 |
| Gm5150 | 0.079983654 | 0.016740032 | 0.027674178 | 0.604897175 | 0.54524732 | NA | | | Dpysl3 | 602.1363725 | 0.120316353 | 0.08213525 | 1.46485649 | 0.142960076 | 0.589284353 |
| Gm5166 | 0 | NA | NA | NA | NA | NA | | | Rnf219 | 678.3980867 | 0.120377278 | 0.082600857 | 1.457336907 | 0.145023404 | 0.590583113 |
| Gm5168 | 0 | NA | NA | NA | NA | NA | | | Rbmxl1 | 1633.381806 | 0.120417265 | 0.058016575 | 2.075566594 | 0.037934054 | 0.352772129 |
| Gm5169 | 0 | NA | NA | NA | NA | NA | | | Kazn | 4201.449838 | 0.120447346 | 0.047467046 | 2.537494041 | 0.011164927 | 0.20051086 |
| Gm5177 | 1.83737654 | 0.049376459 | 0.100508986 | 0.491264122 | 0.623239652 | NA | | | Cxx1c | 4518.893102 | 0.120646975 | 0.05695696 | 2.118213044 | 0.034157029 | 0.337297758 |
| Gm525 | 1.273412036 | -0.047122374 | 0.076601304 | -0.615164125 | 0.538446337 | NA | | | Xpa | 208.8159764 | 0.120654882 | 0.09372806 | 1.287286677 | 0.197994386 | 0.653535282 |
| Gm5294 | 0.178993538 | 0.024685672 | 0.031260828 | 0.789667872 | 0.429721759 | NA | | | Olfr287 | 186.170199 | 0.120692742 | 0.091763686 | 1.315256047 | 0.188423864 | 0.64225749 |
| Gm53 | 0 | NA | NA | NA | NA | NA | | | Pdia6 | 4175.617211 | 0.120737431 | 0.05956015 | 2.027151209 | 0.042646951 | 0.37389754 |
| Gm5334 | 0 | NA | NA | NA | NA | NA | | | Stk40 | 1722.65826 | 0.120741717 | 0.081823875 | 1.47562943 | 0.1400434 | 0.585376897 |
| Gm5346 | 0.116696964 | -0.020477583 | 0.027954224 | -0.732539847 | 0.463839131 | NA | | | Zfp251 | 2234.919922 | 0.120753429 | 0.056898281 | 2.122268496 | 0.033815197 | 0.336326295 |
| Gm5347 | 0 | NA | NA | NA | NA | NA | | | Pmaip1 | 104.2672402 | 0.120765627 | 0.094002385 | 1.284708111 | 0.198894303 | 0.65377551 |
| Gm5382 | 0 | NA | NA | NA | NA | NA | | | Doc2a | 1717.993374 | 0.120789354 | 0.052071041 | 2.319703079 | 0.020356945 | 0.269758295 |
| Gm5409 | 0 | NA | NA | NA | NA | NA | | | Klhl13 | 1985.3741 | 0.120805689 | 0.080890531 | 1.493446603 | 0.135320322 | 0.57866043 |
| Gm5414 | 0 | NA | NA | NA | NA | NA | | | Zfp758 | 227.9333098 | 0.12081965 | 0.094101695 | 1.283926404 | 0.199167708 | 0.65377551 |
| Gm5415 | 0.178993538 | 0.024685672 | 0.031260828 | 0.789667872 | 0.429721759 | NA | | | Ublcp1 | 2051.137826 | 0.120821503 | 0.060543341 | 1.995620029 | 0.045975298 | 0.387151986 |
| Gm5416 | 0 | NA | NA | NA | NA | NA | | | Mob2 | 1007.235113 | 0.120853481 | 0.054293469 | 2.225930364 | 0.026018846 | 0.3021678 |
| Gm5420 | 0.098353386 | -0.020212831 | 0.02786466 | -0.725393061 | 0.468210929 | NA | | | Cplx3 | 925.090099 | 0.120923654 | 0.08244829 | 1.466660549 | 0.14246842 | 0.588697979 |
| Gm5424 | 0.525806514 | 0.006325424 | 0.04539272 | 0.139348856 | 0.889174485 | NA | | | Ubb | 32193.39601 | 0.12099499 | 0.049112827 | 2.463612791 | 0.013754461 | 0.224239397 |
| Gm5434 | 0 | NA | NA | NA | NA | NA | | | Fbxl7 | 57.6289795 | 0.121069854 | 0.095094813 | 1.273148864 | 0.202965222 | 0.656606302 |
| Gm5441 | 0.094392817 | 0.017895942 | 0.027818137 | 0.643319236 | 0.520016977 | NA | | | Rars | 1823.757981 | 0.121115362 | 0.055560922 | 2.179865959 | 0.029267399 | 0.317415456 |
| Gm5458 | 2.847557002 | 0.108818506 | 0.112722406 | 0.965367134 | 0.334360957 | NA | | | Nynrin | 640.9739895 | 0.121156178 | 0.09468047 | 1.279632201 | 0.200674519 | 0.654716629 |
| Gm5460 | 0 | NA | NA | NA | NA | NA | | | Htra1 | 5480.094645 | 0.12118779 | 0.077554292 | 1.562618741 | 0.118142297 | 0.549241002 |
| Gm5464 | 5.830661967 | -0.028250892 | 0.142275042 | -0.198565339 | 0.842602769 | NA | | | Tusc2 | 3378.707365 | 0.121220339 | 0.052188827 | 2.322725899 | 0.020193882 | 0.26886923 |
| Gm5475 | 0.650129455 | 0.071216437 | 0.063544202 | 1.120738548 | 0.262399169 | NA | | | Gpc3 | 206.7921396 | 0.121224817 | 0.09398262 | 1.289864195 | 0.197097814 | 0.653065027 |
| Gm5476 | 0 | NA | NA | NA | NA | NA | | | Bysl | 1211.124985 | 0.121242657 | 0.054216672 | 2.236261515 | 0.025334646 | 0.298582503 |
| Gm5477 | 0 | NA | NA | NA | NA | NA | | | Prkra | 529.9367838 | 0.121275918 | 0.068802956 | 1.762655643 | 0.077958586 | 0.475337606 |
| Gm5478 | 0 | NA | NA | NA | NA | NA | | | Mid1 | 398.7893642 | 0.121329389 | 0.06734212 | 1.801686505 | 0.071594742 | 0.459800193 |
| Gm5480 | 2.052333237 | 0.069070467 | 0.078997461 | 0.874337808 | 0.381934315 | NA | | | Nmnat1 | 135.1929783 | 0.121484617 | 0.090853257 | 1.337152018 | 0.18117302 | 0.634469716 |
| Gm5483 | 0.176054697 | -0.002190495 | 0.031239146 | -0.070120197 | 0.944097991 | NA | | | Vamp1 | 9079.100309 | 0.121508421 | 0.071960017 | 1.688554643 | 0.091304809 | 0.503727415 |
| Gm5485 | 0 | NA | NA | NA | NA | NA | | | Kcnj10 | 10969.15985 | 0.121509345 | 0.076043278 | 1.597897246 | 0.110065847 | 0.535496439 |
| Gm5494 | 0 | NA | NA | NA | NA | NA | | | Trim62 | 1125.869008 | 0.121521421 | 0.062771279 | 1.935939871 | 0.052875066 | 0.406864177 |
| Gm5512 | 0.779778063 | -0.006753362 | 0.062007213 | -0.108912518 | 0.913271878 | NA | | | Txndc12 | 927.9028842 | 0.121550073 | 0.065241388 | 1.863082264 | 0.062450693 | 0.437184949 |
| Gm5523 | 2.538534456 | -0.010241435 | 0.113260938 | -0.090423363 | 0.927950792 | NA | | | Stk24 | 2074.355092 | 0.121615979 | 0.068238516 | 1.782218987 | 0.074713528 | 0.468388672 |
| Gm5531 | 1.678235804 | 0.061666331 | 0.095355726 | 0.646697731 | 0.517827592 | NA | | | Chst8 | 427.7406975 | 0.121617956 | 0.082680988 | 1.470930116 | 0.141310017 | 0.586989325 |
| Gm5538 | 0.105923158 | 0.018524693 | 0.027977518 | 0.662127842 | 0.507889294 | NA | | | Snx30 | 2714.019246 | 0.121677426 | 0.070356966 | 1.729429682 | 0.083732221 | 0.486580734 |
| Gm5544 | 1.970400815 | -0.026780741 | 0.093228206 | -0.287260072 | 0.773913193 | NA | | | Traf3ip2 | 29.39320858 | 0.121705415 | 0.084663499 | 1.437519315 | 0.150570489 | 0.598511432 |
| Gm5547 | 0.644719697 | 0.02588267 | 0.053786651 | 0.481209923 | 0.630367309 | NA | | | Wdr6 | 8302.65968 | 0.121826285 | 0.08561339 | 1.422981666 | 0.15474147 | 0.604402956 |
| Gm5549 | 0 | NA | NA | NA | NA | NA | | | Pcdhb1 | 7.912004206 | 0.121863948 | 0.056946403 | 2.139976201 | 0.03235669 | NA |
| Gm5591 | 0 | NA | NA | NA | NA | NA | | | Rnf125 | 55.41134693 | 0.121941694 | 0.0920367 | 1.324924662 | 0.18519614 | 0.637780733 |
| Gm5592 | 0 | NA | NA | NA | NA | NA | | | Atp6v1h | 7194.111734 | 0.121947221 | 0.061556351 | 1.981066432 | 0.047583824 | 0.392147258 |
| Gm5615 | 0 | NA | NA | NA | NA | NA | | | Gm9866 | 1061.818569 | 0.121983452 | 0.055882113 | 2.182871128 | 0.029045302 | 0.317366934 |
| Gm5622 | 2.363747454 | 0.040331076 | 0.100621713 | 0.400818821 | 0.688553521 | NA | | | Med26 | 385.6557246 | 0.121990549 | 0.07320882 | 1.666336777 | 0.095646356 | 0.514958844 |
| Gm5627 | 0 | NA | NA | NA | NA | NA | | | Atmin | 4935.804709 | 0.122008668 | 0.05901576 | 2.067391293 | 0.038697298 | 0.356201815 |
| Gm5634 | 0 | NA | NA | NA | NA | NA | | | Gm16532 | 406.282521 | 0.12208429 | 0.069764592 | 1.7499463 | 0.08012758 | 0.478984869 |

| | | | | | | | | | | | | | | | |
|---|---|---|---|---|---|---|---|---|---|---|---|---|---|---|---|
| Gm5635 | 0 | NA | NA | NA | NA | NA | | | Slc2a13 | 6957.325846 | 0.122089847 | 0.080426052 | 1.518038538 | 0.129004684 | 0.567618221 |
| Gm5640 | 2.475794889 | -0.075222178 | 0.106993061 | -0.703056604 | 0.482020475 | NA | | | Acvr2a | 2021.480382 | 0.122142698 | 0.076726082 | 1.591931912 | 0.111400004 | 0.53885996 |
| Gm5643 | 6.117196228 | -0.192416509 | 0.125489112 | -1.533332305 | 0.125193999 | NA | | | Zfp92 | 166.895916 | 0.122207419 | 0.091835273 | 1.330724189 | 0.183279781 | 0.63672274 |
| Gm5662 | 0 | NA | NA | NA | NA | NA | | | Prss50 | 29.61765786 | 0.122216668 | 0.088037365 | 1.388236318 | 0.165065097 | 0.615077498 |
| Gm5712 | 0 | NA | NA | NA | NA | NA | | | Tmem163 | 707.7050858 | 0.122310861 | 0.07438433 | 1.644309518 | 0.100112284 | 0.5211685 |
| Gm572 | 2.4021855 | -0.043518809 | 0.112342109 | -0.387377537 | 0.698476731 | NA | | | Ints5 | 977.2759659 | 0.122317579 | 0.066803873 | 1.830995308 | 0.067101243 | 0.447688159 |
| Gm5725 | 0 | NA | NA | NA | NA | NA | | | B630019K06Rik | 1216.700635 | 0.122345576 | 0.081409495 | 1.502841614 | 0.132879892 | 0.575107585 |
| Gm5726 | 0 | NA | NA | NA | NA | NA | | | Tigd2 | 534.3147574 | 0.122408839 | 0.07246933 | 1.689112329 | 0.091197906 | 0.503727415 |
| Gm5728 | 0 | NA | NA | NA | NA | NA | | | Cpa4 | 15.08362295 | 0.122429777 | 0.073813807 | 1.658629761 | 0.097190421 | NA |
| Gm5741 | 1.964853267 | 0.273286686 | 0.102867309 | 2.656691307 | 0.007891166 | NA | | | Slc1a1 | 5784.442155 | 0.122435411 | 0.050941424 | 2.403454793 | 0.016240975 | 0.2424629 |
| Gm5766 | 0.096071043 | -0.020098795 | 0.027825948 | -0.722304036 | 0.470107572 | NA | | | Csrnp3 | 4322.544056 | 0.122476045 | 0.063563399 | 1.92683283 | 0.054000467 | 0.409506392 |
| Gm5771 | 0.159967308 | 0.0240345 | 0.030870969 | 0.778547006 | 0.436246606 | NA | | | Tspan2 | 4472.108502 | 0.122512464 | 0.075366907 | 1.625547181 | 0.104046019 | 0.523645898 |
| Gm5779 | 0.215433205 | -0.027029776 | 0.031922004 | -0.84674434 | 0.397137638 | NA | | | Cend1 | 11363.2381 | 0.122561944 | 0.049090868 | 2.496634266 | 0.012537819 | 0.214455135 |
| Gm5795 | 0 | NA | NA | NA | NA | NA | | | Csdc2 | 5096.27762 | 0.122610181 | 0.06122221 | 2.002707541 | 0.045208689 | 0.383811464 |
| Gm5796 | 4.550372248 | -0.03313484 | 0.138826635 | -0.238677831 | 0.811355408 | NA | | | Tmem231 | 594.0926225 | 0.122692279 | 0.065683372 | 1.867935137 | 0.061771113 | 0.435153822 |
| Gm5797 | 1.07331165 | -0.036005461 | 0.071304466 | -0.5049538 | 0.613591286 | NA | | | Atf4 | 4450.795165 | 0.122778434 | 0.057722867 | 2.12703286 | 0.033417353 | 0.335235132 |
| Gm5800 | 0 | NA | NA | NA | NA | NA | | | Clic4 | 4257.059909 | 0.122834716 | 0.08274661 | 1.484468253 | 0.137684739 | 0.582244112 |
| Gm5801 | 0.107716603 | -0.020477583 | 0.027954224 | -0.732539849 | 0.46383913 | NA | | | Uqcc | 2368.207371 | 0.122847944 | 0.045768987 | 2.684087003 | 0.007272818 | 0.162353883 |
| Gm5803 | 1.760068487 | 0.136834324 | 0.097766945 | 1.399597007 | 0.161634031 | NA | | | Fgf12 | 7081.309505 | 0.12291188 | 0.065373046 | 1.880161416 | 0.060086082 | 0.429321413 |
| Gm5833 | 0 | NA | NA | NA | NA | NA | | | Pdlim2 | 422.6319256 | 0.122993898 | 0.081092002 | 1.516720447 | 0.12933728 | 0.568274875 |
| Gm5868 | 1.537306715 | -0.052823434 | 0.069287568 | -0.762379681 | 0.44583343 | NA | | | Mcin | 10.43921798 | 0.123115489 | 0.065607098 | 1.876557439 | 0.060578775 | NA |
| Gm5878 | 0.680997126 | -0.019373436 | 0.059986914 | -0.322961042 | 0.746724746 | NA | | | Mrpl46 | 757.9914207 | 0.123221788 | 0.063104526 | 1.952661665 | 0.050859699 | 0.403266245 |
| Gm5885 | 0 | NA | NA | NA | NA | NA | | | Psmd12 | 2716.191407 | 0.123303793 | 0.052081917 | 2.367497198 | 0.017908856 | 0.255043174 |
| Gm5886 | 0 | NA | NA | NA | NA | NA | | | Abhd15 | 61.77201023 | 0.123303925 | 0.086534362 | 1.424912859 | 0.154182392 | 0.603517496 |
| Gm5891 | 0 | NA | NA | NA | NA | NA | | | Pcdhb19 | 283.4569432 | 0.123334698 | 0.082281354 | 1.498938594 | 0.133889563 | 0.576260679 |
| Gm590 | 0.318806384 | -0.016982193 | 0.040116674 | -0.423320068 | 0.672061754 | NA | | | Rgs17 | 3403.223187 | 0.123359487 | 0.0691693 | 1.783442749 | 0.07451426 | 0.468021138 |
| Gm5901 | 0.4546761 | 0.062003256 | 0.049007249 | 1.265185398 | 0.205804876 | NA | | | Txnip | 1391.184153 | 0.123395223 | 0.091159987 | 1.353611673 | 0.175860301 | 0.628658305 |
| Gm5916 | 0 | NA | NA | NA | NA | NA | | | Inpp4b | 301.5240164 | 0.123415793 | 0.086539847 | 1.426115226 | 0.153835084 | 0.60340295 |
| Gm5925 | 0 | NA | NA | NA | NA | NA | | | Rap2a | 5366.838403 | 0.123521073 | 0.066424504 | 1.859570877 | 0.062946263 | 0.43959491 |
| Gm5934 | 0 | NA | NA | NA | NA | NA | | | Spred1 | 4980.922216 | 0.123529318 | 0.095063564 | 1.299439163 | 0.193793259 | 0.649958942 |
| Gm5935 | 0 | NA | NA | NA | NA | NA | | | Timm17a | 1814.618548 | 0.123592316 | 0.050651894 | 2.440033473 | 0.014685901 | 0.232209004 |
| Gm5936 | 1.368957027 | -0.018483132 | 0.083396809 | -0.221628774 | 0.824602881 | NA | | | Aass | 233.1911046 | 0.123645688 | 0.092931995 | 1.33049644 | 0.183354758 | 0.63672274 |
| Gm5938 | 0 | NA | NA | NA | NA | NA | | | Htr7 | 466.6932084 | 0.123787969 | 0.088496821 | 1.398784353 | 0.16187766 | 0.613952162 |
| Gm5941 | 0 | NA | NA | NA | NA | NA | | | Blvra | 614.1318862 | 0.123797298 | 0.07533585 | 1.64327206 | 0.100326656 | 0.5211685 |
| Gm595 | 0 | NA | NA | NA | NA | NA | | | Gm15417 | 71.88039497 | 0.123797313 | 0.094905998 | 1.304420321 | 0.19209031 | 0.64725211 |
| Gm597 | 0.199381295 | -0.004055078 | 0.031639813 | -0.128163771 | 0.89801937 | NA | | | Morn2 | 289.4863291 | 0.123895496 | 0.088908722 | 1.393513396 | 0.163464597 | 0.614351408 |
| Gm6026 | 0 | NA | NA | NA | NA | NA | | | Smn1 | 542.9803023 | 0.123942905 | 0.073711794 | 1.681452826 | 0.092674992 | 0.506868248 |
| Gm6034 | 1.095546289 | 0.039378448 | 0.075289599 | 0.523026405 | 0.600955879 | NA | | | Fgf5 | 121.1004977 | 0.124096809 | 0.095082129 | 1.305153872 | 0.191840458 | 0.646997681 |
| Gm6040 | 0 | NA | NA | NA | NA | NA | | | Fam211b | 924.8721535 | 0.124110332 | 0.073907428 | 1.679267367 | 0.093099948 | 0.507646338 |
| Gm6042 | 0 | NA | NA | NA | NA | NA | | | Psmb6 | 2778.661781 | 0.124151088 | 0.060344611 | 2.057368259 | 0.039650816 | 0.36080087 |
| Gm6084 | 0 | NA | NA | NA | NA | NA | | | Vkorc1l1 | 687.2155858 | 0.124219649 | 0.077138219 | 1.610351536 | 0.107321134 | 0.532178904 |
| Gm6086 | 3.953030292 | -0.013715926 | 0.120487487 | -0.113836931 | 0.909367063 | NA | | | Zbtb33 | 1033.829169 | 0.124251556 | 0.086759482 | 1.432138057 | 0.152104315 | 0.60007259 |
| Gm6116 | 0 | NA | NA | NA | NA | NA | | | BC023829 | 1769.759221 | 0.124364938 | 0.05918877 | 2.101157687 | 0.035627126 | 0.342728236 |
| Gm6121 | 0 | NA | NA | NA | NA | NA | | | Gmps | 1644.285673 | 0.124442945 | 0.066081247 | 1.88318094 | 0.059675851 | 0.428234899 |
| Gm614 | 0.61296065 | 0.018665908 | 0.053631683 | 0.348038835 | 0.727811019 | NA | | | Psat1 | 4873.933921 | 0.124484856 | 0.055131187 | 2.257975261 | 0.023947202 | 0.29172307 |
| Gm6150 | 1.559676214 | 0.015996788 | 0.094049439 | 0.170089135 | 0.864940038 | NA | | | Zcchc12 | 3203.941783 | 0.12448603 | 0.094414625 | 1.31850367 | 0.187335097 | 0.641710498 |
| Gm6164 | 0 | NA | NA | NA | NA | NA | | | Gm11627 | 83.0364012 | 0.124513756 | 0.094844149 | 1.312824854 | 0.189241973 | 0.64377726 |
| Gm6194 | 0.521672338 | 0.008656229 | 0.045342019 | 0.190909656 | 0.848596375 | NA | | | Pappa | 32.14503079 | 0.124579326 | 0.082061354 | 1.518124188 | 0.128983095 | 0.567618221 |
| Gm6213 | 0.39969727 | 0.025482209 | 0.048473027 | 0.525698742 | 0.599097538 | NA | | | Nuak1 | 4415.899705 | 0.124597209 | 0.078657181 | 1.584053833 | 0.113181469 | 0.540327972 |
| Gm6249 | 3.350400091 | -0.001580262 | 0.125806947 | -0.012561006 | 0.989978031 | NA | | | Tmem130 | 9335.829822 | 0.124616113 | 0.095019172 | 1.311483881 | 0.189694336 | 0.644323531 |
| Gm6251 | 0.506154836 | -0.058553951 | 0.057703036 | -1.014746452 | 0.310226705 | NA | | | Acyp2 | 366.4468367 | 0.124656627 | 0.081960842 | 1.520929081 | 0.128277635 | 0.566288196 |
| Gm6260 | 1.226944435 | -0.07725345 | 0.082825485 | -0.9327256 | 0.350961664 | NA | | | Nalcn | 4703.648239 | 0.124709038 | 0.06977417 | 1.787323843 | 0.073885163 | 0.466430192 |
| Gm6268 | 0 | NA | NA | NA | NA | NA | | | 2510009E07Rik | 2655.827328 | 0.124728236 | 0.062106655 | 2.008290981 | 0.04461238 | 0.380589751 |
| Gm6289 | 0.105923158 | 0.018524693 | 0.027977518 | 0.662127842 | 0.507889294 | NA | | | Rlim | 2302.878944 | 0.124773414 | 0.06768589 | 1.843418394 | 0.065267947 | 0.444691459 |
| Gm6300 | 0.999400785 | -0.066967927 | 0.073744079 | -0.908112592 | 0.363818738 | NA | | | Rhobtb2 | 6123.58087 | 0.124776599 | 0.06344796 | 1.966597498 | 0.049229632 | 0.397725048 |
| Gm6307 | 0.08731684 | 0.017365056 | 0.027733412 | 0.626142072 | 0.531221759 | NA | | | Aplp1 | 54686.73258 | 0.124802475 | 0.045997968 | 2.713217129 | 0.006663343 | 0.156678744 |
| Gm6329 | 0 | NA | NA | NA | NA | NA | | | Magee1 | 7361.492358 | 0.124820008 | 0.052698062 | 2.368588189 | 0.017856123 | 0.255043174 |
| Gm6367 | 0 | NA | NA | NA | NA | NA | | | Gpr3 | 313.7306394 | 0.124855618 | 0.085640113 | 1.457910469 | 0.144865223 | 0.590269908 |
| Gm6370 | 0 | NA | NA | NA | NA | NA | | | Xkr7 | 147.1795436 | 0.12487623 | 0.095062633 | 1.313620573 | 0.188973922 | 0.643290796 |
| Gm6377 | 0.621055902 | 0.040831999 | 0.04851525 | 0.841632255 | 0.399993829 | NA | | | Pptc7 | 2907.057664 | 0.12488132 | 0.068319236 | 1.827908633 | 0.067563265 | 0.448803776 |
| Gm6406 | 0.08731684 | 0.017365056 | 0.027733412 | 0.626142072 | 0.531221759 | NA | | | Vangl1 | 152.6091285 | 0.124923587 | 0.094135895 | 1.327055822 | 0.184490218 | 0.637780733 |
| Gm6408 | 1.368388281 | 0.045329629 | 0.082036148 | 0.552556767 | 0.580566953 | NA | | | Tnfsf12 | 168.4604228 | 0.124960935 | 0.092808525 | 1.346438112 | 0.178161267 | 0.631366323 |
| Gm6416 | 0.396533849 | -0.033875382 | 0.048794365 | -0.694247834 | 0.4875268 | NA | | | Cmc2 | 444.5011765 | 0.124976111 | 0.071188166 | 1.75557425 | 0.079161132 | 0.477013957 |
| Gm6432 | 0 | NA | NA | NA | NA | NA | | | Mmp15 | 1145.306193 | 0.125058962 | 0.074534927 | 1.677857173 | 0.093374985 | 0.50854554 |
| Gm6455 | 0 | NA | NA | NA | NA | NA | | | Sst | 6090.707934 | 0.125296648 | 0.056299029 | 2.225556135 | 0.026043927 | 0.3021678 |
| Gm6460 | 0 | NA | NA | NA | NA | NA | | | Hexim1 | 1945.487051 | 0.125303935 | 0.066140636 | 1.894507559 | 0.058157672 | 0.422610477 |
| Gm648 | 1.514310546 | -0.053729152 | 0.086623841 | -0.620258251 | 0.535087777 | NA | | | Hint1 | 3375.941827 | 0.125351503 | 0.0548816 | 2.284035151 | 0.022369465 | 0.281750998 |
| Gm6484 | 0.107716603 | -0.020477583 | 0.027954224 | -0.732539849 | 0.46383913 | NA | | | Ints12 | 499.0943925 | 0.125428464 | 0.071824221 | 1.746325429 | 0.08075442 | 0.479417114 |
| Gm6498 | 0.720587189 | 0.082389711 | 0.065783028 | 1.252446319 | 0.210407277 | NA | | | Pdyn | 627.3360027 | 0.125444517 | 0.08424873 | 1.488978146 | 0.136493124 | 0.580415431 |
| Gm6524 | 0 | NA | NA | NA | NA | NA | | | Mtus2 | 2562.305862 | 0.125498427 | 0.056705117 | 2.213176399 | 0.026885487 | 0.306064716 |
| Gm6525 | 0.17833704 | -0.002390466 | 0.031282281 | -0.076415987 | 0.939088151 | NA | | | Sdr42e1 | 369.396636 | 0.125606238 | 0.082049269 | 1.530863578 | 0.125803112 | 0.562787717 |
| Gm6531 | 0.099009884 | 0.018205726 | 0.027896865 | 0.652608315 | 0.514008823 | NA | | | Mrpl51 | 1246.711081 | 0.125691116 | 0.061796107 | 2.033964957 | 0.041955129 | 0.370320688 |
| Gm6567 | 0.183387883 | -0.001444816 | 0.031390672 | -0.046026918 | 0.963288796 | NA | | | Tmem192 | 613.1924042 | 0.125774601 | 0.074566548 | 1.686742961 | 0.091652787 | 0.504543511 |
| Gm6568 | 0.198019769 | 0.025308392 | 0.031628095 | 0.80018706 | 0.423602426 | NA | | | Gpt2 | 2744.899661 | 0.125827957 | 0.078553344 | 1.601815397 | 0.109196437 | 0.535010276 |
| Gm6578 | 0 | NA | NA | NA | NA | NA | | | Vmp1 | 4214.215385 | 0.125916472 | 0.072556731 | 1.735420961 | 0.08266627 | 0.484002515 |
| Gm6583 | 1.648508042 | 0.087776321 | 0.088978852 | 0.986485205 | 0.323895072 | NA | | | Cox6a2 | 186.6024851 | 0.125936645 | 0.087719574 | 1.435673231 | 0.151095346 | 0.598846773 |
| Gm6588 | 0 | NA | NA | NA | NA | NA | | | Lsm4 | 775.0006872 | 0.125951121 | 0.06381771 | 1.973607656 | 0.048426372 | 0.394969953 |
| Gm6592 | 0 | NA | NA | NA | NA | NA | | | Prune | 1424.140766 | 0.126002887 | 0.049711233 | 2.534696455 | 0.01125448 | 0.201333224 |
| Gm6602 | 0.384346493 | 0.039441134 | 0.039210102 | 1.005892153 | 0.314467451 | NA | | | Magee2 | 305.942482 | 0.126059384 | 0.08851629 | 1.424137678 | 0.154406621 | 0.60421231 |
| Gm6607 | 0.601453029 | -0.00149993 | 0.046549396 | -0.032222328 | 0.97429475 | NA | | | Eif4a3 | 1726.310567 | 0.126082455 | 0.053850677 | 2.34133464 | 0.019214936 | 0.263564471 |
| Gm6614 | 0 | NA | NA | NA | NA | NA | | | Kat5 | 2342.018085 | 0.126182756 | 0.04904836 | 2.572619295 | 0.010093216 | 0.192154735 |
| Gm6623 | 1.254307926 | 0.091666933 | 0.083938858 | 1.092067902 | 0.274803256 | NA | | | Rrp7a | 4104.761485 | 0.126190374 | 0.050688518 | 2.48952581 | 0.012791363 | 0.215303986 |
| Gm6634 | 0 | NA | NA | NA | NA | NA | | | Sv2c | 1217.817121 | 0.126197978 | 0.077046176 | 1.637952533 | 0.101431596 | 0.522606076 |
| Gm6639 | 0 | NA | NA | NA | NA | NA | | | Glrb | 8548.445517 | 0.126250392 | 0.072116916 | 1.750634931 | 0.080008814 | 0.478751136 |
| Gm6642 | 0.08731684 | 0.017365056 | 0.027733412 | 0.626142072 | 0.531221759 | NA | | | Atoh8 | 59.00301956 | 0.126296728 | 0.093742759 | 1.34726915 | 0.177893564 | 0.631033434 |
| Gm6644 | 0 | NA | NA | NA | NA | NA | | | Retsat | 465.5941776 | 0.126357775 | 0.085117466 | 1.484510528 | 0.137673532 | 0.582244112 |
| Gm6654 | 0.645748536 | -0.016484187 | 0.054188787 | -0.304199228 | 0.760976113 | NA | | | Tmub1 | 699.3632772 | 0.126372804 | 0.065138461 | 1.94006431 | 0.052371875 | 0.406355487 |
| Gm6696 | 0 | NA | NA | NA | NA | NA | | | Trappc5 | 1267.654494 | 0.126413718 | 0.051500654 | 2.454604156 | 0.01410398 | 0.227385586 |
| Gm6710 | 0 | NA | NA | NA | NA | NA | | | Ndufs8 | 2151.439226 | 0.126497552 | 0.047415976 | 2.66782557 | 0.007634388 | 0.167265646 |
| Gm6756 | 0.094392817 | 0.017895942 | 0.027818137 | 0.643319236 | 0.520016977 | NA | | | Ass1 | 1892.014804 | 0.126612185 | 0.062751916 | 2.017662471 | 0.043626426 | 0.378214766 |
| Gm6760 | 0.099009884 | 0.018205726 | 0.027896865 | 0.652608315 | 0.514008823 | NA | | | Irx1 | 19.36873148 | 0.126711049 | 0.071494935 | 1.77230805 | 0.076343436 | NA |
| Gm6763 | 0 | NA | NA | NA | NA | NA | | | Phlda1 | 746.4759918 | 0.126805689 | 0.080999779 | 1.565506608 | 0.117464169 | 0.548468184 |
| Gm6772 | 0.224413567 | -0.027287719 | 0.032077135 | -0.850690659 | 0.394941214 | NA | | | Kcnj12 | 761.5273563 | 0.126809197 | 0.066628946 | 1.903214817 | 0.057012521 | 0.418421024 |
| Gm6787 | 6.609826932 | -0.066410947 | 0.14908206 | -0.445465716 | 0.65598323 | NA | | | Fam124a | 1174.825987 | 0.126893111 | 0.057396195 | 2.210827925 | 0.027047755 | 0.306595176 |
| Gm6792 | 0.116696964 | -0.020477583 | 0.027954224 | -0.732539847 | 0.463839131 | NA | | | Ahcy | 1464.750788 | 0.126905043 | 0.061627023 | 2.059243452 | 0.039470923 | 0.359835305 |
| Gm6793 | 0 | NA | NA | NA | NA | NA | | | Tram1l1 | 1136.812338 | 0.126916483 | 0.061731742 | 2.055935548 | 0.039788729 | 0.36080087 |
| Gm6812 | 0 | NA | NA | NA | NA | NA | | | Mcf2 | 347.9225612 | 0.126974376 | 0.079909339 | 1.588980443 | 0.112064808 | 0.538992394 |
| Gm6878 | 0.69449026 | -0.032567002 | 0.049908573 | -0.65253322 | 0.51405725 | NA | | | Osbpl8 | 4014.019068 | 0.127062695 | 0.085210658 | 1.491159644 | 0.135919586 | 0.579435507 |
| Gm6880 | 0 | NA | NA | NA | NA | NA | | | Nudt1 | 141.1066652 | 0.1271367 | 0.091560836 | 1.388548928 | 0.164969957 | 0.615077498 |
| Gm6890 | 0.176054697 | -0.002190495 | 0.031239146 | -0.070120197 | 0.944097991 | NA | | | Gucy2g | 362.0691768 | 0.127145556 | 0.077110347 | 1.648878029 | 0.099172625 | 0.520271327 |
| Gm6902 | 0 | NA | NA | NA | NA | NA | | | Tsku | 98.6130307 | 0.127150883 | 0.093834279 | 1.355057928 | 0.175399102 | 0.628167492 |
| Gm6904 | 0.176054697 | -0.002190495 | 0.031239146 | -0.070120197 | 0.944097991 | NA | | | Adamts15 | 294.0478007 | 0.127182917 | 0.082319572 | 1.544990011 | 0.122348679 | 0.556819993 |
| Gm6927 | 0.330336725 | -0.015889236 | 0.040460155 | -0.392713161 | 0.694531349 | NA | | | Tspan13 | 5691.579939 | 0.12722513 | 0.06400229 | 1.987821535 | 0.046831431 | 0.390961706 |
| Gm6936 | 2.865676108 | -0.001987056 | 0.106925254 | -0.018583596 | 0.985173289 | NA | | | Ncam2 | 2660.693913 | 0.127331205 | 0.077928217 | 1.633955062 | 0.102268287 | 0.522856086 |
| Gm6938 | 1.53743108 | 0.000985347 | 0.094385048 | 0.01043965 | 0.991670516 | NA | | | Aldh3a1 | 54.35531189 | 0.127340577 | 0.09187357 | 1.386041466 | 0.165734242 | 0.616168889 |
| Gm6994 | 0.211089781 | -0.002448206 | 0.031868916 | -0.076821119 | 0.93876585 | NA | | | Slc6a4 | 6.531249875 | 0.127359554 | 0.049614594 | 2.566977634 | 0.010258923 | NA |
| Gm7030 | 0 | NA | NA | NA | NA | NA | | | Hs6st1 | 2935.318884 | 0.127365596 | 0.052814984 | 2.411542838 | 0.015885186 | 0.239365359 |
| Gm7056 | 1.62794826 | 0.077392097 | 0.087061263 | 0.888938362 | 0.374036207 | NA | | | Dnajb1 | 2422.213773 | 0.127505846 | 0.06922823 | 1.841818668 | 0.065501678 | 0.445445383 |
| Gm7073 | 0 | NA | NA | NA | NA | NA | | | Ifitm1 | 69.53064478 | 0.127520728 | 0.094160174 | 1.354295795 | 0.175642027 | 0.628459435 |
| Gm7092 | 2.727667465 | -0.206376365 | 0.114401215 | -1.803970045 | 0.071236003 | NA | | | Fam96b | 507.6215908 | 0.127559957 | 0.069195097 | 1.843482599 | 0.06525858 | 0.444691459 |
| Gm7104 | 0 | NA | NA | NA | NA | NA | | | Nme1 | 5252.344733 | 0.12758534 | 0.051833516 | 2.461444806 | 0.013837869 | 0.22534628 |
| Gm7134 | 0 | NA | NA | NA | NA | NA | | | Wdr25 | 321.9795314 | 0.127636762 | 0.07489165 | 1.704285615 | 0.088327741 | 0.496728133 |
| Gm7157 | 0 | NA | NA | NA | NA | NA | | | Mpzl2 | 62.84164535 | 0.127894967 | 0.087300132 | 1.465003142 | 0.142920061 | 0.589284353 |
| Gm7168 | 0.199381295 | -0.004055078 | 0.031639813 | -0.128163771 | 0.89801937 | NA | | | Cdh18 | 998.1466103 | 0.127939492 | 0.077676082 | 1.647089926 | 0.099539563 | 0.520674 |
| Gm7173 | 7.355165769 | -0.168024644 | 0.156136567 | -1.076138967 | 0.281865111 | NA | | | Sema3g | 318.3997811 | 0.127957346 | 0.083343704 | 1.53529709 | 0.124710869 | 0.560776598 |
| Gm7257 | 0 | NA | NA | NA | NA | NA | | | Dos | 11752.11259 | 0.12803654 | 0.057370889 | 2.231733592 | 0.025632576 | 0.300515573 |
| Gm7271 | 1.876384503 | -0.043772021 | 0.089319141 | -0.49006317 | 0.624089199 | NA | | | Wnt2b | 147.142491 | 0.128082735 | 0.092021166 | 1.391883407 | 0.163957707 | 0.614351408 |
| Gm732 | 0 | NA | NA | NA | NA | NA | | | Ptgs2 | 485.1128347 | 0.128482826 | 0.092933228 | 1.382528391 | 0.166809519 | 0.617356019 |
| Gm7325 | 0 | NA | NA | NA | NA | NA | | | Ostc | 1042.831168 | 0.128579288 | 0.058908436 | 2.182697363 | 0.029058104 | 0.317366934 |
| Gm7334 | 0 | NA | NA | NA | NA | NA | | | Lman2 | 3685.155113 | 0.128590487 | 0.048391305 | 2.657305599 | 0.0078768 | 0.169996349 |
| Gm7337 | 0 | NA | NA | NA | NA | NA | | | Fam81a | 6444.989512 | 0.128848253 | 0.053033203 | 2.429577073 | 0.015116451 | 0.233312354 |
| Gm7367 | 0 | NA | NA | NA | NA | NA | | | Gpc4 | 333.9421758 | 0.128848391 | 0.093626653 | 1.3761935 | 0.168761734 | 0.619205089 |
| Gm7444 | 4.761500957 | -0.161747097 | 0.120763461 | -1.339371161 | 0.180449873 | NA | | | Pcdha7 | 122.486808 | 0.128868667 | 0.095079241 | 1.35538174 | 0.175295964 | 0.628089033 |
| Gm7457 | 3.116500362 | 0.000662869 | 0.119346788 | 0.005554141 | 0.99556846 | NA | | | Fhdc1 | 278.1016824 | 0.128897459 | 0.088544018 | 1.455744397 | 0.145463292 | 0.59154529 |
| Gm7534 | 0 | NA | NA | NA | NA | NA | | | Zhx3 | 1376.129605 | 0.128929272 | 0.072413596 | 1.780456686 | 0.075001252 | 0.468589657 |
| Gm7538 | 0.491394591 | 0.007996605 | 0.050513606 | 0.158305954 | 0.874215717 | NA | | | Stambpl1 | 618.4527645 | 0.128954438 | 0.065395133 | 1.971927134 | 0.048617925 | 0.395027926 |
| Gm7550 | 0 | NA | NA | NA | NA | NA | | | Nudt11 | 929.3578584 | 0.128972026 | 0.06795753 | 1.897832756 | 0.057718117 | 0.420257325 |
| Gm757 | 0 | NA | NA | NA | NA | NA | | | Srp54c | 631.7076961 | 0.128984022 | 0.075928149 | 1.698764196 | 0.089363622 | 0.499459783 |
| Gm7609 | 5.468562067 | 0.066313116 | 0.143798188 | 0.461154045 | 0.64468809 | NA | | | Fdft1 | 4056.040943 | 0.129043425 | 0.056960302 | 2.265497564 | 0.023482163 | 0.288824644 |
| Gm7616 | 0.41930074 | -0.004836851 | 0.042752125 | -0.113137094 | 0.909921867 | NA | | | Clec3b | 46.27576837 | 0.129350611 | 0.083554164 | 1.548104896 | 0.121597043 | 0.555970618 |
| Gm766 | 3.5986732 | 0.034657506 | 0.114361721 | 0.303051632 | 0.761850514 | NA | | | Vgf | 9574.497451 | 0.129496584 | 0.076778576 | 1.686623933 | 0.091675687 | 0.504543511 |
| Gm7714 | 0 | NA | NA | NA | NA | NA | | | Impdh2 | 1860.535499 | 0.129620634 | 0.054934014 | 2.359569675 | 0.018296145 | 0.257279572 |
| Gm773 | 0 | NA | NA | NA | NA | NA | | | Naif1 | 113.8994186 | 0.129633102 | 0.09365994 | 1.384082697 | 0.166333134 | 0.616741949 |
| Gm7849 | 0 | NA | NA | NA | NA | NA | | | 2410131K14Rik | 918.8072191 | 0.129647311 | 0.059545195 | 2.177292566 | 0.029458745 | 0.317771905 |
| Gm7861 | 0 | NA | NA | NA | NA | NA | | | Lclat1 | 1904.883151 | 0.129651416 | 0.055941906 | 2.31760814 | 0.020470627 | 0.269900834 |
| Gm7903 | 0 | NA | NA | NA | NA | NA | | | Apmap | 2512.695152 | 0.129669665 | 0.053383445 | 2.429023934 | 0.015139533 | 0.233312354 |
| Gm7904 | 0.20671448 | -0.003314027 | 0.03178208 | -0.10427343 | 0.916952363 | NA | | | Gaa | 16557.79508 | 0.129773688 | 0.083810267 | 1.548422319 | 0.121520651 | 0.555796278 |
| Gm7977 | 0 | NA | NA | NA | NA | NA | | | Smpd1 | 5104.575804 | 0.129832105 | 0.05964356 | 2.176800049 | 0.029495489 | 0.317771905 |
| Gm7978 | 0 | NA | NA | NA | NA | NA | | | Utp3 | 1034.974321 | 0.129848228 | 0.0600589 | 2.162014748 | 0.030617039 | 0.320651128 |
| Gm805 | 0.590147267 | 0.023835975 | 0.058603081 | 0.406735863 | 0.684201998 | NA | | | Arpc3 | 4141.371696 | 0.130046481 | 0.045292591 | 2.871252877 | 0.004088483 | 0.115095555 |
| Gm806 | 0.094392817 | 0.017895942 | 0.027818137 | 0.643319236 | 0.520016977 | NA | | | Micalcl | 115.7806354 | 0.13006439 | 0.094776451 | 1.372328139 | 0.169961309 | 0.620316915 |
| Gm8096 | 0 | NA | NA | NA | NA | NA | | | 2210408I21Rik | 425.569192 | 0.13015327 | 0.090953523 | 1.430986567 | 0.152434065 | 0.600557969 |
| Gm813 | 0.389920616 | 0.025130846 | 0.048220387 | 0.521166418 | 0.602250846 | NA | | | 2210404O07Rik | 20.4099799 | 0.130286001 | 0.081074505 | 1.606991004 | 0.108056342 | NA |
| Gm815 | 0 | NA | NA | NA | NA | NA | | | Inadl | 519.6572493 | 0.130300337 | 0.094880436 | 1.373310897 | 0.169655716 | 0.619945581 |
| Gm8179 | 0.211846316 | 0.025707621 | 0.031860375 | 0.806883811 | 0.419733426 | NA | | | Fam19a1 | 1498.450202 | 0.130317898 | 0.063689135 | 2.046155881 | 0.040741037 | 0.365321333 |
| Gm8221 | 0 | NA | NA | NA | NA | NA | | | Hps6 | 300.3633759 | 0.130347941 | 0.074332432 | 1.753581016 | 0.079502326 | 0.477013957 |
| Gm826 | 2.326774602 | -0.071440128 | 0.101492335 | -0.703896777 | 0.481497059 | NA | | | Vash2 | 121.7421358 | 0.130354643 | 0.094923182 | 1.373264579 | 0.16967011 | 0.619945581 |
| Gm8267 | 1.622612839 | 0.01422999 | 0.092979317 | 0.15304468 | 0.878363041 | NA | | | Pdzd2 | 3666.639281 | 0.1305798 | 0.07012786 | 1.862024598 | 0.062599623 | 0.437594862 |
| Gm829 | 0.107716603 | -0.020477583 | 0.027954224 | -0.732539849 | 0.46383913 | NA | | | Panx1 | 622.0165107 | 0.130647177 | 0.062305301 | 2.096887 | 0.03600358 | 0.344080613 |
| Gm8298 | 0 | NA | NA | NA | NA | NA | | | Dsel | 598.6474598 | 0.130767374 | 0.08820802 | 1.482488488 | 0.138210364 | 0.582904025 |
| Gm8300 | 0 | NA | NA | NA | NA | NA | | | Zfp667 | 748.1717169 | 0.130812224 | 0.081699891 | 1.60113095 | 0.109347918 | 0.535243338 |
| Gm833 | 0.743993901 | 0.030682617 | 0.050374558 | 0.60908956 | 0.542465077 | NA | | | Rsg1 | 43.84270447 | 0.130824187 | 0.093636629 | 1.397147558 | 0.162369203 | 0.614052341 |
| Gm8363 | 0.575591037 | 0.053180578 | 0.045290032 | 1.174222573 | 0.240305887 | NA | | | Ccdc6 | 2635.932676 | 0.1309598 | 0.078696672 | 1.66410848 | 0.096090753 | 0.515631428 |
| Gm8369 | 1.154714898 | 0.047998337 | 0.078758791 | 0.609434669 | 0.542236364 | NA | | | Cyb561 | 3517.665675 | 0.131104256 | 0.052896892 | 2.47848694 | 0.013194094 | 0.219077909 |
| Gm839 | 0 | NA | NA | NA | NA | NA | | | Eefsec | 692.6424699 | 0.131222665 | 0.068361594 | 1.919537824 | 0.054916304 | 0.412110852 |
| Gm8439 | 0.116696964 | -0.020477583 | 0.027954224 | -0.732539847 | 0.463839131 | NA | | | Slc9a7 | 814.2707811 | 0.131522535 | 0.07436037 | 1.768718145 | 0.076940923 | 0.472576677 |
| Gm8453 | 0 | NA | NA | NA | NA | NA | | | Ltbp2 | 26.70398679 | 0.131539749 | 0.085686097 | 1.535135261 | 0.124750606 | 0.560776598 |
| Gm853 | 0.17463368 | 0.024560283 | 0.031186242 | 0.787535833 | 0.430968261 | NA | | | Kcnj5 | 84.50146023 | 0.131575775 | 0.09076892 | 1.449568579 | 0.147178863 | 0.594031723 |
| Gm8579 | 0.81708555 | -0.12266306 | 0.06615006 | -1.854315171 | 0.063694083 | NA | | | Zbtb7c | 309.7536658 | 0.131750239 | 0.082158829 | 1.603604134 | 0.108801339 | 0.534621387 |
| Gm8580 | 3.072772518 | 0.010377089 | 0.120374847 | 0.086206454 | 0.9313023 | NA | | | Msantd4 | 2423.12027 | 0.131807433 | 0.058566181 | 2.25057246 | 0.024412629 | 0.292830597 |
| Gm8615 | 3.091924734 | 0.043490516 | 0.121885067 | 0.356815789 | 0.721229716 | NA | | | Nefl | 12246.3166 | 0.131913578 | 0.05869047 | 2.247614958 | 0.024600753 | 0.293873797 |
| Gm8633 | 0.107716603 | -0.020477583 | 0.027954224 | -0.732539849 | 0.46383913 | NA | | | Tada2b | 1547.942453 | 0.131969682 | 0.060648986 | 2.175958593 | 0.029558357 | 0.317771905 |
| Gm8660 | 0 | NA | NA | NA | NA | NA | | | Smox | 1600.132051 | 0.131972231 | 0.084077375 | 1.569652137 | 0.116496062 | 0.546452317 |
| Gm8677 | 0 | NA | NA | NA | NA | NA | | | Trim67 | 205.4035967 | 0.132002669 | 0.091208109 | 1.447269004 | 0.147821597 | 0.594842535 |
| Gm8693 | 0 | NA | NA | NA | NA | NA | | | Ipmk | 1542.368614 | 0.132027735 | 0.069345938 | 1.903900052 | 0.056923202 | 0.418112798 |
| Gm8709 | 0 | NA | NA | NA | NA | NA | | | Tspyl4 | 24636.84188 | 0.132057056 | 0.05115077 | 2.58172176 | 0.009830881 | 0.189645917 |
| Gm8720 | 0 | NA | NA | NA | NA | NA | | | Selk | 2711.064631 | 0.132076552 | 0.048848884 | 2.703778269 | 0.006855602 | 0.159590502 |
| Gm8764 | 0 | NA | NA | NA | NA | NA | | | 6330409D20Rik | 71.98671376 | 0.132131511 | 0.09496071 | 1.391433489 | 0.164094015 | 0.614351408 |
| Gm8765 | 0 | NA | NA | NA | NA | NA | | | Rin3 | 156.7518132 | 0.132168766 | 0.094890213 | 1.392859831 | 0.163662182 | 0.614351408 |
| Gm8773 | 2.689334559 | -0.061088421 | 0.108456042 | -0.563255123 | 0.573261172 | NA | | | Hdac11 | 9034.083815 | 0.132197945 | 0.066557372 | 1.986225442 | 0.047008297 | 0.39102736 |
| Gm8787 | 7.939486688 | 0.178328192 | 0.154002115 | 1.15795937 | 0.246880617 | NA | | | Amotl1 | 1873.057676 | 0.132251728 | 0.08586296 | 1.540265183 | 0.123495727 | 0.558325219 |
| Gm8801 | 0.567932444 | 0.090023784 | 0.05875239 | 1.532257406 | 0.125458928 | NA | | | Morf4l2 | 7399.576812 | 0.132260293 | 0.050482921 | 2.619901754 | 0.00879551 | 0.179191559 |
| Gm8817 | 0.281977855 | 0.011495295 | 0.038675761 | 0.297222198 | 0.766296876 | NA | | | Cdo1 | 1047.41333 | 0.132329968 | 0.063709616 | 2.077079976 | 0.037794178 | 0.351969169 |
| Gm884 | 1.742690937 | 0.105767348 | 0.087959738 | 1.202451825 | 0.22918852 | NA | | | Hcn1 | 2468.128681 | 0.132332343 | 0.064551828 | 2.05001697 | 0.040362775 | 0.363536189 |
| Gm8882 | 0 | NA | NA | NA | NA | NA | | | Dcbld2 | 1855.127746 | 0.132387358 | 0.050458102 | 2.62370862 | 0.008697816 | 0.178452648 |
| Gm8883 | 0 | NA | NA | NA | NA | NA | | | Hsf5 | 18.94364384 | 0.132464253 | 0.078747579 | 1.682137471 | 0.092542186 | NA |
| Gm8884 | 0 | NA | NA | NA | NA | NA | | | Csrnp1 | 539.1580003 | 0.132489066 | 0.093644125 | 1.414814495 | 0.157122893 | 0.607833353 |
| Gm8898 | 0 | NA | NA | NA | NA | NA | | | Lemd1 | 40.03279641 | 0.132577125 | 0.090521523 | 1.464592302 | 0.143032183 | 0.589285584 |
| Gm8909 | 0.08731684 | 0.017365056 | 0.027733412 | 0.626142072 | 0.531221759 | NA | | | Shc4 | 297.1892486 | 0.132595667 | 0.082691913 | 1.603490125 | 0.108826488 | 0.534621387 |
| Gm8979 | 0.286710141 | 0.009849273 | 0.038825804 | 0.25367852 | 0.79974393 | NA | | | Pspc1 | 1634.801999 | 0.132605756 | 0.055881065 | 2.372999815 | 0.017644273 | 0.253763076 |
| Gm8989 | 0 | NA | NA | NA | NA | NA | | | Rgag1 | 109.057169 | 0.132633037 | 0.094223554 | 1.407642047 | 0.15923709 | 0.610955619 |
| Gm8994 | 0 | NA | NA | NA | NA | NA | | | Cab39l | 1458.728205 | 0.132755597 | 0.083640013 | 1.58722592 | 0.112461486 | 0.539325042 |
| Gm9 | 0 | NA | NA | NA | NA | NA | | | Ddt | 839.765192 | 0.132901376 | 0.060539957 | 2.195267124 | 0.028144442 | 0.312281033 |
| Gm904 | 0.099009884 | 0.018205726 | 0.027896865 | 0.652608315 | 0.514008823 | NA | | | Slc25a4 | 20302.41444 | 0.132912459 | 0.046355066 | 2.86726932 | 0.004140305 | 0.11571384 |
| Gm9041 | 0 | NA | NA | NA | NA | NA | | | Pycrl | 609.7080917 | 0.133023466 | 0.068844369 | 1.932234527 | 0.053330565 | 0.407726207 |
| Gm9047 | 0.626586934 | -0.036676868 | 0.059529235 | -0.616115222 | 0.537818476 | NA | | | Rab37 | 211.8571935 | 0.133036401 | 0.092032667 | 1.44553456 | 0.148307793 | 0.595609345 |
| Gm9054 | 2.778805595 | 0.021395882 | 0.111773788 | 0.191421283 | 0.848195547 | NA | | | Cln8 | 1263.889962 | 0.133081097 | 0.052033539 | 2.557602248 | 0.010539657 | 0.196333074 |
| Gm906 | 0 | NA | NA | NA | NA | NA | | | Pgrmc1 | 9221.06108 | 0.133126374 | 0.056914769 | 2.339047963 | 0.019332951 | 0.263564471 |
| Gm9112 | 0 | NA | NA | NA | NA | NA | | | Mtmr2 | 2525.118719 | 0.133154193 | 0.068551457 | 1.942397719 | 0.052088971 | 0.406272098 |
| Gm9125 | 0 | NA | NA | NA | NA | NA | | | Spry2 | 1994.832337 | 0.133204393 | 0.079042629 | 1.685222207 | 0.091945708 | 0.505527387 |
| Gm9199 | 0.119397641 | -0.020477583 | 0.027954224 | -0.732539845 | 0.463839132 | NA | | | Fbln1 | 262.7160553 | 0.133243209 | 0.094025241 | 1.417100436 | 0.156453565 | 0.60719411 |
| Gm9268 | 0 | NA | NA | NA | NA | NA | | | Sepp1 | 9299.574653 | 0.133246529 | 0.077152828 | 1.727046608 | 0.084159293 | 0.488218009 |
| Gm933 | 0 | NA | NA | NA | NA | NA | | | Dusp4 | 411.700476 | 0.133256718 | 0.083595281 | 1.594069863 | 0.110920388 | 0.537434808 |
| Gm9376 | 0 | NA | NA | NA | NA | NA | | | C1ql1 | 205.7041733 | 0.133341743 | 0.091006546 | 1.465188489 | 0.142869499 | 0.589284353 |
| Gm94 | 0 | NA | NA | NA | NA | NA | | | Taf12 | 515.9449681 | 0.13343298 | 0.074062247 | 1.801632901 | 0.071603181 | 0.459800193 |
| Gm9513 | 0 | NA | NA | NA | NA | NA | | | Phb | 2861.806751 | 0.133593254 | 0.050705374 | 2.634696167 | 0.008421263 | 0.17450394 |
| Gm9573 | 2.274881693 | 0.061073732 | 0.095381592 | 0.64030942 | 0.521971458 | NA | | | Phlda3 | 686.3395049 | 0.133633386 | 0.075890859 | 1.760862745 | 0.078261637 | 0.475461538 |
| Gm9696 | 0 | NA | NA | NA | NA | NA | | | Gtf2b | 1152.208518 | 0.133669175 | 0.060959535 | 2.192752515 | 0.028325215 | 0.31312943 |
| Gm97 | 0 | NA | NA | NA | NA | NA | | | Tcerg1l | 343.1125089 | 0.133816386 | 0.084112713 | 1.590917487 | 0.111628147 | 0.538992394 |
| Gm9705 | 0.320973508 | -0.015129019 | 0.040159764 | -0.376720802 | 0.706381102 | NA | | | B3gnt2 | 1393.508236 | 0.133853025 | 0.083428552 | 1.604403066 | 0.108625236 | 0.534621387 |
| Gm9731 | 0.096071043 | -0.020098795 | 0.027825948 | -0.722304036 | 0.470107572 | NA | | | Slc38a1 | 9125.924335 | 0.133876259 | 0.057822601 | 2.315292929 | 0.020596906 | 0.27051597 |
| Gm9733 | 0 | NA | NA | NA | NA | NA | | | Coq7 | 726.3801508 | 0.133982569 | 0.070458985 | 1.901568264 | 0.05722762 | 0.419207468 |
| Gm9758 | 0.116696964 | -0.020477583 | 0.027954224 | -0.732539847 | 0.463839131 | NA | | | Cct2 | 5554.025709 | 0.134130019 | 0.04830786 | 2.776567169 | 0.005493628 | 0.139755594 |
| Gm9839 | 0 | NA | NA | NA | NA | NA | | | Cd164 | 2461.93788 | 0.134158743 | 0.069309662 | 1.935642726 | 0.052911474 | 0.406864177 |
| Gm9846 | 0 | NA | NA | NA | NA | NA | | | Acsl3 | 8060.371103 | 0.134247157 | 0.051810388 | 2.591124316 | 0.009566293 | 0.186784235 |
| Gm9855 | 0 | NA | NA | NA | NA | NA | | | E130309D02Rik | 844.9959762 | 0.134369525 | 0.059849088 | 2.245139056 | 0.024759207 | 0.294610834 |
| Gm9871 | 1.567935623 | 0.049808998 | 0.08930336 | 0.557750549 | 0.577014735 | NA | | | Slc32a1 | 4616.49664 | 0.134402156 | 0.062147652 | 2.162626463 | 0.030569921 | 0.320602541 |
| Gm9895 | 7.430599815 | 0.18773158 | 0.155673391 | 1.205932359 | 0.227843572 | NA | | | Gdf5 | 17.1008736 | 0.134404831 | 0.072873101 | 1.844368214 | 0.065129496 | NA |
| Gm9920 | 3.84398225 | 0.065978439 | 0.13475695 | 0.48961066 | 0.624409432 | NA | | | Crispld2 | 138.3228603 | 0.134439897 | 0.094761157 | 1.418723679 | 0.155979591 | 0.606466686 |
| Gm9926 | 2.168183447 | -0.258871212 | 0.102067534 | -2.536273797 | 0.01120391 | NA | | | Fam103a1 | 2499.872498 | 0.134459621 | 0.057049926 | 2.356876329 | 0.018429383 | 0.258617131 |
| Gm9961 | 0.096071043 | -0.020098795 | 0.027825948 | -0.722304036 | 0.470107572 | NA | | | Slit3 | 1343.43177 | 0.13467828 | 0.070040539 | 1.922861864 | 0.0544974 | 0.411449707 |
| Gm9992 | 0.331243956 | 0.008596723 | 0.034710864 | 0.247666645 | 0.804392342 | NA | | | Tnmd | 26.7821999 | 0.134746807 | 0.084017963 | 1.603785698 | 0.108761298 | 0.534621387 |
| Gm9994 | 0 | NA | NA | NA | NA | NA | | | E030019B06Rik | 86.68092014 | 0.134762024 | 0.094391139 | 1.427697832 | 0.153378851 | 0.602567676 |
| Gm9999 | 0.159967308 | 0.0240345 | 0.030870969 | 0.778547006 | 0.436246606 | NA | | | Mybpc1 | 210.2743889 | 0.134767168 | 0.093899494 | 1.435227838 | 0.151222183 | 0.598846773 |
| Gmcl1l | 0 | NA | NA | NA | NA | NA | | | Ier3 | 360.4441047 | 0.134882756 | 0.086550769 | 1.558423555 | 0.119132877 | 0.551649399 |
| Gml | 0 | NA | NA | NA | NA | NA | | | Ckmt1 | 8311.91224 | 0.134931033 | 0.055242952 | 2.44250221 | 0.01458584 | 0.231556193 |
| Gnat1 | 5.119184197 | 0.067078729 | 0.141623744 | 0.473640413 | 0.635756351 | NA | | | Ttll12 | 779.7917044 | 0.134947641 | 0.070536131 | 1.913170451 | 0.055726235 | 0.413057568 |
| Gnat2 | 2.442187145 | 0.050275045 | 0.104274793 | 0.482139964 | 0.629706522 | NA | | | D730039F16Rik | 21.61974239 | 0.134973078 | 0.083428562 | 1.617828168 | 0.105699634 | NA |
| Gnat3 | 0 | NA | NA | NA | NA | NA | | | Cdkn2aipnl | 1661.738224 | 0.134983513 | 0.057374278 | 2.352683413 | 0.018638495 | 0.260580156 |
| Gnb3 | 5.962343211 | 0.272265562 | 0.138601293 | 1.964379668 | 0.049486079 | NA | | | Spin2 | 207.8280499 | 0.135074208 | 0.088475556 | 1.52668391 | 0.12683963 | 0.563718247 |
| Gngt1 | 0 | NA | NA | NA | NA | NA | | | F3 | 1652.511105 | 0.13509259 | 0.063692803 | 2.12100245 | 0.033921596 | 0.336326295 |
| Gnrhr | 0 | NA | NA | NA | NA | NA | | | Agt | 1137.312258 | 0.135154798 | 0.077936029 | 1.734176102 | 0.082886842 | 0.484423387 |
| Gp1ba | 0.439846772 | -0.045662679 | 0.041334256 | -1.104717565 | 0.26928199 | NA | | | Rps6ka2 | 4754.851384 | 0.135163872 | 0.06593757 | 2.049876452 | 0.040376489 | 0.363536189 |
| Gp2 | 5.904305298 | -0.208975532 | 0.132330794 | -1.579190494 | 0.11429237 | NA | | | Acly | 9003.713678 | 0.13519143 | 0.04419469 | 3.058997118 | 0.002220793 | 0.079261025 |
| Gp49a | 1.928387987 | 0.153676866 | 0.088963825 | 1.727408476 | 0.084094329 | NA | | | B3galt1 | 1940.860356 | 0.135210093 | 0.068779824 | 1.96583947 | 0.049317157 | 0.398210685 |
| Gp6 | 0.098353386 | -0.020212831 | 0.02786466 | -0.725393061 | 0.468210929 | NA | | | Plcl2 | 3310.397728 | 0.135279814 | 0.073178952 | 1.848616447 | 0.064513216 | 0.443022969 |
| Gpa33 | 0 | NA | NA | NA | NA | NA | | | Grid2 | 430.4439247 | 0.135341764 | 0.085708033 | 1.579102438 | 0.114312563 | 0.543130934 |
| Gpbar1 | 0.17463368 | 0.024560283 | 0.031186242 | 0.787535833 | 0.430968261 | NA | | | Lhfpl3 | 1161.671131 | 0.135390092 | 0.070367579 | 1.924040793 | 0.05434947 | 0.410949742 |
| Gpha2 | 0.547215998 | -0.090429391 | 0.05723373 | -1.58000172 | 0.114106473 | NA | | | Lancl3 | 253.86312 | 0.135438387 | 0.093517287 | 1.448271127 | 0.147541239 | 0.594737171 |
| Gphb5 | 0 | NA | NA | NA | NA | NA | | | Slc6a17 | 28549.08962 | 0.135442958 | 0.042890899 | 3.157848424 | 0.001589382 | 0.06480716 |
| Gpihbp1 | 0.356086128 | 0.051237183 | 0.044718313 | 1.145776311 | 0.251887719 | NA | | | Tac1 | 1346.900866 | 0.135466711 | 0.085088292 | 1.592072271 | 0.111368467 | 0.53885996 |
| Gpr110 | 0.407193159 | 0.02051662 | 0.04329096 | 0.473923893 | 0.63555418 | NA | | | Wwox | 770.6776338 | 0.135622579 | 0.058632636 | 2.31309025 | 0.020717678 | 0.270947196 |
| Gpr113 | 0.094392817 | 0.017895942 | 0.027818137 | 0.643319236 | 0.520016977 | NA | | | Cacybp | 2180.555814 | 0.135676504 | 0.062742347 | 2.16243909 | 0.030584347 | 0.320602541 |
| Gpr119 | 0.38488829 | -0.02558324 | 0.043075085 | -0.593921989 | 0.552564291 | NA | | | Chrac1 | 399.29143 | 0.135790177 | 0.070008072 | 1.939636004 | 0.052423942 | 0.406355487 |
| Gpr128 | 0.929519008 | 0.066206034 | 0.050206807 | 1.318666504 | NA | NA | | | Ttbk2 | 4437.3676 | 0.135849748 | 0.073829298 | 1.840052004 | 0.065760603 | 0.445799444 |

| | | | | | | | | | | | | | |
|---|---|---|---|---|---|---|---|---|---|---|---|---|---|
| Gpr132 | 0.655039105 | 0.023132371 | 0.053842275 | 0.429632135 | 0.667463257 | NA | Lhx6 | 1076.362021 | 0.135918234 | 0.066114226 | 2.055809201 | 0.039800911 | 0.36080087 | |
| Gpr141 | 0.445240244 | -0.001897651 | 0.037393919 | -0.050747581 | 0.959526662 | NA | Tmem121 | 414.6115833 | 0.136010011 | 0.079606748 | 1.708523649 | 0.087539222 | 0.495597906 | |
| Gpr142 | 0.577809731 | -0.005504678 | 0.058021926 | -0.094872373 | 0.924416201 | NA | Edaradd | 14.00363609 | 0.136016051 | 0.069681065 | 1.951980085 | 0.050940569 | NA | |
| Gpr143 | 2.696545174 | 0.098010003 | 0.112966934 | 0.86759904 | 0.385613872 | NA | Zwint | 17308.64718 | 0.136042768 | 0.058246018 | 2.335657821 | 0.019509082 | 0.263578327 | |
| Gpr15 | 0.999636632 | 0.044679399 | 0.071154745 | 0.627918755 | 0.530057164 | NA | Spock3 | 3966.969148 | 0.136213977 | 0.079993997 | 1.702802493 | 0.088605036 | 0.497517107 | |
| Gpr152 | 2.458535501 | 0.05356629 | 0.111667445 | 0.479694778 | 0.631444441 | NA | D830030K20Rik | 111.633145 | 0.136222973 | 0.091443227 | 1.489699965 | 0.136303144 | 0.579855848 | |
| Gpr171 | 0.692149418 | -0.025692469 | 0.061038031 | -0.420925582 | 0.673809424 | NA | Fxyd6 | 4317.128834 | 0.136389561 | 0.062540481 | 2.180820468 | 0.029196698 | 0.317415456 | |
| Gpr174 | 4.064567381 | 0.10918384 | 0.128744314 | 0.848067281 | 0.396400502 | NA | Cyb561d2 | 472.4059869 | 0.136457503 | 0.077762817 | 1.754791157 | 0.079295037 | 0.477013957 | |
| Gpr18 | 0.369409914 | 0.025046824 | 0.047675316 | 0.52536252 | 0.599331203 | NA | Lhfpl5 | 125.6131416 | 0.136543022 | 0.095096513 | 1.435836266 | 0.151048938 | 0.598846773 | |
| Gpr20 | 7.823481412 | -0.207879345 | 0.154900744 | -1.342016442 | 0.179590665 | NA | Zfp948 | 406.5880409 | 0.136592084 | 0.088573178 | 1.542138223 | 0.123040008 | 0.557484892 | |
| Gpr31b | 0.094392817 | 0.017895942 | 0.027818137 | 0.643319236 | 0.520016977 | NA | Usp29 | 2547.031806 | 0.136654112 | 0.085687531 | 1.594795773 | 0.110757913 | 0.537406303 | |
| Gpr33 | 0.079983654 | 0.016740032 | 0.027674178 | 0.604897175 | 0.54524732 | NA | Higd2a | 2175.929412 | 0.136711716 | 0.066321807 | 2.061338817 | 0.039270727 | 0.359615903 | |
| Gpr44 | 3.09601992 | -0.16617475 | 0.125894881 | -1.319948434 | 0.186852235 | NA | Cdh19 | 492.9611507 | 0.136757509 | 0.08576666 | 1.594529954 | 0.110817387 | 0.537406303 | |
| Gpr55 | 6.356275839 | -0.303130177 | 0.142468495 | -2.127699726 | 0.033361988 | NA | Pgr | 1112.645487 | 0.136840012 | 0.079777074 | 1.715279908 | 0.08629392 | 0.491660395 | |
| Gpr65 | 12.8400628 | 0.507655199 | 0.144868566 | 3.504246718 | NA | NA | Fbxo10 | 2024.484886 | 0.136941547 | 0.069128444 | 1.980972514 | 0.047594356 | 0.392147258 | |
| Gpr82 | 0.485069967 | 0.062268854 | 0.049079739 | 1.268728307 | 0.204537983 | NA | Zfp14 | 696.952305 | 0.792505563 | 0.064665398 | 12.25548111 | 1.57E-34 | 7.60E-31 | |
| Gpr87 | 0 | NA | NA | NA | NA | NA | Moxd1 | 415.1310954 | 0.137127705 | 0.075219308 | 1.823038642 | 0.068297539 | 0.451155091 | Gm1082 |
| Gpr97 | 3.673893711 | 0.1026279 | 0.117888708 | 0.870549025 | 0.38400044 | NA | Glra1 | 30.1688648 | 0.13718235 | 0.069880558 | 1.96309752 | 0.049634844 | 0.399665044 | Blvrb |
| Gprc5d | 4.99614278 | 0.030720847 | 0.13167471 | 0.233308637 | 0.815521753 | NA | Cnih3 | 3111.93286 | 0.137223899 | 0.061219537 | 2.24150499 | 0.024993383 | 0.296370511 | Kirrel2 |
| Gprc6a | 0.579680688 | 0.02709434 | 0.040889424 | 0.662624645 | 0.507570982 | NA | Fam20c | 1679.222798 | 0.137239028 | 0.052658711 | 2.606197994 | 0.009155352 | 0.182762906 | Gm6537 |
| Gpx2 | 1.839163825 | 0.064538389 | 0.095235325 | 0.67767279 | 0.497979182 | NA | Fezf2 | 913.969935 | 0.137332451 | 0.0826563 | 1.661487998 | 0.096615477 | 0.516802454 | Plekhf1 |
| Gpx2-ps1 | 7.184999987 | 0.196824021 | 0.134395797 | 1.464510242 | 0.143054587 | NA | Galnt10 | 640.0203449 | 0.13739268 | 0.083307738 | 1.649218706 | 0.099102837 | 0.520271327 | Zfp27 |
| Gpx5 | 0 | NA | NA | NA | NA | NA | Tirap | 115.9880912 | 0.137403724 | 0.092507491 | 1.485325379 | 0.137457652 | 0.582012447 | Hif3a |
| Grhl2 | 3.923822387 | -0.099916815 | 0.132772092 | -0.752543804 | 0.451724098 | NA | Ets2 | 3462.492384 | 0.13745141 | 0.062552321 | 2.197383065 | 0.0279931 | 0.311357668 | Rinl |
| Grifin | 3.414847239 | -0.120549396 | 0.126160407 | -0.955524786 | 0.339312377 | NA | Lyar | 351.7243037 | 0.137458646 | 0.079721802 | 1.72422903 | 0.084666505 | 0.488499377 | Tsc22d3 |
| Grin3b | 5.485007072 | 0.123063841 | 0.137212765 | 0.896883331 | 0.369781176 | NA | Pdxk | 9990.507513 | 0.137531169 | 0.056150413 | 2.449334935 | 0.014312029 | 0.229466376 | Cbln4 |
| Grm6 | 0.477384717 | 0.028178173 | 0.045982715 | 0.612799235 | 0.540009097 | NA | Dclre1b | 376.4343612 | 0.13755605 | 0.074577294 | 1.84447628 | 0.06511376 | 0.444683814 | Adipor2 |
| Grxcr1 | 0.469085263 | 0.061308216 | 0.048717378 | 1.258446538 | 0.20823031 | NA | Cpt2 | 290.0582229 | 0.137729521 | 0.087407932 | 1.575709651 | 0.115092732 | 0.543394118 | Kcna5 |
| Grxcr2 | 6.163260754 | 0.192086484 | 0.118781075 | 1.61714721 | 0.105846508 | NA | Crim1 | 2308.182542 | 0.137775503 | 0.07639758 | 1.803401408 | 0.071325197 | 0.459174089 | Pcdha10 |
| Gsc | 0.17463368 | 0.024560283 | 0.031186242 | 0.787535833 | 0.430968261 | NA | Rps6ka6 | 173.8396539 | 0.137903096 | 0.093471357 | 1.475351391 | 0.140118097 | 0.585376897 | Atp4a |
| Gsc2 | 0.474461318 | 0.009131132 | 0.05516185 | 0.165533453 | 0.868524121 | NA | Oprm1 | 51.94165578 | 0.137947903 | 0.089894285 | 1.534556974 | 0.124892688 | 0.560889589 | Ret |
| Gsdma | 2.521827192 | -0.235250413 | 0.104617938 | -2.248662297 | 0.02453399 | NA | Crym | 1785.395939 | 0.137969767 | 0.083844195 | 1.645549427 | 0.099856559 | 0.521115714 | Arrdc2 |
| Gsdma2 | 0 | NA | NA | NA | NA | NA | Por | 3143.793841 | 0.138022535 | 0.06444901 | 2.141577279 | 0.032227514 | 0.327711599 | Dyrk3 |
| Gsdma3 | 0 | NA | NA | NA | NA | NA | Prkar2a | 2996.000896 | 0.138033059 | 0.057953686 | 2.381782217 | 0.017229083 | 0.250771207 | Fkbp5 |
| Gsdmc | 1.176063034 | -0.050141483 | 0.086599856 | -0.579001921 | 0.562587877 | NA | Rps4y2 | 457.7483298 | 0.138406859 | 0.090486506 | 1.529585624 | 0.12611933 | 0.562787717 | Meig1 |
| Gsdmc2 | 0 | NA | NA | NA | NA | NA | Sertad1 | 378.4443686 | 0.138663635 | 0.092090367 | 1.505734414 | 0.132135366 | 0.573639666 | Ada |
| Gsdmc3 | 0 | NA | NA | NA | NA | NA | Gpr176 | 725.0177437 | 0.138683382 | 0.06648852 | 2.085824473 | 0.036994526 | 0.348173348 | Camk1g |
| Gsdmc4 | 2.780089505 | 0.074454858 | 0.109044961 | 0.682790451 | 0.494739265 | NA | Arhgef3 | 3716.36794 | 0.138810751 | 0.056773765 | 2.444980555 | 0.014485995 | 0.230474881 | Prr5 |
| Gsdmcl-ps | 0 | NA | NA | NA | NA | NA | Dolk | 757.239844 | 0.138861952 | 0.066001057 | 2.103935289 | 0.035384092 | 0.342065168 | Klf15 |
| Gsdmcl1 | 0 | NA | NA | NA | NA | NA | Pdgfra | 2616.831921 | 0.139218861 | 0.064229027 | 2.167538074 | 0.03019385 | 0.319909454 | Ovol2 |
| Gsg1 | 3.269141262 | 0.006816792 | 0.115749724 | 0.05889251 | 0.953037724 | NA | Gm1082 | 32.27045956 | 0.708453774 | 0.082361766 | 8.601731199 | 7.85E-18 | 1.90E-14 | |
| Gsta1 | 0 | NA | NA | NA | NA | NA | Kcnk3 | 1174.965356 | 0.139338445 | 0.0632214 | 2.203975957 | 0.027526035 | 0.308173879 | Tekt4 |
| Gsta2 | 1.303579416 | 0.095315121 | 0.082125978 | 1.160596481 | 0.245806037 | NA | Rims2 | 5585.929981 | 0.139404224 | 0.049699254 | 2.80495606 | 0.005032344 | 0.133393847 | Gpr123 |
| Gstm3 | 0.787779753 | 0.017407436 | 0.055623753 | 0.312949687 | 0.754318887 | NA | Hlf | 10355.61112 | 0.139412654 | 0.075448505 | 1.847785508 | 0.064633378 | 0.443022969 | Ppapdc2 |
| Gstt4 | 0.489742234 | 0.039007481 | 0.051990224 | 0.750284927 | 0.453083119 | NA | Gla | 605.7297034 | 0.139414829 | 0.069401249 | 2.008823061 | 0.044555902 | 0.380589751 | Plin4 |
| Gtf2a1l | 4.33086103 | 0.15690267 | 0.132253029 | 1.186382428 | 0.235471308 | NA | Spsb2 | 89.85255695 | 0.1395133 | 0.094613048 | 1.474567223 | 0.140328932 | 0.585584048 | Rhou |
| Gtsf1 | 2.568829213 | -0.032771883 | 0.11391905 | -0.287676935 | 0.773594047 | NA | 6330403K07Rik | 10108.15504 | 0.139517027 | 0.072224615 | 1.931710211 | 0.053395284 | 0.407791741 | Hs6st2 |
| Gtsf1l | 2.368506006 | -0.015161177 | 0.082039574 | -0.184803214 | 0.853383387 | NA | Ccdc147 | 12.97084452 | 0.139521867 | 0.072608141 | 1.921573313 | 0.054659469 | NA | B3galt5 |
| Guca1a | 3.181988315 | 0.157189115 | 0.118609329 | 1.325267722 | 0.185082371 | NA | Sde2 | 982.9217198 | 0.139593751 | 0.071966454 | 1.939705824 | 0.052415451 | 0.406355487 | Fzd2 |
| Guca2a | 0 | NA | NA | NA | NA | NA | Nrg1 | 233.3843663 | 0.13964104 | 0.085336457 | 1.636358536 | 0.101764572 | 0.522606076 | Arid5b |
| Guca2b | 0.378048898 | -0.001605613 | 0.041752189 | -0.03845579 | 0.96932428 | NA | Arrdc3 | 2271.91487 | 0.139646072 | 0.081380101 | 1.715973202 | 0.086166947 | 0.491660395 | Lfng |
| Gucy2d | 0.759356684 | 0.023263342 | 0.067947642 | 0.342371591 | 0.732071265 | NA | Pifo | 31.1456515 | 0.139663164 | 0.079952536 | 1.746825927 | 0.080667538 | 0.479417114 | Rhbdl3 |
| Gulo | 2.04977914 | -0.006919657 | 0.105252772 | -0.065743226 | 0.947582258 | NA | Asah2 | 796.25584 | 0.139667762 | 0.072119779 | 1.936608279 | 0.052793245 | 0.406864177 | Endod1 |
| Gvin1 | 0 | NA | NA | NA | NA | NA | Slc2a5 | 238.6637701 | 0.139702816 | 0.0856946 | 1.630240606 | 0.103050655 | 0.522975271 | Bcat1 |
| Gykl1 | 0.079983654 | 0.016740032 | 0.027674178 | 0.604897175 | 0.54524732 | NA | Lrrtm2 | 2846.858699 | 0.139824934 | 0.084520841 | 1.654324922 | 0.098061512 | 0.519101888 | Rab3c |
| Gyltl1b | 7.647445691 | 0.212429596 | 0.14326102 | 1.482815046 | 0.138123557 | NA | Neb | 68.51955825 | 0.139839394 | 0.094943505 | 1.472869517 | 0.140786223 | 0.586773565 | Ndufc2 |
| Gypa | 1.66986205 | -0.064037491 | 0.08945648 | -0.715850776 | 0.474083496 | NA | Ptger4 | 45.308755 | 0.139924955 | 0.09056911 | 1.544952302 | 0.122357801 | 0.556819993 | Cables1 |
| Gys2 | 3.691120737 | -0.087120239 | 0.122240439 | -0.712695727 | 0.476034061 | NA | Snrpd1 | 929.8584777 | 0.13994033 | 0.066216283 | 2.113382461 | 0.034568046 | 0.33922827 | Tmeff1 |
| Gzma | 0.546533824 | -0.023452341 | 0.046571786 | -0.50357401 | 0.61456076 | NA | Slc8a1 | 8098.614869 | 0.139943512 | 0.072265973 | 1.936506319 | 0.052805719 | 0.406864177 | Akt2 |
| Gzmb | 0.877155726 | 0.074701388 | 0.061603867 | 1.212608758 | 0.225279439 | NA | St6galnac3 | 398.2283711 | 0.14002181 | 0.082585573 | 1.695475423 | 0.089985269 | 0.500949827 | Bcl2l1 |
| Gzmc | 0.107716603 | -0.020477583 | 0.027954224 | -0.732539849 | 0.46383913 | NA | Hspa13 | 1553.128984 | 0.140101703 | 0.06801494 | 2.059866603 | 0.039411295 | 0.359829336 | Kbtbd4 |
| Gzmd | 0 | NA | NA | NA | NA | NA | Rin2 | 1585.116052 | 0.140182445 | 0.063716682 | 2.200090161 | 0.027800499 | 0.310219758 | Pja1 |
| Gzme | 0 | NA | NA | NA | NA | NA | Pbdc1 | 598.2260074 | 0.140263716 | 0.066918457 | 2.096039308 | 0.036078704 | 0.34411534 | Stip1 |
| Gzmf | 0 | NA | NA | NA | NA | NA | Tox2 | 789.2119309 | 0.14036966 | 0.088983596 | 1.577477944 | 0.114685593 | 0.543130934 | Sccpdh |
| Gzmg | 0 | NA | NA | NA | NA | NA | Cdh4 | 1449.650056 | 0.140465197 | 0.075689453 | 1.855809394 | 0.063480731 | 0.440995266 | Gpi1 |
| Gzmk | 2.414602173 | 0.018978304 | 0.108979724 | 0.174145273 | 0.861751294 | NA | Erp44 | 1278.556697 | 0.140547082 | 0.069662259 | 2.017549868 | 0.043638162 | 0.378214766 | |
| Gzmn | 0 | NA | NA | NA | NA | NA | Gpr179 | 35.56088188 | 0.140586797 | 0.077534069 | 1.813226096 | 0.069796957 | 0.455201066 | |
| H19 | 7.225761096 | -0.030305485 | 0.145582921 | -0.208166486 | 0.835098976 | NA | Carhsp1 | 2585.582094 | 0.140628683 | 0.067746873 | 2.075795926 | 0.037912829 | 0.352772129 | |
| H1fnt | 0 | NA | NA | NA | NA | NA | Pcdh9 | 5884.299793 | 0.140664097 | 0.056923919 | 2.471089463 | 0.013470212 | 0.222603292 | |
| H1foo | 0.308183275 | 0.007047773 | 0.033991235 | 0.207340891 | 0.835743642 | NA | Epm2a | 539.3386658 | 0.140782099 | 0.071420565 | 1.971170336 | 0.048704396 | 0.395027926 | |
| H2-Ab1 | 94.76645758 | 0.108464792 | 0.114756139 | 0.945176378 | NA | NA | Heca | 991.8241885 | 0.140894817 | 0.078746016 | 1.789231043 | 0.073577614 | 0.465500184 | |
| H2-Ea-ps | 0.079983654 | 0.016740032 | 0.027674178 | 0.604897175 | 0.54524732 | NA | Ccdc65 | 292.4224862 | 0.140914083 | 0.084790726 | 1.661904433 | 0.096531938 | 0.516802454 | |
| H2-Eb2 | 0.536442192 | -0.057169132 | 0.05827754 | -0.980980534 | 0.326602341 | NA | Mcee | 533.0077857 | 0.140944342 | 0.065244369 | 2.160252984 | 0.03075309 | 0.321052104 | |
| H2-L | 0 | NA | NA | NA | NA | NA | Tm2d1 | 692.4928063 | 0.140969577 | 0.073259616 | 1.924246723 | 0.054323665 | 0.410949742 | |
| H2-M1 | 0 | NA | NA | NA | NA | NA | Spsb4 | 147.1023634 | 0.141151809 | 0.094153903 | 1.499160463 | 0.133832009 | 0.576183689 | |
| H2-M10.1 | 0 | NA | NA | NA | NA | NA | Slc8a3 | 1504.490468 | 0.141160684 | 0.063634411 | 2.218307402 | 0.02653388 | 0.303967775 | |
| H2-M10.2 | 0 | NA | NA | NA | NA | NA | Tceal1 | 626.8341425 | 0.141182509 | 0.076862704 | 1.836814242 | 0.066237323 | 0.446279967 | |
| H2-M10.3 | 0.099009884 | 0.018205726 | 0.027896865 | 0.652608315 | 0.514008823 | NA | Tmem177 | 474.034511 | 0.141254051 | 0.075564503 | 1.869317534 | 0.06157865 | 0.435077842 | |
| H2-M10.4 | 0 | NA | NA | NA | NA | NA | Synpo2 | 373.0600541 | 0.141316096 | 0.093300558 | 1.514632906 | 0.129865394 | 0.568885619 | |
| H2-M10.5 | 0 | NA | NA | NA | NA | NA | Hspa5 | 11784.71785 | 0.141357417 | 0.075093623 | 1.882415746 | 0.05977959 | 0.428607265 | |
| H2-M10.6 | 0 | NA | NA | NA | NA | NA | P2ry1 | 179.8881599 | 0.14140864 | 0.085710653 | 1.649837394 | 0.098976198 | 0.520271327 | |
| H2-M11 | 0.233393928 | -0.027475306 | 0.032191697 | -0.85349047 | 0.39338737 | NA | Dusp14 | 2265.382841 | 0.141595665 | 0.079750421 | 1.775484867 | 0.075817862 | 0.471258137 | |
| H2-M2 | 0.480100836 | 0.043799064 | 0.056601691 | 0.773811928 | 0.439042018 | NA | Rsph3b | 120.3288833 | 0.141657638 | 0.09305821 | 1.522247609 | 0.12794705 | 0.565771339 | |
| H2-M9 | 0.119397641 | -0.020477583 | 0.027954224 | -0.732539845 | 0.463839132 | NA | Emd | 1228.207221 | 0.141698386 | 0.059699226 | 2.373538071 | 0.017618577 | 0.253730508 | |
| H2-Q10 | 1.376818378 | 0.222349937 | 0.089057002 | 2.496714807 | 0.012534972 | NA | Isoc1 | 764.4282219 | 0.141843142 | 0.073560195 | 1.928259467 | 0.053822861 | 0.408907362 | |
| H2-Q6 | 0 | NA | NA | NA | NA | NA | Bex2 | 6075.911899 | 0.14201137 | 0.055636422 | 2.552489249 | 0.010695621 | 0.19724805 | |
| H2-Q7 | 0 | NA | NA | NA | NA | NA | Gm10033 | 162.0509378 | 0.142043126 | 0.095071508 | 1.494066194 | 0.135158319 | 0.578555714 | |
| H2-Q8 | 0 | NA | NA | NA | NA | NA | Rspo3 | 375.1703646 | 0.14217575 | 0.086658606 | 1.640642006 | 0.10087175 | 0.521542547 | |
| H2-Q9 | 0 | NA | NA | NA | NA | NA | Sgpp2 | 1440.479995 | 0.142298513 | 0.074773869 | 1.903051354 | 0.057033845 | 0.418421024 | |
| H2-T22 | 1.100885548 | -0.034925214 | 0.086748596 | -0.40260264 | 0.687240568 | NA | Oxtr | 483.0012484 | 0.142316487 | 0.084993092 | 1.674447703 | 0.094042644 | 0.51048709 | |
| H2-T3 | 0.370495291 | 0.0385767 | 0.038790203 | 0.994495936 | 0.319981483 | NA | Fut9 | 3594.986023 | 0.142342121 | 0.08179921 | 1.740140532 | 0.081834345 | 0.482437374 | |
| H2-T9 | 0 | NA | NA | NA | NA | NA | Ptprr | 811.9118465 | 0.142398025 | 0.07597629 | 1.874242943 | 0.060896948 | 0.432354503 | |
| H2afb1 | 0.099009884 | 0.018205726 | 0.027896865 | 0.652608315 | 0.514008823 | NA | Polr1c | 728.0568048 | 0.142412541 | 0.063121028 | 2.256182237 | 0.024059221 | 0.29172307 | |
| H2afy3 | 0.08731684 | 0.017365056 | 0.027733412 | 0.626142072 | 0.531221759 | NA | Kcnmb2 | 135.1212676 | 0.142497786 | 0.091059258 | 1.564890704 | 0.117608539 | 0.548537627 | |
| H2bfm | 0 | NA | NA | NA | NA | NA | Tmem14c | 1093.348618 | 0.142547528 | 0.060410406 | 2.359651894 | 0.018292091 | 0.257279572 | |
| H60c | 0.178993538 | 0.024685672 | 0.031260828 | 0.789667872 | 0.429721759 | NA | Zbtb16 | 1879.614671 | 0.142553131 | 0.094410674 | 1.509925988 | 0.13106231 | 0.571029131 | |
| Haao | 6.298059786 | 0.074869019 | 0.145676105 | 0.513941658 | 0.607292781 | NA | Bri3bp | 5750.01452 | 0.142597509 | 0.04528839 | 3.148654849 | 0.001640238 | 0.066000268 | |
| Habp2 | 0.079983654 | 0.016740032 | 0.027674178 | 0.604897175 | 0.54524732 | NA | Fez1 | 8756.207314 | 0.14264745 | 0.047029695 | 3.033135802 | 0.002420267 | 0.083112984 | |
| Hal | 0.721176579 | 0.036635741 | 0.061061229 | 0.599983687 | 0.548517108 | NA | Hmgcr | 4505.599682 | 0.142684088 | 0.058513402 | 2.438485591 | 0.014748947 | 0.232209004 | |
| Hamp | 0.283178451 | 0.028390488 | 0.033368195 | 0.850824814 | 0.394866675 | NA | Htr4 | 155.9868428 | 0.143050476 | 0.093358385 | 1.532272387 | 0.125455232 | 0.562279816 | |
| Hamp2 | 0.505798573 | 0.029734949 | 0.046020057 | 0.646130196 | 0.518195041 | NA | Fastkd5 | 386.1333111 | 0.143114647 | 0.09213733 | 1.553275397 | 0.120357355 | 0.554315455 | |
| Hand1 | 0 | NA | NA | NA | NA | NA | Sowahb | 1002.507321 | 0.143154822 | 0.089302543 | 1.603031857 | 0.108927621 | 0.534621387 | |
| Hand2 | 0 | NA | NA | NA | NA | NA | Kcnk13 | 415.2137043 | 0.143164922 | 0.07946668 | 1.801571709 | 0.071612816 | 0.459800193 | |
| Hao1 | 3.010711308 | 0.088411103 | 0.097130101 | 0.910233815 | 0.362699215 | NA | Fosl2 | 1641.438958 | 0.143312447 | 0.085289223 | 1.680311329 | 0.092896758 | 0.507109472 | |
| Hao2 | 0 | NA | NA | NA | NA | NA | Gng4 | 2375.976669 | 0.143337385 | 0.073528621 | 1.949409413 | 0.051246551 | 0.403266245 | |
| Has2as | 2.140560232 | 0.089250519 | 0.104967094 | 0.850271409 | 0.395174208 | NA | 4921524J17Rik | 271.9149484 | 0.143350856 | 0.089156496 | 1.60785654 | 0.107866602 | 0.533015565 | |
| Havcr1 | 0.275064582 | 0.010932281 | 0.038417727 | 0.284563456 | 0.775978612 | NA | Dbpht2 | 1752.426363 | 0.143359724 | 0.076573315 | 1.872189072 | 0.06118045 | 0.43330435 | |
| Hba-a1 | 0 | NA | NA | NA | NA | NA | Kras | 3479.935955 | 0.143475045 | 0.071957171 | 1.993895029 | 0.046163527 | 0.387613525 | |
| Hba-a2 | 0 | NA | NA | NA | NA | NA | 1700030J22Rik | 395.528747 | 0.143672788 | 0.085731496 | 1.675846039 | 0.093768354 | 0.509378873 | |
| Hba-x | 0 | NA | NA | NA | NA | NA | Map3k19 | 214.3781292 | 0.143693492 | 0.095032578 | 1.512044549 | 0.13052253 | 0.569619261 | |
| Hbb-b1 | 0 | NA | NA | NA | NA | NA | Rims3 | 2333.013331 | 0.143768015 | 0.060890017 | 2.361109778 | 0.018220336 | 0.257188614 | |
| Hbb-b2 | 0 | NA | NA | NA | NA | NA | Tcfl5 | 21.50036819 | 0.143983821 | 0.07908376 | 1.820649652 | 0.06866013 | NA | |
| Hbb-bh1 | 0 | NA | NA | NA | NA | NA | Klhdc7a | 848.7677886 | 0.144042293 | 0.077994892 | 1.846817006 | 0.064773666 | 0.443414795 | |
| Hbb-bh2 | 0 | NA | NA | NA | NA | NA | Ficd | 660.3607536 | 0.144054274 | 0.064823244 | 2.222262666 | 0.02626556 | 0.302739577 | |
| Hbb-y | 1.966967144 | 0.046089237 | 0.090247736 | 0.510696876 | 0.609563329 | NA | Scg2 | 11339.01683 | 0.14408504 | 0.095019999 | 1.516365415 | 0.129426979 | 0.568336245 | |
| Hbq1b | 3.15856341 | 0.03774795 | 0.120402931 | 0.313513551 | 0.753890527 | NA | Nell1 | 2346.114181 | 0.144286204 | 0.066374587 | 2.173816988 | 0.029718882 | 0.317891372 | |
| Hcrt | 2.286357048 | 0.205823382 | 0.106548396 | 1.931736086 | 0.053392088 | NA | Adcy3 | 2953.746953 | 0.144416822 | 0.056795592 | 2.54274703 | 0.010998482 | 0.199361888 | |
| Hdgfl1 | 0.094392817 | 0.017895942 | 0.027818137 | 0.643319236 | 0.520016977 | NA | Triap1 | 533.0247021 | 0.144661306 | 0.072286614 | 2.001218464 | 0.045368852 | 0.383869599 | |
| Hdhd1a | 0 | NA | NA | NA | NA | NA | Rasd1 | 315.0932593 | 0.144711503 | 0.094640999 | 1.52905722 | 0.126250259 | 0.562787717 | |
| Helt | 5.602517392 | -0.02684434 | 0.147888845 | -0.181517001 | 0.85596179 | NA | Hexim2 | 356.6578104 | 0.144739774 | 0.08166438 | 1.772373392 | 0.076332596 | 0.472435997 | |
| Hemgn | 1.339731961 | -0.144898562 | 0.077259302 | -1.875483705 | 0.060726209 | NA | Cacng4 | 503.1229735 | 0.145008624 | 0.066366913 | 2.184953561 | 0.028892252 | 0.316746301 | |
| Hemt1 | 0 | NA | NA | NA | NA | NA | Plch1 | 555.6515981 | 0.14522819 | 0.077212878 | 1.880880413 | 0.059988188 | 0.429169608 | |
| Hephl1 | 5.853980551 | -0.080719501 | 0.058643611 | -1.376441522 | 0.16868498 | NA | Ube2v2 | 2682.990195 | 0.145247756 | 0.069703954 | 2.08378073 | 0.037180117 | 0.348662607 | |
| Herpud1 | 2018.258476 | 0.136115423 | 0.117697413 | 1.156486104 | NA | NA | Marveld2 | 49.32369062 | 0.145509212 | 0.094361903 | 1.542033459 | 0.123065463 | 0.557484892 | |
| Hes2 | 0.494522006 | 0.031199547 | 0.04639328 | 0.672501424 | 0.501264534 | NA | Sgms1 | 1794.030857 | 0.145610949 | 0.058587795 | 2.485346121 | 0.012942553 | 0.216344675 | |
| Hesx1 | 0.486850878 | 0.009782776 | 0.044755885 | 0.218580767 | 0.826976636 | NA | Echdc1 | 1023.396312 | 0.145630072 | 0.066680197 | 2.184007808 | 0.028961674 | 0.317028848 | |
| Hexim1 | 1888.243271 | 0.159838936 | 0.138901855 | 1.150732915 | NA | NA | Herpud1 | 2196.25784 | 0.14566963 | 0.053641621 | 2.715608284 | 0.006615414 | 0.156472283 | |
| Hgd | 1.956849132 | -0.01272358 | 0.100747789 | -0.126291406 | 0.899501259 | NA | Dok4 | 673.9559663 | 0.145685479 | 0.078784731 | 1.849158812 | 0.064434884 | 0.443022969 | |
| Hgfac | 4.516198336 | 0.12697407 | 0.117480587 | 1.080808945 | 0.279782109 | NA | Ccrn4l | 2086.736107 | 0.145688416 | 0.069426554 | 2.098453798 | 0.035865077 | 0.344080613 | |
| Hhla1 | 1.092684631 | 0.036603875 | 0.080081372 | 0.457083514 | 0.64761102 | NA | Sh3bp4 | 947.887031 | 0.14570558 | 0.061497872 | 2.369278426 | 0.017822831 | 0.255043174 | |
| Hibch | 504.4762041 | -0.154015475 | 0.11809773 | -1.304135786 | NA | NA | Zfp503 | 202.8575333 | 0.145801135 | 0.094698902 | 1.539628557 | 0.12365092 | 0.558428989 | |
| Higd1c | 0 | NA | NA | NA | NA | NA | St6galnac5 | 1343.307021 | 0.14584335 | 0.071471795 | 2.040572086 | 0.041293379 | 0.367273584 | |
| Hils1 | 0 | NA | NA | NA | NA | NA | Prph | 41.02283042 | 0.145892721 | 0.085039922 | 1.715579202 | 0.086239088 | 0.491660395 | |
| Hipk1 | 6342.8212 | 0.009305169 | 0.097045344 | 0.095884753 | NA | NA | Lingo4 | 44.95769299 | 0.146024716 | 0.094061696 | 1.552435494 | 0.120558056 | 0.554583368 | |
| Hipk4 | 7.761176972 | -0.056312021 | 0.141812906 | -0.397086714 | 0.691303517 | NA | Rxra | 1934.120258 | 0.14604869 | 0.062051741 | 2.353659814 | 0.018589615 | 0.26039802 | |
| Hist1h1a | 0.735221671 | 0.004778139 | 0.061109148 | 0.078190247 | 0.93767672 | NA | Thsd7b | 328.4955616 | 0.146183171 | 0.073396341 | 1.99169561 | 0.046404466 | 0.388890657 | |
| Hist1h1b | 0.892063337 | 0.057228649 | 0.062091197 | 0.921687 | 0.356691864 | NA | Rit2 | 2713.528529 | 0.146196855 | 0.072979768 | 2.003251854 | 0.045150262 | 0.383539597 | |
| Hist1h1d | 3.359303027 | -0.074191839 | 0.098423094 | -0.753805192 | 0.450966206 | NA | Dnajc12 | 407.9830664 | 0.146279773 | 0.074983571 | 1.950824303 | 0.05107795 | 0.403266245 | |
| Hist1h1t | 0 | NA | NA | NA | NA | NA | Lrp3 | 4102.01723 | 0.146281191 | 0.059122597 | 2.474201048 | 0.013353453 | 0.221177028 | |
| Hist1h2aa | 0 | NA | NA | NA | NA | NA | Spcs1 | 2218.148598 | 0.146326933 | 0.05703671 | 2.565486912 | 0.010303111 | 0.194115419 | |
| Hist1h2ab | 0.411730969 | -0.0302294 | 0.043889081 | -0.688768129 | 0.490969195 | NA | Gpr68 | 621.9785986 | 0.146404181 | 0.089088078 | 1.643364448 | 0.100307551 | 0.5211685 | |
| Hist1h2ad | 0.850310816 | 0.055425758 | 0.053806532 | 1.030093493 | 0.302966119 | NA | Rbm24 | 731.0601767 | 0.14641259 | 0.075246922 | 1.945761849 | 0.051683353 | 0.404936564 | |
| Hist1h2ae | 0.401150917 | -0.033366739 | 0.048906957 | -0.682249333 | 0.495081305 | NA | Zscan2 | 302.1421328 | 0.146438023 | 0.07995813 | 1.831433818 | 0.067035818 | 0.447688159 | |
| Hist1h2af | 0.096071043 | -0.020098795 | 0.027825948 | -0.722304036 | 0.470107572 | NA | Th | 146.8706803 | 0.146442159 | 0.070222227 | 2.08541035 | 0.037032068 | 0.348173348 | |
| Hist1h2ag | 0 | NA | NA | NA | NA | NA | Cdkn1b | 876.8439203 | 0.146478805 | 0.059269668 | 2.471395725 | 0.01345868 | 0.222603292 | |
| Hist1h2ah | 0 | NA | NA | NA | NA | NA | Mtfp1 | 2303.136147 | 0.146833663 | 0.069159347 | 2.123121015 | 0.033743712 | 0.336187357 | |
| Hist1h2ai | 0 | NA | NA | NA | NA | NA | Ccdc85a | 1884.498228 | 0.146833798 | 0.070053197 | 2.096032784 | 0.036079283 | 0.34411534 | |
| Hist1h2ak | 0.116696964 | -0.020477583 | 0.027954224 | -0.732539847 | 0.463839131 | NA | Calb2 | 843.1563154 | 0.146860096 | 0.078094876 | 1.880534335 | 0.060035291 | 0.429169608 | |
| Hist1h2an | 0 | NA | NA | NA | NA | NA | Phf15 | 4326.826621 | 0.146877741 | 0.074325311 | 1.976147008 | 0.04813813 | 0.393789375 | |
| Hist1h2ao | 0 | NA | NA | NA | NA | NA | AI836003 | 1079.522323 | 0.146898108 | 0.070283779 | 2.090071278 | 0.036611397 | 0.346440078 | |
| Hist1h2ap | 0 | NA | NA | NA | NA | NA | Mapk6 | 3899.669533 | 0.1471122 | 0.079614983 | 1.847795401 | 0.064631946 | 0.443022969 | |
| Hist1h2ba | 0 | NA | NA | NA | NA | NA | Khdrbs2 | 182.033972 | 0.147119751 | 0.090497069 | 1.625685248 | 0.10401663 | 0.523645898 | |
| Hist1h2bb | 1.107346981 | -0.060832732 | 0.079704043 | -0.763232701 | 0.44532463 | NA | Proca1 | 127.22638 | 0.147454089 | 0.093616302 | 1.575089863 | 0.115235704 | 0.543478516 | |
| Hist1h2bf | 0.215050351 | -0.027006021 | 0.031907566 | -0.846382992 | 0.397339123 | NA | Brf2 | 553.3077745 | 0.147531987 | 0.065713859 | 2.245066561 | 0.02476386 | 0.294610834 | |
| Hist1h2bh | 1.279162208 | 0.0237592 | 0.074111338 | 0.320587925 | 0.74852269 | NA | Cyp51 | 2341.597749 | 0.147532846 | 0.073801387 | 1.999052487 | 0.045602675 | 0.3851305 | |
| Hist1h2bj | 0.75689519 | -0.059168597 | 0.067594424 | -0.875347296 | 0.381384967 | NA | Ebf1 | 139.0909639 | 0.147584257 | 0.092376972 | 1.597630382 | 0.110125261 | 0.535496439 | |
| Hist1h2bk | 0 | NA | NA | NA | NA | NA | Blvrb | 313.9339086 | 0.547711457 | 0.079836231 | 6.860437294 | 6.87E-12 | 1.11E-08 | |
| Hist1h2bl | 0 | NA | NA | NA | NA | NA | Crb1 | 70.57484767 | 0.147646678 | 0.085436095 | 1.728153402 | 0.083960725 | 0.487260687 | |
| Hist1h2bm | 0 | NA | NA | NA | NA | NA | Hivep3 | 1921.721076 | 0.147675799 | 0.081490234 | 1.81219015 | 0.06995682 | 0.455201066 | |
| Hist1h2bn | 3.186758882 | -0.141833258 | 0.11571051 | -1.225759514 | 0.220289187 | NA | Pvalb | 2358.121772 | 0.147813292 | 0.089260729 | 1.655972272 | 0.097727434 | 0.518456434 | |
| Hist1h2bp | 0.646335094 | -0.079163576 | 0.053841353 | -1.470311779 | 0.141477332 | NA | Oxld1 | 142.2193499 | 0.147941436 | 0.092505135 | 1.599278086 | 0.109758827 | 0.535311472 | |
| Hist1h2bq | 0 | NA | NA | NA | NA | NA | Tmem246 | 3136.660697 | 0.147946331 | 0.048078375 | 3.077190771 | 0.002089615 | 0.076882121 | |
| Hist1h2br | 0 | NA | NA | NA | NA | NA | Dynlt1a | 46.88893786 | 0.147946624 | 0.094292848 | 1.569012147 | 0.116645109 | 0.546752774 | |
| Hist1h3a | 0 | NA | NA | NA | NA | NA | Gap43 | 8915.506367 | 0.148176636 | 0.068153604 | 2.174157013 | 0.029693346 | 0.317852275 | |
| Hist1h3b | 0.303023688 | 0.009739077 | 0.039388711 | 0.247255542 | 0.804710463 | NA | Gpd1 | 4326.337035 | 0.148297041 | 0.068925023 | 2.151570426 | 0.031431204 | 0.32443523 | |
| Hist1h3c | 0.390161855 | -0.000140598 | 0.036193922 | -0.003884577 | 0.996900564 | NA | Pgm2 | 2884.922142 | 0.148336085 | 0.069127657 | 2.145828326 | 0.031886678 | 0.326647307 | |
| Hist1h3d | 2.527979752 | -0.088721967 | 0.106429093 | -0.833625136 | 0.404492257 | NA | Slco3a1 | 1643.647199 | 0.148395705 | 0.054020426 | 2.747029508 | 0.006013772 | 0.147900395 | |
| Hist1h3e | 0.079983654 | 0.016740032 | 0.027674178 | 0.604897175 | 0.54524732 | NA | Wnt4 | 983.8790969 | 0.14853368 | 0.084849471 | 1.750555167 | 0.080022564 | 0.478751136 | |
| Hist1h3f | 0.094392817 | 0.017895942 | 0.027818137 | 0.643319236 | 0.520016977 | NA | Chml | 509.0721088 | 0.148562767 | 0.079953643 | 1.858111302 | 0.06315321 | 0.440342223 | |
| Hist1h3g | 0.517847881 | -0.057108045 | 0.05793189 | -0.985779063 | 0.324241542 | NA | Pard6a | 760.3071007 | 0.148586419 | 0.073361609 | 2.02539749 | 0.042826565 | 0.374984137 | |
| Hist1h3h | 0.116696964 | -0.020477583 | 0.027954224 | -0.732539847 | 0.463839131 | NA | Tubb4b | 11093.65471 | 0.148779234 | 0.057925117 | 2.568475358 | 0.010214697 | 0.193453318 | |
| Hist1h3i | 0.497448118 | -0.02127213 | 0.055798949 | -0.381228152 | 0.703033961 | NA | Wnt3 | 108.4961069 | 0.149123565 | 0.094141346 | 1.58403901 | 0.113184842 | 0.540327972 | |
| Hist1h4a | 0.286053643 | -0.017027416 | 0.03902234 | -0.436350468 | 0.662582471 | NA | Lsm12 | 608.5827429 | 0.149219262 | 0.064825025 | 2.301877432 | 0.021342085 | 0.275079973 | |
| Hist1h4b | 0.496263686 | 0.037917571 | 0.052109423 | 0.727652863 | 0.466826112 | NA | Pnpla2 | 951.4351372 | 0.149283358 | 0.082026224 | 1.819946726 | 0.068767118 | 0.452815575 | |
| Hist1h4c | 1.888179012 | -0.122173901 | 0.088073245 | -1.387185192 | 0.1653853 | NA | Rcan2 | 14593.71055 | 0.149391748 | 0.064800826 | 2.305398801 | 0.021144249 | 0.273744523 | |
| Hist1h4d | 1.393488188 | 0.116951635 | 0.087710608 | 1.333380738 | 0.18240689 | NA | Ruvbl1 | 1452.065254 | 0.149749335 | 0.051055114 | 2.933091792 | 0.003356047 | 0.10135121 | |
| Hist1h4f | 0.392621292 | 0.024891049 | 0.048289092 | 0.515459033 | 0.60623229 | NA | Tubb2a | 12311.54337 | 0.149794923 | 0.058957888 | 2.540710464 | 0.011062749 | 0.199827723 | |
| Hist1h4j | 1.149130049 | -0.015105454 | 0.073409579 | -0.205769517 | 0.836970954 | NA | Ptpro | 1674.014219 | 0.149951638 | 0.088555261 | 1.693311463 | 0.090396197 | 0.502238744 | |
| Hist1h4k | 7.871869586 | -0.063463666 | 0.135785275 | -0.467382534 | 0.640226217 | NA | Itk | 26.07622598 | 0.14995632 | 0.084718865 | 1.770046364 | 0.076719418 | 0.472560915 | |
| Hist1h4m | 0.211846316 | 0.025707621 | 0.031860375 | 0.806883811 | 0.419733426 | NA | Nde1 | 935.6077864 | 0.150006311 | 0.066777325 | 2.246365984 | 0.024680575 | 0.294362343 | |
| Hist1h4n | 0 | NA | NA | NA | NA | NA | Akap12 | 1383.777628 | 0.150082476 | 0.090290213 | 1.662223079 | 0.096468054 | 0.51670168 | |
| Hist2h2aa1 | 0 | NA | NA | NA | NA | NA | Akap2 | 3929.512789 | 0.150097384 | 0.052977909 | 2.833207035 | 0.004608353 | 0.126226882 | |
| Hist2h2aa2 | 0 | NA | NA | NA | NA | NA | Pofut2 | 1122.351609 | 0.150172291 | 0.061685503 | 2.434482719 | 0.014913092 | 0.232332035 | |
| Hist2h2ab | 1.662834137 | 0.028157252 | 0.095539667 | 0.294717924 | 0.768209369 | NA | Zbtb8a | 258.8554085 | 0.150283885 | 0.086039864 | 1.746677394 | 0.080693314 | 0.479417114 | |
| Hist2h2ac | 0.099009884 | 0.018205726 | 0.027896865 | 0.652608315 | 0.514008823 | NA | Olig1 | 3549.355727 | 0.150308014 | 0.071021424 | 2.116375662 | 0.03431287 | 0.338606491 | |
| Hist2h2bb | 0.094392817 | 0.017895942 | 0.027818137 | 0.643319236 | 0.520016977 | NA | S1pr4 | 21.09466036 | 0.15049429 | 0.083492886 | 1.802480404 | 0.071469855 | NA | |
| Hist2h3b | 0 | NA | NA | NA | NA | NA | Zswim3 | 383.3939188 | 0.150544316 | 0.083064924 | 1.81236927 | 0.069929158 | 0.455201066 | |
| Hist2h4 | 4.721943855 | 0.038976049 | 0.137838291 | 0.282766486 | 0.777355847 | NA | 2700094K13Rik | 376.5607273 | 0.15058745 | 0.070334372 | 2.141022179 | 0.03227225 | 0.327711599 | |
| Hist3h2bb-ps | 4.869970795 | 0.030168454 | 0.132748238 | 0.227260677 | 0.820221048 | NA | Cxxc4 | 908.1263559 | 0.150790366 | 0.070998922 | 2.123840202 | 0.033683508 | 0.335817871 | |
| Hmga1-rs1 | 0 | NA | NA | NA | NA | NA | Ramp2 | 574.3895684 | 0.150863206 | 0.077102434 | 1.956659445 | 0.050387521 | 0.402601282 | |
| Hmga2 | 2.380176363 | 0.038319994 | 0.11059335 | 0.346494555 | 0.728971076 | NA | Timm17b | 530.0089425 | 0.150955887 | 0.070699829 | 2.135166234 | 0.032747434 | 0.330481625 | |
| Hmgb1-rs17 | 0.683694727 | 0.038873006 | 0.066191532 | 0.587280648 | 0.557015236 | NA | Emb | 1139.043993 | 0.151047435 | 0.077768871 | 1.942260855 | 0.052105529 | 0.406272098 | |
| Hmgb4 | 0.105923158 | 0.018524693 | 0.027977518 | 0.662127842 | 0.507889294 | NA | Rragb | 2554.29293 | 0.151099464 | 0.054839157 | 2.755320695 | 0.005863463 | 0.14609377 | |
| Hmx1 | 0 | NA | NA | NA | NA | NA | Hsd17b12 | 2902.952278 | 0.151276588 | 0.056667359 | 2.669554225 | 0.007595201 | 0.16691057 | |
| Hmx2 | 0.489484589 | -0.049559653 | 0.043029867 | -1.15175008 | 0.249423787 | NA | Ccnd2 | 3005.335572 | 0.15152151 | 0.083415617 | 1.816464533 | 0.069299144 | 0.454875446 | |
| Hmx3 | 1.269676182 | -0.121350778 | 0.065944701 | -1.840189982 | 0.06574035 | NA | Tango6 | 279.8449221 | 0.151632463 | 0.082395446 | 1.840301508 | 0.065723984 | 0.445799444 | |
| Hnf1a | 0.362381421 | 0.026644729 | 0.047496261 | 0.560985827 | 0.574807199 | NA | Pak3 | 4653.636724 | 0.151701277 | 0.070859366 | 2.140878264 | 0.032283856 | 0.327711599 | |
| Hnf4a | 0.311993147 | -0.014421208 | 0.039860783 | -0.361789373 | 0.717509434 | NA | Nkain4 | 870.3771688 | 0.151726305 | 0.075936442 | 1.998069697 | 0.045709105 | 0.385357198 | |
| Hnf4g | 0.824562923 | -0.076709381 | 0.070664541 | -1.085542766 | 0.277681333 | NA | Zrsr1 | 3815.457116 | 0.151933817 | 0.053190653 | 2.85640065 | 0.004284741 | 0.119462849 | |
| Hormad1 | 0 | NA | NA | NA | NA | NA | Nap1l5 | 11205.89854 | 0.151959432 | 0.077538261 | 1.959799339 | 0.050019249 | 0.401203537 | |
| Hormad2 | 1.272288978 | -0.024683576 | 0.075761823 | -0.325804936 | 0.744571944 | NA | Tnfrsf21 | 3328.765976 | 0.152045401 | 0.058152375 | 2.614603441 | 0.008933111 | 0.180225506 | |
| Hotair | 0 | NA | NA | NA | NA | NA | Cntnap4 | 1127.093597 | 0.152151885 | 0.07200596 | 2.113045709 | 0.034596856 | 0.33922827 | |
| Hoxa1 | 1.25874131 | 0.084597018 | 0.078748822 | 1.07426391 | 0.282704415 | NA | Nap1l2 | 3749.046765 | 0.152168168 | 0.06446452 | 2.360494866 | 0.018250571 | 0.257188614 | |
| Hoxa10 | 0 | NA | NA | NA | NA | NA | Sstr2 | 685.0597589 | 0.152728597 | 0.08563977 | 1.783384025 | 0.074523812 | 0.468021138 | |
| Hoxa11 | 0 | NA | NA | NA | NA | NA | Inhbb | 238.5355568 | 0.152845364 | 0.08963597 | 1.705178877 | 0.088161068 | 0.496559782 | |
| Hoxa11as | 0 | NA | NA | NA | NA | NA | Bet1 | 449.8418768 | 0.152871611 | 0.070130273 | 2.179823406 | 0.029270554 | 0.317415456 | |
| Hoxa13 | 0 | NA | NA | NA | NA | NA | Cntn1 | 14603.99716 | 0.152899968 | 0.050329679 | 3.037968262 | 0.00238179 | 0.082506178 | |
| Hoxa2 | 0.098353386 | -0.020212831 | 0.02786466 | -0.725393061 | 0.468210929 | NA | Ldha | 10780.40961 | 0.153156656 | 0.055843703 | 2.742594923 | 0.006095583 | 0.14881418 | |
| Hoxa3 | 0 | NA | NA | NA | NA | NA | Ube2a | 1198.009111 | 0.153313315 | 0.078637006 | 1.949633169 | 0.051219857 | 0.403266245 | |
| Hoxa4 | 0.176054697 | -0.002190495 | 0.031239146 | -0.070120197 | 0.944097991 | NA | Syt12 | 2056.329535 | 0.153331032 | 0.069534582 | 2.205104671 | 0.02744675 | 0.308173879 | |
| Hoxa5 | 0.192142086 | -0.026384592 | 0.031527717 | -0.836869715 | 0.402665805 | NA | Tnfaip6 | 393.3999466 | 0.153332864 | 0.089031257 | 1.722236321 | 0.085026718 | 0.488760627 | |
| Hoxa6 | 0 | NA | NA | NA | NA | NA | Vma21 | 1310.528026 | 0.153354383 | 0.063298136 | 2.422731401 | 0.015404312 | 0.236036952 | |
| Hoxa7 | 0.815988888 | -0.065845293 | 0.064875033 | -1.014955824 | 0.310126886 | NA | Ankrd9 | 387.5161427 | 0.153510454 | 0.078148017 | 1.964355083 | 0.049488928 | 0.39893239 | |
| Hoxa9 | 0 | NA | NA | NA | NA | NA | Gm4636 | 356.4736005 | 0.153525897 | 0.086198668 | 1.781070418 | 0.074900948 | 0.468589657 | |
| Hoxb1 | 0 | NA | NA | NA | NA | NA | Defb1 | 25.82907545 | 0.153632181 | 0.083381602 | 1.842518939 | 0.065399279 | 0.44524805 | |
| Hoxb13 | 0.186326724 | 0.024939385 | 0.031411053 | 0.793968451 | 0.427213798 | NA | Esrrg | 1486.445535 | 0.153997538 | 0.087152277 | 1.766993849 | 0.077229258 | 0.4735552 | |
| Hoxb2 | 0.183387883 | -0.001444816 | 0.031390672 | -0.046026918 | 0.963288796 | NA | Fam220a | 1479.969226 | 0.154132997 | 0.063713067 | 2.419174043 | 0.015555795 | 0.236858998 | |
| Hoxb3 | 0.200315975 | 0.025370485 | 0.031664208 | 0.801235428 | 0.422995365 | NA | Mobp | 10624.55495 | 0.154309917 | 0.094962105 | 1.624963112 | 0.104170419 | 0.523645898 | |
| Hoxb4 | 1.229976046 | -0.070115861 | 0.075895558 | -0.923846696 | 0.355566139 | NA | Abt1 | 572.9403427 | 0.154315359 | 0.063368732 | 2.435197187 | 0.014883676 | 0.232332035 | |
| Hoxb5 | 0.482559746 | 0.027836625 | 0.0455364 | 0.611304907 | 0.540997743 | NA | Sbk1 | 3940.383251 | 0.15448888 | 0.087598329 | 1.76360534 | 0.077798448 | 0.475337606 | |
| Hoxb6 | 0.096071043 | -0.020098795 | 0.027825948 | -0.722304036 | 0.470107572 | NA | Fsip1 | 19.92612328 | 0.15477322 | 0.080088151 | 1.932535804 | 0.053293407 | NA | |
| Hoxb7 | 0 | NA | NA | NA | NA | NA | Trpc7 | 197.9805688 | 0.154979847 | 0.093950105 | 1.649597379 | 0.099025311 | 0.520271327 | |
| Hoxb8 | 0.964034454 | 0.053194504 | 0.061906427 | 0.859272723 | 0.390190069 | NA | Gtdc2 | 1299.480329 | 0.155063463 | 0.05877434 | 2.63828507 | 0.00833265 | 0.174157793 | |
| Hoxb9 | 0.352791569 | -0.038810225 | 0.038158703 | -1.017074017 | 0.30911822 | NA | Grm4 | 808.2883943 | 0.155443298 | 0.075836792 | 2.049708246 | 0.04039291 | 0.363536189 | |
| Hoxc10 | 0.238795281 | -0.027601606 | 0.032267774 | -0.855392297 | 0.392334007 | NA | Sqle | 2545.213866 | 0.155461133 | 0.047689423 | 3.259866088 | 0.001114648 | 0.051895451 | |
| Hoxc11 | 0 | NA | NA | NA | NA | NA | Pdcl3 | 741.5928016 | 0.155463458 | 0.06209472 | 2.503650216 | 0.012291949 | 0.212038374 | |
| Hoxc12 | 0 | NA | NA | NA | NA | NA | Pepd | 1004.394568 | 0.155998235 | 0.058209101 | 2.679962958 | 0.007363031 | 0.163290664 | |
| Hoxc13 | 0.321356363 | -0.01100297 | 0.034551013 | -0.318455781 | 0.750139232 | NA | Spred2 | 2662.482463 | 0.15600749 | 0.089431356 | 1.74443838 | 0.081082677 | 0.479898997 | |
| Hoxc4 | 0.990867558 | 0.0197887 | 0.077313249 | 0.255954836 | 0.797985711 | NA | Qpct | 417.6700302 | 0.156077729 | 0.081273507 | 1.920401066 | 0.054807259 | 0.412110852 | |
| Hoxc5 | 0.40113891 | -0.035620199 | 0.048927249 | -0.728023742 | 0.466599053 | NA | AW209491 | 843.531026 | 0.156308685 | 0.069412881 | 2.251868554 | 0.02433058 | 0.292457679 | |
| Hoxc6 | 0 | NA | NA | NA | NA | NA | Galntl6 | 904.8298453 | 0.156387786 | 0.078932203 | 1.981292548 | 0.047558476 | 0.392147258 | |
| Hoxc8 | 0.116696964 | -0.020477583 | 0.027954224 | -0.732539847 | 0.463839131 | NA | Gabra1 | 15305.91778 | 0.156388066 | 0.073934544 | 2.115223227 | 0.034410926 | 0.338653871 | |
| Hoxc9 | 0 | NA | NA | NA | NA | NA | Timp3 | 6121.308681 | 0.156391612 | 0.065690418 | 2.380737037 | 0.017278041 | 0.251232055 | |
| Hoxd1 | 2.362826339 | 0.059516032 | 0.105533821 | 0.563952218 | 0.572786651 | NA | D19Bwg1357e | 1422.194531 | 0.156443973 | 0.069698016 | 2.244597214 | 0.024794002 | 0.294650465 | |
| Hoxd10 | 0 | NA | NA | NA | NA | NA | Mrpl19 | 1275.987491 | 0.156457972 | 0.057170694 | 2.736681333 | 0.006206238 | 0.150747361 | |

| | | | | | | | | | | | | | |
|---|---|---|---|---|---|---|---|---|---|---|---|---|---|
| Hoxd11 | 0 | NA | NA | NA | NA | NA | Map3k13 | 1161.403042 | 0.156510071 | 0.084786825 | 1.845924421 | 0.06490318 | 0.444080827 |
| Hoxd12 | 0 | NA | NA | NA | NA | NA | Cdh8 | 1646.775337 | 0.15654372 | 0.074004831 | 2.115317594 | 0.034402888 | 0.338653871 |
| Hoxd13 | 0 | NA | NA | NA | NA | NA | Tmem35 | 2743.610858 | 0.15678765 | 0.053827907 | 2.912757705 | 0.003582525 | 0.106203575 |
| Hoxd3 | 0.301004085 | 0.013136417 | 0.039354285 | 0.333798913 | 0.738531304 | NA | Supt3 | 387.9169392 | 0.156799195 | 0.079383261 | 1.975217361 | 0.048243487 | 0.394364033 |
| Hoxd4 | 0.099009884 | 0.018205726 | 0.027896865 | 0.652608315 | 0.514008823 | NA | Nrsn2 | 4060.956761 | 0.156926509 | 0.078987014 | 1.986738083 | 0.046951429 | 0.39102736 |
| Hoxd8 | 0.079983654 | 0.016740032 | 0.027674178 | 0.604897175 | 0.54524732 | NA | Celf6 | 954.836482 | 0.157256112 | 0.085921816 | 1.830223332 | 0.06721655 | 0.447884225 |
| Hoxd9 | 0 | NA | NA | NA | NA | NA | Prokr2 | 126.60478 | 0.15739267 | 0.095100939 | 1.655006483 | 0.097923183 | 0.518695741 |
| Hp | 6.869078324 | -0.214982673 | 0.130941795 | -1.641818585 | 0.100627606 | NA | Cyp2s1 | 271.8208914 | 0.157448364 | 0.084525571 | 1.862730567 | 0.062500183 | 0.437320645 |
| Hpd | 2.115674275 | 0.049641122 | 0.095095014 | 0.522016028 | 0.601659171 | NA | Lrrc8a | 2487.619988 | 0.157532817 | 0.067742061 | 2.325480138 | 0.020046301 | 0.267640225 |
| Hpse2 | 2.599510969 | -0.011149662 | 0.107279551 | -0.103930919 | 0.91722417 | NA | Cycs | 5004.23945 | 0.157542554 | 0.056964295 | 2.765636846 | 0.005681176 | 0.142529821 |
| Hrasls5 | 0.936130138 | -0.042000539 | 0.065679436 | -0.63947777 | 0.522512171 | NA | Plagl1 | 651.5089354 | 0.157603882 | 0.084492288 | 1.865304936 | 0.062138674 | 0.436531911 |
| Hrg | 1.017409249 | 0.01373711 | 0.070249651 | 0.195547015 | 0.844964735 | NA | Coro2a | 1797.629434 | 0.1577502 | 0.056985986 | 2.76822797 | 0.005636202 | 0.141672083 |
| Hrh4 | 0 | NA | NA | NA | NA | NA | Susd1 | 518.1174264 | 0.157770197 | 0.067982697 | 2.320740498 | 0.020300854 | 0.269537155 |
| Hrnr | 0.570133794 | 0.036094382 | 0.048404861 | 0.745676794 | 0.45586268 | NA | Zfp418 | 169.2569071 | 0.157881779 | 0.091229015 | 1.73060928 | 0.083521475 | 0.486265712 |
| Hs3st6 | 7.689334033 | -0.138802215 | 0.156066491 | -0.889378713 | 0.373799581 | NA | Apoo | 217.9382852 | 0.157978656 | 0.095099879 | 1.661186713 | 0.096675953 | 0.516802454 |
| Hsd11b2 | 0.67038813 | 0.035419061 | 0.061414033 | 0.576725864 | 0.564124657 | NA | Adcyap1 | 526.0617032 | 0.158012918 | 0.094635488 | 1.669700474 | 0.09497864 | 0.513457284 |
| Hsd17b1 | 7.248374535 | 0.160981282 | 0.134076167 | 1.200670373 | 0.229879091 | NA | Tceal8 | 1040.555968 | 0.158203831 | 0.056396552 | 2.805203961 | 0.005028475 | 0.133393847 |
| Hsd17b13 | 5.879659122 | 0.06002237 | 0.135413755 | 0.443251651 | 0.657583723 | NA | Arl5b | 423.9588839 | 0.158251849 | 0.092684791 | 1.707419821 | 0.087744049 | 0.495775125 |
| Hsd17b14 | 6.812879294 | 0.279941837 | 0.146770905 | 1.907338764 | 0.056476734 | NA | Grid1 | 2641.688723 | 0.158371075 | 0.049642557 | 3.190227987 | 0.001421606 | 0.061125282 |
| Hsd17b2 | 5.158410381 | 0.056811922 | 0.131521664 | 0.431958663 | 0.665771457 | NA | Cd200 | 3907.646093 | 0.158452442 | 0.072991664 | 2.170829282 | 0.02994408 | 0.318891276 |
| Hsd17b3 | 0 | NA | NA | NA | NA | NA | Errfi1 | 1385.465743 | 0.158456905 | 0.071051565 | 2.230167689 | 0.025736312 | 0.300614489 |
| Hsd17b6 | 0 | NA | NA | NA | NA | NA | 3632451O06Rik | 1257.520342 | 0.158467356 | 0.069623221 | 2.276070438 | 0.022841794 | 0.284318681 |
| Hsd3b1 | 0 | NA | NA | NA | NA | NA | Lrrc58 | 4148.413384 | 0.158560971 | 0.063796406 | 2.485421701 | 0.012939805 | 0.216344675 |
| Hsd3b2 | 0.098353386 | -0.020212831 | 0.02786466 | -0.725393061 | 0.468210929 | NA | Lrrtm1 | 1827.148698 | 0.15873718 | 0.070045707 | 2.266194283 | 0.023439491 | 0.288788838 |
| Hsd3b3 | 0 | NA | NA | NA | NA | NA | Pcdhb2 | 136.315463 | 0.158812854 | 0.093049458 | 1.706757428 | 0.087867148 | 0.495973914 |
| Hsd3b4 | 0 | NA | NA | NA | NA | NA | Gpr135 | 206.8982888 | 0.158828946 | 0.082646214 | 1.921793363 | 0.054631764 | 0.412035826 |
| Hsd3b5 | 0 | NA | NA | NA | NA | NA | Eya2 | 66.18497213 | 0.158946603 | 0.092218453 | 1.723587826 | 0.084782278 | 0.488499377 |
| Hsd3b6 | 0 | NA | NA | NA | NA | NA | Mafk | 404.2418589 | 0.158993169 | 0.08370473 | 1.899452626 | 0.05750499 | 0.419820382 |
| Hsf3 | 1.66649381 | 0.031307164 | 0.098873152 | 0.316639691 | 0.751517015 | NA | Mum1l1 | 154.8451774 | 0.159159855 | 0.093575587 | 1.700869437 | 0.088967507 | 0.49820432 |
| Hsfy2 | 0 | NA | NA | NA | NA | NA | Sgsm1 | 6394.063931 | 0.159210516 | 0.074285357 | 2.143228788 | 0.032094732 | 0.327711599 |
| Hsh2d | 1.200692057 | 0.095444877 | 0.066470454 | 1.435899284 | 0.151031002 | NA | Sc4mol | 3141.761194 | 0.159538168 | 0.057137278 | 2.792190572 | 0.005235251 | 0.137269422 |
| Hspb9 | 0 | NA | NA | NA | NA | NA | Me2 | 957.4345347 | 0.159546715 | 0.0681725 | 2.340338338 | 0.019266277 | 0.263564471 |
| Htr2b | 3.310832292 | 0.007602766 | 0.119997645 | 0.063357624 | 0.949481731 | NA | Ttc9 | 1451.605218 | 0.159713246 | 0.065171551 | 2.450658958 | 0.014259499 | 0.229129952 |
| Htr3b | 0.794161071 | 0.023670767 | 0.072294609 | 0.327420914 | 0.743349551 | NA | Notch4 | 343.3685974 | 0.159936954 | 0.093757046 | 1.705865971 | 0.088033037 | 0.496223473 |
| Hus1b | 0.429334602 | -0.005411584 | 0.048367196 | -0.111885422 | 0.910914255 | NA | Tinagl1 | 346.5218157 | 0.159978566 | 0.093780076 | 1.705890766 | 0.088028419 | 0.496223473 |
| Hyal4 | 0 | NA | NA | NA | NA | NA | Nxph3 | 1016.169389 | 0.160095178 | 0.082048312 | 1.951230599 | 0.051029621 | 0.403266245 |
| Hyal5 | 0 | NA | NA | NA | NA | NA | Cd24a | 324.9068361 | 0.160546073 | 0.090071848 | 1.782422328 | 0.074680387 | 0.468388672 |
| Hyal6 | 0.105923158 | 0.018524693 | 0.027977518 | 0.662127842 | 0.507889294 | NA | Pcdhb7 | 294.6513207 | 0.160652587 | 0.083438263 | 1.925406664 | 0.054178503 | 0.410426203 |
| I730028E13Rik | 2.004273009 | -0.160314373 | 0.102328518 | -1.566663683 | 0.117193324 | NA | Cbln3 | 56.7736988 | 0.160668326 | 0.072019186 | 2.230910047 | 0.025687088 | 0.300614489 |
| I730030J21Rik | 7.334782236 | 0.115656824 | 0.132390552 | 0.87360331 | 0.382334323 | NA | Hax1 | 1576.723415 | 0.160669573 | 0.061507621 | 2.61218968 | 0.008996432 | 0.181251287 |
| Iapp | 0 | NA | NA | NA | NA | NA | Pinx1 | 166.9244807 | 0.16075009 | 0.087248521 | 1.842439136 | 0.065410942 | 0.44524805 |
| Ibsp | 7.028872326 | 0.085146592 | 0.127034714 | 0.670262395 | 0.502690535 | NA | Zbtb8b | 266.9117646 | 0.160894084 | 0.083110332 | 1.935909545 | 0.052878781 | 0.406864177 |
| Icos | 3.419283575 | -0.07026302 | 0.11921927 | -0.589359586 | 0.555620081 | NA | Slc35f1 | 3725.802227 | 0.160956454 | 0.070614117 | 2.279380687 | 0.022644445 | 0.28302679 |
| Idi2 | 3.00797442 | 0.093299555 | 0.120094986 | 0.776881349 | 0.437228776 | NA | Ankrd13a | 2047.784755 | 0.160972933 | 0.056144546 | 2.867116107 | 0.00414231 | 0.11571384 |
| Ifi202b | 0.803429042 | 0.090552676 | 0.058568157 | 1.546107653 | 0.122078571 | NA | Ramp3 | 491.3450762 | 0.16101906 | 0.08898612 | 1.809485123 | 0.070375669 | 0.456017009 |
| Ifi205 | 0.801425426 | -0.042094233 | 0.068780797 | -0.612005601 | 0.540534052 | NA | Pvrl1 | 2250.657718 | 0.161131714 | 0.080277266 | 2.007189857 | 0.044729451 | 0.380855808 |
| Ifi27l2b | 2.796705782 | -0.154592643 | 0.113204604 | -1.365603845 | 0.172063339 | NA | Pcdh11x | 741.1099835 | 0.16145691 | 0.0855829 | 1.886555723 | 0.059220107 | 0.426913783 |
| Ifi44l | 0.493865508 | 0.010205312 | 0.05056179 | 0.201838429 | 0.840043037 | NA | Hs3st5 | 241.1319307 | 0.161605113 | 0.092615951 | 1.744895024 | 0.081003144 | 0.479898997 |
| Ifitm2 | 259.4752542 | 0.308139079 | 0.162140184 | 1.900448565 | NA | NA | Calca | 20.57864512 | 0.161712199 | 0.083475574 | 1.937239747 | 0.052716044 | NA |
| Ifitm5 | 0.977458471 | 0.100649779 | 0.069659942 | 1.444873139 | 0.148493522 | NA | Pglyrp1 | 179.7573408 | 0.161905502 | 0.094861418 | 1.706758187 | 0.087867007 | 0.495973914 |
| Ifitm6 | 5.42448444 | 0.205485975 | 0.137368359 | 1.49587559 | 0.134686078 | NA | Cited1 | 89.92900043 | 0.16205838 | 0.082498644 | 1.964376279 | 0.049486472 | 0.39893239 |
| Ifna1 | 0 | NA | NA | NA | NA | NA | March3 | 52.63980417 | 0.162107104 | 0.092733566 | 1.748095227 | 0.080447539 | 0.479417114 |
| Ifna11 | 0 | NA | NA | NA | NA | NA | Nfkbia | 986.4424724 | 0.162138807 | 0.089599732 | 1.809590302 | 0.070359345 | 0.456017009 |
| Ifna12 | 0 | NA | NA | NA | NA | NA | Gpr165 | 438.2791147 | 0.162244717 | 0.08860217 | 1.831159631 | 0.06707672 | 0.447688159 |
| Ifna13 | 0 | NA | NA | NA | NA | NA | Cdyl | 351.2901938 | 0.16230647 | 0.073613636 | 2.204842459 | 0.027465151 | 0.308173879 |
| Ifna14 | 0 | NA | NA | NA | NA | NA | Mnt | 2262.889537 | 0.16244334 | 0.070551421 | 2.302481483 | 0.021308034 | 0.275079973 |
| Ifna2 | 0.107716603 | -0.020477583 | 0.027954224 | -0.732539849 | 0.46383913 | NA | Cadps2 | 5217.427467 | 0.162537855 | 0.053926918 | 3.014039409 | 0.002577943 | 0.085301145 |
| Ifna4 | 0 | NA | NA | NA | NA | NA | Fuca2 | 1671.793346 | 0.162669308 | 0.059252635 | 2.745351469 | 0.006044612 | 0.148317631 |
| Ifna5 | 0 | NA | NA | NA | NA | NA | Haus7 | 207.7427599 | 0.162731379 | 0.087799703 | 1.853438827 | 0.063819486 | 0.441368101 |
| Ifna6 | 0 | NA | NA | NA | NA | NA | Ufsp2 | 1673.782011 | 0.16314644 | 0.060543381 | 2.694703166 | 0.007045136 | 0.160386105 |
| Ifna7 | 0 | NA | NA | NA | NA | NA | Klhdc8a | 760.9004114 | 0.163427877 | 0.071516482 | 2.285177811 | 0.022302404 | 0.281708449 |
| Ifna9 | 0 | NA | NA | NA | NA | NA | Btbd10 | 3644.22168 | 0.163523242 | 0.078575839 | 2.081088086 | 0.037425844 | 0.350065015 |
| Ifnab | 0 | NA | NA | NA | NA | NA | Zswim4 | 968.0245772 | 0.163548553 | 0.086035299 | 1.900947103 | 0.057308941 | 0.419380194 |
| Ifnb1 | 0 | NA | NA | NA | NA | NA | Impact | 10597.85286 | 0.163667606 | 0.066801613 | 2.450054708 | 0.014283451 | 0.229261227 |
| Ifne | 0 | NA | NA | NA | NA | NA | Med29 | 694.843713 | 0.163692249 | 0.064682162 | 2.530717015 | 0.011382964 | 0.202439555 |
| Ifng | 0.293623414 | 0.010408489 | 0.039073364 | 0.266383235 | 0.789944076 | NA | Zfp455 | 76.63077951 | 0.163941662 | 0.095101794 | 1.723854566 | 0.084734101 | 0.488499377 |
| Ifnk | 0 | NA | NA | NA | NA | NA | Elavl4 | 3090.182258 | 0.164056883 | 0.062333318 | 2.631929239 | 0.008490154 | 0.175232365 |
| Ifnl2 | 0 | NA | NA | NA | NA | NA | Fam110a | 201.4536555 | 0.164727385 | 0.082659483 | 1.99284315 | 0.046278625 | 0.38813124 |
| Ifnl3 | 0 | NA | NA | NA | NA | NA | Fgf10 | 217.2360493 | 0.164840577 | 0.08886191 | 1.855019514 | 0.06359344 | 0.440996088 |
| Ifnz | 0 | NA | NA | NA | NA | NA | Lims2 | 491.1160907 | 0.164878995 | 0.076349922 | 2.159517523 | 0.030810039 | 0.321052104 |
| Igf2bp1 | 0.948524069 | 0.061319841 | 0.066722407 | 0.919029207 | 0.358080295 | NA | Lrfn1 | 1650.405855 | 0.165100246 | 0.068598288 | 2.406769186 | 0.016094338 | 0.24095394 |
| Igf2bp3 | 1.220477769 | -0.013382445 | 0.077957831 | -0.171662608 | 0.863702784 | NA | Clcn5 | 290.7588735 | 0.165874987 | 0.091385873 | 1.815105356 | 0.06950772 | 0.455055507 |
| Igfals | 2.102523036 | 0.133804318 | 0.096411606 | 1.38784451 | 0.165184398 | NA | Cntn5 | 293.461367 | 0.166123634 | 0.080349291 | 2.067518367 | 0.038685335 | 0.356201815 |
| Igfbp1 | 0.505383842 | 0.011024618 | 0.05572892 | 0.197825804 | 0.843181356 | NA | Rab27b | 1413.125364 | 0.166367728 | 0.068548625 | 2.427003124 | 0.015224125 | 0.233769172 |
| Igfl3 | 0 | NA | NA | NA | NA | NA | Mmp11 | 73.33387872 | 0.166424468 | 0.094885959 | 1.753941988 | 0.079440448 | 0.477013957 |
| Igll1 | 0.669316088 | -0.01851675 | 0.059854163 | -0.309364442 | 0.757044316 | NA | Wisp1 | 189.2163406 | 0.166524244 | 0.095085008 | 1.751319665 | 0.079890862 | 0.478751136 |
| Igsf23 | 0.079983654 | 0.016740032 | 0.027674178 | 0.604897175 | 0.54524732 | NA | Arl4d | 1029.957231 | 0.167092357 | 0.094101521 | 1.775660541 | 0.075788885 | 0.471258137 |
| Igtp | 342.6334407 | 0.368037063 | 0.154679316 | 2.379355383 | NA | NA | Slc10a7 | 203.4516701 | 0.167148559 | 0.084573981 | 1.976359114 | 0.048114119 | 0.393789375 |
| Iigp1 | 113.2435397 | 0.197562664 | 0.118672532 | 1.664771619 | NA | NA | Pcdhgc4 | 562.3440901 | 0.167324962 | 0.07779608 | 2.150814826 | 0.031490818 | 0.32443523 |
| Il10 | 0.319934616 | 0.029547171 | 0.033991092 | 0.869262165 | 0.384703752 | NA | Epb4.1l4a | 152.7379955 | 0.167371032 | 0.094582536 | 1.769576493 | 0.076797718 | 0.472560915 |
| Il11ra2 | 0 | NA | NA | NA | NA | NA | Alox5ap | 130.837655 | 0.167821029 | 0.093715686 | 1.790746411 | 0.073333998 | 0.464565922 |
| Il12b | 0.71227231 | -0.086203635 | 0.065414181 | -1.317812664 | 0.187566367 | NA | Tubb3 | 12967.40978 | 0.167871113 | 0.058072988 | 2.890691851 | 0.003843948 | 0.111261959 |
| Il12rb2 | 7.465627489 | -0.000413354 | 0.120699625 | -0.003424651 | 0.99726753 | NA | Cldn11 | 4344.774523 | 0.168376169 | 0.083657719 | 2.012679428 | 0.044148365 | 0.380406522 |
| Il13 | 0 | NA | NA | NA | NA | NA | Wbp5 | 1685.60334 | 0.168558249 | 0.067963886 | 2.480114952 | 0.013134003 | 0.218789601 |
| Il13ra2 | 7.424376002 | -0.369589113 | 0.140048974 | -2.638999081 | 0.008315121 | NA | Zfhx3 | 280.731514 | 0.168559619 | 0.09479375 | 1.778172281 | 0.075375564 | 0.469714905 |
| Il17a | 0 | NA | NA | NA | NA | NA | Elovl4 | 2000.42257 | 0.168718017 | 0.064168948 | 2.62927819 | 0.008556633 | 0.176053334 |
| Il17b | 6.968546564 | 0.036197373 | 0.151297694 | 0.239246034 | 0.810914809 | NA | H2-Q2 | 27.40243684 | 0.168727181 | 0.06387841 | 2.641380418 | 0.008256895 | 0.174077875 |
| Il17c | 1.051226501 | 0.034639185 | 0.07267005 | 0.476663841 | 0.633601515 | NA | A730020M07Rik | 19.82716672 | 0.168763197 | 0.077901307 | 2.166371832 | 0.030282785 | NA |
| Il17f | 0.298391179 | 0.008861245 | 0.039220874 | 0.225931861 | 0.821254417 | NA | Frmd6 | 1944.620664 | 0.168793287 | 0.084615918 | 1.99481717 | 0.046062824 | 0.387222051 |
| Il17re | 7.915955225 | 0.214883859 | 0.156594606 | 1.372230273 | 0.169991764 | NA | Pcdhgb4 | 196.5609926 | 0.168815591 | 0.087978558 | 1.918826539 | 0.05500629 | 0.41218506 |
| Il18r1 | 2.089358884 | -0.051005697 | 0.089413523 | -0.570447237 | 0.568374399 | NA | Ccdc160 | 85.89560752 | 0.168844396 | 0.094285058 | 1.790786367 | 0.073327584 | 0.464565922 |
| Il18rap | 2.655103376 | -0.088742223 | 0.109395688 | -0.811204029 | 0.417248512 | NA | Bex4 | 428.3235237 | 0.168939747 | 0.085184092 | 1.983231165 | 0.047341614 | 0.391951501 |
| Il19 | 0.57410637 | 0.02462778 | 0.058376336 | 0.421879507 | 0.673112969 | NA | Hspb6 | 266.8516857 | 0.169185664 | 0.088982341 | 1.90133978 | 0.057257522 | 0.419215102 |
| Il1f10 | 0.096071043 | -0.020098795 | 0.027825948 | -0.722304036 | 0.470107572 | NA | Vwc2 | 255.6871392 | 0.169432344 | 0.081678471 | 2.074381932 | 0.038043854 | 0.352889542 |
| Il1f5 | 3.248080251 | 0.065925601 | 0.109009998 | 0.604766556 | 0.545334118 | NA | Rnmtl1 | 409.8530154 | 0.169722874 | 0.073775892 | 2.300519446 | 0.021418809 | 0.275274875 |
| Il1f6 | 0 | NA | NA | NA | NA | NA | Cdr2 | 761.7632911 | 0.1697464 | 0.078580191 | 2.160167834 | 0.030759679 | 0.321052104 |
| Il1f8 | 0 | NA | NA | NA | NA | NA | Zfp64 | 500.9764408 | 0.169780829 | 0.071654092 | 2.369450564 | 0.017814536 | 0.255043174 |
| Il1rn | 3.015185515 | 0.148202274 | 0.099959021 | 1.482630301 | 0.138172662 | NA | Pcdha5 | 186.7102108 | 0.169865228 | 0.095058254 | 1.786959275 | 0.073944071 | 0.466583525 |
| Il2 | 0 | NA | NA | NA | NA | NA | 1700037C18Rik | 137.7539736 | 0.169889228 | 0.092532194 | 1.836001282 | 0.066357467 | 0.446279967 |
| Il20 | 0 | NA | NA | NA | NA | NA | Chac2 | 211.0654914 | 0.169956476 | 0.087181235 | 1.949461674 | 0.051240316 | 0.403266245 |
| Il21 | 2.68038864 | 0.036686079 | 0.089529743 | 0.409764148 | 0.681978968 | NA | Bex1 | 1202.57815 | 0.170028894 | 0.080776933 | 2.10491891 | 0.035298367 | 0.342057427 |
| Il22 | 1.313344777 | 0.030458062 | 0.081741128 | 0.372616127 | 0.709434168 | NA | Galnt15 | 130.5742503 | 0.170051361 | 0.095082635 | 1.788458648 | 0.073702041 | 0.465836525 |
| Il22ra1 | 3.056421495 | 0.079193051 | 0.116960709 | 0.677091061 | 0.49834818 | NA | Hcrtr2 | 81.84969887 | 0.170142566 | 0.094971479 | 1.791512229 | 0.073211134 | 0.464395167 |
| Il22ra2 | 0.119397641 | -0.020477583 | 0.027954224 | -0.732539845 | 0.463839132 | NA | Shq1 | 154.9286031 | 0.170401205 | 0.088764758 | 1.91969436 | 0.054896517 | 0.412110852 |
| Il23a | 3.721289891 | -0.281273911 | 0.130513745 | -2.155128646 | 0.031151769 | NA | Etv1 | 2585.266828 | 0.170504916 | 0.066989071 | 2.545264686 | 0.010919493 | 0.198767619 |
| Il23r | 0.451066898 | 0.006365339 | 0.044041609 | 0.144530111 | 0.885081881 | NA | Mrap2 | 433.3509991 | 0.170568847 | 0.093792512 | 1.818576386 | 0.068976083 | 0.453369495 |
| Il24 | 0 | NA | NA | NA | NA | NA | Fibin | 98.75554232 | 0.171114368 | 0.09457367 | 1.809323549 | 0.070400753 | 0.456017009 |
| Il25 | 2.540226355 | -0.015535638 | 0.103527021 | -0.150063604 | 0.880714435 | NA | Anln | 2165.98366 | 0.171323572 | 0.087909973 | 1.948852508 | 0.051313041 | 0.403340495 |
| Il27 | 1.241576115 | 0.158271463 | 0.083179902 | 1.902760871 | 0.057071755 | NA | Kirrel2 | 538.3397882 | 0.493308697 | 0.094783284 | 5.204595953 | 1.94E-07 | 9.41E-05 |
| Il3 | 0 | NA | NA | NA | NA | NA | Grm8 | 774.8420324 | 0.171600163 | 0.070876214 | 2.421124849 | 0.015472562 | 0.236284305 |
| Il31 | 0.096071043 | -0.020098795 | 0.027825948 | -0.722304036 | 0.470107572 | NA | Ina | 5401.029966 | 0.17199524 | 0.061664167 | 2.789225052 | 0.005283434 | 0.137476727 |
| Il4 | 0 | NA | NA | NA | NA | NA | Pcbd1 | 155.3952881 | 0.172139525 | 0.094370025 | 1.824091117 | 0.068138298 | 0.450923427 |
| Il4i1 | 0.330487422 | -0.017886856 | 0.040493356 | -0.441723219 | 0.658689506 | NA | Doc2b | 2379.760899 | 0.172397807 | 0.076432029 | 2.25557019 | 0.024097562 | 0.29178139 |
| Il5 | 2.601214813 | 0.125010189 | 0.108609286 | 1.151008288 | 0.249728826 | NA | Slc7a1 | 2276.956428 | 0.172614579 | 0.061567055 | 2.803684176 | 0.005052237 | 0.133434166 |
| Il5ra | 0.385382084 | -0.000943527 | 0.041930191 | -0.022502333 | 0.982047251 | NA | Pdlim1 | 395.8872353 | 0.172780455 | 0.083984857 | 2.057281047 | 0.039659199 | 0.36080087 |
| Il6 | 2.260663149 | 0.142455921 | 0.090968713 | 1.565988088 | 0.117351406 | NA | Efna2 | 247.7500587 | 0.172964122 | 0.08049534 | 2.148747021 | 0.031654459 | 0.325415905 |
| Il7 | 3.796125002 | 0.072403111 | 0.130125431 | 0.556410156 | 0.577930498 | NA | Trim16 | 90.07251888 | 0.173027855 | 0.094748808 | 1.826174482 | 0.067823984 | 0.449662797 |
| Il8 | 0 | NA | NA | NA | NA | NA | Gm6537 | 118.5045039 | 0.489308436 | 0.093611714 | 5.227000057 | 1.72E-07 | 8.94E-05 |
| Il9 | 0 | NA | NA | NA | NA | NA | Lad1 | 84.69601444 | 0.17355348 | 0.094897313 | 1.828855577 | 0.067421247 | 0.448803776 |
| Il9r | 0.305736371 | 0.008542183 | 0.033941149 | 0.251676297 | 0.80129128 | NA | Rhcg | 26.75759804 | 0.173573279 | 0.085780277 | 2.023463719 | 0.043025361 | 0.375514735 |
| Iltifb | 1.699873486 | 0.097147322 | 0.096393182 | 1.007823587 | 0.313539161 | NA | Clvs2 | 1132.515432 | 0.173924963 | 0.064010588 | 2.71712804 | 0.006585113 | 0.156299587 |
| Inhbc | 0.107716603 | -0.020477583 | 0.027954224 | -0.732539849 | 0.46383913 | NA | Dtd1 | 1773.65601 | 0.174116289 | 0.054355253 | 3.203301942 | 0.001358615 | 0.059797894 |
| Ins1 | 0 | NA | NA | NA | NA | NA | Tacr3 | 128.6748117 | 0.174187235 | 0.094736274 | 1.838654063 | 0.065966084 | 0.446279967 |
| Ins2 | 0.468677007 | 0.028797021 | 0.045304689 | 0.635630028 | 0.525017593 | NA | Grb10 | 2384.079407 | 0.174757587 | 0.087873174 | 1.988747864 | 0.04672904 | 0.390555831 |
| Insl3 | 1.825258177 | 0.037173765 | 0.093714226 | 0.396671532 | 0.691609696 | NA | Parm1 | 1235.622139 | 0.174773574 | 0.076123414 | 2.295923994 | 0.021680228 | 0.276408712 |
| Insm2 | 3.910246587 | -0.113580811 | 0.130822248 | -0.868207148 | 0.385280942 | NA | Gpr161 | 137.9846312 | 0.174777617 | 0.090505177 | 1.931133925 | 0.053466493 | 0.407906655 |
| Iqcf1 | 0 | NA | NA | NA | NA | NA | C1galt1c1 | 683.1428844 | 0.175147176 | 0.066112153 | 2.649243273 | 0.008067224 | 0.172465045 |
| Iqcf3 | 0.186326724 | 0.024939385 | 0.031411053 | 0.793968451 | 0.427213798 | NA | Acot11 | 2572.180264 | 0.175489478 | 0.070860866 | 2.476535879 | 0.013266429 | 0.21998647 |
| Iqcf4 | 0.211846316 | 0.025707621 | 0.031860375 | 0.806883811 | 0.419733426 | NA | Gpr45 | 422.0817296 | 0.175649553 | 0.069137622 | 2.540578458 | 0.011066926 | 0.199827723 |
| Iqcf5 | 2.293894031 | 0.022767951 | 0.095841592 | 0.237558149 | 0.812223814 | NA | 1110012L19Rik | 339.2860029 | 0.17616862 | 0.07416485 | 2.375365437 | 0.017531583 | 0.253361317 |
| Iqcf6 | 0 | NA | NA | NA | NA | NA | Olfm3 | 1139.923666 | 0.176280591 | 0.078091353 | 2.257363759 | 0.023985354 | 0.29172307 |
| Iqch | 2.970735406 | 0.202007729 | 0.124224906 | 1.626145151 | 0.103918781 | NA | Tshz2 | 1588.398541 | 0.17640354 | 0.091938384 | 1.918714823 | 0.055020434 | 0.41218506 |
| Iqcj | 1.623363065 | -0.102028201 | 0.091583758 | -1.114042529 | 0.265260942 | NA | Cry1 | 817.3984228 | 0.176701404 | 0.08933634 | 1.977934229 | 0.047936127 | 0.393352856 |
| Irg1 | 1.562964034 | 0.074159782 | 0.078666255 | 0.942714026 | 0.345827197 | NA | Usp27x | 722.0606312 | 0.177103991 | 0.08117781 | 2.181679837 | 0.02913317 | 0.317415456 |
| Irgc1 | 4.580136633 | -0.23723407 | 0.128670826 | -1.84372851 | 0.065222716 | NA | Cbln1 | 372.7143689 | 0.177400325 | 0.076003251 | 2.334114963 | 0.019589703 | 0.26397034 |
| Irx2 | 5.247987084 | 0.188455073 | 0.112781148 | 1.670980273 | 0.094725574 | NA | Fam196a | 537.9573829 | 0.177591799 | 0.085753015 | 2.070968574 | 0.038361733 | 0.354869883 |
| Irx3 | 6.141851399 | 0.334152361 | 0.142946491 | 2.337604503 | 0.019407774 | NA | Rara | 529.066335 | 0.177744155 | 0.084523228 | 2.102903062 | 0.035474244 | 0.342120019 |
| Irx4 | 0.094392817 | 0.017895942 | 0.027818137 | 0.643319236 | 0.520016977 | NA | Ngb | 366.2476873 | 0.178191945 | 0.094953731 | 1.876618667 | 0.060570377 | 0.431188114 |
| Irx5 | 3.390072367 | 0.181013927 | 0.111149641 | 1.628560606 | 0.103406069 | NA | Agxt2l1 | 1352.944949 | 0.178486738 | 0.08727911 | 2.045010971 | 0.040853777 | 0.365377968 |
| Isl2 | 0.852951773 | 0.017080043 | 0.056027866 | 0.304849064 | 0.760481112 | NA | Vat1l | 4169.797223 | 0.179281569 | 0.095079546 | 1.885595561 | 0.059349476 | 0.427634171 |
| Isx | 2.802752224 | -0.096197048 | 0.12116548 | -0.793931144 | 0.427235518 | NA | Ky | 184.789662 | 0.179852103 | 0.094685644 | 1.899465393 | 0.057503313 | 0.419820382 |
| Itga2 | 5.231023707 | -0.105884164 | 0.138144283 | -0.766475176 | 0.443393617 | NA | Tmtc4 | 478.2228719 | 0.180086014 | 0.071880768 | 2.505343485 | 0.012233253 | 0.211851282 |
| Itgad | 11.56699328 | 0.276335662 | 0.13545839 | 2.040004035 | NA | NA | Chrna4 | 2120.981049 | 0.180089019 | 0.068724385 | 2.620452967 | 0.008781304 | 0.179191559 |
| Itgae | 7.832432227 | 0.064394042 | 0.150255071 | 0.428564854 | 0.668239927 | NA | Galnt16 | 6798.867056 | 0.180201934 | 0.071474125 | 2.521219177 | 0.011694898 | 0.205417281 |
| Itgam | 1131.312302 | -0.111903595 | 0.166502778 | -0.672082447 | NA | NA | Spryd4 | 286.3731754 | 0.180229432 | 0.07773834 | 2.31841113 | 0.020426988 | 0.269900834 |
| Itgb2l | 0.36509754 | 0.021286825 | 0.04223302 | 0.504032755 | 0.61423836 | NA | Nudt14 | 183.4812961 | 0.180283907 | 0.086662608 | 2.080296353 | 0.037498359 | 0.350065015 |
| Itgb6 | 0.910840021 | -0.026687877 | 0.070723857 | -0.377353252 | 0.705911103 | NA | Plcxd3 | 330.005716 | 0.180754522 | 0.083476903 | 2.165323767 | 0.0303629 | 0.32021294 |
| Itih1 | 0 | NA | NA | NA | NA | NA | Adcy8 | 1126.185955 | 0.180920649 | 0.067500823 | 2.680273227 | 0.007356209 | 0.163290664 |
| Itih4 | 4.566125969 | 0.206095701 | 0.126725605 | 1.626314596 | 0.103882748 | NA | Pcolce2 | 244.8938731 | 0.181005617 | 0.094330852 | 1.918837929 | 0.055004848 | 0.41218506 |
| Itln1 | 0 | NA | NA | NA | NA | NA | Vwc2l | 360.1705871 | 0.181235221 | 0.086710439 | 2.090119984 | 0.036607023 | 0.346440078 |
| Ivl | 0.096071043 | -0.020098795 | 0.027825948 | -0.722304036 | 0.470107572 | NA | Arhgap24 | 286.4299536 | 0.181249386 | 0.094071448 | 1.926720483 | 0.054014474 | 0.409506392 |
| Izumo1 | 1.858657362 | 0.082709532 | 0.099163 | 0.834076536 | 0.404237857 | NA | Bdnf | 930.5590876 | 0.181393304 | 0.09338076 | 1.942512609 | 0.052075075 | 0.406272098 |
| Izumo2 | 0 | NA | NA | NA | NA | NA | Fstl5 | 838.4596628 | 0.181410939 | 0.081641713 | 2.222037377 | 0.02628078 | 0.302739577 |
| Izumo3 | 0 | NA | NA | NA | NA | NA | Mgp | 519.2752032 | 0.181623933 | 0.094964939 | 1.912536728 | 0.055807386 | 0.413057568 |
| Kap | 0 | NA | NA | NA | NA | NA | Plekhf1 | 189.2838919 | 0.418278595 | 0.094469896 | 4.427638995 | 9.53E-06 | 0.001976992 |
| Kcna10 | 4.083566732 | 0.206485294 | 0.124999434 | 1.651889826 | 0.098557013 | NA | Dusp10 | 289.4369783 | 0.182614792 | 0.086104448 | 2.120852025 | 0.033934257 | 0.336326295 |
| Kcna7 | 1.007082075 | 0.091882827 | 0.076574122 | 1.199920073 | 0.230170383 | NA | Rec8 | 225.4806122 | 0.182645382 | 0.095073075 | 1.92110524 | 0.054718441 | 0.412110852 |
| Kcne1 | 0.38827344 | 0.020990736 | 0.04283191 | 0.490072386 | 0.624082678 | NA | Klf13 | 9033.838528 | 0.182713773 | 0.061587985 | 2.966711312 | 0.003010034 | 0.095258736 |
| Kcne2 | 26.11502812 | 0.212993065 | 0.090336557 | 2.357772676 | NA | NA | Hivep1 | 3982.196423 | 0.183029048 | 0.078285542 | 2.337967439 | 0.019388937 | 0.263564471 |
| Kcne3 | 5.558619948 | 0.146664536 | 0.140124964 | 1.046669567 | 0.295252005 | NA | Kctd8 | 275.729378 | 0.183070882 | 0.093700368 | 1.953790437 | 0.050726006 | 0.403266245 |
| Kcnj1 | 0.099009884 | 0.018205726 | 0.027896865 | 0.652608315 | 0.514008823 | NA | Zfp27 | 913.3537307 | 0.395164054 | 0.063899355 | 6.184163444 | 6.24E-10 | 5.67E-07 |
| Kcnj15 | 2.982287336 | -0.067712734 | 0.10854698 | -0.623810392 | 0.532752114 | NA | Klf9 | 7912.969091 | 0.1838971 | 0.078664963 | 2.337725624 | 0.019401486 | 0.263564471 |
| Kcnk15 | 3.251105874 | -0.183021399 | 0.090816472 | -2.015288578 | 0.043874421 | NA | Bhlhe40 | 6666.085036 | 0.18405423 | 0.094942481 | 1.938586693 | 0.052551685 | 0.406864177 |
| Kcnk16 | 0.17463368 | 0.024560283 | 0.031186242 | 0.787535833 | 0.430968261 | NA | Capn11 | 27.47177218 | 0.184091129 | 0.06096146 | 3.019795287 | 0.002529456 | 0.084486004 |
| Kcnk18 | 0 | NA | NA | NA | NA | NA | Fas | 72.79650208 | 0.184121557 | 0.09491974 | 1.93976044 | 0.05240881 | 0.406355487 |
| Kcnk7 | 3.266791574 | 0.049750173 | 0.12016794 | 0.41400538 | 0.678870167 | NA | Rasl11a | 122.3092335 | 0.184195157 | 0.095035661 | 1.938168849 | 0.052602626 | 0.406864177 |
| Kcnmb3 | 0.099009884 | 0.018205726 | 0.027896865 | 0.652608315 | 0.514008823 | NA | Slc25a13 | 159.9185041 | 0.1846618 | 0.094652977 | 1.950934937 | 0.051064786 | 0.403266245 |
| Kcnv2 | 0.531537619 | -0.072805061 | 0.051684011 | -1.408657322 | 0.15893652 | NA | Cthrc1 | 221.2783517 | 0.184877736 | 0.086281781 | 2.142720449 | 0.032135552 | 0.327711599 |
| Kctd19 | 0.602691698 | -0.043058929 | 0.048770142 | -0.882895305 | 0.37729284 | NA | Lrrtm3 | 2193.788201 | 0.185061808 | 0.071072148 | 2.603858364 | 0.009218084 | 0.183045438 |
| Keg1 | 0 | NA | NA | NA | NA | NA | 1810011O10Rik | 39.96145116 | 0.185622125 | 0.09067549 | 2.047103626 | 0.040647911 | 0.364926796 |
| Kel | 0.612869747 | -0.006355786 | 0.053223994 | -0.119415802 | 0.904945943 | NA | Cpm | 688.9688254 | 0.18581537 | 0.085867939 | 2.163966816 | 0.030466896 | 0.320602541 |
| Kera | 0.239950962 | 0.027417169 | 0.032831728 | 0.835081494 | 0.403671825 | NA | Sgcz | 95.79465902 | 0.185879068 | 0.092148734 | 2.017163553 | 0.043678448 | 0.378338186 |
| Khdc1a | 0 | NA | NA | NA | NA | NA | Eva1a | 351.3101482 | 0.186167035 | 0.079548687 | 2.340290474 | 0.019268747 | 0.263564471 |
| Khdc1b | 0 | NA | NA | NA | NA | NA | Pcsk1 | 1446.409785 | 0.186789215 | 0.092499522 | 2.019353302 | 0.043450512 | 0.37703831 |
| Khdc1c | 0 | NA | NA | NA | NA | NA | Catsperg2 | 10.77627923 | 0.18764468 | 0.05678431 | 3.304516349 | 0.000951404 | NA |
| Kif12 | 0.197363271 | -0.000561558 | 0.031650166 | -0.017742642 | 0.985844163 | NA | Slc6a11 | 8761.552864 | 0.187713764 | 0.082941069 | 2.263218529 | 0.023622222 | 0.289811141 |
| Kif2b | 0 | NA | NA | NA | NA | NA | Dpysl5 | 1526.16481 | 0.187848046 | 0.067531836 | 2.78162209 | 0.005408798 | 0.138877279 |
| Kif4-ps | 0 | NA | NA | NA | NA | NA | Fam83d | 66.9597569 | 0.189153526 | 0.090806063 | 2.083049518 | 0.03724671 | 0.348836693 |
| Kir3dl1 | 0 | NA | NA | NA | NA | NA | Lgi2 | 1481.492502 | 0.189172056 | 0.068922514 | 2.744706276 | 0.006056508 | 0.148321973 |
| Kir3dl2 | 5.239499727 | 0.173088669 | 0.128892893 | 1.342887615 | 0.179308367 | NA | Pak4 | 388.7309475 | 0.189298589 | 0.072765142 | 2.601500978 | 0.00928168 | 0.183436305 |
| Kis2 | 0.105923158 | 0.018524693 | 0.027977518 | 0.662127842 | 0.507889294 | NA | Ctgf | 1598.45317 | 0.189357546 | 0.073846481 | 2.564205406 | 0.010341232 | 0.194329551 |
| Kl | 348.806466 | 0.383400764 | 0.150019669 | 2.555669971 | NA | NA | 1700001L19Rik | 361.0556128 | 0.189858647 | 0.082338487 | 2.30583115 | 0.021120069 | 0.273675398 |
| Klb | 6.73275033 | 0.035676134 | 0.148306622 | 0.24055658 | 0.809898806 | NA | Trib2 | 1747.521962 | 0.189934193 | 0.086130007 | 2.205203497 | 0.027439818 | 0.308173879 |
| Klf1 | 4.867912659 | 0.223518709 | 0.142719895 | 1.5661356 | 0.117316875 | NA | Hif3a | 486.7148821 | 0.383189887 | 0.093594459 | 4.094151431 | 4.24E-05 | 0.006093978 |
| Klf17 | 0.639411646 | 0.032045962 | 0.060168178 | 0.532606496 | 0.594306005 | NA | Sdpr | 447.1505565 | 0.190753077 | 0.072128242 | 2.644637829 | 0.008177839 | 0.173925762 |
| Klhl31 | 2.227178724 | 0.106658768 | 0.105957151 | 1.006621709 | 0.314116598 | NA | Scube2 | 64.9935507 | 0.19078153 | 0.092463046 | 2.063327341 | 0.039081538 | 0.358395469 |
| Klhl34 | 6.350984302 | 0.181332575 | 0.136217171 | 1.331202032 | 0.183122543 | NA | Nedd9 | 473.9676271 | 0.190789888 | 0.080443031 | 2.371739175 | 0.017704584 | 0.254345817 |
| Klk1 | 0.233393928 | -0.027475306 | 0.032191697 | -0.85349047 | 0.39338737 | NA | Kcnf1 | 6682.41231 | 0.190931158 | 0.070997969 | 2.689248148 | 0.007161316 | 0.161546928 |
| Klk10 | 3.584123508 | 0.024521382 | 0.112562779 | 0.21784627 | 0.827548892 | NA | Hpcal1 | 2717.607546 | 0.191730348 | 0.088290221 | 2.171592112 | 0.029886443 | 0.3187222 |
| Klk11 | 0.365320263 | 0.05192513 | 0.045006765 | 1.15371833 | 0.248615666 | NA | Mal | 10355.81967 | 0.191782747 | 0.084453908 | 2.270857 | 0.023155636 | 0.286019362 |
| Klk12 | 0 | NA | NA | NA | NA | NA | Rinl | 112.8997985 | 0.362270192 | 0.094948093 | 3.815455177 | 0.000135932 | 0.012666269 |
| Klk13 | 1.002455104 | -0.033529498 | 0.073335503 | -0.457206902 | 0.647522338 | NA | Cdkl4 | 890.9914856 | 0.192237953 | 0.075271793 | 2.553917528 | 0.010651848 | 0.197106686 |
| Klk15 | 0 | NA | NA | NA | NA | NA | Capsl | 50.74347038 | 0.193223315 | 0.092327233 | 2.092809549 | 0.036366159 | 0.345417071 |
| Klk1b1 | 0 | NA | NA | NA | NA | NA | H60b | 29.65253437 | 0.193280477 | 0.080482515 | 2.40152135 | 0.016327056 | 0.242552129 |
| Klk1b11 | 0 | NA | NA | NA | NA | NA | Nxn | 433.9890011 | 0.193309458 | 0.084524497 | 2.287022867 | 0.022194488 | 0.280833744 |
| Klk1b16 | 0 | NA | NA | NA | NA | NA | Zxdb | 175.5375258 | 0.193684788 | 0.091820861 | 2.109376739 | 0.034912076 | 0.340200772 |
| Klk1b21 | 0 | NA | NA | NA | NA | NA | Etv5 | 5824.668028 | 0.194367288 | 0.082085626 | 2.367860226 | 0.017891294 | 0.255043174 |
| Klk1b22 | 0 | NA | NA | NA | NA | NA | Pcdha9 | 151.6822324 | 0.194657032 | 0.092233027 | 2.110491643 | 0.03481603 | 0.339877456 |
| Klk1b24 | 0 | NA | NA | NA | NA | NA | Fzd1 | 454.8447695 | 0.194871038 | 0.07981609 | 2.441500677 | 0.014626361 | 0.231945981 |
| Klk1b26 | 0 | NA | NA | NA | NA | NA | Eif5a2 | 2584.746919 | 0.195276687 | 0.064067699 | 3.047974076 | 0.002303898 | 0.081073853 |
| Klk1b27 | 0.638908139 | -0.060041808 | 0.056221338 | -1.067954093 | 0.28554122 | NA | Kcnk10 | 146.1603196 | 0.195284009 | 0.091106007 | 2.143481153 | 0.032074483 | 0.327711599 |
| Klk1b3 | 0.65992155 | 0.044480374 | 0.050358557 | 0.883273402 | 0.377088571 | NA | Ankrd33b | 4961.022245 | 0.195531919 | 0.086058042 | 2.272093505 | 0.023080863 | 0.286019362 |
| Klk1b4 | 0 | NA | NA | NA | NA | NA | Vasn | 596.3315605 | 0.196827322 | 0.088072947 | 2.234821572 | 0.025429065 | 0.298853235 |
| Klk1b5 | 0 | NA | NA | NA | NA | NA | Ldb2 | 1705.957845 | 0.197337201 | 0.065913869 | 2.99386463 | 0.002754681 | 0.088921114 |
| Klk1b7-ps | 0 | NA | NA | NA | NA | NA | Mest | 1081.937997 | 0.197511043 | 0.078048378 | 2.530623309 | 0.011386006 | 0.202439555 |
| Klk1b8 | 0 | NA | NA | NA | NA | NA | Slc35g2 | 650.1881054 | 0.198152572 | 0.077214171 | 2.566272103 | 0.010279815 | 0.193928045 |
| Klk1b9 | 0 | NA | NA | NA | NA | NA | Mthfd1l | 667.7189084 | 0.198596083 | 0.07824077 | 2.538268522 | 0.011140247 | 0.20051086 |
| Klk4 | 0.218407525 | -0.002234018 | 0.031987735 | -0.069839845 | 0.944321133 | NA | Scml4 | 184.5308144 | 0.198625021 | 0.094593349 | 2.09977787 | 0.035748386 | 0.343212857 |
| Klk5 | 0.116696964 | -0.020477583 | 0.027954224 | -0.732539847 | 0.463839131 | NA | Mgat4c | 164.3738039 | 0.198887906 | 0.090047988 | 2.208687953 | 0.027196353 | 0.307674628 |
| Klk7 | 0.830848704 | 0.060916436 | 0.066285696 | 0.918998219 | 0.358096504 | NA | Spire2 | 672.3543413 | 0.199032689 | 0.0620687 | 3.206651493 | 0.001342896 | 0.059472268 |
| Klk9 | 1.340873711 | 0.000938095 | 0.090559374 | 0.010358894 | 0.991734946 | NA | Tekt5 | 272.7150904 | 0.199223033 | 0.088584326 | 2.248964811 | 0.024514735 | 0.293328698 |
| Klkb1 | 3.41490234 | 0.061658413 | 0.124287637 | 0.496094495 | 0.619827743 | NA | Slc25a34 | 352.5844213 | 0.20025446 | 0.095098459 | 2.105759243 | 0.035225271 | 0.341805132 |
| Klra1 | 0 | NA | NA | NA | NA | NA | Efnb1 | 670.5823622 | 0.200370796 | 0.072280782 | 2.772117165 | 0.005569299 | 0.141186095 |
| Klra10 | 0 | NA | NA | NA | NA | NA | Cbln2 | 1070.749602 | 0.200375869 | 0.079336894 | 2.525632903 | 0.011549009 | 0.203840709 |
| Klra12 | 0 | NA | NA | NA | NA | NA | Cartpt | 277.7391602 | 0.200381385 | 0.095044009 | 2.108301067 | 0.035004956 | 0.34025917 |
| Klra13-ps | 0 | NA | NA | NA | NA | NA | Maff | 50.92857626 | 0.200592653 | 0.082003592 | 2.446144709 | 0.014439303 | 0.230050259 |
| Klra14 | 0.196706773 | -0.026517754 | 0.031609464 | -0.838918169 | 0.401515231 | NA | Slc25a25 | 2961.587398 | 0.200869069 | 0.061886962 | 3.245741298 | 0.001171452 | 0.05383536 |
| Klra15 | 0 | NA | NA | NA | NA | NA | Paqr8 | 6573.153978 | 0.201208448 | 0.067086606 | 2.99923429 | 0.002706591 | 0.087924428 |
| Klra17 | 0 | NA | NA | NA | NA | NA | Gmfg | 51.27470603 | 0.201553347 | 0.093897126 | 2.146533714 | 0.031830422 | 0.326301135 |
| Klra18 | 0 | NA | NA | NA | NA | NA | Nkrf | 1494.21553 | 0.201688986 | 0.074009454 | 2.725178656 | 0.00642667 | 0.154049193 |
| Klra19 | 0 | NA | NA | NA | NA | NA | Jdp2 | 1709.697427 | 0.201702539 | 0.066573137 | 3.029788689 | 0.002447249 | 0.083252314 |
| Klra2 | 3.456128443 | 0.225087158 | 0.125761906 | 1.789788064 | 0.073487989 | NA | Fam107a | 19695.42614 | 0.201872911 | 0.080663348 | 2.502659717 | 0.012326399 | 0.212148434 |
| Klra21 | 0 | NA | NA | NA | NA | NA | Marcksl1 | 1803.716204 | 0.201925094 | 0.083197394 | 2.427060326 | 0.015221725 | 0.233769172 |
| Klra22 | 0 | NA | NA | NA | NA | NA | Tsc22d3 | 3720.830662 | 0.3617828 | 0.087409552 | 4.138938973 | 3.49E-05 | 0.005335105 |
| Klra23 | 0 | NA | NA | NA | NA | NA | Cbln4 | 841.5159451 | 0.361741645 | 0.088886346 | 4.069709916 | 4.71E-05 | 0.006479723 |
| Klra3 | 0 | NA | NA | NA | NA | NA | Rwdd2a | 619.035592 | 0.202669126 | 0.092775775 | 2.184504804 | 0.028925175 | 0.31686809 |
| Klra33 | 0 | NA | NA | NA | NA | NA | Glra2 | 599.0492871 | 0.202671029 | 0.079626053 | 2.545285391 | 0.010918846 | 0.198767619 |
| Klra4 | 0 | NA | NA | NA | NA | NA | Zfp458 | 183.5567779 | 0.203235486 | 0.087266403 | 2.328908723 | 0.019863902 | 0.266183613 |
| Klra5 | 0 | NA | NA | NA | NA | NA | Ddc | 374.3889615 | 0.203319212 | 0.094857728 | 2.143412197 | 0.032080015 | 0.327711599 |
| Klra6 | 0 | NA | NA | NA | NA | NA | Dact1 | 278.615499 | 0.203910149 | 0.089638813 | 2.27479752 | 0.022918079 | 0.285023987 |
| Klra7 | 0 | NA | NA | NA | NA | NA | Adipor2 | 3828.371183 | 0.344487292 | 0.067949921 | 5.069723237 | 3.98E-07 | 0.000170208 |
| Klra8 | 0.079983654 | 0.016740032 | 0.027674178 | 0.604897175 | 0.54524732 | NA | Lrfn3 | 1084.466002 | 0.204962919 | 0.074471748 | 2.752223827 | 0.005919205 | 0.146477633 |
| Klra9 | 0.08731684 | 0.017365056 | 0.027733412 | 0.626142072 | 0.531221759 | NA | Kcna5 | 282.5257998 | 0.342127532 | 0.088865116 | 3.849964392 | 0.000118135 | 0.011600059 |
| Klrb1 | 0.098353386 | -0.020212831 | 0.02786466 | -0.725393061 | 0.468210929 | NA | Mc4r | 74.20819186 | 0.206451598 | 0.091945503 | 2.245369171 | 0.024744443 | 0.294610834 |
| Klrb1-ps1 | 0 | NA | NA | NA | NA | NA | Tmem132c | 441.2681604 | 0.207146877 | 0.080386128 | 2.576898318 | 0.009969127 | 0.19094795 |

| | | | | | | | | | | | | | |
|---|---|---|---|---|---|---|---|---|---|---|---|---|---|
| Klrb1a | 1.517537733 | 0.114815172 | 0.095204409 | 1.205985873 | 0.227822937 | NA | Metrnl | 281.5624441 | 0.207313828 | 0.086054069 | 2.409111278 | 0.015991421 | 0.240716462 |
| Klrb1b | 2.323347446 | 0.06007905 | 0.086588676 | 0.693844191 | 0.487779926 | NA | Kcng1 | 767.4171907 | 0.208006777 | 0.094254941 | 2.206852766 | 0.027324347 | 0.308173879 |
| Klrb1c | 2.511132967 | 0.176184219 | 0.101887222 | 1.729208193 | 0.08377184 | NA | Itgad | 13.78773318 | 0.208233107 | 0.074459594 | 2.796592012 | 0.00516447 | NA |
| Klrb1f | 0.478448042 | -0.035850381 | 0.046036495 | -0.778738272 | 0.436133906 | NA | 2810011L19Rik | 584.6754935 | 0.20829863 | 0.079608252 | 2.616545693 | 0.008882447 | 0.179953183 |
| Klrc1 | 0.096071043 | -0.020098795 | 0.027825948 | -0.722304036 | 0.470107572 | NA | Rrp8 | 515.197705 | 0.208715887 | 0.081003486 | 2.576628452 | 0.009976913 | 0.19094795 |
| Klrc2 | 0 | NA | NA | NA | NA | NA | Slc17a6 | 1486.373668 | 0.208799191 | 0.094908018 | 2.200016333 | 0.027805736 | 0.310219758 |
| Klrc3 | 0 | NA | NA | NA | NA | NA | Sdc4 | 2422.379266 | 0.209963868 | 0.072150305 | 2.910089843 | 0.003613249 | 0.106640918 |
| Klrd1 | 1.248728633 | -0.11829835 | 0.084904083 | -1.393317575 | 0.163523778 | NA | Kcng4 | 472.4891411 | 0.210804095 | 0.087093336 | 2.420438858 | 0.015501786 | 0.236284305 |
| Klre1 | 1.200119047 | -0.119395654 | 0.078267184 | -1.525488039 | 0.127137415 | NA | Ndn | 7794.906231 | 0.212630117 | 0.077959315 | 2.727449792 | 0.006382597 | 0.154009309 |
| Klrg1 | 4.883981353 | 0.018926089 | 0.135593325 | 0.139579796 | 0.888992005 | NA | Gjb6 | 2405.202571 | 0.212850052 | 0.066464573 | 3.202458716 | 0.001362598 | 0.059797894 |
| Klri1 | 0.661489635 | -0.024113131 | 0.059861226 | -0.402817203 | 0.687082704 | NA | Pafah1b3 | 323.2204406 | 0.213364331 | 0.091093199 | 2.342264125 | 0.019167145 | 0.263564471 |
| Klri2 | 3.935442842 | 0.05164906 | 0.13296822 | 0.388431613 | 0.697696654 | NA | Zdbf2 | 1073.958188 | 0.213432504 | 0.094571249 | 2.256843443 | 0.024017859 | 0.29172307 |
| Klrk1 | 1.188686129 | 0.022501498 | 0.075096263 | 0.29963539 | 0.764455287 | NA | Ano2 | 296.1402496 | 0.215207965 | 0.094275919 | 2.282745864 | 0.022445342 | 0.281750998 |
| Kmo | 3.029795415 | -0.081131485 | 0.111186097 | -0.729690915 | 0.465579136 | NA | Socs6 | 458.3522098 | 0.215455111 | 0.072855317 | 2.957301118 | 0.003103449 | 0.097207883 |
| Kncn | 2.897474947 | 0.218710898 | 0.121601032 | 1.798594092 | 0.072082913 | NA | Lypd1 | 3488.447078 | 0.217165001 | 0.074965418 | 2.896869067 | 0.00376907 | 0.109938784 |
| Kng1 | 0 | NA | NA | NA | NA | NA | Ankrd34b | 978.8470846 | 0.219237363 | 0.074561299 | 2.940364039 | 0.003278268 | 0.099891275 |
| Kng2 | 0.176054697 | -0.002190495 | 0.031239146 | -0.070120197 | 0.944097991 | NA | Adra2a | 745.5339729 | 0.219722123 | 0.083145644 | 2.642617381 | 0.008226794 | 0.174077875 |
| Kpna7 | 0 | NA | NA | NA | NA | NA | Vipr2 | 44.254366 | 0.22044043 | 0.087659601 | 2.514732315 | 0.011912279 | 0.207728412 |
| Kprp | 0.292412586 | 0.032260595 | 0.035568585 | 0.906996868 | 0.364408454 | NA | Fam20a | 563.7032857 | 0.221179447 | 0.077068367 | 2.869912197 | 0.004105858 | 0.115138402 |
| Krt13 | 0.195033442 | -0.002424765 | 0.031596358 | -0.076741901 | 0.93882887 | NA | Zfp189 | 436.5412318 | 0.222156217 | 0.094994669 | 2.338617731 | 0.019355226 | 0.263564471 |
| Krt14 | 0.501644273 | 0.06509729 | 0.050114127 | 1.298980819 | 0.193950512 | NA | Bcl2 | 1172.74776 | 0.22253138 | 0.085521606 | 2.602048665 | 0.00926687 | 0.183393127 |
| Krt15 | 1.894032703 | 0.063778806 | 0.073840837 | 0.86373353 | 0.38773429 | NA | Tfap2d | 33.76859537 | 0.223086572 | 0.085440084 | 2.611029406 | 0.009027013 | 0.181364305 |
| Krt16 | 1.016873591 | -0.007411404 | 0.076118005 | -0.097367296 | 0.922434715 | NA | Ankrd55 | 171.9758203 | 0.223153253 | 0.094916391 | 2.351050764 | 0.01872048 | 0.261184487 |
| Krt17 | 3.942048268 | -0.116151483 | 0.131116868 | -0.88586224 | 0.375691766 | NA | Htr2a | 806.714062 | 0.223827884 | 0.084796948 | 2.639574756 | 0.008301011 | 0.174157793 |
| Krt18 | 18.54729659 | 0.509573785 | 0.144505286 | 3.52633319 | NA | NA | Magohb | 99.95561225 | 0.223946188 | 0.09412068 | 2.379351572 | 0.017343126 | 0.251926248 |
| Krt19 | 2.742119319 | -0.119798582 | 0.10298342 | -1.163280286 | 0.244715802 | NA | Ddit4 | 1569.912939 | 0.226492347 | 0.085501758 | 2.648978817 | 0.008073539 | 0.172465045 |
| Krt20 | 4.468513605 | -0.0499318 | 0.135565812 | -0.368321479 | 0.712633539 | NA | Slco4a1 | 110.0727752 | 0.228443397 | 0.093708367 | 2.437812178 | 0.014776449 | 0.232209004 |
| Krt23 | 0.748066559 | 0.059406784 | 0.047704345 | 1.24531181 | 0.213017156 | NA | Cdkn1a | 825.7757488 | 0.229460298 | 0.093466754 | 2.454993765 | 0.014088703 | 0.227385586 |
| Krt24 | 1.105906919 | 0.000490973 | 0.072693744 | 0.006753991 | 0.994611136 | NA | Ppara | 322.431572 | 0.229466169 | 0.076244667 | 3.009602886 | 0.002615895 | 0.086164366 |
| Krt25 | 2.731886909 | -0.173978101 | 0.104059339 | -1.671912429 | 0.094541591 | NA | Cort | 124.9854348 | 0.229716133 | 0.094534389 | 2.429974279 | 0.015099894 | 0.233312354 |
| Krt26 | 2.268360061 | -0.257914606 | 0.105798543 | -2.43778978 | 0.014777365 | NA | Stc1 | 178.4036789 | 0.229916696 | 0.090241206 | 2.547801684 | 0.010840408 | 0.198744008 |
| Krt27 | 0.116696964 | -0.020477583 | 0.027954224 | -0.732539847 | 0.463839131 | NA | Ppp1r3g | 368.5976202 | 0.231572211 | 0.09389517 | 2.466284584 | 0.013652282 | 0.224208584 |
| Krt28 | 3.243874308 | -0.020074229 | 0.11958889 | -0.167860318 | 0.866693169 | NA | Fndc9 | 189.0041547 | 0.232578015 | 0.0947512 | 2.454618134 | 0.014103432 | 0.227385586 |
| Krt31 | 0.280299629 | 0.041967777 | 0.040615819 | 1.033286501 | 0.301469842 | NA | Pcdha10 | 144.3092878 | 0.33458462 | 0.093277255 | 3.586990399 | 0.000334517 | 0.02373572 |
| Krt32 | 1.128431217 | -0.017479721 | 0.059519 | -0.293683037 | 0.769000118 | NA | Ccdc138 | 120.9142237 | 0.233451072 | 0.09128062 | 2.557509704 | 0.010542462 | 0.196333074 |
| Krt33a | 0.185906812 | 0.024895454 | 0.031385107 | 0.793225068 | 0.427646705 | NA | Xdh | 654.2369949 | 0.233866192 | 0.094262749 | 2.481003321 | 0.013101315 | 0.218495643 |
| Krt33b | 0.224413567 | -0.027287719 | 0.032077135 | -0.850690659 | 0.394941214 | NA | Ccdc19 | 85.65490628 | 0.234611037 | 0.095060727 | 2.468012236 | 0.013586568 | 0.223762449 |
| Krt34 | 0 | NA | NA | NA | NA | NA | Impg1 | 69.78512205 | 0.235225257 | 0.088853012 | 2.647352641 | 0.008112471 | 0.173042217 |
| Krt35 | 0.116696964 | -0.020477583 | 0.027954224 | -0.732539847 | 0.463839131 | NA | Trabd2b | 282.7496825 | 0.23595194 | 0.083448785 | 2.827505992 | 0.004691214 | 0.126898642 |
| Krt36 | 0 | NA | NA | NA | NA | NA | Rtp1 | 56.69441495 | 0.237172404 | 0.093361945 | 2.540354151 | 0.011074027 | 0.199827723 |
| Krt39 | 0 | NA | NA | NA | NA | NA | Ntng1 | 943.6368591 | 0.237215927 | 0.086143735 | 2.753722352 | 0.005892173 | 0.14630719 |
| Krt4 | 0.119397641 | -0.020477583 | 0.027954224 | -0.732539845 | 0.463839132 | NA | Atp4a | 36.59909863 | 0.326942083 | 0.090487223 | 3.61312981 | 0.000302523 | 0.022091286 |
| Krt40 | 0 | NA | NA | NA | NA | NA | F2r | 245.4550261 | 0.238379198 | 0.087442447 | 2.726126787 | 0.006408238 | 0.154049193 |
| Krt42 | 0.622661843 | -0.005421728 | 0.053328618 | -0.101666391 | 0.91902148 | NA | Tiparp | 798.917535 | 0.241091836 | 0.093860544 | 2.568617496 | 0.010210509 | 0.193453318 |
| Krt5 | 0.213790458 | -0.002646789 | 0.031909369 | -0.082947074 | 0.933893623 | NA | Slc2a1 | 5177.346636 | 0.241506656 | 0.088357674 | 2.73328445 | 0.006270615 | 0.151811594 |
| Krt6a | 0.363162105 | 0.038102764 | 0.038558083 | 0.988191333 | 0.323058952 | NA | Ret | 307.5358222 | 0.322376917 | 0.095101822 | 3.389808007 | 0.000699416 | 0.039105972 |
| Krt6b | 0 | NA | NA | NA | NA | NA | Pla2g3 | 131.2913536 | 0.243071012 | 0.094885645 | 2.561725876 | 0.010415349 | 0.195217237 |
| Krt7 | 4.331912649 | -0.085230925 | 0.133233088 | -0.639712899 | 0.522359268 | NA | Gpr1 | 35.22336774 | 0.244165747 | 0.09003859 | 2.711789979 | 0.006692098 | 0.156789377 |
| Krt71 | 0.183387883 | -0.001444816 | 0.031390672 | -0.046026918 | 0.963288796 | NA | Chrm2 | 332.8426123 | 0.247278485 | 0.079323527 | 3.117341041 | 0.001824903 | 0.070354982 |
| Krt72 | 0 | NA | NA | NA | NA | NA | Htr2c | 1970.943895 | 0.248577196 | 0.084513524 | 2.941271228 | 0.003268682 | 0.099891275 |
| Krt74 | 0 | NA | NA | NA | NA | NA | Gimap6 | 135.0918472 | 0.249146783 | 0.094420062 | 2.638706002 | 0.008322312 | 0.174157793 |
| Krt75 | 0.533252617 | -0.072906618 | 0.051718649 | -1.409677544 | 0.158634918 | NA | Tmem212 | 50.12157489 | 0.249812112 | 0.089530778 | 2.790237259 | 0.005266943 | 0.137476727 |
| Krt76 | 0.187700257 | -0.003168524 | 0.031449734 | -0.100748845 | 0.919749836 | NA | Vit | 111.1560213 | 0.25015084 | 0.094919265 | 2.635406408 | 0.00840366 | 0.174387945 |
| Krt78 | 1.113675977 | 0.006779323 | 0.078429329 | 0.086438619 | 0.931117748 | NA | Arrdc2 | 663.14207 | 0.317256942 | 0.091217117 | 3.478041758 | 0.000505091 | 0.031489078 |
| Krt79 | 0.185670226 | -0.001645086 | 0.031432796 | -0.052336624 | 0.958260472 | NA | Pcdh17 | 2655.50602 | 0.250933001 | 0.082611902 | 3.037492099 | 0.002385557 | 0.082506178 |
| Krt8 | 10.70122885 | 0.38933577 | 0.129842208 | 2.998530106 | NA | NA | Nlrp6 | 62.92323332 | 0.25171032 | 0.093300904 | 2.69783366 | 0.00697923 | 0.160386105 |
| Krt81 | 0 | NA | NA | NA | NA | NA | Dyrk3 | 366.3415844 | 0.316772986 | 0.087234237 | 3.631291972 | 0.000282006 | 0.021115557 |
| Krt82 | 0.079983654 | 0.016740032 | 0.027674178 | 0.604897175 | 0.54524732 | NA | Fam167a | 109.3146561 | 0.251959026 | 0.094278777 | 2.67248935 | 0.007529076 | 0.165959582 |
| Krt83 | 2.889392607 | -0.217218845 | 0.112683449 | -1.927690773 | 0.0538936 | NA | Fkbp5 | 1931.480402 | 0.315738016 | 0.082254524 | 3.838548923 | 0.000123764 | 0.011827562 |
| Krt84 | 0.094392817 | 0.017895942 | 0.027818137 | 0.643319236 | 0.520016977 | NA | Meig1 | 39.88273091 | 0.308155249 | 0.09319524 | 3.30655568 | 0.000944506 | 0.046581215 |
| Krt85 | 0.467810785 | 0.011930807 | 0.055036386 | 0.216780353 | 0.828379524 | NA | Ada | 29.91823767 | 0.30188568 | 0.083281807 | 3.624869472 | 0.000289107 | 0.02153628 |
| Krt86 | 0.32546763 | -0.049399607 | 0.042801411 | -1.154158374 | 0.248435245 | NA | Irs2 | 1706.125027 | 0.2603382 | 0.088015505 | 2.957867485 | 0.003097753 | 0.097207883 |
| Krtap1-3 | 0 | NA | NA | NA | NA | NA | Paqr5 | 238.1917029 | 0.261305131 | 0.087965018 | 2.970557337 | 0.002972599 | 0.09427942 |
| Krtap1-4 | 0 | NA | NA | NA | NA | NA | Camk1g | 1872.426373 | 0.297280932 | 0.087649508 | 3.391701088 | 0.000694602 | 0.039105972 |
| Krtap1-5 | 0 | NA | NA | NA | NA | NA | Syt9 | 646.4014909 | 0.264435817 | 0.089513364 | 2.954149028 | 0.003135326 | 0.097733347 |
| Krtap10-10 | 0.079983654 | 0.016740032 | 0.027674178 | 0.604897175 | 0.54524732 | NA | Galnt9 | 3416.658703 | 0.265566834 | 0.085300086 | 3.113324342 | 0.001849926 | 0.070715853 |
| Krtap10-4 | 0.116696964 | -0.020477583 | 0.027954224 | -0.732539847 | 0.463839131 | NA | Tppp3 | 1475.472121 | 0.265787757 | 0.086853772 | 3.060175185 | 0.002212076 | 0.079261025 |
| Krtap11-1 | 0.178993538 | 0.024685672 | 0.031260828 | 0.789667872 | 0.429721759 | NA | Prr5 | 281.8088457 | 0.296474453 | 0.090408788 | 3.279265883 | 0.001040775 | 0.049245282 |
| Krtap12-1 | 0 | NA | NA | NA | NA | NA | Pcdha1 | 58.83509195 | 0.266760675 | 0.092740425 | 2.876422808 | 0.004022106 | 0.113889095 |
| Krtap13 | 0 | NA | NA | NA | NA | NA | Fzd4 | 378.239814 | 0.269452118 | 0.084743342 | 3.179625812 | 0.001474653 | 0.062269811 |
| Krtap13-1 | 0 | NA | NA | NA | NA | NA | Htr1b | 276.8622648 | 0.275385781 | 0.094939463 | 2.900646077 | 0.003723942 | 0.109060453 |
| Krtap14 | 0 | NA | NA | NA | NA | NA | Nccrp1 | 13.81151309 | 0.276137572 | 0.064286334 | 4.295431911 | 1.74E-05 | NA |
| Krtap15 | 0 | NA | NA | NA | NA | NA | Klf15 | 1102.528966 | 0.289761012 | 0.079968569 | 3.62343627 | 0.000290715 | 0.021545531 |
| Krtap16-1 | 0 | NA | NA | NA | NA | NA | Emilin2 | 289.8845343 | 0.281732433 | 0.090365875 | 3.117686114 | 0.001822768 | 0.070354982 |
| Krtap16-3 | 0 | NA | NA | NA | NA | NA | Grp | 145.7184483 | 0.28440368 | 0.088936658 | 3.197822886 | 0.001384693 | 0.060402553 |
| Krtap17-1 | 0.912211048 | 0.041259068 | 0.075318195 | 0.547796821 | 0.583831422 | NA | Ovol2 | 249.6968497 | 0.285619939 | 0.08058864 | 3.544171231 | 0.000393849 | 0.02634142 |
| Krtap19-1 | 0 | NA | NA | NA | NA | NA | Rgs16 | 566.9801326 | 0.285470793 | 0.083343517 | 3.425230957 | 0.000614277 | 0.035388683 |
| Krtap19-3 | 0 | NA | NA | NA | NA | NA | Mfsd2a | 972.7537345 | 0.288421608 | 0.09115612 | 3.164039974 | 0.001555954 | 0.064392567 |
| Krtap19-4 | 0 | NA | NA | NA | NA | NA | Map3k6 | 241.8817111 | 0.288892165 | 0.094630947 | 3.052829708 | 0.002266945 | 0.080316218 |
| Krtap19-5 | 0 | NA | NA | NA | NA | NA | Cd59a | 594.4598378 | 0.289009248 | 0.090203248 | 3.203978299 | 0.001355427 | 0.059797894 |
| Krtap19-9b | 0 | NA | NA | NA | NA | NA | Tekt4 | 42.82700988 | 0.277008106 | 0.083244912 | 3.327628056 | 0.000875887 | 0.044373556 |
| Krtap2-4 | 0 | NA | NA | NA | NA | NA | Gpr123 | 3338.540836 | 0.265997824 | 0.068919093 | 3.859566541 | 0.000113588 | 0.011329455 |
| Krtap20-2 | 0 | NA | NA | NA | NA | NA | Ppapdc2 | 2005.274858 | 0.261655261 | 0.065828824 | 3.974782559 | 7.04E-05 | 0.008130208 |
| Krtap21-1 | 0 | NA | NA | NA | NA | NA | Sgk1 | 4723.226973 | 0.298356571 | 0.09337943 | 3.195099517 | 0.001397826 | 0.06073864 |
| Krtap22-2 | 0 | NA | NA | NA | NA | NA | Plin4 | 662.3437959 | 0.257343905 | 0.077920633 | 3.302641367 | 0.000957788 | 0.047002797 |
| Krtap24-1 | 0 | NA | NA | NA | NA | NA | Rhou | 2076.716139 | 0.257145582 | 0.053811914 | 4.778599408 | 1.77E-06 | 0.000545561 |
| Krtap26-1 | 0 | NA | NA | NA | NA | NA | Hs6st2 | 1403.325246 | 0.25377825 | 0.073026783 | 3.475139404 | 0.000510588 | 0.031560853 |
| Krtap27-1 | 0 | NA | NA | NA | NA | NA | B3galt5 | 2862.056054 | 0.251852883 | 0.066802574 | 3.770107454 | 0.000163177 | 0.013732714 |
| Krtap3-1 | 0 | NA | NA | NA | NA | NA | Fzd2 | 553.8499483 | 0.250405675 | 0.076523399 | 3.272275886 | 0.001066854 | 0.049990721 |
| Krtap3-2 | 0 | NA | NA | NA | NA | NA | Arid5b | 1101.463601 | 0.241584838 | 0.061786876 | 3.909970103 | 9.23E-05 | 0.009815749 |
| Krtap3-3 | 0 | NA | NA | NA | NA | NA | Lfng | 646.6805627 | 0.237721572 | 0.07241899 | 3.282586142 | 0.001028596 | 0.048828035 |
| Krtap31-1 | 0 | NA | NA | NA | NA | NA | Rhbdl3 | 969.2947975 | 0.233042119 | 0.070849409 | 3.289259906 | 0.001004512 | 0.048083105 |
| Krtap31-2 | 0 | NA | NA | NA | NA | NA | Endod1 | 5377.441772 | 0.205182169 | 0.062375042 | 3.289491453 | 0.001003686 | 0.048083105 |
| Krtap4-1 | 0 | NA | NA | NA | NA | NA | Bcat1 | 4427.138739 | 0.204071611 | 0.060623612 | 3.366206752 | 0.000762096 | 0.041306721 |
| Krtap4-13 | 0 | NA | NA | NA | NA | NA | Rab3c | 17590.22349 | 0.202413165 | 0.060458431 | 3.34797248 | 0.000814051 | 0.042689184 |
| Krtap4-16 | 0.211846316 | 0.025707621 | 0.031860375 | 0.806883811 | 0.419733426 | NA | Ndufc2 | 1894.914091 | 0.202267486 | 0.060774125 | 3.328184267 | 0.00087414 | 0.044373556 |
| Krtap4-2 | 0 | NA | NA | NA | NA | NA | Cables1 | 1103.82638 | 0.192055811 | 0.056357605 | 3.407806477 | 0.000654873 | 0.037158939 |
| Krtap4-6 | 0 | NA | NA | NA | NA | NA | Tmeff1 | 2931.701249 | 0.190124086 | 0.051944916 | 3.660109584 | 0.000252107 | 0.019720421 |
| Krtap4-7 | 0 | NA | NA | NA | NA | NA | Akt2 | 4959.349386 | 0.183634213 | 0.054628424 | 3.361514027 | 0.000775164 | 0.041703834 |
| Krtap4-8 | 0 | NA | NA | NA | NA | NA | Bcl2l1 | 4645.653376 | 0.182224204 | 0.051151727 | 3.56242523 | 0.000367445 | 0.025538277 |
| Krtap4-9 | 0 | NA | NA | NA | NA | NA | Kbtbd4 | 1330.902444 | 0.173205942 | 0.052649376 | 3.289800468 | 0.001002585 | 0.048083105 |
| Krtap5-1 | 0.105923158 | 0.018524693 | 0.027977518 | 0.662127842 | 0.507889294 | NA | Pja1 | 7886.888833 | 0.171350617 | 0.047790407 | 3.585460475 | 0.000336484 | 0.02373572 |
| Krtap5-2 | 0.211846316 | 0.025707621 | 0.031860375 | 0.806883811 | 0.419733426 | NA | Stip1 | 7185.759144 | 0.147588959 | 0.044766962 | 3.296827638 | 0.000977835 | 0.047664529 |
| Krtap5-3 | 0 | NA | NA | NA | NA | NA | Sccpdh | 2583.297034 | 0.139299815 | 0.042118497 | 3.3073311 | 0.000941895 | 0.046581215 |
| Krtap5-4 | 0 | NA | NA | NA | NA | NA | Gpi1 | 26520.80069 | 0.136990162 | 0.039929322 | 3.430816113 | 0.000601768 | 0.03496515 |
| Krtap5-5 | 0.107716603 | -0.020477583 | 0.027954224 | -0.732539849 | 0.46383913 | NA | Gfra1 | 1810.269635 | 1.49E-05 | 0.054068807 | 0.00027627 | 0.999779569 | 0.999917241 |
| Krtap6-1 | 0 | NA | NA | NA | NA | NA | Rnf183 | 0.976674389 | 3.07E-05 | 0.024493265 | 0.001252459 | 0.999000683 | NA |
| Krtap6-2 | 0 | NA | NA | NA | NA | NA | 4932415M13Rik | 0.714984696 | 3.32E-05 | 0.015894788 | 0.002087505 | 0.998334413 | NA |
| Krtap6-5 | 0 | NA | NA | NA | NA | NA | Il11ra1 | 646.1591078 | 3.36E-05 | 0.082655616 | 0.000406129 | 0.999675956 | 0.999917241 |
| Krtap7-1 | 0.341676965 | 0.050095582 | 0.044235127 | 1.132484193 | 0.257430926 | NA | Ank3 | 10207.45761 | 4.82E-05 | 0.071912155 | 0.000669607 | 0.999465731 | 0.999917241 |
| Krtap8-1 | 0 | NA | NA | NA | NA | NA | Sept1 | 28.48284513 | 6.74E-06 | 0.085629408 | 7.88E-05 | 0.999937155 | 0.999947932 |
| Krtap9-1 | 0 | NA | NA | NA | NA | NA | Ubap2l | 5705.227779 | 7.03E-05 | 0.044657184 | 0.001575318 | 0.998743079 | 0.999637701 |
| Krtap9-3 | 0 | NA | NA | NA | NA | NA | Atic | 1283.481602 | 8.42E-05 | 0.061106417 | 0.001377499 | 0.998900915 | 0.999662491 |
| Krtap9-5 | 0 | NA | NA | NA | NA | NA | Tfcp2l1 | 403.8052919 | 9.08E-05 | 0.078739634 | 0.001153502 | 0.999079639 | 0.9997679 |
| Krtdap | 0.17463368 | 0.024560283 | 0.031186242 | 0.787535833 | 0.430968261 | NA | Opn1mw | 16.8867866 | 9.17E-05 | 0.068435811 | 0.001339296 | 0.998931397 | NA |
| Kynu | 4.014358091 | -0.026246882 | 0.125619344 | -0.208939807 | 0.834495228 | NA | 0610008F07Rik | 0 | NA | NA | NA | NA | NA |
| L1td1 | 4.493681656 | -0.106457821 | 0.123886038 | -0.859320573 | 0.390163677 | NA | 0610031O16Rik | 0 | NA | NA | NA | NA | NA |
| LOC100038947 | 0.203787646 | -0.02671144 | 0.031728022 | -0.841887971 | 0.399850664 | NA | 1100001G20Rik | 0 | NA | NA | NA | NA | NA |
| LOC100043315 | 0.41925633 | -0.060548742 | 0.047272363 | -1.280848629 | 0.20024684 | NA | 1110025L11Rik | 0 | NA | NA | NA | NA | NA |
| LOC100048884 | 0 | NA | NA | NA | NA | NA | 1110028F18Rik | 0 | NA | NA | NA | NA | NA |
| LOC100502924 | 0 | NA | NA | NA | NA | NA | 1600015I10Rik | 0 | NA | NA | NA | NA | NA |
| LOC100503280 | 0.305724365 | 0.009518016 | 0.039476503 | 0.241105846 | 0.809473082 | NA | 1600019K03Rik | 0 | NA | NA | NA | NA | NA |
| LOC100504608 | 1.228183666 | -0.022943163 | 0.073840375 | -0.31071298 | 0.756018829 | NA | 1600025M17Rik | 0 | NA | NA | NA | NA | NA |
| LOC100861615 | 6.091513533 | -0.048265545 | 0.135024574 | -0.357457486 | 0.720749348 | NA | 1700001D01Rik | 0 | NA | NA | NA | NA | NA |
| LOC100861978 | 0 | NA | NA | NA | NA | NA | 1700001F09Rik | 0 | NA | NA | NA | NA | NA |
| LOC171588 | 5.818998857 | -0.020013229 | 0.148150586 | -0.135087074 | 0.89254303 | NA | 1700003C15Rik | 0 | NA | NA | NA | NA | NA |
| LOC547349 | 0.105923158 | 0.018524693 | 0.027977518 | 0.662127842 | 0.507889294 | NA | 1700003F17Rik | 0 | NA | NA | NA | NA | NA |
| LOC666331 | 0 | NA | NA | NA | NA | NA | 1700003G13Rik | 0 | NA | NA | NA | NA | NA |
| Lactbl1 | 0.593755747 | -0.004769304 | 0.058238563 | -0.081892538 | 0.934732168 | NA | 1700003H04Rik | 0 | NA | NA | NA | NA | NA |
| Lalba | 0.798357233 | -0.007830999 | 0.062135221 | -0.126031561 | 0.899706942 | NA | 1700003P14Rik | 0 | NA | NA | NA | NA | NA |
| Lamp3 | 0 | NA | NA | NA | NA | NA | 1700006A11Rik | 0 | NA | NA | NA | NA | NA |
| Lao1 | 6.420233442 | 0.088157996 | 0.116576444 | 0.756224782 | 0.449514437 | NA | 1700006E09Rik | 0 | NA | NA | NA | NA | NA |
| Lbx1 | 0.079983654 | 0.016740032 | 0.027674178 | 0.604897175 | 0.54524732 | NA | 1700006H21Rik | 0 | NA | NA | NA | NA | NA |
| Lbx2 | 1.310361776 | -0.029823595 | 0.087516462 | -0.340776974 | 0.733271486 | NA | 1700008A04Rik | 0 | NA | NA | NA | NA | NA |
| Lce1a1 | 0 | NA | NA | NA | NA | NA | 1700008K24Rik | 0 | NA | NA | NA | NA | NA |
| Lce1a2 | 0 | NA | NA | NA | NA | NA | 1700008P02Rik | 0 | NA | NA | NA | NA | NA |
| Lce1b | 0 | NA | NA | NA | NA | NA | 1700009N14Rik | 0 | NA | NA | NA | NA | NA |
| Lce1c | 0 | NA | NA | NA | NA | NA | 1700010D01Rik | 0 | NA | NA | NA | NA | NA |
| Lce1d | 0 | NA | NA | NA | NA | NA | 1700010I02Rik | 0 | NA | NA | NA | NA | NA |
| Lce1e | 0 | NA | NA | NA | NA | NA | 1700011A15Rik | 0 | NA | NA | NA | NA | NA |
| Lce1f | 0 | NA | NA | NA | NA | NA | 1700011B04Rik | 0 | NA | NA | NA | NA | NA |
| Lce1g | 0.17463368 | 0.024560283 | 0.031186242 | 0.787535833 | 0.430968261 | NA | 1700011M02Rik | 0 | NA | NA | NA | NA | NA |
| Lce1h | 0 | NA | NA | NA | NA | NA | 1700012A03Rik | 0 | NA | NA | NA | NA | NA |
| Lce1i | 0 | NA | NA | NA | NA | NA | 1700012B07Rik | 0 | NA | NA | NA | NA | NA |
| Lce1k | 0 | NA | NA | NA | NA | NA | 1700012I11Rik | 0 | NA | NA | NA | NA | NA |
| Lce1l | 0 | NA | NA | NA | NA | NA | 1700013D24Rik | 0 | NA | NA | NA | NA | NA |
| Lce1m | 0 | NA | NA | NA | NA | NA | 1700013G24Rik | 0 | NA | NA | NA | NA | NA |
| Lce3a | 0.302141032 | -0.047437531 | 0.04196367 | -1.130442867 | 0.25828966 | NA | 1700015E13Rik | 0 | NA | NA | NA | NA | NA |
| Lce3b | 0 | NA | NA | NA | NA | NA | 1700015G11Rik | 0 | NA | NA | NA | NA | NA |
| Lce3c | 0.717316046 | 0.045366621 | 0.050111211 | 0.905318791 | 0.365296524 | NA | 1700016C15Rik | 0 | NA | NA | NA | NA | NA |
| Lce3d | 0 | NA | NA | NA | NA | NA | 1700016L21Rik | 0 | NA | NA | NA | NA | NA |
| Lce3e | 0 | NA | NA | NA | NA | NA | 1700016P04Rik | 0 | NA | NA | NA | NA | NA |
| Lce3f | 1.181425567 | -0.106516642 | 0.066956276 | -1.590838815 | 0.111645855 | NA | 1700017D01Rik | 0 | NA | NA | NA | NA | NA |
| Lce6a | 0 | NA | NA | NA | NA | NA | 1700017N19Rik | 0 | NA | NA | NA | NA | NA |
| Lcn10 | 0 | NA | NA | NA | NA | NA | 1700018B08Rik | 0 | NA | NA | NA | NA | NA |
| Lcn11 | 0.593158268 | 0.071669397 | 0.052434572 | 1.366834783 | 0.171677098 | NA | 1700018C11Rik | 0 | NA | NA | NA | NA | NA |
| Lcn12 | 0 | NA | NA | NA | NA | NA | 1700018G05Rik | 0 | NA | NA | NA | NA | NA |
| Lcn3 | 0.293623414 | 0.010408489 | 0.039073364 | 0.266383235 | 0.789944076 | NA | 1700019B21Rik | 0 | NA | NA | NA | NA | NA |
| Lcn4 | 0 | NA | NA | NA | NA | NA | 1700019E08Rik | 0 | NA | NA | NA | NA | NA |
| Lcn5 | 0 | NA | NA | NA | NA | NA | 1700019M22Rik | 0 | NA | NA | NA | NA | NA |
| Lcn6 | 0 | NA | NA | NA | NA | NA | 1700020A23Rik | 0 | NA | NA | NA | NA | NA |
| Lcn8 | 0.238795281 | -0.027601606 | 0.032267774 | -0.855392297 | 0.392334007 | NA | 1700020M21Rik | 0 | NA | NA | NA | NA | NA |
| Lcn9 | 0 | NA | NA | NA | NA | NA | 1700020N15Rik | 0 | NA | NA | NA | NA | NA |
| Ldhal6b | 7.216227394 | -0.168309871 | 0.154698833 | -1.087984105 | 0.276602124 | NA | 1700022A21Rik | 0 | NA | NA | NA | NA | NA |
| Ldhc | 0.994426385 | 0.049832203 | 0.071188917 | 0.699999454 | 0.483927646 | NA | 1700022A22Rik | 0 | NA | NA | NA | NA | NA |
| Ldlrad1 | 0.239950962 | 0.027417169 | 0.032831728 | 0.835081494 | 0.403671825 | NA | 1700022E09Rik | 0 | NA | NA | NA | NA | NA |
| Ldlrad2 | 2.087130886 | 0.110319212 | 0.101415702 | 1.087792229 | 0.276686841 | NA | 1700022H16Rik | 0 | NA | NA | NA | NA | NA |
| Leap2 | 0.305079873 | -0.015054679 | 0.039642215 | -0.379763828 | 0.704120735 | NA | 1700023C21Rik | 0 | NA | NA | NA | NA | NA |
| Lect2 | 0 | NA | NA | NA | NA | NA | 1700024B18Rik | 0 | NA | NA | NA | NA | NA |
| Lelp1 | 0 | NA | NA | NA | NA | NA | 1700024P03Rik | 0 | NA | NA | NA | NA | NA |
| Lep | 0 | NA | NA | NA | NA | NA | 1700024P04Rik | 0 | NA | NA | NA | NA | NA |
| Lgals2 | 2.826545916 | -0.059761391 | 0.122620378 | -0.487369159 | 0.625996745 | NA | 1700025B11Rik | 0 | NA | NA | NA | NA | NA |
| Lgals4 | 7.737277884 | 0.425836631 | 0.156769947 | 2.716315469 | 0.006601298 | NA | 1700025F22Rik | 0 | NA | NA | NA | NA | NA |
| Lgals6 | 0.60415196 | 0.020563279 | 0.052843793 | 0.389133293 | 0.697177546 | NA | 1700025K24Rik | 0 | NA | NA | NA | NA | NA |
| Lgals7 | 3.880974019 | 0.009929444 | 0.131055042 | 0.075765448 | 0.939605706 | NA | 1700025M24Rik | 0 | NA | NA | NA | NA | NA |
| Lgsn | 0.473045832 | 0.03947874 | 0.051663738 | 0.764147964 | 0.444779072 | NA | 1700026F02Rik | 0 | NA | NA | NA | NA | NA |
| Lhb | 2.559019543 | 0.252937408 | 0.107532361 | 2.352198041 | 0.018662836 | NA | 1700027F09Rik | 0 | NA | NA | NA | NA | NA |
| Lhfpl1 | 4.535166914 | -0.100523214 | 0.120567251 | -0.833752223 | 0.404420623 | NA | 1700028D13Rik | 0 | NA | NA | NA | NA | NA |
| Lhx1 | 5.917210998 | -0.18460631 | 0.124027708 | -1.488427972 | 0.136638064 | NA | 1700028I16Rik | 0 | NA | NA | NA | NA | NA |
| Lhx3 | 5.555580945 | 0.322154567 | 0.119185858 | 2.702959659 | 0.006872509 | NA | 1700028M03Rik | 0 | NA | NA | NA | NA | NA |
| Lhx4 | 0.426277686 | 0.058476664 | 0.047630606 | 1.227711936 | 0.219555124 | NA | 1700028P15Rik | 0 | NA | NA | NA | NA | NA |
| Lilra6 | 0.990536273 | -0.061771697 | 0.067614392 | -0.913587997 | 0.360933376 | NA | 1700029B22Rik | 0 | NA | NA | NA | NA | NA |
| Lim2 | 0 | NA | NA | NA | NA | NA | 1700029F12Rik | 0 | NA | NA | NA | NA | NA |
| Lin28a | 0.514098028 | 0.006684456 | 0.05090618 | 0.131309328 | 0.895530613 | NA | 1700029J03Rik | 0 | NA | NA | NA | NA | NA |
| Lipf | 0.338960846 | 0.036479516 | 0.037750094 | 0.966342396 | 0.333872878 | NA | 1700029N11Rik | 0 | NA | NA | NA | NA | NA |
| Lipi | 0.300347587 | -0.013446771 | 0.039455705 | -0.340806761 | 0.73324906 | NA | 1700029P11Rik | 0 | NA | NA | NA | NA | NA |
| Lipk | 0.238795281 | -0.027601606 | 0.032267774 | -0.855392297 | 0.392334007 | NA | 1700030N03Rik | 0 | NA | NA | NA | NA | NA |
| Lipm | 3.142511058 | 0.230372281 | 0.109191951 | 2.109791775 | 0.034876295 | NA | 1700030O20Rik | 0 | NA | NA | NA | NA | NA |
| Lipn | 0 | NA | NA | NA | NA | NA | 1700031A10Rik | 0 | NA | NA | NA | NA | NA |
| Lman1l | 1.393655212 | 0.138459737 | 0.077590734 | 1.784488049 | 0.074344395 | NA | 1700031F05Rik | 0 | NA | NA | NA | NA | NA |
| Lmod2 | 0.094392817 | 0.017895942 | 0.027818137 | 0.643319236 | 0.520016977 | NA | 1700031P21Rik | 0 | NA | NA | NA | NA | NA |
| Lmod3 | 1.843987541 | -0.004747097 | 0.087084654 | -0.054511294 | 0.956527811 | NA | 1700034E13Rik | 0 | NA | NA | NA | NA | NA |
| Lmx1a | 5.016530087 | 0.264831897 | 0.10694634 | 2.476306323 | NA | NA | 1700034K08Rik | 0 | NA | NA | NA | NA | NA |
| Lmx1b | 0.819267527 | -0.011522036 | 0.055657027 | -0.207018536 | 0.835995383 | NA | 1700034O15Rik | 0 | NA | NA | NA | NA | NA |
| Loxl4 | 6.44770556 | 0.002642171 | 0.124206831 | 0.021272352 | 0.983028399 | NA | 1700041C23Rik | 0 | NA | NA | NA | NA | NA |
| Lpcat2b | 1.298488313 | -0.022962054 | 0.080392725 | -0.285623537 | 0.775166474 | NA | 1700041M19Rik | 0 | NA | NA | NA | NA | NA |
| Lrat | 5.053319707 | -0.021652086 | 0.106831773 | -0.202674591 | 0.839389384 | NA | 1700042G07Rik | 0 | NA | NA | NA | NA | NA |
| Lrch4 | 0 | NA | NA | NA | NA | NA | 1700044C05Rik | 0 | NA | NA | NA | NA | NA |
| Lrcol1 | 0.107716603 | -0.020477583 | 0.027954224 | -0.732539849 | 0.46383913 | NA | 1700044K03Rik | 0 | NA | NA | NA | NA | NA |
| Lrg1 | 49.01134731 | 0.271078349 | 0.16780799 | 1.615407876 | NA | NA | 1700046C09Rik | 0 | NA | NA | NA | NA | NA |
| Lrit1 | 0.515459553 | -0.019052845 | 0.045554446 | -0.418243369 | 0.67576919 | NA | 1700047G03Rik | 0 | NA | NA | NA | NA | NA |
| Lrit2 | 0.293623414 | 0.010408489 | 0.039073364 | 0.266383235 | 0.789944076 | NA | 1700047I17Rik2 | 0 | NA | NA | NA | NA | NA |
| Lrit3 | 2.173935032 | 0.14230611 | 0.107900473 | 1.318864558 | 0.187214397 | NA | 1700047L14Rik | 0 | NA | NA | NA | NA | NA |
| Lrr1 | 1.045045392 | -0.104975171 | 0.076446864 | -1.37317825 | 0.169696939 | NA | 1700048M11Rik | 0 | NA | NA | NA | NA | NA |
| Lrrc10 | 1.328786961 | 0.04053153 | 0.081204073 | 0.499131732 | 0.617686584 | NA | 1700049E15Rik | 0 | NA | NA | NA | NA | NA |
| Lrrc15 | 1.154903561 | -0.039247414 | 0.080272433 | -0.488927671 | 0.624892905 | NA | 1700049E22Rik | 0 | NA | NA | NA | NA | NA |
| Lrrc19 | 0 | NA | NA | NA | NA | NA | 1700049L16Rik | 0 | NA | NA | NA | NA | NA |
| Lrrc26 | 0.287139021 | 0.008693067 | 0.033441491 | 0.259948559 | 0.794903454 | NA | 1700052I22Rik | 0 | NA | NA | NA | NA | NA |
| Lrrc30 | 0 | NA | NA | NA | NA | NA | 1700054A03Rik | 0 | NA | NA | NA | NA | NA |
| Lrrc34 | 4.067439929 | 0.197196167 | 0.127441899 | 1.547341726 | 0.121780865 | NA | 1700054K19Rik | 0 | NA | NA | NA | NA | NA |
| Lrrc52 | 0.603663061 | -0.004610385 | 0.052400067 | -0.087984343 | 0.92988912 | NA | 1700054O13Rik | 0 | NA | NA | NA | NA | NA |
| Lrrc58 | 4206.490262 | -0.008980542 | 0.135290053 | -0.066379912 | NA | NA | 1700055C04Rik | 0 | NA | NA | NA | NA | NA |
| Lrrc63 | 0.903760442 | 0.00255845 | 0.070410694 | 0.036336097 | 0.971014367 | NA | 1700057G04Rik | 0 | NA | NA | NA | NA | NA |
| Lrrc66 | 1.166819058 | 0.045440474 | 0.077118192 | 0.58923158 | 0.555705935 | NA | 1700061F12Rik | 0 | NA | NA | NA | NA | NA |
| Lrrc69 | 2.091909967 | 0.054846023 | 0.107070421 | 0.512242529 | 0.608481282 | NA | 1700063A18Rik | 0 | NA | NA | NA | NA | NA |
| Lrrc72 | 0.211846316 | 0.025707621 | 0.031860375 | 0.806883811 | 0.419733426 | NA | 1700064J06Rik | 0 | NA | NA | NA | NA | NA |
| Lrrc8e | 0.789559445 | 0.014020811 | 0.056659565 | 0.247457092 | 0.804554494 | NA | 1700065I16Rik | 0 | NA | NA | NA | NA | NA |
| Lrriq4 | 0.275017096 | 0.008844542 | 0.038389071 | 0.2303922 | 0.817787022 | NA | 1700065O20Rik | 0 | NA | NA | NA | NA | NA |
| Lrtm1 | 0.295716657 | -0.010253415 | 0.03372058 | -0.304069933 | 0.761074613 | NA | 1700066N21Rik | 0 | NA | NA | NA | NA | NA |
| Lta | 5.73735428 | -0.10244084 | 0.13750598 | -0.744991892 | 0.456276622 | NA | 1700066O22Rik | 0 | NA | NA | NA | NA | NA |
| Ltb4r1 | 4.365710893 | 0.055336194 | 0.135407208 | 0.408665052 | 0.682785484 | NA | 1700067G17Rik | 0 | NA | NA | NA | NA | NA |
| Ltb4r2 | 2.065653484 | 0.051845857 | 0.096794205 | 0.535629757 | 0.592214459 | NA | 1700069P05Rik | 0 | NA | NA | NA | NA | NA |
| Luzp4 | 0 | NA | NA | NA | NA | NA | 1700072B07Rik | 0 | NA | NA | NA | NA | NA |
| Ly6c2 | 1.112398198 | -0.030243582 | 0.06703571 | -0.451156285 | 0.651876914 | NA | 1700072O05Rik | 0 | NA | NA | NA | NA | NA |
| Ly6f | 0 | NA | NA | NA | NA | NA | 1700074P13Rik | 0 | NA | NA | NA | NA | NA |
| Ly6g5c | 0 | NA | NA | NA | NA | NA | 1700080E11Rik | 0 | NA | NA | NA | NA | NA |
| Ly6g6c | 0 | NA | NA | NA | NA | NA | 1700080N15Rik | 0 | NA | NA | NA | NA | NA |
| Ly6i | 0.484297991 | 0.036542974 | 0.051863038 | 0.704605351 | 0.481055868 | NA | 1700081H04Rik | 0 | NA | NA | NA | NA | NA |
| Ly6k | 0.982137931 | 0.021872688 | 0.07729474 | 0.282977698 | 0.777193933 | NA | 1700081H22Rik | 0 | NA | NA | NA | NA | NA |
| Ly9 | 43.63071773 | 0.8464765 | 0.167386621 | 5.057014072 | NA | NA | 1700084F23Rik | 0 | NA | NA | NA | NA | NA |
| Lyg1 | 0 | NA | NA | NA | NA | NA | 1700084J12Rik | 0 | NA | NA | NA | NA | NA |
| Lyg2 | 2.377896396 | 0.067007288 | 0.102529546 | 0.653541248 | 0.513407407 | NA | 1700085B03Rik | 0 | NA | NA | NA | NA | NA |
| Lypd2 | 4.849282279 | -0.099581706 | 0.134744639 | -0.73904021 | 0.459882581 | NA | 1700092C10Rik | 0 | NA | NA | NA | NA | NA |
| Lypd3 | 0.362333936 | 0.019030129 | 0.042103636 | 0.451983023 | 0.651281213 | NA | 1700092E19Rik | 0 | NA | NA | NA | NA | NA |
| Lypd4 | 0 | NA | NA | NA | NA | NA | 1700094M24Rik | 0 | NA | NA | NA | NA | NA |

| | | | | | | | | | | | | | | | |
|---|---|---|---|---|---|---|---|---|---|---|---|---|---|---|---|
| Lypd5 | 0 | NA | NA | NA | NA | NA | | | 1700095A21Rik | 0 | NA | NA | NA | NA | NA |
| Lypd8 | 0.287795519 | 0.031925978 | 0.035387605 | 0.902179657 | 0.366961439 | NA | | | 1700100L14Rik | 0 | NA | NA | NA | NA | NA |
| Lyzl1 | 0 | NA | NA | NA | NA | NA | | | 1700101O22Rik | 0 | NA | NA | NA | NA | NA |
| Lyzl6 | 0.907724086 | -0.087049886 | 0.072820354 | -1.19540596 | 0.231928458 | NA | | | 1700106J16Rik | 0 | NA | NA | NA | NA | NA |
| M1ap | 1.325353327 | 0.073222765 | 0.086063402 | 0.850800263 | 0.394880316 | NA | | | 1700108J01Rik | 0 | NA | NA | NA | NA | NA |
| Mab21l3 | 3.198626589 | 0.014856769 | 0.105264105 | 0.141138032 | 0.887760899 | NA | | | 1700109F18Rik | 0 | NA | NA | NA | NA | NA |
| Macc1 | 4.742395832 | -0.142333648 | 0.13380872 | -1.063709807 | 0.287460181 | NA | | | 1700109G15Rik | 0 | NA | NA | NA | NA | NA |
| Madcam1 | 2.160123571 | 0.099281291 | 0.100597065 | 0.986920356 | 0.323681684 | NA | | | 1700109I08Rik | 0 | NA | NA | NA | NA | NA |
| Mael | 2.721507992 | 0.073588115 | 0.09825567 | 0.748945223 | 0.453890222 | NA | | | 1700110I01Rik | 0 | NA | NA | NA | NA | NA |
| Maff | 69.17769177 | 0.257166854 | 0.117863747 | 2.181899534 | NA | NA | | | 1700111N16Rik | 0 | NA | NA | NA | NA | NA |
| Magea1 | 0 | NA | NA | NA | NA | NA | | | 1700112H15Rik | 0 | NA | NA | NA | NA | NA |
| Magea10 | 0.17833704 | -0.002390466 | 0.031282281 | -0.076415987 | 0.939088151 | NA | | | 1700112J05Rik | 0 | NA | NA | NA | NA | NA |
| Magea2 | 0 | NA | NA | NA | NA | NA | | | 1700120E14Rik | 0 | NA | NA | NA | NA | NA |
| Magea3 | 0 | NA | NA | NA | NA | NA | | | 1700121L16Rik | 0 | NA | NA | NA | NA | NA |
| Magea4 | 0 | NA | NA | NA | NA | NA | | | 1700123O12Rik | 0 | NA | NA | NA | NA | NA |
| Magea5 | 0 | NA | NA | NA | NA | NA | | | 1700126H18Rik | 0 | NA | NA | NA | NA | NA |
| Magea6 | 0 | NA | NA | NA | NA | NA | | | 1700128A07Rik | 0 | NA | NA | NA | NA | NA |
| Magea8 | 0 | NA | NA | NA | NA | NA | | | 1700128F08Rik | 0 | NA | NA | NA | NA | NA |
| Mageb1 | 0 | NA | NA | NA | NA | NA | | | 1700129C05Rik | 0 | NA | NA | NA | NA | NA |
| Mageb16 | 0 | NA | NA | NA | NA | NA | | | 1810006J02Rik | 0 | NA | NA | NA | NA | NA |
| Mageb16-ps1 | 0.258320694 | 0.006870708 | 0.033097595 | 0.207589347 | 0.835549623 | NA | | | 1810007I06Rik | 0 | NA | NA | NA | NA | NA |
| Mageb18 | 0.413070119 | -0.002318447 | 0.042591244 | -0.054434824 | 0.956588734 | NA | | | 1810008I18Rik | 0 | NA | NA | NA | NA | NA |
| Mageb2 | 0 | NA | NA | NA | NA | NA | | | 1810009J06Rik | 0 | NA | NA | NA | NA | NA |
| Mageb3 | 3.848484583 | 0.123354358 | 0.116432672 | 1.059447962 | 0.289395816 | NA | | | 1810013A23Rik | 0 | NA | NA | NA | NA | NA |
| Mageb4 | 0.188785634 | 0.025025111 | 0.031461603 | 0.79541754 | 0.42637066 | NA | | | 1810018F18Rik | 0 | NA | NA | NA | NA | NA |
| Mageb5 | 0 | NA | NA | NA | NA | NA | | | 1810035I16Rik | 0 | NA | NA | NA | NA | NA |
| Mansc4 | 6.531528774 | 0.118896125 | 0.134343551 | 0.885015499 | 0.376148272 | NA | | | 1810053B23Rik | 0 | NA | NA | NA | NA | NA |
| 1-Mar | 3.674522383 | 0.051839156 | 0.132232611 | 0.392030044 | 0.695036015 | NA | | | 1810065E05Rik | 0 | NA | NA | NA | NA | NA |
| Marcksl1-ps4 | 0.793177326 | -0.012109046 | 0.055799279 | -0.217010791 | 0.828199935 | NA | | | 2010003K11Rik | 0 | NA | NA | NA | NA | NA |
| Marco | 0.19442443 | -0.026451352 | 0.031568725 | -0.837897386 | 0.402088337 | NA | | | 2010009K17Rik | 0 | NA | NA | NA | NA | NA |
| Mat1a | 1.361257865 | -0.000613543 | 0.081854933 | -0.007495494 | 0.994019517 | NA | | | 2010310C07Rik | 0 | NA | NA | NA | NA | NA |
| Matn3 | 2.865118358 | -0.089187609 | 0.118298064 | -0.753922813 | 0.450895572 | NA | | | 2210019I11Rik | 0 | NA | NA | NA | NA | NA |
| Mb | 5.31222965 | 0.13556448 | 0.121250778 | 1.118050393 | 0.263545471 | NA | | | 2210409E12Rik | 0 | NA | NA | NA | NA | NA |
| Mbd3l1 | 0 | NA | NA | NA | NA | NA | | | 2210414B05Rik | 0 | NA | NA | NA | NA | NA |
| Mbd3l2 | 0.099009884 | 0.018205726 | 0.027896865 | 0.652608315 | 0.514008823 | NA | | | 2210420H20Rik | 0 | NA | NA | NA | NA | NA |
| Mbl1 | 0 | NA | NA | NA | NA | NA | | | 2300003K06Rik | 0 | NA | NA | NA | NA | NA |
| Mbl2 | 0.681282946 | 0.008839375 | 0.054361526 | 0.162603511 | 0.870830622 | NA | | | 2310005A03Rik | 0 | NA | NA | NA | NA | NA |
| Mbnl3 | 3.407290376 | -0.09714654 | 0.106540491 | -0.911827407 | 0.361859581 | NA | | | 2310005E17Rik | 0 | NA | NA | NA | NA | NA |
| Mboat4 | 5.240030806 | 0.305571521 | 0.136591577 | 2.237118329 | 0.025278607 | NA | | | 2310020H05Rik | 0 | NA | NA | NA | NA | NA |
| Mc1r | 1.717398057 | 0.107937639 | 0.088991717 | 1.21289534 | 0.225169837 | NA | | | 2310034C09Rik | 0 | NA | NA | NA | NA | NA |
| Mc2r | 0.831689407 | 0.122187568 | 0.067723391 | 1.804215155 | 0.071197584 | NA | | | 2310043O21Rik | 0 | NA | NA | NA | NA | NA |
| Mcm7 | 492.3551671 | 0.026997463 | 0.11970892 | 0.225525907 | NA | NA | | | 2310050C09Rik | 0 | NA | NA | NA | NA | NA |
| Mcoln2 | 0.962157358 | -0.03636776 | 0.072161417 | -0.503977905 | 0.614276904 | NA | | | 2310057N15Rik | 0 | NA | NA | NA | NA | NA |
| Mcoln3 | 0.760998328 | 0.011477899 | 0.05537348 | 0.207281514 | 0.83579001 | NA | | | 2310061N02Rik | 0 | NA | NA | NA | NA | NA |
| Mcpt-ps1 | 0.21363976 | -0.000753674 | 0.031922632 | -0.02360938 | 0.98116419 | NA | | | 2310079G19Rik | 0 | NA | NA | NA | NA | NA |
| Mcpt1 | 0 | NA | NA | NA | NA | NA | | | 2410012E07Rik | 0 | NA | NA | NA | NA | NA |
| Mcpt2 | 0 | NA | NA | NA | NA | NA | | | 2410114N07Rik | 0 | NA | NA | NA | NA | NA |
| Mcpt4 | 0.670575502 | -0.080313968 | 0.054217216 | -1.481336999 | 0.138516794 | NA | | | 2410141K09Rik | 0 | NA | NA | NA | NA | NA |
| Mcpt8 | 0 | NA | NA | NA | NA | NA | | | 2600006L11Rik | 0 | NA | NA | NA | NA | NA |
| Mcpt9 | 0 | NA | NA | NA | NA | NA | | | 2700086A05Rik | 0 | NA | NA | NA | NA | NA |
| Mef2b | 6.58410113 | 0.046991247 | 0.134334315 | 0.349808215 | 0.726482634 | NA | | | 2700089I24Rik | 0 | NA | NA | NA | NA | NA |
| Mefv | 0.958981039 | 0.044863195 | 0.072618989 | 0.617788754 | 0.536714597 | NA | | | 2810007J24Rik | 0 | NA | NA | NA | NA | NA |
| Meiob | 0 | NA | NA | NA | NA | NA | | | 2900057B20Rik | 0 | NA | NA | NA | NA | NA |
| Meox2 | 1.368920632 | -0.072981926 | 0.07790776 | -0.936773509 | 0.348875075 | NA | | | 3300005D01Rik | 0 | NA | NA | NA | NA | NA |
| Mep1a | 1.300725613 | 0.045436634 | 0.070135083 | 0.64784458 | 0.517085479 | NA | | | 3830417A13Rik | 0 | NA | NA | NA | NA | NA |
| Mep1b | 0.215706849 | -0.002035226 | 0.0319478 | -0.063704747 | 0.949205325 | NA | | | 4921509C19Rik | 0 | NA | NA | NA | NA | NA |
| Mepe | 0.192746203 | -0.000975302 | 0.031567365 | -0.030895906 | 0.975352555 | NA | | | 4921511C20Rik | 0 | NA | NA | NA | NA | NA |
| Mesp1 | 0.686827323 | 0.040767534 | 0.066213896 | 0.615694537 | 0.538096144 | NA | | | 4921511H03Rik | 0 | NA | NA | NA | NA | NA |
| Mettl21c | 1.186620711 | 0.016888272 | 0.076282014 | 0.221392583 | 0.824786767 | NA | | | 4921511M17Rik | 0 | NA | NA | NA | NA | NA |
| Mettl21e | 1.304414302 | -0.084831958 | 0.067412641 | -1.258398364 | 0.208247723 | NA | | | 4921513I03Rik | 0 | NA | NA | NA | NA | NA |
| Mettl7a2 | 0 | NA | NA | NA | NA | NA | | | 4921524L21Rik | 0 | NA | NA | NA | NA | NA |
| Mettl7a3 | 5.440438738 | 0.081190936 | 0.12005675 | 0.676271311 | 0.498868404 | NA | | | 4921533I20Rik | 0 | NA | NA | NA | NA | NA |
| Mettl7b | 0.609883421 | -0.040524199 | 0.059374544 | -0.682518065 | 0.494911424 | NA | | | 4921539E11Rik | 0 | NA | NA | NA | NA | NA |
| Mfap5 | 5.888324268 | -0.088737789 | 0.137057818 | -0.647447843 | 0.51734214 | NA | | | 4922502D21Rik | 0 | NA | NA | NA | NA | NA |
| Mfrp | 0 | NA | NA | NA | NA | NA | | | 4922502H24Rik | 0 | NA | NA | NA | NA | NA |
| Mfsd6l | 1.847446006 | 0.041419523 | 0.093546088 | 0.442771303 | 0.657931162 | NA | | | 4930401O10Rik | 0 | NA | NA | NA | NA | NA |
| Mgam | 4.339769707 | -0.286992334 | 0.103171518 | -2.781701188 | 0.00540748 | NA | | | 4930401O12Rik | 0 | NA | NA | NA | NA | NA |
| Mgarp | 2.090185046 | 0.056731167 | 0.100188161 | 0.566246213 | 0.571226419 | NA | | | 4930402F11Rik | 0 | NA | NA | NA | NA | NA |
| Mgst2 | 1.910121513 | 0.051605332 | 0.095108525 | 0.542594181 | 0.587409246 | NA | | | 4930402K13Rik | 0 | NA | NA | NA | NA | NA |
| Mia2 | 0 | NA | NA | NA | NA | NA | | | 4930404A05Rik | 0 | NA | NA | NA | NA | NA |
| Mill1 | 0 | NA | NA | NA | NA | NA | | | 4930404A10Rik | 0 | NA | NA | NA | NA | NA |
| Miox | 0 | NA | NA | NA | NA | NA | | | 4930405A10Rik | 0 | NA | NA | NA | NA | NA |
| Mip | 0 | NA | NA | NA | NA | NA | | | 4930405D11Rik | 0 | NA | NA | NA | NA | NA |
| Mir100 | 0.099009884 | 0.018205726 | 0.027896865 | 0.652608315 | 0.514008823 | NA | | | 4930405J17Rik | 0 | NA | NA | NA | NA | NA |
| Mir101a | 0 | NA | NA | NA | NA | NA | | | 4930406D18Rik | 0 | NA | NA | NA | NA | NA |
| Mir101b | 0.188785634 | 0.025025111 | 0.031461603 | 0.79541754 | 0.42637066 | NA | | | 4930408F14Rik | 0 | NA | NA | NA | NA | NA |
| Mir101c | 0 | NA | NA | NA | NA | NA | | | 4930412B13Rik | 0 | NA | NA | NA | NA | NA |
| Mir103-1 | 0 | NA | NA | NA | NA | NA | | | 4930413E15Rik | 0 | NA | NA | NA | NA | NA |
| Mir103-2 | 1.920915016 | 0.167017145 | 0.088579629 | 1.885502865 | 0.059361978 | NA | | | 4930413M19Rik | 0 | NA | NA | NA | NA | NA |
| Mir105 | 0 | NA | NA | NA | NA | NA | | | 4930415L06Rik | 0 | NA | NA | NA | NA | NA |
| Mir106a | 0 | NA | NA | NA | NA | NA | | | 4930423M02Rik | 0 | NA | NA | NA | NA | NA |
| Mir106b | 0.909184964 | 0.024388887 | 0.058249921 | 0.418693912 | 0.675439846 | NA | | | 4930428D18Rik | 0 | NA | NA | NA | NA | NA |
| Mir107 | 0 | NA | NA | NA | NA | NA | | | 4930428O21Rik | 0 | NA | NA | NA | NA | NA |
| Mir10a | 0 | NA | NA | NA | NA | NA | | | 4930429F11Rik | 0 | NA | NA | NA | NA | NA |
| Mir10b | 0 | NA | NA | NA | NA | NA | | | 4930430A15Rik | 0 | NA | NA | NA | NA | NA |
| Mir1186 | 0 | NA | NA | NA | NA | NA | | | 4930430D24Rik | 0 | NA | NA | NA | NA | NA |
| Mir1186b | 0 | NA | NA | NA | NA | NA | | | 4930430J02Rik | 0 | NA | NA | NA | NA | NA |
| Mir1187 | 0 | NA | NA | NA | NA | NA | | | 4930432K09Rik | 0 | NA | NA | NA | NA | NA |
| Mir1190 | 0 | NA | NA | NA | NA | NA | | | 4930433B08Rik | 0 | NA | NA | NA | NA | NA |
| Mir1192 | 0.383703858 | 0.028442486 | 0.048086527 | 0.591485549 | 0.554195135 | NA | | | 4930433N12Rik | 0 | NA | NA | NA | NA | NA |
| Mir1193 | 0.099009884 | 0.018205726 | 0.027896865 | 0.652608315 | 0.514008823 | NA | | | 4930435E12Rik | 0 | NA | NA | NA | NA | NA |
| Mir1196 | 0 | NA | NA | NA | NA | NA | | | 4930438E09Rik | 0 | NA | NA | NA | NA | NA |
| Mir1197 | 0 | NA | NA | NA | NA | NA | | | 4930440C22Rik | 0 | NA | NA | NA | NA | NA |
| Mir1199 | 0.116696964 | -0.020477583 | 0.027954224 | -0.732539847 | 0.463839131 | NA | | | 4930442J19Rik | 0 | NA | NA | NA | NA | NA |
| Mir1224 | 0 | NA | NA | NA | NA | NA | | | 4930442L01Rik | 0 | NA | NA | NA | NA | NA |
| Mir122a | 0 | NA | NA | NA | NA | NA | | | 4930444F02Rik | 0 | NA | NA | NA | NA | NA |
| Mir1247 | 0 | NA | NA | NA | NA | NA | | | 4930447A16Rik | 0 | NA | NA | NA | NA | NA |
| Mir124a-1 | 0 | NA | NA | NA | NA | NA | | | 4930447J18Rik | 0 | NA | NA | NA | NA | NA |
| Mir124a-2 | 6.188467794 | 0.159161527 | 0.132752934 | 1.198930394 | 0.230555012 | NA | | | 4930448I06Rik | 0 | NA | NA | NA | NA | NA |
| Mir124a-3 | 0.398821088 | -0.027919467 | 0.043501191 | -0.641809245 | 0.520997046 | NA | | | 4930448K20Rik | 0 | NA | NA | NA | NA | NA |
| Mir1251 | 0.119397641 | -0.020477583 | 0.027954224 | -0.732539845 | 0.463839132 | NA | | | 4930449E18Rik | 0 | NA | NA | NA | NA | NA |
| Mir125a | 4.436323189 | 0.310515415 | 0.135676455 | 2.288646281 | 0.022099913 | NA | | | 4930449I24Rik | 0 | NA | NA | NA | NA | NA |
| Mir125b-1 | 0 | NA | NA | NA | NA | NA | | | 4930453L07Rik | 0 | NA | NA | NA | NA | NA |
| Mir125b-2 | 0 | NA | NA | NA | NA | NA | | | 4930455J16Rik | 0 | NA | NA | NA | NA | NA |
| Mir126 | 1.690534935 | 0.064467984 | 0.088732415 | 0.726543777 | 0.467505481 | NA | | | 4930459L07Rik | 0 | NA | NA | NA | NA | NA |
| Mir1264 | 0 | NA | NA | NA | NA | NA | | | 4930465A12Rik | 0 | NA | NA | NA | NA | NA |
| Mir127 | 0 | NA | NA | NA | NA | NA | | | 4930467E23Rik | 0 | NA | NA | NA | NA | NA |
| Mir128-1 | 6.461650651 | 0.179996805 | 0.137606822 | 1.308051461 | 0.190855855 | NA | | | 4930470P17Rik | 0 | NA | NA | NA | NA | NA |
| Mir128-2 | 0 | NA | NA | NA | NA | NA | | | 4930474H20Rik | 0 | NA | NA | NA | NA | NA |
| Mir129-1 | 5.202608281 | 0.20374429 | 0.134992744 | 1.509298087 | 0.131222623 | NA | | | 4930474M22Rik | 0 | NA | NA | NA | NA | NA |
| Mir129-2 | 0 | NA | NA | NA | NA | NA | | | 4930474N05Rik | 0 | NA | NA | NA | NA | NA |
| Mir1298 | 0 | NA | NA | NA | NA | NA | | | 4930478L05Rik | 0 | NA | NA | NA | NA | NA |
| Mir1306 | 0 | NA | NA | NA | NA | NA | | | 4930478P22Rik | 0 | NA | NA | NA | NA | NA |
| Mir130a | 0 | NA | NA | NA | NA | NA | | | 4930480E11Rik | 0 | NA | NA | NA | NA | NA |
| Mir130b | 0 | NA | NA | NA | NA | NA | | | 4930480K15Rik | 0 | NA | NA | NA | NA | NA |
| Mir133a-1 | 0 | NA | NA | NA | NA | NA | | | 4930480M12Rik | 0 | NA | NA | NA | NA | NA |
| Mir133a-2 | 0 | NA | NA | NA | NA | NA | | | 4930486F22Rik | 0 | NA | NA | NA | NA | NA |
| Mir133b | 0 | NA | NA | NA | NA | NA | | | 4930487D11Rik | 0 | NA | NA | NA | NA | NA |
| Mir134 | 5.490815863 | -0.122742067 | 0.141703851 | -0.866187235 | 0.386387492 | NA | | | 4930488B22Rik | 0 | NA | NA | NA | NA | NA |
| Mir135a-1 | 0.105923158 | 0.018524693 | 0.027977518 | 0.662127842 | 0.507889294 | NA | | | 4930503E24Rik | 0 | NA | NA | NA | NA | NA |
| Mir135a-2 | 0.416779358 | 0.041366775 | 0.04012708 | 1.03089422 | 0.302590428 | NA | | | 4930504O13Rik | 0 | NA | NA | NA | NA | NA |
| Mir135b | 0.222620122 | -0.001447027 | 0.032059667 | -0.04513545 | 0.963999345 | NA | | | 4930505G20Rik | 0 | NA | NA | NA | NA | NA |
| Mir136 | 0.525223898 | 0.006204767 | 0.045384814 | 0.136714611 | 0.89125638 | NA | | | 4930509J09Rik | 0 | NA | NA | NA | NA | NA |
| Mir137 | 1.72245743 | 0.012298194 | 0.092857863 | 0.132441065 | 0.894635436 | NA | | | 4930509K18Rik | 0 | NA | NA | NA | NA | NA |
| Mir138-1 | 4.336868572 | 0.16737562 | 0.128531445 | 1.302215339 | 0.192842783 | NA | | | 4930513O06Rik | 0 | NA | NA | NA | NA | NA |
| Mir138-2 | 7.888519922 | -0.090004968 | 0.158504068 | -0.567840113 | 0.570143539 | NA | | | 4930515B02Rik | 0 | NA | NA | NA | NA | NA |
| Mir139 | 0 | NA | NA | NA | NA | NA | | | 4930515G16Rik | 0 | NA | NA | NA | NA | NA |
| Mir140 | 0.813705528 | 0.030122804 | 0.06445444 | 0.467350329 | 0.640249254 | NA | | | 4930515L19Rik | 0 | NA | NA | NA | NA | NA |
| Mir141 | 0 | NA | NA | NA | NA | NA | | | 4930517E11Rik | 0 | NA | NA | NA | NA | NA |
| Mir142 | 0.598714191 | 0.017606709 | 0.053437106 | 0.3294847 | 0.741789357 | NA | | | 4930518P08Rik | 0 | NA | NA | NA | NA | NA |
| Mir143 | 0.212768007 | -0.02694223 | 0.031868764 | -0.845411806 | 0.397880955 | NA | | | 4930519F24Rik | 0 | NA | NA | NA | NA | NA |
| Mir144 | 0 | NA | NA | NA | NA | NA | | | 4930519H02Rik | 0 | NA | NA | NA | NA | NA |
| Mir145 | 1.422393584 | -0.006857031 | 0.075784459 | -0.090480703 | 0.927905228 | NA | | | 4930521E06Rik | 0 | NA | NA | NA | NA | NA |
| Mir146 | 0 | NA | NA | NA | NA | NA | | | 4930522H14Rik | 0 | NA | NA | NA | NA | NA |
| Mir146b | 0.119397641 | -0.020477583 | 0.027954224 | -0.732539845 | 0.463839132 | NA | | | 4930522O17Rik | 0 | NA | NA | NA | NA | NA |
| Mir147 | 0 | NA | NA | NA | NA | NA | | | 4930523O13Rik | 0 | NA | NA | NA | NA | NA |
| Mir148a | 0 | NA | NA | NA | NA | NA | | | 4930524C18Rik | 0 | NA | NA | NA | NA | NA |
| Mir148b | 0 | NA | NA | NA | NA | NA | | | 4930524N10Rik | 0 | NA | NA | NA | NA | NA |
| Mir149 | 2.972681264 | 0.073213464 | 0.110445894 | 0.662889864 | 0.507401094 | NA | | | 4930524O05Rik | 0 | NA | NA | NA | NA | NA |
| Mir150 | 0 | NA | NA | NA | NA | NA | | | 4930525M21Rik | 0 | NA | NA | NA | NA | NA |
| Mir152 | 0 | NA | NA | NA | NA | NA | | | 4930526D03Rik | 0 | NA | NA | NA | NA | NA |
| Mir153 | 0.298180463 | -0.015304498 | 0.039412854 | -0.388312359 | 0.697784893 | NA | | | 4930527G23Rik | 0 | NA | NA | NA | NA | NA |
| Mir154 | 1.201797363 | 0.081344257 | 0.081416452 | 0.999113264 | 0.317739826 | NA | | | 4930529C04Rik | 0 | NA | NA | NA | NA | NA |
| Mir155 | 0.119397641 | -0.020477583 | 0.027954224 | -0.732539845 | 0.463839132 | NA | | | 4930529F24Rik | 0 | NA | NA | NA | NA | NA |
| Mir15a | 0.321391842 | -0.01514286 | 0.04017217 | -0.376949019 | 0.706211492 | NA | | | 4930529L06Rik | 0 | NA | NA | NA | NA | NA |
| Mir15b | 0 | NA | NA | NA | NA | NA | | | 4930533B01Rik | 0 | NA | NA | NA | NA | NA |
| Mir16-1 | 0.303286429 | 0.012912943 | 0.039427608 | 0.32751017 | 0.743282052 | NA | | | 4930533P14Rik | 0 | NA | NA | NA | NA | NA |
| Mir16-2 | 0 | NA | NA | NA | NA | NA | | | 4930539C22Rik | 0 | NA | NA | NA | NA | NA |
| Mir17 | 0 | NA | NA | NA | NA | NA | | | 4930539N22Rik | 0 | NA | NA | NA | NA | NA |
| Mir18 | 0 | NA | NA | NA | NA | NA | | | 4930540M03Rik | 0 | NA | NA | NA | NA | NA |
| Mir181a-1 | 0 | NA | NA | NA | NA | NA | | | 4930542C21Rik | 0 | NA | NA | NA | NA | NA |
| Mir181a-2 | 0.215468684 | -0.027008432 | 0.031909031 | -0.846419662 | 0.397318674 | NA | | | 4930542H20Rik | 0 | NA | NA | NA | NA | NA |
| Mir181b-1 | 0.599170685 | -0.002006017 | 0.052323611 | -0.038338665 | 0.969417664 | NA | | | 4930544G11Rik | 0 | NA | NA | NA | NA | NA |
| Mir181b-2 | 0 | NA | NA | NA | NA | NA | | | 4930544M13Rik | 0 | NA | NA | NA | NA | NA |
| Mir181c | 0.859830262 | 0.09472059 | 0.069693768 | 1.359096978 | 0.17411586 | NA | | | 4930545E07Rik | 0 | NA | NA | NA | NA | NA |
| Mir181d | 1.161288641 | 0.052504258 | 0.086764318 | 0.605136528 | 0.545088285 | NA | | | 4930547C10Rik | 0 | NA | NA | NA | NA | NA |
| Mir182 | 0 | NA | NA | NA | NA | NA | | | 4930547E08Rik | 0 | NA | NA | NA | NA | NA |
| Mir183 | 0 | NA | NA | NA | NA | NA | | | 4930547E14Rik | 0 | NA | NA | NA | NA | NA |
| Mir184 | 0 | NA | NA | NA | NA | NA | | | 4930548G14Rik | 0 | NA | NA | NA | NA | NA |
| Mir1843 | 0 | NA | NA | NA | NA | NA | | | 4930548J01Rik | 0 | NA | NA | NA | NA | NA |
| Mir1843b | 0 | NA | NA | NA | NA | NA | | | 4930548K13Rik | 0 | NA | NA | NA | NA | NA |
| Mir185 | 0.116696964 | -0.020477583 | 0.027954224 | -0.732539847 | 0.463839131 | NA | | | 4930549C01Rik | 0 | NA | NA | NA | NA | NA |
| Mir187 | 0 | NA | NA | NA | NA | NA | | | 4930552N02Rik | 0 | NA | NA | NA | NA | NA |
| Mir188 | 0 | NA | NA | NA | NA | NA | | | 4930554C24Rik | 0 | NA | NA | NA | NA | NA |
| Mir1892 | 0 | NA | NA | NA | NA | NA | | | 4930555B11Rik | 0 | NA | NA | NA | NA | NA |
| Mir1893 | 0 | NA | NA | NA | NA | NA | | | 4930556G01Rik | 0 | NA | NA | NA | NA | NA |
| Mir1894 | 3.582350746 | -0.174863184 | 0.118121118 | -1.480371907 | 0.138774023 | NA | | | 4930556J02Rik | 0 | NA | NA | NA | NA | NA |
| Mir1895 | 1.373629463 | 0.032823391 | 0.073130882 | 0.448830776 | 0.653553736 | NA | | | 4930556N09Rik | 0 | NA | NA | NA | NA | NA |
| Mir1896 | 1.097917236 | -0.026940225 | 0.074160089 | -0.363271205 | 0.716402301 | NA | | | 4930557A04Rik | 0 | NA | NA | NA | NA | NA |
| Mir1897 | 0 | NA | NA | NA | NA | NA | | | 4930557J02Rik | 0 | NA | NA | NA | NA | NA |
| Mir1898 | 6.583395593 | 0.084719094 | 0.148226174 | 0.571552861 | 0.567624939 | NA | | | 4930558C23Rik | 0 | NA | NA | NA | NA | NA |
| Mir1899 | 0 | NA | NA | NA | NA | NA | | | 4930558G05Rik | 0 | NA | NA | NA | NA | NA |
| Mir18b | 0 | NA | NA | NA | NA | NA | | | 4930563F08Rik | 0 | NA | NA | NA | NA | NA |
| Mir190 | 0 | NA | NA | NA | NA | NA | | | 4930563M20Rik | 0 | NA | NA | NA | NA | NA |
| Mir1900 | 0 | NA | NA | NA | NA | NA | | | 4930564C03Rik | 0 | NA | NA | NA | NA | NA |
| Mir1901 | 6.378478339 | 0.020424364 | 0.153466177 | 0.133087073 | 0.894124518 | NA | | | 4930564D02Rik | 0 | NA | NA | NA | NA | NA |
| Mir1902 | 0 | NA | NA | NA | NA | NA | | | 4930564G21Rik | 0 | NA | NA | NA | NA | NA |
| Mir1903 | 0 | NA | NA | NA | NA | NA | | | 4930564K09Rik | 0 | NA | NA | NA | NA | NA |
| Mir1904 | 0 | NA | NA | NA | NA | NA | | | 4930565D16Rik | 0 | NA | NA | NA | NA | NA |
| Mir1905 | 1.115578782 | 0.005731202 | 0.083430484 | 0.068694338 | 0.945232925 | NA | | | 4930565N06Rik | 0 | NA | NA | NA | NA | NA |
| Mir1906-1 | 0 | NA | NA | NA | NA | NA | | | 4930567H17Rik | 0 | NA | NA | NA | NA | NA |
| Mir1906-2 | 0 | NA | NA | NA | NA | NA | | | 4930567J20Rik | 0 | NA | NA | NA | NA | NA |
| Mir1907 | 0.518444887 | -0.021172618 | 0.051250408 | -0.413120974 | 0.679517983 | NA | | | 4930567K20Rik | 0 | NA | NA | NA | NA | NA |
| Mir190b | 0 | NA | NA | NA | NA | NA | | | 4930568D16Rik | 0 | NA | NA | NA | NA | NA |
| Mir191 | 4.417109132 | 0.032452265 | 0.123406214 | 0.262971074 | 0.792572875 | NA | | | 4930568E12Rik | 0 | NA | NA | NA | NA | NA |
| Mir1912 | 0 | NA | NA | NA | NA | NA | | | 4930568K20Rik | 0 | NA | NA | NA | NA | NA |
| Mir192 | 0.099009884 | 0.018205726 | 0.027896865 | 0.652608315 | 0.514008823 | NA | | | 4930571K23Rik | 0 | NA | NA | NA | NA | NA |
| Mir1929 | 0.527163612 | -0.038923887 | 0.04705911 | -0.827127563 | 0.408164763 | NA | | | 4930572K03Rik | 0 | NA | NA | NA | NA | NA |
| Mir193 | 0 | NA | NA | NA | NA | NA | | | 4930573O16Rik | 0 | NA | NA | NA | NA | NA |
| Mir1930 | 0.707852793 | 0.070978149 | 0.063325329 | 1.120849269 | 0.262352028 | NA | | | 4930578I06Rik | 0 | NA | NA | NA | NA | NA |
| Mir1931 | 0.167300494 | 0.024299763 | 0.031030539 | 0.783091891 | 0.433573153 | NA | | | 4930578N18Rik | 0 | NA | NA | NA | NA | NA |
| Mir1932 | 0.096071043 | -0.020098795 | 0.027825948 | -0.722304036 | 0.470107572 | NA | | | 4930579F01Rik | 0 | NA | NA | NA | NA | NA |
| Mir1933 | 0 | NA | NA | NA | NA | NA | | | 4930583P06Rik | 0 | NA | NA | NA | NA | NA |
| Mir1934 | 0.659372557 | 0.083679911 | 0.06789324 | 1.23252199 | 0.217754157 | NA | | | 4930584F24Rik | 0 | NA | NA | NA | NA | NA |
| Mir1936 | 0 | NA | NA | NA | NA | NA | | | 4930591A17Rik | 0 | NA | NA | NA | NA | NA |
| Mir1938 | 0.392396713 | -0.003471664 | 0.047500251 | -0.073087276 | 0.941736667 | NA | | | 4930592I03Rik | 0 | NA | NA | NA | NA | NA |
| Mir193b | 0.296502237 | 0.007885408 | 0.033590428 | 0.234751624 | 0.814401517 | NA | | | 4930595M18Rik | 0 | NA | NA | NA | NA | NA |
| Mir194-1 | 0 | NA | NA | NA | NA | NA | | | 4930596D02Rik | 0 | NA | NA | NA | NA | NA |
| Mir194-2 | 0 | NA | NA | NA | NA | NA | | | 4930597G03Rik | 0 | NA | NA | NA | NA | NA |
| Mir1940 | 0 | NA | NA | NA | NA | NA | | | 4930598F16Rik | 0 | NA | NA | NA | NA | NA |
| Mir1941 | 0 | NA | NA | NA | NA | NA | | | 4931403G20Rik | 0 | NA | NA | NA | NA | NA |
| Mir1942 | 0 | NA | NA | NA | NA | NA | | | 4931406B18Rik | 0 | NA | NA | NA | NA | NA |
| Mir1943 | 5.355910632 | 0.236561267 | 0.11430196 | 2.069616889 | 0.038488235 | NA | | | 4931408C20Rik | 0 | NA | NA | NA | NA | NA |
| Mir1945 | 0 | NA | NA | NA | NA | NA | | | 4931417E11Rik | 0 | NA | NA | NA | NA | NA |
| Mir1946a | 0 | NA | NA | NA | NA | NA | | | 4931420L22Rik | 0 | NA | NA | NA | NA | NA |
| Mir1947 | 0.19046386 | -0.000774927 | 0.031526133 | -0.024580468 | 0.980389599 | NA | | | 4931431F19Rik | 0 | NA | NA | NA | NA | NA |
| Mir1948 | 0.094392817 | 0.017895942 | 0.027818137 | 0.643319236 | 0.520016977 | NA | | | 4932411N23Rik | 0 | NA | NA | NA | NA | NA |
| Mir1949 | 3.481742175 | 0.174092853 | 0.131864775 | 1.32023774 | 0.186755655 | NA | | | 4932413F04Rik | 0 | NA | NA | NA | NA | NA |
| Mir195 | 0.079983654 | 0.016740032 | 0.027674178 | 0.604897175 | 0.54524732 | NA | | | 4932414J04Rik | 0 | NA | NA | NA | NA | NA |
| Mir1951 | 0 | NA | NA | NA | NA | NA | | | 4932414N04Rik | 0 | NA | NA | NA | NA | NA |
| Mir1952 | 0 | NA | NA | NA | NA | NA | | | 4932416K20Rik | 0 | NA | NA | NA | NA | NA |
| Mir1953 | 0 | NA | NA | NA | NA | NA | | | 4932429P05Rik | 0 | NA | NA | NA | NA | NA |
| Mir1955 | 0.098353386 | -0.020212831 | 0.02786466 | -0.725393061 | 0.468210929 | NA | | | 4933400A11Rik | 0 | NA | NA | NA | NA | NA |
| Mir1956 | 0 | NA | NA | NA | NA | NA | | | 4933400L20Rik | 0 | NA | NA | NA | NA | NA |
| Mir1960 | 0 | NA | NA | NA | NA | NA | | | 4933401B06Rik | 0 | NA | NA | NA | NA | NA |
| Mir1961 | 0 | NA | NA | NA | NA | NA | | | 4933401P06Rik | 0 | NA | NA | NA | NA | NA |
| Mir1962 | 0 | NA | NA | NA | NA | NA | | | 4933402E13Rik | 0 | NA | NA | NA | NA | NA |
| Mir1963 | 0 | NA | NA | NA | NA | NA | | | 4933402J10Rik | 0 | NA | NA | NA | NA | NA |
| Mir1964 | 1.098050894 | -0.021040418 | 0.072437663 | -0.290462406 | 0.771462507 | NA | | | 4933402N22Rik | 0 | NA | NA | NA | NA | NA |
| Mir1965 | 0 | NA | NA | NA | NA | NA | | | 4933402P03Rik | 0 | NA | NA | NA | NA | NA |
| Mir1966 | 0 | NA | NA | NA | NA | NA | | | 4933404G15Rik | 0 | NA | NA | NA | NA | NA |
| Mir1967 | 0.185670226 | -0.001645086 | 0.031432796 | -0.052336624 | 0.958260472 | NA | | | 4933405E24Rik | 0 | NA | NA | NA | NA | NA |
| Mir1968 | 2.217277567 | 0.039560172 | 0.105632057 | 0.374509154 | 0.708025547 | NA | | | 4933406D12Rik | 0 | NA | NA | NA | NA | NA |
| Mir1969 | 1.549136228 | 0.168408375 | 0.091425718 | 1.842024088 | 0.065471627 | NA | | | 4933406G16Rik | 0 | NA | NA | NA | NA | NA |
| Mir196a-1 | 0 | NA | NA | NA | NA | NA | | | 4933406J08Rik | 0 | NA | NA | NA | NA | NA |
| Mir196a-2 | 0 | NA | NA | NA | NA | NA | | | 4933406J10Rik | 0 | NA | NA | NA | NA | NA |
| Mir196b | 0 | NA | NA | NA | NA | NA | | | 4933406K04Rik | 0 | NA | NA | NA | NA | NA |
| Mir1971 | 0 | NA | NA | NA | NA | NA | | | 4933407G14Rik | 0 | NA | NA | NA | NA | NA |
| Mir1981 | 0 | NA | NA | NA | NA | NA | | | 4933409G03Rik | 0 | NA | NA | NA | NA | NA |
| Mir1982 | 0 | NA | NA | NA | NA | NA | | | 4933411G06Rik | 0 | NA | NA | NA | NA | NA |
| Mir1983 | 0.186326724 | 0.024939385 | 0.031411053 | 0.793968451 | 0.427213798 | NA | | | 4933412E24Rik | 0 | NA | NA | NA | NA | NA |
| Mir199a-1 | 0.198019769 | 0.025308392 | 0.031628095 | 0.80018706 | 0.423602426 | NA | | | 4933413J09Rik | 0 | NA | NA | NA | NA | NA |
| Mir199a-2 | 0 | NA | NA | NA | NA | NA | | | 4933413L06Rik | 0 | NA | NA | NA | NA | NA |
| Mir199b | 0 | NA | NA | NA | NA | NA | | | 4933416C03Rik | 0 | NA | NA | NA | NA | NA |
| Mir19a | 0 | NA | NA | NA | NA | NA | | | 4933416E03Rik | 0 | NA | NA | NA | NA | NA |

| | | | | | | | | | | | | | | | |
|---|---|---|---|---|---|---|---|---|---|---|---|---|---|---|---|
| Mir19b-1 | 0 | NA | NA | NA | NA | NA | | | 4933416M06Rik | 0 | NA | NA | NA | NA | NA |
| Mir19b-2 | 0 | NA | NA | NA | NA | NA | | | 4933416M07Rik | 0 | NA | NA | NA | NA | NA |
| Mir1a-1 | 0 | NA | NA | NA | NA | NA | | | 4933417A18Rik | 0 | NA | NA | NA | NA | NA |
| Mir1a-2 | 0 | NA | NA | NA | NA | NA | | | 4933417G07Rik | 0 | NA | NA | NA | NA | NA |
| Mir1b | 0.096071043 | -0.020098795 | 0.027825948 | -0.722304036 | 0.470107572 | NA | | | 4933421I07Rik | 0 | NA | NA | NA | NA | NA |
| Mir200a | 0 | NA | NA | NA | NA | NA | | | 4933425B07Rik | 0 | NA | NA | NA | NA | NA |
| Mir200b | 0 | NA | NA | NA | NA | NA | | | 4933429K18Rik | 0 | NA | NA | NA | NA | NA |
| Mir200c | 0 | NA | NA | NA | NA | NA | | | 4933430N04Rik | 0 | NA | NA | NA | NA | NA |
| Mir201 | 0 | NA | NA | NA | NA | NA | | | 4933431G14Rik | 0 | NA | NA | NA | NA | NA |
| Mir202 | 0 | NA | NA | NA | NA | NA | | | 4933432G23Rik | 0 | NA | NA | NA | NA | NA |
| Mir203 | 0 | NA | NA | NA | NA | NA | | | 4933433F19Rik | 0 | NA | NA | NA | NA | NA |
| Mir204 | 0 | NA | NA | NA | NA | NA | | | 4933436I01Rik | 0 | NA | NA | NA | NA | NA |
| Mir205 | 0 | NA | NA | NA | NA | NA | | | 4933438B17Rik | 0 | NA | NA | NA | NA | NA |
| Mir206 | 0 | NA | NA | NA | NA | NA | | | 4933440J02Rik | 0 | NA | NA | NA | NA | NA |
| Mir207 | 0 | NA | NA | NA | NA | NA | | | 5330439B14Rik | 0 | NA | NA | NA | NA | NA |
| Mir208a | 0 | NA | NA | NA | NA | NA | | | 5430425J12Rik | 0 | NA | NA | NA | NA | NA |
| Mir208b | 0 | NA | NA | NA | NA | NA | | | 5430427M07Rik | 0 | NA | NA | NA | NA | NA |
| Mir20a | 0 | NA | NA | NA | NA | NA | | | 5430428K19Rik | 0 | NA | NA | NA | NA | NA |
| Mir20b | 0 | NA | NA | NA | NA | NA | | | 5430434I15Rik | 0 | NA | NA | NA | NA | NA |
| Mir21 | 0 | NA | NA | NA | NA | NA | | | 5530400C23Rik | 0 | NA | NA | NA | NA | NA |
| Mir210 | 0 | NA | NA | NA | NA | NA | | | 5730416F02Rik | 0 | NA | NA | NA | NA | NA |
| Mir211 | 0 | NA | NA | NA | NA | NA | | | 5730457N03Rik | 0 | NA | NA | NA | NA | NA |
| Mir2136 | 0 | NA | NA | NA | NA | NA | | | 5730460C07Rik | 0 | NA | NA | NA | NA | NA |
| Mir2137 | 0 | NA | NA | NA | NA | NA | | | 5830428M24Rik | 0 | NA | NA | NA | NA | NA |
| Mir214 | 0 | NA | NA | NA | NA | NA | | | 6030440G07Rik | 0 | NA | NA | NA | NA | NA |
| Mir215 | 0 | NA | NA | NA | NA | NA | | | 6330407A03Rik | 0 | NA | NA | NA | NA | NA |
| Mir216a | 0 | NA | NA | NA | NA | NA | | | 6330415B21Rik | 0 | NA | NA | NA | NA | NA |
| Mir216b | 0 | NA | NA | NA | NA | NA | | | 6430706D22Rik | 0 | NA | NA | NA | NA | NA |
| Mir217 | 0 | NA | NA | NA | NA | NA | | | 7420426K07Rik | 0 | NA | NA | NA | NA | NA |
| Mir218-1 | 1.62510428 | 0.082863464 | 0.092234695 | 0.898397979 | 0.368973413 | NA | | | 7420700N18Rik | 0 | NA | NA | NA | NA | NA |
| Mir218-2 | 0.200315975 | 0.025370485 | 0.031664208 | 0.801235428 | 0.422995365 | NA | | | 7420701I03Rik | 0 | NA | NA | NA | NA | NA |
| Mir219-1 | 0 | NA | NA | NA | NA | NA | | | 8030411F24Rik | 0 | NA | NA | NA | NA | NA |
| Mir219-2 | 1.863088764 | 0.058447026 | 0.102840709 | 0.568325775 | 0.56981378 | NA | | | 8430403D17Rik | 0 | NA | NA | NA | NA | NA |
| Mir22 | 0 | NA | NA | NA | NA | NA | | | 8430422H06Rik | 0 | NA | NA | NA | NA | NA |
| Mir221 | 1.732378575 | 0.016218783 | 0.08655574 | 0.187379638 | 0.851362987 | NA | | | 8430423G03Rik | 0 | NA | NA | NA | NA | NA |
| Mir222 | 0.331747315 | -0.03734372 | 0.03744515 | -0.99729124 | 0.318623164 | NA | | | 8430437L04Rik | 0 | NA | NA | NA | NA | NA |
| Mir223 | 0 | NA | NA | NA | NA | NA | | | 9130015L21Rik | 0 | NA | NA | NA | NA | NA |
| Mir224 | 0 | NA | NA | NA | NA | NA | | | 9130204L05Rik | 0 | NA | NA | NA | NA | NA |
| Mir23a | 0.389426822 | -0.00542245 | 0.0474348 | -0.114313744 | 0.908989088 | NA | | | 9130209A04Rik | 0 | NA | NA | NA | NA | NA |
| Mir24-2 | 0.898641866 | 0.005814992 | 0.070393532 | 0.082606915 | 0.934164103 | NA | | | 9130230L23Rik | 0 | NA | NA | NA | NA | NA |
| Mir25 | 3.069236884 | 0.051379836 | 0.102669605 | 0.500438625 | 0.616766262 | NA | | | 9230105E05Rik | 0 | NA | NA | NA | NA | NA |
| Mir26a-1 | 1.967147038 | 0.165729845 | 0.101439177 | 1.633785391 | 0.102303921 | NA | | | 9230110F15Rik | 0 | NA | NA | NA | NA | NA |
| Mir26a-2 | 0.176054697 | -0.002190495 | 0.031239146 | -0.070120197 | 0.944097991 | NA | | | 9330178D15Rik | 0 | NA | NA | NA | NA | NA |
| Mir26b | 0.116696964 | -0.020477583 | 0.027954224 | -0.732539847 | 0.463839131 | NA | | | 9330182O14Rik | 0 | NA | NA | NA | NA | NA |
| Mir27a | 1.218550218 | 0.080075385 | 0.079375039 | 1.008823258 | 0.313059405 | NA | | | 9430019J16Rik | 0 | NA | NA | NA | NA | NA |
| Mir2861 | 0.927827864 | 0.009771901 | 0.056235838 | 0.173766435 | 0.862049024 | NA | | | 9430069I07Rik | 0 | NA | NA | NA | NA | NA |
| Mir28b | 0 | NA | NA | NA | NA | NA | | | 9530002B09Rik | 0 | NA | NA | NA | NA | NA |
| Mir290 | 0 | NA | NA | NA | NA | NA | | | 9530026F06Rik | 0 | NA | NA | NA | NA | NA |
| Mir291a | 0 | NA | NA | NA | NA | NA | | | A430089I19Rik | 0 | NA | NA | NA | NA | NA |
| Mir291b | 0 | NA | NA | NA | NA | NA | | | A530006G24Rik | 0 | NA | NA | NA | NA | NA |
| Mir292 | 0 | NA | NA | NA | NA | NA | | | A530046M15Rik | 0 | NA | NA | NA | NA | NA |
| Mir293 | 0 | NA | NA | NA | NA | NA | | | A630010A05Rik | 0 | NA | NA | NA | NA | NA |
| Mir294 | 0 | NA | NA | NA | NA | NA | | | A630012P03Rik | 0 | NA | NA | NA | NA | NA |
| Mir295 | 0 | NA | NA | NA | NA | NA | | | A630023A22Rik | 0 | NA | NA | NA | NA | NA |
| Mir296 | 0.238795281 | -0.027601606 | 0.032267774 | -0.855392297 | 0.392334007 | NA | | | A630073D07Rik | 0 | NA | NA | NA | NA | NA |
| Mir297-1 | 0 | NA | NA | NA | NA | NA | | | A630095E13Rik | 0 | NA | NA | NA | NA | NA |
| Mir297a-3 | 0 | NA | NA | NA | NA | NA | | | A730008H23Rik | 0 | NA | NA | NA | NA | NA |
| Mir297b | 0 | NA | NA | NA | NA | NA | | | A730082K24Rik | 0 | NA | NA | NA | NA | NA |
| Mir297c | 0 | NA | NA | NA | NA | NA | | | AA545190 | 0 | NA | NA | NA | NA | NA |
| Mir298 | 0 | NA | NA | NA | NA | NA | | | AF366264 | 0 | NA | NA | NA | NA | NA |
| Mir299 | 0 | NA | NA | NA | NA | NA | | | AI646519 | 0 | NA | NA | NA | NA | NA |
| Mir29a | 0 | NA | NA | NA | NA | NA | | | AU016765 | 0 | NA | NA | NA | NA | NA |
| Mir29b-1 | 0 | NA | NA | NA | NA | NA | | | AU018091 | 0 | NA | NA | NA | NA | NA |
| Mir29b-2 | 0 | NA | NA | NA | NA | NA | | | AU018829 | 0 | NA | NA | NA | NA | NA |
| Mir29c | 0 | NA | NA | NA | NA | NA | | | AV320801 | 0 | NA | NA | NA | NA | NA |
| Mir300 | 0.201994201 | 0.000227241 | 0.031727614 | 0.00716224 | 0.994285408 | NA | | | AY761184 | 0 | NA | NA | NA | NA | NA |
| Mir301 | 0.096071043 | -0.020098795 | 0.027825948 | -0.722304036 | 0.470107572 | NA | | | AY761185 | 0 | NA | NA | NA | NA | NA |
| Mir301b | 0 | NA | NA | NA | NA | NA | | | Aadac | 0 | NA | NA | NA | NA | NA |
| Mir302a | 0 | NA | NA | NA | NA | NA | | | Aadacl2 | 0 | NA | NA | NA | NA | NA |
| Mir302b | 0 | NA | NA | NA | NA | NA | | | Abcb5 | 0 | NA | NA | NA | NA | NA |
| Mir302c | 0 | NA | NA | NA | NA | NA | | | Abhd16b | 0 | NA | NA | NA | NA | NA |
| Mir302d | 0 | NA | NA | NA | NA | NA | | | Ace3 | 0 | NA | NA | NA | NA | NA |
| Mir3057 | 0.791376948 | 0.082635319 | 0.066510581 | 1.24243869 | 0.214074748 | NA | | | Acpt | 0 | NA | NA | NA | NA | NA |
| Mir3058 | 0.676440077 | 0.007297889 | 0.060582559 | 0.120461886 | 0.90411727 | NA | | | Acrv1 | 0 | NA | NA | NA | NA | NA |
| Mir3059 | 0 | NA | NA | NA | NA | NA | | | Acsm1 | 0 | NA | NA | NA | NA | NA |
| Mir3062 | 0 | NA | NA | NA | NA | NA | | | Actl11 | 0 | NA | NA | NA | NA | NA |
| Mir3063 | 0 | NA | NA | NA | NA | NA | | | Actl7a | 0 | NA | NA | NA | NA | NA |
| Mir3065 | 0 | NA | NA | NA | NA | NA | | | Actrt1 | 0 | NA | NA | NA | NA | NA |
| Mir3066 | 0 | NA | NA | NA | NA | NA | | | Actrt2 | 0 | NA | NA | NA | NA | NA |
| Mir3067 | 0 | NA | NA | NA | NA | NA | | | Adam20 | 0 | NA | NA | NA | NA | NA |
| Mir3068 | 3.272901237 | 0.09265432 | 0.107422202 | 0.862524867 | 0.388398755 | NA | | | Adam24 | 0 | NA | NA | NA | NA | NA |
| Mir3069 | 0 | NA | NA | NA | NA | NA | | | Adam26b | 0 | NA | NA | NA | NA | NA |
| Mir3070a | 4.880662248 | 0.266296126 | 0.130175445 | 2.045670945 | 0.040788756 | NA | | | Adam29 | 0 | NA | NA | NA | NA | NA |
| Mir3070b | 6.499130046 | 0.18487701 | 0.149589309 | 1.235897218 | 0.216496771 | NA | | | Adam30 | 0 | NA | NA | NA | NA | NA |
| Mir3071 | 0 | NA | NA | NA | NA | NA | | | Adam39 | 0 | NA | NA | NA | NA | NA |
| Mir3073 | 0 | NA | NA | NA | NA | NA | | | Adam6a | 0 | NA | NA | NA | NA | NA |
| Mir3074-1 | 0.17833704 | -0.002390466 | 0.031282281 | -0.076415987 | 0.939088151 | NA | | | Adam6b | 0 | NA | NA | NA | NA | NA |
| Mir3074-2 | 0.217751027 | -0.027071875 | 0.031947577 | -0.847384294 | 0.396780956 | NA | | | Adh4 | 0 | NA | NA | NA | NA | NA |
| Mir3075 | 0 | NA | NA | NA | NA | NA | | | Adh6-ps1 | 0 | NA | NA | NA | NA | NA |
| Mir3076 | 0.096071043 | -0.020098795 | 0.027825948 | -0.722304036 | 0.470107572 | NA | | | Adh6a | 0 | NA | NA | NA | NA | NA |
| Mir3077 | 0 | NA | NA | NA | NA | NA | | | Adm2 | 0 | NA | NA | NA | NA | NA |
| Mir3078 | 0 | NA | NA | NA | NA | NA | | | Afm | 0 | NA | NA | NA | NA | NA |
| Mir3081 | 0 | NA | NA | NA | NA | NA | | | Agxt | 0 | NA | NA | NA | NA | NA |
| Mir3083 | 0 | NA | NA | NA | NA | NA | | | Aicda | 0 | NA | NA | NA | NA | NA |
| Mir3085 | 0 | NA | NA | NA | NA | NA | | | Akap4 | 0 | NA | NA | NA | NA | NA |
| Mir3086 | 0 | NA | NA | NA | NA | NA | | | Akr1c20 | 0 | NA | NA | NA | NA | NA |
| Mir3087 | 0 | NA | NA | NA | NA | NA | | | Akr1c6 | 0 | NA | NA | NA | NA | NA |
| Mir3088 | 0 | NA | NA | NA | NA | NA | | | Alpi | 0 | NA | NA | NA | NA | NA |
| Mir3089 | 0 | NA | NA | NA | NA | NA | | | Alppl2 | 0 | NA | NA | NA | NA | NA |
| Mir3091 | 2.100454759 | -0.029999523 | 0.092428139 | -0.324571322 | 0.745505535 | NA | | | Ambp | 0 | NA | NA | NA | NA | NA |
| Mir3092 | 0 | NA | NA | NA | NA | NA | | | Amd1 | 0 | NA | NA | NA | NA | NA |
| Mir3093 | 2.001486636 | 0.026368064 | 0.089586297 | 0.294331446 | 0.768504646 | NA | | | Amd2 | 0 | NA | NA | NA | NA | NA |
| Mir3094 | 0.316104413 | -0.036204367 | 0.036880698 | -0.981661674 | 0.326266553 | NA | | | Amtn | 0 | NA | NA | NA | NA | NA |
| Mir3095 | 0 | NA | NA | NA | NA | NA | | | Amy2a2 | 0 | NA | NA | NA | NA | NA |
| Mir3096 | 0 | NA | NA | NA | NA | NA | | | Amy2a3 | 0 | NA | NA | NA | NA | NA |
| Mir3096b | 0 | NA | NA | NA | NA | NA | | | Amy2a4 | 0 | NA | NA | NA | NA | NA |
| Mir3097 | 0.706253102 | 0.062939234 | 0.058126208 | 1.082803018 | 0.278895866 | NA | | | Amy2a5 | 0 | NA | NA | NA | NA | NA |
| Mir3099 | 0 | NA | NA | NA | NA | NA | | | Ang2 | 0 | NA | NA | NA | NA | NA |
| Mir30a | 0 | NA | NA | NA | NA | NA | | | Ang3 | 0 | NA | NA | NA | NA | NA |
| Mir30b | 0.383703858 | 0.028442486 | 0.048086527 | 0.591485549 | 0.554195135 | NA | | | Ang4 | 0 | NA | NA | NA | NA | NA |
| Mir30c-1 | 0 | NA | NA | NA | NA | NA | | | Ang5 | 0 | NA | NA | NA | NA | NA |
| Mir30c-2 | 0.079983654 | 0.016740032 | 0.027674178 | 0.604897175 | 0.54524732 | NA | | | Ang6 | 0 | NA | NA | NA | NA | NA |
| Mir30d | 0.327267802 | 0.035667551 | 0.037337881 | 0.955264473 | 0.339443969 | NA | | | Ankrd36 | 0 | NA | NA | NA | NA | NA |
| Mir30e | 0 | NA | NA | NA | NA | NA | | | Ankrd53 | 0 | NA | NA | NA | NA | NA |
| Mir31 | 0.309826022 | -0.01193339 | 0.034225532 | -0.348669245 | 0.727337636 | NA | | | Antxrl | 0 | NA | NA | NA | NA | NA |
| Mir3100 | 0.70093952 | 0.072751279 | 0.064177266 | 1.133598915 | 0.256962828 | NA | | | Apcs | 0 | NA | NA | NA | NA | NA |
| Mir3102 | 0 | NA | NA | NA | NA | NA | | | Apoa4 | 0 | NA | NA | NA | NA | NA |
| Mir3103 | 0 | NA | NA | NA | NA | NA | | | Apol10a | 0 | NA | NA | NA | NA | NA |
| Mir3104 | 0.588203125 | 0.0520691 | 0.055442676 | 0.939151999 | 0.34765271 | NA | | | Apol11a | 0 | NA | NA | NA | NA | NA |
| Mir3106 | 0 | NA | NA | NA | NA | NA | | | Apol7e | 0 | NA | NA | NA | NA | NA |
| Mir3107 | 0 | NA | NA | NA | NA | NA | | | Apoo-ps | 0 | NA | NA | NA | NA | NA |
| Mir3108 | 0 | NA | NA | NA | NA | NA | | | Arl14 | 0 | NA | NA | NA | NA | NA |
| Mir3109 | 0 | NA | NA | NA | NA | NA | | | Arrdc5 | 0 | NA | NA | NA | NA | NA |
| Mir3110 | 0 | NA | NA | NA | NA | NA | | | Art2a-ps | 0 | NA | NA | NA | NA | NA |
| Mir3112 | 4.820484465 | 0.195429931 | 0.137986327 | 1.416299248 | 0.156687908 | NA | | | Art2b | 0 | NA | NA | NA | NA | NA |
| Mir32 | 0.107716603 | -0.020477583 | 0.027954224 | -0.732539849 | 0.46383913 | NA | | | Asb9 | 0 | NA | NA | NA | NA | NA |
| Mir320 | 1.168957922 | 0.046872974 | 0.075889022 | 0.617651578 | 0.536805037 | NA | | | Atoh1 | 0 | NA | NA | NA | NA | NA |
| Mir322 | 0 | NA | NA | NA | NA | NA | | | Atp4b | 0 | NA | NA | NA | NA | NA |
| Mir323 | 0.198019769 | 0.025308392 | 0.031628095 | 0.80018706 | 0.423602426 | NA | | | Atp6v0c-ps2 | 0 | NA | NA | NA | NA | NA |
| Mir324 | 0.671194683 | 0.056551227 | 0.056497169 | 1.000956827 | 0.316847681 | NA | | | Avpr2 | 0 | NA | NA | NA | NA | NA |
| Mir326 | 0 | NA | NA | NA | NA | NA | | | B020004C17Rik | 0 | NA | NA | NA | NA | NA |
| Mir328 | 0.389457872 | -0.034711543 | 0.048622122 | -0.713904319 | 0.475286347 | NA | | | B020014A21Rik | 0 | NA | NA | NA | NA | NA |
| Mir329 | 0 | NA | NA | NA | NA | NA | | | BB014433 | 0 | NA | NA | NA | NA | NA |
| Mir33 | 0 | NA | NA | NA | NA | NA | | | BB283400 | 0 | NA | NA | NA | NA | NA |
| Mir330 | 0.203787646 | -0.02671144 | 0.031728022 | -0.841887971 | 0.399850664 | NA | | | BB287469 | 0 | NA | NA | NA | NA | NA |
| Mir335 | 0.681939444 | 0.039928515 | 0.061568448 | 0.648522352 | 0.516647159 | NA | | | BC021614 | 0 | NA | NA | NA | NA | NA |
| Mir337 | 1.634991426 | 0.155901605 | 0.086358302 | 1.805287994 | 0.071029627 | NA | | | BC024386 | 0 | NA | NA | NA | NA | NA |
| Mir338 | 3.325487554 | 0.251093364 | 0.116348254 | 2.158118885 | 0.03091859 | NA | | | BC030870 | 0 | NA | NA | NA | NA | NA |
| Mir339 | 0.422001417 | -0.006176072 | 0.048208622 | -0.128111359 | 0.898060847 | NA | | | BC048502 | 0 | NA | NA | NA | NA | NA |
| Mir340 | 0 | NA | NA | NA | NA | NA | | | BC048562 | 0 | NA | NA | NA | NA | NA |
| Mir344 | 0.192142086 | -0.026384592 | 0.031527717 | -0.836869715 | 0.402665805 | NA | | | BC048602 | 0 | NA | NA | NA | NA | NA |
| Mir344-2 | 6.856383389 | 0.192744819 | 0.143224784 | 1.34575046 | 0.178383007 | NA | | | BC048671 | 0 | NA | NA | NA | NA | NA |
| Mir344b | 0.811177505 | -0.011323829 | 0.062251274 | -0.181905174 | 0.855657144 | NA | | | BC048679 | 0 | NA | NA | NA | NA | NA |
| Mir344c | 0 | NA | NA | NA | NA | NA | | | BC049730 | 0 | NA | NA | NA | NA | NA |
| Mir344d-1 | 0.266310378 | 0.0407635 | 0.040043874 | 1.017970943 | 0.308691765 | NA | | | BC061212 | 0 | NA | NA | NA | NA | NA |
| Mir344d-2 | 0.105923158 | 0.018524693 | 0.027977518 | 0.662127842 | 0.507889294 | NA | | | BC061237 | 0 | NA | NA | NA | NA | NA |
| Mir344d-3 | 0.272987066 | 0.007834485 | 0.033279985 | 0.2354113 | 0.813889516 | NA | | | BC080695 | 0 | NA | NA | NA | NA | NA |
| Mir344e | 0.08731684 | 0.017365056 | 0.027733412 | 0.626142072 | 0.531221759 | NA | | | BC089597 | 0 | NA | NA | NA | NA | NA |
| Mir344f | 0.20671448 | -0.003314027 | 0.03178208 | -0.10427343 | 0.916952363 | NA | | | BC090627 | 0 | NA | NA | NA | NA | NA |
| Mir344g | 1.23487479 | -0.007358988 | 0.080815105 | -0.091059565 | 0.927445261 | NA | | | BC096441 | 0 | NA | NA | NA | NA | NA |
| Mir345 | 2.224772481 | 0.004362455 | 0.102982554 | 0.042361109 | 0.966210831 | NA | | | BC100530 | 0 | NA | NA | NA | NA | NA |
| Mir346 | 0 | NA | NA | NA | NA | NA | | | BC117090 | 0 | NA | NA | NA | NA | NA |
| Mir3471-1 | 0 | NA | NA | NA | NA | NA | | | Banf2 | 0 | NA | NA | NA | NA | NA |
| Mir3473 | 0 | NA | NA | NA | NA | NA | | | Barx1 | 0 | NA | NA | NA | NA | NA |
| Mir3473c | 0.193239998 | 0.025145508 | 0.031532422 | 0.797449311 | 0.42519013 | NA | | | Bcl2a1c | 0 | NA | NA | NA | NA | NA |
| Mir3473d | 0.988148424 | -0.001676897 | 0.057931531 | -0.028946194 | 0.976907504 | NA | | | Bex6 | 0 | NA | NA | NA | NA | NA |
| Mir3474 | 0 | NA | NA | NA | NA | NA | | | Bmp10 | 0 | NA | NA | NA | NA | NA |
| Mir3475 | 0 | NA | NA | NA | NA | NA | | | Boll | 0 | NA | NA | NA | NA | NA |
| Mir34a | 0.20671448 | -0.003314027 | 0.03178208 | -0.10427343 | 0.916952363 | NA | | | Bpifa3 | 0 | NA | NA | NA | NA | NA |
| Mir34b | 0 | NA | NA | NA | NA | NA | | | Bpifb2 | 0 | NA | NA | NA | NA | NA |
| Mir34c | 0 | NA | NA | NA | NA | NA | | | Bsph2 | 0 | NA | NA | NA | NA | NA |
| Mir350 | 0 | NA | NA | NA | NA | NA | | | Btg3 | 0 | NA | NA | NA | NA | NA |
| Mir351 | 0 | NA | NA | NA | NA | NA | | | Btnl1 | 0 | NA | NA | NA | NA | NA |
| Mir3572 | 0 | NA | NA | NA | NA | NA | | | Btnl4 | 0 | NA | NA | NA | NA | NA |
| Mir362 | 0 | NA | NA | NA | NA | NA | | | C130079G13Rik | 0 | NA | NA | NA | NA | NA |
| Mir363 | 0.08731684 | 0.017365056 | 0.027733412 | 0.626142072 | 0.531221759 | NA | | | C1qtnf5 | 0 | NA | NA | NA | NA | NA |
| Mir365-1 | 0.984004379 | -0.031039372 | 0.065916481 | -0.4708894 | 0.637719718 | NA | | | C230079O03Rik | 0 | NA | NA | NA | NA | NA |
| Mir365-2 | 0 | NA | NA | NA | NA | NA | | | C330008G21Rik | 0 | NA | NA | NA | NA | NA |
| Mir367 | 0 | NA | NA | NA | NA | NA | | | C330011F03Rik | 0 | NA | NA | NA | NA | NA |
| Mir369 | 3.201080901 | 0.165684723 | 0.118335717 | 1.400124379 | 0.161476076 | NA | | | C330046G13Rik | 0 | NA | NA | NA | NA | NA |
| Mir370 | 2.605066898 | -0.025067966 | 0.111088766 | -0.225657072 | 0.821468148 | NA | | | C430002E04Rik | 0 | NA | NA | NA | NA | NA |
| Mir374c | 0 | NA | NA | NA | NA | NA | | | C630028M04Rik | 0 | NA | NA | NA | NA | NA |
| Mir375 | 0 | NA | NA | NA | NA | NA | | | C730027H18Rik | 0 | NA | NA | NA | NA | NA |
| Mir376a | 0 | NA | NA | NA | NA | NA | | | C87414 | 0 | NA | NA | NA | NA | NA |
| Mir376b | 0 | NA | NA | NA | NA | NA | | | C87977 | 0 | NA | NA | NA | NA | NA |
| Mir376c | 0.119397641 | -0.020477583 | 0.027954224 | -0.732539845 | 0.463839132 | NA | | | Calhm1 | 0 | NA | NA | NA | NA | NA |
| Mir378b | 0 | NA | NA | NA | NA | NA | | | Calm4 | 0 | NA | NA | NA | NA | NA |
| Mir379 | 0 | NA | NA | NA | NA | NA | | | Capza3 | 0 | NA | NA | NA | NA | NA |
| Mir380 | 0 | NA | NA | NA | NA | NA | | | Catsper1 | 0 | NA | NA | NA | NA | NA |
| Mir381 | 0.096071043 | -0.020098795 | 0.027825948 | -0.722304036 | 0.470107572 | NA | | | Catsperb | 0 | NA | NA | NA | NA | NA |
| Mir382 | 7.259339403 | 0.137739379 | 0.146864577 | 0.937866586 | 0.348312977 | NA | | | Ccdc178 | 0 | NA | NA | NA | NA | NA |
| Mir383 | 0 | NA | NA | NA | NA | NA | | | Ccdc7 | 0 | NA | NA | NA | NA | NA |
| Mir384 | 0.794367966 | -0.094767332 | 0.066368305 | -1.427900438 | 0.153320518 | NA | | | Ccdc70 | 0 | NA | NA | NA | NA | NA |
| Mir3960 | 1.285712233 | 0.048271091 | 0.053775144 | 0.89764688 | 0.369373837 | NA | | | Ccl1 | 0 | NA | NA | NA | NA | NA |
| Mir3966 | 0 | NA | NA | NA | NA | NA | | | Ccl20 | 0 | NA | NA | NA | NA | NA |
| Mir3967 | 0 | NA | NA | NA | NA | NA | | | Ccl21a | 0 | NA | NA | NA | NA | NA |
| Mir3968 | 0 | NA | NA | NA | NA | NA | | | Ccl21b | 0 | NA | NA | NA | NA | NA |
| Mir3969 | 0 | NA | NA | NA | NA | NA | | | Ccl21c | 0 | NA | NA | NA | NA | NA |
| Mir3971 | 0.096071043 | -0.020098795 | 0.027825948 | -0.722304036 | 0.470107572 | NA | | | Ccl26 | 0 | NA | NA | NA | NA | NA |
| Mir409 | 7.371817728 | 0.159969339 | 0.123064958 | 1.299877251 | 0.193643043 | NA | | | Ccl27b | 0 | NA | NA | NA | NA | NA |
| Mir410 | 0 | NA | NA | NA | NA | NA | | | Ccr8 | 0 | NA | NA | NA | NA | NA |
| Mir411 | 0 | NA | NA | NA | NA | NA | | | Cd1d2 | 0 | NA | NA | NA | NA | NA |
| Mir412 | 0.094392817 | 0.017895942 | 0.027818137 | 0.643319236 | 0.520016977 | NA | | | Cd200r3 | 0 | NA | NA | NA | NA | NA |
| Mir423 | 0.681390527 | -0.014222642 | 0.054510777 | -0.26091431 | 0.794158589 | NA | | | Cd5l | 0 | NA | NA | NA | NA | NA |
| Mir429 | 0 | NA | NA | NA | NA | NA | | | Cdx1 | 0 | NA | NA | NA | NA | NA |
| Mir431 | 3.907237645 | 0.135417483 | 0.122655561 | 1.10404683 | 0.269572824 | NA | | | Cdx2 | 0 | NA | NA | NA | NA | NA |
| Mir432 | 1.196030218 | 0.044886332 | 0.075085309 | 0.597804446 | 0.549970424 | NA | | | Cdx4 | 0 | NA | NA | NA | NA | NA |
| Mir433 | 0 | NA | NA | NA | NA | NA | | | Ceacam11 | 0 | NA | NA | NA | NA | NA |
| Mir434 | 3.581388194 | 0.238684672 | 0.128963735 | 1.850789073 | 0.064199904 | NA | | | Ceacam12 | 0 | NA | NA | NA | NA | NA |
| Mir448 | 0 | NA | NA | NA | NA | NA | | | Ceacam13 | 0 | NA | NA | NA | NA | NA |
| Mir449a | 0 | NA | NA | NA | NA | NA | | | Ceacam14 | 0 | NA | NA | NA | NA | NA |
| Mir449b | 0 | NA | NA | NA | NA | NA | | | Ceacam19 | 0 | NA | NA | NA | NA | NA |
| Mir449c | 0 | NA | NA | NA | NA | NA | | | Ceacam20 | 0 | NA | NA | NA | NA | NA |
| Mir450-1 | 0 | NA | NA | NA | NA | NA | | | Ceacam5 | 0 | NA | NA | NA | NA | NA |
| Mir450-2 | 0 | NA | NA | NA | NA | NA | | | Cela3b | 0 | NA | NA | NA | NA | NA |
| Mir450b | 0 | NA | NA | NA | NA | NA | | | Cers1 | 0 | NA | NA | NA | NA | NA |
| Mir451 | 0 | NA | NA | NA | NA | NA | | | Ces1b | 0 | NA | NA | NA | NA | NA |
| Mir452 | 0 | NA | NA | NA | NA | NA | | | Ces1c | 0 | NA | NA | NA | NA | NA |
| Mir455 | 0 | NA | NA | NA | NA | NA | | | Ces2a | 0 | NA | NA | NA | NA | NA |
| Mir463 | 0 | NA | NA | NA | NA | NA | | | Ces2c | 0 | NA | NA | NA | NA | NA |
| Mir465 | 0 | NA | NA | NA | NA | NA | | | Ces2d-ps | 0 | NA | NA | NA | NA | NA |
| Mir465b-1 | 0 | NA | NA | NA | NA | NA | | | Ces2f | 0 | NA | NA | NA | NA | NA |
| Mir465b-2 | 0 | NA | NA | NA | NA | NA | | | Ces2h | 0 | NA | NA | NA | NA | NA |
| Mir465c-1 | 0 | NA | NA | NA | NA | NA | | | Ces3a | 0 | NA | NA | NA | NA | NA |
| Mir465c-2 | 0 | NA | NA | NA | NA | NA | | | Ces3b | 0 | NA | NA | NA | NA | NA |
| Mir466 | 0 | NA | NA | NA | NA | NA | | | Ces4a | 0 | NA | NA | NA | NA | NA |
| Mir4660 | 0 | NA | NA | NA | NA | NA | | | Cetn1 | 0 | NA | NA | NA | NA | NA |
| Mir466g | 0 | NA | NA | NA | NA | NA | | | Cfhr1 | 0 | NA | NA | NA | NA | NA |
| Mir466i | 0 | NA | NA | NA | NA | NA | | | Chi3l4 | 0 | NA | NA | NA | NA | NA |
| Mir467a-1 | 0 | NA | NA | NA | NA | NA | | | Chrnd | 0 | NA | NA | NA | NA | NA |
| Mir467a-10 | 0 | NA | NA | NA | NA | NA | | | Chrng | 0 | NA | NA | NA | NA | NA |
| Mir467a-2 | 0 | NA | NA | NA | NA | NA | | | Cib4 | 0 | NA | NA | NA | NA | NA |
| Mir467a-3 | 0 | NA | NA | NA | NA | NA | | | Cldn17 | 0 | NA | NA | NA | NA | NA |
| Mir467a-4 | 0 | NA | NA | NA | NA | NA | | | Cldn24 | 0 | NA | NA | NA | NA | NA |
| Mir467a-5 | 0 | NA | NA | NA | NA | NA | | | Cldn8 | 0 | NA | NA | NA | NA | NA |
| Mir467a-6 | 0 | NA | NA | NA | NA | NA | | | Clec2g | 0 | NA | NA | NA | NA | NA |
| Mir467a-7 | 0 | NA | NA | NA | NA | NA | | | Clec2h | 0 | NA | NA | NA | NA | NA |
| Mir467a-8 | 0 | NA | NA | NA | NA | NA | | | Clec4b2 | 0 | NA | NA | NA | NA | NA |
| Mir467a-9 | 0 | NA | NA | NA | NA | NA | | | Clpsl2 | 0 | NA | NA | NA | NA | NA |
| Mir467b | 0 | NA | NA | NA | NA | NA | | | Cma2 | 0 | NA | NA | NA | NA | NA |
| Mir467d | 0 | NA | NA | NA | NA | NA | | | Cmtm2b | 0 | NA | NA | NA | NA | NA |
| Mir467f | 0 | NA | NA | NA | NA | NA | | | Cnga1 | 0 | NA | NA | NA | NA | NA |
| Mir468 | 0.62501449 | 0.025957127 | 0.053549922 | 0.48472764 | 0.627869557 | NA | | | Colec10 | 0 | NA | NA | NA | NA | NA |
| Mir470 | 0 | NA | NA | NA | NA | NA | | | Cox8c | 0 | NA | NA | NA | NA | NA |
| Mir471 | 0 | NA | NA | NA | NA | NA | | | Cphx1 | 0 | NA | NA | NA | NA | NA |

| | | | | | | | | | | | | | | | |
|---|---|---|---|---|---|---|---|---|---|---|---|---|---|---|---|
| Mir483 | 0 | NA | NA | NA | NA | NA | | | Cphx2 | 0 | NA | NA | NA | NA | NA |
| Mir484 | 0 | NA | NA | NA | NA | NA | | | Cpn2 | 0 | NA | NA | NA | NA | NA |
| Mir485 | 0.590545564 | 0.020004006 | 0.053347245 | 0.374977293 | 0.707677354 | NA | | | Cpxcr1 | 0 | NA | NA | NA | NA | NA |
| Mir486 | 0.94480822 | 0.019528546 | 0.071623939 | 0.272653891 | 0.785119282 | NA | | | Crisp3 | 0 | NA | NA | NA | NA | NA |
| Mir487b | 0.222620122 | -0.001447027 | 0.032059667 | -0.04513545 | 0.963999345 | NA | | | Crisp4 | 0 | NA | NA | NA | NA | NA |
| Mir488 | 0 | NA | NA | NA | NA | NA | | | Crnn | 0 | NA | NA | NA | NA | NA |
| Mir489 | 0 | NA | NA | NA | NA | NA | | | Crp | 0 | NA | NA | NA | NA | NA |
| Mir490 | 0 | NA | NA | NA | NA | NA | | | Crygc | 0 | NA | NA | NA | NA | NA |
| Mir491 | 2.293765607 | -0.007964255 | 0.104865042 | -0.075947666 | 0.939460735 | NA | | | Crygd | 0 | NA | NA | NA | NA | NA |
| Mir493 | 0.635690241 | -0.003851216 | 0.053467645 | -0.072028909 | 0.942578901 | NA | | | Crygf | 0 | NA | NA | NA | NA | NA |
| Mir494 | 0.256038351 | 0.007085343 | 0.03307033 | 0.214250763 | 0.830351512 | NA | | | Csl | 0 | NA | NA | NA | NA | NA |
| Mir495 | 0 | NA | NA | NA | NA | NA | | | Csn1s2a | 0 | NA | NA | NA | NA | NA |
| Mir496 | 4.52837712 | 0.163733886 | 0.122973017 | 1.33146189 | 0.183037077 | NA | | | Csn3 | 0 | NA | NA | NA | NA | NA |
| Mir497 | 0 | NA | NA | NA | NA | NA | | | Cst10 | 0 | NA | NA | NA | NA | NA |
| Mir499 | 0 | NA | NA | NA | NA | NA | | | Cst13 | 0 | NA | NA | NA | NA | NA |
| Mir500 | 0 | NA | NA | NA | NA | NA | | | Cst8 | 0 | NA | NA | NA | NA | NA |
| Mir501 | 0 | NA | NA | NA | NA | NA | | | Cst9 | 0 | NA | NA | NA | NA | NA |
| Mir503 | 0 | NA | NA | NA | NA | NA | | | Csta | 0 | NA | NA | NA | NA | NA |
| Mir504 | 0 | NA | NA | NA | NA | NA | | | Ctag2 | 0 | NA | NA | NA | NA | NA |
| Mir5046 | 0 | NA | NA | NA | NA | NA | | | Ctf2 | 0 | NA | NA | NA | NA | NA |
| Mir505 | 0 | NA | NA | NA | NA | NA | | | Ctrc | 0 | NA | NA | NA | NA | NA |
| Mir509 | 0 | NA | NA | NA | NA | NA | | | Cts3 | 0 | NA | NA | NA | NA | NA |
| Mir5098 | 0 | NA | NA | NA | NA | NA | | | Cts6 | 0 | NA | NA | NA | NA | NA |
| Mir5100 | 0.502431138 | -0.017833366 | 0.050924809 | -0.350190145 | 0.726196002 | NA | | | Cts7 | 0 | NA | NA | NA | NA | NA |
| Mir5101 | 0 | NA | NA | NA | NA | NA | | | Cts8 | 0 | NA | NA | NA | NA | NA |
| Mir5103 | 3.393330191 | -0.011278602 | 0.127968635 | -0.088135673 | 0.929768843 | NA | | | Cts8-ps | 0 | NA | NA | NA | NA | NA |
| Mir5104 | 0 | NA | NA | NA | NA | NA | | | Ctsj | 0 | NA | NA | NA | NA | NA |
| Mir5106 | 0 | NA | NA | NA | NA | NA | | | Ctsm | 0 | NA | NA | NA | NA | NA |
| Mir5107 | 0.079983654 | 0.016740032 | 0.027674178 | 0.604897175 | 0.54524732 | NA | | | Ctsq | 0 | NA | NA | NA | NA | NA |
| Mir5108 | 0 | NA | NA | NA | NA | NA | | | Ctsr | 0 | NA | NA | NA | NA | NA |
| Mir5109 | 0 | NA | NA | NA | NA | NA | | | Cxcl2 | 0 | NA | NA | NA | NA | NA |
| Mir511 | 0 | NA | NA | NA | NA | NA | | | Cxcl3 | 0 | NA | NA | NA | NA | NA |
| Mir5112 | 0.813522201 | 0.05673995 | 0.064692596 | 0.877070229 | 0.380448494 | NA | | | Cxcr1 | 0 | NA | NA | NA | NA | NA |
| Mir5113 | 0 | NA | NA | NA | NA | NA | | | Cxcr5 | 0 | NA | NA | NA | NA | NA |
| Mir5114 | 0 | NA | NA | NA | NA | NA | | | Cylc1 | 0 | NA | NA | NA | NA | NA |
| Mir5116 | 0 | NA | NA | NA | NA | NA | | | Cylc2 | 0 | NA | NA | NA | NA | NA |
| Mir5117 | 0 | NA | NA | NA | NA | NA | | | Cyp2a12 | 0 | NA | NA | NA | NA | NA |
| Mir5119 | 3.771806447 | 0.167827703 | 0.123225056 | 1.361960864 | 0.173210236 | NA | | | Cyp2a22 | 0 | NA | NA | NA | NA | NA |
| Mir5121 | 0 | NA | NA | NA | NA | NA | | | Cyp2b13 | 0 | NA | NA | NA | NA | NA |
| Mir5122 | 0 | NA | NA | NA | NA | NA | | | Cyp2b19 | 0 | NA | NA | NA | NA | NA |
| Mir5123 | 0.094392817 | 0.017895942 | 0.027818137 | 0.643319236 | 0.520016977 | NA | | | Cyp2b9 | 0 | NA | NA | NA | NA | NA |
| Mir5126 | 0 | NA | NA | NA | NA | NA | | | Cyp2c29 | 0 | NA | NA | NA | NA | NA |
| Mir5127 | 0 | NA | NA | NA | NA | NA | | | Cyp2c37 | 0 | NA | NA | NA | NA | NA |
| Mir5128 | 0.197363271 | -0.000561558 | 0.031650166 | -0.017742642 | 0.985844163 | NA | | | Cyp2c38 | 0 | NA | NA | NA | NA | NA |
| Mir5129 | 0 | NA | NA | NA | NA | NA | | | Cyp2c39 | 0 | NA | NA | NA | NA | NA |
| Mir5130 | 0 | NA | NA | NA | NA | NA | | | Cyp2c40 | 0 | NA | NA | NA | NA | NA |
| Mir5131 | 0 | NA | NA | NA | NA | NA | | | Cyp2c50 | 0 | NA | NA | NA | NA | NA |
| Mir5132 | 0 | NA | NA | NA | NA | NA | | | Cyp2c53-ps | 0 | NA | NA | NA | NA | NA |
| Mir5134 | 0 | NA | NA | NA | NA | NA | | | Cyp2c54 | 0 | NA | NA | NA | NA | NA |
| Mir5135 | 0 | NA | NA | NA | NA | NA | | | Cyp2c66 | 0 | NA | NA | NA | NA | NA |
| Mir5136 | 0 | NA | NA | NA | NA | NA | | | Cyp2c67 | 0 | NA | NA | NA | NA | NA |
| Mir532 | 0 | NA | NA | NA | NA | NA | | | Cyp2c68 | 0 | NA | NA | NA | NA | NA |
| Mir539 | 0 | NA | NA | NA | NA | NA | | | Cyp2c69 | 0 | NA | NA | NA | NA | NA |
| Mir540 | 1.078837746 | 0.02419156 | 0.065681076 | 0.368318573 | 0.712635705 | NA | | | Cyp2d10 | 0 | NA | NA | NA | NA | NA |
| Mir542 | 0 | NA | NA | NA | NA | NA | | | Cyp2d11 | 0 | NA | NA | NA | NA | NA |
| Mir543 | 0.238795281 | -0.027601606 | 0.032267774 | -0.855392297 | 0.392334007 | NA | | | Cyp2d12 | 0 | NA | NA | NA | NA | NA |
| Mir546 | 0 | NA | NA | NA | NA | NA | | | Cyp2d13 | 0 | NA | NA | NA | NA | NA |
| Mir547 | 0 | NA | NA | NA | NA | NA | | | Cyp2d34 | 0 | NA | NA | NA | NA | NA |
| Mir551b | 0 | NA | NA | NA | NA | NA | | | Cyp2d37-ps | 0 | NA | NA | NA | NA | NA |
| Mir574 | 0 | NA | NA | NA | NA | NA | | | Cyp2d40 | 0 | NA | NA | NA | NA | NA |
| Mir582 | 0 | NA | NA | NA | NA | NA | | | Cyp2d9 | 0 | NA | NA | NA | NA | NA |
| Mir592 | 0 | NA | NA | NA | NA | NA | | | Cyp2j11 | 0 | NA | NA | NA | NA | NA |
| Mir598 | 0.096071043 | -0.020098795 | 0.027825948 | -0.722304036 | 0.470107572 | NA | | | Cyp2j5 | 0 | NA | NA | NA | NA | NA |
| Mir599 | 0 | NA | NA | NA | NA | NA | | | Cyp2t4 | 0 | NA | NA | NA | NA | NA |
| Mir615 | 0 | NA | NA | NA | NA | NA | | | Cyp3a11 | 0 | NA | NA | NA | NA | NA |
| Mir653 | 0 | NA | NA | NA | NA | NA | | | Cyp3a16 | 0 | NA | NA | NA | NA | NA |
| Mir654 | 0.176054697 | -0.002190495 | 0.031239146 | -0.070120197 | 0.944097991 | NA | | | Cyp3a25 | 0 | NA | NA | NA | NA | NA |
| Mir664 | 0 | NA | NA | NA | NA | NA | | | Cyp3a41a | 0 | NA | NA | NA | NA | NA |
| Mir665 | 1.005556649 | -0.004527101 | 0.071578148 | -0.063246976 | 0.949569839 | NA | | | Cyp3a41b | 0 | NA | NA | NA | NA | NA |
| Mir666 | 0 | NA | NA | NA | NA | NA | | | Cyp3a44 | 0 | NA | NA | NA | NA | NA |
| Mir667 | 0 | NA | NA | NA | NA | NA | | | Cyp3a59 | 0 | NA | NA | NA | NA | NA |
| Mir668 | 5.45249312 | 0.092449238 | 0.139097008 | 0.664638584 | 0.506281684 | NA | | | Cyp4a10 | 0 | NA | NA | NA | NA | NA |
| Mir669a-1 | 0 | NA | NA | NA | NA | NA | | | Cyp4a12a | 0 | NA | NA | NA | NA | NA |
| Mir669a-10 | 0 | NA | NA | NA | NA | NA | | | Cyp4a14 | 0 | NA | NA | NA | NA | NA |
| Mir669a-11 | 0 | NA | NA | NA | NA | NA | | | Cyp4a29-ps | 0 | NA | NA | NA | NA | NA |
| Mir669a-12 | 0 | NA | NA | NA | NA | NA | | | Cyp4a30b-ps | 0 | NA | NA | NA | NA | NA |
| Mir669a-2 | 0 | NA | NA | NA | NA | NA | | | Cyp4a31 | 0 | NA | NA | NA | NA | NA |
| Mir669a-3 | 0 | NA | NA | NA | NA | NA | | | Cyp4a32 | 0 | NA | NA | NA | NA | NA |
| Mir669a-4 | 0 | NA | NA | NA | NA | NA | | | Cyp7a1 | 0 | NA | NA | NA | NA | NA |
| Mir669a-5 | 0 | NA | NA | NA | NA | NA | | | Cyp8b1 | 0 | NA | NA | NA | NA | NA |
| Mir669a-6 | 0 | NA | NA | NA | NA | NA | | | Cypt1 | 0 | NA | NA | NA | NA | NA |
| Mir669a-7 | 0 | NA | NA | NA | NA | NA | | | Cypt10 | 0 | NA | NA | NA | NA | NA |
| Mir669a-8 | 0 | NA | NA | NA | NA | NA | | | Cypt14 | 0 | NA | NA | NA | NA | NA |
| Mir669a-9 | 0 | NA | NA | NA | NA | NA | | | Cypt15 | 0 | NA | NA | NA | NA | NA |
| Mir669b | 0 | NA | NA | NA | NA | NA | | | Cypt2 | 0 | NA | NA | NA | NA | NA |
| Mir669c | 0 | NA | NA | NA | NA | NA | | | Cypt3 | 0 | NA | NA | NA | NA | NA |
| Mir669e | 0 | NA | NA | NA | NA | NA | | | Cypt4 | 0 | NA | NA | NA | NA | NA |
| Mir669g | 0 | NA | NA | NA | NA | NA | | | Cypt7 | 0 | NA | NA | NA | NA | NA |
| Mir669h | 0 | NA | NA | NA | NA | NA | | | Cypt8 | 0 | NA | NA | NA | NA | NA |
| Mir669i | 0 | NA | NA | NA | NA | NA | | | Cypt9 | 0 | NA | NA | NA | NA | NA |
| Mir669j | 0 | NA | NA | NA | NA | NA | | | D030024E09Rik | 0 | NA | NA | NA | NA | NA |
| Mir669k | 0 | NA | NA | NA | NA | NA | | | D130009I18Rik | 0 | NA | NA | NA | NA | NA |
| Mir669m-1 | 0 | NA | NA | NA | NA | NA | | | D130058E03 | 0 | NA | NA | NA | NA | NA |
| Mir669m-2 | 0 | NA | NA | NA | NA | NA | | | D1Pas1 | 0 | NA | NA | NA | NA | NA |
| Mir669p-1 | 0 | NA | NA | NA | NA | NA | | | D530049I02Rik | 0 | NA | NA | NA | NA | NA |
| Mir669p-2 | 0 | NA | NA | NA | NA | NA | | | D5Ertd577e | 0 | NA | NA | NA | NA | NA |
| Mir670 | 1.415539497 | -0.013598678 | 0.061933765 | -0.219568086 | 0.826207549 | NA | | | D730050B12Rik | 0 | NA | NA | NA | NA | NA |
| Mir671 | 0 | NA | NA | NA | NA | NA | | | D7Ertd143e | 0 | NA | NA | NA | NA | NA |
| Mir672 | 0 | NA | NA | NA | NA | NA | | | D830005E20Rik | 0 | NA | NA | NA | NA | NA |
| Mir673 | 0 | NA | NA | NA | NA | NA | | | DXBay18 | 0 | NA | NA | NA | NA | NA |
| Mir674 | 2.601193737 | 0.090443513 | 0.115514443 | 0.782962812 | 0.43364895 | NA | | | Daf2 | 0 | NA | NA | NA | NA | NA |
| Mir675 | 0 | NA | NA | NA | NA | NA | | | Dbx1 | 0 | NA | NA | NA | NA | NA |
| Mir676 | 0 | NA | NA | NA | NA | NA | | | Dcpp1 | 0 | NA | NA | NA | NA | NA |
| Mir677 | 7.452538808 | 0.21603724 | 0.150460123 | 1.435843828 | 0.151046785 | NA | | | Dcpp2 | 0 | NA | NA | NA | NA | NA |
| Mir678 | 0 | NA | NA | NA | NA | NA | | | Dcpp3 | 0 | NA | NA | NA | NA | NA |
| Mir679 | 0 | NA | NA | NA | NA | NA | | | Ddi1 | 0 | NA | NA | NA | NA | NA |
| Mir680-1 | 0 | NA | NA | NA | NA | NA | | | Defa-ps1 | 0 | NA | NA | NA | NA | NA |
| Mir680-2 | 0 | NA | NA | NA | NA | NA | | | Defa-ps12 | 0 | NA | NA | NA | NA | NA |
| Mir680-3 | 0 | NA | NA | NA | NA | NA | | | Defa-ps13 | 0 | NA | NA | NA | NA | NA |
| Mir681 | 0.422179562 | -0.00343537 | 0.036938882 | -0.093001469 | 0.925902394 | NA | | | Defa-rs1 | 0 | NA | NA | NA | NA | NA |
| Mir683-1 | 0 | NA | NA | NA | NA | NA | | | Defa-rs7 | 0 | NA | NA | NA | NA | NA |
| Mir683-2 | 0 | NA | NA | NA | NA | NA | | | Defa17 | 0 | NA | NA | NA | NA | NA |
| Mir684-1 | 0.08731684 | 0.017365056 | 0.027733412 | 0.626142072 | 0.531221759 | NA | | | Defa20 | 0 | NA | NA | NA | NA | NA |
| Mir684-2 | 0 | NA | NA | NA | NA | NA | | | Defa21 | 0 | NA | NA | NA | NA | NA |
| Mir686 | 0 | NA | NA | NA | NA | NA | | | Defa22 | 0 | NA | NA | NA | NA | NA |
| Mir687 | 0 | NA | NA | NA | NA | NA | | | Defa23 | 0 | NA | NA | NA | NA | NA |
| Mir688 | 1.329130058 | 0.089015312 | 0.091802701 | 0.969637186 | 0.33222737 | NA | | | Defa24 | 0 | NA | NA | NA | NA | NA |
| Mir690 | 0 | NA | NA | NA | NA | NA | | | Defa25 | 0 | NA | NA | NA | NA | NA |
| Mir691 | 0 | NA | NA | NA | NA | NA | | | Defa26 | 0 | NA | NA | NA | NA | NA |
| Mir692-1 | 0 | NA | NA | NA | NA | NA | | | Defa3 | 0 | NA | NA | NA | NA | NA |
| Mir693 | 0.297029653 | 0.02866449 | 0.033517577 | 0.855207712 | 0.392436167 | NA | | | Defa4 | 0 | NA | NA | NA | NA | NA |
| Mir694 | 0 | NA | NA | NA | NA | NA | | | Defa5 | 0 | NA | NA | NA | NA | NA |
| Mir695 | 0 | NA | NA | NA | NA | NA | | | Defa6 | 0 | NA | NA | NA | NA | NA |
| Mir697 | 4.83199621 | 0.242706135 | 0.128600216 | 1.887291823 | 0.059121086 | NA | | | Defb13 | 0 | NA | NA | NA | NA | NA |
| Mir698 | 0 | NA | NA | NA | NA | NA | | | Defb15 | 0 | NA | NA | NA | NA | NA |
| Mir7-1 | 6.288786782 | 0.176047001 | 0.141922735 | 1.240442562 | 0.214811747 | NA | | | Defb18 | 0 | NA | NA | NA | NA | NA |
| Mir7-2 | 0 | NA | NA | NA | NA | NA | | | Defb19 | 0 | NA | NA | NA | NA | NA |
| Mir700 | 0.299325859 | 0.043567584 | 0.041359933 | 1.053376566 | 0.29216844 | NA | | | Defb2 | 0 | NA | NA | NA | NA | NA |
| Mir701 | 0 | NA | NA | NA | NA | NA | | | Defb21 | 0 | NA | NA | NA | NA | NA |
| Mir702 | 0 | NA | NA | NA | NA | NA | | | Defb22 | 0 | NA | NA | NA | NA | NA |
| Mir703 | 7.726868215 | 0.02662373 | 0.152955264 | 0.174062199 | 0.86181658 | NA | | | Defb23 | 0 | NA | NA | NA | NA | NA |
| Mir704 | 0 | NA | NA | NA | NA | NA | | | Defb26 | 0 | NA | NA | NA | NA | NA |
| Mir705 | 0 | NA | NA | NA | NA | NA | | | Defb29 | 0 | NA | NA | NA | NA | NA |
| Mir706 | 0 | NA | NA | NA | NA | NA | | | Defb3 | 0 | NA | NA | NA | NA | NA |
| Mir708 | 0 | NA | NA | NA | NA | NA | | | Defb34 | 0 | NA | NA | NA | NA | NA |
| Mir709 | 3.257913912 | -0.000278494 | 0.107607283 | -0.002588055 | 0.997935034 | NA | | | Defb35 | 0 | NA | NA | NA | NA | NA |
| Mir710 | 0 | NA | NA | NA | NA | NA | | | Defb36 | 0 | NA | NA | NA | NA | NA |
| Mir711 | 0.079983654 | 0.016740032 | 0.027674178 | 0.604897175 | 0.54524732 | NA | | | Defb37 | 0 | NA | NA | NA | NA | NA |
| Mir713 | 0 | NA | NA | NA | NA | NA | | | Defb38 | 0 | NA | NA | NA | NA | NA |
| Mir717 | 0 | NA | NA | NA | NA | NA | | | Defb39 | 0 | NA | NA | NA | NA | NA |
| Mir718 | 0 | NA | NA | NA | NA | NA | | | Defb4 | 0 | NA | NA | NA | NA | NA |
| Mir719 | 0 | NA | NA | NA | NA | NA | | | Defb40 | 0 | NA | NA | NA | NA | NA |
| Mir721 | 0 | NA | NA | NA | NA | NA | | | Defb43 | 0 | NA | NA | NA | NA | NA |
| Mir741 | 0 | NA | NA | NA | NA | NA | | | Defb44-ps | 0 | NA | NA | NA | NA | NA |
| Mir742 | 0 | NA | NA | NA | NA | NA | | | Defb45 | 0 | NA | NA | NA | NA | NA |
| Mir743 | 0 | NA | NA | NA | NA | NA | | | Defb46 | 0 | NA | NA | NA | NA | NA |
| Mir743b | 0 | NA | NA | NA | NA | NA | | | Defb48 | 0 | NA | NA | NA | NA | NA |
| Mir744 | 1.087655921 | -0.092254674 | 0.079640209 | -1.158393175 | 0.246703623 | NA | | | Defb5 | 0 | NA | NA | NA | NA | NA |
| Mir758 | 0 | NA | NA | NA | NA | NA | | | Defb50 | 0 | NA | NA | NA | NA | NA |
| Mir759 | 0 | NA | NA | NA | NA | NA | | | Defb7 | 0 | NA | NA | NA | NA | NA |
| Mir760 | 2.69608766 | 0.077396299 | 0.110639458 | 0.699536143 | 0.484217034 | NA | | | Defb8 | 0 | NA | NA | NA | NA | NA |
| Mir761 | 6.735877293 | 0.133968576 | 0.146888601 | 0.912042016 | 0.3617466 | NA | | | Defb9 | 0 | NA | NA | NA | NA | NA |
| Mir762 | 0.196706773 | -0.026517754 | 0.031609464 | -0.838918169 | 0.401515231 | NA | | | Dio1 | 0 | NA | NA | NA | NA | NA |
| Mir763 | 0 | NA | NA | NA | NA | NA | | | Dkk4 | 0 | NA | NA | NA | NA | NA |
| Mir764 | 0 | NA | NA | NA | NA | NA | | | Dmrt1 | 0 | NA | NA | NA | NA | NA |
| Mir767 | 0 | NA | NA | NA | NA | NA | | | Dmrtc1b | 0 | NA | NA | NA | NA | NA |
| Mir770 | 0 | NA | NA | NA | NA | NA | | | Dmrtc1c1 | 0 | NA | NA | NA | NA | NA |
| Mir7b | 0.412413621 | -0.026778838 | 0.043795168 | -0.611456443 | 0.540897446 | NA | | | Dmrtc1c2 | 0 | NA | NA | NA | NA | NA |
| Mir802 | 0 | NA | NA | NA | NA | NA | | | Dnajb8 | 0 | NA | NA | NA | NA | NA |
| Mir804 | 0.581676142 | 0.059713211 | 0.060372883 | 0.989073377 | 0.322627242 | NA | | | Doxl2 | 0 | NA | NA | NA | NA | NA |
| Mir871 | 0 | NA | NA | NA | NA | NA | | | Dpep3 | 0 | NA | NA | NA | NA | NA |
| Mir872 | 0.079983654 | 0.016740032 | 0.027674178 | 0.604897175 | 0.54524732 | NA | | | Dppa3 | 0 | NA | NA | NA | NA | NA |
| Mir873 | 0.389505357 | -0.025109253 | 0.043180094 | -0.581500646 | 0.560903083 | NA | | | Dppa4 | 0 | NA | NA | NA | NA | NA |
| Mir874 | 0.87355605 | 0.004156921 | 0.070283927 | 0.059144693 | 0.952836861 | NA | | | Dsg4 | 0 | NA | NA | NA | NA | NA |
| Mir875 | 0 | NA | NA | NA | NA | NA | | | Dusp21 | 0 | NA | NA | NA | NA | NA |
| Mir876 | 0 | NA | NA | NA | NA | NA | | | Duxbl1 | 0 | NA | NA | NA | NA | NA |
| Mir877 | 0 | NA | NA | NA | NA | NA | | | Duxbl2 | 0 | NA | NA | NA | NA | NA |
| Mir878 | 0 | NA | NA | NA | NA | NA | | | Duxbl3 | 0 | NA | NA | NA | NA | NA |
| Mir879 | 0 | NA | NA | NA | NA | NA | | | Dynap | 0 | NA | NA | NA | NA | NA |
| Mir880 | 0 | NA | NA | NA | NA | NA | | | Dynlt1c | 0 | NA | NA | NA | NA | NA |
| Mir881 | 0 | NA | NA | NA | NA | NA | | | Dynlt1f | 0 | NA | NA | NA | NA | NA |
| Mir882 | 0 | NA | NA | NA | NA | NA | | | E030018B13Rik | 0 | NA | NA | NA | NA | NA |
| Mir883a | 0 | NA | NA | NA | NA | NA | | | E330014E10Rik | 0 | NA | NA | NA | NA | NA |
| Mir883b | 0 | NA | NA | NA | NA | NA | | | EU599041 | 0 | NA | NA | NA | NA | NA |
| Mir9-1 | 4.354946689 | 0.154274195 | 0.13631842 | 1.131719364 | 0.257752438 | NA | | | Ear10 | 0 | NA | NA | NA | NA | NA |
| Mir9-3 | 0.574762868 | 0.049560631 | 0.055201391 | 0.897814902 | 0.369284239 | NA | | | Ear12 | 0 | NA | NA | NA | NA | NA |
| Mir92-1 | 0 | NA | NA | NA | NA | NA | | | Ear14 | 0 | NA | NA | NA | NA | NA |
| Mir92-2 | 0 | NA | NA | NA | NA | NA | | | Ear3 | 0 | NA | NA | NA | NA | NA |
| Mir92b | 0 | NA | NA | NA | NA | NA | | | Ear4 | 0 | NA | NA | NA | NA | NA |
| Mir93 | 0.685873859 | 0.036908102 | 0.061591586 | 0.599239353 | 0.549013284 | NA | | | Ear5 | 0 | NA | NA | NA | NA | NA |
| Mir96 | 0 | NA | NA | NA | NA | NA | | | Ear6 | 0 | NA | NA | NA | NA | NA |
| Mir98 | 0 | NA | NA | NA | NA | NA | | | Ear7 | 0 | NA | NA | NA | NA | NA |
| Mir99a | 1.058404621 | 0.052246972 | 0.067612117 | 0.772745691 | 0.4396729 | NA | | | Eddm3b | 0 | NA | NA | NA | NA | NA |
| Mir99b | 0.29182997 | 0.032201186 | 0.035537591 | 0.906116166 | 0.36487437 | NA | | | Egfbp2 | 0 | NA | NA | NA | NA | NA |
| Mira | 0.957081083 | 0.088239072 | 0.078769759 | 1.120215085 | 0.262622117 | NA | | | Eif3j1 | 0 | NA | NA | NA | NA | NA |
| Mirlet7a-1 | 1.968685763 | -0.017095067 | 0.101609264 | -0.168243196 | 0.866391959 | NA | | | Eif3j2 | 0 | NA | NA | NA | NA | NA |
| Mirlet7a-2 | 0 | NA | NA | NA | NA | NA | | | Enam | 0 | NA | NA | NA | NA | NA |
| Mirlet7c-1 | 0.079983654 | 0.016740032 | 0.027674178 | 0.604897175 | 0.54524732 | NA | | | Enpp7 | 0 | NA | NA | NA | NA | NA |
| Mirlet7e | 1.250852019 | 0.088818589 | 0.081499644 | 1.089803398 | 0.275799757 | NA | | | Eras | 0 | NA | NA | NA | NA | NA |
| Mirlet7f-1 | 2.284652334 | 0.146834136 | 0.097290874 | 1.50922826 | 0.131240461 | NA | | | Esx1 | 0 | NA | NA | NA | NA | NA |
| Mirlet7f-2 | 0 | NA | NA | NA | NA | NA | | | Etd | 0 | NA | NA | NA | NA | NA |
| Mirlet7g | 0 | NA | NA | NA | NA | NA | | | Etv2 | 0 | NA | NA | NA | NA | NA |
| Mirlet7i | 1.93374864 | -0.039596899 | 0.087544269 | -0.45230715 | 0.651047726 | NA | | | Evx2 | 0 | NA | NA | NA | NA | NA |
| Mixl1 | 0 | NA | NA | NA | NA | NA | | | F13b | 0 | NA | NA | NA | NA | NA |
| Mlana | 1.605273873 | -0.035104211 | 0.061353749 | -0.572160804 | 0.567213039 | NA | | | F630111L10Rik | 0 | NA | NA | NA | NA | NA |
| Mmp10 | 0.079983654 | 0.016740032 | 0.027674178 | 0.604897175 | 0.54524732 | NA | | | F830002L21Rik | 0 | NA | NA | NA | NA | NA |
| Mmp12 | 2.696035589 | 0.289662924 | 0.117370765 | 2.467930788 | 0.013589659 | NA | | | Fabp1 | 0 | NA | NA | NA | NA | NA |
| Mmp13 | 1.151436786 | 0.019515823 | 0.079789089 | 0.244592625 | 0.806771875 | NA | | | Fabp2 | 0 | NA | NA | NA | NA | NA |
| Mmp1a | 0 | NA | NA | NA | NA | NA | | | Fabp6 | 0 | NA | NA | NA | NA | NA |
| Mmp1b | 0 | NA | NA | NA | NA | NA | | | Fabp9 | 0 | NA | NA | NA | NA | NA |
| Mmp20 | 0.098353386 | -0.020212831 | 0.02786466 | -0.725393061 | 0.468210929 | NA | | | Fam115e | 0 | NA | NA | NA | NA | NA |
| Mmp21 | 3.056002172 | 0.345337112 | 0.123704552 | 2.791628179 | 0.005244358 | NA | | | Fam122c | 0 | NA | NA | NA | NA | NA |
| Mmp27 | 0 | NA | NA | NA | NA | NA | | | Fam162b | 0 | NA | NA | NA | NA | NA |
| Mmp3 | 0.456546308 | -0.046631236 | 0.041759841 | -1.116652618 | 0.264142884 | NA | | | Fam177a | 0 | NA | NA | NA | NA | NA |
| Mmp7 | 0 | NA | NA | NA | NA | NA | | | Fam24a | 0 | NA | NA | NA | NA | NA |
| Mmp8 | 2.287495582 | -0.025068287 | 0.091182534 | -0.274924221 | 0.783374459 | NA | | | Fam47c | 0 | NA | NA | NA | NA | NA |
| Mmrn1 | 0.947272025 | 0.019086986 | 0.072770315 | 0.262290825 | 0.793097235 | NA | | | Fam50b | 0 | NA | NA | NA | NA | NA |
| Mnda | 5.093501151 | 0.270803598 | 0.126623554 | 2.138651066 | 0.032463938 | NA | | | Fam71a | 0 | NA | NA | NA | NA | NA |
| Mnx1 | 0.098353386 | -0.020212831 | 0.02786466 | -0.725393061 | 0.468210929 | NA | | | Fam71f1 | 0 | NA | NA | NA | NA | NA |
| Moap1 | 3.153424689 | -0.001762253 | 0.11992441 | -0.014694701 | 0.988275747 | NA | | | Fam83c | 0 | NA | NA | NA | NA | NA |
| Mogat1 | 3.653524889 | -0.057343705 | 0.122701267 | -0.467344033 | 0.640253757 | NA | | | Fbxw14 | 0 | NA | NA | NA | NA | NA |
| Mogat2 | 0.745606615 | -0.061062738 | 0.067492909 | -0.904728193 | 0.365609401 | NA | | | Fbxw16 | 0 | NA | NA | NA | NA | NA |
| Morc1 | 1.391559479 | 0.015745358 | 0.061596964 | 0.255619067 | 0.798244994 | NA | | | Fbxw20 | 0 | NA | NA | NA | NA | NA |
| Morn3 | 4.154172981 | 0.041431942 | 0.128366215 | 0.322763604 | 0.746874279 | NA | | | Fbxw21 | 0 | NA | NA | NA | NA | NA |
| Mos | 3.193792803 | 0.102481655 | 0.121620599 | 0.842634027 | 0.399433154 | NA | | | Fbxw22 | 0 | NA | NA | NA | NA | NA |
| Mospd4 | 1.28926284 | -0.115640977 | 0.084031933 | -1.37615514 | 0.168773607 | NA | | | Fbxw26 | 0 | NA | NA | NA | NA | NA |
| Mov10l1 | 0.195033442 | -0.002424765 | 0.031596358 | -0.076741901 | 0.93882887 | NA | | | Fbxw28 | 0 | NA | NA | NA | NA | NA |
| Moxd2 | 0.098353386 | -0.020212831 | 0.02786466 | -0.725393061 | 0.468210929 | NA | | | Fcrl5 | 0 | NA | NA | NA | NA | NA |
| Mpo | 3.785362582 | -0.217414719 | 0.12679216 | -1.714733145 | 0.086394164 | NA | | | Ffar1 | 0 | NA | NA | NA | NA | NA |
| Mptx1 | 0 | NA | NA | NA | NA | NA | | | Fgf4 | 0 | NA | NA | NA | NA | NA |
| Mptx2 | 0 | NA | NA | NA | NA | NA | | | Fgf6 | 0 | NA | NA | NA | NA | NA |
| Mpz | 7.078823613 | -0.169653088 | 0.131092866 | -1.294144324 | 0.195615574 | NA | | | Fgg | 0 | NA | NA | NA | NA | NA |
| Mrgpra1 | 0 | NA | NA | NA | NA | NA | | | Fitm1 | 0 | NA | NA | NA | NA | NA |
| Mrgpra2a | 0 | NA | NA | NA | NA | NA | | | Flg2 | 0 | NA | NA | NA | NA | NA |
| Mrgpra2b | 0.079983654 | 0.016740032 | 0.027674178 | 0.604897175 | 0.54524732 | NA | | | Fmo3 | 0 | NA | NA | NA | NA | NA |
| Mrgpra3 | 0 | NA | NA | NA | NA | NA | | | Fnd3c2 | 0 | NA | NA | NA | NA | NA |
| Mrgpra4 | 0 | NA | NA | NA | NA | NA | | | Foxa3 | 0 | NA | NA | NA | NA | NA |
| Mrgpra6 | 0 | NA | NA | NA | NA | NA | | | Foxe1 | 0 | NA | NA | NA | NA | NA |
| Mrgpra9 | 0 | NA | NA | NA | NA | NA | | | Foxe3 | 0 | NA | NA | NA | NA | NA |
| Mrgprb1 | 0.119397641 | -0.020477583 | 0.027954224 | -0.732539845 | 0.463839132 | NA | | | Foxi3 | 0 | NA | NA | NA | NA | NA |
| Mrgprb2 | 0 | NA | NA | NA | NA | NA | | | Foxr1 | 0 | NA | NA | NA | NA | NA |
| Mrgprb3 | 0 | NA | NA | NA | NA | NA | | | Fpr-rs3 | 0 | NA | NA | NA | NA | NA |
| Mrgprb4 | 0 | NA | NA | NA | NA | NA | | | Fpr-rs4 | 0 | NA | NA | NA | NA | NA |
| Mrgprb5 | 0 | NA | NA | NA | NA | NA | | | Fpr-rs6 | 0 | NA | NA | NA | NA | NA |
| Mrgprb8 | 0 | NA | NA | NA | NA | NA | | | Fpr3 | 0 | NA | NA | NA | NA | NA |

| | | | | | | | | | | | | | | | |
|---|---|---|---|---|---|---|---|---|---|---|---|---|---|---|---|
| Mrgprd | 0.196706773 | -0.026517754 | 0.031609464 | -0.838918169 | 0.401515231 | NA | | | Fthl17 | 0 | NA | NA | NA | NA | NA |
| Mrgprg | 0 | NA | NA | NA | NA | NA | | | Ftmt | 0 | NA | NA | NA | NA | NA |
| Mrgprh | 5.56264731 | -0.119494366 | 0.146952059 | -0.813152035 | 0.416130896 | NA | | | G630055G22Rik | 0 | NA | NA | NA | NA | NA |
| Mrgprx1 | 0 | NA | NA | NA | NA | NA | | | G630071F17Rik | 0 | NA | NA | NA | NA | NA |
| Mrgprx2 | 0 | NA | NA | NA | NA | NA | | | G6b | 0 | NA | NA | NA | NA | NA |
| Mroh2a | 1.308692255 | 0.012060884 | 0.082918141 | 0.145455316 | 0.884351393 | NA | | | G6pd2 | 0 | NA | NA | NA | NA | NA |
| Mroh2b | 0.408027608 | -0.004440697 | 0.036642628 | -0.121189371 | 0.903541043 | NA | | | Gata1 | 0 | NA | NA | NA | NA | NA |
| Mroh4 | 3.747974313 | 0.275704126 | 0.122883014 | 2.243630889 | 0.02485616 | NA | | | Gata4 | 0 | NA | NA | NA | NA | NA |
| Mroh9 | 0.098353386 | -0.020212831 | 0.02786466 | -0.725393061 | 0.468210929 | NA | | | Gata5 | 0 | NA | NA | NA | NA | NA |
| Mrps6 | 615.8638723 | 0.063948532 | 0.126733364 | 0.504591134 | NA | NA | | | Gbp1 | 0 | NA | NA | NA | NA | NA |
| Ms4a1 | 0.828011001 | -0.008119904 | 0.062313387 | -0.130307538 | 0.896323116 | NA | | | Gcnt3 | 0 | NA | NA | NA | NA | NA |
| Ms4a10 | 0 | NA | NA | NA | NA | NA | | | Gcnt7 | 0 | NA | NA | NA | NA | NA |
| Ms4a13 | 0 | NA | NA | NA | NA | NA | | | Gdf1 | 0 | NA | NA | NA | NA | NA |
| Ms4a18 | 0 | NA | NA | NA | NA | NA | | | Gfi1b | 0 | NA | NA | NA | NA | NA |
| Ms4a2 | 0.58646488 | 0.050588692 | 0.054651257 | 0.925663823 | 0.354620714 | NA | | | Gif | 0 | NA | NA | NA | NA | NA |
| Ms4a3 | 0.849765129 | -0.040631658 | 0.062482766 | -0.65028583 | 0.515507608 | NA | | | Gip | 0 | NA | NA | NA | NA | NA |
| Ms4a4b | 0.698099462 | 0.00568318 | 0.054502766 | 0.104273234 | 0.916952518 | NA | | | Gkn2 | 0 | NA | NA | NA | NA | NA |
| Ms4a4c | 2.191486038 | 0.112669057 | 0.097010559 | 1.16141025 | 0.245475102 | NA | | | Glipr1l2 | 0 | NA | NA | NA | NA | NA |
| Ms4a4d | 0.68531206 | 0.075058946 | 0.063967246 | 1.17339654 | 0.240636824 | NA | | | Glt6d1 | 0 | NA | NA | NA | NA | NA |
| Ms4a5 | 0 | NA | NA | NA | NA | NA | | | Gm10024 | 0 | NA | NA | NA | NA | NA |
| Ms4a8a | 0.838007076 | 0.022662155 | 0.061762684 | 0.366923086 | 0.713676391 | NA | | | Gm10052 | 0 | NA | NA | NA | NA | NA |
| Msgn1 | 0 | NA | NA | NA | NA | NA | | | Gm10057 | 0 | NA | NA | NA | NA | NA |
| Msh4 | 0.757306031 | 0.042257997 | 0.058686886 | 0.720058609 | 0.471488911 | NA | | | Gm10058 | 0 | NA | NA | NA | NA | NA |
| Msmb | 0 | NA | NA | NA | NA | NA | | | Gm10081 | 0 | NA | NA | NA | NA | NA |
| Msmp | 0.897514379 | 0.004682035 | 0.069324716 | 0.067537749 | 0.946153612 | NA | | | Gm10094 | 0 | NA | NA | NA | NA | NA |
| Mst1 | 0.594987921 | 0.017868075 | 0.05269327 | 0.339095951 | 0.734537448 | NA | | | Gm10096 | 0 | NA | NA | NA | NA | NA |
| Msx1as | 0.185906812 | 0.024895454 | 0.031385107 | 0.793225068 | 0.427646705 | NA | | | Gm10100 | 0 | NA | NA | NA | NA | NA |
| Msx3 | 3.894539214 | -0.350889337 | 0.126935976 | -2.764301712 | 0.005704477 | NA | | | Gm10104 | 0 | NA | NA | NA | NA | NA |
| Mt4 | 0 | NA | NA | NA | NA | NA | | | Gm10142 | 0 | NA | NA | NA | NA | NA |
| Mtap7d3 | 2.627510154 | 0.073243988 | 0.112918892 | 0.648642458 | 0.516569505 | NA | | | Gm10147 | 0 | NA | NA | NA | NA | NA |
| Mtl5 | 4.998152067 | 0.213281802 | 0.1428941 | 1.492586483 | 0.135545464 | NA | | | Gm10228 | 0 | NA | NA | NA | NA | NA |
| Mtnr1a | 0.638084606 | 0.0288786 | 0.053770188 | 0.537074568 | 0.591216107 | NA | | | Gm10229 | 0 | NA | NA | NA | NA | NA |
| Mtnr1b | 0 | NA | NA | NA | NA | NA | | | Gm10230 | 0 | NA | NA | NA | NA | NA |
| Muc13 | 1.682814255 | 0.013843078 | 0.087901789 | 0.157483463 | 0.87486384 | NA | | | Gm10256 | 0 | NA | NA | NA | NA | NA |
| Muc15 | 5.511925791 | 0.004302542 | 0.147961621 | 0.029078767 | 0.97680177 | NA | | | Gm10267 | 0 | NA | NA | NA | NA | NA |
| Muc19 | 0.498271657 | 0.024449561 | 0.039505462 | 0.618890637 | 0.535988407 | NA | | | Gm10272 | 0 | NA | NA | NA | NA | NA |
| Muc20 | 0.633091198 | -0.008942341 | 0.058752393 | -0.152203852 | 0.879026156 | NA | | | Gm10318 | 0 | NA | NA | NA | NA | NA |
| Muc4 | 0.699494611 | -0.026379251 | 0.065754389 | -0.401178553 | 0.68828867 | NA | | | Gm10319 | 0 | NA | NA | NA | NA | NA |
| Muc5ac | 0.894578973 | -0.028540078 | 0.073878633 | -0.386310311 | 0.699266865 | NA | | | Gm10352 | 0 | NA | NA | NA | NA | NA |
| Muc5b | 0.079983654 | 0.016740032 | 0.027674178 | 0.604897175 | 0.54524732 | NA | | | Gm10375 | 0 | NA | NA | NA | NA | NA |
| Muc6 | 0.181709657 | 0.024794574 | 0.031325423 | 0.791516008 | 0.428642938 | NA | | | Gm10377 | 0 | NA | NA | NA | NA | NA |
| Mug-ps1 | 0 | NA | NA | NA | NA | NA | | | Gm10400 | 0 | NA | NA | NA | NA | NA |
| Mug1 | 0 | NA | NA | NA | NA | NA | | | Gm10409 | 0 | NA | NA | NA | NA | NA |
| Mug2 | 0.08731684 | 0.017365056 | 0.027733412 | 0.626142072 | 0.531221759 | NA | | | Gm10413 | 0 | NA | NA | NA | NA | NA |
| Mup1 | 0.098353386 | -0.020212831 | 0.02786466 | -0.725393061 | 0.468210929 | NA | | | Gm10436 | 0 | NA | NA | NA | NA | NA |
| Mup10 | 0 | NA | NA | NA | NA | NA | | | Gm10439 | 0 | NA | NA | NA | NA | NA |
| Mup11 | 0.08731684 | 0.017365056 | 0.027733412 | 0.626142072 | 0.531221759 | NA | | | Gm10445 | 0 | NA | NA | NA | NA | NA |
| Mup12 | 0 | NA | NA | NA | NA | NA | | | Gm10486 | 0 | NA | NA | NA | NA | NA |
| Mup13 | 0 | NA | NA | NA | NA | NA | | | Gm10487 | 0 | NA | NA | NA | NA | NA |
| Mup14 | 0 | NA | NA | NA | NA | NA | | | Gm10488 | 0 | NA | NA | NA | NA | NA |
| Mup15 | 0 | NA | NA | NA | NA | NA | | | Gm10512 | 0 | NA | NA | NA | NA | NA |
| Mup16 | 0 | NA | NA | NA | NA | NA | | | Gm10538 | 0 | NA | NA | NA | NA | NA |
| Mup17 | 0 | NA | NA | NA | NA | NA | | | Gm10578 | 0 | NA | NA | NA | NA | NA |
| Mup19 | 0 | NA | NA | NA | NA | NA | | | Gm10591 | 0 | NA | NA | NA | NA | NA |
| Mup2 | 0 | NA | NA | NA | NA | NA | | | Gm10619 | 0 | NA | NA | NA | NA | NA |
| Mup20 | 0.119397641 | -0.020477583 | 0.027954224 | -0.732539845 | 0.463839132 | NA | | | Gm10662 | 0 | NA | NA | NA | NA | NA |
| Mup21 | 0 | NA | NA | NA | NA | NA | | | Gm10665 | 0 | NA | NA | NA | NA | NA |
| Mup3 | 0.480379018 | 0.040041947 | 0.051818222 | 0.772738725 | 0.439677024 | NA | | | Gm10666 | 0 | NA | NA | NA | NA | NA |
| Mup4 | 1.151479737 | -0.029335583 | 0.052330001 | -0.56058823 | 0.575078277 | NA | | | Gm10670 | 0 | NA | NA | NA | NA | NA |
| Mup5 | 6.084986267 | 0.067297747 | 0.135854788 | 0.495365297 | 0.620342286 | NA | | | Gm10681 | 0 | NA | NA | NA | NA | NA |
| Mup6 | 0.116696964 | -0.020477583 | 0.027954224 | -0.732539847 | 0.463839131 | NA | | | Gm10696 | 0 | NA | NA | NA | NA | NA |
| Mup7 | 0 | NA | NA | NA | NA | NA | | | Gm10697 | 0 | NA | NA | NA | NA | NA |
| Mup8 | 0 | NA | NA | NA | NA | NA | | | Gm10745 | 0 | NA | NA | NA | NA | NA |
| Mup9 | 0 | NA | NA | NA | NA | NA | | | Gm10782 | 0 | NA | NA | NA | NA | NA |
| Mybphl | 1.896426157 | 0.002717667 | 0.090605352 | 0.029994549 | 0.9760714 | NA | | | Gm10787 | 0 | NA | NA | NA | NA | NA |
| Mycs | 0.453998628 | 0.026589972 | 0.045454031 | 0.584986 | 0.55855713 | NA | | | Gm10804 | 0 | NA | NA | NA | NA | NA |
| Myf5 | 0 | NA | NA | NA | NA | NA | | | Gm10823 | 0 | NA | NA | NA | NA | NA |
| Myf6 | 0 | NA | NA | NA | NA | NA | | | Gm10921 | 0 | NA | NA | NA | NA | NA |
| Myh6 | 1.403832196 | -0.044467476 | 0.085158715 | -0.522171763 | 0.601550744 | NA | | | Gm10922 | 0 | NA | NA | NA | NA | NA |
| Myl1 | 6.913453372 | 0.158164195 | 0.151182927 | 1.046177625 | 0.295479031 | NA | | | Gm11194 | 0 | NA | NA | NA | NA | NA |
| Myl10 | 0.08731684 | 0.017365056 | 0.027733412 | 0.626142072 | 0.531221759 | NA | | | Gm11213 | 0 | NA | NA | NA | NA | NA |
| Myl3 | 2.212555087 | 0.002935246 | 0.090917319 | 0.032284788 | 0.97424494 | NA | | | Gm1140 | 0 | NA | NA | NA | NA | NA |
| Myl7 | 0.59419716 | 0.024548688 | 0.058662177 | 0.418475568 | 0.675599446 | NA | | | Gm11435 | 0 | NA | NA | NA | NA | NA |
| Mylk2 | 0.69074833 | 0.079811121 | 0.055198329 | 1.445897412 | 0.148205978 | NA | | | Gm11487 | 0 | NA | NA | NA | NA | NA |
| Myo3a | 1.316026059 | 0.023699925 | 0.077516712 | 0.305739554 | 0.759802955 | NA | | | Gm11541 | 0 | NA | NA | NA | NA | NA |
| Myo7b | 4.585168708 | 0.099602232 | 0.132212434 | 0.753349961 | 0.451239644 | NA | | | Gm11544 | 0 | NA | NA | NA | NA | NA |
| Myod1 | 3.454667478 | 0.054640306 | 0.11792442 | 0.463350219 | 0.643113355 | NA | | | Gm11548 | 0 | NA | NA | NA | NA | NA |
| Myog | 0.535761119 | -0.039416426 | 0.047223402 | -0.834679942 | 0.403897938 | NA | | | Gm11554 | 0 | NA | NA | NA | NA | NA |
| Myom2 | 6.623072989 | -0.070833386 | 0.152056726 | -0.465835267 | 0.641333419 | NA | | | Gm11559 | 0 | NA | NA | NA | NA | NA |
| Myot | 2.399949825 | -0.322176302 | 0.110278779 | -2.921471435 | 0.003483822 | NA | | | Gm11562 | 0 | NA | NA | NA | NA | NA |
| Myoz1 | 1.091107073 | -0.088395639 | 0.080114811 | -1.103362008 | 0.269869988 | NA | | | Gm11563 | 0 | NA | NA | NA | NA | NA |
| Myoz2 | 2.173228614 | 0.010710975 | 0.10458328 | 0.102415745 | 0.918426686 | NA | | | Gm11564 | 0 | NA | NA | NA | NA | NA |
| Myoz3 | 0.863460983 | -0.006944181 | 0.062492514 | -0.111120205 | 0.911521027 | NA | | | Gm11565 | 0 | NA | NA | NA | NA | NA |
| Myrfl | 5.680828004 | -0.212031363 | 0.143188575 | -1.480784078 | 0.138664121 | NA | | | Gm11567 | 0 | NA | NA | NA | NA | NA |
| Mzb1 | 3.839294082 | 0.012605687 | 0.119707059 | 0.105304458 | 0.916134226 | NA | | | Gm11568 | 0 | NA | NA | NA | NA | NA |
| Naa11 | 0.789799633 | 0.020365278 | 0.068182092 | 0.298689542 | 0.765176937 | NA | | | Gm11569 | 0 | NA | NA | NA | NA | NA |
| Naaladl1 | 6.947671097 | -0.206554886 | 0.150085579 | -1.376247385 | 0.168745056 | NA | | | Gm11595 | 0 | NA | NA | NA | NA | NA |
| Nags | 3.153450993 | -0.008749018 | 0.125862701 | -0.069512399 | 0.944581763 | NA | | | Gm11596 | 0 | NA | NA | NA | NA | NA |
| Naip7 | 0.334447991 | -0.050120769 | 0.043104878 | -1.162763289 | 0.244925556 | NA | | | Gm11710 | 0 | NA | NA | NA | NA | NA |
| Nanog | 1.060817981 | 0.072334201 | 0.068997377 | 1.048361611 | 0.294472033 | NA | | | Gm11711 | 0 | NA | NA | NA | NA | NA |
| Napsa | 4.658449301 | -0.41834785 | 0.140281831 | -2.982195533 | 0.002861892 | NA | | | Gm11756 | 0 | NA | NA | NA | NA | NA |
| Nat3 | 0 | NA | NA | NA | NA | NA | | | Gm11757 | 0 | NA | NA | NA | NA | NA |
| Nccrp1 | 4.830353489 | -0.190264658 | 0.120531099 | -1.578552428 | 0.114438754 | NA | | | Gm11758 | 0 | NA | NA | NA | NA | NA |
| Ncr1 | 0.472604946 | 0.035675895 | 0.051617383 | 0.691160469 | 0.489464703 | NA | | | Gm11937 | 0 | NA | NA | NA | NA | NA |
| Nctc1 | 0 | NA | NA | NA | NA | NA | | | Gm11938 | 0 | NA | NA | NA | NA | NA |
| Ndufs5 | 0 | NA | NA | NA | NA | NA | | | Gm11981 | 0 | NA | NA | NA | NA | NA |
| Neil3 | 5.018182292 | -0.212206768 | 0.13556121 | -1.565394466 | 0.117490445 | NA | | | Gm11985 | 0 | NA | NA | NA | NA | NA |
| Nepn | 1.19780767 | 0.01177341 | 0.067964026 | 0.173230031 | 0.862470619 | NA | | | Gm12 | 0 | NA | NA | NA | NA | NA |
| Nespas | 2.015159464 | 0.063619504 | 0.078921413 | 0.806112077 | 0.420178229 | NA | | | Gm12130 | 0 | NA | NA | NA | NA | NA |
| Neurog1 | 3.385506913 | -0.031957493 | 0.117260483 | -0.272534212 | 0.785211289 | NA | | | Gm12159 | 0 | NA | NA | NA | NA | NA |
| Neurog2 | 7.943221544 | 0.06639644 | 0.154118602 | 0.430813927 | 0.666603672 | NA | | | Gm12169 | 0 | NA | NA | NA | NA | NA |
| Neurog3 | 2.230123776 | -0.011449681 | 0.09433143 | -0.12137716 | 0.903392307 | NA | | | Gm12191 | 0 | NA | NA | NA | NA | NA |
| Nfe2 | 2.013152324 | 0.001533329 | 0.094740235 | 0.016184557 | 0.987087155 | NA | | | Gm12228 | 0 | NA | NA | NA | NA | NA |
| Nipal1 | 0.378164117 | -0.004452483 | 0.047147495 | -0.094437328 | 0.924761765 | NA | | | Gm12253 | 0 | NA | NA | NA | NA | NA |
| Nipsnap3a | 0 | NA | NA | NA | NA | NA | | | Gm12298 | 0 | NA | NA | NA | NA | NA |
| Nkapl | 7.805105387 | 0.041705858 | 0.154498785 | 0.269942955 | 0.78720414 | NA | | | Gm12505 | 0 | NA | NA | NA | NA | NA |
| Nkg7 | 1.097492992 | 0.10270995 | 0.078095624 | 1.315181875 | 0.188448785 | NA | | | Gm12597 | 0 | NA | NA | NA | NA | NA |
| Nkx1-2 | 1.490843325 | 0.055405805 | 0.081854543 | 0.676881241 | 0.498481307 | NA | | | Gm12669 | 0 | NA | NA | NA | NA | NA |
| Nkx2-3 | 0.28821313 | -0.029561117 | 0.033427731 | -0.884329147 | 0.376518559 | NA | | | Gm12789 | 0 | NA | NA | NA | NA | NA |
| Nkx2-4 | 0 | NA | NA | NA | NA | NA | | | Gm12886 | 0 | NA | NA | NA | NA | NA |
| Nkx2-5 | 0 | NA | NA | NA | NA | NA | | | Gm12887 | 0 | NA | NA | NA | NA | NA |
| Nkx2-6 | 0 | NA | NA | NA | NA | NA | | | Gm12888 | 0 | NA | NA | NA | NA | NA |
| Nkx3-2 | 0 | NA | NA | NA | NA | NA | | | Gm13011 | 0 | NA | NA | NA | NA | NA |
| Nkx6-1 | 3.139826244 | 0.08464138 | 0.108929772 | 0.777027055 | 0.437142809 | NA | | | Gm13023 | 0 | NA | NA | NA | NA | NA |
| Nkx6-3 | 1.619407367 | 0.059613199 | 0.095760045 | 0.622526851 | 0.533595494 | NA | | | Gm13037 | 0 | NA | NA | NA | NA | NA |
| Nlrc5 | 74.26540573 | 0.334813377 | 0.144531556 | 2.316541709 | NA | NA | | | Gm13040 | 0 | NA | NA | NA | NA | NA |
| Nlrp12 | 1.391324294 | 0.029897338 | 0.087895529 | 0.340146295 | 0.73374636 | NA | | | Gm13043 | 0 | NA | NA | NA | NA | NA |
| Nlrp14 | 0 | NA | NA | NA | NA | NA | | | Gm13057 | 0 | NA | NA | NA | NA | NA |
| Nlrp2 | 2.613462047 | -0.065946138 | 0.114997382 | -0.573457735 | 0.566334808 | NA | | | Gm13078 | 0 | NA | NA | NA | NA | NA |
| Nlrp3 | 53.20753095 | 0.210094283 | 0.164305906 | 1.278677605 | NA | NA | | | Gm13083 | 0 | NA | NA | NA | NA | NA |
| Nlrp4a | 0 | NA | NA | NA | NA | NA | | | Gm13084 | 0 | NA | NA | NA | NA | NA |
| Nlrp4b | 0 | NA | NA | NA | NA | NA | | | Gm13088 | 0 | NA | NA | NA | NA | NA |
| Nlrp4c | 0.08731684 | 0.017365056 | 0.027733412 | 0.626142072 | 0.531221759 | NA | | | Gm13102 | 0 | NA | NA | NA | NA | NA |
| Nlrp4e | 0 | NA | NA | NA | NA | NA | | | Gm13103 | 0 | NA | NA | NA | NA | NA |
| Nlrp4f | 4.652014542 | 0.111635616 | 0.134249304 | 0.831554525 | 0.40566044 | NA | | | Gm13109 | 0 | NA | NA | NA | NA | NA |
| Nlrp5 | 0.167300494 | 0.024299763 | 0.031030539 | 0.783091891 | 0.433573153 | NA | | | Gm13119 | 0 | NA | NA | NA | NA | NA |
| Nlrp9a | 0 | NA | NA | NA | NA | NA | | | Gm13124 | 0 | NA | NA | NA | NA | NA |
| Nlrp9b | 0.099009884 | 0.018205726 | 0.027896865 | 0.652608315 | 0.514008823 | NA | | | Gm13125 | 0 | NA | NA | NA | NA | NA |
| Nlrp9c | 0 | NA | NA | NA | NA | NA | | | Gm13128 | 0 | NA | NA | NA | NA | NA |
| Nme8 | 0.195080928 | -0.000361156 | 0.031609476 | -0.011425558 | 0.990883922 | NA | | | Gm13177 | 0 | NA | NA | NA | NA | NA |
| Nme9 | 5.450462961 | 0.225083681 | 0.119367065 | 1.885643088 | 0.059343067 | NA | | | Gm1322 | 0 | NA | NA | NA | NA | NA |
| Nmrk2 | 3.515925956 | 0.172381982 | 0.107719335 | 1.600288213 | 0.10953466 | NA | | | Gm13271 | 0 | NA | NA | NA | NA | NA |
| Nms | 1.56081153 | -0.040459704 | 0.086896442 | -0.465608296 | 0.641495904 | NA | | | Gm13272 | 0 | NA | NA | NA | NA | NA |
| Nmu | 0.698265872 | -0.052029371 | 0.062441752 | -0.833246491 | 0.404705728 | NA | | | Gm13275 | 0 | NA | NA | NA | NA | NA |
| Nmur1 | 4.628174789 | -0.19431463 | 0.124295954 | -1.563322252 | 0.117976817 | NA | | | Gm13276 | 0 | NA | NA | NA | NA | NA |
| Nmur2 | 1.441198444 | -0.071728574 | 0.083144879 | -0.862693826 | 0.388305828 | NA | | | Gm13277 | 0 | NA | NA | NA | NA | NA |
| Nobox | 0.609036663 | 0.027795583 | 0.058866341 | 0.472181252 | 0.63679742 | NA | | | Gm13278 | 0 | NA | NA | NA | NA | NA |
| Noto | 0 | NA | NA | NA | NA | NA | | | Gm13279 | 0 | NA | NA | NA | NA | NA |
| Nox1 | 1.646020596 | -0.126685703 | 0.099899233 | -1.26813489 | 0.204749784 | NA | | | Gm13280 | 0 | NA | NA | NA | NA | NA |
| Nox3 | 0.592096564 | -0.004291432 | 0.052941551 | -0.081059815 | 0.935394384 | NA | | | Gm13283 | 0 | NA | NA | NA | NA | NA |
| Nox4 | 4.143342874 | -0.018443496 | 0.12100948 | -0.152413645 | 0.878860696 | NA | | | Gm13285 | 0 | NA | NA | NA | NA | NA |
| Noxa1 | 0.079983654 | 0.016740032 | 0.027674178 | 0.604897175 | 0.54524732 | NA | | | Gm13286 | 0 | NA | NA | NA | NA | NA |
| Npb | 4.868376212 | 0.13055971 | 0.141309944 | 0.923924435 | 0.35552566 | NA | | | Gm13287 | 0 | NA | NA | NA | NA | NA |
| Npcd | 0 | NA | NA | NA | NA | NA | | | Gm13288 | 0 | NA | NA | NA | NA | NA |
| Npm2 | 7.918720856 | 0.251925001 | 0.155487655 | 1.620225092 | 0.105183933 | NA | | | Gm13289 | 0 | NA | NA | NA | NA | NA |
| Npm3-ps1 | 0.832578476 | -0.037757824 | 0.068913294 | -0.547903343 | 0.583758273 | NA | | | Gm13290 | 0 | NA | NA | NA | NA | NA |
| Nppb | 0.215468684 | -0.027008432 | 0.031909031 | -0.846419662 | 0.397318674 | NA | | | Gm13304 | 0 | NA | NA | NA | NA | NA |
| Nps | 0.556402103 | 0.086806466 | 0.057717414 | 1.503990893 | 0.132583712 | NA | | | Gm13305 | 0 | NA | NA | NA | NA | NA |
| Npvf | 1.236846178 | 0.019351132 | 0.073871169 | 0.261957839 | 0.793353947 | NA | | | Gm13306 | 0 | NA | NA | NA | NA | NA |
| Npw | 1.693065885 | -0.004491479 | 0.082043825 | -0.054744878 | 0.956341715 | NA | | | Gm13308 | 0 | NA | NA | NA | NA | NA |
| Npy6r | 0.513115595 | -0.045873581 | 0.053031465 | -0.86502571 | 0.387024678 | NA | | | Gm13363 | 0 | NA | NA | NA | NA | NA |
| Nr0b1 | 0.294043327 | 0.010508917 | 0.039092051 | 0.2688249 | 0.788064431 | NA | | | Gm13539 | 0 | NA | NA | NA | NA | NA |
| Nr0b2 | 0.094392817 | 0.017895942 | 0.027818137 | 0.643319236 | 0.520016977 | NA | | | Gm13718 | 0 | NA | NA | NA | NA | NA |
| Nr1h4 | 4.708145276 | -0.080325303 | 0.139081023 | -0.577543231 | 0.563572543 | NA | | | Gm13752 | 0 | NA | NA | NA | NA | NA |
| Nr1h5 | 0.895456348 | 0.009786724 | 0.056538074 | 0.17309971 | 0.862573053 | NA | | | Gm13769 | 0 | NA | NA | NA | NA | NA |
| Nr1i2 | 0.47910454 | -0.016023567 | 0.050473258 | -0.317466479 | 0.750889671 | NA | | | Gm13807 | 0 | NA | NA | NA | NA | NA |
| Nr1i3 | 3.697592198 | -0.084714827 | 0.126150329 | -0.671538691 | 0.50187742 | NA | | | Gm13871 | 0 | NA | NA | NA | NA | NA |
| Nr2e3 | 0.197363271 | -0.000561558 | 0.031650166 | -0.017742642 | 0.985844163 | NA | | | Gm13939 | 0 | NA | NA | NA | NA | NA |
| Nr5a1 | 2.325661084 | 0.022814063 | 0.099116043 | 0.230175278 | 0.817955571 | NA | | | Gm14288 | 0 | NA | NA | NA | NA | NA |
| Nr5a2 | 1.262916509 | -0.002348899 | 0.080299129 | -0.029251859 | 0.976663722 | NA | | | Gm14305 | 0 | NA | NA | NA | NA | NA |
| Nrk | 1.510314622 | -0.114129947 | 0.068192908 | -1.673633672 | 0.094202617 | NA | | | Gm14306 | 0 | NA | NA | NA | NA | NA |
| Nron | 0.623077958 | -0.025656387 | 0.047703389 | -0.537831528 | 0.590693363 | NA | | | Gm14345 | 0 | NA | NA | NA | NA | NA |
| Nrp | 4.814916409 | -0.049398025 | 0.144472626 | -0.34191962 | 0.732411384 | NA | | | Gm14346 | 0 | NA | NA | NA | NA | NA |
| Nt5c1b | 0.345637534 | 0.019330126 | 0.041637051 | 0.464253003 | 0.642466491 | NA | | | Gm14347 | 0 | NA | NA | NA | NA | NA |
| Ntf5 | 4.669376945 | 0.011842635 | 0.12996755 | 0.091119935 | 0.927397292 | NA | | | Gm14351 | 0 | NA | NA | NA | NA | NA |
| Ntrk2 | 24371.76408 | -0.034132687 | 0.095398685 | -0.357789908 | NA | NA | | | Gm14354 | 0 | NA | NA | NA | NA | NA |
| Nuggc | 2.196805493 | 0.065764734 | 0.105089153 | 0.625799448 | 0.531446494 | NA | | | Gm14374 | 0 | NA | NA | NA | NA | NA |
| Nup62-il4i1 | 3.424502979 | 0.13945278 | 0.114487382 | 1.218062442 | 0.223200245 | NA | | | Gm14391 | 0 | NA | NA | NA | NA | NA |
| Nup62cl | 5.384119915 | 0.217158713 | 0.140382346 | 1.546908989 | 0.121885193 | NA | | | Gm14430 | 0 | NA | NA | NA | NA | NA |
| Nutf2-ps1 | 0 | NA | NA | NA | NA | NA | | | Gm14431 | 0 | NA | NA | NA | NA | NA |
| Nxf2 | 2.731299807 | -0.210425303 | 0.114537431 | -1.837174986 | 0.066184067 | NA | | | Gm14434 | 0 | NA | NA | NA | NA | NA |
| Nxf3 | 4.464267893 | 0.280771482 | 0.126178935 | 2.225185069 | 0.026068817 | NA | | | Gm14436 | 0 | NA | NA | NA | NA | NA |
| Nxnl1 | 4.194395197 | 0.014482128 | 0.128491486 | 0.112708856 | 0.91026138 | NA | | | Gm14440 | 0 | NA | NA | NA | NA | NA |
| Nxnl2 | 4.299733494 | 0.067201784 | 0.125121139 | 0.537093769 | 0.591202845 | NA | | | Gm14458 | 0 | NA | NA | NA | NA | NA |
| Nxpe5 | 0.482637559 | -0.049175986 | 0.042867026 | -1.14717514 | 0.251309244 | NA | | | Gm14459 | 0 | NA | NA | NA | NA | NA |
| OTTMUSG00000016609 | 0 | NA | NA | NA | NA | NA | | | Gm14474 | 0 | NA | NA | NA | NA | NA |
| Oas1d | 0.748706893 | 0.039722983 | 0.056595821 | 0.701871312 | 0.482759421 | NA | | | Gm14475 | 0 | NA | NA | NA | NA | NA |
| Oas1e | 0.384284173 | -0.031469349 | 0.034522933 | -0.911549113 | 0.362006121 | NA | | | Gm14476 | 0 | NA | NA | NA | NA | NA |
| Oas1f | 0.19046386 | -0.000774927 | 0.031526133 | -0.024580468 | 0.980389599 | NA | | | Gm14477 | 0 | NA | NA | NA | NA | NA |
| Oas1g | 3.14379104 | 0.04901949 | 0.09227846 | 0.531212698 | 0.595271395 | NA | | | Gm14478 | 0 | NA | NA | NA | NA | NA |
| Oas1h | 0.575354256 | -0.004736431 | 0.046231168 | -0.102451031 | 0.91839868 | NA | | | Gm14479 | 0 | NA | NA | NA | NA | NA |
| Oas3 | 3.864437366 | 0.060347289 | 0.120403974 | 0.501206783 | 0.616225601 | NA | | | Gm14482 | 0 | NA | NA | NA | NA | NA |
| Oaz1 | 0 | NA | NA | NA | NA | NA | | | Gm14483 | 0 | NA | NA | NA | NA | NA |
| Oaz1-ps | 0 | NA | NA | NA | NA | NA | | | Gm14484 | 0 | NA | NA | NA | NA | NA |
| Oaz3 | 3.082645033 | 0.125802511 | 0.103314688 | 1.217663369 | 0.223351923 | NA | | | Gm14496 | 0 | NA | NA | NA | NA | NA |
| Obox1 | 0 | NA | NA | NA | NA | NA | | | Gm14511 | 0 | NA | NA | NA | NA | NA |
| Obox2 | 0 | NA | NA | NA | NA | NA | | | Gm14525 | 0 | NA | NA | NA | NA | NA |
| Obox3 | 0 | NA | NA | NA | NA | NA | | | Gm14548 | 0 | NA | NA | NA | NA | NA |
| Obox5 | 0.393053211 | 0.021363521 | 0.042959842 | 0.497290494 | 0.618984214 | NA | | | Gm14625 | 0 | NA | NA | NA | NA | NA |
| Obox6 | 0 | NA | NA | NA | NA | NA | | | Gm14632 | 0 | NA | NA | NA | NA | NA |
| Obp1a | 1.264166687 | 0.110742242 | 0.064169661 | 1.725772584 | 0.084388334 | NA | | | Gm14635 | 0 | NA | NA | NA | NA | NA |
| Obp2a | 0.914950978 | 0.096606065 | 0.069296257 | 1.39410221 | 0.163286742 | NA | | | Gm14685 | 0 | NA | NA | NA | NA | NA |
| Obp2b | 1.992791227 | 0.094284877 | 0.089429586 | 1.05429178 | 0.291749351 | NA | | | Gm14692 | 0 | NA | NA | NA | NA | NA |
| Oc90 | 2.799342917 | 0.002494517 | 0.12108851 | 0.020600777 | 0.983564121 | NA | | | Gm14718 | 0 | NA | NA | NA | NA | NA |
| Oca2 | 16.29520498 | 0.102392568 | 0.148485906 | 0.68957769 | NA | NA | | | Gm14725 | 0 | NA | NA | NA | NA | NA |
| Ociad2 | 3605.454997 | -0.003095451 | 0.103416732 | -0.029931821 | NA | NA | | | Gm14781 | 0 | NA | NA | NA | NA | NA |
| Ocstamp | 0.582208971 | -0.023855297 | 0.047658271 | -0.500548931 | 0.616688612 | NA | | | Gm14819 | 0 | NA | NA | NA | NA | NA |
| Odam | 0 | NA | NA | NA | NA | NA | | | Gm14850 | 0 | NA | NA | NA | NA | NA |
| Odf1 | 1.331330041 | 0.041256347 | 0.071099425 | 0.580262741 | 0.561737449 | NA | | | Gm14851 | 0 | NA | NA | NA | NA | NA |
| Odf3 | 2.868109075 | 0.091449229 | 0.112579873 | 0.81230531 | 0.416616464 | NA | | | Gm14920 | 0 | NA | NA | NA | NA | NA |
| Odf3l1 | 1.819315222 | 0.04220695 | 0.094135522 | 0.448363688 | 0.653890744 | NA | | | Gm15023 | 0 | NA | NA | NA | NA | NA |
| Odf3l2 | 7.508541522 | -0.01851026 | 0.156537286 | -0.118248251 | 0.905870959 | NA | | | Gm15056 | 0 | NA | NA | NA | NA | NA |
| Ofcc1 | 0.116696964 | -0.020477583 | 0.027954224 | -0.732539847 | 0.463839131 | NA | | | Gm15080 | 0 | NA | NA | NA | NA | NA |
| Oit1 | 0.098353386 | -0.020212831 | 0.02786466 | -0.725393061 | 0.468210929 | NA | | | Gm15085 | 0 | NA | NA | NA | NA | NA |
| Oit3 | 5.117316794 | 0.212135524 | 0.124677522 | 1.701473695 | 0.088854073 | NA | | | Gm15091 | 0 | NA | NA | NA | NA | NA |
| Olah | 1.314458349 | 0.119766747 | 0.080139039 | 1.49448693 | 0.135048395 | NA | | | Gm15093 | 0 | NA | NA | NA | NA | NA |
| Olfr1 | 0 | NA | NA | NA | NA | NA | | | Gm15097 | 0 | NA | NA | NA | NA | NA |
| Olfr10 | 0.188785634 | 0.025025111 | 0.031461603 | 0.79541754 | 0.42637066 | NA | | | Gm15107 | 0 | NA | NA | NA | NA | NA |
| Olfr100 | 0 | NA | NA | NA | NA | NA | | | Gm15114 | 0 | NA | NA | NA | NA | NA |
| Olfr1000 | 0 | NA | NA | NA | NA | NA | | | Gm15127 | 0 | NA | NA | NA | NA | NA |
| Olfr1002 | 0 | NA | NA | NA | NA | NA | | | Gm15140 | 0 | NA | NA | NA | NA | NA |
| Olfr1006 | 0 | NA | NA | NA | NA | NA | | | Gm15217 | 0 | NA | NA | NA | NA | NA |
| Olfr1008 | 0 | NA | NA | NA | NA | NA | | | Gm1527 | 0 | NA | NA | NA | NA | NA |
| Olfr1009 | 0 | NA | NA | NA | NA | NA | | | Gm15284 | 0 | NA | NA | NA | NA | NA |
| Olfr101 | 0 | NA | NA | NA | NA | NA | | | Gm15292 | 0 | NA | NA | NA | NA | NA |
| Olfr1010 | 0 | NA | NA | NA | NA | NA | | | Gm15293 | 0 | NA | NA | NA | NA | NA |
| Olfr1012 | 0 | NA | NA | NA | NA | NA | | | Gm15299 | 0 | NA | NA | NA | NA | NA |
| Olfr1013 | 0 | NA | NA | NA | NA | NA | | | Gm15308 | 0 | NA | NA | NA | NA | NA |
| Olfr1014 | 0 | NA | NA | NA | NA | NA | | | Gm15315 | 0 | NA | NA | NA | NA | NA |
| Olfr1015 | 0 | NA | NA | NA | NA | NA | | | Gm15319 | 0 | NA | NA | NA | NA | NA |
| Olfr1016 | 0 | NA | NA | NA | NA | NA | | | Gm15386 | 0 | NA | NA | NA | NA | NA |
| Olfr1018 | 0 | NA | NA | NA | NA | NA | | | Gm15412 | 0 | NA | NA | NA | NA | NA |
| Olfr1019 | 0 | NA | NA | NA | NA | NA | | | Gm156 | 0 | NA | NA | NA | NA | NA |
| Olfr102 | 0 | NA | NA | NA | NA | NA | | | Gm15698 | 0 | NA | NA | NA | NA | NA |
| Olfr1020 | 0 | NA | NA | NA | NA | NA | | | Gm15713 | 0 | NA | NA | NA | NA | NA |
| Olfr1022 | 0 | NA | NA | NA | NA | NA | | | Gm15772 | 0 | NA | NA | NA | NA | NA |
| Olfr1023 | 0 | NA | NA | NA | NA | NA | | | Gm15881 | 0 | NA | NA | NA | NA | NA |
| Olfr1024 | 0 | NA | NA | NA | NA | NA | | | Gm16063 | 0 | NA | NA | NA | NA | NA |
| Olfr1026 | 0 | NA | NA | NA | NA | NA | | | Gm16157 | 0 | NA | NA | NA | NA | NA |
| Olfr1028 | 0 | NA | NA | NA | NA | NA | | | Gm1631 | 0 | NA | NA | NA | NA | NA |
| Olfr1029 | 0.098353386 | -0.020212831 | 0.02786466 | -0.725393061 | 0.468210929 | NA | | | Gm16336 | 0 | NA | NA | NA | NA | NA |
| Olfr103 | 0 | NA | NA | NA | NA | NA | | | Gm16367 | 0 | NA | NA | NA | NA | NA |
| Olfr1030 | 0 | NA | NA | NA | NA | NA | | | Gm16381 | 0 | NA | NA | NA | NA | NA |
| Olfr1031 | 0 | NA | NA | NA | NA | NA | | | Gm16390 | 0 | NA | NA | NA | NA | NA |
| Olfr1032 | 0 | NA | NA | NA | NA | NA | | | Gm16404 | 0 | NA | NA | NA | NA | NA |

| | | | | | | | | | | | | | | | |
|---|---|---|---|---|---|---|---|---|---|---|---|---|---|---|---|
| Olfr1033 | 5.192019563 | -0.140129054 | 0.131726217 | -1.063790171 | 0.287423765 | NA | | | Gm16405 | 0 | NA | NA | NA | NA | NA |
| Olfr1034 | 0.188785634 | 0.025025111 | 0.031461603 | 0.79541754 | 0.42637066 | NA | | | Gm16430 | 0 | NA | NA | NA | NA | NA |
| Olfr1036 | 0 | NA | NA | NA | NA | NA | | | Gm16442 | 0 | NA | NA | NA | NA | NA |
| Olfr1037 | 0 | NA | NA | NA | NA | NA | | | Gm16445 | 0 | NA | NA | NA | NA | NA |
| Olfr1038-ps | 0 | NA | NA | NA | NA | NA | | | Gm16451 | 0 | NA | NA | NA | NA | NA |
| Olfr1039 | 0 | NA | NA | NA | NA | NA | | | Gm16501 | 0 | NA | NA | NA | NA | NA |
| Olfr1040 | 0 | NA | NA | NA | NA | NA | | | Gm16523 | 0 | NA | NA | NA | NA | NA |
| Olfr1042 | 0 | NA | NA | NA | NA | NA | | | Gm17252 | 0 | NA | NA | NA | NA | NA |
| Olfr1043 | 0 | NA | NA | NA | NA | NA | | | Gm17365 | 0 | NA | NA | NA | NA | NA |
| Olfr1044 | 0 | NA | NA | NA | NA | NA | | | Gm17495 | 0 | NA | NA | NA | NA | NA |
| Olfr1045 | 0 | NA | NA | NA | NA | NA | | | Gm17689 | 0 | NA | NA | NA | NA | NA |
| Olfr1046 | 0 | NA | NA | NA | NA | NA | | | Gm17757 | 0 | NA | NA | NA | NA | NA |
| Olfr1047 | 0 | NA | NA | NA | NA | NA | | | Gm18853 | 0 | NA | NA | NA | NA | NA |
| Olfr1048 | 0 | NA | NA | NA | NA | NA | | | Gm19277 | 0 | NA | NA | NA | NA | NA |
| Olfr1049 | 0 | NA | NA | NA | NA | NA | | | Gm19395 | 0 | NA | NA | NA | NA | NA |
| Olfr1051 | 0 | NA | NA | NA | NA | NA | | | Gm19402 | 0 | NA | NA | NA | NA | NA |
| Olfr1052 | 0 | NA | NA | NA | NA | NA | | | Gm1965 | 0 | NA | NA | NA | NA | NA |
| Olfr1053 | 0 | NA | NA | NA | NA | NA | | | Gm19668 | 0 | NA | NA | NA | NA | NA |
| Olfr1054 | 0 | NA | NA | NA | NA | NA | | | Gm19784 | 0 | NA | NA | NA | NA | NA |
| Olfr1055 | 0 | NA | NA | NA | NA | NA | | | Gm1987 | 0 | NA | NA | NA | NA | NA |
| Olfr1056 | 0 | NA | NA | NA | NA | NA | | | Gm1993 | 0 | NA | NA | NA | NA | NA |
| Olfr1057 | 0 | NA | NA | NA | NA | NA | | | Gm1995 | 0 | NA | NA | NA | NA | NA |
| Olfr1058 | 0 | NA | NA | NA | NA | NA | | | Gm2002 | 0 | NA | NA | NA | NA | NA |
| Olfr1061 | 0 | NA | NA | NA | NA | NA | | | Gm2012 | 0 | NA | NA | NA | NA | NA |
| Olfr1062 | 0 | NA | NA | NA | NA | NA | | | Gm2016 | 0 | NA | NA | NA | NA | NA |
| Olfr1065 | 0.098353386 | -0.020212831 | 0.02786466 | -0.725393061 | 0.468210929 | NA | | | Gm20172 | 0 | NA | NA | NA | NA | NA |
| Olfr1066 | 0 | NA | NA | NA | NA | NA | | | Gm2022 | 0 | NA | NA | NA | NA | NA |
| Olfr107 | 0 | NA | NA | NA | NA | NA | | | Gm20268 | 0 | NA | NA | NA | NA | NA |
| Olfr1076 | 0 | NA | NA | NA | NA | NA | | | Gm2030 | 0 | NA | NA | NA | NA | NA |
| Olfr1077-ps1 | 0 | NA | NA | NA | NA | NA | | | Gm20356 | 0 | NA | NA | NA | NA | NA |
| Olfr1079 | 0 | NA | NA | NA | NA | NA | | | Gm20408 | 0 | NA | NA | NA | NA | NA |
| Olfr108 | 0 | NA | NA | NA | NA | NA | | | Gm20410 | 0 | NA | NA | NA | NA | NA |
| Olfr1080 | 0 | NA | NA | NA | NA | NA | | | Gm2042 | 0 | NA | NA | NA | NA | NA |
| Olfr1082 | 0 | NA | NA | NA | NA | NA | | | Gm20516 | 0 | NA | NA | NA | NA | NA |
| Olfr1084 | 0 | NA | NA | NA | NA | NA | | | Gm20556 | 0 | NA | NA | NA | NA | NA |
| Olfr1085 | 0 | NA | NA | NA | NA | NA | | | Gm20579 | 0 | NA | NA | NA | NA | NA |
| Olfr1086 | 0 | NA | NA | NA | NA | NA | | | Gm20580 | 0 | NA | NA | NA | NA | NA |
| Olfr1087 | 0 | NA | NA | NA | NA | NA | | | Gm20581 | 0 | NA | NA | NA | NA | NA |
| Olfr1089 | 0 | NA | NA | NA | NA | NA | | | Gm20604 | 0 | NA | NA | NA | NA | NA |
| Olfr109 | 0 | NA | NA | NA | NA | NA | | | Gm20740 | 0 | NA | NA | NA | NA | NA |
| Olfr1090 | 0 | NA | NA | NA | NA | NA | | | Gm20741 | 0 | NA | NA | NA | NA | NA |
| Olfr1093 | 0 | NA | NA | NA | NA | NA | | | Gm20759 | 0 | NA | NA | NA | NA | NA |
| Olfr1094 | 0 | NA | NA | NA | NA | NA | | | Gm20765 | 0 | NA | NA | NA | NA | NA |
| Olfr1095 | 0 | NA | NA | NA | NA | NA | | | Gm2083 | 0 | NA | NA | NA | NA | NA |
| Olfr1097 | 0 | NA | NA | NA | NA | NA | | | Gm20878 | 0 | NA | NA | NA | NA | NA |
| Olfr1098 | 0 | NA | NA | NA | NA | NA | | | Gm21002 | 0 | NA | NA | NA | NA | NA |
| Olfr1099 | 0 | NA | NA | NA | NA | NA | | | Gm21119 | 0 | NA | NA | NA | NA | NA |
| Olfr11 | 0 | NA | NA | NA | NA | NA | | | Gm21276 | 0 | NA | NA | NA | NA | NA |
| Olfr110 | 1.747658581 | -0.023432014 | 0.068517071 | -0.341987971 | 0.732359945 | NA | | | Gm21293 | 0 | NA | NA | NA | NA | NA |
| Olfr1100 | 0 | NA | NA | NA | NA | NA | | | Gm21304 | 0 | NA | NA | NA | NA | NA |
| Olfr1101 | 0 | NA | NA | NA | NA | NA | | | Gm21312 | 0 | NA | NA | NA | NA | NA |
| Olfr1102 | 0 | NA | NA | NA | NA | NA | | | Gm21319 | 0 | NA | NA | NA | NA | NA |
| Olfr1104 | 0 | NA | NA | NA | NA | NA | | | Gm21498 | 0 | NA | NA | NA | NA | NA |
| Olfr1105 | 0 | NA | NA | NA | NA | NA | | | Gm21541 | 0 | NA | NA | NA | NA | NA |
| Olfr1106 | 0 | NA | NA | NA | NA | NA | | | Gm21586 | 0 | NA | NA | NA | NA | NA |
| Olfr1107 | 0 | NA | NA | NA | NA | NA | | | Gm21637 | 0 | NA | NA | NA | NA | NA |
| Olfr1109 | 0 | NA | NA | NA | NA | NA | | | Gm21677 | 0 | NA | NA | NA | NA | NA |
| Olfr111 | 3.236110612 | 0.119609163 | 0.086359032 | 1.385022039 | 0.166045728 | NA | | | Gm21693 | 0 | NA | NA | NA | NA | NA |
| Olfr1110 | 0 | NA | NA | NA | NA | NA | | | Gm21704 | 0 | NA | NA | NA | NA | NA |
| Olfr1111 | 0 | NA | NA | NA | NA | NA | | | Gm21708 | 0 | NA | NA | NA | NA | NA |
| Olfr1112 | 0 | NA | NA | NA | NA | NA | | | Gm2176 | 0 | NA | NA | NA | NA | NA |
| Olfr1113 | 0 | NA | NA | NA | NA | NA | | | Gm21945 | 0 | NA | NA | NA | NA | NA |
| Olfr1115 | 0 | NA | NA | NA | NA | NA | | | Gm21946 | 0 | NA | NA | NA | NA | NA |
| Olfr1116-ps | 0 | NA | NA | NA | NA | NA | | | Gm21947 | 0 | NA | NA | NA | NA | NA |
| Olfr1118 | 0 | NA | NA | NA | NA | NA | | | Gm21949 | 0 | NA | NA | NA | NA | NA |
| Olfr112 | 0 | NA | NA | NA | NA | NA | | | Gm21950 | 0 | NA | NA | NA | NA | NA |
| Olfr1120 | 0 | NA | NA | NA | NA | NA | | | Gm21951 | 0 | NA | NA | NA | NA | NA |
| Olfr1121 | 0 | NA | NA | NA | NA | NA | | | Gm2506 | 0 | NA | NA | NA | NA | NA |
| Olfr1122 | 0 | NA | NA | NA | NA | NA | | | Gm2663 | 0 | NA | NA | NA | NA | NA |
| Olfr1123 | 0 | NA | NA | NA | NA | NA | | | Gm2696 | 0 | NA | NA | NA | NA | NA |
| Olfr1124 | 0 | NA | NA | NA | NA | NA | | | Gm2799 | 0 | NA | NA | NA | NA | NA |
| Olfr1126 | 0.099009884 | 0.018205726 | 0.027896865 | 0.652608315 | 0.514008823 | NA | | | Gm2825 | 0 | NA | NA | NA | NA | NA |
| Olfr1128 | 0 | NA | NA | NA | NA | NA | | | Gm2837 | 0 | NA | NA | NA | NA | NA |
| Olfr1129 | 0 | NA | NA | NA | NA | NA | | | Gm2863 | 0 | NA | NA | NA | NA | NA |
| Olfr113 | 0 | NA | NA | NA | NA | NA | | | Gm2913 | 0 | NA | NA | NA | NA | NA |
| Olfr1130 | 0 | NA | NA | NA | NA | NA | | | Gm2933 | 0 | NA | NA | NA | NA | NA |
| Olfr1131 | 0 | NA | NA | NA | NA | NA | | | Gm3020 | 0 | NA | NA | NA | NA | NA |
| Olfr1132 | 0 | NA | NA | NA | NA | NA | | | Gm3139 | 0 | NA | NA | NA | NA | NA |
| Olfr1133 | 0 | NA | NA | NA | NA | NA | | | Gm3238 | 0 | NA | NA | NA | NA | NA |
| Olfr1134 | 0 | NA | NA | NA | NA | NA | | | Gm3259 | 0 | NA | NA | NA | NA | NA |
| Olfr1135 | 0 | NA | NA | NA | NA | NA | | | Gm3285 | 0 | NA | NA | NA | NA | NA |
| Olfr1136 | 0 | NA | NA | NA | NA | NA | | | Gm3286 | 0 | NA | NA | NA | NA | NA |
| Olfr1137 | 0 | NA | NA | NA | NA | NA | | | Gm3317 | 0 | NA | NA | NA | NA | NA |
| Olfr1138 | 0 | NA | NA | NA | NA | NA | | | Gm3376 | 0 | NA | NA | NA | NA | NA |
| Olfr114 | 0 | NA | NA | NA | NA | NA | | | Gm3383 | 0 | NA | NA | NA | NA | NA |
| Olfr1140 | 0.079983654 | 0.016740032 | 0.027674178 | 0.604897175 | 0.54524732 | NA | | | Gm3402 | 0 | NA | NA | NA | NA | NA |
| Olfr1141 | 0 | NA | NA | NA | NA | NA | | | Gm3404 | 0 | NA | NA | NA | NA | NA |
| Olfr1143 | 0 | NA | NA | NA | NA | NA | | | Gm3409 | 0 | NA | NA | NA | NA | NA |
| Olfr1145 | 0 | NA | NA | NA | NA | NA | | | Gm3415 | 0 | NA | NA | NA | NA | NA |
| Olfr1148 | 0 | NA | NA | NA | NA | NA | | | Gm3417 | 0 | NA | NA | NA | NA | NA |
| Olfr115 | 0 | NA | NA | NA | NA | NA | | | Gm3428 | 0 | NA | NA | NA | NA | NA |
| Olfr1151 | 0 | NA | NA | NA | NA | NA | | | Gm3434 | 0 | NA | NA | NA | NA | NA |
| Olfr1152 | 0 | NA | NA | NA | NA | NA | | | Gm3448 | 0 | NA | NA | NA | NA | NA |
| Olfr1153 | 0 | NA | NA | NA | NA | NA | | | Gm3494 | 0 | NA | NA | NA | NA | NA |
| Olfr1154 | 0 | NA | NA | NA | NA | NA | | | Gm3500 | 0 | NA | NA | NA | NA | NA |
| Olfr1155 | 0 | NA | NA | NA | NA | NA | | | Gm362 | 0 | NA | NA | NA | NA | NA |
| Olfr1156 | 0 | NA | NA | NA | NA | NA | | | Gm364 | 0 | NA | NA | NA | NA | NA |
| Olfr1157 | 0 | NA | NA | NA | NA | NA | | | Gm3646 | 0 | NA | NA | NA | NA | NA |
| Olfr1158 | 0 | NA | NA | NA | NA | NA | | | Gm3701 | 0 | NA | NA | NA | NA | NA |
| Olfr116 | 0 | NA | NA | NA | NA | NA | | | Gm3706 | 0 | NA | NA | NA | NA | NA |
| Olfr1160 | 0 | NA | NA | NA | NA | NA | | | Gm3750 | 0 | NA | NA | NA | NA | NA |
| Olfr1161 | 0 | NA | NA | NA | NA | NA | | | Gm3763 | 0 | NA | NA | NA | NA | NA |
| Olfr1162 | 0 | NA | NA | NA | NA | NA | | | Gm382 | 0 | NA | NA | NA | NA | NA |
| Olfr1163 | 0 | NA | NA | NA | NA | NA | | | Gm3833 | 0 | NA | NA | NA | NA | NA |
| Olfr1164 | 0 | NA | NA | NA | NA | NA | | | Gm4027 | 0 | NA | NA | NA | NA | NA |
| Olfr1166 | 0 | NA | NA | NA | NA | NA | | | Gm4064 | 0 | NA | NA | NA | NA | NA |
| Olfr1167 | 0 | NA | NA | NA | NA | NA | | | Gm4070 | 0 | NA | NA | NA | NA | NA |
| Olfr1168 | 0 | NA | NA | NA | NA | NA | | | Gm41 | 0 | NA | NA | NA | NA | NA |
| Olfr117 | 0 | NA | NA | NA | NA | NA | | | Gm4133 | 0 | NA | NA | NA | NA | NA |
| Olfr1170 | 0 | NA | NA | NA | NA | NA | | | Gm4141 | 0 | NA | NA | NA | NA | NA |
| Olfr1173 | 0 | NA | NA | NA | NA | NA | | | Gm4172 | 0 | NA | NA | NA | NA | NA |
| Olfr1176 | 0 | NA | NA | NA | NA | NA | | | Gm4175 | 0 | NA | NA | NA | NA | NA |
| Olfr1178 | 0 | NA | NA | NA | NA | NA | | | Gm4177 | 0 | NA | NA | NA | NA | NA |
| Olfr1179 | 0 | NA | NA | NA | NA | NA | | | Gm4179 | 0 | NA | NA | NA | NA | NA |
| Olfr118 | 0 | NA | NA | NA | NA | NA | | | Gm4187 | 0 | NA | NA | NA | NA | NA |
| Olfr1180 | 0 | NA | NA | NA | NA | NA | | | Gm4201 | 0 | NA | NA | NA | NA | NA |
| Olfr1181 | 0 | NA | NA | NA | NA | NA | | | Gm4214 | 0 | NA | NA | NA | NA | NA |
| Olfr1182 | 0 | NA | NA | NA | NA | NA | | | Gm4216 | 0 | NA | NA | NA | NA | NA |
| Olfr1183 | 0 | NA | NA | NA | NA | NA | | | Gm4220 | 0 | NA | NA | NA | NA | NA |
| Olfr1184 | 0 | NA | NA | NA | NA | NA | | | Gm4224 | 0 | NA | NA | NA | NA | NA |
| Olfr1186 | 0 | NA | NA | NA | NA | NA | | | Gm4278 | 0 | NA | NA | NA | NA | NA |
| Olfr1188 | 0 | NA | NA | NA | NA | NA | | | Gm428 | 0 | NA | NA | NA | NA | NA |
| Olfr1189 | 0 | NA | NA | NA | NA | NA | | | Gm4297 | 0 | NA | NA | NA | NA | NA |
| Olfr119 | 0.098353386 | -0.020212831 | 0.02786466 | -0.725393061 | 0.468210929 | NA | | | Gm4301 | 0 | NA | NA | NA | NA | NA |
| Olfr1193 | 0.233393928 | -0.027475306 | 0.032191697 | -0.85349047 | 0.39338737 | NA | | | Gm4302 | 0 | NA | NA | NA | NA | NA |
| Olfr1195 | 0.098353386 | -0.020212831 | 0.02786466 | -0.725393061 | 0.468210929 | NA | | | Gm4303 | 0 | NA | NA | NA | NA | NA |
| Olfr1196 | 0 | NA | NA | NA | NA | NA | | | Gm4305 | 0 | NA | NA | NA | NA | NA |
| Olfr1197 | 0 | NA | NA | NA | NA | NA | | | Gm4307 | 0 | NA | NA | NA | NA | NA |
| Olfr1198 | 0 | NA | NA | NA | NA | NA | | | Gm4312 | 0 | NA | NA | NA | NA | NA |
| Olfr1199 | 0 | NA | NA | NA | NA | NA | | | Gm4340 | 0 | NA | NA | NA | NA | NA |
| Olfr12 | 0.215050351 | -0.027006021 | 0.031907566 | -0.846382992 | 0.397339123 | NA | | | Gm4345 | 0 | NA | NA | NA | NA | NA |
| Olfr120 | 0 | NA | NA | NA | NA | NA | | | Gm436 | 0 | NA | NA | NA | NA | NA |
| Olfr1200 | 0 | NA | NA | NA | NA | NA | | | Gm438 | 0 | NA | NA | NA | NA | NA |
| Olfr1201 | 0 | NA | NA | NA | NA | NA | | | Gm44 | 0 | NA | NA | NA | NA | NA |
| Olfr1202 | 0 | NA | NA | NA | NA | NA | | | Gm4498 | 0 | NA | NA | NA | NA | NA |
| Olfr1204 | 0 | NA | NA | NA | NA | NA | | | Gm4541 | 0 | NA | NA | NA | NA | NA |
| Olfr1205 | 0.098353386 | -0.020212831 | 0.02786466 | -0.725393061 | 0.468210929 | NA | | | Gm4559 | 0 | NA | NA | NA | NA | NA |
| Olfr1206 | 0 | NA | NA | NA | NA | NA | | | Gm4567 | 0 | NA | NA | NA | NA | NA |
| Olfr1208 | 0.08731684 | 0.017365056 | 0.027733412 | 0.626142072 | 0.531221759 | NA | | | Gm4710 | 0 | NA | NA | NA | NA | NA |
| Olfr1209 | 0 | NA | NA | NA | NA | NA | | | Gm4719 | 0 | NA | NA | NA | NA | NA |
| Olfr121 | 0.08731684 | 0.017365056 | 0.027733412 | 0.626142072 | 0.531221759 | NA | | | Gm4724 | 0 | NA | NA | NA | NA | NA |
| Olfr1211 | 0.099009884 | 0.018205726 | 0.027896865 | 0.652608315 | 0.514008823 | NA | | | Gm4736 | 0 | NA | NA | NA | NA | NA |
| Olfr1212 | 0 | NA | NA | NA | NA | NA | | | Gm4763 | 0 | NA | NA | NA | NA | NA |
| Olfr1213 | 0 | NA | NA | NA | NA | NA | | | Gm4776 | 0 | NA | NA | NA | NA | NA |
| Olfr1214 | 0 | NA | NA | NA | NA | NA | | | Gm4794 | 0 | NA | NA | NA | NA | NA |
| Olfr1215 | 0 | NA | NA | NA | NA | NA | | | Gm4836 | 0 | NA | NA | NA | NA | NA |
| Olfr1216 | 0 | NA | NA | NA | NA | NA | | | Gm4846 | 0 | NA | NA | NA | NA | NA |
| Olfr1217 | 0 | NA | NA | NA | NA | NA | | | Gm4850 | 0 | NA | NA | NA | NA | NA |
| Olfr1218 | 0 | NA | NA | NA | NA | NA | | | Gm4858 | 0 | NA | NA | NA | NA | NA |
| Olfr1219 | 0 | NA | NA | NA | NA | NA | | | Gm4871 | 0 | NA | NA | NA | NA | NA |
| Olfr122 | 0 | NA | NA | NA | NA | NA | | | Gm4872 | 0 | NA | NA | NA | NA | NA |
| Olfr1220 | 0 | NA | NA | NA | NA | NA | | | Gm4884 | 0 | NA | NA | NA | NA | NA |
| Olfr1221 | 0 | NA | NA | NA | NA | NA | | | Gm4906 | 0 | NA | NA | NA | NA | NA |
| Olfr1222 | 0 | NA | NA | NA | NA | NA | | | Gm4907 | 0 | NA | NA | NA | NA | NA |
| Olfr1223 | 0.098353386 | -0.020212831 | 0.02786466 | -0.725393061 | 0.468210929 | NA | | | Gm4922 | 0 | NA | NA | NA | NA | NA |
| Olfr1225 | 0 | NA | NA | NA | NA | NA | | | Gm4937 | 0 | NA | NA | NA | NA | NA |
| Olfr1226 | 0.098353386 | -0.020212831 | 0.02786466 | -0.725393061 | 0.468210929 | NA | | | Gm4971 | 0 | NA | NA | NA | NA | NA |
| Olfr1228 | 0.295060159 | -0.029687054 | 0.033501022 | -0.886153701 | 0.375534709 | NA | | | Gm4984 | 0 | NA | NA | NA | NA | NA |
| Olfr1229 | 0 | NA | NA | NA | NA | NA | | | Gm5039 | 0 | NA | NA | NA | NA | NA |
| Olfr123 | 0.105923158 | 0.018524693 | 0.027977518 | 0.662127842 | 0.507889294 | NA | | | Gm5071 | 0 | NA | NA | NA | NA | NA |
| Olfr1230 | 0 | NA | NA | NA | NA | NA | | | Gm5072 | 0 | NA | NA | NA | NA | NA |
| Olfr1231 | 0 | NA | NA | NA | NA | NA | | | Gm5082 | 0 | NA | NA | NA | NA | NA |
| Olfr1232 | 0 | NA | NA | NA | NA | NA | | | Gm5091 | 0 | NA | NA | NA | NA | NA |
| Olfr1233 | 0 | NA | NA | NA | NA | NA | | | Gm5095 | 0 | NA | NA | NA | NA | NA |
| Olfr1234 | 0 | NA | NA | NA | NA | NA | | | Gm5114 | 0 | NA | NA | NA | NA | NA |
| Olfr1238 | 0 | NA | NA | NA | NA | NA | | | Gm5128 | 0 | NA | NA | NA | NA | NA |
| Olfr1239 | 0 | NA | NA | NA | NA | NA | | | Gm5136 | 0 | NA | NA | NA | NA | NA |
| Olfr124 | 0 | NA | NA | NA | NA | NA | | | Gm5142 | 0 | NA | NA | NA | NA | NA |
| Olfr1240 | 0 | NA | NA | NA | NA | NA | | | Gm5166 | 0 | NA | NA | NA | NA | NA |
| Olfr1241 | 0 | NA | NA | NA | NA | NA | | | Gm5168 | 0 | NA | NA | NA | NA | NA |
| Olfr1242 | 0 | NA | NA | NA | NA | NA | | | Gm5169 | 0 | NA | NA | NA | NA | NA |
| Olfr1243 | 0 | NA | NA | NA | NA | NA | | | Gm53 | 0 | NA | NA | NA | NA | NA |
| Olfr1245 | 0 | NA | NA | NA | NA | NA | | | Gm5334 | 0 | NA | NA | NA | NA | NA |
| Olfr1246 | 0 | NA | NA | NA | NA | NA | | | Gm5346 | 0 | NA | NA | NA | NA | NA |
| Olfr1247 | 0 | NA | NA | NA | NA | NA | | | Gm5347 | 0 | NA | NA | NA | NA | NA |
| Olfr1248 | 0 | NA | NA | NA | NA | NA | | | Gm5382 | 0 | NA | NA | NA | NA | NA |
| Olfr1249 | 0 | NA | NA | NA | NA | NA | | | Gm5409 | 0 | NA | NA | NA | NA | NA |
| Olfr125 | 0 | NA | NA | NA | NA | NA | | | Gm5414 | 0 | NA | NA | NA | NA | NA |
| Olfr1250 | 0 | NA | NA | NA | NA | NA | | | Gm5416 | 0 | NA | NA | NA | NA | NA |
| Olfr1251 | 0 | NA | NA | NA | NA | NA | | | Gm5441 | 0 | NA | NA | NA | NA | NA |
| Olfr1252 | 0.192142086 | -0.026384592 | 0.031527717 | -0.836869715 | 0.402665805 | NA | | | Gm5460 | 0 | NA | NA | NA | NA | NA |
| Olfr1253 | 0 | NA | NA | NA | NA | NA | | | Gm5476 | 0 | NA | NA | NA | NA | NA |
| Olfr1254 | 0 | NA | NA | NA | NA | NA | | | Gm5477 | 0 | NA | NA | NA | NA | NA |
| Olfr1255 | 0 | NA | NA | NA | NA | NA | | | Gm5478 | 0 | NA | NA | NA | NA | NA |
| Olfr1256 | 0 | NA | NA | NA | NA | NA | | | Gm5485 | 0 | NA | NA | NA | NA | NA |
| Olfr1257 | 0 | NA | NA | NA | NA | NA | | | Gm5549 | 0 | NA | NA | NA | NA | NA |
| Olfr1258 | 0 | NA | NA | NA | NA | NA | | | Gm5591 | 0 | NA | NA | NA | NA | NA |
| Olfr1259 | 0.196706773 | -0.026517754 | 0.031609464 | -0.838918169 | 0.401515231 | NA | | | Gm5592 | 0 | NA | NA | NA | NA | NA |
| Olfr126 | 0 | NA | NA | NA | NA | NA | | | Gm5615 | 0 | NA | NA | NA | NA | NA |
| Olfr1260 | 0 | NA | NA | NA | NA | NA | | | Gm5627 | 0 | NA | NA | NA | NA | NA |
| Olfr1261 | 0.196706773 | -0.026517754 | 0.031609464 | -0.838918169 | 0.401515231 | NA | | | Gm5634 | 0 | NA | NA | NA | NA | NA |
| Olfr1262 | 0 | NA | NA | NA | NA | NA | | | Gm5635 | 0 | NA | NA | NA | NA | NA |
| Olfr1263 | 0 | NA | NA | NA | NA | NA | | | Gm5662 | 0 | NA | NA | NA | NA | NA |
| Olfr1264 | 0 | NA | NA | NA | NA | NA | | | Gm5712 | 0 | NA | NA | NA | NA | NA |
| Olfr1265 | 0 | NA | NA | NA | NA | NA | | | Gm5725 | 0 | NA | NA | NA | NA | NA |
| Olfr1269 | 0 | NA | NA | NA | NA | NA | | | Gm5726 | 0 | NA | NA | NA | NA | NA |
| Olfr127 | 0 | NA | NA | NA | NA | NA | | | Gm5728 | 0 | NA | NA | NA | NA | NA |
| Olfr1270 | 0.107716603 | -0.020477583 | 0.027954224 | -0.732539849 | 0.46383913 | NA | | | Gm5766 | 0 | NA | NA | NA | NA | NA |
| Olfr1271 | 0 | NA | NA | NA | NA | NA | | | Gm5795 | 0 | NA | NA | NA | NA | NA |
| Olfr1272 | 0 | NA | NA | NA | NA | NA | | | Gm5800 | 0 | NA | NA | NA | NA | NA |
| Olfr1273-ps | 0 | NA | NA | NA | NA | NA | | | Gm5833 | 0 | NA | NA | NA | NA | NA |
| Olfr1274-ps | 0 | NA | NA | NA | NA | NA | | | Gm5885 | 0 | NA | NA | NA | NA | NA |
| Olfr1275 | 0 | NA | NA | NA | NA | NA | | | Gm5886 | 0 | NA | NA | NA | NA | NA |
| Olfr1276 | 0 | NA | NA | NA | NA | NA | | | Gm5891 | 0 | NA | NA | NA | NA | NA |
| Olfr1277 | 0 | NA | NA | NA | NA | NA | | | Gm590 | 0 | NA | NA | NA | NA | NA |
| Olfr1278 | 0.099009884 | 0.018205726 | 0.027896865 | 0.652608315 | 0.514008823 | NA | | | Gm5925 | 0 | NA | NA | NA | NA | NA |
| Olfr1279 | 0 | NA | NA | NA | NA | NA | | | Gm5934 | 0 | NA | NA | NA | NA | NA |
| Olfr128 | 0 | NA | NA | NA | NA | NA | | | Gm5935 | 0 | NA | NA | NA | NA | NA |
| Olfr1280 | 0 | NA | NA | NA | NA | NA | | | Gm5941 | 0 | NA | NA | NA | NA | NA |
| Olfr1281 | 0.099009884 | 0.018205726 | 0.027896865 | 0.652608315 | 0.514008823 | NA | | | Gm595 | 0 | NA | NA | NA | NA | NA |
| Olfr1282 | 0 | NA | NA | NA | NA | NA | | | Gm6026 | 0 | NA | NA | NA | NA | NA |
| Olfr1283 | 0 | NA | NA | NA | NA | NA | | | Gm6040 | 0 | NA | NA | NA | NA | NA |
| Olfr1284 | 0 | NA | NA | NA | NA | NA | | | Gm6042 | 0 | NA | NA | NA | NA | NA |
| Olfr1286 | 0 | NA | NA | NA | NA | NA | | | Gm6084 | 0 | NA | NA | NA | NA | NA |
| Olfr1287 | 0 | NA | NA | NA | NA | NA | | | Gm6116 | 0 | NA | NA | NA | NA | NA |
| Olfr1288 | 0 | NA | NA | NA | NA | NA | | | Gm6121 | 0 | NA | NA | NA | NA | NA |
| Olfr1289 | 0 | NA | NA | NA | NA | NA | | | Gm6164 | 0 | NA | NA | NA | NA | NA |
| Olfr129 | 0 | NA | NA | NA | NA | NA | | | Gm6268 | 0 | NA | NA | NA | NA | NA |
| Olfr1290 | 0.096071043 | -0.020098795 | 0.027825948 | -0.722304036 | 0.470107572 | NA | | | Gm6329 | 0 | NA | NA | NA | NA | NA |
| Olfr1294 | 0 | NA | NA | NA | NA | NA | | | Gm6367 | 0 | NA | NA | NA | NA | NA |
| Olfr1295 | 0 | NA | NA | NA | NA | NA | | | Gm6370 | 0 | NA | NA | NA | NA | NA |
| Olfr1297 | 0 | NA | NA | NA | NA | NA | | | Gm6432 | 0 | NA | NA | NA | NA | NA |
| Olfr1298 | 0 | NA | NA | NA | NA | NA | | | Gm6455 | 0 | NA | NA | NA | NA | NA |
| Olfr1299 | 0 | NA | NA | NA | NA | NA | | | Gm6460 | 0 | NA | NA | NA | NA | NA |
| Olfr13 | 0 | NA | NA | NA | NA | NA | | | Gm6484 | 0 | NA | NA | NA | NA | NA |
| Olfr130 | 0 | NA | NA | NA | NA | NA | | | Gm6524 | 0 | NA | NA | NA | NA | NA |
| Olfr1300-ps1 | 0 | NA | NA | NA | NA | NA | | | Gm6578 | 0 | NA | NA | NA | NA | NA |
| Olfr1301 | 0 | NA | NA | NA | NA | NA | | | Gm6588 | 0 | NA | NA | NA | NA | NA |
| Olfr1302 | 0 | NA | NA | NA | NA | NA | | | Gm6592 | 0 | NA | NA | NA | NA | NA |
| Olfr1303 | 0 | NA | NA | NA | NA | NA | | | Gm6614 | 0 | NA | NA | NA | NA | NA |
| Olfr1305 | 0 | NA | NA | NA | NA | NA | | | Gm6634 | 0 | NA | NA | NA | NA | NA |
| Olfr1306 | 0 | NA | NA | NA | NA | NA | | | Gm6639 | 0 | NA | NA | NA | NA | NA |
| Olfr1307 | 0 | NA | NA | NA | NA | NA | | | Gm6644 | 0 | NA | NA | NA | NA | NA |
| Olfr1308 | 0 | NA | NA | NA | NA | NA | | | Gm6696 | 0 | NA | NA | NA | NA | NA |
| Olfr1309 | 0 | NA | NA | NA | NA | NA | | | Gm6710 | 0 | NA | NA | NA | NA | NA |

| | | | | | | | | | | | | | | | |
|---|---|---|---|---|---|---|---|---|---|---|---|---|---|---|---|
| Olfr131 | 0 | NA | NA | NA | NA | NA | | | Gm6763 | 0 | NA | NA | NA | NA | NA |
| Olfr1310 | 0 | NA | NA | NA | NA | NA | | | Gm6772 | 0 | NA | NA | NA | NA | NA |
| Olfr1311 | 0 | NA | NA | NA | NA | NA | | | Gm6793 | 0 | NA | NA | NA | NA | NA |
| Olfr1312 | 0 | NA | NA | NA | NA | NA | | | Gm6812 | 0 | NA | NA | NA | NA | NA |
| Olfr1313 | 0 | NA | NA | NA | NA | NA | | | Gm6902 | 0 | NA | NA | NA | NA | NA |
| Olfr1314 | 0 | NA | NA | NA | NA | NA | | | Gm6994 | 0 | NA | NA | NA | NA | NA |
| Olfr1316 | 0 | NA | NA | NA | NA | NA | | | Gm7030 | 0 | NA | NA | NA | NA | NA |
| Olfr1317 | 0 | NA | NA | NA | NA | NA | | | Gm7073 | 0 | NA | NA | NA | NA | NA |
| Olfr1318 | 0 | NA | NA | NA | NA | NA | | | Gm7104 | 0 | NA | NA | NA | NA | NA |
| Olfr132 | 0.098353386 | -0.020212831 | 0.02786466 | -0.725393061 | 0.468210929 | NA | | | Gm7134 | 0 | NA | NA | NA | NA | NA |
| Olfr1320 | 0 | NA | NA | NA | NA | NA | | | Gm7157 | 0 | NA | NA | NA | NA | NA |
| Olfr1321 | 0 | NA | NA | NA | NA | NA | | | Gm7257 | 0 | NA | NA | NA | NA | NA |
| Olfr1322 | 0 | NA | NA | NA | NA | NA | | | Gm732 | 0 | NA | NA | NA | NA | NA |
| Olfr1323 | 0 | NA | NA | NA | NA | NA | | | Gm7325 | 0 | NA | NA | NA | NA | NA |
| Olfr1324 | 0 | NA | NA | NA | NA | NA | | | Gm7334 | 0 | NA | NA | NA | NA | NA |
| Olfr1325 | 0 | NA | NA | NA | NA | NA | | | Gm7337 | 0 | NA | NA | NA | NA | NA |
| Olfr1328 | 0 | NA | NA | NA | NA | NA | | | Gm7367 | 0 | NA | NA | NA | NA | NA |
| Olfr1329 | 0 | NA | NA | NA | NA | NA | | | Gm7534 | 0 | NA | NA | NA | NA | NA |
| Olfr133 | 0 | NA | NA | NA | NA | NA | | | Gm7550 | 0 | NA | NA | NA | NA | NA |
| Olfr1330 | 0 | NA | NA | NA | NA | NA | | | Gm757 | 0 | NA | NA | NA | NA | NA |
| Olfr1331 | 0 | NA | NA | NA | NA | NA | | | Gm773 | 0 | NA | NA | NA | NA | NA |
| Olfr1333 | 0 | NA | NA | NA | NA | NA | | | Gm7849 | 0 | NA | NA | NA | NA | NA |
| Olfr1335 | 0 | NA | NA | NA | NA | NA | | | Gm7861 | 0 | NA | NA | NA | NA | NA |
| Olfr1336 | 0 | NA | NA | NA | NA | NA | | | Gm7903 | 0 | NA | NA | NA | NA | NA |
| Olfr1337 | 0 | NA | NA | NA | NA | NA | | | Gm7977 | 0 | NA | NA | NA | NA | NA |
| Olfr1338 | 0 | NA | NA | NA | NA | NA | | | Gm7978 | 0 | NA | NA | NA | NA | NA |
| Olfr1339 | 0.233393928 | -0.027475306 | 0.032191697 | -0.85349047 | 0.39338737 | NA | | | Gm806 | 0 | NA | NA | NA | NA | NA |
| Olfr134 | 0 | NA | NA | NA | NA | NA | | | Gm8096 | 0 | NA | NA | NA | NA | NA |
| Olfr1340 | 0 | NA | NA | NA | NA | NA | | | Gm815 | 0 | NA | NA | NA | NA | NA |
| Olfr1341 | 0 | NA | NA | NA | NA | NA | | | Gm8221 | 0 | NA | NA | NA | NA | NA |
| Olfr1342 | 0.08731684 | 0.017365056 | 0.027733412 | 0.626142072 | 0.531221759 | NA | | | Gm829 | 0 | NA | NA | NA | NA | NA |
| Olfr1346 | 0.29403132 | 0.006032586 | 0.033500653 | 0.180073692 | 0.857094716 | NA | | | Gm8298 | 0 | NA | NA | NA | NA | NA |
| Olfr1347 | 0.096071043 | -0.020098795 | 0.027825948 | -0.722304036 | 0.470107572 | NA | | | Gm8300 | 0 | NA | NA | NA | NA | NA |
| Olfr1348 | 0.094392817 | 0.017895942 | 0.027818137 | 0.643319236 | 0.520016977 | NA | | | Gm839 | 0 | NA | NA | NA | NA | NA |
| Olfr1349 | 0.212768007 | -0.02694223 | 0.031868764 | -0.845411806 | 0.397880955 | NA | | | Gm8439 | 0 | NA | NA | NA | NA | NA |
| Olfr135 | 0 | NA | NA | NA | NA | NA | | | Gm8453 | 0 | NA | NA | NA | NA | NA |
| Olfr1350 | 0.099009884 | 0.018205726 | 0.027896865 | 0.652608315 | 0.514008823 | NA | | | Gm8579 | 0 | NA | NA | NA | NA | NA |
| Olfr1351 | 0 | NA | NA | NA | NA | NA | | | Gm8633 | 0 | NA | NA | NA | NA | NA |
| Olfr1352 | 0 | NA | NA | NA | NA | NA | | | Gm8660 | 0 | NA | NA | NA | NA | NA |
| Olfr1353 | 0 | NA | NA | NA | NA | NA | | | Gm8677 | 0 | NA | NA | NA | NA | NA |
| Olfr1354 | 0 | NA | NA | NA | NA | NA | | | Gm8693 | 0 | NA | NA | NA | NA | NA |
| Olfr1355 | 0 | NA | NA | NA | NA | NA | | | Gm8709 | 0 | NA | NA | NA | NA | NA |
| Olfr1356 | 0 | NA | NA | NA | NA | NA | | | Gm8720 | 0 | NA | NA | NA | NA | NA |
| Olfr1357 | 0 | NA | NA | NA | NA | NA | | | Gm8764 | 0 | NA | NA | NA | NA | NA |
| Olfr1359 | 0 | NA | NA | NA | NA | NA | | | Gm8765 | 0 | NA | NA | NA | NA | NA |
| Olfr136 | 0.098353386 | -0.020212831 | 0.02786466 | -0.725393061 | 0.468210929 | NA | | | Gm8882 | 0 | NA | NA | NA | NA | NA |
| Olfr1360 | 0 | NA | NA | NA | NA | NA | | | Gm8883 | 0 | NA | NA | NA | NA | NA |
| Olfr1361 | 0 | NA | NA | NA | NA | NA | | | Gm8884 | 0 | NA | NA | NA | NA | NA |
| Olfr1362 | 0 | NA | NA | NA | NA | NA | | | Gm8898 | 0 | NA | NA | NA | NA | NA |
| Olfr1364 | 0 | NA | NA | NA | NA | NA | | | Gm8989 | 0 | NA | NA | NA | NA | NA |
| Olfr1366 | 0 | NA | NA | NA | NA | NA | | | Gm8994 | 0 | NA | NA | NA | NA | NA |
| Olfr1367 | 0 | NA | NA | NA | NA | NA | | | Gm9 | 0 | NA | NA | NA | NA | NA |
| Olfr1368 | 0 | NA | NA | NA | NA | NA | | | Gm9041 | 0 | NA | NA | NA | NA | NA |
| Olfr137 | 0 | NA | NA | NA | NA | NA | | | Gm906 | 0 | NA | NA | NA | NA | NA |
| Olfr1370 | 0 | NA | NA | NA | NA | NA | | | Gm9112 | 0 | NA | NA | NA | NA | NA |
| Olfr1371 | 0.196680618 | -0.003857632 | 0.031597546 | -0.122086447 | 0.902830556 | NA | | | Gm9125 | 0 | NA | NA | NA | NA | NA |
| Olfr1372-ps1 | 7.366762932 | 0.011213609 | 0.149911666 | 0.074801442 | 0.940372694 | NA | | | Gm9199 | 0 | NA | NA | NA | NA | NA |
| Olfr1373 | 0 | NA | NA | NA | NA | NA | | | Gm9268 | 0 | NA | NA | NA | NA | NA |
| Olfr1377 | 0.406008005 | -0.00339866 | 0.047828467 | -0.071059364 | 0.943350509 | NA | | | Gm933 | 0 | NA | NA | NA | NA | NA |
| Olfr1378 | 0.612124479 | 0.05749822 | 0.056841187 | 1.011559104 | 0.311748908 | NA | | | Gm9376 | 0 | NA | NA | NA | NA | NA |
| Olfr138 | 0 | NA | NA | NA | NA | NA | | | Gm94 | 0 | NA | NA | NA | NA | NA |
| Olfr1380 | 0 | NA | NA | NA | NA | NA | | | Gm9513 | 0 | NA | NA | NA | NA | NA |
| Olfr1381 | 0 | NA | NA | NA | NA | NA | | | Gm9696 | 0 | NA | NA | NA | NA | NA |
| Olfr1382 | 0 | NA | NA | NA | NA | NA | | | Gm97 | 0 | NA | NA | NA | NA | NA |
| Olfr1383 | 0 | NA | NA | NA | NA | NA | | | Gm9731 | 0 | NA | NA | NA | NA | NA |
| Olfr1384 | 0 | NA | NA | NA | NA | NA | | | Gm9733 | 0 | NA | NA | NA | NA | NA |
| Olfr1385 | 0 | NA | NA | NA | NA | NA | | | Gm9839 | 0 | NA | NA | NA | NA | NA |
| Olfr1386 | 0 | NA | NA | NA | NA | NA | | | Gm9846 | 0 | NA | NA | NA | NA | NA |
| Olfr1387 | 0 | NA | NA | NA | NA | NA | | | Gm9855 | 0 | NA | NA | NA | NA | NA |
| Olfr1388 | 0 | NA | NA | NA | NA | NA | | | Gm9961 | 0 | NA | NA | NA | NA | NA |
| Olfr1389 | 0.198019769 | 0.025308392 | 0.031628095 | 0.80018706 | 0.423602426 | NA | | | Gmcl1l | 0 | NA | NA | NA | NA | NA |
| Olfr139 | 0 | NA | NA | NA | NA | NA | | | Gnat3 | 0 | NA | NA | NA | NA | NA |
| Olfr1390 | 0.192142086 | -0.026384592 | 0.031527717 | -0.836869715 | 0.402665805 | NA | | | Gngt1 | 0 | NA | NA | NA | NA | NA |
| Olfr1391 | 0.096071043 | -0.020098795 | 0.027825948 | -0.722304036 | 0.470107572 | NA | | | Gnrhr | 0 | NA | NA | NA | NA | NA |
| Olfr1392 | 2.880441076 | -0.095750908 | 0.116126803 | -0.824537533 | 0.409634198 | NA | | | Gpa33 | 0 | NA | NA | NA | NA | NA |
| Olfr1394 | 0 | NA | NA | NA | NA | NA | | | Gpr113 | 0 | NA | NA | NA | NA | NA |
| Olfr1395 | 0.079983654 | 0.016740032 | 0.027674178 | 0.604897175 | 0.54524732 | NA | | | Gpr87 | 0 | NA | NA | NA | NA | NA |
| Olfr1396 | 0.284680111 | 0.011373957 | 0.038774397 | 0.293336791 | 0.769264735 | NA | | | Gpx5 | 0 | NA | NA | NA | NA | NA |
| Olfr140 | 0 | NA | NA | NA | NA | NA | | | Gsdmc2 | 0 | NA | NA | NA | NA | NA |
| Olfr1402 | 0 | NA | NA | NA | NA | NA | | | Gsdmc3 | 0 | NA | NA | NA | NA | NA |
| Olfr1404 | 0 | NA | NA | NA | NA | NA | | | Gsdmcl-ps | 0 | NA | NA | NA | NA | NA |
| Olfr1406 | 0 | NA | NA | NA | NA | NA | | | Gsdmcl1 | 0 | NA | NA | NA | NA | NA |
| Olfr1408 | 0.116696964 | -0.020477583 | 0.027954224 | -0.732539847 | 0.463839131 | NA | | | Gvin1 | 0 | NA | NA | NA | NA | NA |
| Olfr141 | 0 | NA | NA | NA | NA | NA | | | Gzmd | 0 | NA | NA | NA | NA | NA |
| Olfr1410 | 0 | NA | NA | NA | NA | NA | | | Gzme | 0 | NA | NA | NA | NA | NA |
| Olfr1411 | 0 | NA | NA | NA | NA | NA | | | Gzmf | 0 | NA | NA | NA | NA | NA |
| Olfr1412 | 0 | NA | NA | NA | NA | NA | | | Gzmn | 0 | NA | NA | NA | NA | NA |
| Olfr1413 | 0 | NA | NA | NA | NA | NA | | | H1fnt | 0 | NA | NA | NA | NA | NA |
| Olfr1414 | 0.096071043 | -0.020098795 | 0.027825948 | -0.722304036 | 0.470107572 | NA | | | H1foo | 0 | NA | NA | NA | NA | NA |
| Olfr1415 | 0 | NA | NA | NA | NA | NA | | | H2-L | 0 | NA | NA | NA | NA | NA |
| Olfr1416 | 0 | NA | NA | NA | NA | NA | | | H2-M1 | 0 | NA | NA | NA | NA | NA |
| Olfr1417 | 0 | NA | NA | NA | NA | NA | | | H2-M10.1 | 0 | NA | NA | NA | NA | NA |
| Olfr1418 | 0 | NA | NA | NA | NA | NA | | | H2-M10.2 | 0 | NA | NA | NA | NA | NA |
| Olfr1419 | 0.263371537 | 0.009921338 | 0.037961873 | 0.261350055 | 0.793822569 | NA | | | H2-M10.4 | 0 | NA | NA | NA | NA | NA |
| Olfr142 | 0 | NA | NA | NA | NA | NA | | | H2-M10.5 | 0 | NA | NA | NA | NA | NA |
| Olfr1420 | 0 | NA | NA | NA | NA | NA | | | H2-M10.6 | 0 | NA | NA | NA | NA | NA |
| Olfr1423 | 0 | NA | NA | NA | NA | NA | | | H2-Q6 | 0 | NA | NA | NA | NA | NA |
| Olfr1424 | 0 | NA | NA | NA | NA | NA | | | H2-Q7 | 0 | NA | NA | NA | NA | NA |
| Olfr1425 | 0 | NA | NA | NA | NA | NA | | | H2-Q8 | 0 | NA | NA | NA | NA | NA |
| Olfr1426 | 0 | NA | NA | NA | NA | NA | | | H2-Q9 | 0 | NA | NA | NA | NA | NA |
| Olfr1427 | 0 | NA | NA | NA | NA | NA | | | H2-T9 | 0 | NA | NA | NA | NA | NA |
| Olfr1428 | 0.096071043 | -0.020098795 | 0.027825948 | -0.722304036 | 0.470107572 | NA | | | H2bfm | 0 | NA | NA | NA | NA | NA |
| Olfr143 | 0 | NA | NA | NA | NA | NA | | | Hand2 | 0 | NA | NA | NA | NA | NA |
| Olfr1431 | 0.107716603 | -0.020477583 | 0.027954224 | -0.732539849 | 0.46383913 | NA | | | Hao2 | 0 | NA | NA | NA | NA | NA |
| Olfr1433 | 0 | NA | NA | NA | NA | NA | | | Hba-a1 | 0 | NA | NA | NA | NA | NA |
| Olfr1434 | 0 | NA | NA | NA | NA | NA | | | Hba-a2 | 0 | NA | NA | NA | NA | NA |
| Olfr1436 | 0 | NA | NA | NA | NA | NA | | | Hba-x | 0 | NA | NA | NA | NA | NA |
| Olfr1437 | 0.08731684 | 0.017365056 | 0.027733412 | 0.626142072 | 0.531221759 | NA | | | Hbb-b1 | 0 | NA | NA | NA | NA | NA |
| Olfr1440 | 0 | NA | NA | NA | NA | NA | | | Hbb-b2 | 0 | NA | NA | NA | NA | NA |
| Olfr1441 | 0 | NA | NA | NA | NA | NA | | | Hbb-bh1 | 0 | NA | NA | NA | NA | NA |
| Olfr1442 | 0 | NA | NA | NA | NA | NA | | | Hbb-bh2 | 0 | NA | NA | NA | NA | NA |
| Olfr1443 | 0 | NA | NA | NA | NA | NA | | | Hemt1 | 0 | NA | NA | NA | NA | NA |
| Olfr1444 | 0.099009884 | 0.018205726 | 0.027896865 | 0.652608315 | 0.514008823 | NA | | | Higd1c | 0 | NA | NA | NA | NA | NA |
| Olfr1445 | 0.094392817 | 0.017895942 | 0.027818137 | 0.643319236 | 0.520016977 | NA | | | Hils1 | 0 | NA | NA | NA | NA | NA |
| Olfr1446 | 0 | NA | NA | NA | NA | NA | | | Hist1h1t | 0 | NA | NA | NA | NA | NA |
| Olfr1447 | 0 | NA | NA | NA | NA | NA | | | Hist1h2aa | 0 | NA | NA | NA | NA | NA |
| Olfr1448 | 0 | NA | NA | NA | NA | NA | | | Hist1h2ag | 0 | NA | NA | NA | NA | NA |
| Olfr1449 | 0 | NA | NA | NA | NA | NA | | | Hist1h2ah | 0 | NA | NA | NA | NA | NA |
| Olfr145 | 0 | NA | NA | NA | NA | NA | | | Hist1h2an | 0 | NA | NA | NA | NA | NA |
| Olfr1450 | 0 | NA | NA | NA | NA | NA | | | Hist1h2ao | 0 | NA | NA | NA | NA | NA |
| Olfr1451 | 0 | NA | NA | NA | NA | NA | | | Hist1h2ap | 0 | NA | NA | NA | NA | NA |
| Olfr1453 | 0 | NA | NA | NA | NA | NA | | | Hist1h2ba | 0 | NA | NA | NA | NA | NA |
| Olfr1454 | 0 | NA | NA | NA | NA | NA | | | Hist1h2bf | 0 | NA | NA | NA | NA | NA |
| Olfr1457 | 0 | NA | NA | NA | NA | NA | | | Hist1h2bk | 0 | NA | NA | NA | NA | NA |
| Olfr1459 | 0 | NA | NA | NA | NA | NA | | | Hist1h2bl | 0 | NA | NA | NA | NA | NA |
| Olfr146 | 0 | NA | NA | NA | NA | NA | | | Hist1h2bm | 0 | NA | NA | NA | NA | NA |
| Olfr1461 | 0 | NA | NA | NA | NA | NA | | | Hist1h2bq | 0 | NA | NA | NA | NA | NA |
| Olfr1462 | 0 | NA | NA | NA | NA | NA | | | Hist1h2br | 0 | NA | NA | NA | NA | NA |
| Olfr1463 | 0 | NA | NA | NA | NA | NA | | | Hist1h3a | 0 | NA | NA | NA | NA | NA |
| Olfr1465 | 0 | NA | NA | NA | NA | NA | | | Hist1h3h | 0 | NA | NA | NA | NA | NA |
| Olfr1466 | 0 | NA | NA | NA | NA | NA | | | Hist1h4n | 0 | NA | NA | NA | NA | NA |
| Olfr1467 | 0 | NA | NA | NA | NA | NA | | | Hist2h2aa1 | 0 | NA | NA | NA | NA | NA |
| Olfr1469 | 0 | NA | NA | NA | NA | NA | | | Hist2h2aa2 | 0 | NA | NA | NA | NA | NA |
| Olfr147 | 0 | NA | NA | NA | NA | NA | | | Hist2h3b | 0 | NA | NA | NA | NA | NA |
| Olfr1471 | 0 | NA | NA | NA | NA | NA | | | Hmga1-rs1 | 0 | NA | NA | NA | NA | NA |
| Olfr1472 | 0 | NA | NA | NA | NA | NA | | | Hmx1 | 0 | NA | NA | NA | NA | NA |
| Olfr1474 | 0 | NA | NA | NA | NA | NA | | | Hotair | 0 | NA | NA | NA | NA | NA |
| Olfr1475 | 0 | NA | NA | NA | NA | NA | | | Hoxa11 | 0 | NA | NA | NA | NA | NA |
| Olfr1477 | 0 | NA | NA | NA | NA | NA | | | Hoxa11as | 0 | NA | NA | NA | NA | NA |
| Olfr148 | 0 | NA | NA | NA | NA | NA | | | Hoxa13 | 0 | NA | NA | NA | NA | NA |
| Olfr1480 | 0 | NA | NA | NA | NA | NA | | | Hoxa3 | 0 | NA | NA | NA | NA | NA |
| Olfr1484 | 0 | NA | NA | NA | NA | NA | | | Hoxa6 | 0 | NA | NA | NA | NA | NA |
| Olfr1487 | 0.098353386 | -0.020212831 | 0.02786466 | -0.725393061 | 0.468210929 | NA | | | Hoxa9 | 0 | NA | NA | NA | NA | NA |
| Olfr1489 | 0 | NA | NA | NA | NA | NA | | | Hoxb1 | 0 | NA | NA | NA | NA | NA |
| Olfr149 | 0 | NA | NA | NA | NA | NA | | | Hoxc10 | 0 | NA | NA | NA | NA | NA |
| Olfr1490 | 0 | NA | NA | NA | NA | NA | | | Hoxc11 | 0 | NA | NA | NA | NA | NA |
| Olfr1491 | 0 | NA | NA | NA | NA | NA | | | Hoxc12 | 0 | NA | NA | NA | NA | NA |
| Olfr1494 | 0 | NA | NA | NA | NA | NA | | | Hoxc8 | 0 | NA | NA | NA | NA | NA |
| Olfr1495 | 0 | NA | NA | NA | NA | NA | | | Hoxc9 | 0 | NA | NA | NA | NA | NA |
| Olfr1496 | 0 | NA | NA | NA | NA | NA | | | Hoxd10 | 0 | NA | NA | NA | NA | NA |
| Olfr1497 | 0 | NA | NA | NA | NA | NA | | | Hoxd11 | 0 | NA | NA | NA | NA | NA |
| Olfr1499 | 0 | NA | NA | NA | NA | NA | | | Hoxd12 | 0 | NA | NA | NA | NA | NA |
| Olfr15 | 0.098353386 | -0.020212831 | 0.02786466 | -0.725393061 | 0.468210929 | NA | | | Hoxd13 | 0 | NA | NA | NA | NA | NA |
| Olfr150 | 0.079983654 | 0.016740032 | 0.027674178 | 0.604897175 | 0.54524732 | NA | | | Hoxd9 | 0 | NA | NA | NA | NA | NA |
| Olfr1500 | 0 | NA | NA | NA | NA | NA | | | Hrh4 | 0 | NA | NA | NA | NA | NA |
| Olfr1501 | 0 | NA | NA | NA | NA | NA | | | Hsd17b3 | 0 | NA | NA | NA | NA | NA |
| Olfr1502 | 0 | NA | NA | NA | NA | NA | | | Hsd17b6 | 0 | NA | NA | NA | NA | NA |
| Olfr1504 | 0 | NA | NA | NA | NA | NA | | | Hsd3b1 | 0 | NA | NA | NA | NA | NA |
| Olfr1505 | 0 | NA | NA | NA | NA | NA | | | Hsd3b2 | 0 | NA | NA | NA | NA | NA |
| Olfr1506 | 0 | NA | NA | NA | NA | NA | | | Hsd3b3 | 0 | NA | NA | NA | NA | NA |
| Olfr1507 | 2.68215537 | -0.062068599 | 0.070310921 | -0.882773226 | 0.377358809 | NA | | | Hsd3b4 | 0 | NA | NA | NA | NA | NA |
| Olfr1508 | 0.096071043 | -0.020098795 | 0.027825948 | -0.722304036 | 0.470107572 | NA | | | Hsd3b5 | 0 | NA | NA | NA | NA | NA |
| Olfr1509 | 0 | NA | NA | NA | NA | NA | | | Hsd3b6 | 0 | NA | NA | NA | NA | NA |
| Olfr151 | 0 | NA | NA | NA | NA | NA | | | Hsfy2 | 0 | NA | NA | NA | NA | NA |
| Olfr1510 | 0 | NA | NA | NA | NA | NA | | | Hspb9 | 0 | NA | NA | NA | NA | NA |
| Olfr1511 | 0 | NA | NA | NA | NA | NA | | | Hyal4 | 0 | NA | NA | NA | NA | NA |
| Olfr1512 | 0 | NA | NA | NA | NA | NA | | | Hyal5 | 0 | NA | NA | NA | NA | NA |
| Olfr1513 | 0 | NA | NA | NA | NA | NA | | | Ifna1 | 0 | NA | NA | NA | NA | NA |
| Olfr152 | 0 | NA | NA | NA | NA | NA | | | Ifna11 | 0 | NA | NA | NA | NA | NA |
| Olfr153 | 0.49741106 | -0.017678863 | 0.050840067 | -0.347734855 | 0.728039318 | NA | | | Ifna12 | 0 | NA | NA | NA | NA | NA |
| Olfr1532-ps1 | 0 | NA | NA | NA | NA | NA | | | Ifna13 | 0 | NA | NA | NA | NA | NA |
| Olfr1535 | 0 | NA | NA | NA | NA | NA | | | Ifna14 | 0 | NA | NA | NA | NA | NA |
| Olfr1537 | 0 | NA | NA | NA | NA | NA | | | Ifna2 | 0 | NA | NA | NA | NA | NA |
| Olfr154 | 0 | NA | NA | NA | NA | NA | | | Ifna4 | 0 | NA | NA | NA | NA | NA |
| Olfr155 | 0 | NA | NA | NA | NA | NA | | | Ifna5 | 0 | NA | NA | NA | NA | NA |
| Olfr156 | 0.107716603 | -0.020477583 | 0.027954224 | -0.732539849 | 0.46383913 | NA | | | Ifna6 | 0 | NA | NA | NA | NA | NA |
| Olfr157 | 0 | NA | NA | NA | NA | NA | | | Ifna7 | 0 | NA | NA | NA | NA | NA |
| Olfr159 | 0 | NA | NA | NA | NA | NA | | | Ifna9 | 0 | NA | NA | NA | NA | NA |
| Olfr16 | 0.094392817 | 0.017895942 | 0.027818137 | 0.643319236 | 0.520016977 | NA | | | Ifnab | 0 | NA | NA | NA | NA | NA |
| Olfr160 | 0 | NA | NA | NA | NA | NA | | | Ifnb1 | 0 | NA | NA | NA | NA | NA |
| Olfr161 | 0 | NA | NA | NA | NA | NA | | | Ifnl2 | 0 | NA | NA | NA | NA | NA |
| Olfr164 | 0 | NA | NA | NA | NA | NA | | | Ifnl3 | 0 | NA | NA | NA | NA | NA |
| Olfr165 | 0 | NA | NA | NA | NA | NA | | | Igfl3 | 0 | NA | NA | NA | NA | NA |
| Olfr166 | 0.116696964 | -0.020477583 | 0.027954224 | -0.732539847 | 0.463839131 | NA | | | Il11ra2 | 0 | NA | NA | NA | NA | NA |
| Olfr167 | 0 | NA | NA | NA | NA | NA | | | Il17a | 0 | NA | NA | NA | NA | NA |
| Olfr168 | 0.116696964 | -0.020477583 | 0.027954224 | -0.732539847 | 0.463839131 | NA | | | Il1f10 | 0 | NA | NA | NA | NA | NA |
| Olfr169 | 0 | NA | NA | NA | NA | NA | | | Il1f6 | 0 | NA | NA | NA | NA | NA |
| Olfr17 | 0 | NA | NA | NA | NA | NA | | | Il2 | 0 | NA | NA | NA | NA | NA |
| Olfr170 | 0 | NA | NA | NA | NA | NA | | | Il20 | 0 | NA | NA | NA | NA | NA |
| Olfr171 | 0 | NA | NA | NA | NA | NA | | | Il22ra2 | 0 | NA | NA | NA | NA | NA |
| Olfr172 | 0 | NA | NA | NA | NA | NA | | | Il24 | 0 | NA | NA | NA | NA | NA |
| Olfr173 | 0 | NA | NA | NA | NA | NA | | | Il31 | 0 | NA | NA | NA | NA | NA |
| Olfr175-ps1 | 0 | NA | NA | NA | NA | NA | | | Il8 | 0 | NA | NA | NA | NA | NA |
| Olfr176 | 0 | NA | NA | NA | NA | NA | | | Il9 | 0 | NA | NA | NA | NA | NA |
| Olfr177 | 0 | NA | NA | NA | NA | NA | | | Inhbc | 0 | NA | NA | NA | NA | NA |
| Olfr178 | 0 | NA | NA | NA | NA | NA | | | Ins1 | 0 | NA | NA | NA | NA | NA |
| Olfr179 | 0 | NA | NA | NA | NA | NA | | | Iqcf1 | 0 | NA | NA | NA | NA | NA |
| Olfr18 | 0.478259379 | 0.031371149 | 0.045569182 | 0.688429064 | 0.491182626 | NA | | | Iqcf6 | 0 | NA | NA | NA | NA | NA |
| Olfr180 | 0 | NA | NA | NA | NA | NA | | | Itln1 | 0 | NA | NA | NA | NA | NA |
| Olfr181 | 0 | NA | NA | NA | NA | NA | | | Ivl | 0 | NA | NA | NA | NA | NA |
| Olfr183 | 0.188785634 | 0.025025111 | 0.031461603 | 0.79541754 | 0.42637066 | NA | | | Izumo2 | 0 | NA | NA | NA | NA | NA |
| Olfr186 | 0 | NA | NA | NA | NA | NA | | | Izumo3 | 0 | NA | NA | NA | NA | NA |
| Olfr187 | 0.233393928 | -0.027475306 | 0.032191697 | -0.85349047 | 0.39338737 | NA | | | Kap | 0 | NA | NA | NA | NA | NA |
| Olfr19 | 0 | NA | NA | NA | NA | NA | | | Kdm5d | 0 | NA | NA | NA | NA | NA |
| Olfr190 | 0 | NA | NA | NA | NA | NA | | | Keg1 | 0 | NA | NA | NA | NA | NA |
| Olfr191 | 0.098353386 | -0.020212831 | 0.02786466 | -0.725393061 | 0.468210929 | NA | | | Khdc1a | 0 | NA | NA | NA | NA | NA |
| Olfr192 | 0 | NA | NA | NA | NA | NA | | | Khdc1c | 0 | NA | NA | NA | NA | NA |
| Olfr193 | 0 | NA | NA | NA | NA | NA | | | Kif2b | 0 | NA | NA | NA | NA | NA |
| Olfr194 | 0 | NA | NA | NA | NA | NA | | | Kif4-ps | 0 | NA | NA | NA | NA | NA |
| Olfr195 | 0 | NA | NA | NA | NA | NA | | | Kir3dl1 | 0 | NA | NA | NA | NA | NA |
| Olfr196 | 0 | NA | NA | NA | NA | NA | | | Klk12 | 0 | NA | NA | NA | NA | NA |
| Olfr197 | 0 | NA | NA | NA | NA | NA | | | Klk15 | 0 | NA | NA | NA | NA | NA |
| Olfr198 | 0 | NA | NA | NA | NA | NA | | | Klk1b1 | 0 | NA | NA | NA | NA | NA |
| Olfr199 | 0 | NA | NA | NA | NA | NA | | | Klk1b11 | 0 | NA | NA | NA | NA | NA |
| Olfr2 | 0 | NA | NA | NA | NA | NA | | | Klk1b16 | 0 | NA | NA | NA | NA | NA |
| Olfr20 | 0.116696964 | -0.020477583 | 0.027954224 | -0.732539847 | 0.463839131 | NA | | | Klk1b22 | 0 | NA | NA | NA | NA | NA |
| Olfr201 | 0 | NA | NA | NA | NA | NA | | | Klk1b24 | 0 | NA | NA | NA | NA | NA |
| Olfr202 | 0 | NA | NA | NA | NA | NA | | | Klk1b26 | 0 | NA | NA | NA | NA | NA |
| Olfr203 | 0 | NA | NA | NA | NA | NA | | | Klk1b27 | 0 | NA | NA | NA | NA | NA |
| Olfr204 | 0 | NA | NA | NA | NA | NA | | | Klk1b5 | 0 | NA | NA | NA | NA | NA |
| Olfr205 | 0 | NA | NA | NA | NA | NA | | | Klk1b7-ps | 0 | NA | NA | NA | NA | NA |
| Olfr206 | 0 | NA | NA | NA | NA | NA | | | Klk1b8 | 0 | NA | NA | NA | NA | NA |
| Olfr207 | 0 | NA | NA | NA | NA | NA | | | Klk1b9 | 0 | NA | NA | NA | NA | NA |
| Olfr208 | 0 | NA | NA | NA | NA | NA | | | Klk5 | 0 | NA | NA | NA | NA | NA |
| Olfr209 | 0 | NA | NA | NA | NA | NA | | | Klra1 | 0 | NA | NA | NA | NA | NA |
| Olfr211 | 0 | NA | NA | NA | NA | NA | | | Klra10 | 0 | NA | NA | NA | NA | NA |
| Olfr212 | 3.176389577 | 0.095130843 | 0.092764089 | 1.025513684 | 0.305120872 | NA | | | Klra12 | 0 | NA | NA | NA | NA | NA |
| Olfr213 | 0 | NA | NA | NA | NA | NA | | | Klra13-ps | 0 | NA | NA | NA | NA | NA |
| Olfr214 | 0.079983654 | 0.016740032 | 0.027674178 | 0.604897175 | 0.54524732 | NA | | | Klra14 | 0 | NA | NA | NA | NA | NA |
| Olfr215 | 0 | NA | NA | NA | NA | NA | | | Klra15 | 0 | NA | NA | NA | NA | NA |
| Olfr218 | 0 | NA | NA | NA | NA | NA | | | Klra17 | 0 | NA | NA | NA | NA | NA |
| Olfr220 | 0 | NA | NA | NA | NA | NA | | | Klra18 | 0 | NA | NA | NA | NA | NA |
| Olfr221 | 0.979208136 | 0.017547049 | 0.071719569 | 0.244661946 | 0.806718195 | NA | | | Klra19 | 0 | NA | NA | NA | NA | NA |
| Olfr222 | 0.290495473 | -0.034250308 | 0.035891155 | -0.954282697 | 0.339940564 | NA | | | Klra21 | 0 | NA | NA | NA | NA | NA |
| Olfr223 | 0 | NA | NA | NA | NA | NA | | | Klra22 | 0 | NA | NA | NA | NA | NA |

| | | | | | | | | | | | | | | | |
|---|---|---|---|---|---|---|---|---|---|---|---|---|---|---|---|
| Olfr224 | 0.192142086 | -0.026384592 | 0.031527717 | -0.836869715 | 0.402665805 | NA | | | Klra23 | 0 | NA | NA | NA | NA | NA |
| Olfr225 | 0 | NA | NA | NA | NA | NA | | | Klra3 | 0 | NA | NA | NA | NA | NA |
| Olfr228 | 0 | NA | NA | NA | NA | NA | | | Klra33 | 0 | NA | NA | NA | NA | NA |
| Olfr229 | 0 | NA | NA | NA | NA | NA | | | Klra4 | 0 | NA | NA | NA | NA | NA |
| Olfr23 | 0 | NA | NA | NA | NA | NA | | | Klra5 | 0 | NA | NA | NA | NA | NA |
| Olfr231 | 0 | NA | NA | NA | NA | NA | | | Klra6 | 0 | NA | NA | NA | NA | NA |
| Olfr235 | 0 | NA | NA | NA | NA | NA | | | Klra7 | 0 | NA | NA | NA | NA | NA |
| Olfr237-ps1 | 0 | NA | NA | NA | NA | NA | | | Klrb1-ps1 | 0 | NA | NA | NA | NA | NA |
| Olfr239 | 0 | NA | NA | NA | NA | NA | | | Klrc2 | 0 | NA | NA | NA | NA | NA |
| Olfr24 | 0 | NA | NA | NA | NA | NA | | | Klrc3 | 0 | NA | NA | NA | NA | NA |
| Olfr242 | 0 | NA | NA | NA | NA | NA | | | Kng1 | 0 | NA | NA | NA | NA | NA |
| Olfr243 | 0 | NA | NA | NA | NA | NA | | | Krt35 | 0 | NA | NA | NA | NA | NA |
| Olfr247 | 0 | NA | NA | NA | NA | NA | | | Krt36 | 0 | NA | NA | NA | NA | NA |
| Olfr248 | 0 | NA | NA | NA | NA | NA | | | Krt39 | 0 | NA | NA | NA | NA | NA |
| Olfr25 | 0 | NA | NA | NA | NA | NA | | | Krt4 | 0 | NA | NA | NA | NA | NA |
| Olfr250 | 0 | NA | NA | NA | NA | NA | | | Krt40 | 0 | NA | NA | NA | NA | NA |
| Olfr251 | 0 | NA | NA | NA | NA | NA | | | Krt5 | 0 | NA | NA | NA | NA | NA |
| Olfr257 | 0 | NA | NA | NA | NA | NA | | | Krt6b | 0 | NA | NA | NA | NA | NA |
| Olfr259 | 0 | NA | NA | NA | NA | NA | | | Krt72 | 0 | NA | NA | NA | NA | NA |
| Olfr26 | 0 | NA | NA | NA | NA | NA | | | Krt81 | 0 | NA | NA | NA | NA | NA |
| Olfr262 | 0 | NA | NA | NA | NA | NA | | | Krtap1-3 | 0 | NA | NA | NA | NA | NA |
| Olfr263 | 0 | NA | NA | NA | NA | NA | | | Krtap1-4 | 0 | NA | NA | NA | NA | NA |
| Olfr266 | 0.098353386 | -0.020212831 | 0.02786466 | -0.725393061 | 0.468210929 | NA | | | Krtap1-5 | 0 | NA | NA | NA | NA | NA |
| Olfr267 | 0.587805432 | 0.015924212 | 0.053274974 | 0.29890606 | 0.765011723 | NA | | | Krtap10-4 | 0 | NA | NA | NA | NA | NA |
| Olfr27 | 0 | NA | NA | NA | NA | NA | | | Krtap12-1 | 0 | NA | NA | NA | NA | NA |
| Olfr270 | 0 | NA | NA | NA | NA | NA | | | Krtap13 | 0 | NA | NA | NA | NA | NA |
| Olfr272 | 0.238795281 | -0.027601606 | 0.032267774 | -0.855392297 | 0.392334007 | NA | | | Krtap14 | 0 | NA | NA | NA | NA | NA |
| Olfr273 | 0 | NA | NA | NA | NA | NA | | | Krtap15 | 0 | NA | NA | NA | NA | NA |
| Olfr275 | 0 | NA | NA | NA | NA | NA | | | Krtap16-1 | 0 | NA | NA | NA | NA | NA |
| Olfr279 | 0 | NA | NA | NA | NA | NA | | | Krtap16-3 | 0 | NA | NA | NA | NA | NA |
| Olfr281 | 0.098353386 | -0.020212831 | 0.02786466 | -0.725393061 | 0.468210929 | NA | | | Krtap19-1 | 0 | NA | NA | NA | NA | NA |
| Olfr282 | 0 | NA | NA | NA | NA | NA | | | Krtap19-3 | 0 | NA | NA | NA | NA | NA |
| Olfr283 | 0.577209771 | 0.036509886 | 0.048527698 | 0.752351489 | 0.451839712 | NA | | | Krtap19-4 | 0 | NA | NA | NA | NA | NA |
| Olfr284 | 0 | NA | NA | NA | NA | NA | | | Krtap19-5 | 0 | NA | NA | NA | NA | NA |
| Olfr285 | 0 | NA | NA | NA | NA | NA | | | Krtap19-9b | 0 | NA | NA | NA | NA | NA |
| Olfr286 | 0.159967308 | 0.0240345 | 0.030870969 | 0.778547006 | 0.436246606 | NA | | | Krtap2-4 | 0 | NA | NA | NA | NA | NA |
| Olfr288 | 7.316914825 | 0.078441901 | 0.128301048 | 0.6113894 | 0.540941818 | NA | | | Krtap20-2 | 0 | NA | NA | NA | NA | NA |
| Olfr29-ps1 | 0 | NA | NA | NA | NA | NA | | | Krtap21-1 | 0 | NA | NA | NA | NA | NA |
| Olfr290 | 0 | NA | NA | NA | NA | NA | | | Krtap22-2 | 0 | NA | NA | NA | NA | NA |
| Olfr291 | 0 | NA | NA | NA | NA | NA | | | Krtap24-1 | 0 | NA | NA | NA | NA | NA |
| Olfr292 | 0 | NA | NA | NA | NA | NA | | | Krtap26-1 | 0 | NA | NA | NA | NA | NA |
| Olfr293 | 0 | NA | NA | NA | NA | NA | | | Krtap27-1 | 0 | NA | NA | NA | NA | NA |
| Olfr294 | 0 | NA | NA | NA | NA | NA | | | Krtap3-1 | 0 | NA | NA | NA | NA | NA |
| Olfr295 | 0 | NA | NA | NA | NA | NA | | | Krtap3-2 | 0 | NA | NA | NA | NA | NA |
| Olfr297 | 0 | NA | NA | NA | NA | NA | | | Krtap3-3 | 0 | NA | NA | NA | NA | NA |
| Olfr298 | 0 | NA | NA | NA | NA | NA | | | Krtap31-1 | 0 | NA | NA | NA | NA | NA |
| Olfr299 | 0 | NA | NA | NA | NA | NA | | | Krtap31-2 | 0 | NA | NA | NA | NA | NA |
| Olfr3 | 0.099009884 | 0.018205726 | 0.027896865 | 0.652608315 | 0.514008823 | NA | | | Krtap4-1 | 0 | NA | NA | NA | NA | NA |
| Olfr30 | 0 | NA | NA | NA | NA | NA | | | Krtap4-13 | 0 | NA | NA | NA | NA | NA |
| Olfr301 | 0 | NA | NA | NA | NA | NA | | | Krtap4-2 | 0 | NA | NA | NA | NA | NA |
| Olfr303 | 0 | NA | NA | NA | NA | NA | | | Krtap4-6 | 0 | NA | NA | NA | NA | NA |
| Olfr304 | 0 | NA | NA | NA | NA | NA | | | Krtap4-7 | 0 | NA | NA | NA | NA | NA |
| Olfr305 | 0 | NA | NA | NA | NA | NA | | | Krtap4-8 | 0 | NA | NA | NA | NA | NA |
| Olfr307 | 0 | NA | NA | NA | NA | NA | | | Krtap4-9 | 0 | NA | NA | NA | NA | NA |
| Olfr308 | 0 | NA | NA | NA | NA | NA | | | Krtap5-3 | 0 | NA | NA | NA | NA | NA |
| Olfr309 | 0.290495473 | -0.034250308 | 0.035891155 | -0.954282697 | 0.339940564 | NA | | | Krtap5-4 | 0 | NA | NA | NA | NA | NA |
| Olfr31 | 0.094392817 | 0.017895942 | 0.027818137 | 0.643319236 | 0.520016977 | NA | | | Krtap5-5 | 0 | NA | NA | NA | NA | NA |
| Olfr310 | 0 | NA | NA | NA | NA | NA | | | Krtap6-1 | 0 | NA | NA | NA | NA | NA |
| Olfr311 | 0 | NA | NA | NA | NA | NA | | | Krtap6-2 | 0 | NA | NA | NA | NA | NA |
| Olfr312 | 0 | NA | NA | NA | NA | NA | | | Krtap6-5 | 0 | NA | NA | NA | NA | NA |
| Olfr313 | 0 | NA | NA | NA | NA | NA | | | Krtap8-1 | 0 | NA | NA | NA | NA | NA |
| Olfr314 | 0 | NA | NA | NA | NA | NA | | | Krtap9-1 | 0 | NA | NA | NA | NA | NA |
| Olfr315 | 0 | NA | NA | NA | NA | NA | | | Krtap9-3 | 0 | NA | NA | NA | NA | NA |
| Olfr317 | 1.721536622 | 0.125837949 | 0.096092464 | 1.309550652 | 0.190347892 | NA | | | Krtap9-5 | 0 | NA | NA | NA | NA | NA |
| Olfr318 | 0 | NA | NA | NA | NA | NA | | | LOC100502924 | 0 | NA | NA | NA | NA | NA |
| Olfr319 | 0 | NA | NA | NA | NA | NA | | | LOC100861978 | 0 | NA | NA | NA | NA | NA |
| Olfr32 | 0.775415375 | -0.069517603 | 0.050698856 | -1.371186808 | 0.17031673 | NA | | | LOC666331 | 0 | NA | NA | NA | NA | NA |
| Olfr320 | 0.079983654 | 0.016740032 | 0.027674178 | 0.604897175 | 0.54524732 | NA | | | Lce1a1 | 0 | NA | NA | NA | NA | NA |
| Olfr323 | 0 | NA | NA | NA | NA | NA | | | Lce1a2 | 0 | NA | NA | NA | NA | NA |
| Olfr324 | 1.583113224 | 0.018576751 | 0.093897186 | 0.197841397 | 0.843169156 | NA | | | Lce1b | 0 | NA | NA | NA | NA | NA |
| Olfr325 | 0 | NA | NA | NA | NA | NA | | | Lce1c | 0 | NA | NA | NA | NA | NA |
| Olfr328 | 0.105923158 | 0.018524693 | 0.027977518 | 0.662127842 | 0.507889294 | NA | | | Lce1d | 0 | NA | NA | NA | NA | NA |
| Olfr329-ps | 0.167300494 | 0.024299763 | 0.031030539 | 0.783091891 | 0.433573153 | NA | | | Lce1e | 0 | NA | NA | NA | NA | NA |
| Olfr33 | 0.298065244 | -0.009392943 | 0.033782767 | -0.278039479 | 0.780982053 | NA | | | Lce1f | 0 | NA | NA | NA | NA | NA |
| Olfr330 | 0 | NA | NA | NA | NA | NA | | | Lce1h | 0 | NA | NA | NA | NA | NA |
| Olfr331 | 0 | NA | NA | NA | NA | NA | | | Lce1i | 0 | NA | NA | NA | NA | NA |
| Olfr332 | 0 | NA | NA | NA | NA | NA | | | Lce1k | 0 | NA | NA | NA | NA | NA |
| Olfr338 | 0 | NA | NA | NA | NA | NA | | | Lce1l | 0 | NA | NA | NA | NA | NA |
| Olfr339 | 0 | NA | NA | NA | NA | NA | | | Lce1m | 0 | NA | NA | NA | NA | NA |
| Olfr340 | 0 | NA | NA | NA | NA | NA | | | Lce3b | 0 | NA | NA | NA | NA | NA |
| Olfr341 | 0 | NA | NA | NA | NA | NA | | | Lce3e | 0 | NA | NA | NA | NA | NA |
| Olfr342 | 0.098353386 | -0.020212831 | 0.02786466 | -0.725393061 | 0.468210929 | NA | | | Lce6a | 0 | NA | NA | NA | NA | NA |
| Olfr344 | 0 | NA | NA | NA | NA | NA | | | Lcn10 | 0 | NA | NA | NA | NA | NA |
| Olfr345 | 0 | NA | NA | NA | NA | NA | | | Lcn4 | 0 | NA | NA | NA | NA | NA |
| Olfr346 | 0 | NA | NA | NA | NA | NA | | | Lcn5 | 0 | NA | NA | NA | NA | NA |
| Olfr347 | 0 | NA | NA | NA | NA | NA | | | Lcn6 | 0 | NA | NA | NA | NA | NA |
| Olfr348 | 0 | NA | NA | NA | NA | NA | | | Lcn9 | 0 | NA | NA | NA | NA | NA |
| Olfr350 | 0 | NA | NA | NA | NA | NA | | | Lelp1 | 0 | NA | NA | NA | NA | NA |
| Olfr351 | 0 | NA | NA | NA | NA | NA | | | Lep | 0 | NA | NA | NA | NA | NA |
| Olfr352 | 0 | NA | NA | NA | NA | NA | | | Lim2 | 0 | NA | NA | NA | NA | NA |
| Olfr353 | 0 | NA | NA | NA | NA | NA | | | Lrch4 | 0 | NA | NA | NA | NA | NA |
| Olfr354 | 0 | NA | NA | NA | NA | NA | | | Lrcol1 | 0 | NA | NA | NA | NA | NA |
| Olfr355 | 0 | NA | NA | NA | NA | NA | | | Lrrc19 | 0 | NA | NA | NA | NA | NA |
| Olfr356 | 0 | NA | NA | NA | NA | NA | | | Lrrc26 | 0 | NA | NA | NA | NA | NA |
| Olfr357 | 0 | NA | NA | NA | NA | NA | | | Lrrc52 | 0 | NA | NA | NA | NA | NA |
| Olfr358 | 0 | NA | NA | NA | NA | NA | | | Luzp4 | 0 | NA | NA | NA | NA | NA |
| Olfr360 | 0 | NA | NA | NA | NA | NA | | | Ly6f | 0 | NA | NA | NA | NA | NA |
| Olfr361 | 0 | NA | NA | NA | NA | NA | | | Ly6g5c | 0 | NA | NA | NA | NA | NA |
| Olfr362 | 0 | NA | NA | NA | NA | NA | | | Lyg1 | 0 | NA | NA | NA | NA | NA |
| Olfr365 | 0 | NA | NA | NA | NA | NA | | | Lypd4 | 0 | NA | NA | NA | NA | NA |
| Olfr366 | 0 | NA | NA | NA | NA | NA | | | Lypd5 | 0 | NA | NA | NA | NA | NA |
| Olfr367-ps | 0 | NA | NA | NA | NA | NA | | | Lyzl1 | 0 | NA | NA | NA | NA | NA |
| Olfr368 | 0 | NA | NA | NA | NA | NA | | | Magea1 | 0 | NA | NA | NA | NA | NA |
| Olfr370 | 0 | NA | NA | NA | NA | NA | | | Magea2 | 0 | NA | NA | NA | NA | NA |
| Olfr371 | 0 | NA | NA | NA | NA | NA | | | Magea3 | 0 | NA | NA | NA | NA | NA |
| Olfr372 | 0 | NA | NA | NA | NA | NA | | | Magea4 | 0 | NA | NA | NA | NA | NA |
| Olfr373 | 0 | NA | NA | NA | NA | NA | | | Magea5 | 0 | NA | NA | NA | NA | NA |
| Olfr374 | 0 | NA | NA | NA | NA | NA | | | Magea6 | 0 | NA | NA | NA | NA | NA |
| Olfr376 | 0 | NA | NA | NA | NA | NA | | | Magea8 | 0 | NA | NA | NA | NA | NA |
| Olfr378 | 0 | NA | NA | NA | NA | NA | | | Mageb1 | 0 | NA | NA | NA | NA | NA |
| Olfr38 | 0 | NA | NA | NA | NA | NA | | | Mageb16 | 0 | NA | NA | NA | NA | NA |
| Olfr380 | 0 | NA | NA | NA | NA | NA | | | Mageb2 | 0 | NA | NA | NA | NA | NA |
| Olfr381 | 0 | NA | NA | NA | NA | NA | | | Mageb4 | 0 | NA | NA | NA | NA | NA |
| Olfr382 | 0 | NA | NA | NA | NA | NA | | | Mageb5 | 0 | NA | NA | NA | NA | NA |
| Olfr384 | 0 | NA | NA | NA | NA | NA | | | Marco | 0 | NA | NA | NA | NA | NA |
| Olfr385 | 0 | NA | NA | NA | NA | NA | | | Mbd3l1 | 0 | NA | NA | NA | NA | NA |
| Olfr389 | 0 | NA | NA | NA | NA | NA | | | Mbl1 | 0 | NA | NA | NA | NA | NA |
| Olfr39 | 0 | NA | NA | NA | NA | NA | | | Mcpt1 | 0 | NA | NA | NA | NA | NA |
| Olfr390 | 0 | NA | NA | NA | NA | NA | | | Mcpt2 | 0 | NA | NA | NA | NA | NA |
| Olfr391-ps | 0 | NA | NA | NA | NA | NA | | | Mcpt9 | 0 | NA | NA | NA | NA | NA |
| Olfr392 | 0 | NA | NA | NA | NA | NA | | | Mfrp | 0 | NA | NA | NA | NA | NA |
| Olfr393 | 0 | NA | NA | NA | NA | NA | | | Mia2 | 0 | NA | NA | NA | NA | NA |
| Olfr394 | 0 | NA | NA | NA | NA | NA | | | Mill1 | 0 | NA | NA | NA | NA | NA |
| Olfr395 | 0 | NA | NA | NA | NA | NA | | | Miox | 0 | NA | NA | NA | NA | NA |
| Olfr397 | 0 | NA | NA | NA | NA | NA | | | Mip | 0 | NA | NA | NA | NA | NA |
| Olfr398 | 0 | NA | NA | NA | NA | NA | | | Mir101a | 0 | NA | NA | NA | NA | NA |
| Olfr399 | 0 | NA | NA | NA | NA | NA | | | Mir101b | 0 | NA | NA | NA | NA | NA |
| Olfr401 | 0 | NA | NA | NA | NA | NA | | | Mir101c | 0 | NA | NA | NA | NA | NA |
| Olfr402 | 0 | NA | NA | NA | NA | NA | | | Mir103-1 | 0 | NA | NA | NA | NA | NA |
| Olfr403 | 0 | NA | NA | NA | NA | NA | | | Mir105 | 0 | NA | NA | NA | NA | NA |
| Olfr406 | 0 | NA | NA | NA | NA | NA | | | Mir106a | 0 | NA | NA | NA | NA | NA |
| Olfr410 | 0 | NA | NA | NA | NA | NA | | | Mir107 | 0 | NA | NA | NA | NA | NA |
| Olfr411 | 0 | NA | NA | NA | NA | NA | | | Mir10a | 0 | NA | NA | NA | NA | NA |
| Olfr412 | 0 | NA | NA | NA | NA | NA | | | Mir10b | 0 | NA | NA | NA | NA | NA |
| Olfr414 | 0.098353386 | -0.020212831 | 0.02786466 | -0.725393061 | 0.468210929 | NA | | | Mir1186 | 0 | NA | NA | NA | NA | NA |
| Olfr417 | 0 | NA | NA | NA | NA | NA | | | Mir1186b | 0 | NA | NA | NA | NA | NA |
| Olfr418-ps1 | 0 | NA | NA | NA | NA | NA | | | Mir1187 | 0 | NA | NA | NA | NA | NA |
| Olfr419 | 0 | NA | NA | NA | NA | NA | | | Mir1190 | 0 | NA | NA | NA | NA | NA |
| Olfr420 | 0 | NA | NA | NA | NA | NA | | | Mir1196 | 0 | NA | NA | NA | NA | NA |
| Olfr421-ps1 | 0 | NA | NA | NA | NA | NA | | | Mir1197 | 0 | NA | NA | NA | NA | NA |
| Olfr424 | 0 | NA | NA | NA | NA | NA | | | Mir1199 | 0 | NA | NA | NA | NA | NA |
| Olfr426 | 0 | NA | NA | NA | NA | NA | | | Mir1224 | 0 | NA | NA | NA | NA | NA |
| Olfr427 | 0 | NA | NA | NA | NA | NA | | | Mir122a | 0 | NA | NA | NA | NA | NA |
| Olfr429 | 0 | NA | NA | NA | NA | NA | | | Mir1247 | 0 | NA | NA | NA | NA | NA |
| Olfr43 | 0.096071043 | -0.020098795 | 0.027825948 | -0.722304036 | 0.470107572 | NA | | | Mir124a-1 | 0 | NA | NA | NA | NA | NA |
| Olfr430 | 0 | NA | NA | NA | NA | NA | | | Mir125b-1 | 0 | NA | NA | NA | NA | NA |
| Olfr432 | 0 | NA | NA | NA | NA | NA | | | Mir125b-2 | 0 | NA | NA | NA | NA | NA |
| Olfr433 | 0 | NA | NA | NA | NA | NA | | | Mir1264 | 0 | NA | NA | NA | NA | NA |
| Olfr434 | 0 | NA | NA | NA | NA | NA | | | Mir127 | 0 | NA | NA | NA | NA | NA |
| Olfr435 | 0 | NA | NA | NA | NA | NA | | | Mir128-2 | 0 | NA | NA | NA | NA | NA |
| Olfr437 | 0.17463368 | 0.024560283 | 0.031186242 | 0.787535833 | 0.430968261 | NA | | | Mir129-2 | 0 | NA | NA | NA | NA | NA |
| Olfr44 | 0 | NA | NA | NA | NA | NA | | | Mir1298 | 0 | NA | NA | NA | NA | NA |
| Olfr441 | 0 | NA | NA | NA | NA | NA | | | Mir1306 | 0 | NA | NA | NA | NA | NA |
| Olfr444 | 0 | NA | NA | NA | NA | NA | | | Mir130a | 0 | NA | NA | NA | NA | NA |
| Olfr446 | 0.19442443 | -0.026451352 | 0.031568725 | -0.837897386 | 0.402088337 | NA | | | Mir130b | 0 | NA | NA | NA | NA | NA |
| Olfr447 | 0 | NA | NA | NA | NA | NA | | | Mir133a-2 | 0 | NA | NA | NA | NA | NA |
| Olfr448 | 0 | NA | NA | NA | NA | NA | | | Mir133b | 0 | NA | NA | NA | NA | NA |
| Olfr449 | 0.119397641 | -0.020477583 | 0.027954224 | -0.732539845 | 0.463839132 | NA | | | Mir139 | 0 | NA | NA | NA | NA | NA |
| Olfr45 | 0 | NA | NA | NA | NA | NA | | | Mir141 | 0 | NA | NA | NA | NA | NA |
| Olfr450 | 0 | NA | NA | NA | NA | NA | | | Mir143 | 0 | NA | NA | NA | NA | NA |
| Olfr452 | 0 | NA | NA | NA | NA | NA | | | Mir144 | 0 | NA | NA | NA | NA | NA |
| Olfr453 | 0 | NA | NA | NA | NA | NA | | | Mir146 | 0 | NA | NA | NA | NA | NA |
| Olfr455 | 0 | NA | NA | NA | NA | NA | | | Mir146b | 0 | NA | NA | NA | NA | NA |
| Olfr456 | 0 | NA | NA | NA | NA | NA | | | Mir147 | 0 | NA | NA | NA | NA | NA |
| Olfr457 | 0 | NA | NA | NA | NA | NA | | | Mir148a | 0 | NA | NA | NA | NA | NA |
| Olfr458 | 0 | NA | NA | NA | NA | NA | | | Mir148b | 0 | NA | NA | NA | NA | NA |
| Olfr459 | 0 | NA | NA | NA | NA | NA | | | Mir150 | 0 | NA | NA | NA | NA | NA |
| Olfr46 | 0.099009884 | 0.018205726 | 0.027896865 | 0.652608315 | 0.514008823 | NA | | | Mir152 | 0 | NA | NA | NA | NA | NA |
| Olfr460 | 0.08731684 | 0.017365056 | 0.027733412 | 0.626142072 | 0.531221759 | NA | | | Mir153 | 0 | NA | NA | NA | NA | NA |
| Olfr461 | 0 | NA | NA | NA | NA | NA | | | Mir155 | 0 | NA | NA | NA | NA | NA |
| Olfr462 | 0 | NA | NA | NA | NA | NA | | | Mir15b | 0 | NA | NA | NA | NA | NA |
| Olfr463 | 0.346511884 | -0.038380278 | 0.037950969 | -1.011312187 | 0.311867035 | NA | | | Mir16-2 | 0 | NA | NA | NA | NA | NA |
| Olfr466 | 0 | NA | NA | NA | NA | NA | | | Mir17 | 0 | NA | NA | NA | NA | NA |
| Olfr467 | 0 | NA | NA | NA | NA | NA | | | Mir18 | 0 | NA | NA | NA | NA | NA |
| Olfr469 | 0 | NA | NA | NA | NA | NA | | | Mir181a-2 | 0 | NA | NA | NA | NA | NA |
| Olfr47 | 0 | NA | NA | NA | NA | NA | | | Mir181b-2 | 0 | NA | NA | NA | NA | NA |
| Olfr470 | 0 | NA | NA | NA | NA | NA | | | Mir182 | 0 | NA | NA | NA | NA | NA |
| Olfr472 | 0 | NA | NA | NA | NA | NA | | | Mir183 | 0 | NA | NA | NA | NA | NA |
| Olfr473 | 0 | NA | NA | NA | NA | NA | | | Mir184 | 0 | NA | NA | NA | NA | NA |
| Olfr474 | 0 | NA | NA | NA | NA | NA | | | Mir1843 | 0 | NA | NA | NA | NA | NA |
| Olfr476 | 0 | NA | NA | NA | NA | NA | | | Mir1843b | 0 | NA | NA | NA | NA | NA |
| Olfr477 | 0 | NA | NA | NA | NA | NA | | | Mir185 | 0 | NA | NA | NA | NA | NA |
| Olfr478 | 0 | NA | NA | NA | NA | NA | | | Mir187 | 0 | NA | NA | NA | NA | NA |
| Olfr479 | 0 | NA | NA | NA | NA | NA | | | Mir188 | 0 | NA | NA | NA | NA | NA |
| Olfr48 | 0.098353386 | -0.020212831 | 0.02786466 | -0.725393061 | 0.468210929 | NA | | | Mir1892 | 0 | NA | NA | NA | NA | NA |
| Olfr480 | 0 | NA | NA | NA | NA | NA | | | Mir1893 | 0 | NA | NA | NA | NA | NA |
| Olfr481 | 0 | NA | NA | NA | NA | NA | | | Mir1897 | 0 | NA | NA | NA | NA | NA |
| Olfr482 | 0 | NA | NA | NA | NA | NA | | | Mir1899 | 0 | NA | NA | NA | NA | NA |
| Olfr483 | 0 | NA | NA | NA | NA | NA | | | Mir18b | 0 | NA | NA | NA | NA | NA |
| Olfr484 | 0 | NA | NA | NA | NA | NA | | | Mir1900 | 0 | NA | NA | NA | NA | NA |
| Olfr485 | 0 | NA | NA | NA | NA | NA | | | Mir1902 | 0 | NA | NA | NA | NA | NA |
| Olfr486 | 0 | NA | NA | NA | NA | NA | | | Mir1903 | 0 | NA | NA | NA | NA | NA |
| Olfr487 | 0 | NA | NA | NA | NA | NA | | | Mir1904 | 0 | NA | NA | NA | NA | NA |
| Olfr488 | 0 | NA | NA | NA | NA | NA | | | Mir1906-1 | 0 | NA | NA | NA | NA | NA |
| Olfr49 | 0 | NA | NA | NA | NA | NA | | | Mir1906-2 | 0 | NA | NA | NA | NA | NA |
| Olfr490 | 0 | NA | NA | NA | NA | NA | | | Mir190b | 0 | NA | NA | NA | NA | NA |
| Olfr491 | 0 | NA | NA | NA | NA | NA | | | Mir1912 | 0 | NA | NA | NA | NA | NA |
| Olfr492 | 0 | NA | NA | NA | NA | NA | | | Mir193 | 0 | NA | NA | NA | NA | NA |
| Olfr493 | 0 | NA | NA | NA | NA | NA | | | Mir1932 | 0 | NA | NA | NA | NA | NA |
| Olfr494 | 0 | NA | NA | NA | NA | NA | | | Mir1933 | 0 | NA | NA | NA | NA | NA |
| Olfr495 | 0 | NA | NA | NA | NA | NA | | | Mir1936 | 0 | NA | NA | NA | NA | NA |
| Olfr497 | 0 | NA | NA | NA | NA | NA | | | Mir193b | 0 | NA | NA | NA | NA | NA |
| Olfr498 | 0 | NA | NA | NA | NA | NA | | | Mir194-1 | 0 | NA | NA | NA | NA | NA |
| Olfr5 | 0 | NA | NA | NA | NA | NA | | | Mir194-2 | 0 | NA | NA | NA | NA | NA |
| Olfr50 | 0 | NA | NA | NA | NA | NA | | | Mir1941 | 0 | NA | NA | NA | NA | NA |
| Olfr502 | 0 | NA | NA | NA | NA | NA | | | Mir1942 | 0 | NA | NA | NA | NA | NA |
| Olfr503 | 0 | NA | NA | NA | NA | NA | | | Mir1945 | 0 | NA | NA | NA | NA | NA |
| Olfr504 | 0 | NA | NA | NA | NA | NA | | | Mir1946a | 0 | NA | NA | NA | NA | NA |
| Olfr506 | 0 | NA | NA | NA | NA | NA | | | Mir1948 | 0 | NA | NA | NA | NA | NA |
| Olfr507 | 0 | NA | NA | NA | NA | NA | | | Mir1951 | 0 | NA | NA | NA | NA | NA |
| Olfr508 | 0 | NA | NA | NA | NA | NA | | | Mir1952 | 0 | NA | NA | NA | NA | NA |
| Olfr509 | 0 | NA | NA | NA | NA | NA | | | Mir1953 | 0 | NA | NA | NA | NA | NA |
| Olfr51 | 0 | NA | NA | NA | NA | NA | | | Mir1955 | 0 | NA | NA | NA | NA | NA |
| Olfr510 | 0 | NA | NA | NA | NA | NA | | | Mir1960 | 0 | NA | NA | NA | NA | NA |
| Olfr512 | 0 | NA | NA | NA | NA | NA | | | Mir1961 | 0 | NA | NA | NA | NA | NA |
| Olfr513 | 0 | NA | NA | NA | NA | NA | | | Mir1962 | 0 | NA | NA | NA | NA | NA |
| Olfr514 | 0 | NA | NA | NA | NA | NA | | | Mir1963 | 0 | NA | NA | NA | NA | NA |
| Olfr516 | 0 | NA | NA | NA | NA | NA | | | Mir1965 | 0 | NA | NA | NA | NA | NA |
| Olfr517 | 0 | NA | NA | NA | NA | NA | | | Mir1966 | 0 | NA | NA | NA | NA | NA |
| Olfr518 | 0 | NA | NA | NA | NA | NA | | | Mir196a-1 | 0 | NA | NA | NA | NA | NA |
| Olfr519 | 0 | NA | NA | NA | NA | NA | | | Mir196a-2 | 0 | NA | NA | NA | NA | NA |
| Olfr52 | 0 | NA | NA | NA | NA | NA | | | Mir196b | 0 | NA | NA | NA | NA | NA |
| Olfr520 | 0.185670226 | -0.001645086 | 0.031432796 | -0.052336624 | 0.958260472 | NA | | | Mir1971 | 0 | NA | NA | NA | NA | NA |
| Olfr521 | 0 | NA | NA | NA | NA | NA | | | Mir1981 | 0 | NA | NA | NA | NA | NA |
| Olfr522 | 0.096071043 | -0.020098795 | 0.027825948 | -0.722304036 | 0.470107572 | NA | | | Mir1982 | 0 | NA | NA | NA | NA | NA |
| Olfr523 | 0.215050351 | -0.027006021 | 0.031907566 | -0.846382992 | 0.397339123 | NA | | | Mir199a-2 | 0 | NA | NA | NA | NA | NA |
| Olfr524 | 0 | NA | NA | NA | NA | NA | | | Mir199b | 0 | NA | NA | NA | NA | NA |
| Olfr525 | 0 | NA | NA | NA | NA | NA | | | Mir19a | 0 | NA | NA | NA | NA | NA |
| Olfr527 | 0 | NA | NA | NA | NA | NA | | | Mir19b-1 | 0 | NA | NA | NA | NA | NA |
| Olfr53 | 0 | NA | NA | NA | NA | NA | | | Mir19b-2 | 0 | NA | NA | NA | NA | NA |
| Olfr530 | 0 | NA | NA | NA | NA | NA | | | Mir1a-1 | 0 | NA | NA | NA | NA | NA |
| Olfr531 | 0 | NA | NA | NA | NA | NA | | | Mir1a-2 | 0 | NA | NA | NA | NA | NA |
| Olfr532 | 0.079983654 | 0.016740032 | 0.027674178 | 0.604897175 | 0.54524732 | NA | | | Mir200a | 0 | NA | NA | NA | NA | NA |
| Olfr533 | 0.116696964 | -0.020477583 | 0.027954224 | -0.732539847 | 0.463839131 | NA | | | Mir200b | 0 | NA | NA | NA | NA | NA |
| Olfr535 | 0 | NA | NA | NA | NA | NA | | | Mir200c | 0 | NA | NA | NA | NA | NA |

| | | | | | | | | | | | | | | | |
|---|---|---|---|---|---|---|---|---|---|---|---|---|---|---|---|
| Olfr536 | 0.215706849 | -0.002035226 | 0.0319478 | -0.063704747 | 0.949205325 | NA | | | Mir201 | 0 | NA | NA | NA | NA | NA |
| Olfr538 | 0 | NA | NA | NA | NA | NA | | | Mir202 | 0 | NA | NA | NA | NA | NA |
| Olfr539 | 0 | NA | NA | NA | NA | NA | | | Mir203 | 0 | NA | NA | NA | NA | NA |
| Olfr54 | 0 | NA | NA | NA | NA | NA | | | Mir204 | 0 | NA | NA | NA | NA | NA |
| Olfr541 | 0 | NA | NA | NA | NA | NA | | | Mir205 | 0 | NA | NA | NA | NA | NA |
| Olfr543 | 0 | NA | NA | NA | NA | NA | | | Mir206 | 0 | NA | NA | NA | NA | NA |
| Olfr544 | 0 | NA | NA | NA | NA | NA | | | Mir207 | 0 | NA | NA | NA | NA | NA |
| Olfr545 | 0 | NA | NA | NA | NA | NA | | | Mir208a | 0 | NA | NA | NA | NA | NA |
| Olfr547 | 0 | NA | NA | NA | NA | NA | | | Mir208b | 0 | NA | NA | NA | NA | NA |
| Olfr549 | 0 | NA | NA | NA | NA | NA | | | Mir20a | 0 | NA | NA | NA | NA | NA |
| Olfr55 | 0 | NA | NA | NA | NA | NA | | | Mir20b | 0 | NA | NA | NA | NA | NA |
| Olfr550 | 2.860648009 | 0.150094874 | 0.097489485 | 1.53960065 | 0.123657727 | NA | | | Mir21 | 0 | NA | NA | NA | NA | NA |
| Olfr551 | 0 | NA | NA | NA | NA | NA | | | Mir211 | 0 | NA | NA | NA | NA | NA |
| Olfr552 | 0 | NA | NA | NA | NA | NA | | | Mir2136 | 0 | NA | NA | NA | NA | NA |
| Olfr553 | 0 | NA | NA | NA | NA | NA | | | Mir2137 | 0 | NA | NA | NA | NA | NA |
| Olfr554 | 0 | NA | NA | NA | NA | NA | | | Mir214 | 0 | NA | NA | NA | NA | NA |
| Olfr555 | 0 | NA | NA | NA | NA | NA | | | Mir215 | 0 | NA | NA | NA | NA | NA |
| Olfr556 | 0 | NA | NA | NA | NA | NA | | | Mir216a | 0 | NA | NA | NA | NA | NA |
| Olfr557 | 0 | NA | NA | NA | NA | NA | | | Mir216b | 0 | NA | NA | NA | NA | NA |
| Olfr559 | 0 | NA | NA | NA | NA | NA | | | Mir217 | 0 | NA | NA | NA | NA | NA |
| Olfr56 | 1.12784763 | -0.033726047 | 0.068604092 | -0.491604007 | 0.62299931 | NA | | | Mir219-1 | 0 | NA | NA | NA | NA | NA |
| Olfr560 | 0.669314316 | -0.102290746 | 0.060715769 | -1.684747598 | 0.092037279 | NA | | | Mir22 | 0 | NA | NA | NA | NA | NA |
| Olfr561 | 0.292412586 | 0.032260595 | 0.035568585 | 0.906996868 | 0.364408454 | NA | | | Mir222 | 0 | NA | NA | NA | NA | NA |
| Olfr564 | 0.510820111 | 0.029651871 | 0.046114588 | 0.643004131 | 0.520221419 | NA | | | Mir223 | 0 | NA | NA | NA | NA | NA |
| Olfr566 | 0 | NA | NA | NA | NA | NA | | | Mir28b | 0 | NA | NA | NA | NA | NA |
| Olfr568 | 0 | NA | NA | NA | NA | NA | | | Mir290 | 0 | NA | NA | NA | NA | NA |
| Olfr569 | 0 | NA | NA | NA | NA | NA | | | Mir291a | 0 | NA | NA | NA | NA | NA |
| Olfr57 | 0 | NA | NA | NA | NA | NA | | | Mir291b | 0 | NA | NA | NA | NA | NA |
| Olfr570 | 0.254617334 | 0.029389883 | 0.033981526 | 0.864878265 | 0.387105608 | NA | | | Mir292 | 0 | NA | NA | NA | NA | NA |
| Olfr571 | 0 | NA | NA | NA | NA | NA | | | Mir293 | 0 | NA | NA | NA | NA | NA |
| Olfr572 | 0 | NA | NA | NA | NA | NA | | | Mir294 | 0 | NA | NA | NA | NA | NA |
| Olfr574 | 0.195033442 | -0.002424765 | 0.031596358 | -0.076741901 | 0.93882887 | NA | | | Mir295 | 0 | NA | NA | NA | NA | NA |
| Olfr575 | 0 | NA | NA | NA | NA | NA | | | Mir296 | 0 | NA | NA | NA | NA | NA |
| Olfr576 | 0 | NA | NA | NA | NA | NA | | | Mir297-1 | 0 | NA | NA | NA | NA | NA |
| Olfr577 | 0 | NA | NA | NA | NA | NA | | | Mir297a-3 | 0 | NA | NA | NA | NA | NA |
| Olfr578 | 0 | NA | NA | NA | NA | NA | | | Mir297b | 0 | NA | NA | NA | NA | NA |
| Olfr58 | 0.08731684 | 0.017365056 | 0.027733412 | 0.626142072 | 0.531221759 | NA | | | Mir297c | 0 | NA | NA | NA | NA | NA |
| Olfr582 | 0 | NA | NA | NA | NA | NA | | | Mir298 | 0 | NA | NA | NA | NA | NA |
| Olfr583 | 0 | NA | NA | NA | NA | NA | | | Mir299 | 0 | NA | NA | NA | NA | NA |
| Olfr584 | 0 | NA | NA | NA | NA | NA | | | Mir29a | 0 | NA | NA | NA | NA | NA |
| Olfr585 | 0 | NA | NA | NA | NA | NA | | | Mir29b-1 | 0 | NA | NA | NA | NA | NA |
| Olfr586 | 0 | NA | NA | NA | NA | NA | | | Mir29b-2 | 0 | NA | NA | NA | NA | NA |
| Olfr589 | 0 | NA | NA | NA | NA | NA | | | Mir29c | 0 | NA | NA | NA | NA | NA |
| Olfr59 | 0 | NA | NA | NA | NA | NA | | | Mir301 | 0 | NA | NA | NA | NA | NA |
| Olfr591 | 0 | NA | NA | NA | NA | NA | | | Mir301b | 0 | NA | NA | NA | NA | NA |
| Olfr592 | 0 | NA | NA | NA | NA | NA | | | Mir302a | 0 | NA | NA | NA | NA | NA |
| Olfr593 | 0 | NA | NA | NA | NA | NA | | | Mir302b | 0 | NA | NA | NA | NA | NA |
| Olfr594 | 0 | NA | NA | NA | NA | NA | | | Mir302c | 0 | NA | NA | NA | NA | NA |
| Olfr596 | 0 | NA | NA | NA | NA | NA | | | Mir302d | 0 | NA | NA | NA | NA | NA |
| Olfr597 | 0 | NA | NA | NA | NA | NA | | | Mir3059 | 0 | NA | NA | NA | NA | NA |
| Olfr598 | 0 | NA | NA | NA | NA | NA | | | Mir3062 | 0 | NA | NA | NA | NA | NA |
| Olfr599 | 0 | NA | NA | NA | NA | NA | | | Mir3063 | 0 | NA | NA | NA | NA | NA |
| Olfr6 | 0.098353386 | -0.020212831 | 0.02786466 | -0.725393061 | 0.468210929 | NA | | | Mir3065 | 0 | NA | NA | NA | NA | NA |
| Olfr60 | 0 | NA | NA | NA | NA | NA | | | Mir3066 | 0 | NA | NA | NA | NA | NA |
| Olfr600 | 0 | NA | NA | NA | NA | NA | | | Mir3067 | 0 | NA | NA | NA | NA | NA |
| Olfr601 | 0 | NA | NA | NA | NA | NA | | | Mir3069 | 0 | NA | NA | NA | NA | NA |
| Olfr603 | 0 | NA | NA | NA | NA | NA | | | Mir3071 | 0 | NA | NA | NA | NA | NA |
| Olfr605 | 0 | NA | NA | NA | NA | NA | | | Mir3073 | 0 | NA | NA | NA | NA | NA |
| Olfr606 | 0 | NA | NA | NA | NA | NA | | | Mir3074-2 | 0 | NA | NA | NA | NA | NA |
| Olfr608 | 0 | NA | NA | NA | NA | NA | | | Mir3075 | 0 | NA | NA | NA | NA | NA |
| Olfr609 | 0.198019769 | 0.025308392 | 0.031628095 | 0.80018706 | 0.423602426 | NA | | | Mir3076 | 0 | NA | NA | NA | NA | NA |
| Olfr61 | 0.098353386 | -0.020212831 | 0.02786466 | -0.725393061 | 0.468210929 | NA | | | Mir3077 | 0 | NA | NA | NA | NA | NA |
| Olfr610 | 0.094392817 | 0.017895942 | 0.027818137 | 0.643319236 | 0.520016977 | NA | | | Mir3078 | 0 | NA | NA | NA | NA | NA |
| Olfr611 | 0 | NA | NA | NA | NA | NA | | | Mir3081 | 0 | NA | NA | NA | NA | NA |
| Olfr612 | 0 | NA | NA | NA | NA | NA | | | Mir3083 | 0 | NA | NA | NA | NA | NA |
| Olfr615 | 0 | NA | NA | NA | NA | NA | | | Mir3085 | 0 | NA | NA | NA | NA | NA |
| Olfr616 | 0 | NA | NA | NA | NA | NA | | | Mir3086 | 0 | NA | NA | NA | NA | NA |
| Olfr617 | 0 | NA | NA | NA | NA | NA | | | Mir3087 | 0 | NA | NA | NA | NA | NA |
| Olfr618 | 0 | NA | NA | NA | NA | NA | | | Mir3088 | 0 | NA | NA | NA | NA | NA |
| Olfr619 | 0 | NA | NA | NA | NA | NA | | | Mir3089 | 0 | NA | NA | NA | NA | NA |
| Olfr62 | 0 | NA | NA | NA | NA | NA | | | Mir3092 | 0 | NA | NA | NA | NA | NA |
| Olfr620 | 0 | NA | NA | NA | NA | NA | | | Mir3094 | 0 | NA | NA | NA | NA | NA |
| Olfr622 | 0.159967308 | 0.0240345 | 0.030870969 | 0.778547006 | 0.436246606 | NA | | | Mir3095 | 0 | NA | NA | NA | NA | NA |
| Olfr623 | 0 | NA | NA | NA | NA | NA | | | Mir3096 | 0 | NA | NA | NA | NA | NA |
| Olfr624 | 0 | NA | NA | NA | NA | NA | | | Mir3096b | 0 | NA | NA | NA | NA | NA |
| Olfr628 | 0 | NA | NA | NA | NA | NA | | | Mir3099 | 0 | NA | NA | NA | NA | NA |
| Olfr629 | 0 | NA | NA | NA | NA | NA | | | Mir30a | 0 | NA | NA | NA | NA | NA |
| Olfr63 | 0 | NA | NA | NA | NA | NA | | | Mir31 | 0 | NA | NA | NA | NA | NA |
| Olfr630 | 0 | NA | NA | NA | NA | NA | | | Mir3102 | 0 | NA | NA | NA | NA | NA |
| Olfr631 | 0 | NA | NA | NA | NA | NA | | | Mir3103 | 0 | NA | NA | NA | NA | NA |
| Olfr632 | 0 | NA | NA | NA | NA | NA | | | Mir3106 | 0 | NA | NA | NA | NA | NA |
| Olfr633 | 0 | NA | NA | NA | NA | NA | | | Mir3107 | 0 | NA | NA | NA | NA | NA |
| Olfr635 | 0 | NA | NA | NA | NA | NA | | | Mir3108 | 0 | NA | NA | NA | NA | NA |
| Olfr638 | 0 | NA | NA | NA | NA | NA | | | Mir3109 | 0 | NA | NA | NA | NA | NA |
| Olfr639 | 0 | NA | NA | NA | NA | NA | | | Mir3110 | 0 | NA | NA | NA | NA | NA |
| Olfr64 | 0 | NA | NA | NA | NA | NA | | | Mir322 | 0 | NA | NA | NA | NA | NA |
| Olfr640 | 0.098353386 | -0.020212831 | 0.02786466 | -0.725393061 | 0.468210929 | NA | | | Mir326 | 0 | NA | NA | NA | NA | NA |
| Olfr641 | 0 | NA | NA | NA | NA | NA | | | Mir329 | 0 | NA | NA | NA | NA | NA |
| Olfr642 | 0 | NA | NA | NA | NA | NA | | | Mir33 | 0 | NA | NA | NA | NA | NA |
| Olfr643 | 0 | NA | NA | NA | NA | NA | | | Mir344 | 0 | NA | NA | NA | NA | NA |
| Olfr644 | 0 | NA | NA | NA | NA | NA | | | Mir344c | 0 | NA | NA | NA | NA | NA |
| Olfr645 | 0 | NA | NA | NA | NA | NA | | | Mir346 | 0 | NA | NA | NA | NA | NA |
| Olfr646 | 0 | NA | NA | NA | NA | NA | | | Mir3471-1 | 0 | NA | NA | NA | NA | NA |
| Olfr648 | 0 | NA | NA | NA | NA | NA | | | Mir3473 | 0 | NA | NA | NA | NA | NA |
| Olfr649 | 0 | NA | NA | NA | NA | NA | | | Mir3474 | 0 | NA | NA | NA | NA | NA |
| Olfr65 | 0 | NA | NA | NA | NA | NA | | | Mir3475 | 0 | NA | NA | NA | NA | NA |
| Olfr651 | 0 | NA | NA | NA | NA | NA | | | Mir34b | 0 | NA | NA | NA | NA | NA |
| Olfr652 | 0 | NA | NA | NA | NA | NA | | | Mir34c | 0 | NA | NA | NA | NA | NA |
| Olfr653 | 0 | NA | NA | NA | NA | NA | | | Mir350 | 0 | NA | NA | NA | NA | NA |
| Olfr654 | 0 | NA | NA | NA | NA | NA | | | Mir351 | 0 | NA | NA | NA | NA | NA |
| Olfr655 | 0 | NA | NA | NA | NA | NA | | | Mir3572 | 0 | NA | NA | NA | NA | NA |
| Olfr656 | 0 | NA | NA | NA | NA | NA | | | Mir362 | 0 | NA | NA | NA | NA | NA |
| Olfr657 | 0 | NA | NA | NA | NA | NA | | | Mir365-2 | 0 | NA | NA | NA | NA | NA |
| Olfr658 | 2.070826888 | 0.058282488 | 0.102596741 | 0.568073485 | 0.569985071 | NA | | | Mir367 | 0 | NA | NA | NA | NA | NA |
| Olfr659 | 0 | NA | NA | NA | NA | NA | | | Mir374c | 0 | NA | NA | NA | NA | NA |
| Olfr66 | 0 | NA | NA | NA | NA | NA | | | Mir375 | 0 | NA | NA | NA | NA | NA |
| Olfr661 | 0 | NA | NA | NA | NA | NA | | | Mir376a | 0 | NA | NA | NA | NA | NA |
| Olfr663 | 0 | NA | NA | NA | NA | NA | | | Mir376b | 0 | NA | NA | NA | NA | NA |
| Olfr665 | 0 | NA | NA | NA | NA | NA | | | Mir378b | 0 | NA | NA | NA | NA | NA |
| Olfr666 | 0 | NA | NA | NA | NA | NA | | | Mir379 | 0 | NA | NA | NA | NA | NA |
| Olfr667 | 0 | NA | NA | NA | NA | NA | | | Mir380 | 0 | NA | NA | NA | NA | NA |
| Olfr668 | 0 | NA | NA | NA | NA | NA | | | Mir383 | 0 | NA | NA | NA | NA | NA |
| Olfr669 | 0 | NA | NA | NA | NA | NA | | | Mir3966 | 0 | NA | NA | NA | NA | NA |
| Olfr67 | 0 | NA | NA | NA | NA | NA | | | Mir3967 | 0 | NA | NA | NA | NA | NA |
| Olfr670 | 0 | NA | NA | NA | NA | NA | | | Mir3968 | 0 | NA | NA | NA | NA | NA |
| Olfr671 | 0 | NA | NA | NA | NA | NA | | | Mir3969 | 0 | NA | NA | NA | NA | NA |
| Olfr672 | 0 | NA | NA | NA | NA | NA | | | Mir410 | 0 | NA | NA | NA | NA | NA |
| Olfr675 | 0 | NA | NA | NA | NA | NA | | | Mir411 | 0 | NA | NA | NA | NA | NA |
| Olfr676 | 0 | NA | NA | NA | NA | NA | | | Mir423 | 0 | NA | NA | NA | NA | NA |
| Olfr677 | 0 | NA | NA | NA | NA | NA | | | Mir429 | 0 | NA | NA | NA | NA | NA |
| Olfr678 | 0 | NA | NA | NA | NA | NA | | | Mir433 | 0 | NA | NA | NA | NA | NA |
| Olfr679 | 0.094392817 | 0.017895942 | 0.027818137 | 0.643319236 | 0.520016977 | NA | | | Mir448 | 0 | NA | NA | NA | NA | NA |
| Olfr68 | 0 | NA | NA | NA | NA | NA | | | Mir449a | 0 | NA | NA | NA | NA | NA |
| Olfr681 | 0.099009884 | 0.018205726 | 0.027896865 | 0.652608315 | 0.514008823 | NA | | | Mir449b | 0 | NA | NA | NA | NA | NA |
| Olfr683 | 0 | NA | NA | NA | NA | NA | | | Mir449c | 0 | NA | NA | NA | NA | NA |
| Olfr684 | 0 | NA | NA | NA | NA | NA | | | Mir450-1 | 0 | NA | NA | NA | NA | NA |
| Olfr685 | 0 | NA | NA | NA | NA | NA | | | Mir450-2 | 0 | NA | NA | NA | NA | NA |
| Olfr686 | 0 | NA | NA | NA | NA | NA | | | Mir450b | 0 | NA | NA | NA | NA | NA |
| Olfr688 | 0 | NA | NA | NA | NA | NA | | | Mir451 | 0 | NA | NA | NA | NA | NA |
| Olfr689 | 0 | NA | NA | NA | NA | NA | | | Mir452 | 0 | NA | NA | NA | NA | NA |
| Olfr69 | 0 | NA | NA | NA | NA | NA | | | Mir455 | 0 | NA | NA | NA | NA | NA |
| Olfr690 | 1.599022041 | 0.012544975 | 0.091741038 | 0.136743326 | 0.891233682 | NA | | | Mir463 | 0 | NA | NA | NA | NA | NA |
| Olfr691 | 1.723081843 | -0.107161713 | 0.100602915 | -1.065194912 | 0.286787736 | NA | | | Mir465 | 0 | NA | NA | NA | NA | NA |
| Olfr692 | 0 | NA | NA | NA | NA | NA | | | Mir465b-1 | 0 | NA | NA | NA | NA | NA |
| Olfr693 | 0 | NA | NA | NA | NA | NA | | | Mir465b-2 | 0 | NA | NA | NA | NA | NA |
| Olfr694 | 0 | NA | NA | NA | NA | NA | | | Mir465c-1 | 0 | NA | NA | NA | NA | NA |
| Olfr695 | 0 | NA | NA | NA | NA | NA | | | Mir465c-2 | 0 | NA | NA | NA | NA | NA |
| Olfr697 | 0.695061837 | 0.009378663 | 0.060757132 | 0.154363163 | 0.877323397 | NA | | | Mir466 | 0 | NA | NA | NA | NA | NA |
| Olfr698 | 0 | NA | NA | NA | NA | NA | | | Mir4660 | 0 | NA | NA | NA | NA | NA |
| Olfr699 | 0 | NA | NA | NA | NA | NA | | | Mir466g | 0 | NA | NA | NA | NA | NA |
| Olfr70 | 0 | NA | NA | NA | NA | NA | | | Mir466i | 0 | NA | NA | NA | NA | NA |
| Olfr700 | 0.119397641 | -0.020477583 | 0.027954224 | -0.732539845 | 0.463839132 | NA | | | Mir467a-1 | 0 | NA | NA | NA | NA | NA |
| Olfr701 | 0 | NA | NA | NA | NA | NA | | | Mir467a-10 | 0 | NA | NA | NA | NA | NA |
| Olfr702 | 0 | NA | NA | NA | NA | NA | | | Mir467a-2 | 0 | NA | NA | NA | NA | NA |
| Olfr703 | 0 | NA | NA | NA | NA | NA | | | Mir467a-3 | 0 | NA | NA | NA | NA | NA |
| Olfr704 | 0 | NA | NA | NA | NA | NA | | | Mir467a-4 | 0 | NA | NA | NA | NA | NA |
| Olfr705 | 0 | NA | NA | NA | NA | NA | | | Mir467a-5 | 0 | NA | NA | NA | NA | NA |
| Olfr706 | 0 | NA | NA | NA | NA | NA | | | Mir467a-6 | 0 | NA | NA | NA | NA | NA |
| Olfr707 | 0.08731684 | 0.017365056 | 0.027733412 | 0.626142072 | 0.531221759 | NA | | | Mir467a-7 | 0 | NA | NA | NA | NA | NA |
| Olfr71 | 0 | NA | NA | NA | NA | NA | | | Mir467a-8 | 0 | NA | NA | NA | NA | NA |
| Olfr710 | 0 | NA | NA | NA | NA | NA | | | Mir467a-9 | 0 | NA | NA | NA | NA | NA |
| Olfr711 | 0 | NA | NA | NA | NA | NA | | | Mir467b | 0 | NA | NA | NA | NA | NA |
| Olfr713 | 0 | NA | NA | NA | NA | NA | | | Mir467d | 0 | NA | NA | NA | NA | NA |
| Olfr714 | 0 | NA | NA | NA | NA | NA | | | Mir467f | 0 | NA | NA | NA | NA | NA |
| Olfr715 | 0 | NA | NA | NA | NA | NA | | | Mir470 | 0 | NA | NA | NA | NA | NA |
| Olfr716 | 0 | NA | NA | NA | NA | NA | | | Mir471 | 0 | NA | NA | NA | NA | NA |
| Olfr720 | 0 | NA | NA | NA | NA | NA | | | Mir483 | 0 | NA | NA | NA | NA | NA |
| Olfr722 | 0 | NA | NA | NA | NA | NA | | | Mir488 | 0 | NA | NA | NA | NA | NA |
| Olfr723 | 0 | NA | NA | NA | NA | NA | | | Mir489 | 0 | NA | NA | NA | NA | NA |
| Olfr724 | 0 | NA | NA | NA | NA | NA | | | Mir490 | 0 | NA | NA | NA | NA | NA |
| Olfr725 | 0 | NA | NA | NA | NA | NA | | | Mir495 | 0 | NA | NA | NA | NA | NA |
| Olfr726 | 0 | NA | NA | NA | NA | NA | | | Mir497 | 0 | NA | NA | NA | NA | NA |
| Olfr727 | 0.274408084 | -0.016030385 | 0.038579263 | -0.415518171 | 0.677762616 | NA | | | Mir499 | 0 | NA | NA | NA | NA | NA |
| Olfr728 | 0 | NA | NA | NA | NA | NA | | | Mir500 | 0 | NA | NA | NA | NA | NA |
| Olfr729 | 0 | NA | NA | NA | NA | NA | | | Mir501 | 0 | NA | NA | NA | NA | NA |
| Olfr73 | 0 | NA | NA | NA | NA | NA | | | Mir503 | 0 | NA | NA | NA | NA | NA |
| Olfr730 | 0 | NA | NA | NA | NA | NA | | | Mir504 | 0 | NA | NA | NA | NA | NA |
| Olfr731 | 0 | NA | NA | NA | NA | NA | | | Mir5046 | 0 | NA | NA | NA | NA | NA |
| Olfr732 | 0 | NA | NA | NA | NA | NA | | | Mir505 | 0 | NA | NA | NA | NA | NA |
| Olfr733 | 0 | NA | NA | NA | NA | NA | | | Mir509 | 0 | NA | NA | NA | NA | NA |
| Olfr734 | 0 | NA | NA | NA | NA | NA | | | Mir5098 | 0 | NA | NA | NA | NA | NA |
| Olfr735 | 0 | NA | NA | NA | NA | NA | | | Mir5101 | 0 | NA | NA | NA | NA | NA |
| Olfr736 | 0 | NA | NA | NA | NA | NA | | | Mir5104 | 0 | NA | NA | NA | NA | NA |
| Olfr738 | 0 | NA | NA | NA | NA | NA | | | Mir5106 | 0 | NA | NA | NA | NA | NA |
| Olfr739 | 0 | NA | NA | NA | NA | NA | | | Mir5108 | 0 | NA | NA | NA | NA | NA |
| Olfr74 | 0 | NA | NA | NA | NA | NA | | | Mir5109 | 0 | NA | NA | NA | NA | NA |
| Olfr740 | 0.096071043 | -0.020098795 | 0.027825948 | -0.722304036 | 0.470107572 | NA | | | Mir511 | 0 | NA | NA | NA | NA | NA |
| Olfr741 | 0.222620122 | -0.001447027 | 0.032059667 | -0.04513545 | 0.963999345 | NA | | | Mir5113 | 0 | NA | NA | NA | NA | NA |
| Olfr742 | 0.159967308 | 0.0240345 | 0.030870969 | 0.778547006 | 0.436246606 | NA | | | Mir5114 | 0 | NA | NA | NA | NA | NA |
| Olfr743 | 0 | NA | NA | NA | NA | NA | | | Mir5116 | 0 | NA | NA | NA | NA | NA |
| Olfr744 | 0 | NA | NA | NA | NA | NA | | | Mir5117 | 0 | NA | NA | NA | NA | NA |
| Olfr745 | 0 | NA | NA | NA | NA | NA | | | Mir5121 | 0 | NA | NA | NA | NA | NA |
| Olfr746 | 0 | NA | NA | NA | NA | NA | | | Mir5122 | 0 | NA | NA | NA | NA | NA |
| Olfr747 | 0 | NA | NA | NA | NA | NA | | | Mir5123 | 0 | NA | NA | NA | NA | NA |
| Olfr748 | 0 | NA | NA | NA | NA | NA | | | Mir5126 | 0 | NA | NA | NA | NA | NA |
| Olfr749 | 0 | NA | NA | NA | NA | NA | | | Mir5127 | 0 | NA | NA | NA | NA | NA |
| Olfr75-ps1 | 0 | NA | NA | NA | NA | NA | | | Mir5129 | 0 | NA | NA | NA | NA | NA |
| Olfr750 | 0.116696964 | -0.020477583 | 0.027954224 | -0.732539847 | 0.463839131 | NA | | | Mir5130 | 0 | NA | NA | NA | NA | NA |
| Olfr76 | 0 | NA | NA | NA | NA | NA | | | Mir5131 | 0 | NA | NA | NA | NA | NA |
| Olfr761 | 0 | NA | NA | NA | NA | NA | | | Mir5132 | 0 | NA | NA | NA | NA | NA |
| Olfr763 | 0 | NA | NA | NA | NA | NA | | | Mir5134 | 0 | NA | NA | NA | NA | NA |
| Olfr765 | 0 | NA | NA | NA | NA | NA | | | Mir5135 | 0 | NA | NA | NA | NA | NA |
| Olfr767 | 0 | NA | NA | NA | NA | NA | | | Mir5136 | 0 | NA | NA | NA | NA | NA |
| Olfr768 | 0 | NA | NA | NA | NA | NA | | | Mir532 | 0 | NA | NA | NA | NA | NA |
| Olfr769 | 0 | NA | NA | NA | NA | NA | | | Mir539 | 0 | NA | NA | NA | NA | NA |
| Olfr77 | 0 | NA | NA | NA | NA | NA | | | Mir542 | 0 | NA | NA | NA | NA | NA |
| Olfr770 | 0 | NA | NA | NA | NA | NA | | | Mir543 | 0 | NA | NA | NA | NA | NA |
| Olfr771 | 0 | NA | NA | NA | NA | NA | | | Mir546 | 0 | NA | NA | NA | NA | NA |
| Olfr772 | 0 | NA | NA | NA | NA | NA | | | Mir547 | 0 | NA | NA | NA | NA | NA |
| Olfr773 | 0 | NA | NA | NA | NA | NA | | | Mir551b | 0 | NA | NA | NA | NA | NA |
| Olfr774 | 0 | NA | NA | NA | NA | NA | | | Mir574 | 0 | NA | NA | NA | NA | NA |
| Olfr775 | 0 | NA | NA | NA | NA | NA | | | Mir582 | 0 | NA | NA | NA | NA | NA |
| Olfr776 | 0 | NA | NA | NA | NA | NA | | | Mir598 | 0 | NA | NA | NA | NA | NA |
| Olfr777 | 0 | NA | NA | NA | NA | NA | | | Mir599 | 0 | NA | NA | NA | NA | NA |
| Olfr780 | 0 | NA | NA | NA | NA | NA | | | Mir615 | 0 | NA | NA | NA | NA | NA |
| Olfr781 | 0 | NA | NA | NA | NA | NA | | | Mir653 | 0 | NA | NA | NA | NA | NA |
| Olfr782 | 0 | NA | NA | NA | NA | NA | | | Mir664 | 0 | NA | NA | NA | NA | NA |
| Olfr784 | 0 | NA | NA | NA | NA | NA | | | Mir666 | 0 | NA | NA | NA | NA | NA |
| Olfr786 | 0 | NA | NA | NA | NA | NA | | | Mir669a-1 | 0 | NA | NA | NA | NA | NA |
| Olfr787 | 0 | NA | NA | NA | NA | NA | | | Mir669a-10 | 0 | NA | NA | NA | NA | NA |
| Olfr788 | 0.096071043 | -0.020098795 | 0.027825948 | -0.722304036 | 0.470107572 | NA | | | Mir669a-11 | 0 | NA | NA | NA | NA | NA |
| Olfr790 | 0 | NA | NA | NA | NA | NA | | | Mir669a-12 | 0 | NA | NA | NA | NA | NA |
| Olfr791 | 0 | NA | NA | NA | NA | NA | | | Mir669a-3 | 0 | NA | NA | NA | NA | NA |
| Olfr792 | 0 | NA | NA | NA | NA | NA | | | Mir669a-4 | 0 | NA | NA | NA | NA | NA |
| Olfr794 | 0 | NA | NA | NA | NA | NA | | | Mir669a-5 | 0 | NA | NA | NA | NA | NA |
| Olfr796 | 0.098353386 | -0.020212831 | 0.02786466 | -0.725393061 | 0.468210929 | NA | | | Mir669a-6 | 0 | NA | NA | NA | NA | NA |
| Olfr798 | 0 | NA | NA | NA | NA | NA | | | Mir669a-7 | 0 | NA | NA | NA | NA | NA |
| Olfr799 | 0 | NA | NA | NA | NA | NA | | | Mir669a-8 | 0 | NA | NA | NA | NA | NA |
| Olfr8 | 0 | NA | NA | NA | NA | NA | | | Mir669a-9 | 0 | NA | NA | NA | NA | NA |
| Olfr800 | 0 | NA | NA | NA | NA | NA | | | Mir669b | 0 | NA | NA | NA | NA | NA |
| Olfr801 | 0 | NA | NA | NA | NA | NA | | | Mir669c | 0 | NA | NA | NA | NA | NA |
| Olfr802 | 0 | NA | NA | NA | NA | NA | | | Mir669e | 0 | NA | NA | NA | NA | NA |
| Olfr803 | 0 | NA | NA | NA | NA | NA | | | Mir669g | 0 | NA | NA | NA | NA | NA |
| Olfr804 | 0 | NA | NA | NA | NA | NA | | | Mir669h | 0 | NA | NA | NA | NA | NA |
| Olfr805 | 0 | NA | NA | NA | NA | NA | | | Mir669i | 0 | NA | NA | NA | NA | NA |
| Olfr806 | 0 | NA | NA | NA | NA | NA | | | Mir669j | 0 | NA | NA | NA | NA | NA |
| Olfr807 | 0 | NA | NA | NA | NA | NA | | | Mir669k | 0 | NA | NA | NA | NA | NA |
| Olfr808 | 0 | NA | NA | NA | NA | NA | | | Mir669m-1 | 0 | NA | NA | NA | NA | NA |
| Olfr809 | 0 | NA | NA | NA | NA | NA | | | Mir669m-2 | 0 | NA | NA | NA | NA | NA |
| Olfr810 | 0 | NA | NA | NA | NA | NA | | | Mir669p-1 | 0 | NA | NA | NA | NA | NA |
| Olfr811 | 0 | NA | NA | NA | NA | NA | | | Mir669p-2 | 0 | NA | NA | NA | NA | NA |
| Olfr812 | 0 | NA | NA | NA | NA | NA | | | Mir671 | 0 | NA | NA | NA | NA | NA |
| Olfr813 | 0 | NA | NA | NA | NA | NA | | | Mir672 | 0 | NA | NA | NA | NA | NA |
| Olfr814 | 0 | NA | NA | NA | NA | NA | | | Mir673 | 0 | NA | NA | NA | NA | NA |
| Olfr815 | 0 | NA | NA | NA | NA | NA | | | Mir675 | 0 | NA | NA | NA | NA | NA |

| | | | | | | | | | | | | | |
|---|---|---|---|---|---|---|---|---|---|---|---|---|---|
| Olfr816 | 0 | NA | NA | NA | NA | NA | Mir676 | 0 | NA | NA | NA | NA | NA |
| Olfr818 | 0 | NA | NA | NA | NA | NA | Mir678 | 0 | NA | NA | NA | NA | NA |
| Olfr819 | 0 | NA | NA | NA | NA | NA | Mir679 | 0 | NA | NA | NA | NA | NA |
| Olfr820 | 0 | NA | NA | NA | NA | NA | Mir680-1 | 0 | NA | NA | NA | NA | NA |
| Olfr821 | 0 | NA | NA | NA | NA | NA | Mir680-2 | 0 | NA | NA | NA | NA | NA |
| Olfr822 | 0 | NA | NA | NA | NA | NA | Mir680-3 | 0 | NA | NA | NA | NA | NA |
| Olfr823 | 0 | NA | NA | NA | NA | NA | Mir681 | 0 | NA | NA | NA | NA | NA |
| Olfr824 | 0 | NA | NA | NA | NA | NA | Mir683-1 | 0 | NA | NA | NA | NA | NA |
| Olfr825 | 0 | NA | NA | NA | NA | NA | Mir683-2 | 0 | NA | NA | NA | NA | NA |
| Olfr826 | 0 | NA | NA | NA | NA | NA | Mir684-2 | 0 | NA | NA | NA | NA | NA |
| Olfr827 | 0 | NA | NA | NA | NA | NA | Mir686 | 0 | NA | NA | NA | NA | NA |
| Olfr828 | 0 | NA | NA | NA | NA | NA | Mir687 | 0 | NA | NA | NA | NA | NA |
| Olfr829 | 0 | NA | NA | NA | NA | NA | Mir690 | 0 | NA | NA | NA | NA | NA |
| Olfr830 | 0 | NA | NA | NA | NA | NA | Mir691 | 0 | NA | NA | NA | NA | NA |
| Olfr832 | 0 | NA | NA | NA | NA | NA | Mir692-1 | 0 | NA | NA | NA | NA | NA |
| Olfr834 | 0 | NA | NA | NA | NA | NA | Mir694 | 0 | NA | NA | NA | NA | NA |
| Olfr835 | 0 | NA | NA | NA | NA | NA | Mir695 | 0 | NA | NA | NA | NA | NA |
| Olfr836 | 0.099009884 | 0.018205726 | 0.027896865 | 0.652608315 | 0.514008823 | NA | Mir698 | 0 | NA | NA | NA | NA | NA |
| Olfr837 | 0 | NA | NA | NA | NA | NA | Mir7-2 | 0 | NA | NA | NA | NA | NA |
| Olfr843 | 0 | NA | NA | NA | NA | NA | Mir701 | 0 | NA | NA | NA | NA | NA |
| Olfr845 | 0 | NA | NA | NA | NA | NA | Mir702 | 0 | NA | NA | NA | NA | NA |
| Olfr846 | 0 | NA | NA | NA | NA | NA | Mir705 | 0 | NA | NA | NA | NA | NA |
| Olfr847 | 0 | NA | NA | NA | NA | NA | Mir706 | 0 | NA | NA | NA | NA | NA |
| Olfr849 | 0.098353386 | -0.020212831 | 0.02786466 | -0.725393061 | 0.468210929 | NA | Mir710 | 0 | NA | NA | NA | NA | NA |
| Olfr850 | 0 | NA | NA | NA | NA | NA | Mir713 | 0 | NA | NA | NA | NA | NA |
| Olfr851 | 0 | NA | NA | NA | NA | NA | Mir717 | 0 | NA | NA | NA | NA | NA |
| Olfr853 | 0 | NA | NA | NA | NA | NA | Mir718 | 0 | NA | NA | NA | NA | NA |
| Olfr854 | 0 | NA | NA | NA | NA | NA | Mir719 | 0 | NA | NA | NA | NA | NA |
| Olfr855 | 0 | NA | NA | NA | NA | NA | Mir721 | 0 | NA | NA | NA | NA | NA |
| Olfr857 | 0.299325859 | 0.043567584 | 0.041359933 | 1.053376566 | 0.29216844 | NA | Mir741 | 0 | NA | NA | NA | NA | NA |
| Olfr859 | 0.500059445 | -0.006790017 | 0.038562195 | -0.176079638 | 0.860231377 | NA | Mir742 | 0 | NA | NA | NA | NA | NA |
| Olfr860 | 0 | NA | NA | NA | NA | NA | Mir743 | 0 | NA | NA | NA | NA | NA |
| Olfr862 | 0 | NA | NA | NA | NA | NA | Mir743b | 0 | NA | NA | NA | NA | NA |
| Olfr866 | 0 | NA | NA | NA | NA | NA | Mir758 | 0 | NA | NA | NA | NA | NA |
| Olfr867 | 0 | NA | NA | NA | NA | NA | Mir759 | 0 | NA | NA | NA | NA | NA |
| Olfr868 | 0 | NA | NA | NA | NA | NA | Mir763 | 0 | NA | NA | NA | NA | NA |
| Olfr869 | 0 | NA | NA | NA | NA | NA | Mir764 | 0 | NA | NA | NA | NA | NA |
| Olfr870 | 0 | NA | NA | NA | NA | NA | Mir767 | 0 | NA | NA | NA | NA | NA |
| Olfr871 | 0 | NA | NA | NA | NA | NA | Mir770 | 0 | NA | NA | NA | NA | NA |
| Olfr872 | 0 | NA | NA | NA | NA | NA | Mir802 | 0 | NA | NA | NA | NA | NA |
| Olfr873 | 0 | NA | NA | NA | NA | NA | Mir871 | 0 | NA | NA | NA | NA | NA |
| Olfr874 | 0.099009884 | 0.018205726 | 0.027896865 | 0.652608315 | 0.514008823 | NA | Mir875 | 0 | NA | NA | NA | NA | NA |
| Olfr875 | 0 | NA | NA | NA | NA | NA | Mir876 | 0 | NA | NA | NA | NA | NA |
| Olfr876 | 0 | NA | NA | NA | NA | NA | Mir877 | 0 | NA | NA | NA | NA | NA |
| Olfr877 | 0.520888355 | -0.021443664 | 0.051299307 | -0.418010783 | 0.675939233 | NA | Mir878 | 0 | NA | NA | NA | NA | NA |
| Olfr878 | 0.831003462 | -0.009820528 | 0.06234679 | -0.157514569 | 0.874839327 | NA | Mir879 | 0 | NA | NA | NA | NA | NA |
| Olfr881 | 0.375043813 | -0.019924845 | 0.036566726 | -0.544890048 | 0.585829141 | NA | Mir880 | 0 | NA | NA | NA | NA | NA |
| Olfr883 | 0.105923158 | 0.018524693 | 0.027977518 | 0.662127842 | 0.507889294 | NA | Mir881 | 0 | NA | NA | NA | NA | NA |
| Olfr884 | 0 | NA | NA | NA | NA | NA | Mir882 | 0 | NA | NA | NA | NA | NA |
| Olfr885 | 0 | NA | NA | NA | NA | NA | Mir883a | 0 | NA | NA | NA | NA | NA |
| Olfr887 | 0 | NA | NA | NA | NA | NA | Mir883b | 0 | NA | NA | NA | NA | NA |
| Olfr888 | 0 | NA | NA | NA | NA | NA | Mir92-1 | 0 | NA | NA | NA | NA | NA |
| Olfr889 | 0 | NA | NA | NA | NA | NA | Mir92-2 | 0 | NA | NA | NA | NA | NA |
| Olfr890 | 0 | NA | NA | NA | NA | NA | Mir92b | 0 | NA | NA | NA | NA | NA |
| Olfr891 | 0 | NA | NA | NA | NA | NA | Mir96 | 0 | NA | NA | NA | NA | NA |
| Olfr893 | 0 | NA | NA | NA | NA | NA | Mir98 | 0 | NA | NA | NA | NA | NA |
| Olfr894 | 0 | NA | NA | NA | NA | NA | Mirlet7a-2 | 0 | NA | NA | NA | NA | NA |
| Olfr895 | 0 | NA | NA | NA | NA | NA | Mirlet7f-2 | 0 | NA | NA | NA | NA | NA |
| Olfr898 | 0.201994201 | 0.000227241 | 0.031727614 | 0.00716224 | 0.994285408 | NA | Mixl1 | 0 | NA | NA | NA | NA | NA |
| Olfr899 | 0 | NA | NA | NA | NA | NA | Mmp1a | 0 | NA | NA | NA | NA | NA |
| Olfr9 | 0 | NA | NA | NA | NA | NA | Mmp1b | 0 | NA | NA | NA | NA | NA |
| Olfr90 | 0 | NA | NA | NA | NA | NA | Mmp20 | 0 | NA | NA | NA | NA | NA |
| Olfr900 | 0 | NA | NA | NA | NA | NA | Mmp27 | 0 | NA | NA | NA | NA | NA |
| Olfr901 | 0.098353386 | -0.020212831 | 0.02786466 | -0.725393061 | 0.468210929 | NA | Mmp3 | 0 | NA | NA | NA | NA | NA |
| Olfr902 | 0 | NA | NA | NA | NA | NA | Mmp7 | 0 | NA | NA | NA | NA | NA |
| Olfr904 | 0 | NA | NA | NA | NA | NA | Mnx1 | 0 | NA | NA | NA | NA | NA |
| Olfr905 | 0 | NA | NA | NA | NA | NA | Moxd2 | 0 | NA | NA | NA | NA | NA |
| Olfr906 | 0 | NA | NA | NA | NA | NA | Mptx1 | 0 | NA | NA | NA | NA | NA |
| Olfr907 | 0 | NA | NA | NA | NA | NA | Mrgpra2a | 0 | NA | NA | NA | NA | NA |
| Olfr908 | 0 | NA | NA | NA | NA | NA | Mrgpra3 | 0 | NA | NA | NA | NA | NA |
| Olfr91 | 0 | NA | NA | NA | NA | NA | Mrgpra4 | 0 | NA | NA | NA | NA | NA |
| Olfr910 | 0 | NA | NA | NA | NA | NA | Mrgpra6 | 0 | NA | NA | NA | NA | NA |
| Olfr911-ps1 | 0 | NA | NA | NA | NA | NA | Mrgpra9 | 0 | NA | NA | NA | NA | NA |
| Olfr912 | 0 | NA | NA | NA | NA | NA | Mrgprb2 | 0 | NA | NA | NA | NA | NA |
| Olfr913 | 0 | NA | NA | NA | NA | NA | Mrgprb3 | 0 | NA | NA | NA | NA | NA |
| Olfr914 | 0 | NA | NA | NA | NA | NA | Mrgprb4 | 0 | NA | NA | NA | NA | NA |
| Olfr915 | 0.096071043 | -0.020098795 | 0.027825948 | -0.722304036 | 0.470107572 | NA | Mrgprb5 | 0 | NA | NA | NA | NA | NA |
| Olfr916 | 0 | NA | NA | NA | NA | NA | Mrgprb8 | 0 | NA | NA | NA | NA | NA |
| Olfr917 | 0 | NA | NA | NA | NA | NA | Mrgprd | 0 | NA | NA | NA | NA | NA |
| Olfr918 | 0 | NA | NA | NA | NA | NA | Mrgprg | 0 | NA | NA | NA | NA | NA |
| Olfr919 | 0 | NA | NA | NA | NA | NA | Mrgprx1 | 0 | NA | NA | NA | NA | NA |
| Olfr92 | 0 | NA | NA | NA | NA | NA | Mrgprx2 | 0 | NA | NA | NA | NA | NA |
| Olfr920 | 3.481590909 | 0.274191145 | 0.115275251 | 2.378577742 | 0.017379572 | NA | Mroh9 | 0 | NA | NA | NA | NA | NA |
| Olfr921 | 0.295060159 | -0.029687054 | 0.033501022 | -0.886153701 | 0.375534709 | NA | Ms4a10 | 0 | NA | NA | NA | NA | NA |
| Olfr922 | 0 | NA | NA | NA | NA | NA | Ms4a13 | 0 | NA | NA | NA | NA | NA |
| Olfr923 | 0 | NA | NA | NA | NA | NA | Ms4a18 | 0 | NA | NA | NA | NA | NA |
| Olfr924 | 0 | NA | NA | NA | NA | NA | Ms4a5 | 0 | NA | NA | NA | NA | NA |
| Olfr926 | 0 | NA | NA | NA | NA | NA | Msgn1 | 0 | NA | NA | NA | NA | NA |
| Olfr93 | 0.098353386 | -0.020212831 | 0.02786466 | -0.725393061 | 0.468210929 | NA | Msmb | 0 | NA | NA | NA | NA | NA |
| Olfr930 | 0 | NA | NA | NA | NA | NA | Mt4 | 0 | NA | NA | NA | NA | NA |
| Olfr933 | 0 | NA | NA | NA | NA | NA | Mtnr1b | 0 | NA | NA | NA | NA | NA |
| Olfr934 | 0 | NA | NA | NA | NA | NA | Mug-ps1 | 0 | NA | NA | NA | NA | NA |
| Olfr935 | 0 | NA | NA | NA | NA | NA | Mug1 | 0 | NA | NA | NA | NA | NA |
| Olfr936 | 0 | NA | NA | NA | NA | NA | Mup1 | 0 | NA | NA | NA | NA | NA |
| Olfr937 | 0 | NA | NA | NA | NA | NA | Mup10 | 0 | NA | NA | NA | NA | NA |
| Olfr938 | 0 | NA | NA | NA | NA | NA | Mup12 | 0 | NA | NA | NA | NA | NA |
| Olfr94 | 0 | NA | NA | NA | NA | NA | Mup13 | 0 | NA | NA | NA | NA | NA |
| Olfr943 | 0 | NA | NA | NA | NA | NA | Mup14 | 0 | NA | NA | NA | NA | NA |
| Olfr944 | 0 | NA | NA | NA | NA | NA | Mup15 | 0 | NA | NA | NA | NA | NA |
| Olfr945 | 0 | NA | NA | NA | NA | NA | Mup16 | 0 | NA | NA | NA | NA | NA |
| Olfr947-ps1 | 0.098353386 | -0.020212831 | 0.02786466 | -0.725393061 | 0.468210929 | NA | Mup17 | 0 | NA | NA | NA | NA | NA |
| Olfr948 | 0 | NA | NA | NA | NA | NA | Mup19 | 0 | NA | NA | NA | NA | NA |
| Olfr95 | 0.187700257 | -0.003168524 | 0.031449734 | -0.100748845 | 0.919749836 | NA | Mup2 | 0 | NA | NA | NA | NA | NA |
| Olfr951 | 0 | NA | NA | NA | NA | NA | Mup20 | 0 | NA | NA | NA | NA | NA |
| Olfr952 | 0 | NA | NA | NA | NA | NA | Mup21 | 0 | NA | NA | NA | NA | NA |
| Olfr954 | 0 | NA | NA | NA | NA | NA | Mup7 | 0 | NA | NA | NA | NA | NA |
| Olfr955 | 0 | NA | NA | NA | NA | NA | Mup8 | 0 | NA | NA | NA | NA | NA |
| Olfr957 | 0 | NA | NA | NA | NA | NA | Mup9 | 0 | NA | NA | NA | NA | NA |
| Olfr958 | 0 | NA | NA | NA | NA | NA | Myf5 | 0 | NA | NA | NA | NA | NA |
| Olfr959 | 0 | NA | NA | NA | NA | NA | Myf6 | 0 | NA | NA | NA | NA | NA |
| Olfr96 | 0 | NA | NA | NA | NA | NA | Nat3 | 0 | NA | NA | NA | NA | NA |
| Olfr960 | 0 | NA | NA | NA | NA | NA | Nctc1 | 0 | NA | NA | NA | NA | NA |
| Olfr961 | 0 | NA | NA | NA | NA | NA | Ndufs5 | 0 | NA | NA | NA | NA | NA |
| Olfr963 | 0 | NA | NA | NA | NA | NA | Nipsnap3a | 0 | NA | NA | NA | NA | NA |
| Olfr965 | 0 | NA | NA | NA | NA | NA | Nkx2-3 | 0 | NA | NA | NA | NA | NA |
| Olfr967 | 0 | NA | NA | NA | NA | NA | Nkx2-5 | 0 | NA | NA | NA | NA | NA |
| Olfr968 | 0 | NA | NA | NA | NA | NA | Nkx2-6 | 0 | NA | NA | NA | NA | NA |
| Olfr969 | 0 | NA | NA | NA | NA | NA | Nkx3-2 | 0 | NA | NA | NA | NA | NA |
| Olfr97 | 0 | NA | NA | NA | NA | NA | Nlrp14 | 0 | NA | NA | NA | NA | NA |
| Olfr970 | 0.431413697 | -0.00280547 | 0.037124934 | -0.075568363 | 0.939762507 | NA | Nlrp4a | 0 | NA | NA | NA | NA | NA |
| Olfr971 | 0 | NA | NA | NA | NA | NA | Nlrp4b | 0 | NA | NA | NA | NA | NA |
| Olfr972 | 0 | NA | NA | NA | NA | NA | Nlrp4e | 0 | NA | NA | NA | NA | NA |
| Olfr974 | 0 | NA | NA | NA | NA | NA | Nlrp9a | 0 | NA | NA | NA | NA | NA |
| Olfr975 | 0 | NA | NA | NA | NA | NA | Nlrp9c | 0 | NA | NA | NA | NA | NA |
| Olfr976 | 0 | NA | NA | NA | NA | NA | Noto | 0 | NA | NA | NA | NA | NA |
| Olfr978 | 0 | NA | NA | NA | NA | NA | Npcd | 0 | NA | NA | NA | NA | NA |
| Olfr979 | 0 | NA | NA | NA | NA | NA | Nutf2-ps1 | 0 | NA | NA | NA | NA | NA |
| Olfr98 | 0 | NA | NA | NA | NA | NA | OTTMUSG00000016609 | 0 | NA | NA | NA | NA | NA |
| Olfr980 | 0 | NA | NA | NA | NA | NA | Oas1e | 0 | NA | NA | NA | NA | NA |
| Olfr981 | 0 | NA | NA | NA | NA | NA | Oaz1 | 0 | NA | NA | NA | NA | NA |
| Olfr982 | 0 | NA | NA | NA | NA | NA | Oaz1-ps | 0 | NA | NA | NA | NA | NA |
| Olfr983 | 0.096071043 | -0.020098795 | 0.027825948 | -0.722304036 | 0.470107572 | NA | Obox1 | 0 | NA | NA | NA | NA | NA |
| Olfr984 | 0 | NA | NA | NA | NA | NA | Obox2 | 0 | NA | NA | NA | NA | NA |
| Olfr985 | 0.107716603 | -0.020477583 | 0.027954224 | -0.732539849 | 0.46383913 | NA | Obox3 | 0 | NA | NA | NA | NA | NA |
| Olfr986 | 0 | NA | NA | NA | NA | NA | Obox6 | 0 | NA | NA | NA | NA | NA |
| Olfr987 | 0 | NA | NA | NA | NA | NA | Odam | 0 | NA | NA | NA | NA | NA |
| Olfr988 | 0 | NA | NA | NA | NA | NA | Oit1 | 0 | NA | NA | NA | NA | NA |
| Olfr99 | 0 | NA | NA | NA | NA | NA | Olfr1 | 0 | NA | NA | NA | NA | NA |
| Olfr992 | 0 | NA | NA | NA | NA | NA | Olfr100 | 0 | NA | NA | NA | NA | NA |
| Olfr993 | 0 | NA | NA | NA | NA | NA | Olfr1000 | 0 | NA | NA | NA | NA | NA |
| Olfr994 | 0.116696964 | -0.020477583 | 0.027954224 | -0.732539847 | 0.463839131 | NA | Olfr1002 | 0 | NA | NA | NA | NA | NA |
| Olfr995 | 0 | NA | NA | NA | NA | NA | Olfr1006 | 0 | NA | NA | NA | NA | NA |
| Olfr996 | 0 | NA | NA | NA | NA | NA | Olfr1008 | 0 | NA | NA | NA | NA | NA |
| Olfr998 | 0 | NA | NA | NA | NA | NA | Olfr1009 | 0 | NA | NA | NA | NA | NA |
| Olig3 | 1.495177282 | 0.133257063 | 0.069937737 | 1.905367104 | 0.056732368 | NA | Olfr101 | 0 | NA | NA | NA | NA | NA |
| Olr1 | 2.329184167 | 0.193705392 | 0.109915037 | 1.762319309 | 0.078015363 | NA | Olfr1010 | 0 | NA | NA | NA | NA | NA |
| Omt2a | 0 | NA | NA | NA | NA | NA | Olfr1012 | 0 | NA | NA | NA | NA | NA |
| Omt2b | 0 | NA | NA | NA | NA | NA | Olfr1013 | 0 | NA | NA | NA | NA | NA |
| Onecut1 | 3.472786976 | 0.220068871 | 0.112243424 | 1.960639328 | 0.049921111 | NA | Olfr1014 | 0 | NA | NA | NA | NA | NA |
| Onecut3 | 2.239489832 | -0.17551623 | 0.081101217 | -2.164162712 | 0.030451864 | NA | Olfr1015 | 0 | NA | NA | NA | NA | NA |
| Ooep | 5.206067247 | 0.001695608 | 0.127415209 | 0.013307738 | 0.989382275 | NA | Olfr1016 | 0 | NA | NA | NA | NA | NA |
| Oog1 | 0 | NA | NA | NA | NA | NA | Olfr1018 | 0 | NA | NA | NA | NA | NA |
| Oog2 | 0 | NA | NA | NA | NA | NA | Olfr1019 | 0 | NA | NA | NA | NA | NA |
| Oog3 | 0 | NA | NA | NA | NA | NA | Olfr102 | 0 | NA | NA | NA | NA | NA |
| Oog4 | 0 | NA | NA | NA | NA | NA | Olfr1020 | 0 | NA | NA | NA | NA | NA |
| Oosp1 | 0.206069989 | -0.026776453 | 0.031767726 | -0.842882251 | 0.399294301 | NA | Olfr1022 | 0 | NA | NA | NA | NA | NA |
| Opn1sw | 2.057131987 | 0.017078223 | 0.104042247 | 0.164147005 | 0.869615419 | NA | Olfr1023 | 0 | NA | NA | NA | NA | NA |
| Opn4 | 1.291219282 | 0.041594807 | 0.065863348 | 0.631531924 | 0.527692782 | NA | Olfr1024 | 0 | NA | NA | NA | NA | NA |
| Opn5 | 1.030044237 | -0.128301444 | 0.074244241 | -1.728099602 | 0.083970369 | NA | Olfr1026 | 0 | NA | NA | NA | NA | NA |
| Orc1 | 4.243598391 | 0.064110754 | 0.139122119 | 0.460823586 | 0.644925179 | NA | Olfr1028 | 0 | NA | NA | NA | NA | NA |
| Orm1 | 0.08731684 | 0.017365056 | 0.027733412 | 0.626142072 | 0.531221759 | NA | Olfr103 | 0 | NA | NA | NA | NA | NA |
| Orm2 | 0 | NA | NA | NA | NA | NA | Olfr1031 | 0 | NA | NA | NA | NA | NA |
| Orm3 | 0.079983654 | 0.016740032 | 0.027674178 | 0.604897175 | 0.54524732 | NA | Olfr1032 | 0 | NA | NA | NA | NA | NA |
| Osm | 5.813465361 | 0.176672858 | 0.148990891 | 1.185796377 | 0.235702723 | NA | Olfr1036 | 0 | NA | NA | NA | NA | NA |
| Osr2 | 0.698905935 | -0.042900743 | 0.056927169 | -0.753607537 | 0.451084917 | NA | Olfr1037 | 0 | NA | NA | NA | NA | NA |
| Ostn | 0.654429243 | -0.028855515 | 0.054562931 | -0.528848329 | 0.596910669 | NA | Olfr1038-ps | 0 | NA | NA | NA | NA | NA |
| Otc | 0.820565582 | -0.063884022 | 0.064857528 | -0.98499009 | 0.324628939 | NA | Olfr1039 | 0 | NA | NA | NA | NA | NA |
| Otoa | 5.009308354 | 0.127446385 | 0.137663904 | 0.925779245 | 0.354560715 | NA | Olfr1040 | 0 | NA | NA | NA | NA | NA |
| Otog | 5.739167022 | 0.202669602 | 0.145976704 | 1.388369492 | 0.165024561 | NA | Olfr1042 | 0 | NA | NA | NA | NA | NA |
| Otol1 | 0 | NA | NA | NA | NA | NA | Olfr1043 | 0 | NA | NA | NA | NA | NA |
| Otop1 | 1.24152155 | 0.028934276 | 0.045989539 | 0.629149084 | 0.529251459 | NA | Olfr1044 | 0 | NA | NA | NA | NA | NA |
| Otop3 | 5.621215564 | -0.09303327 | 0.115766404 | -0.803629259 | 0.42161112 | NA | Olfr1045 | 0 | NA | NA | NA | NA | NA |
| Otor | 0.39538304 | 0.017028994 | 0.036795472 | 0.462801355 | 0.64350676 | NA | Olfr1046 | 0 | NA | NA | NA | NA | NA |
| Otos | 2.505579875 | -0.008737251 | 0.110786601 | -0.07886559 | 0.937139533 | NA | Olfr1047 | 0 | NA | NA | NA | NA | NA |
| Otp | 6.230244395 | 0.303533233 | 0.119398721 | 2.542181609 | 0.011016292 | NA | Olfr1048 | 0 | NA | NA | NA | NA | NA |
| Ott | 0 | NA | NA | NA | NA | NA | Olfr1049 | 0 | NA | NA | NA | NA | NA |
| Otud6a | 0 | NA | NA | NA | NA | NA | Olfr1051 | 0 | NA | NA | NA | NA | NA |
| Otx2os1 | 0.273223652 | 0.041357205 | 0.040327385 | 1.025536483 | 0.30511012 | NA | Olfr1052 | 0 | NA | NA | NA | NA | NA |
| Ovch2 | 0 | NA | NA | NA | NA | NA | Olfr1053 | 0 | NA | NA | NA | NA | NA |
| Ovol1 | 0.406270745 | 0.000396321 | 0.042421082 | 0.009342549 | 0.992545833 | NA | Olfr1054 | 0 | NA | NA | NA | NA | NA |
| Oxct2a | 0.838197859 | 0.08496648 | 0.066223674 | 1.283022739 | 0.19948411 | NA | Olfr1055 | 0 | NA | NA | NA | NA | NA |
| Oxct2b | 3.565518149 | -0.121942891 | 0.121289381 | -1.00538803 | 0.314710041 | NA | Olfr1056 | 0 | NA | NA | NA | NA | NA |
| P2rx2 | 3.147652957 | -0.067471213 | 0.126862922 | -0.53184344 | 0.594834434 | NA | Olfr1057 | 0 | NA | NA | NA | NA | NA |
| P2ry10 | 0.105923158 | 0.018524693 | 0.027977518 | 0.662127842 | 0.507889294 | NA | Olfr1058 | 0 | NA | NA | NA | NA | NA |
| P2ry4 | 0.284916696 | 0.042355674 | 0.040798016 | 1.038179738 | 0.299186384 | NA | Olfr1061 | 0 | NA | NA | NA | NA | NA |
| P4hb | 4610.350828 | -0.056078986 | 0.065364279 | -0.857945462 | NA | NA | Olfr1062 | 0 | NA | NA | NA | NA | NA |
| Pabpc2 | 0 | NA | NA | NA | NA | NA | Olfr1065 | 0 | NA | NA | NA | NA | NA |
| Pabpc6 | 0.098353386 | -0.020212831 | 0.02786466 | -0.725393061 | 0.468210929 | NA | Olfr1066 | 0 | NA | NA | NA | NA | NA |
| Pabpn1l | 0.474724059 | 0.012514508 | 0.055169993 | 0.226835404 | 0.820551733 | NA | Olfr107 | 0 | NA | NA | NA | NA | NA |
| Padi1 | 4.905138669 | -0.245018244 | 0.115233128 | -2.126283034 | 0.0334797 | NA | Olfr1076 | 0 | NA | NA | NA | NA | NA |
| Padi3 | 1.366927568 | -0.06609889 | 0.080328433 | -0.822857947 | 0.41058878 | NA | Olfr1077-ps1 | 0 | NA | NA | NA | NA | NA |
| Padi4 | 2.314169231 | -0.052098663 | 0.100979433 | -0.515933412 | 0.605900917 | NA | Olfr108 | 0 | NA | NA | NA | NA | NA |
| Pah | 3.588377053 | 0.128634487 | 0.125737589 | 1.023039238 | 0.306289294 | NA | Olfr1080 | 0 | NA | NA | NA | NA | NA |
| Panx3 | 0.094392817 | 0.017895942 | 0.027818137 | 0.643319236 | 0.520016977 | NA | Olfr1082 | 0 | NA | NA | NA | NA | NA |
| Papolb | 6.672834408 | -0.203629957 | 0.152389182 | -1.336249424 | 0.181467761 | NA | Olfr1084 | 0 | NA | NA | NA | NA | NA |
| Papss2 | 731.0706084 | -0.159608637 | 0.121798581 | -1.310431009 | NA | NA | Olfr1085 | 0 | NA | NA | NA | NA | NA |
| Pate2 | 6.935396093 | -0.033564293 | 0.152523321 | -0.220060074 | 0.825824369 | NA | Olfr1086 | 0 | NA | NA | NA | NA | NA |
| Pate4 | 0.096071043 | -0.020098795 | 0.027825948 | -0.722304036 | 0.470107572 | NA | Olfr1087 | 0 | NA | NA | NA | NA | NA |
| Patl2 | 3.437888617 | -0.10195791 | 0.129845326 | -0.785225873 | 0.432321148 | NA | Olfr1089 | 0 | NA | NA | NA | NA | NA |
| Pax1 | 2.52653858 | 0.128005298 | 0.100384222 | 1.275153561 | 0.202254895 | NA | Olfr109 | 0 | NA | NA | NA | NA | NA |
| Pax2 | 5.556533266 | 0.171593911 | 0.137104289 | 1.251557565 | 0.210731126 | NA | Olfr1090 | 0 | NA | NA | NA | NA | NA |
| Pax3 | 0.516243536 | 0.008771227 | 0.050948872 | 0.172157442 | 0.863313755 | NA | Olfr1093 | 0 | NA | NA | NA | NA | NA |
| Pax4 | 0.721075248 | 0.006675597 | 0.060994402 | 0.109446069 | 0.912848696 | NA | Olfr1094 | 0 | NA | NA | NA | NA | NA |
| Pax5 | 0.637093824 | 0.026258985 | 0.054685418 | 0.480182574 | 0.631097576 | NA | Olfr1095 | 0 | NA | NA | NA | NA | NA |
| Pax6os1 | 7.078914153 | -0.034704897 | 0.147094445 | -0.235936147 | 0.813482219 | NA | Olfr1097 | 0 | NA | NA | NA | NA | NA |
| Pax7 | 4.52595865 | 0.175388583 | 0.082573036 | 2.124041836 | 0.033666645 | NA | Olfr1098 | 0 | NA | NA | NA | NA | NA |
| Pax8 | 0.580633354 | 0.022423036 | 0.053248904 | 0.421098549 | 0.673683121 | NA | Olfr1099 | 0 | NA | NA | NA | NA | NA |
| Pax9 | 0 | NA | NA | NA | NA | NA | Olfr11 | 0 | NA | NA | NA | NA | NA |
| Pbp2 | 5.181955143 | -0.216329473 | 0.13401829 | -1.614178726 | 0.106488665 | NA | Olfr1100 | 0 | NA | NA | NA | NA | NA |
| Pbsn | 0 | NA | NA | NA | NA | NA | Olfr1101 | 0 | NA | NA | NA | NA | NA |
| Pcdha4 | 0.08731684 | 0.017365056 | 0.027733412 | 0.626142072 | 0.531221759 | NA | Olfr1102 | 0 | NA | NA | NA | NA | NA |
| Pcdha4-g | 0 | NA | NA | NA | NA | NA | Olfr1104 | 0 | NA | NA | NA | NA | NA |
| Pcdhb1 | 6.652721741 | -0.066901165 | 0.136739181 | -0.48926112 | 0.624656843 | NA | Olfr1105 | 0 | NA | NA | NA | NA | NA |
| Pck1 | 1.585148945 | -0.004545358 | 0.0723927 | -0.062787513 | 0.949935709 | NA | Olfr1106 | 0 | NA | NA | NA | NA | NA |
| Pcnt | 1554.201552 | 0.130183332 | 0.118868667 | 1.095186277 | NA | NA | Olfr1107 | 0 | NA | NA | NA | NA | NA |
| Pcp2 | 6.44973494 | -0.096827672 | 0.137231645 | -0.705578304 | 0.480450421 | NA | Olfr1109 | 0 | NA | NA | NA | NA | NA |
| Pdc | 0.298669361 | 0.012533675 | 0.03926893 | 0.319175364 | 0.749593537 | NA | Olfr1110 | 0 | NA | NA | NA | NA | NA |
| Pdcd1lg2 | 4.391468329 | 0.02915534 | 0.134584049 | 0.216632954 | 0.828494402 | NA | Olfr1111 | 0 | NA | NA | NA | NA | NA |
| Pdcl2 | 0.206069989 | -0.026776453 | 0.031767726 | -0.842882251 | 0.399294301 | NA | Olfr1112 | 0 | NA | NA | NA | NA | NA |
| Pde6a | 6.060335561 | -0.171546757 | 0.135696943 | -1.264190284 | 0.206161737 | NA | Olfr1113 | 0 | NA | NA | NA | NA | NA |
| Pde6c | 7.108978609 | 0.117915475 | 0.146447249 | 0.805173707 | 0.420719447 | NA | Olfr1115 | 0 | NA | NA | NA | NA | NA |
| Pde6g | 5.982113432 | 0.068252227 | 0.129232659 | 0.528134512 | 0.597405979 | NA | Olfr1116-ps | 0 | NA | NA | NA | NA | NA |
| Pde6h | 4.163190705 | 0.054426054 | 0.135918579 | 0.400431303 | 0.688838872 | NA | Olfr1118 | 0 | NA | NA | NA | NA | NA |
| Pdha2 | 0 | NA | NA | NA | NA | NA | Olfr112 | 0 | NA | NA | NA | NA | NA |
| Pdilt | 1.620847932 | 0.034163249 | 0.084139753 | 0.406029824 | 0.684720687 | NA | Olfr1120 | 0 | NA | NA | NA | NA | NA |
| Pdx1 | 0 | NA | NA | NA | NA | NA | Olfr1121 | 0 | NA | NA | NA | NA | NA |
| Pdxk-ps | 2.905076689 | 0.015946435 | 0.110471739 | 0.144348549 | 0.885225244 | NA | Olfr1122 | 0 | NA | NA | NA | NA | NA |
| Pdzd9 | 2.126268668 | -0.040047852 | 0.109038784 | -0.367280803 | 0.713409572 | NA | Olfr1123 | 0 | NA | NA | NA | NA | NA |
| Pea15b | 0 | NA | NA | NA | NA | NA | Olfr1124 | 0 | NA | NA | NA | NA | NA |
| Pebp4 | 3.5279941 | -0.032343569 | 0.11958746 | -0.270459539 | 0.78680674 | NA | Olfr1128 | 0 | NA | NA | NA | NA | NA |
| Peg12 | 3.499186594 | -0.204394151 | 0.13043656 | -1.567000475 | 0.11711458 | NA | Olfr1129 | 0 | NA | NA | NA | NA | NA |
| Peg3as | 6.231941421 | 0.154966317 | 0.098504051 | 1.573197401 | NA | NA | Olfr113 | 0 | NA | NA | NA | NA | NA |
| Pet117 | 0 | NA | NA | NA | NA | NA | Olfr1130 | 0 | NA | NA | NA | NA | NA |
| Pet2 | 0 | NA | NA | NA | NA | NA | Olfr1132 | 0 | NA | NA | NA | NA | NA |
| Pfn3 | 1.414062082 | -0.006063038 | 0.080297207 | -0.07550746 | 0.939810962 | NA | Olfr1133 | 0 | NA | NA | NA | NA | NA |

| | | | | | | | | | | | | | | | |
|---|---|---|---|---|---|---|---|---|---|---|---|---|---|---|---|
| Pfpl | 0.57091032 | -0.005729398 | 0.057927547 | -0.098906271 | 0.92121269 | NA | | | Olfr1134 | 0 | NA | NA | NA | NA | NA |
| Pga5 | 0.096071043 | -0.020098795 | 0.027825948 | -0.722304036 | 0.470107572 | NA | | | Olfr1135 | 0 | NA | NA | NA | NA | NA |
| Pgc | 0.784043763 | 0.094117759 | 0.070872186 | 1.327992891 | 0.184180455 | NA | | | Olfr1136 | 0 | NA | NA | NA | NA | NA |
| Pgk2 | 0 | NA | NA | NA | NA | NA | | | Olfr1137 | 0 | NA | NA | NA | NA | NA |
| Pglyrp2 | 0.310199702 | 0.010205801 | 0.034113052 | 0.299175856 | 0.76480587 | NA | | | Olfr1138 | 0 | NA | NA | NA | NA | NA |
| Pglyrp3 | 0 | NA | NA | NA | NA | NA | | | Olfr114 | 0 | NA | NA | NA | NA | NA |
| Pglyrp4 | 0 | NA | NA | NA | NA | NA | | | Olfr1141 | 0 | NA | NA | NA | NA | NA |
| Phf11a | 3.333410359 | 0.10281034 | 0.119701353 | 0.858890377 | 0.390400999 | NA | | | Olfr1143 | 0 | NA | NA | NA | NA | NA |
| Phf2 | 2545.963116 | 0.026272047 | 0.097650381 | 0.269041926 | NA | NA | | | Olfr1145 | 0 | NA | NA | NA | NA | NA |
| Phgr1 | 0 | NA | NA | NA | NA | NA | | | Olfr1148 | 0 | NA | NA | NA | NA | NA |
| Phlda2 | 0.740683258 | -0.038907871 | 0.057196921 | -0.680244161 | 0.496349875 | NA | | | Olfr115 | 0 | NA | NA | NA | NA | NA |
| Phldb3 | 4.528429979 | 0.1785205 | 0.130187157 | 1.37126045 | 0.17029378 | NA | | | Olfr1151 | 0 | NA | NA | NA | NA | NA |
| Phox2a | 0.266310378 | 0.0407635 | 0.040043874 | 1.017970943 | 0.308691765 | NA | | | Olfr1152 | 0 | NA | NA | NA | NA | NA |
| Phox2b | 0.098353386 | -0.020212831 | 0.02786466 | -0.725393061 | 0.468210929 | NA | | | Olfr1153 | 0 | NA | NA | NA | NA | NA |
| Phxr4 | 0 | NA | NA | NA | NA | NA | | | Olfr1154 | 0 | NA | NA | NA | NA | NA |
| Pigr | 3.885978194 | -0.070447289 | 0.116903789 | -0.602609111 | 0.546768758 | NA | | | Olfr1155 | 0 | NA | NA | NA | NA | NA |
| Pih1d3 | 0.196706773 | -0.026517754 | 0.031609464 | -0.838918169 | 0.401515231 | NA | | | Olfr1156 | 0 | NA | NA | NA | NA | NA |
| Pik3c2g | 7.169320049 | 0.004016622 | 0.138447562 | 0.029011863 | 0.97685513 | NA | | | Olfr1157 | 0 | NA | NA | NA | NA | NA |
| Pilrb1 | 0.932836247 | -0.05196266 | 0.070817098 | -0.733758678 | 0.463095829 | NA | | | Olfr1158 | 0 | NA | NA | NA | NA | NA |
| Pilrb2 | 2.07725609 | 0.103125854 | 0.097169719 | 1.061296197 | 0.288555313 | NA | | | Olfr116 | 0 | NA | NA | NA | NA | NA |
| Pin1-ps1 | 0 | NA | NA | NA | NA | NA | | | Olfr1160 | 0 | NA | NA | NA | NA | NA |
| Pinc | 0.40343326 | -0.026145848 | 0.043560066 | -0.600225165 | 0.548356185 | NA | | | Olfr1161 | 0 | NA | NA | NA | NA | NA |
| Pinlyp | 0.876573426 | 0.005823502 | 0.0575016 | 0.101275475 | 0.919331784 | NA | | | Olfr1162 | 0 | NA | NA | NA | NA | NA |
| Pip | 0 | NA | NA | NA | NA | NA | | | Olfr1163 | 0 | NA | NA | NA | NA | NA |
| Pira1 | 0 | NA | NA | NA | NA | NA | | | Olfr1164 | 0 | NA | NA | NA | NA | NA |
| Pira11 | 0 | NA | NA | NA | NA | NA | | | Olfr1166 | 0 | NA | NA | NA | NA | NA |
| Pira2 | 0.08731684 | 0.017365056 | 0.027733412 | 0.626142072 | 0.531221759 | NA | | | Olfr1167 | 0 | NA | NA | NA | NA | NA |
| Pira4 | 0 | NA | NA | NA | NA | NA | | | Olfr1168 | 0 | NA | NA | NA | NA | NA |
| Pira6 | 1.055108535 | 0.037618857 | 0.080056788 | 0.469902153 | 0.638424926 | NA | | | Olfr117 | 0 | NA | NA | NA | NA | NA |
| Pira7 | 0 | NA | NA | NA | NA | NA | | | Olfr1170 | 0 | NA | NA | NA | NA | NA |
| Pisd-ps1 | 0.094392817 | 0.017895942 | 0.027818137 | 0.643319236 | 0.520016977 | NA | | | Olfr1173 | 0 | NA | NA | NA | NA | NA |
| Pisd-ps3 | 0 | NA | NA | NA | NA | NA | | | Olfr1176 | 0 | NA | NA | NA | NA | NA |
| Pitx1 | 0.159967308 | 0.0240345 | 0.030870969 | 0.778547006 | 0.436246606 | NA | | | Olfr1178 | 0 | NA | NA | NA | NA | NA |
| Pitx2 | 1.539596081 | 0.097937816 | 0.069972898 | 1.399653567 | 0.161617085 | NA | | | Olfr1179 | 0 | NA | NA | NA | NA | NA |
| Pitx3 | 1.584338663 | 0.042659999 | 0.094421335 | 0.451804664 | 0.651409709 | NA | | | Olfr118 | 0 | NA | NA | NA | NA | NA |
| Piwil1 | 7.162601587 | -0.106265545 | 0.145177836 | -0.731968106 | 0.464188035 | NA | | | Olfr1180 | 0 | NA | NA | NA | NA | NA |
| Piwil4 | 0.956200006 | 0.0198166 | 0.077208804 | 0.256662444 | 0.797439366 | NA | | | Olfr1181 | 0 | NA | NA | NA | NA | NA |
| Pkhd1 | 3.419336713 | 0.122665716 | 0.117696724 | 1.042218617 | 0.297310334 | NA | | | Olfr1182 | 0 | NA | NA | NA | NA | NA |
| Pklr | 2.107931662 | 0.125979004 | 0.105215142 | 1.1973467 | 0.231171449 | NA | | | Olfr1183 | 0 | NA | NA | NA | NA | NA |
| Pkp3 | 2.513033096 | -0.038644634 | 0.113222479 | -0.341315911 | 0.732865771 | NA | | | Olfr1184 | 0 | NA | NA | NA | NA | NA |
| Pla2g10 | 0.107716603 | -0.020477583 | 0.027954224 | -0.732539849 | 0.46383913 | NA | | | Olfr1186 | 0 | NA | NA | NA | NA | NA |
| Pla2g12b | 0 | NA | NA | NA | NA | NA | | | Olfr1188 | 0 | NA | NA | NA | NA | NA |
| Pla2g1b | 0 | NA | NA | NA | NA | NA | | | Olfr1189 | 0 | NA | NA | NA | NA | NA |
| Pla2g2a | 0.105923158 | 0.018524693 | 0.027977518 | 0.662127842 | 0.507889294 | NA | | | Olfr119 | 0 | NA | NA | NA | NA | NA |
| Pla2g2c | 6.13330658 | 0.093089917 | 0.146138441 | 0.636998152 | 0.524126048 | NA | | | Olfr1193 | 0 | NA | NA | NA | NA | NA |
| Pla2g2e | 0.29109959 | -0.010630118 | 0.033578644 | -0.316573778 | 0.751567036 | NA | | | Olfr1195 | 0 | NA | NA | NA | NA | NA |
| Pla2g2f | 5.340863401 | 0.15590213 | 0.141126347 | 1.104698963 | 0.269290053 | NA | | | Olfr1196 | 0 | NA | NA | NA | NA | NA |
| Pla2g3 | 119.7263356 | 0.381232508 | 0.15684787 | 2.430587733 | NA | NA | | | Olfr1197 | 0 | NA | NA | NA | NA | NA |
| Pla2g4c | 6.523544008 | 0.100244289 | 0.146846293 | 0.682647735 | 0.494829463 | NA | | | Olfr1198 | 0 | NA | NA | NA | NA | NA |
| Pla2g4d | 2.085772536 | 0.117154982 | 0.095012777 | 1.233044501 | 0.217559162 | NA | | | Olfr1199 | 0 | NA | NA | NA | NA | NA |
| Pla2g4f | 2.029914565 | -0.058556496 | 0.094340198 | -0.620695075 | 0.534800271 | NA | | | Olfr120 | 0 | NA | NA | NA | NA | NA |
| Plac1 | 0.206726487 | -0.001342295 | 0.031808201 | -0.04219965 | 0.966339542 | NA | | | Olfr1200 | 0 | NA | NA | NA | NA | NA |
| Plac1l | 0.407192437 | -0.059678651 | 0.046941173 | -1.2713498 | 0.203604233 | NA | | | Olfr1201 | 0 | NA | NA | NA | NA | NA |
| Plac8 | 1.839029156 | 0.020541468 | 0.096883849 | 0.212021595 | 0.832090187 | NA | | | Olfr1202 | 0 | NA | NA | NA | NA | NA |
| Plac8l1 | 1.457471117 | 0.027500759 | 0.085429105 | 0.321913228 | 0.747518431 | NA | | | Olfr1204 | 0 | NA | NA | NA | NA | NA |
| Plac9a | 0 | NA | NA | NA | NA | NA | | | Olfr1205 | 0 | NA | NA | NA | NA | NA |
| Plac9b | 0 | NA | NA | NA | NA | NA | | | Olfr1206 | 0 | NA | NA | NA | NA | NA |
| Plaur | 55.7093842 | 0.254864934 | 0.161760859 | 1.575566151 | NA | NA | | | Olfr1209 | 0 | NA | NA | NA | NA | NA |
| Plbd1 | 5.892752642 | -0.185260901 | 0.144428705 | -1.282715241 | 0.199591858 | NA | | | Olfr1212 | 0 | NA | NA | NA | NA | NA |
| Pldi | 0 | NA | NA | NA | NA | NA | | | Olfr1213 | 0 | NA | NA | NA | NA | NA |
| Plg | 3.298327254 | 0.142602649 | 0.126589991 | 1.126492292 | 0.259957201 | NA | | | Olfr1214 | 0 | NA | NA | NA | NA | NA |
| Plin1 | 1.10321352 | -0.016789173 | 0.066089342 | -0.254037532 | 0.799466563 | NA | | | Olfr1215 | 0 | NA | NA | NA | NA | NA |
| Plscr5 | 1.185070533 | -0.13680812 | 0.084077097 | -1.627174639 | 0.103700013 | NA | | | Olfr1216 | 0 | NA | NA | NA | NA | NA |
| Pmp2 | 1.392519306 | -0.014294857 | 0.082638399 | -0.172980812 | 0.86266651 | NA | | | Olfr1217 | 0 | NA | NA | NA | NA | NA |
| Pnliprp1 | 0.796903708 | -0.040192877 | 0.050691198 | -0.792896579 | 0.427838081 | NA | | | Olfr1218 | 0 | NA | NA | NA | NA | NA |
| Pnliprp2 | 3.244481446 | -0.137778797 | 0.121076856 | -1.13794495 | 0.25514347 | NA | | | Olfr1219 | 0 | NA | NA | NA | NA | NA |
| Pnma5 | 5.469156186 | -0.062300479 | 0.132487665 | -0.470236068 | 0.638186368 | NA | | | Olfr122 | 0 | NA | NA | NA | NA | NA |
| Pnpla2 | 849.433174 | 0.395853393 | 0.167492267 | 2.363412946 | NA | NA | | | Olfr1220 | 0 | NA | NA | NA | NA | NA |
| Pnpla5 | 0.394700387 | 0.01999451 | 0.042974513 | 0.465264371 | 0.641742146 | NA | | | Olfr1221 | 0 | NA | NA | NA | NA | NA |
| Podnl1 | 1.495842343 | -0.0117407 | 0.074261612 | -0.158099182 | 0.874378645 | NA | | | Olfr1222 | 0 | NA | NA | NA | NA | NA |
| Pof1b | 0.384380875 | -0.006409867 | 0.041941234 | -0.152829724 | 0.878532557 | NA | | | Olfr1223 | 0 | NA | NA | NA | NA | NA |
| Pom121l12 | 0 | NA | NA | NA | NA | NA | | | Olfr1225 | 0 | NA | NA | NA | NA | NA |
| Pom121l2 | 1.548674272 | 0.114262852 | 0.082902948 | 1.378272486 | 0.168119175 | NA | | | Olfr1226 | 0 | NA | NA | NA | NA | NA |
| Pon1 | 0.949583441 | 0.083272253 | 0.063202749 | 1.31754164 | 0.187657133 | NA | | | Olfr1228 | 0 | NA | NA | NA | NA | NA |
| Poteg | 0 | NA | NA | NA | NA | NA | | | Olfr1229 | 0 | NA | NA | NA | NA | NA |
| Pou1f1 | 0.516054436 | -0.019319368 | 0.056115458 | -0.344278905 | 0.730636547 | NA | | | Olfr1230 | 0 | NA | NA | NA | NA | NA |
| Pou2af1 | 3.088322072 | -0.118224156 | 0.116071805 | -1.018543273 | 0.308419847 | NA | | | Olfr1231 | 0 | NA | NA | NA | NA | NA |
| Pou4f1 | 3.232891575 | 0.254519548 | 0.117724772 | 2.16198803 | 0.030619099 | NA | | | Olfr1232 | 0 | NA | NA | NA | NA | NA |
| Pou4f2 | 0.587618527 | 0.02809576 | 0.041193526 | 0.68204309 | 0.495211705 | NA | | | Olfr1233 | 0 | NA | NA | NA | NA | NA |
| Pou4f3 | 0.376356809 | 0.021703132 | 0.042541541 | 0.510163288 | 0.60993707 | NA | | | Olfr1234 | 0 | NA | NA | NA | NA | NA |
| Pou5f1 | 0 | NA | NA | NA | NA | NA | | | Olfr1238 | 0 | NA | NA | NA | NA | NA |
| Pou5f2 | 0 | NA | NA | NA | NA | NA | | | Olfr1239 | 0 | NA | NA | NA | NA | NA |
| Pp2d1 | 5.414729412 | -0.026397436 | 0.137055751 | -0.192603638 | 0.847269395 | NA | | | Olfr124 | 0 | NA | NA | NA | NA | NA |
| Ppbp | 1.959774784 | -0.166789816 | 0.090069715 | -1.851785756 | 0.064056594 | NA | | | Olfr1240 | 0 | NA | NA | NA | NA | NA |
| Ppifos | 0.669327356 | 0.044675377 | 0.049579597 | 0.901083916 | 0.367543704 | NA | | | Olfr1241 | 0 | NA | NA | NA | NA | NA |
| Ppm1n | 1.551335448 | 0.050106203 | 0.096567402 | 0.518872845 | 0.603849415 | NA | | | Olfr1242 | 0 | NA | NA | NA | NA | NA |
| Ppp1r14d | 0.608846751 | -0.058874571 | 0.04678185 | -1.258491744 | 0.208213971 | NA | | | Olfr1243 | 0 | NA | NA | NA | NA | NA |
| Ppp1r1c | 3.493365301 | -0.244963968 | 0.112490805 | -2.177635481 | 0.029433186 | NA | | | Olfr1245 | 0 | NA | NA | NA | NA | NA |
| Ppp1r2-ps3 | 1.174269328 | -0.001670633 | 0.060912306 | -0.027426856 | 0.978119278 | NA | | | Olfr1246 | 0 | NA | NA | NA | NA | NA |
| Ppp1r2-ps7 | 0 | NA | NA | NA | NA | NA | | | Olfr1247 | 0 | NA | NA | NA | NA | NA |
| Ppp1r2-ps9 | 0 | NA | NA | NA | NA | NA | | | Olfr1248 | 0 | NA | NA | NA | NA | NA |
| Ppp1r27 | 0 | NA | NA | NA | NA | NA | | | Olfr1249 | 0 | NA | NA | NA | NA | NA |
| Ppp1r3a | 1.744453034 | 0.119676409 | 0.090430701 | 1.323404638 | 0.18570085 | NA | | | Olfr125 | 0 | NA | NA | NA | NA | NA |
| Ppp3r2 | 5.521500261 | -0.120183725 | 0.14070536 | -0.854151718 | 0.393020931 | NA | | | Olfr1250 | 0 | NA | NA | NA | NA | NA |
| Ppy | 0.107716603 | -0.020477583 | 0.027954224 | -0.732539849 | 0.46383913 | NA | | | Olfr1251 | 0 | NA | NA | NA | NA | NA |
| Ppyr1 | 0.573420282 | 0.014545625 | 0.047116924 | 0.308713384 | 0.757539562 | NA | | | Olfr1252 | 0 | NA | NA | NA | NA | NA |
| Pqbp1 | 2003.471158 | 0.092169247 | 0.075802169 | 1.215918333 | NA | NA | | | Olfr1253 | 0 | NA | NA | NA | NA | NA |
| Prame | 0.247284148 | 0.028796741 | 0.033643376 | 0.855940884 | 0.392030479 | NA | | | Olfr1254 | 0 | NA | NA | NA | NA | NA |
| Pramef12 | 0 | NA | NA | NA | NA | NA | | | Olfr1255 | 0 | NA | NA | NA | NA | NA |
| Pramef17 | 0 | NA | NA | NA | NA | NA | | | Olfr1256 | 0 | NA | NA | NA | NA | NA |
| Pramef6 | 0 | NA | NA | NA | NA | NA | | | Olfr1257 | 0 | NA | NA | NA | NA | NA |
| Pramel1 | 1.116174192 | 0.037311252 | 0.077241891 | 0.483044253 | 0.629064315 | NA | | | Olfr1258 | 0 | NA | NA | NA | NA | NA |
| Pramel3 | 0.079983654 | 0.016740032 | 0.027674178 | 0.604897175 | 0.54524732 | NA | | | Olfr1259 | 0 | NA | NA | NA | NA | NA |
| Pramel4 | 0 | NA | NA | NA | NA | NA | | | Olfr126 | 0 | NA | NA | NA | NA | NA |
| Pramel5 | 0 | NA | NA | NA | NA | NA | | | Olfr1260 | 0 | NA | NA | NA | NA | NA |
| Pramel6 | 0 | NA | NA | NA | NA | NA | | | Olfr1261 | 0 | NA | NA | NA | NA | NA |
| Pramel7 | 0 | NA | NA | NA | NA | NA | | | Olfr1262 | 0 | NA | NA | NA | NA | NA |
| Prap1 | 0.119397641 | -0.020477583 | 0.027954224 | -0.732539845 | 0.463839132 | NA | | | Olfr1263 | 0 | NA | NA | NA | NA | NA |
| Prb1 | 0 | NA | NA | NA | NA | NA | | | Olfr1264 | 0 | NA | NA | NA | NA | NA |
| Prdm13 | 0.734759766 | 0.030213253 | 0.050271506 | 0.601001553 | 0.547838952 | NA | | | Olfr1265 | 0 | NA | NA | NA | NA | NA |
| Prdm14 | 0.445240244 | -0.001897651 | 0.037393919 | -0.050747581 | 0.959526662 | NA | | | Olfr1269 | 0 | NA | NA | NA | NA | NA |
| Prdx6b | 0 | NA | NA | NA | NA | NA | | | Olfr127 | 0 | NA | NA | NA | NA | NA |
| Prelid2 | 6.692044179 | -0.015802065 | 0.133551379 | -0.118321988 | 0.905812535 | NA | | | Olfr1270 | 0 | NA | NA | NA | NA | NA |
| Prf1 | 0.323149808 | -0.030167563 | 0.033779321 | -0.893077861 | 0.371815478 | NA | | | Olfr1271 | 0 | NA | NA | NA | NA | NA |
| Prg2 | 1.141916517 | -0.093944404 | 0.069903023 | -1.34392476 | 0.178972718 | NA | | | Olfr1272 | 0 | NA | NA | NA | NA | NA |
| Prg3 | 0.096071043 | -0.020098795 | 0.027825948 | -0.722304036 | 0.470107572 | NA | | | Olfr1273-ps | 0 | NA | NA | NA | NA | NA |
| Prh1 | 0 | NA | NA | NA | NA | NA | | | Olfr1274-ps | 0 | NA | NA | NA | NA | NA |
| Prhoxnb | 0 | NA | NA | NA | NA | NA | | | Olfr1275 | 0 | NA | NA | NA | NA | NA |
| Prkag3 | 3.219630112 | -0.001288579 | 0.108444624 | -0.011882372 | 0.990519462 | NA | | | Olfr1276 | 0 | NA | NA | NA | NA | NA |
| Prl2a1 | 0 | NA | NA | NA | NA | NA | | | Olfr1277 | 0 | NA | NA | NA | NA | NA |
| Prl2b1 | 0 | NA | NA | NA | NA | NA | | | Olfr1279 | 0 | NA | NA | NA | NA | NA |
| Prl2c1 | 0 | NA | NA | NA | NA | NA | | | Olfr128 | 0 | NA | NA | NA | NA | NA |
| Prl2c2 | 0 | NA | NA | NA | NA | NA | | | Olfr1280 | 0 | NA | NA | NA | NA | NA |
| Prl2c3 | 0 | NA | NA | NA | NA | NA | | | Olfr1282 | 0 | NA | NA | NA | NA | NA |
| Prl2c4 | 0 | NA | NA | NA | NA | NA | | | Olfr1283 | 0 | NA | NA | NA | NA | NA |
| Prl2c5 | 0 | NA | NA | NA | NA | NA | | | Olfr1284 | 0 | NA | NA | NA | NA | NA |
| Prl3a1 | 0 | NA | NA | NA | NA | NA | | | Olfr1286 | 0 | NA | NA | NA | NA | NA |
| Prl3b1 | 0 | NA | NA | NA | NA | NA | | | Olfr1287 | 0 | NA | NA | NA | NA | NA |
| Prl3c1 | 0 | NA | NA | NA | NA | NA | | | Olfr1288 | 0 | NA | NA | NA | NA | NA |
| Prl3d1 | 0 | NA | NA | NA | NA | NA | | | Olfr1289 | 0 | NA | NA | NA | NA | NA |
| Prl3d2 | 0 | NA | NA | NA | NA | NA | | | Olfr129 | 0 | NA | NA | NA | NA | NA |
| Prl3d3 | 0 | NA | NA | NA | NA | NA | | | Olfr1290 | 0 | NA | NA | NA | NA | NA |
| Prl4a1 | 0 | NA | NA | NA | NA | NA | | | Olfr1294 | 0 | NA | NA | NA | NA | NA |
| Prl5a1 | 0 | NA | NA | NA | NA | NA | | | Olfr1295 | 0 | NA | NA | NA | NA | NA |
| Prl6a1 | 0 | NA | NA | NA | NA | NA | | | Olfr1297 | 0 | NA | NA | NA | NA | NA |
| Prl7a1 | 0 | NA | NA | NA | NA | NA | | | Olfr1298 | 0 | NA | NA | NA | NA | NA |
| Prl7a2 | 0 | NA | NA | NA | NA | NA | | | Olfr1299 | 0 | NA | NA | NA | NA | NA |
| Prl7b1 | 0 | NA | NA | NA | NA | NA | | | Olfr13 | 0 | NA | NA | NA | NA | NA |
| Prl7c1 | 0 | NA | NA | NA | NA | NA | | | Olfr130 | 0 | NA | NA | NA | NA | NA |
| Prl7d1 | 0 | NA | NA | NA | NA | NA | | | Olfr1300-ps1 | 0 | NA | NA | NA | NA | NA |
| Prl8a1 | 0 | NA | NA | NA | NA | NA | | | Olfr1301 | 0 | NA | NA | NA | NA | NA |
| Prl8a2 | 0 | NA | NA | NA | NA | NA | | | Olfr1302 | 0 | NA | NA | NA | NA | NA |
| Prl8a6 | 0 | NA | NA | NA | NA | NA | | | Olfr1303 | 0 | NA | NA | NA | NA | NA |
| Prl8a8 | 0 | NA | NA | NA | NA | NA | | | Olfr1305 | 0 | NA | NA | NA | NA | NA |
| Prl8a9 | 0 | NA | NA | NA | NA | NA | | | Olfr1306 | 0 | NA | NA | NA | NA | NA |
| Prlh | 0 | NA | NA | NA | NA | NA | | | Olfr1307 | 0 | NA | NA | NA | NA | NA |
| Prm1 | 0 | NA | NA | NA | NA | NA | | | Olfr1308 | 0 | NA | NA | NA | NA | NA |
| Prm2 | 0.094392817 | 0.017895942 | 0.027818137 | 0.643319236 | 0.520016977 | NA | | | Olfr1309 | 0 | NA | NA | NA | NA | NA |
| Prm3 | 0 | NA | NA | NA | NA | NA | | | Olfr131 | 0 | NA | NA | NA | NA | NA |
| Prnd | 7.866505926 | -0.00404365 | 0.148905287 | -0.027155852 | 0.978335428 | NA | | | Olfr1310 | 0 | NA | NA | NA | NA | NA |
| Proc | 0 | NA | NA | NA | NA | NA | | | Olfr1311 | 0 | NA | NA | NA | NA | NA |
| Prodh2 | 0.415899877 | -0.004022046 | 0.042665509 | -0.094269271 | 0.924895259 | NA | | | Olfr1312 | 0 | NA | NA | NA | NA | NA |
| Prok1 | 0.614929368 | -0.038004199 | 0.059400239 | -0.639798746 | 0.522303448 | NA | | | Olfr1313 | 0 | NA | NA | NA | NA | NA |
| Prokr1 | 7.357789906 | -0.019095274 | 0.148106986 | -0.12892892 | 0.897413893 | NA | | | Olfr1314 | 0 | NA | NA | NA | NA | NA |
| Prol1 | 0 | NA | NA | NA | NA | NA | | | Olfr1316 | 0 | NA | NA | NA | NA | NA |
| Prop1 | 0.403421253 | -0.028229169 | 0.04362283 | -0.647119161 | 0.517554825 | NA | | | Olfr1317 | 0 | NA | NA | NA | NA | NA |
| Proz | 1.66627929 | 0.075725797 | 0.085845564 | 0.882116601 | 0.377713756 | NA | | | Olfr1318 | 0 | NA | NA | NA | NA | NA |
| Prp2 | 0 | NA | NA | NA | NA | NA | | | Olfr132 | 0 | NA | NA | NA | NA | NA |
| Prph2 | 1.475740698 | 0.033545409 | 0.097006331 | 0.345806383 | 0.729488228 | NA | | | Olfr1320 | 0 | NA | NA | NA | NA | NA |
| Prpmp5 | 0 | NA | NA | NA | NA | NA | | | Olfr1321 | 0 | NA | NA | NA | NA | NA |
| Prps1l1 | 0 | NA | NA | NA | NA | NA | | | Olfr1322 | 0 | NA | NA | NA | NA | NA |
| Prr15l | 0.282093074 | 0.009461846 | 0.038656194 | 0.244769212 | 0.806635134 | NA | | | Olfr1323 | 0 | NA | NA | NA | NA | NA |
| Prr19 | 4.462012358 | 0.208689708 | 0.138380097 | 1.508090485 | 0.13153137 | NA | | | Olfr1324 | 0 | NA | NA | NA | NA | NA |
| Prr23a | 0.174376471 | 0.024538273 | 0.031173126 | 0.787161136 | 0.431187544 | NA | | | Olfr1325 | 0 | NA | NA | NA | NA | NA |
| Prr9 | 0.17833704 | -0.002390466 | 0.031282281 | -0.076415987 | 0.939088151 | NA | | | Olfr1328 | 0 | NA | NA | NA | NA | NA |
| Prrxl1 | 0.651198668 | 0.010138487 | 0.065005828 | 0.155962742 | 0.876062389 | NA | | | Olfr1329 | 0 | NA | NA | NA | NA | NA |
| Prss1 | 0 | NA | NA | NA | NA | NA | | | Olfr133 | 0 | NA | NA | NA | NA | NA |
| Prss2 | 0 | NA | NA | NA | NA | NA | | | Olfr1330 | 0 | NA | NA | NA | NA | NA |
| Prss21 | 0 | NA | NA | NA | NA | NA | | | Olfr1331 | 0 | NA | NA | NA | NA | NA |
| Prss27 | 0.258977192 | 0.029727007 | 0.03417262 | 0.869907163 | 0.38435114 | NA | | | Olfr1333 | 0 | NA | NA | NA | NA | NA |
| Prss28 | 0 | NA | NA | NA | NA | NA | | | Olfr1336 | 0 | NA | NA | NA | NA | NA |
| Prss29 | 0.08731684 | 0.017365056 | 0.027733412 | 0.626142072 | 0.531221759 | NA | | | Olfr1337 | 0 | NA | NA | NA | NA | NA |
| Prss3 | 0 | NA | NA | NA | NA | NA | | | Olfr1338 | 0 | NA | NA | NA | NA | NA |
| Prss32 | 1.242096834 | 0.027552611 | 0.083794036 | 0.328813513 | 0.742296649 | NA | | | Olfr1339 | 0 | NA | NA | NA | NA | NA |
| Prss33 | 0.985455347 | 0.079724183 | 0.073074396 | 1.091000244 | 0.275272775 | NA | | | Olfr134 | 0 | NA | NA | NA | NA | NA |
| Prss34 | 0.119397641 | -0.020477583 | 0.027954224 | -0.732539845 | 0.463839132 | NA | | | Olfr1341 | 0 | NA | NA | NA | NA | NA |
| Prss37 | 0 | NA | NA | NA | NA | NA | | | Olfr1348 | 0 | NA | NA | NA | NA | NA |
| Prss38 | 1.430750791 | 0.046143255 | 0.087228565 | 0.528992478 | 0.596810668 | NA | | | Olfr1349 | 0 | NA | NA | NA | NA | NA |
| Prss39 | 0 | NA | NA | NA | NA | NA | | | Olfr135 | 0 | NA | NA | NA | NA | NA |
| Prss40 | 0 | NA | NA | NA | NA | NA | | | Olfr1351 | 0 | NA | NA | NA | NA | NA |
| Prss41 | 4.056226237 | 0.027637124 | 0.136255618 | 0.202832913 | 0.839265632 | NA | | | Olfr1352 | 0 | NA | NA | NA | NA | NA |
| Prss42 | 0.099009884 | 0.018205726 | 0.027896865 | 0.652608315 | 0.514008823 | NA | | | Olfr1353 | 0 | NA | NA | NA | NA | NA |
| Prss43 | 5.199971063 | 0.076795151 | 0.134078207 | 0.572763858 | 0.566804593 | NA | | | Olfr1354 | 0 | NA | NA | NA | NA | NA |
| Prss44 | 0.196706773 | -0.026517754 | 0.031609464 | -0.838918169 | 0.401515231 | NA | | | Olfr1355 | 0 | NA | NA | NA | NA | NA |
| Prss46 | 0.185670226 | -0.001645086 | 0.031432796 | -0.052336624 | 0.958260472 | NA | | | Olfr1356 | 0 | NA | NA | NA | NA | NA |
| Prss48 | 0 | NA | NA | NA | NA | NA | | | Olfr1357 | 0 | NA | NA | NA | NA | NA |
| Prss51 | 0 | NA | NA | NA | NA | NA | | | Olfr1359 | 0 | NA | NA | NA | NA | NA |
| Prss52 | 0.119397641 | -0.020477583 | 0.027954224 | -0.732539845 | 0.463839132 | NA | | | Olfr136 | 0 | NA | NA | NA | NA | NA |
| Prss54 | 4.806953644 | 0.385914742 | 0.142130464 | 2.715214822 | 0.006623279 | NA | | | Olfr1360 | 0 | NA | NA | NA | NA | NA |
| Prss55 | 1.471134817 | 0.035870757 | 0.08525442 | 0.420749531 | 0.673937989 | NA | | | Olfr1361 | 0 | NA | NA | NA | NA | NA |
| Prss58 | 0 | NA | NA | NA | NA | NA | | | Olfr1362 | 0 | NA | NA | NA | NA | NA |
| Prss8 | 2.008251573 | 0.049147042 | 0.101145204 | 0.485905811 | 0.627033948 | NA | | | Olfr1364 | 0 | NA | NA | NA | NA | NA |
| Psapl1 | 0.803193456 | -0.004987786 | 0.068001862 | -0.07334778 | 0.941529371 | NA | | | Olfr1366 | 0 | NA | NA | NA | NA | NA |
| Psca | 2.727767399 | 0.033442762 | 0.092677388 | 0.360851366 | 0.718210562 | NA | | | Olfr1367 | 0 | NA | NA | NA | NA | NA |
| Psg-ps1 | 0 | NA | NA | NA | NA | NA | | | Olfr1368 | 0 | NA | NA | NA | NA | NA |
| Psg17 | 1.455416154 | 0.025154363 | 0.077252925 | 0.325610491 | 0.744719073 | NA | | | Olfr137 | 0 | NA | NA | NA | NA | NA |
| Psg18 | 0 | NA | NA | NA | NA | NA | | | Olfr1370 | 0 | NA | NA | NA | NA | NA |
| Psg19 | 0 | NA | NA | NA | NA | NA | | | Olfr1373 | 0 | NA | NA | NA | NA | NA |
| Psg20 | 0 | NA | NA | NA | NA | NA | | | Olfr138 | 0 | NA | NA | NA | NA | NA |
| Psg21 | 0 | NA | NA | NA | NA | NA | | | Olfr1380 | 0 | NA | NA | NA | NA | NA |
| Psg22 | 0.094392817 | 0.017895942 | 0.027818137 | 0.643319236 | 0.520016977 | NA | | | Olfr1381 | 0 | NA | NA | NA | NA | NA |
| Psg25 | 0 | NA | NA | NA | NA | NA | | | Olfr1382 | 0 | NA | NA | NA | NA | NA |
| Psg26 | 0 | NA | NA | NA | NA | NA | | | Olfr1383 | 0 | NA | NA | NA | NA | NA |
| Psg27 | 0 | NA | NA | NA | NA | NA | | | Olfr1384 | 0 | NA | NA | NA | NA | NA |
| Psg28 | 0 | NA | NA | NA | NA | NA | | | Olfr1385 | 0 | NA | NA | NA | NA | NA |
| Psg29 | 3.212231462 | 0.057291191 | 0.125186582 | 0.457646421 | 0.647206487 | NA | | | Olfr1386 | 0 | NA | NA | NA | NA | NA |
| Psmb11 | 6.886808652 | 0.039236154 | 0.149717425 | 0.26206805 | 0.793268979 | NA | | | Olfr1387 | 0 | NA | NA | NA | NA | NA |
| Psmb2 | 1840.493634 | -0.021000219 | 0.080518345 | -0.26081285 | NA | NA | | | Olfr1388 | 0 | NA | NA | NA | NA | NA |
| Psors1c2 | 4.538298468 | 0.233022611 | 0.141324948 | 1.648842717 | 0.099179861 | NA | | | Olfr139 | 0 | NA | NA | NA | NA | NA |
| Pspn | 0.481615831 | -0.011066428 | 0.044268784 | -0.249982642 | 0.802600772 | NA | | | Olfr1390 | 0 | NA | NA | NA | NA | NA |
| Ptchd3 | 0 | NA | NA | NA | NA | NA | | | Olfr1391 | 0 | NA | NA | NA | NA | NA |
| Ptf1a | 0.381531838 | 0.016204264 | 0.036434783 | 0.444747101 | 0.656502526 | NA | | | Olfr1394 | 0 | NA | NA | NA | NA | NA |
| Ptgir | 5.488531867 | -0.101224414 | 0.138770167 | -0.729439306 | 0.465732982 | NA | | | Olfr140 | 0 | NA | NA | NA | NA | NA |
| Pth | 0.20671448 | -0.003314027 | 0.03178208 | -0.10427343 | 0.916952363 | NA | | | Olfr1402 | 0 | NA | NA | NA | NA | NA |
| Pth2 | 0.731905974 | -0.048379781 | 0.062701219 | -0.771592351 | 0.440355906 | NA | | | Olfr1404 | 0 | NA | NA | NA | NA | NA |
| Ptk6 | 2.504663572 | 0.060484835 | 0.109573857 | 0.552000604 | 0.580947939 | NA | | | Olfr1406 | 0 | NA | NA | NA | NA | NA |
| Ptpn11 | 8646.390569 | 0.016434726 | 0.100931217 | 0.162830953 | NA | NA | | | Olfr1408 | 0 | NA | NA | NA | NA | NA |
| Ptprcap | 4.998701277 | -0.029774673 | 0.145182462 | -0.205084505 | 0.837506103 | NA | | | Olfr141 | 0 | NA | NA | NA | NA | NA |
| Ptprq | 6.32643376 | -0.159113074 | 0.149571784 | -1.063790715 | 0.287423519 | NA | | | Olfr1410 | 0 | NA | NA | NA | NA | NA |
| Ptx3 | 5.661119263 | -0.018929939 | 0.127926135 | -0.147975538 | 0.882362074 | NA | | | Olfr1411 | 0 | NA | NA | NA | NA | NA |
| Ptx4 | 4.32173572 | 0.270517178 | 0.124909831 | 2.165699663 | 0.030334145 | NA | | | Olfr1412 | 0 | NA | NA | NA | NA | NA |
| Pxt1 | 1.172185394 | 0.038541959 | 0.075817592 | 0.508351143 | 0.611207109 | NA | | | Olfr1413 | 0 | NA | NA | NA | NA | NA |
| Pydc3 | 1.79767076 | 0.152674599 | 0.085799484 | 1.77943494 | 0.075168482 | NA | | | Olfr1414 | 0 | NA | NA | NA | NA | NA |
| Pydc4 | 0.92872216 | 0.017464004 | 0.065307653 | 0.267411287 | 0.78915251 | NA | | | Olfr1415 | 0 | NA | NA | NA | NA | NA |
| Pyy | 0 | NA | NA | NA | NA | NA | | | Olfr1416 | 0 | NA | NA | NA | NA | NA |
| Pzp | 0 | NA | NA | NA | NA | NA | | | Olfr1417 | 0 | NA | NA | NA | NA | NA |
| Qrfp | 0.105923158 | 0.018524693 | 0.027977518 | 0.662127842 | 0.507889294 | NA | | | Olfr1418 | 0 | NA | NA | NA | NA | NA |
| Qrich2 | 0.922304787 | 0.065687929 | 0.067719607 | 0.969998679 | 0.332047151 | NA | | | Olfr142 | 0 | NA | NA | NA | NA | NA |
| R3hdml | 0.682018854 | -0.0685167 | 0.057665156 | -1.188181988 | 0.234761718 | NA | | | Olfr1420 | 0 | NA | NA | NA | NA | NA |
| Rab17 | 1.031810426 | -0.040174341 | 0.05338871 | -0.752487574 | 0.4517579 | NA | | | Olfr1423 | 0 | NA | NA | NA | NA | NA |
| Rab19 | 2.959517648 | -0.064979996 | 0.123729994 | -0.525175778 | 0.599461002 | NA | | | Olfr1424 | 0 | NA | NA | NA | NA | NA |
| Rab25 | 1.493587948 | -3.10E-05 | 0.090496353 | -0.000342893 | 0.999726411 | NA | | | Olfr1425 | 0 | NA | NA | NA | NA | NA |
| Rab44 | 2.617581641 | 0.01035885 | 0.103891834 | 0.099708028 | 0.920576127 | NA | | | Olfr1426 | 0 | NA | NA | NA | NA | NA |
| Rad51b | 7.232306618 | 0.070867641 | 0.154712864 | 0.458059138 | 0.646909955 | NA | | | Olfr1427 | 0 | NA | NA | NA | NA | NA |
| Raet1a | 0.105923158 | 0.018524693 | 0.027977518 | 0.662127842 | 0.507889294 | NA | | | Olfr143 | 0 | NA | NA | NA | NA | NA |

| | | | | | | | | | | | | | | | |
|---|---|---|---|---|---|---|---|---|---|---|---|---|---|---|---|
| Raet1b | 0 | NA | NA | NA | NA | NA | | | Olfr1431 | 0 | NA | NA | NA | NA | NA |
| Raet1c | 0 | NA | NA | NA | NA | NA | | | Olfr1433 | 0 | NA | NA | NA | NA | NA |
| Raet1d | 0.307160824 | -0.016020939 | 0.039725656 | -0.403289483 | 0.686735278 | NA | | | Olfr1434 | 0 | NA | NA | NA | NA | NA |
| Raet1e | 0 | NA | NA | NA | NA | NA | | | Olfr1436 | 0 | NA | NA | NA | NA | NA |
| Rag2 | 0.116696964 | -0.020477583 | 0.027954224 | -0.732539847 | 0.463839131 | NA | | | Olfr1440 | 0 | NA | NA | NA | NA | NA |
| Rasef | 3.455393393 | -0.069929101 | 0.125673393 | -0.556435212 | 0.577913373 | NA | | | Olfr1441 | 0 | NA | NA | NA | NA | NA |
| Rasl2-9 | 3.856998568 | -0.089202206 | 0.122030712 | -0.730981611 | 0.464790385 | NA | | | Olfr1442 | 0 | NA | NA | NA | NA | NA |
| Raver1-fdx1l | 0 | NA | NA | NA | NA | NA | | | Olfr1443 | 0 | NA | NA | NA | NA | NA |
| Rax | 2.36282512 | 0.180793304 | 0.098567262 | 1.834212496 | 0.066622457 | NA | | | Olfr1445 | 0 | NA | NA | NA | NA | NA |
| Rbbp8nl | 0 | NA | NA | NA | NA | NA | | | Olfr1446 | 0 | NA | NA | NA | NA | NA |
| Rbm44 | 1.102006092 | -0.063204017 | 0.070650799 | -0.894597346 | 0.371002371 | NA | | | Olfr1447 | 0 | NA | NA | NA | NA | NA |
| Rbmxl2 | 0.099009884 | 0.018205726 | 0.027896865 | 0.652608315 | 0.514008823 | NA | | | Olfr1448 | 0 | NA | NA | NA | NA | NA |
| Rbmy | 0 | NA | NA | NA | NA | NA | | | Olfr1449 | 0 | NA | NA | NA | NA | NA |
| Rbp2 | 0.991301619 | -0.011024931 | 0.077182441 | -0.142842476 | 0.88641459 | NA | | | Olfr145 | 0 | NA | NA | NA | NA | NA |
| Rbp3 | 3.562431764 | 0.214066409 | 0.117492438 | 1.82195904 | 0.068461201 | NA | | | Olfr1450 | 0 | NA | NA | NA | NA | NA |
| Rbp7 | 2.043720657 | -0.027469301 | 0.098803711 | -0.278018918 | 0.780997837 | NA | | | Olfr1451 | 0 | NA | NA | NA | NA | NA |
| Rcvrn | 6.934324495 | 0.106488628 | 0.137276122 | 0.77572579 | 0.437910909 | NA | | | Olfr1453 | 0 | NA | NA | NA | NA | NA |
| Rdh1 | 4.02785589 | 0.134807819 | 0.120160741 | 1.121895709 | 0.261906786 | NA | | | Olfr1454 | 0 | NA | NA | NA | NA | NA |
| Rdh16 | 5.69937055 | 0.241813316 | 0.140816375 | 1.717224401 | 0.085938179 | NA | | | Olfr1457 | 0 | NA | NA | NA | NA | NA |
| Rdh18-ps | 6.089229591 | 0.105723181 | 0.132679412 | 0.796831845 | 0.425548698 | NA | | | Olfr1459 | 0 | NA | NA | NA | NA | NA |
| Rdh19 | 0.08731684 | 0.017365056 | 0.027733412 | 0.626142072 | 0.531221759 | NA | | | Olfr146 | 0 | NA | NA | NA | NA | NA |
| Rdh5 | 91.15875645 | 0.306568967 | 0.166926322 | 1.836552579 | NA | NA | | | Olfr1461 | 0 | NA | NA | NA | NA | NA |
| Rdh7 | 0 | NA | NA | NA | NA | NA | | | Olfr1462 | 0 | NA | NA | NA | NA | NA |
| Rdh8 | 0.283178451 | 0.028390488 | 0.033368195 | 0.850824814 | 0.394866675 | NA | | | Olfr1463 | 0 | NA | NA | NA | NA | NA |
| Reg1 | 0 | NA | NA | NA | NA | NA | | | Olfr1465 | 0 | NA | NA | NA | NA | NA |
| Reg2 | 0 | NA | NA | NA | NA | NA | | | Olfr1466 | 0 | NA | NA | NA | NA | NA |
| Reg3a | 0.357330579 | 0.020235566 | 0.041992186 | 0.481888834 | 0.629884919 | NA | | | Olfr1467 | 0 | NA | NA | NA | NA | NA |
| Reg3b | 0.79644831 | 0.019239514 | 0.068219575 | 0.282023363 | 0.777925596 | NA | | | Olfr1469 | 0 | NA | NA | NA | NA | NA |
| Reg3d | 0 | NA | NA | NA | NA | NA | | | Olfr147 | 0 | NA | NA | NA | NA | NA |
| Reg3g | 0.383007342 | 0.024584967 | 0.04803763 | 0.511785598 | 0.608801071 | NA | | | Olfr1471 | 0 | NA | NA | NA | NA | NA |
| Reg4 | 0.201994201 | 0.000227241 | 0.031727614 | 0.00716224 | 0.994285408 | NA | | | Olfr1472 | 0 | NA | NA | NA | NA | NA |
| Ren1 | 0 | NA | NA | NA | NA | NA | | | Olfr1474 | 0 | NA | NA | NA | NA | NA |
| Ren2 | 0 | NA | NA | NA | NA | NA | | | Olfr1475 | 0 | NA | NA | NA | NA | NA |
| Rergl | 1.981022479 | -0.21192552 | 0.094872729 | -2.233787539 | 0.025497056 | NA | | | Olfr1477 | 0 | NA | NA | NA | NA | NA |
| Retnla | 2.229168003 | -0.225033942 | 0.087173379 | -2.581452553 | 0.009838551 | NA | | | Olfr148 | 0 | NA | NA | NA | NA | NA |
| Retnlb | 0 | NA | NA | NA | NA | NA | | | Olfr1480 | 0 | NA | NA | NA | NA | NA |
| Retnlg | 3.754170542 | -0.033909857 | 0.118334712 | -0.28655883 | 0.774450142 | NA | | | Olfr1484 | 0 | NA | NA | NA | NA | NA |
| Rex2 | 0 | NA | NA | NA | NA | NA | | | Olfr1487 | 0 | NA | NA | NA | NA | NA |
| Rfpl3s | 0 | NA | NA | NA | NA | NA | | | Olfr1489 | 0 | NA | NA | NA | NA | NA |
| Rfpl4 | 0 | NA | NA | NA | NA | NA | | | Olfr149 | 0 | NA | NA | NA | NA | NA |
| Rfpl4b | 0.119397641 | -0.020477583 | 0.027954224 | -0.732539845 | 0.463839132 | NA | | | Olfr1490 | 0 | NA | NA | NA | NA | NA |
| Rfx6 | 0.19046386 | -0.000774927 | 0.031526133 | -0.024580468 | 0.980389599 | NA | | | Olfr1491 | 0 | NA | NA | NA | NA | NA |
| Rgr | 0.5855302 | 0.02347199 | 0.058537891 | 0.400970889 | 0.688441557 | NA | | | Olfr1494 | 0 | NA | NA | NA | NA | NA |
| Rgs1 | 6.11725672 | 0.096417133 | 0.138892139 | 0.694187115 | 0.487564872 | NA | | | Olfr1495 | 0 | NA | NA | NA | NA | NA |
| Rgs18 | 4.329224149 | -0.117003015 | 0.130286346 | -0.898045103 | 0.369161504 | NA | | | Olfr1496 | 0 | NA | NA | NA | NA | NA |
| Rhag | 0.296387018 | 0.012756677 | 0.039194327 | 0.325472542 | 0.74482346 | NA | | | Olfr1497 | 0 | NA | NA | NA | NA | NA |
| Rhbg | 0.686557386 | -0.05415787 | 0.062368571 | -0.868351935 | 0.385201699 | NA | | | Olfr1499 | 0 | NA | NA | NA | NA | NA |
| Rhd | 3.913436963 | 0.107411919 | 0.122906442 | 0.87393237 | 0.382155085 | NA | | | Olfr15 | 0 | NA | NA | NA | NA | NA |
| Rhou | 1799.611557 | 0.154249562 | 0.12017974 | 1.28349056 | NA | NA | | | Olfr1500 | 0 | NA | NA | NA | NA | NA |
| Rhox1 | 0 | NA | NA | NA | NA | NA | | | Olfr1501 | 0 | NA | NA | NA | NA | NA |
| Rhox10 | 0 | NA | NA | NA | NA | NA | | | Olfr1502 | 0 | NA | NA | NA | NA | NA |
| Rhox11 | 0 | NA | NA | NA | NA | NA | | | Olfr1504 | 0 | NA | NA | NA | NA | NA |
| Rhox12 | 0 | NA | NA | NA | NA | NA | | | Olfr1505 | 0 | NA | NA | NA | NA | NA |
| Rhox13 | 0 | NA | NA | NA | NA | NA | | | Olfr1506 | 0 | NA | NA | NA | NA | NA |
| Rhox2a | 0 | NA | NA | NA | NA | NA | | | Olfr1509 | 0 | NA | NA | NA | NA | NA |
| Rhox2b | 0 | NA | NA | NA | NA | NA | | | Olfr151 | 0 | NA | NA | NA | NA | NA |
| Rhox2c | 0 | NA | NA | NA | NA | NA | | | Olfr1510 | 0 | NA | NA | NA | NA | NA |
| Rhox2d | 0 | NA | NA | NA | NA | NA | | | Olfr1511 | 0 | NA | NA | NA | NA | NA |
| Rhox2e | 0 | NA | NA | NA | NA | NA | | | Olfr1512 | 0 | NA | NA | NA | NA | NA |
| Rhox2f | 0 | NA | NA | NA | NA | NA | | | Olfr1513 | 0 | NA | NA | NA | NA | NA |
| Rhox2g | 0 | NA | NA | NA | NA | NA | | | Olfr152 | 0 | NA | NA | NA | NA | NA |
| Rhox2h | 0 | NA | NA | NA | NA | NA | | | Olfr1532-ps1 | 0 | NA | NA | NA | NA | NA |
| Rhox3a | 0 | NA | NA | NA | NA | NA | | | Olfr1535 | 0 | NA | NA | NA | NA | NA |
| Rhox3c | 0 | NA | NA | NA | NA | NA | | | Olfr1537 | 0 | NA | NA | NA | NA | NA |
| Rhox3e | 0 | NA | NA | NA | NA | NA | | | Olfr154 | 0 | NA | NA | NA | NA | NA |
| Rhox3f | 0 | NA | NA | NA | NA | NA | | | Olfr155 | 0 | NA | NA | NA | NA | NA |
| Rhox3g-ps | 0 | NA | NA | NA | NA | NA | | | Olfr156 | 0 | NA | NA | NA | NA | NA |
| Rhox3h | 0 | NA | NA | NA | NA | NA | | | Olfr157 | 0 | NA | NA | NA | NA | NA |
| Rhox4a | 0.174376471 | 0.024538273 | 0.031173126 | 0.787161136 | 0.431187544 | NA | | | Olfr159 | 0 | NA | NA | NA | NA | NA |
| Rhox4b | 0 | NA | NA | NA | NA | NA | | | Olfr16 | 0 | NA | NA | NA | NA | NA |
| Rhox4c | 0 | NA | NA | NA | NA | NA | | | Olfr160 | 0 | NA | NA | NA | NA | NA |
| Rhox4d | 0 | NA | NA | NA | NA | NA | | | Olfr161 | 0 | NA | NA | NA | NA | NA |
| Rhox4e | 0 | NA | NA | NA | NA | NA | | | Olfr164 | 0 | NA | NA | NA | NA | NA |
| Rhox4f | 0 | NA | NA | NA | NA | NA | | | Olfr165 | 0 | NA | NA | NA | NA | NA |
| Rhox4g | 0 | NA | NA | NA | NA | NA | | | Olfr167 | 0 | NA | NA | NA | NA | NA |
| Rhox6 | 0 | NA | NA | NA | NA | NA | | | Olfr168 | 0 | NA | NA | NA | NA | NA |
| Rhox7 | 0 | NA | NA | NA | NA | NA | | | Olfr169 | 0 | NA | NA | NA | NA | NA |
| Rhox8 | 0.099009884 | 0.018205726 | 0.027896865 | 0.652608315 | 0.514008823 | NA | | | Olfr17 | 0 | NA | NA | NA | NA | NA |
| Rhox9 | 0 | NA | NA | NA | NA | NA | | | Olfr170 | 0 | NA | NA | NA | NA | NA |
| Ribc2 | 6.357543594 | 0.015626106 | 0.135185025 | 0.115590513 | 0.907977082 | NA | | | Olfr171 | 0 | NA | NA | NA | NA | NA |
| Ripk3 | 7.602369188 | 0.043477767 | 0.151801714 | 0.286411571 | 0.774562913 | NA | | | Olfr172 | 0 | NA | NA | NA | NA | NA |
| Ripply1 | 0.462846809 | -0.009373068 | 0.037958243 | -0.246931029 | 0.804961602 | NA | | | Olfr173 | 0 | NA | NA | NA | NA | NA |
| Rln3 | 0.331747315 | -0.03734372 | 0.03744515 | -0.99729124 | 0.318623164 | NA | | | Olfr175-ps1 | 0 | NA | NA | NA | NA | NA |
| Rmi2 | 7.68667395 | -0.104345867 | 0.146692338 | -0.711324593 | 0.476883115 | NA | | | Olfr176 | 0 | NA | NA | NA | NA | NA |
| Rmnd1 | 0.440108409 | -0.030798726 | 0.044545461 | -0.691399874 | 0.489314283 | NA | | | Olfr177 | 0 | NA | NA | NA | NA | NA |
| Rmrp | 3.751697272 | -0.24950485 | 0.123210895 | -2.025022636 | 0.04286504 | NA | | | Olfr178 | 0 | NA | NA | NA | NA | NA |
| Rn4.5s | 0.284023613 | -0.011266173 | 0.033405616 | -0.337253873 | 0.735925529 | NA | | | Olfr179 | 0 | NA | NA | NA | NA | NA |
| Rnase10 | 1.441550775 | -0.032974422 | 0.091699494 | -0.359592186 | 0.719152128 | NA | | | Olfr180 | 0 | NA | NA | NA | NA | NA |
| Rnase11 | 0.119397641 | -0.020477583 | 0.027954224 | -0.732539845 | 0.463839132 | NA | | | Olfr181 | 0 | NA | NA | NA | NA | NA |
| Rnase12 | 0.412784469 | -0.036563067 | 0.049225712 | -0.742763586 | 0.457624825 | NA | | | Olfr183 | 0 | NA | NA | NA | NA | NA |
| Rnase13 | 1.366069526 | 0.075899547 | 0.086602413 | 0.876413765 | 0.380805138 | NA | | | Olfr186 | 0 | NA | NA | NA | NA | NA |
| Rnase9 | 0 | NA | NA | NA | NA | NA | | | Olfr187 | 0 | NA | NA | NA | NA | NA |
| Rnaset2a | 0 | NA | NA | NA | NA | NA | | | Olfr190 | 0 | NA | NA | NA | NA | NA |
| Rnf125 | 56.35163046 | 0.178217722 | 0.168039044 | 1.060573292 | NA | NA | | | Olfr191 | 0 | NA | NA | NA | NA | NA |
| Rnf133 | 0.079983654 | 0.016740032 | 0.027674178 | 0.604897175 | 0.54524732 | NA | | | Olfr192 | 0 | NA | NA | NA | NA | NA |
| Rnf148 | 0 | NA | NA | NA | NA | NA | | | Olfr193 | 0 | NA | NA | NA | NA | NA |
| Rnf151 | 1.976521841 | 0.040251082 | 0.100397971 | 0.400915297 | 0.688482487 | NA | | | Olfr194 | 0 | NA | NA | NA | NA | NA |
| Rnf183 | 0.472604946 | 0.035675895 | 0.051617383 | 0.691160469 | 0.489464703 | NA | | | Olfr195 | 0 | NA | NA | NA | NA | NA |
| Rnf186 | 0.204013804 | -0.003115899 | 0.031740777 | -0.098167077 | 0.921799625 | NA | | | Olfr196 | 0 | NA | NA | NA | NA | NA |
| Rnf222 | 1.535178008 | 0.014924316 | 0.078164365 | 0.190935034 | 0.848576492 | NA | | | Olfr197 | 0 | NA | NA | NA | NA | NA |
| Rnu11 | 0 | NA | NA | NA | NA | NA | | | Olfr198 | 0 | NA | NA | NA | NA | NA |
| Rnu12 | 0.211846316 | 0.025707621 | 0.031860375 | 0.806883811 | 0.419733426 | NA | | | Olfr199 | 0 | NA | NA | NA | NA | NA |
| Rnu73b | 4.901059966 | 0.180714247 | 0.135132054 | 1.337315922 | 0.181119536 | NA | | | Olfr2 | 0 | NA | NA | NA | NA | NA |
| Ropn1 | 2.88413121 | -0.113528915 | 0.114095572 | -0.995033484 | 0.319719979 | NA | | | Olfr201 | 0 | NA | NA | NA | NA | NA |
| Ros1 | 0.079983654 | 0.016740032 | 0.027674178 | 0.604897175 | 0.54524732 | NA | | | Olfr202 | 0 | NA | NA | NA | NA | NA |
| Rp1 | 1.924705671 | 0.029416796 | 0.095110125 | 0.309291945 | 0.757099458 | NA | | | Olfr203 | 0 | NA | NA | NA | NA | NA |
| Rp1l1 | 0.159967308 | 0.0240345 | 0.030870969 | 0.778547006 | 0.436246606 | NA | | | Olfr204 | 0 | NA | NA | NA | NA | NA |
| Rpl10l | 0 | NA | NA | NA | NA | NA | | | Olfr205 | 0 | NA | NA | NA | NA | NA |
| Rpl26 | 0 | NA | NA | NA | NA | NA | | | Olfr206 | 0 | NA | NA | NA | NA | NA |
| Rpl31-ps12 | 2.541509973 | -0.092173001 | 0.102889947 | -0.895840684 | 0.370337857 | NA | | | Olfr207 | 0 | NA | NA | NA | NA | NA |
| Rpl34 | 7.220905668 | -0.044252212 | 0.1528405 | -0.289531975 | 0.772174314 | NA | | | Olfr208 | 0 | NA | NA | NA | NA | NA |
| Rpl34-ps1 | 0 | NA | NA | NA | NA | NA | | | Olfr209 | 0 | NA | NA | NA | NA | NA |
| Rpl39l | 1.060129594 | -0.013374988 | 0.071765407 | -0.186370961 | 0.852153863 | NA | | | Olfr211 | 0 | NA | NA | NA | NA | NA |
| Rplp2-ps1 | 0 | NA | NA | NA | NA | NA | | | Olfr213 | 0 | NA | NA | NA | NA | NA |
| Rpph1 | 6.871012262 | -0.10230378 | 0.151463495 | -0.675435222 | 0.499399294 | NA | | | Olfr215 | 0 | NA | NA | NA | NA | NA |
| Rprl1 | 0 | NA | NA | NA | NA | NA | | | Olfr218 | 0 | NA | NA | NA | NA | NA |
| Rprl2 | 0 | NA | NA | NA | NA | NA | | | Olfr220 | 0 | NA | NA | NA | NA | NA |
| Rprl3 | 0 | NA | NA | NA | NA | NA | | | Olfr222 | 0 | NA | NA | NA | NA | NA |
| Rptn | 0.745603014 | -0.017866445 | 0.055148533 | -0.323969547 | 0.74596109 | NA | | | Olfr223 | 0 | NA | NA | NA | NA | NA |
| Rrh | 5.27185439 | 0.443849627 | 0.138547587 | 3.203589726 | 0.001357257 | NA | | | Olfr224 | 0 | NA | NA | NA | NA | NA |
| Rsc1a1 | 0.08731684 | 0.017365056 | 0.027733412 | 0.626142072 | 0.531221759 | NA | | | Olfr225 | 0 | NA | NA | NA | NA | NA |
| Rsph6a | 6.251109469 | 0.08820279 | 0.136193845 | 0.647626847 | 0.517226329 | NA | | | Olfr228 | 0 | NA | NA | NA | NA | NA |
| Rspo4 | 8.088924366 | -0.116627937 | 0.154057832 | -0.757039972 | 0.449025915 | NA | | | Olfr229 | 0 | NA | NA | NA | NA | NA |
| Rtdr1 | 5.549808354 | 0.098825807 | 0.13748284 | 0.718822851 | 0.472250076 | NA | | | Olfr23 | 0 | NA | NA | NA | NA | NA |
| Rtp1 | 51.04770619 | 0.597057864 | 0.155319259 | 3.844068461 | NA | NA | | | Olfr231 | 0 | NA | NA | NA | NA | NA |
| Rtp2 | 4.043698216 | -0.114263572 | 0.123126775 | -0.928015634 | 0.35339946 | NA | | | Olfr237-ps1 | 0 | NA | NA | NA | NA | NA |
| Rubie | 1.345916069 | 0.052699266 | 0.07091647 | 0.743117448 | 0.457410577 | NA | | | Olfr239 | 0 | NA | NA | NA | NA | NA |
| Rufy4 | 2.09458199 | -0.114836471 | 0.093463254 | -1.228680432 | 0.219191646 | NA | | | Olfr24 | 0 | NA | NA | NA | NA | NA |
| Runx3 | 0.842833339 | 0.002721243 | 0.069092262 | 0.039385634 | 0.968582934 | NA | | | Olfr242 | 0 | NA | NA | NA | NA | NA |
| Rxfp4 | 0 | NA | NA | NA | NA | NA | | | Olfr243 | 0 | NA | NA | NA | NA | NA |
| S100a14 | 1.85227154 | 0.054955778 | 0.096856315 | 0.567394886 | 0.570445924 | NA | | | Olfr247 | 0 | NA | NA | NA | NA | NA |
| S100a2 | 0.099009884 | 0.018205726 | 0.027896865 | 0.652608315 | 0.514008823 | NA | | | Olfr248 | 0 | NA | NA | NA | NA | NA |
| S100g | 0.094392817 | 0.017895942 | 0.027818137 | 0.643319236 | 0.520016977 | NA | | | Olfr25 | 0 | NA | NA | NA | NA | NA |
| S100z | 0.400720577 | -0.028041581 | 0.043551246 | -0.643875506 | 0.519656166 | NA | | | Olfr250 | 0 | NA | NA | NA | NA | NA |
| Saa1 | 0.842506355 | -0.012771225 | 0.067335073 | -0.189666756 | 0.849570272 | NA | | | Olfr251 | 0 | NA | NA | NA | NA | NA |
| Saa2 | 1.548906565 | 0.122461294 | 0.087530823 | 1.399064815 | 0.161793548 | NA | | | Olfr257 | 0 | NA | NA | NA | NA | NA |
| Saa3 | 0.233393928 | -0.027475306 | 0.032191697 | -0.85349047 | 0.39338737 | NA | | | Olfr259 | 0 | NA | NA | NA | NA | NA |
| Saa4 | 0 | NA | NA | NA | NA | NA | | | Olfr26 | 0 | NA | NA | NA | NA | NA |
| Sall4 | 5.625156988 | 0.100267113 | 0.150266578 | 0.667261572 | 0.504605068 | NA | | | Olfr262 | 0 | NA | NA | NA | NA | NA |
| Samd7 | 1.823176112 | -0.07817623 | 0.097817348 | -0.799206193 | 0.424170861 | NA | | | Olfr263 | 0 | NA | NA | NA | NA | NA |
| Samt1 | 0 | NA | NA | NA | NA | NA | | | Olfr266 | 0 | NA | NA | NA | NA | NA |
| Samt2 | 0 | NA | NA | NA | NA | NA | | | Olfr27 | 0 | NA | NA | NA | NA | NA |
| Samt3 | 0 | NA | NA | NA | NA | NA | | | Olfr270 | 0 | NA | NA | NA | NA | NA |
| Samt4 | 0 | NA | NA | NA | NA | NA | | | Olfr272 | 0 | NA | NA | NA | NA | NA |
| Sap25 | 0 | NA | NA | NA | NA | NA | | | Olfr273 | 0 | NA | NA | NA | NA | NA |
| Sapcd1 | 7.798835615 | 0.12963868 | 0.140046685 | 0.925681896 | 0.354611319 | NA | | | Olfr275 | 0 | NA | NA | NA | NA | NA |
| Sat1 | 1335.670063 | 0.349017909 | 0.168019537 | 2.07724598 | NA | NA | | | Olfr282 | 0 | NA | NA | NA | NA | NA |
| Satl1 | 0 | NA | NA | NA | NA | NA | | | Olfr284 | 0 | NA | NA | NA | NA | NA |
| Sbk2 | 4.261273317 | 0.00978297 | 0.126437501 | 0.077373956 | 0.938326059 | NA | | | Olfr285 | 0 | NA | NA | NA | NA | NA |
| Sbp | 0 | NA | NA | NA | NA | NA | | | Olfr29-ps1 | 0 | NA | NA | NA | NA | NA |
| Sbpl | 0.096071043 | -0.020098795 | 0.027825948 | -0.722304036 | 0.470107572 | NA | | | Olfr290 | 0 | NA | NA | NA | NA | NA |
| Scarna10 | 0.743813171 | -0.024893294 | 0.066152898 | -0.376299372 | 0.706694346 | NA | | | Olfr291 | 0 | NA | NA | NA | NA | NA |
| Scarna13 | 3.647477527 | 0.087140605 | 0.124696778 | 0.698820024 | 0.484664513 | NA | | | Olfr292 | 0 | NA | NA | NA | NA | NA |
| Scarna17 | 2.04761523 | -0.070980183 | 0.088615341 | -0.800992045 | 0.423136251 | NA | | | Olfr293 | 0 | NA | NA | NA | NA | NA |
| Scarna2 | 0.098353386 | -0.020212831 | 0.02786466 | -0.725393061 | 0.468210929 | NA | | | Olfr294 | 0 | NA | NA | NA | NA | NA |
| Scarna3a | 0 | NA | NA | NA | NA | NA | | | Olfr295 | 0 | NA | NA | NA | NA | NA |
| Scarna3b | 0 | NA | NA | NA | NA | NA | | | Olfr297 | 0 | NA | NA | NA | NA | NA |
| Scarna6 | 2.707959894 | 0.110666663 | 0.11640052 | 0.950740281 | 0.341736235 | NA | | | Olfr298 | 0 | NA | NA | NA | NA | NA |
| Scarna8 | 0.099009884 | 0.018205726 | 0.027896865 | 0.652608315 | 0.514008823 | NA | | | Olfr299 | 0 | NA | NA | NA | NA | NA |
| Scarna9 | 0 | NA | NA | NA | NA | NA | | | Olfr30 | 0 | NA | NA | NA | NA | NA |
| Sccpdh | 2502.747246 | -0.404563707 | 0.071706625 | -5.641929248 | NA | NA | | | Olfr301 | 0 | NA | NA | NA | NA | NA |
| Scgb1a1 | 0 | NA | NA | NA | NA | NA | | | Olfr303 | 0 | NA | NA | NA | NA | NA |
| Scgb1b2 | 0 | NA | NA | NA | NA | NA | | | Olfr304 | 0 | NA | NA | NA | NA | NA |
| Scgb1b20 | 0 | NA | NA | NA | NA | NA | | | Olfr305 | 0 | NA | NA | NA | NA | NA |
| Scgb1b24 | 0 | NA | NA | NA | NA | NA | | | Olfr307 | 0 | NA | NA | NA | NA | NA |
| Scgb1b27 | 0 | NA | NA | NA | NA | NA | | | Olfr308 | 0 | NA | NA | NA | NA | NA |
| Scgb1b29 | 0 | NA | NA | NA | NA | NA | | | Olfr309 | 0 | NA | NA | NA | NA | NA |
| Scgb1b3 | 0 | NA | NA | NA | NA | NA | | | Olfr31 | 0 | NA | NA | NA | NA | NA |
| Scgb1b30 | 0 | NA | NA | NA | NA | NA | | | Olfr310 | 0 | NA | NA | NA | NA | NA |
| Scgb1b7 | 0 | NA | NA | NA | NA | NA | | | Olfr311 | 0 | NA | NA | NA | NA | NA |
| Scgb1c1 | 2.319617599 | 0.129513312 | 0.100420196 | 1.289713797 | 0.197150048 | NA | | | Olfr312 | 0 | NA | NA | NA | NA | NA |
| Scgb2b19 | 0.107716603 | -0.020477583 | 0.027954224 | -0.732539849 | 0.46383913 | NA | | | Olfr313 | 0 | NA | NA | NA | NA | NA |
| Scgb2b2 | 0 | NA | NA | NA | NA | NA | | | Olfr314 | 0 | NA | NA | NA | NA | NA |
| Scgb2b20 | 0 | NA | NA | NA | NA | NA | | | Olfr315 | 0 | NA | NA | NA | NA | NA |
| Scgb2b23-ps | 0 | NA | NA | NA | NA | NA | | | Olfr318 | 0 | NA | NA | NA | NA | NA |
| Scgb2b24 | 0 | NA | NA | NA | NA | NA | | | Olfr319 | 0 | NA | NA | NA | NA | NA |
| Scgb2b26 | 0 | NA | NA | NA | NA | NA | | | Olfr32 | 0 | NA | NA | NA | NA | NA |
| Scgb2b27 | 0 | NA | NA | NA | NA | NA | | | Olfr323 | 0 | NA | NA | NA | NA | NA |
| Scgb2b3 | 0 | NA | NA | NA | NA | NA | | | Olfr325 | 0 | NA | NA | NA | NA | NA |
| Scgb2b7 | 0 | NA | NA | NA | NA | NA | | | Olfr330 | 0 | NA | NA | NA | NA | NA |
| Scgb3a1 | 1.612864918 | 0.084380987 | 0.098480193 | 0.856832057 | 0.391537706 | NA | | | Olfr331 | 0 | NA | NA | NA | NA | NA |
| Scgb3a2 | 0.08731684 | 0.017365056 | 0.027733412 | 0.626142072 | 0.531221759 | NA | | | Olfr332 | 0 | NA | NA | NA | NA | NA |
| Scimp | 1.164911991 | 0.079176849 | 0.08200121 | 0.965557078 | 0.334265862 | NA | | | Olfr338 | 0 | NA | NA | NA | NA | NA |
| Scin | 6.758425107 | 0.05192602 | 0.137656258 | 0.377215107 | 0.706013754 | NA | | | Olfr339 | 0 | NA | NA | NA | NA | NA |
| Scn10a | 40.82910494 | 0.315661711 | 0.125490269 | 2.515427796 | NA | NA | | | Olfr340 | 0 | NA | NA | NA | NA | NA |
| Scn11a | 16.77910324 | 0.403017476 | 0.15671383 | 2.571677794 | NA | NA | | | Olfr341 | 0 | NA | NA | NA | NA | NA |
| Scn4a | 0.671056634 | 0.056541089 | 0.056494479 | 1.000825029 | 0.316911407 | NA | | | Olfr342 | 0 | NA | NA | NA | NA | NA |
| Scnn1b | 0.919709443 | -0.056949659 | 0.070846114 | -0.803850137 | 0.42148353 | NA | | | Olfr344 | 0 | NA | NA | NA | NA | NA |
| Scnn1g | 6.319470388 | 0.070038414 | 0.13893941 | 0.504093217 | 0.614195873 | NA | | | Olfr345 | 0 | NA | NA | NA | NA | NA |
| Scp2d1 | 0 | NA | NA | NA | NA | NA | | | Olfr346 | 0 | NA | NA | NA | NA | NA |
| Sct | 1.116057616 | -0.017872692 | 0.060353447 | -0.296133734 | 0.767127953 | NA | | | Olfr347 | 0 | NA | NA | NA | NA | NA |
| Sdhc | 2822.274744 | -0.132054187 | 0.065111067 | -2.02813736 | NA | NA | | | Olfr348 | 0 | NA | NA | NA | NA | NA |
| Sdr16c5 | 0 | NA | NA | NA | NA | NA | | | Olfr350 | 0 | NA | NA | NA | NA | NA |
| Sdr16c6 | 0 | NA | NA | NA | NA | NA | | | Olfr351 | 0 | NA | NA | NA | NA | NA |
| Sdr9c7 | 6.742021621 | 0.010525215 | 0.141459066 | 0.074404668 | 0.940688395 | NA | | | Olfr352 | 0 | NA | NA | NA | NA | NA |
| Sds | 4.559517294 | 0.065796938 | 0.123047332 | 0.534728688 | 0.592837481 | NA | | | Olfr353 | 0 | NA | NA | NA | NA | NA |
| Sectm1a | 0.096071043 | -0.020098795 | 0.027825948 | -0.722304036 | 0.470107572 | NA | | | Olfr354 | 0 | NA | NA | NA | NA | NA |
| Sectm1b | 0 | NA | NA | NA | NA | NA | | | Olfr355 | 0 | NA | NA | NA | NA | NA |
| Sel1l2 | 0.098353386 | -0.020212831 | 0.02786466 | -0.725393061 | 0.468210929 | NA | | | Olfr356 | 0 | NA | NA | NA | NA | NA |
| Sele | 1.286676614 | -0.024345638 | 0.085698685 | -0.284084155 | 0.776345889 | NA | | | Olfr357 | 0 | NA | NA | NA | NA | NA |
| Selenbp2 | 0.789636402 | -0.004616457 | 0.062054785 | -0.074393247 | 0.940697482 | NA | | | Olfr358 | 0 | NA | NA | NA | NA | NA |
| Sell | 2.355369266 | -0.081621642 | 0.101007089 | -0.808078355 | 0.419045477 | NA | | | Olfr360 | 0 | NA | NA | NA | NA | NA |
| Selp | 0.626781319 | -0.049557557 | 0.050238808 | -0.986439744 | 0.32391737 | NA | | | Olfr361 | 0 | NA | NA | NA | NA | NA |
| 12-Sep | 0 | NA | NA | NA | NA | NA | | | Olfr362 | 0 | NA | NA | NA | NA | NA |
| 14-Sep | 0.97104811 | -0.031165945 | 0.065495028 | -0.475852076 | 0.634179766 | NA | | | Olfr365 | 0 | NA | NA | NA | NA | NA |
| Serpina10 | 0.650108395 | -0.016118184 | 0.054231577 | -0.297210319 | 0.766305944 | NA | | | Olfr366 | 0 | NA | NA | NA | NA | NA |
| Serpina11 | 0.096071043 | -0.020098795 | 0.027825948 | -0.722304036 | 0.470107572 | NA | | | Olfr367-ps | 0 | NA | NA | NA | NA | NA |
| Serpina12 | 0.606904295 | -0.042153114 | 0.04878701 | -0.864023309 | 0.387575086 | NA | | | Olfr368 | 0 | NA | NA | NA | NA | NA |
| Serpina1a | 0.119397641 | -0.020477583 | 0.027954224 | -0.732539845 | 0.463839132 | NA | | | Olfr370 | 0 | NA | NA | NA | NA | NA |
| Serpina1b | 0.676114142 | -0.009201815 | 0.047624202 | -0.193217197 | 0.846788871 | NA | | | Olfr371 | 0 | NA | NA | NA | NA | NA |
| Serpina1c | 0.098353386 | -0.020212831 | 0.02786466 | -0.725393061 | 0.468210929 | NA | | | Olfr372 | 0 | NA | NA | NA | NA | NA |
| Serpina1d | 0.38092772 | -0.000766687 | 0.035977 | -0.021310467 | 0.982997994 | NA | | | Olfr373 | 0 | NA | NA | NA | NA | NA |
| Serpina1e | 0.08731684 | 0.017365056 | 0.027733412 | 0.626142072 | 0.531221759 | NA | | | Olfr374 | 0 | NA | NA | NA | NA | NA |
| Serpina1f | 0 | NA | NA | NA | NA | NA | | | Olfr376 | 0 | NA | NA | NA | NA | NA |
| Serpina3a | 0 | NA | NA | NA | NA | NA | | | Olfr378 | 0 | NA | NA | NA | NA | NA |
| Serpina3b | 0.413840517 | 0.018406376 | 0.037259486 | 0.494005092 | 0.621302583 | NA | | | Olfr38 | 0 | NA | NA | NA | NA | NA |
| Serpina3c | 0.986744043 | -0.004546526 | 0.066284846 | -0.068590737 | 0.945315393 | NA | | | Olfr380 | 0 | NA | NA | NA | NA | NA |
| Serpina3f | 4.307982241 | 0.119560657 | 0.090160649 | 1.326084693 | NA | NA | | | Olfr381 | 0 | NA | NA | NA | NA | NA |
| Serpina3h | 1.700969836 | -0.033251273 | 0.091200833 | -0.364593967 | 0.715414519 | NA | | | Olfr382 | 0 | NA | NA | NA | NA | NA |
| Serpina3i | 1.437030694 | 0.085119104 | 0.08647326 | 0.984340174 | 0.324948283 | NA | | | Olfr384 | 0 | NA | NA | NA | NA | NA |
| Serpina3j | 0 | NA | NA | NA | NA | NA | | | Olfr385 | 0 | NA | NA | NA | NA | NA |
| Serpina3k | 0 | NA | NA | NA | NA | NA | | | Olfr389 | 0 | NA | NA | NA | NA | NA |
| Serpina3m | 0.188785634 | 0.025025111 | 0.031461603 | 0.79541754 | 0.42637066 | NA | | | Olfr39 | 0 | NA | NA | NA | NA | NA |
| Serpina4-ps1 | 0.096071043 | -0.020098795 | 0.027825948 | -0.722304036 | 0.470107572 | NA | | | Olfr390 | 0 | NA | NA | NA | NA | NA |
| Serpina5 | 0.079983654 | 0.016740032 | 0.027674178 | 0.604897175 | 0.54524732 | NA | | | Olfr391-ps | 0 | NA | NA | NA | NA | NA |
| Serpina6 | 0 | NA | NA | NA | NA | NA | | | Olfr392 | 0 | NA | NA | NA | NA | NA |
| Serpina7 | 0.098353386 | -0.020212831 | 0.02786466 | -0.725393061 | 0.468210929 | NA | | | Olfr393 | 0 | NA | NA | NA | NA | NA |
| Serpinb10 | 0.725213963 | -0.019866562 | 0.06124627 | -0.324371786 | 0.745656578 | NA | | | Olfr394 | 0 | NA | NA | NA | NA | NA |
| Serpinb11 | 2.096562876 | -0.010921561 | 0.099007949 | -0.110309938 | 0.912163576 | NA | | | Olfr395 | 0 | NA | NA | NA | NA | NA |
| Serpinb12 | 0.079983654 | 0.016740032 | 0.027674178 | 0.604897175 | 0.54524732 | NA | | | Olfr397 | 0 | NA | NA | NA | NA | NA |
| Serpinb13 | 0 | NA | NA | NA | NA | NA | | | Olfr398 | 0 | NA | NA | NA | NA | NA |
| Serpinb1c | 0.341110531 | -0.038008058 | 0.037770309 | -1.006294602 | 0.314273876 | NA | | | Olfr399 | 0 | NA | NA | NA | NA | NA |
| Serpinb2 | 2.01169938 | -0.025748391 | 0.084858552 | -0.303427176 | 0.761564337 | NA | | | Olfr401 | 0 | NA | NA | NA | NA | NA |
| Serpinb3a | 0.192142086 | -0.026384592 | 0.031527717 | -0.836869715 | 0.402665805 | NA | | | Olfr402 | 0 | NA | NA | NA | NA | NA |
| Serpinb3b | 0.519763674 | 0.02504685 | 0.039752317 | 0.630072711 | 0.528647013 | NA | | | Olfr403 | 0 | NA | NA | NA | NA | NA |

| | | | | | | | | | | | | | | | |
|---|---|---|---|---|---|---|---|---|---|---|---|---|---|---|---|
| Serpinb3c | 0.211846316 | 0.025707621 | 0.031860375 | 0.806883811 | 0.419733426 | NA | | | Olfr406 | 0 | NA | NA | NA | NA | NA |
| Serpinb3d | 0 | NA | NA | NA | NA | NA | | | Olfr410 | 0 | NA | NA | NA | NA | NA |
| Serpinb5 | 4.934441871 | -0.232121113 | 0.139900385 | -1.659188523 | 0.09707781 | NA | | | Olfr412 | 0 | NA | NA | NA | NA | NA |
| Serpinb6c | 5.590896087 | 0.015753888 | 0.142647351 | 0.110439399 | 0.912060908 | NA | | | Olfr414 | 0 | NA | NA | NA | NA | NA |
| Serpinb6d | 0 | NA | NA | NA | NA | NA | | | Olfr417 | 0 | NA | NA | NA | NA | NA |
| Serpinb6e | 0.898256634 | 0.026555098 | 0.057925231 | 0.458437501 | 0.646638154 | NA | | | Olfr418-ps1 | 0 | NA | NA | NA | NA | NA |
| Serpinb7 | 0.649309414 | -0.061653223 | 0.057177405 | -1.078279488 | 0.280909048 | NA | | | Olfr419 | 0 | NA | NA | NA | NA | NA |
| Serpinb9b | 1.33413296 | 0.004628374 | 0.074655478 | 0.061996436 | 0.950565671 | NA | | | Olfr420 | 0 | NA | NA | NA | NA | NA |
| Serpinb9c | 0.094392817 | 0.017895942 | 0.027818137 | 0.643319236 | 0.520016977 | NA | | | Olfr421-ps1 | 0 | NA | NA | NA | NA | NA |
| Serpinb9d | 0 | NA | NA | NA | NA | NA | | | Olfr424 | 0 | NA | NA | NA | NA | NA |
| Serpinb9e | 0.098353386 | -0.020212831 | 0.02786466 | -0.725393061 | 0.468210929 | NA | | | Olfr426 | 0 | NA | NA | NA | NA | NA |
| Serpinb9f | 0 | NA | NA | NA | NA | NA | | | Olfr427 | 0 | NA | NA | NA | NA | NA |
| Serpinb9g | 0 | NA | NA | NA | NA | NA | | | Olfr429 | 0 | NA | NA | NA | NA | NA |
| Serpinc1 | 0.569860421 | -0.032391892 | 0.053568423 | -0.604682578 | 0.545389926 | NA | | | Olfr43 | 0 | NA | NA | NA | NA | NA |
| Serpine3 | 0.098353386 | -0.020212831 | 0.02786466 | -0.725393061 | 0.468210929 | NA | | | Olfr430 | 0 | NA | NA | NA | NA | NA |
| Serpinf2 | 5.348155506 | 0.082448973 | 0.115921372 | 0.711249119 | 0.476929876 | NA | | | Olfr432 | 0 | NA | NA | NA | NA | NA |
| Serpini2 | 0 | NA | NA | NA | NA | NA | | | Olfr433 | 0 | NA | NA | NA | NA | NA |
| Sfta2 | 0.376370672 | 0.027798864 | 0.047885955 | 0.580522285 | 0.561562463 | NA | | | Olfr434 | 0 | NA | NA | NA | NA | NA |
| Sftpa1 | 0.297029653 | 0.02866449 | 0.033517577 | 0.855207712 | 0.392436167 | NA | | | Olfr435 | 0 | NA | NA | NA | NA | NA |
| Sftpb | 0.185670226 | -0.001645086 | 0.031432796 | -0.052336624 | 0.958260472 | NA | | | Olfr44 | 0 | NA | NA | NA | NA | NA |
| Sftpd | 0 | NA | NA | NA | NA | NA | | | Olfr441 | 0 | NA | NA | NA | NA | NA |
| Sgcg | 3.714718138 | -0.005543271 | 0.109092518 | -0.050812566 | 0.959474878 | NA | | | Olfr444 | 0 | NA | NA | NA | NA | NA |
| Sh2d1a | 0.551455211 | -0.006423304 | 0.04591486 | -0.139895971 | 0.888742185 | NA | | | Olfr447 | 0 | NA | NA | NA | NA | NA |
| Sh2d1b1 | 1.607386356 | -0.052718474 | 0.092387153 | -0.570625599 | 0.568253462 | NA | | | Olfr448 | 0 | NA | NA | NA | NA | NA |
| Sh2d1b2 | 0 | NA | NA | NA | NA | NA | | | Olfr449 | 0 | NA | NA | NA | NA | NA |
| Sh2d2a | 0.610399768 | -0.006711683 | 0.062901991 | -0.106700643 | 0.915026472 | NA | | | Olfr45 | 0 | NA | NA | NA | NA | NA |
| Sh2d7 | 3.465215393 | 0.089890027 | 0.121590735 | 0.739283524 | 0.459734852 | NA | | | Olfr450 | 0 | NA | NA | NA | NA | NA |
| Shbg | 1.281199613 | 0.033476079 | 0.077403559 | 0.432487592 | 0.665387069 | NA | | | Olfr452 | 0 | NA | NA | NA | NA | NA |
| Shox2 | 6.221846719 | 0.259673605 | 0.104389761 | 2.487539038 | 0.012863034 | NA | | | Olfr453 | 0 | NA | NA | NA | NA | NA |
| Siglec15 | 0.932205047 | -0.017555772 | 0.06557263 | -0.267730174 | 0.788907022 | NA | | | Olfr455 | 0 | NA | NA | NA | NA | NA |
| Siglecg | 2.662740426 | 0.018562599 | 0.11140283 | 0.166625918 | 0.867664399 | NA | | | Olfr456 | 0 | NA | NA | NA | NA | NA |
| Sim2 | 4.214574456 | -0.077544818 | 0.129059375 | -0.600846068 | 0.547942517 | NA | | | Olfr457 | 0 | NA | NA | NA | NA | NA |
| Sirpb1a | 0.582088857 | -0.027172096 | 0.053599983 | -0.506942252 | 0.612195339 | NA | | | Olfr458 | 0 | NA | NA | NA | NA | NA |
| Sirpb1b | 0.299858689 | -0.034980473 | 0.036264327 | -0.964597324 | 0.33474654 | NA | | | Olfr461 | 0 | NA | NA | NA | NA | NA |
| Sis | 4.599129682 | 0.152244774 | 0.135595123 | 1.122789455 | 0.261526925 | NA | | | Olfr462 | 0 | NA | NA | NA | NA | NA |
| Sit1 | 0.286710141 | 0.009849273 | 0.038825804 | 0.25367852 | 0.79974393 | NA | | | Olfr463 | 0 | NA | NA | NA | NA | NA |
| Six6 | 1.810013564 | -0.049986132 | 0.089873706 | -0.556181931 | 0.57808649 | NA | | | Olfr466 | 0 | NA | NA | NA | NA | NA |
| Ska1 | 2.185053297 | -0.081721159 | 0.091527124 | -0.892862748 | 0.371930678 | NA | | | Olfr467 | 0 | NA | NA | NA | NA | NA |
| Skint1 | 0 | NA | NA | NA | NA | NA | | | Olfr469 | 0 | NA | NA | NA | NA | NA |
| Skint10 | 0.318806384 | -0.016982193 | 0.040116674 | -0.423320068 | 0.672061754 | NA | | | Olfr47 | 0 | NA | NA | NA | NA | NA |
| Skint11 | 0 | NA | NA | NA | NA | NA | | | Olfr470 | 0 | NA | NA | NA | NA | NA |
| Skint2 | 0 | NA | NA | NA | NA | NA | | | Olfr472 | 0 | NA | NA | NA | NA | NA |
| Skint3 | 1.581200634 | -0.05378995 | 0.084321719 | -0.637913348 | 0.523530089 | NA | | | Olfr473 | 0 | NA | NA | NA | NA | NA |
| Skint4 | 0.107716603 | -0.020477583 | 0.027954224 | -0.732539849 | 0.46383913 | NA | | | Olfr474 | 0 | NA | NA | NA | NA | NA |
| Skint5 | 0.107716603 | -0.020477583 | 0.027954224 | -0.732539849 | 0.46383913 | NA | | | Olfr476 | 0 | NA | NA | NA | NA | NA |
| Skint6 | 0 | NA | NA | NA | NA | NA | | | Olfr477 | 0 | NA | NA | NA | NA | NA |
| Skint7 | 0 | NA | NA | NA | NA | NA | | | Olfr478 | 0 | NA | NA | NA | NA | NA |
| Skint8 | 0 | NA | NA | NA | NA | NA | | | Olfr479 | 0 | NA | NA | NA | NA | NA |
| Skint9 | 0 | NA | NA | NA | NA | NA | | | Olfr48 | 0 | NA | NA | NA | NA | NA |
| Skor2 | 0.499336704 | -0.024198502 | 0.039525929 | -0.612218432 | 0.540393248 | NA | | | Olfr480 | 0 | NA | NA | NA | NA | NA |
| Slamf7 | 4.005037745 | -0.019113612 | 0.111189675 | -0.171900959 | 0.863515392 | NA | | | Olfr481 | 0 | NA | NA | NA | NA | NA |
| Slc10a1 | 0.096071043 | -0.020098795 | 0.027825948 | -0.722304036 | 0.470107572 | NA | | | Olfr482 | 0 | NA | NA | NA | NA | NA |
| Slc10a2 | 0 | NA | NA | NA | NA | NA | | | Olfr483 | 0 | NA | NA | NA | NA | NA |
| Slc10a5 | 0.159967308 | 0.0240345 | 0.030870969 | 0.778547006 | 0.436246606 | NA | | | Olfr484 | 0 | NA | NA | NA | NA | NA |
| Slc10a6 | 1.560736871 | 0.063371788 | 0.0726491 | 0.872299694 | 0.383044904 | NA | | | Olfr485 | 0 | NA | NA | NA | NA | NA |
| Slc12a1 | 3.49259856 | 0.19811565 | 0.106583684 | 1.858780267 | 0.063058291 | NA | | | Olfr486 | 0 | NA | NA | NA | NA | NA |
| Slc12a3 | 7.353508885 | 0.111521512 | 0.133253346 | 0.836913415 | 0.402641239 | NA | | | Olfr487 | 0 | NA | NA | NA | NA | NA |
| Slc13a1 | 4.86241085 | 0.079856996 | 0.131174675 | 0.608783641 | 0.542667858 | NA | | | Olfr488 | 0 | NA | NA | NA | NA | NA |
| Slc13a2 | 0.427314999 | -0.00123242 | 0.042901703 | -0.02872661 | 0.977082634 | NA | | | Olfr49 | 0 | NA | NA | NA | NA | NA |
| Slc15a1 | 2.06578724 | 0.083811144 | 0.090251847 | 0.928636337 | 0.353077584 | NA | | | Olfr490 | 0 | NA | NA | NA | NA | NA |
| Slc15a5 | 2.432967051 | -0.124535988 | 0.102797106 | -1.211473678 | 0.225713917 | NA | | | Olfr491 | 0 | NA | NA | NA | NA | NA |
| Slc16a5 | 1.269995199 | -0.081054214 | 0.078926637 | -1.026956382 | 0.304441003 | NA | | | Olfr492 | 0 | NA | NA | NA | NA | NA |
| Slc16a8 | 6.573843282 | 0.326076484 | 0.107915393 | 3.021593808 | NA | NA | | | Olfr493 | 0 | NA | NA | NA | NA | NA |
| Slc17a1 | 0.099009884 | 0.018205726 | 0.027896865 | 0.652608315 | 0.514008823 | NA | | | Olfr494 | 0 | NA | NA | NA | NA | NA |
| Slc17a2 | 0.600329945 | 0.053285905 | 0.055639064 | 0.957706721 | 0.338210667 | NA | | | Olfr495 | 0 | NA | NA | NA | NA | NA |
| Slc17a3 | 0.816087363 | 0.053149322 | 0.064635028 | 0.82229905 | 0.410906717 | NA | | | Olfr497 | 0 | NA | NA | NA | NA | NA |
| Slc17a4 | 0 | NA | NA | NA | NA | NA | | | Olfr498 | 0 | NA | NA | NA | NA | NA |
| Slc18a1 | 5.787397128 | -0.130868944 | 0.148436108 | -0.881651674 | 0.377965202 | NA | | | Olfr50 | 0 | NA | NA | NA | NA | NA |
| Slc1a7 | 3.047235252 | 0.010976726 | 0.122253463 | 0.08978663 | 0.928456773 | NA | | | Olfr502 | 0 | NA | NA | NA | NA | NA |
| Slc22a1 | 7.124424939 | -0.00445354 | 0.148188891 | -0.03005313 | 0.976024681 | NA | | | Olfr503 | 0 | NA | NA | NA | NA | NA |
| Slc22a12 | 7.10112088 | 0.112817447 | 0.147475559 | 0.764990805 | 0.44427702 | NA | | | Olfr504 | 0 | NA | NA | NA | NA | NA |
| Slc22a13 | 0.831476197 | -0.075869228 | 0.070717418 | -1.072850644 | 0.283338134 | NA | | | Olfr506 | 0 | NA | NA | NA | NA | NA |
| Slc22a14 | 0.56700977 | 0.058647955 | 0.060141457 | 0.97516684 | 0.329477508 | NA | | | Olfr507 | 0 | NA | NA | NA | NA | NA |
| Slc22a16 | 0 | NA | NA | NA | NA | NA | | | Olfr508 | 0 | NA | NA | NA | NA | NA |
| Slc22a19 | 0 | NA | NA | NA | NA | NA | | | Olfr509 | 0 | NA | NA | NA | NA | NA |
| Slc22a20 | 0.284916696 | 0.042355674 | 0.040798016 | 1.038179738 | 0.299186384 | NA | | | Olfr51 | 0 | NA | NA | NA | NA | NA |
| Slc22a22 | 0 | NA | NA | NA | NA | NA | | | Olfr510 | 0 | NA | NA | NA | NA | NA |
| Slc22a26 | 0 | NA | NA | NA | NA | NA | | | Olfr512 | 0 | NA | NA | NA | NA | NA |
| Slc22a27 | 0 | NA | NA | NA | NA | NA | | | Olfr513 | 0 | NA | NA | NA | NA | NA |
| Slc22a28 | 0 | NA | NA | NA | NA | NA | | | Olfr514 | 0 | NA | NA | NA | NA | NA |
| Slc22a29 | 1.288823074 | 0.094167423 | 0.072361247 | 1.301351585 | 0.193138137 | NA | | | Olfr516 | 0 | NA | NA | NA | NA | NA |
| Slc22a30 | 0.218407525 | -0.002234018 | 0.031987735 | -0.069839845 | 0.944321133 | NA | | | Olfr517 | 0 | NA | NA | NA | NA | NA |
| Slc23a1 | 6.475352322 | 0.04140952 | 0.152293342 | 0.271906303 | 0.785694064 | NA | | | Olfr518 | 0 | NA | NA | NA | NA | NA |
| Slc24a1 | 7.99271006 | -0.135920025 | 0.156082007 | -0.870824429 | 0.383850024 | NA | | | Olfr519 | 0 | NA | NA | NA | NA | NA |
| Slc25a2 | 5.289379588 | -0.123422725 | 0.137415725 | -0.898170313 | 0.369094758 | NA | | | Olfr52 | 0 | NA | NA | NA | NA | NA |
| Slc25a31 | 1.120484986 | 0.025598664 | 0.065749371 | 0.389337017 | 0.697026856 | NA | | | Olfr521 | 0 | NA | NA | NA | NA | NA |
| Slc25a41 | 3.529875707 | 0.169315375 | 0.112299314 | 1.507715135 | 0.13162745 | NA | | | Olfr522 | 0 | NA | NA | NA | NA | NA |
| Slc25a43 | 6.904239365 | 0.056192651 | 0.147297934 | 0.381489742 | 0.702839881 | NA | | | Olfr524 | 0 | NA | NA | NA | NA | NA |
| Slc25a48 | 6.14951664 | -0.118222169 | 0.138160763 | -0.855685553 | 0.392171733 | NA | | | Olfr525 | 0 | NA | NA | NA | NA | NA |
| Slc25a54 | 0 | NA | NA | NA | NA | NA | | | Olfr527 | 0 | NA | NA | NA | NA | NA |
| Slc26a3 | 5.006784055 | 0.026397753 | 0.138252049 | 0.190939327 | 0.848573129 | NA | | | Olfr530 | 0 | NA | NA | NA | NA | NA |
| Slc26a9 | 0.869904894 | -0.055282602 | 0.070581208 | -0.78324817 | 0.433481393 | NA | | | Olfr531 | 0 | NA | NA | NA | NA | NA |
| Slc27a6 | 7.372042461 | -0.334043524 | 0.149942228 | -2.227814868 | 0.025892863 | NA | | | Olfr533 | 0 | NA | NA | NA | NA | NA |
| Slc28a2 | 3.445389332 | 0.018368091 | 0.126347099 | 0.145378018 | 0.884412419 | NA | | | Olfr535 | 0 | NA | NA | NA | NA | NA |
| Slc28a3 | 2.271840048 | 0.068000825 | 0.091578763 | 0.742539238 | 0.457760688 | NA | | | Olfr538 | 0 | NA | NA | NA | NA | NA |
| Slc2a12 | 130.0413045 | 0.240041497 | 0.165165798 | 1.453336588 | NA | NA | | | Olfr539 | 0 | NA | NA | NA | NA | NA |
| Slc2a2 | 1.415985396 | -0.006921476 | 0.076339664 | -0.090666837 | 0.927757322 | NA | | | Olfr541 | 0 | NA | NA | NA | NA | NA |
| Slc2a5 | 204.357231 | -0.218856896 | 0.166955716 | -1.310867944 | NA | NA | | | Olfr543 | 0 | NA | NA | NA | NA | NA |
| Slc2a7 | 0 | NA | NA | NA | NA | NA | | | Olfr544 | 0 | NA | NA | NA | NA | NA |
| Slc30a2 | 3.854484238 | -0.122897272 | 0.120445105 | -1.020359213 | 0.30755813 | NA | | | Olfr547 | 0 | NA | NA | NA | NA | NA |
| Slc30a8 | 1.516755232 | -0.087377064 | 0.087474128 | -0.99889037 | 0.317847802 | NA | | | Olfr549 | 0 | NA | NA | NA | NA | NA |
| Slc34a1 | 1.880407214 | 0.061162988 | 0.102707221 | 0.595508155 | 0.55150385 | NA | | | Olfr55 | 0 | NA | NA | NA | NA | NA |
| Slc34a2 | 2.449511793 | -0.144336872 | 0.105649065 | -1.366191658 | 0.171878815 | NA | | | Olfr551 | 0 | NA | NA | NA | NA | NA |
| Slc34a3 | 6.109588583 | 0.228385488 | 0.131853278 | 1.732118389 | 0.083252486 | NA | | | Olfr552 | 0 | NA | NA | NA | NA | NA |
| Slc35g3 | 2.437441128 | 0.13187174 | 0.099519648 | 1.325082454 | 0.185143805 | NA | | | Olfr553 | 0 | NA | NA | NA | NA | NA |
| Slc36a3 | 0.17463368 | 0.024560283 | 0.031186242 | 0.787535833 | 0.430968261 | NA | | | Olfr554 | 0 | NA | NA | NA | NA | NA |
| Slc38a2 | 5998.012261 | 0.24738027 | 0.166917212 | 1.482053686 | NA | NA | | | Olfr555 | 0 | NA | NA | NA | NA | NA |
| Slc39a14 | 1535.394057 | 0.06927232 | 0.085215767 | 0.812904963 | NA | NA | | | Olfr556 | 0 | NA | NA | NA | NA | NA |
| Slc44a3 | 4.487310293 | 0.161364895 | 0.130852994 | 1.233176944 | 0.217509755 | NA | | | Olfr557 | 0 | NA | NA | NA | NA | NA |
| Slc45a2 | 2.017695165 | 0.01360999 | 0.094633027 | 0.143818603 | 0.885643713 | NA | | | Olfr559 | 0 | NA | NA | NA | NA | NA |
| Slc46a2 | 0 | NA | NA | NA | NA | NA | | | Olfr566 | 0 | NA | NA | NA | NA | NA |
| Slc47a2 | 3.802909098 | 0.076984124 | 0.127448864 | 0.604039314 | 0.545817504 | NA | | | Olfr568 | 0 | NA | NA | NA | NA | NA |
| Slc4a1 | 6.459954439 | -0.196756786 | 0.129307359 | -1.521620947 | 0.128104086 | NA | | | Olfr569 | 0 | NA | NA | NA | NA | NA |
| Slc4a5 | 49.33279172 | 0.303545429 | 0.103950937 | 2.920083635 | NA | NA | | | Olfr57 | 0 | NA | NA | NA | NA | NA |
| Slc4a9 | 5.511057126 | 0.123489862 | 0.12628237 | 0.977886794 | 0.328130317 | NA | | | Olfr571 | 0 | NA | NA | NA | NA | NA |
| Slc51a | 0.079983654 | 0.016740032 | 0.027674178 | 0.604897175 | 0.54524732 | NA | | | Olfr572 | 0 | NA | NA | NA | NA | NA |
| Slc51b | 0 | NA | NA | NA | NA | NA | | | Olfr575 | 0 | NA | NA | NA | NA | NA |
| Slc5a1 | 1.776605274 | 0.044468868 | 0.099695053 | 0.44604889 | 0.655561931 | NA | | | Olfr576 | 0 | NA | NA | NA | NA | NA |
| Slc5a10 | 4.65089161 | 0.130441909 | 0.128306044 | 1.016646643 | 0.309321557 | NA | | | Olfr578 | 0 | NA | NA | NA | NA | NA |
| Slc5a12 | 1.361654606 | -0.017884823 | 0.082788648 | -0.216029898 | 0.828964443 | NA | | | Olfr582 | 0 | NA | NA | NA | NA | NA |
| Slc5a2 | 0.357764354 | 0.026260735 | 0.047363929 | 0.554445872 | 0.57927374 | NA | | | Olfr583 | 0 | NA | NA | NA | NA | NA |
| Slc5a4a | 1.07156002 | 0.007911469 | 0.078408129 | 0.10090113 | 0.919628946 | NA | | | Olfr584 | 0 | NA | NA | NA | NA | NA |
| Slc5a4b | 1.63457504 | -0.055961005 | 0.095701821 | -0.584743374 | 0.558720284 | NA | | | Olfr585 | 0 | NA | NA | NA | NA | NA |
| Slc5a8 | 3.000929632 | -0.014338427 | 0.115992831 | -0.123614772 | 0.901620296 | NA | | | Olfr586 | 0 | NA | NA | NA | NA | NA |
| Slc5a9 | 0.983728672 | -0.024520894 | 0.065357323 | -0.375182046 | 0.707525081 | NA | | | Olfr589 | 0 | NA | NA | NA | NA | NA |
| Slc6a14 | 1.598951379 | -0.127632625 | 0.077648368 | -1.643725785 | 0.100232857 | NA | | | Olfr59 | 0 | NA | NA | NA | NA | NA |
| Slc6a18 | 0.334830846 | -0.037566726 | 0.037554766 | -1.000318476 | 0.317156409 | NA | | | Olfr591 | 0 | NA | NA | NA | NA | NA |
| Slc6a2 | 2.51436945 | 0.119831659 | 0.103380424 | 1.159132978 | 0.246401986 | NA | | | Olfr592 | 0 | NA | NA | NA | NA | NA |
| Slc6a20b | 1.580178914 | 0.056601605 | 0.074818355 | 0.756520312 | 0.449337299 | NA | | | Olfr593 | 0 | NA | NA | NA | NA | NA |
| Slc6a4 | 8.2200236 | -0.236383931 | 0.116077468 | -2.036432522 | NA | NA | | | Olfr596 | 0 | NA | NA | NA | NA | NA |
| Slc6a5 | 9.398780647 | 0.052891473 | 0.047345844 | 1.117130217 | NA | NA | | | Olfr597 | 0 | NA | NA | NA | NA | NA |
| Slc7a12 | 0 | NA | NA | NA | NA | NA | | | Olfr598 | 0 | NA | NA | NA | NA | NA |
| Slc7a13 | 0.096071043 | -0.020098795 | 0.027825948 | -0.722304036 | 0.470107572 | NA | | | Olfr599 | 0 | NA | NA | NA | NA | NA |
| Slc7a15 | 0.412796476 | -0.02682719 | 0.043806178 | -0.612406537 | 0.540268818 | NA | | | Olfr6 | 0 | NA | NA | NA | NA | NA |
| Slc7a9 | 0.369714607 | 0.021630852 | 0.042365317 | 0.510579254 | 0.609645706 | NA | | | Olfr60 | 0 | NA | NA | NA | NA | NA |
| Slc9b1 | 2.688258334 | 0.11790608 | 0.114244178 | 1.032053288 | 0.302047155 | NA | | | Olfr600 | 0 | NA | NA | NA | NA | NA |
| Slc9c1 | 0 | NA | NA | NA | NA | NA | | | Olfr601 | 0 | NA | NA | NA | NA | NA |
| Slco1a1 | 0 | NA | NA | NA | NA | NA | | | Olfr603 | 0 | NA | NA | NA | NA | NA |
| Slco1a5 | 5.505525164 | 0.190158591 | 0.118893587 | 1.599401576 | 0.109731402 | NA | | | Olfr605 | 0 | NA | NA | NA | NA | NA |
| Slco1a6 | 0.096071043 | -0.020098795 | 0.027825948 | -0.722304036 | 0.470107572 | NA | | | Olfr606 | 0 | NA | NA | NA | NA | NA |
| Slco1b2 | 4.756397274 | 0.232516176 | 0.126333168 | 1.840499847 | 0.065694887 | NA | | | Olfr608 | 0 | NA | NA | NA | NA | NA |
| Slco6b1 | 0 | NA | NA | NA | NA | NA | | | Olfr61 | 0 | NA | NA | NA | NA | NA |
| Slco6c1 | 0.951824705 | 0.018973528 | 0.077202833 | 0.245762072 | 0.805866419 | NA | | | Olfr610 | 0 | NA | NA | NA | NA | NA |
| Slco6d1 | 0.079983654 | 0.016740032 | 0.027674178 | 0.604897175 | 0.54524732 | NA | | | Olfr611 | 0 | NA | NA | NA | NA | NA |
| Slfn1 | 1.843410828 | -0.010768272 | 0.084241161 | -0.127826724 | 0.898286101 | NA | | | Olfr612 | 0 | NA | NA | NA | NA | NA |
| Slfn10-ps | 5.851185142 | 0.055919566 | 0.132030294 | 0.423535875 | 0.67190433 | NA | | | Olfr615 | 0 | NA | NA | NA | NA | NA |
| Slfn14-ps | 0.828450308 | -0.004796219 | 0.063212146 | -0.075874957 | 0.939518581 | NA | | | Olfr616 | 0 | NA | NA | NA | NA | NA |
| Slfn2 | 90.02144944 | 0.474883087 | 0.163292962 | 2.908166286 | NA | NA | | | Olfr617 | 0 | NA | NA | NA | NA | NA |
| Slfn4 | 1.779416325 | -0.11518335 | 0.077243412 | -1.491173767 | 0.135915879 | NA | | | Olfr618 | 0 | NA | NA | NA | NA | NA |
| Slfnl1 | 1.853738839 | -0.020551781 | 0.082403985 | -0.249402763 | 0.803049248 | NA | | | Olfr619 | 0 | NA | NA | NA | NA | NA |
| Sln | 7.139820866 | 0.053572088 | 0.148022269 | 0.361919112 | 0.717412477 | NA | | | Olfr620 | 0 | NA | NA | NA | NA | NA |
| Slpi | 1.524041911 | -0.026465255 | 0.085757308 | -0.308606411 | 0.757620943 | NA | | | Olfr623 | 0 | NA | NA | NA | NA | NA |
| Slurp1 | 0 | NA | NA | NA | NA | NA | | | Olfr624 | 0 | NA | NA | NA | NA | NA |
| Slx | 0 | NA | NA | NA | NA | NA | | | Olfr628 | 0 | NA | NA | NA | NA | NA |
| Slxl1 | 0 | NA | NA | NA | NA | NA | | | Olfr629 | 0 | NA | NA | NA | NA | NA |
| Smarca5-ps | 6.142828382 | -0.140467067 | 0.14601476 | -0.962005944 | 0.336046618 | NA | | | Olfr63 | 0 | NA | NA | NA | NA | NA |
| Smc1b | 3.729585714 | 0.190189383 | 0.122260053 | 1.555613457 | 0.119800037 | NA | | | Olfr630 | 0 | NA | NA | NA | NA | NA |
| Smcp | 0 | NA | NA | NA | NA | NA | | | Olfr631 | 0 | NA | NA | NA | NA | NA |
| Smgc | 0 | NA | NA | NA | NA | NA | | | Olfr632 | 0 | NA | NA | NA | NA | NA |
| Smim6 | 0.73395846 | 0.057529561 | 0.068965195 | 0.834182529 | 0.404178135 | NA | | | Olfr633 | 0 | NA | NA | NA | NA | NA |
| Smim9 | 2.064742545 | 0.007331969 | 0.090417284 | 0.081090348 | 0.935370101 | NA | | | Olfr635 | 0 | NA | NA | NA | NA | NA |
| Smlr1 | 1.861110957 | -0.034383872 | 0.092834115 | -0.3703797 | 0.711099597 | NA | | | Olfr638 | 0 | NA | NA | NA | NA | NA |
| Smok2a | 0.203787646 | -0.02671144 | 0.031728022 | -0.841887971 | 0.399850664 | NA | | | Olfr639 | 0 | NA | NA | NA | NA | NA |
| Smok2b | 0 | NA | NA | NA | NA | NA | | | Olfr64 | 0 | NA | NA | NA | NA | NA |
| Smok3a | 0 | NA | NA | NA | NA | NA | | | Olfr640 | 0 | NA | NA | NA | NA | NA |
| Smok3b | 0 | NA | NA | NA | NA | NA | | | Olfr641 | 0 | NA | NA | NA | NA | NA |
| Smok4a | 0 | NA | NA | NA | NA | NA | | | Olfr642 | 0 | NA | NA | NA | NA | NA |
| Smpx | 2.002337781 | -0.078651516 | 0.090071661 | -0.873210455 | 0.382548377 | NA | | | Olfr643 | 0 | NA | NA | NA | NA | NA |
| Smr2 | 0 | NA | NA | NA | NA | NA | | | Olfr644 | 0 | NA | NA | NA | NA | NA |
| Smr3a | 0.498722596 | 0.038106329 | 0.05215842 | 0.730588245 | 0.465030694 | NA | | | Olfr645 | 0 | NA | NA | NA | NA | NA |
| Smtnl1 | 0.297029653 | 0.02866449 | 0.033517577 | 0.855207712 | 0.392436167 | NA | | | Olfr646 | 0 | NA | NA | NA | NA | NA |
| Snhg9 | 6.228410644 | 0.120158539 | 0.144461314 | 0.831769668 | 0.405538968 | NA | | | Olfr648 | 0 | NA | NA | NA | NA | NA |
| Snora15 | 0.176054697 | -0.002190495 | 0.031239146 | -0.070120197 | 0.944097991 | NA | | | Olfr649 | 0 | NA | NA | NA | NA | NA |
| Snora17 | 7.419739404 | 0.163646196 | 0.148316452 | 1.103358354 | 0.269871574 | NA | | | Olfr65 | 0 | NA | NA | NA | NA | NA |
| Snora19 | 0 | NA | NA | NA | NA | NA | | | Olfr651 | 0 | NA | NA | NA | NA | NA |
| Snora20 | 0.776311287 | 0.056253754 | 0.069333946 | 0.811345061 | 0.41716754 | NA | | | Olfr652 | 0 | NA | NA | NA | NA | NA |
| Snora21 | 0.756517709 | -0.003487157 | 0.060621506 | -0.057523435 | 0.954128239 | NA | | | Olfr653 | 0 | NA | NA | NA | NA | NA |
| Snora23 | 4.454482807 | 0.052035229 | 0.127935034 | 0.406731663 | 0.684205083 | NA | | | Olfr654 | 0 | NA | NA | NA | NA | NA |
| Snora24 | 0 | NA | NA | NA | NA | NA | | | Olfr655 | 0 | NA | NA | NA | NA | NA |
| Snora26 | 0 | NA | NA | NA | NA | NA | | | Olfr656 | 0 | NA | NA | NA | NA | NA |
| Snora2b | 4.287580726 | 0.091161089 | 0.124264218 | 0.733606909 | 0.463188349 | NA | | | Olfr657 | 0 | NA | NA | NA | NA | NA |
| Snora30 | 5.453059429 | 0.04289294 | 0.143507137 | 0.298890639 | 0.76502349 | NA | | | Olfr659 | 0 | NA | NA | NA | NA | NA |
| Snora33 | 4.880171593 | 0.107710074 | 0.138614036 | 0.777050271 | 0.437129112 | NA | | | Olfr66 | 0 | NA | NA | NA | NA | NA |
| Snora35 | 0 | NA | NA | NA | NA | NA | | | Olfr663 | 0 | NA | NA | NA | NA | NA |
| Snora36b | 0.098353386 | -0.020212831 | 0.02786466 | -0.725393061 | 0.468210929 | NA | | | Olfr665 | 0 | NA | NA | NA | NA | NA |
| Snora44 | 7.757554327 | 0.236738768 | 0.152725308 | 1.550095202 | 0.121118668 | NA | | | Olfr666 | 0 | NA | NA | NA | NA | NA |
| Snora47 | 1.661032965 | 0.030399447 | 0.102875619 | 0.295497095 | 0.767614171 | NA | | | Olfr667 | 0 | NA | NA | NA | NA | NA |
| Snora5c | 4.709046317 | 0.12076528 | 0.124500499 | 0.969998363 | 0.332047308 | NA | | | Olfr668 | 0 | NA | NA | NA | NA | NA |
| Snora62 | 1.647731421 | -0.020662579 | 0.097009475 | -0.212995475 | 0.831330492 | NA | | | Olfr669 | 0 | NA | NA | NA | NA | NA |
| Snora68 | 5.229687796 | 0.143930448 | 0.137887133 | 1.043827977 | 0.296564985 | NA | | | Olfr67 | 0 | NA | NA | NA | NA | NA |
| Snora69 | 1.614440914 | 0.111076425 | 0.082010488 | 1.354417306 | 0.175603279 | NA | | | Olfr670 | 0 | NA | NA | NA | NA | NA |
| Snora74a | 2.927036111 | 0.079155726 | 0.112849439 | 0.701427737 | 0.483036117 | NA | | | Olfr671 | 0 | NA | NA | NA | NA | NA |
| Snora75 | 5.714637261 | 0.140444187 | 0.136321732 | 1.03024063 | 0.302897061 | NA | | | Olfr672 | 0 | NA | NA | NA | NA | NA |
| Snora78 | 4.210737126 | 0.08472242 | 0.12842511 | 0.659702924 | 0.50944449 | NA | | | Olfr675 | 0 | NA | NA | NA | NA | NA |
| Snord11 | 0 | NA | NA | NA | NA | NA | | | Olfr676 | 0 | NA | NA | NA | NA | NA |
| Snord110 | 0.836563683 | 0.120566208 | 0.067280645 | 1.791989478 | 0.073134651 | NA | | | Olfr677 | 0 | NA | NA | NA | NA | NA |
| Snord111 | 0.17463368 | 0.024560283 | 0.031186242 | 0.787535833 | 0.430968261 | NA | | | Olfr678 | 0 | NA | NA | NA | NA | NA |
| Snord116 | 0 | NA | NA | NA | NA | NA | | | Olfr679 | 0 | NA | NA | NA | NA | NA |
| Snord116l1 | 0 | NA | NA | NA | NA | NA | | | Olfr68 | 0 | NA | NA | NA | NA | NA |
| Snord116l2 | 0 | NA | NA | NA | NA | NA | | | Olfr684 | 0 | NA | NA | NA | NA | NA |
| Snord123 | 0.079983654 | 0.016740032 | 0.027674178 | 0.604897175 | 0.54524732 | NA | | | Olfr685 | 0 | NA | NA | NA | NA | NA |
| Snord15a | 0.185670226 | -0.001645086 | 0.031432796 | -0.052336624 | 0.958260472 | NA | | | Olfr686 | 0 | NA | NA | NA | NA | NA |
| Snord15b | 4.508321953 | 0.021899669 | 0.129521653 | 0.169081139 | 0.86573282 | NA | | | Olfr688 | 0 | NA | NA | NA | NA | NA |
| Snord16a | 4.669576605 | 0.108281837 | 0.123963081 | 0.873500687 | 0.382390232 | NA | | | Olfr689 | 0 | NA | NA | NA | NA | NA |
| Snord17 | 0.856084225 | 0.046257005 | 0.078427052 | 0.589809308 | 0.555318502 | NA | | | Olfr69 | 0 | NA | NA | NA | NA | NA |
| Snord1a | 0 | NA | NA | NA | NA | NA | | | Olfr692 | 0 | NA | NA | NA | NA | NA |
| Snord1b | 0.944323739 | 0.070449576 | 0.067910535 | 1.037388016 | 0.299555061 | NA | | | Olfr693 | 0 | NA | NA | NA | NA | NA |
| Snord1c | 1.966396074 | -0.024856255 | 0.087490008 | -0.28410393 | 0.776330735 | NA | | | Olfr694 | 0 | NA | NA | NA | NA | NA |
| Snord23 | 1.729293513 | 0.137780731 | 0.089556602 | 1.538476528 | 0.123932143 | NA | | | Olfr695 | 0 | NA | NA | NA | NA | NA |
| Snord33 | 0.079983654 | 0.016740032 | 0.027674178 | 0.604897175 | 0.54524732 | NA | | | Olfr698 | 0 | NA | NA | NA | NA | NA |
| Snord34 | 2.761218143 | 0.148125241 | 0.114082284 | 1.298407041 | 0.194147501 | NA | | | Olfr699 | 0 | NA | NA | NA | NA | NA |
| Snord35a | 3.313797802 | 0.112011181 | 0.107819687 | 1.03887503 | 0.298862861 | NA | | | Olfr70 | 0 | NA | NA | NA | NA | NA |
| Snord35b | 2.7360309 | -0.033000222 | 0.111615197 | -0.29566065 | 0.767489251 | NA | | | Olfr702 | 0 | NA | NA | NA | NA | NA |
| Snord38a | 0.97346973 | 0.040727718 | 0.065993421 | 0.61714815 | 0.537137012 | NA | | | Olfr703 | 0 | NA | NA | NA | NA | NA |
| Snord42a | 1.601975893 | 0.101605255 | 0.089815358 | 1.131268163 | 0.25794224 | NA | | | Olfr704 | 0 | NA | NA | NA | NA | NA |
| Snord43 | 1.06343616 | 0.106829216 | 0.082129948 | 1.300734002 | 0.193349519 | NA | | | Olfr705 | 0 | NA | NA | NA | NA | NA |
| Snord45b | 7.299069619 | 0.155278879 | 0.139179939 | 1.115669977 | 0.264563426 | NA | | | Olfr706 | 0 | NA | NA | NA | NA | NA |
| Snord45c | 0 | NA | NA | NA | NA | NA | | | Olfr71 | 0 | NA | NA | NA | NA | NA |
| Snord47 | 1.659266406 | 0.051411949 | 0.083406692 | 0.616400772 | 0.537630044 | NA | | | Olfr710 | 0 | NA | NA | NA | NA | NA |
| Snord49b | 2.026158894 | 0.010474814 | 0.104496421 | 0.100240892 | 0.920153082 | NA | | | Olfr711 | 0 | NA | NA | NA | NA | NA |
| Snord4a | 5.077063052 | -0.0371771 | 0.140092135 | -0.265376069 | 0.79071977 | NA | | | Olfr713 | 0 | NA | NA | NA | NA | NA |
| Snord55 | 1.899069637 | -0.072711789 | 0.092540098 | -0.785732788 | 0.43202405 | NA | | | Olfr714 | 0 | NA | NA | NA | NA | NA |
| Snord57 | 4.032632855 | 0.095197874 | 0.124156007 | 0.766760116 | 0.443224154 | NA | | | Olfr715 | 0 | NA | NA | NA | NA | NA |
| Snord58b | 4.051392012 | 0.111357297 | 0.130557126 | 0.852939244 | 0.393692997 | NA | | | Olfr716 | 0 | NA | NA | NA | NA | NA |
| Snord61 | 0 | NA | NA | NA | NA | NA | | | Olfr720 | 0 | NA | NA | NA | NA | NA |
| Snord64 | 2.774180084 | 0.094359898 | 0.117405762 | 0.803707559 | 0.421565888 | NA | | | Olfr722 | 0 | NA | NA | NA | NA | NA |
| Snord65 | 5.091137723 | 0.093665193 | 0.135875982 | 0.689343261 | 0.49060728 | NA | | | Olfr723 | 0 | NA | NA | NA | NA | NA |
| Snord66 | 2.34916592 | 0.050203948 | 0.100213218 | 0.500971322 | 0.616391306 | NA | | | Olfr724 | 0 | NA | NA | NA | NA | NA |
| Snord67 | 0 | NA | NA | NA | NA | NA | | | Olfr725 | 0 | NA | NA | NA | NA | NA |
| Snord68 | 5.346287061 | 0.215082612 | 0.133920108 | 1.606051659 | 0.10826256 | NA | | | Olfr726 | 0 | NA | NA | NA | NA | NA |
| Snord7 | 0 | NA | NA | NA | NA | NA | | | Olfr728 | 0 | NA | NA | NA | NA | NA |
| Snord70 | 0 | NA | NA | NA | NA | NA | | | Olfr729 | 0 | NA | NA | NA | NA | NA |
| Snord71 | 1.914277662 | -0.041711868 | 0.106185353 | -0.392821299 | 0.694451472 | NA | | | Olfr73 | 0 | NA | NA | NA | NA | NA |
| Snord72 | 2.028357065 | -0.093775826 | 0.088327722 | -1.061680569 | 0.288380723 | NA | | | Olfr730 | 0 | NA | NA | NA | NA | NA |
| Snord73a | 1.37279849 | 0.010632801 | 0.082723032 | 0.128534959 | 0.897725635 | NA | | | Olfr731 | 0 | NA | NA | NA | NA | NA |

| | | | | | | | | | | | | | |
|---|---|---|---|---|---|---|---|---|---|---|---|---|---|
| Snord8 | 0.921924053 | -0.061698577 | 0.076084357 | -0.810923289 | 0.417409725 | NA | Olfr732 | 0 | NA | NA | NA | NA | NA |
| Snord82 | 0 | NA | NA | NA | NA | NA | Olfr733 | 0 | NA | NA | NA | NA | NA |
| Snord83b | 4.547536993 | 0.227758917 | 0.140969012 | 1.615666541 | 0.106166429 | NA | Olfr734 | 0 | NA | NA | NA | NA | NA |
| Snord85 | 0.29403132 | 0.006032586 | 0.033500653 | 0.180073692 | 0.857094716 | NA | Olfr735 | 0 | NA | NA | NA | NA | NA |
| Snord87 | 1.214169242 | 0.070065494 | 0.07329384 | 0.955953374 | 0.339095791 | NA | Olfr736 | 0 | NA | NA | NA | NA | NA |
| Snord88a | 1.991792652 | 0.069812155 | 0.093519149 | 0.746501171 | 0.455364725 | NA | Olfr738 | 0 | NA | NA | NA | NA | NA |
| Snord89 | 5.937936274 | 0.329545783 | 0.147550232 | 2.233448083 | 0.02551941 | NA | Olfr739 | 0 | NA | NA | NA | NA | NA |
| Snord90 | 0 | NA | NA | NA | NA | NA | Olfr74 | 0 | NA | NA | NA | NA | NA |
| Snord91a | 0.855062497 | 0.11313603 | 0.075230074 | 1.503867056 | 0.132615601 | NA | Olfr740 | 0 | NA | NA | NA | NA | NA |
| Snord92 | 0 | NA | NA | NA | NA | NA | Olfr743 | 0 | NA | NA | NA | NA | NA |
| Snord93 | 0 | NA | NA | NA | NA | NA | Olfr744 | 0 | NA | NA | NA | NA | NA |
| Snord95 | 5.746609284 | 0.198417805 | 0.141353351 | 1.403700749 | 0.160407979 | NA | Olfr745 | 0 | NA | NA | NA | NA | NA |
| Sntn | 7.934930054 | 0.172702571 | 0.142813564 | 1.209286894 | 0.226552644 | NA | Olfr746 | 0 | NA | NA | NA | NA | NA |
| Snurf | 0 | NA | NA | NA | NA | NA | Olfr747 | 0 | NA | NA | NA | NA | NA |
| Soat2 | 7.165783843 | 0.234555454 | 0.151916526 | 1.54397589 | 0.122594173 | NA | Olfr748 | 0 | NA | NA | NA | NA | NA |
| Sohlh1 | 2.407972676 | 0.13330053 | 0.112137835 | 1.188720381 | 0.234549718 | NA | Olfr749 | 0 | NA | NA | NA | NA | NA |
| Sohlh2 | 0.570960929 | -0.072956975 | 0.051753376 | -1.409704638 | 0.158626915 | NA | Olfr75-ps1 | 0 | NA | NA | NA | NA | NA |
| Sorbs1 | 6155.148757 | 0.252065664 | 0.102485928 | 2.459514869 | NA | NA | Olfr750 | 0 | NA | NA | NA | NA | NA |
| Sost | 4.074268789 | -0.034610803 | 0.110089933 | -0.314386632 | 0.753227408 | NA | Olfr76 | 0 | NA | NA | NA | NA | NA |
| Sostdc1 | 49.54682155 | 0.385153684 | 0.157084879 | 2.451882618 | NA | NA | Olfr761 | 0 | NA | NA | NA | NA | NA |
| Sowahd | 0.528070844 | -0.018641267 | 0.051372038 | -0.362867963 | 0.71670352 | NA | Olfr763 | 0 | NA | NA | NA | NA | NA |
| Sox14 | 1.838380592 | 0.081284516 | 0.062738168 | 1.295615074 | NA | NA | Olfr765 | 0 | NA | NA | NA | NA | NA |
| Sox15 | 2.038582862 | -0.089736471 | 0.105899275 | -0.847375685 | 0.396785753 | NA | Olfr767 | 0 | NA | NA | NA | NA | NA |
| Sox30 | 4.138357827 | 0.087630711 | 0.127423461 | 0.687712532 | 0.491633826 | NA | Olfr768 | 0 | NA | NA | NA | NA | NA |
| Spaca1 | 3.615942245 | -0.296482065 | 0.114712299 | -2.584570858 | 0.009750027 | NA | Olfr769 | 0 | NA | NA | NA | NA | NA |
| Spaca3 | 0.107716603 | -0.020477583 | 0.027954224 | -0.732539849 | 0.46383913 | NA | Olfr77 | 0 | NA | NA | NA | NA | NA |
| Spaca4 | 0 | NA | NA | NA | NA | NA | Olfr770 | 0 | NA | NA | NA | NA | NA |
| Spaca5 | 0.303023688 | 0.009739077 | 0.039388711 | 0.247255542 | 0.804710463 | NA | Olfr771 | 0 | NA | NA | NA | NA | NA |
| Spaca7 | 0 | NA | NA | NA | NA | NA | Olfr772 | 0 | NA | NA | NA | NA | NA |
| Spag11a | 0 | NA | NA | NA | NA | NA | Olfr773 | 0 | NA | NA | NA | NA | NA |
| Spag11b | 0.105923158 | 0.018524693 | 0.027977518 | 0.662127842 | 0.507889294 | NA | Olfr774 | 0 | NA | NA | NA | NA | NA |
| Spam1 | 0 | NA | NA | NA | NA | NA | Olfr775 | 0 | NA | NA | NA | NA | NA |
| Spata16 | 3.264970887 | -0.101572799 | 0.120909337 | -0.84007407 | 0.400866858 | NA | Olfr776 | 0 | NA | NA | NA | NA | NA |
| Spata19 | 5.673169361 | 0.135287959 | 0.147839571 | 0.915099783 | 0.360139249 | NA | Olfr777 | 0 | NA | NA | NA | NA | NA |
| Spata20 | 4.280226671 | 0.07261472 | 0.129458911 | 0.560909398 | 0.574859303 | NA | Olfr780 | 0 | NA | NA | NA | NA | NA |
| Spata22 | 0.745115121 | 0.01765305 | 0.062917093 | 0.280576371 | 0.779035342 | NA | Olfr781 | 0 | NA | NA | NA | NA | NA |
| Spata25 | 2.817128849 | -0.047127984 | 0.110879361 | -0.425038376 | 0.6708087 | NA | Olfr782 | 0 | NA | NA | NA | NA | NA |
| Spata31 | 0 | NA | NA | NA | NA | NA | Olfr784 | 0 | NA | NA | NA | NA | NA |
| Spata31d1a | 1.725001558 | 0.047203527 | 0.086973592 | 0.54273402 | 0.587312948 | NA | Olfr786 | 0 | NA | NA | NA | NA | NA |
| Spata31d1b | 2.590111147 | 0.079996456 | 0.072614066 | 1.101666118 | 0.270606852 | NA | Olfr787 | 0 | NA | NA | NA | NA | NA |
| Spata31d1c | 0.193239998 | 0.025145508 | 0.031532422 | 0.797449311 | 0.42519013 | NA | Olfr788 | 0 | NA | NA | NA | NA | NA |
| Spata31d1d | 2.074998987 | 0.041834143 | 0.097289076 | 0.42999836 | 0.667196834 | NA | Olfr790 | 0 | NA | NA | NA | NA | NA |
| Spata4 | 1.825730401 | -0.045155613 | 0.094649565 | -0.477082103 | 0.633303659 | NA | Olfr791 | 0 | NA | NA | NA | NA | NA |
| Spata9 | 3.203432809 | 0.007843334 | 0.124095971 | 0.063203774 | 0.949604239 | NA | Olfr792 | 0 | NA | NA | NA | NA | NA |
| Spatc1 | 0.913368603 | -0.058414623 | 0.059789942 | -0.976997495 | 0.328570393 | NA | Olfr794 | 0 | NA | NA | NA | NA | NA |
| Spatc1l | 1.20437504 | 0.045352351 | 0.06979633 | 0.64978132 | 0.515833487 | NA | Olfr796 | 0 | NA | NA | NA | NA | NA |
| Spdef | 7.113733933 | 0.199837831 | 0.136978785 | 1.458896217 | 0.144593675 | NA | Olfr798 | 0 | NA | NA | NA | NA | NA |
| Spdyb | 0.186326724 | 0.024939385 | 0.031411053 | 0.793968451 | 0.427213798 | NA | Olfr799 | 0 | NA | NA | NA | NA | NA |
| Speer1-ps1 | 0 | NA | NA | NA | NA | NA | Olfr8 | 0 | NA | NA | NA | NA | NA |
| Speer2 | 0.714780812 | -0.040502242 | 0.056225019 | -0.720359779 | 0.471303507 | NA | Olfr800 | 0 | NA | NA | NA | NA | NA |
| Speer3 | 0 | NA | NA | NA | NA | NA | Olfr801 | 0 | NA | NA | NA | NA | NA |
| Speer4a | 1.675574735 | 0.048176622 | 0.088345027 | 0.545323528 | 0.585531025 | NA | Olfr802 | 0 | NA | NA | NA | NA | NA |
| Speer4b | 2.754380874 | -0.022620124 | 0.111163425 | -0.203485315 | 0.838755723 | NA | Olfr803 | 0 | NA | NA | NA | NA | NA |
| Speer4d | 0.38488829 | -0.02558324 | 0.043075085 | -0.593921989 | 0.552564291 | NA | Olfr804 | 0 | NA | NA | NA | NA | NA |
| Speer4e | 0.688668513 | 0.012091754 | 0.060689169 | 0.199240727 | 0.842074443 | NA | Olfr805 | 0 | NA | NA | NA | NA | NA |
| Speer4f | 0 | NA | NA | NA | NA | NA | Olfr806 | 0 | NA | NA | NA | NA | NA |
| Speer5-ps1 | 0 | NA | NA | NA | NA | NA | Olfr807 | 0 | NA | NA | NA | NA | NA |
| Speer6-ps1 | 0.507592018 | 0.0062035 | 0.044582043 | 0.139147943 | 0.889333244 | NA | Olfr808 | 0 | NA | NA | NA | NA | NA |
| Speer7-ps1 | 4.087694419 | -0.202015384 | 0.130756363 | -1.544975555 | 0.122352176 | NA | Olfr809 | 0 | NA | NA | NA | NA | NA |
| Speer8-ps1 | 0.302629931 | -0.009711995 | 0.033929987 | -0.286236321 | 0.774697127 | NA | Olfr810 | 0 | NA | NA | NA | NA | NA |
| Speer9-ps1 | 0 | NA | NA | NA | NA | NA | Olfr811 | 0 | NA | NA | NA | NA | NA |
| Spem1 | 4.868364845 | 0.031948161 | 0.129093631 | 0.247480533 | 0.804536356 | NA | Olfr812 | 0 | NA | NA | NA | NA | NA |
| Spert | 0 | NA | NA | NA | NA | NA | Olfr813 | 0 | NA | NA | NA | NA | NA |
| Spesp1 | 1.511865256 | 0.02825672 | 0.085173065 | 0.331756519 | 0.740073119 | NA | Olfr814 | 0 | NA | NA | NA | NA | NA |
| Spib | 1.633440668 | -0.056844572 | 0.090236294 | -0.629952426 | 0.528725711 | NA | Olfr815 | 0 | NA | NA | NA | NA | NA |
| Spic | 1.367443798 | -0.0486333 | 0.078163966 | -0.622195902 | 0.533813061 | NA | Olfr816 | 0 | NA | NA | NA | NA | NA |
| Spink11 | 0.079983654 | 0.016740032 | 0.027674178 | 0.604897175 | 0.54524732 | NA | Olfr818 | 0 | NA | NA | NA | NA | NA |
| Spink12 | 0.192746203 | -0.000975302 | 0.031567365 | -0.030895906 | 0.975352555 | NA | Olfr819 | 0 | NA | NA | NA | NA | NA |
| Spink13 | 1.634970793 | -0.19230465 | 0.091671591 | -2.09775622 | 0.035926686 | NA | Olfr820 | 0 | NA | NA | NA | NA | NA |
| Spink14 | 0 | NA | NA | NA | NA | NA | Olfr821 | 0 | NA | NA | NA | NA | NA |
| Spink2 | 0.17833704 | -0.002390466 | 0.031282281 | -0.076415987 | 0.939088151 | NA | Olfr822 | 0 | NA | NA | NA | NA | NA |
| Spink3 | 0 | NA | NA | NA | NA | NA | Olfr823 | 0 | NA | NA | NA | NA | NA |
| Spink4 | 0.289473745 | 0.012204471 | 0.038954522 | 0.313300481 | 0.754052385 | NA | Olfr824 | 0 | NA | NA | NA | NA | NA |
| Spink5 | 0.472189209 | -0.048684432 | 0.042637861 | -1.141812248 | 0.25353207 | NA | Olfr825 | 0 | NA | NA | NA | NA | NA |
| Spink6 | 0 | NA | NA | NA | NA | NA | Olfr826 | 0 | NA | NA | NA | NA | NA |
| Spink7 | 0 | NA | NA | NA | NA | NA | Olfr827 | 0 | NA | NA | NA | NA | NA |
| Spinkl | 0 | NA | NA | NA | NA | NA | Olfr828 | 0 | NA | NA | NA | NA | NA |
| Spint3 | 0 | NA | NA | NA | NA | NA | Olfr829 | 0 | NA | NA | NA | NA | NA |
| Spint4 | 0.458631138 | 0.026681677 | 0.045037875 | 0.592427534 | 0.553564333 | NA | Olfr830 | 0 | NA | NA | NA | NA | NA |
| Spint5 | 0 | NA | NA | NA | NA | NA | Olfr832 | 0 | NA | NA | NA | NA | NA |
| Spire1 | 8371.158319 | 0.165101596 | 0.112973923 | 1.461413315 | NA | NA | Olfr834 | 0 | NA | NA | NA | NA | NA |
| Spn | 6.57032193 | -0.126585441 | 0.153581259 | -0.824224529 | 0.409811992 | NA | Olfr835 | 0 | NA | NA | NA | NA | NA |
| Spn-ps | 0.518310624 | 0.007452973 | 0.050979418 | 0.146195733 | 0.883766874 | NA | Olfr837 | 0 | NA | NA | NA | NA | NA |
| Spns3 | 7.270725561 | 0.056149556 | 0.155121155 | 0.361972269 | 0.717372753 | NA | Olfr843 | 0 | NA | NA | NA | NA | NA |
| Spon2 | 3.084358915 | -0.086554282 | 0.112880987 | -0.766774676 | 0.443215496 | NA | Olfr845 | 0 | NA | NA | NA | NA | NA |
| Spp2 | 3.712950175 | -0.240209296 | 0.127036008 | -1.890875661 | 0.058640944 | NA | Olfr846 | 0 | NA | NA | NA | NA | NA |
| Sppl2c | 0 | NA | NA | NA | NA | NA | Olfr847 | 0 | NA | NA | NA | NA | NA |
| Sprr1a | 4.524972019 | -0.15121316 | 0.133439168 | -1.133199209 | 0.257130606 | NA | Olfr849 | 0 | NA | NA | NA | NA | NA |
| Sprr1b | 0 | NA | NA | NA | NA | NA | Olfr850 | 0 | NA | NA | NA | NA | NA |
| Sprr2a1 | 0 | NA | NA | NA | NA | NA | Olfr851 | 0 | NA | NA | NA | NA | NA |
| Sprr2a2 | 0 | NA | NA | NA | NA | NA | Olfr853 | 0 | NA | NA | NA | NA | NA |
| Sprr2b | 0.096071043 | -0.020098795 | 0.027825948 | -0.722304036 | 0.470107572 | NA | Olfr854 | 0 | NA | NA | NA | NA | NA |
| Sprr2d | 0 | NA | NA | NA | NA | NA | Olfr855 | 0 | NA | NA | NA | NA | NA |
| Sprr2e | 0 | NA | NA | NA | NA | NA | Olfr860 | 0 | NA | NA | NA | NA | NA |
| Sprr2f | 0 | NA | NA | NA | NA | NA | Olfr862 | 0 | NA | NA | NA | NA | NA |
| Sprr2g | 0 | NA | NA | NA | NA | NA | Olfr866 | 0 | NA | NA | NA | NA | NA |
| Sprr2h | 0 | NA | NA | NA | NA | NA | Olfr867 | 0 | NA | NA | NA | NA | NA |
| Sprr2i | 0 | NA | NA | NA | NA | NA | Olfr868 | 0 | NA | NA | NA | NA | NA |
| Sprr2j-ps | 0 | NA | NA | NA | NA | NA | Olfr869 | 0 | NA | NA | NA | NA | NA |
| Sprr2k | 0 | NA | NA | NA | NA | NA | Olfr870 | 0 | NA | NA | NA | NA | NA |
| Sprr3 | 0 | NA | NA | NA | NA | NA | Olfr871 | 0 | NA | NA | NA | NA | NA |
| Sprr4 | 0.323149808 | -0.030167563 | 0.033779321 | -0.893077861 | 0.371815478 | NA | Olfr872 | 0 | NA | NA | NA | NA | NA |
| Spta1 | 0.583851251 | -0.037236951 | 0.058974167 | -0.631411228 | 0.527771676 | NA | Olfr873 | 0 | NA | NA | NA | NA | NA |
| Sptlc3 | 2.853528592 | 0.009473679 | 0.113370541 | 0.083563854 | 0.933403207 | NA | Olfr884 | 0 | NA | NA | NA | NA | NA |
| Spz1 | 0 | NA | NA | NA | NA | NA | Olfr885 | 0 | NA | NA | NA | NA | NA |
| Srd5a2 | 1.432144443 | -0.05491518 | 0.086347828 | -0.63597639 | 0.524791811 | NA | Olfr887 | 0 | NA | NA | NA | NA | NA |
| Sry | 0 | NA | NA | NA | NA | NA | Olfr888 | 0 | NA | NA | NA | NA | NA |
| Ssu2 | 4.484801144 | 0.02334115 | 0.135817984 | 0.171856106 | 0.863550656 | NA | Olfr889 | 0 | NA | NA | NA | NA | NA |
| Ssx9 | 0 | NA | NA | NA | NA | NA | Olfr890 | 0 | NA | NA | NA | NA | NA |
| Ssxb1 | 0 | NA | NA | NA | NA | NA | Olfr891 | 0 | NA | NA | NA | NA | NA |
| Ssxb10 | 0 | NA | NA | NA | NA | NA | Olfr893 | 0 | NA | NA | NA | NA | NA |
| Ssxb2 | 0 | NA | NA | NA | NA | NA | Olfr894 | 0 | NA | NA | NA | NA | NA |
| Ssxb3 | 0 | NA | NA | NA | NA | NA | Olfr895 | 0 | NA | NA | NA | NA | NA |
| Ssxb5 | 0 | NA | NA | NA | NA | NA | Olfr899 | 0 | NA | NA | NA | NA | NA |
| Ssxb6 | 0.08731684 | 0.017365056 | 0.027733412 | 0.626142072 | 0.531221759 | NA | Olfr9 | 0 | NA | NA | NA | NA | NA |
| Ssxb8 | 0.192142086 | -0.026384592 | 0.031527717 | -0.836869715 | 0.402665805 | NA | Olfr90 | 0 | NA | NA | NA | NA | NA |
| Ssxb9 | 0.225320798 | -0.001646023 | 0.032098875 | -0.051279768 | 0.95910259 | NA | Olfr900 | 0 | NA | NA | NA | NA | NA |
| St6galnac1 | 0.167300494 | 0.024299763 | 0.031030539 | 0.783091891 | 0.433573153 | NA | Olfr901 | 0 | NA | NA | NA | NA | NA |
| Stap1 | 1.094190362 | 0.00690199 | 0.071121416 | 0.097045169 | 0.922690524 | NA | Olfr902 | 0 | NA | NA | NA | NA | NA |
| Steap1 | 10.8170662 | 0.206857191 | 0.121166028 | 1.707221036 | NA | NA | Olfr904 | 0 | NA | NA | NA | NA | NA |
| Stfa1 | 0 | NA | NA | NA | NA | NA | Olfr905 | 0 | NA | NA | NA | NA | NA |
| Stfa2 | 0 | NA | NA | NA | NA | NA | Olfr906 | 0 | NA | NA | NA | NA | NA |
| Stfa2l1 | 0.188785634 | 0.025025111 | 0.031461603 | 0.79541754 | 0.42637066 | NA | Olfr907 | 0 | NA | NA | NA | NA | NA |
| Stfa3 | 0 | NA | NA | NA | NA | NA | Olfr908 | 0 | NA | NA | NA | NA | NA |
| Stk31 | 1.559446096 | 0.112133069 | 0.092909366 | 1.206908134 | 0.227467525 | NA | Olfr91 | 0 | NA | NA | NA | NA | NA |
| Stmn1-rs1 | 0.079983654 | 0.016740032 | 0.027674178 | 0.604897175 | 0.54524732 | NA | Olfr910 | 0 | NA | NA | NA | NA | NA |
| Stmnd1 | 6.707037581 | 0.368062238 | 0.150432828 | 2.446688291 | 0.014417547 | NA | Olfr911-ps1 | 0 | NA | NA | NA | NA | NA |
| Ston1 | 177.4926521 | 0.163835912 | 0.157622854 | 1.039417238 | NA | NA | Olfr912 | 0 | NA | NA | NA | NA | NA |
| Stpg2 | 2.114023909 | 0.045308749 | 0.097303285 | 0.4656446 | 0.641469913 | NA | Olfr913 | 0 | NA | NA | NA | NA | NA |
| Stra8 | 0.116696964 | -0.020477583 | 0.027954224 | -0.732539847 | 0.463839131 | NA | Olfr914 | 0 | NA | NA | NA | NA | NA |
| Stx19 | 0.192142086 | -0.026384592 | 0.031527717 | -0.836869715 | 0.402665805 | NA | Olfr915 | 0 | NA | NA | NA | NA | NA |
| Styxl1 | 5.612173477 | 0.051955929 | 0.138398518 | 0.375408132 | 0.707356958 | NA | Olfr916 | 0 | NA | NA | NA | NA | NA |
| Sucnr1 | 0.311993147 | -0.014421208 | 0.039860783 | -0.361789373 | 0.717509434 | NA | Olfr917 | 0 | NA | NA | NA | NA | NA |
| Sult1b1 | 0 | NA | NA | NA | NA | NA | Olfr918 | 0 | NA | NA | NA | NA | NA |
| Sult1c1 | 4.124745454 | 0.239600029 | 0.107349233 | 2.23196778 | 0.025617093 | NA | Olfr919 | 0 | NA | NA | NA | NA | NA |
| Sult1c2 | 3.003228312 | 0.177472826 | 0.086205761 | 2.058711889 | 0.039521847 | NA | Olfr92 | 0 | NA | NA | NA | NA | NA |
| Sult1d1 | 0.898933214 | -0.093259539 | 0.064431186 | -1.447428561 | 0.147776931 | NA | Olfr922 | 0 | NA | NA | NA | NA | NA |
| Sult1e1 | 0 | NA | NA | NA | NA | NA | Olfr923 | 0 | NA | NA | NA | NA | NA |
| Sult2a1 | 0 | NA | NA | NA | NA | NA | Olfr924 | 0 | NA | NA | NA | NA | NA |
| Sult2a2 | 0 | NA | NA | NA | NA | NA | Olfr926 | 0 | NA | NA | NA | NA | NA |
| Sult2a3 | 0 | NA | NA | NA | NA | NA | Olfr93 | 0 | NA | NA | NA | NA | NA |
| Sult2a4 | 0 | NA | NA | NA | NA | NA | Olfr930 | 0 | NA | NA | NA | NA | NA |
| Sult2a5 | 0.707463931 | 0.031633586 | 0.054543609 | 0.579968699 | 0.561935726 | NA | Olfr933 | 0 | NA | NA | NA | NA | NA |
| Sult2a6 | 0 | NA | NA | NA | NA | NA | Olfr934 | 0 | NA | NA | NA | NA | NA |
| Sult2a7 | 0 | NA | NA | NA | NA | NA | Olfr935 | 0 | NA | NA | NA | NA | NA |
| Sult3a1 | 0.105923158 | 0.018524693 | 0.027977518 | 0.662127842 | 0.507889294 | NA | Olfr936 | 0 | NA | NA | NA | NA | NA |
| Sult5a1 | 5.699542381 | 0.026347987 | 0.145060757 | 0.18163415 | 0.855869847 | NA | Olfr937 | 0 | NA | NA | NA | NA | NA |
| Sult6b1 | 1.912376999 | -0.065649506 | 0.100216088 | -0.655079509 | 0.512416564 | NA | Olfr938 | 0 | NA | NA | NA | NA | NA |
| Sun3 | 2.050813791 | 0.090764811 | 0.10590399 | 0.857048079 | 0.391418314 | NA | Olfr94 | 0 | NA | NA | NA | NA | NA |
| Sun5 | 0.096071043 | -0.020098795 | 0.027825948 | -0.722304036 | 0.470107572 | NA | Olfr943 | 0 | NA | NA | NA | NA | NA |
| Sva | 0 | NA | NA | NA | NA | NA | Olfr944 | 0 | NA | NA | NA | NA | NA |
| Sval1 | 0 | NA | NA | NA | NA | NA | Olfr945 | 0 | NA | NA | NA | NA | NA |
| Sval2 | 0 | NA | NA | NA | NA | NA | Olfr947-ps1 | 0 | NA | NA | NA | NA | NA |
| Sval3 | 0 | NA | NA | NA | NA | NA | Olfr948 | 0 | NA | NA | NA | NA | NA |
| Svopl | 0.957201742 | 0.038953393 | 0.065902215 | 0.591078666 | 0.554467713 | NA | Olfr951 | 0 | NA | NA | NA | NA | NA |
| Svs1 | 0 | NA | NA | NA | NA | NA | Olfr952 | 0 | NA | NA | NA | NA | NA |
| Svs2 | 0 | NA | NA | NA | NA | NA | Olfr954 | 0 | NA | NA | NA | NA | NA |
| Svs3a | 0.094392817 | 0.017895942 | 0.027818137 | 0.643319236 | 0.520016977 | NA | Olfr955 | 0 | NA | NA | NA | NA | NA |
| Svs3b | 0 | NA | NA | NA | NA | NA | Olfr957 | 0 | NA | NA | NA | NA | NA |
| Svs4 | 0 | NA | NA | NA | NA | NA | Olfr958 | 0 | NA | NA | NA | NA | NA |
| Svs5 | 0 | NA | NA | NA | NA | NA | Olfr959 | 0 | NA | NA | NA | NA | NA |
| Svs6 | 0 | NA | NA | NA | NA | NA | Olfr96 | 0 | NA | NA | NA | NA | NA |
| Syce1 | 5.775800117 | 0.047463816 | 0.142759344 | 0.332474325 | 0.739531124 | NA | Olfr960 | 0 | NA | NA | NA | NA | NA |
| Syce1l | 4.444955211 | 0.146339423 | 0.128739573 | 1.136708939 | 0.255659981 | NA | Olfr961 | 0 | NA | NA | NA | NA | NA |
| Syce3 | 0.314431083 | -0.017921647 | 0.039992366 | -0.448126696 | 0.654061762 | NA | Olfr963 | 0 | NA | NA | NA | NA | NA |
| Sycn | 0 | NA | NA | NA | NA | NA | Olfr965 | 0 | NA | NA | NA | NA | NA |
| Sycp1 | 0.37239624 | 0.052452486 | 0.04522608 | 1.159784039 | 0.246136744 | NA | Olfr967 | 0 | NA | NA | NA | NA | NA |
| Sycp1-ps1 | 0 | NA | NA | NA | NA | NA | Olfr968 | 0 | NA | NA | NA | NA | NA |
| Sycp3 | 6.175903059 | 0.070594094 | 0.138200365 | 0.510809751 | 0.60948428 | NA | Olfr969 | 0 | NA | NA | NA | NA | NA |
| Synb | 2.827455305 | 0.171654527 | 0.099670056 | 1.722227658 | 0.085028286 | NA | Olfr97 | 0 | NA | NA | NA | NA | NA |
| Syne1 | 19737.15193 | -0.169729761 | 0.097243582 | -1.745408354 | NA | NA | Olfr971 | 0 | NA | NA | NA | NA | NA |
| Syngr4 | 2.701203929 | 0.102966466 | 0.114471088 | 0.899497578 | 0.368387685 | NA | Olfr972 | 0 | NA | NA | NA | NA | NA |
| Syt8 | 0.607073637 | 0.043174474 | 0.049620922 | 0.870086075 | 0.384253367 | NA | Olfr974 | 0 | NA | NA | NA | NA | NA |
| T | 2.95375557 | 0.132193287 | 0.110911914 | 1.191876349 | 0.233309733 | NA | Olfr975 | 0 | NA | NA | NA | NA | NA |
| Taar1 | 0 | NA | NA | NA | NA | NA | Olfr976 | 0 | NA | NA | NA | NA | NA |
| Taar2 | 0.116696964 | -0.020477583 | 0.027954224 | -0.732539847 | 0.463839131 | NA | Olfr978 | 0 | NA | NA | NA | NA | NA |
| Taar3 | 0.305736371 | 0.008542183 | 0.033941149 | 0.251676297 | 0.80129128 | NA | Olfr979 | 0 | NA | NA | NA | NA | NA |
| Taar4 | 0 | NA | NA | NA | NA | NA | Olfr98 | 0 | NA | NA | NA | NA | NA |
| Taar5 | 0.094392817 | 0.017895942 | 0.027818137 | 0.643319236 | 0.520016977 | NA | Olfr980 | 0 | NA | NA | NA | NA | NA |
| Taar6 | 0 | NA | NA | NA | NA | NA | Olfr981 | 0 | NA | NA | NA | NA | NA |
| Taar7a | 0 | NA | NA | NA | NA | NA | Olfr982 | 0 | NA | NA | NA | NA | NA |
| Taar7b | 0 | NA | NA | NA | NA | NA | Olfr983 | 0 | NA | NA | NA | NA | NA |
| Taar7d | 0 | NA | NA | NA | NA | NA | Olfr984 | 0 | NA | NA | NA | NA | NA |
| Taar7e | 0 | NA | NA | NA | NA | NA | Olfr985 | 0 | NA | NA | NA | NA | NA |
| Taar7f | 0 | NA | NA | NA | NA | NA | Olfr986 | 0 | NA | NA | NA | NA | NA |
| Taar8a | 0 | NA | NA | NA | NA | NA | Olfr987 | 0 | NA | NA | NA | NA | NA |
| Taar8b | 0 | NA | NA | NA | NA | NA | Olfr988 | 0 | NA | NA | NA | NA | NA |
| Taar8c | 0 | NA | NA | NA | NA | NA | Olfr99 | 0 | NA | NA | NA | NA | NA |
| Taar9 | 0 | NA | NA | NA | NA | NA | Olfr992 | 0 | NA | NA | NA | NA | NA |
| Tac4 | 0.783678328 | -0.040057892 | 0.068613333 | -0.583820817 | 0.559340871 | NA | Olfr993 | 0 | NA | NA | NA | NA | NA |
| Tacstd2 | 5.654824625 | -0.017623005 | 0.145771708 | -0.120894548 | 0.903774561 | NA | Olfr994 | 0 | NA | NA | NA | NA | NA |
| Taf7l | 3.465168167 | -0.270614826 | 0.115410054 | -2.344811532 | 0.0190367 | NA | Olfr995 | 0 | NA | NA | NA | NA | NA |
| Tal2 | 3.541156078 | 0.236510616 | 0.121988916 | 1.938787752 | 0.052527188 | NA | Olfr996 | 0 | NA | NA | NA | NA | NA |
| Tap1 | 175.3532398 | 0.407157652 | 0.164556707 | 2.474269568 | NA | NA | Olfr998 | 0 | NA | NA | NA | NA | NA |
| Tarm1 | 0.858186872 | 0.089817948 | 0.05829794 | 1.54067106 | 0.123396864 | NA | Omt2a | 0 | NA | NA | NA | NA | NA |
| Tas1r1 | 4.508613947 | 0.036943638 | 0.128455738 | 0.28759819 | 0.773654331 | NA | Omt2b | 0 | NA | NA | NA | NA | NA |
| Tas1r2 | 0.181709657 | 0.024794574 | 0.031325423 | 0.791516008 | 0.428642938 | NA | Oog1 | 0 | NA | NA | NA | NA | NA |
| Tas2r102 | 0 | NA | NA | NA | NA | NA | Oog2 | 0 | NA | NA | NA | NA | NA |
| Tas2r103 | 0.079983654 | 0.016740032 | 0.027674178 | 0.604897175 | 0.54524732 | NA | Oog3 | 0 | NA | NA | NA | NA | NA |
| Tas2r104 | 0 | NA | NA | NA | NA | NA | Oog4 | 0 | NA | NA | NA | NA | NA |
| Tas2r105 | 0 | NA | NA | NA | NA | NA | Orm2 | 0 | NA | NA | NA | NA | NA |
| Tas2r106 | 0 | NA | NA | NA | NA | NA | Ott | 0 | NA | NA | NA | NA | NA |
| Tas2r107 | 0 | NA | NA | NA | NA | NA | Otud6a | 0 | NA | NA | NA | NA | NA |
| Tas2r108 | 0.631566968 | 0.019997227 | 0.053874878 | 0.371179071 | 0.71050416 | NA | Ovch2 | 0 | NA | NA | NA | NA | NA |
| Tas2r109 | 0 | NA | NA | NA | NA | NA | Pabpc2 | 0 | NA | NA | NA | NA | NA |
| Tas2r110 | 0 | NA | NA | NA | NA | NA | Pabpc6 | 0 | NA | NA | NA | NA | NA |
| Tas2r113 | 0 | NA | NA | NA | NA | NA | Panx3 | 0 | NA | NA | NA | NA | NA |
| Tas2r114 | 0 | NA | NA | NA | NA | NA | Pate4 | 0 | NA | NA | NA | NA | NA |
| Tas2r115 | 0 | NA | NA | NA | NA | NA | Pbsn | 0 | NA | NA | NA | NA | NA |
| Tas2r116 | 0 | NA | NA | NA | NA | NA | Pcdha4-g | 0 | NA | NA | NA | NA | NA |
| Tas2r117 | 0 | NA | NA | NA | NA | NA | Pdha2 | 0 | NA | NA | NA | NA | NA |
| Tas2r118 | 0 | NA | NA | NA | NA | NA | Pdx1 | 0 | NA | NA | NA | NA | NA |
| Tas2r119 | 0 | NA | NA | NA | NA | NA | Pea15b | 0 | NA | NA | NA | NA | NA |
| Tas2r120 | 0 | NA | NA | NA | NA | NA | Pet117 | 0 | NA | NA | NA | NA | NA |
| Tas2r121 | 0 | NA | NA | NA | NA | NA | Pet2 | 0 | NA | NA | NA | NA | NA |
| Tas2r122 | 0 | NA | NA | NA | NA | NA | Pga5 | 0 | NA | NA | NA | NA | NA |
| Tas2r123 | 0 | NA | NA | NA | NA | NA | Pgk2 | 0 | NA | NA | NA | NA | NA |
| Tas2r124 | 0 | NA | NA | NA | NA | NA | Pglyrp4 | 0 | NA | NA | NA | NA | NA |
| Tas2r125 | 0 | NA | NA | NA | NA | NA | Phgr1 | 0 | NA | NA | NA | NA | NA |
| Tas2r126 | 0 | NA | NA | NA | NA | NA | Phox2b | 0 | NA | NA | NA | NA | NA |
| Tas2r129 | 0 | NA | NA | NA | NA | NA | Phxr4 | 0 | NA | NA | NA | NA | NA |
| Tas2r130 | 0 | NA | NA | NA | NA | NA | Pih1d3 | 0 | NA | NA | NA | NA | NA |
| Tas2r131 | 0 | NA | NA | NA | NA | NA | Pira1 | 0 | NA | NA | NA | NA | NA |
| Tas2r134 | 0 | NA | NA | NA | NA | NA | Pira11 | 0 | NA | NA | NA | NA | NA |
| Tas2r135 | 0.254360125 | 0.029364299 | 0.033967282 | 0.864487756 | 0.387320003 | NA | Pira4 | 0 | NA | NA | NA | NA | NA |
| Tas2r136 | 0 | NA | NA | NA | NA | NA | Pira7 | 0 | NA | NA | NA | NA | NA |
| Tas2r137 | 0.725227826 | -0.022946998 | 0.065969169 | -0.347844275 | 0.727957137 | NA | Pisd-ps1 | 0 | NA | NA | NA | NA | NA |
| Tas2r138 | 0 | NA | NA | NA | NA | NA | Pisd-ps3 | 0 | NA | NA | NA | NA | NA |
| Tas2r139 | 0 | NA | NA | NA | NA | NA | Pla2g10 | 0 | NA | NA | NA | NA | NA |
| Tas2r140 | 0 | NA | NA | NA | NA | NA | Pla2g12b | 0 | NA | NA | NA | NA | NA |
| Tas2r143 | 0.293774112 | 0.008475859 | 0.039057974 | 0.217007122 | 0.828202795 | NA | Pla2g1b | 0 | NA | NA | NA | NA | NA |
| Tas2r144 | 0 | NA | NA | NA | NA | NA | Pla2g2e | 0 | NA | NA | NA | NA | NA |
| Tat | 1.611244111 | 0.079196092 | 0.091588384 | 0.864695809 | 0.38720577 | NA | Plac9a | 0 | NA | NA | NA | NA | NA |
| Tbc1d21 | 0 | NA | NA | NA | NA | NA | Plac9b | 0 | NA | NA | NA | NA | NA |
| Tbpl2 | 0 | NA | NA | NA | NA | NA | Pom121l12 | 0 | NA | NA | NA | NA | NA |

| | | | | | | | | | | | | | | | |
|---|---|---|---|---|---|---|---|---|---|---|---|---|---|---|---|
| Tbrg3 | 0.494745862 | -0.017414199 | 0.05079001 | -0.342866614 | 0.73169881 | NA | | | Poteg | 0 | NA | NA | NA | NA | NA |
| Tbx10 | 0.47124342 | 0.077409868 | 0.054708132 | 1.414960895 | 0.157079962 | NA | | | Pou5f2 | 0 | NA | NA | NA | NA | NA |
| Tbx19 | 1.19880358 | -0.039255453 | 0.076895544 | -0.510503613 | 0.609698684 | NA | | | Ppp1r2-ps7 | 0 | NA | NA | NA | NA | NA |
| Tbx20 | 0.656506615 | -0.019769438 | 0.054376061 | -0.363568787 | 0.716180038 | NA | | | Ppp1r2-ps9 | 0 | NA | NA | NA | NA | NA |
| Tbx22 | 6.96165136 | -0.312816501 | 0.152173296 | -2.055659632 | 0.039815336 | NA | | | Ppp1r27 | 0 | NA | NA | NA | NA | NA |
| Tbx5 | 0.079983654 | 0.016740032 | 0.027674178 | 0.604897175 | 0.54524732 | NA | | | Ppy | 0 | NA | NA | NA | NA | NA |
| Tc2n | 4.428694762 | 0.144517302 | 0.086317465 | 1.674253308 | 0.094080827 | NA | | | Pramef12 | 0 | NA | NA | NA | NA | NA |
| Tcam1 | 7.072622623 | 0.138219208 | 0.146330259 | 0.944570241 | 0.344878326 | NA | | | Pramef17 | 0 | NA | NA | NA | NA | NA |
| Tcf21 | 0.08731684 | 0.017365056 | 0.027733412 | 0.626142072 | 0.531221759 | NA | | | Pramef6 | 0 | NA | NA | NA | NA | NA |
| Tchhl1 | 0.487806363 | -0.025218575 | 0.039384805 | -0.640312313 | 0.521969578 | NA | | | Pramel4 | 0 | NA | NA | NA | NA | NA |
| Tcl1 | 0 | NA | NA | NA | NA | NA | | | Pramel5 | 0 | NA | NA | NA | NA | NA |
| Tcl1b1 | 0.159967308 | 0.0240345 | 0.030870969 | 0.778547006 | 0.436246606 | NA | | | Pramel6 | 0 | NA | NA | NA | NA | NA |
| Tcl1b2 | 0.904023182 | 0.00650544 | 0.074708519 | 0.087077624 | 0.930609811 | NA | | | Pramel7 | 0 | NA | NA | NA | NA | NA |
| Tcl1b3 | 0 | NA | NA | NA | NA | NA | | | Prb1 | 0 | NA | NA | NA | NA | NA |
| Tcl1b4 | 0 | NA | NA | NA | NA | NA | | | Prg3 | 0 | NA | NA | NA | NA | NA |
| Tcl1b5 | 0 | NA | NA | NA | NA | NA | | | Prh1 | 0 | NA | NA | NA | NA | NA |
| Tcp10a | 0.406927243 | 0.024331845 | 0.043361066 | 0.561144991 | 0.574698699 | NA | | | Prhoxnb | 0 | NA | NA | NA | NA | NA |
| Tcp10b | 4.719536598 | 0.079476574 | 0.120586515 | 0.659083432 | 0.509842194 | NA | | | Prl2a1 | 0 | NA | NA | NA | NA | NA |
| Tcp10c | 0.487454995 | 0.031555684 | 0.045749554 | 0.689748452 | 0.49035239 | NA | | | Prl2b1 | 0 | NA | NA | NA | NA | NA |
| Tcte3 | 0 | NA | NA | NA | NA | NA | | | Prl2c1 | 0 | NA | NA | NA | NA | NA |
| Tdg | 2.028800312 | 0.008983561 | 0.08875971 | 0.101212147 | 0.919382055 | NA | | | Prl2c2 | 0 | NA | NA | NA | NA | NA |
| Tdh | 1.156995207 | -0.018693609 | 0.079249747 | -0.23588226 | 0.813524034 | NA | | | Prl2c3 | 0 | NA | NA | NA | NA | NA |
| Tdpoz1 | 0 | NA | NA | NA | NA | NA | | | Prl2c4 | 0 | NA | NA | NA | NA | NA |
| Tdpoz2 | 0 | NA | NA | NA | NA | NA | | | Prl2c5 | 0 | NA | NA | NA | NA | NA |
| Tdpoz3 | 0.096071043 | -0.020098795 | 0.027825948 | -0.722304036 | 0.470107572 | NA | | | Prl3a1 | 0 | NA | NA | NA | NA | NA |
| Tdpoz4 | 0 | NA | NA | NA | NA | NA | | | Prl3b1 | 0 | NA | NA | NA | NA | NA |
| Tdpoz5 | 0 | NA | NA | NA | NA | NA | | | Prl3c1 | 0 | NA | NA | NA | NA | NA |
| Tecrl | 6.731038187 | 0.354808408 | 0.14026961 | 2.529474539 | 0.011423346 | NA | | | Prl3d1 | 0 | NA | NA | NA | NA | NA |
| Tectb | 3.84552122 | 0.021939578 | 0.126013754 | 0.174104628 | 0.861783236 | NA | | | Prl3d2 | 0 | NA | NA | NA | NA | NA |
| Teddm1 | 0 | NA | NA | NA | NA | NA | | | Prl3d3 | 0 | NA | NA | NA | NA | NA |
| Tekt3 | 2.135990805 | 0.130476131 | 0.106453419 | 1.225664071 | 0.220325116 | NA | | | Prl4a1 | 0 | NA | NA | NA | NA | NA |
| Tepp | 3.447898974 | 0.095804654 | 0.109689984 | 0.873412965 | 0.382438027 | NA | | | Prl5a1 | 0 | NA | NA | NA | NA | NA |
| Terc | 0.729646711 | -0.019163599 | 0.061280247 | -0.312720663 | 0.754492896 | NA | | | Prl6a1 | 0 | NA | NA | NA | NA | NA |
| Tescl | 0.105923158 | 0.018524693 | 0.027977518 | 0.662127842 | 0.507889294 | NA | | | Prl7a1 | 0 | NA | NA | NA | NA | NA |
| Tespa1 | 0.284260198 | 0.011273774 | 0.038754816 | 0.290899955 | 0.771127836 | NA | | | Prl7a2 | 0 | NA | NA | NA | NA | NA |
| Tex101 | 0 | NA | NA | NA | NA | NA | | | Prl7b1 | 0 | NA | NA | NA | NA | NA |
| Tex12 | 0 | NA | NA | NA | NA | NA | | | Prl7c1 | 0 | NA | NA | NA | NA | NA |
| Tex13 | 0.098353386 | -0.020212831 | 0.02786466 | -0.725393061 | 0.468210929 | NA | | | Prl7d1 | 0 | NA | NA | NA | NA | NA |
| Tex13a | 0 | NA | NA | NA | NA | NA | | | Prl8a1 | 0 | NA | NA | NA | NA | NA |
| Tex16 | 0.272987066 | 0.007834485 | 0.033279985 | 0.2354113 | 0.813889516 | NA | | | Prl8a2 | 0 | NA | NA | NA | NA | NA |
| Tex19.1 | 0 | NA | NA | NA | NA | NA | | | Prl8a6 | 0 | NA | NA | NA | NA | NA |
| Tex19.2 | 0.099009884 | 0.018205726 | 0.027896865 | 0.652608315 | 0.514008823 | NA | | | Prl8a8 | 0 | NA | NA | NA | NA | NA |
| Tex21 | 0.320973508 | -0.015129019 | 0.040159764 | -0.376720802 | 0.706381102 | NA | | | Prl8a9 | 0 | NA | NA | NA | NA | NA |
| Tex22 | 0.913014257 | -0.020655493 | 0.069611272 | -0.296726267 | 0.766675501 | NA | | | Prlh | 0 | NA | NA | NA | NA | NA |
| Tex24 | 0.105923158 | 0.018524693 | 0.027977518 | 0.662127842 | 0.507889294 | NA | | | Prm1 | 0 | NA | NA | NA | NA | NA |
| Tex28 | 0 | NA | NA | NA | NA | NA | | | Prm3 | 0 | NA | NA | NA | NA | NA |
| Tex29 | 0.573334654 | -0.00282487 | 0.052673623 | -0.053629686 | 0.957230204 | NA | | | Prodh2 | 0 | NA | NA | NA | NA | NA |
| Tex33 | 1.096528261 | 0.001265473 | 0.079649808 | 0.015887959 | 0.987323776 | NA | | | Prol1 | 0 | NA | NA | NA | NA | NA |
| Tex35 | 1.371065569 | -0.028309733 | 0.086162026 | -0.328563914 | 0.742485327 | NA | | | Prpmp5 | 0 | NA | NA | NA | NA | NA |
| Tex36 | 0.453998628 | 0.026589972 | 0.045454031 | 0.584986 | 0.55855713 | NA | | | Prps1l1 | 0 | NA | NA | NA | NA | NA |
| Tex37 | 0 | NA | NA | NA | NA | NA | | | Prss1 | 0 | NA | NA | NA | NA | NA |
| Tex38 | 4.355556834 | -0.059106171 | 0.136627621 | -0.432607776 | 0.665299739 | NA | | | Prss2 | 0 | NA | NA | NA | NA | NA |
| Tfap2a | 6.742375519 | -0.037887232 | 0.151565133 | -0.24997327 | 0.80260802 | NA | | | Prss21 | 0 | NA | NA | NA | NA | NA |
| Tfap2c | 7.507192167 | -0.203723485 | 0.140626394 | -1.448685976 | 0.147425299 | NA | | | Prss28 | 0 | NA | NA | NA | NA | NA |
| Tfap2e | 0.272729858 | 0.010322181 | 0.038323464 | 0.26934363 | 0.78766526 | NA | | | Prss37 | 0 | NA | NA | NA | NA | NA |
| Tfec | 1.883608677 | -0.038493707 | 0.105120215 | -0.36618748 | 0.714225184 | NA | | | Prss39 | 0 | NA | NA | NA | NA | NA |
| Tff1 | 0 | NA | NA | NA | NA | NA | | | Prss40 | 0 | NA | NA | NA | NA | NA |
| Tff2 | 0 | NA | NA | NA | NA | NA | | | Prss48 | 0 | NA | NA | NA | NA | NA |
| Tff3 | 0 | NA | NA | NA | NA | NA | | | Prss51 | 0 | NA | NA | NA | NA | NA |
| Tfpi2 | 1.029540879 | -0.073577205 | 0.078163571 | -0.941323481 | 0.346539113 | NA | | | Prss52 | 0 | NA | NA | NA | NA | NA |
| Tg | 7.738540515 | 0.082199922 | 0.150480033 | 0.546251359 | 0.584893167 | NA | | | Prss58 | 0 | NA | NA | NA | NA | NA |
| Tgif2lx1 | 0 | NA | NA | NA | NA | NA | | | Psg-ps1 | 0 | NA | NA | NA | NA | NA |
| Tgif2lx2 | 0 | NA | NA | NA | NA | NA | | | Psg18 | 0 | NA | NA | NA | NA | NA |
| Tgm1 | 4.589164824 | 0.094308232 | 0.124129342 | 0.759757765 | 0.447399393 | NA | | | Psg20 | 0 | NA | NA | NA | NA | NA |
| Tgm5 | 0.505937128 | 0.03385261 | 0.046683903 | 0.725145244 | 0.46836293 | NA | | | Psg21 | 0 | NA | NA | NA | NA | NA |
| Tgm6 | 7.479841179 | -0.135278741 | 0.1533584 | -0.882108458 | 0.377718159 | NA | | | Psg25 | 0 | NA | NA | NA | NA | NA |
| Tgoln2 | 0 | NA | NA | NA | NA | NA | | | Psg26 | 0 | NA | NA | NA | NA | NA |
| Tgtp1 | 0 | NA | NA | NA | NA | NA | | | Psg27 | 0 | NA | NA | NA | NA | NA |
| Them5 | 0.463017151 | 0.009204469 | 0.04998486 | 0.184145136 | 0.853899599 | NA | | | Psg28 | 0 | NA | NA | NA | NA | NA |
| Themis | 0.674726373 | 0.035983769 | 0.061458617 | 0.585495914 | 0.558214313 | NA | | | Ptchd3 | 0 | NA | NA | NA | NA | NA |
| Themis3 | 0.63671097 | 0.032300813 | 0.060136437 | 0.537125483 | 0.591180939 | NA | | | Raet1b | 0 | NA | NA | NA | NA | NA |
| Tigd4 | 2.461664092 | 0.017037577 | 0.108471884 | 0.157069061 | 0.875190421 | NA | | | Raet1c | 0 | NA | NA | NA | NA | NA |
| Tigit | 0.400129188 | 0.02188733 | 0.04314476 | 0.507299852 | 0.611944442 | NA | | | Raet1e | 0 | NA | NA | NA | NA | NA |
| Timd2 | 0.400720577 | -0.028041581 | 0.043551246 | -0.643875506 | 0.519656166 | NA | | | Rag2 | 0 | NA | NA | NA | NA | NA |
| Timd4 | 2.710522476 | 0.14887368 | 0.112577536 | 1.322410185 | 0.186031599 | NA | | | Raver1-fdx1l | 0 | NA | NA | NA | NA | NA |
| Timm8a2 | 0.176054697 | -0.002190495 | 0.031239146 | -0.070120197 | 0.944097991 | NA | | | Rbbp8nl | 0 | NA | NA | NA | NA | NA |
| Tinag | 1.018815137 | 0.073477377 | 0.081170573 | 0.905221854 | 0.365347867 | NA | | | Rbmy | 0 | NA | NA | NA | NA | NA |
| Tktl1 | 3.655482423 | 0.05044423 | 0.125631359 | 0.401525783 | 0.68803306 | NA | | | Rdh7 | 0 | NA | NA | NA | NA | NA |
| Tktl2 | 6.943658428 | -0.559927057 | 0.146636401 | -3.818472457 | 0.000134281 | NA | | | Rdh8 | 0 | NA | NA | NA | NA | NA |
| Tlr11 | 0.201994201 | 0.000227241 | 0.031727614 | 0.00716224 | 0.994285408 | NA | | | Reg1 | 0 | NA | NA | NA | NA | NA |
| Tlr8 | 6.668945966 | -0.181774026 | 0.140074093 | -1.297699115 | 0.194390748 | NA | | | Reg2 | 0 | NA | NA | NA | NA | NA |
| Tlx1 | 0 | NA | NA | NA | NA | NA | | | Reg3d | 0 | NA | NA | NA | NA | NA |
| Tlx2 | 0 | NA | NA | NA | NA | NA | | | Ren1 | 0 | NA | NA | NA | NA | NA |
| Tlx3 | 0 | NA | NA | NA | NA | NA | | | Ren2 | 0 | NA | NA | NA | NA | NA |
| Tm4sf19 | 1.642775276 | 0.067198221 | 0.100893972 | 0.666028104 | 0.505393137 | NA | | | Retnlb | 0 | NA | NA | NA | NA | NA |
| Tm4sf20 | 0 | NA | NA | NA | NA | NA | | | Rex2 | 0 | NA | NA | NA | NA | NA |
| Tm4sf4 | 0.55073328 | 0.049633073 | 0.054814118 | 0.905479733 | 0.365211292 | NA | | | Rfpl3s | 0 | NA | NA | NA | NA | NA |
| Tmc1 | 7.74719123 | 0.208245389 | 0.146201061 | 1.424376731 | 0.154337446 | NA | | | Rfpl4 | 0 | NA | NA | NA | NA | NA |
| Tmc2 | 0.291593384 | 0.011931959 | 0.039023223 | 0.305765587 | 0.759783132 | NA | | | Rfpl4b | 0 | NA | NA | NA | NA | NA |
| Tmc5 | 5.999963146 | 0.112682217 | 0.126481366 | 0.890899749 | 0.372982955 | NA | | | Rhox1 | 0 | NA | NA | NA | NA | NA |
| Tmco2 | 0.837310763 | -0.033554311 | 0.064018926 | -0.524131119 | 0.600187346 | NA | | | Rhox10 | 0 | NA | NA | NA | NA | NA |
| Tmco5b | 2.087200257 | 0.036995967 | 0.104147049 | 0.355228187 | 0.72241865 | NA | | | Rhox11 | 0 | NA | NA | NA | NA | NA |
| Tmed11 | 0 | NA | NA | NA | NA | NA | | | Rhox12 | 0 | NA | NA | NA | NA | NA |
| Tmed6 | 0.105923158 | 0.018524693 | 0.027977518 | 0.662127842 | 0.507889294 | NA | | | Rhox13 | 0 | NA | NA | NA | NA | NA |
| Tmem102 | 3.998689575 | 0.03048839 | 0.107871677 | 0.282635732 | 0.777456087 | NA | | | Rhox2a | 0 | NA | NA | NA | NA | NA |
| Tmem139 | 0 | NA | NA | NA | NA | NA | | | Rhox2b | 0 | NA | NA | NA | NA | NA |
| Tmem171 | 4.538185253 | 0.227651133 | 0.129760802 | 1.754390608 | 0.0793636 | NA | | | Rhox2c | 0 | NA | NA | NA | NA | NA |
| Tmem181c-ps | 0.206069989 | -0.026776453 | 0.031767726 | -0.842882251 | 0.399294301 | NA | | | Rhox2d | 0 | NA | NA | NA | NA | NA |
| Tmem182 | 3.632067248 | -0.316780394 | 0.122653805 | -2.582719657 | 0.009802494 | NA | | | Rhox2e | 0 | NA | NA | NA | NA | NA |
| Tmem184a | 4.528773212 | 0.212630843 | 0.110302079 | 1.927713838 | 0.05389073 | NA | | | Rhox2f | 0 | NA | NA | NA | NA | NA |
| Tmem190 | 0 | NA | NA | NA | NA | NA | | | Rhox2g | 0 | NA | NA | NA | NA | NA |
| Tmem207 | 0 | NA | NA | NA | NA | NA | | | Rhox2h | 0 | NA | NA | NA | NA | NA |
| Tmem210 | 0.283178451 | 0.028390488 | 0.033368195 | 0.850824814 | 0.394866675 | NA | | | Rhox3a | 0 | NA | NA | NA | NA | NA |
| Tmem211 | 1.154859085 | 0.086358106 | 0.072904567 | 1.184536298 | 0.236200836 | NA | | | Rhox3c | 0 | NA | NA | NA | NA | NA |
| Tmem213 | 0.116696964 | -0.020477583 | 0.027954224 | -0.732539847 | 0.463839131 | NA | | | Rhox3e | 0 | NA | NA | NA | NA | NA |
| Tmem217 | 1.887823986 | 0.002348191 | 0.084237717 | 0.027875769 | 0.977761235 | NA | | | Rhox3f | 0 | NA | NA | NA | NA | NA |
| Tmem221 | 7.859498067 | 0.085292255 | 0.156803724 | 0.543942789 | 0.586480842 | NA | | | Rhox3g-ps | 0 | NA | NA | NA | NA | NA |
| Tmem225 | 0 | NA | NA | NA | NA | NA | | | Rhox3h | 0 | NA | NA | NA | NA | NA |
| Tmem233 | 0 | NA | NA | NA | NA | NA | | | Rhox4b | 0 | NA | NA | NA | NA | NA |
| Tmem235 | 0.99394839 | 0.049346079 | 0.073902494 | 0.667718726 | 0.504313154 | NA | | | Rhox4c | 0 | NA | NA | NA | NA | NA |
| Tmem239 | 0.094392817 | 0.017895942 | 0.027818137 | 0.643319236 | 0.520016977 | NA | | | Rhox4d | 0 | NA | NA | NA | NA | NA |
| Tmem247 | 0 | NA | NA | NA | NA | NA | | | Rhox4e | 0 | NA | NA | NA | NA | NA |
| Tmem252 | 47.3771998 | 0.204360485 | 0.13008139 | 1.571020154 | NA | NA | | | Rhox4f | 0 | NA | NA | NA | NA | NA |
| Tmem253 | 4.460908769 | 0.063004086 | 0.110064042 | 0.572431143 | 0.567029923 | NA | | | Rhox6 | 0 | NA | NA | NA | NA | NA |
| Tmem254a | 0 | NA | NA | NA | NA | NA | | | Rhox7 | 0 | NA | NA | NA | NA | NA |
| Tmem254b | 0 | NA | NA | NA | NA | NA | | | Rhox9 | 0 | NA | NA | NA | NA | NA |
| Tmem254c | 0 | NA | NA | NA | NA | NA | | | Rnase11 | 0 | NA | NA | NA | NA | NA |
| Tmem27 | 4.323201814 | 0.210631168 | 0.120882324 | 1.742448037 | 0.081430077 | NA | | | Rnase9 | 0 | NA | NA | NA | NA | NA |
| Tmem30b | 3.576165431 | 0.007358545 | 0.115199793 | 0.06387637 | 0.949068668 | NA | | | Rnaset2a | 0 | NA | NA | NA | NA | NA |
| Tmem30c | 7.202564556 | 0.17144171 | 0.149584489 | 1.14611957 | 0.251745681 | NA | | | Rpl10l | 0 | NA | NA | NA | NA | NA |
| Tmem45b | 1.274413075 | -0.036644497 | 0.07236703 | -0.506370065 | 0.612596885 | NA | | | Rpl26 | 0 | NA | NA | NA | NA | NA |
| Tmem51as1 | 3.567674565 | 0.150962731 | 0.130016648 | 1.161103083 | 0.245599981 | NA | | | Rpl34-ps1 | 0 | NA | NA | NA | NA | NA |
| Tmem52b | 0.099009884 | 0.018205726 | 0.027896865 | 0.652608315 | 0.514008823 | NA | | | Rplp2-ps1 | 0 | NA | NA | NA | NA | NA |
| Tmem54 | 2.143751428 | -0.021525632 | 0.102567875 | -0.209867196 | 0.833771326 | NA | | | Rprl1 | 0 | NA | NA | NA | NA | NA |
| Tmem72 | 24.87395896 | 0.215595491 | 0.110048562 | 1.959094128 | NA | NA | | | Rprl2 | 0 | NA | NA | NA | NA | NA |
| Tmem89 | 1.997515811 | -0.080396475 | 0.102131164 | -0.78718847 | 0.431171546 | NA | | | Rprl3 | 0 | NA | NA | NA | NA | NA |
| Tmem92 | 0.079983654 | 0.016740032 | 0.027674178 | 0.604897175 | 0.54524732 | NA | | | Rxfp4 | 0 | NA | NA | NA | NA | NA |
| Tmem95 | 1.153741678 | -0.001538765 | 0.058017103 | -0.026522614 | 0.978840496 | NA | | | S100g | 0 | NA | NA | NA | NA | NA |
| Tmigd1 | 0.119397641 | -0.020477583 | 0.027954224 | -0.732539845 | 0.463839132 | NA | | | Saa4 | 0 | NA | NA | NA | NA | NA |
| Tmod4 | 7.563143088 | 0.054579957 | 0.142124937 | 0.384028009 | 0.700957688 | NA | | | Samt1 | 0 | NA | NA | NA | NA | NA |
| Tmprss11a | 5.172311587 | 0.185875555 | 0.102622617 | 1.811253316 | 0.070101649 | NA | | | Samt2 | 0 | NA | NA | NA | NA | NA |
| Tmprss11bnl | 0 | NA | NA | NA | NA | NA | | | Samt3 | 0 | NA | NA | NA | NA | NA |
| Tmprss11c | 0 | NA | NA | NA | NA | NA | | | Samt4 | 0 | NA | NA | NA | NA | NA |
| Tmprss11d | 0.19442443 | -0.026451352 | 0.031568725 | -0.837897386 | 0.402088337 | NA | | | Sap25 | 0 | NA | NA | NA | NA | NA |
| Tmprss11e | 0.116696964 | -0.020477583 | 0.027954224 | -0.732539847 | 0.463839131 | NA | | | Satl1 | 0 | NA | NA | NA | NA | NA |
| Tmprss11f | 0.176054697 | -0.002190495 | 0.031239146 | -0.070120197 | 0.944097991 | NA | | | Sbp | 0 | NA | NA | NA | NA | NA |
| Tmprss11g | 0 | NA | NA | NA | NA | NA | | | Sbpl | 0 | NA | NA | NA | NA | NA |
| Tmprss12 | 0 | NA | NA | NA | NA | NA | | | Scarna2 | 0 | NA | NA | NA | NA | NA |
| Tmprss13 | 0.512850401 | 0.033489004 | 0.046267694 | 0.723809661 | 0.469182597 | NA | | | Scarna3a | 0 | NA | NA | NA | NA | NA |
| Tmprss15 | 0.871501254 | -0.017336007 | 0.064315405 | -0.269546729 | 0.787508988 | NA | | | Scarna3b | 0 | NA | NA | NA | NA | NA |
| Tmprss2 | 1.442318935 | 0.123906339 | 0.090922286 | 1.362771921 | 0.172954403 | NA | | | Scgb1a1 | 0 | NA | NA | NA | NA | NA |
| Tmprss3 | 1.795150573 | 0.108706306 | 0.096860485 | 1.122297767 | 0.261735855 | NA | | | Scgb1b20 | 0 | NA | NA | NA | NA | NA |
| Tmsb15a | 0 | NA | NA | NA | NA | NA | | | Scgb1b24 | 0 | NA | NA | NA | NA | NA |
| Tmsb15b2 | 1.342234934 | 0.003684634 | 0.084180385 | 0.043770692 | 0.965087189 | NA | | | Scgb1b27 | 0 | NA | NA | NA | NA | NA |
| Tmsb15l | 0 | NA | NA | NA | NA | NA | | | Scgb1b29 | 0 | NA | NA | NA | NA | NA |
| Tnf | 2.607046118 | 0.104075028 | 0.112160005 | 0.927915685 | 0.353451308 | NA | | | Scgb1b3 | 0 | NA | NA | NA | NA | NA |
| Tnfrsf13b | 4.072251826 | 0.03083063 | 0.131797657 | 0.233923957 | 0.815044014 | NA | | | Scgb1b30 | 0 | NA | NA | NA | NA | NA |
| Tnfrsf17 | 7.474252658 | -0.267831735 | 0.154031663 | -1.738809612 | 0.082068257 | NA | | | Scgb1b7 | 0 | NA | NA | NA | NA | NA |
| Tnfrsf18 | 2.22352369 | -0.044768391 | 0.100717773 | -0.444493456 | 0.656685859 | NA | | | Scgb2b19 | 0 | NA | NA | NA | NA | NA |
| Tnfrsf26 | 1.509362283 | -0.131811912 | 0.085485001 | -1.541930285 | 0.123090536 | NA | | | Scgb2b20 | 0 | NA | NA | NA | NA | NA |
| Tnfsf11 | 0.953514938 | -0.009882079 | 0.072650881 | -0.136021457 | 0.891804318 | NA | | | Scgb2b23-ps | 0 | NA | NA | NA | NA | NA |
| Tnfsf14 | 0.280299629 | 0.041967777 | 0.040615819 | 1.033286501 | 0.301469842 | NA | | | Scgb2b24 | 0 | NA | NA | NA | NA | NA |
| Tnfsf15 | 1.036246275 | -0.043177279 | 0.073581656 | -0.586794062 | 0.557342025 | NA | | | Scgb2b26 | 0 | NA | NA | NA | NA | NA |
| Tnfsf18 | 0.895316464 | 0.045505672 | 0.078674053 | 0.57840762 | 0.562988951 | NA | | | Scgb2b27 | 0 | NA | NA | NA | NA | NA |
| Tnfsf4 | 0 | NA | NA | NA | NA | NA | | | Scgb2b3 | 0 | NA | NA | NA | NA | NA |
| Tnfsf8 | 3.460287471 | 0.198018841 | 0.119834684 | 1.652433447 | 0.098446222 | NA | | | Scgb2b7 | 0 | NA | NA | NA | NA | NA |
| Tnip3 | 2.977704263 | 0.030418475 | 0.119087463 | 0.255429701 | 0.798391234 | NA | | | Sdr16c5 | 0 | NA | NA | NA | NA | NA |
| Tnk1 | 6.239772721 | 0.102355081 | 0.144356066 | 0.709045932 | 0.478295974 | NA | | | Sdr16c6 | 0 | NA | NA | NA | NA | NA |
| Tnnc2 | 4.409021046 | 0.169022739 | 0.137940729 | 1.225328738 | 0.220451385 | NA | | | Sectm1b | 0 | NA | NA | NA | NA | NA |
| Tnnt3 | 2.97399399 | -0.03655763 | 0.10425375 | -0.350660099 | 0.725843363 | NA | | | Sel1l2 | 0 | NA | NA | NA | NA | NA |
| Tnp1 | 0 | NA | NA | NA | NA | NA | | | Sept12 | 0 | NA | NA | NA | NA | NA |
| Tnp2 | 0 | NA | NA | NA | NA | NA | | | Serpina1a | 0 | NA | NA | NA | NA | NA |
| Tomm20l | 0.116696964 | -0.020477583 | 0.027954224 | -0.732539847 | 0.463839131 | NA | | | Serpina1c | 0 | NA | NA | NA | NA | NA |
| Tomt | 2.604535377 | 0.178115516 | 0.106927723 | 1.665756194 | 0.095761985 | NA | | | Serpina1f | 0 | NA | NA | NA | NA | NA |
| Topaz1 | 0.754584575 | 0.085968027 | 0.057162772 | 1.503916326 | 0.132602913 | NA | | | Serpina3a | 0 | NA | NA | NA | NA | NA |
| Tpbpa | 0 | NA | NA | NA | NA | NA | | | Serpina3j | 0 | NA | NA | NA | NA | NA |
| Tpbpb | 0 | NA | NA | NA | NA | NA | | | Serpina3k | 0 | NA | NA | NA | NA | NA |
| Tppp2 | 0 | NA | NA | NA | NA | NA | | | Serpina4-ps1 | 0 | NA | NA | NA | NA | NA |
| Tprg | 0.427430218 | -0.003973439 | 0.048318641 | -0.082234071 | 0.93446058 | NA | | | Serpina6 | 0 | NA | NA | NA | NA | NA |
| Tpsab1 | 0 | NA | NA | NA | NA | NA | | | Serpinb13 | 0 | NA | NA | NA | NA | NA |
| Tpsb2 | 1.029588275 | -0.068296389 | 0.07386073 | -0.924664408 | 0.355140499 | NA | | | Serpinb3a | 0 | NA | NA | NA | NA | NA |
| Tpsg1 | 0.215433205 | -0.027029776 | 0.031922004 | -0.84674434 | 0.397137638 | NA | | | Serpinb6d | 0 | NA | NA | NA | NA | NA |
| Tpte | 0.523233625 | -0.018953331 | 0.051293849 | -0.369504955 | 0.71175138 | NA | | | Serpinb9d | 0 | NA | NA | NA | NA | NA |
| Trap1a | 0.079983654 | 0.016740032 | 0.027674178 | 0.604897175 | 0.54524732 | NA | | | Serpinb9f | 0 | NA | NA | NA | NA | NA |
| Trat1 | 4.656008607 | -0.029134172 | 0.140489697 | -0.207375865 | 0.83571633 | NA | | | Serpinb9g | 0 | NA | NA | NA | NA | NA |
| Trcg1 | 0 | NA | NA | NA | NA | NA | | | Serpini2 | 0 | NA | NA | NA | NA | NA |
| Treh | 1.119991334 | -0.008487325 | 0.066260281 | -0.128090685 | 0.898077207 | NA | | | Sftpd | 0 | NA | NA | NA | NA | NA |
| Trem1 | 0.079983654 | 0.016740032 | 0.027674178 | 0.604897175 | 0.54524732 | NA | | | Sh2d1b2 | 0 | NA | NA | NA | NA | NA |
| Trem3 | 1.974725285 | -0.1263264 | 0.101647741 | -1.242786103 | 0.213946665 | NA | | | Sirpb1b | 0 | NA | NA | NA | NA | NA |
| Treml1 | 3.581138038 | 0.081416692 | 0.120772832 | 0.674130851 | 0.500228127 | NA | | | Skint1 | 0 | NA | NA | NA | NA | NA |
| Treml4 | 2.56864326 | 0.116636441 | 0.109414127 | 1.066008965 | 0.286419589 | NA | | | Skint10 | 0 | NA | NA | NA | NA | NA |
| Trex2 | 0 | NA | NA | NA | NA | NA | | | Skint11 | 0 | NA | NA | NA | NA | NA |
| Trim10 | 1.496180113 | -0.092272274 | 0.090491902 | -1.019674371 | 0.30788292 | NA | | | Skint2 | 0 | NA | NA | NA | NA | NA |
| Trim15 | 2.241585677 | 0.049665234 | 0.106436454 | 0.466618646 | 0.640772744 | NA | | | Skint6 | 0 | NA | NA | NA | NA | NA |
| Trim30b | 3.204279456 | 0.174378247 | 0.114578223 | 1.521914401 | 0.12803053 | NA | | | Skint8 | 0 | NA | NA | NA | NA | NA |
| Trim30e-ps1 | 3.224145527 | 0.026776696 | 0.113177776 | 0.236589702 | 0.81297511 | NA | | | Skint9 | 0 | NA | NA | NA | NA | NA |
| Trim31 | 0.238795281 | -0.027601606 | 0.032267774 | -0.855392297 | 0.392334007 | NA | | | Slc10a1 | 0 | NA | NA | NA | NA | NA |
| Trim34b | 1.334494249 | 0.003706998 | 0.081910033 | 0.045256947 | 0.963902504 | NA | | | Slc10a2 | 0 | NA | NA | NA | NA | NA |
| Trim38 | 1.494402856 | 0.00830225 | 0.084687017 | 0.098034505 | 0.921904895 | NA | | | Slc17a4 | 0 | NA | NA | NA | NA | NA |
| Trim40 | 5.231497624 | -0.138444575 | 0.141706871 | -0.976978558 | 0.328579768 | NA | | | Slc22a16 | 0 | NA | NA | NA | NA | NA |
| Trim42 | 0.225320798 | -0.001646023 | 0.032098875 | -0.051279768 | 0.95910259 | NA | | | Slc22a19 | 0 | NA | NA | NA | NA | NA |
| Trim43b | 2.192713984 | 0.070831641 | 0.087001845 | 0.814139527 | 0.415565024 | NA | | | Slc22a22 | 0 | NA | NA | NA | NA | NA |
| Trim43c | 0.484476137 | 0.03706576 | 0.05187087 | 0.714577567 | 0.474870111 | NA | | | Slc22a27 | 0 | NA | NA | NA | NA | NA |
| Trim50 | 0.451324107 | 0.006406988 | 0.044046195 | 0.145460652 | 0.884347181 | NA | | | Slc22a28 | 0 | NA | NA | NA | NA | NA |
| Trim52 | 0 | NA | NA | NA | NA | NA | | | Slc25a54 | 0 | NA | NA | NA | NA | NA |
| Trim55 | 0.536358603 | 0.018999954 | 0.051864556 | 0.366337928 | 0.714112931 | NA | | | Slc2a7 | 0 | NA | NA | NA | NA | NA |
| Trim58 | 4.134495169 | -0.159624254 | 0.128512812 | -1.242088251 | 0.214204003 | NA | | | Slc46a2 | 0 | NA | NA | NA | NA | NA |
| Trim60 | 0.079983654 | 0.016740032 | 0.027674178 | 0.604897175 | 0.54524732 | NA | | | Slc51b | 0 | NA | NA | NA | NA | NA |
| Trim61 | 0.755112314 | -0.0351974 | 0.062581738 | -0.562422859 | 0.573827948 | NA | | | Slc7a12 | 0 | NA | NA | NA | NA | NA |
| Trim63 | 5.998641539 | -0.103461712 | 0.145400001 | -0.711566103 | 0.476733503 | NA | | | Slc9c1 | 0 | NA | NA | NA | NA | NA |
| Trim69 | 1.187744795 | 0.035950389 | 0.074039548 | 0.48555658 | 0.627281588 | NA | | | Slco1a1 | 0 | NA | NA | NA | NA | NA |
| Trim75 | 0.388436144 | 0.026670728 | 0.048195357 | 0.553387917 | 0.579997811 | NA | | | Slco1a6 | 0 | NA | NA | NA | NA | NA |
| Triml1 | 0 | NA | NA | NA | NA | NA | | | Slco6b1 | 0 | NA | NA | NA | NA | NA |
| Triml2 | 1.481389728 | -0.116726324 | 0.091785309 | -1.271732097 | 0.203468322 | NA | | | Slurp1 | 0 | NA | NA | NA | NA | NA |
| Troap | 7.196862209 | -0.097420481 | 0.141967602 | -0.686216293 | 0.492576725 | NA | | | Slx | 0 | NA | NA | NA | NA | NA |
| Trp53tg5 | 0.373757766 | 0.017971539 | 0.042393413 | 0.423922914 | 0.671622033 | NA | | | Slxl1 | 0 | NA | NA | NA | NA | NA |
| Trpd52l3 | 0 | NA | NA | NA | NA | NA | | | Smcp | 0 | NA | NA | NA | NA | NA |
| Trpm5 | 0.961060696 | 0.01515042 | 0.07168587 | 0.211344577 | 0.832618401 | NA | | | Smgc | 0 | NA | NA | NA | NA | NA |
| Trpm8 | 5.072425431 | 0.007314258 | 0.130694487 | 0.055964546 | 0.955370051 | NA | | | Smok2a | 0 | NA | NA | NA | NA | NA |
| Trpv1 | 4.831119748 | 0.266247382 | 0.138219288 | 1.926267933 | 0.054070927 | NA | | | Smok2b | 0 | NA | NA | NA | NA | NA |
| Trpv5 | 0 | NA | NA | NA | NA | NA | | | Smok3a | 0 | NA | NA | NA | NA | NA |
| Try10 | 0.08731684 | 0.017365056 | 0.027733412 | 0.626142072 | 0.531221759 | NA | | | Smok3b | 0 | NA | NA | NA | NA | NA |
| Try4 | 0 | NA | NA | NA | NA | NA | | | Smok4a | 0 | NA | NA | NA | NA | NA |
| Try5 | 0 | NA | NA | NA | NA | NA | | | Smr2 | 0 | NA | NA | NA | NA | NA |
| Tsc1 | 3905.656923 | 0.187964682 | 0.099547126 | 1.888197976 | NA | NA | | | Snora19 | 0 | NA | NA | NA | NA | NA |
| Tsga13 | 3.634188703 | 0.122436544 | 0.127197137 | 0.962573116 | 0.335761794 | NA | | | Snora24 | 0 | NA | NA | NA | NA | NA |
| Tsga8 | 0 | NA | NA | NA | NA | NA | | | Snora26 | 0 | NA | NA | NA | NA | NA |
| Tshb | 1.259273622 | 0.134503086 | 0.085858944 | 1.566558828 | 0.117217848 | NA | | | Snora35 | 0 | NA | NA | NA | NA | NA |
| Tsix | 5.271486292 | 0.212349726 | 0.13748138 | 1.544570806 | 0.122450112 | NA | | | Snora36b | 0 | NA | NA | NA | NA | NA |
| Tslp | 5.126907474 | -0.201091064 | 0.141880112 | -1.417330883 | 0.15638621 | NA | | | Snord11 | 0 | NA | NA | NA | NA | NA |
| Tspan1 | 4.514927444 | -0.191255442 | 0.136863564 | -1.39741679 | 0.162288273 | NA | | | Snord116 | 0 | NA | NA | NA | NA | NA |
| Tspan10 | 2.59523789 | -0.088338573 | 0.099418998 | -0.888548208 | 0.374245936 | NA | | | Snord116l1 | 0 | NA | NA | NA | NA | NA |
| Tspan32 | 6.197854676 | 0.050745529 | 0.151579969 | 0.334777273 | 0.737793106 | NA | | | Snord116l2 | 0 | NA | NA | NA | NA | NA |
| Tspo2 | 1.044963501 | 0.077922586 | 0.069093725 | 1.127780945 | 0.259412443 | NA | | | Snord1a | 0 | NA | NA | NA | NA | NA |
| Tspy-ps | 0.215050351 | -0.027006021 | 0.031907566 | -0.846382992 | 0.397339123 | NA | | | Snord45c | 0 | NA | NA | NA | NA | NA |
| Tssk1 | 2.886031018 | 0.103274677 | 0.115846973 | 0.89147497 | 0.372674414 | NA | | | Snord61 | 0 | NA | NA | NA | NA | NA |
| Tssk2 | 0 | NA | NA | NA | NA | NA | | | Snord67 | 0 | NA | NA | NA | NA | NA |
| Tssk3 | 1.824711376 | 0.081781891 | 0.097352299 | 0.840061223 | 0.40087406 | NA | | | Snord7 | 0 | NA | NA | NA | NA | NA |
| Tssk5 | 0.233393928 | -0.027475306 | 0.032191697 | -0.85349047 | 0.39338737 | NA | | | Snord70 | 0 | NA | NA | NA | NA | NA |
| Tssk6 | 2.3286089 | -0.077148753 | 0.116204333 | -0.663905995 | 0.506750479 | NA | | | Snord82 | 0 | NA | NA | NA | NA | NA |
| Tsx | 2.84739085 | -0.031909003 | 0.106203304 | -0.300452074 | 0.763832348 | NA | | | Snord90 | 0 | NA | NA | NA | NA | NA |
| Ttc23l | 0.099009884 | 0.018205726 | 0.027896865 | 0.652608315 | 0.514008823 | NA | | | Snord92 | 0 | NA | NA | NA | NA | NA |
| Ttc24 | 0.588839167 | -0.044984229 | 0.06340616 | -0.709461493 | 0.47803814 | NA | | | Snord93 | 0 | NA | NA | NA | NA | NA |
| Ttc29 | 7.994592152 | 0.177986267 | 0.142321928 | 1.250589206 | 0.211084391 | NA | | | Snurf | 0 | NA | NA | NA | NA | NA |
| Ttc30a2 | 2.517119421 | -0.12207371 | 0.093403509 | -1.306949929 | 0.191229717 | NA | | | Spaca4 | 0 | NA | NA | NA | NA | NA |

| | | | | | | | | | | | | | | | |
|---|---|---|---|---|---|---|---|---|---|---|---|---|---|---|---|
| Ttc36 | 1.787386729 | -0.123140353 | 0.095388043 | -1.290941185 | 0.196724073 | NA | | | Spaca7 | 0 | NA | NA | NA | NA | NA |
| Ttc39d | 2.53430735 | -0.142404698 | 0.107101953 | -1.329618124 | 0.183644124 | NA | | | Spag11a | 0 | NA | NA | NA | NA | NA |
| Ttll10 | 3.342846972 | 0.231011973 | 0.122705588 | 1.882652425 | 0.059747487 | NA | | | Spam1 | 0 | NA | NA | NA | NA | NA |
| Ttll2 | 0.834746954 | -0.043086269 | 0.051353324 | -0.839016171 | 0.401460235 | NA | | | Spata31 | 0 | NA | NA | NA | NA | NA |
| Ttr | 3242.384781 | 0.264883478 | 0.09876165 | 2.682047932 | NA | NA | | | Speer1-ps1 | 0 | NA | NA | NA | NA | NA |
| Tuba3a | 0.099009884 | 0.018205726 | 0.027896865 | 0.652608315 | 0.514008823 | NA | | | Speer3 | 0 | NA | NA | NA | NA | NA |
| Tuba3b | 0 | NA | NA | NA | NA | NA | | | Speer4f | 0 | NA | NA | NA | NA | NA |
| Tubal3 | 0 | NA | NA | NA | NA | NA | | | Speer6-ps1 | 0 | NA | NA | NA | NA | NA |
| Tubb1 | 1.56694682 | -0.108455904 | 0.092576313 | -1.171529743 | 0.241385909 | NA | | | Spert | 0 | NA | NA | NA | NA | NA |
| Tubb2a-ps2 | 0.280719541 | 0.042027364 | 0.040642707 | 1.034069012 | 0.301103902 | NA | | | Spink14 | 0 | NA | NA | NA | NA | NA |
| Tulp1 | 1.443701084 | 0.038351081 | 0.076630828 | 0.500465438 | 0.616747387 | NA | | | Spink3 | 0 | NA | NA | NA | NA | NA |
| Tulp2 | 2.623988091 | -0.046266979 | 0.11508258 | -0.402032863 | 0.687659841 | NA | | | Spink6 | 0 | NA | NA | NA | NA | NA |
| Txk | 5.499631949 | 0.067900859 | 0.13909452 | 0.488163436 | 0.625434081 | NA | | | Spink7 | 0 | NA | NA | NA | NA | NA |
| Txndc2 | 2.185929593 | -0.022596257 | 0.0896969 | -0.251917929 | 0.801104501 | NA | | | Spinkl | 0 | NA | NA | NA | NA | NA |
| Txndc8 | 0 | NA | NA | NA | NA | NA | | | Spint3 | 0 | NA | NA | NA | NA | NA |
| Tyms-ps | 0.105923158 | 0.018524693 | 0.027977518 | 0.662127842 | 0.507889294 | NA | | | Spint5 | 0 | NA | NA | NA | NA | NA |
| Tyr | 4.464869866 | -0.147637378 | 0.137195215 | -1.076111708 | 0.2818773 | NA | | | Sppl2c | 0 | NA | NA | NA | NA | NA |
| U05342 | 0 | NA | NA | NA | NA | NA | | | Sprr1b | 0 | NA | NA | NA | NA | NA |
| U90926 | 0.291330644 | 0.006248311 | 0.033473087 | 0.186666694 | 0.851921971 | NA | | | Sprr2a1 | 0 | NA | NA | NA | NA | NA |
| Ubald2 | 0.305482598 | 0.007240588 | 0.03390939 | 0.21352751 | 0.830915534 | NA | | | Sprr2a2 | 0 | NA | NA | NA | NA | NA |
| Ubap1 | 1953.538303 | 0.116286704 | 0.102450721 | 1.13505013 | NA | NA | | | Sprr2b | 0 | NA | NA | NA | NA | NA |
| Ubash3a | 1.943408164 | 0.070357288 | 0.070260689 | 1.001374872 | 0.316645608 | NA | | | Sprr2d | 0 | NA | NA | NA | NA | NA |
| Ubb | 29757.92344 | -0.021607675 | 0.076180919 | -0.283636317 | NA | NA | | | Sprr2e | 0 | NA | NA | NA | NA | NA |
| Ubd | 0.239950962 | 0.027417169 | 0.032831728 | 0.835081494 | 0.403671825 | NA | | | Sprr2f | 0 | NA | NA | NA | NA | NA |
| Ube2d2b | 0.59705218 | -0.055901067 | 0.045635394 | -1.224949792 | 0.220594138 | NA | | | Sprr2h | 0 | NA | NA | NA | NA | NA |
| Ube2dnl1 | 0 | NA | NA | NA | NA | NA | | | Sprr2i | 0 | NA | NA | NA | NA | NA |
| Ube2dnl2 | 0 | NA | NA | NA | NA | NA | | | Sprr2j-ps | 0 | NA | NA | NA | NA | NA |
| Ube2u | 0.886445463 | 0.057284504 | 0.065158193 | 0.879160416 | 0.379314303 | NA | | | Sprr2k | 0 | NA | NA | NA | NA | NA |
| Ubl4b | 0.963937922 | 0.019238703 | 0.06542135 | 0.294073771 | 0.768701533 | NA | | | Sprr3 | 0 | NA | NA | NA | NA | NA |
| Ubqln3 | 0.08731684 | 0.017365056 | 0.027733412 | 0.626142072 | 0.531221759 | NA | | | Sprr4 | 0 | NA | NA | NA | NA | NA |
| Ubqlnl | 0.096071043 | -0.020098795 | 0.027825948 | -0.722304036 | 0.470107572 | NA | | | Spz1 | 0 | NA | NA | NA | NA | NA |
| Ubtfl1 | 0 | NA | NA | NA | NA | NA | | | Sry | 0 | NA | NA | NA | NA | NA |
| Uchl4 | 0 | NA | NA | NA | NA | NA | | | Ssx9 | 0 | NA | NA | NA | NA | NA |
| Ucn | 7.068471288 | -0.019200273 | 0.150350244 | -0.127703638 | 0.898383511 | NA | | | Ssxb1 | 0 | NA | NA | NA | NA | NA |
| Ucn2 | 1.776156778 | 0.013873087 | 0.084654647 | 0.163878627 | 0.869826693 | NA | | | Ssxb10 | 0 | NA | NA | NA | NA | NA |
| Ucp1 | 2.024471489 | -0.032467156 | 0.089660064 | -0.3621139 | 0.717266917 | NA | | | Ssxb3 | 0 | NA | NA | NA | NA | NA |
| Ugt1a1 | 0.284856677 | 0.008904946 | 0.033416532 | 0.266483248 | 0.789867059 | NA | | | Ssxb5 | 0 | NA | NA | NA | NA | NA |
| Ugt1a10 | 0 | NA | NA | NA | NA | NA | | | Ssxb8 | 0 | NA | NA | NA | NA | NA |
| Ugt1a2 | 0 | NA | NA | NA | NA | NA | | | Stfa1 | 0 | NA | NA | NA | NA | NA |
| Ugt1a5 | 0 | NA | NA | NA | NA | NA | | | Stfa2 | 0 | NA | NA | NA | NA | NA |
| Ugt1a6a | 56.13477206 | 0.811262311 | 0.166727719 | 4.865791446 | NA | NA | | | Stfa2l1 | 0 | NA | NA | NA | NA | NA |
| Ugt1a6b | 0 | NA | NA | NA | NA | NA | | | Stfa3 | 0 | NA | NA | NA | NA | NA |
| Ugt1a9 | 0 | NA | NA | NA | NA | NA | | | Stra8 | 0 | NA | NA | NA | NA | NA |
| Ugt2a1 | 1.36967311 | -0.06198634 | 0.069773716 | -0.888390979 | 0.374330476 | NA | | | Stx19 | 0 | NA | NA | NA | NA | NA |
| Ugt2a2 | 0 | NA | NA | NA | NA | NA | | | Sult1e1 | 0 | NA | NA | NA | NA | NA |
| Ugt2a3 | 0 | NA | NA | NA | NA | NA | | | Sult2a1 | 0 | NA | NA | NA | NA | NA |
| Ugt2b1 | 0 | NA | NA | NA | NA | NA | | | Sult2a2 | 0 | NA | NA | NA | NA | NA |
| Ugt2b34 | 0 | NA | NA | NA | NA | NA | | | Sult2a3 | 0 | NA | NA | NA | NA | NA |
| Ugt2b35 | 0 | NA | NA | NA | NA | NA | | | Sult2a4 | 0 | NA | NA | NA | NA | NA |
| Ugt2b36 | 0 | NA | NA | NA | NA | NA | | | Sult2a6 | 0 | NA | NA | NA | NA | NA |
| Ugt2b37 | 0 | NA | NA | NA | NA | NA | | | Sult2a7 | 0 | NA | NA | NA | NA | NA |
| Ugt2b38 | 0 | NA | NA | NA | NA | NA | | | Sun5 | 0 | NA | NA | NA | NA | NA |
| Ugt2b5 | 0.284916696 | 0.042355674 | 0.040798016 | 1.038179738 | 0.299186384 | NA | | | Sva | 0 | NA | NA | NA | NA | NA |
| Ugt3a1 | 0.458657292 | 0.005116849 | 0.037753393 | 0.135533495 | 0.892190083 | NA | | | Sval1 | 0 | NA | NA | NA | NA | NA |
| Ugt3a2 | 0 | NA | NA | NA | NA | NA | | | Sval3 | 0 | NA | NA | NA | NA | NA |
| Umod | 0 | NA | NA | NA | NA | NA | | | Svs1 | 0 | NA | NA | NA | NA | NA |
| Unc5cl | 5.534235381 | -0.059142708 | 0.146730175 | -0.403071203 | 0.686895844 | NA | | | Svs2 | 0 | NA | NA | NA | NA | NA |
| Unc93a | 0.508233334 | 0.033209451 | 0.046178565 | 0.719152935 | 0.472046695 | NA | | | Svs3a | 0 | NA | NA | NA | NA | NA |
| Uncx | 4.103913735 | 0.121382362 | 0.126181977 | 0.961962753 | 0.336068314 | NA | | | Svs3b | 0 | NA | NA | NA | NA | NA |
| Upk2 | 0.481360729 | 0.009303061 | 0.05529186 | 0.168253724 | 0.866383678 | NA | | | Svs4 | 0 | NA | NA | NA | NA | NA |
| Upk3a | 2.505005272 | -0.068714024 | 0.099994133 | -0.687180562 | 0.491968952 | NA | | | Svs5 | 0 | NA | NA | NA | NA | NA |
| Upk3b | 0.178993538 | 0.024685672 | 0.031260828 | 0.789667872 | 0.429721759 | NA | | | Svs6 | 0 | NA | NA | NA | NA | NA |
| Upk3bl | 2.866502003 | 0.097853592 | 0.1187077 | 0.824323884 | 0.409755551 | NA | | | Sycn | 0 | NA | NA | NA | NA | NA |
| Uroc1 | 0 | NA | NA | NA | NA | NA | | | Sycp1-ps1 | 0 | NA | NA | NA | NA | NA |
| Ush1c | 6.951718787 | 0.176395112 | 0.126009543 | 1.399855189 | 0.161556687 | NA | | | Taar1 | 0 | NA | NA | NA | NA | NA |
| Ush2a | 3.868210486 | 0.043616446 | 0.109289363 | 0.399091406 | 0.689825855 | NA | | | Taar2 | 0 | NA | NA | NA | NA | NA |
| Usp17la | 3.144878331 | 0.185500021 | 0.123682925 | 1.499802997 | 0.133665441 | NA | | | Taar4 | 0 | NA | NA | NA | NA | NA |
| Usp17lb | 0 | NA | NA | NA | NA | NA | | | Taar5 | 0 | NA | NA | NA | NA | NA |
| Usp17lc | 0 | NA | NA | NA | NA | NA | | | Taar6 | 0 | NA | NA | NA | NA | NA |
| Usp17ld | 0.099009884 | 0.018205726 | 0.027896865 | 0.652608315 | 0.514008823 | NA | | | Taar7a | 0 | NA | NA | NA | NA | NA |
| Usp17le | 0 | NA | NA | NA | NA | NA | | | Taar7b | 0 | NA | NA | NA | NA | NA |
| Usp26 | 0 | NA | NA | NA | NA | NA | | | Taar7d | 0 | NA | NA | NA | NA | NA |
| Usp44 | 0.480966971 | -0.012302917 | 0.044293682 | -0.277757829 | 0.781198266 | NA | | | Taar7e | 0 | NA | NA | NA | NA | NA |
| Usp50 | 2.553787272 | 0.13238994 | 0.087912352 | 1.505931046 | 0.132084876 | NA | | | Taar7f | 0 | NA | NA | NA | NA | NA |
| Usp53 | 924.941957 | 0.237369371 | 0.166002668 | 1.429912988 | NA | NA | | | Taar8a | 0 | NA | NA | NA | NA | NA |
| Usp9y | 0 | NA | NA | NA | NA | NA | | | Taar8b | 0 | NA | NA | NA | NA | NA |
| Utf1 | 0.465691673 | 0.035179317 | 0.051468901 | 0.683506283 | 0.494286982 | NA | | | Taar8c | 0 | NA | NA | NA | NA | NA |
| Uts2 | 0.200315975 | 0.025370485 | 0.031664208 | 0.801235428 | 0.422995365 | NA | | | Taar9 | 0 | NA | NA | NA | NA | NA |
| Uts2d | 0.099009884 | 0.018205726 | 0.027896865 | 0.652608315 | 0.514008823 | NA | | | Tas2r102 | 0 | NA | NA | NA | NA | NA |
| Uts2r | 6.559290417 | 0.091557576 | 0.14883831 | 0.615147914 | 0.538457042 | NA | | | Tas2r104 | 0 | NA | NA | NA | NA | NA |
| V1ra8 | 0 | NA | NA | NA | NA | NA | | | Tas2r105 | 0 | NA | NA | NA | NA | NA |
| V1rd18 | 0 | NA | NA | NA | NA | NA | | | Tas2r106 | 0 | NA | NA | NA | NA | NA |
| V1rd19 | 0 | NA | NA | NA | NA | NA | | | Tas2r107 | 0 | NA | NA | NA | NA | NA |
| Vaultrc5 | 1.329536028 | -0.091579739 | 0.08414314 | -1.088380347 | 0.276427234 | NA | | | Tas2r109 | 0 | NA | NA | NA | NA | NA |
| Vax2 | 0 | NA | NA | NA | NA | NA | | | Tas2r110 | 0 | NA | NA | NA | NA | NA |
| Vax2os | 1.1654809 | 0.04742536 | 0.081762314 | 0.580039354 | 0.561888079 | NA | | | Tas2r113 | 0 | NA | NA | NA | NA | NA |
| Veph1 | 4.757637665 | -0.114894081 | 0.13399422 | -0.857455502 | 0.391193197 | NA | | | Tas2r114 | 0 | NA | NA | NA | NA | NA |
| Vgll1 | 0 | NA | NA | NA | NA | NA | | | Tas2r115 | 0 | NA | NA | NA | NA | NA |
| Vgll2 | 3.827524474 | 0.066210044 | 0.125171202 | 0.528955889 | 0.59683605 | NA | | | Tas2r116 | 0 | NA | NA | NA | NA | NA |
| Vmn1r-ps103 | 0 | NA | NA | NA | NA | NA | | | Tas2r117 | 0 | NA | NA | NA | NA | NA |
| Vmn1r-ps79 | 0 | NA | NA | NA | NA | NA | | | Tas2r118 | 0 | NA | NA | NA | NA | NA |
| Vmn1r1 | 0 | NA | NA | NA | NA | NA | | | Tas2r119 | 0 | NA | NA | NA | NA | NA |
| Vmn1r10 | 0 | NA | NA | NA | NA | NA | | | Tas2r120 | 0 | NA | NA | NA | NA | NA |
| Vmn1r100 | 0 | NA | NA | NA | NA | NA | | | Tas2r121 | 0 | NA | NA | NA | NA | NA |
| Vmn1r101 | 0 | NA | NA | NA | NA | NA | | | Tas2r122 | 0 | NA | NA | NA | NA | NA |
| Vmn1r103 | 0 | NA | NA | NA | NA | NA | | | Tas2r123 | 0 | NA | NA | NA | NA | NA |
| Vmn1r104 | 0 | NA | NA | NA | NA | NA | | | Tas2r124 | 0 | NA | NA | NA | NA | NA |
| Vmn1r107 | 0 | NA | NA | NA | NA | NA | | | Tas2r125 | 0 | NA | NA | NA | NA | NA |
| Vmn1r11 | 0 | NA | NA | NA | NA | NA | | | Tas2r126 | 0 | NA | NA | NA | NA | NA |
| Vmn1r111 | 0 | NA | NA | NA | NA | NA | | | Tas2r129 | 0 | NA | NA | NA | NA | NA |
| Vmn1r112 | 0 | NA | NA | NA | NA | NA | | | Tas2r130 | 0 | NA | NA | NA | NA | NA |
| Vmn1r113 | 0 | NA | NA | NA | NA | NA | | | Tas2r131 | 0 | NA | NA | NA | NA | NA |
| Vmn1r114 | 0 | NA | NA | NA | NA | NA | | | Tas2r134 | 0 | NA | NA | NA | NA | NA |
| Vmn1r115 | 0 | NA | NA | NA | NA | NA | | | Tas2r136 | 0 | NA | NA | NA | NA | NA |
| Vmn1r116 | 0 | NA | NA | NA | NA | NA | | | Tas2r138 | 0 | NA | NA | NA | NA | NA |
| Vmn1r117 | 0 | NA | NA | NA | NA | NA | | | Tas2r139 | 0 | NA | NA | NA | NA | NA |
| Vmn1r118 | 0 | NA | NA | NA | NA | NA | | | Tas2r140 | 0 | NA | NA | NA | NA | NA |
| Vmn1r119 | 0 | NA | NA | NA | NA | NA | | | Tas2r144 | 0 | NA | NA | NA | NA | NA |
| Vmn1r12 | 0 | NA | NA | NA | NA | NA | | | Tbc1d21 | 0 | NA | NA | NA | NA | NA |
| Vmn1r120 | 0 | NA | NA | NA | NA | NA | | | Tcl1 | 0 | NA | NA | NA | NA | NA |
| Vmn1r121 | 0 | NA | NA | NA | NA | NA | | | Tcl1b3 | 0 | NA | NA | NA | NA | NA |
| Vmn1r122 | 0 | NA | NA | NA | NA | NA | | | Tcl1b4 | 0 | NA | NA | NA | NA | NA |
| Vmn1r123 | 0 | NA | NA | NA | NA | NA | | | Tcl1b5 | 0 | NA | NA | NA | NA | NA |
| Vmn1r124 | 0 | NA | NA | NA | NA | NA | | | Tcte3 | 0 | NA | NA | NA | NA | NA |
| Vmn1r125 | 0 | NA | NA | NA | NA | NA | | | Tdpoz1 | 0 | NA | NA | NA | NA | NA |
| Vmn1r126 | 0 | NA | NA | NA | NA | NA | | | Tdpoz2 | 0 | NA | NA | NA | NA | NA |
| Vmn1r127 | 0 | NA | NA | NA | NA | NA | | | Tdpoz3 | 0 | NA | NA | NA | NA | NA |
| Vmn1r128 | 0 | NA | NA | NA | NA | NA | | | Tdpoz4 | 0 | NA | NA | NA | NA | NA |
| Vmn1r129 | 0 | NA | NA | NA | NA | NA | | | Tdpoz5 | 0 | NA | NA | NA | NA | NA |
| Vmn1r13 | 0 | NA | NA | NA | NA | NA | | | Teddm1 | 0 | NA | NA | NA | NA | NA |
| Vmn1r130 | 0 | NA | NA | NA | NA | NA | | | Tex101 | 0 | NA | NA | NA | NA | NA |
| Vmn1r132 | 0 | NA | NA | NA | NA | NA | | | Tex12 | 0 | NA | NA | NA | NA | NA |
| Vmn1r135 | 0 | NA | NA | NA | NA | NA | | | Tex13 | 0 | NA | NA | NA | NA | NA |
| Vmn1r137 | 0 | NA | NA | NA | NA | NA | | | Tex13a | 0 | NA | NA | NA | NA | NA |
| Vmn1r139 | 0 | NA | NA | NA | NA | NA | | | Tex19.1 | 0 | NA | NA | NA | NA | NA |
| Vmn1r14 | 0 | NA | NA | NA | NA | NA | | | Tex28 | 0 | NA | NA | NA | NA | NA |
| Vmn1r142 | 0 | NA | NA | NA | NA | NA | | | Tex37 | 0 | NA | NA | NA | NA | NA |
| Vmn1r143 | 0 | NA | NA | NA | NA | NA | | | Tff1 | 0 | NA | NA | NA | NA | NA |
| Vmn1r148 | 0 | NA | NA | NA | NA | NA | | | Tff2 | 0 | NA | NA | NA | NA | NA |
| Vmn1r149 | 0 | NA | NA | NA | NA | NA | | | Tff3 | 0 | NA | NA | NA | NA | NA |
| Vmn1r15 | 0 | NA | NA | NA | NA | NA | | | Tgif2lx1 | 0 | NA | NA | NA | NA | NA |
| Vmn1r151 | 0 | NA | NA | NA | NA | NA | | | Tgif2lx2 | 0 | NA | NA | NA | NA | NA |
| Vmn1r152 | 0 | NA | NA | NA | NA | NA | | | Tgoln2 | 0 | NA | NA | NA | NA | NA |
| Vmn1r155 | 0 | NA | NA | NA | NA | NA | | | Tgtp1 | 0 | NA | NA | NA | NA | NA |
| Vmn1r157 | 0 | NA | NA | NA | NA | NA | | | Tlx1 | 0 | NA | NA | NA | NA | NA |
| Vmn1r158 | 0 | NA | NA | NA | NA | NA | | | Tm4sf20 | 0 | NA | NA | NA | NA | NA |
| Vmn1r159 | 0 | NA | NA | NA | NA | NA | | | Tmed11 | 0 | NA | NA | NA | NA | NA |
| Vmn1r16 | 0 | NA | NA | NA | NA | NA | | | Tmem190 | 0 | NA | NA | NA | NA | NA |
| Vmn1r160 | 0 | NA | NA | NA | NA | NA | | | Tmem225 | 0 | NA | NA | NA | NA | NA |
| Vmn1r163 | 0 | NA | NA | NA | NA | NA | | | Tmem247 | 0 | NA | NA | NA | NA | NA |
| Vmn1r165 | 0 | NA | NA | NA | NA | NA | | | Tmem254a | 0 | NA | NA | NA | NA | NA |
| Vmn1r167 | 0 | NA | NA | NA | NA | NA | | | Tmem254b | 0 | NA | NA | NA | NA | NA |
| Vmn1r168 | 0 | NA | NA | NA | NA | NA | | | Tmem254c | 0 | NA | NA | NA | NA | NA |
| Vmn1r169 | 0 | NA | NA | NA | NA | NA | | | Tmprss11c | 0 | NA | NA | NA | NA | NA |
| Vmn1r17 | 0 | NA | NA | NA | NA | NA | | | Tmprss11e | 0 | NA | NA | NA | NA | NA |
| Vmn1r170 | 0 | NA | NA | NA | NA | NA | | | Tmprss11g | 0 | NA | NA | NA | NA | NA |
| Vmn1r171 | 0 | NA | NA | NA | NA | NA | | | Tmprss12 | 0 | NA | NA | NA | NA | NA |
| Vmn1r172 | 0 | NA | NA | NA | NA | NA | | | Tmsb15a | 0 | NA | NA | NA | NA | NA |
| Vmn1r173 | 0 | NA | NA | NA | NA | NA | | | Tmsb15l | 0 | NA | NA | NA | NA | NA |
| Vmn1r174 | 0 | NA | NA | NA | NA | NA | | | Tnp1 | 0 | NA | NA | NA | NA | NA |
| Vmn1r175 | 0 | NA | NA | NA | NA | NA | | | Tnp2 | 0 | NA | NA | NA | NA | NA |
| Vmn1r176 | 0 | NA | NA | NA | NA | NA | | | Tomm20l | 0 | NA | NA | NA | NA | NA |
| Vmn1r177 | 0 | NA | NA | NA | NA | NA | | | Tpbpa | 0 | NA | NA | NA | NA | NA |
| Vmn1r178 | 0 | NA | NA | NA | NA | NA | | | Tpbpb | 0 | NA | NA | NA | NA | NA |
| Vmn1r179 | 0 | NA | NA | NA | NA | NA | | | Tppp2 | 0 | NA | NA | NA | NA | NA |
| Vmn1r18 | 0 | NA | NA | NA | NA | NA | | | Tpsab1 | 0 | NA | NA | NA | NA | NA |
| Vmn1r180 | 0 | NA | NA | NA | NA | NA | | | Tpsg1 | 0 | NA | NA | NA | NA | NA |
| Vmn1r181 | 0 | NA | NA | NA | NA | NA | | | Trcg1 | 0 | NA | NA | NA | NA | NA |
| Vmn1r183 | 0 | NA | NA | NA | NA | NA | | | Trex2 | 0 | NA | NA | NA | NA | NA |
| Vmn1r184 | 0 | NA | NA | NA | NA | NA | | | Trim52 | 0 | NA | NA | NA | NA | NA |
| Vmn1r185 | 0 | NA | NA | NA | NA | NA | | | Triml1 | 0 | NA | NA | NA | NA | NA |
| Vmn1r186 | 0 | NA | NA | NA | NA | NA | | | Trpd52l3 | 0 | NA | NA | NA | NA | NA |
| Vmn1r187 | 0 | NA | NA | NA | NA | NA | | | Try4 | 0 | NA | NA | NA | NA | NA |
| Vmn1r188 | 0 | NA | NA | NA | NA | NA | | | Try5 | 0 | NA | NA | NA | NA | NA |
| Vmn1r189 | 0 | NA | NA | NA | NA | NA | | | Tsga8 | 0 | NA | NA | NA | NA | NA |
| Vmn1r19 | 0 | NA | NA | NA | NA | NA | | | Tspy-ps | 0 | NA | NA | NA | NA | NA |
| Vmn1r191 | 0 | NA | NA | NA | NA | NA | | | Tssk2 | 0 | NA | NA | NA | NA | NA |
| Vmn1r192 | 0 | NA | NA | NA | NA | NA | | | Tuba3b | 0 | NA | NA | NA | NA | NA |
| Vmn1r193 | 0 | NA | NA | NA | NA | NA | | | Txndc8 | 0 | NA | NA | NA | NA | NA |
| Vmn1r194 | 0 | NA | NA | NA | NA | NA | | | U05342 | 0 | NA | NA | NA | NA | NA |
| Vmn1r195 | 0 | NA | NA | NA | NA | NA | | | Ubald2 | 0 | NA | NA | NA | NA | NA |
| Vmn1r196 | 0 | NA | NA | NA | NA | NA | | | Ube2d2b | 0 | NA | NA | NA | NA | NA |
| Vmn1r197 | 0 | NA | NA | NA | NA | NA | | | Ube2dnl1 | 0 | NA | NA | NA | NA | NA |
| Vmn1r198 | 0 | NA | NA | NA | NA | NA | | | Ube2dnl2 | 0 | NA | NA | NA | NA | NA |
| Vmn1r199 | 0 | NA | NA | NA | NA | NA | | | Ubqlnl | 0 | NA | NA | NA | NA | NA |
| Vmn1r2 | 0 | NA | NA | NA | NA | NA | | | Ubtfl1 | 0 | NA | NA | NA | NA | NA |
| Vmn1r20 | 0 | NA | NA | NA | NA | NA | | | Uchl4 | 0 | NA | NA | NA | NA | NA |
| Vmn1r200 | 0.486850878 | 0.009782776 | 0.044755885 | 0.218580767 | 0.826976636 | NA | | | Ugt1a10 | 0 | NA | NA | NA | NA | NA |
| Vmn1r201 | 0 | NA | NA | NA | NA | NA | | | Ugt1a2 | 0 | NA | NA | NA | NA | NA |
| Vmn1r202 | 0 | NA | NA | NA | NA | NA | | | Ugt1a5 | 0 | NA | NA | NA | NA | NA |
| Vmn1r203 | 0.116696964 | -0.020477583 | 0.027954224 | -0.732539847 | 0.463839131 | NA | | | Ugt1a6b | 0 | NA | NA | NA | NA | NA |
| Vmn1r204 | 0 | NA | NA | NA | NA | NA | | | Ugt1a9 | 0 | NA | NA | NA | NA | NA |
| Vmn1r205 | 0 | NA | NA | NA | NA | NA | | | Ugt2a3 | 0 | NA | NA | NA | NA | NA |
| Vmn1r206 | 3.545874401 | -0.118193148 | 0.108800615 | -1.086327933 | 0.277333933 | NA | | | Ugt2b1 | 0 | NA | NA | NA | NA | NA |
| Vmn1r207-ps | 0.39257328 | -0.00221316 | 0.042111238 | -0.052555094 | 0.958086398 | NA | | | Ugt2b34 | 0 | NA | NA | NA | NA | NA |
| Vmn1r208 | 0 | NA | NA | NA | NA | NA | | | Ugt2b35 | 0 | NA | NA | NA | NA | NA |
| Vmn1r209 | 0 | NA | NA | NA | NA | NA | | | Ugt2b36 | 0 | NA | NA | NA | NA | NA |
| Vmn1r21 | 0 | NA | NA | NA | NA | NA | | | Ugt2b37 | 0 | NA | NA | NA | NA | NA |
| Vmn1r210 | 0 | NA | NA | NA | NA | NA | | | Ugt2b38 | 0 | NA | NA | NA | NA | NA |
| Vmn1r211 | 0.094392817 | 0.017895942 | 0.027818137 | 0.643319236 | 0.520016977 | NA | | | Ugt3a2 | 0 | NA | NA | NA | NA | NA |
| Vmn1r212 | 0 | NA | NA | NA | NA | NA | | | Umod | 0 | NA | NA | NA | NA | NA |
| Vmn1r213 | 0 | NA | NA | NA | NA | NA | | | Usp17lb | 0 | NA | NA | NA | NA | NA |
| Vmn1r214 | 0 | NA | NA | NA | NA | NA | | | Usp17lc | 0 | NA | NA | NA | NA | NA |
| Vmn1r215 | 0 | NA | NA | NA | NA | NA | | | Usp17le | 0 | NA | NA | NA | NA | NA |
| Vmn1r216 | 0 | NA | NA | NA | NA | NA | | | Usp9y | 0 | NA | NA | NA | NA | NA |
| Vmn1r217 | 0 | NA | NA | NA | NA | NA | | | V1ra8 | 0 | NA | NA | NA | NA | NA |
| Vmn1r218 | 0 | NA | NA | NA | NA | NA | | | V1rd18 | 0 | NA | NA | NA | NA | NA |
| Vmn1r219 | 0 | NA | NA | NA | NA | NA | | | V1rd19 | 0 | NA | NA | NA | NA | NA |
| Vmn1r22 | 0 | NA | NA | NA | NA | NA | | | Vax2 | 0 | NA | NA | NA | NA | NA |
| Vmn1r220 | 0 | NA | NA | NA | NA | NA | | | Vgll1 | 0 | NA | NA | NA | NA | NA |
| Vmn1r221 | 0 | NA | NA | NA | NA | NA | | | Vmn1r-ps103 | 0 | NA | NA | NA | NA | NA |
| Vmn1r222 | 0.295034005 | -0.017733924 | 0.039348288 | -0.45069111 | 0.652212191 | NA | | | Vmn1r-ps79 | 0 | NA | NA | NA | NA | NA |
| Vmn1r223 | 0.079983654 | 0.016740032 | 0.027674178 | 0.604897175 | 0.54524732 | NA | | | Vmn1r1 | 0 | NA | NA | NA | NA | NA |
| Vmn1r224 | 0 | NA | NA | NA | NA | NA | | | Vmn1r10 | 0 | NA | NA | NA | NA | NA |
| Vmn1r225 | 0 | NA | NA | NA | NA | NA | | | Vmn1r100 | 0 | NA | NA | NA | NA | NA |
| Vmn1r226 | 0 | NA | NA | NA | NA | NA | | | Vmn1r101 | 0 | NA | NA | NA | NA | NA |
| Vmn1r227 | 0 | NA | NA | NA | NA | NA | | | Vmn1r103 | 0 | NA | NA | NA | NA | NA |
| Vmn1r228 | 0 | NA | NA | NA | NA | NA | | | Vmn1r104 | 0 | NA | NA | NA | NA | NA |
| Vmn1r229 | 0 | NA | NA | NA | NA | NA | | | Vmn1r107 | 0 | NA | NA | NA | NA | NA |
| Vmn1r23 | 0 | NA | NA | NA | NA | NA | | | Vmn1r11 | 0 | NA | NA | NA | NA | NA |
| Vmn1r230 | 0 | NA | NA | NA | NA | NA | | | Vmn1r111 | 0 | NA | NA | NA | NA | NA |
| Vmn1r231 | 0 | NA | NA | NA | NA | NA | | | Vmn1r112 | 0 | NA | NA | NA | NA | NA |
| Vmn1r232 | 0.099009884 | 0.018205726 | 0.027896865 | 0.652608315 | 0.514008823 | NA | | | Vmn1r113 | 0 | NA | NA | NA | NA | NA |
| Vmn1r233 | 0 | NA | NA | NA | NA | NA | | | Vmn1r114 | 0 | NA | NA | NA | NA | NA |
| Vmn1r234 | 0 | NA | NA | NA | NA | NA | | | Vmn1r115 | 0 | NA | NA | NA | NA | NA |
| Vmn1r235 | 0 | NA | NA | NA | NA | NA | | | Vmn1r116 | 0 | NA | NA | NA | NA | NA |
| Vmn1r236 | 0 | NA | NA | NA | NA | NA | | | Vmn1r117 | 0 | NA | NA | NA | NA | NA |
| Vmn1r237 | 0 | NA | NA | NA | NA | NA | | | Vmn1r118 | 0 | NA | NA | NA | NA | NA |
| Vmn1r238 | 0 | NA | NA | NA | NA | NA | | | Vmn1r119 | 0 | NA | NA | NA | NA | NA |
| Vmn1r24 | 0 | NA | NA | NA | NA | NA | | | Vmn1r12 | 0 | NA | NA | NA | NA | NA |
| Vmn1r25 | 0 | NA | NA | NA | NA | NA | | | Vmn1r120 | 0 | NA | NA | NA | NA | NA |
| Vmn1r26 | 0 | NA | NA | NA | NA | NA | | | Vmn1r121 | 0 | NA | NA | NA | NA | NA |
| Vmn1r27 | 0 | NA | NA | NA | NA | NA | | | Vmn1r122 | 0 | NA | NA | NA | NA | NA |
| Vmn1r28 | 0 | NA | NA | NA | NA | NA | | | Vmn1r123 | 0 | NA | NA | NA | NA | NA |
| Vmn1r29 | 0 | NA | NA | NA | NA | NA | | | Vmn1r124 | 0 | NA | NA | NA | NA | NA |
| Vmn1r3 | 0 | NA | NA | NA | NA | NA | | | Vmn1r125 | 0 | NA | NA | NA | NA | NA |
| Vmn1r30 | 0 | NA | NA | NA | NA | NA | | | Vmn1r126 | 0 | NA | NA | NA | NA | NA |
| Vmn1r31 | 0 | NA | NA | NA | NA | NA | | | Vmn1r127 | 0 | NA | NA | NA | NA | NA |
| Vmn1r32 | 0.28821313 | -0.029561117 | 0.033427731 | -0.884329147 | 0.376518559 | NA | | | Vmn1r128 | 0 | NA | NA | NA | NA | NA |
| Vmn1r33 | 0 | NA | NA | NA | NA | NA | | | Vmn1r129 | 0 | NA | NA | NA | NA | NA |
| Vmn1r34 | 0 | NA | NA | NA | NA | NA | | | Vmn1r13 | 0 | NA | NA | NA | NA | NA |
| Vmn1r35 | 0 | NA | NA | NA | NA | NA | | | Vmn1r130 | 0 | NA | NA | NA | NA | NA |
| Vmn1r36 | 0 | NA | NA | NA | NA | NA | | | Vmn1r132 | 0 | NA | NA | NA | NA | NA |
| Vmn1r37 | 0 | NA | NA | NA | NA | NA | | | Vmn1r135 | 0 | NA | NA | NA | NA | NA |
| Vmn1r38 | 0 | NA | NA | NA | NA | NA | | | Vmn1r137 | 0 | NA | NA | NA | NA | NA |

| | | | | | | |
|---|---|---|---|---|---|---|
| Vmn1r39 | 0 | NA | NA | NA | NA | NA |
| Vmn1r4 | 0.107716603 | -0.020477583 | 0.027954224 | -0.732539849 | 0.46383913 | NA |
| Vmn1r40 | 0 | NA | NA | NA | NA | NA |
| Vmn1r41 | 0 | NA | NA | NA | NA | NA |
| Vmn1r42 | 0 | NA | NA | NA | NA | NA |
| Vmn1r43 | 0 | NA | NA | NA | NA | NA |
| Vmn1r44 | 0 | NA | NA | NA | NA | NA |
| Vmn1r45 | 0 | NA | NA | NA | NA | NA |
| Vmn1r46 | 0 | NA | NA | NA | NA | NA |
| Vmn1r47 | 0 | NA | NA | NA | NA | NA |
| Vmn1r48 | 0 | NA | NA | NA | NA | NA |
| Vmn1r49 | 0 | NA | NA | NA | NA | NA |
| Vmn1r5 | 0 | NA | NA | NA | NA | NA |
| Vmn1r50 | 0 | NA | NA | NA | NA | NA |
| Vmn1r51 | 0.098353386 | -0.020212831 | 0.02786466 | -0.725393061 | 0.468210929 | NA |
| Vmn1r52 | 0 | NA | NA | NA | NA | NA |
| Vmn1r53 | 0 | NA | NA | NA | NA | NA |
| Vmn1r54 | 0 | NA | NA | NA | NA | NA |
| Vmn1r55 | 0 | NA | NA | NA | NA | NA |
| Vmn1r56 | 0 | NA | NA | NA | NA | NA |
| Vmn1r57 | 0 | NA | NA | NA | NA | NA |
| Vmn1r59 | 0 | NA | NA | NA | NA | NA |
| Vmn1r6 | 0 | NA | NA | NA | NA | NA |
| Vmn1r60 | 0 | NA | NA | NA | NA | NA |
| Vmn1r61 | 0 | NA | NA | NA | NA | NA |
| Vmn1r62 | 0 | NA | NA | NA | NA | NA |
| Vmn1r63 | 0 | NA | NA | NA | NA | NA |
| Vmn1r64 | 0 | NA | NA | NA | NA | NA |
| Vmn1r65 | 0.927369673 | 0.009575659 | 0.064866054 | 0.147622033 | 0.882641066 | NA |
| Vmn1r66 | 0 | NA | NA | NA | NA | NA |
| Vmn1r67 | 0 | NA | NA | NA | NA | NA |
| Vmn1r68 | 0 | NA | NA | NA | NA | NA |
| Vmn1r69 | 0 | NA | NA | NA | NA | NA |
| Vmn1r7 | 0 | NA | NA | NA | NA | NA |
| Vmn1r70 | 0 | NA | NA | NA | NA | NA |
| Vmn1r71 | 0 | NA | NA | NA | NA | NA |
| Vmn1r72 | 0 | NA | NA | NA | NA | NA |
| Vmn1r73 | 0 | NA | NA | NA | NA | NA |
| Vmn1r74 | 0 | NA | NA | NA | NA | NA |
| Vmn1r75 | 0 | NA | NA | NA | NA | NA |
| Vmn1r76 | 0 | NA | NA | NA | NA | NA |
| Vmn1r77 | 0 | NA | NA | NA | NA | NA |
| Vmn1r78 | 0.116696964 | -0.020477583 | 0.027954224 | -0.732539847 | 0.463839131 | NA |
| Vmn1r79 | 0 | NA | NA | NA | NA | NA |
| Vmn1r8 | 0 | NA | NA | NA | NA | NA |
| Vmn1r80 | 0 | NA | NA | NA | NA | NA |
| Vmn1r81 | 0 | NA | NA | NA | NA | NA |
| Vmn1r82 | 0 | NA | NA | NA | NA | NA |
| Vmn1r83 | 0 | NA | NA | NA | NA | NA |
| Vmn1r84 | 0 | NA | NA | NA | NA | NA |
| Vmn1r85 | 0 | NA | NA | NA | NA | NA |
| Vmn1r86 | 0 | NA | NA | NA | NA | NA |
| Vmn1r87 | 0 | NA | NA | NA | NA | NA |
| Vmn1r88 | 0 | NA | NA | NA | NA | NA |
| Vmn1r89 | 0 | NA | NA | NA | NA | NA |
| Vmn1r9 | 0 | NA | NA | NA | NA | NA |
| Vmn1r90 | 0 | NA | NA | NA | NA | NA |
| Vmn1r91 | 0 | NA | NA | NA | NA | NA |
| Vmn1r93 | 0 | NA | NA | NA | NA | NA |
| Vmn1r94 | 0 | NA | NA | NA | NA | NA |
| Vmn1r95 | 0 | NA | NA | NA | NA | NA |
| Vmn2r-ps11 | 0.864449938 | -0.020465133 | 0.056685193 | -0.361031362 | 0.718076002 | NA |
| Vmn2r-ps54 | 1.387477171 | -0.021122606 | 0.087481186 | -0.241453125 | 0.809203943 | NA |
| Vmn2r-ps60 | 0 | NA | NA | NA | NA | NA |
| Vmn2r10 | 0 | NA | NA | NA | NA | NA |
| Vmn2r100 | 0 | NA | NA | NA | NA | NA |
| Vmn2r101 | 0 | NA | NA | NA | NA | NA |
| Vmn2r102 | 0 | NA | NA | NA | NA | NA |
| Vmn2r103 | 0 | NA | NA | NA | NA | NA |
| Vmn2r104 | 0 | NA | NA | NA | NA | NA |
| Vmn2r105 | 0 | NA | NA | NA | NA | NA |
| Vmn2r106 | 0 | NA | NA | NA | NA | NA |
| Vmn2r107 | 0 | NA | NA | NA | NA | NA |
| Vmn2r108 | 0 | NA | NA | NA | NA | NA |
| Vmn2r109 | 0.099009884 | 0.018205726 | 0.027896865 | 0.652608315 | 0.514008823 | NA |
| Vmn2r11 | 0 | NA | NA | NA | NA | NA |
| Vmn2r110 | 0 | NA | NA | NA | NA | NA |
| Vmn2r111 | 0 | NA | NA | NA | NA | NA |
| Vmn2r112 | 0 | NA | NA | NA | NA | NA |
| Vmn2r113 | 0 | NA | NA | NA | NA | NA |
| Vmn2r114 | 0 | NA | NA | NA | NA | NA |
| Vmn2r115 | 0 | NA | NA | NA | NA | NA |
| Vmn2r116 | 0 | NA | NA | NA | NA | NA |
| Vmn2r117 | 0 | NA | NA | NA | NA | NA |
| Vmn2r118 | 0.116696964 | -0.020477583 | 0.027954224 | -0.732539847 | 0.463839131 | NA |
| Vmn2r12 | 0 | NA | NA | NA | NA | NA |
| Vmn2r120 | 0 | NA | NA | NA | NA | NA |
| Vmn2r121 | 0 | NA | NA | NA | NA | NA |
| Vmn2r124 | 0 | NA | NA | NA | NA | NA |
| Vmn2r13 | 0.159967308 | 0.0240345 | 0.030870969 | 0.778547006 | 0.436246606 | NA |
| Vmn2r14 | 0 | NA | NA | NA | NA | NA |
| Vmn2r15 | 0.107716603 | -0.020477583 | 0.027954224 | -0.732539849 | 0.46383913 | NA |
| Vmn2r16 | 0 | NA | NA | NA | NA | NA |
| Vmn2r17 | 0 | NA | NA | NA | NA | NA |
| Vmn2r18 | 0.287795519 | 0.031925978 | 0.035387605 | 0.902179657 | 0.366961439 | NA |
| Vmn2r19 | 0 | NA | NA | NA | NA | NA |
| Vmn2r2 | 0.279458926 | -0.011015785 | 0.033355472 | -0.330254219 | 0.741207882 | NA |
| Vmn2r20 | 0 | NA | NA | NA | NA | NA |
| Vmn2r21 | 0 | NA | NA | NA | NA | NA |
| Vmn2r22 | 0 | NA | NA | NA | NA | NA |
| Vmn2r23 | 0 | NA | NA | NA | NA | NA |
| Vmn2r24 | 0 | NA | NA | NA | NA | NA |
| Vmn2r25 | 0 | NA | NA | NA | NA | NA |
| Vmn2r26 | 0 | NA | NA | NA | NA | NA |
| Vmn2r27 | 1.71778417 | -0.096222303 | 0.083292515 | -1.155233491 | 0.247994823 | NA |
| Vmn2r28 | 1.080958715 | 0.007275374 | 0.079606288 | 0.091391953 | 0.927181156 | NA |
| Vmn2r3 | 0 | NA | NA | NA | NA | NA |
| Vmn2r30 | 0.239950962 | 0.027417169 | 0.032831728 | 0.835081494 | 0.403671825 | NA |
| Vmn2r31 | 0 | NA | NA | NA | NA | NA |
| Vmn2r32 | 0 | NA | NA | NA | NA | NA |
| Vmn2r33 | 0 | NA | NA | NA | NA | NA |
| Vmn2r34 | 0 | NA | NA | NA | NA | NA |
| Vmn2r35 | 0 | NA | NA | NA | NA | NA |
| Vmn2r36 | 0 | NA | NA | NA | NA | NA |
| Vmn2r37 | 0.099009884 | 0.018205726 | 0.027896865 | 0.652608315 | 0.514008823 | NA |
| Vmn2r38 | 0 | NA | NA | NA | NA | NA |
| Vmn2r39 | 0 | NA | NA | NA | NA | NA |
| Vmn2r4 | 0 | NA | NA | NA | NA | NA |
| Vmn2r40 | 0 | NA | NA | NA | NA | NA |
| Vmn2r41 | 0 | NA | NA | NA | NA | NA |
| Vmn2r42 | 0 | NA | NA | NA | NA | NA |
| Vmn2r43 | 0 | NA | NA | NA | NA | NA |
| Vmn2r44 | 0 | NA | NA | NA | NA | NA |
| Vmn2r45 | 0 | NA | NA | NA | NA | NA |
| Vmn2r46 | 0 | NA | NA | NA | NA | NA |
| Vmn2r47 | 0 | NA | NA | NA | NA | NA |
| Vmn2r48 | 0 | NA | NA | NA | NA | NA |
| Vmn2r49 | 0 | NA | NA | NA | NA | NA |
| Vmn2r5 | 0.492397275 | -0.018455921 | 0.05075977 | -0.363593463 | 0.716161609 | NA |
| Vmn2r50 | 0 | NA | NA | NA | NA | NA |
| Vmn2r51 | 0.08731684 | 0.017365056 | 0.027733412 | 0.626142072 | 0.531221759 | NA |
| Vmn2r52 | 0 | NA | NA | NA | NA | NA |
| Vmn2r53 | 0 | NA | NA | NA | NA | NA |
| Vmn2r54 | 0.715075039 | 0.032315171 | 0.056394524 | 0.573019664 | 0.56663138 | NA |
| Vmn2r55 | 0.215050351 | -0.027006021 | 0.031907566 | -0.846382992 | 0.397339123 | NA |
| Vmn2r56 | 0.313786592 | -0.036039695 | 0.036798757 | -0.979372609 | 0.327395906 | NA |
| Vmn2r58 | 0 | NA | NA | NA | NA | NA |
| Vmn2r59 | 0 | NA | NA | NA | NA | NA |
| Vmn2r6 | 0 | NA | NA | NA | NA | NA |
| Vmn2r60 | 0 | NA | NA | NA | NA | NA |
| Vmn2r61 | 0 | NA | NA | NA | NA | NA |
| Vmn2r62 | 0 | NA | NA | NA | NA | NA |
| Vmn2r63 | 0 | NA | NA | NA | NA | NA |
| Vmn2r65 | 0 | NA | NA | NA | NA | NA |
| Vmn2r66 | 0 | NA | NA | NA | NA | NA |
| Vmn2r67 | 0 | NA | NA | NA | NA | NA |
| Vmn2r68 | 0 | NA | NA | NA | NA | NA |
| Vmn2r69 | 0 | NA | NA | NA | NA | NA |
| Vmn2r7 | 0 | NA | NA | NA | NA | NA |
| Vmn2r70 | 0 | NA | NA | NA | NA | NA |
| Vmn2r71 | 0 | NA | NA | NA | NA | NA |
| Vmn2r72 | 0 | NA | NA | NA | NA | NA |
| Vmn2r73 | 0 | NA | NA | NA | NA | NA |
| Vmn2r74 | 0 | NA | NA | NA | NA | NA |
| Vmn2r75 | 0 | NA | NA | NA | NA | NA |
| Vmn2r76 | 0 | NA | NA | NA | NA | NA |
| Vmn2r77 | 0.098353386 | -0.020212831 | 0.02786466 | -0.725393061 | 0.468210929 | NA |
| Vmn2r78 | 0.32854328 | 0.00879128 | 0.034635881 | 0.253820007 | 0.799634616 | NA |
| Vmn2r79 | 0 | NA | NA | NA | NA | NA |
| Vmn2r8 | 0 | NA | NA | NA | NA | NA |
| Vmn2r80 | 0 | NA | NA | NA | NA | NA |
| Vmn2r81 | 0 | NA | NA | NA | NA | NA |
| Vmn2r82 | 0 | NA | NA | NA | NA | NA |
| Vmn2r83 | 0 | NA | NA | NA | NA | NA |
| Vmn2r85 | 6.843460148 | -0.070462639 | 0.139289425 | -0.505872138 | 0.612946413 | NA |
| Vmn2r86 | 2.533008175 | -0.032033396 | 0.105125763 | -0.30471499 | 0.760583232 | NA |
| Vmn2r88 | 0 | NA | NA | NA | NA | NA |
| Vmn2r89 | 0 | NA | NA | NA | NA | NA |
| Vmn2r9 | 0 | NA | NA | NA | NA | NA |
| Vmn2r90 | 0 | NA | NA | NA | NA | NA |
| Vmn2r91 | 0 | NA | NA | NA | NA | NA |
| Vmn2r92 | 0 | NA | NA | NA | NA | NA |
| Vmn2r93 | 0 | NA | NA | NA | NA | NA |
| Vmn2r94 | 0 | NA | NA | NA | NA | NA |
| Vmn2r95 | 0 | NA | NA | NA | NA | NA |
| Vmn2r96 | 0 | NA | NA | NA | NA | NA |
| Vmn2r97 | 0 | NA | NA | NA | NA | NA |
| Vmn2r98 | 0 | NA | NA | NA | NA | NA |
| Vmn2r99 | 0 | NA | NA | NA | NA | NA |
| Vmo1 | 3.032667467 | -0.050000583 | 0.121282702 | -0.412264755 | 0.680145379 | NA |
| Vnn3 | 3.56905524 | -0.106108741 | 0.118010439 | -0.899147072 | 0.368574328 | NA |
| Vpreb1 | 1.717552634 | -0.10940848 | 0.103708828 | -1.054958217 | 0.291444435 | NA |
| Vpreb2 | 0.212768007 | -0.02694223 | 0.031868764 | -0.845411806 | 0.397880955 | NA |
| Vpreb3 | 1.674477876 | -0.028956127 | 0.096876275 | -0.298898024 | 0.765017855 | NA |
| Vrtn | 4.096327904 | -0.199447721 | 0.128421976 | -1.55306535 | 0.120407523 | NA |
| Vsig1 | 0.233393928 | -0.027475306 | 0.032191697 | -0.85349047 | 0.39338737 | NA |
| Vsig4 | 0 | NA | NA | NA | NA | NA |
| Vsig8 | 7.590406203 | 0.020972284 | 0.15625742 | 0.134216246 | 0.89323158 | NA |
| Vsx1 | 6.03869358 | -0.068781925 | 0.118307491 | -0.581382669 | 0.560982575 | NA |
| Vwa2 | 0.589190971 | -0.008579232 | 0.052937852 | -0.162062328 | 0.871256772 | NA |
| Vwde | 0.658277769 | -0.062529329 | 0.056532976 | -1.106068236 | 0.268696986 | NA |
| Wap | 0 | NA | NA | NA | NA | NA |
| Wbp2nl | 1.525674379 | 0.031567476 | 0.086285278 | 0.36585008 | 0.714476948 | NA |
| Wbscr28 | 0 | NA | NA | NA | NA | NA |
| Wdr20b | 0.233393928 | -0.027475306 | 0.032191697 | -0.85349047 | 0.39338737 | NA |
| Wdr38 | 5.02981735 | -0.002470608 | 0.120626277 | -0.020481509 | 0.983659263 | NA |
| Wdr61 | 1237.717804 | -0.009051068 | 0.078275304 | -0.115631212 | NA | NA |
| Wdr64 | 0.492768124 | -0.024633558 | 0.055739517 | -0.44194064 | 0.658532161 | NA |
| Wdr72 | 4.569572229 | 0.167982415 | 0.083677933 | 2.007487623 | NA | NA |
| Wee2 | 6.406454477 | 0.172377554 | 0.144930786 | 1.189378448 | 0.234290778 | NA |
| Wfdc10 | 0 | NA | NA | NA | NA | NA |
| Wfdc11 | 0 | NA | NA | NA | NA | NA |
| Wfdc12 | 0.603371276 | -0.005189871 | 0.062807235 | -0.082631732 | 0.934144369 | NA |
| Wfdc13 | 0 | NA | NA | NA | NA | NA |
| Wfdc15a | 0.280299629 | 0.041967777 | 0.040615819 | 1.033286501 | 0.301469842 | NA |
| Wfdc15b | 0.098353386 | -0.020212831 | 0.02786466 | -0.725393061 | 0.468210929 | NA |
| Wfdc16 | 0 | NA | NA | NA | NA | NA |
| Wfdc17 | 7.863379813 | -0.357414337 | 0.142113425 | -2.514993479 | 0.011903459 | NA |
| Wfdc2 | 7.400308631 | 0.323755539 | 0.13551065 | 2.389151985 | NA | NA |
| Wfdc5 | 0 | NA | NA | NA | NA | NA |
| Wfdc6a | 0.818336458 | 0.030833894 | 0.064495082 | 0.478081325 | 0.632592323 | NA |
| Wfdc6b | 0 | NA | NA | NA | NA | NA |
| Wfdc8 | 0 | NA | NA | NA | NA | NA |
| Wfdc9 | 0 | NA | NA | NA | NA | NA |
| Wfikkn1 | 7.145698986 | 0.082598747 | 0.141746252 | 0.582722615 | 0.560080047 | NA |
| Wisp2 | 7.165727003 | 0.110055098 | 0.138609263 | 0.793995262 | 0.427198189 | NA |
| Wisp3 | 4.077437145 | 0.056489555 | 0.122511343 | 0.461096523 | 0.644729356 | NA |
| Wnt1 | 3.810729552 | -0.106960266 | 0.12112162 | -0.88308153 | 0.377192223 | NA |
| Wnt3 | 101.8637554 | 0.341800524 | 0.167404788 | 2.041760736 | NA | NA |
| Wnt3a | 0.791119018 | -0.005831258 | 0.067937947 | -0.085832117 | 0.931599875 | NA |
| Wnt8a | 0.17463368 | 0.024560283 | 0.031186242 | 0.787535833 | 0.430968261 | NA |
| Wt1 | 4.825393899 | -0.087355315 | 0.131370515 | -0.664953737 | 0.506080083 | NA |
| Xcl1 | 0.414584642 | 0.043381126 | 0.041065838 | 1.056379903 | 0.290794684 | NA |
| Xcr1 | 2.082470675 | -0.017091836 | 0.099124538 | -0.172427904 | 0.863101136 | NA |
| Xirp1 | 6.475558845 | -0.315428202 | 0.149592908 | -2.108577251 | 0.034981088 | NA |
| Xkr5 | 7.521844008 | 0.031874979 | 0.141376145 | 0.225462215 | 0.821619717 | NA |
| Xkr9 | 0 | NA | NA | NA | NA | NA |
| Xlr3c | 0.303023688 | 0.009739077 | 0.039388711 | 0.247255542 | 0.804710463 | NA |
| Xlr4a | 2.900418569 | 0.142513721 | 0.096346356 | 1.479181231 | 0.139091884 | NA |
| Xlr4b | 4.791741083 | 0.143096307 | 0.115648122 | 1.237342246 | 0.21596005 | NA |
| Xlr4c | 0.696916283 | 0.076469279 | 0.054048147 | 1.414836276 | 0.157116505 | NA |
| Xlr5a | 0.119397641 | -0.020477583 | 0.027954224 | -0.732539845 | 0.463839132 | NA |
| Xlr5b | 0 | NA | NA | NA | NA | NA |
| Xlr5c | 0 | NA | NA | NA | NA | NA |
| Yipf7 | 0.550767766 | 0.020121298 | 0.052098809 | 0.386214157 | 0.69933807 | NA |

| | | | | | | |
|---|---|---|---|---|---|---|
| Vmn1r139 | 0 | NA | NA | NA | NA | NA |
| Vmn1r14 | 0 | NA | NA | NA | NA | NA |
| Vmn1r142 | 0 | NA | NA | NA | NA | NA |
| Vmn1r143 | 0 | NA | NA | NA | NA | NA |
| Vmn1r148 | 0 | NA | NA | NA | NA | NA |
| Vmn1r149 | 0 | NA | NA | NA | NA | NA |
| Vmn1r151 | 0 | NA | NA | NA | NA | NA |
| Vmn1r152 | 0 | NA | NA | NA | NA | NA |
| Vmn1r155 | 0 | NA | NA | NA | NA | NA |
| Vmn1r157 | 0 | NA | NA | NA | NA | NA |
| Vmn1r158 | 0 | NA | NA | NA | NA | NA |
| Vmn1r159 | 0 | NA | NA | NA | NA | NA |
| Vmn1r16 | 0 | NA | NA | NA | NA | NA |
| Vmn1r160 | 0 | NA | NA | NA | NA | NA |
| Vmn1r163 | 0 | NA | NA | NA | NA | NA |
| Vmn1r165 | 0 | NA | NA | NA | NA | NA |
| Vmn1r167 | 0 | NA | NA | NA | NA | NA |
| Vmn1r168 | 0 | NA | NA | NA | NA | NA |
| Vmn1r169 | 0 | NA | NA | NA | NA | NA |
| Vmn1r17 | 0 | NA | NA | NA | NA | NA |
| Vmn1r170 | 0 | NA | NA | NA | NA | NA |
| Vmn1r171 | 0 | NA | NA | NA | NA | NA |
| Vmn1r172 | 0 | NA | NA | NA | NA | NA |
| Vmn1r173 | 0 | NA | NA | NA | NA | NA |
| Vmn1r174 | 0 | NA | NA | NA | NA | NA |
| Vmn1r175 | 0 | NA | NA | NA | NA | NA |
| Vmn1r176 | 0 | NA | NA | NA | NA | NA |
| Vmn1r177 | 0 | NA | NA | NA | NA | NA |
| Vmn1r178 | 0 | NA | NA | NA | NA | NA |
| Vmn1r179 | 0 | NA | NA | NA | NA | NA |
| Vmn1r18 | 0 | NA | NA | NA | NA | NA |
| Vmn1r180 | 0 | NA | NA | NA | NA | NA |
| Vmn1r181 | 0 | NA | NA | NA | NA | NA |
| Vmn1r183 | 0 | NA | NA | NA | NA | NA |
| Vmn1r184 | 0 | NA | NA | NA | NA | NA |
| Vmn1r185 | 0 | NA | NA | NA | NA | NA |
| Vmn1r186 | 0 | NA | NA | NA | NA | NA |
| Vmn1r187 | 0 | NA | NA | NA | NA | NA |
| Vmn1r188 | 0 | NA | NA | NA | NA | NA |
| Vmn1r189 | 0 | NA | NA | NA | NA | NA |
| Vmn1r19 | 0 | NA | NA | NA | NA | NA |
| Vmn1r191 | 0 | NA | NA | NA | NA | NA |
| Vmn1r192 | 0 | NA | NA | NA | NA | NA |
| Vmn1r193 | 0 | NA | NA | NA | NA | NA |
| Vmn1r194 | 0 | NA | NA | NA | NA | NA |
| Vmn1r195 | 0 | NA | NA | NA | NA | NA |
| Vmn1r196 | 0 | NA | NA | NA | NA | NA |
| Vmn1r197 | 0 | NA | NA | NA | NA | NA |
| Vmn1r198 | 0 | NA | NA | NA | NA | NA |
| Vmn1r199 | 0 | NA | NA | NA | NA | NA |
| Vmn1r2 | 0 | NA | NA | NA | NA | NA |
| Vmn1r20 | 0 | NA | NA | NA | NA | NA |
| Vmn1r201 | 0 | NA | NA | NA | NA | NA |
| Vmn1r202 | 0 | NA | NA | NA | NA | NA |
| Vmn1r203 | 0 | NA | NA | NA | NA | NA |
| Vmn1r204 | 0 | NA | NA | NA | NA | NA |
| Vmn1r205 | 0 | NA | NA | NA | NA | NA |
| Vmn1r209 | 0 | NA | NA | NA | NA | NA |
| Vmn1r21 | 0 | NA | NA | NA | NA | NA |
| Vmn1r210 | 0 | NA | NA | NA | NA | NA |
| Vmn1r211 | 0 | NA | NA | NA | NA | NA |
| Vmn1r212 | 0 | NA | NA | NA | NA | NA |
| Vmn1r213 | 0 | NA | NA | NA | NA | NA |
| Vmn1r214 | 0 | NA | NA | NA | NA | NA |
| Vmn1r215 | 0 | NA | NA | NA | NA | NA |
| Vmn1r216 | 0 | NA | NA | NA | NA | NA |
| Vmn1r217 | 0 | NA | NA | NA | NA | NA |
| Vmn1r218 | 0 | NA | NA | NA | NA | NA |
| Vmn1r219 | 0 | NA | NA | NA | NA | NA |
| Vmn1r22 | 0 | NA | NA | NA | NA | NA |
| Vmn1r220 | 0 | NA | NA | NA | NA | NA |
| Vmn1r221 | 0 | NA | NA | NA | NA | NA |
| Vmn1r224 | 0 | NA | NA | NA | NA | NA |
| Vmn1r225 | 0 | NA | NA | NA | NA | NA |
| Vmn1r226 | 0 | NA | NA | NA | NA | NA |
| Vmn1r227 | 0 | NA | NA | NA | NA | NA |
| Vmn1r228 | 0 | NA | NA | NA | NA | NA |
| Vmn1r229 | 0 | NA | NA | NA | NA | NA |
| Vmn1r23 | 0 | NA | NA | NA | NA | NA |
| Vmn1r230 | 0 | NA | NA | NA | NA | NA |
| Vmn1r231 | 0 | NA | NA | NA | NA | NA |
| Vmn1r233 | 0 | NA | NA | NA | NA | NA |
| Vmn1r234 | 0 | NA | NA | NA | NA | NA |
| Vmn1r235 | 0 | NA | NA | NA | NA | NA |
| Vmn1r236 | 0 | NA | NA | NA | NA | NA |
| Vmn1r237 | 0 | NA | NA | NA | NA | NA |
| Vmn1r238 | 0 | NA | NA | NA | NA | NA |
| Vmn1r24 | 0 | NA | NA | NA | NA | NA |
| Vmn1r25 | 0 | NA | NA | NA | NA | NA |
| Vmn1r26 | 0 | NA | NA | NA | NA | NA |
| Vmn1r27 | 0 | NA | NA | NA | NA | NA |
| Vmn1r28 | 0 | NA | NA | NA | NA | NA |
| Vmn1r29 | 0 | NA | NA | NA | NA | NA |
| Vmn1r3 | 0 | NA | NA | NA | NA | NA |
| Vmn1r30 | 0 | NA | NA | NA | NA | NA |
| Vmn1r32 | 0 | NA | NA | NA | NA | NA |
| Vmn1r33 | 0 | NA | NA | NA | NA | NA |
| Vmn1r34 | 0 | NA | NA | NA | NA | NA |
| Vmn1r35 | 0 | NA | NA | NA | NA | NA |
| Vmn1r36 | 0 | NA | NA | NA | NA | NA |
| Vmn1r37 | 0 | NA | NA | NA | NA | NA |
| Vmn1r38 | 0 | NA | NA | NA | NA | NA |
| Vmn1r39 | 0 | NA | NA | NA | NA | NA |
| Vmn1r40 | 0 | NA | NA | NA | NA | NA |
| Vmn1r41 | 0 | NA | NA | NA | NA | NA |
| Vmn1r42 | 0 | NA | NA | NA | NA | NA |
| Vmn1r43 | 0 | NA | NA | NA | NA | NA |
| Vmn1r44 | 0 | NA | NA | NA | NA | NA |
| Vmn1r45 | 0 | NA | NA | NA | NA | NA |
| Vmn1r46 | 0 | NA | NA | NA | NA | NA |
| Vmn1r47 | 0 | NA | NA | NA | NA | NA |
| Vmn1r48 | 0 | NA | NA | NA | NA | NA |
| Vmn1r49 | 0 | NA | NA | NA | NA | NA |
| Vmn1r50 | 0 | NA | NA | NA | NA | NA |
| Vmn1r51 | 0 | NA | NA | NA | NA | NA |
| Vmn1r52 | 0 | NA | NA | NA | NA | NA |
| Vmn1r54 | 0 | NA | NA | NA | NA | NA |
| Vmn1r55 | 0 | NA | NA | NA | NA | NA |
| Vmn1r56 | 0 | NA | NA | NA | NA | NA |
| Vmn1r57 | 0 | NA | NA | NA | NA | NA |
| Vmn1r59 | 0 | NA | NA | NA | NA | NA |
| Vmn1r6 | 0 | NA | NA | NA | NA | NA |
| Vmn1r60 | 0 | NA | NA | NA | NA | NA |
| Vmn1r61 | 0 | NA | NA | NA | NA | NA |
| Vmn1r62 | 0 | NA | NA | NA | NA | NA |
| Vmn1r63 | 0 | NA | NA | NA | NA | NA |
| Vmn1r64 | 0 | NA | NA | NA | NA | NA |
| Vmn1r66 | 0 | NA | NA | NA | NA | NA |
| Vmn1r67 | 0 | NA | NA | NA | NA | NA |
| Vmn1r68 | 0 | NA | NA | NA | NA | NA |
| Vmn1r69 | 0 | NA | NA | NA | NA | NA |
| Vmn1r7 | 0 | NA | NA | NA | NA | NA |
| Vmn1r70 | 0 | NA | NA | NA | NA | NA |
| Vmn1r71 | 0 | NA | NA | NA | NA | NA |
| Vmn1r72 | 0 | NA | NA | NA | NA | NA |
| Vmn1r73 | 0 | NA | NA | NA | NA | NA |
| Vmn1r74 | 0 | NA | NA | NA | NA | NA |
| Vmn1r75 | 0 | NA | NA | NA | NA | NA |
| Vmn1r76 | 0 | NA | NA | NA | NA | NA |
| Vmn1r77 | 0 | NA | NA | NA | NA | NA |
| Vmn1r78 | 0 | NA | NA | NA | NA | NA |
| Vmn1r79 | 0 | NA | NA | NA | NA | NA |
| Vmn1r8 | 0 | NA | NA | NA | NA | NA |
| Vmn1r80 | 0 | NA | NA | NA | NA | NA |
| Vmn1r81 | 0 | NA | NA | NA | NA | NA |
| Vmn1r82 | 0 | NA | NA | NA | NA | NA |
| Vmn1r83 | 0 | NA | NA | NA | NA | NA |
| Vmn1r84 | 0 | NA | NA | NA | NA | NA |
| Vmn1r85 | 0 | NA | NA | NA | NA | NA |
| Vmn1r86 | 0 | NA | NA | NA | NA | NA |
| Vmn1r87 | 0 | NA | NA | NA | NA | NA |
| Vmn1r88 | 0 | NA | NA | NA | NA | NA |
| Vmn1r89 | 0 | NA | NA | NA | NA | NA |
| Vmn1r9 | 0 | NA | NA | NA | NA | NA |
| Vmn1r90 | 0 | NA | NA | NA | NA | NA |
| Vmn1r91 | 0 | NA | NA | NA | NA | NA |
| Vmn1r93 | 0 | NA | NA | NA | NA | NA |
| Vmn1r94 | 0 | NA | NA | NA | NA | NA |
| Vmn1r95 | 0 | NA | NA | NA | NA | NA |
| Vmn2r-ps60 | 0 | NA | NA | NA | NA | NA |
| Vmn2r10 | 0 | NA | NA | NA | NA | NA |
| Vmn2r100 | 0 | NA | NA | NA | NA | NA |
| Vmn2r101 | 0 | NA | NA | NA | NA | NA |
| Vmn2r102 | 0 | NA | NA | NA | NA | NA |
| Vmn2r103 | 0 | NA | NA | NA | NA | NA |
| Vmn2r104 | 0 | NA | NA | NA | NA | NA |
| Vmn2r105 | 0 | NA | NA | NA | NA | NA |
| Vmn2r106 | 0 | NA | NA | NA | NA | NA |
| Vmn2r107 | 0 | NA | NA | NA | NA | NA |
| Vmn2r108 | 0 | NA | NA | NA | NA | NA |
| Vmn2r11 | 0 | NA | NA | NA | NA | NA |
| Vmn2r110 | 0 | NA | NA | NA | NA | NA |
| Vmn2r112 | 0 | NA | NA | NA | NA | NA |
| Vmn2r113 | 0 | NA | NA | NA | NA | NA |
| Vmn2r114 | 0 | NA | NA | NA | NA | NA |
| Vmn2r115 | 0 | NA | NA | NA | NA | NA |
| Vmn2r116 | 0 | NA | NA | NA | NA | NA |
| Vmn2r117 | 0 | NA | NA | NA | NA | NA |
| Vmn2r118 | 0 | NA | NA | NA | NA | NA |
| Vmn2r12 | 0 | NA | NA | NA | NA | NA |
| Vmn2r120 | 0 | NA | NA | NA | NA | NA |
| Vmn2r121 | 0 | NA | NA | NA | NA | NA |
| Vmn2r124 | 0 | NA | NA | NA | NA | NA |
| Vmn2r14 | 0 | NA | NA | NA | NA | NA |
| Vmn2r16 | 0 | NA | NA | NA | NA | NA |
| Vmn2r17 | 0 | NA | NA | NA | NA | NA |
| Vmn2r19 | 0 | NA | NA | NA | NA | NA |
| Vmn2r20 | 0 | NA | NA | NA | NA | NA |
| Vmn2r21 | 0 | NA | NA | NA | NA | NA |
| Vmn2r22 | 0 | NA | NA | NA | NA | NA |
| Vmn2r23 | 0 | NA | NA | NA | NA | NA |
| Vmn2r24 | 0 | NA | NA | NA | NA | NA |
| Vmn2r25 | 0 | NA | NA | NA | NA | NA |
| Vmn2r26 | 0 | NA | NA | NA | NA | NA |
| Vmn2r3 | 0 | NA | NA | NA | NA | NA |
| Vmn2r31 | 0 | NA | NA | NA | NA | NA |
| Vmn2r32 | 0 | NA | NA | NA | NA | NA |
| Vmn2r33 | 0 | NA | NA | NA | NA | NA |
| Vmn2r34 | 0 | NA | NA | NA | NA | NA |
| Vmn2r35 | 0 | NA | NA | NA | NA | NA |
| Vmn2r36 | 0 | NA | NA | NA | NA | NA |
| Vmn2r38 | 0 | NA | NA | NA | NA | NA |
| Vmn2r39 | 0 | NA | NA | NA | NA | NA |
| Vmn2r4 | 0 | NA | NA | NA | NA | NA |
| Vmn2r40 | 0 | NA | NA | NA | NA | NA |
| Vmn2r41 | 0 | NA | NA | NA | NA | NA |
| Vmn2r42 | 0 | NA | NA | NA | NA | NA |
| Vmn2r43 | 0 | NA | NA | NA | NA | NA |
| Vmn2r44 | 0 | NA | NA | NA | NA | NA |
| Vmn2r45 | 0 | NA | NA | NA | NA | NA |
| Vmn2r46 | 0 | NA | NA | NA | NA | NA |
| Vmn2r47 | 0 | NA | NA | NA | NA | NA |
| Vmn2r48 | 0 | NA | NA | NA | NA | NA |
| Vmn2r49 | 0 | NA | NA | NA | NA | NA |
| Vmn2r50 | 0 | NA | NA | NA | NA | NA |
| Vmn2r52 | 0 | NA | NA | NA | NA | NA |
| Vmn2r53 | 0 | NA | NA | NA | NA | NA |
| Vmn2r56 | 0 | NA | NA | NA | NA | NA |
| Vmn2r58 | 0 | NA | NA | NA | NA | NA |
| Vmn2r59 | 0 | NA | NA | NA | NA | NA |
| Vmn2r6 | 0 | NA | NA | NA | NA | NA |
| Vmn2r60 | 0 | NA | NA | NA | NA | NA |
| Vmn2r61 | 0 | NA | NA | NA | NA | NA |
| Vmn2r62 | 0 | NA | NA | NA | NA | NA |
| Vmn2r63 | 0 | NA | NA | NA | NA | NA |
| Vmn2r65 | 0 | NA | NA | NA | NA | NA |
| Vmn2r66 | 0 | NA | NA | NA | NA | NA |
| Vmn2r67 | 0 | NA | NA | NA | NA | NA |
| Vmn2r68 | 0 | NA | NA | NA | NA | NA |
| Vmn2r69 | 0 | NA | NA | NA | NA | NA |
| Vmn2r7 | 0 | NA | NA | NA | NA | NA |
| Vmn2r70 | 0 | NA | NA | NA | NA | NA |
| Vmn2r71 | 0 | NA | NA | NA | NA | NA |
| Vmn2r72 | 0 | NA | NA | NA | NA | NA |
| Vmn2r74 | 0 | NA | NA | NA | NA | NA |

| | | | | | | | | | | | | | | | |
|---|---|---|---|---|---|---|---|---|---|---|---|---|---|---|---|
| Zbtb32 | 1.152464342 | 0.040710781 | 0.060016863 | 0.678322367 | 0.49756732 | NA | | | Vmn2r75 | 0 | NA | NA | NA | NA | NA |
| Zc2hc1b | 0 | NA | NA | NA | NA | NA | | | Vmn2r76 | 0 | NA | NA | NA | NA | NA |
| Zc3h12d | 2.241074628 | 0.00248366 | 0.097761553 | 0.025405284 | 0.979731696 | NA | | | Vmn2r79 | 0 | NA | NA | NA | NA | NA |
| Zcchc13 | 0.174376471 | 0.024538273 | 0.031173126 | 0.787161136 | 0.431187544 | NA | | | Vmn2r8 | 0 | NA | NA | NA | NA | NA |
| Zcchc5 | 4.805242689 | 0.139991527 | 0.104361919 | 1.34140431 | 0.179789219 | NA | | | Vmn2r80 | 0 | NA | NA | NA | NA | NA |
| Zdhhc11 | 0.188785634 | 0.025025111 | 0.031461603 | 0.79541754 | 0.42637066 | NA | | | Vmn2r81 | 0 | NA | NA | NA | NA | NA |
| Zdhhc19 | 0.4144578 | -0.044107517 | 0.040639925 | -1.085324756 | 0.277777845 | NA | | | Vmn2r82 | 0 | NA | NA | NA | NA | NA |
| Zdhhc25 | 0 | NA | NA | NA | NA | NA | | | Vmn2r83 | 0 | NA | NA | NA | NA | NA |
| Zfa-ps | 0.706356315 | 0.072659511 | 0.06424429 | 1.130987858 | 0.258060202 | NA | | | Vmn2r88 | 0 | NA | NA | NA | NA | NA |
| Zfp345 | 0.08731684 | 0.017365056 | 0.027733412 | 0.626142072 | 0.531221759 | NA | | | Vmn2r89 | 0 | NA | NA | NA | NA | NA |
| Zfp36l3 | 1.492030096 | 0.035174536 | 0.094780073 | 0.371117415 | 0.710550081 | NA | | | Vmn2r9 | 0 | NA | NA | NA | NA | NA |
| Zfp42 | 0.079983654 | 0.016740032 | 0.027674178 | 0.604897175 | 0.54524732 | NA | | | Vmn2r90 | 0 | NA | NA | NA | NA | NA |
| Zfp534 | 2.000606159 | 0.054859949 | 0.102743157 | 0.533952343 | 0.593374507 | NA | | | Vmn2r91 | 0 | NA | NA | NA | NA | NA |
| Zfp572 | 0.17463368 | 0.024560283 | 0.031186242 | 0.787535833 | 0.430968261 | NA | | | Vmn2r92 | 0 | NA | NA | NA | NA | NA |
| Zfp600 | 0.183387883 | -0.001444816 | 0.031390672 | -0.046026918 | 0.963288796 | NA | | | Vmn2r93 | 0 | NA | NA | NA | NA | NA |
| Zfp616 | 0 | NA | NA | NA | NA | NA | | | Vmn2r94 | 0 | NA | NA | NA | NA | NA |
| Zfp663 | 2.048067755 | -0.192161354 | 0.095886602 | -2.004048002 | 0.045064919 | NA | | | Vmn2r95 | 0 | NA | NA | NA | NA | NA |
| Zfp735 | 0 | NA | NA | NA | NA | NA | | | Vmn2r96 | 0 | NA | NA | NA | NA | NA |
| Zfp750 | 6.930758443 | 0.164315773 | 0.145321112 | 1.130708199 | 0.258177929 | NA | | | Vmn2r97 | 0 | NA | NA | NA | NA | NA |
| Zfp819 | 4.045034739 | 0.172203602 | 0.129765696 | 1.327034857 | 0.184497152 | NA | | | Vmn2r98 | 0 | NA | NA | NA | NA | NA |
| Zfp872 | 2.2216008 | 0.06052693 | 0.079721162 | 0.759232918 | 0.447713238 | NA | | | Vmn2r99 | 0 | NA | NA | NA | NA | NA |
| Zfp936 | 0.293434314 | -0.014075032 | 0.039227749 | -0.358802961 | 0.719742498 | NA | | | Vpreb2 | 0 | NA | NA | NA | NA | NA |
| Zfp957 | 0.17463368 | 0.024560283 | 0.031186242 | 0.787535833 | 0.430968261 | NA | | | Vsig4 | 0 | NA | NA | NA | NA | NA |
| Zfy1 | 0 | NA | NA | NA | NA | NA | | | Wap | 0 | NA | NA | NA | NA | NA |
| Zfy2 | 0 | NA | NA | NA | NA | NA | | | Wfdc11 | 0 | NA | NA | NA | NA | NA |
| Zg16 | 0 | NA | NA | NA | NA | NA | | | Wfdc13 | 0 | NA | NA | NA | NA | NA |
| Zim3 | 4.2945162 | 0.135609547 | 0.100124632 | 1.354407449 | 0.175606422 | NA | | | Wfdc15b | 0 | NA | NA | NA | NA | NA |
| Znrf4 | 0 | NA | NA | NA | NA | NA | | | Wfdc16 | 0 | NA | NA | NA | NA | NA |
| Zp1 | 4.558993664 | -0.002736762 | 0.136103234 | -0.020107986 | 0.983957229 | NA | | | Wfdc6b | 0 | NA | NA | NA | NA | NA |
| Zp3 | 0.930606196 | -0.065800364 | 0.076152546 | -0.864059935 | 0.387554967 | NA | | | Xlr5b | 0 | NA | NA | NA | NA | NA |
| Zp3r | 3.360176357 | 0.036943991 | 0.120062992 | 0.307705068 | 0.758306763 | NA | | | Xlr5c | 0 | NA | NA | NA | NA | NA |
| Zp4-ps | 0 | NA | NA | NA | NA | NA | | | Zc2hc1b | 0 | NA | NA | NA | NA | NA |
| Zpbp2 | 1.182448374 | 0.052326943 | 0.085574332 | 0.611479415 | 0.540882242 | NA | | | Zdhhc25 | 0 | NA | NA | NA | NA | NA |
| Zpld1 | 6.036963971 | 0.025764405 | 0.135165346 | 0.19061398 | 0.848828037 | NA | | | Zfp616 | 0 | NA | NA | NA | NA | NA |
| Zscan10 | 1.096775958 | 0.146845934 | 0.073590533 | 1.995445999 | 0.045994258 | NA | | | Zfp735 | 0 | NA | NA | NA | NA | NA |
| Zscan4a | 0 | NA | NA | NA | NA | NA | | | Zfy1 | 0 | NA | NA | NA | NA | NA |
| Zscan4b | 0 | NA | NA | NA | NA | NA | | | Zfy2 | 0 | NA | NA | NA | NA | NA |
| Zscan4c | 0 | NA | NA | NA | NA | NA | | | Znrf4 | 0 | NA | NA | NA | NA | NA |
| Zscan4d | 0 | NA | NA | NA | NA | NA | | | Zp4-ps | 0 | NA | NA | NA | NA | NA |
| Zscan4e | 0 | NA | NA | NA | NA | NA | | | Zscan4a | 0 | NA | NA | NA | NA | NA |
| Zscan4f | 0 | NA | NA | NA | NA | NA | | | Zscan4b | 0 | NA | NA | NA | NA | NA |
| Zscan5b | 0.256038351 | 0.007085343 | 0.03307033 | 0.214250763 | 0.830351512 | NA | | | Zscan4c | 0 | NA | NA | NA | NA | NA |
| Zswim2 | 0.686664619 | 0.040716927 | 0.066216252 | 0.614908356 | 0.538615244 | NA | | | Zscan4d | 0 | NA | NA | NA | NA | NA |
| Zyg11a | 0.270704723 | 0.00804778 | 0.033253949 | 0.242009739 | 0.808772617 | NA | | | Zscan4e | 0 | NA | NA | NA | NA | NA |
| a | 0.521693954 | 0.010491765 | 0.051043378 | 0.205546051 | 0.837145523 | NA | | | Zscan4f | 0 | NA | NA | NA | NA | NA |

sFigure2AB_DEGs_of_bulkseq

DAP12+/+ Tau+ vs DAP12+/+ Tau-

| | baseMean | log2FoldChange | lfcSE | stat | pvalue | padj |
|---|---|---|---|---|---|---|
| Xist | 9788.905385 | 8.703279608 | 0.126159893 | 68.98610486 | 0 | 0 |
| Slc25a20 | 872.0526975 | 1.445819194 | 0.063000299 | 22.94940208 | 1.49E-116 | 3.31E-113 |
| Tspan11 | 208.9624499 | 1.375178007 | 0.107852361 | 12.75056011 | 3.10E-37 | 5.33E-34 |
| Wnt2b | 77.90124492 | 1.329179115 | 0.153024737 | 8.686040816 | 3.75E-18 | 3.06E-15 |
| C4b | 1933.281053 | 1.310089731 | 0.114562669 | 11.43557272 | 2.78E-30 | 4.30E-27 |
| Rspo1 | 356.2679796 | 1.214775244 | 0.148470213 | 8.181945875 | 2.79E-16 | 1.88E-13 |
| Gm765 | 60.65268662 | 1.19169213 | 0.168110463 | 7.088744567 | 1.35E-12 | 3.88E-10 |
| Ntn5 | 150.9054749 | 1.118404744 | 0.16373717 | 6.830487818 | 8.46E-12 | 2.22E-09 |
| Clec7a | 69.16320609 | 1.107588906 | 0.163720099 | 6.76513705 | 1.33E-11 | 3.26E-09 |
| Myo5c | 158.7856697 | 1.081513555 | 0.151685928 | 7.129953129 | 1.00E-12 | 2.94E-10 |
| Cd52 | 80.47788727 | 1.07457152 | 0.162912389 | 6.596008629 | 4.22E-11 | 9.26E-09 |
| Adam33 | 276.5512317 | 1.065251902 | 0.157074763 | 6.781814493 | 1.19E-11 | 2.99E-09 |
| Nr2f2 | 651.2817828 | 1.044731182 | 0.133571384 | 7.821519505 | 5.22E-15 | 2.96E-12 |
| Igfn1 | 1449.509837 | 1.007539751 | 0.159582665 | 6.313591443 | 2.73E-10 | 4.59E-08 |
| Gpr114 | 11.3145576 | 0.998030924 | 0.151323956 | 6.595326673 | 4.24E-11 | 9.26E-09 |
| Gfap | 9181.31116 | 0.990633417 | 0.157642669 | 6.284043668 | 3.30E-10 | 5.44E-08 |
| Entpd4 | 5040.613621 | 0.958156073 | 0.073369172 | 13.05938241 | 5.62E-39 | 1.09E-35 |
| Scnn1a | 189.9552663 | 0.931182536 | 0.121517073 | 7.66297703 | 1.82E-14 | 8.04E-12 |
| Crb1 | 33.17447847 | 0.926287801 | 0.166155692 | 5.574818343 | 2.48E-08 | 2.30E-06 |
| Ccl6 | 85.51648389 | 0.922088455 | 0.167185195 | 5.515371459 | 3.48E-08 | 3.08E-06 |
| Sec1 | 46.14779074 | 0.906037211 | 0.167815582 | 5.399005272 | 6.70E-08 | 5.35E-06 |
| Sytl1 | 49.0441286 | 0.877653634 | 0.165681133 | 5.297245486 | 1.18E-07 | 8.51E-06 |
| Cyp39a1 | 136.7727229 | 0.875057103 | 0.130831669 | 6.688419644 | 2.26E-11 | 5.14E-09 |
| Miat | 16854.34769 | 0.873351165 | 0.155958359 | 5.599899691 | 2.14E-08 | 2.03E-06 |
| Thy1 | 37702.74949 | 0.870839947 | 0.079623348 | 10.93699238 | 7.67E-28 | 9.90E-25 |
| Clec18a | 204.5870021 | 0.870597815 | 0.155166843 | 5.610720666 | 2.01E-08 | 1.94E-06 |
| A330049N07Rik | 62.36615135 | 0.868878623 | 0.152994381 | 5.679153836 | 1.35E-08 | 1.38E-06 |
| Lag3 | 407.474968 | 0.862273253 | 0.158321784 | 5.446333617 | 5.14E-08 | 4.24E-06 |
| | 446.5826992 | 0.858681124 | 0.116028675 | 7.400594041 | 1.36E-13 | 5.25E-11 |
| Barx2 | 133.4700717 | 0.85692625 | 0.126382557 | 6.780415499 | 1.20E-11 | 2.99E-09 |
| Itgax | 89.51715568 | 0.845569665 | 0.151973521 | 5.563927566 | 2.64E-08 | 2.42E-06 |
| Tmem215 | 450.7499935 | 0.832106122 | 0.143042301 | 5.817203131 | 5.98E-09 | 6.92E-07 |
| Cxcl10 | 24.64422152 | 0.828484152 | 0.160237074 | 5.170364943 | 2.34E-07 | 1.50E-05 |
| Cldn22 | 11.60959755 | 0.827013419 | 0.158006096 | 5.234060204 | 1.66E-07 | 1.13E-05 |
| Mir27b | 19.09149771 | 0.81414511 | 0.167876699 | 4.849661183 | 1.24E-06 | 5.88E-05 |
| Endou | 142.3765054 | 0.808710506 | 0.15334902 | 5.273659449 | 1.34E-07 | 9.46E-06 |
| Ppl | 72.9663435 | 0.808626529 | 0.15197777 | 5.320689538 | 1.03E-07 | 7.66E-06 |
| Thbs3 | 538.984351 | 0.808257379 | 0.123621781 | 6.538147008 | 6.23E-11 | 1.30E-08 |
| Coro6 | 1141.994843 | 0.803160271 | 0.138197986 | 5.811664096 | 6.19E-09 | 7.00E-07 |
| Kcnab3 | 3234.69678 | 0.801421739 | 0.11213476 | 7.146951909 | 8.87E-13 | 2.64E-10 |
| Slamf9 | 47.10396413 | 0.789940209 | 0.164702297 | 4.796169948 | 1.62E-06 | 7.46E-05 |
| C920025E04Rik | 37.70378763 | 0.785211291 | 0.167428048 | 4.689843182 | 2.73E-06 | 0.000114804 |
| Tmem145 | 1516.001754 | 0.779047173 | 0.116817463 | 6.66892737 | 2.58E-11 | 5.79E-09 |
| Zan | 25.336803 | 0.776439059 | 0.167533641 | 4.634526263 | 3.58E-06 | 0.000140331 |
| Cd22 | 30.35036383 | 0.77279636 | 0.168101702 | 4.597195337 | 4.28E-06 | 0.00016222 |
| Ap1m2 | 36.06865584 | 0.757293797 | 0.163898605 | 4.620501775 | 3.83E-06 | 0.000148283 |
| H2-Bl | 23.35922168 | 0.751782931 | 0.16559525 | 4.539882229 | 5.63E-06 | 0.000201873 |
| Aifm3 | 3496.945545 | 0.745002989 | 0.095699294 | 7.784832698 | 6.98E-15 | 3.73E-12 |
| Col5a1 | 1179.554475 | 0.74149563 | 0.165678381 | 4.47551228 | 7.62E-06 | 0.000254543 |
| Pnpla3 | 472.15872 | 0.741295259 | 0.133004398 | 5.573464254 | 2.50E-08 | 2.30E-06 |
| Fmo2 | 121.9446464 | 0.739835358 | 0.155269311 | 4.764852442 | 1.89E-06 | 8.51E-05 |
| C1ra | 135.4376341 | 0.736809088 | 0.160800145 | 4.58214193 | 4.60E-06 | 0.000173081 |
| Obscn | 273.2247426 | 0.735838921 | 0.11639606 | 6.321854219 | 2.58E-10 | 4.40E-08 |
| Gm3893 | 469.6865392 | 0.7355971 | 0.163005489 | 4.512713679 | 6.40E-06 | 0.000222847 |
| Casr | 25.82938339 | 0.735098739 | 0.165662058 | 4.437339179 | 9.11E-06 | 0.000292771 |
| 5830416P10Rik | 43.74549909 | 0.723926795 | 0.164941543 | 4.388990089 | 1.14E-05 | 0.000353593 |
| Lgals3 | 25.58904438 | 0.722818856 | 0.167068041 | 4.326493872 | 1.52E-05 | 0.000445802 |
| Thpo | 233.75172 | 0.720308059 | 0.146402317 | 4.920059153 | 8.65E-07 | 4.33E-05 |
| Ncrna00085 | 411.7530876 | 0.719298456 | 0.137304392 | 5.238714101 | 1.62E-07 | 1.11E-05 |
| Ccl9 | 67.04289552 | 0.715476926 | 0.154999199 | 4.616004021 | 3.91E-06 | 0.000150403 |
| Chrd | 1265.389473 | 0.713978323 | 0.109066891 | 6.546242576 | 5.90E-11 | 1.25E-08 |
| Lgals3bp | 855.1685834 | 0.709110844 | 0.13450516 | 5.271997338 | 1.35E-07 | 9.50E-06 |
| Krt2 | 29.95118576 | 0.704135642 | 0.148487189 | 4.74206325 | 2.12E-06 | 9.29E-05 |
| Stx3 | 1104.88517 | 0.70410261 | 0.096457325 | 7.299628209 | 2.89E-13 | 9.37E-11 |
| Gm5577 | 181.2691896 | 0.703416888 | 0.147188577 | 4.779018194 | 1.76E-06 | 7.98E-05 |
| Nr4a2 | 1623.977445 | 0.699106764 | 0.086432638 | 8.088458038 | 6.04E-16 | 3.90E-13 |
| Vwa5b2 | 2339.746938 | 0.69844737 | 0.108719406 | 6.424311883 | 1.32E-10 | 2.53E-08 |
| Gm6537 | 63.20351091 | 0.69558237 | 0.153572187 | 4.529351209 | 5.92E-06 | 0.000208816 |
| C1ql2 | 88.92896909 | 0.693525747 | 0.16803255 | 4.127329774 | 3.67E-05 | 0.00089548 |
| Oas1b | 28.63052397 | 0.69301528 | 0.167831421 | 4.129234428 | 3.64E-05 | 0.0008909 |
| Arntl2 | 162.7786887 | 0.691742609 | 0.118471689 | 5.838885336 | 5.26E-09 | 6.17E-07 |
| Rnf207 | 283.6688619 | 0.69161035 | 0.138242145 | 5.00289076 | 5.65E-07 | 3.10E-05 |
| Gm129 | 582.3494173 | 0.688767371 | 0.096355988 | 7.148153293 | 8.80E-13 | 2.64E-10 |
| Spag5 | 517.0595542 | 0.688255661 | 0.137033559 | 5.022533642 | 5.10E-07 | 2.84E-05 |
| Ccl17 | 46.85776714 | 0.686370154 | 0.166286631 | 4.1276328 | 3.67E-05 | 0.00089548 |
| Rnpc3 | 774.9279501 | 0.68345431 | 0.112271155 | 6.087532549 | 1.15E-09 | 1.54E-07 |
| Ildr1 | 14.60061707 | 0.683200407 | 0.166648258 | 4.099655252 | 4.14E-05 | 0.000990864 |
| Mslnl | 18.16827862 | 0.683047468 | 0.166031507 | 4.113962958 | 3.89E-05 | 0.000944512 |
| Kcns1 | 673.7610618 | 0.68286652 | 0.138232449 | 4.939987123 | 7.81E-07 | 4.04E-05 |
| Ccdc108 | 321.2306714 | 0.681796538 | 0.157066532 | 4.340813587 | 1.42E-05 | 0.000424608 |
| Zfp185 | 70.88662 | 0.680386425 | 0.165938367 | 4.100235751 | 4.13E-05 | 0.000989912 |
| Cdc25b | 817.8909763 | 0.678073689 | 0.097581754 | 6.948775353 | 3.68E-12 | 1.02E-09 |
| Agmat | 63.00777509 | 0.677291518 | 0.153191999 | 4.421193802 | 9.82E-06 | 0.000309115 |
| Mir24-1 | 29.71920695 | 0.675921481 | 0.167765917 | 4.028955889 | 5.60E-05 | 0.001258048 |
| Plekha4 | 85.21933627 | 0.6723507 | 0.144425374 | 4.655350249 | 3.23E-06 | 0.000130501 |
| Slc23a3 | 46.00813392 | 0.669340157 | 0.166061419 | 4.030678308 | 5.56E-05 | 0.001250677 |
| Vsig2 | 62.16208071 | 0.665546841 | 0.152709347 | 4.358258717 | 1.31E-05 | 0.000398291 |
| Plxdc1 | 1523.841173 | 0.66477181 | 0.118487708 | 5.61047068 | 2.02E-08 | 1.94E-06 |
| BC024139 | 40.83153772 | 0.660358548 | 0.164921055 | 4.004088803 | 6.23E-05 | 0.001364151 |
| Siglec5 | 32.31699214 | 0.656799616 | 0.164795556 | 3.98554204 | 6.73E-05 | 0.001446814 |
| Vill | 288.3142611 | 0.656727208 | 0.106879005 | 6.144585747 | 8.02E-10 | 1.12E-07 |
| Col23a1 | 646.4326187 | 0.656065592 | 0.141480978 | 4.637129323 | 3.53E-06 | 0.000139281 |
| Psrc1 | 319.8846057 | 0.655551221 | 0.126468183 | 5.183526855 | 2.18E-07 | 1.42E-05 |
| Tdrd5 | 48.3000345 | 0.653685879 | 0.160102213 | 4.082928453 | 4.45E-05 | 0.001041174 |
| Tmc8 | 23.44996665 | 0.653027827 | 0.166773954 | 3.915646377 | 9.02E-05 | 0.0018119 |
| Hnf1b | 16.16815233 | 0.652764166 | 0.166638127 | 3.917255792 | 8.96E-05 | 0.001802188 |
| Bmp1 | 2709.289137 | 0.652113949 | 0.108619921 | 6.003631224 | 1.93E-09 | 2.45E-07 |
| Vamp1 | 6675.022313 | 0.651168277 | 0.101469624 | 6.417371544 | 1.39E-10 | 2.61E-08 |
| Arhgef37 | 106.9466907 | 0.64857888 | 0.163450613 | 3.9680419 | 7.25E-05 | 0.001531763 |
| C3ar1 | 128.7260767 | 0.647309963 | 0.129734365 | 4.989502709 | 6.05E-07 | 3.28E-05 |
| BC061194 | 114.0649251 | 0.647240143 | 0.121346098 | 5.333835657 | 9.62E-08 | 7.23E-06 |
| Slc2a4 | 86.66880659 | 0.647084388 | 0.168121297 | 3.848913851 | 0.000118643 | 0.002230886 |
| Tns4 | 15.05116877 | 0.643322577 | 0.165776906 | 3.880652587 | 0.000104177 | 0.002030329 |
| Dnahc9 | 236.1863733 | 0.642961656 | 0.123857143 | 5.191155237 | 2.09E-07 | 1.38E-05 |
| E130102H24Rik | 136.2400793 | 0.641203881 | 0.148826725 | 4.308392074 | 1.64E-05 | 0.000478933 |
| Phyhd1 | 512.6650775 | 0.641193533 | 0.149444719 | 4.290506467 | 1.78E-05 | 0.000509604 |
| Lhx9 | 40.80891809 | 0.640377013 | 0.157658748 | 4.061791818 | 4.87E-05 | 0.001126148 |
| Eno3 | 254.4885681 | 0.640140949 | 0.121543379 | 5.266769408 | 1.39E-07 | 9.69E-06 |
| Cd68 | 429.4261248 | 0.639284216 | 0.124790287 | 5.12286837 | 3.01E-07 | 1.86E-05 |
| Proser2 | 107.7436 | 0.639061454 | 0.126375446 | 5.056848274 | 4.26E-07 | 2.46E-05 |
| Capg | 88.0298116 | 0.638389445 | 0.14043478 | 4.54580728 | 5.47E-06 | 0.000198718 |
| Flt3 | 274.0022319 | 0.638340438 | 0.115336276 | 5.534602464 | 3.12E-08 | 2.79E-06 |
| Sfrs18 | 5999.036191 | 0.638272853 | 0.122117262 | 5.226720985 | 1.73E-07 | 1.17E-05 |
| Ucn3 | 16.4104385 | 0.638030396 | 0.160218136 | 3.982260761 | 6.83E-05 | 0.001464909 |
| Tmc6 | 184.8515502 | 0.637606888 | 0.126603053 | 5.036267886 | 4.75E-07 | 2.68E-05 |
| Snhg11 | 57641.62633 | 0.635492944 | 0.159991209 | 3.972049128 | 7.13E-05 | 0.001512406 |
| Cd14 | 97.32257121 | 0.6321603 | 0.130408646 | 4.847533663 | 1.25E-06 | 5.91E-05 |
| Aloxe3 | 206.2570327 | 0.6321384 | 0.145349035 | 4.349106274 | 1.37E-05 | 0.000411358 |
| Kazald1 | 161.9342514 | 0.629186885 | 0.133488978 | 4.713399517 | 2.44E-06 | 0.00010485 |
| Ifit3 | 374.1335853 | 0.627249931 | 0.14617201 | 4.291176741 | 1.78E-05 | 0.000509164 |
| Ifi204 | 25.57338141 | 0.626309725 | 0.15667187 | 3.997588879 | 6.40E-05 | 0.001387951 |
| Pvalb | 1580.452374 | 0.625764924 | 0.137677142 | 4.545162085 | 5.49E-06 | 0.000198718 |
| Shmt1 | 68.83104912 | 0.62358858 | 0.158745107 | 3.928238104 | 8.56E-05 | 0.001745007 |
| Gpr84 | 42.23628281 | 0.620402895 | 0.165985651 | 3.737689934 | 0.000185719 | 0.003222314 |
| St14 | 36.89020823 | 0.620137979 | 0.167401967 | 3.704484417 | 0.000211821 | 0.003548499 |
| Il16 | 158.1856135 | 0.619670187 | 0.163541933 | 3.789059941 | 0.000151218 | 0.002724393 |
| Gm13629 | 161.09298 | 0.615423522 | 0.132135136 | 4.657531217 | 3.20E-06 | 0.000129463 |
| Syt2 | 1570.135915 | 0.615048048 | 0.111094675 | 5.5362514 | 3.09E-08 | 2.78E-06 |
| 5930438M14Rik | 17.45696186 | 0.613516477 | 0.155833964 | 3.936988205 | 8.25E-05 | 0.001691032 |
| Npc1l1 | 78.63167845 | 0.611107303 | 0.149289547 | 4.093436645 | 4.25E-05 | 0.001010024 |
| Mylk3 | 141.5205768 | 0.610234573 | 0.136596792 | 4.467415121 | 7.92E-06 | 0.000261549 |
| Hdc | 64.38314051 | 0.607773155 | 0.153306357 | 3.964435439 | 7.36E-05 | 0.001544568 |
| Six4 | 61.13542265 | 0.607204373 | 0.154777094 | 3.923089368 | 8.74E-05 | 0.001768185 |
| I830012O16Rik | 207.5166096 | 0.606921303 | 0.141195899 | 4.298434365 | 1.72E-05 | 0.000495373 |
| A930011O12Rik | 6007.734216 | 0.606534719 | 0.157869559 | 3.841999207 | 0.000122036 | 0.002280856 |
| Pparg | 162.7000464 | 0.606513618 | 0.127443221 | 4.759088908 | 1.94E-06 | 8.68E-05 |
| T2 | 28.84656258 | 0.602333471 | 0.166790932 | 3.611308271 | 0.000304656 | 0.004725069 |
| Kndc1 | 4882.897485 | 0.602118247 | 0.089168546 | 6.752585668 | 1.45E-11 | 3.46E-09 |
| Emilin2 | 208.3914897 | 0.601685066 | 0.154238834 | 3.900995943 | 9.58E-05 | 0.001895646 |
| Cbln4 | 572.1106744 | 0.601274644 | 0.148253734 | 4.055713328 | 5.00E-05 | 0.001145662 |
| Tex40 | 206.3128231 | 0.600476649 | 0.146957363 | 4.086060327 | 4.39E-05 | 0.001031585 |
| 5530601H04Rik | 333.8879106 | 0.599554699 | 0.097243256 | 6.165514455 | 7.03E-10 | 1.03E-07 |
| Podn | 206.8579342 | 0.59878885 | 0.141278473 | 4.238358726 | 2.25E-05 | 0.000613905 |
| Eif2s3x | 6066.41865 | 0.592400434 | 0.060726273 | 9.755257506 | 1.75E-22 | 1.81E-19 |
| Gsta3 | 55.33800495 | 0.590936087 | 0.162360854 | 3.639646329 | 0.000273013 | 0.004325214 |
| A930013F10Rik | 293.0005318 | 0.590065387 | 0.160459347 | 3.677351282 | 0.000235668 | 0.003831528 |
| Hpn | 42.1963727 | 0.589751697 | 0.16672172 | 3.537341726 | 0.000404176 | 0.005876062 |
| Myh7b | 303.2394482 | 0.589056212 | 0.092248831 | 6.385514096 | 1.71E-10 | 3.11E-08 |
| Pcolce2 | 167.7500345 | 0.588346204 | 0.13816828 | 4.25818576 | 2.06E-05 | 0.000572258 |
| Cst7 | 45.35413944 | 0.586135197 | 0.131382376 | 4.4612924 | 8.15E-06 | 0.000266837 |
| Cchcr1 | 200.8938411 | 0.586079184 | 0.137577279 | 4.259999808 | 2.04E-05 | 0.000569675 |
| Plcz1 | 36.57405176 | 0.584331196 | 0.168010054 | 3.477953753 | 0.000505257 | 0.006882848 |
| Xaf1 | 204.349273 | 0.583772651 | 0.140600049 | 4.15200887 | 3.30E-05 | 0.000827609 |
| Klf14 | 19.25838698 | 0.58086797 | 0.167768834 | 3.462311544 | 0.000535557 | 0.007178129 |
| 2010111I01Rik | 631.3388566 | 0.579226212 | 0.114069079 | 5.077854732 | 3.82E-07 | 2.25E-05 |
| Prrg4 | 26.58797669 | 0.57670018 | 0.166887056 | 3.455631576 | 0.000549005 | 0.007320384 |
| Ankrd35 | 555.7922512 | 0.576004171 | 0.095004608 | 6.062907696 | 1.34E-09 | 1.76E-07 |
| Bst2 | 133.3812876 | 0.575379262 | 0.165055871 | 3.48596665 | 0.000490362 | 0.006749647 |
| Ap1g2 | 260.8641313 | 0.574648173 | 0.131629733 | 4.365641102 | 1.27E-05 | 0.000388676 |
| Nudt17 | 214.4373674 | 0.574134011 | 0.117377266 | 4.89135614 | 1.00E-06 | 4.88E-05 |
| Krt12 | 436.6734184 | 0.573160843 | 0.104139081 | 5.503801615 | 3.72E-08 | 3.25E-06 |
| Gm3002 | 494.5042043 | 0.572045443 | 0.145621492 | 3.92830369 | 8.55E-05 | 0.001745007 |
| Atf3 | 52.25598925 | 0.571528818 | 0.168120904 | 3.399510731 | 0.000675065 | 0.008462348 |
| Plb1 | 48.81217099 | 0.57120599 | 0.163632616 | 3.490783215 | 0.000481607 | 0.006658136 |
| Kcnc3 | 5344.180163 | 0.570918041 | 0.090143566 | 6.333430868 | 2.40E-10 | 4.13E-08 |
| Amy1 | 699.7132709 | 0.57065479 | 0.14427671 | 3.955280025 | 7.64E-05 | 0.00159628 |
| Pdcd1 | 8.41787075 | 0.570292581 | 0.135497191 | 4.208888584 | 2.57E-05 | 0.000679696 |
| Il18bp | 142.584881 | 0.569456576 | 0.158984556 | 3.581835802 | 0.000341188 | 0.005167517 |
| Aldob | 49.26684737 | 0.569412404 | 0.1666312 | 3.417201609 | 0.000632684 | 0.008095075 |
| Mapk13 | 24.59753223 | 0.569140682 | 0.166422005 | 3.419864339 | 0.000626524 | 0.008054403 |
| Ifi30 | 113.1439711 | 0.567974349 | 0.138705738 | 4.094815075 | 4.23E-05 | 0.001005576 |
| Snapc4 | 1839.848641 | 0.565921866 | 0.101054208 | 5.600181114 | 2.14E-08 | 2.03E-06 |
| Akr1b8 | 32.90242689 | 0.565121734 | 0.16721155 | 3.379681221 | 0.000725699 | 0.0089238 |
| Nxf7 | 91.38510561 | 0.564298089 | 0.161314342 | 3.498127206 | 0.000468538 | 0.006510782 |
| B230312C02Rik | 25.57658329 | 0.564086595 | 0.16809139 | 3.355832779 | 0.000791264 | 0.009492476 |
| Rtp4 | 88.21087806 | 0.563871015 | 0.158718159 | 3.552655975 | 0.000381363 | 0.005638202 |
| Cpeb1 | 2576.114924 | 0.562557535 | 0.089226242 | 6.304844023 | 2.88E-10 | 4.81E-08 |
| Uggt2 | 1071.344552 | 0.562437283 | 0.090183341 | 6.236598431 | 4.47E-10 | 7.00E-08 |
| Mafb | 843.6973662 | 0.562398299 | 0.099433559 | 5.656020989 | 1.55E-08 | 1.56E-06 |
| Hspb6 | 190.731772 | 0.560143428 | 0.125651625 | 4.457908359 | 8.28E-06 | 0.000269398 |
| Casp1 | 96.12265149 | 0.559826726 | 0.158548855 | 3.530941469 | 0.000414083 | 0.005968193 |
| Vmn2r84 | 21.89585925 | 0.55928314 | 0.162672249 | 3.438098043 | 0.000585816 | 0.007673777 |
| Myo19 | 356.7218086 | 0.558430401 | 0.132652582 | 4.209721317 | 2.56E-05 | 0.000678356 |
| Tbc1d10c | 47.97776154 | 0.55737192 | 0.15237634 | 3.657863946 | 0.000254326 | 0.004091928 |
| Spata32 | 28.38503249 | 0.552239221 | 0.162390024 | 3.400696708 | 0.000672144 | 0.008432544 |
| Cpne9 | 1962.2034 | 0.552008186 | 0.120911553 | 4.565388273 | 4.99E-06 | 0.000184364 |
| Parp3 | 215.7890304 | 0.551963467 | 0.134055047 | 4.117438925 | 3.83E-05 | 0.000931844 |
| Rorc | 95.43351474 | 0.551602804 | 0.129582629 | 4.256765031 | 2.07E-05 | 0.000573848 |
| D4Wsu53e | 5896.814657 | 0.551128304 | 0.106799626 | 5.160395456 | 2.46E-07 | 1.56E-05 |
| Npff | 34.08243052 | 0.550987823 | 0.158224005 | 3.482327619 | 0.000497075 | 0.006796205 |
| Pagr1a | 1471.115104 | 0.550577507 | 0.081945903 | 6.718792389 | 1.83E-11 | 4.24E-09 |
| Cerkl | 46.25713664 | 0.55052048 | 0.157939409 | 3.485643543 | 0.000490955 | 0.006749647 |
| Esyt1 | 596.2409955 | 0.55043157 | 0.106174214 | 5.184230248 | 2.17E-07 | 1.42E-05 |
| 4930570G19Rik | 287.7082891 | 0.549468517 | 0.12580114 | 4.367754674 | 1.26E-05 | 0.000386674 |
| Abhd14b | 557.6240189 | 0.549429402 | 0.084519451 | 6.500626688 | 8.00E-11 | 1.65E-08 |
| Cd84 | 164.8309008 | 0.548032146 | 0.12855625 | 4.2629755 | 2.02E-05 | 0.000563151 |
| Il12rb1 | 36.82283005 | 0.547840569 | 0.163880434 | 3.342928467 | 0.000828993 | 0.009842462 |
| Crhr2 | 67.63715087 | 0.54778498 | 0.165427121 | 3.311337191 | 0.000928513 | 0.010720099 |
| Stc2 | 33.38240404 | 0.547734858 | 0.162170251 | 3.37752981 | 0.0007314 | 0.008972538 |
| Evc | 391.0452063 | 0.546734199 | 0.1118509 | 4.888062606 | 1.02E-06 | 4.95E-05 |
| Col19a1 | 1105.15736 | 0.54631796 | 0.106679956 | 5.121092838 | 3.04E-07 | 1.87E-05 |
| Lilrb4 | 18.24260137 | 0.54491141 | 0.158174217 | 3.445007784 | 0.000571043 | 0.007510814 |
| Xylb | 299.337968 | 0.544762963 | 0.111693663 | 4.877295171 | 1.08E-06 | 5.16E-05 |
| Emx1 | 445.9827917 | 0.544150679 | 0.112441129 | 4.839427388 | 1.30E-06 | 6.10E-05 |
| Met | 263.0066217 | 0.543404877 | 0.130774315 | 4.15528751 | 3.25E-05 | 0.000822495 |
| Abcc8 | 1206.904999 | 0.54237955 | 0.119633736 | 4.533667257 | 5.80E-06 | 0.000206 |
| Fcgr4 | 18.21659617 | 0.541197242 | 0.14928507 | 3.625260337 | 0.000288671 | 0.00451328 |
| C3 | 41.06919178 | 0.541143698 | 0.16673764 | 3.245480131 | 0.001172527 | 0.012845805 |
| Acr | 39.6630239 | 0.540924191 | 0.167977053 | 3.220226707 | 0.001280893 | 0.013715376 |
| Serpina3n | 821.3954831 | 0.53937954 | 0.163707744 | 3.29477107 | 0.000985019 | 0.011255076 |
| Miip | 522.0950795 | 0.538604951 | 0.113464141 | 4.746917819 | 2.07E-06 | 9.12E-05 |
| Luc7l3 | 4668.850558 | 0.538298836 | 0.107401245 | 5.012035354 | 5.39E-07 | 2.99E-05 |
| Mir3072 | 34.80119347 | 0.538047947 | 0.168120306 | 3.200374545 | 0.001372491 | 0.014433037 |
| Hdac7 | 1242.897745 | 0.538011846 | 0.129227496 | 4.163292359 | 3.14E-05 | 0.000798084 |
| Col5a3 | 202.0086317 | 0.537715696 | 0.162112967 | 3.316919707 | 0.000910158 | 0.010547482 |
| Syne4 | 85.68276377 | 0.537357883 | 0.160733436 | 3.3431618 | 0.000828296 | 0.009842462 |
| Srpk3 | 215.437209 | 0.536711183 | 0.144951976 | 3.702682767 | 0.000213332 | 0.003556118 |
| Clk1 | 2833.366666 | 0.534807707 | 0.137186122 | 3.8984097 | 9.68E-05 | 0.001913558 |
| Catsper2 | 93.83404998 | 0.533968231 | 0.151466892 | 3.525313193 | 0.000422982 | 0.006043541 |
| Shisa6 | 968.6956184 | 0.532741747 | 0.132302384 | 4.026698007 | 5.66E-05 | 0.001268347 |
| Pbld2 | 41.93246654 | 0.532493738 | 0.155085353 | 3.433552731 | 0.000595726 | 0.00777606 |
| Sh2b2 | 199.8019305 | 0.532274438 | 0.139943963 | 3.803482681 | 0.000142676 | 0.002600731 |
| Ovgp1 | 285.7400495 | 0.530035867 | 0.167759261 | 3.159502877 | 0.001580385 | 0.016067252 |
| Ttc14 | 3698.981742 | 0.529844484 | 0.164163585 | 3.227539684 | 0.001248597 | 0.013490771 |
| Meg3 | 106402.8544 | 0.529597234 | 0.16278939 | 3.253266294 | 0.001140865 | 0.012610872 |
| Ptprc | 162.9082999 | 0.528853482 | 0.127446535 | 4.149610524 | 3.33E-05 | 0.000833626 |
| Hr | 1742.726301 | 0.528200436 | 0.103777027 | 5.089762665 | 3.59E-07 | 2.15E-05 |
| Eqtn | 30.09263517 | 0.52756931 | 0.166071736 | 3.176755564 | 0.001489325 | 0.01539287 |
| Pnpla7 | 679.3112604 | 0.527251026 | 0.080060702 | 6.585640835 | 4.53E-11 | 9.75E-09 |
| Efcab12 | 208.3753231 | 0.52700164 | 0.13504409 | 3.902441338 | 9.52E-05 | 0.001886767 |
| C4a | 90.02179343 | 0.526973429 | 0.163552929 | 3.222036019 | 0.001272831 | 0.013683744 |
| Asprv1 | 40.53947689 | 0.526915855 | 0.158213556 | 3.330409021 | 0.000867185 | 0.010225679 |

DAP12-/- Tau+ vs DAP12+/+ Tau+

| gene | baseMean | log2FoldChange | lfcSE | stat | pvalue | padj | refseq_id |
|---|---|---|---|---|---|---|---|
| Tyrobp | 359.6387953 | -2.40291984988619 | 0.0950231415271382 | -25.2877331907609 | 4.3583983657463e-141 | 6.33100946608307e-137 | NM_011662 |
| Nudt19 | 1406.974187 | -1.21111083377982 | 0.061162364308091 | -19.8015699275315 | 2.88559298831442e-87 | 2.09580618741277e-83 | NM_033080 |
| Zfp14 | 696.952305 | 0.792505562773603 | 0.064665397941985 | 12.2554811072933 | 1.57014718313813e-34 | 7.60265266075481e-31 | NM_011748,NM_178733 |
| C4b | 2177.430204 | -0.787387232403712 | 0.0854832440364647 | -9.21101253560095 | 3.23062913518304e-20 | 1.17320297044172e-16 | NM_009780 |
| Sbsn | 264.8129095 | -0.72997241278742 | 0.0840460969012288 | -8.68538147161414 | 3.7747346009186e-18 | 1.09663589625887e-14 | NM_172205,NM_001083903 |
| Gm1082 | 32.27045956 | 0.708453773923395 | 0.082361766201253 | 8.60173119882193 | 7.85225292191983e-18 | 1.90103043239679e-14 | NR_036630 |
| Mrps12 | 803.7912892 | -0.492843662206338 | 0.0650271417834732 | -7.57904543686396 | 3.48106592679104e-14 | 7.22370909322381e-11 | NM_011885 |
| Lag3 | 504.3588427 | -0.664146063304481 | 0.0940716433754775 | -7.06000277526363 | 1.66499278885148e-12 | 3.02321065635707e-09 | NM_008479 |
| Blvrb | 313.9339086 | 0.547711457290422 | 0.0798362311007341 | 6.86043729443268 | 6.86500730170466e-12 | 1.10801217849513e-08 | NM_144923 |
| Cd52 | 94.61086518 | -0.618505511041228 | 0.0935981136669403 | -6.60809803541682 | 3.89288876346661e-11 | 5.65481021781196e-08 | NM_013706 |
| Gpatch1 | 977.1291577 | -0.349113433763029 | 0.0538139971793471 | -6.48740944850336 | 8.73247898706909e-11 | 1.15316354332878e-07 | NM_026181 |
| Cep89 | 462.1990602 | -0.487892182972809 | 0.0754168720215634 | -6.46927099858119 | 9.84768472263976e-11 | 1.19206223567554e-07 | NM_028120 |
| Ryr1 | 814.1892978 | -0.527323355573007 | 0.0821114342885364 | -6.42204535996794 | 1.34455341043115e-10 | 1.50238329537869e-07 | NM_009109 |
| Itgax | 102.6554611 | -0.408936713221158 | 0.0645775440870206 | -6.33249094561571 | 2.41234448026495e-10 | 2.50297970859491e-07 | NM_021334 |
| Zkscan3 | 1163.650182 | -0.446284155141329 | 0.0707188317154037 | -6.31068336843185 | 2.77806041315346e-10 | 2.69027370409781e-07 | NM_001145778,NM_023685 |
| Zfp27 | 913.3537307 | 0.395164053560711 | 0.0638993547192485 | 6.18416344416817 | 6.24326565780827e-10 | 5.66810480908269e-07 | NM_011754,NM_001037707 |
| Col11a1 | 849.3852423 | -0.512153256127347 | 0.08582477354343 | -5.9674291580645 | 2.41020875738938e-09 | 2.05945249469636e-06 | NM_007729 |
| Dmkn | 74.88580842 | -0.562152356119932 | 0.0949489192255246 | -5.92057667117512 | 3.20814685653742e-09 | 2.5889745132257e-06 | NM_028618,NM_001166173,NM_001166174,NM_172899 |
| Ech1 | 1234.130815 | -0.339138321793787 | 0.0581011559383374 | -5.83703226410355 | 5.31388089174296e-09 | 4.06260178070833e-06 | NM_016772 |
| Fbxo17 | 72.42112456 | -0.534556475552189 | 0.0949609690681557 | -5.62922304603405 | 1.81023184496976e-08 | 1.31477138900154e-05 | NM_015796 |
| Klhl6 | 80.66227421 | -0.529154941054153 | 0.0950638641606262 | -5.56631003511553 | 2.60190139380268e-08 | 1.79977236411322e-05 | NM_183390 |
| Sfrs18 | 7556.932928 | -0.360726385032001 | 0.0665081627800156 | -5.42379115515715 | 5.83480505378186e-08 | 3.85256264596524e-05 | NM_025669 |
| Zkscan2 | 845.8572196 | -0.394282698836982 | 0.0731518421161114 | -5.38992166747012 | 7.04884015502599e-08 | 4.45180226486555e-05 | NM_001081329 |
| Snhg11 | 73090.53635 | -0.450347664499575 | 0.0850995125794518 | -5.29201226715741 | 1.20977775283854e-07 | 7.32217984905527e-05 | NM_175692 |
| Gbp9 | 153.547225 | -0.496320855107311 | 0.0947602061557952 | -5.2376506472697 | 1.62633537880121e-07 | 8.88165337677487e-05 | NM_172777 |
| Kctd15 | 410.9900555 | -0.358287522600791 | 0.0683691461859688 | -5.24048554923041 | 1.60154629863265e-07 | 8.88165337677487e-05 | NM_146188 |
| Robo3 | 1385.509849 | -0.488242925544516 | 0.0932671507129933 | -5.2348862575095 | 1.65086493992098e-07 | 8.88165337677487e-05 | NM_001164767 |
| Gm6537 | 118.5045039 | 0.489308436225462 | 0.0936117143472559 | 5.22700005696248 | 1.72282447302814e-07 | 8.93776724828815e-05 | NM_001195091 |
| Irf8 | 278.0190289 | -0.490156352196579 | 0.0941089206478175 | -5.20839415458695 | 1.90481958797656e-07 | 9.4137751627637e-05 | NM_008320 |
| Kirrel2 | 538.3397882 | 0.493308696612428 | 0.0947832840524713 | 5.20459595322036 | 1.9441914834291e-07 | 9.4137751627637e-05 | NM_172898 |
| Ttc14 | 4985.536852 | -0.374806833004513 | 0.0721524364837758 | -5.19465247842035 | 2.05102611081286e-07 | 9.61071138247342e-05 | NM_027619,NM_025978 |
| Myo1f | 177.6744227 | -0.481449214893432 | 0.0936206547462954 | -5.14255338416636 | 2.7102939302578e-07 | 0.000123030405096664 | NM_053214 |
| Slc26a10 | 206.126767 | -0.429691495323828 | 0.0840312475438696 | -5.11347276023127 | 3.16289384407524e-07 | 0.000139224836300112 | NM_177615 |
| Adipor2 | 3828.371184 | 0.344487291662744 | 0.0679499206515552 | 5.06972323675347 | 3.98394604264493e-07 | 0.000170208235927824 | NM_197985 |
| Ddx26b | 2180.945356 | -0.318737944656715 | 0.063189679179307 | -5.04414563891462 | 4.55552231685146e-07 | 0.000189067191927384 | NM_172779 |
| Mpeg1 | 1997.379678 | -0.44172349723431 | 0.0887329263362837 | -4.97812385404993 | 6.42035694482932e-07 | 0.000259061402723863 | NM_010821 |
| Gm5577 | 221.6962085 | -0.435279408315105 | 0.0888094939418842 | -4.90127112535903 | 9.5218537490163e-07 | 0.000373822831238407 | NR_026990 |
| Atp11b | 3806.078508 | -0.290789676879999 | 0.0594498093488674 | -4.8913475091832 | 1.00147967186867e-06 | 0.000376211074681689 | NM_029570 |
| Luc7l3 | 5691.921881 | -0.302664573205028 | 0.0618988105223623 | -4.88966701865271 | 1.01006690848037e-06 | 0.000376211074681689 | NM_026313 |
| Cd14 | 98.2637337 | -0.461192412574898 | 0.0944200313621335 | -4.88447637563332 | 1.03704075079701e-06 | 0.000376601348651933 | NM_009841 |
| Cd33 | 276.1730083 | -0.417140194314924 | 0.0856568250599698 | -4.86990025631789 | 1.11654595738449e-06 | 0.000395584062852855 | NM_021293,NM_001111058 |
| Fcgr1 | 152.2107014 | -0.449813291572231 | 0.0926511994515923 | -4.85491061351285 | 1.20441058917874e-06 | 0.000409026548960365 | NM_010186 |
| Stx3 | 1309.629746 | -0.303428616949977 | 0.0625128318200339 | -4.85386132919891 | 1.2108041859628e-06 | 0.000409026548960365 | NM_152220,NM_011502,NM_001025307 |
| Nckap1l | 737.8115118 | -0.286273931965638 | 0.0591523204012365 | -4.83960612236023 | 1.30096707895454e-06 | 0.000429496540656672 | NM_153505 |
| Emr1 | 327.1843711 | -0.396968152019382 | 0.0822545373636002 | -4.82609427689821 | 1.39236616524139e-06 | 0.000449455798139921 | NM_010130 |
| A930014F10Rik | 374.1246032 | -0.426274782844462 | 0.08911779547584 | -4.78327342556432 | 1.72463271384661e-06 | 0.000544609017420344 | NR_027886 |
| Rhou | 2076.716139 | 0.257145581601538 | 0.0538119142554578 | 4.77859940794537 | 1.76520485272119e-06 | 0.00054556097214102 | NM_133955 |
| Ovgp1 | 266.9610844 | -0.445222446229205 | 0.0936929426885928 | -4.75193150575909 | 2.01482636796523e-06 | 0.000609736911448431 | NM_007696 |
| Lair1 | 380.3287927 | -0.393355671890463 | 0.082910601491119 | -4.74433504058704 | 2.09192368407989e-06 | 0.000620148641529481 | NM_001113474,NM_178611 |
| Col19a1 | 1298.402088 | -0.370690028018518 | 0.0787038575504339 | -4.70993467862713 | 2.47796196144481e-06 | 0.000719897509038945 | NM_007733 |
| Ptpn6 | 220.8980729 | -0.434744984448747 | 0.0934325621495491 | -4.65303502811889 | 3.27084658196932e-06 | 0.000931614067640908 | NM_013545,NM_001077705 |
| Ets1 | 514.946524 | -0.302135644808435 | 0.0650171380304771 | -4.64701544793939 | 3.36771954914958e-06 | 0.000940759503287438 | NM_001038642,NM_011808 |
| Aif1 | 230.4358008 | -0.375054553354123 | 0.0808969503070345 | -4.63620138868832 | 3.54870235380025e-06 | 0.000972612271534009 | NM_019467 |
| Gm3002 | 637.324772 | -0.325282282362096 | 0.0706175175836034 | -4.6062548428865 | 4.09985897301741e-06 | 0.00110286206374168 | NR_033388 |
| Ly86 | 561.0912527 | -0.410913160123243 | 0.0895195642621519 | -4.59020509661901 | 4.42810683862875e-06 | 0.00114861928460574 | NM_010745 |
| Siglec5 | 36.62049432 | -0.358877236456539 | 0.0781648864715564 | -4.59128456083834 | 4.40526221126048e-06 | 0.00114861928460574 | NM_001271019,NM_145581 |
| Cep70 | 745.9681042 | -0.303987948334176 | 0.0664866975853687 | -4.57216194177575 | 4.82717629216256e-06 | 0.00121391585400917 | NM_023873 |
| Haus5 | 232.0451918 | -0.381941079782139 | 0.0836819969632151 | -4.5641965258106 | 5.01410926893502e-06 | 0.00121391585400917 | NM_027999 |
| Malat1 | 8083.783777 | -0.398032461620918 | 0.0870830345401398 | -4.57072337594586 | 4.86043523190145e-06 | 0.00121391585400917 | NR_002847 |
| Tie1 | 491.7339399 | -0.294713484941079 | 0.0645645127121327 | -4.56463578150258 | 5.00362266780618e-06 | 0.00121391585400917 | NM_011587 |
| Naip5 | 88.86037488 | -0.429307414298131 | 0.0950777146047279 | -4.51533165351013 | 6.32177500970931e-06 | 0.00150541153755799 | NM_010870 |
| Itgam | 1199.284169 | -0.271064873261054 | 0.0602682699096902 | -4.4976382044 | 6.87124947508523e-06 | 0.00160986725604981 | NM_001082960,NM_008401 |
| Miat | 23357.0276 | -0.371041217685309 | 0.0826254099425863 | -4.49064298674142 | 7.1008486861765e-06 | 0.00163725282564127 | NR_033657,NR_003718 |
| Uvssa | 535.0820942 | -0.32378369868314 | 0.0722068655206848 | -4.48411236727049 | 7.32180738647817e-06 | 0.00166182147024972 | NM_001081101 |
| Dock2 | 244.9161645 | -0.389378356606729 | 0.0869098179838523 | -4.48025741670607 | 7.45530698972011e-06 | 0.00166608906665653 | NM_033374 |
| I830012O16Rik | 248.0010337 | -0.413135649963232 | 0.0927164524126037 | -4.45590441839502 | 8.35402489397793e-06 | 0.0018112024717899 | NM_001005858 |
| Leng8 | 11174.74125 | -0.399388558650658 | 0.0896283402294687 | -4.45605215264613 | 8.34827302204529e-06 | 0.0018112024717899 | NM_172736 |
| Apobec3 | 130.5387185 | -0.413054695508539 | 0.0927883860883762 | -4.45157754026593 | 8.52417411773162e-06 | 0.00182091401814955 | NM_030255,NM_001160415 |
| Alkbh6 | 956.9097166 | -0.250987433077155 | 0.0564566548627971 | -4.44566603684036 | 8.76199762730561e-06 | 0.00184459097875712 | NM_198027 |
| Plekhf1 | 189.2838919 | 0.418278595141035 | 0.0944698959444676 | 4.42763899503936 | 9.52701791888276e-06 | 0.00197699231842416 | NM_024413 |
| Il18bp | 169.3893851 | -0.412845495861055 | 0.093472143860561 | -4.41677572386621 | 1.00184083374961e-05 | 0.00199352602069135 | NM_010531 |
| Meg3 | 133078.2528 | -0.392067205483107 | 0.0887664267492486 | -4.41684113961955 | 1.00153781373426e-05 | 0.00199352602069135 | NR_046475,NR_027651,NR_027652,NR_003633 |
| Zfc3h1 | 2588.245007 | -0.287546225809227 | 0.0650332093015781 | -4.4215290756418 | 9.80048732067536e-06 | 0.00199352602069135 | NM_001033261 |
| Cd84 | 177.1092639 | -0.413512276350292 | 0.0940141005397917 | -4.39840698337876 | 1.0904834481003e-05 | 0.00214058953609526 | NM_013489,NM_001252472 |
| Phf11d | 27.76999338 | -0.349630302831045 | 0.0798333226750422 | -4.37950333414279 | 1.1895010182116e-05 | 0.00229510566604579 | NM_199015 |
| Srsf11 | 3374.870484 | -0.269779165643664 | 0.0616293986893178 | -4.37744276889115 | 1.20079877887567e-05 | 0.00229510566604579 | NM_026989,NM_001093753,NM_001093752 |
| Zfp217 | 175.1211143 | -0.371529203787759 | 0.0849785518493185 | -4.37203501004046 | 1.23093780209876e-05 | 0.00232215617055669 | NM_001033299,NM_001159683 |
| Cxcl16 | 75.5319207 | -0.403461317876058 | 0.0923897247536674 | -4.36695010134277 | 1.25993496751834e-05 | 0.00234638658181684 | NM_023158 |
| Man2b1 | 2146.94953 | -0.363986953895851 | 0.0836696075875981 | -4.35028876542505 | 1.35958370835475e-05 | 0.00249049020409843 | NM_010764 |
| Nvl | 1987.591374 | -0.284658404941717 | 0.065486480814993 | -4.34682703054254 | 1.38121054614068e-05 | 0.00249049020409843 | NM_026171 |
| Spag5 | 649.0770472 | -0.355277427672769 | 0.0817550560522105 | -4.34563248841615 | 1.3887491844415e-05 | 0.00249049020409843 | NM_017407 |
| Col11a2 | 1033.387715 | -0.336296843011168 | 0.0774469036282748 | -4.34228906846046 | 1.41005838723082e-05 | 0.00249786684547743 | NM_009926 |
| Lcp1 | 560.9722753 | -0.333411231369673 | 0.07683965144464 | -4.33905184499545 | 1.43098764143551e-05 | 0.00250440078066171 | NM_008879,NM_001247984 |
| Cep110 | 638.3545329 | -0.340343939095525 | 0.0786228423742528 | -4.32881753976094 | 1.4991206699607e-05 | 0.00259240795855347 | NM_030000,NM_012018 |
| Gm15328 | 107.6535738 | -0.39409344281951 | 0.0918949905852544 | -4.28851932308426 | 1.79868153581115e-05 | 0.00303809860339451 | NR_045399 |
| Srrm2 | 30891.1576 | -0.315263770065499 | 0.0734691753382848 | -4.29110261022918 | 1.7778813366539e-05 | 0.00303809860339451 | NM_175229 |
| Chuk | 1305.646425 | -0.315493878921368 | 0.0736277525184144 | -4.28498586647015 | 1.82750805734605e-05 | 0.00305130828057572 | NM_007700,NM_001162410 |
| Cx3cr1 | 2682.254466 | -0.284030971208205 | 0.0665323205346839 | -4.26906755882859 | 1.96291813726347e-05 | 0.00324015327976013 | NM_009987 |
| Ogt | 12777.40456 | -0.254510356346119 | 0.0598281641177064 | -4.25402250093105 | 2.09964189083706e-05 | 0.00342689866362912 | NM_139144 |
| Ccl6 | 107.509393 | -0.37402255046444 | 0.0884757904431775 | -4.22739993156265 | 2.3640725657356e-05 | 0.00377368330657971 | NM_009139 |
| Slc11a1 | 135.5536509 | -0.402194972775397 | 0.0950959733482861 | -4.22935860073034 | 2.34358523980207e-05 | 0.00377368330657971 | NM_013612 |
| Zfp36 | 200.9219435 | -0.360842275709323 | 0.0855791768222731 | -4.21647285131877 | 2.48153146768934e-05 | 0.00391812240213645 | NM_011756 |
| Parp14 | 203.9879468 | -0.395434676740447 | 0.0945620931552548 | -4.18174623198337 | 2.8927873935482e-05 | 0.00451834727727755 | NM_001039530 |
| Sall1 | 1219.244421 | -0.211152761774178 | 0.0506296311365157 | -4.17053723351912 | 3.03882397959333e-05 | 0.00469595288591199 | NM_021390 |
| Tsc22d3 | 3720.830662 | 0.361782799565582 | 0.0874095515526477 | 4.13893897336467 | 3.4891573895351e-05 | 0.00533510528846178 | NM_001077364,NM_010286 |
| Rbm5 | 5107.812644 | -0.196961597269706 | 0.0476364659354664 | -4.13468113979176 | 3.5544780279529e-05 | 0.00537836956604623 | NM_148930 |
| Ctsc | 306.293372 | -0.381779572431914 | 0.0925392775008706 | -4.12559491215309 | 3.69777642013678e-05 | 0.00553751549267081 | NM_009982 |
| 6820431F20Rik | 2800.15088 | -0.196984418552859 | 0.0478965148559094 | -4.11270880867765 | 3.91043501542069e-05 | 0.00579622235040826 | NR_030708 |
| Gbp4 | 131.6410243 | -0.373560214612436 | 0.0909197595796239 | -4.10868018502939 | 3.97926675922049e-05 | 0.0058036203701741 | NM_008620,NM_001256005 |
| Unc93b1 | 693.6997405 | -0.335496374388954 | 0.0816740123687581 | -4.10774938880422 | 3.99533276206396e-05 | 0.0058036203701741 | NM_019449,NM_001161428 |
| Hif3a | 486.7148821 | 0.383189886986828 | 0.093594458702543 | 4.0941514305314 | 4.23717328124341e-05 | 0.00609397812706354 | NM_016868,NM_001162950 |
| Fmnl3 | 666.2656188 | -0.286021516029054 | 0.0701188745071539 | -4.07909450970827 | 4.52114552734219e-05 | 0.00643864313040909 | NM_011711 |
| Cbln4 | 841.5159451 | 0.361741645349437 | 0.0888863463938618 | 4.06970991637494 | 4.70717107193091e-05 | 0.00647972265111568 | NM_175631 |
| Cyth4 | 504.1714374 | -0.324515540291918 | 0.0796851130664544 | -4.07247386373518 | 4.65164213106404e-05 | 0.00647972265111568 | NM_028195 |
| Lacc1 | 219.6643466 | -0.334321224153048 | 0.082157253077911 | -4.06928434956322 | 4.71577653677257e-05 | 0.00647972265111568 | NM_172488 |
| Trank1 | 19038.44546 | -0.265062349736 | 0.065147328343461 | -4.06866031924707 | 4.72842214662166e-05 | 0.00647972265111568 | NM_001164659 |
| Irf9 | 529.277702 | -0.333430950748619 | 0.0820952063295775 | -4.06151547277968 | 4.87551862043741e-05 | 0.00656453901416076 | NM_001159418,NM_008394,NM_001159417 |
| Smc5 | 1223.408165 | -0.230696231877219 | 0.0568039963821783 | -4.0612676320358 | 4.88069815179239e-05 | 0.00656453901416076 | NM_001252684,NM_153808,NM_001252685 |
| Med12 | 1864.223624 | -0.281606891675423 | 0.0694353847254249 | -4.05566834243101 | 4.9991152182501e-05 | 0.00666212363855972 | NM_021521 |
| Lgi4 | 1540.32414 | -0.272967291177452 | 0.0673613687610468 | -4.05228243128131 | 5.07203838010502e-05 | 0.00669785722812778 | NM_144556 |
| Slc14a1 | 550.7817664 | -0.319536826856313 | 0.0789825684867271 | -4.04566264940623 | 5.21753256902352e-05 | 0.00682791694573295 | NM_028122,NM_001171011,NM_001171010 |
| Uggt2 | 1319.686242 | -0.256895802558219 | 0.0635649506621038 | -4.04146939283908 | 5.31173148365564e-05 | 0.00688912602960551 | NM_001081252 |
| Hdac7 | 1532.395537 | -0.344945448252702 | 0.0854807147945204 | -4.03535989470708 | 5.45186646837361e-05 | 0.0070083019751854 | NM_019572,NM_001204278,NM_001204281,NM_001204280,NM_001204276,NM_001204277,NM_001204275,NM_001204279 |
| Cd86 | 111.5448965 | -0.382234780686972 | 0.0949649301770866 | -4.02500986389604 | 5.69729273458561e-05 | 0.00717959921740751 | NM_019388 |
| Ddx58 | 254.1809065 | -0.354680512037549 | 0.088186844308658 | -4.02192089781726 | 5.7725445288841e-05 | 0.00717959921740751 | NM_172689 |
| Plek | 762.7550869 | -0.347521244303612 | 0.0864157856195978 | -4.02150187968435 | 5.7828246484695e-05 | 0.00717959921740751 | NM_019549 |
| Zcchc7 | 841.2395217 | -0.261415531916062 | 0.0649523167626359 | -4.02472990873272 | 5.70407437852471e-05 | 0.00717959921740751 | NM_138590 |
| Aspg | 121.0110147 | -0.372163273868755 | 0.0926307050520943 | -4.01770960999871 | 5.87665540756157e-05 | 0.00722952240986051 | NM_001081169 |
| Pnn | 3176.347877 | -0.256077572336065 | 0.0637663269515498 | -4.01587459366502 | 5.92257446491395e-05 | 0.00722952240986051 | NM_008891 |
| Ifi44 | 51.24543977 | -0.274185060656044 | 0.0683498672083733 | -4.01149368469372 | 6.03357913144008e-05 | 0.00730364753860822 | NM_133871 |
| Tnfsf10 | 121.9224566 | -0.379715936139552 | 0.0949082835290583 | -4.00087244253336 | 6.31093723225072e-05 | 0.00757625406906396 | NM_009425 |
| 4831440E17Rik | 178.1454702 | -0.354931177477553 | 0.0891007004805977 | -3.98348358164527 | 6.79123654594483e-05 | 0.00808602475954054 | NR_030700 |
| Akap8 | 3390.384067 | -0.264554828337596 | 0.0664710481852223 | -3.98000085090296 | 6.89150215579065e-05 | 0.00813020773651803 | NM_019774 |
| Ppapdc2 | 2005.274858 | 0.261655261217912 | 0.0658288239282835 | 3.9747825588221 | 7.04435852246396e-05 | 0.00813020773651803 | NM_028922 |
| Ptprc | 183.2461383 | -0.365003777163134 | 0.0918360103237697 | -3.97451692289664 | 7.052224802432e-05 | 0.00813020773651803 | NM_011210,NM_001268286,NM_001111316 |
| Trim30d | 71.96198326 | -0.374869548785096 | 0.094314340910117 | -3.97468237765742 | 7.04732421019063e-05 | 0.00813020773651803 | NM_199146,NM_001167828 |
| Slc15a3 | 106.1324819 | -0.37249978586481 | 0.0942044174470841 | -3.95416474045973 | 7.68024800394943e-05 | 0.00871588144573199 | NM_023044 |
| Tnip2 | 571.8799493 | -0.281547766641211 | 0.0711912605886249 | -3.9548079962809 | 7.65961400050452e-05 | 0.00871588144573199 | NM_139064 |
| Ddx17 | 18592.86926 | -0.195972766325697 | 0.04962026439271 | -3.94945026440625 | 7.83308839287585e-05 | 0.0088204218600709 | NM_001040187,NM_199079,NM_199080,NM_152806 |
| Fes | 165.0930382 | -0.341573725403749 | 0.0867531471185191 | -3.93730644649817 | 8.24013484649613e-05 | 0.00920739990616944 | NM_010194 |
| Col5a1 | 1668.118593 | -0.348921463074035 | 0.0887369037280338 | -3.93208967650517 | 8.42106382157331e-05 | 0.00933773840245603 | NM_015734 |
| Ccdc39 | 737.3523386 | -0.279633731692804 | 0.0712245756070935 | -3.92608491253621 | 8.63396965625416e-05 | 0.00950129009710514 | NM_026222 |
| Parp12 | 508.2499547 | -0.300342063447896 | 0.0766407169323352 | -3.91883160113263 | 8.89792520990802e-05 | 0.00971813996985894 | NM_172893 |
| Tia1 | 2489.824 | -0.274761543483474 | 0.0701983878737522 | -3.91407198663333 | 9.07525602450749e-05 | 0.00978668659253215 | NM_011585,NM_001164079,NM_001164078 |
| Wsb1 | 3248.782231 | -0.254632595578902 | 0.0650645851081971 | -3.91353599128418 | 9.0954336361823e-05 | 0.00978668659253215 | NM_001042565,NM_019653 |
| Arid5b | 1101.463601 | 0.241584838423449 | 0.0617868761253579 | 3.90997010325143 | 9.23075451775391e-05 | 0.00981574909690799 | NM_023598 |
| Flna | 4352.157067 | -0.28175526959398 | 0.0720736494939614 | -3.90926880421777 | 9.25759070822246e-05 | 0.00981574909690799 | NM_010227 |
| Rnf207 | 354.4363388 | -0.340600586605238 | 0.0874478424059431 | -3.89489983096587 | 9.82393191180473e-05 | 0.010306487659487 | NM_001033489 |
| Xaf1 | 257.4136361 | -0.340503168559174 | 0.0874440611642593 | -3.89395419226419 | 9.862328133347581e-05 | 0.010306487659487 | NM_001037713 |
| Tatdn3 | 518.9302249 | -0.299045015454035 | 0.0770383119101615 | -3.88177009645237 | 0.000103698891199207 | 0.0107595006682834 | NM_026895,NM_001163421,NM_001163422 |
| Guf1 | 1262.572621 | -0.249742998267656 | 0.0643733431984057 | -3.87960273397516 | 0.000104627204617116 | 0.0107788281862995 | NM_172711 |
| Dclre1c | 630.0777169 | -0.304277804416391 | 0.0785872663015364 | -3.8718461493353 | 0.000108014125501691 | 0.0109721201890738 | NM_146114,NM_001110214,NM_175683 |
| Ncf2 | 204.4404366 | -0.356569866438547 | 0.0920704420821399 | -3.87279411692665 | 0.000107594713741344 | 0.0109721201890738 | NM_010877 |
| Csf3r | 361.4791217 | -0.357082923521769 | 0.0925335190559608 | -3.85895756656373 | 0.000113871705329449 | 0.0113294547370929 | NM_001252651,NR_045561,NM_007782 |
| Gpr123 | 3338.540836 | 0.265997823698726 | 0.0689190925673221 | 3.85956654085212 | 0.00011358832543268 | 0.0113294547370929 | NM_177469 |
| Pfkfb2 | 2328.368369 | -0.203812067268248 | 0.0527858647282346 | -3.86111070298013 | 0.000112872744361306 | 0.0113294547370929 | NM_008825,NM_001162416,NM_001162415,NR_027859 |
| Arglu1 | 3327.755096 | -0.281284147385129 | 0.0730949272722345 | -3.84820339635219 | 0.000118987245550324 | 0.0116000585829799 | NM_176849 |
| Dusp11 | 2594.484577 | -0.280606459983912 | 0.0728928528853638 | -3.84957439414825 | 0.000118323242849982 | 0.0116000585829799 | NM_028099 |
| Kcna5 | 282.5257998 | 0.342127531696297 | 0.08886511585324 | 3.84996439166656 | 0.000118134998246325 | 0.0116000585829799 | NM_145983 |
| 6720401G13Rik | 3516.251139 | -0.275941128767798 | 0.0718021928942127 | -3.84307383444885 | 0.000121502895300666 | 0.0116884175969369 | NR_015505,NR_026976 |
| Odf2l | 439.5629013 | -0.260288302505576 | 0.0677049876789251 | -3.84444797095204 | 0.000120824115424549 | 0.0116884175969369 | NM_001162538,NM_001162539,NM_025714 |
| Fkbp5 | 1931.480402 | 0.315738016015486 | 0.0822545244022119 | 3.8385489225076 | 0.000123763557439529 | 0.0118275620747803 | NM_010220 |
| Lyn | 409.3667518 | -0.309025703263271 | 0.0805627707499044 | -3.83583757592694 | 0.000125137092179033 | 0.0118806627515858 | NM_010747,NM_001111096 |
| Parp9 | 177.7816606 | -0.356530444604407 | 0.0931422898626516 | -3.82780415995946 | 0.000129291562645044 | 0.0121953846687137 | NM_030253 |
| Rinl | 112.8997985 | 0.362270192444225 | 0.0949480928709928 | 3.81545517650835 | 0.000135932130350605 | 0.0126662689125986 | NM_177158 |
| Srek1 | 1992.847244 | -0.243360761731765 | 0.0637857913477705 | -3.81528168875295 | 0.000136027671097713 | 0.0126662689125986 | NM_172592 |
| Arhgap30 | 168.4926081 | -0.328316007773427 | 0.086161679084925 | -3.8104643649043 | 0.000138706005777227 | 0.0127521736703798 | NM_001005508 |
| Szt2 | 2505.216463 | -0.263936531992258 | 0.0692574188259675 | -3.81094959163129 | 0.000138433996416022 | 0.0127521736703798 | NM_198170 |
| Acp6 | 469.0758991 | -0.263966462746121 | 0.0694792151702979 | -3.79921480257258 | 0.000145155244416851 | 0.0128694201397073 | NM_019800 |
| Ccl12 | 24.89050196 | -0.254399683979627 | 0.066972986167609 | -3.79854174850375 | 0.000145549915126239 | 0.0128694201397073 | NM_011331 |
| Cd48 | 39.5085305 | -0.337126237211923 | 0.0885948692022656 | -3.80525689859364 | 0.000141657086084257 | 0.0128694201397073 | NM_007649 |
| Csf1r | 5157.77762 | -0.246104524602814 | 0.0648005321253638 | -3.79787814283218 | 0.000145940034540227 | 0.0128694201397073 | NM_001037859 |
| Luc7l2 | 5384.002953 | -0.172945470626369 | 0.045537164087001 | -3.79789725807141 | 0.000145928783345552 | 0.0128694201397073 | NM_138680,NM_001170849,NM_001170848 |
| Slamf9 | 57.17082679 | -0.339481901492477 | 0.0895023220692444 | -3.79299546250692 | 0.000148840877286991 | 0.0128694201397073 | NM_029612 |
| Srrm1 | 5284.812112 | -0.208995694962629 | 0.0550427557558579 | -3.79697004796834 | 0.000146475479636164 | 0.0128694201397073 | NM_001130477,NM_016799 |
| Tmem181b-ps | 10365.76073 | -0.256282077738801 | 0.0675193867846039 | -3.79568135825013 | 0.00014723851294941 | 0.0128694201397073 | NR_033520 |
| Trim30a | 123.7714625 | -0.360269648413501 | 0.0949566487428986 | -3.79404342068721 | 0.000148213738026519 | 0.0128694201397073 | NM_009099 |
| Zc3h7a | 1472.437608 | -0.204007135889026 | 0.0537210199838321 | -3.79752908545713 | 0.000146145632771576 | 0.0128694201397073 | NM_145931,NR_027502 |
| Ifit1 | 155.5557556 | -0.347333644409904 | 0.0917058396843256 | -3.78747575514834 | 0.000152185529622826 | 0.0130807514988235 | NM_008331 |
| Mlxipl | 161.7954127 | -0.355793663042907 | 0.094065817943644 | -3.78239057312048 | 0.000155329384081189 | 0.013272439018608 | NM_021455 |
| Dennd6b | 3614.624161 | -0.300625141067903 | 0.0796335213657825 | -3.77510796850279 | 0.000159938374431494 | 0.0135863440174964 | NM_027081 |
| B3galt5 | 2862.056054 | 0.25185288273177 | 0.0668025741390894 | 3.77010745435329 | 0.000163177275117944 | 0.0137327135474272 | NM_001122993,NM_033149 |
| Csf2ra | 781.1309724 | -0.322208983960467 | 0.0854771252151337 | -3.76953463455297 | 0.000163552212839385 | 0.0137327135474272 | NM_009970 |
| Lpcat2 | 339.7725945 | -0.309336153592072 | 0.0822308722551156 | -3.76180070949969 | 0.000168694442486002 | 0.0140830774227107 | NM_173014 |
| Ankrd16 | 744.2242263 | -0.266965577032735 | 0.071210012662461 | -3.74898932118107 | 0.000177548644587868 | 0.0147375520644764 | NM_177268 |
| Tmc4 | 184.108921 | -0.353452527424893 | 0.0943836000725785 | -3.74485108803963 | 0.000180500803212464 | 0.0148974697015014 | NM_181820 |
| Rbm33 | 3862.600328 | -0.230339859600552 | 0.0618069339327472 | -3.7267640529004 | 0.000193953866197115 | 0.0159173664428208 | NM_028234 |
| Ncrna00085 | 430.7779093 | -0.336751891857786 | 0.0906595881666798 | -3.71446527242832 | 0.000203633830775916 | 0.0166178934036571 | NM_001162909 |
| Ubp1 | 4011.714008 | -0.211473526505821 | 0.0571382875635682 | -3.70108268069024 | 0.000214681504210824 | 0.0174215839674102 | NM_013699,NM_001083319 |
| Hk2 | 209.8444779 | -0.333115117512698 | 0.0902295847549638 | -3.69186136029926 | 0.000222618813456962 | 0.0179653382459768 | NM_013820 |
| Thpo | 289.4104594 | -0.345231240763803 | 0.0937262730435432 | -3.68339879046952 | 0.000230144684996509 | 0.0184700646091673 | NM_001173505,NM_009379 |
| Catsper2 | 114.7450725 | -0.344185124849999 | 0.0935643447716709 | -3.67859279824947 | 0.00023452435992153 | 0.0186158516514762 | NM_153075 |
| Was | 68.80934543 | -0.345541391580127 | 0.0939205143720342 | -3.67908325343474 | 0.000234073853040473 | 0.0186158516514762 | NM_009515 |
| Eml5 | 3035.789718 | -0.274741017257679 | 0.074902650894644 | -3.66797455064872 | 0.000244479511177166 | 0.0193005944530408 | NM_001081191 |
| Csf2rb2 | 52.90440529 | -0.341395936465361 | 0.093285274374327 | -3.65969804725489 | 0.000252512621224766 | 0.0197204211608116 | NM_007781 |
| Tmeff1 | 2931.701249 | 0.190124085539912 | 0.0519449161806608 | 3.66010958347971 | 0.00025210740786286 | 0.0197204211608116 | NM_021436 |
| Samsn1 | 56.79513362 | -0.341778385135851 | 0.0935241973862822 | -3.65443804584824 | 0.000257745890258304 | 0.020021480224022 | NM_023380 |
| Sp100 | 142.738279 | -0.337955746115234 | 0.0925327638203695 | -3.65228198275041 | 0.000259920241933189 | 0.0200829863527739 | NM_013673 |
| Cdc25b | 986.7786974 | -0.291871678744864 | 0.0800565638143818 | -3.64581821699959 | 0.000266542352471046 | 0.0204856836613461 | NM_001111075,NM_023117 |
| Akap8l | 5134.106955 | -0.276626134593611 | 0.0759796283287298 | -3.64079346896479 | 0.000271799104840704 | 0.0205633010256045 | NM_017476 |
| Gabpb2 | 1054.44523 | -0.247381649458765 | 0.0679334782608882 | -3.64152779736573 | 0.000271024854572862 | 0.0205633010256045 | NM_172512,NM_029885 |
| Hvcn1 | 178.7917752 | -0.345656851520226 | 0.0948712483125453 | -3.64343104647984 | 0.000269027746652754 | 0.0205633010256045 | NM_001042489,NM_028752 |
| Ifi204 | 27.12459308 | -0.232358994903303 | 0.0639354510905792 | -3.63427474022375 | 0.000278763688161972 | 0.0209809395556519 | NM_008329 |
| Dyrk3 | 366.3415844 | 0.316772986021472 | 0.0872342374138302 | 3.63129197219588 | 0.000282005922585724 | 0.0211155568633001 | NM_145508 |
| Ada | 29.91823767 | 0.301885680015381 | 0.083281807067605 | 3.62486947203634 | 0.000289107432619888 | 0.0215362798268538 | NM_007398,NM_001272052 |
| Klf15 | 1102.528966 | 0.289761011681878 | 0.079968568529661 | 3.6234362701441 | 0.000290714858351841 | 0.0215455307776472 | NM_023184 |
| Atp4a | 36.59909863 | 0.3269420833156 | 0.0904872231311212 | 3.61312980996048 | 0.000302523153534167 | 0.0220912857015525 | NM_018731 |
| Ifi203 | 49.80148407 | -0.315950437757333 | 0.0874220645538546 | -3.61408117469816 | 0.000301414624109066 | 0.0220912857015525 | NM_008328,NM_001045481 |
| Pion | 276.7235449 | -0.290858432737078 | 0.0805026631348074 | -3.61302870502576 | 0.000302641185089422 | 0.0220912857015525 | NM_175437 |
| A230057D06Rik | 249.4336797 | -0.337734990420834 | 0.0940207796993929 | -3.59213135118272 | 0.000327984449297147 | 0.0235856540123285 | NR_015533 |
| Iqgap1 | 1261.324977 | -0.196294170222821 | 0.054644859092475 | -3.59218001442065 | 0.000327923191166878 | 0.0235856540123285 | NM_016721 |
| Pabpn1 | 5687.946795 | -0.259533746183652 | 0.0722171264100187 | -3.59379774695172 | 0.000325892845265622 | 0.0235856540123285 | NM_019402 |
| Invs | 617.0892343 | -0.26642355955181 | 0.074301179265358 | -3.58572450916706 | 0.00033614367182307 | 0.0237357198071778 | NM_010569 |
| Pcdha10 | 144.3092878 | 0.334584619642469 | 0.0932772554228303 | 3.58699039895397 | 0.000334516635606054 | 0.0237357198071778 | NM_009961 |
| Pja1 | 7886.888833 | 0.171350616650478 | 0.0477904073547458 | 3.58546047491395 | 0.000336483964493482 | 0.0237357198071778 | NM_008853,NM_001083110 |
| Rbm25 | 4392.493867 | -0.207559616732852 | 0.0578907926990547 | -3.58536480596925 | 0.000336607344091878 | 0.0237357198071778 | NM_027349 |
| Edem1 | 1114.190989 | -0.19195429225525 | 0.0535585045933147 | -3.58401142288821 | 0.000338357277676387 | 0.0237438541812908 | NM_138677 |
| Ifit3 | 441.0825018 | -0.339259462586775 | 0.0948429870194766 | -3.57706429593056 | 0.000347474728056119 | 0.0242664322103038 | NM_010501 |
| Bcl2l1 | 4645.653376 | 0.182224204474796 | 0.0511517274671908 | 3.56242522975742 | 0.000367444578623704 | 0.0255382772683633 | NM_009743 |
| Unc13b | 2407.33301 | -0.229044652740082 | 0.0643305384060807 | -3.56043425743242 | 0.000370242000044557 | 0.0256101680602249 | NM_021468,NM_001081413 |
| Ptprv | 393.0429352 | -0.306467014850702 | 0.0861743776982923 | -3.55635889734746 | 0.000376030296364512 | 0.0258872800236536 | NM_007955 |
| Whamm | 762.3049476 | -0.238621138714453 | 0.0671507671641691 | -3.55351321796631 | 0.000380122094450792 | 0.0260455355848689 | NM_001004185 |
| Slco1a4 | 2061.883328 | -0.216785385597485 | 0.0610418629785859 | -3.55142151663253 | 0.000383156245353607 | 0.0261301766197488 | NM_030687 |
| Arhgef1 | 1855.373335 | -0.293716826445807 | 0.0830021926812764 | -3.53866346126134 | 0.000402158205049596 | 0.0263414195588594 | NM_001130151,NM_001130150,NM_001130153,NM_001130152,NM_008488 |
| Capg | 96.58287031 | -0.337347480700783 | 0.0950875600082959 | -3.54775620145633 | 0.000388527675600156 | 0.0263414195588594 | NM_001271395,NM_007599,NM_001042534,NM_001271415 |
| Col6a2 | 1235.455651 | -0.308965185584797 | 0.0873610845629867 | -3.53664548843869 | 0.000405243229993332 | 0.0263414195588594 | NM_146007 |
| Ctss | 3301.869585 | -0.289898851884558 | 0.0817510952194856 | -3.54611581785219 | 0.000390954340428335 | 0.0263414195588594 | NM_021281,NM_001267695 |
| Ddb2 | 496.9790336 | -0.248035061538319 | 0.0701894058318974 | -3.53379628447574 | 0.000409636676944269 | 0.0263414195588594 | NM_028119 |
| Gbp5 | 158.3434278 | -0.335275917696126 | 0.0945677509316987 | -3.54535150083328 | 0.000392089845997165 | 0.0263414195588594 | NM_153564 |
| Nxf1 | 2696.109198 | -0.209845469142208 | 0.0593523932274549 | -3.53558563911689 | 0.000406872341166148 | 0.0263414195588594 | NM_016813 |
| Ovol2 | 249.6968497 | 0.285619939120328 | 0.0805886399226145 | 3.54417123051829 | 0.000393849366834682 | 0.0263414195588594 | NM_026924,NM_152947 |
| Phkg1 | 631.6367996 | -0.243482630644716 | 0.0688258805451164 | -3.53766096003827 | 0.000403688050311786 | 0.0263414195588594 | NM_011079 |
| Pnpla3 | 611.3469588 | -0.311155996673145 | 0.0880545557958771 | -3.53367289018468 | 0.000409827951280615 | 0.0263414195588594 | NM_054088 |
| Tardbp | 7760.601967 | -0.165372985147777 | 0.0467831289027335 | -3.5348850969674 | 0.000407952511627526 | 0.0263414195588594 | NR_027864,NM_001008546,NM_001008545,NM_001003899,NM_001003898,NM_145556 |
| Tbce | 1843.727523 | -0.214668307529977 | 0.06059141306388 | -3.54288333404037 | 0.00039577775050413 | 0.0263414195588594 | NM_178337 |
| Vmn2r84 | 31.66865262 | -0.32182542708426 | 0.0909550326237678 | -3.53829159091701 | 0.00040272505671031 | 0.0263414195588594 | NM_001081448 |
| Ccdc159 | 477.4810088 | -0.304130774877352 | 0.0865070822306865 | -3.51567486655408 | 0.000438637793691957 | 0.0280689541461206 | NM_025977,NM_001164614 |
| Phka2 | 1662.411297 | -0.233901889806954 | 0.0666770895889589 | -3.50797989607642 | 0.000451523169876767 | 0.0287667787966225 | NM_172783,NM_001177879,NM_001177878 |
| Bin2 | 266.5983599 | -0.298765056826683 | 0.0852715923090951 | -3.50368802477278 | 0.000458862587908442 | 0.029106715947415 | NM_001270537 |
| Zfp932 | 1563.847263 | -0.255446771079409 | 0.0730086183959264 | -3.49885776079365 | 0.000467255772823765 | 0.0295102493740783 | NM_145563 |
| Zfp280c | 893.3975499 | -0.230462155843629 | 0.0660579999803195 | -3.48878494523434 | 0.000485221353561154 | 0.0305122310901702 | NM_001166650,NM_153532,NM_001166648,NM_001166649 |
| Lgals9 | 344.8743039 | -0.31108912834581 | 0.089248872746691 | -3.4856364990598 | 0.000490967665682762 | 0.0307405013435681 | NM_001159301,NM_010708 |

| | | | | | | | | | | | | | | | |
|---|---|---|---|---|---|---|---|---|---|---|---|---|---|---|---|
| Gnmt | 16.5160432 | 0.526301318 | 0.16170062 | 3.254788496 | 0.001134768 | 0.012585613 | | Arrdc2 | 663.14207 | 0.317256941984492 | 0.0912171170136007 | 3.47804175763621 | 0.000505091233676747 | 0.0314890783707658 | NM_027560 |
| Irf9 | 457.3179858 | 0.526288072 | 0.116653649 | 4.511544031 | 6.44E-06 | 0.000222978 | | Hs6st2 | 1403.325246 | 0.253778250384948 | 0.0730267827786929 | 3.47513940404606 | 0.000510587938418652 | 0.0315608527381674 | NM_001077202,NM_015819 |
| 4933406C10Rik | 52.29709545 | 0.525632672 | 0.166304815 | 3.160658169 | 0.001574131 | 0.016024696 | | Sfi1 | 933.6173876 | -0.278193583457601 | 0.0800485528501741 | -3.47531059029003 | 0.000510262191206014 | 0.0315608527381674 | NM_030207 |
| Uvssa | 459.0080038 | 0.525487171 | 0.106474448 | 4.935335936 | 8.00E-07 | 4.09E-05 | | Bmp1 | 3285.541053 | -0.292467474929143 | 0.0841928297330426 | -3.47378126921847 | 0.000513179186825654 | 0.0315866138467349 | NR_033241,NM_009755 |
| Cyp26b1 | 595.3830254 | 0.525322735 | 0.123661523 | 4.248069411 | 2.16E-05 | 0.000592345 | | Ap1g2 | 284.3237942 | -0.315949893132479 | 0.0911644370466595 | -3.46571430009237 | 0.000528824898310523 | 0.0322760944237759 | NM_007455 |
| Mpl | 31.11447344 | 0.524515976 | 0.167727666 | 3.127188195 | 0.001764869 | 0.017394963 | | Nsun7 | 165.5883214 | -0.295494147526331 | 0.0852421717624451 | -3.4665253291507 | 0.000527232056778496 | 0.0322760944237759 | NM_027602 |
| Oasl2 | 242.1392103 | 0.52342252 | 0.162639367 | 3.218301516 | 0.001289522 | 0.013779208 | | Itpripl2 | 384.0352659 | -0.238466898691094 | 0.0689982399399064 | -3.45613016938962 | 0.000547990618248956 | 0.033305906781106 | NM_001033380 |
| Pmch | 32.05889035 | 0.522935987 | 0.148988516 | 3.509908029 | 0.000448262 | 0.006323296 | | Eif2ak2 | 335.0410949 | -0.299821310700744 | 0.086796366769983 | -3.45430715429936 | 0.000551708630609579 | 0.0333139411343229 | NM_011163 |
| Adam8 | 113.0969699 | 0.522741086 | 0.153099885 | 3.414379349 | 0.000639275 | 0.008138806 | | Qtrtd1 | 567.4114928 | -0.233788205762582 | 0.0676897812355287 | -3.45381830898683 | 0.000552709611274392 | 0.0333139411343229 | NM_029128 |
| Cd109 | 81.48102203 | 0.522338637 | 0.167612776 | 3.116341429 | 0.001831101 | 0.017883806 | | Prpf39 | 1383.500281 | -0.228811843352954 | 0.066289217964449 | -3.45172036085365 | 0.000557024699909539 | 0.033435292475453 | NM_177806 |
| P2rx6 | 86.4158704 | 0.521200156 | 0.151499789 | 3.440269835 | 0.000581134 | 0.00762413 | | Dock6 | 857.7042041 | -0.286578923565672 | 0.0830918331804808 | -3.4489421233878 | 0.000562787313807457 | 0.0335899904504301 | NM_177030 |
| Tnip2 | 474.5824552 | 0.52102588 | 0.098551445 | 5.286841602 | 1.24E-07 | 8.93E-06 | | Ifitm3 | 328.5259502 | -0.304029191571984 | 0.0882205483759085 | -3.44624010130285 | 0.000568445058858605 | 0.0335899904504301 | NM_025378 |
| Col6a2 | 1063.045587 | 0.520976416 | 0.106699367 | 4.882657024 | 1.05E-06 | 5.05E-05 | | Itgb2 | 311.0803894 | -0.32442871722943 | 0.0941283902140459 | -3.446661697833 | 0.000567558807726825 | 0.0335899904504301 | NM_008404 |
| Mx1 | 17.65054141 | 0.520581474 | 0.163624021 | 3.181571197 | 0.001464785 | 0.015180857 | | Sipa1l3 | 2410.47067 | -0.234303754609612 | 0.0679920381522775 | -3.4460469339786 | 0.000568851552444293 | 0.0335899904504301 | NM_001081028 |
| Ftx | 809.9636154 | 0.520100565 | 0.145803079 | 3.567143903 | 0.000360893 | 0.005392171 | | D4Wsu53e | 7279.580763 | -0.263531718701014 | 0.0765341371504847 | -3.44332252917214 | 0.000574613588080914 | 0.0337928622690824 | NM_023665 |
| Robo3 | 988.5187055 | 0.518671828 | 0.165098148 | 3.141596879 | 0.001680292 | 0.01680726 | | C1qc | 1930.48976 | -0.302331526030363 | 0.0880408725432333 | -3.43399056934494 | 0.000594764888437336 | 0.0348369143929062 | NM_007574 |
| Atp2c2 | 43.43024302 | 0.517888136 | 0.168028324 | 3.082147832 | 0.002055127 | 0.019478131 | | Gpi1 | 26520.80069 | 0.136990162411271 | 0.0399293223256357 | 3.43081611288252 | 0.000601768377973087 | 0.0349651498337483 | NM_008155 |
| Col24a1 | 404.9902976 | 0.515780321 | 0.16040753 | 3.215437084 | 0.00130246 | 0.013850598 | | Slc4a7 | 722.2505516 | -0.231252390265638 | 0.0673874038454647 | -3.43168570191478 | 0.000599842295149483 | 0.0349651498337483 | NM_001033270 |
| Clk4 | 1456.694376 | 0.514390789 | 0.118380271 | 4.345240858 | 1.39E-05 | 0.000416938 | | Cdk2 | 89.54156948 | -0.321140693258953 | 0.0937826431167042 | -3.4243084070399 | 0.000616366287836454 | 0.0353886825972819 | NM_183417,NM_016756 |
| Ccdc11 | 11.27390938 | 0.514226207 | 0.163423258 | 3.146591337 | 0.001651857 | 0.016576338 | | Pld4 | 485.704191 | -0.296851407005403 | 0.0866693508364668 | -3.42510246287088 | 0.000614567597074137 | 0.0353886825972819 | NM_178911 |
| Ddx26b | 1788.152824 | 0.513122808 | 0.114839497 | 4.468173577 | 7.89E-06 | 0.000261181 | | Rgs16 | 566.9801326 | 0.285470793030238 | 0.0833435165769155 | 3.42523095682895 | 0.000614278992839568 | 0.0353886825972819 | NM_011267 |
| Casc1 | 26.31824637 | 0.512406761 | 0.167908603 | 3.051700459 | 0.002275491 | 0.021036068 | | Col4a1 | 2174.644013 | -0.285208608729477 | 0.0833991608513906 | -3.41980189989791 | 0.000626667531547165 | 0.0357757040052952 | NM_009931 |
| Cep110 | 550.9013864 | 0.510986903 | 0.105475049 | 4.844623507 | 1.27E-06 | 5.97E-05 | | Safb2 | 2395.916686 | -0.252803115084196 | 0.073936122384503 | -3.41920981153839 | 0.000628032804719143 | 0.0357757040052952 | NM_001029979 |
| 1700012B15Rik | 372.9530798 | 0.510193113 | 0.098529768 | 5.178060611 | 2.24E-07 | 1.45E-05 | | Cables1 | 1103.82638 | 0.192055810700644 | 0.0563576048056744 | 3.40780647727788 | 0.000654873212512135 | 0.0371589386130909 | NM_022021,NM_001146287 |
| Rsad1 | 634.1711095 | 0.509872078 | 0.09648567 | 5.284433188 | 1.26E-07 | 9.00E-06 | | Camk1g | 1872.426373 | 0.297280932474083 | 0.0876495082518948 | 3.39170108769728 | 0.000694601724444 77 | 0.0391059723208814 | NM_144817 |
| Neurod1 | 222.1712959 | 0.508871435 | 0.107637183 | 4.727654724 | 2.27E-06 | 9.94E-05 | | Lat2 | 91.19534975 | -0.322318081926156 | 0.0950777180174761 | -3.39004856918117 | 0.000698802174786669 | 0.0391059723208814 | NM_020044,NM_022964 |
| E030003E18Rik | 239.7697447 | 0.507774735 | 0.143032662 | 3.550061423 | 0.000385141 | 0.005672413 | | Ret | 307.5358222 | 0.32237691694505 | 0.095101821780405 | 3.38980800693214 | 0.000699415890486618 | 0.0391059723208814 | NM_001080780,NM_009050 |
| Cfp | 305.9397896 | 0.507097738 | 0.155441238 | 3.262311507 | 0.001105076 | 0.012318024 | | Rgs11 | 1593.992311 | -0.292876981807339 | 0.0864046720341867 | -3.38959659136497 | 0.000699955445644304 | 0.0391059723208814 | NM_001081069 |
| Gdpd2 | 231.9702453 | 0.506007696 | 0.159293062 | 3.176583388 | 0.001490209 | 0.01539287 | | 2610005L07Rik | 718.1818116 | -0.313041650088178 | 0.0924122031369021 | -3.38744927035696 | 0.000705457620864402 | 0.0392623655198326 | NR_028428 |
| Tlr2 | 77.92161558 | 0.505953899 | 0.161640756 | 3.130113411 | 0.001747388 | 0.017321839 | | Trub1 | 1163.165969 | -0.195856000204069 | 0.057861913288443 | -3.38488634531875 | 0.000712077319057088 | 0.0394795234222262 | NM_028115,NM_029839 |
| Boc | 889.1728922 | 0.505927061 | 0.084621444 | 5.978709843 | 2.25E-09 | 2.83E-07 | | Liph | 54.73469408 | -0.310966335368875 | 0.0920156878631142 | -3.37949259077952 | 0.000726197647591863 | 0.0400034927562411 | NM_153404,NM_001083894 |
| Sh3bp1 | 595.5107772 | 0.505569578 | 0.100054185 | 5.052957813 | 4.35E-07 | 2.50E-05 | | Myo9b | 2648.749946 | -0.212887471793769 | 0.0629998269786894 | -3.37917549941496 | 0.000727035803913511 | 0.0400034927562411 | NM_001142323,NM_001142322,NM_015742 |
| Rwdd3 | 128.5576472 | 0.505223793 | 0.117737359 | 4.291108594 | 1.78E-05 | 0.000509164 | | Plekha5 | 2347.283512 | -0.179025446411656 | 0.0530106135655714 | -3.37719112393275 | 0.000732301475193939 | 0.0401411744480994 | NM_144920 |
| Ssh3 | 1564.140088 | 0.50491102 | 0.121315544 | 4.161964776 | 3.16E-05 | 0.000801423 | | Heatr6 | 928.7800134 | -0.204833853459715 | 0.0606758691834425 | -3.3758712271757 | 0.000735823502673543 | 0.0401826002550221 | NM_145432 |
| Mir186 | 30.42506888 | 0.504343433 | 0.167340389 | 3.013877503 | 0.002579319 | 0.02290199 | | Jag1 | 337.4349971 | -0.284412137380215 | 0.0842819538527479 | -3.37453184672393 | 0.000739413599427918 | 0.0402274230160672 | NM_013822 |
| Nr1d1 | 5181.075221 | 0.503915447 | 0.085213962 | 5.913531513 | 3.35E-09 | 4.18E-07 | | Bcat1 | 4427.138739 | 0.204071610875366 | 0.0606236116454682 | 3.36620675239202 | 0.000762095643493664 | 0.0413067213335409 | NM_007532,NM_001024468 |
| Wdr96 | 44.31362833 | 0.503692851 | 0.163060594 | 3.088991886 | 0.002008369 | 0.019184755 | | Akt2 | 4959.349386 | 0.183634212926778 | 0.0546284238177213 | 3.36151402682805 | 0.000775164193599843 | 0.0417038336156716 | NM_001110208,NM_007434 |
| Hmox1 | 233.4038846 | 0.502851508 | 0.121319761 | 4.144844199 | 3.40E-05 | 0.000847051 | | Fam227a | 302.2116472 | -0.285192236301888 | 0.08483517979221 | -3.36172136371279 | 0.000774582425384435 | 0.0417038336156716 | NM_029407 |
| Ankrd16 | 646.0294911 | 0.502361221 | 0.089646167 | 5.603822636 | 2.10E-08 | 2.01E-06 | | Col4a2 | 2917.356857 | -0.291063731943816 | 0.0866977118070973 | -3.35722507407617 | 0.000787289958395639 | 0.0420822903534047 | NM_009932 |
| Alox8 | 246.7224836 | 0.50210292 | 0.121299727 | 4.13935739 | 3.48E-05 | 0.000857909 | | Slc38a6 | 331.5049198 | -0.287498676832012 | 0.0856663405159307 | -3.35602845995911 | 0.000790704314451817 | 0.0420822903534047 | NM_001037717 |
| Treml2 | 27.54744045 | 0.501679346 | 0.165546968 | 3.030435117 | 0.002442016 | 0.022066759 | | Tlr2 | 78.64670622 | -0.296160878210996 | 0.0882491332932993 | -3.35596359033572 | 0.000790889802180881 | 0.0420822903534047 | NM_011905 |
| Gdap10 | 63.09027955 | 0.501216382 | 0.15520918 | 3.229295976 | 0.001240954 | 0.013431707 | | Gigyf1 | 3159.268209 | -0.246002023972388 | 0.0733693456669268 | -3.35292650815175 | 0.000799619356518174 | 0.0423914991707408 | NM_031408 |
| Hook1 | 2087.80065 | 0.500913073 | 0.079815354 | 6.275898628 | 3.48E-10 | 5.61E-08 | | Ccdc108 | 434.2583441 | -0.311297369264211 | 0.092958049298269 | -3.34877261729238 | 0.000811703816587145 | 0.0426891841869858 | NM_001039495 |
| Irf8 | 253.1120082 | 0.499735003 | 0.142764546 | 3.500413913 | 0.000464536 | 0.006460974 | | Rab3c | 17590.22349 | 0.202413164851978 | 0.0604584314950097 | 3.3479724803758 | 0.000814050944499178 | 0.0426891841869858 | NM_023852 |
| Rnf112 | 13465.15019 | 0.499567263 | 0.098013838 | 5.096905413 | 3.45E-07 | 2.09E-05 | | Zfp445 | 8652.042808 | -0.220750123804074 | 0.0659314140682917 | -3.34817820796959 | 0.000813446860884838 | 0.0426891841869858 | NM_173364 |
| B3galnt2 | 683.8561278 | 0.498581888 | 0.095613863 | 5.214326399 | 1.84E-07 | 1.24E-05 | | Irf1 | 323.049198 | -0.317631308638647 | 0.0949817874840284 | -3.34412856456358 | 0.000825414765857268 | 0.0431294060749737 | NM_001159396,NM_001159393,NM_008390 |
| Naip5 | 77.38687791 | 0.498421629 | 0.16350802 | 3.048300811 | 0.002301394 | 0.021168869 | | I830077J02Rik | 30.73612512 | -0.2921725715703 | 0.0874218651525185 | -3.3420994971975 | 0.000831472502398355 | 0.043290213511966 | NM_001033780 |
| Frem3 | 55.33041029 | 0.497691171 | 0.159946423 | 3.111617759 | 0.00186069 | 0.018120387 | | Ccdc84 | 327.9165278 | -0.29683257933338 | 0.0889339186578481 | -3.33767570138646 | 0.000844822894123315 | 0.0438282048572688 | NM_201372 |
| Mirg | 554.1907585 | 0.497010713 | 0.147457804 | 3.370528383 | 0.000750242 | 0.009126188 | | Stat2 | 1401.135806 | -0.257316013764241 | 0.077159072452693 | -3.33487697019692 | 0.000853371459752586 | 0.0441141417237227 | NM_019963 |
| Lrrc29 | 42.95972806 | 0.49657143 | 0.165265519 | 3.004688654 | 0.002658528 | 0.023324597 | | AF251705 | 110.3470426 | -0.316290929619203 | 0.0950256214016915 | -3.328480424193 | 0.000873211414723407 | 0.0443735562406493 | NM_134158 |
| Lrrc23 | 61.69704346 | 0.496427525 | 0.166764377 | 2.976819954 | 0.00291255 | 0.024822359 | | Flnb | 3768.40108 | -0.260646137265489 | 0.0783133677645574 | -3.32748099138164 | 0.000876349676902056 | 0.0443735562406493 | NM_134080 |
| Ccdc135 | 62.59120814 | 0.49629014 | 0.166608992 | 2.978771646 | 0.002894064 | 0.024719197 | | Mov10 | 381.4679661 | -0.266895198731244 | 0.0802122084189463 | -3.32736380149586 | 0.000876718342356212 | 0.0443735562406493 | NM_001163441,NM_001163440,NM_008619 |
| Flnb | 3420.57541 | 0.496075006 | 0.099803472 | 4.970518523 | 6.68E-07 | 3.53E-05 | | Ndufc2 | 1894.914091 | 0.202267485617636 | 0.0607741498427445 | 3.32818426685501 | 0.000874140273510889 | 0.0443735562406493 | NM_024220 |
| Myo1h | 66.98676742 | 0.495256245 | 0.165662142 | 2.989555959 | 0.002793833 | 0.024150338 | | Tekt4 | 42.82700988 | 0.277008105853229 | 0.0832449123544289 | 3.32762805580024 | 0.000875887233075155 | 0.0443735562406493 | NM_027951 |
| Ace | 1023.649079 | 0.495163135 | 0.133745122 | 3.702289306 | 0.000213663 | 0.003556118 | | Usp48 | 4073.196016 | -0.175246407675894 | 0.0526360981247025 | -3.32939586936536 | 0.000870346023535921 | 0.0443735562406493 | NM_130879 |
| Igflr1 | 213.4126547 | 0.494671856 | 0.146725447 | 3.371411489 | 0.000747841 | 0.009109311 | | Ccdc142 | 374.4284979 | -0.290483565473538 | 0.0873748778096516 | -3.32456219975326 | 0.000885562067149501 | 0.044665536761853 | NM_001081266 |
| Mroh8 | 49.45192197 | 0.494563044 | 0.153448469 | 3.222991053 | 0.001268595 | 0.013659217 | | H2-Q4 | 430.3689413 | -0.296610198920395 | 0.0893059831799046 | -3.32128032565506 | 0.000896054891656238 | 0.0450383855155658 | NM_001143689 |
| Fermt1 | 11.79605464 | 0.494441377 | 0.154771501 | 3.194653887 | 0.001399986 | 0.014666252 | | Ipw | 443.7409427 | -0.306254519178604 | 0.0922760019531972 | -3.3188967087449 | 0.000903738516820201 | 0.0452679506735525 | NR_015351 |
| Espn | 31.53034311 | 0.493933235 | 0.165842309 | 2.978330665 | 0.002898231 | 0.024741154 | | 4930570G19Rik | 315.11887 | -0.276880881995891 | 0.0835847658045634 | -3.31257591417185 | 0.000924410345573392 | 0.0461442772501687 | NR_040398,NR_040399,NR_040400 |
| Whamm | 661.5184479 | 0.492955525 | 0.107616891 | 4.580651963 | 4.64E-06 | 0.000173896 | | Ctsh | 461.9397725 | -0.27856555067176 | 0.0841720870260237 | -3.30947605744284 | 0.000934707599698521 | 0.0464985020315778 | NM_007801 |
| 5830418P13Rik | 18.02715787 | 0.492797883 | 0.167719583 | 2.938225069 | 0.003300973 | 0.027132771 | | Anxa3 | 446.8128622 | -0.273900667322167 | 0.0828466699262712 | -3.30611559361315 | 0.000945990530119 42 | 0.0465812150525922 | NM_013470 |
| Mybpc1 | 171.7075507 | 0.492514362 | 0.159582983 | 3.086249133 | 0.002026833 | 0.019325387 | | Meig1 | 39.88273091 | 0.308155576293 5102 | 0.093195240325982 | 3.30655567957102 | 0.000944505773118318 | 0.0465812150525922 | NM_008579 |
| Cdk15 | 18.35479181 | 0.491527816 | 0.16804288 | 2.925014231 | 0.003444405 | 0.027955792 | | Sccpdh | 2583.297034 | 0.139299814586236 | 0.0421184968648936 | 3.30733110046822 | 0.000941894918575007 | 0.0465812150525922 | NM_178653 |
| Efcab4b | 31.26931778 | 0.491000666 | 0.162028958 | 3.030326626 | 0.002442894 | 0.022066759 | | Plin4 | 662.3437959 | 0.257343904694137 | 0.0779206326426512 | 3.30264136681657 | 0.000957787966998564 | 0.0470027973264228 | NM_020568 |
| Oscar | 72.82397407 | 0.489328862 | 0.157936155 | 3.098270079 | 0.001946539 | 0.018724864 | | 1700024P16Rik | 48.93689332 | -0.30228205227117 | 0.0916391040829591 | -3.29861422474755 | 0.000971633403023798 | 0.0475217064388003 | NM_001162980 |
| 4930451C15Rik | 44.73283795 | 0.489239209 | 0.158735421 | 3.082104839 | 0.002055424 | 0.019478131 | | Stip1 | 7185.759144 | 0.147588958825708 | 0.0447669623722212 | 3.29682763817118 | 0.000977834898244396 | 0.0476645293016502 | NM_016737 |
| Nek5 | 13.9735857 | 0.488309147 | 0.157858604 | 3.103950425 | 0.001909553 | 0.018503201 | | Csf2rb | 75.07014121 | -0.312024939060842354 | 0.0948239060842354 | -3.29057247684639 | 0.000999837415819347 | 0.0480831047618658 | NM_007780 |
| Lilra5 | 13.91655782 | 0.48700346 | 0.164229125 | 2.965390333 | 0.003022991 | 0.025445766 | | Endod1 | 5377.441772 | 0.205182169102121 | 0.0623750424708765 | 3.28949145321901 | 0.00100368604734771 | 0.0480831047618658 | NM_028013 |
| Arrdc1 | 272.4660883 | 0.486139135 | 0.115663437 | 4.203049354 | 2.63E-05 | 0.000692746 | | Izumo4 | 970.2414993 | -0.268341700484965 | 0.0815934873250477 | -3.28876423178713 | 0.00100628279275833 | 0.0480831047618658 | NM_027829 |
| Fgf5 | 86.1659671 | 0.485459658 | 0.150935112 | 3.216346756 | 0.001298339 | 0.013849883 | | Kbtbd4 | 1330.902444 | 0.173205942008107 | 0.0526493760673915 | 3.2898004676523 | 0.00100258450492576 | 0.0480831047618658 | NM_025991 |
| Slc17a6 | 915.9812025 | 0.485141401 | 0.130313757 | 3.722871713 | 0.00019697 | 0.003364771 | | Rhbdl3 | 969.2947975 | 0.233042117878396 | 0.0708494085172687 | 3.28925990569837 | 0.00100451217796487 | 0.0480831047618658 | NM_139228 |
| Paxbp1 | 1499.775334 | 0.484917583 | 0.098464877 | 4.924777208 | 8.45E-07 | 4.25E-05 | | Slc6a6 | 7760.754252 | -0.168449042061349 | 0.0511656409732563 | -3.29222968494415 | 0.000993963985589788 | 0.0480831047618658 | NM_009320 |
| Aldh1a1 | 1796.04932 | 0.484897496 | 0.108134416 | 4.484210615 | 7.32E-06 | 0.000246462 | | Tnfrsf14 | 22.64225463 | -0.265237027914364 | 0.0807299963494962 | -3.28548296677854 | 0.00101807697722253 | 0.0484871677742112 | NM_178931 |
| Otx1 | 194.3325165 | 0.484569328 | 0.124725842 | 3.885075618 | 0.000102298 | 0.001996229 | | Lfng | 646.6805627 | 0.23772157242522 | 0.0724189982285122 | 3.28258614195177 | 0.00102859553516633 | 0.0488280351105427 | NM_008494 |
| Lrrc45 | 1936.627398 | 0.483971482 | 0.1350818 | 3.582803034 | 0.000339927 | 0.005158499 | | Prr5 | 281.8088457 | 0.296474453211028 | 0.0904087877660761 | 3.27926588262777 | 0.00104077527073779 | 0.0492452820284598 | NM_146061 |
| Vasn | 427.0462911 | 0.48298555 | 0.13936595 | 3.465592196 | 0.000529065 | 0.007115035 | | C1s | 86.01757752 | -0.3117535650005 | 0.0951019176238176 | -3.27809967232905 | 0.00104508486401297 | 0.0492886452423779 | NM_144938,NM_001097617 |
| Ifi44 | 46.20666261 | 0.482753037 | 0.158098913 | 3.053487381 | 0.002261982 | 0.020948688 | | Rasal3 | 164.7337673 | -0.309976523505615 | 0.094654557312099 | -3.2748187969814 | 0.00105729768127521 | 0.0497032560459668 | NM_178785 |
| Cd72 | 34.16620951 | 0.482498999 | 0.167951457 | 2.872847956 | 0.004067898 | 0.031404095 | | Fzd2 | 553.8499483 | 0.250405674735271 | 0.0765233994644902 | 3.27227588613199 | 0.00106685416524709 | 0.0499907213044902 | NM_020510 |
| Dna2 | 62.42824683 | 0.482103095 | 0.151979033 | 3.172168454 | 0.001513052 | 0.015566554 | | Snrnp70 | 6880.890335 | -0.250763747663968 | 0.076864112591225 | -3.26242949030801 | 0.00110461648219437 | 0.0515937589078953 | NM_009224 |
| Rbm38 | 73.19605725 | 0.481685913 | 0.144085832 | 3.343048421 | 0.000828634 | 0.009842462 | | Sqle | 2545.213866 | 0.155461133444629 | 0.0476894231988727 | 3.25986608804913 | 0.00111464827666638 | 0.0518954514963327 | NM_009270 |
| Relt | 727.0285178 | 0.481383405 | 0.142614851 | 3.375408662 | 0.00073706 2 | 0.009013442 | | Stx2 | 647.6704684 | -0.255113444292275 | 0.0783207770183716 | -3.25728949585499 | 0.00112481652166941 | 0.0522015488618846 | NM_007941 |
| Dusp9 | 31.11231638 | 0.481132063 | 0.163218104 | 2.947786124 | 0.003200584 | 0.026528802 | | Asun | 1070.292666 | -0.206382874630482 | 0.0634233848065917 | -3.25402705875224 | 0.00113781441483243 | 0.0526365993307511 | NM_138757 |
| Ccdc153 | 66.79771211 | 0.480928647 | 0.159037758 | 3.023990356 | 0.002494644 | 0.022341444 | | Col15a1 | 316.2576924 | -0.303153236667415 | 0.0934240143283328 | -3.24491769937735 | 0.00117484573305966 | 0.0538353599950303 | NM_009928 |
| Isg15 | 78.27792383 | 0.479806917 | 0.16683061 | 2.876012482 | 0.004027338 | 0.031168618 | | Glt8d1 | 1211.216073 | -0.215528425703117 | 0.0664181393351881 | -3.24502354116583 | 0.00117440911440876 | 0.0538353599950303 | NM_029626,NM_001165930 |
| B2m | 3100.791312 | 0.478939498 | 0.084022131 | 5.700158894 | 1.20E-08 | 1.23E-06 | | Slc25a25 | 2961.587398 | 0.200869069174516 | 0.0618869622527785 | 3.24574129772378 | 0.0011714524086011 | 0.0538353599950303 | NM_146118,NM_001164358,NM_001164357 |
| Fam221a | 29.11745327 | 0.47893661 | 0.166399963 | 2.878225454 | 0.003999192 | 0.031028284 | | Cst7 | 58.57180908 | -0.144851779862425 | 0.0447166722528648 | -3.23932378159345 | 0.00119813477104191 | 0.0547298920885369 | NM_009977 |
| Rassf3 | 2023.315597 | 0.478801562 | 0.096492857 | 4.962041502 | 6.98E-07 | 3.66E-05 | | Atxn2l | 6339.308554 | -0.237201140014202 | 0.0734024085513038 | -3.23151712179054 | 0.00123134946639344 | 0.0560707910621663 | NM_183020 |
| Gm14872 | 186.5972643 | 0.478577538 | 0.113034974 | 4.233889039 | 2.30E-05 | 0.000623245 | | Ophn1 | 1889.627723 | -0.151183030748925 | 0.0468436777490895 | -3.22739456023748 | 0.00124923071849813 | 0.0567072669278247 | NM_052976 |
| Pias3 | 953.0465659 | 0.478528523 | 0.091924722 | 5.205656451 | 1.93E-07 | 1.29E-05 | | Stxbp2 | 1700.902879 | -0.268101621746094 | 0.0831963421531723 | -3.2225169377338 | 0.00127069652690697 | 0.0575019867596594 | NM_011503 |
| Ifit1 | 136.2514187 | 0.477810762 | 0.166849983 | 2.863714784 | 0.004187048 | 0.032100014 | | H2-T24 | 85.66416614 | -0.306313863596242 | 0.0950953599009273 | -3.22112191336126 | 0.00127689818299898 | 0.0576031770380224 | NM_008207 |
| Ttc21a | 68.87641437 | 0.477126829 | 0.150794935 | 3.164077285 | 0.001555755 | 0.015900306 | | Hpgds | 256.6455066 | -0.279859498309424 | 0.0869254444969694 | -3.21991331595856 | 0.00128229364461916 | 0.057667484463585 | NM_019455 |
| B230216G23Rik | 53.79093156 | 0.477089856 | 0.157925875 | 3.020973319 | 0.002519636 | 0.02250152 | | Ints2 | 908.0333464 | -0.201256431123288 | 0.0625409609538192 | -3.21799390437728 | 0.00129090561260567 | 0.0578756016318207 | NM_027421 |
| Morn1 | 148.9727993 | 0.477029221 | 0.124191597 | 3.841074859 | 0.000122497 | 0.00228670 4 | | Atad2 | 475.1802548 | -0.269217618080475 | 0.083757321001525 | -3.21425774918915 | 0.00130782218492231 | 0.0584536155636354 | NM_027435 |
| Trpm4 | 505.6135134 | 0.477006058 | 0.116041976 | 4.11063369 | 3.95E-05 | 0.000953751 | | Fyco1 | 2996.189938 | -0.220700432594015 | 0.0687521233461208 | -3.21008896674996 | 0.00132693895271826 | 0.0591261203287898 | NM_148925,NM_001110253 |
| Slc39a2 | 234.4910531 | 0.47695963 | 0.164503252 | 2.899393322 | 0.003738855 | 0.029707604 | | Plcb2 | 175.4814559 | -0.28428612209096 | 0.0885920499382662 | -3.20893564247508 | 0.00133227309591437 | 0.0591822599120861 | NM_177568 |
| Lime1 | 1632.314876 | 0.476643487 | 0.130300977 | 3.658019291 | 0.000254172 | 0.004091928 | | Spire2 | 672.3543413 | 0.199102688785663 | 0.0620686997741198 | 3.2066514927811 | 0.00134289576158971 | 0.0594722677830857 | NM_172287 |
| Ccl3 | 19.06722801 | 0.476061748 | 0.147295898 | 3.232009549 | 0.001229229 | 0.013327979 | | Cd59a | 594.4598378 | 0.289092824779309 | 0.0902032479863904 | 3.20397829863904 | 0.00135542692936148 | 0.0597978938467153 | NM_001111060,NM_007652 |
| Ggnbp1 | 94.12601739 | 0.475929174 | 0.144261961 | 3.299062143 | 0.000970084 | 0.011141947 | | Dtd1 | 1773.65601 | 0.174116288813117 | 0.0543552534098841 | 3.20330194213491 | 0.0013586145479800 1 | 0.0597978938467153 | NM_025314 |
| Sfi1 | 741.8557003 | 0.475739353 | 0.134966108 | 3.524880143 | 0.000423674 | 0.006044578 | | Gjb6 | 2405.202571 | 0.21285005190596 | 0.0664645732569822 | 3.20245871560756 | 0.00136259829707165 | 0.0597978938467153 | NM_001010937,NM_001271663 |
| Ak7 | 72.38386989 | 0.475048517 | 0.164029493 | 2.896116479 | 0.003778121 | 0.029958144 | | Zfyve16 | 778.9209442 | -0.214189549837214 | 0.0669428002945887 | -3.19959052944679 | 0.00137622958330418 | 0.060214189539387 | NM_173392 |
| Col26a1 | 211.9286931 | 0.475028497 | 0.144104859 | 3.296408611 | 0.000979295 | 0.01121448 | | Grp | 145.7184483 | 0.28440367967149 | 0.0889365779967352 | 3.19782288560353 | 0.00138469297773105 | 0.0604025531367005 | NM_175012 |
| 4930422G04Rik | 202.0971971 | 0.474853921 | 0.101221282 | 4.691245893 | 2.72E-06 | 0.000114641 | | Kcnt1 | 8027.833192 | -0.202669937022182 | 0.0634435110355113 | -3.19449434172616 | 0.00140076031022285 | 0.0607386396008869 | NM_175462,NM_001145403 |
| Leng8 | 9341.863843 | 0.473997794 | 0.157360662 | 3.01217463 | 0.002593833 | 0.02297819 | | Sgk1 | 4723.226973 | 0.298356570788585 | 0.0933794297544911 | 3.19509951685303 | 0.00139782632113594 | 0.0607386396008869 | NM_001161845,NM_001161847,NM_001161850,NM_001161849,NM_001161848,NM_011361 |
| Skp2 | 252.5865249 | 0.473736265 | 0.120315077 | 3.937463847 | 8.23E-05 | 0.00168992 | | Col7a1 | 192.0368488 | -0.297006704412492 | 0.093072926424288 | -3.19111814598521 | 0.00141723331342231 7 | 0.0611252816644653 | NM_007738 |
| Tcea2 | 1135.620647 | 0.472711561 | 0.092883753 | 5.089281458 | 3.59E-07 | 2.15E-05 | | Grid1 | 2641.688723 | 0.158371074588595 | 0.0496425569640814 | 3.19022798731305 | 0.00142160596848556 | 0.0611252816644653 | NM_008166 |
| Cdan1 | 1123.913599 | 0.471901744 | 0.093743464 | 5.033969544 | 4.80E-07 | 2.69E-05 | | Rufy2 | 2620.854008 | -0.188261586264075 | 0.0590190566359706 | -3.18984404351415 | 0.00142349589887749 | 0.0611252816644653 | NM_027425 |
| Zkscan3 | 1011.359314 | 0.471207977 | 0.110462627 | 4.265768359 | 1.99E-05 | 0.000557702 | | Smpd4 | 2883.067784 | -0.272007553225405 | 0.0852893354025672 | -3.18923288523148 | 0.00142650905164903 | 0.0611252816644653 | NM_029945,NM_001164609,NM_001164611,NM_001164610 |
| Dcdc2b | 97.6171826 | 0.470001665 | 0.165819006 | 2.834425768 | 0.004590812 | 0.034079589 | | Malt1 | 316.9152143 | -0.241608789225767 | 0.0757872644577828 | -3.18798667499544 | 0.00143267138495667 | 0.0612087780525898 | NM_172833 |
| Cep70 | 624.5507929 | 0.4698769 | 0.112736792 | 4.167910871 | 3.07E-05 | 0.000784665 | | Tnfrsf4 | 39.93369917 | -0.288493074879451 | 0.0905269295276205 | -3.18679067527006 | 0.00143860850527436 | 0.0612821908959494 | NM_011659 |
| Hes3 | 9.773242983 | 0.469795598 | 0.149405714 | 3.144428588 | 0.001664115 | 0.016666969 | | Ccdc62 | 109.6592516 | -0.301092508589765 | 0.0946170530211899 | -3.18222243216911 | 0.00146149528436345 | 0.0620754891832264 | NM_001134767 |
| Wdr62 | 69.84615674 | 0.46958471 | 0.149224588 | 3.14683202 | 0.001650498 | 0.016573434 | | Ltbp4 | 2859.080522 | -0.232265591650228 | 0.0730116084661965 | -3.18123880855196 | 0.00146647057313955 | 0.0621048150012394 | NM_001113549,NM_175641 |
| Satb1 | 3895.927084 | 0.46950607 | 0.108996354 | 4.307539241 | 1.65E-05 | 0.000478982 | | Fzd4 | 378.239814 | 0.269452117833335 | 0.0847433420572224 | 3.17962581238995 | 0.00147465338867666 | 0.0622698114067359 | NM_008055 |
| Wdr90 | 923.9693233 | 0.469363803 | 0.110631795 | 4.242576056 | 2.21E-05 | 0.000604892 | | 4930503L19Rik | 353.4793158 | -0.234809429762248 | 0.0739646201283973 | -3.17461820738936 | 0.00150033766664016 | 0.0630177839423566 | NM_172967 |
| Slc11a1 | 141.381587 | 0.469101562 | 0.148179156 | 3.165772947 | 0.001546715 | 0.015860224 | | Pxn | 1335.464942 | -0.247192801347868 | 0.0778703521347474 | -3.17448177260859 | 0.00150104318078311 | 0.0630177839423566 | NM_133915,NM_011223 |
| Wnt9b | 17.26626721 | 0.468732391 | 0.165463809 | 2.832839362 | 0.004613656 | 0.03418651 | | Fcgr3 | 628.8817501 | -0.256009566554254 | 0.0806798196554624 | -3.17315491836404 | 0.00150792039773863 | 0.0631240682926552 | NM_010188 |
| BC051142 | 77.83953271 | 0.468026018 | 0.157832079 | 2.965341527 | 0.003023471 | 0.025445766 | | Rbms1 | 633.3415844 | -0.224053444405106 | 0.0706603826005967 | -3.17084957877392 | 0.00151993826185045 | 0.0634443198045935 | NM_001141932,NM_020296,NM_001141931 |
| Ccdc84 | 290.6932066 | 0.467663696 | 0.14349943 | 3.258993397 | 0.001118083 | 0.012436162 | | Aanat | 24.11580607 | -0.264234752912689 | 0.0834001040361022 | -3.16827845680273 | 0.00153344571024 75 | 0.0638247346333958 | NM_009591,NR_033223 |
| Arl11 | 25.4035542 | 0.467547731 | 0.166893923 | 2.801465642 | 0.005087106 | 0.036290258 | | 5830418K08Rik | 1149.619542 | -0.197426923860297 | 0.0623692167214539 | -3.16545498925193 | 0.00154840824442136 | 0.0642633661670419 | NM_176976 |
| E230016M11Rik | 41.37623989 | 0.467542876 | 0.160631853 | 2.910648592 | 0.003606794 | 0.028895382 | | Mfsd2a | 972.7537345 | 0.288421608391301 | 0.091156120257681 | 3.16403997423309 | 0.001555954230448 9 | 0.0643925673831929 | NM_029662 |
| Hsf4 | 654.2707419 | 0.467490676 | 0.090715709 | 5.153359676 | 2.56E-07 | 1.62E-05 | | Tyk2 | 837.052247 | -0.24935169529593 | 0.0788406478773279 | -3.16273019948042 | 0.00156297109970876 | 0.0644317343827717 | NM_001205312,NM_018793 |
| Tlr1 | 33.11334525 | 0.467422005 | 0.164705635 | 2.837923576 | 0.004540805 | 0.033817188 | | Uba7 | 186.6273189 | -0.298326124839313 | 0.0943410548635429 | -3.16220891605271 | 0.00156577187368294 | 0.0644317343827717 | NM_023738 |
| Cdsn | 75.82981681 | 0.467340099 | 0.158737931 | 2.94409847 | 0.003238969 | 0.026736596 | | Atr | 697.4807259 | -0.196796661666496 | 0.0622878661076845 | -3.15947027830862 | 0.00158056220798284 | 0.0648071601733144 | NM_019864 |
| Fgr | 14.57654321 | 0.466914266 | 0.151354355 | 3.084908029 | 0.00203615 1 | 0.019331788 | | Gm15446 | 180.683477 | -0.279901789848136 | 0.0886664348440263 | -3.15679535712147 | 0.0015951325059645 | 0.0648071601733144 | NR_040366 |
| Dnase1l2 | 119.7554382 | 0.466911385 | 0.167350681 | 2.790017843 | 0.005270514 | 0.037152566 | | Pdlim7 | 1719.583252 | -0.249811814862209 | 0.0791441061549545 | -3.15641716103418 | 0.00159720248809352 | 0.0648071601733144 | NM_001114087,NM_001114088,NM_026131 |
| Dennd6b | 3013.996015 | 0.466728347 | 0.125203686 | 3.72775246 | 0.000193195 | 0.003311243 | | Slc6a17 | 28549.08962 | 0.135442958404498 | 0.0428908991908119 | 3.15784842378666 | 0.0015893817568811 2 | 0.0648071601733144 | NM_172271 |
| Dsp | 39.76159768 | 0.466514801 | 0.149612224 | 3.118159652 | 0.001819842 | 0.017812149 | | Taf1d | 477.5356907 | -0.226145166640747 | 0.0716026862314787 | -3.15833317724655 | 0.00158674117854943 | 0.0648071601733144 | NR_028401,NM_029248,NM_027261 |
| Arrdc2 | 523.6972478 | 0.46592091 | 0.165794666 | 2.810228588 | 0.004950632 | 0.035738209 | | BC018507 | 6497.022868 | -0.173471729024035 | 0.055052632589312 | -3.15101605254955 | 0.00162703547694061 | 0.0656508814945535 | NM_144837 |
| Syna | 13.95154812 | 0.465681156 | 0.150933715 | 3.085335548 | 0.002033226 | 0.019331788 | | Oas2 | 39.36332655 | -0.252492448025565 | 0.0801187671960773 | -3.15147694930992 | 0.00162446984314386 | 0.0656508814945535 | NM_145227 |
| Gpr64 | 67.06178327 | 0.465566784 | 0.144035817 | 3.232298694 | 0.001227986 | 0.013323823 | | Bri3bp | 5750.01452 | 0.142597509146881 | 0.045288390115569 | 3.1486548491345 | 0.00164023797060652 | 0.0660002680360952 | NM_029752 |
| Slc45a3 | 46.75169494 | 0.464440564 | 0.167981038 | 2.764839238 | 0.005695086 | 0.039424659 | | Hook1 | 2559.032066 | -0.196058391285243 | 0.0622984317119734 | -3.14708362479228 | 0.00164907790652434 | 0.0661726675975625 | NM_030014 |
| Apol7a | 10.63112137 | 0.464384891 | 0.162400881 | 2.859497369 | 0.004243129 | 0.032353861 | | Adam33 | 409.5614273 | -0.298441982890912 | 0.0950875631102582 | -3.13860165440201 | 0.00169756042985064 | 0.0673736688634163 | NM_001163529,NM_033615 |
| 4933439C10Rik | 776.901937 | 0.463497599 | 0.114283089 | 4.055697148 | 5.00E-05 | 0.001145662 | | Atxn7l2 | 934.6593169 | -0.256599871128324 | 0.0817440454411168 | -3.13906498930448 | 0.00169487854064861 | 0.0673736688634163 | NM_175183 |
| Gm20751 | 29.27064596 | 0.462648674 | 0.168073602 | 2.752655193 | 0.005911412 | 0.040384223 | | C330005M16Rik | 68.79526308 | -0.294687069731102 | 0.093831236638967 | -3.14060733170303 | 0.0016859791610522 | 0.0673736688634163 | NM_175319 |
| Ceacam2 | 11.27049967 | 0.462617616 | 0.158145267 | 2.925270065 | 0.003441574 | 0.027955792 | | Cep95 | 391.5742724 | -0.270339830743639 | 0.0861068392312944 | -3.13958604400135 | 0.00169186721174308 | 0.0673736688634163 | NM_177088,NM_001166685 |
| Ddx3x | 10048.43464 | 0.461210136 | 0.101412068 | 4.547882188 | 5.42E-06 | 0.000198486 | | Ankzf1 | 1170.269729 | -0.240339318580429 | 0.0766241225763004 | -3.1366012490532 | 0.00170918410292144 | 0.0676501587984654 | NR_051998,NM_026187,NM_001267620,NR_051999 |
| Car14 | 494.2670555 | 0.459803102 | 0.152395429 | 3.017171225 | 0.002551456 | 0.022693609 | | Crebzf | 1853.913958 | -0.195696464771274 | 0.0624657955694171 | -3.13285795830135 | 0.00173113195633309 | 0.0683326706459089 | NR_073437,NR_073436,NM_145151,NR_073438 |
| Car4 | 1035.075922 | 0.459795566 | 0.099946695 | 4.600407896 | 4.22E-06 | 0.000160129 | | Flt3l | 125.3329142 | -0.287470513478559 | 0.0917843597267469 | -3.13202068777506 | 0.00173607641760689 | 0.0683421302073589 | NM_013520 |
| Mfsd7a | 35.74702236 | 0.459121206 | 0.162606118 | 2.823517422 | 0.004749984 | 0.034731599 | | Arhgef2 | 6865.59606 | -0.182005591766566 | 0.058159252567331 | -3.12943484883782 | 0.00175142906939787 | 0.0687037740446095 | NM_008487,NM_001198913,NM_001198911,NM_001198912 |
| Myo1f | 165.9635705 | 0.458657398 | 0.136283283 | 3.366829462 | 0.000760377 | 0.009204125 | | Clasrp | 1776.435166 | -0.263060679112077 | 0.0840879323807331 | -3.12839978750337 | 0.00175760938678861 | 0.0687037740446095 | NM_016680 |
| Hspa1l | 14.96772533 | 0.458436397 | 0.165849496 | 2.764171189 | 0.005706759 | 0.039425264 | | Tctn3 | 1169.159403 | -0.182199115498841 | 0.0582460899969175 | -3.1280917827872 | 0.00175945228862693 | 0.0687037740446095 | NM_026260 |
| 5730480H06Rik | 21.571123 | 0.457622664 | 0.167833468 | 2.726647257 | 0.00639814 | 0.042690885 | | Celsr3 | 5384.476084 | -0.24729735869306 | 0.0790905769397021 | -3.12676164783356 | 0.00176743155161501 | 0.0688303236427871 | NM_080437 |
| Tacc3 | 210.6457061 | 0.45707704 | 0.11049967 | 4.136456147 | 3.53E-05 | 0.00086607 | | Slc2a9 | 34.33935671 | -0.284638588783877 | 0.0910383414748634 | -3.12653203321267 | 0.00177785698263909 | 0.0690512046251751 | NM_145559,NM_001102414,NM_001012363,NM_001102415 |
| Piwil2 | 42.01146632 | 0.456232624 | 0.167892392 | 2.717410954 | 0.006579486 | 0.043453773 | | Chrm2 | 332.8426123 | 0.24727848501107 | 0.0793235266130817 | 3.1173410407889 | 0.00182490330042298 | 0.0703549815492015 | NM_203491 |
| Pcgf6 | 428.3916584 | 0.456061467 | 0.114091037 | 3.997347032 | 6.41E-05 | 0.001387951 | | Dhtkd1 | 247.3277377 | -0.281664769972011 | 0.0903818419006421 | -3.11639014748851 | 0.00183079877627527 | 0.0703549815492015 | NM_001081131 |
| Il2rb | 12.36692174 | 0.455863515 | 0.162501863 | 2.805281777 | 0.005027261 | 0.036077991 | | Emilin2 | 289.8845343 | 0.281732432609342 | 0.0903658746560048 | 3.11768611416435 | 0.00182276818583884 | 0.0703549815492015 | NM_145158 |
| Ppil6 | 80.39851164 | 0.45575248 | 0.140678881 | 3.23966524 | 0.001196701 | 0.013057525 | | Fgd2 | 232.4174123 | -0.257720817135 95873 | 0.0826973319902504 | -3.11643444103199 | 0.00183052377116018 | 0.0703549815492015 | NM_001159538,NM_013710 |
| Muc1 | 23.29115415 | 0.455382881 | 0.168012479 | 2.710411057 | 0.006719987 | 0.044107886 | | Galnt9 | 3416.658703 | 0.265566834453497 | 0.0853000861154193 | 3.11332434171472 | 0.00184992594828 66 | 0.0707158534863452 | NM_198306,NM_001122639 |
| Slc50a1 | 909.4419845 | 0.454835969 | 0.084753436 | 5.366578517 | 8.02E-08 | 6.25E-06 | | Oas1b | 30.76809102 | -0.258322770638813 | 0.0829723908666137 | -3.11335816577942 | 0.00184971392638729 | 0.0707158534863452 | NM_001083925,NR_003507 |
| Dnaaf3 | 43.07708059 | 0.454770008 | 0.157518664 | 2.887086495 | 0.003888273 | 0.030510193 | | AW549542 | 51.78402886 | -0.29104774715608 | 0.0935612087374049 | -3.11077369653222 | 0.00186597882506 07 | 0.0709560429655281 | NR_045702 |
| Ccdc81 | 33.80986404 | 0.454635054 | 0.16508877 | 2.753882379 | 0.005889293 | 0.040250862 | | Dot1l | 2620.972567 | -0.255160105556612 | 0.0820208210650385 | -3.1109192427406 | 0.00186505937597139 | 0.0709560429655281 | NM_199322 |
| 1110059M19Rik | 16.77721501 | 0.453733566 | 0.125972396 | 3.601849134 | 0.000315962 | 0.004866313 | | 2810459M11Rik | 465.0791344 | -0.251800456147264 | 0.0809915756261029 | -3.10897095408662 | 0.00187740173802053 | 0.0712040147427838 | NM_001144992,NM_001144993 |
| Ly86 | 461.6558974 | 0.453065112 | 0.150626978 | 3.007861654 | 0.002630929 | 0.023198299 | | Nktr | 4630.989869 | -0.219016462481857 | 0.0704732875741448 | -3.10779403119842 | 0.00188489382112208 | 0.0713019990771336 | NM_010918 |
| Exoc3l4 | 56.36822769 | 0.452245602 | 0.157789629 | 2.866148243 | 0.004154996 | 0.031917458 | | Pde7a | 897.9388442 | -0.188028052463131 | 0.0605403691085562 | -3.10582926702539 | 0.00189746237827304 | 0.0715910091085562 | NM_008802,NM_001122759 |
| Kcnk4 | 445.133316 | 0.452201891 | 0.132699223 | 3.407720708 | 0.000655079 | 0.008270829 | | Cacna1h | 2892.532099 | -0.181977546944737 | 0.0586251663997329 | -3.10408580463477 | 0.00190867966584497 | 0.0718276705338448 | NM_021415,NM_001163691 |
| Mei1 | 242.1362796 | 0.452147357 | 0.106687102 | 4.238069551 | 2.25E-05 | 0.000613905 | | 0610030E20Rik | 875.410351 | -0.199238924127083 | 0.0642570567410706 | -3.10065437524816 | 0.00193093530902616 | 0.0724774322969353 | NM_026696 |
| Usb1 | 329.0973659 | 0.451582228 | 0.106955769 | 4.222139961 | 2.42E-05 | 0.000648694 | | Adamts3 | 1227.458387 | -0.224461140104771 | 0.0724362089953347 | -3.09874223427716 | 0.00194344025204628 | 0.0727587966526397 | NM_177872,NM_001081401 |
| Ccdc57 | 341.7458459 | 0.451573212 | 0.125098281 | 3.609747528 | 0.000306495 | 0.004744092 | | Apcdd1 | 878.0516347 | -0.20394413503461 | 0.0658688273960784 | -3.09621626946941 | 0.00196007344783963 | 0.073192871216757 | NM_133237 |
| Zc3h12a | 38.47160976 | 0.451382516 | 0.153357731 | 2.943138774 | 0.003249027 | 0.026805341 | | Rad52 | 521.4349738 | -0.20261415119509 | 0.0655149392980916 | -3.09263789852195 | 0.00198386038727352 | 0.0738911691936799 | NM_011236,NM_001166383,NM_001166382,NM_001166381 |
| Pvrl4 | 237.9846808 | 0.451288363 | 0.11878786 | 3.799111822 | 0.000145216 | 0.00264081 | | 2610203C20Rik | 752.5465455 | -0.211959880336184 | 0.0686143588083183 | -3.0891475781085 | 0.00200731700941806 | 0.0743833848949152 | NM_015483 |
| 1700112E06Rik | 8.650158005 | 0.450717204 | 0.14643425 | 3.077949358 | 0.002084303 | 0.019715256 | | Abcc8 | 1566.672775 | -0.208419439245935 | 0.0674673851083157 | -3.08918803764579 | 0.00200704365000801 | 0.0743833848949152 | NM_011510 |
| Efhc1 | 52.1398574 | 0.450053072 | 0.167406898 | 2.688378295 | 0.00718 | 0.04643027 | | Xlr3b | 43.83685867 | -0.233887546295925 | 0.0758612604381539 | -3.08309596947182 | 0.00204859078230285 | 0.0757196684573314 | NM_001081643 |
| Snord2 | 25.57595321 | 0.450003461 | 0.165440366 | 2.72003424 | 0.006527516 | 0.043239925 | | Tlr9 | 59.26938402 | -0.290225602282864 | 0.0943195349485353 | -3.07704657832784 | 0.00209062630210535 | 0.0768821206693224 | NM_031178 |
| Myh3 | 120.1901526 | 0.449980489 | 0.150268575 | 2.994508252 | 0.002748876 | 0.023900723 | | Tmem246 | 3136.660697 | 0.147946330637354 | 0.0480783746109177 | 3.0771907709991 | 0.00208961521167163 | 0.0768821206693224 | NM_025944 |
| 9830166K06Rik | 60.21086627 | 0.449650747 | 0.161015997 | 2.792584321 | 0.005228884 | 0.036943149 | | 4930422G04Rik | 225.466564 | -0.265571607817041 | 0.0863896395684543 | -3.07375828352419 | 0.00211380623202503 | 0.0770941310869857 | NM_197997 |
| Neil1 | 250.0495939 | 0.449614886 | 0.152062163 | 2.956961012 | 0.003106874 | 0.025894516 | | Bzrap1 | 15593.29937 | -0.231570457430713 | 0.0753412802492203 | -3.07361988972018 | 0.00211478695564581 | 0.0770941310869857 | NM_172449 |
| Apoc1 | 32.9167387 | 0.449418026 | 0.159777274 | 2.812778156 | 0.004911552 | 0.035593822 | | Cd180 | 176.9553587 | -0.234864180975054 | 0.0764141857498407 | -3.07356780250116 | 0.00211515615670 45246 | 0.0770941310869857 | NM_008533 |
| Rgs22 | 40.47606263 | 0.449242999 | 0.166123384 | 2.704110335 | 0.006848754 | 0.044661027 | | Pcgf2 | 1083.254423 | -0.251729641724106 | 0.0819360189546678 | -3.07247333092147 | 0.00212292802112036 | 0.0770941310869857 | NM_009545,NM_001163308,NM_001163307 |
| Ccdc37 | 103.2056304 | 0.449151474 | 0.160070435 | 2.805961481 | 0.005016669 | 0.03601866 | | Zfp566 | 166.8703862 | -0.271485822730617 | 0.088352925490742 | -3.07274288002003 | 0.0021210115243021 | 0.0770941310869857 | NM_152814 |
| Icosl | 183.8017346 | 0.449083539 | 0.12699202 | 3.536313072 | 0.000405754 | 0.005884268 | | D11Wsu47e | 324.6479892 | -0.236410926783259 | 0.0770043151937618 | -3.07009972348161 | 0.00213987308143116 | 0.0775157016979277 | NM_177777 |
| Igsf6 | 43.74293926 | 0.448869066 | 0.162898542 | 2.75551309 | 0.005860016 | 0.040157049 | | Gbp7 | 262.0714339 | -0.287393344983168 | 0.0936434016337033 | -3.06901863846573 | 0.00214763192062735 | 0.0776032370125197 | NM_001083312,NM_145545 |
| Tnfrsf4 | 33.97468857 | 0.448361619 | 0.167335646 | 2.679414871 | 0.007375095 | 0.047316657 | | Rtel1 | 898.9632671 | -0.2539862968416 | 0.0829367144042405 | -3.06241088360036 | 0.00219561841703308 | 0.0791403303370284 | NM_001001882,NM_001166665,NM_001166666,NM_001166667,NM_001166668,NR_030710 |
| 2900005J15Rik | 74.23679272 | 0.448278439 | 0.158933802 | 2.820535551 | 0.004794356 | 0.035022986 | | Acly | 9003.713678 | 0.135191429675624 | 0.0441946901144749 | 3.05899711765023 | 0.00222079292863955 | 0.0792610250055552 | NM_134037,NM_001199296 |
| Hapln4 | 4844.91108 | 0.448028645 | 0.097185928 | 4.610015607 | 4.03E-06 | 0.000154037 | | Fancg | 595.5571862 | -0.204286708616769 | 0.0667737589769862 | -3.0593860784021 | 0.00221791122365678 | 0.0792610250055552 | NM_053081,NM_001163233 |
| Gm216 | 90.47575467 | 0.447978306 | 0.153937817 | 2.910125101 | 0.003612841 | 0.028913926 | | Slc20a1 | 7246.849784 | -0.158315170419232 | 0.0517313077223994 | -3.06034618859407 | 0.0022108128412 0767 | 0.0792610250055552 | NM_015747,NM_001159593 |
| Ppp1r36 | 37.06551535 | 0.447799976 | 0.156900116 | 2.854044893 | 0.004316643 | 0.032801407 | | Tppp3 | 1475.472121 | 0.265787756724805 | 0.0868537716367364 | 3.06017518544221 | 0.00221207559450475 | 0.0792610250055552 | NM_026481 |
| Pip5k1b | 695.5536278 | 0.447278948 | 0.115021403 | 3.888658439 | 0.0001008 | 0.001974454 | | Pou2f2 | 228.4269991 | -0.266457315979886 | 0.0871457694374593 | -3.0576047202199 | 0.00222311366878155 45 | 0.0794350282529587 | NM_001163555,NM_001163554,NM_001163556,NM_011138 |
| Casq1 | 19.50276899 | 0.447098517 | 0.168047709 | 2.660545152 | 0.007801427 | 0.04927652 | | Lrrc45 | 2405.997923 | -0.264621232109473 | 0.0866726104977221 | -3.05311251839528 | 0.00226481000545273 | 0.0803162180514787 | NM_153545 |
| Msantd2 | 321.3338878 | 0.446856602 | 0.122915495 | 3.635478188 | 0.000277465 | 0.004355674 | | Map3k6 | 241.8817111 | 0.288892164855694 | 0.0946309465213631 | 3.05282970820202 | 0.0022669454358465 | 0.0803162180514787 | NM_016693 |
| Ankzf1 | 944.4471104 | 0.446489849 | 0.121097487 | 3.687028203 | 0.000226888 | 0.003716074 | | Rgs5 | 1569.179504 | -0.204543106370544 | 0.0670606844598064 | -3.05011143337185 | 0.00228756473704503 | 0.0808495507793581 | NM_009063 |
| Zfp57 | 301.2274226 | 0.445814719 | 0.145327214 | 3.067661626 | 0.002157408 | 0.020258711 | | Eif5a2 | 2584.746919 | 0.19527668714158 | 0.0640676994796724 | 3.04797407629 | 0.00230389801636474 | 0.0810738528223631 | NM_177586 |
| Gm609 | 9.178286176 | 0.445636307 | 0.155687814 | 2.862371146 | 0.004204842 | 0.03220455 8 | | Tmem145 | 2051.42566 | -0.231667039339356 | 0.0760107164035446 | -3.0478207576302 | 0.00230507374470852 | 0.0810738528223631 | NM_183311 |
| Cd46 | 40.01352506 | 0.44495608 | 0.163594698 | 2.71986859 | 0.006530786 | 0.043239925 | | Hmha1 | 351.1610005 | -0.250692841050755 | 0.0822899500835684 | -3.04569198056666 | 0.00232145521720 24 | 0.0814529827317449 | NM_001142701,NM_027521 |
| Hapln2 | 790.3090677 | 0.444185585 | 0.120377161 | 3.695149328 | 0.000219758 | 0.003630313 | | Dock8 | 268.5886384 | -0.237042792877287 | 0.0778874182605522 | -3.04340287778343 | 0.00233918934931513 | 0.0814845671178694 | NM_028785 |
| Abcb1b | 124.479352 | 0.444411475 | 0.123186869 | 3.607620509 | 0.000309018 | 0.004773604 | | Tln1 | 2302.005325 | -0.193798895838284 | 0.0636720660810797 | -3.04370358567447 | 0.00233685264885811 | 0.0814845671178694 | NM_011602 |
| Plbd2 | 3043.140017 | 0.444376299 | 0.0799921 | 5.555252352 | 2.77E-08 | 2.53E-06 | | Tlr1 | 35.53224131 | -0.266601471905186 | 0.087566283679197 | -3.0445676201344 | 0.00233015041573883 | 0.0814845671178694 | NM_001276445,NM_030682 |
| 5031434O11Rik | 70.85937099 | 0.444121975 | 0.155659777 | 2.85315824 | 0.004328706 | 0.032867482 | | Slco2b1 | 1063.35345 | -0.17575111050131 | 0.0577849304142093 | -3.04146962264219 | 0.00235426319589652 | 0.0818134621617053 | NM_001252530,NM_001252531,NM_175316 |
| Prpf39 | 1175.645894 | 0.443079558 | 0.111593199 | 3.970488908 | 7.17E-05 | 0.001518185 | | Cntn1 | 14603.99716 | 0.152899967614029 | 0.0503029790525842 | 3.03796826231059 | 0.00238179038111873 | 0.0825061784356232 | NM_001159648,NM_007727,NM_001159647 |
| Tanc1 | 1576.170336 | 0.442933464 | 0.12597707 | 3.515984792 | 0.000438126 | 0.006199384 | | Pcdh17 | 2655.50602 | 0.250933000 1083121 | 0.0826119024866177 | 3.03749209895958 | 0.00238555658426007 | 0.0825061784356232 | NM_001013753 |
| Gpat2 | 10.94092258 | 0.442765655 | 0.159509327 | 2.775797899 | 0.005506643 | 0.038519152 | | Helq | 560.64088 | -0.240768187907593 | 0.0792962203933175 | -3.03631354323519 | 0.00239490180214471 | 0.0826326450782757 | NM_001081107 |
| Cabp7 | 277.4505003 | 0.442746738 | 0.151037819 | 2.931363416 | 0.003374777 | 0.027563943 | | Pot1b | 278.3682234 | -0.252954630741029 | 0.0833577334287148 | -3.03456704418851 | 0.00240881209864527 | 0.0829156505803821 | NM_028370 |
| Stxbp2 | 1447.003217 | 0.442260263 | 0.109109186 | 4.053373317 | 5.05E-05 | 0.001155398 | | Fez1 | 8756.207314 | 0.142647450238279 | 0.047029694537448 | 3.0331358015667 | 0.00242026657305472 | 0.0831129840193685 | NM_183171 |
| BC048546 | 258.7041318 | 0.442013699 | 0.098706759 | 4.478048971 | 7.53E-06 | 0.000252627 | | Arl5c | 73.40569053 | -0.288340421659046 | 0.0950964972628997 | -3.03208246316279 | 0.0024287284396175 | 0.0832068615893485 | NM_207231 |
| Med12 | 1713.181044 | 0.441762167 | 0.103751948 | 4.259364574 | 2.05E-05 | 0.000570317 | | Cnot10 | 2323.337586 | -0.179810469172265 | 0.059344496230458 | -3.02994347570144 | 0.00244599511291821 | 0.0832523144918091 | NM_153585 |
| Plcd3 | 539.3248578 | 0.441747606 | 0.098310492 | 4.493392285 | 7.01E-06 | 0.000238701 | | Efcab12 | 264.9584202 | -0.255274517198827 | 0.084250693351242 | -3.0299396603727 | 0.00244602601173234 | 0.0832523144918091 | NM_001110506 |
| Sun2 | 2983.500869 | 0.441292907 | 0.081632014 | 5.405880453 | 6.45E-08 | 5.20E-06 | | Jdp2 | 1709.697427 | 0.201702538753366 | 0.0665731374164004 | 3.02978868792691 | 0.00244724895277451 | 0.0832523144918091 | NM_001205053,NM_001205052,NM_030887 |
| Wdr63 | 14.06777705 | 0.441078577 | 0.157117128 | 2.807323315 | 0.004995508 | 0.03594248 | | Pus10 | 439.3096895 | -0.224879157750362 | 0.0743040786351324 | -3.02647125004569 | 0.00247426349394876 | 0.0839746530679899 | NM_001033654,NM_028956,NM_028304 |
| Sfxn2 | 221.7630986 | 0.439940641 | 0.117303182 | 3.750457877 | 0.000176512 | 0.003097255 | | Arhgap29 | 940.6322137 | -0.221138645860327 | 0.0731513527271909 | -3.02302991315615 | 0.00250257504635628 | 0.0844860040817242 | NM_172525 |
| Tekt4 | 24.96881038 | 0.438625361 | 0.161052983 | 2.723484862 | 0.006459718 | 0.042939425 | | Capn11 | 27.47177218 | 0.184091128883712 | 0.0609614597677181 | 3.01979528681164 | 0.00252945588830715 | 0.0844860040817242 | NM_001013767 |
| Prdm8 | 849.4752006 | 0.438233051 | 0.114953388 | 3.812267383 | 0.000137698 | 0.002515908 | | Fnbp4 | 2009.572349 | -0.177063919684953 | 0.0585732106700425 | -3.02295055468955 | 0.00250323140285992 | 0.0844860040817242 | NM_018828 |
| Akap8l | 4295.475447 | 0.437891996 | 0.116826733 | 3.748249508 | 0.000178073 | 0.003110557 | | Mbip | 526.3336678 | -0.221330379656085 | 0.0732669852183533 | -3.02087466582599 | 0.00252045664359911 | 0.0844860040817242 | NM_145442 |
| Slamf8 | 15.56503004 | 0.437497472 | 0.15455723 | 2.830650319 | 0.004645348 | 0.034306491 | | Olfr856-ps1 | 153.0915155 | -0.274381881101597 | 0.0908022899146933 | -3.02175067786696 | 0.00251317453120635 | 0.0844860040817242 | NR_033621 |
| Aim2 | 71.15896317 | 0.436468281 | 0.150967624 | 2.891138180 | 0.003838491 | 0.030256816 | | Pagr1a | 1737.824094 | -0.218892180973938 | 0.0724713990468379 | -3.02039402927035 | 0.00252446030366614 | 0.0844860040817242 | NM_030240 |
| Abca4 | 167.8939466 | 0.436262523 | 0.163772426 | 2.663833796 | 0.007725572 | 0.048957062 | | Thoc1 | 1075.572384 | -0.209876517368744 | 0.0695018659531879 | -3.01972493098393 | 0.00253004349274061 | 0.0844860040817242 | NM_153552 |
| Net1 | 1638.574971 | 0.43600555 | 0.124703849 | 3.49632793 | 0.000471709 | 0.006531924 | | Gpr84 | 47.82370797 | -0.239594798792728 | 0.0794128129706809 | -3.01707986192589 | 0.00255222562144038 | 0.0848366804966658 | NM_030720 |
| Rsph4a | 168.3673849 | 0.435846851 | 0.098640488 | 4.418539065 | 9.94E-06 | 0.000311669 | | Usp18 | 62.80808088 | -0.272928105747623 | 0.0904479305698581 | -3.01751631052327 | 0.00254855325798313 | 0.0848366804966658 | NM_011909 |
| Sfswap | 2048.062988 | 0.434417132 | 0.092383846 | 4.70230622 | 2.57E-06 | 0.000109196 | | Sfswap | 2411.193226 | -0.205535531968651 | 0.0681707404293888 | -3.01501099553913 | 0.00256969939624298 | 0.0852224964151269 | NM_172276 |
| Eif4ebp3 | 180.6790243 | 0.43413569 | 0.127218902 | 3.412509331 | 0.000643677 | 0.008181407 | | Cadps2 | 5217.427467 | 0.169320762 | 0.0539269176389946 | 3.01403940851479 | 0.00257794317407511 1 | 0.0853011447523153 | NM_153163,NM_001252109,NM_001252108,NM_001252110,NM_001252107,NM_001252106,NM_001252105 |
| Syde1 | 485.026951 | 0.434114466 | 0.09981828 | 4.349047762 | 1.37E-05 | 0.000411358 | | Tgfbr2 | 1031.626482 | -0.236876174504284 | 0.0786662005706524 | -3.01115565243981 | 0.00260255404912453 | 0.0859197729945068 | NM_029575,NM_009371 |
| Grid2ip | 111.1608945 | 0.434095213 | 0.150574921 | 2.882918417 | 0.003940095 | 0.030816672 | | Ppara | 322.431572 | 0.229466169067326 | 0.0762446667339537 | 3.00960288629786 | 0.00261589463571332 | 0.0861643661641082 | NM_011144,NM_001113418 |
| Ccdc176 | 158.4141381 | 0.433570523 | 0.133690279 | 3.243096853 | 0.00118238 | 0.012919462 | | Sfxn2 | 252.7220463 | -0.254849121289664 | 0.0847105531043449 | -3.00846957020509 | 0.00262567095220725 | 0.0862907155017252 | NM_053196 |
| Renbp | 136.9093117 | 0.43295196 | 0.111793981 | 3.87276628 | 0.000107607 | 0.002067036 | | Arntl2 | 199.7486263 | -0.264138105187622 | 0.0878916366425031 | -3.00527007207746 | 0.00265345128336394 | 0.0870068472734642 | NM_172309 |
| Cyp27a1 | 227.0613703 | 0.43285843 | 0.13077698 | 3.309897749 | 0.000933301 | 0.010767356 | | Ccnl2 | 2226.655765 | -0.220970762728219 | 0.073607622021955 | -3.00200925744225 | 0.00268204019434553 | 0.0875069574296561 | NM_207678 |
| Gm14446 | 131.7411296 | 0.432347412 | 0.132918517 | 3.252725209 | 0.00114304 | 0.012623135 | | Fam193b | 3745.83994 | -0.250332016540913 | 0.083403076645635 | -3.00147220712887 | 0.00268677564461 15 | 0.0875069574296561 | NM_145382 |
| Apobec3 | 120.3943397 | 0.431709652 | 0.132312316 | 3.262807174 | 0.001103143 | 0.012305329 | | Rrnad1 | 958.8813731 | -0.22697118720271 | 0.0755937711947941 | -3.00251176274615 | 0.002677616253728 | 0.0875069574296561 | NM_153562 |
| Cdr2l | 1028.27692 | 0.431347814 | 0.113125988 | 3.812986064 | 0.000137298 | 0.002511562 | | Irf7 | 125.0842672 | -0.26521948539306 | 0.0884565457173811 | -2.99830253648426 | 0.00271488023264197 | 0.0879244280435542 | NM_001252600,NM_001252601,NM_016850 |
| Il12a | 79.00272076 | 0.430821237 | 0.149071202 | 2.890036652 | 0.003851969 | 0.030341845 | | Naa40 | 1537.459359 | -0.203925422987547 | 0.068020934390441 | -2.99798023495427 | 0.0027177521817125 | 0.0879244280435542 | NM_027643 |
| Gm581 | 15.94165174 | 0.430598755 | 0.149784135 | 2.874795481 | 0.004042893 | 0.031273381 | | Paqr8 | 6573.153978 | 0.201208447876299 | 0.0670866055775105 | 2.99923429042519 | 0.00270659088142094 | 0.0879244280435542 | NM_028829 |
| 2810098E08Rik | 122.5407629 | 0.429492277 | 0.125402147 | 3.424919644 | 0.000614981 | 0.007960334 | | Ldb2 | 1705.957845 | 0.197337200782729 | 0.0659138689114515 | 2.99386463033799 | 0.00275468133977022 | 0.0889211136477828 | NM_010698,NM_001077398 |
| Snrnp70 | 5977.715205 | 0.429189101 | 0.097328823 | 4.409681425 | 1.04E-05 | 0.000323384 | | Hexb | 5099.463387 | -0.230381287638673 | 0.0769774647645052 | -2.99284067542978 | 0.00276393997553816 | 0.0890221155983756 | NM_010422 |
| Mthfr | 1351.57992 | 0.429180644 | 0.084442751 | 5.082504277 | 3.72E-07 | 2.20E-05 | | Ccdc57 | 409.207547 | -0.236815458406939 | 0.0792240191833455 | -2.98918763334747 | 0.00279720297914062 | 0.0898941824668067 | NM_027745 |
| Helq | 481.3091982 | 0.429020795 | 0.109245105 | 3.927139769 | 8.60E-05 | 0.001750191 | | Abca9 | 445.012748 | -0.217885847279782 | 0.0729883557232496 | -2.98521366247181 | 0.00283380308380171 | 0.0908670635596945 | NM_147220 |
| Cd6 | 107.2496666 | 0.428775579 | 0.156264713 | 2.743905283 | 0.006071304 | 0.041101633 | | Hps1 | 583.6512228 | -0.261323615553555 | 0.0875588909340628 | -2.98454688913714 | 0.00283998670357299 | 0.0908670635596945 | NM_019424 |
| Hdac10 | 537.4239533 | 0.428547335 | 0.113627166 | 3.771521797 | 0.000162255 | 0.002886313 | | Rras2 | 417.6854411 | -0.249427801149044 | 0.0837010487124145 | -2.97998419421182 | 0.00288263269438376 | 0.0920253020826503 | NM_025846 |
| Zc2hc1c | 121.000458 | 0.428206817 | 0.153764508 | 2.784822209 | 0.005355708 | 0.037599157 | | Srpk3 | 271.4553314 | -0.28024411214825 | 0.0940630052975305 | -2.97932339352554 | 0.00288888570666176 9 | 0.0920253020826503 | NM_019684 |
| Tgm3 | 159.9422721 | 0.427301187 | 0.146006058 | 2.92659903 | 0.003426904 | 0.027877541 | | Paqr5 | 238.1917029 | 0.261305130516046 | 0.0879650183116 69 | 2.97055733667008 | 0.00297259910157735 | 0.094279420413783 | NM_028748 |
| Pomc | 104.2136796 | 0.427262683 | 0.141584063 | 3.017731465 | 0.002546745 | 0.022677737 | | Psmd7 | 3695.8742 | 0.116378122134907 | 0.0391699941040668 | 2.97110389717481 | 0.00296731376405136 | 0.094279420413783 | NM_010817 |
| G530011O06Rik | 121.7452738 | 0.426995434 | 0.138849729 | 3.075234194 | 0.002103373 | 0.019861191 | | Klf13 | 9033.838528 | 0.182713772897145 | 0.0615879853761793 | 2.96671131197311 | 0.00301003439502185 | 0.0952587355601033 | NM_021366 |
| Tonsl | 275.5385023 | 0.426861036 | 0.108629658 | 3.929507328 | 8.51E-05 | 0.001742207 | | Ccnt2 | 2045.763996 | -0.196134327295929 | 0.066325717113645 | -2.95713865777822 | 0.00310508451287732 | 0.0972078828320172 | NM_028399 |
| C1s | 76.64378713 | 0.426434273 | 0.145497406 | 2.930871991 | 0.00338012 | 0.027584543 | | Irs2 | 1706.125027 | 0.260338200209641 | 0.0880155049440581 | 2.95786748454275 | 0.0030977526276203 | 0.0972078828320172 | NM_001081212 |
| Cdhr3 | 79.0877983 | 0.426367305 | 0.155338658 | 2.744759808 | 0.00605552 | 0.041042969 | | Mirg | 688.173883 | -0.257060315034702 | 0.0868905388615 | -2.95843849517893 | 0.00309201938063865 | 0.0972078828320172 | NR_028265 |
| Tle6 | 155.1927921 | 0.425930202 | 0.141854745 | 3.002579873 | 0.002677017 | 0.023446978 | | Slfn8 | 59.03905568 | -0.270452452267259 | 0.0914129085226326 | -2.95858504008093 | 0.003090594678751 5 | 0.0972078828320172 | NM_001167743,NM_181545 |
| Slc38a6 | 274.2003973 | 0.425211586 | 0.134450186 | 3.162595757 | 0.001563693 | 0.015960382 | | Socs6 | 458.3522098 | 0.215455111119837 | 0.0728553172325735 | 2.95730111825153 | 0.00310344881682744 | 0.0972078828320172 | NM_018821 |
| Anxa11 | 1782.042492 | 0.424907234 | 0.120535259 | 3.525169631 | 0.000423212 | 0.006043541 | | Nsun5 | 412.8039845 | -0.250481611367662 | 0.0847439068391834 | -2.95574774293797 | 0.00311912082985894 | 0.0974373100527546 | NM_145414 |
| Klhl38 | 43.85321957 | 0.42481891 | 0.154242799 | 2.754221962 | 0.005883185 | 0.040226861 | | Syt9 | 646.4014909 | 0.264435816962151 | 0.0895133639085611 | 2.95414902776166 | 0.00313532559043092 | 0.097733346623604 | NM_021889 |
| Rorb | 1344.004057 | 0.424654058 | 0.103778147 | 4.091941022 | 4.28E-05 | 0.001011906 | | Amy1 | 848.0352953 | -0.220712350247257 | 0.074791007734142 | -2.95105463790271 | 0.00316690890505578 | 0.0985064641431267 | NM_001110505,NM_007446 |

| | | | | | | | | | | | | | | | |
|---|---|---|---|---|---|---|---|---|---|---|---|---|---|---|---|
| Fam160a1 | 110.5692327 | 0.424522703 | 0.126933764 | 3.344442716 | 0.000824481 | 0.009818987 | | Cwf19l2 | 532.4078865 | -0.199737246079508 | 0.0677277180309255 | -2.9491215580662 | 0.00318679018134426 | 0.0987021624183512 | NM_027545 |
| Rnf39 | 141.4456213 | 0.424502989 | 0.154459052 | 2.748320559 | 0.005990141 | 0.040814094 | | Rnpc3 | 999.3496069 | -0.195531265212536 | 0.0662956494032731 | -2.94938305865485 | 0.00318409048770374 | 0.0987021624183512 | NM_026043,NM_001038696 |
| Cxcl16 | 73.41352993 | 0.424462181 | 0.154554809 | 2.746353765 | 0.006026174 | 0.040952059 | | Pcgf6 | 512.1137082 | -0.248115471603892 | 0.084178058314055 | -2.94750765904118 | 0.00320346816600758 | 0.099007613998779 | NM_027654 |
| 1500015A07Rik | 123.8775294 | 0.424236261 | 0.121731144 | 3.485026479 | 0.000492088 | 0.00675805 | | Wdr60 | 883.1989925 | -0.21401214531079 6 | 0.07267512664255 | -2.94477842967384 | 0.00323186013085575 | 0.0994618649593444 | NM_146039 |
| Npy2r | 337.064204 | 0.423848103 | 0.146441847 | 2.894309998 | 0.003799928 | 0.030084867 | | Zbed6 | 1549.130725 | -0.24785222741948 | 0.084165250860232 3 | -2.94482842038384 | 0.00323133802648635 | 0.0994618649593444 | NM_001166552 |
| Odf2 | 2782.072658 | 0.423824338 | 0.105568589 | 4.014682238 | 5.95E-05 | 0.001321339 | | Ercc2 | 1409.594554 | -0.19802135503014 8 | 0.0672894339500054 | -2.94282985315754 | 0.00325227110780734 | 0.0998784146131276 | NM_007949 |
| Crebzf | 1541.389889 | 0.422939118 | 0.094547822 | 4.473282498 | 7.70E-06 | 0.00025648 | | Ankrd34b | 978.8470846 | 0.21923736332571 9 | 0.07456129936659 8 | 2.94036403855823 | 0.00327826838445948 | 0.0998912747499865 | NM_175455 |
| Jpx | 131.3863067 | 0.422480347 | 0.124085338 | 3.404756386 | 0.000662231 | 0.008335181 | | Ap4e1 | 689.8798976 | -0.211593883579215 | 0.0719521859631579 | -2.94075684771493 | 0.00327411433046226 | 0.0998912747499865 | NM_175550 |
| Rims3 | 1776.609993 | 0.422387531 | 0.102889699 | 4.105246042 | 4.04E-05 | 0.0009702 | | Htr2c | 1970.943895 | 0.24857719613922 5 | 0.0845135238746263 | 2.94127212788044 | 0.00326868188409722 | 0.0998912747499865 | NM_008312 |
| Agxt2l2 | 92.79025906 | 0.422221749 | 0.136641225 | 3.090002658 | 0.002001547 | 0.019149082 | | Lrrcc1 | 737.6794115 | -0.21410964677876 5 | 0.0728379822420711 | -2.93952287208438 | 0.00328707347724725 | 0.0998912747499865 | NM_028915,NM_001163579,NM_001163580 |
| 4933421O10Rik | 333.1524645 | 0.421850902 | 0.092811964 | 4.545221162 | 5.49E-06 | 0.000198718 | | Ncapd2 | 1073.102104 | -0.17303532818610 2 | 0.0588537723429414 | -2.94008899771024 | 0.00328117993022425 | 0.0998912747499865 | NM_146171 |
| Hjurp | 181.0217292 | 0.421656425 | 0.123577783 | 3.412073067 | 0.000644708 | 0.008187795 | | Neurl3 | 42.74228278 | -0.267719190770741 | 0.0911753730636077 | -2.93631034099493 | 0.00332141853368793 | 0.10072427060616 | NM_153408 |
| Ncf2 | 186.8801818 | 0.421386171 | 0.123193996 | 3.420508981 | 0.000625041 | 0.008054403 | | Jak3 | 263.8473183 | -0.261065993364878 | 0.0889770140496 21 | -2.9340608853975 | 0.00334558573678814 | 0.101245788359551 | NM_001190830,NM_010589 |
| Cdt1 | 58.6889381 | 0.421190729 | 0.156836211 | 2.685545181 | 0.007241158 | 0.046631134 | | Ruvbl1 | 1452.065254 | 0.149749335352859 | 0.05105511384888 5 | 2.93309179166838 | 0.00335604653966832 | 0.101351210052436 | NM_019685 |
| Rxfp2 | 49.83036557 | 0.421030845 | 0.157731743 | 2.669284165 | 0.007601311 | 0.048288115 | | Cfp | 385.9824354 | -0.275665398151484 | 0.0940982639419279 | -2.92954818296764 | 0.00339455180831054 | 0.102281257583347 | NM_008823 |
| Niacr1 | 8.704669408 | 0.42011272 | 0.142905427 | 2.939795433 | 0.00328429 | 0.027038677 | | Tmem80 | 922.3945185 | -0.185069220981521 | 0.0631858738106014 | -2.92896512812757 | 0.00340092574781471 | 0.102281257583347 | NM_027797,NR_024509,NM_001141950 |
| Tlr7 | 183.9730582 | 0.418724091 | 0.136888965 | 3.058859347 | 0.002221814 | 0.020687976 | | Myo5c | 227.4913003 | -0.27798413375891 | 0.0950889809719 27 | -2.92341313572459 | 0.00346216810593996 | 0.103907962617529 | NM_001081322 |
| Esrrb | 106.1518655 | 0.418325939 | 0.132399224 | 3.159579992 | 0.001579967 | 0.016067252 | | Akap13 | 1848.860655 | -0.205917301972427 | 0.0705198812992047 | -2.91999834454743 | 0.00350043354090778 | 0.104182436347003 | NM_029332 |
| Kdm6a | 1413.707683 | 0.417917006 | 0.083617184 | 4.997979924 | 5.79E-07 | 3.15E-05 | | Rxfp1 | 308.2707309 | -0.244209684090451 | 0.0836509352644957 | -2.91938976316624 | 0.00350717412733611 | 0.104182436347003 | NM_212452 |
| Cadps2 | 3889.122079 | 0.41755448 | 0.071398308 | 5.848240576 | 4.97E-09 | 5.92E-07 | | Taz | 1709.933222 | -0.204629587550372 | 0.0700778073636558 | -2.92003410564039 | 0.00349993080299256 | 0.104182436347003 | NM_001242616,NM_181516,NM_001173547,NM_001242615 |
| Slc26a11 | 164.1703546 | 0.417296465 | 0.108877041 | 3.832731498 | 0.000126728 | 0.002347184 | | Tcirg1 | 612.4305361 | -0.228952355369373 | 0.0783868950244968 | -2.92079888223437 | 0.00349135129696789 | 0.104182436347003 | NM_016921,NM_001136091,NM_001167784 |
| Rgl2 | 1786.332434 | 0.416900757 | 0.114630883 | 3.636897378 | 0.000275942 | 0.00434933 | | Tle2 | 1388.643769 | -0.238534768138244 | 0.0817037577284471 | -2.91950792436065 | 0.0035058448072173 | 0.104182436347003 | NM_019725,NM_001252401 |
| Kcnq4 | 214.2834339 | 0.416821847 | 0.143003149 | 2.914773916 | 0.003559463 | 0.028648293 | | Tmem35 | 2743.610858 | 0.156787649579883 | 0.0538279065634654 | 2.91275770487235 | 0.003582524575603 | 0.106203575480019 | NM_026239 |
| Pamr1 | 1324.375835 | 0.41595314 | 0.141470197 | 2.940217427 | 0.00327982 | 0.027016232 | | Arhgap9 | 34.16223775 | -0.26426759689227 | 0.0908259317346227 | -2.90960512978181 | 0.00361885654333388 | 0.106640918248526 | NM_146011 |
| Osmr | 236.4491491 | 0.415561741 | 0.152611499 | 2.723004126 | 0.006469125 | 0.042962976 | | Sdc4 | 2422.379266 | 0.209963868349864 | 0.0721503045078331 | 2.91008984344714 | 0.00361324873002733 | 0.106640918248526 | NM_011521 |
| Parp12 | 419.7776746 | 0.415409769 | 0.123159648 | 3.37293728 | 0.000743709 | 0.009066516 | | Vwa5b2 | 2981.755551 | -0.244183351480645 | 0.0839243008648169 | -2.90956670433239 | 0.00361930143856006 | 0.106640918248526 | NM_001144953,NM_182636 |
| Eya4 | 175.7902938 | 0.415135175 | 0.113460164 | 3.6588628 | 0.000253337 | 0.004084498 | | Zmym6 | 1757.103646 | -0.199791302994539 | 0.068737600401792 | -2.90657954055274 | 0.00365403986361576 | 0.107446524410693 | NM_177462 |
| Carns1 | 487.1543612 | 0.415064307 | 0.114101138 | 3.6376877 | 0.000275097 | 0.004344901 | | Usp28 | 1248.877818 | -0.211795436247524 | 0.0729234780451733 | -2.9043518209091 | 0.00368014360194664 | 0.10799548679167 | NM_175482 |
| 2610005L07Rik | 599.4656951 | 0.413935451 | 0.150693211 | 2.746875238 | 0.006016602 | 0.0409224 | | Htr1b | 276.8622648 | 0.275385781363288 | 0.0949394631701783 | 2.90066076654424 | 0.00372394221802523 | 0.109060452941602 | NM_010482 |
| Gm10865 | 9.873488124 | 0.413572823 | 0.154621173 | 2.674748962 | 0.007478523 | 0.047684045 | | Arnt | 1453.547339 | -0.200785946378931 | 0.0692811515987649 | -2.89813234545758 | 0.00375392145843145 | 0.109717229587878 | NM_001037737,NM_009709 |
| 4930563E22Rik | 98.28359016 | 0.413304426 | 0.124492847 | 3.319905016 | 0.000900481 | 0.010498156 | | Lypd1 | 3488.447078 | 0.21716500069043 4 | 0.0749654180746706 | 2.89686906666915 | 0.00376907023541278 | 0.109938783613667 | NM_145100 |
| Guf1 | 1073.655258 | 0.412992843 | 0.104141767 | 3.965679217 | 7.32E-05 | 0.001540028 | | Oasl2 | 282.4421304 | -0.195525909440766 | 0.0675360802757824 | -2.8951326203247 | 0.00378998367114335 | 0.110327260134325 | NM_011854 |
| Klf15 | 871.8461861 | 0.412893692 | 0.141516729 | 2.917631692 | 0.003527007 | 0.02846221 | | Il10rb | 391.1982218 | -0.217360849596926 | 0.0751585910093963 | -2.89202932995047 | 0.0038276221253339 | 0.111200077985201 | NM_008349 |
| Taf1d | 416.5507127 | 0.412550898 | 0.096267806 | 4.285450297 | 1.82E-05 | 0.000518465 | | Cd9 | 1411.09227 | -0.250557981246262 | 0.0866990281112114 | -2.88997450957425 | 0.00385273065477369 | 0.111261959227123 | NM_007657 |
| Csf2rb | 74.08500698 | 0.412294298 | 0.135251349 | 3.048356266 | 0.00230097 | 0.021168869 | | Spata32 | 31.85378847 | -0.254335598708236 | 0.0879878269041465 | -2.89057711341488 | 0.00384535177654535 | 0.111261959227123 | NM_177801 |
| Asun | 927.1327229 | 0.412202871 | 0.081087982 | 5.083402746 | 3.71E-07 | 2.20E-05 | | Tubb3 | 12967.40978 | 0.167871113092712 | 0.0580729879652586 | 2.89069185131542 | 0.00384394826801099 | 0.111261959227123 | NM_023279 |
| Azi1 | 902.9673431 | 0.412059687 | 0.091215177 | 4.517446547 | 6.26E-06 | 0.000218909 | | Ppox | 944.2365591 | -0.231481345495756 | 0.0801702243930679 | -2.88737305212996 | 0.00388473333750751 | 0.11196356440602 | NM_008911 |
| Nr2f1 | 1976.305543 | 0.411975361 | 0.065625128 | 6.277707565 | 3.44E-10 | 5.60E-08 | | Fcer1g | 369.1126506 | -0.247133576028749 | 0.0856761063278245 | -2.88450988987683 | 0.00392023445020172 | 0.112763021036891 | NM_010185 |
| Otx2 | 34.56064157 | 0.411885017 | 0.154112164 | 2.672631462 | 0.007525888 | 0.047926884 | | Ccdc176 | 185.7948878 | -0.261700868781247 | 0.0908118815181974 | -2.88179106528925 | 0.00395421837084179 | 0.113403194240242 | NM_028377 |
| Dpp7 | 782.6560436 | 0.411862974 | 0.086146189 | 4.780977322 | 1.74E-06 | 7.93E-05 | | Treml2 | 27.30688501 | -0.253273196053551 | 0.0878968654009313 | -2.88148155111419 | 0.00395810405340788 | 0.113403194240242 | NM_001033405 |
| Pisd-ps2 | 891.1261271 | 0.411808925 | 0.141550129 | 2.909279768 | 0.003622625 | 0.028942617 | | Atp5a1 | 35358.49742 | 0.116965914912102 | 0.0406528706961452 | 2.8771870942731 | 0.0040123762217545 | 0.113864147262278 | NM_007505 |
| A230057D06Rik | 222.7510123 | 0.411210893 | 0.146185577 | 2.812937507 | 0.004909119 | 0.035592836 | | Col6a1 | 4211.869628 | -0.273388479162507 | 0.0949935569581992 | -2.87796865299831 | 0.00400244908677724 | 0.113864147262278 | NM_009933 |
| Pole | 84.27169108 | 0.410610961 | 0.14566909 | 2.818792657 | 0.004820465 | 0.035114376 | | Ftx | 1032.960593 | -0.242628039312235 | 0.0843005627029333 | -2.87813072099212 | 0.00400039333909594 | 0.113864147262278 | NR_028381,NR_028380 |
| Efna5 | 1114.893734 | 0.410021757 | 0.088567333 | 4.629491988 | 3.67E-06 | 0.000143423 | | Myo19 | 446.3690144 | -0.261027587046277 | 0.0907256919336715 | -2.87710770216127 | 0.00401338588725639 | 0.113864147262278 | NM_025414 |
| Rgsl1 | 97.99706985 | 0.409837516 | 0.150586721 | 2.721604633 | 0.006496582 | 0.043126835 | | Stk38 | 1200.190107 | -0.208648721419015 | 0.0725003313676053 | -2.87790024518762 | 0.00400331709187762 | 0.113864147262278 | NM_134115 |
| H2-D1 | 646.1180434 | 0.409644746 | 0.103068222 | 3.974500954 | 7.05E-05 | 0.001501023 | | Pcdha1 | 58.83509195 | 0.266760675142562 | 0.092740425947016 | 2.87642280819196 | 0.00402210557702094 | 0.113889094759856 | NM_054072 |
| Ttc39a | 112.1888321 | 0.40828564 | 0.141939095 | 2.876484734 | 0.004021316 | 0.031153139 | | Clk4 | 1725.260698 | -0.250332650383412 | 0.0870957543059723 | -2.87422334622626 | 0.00405022433117872 | 0.114239919679033 | NM_007714 |
| Ctnnal1 | 644.0477964 | 0.407932335 | 0.106527333 | 3.829367767 | 0.000128473 | 0.002372538 | | Sox9 | 803.0536186 | -0.240774593573728 | 0.0837557213279459 | -2.87472413533368 | 0.00404380641071945 | 0.114239919679033 | NM_011448 |
| Slc26a4 | 119.7147907 | 0.407358269 | 0.131481796 | 3.098210408 | 0.001946931 | 0.018724864 | | Arpc3 | 4141.371696 | 0.130046481227915 | 0.0452925906527371 | 2.87125287720888 | 0.00408848315884041 | 0.115095554971542 | NM_019824 |
| Cep164 | 638.570317 | 0.407302036 | 0.089816394 | 4.534829548 | 5.77E-06 | 0.00020534 | | Fam20a | 563.7032857 | 0.221179446527361 | 0.0770683670115584 | 2.86991219749329 | 0.00410585792881515 | 0.115138402073299 | NM_153782 |
| Srrm2 | 26989.0295 | 0.407065876 | 0.115031785 | 3.538725205 | 0.000402064 | 0.005876062 | | Gm17296 | 287.682353 | -0.24502088347525 | 0.0853656304170665 | -2.87025214103339 | 0.00410144604221568 | 0.115138402073299 | NM_001159907 |
| Tap2 | 404.9578748 | 0.406917864 | 0.120374429 | 3.380434427 | 0.000723713 | 0.008920617 | | Ankrd13a | 2047.784755 | 0.160972933293658 | 0.0561445463886215 | 2.86711610740311 | 0.00414231012120673 | 0.11571384003971 | NM_026718 |
| Il4ra | 312.3189014 | 0.406074535 | 0.113392784 | 3.58113207 | 0.000342109 | 0.005171348 | | Slc25a4 | 20302.41444 | 0.132912458918202 | 0.046355066118269 | 2.86726932023122 | 0.00414030513012341 | 0.11571384003971 | NM_007450 |
| Osbp2 | 4214.553581 | 0.405888975 | 0.079155281 | 5.127756096 | 2.93E-07 | 1.82E-05 | | Zrsr1 | 3815.457116 | 0.151933816708039 | 0.0531906533214427 | 2.85640064975081 | 0.00428474076249817 | 0.119462848975141 | NM_011663 |
| Kcnt1 | 6820.849648 | 0.405563177 | 0.076972785 | 5.268916491 | 1.37E-07 | 9.62E-06 | | Pald1 | 564.9280957 | -0.201630709163193 | 0.0706596100465732 | -2.85354970159465 | 0.00432337644808843 | 0.120309130814047 | NM_013753 |
| Ppm1m | 632.7102671 | 0.404761785 | 0.08570283 | 4.722852007 | 2.33E-06 | 0.000100932 | | A330049N07Rik | 85.79137884 | -0.271157841767178 | 0.0950517514098033 | -2.85273903684442 | 0.00433442000350337 | 0.120385822124073 | NR_040646 |
| Ak8 | 73.35274009 | 0.403538322 | 0.14253715 | 2.831109795 | 0.004638679 | 0.034306301 | | Zfp420 | 504.7053987 | -0.207373397644368 | 0.0727778755814899 | -2.84940163459663 | 0.00438015477803949 | 0.121423908980537 | NM_172740 |
| Pgm1 | 282.1419386 | 0.402899547 | 0.095536395 | 4.217236257 | 2.47E-05 | 0.000661813 | | Bcl2a1b | 23.82138782 | -0.22267448621083 5 | 0.0781816025890235 | -2.84816988699203 | 0.00439714449367609 | 0.121662706505026 | NM_007534 |
| Phka2 | 1480.198793 | 0.401821179 | 0.081290952 | 4.943000072 | 7.69E-07 | 4.00E-05 | | Pan2 | 1140.4154 | -0.20481336068634 | 0.0720776203587686 | -2.84156662868274 | 0.00448924701524594 | 0.12397490901799 | NM_001252327,NM_001252326,NM_133992 |
| Gata2 | 178.5816356 | 0.401768888 | 0.127322477 | 3.155522072 | 0.001602111 | 0.01619582 | | Abcc5 | 6775.532219 | -0.173838668073934 | 0.0612633289524393 | -2.83756483766807 | 0.00454591128660567 | 0.125301531971981 | NM_176839,NM_013790 |
| Scn1b | 14137.37845 | 0.401690107 | 0.104320172 | 3.850550677 | 0.000117853 | 0.002218721 | | Psmc1 | 5905.22531 | 0.119073320831618 | 0.0419843664869668 | 2.83613475193397 | 0.00456631752268221 | 0.12562562184561 | NM_008947 |
| 4831440E17Rik | 156.4821858 | 0.400932099 | 0.147641208 | 2.71558398 | 0.006615899 | 0.043619892 | | Akap2 | 3929.512789 | 0.150097383828118 | 0.0529779087638069 | 2.83320703535698 | 0.0046083528207409 | 0.126226881930386 | NM_001035533,NM_001035532,NM_009649 |
| Arglu1 | 2910.74368 | 0.40077953 | 0.107168628 | 3.739709423 | 0.000184233 | 0.003203712 | | B3galnt2 | 820.925279 | -0.192447514155026 | 0.0679419745322268 | -2.83252754780836 | 0.00461815867073482 | 0.126226881930386 | NM_178640 |
| Zbed6 | 1317.7453 | 0.400745696 | 0.115229356 | 3.477809036 | 0.00050553 | 0.006882848 | | Gm17821 | 372.011346 | -0.218669063142929 | 0.0771695987712363 | -2.83361669134963 | 0.0046024500900783 | 0.126226881930386 | NR_033146 |
| Nrbp2 | 5296.305868 | 0.399380579 | 0.111883609 | 3.569607577 | 0.000357516 | 0.005357214 | | Slc7a6 | 1059.729648 | -0.184648787415884 | 0.0651963009132344 | -2.83219729999132 | 0.00462293137732103 | 0.126226881930386 | NM_178798 |
| Olfml2b | 250.5628013 | 0.39911716 | 0.148895232 | 2.68052344 | 0.007350712 | 0.047199306 | | Bbc3 | 153.8244273 | -0.266823265836459 | 0.0942349001457954 | -2.8314697147622 | 0.00463346212618758 | 0.126277055994373 | NM_133234 |
| Frmd5 | 1190.066043 | 0.398958307 | 0.098274685 | 4.059624391 | 4.92E-05 | 0.001131652 | | Clk1 | 3345.062499 | -0.25157162698148 | 0.0889136073797588 | -2.82939399713401 | 0.00466362463967968 | 0.126643249817882 | NM_001042634,NR_027854,NR_027853 |
| Mypn | 133.2386028 | 0.397785758 | 0.140799648 | 2.825190001 | 0.004725259 | 0.034663987 | | Ikzf1 | 214.4058847 | -0.241613854469113 | 0.0853956784380779 | -2.82934521849735 | 0.00466433558120383 | 0.126643249817882 | NM_001025597,NM_009578 |
| Slc4a2 | 1164.158778 | 0.397675655 | 0.097579421 | 4.075404935 | 4.59E-05 | 0.001068629 | | Crtc2 | 1115.700987 | -0.220071178052152 | 0.0778143432877394 | -2.82815697921371 | 0.00468168433309127 | 0.126877139221052 | NM_028881 |
| Fam227a | 266.1560058 | 0.39734091 | 0.132586973 | 2.996832198 | 0.002728008 | 0.023786022 | | Trabd2b | 282.7496825 | 0.235951940261696 | 0.0834487852180337 | 2.82750599239048 | 0.00469121374223417 | 0.126898642122334 | NM_001085549 |
| Qrfpr | 81.90504758 | 0.396995821 | 0.143195914 | 2.772396292 | 0.005564525 | 0.038738759 | | Trpm4 | 614.9359421 | -0.234904405914495 | 0.0831235540652542 | -2.82596682199234 | 0.00471381462186543 | 0.127272994790367 | NM_175130 |
| Pde4c | 114.3133344 | 0.396739469 | 0.143036247 | 2.773698817 | 0.005542296 | 0.038692429 | | Rccd1 | 394.8716509 | -0.227191000378722 | 0.0804466158519905 | -2.82412128804421 | 0.00474104400354334 | 0.127298346017506 | NM_173445 |
| Gm10560 | 137.6570289 | 0.396621111 | 0.138583945 | 2.861955706 | 0.004210357 | 0.032225775 | | Slc7a11 | 1693.821151 | -0.23467915971297 | 0.0830818808550955 | -2.82467316937947 | 0.00473288655098366 | 0.127298346017506 | NM_011990 |
| Akap8 | 2964.72911 | 0.396454666 | 0.10146692 | 3.907230708 | 9.34E-05 | 0.00185928 | | Snap23 | 432.6682099 | -0.21170887889447 | 0.074955204633805 | -2.82447202871072 | 0.00473585817313703 | 0.127298346017506 | NM_001177792,NM_001177793,NM_009222 |
| Ndor1 | 1375.873407 | 0.396379179 | 0.084893467 | 4.669136417 | 3.02E-06 | 0.000123328 | | Dvl2 | 811.5811287 | -0.230042791973391 | 0.0816400309352818 | -2.81776953460187 | 0.00483585069796827 | 0.129604367599054 | NM_007888 |
| Ccnl2 | 1906.477927 | 0.396049701 | 0.119327502 | 3.319014425 | 0.000903358 | 0.010507975 | | Coro6 | 1507.263176 | -0.247682901678629 | 0.0879696020444909 | -2.81555100764651 | 0.00486936664139565 | 0.130262283301866 | NM_139130,NM_139129,NM_139128 |
| Ccl4 | 8.703824082 | 0.395639078 | 0.129547293 | 3.054012704 | 0.002258025 | 0.020941771 | | Il1a | 38.04261118 | -0.252822232408177 | 0.0898336523836446 | -2.81433767524526 | 0.00488778557531908 | 0.130514656740965 | NM_010554 |
| Slc16a10 | 102.7778088 | 0.395466299 | 0.14192466 | 2.786452332 | 0.005328845 | 0.037473304 | | Gbp2 | 238.2975561 | -0.191456036788644 | 0.0682254976307037 | -2.80622411616507 | 0.00501258153888782 | 0.1333938471437 | NM_010260 |
| 2810403A07Rik | 2212.842361 | 0.395099973 | 0.109819306 | 3.597727821 | 0.000321009 | 0.004932302 | | Rims2 | 5585.929981 | 0.139404224095849 | 0.0496992541487789 | 2.80495605987428 | 0.00503234395117359 | 0.1333938471437 | NM_001256384,NM_053271,NM_001256383,NM_001256382 |
| Fcgr2b | 150.5227812 | 0.393709344 | 0.136328172 | 2.887952942 | 0.003877579 | 0.030481586 | | Tceal8 | 1040.555968 | 0.158203830660639 | 0.0563965518560703 | 2.80520396112853 | 0.00502847492735951 | 0.1333938471437 | NM_025703,NM_001168578 |
| Ctss | 2868.987884 | 0.393089456 | 0.101346622 | 3.878663618 | 0.000105032 | 0.002036751 | | Usp40 | 867.2412304 | -0.188617971674478 | 0.0672307525756339 | -2.80553116614729 | 0.00502337231805326 | 0.1333938471437 | NM_001033291,NM_001198573 |
| Trip6 | 196.8315502 | 0.392629641 | 0.127649664 | 3.075837632 | 0.002099121 | 0.019843675 | | Shkbp1 | 577.3892483 | -0.254112675490399 | 0.0906234567269413 | -2.80404968722468 | 0.0050465127351462 | 0.133434165580226 | NM_138676 |
| Nfkbiz | 170.9904054 | 0.392587452 | 0.121574444 | 3.229193898 | 0.001241397 | 0.013431707 | | Slc7a1 | 2276.956428 | 0.172614578794815 | 0.0615670553321888 | 2.80368417595193 | 0.00505223675265899 | 0.133434165580226 | NM_007513 |
| 1700094D03Rik | 260.8815311 | 0.392473969 | 0.135753479 | 2.89107854 | 0.003839222 | 0.030256816 | | Neil1 | 324.7003463 | -0.244706814883486 | 0.0873662418632561 | -2.80093099651117 | 0.00509554134079248 | 0.134333636145829 | NM_028347 |
| Ppip5k2 | 1173.157462 | 0.391262511 | 0.070977746 | 5.5124674 | 3.54E-08 | 3.11E-06 | | Pik3ap1 | 130.5533675 | -0.26271973770087 | 0.0938668944277552 | -2.79885405075453 | 0.00512843120435963 | 0.134955782019072 | NM_031376 |
| Atf7 | 313.9838461 | 0.3910663 | 0.094367629 | 4.14407252 | 3.41E-05 | 0.000847891 | | C3ar1 | 141.5368275 | -0.263384807582003 | 0.0941927358210394 | -2.79623269550657 | 0.00517021601420903 | 0.135809326984449 | NM_009779 |
| Ntsr1 | 361.6443807 | 0.390801292 | 0.133669808 | 2.923631741 | 0.003459738 | 0.028006885 | | Sc4mol | 3141.761194 | 0.159538167772718 | 0.0571372775790717 | 2.79219057211702 | 0.00523525126084957 | 0.137269422048919 | NM_025436 |
| Gm6548 | 132.9833753 | 0.390653733 | 0.126509694 | 3.08793516 | 0.002015524 | 0.019241242 | | Osmr | 216.6755115 | -0.263527620571973 | 0.0944142455228805 | -2.79118494367578 | 0.00525154558123616 | 0.13744856056403 | NM_011019 |
| Pstpip2 | 363.3532419 | 0.390492311 | 0.129015807 | 3.026701297 | 0.002472381 | 0.022169068 | | Clec5a | 105.5540405 | -0.263355222564589 | 0.0944415139269505 | -2.78855358850231 | 0.00529439875079369 | 0.137476726802077 | NM_001038604,NM_021364 |
| Dgka | 438.2916731 | 0.389287384 | 0.087395892 | 4.454298448 | 8.42E-06 | 0.000273396 | | Ina | 5401.029966 | 0.171995239934191 | 0.0616641671999855 | 2.78922505150173 | 0.00528343366870643 | 0.137476726802077 | NM_146100 |
| Ciz1 | 1788.654606 | 0.389287067 | 0.075708734 | 5.141904338 | 2.72E-07 | 1.70E-05 | | Odf2 | 3371.3231 | -0.219014522677695 | 0.078582316860336 | -2.78707133396116 | 0.00531867697084034 | 0.137476726802077 | NM_001113214,NM_001177661,NM_001177659,NM_001113213,NM_013615 |
| Tle2 | 1111.733298 | 0.388856397 | 0.140206334 | 2.773458135 | 0.005546398 | 0.038692429 | | Pask | 174.7327919 | -0.251374593783457 | 0.0902097880456623 | -2.78655564134811 | 0.00532714717736701 | 0.137476726802077 | NM_080850 |
| Trmt13 | 212.5896922 | 0.388451197 | 0.109934496 | 3.533478683 | 0.000410129 | 0.005927744 | | Pisd-ps2 | 1040.580608 | -0.256181034747458 | 0.0919370406189498 | -2.78648336973612 | 0.00532833520511973 | 0.137476726802077 | NR_003519 |
| Ankrd24 | 1097.561926 | 0.388268049 | 0.142556471 | 2.723608738 | 0.006457295 | 0.042939425 | | Prpf4b | 3593.285223 | -0.17198902874179 | 0.0616879284721704 | -2.78804999618978 | 0.00530263596589264 | 0.137476726802077 | NM_013830 |
| Anxa4 | 148.4624249 | 0.386832245 | 0.12062636 | 3.206863288 | 0.001341908 | 0.014201854 | | Tmem212 | 50.12157489 | 0.249812111518682 | 0.0895307776065749 | 2.79023725915161 | 0.00526694294548437 | 0.137476726802077 | NM_001164437 |
| Odf2l | 365.1519257 | 0.386517432 | 0.114760746 | 3.368028227 | 0.000757078 | 0.009189519 | | Tyms | 156.8724868 | -0.247260952925286 | 0.0886627919038674 | -2.78877923439833 | 0.00529071163168682 | 0.137476726802077 | NM_021288,NR_033402 |
| Ctps2 | 1573.638203 | 0.385517154 | 0.066985222 | 5.755256781 | 8.65E-09 | 9.39E-07 | | Nek1 | 1612.923193 | -0.167701028505582 | 0.0602258062870149 | -2.78453770641738 | 0.00536040881181015 | 0.13805903971694 | NM_175089 |
| Plekhh3 | 409.0509653 | 0.385244042 | 0.100568286 | 3.830671236 | 0.000127794 | 0.00236282 | | Atp13a5 | 563.6572922 | -0.216855842429749 | 0.0779661185297301 | -2.78141129145806 | 0.00541231199441578 | 0.138877278577871 | NM_175650 |
| Gpr115 | 95.86240275 | 0.385223841 | 0.13501968 | 2.853094008 | 0.004329581 | 0.032867482 | | Cxcl10 | 27.56363512 | -0.166164706230147 | 0.0597889977122673 | -2.77918534493268 | 0.0054495421168516 | 0.138877278577871 | NM_021274 |
| Fcgr3 | 627.8433476 | 0.384514396 | 0.125639206 | 3.06046502 | 0.002209936 | 0.020614536 | | Dpysl5 | 1526.16481 | 0.187848045770875 | 0.0675318356276048 | 2.78162209016112 | 0.00540879820600967 | 0.138877278577871 | NM_023047 |
| Ifit2 | 528.9881618 | 0.383933555 | 0.112671032 | 3.407562246 | 0.00065546 | 0.008270829 | | Ifi27l2a | 25.45252315 | -0.214504387660046 | 0.0771593843565494 | -2.78001683721105 | 0.00543560800755315 | 0.138877278577871 | NM_029803 |
| Atxn7l2 | 803.2788139 | 0.383700209 | 0.123955055 | 3.095478496 | 0.001964956 | 0.018886496 | | Mettl17 | 771.604056 | -0.222824141671736 | 0.0801322906308801 | -2.78070350812944 | 0.00542412506590018 | 0.138877278577871 | NM_001029990 |
| C77080 | 1044.137935 | 0.383297434 | 0.131411854 | 2.916764537 | 0.003536827 | 0.028526596 | | Snapc4 | 2223.079437 | -0.227826905061158 | 0.0819638317361459 | -2.77960290820185 | 0.0054425405853115 | 0.138877278577871 | NM_172339,NR_033572 |
| Pxmp2 | 119.1799182 | 0.383139418 | 0.121312385 | 3.158287723 | 0.001586988 | 0.016102684 | | Cct2 | 5554.025709 | 0.134130018923662 | 0.0483078602972992 | 2.77656716936313 | 0.00549362825134414 | 0.139755593658538 | NM_007636 |
| Tmem181a | 1656.620027 | 0.382756788 | 0.081951111 | 4.670550316 | 3.00E-06 | 0.000122805 | | Got1 | 15572.72228 | 0.109737767609636 | 0.0395721609463548 | 2.77310525847703 | 0.0055524157944638 | 0.141004181521646 | NM_010324 |
| 2610015P09Rik | 203.1639337 | 0.382747078 | 0.134749218 | 2.840440058 | 0.004505134 | 0.03360928 | | Efnb1 | 670.5823622 | 0.200370795817635 | 0.0722807817491037 | 2.7721171654334 | 0.00556929866145333 | 0.141186094862602 | NM_010110 |
| Arid3b | 295.5686109 | 0.382067784 | 0.105966617 | 3.60554857 | 0.000311494 | 0.004807063 | | Mrgbp | 320.6562358 | -0.233356902045854 | 0.0842311474854866 | -2.77043479772209 | 0.00559815072693648 | 0.14167027431965 | NM_028479 |
| Pcdhb3 | 236.6135549 | 0.381547815 | 0.095716577 | 3.986225024 | 6.71E-05 | 0.001444662 | | Col12a1 | 509.5265694 | -0.236981174301262 | 0.0856093818550793 | -2.76816826808103 | 0.00563723420152984 | 0.141672083064745 | NM_007730 |
| Srsf11 | 2978.152753 | 0.381315642 | 0.087457831 | 4.359994268 | 1.30E-05 | 0.000395921 | | Coro2a | 1797.629434 | 0.157750200203652 | 0.0569859859536383 | 2.76822796980281 | 0.00563620157137483 | 0.141672083064745 | NM_001164804,NM_178893 |
| Mlip | 866.6974943 | 0.379839657 | 0.114788737 | 3.309032468 | 0.00093619 | 0.010792652 | | Lhfpl2 | 756.91459 | -0.178209575358282 | 0.0643640767241498 | -2.76877389420264 | 0.00562676690772323 | 0.141672083064745 | NM_172589 |
| C1qc | 1710.56872 | 0.378925295 | 0.104071683 | 3.641002863 | 0.000271578 | 0.004310233 | | Thbs3 | 663.4589144 | -0.251884612797945 | 0.090940504657933 | -2.76977364206844 | 0.00560952619791372 | 0.141672083064745 | NM_013691 |
| Pcsk4 | 157.2358002 | 0.376972912 | 0.122469657 | 3.078092317 | 0.002083304 | 0.019715626 | | Cycs | 5004.23945 | 0.157542553894138 | 0.0569642952621857 | 2.76563684618633 | 0.00568117626301187 | 0.142529821064785 | NM_007808 |
| Ddit4l | 1039.655694 | 0.376904493 | 0.122922457 | 3.066197184 | 0.002168003 | 0.020333557 | | Catsperg1 | 39.27333461 | -0.240884246848345 | 0.0871344827599963 | -2.76451112370562 | 0.00570081634035055 | 0.142775962344711 | NM_001164658 |
| Heph | 310.0129627 | 0.37662199 | 0.13927249 | 2.7042095 | 0.006846711 | 0.044661027 | | Mbd6 | 1973.830687 | -0.243737232731064 | 0.0882106549938955 | -2.76312674828071 | 0.00572505299112222 | 0.143136178569778 | NM_033072 |
| Fam21 | 4914.122686 | 0.376505101 | 0.086780986 | 4.338566756 | 1.43E-05 | 0.000428145 | | Tmem67 | 485.8695036 | -0.188069427024363 | 0.0680816135387934 | -2.76241142430034 | 0.005737612748404 | 0.143203716122537 | NM_177861 |
| Sytl2 | 3140.953628 | 0.3757349 | 0.104481081 | 3.596200363 | 0.000322899 | 0.004948565 | | Rragb | 2554.29293 | 0.151099464311125 | 0.0548391570496329 | 2.75532069492553 | 0.00586346328912763 | 0.146093769704748 | NM_001004154 |
| Pnn | 2791.000645 | 0.375389536 | 0.088341018 | 4.249323164 | 2.14E-05 | 0.000590086 | | Adamts16 | 174.0048673 | -0.256181527348524 | 0.0930284837568024 | -2.75379665456271 | 0.00589083562214246 | 0.14630718973507 | NM_172053 |
| Neu2 | 207.7879031 | 0.37515095 | 0.126427292 | 2.967325678 | 0.003004026 | 0.025392458 | | Ntng1 | 943.6368591 | 0.237215927288398 | 0.0861437345491486 | 2.75372235171737 | 0.00589217306863665 | 0.14630718973507 | NM_133488,NM_001163348,NM_001163349,NM_030699,NM_001163351,NM_001163350 |
| Csf1 | 1005.125789 | 0.375019592 | 0.068799224 | 5.450927629 | 5.01E-08 | 4.17E-06 | | Capn3 | 227.9967817 | -0.24606950632655 | 0.0893933544851307 | -2.75265994596366 | 0.00591132626091335 | 0.146477632799513 | NM_001109761,NM_007601,NM_001177799 |
| Cc2d2a | 1088.406195 | 0.374362934 | 0.082939001 | 4.513714064 | 6.37E-06 | 0.000222297 | | Lrfn3 | 1084.466002 | 0.204962919499279 | 0.0744717480812935 | 2.75222382688723 | 0.0059192049052261 | 0.146477632799513 | NM_175478 |
| Ccdc136 | 2760.632067 | 0.373653732 | 0.097643577 | 3.826710836 | 0.000129867 | 0.002392578 | | Lpl | 1062.026025 | -0.219505508906928 | 0.079772340179515 | -2.75164936133208 | 0.00592959726987366 | 0.146485255003716 | NM_008509 |
| Ucp2 | 845.7712423 | 0.373466475 | 0.126801358 | 2.945287671 | 0.003226545 | 0.026676676 | | Fcgr2b | 159.8117014 | -0.261007001802392 | 0.0949469952819133 | -2.74897589994733 | 0.00597817815433437 | 0.14743466191827 | NM_001077189,NM_010187 |
| Zbp1 | 16.0414848 | 0.373138245 | 0.131747341 | 2.832226011 | 0.004622516 | 0.034225481 | | Cpeb1 | 2996.80889 | -0.186040469611754 | 0.0677291459163909 | -2.7468302913699 | 0.00601742624127163 | 0.147900395229631 | NM_001252525,NM_001252526,NM_007755 |
| Stat6 | 677.9555643 | 0.373091623 | 0.070977141 | 5.256503981 | 1.47E-07 | 1.02E-05 | | Slco3a1 | 1643.647199 | 0.148395705200987 | 0.0540204263386046 | 2.747029508261 | 0.00601377235673895 | 0.147900395229631 | NM_001038643,NM_023908 |
| Slc14a1 | 460.2558083 | 0.372643575 | 0.12888889 | 2.891200127 | 0.003837736 | 0.030256816 | | Fuca2 | 1671.793346 | 0.162669307769407 | 0.0592526347094192 | 2.74535146946888 | 0.00604461225713882 | 0.148317631160808 | NM_025799 |
| Camk2n2 | 2940.361701 | 0.372580667 | 0.091083597 | 4.090535266 | 4.30E-05 | 0.00101496 | | Lgi2 | 1481.492502 | 0.189172055906101 | 0.0689225137064998 | 2.74470627568333 | 0.00605650784685628 | 0.148321973455592 | NM_144945 |
| Cast | 469.7537172 | 0.371928365 | 0.099962635 | 3.720673862 | 0.000198692 | 0.003386724 | | Setd4 | 246.8814842 | -0.243466157462494 | 0.0887191368224269 | -2.74423496646272 | 0.00606521081045171 | 0.148321973455592 | NM_145482 |
| Cpsf4 | 587.304177 | 0.37187798 | 0.096485577 | 3.854233888 | 0.000116093 | 0.002190912 | | Ldha | 10780.40961 | 0.153156656007092 | 0.0558437028743867 | 2.74259492339894 | 0.00609558290904747 | 0.148814180398023 | NM_010699,NM_001136069 |
| Hpcal1 | 1911.672466 | 0.371098077 | 0.106309882 | 3.490720438 | 0.00048172 | 0.006658136 | | Phldb2 | 303.21987 | -0.230633724663653 | 0.0841503170111308 | -2.74073506619287 | 0.00613019147102952 | 0.149407988772105 | NM_001252442,NM_153412 |
| Kifc5b | 71.98163534 | 0.371066222 | 0.138547338 | 2.678263089 | 0.007400506 | 0.047350844 | | Ncapd3 | 1276.259089 | -0.153843760027142 | 0.056196321475284 | -2.73761264062105 | 0.00618869224562727 | 0.150581144991594 | NM_178113 |
| Man2c1 | 2203.38762 | 0.370688366 | 0.107201792 | 3.457856074 | 0.000544492 | 0.007285287 | | Mrpl19 | 1275.987491 | 0.156457972082134 | 0.0571706943721589 | 2.73668133298598 | 0.00620623799511364 | 0.150747361160254 | NM_026490 |
| Unc5d | 2101.251659 | 0.369527162 | 0.096597572 | 3.825429094 | 0.000130545 | 0.002399355 | | Styx | 385.9951812 | -0.2299131718872 | 0.0840280027047355 | -2.73614943217308 | 0.0062162790399967 | 0.150747361160254 | NM_019637 |
| 9030624G23Rik | 163.964947 | 0.368922236 | 0.111909717 | 3.296605931 | 0.000978607 | 0.01121448 | | Slc2a1 | 5177.346636 | 0.241506655983886 | 0.0883576738693086 | 2.73328444953296 | 0.00627061520675141 | 0.151811594155452 | NM_011400 |
| Pcgf2 | 970.1732607 | 0.368507306 | 0.101339147 | 3.63637664 | 0.0002765 | 0.00434933 | | Khdrbs3 | 5337.640607 | 0.101681230153625 | 0.0372171635925863 | 2.73210584414015 | 0.00629309203194313 | 0.152102254336116 | NM_010158 |
| Tbc1d2 | 156.2386477 | 0.368254297 | 0.134210284 | 2.743860501 | 0.006072134 | 0.041101633 | | Ndn | 7794.906231 | 0.212630116857962 | 0.077959314770763 | 2.72744979202543 | 0.00638259700506754 | 0.154009309128922 | NM_010882 |
| Ccdc39 | 605.5723044 | 0.368186139 | 0.129721476 | 2.83828206 | 0.004535708 | 0.033802916 | | F2r | 245.4550261 | 0.238379197546595 | 0.0874424471812655 | 2.72612678659876 | 0.00640823776307845 | 0.154049192620522 | NM_010169 |
| Anks6 | 444.6349305 | 0.367987208 | 0.09068614 | 4.057810898 | 4.95E-05 | 0.001138714 | | Micu1 | 3702.047633 | 0.119606053693764 | 0.043881282873193 | 2.72567358705986 | 0.00641704237177439 | 0.154049192620522 | NM_144822 |
| 4833420G17Rik | 1280.317341 | 0.366745782 | 0.087974402 | 4.168778337 | 3.06E-05 | 0.000782975 | | Nkrf | 1494.21553 | 0.20168898554872 | 0.0740094544275907 | 2.72517865600601 | 0.00642667015889 | 0.154049192620522 | NM_029891 |
| Kcnh4 | 1011.259504 | 0.366222177 | 0.10525568 | 3.479357868 | 0.000502617 | 0.006855234 | | Spint1 | 162.9265907 | -0.257305744277092 | 0.0943650272800019 | -2.72670661678091 | 0.00639698886704907 | 0.154049192620522 | NM_016907 |
| Dvl2 | 691.8969593 | 0.365570042 | 0.10401941 | 3.514440655 | 0.000440681 | 0.006229848 | | Ppfibp1 | 1394.768536 | -0.144279683068996 | 0.0529927801533132 | -2.72262905723347 | 0.00647647317555237 | 0.154987231215937 | NM_001170433,NM_026221 |
| Apol8 | 222.9614253 | 0.365031068 | 0.098706634 | 3.698141186 | 0.000217184 | 0.003602837 | | Epb4.1l2 | 4107.371655 | -0.130797889316294 | 0.0480769836048633 | -2.72059267260401 | 0.00651650033374341 | 0.155688624749929 | NM_013511,NM_001199265 |
| Slc38a7 | 1215.778497 | 0.364952929 | 0.066647459 | 5.475871585 | 4.35E-08 | 3.69E-06 | | Parvg | 313.3323539 | -0.249760823627063 | 0.0918409504862951 | -2.71949301814263 | 0.00653820752745024 | 0.155695086137282 | NM_001162500,NM_022321 |
| Rhbdf1 | 444.8602606 | 0.364897929 | 0.101014648 | 3.61232687 | 0.000303462 | 0.004711258 | | Tstd2 | 723.0854299 | -0.168853980052879 | 0.06208167594442 | -2.7198682619981 | 0.00653079290995656 | 0.155695086137282 | NM_173033 |
| Ptpn2 | 1044.223472 | 0.364383735 | 0.107556114 | 3.387847719 | 0.000704434 | 0.008745589 | | Ccr5 | 217.0915648 | -0.224377169589513 | 0.0825330720369988 | -2.71863344053069 | 0.00655522086403208 | 0.15584474348761 | NM_009917 |
| Dusp15 | 622.4026598 | 0.363932292 | 0.121204463 | 3.002631117 | 0.002676566 | 0.023446978 | | Clvs2 | 1132.515432 | 0.173924962502864 | 0.0640105876397119 | 2.71712803953328 | 0.00658511271729295 | 0.156299587142806 | NM_175448 |
| Adamts17 | 194.9772686 | 0.363473049 | 0.132883709 | 2.735271702 | 0.006232881 | 0.041951456 | | C2cd5 | 2440.098642 | -0.14374053116368 | 0.0529403057176729 | -2.71514357945416 | 0.0066247042383919 | 0.156472282547774 | NM_029081,NM_001109688,NM_029897 |
| 9330151L19Rik | 740.6043178 | 0.363002254 | 0.081927183 | 4.430791339 | 9.39E-06 | 0.000297485 | | Exd2 | 3909.343377 | -0.169342144062776 | 0.0623583545421237 | -2.71562880878109 | 0.00661500381787822 | 0.156472282547774 | NM_133798 |
| Gnl3 | 885.9117574 | 0.362312976 | 0.08953426 | 4.04664066 | 5.20E-05 | 0.001183876 | | Herpud1 | 2196.25784 | 0.145669629750614 | 0.0536416207733049 | 2.71560828421328 | 0.00661541387418338 | 0.156472282547774 | NM_022331 |
| Kcnj9 | 5660.677355 | 0.362271942 | 0.096245895 | 3.764024881 | 0.0001672 | 0.002947213 | | Aplp1 | 54686.73258 | 0.124802475337014 | 0.0459979682398231 | 2.71321712920713 | 0.00666334301981252 | 0.156678743676988 | NM_007467 |
| Ccdc22 | 1247.335818 | 0.362124209 | 0.071704768 | 5.050210993 | 4.41E-07 | 2.52E-05 | | Appl2 | 3560.226216 | -0.162180965683588 | 0.0597764167276138 | -2.71312421304903 | 0.00666521174908233 | 0.156678743676988 | NM_145220 |
| Krba1 | 1777.096354 | 0.361598954 | 0.085273431 | 4.240464453 | 2.23E-05 | 0.000609534 | | Ccl9 | 83.63069379 | -0.250637718200275 | 0.0923807451422968 | -2.71309478846712 | 0.00666580363433694 | 0.156678743676988 | NM_011338 |
| Rgs16 | 397.1625174 | 0.360849725 | 0.086803831 | 4.157071408 | 3.22E-05 | 0.000817437 | | 2010015L04Rik | 776.2323578 | -0.164275102019275 | 0.06057739090263 | -2.71182201100944 | 0.00669145124863266 | 0.156789376562465 | NM_001166029,NM_177674 |
| Neurod2 | 2371.290604 | 0.359629948 | 0.101926648 | 3.528321143 | 0.000418204 | 0.006005245 | | Gpr1 | 35.22336774 | 0.244165747015605 | 0.0900385903290677 | 2.71178997942152 | 0.0066920978568586 | 0.156789376562465 | NM_146250 |
| Letm2 | 393.1244131 | 0.359541351 | 0.094616177 | 3.799998716 | 0.000144697 | 0.002634469 | | Sned1 | 271.1566463 | -0.246544824790295 | 0.090979715179922 | -2.70988785030516 | 0.0067305961889707 | 0.157437423898532 | NM_172463 |
| Tmem181b-ps | 8877.054085 | 0.359107604 | 0.114501339 | 3.136274265 | 0.001711091 | 0.017022514 | | Gdap10 | 76.15709017 | -0.257356167060235 | 0.0950132195041847 | -2.7086353709854 | 0.00675605444636529 | 0.157778853517528 | NR_045032 |
| Car7 | 613.3628753 | 0.359092223 | 0.119930463 | 2.994170246 | 0.002751923 | 0.023913798 | | S100a6 | 453.5323448 | -0.213122551515852 | 0.0787528791172629 | -2.70621917452076 | 0.00680541140898465 | 0.15867641432891 | NM_011313 |
| Rrnad1 | 824.4262619 | 0.358473555 | 0.10800452 | 3.31906066 | 0.000903208 | 0.010507975 | | Selk | 2711.064631 | 0.132076555767468 | 0.0488488842806818 | 2.70377826868197 | 0.00685560190409936 | 0.159590502017544 | NM_019979 |
| Sirt4 | 484.8012323 | 0.358470251 | 0.126398203 | 2.836039147 | 0.004567685 | 0.033967062 | | Cct7 | 7176.798944 | 0.111114478634707 | 0.0411134974907722 | 2.70262773580978 | 0.00687937456552209 | 0.159632260285581 | NM_007638 |
| D930048N14Rik | 212.2627533 | 0.358134651 | 0.122139116 | 2.932186368 | 0.003365847 | 0.027505503 | | Rtp4 | 98.62329338 | -0.246128212842133 | 0.0910674157992305 | -2.70270338388379 | 0.0068778092301983 | 0.159632260285581 | NM_023386 |
| Rbm5 | 4432.782038 | 0.357669824 | 0.066206006 | 5.402377324 | 6.58E-08 | 5.28E-06 | | E230029C05Rik | 28.90041993 | -0.224838886313471 | 0.0832296525400813 | -2.70142766972618 | 0.00690424958524968 | 0.159953954506119 | NR_015614 |
| Fcer1g | 321.350714 | 0.357325884 | 0.102884591 | 3.473074842 | 0.000514532 | 0.006962582 | | Akna | 261.7962092 | -0.238710088049627 | 0.0885901926019481 | -2.69454305311419 | 0.00704852199803134 | 0.160386104562512 | NM_001045514 |
| 9430008C03Rik | 289.9068602 | 0.357255213 | 0.127117382 | 2.810435586 | 0.004947449 | 0.035738209 | | Dapp1 | 235.0321283 | -0.230705497839243 | 0.0855473375040325 | -2.69681681008912 | 0.00700057703987339 | 0.160386104562512 | NM_011932 |
| Lgi3 | 4565.731009 | 0.357144893 | 0.084502183 | 4.226457606 | 2.37E-05 | 0.000639697 | | Dennd1c | 68.41112044 | -0.256670651941208 | 0.0950770066825608 | -2.69960804296426 | 0.00694212104745929 | 0.160386104562512 | NM_153551 |
| Ctns | 694.6977618 | 0.357028161 | 0.083472835 | 4.277177855 | 1.89E-05 | 0.000533212 | | Krba1 | 2082.149685 | -0.186643224550221 | 0.069263315504307 | -2.69469087844365 | 0.00704539598085279 | 0.160386104562512 | NM_133922 |
| Parvg | 262.5668192 | 0.356994034 | 0.133330038 | 2.677521438 | 0.007416911 | 0.047350844 | | Laptm5 | 2213.804705 | -0.216616818872849 | 0.0803955784216156 | -2.69438721787476 | 0.00705181874680296 | 0.160386104562512 | NM_010686 |
| Arhgef1 | 1649.310176 | 0.356879914 | 0.115621561 | 3.086620783 | 0.002024457 | 0.019314614 | | Mdc1 | 1113.266169 | -0.160616846357408 | 0.0596436251497111 | -2.69369686960059 | 0.00706643996420267 | 0.160386104562512 | NM_001010833 |
| Sec14l5 | 377.5058558 | 0.356797305 | 0.120089584 | 2.971092856 | 0.00296742 | 0.025179196 | | Ndor1 | 1598.914242 | -0.204024599733246 | 0.075729225392485 | -2.69412236181135 | 0.00705742504559991 | 0.160386104562512 | NM_001252542,NM_001252541,NM_001082476 |
| Cyp4f13 | 287.6000724 | 0.356327703 | 0.127470818 | 2.795366877 | 0.005184085 | 0.03674392 | | Nlrp6 | 62.92323332 | 0.251710320199542 | 0.0933009043334253 | 2.69783365978979 | 0.00697923045421892 | 0.160386104562512 | NM_001081389 |
| Gm166 | 183.7862171 | 0.356006853 | 0.110238709 | 3.229417846 | 0.001240425 | 0.013431707 | | Rasgrp3 | 857.4523741 | -0.211589253264522 | 0.0785041079684017 | -2.69526345487154 | 0.00703329962795801 | 0.160386104562512 | NM_001166493,NM_207246 |
| H2-Q4 | 388.8534656 | 0.35592999 | 0.115381615 | 3.084806793 | 0.002036844 | 0.019331788 | | Rgl2 | 2142.26731 | -0.229498649879004 | 0.0850843807615208 | -2.6973064600688 | 0.00699029057658526 | 0.160386104562512 | NM_009059 |
| Cep44 | 266.9074885 | 0.355404659 | 0.104782226 | 3.391841079 | 0.000694247 | 0.008660759 | | Rps6kb2 | 1092.403585 | -0.23733434474357 | 0.0880374316643549 | -2.69583449058821 | 0.00702125440243711 | 0.160386104562512 | NM_021485 |
| Ddb2 | 426.7535829 | 0.355076098 | 0.100790093 | 3.522926572 | 0.00042681 | 0.006083705 | | Ufsp2 | 1673.782011 | 0.163146439572099 | 0.0605433806811498 | 2.69470316550881 | 0.00704513620607872 | 0.160386104562512 | NM_138668 |
| Rufy2 | 2253.489513 | 0.354842098 | 0.080081661 | 4.431003201 | 9.38E-06 | 0.000297485 | | Yeats2 | 1529.285586 | -0.161181140887953 | 0.0598295924639115 | -2.69400365688895 | 0.00705993901181634 | 0.160386104562512 | NM_001033237,NM_001145931,NM_001145930 |
| Itga7 | 537.0391671 | 0.3545551 | 0.117092114 | 3.028001519 | 0.002461768 | 0.022124502 | | B930059L03Rik | 34.65876254 | -0.246100397666249 | 0.0914131683818555 | -2.69217665269216 | 0.00709873337974223 | 0.160867708383987 | NR_033340 |
| Napepld | 709.4210356 | 0.353240419 | 0.090384861 | 3.908181232 | 9.30E-05 | 0.001854365 | | Flywch1 | 9607.769639 | 0.108799708686686 | 0.0404734041260343 | 2.68817785496579 | 0.00718431195826183 | 0.161546928027417 | NM_153791 |
| Sh2d3c | 806.80103 | 0.353164631 | 0.081838263 | 4.315397413 | 1.59E-05 | 0.000465744 | | Gcfc2 | 279.010927 | -0.212427034281832 | 0.0789821863614774 | -2.68955626664497 | 0.00715470843855136 | 0.161546928027417 | NM_177884 |
| Csf3r | 333.1659592 | 0.352781434 | 0.123881412 | 2.847735011 | 0.004403157 | 0.033182157 | | Kcnf1 | 6682.41231 | 0.190931157652598 | 0.0709979693830008 | 2.68924814768453 | 0.00716131624998447 | 0.161546928027417 | NM_201531 |
| Fah | 808.7980251 | 0.352217265 | 0.096741286 | 3.64081646 | 0.000271775 | 0.004310233 | | Msn | 1551.033671 | -0.175934950466129 | 0.0654363087848129 | -2.68864417528027 | 0.00717428472639228 | 0.161546928027417 | NM_010833 |
| Epb4.1l4b | 414.2891405 | 0.352062702 | 0.116893856 | 3.011815283 | 0.002596906 | 0.022992231 | | Mysm1 | 1233.874185 | -0.163924490550425 | 0.0609660734468831 | -2.68878215837936 | 0.00717132010120282 | 0.161546928027417 | NM_177239 |
| Clcn2 | 2488.70766 | 0.35193773 | 0.079367625 | 4.434273153 | 9.24E-06 | 0.000295885 | | 4933427D14Rik | 1608.487505 | -0.17222421642084 7 | 0.064154959543081 | -2.68450354652933 | 0.00726376126580921 | 0.162353882907151 | NM_028963 |
| Fam161a | 90.70667909 | 0.351657004 | 0.130943181 | 2.685569425 | 0.007240633 | 0.046631134 | | Cchcr1 | 269.7157 | -0.226643634569063 | 0.0844015064678308 | -2.68530319012075 | 0.00724640381454586 | 0.162353882907151 | NM_146248 |
| Prpf4b | 3170.185032 | 0.351609355 | 0.076831034 | 4.576397556 | 4.73E-06 | 0.000176613 | | L3mbtl1 | 424.6778311 | -0.249536840102459 | 0.0929699638243997 | -2.68405869850375 | 0.00727343349524298 | 0.162353882907151 | NM_001081338 |
| Tardbp | 6429.134021 | 0.350995277 | 0.086632344 | 4.051550036 | 5.09E-05 | 0.001162723 | | Mccc1 | 968.5752052 | -0.192054160467708 | 0.0715568824036612 | -2.68393694661412 | 0.00727608273251793 | 0.162353882907151 | NM_023644 |
| Stat2 | 1223.068967 | 0.350726365 | 0.100142316 | 3.502279359 | 0.000461296 | 0.006438851 | | Uqcc | 2368.207371 | 0.122847943615577 | 0.0457689871724692 | 2.68408700311971 | 0.00727281773030783 | 0.162353882907151 | NM_018888,NR_024487 |
| Rtel1 | 746.5488996 | 0.350557246 | 0.130086218 | 2.694806949 | 0.007042942 | 0.045660528 | | Zmym3 | 6877.608952 | -0.162001466788397 | 0.0603872471680543 | -2.68270991617743 | 0.00730283044534142 | 0.162700789952499 | NM_001177987,NM_019831,NM_001177985,NM_001177988,NM_001177986 |
| Inpp5e | 1351.246761 | 0.350212131 | 0.065619225 | 5.337035433 | 9.45E-08 | 7.18E-06 | | Plekha2 | 1354.776382 | -0.169606980123603 | 0.0632597334066227 | -2.68112069068326 | 0.00733760467264241 | 0.163225184494339 | NM_031257 |
| Safb2 | 2111.924117 | 0.350015125 | 0.097213311 | 3.600485575 | 0.000317623 | 0.004887048 | | Adcy8 | 1126.185955 | 0.180920648730264 | 0.0675008230260734 | 2.68027322660021 | 0.00735620890682269 | 0.163290663754399 | NM_009623 |
| Ogt | 11070.76687 | 0.34930878 | 0.087057659 | 4.012384256 | 6.01E-05 | 0.00132634 | | Pepd | 1004.394568 | 0.155998234721144 | 0.058209101083068 | 2.67996295800076 | 0.00736303075582619 | 0.163290663754399 | NM_008820 |
| Map4k2 | 2475.690992 | 0.349150339 | 0.090162815 | 3.872442752 | 0.00010775 | 0.002067036 | | Rfxank | 710.6413315 | -0.166764078674895 | 0.0622786876096799 | -2.6777070146381 | 0.00741280287753129 | 0.16389402526487 | NM_001025589,NM_011266 |
| Usp54 | 3702.560475 | 0.348788941 | 0.129108445 | 2.701519193 | 0.00690235 | 0.044916004 | | Tollip | 7439.439908 | 0.107136865101796 | 0.0400083816378154 | 2.67786050612284 | 0.00740940690285606 | 0.16389402526487 | NM_023764 |
| Amt | 679.7474495 | 0.347780219 | 0.10655192 | 3.263950762 | 0.001098703 | 0.012273466 | | H2-Oa | 25.35050634 | -0.194900367797611 | 0.0728753393255026 | -2.67443513267328 | 0.00748552559533799 | 0.165250372033252 | NM_008206 |
| Pxn | 1177.552426 | 0.347745972 | 0.102770294 | 3.383720695 | 0.000715107 | 0.008835621 | | Fam167a | 109.3146561 | 0.25195902613943 | 0.09427877651005 | 2.67248934984399 | 0.0075290764399598 | 0.165959581740297 | NM_177628 |
| Srrm1 | 4548.41591 | 0.347485318 | 0.091854072 | 3.78301485 | 0.00015494 | 0.00277531 | | Zfp82 | 208.193946 | -0.229887231076008 | 0.0860844761182024 | -2.67048417371269 | 0.00757419420616954 | 0.166701128846695 | NM_177889,NM_001252519,NM_001253385 |
| Ppp1r13l | 149.5011414 | 0.34724911 | 0.120248328 | 2.887766635 | 0.003879876 | 0.030484179 | | Hsd17b12 | 2902.952278 | 0.151276588257169 | 0.0566673592280196 | 2.66955422518382 | 0.00759520080975651 | 0.16691057029126 | NM_019657 |
| Havcr2 | 188.1072072 | 0.347015228 | 0.116411771 | 2.980929039 | 0.002873754 | 0.024599967 | | Ltk | 1366.09663 | -0.23768148146322 | 0.0890558273206901 | -2.66890431108263 | 0.00760991270717094 | 0.166981256773965 | NM_206942,NM_206941,NM_203345,NM_008523 |
| Bhlhe41 | 1655.531812 | 0.345901421 | 0.078808405 | 4.389143776 | 1.14E-05 | 0.000353593 | | Ndufs8 | 2151.439226 | 0.126497552018356 | 0.0474159755628238 | 2.66782556969124 | 0.00763438822825748 | 0.167265646159379 | NM_144870,NM_001271443,NM_001271444 |
| Mtmr11 | 357.41333 | 0.345861613 | 0.099931997 | 3.460969704 | 0.000538233 | 0.007207768 | | Cd68 | 508.8635483 | -0.248952969838486 | 0.0933372162483421 | -2.66724228389346 | 0.00764765174805331 | 0.167303899536479 | NM_009853 |
| Atp11b | 3451.526136 | 0.345730384 | 0.083639093 | 4.133597948 | 3.57E-05 | 0.000875529 | | Lama4 | 382.7683066 | -0.249453808145412 | 0.0935526213395466 | -2.6664544998694 | 0.00766559820308059 | 0.167444330072103 | NM_010681 |
| Nsun5 | 372.968735 | 0.34513212 | 0.093708652 | 3.683033685 | 0.000230475 | 0.003762882 | | Tgfbr1 | 1246.408669 | -0.190365135514971 | 0.0714228186679315 | -2.66532655900969 | 0.00769135946312872 | 0.167597786315907 | NM_009370 |
| Ypel4 | 1336.54292 | 0.344873913 | 0.07785224 | 4.429852149 | 9.43E-06 | 0.000298171 | | Wdr90 | 1070.92114 | -0.242981421176826 | 0.0911703343198253 | -2.6651368889846 | 0.0076956989861428 | 0.167597786315907 | NM_001163766 |
| Fancm | 553.0535076 | 0.344632378 | 0.104006582 | 3.313563148 | 0.000921153 | 0.010651004 | | H2-DMa | 518.8129695 | -0.237110889232144 | 0.0891267271555747 | -2.66037917916873 | 0.00780527252567569 | 0.169708597463407 | NM_010386 |
| Wdr67 | 291.2082765 | 0.344571742 | 0.092531309 | 3.723839482 | 0.000196216 | 0.003355592 | | Traf7 | 2170.505058 | -0.183836022538032 | 0.0691134406456929 | -2.65991709891075 | 0.00781598868945471 | 0.169708597463407 | NM_153792,NM_001172114,NM_001172113 |
| Wdr60 | 797.2392328 | 0.34450001 | 0.086051712 | 4.003406834 | 6.24E-05 | 0.001364451 | | Col26a1 | 272.5388456 | -0.242653812600744 | 0.0912698882351882 | -2.65864040476815 | 0.00784566522351762 | 0.169996348783533 | NM_024474 |
| 2010015L04Rik | 691.6729612 | 0.344360354 | 0.071396396 | 4.82321761 | 1.41E-06 | 6.55E-05 | | Icam1 | 131.7586345 | -0.245009191102163 | 0.0922133834054743 | -2.65698082050439 | 0.00788439284672517 | 0.169996348783533 | NM_010493 |
| Fam214a | 1950.276933 | 0.34423913 | 0.117253706 | 2.935848605 | 0.003326366 | 0.027283599 | | Lman2 | 3685.155113 | 0.128590486722078 | 0.0483913053813176 | 2.65730559878062 | 0.00787680045711714 | 0.169996348783533 | NM_025828 |
| Cln3 | 541.7537533 | 0.344016268 | 0.099687509 | 3.450946587 | 0.000558624 | 0.007404262 | | Pxdn | 3387.253934 | -0.221808486567655 | 0.0834859201691363 | -2.65683705849187 | 0.00788775568498563 | 0.169996348783533 | NM_181395 |
| Ppox | 781.6305115 | 0.343877204 | 0.126647787 | 2.71522474 | 0.006623081 | 0.043630108 | | Surf4 | 3165.405312 | 0.115184931001777 | 0.0433436214948555 | 2.65748285512894 | 0.0078726594724703 | 0.169996348783533 | NM_011512 |
| A930017M01Rik | 477.1660834 | 0.343538779 | 0.093780295 | 3.663229868 | 0.000249055 | 0.004028033 | | B2m | 3685.558993 | -0.206919632472882 | 0.0779522304180923 | -2.6544414619451 | 0.00794398215896907 | 0.170584808343056 | NM_009735 |
| Fbxo27 | 483.2581327 | 0.342401876 | 0.094370067 | 3.628288988 | 0.000285306 | 0.00446518 | | Lpar6 | 139.6021702 | -0.236635116337792 | 0.0891558532442912 | -2.65417364903009 | 0.00795029018644149 | 0.170584808343056 | NM_175116 |
| Tuft1 | 90.24635769 | 0.342363264 | 0.121823713 | 2.8103171 | 0.004949271 | 0.035738209 | | Mcmdc2 | 96.01463404 | -0.252422979143915 | 0.0950720036191914 | -2.65507162502843 | 0.00792915706559708 | 0.170584808343056 | NM_177722 |
| Mettl17 | 661.8600698 | 0.342293597 | 0.114670978 | 2.98500635 | 0.002835724 | 0.024368671 | | Wtip | 183.7298073 | -0.232780864042622 | 0.0878372410073814 | -2.6501386128813 | 0.00804587531115663 | 0.172381098480621 | NM_207212 |
| Nt5dc3 | 6242.31945 | 0.342147904 | 0.10238873 | 3.341655918 | 0.000832802 | 0.009880121 | | C1galt1c1 | 683.1428844 | 0.175147176020373 | 0.0661121527900934 | 2.64924327266223 | 0.00806722381333074 | 0.172465044500895 | NM_021550 |
| Jun | 1733.054572 | 0.341968254 | 0.085701082 | 3.990244276 | 6.60E-05 | 0.001424354 | | Ddit4 | 1569.912939 | 0.226492346878581 | 0.0855017584199036 | 2.64897881709363 | 0.00807353918908222 | 0.172465044500895 | NM_029083 |
| Dmpk | 531.2922133 | 0.341531986 | 0.111442362 | 3.06465136 | 0.002179239 | 0.020414223 | | Impg1 | 69.78512205 | 0.235225257224766 | 0.0888530124803862 | 2.6473526407075 | 0.00811247074814978 | 0.173042217456129 | NM_022016 |
| Ltk | 1168.342884 | 0.34128215 | 0.12664514 | 2.694790725 | 0.007043285 | 0.045660528 | | Tbc1d10c | 54.36077467 | -0.246124184843099 | 0.0929982999192991 | -2.6465449912168 | 0.00813186870918369 | 0.173201649368918 | NM_178650 |
| 4732491K20Rik | 148.1657636 | 0.341199546 | 0.118832041 | 2.87127566 | 0.004088188 | 0.031506284 | | Sdpr | 447.1505565 | 0.190753076514841 | 0.0721282416869109 | 2.64463782914393 | 0.00817783940711923 | 0.173925761680548 | NM_138741 |
| Zcchc7 | 763.5175243 | 0.341078071 | 0.111068382 | 3.070883581 | 0.002134263 | 0.020096898 | | Adamts10 | 965.1935216 | -0.231492442591364 | 0.0876136764541716 | -2.6421952822908 | 0.00823705477841524 | 0.174077875368871 | NR_037707,NM_172619,NR_037708 |
| Mlh1 | 431.2374349 | 0.340751541 | 0.105953683 | 3.216042439 | 0.001299716 | 0.013849883 | | Adra2a | 745.5339729 | 0.219722122712631 | 0.0831456435284489 | 2.64261738063824 | 0.00822679438258255 | 0.174077875368871 | NM_007417 |
| Gtdc1 | 2151.831098 | 0.340440626 | 0.069206793 | 4.919179345 | 8.69E-07 | 4.33E-05 | | Fam178a | 2663.883213 | -0.183424259886399 | 0.0694424813095776 | -2.64138401202407 | 0.00825680735200509 | 0.174077875368871 | NM_001081225 |
| Gba | 1378.29634 | 0.339685828 | 0.077423396 | 4.387379621 | 1.15E-05 | 0.000354799 | | Fyb | 175.049246 | -0.234258898456402 | 0.0886667167220825 | -2.64201616025396 | 0.00824141234891732 | 0.174077875368871 | NM_011815 |
| Usp35 | 936.0955222 | 0.33959468 | 0.073258454 | 4.635569819 | 3.56E-06 | 0.000139979 | | H2-Q2 | 27.40243684 | 0.167727180900337 | 0.0638784098494277 | 2.64138041786036 | 0.00825689495588271 | 0.174077875368871 | NM_010392 |
| Gigyf1 | 2770.800586 | 0.339511098 | 0.095739763 | 3.546186948 | 0.000390849 | 0.005732886 | | R3hdm1 | 13183.71669 | -0.143698738705889 | 0.0543978322827629 | -2.64162619493614 | 0.00825090631952793 | 0.174077875368871 | NM_181750 |
| Crtc2 | 995.1795021 | 0.339118138 | 0.091343487 | 3.712559597 | 0.000205174 | 0.003466698 | | Adam17 | 729.775038 | -0.16604311623491 | 0.0629053168547562 | -2.63957205109214 | 0.00830107757449138 | 0.174157793009078 | NM_009615 |
| Nphp4 | 709.628538 | 0.338924177 | 0.101211468 | 3.348673648 | 0.000811994 | 0.009692629 | | Cttnbp2 | 4874.898029 | -0.150163548727104 | 0.056905982253593 | -2.63880075135023 | 0.00831998648815606 | 0.174157793009078 | NM_080285 |
| Prrt2 | 11461.04187 | 0.338893826 | 0.090389591 | 3.749257228 | 0.000177359 | 0.003101582 | | D930015E06Rik | 695.2503505 | -0.161150941266734 | 0.0611155153073034 | -2.63682537006239 | 0.00836859008125682 | 0.174157793009078 | NM_172681 |
| Pcdhga1 | 235.6751227 | 0.338581142 | 0.119396448 | 2.835772323 | 0.004571502 | 0.033971634 | | Gimap6 | 135.0918472 | 0.249146782964004 | 0.0944200615054026 | 2.63870600158154 | 0.00832231199581084 | 0.174157793009078 | NM_153175 |
| Nvl | 1678.933046 | 0.338191342 | 0.118745577 | 2.848033158 | 0.004399034 | 0.033182157 | | Gtdc2 | 1299.480329 | 0.155063463309021 | 0.0587743398529127 | 2.63828506959123 | 0.00833265024595577 | 0.174157793009078 | NM_153540 |
| Mapk1ip1 | 1139.927569 | 0.33815085 | 0.080959079 | 4.176811942 | 2.96E-05 | 0.0007596 | | Hcls1 | 241.0245947 | -0.234810946623198 | 0.0890463188163878 | -2.63695287738256 | 0.00836544515544054 | 0.174157793009078 | NM_008225 |
| Vopp1 | 6048.025157 | 0.338063605 | 0.115613605 | 2.924081511 | 0.003454743 | 0.027983052 | | Htr2a | 806.714062 | 0.223827883947037 | 0.0847969482280352 | 2.63957475621789 | 0.00830101132429382 | 0.174157793009078 | NM_172812 |
| Tprn | 705.7493628 | 0.337821789 | 0.0982861 | 3.4371268 | 0.00058792 | 0.007693612 | | Snord2 | 32.56535568 | -0.216252389102858 | 0.0819102364770504 | -2.64011433007458 | 0.00828780628148631 | 0.174157793009078 | NR_030705 |
| Usp40 | 771.5502997 | 0.337582827 | 0.104012551 | 3.245597021 | 0.001172046 | 0.012845805 | | Tmc8 | 27.27931103 | -0.216489736942475 | 0.0821009413815551 | -2.63687277270504 | 0.0083674207872957 | 0.174157793009078 | NM_001195088,NM_001195089,NM_001195090,NM_181856 |
| Rbm25 | 3887.998302 | 0.337021307 | 0.073849074 | 4.563649715 | 5.03E-06 | 0.000185456 | | Itgb3 | 238.0905467 | -0.217909036018563 | 0.0826675006760646 | -2.63596980961656 | 0.00838971956606611 | 0.174347734501683 | NM_016780 |
| Rbm12b1 | 233.9514309 | 0.336852811 | 0.110031481 | 3.061422126 | 0.002202883 | 0.020597302 | | Vit | 111.1560213 | 0.250150839660411 | 0.0949192651786724 | 2.6354064076406 | 0.00840365975816526 | 0.174387945210155 | NM_028813,NM_001197028 |
| A230050P20Rik | 1991.438953 | 0.336448249 | 0.081271092 | 4.139826856 | 3.48E-05 | 0.000857519 | | Phb | 2861.806751 | 0.133593254184294 | 0.0507053738712183 | 2.63469616699 | 0.00842126268148572 | 0.174503939673697 | NM_008831 |
| Farsa | 1127.283223 | 0.336445413 | 0.083847012 | 4.012610665 | 6.01E-05 | 0.00132634 | | Asap3 | 246.0337934 | -0.221788456941038 | 0.0842715366685236 | -2.63183116991717 | 0.00849260534469894 | 0.175232365393603 | NM_001008232 |

| | | | | | | | | | | | | | | | |
|---|---|---|---|---|---|---|---|---|---|---|---|---|---|---|---|
| Pygm | 1424.660383 | 0.336149771 | 0.083783852 | 4.012106884 | 6.02E-05 | 0.00132634 | | Elavl4 | 3090.182258 | 0.164056882654989 | 0.0623333181600327 | 2.63192923941244 | 0.00849015436381142 | 0.175232365393603 | NM_001038698,NM_001163399,NM_001163397,NM_010488 |
| Cc2d1b | 1044.503122 | 0.33605806 | 0.111968883 | 3.001352248 | 0.002687834 | 0.023503194 | | Srsf5 | 8446.686554 | -0.172021579831034 | 0.0653386773285614 | -2.63276801527535 | 0.00846921726399612 | 0.175232365393603 | NM_001079694,NM_009159,NM_001079695 |
| Cmbl | 292.6751902 | 0.335857891 | 0.103799823 | 3.235630669 | 0.001213743 | 0.01322485 | | Podn | 269.4063983 | -0.243540337822962 | 0.0925637651542154 | -2.63105479144256 | 0.00851203115951531 | 0.175384063295205 | NM_172874 |
| Clasrp | 1568.212077 | 0.33559437 | 0.118531473 | 2.831268037 | 0.004636385 | 0.034305706 | | Elovl4 | 2000.42257 | 0.168718016621104 | 0.0641689484551568 | 2.62927818957496 | 0.00855663318610732 | 0.17605333379801 | NM_148941 |
| Ccnl1 | 1004.294713 | 0.335005052 | 0.118017966 | 2.838593666 | 0.004531282 | 0.033786179 | | Mpzl1 | 674.6399346 | -0.185393591062134 | 0.0705619517958482 | -2.62738751329498 | 0.00860432847380069 | 0.1767842650784 | NM_001001880,NM_001083897 |
| Mvp | 298.5631138 | 0.334685876 | 0.089969009 | 3.720012927 | 0.000199213 | 0.003388145 | | Dcbld2 | 1855.127746 | 0.132387358323812 | 0.0504581024442594 | 2.62370861984077 | 0.00869781599273803 | 0.178452648461176 | NM_028523 |
| Pask | 159.3746101 | 0.333820147 | 0.111594357 | 2.991371205 | 0.002777277 | 0.024026312 | | Lig3 | 1374.209589 | -0.176025726822437 | 0.0671292561221341 | -2.62219093419087 | 0.00873664696956992 | 0.178996521692486 | NM_010716 |
| Masp1 | 346.0463744 | 0.333689261 | 0.085839996 | 3.887340144 | 0.000101349 | 0.001982698 | | Arhgap4 | 202.3418634 | -0.240171574710143 | 0.0916205229533253 | -2.62137312068412 | 0.00875763542688803 | 0.179173820015458 | NM_138630,NM_001162423,NM_001162424 |
| Pabpn1 | 5170.477244 | 0.333579953 | 0.097777345 | 3.411628259 | 0.000645761 | 0.00819445 | | Chrna4 | 2120.981049 | 0.18008901898813 | 0.0687243851644265 | 2.62045296669091 | 0.00878130421689825 | 0.179191559025074 | NM_015730 |
| Rbm33 | 3309.389709 | 0.333386399 | 0.105308453 | 3.165808527 | 0.001546526 | 0.015860224 | | Morf4l2 | 7399.576812 | 0.13226029302779 | 0.0504829209108582 | 2.6199017537304 | 0.00879551022889149 | 0.179191559025074 | NM_019768,NM_001168230,NM_001168226,NM_001168227,NM_001168225,NM_001168228,NM_001168229 |
| Grn | 2735.47283 | 0.332777116 | 0.080333943 | 4.142422302 | 3.44E-05 | 0.000849221 | | Ptpn11 | 9081.521826 | 0.113885046230864 | 0.0434688713807692 | 2.61992185702909 | 0.00879499176058099 | 0.179191559025074 | NM_011202,NM_001109992 |
| Pola2 | 232.2705326 | 0.332578668 | 0.10451109 | 3.182233263 | 0.001461441 | 0.015166484 | | Atcay | 11732.74833 | 0.107221737837083 | 0.0409383090792444 | 2.61910519141212 | 0.00881607572956645 | 0.17935898606118 | NM_178662 |
| C1qb | 2030.231878 | 0.332517819 | 0.124008583 | 2.681409705 | 0.00733127 | 0.047113518 | | Cdan1 | 1295.455413 | -0.181282285810689 | 0.0692326350373478 | -2.61845133747827 | 0.00883298890511279 | 0.17945174382611 | NM_026891 |
| 9430015G10Rik | 830.7677007 | 0.332331743 | 0.084706638 | 3.923325875 | 8.73E-05 | 0.001768185 | | 2810011L19Rik | 584.6754935 | 0.208298629724998 | 0.079608252314761 | 2.61654569304463 | 0.00888244749788038 | 0.179953183199736 | NR_045047 |
| Slc27a2 | 225.1095458 | 0.331923287 | 0.113817529 | 2.91627563 | 0.003542374 | 0.028541624 | | Mndal | 41.64685387 | -0.229408151404135 | 0.0876746820393392 | -2.61658378528479 | 0.00888145644390281 | 0.179953183199736 | NM_001170853 |
| Aasdh | 497.0489729 | 0.331884153 | 0.114902483 | 2.888398442 | 0.00387209 | 0.03046936 | | Ovca2 | 483.4350975 | -0.185313075505471 | 0.0708410132449854 | -2.61590097341796 | 0.00889923629824056 | 0.180042209565797 | NM_027136 |
| Zgpat | 1807.58806 | 0.331651004 | 0.11686739 | 2.837840425 | 0.004541988 | 0.033817188 | | Lmbr1l | 353.0906235 | -0.219369007413963 | 0.0838764809761473 | -2.61538162856816 | 0.00891278088705748 | 0.180065445292624 | NM_029098 |
| Fam19a2 | 1935.416214 | 0.331614507 | 0.122967551 | 2.696764338 | 0.00700168 | 0.045466904 | | Tnfrsf21 | 3328.765976 | 0.152045400939787 | 0.0581523754510158 | 2.61460344071174 | 0.00893311060537999 | 0.180225506463541 | NM_178589 |
| Dcaf15 | 833.9241622 | 0.330982057 | 0.084972005 | 3.895189463 | 9.81E-05 | 0.001929318 | | Hax1 | 1576.723415 | 0.160669572555064 | 0.0615076208902879 | 2.61218968039185 | 0.00899643245089588 | 0.181251286798493 | NM_011826 |
| Mcoln1 | 2340.258499 | 0.330149644 | 0.073185026 | 4.511163822 | 6.45E-06 | 0.000222978 | | Csf1 | 1154.344337 | -0.176144659005512 | 0.0674518868551786 | -2.61141188509226 | 0.00901692211375559 | 0.181364305361562 | NM_007778,NM_001113530,NM_001113529 |
| Pan2 | 975.5096009 | 0.32961323 | 0.093179539 | 3.537399244 | 0.000404088 | 0.005876062 | | Tfap2d | 33.76859537 | 0.223086572443743 | 0.0854400842541363 | 2.6110294060594 | 0.00902701313344414 | 0.181364305361562 | NM_153154 |
| Dhx33 | 1744.173362 | 0.329472698 | 0.088666355 | 3.715870577 | 0.000202505 | 0.003440368 | | H2-D1 | 733.6634636 | -0.220461659328093 | 0.0844510369618186 | -2.61052637432702 | 0.00904030007037722 | 0.181380385113673 | NM_010380 |
| Ifih1 | 237.68493 | 0.329447824 | 0.110960888 | 2.969044597 | 0.002987272 | 0.025306068 | | Plau | 174.3059733 | -0.247775110724237 | 0.0949608298012871 | -2.60923489445833 | 0.00907449285388339 | 0.181815287166221 | NM_008873 |
| Las1l | 2084.734115 | 0.329356397 | 0.066966163 | 4.918250974 | 8.73E-07 | 4.34E-05 | | Arfrp1 | 1299.37266 | -0.141653920423339 | 0.0543078635734639 | -2.60835008233603 | 0.00909798544369526 | 0.18203489883625 | NM_001165995,NR_028585,NM_001165992,NM_001165991,NM_029702 |
| Nxf1 | 2383.938123 | 0.329312612 | 0.083159437 | 3.960014946 | 7.49E-05 | 0.001571311 | | A330021E22Rik | 489.5540278 | -0.218049769641371 | 0.0837010567644211 | -2.60510175224046 | 0.00918469789765361 | 0.182762906385365 | NM_172447 |
| Rhebl1 | 205.4009882 | 0.329018655 | 0.118082307 | 2.786350162 | 0.005330526 | 0.037473304 | | Aasdh | 595.634955 | -0.20282538607813 | 0.0778543680081057 | -2.60518955156135 | 0.00918234447555438 | 0.182762906385365 | NM_173765 |
| Whrn | 1215.662567 | 0.328996266 | 0.090363652 | 3.64080311 | 0.000271789 | 0.004310233 | | Ddx60 | 56.81493954 | -0.212843368558752 | 0.0816822557483418 | -2.6057479266306 | 0.00916739006987004 | 0.182762906385365 | NM_001081215 |
| Pnma3 | 918.0572535 | 0.328930289 | 0.103069851 | 3.191333693 | 0.001416176 | 0.014775915 | | Fam20c | 1679.222798 | 0.137239027808113 | 0.0526587113135299 | 2.60619799430699 | 0.00915535218048423 | 0.182762906385365 | NM_030565 |
| Kcns3 | 225.534418 | 0.328604135 | 0.112801713 | 2.913112973 | 0.003578451 | 0.028727732 | | Cep164 | 730.9754028 | -0.19949893567769 | 0.0766195618164218 | -2.60363479048158 | 0.00922409888319245 | 0.183045437673844 | NM_001081373 |
| Cables1 | 840.2816142 | 0.328589092 | 0.066537344 | 4.938416111 | 7.88E-07 | 4.05E-05 | | Lrrtm3 | 2193.788201 | 0.185061808289509 | 0.071072148478862 | 2.60385836435703 | 0.00921808420746073 | 0.183045437673844 | NM_178678 |
| Gm4285 | 124.083663 | 0.327332358 | 0.113805615 | 2.876240853 | 0.004024425 | 0.03116164 | | Bcl2 | 1172.74776 | 0.222531379930503 | 0.0855216056891418 | 2.60204866521532 | 0.00926687013132165 | 0.183393127421769 | NM_177410,NM_009741 |
| Atp1b2 | 19777.22233 | 0.326764687 | 0.063005909 | 5.186254615 | 2.15E-07 | 1.41E-05 | | Tnfrsf11a | 258.29117 | -0.222948261202114 | 0.0856728571520919 | -2.60232083548145 | 0.00925951826813014 | 0.183393127421769 | NM_009399 |
| Cwc27 | 617.0499789 | 0.326582686 | 0.103826399 | 3.145468676 | 0.00165821 | 0.016629319 | | Pak4 | 388.7309475 | 0.189298589030819 | 0.0727651423773033 | 2.60150097761459 | 0.00928168005776432 | 0.183436305468142 | NM_027470 |
| Bzrap1 | 14045.0598 | 0.325826249 | 0.084855594 | 3.839773333 | 0.000123148 | 0.002296094 | | AW011738 | 159.228046 | -0.225488905106174 | 0.0867422339904794 | -2.5995284503616 | 0.0093351939914041 | 0.183993253621623 | NR_030671 |
| Dot1l | 2348.723724 | 0.325487009 | 0.116396193 | 2.796371595 | 0.005167994 | 0.036680212 | | Eqtn | 32.25223171 | -0.231265721238932 | 0.0889608000825708 | -2.59963625579219 | 0.00933226217407758 | 0.183993253621623 | NM_027089 |
| Pisd | 2971.114649 | 0.325221842 | 0.082882631 | 3.923884141 | 8.71E-05 | 0.001768185 | | AF357359 | 237.315513 | -0.247114818808857 | 0.0951027395359408 | -2.59839853209979 | 0.00936597210726704 | 0.184100285291152 | NR_028434 |
| Sidt1 | 2371.647412 | 0.324911083 | 0.094011485 | 3.456078639 | 0.000548095 | 0.007319044 | | Dennd2c | 34.16499045 | -0.224632802679143 | 0.0864477318067947 | -2.59848116294344 | 0.00936371824152626 | 0.184100285291152 | NM_177857 |
| Eef2k | 1782.81568 | 0.32480299 | 0.094957472 | 3.420510093 | 0.000625038 | 0.008054403 | | Npr1 | 157.8236296 | -0.245299624719702 | 0.0944233657196344 | -2.59786995358813 | 0.00938040124033182 | 0.184134741104135 | NM_008727 |
| Thoc1 | 919.6803635 | 0.324104407 | 0.098472957 | 3.291303709 | 0.000997242 | 0.011344541 | | Ifih1 | 268.2169271 | -0.22895912365163 | 0.0882158070319447 | -2.59544328114256 | 0.00944689934832794 | 0.185189824472081 | NM_001164477,NM_027835 |
| Abca8b | 931.8301559 | 0.324079243 | 0.090537872 | 3.579488174 | 0.000344268 | 0.005197309 | | Snap47 | 15247.82908 | 0.11577271860494 | 0.0446479295101073 | 2.59301427580715 | 0.00951388213720864 | 0.186251552459694 | NM_144521 |
| Slc7a6 | 999.199062 | 0.323996816 | 0.09444127 | 3.430669829 | 0.000602093 | 0.007819638 | | Acsl3 | 8060.371103 | 0.134247157114844 | 0.0518103883616516 | 2.59112431618461 | 0.00956629272484052 | 0.186784234661884 | NM_001033606,NM_028817,NM_001136222 |
| Tbce | 1529.441307 | 0.323969718 | 0.111236698 | 2.912435594 | 0.003586221 | 0.02876272 | | Parp4 | 398.7614629 | -0.209758747673431 | 0.0809533723953657 | -2.5911057373743 | 0.00956680921027411 | 0.186784234661884 | NM_001145978 |
| Rere | 6324.29544 | 0.323921212 | 0.109896846 | 2.947502346 | 0.003203523 | 0.026528802 | | Cldn5 | 726.5244789 | -0.213929361147696 | 0.0826354229809721 | -2.58883361917269 | 0.00963016119348032 | 0.187266026099726 | NM_013805 |
| Cntrob | 690.0697006 | 0.322643718 | 0.108224043 | 2.981257296 | 0.002870675 | 0.024587196 | | Idua | 867.2895184 | -0.2054047689351 | 0.079315844461998 | -2.58970663841969 | 0.00960577525849333 | 0.187266026099726 | NM_008325 |
| Acox3 | 1582.79283 | 0.322564717 | 0.072569368 | 4.44491564 | 8.79E-06 | 0.000284411 | | Scaf4 | 1443.686162 | -0.159769026243615 | 0.0617088763125222 | -2.58907690093839 | 0.00962336009395939 | 0.187266026099726 | NM_178923 |
| Tmem108 | 727.467146 | 0.322520216 | 0.085289718 | 3.781466558 | 0.000155907 | 0.002789406 | | Suv420h2 | 505.0905168 | -0.240709548948796 | 0.093023035347344 | -2.58763378933416 | 0.00966376608603879 | 0.187668270275133 | NM_146177,NM_001115018 |
| Sbsn | 305.8128012 | 0.32222278 | 0.105227154 | 3.062163779 | 0.002197432 | 0.020565282 | | 2810055G20Rik | 34.75497903 | -0.229524585561762 | 0.088746132419903 | -2.58630522033078 | 0.00970109831881212 | 0.188141727875921 | NR_015543 |
| Fam13a | 776.0971491 | 0.322103758 | 0.076230832 | 4.22537377 | 2.39E-05 | 0.000641668 | | Slc39a2 | 289.17254 | -0.244350696163768 | 0.0945079098923424 | -2.58550523910769 | 0.00972363950485547 | 0.188327449930041 | NM_001039676 |
| 3010026O09Rik | 345.4311167 | 0.320641882 | 0.098773127 | 3.246246147 | 0.001169377 | 0.012845805 | | Mapk1ip1 | 1277.521033 | -0.154339354385935 | 0.0597077656916259 | -2.58491257541028 | 0.0097403691666759 | 0.18840026960631 | NM_027115,NM_001045483 |
| Adamts10 | 842.2451344 | 0.320253927 | 0.119564443 | 2.678504724 | 0.007395169 | 0.047350844 | | Irf5 | 184.8896405 | -0.22960666926343 | 0.0888637595303101 | -2.58296074136483 | 0.00979564688848909 | 0.18921750891249 | NM_001252382,NM_012057 |
| Prr14 | 906.9558508 | 0.319805576 | 0.073391977 | 4.357500502 | 1.32E-05 | 0.000398891 | | Tspyl4 | 24636.84188 | 0.132057056138511 | 0.0511507700720419 | 2.58172176005403 | 0.00983088085728122 | 0.189645916776716 | NM_030203 |
| Prkab2 | 2010.745757 | 0.319428643 | 0.055203199 | 5.786415438 | 7.19E-09 | 7.96E-07 | | Sfxn4 | 526.9561023 | -0.181303428024202 | 0.0702414751491694 | -2.58114493807564 | 0.00984732292817278 | 0.189711157632146 | NM_053198 |
| Chfr | 1279.736261 | 0.318830516 | 0.076768597 | 4.153137207 | 3.28E-05 | 0.000825264 | | Sfpi1 | 113.9206449 | -0.240239353806397 | 0.0931059463469824 | -2.58027938313505 | 0.00987204118334489 | 0.189935457257308 | NM_011355 |
| Gstm1 | 6963.73249 | 0.318698504 | 0.066819478 | 4.769544923 | 1.85E-06 | 8.34E-05 | | Polk | 555.5426304 | -0.18013709010857 | 0.0699122530418179 | -2.57661686286696 | 0.00997724727773617 | 0.190947949876674 | NM_012048 |
| Mfsd10 | 619.6289312 | 0.318677284 | 0.094875376 | 3.358904032 | 0.000782522 | 0.009413355 | | Rrp8 | 515.197705 | 0.208715886943134 | 0.0810034860826353 | 2.5766284518942 | 0.00997691281332157 | 0.190947949876674 | NM_133951,NM_025897 |
| Macrod1 | 270.6548945 | 0.317852033 | 0.102867689 | 3.089911281 | 0.002002163 | 0.019149082 | | Tmem132c | 441.2681604 | 0.207146876910913 | 0.0803861275579715 | 2.57689831820206 | 0.00996912717672998 | 0.190947949876674 | NM_175432 |
| Eif2ak2 | 313.3758622 | 0.317807974 | 0.107486007 | 2.956738128 | 0.003109121 | 0.025899308 | | Trmt13 | 250.2192641 | -0.216400789975762 | 0.0839785939277881 | -2.57685655182369 | 0.00997033178166787 | 0.190947949876674 | NM_030016 |
| Vwa5b1 | 165.1069862 | 0.317475661 | 0.114426745 | 2.774488252 | 0.005528863 | 0.03865614 | | Dna2 | 68.11805203 | -0.244929532179148 | 0.0950887744738586 | -2.57579860014373 | 0.01000088796181 | 0.191148550701648 | NM_177372 |
| Gm15706 | 164.2655142 | 0.317192673 | 0.111958868 | 2.833117906 | 0.004609638 | 0.034173077 | | Phf20l1 | 1698.364099 | -0.171665462763073 | 0.0666812481725068 | -2.57441885789193 | 0.0100408635350084 | 0.191660425373892 | NM_001081409 |
| Pcdhb5 | 234.8292852 | 0.316416642 | 0.109839831 | 2.880709467 | 0.003967812 | 0.030908637 | | Ankhd1 | 3049.379164 | -0.162893841988444 | 0.0632976563526126 | -2.57345771352119 | 0.0100687950142761 | 0.191941360075295 | NM_175375 |
| Slc20a1 | 6211.789557 | 0.316395899 | 0.067609707 | 4.67974074 | 2.87E-06 | 0.000119315 | | Kat5 | 2342.018085 | 0.126182756397307 | 0.049048359631663 | 2.57261929542391 | 0.0100932164714458 | 0.19215473455337 | NM_001199248,NM_001199249,NM_178637,NR_037603,NM_001199247 |
| Faah | 3515.38889 | 0.3143779 | 0.081520087 | 3.856447059 | 0.000115047 | 0.002173827 | | Tial1 | 3403.502488 | -0.150979010564211 | 0.0587449050381443 | -2.57007838324323 | 0.0101675506721964 | 0.193316545895713 | NM_009383 |
| Ccdc141 | 853.3714656 | 0.31429999 | 0.094336286 | 3.331697734 | 0.00086318 | 0.010193677 | | Pcsk1n | 8130.877654 | -0.228102642157346 | 0.0888017261687858 | -2.56867351569035 | 0.0102088588723866 | 0.193453318025817 | NM_013892 |
| Zfp280c | 753.4024207 | 0.314246635 | 0.113450273 | 2.76990637 | 0.005607241 | 0.038941547 | | Tiparp | 798.917535 | 0.241091835798667 | 0.0938605441301587 | 2.56861749559366 | 0.0102105091594934 | 0.193453318025817 | NM_178892 |
| Sesn2 | 426.7326696 | 0.314139299 | 0.110059658 | 2.854263808 | 0.004313669 | 0.032794894 | | Tubb4b | 11093.65471 | 0.148779234464113 | 0.05792511654848 | 2.5684753580011 | 0.0102146974339668 | 0.193453318025817 | NM_146116 |
| Smyd4 | 500.8306239 | 0.313266091 | 0.07139473 | 4.387804109 | 1.15E-05 | 0.000354799 | | Abcg1 | 3870.052705 | 0.119836035975205 | 0.0466762775435545 | 2.56738631017404 | 0.0102468384695175 | 0.193809343239858 | NM_009593 |
| Gm10845 | 360.462726 | 0.313211143 | 0.10826656 | 2.892962901 | 0.003816264 | 0.030167954 | | Slc35g2 | 650.1881054 | 0.198152572258036 | 0.0772141707056966 | 2.56627210325549 | 0.0102798151610667 | 0.193928045493058 | NM_001101483 |
| Atxn7 | 1165.622131 | 0.313196361 | 0.095145234 | 3.291771423 | 0.000995585 | 0.011334014 | | Tmem170 | 107.0557786 | -0.240591487162773 | 0.0937375596597372 | -2.5666497830337 | 0.0102686265713554 | 0.193928045493058 | NM_025781 |
| Cnot10 | 2009.417662 | 0.313147417 | 0.07755455 | 4.037769754 | 5.40E-05 | 0.001224134 | | Spcs1 | 2218.148598 | 0.14632693259178 | 0.0570367098440805 | 2.56548691170633 | 0.0103031108348087 | 0.194115418918848 | NM_026911 |
| Scrt1 | 5623.778362 | 0.313076549 | 0.0749237 | 4.17860499 | 2.93E-05 | 0.000757405 | | Ctgf | 1598.45317 | 0.189357545835629 | 0.0738464810219914 | 2.56420540579637 | 0.0103412324815825 | 0.194325551135145 | NM_010217 |
| Stx2 | 578.2145827 | 0.312874178 | 0.098696231 | 3.170072205 | 0.001524011 | 0.015659114 | | Cxadr | 979.969776 | -0.1693231813104 | 0.0660295245664625 | -2.5643556033781 | 0.0103367579886312 | 0.194325551135145 | NM_001025192,NM_009988,NM_001276263 |
| Cecr5 | 890.9432578 | 0.312371589 | 0.094462208 | 3.306841935 | 0.000943541 | 0.010861238 | | Tapbp | 1258.598886 | -0.204879883981692 | 0.0799441712621138 | -2.56278701432716 | 0.0103835724468449 | 0.194873092200089 | NM_009318,NM_001025313 |
| Timeless | 204.7374346 | 0.312166634 | 0.112923207 | 2.764415229 | 0.005702492 | 0.039425264 | | Pla2g3 | 131.2913536 | 0.243071011905145 | 0.0948856449315059 | 2.56172587624407 | 0.0104153489504771 | 0.195217237231782 | NM_172791 |
| Taz | 1448.989399 | 0.311874921 | 0.098230626 | 3.17492552 | 0.00149875 | 0.015450184 | | Il2rg | 45.16385225 | -0.238677773968298 | 0.0931905204322824 | -2.56118082462835 | 0.0104317045052462 | 0.195278129437121 | NM_013563 |
| Lypd1 | 2570.951793 | 0.311779271 | 0.08858678 | 3.519478525 | 0.000432396 | 0.006144506 | | Brd9 | 2820.315329 | -0.134242730695785 | 0.0524629705798485 | -2.55880917935189 | 0.0105031376480485 | 0.196333074261771 | NM_001024508 |
| Tspyl2 | 4051.587203 | 0.310947584 | 0.093636655 | 3.320718175 | 0.000897862 | 0.010498156 | | Ccdc138 | 120.9142237 | 0.233451072345443 | 0.0912806203552995 | 2.55750970399588 | 0.0105424616497444 | 0.196333074261771 | NM_001162956 |
| Sdccag8 | 611.577516 | 0.310892893 | 0.08049897 | 3.862072935 | 0.000112429 | 0.002134773 | | Cln8 | 1263.889962 | 0.133081097383022 | 0.0520335394248347 | 2.55760224759004 | 0.0105396568218329 | 0.196333074261771 | NM_012000 |
| Atad2 | 428.2355735 | 0.310825511 | 0.099693905 | 3.117798546 | 0.001822073 | 0.017822727 | | Col16a1 | 1204.200657 | -0.230809435684668 | 0.09024663915305 | -2.55754051176617 | 0.0105415278458581 | 0.196333074261771 | NM_028266 |
| Mbip | 451.5009074 | 0.310688757 | 0.111249363 | 2.792723905 | 0.005226628 | 0.036943149 | | Cct5 | 5809.621035 | 0.119480638713634 | 0.0467400621640128 | 2.5562789859879 | 0.0105798256497997 | 0.196611595141644 | NM_007637 |
| Zfp112 | 399.7055917 | 0.310682449 | 0.083253913 | 3.731745918 | 0.000190157 | 0.003277304 | | Miip | 630.3988449 | -0.213315213972722 | 0.0834525524843871 | -2.55612569804417 | 0.0105844876360158 | 0.196611595141644 | NM_001025365 |
| Tenc1 | 1143.793915 | 0.310077897 | 0.099881346 | 3.104462524 | 0.001906251 | 0.018494333 | | Tmem136 | 923.6650443 | -0.168707112664516 | 0.0660159682387724 | -2.55555007622279 | 0.01060201048974 | 0.196685573913108 | NM_001034863 |
| Inpp5k | 1800.335274 | 0.309842966 | 0.088483931 | 3.501686254 | 0.000462324 | 0.006438851 | | Zfp641 | 611.074131 | -0.17352876915599 | 0.0679331053322275 | -2.55440654902122 | 0.0106368978523489 | 0.197081094646964 | NM_173769 |
| Map7 | 3871.371567 | 0.30952292 | 0.072751631 | 4.254515168 | 2.10E-05 | 0.000578615 | | Cdkl4 | 890.9914856 | 0.192237952728231 | 0.0752717934929295 | 2.55391752750369 | 0.0106518483312847 | 0.197106686446168 | NM_001033443 |
| Zfpm2 | 525.7586171 | 0.309449321 | 0.108514572 | 2.851684473 | 0.004348824 | 0.032932886 | | Gpr183 | 59.33522384 | -0.238754698188752 | 0.0935079729145274 | -2.55330845859518 | 0.0106704950554955 | 0.197200523124845 | NM_183031 |
| Ncapd3 | 1132.415469 | 0.309225511 | 0.068658795 | 4.503800432 | 6.67E-06 | 0.000228302 | | Bex2 | 6075.911899 | 0.142011370014594 | 0.0556364223959585 | 2.55248924892967 | 0.0106956210406515 | 0.197248049705551 | NM_009749 |
| Oplah | 1116.562181 | 0.308660012 | 0.076529015 | 4.033241649 | 5.50E-05 | 0.001241224 | | Tom1l1 | 752.4558003 | -0.155475894794892 | 0.0609150564217845 | -2.55233933821476 | 0.0107002246432586 | 0.197248049705551 | NM_028011 |
| Dpysl4 | 1914.72139 | 0.308419097 | 0.084639438 | 3.643917101 | 0.00026852 | 0.004275897 | | Zfp692 | 1310.332432 | -0.224322312497821 | 0.0879243375931169 | -2.55131080470468 | 0.0107318573808081 | 0.197580431322709 | NM_182996,NM_001040686 |
| Rabep2 | 373.6232332 | 0.308278704 | 0.097661614 | 3.156600555 | 0.001596198 | 0.016185535 | | Cc2d2a | 1272.458544 | -0.174705102822077 | 0.0685310209005468 | -2.54928498840856 | 0.010794404983846 | 0.198480413664997 | NM_172274 |
| Cplx3 | 699.8939804 | 0.307959893 | 0.114865188 | 2.681055061 | 0.007339044 | 0.047143925 | | Stc1 | 178.4036789 | 0.229916695665415 | 0.0902412056166307 | 2.54780168432328 | 0.0108404076774304 | 0.198744007962872 | NM_009285 |
| Kctd6 | 1089.93197 | 0.307583891 | 0.07545159 | 4.076572715 | 4.57E-05 | 0.001064877 | | Suclg1 | 3490.521047 | 0.117930464522104 | 0.0462884410784745 | 2.54773031397131 | 0.0108426255210913 | 0.198744007962872 | NM_019879 |
| Fbxo32 | 1311.789123 | 0.307430398 | 0.090675702 | 3.390438557 | 0.000697809 | 0.008684218 | | Ulk3 | 1039.091676 | -0.205267437173875 | 0.0805760314764495 | -2.54749996261445 | 0.0108497864735342 | 0.198744007962872 | NM_027895 |
| Atg4a | 215.4847605 | 0.307263807 | 0.109932165 | 2.795030989 | 0.005189474 | 0.036748497 | | 9930111J21Rik1 | 92.70673649 | -0.241698627604584 | 0.0948977859204576 | -2.54693642491484 | 0.0108673229431495 | 0.198764661801193 | NM_001114679 |
| Pcdhga2 | 477.9428513 | 0.30644184 | 0.088917391 | 3.446365652 | 0.000568181 | 0.007492252 | | Mllt6 | 8366.595943 | -0.133540822529562 | 0.052439183474973 | -2.54658470403711 | 0.0108782807470707 | 0.198764661801193 | NM_139311 |
| 4930503L19Rik | 319.053047 | 0.306301351 | 0.101508592 | 3.017491869 | 0.002548759 | 0.022682636 | | Etv1 | 2585.266828 | 0.170504915943611 | 0.0669890706682616 | 2.54526468635418 | 0.010919493311848 | 0.198767618857023 | NM_007960,NM_001163154 |
| Fam57b | 2112.631604 | 0.304689943 | 0.071314729 | 4.272468652 | 1.93E-05 | 0.000543614 | | Glra2 | 599.0492871 | 0.202671029438974 | 0.0796260529928753 | 2.54528539116598 | 0.0109188458126583 | 0.198767618857023 | NM_183427 |
| Map2k3 | 503.3716609 | 0.304415422 | 0.107054515 | 2.843555207 | 0.004461328 | 0.033425443 | | Zfp488 | 532.6969622 | -0.204584342728381 | 0.0803638280687942 | -2.54572669874872 | 0.0109050529635373 | 0.198767618857023 | NM_001013777 |
| Ctsz | 1526.115012 | 0.304061443 | 0.114229987 | 2.661835572 | 0.007771583 | 0.049168195 | | Znf512b | 3709.018787 | -0.202743278864607 | 0.079674015769395 | -2.54465997360317 | 0.0109384194824815 | 0.198862930416178 | NM_001164597 |
| Srrt | 4418.946013 | 0.303729867 | 0.092147405 | 3.296130466 | 0.000980265 | 0.011217299 | | Adcy3 | 2953.746953 | 0.144416822171359 | 0.0567955917258975 | 2.54274702988099 | 0.0109984824985838 | 0.1993618877681 | NM_138305,NM_001159536,NM_001159537 |
| Atg16l2 | 576.6098939 | 0.303710352 | 0.108434572 | 2.800862744 | 0.005096619 | 0.036290258 | | Mdh1 | 29272.27832 | 0.110194177989565 | 0.0433412973211189 | 2.54247530186115 | 0.0110070380001388 | 0.1993618877681 | NM_008618 |
| Sun1 | 4086.254053 | 0.303509694 | 0.066602029 | 4.55706381 | 5.19E-06 | 0.000190457 | | Son | 13919.88995 | -0.123258256254358 | 0.0484762479406249 | -2.54265256875468 | 0.011001456049258 | 0.1993618877681 | NM_178880,NM_019973 |
| Nfic | 5115.850644 | 0.303190005 | 0.085851954 | 3.531544616 | 0.00041314 | 0.005960143 | | Gpr45 | 422.0817296 | 0.175649552847853 | 0.0691376219012825 | 2.54057845811724 | 0.0110669260282996 | 0.19982772316468 | NM_053107 |
| Atxn2l | 5603.951341 | 0.303033157 | 0.099477027 | 3.046262711 | 0.002317053 | 0.021280627 | | Rtp1 | 56.69441495 | 0.237172403722905 | 0.0933619446878511 | 2.54035415088957 | 0.0110740270650948 | 0.19982772316468 | NM_001004151 |
| Prox1 | 382.9755278 | 0.303002734 | 0.108441916 | 2.79414775 | 0.00520367 | 0.036815369 | | Tubb2a | 12311.54337 | 0.149794923109653 | 0.0589578880564827 | 2.54071046181693 | 0.0110627489308557 | 0.19982772316468 | NM_009450 |
| Mcm8 | 190.3674274 | 0.302922841 | 0.098843651 | 3.064666652 | 0.002179127 | 0.020414223 | | Fam76b | 584.9143322 | -0.207549323693193 | 0.0817952242991765 | -2.53742593741237 | 0.0111670994056471 | 0.200510860279888 | NM_176836 |
| Pfkfb4 | 576.0966674 | 0.302913637 | 0.09545362 | 3.173411733 | 0.001506587 | 0.015520651 | | Kazn | 4201.449838 | 0.12044734622911 | 0.0474670459460258 | 2.53749404094094 | 0.011164926889357 | 0.200510860279888 | NM_001109685,NM_144531,NM_001109684 |
| Prpf38b | 1486.065958 | 0.301509747 | 0.088277439 | 3.415479073 | 0.000636699 | 0.008131449 | | Mthfd1l | 667.7189084 | 0.198596083172778 | 0.0782407698243103 | 2.53826852182092 | 0.0111402471871161 | 0.200510860279888 | NM_172308,NM_001170785,NM_001170786 |
| Gnl2 | 1900.318567 | 0.301496283 | 0.070129974 | 4.29910732 | 1.71E-05 | 0.000494791 | | Rmdn1 | 273.8044857 | -0.211715955829273 | 0.0834014387413361 | -2.53851682925873 | 0.011132344859279 | 0.200510860279888 | NM_025476 |
| Wbp2 | 15954.24471 | 0.301228942 | 0.100537949 | 2.996171545 | 0.002733926 | 0.023821531 | | Fgfbp1 | 52.8466015 | -0.228189630265103 | 0.0899645073733964 | -2.53644061338552 | 0.0111985734807289 | 0.200580121308345 | NM_001271616,NM_008009 |
| Ptpn3 | 2925.247553 | 0.30054298 | 0.105809267 | 2.8404221 | 0.004505388 | 0.03360928 | | Parp10 | 124.8363507 | -0.237077711687987 | 0.0934654699179106 | -2.53652725328625 | 0.0111958027980785 | 0.200580121308345 | NM_001163576,NM_001163575 |
| Plagl1 | 520.9508103 | 0.298715433 | 0.106742185 | 2.798475902 | 0.00513444 | 0.036475477 | | Prune | 1424.140766 | 0.126002886757443 | 0.0497112332791203 | 2.53469645482255 | 0.011254480109702 | 0.201333224228487 | NM_173347 |
| Acp6 | 430.7721388 | 0.298687608 | 0.084503112 | 3.534634431 | 0.00040834 | 0.00590739 | | Med29 | 694.843713 | 0.163692248844782 | 0.0646821623451437 | 2.53071701547825 | 0.0113829644121612 | 0.202439555379673 | NM_026042 |
| Mdm1 | 215.0303331 | 0.298561027 | 0.112174952 | 2.661565896 | 0.007777812 | 0.049187515 | | Mest | 1081.937997 | 0.197511043419977 | 0.0780483775476638 | 2.5306233085929 | 0.0113860055586667 | 0.202439555379673 | NM_001252293,NM_001252292,NM_008590 |
| Adam11 | 6054.975929 | 0.298280293 | 0.077255317 | 3.86096784 | 0.000112939 | 0.002141828 | | Rab3a | 21820.18913 | 0.0968810200010232 | 0.0382586486356556 | 2.53226455862672 | 0.0113328449062708 | 0.202439555379673 | NM_009001,NM_001166399 |
| Zranb2 | 4255.729807 | 0.298238338 | 0.069633816 | 4.282952662 | 1.84E-05 | 0.000521593 | | Stxbp3b | 36.01320272 | -0.23207507158725 | 0.0916670455857141 | -2.53171758095642 | 0.0113505371671504 | 0.202439555379673 | NR_073559 |
| Mcf2 | 260.0971539 | 0.298151199 | 0.106433974 | 2.801278446 | 0.005090058 | 0.036290258 | | Tlr12 | 40.80684099 | -0.221782701321144 | 0.0876286814171621 | -2.53093733392304 | 0.011375817077302 | 0.202439555379673 | NM_205823 |
| Trub1 | 1022.216099 | 0.297550221 | 0.073765009 | 4.033758344 | 5.49E-05 | 0.001241224 | | Rnasel | 627.7691771 | -0.165766578790182 | 0.0655598429324179 | -2.52847736320939 | 0.0114558473600921 | 0.203432321213567 | NM_011882 |
| 4933427D14Rik | 1422.16298 | 0.297496851 | 0.074377479 | 3.999824334 | 6.34E-05 | 0.001377705 | | 1700029J07Rik | 177.8657883 | -0.227584126000539 | 0.0900254858185655 | -2.52799664374158 | 0.0114715448590773 | 0.203462345082976 | NM_001033148 |
| Dhodh | 613.2768401 | 0.297421919 | 0.089532647 | 3.321938192 | 0.000893945 | 0.010481817 | | 5930403L14Rik | 1978.88434 | -0.186726343419741 | 0.0738916478070535 | -2.52702908869109 | 0.0115031974667498 | 0.203514823003249 | NR_045643 |
| Sdccag3 | 3082.901601 | 0.297056165 | 0.083610307 | 3.552865389 | 0.000381059 | 0.005638202 | | H2-K1 | 1224.405121 | -0.228149208189673 | 0.0902728377618919 | -2.52732952509425 | 0.0114933606993789 | 0.203514823003249 | NM_001001892 |
| Ptpn21 | 408.9176946 | 0.296908519 | 0.105514815 | 2.813903605 | 0.00489439 | 0.03550266 | | Ppip5k2 | 1359.53103 | -0.143015883511957 | 0.0566035907939146 | -2.5266221012843 | 0.0115165348002664 | 0.203514823003249 | NM_173760 |
| Slc25a28 | 1085.443518 | 0.296551244 | 0.084785871 | 3.497649341 | 0.000469378 | 0.006516613 | | Cbln2 | 1070.749602 | 0.200375869160957 | 0.0793368936908674 | 2.52563290342211 | 0.0115490089104512 | 0.203840708910345 | NM_172633 |
| Clk2 | 1090.389031 | 0.296430327 | 0.07725432 | 3.837071226 | 0.00012451 | 0.002318706 | | 1700109H08Rik | 25.57412999 | -0.21585003425603 | 0.0854876019399223 | -2.52492793525454 | 0.0115722016855024 | 0.204002186509232 | NM_029843 |
| Cblb | 927.0625301 | 0.296321232 | 0.06454882 | 4.590652959 | 4.42E-06 | 0.000166574 | | Sestd1 | 3113.668229 | -0.137959543229048 | 0.0546591454553016 | -2.52399744909908 | 0.0116028773407017 | 0.204295025758828 | NM_175465 |
| Myh7 | 541.9250231 | 0.296076649 | 0.0866067 | 3.418634455 | 0.000629362 | 0.008072299 | | Slc7a6os | 695.002622 | -0.152338855419265 | 0.0603972914505974 | -2.52227958837976 | 0.0116596994414235 | 0.205046966205954 | NM_001007567 |
| Hook2 | 499.4074124 | 0.296071548 | 0.103699038 | 2.855104096 | 0.004302273 | 0.032740378 | | Galnt16 | 6798.867056 | 0.180201934184086 | 0.0714741248250017 | 2.52121917722385 | 0.0116948982036672 | 0.205417280902624 | NM_001081421 |
| Ptpn23 | 2885.085132 | 0.296010653 | 0.069040502 | 4.287492773 | 1.81E-05 | 0.000515614 | | Gtf2ird1 | 1231.874313 | -0.142882970426271 | 0.0566932763377819 | -2.52028070445189 | 0.011726127997042 | 0.205471748137973 | [illegible]4936,NM_001081468,NM_001081469,NM_001081466,NM_001081470,NM_001081464,NM_001081465,NM_001081462,NM_001081463,NM_001081467,N[illegible] |
| Cux2 | 3319.031458 | 0.295396006 | 0.1025118 | 2.881580517 | 0.003956861 | 0.030869893 | | Serpina3n | 778.4488747 | -0.232819203948119 | 0.0923784604410106 | -2.52027586124136 | 0.0117262893574542 | 0.205471748137973 | NM_009252 |
| Zswim8 | 5726.546978 | 0.295086964 | 0.069179119 | 4.265549615 | 1.99E-05 | 0.000557702 | | Col6a4 | 215.9328858 | -0.237808469351963 | 0.0943837387762364 | -2.51959153595045 | 0.0117491087153254 | 0.205623558070863 | NM_026763 |
| Zfp566 | 162.5020211 | 0.294296404 | 0.104813665 | 2.807805675 | 0.004988032 | 0.0359129 | | Col22a1 | 103.2584293 | -0.237727008230121 | 0.0944157734872406 | -2.51787386206445 | 0.0118065594827552 | 0.206380364676898 | NM_027174 |
| Mroh1 | 3300.376229 | 0.293868978 | 0.062650901 | 4.690578628 | 2.72E-06 | 0.000114703 | | Dgkq | 2219.894605 | -0.151635507887017 | 0.0602407565744413 | -2.51715809212384 | 0.011830573175944 | 0.206551569655964 | NM_199011 |
| Unc13b | 2028.390901 | 0.29371934 | 0.104841628 | 2.801552638 | 0.005085734 | 0.036290258 | | Vipr2 | 44.254366 | 0.220440430481283 | 0.0876596006618747 | 2.51473231473615 | 0.0119122791519636 | 0.207728411718394 | NM_009511 |
| Sorbs3 | 1293.567553 | 0.293297601 | 0.084119 | 3.486698609 | 0.000489022 | 0.006741022 | | Slc25a12 | 6993.898921 | 0.11023740650932 | 0.0438620925373188 | 2.51327285435657 | 0.0119616779782895 | 0.208339729391647 | NM_172436 |
| Slc25a27 | 1831.970394 | 0.293018547 | 0.070120411 | 4.178791063 | 2.93E-05 | 0.000757405 | | Rnf112 | 16509.88398 | -0.207493422376769 | 0.0826553749613161 | -2.51034397307953 | 0.0120613610234365 | 0.209573361514878 | NM_009548 |
| Fam5c | 1270.646754 | 0.292953368 | 0.078442276 | 3.734636266 | 0.000187987 | 0.003250786 | | Samd9l | 273.8234423 | -0.234328700124706 | 0.0933330296612829 | -2.51067281299143 | 0.0120501325422011 | 0.209573361514878 | NM_010156 |
| Pnpt1 | 772.7019076 | 0.292796823 | 0.085747452 | 3.414641661 | 0.00063866 | 0.008137659 | | Trem2 | 583.7764962 | -0.236312408134837 | 0.0942390583104879 | -2.50758456601149 | 0.0121559490197853 | 0.210964534601436 | NM_001272078,NM_031254 |
| Tmem8 | 698.9716938 | 0.292489412 | 0.100501539 | 2.910297851 | 0.003610845 | 0.028912882 | | Fam21 | 5475.245017 | -0.122405522328862 | 0.0488639728156652 | -2.50502599922904 | 0.0122442392566056 | 0.211851282043827 | NM_026585 |
| Fam214b | 1575.650912 | 0.292017042 | 0.076876462 | 3.798523426 | 0.000145561 | 0.002643982 | | Prpf40b | 2943.474344 | -0.174898222149459 | 0.0698242037285692 | -2.50483661552875 | 0.0122507969789904 | 0.211851282043827 | NM_018786 |
| Asl | 496.4649404 | 0.291931987 | 0.099247199 | 2.941463221 | 0.003266656 | 0.026936441 | | Tmtc4 | 478.2228719 | 0.180086013516246 | 0.0718807678737839 | 2.50534348537373 | 0.0122332527532521 | 0.211851282043827 | NM_028651 |
| Ankrd27 | 1449.431275 | 0.291888486 | 0.066315099 | 4.401538884 | 1.07E-05 | 0.000335088 | | Mdm4 | 1629.448363 | -0.176013431483021 | 0.0702886716541163 | -2.50415077338729 | 0.0122745714392142 | 0.212010017510137 | NM_008575 |
| Myd88 | 236.0652912 | 0.291814065 | 0.105647672 | 2.76214382 | 0.005742318 | 0.039586803 | | Pdcl3 | 741.5928016 | 0.155463458380296 | 0.0620947196881523 | 2.50365021633165 | 0.0122919488554867 | 0.212038373702268 | NM_026850 |
| Uimc1 | 538.9098145 | 0.290979352 | 0.10297029 | 2.825857367 | 0.004715426 | 0.034613354 | | Zfp828 | 1932.18921 | 0.115501067978614 | 0.0461402055329969 | 2.50326297085985 | 0.0123054074783844 | 0.212038373702268 | NM_181854 |
| Rad52 | 455.3295744 | 0.290381844 | 0.094902277 | 3.059798485 | 0.00221486 | 0.020635619 | | Fam107a | 19695.42614 | 0.201872911197323 | 0.0806633477958287 | 2.50265971737615 | 0.0123263994164315 | 0.212148433558156 | NM_183187 |
| Sord | 908.8580345 | 0.290340785 | 0.097594791 | 2.9749619 | 0.002930249 | 0.024959475 | | Myct1 | 29.15773614 | -0.220564833477087 | 0.0881764596008266 | -2.50140269266869 | 0.0123702431484935 | 0.212148939757988 | NM_026793 |
| Mbd4 | 357.6312976 | 0.290268302 | 0.084494207 | 3.435363345 | 0.00059176 | 0.007730798 | | Sirt4 | 570.7969222 | -0.232741475497029 | 0.0930401586719446 | -2.50151632176021 | 0.0123662742118159 | 0.212148939757988 | NM_133760,NM_001167691 |
| Gabpb2 | 928.7104098 | 0.28907963 | 0.103185531 | 2.801551987 | 0.005085745 | 0.036290258 | | Ypel4 | 1531.213365 | -0.161052736365865 | 0.0643724073344955 | -2.5018908416613 | 0.0123532006342365 | 0.212148939757988 | NM_001005342 |
| Snrnp48 | 371.5549915 | 0.288690963 | 0.094809093 | 3.04497125 | 0.002327026 | 0.021346912 | | Col24a1 | 608.4075453 | -0.230514783182609 | 0.0922248353288862 | -2.49948706723696 | 0.0124373238641404 | 0.213047837795404 | NM_027770 |
| Fyco1 | 2632.760082 | 0.288383837 | 0.091294063 | 3.158845455 | 0.001583955 | 0.016082433 | | Atp8b1 | 199.4570568 | -0.23108953358391 | 0.0925721553679726 | -2.49631795506255 | 0.0125490062630294 | 0.214455135266783 | NM_001001488 |
| Rnft1 | 535.2317867 | 0.288129581 | 0.084367296 | 3.415180947 | 0.000637397 | 0.008131449 | | Cend1 | 11363.2381 | 0.122561943574198 | 0.0490908681584955 | 2.49663426563353 | 0.0125378194381603 | 0.214455135266783 | NM_021316 |
| Snap23 | 381.6245255 | 0.288024505 | 0.092143945 | 3.125810438 | 0.001773158 | 0.017434924 | | 6430411K18Rik | 104.4736144 | -0.237247610954777 | 0.0950934569054425 | -2.49488890903069 | 0.0125996569724658 | 0.21506770526679 | NR_002848 |
| Slc31a2 | 346.4913312 | 0.287888504 | 0.082974544 | 3.469600301 | 0.000521233 | 0.007034835 | | Slc12a9 | 1238.569545 | -0.206061651444183 | 0.082615752732748 | -2.49421744192984 | 0.0126235187042254 | 0.21522210410514 | NR_031406 |
| Scaf4 | 1297.230621 | 0.287589093 | 0.06760416 | 4.254014764 | 2.10E-05 | 0.000578879 | | Clk2 | 1253.894316 | -0.143030579963975 | 0.0573746154024087 | -2.49292442242619 | 0.0126695810801645 | 0.215249514351426 | NM_001163432,NM_007712 |
| Flcn | 2118.630784 | 0.287512686 | 0.095970467 | 2.995845436 | 0.002736851 | 0.023822906 | | Rabl2 | 1043.933166 | -0.165962034813174 | 0.0665675228152327 | -2.49313821206513 | 0.0126619548120493 | 0.215249514351426 | NM_026817 |
| Pml | 680.1410121 | 0.287419935 | 0.073617661 | 3.904225305 | 9.45E-05 | 0.001877708 | | Xlr | 29.22583788 | -0.208780581304288 | 0.0837368093508842 | -2.49329515804012 | 0.0126563588483429 | 0.215249514351426 | NM_011725 |
| Inpp5b | 1521.8713 | 0.28716107 | 0.059963241 | 4.788951769 | 1.68E-06 | 7.69E-05 | | 2810410L24Rik | 141.1012059 | -0.232805697290284 | 0.0934437843261994 | -2.49139842707559 | 0.0127241343162877 | 0.215303986117625 | NR_030682 |
| Zfc3h1 | 2436.287963 | 0.287098105 | 0.095853903 | 2.995163422 | 0.002742978 | 0.023862834 | | Bsdc1 | 2859.334589 | 0.11001850207687 | 0.0441863961126661 | 2.48987271549246 | 0.0127788851485462 | 0.215303986117625 | NM_133889 |
| Ldb3 | 313.817871 | 0.286870609 | 0.10633016 | 2.697923228 | 0.006977353 | 0.04536597 | | Gpatch8 | 3367.737096 | -0.111263729950518 | 0.0446732556293484 | -2.49061162843531 | 0.012752342950697 | 0.215303986117625 | NM_001159492 |
| Abcc5 | 6156.817105 | 0.286391772 | 0.063040982 | 4.542945943 | 5.55E-06 | 0.000199939 | | Nod1 | 274.9060099 | -0.207558605707171 | 0.0833199676986449 | -2.49110281052889 | 0.0127347264011134 | 0.215303986117625 | NM_172729,NM_001171007 |
| Lrrc27 | 297.6956938 | 0.28615807 | 0.091572813 | 3.124923863 | 0.001778511 | 0.017473841 | | Pnpt1 | 865.8025638 | -0.164388554577222 | 0.0659891586526233 | -2.49114487794257 | 0.0127332186291152 | 0.215303986117625 | NM_027869 |
| Smc2 | 386.6074948 | 0.286138802 | 0.07854247 | 3.643109309 | 0.000269364 | 0.00428494 | | Rrp7a | 4104.761485 | 0.126190374294376 | 0.050688518195083 | 2.48952580954749 | 0.0127913630744534 | 0.215303986117625 | NM_029101 |
| Fam46a | 251.8735383 | 0.286048526 | 0.101788157 | 2.810233855 | 0.004950551 | 0.035738209 | | Tmem150a | 378.3251875 | -0.218160439080344 | 0.0876171523328741 | -2.48992843606138 | 0.0127768819291821 | 0.215303986117625 | NM_144916 |
| Top3a | 254.2554316 | 0.285927756 | 0.101152764 | 2.826692471 | 0.004703147 | 0.034552186 | | Zfp383 | 537.2766942 | -0.166118319357383 | 0.0666720303191118 | -2.49157433127943 | 0.0127178352840044 | 0.215303986117625 | NM_001243908 |
| Alkbh3 | 292.483378 | 0.285695303 | 0.103363453 | 2.763987591 | 0.005709971 | 0.039425264 | | Rbl1 | 89.85241024 | -0.232066071580048 | 0.0932864117398886 | -2.48767282663975 | 0.0128581961505697 | 0.216178422781454 | NM_001139516,NM_011249 |
| Camk2g | 11881.57758 | 0.285690176 | 0.079894857 | 3.575826859 | 0.000349123 | 0.005251754 | | Blnk | 366.0139861 | -0.199827126799855 | 0.0803566282714951 | -2.48675350245798 | 0.0128914687088673 | 0.216344674793938 | NM_008528 |
| Ell2 | 1036.079926 | 0.285578093 | 0.077598636 | 3.680194734 | 0.000233056 | 0.003801019 | | Lrrc58 | 4148.413384 | 0.158560971227824 | 0.0637964057129028 | 2.48542170136327 | 0.0129398049087395 | 0.216344674793938 | NM_177093 |
| Tox3 | 471.1725551 | 0.285390303 | 0.101587887 | 2.809294601 | 0.004965019 | 0.035795744 | | Pik3r6 | 168.7819858 | -0.231730528373913 | 0.0932058980231355 | -2.48622172296857 | 0.0129107498465168 | 0.216344674793938 | NM_001004435,NM_001081566 |
| Clk3 | 1896.046402 | 0.285251644 | 0.072964008 | 3.909484282 | 9.25E-05 | 0.001849151 | | Rnf220 | 6073.003461 | 0.110911411119875 | 0.044625594460061 | 2.48537666471061 | 0.0129414422614881 | 0.216344674793938 | NM_025739 |
| Myrf | 2570.106459 | 0.284740795 | 0.099634379 | 2.85785688 | 0.004265127 | 0.032489617 | | Sgms1 | 1794.030857 | 0.145610949250573 | 0.0585877950946565 | 2.48534612055837 | 0.0129425528291293 | 0.216344674793938 | NM_001168525,NM_144792,NM_001168526 |
| Mvb12a | 487.5058516 | 0.2847014 | 0.087737419 | 3.244925626 | 0.001174813 | 0.012845838 | | Hdac10 | 655.8335272 | -0.205600465182423 | 0.082746599904188 | -2.48469985981886 | 0.0129660702662329 | 0.216488662858965 | NM_199198,NR_028447,NR_028448,NR_028449 |
| Ly6e | 7638.14226 | 0.284524575 | 0.097600639 | 2.915191723 | 0.003554701 | 0.028626059 | | Xdh | 654.2369949 | 0.233866192288286 | 0.0942627485962902 | 2.48100332072738 | 0.0131013152540435 | 0.218495643375702 | NM_011723 |
| Pcsk7 | 1037.497521 | 0.284275344 | 0.098961071 | 2.872597694 | 0.004071122 | 0.031413328 | | Wbp5 | 1685.60334 | 0.168558249247264 | 0.0679638857491519 | 2.48011495207022 | 0.0131340032793912 | 0.2187896005005 | NM_011712 |
| Plxnd1 | 5240.027227 | 0.28420993 | 0.097586968 | 2.912375857 | 0.003586907 | 0.02876272 | | Cyb561 | 3517.665675 | 0.131104255535588 | 0.0528968918166162 | 2.47848694002858 | 0.0131940941284682 | 0.219077908902302 | NM_007805 |
| Zfp940 | 643.8843164 | 0.283847092 | 0.106105275 | 2.675145907 | 0.007469673 | 0.047647229 | | Psmb8 | 153.3258806 | -0.189638233831947 | 0.0765157658191041 | -2.47842038567951 | 0.0131965558508546 | 0.219077908902302 | NM_010724 |
| Setd1a | 2831.862923 | 0.283765665 | 0.077139908 | 3.678584454 | 0.000234532 | 0.003821072 | | Slc24a6 | 241.0781078 | -0.217430530470744 | 0.0877071837079526 | -2.47905041843259 | 0.0131732683693247 | 0.219077908902302 | NM_133221,NM_001177594,NM_001177595 |
| Prpf40b | 2541.661095 | 0.283651333 | 0.080610625 | 3.518783456 | 0.00043353 | 0.006145581 | | Acot11 | 2572.180264 | 0.17548947757476 | 0.0708608662015421 | 2.47653587913579 | 0.0132664290210593 | 0.219986470273866 | NM_025590 |
| Pnpla6 | 2151.687758 | 0.283483454 | 0.06825944 | 4.153029275 | 3.28E-05 | 0.000825264 | | Lrp3 | 4102.01723 | 0.14628119078231 | 0.059122596718256 | 2.47420104836398 | 0.0133534526770153 | 0.221177028034577 | NM_001024707 |
| Clu | 28301.34007 | 0.283418634 | 0.074063093 | 3.826718865 | 0.000129863 | 0.002392578 | | Cdkn1b | 876.8439203 | 0.146478804640728 | 0.059269668216112 | 2.4713957248188 | 0.0134586795328519 | 0.222603291848492 | NM_009875 |
| Fnbp4 | 1797.976227 | 0.283207434 | 0.087136678 | 3.250151829 | 0.001153434 | 0.012719792 | | Pcdh9 | 5884.299793 | 0.140664096609511 | 0.056923919082242 | 2.47108946252073 | 0.0134702115885188 | 0.222603291848492 | NM_001271799,NM_001271800,NM_001081377,NM_001271798 |
| Cep135 | 380.2416604 | 0.283107777 | 0.105223732 | 2.690531617 | 0.007133827 | 0.046162698 | | 1700094D03Rik | 293.0949867 | -0.224022858790624 | 0.090682755683511 | -2.47040197556941 | 0.0134961301505629 | 0.222778166553496 | NM_028567 |
| Rltpr | 1889.519764 | 0.283093426 | 0.102761228 | 2.754866137 | 0.005871615 | 0.040200975 | | Clstn3 | 20819.20551 | 0.0924105437742652 | 0.0374228888584718 | 2.46935890288291 | 0.0135355386032673 | 0.223175066686789 | NM_153508 |
| Pus10 | 394.9943801 | 0.2820996 | 0.100209424 | 2.815100514 | 0.004876198 | 0.035420446 | | Ccdc19 | 85.65490628 | 0.234611037282633 | 0.0950607269586918 | 2.46801223584774 | 0.0135865675353717 | 0.223762449000918 | NM_027972 |
| Eml3 | 431.2331535 | 0.282017316 | 0.104668539 | 2.694384766 | 0.007051871 | 0.045697065 | | 2900005J15Rik | 91.97407777 | -0.233283552729843 | 0.0946085950740381 | -2.46577546730592 | 0.0136717005199254 | 0.224208584142078 | NR_027851 |
| Adcy6 | 2340.931852 | 0.28173465 | 0.079546519 | 3.541759647 | 0.000397467 | 0.005815261 | | Cd2ap | 1016.826608 | -0.157584101106433 | 0.0639494619067588 | -2.46419745229753 | 0.0137320440094392 | 0.224208584142078 | NM_009847 |
| Gns | 4767.12951 | 0.280845004 | 0.062319401 | 4.506542083 | 6.59E-06 | 0.000227381 | | Chp1 | 7148.898005 | 0.106032622469175 | 0.0430156147619043 | 2.46497982316599 | 0.0137020967120036 | 0.224208584142078 | NM_019769 |
| Scpep1 | 642.2410366 | 0.28051406 | 0.100521574 | 2.790585625 | 0.005261278 | 0.037121241 | | Gbp3 | 254.1353809 | -0.202593068810058 | 0.0822190584069491 | -2.46406457013055 | 0.0137371361618098 | 0.224208584142078 | NM_018734 |
| Gm19757 | 152.1302568 | 0.280503242 | 0.105548652 | 2.657572932 | 0.007870556 | 0.049612039 | | Gm10560 | 161.6435471 | -0.231129145140619 | 0.0937736600755475 | -2.4647555076171 | 0.0137106770741003 | 0.224208584142078 | NR_040563 |
| Arhgap17 | 487.2926756 | 0.280231185 | 0.077659416 | 3.608463734 | 0.000308016 | 0.004762867 | | Memo1 | 713.8941971 | -0.154628058831192 | 0.0627317907691122 | -2.46490745657667 | 0.013704864309998 | 0.224208584142078 | NM_133771 |
| Mitd1 | 194.336654 | 0.280161626 | 0.099339725 | 2.820237583 | 0.004798811 | 0.035023534 | | Ppp1r3g | 368.5976202 | 0.231572211250573 | 0.0938951703754798 | 2.4662845844417 | 0.0136522818537534 | 0.224208584142078 | NM_029628 |
| Sbno2 | 741.4026484 | 0.280136684 | 0.096458033 | 2.904233843 | 0.003681531 | 0.029312249 | | Vav1 | 196.5613509 | -0.228107415525435 | 0.0925372787691401 | -2.46503267180043 | 0.0137000758733071 | 0.224208584142078 | NM_011691,NM_001163815,NM_001163816 |
| Zmat4 | 2075.561661 | 0.280077682 | 0.080432483 | 3.482146405 | 0.000497412 | 0.006796205 | | Ubb | 32193.39601 | 0.120994989987554 | 0.0491128274756549 | 2.463612791333 | 0.0137544611669583 | 0.224239397206774 | NM_011664 |
| Riok1 | 492.5142375 | 0.279662668 | 0.095842684 | 2.917934424 | 0.003523584 | 0.02844941 | | Nme1 | 5252.344733 | 0.127585339528984 | 0.0518335163224735 | 2.46144480600609 | 0.0138378687468698 | 0.225346279615505 | NM_008704 |
| Slc5a5 | 857.4069729 | 0.278953566 | 0.100320731 | 2.780617351 | 0.005425565 | 0.038020669 | | Cdc37 | 4498.253983 | 0.104944536559899 | 0.0426715095124146 | 2.45935842811858 | 0.0139185580712344 | 0.226406466453249 | NM_016742 |
| Calcoco1 | 3201.173313 | 0.278916813 | 0.078427951 | 3.55634452 | 0.000376051 | 0.005580969 | | 9930012K11Rik | 68.57147636 | -0.231893752930094 | 0.0944272728881307 | -2.45579212273579 | 0.0140574447906992 | 0.227385585865846 | NM_001004155,NM_001112735 |
| Celsr3 | 4967.381609 | 0.278794021 | 0.097946535 | 2.846389821 | 0.004421803 | 0.033297621 | | Cdkn1a | 825.7757488 | 0.229460298426282 | 0.0934667540536047 | 2.45499376489187 | 0.0140887032003731 | 0.227385585865846 | NM_001111099,NM_007669 |
| Hectd3 | 2374.141254 | 0.278637484 | 0.060405345 | 4.612795149 | 3.97E-06 | 0.000152367 | | Fndc9 | 189.0041547 | 0.232578014916923 | 0.0947512004881915 | 2.4546181337925 | 0.0141034316370885 | 0.227385585865846 | NM_177075 |
| Bms1 | 1511.386886 | 0.278424845 | 0.0658206 | 4.230056322 | 2.34E-05 | 0.000631746 | | Gabarapl1 | 26218.28507 | 0.0838026628260577 | 0.0341202482108208 | 2.45609769038787 | 0.0140454969905195 | 0.227385585865846 | NM_020590 |
| Utp15 | 1158.978164 | 0.278360302 | 0.069710482 | 3.993091046 | 6.52E-05 | 0.001409317 | | Gm684 | 52.85048096 | -0.232673301993192 | 0.0947827453548916 | -2.45480652751724 | 0.014096043050955 | 0.227385585865846 | NM_001195681 |
| Scn1a | 6633.89606 | 0.278343336 | 0.096943473 | 2.871192114 | 0.004089269 | 0.031506284 | | Gramd1a | 3939.658061 | -0.158072220165612 | 0.0643532501614353 | -2.45632069505542 | 0.0140367830895723 | 0.227385585865846 | NM_027898 |
| Plekhm2 | 5101.002108 | 0.27827922 | 0.080065449 | 3.475646801 | 0.000509623 | 0.006914272 | | Kdelc1 | 386.811152 | -0.191987741400916 | 0.0781948765131 78 | -2.45524706939797 | 0.0140787788350636 | 0.227385585865846 | NM_023645 |
| Cux1 | 3137.840507 | 0.278182169 | 0.067072418 | 4.147489786 | 3.36E-05 | 0.000840026 | | Trappc5 | 1267.654494 | 0.126413718254216 | 0.0515006535509677 | 2.4546041562193 | 0.0141039799576709 | 0.227385585865846 | NM_025701 |
| Fubp3 | 1172.234895 | 0.277395808 | 0.07830315 | 3.542588126 | 0.000396221 | 0.005802503 | | Psma2 | 2835.465863 | 0.120125758999444 | 0.0489725669383653 | 2.45291938955596 | 0.0141702089406574 | 0.228200061055421 | NM_008944 |
| AI854703 | 1146.145472 | 0.277139183 | 0.061717627 | 4.490438074 | 7.11E-06 | 0.000241505 | | Zfp329 | 1325.573161 | -0.17299357660309 | 0.0705511373944224 | -2.45203100888301 | 0.0142052419887694 | 0.228510902689772 | NM_026046 |
| Adora1 | 10313.67738 | 0.277076233 | 0.099042012 | 2.79756266 | 0.005148978 | 0.036561991 | | Ttc9 | 1451.605218 | 0.15971324583705 | 0.0651715512382245 | 2.45065895782108 | 0.0142594986054423 | 0.229129952148955 | NM_001033149 |
| Fancg | 507.5489224 | 0.277050229 | 0.10044594 | 2.75820237 | 0.00581202 | 0.03995879 | | Impact | 10597.85286 | 0.163667605657059 | 0.066801612673756 | 2.45005470835524 | 0.0142834510779396 | 0.229261226915084 | NM_008378 |
| Rbms1 | 569.3201675 | 0.276738619 | 0.079031736 | 3.50161382 | 0.000462449 | 0.006438851 | | Pdxk | 9990.507513 | 0.137531168751589 | 0.0561504132390104 | 2.44933493483248 | 0.0143120292441382 | 0.229466376159329 | NM_172134 |
| Dennd3 | 326.8464269 | 0.275588881 | 0.099717699 | 2.76369072 | 0.005715168 | 0.039443571 | | App | 72861.48143 | 0.0999327579512934 | 0.0408153930090936 | 2.44840856803776 | 0.0143488842945089 | 0.229550543240128 | NM_001198823,NM_001198825,NM_001198826,NM_007471,NM_001198824 |
| Gnpda1 | 743.8797548 | 0.275482641 | 0.063318726 | 4.350729349 | 1.36E-05 | 0.000409807 | | Ctso | 1475.274626 | -0.138193329543747 | 0.056440720268509 | -2.4484685681953 | 0.0143464946846231 | 0.229550543240128 | NM_177662 |
| Pdk2 | 4565.772657 | 0.27508552 | 0.082922403 | 3.317384809 | 0.000908644 | 0.010547482 | | A930011O12Rik | 7787.879579 | -0.232293334019196 | 0.0949084015115133 | -2.4475529070102 | 0.01438300069 6174 | 0.229590624299586 | NR_040709 |
| Chka | 1678.053777 | 0.275015342 | 0.096159892 | 2.859979723 | 0.004236681 | 0.032353861 | | Gm10845 | 430.7382562 | -0.179748508430833 | 0.0734368172851185 | -2.44766201853437 | 0.0143786462909813 | 0.229590624299586 | NR_033535 |
| Lrrc8a | 2055.320247 | 0.274907997 | 0.095482478 | 2.879146041 | 0.003987536 | 0.031002229 | | Ints10 | 2119.56311 | -0.125546892429725 | 0.051326575366341 | -2.44604070179318 | 0.0144434694094028 | 0.230050259474765 | NM_027590 |
| Per3 | 2035.965649 | 0.274901301 | 0.077834744 | 3.531858506 | 0.00041265 | 0.005958621 | | Maff | 50.92857626 | 0.200592653398461 | 0.0820035922969589 | 2.44614470878369 | 0.0144393033025076 | 0.230050259474765 | NM_010755 |
| Polk | 494.6675454 | 0.27474836 | 0.078342053 | 3.507035486 | 0.000453129 | 0.006353644 | | Arhgef3 | 3716.36794 | 0.138810750509548 | 0.0567737646043801 | 2.44498055531161 | 0.0144859951745556 | 0.230474880509961 | NM_027871 |
| Foxo3 | 1451.09523 | 0.274144117 | 0.077859274 | 3.521020712 | 0.000429889 | 0.006116346 | | Nmnat2 | 8385.882319 | 0.106531918369798 | 0.0436055801789804 | 2.44307994372589 | 0.0145625110671971 | 0.231438769980422 | NM_175460 |
| Lnx1 | 908.489144 | 0.273969318 | 0.079373754 | 3.451636123 | 0.000557199 | 0.007397802 | | Ckmt1 | 8311.91224 | 0.134931033272398 | 0.0552429523856512 | 2.44250220970168 | 0.0145858403231034 | 0.231556192932678 | NM_009897 |
| Zfp108 | 243.448303 | 0.273240983 | 0.099526877 | 2.745398932 | 0.006043738 | 0.041016941 | | Fzd1 | 454.8447695 | 0.194871037629707 | 0.0798160899479996 | 2.44150067682576 | 0.0146263609159641 | 0.231945981075649 | NM_021457 |
| Sltm | 2513.626454 | 0.272462165 | 0.092528366 | 2.944633906 | 0.00323337 | 0.026704603 | | Wbp7 | 3108.802392 | -0.140075879080046 | 0.0573870190407877 | -2.44089833243451 | 0.0146507786719874 | 0.232079837501951 | NM_029274 |
| Zscan26 | 3193.936833 | 0.272009692 | 0.085462884 | 3.182781575 | 0.001458676 | 0.01514794 | | A230050P20Rik | 2277.398145 | -0.170216974589509 | 0.0698644868008088 | -2.43638767539817 | 0.0148347759744729 | 0.232209004100424 | NM_175687 |
| Zdhhc4 | 706.8152557 | 0.271803363 | 0.087558423 | 3.104251451 | 0.001907611 | 0.018495951 | | Arap1 | 736.0495003 | -0.173564930055309 | 0.0711357896157717 | -2.43991007891796 | 0.0146909182072748 | 0.232209004100424 | NM_001040111,NM_198096,NM_001040112,NM_027180 |
| Nckipsd | 3224.885303 | 0.271550756 | 0.090981177 | 2.9846916 | 0.002838644 | 0.024380235 | | Azi1 | 1036.396064 | -0.188090803476646 | 0.0771955997292233 | -2.43654825063095 | 0.014828191048074 | 0.232209004100424 | NM_009734 |
| Dcdc2a | 376.6574047 | 0.271538895 | 0.082557734 | 3.289078852 | 0.001005159 | 0.011417835 | | Hmgcr | 4505.599682 | 0.142684088441101 | 0.0585134023293 | 2.43848559067045 | 0.0147489466876429 | 0.232209004100424 | NM_008255 |
| Osbpl7 | 642.3745631 | 0.271270809 | 0.09077392 | 2.988422318 | 0.002804218 | 0.02420532 | | Hps5 | 728.0910038 | -0.168975413120628 | 0.0692751196582456 | -2.43919337785677 | 0.0147200888045471 | 0.232209004100424 | NM_001005248,NM_001005247,NM_001167864 |
| Zfp276 | 839.1398312 | 0.270661147 | 0.099993027 | 2.706800205 | 0.006793513 | 0.044436233 | | P2ry6 | 98.37981341 | -0.230946789702426 | 0.0947740594351444 | -2.43681436754819 | 0.0148172837020257 | 0.232209004100424 | NM_183168 |
| Sel1l3 | 2790.240901 | 0.270621442 | 0.097823558 | 2.766424032 | 0.005667479 | 0.039254321 | | Papln | 199.4567752 | -0.223389494152351 | 0.0916268005508622 | -2.43803660947811 | 0.0147672783746295 | 0.232209004100424 | NM_130887,NM_001205343 |
| Prodh | 983.0157978 | 0.269619886 | 0.088297107 | 3.053552868 | 0.002261489 | 0.020948688 | | Sbno2 | 816.18807 | -0.202640475160378 | 0.0831612526893932 | -2.43671744480858 | 0.0148212554598392 | 0.232209004100424 | NM_183426 |
| Sema6c | 689.6047835 | 0.269527297 | 0.091112643 | 2.958176706 | 0.003094647 | 0.025824561 | | Slco4a1 | 110.0727752 | 0.228443397066008 | 0.0937083665005285 | 2.43781217884281 | 0.0147764493163003 | 0.232209004100424 | NM_148933 |
| Zfp956 | 802.9712627 | 0.269472669 | 0.079610977 | 3.384868237 | 0.000712124 | 0.008819867 | | Timm17a | 1814.618548 | 0.123592315703803 | 0.0506518935372249 | 2.4400334730423 | 0.014685901062448 | 0.232209004100424 | NM_011590 |
| Grhl1 | 514.0701098 | 0.269261792 | 0.094136075 | 2.860346477 | 0.004231784 | 0.032346946 | | Trim25 | 370.2814394 | -0.20387185141584 | 0.0836580403777833 | -2.43696661426917 | 0.0148110467388768 | 0.232209004100424 | NM_009546 |
| Elmod3 | 804.7182459 | 0.269060857 | 0.075817992 | 3.548773202 | 0.00038703 | 0.005694822 | | Abt1 | 572.9403427 | 0.154315358886001 | 0.0633687323962503 | 2.43519718717196 | 0.0148836763624532 | 0.232332035333844 | NM_013924 |
| Rnf31 | 1005.021949 | 0.26904605 | 0.063904177 | 4.210148134 | 2.55E-05 | 0.000678356 | | Eya4 | 201.5054874 | -0.218537445281557 | 0.0897760200644587 | -2.43425187622094 | 0.0149226069782787 | 0.232332035333844 | NM_010167 |
| Tesk2 | 243.9767993 | 0.268669483 | 0.093248934 | 2.88120701 | 0.003961554 | 0.030875408 | | Man2c1 | 2667.685955 | -0.193865243484176 | 0.0796378006148972 | -2.4343369855434 | 0.0149190982609492 | 0.232332035333844 | NM_028636 |
| Fbrs | 1567.645785 | 0.268057581 | 0.075947793 | 3.529497964 | 0.000416349 | 0.005984147 | | Pofut2 | 1122.351609 | 0.150172291005686 | 0.0616855029790754 | 2.43448271884281 | 0.0149130919471109 | 0.232332035333844 | NM_030262 |
| Zfp335 | 1569.866418 | 0.267800344 | 0.090032125 | 2.974497638 | 0.002934687 | 0.02498354 | | Sh3tc1 | 115.1227563 | -0.227493421693694 | 0.0934421239032251 | -2.43459172577564 | 0.0149086006821503 | 0.232332035333844 | NM_194344 |
| Kctd18 | 345.9035113 | 0.267582382 | 0.09499846 | 2.816702305 | 0.004851948 | 0.035293932 | | Sdha | 18069.20178 | 0.0910793442072012 | 0.0374257716409861 | 2.43359963505622 | 0.0149495204191035 | 0.232501856111239 | NM_023281 |
| Ndst2 | 694.5783561 | 0.267484081 | 0.080320982 | 3.330189386 | 0.000867869 | 0.010225679 | | Gm20751 | 31.65224638 | -0.2164312247372 | 0.0889637160866868 | -2.43280332991383 | 0.0149824363416038 | 0.232764567163783 | NR_046022 |
| Trpc2 | 511.711799 | 0.266880067 | 0.095283401 | 2.800908284 | 0.0050959 | 0.036290258 | | Slc2a4rg-ps | 404.0839001 | -0.230431902421616 | 0.0947495885777854 | -2.43200953038906 | 0.0150153122196047 | 0.233026095408096 | NR_045164 |
| Gmppa | 1152.552036 | 0.266587613 | 0.07484588 | 3.561820807 | 0.000368292 | 0.005486838 | | Apmap | 2512.695152 | 0.129669665364447 | 0.0533834449043399 | 2.42902393423293 | 0.0151395328671691 | 0.233312353578214 | NM_027977 |

| | | | | | | | | | | | | | | | |
|---|---|---|---|---|---|---|---|---|---|---|---|---|---|---|---|
| 4-Sep | 6547.743147 | 0.266501629 | 0.09947411 | 2.679105445 | 0.007381914 | 0.047340803 | | Cacna1d | 2261.665325 | -0.130019829228953 | 0.0535155145551369 | -2.42957262599042 | 0.0151166359915816 | 0.233312353578214 | NM_028981,NM_001083616 |
| 2700050L05Rik | 992.9651654 | 0.266220739 | 0.076558811 | 3.477336387 | 0.000506422 | 0.00688894 | | Cort | 124.9854348 | 0.22971613259231 | 0.0945343885377591 | 2.42997427862514 | 0.0150998943891049 | 0.233312353578214 | NM_007745 |
| Cwf19l2 | 474.4366147 | 0.265974466 | 0.08840914 | 3.008449888 | 0.002625841 | 0.023191304 | | Fam81a | 6444.989512 | 0.128848253446259 | 0.0530332027341605 | 2.42957707253957 | 0.0151164505619855 | 0.233312353578214 | NM_029784 |
| Grina | 14385.91329 | 0.265427305 | 0.072165788 | 3.678021277 | 0.00023505 | 0.003825494 | | Fus | 8774.019803 | -0.121272886062567 | 0.0499298678075728 | -2.4288645531758 | 0.0151461895514426 | 0.233312353578214 | NM_139149 |
| Hmha1 | 319.927829 | 0.265395447 | 0.097393904 | 2.724969805 | 0.006430737 | 0.042836559 | | Pla2g4e | 570.7372835 | -0.206775557898807 | 0.0850934892504967 | -2.42998095060017 | 0.0150996164271517 | 0.233312353578214 | NM_177845 |
| Pex5l | 7514.827652 | 0.264905892 | 0.094115201 | 2.814698256 | 0.004882305 | 0.035431587 | | Zwilch | 136.9667513 | -0.220188629008889 | 0.0905783341282392 | -2.43091939289974 | 0.0150605647360531 | 0.233312353578214 | NM_026507 |
| Szt2 | 2301.531666 | 0.26454484 | 0.091117031 | 2.903352296 | 0.003691911 | 0.029379797 | | Marcksl1 | 1803.716204 | 0.201925093704171 | 0.0831973937802966 | 2.42706032640163 | 0.0152217245348858 | 0.233769171552376 | NM_010807 |
| Anks3 | 1157.292551 | 0.264533166 | 0.073178985 | 3.614878874 | 0.000300488 | 0.004669772 | | Peg13 | 18894.70533 | 0.107051104656337 | 0.0440962523891541 | 2.42766899353713 | 0.0151962053461304 | 0.233769171552376 | NR_002864 |
| Bbs2 | 1120.689392 | 0.263800713 | 0.090696917 | 2.908596238 | 0.003630554 | 0.028965914 | | Rab27b | 1413.125364 | 0.166367728051862 | 0.0685486254251205 | 2.42700312398816 | 0.0152241247617064 | 0.233769171552376 | NM_001082553,NM_030554 |
| Hirip3 | 1041.647189 | 0.263528368 | 0.069536504 | 3.789784556 | 0.000150778 | 0.002722791 | | Neat1 | 6240.970233 | -0.230145222804651 | 0.0948905372001729 | -2.42537590781214 | 0.0152925428589477 | 0.234571782015918 | NR_003513 |
| Mxra7 | 487.0823002 | 0.263473212 | 0.098190045 | 2.683298623 | 0.007289986 | 0.046926069 | | Vma21 | 1310.528026 | 0.153354382708423 | 0.0632981364125107 | 2.42273140095342 | 0.0154043115964248 | 0.236036951740154 | NM_001081356 |
| Sirt6 | 499.704828 | 0.263342774 | 0.089866252 | 2.930385644 | 0.003385416 | 0.027607174 | | Crhr1 | 1077.973133 | -0.157173479319266 | 0.0649238454355331 | -2.42088986357615 | 0.0154825673221075 | 0.236284305283788 | NM_007762 |
| Srek1 | 1838.959289 | 0.262677687 | 0.091166617 | 2.881292462 | 0.00396048 | 0.030875408 | | Grm8 | 774.8420324 | 0.171600163280167 | 0.0708762141593344 | 2.42112484866077 | 0.0154725622478326 | 0.236284305283788 | NM_008174 |
| Zfp316 | 1198.474351 | 0.262461325 | 0.078212821 | 3.355732739 | 0.00079155 | 0.009492476 | | Hk3 | 85.34104637 | -0.223004718997832 | 0.0921217278611496 | -2.42076135755894 | 0.0154880411932042 | 0.236284305283788 | NM_001033245,NM_001206392,NM_001206391,NM_001206390 |
| Slc37a4 | 784.5343816 | 0.261652654 | 0.070185919 | 3.727993573 | 0.00019301 | 0.003311243 | | Kcng4 | 472.4891411 | 0.21080409529976 | 0.0870933362430505 | 2.42043885781882 | 0.0155017859655411 | 0.236284305283788 | NM_025734 |
| Chpf2 | 2194.501165 | 0.261371227 | 0.054958612 | 4.755782874 | 1.98E-06 | 8.78E-05 | | Tcea2 | 1381.156586 | -0.17902568153615 | 0.0739380505718984 | -2.42129296284412 | 0.0154654078604979 | 0.236284305283788 | NM_009326 |
| Btbd11 | 1159.746864 | 0.261300517 | 0.080296082 | 3.254212532 | 0.001137072 | 0.012602139 | | Fam220a | 1479.969226 | 0.154132997205591 | 0.0637130667302684 | 2.41917404255737 | 0.0155557953973768 | 0.236858997843077 | NM_026050,NM_133703 |
| Lgals8 | 1017.492143 | 0.261051122 | 0.074987893 | 3.481243576 | 0.000499091 | 0.006813147 | | Zfp579 | 433.8169117 | -0.206851424395469 | 0.0855343185719519 | -2.4183442137493 | 0.0155913201727427 | 0.237151326522786 | NM_026741 |
| B3gat2 | 446.4675839 | 0.260817909 | 0.089071412 | 2.928188767 | 0.00340943 | 0.027759173 | | Gm266 | 58.03992489 | -0.224010237939942 | 0.0926960881017102 | -2.41660940097222 | 0.0156658177775658 | 0.237621413155408 | NM_001033248 |
| Taf1b | 675.1268158 | 0.260624888 | 0.088255243 | 2.953081076 | 0.003146193 | 0.026194045 | | Nphp3 | 455.3317195 | -0.192237062040117 | 0.0795602216779959 | -2.41624593277477 | 0.015681465715434 | 0.237621413155408 | NM_172460,NM_028721 |
| Utp14a | 597.7831145 | 0.259846116 | 0.079033656 | 3.287790672 | 0.001009769 | 0.011447859 | | Ubxn11 | 484.5464336 | -0.208141168379603 | 0.0861301961110128 | -2.41658765192327 | 0.0156667537253957 | 0.237621413155408 | NM_026257,NR_045571 |
| Cnot3 | 1583.700684 | 0.259822933 | 0.067178437 | 3.867653738 | 0.000109888 | 0.002101972 | | Uqcrc2 | 8409.644389 | 0.116935101667449 | 0.0483982438528174 | 2.41610216319123 | 0.0156876590400686 | 0.237621413155408 | NM_025899 |
| Surf2 | 827.2797765 | 0.259701239 | 0.078220673 | 3.320109991 | 0.00089982 | 0.010498156 | | Ttc8 | 761.5016844 | -0.161267057453519 | 0.0668037882597794 | -2.41404060539803 | 0.0157767040341802 | 0.23872125291719 | NM_198311,NM_029553 |
| Ubp1 | 3567.846609 | 0.259582492 | 0.087962283 | 2.951065877 | 0.003166794 | 0.026337252 | | Tra2b | 2162.143002 | 0.117591032106705 | 0.0487277354118843 | 2.41322587870615 | 0.0158120169497384 | 0.239006616245474 | NM_009186 |
| Trp53 | 801.2736227 | 0.259267504 | 0.075803784 | 3.420244871 | 0.000625648 | 0.008054403 | | Rab32 | 83.58630381 | -0.228217974572208 | 0.0945958772093685 | -2.4125573048717 | 0.0158410470333641 | 0.239084041897695 | NM_026405 |
| Apbb3 | 1424.740789 | 0.25915149 | 0.059365892 | 4.365326323 | 1.27E-05 | 0.000388676 | | Slc25a28 | 1174.424189 | -0.143636418257092 | 0.059542113464569 | -2.41235001412176 | 0.0158500573005288 | 0.239084041897695 | NM_145156 |
| Trex1 | 383.859503 | 0.258539066 | 0.084582579 | 3.056646766 | 0.002238279 | 0.020803774 | | Hs6st1 | 2935.318884 | 0.127365595754242 | 0.0528149837319965 | 2.41154283792917 | 0.0158851856320913 | 0.239365359431285 | NM_015818 |
| Wdr19 | 906.1033872 | 0.258475462 | 0.070119223 | 3.686228287 | 0.000227602 | 0.003723832 | | Metrnl | 281.5624441 | 0.207313827987339 | 0.0860540689496982 | 2.40911127756808 | 0.0159914213122502 | 0.24071646215725 | NM_144797 |
| Rfxank | 641.5151331 | 0.258362765 | 0.075857229 | 3.405908271 | 0.000659443 | 0.0083136 | | Agrn | 9509.708624 | -0.173205640551497 | 0.0719313146751897 | -2.40793097322937 | 0.0160432140280968 | 0.240747651830718 | NM_021604 |
| Plekha6 | 5032.208944 | 0.257134812 | 0.052212076 | 4.924814984 | 8.44E-07 | 4.25E-05 | | Hars2 | 1117.019929 | -0.153734277595111 | 0.0638402400497382 | -2.4081093284633 | 0.0160353782092006 | 0.240747651830718 | NM_080636 |
| Polrmt | 1180.968823 | 0.257033945 | 0.071465002 | 3.596640831 | 0.000322353 | 0.004945088 | | Trp53 | 878.1366384 | -0.154266130244408 | 0.0640595866167295 | -2.40816618389044 | 0.0160328810429193 | 0.240747651830718 | NM_011640,NM_001127233 |
| Lrrc57 | 1263.270748 | 0.256485513 | 0.065293896 | 3.928169816 | 8.56E-05 | 0.001745007 | | Fgfrl1 | 675.2921841 | -0.177195468657984 | 0.0736213422076886 | -2.4068492008487 | 0.0160908125369341 | 0.240953940414456 | NM_054071,NM_001164259 |
| 4922501C03Rik | 415.6370696 | 0.256438059 | 0.091484063 | 2.80308996 | 0.005061555 | 0.036240172 | | Igflr1 | 263.4690097 | -0.22881764403575 | 0.0950836342231661 | -2.40648820278265 | 0.0161067242284481 | 0.240953940414456 | NM_145580 |
| Rhno1 | 429.9475416 | 0.256332015 | 0.093782155 | 2.733270685 | 0.006270877 | 0.042079243 | | Lrfn1 | 1650.405855 | 0.165100245684805 | 0.0685982879617043 | 2.40676918609068 | 0.0160943381505437 | 0.240953940414456 | NM_001141921,NM_030562 |
| Polr3e | 1272.856745 | 0.256065996 | 0.067459418 | 3.795852445 | 0.000147137 | 0.002669486 | | Nptxr | 105.2976328 | -0.120112211523121 | 0.0499500609995237 | -2.40464594276003 | 0.0161881409945462 | 0.241922773751829 | NM_030689 |
| Rffl | 562.5782036 | 0.256006693 | 0.082037893 | 3.1205908 | 0.001804886 | 0.017699311 | | Slc1a1 | 5784.442155 | 0.122435410574295 | 0.0509414243644724 | 2.40345479345655 | 0.0162409749333699 | 0.242462900187186 | NM_009199 |
| Slc6a8 | 4564.88945 | 0.255823656 | 0.064821429 | 3.946590783 | 7.93E-05 | 0.001635816 | | Cntrob | 818.9833046 | -0.187717422781081 | 0.0781453600563863 | -2.40215698853563 | 0.0162987120653301 | 0.242552129102769 | NM_172560 |
| Adam15 | 4099.675672 | 0.255821041 | 0.067950603 | 3.764808986 | 0.000166676 | 0.002941327 | | Glb1l | 436.6549673 | -0.204156430584957 | 0.0849781400235192 | -2.40245821488271 | 0.0162852949691972 | 0.242552129102769 | NM_029010 |
| Arhgap15 | 418.2416294 | 0.255660185 | 0.096133849 | 2.659419012 | 0.007827555 | 0.049401276 | | H60b | 29.65253437 | 0.193280477497781 | 0.0804825147650301 | 2.40152134984929 | 0.0163270562909889 | 0.242552129102769 | NM_001177775 |
| Slc12a9 | 1121.625281 | 0.255157079 | 0.086137375 | 2.962210985 | 0.003054384 | 0.025594714 | | Hpgd | 189.6485636 | -0.21865113234444 | 0.0910497993360331 | -2.40144551595852 | 0.0163304407450439 | 0.242552129102769 | NM_008278 |
| Mtcp1 | 241.7527577 | 0.254776735 | 0.094540185 | 2.69490414 | 0.007040888 | 0.045660528 | | Inha | 1047.304125 | -0.210517445625783 | 0.0876216745002263 | -2.40257272902539 | 0.0162801968754964 | 0.242552129102769 | NM_010564 |
| Tmem25 | 2406.135191 | 0.254759116 | 0.085219172 | 2.989457786 | 0.002794731 | 0.024150338 | | Plb1 | 62.90552893 | -0.226440754307499 | 0.0943282466375752 | -2.40056146890466 | 0.0163699411216722 | 0.24289046448765 | NM_172147,NM_001081407,NM_030072 |
| Rcan2 | 11886.4631 | 0.254635249 | 0.077586453 | 3.28195503 | 0.0010309 | 0.011633483 | | Mafb | 1002.130043 | -0.165944027433641 | 0.0691632641325842 | -2.39930878790698 | 0.0164260562801109 | 0.243474381147848 | NM_010658 |
| Rhbdd3 | 559.7434643 | 0.253408777 | 0.09045072 | 2.80162257 | 0.005084632 | 0.036290258 | | Elk3 | 355.247573 | -0.182201671376349 | 0.0760083034781584 | -2.39712851147514 | 0.0165241271318068 | 0.244678359548038 | NM_013508 |
| Lypd6b | 574.2849864 | 0.253148173 | 0.072175624 | 3.507391516 | 0.000452523 | 0.006350895 | | Clec14a | 166.5784525 | -0.224624513420812 | 0.0937673435065386 | -2.39555163898985 | 0.0165953764311463 | 0.245483134459095 | NM_025809 |
| Vars | 1679.577633 | 0.252971891 | 0.071339678 | 3.546019507 | 0.000391097 | 0.005732886 | | C230091D08Rik | 65.39358164 | -0.226808865219344 | 0.0947301537353632 | -2.39426383328652 | 0.0166537645535 77 | 0.245893369754036 | NR_015479 |
| Znf512b | 3330.081277 | 0.252897834 | 0.091631693 | 2.759938464 | 0.005781225 | 0.039775444 | | D330050I16Rik | 40.57469832 | -0.218291029214152 | 0.0911751879596626 | -2.39419335566084 | 0.016656965154755 | 0.245893369754036 | NR_033224 |
| Hrsp12 | 1043.445333 | 0.252038796 | 0.089802075 | 2.806603254 | 0.005006686 | 0.035980333 | | Heatr8 | 314.0527982 | -0.212493202769932 | 0.0887853172052848 | -2.39333720325194 | 0.0166958887681005 | 0.245968032703273 | NM_001126487 |
| Ell | 865.5372805 | 0.251905219 | 0.071585962 | 3.518919238 | 0.000433309 | 0.006145581 | | Isg15 | 83.97133234 | -0.142212549371526 | 0.0594144609265162 | -2.39356794884354 | 0.0166853904316691 | 0.245968032703273 | NM_015783 |
| Rela | 820.8787464 | 0.251731491 | 0.078131853 | 3.221880457 | 0.001273523 | 0.013683744 | | A330076H08Rik | 85.12221156 | -0.226481196584013 | 0.0947083063595737 | -2.39135515446918 | 0.0167863060687754 | 0.246336260957249 | NR_015599 |
| Exoc1 | 2049.673333 | 0.251699664 | 0.050887073 | 4.946239804 | 7.57E-07 | 3.95E-05 | | Cnga4 | 57.01044436 | -0.22408289428188 | 0.0937074669245425 | -2.39130243977593 | 0.0167887166699488 | 0.246336260957249 | NM_001033317 |
| Cnppd1 | 1165.702659 | 0.251658223 | 0.082404299 | 3.053945333 | 0.002258532 | 0.020941771 | | Ggta1 | 203.2759328 | -0.223510585522334 | 0.0934440243414502 | -2.39191951649698 | 0.0167605172742569 | 0.246336260957249 | NM_001145821,NM_010283 |
| Zfp423 | 640.5460814 | 0.251462017 | 0.085007737 | 2.958107399 | 0.003095343 | 0.025824561 | | Sema4a | 2325.548078 | -0.137351477917011 | 0.0574359584565159 | -2.39138479809645 | 0.0167849506226603 | 0.246336260957249 | NM_013658,NM_001163490,NM_001163491,NM_001163489 |
| Fam160b2 | 2431.822796 | 0.251254355 | 0.055083128 | 4.561366868 | 5.08E-06 | 0.000187038 | | Psmc5 | 6209.356671 | 0.107748709786087 | 0.0450698691611506 | 2.39070385140953 | 0.0168161109670625 | 0.246489230986427 | NM_008950 |
| Acd | 1247.255523 | 0.250851148 | 0.079497429 | 3.155462404 | 0.001602439 | 0.01619582 | | Ahsa1 | 4364.075358 | 0.111837773387369 | 0.0467946298409496 | 2.38997025443933 | 0.0168497374160869 | 0.246536560014131 | NM_146036 |
| Rnaseh1 | 307.5830117 | 0.250480512 | 0.091588195 | 2.734855861 | 0.00624076 | 0.041968026 | | Snhg5 | 263.0220603 | -0.210020117505493 | 0.0878784620863584 | -2.38989295578597 | 0.0168532840488801 | 0.246536560014131 | NR_040721 |
| Msn | 1403.761535 | 0.250469502 | 0.055618761 | 4.503327627 | 6.69E-06 | 0.000228307 | | Lime1 | 2046.85709 | -0.209687808660029 | 0.0878436702198839 | -2.3870565532514 | 0.0169838784327006 | 0.248007351244694 | NM_023684 |
| Katna1 | 359.0367202 | 0.25033736 | 0.091623123 | 2.732250894 | 0.006290322 | 0.042154952 | | Pamr1 | 1809.134718 | -0.148248779758088 | 0.0621075714798809 | -2.38696790464769 | 0.0169879742866908 | 0.248007351244694 | NM_173749 |
| Cars | 1454.770105 | 0.250220057 | 0.04621781 | 5.413931438 | 6.17E-08 | 5.05E-06 | | Fgd3 | 300.4001706 | -0.193121482618384 | 0.0809907499265056 | -2.38448813961632 | 0.0171028992980936 | 0.249184268008132 | NM_015759 |
| Smarcd2 | 559.1782418 | 0.25008658 | 0.073908746 | 3.383721059 | 0.000715106 | 0.008835621 | | Stx1b | 10289.50076 | 0.0868581412984756 | 0.0364253826876408 | 2.38454986302578 | 0.0171000304617826 | 0.249184268008132 | NM_024414 |
| Sec24a | 666.7153159 | 0.249933717 | 0.074936096 | 3.335291402 | 0.000852101 | 0.010070516 | | Prkar2a | 2996.000896 | 0.138033058680925 | 0.0579536859826539 | 2.38178221696269 | 0.0172290833614123 | 0.250771207322521 | NM_008924 |
| Wdr73 | 1030.124251 | 0.249297798 | 0.071772172 | 3.473460412 | 0.000513793 | 0.006962544 | | Timp3 | 6121.308681 | 0.156391611812616 | 0.0656904182945016 | 2.38073703704374 | 0.0172780409544488 | 0.251232054959283 | NM_011595 |
| Lrrfip1 | 1979.103503 | 0.249128055 | 0.067250966 | 3.70445317 | 0.000211847 | 0.003548499 | | Magohb | 99.95561225 | 0.223946187605137 | 0.0941206798608458 | 2.37935157221807 | 0.0173431259832835 | 0.251926248033176 | NM_025564 |
| Cables2 | 1467.331658 | 0.248956697 | 0.061931072 | 4.019899704 | 5.82E-05 | 0.001299865 | | Paxbp1 | 1825.068172 | -0.160722067242122 | 0.0675647865490226 | -2.37878450375193 | 0.0173698271268079 | 0.252062046797214 | NM_026110 |
| Zfp414 | 798.413644 | 0.248908362 | 0.085903554 | 2.897532778 | 0.003761104 | 0.029853766 | | Ctns | 794.2379829 | -0.152447081474333 | 0.0641048771535257 | -2.37808866101147 | 0.0174026410246478 | 0.252286191141751 | NM_031251 |
| C1qtnf1 | 326.247084 | 0.248296434 | 0.087234205 | 2.84631969 | 0.004422777 | 0.033297621 | | Zfhx2as | 248.3904996 | -0.224642110202014 | 0.0944858140706467 | -2.37752209060769 | 0.0174293989541194 | 0.252422182659559 | NR_004444 |
| Hps5 | 638.7026078 | 0.247686739 | 0.092061568 | 2.690446686 | 0.007135644 | 0.046162698 | | Cecr5 | 1046.903996 | -0.175361222825054 | 0.0737831118635551 | -2.37671220955473 | 0.0174677105720549 | 0.252725063515608 | NM_144815 |
| Timm44 | 1550.311694 | 0.247430173 | 0.076150048 | 3.249245133 | 0.001157117 | 0.012746794 | | 1110012L19Rik | 339.2860029 | 0.176168620142227 | 0.0741648495048883 | 2.37536543683832 | 0.0175315834727894 | 0.253361317169201 | NM_026787 |
| Zfp598 | 1537.487058 | 0.247382855 | 0.091273323 | 2.710352242 | 0.006721179 | 0.044107886 | | Rsad2 | 73.10873353 | -0.21840286174357 | 0.0919571617005049 | -2.37505005270699 | 0.0175465706369418 | 0.253361317169201 | NM_021384 |
| Gpr137b-ps | 421.3135084 | 0.247311136 | 0.084101142 | 2.940639462 | 0.003275355 | 0.026993805 | | Leprel2 | 834.301798 | -0.205965695622991 | 0.0867365200470748 | -2.37461331756458 | 0.0175673429884442 | 0.253409358738968 | NM_013534 |
| Rbmx | 2110.804637 | 0.246918016 | 0.088030362 | 2.804918768 | 0.005032926 | 0.036101925 | | Emd | 1228.207221 | 0.141698385578521 | 0.059699225933081 | 2.37353807128682 | 0.0176185766198183 | 0.253730507684552 | NM_007927 |
| Sez6l2 | 15571.27814 | 0.24669124 | 0.057203884 | 4.312491053 | 1.61E-05 | 0.000471022 | | Zgpat | 2154.25622 | -0.186106601408897 | 0.0784130682470958 | -2.37341307474969 | 0.0176245409785015 | 0.253730507684552 | NM_001048148,NM_144894 |
| Tbc1d30 | 4777.718345 | 0.24652925 | 0.058546292 | 4.210843075 | 2.54E-05 | 0.000678356 | | Pspc1 | 1634.801999 | 0.132605756484295 | 0.0558810648268822 | 2.37299981478706 | 0.0176442727698685 | 0.253763075500108 | NM_025682 |
| Cbx7 | 2483.46923 | 0.246002687 | 0.085689616 | 2.870857627 | 0.004093599 | 0.031523964 | | Cog7 | 2543.80885 | 0.104788526496933 | 0.0441880646563142 | 2.37142149836064 | 0.0177198104369128 | 0.254345816607307 | NM_001033318 |
| Trhde | 1056.963841 | 0.245531176 | 0.087230119 | 2.814752272 | 0.004881485 | 0.035431587 | | Nedd9 | 473.9676271 | 0.1907898881526 | 0.080443031076714 | 2.37173917490323 | 0.0177045838091194 | 0.254345816607307 | NM_017464,NM_001111324 |
| Sf3a2 | 1214.065511 | 0.24526687 | 0.080400819 | 3.050551878 | 0.002284212 | 0.021078966 | | Atp5b | 45294.39078 | 0.0904544226891126 | 0.0382025747088431 | 2.36775723569684 | 0.0178962744133374 | 0.255043173831708 | NM_016774 |
| Gatad2a | 1263.692408 | 0.245047978 | 0.082698971 | 2.963132121 | 0.003045258 | 0.025559714 | | Etv5 | 5824.668028 | 0.194367288064697 | 0.0820856256253319 | 2.36786022624544 | 0.0178912936319742 | 0.255043173831708 | NM_023794 |
| Tmem209 | 1177.039335 | 0.244813797 | 0.077653752 | 3.152633191 | 0.00161805 | 0.016318843 | | Hspa8 | 89369.25306 | 0.118776390537503 | 0.0501520727395084 | 2.36832465837318 | 0.0178688480632672 | 0.255043173831708 | NM_031165 |
| Arsa | 1495.119009 | 0.244375212 | 0.071716055 | 3.407538415 | 0.000655517 | 0.008270829 | | Kif19a | 106.3604411 | -0.220166404755355 | 0.0929796429696408 | -2.36789901233803 | 0.0178894181916977 | 0.255043173831708 | NM_001102615 |
| Tarbp2 | 596.6402171 | 0.244276449 | 0.090406132 | 2.701989845 | 0.006892587 | 0.04487205 | | Magee1 | 7361.492358 | 0.12482000768647 | 0.0526980622001703 | 2.36858818854381 | 0.0178561228718269 | 0.255043173831708 | NM_053201 |
| Brsk2 | 8256.83612 | 0.243474526 | 0.090319628 | 2.695698942 | 0.007024112 | 0.045593461 | | Psmd12 | 2716.191407 | 0.123303793031325 | 0.0520819172051094 | 2.36749719764979 | 0.0179088556593929 | 0.255043173831708 | NM_025894 |
| Ctdp1 | 671.3858532 | 0.242888522 | 0.085145105 | 2.852642235 | 0.00433574 | 0.032878466 | | Sh3bp4 | 947.887031 | 0.145705580356831 | 0.0614978715821176 | 2.36927842555123 | 0.0178228307342628 | 0.255043173831708 | NM_133816 |
| Zfp740 | 2700.0881 | 0.242882392 | 0.079665021 | 3.048795925 | 0.002297605 | 0.021168869 | | Zfp64 | 500.9764408 | 0.169780828692039 | 0.0716540199553732 | 2.36945056400968 | 0.0178145364739877 | 0.255043173831708 | NM_009564 |
| Camta2 | 13166.71322 | 0.242246741 | 0.085970848 | 2.817777741 | 0.004835727 | 0.035192464 | | Zcchc8 | 1090.380134 | -0.135166485509343 | 0.0571240778257349 | -2.36619111684722 | 0.0179721641542619 | 0.255694080807844 | NM_027494 |
| Gng2 | 7105.148466 | 0.242130475 | 0.062533033 | 3.872041123 | 0.000107928 | 0.002067036 | | 9430008C03Rik | 354.7296345 | -0.187910958397108 | 0.0794795617311152 | -2.36426767214474 | 0.018065754504711 | 0.25627260735882 | NR_015463,NR_027887 |
| Tub | 9202.683926 | 0.241341825 | 0.064192826 | 3.759638582 | 0.000170159 | 0.002992558 | | Mylip | 581.7375014 | -0.171020023600475 | 0.0723188029442048 | -2.36480716823294 | 0.0180394609029368 | 0.25627260735882 | NM_153789 |
| Rhov | 270.874117 | 0.241235283 | 0.088902058 | 2.713494937 | 0.006657759 | 0.043784088 | | Ptpn2 | 1278.009567 | -0.167783785263027 | 0.0709597734520322 | -2.36449155769144 | 0.0180548388481074 | 0.25627260735882 | NM_001127177,NM_008977 |
| Stx17 | 1127.471374 | 0.241221591 | 0.07763604 | 3.107082619 | 0.001889436 | 0.018346874 | | Atf7 | 345.4430936 | -0.190178972276744 | 0.0804924367314133 | -2.36269368899008 | 0.0181426581581016 | 0.256569933674424 | NM_146065 |
| Atg4c | 1520.340315 | 0.240916624 | 0.071714122 | 3.359402797 | 0.000781111 | 0.009403681 | | Fkbp4 | 4651.812433 | 0.119294613124948 | 0.0504973541576204 | 2.36239333951213 | 0.0181573655388482 | 0.256569933674424 | NM_010219 |
| Ercc2 | 1265.212997 | 0.240822727 | 0.073099686 | 3.294442686 | 0.000986171 | 0.011259931 | | Gm4951 | 23.85016393 | -0.122136413858376 | 0.05168779370453 | -2.36296435008547 | 0.0181294134823545 | 0.256569933674424 | NM_001033767 |
| Cstf2 | 3899.973998 | 0.240490761 | 0.068994133 | 3.485669724 | 0.000490907 | 0.006749647 | | Gpd1l | 6436.086605 | 0.103736099569655 | 0.0439014748729581 | 2.36292971636708 | 0.0181311077948209 | 0.256569933674424 | NM_175380 |
| Cyth1 | 3261.713684 | 0.240275061 | 0.066457323 | 3.615479101 | 0.000299793 | 0.004663642 | | Fzd6 | 421.1218243 | -0.185467300143433 | 0.0785738626358809 | -2.36041978746682 | 0.0182542655057979 | 0.257188613712144 | NM_001162494,NM_008056 |
| Kcnk9 | 560.879223 | 0.239946381 | 0.085429878 | 2.808693941 | 0.004974291 | 0.035830621 | | Nap1l2 | 3749.046765 | 0.152168167523427 | 0.0644645195816889 | 2.3604948661814 | 0.0182505709277829 | 0.257188613712144 | NM_008671 |
| Map2k7 | 3612.361853 | 0.239217009 | 0.06013337 | 3.978107512 | 6.95E-05 | 0.001482515 | | Rims3 | 2333.013331 | 0.143768015118026 | 0.0608900172418028 | 2.36110977842399 | 0.0182203361005502 | 0.257188613712144 | NM_182929 |
| Mccc1 | 851.0861434 | 0.23914193 | 0.090030254 | 2.656239655 | 0.007901744 | 0.049768137 | | Impdh2 | 1860.535499 | 0.129620633526751 | 0.0549340139783278 | 2.35956967531788 | 0.0182961447148413 | 0.257279572243742 | NM_011830 |
| Dgkq | 1906.872318 | 0.238881648 | 0.089903271 | 2.657096295 | 0.007881693 | 0.049662036 | | Tmem14c | 1093.348618 | 0.142547527818846 | 0.0604104055385483 | 2.35965189354482 | 0.0182920907144763 | 0.257279572243742 | NM_025387 |
| Slc9a8 | 1546.263877 | 0.238460248 | 0.074351435 | 3.20720436 | 0.001340317 | 0.014194722 | | Pkn3 | 64.49379296 | -0.224212153632494 | 0.0951027260736153 | -2.3575786193441 | 0.0183945597721605 | 0.25841332229246 | NM_153805 |
| Gls2 | 932.0295382 | 0.237450278 | 0.082450957 | 2.879897153 | 0.003978049 | 0.030957254 | | Eps15l1 | 2490.98124 | -0.107191815841494 | 0.045486402993002 | -2.35656831027033 | 0.018444674892138 | 0.258617130775286 | NM_007944,NM_001122832 |
| Fan1 | 408.7100891 | 0.237423189 | 0.086461258 | 2.746006629 | 0.006032554 | 0.04097694 | | Fam103a1 | 2499.872498 | 0.134459620806814 | 0.0570499262653145 | 2.35687632936632 | 0.0184293833419582 | 0.258617130775286 | NM_025997 |
| Vps33b | 799.5557611 | 0.236931992 | 0.065139876 | 3.637280355 | 0.000275532 | 0.004347345 | | Rxra | 1934.120258 | 0.146048689746017 | 0.0620517412296862 | 2.35365981440253 | 0.0185896149799023 | 0.260398020441717 | NM_011305 |
| Trpm7 | 1925.682531 | 0.236438089 | 0.077825514 | 3.038053689 | 0.002381115 | 0.021727326 | | Ctnnal1 | 770.8452213 | -0.189853606250775 | 0.0806801310153125 | -2.35316432759315 | 0.0186144058448991 | 0.260494084106941 | NM_018761 |
| Kdm5c | 2944.571708 | 0.236430452 | 0.072172651 | 3.275900894 | 0.001053255 | 0.011851224 | | Cdkn2aipnl | 1661.738224 | 0.134983512626706 | 0.0573742781978369 | 2.3526834126132 | 0.0186384952909033 | 0.260580156492456 | NM_029976 |
| Zc3h7a | 1314.597311 | 0.236248833 | 0.067355606 | 3.507485808 | 0.000452362 | 0.006350895 | | Celf3 | 5991.565073 | -0.172116957670502 | 0.0731865070676974 | -2.35175805714152 | 0.0186849238874228 | 0.260978081142984 | NM_172434 |
| Proser1 | 940.6849074 | 0.236081856 | 0.070936165 | 3.32808881 | 0.00087444 | 0.01028745 | | Ankrd55 | 171.9758203 | 0.223153252746354 | 0.0949163906462407 | 2.35105076401461 | 0.0187204796619616 | 0.261184487217208 | NM_001168403,NM_001168405,NM_001168404,NM_029898 |
| Cep85l | 382.2132178 | 0.2356606 | 0.086770658 | 2.715901957 | 0.006609549 | 0.043599054 | | Psmd2 | 9939.602825 | 0.101879367248479 | 0.0433391073038569 | 2.35074909444228 | 0.0187356626518196 | 0.261184487217208 | NR_027485,NM_134101 |
| Mfsd11 | 1225.161794 | 0.235396574 | 0.086302759 | 2.727567202 | 0.006380326 | 0.042645781 | | Cmtm6 | 1212.866882 | -0.147202535451953 | 0.0626415042371589 | -2.34992018861246 | 0.0187774368494375 | 0.261433927127791 | NM_026036 |
| Safb | 3661.919541 | 0.235238504 | 0.069361439 | 3.391488212 | 0.000695142 | 0.008664942 | | Mms19 | 1677.284714 | -0.144872720530506 | 0.0616563590027473 | -2.34968011205545 | 0.0187895511442526 | 0.261433927127791 | NM_028152 |
| Gpatch1 | 978.0450182 | 0.235212516 | 0.065509236 | 3.590524512 | 0.000330013 | 0.005042628 | | Nfrkb | 1409.710376 | -0.148193526093551 | 0.0630897912579156 | -2.34893036002806 | 0.0188274277526501 | 0.261710254100474 | NM_172766 |
| Tstd2 | 645.6827232 | 0.234716636 | 0.069733785 | 3.36589553 | 0.000762956 | 0.009220936 | | Anks6 | 524.2733561 | -0.179462018700703 | 0.076429152546361 | -2.34808332582052 | 0.0188702992697027 | 0.261805126257594 | NM_001024136 |
| Gtpbp2 | 2522.137033 | 0.234139806 | 0.07417936 | 3.156401012 | 0.001597291 | 0.016186021 | | Mtf2 | 1398.251006 | -0.172207721065448 | 0.0733299161202982 | -2.34839653686416 | 0.0188544365687338 | 0.261805126257594 | NM_013827,NM_001253878,NM_001253879,NM_001253880,NR_045608,NM_001253877 |
| Mtfmt | 384.2911971 | 0.233899175 | 0.083234456 | 2.810124401 | 0.004952235 | 0.035738209 | | Slc40a1 | 453.1770367 | -0.174174253586983 | 0.0742287383530572 | -2.34645310497601 | 0.0189530510792179 | 0.262451877956834 | NM_016917 |
| Ddx17 | 16513.67643 | 0.233648534 | 0.082148333 | 2.844227318 | 0.004451927 | 0.033404246 | | Ticam2 | 32.49029428 | -0.205288105839418 | 0.0874869417034261 | -2.34649996722172 | 0.0189506678772784 | 0.262451877956834 | NM_173394 |
| Sprtn | 370.1995883 | 0.232885513 | 0.082270359 | 2.830734143 | 0.004644131 | 0.034306491 | | Col5a3 | 202.3335351 | -0.215704427654255 | 0.0919666941798008 | -2.34546244787857 | 0.0190034927858221 | 0.26289974876843 | NM_016919 |
| Dus3l | 2188.34161 | 0.232743448 | 0.083147916 | 2.799149493 | 0.005123741 | 0.036432879 | | Atg16l2 | 655.6892323 | -0.211363081123276 | 0.0901509833288843 | -2.34454548712122 | 0.0190502866308563 | 0.263085246439164 | NM_001111111 |
| Plcb4 | 2109.860872 | 0.23238591 | 0.077329999 | 3.005119805 | 0.002654763 | 0.023304754 | | Vegfc | 117.5595011 | -0.218659147211809 | 0.0932651245566274 | -2.34448995003536 | 0.0190531240020653 | 0.263085246439164 | NM_009506 |
| St5 | 707.499199 | 0.232323052 | 0.085316581 | 2.723070358 | 0.006467828 | 0.042962976 | | Skp2 | 302.0220928 | -0.206651106113488 | 0.0881762481222983 | -2.34361418765367 | 0.0190979152848876 | 0.263453292904347 | NM_145468,NM_013787 |
| Dnm1 | 49898.31488 | 0.232188091 | 0.075826825 | 3.062083799 | 0.002198019 | 0.020565282 | | 9430015G10Rik | 974.8670684 | -0.159464624793503 | 0.0681511340676647 | -2.33986751614693 | 0.0192905815099179 | 0.263564471397747 | NM_177205,NM_145557 |
| Ttc13 | 995.6755259 | 0.232179889 | 0.06217734 | 3.734156014 | 0.000188346 | 0.003253321 | | Abhd14b | 652.3893308 | -0.207141516832843 | 0.088606061562451 | -2.33778043149843 | 0.0193986412404308 | 0.263564471397747 | NM_029631 |
| Xrcc1 | 933.5953986 | 0.232040905 | 0.084545205 | 2.744577955 | 0.006058876 | 0.041047761 | | Adap2 | 213.3738463 | -0.211128764792879 | 0.0901588919090056 | -2.34170090122984 | 0.0191940345608524 | 0.263564471397747 | NM_172133 |
| Wrn | 463.7214933 | 0.231922385 | 0.07392145 | 3.137416595 | 0.001704438 | 0.017004866 | | Atp6v0d1 | 9892.060536 | 0.0998818140678757 | 0.0426343353882061 | 2.342755273617 | 0.0191419342377346 | 0.263564471397747 | NM_013477 |
| Ogdhl | 4711.592687 | 0.23130974 | 0.072653394 | 3.183743083 | 0.00145384 | 0.015115049 | | C1ra | 173.9585403 | -0.217801430075417 | 0.0931816381275704 | -2.3373857173152 | 0.0194191371078739 | 0.263564471397747 | NM_023143 |
| Edc4 | 2919.170175 | 0.230998986 | 0.062864633 | 3.674545978 | 0.000238273 | 0.003869809 | | Cyfip1 | 3776.551507 | -0.101067798948432 | 0.0431510591091846 | -2.34218582428537 | 0.0191711671258457 | 0.263564471397747 | NM_011370,NM_001164662,NM_001164661 |
| Zfp207 | 3445.073966 | 0.230304199 | 0.074268152 | 3.100981933 | 0.001928801 | 0.01865818 | | Eif4a3 | 1726.310567 | 0.126008245477097 | 0.0538506767061797 | 2.34133464021084 | 0.0192149357273211 | 0.263564471397747 | NM_138669 |
| Ganc | 367.6424506 | 0.228980041 | 0.084674312 | 2.704244501 | 0.00684599 | 0.044661027 | | Eva1a | 351.3101482 | 0.186167034670445 | 0.0795486871099236 | 2.34029047409912 | 0.0192687468939404 | 0.263564471397747 | NM_145570 |
| Zfyve27 | 3183.596224 | 0.228932052 | 0.062710735 | 3.650603863 | 0.000261624 | 0.004183292 | | Fbxo7 | 1136.231077 | 0.1168267832121 | 0.0499937594089406 | 2.33682732791661 | 0.0194481642643562 | 0.263564471397747 | NM_153195 |
| Sugp1 | 1827.14392 | 0.228542198 | 0.068476659 | 3.337519681 | 0.000845297 | 0.009997738 | | Hivep1 | 3982.196423 | 0.18302904816052 | 0.0782855419988437 | 2.33796743929068 | 0.0193889373035152 | 0.263564471397747 | NM_007772 |
| Gys1 | 730.880899 | 0.228230043 | 0.085740874 | 2.661858129 | 0.007771062 | 0.049168195 | | Klf9 | 7912.969091 | 0.18389710023636 | 0.078664963227249 | 2.3377256238602 | 0.0194014860428215 | 0.263564471397747 | NM_010638 |
| Wash | 1139.960495 | 0.22803636 | 0.081683285 | 2.79171387 | 0.00524297 | 0.037025785 | | Me2 | 957.4345347 | 0.159546714521197 | 0.0681724996551669 | 2.34033833772009 | 0.0192662773620224 | 0.263564471397747 | NM_145494 |
| Zc3h3 | 897.6908638 | 0.227819291 | 0.076392928 | 2.982203927 | 0.002861813 | 0.024524851 | | Pafah1b3 | 323.2204406 | 0.213364331003995 | 0.0910931985470908 | 2.34226412517172 | 0.0191671452088433 | 0.263564471397747 | NM_008776 |
| Iffo1 | 1770.609293 | 0.22770364 | 0.080847799 | 2.816448229 | 0.004855787 | 0.035305284 | | Pgrmc1 | 9221.06108 | 0.133126373528404 | 0.0569147685832168 | 2.33904796316189 | 0.0193329513405019 | 0.263564471397747 | NM_016783 |
| Ddx55 | 766.3982168 | 0.227493134 | 0.080543267 | 2.824483568 | 0.004735688 | 0.03467463 | | Ssh3 | 1986.830878 | -0.201229728905355 | 0.0861141737160599 | -2.33677825869711 | 0.0194507168758354 | 0.263564471397747 | NM_198113 |
| Kremen1 | 1244.462007 | 0.227298875 | 0.083946491 | 2.707663792 | 0.006775863 | 0.044353703 | | Tnfrsf1b | 251.3004539 | -0.189802527858741 | 0.0810900119560773 | -2.34064002804128 | 0.019250717962584 | 0.263564471397747 | NM_011610 |
| Ggt7 | 5111.810648 | 0.226939944 | 0.070575087 | 3.215581472 | 0.001301805 | 0.013850598 | | Vill | 370.5896287 | -0.199560075739206 | 0.0853114349212105 | -2.3391942419385 | 0.0193253829599485 | 0.263564471397747 | NM_011700,NM_001164567 |
| Exosc8 | 446.6885192 | 0.226851198 | 0.079861179 | 2.840569117 | 0.004503311 | 0.03360928 | | Ywhag | 50321.20208 | 0.0901598368503808 | 0.0385451869344029 | 2.33906861066252 | 0.0193318828936115 | 0.263564471397747 | NM_018871 |
| Phf20l1 | 1521.505311 | 0.226698132 | 0.070068523 | 3.235377646 | 0.001214819 | 0.013227274 | | Zfp189 | 436.5412318 | 0.222156216902714 | 0.0949946688371481 | 2.33861773110196 | 0.0193552263160534 | 0.263564471397747 | NM_145547 |
| Trim3 | 4643.726712 | 0.226311266 | 0.081676089 | 2.77083869 | 0.005591212 | 0.038865068 | | Cers4 | 2097.65404 | -0.142417984002408 | 0.0609831595729562 | -2.33536577966297 | 0.0195243205097464 | 0.263578326881576 | NM_026058 |
| 5031439G07Rik | 5181.112955 | 0.225677148 | 0.065687217 | 3.435632641 | 0.000591172 | 0.007729636 | | Stat1 | 1443.514868 | -0.206146703154507 | 0.088245181603888 | -2.3360675269484 | 0.0194877224237643 | 0.263578326881576 | NM_001205314,NM_009283,NM_001205313 |
| Hdgfrp2 | 2852.160487 | 0.225591066 | 0.057950236 | 3.892841224 | 9.91E-05 | 0.001945626 | | Vdac1 | 18000.96992 | 0.0951004423367408 | 0.0407125440919075 | 2.33590026017667 | 0.0194964404097862 | 0.263578326881576 | NM_011694 |
| Ezh1 | 3022.571278 | 0.225525074 | 0.080884039 | 2.788251876 | 0.005299332 | 0.037321753 | | Zwint | 17308.64718 | 0.136042767822794 | 0.0582460181346124 | 2.33565782142199 | 0.0195090824263595 | 0.263578326881576 | NM_025635 |
| Scrib | 1988.240606 | 0.225349352 | 0.080461449 | 2.800712074 | 0.005098999 | 0.036290258 | | Cbln1 | 372.7143689 | 0.177400324800057 | 0.0760032507578857 | 2.33411496259785 | 0.0195897030272372 | 0.263970339678708 | NM_019626 |
| F3 | 1375.846653 | 0.224962826 | 0.081913394 | 2.746349731 | 0.006026249 | 0.040952059 | | Snx1 | 2966.863467 | 0.10132060232511 | 0.0434055277306116 | 2.3342787802037 | 0.0195811291094064 | 0.263970339678708 | NM_019727 |
| B930041F14Rik | 1093.17697 | 0.224360183 | 0.080032062 | 2.803378746 | 0.005057024 | 0.036224473 | | Gm216 | 116.3069484 | -0.216234035043704 | 0.0926691113695483 | -2.33339925081832 | 0.0196272005611713 | 0.264230505423146 | NM_001033345 |
| Ik | 4229.998476 | 0.224063645 | 0.075462476 | 2.969206099 | 0.002985703 | 0.025306068 | | D930016D06Rik | 481.674099 | -0.175071852477182 | 0.0750593444430657 | -2.33244579707166 | 0.0196772512355178 | 0.264659029117714 | NR_030673 |
| Htra2 | 905.5619878 | 0.22396659 | 0.07662938 | 2.922724792 | 0.003469831 | 0.028073919 | | Nbeal1 | 611.8424663 | -0.201808356797679 | 0.0865745500187351 | -2.33103558440681 | 0.0197514833962058 | 0.265411700104797 | NM_173444 |
| Tnrc6a | 3577.033996 | 0.223813904 | 0.07845248 | 2.852859521 | 0.004332777 | 0.032875635 | | Apobec1 | 73.84144659 | -0.220751979315239 | 0.0947296632489218 | -2.33033636713315 | 0.019788380081137 | 0.26566174589519 | NM_031159,NM_001134391 |
| Ift140 | 1236.615601 | 0.223714689 | 0.064550668 | 3.465722299 | 0.000528809 | 0.007115035 | | Aox1 | 292.4837882 | -0.218104120217424 | 0.0936292458271951 | -2.32944437702685 | 0.01983553645875 | 0.266048940535367 | NM_009676 |
| Manba | 471.8620065 | 0.223542534 | 0.074848785 | 2.986588656 | 0.00282109 | 0.024320521 | | Zfp458 | 183.5567779 | 0.203235486220101 | 0.087266402582768 | 2.3289087232322 | 0.0198639017156113 | 0.266183612842223 | NM_001001152 |
| Spns1 | 2129.241976 | 0.222958604 | 0.076140246 | 2.928262173 | 0.003408625 | 0.027759173 | | Prkar1a | 28417.63793 | 0.110723876581267 | 0.0475570122046678 | 2.32823450103957 | 0.0198996551229201 | 0.266416949599573 | NM_021880 |
| Tpra1 | 969.257986 | 0.222889506 | 0.082956033 | 2.686839 | 0.007213171 | 0.046567031 | | Inppl1 | 1724.954143 | -0.128860974698974 | 0.0553931206078788 | -2.32629924591477 | 0.0200025922340828 | 0.267548485075771 | NM_010567,NM_001122739 |
| Mat2a | 7325.374297 | 0.222587644 | 0.073450778 | 3.030432743 | 0.002442036 | 0.022066759 | | Lrrc8a | 2487.619988 | 0.157532817252428 | 0.067742060954155 | 2.3254801379462 | 0.0200463007627051 | 0.26764022507266 | NM_177725 |
| Wipf1 | 531.6603312 | 0.222518892 | 0.069193912 | 3.21587384 | 0.00130048 | 0.013849883 | | Mybl2 | 37.75923922 | -0.204187475629373 | 0.0877972477914434 | -2.32567057357432 | 0.0200361314714927 | 0.26764022507266 | NM_008652 |
| Rps6kb1 | 1808.931475 | 0.222070932 | 0.07640764 | 2.906396933 | 0.003656173 | 0.0291553 | | Mical1 | 506.4218499 | -0.191568802245502 | 0.082393571517239 | -2.32504549466484 | 0.0200695276548467 | 0.267704277974567 | NM_001164433,NM_138315 |
| Kdsr | 1281.022261 | 0.221775382 | 0.073892244 | 3.00133505 | 0.002687986 | 0.023503194 | | Tusc2 | 3378.707365 | 0.121220339203136 | 0.052188826604052 | 2.32272589922005 | 0.0201938820282653 | 0.268869230378168 | NM_019742 |
| Setdb1 | 1855.798707 | 0.221693616 | 0.078988609 | 2.806652981 | 0.005005914 | 0.035980333 | | Txn1 | 1573.28178 | 0.118309257884446 | 0.0509321648993186 | 2.32287903171437 | 0.0201856518594437 | 0.268869230378168 | NM_011660 |
| Srsf2 | 3193.134811 | 0.221522957 | 0.078272233 | 2.830160163 | 0.004652471 | 0.034310034 | | Zkscan7 | 136.0297037 | -0.213781002949407 | 0.0920629158290517 | -2.32211853192191 | 0.0202265540776427 | 0.269057623197654 | NM_001177505 |
| Coro2b | 8388.601296 | 0.22104682 | 0.064835704 | 3.409337861 | 0.000651208 | 0.008237633 | | Naip2 | 76.808045 | -0.216174550210054 | 0.0931176057289623 | -2.32152178439031 | 0.0202586997687774 | 0.269238675975536 | NM_010872,NM_001126182 |
| Serac1 | 1140.182458 | 0.220950735 | 0.064392388 | 3.431317635 | 0.000600657 | 0.007807531 | | Ppm1g | 2862.781001 | 0.106067153235933 | 0.0457103624232901 | 2.32041811993795 | 0.0203182696056451 | 0.269537154604201 | NM_008014 |
| Pusl1 | 564.044989 | 0.220525598 | 0.081233306 | 2.71471898 | 0.006633203 | 0.043659664 | | Susd1 | 518.1174264 | 0.157770197490881 | 0.0679826967307284 | 2.32074049836226 | 0.020300853583637 | 0.269537154604201 | NM_001163288 |
| Plod3 | 1106.202768 | 0.220450842 | 0.077470988 | 2.845592247 | 0.004432892 | 0.033344824 | | Bst2 | 159.5387085 | -0.177382465030552 | 0.0764800598762237 | -2.31932957842385 | 0.0203771728130579 | 0.269758294844772 | NM_198095 |
| Paf1 | 2096.52074 | 0.220235406 | 0.080999456 | 2.718973891 | 0.006548478 | 0.043304359 | | Crebl2 | 1521.161969 | 0.118802798117096 | 0.0512283981188092 | 2.31908087076172 | 0.0203906517788489 | 0.269758294844772 | NM_177687 |
| Eif2ak4 | 1107.369832 | 0.220100668 | 0.069767968 | 3.154752443 | 0.001606343 | 0.016224697 | | Doc2a | 1717.993374 | 0.1207893536331 | 0.0520710407896435 | 2.31970307874322 | 0.0203569451822159 | 0.269758294844772 | NM_010069 |
| Tcerg1 | 2513.03226 | 0.220091098 | 0.059400563 | 3.705202246 | 0.000211222 | 0.003545694 | | Fam228a | 63.44013144 | -0.219266962824665 | 0.0946289224580061 | -2.31712416383025 | 0.0204969690669288 | 0.269900833579386 | NM_029107 |
| Abca7 | 1145.326111 | 0.219920398 | 0.081860181 | 2.686536907 | 0.007219697 | 0.046578655 | | Gna15 | 79.21182506 | -0.217858408009705 | 0.09404610986719 | -2.31650632139235 | 0.0205306396904878 | 0.269900833579386 | NM_010304 |
| Rbm39 | 5548.235615 | 0.219635653 | 0.072526501 | 3.028350344 | 0.002458928 | 0.022123355 | | Lclat1 | 1904.883151 | 0.129651416124231 | 0.0559419057389788 | 2.31760814029425 | 0.0204706273987698 | 0.269900833579386 | NM_001177968,NM_001081071,NM_001177967 |
| Atp13a1 | 2066.6212 | 0.219274921 | 0.073294217 | 2.99170833 | 0.002774212 | 0.024026036 | | Lrrfip1 | 2215.43964 | -0.115284694660585 | 0.0497669569760079 | -2.31649073332256 | 0.0205314898186164 | 0.269900833579386 | NM_001111312,NM_008515,NM_001111311 |
| Bnip2 | 1212.504127 | 0.218505437 | 0.070213427 | 3.112017843 | 0.001858133 | 0.01810686 | | Myef2 | 2318.968179 | -0.110753340161021 | 0.0477773350344811 | -2.31811464748066 | 0.0204430910615634 | 0.269900833579386 | NM_001162417,NM_010852,NM_001162418 |
| Bak1 | 434.0666152 | 0.218256932 | 0.073438168 | 2.971982257 | 0.002958838 | 0.025129358 | | Nbeal2 | 664.2823081 | -0.198671111412289 | 0.085752901721602 | -2.3167858745734 | 0.0205153988814316 | 0.269900833579386 | NM_183276 |
| Edem1 | 1032.399806 | 0.218055878 | 0.057862968 | 3.768487576 | 0.00016424 | 0.00291159 | | Spryd4 | 286.3731754 | 0.180229431729016 | 0.0777383396012434 | 2.31841112961117 | 0.0204269877618569 | 0.269900833579386 | NM_025716 |
| Eps15l1 | 2228.235249 | 0.217315122 | 0.060660209 | 3.582498747 | 0.000340323 | 0.00515946 | | Slc38a1 | 9125.924335 | 0.133876259439843 | 0.0578226010846248 | 2.31529292921139 | 0.0205969064594586 | 0.270515970370792 | NM_001166456,NM_134086,NM_001166458 |
| Yeats2 | 1368.08169 | 0.217197597 | 0.064101941 | 3.388315421 | 0.000703233 | 0.008740529 | | Col4a5 | 188.6446047 | -0.219649004583552 | 0.0948935240627334 | -2.31468908709032 | 0.0206299534268697 | 0.270705242528192 | NM_007736,NM_001163155 |
| Msh2 | 1276.275588 | 0.216356276 | 0.077517671 | 2.791057498 | 0.005253614 | 0.037084052 | | Caprin2 | 309.0361386 | -0.188744373812994 | 0.0815661071220076 | -2.31400492769218 | 0.02066745184038 | 0.270708210489955 | NM_181541 |
| Kcnc1 | 8453.316143 | 0.21608876 | 0.053106579 | 4.068964044 | 4.72E-05 | 0.001095311 | | Sh3bp1 | 682.4073089 | -0.193312918570548 | 0.0835281803913585 | -2.31434370609787 | 0.0206488761546491 | 0.270708210489955 | NM_009164 |
| Zfp385a | 2832.997442 | 0.215414447 | 0.066069187 | 3.260437353 | 0.001112405 | 0.012381903 | | Clec2d | 152.6503099 | -0.21729818880316 | 0.0939467400254929 | -2.31299339119372 | 0.0207230025000905 | 0.270947195604243 | NM_053109 |
| Usp36 | 1634.107729 | 0.215036903 | 0.067326259 | 3.19395294 | 0.00140339 | 0.014691975 | | Wwox | 770.6776338 | 0.135622578790947 | 0.0586326360521643 | 2.31309025011746 | 0.020717677658609 | 0.270947195604243 | NM_019573 |
| Hs1bp3 | 669.1468416 | 0.215010552 | 0.078828054 | 2.727589243 | 0.0063799 | 0.042645781 | | Pros1 | 612.7332853 | -0.163731236009667 | 0.0708151213316965 | -2.31209426645975 | 0.0207724890585292 | 0.271349978474995 | NM_011173 |
| Slc9a1 | 2588.288683 | 0.214714003 | 0.067898202 | 3.162292891 | 0.00156532 | 0.015966474 | | Proser1 | 1061.334287 | -0.147291777555909 | 0.0637145167452534 | -2.31174597376009 | 0.0207916862610361 | 0.271356724732983 | NM_173382 |
| Rufy1 | 1369.765958 | 0.214346916 | 0.060603036 | 3.536900636 | 0.000404852 | 0.00587889 | | Tbx18 | 104.8411077 | -0.219614058711921 | 0.0950347994507113 | -2.31088043518018 | 0.0208394600130306 | 0.271736082719285 | NM_023814 |
| Jag2 | 1352.821726 | 0.213584973 | 0.071735592 | 2.977391939 | 0.002907121 | 0.024789727 | | Psmb4 | 3719.480736 | 0.102231238825767 | 0.044261135230095 | 2.3097291177533 | 0.0209031557069624 | 0.272322188160839 | NM_008945 |
| Usp48 | 3692.080673 | 0.213378775 | 0.07539388 | 2.830186946 | 0.004652081 | 0.034310034 | | Kif21b | 4919.488407 | -0.147054036041453 | 0.0637051292025613 | -2.30835472567765 | 0.0209794149411405 | 0.273070771895168 | NM_001039472 |
| Rbfox3 | 11023.49346 | 0.213232468 | 0.074568518 | 2.859550813 | 0.004242414 | 0.032353861 | | Anks3 | 1304.288553 | -0.170196079560576 | 0.0737484420001233 | -2.30779220475318 | 0.0210106967076328 | 0.27323310687115 | NM_028301 |
| Nckap5l | 954.9283759 | 0.212273195 | 0.070586979 | 3.007257129 | 0.002636167 | 0.023198299 | | Clcn1 | 95.75726186 | -0.219363562546811 | 0.0950832353133283 | -2.3070687679478 | 0.0210509867444707 | 0.273267768945649 | NM_013491 |
| Sparcl1 | 82107.6409 | 0.212215797 | 0.054732908 | 3.877298052 | 0.000105623 | 0.002043399 | | Trpm7 | 2099.75254 | -0.150879069600304 | 0.0653896007394452 | -2.30738631057719 | 0.021033293705537 | 0.273267768945649 | NM_021450,NM_001164325 |
| Dock9 | 5818.375937 | 0.212110072 | 0.067574082 | 3.138926435 | 0.00169568 | 0.016939309 | | 1700001L19Rik | 361.0556128 | 0.189858647297695 | 0.0823384866085748 | 2.30583115038604 | 0.0211200689172111 | 0.273675397940596 | NM_027035 |
| Dennd4b | 2263.169587 | 0.211943385 | 0.070554786 | 3.003954747 | 0.00266495 | 0.023367702 | | Rbm28 | 2102.872684 | -0.150913077984973 | 0.0654484138948129 | -2.3058324717772 | 0.0211199950537075 | 0.273675397940596 | NM_133925 |
| Slc11a2 | 2021.013837 | 0.211377605 | 0.054981145 | 3.844547172 | 0.000120775 | 0.002260014 | | Rcan2 | 14593.71055 | 0.14939174762144 | 0.0648008264577639 | 2.30539880102935 | 0.0211442485928418 | 0.273744523226043 | NM_207649,NM_030598 |
| Git2 | 1332.932688 | 0.211367968 | 0.072005818 | 2.935429014 | 0.003330868 | 0.027292548 | | Hdhd2 | 3031.820926 | 0.114299989592774 | 0.0495969119453027 | 2.30457875520211 | 0.0211901769330457 | 0.274094844282655 | NM_001039201,NM_001039202,NM_029826 |
| Otub1 | 6737.247127 | 0.21135559 | 0.074954729 | 2.819776574 | 0.00480571 | 0.035039843 | | Hist4h4 | 36.84747879 | -0.207323973016867 | 0.0899752679292795 | -2.30423290520038 | 0.0212095730078076 | 0.274101652590225 | NM_175652 |
| Bin1 | 8671.158807 | 0.211301844 | 0.072497091 | 2.914625151 | 0.00356116 | 0.028648293 | | 4930447C04Rik | 81.6116268 | -0.21790048085329 | 0.0946597047189579 | -2.30193493102721 | 0.0213388419486623 | 0.27507997339773 | NM_029444 |
| Jakmip3 | 1361.412843 | 0.211097364 | 0.074094036 | 2.84904665 | 0.004385045 | 0.033093954 | | Lsm12 | 608.5827429 | 0.149219262213013 | 0.0648250250500808 | 2.30187743233008 | 0.0213420852278151 | 0.27507997339773 | NM_172947 |
| Rgs7 | 3215.845546 | 0.210484484 | 0.069967202 | 3.008330731 | 0.002626871 | 0.023191304 | | Mnt | 2262.889537 | 0.162443340018237 | 0.0705514207970538 | 2.30248148347737 | 0.0213080344641594 | 0.27507997339773 | NM_010813 |
| Pxk | 1437.086382 | 0.210355912 | 0.076335768 | 2.755666405 | 0.00585727 | 0.040157049 | | Hgf | 279.7099871 | -0.195464787567965 | 0.0849281774542776 | -2.30153046288078 | 0.0213616655456853 | 0.275088256840979 | NM_010427 |
| Snx32 | 2891.815261 | 0.210202736 | 0.063835362 | 3.292888609 | 0.000991638 | 0.011314015 | | Fancm | 612.4370763 | -0.173328744158396 | 0.0753229644799132 | -2.30114076570392 | 0.0213836757425026 | 0.275127780948355 | NM_178912 |
| Osbpl5 | 1134.127178 | 0.209682546 | 0.069256678 | 3.027614856 | 0.00246492 | 0.022127158 | | Mtmr11 | 396.5878766 | -0.187095526456687 | 0.0813363831231806 | -2.30026857935567 | 0.0214330086272222 | 0.27527487472947 | NM_181409 |
| Arhgef10 | 1900.040938 | 0.209320979 | 0.074228073 | 2.819970525 | 0.004802806 | 0.035035159 | | Rnmtl1 | 409.8530154 | 0.169722874449405 | 0.0737758921040317 | 2.3005194462451 | 0.0214188088705454 | 0.27527487472947 | NM_183263 |
| Zfp263 | 793.5984477 | 0.209028455 | 0.07725144 | 2.70581953 | 0.006813606 | 0.044544311 | | 4930470H14Rik | 73.68447443 | -0.218562036291067 | 0.0950501957097076 | -2.29943804596233 | 0.0214800776115713 | 0.275635695570393 | NR_045764 |
| Cyth3 | 1471.431858 | 0.207874083 | 0.053662351 | 3.873741596 | 0.000107177 | 0.002065424 | | Psmd1 | 6504.291334 | 0.108402943605703 | 0.0471532595742277 | 2.29894909884347 | 0.0215078298774626 | 0.275709409060662 | NM_027357 |
| Dnajb2 | 3706.13127 | 0.207468747 | 0.066246137 | 3.131786349 | 0.001737463 | 0.017245513 | | Vdac2 | 7090.679269 | 0.11203403689878 | 0.0487386708700843 | 2.29866828329014 | 0.0215237828634718 | 0.275709409060662 | NM_011695 |
| Slc17a5 | 784.0980704 | 0.206661443 | 0.07717882 | 2.677696326 | 0.007413039 | 0.047350844 | | Psmb2 | 1974.985633 | 0.108862675358161 | 0.0473818902042648 | 2.29755872737137 | 0.0215869169717256 | 0.276274498617873 | NM_011970 |
| Nop56 | 2725.424052 | 0.206619088 | 0.073828409 | 2.798639306 | 0.005131843 | 0.036473748 | | Ccdc3 | 664.3007976 | -0.201469979925111 | 0.0877018734244453 | -2.2972140965572 | 0.0216065593668299 | 0.276282465988179 | NM_028804 |
| Gps2 | 733.7424196 | 0.206245491 | 0.07626673 | 2.704255553 | 0.006845556 | 0.044661027 | | Cabin1 | 4309.508141 | -0.126579350161276 | 0.0551373760586636 | -2.29570863195596 | 0.0216925465701745 | 0.276408711823118 | NM_172549 |
| Atrip | 764.6469971 | 0.206079681 | 0.069092446 | 2.982665886 | 0.002857497 | 0.024507848 | | Fto | 9756.982879 | 0.101122788567881 | 0.0440400886795041 | 2.29615315499904 | 0.021667125935251 | 0.276408711823118 | NM_011936 |
| Pde7a | 829.317474 | 0.205904682 | 0.074217274 | 2.774349846 | 0.005531216 | 0.03865614 | | Parm1 | 1235.622139 | 0.174773573516874 | 0.0761234143659721 | 2.29592399359059 | 0.0216802275919144 | 0.276408711823118 | NM_145562 |
| Map4k5 | 1775.033601 | 0.205323655 | 0.065572567 | 3.131243188 | 0.00174068 | 0.017266382 | | Parp3 | 230.876434 | -0.215076579321417 | 0.0936714305295365 | -2.29607446054322 | 0.02167162429147 | 0.276408711823118 | NM_145619 |
| Ercc5 | 965.5107105 | 0.204674286 | 0.076989841 | 2.658458351 | 0.007849905 | 0.049502007 | | Exoc6b | 5186.665773 | 0.114649083762484 | 0.0499549625147763 | 2.29504893990406 | 0.0217303197747434 | 0.276647348858828 | NM_177077 |
| Brd8 | 2330.470484 | 0.204309109 | 0.063531991 | 3.215846156 | 0.001300605 | 0.013849883 | | 4833420G17Rik | 1452.447021 | -0.139671634814936 | 0.0609142608512738 | -2.29292177009179 | 0.0218525092230192 | 0.277716140834276 | NM_001113550,NM_026127 |
| Ints6 | 789.9246417 | 0.203931576 | 0.074510615 | 2.736946628 | 0.006201235 | 0.041781366 | | Gne | 1724.886544 | 0.120249072220107 | 0.0524360687038712 | 2.29325109971925 | 0.0218335527563188 | 0.277716140834276 | NM_015828,NM_001190414 |
| Kri1 | 1271.519384 | 0.203730929 | 0.065743231 | 3.09888827 | 0.001942483 | 0.018716931 | | Blmh | 2373.356012 | 0.113178973982991 | 0.0494123141103711 | 2.29050138656097 | 0.0219922682909192 | 0.278517601738354 | NM_178645 |
| Mms19 | 1492.99769 | 0.203594273 | 0.074832328 | 2.720672702 | 0.006514923 | 0.043211566 | | Dgka | 506.0286943 | -0.153130675871447 | 0.0668446509736596 | -2.29084412351541 | 0.0219724306519923 | 0.278517601738354 | NM_016811 |
| 9930104L06Rik | 705.0685561 | 0.20319383 | 0.073288245 | 2.77252962 | 0.005562246 | 0.038738759 | | Hhex | 75.35210837 | -0.217404427698969 | 0.0948981077961946 | -2.29092479025896 | 0.0219677639190551 | 0.278517601738354 | NM_008245 |
| Luc7l2 | 4917.28795 | 0.202934333 | 0.058789306 | 3.451891982 | 0.000556671 | 0.00739713 | | Rasl12 | 130.3664543 | -0.210248178433477 | 0.0917900265384337 | -2.29053401946063 | 0.0219903788254942 | 0.278517601738354 | NM_001162923,NM_001033158 |
| Khnyn | 704.1481221 | 0.202875743 | 0.071329134 | 2.8442199 | 0.004452031 | 0.033404246 | | Nxn | 433.9890011 | 0.193309458487747 | 0.0845244974429897 | 2.287022867165 | 0.0221944884031933 | 0.280833744376991 | NM_008750 |
| Edil3 | 4441.239623 | 0.20287005 | 0.072183055 | 2.810494094 | 0.00494655 | 0.035738209 | | Ccdc88c | 1421.006506 | -0.151516004575593 | 0.0662765938944375 | -2.28611634473735 | 0.0222474530305056 | 0.28125892316895 | NM_026681 |
| Atg13 | 4342.891724 | 0.202598679 | 0.064018299 | 3.164699514 | 0.001552432 | 0.015897805 | | Klhdc8a | 760.9004114 | 0.163427876747652 | 0.0715164815300133 | 2.28517781148205 | 0.0223024036969777 | 0.281708448784607 | NM_144810 |
| Cyb561d1 | 1229.815026 | 0.202580603 | 0.071057204 | 2.850950953 | 0.004358869 | 0.032960625 | | 2410089E03Rik | 2455.684771 | -0.150807667696275 | 0.0660093594220168 | -2.28464067848498 | 0.0223339056422888 | 0.281750997682713 | NM_001162906 |
| Ephx1 | 1009.826587 | 0.202508578 | 0.073520981 | 2.754432485 | 0.005879402 | 0.040218743 | | Ano2 | 296.1402496 | 0.215207965138431 | 0.0942759194263566 | 2.282745863927 | 0.0224453424912712 | 0.281750997682713 | NM_153589 |
| Atp8b2 | 3769.512524 | 0.202137081 | 0.061289643 | 3.298062623 | 0.000973544 | 0.011167224 | | Fdps | 2348.381945 | 0.115171196108979 | 0.0504328687369861 | 2.28365347824276 | 0.0223919041788957 | 0.281750997682713 | NM_001253751,NM_134469 |
| Diexf | 728.5598181 | 0.201275567 | 0.067345577 | 2.988697625 | 0.002801692 | 0.024197003 | | Hint1 | 3375.941827 | 0.125351502996424 | 0.0548815997505039 | 2.284035151422 | 0.0223694651694703 | 0.281750997682713 | NM_008248 |
| Trmt1l | 1067.310916 | 0.200470394 | 0.063278687 | 3.168055537 | 0.001534622 | 0.01575708 | | Myl12b | 7901.317927 | 0.115397732859671 | 0.0505173641585603 | 2.28431817023288 | 0.0223528387906862 | 0.281750997682713 | NM_023402 |
| Pprc1 | 1061.371014 | 0.199223196 | 0.074336381 | 2.680022809 | 0.007361714 | 0.047250374 | | Rsad1 | 808.3126686 | -0.154058923342191 | 0.0675026389049614 | -2.28226519498141 | 0.0224736880883492 | 0.281750997682713 | NM_001013381 |
| Fam189b | 1975.149822 | 0.198671218 | 0.066364695 | 2.993628144 | 0.002756817 | 0.023916083 | | Sult4a1 | 18853.22657 | 0.1014906278716 | 0.0444714489813992 | 2.28215248650995 | 0.0224803391377024 | 0.281750997682713 | NM_013873 |
| Gpr108 | 946.1526402 | 0.198046143 | 0.064502297 | 3.070373482 | 0.002137912 | 0.020112213 | | Tbata | 106.2597532 | -0.215486177593611 | 0.0943845535621124 | -2.28306613170347 | 0.0224264731921677 | 0.281750997682713 | NM_001017441,NM_001017419,NM_001017409,NM_001017407,NM_001017433,NM_023064 |
| Repin1 | 1517.461449 | 0.197788166 | 0.071470793 | 2.767398519 | 0.005650563 | 0.039197474 | | Tpcn2 | 141.3275075 | -0.207497647278327 | 0.0908704246281269 | -2.28344533578972 | 0.0224041493593042 | 0.281750997682713 | NM_146206 |
| Zfp286 | 583.5372339 | 0.197153628 | 0.069404051 | 2.840664574 | 0.004501964 | 0.03360928 | | Swi5 | 3509.847136 | 0.111064569593538 | 0.0486794417566019 | 2.28154977924487 | 0.0225159345951296 | 0.281953849938666 | NM_175190 |
| Clcn7 | 2531.298654 | 0.196876359 | 0.058631858 | 3.35783934 | 0.000785543 | 0.009435036 | | 9530068E07Rik | 3415.271996 | 0.0968965268168549 | 0.0425078557418812 | 2.27949693358413 | 0.0226375420448935 | 0.283026790342007 | NM_153117 |
| Ubl7 | 2630.996206 | 0.196757964 | 0.07346692 | 2.678184474 | 0.007402244 | 0.047350844 | | Sft2d3 | 177.8859836 | -0.207180875572633 | 0.0909039818723321 | -2.27911771635706 | 0.0226600686470986 | 0.283026790342007 | NM_026006 |
| Top3b | 1329.62685 | 0.196333337 | 0.05818472 | 3.374310971 | 0.000740007 | 0.009042328 | | Slc35f1 | 3725.802227 | 0.160956453525703 | 0.0706141165512588 | 2.27938068741347 | 0.0226444453348806 | 0.283026790342007 | NM_178675 |
| Ccng1 | 2917.240865 | 0.19538184 | 0.064287684 | 3.039179972 | 0.002372231 | 0.021674644 | | Hook2 | 562.3414342 | -0.176754714157023 | 0.0775885437574122 | -2.27810325593509 | 0.0227204263212058 | 0.283293487332048 | NM_001167991,NM_133255 |
| Map3k11 | 992.9405756 | 0.194722724 | 0.072895191 | 2.671269821 | 0.007556488 | 0.048062489 | | Uba2 | 3645.001772 | 0.118851972152306 | 0.0521710499594752 | 2.27812114658659 | 0.0227193606664969 | 0.283293487332048 | NM_016682 |
| Lonp1 | 2678.236127 | 0.194690895 | 0.065065299 | 2.992238511 | 0.002769398 | 0.023998352 | | Ppp2r5d | 4937.991577 | 0.0879127392794826 | 0.0386146566443189 | 2.27666764175195 | 0.0228060802251885 | 0.284117599786525 | NM_009358 |
| Sf3b1 | 10501.41869 | 0.194511355 | 0.068711812 | 2.830828473 | 0.004642761 | 0.034306491 | | 3632451O06Rik | 1257.520342 | 0.158467355981417 | 0.0696232213847145 | 2.27607043784687 | 0.0228417941017134 | 0.284318681338036 | NM_026142 |
| Zfp239 | 1763.672314 | 0.194291565 | 0.068892145 | 2.820228125 | 0.004798952 | 0.035023534 | | Dact1 | 278.615499 | 0.203910148739424 | 0.0896388126732527 | 2.27479751971624 | 0.0229180791010482 | 0.285023987176221 | NM_021532,NM_001190466 |
| Coq10a | 1147.109385 | 0.192805137 | 0.06648287 | 2.900072399 | 0.003730765 | 0.029658527 | | Arhgap17 | 553.3169907 | -0.156086245198935 | 0.068650558291038 | -2.27363402548339 | 0.0229879999042352 | 0.285162809961452 | NM_001122640,NM_001122642,NM_001122643,NM_144529,NM_001122641 |
| Nprl3 | 968.6154974 | 0.192647555 | 0.062355401 | 3.089508732 | 0.002004878 | 0.019163221 | | Lrrk1 | 249.1810065 | -0.192846503604101 | 0.0848187052116459 | -2.27363177877918 | 0.0229881351001556 | 0.285162809961452 | NM_146191 |
| Map2k5 | 1149.534504 | 0.191064593 | 0.069174379 | 2.762071676 | 0.005743587 | 0.039586803 | | Mapk7 | 811.9406836 | -0.182134111390077 | 0.0801019127567809 | -2.27377980277579 | 0.0229792291996112 | 0.285162809961452 | NM_011841 |
| Rsrc2 | 2248.694519 | 0.188901973 | 0.066869686 | 2.824926866 | 0.004729141 | 0.034663987 | | Adam1a | 223.7027506 | -0.206436075486228 | 0.0909024453339555 | -2.27096284074457 | 0.023149227527555 | 0.286019361930858 | NM_172126 |

| | | | | | | | | | | | | | | |
|---|---|---|---|---|---|---|---|---|---|---|---|---|---|---|
| Sart1 | 2107.474655 | 0.188402257 | 0.06833298 | 2.757120451 | 0.005831287 | 0.040013267 | | Ankrd33b | 4961.022245 | 0.195531919271738 | 0.0860580424331068 | 2.27209350507509 | 0.0230808633210573 | 0.286019361930858 | NM_027496,NM_001164441,NM_026153 |
| Map3k3 | 1139.059462 | 0.187688536 | 0.070081808 | 2.678134922 | 0.007403339 | 0.047350844 | | Chst10 | 3375.405718 | 0.110029237876549 | 0.0484504505533787 | 2.2709641834048 | 0.0231491462411174 | 0.286019361930858 | NM_145142 |
| Nxpe3 | 1012.80732 | 0.187608551 | 0.062665359 | 2.993815961 | 0.002755121 | 0.023914757 | | Mal | 10355.81967 | 0.19178274711993 | 0.0844539075597726 | 2.27085699953191 | 0.0231556360753607 | 0.286019361930858 | NM_001171187,NM_010762 |
| Vps16 | 2089.322466 | 0.187293614 | 0.064380431 | 2.909169947 | 0.003623898 | 0.028942617 | | Sqstm1 | 18338.90914 | 0.0892914936536058 | 0.0393110270262671 | 2.27141085868712 | 0.0231221176780856 | 0.286019361930858 | NM_011018 |
| Cog2 | 1021.066288 | 0.187098248 | 0.063234034 | 2.958821929 | 0.003088175 | 0.025822009 | | Cmas | 3313.379544 | 0.109412609212532 | 0.0481897963580788 | 2.27045178609038 | 0.023180185463586 | 0.286020751576722 | NM_009908 |
| Flii | 2452.874925 | 0.18675676 | 0.049972148 | 3.737217002 | 0.000186068 | 0.003224768 | | Lepre1 | 327.4922715 | -0.200291232666334 | 0.0882260436788865 | -2.27020530803045 | 0.0231951291034957 | 0.286020751576722 | NM_019782,NM_019783,NM_001042411 |
| Vac14 | 1119.321733 | 0.186673281 | 0.069592709 | 2.682368353 | 0.007310292 | 0.047017709 | | Lrrtm1 | 1827.148698 | 0.158737180117791 | 0.0700457067237671 | 2.26619428288144 | 0.0234394905348833 | 0.288788837582455 | NM_028880 |
| Hnrnpu | 13084.91441 | 0.186524227 | 0.055542552 | 3.358222146 | 0.000784455 | 0.009429288 | | Atxn7 | 1333.903391 | -0.16626550714399 | 0.0733821166666074 | -2.26574967712329 | 0.0234667140772485 | 0.288824643867809 | NM_139227 |
| Sncb | 13839.00737 | 0.185278921 | 0.067874788 | 2.729716375 | 0.006338884 | 0.042445806 | | Fdft1 | 4056.040943 | 0.12904342505822 | 0.0569603018297643 | 2.26549756431925 | 0.0234821633214844 | 0.288824643867809 | NM_010191 |
| Sgpl1 | 1627.701312 | 0.185059172 | 0.052154185 | 3.548309155 | 0.000387713 | 0.005699453 | | Tifab | 165.4027612 | -0.214071999466387 | 0.0945266653036228 | -2.26467313512839 | 0.0235327452415037 | 0.289201909795332 | NM_145976,NM_001168615 |
| BC017158 | 831.5444776 | 0.18504505 | 0.065066489 | 2.843937849 | 0.004455974 | 0.033417647 | | Chi3l1 | 386.9040664 | -0.191388898523191 | 0.084555101979239 | -2.26348137537796 | 0.0236060314367061 | 0.289811140622062 | NM_007695 |
| Syt3 | 2067.102553 | 0.184685935 | 0.064126723 | 2.880015188 | 0.00397656 | 0.030957254 | | Slc6a11 | 8761.552864 | 0.187713763637335 | 0.0829410687617876 | 2.26321852900717 | 0.023622221567983 | 0.289811140622062 | NM_172890 |
| Gpr61 | 803.4361203 | 0.184450127 | 0.064255099 | 2.870591284 | 0.004097049 | 0.031534861 | | Lmtk3 | 9300.216162 | -0.162985823834281 | 0.0720499478806021 | -2.2621227166517 | 0.0236898224428042 | 0.290395241184957 | NM_001005511 |
| Tspan33 | 824.9971622 | 0.184255457 | 0.063123061 | 2.918987976 | 0.003511698 | 0.028383017 | | Cfh | 1271.577537 | -0.17498777162115 | 0.0773863219756418 | -2.26122352314718 | 0.0237454191730416 | 0.29083133128803 | NM_009888 |
| Fbf1 | 2954.201891 | 0.181905349 | 0.068067721 | 2.672417204 | 0.007530696 | 0.047937797 | | Rfx1 | 1109.95299 | -0.188993009366658 | 0.0836133364403252 | -2.26032134839569 | 0.023801313954603 | 0.291270334039228 | NM_009055 |
| Ints1 | 4194.901813 | 0.181904089 | 0.065613991 | 2.772336905 | 0.00556554 | 0.038738759 | | Alkbh3 | 320.7247417 | -0.188582762373912 | 0.0834645111285825 | -2.25943649371394 | 0.0238562464921633 | 0.291530238803462 | NM_026944 |
| Ddx23 | 2744.353771 | 0.18174851 | 0.065025335 | 2.795041497 | 0.005189305 | 0.036748497 | | Cacna1a | 6389.351803 | -0.105253192807758 | 0.0465877020907693 | -2.25924843004034 | 0.0238679358058957 | 0.291530238803462 | NM_001252059,NM_001252061,NM_001252060,NM_007578 |
| Tada2a | 983.1500646 | 0.181732361 | 0.064478002 | 2.818517262 | 0.004824602 | 0.035127997 | | Cep44 | 320.4946813 | -0.162174758295644 | 0.07179018950941 | -2.25901003192625 | 0.0238827608547514 | 0.291530238803462 | NM_001009951 |
| Rbbp5 | 2289.22541 | 0.181660662 | 0.058467905 | 3.107015059 | 0.001889868 | 0.018346874 | | Cspp1 | 726.1325055 | -0.174302495719233 | 0.0771785817507945 | -2.25843092429511 | 0.0239188065684291 | 0.291723069562603 | NM_026493 |
| Rnh1 | 1210.727884 | 0.180503103 | 0.067880607 | 2.659126233 | 0.00783436 | 0.049424096 | | Dync1i1 | 5419.843574 | 0.0987099601993571 | 0.0437390104490929 | 2.25679454532342 | 0.0240209157460705 | 0.291723069562603 | NM_010063,NM_001191023,NM_001191025,NM_001191026,NM_001191027 |
| Ctsa | 4034.581419 | 0.180425617 | 0.058670974 | 3.075210864 | 0.002103538 | 0.019861191 | | Gpatch2l | 1684.598212 | 0.117883938225706 | 0.0522491873333199 | 2.25618717466261 | 0.0240589114256571 | 0.291723069562603 | NM_027405 |
| Polr1b | 781.07155 | 0.180254703 | 0.06468694 | 2.78657026 | 0.005326907 | 0.037473304 | | Lca5 | 385.7175269 | -0.175438018102867 | 0.0777087552796914 | -2.25763515927421 | 0.0239684147657675 | 0.291723069562603 | NM_027448,NM_029434 |
| Arl4c | 2056.746766 | 0.180074452 | 0.056243338 | 3.201702784 | 0.001366179 | 0.01439971 | | Olfm3 | 1139.923666 | 0.176280590847046 | 0.0780913532915972 | 2.2573637594524 | 0.0239853542941139 | 0.291723069562603 | NM_153157,NM_153458 |
| Mbd1 | 1067.608251 | 0.177968009 | 0.058384587 | 3.04820189 | 0.002302152 | 0.021168869 | | Polr1c | 728.0568048 | 0.142412541035033 | 0.0631210275022049 | 2.25618223705339 | 0.0240592205243012 | 0.291723069562603 | NM_009085 |
| Taok3 | 2185.693089 | 0.176669363 | 0.052239749 | 3.381895329 | 0.000719876 | 0.008887453 | | Psat1 | 4873.933921 | 0.124484856000736 | 0.0551311868366505 | 2.25797526125412 | 0.0239472018125289 | 0.291723069562603 | NM_001205339,NM_177420 |
| Dpf2 | 2088.632595 | 0.175884713 | 0.064207789 | 2.739304932 | 0.006156924 | 0.041559293 | | Zdbf2 | 1073.958188 | 0.213432503822661 | 0.0945712492676183 | 2.25684344317678 | 0.0240178590758564 | 0.291723069562603 | NM_028673,NM_001267872 |
| Tbcd | 1839.935948 | 0.175554281 | 0.059463888 | 2.952283913 | 0.003154328 | 0.026247665 | | Cnot8 | 2239.411061 | 0.117501749372817 | 0.052101959888551 | 2.25522705142301 | 0.0241190807803996 | 0.291781390175991 | NM_026949 |
| Prpf6 | 2552.293501 | 0.17504458 | 0.063342568 | 2.763458865 | 0.00571923 | 0.03945403 | | Doc2b | 2379.760899 | 0.172397806674061 | 0.0764320292221882 | 2.25555018999064 | 0.0240975618818554 | 0.291781390175991 | NM_007873 |
| Tubgcp2 | 1570.010678 | 0.174888828 | 0.06282501 | 2.783745323 | 0.005373521 | 0.037690055 | | Trpc2 | 578.9649251 | -0.176872937326042 | 0.0784362359476513 | -2.2549901227296 | 0.0241339487693812 | 0.291781390175991 | NM_001109897,NM_011644 |
| Traf3ip1 | 659.0589943 | 0.174314584 | 0.062917765 | 2.770514549 | 0.00559678 | 0.038886327 | | Ttll4 | 500.3312232 | -0.158390607122893 | 0.0702452194021494 | -2.25482400754017 | 0.0241443777358902 | 0.291781390175991 | NM_001014974 |
| Kat2a | 3135.53649 | 0.173507368 | 0.055508747 | 3.125766245 | 0.001773425 | 0.017434924 | | Eif2ak4 | 1228.65505 | -0.144079684367198 | 0.0639077411148974 | -2.25449502444724 | 0.0241650433312874 | 0.291788378578787 | NM_001177806,NM_013719 |
| Reps1 | 1495.331577 | 0.172439414 | 0.062371376 | 2.76472038 | 0.005697161 | 0.039424659 | | Atp6ap1 | 17447.159 | 0.0906035552440897 | 0.0402071263133576 | 2.25342031504473 | 0.0242326598177561 | 0.291915574352588 | NM_018794 |
| Capn2 | 3133.250182 | 0.171004796 | 0.04273218 | 4.001780254 | 6.29E-05 | 0.001370002 | | Rreb1 | 2620.314921 | -0.214050944592467 | 0.0949915011654442 | -2.25336942743604 | 0.0242358655286535 | 0.291915574352588 | NR_033615,NM_026830,NR_033218,NM_001039188,NM_001177868,NM_001177869 |
| Golga1 | 1499.963615 | 0.170413094 | 0.06096032 | 2.795475718 | 0.005182339 | 0.03674392 | | Tap2 | 456.2824726 | -0.20589870155529 | 0.0913709376677465 | -2.25343754601713 | 0.0242315744203855 | 0.291915574352588 | NM_011530 |
| Ddx46 | 2850.474865 | 0.169291541 | 0.05382713 | 3.145096913 | 0.001660318 | 0.016639696 | | Nek9 | 3330.294899 | -0.097081969343639 | 0.0430993601633419 | -2.25251532681016 | 0.0242897252605543 | 0.292321913119148 | NM_145138 |
| Ccpg1 | 3902.468409 | 0.169286814 | 0.051509306 | 3.286528709 | 0.001014304 | 0.011471264 | | AW209491 | 843.531026 | 0.156308684772274 | 0.0694128813516508 | 2.25186855420109 | 0.0243305798526391 | 0.292457679088999 | NM_001104646,NM_134067 |
| Bcl9 | 2155.561692 | 0.169048826 | 0.061811402 | 2.73491331 | 0.006239671 | 0.041968026 | | Foxn2 | 626.6196937 | -0.172159176831362 | 0.0764682498646888 | -2.25138115670228 | 0.0243614065604907 | 0.292457679088999 | NM_180974 |
| Sars | 6397.6773 | 0.168967269 | 0.0561883 | 3.007161085 | 0.002637 | 0.023198299 | | Pnkd | 8518.101533 | 0.100669598372708 | 0.044708330659732 | 2.25169664997979 | 0.0243414485154558 | 0.292457679088999 | NM_025580,NM_019999,NM_001039509 |
| Fam222b | 1476.115515 | 0.168436902 | 0.056014286 | 3.00703469 | 0.002638097 | 0.023198299 | | Msantd4 | 2423.12027 | 0.131807433238854 | 0.058566180632658 | 2.25057246033481 | 0.0244126293218071 | 0.292830597463724 | NM_145609 |
| Ctnna1 | 3137.489736 | 0.167774198 | 0.04876359 | 3.440562911 | 0.000580505 | 0.007622332 | | Srsf6 | 3817.196327 | -0.140970031701397 | 0.0626575842669863 | -2.2498478572158 | 0.024458604881296 | 0.293140011968405 | NM_026499 |
| Alas1 | 2168.222391 | 0.167597087 | 0.06113985 | 2.741208682 | 0.006121362 | 0.041362572 | | Atg4a | 247.3572108 | -0.189698188357388 | 0.0843490969069473 | -2.24896525645865 | 0.0245147066199922 | 0.293328698261653 | NM_174875 |
| Cherp | 2489.767726 | 0.16728869 | 0.0545513 | 3.06663065 | 0.002164862 | 0.020316395 | | Tekt5 | 272.7150904 | 0.199223032887911 | 0.0885843263871259 | 2.24896481141911 | 0.024514734936641 | 0.293328698261653 | NM_001099275 |
| Plcg1 | 3259.412552 | 0.166452245 | 0.058904002 | 2.825822356 | 0.004715941 | 0.034613354 | | Vars | 1939.050705 | -0.142499416535656 | 0.0633775121860071 | -2.24842237602248 | 0.0245492696931388 | 0.293500157664637 | NM_011690 |
| B4galnt4 | 2923.630059 | 0.166291729 | 0.060838187 | 2.733344565 | 0.006269471 | 0.042079243 | | Nefl | 12246.3166 | 0.131913578248263 | 0.0586904699840632 | 2.24761495834132 | 0.0246007529116555 | 0.293873796706174 | NM_010910 |
| Fam98a | 1312.567607 | 0.165852453 | 0.061062501 | 2.716109743 | 0.006605402 | 0.043599054 | | Drp2 | 4677.431162 | -0.103591023884287 | 0.0461154505615792 | -2.24634092528184 | 0.0246821790857869 | 0.294362342693054 | NM_010078 |
| Phf12 | 2268.931824 | 0.165792252 | 0.048891151 | 3.391048273 | 0.000696258 | 0.008671887 | | Nde1 | 935.6077864 | 0.150006311498153 | 0.0667773250405332 | 2.2463659843682 | 0.0246805752578762 | 0.294362342693054 | NM_001114085,NM_023317 |
| Gpbp1l1 | 1404.311263 | 0.164923375 | 0.061479355 | 2.682581426 | 0.007305636 | 0.04700728 | | Brf2 | 553.3077745 | 0.147531987055649 | 0.0657138588175046 | 2.24506656145949 | 0.0247638598589575 | 0.29461083399772 | NM_025686 |
| Riok3 | 2221.700274 | 0.163690757 | 0.056779411 | 2.882924527 | 0.003940018 | 0.030816672 | | E130309D02Rik | 844.9959762 | 0.134369524540739 | 0.0598490878308887 | 2.24513905576 | 0.0247592070399093 | 0.29461083399772 | NM_172726 |
| Elp2 | 3765.13999 | 0.162250292 | 0.042315519 | 3.8342976 | 0.000125924 | 0.002336599 | | Mc4r | 74.20819186 | 0.206451598022166 | 0.0919455030658082 | 2.24536917128401 | 0.0247444428137955 | 0.29461083399772 | NM_016977 |
| Tmem110 | 892.5459094 | 0.160736471 | 0.05979372 | 2.688183179 | 0.007184197 | 0.046438029 | | D19Bwg1357e | 1422.194531 | 0.156443973355797 | 0.0696980163656376 | 2.2445972140023 | 0.0247940017798465 | 0.294650465081573 | NM_177474 |
| Extl2 | 3119.696627 | 0.160724683 | 0.055972431 | 2.871497292 | 0.004085323 | 0.031506284 | | Hpn | 51.73890666 | -0.198226082942747 | 0.088320963849966 | -2.24438314871066 | 0.024807759795867 | 0.294650465081573 | NM_008281,NM_001276269,NM_001110252 |
| Polg | 2669.093502 | 0.160013919 | 0.055602224 | 2.877833091 | 0.004004169 | 0.031051351 | | Mir682 | 56.29074 | -0.19557706732883 | 0.087162127905153 | -2.2438308016258 | 0.0248432897907587 | 0.294831395016798 | NR_030451 |
| Stx5a | 1904.820363 | 0.159100409 | 0.057561117 | 2.764025726 | 0.005709304 | 0.039425264 | | Cnih3 | 3111.93286 | 0.137223898632591 | 0.0612195374215345 | 2.24150499027328 | 0.0249933826457418 | 0.296370511275139 | NM_028408,NM_001160211,NM_001160212 |
| Sgsm3 | 2538.365112 | 0.158292349 | 0.057327407 | 2.76119848 | 0.005758967 | 0.039657524 | | Sdccag3 | 3648.232078 | -0.137490963264502 | 0.0613762697025129 | -2.24013228452809 | 0.0250823361623317 | 0.297182720305082 | NM_001085408,NM_026563,NM_001085407 |
| Lrsam1 | 1508.892021 | 0.158049562 | 0.058249042 | 2.713341841 | 0.006660836 | 0.043785739 | | Urb1 | 709.3889534 | -0.145764404901637 | 0.0650978626187452 | -2.23915807736004 | 0.0251456325029371 | 0.297689859606898 | NM_029497 |
| Rae1 | 1392.415562 | 0.156708508 | 0.056544557 | 2.771416322 | 0.005581302 | 0.038831022 | | Tnfaip2 | 39.63919234 | -0.20289818484153 | 0.0906678218383077 | -2.23781911518033 | 0.0252328533565505 | 0.298479175779521 | NM_009396 |
| Ncstn | 2680.334264 | 0.154969648 | 0.054689357 | 2.833634472 | 0.004602194 | 0.034134224 | | A230073K19Rik | 115.0779278 | -0.212610315903807 | 0.0950736147473381 | -2.2362704570435 | 0.0253340604124521 | 0.298582503407544 | NR_033229 |
| Tspan9 | 1842.380723 | 0.154198765 | 0.057815768 | 2.667071103 | 0.007651548 | 0.048587331 | | Bysl | 1211.124985 | 0.121242657099355 | 0.054216672005734 | 2.23626151539755 | 0.025334645780522 | 0.298582503407544 | NM_016859 |
| Usp8 | 2877.496576 | 0.151044523 | 0.056225142 | 2.686423129 | 0.007222156 | 0.046578655 | | Diras1 | 5451.239281 | 0.114872362332292 | 0.0513727352567424 | 2.23605696208702 | 0.0253480401336867 | 0.298582503407544 | NM_145217 |
| Rnps1 | 5042.659086 | 0.149000094 | 0.054172458 | 2.750476898 | 0.005950859 | 0.040564493 | | Sdhd | 3060.881682 | 0.100173231665248 | 0.0447889050908404 | 2.23656352978663 | 0.0253148807777072 | 0.298582503407544 | NM_025848 |
| Trim66 | 2087.310147 | 0.147329434 | 0.054035701 | 2.726520283 | 0.006400602 | 0.042690885 | | Tnfrsf19 | 2793.089312 | -0.114648985848099 | 0.0512723425458701 | -2.23607855922577 | 0.0253466256423699 | 0.298582503407544 | NM_001164155,NM_013869 |
| Cog1 | 1949.613719 | 0.144457439 | 0.048251005 | 2.993874225 | 0.002754595 | 0.023914757 | | Tprkb | 2552.806821 | -0.105047612976777 | 0.0469870355127281 | -2.23567228343913 | 0.0253732458717942 | 0.298582503407544 | NM_001170488,NM_176842 |
| Fbxo18 | 3453.87785 | 0.143373486 | 0.050858847 | 2.819047148 | 0.004816644 | 0.035103051 | | Txnl4a | 4564.104754 | 0.104134094332023 | 0.0465823110918756 | 2.23548578615252 | 0.0253854737510888 | 0.298582503407544 | NM_178604,NM_025299,NM_001038608,NM_001042408 |
| Cnnm1 | 5322.957827 | 0.13845579 | 0.051127396 | 2.708054774 | 0.006767885 | 0.044320208 | | Vasn | 596.3315605 | 0.1968273219599 | 0.0880729470277869 | 2.23482157237002 | 0.025429065016475 | 0.298853234975175 | NM_139307 |
| Cyfip1 | 3412.518478 | 0.135210249 | 0.047704999 | 2.834299395 | 0.004592628 | 0.034079589 | | Hars | 3971.040918 | 0.101888347446964 | 0.045642752174052 | 2.23230069603226 | 0.02559509591497 | 0.300343543088629 | NM_008214 |
| Atp6v0a1 | 21809.71969 | 0.131881818 | 0.049324373 | 2.673765712 | 0.007500483 | 0.047804396 | | Slc39a9 | 2762.851126 | 0.0973356987675513 | 0.043603939742341 | 2.23226844506977 | 0.0255972261010411 | 0.300343543088629 | NM_026244 |
| Map7d1 | 9546.742919 | 0.129602514 | 0.044787224 | 2.893738519 | 0.00380685 | 0.030108902 | | Dos | 11752.11259 | 0.128036540305603 | 0.0573708890555179 | 2.23173359195639 | 0.025632575683345 | 0.300515572539362 | NR_036582,NM_001195268 |
| Sfxn3 | 5209.780514 | 0.119885872 | 0.04218895 | 2.84164156 | 0.004488192 | 0.03354561 | | Cbln3 | 56.7736988 | 0.160668325942996 | 0.0720191861468719 | 2.23091004687749 | 0.0256870881007295 | 0.300614489339653 | NM_019820 |
| Bscl2 | 2783.167985 | 0.119041387 | 0.043475584 | 2.738120469 | 0.006179144 | 0.041680301 | | Errfi1 | 1385.465743 | 0.158456905176034 | 0.0710515652996356 | 2.23016768888619 | 0.0257363124553163 | 0.300614489339653 | NM_133753 |
| Vdac3 | 5158.700146 | -0.108268353 | 0.04065859 | -2.662865422 | 0.007747839 | 0.049078096 | | Glipr1 | 42.65719365 | -0.199103502949831 | 0.0892947786143205 | -2.22973286948605 | 0.025765182378347 | 0.300614489339653 | NM_028608 |
| Cs | 11988.39732 | -0.11238592 | 0.040034292 | -2.807241312 | 0.004996779 | 0.03594248 | | Slc16a1 | 1317.211597 | -0.14793898238316 | 0.0663367326058375 | -2.23012163204043 | 0.0257393690836453 | 0.300614489339653 | NM_009196 |
| Slc30a9 | 6116.525227 | -0.114094066 | 0.03984955 | -2.86312055 | 0.004194909 | 0.032144374 | | Taf4a | 942.6251733 | -0.149210513841209 | 0.0669168723081405 | -2.22978912035997 | 0.0257614460134806 | 0.300614489339653 | NM_001081092 |
| Eef1a1 | 84643.77661 | -0.117467409 | 0.038991463 | -3.01264428 | 0.002589823 | 0.022955788 | | Zfp36l1 | 889.3321464 | -0.188768984939004 | 0.084607183631873 | -2.231122427622 | 0.0256730205249674 | 0.300614489339653 | NM_007564 |
| Cyb5b | 7442.71167 | -0.117754099 | 0.042299577 | -2.783812697 | 0.005372405 | 0.037690055 | | 2700038G22Rik | 28.72045214 | -0.193731452797607 | 0.0869455110128037 | -2.22821331455603 | 0.0258662936154695 | 0.301310169252854 | NR_045040,NR_045042 |
| Kidins220 | 15379.33307 | -0.122269131 | 0.039443024 | -3.099892401 | 0.00193591 | 0.018676826 | | Hck | 108.2362102 | -0.211216857989187 | 0.0947852615312001 | -2.2283723711588 | 0.0258556939304462 | 0.301310169252854 | NM_001172117,NM_010407 |
| Eif5a | 9717.907832 | -0.122423501 | 0.045382249 | -2.697607643 | 0.00698397 | 0.045389947 | | Ano8 | 2342.560518 | -0.150148298593876 | 0.0674611114459958 | -2.22570152456016 | 0.0260341805824954 | 0.302167799626426 | NM_001164679 |
| Dennd5a | 14294.85777 | -0.123288321 | 0.042713811 | -2.88638072 | 0.003897004 | 0.030556774 | | Mob2 | 1007.235113 | 0.120853481282754 | 0.0542934690363681 | 2.22593036377545 | 0.0260188461622999 | 0.302167799626426 | NM_028308 |
| Rer1 | 2177.562415 | -0.123859907 | 0.045786633 | -2.705154272 | 0.006827267 | 0.044601532 | | Slc26a4 | 132.5080571 | -0.211701074729654 | 0.0950808938124713 | -2.22653643903678 | 0.0259782710329584 | 0.302167799626426 | NM_011867 |
| Nap1l4 | 4356.610403 | -0.124575023 | 0.042517077 | -2.92999969 | 0.003389623 | 0.027626946 | | Sst | 6090.707934 | 0.125296648329687 | 0.0562990285146493 | 2.22555613543285 | 0.026043927105348 | 0.302167799626426 | NM_009215 |
| Sel1l | 8883.508886 | -0.129959881 | 0.045143294 | -2.878830265 | 0.003991531 | 0.031015437 | | Tek | 684.1709782 | -0.165094294726064 | 0.0741792777164355 | -2.22561205512365 | 0.0260401780161354 | 0.302167799626426 | NM_013690 |
| Pdha1 | 9333.704875 | -0.133561824 | 0.043812584 | -3.048480854 | 0.002300016 | 0.021168869 | | Derl1 | 2920.692473 | 0.118152282607834 | 0.0531050348462987 | 2.2248791089169 | 0.0260893548225038 | 0.302212095814745 | NM_024207 |
| Gzf1 | 1823.659254 | -0.133658747 | 0.043984491 | -3.038769883 | 0.002375462 | 0.021688516 | | Eogt | 1201.405251 | -0.135243942021689 | 0.0607848049906384 | -2.22496299626467 | 0.0260837221593329 | 0.302212095814745 | NM_175313 |
| Glud1 | 9953.459522 | -0.134578907 | 0.0451658 | -2.979664007 | 0.002885647 | 0.024666584 | | 4933432I09Rik | 53.89153596 | -0.209149133866243 | 0.0940619658935719 | -2.2235250122551 | 0.0261804187160006 | 0.30237340574348 | NR_015575,NR_027843 |
| Ubl4 | 3072.112219 | -0.135024756 | 0.049012747 | -2.754890621 | 0.005871176 | 0.040200975 | | Adora3 | 45.92865487 | -0.205179531371641 | 0.0922691642491405 | -2.22370640333997 | 0.0261682041321549 | 0.30237340574348 | NM_001174169,NM_027025,NM_009631 |
| Ndufb10 | 3183.87443 | -0.137191646 | 0.048118945 | -2.851094196 | 0.004356906 | 0.032960625 | | Psen1 | 2859.812873 | 0.112926338976689 | 0.0507798796366235 | 2.22384022539597 | 0.0261591959278584 | 0.30237340574348 | NM_008943 |
| Micu1 | 3353.105412 | -0.138087299 | 0.046090025 | -2.996034398 | 0.002735156 | 0.023821531 | | Tmed10 | 6588.343332 | 0.115191705057669 | 0.0518152132402119 | 2.22312517606842 | 0.0262073604454799 | 0.30237340574348 | NM_026775 |
| Btbd1 | 4944.287773 | -0.139264955 | 0.051511667 | -2.703561416 | 0.006860077 | 0.044716042 | | Uimc1 | 599.1247168 | -0.165737801360453 | 0.0745467759271036 | -2.2232725600705 | 0.0261974266414708 | 0.30237340574348 | NM_011307 |
| 15-Sep | 4544.817021 | -0.139769007 | 0.041841941 | -3.340404518 | 0.000836564 | 0.009917161 | | Ficd | 660.3607536 | 0.144054274210489 | 0.0648232436198136 | 2.22226266638805 | 0.0262655596105191 | 0.302739576664324 | NM_001010825 |
| Prmt1 | 2165.964715 | -0.141638883 | 0.044760607 | -3.164364659 | 0.001554219 | 0.01589782 | | Fstl5 | 838.4596628 | 0.181410938794348 | 0.0816417134299194 | 2.22203737765657 | 0.0262807797173147 | 0.302739576664324 | NM_178673,NM_001253719 |
| Rpl5 | 9176.039278 | -0.141733471 | 0.051387301 | -2.758141983 | 0.005813094 | 0.03995879 | | Gm5086 | 39.34251437 | -0.206708185005937 | 0.0930724032404924 | -2.22093958906184 | 0.026355053357713 | 0.303114414152129 | NR_046157 |
| Prune | 1304.968703 | -0.141848233 | 0.051819735 | -2.737339977 | 0.006193825 | 0.041761148 | | Neu2 | 249.0108219 | -0.18892726790687 | 0.0850618255348799 | -2.22105823286616 | 0.0263470174791771 | 0.303114414152129 | NM_001160163,NM_001160165,NM_001160164,NM_015750 |
| 9530068E07Rik | 3226.008428 | -0.146031171 | 0.048211443 | -3.028973245 | 0.002453864 | 0.02211392 | | Ccm2l | 152.6454866 | -0.208976771228718 | 0.0941824142481611 | -2.21885128871389 | 0.026496843382527 | 0.303950051694259 | NM_145536 |
| Ptp4a2 | 4725.170305 | -0.146884653 | 0.05325766 | -2.758000516 | 0.005815611 | 0.03995879 | | Ndufa10 | 4725.745646 | 0.104323412735534 | 0.0470122219639903 | 2.2190700285437 | 0.0264819606971096 | 0.303950051694259 | NM_024197 |
| Dzank1 | 13288.40347 | -0.14785555 | 0.053434991 | -2.767017399 | 0.005657174 | 0.039218008 | | Nfxl1 | 442.8028178 | -0.152549932565679 | 0.0687583909939936 | -2.2186373235378 | 0.026511408198859 | 0.303950051694259 | NM_133921 |
| Pdgfra | 2428.598642 | -0.148240114 | 0.055817676 | -2.655791601 | 0.007912249 | 0.049814056 | | Tsc22d1 | 16691.0519 | 0.109758372491209 | 0.0494530063616006 | 2.21944792776915 | 0.026456266092682 | 0.303950051694259 | NM_009366,NM_207652,NM_001177751 |
| Dcaf12 | 1994.375653 | -0.148411725 | 0.052331936 | -2.835968527 | 0.004568695 | 0.033967062 | | Slc8a3 | 1504.490468 | 0.141160684115278 | 0.063634410638565 | 2.21830740158893 | 0.0265338798588946 | 0.303967775102763 | NM_001167920,NM_080440 |
| Gabrb3 | 11480.14202 | -0.149244718 | 0.055279254 | -2.69983232 | 0.006937443 | 0.045125418 | | Chrd | 1632.355855 | -0.186779928407619 | 0.0842358103399169 | -2.21734589664308 | 0.0265994638077841 | 0.304478968693358 | NM_009893 |
| Pnpla8 | 4217.219252 | -0.149536597 | 0.055245478 | -2.706766311 | 0.006794206 | 0.044436233 | | Fhod1 | 186.0566558 | -0.188779025042632 | 0.0851796276472373 | -2.21624618769691 | 0.0266746462203151 | 0.305080385907289 | NM_177699 |
| Pgam1 | 20595.79169 | -0.149886192 | 0.045911652 | -3.264665639 | 0.001095934 | 0.012251368 | | Igsf9 | 254.2666241 | -0.20303859017664 | 0.091636075000696 | -2.21570587975427 | 0.0267116519606601 | 0.305080385907289 | NM_001145800,NM_033608 |
| Sec62 | 5581.05613 | -0.150248517 | 0.050316642 | -2.986060128 | 0.002825971 | 0.02433885 | | Intu | 639.5592727 | -0.150981829173771 | 0.0681431459248956 | -2.21565686650535 | 0.0267150110749051 | 0.305080385907289 | NM_175515 |
| Dtd1 | 1597.265016 | -0.150288047 | 0.051190033 | -2.935884959 | 0.003325976 | 0.027283599 | | Gm6548 | 154.3783314 | -0.204521327800267 | 0.0923357425998508 | -2.21497463540838 | 0.026761805554436 | 0.305374695588168 | NR_003363 |
| Gnb2l1 | 5682.51273 | -0.150575168 | 0.056234407 | -2.677634161 | 0.007414415 | 0.047350844 | | Tec | 86.88546946 | -0.209818665081654 | 0.0947674061771687 | -2.21403828115118 | 0.0268261456654921 | 0.305868596496811 | NM_001113464,NM_001113460,NM_001113461 |
| Eef1b2 | 4286.864517 | -0.150755237 | 0.047469155 | -3.175856752 | 0.001493947 | 0.015410929 | | Mtus2 | 2562.305862 | 0.125498426590993 | 0.0567051169634189 | 2.21317639944123 | 0.0268854865234662 | 0.306064715705227 | NM_029920 |
| Cmas | 3069.390949 | -0.151392975 | 0.040752296 | -3.714955689 | 0.000203239 | 0.003445285 | | Ppia | 31521.712 | 0.109906971917046 | 0.0496542620784278 | 2.21344487495254 | 0.0268669897487922 | 0.306064715705227 | NM_008907 |
| Gnb1 | 25047.69788 | -0.15192344 | 0.051223002 | -2.965922227 | 0.003017768 | 0.025425392 | | Eif4h | 14478.46463 | 0.0880274091541093 | 0.0398193416755632 | 2.2106696256139 | 0.027058723391516 | 0.306595176275478 | NM_033561 |
| Arl2bp | 4816.860739 | -0.152453549 | 0.048764719 | -3.126308335 | 0.001770158 | 0.017424927 | | Fam124a | 1174.825987 | 0.1268931109542 | 0.0573961951125318 | 2.21082792518584 | 0.0270477552719869 | 0.306595176275478 | NM_001243857 |
| Selk | 2487.57478 | -0.152474639 | 0.050667885 | -3.009295488 | 0.002618543 | 0.023157366 | | Il10ra | 324.1009558 | -0.183473179500194 | 0.0829550885065656 | -2.21171700016537 | 0.02698622511 5946 | 0.306595176275478 | NM_008348 |
| Rpl10a | 4272.874724 | -0.152653455 | 0.049082012 | -3.110171094 | 0.00186979 | 0.018186144 | | Ptpn21 | 451.7187563 | -0.186798754578929 | 0.0844890907774954 | -2.21092158596983 | 0.0270412675943425 | 0.306595176275478 | NM_011877,NM_001146199 |
| Sae1 | 3527.91207 | -0.152916694 | 0.040961443 | -3.733186207 | 0.000189073 | 0.003262241 | | Setd1a | 3095.27776 | -0.126177285950375 | 0.0570458972959399 | -2.21185557474534 | 0.0269766456865899 | 0.306595176275478 | NM_178029 |
| Naa20 | 1495.727797 | -0.153546228 | 0.052357754 | -2.932635882 | 0.003360978 | 0.027494717 | | Tbxas1 | 110.8418046 | -0.20674423361559 | 0.0934935956126592 | -2.21131974078871 | 0.0270137032710879 | 0.306595176275478 | NM_011539 |
| Calu | 3614.309446 | -0.154439632 | 0.045408354 | -3.401128142 | 0.000671084 | 0.008426069 | | Arhgap11a | 147.2337024 | -0.202581169066199 | 0.0916627826849162 | -2.21007003204949 | 0.0271003023131965 | 0.306826961341771 | NM_181416 |
| Vdac1 | 16919.7671 | -0.154680132 | 0.045579338 | -3.393645881 | 0.000689688 | 0.008617765 | | Mgat4c | 164.3738039 | 0.198887905690392 | 0.0900479877183579 | 2.20868795327722 | 0.027196352918496 | 0.307674628110648 | NM_001162369,NM_001205098,NM_026243,NM_001162368 |
| Mtmr9 | 2721.325589 | -0.155388337 | 0.056781581 | -2.736597574 | 0.006207818 | 0.041800928 | | Cct8 | 6968.949747 | 0.100301652518221 | 0.0454489728141243 | 2.20690691797218 | 0.027320563021062 | 0.308173878949826 | NM_009840 |
| Nfyb | 1592.028691 | -0.155810118 | 0.049999522 | -3.116232153 | 0.00183178 | 0.017883806 | | Cdyl | 351.2901938 | 0.162306469710293 | 0.0736136357610116 | 2.20484245931317 | 0.0274651514207864 | 0.308173878949826 | NM_001123386,NM_009881 |
| Sod2 | 4831.109154 | -0.156084326 | 0.04612495 | -3.383945703 | 0.000714521 | 0.008835621 | | Celf5 | 11722.22512 | -0.133613092195695 | 0.0606004652199994 | -2.20481957870515 | 0.0274667576004456 | 0.308173878949826 | NM_176954 |
| Cadm1 | 5939.79473 | -0.156448554 | 0.051115875 | -3.060664674 | 0.002208463 | 0.020613206 | | Cerkl | 59.36915581 | -0.209478529273166 | 0.0950527695606418 | -2.2038130003095 | 0.0275374979262615 | 0.308173878949826 | NM_001048176 |
| Kctd20 | 1812.647683 | -0.157920407 | 0.05200303 | -3.036753923 | 0.002391406 | 0.021769943 | | Foxq1 | 141.4822234 | -0.207101288651985 | 0.0939568450682095 | -2.20421714353687 | 0.0275090766851152 | 0.308173878949826 | NM_008239 |
| Atpif1 | 3785.672016 | -0.158281607 | 0.055641298 | -2.844678568 | 0.004445626 | 0.033388526 | | Gm14827 | 235.0710097 | -0.209707123455431 | 0.0950389166208323 | -2.20653949888843 | 0.0273462477888711 | 0.308173878949826 | NR_045323 |
| Marf1 | 6931.181184 | -0.158603522 | 0.053962942 | -2.939119231 | 0.003291464 | 0.02708335 | | Kcng1 | 767.4171907 | 0.208006777218178 | 0.0942549410012074 | 2.20685276558089 | 0.0273243472852385 | 0.308173878949826 | NM_001081134 |
| Ctnnb1 | 18229.57502 | -0.159041907 | 0.053017948 | -2.999774868 | 0.002701792 | 0.023570703 | | Kcnk3 | 1174.965356 | 0.139338444898092 | 0.0632213996839572 | 2.20397595742332 | 0.0275260349741962 | 0.308173878949826 | NM_010608 |
| Cbx5 | 8808.556698 | -0.160239879 | 0.056376797 | -2.842301949 | 0.004478905 | 0.03349786 | | Lcp2 | 142.8045705 | -0.209352504411245 | 0.0949474406896461 | -2.20493046353459 | 0.0274589744296687 | 0.308173878949826 | NM_010696 |
| Chchd3 | 1545.551948 | -0.160376824 | 0.060058477 | -2.670344501 | 0.007577346 | 0.048155619 | | Mir186 | 41.80427735 | -0.200285778809059 | 0.0908548520452025 | -2.20445880765302 | 0.0274920938265833 | 0.308173878949826 | NR_029572 |
| Atp5h | 5903.154605 | -0.16039856 | 0.056061667 | -2.861109369 | 0.004221614 | 0.032285136 | | Nsg1 | 12026.7152 | 0.1031488498486 | 0.0468002112622411 | 2.20402530387339 | 0.0275225645907628 | 0.308173878949826 | NM_010942 |
| Rps2 | 10086.45592 | -0.16069142 | 0.056777483 | -2.83019625 | 0.004651946 | 0.034310034 | | Prkdc | 1161.016933 | -0.175647702974648 | 0.0795933722981339 | -2.20681317932757 | 0.0273271139282692 | 0.308173878949826 | NM_011159 |
| Nhp2l1 | 3830.890707 | -0.161209585 | 0.039086195 | -4.124463514 | 3.72E-05 | 0.000905278 | | Syt12 | 2056.329535 | 0.153331032418511 | 0.0695345823740119 | 2.20510467142486 | 0.0274467503619344 | 0.308173878949826 | NM_134164 |
| Ddost | 3363.055252 | -0.162000504 | 0.054605481 | -2.96674438 | 0.003009711 | 0.025407408 | | Trib2 | 1747.521962 | 0.189934193196756 | 0.0861300072443744 | 2.20520349728824 | 0.0274398178950155 | 0.308173878949826 | NM_144551 |
| Cyc1 | 4938.060661 | -0.162019076 | 0.050089561 | -3.234587638 | 0.001218185 | 0.013245307 | | Slc25a45 | 158.554909 | -0.192352993613341 | 0.0872999866527304 | -2.20335650655366 | 0.0275696311601973 | 0.308295967846825 | NM_134154 |
| Ndufb11 | 2399.993251 | -0.162055629 | 0.04980939 | -3.25351564 | 0.001139865 | 0.012610872 | | Itga7 | 621.2869759 | -0.193979388769932 | 0.0880737748957957 | -2.2024647972616 | 0.0276324931244266 | 0.308761227019554 | NM_008398 |
| Srr | 4327.800748 | -0.162653845 | 0.048123093 | -3.379954112 | 0.000724979 | 0.0089238 | | Rin2 | 1585.116052 | 0.140182444594734 | 0.0637166817516594 | 2.20009016070713 | 0.0278004988326283 | 0.310219757573383 | NM_028724 |
| D5Ertd579e | 6460.136281 | -0.162671354 | 0.059250005 | -2.745507867 | 0.006041732 | 0.041016941 | | Slc17a6 | 1486.373668 | 0.208799190816039 | 0.0949080184846701 | 2.20001633318017 | 0.0278057362219844 | 0.310219757573383 | NM_080853 |
| Myl6 | 8423.646643 | -0.163141867 | 0.059765498 | -2.729699813 | 0.006339202 | 0.042445806 | | Tfg | 1871.175527 | 0.119921290327293 | 0.0545282430428069 | 2.19925095024884 | 0.0278600833046097 | 0.310587544192448 | NM_001252443,NM_019678 |
| Tex2 | 4719.898534 | -0.164413376 | 0.047950375 | -3.428823594 | 0.000606203 | 0.007866428 | | Abi3 | 222.9738028 | -0.202860336387044 | 0.0923185810010009 | -2.19739443769012 | 0.0279922881928286 | 0.311357667645585 | NM_025659,NM_001163464 |
| Rps5 | 4994.792777 | -0.165104056 | 0.042729633 | -3.863923986 | 0.00011158 | 0.002126409 | | Apbb1ip | 271.3074809 | -0.200174685808809 | 0.0910970685751798 | -2.1973779062233 | 0.0279934678469733 | 0.311357667645585 | NM_019456 |
| Pik3r3 | 2696.175112 | -0.165537098 | 0.060867147 | -2.719646064 | 0.006535183 | 0.043239925 | | Ets2 | 3462.492384 | 0.137451409907424 | 0.0625523205690577 | 2.19738306520024 | 0.0279930997075555 | 0.311357667645585 | NM_011809 |
| Arf2 | 1918.690885 | -0.166330681 | 0.062116932 | -2.677702751 | 0.007412897 | 0.047350844 | | Clec18a | 293.4421732 | -0.208777789274181 | 0.0950798617396368 | -2.19581502806442 | 0.0281051855734773 | 0.312140588068853 | NM_181549 |
| Uqcrfs1 | 3208.467197 | -0.166462621 | 0.049486367 | -3.36380771 | 0.000768751 | 0.009269281 | | Echdc2 | 374.106329 | -0.185271251871793 | 0.0843755932905949 | -2.19579198977772 | 0.0281068352742709 | 0.312140588068853 | NM_001254754,NM_026728 |
| Cask | 3579.618907 | -0.166598288 | 0.050783928 | -3.280531769 | 0.001036116 | 0.011675331 | | Ddt | 839.765192 | 0.132901376391336 | 0.0605399565948996 | 2.19526712383756 | 0.0281444417537188 | 0.312281032999026 | NM_010027 |
| Ppp1r11 | 2036.17777 | -0.166935923 | 0.062001262 | -2.692460097 | 0.007092702 | 0.045942445 | | Nap1l4 | 4782.192198 | 0.108525945318892 | 0.049441992446271 | 2.19501561222128 | 0.0281624778485972 | 0.312281032999026 | NM_008672 |
| Cops6 | 3183.945707 | -0.167164286 | 0.050323197 | -3.321813737 | 0.000894344 | 0.010481817 | | Nfe2l1 | 17287.5355 | 0.0771199151229936 | 0.0351462551000172 | 2.19425696716564 | 0.0282169412279591 | 0.312646291592169 | NM_001130450,NM_001130451,NM_001130452,NM_001130453,NM_001130454,NM_008686 |
| Ddx6 | 7248.125674 | -0.167303454 | 0.061298893 | -2.729306272 | 0.006346773 | 0.042474471 | | 1700020D05Rik | 41.95066779 | -0.197997201432369 | 0.0902682621916105 | -2.19343096482886 | 0.0282763433936249 | 0.312827238488801 | NM_023781 |
| Atg12 | 2453.663869 | -0.168047612 | 0.053698628 | -3.129458193 | 0.00175129 | 0.01733833 | | Nsmaf | 1411.451093 | -0.140609142038449 | 0.0640993639576742 | -2.19361212587531 | 0.0282633059347926 | 0.312827238488801 | NM_010945 |
| Ran | 7038.030763 | -0.16814012 | 0.061347038 | -2.740802578 | 0.006128932 | 0.041395673 | | Gtf2b | 1152.208518 | 0.13366917474774 | 0.0609595354748413 | 2.19275251536171 | 0.0283252148264212 | 0.313129429656464 | NM_145546 |
| Pgam5 | 1901.74222 | -0.168468658 | 0.051785458 | -3.253204003 | 0.001141115 | 0.012610872 | | Eif5a | 10577.08237 | 0.0883291967305098 | 0.040300559514682 | 2.19176105230823 | 0.0283967647604605 | 0.313443316801253 | NM_001166589,NM_001166596,NM_001166594,NM_001166595,NM_001166592,NM_001166593,NM_001166590,NM_001166591,NM_181582 |
| Fau | 4785.712144 | -0.16925374 | 0.059540577 | -2.842662089 | 0.004473848 | 0.03349786 | | Sf3b4 | 1873.634876 | 0.118552846276399 | 0.0540895964712711 | 2.19178648040695 | 0.0283949277723439 | 0.313443316801253 | NM_153053 |
| Mlc1 | 6892.100298 | -0.169439283 | 0.05519258 | -3.069964886 | 0.002140839 | 0.020127528 | | Ctsz | 1774.930645 | -0.194652182658919 | 0.0888292855342813 | -2.19130640855822 | 0.028429626622589 | 0.313543192127037 | NM_022325 |
| Astn1 | 13183.82223 | -0.169751503 | 0.061717155 | -2.75047517 | 0.00595089 | 0.040564493 | | E4f1 | 1087.674524 | -0.169716810702095 | 0.0774595177321033 | -2.19103882480998 | 0.0284489830113888 | 0.313543192127037 | NM_007893 |
| 2900097C17Rik | 65811.20869 | -0.169824511 | 0.057641696 | -2.94620947 | 0.003216945 | 0.026625716 | | Spata5l1 | 62.96821866 | -0.207869769449886 | 0.0949050195007144 | -2.19029267939112 | 0.0285030174194206 | 0.313900554233892 | NM_001033256 |
| Map4k4 | 4532.113062 | -0.169898088 | 0.058387881 | -2.90981769 | 0.003616396 | 0.028927437 | | Nudt3 | 9080.612077 | 0.0948766243006172 | 0.0433614235534981 | 2.1880421933925 | 0.0286665290113791 | 0.315230284167518 | NM_019837 |
| Rab1 | 6328.232708 | -0.170081229 | 0.058166132 | -2.92405944 | 0.003454988 | 0.027983052 | | Rasgrp4 | 38.86759462 | -0.177846418981633 | 0.0812813969882811 | -2.18803349316541 | 0.0286671627003505 | 0.315230284167518 | NM_001174155,NR_045676,NM_145149 |
| Slmo2 | 1491.282818 | -0.170175402 | 0.062659325 | -2.715883115 | 0.006609925 | 0.043599054 | | Clcn2 | 2886.270063 | -0.139206499456654 | 0.0636372353623519 | -2.18750074015644 | 0.0287059892442062 | 0.31541845670298 | NM_009900 |
| Rala | 2160.378613 | -0.1702305 | 0.055615487 | -3.060847062 | 0.002207118 | 0.020613072 | | Pgam1 | 22164.98758 | 0.0686668251553126 | 0.0314134857038364 | 2.18590276172143 | 0.0288227201583828 | 0.316461702963468 | NM_023418 |
| Ndufb3 | 1582.583249 | -0.171708456 | 0.054351536 | -3.159219911 | 0.001581921 | 0.016072315 | | Cacng4 | 503.1229735 | 0.145008623819187 | 0.0663669134288719 | 2.18495356085225 | 0.0288922517430721 | 0.316746300996125 | NM_019431 |
| Ndufa11 | 2031.963172 | -0.172714373 | 0.056515942 | -3.05602928 | 0.00224294 | 0.020821685 | | Tmc6 | 243.9814104 | -0.194317328465381 | 0.0889224613934947 | -2.18524459872404 | 0.0288709170811842 | 0.316746300996125 | NM_145439,NM_181321 |
| Prps1 | 2908.057748 | -0.17280073 5 | 0.061667202 | -2.802149772 | 0.00507633 | 0.036290258 | | Rwdd2a | 619.035592 | 0.202669125648707 | 0.0927757747691088 | 2.1845048036849 | 0.0289251746867376 | 0.316868090120324 | NM_001145968,NM_001145969,NM_027100 |
| Rplp2 | 3337.463718 | -0.172917251 | 0.058283615 | -2.967842824 | 0.002998976 | 0.025363614 | | Echdc1 | 1023.396312 | 0.145630071552952 | 0.0666801972998384 | 2.18400780816683 | 0.0289616743081865 | 0.317028847777481 | NM_025855,NM_001110195 |
| Trappc4 | 1529.869427 | -0.173523138 | 0.050739023 | -3.419914865 | 0.000626407 | 0.008054403 | | Cdip1 | 11040.3728 | 0.0775425281296277 | 0.0355138027673308 | 2.18344762000422 | 0.029002862353994 | 0.317240646501594 | NM_025670 |
| Dnajc27 | 6705.705458 | -0.173829494 | 0.041722581 | -4.16631691 | 3.10E-05 | 0.00078887 | | Gm9866 | 1061.818569 | 0.121983451788222 | 0.0558821133487133 | 2.18287112777975 | 0.0290453017942985 | 0.317366933890266 | NR_045870,NR_045869,NR_045868 |
| Sdhd | 2890.363856 | -0.174076958 | 0.047063021 | -3.698805458 | 0.000216617 | 0.003599861 | | Ostc | 1042.831168 | 0.128579288405629 | 0.0589084362147897 | 2.18269736335909 | 0.0290581042320015 | 0.317366933890266 | NM_025509 |
| Chmp5 | 2312.633652 | -0.174497479 | 0.060188833 | -2.899167021 | 0.003741555 | 0.029713817 | | Bet1 | 449.8418768 | 0.152871611365647 | 0.0701302733795123 | 2.17982340576911 | 0.0292705542641943 | 0.317415456361503 | NM_009748 |
| Rftn2 | 1203.996563 | -0.175011445 | 0.061521999 | -2.844696972 | 0.004445369 | 0.033388526 | | Dlg3 | 9685.949002 | 0.0985441719867724 | 0.0451955558854552 | 2.18039517505936 | 0.0292281814911541 | 0.317415456361503 | NM_016747,NM_001177780,NM_001177778,NM_001177779 |
| Hibadh | 1929.701794 | -0.175085094 | 0.060445635 | -2.896571328 | 0.003772649 | 0.029930066 | | Fxyd6 | 4317.128834 | 0.136389561241338 | 0.0625404810901621 | 2.18082046802151 | 0.0291966980511285 | 0.317415456361503 | NM_022004 |
| Ndufb5 | 3246.140297 | -0.175949887 | 0.059394782 | -2.962379527 | 0.003052712 | 0.025594549 | | Gcnt1 | 145.7735044 | -0.20110066844518 | 0.0922107960592214 | -2.18087986482651 | 0.0291923033683057 | 0.317415456361503 | NM_010265,NM_173442,NM_001136484 |
| Dmxl2 | 10066.71195 | -0.176039862 | 0.050033502 | -3.518439782 | 0.000434092 | 0.006147922 | | Itga2b | 95.86429326 | -0.205915027589586 | 0.0944702272301058 | -2.1796817222428 | 0.0292810623381808 | 0.317415456361503 | NM_010575 |
| Rwdd1 | 1366.11154 | -0.176187197 | 0.064046844 | -2.750911436 | 0.005942971 | 0.040546187 | | Phf7 | 171.5324099 | -0.186896193181977 | 0.0857066002629036 | -2.18065111215095 | 0.02920923157047 | 0.317415456361503 | NM_027949 |
| Apba1 | 9000.174827 | -0.176506377 | 0.057849693 | -3.051120372 | 0.002279892 | 0.021064186 | | Rars | 1823.757981 | 0.121115361725685 | 0.0555609216381055 | 2.17986595893002 | 0.0292673989080653 | 0.317415456361503 | NM_025936 |
| Dmd | 2511.592565 | -0.176814466 | 0.050495806 | -3.501567349 | 0.00046253 | 0.006438851 | | Slc6a1 | 20951.09253 | 0.0862153837220029 | 0.0395188699073391 | 2.18162573788558 | 0.0291371657469514 | 0.317415456361503 | NM_178703 |
| Slc25a5 | 7442.673467 | -0.17718014 8 | 0.051149805 | -3.46394575 | 0.000532314 | 0.007140841 | | Usp27x | 722.0606312 | 0.177103991243471 | 0.0811778099870783 | 2.18167983678868 | 0.029133170049617 | 0.317415456361503 | NM_019461 |
| St3gal2 | 5037.027951 | -0.177220494 | 0.058845135 | -3.011642256 | 0.002598386 | 0.022992231 | | Zfp446 | 495.4911271 | -0.163226276106371 | 0.0748411871169028 | -2.18096856015137 | 0.0291857419904567 | 0.317415456361503 | NM_175558,NM_001168561,NM_001168562 |
| Gcsh | 1586.700514 | -0.177978017 | 0.05679425 | -3.133733009 | 0.001725978 | 0.017142504 | | 2410131K14Rik | 918.8072191 | 0.129647311129433 | 0.0595451953198116 | 2.17729256631219 | 0.0294587454751855 | 0.317771904512911 | NM_001081236 |
| Bzw2 | 1276.880679 | -0.178740258 | 0.059927956 | -2.982585595 | 0.002858247 | 0.024507848 | | 8430429K09Rik | 197.3866758 | -0.190307523414273 | 0.0874110897174777 | -2.17715537043832 | 0.0294689769239283 | 0.317771904512911 | NR_045276,NR_045275,NR_028317 |
| 0610009D07Rik | 1082.925922 | -0.178807012 | 0.057821125 | -3.092416688 | 0.00198534 | 0.019046966 | | Clcn6 | 2810.800385 | 0.106949603502359 | 0.0491776973051952 | 2.17475826162892 | 0.0296482363629161 | 0.317771904512911 | NM_011929 |
| 6-Mar | 11759.12775 | -0.17935654 | 0.065420174 | -2.741609043 | 0.006113906 | 0.04133022 | | Ddx11 | 80.37712385 | -0.20677533349742 | 0.0950133724341761 | -2.1762761198764 | 0.0295346200546624 | 0.317771904512911 | NM_001003919 |
| Rprd1a | 5401.404916 | -0.180207702 | 0.05911064 | -3.048650849 | 0.002298715 | 0.021168869 | | Ddx46 | 3123.737039 | -0.113190752752808 | 0.0519619605427817 | -2.17833876109457 | 0.0293808254385716 | 0.317771904512911 | NM_145975 |
| 2410015M20Rik | 1190.176276 | -0.180393732 | 0.055056287 | -3.276532826 | 0.001050901 | 0.011833328 | | Dtl | 411.1343753 | -0.187625457365346 | 0.0862683650802435 | -2.17490452254224 | 0.0296372719486384 | 0.317771904512911 | NM_029766 |
| Slc1a3 | 24809.19634 | -0.180875682 | 0.056931553 | -3.177072708 | 0.001487667 | 0.015387438 | | Eif4ebp3 | 202.8462281 | -0.191982509320907 | 0.0882486122863812 | -2.17547340798847 | 0.0295946587398994 | 0.317771904512911 | NM_201256 |
| Spcs1 | 2033.689478 | -0.180892589 | 0.065104782 | -2.778483901 | 0.005461322 | 0.038266657 | | Gm6297 | 74.88183417 | -0.20311914747936 | 0.0933749783711273 | -2.17530596550229 | 0.029607195792069 | 0.317771904512911 | NR_077226,NR_077221 |
| Nsg2 | 26465.299 | -0.181230185 | 0.063371631 | -2.859799891 | 0.004239084 | 0.032353861 | | Igfn1 | 2250.28956 | -0.207100528947312 | 0.0950952233938272 | -2.17782262406205 | 0.0294192448652341 | 0.317771904512911 | NM_177642 |
| Myh10 | 14401.10043 | -0.181637624 | 0.063287904 | -2.870021148 | 0.004104443 | 0.031576091 | | Nanos3 | 23.51254187 | -0.182045073018355 | 0.0837162594928668 | -2.1745485777929 | 0.0296639613465171 | 0.317771904512911 | NM_194059 |
| Abcd3 | 2862.01866 | -0.181875 | 0.063989807 | -2.842249523 | 0.004479642 | 0.03349786 | | Rraga | 3550.711971 | 0.0976956078374911 | 0.0448684478047903 | 2.177378817795 | 0.0294523147935685 | 0.317771904512911 | NM_178376 |
| Prmt2 | 1828.1902 | -0.183112882 | 0.062829272 | -2.91445177 | 0.003563139 | 0.028649336 | | Slc25a42 | 1547.53034 | -0.139127036628652 | 0.0639356593598869 | -2.17604757691668 | 0.0295517032057363 | 0.317771904512911 | NM_001007570 |
| Slc38a10 | 3182.244039 | -0.183482577 | 0.060554903 | -3.030020175 | 0.002445374 | 0.022066759 | | Smpd1 | 5104.575804 | 0.129832104828353 | 0.0596435602166293 | 2.17680004944029 | 0.0294954893701897 | 0.317771904512911 | NM_011421 |
| Csrnp3 | 4086.192875 | -0.183588217 | 0.067185468 | -2.732558445 | 0.006284452 | 0.042133838 | | Syt5 | 6086.151249 | 0.100197276021042 | 0.0460198534517348 | 2.17726195338991 | 0.0294610281838093 | 0.317771904512911 | NM_016908 |
| Itgav | 3728.42287 | -0.184214363 | 0.059227433 | -3.110287841 | 0.001869051 | 0.018186144 | | Tada2b | 1547.942453 | 0.131969682344703 | 0.0606489860512773 | 2.17595859282999 | 0.0295583568950742 | 0.317771904512911 | NM_001170454 |
| Tusc3 | 6600.70704 | -0.185119844 | 0.063710266 | -2.905651741 | 0.003664891 | 0.029209537 | | Vwa5a | 794.5075276 | -0.156421099601905 | 0.0718220474136245 | -2.17789808609984 | 0.0294136250403488 | 0.317771904512911 | NM_172767,NM_001145957 |
| Armcx2 | 3896.06452 | -0.185166701 | 0.057485056 | -3.221127613 | 0.001276873 | 0.013691257 | | Gap43 | 8915.506367 | 0.148176636275692 | 0.0681536040953542 | 2.17415701256791 | 0.0296933455060148 | 0.317852274738667 | NM_008083 |
| Tomm22 | 2047.183508 | -0.185292339 | 0.055303652 | -3.35045394 | 0.000806792 | 0.009645401 | | Nell1 | 2346.114181 | 0.144286204356531 | 0.0663745867859723 | 2.17381698844753 | 0.0297188821766565 | 0.317891371500819 | NM_001037906 |
| Selrc1 | 493.0000477 | -0.18536028 | 0.068601535 | -2.701984426 | 0.006892699 | 0.04487205 | | BC004004 | 2301.903003 | 0.113846595331747 | 0.0523868048418212 | 2.1731921936354 | 0.029765855079468 | 0.318159537074578 | NM_030561 |
| Uchl1 | 17963.70546 | -0.18552021 | 0.058093683 | -3.196437364 | 0.00139136 | 0.014585749 | | Irak1 | 2519.299127 | -0.126488125720303 | 0.0582286831603902 | -2.17226491919615 | 0.0298356864723526 | 0.318437312048048 | NM_001177976,NM_001177974,NM_001177975,NM_001177973,NM_008363 |
| Swi5 | 3323.586994 | -0.186061444 | 0.048362528 | -3.84722333 | 0.000119464 | 0.002238182 | | Ubqln1 | 7937.754046 | 0.097881486068935 | 0.045059230553791 | 2.1722848984757 | 0.0298341803848629 | 0.318437312048048 | NM_152234,NM_026842 |
| Sms | 4538.576634 | -0.186310041 | 0.065783507 | -2.832169474 | 0.004623334 | 0.034225481 | | Hpcal1 | 2717.607546 | 0.191730347951588 | 0.0882902212228748 | 2.17159211174242 | 0.0298864425398099 | 0.318722199578544 | NM_016677 |
| Mrps14 | 891.8300304 | -0.187807754 | 0.067202601 | -2.794650066 | 0.005195592 | 0.036775013 | | Wdr67 | 328.5304268 | -0.166237099647477 | 0.0765600464261495 | -2.17132966093262 | 0.0299062617393333 | 0.318722199578544 | NM_001167679,NM_001081396 |
| Chst10 | 3130.053675 | -0.187883465 | 0.046815333 | -4.013289125 | 5.99E-05 | 0.001325368 | | Cd200 | 3907.646093 | 0.158452441543033 | 0.0729916640063149 | 2.1708292816742 | 0.0299440796182679 | 0.318891276052023 | NM_010818 |
| Pin1 | 2505.096558 | -0.188059039 | 0.068981988 | -2.726728631 | 0.006396597 | 0.042690885 | | Cct3 | 4571.027001 | 0.0893725195874417 | 0.0411821001357843 | 2.17017877409762 | 0.0299933054140422 | 0.318947843663526 | NM_009836 |
| B230219D22Rik | 4196.938807 | -0.1882059 | 0.066628513 | -2.824705091 | 0.004732415 | 0.034668577 | | Oacyl | 78.12565905 | -0.206358304973001 | 0.095079444112044 | -2.17037769730567 | 0.0299782449399163 | 0.318947843663526 | NM_177028 |
| Chchd4 | 1146.835173 | -0.188392155 | 0.058507497 | -3.219966056 | 0.001282058 | 0.013718372 | | Cecr2 | 180.2323834 | -0.203792904602799 | 0.0939518975159304 | -2.16911962388247 | 0.0300736032104892 | 0.319144111214184 | NM_001128151 |
| Armc1 | 2980.334526 | -0.188425044 | 0.065547982 | -2.874612422 | 0.004045237 | 0.031275901 | | Top3a | 274.3791095 | -0.172466347011585 | 0.0795045120052563 | -2.16926489656566 | 0.0300625786707906 | 0.319144111214184 | NM_009410 |
| Dnajb11 | 1876.55572 | -0.18874525 | 0.068044909 | -2.773833534 | 0.005540002 | 0.038692429 | | Vcl | 1860.450452 | -0.13430613780614 | 0.0619188802692336 | -2.16906599767558 | 0.0300776737059216 | 0.319144111214184 | NM_009502 |
| Pdxp | 6085.790717 | -0.188768237 | 0.05921312 | -3.187946115 | 0.001432872 | 0.014919976 | | Setd6 | 448.704934 | -0.156098683804168 | 0.0719835401667072 | -2.16853302078027 | 0.0301181550452425 | 0.31934037969868 | NM_001035123 |
| Guk1 | 2564.350396 | -0.189284634 | 0.068257523 | -2.773095573 | 0.005552581 | 0.038718133 | | Pdgfra | 2616.831921 | 0.1392188609858 | 0.0642290267620579 | 2.16753807435925 | 0.030193849781192 | 0.319909454355649 | NM_001083316,NM_011058 |
| Tmem242 | 1239.410281 | -0.189468682 | 0.055938231 | -3.38710535 | 0.000706343 | 0.008762267 | | Sptbn4 | 8726.290236 | -0.143153841706048 | 0.066064370031977 | -2.16688423179934 | 0.0302436825716648 | 0.32020388705248 | NM_001199235,NM_001199234,NM_001199236,NM_032610 |
| Smarca2 | 11947.44592 | -0.189732511 | 0.070201056 | -2.702701652 | 0.006877845 | 0.044813007 | | Exosc8 | 498.4980204 | -0.139080641723347 | 0.0642243518906259 | -2.16554371712775 | 0.0303460712695935 | 0.320212939565955 | NM_027148,NM_001163570 |
| Dync1h1 | 25479.74989 | -0.190341877 | 0.071548914 | -2.660304202 | 0.00780701 | 0.049291695 | | Plcxd3 | 330.005716 | 0.180754522182818 | 0.0834769030579053 | 2.16532376695186 | 0.0303628995033528 | 0.320212939565955 | NM_177355 |
| Rps20 | 3786.829768 | -0.190355044 | 0.069346441 | -2.744986493 | 0.006051339 | 0.041042969 | | Tctn1 | 471.5544748 | -0.184190967731913 | 0.0850682296129069 | -2.16521454096379 | 0.0303712592858996 | 0.320212939565955 | NM_001039153 |
| Timm22 | 955.6283569 | -0.191270003 | 0.062005768 | -3.084713066 | 0.002037486 | 0.019331788 | | Tlr7 | 190.5483933 | -0.186789163865964 | 0.0862286370008432 | -2.16620800655979 | 0.0302952955877102 | 0.320212939565955 | NM_133211 |
| Vta1 | 1335.944826 | -0.1913323 | 0.070485866 | -2.714477539 | 0.00663804 | 0.043672949 | | Zcchc18 | 12950.05034 | 0.090366742224351 | 0.0417370947166857 | 2.16514213166409 | 0.0303768023352531 | 0.320212939565955 | NM_001035509,NM_001035510,NM_025893 |
| Higd1a | 2675.824238 | -0.191360804 | 0.051424859 | -3.721173136 | 0.000198299 | 0.003383757 | | Zfp771 | 392.0822371 | -0.205354874683507 | 0.0948412761160126 | -2.16524790780241 | 0.0303687052927611 | 0.320212939565955 | NM_177362 |
| Rit1 | 1031.572724 | -0.191661857 | 0.063226582 | -3.031349346 | 0.002434634 | 0.02204625 | | Cacybp | 2180.555814 | 0.135676503805354 | 0.0627423470393226 | 2.16243908950873 | 0.0305843472985887 | 0.3206025414857 | NM_009786 |
| Ext1 | 1673.055144 | -0.191677666 | 0.067782414 | -2.827837707 | 0.004686356 | 0.03444516 | | Cep192 | 412.8996416 | -0.164056633342788 | 0.0758039476499072 | -2.16422281990467 | 0.0304472527389902 | 0.3206025414857 | NM_027556 |
| Ep400 | 4083.179035 | -0.192000293 | 0.066381874 | -2.892360242 | 0.003823593 | 0.030195078 | | Cpm | 688.9688254 | 0.185815369537122 | 0.0858679385264079 | 2.16396681608907 | 0.0304668962649407 | 0.3206025414857 | NM_027468 |
| Ndufa13 | 3499.334994 | -0.1921289 | 0.055696498 | -3.449568733 | 0.000561483 | 0.007429218 | | Mfsd2b | 37.32123703 | -0.193300458007374 | 0.0893646650525307 | -2.16305245360398 | 0.030537145462842 | 0.3206025414857 | NM_001033488 |
| Zfp330 | 793.1130917 | -0.192339612 | 0.060308318 | -3.189271711 | 0.001426317 | 0.01486171 | | Necap1 | 8477.885494 | 0.119648888311049 | 0.0553242048819021 | 2.16268608950563 | 0.0305653317212347 | 0.3206025414857 | NM_026267 |
| H3f3b | 5479.448395 | -0.192351253 | 0.061333172 | -3.136169954 | 0.0017117 | 0.017022514 | | Oxct1 | 12418.4517 | 0.111358768417487 | 0.0514877228071304 | 2.1628217824787 | 0.0305548895645614 | 0.3206025414857 | NM_024188 |
| Thnsl1 | 863.3923196 | -0.192577231 | 0.068096547 | -2.828002861 | 0.004683939 | 0.034443734 | | Slc32a1 | 4616.49664 | 0.134402156153768 | 0.062147651689322 | 2.16262646295388 | 0.0305699212095398 | 0.3206025414857 | NM_009508 |
| Eif2s1 | 1434.508531 | -0.192898108 | 0.069250112 | -2.785527743 | 0.005344067 | 0.03755146 | | Usp49 | 720.1265382 | -0.135076883190008 | 0.062467301828104 | -2.16236141528426 | 0.0305903292371733 | 0.3206025414857 | NM_198421 |
| Arl15 | 1666.808741 | -0.193050821 | 0.070981634 | -2.71972917 | 0.00653354 | 0.043239925 | | Utp3 | 1034.974321 | 0.129848228326194 | 0.0600589003668049 | 2.1620147477419 | 0.030617039456683 | 0.32065112844108 | NM_023054 |
| Timm17b | 477.2129063 | -0.193102696 | 0.071046698 | -2.71796863 | 0.006568407 | 0.043399101 | | Trim12a | 95.65072675 | -0.203610996087144 | 0.0942045895195219 | -2.16137023817666 | 0.0306667511984209 | 0.320940365928142 | NM_023835 |
| Csgalnact1 | 877.9778584 | -0.193183962 | 0.070988587 | -2.721338299 | 0.006501819 | 0.043143116 | | Cdr2 | 761.7632911 | 0.169746400252835 | 0.078580190649978 | 2.16016783426935 | 0.0307596792488519 | 0.321052103585412 | NM_007672 |
| Eef1e1 | 557.9752635 | -0.193248453 | 0.065598039 | -2.945948636 | 0.003219659 | 0.026633951 | | Gfap | 10952.7927 | -0.174500775655028 | 0.080752631543366 | -2.16092989565692 | 0.0307007551502852 | 0.321052103585412 | NM_010277,NM_001131020 |
| Stard10 | 1868.215247 | -0.193401406 | 0.061796075 | -3.129671365 | 0.00175002 | 0.017336832 | | Lama5 | 654.8648405 | -0.165955280875995 | 0.0768388608347412 | -2.15978320179574 | 0.0307894566517857 | 0.321052103585412 | NM_001081171 |
| Acly | 8471.035684 | -0.19397627 | 0.050188938 | -3.864920757 | 0.000111125 | 0.002123027 | | Lims2 | 491.1160907 | 0.164878995343821 | 0.0763499224346092 | 2.15951752256243 | 0.0308100394050712 | 0.321052103585412 | NM_144862 |
| Fam63b | 3500.665236 | -0.194127117 | 0.064869168 | -2.992594505 | 0.00276617 | 0.023983794 | | Mcee | 533.0077857 | 0.14094434204396 | 0.0652443686382015 | 2.16025298406267 | 0.0307530904825564 | 0.321052103585412 | NM_028626 |
| Rasal2 | 4655.799076 | -0.194900065 | 0.064017893 | -3.044462353 | 0.002330966 | 0.021370407 | | Trim34a | 68.54565458 | -0.203427674818536 | 0.0941853119302168 | -2.1598662323193 | 0.0307830265167548 | 0.321052103585412 | NM_030684 |
| 2810407C02Rik | 5665.606864 | -0.195349578 | 0.071941356 | -2.715400262 | 0.006619571 | 0.043625536 | | Sox12 | 898.0973168 | -0.166631444430177 | 0.0772855178581011 | -2.15605004725617 | 0.0310797574346492 | 0.323630506448541 | NM_011438 |
| Fam160a2 | 3704.208157 | -0.195500601 | 0.051661506 | -3.784260595 | 0.000154166 | 0.002767775 | | Ciz1 | 2131.239677 | -0.137754702638792 | 0.0639150032870577 | -2.15527959875246 | 0.0311399612748276 | 0.323885538359364 | NM_028412,NM_001252538,NM_001252536,NM_001252537,NM_001252534 |
| Ptcd2 | 1077.061176 | -0.19563934 | 0.058001811 | -3.372986732 | 0.000743575 | 0.009066516 | | Cpsf7 | 3788.570747 | -0.0973316340326714 | 0.0451620117264659 | -2.15516604136642 | 0.0311488432526526 | 0.323885538359364 | NM_172302,NM_001164272 |
| Wnk1 | 10212.70462 | -0.195668199 | 0.049876691 | -3.923038868 | 8.74E-05 | 0.001768185 | | A130049A11Rik | 66.55608356 | -0.204629671986928 | 0.0950776292438807 | -2.15223784621342 | 0.0313786262650746 | 0.324435230190967 | NR_030676 |
| Pdzd8 | 2967.283641 | -0.196820057 | 0.06294932 | -3.126643081 | 0.001768144 | 0.017416166 | | A130077B15Rik | 377.0621112 | -0.171451767006014 | 0.0796380327929415 | -2.15288802338686 | 0.0313274799720318 | 0.324435230190967 | NR_040616 |
| Rdh14 | 793.9857803 | -0.196907322 | 0.065345608 | -3.013321454 | 0.00258405 | 0.022923025 | | Arl11 | 29.36215852 | -0.180417586341742 | 0.0838126903459151 | -2.15262850526711 | 0.0313478864237397 | 0.324435230190967 | NM_177337 |
| Nudt19 | 2127.233966 | -0.196907481 | 0.057134781 | -3.446368019 | 0.000568176 | 0.007492252 | | Cadm1 | 6397.831807 | 0.119254930526819 | 0.0554022999967624 | 2.15252670978981 | 0.0313558939280957 | 0.324435230190967 | NM_018770,NM_207676,NM_207675,NM_001025600 |
| Nme2 | 2158.006305 | -0.197883874 | 0.048114049 | -4.112808571 | 3.91E-05 | 0.000946283 | | Clns1a | 2504.730721 | 0.10222354341208 | 0.0475406656376152 | 2.15023374286116 | 0.031536730347628 | 0.324435230190967 | NM_023671 |
| Cpne5 | 5519.813838 | -0.19807441 | 0.0618166 | -3.204226834 | 0.001354257 | 0.014303247 | | Dnahc7b | 189.5309022 | -0.192082838588045 | 0.0893227519906612 | -2.15043574349487 | 0.0315207635768828 | 0.324435230190967 | NM_001160386 |
| Jkamp | 1569.531137 | -0.199050378 | 0.064183515 | -3.101269517 | 0.001926928 | 0.01865818 | | Gpd1 | 4326.337035 | 0.148297040797605 | 0.0689250228734592 | 2.15157042558936 | 0.0314312035130218 | 0.324435230190967 | NM_010271 |
| Vps29 | 2268.644399 | -0.199099184 | 0.051222525 | -3.886945893 | 0.000101513 | 0.001983415 | | Klhl17 | 2045.542088 | -0.175578646486126 | 0.0816359094873362 | -2.15075262330927 | 0.0314957303614768 | 0.324435230190967 | NM_198305 |
| Ica1l | 1544.785964 | -0.199305655 | 0.050821683 | -3.921665771 | 8.79E-05 | 0.001774122 | | Map4k1 | 78.6395891 | -0.204791479412006 | 0.0951025017659897 | -2.15337636349376 | 0.0312891116694262 | 0.324435230190967 | NM_008279 |
| 2510009E07Rik | 2462.912785 | -0.199470764 | 0.053777589 | -3.709180102 | 0.000207931 | 0.003505648 | | Metap1d | 534.6563802 | -0.168855541253136 | 0.0784898822797294 | -2.15130328073819 | 0.0314522694930125 | 0.324435230190967 | NM_025633 |

| | | | | | | | | | | | | | | | |
|---|---|---|---|---|---|---|---|---|---|---|---|---|---|---|---|
| Chga | 6075.46708 | -0.199622942 | 0.061472717 | -3.247342097 | 0.001164883 | 0.012809577 | | Pcdhgc4 | 562.3440901 | 0.167324961808328 | 0.0777960797767944 | 2.15081482625348 | 0.0314908183886156 | 0.324435230190967 | NM_033582 |
| Ndufb7 | 1592.123223 | -0.199745477 | 0.060163829 | -3.320025995 | 0.000900091 | 0.010498156 | | Rarg | 437.2887718 | -0.163451689413256 | 0.0759512589804425 | -2.15206030298122 | 0.0313926051779847 | 0.324435230190967 | NM_011244,NM_001042727 |
| Mea1 | 2019.598271 | -0.199805989 | 0.074623723 | -2.677513025 | 0.007417097 | 0.047350844 | | Supt20 | 880.2275896 | -0.157152885878135 | 0.0730267852059358 | -2.15198964920834 | 0.0313981696071823 | 0.324435230190967 | NM_019995 |
| Ppm1l | 2850.436383 | -0.200520314 | 0.059349513 | -3.37863453 | 0.000728468 | 0.008943645 | | Tuba1a | 26588.13307 | 0.106437886853957 | 0.0494254474959744 | 2.15350375659473 | 0.031279109178117 | 0.324435230190967 | NM_011653 |
| Zfp407 | 723.649761 | -0.201442276 | 0.067761555 | -2.972810712 | 0.002950864 | 0.025079913 | | Zfp810 | 652.4624069 | -0.157573516190012 | 0.0732538019926237 | -2.1510626329795 | 0.0314712563856765 | 0.324435230190967 | NM_145612 |
| Pcdhb20 | 424.9154934 | -0.201732851 | 0.073993086 | -2.726374331 | 0.006403433 | 0.042691391 | | Efna2 | 247.7500587 | 0.17296412185157 | 0.0804953399307314 | 2.14874702064017 | 0.0316544591452691 | 0.325415904843722 | NM_007909 |
| Cnep1r1 | 886.2493507 | -0.202233766 | 0.072839997 | -2.776410951 | 0.005496269 | 0.038463952 | | 9030624G23Rik | 189.6497607 | -0.189367279982431 | 0.0881928621110469 | -2.14719508415536 | 0.031777753911794 | 0.326301135408255 | NM_001256489 |
| Rpl15 | 6983.480644 | -0.202484105 | 0.038270184 | -5.290910092 | 1.22E-07 | 8.77E-06 | | Etl4 | 5421.192464 | 0.101299105087887 | 0.04718149486047 | 2.1470092329092 | 0.0317925465762772 | 0.326301135408255 | NM_001177631,NM_178059,NM_029895,NM_001081006,NM_001177630 |
| Lsm6 | 1233.133057 | -0.20289465 | 0.059195937 | -3.427509709 | 0.000609145 | 0.007897979 | | Gmfg | 51.27470603 | 0.201553347325034 | 0.0938971263139516 | 2.14653371447308 | 0.0318304219243768 | 0.326301135408255 | NM_022024,NM_001039192 |
| Bloc1s1 | 978.8985218 | -0.203205424 | 0.073372609 | -2.769499779 | 0.005614244 | 0.038972715 | | Gpam | 2266.059379 | -0.112836702098551 | 0.0525647290872787 | -2.14662386847265 | 0.0318232381299945 | 0.326301135408255 | NM_008149 |
| Cnih3 | 2825.385613 | -0.203378977 | 0.076429609 | -2.660997231 | 0.00779096 | 0.049250563 | | Pgm2 | 2884.922142 | 0.148336084898309 | 0.06912765717065 | 2.14582832645585 | 0.031886677787349 | 0.326647307150234 | NM_028132 |
| Pdpk1 | 7481.442386 | -0.203620416 | 0.072554867 | -2.80643359 | 0.005009324 | 0.035982597 | | Epsti1 | 34.6570421 | -0.183380488590577 | 0.0855071387874364 | -2.14462197181507 | 0.0319830841105899 | 0.327404002671198 | NM_029495,NM_178825 |
| Gabra3 | 6979.437762 | -0.204170357 | 0.066177523 | -3.085191902 | 0.002034209 | 0.019331788 | | 2700094K13Rik | 376.5607273 | 0.150587450438526 | 0.0703343720260596 | 2.1410221787824 | 0.032272249554709 | 0.3277115992225 | NM_001033166,NM_001037279 |
| Rps21 | 4196.097987 | -0.204324777 | 0.075346081 | -2.711816922 | 0.006691554 | 0.043969015 | | Ajuba | 82.2640887 | -0.203623133349868 | 0.0950899279844774 | -2.14137435652604 | 0.0322438611817966 | 0.3277115992225 | NM_010590 |
| Hsd17b12 | 2612.578033 | -0.204361656 | 0.07006929 | -2.916565248 | 0.003539087 | 0.028529975 | | Ccdc92 | 3704.111099 | 0.090930693872558 | 0.0424619608611243 | 2.14146242963096 | 0.0322367651232987 | 0.3277115992225 | NM_144819 |
| Tmem70 | 1094.997445 | -0.204922764 | 0.073378945 | -2.792664351 | 0.00522759 | 0.036943149 | | Cthrc1 | 221.2783517 | 0.184877735831414 | 0.0862817806783535 | 2.142720448951 | 0.0321355523657402 | 0.3277115992225 | NM_026778 |
| H2afv | 934.5230211 | -0.205094048 | 0.066156461 | -3.100136343 | 0.001934316 | 0.018673077 | | Ctif | 5065.395647 | 0.103529873810081 | 0.0483572909052523 | 2.14093618298283 | 0.0322791847636017 | 0.3277115992225 | NM_201354 |
| Ndufb8 | 3688.617777 | -0.205199265 | 0.070545712 | -2.908741834 | 0.003628864 | 0.028965914 | | Ddc | 374.3889615 | 0.203319211906005 | 0.0948577283368853 | 2.14341219709501 | 0.0320800145572887 | 0.3277115992225 | NM_001190448,NM_016672 |
| Atp5g3 | 6698.707539 | -0.205641521 | 0.053048242 | -3.876500175 | 0.00010597 | 0.002047251 | | Epm2aip1 | 5918.878465 | 0.117895289390342 | 0.0550667741503741 | 2.14095143958131 | 0.032277954288271 | 0.3277115992225 | NM_175266 |
| Pds5b | 4342.271676 | -0.205648899 | 0.071207407 | -2.88802678 | 0.003876669 | 0.030481586 | | Kcnk10 | 146.1603196 | 0.195284009250335 | 0.0911060071332762 | 2.14348115338498 | 0.0320744828325845 | 0.3277115992225 | NM_029911 |
| Rps16 | 4309.207592 | -0.205712683 | 0.060236674 | -3.415073725 | 0.000637647 | 0.008131449 | | Ncald | 22907.36797 | 0.093266254825551 | 0.0435594058977476 | 2.1411277978512 | 0.0322637335565831 | 0.3277115992225 | NM_134094,NM_001170868,NM_001170867,NM_001170866 |
| Ralyl | 2226.841434 | -0.205816204 | 0.067428997 | -3.052339705 | 0.00227065 | 0.021003849 | | Pak3 | 4653.636724 | 0.151701277487412 | 0.0708593664821371 | 2.14087826378824 | 0.0322838564289823 | 0.3277115992225 | NM_001195046,NM_008778,NM_001195047,NM_001195048,NM_001195049 |
| Serp2 | 1825.538046 | -0.205860799 | 0.063267164 | -3.253833201 | 0.001138591 | 0.012609958 | | Por | 3143.793841 | 0.138022534510011 | 0.0644490095512138 | 2.14157727901671 | 0.0322275137076125 | 0.3277115992225 | NM_008898 |
| Snx18 | 957.1119867 | -0.205892181 | 0.075997035 | -2.709213344 | 0.006744296 | 0.044184405 | | Sgsm1 | 6394.063931 | 0.159210515996949 | 0.0742853571537298 | 2.14322878824519 | 0.0320947317310968 | 0.3277115992225 | NM_172718,NM_001162965,NM_001254731,NR_027934,NR_027936 |
| Gde1 | 6906.107748 | -0.20627582 | 0.058793548 | -3.508477127 | 0.00045068 | 0.006342267 | | Kcnh4 | 1189.025269 | -0.186717761853611 | 0.0872891328425231 | -2.13907225072869 | 0.0324298173684666 | 0.32896335690946 | NM_001081194 |
| Mzt1 | 1860.166132 | -0.206585931 | 0.05988008 | -3.449994211 | 0.000560599 | 0.007423858 | | Ehd1 | 1595.309918 | 0.108559965670046 | 0.0507668852338979 | 2.13840114810822 | 0.0324841993806944 | 0.329285052480088 | NM_010119 |
| Fam174b | 1377.833796 | -0.206765576 | 0.076066606 | -2.718217474 | 0.006563468 | 0.043384973 | | Elp5 | 2294.160593 | 0.0955150203585465 | 0.0446945275715575 | 2.13706298171792 | 0.0325928693314006 | 0.329798129316311 | NM_018740,NM_001253700 |
| Fxyd6 | 3992.991095 | -0.206808161 | 0.056036932 | -3.690568969 | 0.000223753 | 0.003680286 | | Iqce | 1346.916099 | -0.15997396612923 | 0.0748626165206538 | -2.13690054615036 | 0.0326060815828254 | 0.329798129316311 | NM_028833 |
| Dolk | 684.4458121 | -0.206886123 | 0.072527466 | -2.85252104 | 0.004337394 | 0.032878466 | | Trim72 | 37.78313842 | -0.193915916579868 | 0.090730297172981 | -2.13727853453582 | 0.0325753436869799 | 0.329798129316311 | NM_001079932 |
| Psmb5 | 1781.380598 | -0.206934627 | 0.069826906 | -2.963537099 | 0.003041254 | 0.02553994 | | Zfp57 | 385.51964 | -0.190555929752578 | 0.0891840079537437 | -2.13666030631201 | 0.0326256307192302 | 0.329798129316311 | NM_001168502,NM_001168501,NR_033137,NM_001013745 |
| Pnma2 | 2969.016783 | -0.207968869 | 0.054873099 | -3.789996804 | 0.000150649 | 0.002722791 | | Gabarapl2 | 5004.294183 | 0.0887341076678986 | 0.0415477669850843 | 2.13571303843488 | 0.0327028109968659 | 0.330348423185309 | NM_026693 |
| Mrps21 | 925.033669 | -0.208357331 | 0.075561317 | -2.75746031 | 0.005825228 | 0.039989406 | | Timm17b | 530.0089425 | 0.150955886677042 | 0.0706998285448636 | 2.1351662342611 | 0.0327474339472923 | 0.330481624984955 | NM_011591 |
| Cox7b | 3671.480743 | -0.208374788 | 0.076444561 | -2.725828834 | 0.006414025 | 0.042743615 | | Ubl4 | 3346.348793 | 0.108065569624352 | 0.050616333863144 | 2.13499401036311 | 0.0327614993789299 | 0.330481624984955 | NR_045779,NM_145405 |
| Nme1 | 4792.757464 | -0.208634707 | 0.062062573 | -3.361683151 | 0.00077469 | 0.009333624 | | Aqp9 | 61.27155413 | -0.200352556265054 | 0.0939031681967038 | -2.13360805724217 | 0.032874877885438 | 0.331395195117192 | NR_073483,NM_001271843,NM_022026 |
| Usp31 | 5730.814556 | -0.208769454 | 0.065379466 | -3.193196044 | 0.001407074 | 0.014710663 | | Hfm1 | 74.02917977 | -0.202136160910412 | 0.0947990932804561 | -2.1322583783835 | 0.0329856116520814 | 0.332280856350995 | NM_177873,NM_001252516 |
| Fam49a | 10302.60065 | -0.208938773 | 0.062099451 | -3.364583265 | 0.000766593 | 0.009250467 | | E230016M11Rik | 52.22805279 | -0.20207704630652 | 0.0948161564371649 | -2.13125119072336 | 0.0330684537861432 | 0.332884518154897 | NR_040278 |
| Ppdpf | 1215.558062 | -0.209536907 | 0.060465683 | -3.465385584 | 0.000529472 | 0.007115035 | | Pnmal1 | 3494.502714 | 0.120112201156052 | 0.0564205347791259 | 2.12887385109456 | 0.0332646987407386 | 0.334628125975047 | NM_001007569 |
| Prdx3 | 2166.488645 | -0.209842377 | 0.07492398 | -2.800737204 | 0.005098602 | 0.036290258 | | Zfp207 | 3801.569487 | -0.110057642749161 | 0.0517055660484116 | -2.12854536097841 | 0.0332918931963964 | 0.334669924270487 | NM_011751,NM_001130169,NM_001130171,NM_001130170,NR_045038 |
| Mrpl13 | 874.7496095 | -0.210444558 | 0.071804378 | -2.930804005 | 0.00338086 | 0.027584543 | | Cr1l | 623.300326 | -0.147200571704973 | 0.0691751399950303 | -2.12794035133934 | 0.033342029444709 | 0.334860975734621 | NM_013499 |
| Myo10 | 1760.663839 | -0.21050183 | 0.075467425 | -2.789307178 | 0.005282094 | 0.037217264 | | Sec62 | 6059.649482 | 0.120024725015107 | 0.0564089633833646 | 2.12775980652931 | 0.0333570034343932 | 0.334860975734621 | NM_027016 |
| Sephs2 | 1151.085914 | -0.210865158 | 0.058824576 | -3.584643934 | 0.000337538 | 0.005132306 | | Atf4 | 4450.795165 | 0.122778434124039 | 0.0577228666559636 | 2.12703285953929 | 0.0334173530576511 | 0.335235131571436 | NM_009716 |
| AI597468 | 1112.933861 | -0.211280938 | 0.076514875 | -2.761305399 | 0.005757081 | 0.039657524 | | 4933439C10Rik | 940.3748484 | -0.18400155002432 | 0.0865908051169487 | -2.12495483528314 | 0.0335903817107218 | 0.335456506554182 | NR_015585 |
| D630045J12Rik | 3072.036594 | -0.211491029 | 0.076855296 | -2.751808137 | 0.005926723 | 0.040455567 | | Atp1b1 | 59295.23158 | 0.0936515731931258 | 0.0440688878831447 | 2.12511768941066 | 0.0335767938884554 | 0.335456506554182 | NM_009721 |
| Slc1a2 | 125043.0129 | -0.211759369 | 0.044956061 | -4.710363019 | 2.47E-06 | 0.00010613 | | Gfpt2 | 339.3242941 | -0.19697219049893 | 0.0926674025306905 | -2.1255822988423 | 0.0335380547855501 | 0.335456506554182 | NM_013529 |
| Scn2a1 | 13122.46649 | -0.211823352 | 0.070172149 | -3.018624254 | 0.002539252 | 0.022624024 | | Inpp5d | 792.7526384 | -0.158089222277192 | 0.0744077382179112 | -2.12463415853618 | 0.0336171512977914 | 0.335456506554182 | NM_001110193,NM_001110192,NM_010566 |
| Casd1 | 2507.605979 | -0.212233012 | 0.069921582 | -3.035300506 | 0.002402961 | 0.021823847 | | Oas1a | 23.6211571 | -0.159783266104051 | 0.0751573911850483 | -2.1259820702217 | 0.0335047524825095 | 0.335456506554182 | NM_145211 |
| Tceb2 | 3040.666416 | -0.212528385 | 0.056299188 | -3.774981348 | 0.00016002 | 0.002856387 | | Rgl3 | 159.4592501 | -0.193335533093536 | 0.09090474473623 | -2.12595693671382 | 0.0335068453547782 | 0.335456506554182 | NM_023622 |
| Eif5a2 | 2333.190621 | -0.212749616 | 0.070910817 | -3.000242079 | 0.002697651 | 0.023561109 | | Runx1 | 57.97645615 | -0.198564180358849 | 0.093441091769515 | -2.12501990954432 | 0.033584951640332 | 0.335456506554182 | NM_001111021,NM_001111022,NM_001111023,NM_009821 |
| Notch1 | 1433.46087 | -0.212755381 | 0.073014034 | -2.913897069 | 0.003569476 | 0.028685403 | | Tmem181a | 2008.952096 | -0.114449133884511 | 0.0538698203003439 | -2.12455013301353 | 0.0336241686316184 | 0.335456506554182 | NM_001033178 |
| Snrpd2 | 1338.07867 | -0.212806234 | 0.066437407 | -3.203108658 | 0.001359527 | 0.014339352 | | Cxxc4 | 908.1263559 | 0.150790365716268 | 0.0709989224121533 | 2.1238402019811 | 0.0336835080607312 | 0.335817871029637 | NM_001004367 |
| 1700025G04Rik | 4420.121962 | -0.212982072 | 0.052463671 | -4.059610584 | 4.92E-05 | 0.001131652 | | Mtfp1 | 2303.136147 | 0.146833662696645 | 0.0691593468485835 | 2.12312101527101 | 0.0337437124210128 | 0.336187357083424 | NM_026443 |
| Cplx2 | 59873.1095 | -0.213256739 | 0.075834503 | -2.812133415 | 0.004921409 | 0.035648576 | | Dusp10 | 289.4369783 | 0.182614792369608 | 0.0861044477387703 | 2.12085202524773 | 0.0339342569833958 | 0.33632629510183 | NM_022019 |
| Elovl5 | 2724.835622 | -0.213549169 | 0.073965802 | -2.887133812 | 0.003887689 | 0.030510193 | | F3 | 1652.511105 | 0.135092590239776 | 0.0636928025521513 | 2.12100244967495 | 0.0339215962760107 | 0.33632629510183 | NM_010171 |
| Gria3 | 9985.413447 | -0.213633771 | 0.062662007 | -3.409303048 | 0.000651291 | 0.008237633 | | Pold1 | 307.674381 | -0.174453100401299 | 0.0822236201012613 | -2.12169082541554 | 0.0338637095696559 | 0.33632629510183 | NM_011131 |
| Akap11 | 12092.52585 | -0.214723663 | 0.077859925 | -2.757820065 | 0.005818822 | 0.039963132 | | Ptpn23 | 3290.752167 | -0.141695308408648 | 0.0667775282415797 | -2.12190166671873 | 0.0338460468608613 | 0.33632629510183 | NM_001081043 |
| Kcnd2 | 4019.492144 | -0.214856286 | 0.079300294 | -2.709400871 | 0.006740484 | 0.044184405 | | Rasgef1b | 2730.00515 | -0.16779774279494 | 0.0791203356024442 | -2.12079915970395 | 0.0339393442679035 | 0.33632629510183 | NM_181318,NM_145839 |
| Gabarapl2 | 4780.507071 | -0.214901601 | 0.042361046 | -5.073094771 | 3.91E-07 | 2.30E-05 | | Uqcrc1 | 8886.647328 | 0.108212085687741 | 0.0510253962648816 | 2.12074954006178 | 0.0339428850763653 | 0.33632629510183 | NM_025407 |
| Ndufa8 | 2964.542542 | -0.215202682 | 0.055930349 | -3.847690655 | 0.000119236 | 0.002236622 | | Zc3hav1 | 203.5515442 | -0.191661044319921 | 0.0903011816328142 | -2.12246441136573 | 0.033798758201487 | 0.33632629510183 | NM_028421,NM_028864 |
| Reps2 | 8345.854678 | -0.215274667 | 0.070885114 | -3.036951687 | 0.002389838 | 0.021768456 | | Zfp251 | 2234.919922 | 0.120753429274498 | 0.056898281010419 | 2.1222684961675 | 0.033815197436795 | 0.33632629510183 | NM_001007568 |
| Zeb1 | 3265.258254 | -0.215305155 | 0.070923651 | -3.035731391 | 0.00239953 | 0.021823847 | | Pycard | 88.71190982 | -0.201401005397379 | 0.0950450304487097 | -2.11900616419986 | 0.0340899460102727 | 0.336970371935979 | NM_023258 |
| Gatad1 | 2382.462965 | -0.215418793 | 0.060460136 | -3.562988886 | 0.000366656 | 0.00546773 | | Ran | 7704.165734 | 0.102814795093652 | 0.048523198243887 | 2.11887919211105 | 0.0341006778704316 | 0.336970371935979 | NM_009391 |
| Erh | 1568.037313 | -0.215457421 | 0.04824643 | -4.465769176 | 7.98E-06 | 0.000262449 | | Sdccag8 | 688.4419746 | -0.140381313545771 | 0.066239952417488 | -2.11928463747974 | 0.0340664191641937 | 0.336970371935979 | NM_029756 |
| Atp5g1 | 4323.655338 | -0.21583791 | 0.065008111 | -3.320168934 | 0.00089963 | 0.010498156 | | Uckl1 | 794.0249261 | -0.138555961792165 | 0.065380674857924 | -2.11921889905932 | 0.0340719718301202 | 0.336970371935979 | NM_026765 |
| Celf4 | 16304.22495 | -0.215866214 | 0.058419172 | -3.695126245 | 0.000219778 | 0.003630313 | | Cxx1c | 4518.893102 | 0.120646975371367 | 0.056956959870782 | 2.11821304446513 | 0.0341570289399791 | 0.337297758247543 | NM_028375 |
| Tenm2 | 2511.105801 | -0.215939245 | 0.058987574 | -3.660758163 | 0.00025147 | 0.004058622 | | Olig1 | 3549.355727 | 0.150308014125198 | 0.0710214244181941 | 2.11637566208391 | 0.0343128703155852 | 0.338606490627846 | NM_016968 |
| Tmem183a | 2103.510793 | -0.216436312 | 0.058608487 | -3.692917548 | 0.000221696 | 0.003650326 | | Prrt4 | 159.4053116 | -0.189061827899375 | 0.0893465108005861 | -2.11605160856639 | 0.034340418517496 | 0.338648281999421 | NM_001101443 |
| Dync2h1 | 1764.679479 | -0.216526163 | 0.059801156 | -3.620768867 | 0.000293729 | 0.004573904 | | Cdh8 | 1646.775337 | 0.156543720436591 | 0.0740048306993626 | 2.11531759423294 | 0.0344028879036614 | 0.338653870994267 | NM_001039154,NM_007667 |
| Txn1 | 1471.276216 | -0.216831216 | 0.055450046 | -3.910388399 | 9.21E-05 | 0.001844623 | | Gabra1 | 15305.91778 | 0.156388065597966 | 0.0739345443883598 | 2.11522322740637 | 0.0344109261728995 | 0.338653870994267 | NM_010250 |
| Tanc2 | 8753.849108 | -0.217239088 | 0.072226312 | -3.007755512 | 0.002631848 | 0.023198299 | | Npr2 | 2354.322058 | -0.14313290961439 | 0.0676656480996391 | -2.11529651505922 | 0.034404683311501 | 0.338653870994267 | NM_173788 |
| Spryd7 | 1303.369018 | -0.217324671 | 0.058415596 | -3.720319313 | 0.000198971 | 0.003387755 | | Gmcl1 | 1064.419704 | -0.13081366417024 | 0.0618685546350733 | -2.11438047877202 | 0.0344827836048952 | 0.339131289536024 | NM_011818 |
| Hsbp1 | 6932.186699 | -0.217485076 | 0.040683298 | -5.345807427 | 9.00E-08 | 6.94E-06 | | BC031181 | 3192.022316 | 0.0941771715186015 | 0.0445797745140438 | 2.11255378801732 | 0.0346389775925666 | 0.339228269500411 | NM_001001181 |
| Mfap3l | 2910.079189 | -0.217722191 | 0.058908813 | -3.695918839 | 0.000219093 | 0.003626738 | | Bnip2 | 1334.38692 | -0.120331908911703 | 0.0569820721055659 | -2.11175031839443 | 0.0347078706197565 | 0.339228269500411 | NM_001008238,NM_016787 |
| Nab1 | 1102.338196 | -0.218135486 | 0.076507496 | -2.851164893 | 0.004355937 | 0.032960625 | | Ccnl1 | 1185.351787 | -0.146701415355601 | 0.0694488704495567 | -2.11236575060145 | 0.0346550902715673 | 0.339228269500411 | NM_019937 |
| Cacybp | 2041.666175 | -0.21835868 | 0.071768822 | -3.042528402 | 0.002345997 | 0.021482786 | | Cntnap4 | 1127.093597 | 0.152151884789578 | 0.0720059600061947 | 2.11304570866756 | 0.0345968558222178 | 0.339228269500411 | NM_130457 |
| Ndn | 6907.885412 | -0.218576506 | 0.062972967 | -3.470957703 | 0.000518606 | 0.007005472 | | Coro1c | 4056.083877 | 0.0948908552886638 | 0.0449314713110047 | 2.1119018033452 | 0.0346948726875973 | 0.339228269500411 | NM_011779 |
| Phactr1 | 25008.58176 | -0.218679675 | 0.060110038 | -3.637989277 | 0.000274775 | 0.004344247 | | Gpr124 | 706.0051939 | -0.147451657228054 | 0.0698192929993269 | -2.11190418713457 | 0.034694668183463 | 0.339228269500411 | NM_054044 |
| Bsg | 13114.69301 | -0.218798481 | 0.077189634 | -2.83455782 | 0.004588915 | 0.034079589 | | Ndufb3 | 1683.00762 | 0.115513527293537 | 0.0547024968226229 | 2.11166827847179 | 0.034714911664484 | 0.339228269500411 | NM_025597 |
| Ssbp4 | 2733.23546 | -0.218877457 | 0.073783698 | -2.966474463 | 0.003012354 | 0.025407408 | | Nlrp1a | 106.0997897 | -0.198832714727054 | 0.0941649184547595 | -2.11153705636756 | 0.034726176287148 | 0.339228269500411 | NM_001004142 |
| Tcf20 | 5220.32896 | -0.218901123 | 0.080375858 | -2.723468574 | 0.006460036 | 0.042939425 | | Plekhb2 | 12739.84243 | 0.0928029586814262 | 0.0439216489322209 | 2.11292064249769 | 0.0346075607337265 | 0.339228269500411 | NM_145516,NM_175421 |
| Rps12 | 4654.99579 | -0.218946756 | 0.069381305 | -3.155702481 | 0.001601121 | 0.01619582 | | Snrpd1 | 929.8584777 | 0.139940330440049 | 0.0662162826608697 | 2.11338246148249 | 0.0345680458681889 | 0.339228269500411 | NM_009226 |
| Nedd8 | 2698.066681 | -0.219274063 | 0.059122898 | -3.708784087 | 0.000208257 | 0.003507318 | | Pcdha9 | 151.6822324 | 0.194657032038282 | 0.0922330266754301 | 2.11049164333818 | 0.0348160302162502 | 0.339877456264281 | NM_138661 |
| Sorcs2 | 2631.138908 | -0.219342691 | 0.075284923 | -2.913500879 | 0.003574008 | 0.028706939 | | Rad51ap1 | 61.3464051 | -0.200676561964715 | 0.0951006477654471 | -2.11014926480476 | 0.0348455010003359 | 0.339936700826648 | NM_009013 |
| Mrpl12 | 1528.721775 | -0.219477515 | 0.069962563 | -3.137070821 | 0.001706449 | 0.017013978 | | Irgm1 | 578.2573813 | -0.16091116878312 | 0.0762868256605051 | -2.10929170783975 | 0.0349194101104231 | 0.34020077214219 | NM_008326 |
| Rbm12 | 787.1576654 | -0.21949251 | 0.081139868 | -2.705112976 | 0.006828116 | 0.044601532 | | Zxdb | 175.5375258 | 0.193684787968742 | 0.0918208608220939 | 2.10937673895274 | 0.0349120756749867 | 0.34020077214219 | NM_001081473 |
| Vkorc1l1 | 638.1950196 | -0.219566855 | 0.068484658 | -3.206073629 | 0.001345596 | 0.014231165 | | Cartpt | 277.7391602 | 0.200381385078146 | 0.09504400874952 | 2.10830106720597 | 0.0350049557068181 | 0.340259169885816 | NM_013732,NM_001081493 |
| Ndufa2 | 1610.729781 | -0.219957898 | 0.077130134 | -2.851776436 | 0.004347567 | 0.032932886 | | Gphn | 3130.321217 | 0.096735731068899 | 0.0458797296917123 | 2.10846340462144 | 0.0349909250065439 | 0.340259169885816 | NM_145965,NM_172952 |
| Pak6 | 3889.667023 | -0.220036479 | 0.065606653 | -3.35387447 | 0.000796885 | 0.009541682 | | Slc25a5 | 7927.026801 | 0.115919010531997 | 0.0549704377694362 | 2.10875181708028 | 0.0349660095779115 | 0.340259169885816 | NM_007451 |
| Pgrmc1 | 8558.057799 | -0.220175693 | 0.064397441 | -3.419013057 | 0.000628487 | 0.008067754 | | Zswim8 | 6371.789537 | -0.12134635420469 | 0.0575609306317734 | -2.10813746165463 | 0.0350191008522164 | 0.340259169885816 | NM_027996,NM_001252082,NM_001252081 |
| Polr1d | 1392.615692 | -0.220204446 | 0.062242706 | -3.537835325 | 0.000403422 | 0.005876062 | | Sat1 | 1244.529383 | -0.142653148131404 | 0.0677067723988433 | -2.10692583736042 | 0.0351240085962053 | 0.341050366890694 | NM_009121 |
| Rps18 | 4385.517557 | -0.22023814 | 0.070397349 | -3.128500457 | 0.001757007 | 0.01736241 | | Slc25a34 | 352.5844213 | 0.200254459983347 | 0.0950984594380637 | 2.10575924370642 | 0.0352252707764342 | 0.341805132463917 | NM_001013780 |
| Kcnk2 | 2894.325898 | -0.220496965 | 0.073441897 | -3.002332084 | 0.002679197 | 0.023452816 | | C1qb | 2328.58751 | -0.200067073559943 | 0.0950271512442529 | -2.10536475593013 | 0.0352593326510598 | 0.341907253731172 | NM_009777 |
| Ldoc1l | 3210.175525 | -0.220756163 | 0.074628477 | -2.958068735 | 0.003095731 | 0.025824561 | | Bex1 | 1202.57815 | 0.17002889402529 | 0.0807769331117999 | 2.10491891032754 | 0.0352983673000065 | 0.342057427218075 | NM_009052 |
| Chd9 | 2765.180247 | -0.220845909 | 0.082681786 | -2.671034557 | 0.007561786 | 0.048076452 | | 2810403A07Rik | 2375.268501 | -0.145080008102687 | 0.06895997881179 | -2.10382907017197 | 0.0353933600136156 | 0.342065168035782 | NM_028814 |
| Dnalc1 | 5506.664735 | -0.221324696 | 0.060005543 | -3.688404187 | 0.000225665 | 0.00370387 | | Dolk | 757.239844 | 0.138861955345178 | 0.0660010567297163 | 2.10393252870057 | 0.0353840922036781 | 0.342065168035782 | NM_177648 |
| Rap2b | 2194.056811 | -0.221397533 | 0.075031754 | -2.95071871 | 0.003170364 | 0.026352802 | | Lct | 84.89152746 | -0.12370897007432 | 0.0587989142503026 | -2.10393289827939 | 0.0353843007506218 | 0.342065168035782 | NM_001081078 |
| Tmsb4x | 16876.94959 | -0.221637628 | 0.069301923 | -3.198145429 | 0.001383145 | 0.014519275 | | Ndufa9 | 3857.643713 | 0.0934282526468877 | 0.0444039827554091 | 2.10405118751441 | 0.0353739821292765 | 0.342065168035782 | NM_025358 |
| Vps13a | 2429.544911 | -0.222028936 | 0.070217603 | -3.162012466 | 0.001566829 | 0.015971343 | | Adamts20 | 471.1894728 | -0.160213597854069 | 0.0761696542175658 | -2.10337830071338 | 0.0354327137323392 | 0.342120019231982 | NM_001164785,NM_177431,NM_001164786 |
| Tubb4b | 10125.50264 | -0.222450048 | 0.077982929 | -2.849919232 | 0.004373023 | 0.033045155 | | Fam46c | 64.67680988 | -0.195274430804268 | 0.0928696577557134 | -2.10267201929313 | 0.0354944495994744 | 0.342120019231982 | NM_001142952 |
| Fopnl | 1211.32893 | -0.222637633 | 0.058836911 | -3.783978927 | 0.000154341 | 0.002767775 | | Maml2 | 412.1795482 | -0.170842660552549 | 0.0812477213587711 | -2.10273787000312 | 0.0354886897299875 | 0.342120019231982 | NM_173776,NM_001013813 |
| Tram1l1 | 1079.409376 | -0.223311174 | 0.067762891 | -3.295478845 | 0.000982541 | 0.011235048 | | Rara | 529.066335 | 0.177744154983004 | 0.0845232279933942 | 2.10290306230253 | 0.0354742441000684 | 0.342120019231982 | NM_009024,NM_001176528,NM_001177303,NM_001177302 |
| Mid2 | 1235.345003 | -0.223400024 | 0.079097959 | -2.824346251 | 0.004737717 | 0.03467463 | | Slc16a11 | 525.3114085 | -0.184998524027983 | 0.0879932751737907 | -2.10241661834501 | 0.0355167967094746 | 0.342120019231982 | NM_153081 |
| Cdh10 | 1872.274555 | -0.223563643 | 0.06134207 | -3.644540265 | 0.00026787 | 0.004269939 | | BC023829 | 1769.759221 | 0.124364938228667 | 0.0591887695889229 | 2.10115768738571 | 0.0356271263316071 | 0.34272823648538 | NM_001033328 |
| Gja1 | 6786.808247 | -0.223844228 | 0.07758358 | -2.885201062 | 0.003911638 | 0.030656 | | Vim | 1457.748291 | -0.135949607552668 | 0.0646954480387345 | -2.10137825262872 | 0.0356077543700033 | 0.34272823648538 | NM_011701 |
| Sptbn1 | 40054.53704 | -0.224036527 | 0.038236364 | -5.859252989 | 4.65E-09 | 5.58E-07 | | Adam8 | 147.43239 | -0.199509586051752 | 0.0949737739641746 | -2.10068082718298 | 0.0356689934681632 | 0.342903904115512 | NM_007403 |
| Grid1 | 2481.064177 | -0.22453446 | 0.057910568 | -3.877262293 | 0.000105638 | 0.002043399 | | Hexdc | 1159.918226 | -0.160544964427628 | 0.0764378323272361 | -2.100333794924 | 0.0356994884129007 | 0.342970085109653 | NM_001146073,NM_001001333 |
| Dgat2 | 2655.805663 | -0.224555094 | 0.060203092 | -3.729959477 | 0.000191511 | 0.003296961 | | Scml4 | 184.5308144 | 0.198625020594816 | 0.094593348852436 | 2.09977787026726 | 0.0357483858124867 | 0.343212856782671 | NM_172938 |
| Gpr125 | 865.8734092 | -0.224567457 | 0.074366457 | -3.019741237 | 0.002529907 | 0.022566715 | | Ccrn4l | 2086.736107 | 0.145688415654152 | 0.0694265538617838 | 2.09845379830017 | 0.0358650771542335 | 0.344080613138572 | NM_009834 |
| Mettl9 | 1995.363653 | -0.224583753 | 0.072790646 | -3.085338089 | 0.002033209 | 0.019331788 | | Cd274 | 182.7608905 | -0.146814934440827 | 0.0700249718520427 | -2.09660826070784 | 0.036028267422812 | 0.344080613138572 | NM_021893 |
| Ndufb9 | 4651.72367 | -0.224630582 | 0.075247374 | -2.985228169 | 0.002833669 | 0.024364519 | | E130102H24Rik | 161.193477 | -0.192134137886604 | 0.0916225803990849 | -2.09701731876265 | 0.0359920422657945 | 0.344080613138572 | NR_040708 |
| Hbegf | 540.6838468 | -0.224818615 | 0.079487932 | -2.828336467 | 0.00467906 | 0.034424195 | | Eno3 | 330.4129066 | -0.183102641154321 | 0.0872842174504091 | -2.09777490711138 | 0.0359250341553195 | 0.344080613138572 | NM_001136062,NM_001276285,NM_007933 |
| Cenpv | 534.775648 | -0.225009681 | 0.077263234 | -2.912247782 | 0.003588379 | 0.02876272 | | Panx1 | 622.0165107 | 0.13064717673634 | 0.0623053014952659 | 2.09688699999737 | 0.0360035795968905 | 0.344080613138572 | NM_019482 |
| 2610507B11Rik | 14041.9846 | -0.225524877 | 0.040321262 | -5.593199863 | 2.23E-08 | 2.09E-06 | | Stx7 | 3956.888454 | 0.113819024444617 | 0.054276233447617 | 2.09703248023771 | 0.0359907002006132 | 0.344080613138572 | NM_016797 |
| Atxn7l3b | 8179.446081 | -0.225679697 | 0.048127415 | -4.689212912 | 2.74E-06 | 0.000114847 | | Tm6sf1 | 565.7723281 | -0.140382079340083 | 0.0669490253474573 | -2.09685023212088 | 0.0360068352867559 | 0.344080613138572 | NM_145375 |
| Tulp4 | 8180.879394 | -0.225774764 | 0.07597062 | -2.97186946 | 0.002959925 | 0.025129358 | | Zfp160 | 557.5127804 | -0.142373319220498 | 0.067903938188348 | -2.09668721754534 | 0.0360212727829647 | 0.344080613138572 | NM_145483 |
| Ndufa7 | 1711.799455 | -0.225939321 | 0.060406827 | -3.740294499 | 0.000183805 | 0.003199855 | | Ccdc85a | 1884.498228 | 0.146833797760824 | 0.070053197116684 | 2.09603278371793 | 0.0360792828495916 | 0.34411533990359 | NM_181577,NM_001166661,NM_001166662 |
| Lphn3 | 3829.17901 | -0.225981055 | 0.075668523 | -2.986460503 | 0.002822273 | 0.024320521 | | Pbdc1 | 598.2260074 | 0.140263715657917 | 0.0669184566979721 | 2.09603930782473 | 0.0360787041494637 | 0.34411533990359 | NM_026312 |
| Timmdc1 | 640.732965 | -0.226316892 | 0.084250736 | -2.686230447 | 0.007226323 | 0.046578655 | | Tulp3 | 857.8146805 | -0.153879674206305 | 0.0734382286999045 | -2.09536200600799 | 0.0361388243006149 | 0.344457061542475 | NM_011657 |
| Oxct1 | 11269.40151 | -0.226410841 | 0.074777728 | -3.027784426 | 0.002463538 | 0.022127158 | | Oscar | 82.91914014 | -0.197397414879488 | 0.0942257376581487 | -2.09494157101371 | 0.0361761868197531 | 0.344587075241792 | NM_175632 |
| Phactr3 | 5455.548194 | -0.226497465 | 0.054664533 | -4.143408011 | 3.42E-05 | 0.000848284 | | Nxf7 | 108.8456193 | -0.19055119888248 | 0.0905458652923512 | -2.09430593614961 | 0.0362327358840114 | 0.344899555341514 | NM_130888 |
| Cdc34 | 1222.231111 | -0.226510309 | 0.079691821 | -2.842328169 | 0.004478537 | 0.03349786 | | Clptm1l | 4368.264133 | 0.0936531783209097 | 0.044729363683682 | 2.09377399113495 | 0.0362801181077002 | 0.345124424120795 | NM_146047 |
| Rpsa | 12881.31304 | -0.226776535 | 0.05211697 | -4.35129927 | 1.35E-05 | 0.000409541 | | Sgpl1 | 1804.667408 | -0.105052342906553 | 0.0501801119714734 | -2.09350554989342 | 0.036304049162116 | 0.34512605898488 | NM_009163 |
| Rgs8 | 5912.537735 | -0.22720042 | 0.060931365 | -3.728792537 | 0.000192399 | 0.003308587 | | Capsl | 50.74347038 | 0.193223315445889 | 0.0923272333033556 | 2.09280954852209 | 0.0363661591047738 | 0.345417070841624 | NM_029341 |
| 2610002M06Rik | 2413.129244 | -0.227289666 | 0.085230528 | -2.666763546 | 0.007658553 | 0.048611891 | | Itga5 | 328.2394456 | -0.177195663794297 | 0.0846760705748269 | -2.09262974287065 | 0.036382219357544 | 0.345417070841624 | NM_010577 |
| Dock3 | 14466.78316 | -0.227301123 | 0.073307809 | -3.100639985 | 0.001931029 | 0.018662475 | | Nphp4 | 818.3632044 | -0.177928051462726 | 0.085045495872954 | -2.092151378934 | 0.0364249762909185 | 0.345597129720367 | NM_153424 |
| Lix1l | 855.7449009 | -0.227305766 | 0.078871754 | -2.881966667 | 0.003952015 | 0.030863169 | | Plcg1 | 3640.326508 | -0.107222893924942 | 0.0512654179291039 | -2.09152481841117 | 0.0364810440398236 | 0.345903163004228 | NM_021280 |
| Cntn1 | 13563.94287 | -0.227370013 | 0.054740566 | -4.153592632 | 3.27E-05 | 0.000825264 | | AI836003 | 1079.522323 | 0.14689810839092 | 0.0702837792843731 | 2.09007127799089 | 0.0366113973107542 | 0.346440077950594 | NM_177716 |
| Samd4 | 1406.665266 | -0.227541528 | 0.085331812 | -2.666549824 | 0.007663425 | 0.048622892 | | Ptprn2 | 18312.04008 | 0.107926189309728 | 0.0516435244667582 | 2.08983004982933 | 0.0366330689613185 | 0.346440077950594 | NM_011215 |
| Ndufs6 | 1342.603392 | -0.227597163 | 0.065461304 | -3.476819856 | 0.000507399 | 0.00689617 | | Rnf34 | 2285.808367 | 0.111591535008973 | 0.0533887819830832 | 2.09016821667763 | 0.0366026918909618 | 0.346440077950594 | NM_030564 |
| Ppp1r12b | 2078.398659 | -0.227824161 | 0.076811672 | -2.966009669 | 0.00301691 | 0.025425392 | | Vwc2l | 360.1705871 | 0.181235220748541 | 0.0867104387064944 | 2.0901199838465 | 0.0366070229603241 | 0.346440077950594 | NM_177164 |
| Pard3 | 967.6525454 | -0.228014359 | 0.071295441 | -3.198161838 | 0.001383066 | 0.014519275 | | Gnl1 | 8222.838656 | 0.101653183056927 | 0.0486895243412833 | 2.08778344894893 | 0.0368173732256271 | 0.347730275346373 | NM_008136 |
| Prdx4 | 417.3336089 | -0.228426108 | 0.080882368 | -2.824176817 | 0.004740223 | 0.034676586 | | Tmeff2 | 2822.970069 | 0.0985682380835762 | 0.0472071249351911 | 2.08799494184187 | 0.0367982909605039 | 0.347730275346373 | NM_019790 |
| Pdlim5 | 1232.861789 | -0.228826346 | 0.075552116 | -3.028721883 | 0.002455907 | 0.02211392 | | Bbs2 | 1323.211703 | -0.125449099124803 | 0.0601296028535264 | -2.08631178606638 | 0.0369503897452391 | 0.348173347765048 | NM_026116 |
| Agps | 1742.359557 | -0.229246093 | 0.073092508 | -3.136382913 | 0.001710457 | 0.017022514 | | Fdxr | 457.8959162 | -0.182503572354367 | 0.0875117811144366 | -2.08547432163119 | 0.0370262666707596 | 0.348173347765048 | NM_007997 |
| Mrps25 | 1228.800872 | -0.23041079 | 0.078354106 | -2.935015796 | 0.003335307 | 0.027312724 | | Gpr176 | 725.0177437 | 0.138683381529346 | 0.0664885196744895 | 2.08582447328206 | 0.0369945256997023 | 0.348173347765048 | NM_201367 |
| Ndufa3 | 2112.850974 | -0.230454898 | 0.081440128 | -2.829746264 | 0.004658493 | 0.034336846 | | Naprt1 | 190.6936525 | -0.193114532686926 | 0.0925889239387344 | -2.08573156752109 | 0.0370029245158 | 0.348173347765048 | NM_172607 |
| Tmem35 | 2484.165498 | -0.230491158 | 0.086504925 | -2.664485971 | 0.007710608 | 0.048882228 | | Prdx5 | 5559.963931 | 0.114830698216224 | 0.0550559665592124 | 2.08570851431196 | 0.0370050347218287 | 0.348173347765048 | NM_012021 |
| Akap6 | 10151.29268 | -0.230590514 | 0.073098293 | -3.154526679 | 0.001607587 | 0.016226679 | | Th | 146.8706803 | 0.146442159265506 | 0.0702222272190127 | 2.08541034977109 | 0.0370320681741015 | 0.348173347765048 | NM_009377 |
| Cdkl5 | 2562.332265 | -0.230651214 | 0.080684731 | -2.858672407 | 0.004254179 | 0.032422156 | | Uxs1 | 1528.570298 | 0.11620746365966 | 0.0556994304712436 | 2.08633127262684 | 0.0369486257788456 | 0.348173347765048 | NM_026430 |
| Abcb7 | 1372.37207 | -0.230705519 | 0.076906676 | -2.999811328 | 0.002701469 | 0.023570703 | | Dtx3 | 8256.649771 | -0.141922472777307 | 0.068065034897054 | -2.08510033810986 | 0.0370601935900499 | 0.348202097803143 | NM_030714 |
| Ccm2 | 1996.642015 | -0.230873284 | 0.064397371 | -3.585135237 | 0.000336904 | 0.005132306 | | Iqcc | 535.0023727 | -0.151845837824364 | 0.0728330372638033 | -2.08484835356205 | 0.0370830679678826 | 0.348202097803143 | NM_198026 |
| Ppp1ca | 8033.232645 | -0.231475613 | 0.061089301 | -3.789135093 | 0.000151173 | 0.002724393 | | Cd46 | 49.92900458 | -0.192766336219736 | 0.0925056262246371 | -2.08380150253388 | 0.0371782266192923 | 0.348662607299545 | NM_010778 |
| Cdk14 | 8483.781205 | -0.231623159 | 0.053464618 | -4.332269974 | 1.48E-05 | 0.000438887 | | Ube2v2 | 2682.990195 | 0.145247756217984 | 0.0697039540328066 | 2.08378072999448 | 0.0371801169425165 | 0.348662607299545 | NM_023585,NM_001159351 |
| Zfp827 | 2244.603061 | -0.232856594 | 0.06542321 | -3.559235215 | 0.000371936 | 0.0055305 | | Fam83d | 66.9597569 | 0.189153525912056 | 0.0908060630536714 | 2.08304915840336 | 0.0372467101314539 | 0.348836693339458 | NM_027975 |
| Syt5 | 5924.558638 | -0.232875573 | 0.057564153 | -4.045496388 | 5.22E-05 | 0.001187927 | | Strbp | 7467.188516 | -0.123297211359559 | 0.0591868949177772 | -2.08318431860389 | 0.0372344259264896 | 0.348836693339458 | NM_009261 |
| Lin7a | 2860.656039 | -0.233191432 | 0.07627387 | -3.057291192 | 0.002233472 | 0.020771557 | | AU022754 | 70.13948773 | -0.196986280786202 | 0.094656093419603 | -2.08107342770821 | 0.0374271856210415 | 0.350065015367925 | NR_040433 |
| Ube2e3 | 2025.595618 | -0.233193243 | 0.072121467 | -3.233340246 | 0.001223517 | 0.013284639 | | Btbd10 | 3644.22168 | 0.163523242490724 | 0.0785758390391273 | 2.08108808624108 | 0.0374258441286719 | 0.350065015367925 | NM_133700 |
| Nacc2 | 6513.112437 | -0.233388331 | 0.07461162 | -3.128042666 | 0.001759746 | 0.017366567 | | Ccdc90a | 1054.148433 | -0.125020631406176 | 0.0600919378665845 | -2.0804892610344 | 0.0374806796230499 | 0.350065015367925 | NM_001081059 |
| Tmem164 | 1432.32013 | -0.233571301 | 0.080665203 | -2.895564544 | 0.003784772 | 0.029995527 | | Cntfr | 3163.08383 | 0.103310282433843 | 0.0496559100938386 | 2.08052339064191 | 0.0374775524774973 | 0.350065015367925 | NM_016673,NM_001136056,NM_001146080 |
| Tmem126a | 547.3942268 | -0.233701081 | 0.070403377 | -3.319458397 | 0.000901922 | 0.010507057 | | Nudt14 | 183.4812961 | 0.180283906530583 | 0.0866626075826412 | 2.08029635340322 | 0.0374983590742456 | 0.350065015367925 | NM_025399 |
| Rps3 | 8657.996085 | -0.233787144 | 0.060606091 | -3.857485947 | 0.000114559 | 0.002167254 | | Prelid1 | 3961.159083 | 0.097197526054371 | 0.0467624778689972 | 2.07853669188928 | 0.0376599550570333 | 0.351347788797987 | NM_025596 |
| Rpl36 | 1896.0831 | -0.233950509 | 0.065458725 | -3.574015673 | 0.000351548 | 0.005283106 | | Dph1 | 576.7672237 | -0.156393054351007 | 0.0752556781323038 | -2.07815620339052 | 0.037694974472642 | 0.351448779967649 | NM_144491 |
| Scarb1 | 1118.38109 | -0.233970993 | 0.068449074 | -3.418176164 | 0.000630423 | 0.008079216 | | Cdo1 | 1047.41333 | 0.13232996808019 | 0.0637096161841529 | 2.07707997639932 | 0.0377941783421357 | 0.351969169482281 | NM_033037 |
| Parvb | 1016.349827 | -0.234138962 | 0.068102163 | -3.438054726 | 0.000585909 | 0.007673777 | | Gpr34 | 424.3215779 | -0.184990172687432 | 0.08906497071337 | -2.07702502123725 | 0.0377992499237476 | 0.351969169482281 | NM_011823 |
| Plcl2 | 3234.985032 | -0.234156093 | 0.078438082 | -2.985234828 | 0.002833607 | 0.024364519 | | Carhsp1 | 2585.582094 | 0.140628683481077 | 0.0677468732392931 | 2.07579592617291 | 0.0379128293162364 | 0.352772128684868 | NM_025821 |
| Mrpl53 | 632.3209522 | -0.234259043 | 0.082809834 | -2.828879505 | 0.004671128 | 0.034398508 | | Rbmxl1 | 1633.381806 | 0.120417264791129 | 0.0580165749147282 | 2.0755665939969 | 0.0379340537660583 | 0.352772128684868 | NM_001252089,NM_009033 |
| Rap1b | 1750.435981 | -0.234329511 | 0.05597616 | -4.186237723 | 2.84E-05 | 0.000738546 | | Cyba | 182.3276758 | -0.196192516878516 | 0.0945658272226478 | -2.07466610974169 | 0.0380174904027793 | 0.352889542159351 | NM_007806 |
| Mdga2 | 1494.131084 | -0.234339827 | 0.078938255 | -2.968647175 | 0.002991138 | 0.025311139 | | Dhx58 | 69.97524385 | -0.195533687694617 | 0.0942590270222238 | -2.07442930265465 | 0.0380394582609625 | 0.352889542159351 | NM_030150 |
| 6330403K07Rik | 9439.219227 | -0.23449348 | 0.051586152 | -4.545667245 | 5.48E-06 | 0.000198718 | | Ptafr | 35.39846019 | -0.186153429390645 | 0.0897251664514533 | -2.07470698303319 | 0.0380136998064681 | 0.352889542159351 | NM_001081211 |
| Mtx2 | 1111.998283 | -0.234640591 | 0.085408215 | -2.747283637 | 0.006009114 | 0.040889423 | | Vwc2 | 255.6871392 | 0.169432344474326 | 0.0816784710003442 | 2.07438193197338 | 0.0380438539874394 | 0.352889542159351 | NM_177033 |
| Cldn12 | 1840.913707 | -0.235605854 | 0.069772714 | -3.376762073 | 0.000733445 | 0.008983392 | | Glis2 | 506.0661707 | -0.144890555010149 | 0.0698735717272775 | -2.07361025676015 | 0.0381155218612221 | 0.35332869850422 | NM_031184 |
| 6030419C18Rik | 6021.93407 | -0.235732989 | 0.063666237 | -3.702637394 | 0.00021337 | 0.003556118 | | Ctps2 | 1850.946946 | -0.125422053362189 | 0.0605158758546595 | -2.07254793210651 | 0.0382143710350596 | 0.354019103096477 | NM_001168571,NM_018737,NM_001168569,NM_001168568,NR_033143 |
| Arhgef12 | 11573.19938 | -0.235880654 | 0.07204004 | -3.274299308 | 0.001059244 | 0.011909956 | | Fam196a | 537.9573829 | 0.177591798842685 | 0.0857530148207216 | 2.07096857427071 | 0.0383617329356553 | 0.354869882880824 | NM_001143802 |
| Rnf11 | 6279.563841 | -0.235922382 | 0.071738587 | -3.288639933 | 0.001006727 | 0.011427276 | | Nfatc2ip | 239.3431904 | -0.181783216022342 | 0.0877849628581824 | -2.07077852634073 | 0.0383794978662931 | 0.354869882880824 | NM_010900 |
| Kitl | 1797.266147 | -0.235945227 | 0.082112454 | -2.873440193 | 0.004060279 | 0.031376544 | | Stmn3 | 13586.52853 | 0.0811599538014797 | 0.0391833091494322 | 2.07128891263274 | 0.038331804790597 | 0.354869882880824 | NM_009133 |
| Vti1b | 4628.969292 | -0.235995426 | 0.065860405 | -3.583267148 | 0.000339323 | 0.005154387 | | Cc2d1b | 1272.445497 | -0.171898972418657 | 0.0830508103204578 | -2.06980487915015 | 0.0384706202902521 | 0.35548615161336 | NM_177045 |
| Usp46 | 4307.606762 | -0.236362944 | 0.072023566 | -3.28174451 | 0.00103167 | 0.011633698 | | Fbrs | 1756.702395 | -0.137042144345446 | 0.0662338014468021 | -2.0690665695146 | 0.0385398402952431 | 0.355824003198916 | NM_010183 |
| Mrps36 | 631.8951958 | -0.236426971 | 0.086703569 | -2.72684244 | 0.006394356 | 0.042690885 | | Lrrc17 | 205.9323374 | -0.109656695215763 | 0.0530092911563421 | -2.06863160822787 | 0.038580669491828 | 0.355824003198916 | NM_028977 |
| Slc9a3r1 | 1559.369133 | -0.236507966 | 0.055072315 | -4.294498311 | 1.75E-05 | 0.000503307 | | Pgk1 | 18657.599 | 0.0921820146197066 | 0.0445566178185157 | 2.0688763796239 | 0.0385579404586884 | 0.355824003198916 | NM_008828 |
| Fam171a2 | 2705.365274 | -0.236738197 | 0.087354506 | -2.710085692 | 0.006726583 | 0.044124332 | | Gsdmd | 118.1768303 | -0.194665834093636 | 0.0941294710279232 | -2.06806476469585 | 0.0386339416988062 | 0.356089236749276 | NM_026960 |
| 3632451O06Rik | 1121.995827 | -0.237017373 | 0.086014498 | -2.755551419 | 0.005859329 | 0.040157049 | | Atmin | 4935.804709 | 0.122008668332279 | 0.0590157599904313 | 2.06739129263403 | 0.0386972980673319 | 0.356201814759405 | NM_177700 |
| Ank2 | 20264.45172 | -0.237252924 | 0.052314343 | -4.535141036 | 5.76E-06 | 0.00020534 | | Cntn5 | 293.461367 | 0.166123634268754 | 0.0803492906669091 | 2.06751836749158 | 0.038685335286046 | 0.356201814759405 | NM_001033359,NM_001170787 |
| Ostc | 987.4976411 | -0.237497487 | 0.059095764 | -4.018858051 | 5.85E-05 | 0.001303745 | | Rab1 | 6910.695407 | 0.0972561247214439 | 0.0470483391939277 | 2.06715319579222 | 0.0387197208801529 | 0.356201814759405 | NM_008996 |
| Smarcd3 | 2698.468309 | -0.237531971 | 0.057792087 | -4.110123515 | 3.95E-05 | 0.000954418 | | Gm16039 | 1228.728655 | 0.118779939191967 | 0.0575061522230872 | 2.06551707252429 | 0.0388741018581023 | 0.357169641739908 | NR_033518 |
| C1d | 879.9517057 | -0.238383592 | 0.070675629 | -3.372924964 | 0.000743742 | 0.009066516 | | Herc6 | 747.8590428 | -0.176539826983336 | 0.0854677860278955 | -2.06557255675465 | 0.0388689533872299 | 0.357169641739908 | NM_025992 |
| Wrb | 2033.983469 | -0.238513172 | 0.077307572 | -3.085249815 | 0.002033812 | 0.019331788 | | Slc25a44 | 5141.893077 | 0.0989747255128299 | 0.047951925921021 | 2.06404067431714 | 0.0390128600603345 | 0.358227137317585 | NM_178696,NR_027357,NM_001145876,NM_001145877 |
| Ksr2 | 988.2365406 | -0.238530007 | 0.074926563 | -3.183828231 | 0.001453412 | 0.015115049 | | Kctd3 | 3654.181805 | 0.107797544679305 | 0.0522378468900397 | 2.06359088471273 | 0.0390565225817677 | 0.358392322819178 | NM_172650 |
| Vps13b | 3106.92331 | -0.238990823 | 0.077269009 | -3.092971223 | 0.001981634 | 0.01902319 | | Scube2 | 64.9935507 | 0.190781529857172 | 0.0924630455148513 | 2.06332734115414 | 0.0390815380800349 | 0.358395468529411 | NM_020052 |
| Glrx5 | 920.7001175 | -0.239722055 | 0.074559766 | -3.209130964 | 0.001331368 | 0.014119248 | | Trip6 | 231.3317201 | -0.190577775604811 | 0.0924339436750437 | -2.06177263489695 | 0.0392293875487924 | 0.359524342923507 | NM_011639 |
| Mrps33 | 1625.72663 | -0.239272344 | 0.068839475 | -3.475801396 | 0.000509329 | 0.006914272 | | Higd2a | 2175.929412 | 0.13671171566948 | 0.0663218072475525 | 2.06133881664579 | 0.0392707274571667 | 0.359615903073255 | NM_025933 |
| Ptprg | 2958.695308 | -0.239416329 | 0.06062175 | -3.949347041 | 7.84E-05 | 0.001629775 | | Itgb5 | 2252.775838 | -0.154246572897843 | 0.0748352607157927 | -2.0611483268189 | 0.0392888915170905 | 0.359615903073255 | NM_001145884,NM_010580 |
| Rab15 | 1335.71095 | -0.239502442 | 0.082754061 | -2.894142279 | 0.003801898 | 0.03008509 | | Elmo1 | 7226.94836 | 0.116692206799986 | 0.0566274281741931 | 2.06070115776804 | 0.0393315591230927 | 0.359779740442094 | NM_080288,NM_038122 |
| 1110001A16Rik | 387.6976394 | -0.239689196 | 0.087245107 | -2.747308178 | 0.006008665 | 0.040889423 | | 5430416N02Rik | 122.9885316 | -0.189977873766506 | 0.0917321403758234 | -2.0601053566642 | 0.0393884699041617 | 0.359829336474221 | NR_034038 |
| Ptprz1 | 8916.53016 | -0.239813906 | 0.079142615 | -3.030148884 | 0.002444332 | 0.022066759 | | H2-T23 | 868.6070342 | -0.192146794548977 | 0.0932678726792996 | -2.06016057865575 | 0.0393831921746041 | 0.359829336474221 | NM_010398 |
| Rps6ka3 | 1147.772766 | -0.239865289 | 0.080903381 | -2.964836407 | 0.003028439 | 0.025459923 | | Hspa13 | 1553.128984 | 0.140101702831855 | 0.0680149397211396 | 2.05986660292975 | 0.0394112952175744 | 0.359829336474221 | NM_030201,NR_027492 |
| Timm10 | 558.5059313 | -0.239869575 | 0.083816286 | -2.861849242 | 0.004211772 | 0.032225775 | | Ahcy | 1464.750788 | 0.126905043090237 | 0.0616270227614876 | 2.05924345203876 | 0.0394709225421042 | 0.359835304789816 | NM_016661 |
| Cnn2 | 277.2405564 | -0.240031848 | 0.083765573 | -2.86551907 | 0.004163262 | 0.031965105 | | Cep63 | 1066.140014 | -0.111386483523504 | 0.0540950393708113 | -2.05908868574752 | 0.0394857435019034 | 0.359835304789816 | NM_001081122 |
| Klf16 | 2085.817085 | -0.240216806 | 0.086693292 | -2.770881127 | 0.005590483 | 0.038865068 | | Etnk2 | 357.5770794 | -0.158869392422831 | 0.0771554003902616 | -2.05908324782516 | 0.0394862643422117 | 0.359835304789816 | NM_175443 |
| Ranbp2 | 6090.810573 | -0.240574418 | 0.07319485 | -3.286767024 | 0.001013446 | 0.011471264 | | Acat1 | 3558.343485 | 0.10935330548246 | 0.0531684912545684 | 2.05673140053724 | 0.0397120699516144 | 0.360800870379348 | NM_144784 |
| Rheb | 3197.361137 | -0.240612902 | 0.077204443 | -3.116568068 | 0.001829694 | 0.017883806 | | Eif3d | 2836.794225 | 0.107840286737068 | 0.0524489048431004 | 2.05610178248086 | 0.0397727064792401 | 0.360800870379348 | NM_018749 |
| St8sia3 | 7716.409767 | -0.240626298 | 0.075397308 | -3.191444173 | 0.001415635 | 0.014775915 | | Lefty1 | 32.74894581 | -0.161816336029645 | 0.0787176454625693 | -2.05565518479068 | 0.0398157645062712 | 0.360800870379348 | NM_010094 |
| Coa5 | 4771.896106 | -0.240648456 | 0.074540909 | -3.228407839 | 0.001244814 | 0.013459276 | | Lhx6 | 1076.362021 | 0.13591823351148 | 0.0661142256907373 | 2.05580920129452 | 0.0398009107765961 | 0.360800870379348 | NM_001083125,NM_001083127,NM_001083126,NM_008500 |
| Fdps | 2137.145413 | -0.240917044 | 0.083577252 | -2.882567178 | 0.00394449 | 0.030835485 | | Pdgfrb | 1131.761926 | -0.154959583921327 | 0.0753337095103346 | -2.05697535576778 | 0.0396885964904032 | 0.360800870379348 | NM_008809,NM_001146268 |
| Reep1 | 5450.95174 | -0.241314839 | 0.063098246 | -3.824430209 | 0.000131075 | 0.002406249 | | Pdlim1 | 395.8872353 | 0.172780455078179 | 0.083984857255639 | 2.0572810471328 | 0.0396591993788437 | 0.360800870379348 | NM_016861 |
| Anks1b | 12133.90324 | -0.241680097 | 0.059198954 | -4.082506195 | 4.46E-05 | 0.001041174 | | Psmb6 | 2778.661781 | 0.124151088115776 | 0.0603446114139549 | 2.05736825885113 | 0.0396508159668489 | 0.360800870379348 | NM_008946 |
| Maob | 1130.785681 | -0.241974325 | 0.067793425 | -3.569289013 | 0.000357951 | 0.005358549 | | Taldo1 | 2899.631857 | 0.093896890191344 | 0.0456635763563576 | 2.05627628742488 | 0.0397558925939326 | 0.360800870379348 | NM_011528 |
| Ndufs4 | 2085.278257 | -0.242494368 | 0.061522025 | -3.941586271 | 8.09E-05 | 0.001667758 | | Tram1l1 | 1136.812338 | 0.126916483374877 | 0.061731742263232 | 2.05593554825796 | 0.039788729076428 | 0.360800870379348 | NM_146140 |
| Usp9x | 11207.9479 | -0.242559842 | 0.074512608 | -3.2552859 | 0.001132783 | 0.01257259 | | Casp2 | 509.3200995 | -0.146154241214898 | 0.0711339852379869 | -2.05463310857563 | 0.0399144552734956 | 0.361071695152331 | NM_007610 |
| Zyg11b | 10679.49348 | -0.242698978 | 0.063472132 | -3.82370926 | 0.000131459 | 0.002410443 | | Myo1a | 52.3338673 | -0.187757370753571 | 0.091385085481749 | -2.05457345434195 | 0.0399202218377147 | 0.361071695152331 | NM_001081219 |
| Rps19 | 4419.021782 | -0.242196363 | 0.068063657 | -3.57307223 | 0.000352817 | 0.005291919 | | Sash3 | 97.90768692 | -0.192199688634806 | 0.093527036692708 | -2.05501740920432 | 0.0398773232271972 | 0.361071695152331 | NM_028773 |
| Ndufs8 | 2008.72814 | -0.243402428 | 0.064276381 | -3.786809755 | 0.000152594 | 0.002742795 | | Dnajc27 | 7208.782797 | 0.0970137627478477 | 0.0472378220487975 | 2.05373064506723 | 0.0400017688023531 | 0.361359622203346 | NM_153082 |
| Zfp428 | 721.9231047 | -0.243737071 | 0.080292652 | -3.035608667 | 0.002400507 | 0.021823847 | | Slc6a15 | 2757.264872 | 0.113761989444331 | 0.0553896457444899 | 2.05384937661047 | 0.0399902722533018 | 0.361359622203346 | NM_001252330,NM_175328 |
| Ndrg4 | 80486.46653 | -0.244143841 | 0.05199144 | -4.695846913 | 2.66E-06 | 0.000112397 | | Fuk | 774.5914343 | -0.162273695716154 | 0.0791100382784888 | -2.05124026289693 | 0.0402435547130086 | 0.363317531149387 | NM_172283,NR_052009 |
| Prrc2b | 26769.60951 | -0.244246662 | 0.058784704 | -4.154901627 | 3.25E-05 | 0.000822541 | | Dnahc7a | 42.24349653 | -0.189703865029668 | 0.0925073196953301 | -2.05069032001415 | 0.0402971142458056 | 0.363536188740086 | NM_001252070 |
| Triap1 | 491.408948 | -0.244424523 | 0.083522344 | -2.926456681 | 0.003428472 | 0.027877541 | | Grm4 | 808.2883943 | 0.155443298360655 | 0.0758302992225336 | 2.0497082459992 | 0.0403929098600095 | 0.363536188740086 | NM_001013385 |
| Scoc | 4040.728539 | -0.244469075 | 0.088979296 | -2.747482698 | 0.006005468 | 0.040889423 | | Hcn1 | 2468.128681 | 0.132332342518868 | 0.0645518278313195 | 2.05001697031207 | 0.0403627748086507 | 0.363536188740086 | NM_010408 |
| Irf2bpl | 2676.30094 | -0.245087378 | 0.08059155 | -3.041105136 | 0.002357115 | 0.021559115 | | Rad9b | 105.3313116 | -0.19077321222719 | 0.0930499882054634 | -2.0502228523226 | 0.0403426889489923 | 0.363536188740086 | NM_144912 |
| Agap2 | 43934.69301 | -0.245474386 | 0.081628549 | -3.007212417 | 0.002636555 | 0.023198299 | | Rps6ka2 | 4754.851384 | 0.135163871508007 | 0.0659375697494781 | 2.04987645164275 | 0.0403764886857721 | 0.363536188740086 | NM_011299 |
| Rpl13a | 8858.022542 | -0.245600197 | 0.072669462 | -3.379689202 | 0.000725678 | 0.0089238 | | Ldoc1l | 3538.025691 | 0.11983858127406 | 0.0584783200746387 | 2.04928221469263 | 0.0404345267525327 | 0.363685409044762 | NM_177630 |
| Nrxn3 | 8428.703657 | -0.245780785 | 0.062380551 | -3.940022681 | 8.15E-05 | 0.001676437 | | Ttc13 | 1128.073662 | -0.112722356364205 | 0.055038894538203 | -2.04804906257634 | 0.0405551922540161 | 0.364545001659553 | NM_145607 |
| Mcee | 490.6396541 | -0.245871982 | 0.075255651 | -3.267142788 | 0.001086389 | 0.012179817 | | Ascc2 | 758.425544 | -0.149225281318648 | 0.0728796393359536 | -2.04755790064717 | 0.0406033380102439 | 0.364752064277553 | NM_029291 |
| Xiap | 3660.133863 | -0.245992315 | 0.074158935 | -3.317096089 | 0.000909583 | 0.010547482 | | 1810011O10Rik | 39.96145116 | 0.185622125280774 | 0.0906754904414279 | 2.04710362609703 | 0.0406479110327166 | 0.364926795835131 | NM_026931 |
| Plekha2 | 1474.362606 | -0.246258442 | 0.0876375 | -2.809966516 | 0.004954665 | 0.0357391 | | Fam19a1 | 1498.450202 | 0.130317898470611 | 0.0636891351710284 | 2.04615588075832 | 0.0407410365218388 | 0.365321333377494 | NM_182808 |
| Pdia6 | 3862.583827 | -0.24665017 | 0.081533212 | -3.02514966 | 0.002485101 | 0.022269612 | | Ndnf | 311.3397539 | -0.185450972585877 | 0.0906343480687595 | -2.04614449750535 | 0.040742156138754 | 0.365321333377494 | NM_172399 |
| Panx1 | 577.5617808 | -0.247039215 | 0.066626155 | -3.707841404 | 0.000209033 | 0.003516574 | | Agxt2l1 | 1352.944949 | 0.178486738246203 | 0.0872791107086733 | 2.04501097086733 | 0.0408537765193265 | 0.365377968159263 | NM_001163587,NM_027907 |
| Mmd | 7643.273875 | -0.247054285 | 0.067546034 | -3.657569064 | 0.000254619 | 0.004092388 | | Inpp5b | 1746.904207 | -0.113689005019821 | 0.0555928481194044 | -2.04502933139234 | 0.0408519664630834 | 0.365377968159263 | NM_008385 |
| Klf7 | 350.4906268 | -0.247689514 | 0.086052751 | -2.878345086 | 0.003997676 | 0.031028284 | | Lgals3bp | 1014.970777 | -0.181401396315648 | 0.0887027392268719 | -2.04504841560399 | 0.0408500851348619 | 0.365377968159263 | NM_011150 |
| Mrps24 | 516.7276681 | -0.248268999 | 0.087252614 | -2.845404699 | 0.004435503 | 0.033344824 | | Mrpl17 | 1032.48986 | -0.113248283573877 | 0.0553620667325075 | -2.04559349492962 | 0.0407963820139465 | 0.365377968159263 | NM_025301 |
| Herc1 | 10018.44547 | -0.248472068 | 0.074668033 | -3.327690028 | 0.000875692 | 0.010294369 | | Snrk | 4118.879577 | 0.108021429221935 | 0.0528272920528854 | 2.04480345337018 | 0.0408742391751895 | 0.365377968159263 | NM_133741,NM_001164572 |
| Neurl1b | 2294.944542 | -0.24870611 | 0.090379803 | -2.751788573 | 0.005927077 | 0.040455567 | | Strada | 1074.319836 | -0.116533659897376 | 0.056999140228645 | -2.04448101199273 | 0.0409060513501108 | 0.365437455050252 | NM_001252448,NM_001252449,NM_028126 |
| Vstm2b | 980.1260943 | -0.249224802 | 0.066168002 | -3.766545677 | 0.000165522 | 0.002924282 | | Dnase2a | 169.9712535 | -0.191508410664345 | 0.0936919400570897 | -2.04402225578478 | 0.0409513485507939 | 0.365617264320118 | NM_010062 |
| Hip1 | 2992.186094 | -0.249357203 | 0.066407034 | -3.754981785 | 0.000173354 | 0.003045293 | | Ccdc171 | 159.946862 | -0.187177438109409 | 0.0916137560778598 | -2.04311498756074 | 0.0410410566030299 | 0.366193113154553 | NM_001081012 |
| Ncam2 | 2470.067898 | -0.249430728 | 0.083544124 | -2.985616664 | 0.002830072 | 0.024360628 | | Prmt1 | 2351.33227 | 0.0919603950115058 | 0.0450176385650145 | 2.04276363538475 | 0.0410758419842749 | 0.36627850255591 | NM_019830,NM_001252476,NR_045521,NM_001252477 |
| Nav3 | 2897.819276 | -0.250342557 | 0.058801309 | -4.257413713 | 2.07E-05 | 0.000573164 | | Srpr | 3279.507345 | 0.101051311254876 | 0.0494842127486992 | 2.04209192471254 | 0.0411424137659459 | 0.366647056665111 | NM_026130 |
| Bola2 | 553.6563987 | -0.250780031 | 0.079277396 | -3.163323269 | 0.00155979 | 0.015931042 | | Gng2 | 8707.815231 | 0.0946609803947517 | 0.0463707374150202 | 2.04139476039666 | 0.041211604842542 | 0.367038486782811 | NM_001038637,NM_010315 |
| Hey1 | 1812.835411 | -0.250895648 | 0.07188967 | -3.490009733 | 0.000483003 | 0.006669918 | | Mrps23 | 1241.200586 | 0.101677089339582 | 0.0498148018324639 | 2.04110196566756 | 0.0412406930543356 | 0.367072492222597 | NM_024174 |
| Tmem14c | 1014.624113 | -0.250994079 | 0.05597745 | -4.48385068 | 7.33E-06 | 0.000246462 | | Ndufb8 | 3769.9787 | 0.106734566597033 | 0.0523114724047713 | 2.04036632291959 | 0.0413138535353968 | 0.367273584121894 | NM_026061 |
| Rasa2 | 1346.518983 | -0.25112485 | 0.093455384 | -2.687109507 | 0.007207332 | 0.046548729 | | St6galnac5 | 1343.307021 | 0.145843350124221 | 0.0714717951563918 | 2.04057208588496 | 0.0412933791171068 | 0.367273584121894 | NM_012028 |
| Fam162a | 708.043623 | -0.251302822 | 0.081052638 | -3.100489133 | 0.001932013 | 0.018662475 | | Ushbp1 | 213.0383837 | -0.182765131806274 | 0.0896233368982542 | -2.03925827950102 | 0.0414242571362733 | 0.368029822117129 | NM_181418 |
| Gpx1 | 1960.649955 | -0.251308366 | 0.067647743 | -3.714955649 | 0.000203239 | 0.003445285 | | B3gnt1 | 4438.706786 | 0.089345436311407 | 0.0438273563959521 | 2.03857689941935 | 0.0414922726970153 | 0.368048911302019 | NM_175383 |
| Sar1b | 1243.417941 | -0.251488028 | 0.058417271 | -4.305028701 | 1.67E-05 | 0.00048354 | | C920006O11Rik | 113.6099251 | -0.193746269661137 | 0.0950458861575173 | -2.03844982138465 | 0.0415049681212674 | 0.368048911302019 | NR_040401 |
| Prex1 | 7537.498007 | -0.251518495 | 0.078509773 | -3.203658414 | 0.001356934 | 0.014321751 | | Fam228b | 97.43256362 | -0.192029574202642 | 0.0941977357258107 | -2.03857951279847 | 0.0414920116481952 | 0.368048911302019 | NM_175431 |
| Coa3 | 1872.314449 | -0.251549445 | 0.085783011 | -2.932392359 | 0.003363615 | 0.027501768 | | Gal3st4 | 182.0294937 | -0.182191483397923 | 0.089387466087142 | -2.03822181535281 | 0.0415277547586403 | 0.368048911302019 | NM_001033416 |
| Tbc1d9 | 5213.426491 | -0.251576425 | 0.05113548 | -4.919801768 | 8.66E-07 | 4.33E-05 | | Aspm | 26.31198451 | -0.178335562938341 | 0.0875191395115796 | -2.03767500381725 | 0.0415824455872182 | 0.368308905243861 | NM_009791 |
| Stxbp6 | 2729.466581 | -0.252178537 | 0.09387931 | -2.686199295 | 0.007226997 | 0.046578655 | | Recql5 | 1077.131336 | -0.145304500880961 | 0.0713232837460339 | -2.03726599855326 | 0.041623393206609 | 0.368446928530897 | NM_130454 |
| Nol4 | 3841.267403 | -0.252484477 | 0.074723268 | -3.378927099 | 0.000727693 | 0.008941218 | | Ccdc32 | 1671.534796 | 0.106937067048294 | 0.0525179554805446 | 2.03620011612808 | 0.0417302646155636 | 0.369167980393226 | NM_199310 |

| | | | | | | | | | | | | | | | |
|---|---|---|---|---|---|---|---|---|---|---|---|---|---|---|---|
| Rnf217 | 440.930607 | -0.253135429 | 0.091024327 | -2.780964572 | 0.005419765 | 0.037997213 | | Mir24-1 | 36.55807357 | -0.168218426254111 | 0.082628628708517 | -2.03583708072323 | 0.0417667175937654 | 0.369265575025585 | NR_029575 |
| Pdgfrb | 1216.985982 | -0.253637801 | 0.069759776 | -3.635874637 | 0.000277039 | 0.00435339 | | 2810049E08Rik | 143.0393204 | -0.190236204829117 | 0.0935557343533787 | -2.03339972845018 | 0.042012154765159 | 0.370320688347857 | NR_036594 |
| Tle4 | 2542.709594 | -0.253648161 | 0.077877458 | -3.257016432 | 0.001125899 | 0.012514119 | | Arhgef5 | 164.3021736 | -0.180638691610194 | 0.0888363939136192 | -2.0333861343566 | 0.0420135270822847 | 0.370320688347857 | NM_133674 |
| Cox6b1 | 5843.290998 | -0.25440655 | 0.06460162 | -3.938083146 | 8.21E-05 | 0.001687801 | | Car7 | 742.2795394 | -0.17458915601344 | 0.0858279841196607 | -2.03417519127595 | 0.0419339349461361 | 0.370320688347857 | NM_053070 |
| Cspg4 | 818.8540317 | -0.254689724 | 0.090044956 | -2.828472971 | 0.004677065 | 0.034424195 | | Hps4 | 842.5544044 | -0.158993805096976 | 0.0781769855667585 | -2.03376740538563 | 0.0419750523426961 | 0.370320688347857 | NM_138646 |
| Pde7b | 3354.858462 | -0.254813122 | 0.090052516 | -2.829605802 | 0.004660539 | 0.034336846 | | Mrpl51 | 1246.711081 | 0.125691115801428 | 0.0617961068582388 | 2.03396495655891 | 0.0419551288317801 | 0.370320688347857 | NM_025595 |
| Ttc28 | 824.6842737 | -0.254936955 | 0.085615918 | -2.977681736 | 0.002904374 | 0.02477994 | | Epb4.1l3 | 8982.149257 | 0.115838306000467 | 0.0570126466388168 | 2.03180018521714 | 0.0421738884524697 | 0.371439588301649 | NM_013813 |
| Fam174a | 826.2524668 | -0.25520065 | 0.089240889 | -2.859682961 | 0.004240647 | 0.032353861 | | Etf1 | 3511.59025 | 0.114983973188798 | 0.05659703914151 | 2.03162523928686 | 0.0421916095757759 | 0.371439588301649 | NM_144866 |
| Neto1 | 2877.765954 | -0.255318918 | 0.068808088 | -3.710594593 | 0.000206773 | 0.003489914 | | Actr1a | 9762.794698 | 0.084155731897865 | 0.041464308068113 | 2.02959450715114 | 0.0423977737101683 | 0.373028504490554 | NM_016860 |
| Hmgn2 | 2596.161774 | -0.255446562 | 0.091156224 | -2.802294252 | 0.005074057 | 0.036290258 | | Zfyve28 | 2261.633213 | -0.125102989130259 | 0.0616565682390178 | -2.029029391407 | 0.042455296761613 | 0.373308499248905 | NM_001015039 |
| Ephb4 | 280.9890224 | -0.255593617 | 0.087476461 | -2.921855952 | 0.003479524 | 0.028137656 | | Mapkbp1 | 3426.56809 | -0.104276283436497 | 0.0514011033648241 | -2.02867791954555 | 0.0424911063160311 | 0.373397344432346 | NM_011941 |
| Psd3 | 22599.90361 | -0.255948166 | 0.084333952 | -3.034936207 | 0.002405866 | 0.021837424 | | Dpcd | 1510.483475 | 0.110960488225256 | 0.0547382198191979 | 2.02711174370965 | 0.0426509861339681 | 0.37389754048402 | NM_172639 |
| Immp1l | 466.715662 | -0.255978324 | 0.090016465 | -2.843683359 | 0.004459534 | 0.033425443 | | Pdia6 | 4175.617211 | 0.120737430511719 | 0.0595601502145485 | 2.0271512089341 | 0.0426469511693741 | 0.37389754048402 | NM_027959 |
| Ctxn1 | 12894.94456 | -0.256038227 | 0.083861183 | -3.05311965 | 0.002264756 | 0.020961847 | | Rab3gap1 | 4804.65865 | 0.105674196502195 | 0.0521218750021575 | 2.0274442640028 | 0.0426169990183807 | 0.37389754048402 | NM_178690 |
| Dnajb1 | 2399.650821 | -0.256133902 | 0.086340075 | -2.96657029 | 0.003011415 | 0.025407408 | | Tjap1 | 740.5990857 | -0.180064665041334 | 0.0888259986299812 | -2.02716172988296 | 0.0426458755513843 | 0.37389754048402 | NM_028751,NM_001252475,NM_001252474,NM_001252473 |
| Myo5a | 24699.48704 | -0.256323946 | 0.070488653 | -3.636385934 | 0.00027649 | 0.00434933 | | Ifi27l1 | 1011.166257 | -0.16384537848963 | 0.0808937603272812 | -2.0254390181238 | 0.0428223043452901 | 0.374984137018746 | NM_194067,NM_194069,NM_026790,NM_194068,NM_194066 |
| Sec22a | 583.8708305 | -0.256368425 | 0.068155344 | -3.761530789 | 0.000168877 | 0.00297338 | | Pard6a | 760.3071007 | 0.148586419479644 | 0.0733616093769667 | 2.02539748979792 | 0.0428265650085433 | 0.374984137018746 | NM_001047436,NM_001047435,NM_019695 |
| Serpini1 | 10366.11932 | -0.256457561 | 0.079667928 | -3.219081593 | 0.001286019 | 0.01375126 | | Hinfp | 478.2912574 | -0.15702016630665 | 0.0775395073735222 | -2.02503435507082 | 0.0428638366521537 | 0.375084392294689 | NM_172162 |
| Pcnx | 6221.284507 | -0.25696324 | 0.086774327 | -2.961281836 | 0.003063615 | 0.025630477 | | Alpk1 | 217.4308053 | -0.173600071368749 | 0.0857494673433946 | -2.02450320389216 | 0.0429184026568867 | 0.375335771820551 | NM_027808 |
| Snrpg | 524.2462803 | -0.257041309 | 0.089277647 | -2.87912279 | 0.00398783 | 0.031002229 | | Mthfsd | 1025.394485 | -0.13439918908615 | 0.0663998098627699 | -2.02408996899111 | 0.0429608955369259 | 0.375481328862446 | NM_001166482,NM_172761 |
| Cycs | 4581.692427 | -0.257106475 | 0.07550611 | -3.405108218 | 0.000661378 | 0.008331217 | | Afap1l2 | 412.699764 | -0.146246841042866 | 0.0722954956399676 | -2.02290391328357 | 0.0430830550216356 | 0.375514734922447 | NM_001177796,NM_001177797,NM_146102 |
| Timm8b | 1875.278519 | -0.257265652 | 0.083685655 | -3.07419059 | 0.002110745 | 0.019904984 | | Rab34 | 570.2978219 | -0.169308763001837 | 0.0836916103444399 | -2.02300759066569 | 0.0430723649262888 | 0.375514734922447 | NM_001159482,NM_033475 |
| Ehbp1l1 | 958.3001575 | -0.257320114 | 0.082407978 | -3.122514593 | 0.001793132 | 0.017606331 | | Rhcg | 26.75759804 | 0.173573278637882 | 0.08578027716146 | 2.02346371895224 | 0.0430253605120709 | 0.375514734922447 | NM_019799 |
| Myo9a | 5171.231745 | -0.257835734 | 0.066035379 | -3.90450903 | 9.44E-05 | 0.001877708 | | Slc35d1 | 369.1764293 | -0.154649185696171 | 0.0764396885676614 | -2.02315300590585 | 0.0430573750497118 | 0.375514734922447 | NM_177732 |
| Mn1 | 1909.381199 | -0.257860008 | 0.079740182 | -3.233752428 | 0.001221753 | 0.013274785 | | Tmem173 | 185.8008231 | -0.186258638678791 | 0.0920797009555571 | -2.02279804067446 | 0.0430939737791353 | 0.375514734922447 | NM_028261 |
| Tm2d1 | 637.6335583 | -0.258160068 | 0.094925485 | -2.71960757 | 0.006535943 | 0.043239925 | | Egflam | 167.262533 | -0.188388906472289 | 0.0931651929544491 | -2.02209538238596 | 0.0431664989769812 | 0.37569596413399 | NM_178748 |
| Atp8a1 | 11031.65112 | -0.258356426 | 0.059717567 | -4.32630526 | 1.52E-05 | 0.000445802 | | Gm14872 | 213.4711444 | -0.17698032133371 | 0.0875228452084642 | -2.02210429645167 | 0.0431655782620271 | 0.37569596413399 | NR_045945 |
| Grin3a | 2151.726381 | -0.258530489 | 0.091621142 | -2.821733972 | 0.004776478 | 0.034908845 | | Gas5 | 3082.567876 | -0.14555467045708 | 0.0719914402498595 | -2.02183301170119 | 0.0431936061229047 | 0.375706779964858 | NR_002840 |
| Tomm70a | 8206.006034 | -0.25864699 | 0.082297445 | -3.142831334 | 0.001673222 | 0.016747355 | | Dffb | 267.0915877 | -0.176227027241436 | 0.0872600108099883 | -2.01956228982342 | 0.0434288100422129 | 0.377038309549304 | NM_007859 |
| Atp2b4 | 13680.29093 | -0.259226925 | 0.075225176 | -3.446012875 | 0.000568923 | 0.007495661 | | Fcho1 | 4313.482876 | -0.155070520170282 | 0.0767700049047653 | -2.01993630666886 | 0.0433899946882478 | 0.377038309549304 | NM_028715,NR_028267 |
| Echdc1 | 922.9944129 | -0.259822042 | 0.084354774 | -3.080110705 | 0.002069237 | 0.019597037 | | Pcsk1 | 1446.409785 | 0.186789214725549 | 0.0924995217740901 | 2.01935330197426 | 0.0434505115093993 | 0.377038309549304 | NM_013628 |
| Fem1b | 6727.994771 | -0.260497003 | 0.057362315 | -4.541256842 | 5.59E-06 | 0.000201026 | | Slc44a2 | 2077.922932 | -0.125731094349449 | 0.0622528038183784 | -2.0196856468709 | 0.0434160048447807 | 0.377038309549304 | NM_001199186,NM_152808 |
| Map2k1 | 11514.50852 | -0.260840114 | 0.059022831 | -4.419308754 | 9.90E-06 | 0.000311191 | | Ass1 | 1892.014804 | 0.126612185413914 | 0.0627519157644857 | 2.0176624708814 | 0.0436264258943869 | 0.378214765635393 | NM_007494 |
| Purg | 511.0060247 | -0.260840317 | 0.090433025 | -2.88434802 | 0.00392225 | 0.030723633 | | Erp44 | 1278.556697 | 0.140547082277688 | 0.0696622593968792 | 2.01754986838662 | 0.043638162412565 | 0.378214765635393 | NM_029572 |
| Fasn | 13324.23434 | -0.260847438 | 0.0532988 | -4.894058405 | 9.88E-07 | 4.83E-05 | | Sgcz | 95.79465902 | 0.185879068295346 | 0.092148734302174 | 2.01716355306424 | 0.0436784481924657 | 0.378338186311125 | NM_145841 |
| Il34 | 2084.273178 | -0.261263587 | 0.081090389 | -3.221881061 | 0.00127352 | 0.013683744 | | Agap1 | 6162.315011 | 0.116238258658787 | 0.0576459305182581 | 2.01641742294316 | 0.0437563451419156 | 0.378787049780373 | NM_178119,NM_001037136 |
| Grik2 | 1937.132698 | -0.261445858 | 0.086555275 | -3.020565269 | 0.002523033 | 0.022518362 | | Ppp1r14a | 248.8467534 | -0.191184615160796 | 0.0948347594569567 | -2.01597617008319 | 0.043802467692673 | 0.378860464840841 | NM_026731 |
| Tpbg | 587.2474673 | -0.26158311 | 0.079197772 | -3.302909966 | 0.000956871 | 0.010998338 | | Prps1 | 3155.805415 | 0.106823562173842 | 0.0529921540432939 | 2.01583732728751 | 0.0438169889117867 | 0.378860464840841 | NM_021463 |
| Stox2 | 6053.782691 | -0.261765874 | 0.074159431 | -3.529771863 | 0.000415918 | 0.005984147 | | Prdx2 | 7445.240719 | 0.10102959197339 | 0.0501349451881189 | 2.01515313508973 | 0.0438886062402746 | 0.379253952555758 | NM_011563 |
| Gda | 10924.00403 | -0.261820251 | 0.093831009 | -2.790338218 | 0.005265301 | 0.037132712 | | Ppp4r1 | 1522.880075 | -0.109289059758813 | 0.0542584440980067 | -2.01423136206052 | 0.0439852484257417 | 0.379863090744545 | NM_001114131,NM_146081 |
| Chd7 | 598.8044073 | -0.261830215 | 0.088302556 | -2.965148754 | 0.003025366 | 0.025447894 | | 1700049G17Rik | 244.6542665 | -0.158080089422423 | 0.0785162418166684 | -2.01334253607721 | 0.0440786063992795 | 0.380217242610412 | NM_028538 |
| Lrrn3 | 2645.202435 | -0.261888786 | 0.083367887 | -3.141362879 | 0.001681635 | 0.016809844 | | Cops7a | 5543.562455 | 0.0892387509193204 | 0.0443224202161596 | 2.01339977564638 | 0.0440725891972622 | 0.380217242610412 | NM_001164089,NM_012003 |
| Grik3 | 5849.979912 | -0.261916832 | 0.069139414 | -3.788241998 | 0.000151717 | 0.002730202 | | Cldn11 | 4344.774523 | 0.168376169054167 | 0.0836577185032074 | 2.01267942835079 | 0.0441483649187573 | 0.380406521556425 | NM_008770 |
| Rpl9 | 9690.250307 | -0.26200314 | 0.057817727 | -4.531536517 | 5.86E-06 | 0.000207139 | | Gm996 | 5479.062405 | -0.128276865371724 | 0.063735746806139 | -2.0126361076759 | 0.0441529254677222 | 0.380406521556425 | NM_001005424 |
| Garem | 1406.090467 | -0.262012659 | 0.077827482 | -3.36658277 | 0.000761057 | 0.00920517 | | Dock7 | 1896.403227 | -0.128485938284476 | 0.063847969122628 | -2.01237314279648 | 0.0441806174163521 | 0.380419471600433 | NM_026082 |
| Nbea | 12146.71499 | -0.262750599 | 0.080444539 | -3.266232874 | 0.001089886 | 0.012203413 | | 2510009E07Rik | 2655.827328 | 0.124728235820182 | 0.0621066553706117 | 2.00829098066682 | 0.0446123797661593 | 0.380589750971131 | NM_001001881 |
| Rplp1 | 11093.3193 | -0.26287478 | 0.070775833 | -3.714188445 | 0.000203857 | 0.003448207 | | Ap2b1 | 19084.76379 | 0.0887033513694498 | 0.0441551675930214 | 2.00890079700364 | 0.0445476553958621 | 0.380589750971131 | NM_027915,NM_001035854 |
| Hn1 | 2903.057988 | -0.263207859 | 0.086905359 | -3.028672347 | 0.002456309 | 0.02211392 | | Casp12 | 41.12608815 | -0.185132061791262 | 0.0921983270525266 | -2.00797636692247 | 0.044645803087898 | 0.380589750971131 | NM_009808 |
| Gp1bb | 330.1562422 | -0.263473495 | 0.095208419 | -2.767334012 | 0.005651682 | 0.039197474 | | Cst3 | 30741.28026 | -0.12013969484743 | 0.0597809169741526 | -2.00966631039425 | 0.0444665178972288 | 0.380589750971131 | NM_009976 |
| Fam171a1 | 4272.369906 | -0.263495645 | 0.085218378 | -3.092004928 | 0.001988095 | 0.019061604 | | D630041G03Rik | 191.1480397 | -0.184944936007499 | 0.0920817231101435 | -2.00848691532713 | 0.0445915751101793 | 0.380589750971131 | NR_028416 |
| Uqcrq | 2187.379794 | -0.263505744 | 0.070759396 | -3.723968239 | 0.000196116 | 0.003355592 | | Farsb | 2339.58868 | 0.0980762325460981 | 0.0488092781859009 | 2.00937682734322 | 0.0444971858105626 | 0.380589750971131 | NM_011811 |
| Lrp1 | 17897.30816 | -0.264025808 | 0.08267981 | -3.193352878 | 0.00140631 | 0.014710663 | | Focad | 1670.183517 | -0.115620801576587 | 0.0575537914430727 | -2.00891720037157 | 0.0445459154784624 | 0.380589750971131 | NM_001081184 |
| Scn3b | 5980.761202 | -0.264448407 | 0.091769545 | -2.881657611 | 0.003955893 | 0.030869893 | | Gla | 605.7297034 | 0.139414828989459 | 0.069401248782389 | 2.0088230606138 | 0.0445559017320972 | 0.380589750971131 | NM_013463 |
| Pde1b | 16534.44502 | -0.264751376 | 0.087267495 | -3.033791387 | 0.002415014 | 0.021882001 | | Glis3 | 121.7348104 | -0.184881195058173 | 0.0920312466376924 | -2.00889591103781 | 0.0445481736649627 | 0.380589750971131 | NM_175459 |
| Htt | 8429.087796 | -0.264822722 | 0.056627418 | -4.676581233 | 2.92E-06 | 0.000120521 | | Gps1 | 5265.501852 | 0.0872950745406443 | 0.0434730199117026 | 2.00802876630029 | 0.0446402349192485 | 0.380589750971131 | NM_001177874,NM_145370 |
| Cox4i1 | 13251.02491 | -0.265134656 | 0.061905519 | -4.282892063 | 1.84E-05 | 0.000521593 | | Ifit2 | 608.3127271 | -0.173821220424263 | 0.0864283586240232 | -2.01115956835896 | 0.0443086048729896 | 0.380589750971131 | NM_008332 |
| Lamb1 | 819.589313 | -0.265606827 | 0.086763983 | -3.061256749 | 0.0022041 | 0.020597302 | | Ldhb | 20367.79021 | 0.0938053279997163 | 0.0466593862247773 | 2.01042781719883 | 0.0443859288113403 | 0.380589750971131 | NM_008492 |
| Lsmd1 | 707.218225 | -0.265951943 | 0.065065234 | -4.087466144 | 4.36E-05 | 0.001026914 | | Nsdhl | 1044.978714 | 0.113383854130858 | 0.0564182096373907 | 2.00970315895515 | 0.0444626154305065 | 0.380589750971131 | NM_010941 |
| Nf1 | 4761.694716 | -0.266125825 | 0.075926724 | -3.505035015 | 0.000456547 | 0.006390012 | | Rgs9 | 4569.625627 | -0.163973127336202 | 0.0816001454692748 | -2.00947101740086 | 0.0444872053321194 | 0.380589750971131 | NM_011268,NM_001165934 |
| Cox7c | 3237.854925 | -0.266505499 | 0.065933018 | -4.042064327 | 5.30E-05 | 0.001203686 | | Tmem127 | 6313.310676 | 0.085145459118721 | 0.0423291342711488 | 2.01150958045356 | 0.0442716594049872 | 0.380589750971131 | NM_175145 |
| Thsd7a | 2015.923869 | -0.266567494 | 0.088155704 | -3.023825823 | 0.002496001 | 0.022341444 | | Tpi1 | 24122.11141 | 0.0939869012656694 | 0.0467276792158591 | 2.0113753313426 | 0.0442858269726367 | 0.380589750971131 | NM_009415 |
| Fbll1 | 840.1832984 | -0.266729676 | 0.070785028 | -3.768165136 | 0.000164452 | 0.00291202 | | Zbtb34 | 1007.114384 | -0.157198387584369 | 0.0781526901601076 | -2.01142644306069 | 0.0442804326034257 | 0.380589750971131 | NM_001276332,NM_001085507 |
| Flrt1 | 1367.906598 | -0.266965471 | 0.05924832 | -4.505874111 | 6.61E-06 | 0.000227591 | | C030023E24Rik | 106.5632592 | -0.18951174640526 | 0.09441079515656 | -2.00731013959787 | 0.044716650180388 | 0.380855808258635 | NR_033502 |
| Tmem100 | 403.7641217 | -0.267300509 | 0.093938258 | -2.845491448 | 0.004434295 | 0.033344824 | | Pvrl1 | 2250.657718 | 0.161131713740683 | 0.0802772658518534 | 2.00718985669046 | 0.0447294512521845 | 0.380855808258635 | NM_021424 |
| Nat14 | 2137.580978 | -0.267308243 | 0.08989664 | -2.973506481 | 0.002944182 | 0.025050606 | | Letm1 | 3511.298388 | 0.0869881857695057 | 0.0433533675443341 | 2.00649201427201 | 0.0448037799228505 | 0.381265206303061 | NM_019694 |
| 1700086L19Rik | 799.2459527 | -0.267366468 | 0.096639323 | -2.766642596 | 0.005663681 | 0.03924556 | | Atp9a | 18025.22499 | 0.10655357521813 | 0.0531681745324833 | 2.00408564249334 | 0.0450608871283014 | 0.383004357182976 | NM_015731 |
| Mpc2 | 1622.013928 | -0.267468917 | 0.087819195 | -3.045677171 | 0.00232157 | 0.021309478 | | Itga6 | 910.3856358 | -0.150559472231884 | 0.0751208121014475 | -2.00423115805191 | 0.04504530435208 | 0.383004357182976 | NM_008397 |
| Amotl1 | 1803.631944 | -0.267545407 | 0.095518402 | -2.800982853 | 0.005094723 | 0.036290258 | | Rit2 | 2713.528529 | 0.146196854788899 | 0.0729797676074941 | 2.00325185433841 | 0.0451502623844685 | 0.383539597308064 | NM_009065 |
| Nrp1 | 1207.407396 | -0.267888144 | 0.095525195 | -2.804371611 | 0.005041476 | 0.036129802 | | Csdc2 | 5096.27762 | 0.122610181082893 | 0.0612222097289477 | 2.00270754070673 | 0.0452086889373856 | 0.383811464350943 | NM_145473 |
| Wnk3 | 1689.833754 | -0.268498262 | 0.067873015 | -3.955891179 | 7.62E-05 | 0.001594353 | | Arrdc4 | 332.8258418 | -0.149305397281759 | 0.0745954405655913 | -2.00153516286931 | 0.0453347483306578 | 0.383869598536285 | NM_001042592,NM_025549 |
| Tspan13 | 5489.225108 | -0.268533796 | 0.067844235 | -3.958093086 | 7.56E-05 | 0.001581864 | | Astn2 | 2207.425747 | 0.117233846304634 | 0.058557798415151 | 2.00201936339022 | 0.0452826490213584 | 0.383869598536285 | NM_207109,NM_019514 |
| Kcnh7 | 1223.233201 | -0.268684675 | 0.090633627 | -2.964514207 | 0.003031612 | 0.025472778 | | C130074G19Rik | 3012.863975 | -0.117002459810764 | 0.0584482452773465 | -2.00181304426793 | 0.0453048425005709 | 0.383869598536285 | NM_178692 |
| Gbas | 2749.445477 | -0.268878589 | 0.05943835 | -4.523655018 | 6.08E-06 | 0.000213063 | | Crocc | 2712.846777 | -0.172892935511634 | 0.0863959349592553 | -2.00116979569897 | 0.0453740947739777 | 0.383869598536285 | NM_001145958,NM_172122 |
| Rprml | 1218.404517 | -0.268911028 | 0.098726761 | -2.723790648 | 0.00645374 | 0.042939425 | | Psrc1 | 414.1945429 | -0.168403618640825 | 0.0841028638267444 | -2.00235296372004 | 0.0452467834577671 | 0.383869598536285 | NM_001190161,NM_019976 |
| Chd5 | 12626.58739 | -0.269067142 | 0.052298929 | -5.144792559 | 2.68E-07 | 1.68E-05 | | Triap1 | 533.0247021 | 0.144661305673421 | 0.0722866135243148 | 2.00121846384133 | 0.0453688520336227 | 0.383869598536285 | NM_026933 |
| Zfp41 | 286.9877087 | -0.269119506 | 0.094284481 | -2.854335121 | 0.004312701 | 0.032794894 | | Abca1 | 1691.996749 | -0.151281038766967 | 0.0756135946430575 | -2.00071216665609 | 0.0454234175111204 | 0.384063191365853 | NM_013454 |
| Plxna4 | 4845.4661 | -0.269289805 | 0.072736408 | -3.702269752 | 0.000213679 | 0.003556118 | | F11r | 435.0343667 | -0.175751734348989 | 0.0878530462428065 | -2.00010974990382 | 0.0454884141901696 | 0.384389007868763 | NM_172647 |
| BC030500 | 275.6457558 | -0.269583297 | 0.090476538 | -2.979593435 | 0.002886312 | 0.024666584 | | Cyp51 | 2341.597749 | 0.147532846107249 | 0.073801386952723 | 1.99905248666612 | 0.0456026752078384 | 0.385130500040151 | NM_020010 |
| Dlg2 | 18956.97038 | -0.269927555 | 0.054682573 | -4.936262913 | 7.96E-07 | 4.09E-05 | | Cd37 | 154.8703809 | -0.189476018198741 | 0.0948261966900186 | -1.99814001628819 | 0.0457014824482321 | 0.385357198146082 | NM_007645 |
| BC052040 | 305.307873 | -0.269987223 | 0.099613425 | -2.710349771 | 0.006721229 | 0.044107886 | | Nkain4 | 870.3771688 | 0.151726304561525 | 0.0759364424689 | 1.99806969655795 | 0.045709104530201 | 0.385357198146082 | NM_001141933,NM_021426 |
| Actn2 | 1104.620041 | -0.270039436 | 0.071646334 | -3.769061432 | 0.000163863 | 0.002908232 | | Spag1 | 489.2889648 | -0.145671782662973 | 0.0729036514621483 | -1.99814110461403 | 0.045701364491054 | 0.385357198146082 | NM_012031 |
| H3f3a | 4003.980176 | -0.270058045 | 0.059104089 | -4.569193933 | 4.90E-06 | 0.000181916 | | Raver1 | 631.1233367 | -0.117723975697411 | 0.0589286092252151 | -1.99773891230811 | 0.0457449731999609 | 0.385435893679021 | NM_027911 |
| Srgap1 | 1201.124009 | -0.270475862 | 0.101242399 | -2.67156709 | 0.007549798 | 0.048039658 | | Ublcp1 | 2051.137826 | 0.120821503497036 | 0.0605433407959296 | 1.99560200870445 | 0.0459752977448466 | 0.387151985531387 | NM_024475 |
| 4-Mar | 1093.407799 | -0.270639012 | 0.064561249 | -4.191973002 | 2.77E-05 | 0.000723766 | | Frmd6 | 1944.620664 | 0.168793286802801 | 0.0846159183411482 | 1.99481717047935 | 0.0460628239502763 | 0.387222050524594 | NM_028127 |
| Itgb1bp1 | 827.2080644 | -0.270676359 | 0.068571471 | -3.947361123 | 7.90E-05 | 0.001635816 | | Tmx2 | 9177.878122 | 0.0888700680463091 | 0.0445506395351597 | 1.99481015252704 | 0.0460635896534833 | 0.387222050524594 | NM_025868 |
| D430041D05Rik | 8895.851625 | -0.270948336 | 0.08348969 | -3.245290967 | 0.001173307 | 0.012845805 | | Zscan26 | 3710.197062 | -0.11234068519761 | 0.0563102355468937 | -1.99503134921279 | 0.0460394608440483 | 0.387222050524594 | NM_001013786 |
| Snx10 | 3560.434107 | -0.271057302 | 0.073967226 | -3.66455952 | 0.000247765 | 0.004011353 | | Cpsf1 | 2820.870974 | -0.132971281404557 | 0.0666749323998048 | -1.99432195307252 | 0.0461168816333376 | 0.387445819899284 | NM_001164173,NM_053193 |
| Nhp2 | 826.8979944 | -0.271121549 | 0.074794609 | -3.624880912 | 0.000289095 | 0.004515355 | | Kras | 3479.935955 | 0.143475045416119 | 0.0719571709202257 | 1.9938950292415 | 0.0461635272960704 | 0.387613524568045 | NM_021284 |
| Ccnd2 | 2894.115089 | -0.271326709 | 0.090941152 | -2.98354158 | 0.002849333 | 0.024458487 | | Cdca7l | 41.84915241 | -0.183152793727661 | 0.0918789682802476 | -1.99341369582004 | 0.0462161653876057 | 0.387831322022161 | NM_146040 |
| Bsn | 21922.82964 | -0.271897445 | 0.09446213 | -2.878375136 | 0.003997295 | 0.031028284 | | Fam110a | 201.4536555 | 0.164727384738084 | 0.0826594831169527 | 1.99284315031363 | 0.046278625063587 | 0.388131239996342 | NM_028666,NM_146127 |
| Tubb5 | 17037.70134 | -0.271996295 | 0.064052323 | -4.246470398 | 2.17E-05 | 0.00059553 | | Scand1 | 236.375585 | -0.185835448302234 | 0.0933243537071207 | -1.99128567110617 | 0.046449489853212 | 0.388890656834443 | NM_020255 |
| Neurl1a | 8091.010445 | -0.272028996 | 0.087343759 | -3.114464041 | 0.001842794 | 0.017980008 | | Srrm3 | 1831.392349 | -0.12930899892842 | 0.0649349864986685 | -1.99136098890344 | 0.0464412148376319 | 0.388890656834443 | NM_021403 |
| Phlda3 | 625.2975953 | -0.272084803 | 0.086806275 | -3.134390958 | 0.001722112 | 0.01711508 | | Thsd7b | 328.4955616 | 0.146183170858154 | 0.0733963413652584 | 1.99169560960363 | 0.046404465734163 | 0.388890656834443 | NM_172485 |
| Arl6 | 769.0131891 | -0.272318536 | 0.088580583 | -3.074246379 | 0.002110351 | 0.019904984 | | Nr6a1 | 505.6576918 | -0.139124816236164 | 0.0698904236634208 | -1.99061343376831 | 0.0465234022158348 | 0.389285104024894 | NM_001159549,NM_001159548,NM_010264 |
| Dand5 | 318.7387713 | -0.27239276 | 0.090532554 | -3.008782466 | 0.002622968 | 0.023183271 | | Bmpr1b | 953.1869614 | -0.10802146490126 | 0.0543054374925074 | -1.98914638918367 | 0.0466850472931726 | 0.390412778918034 | NM_007560 |
| Commd2 | 373.6835996 | -0.272597141 | 0.084520064 | -3.225235876 | 0.001258689 | 0.013590335 | | Grb10 | 2384.079407 | 0.17475758730932 | 0.087873174087599 | 1.98874786445187 | 0.0467290399569988 | 0.390555831079036 | NM_010345,NM_001177629 |
| Elmod1 | 10113.68959 | -0.273152044 | 0.06593415 | -4.142800745 | 3.43E-05 | 0.000849175 | | Hnrpdl | 6252.706877 | -0.122161204297497 | 0.0614459995657416 | -1.98810671420188 | 0.0467998889663926 | 0.390923051826233 | NM_016690 |
| Scd1 | 7879.33641 | -0.273542398 | 0.089668927 | -3.050581811 | 0.002283984 | 0.021078966 | | Tspan13 | 5691.579939 | 0.127225130335925 | 0.0640022899853127 | 1.9878215352157 | 0.046831431123805 | 0.390961706037006 | NM_025359 |
| Tmem150c | 1556.800868 | -0.273919926 | 0.072614557 | -3.772245347 | 0.000161785 | 0.002881262 | | Acot7 | 9027.936893 | 0.117086192991959 | 0.0589226085464753 | 1.98711828753487 | 0.0469092901265209 | 0.39102735987968 | NM_001146057,NM_133348,NM_001146058 |
| Cox7a2 | 4106.934733 | -0.273978707 | 0.072280504 | -3.790492456 | 0.000150349 | 0.002721919 | | Gm15706 | 181.4353073 | -0.18002050172499 | 0.0906424645434484 | -1.9860503863363 | 0.0470277294306623 | 0.39102735987968 | NR_045598 |
| Btbd3 | 7288.471037 | -0.274244277 | 0.072549045 | -3.780122507 | 0.000156751 | 0.002801273 | | Hdac11 | 9034.083815 | 0.13219794499084 | 0.0665573716820222 | 1.98622544205043 | 0.0470082970450883 | 0.39102735987968 | NM_144919 |
| Vhl | 2041.025207 | -0.274321206 | 0.067188994 | -4.082829508 | 4.45E-05 | 0.001041174 | | Igsf6 | 37.00315562 | -0.157980017822025 | 0.0795392775431056 | -1.98618874475455 | 0.0470123701370599 | 0.39102735987968 | NM_030691 |
| Vps13c | 2727.000421 | -0.274972029 | 0.082233655 | -3.343789485 | 0.000826424 | 0.009834575 | | Nrsn2 | 4060.956761 | 0.156926508931522 | 0.0789870140896028 | 1.98673808271198 | 0.0469514292654161 | 0.39102735987968 | NM_001009948 |
| Nfya | 1070.05673 | -0.275564693 | 0.075232807 | -3.662826158 | 0.000249448 | 0.004030182 | | Timm23 | 2311.203929 | 0.10130454432783 | 0.0509787628814139 | 1.98719110864819 | 0.046901222795635 | 0.39102735987968 | NM_016897 |
| Nhsl2 | 2959.406824 | -0.275762643 | 0.102232764 | -2.697399868 | 0.00698833 | 0.045399238 | | Zfp36l2 | 879.1711633 | -0.181295199515267 | 0.0912643785691527 | -1.98648369010584 | 0.0469796420721809 | 0.39102735987968 | NM_001001806 |
| Sema5b | 883.0362923 | -0.275899436 | 0.093691818 | -2.944754861 | 0.003232106 | 0.026704603 | | Maea | 3413.970505 | 0.0868046961393726 | 0.0437191378221786 | 1.98550795975068 | 0.0470879854197643 | 0.391304391423053 | NM_021500 |
| Arxes1 | 440.049255 | -0.276369411 | 0.0940584 | -2.938274618 | 0.003300445 | 0.027132771 | | 5430435G22Rik | 68.52469412 | -0.186664775474592 | 0.0940734691780317 | -1.98424462396867 | 0.0472285761213711 | 0.391827401706351 | NM_145509 |
| Tnfrsf19 | 2848.21442 | -0.276392381 | 0.085751347 | -3.223184139 | 0.00126774 | 0.013659217 | | Gm14393 | 22.48566219 | -0.162432295256122 | 0.0818622335258271 | -1.98421528780882 | 0.0472318449943426 | 0.391827401706351 | NM_001085546 |
| Ubxn2b | 1687.276205 | -0.276527401 | 0.061974402 | -4.461961597 | 8.12E-06 | 0.000266591 | | Pkp4 | 10054.8987 | 0.0989638339121234 | 0.0498716057043436 | 1.98437231996932 | 0.0472143494134498 | 0.391827401706351 | NM_175464,NM_026361 |
| Fzd3 | 5088.22638 | -0.277268296 | 0.081139209 | -3.417192489 | 0.000632705 | 0.008095075 | | Ano6 | 1254.364598 | -0.11447030184317 | 0.0577433206676774 | -1.98239901203406 | 0.0474346000911549 | 0.391951500592767 | NM_001253813,NM_175344 |
| Lsm4 | 726.2675527 | -0.277768222 | 0.081626649 | -3.402911005 | 0.00066672 | 0.008378069 | | Bex4 | 428.3235237 | 0.168939746710473 | 0.0851840923454387 | 1.98323116510285 | 0.0473416142445034 | 0.391951500592767 | NM_212457 |
| Rnf150 | 4078.154573 | -0.277907678 | 0.077296743 | -3.595334903 | 0.000323975 | 0.00496014 | | Khdrbs1 | 4457.945494 | 0.0958097179031725 | 0.0483096577889033 | 1.98324149431628 | 0.0473404610088385 | 0.391951500592767 | NR_045036,NM_011317 |
| Tubb2a | 11406.40029 | -0.278208067 | 0.066191991 | -4.203047241 | 2.63E-05 | 0.000692746 | | Klhdc10 | 5028.887036 | 0.0977474650802344 | 0.0493024589370583 | 1.98261196677885 | 0.0474107896468089 | 0.391951500592767 | NM_029742 |
| Myeov2 | 1382.767798 | -0.279163817 | 0.082894727 | -3.367690894 | 0.000758005 | 0.009189776 | | Mphosph9 | 1078.799594 | -0.120028606287568 | 0.0605474438297825 | -1.98238932472534 | 0.0474356834670305 | 0.391951500592767 | NM_001081323 |
| Ttbk2 | 4499.553625 | -0.27924986 | 0.091303711 | -3.058472168 | 0.002224687 | 0.020702287 | | Pctp | 83.15387197 | -0.188071740087215 | 0.0948341402919776 | -1.9831649183319 | 0.0473490111233586 | 0.391951500592767 | NM_008796 |
| Ets1 | 563.7972172 | -0.279800018 | 0.093075585 | -3.006159114 | 0.002645705 | 0.02323841 | | Sirt7 | 1218.627001 | -0.162864329443113 | 0.0821287961507893 | -1.98303563520026 | 0.0473634492171193 | 0.391951500592767 | NM_153056 |
| Romo1 | 786.8249378 | -0.279890598 | 0.089226121 | -3.136868372 | 0.001707628 | 0.017014781 | | Atp6v1h | 7194.111734 | 0.121947221210307 | 0.0615563512829358 | 1.98106643211831 | 0.0475838241163537 | 0.392147257933715 | NM_133826 |
| Tuba1a | 24555.33983 | -0.280077973 | 0.085750789 | -3.266185367 | 0.001090069 | 0.012203413 | | Fbxo10 | 2024.488886 | 0.136941546727485 | 0.0691284436018704 | 1.98097251423985 | 0.0475943560331226 | 0.392147257933715 | NM_001024142 |
| Ndufa1 | 1383.852938 | -0.280154 | 0.075745487 | -3.698622987 | 0.000216772 | 0.003599861 | | Galntl6 | 904.8298453 | 0.156387786289676 | 0.0789322033537996 | 1.98129254784256 | 0.0475584756180379 | 0.392147257933715 | NM_175032 |
| Sc5d | 1706.085197 | -0.280380883 | 0.078461643 | -3.573477072 | 0.000352272 | 0.00528886 | | Greb1l | 458.1728604 | -0.14609682371191 | 0.0737447997508515 | -1.9811135728282 | 0.047578538512846 | 0.392147257933715 | NM_001083628 |
| Slc4a8 | 5343.381303 | -0.280504596 | 0.052758348 | -5.316781226 | 1.06E-07 | 7.79E-06 | | Morc2a | 2446.305072 | -0.0936739110188604 | 0.0472723874882745 | -1.9815777454036 | 0.0475265199945643 | 0.392147257933715 | NM_001159288,NM_198162 |
| Dpy19l3 | 2786.64691 | -0.280828977 | 0.066832451 | -4.201985292 | 2.65E-05 | 0.000694825 | | Map2k4 | 9630.538481 | 0.104677540031595 | 0.0528825026908457 | 1.97943619732875 | 0.0477669164648648 | 0.392678114639856 | NM_009157 |
| Runx1t1 | 821.449698 | -0.281049249 | 0.087820928 | -3.200253689 | 0.001373067 | 0.014433037 | | Pkn1 | 1791.118127 | -0.140853633757504 | 0.0711456124629845 | -1.9797936778011 | 0.0477267170777253 | 0.392678114639856 | NM_001199593,NM_177262 |
| Gnb4 | 1289.234291 | -0.282679962 | 0.103197261 | -2.739219623 | 0.006158522 | 0.041559293 | | Ube2n | 7704.799102 | 0.102128399508613 | 0.0515837091611482 | 1.97985761724815 | 0.0477195299590829 | 0.392678114639856 | NM_080560 |
| Nwd1 | 3589.869655 | -0.283242048 | 0.064875998 | -4.365898867 | 1.27E-05 | 0.000386676 | | Wdr61 | 1339.475806 | 0.10757811128181 | 0.0543414624848063 | 1.97966904758753 | 0.0477407287620543 | 0.392678114639856 | NM_001025375,NM_001025376,NM_023191 |
| Ntn3 | 382.665928 | -0.283600438 | 0.099587534 | -2.84775038 | 0.004402944 | 0.033182157 | | Gli3 | 499.551502 | -0.15598673794421 | 0.0788249687556608 | -1.97890009227575 | 0.0478272558803019 | 0.392951764093475 | NM_008130 |
| Htr1a | 704.2859663 | -0.283939262 | 0.09963389 | -2.849826124 | 0.004374314 | 0.033045155 | | Cry1 | 817.3984228 | 0.176701404180443 | 0.0893363397032248 | 1.97793422886415 | 0.0479361269089619 | 0.393352855610125 | NM_007771 |
| Sec61b | 400.1800827 | -0.284066244 | 0.089296722 | -3.181149721 | 0.001466918 | 0.015152799 | | Lonp2 | 3198.504055 | 0.11876960032536 | 0.0600520421069494 | 1.97777877662638 | 0.0479537703263976 | 0.393352855610125 | NM_001168591,NM_025827 |
| Rragd | 7845.2649 | -0.284243947 | 0.061171156 | -4.64669896 | 3.37E-06 | 0.00013469 | | Prkaca | 8825.434052 | 0.0967111078812492 | 0.0488996480740893 | 1.97774650105292 | 0.0479573115300517 | 0.393352855610125 | NM_008854 |
| Msantd3 | 375.6812436 | -0.284397132 | 0.088306674 | -3.220562136 | 0.001279395 | 0.01370881 | | Timm21 | 361.947193 | -0.149575937688235 | 0.0756563535323263 | -1.97704397191605 | 0.0480366600363737 | 0.393773739320375 | NM_025969 |
| Fam84a | 2741.249843 | -0.284790976 | 0.077687553 | -3.665850775 | 0.000246518 | 0.003995339 | | Utp20 | 1149.436534 | -0.128035138308649 | 0.0647684826335989 | -1.97681328007311 | 0.0480628417881747 | 0.393773739320375 | NM_175158 |
| Hmgn1 | 1661.716221 | -0.284813726 | 0.083180704 | -3.424036014 | 0.000616984 | 0.007974295 | | Phf15 | 4326.826621 | 0.146877740698459 | 0.074325310871444 | 1.97614700801594 | 0.04813812966166 | 0.393789374982573 | NM_199299 |
| Enho | 1337.560229 | -0.285001367 | 0.084612906 | -3.368296655 | 0.000756342 | 0.009189519 | | Polr2m | 7275.627501 | 0.0982140494420878 | 0.0497015342601191 | 1.97607681340524 | 0.0481460780647839 | 0.393789374982573 | NM_001164793,NM_178602 |
| Med21 | 517.342109 | -0.285119741 | 0.08431539 | -3.381585981 | 0.000720687 | 0.008890383 | | Slc10a7 | 203.4516701 | 0.167148558780791 | 0.0845739813061521 | 1.97635911422598 | 0.0481141187660292 | 0.393789374982573 | NM_029736 |
| Gpr124 | 755.0101673 | -0.286131541 | 0.082866592 | -3.452917943 | 0.000554558 | 0.007375379 | | Supt3 | 387.9169392 | 0.156799195157031 | 0.0793832609469984 | 1.97521736051786 | 0.0482434866197324 | 0.394364032998443 | NM_178652 |
| Map1a | 62042.02897 | -0.28625253 | 0.071398196 | -4.009240389 | 6.09E-05 | 0.001338734 | | 9330151L19Rik | 849.1006789 | -0.135156554098091 | 0.0684758068199596 | -1.97378549264052 | 0.0484061391211639 | 0.394969953195978 | NR_033222 |
| Robo2 | 4828.820806 | -0.286324958 | 0.064580302 | -4.433626783 | 9.27E-06 | 0.000296018 | | Fam222a | 416.2719678 | -0.171593014531907 | 0.0869369908315891 | -1.97376298504417 | 0.0484086995390332 | 0.394969953195978 | NM_001004180 |
| Homer1 | 6414.64479 | -0.28665955 | 0.107110646 | -2.676293721 | 0.007444137 | 0.047503899 | | Lsm4 | 775.0006872 | 0.125951120720373 | 0.0638177092226892 | 1.97360765640628 | 0.0484263724798319 | 0.394969953195978 | NM_015816 |
| Erc1 | 2660.048639 | -0.288362174 | 0.055328716 | -5.21179953 | 1.87E-07 | 1.25E-05 | | Rian | 24082.87652 | -0.0967784789122343 | 0.0490298589662839 | -1.97386818874567 | 0.0483967327715287 | 0.394969953195978 | NR_028261 |
| Med13 | 2820.827253 | -0.288401508 | 0.083728069 | -3.444502081 | 0.000572112 | 0.007518496 | | A930024E05Rik | 102.1592212 | -0.186998332233444 | 0.0948572153344447 | -1.97136645403653 | 0.0486819752176774 | 0.39502792612861 | NR_045820 |
| Cdk18 | 1640.290027 | -0.288499536 | 0.100725651 | -2.86421118 | 0.004180492 | 0.032081497 | | Cops3 | 2454.449595 | 0.0995797104730147 | 0.050478313437762 | 1.97272261474807 | 0.0485271739691969 | 0.39502792612861 | NM_011991 |
| Fads1 | 9815.851253 | -0.288719931 | 0.088482758 | -3.26300781 | 0.001102365 | 0.012305329 | | Epm2a | 539.3386658 | 0.140782099331848 | 0.0714205651293827 | 1.97117033555829 | 0.0487043957703287 | 0.39502792612861 | NM_010146 |
| Wwtr1 | 667.5652034 | -0.288955195 | 0.095621116 | -3.021876402 | 0.002512131 | 0.022459872 | | Fancd2 | 197.18293 | -0.176939568565752 | 0.0897446758743951 | -1.97158847409948 | 0.0486566040167485 | 0.39502792612861 | NM_001033244 |
| Ptar1 | 245.9913229 | -0.290153362 | 0.100678613 | -2.881976145 | 0.003951897 | 0.030863169 | | Maged1 | 22501.86593 | 0.118479977823594 | 0.0600629328284844 | 1.97259727829018 | 0.0485414633520169 | 0.39502792612861 | NM_019791 |
| 1110001J03Rik | 383.9704642 | -0.290169745 | 0.090193704 | -3.217184033 | 0.001294555 | 0.01382346 | | Mlh1 | 522.1232483 | -0.140302570129102 | 0.0711061216695422 | -1.97314333611307 | 0.0484792341153842 | 0.39502792612861 | NM_026810 |
| Cd97 | 502.0610453 | -0.290193783 | 0.105721558 | -2.744887485 | 0.006053165 | 0.041042969 | | Orc5 | 969.218877 | 0.112025446169759 | 0.0568322057805872 | 1.97116132712246 | 0.0487054258361794 | 0.39502792612861 | NM_011959 |
| Enox2 | 391.4511916 | -0.290667272 | 0.10600588 | -2.741991977 | 0.006106783 | 0.041300088 | | Stambpl1 | 618.4527645 | 0.128954437599773 | 0.0653951331967988 | 1.97192713426701 | 0.0486179252266285 | 0.39502792612861 | NM_029682 |
| Dpysl5 | 1463.99801 | -0.290968022 | 0.083004783 | -3.505436811 | 0.000455859 | 0.006386143 | | Stk10 | 377.0270149 | -0.146852443798202 | 0.0744680136549023 | -1.97202042314088 | 0.0486072751274343 | 0.39502792612861 | NM_009288 |
| Chml | 479.0348954 | -0.291022253 | 0.095916548 | -3.034119353 | 0.00241239 | 0.021882001 | | Ybx2 | 156.4934039 | -0.186773116949506 | 0.0947334413885627 | -1.9715647844295 | 0.048659310609435 | 0.39502792612861 | NM_016875 |
| Kcna6 | 5070.068811 | -0.291298993 | 0.099803571 | -2.918723155 | 0.003514682 | 0.028392328 | | Aco2 | 23504.93539 | 0.0926168402736016 | 0.0470299810948742 | 1.96931485230165 | 0.0489169466892497 | 0.396522080138416 | NM_080633 |
| Slc32a1 | 4455.48704 | -0.291850313 | 0.094609159 | -3.08479977 | 0.002036892 | 0.019331788 | | Dgat2l6 | 26.56374978 | -0.167055509783042 | 0.0848427788441002 | -1.96900092216461 | 0.0489529851864724 | 0.396592896162129 | NM_001114084 |
| Tomm7 | 1365.140749 | -0.291889305 | 0.070329936 | -4.150285398 | 3.32E-05 | 0.000832517 | | Chd4 | 8757.919544 | -0.0923798248063342 | 0.0469387597138775 | -1.96809258211017 | 0.0490573862023791 | 0.397217163866087 | NM_145979 |
| Lrrk2 | 3437.238212 | -0.291960367 | 0.088103951 | -3.313816938 | 0.000920317 | 0.010649289 | | 2900076A07Rik | 130.4427097 | -0.185708513791088 | 0.0943712703174149 | -1.96785010063405 | 0.049085287644447 | 0.397221664804032 | NR_045299 |
| Lzic | 359.680208 | -0.292329302 | 0.076274447 | -3.832598117 | 0.000126797 | 0.002347184 | | Sp110 | 80.06631555 | -0.186930408993154 | 0.0950121307676966 | -1.96743728914149 | 0.0491328189646766 | 0.39738492665974 | NM_175397,NM_030194 |
| Mycbp2 | 15363.72493 | -0.292445904 | 0.056777002 | -5.150780981 | 2.59E-07 | 1.63E-05 | | Ganc | 420.5319029 | -0.132282745463315 | 0.0672616328184742 | -1.96668947690164 | 0.0492190207999245 | 0.397725047787054 | NM_172672 |
| Ccdc90b | 411.1475481 | -0.292895391 | 0.10421604 | -2.810463639 | 0.004947018 | 0.035738209 | | Rhobtb2 | 6123.58087 | 0.124776599228589 | 0.0634479599039356 | 1.96659749844612 | 0.0492296321025144 | 0.397725047787054 | NM_153514 |
| Atp5k | 2060.648747 | -0.293022495 | 0.090184517 | -3.249144162 | 0.001157528 | 0.012746794 | | B3galt1 | 1940.860356 | 0.135210092681368 | 0.068779823976515 | 1.96583946954466 | 0.0493171569449005 | 0.398210684703516 | NM_020283 |
| Phlda1 | 717.2971049 | -0.293340213 | 0.089662523 | -3.27160338 | 0.001069395 | 0.012015376 | | Ankrd9 | 387.5161427 | 0.153510453544493 | 0.0781480165484132 | 1.96435508314395 | 0.0494889279158074 | 0.398932390069377 | NM_175207 |
| Cnih2 | 5572.093156 | -0.293638488 | 0.075367597 | -3.896084015 | 9.78E-05 | 0.001924652 | | Cited1 | 89.92900043 | 0.162058379953489 | 0.0824986443158038 | 1.96437627911959 | 0.0494864716217875 | 0.398932390069377 | NM_001276474,NM_001276466,NM_001276473,NM_007709 |
| Chac2 | 197.785737 | -0.293843382 | 0.09694021 | -3.031181611 | 0.002435987 | 0.02204625 | | Spock2 | 32981.30078 | 0.10106282125788 | 0.0514419504181356 | 1.96459932868818 | 0.0494606297396376 | 0.398932390069377 | NM_052994 |
| Calr | 12996.75824 | -0.29406075 | 0.08932617 | -3.291988775 | 0.000994816 | 0.011333586 | | Glra1 | 30.1688648 | 0.137182349754357 | 0.0698805578256157 | 1.96309751986654 | 0.0496348436894916 | 0.399665044031904 | NM_020492 |
| Arhgef19 | 604.0598838 | -0.294433515 | 0.108094458 | -2.723853937 | 0.006452504 | 0.042939425 | | Park2 | 246.7325046 | -0.177544472133543 | 0.0904376406693626 | -1.96317610492685 | 0.0496257148627095 | 0.399665044031904 | NM_016694 |
| Arhgap32 | 20321.43569 | -0.295181582 | 0.074326588 | -3.971413036 | 7.14E-05 | 0.001514376 | | 2510002D24Rik | 444.338675 | -0.138102933877017 | 0.0703657631979272 | -1.96264387111892 | 0.0496875692946082 | 0.399757903930456 | NM_001033164 |
| Itga1 | 320.3860742 | -0.295300062 | 0.105415343 | -2.801300594 | 0.005089709 | 0.036290258 | | Rnf31 | 1147.52283 | -0.125945856550715 | 0.0641860881170668 | -1.96219866711625 | 0.0497393590657907 | 0.399757903930456 | NM_194346 |
| St6galnac5 | 1318.961732 | -0.296393559 | 0.08324336 | -3.560566993 | 0.000370055 | 0.00550781 | | Rrp1 | 12144.12895 | 0.0825110831957494 | 0.042053468852665 | 1.96205177472584 | 0.0497564567194178 | 0.399757903930456 | NM_010925 |
| Tnr | 3651.991982 | -0.296721095 | 0.088129551 | -3.366874017 | 0.000760254 | 0.009204125 | | Tnfrsf22 | 38.27127116 | -0.181476104393825 | 0.0924710867168853 | -1.96251726714798 | 0.0497022923002398 | 0.399757903930456 | NM_023680 |
| Arhgap31 | 1703.422183 | -0.296842791 | 0.07865307 | -3.774077606 | 0.000160601 | 0.002863462 | | Plgrkt | 381.377328 | -0.141874715448794 | 0.0723372607437501 | -1.96129510559401 | 0.0498446079132661 | 0.400244762049808 | NM_026362 |
| Ndufb6 | 1608.890521 | -0.296903255 | 0.054284398 | -5.46940312 | 4.52E-08 | 3.78E-06 | | Evi2b | 30.26041139 | -0.1718982397148 | 0.0876699533811763 | -1.96074291231126 | 0.0499090204988063 | 0.400540570036276 | NM_001077496,NM_146023 |
| Sema6a | 1194.054276 | -0.297070256 | 0.085246547 | -3.484836253 | 0.000492438 | 0.00675805 | | Nap1l5 | 11205.89854 | 0.151959432212065 | 0.0775382607902746 | 1.95979933858828 | 0.0500192485937976 | 0.401203536760631 | NM_021432 |
| Brk1 | 5480.3681 | -0.297463365 | 0.063001046 | -4.72156232 | 2.34E-06 | 0.000101291 | | Mapkapk3 | 140.267034 | -0.181310884047967 | 0.0925412554202326 | -1.95924383373263 | 0.050084237956816 | 0.401503112892224 | NM_178907 |
| Ston2 | 1065.013565 | -0.297793201 | 0.079589097 | -3.741633129 | 0.000182828 | 0.003186436 | | Ormdl3 | 2378.949166 | 0.0943183614362659 | 0.0481538679293298 | 1.95868713131594 | 0.0501494384243325 | 0.401582548264528 | NM_025661 |
| Erbb4 | 1115.74852 | -0.298877352 | 0.091519954 | -3.265706976 | 0.001091912 | 0.012215221 | | Slc25a37 | 1558.05316 | -0.128737632895986 | 0.0657206205783868 | -1.95886210085976 | 0.0501289384921879 | 0.401582548264528 | NM_026331 |
| Nfia | 1208.783073 | -0.299005118 | 0.080940377 | -3.694140421 | 0.000220632 | 0.00363667 | | Hnrnpl | 5843.999863 | -0.109009175197364 | 0.0556735975082965 | -1.95800487261704 | 0.0502294409400353 | 0.402001575258927 | NM_177301 |
| Utrn | 3640.423923 | -0.299095429 | 0.054907726 | -5.44723756 | 5.12E-08 | 4.24E-06 | | Ccdc130 | 492.7954894 | -0.129591051944963 | 0.0662051987689449 | -1.95741504224213 | 0.0502986913832398 | 0.402334136031356 | NM_026350 |
| Acvr1c | 1099.384348 | -0.299316444 | 0.095362582 | -3.138720022 | 0.001696875 | 0.016940322 | | Iffo1 | 2039.342718 | -0.139383014519703 | 0.071230809721851 | -1.95677986904795 | 0.0503733548702751 | 0.402601281721812 | NM_001039669,NM_178787 |
| Uqcr10 | 2188.451836 | -0.299482841 | 0.066999488 | -4.469927305 | 7.82E-06 | 0.000259603 | | Ramp2 | 574.3895684 | 0.150863205934168 | 0.0771024341220435 | 1.95665944469887 | 0.0503875210085539 | 0.402601281721812 | NM_019444 |
| Prex2 | 2795.441201 | -0.300043865 | 0.063052733 | -4.758617939 | 1.95E-06 | 8.68E-05 | | Aifm3 | 4608.448095 | -0.160236135429282 | 0.081950265795582 | -1.9552851216978 | 0.0505494263106599 | 0.403264376061763 | NM_175178 |
| Syne2 | 1000.398004 | -0.300064439 | 0.087426316 | -3.432198137 | 0.00059871 | 0.007801861 | | C1qa | 2069.417702 | -0.184425404264298 | 0.0943240133809877 | -1.95523279442508 | 0.0505555994502315 | 0.403264376061763 | NM_007572 |
| Adra1a | 424.2902146 | -0.300085241 | 0.109953862 | -2.729192358 | 0.006348966 | 0.042474471 | | Sec24a | 766.4769622 | -0.126922300156681 | 0.0649152881889427 | -1.95519890148621 | 0.050559598196349 | 0.403264376061763 | NM_175255 |
| Ddn | 44170.29246 | -0.300313971 | 0.084932897 | -3.535896948 | 0.000406393 | 0.005884725 | | Trh | 400.3082846 | 0.099328613728465 | 0.0508071415652684 | 1.95502193228045 | 0.0505815567385745 | 0.403264376061763 | NM_009426 |
| Idh1 | 1481.005645 | -0.301045146 | 0.076280775 | -3.946540226 | 7.93E-05 | 0.001635816 | | Acr | 41.00105014 | -0.17857058240015 | 0.0914342608656851 | -1.95299421365112 | 0.0508202808409687 | 0.40326624518384 | NM_013455,NM_001205049 |
| Eif2c2 | 2985.480442 | -0.301175767 | 0.101056707 | -2.980265008 | 0.002879991 | 0.024639747 | | Alg2 | 6092.892262 | 0.110989965691425 | 0.056917102010887 | 1.95002840570125 | 0.0511727334782108 | 0.40326624518384 | NM_019998 |
| Setbp1 | 1681.934272 | -0.301815316 | 0.075413627 | -4.002132352 | 6.28E-05 | 0.001369891 | | Cadm4 | 9944.230764 | 0.0986332573117213 | 0.0505450505487855 | 1.95139298983432 | 0.0510103147490217 | 0.40326624518384 | NM_153112 |
| Ppapdc2 | 2006.220703 | -0.303022204 | 0.111847721 | -2.709238952 | 0.006743775 | 0.044184405 | | Catsperd | 50.7968235 | -0.180130735234449 | 0.0921579206336484 | -1.95458766860111 | 0.0506317579782905 | 0.40326624518384 | NM_175350 |
| Cacng4 | 490.6135266 | -0.303105591 | 0.090510684 | -3.348837687 | 0.000811513 | 0.009692629 | | Chac2 | 211.0654914 | 0.169956475752551 | 0.0871812347152806 | 1.94946167380631 | 0.0512403155115214 | 0.40326624518384 | NM_026527 |
| Churc1 | 671.2397872 | -0.303132204 | 0.072003783 | -4.209948281 | 2.55E-05 | 0.000678356 | | Cln3 | 620.3859892 | -0.147367553780382 | 0.0755453847428627 | -1.95071551070901 | 0.0510908972162933 | 0.40326624518384 | NM_009907,NM_001146311 |
| Pgm2l1 | 13559.50154 | -0.303375422 | 0.097205401 | -3.120972896 | 0.001802546 | 0.017677556 | | Cpne9 | 2618.68375 | -0.125028017555012 | 0.0641269362801789 | -1.94969578787841 | 0.0512123886634079 | 0.40326624518384 | NM_170673 |
| Tyro3 | 10902.4078 | -0.30356858 | 0.085064771 | -3.568675673 | 0.00035879 | 0.005365923 | | Dnajc12 | 407.9830664 | 0.146279773336942 | 0.074983571357025 | 1.95082430310566 | 0.051077949771154 | 0.40326624518384 | NM_001253685,NM_013888 |
| Spon1 | 3364.181049 | -0.303576995 | 0.078045351 | -3.88975115 | 0.000100347 | 0.001968073 | | Eif2c4 | 704.8700998 | -0.143939125022702 | 0.0736686158328342 | -1.95387307601005 | 0.0507162293863465 | 0.40326624518384 | NM_153177 |
| Tiam2 | 2926.008248 | -0.303600583 | 0.087409564 | -3.473310814 | 0.00051408 | 0.006962544 | | Gng4 | 2375.976669 | 0.143337385350529 | 0.0735286207188909 | 1.94940941294854 | 0.0512465513113551 | 0.40326624518384 | NM_010317 |
| Anpep | 202.6453715 | -0.303631148 | 0.102274403 | -2.968789265 | 0.002989756 | 0.025311139 | | Gsr | 1006.611931 | 0.107749123101535 | 0.055270212179296 | 1.94949718578965 | 0.0512360785608786 | 0.40326624518384 | NM_010344 |
| Prrx1 | 259.6911978 | -0.303673875 | 0.100739229 | -3.014455029 | 0.002574414 | 0.02287154 | | Hmga2-ps1 | 34.24473153 | -0.167823475010228 | 0.0860053256731395 | -1.95131491796259 | 0.0510195955453674 | 0.40326624518384 | NR_037996 |
| Gm15800 | 13692.64898 | -0.303677341 | 0.077360694 | -3.925473312 | 8.66E-05 | 0.00176004 | | Kctd8 | 275.729378 | 0.183070882307586 | 0.093700367655493 | 1.95379043741515 | 0.0507260056385804 | 0.40326624518384 | NM_175519 |
| Rgs9 | 5036.503318 | -0.303951116 | 0.099728576 | -3.047783588 | 0.002305359 | 0.021185783 | | Lin28b | 225.3611139 | -0.177334495280095 | 0.0909727872371953 | -1.94931364274601 | 0.0512579803236833 | 0.40326624518384 | NM_001031772 |
| Mcm5 | 254.5995703 | -0.30434921 | 0.113625859 | -2.678520663 | 0.007394817 | 0.047350844 | | Mrpl46 | 757.9914207 | 0.123221788124101 | 0.0631045256498383 | 1.9526616649945 | 0.0508596989907713 | 0.40326624518384 | NM_023331 |
| Chd3 | 25957.77394 | -0.304549194 | 0.061972413 | -4.914270364 | 8.91E-07 | 4.41E-05 | | Nedd8 | 2864.381167 | 0.0991613904182996 | 0.0508261580585952 | 1.95116401562442 | 0.0510375380737488 | 0.40326624518384 | NM_008683 |
| Prkcdbp | 472.2286655 | -0.304789298 | 0.101586772 | -3.000285288 | 0.002697268 | 0.023561109 | | Nudt13 | 438.5675273 | -0.152633050594484 | 0.078111809579123 | -1.95403296142915 | 0.0506973192162297 | 0.40326624518384 | NM_026341 |
| Ehd2 | 341.660789 | -0.304991899 | 0.08166583 | -3.734632941 | 0.000187989 | 0.003250786 | | Nxph3 | 1016.169389 | 0.160095177533699 | 0.0820483123067737 | 1.95123059856628 | 0.0510296206046319 | 0.40326624518384 | NM_130858 |
| Unc80 | 13103.75364 | -0.305116879 | 0.067731317 | -4.504812452 | 6.64E-06 | 0.000228224 | | Pigk | 3383.311846 | 0.105405528217959 | 0.0540044713778647 | 1.95179261140148 | 0.0509628318581715 | 0.40326624518384 | NM_178016,NM_025662 |
| Wfdc1 | 211.4390817 | -0.305144084 | 0.09943757 | -3.068700118 | 0.002149923 | 0.02020067 | | Scp2 | 4260.485143 | 0.0823932381546238 | 0.0422166947949484 | 1.95167429745075 | 0.0509768860204731 | 0.40326624518384 | NM_011327 |
| Slc16a2 | 1908.756794 | -0.305210956 | 0.064583512 | -4.725833999 | 2.29E-06 | 0.000100023 | | Slc11a2 | 2260.657203 | -0.100330979346203 | 0.0514738499828526 | -1.94916407806345 | 0.0512758333233204 | 0.40326624518384 | NM_001146161,NM_008732 |
| Wdfy3 | 9320.76383 | -0.306152062 | 0.095253058 | -3.214091698 | 0.001308579 | 0.013906117 | | Slc25a13 | 159.9185041 | 0.184661799629249 | 0.0946529769726449 | 1.95093493660128 | 0.0510647860335219 | 0.40326624518384 | NM_001177572,NM_015829 |
| Zswim6 | 1342.007494 | -0.306214515 | 0.076478672 | -4.003920394 | 6.23E-05 | 0.001364151 | | Spock1 | 13826.20038 | 0.0781462387596139 | 0.040015106671727 | 1.95291841655464 | 0.0508292630823722 | 0.40326624518384 | NM_001166463,NM_009262,NM_001166466,NM_001166464,NM_001166465 |
| Ptprr | 773.5433013 | -0.306288759 | 0.088890716 | -3.44567771 | 0.000569629 | 0.007498586 | | Ube2a | 1198.009111 | 0.153313315275544 | 0.0786370060213695 | 1.94963316932337 | 0.0512198570164525 | 0.40326624518384 | NM_019668 |
| Fam124a | 1172.750571 | -0.307017094 | 0.09285312 | -3.306481179 | 0.000944757 | 0.01086716 | | Zc3h15 | 4085.223746 | 0.109858879936254 | 0.0563035243777853 | 1.95119011021625 | 0.0510344350074061 | 0.40326624518384 | NM_026934 |
| Birc6 | 5240.795754 | -0.307880226 | 0.068356539 | -4.504034756 | 6.67E-06 | 0.000228302 | | Anln | 2165.98366 | 0.171323571956838 | 0.0879099733142224 | 1.94885250783167 | 0.0513130410621122 | 0.403340494842122 | NM_028390 |
| Acyp2 | 354.3342021 | -0.308101024 | 0.085913691 | -3.58616911 | 0.000335571 | 0.005117464 | | Tmem11 | 786.1756962 | 0.115017818660965 | 0.0590281733783998 | 1.94852410430598 | 0.05135228349827 | 0.403430649051309 | NM_001168507,NM_173453 |
| Ptn | 7885.585572 | -0.308298955 | 0.095306106 | -3.234828994 | 0.001217156 | 0.01324341 | | Txn2 | 4295.450081 | 0.100007885345049 | 0.051357513636632 | 1.94728829850743 | 0.0515001807880974 | 0.404373851961029 | NM_019913 |
| Ncrna00086 | 942.476719 | -0.308533352 | 0.063013507 | -4.896305015 | 9.77E-07 | 4.79E-05 | | Timm44 | 1702.970899 | -0.106364793644413 | 0.05463563906099 | -1.94680240722869 | 0.0515584282642897 | 0.404394022120449 | NM_011592 |
| Uqcrh | 4551.083169 | -0.308537056 | 0.071623659 | -4.307753362 | 1.65E-05 | 0.000478982 | | Xylt2 | 1102.917814 | 0.113659079073043 | 0.05838037619397 | 1.94687130304553 | 0.0515501658466237 | 0.404394022120449 | NM_145828 |
| Zfp462 | 2122.452167 | -0.3085816 | 0.091969098 | -3.355274843 | 0.000792862 | 0.00950085 | | Ppa1 | 2733.588404 | 0.109937422235474 | 0.0565003178135586 | 1.94578413874147 | 0.051680674693353 | 0.404936564445008 | NM_026438 |
| Hspa5 | 11307.71383 | -0.308810107 | 0.090935049 | -3.395941502 | 0.00068393 | 0.00855271 | | Rbm24 | 731.0601767 | 0.146412590054737 | 0.0752469219765964 | 1.94576184923916 | 0.0516833533306516 | 0.404936564445008 | NM_001081425 |
| Ehhadh | 166.7226753 | -0.309804734 | 0.102814102 | -3.013251376 | 0.002584647 | 0.022923025 | | Dera | 63.95677675 | -0.182541808799354 | 0.0938497143150311 | -1.94504384090723 | 0.051769702032053 | 0.405394442974449 | NM_172733 |
| Nanos1 | 1163.764078 | -0.310231298 | 0.077104969 | -4.023492907 | 5.73E-05 | 0.001283882 | | Cdon | 520.0464829 | -0.137535376721165 | 0.0707418574366153 | -1.94418667681149 | 0.0518729438369494 | 0.405765418511323 | NM_021339 |
| Mageh1 | 887.2726559 | -0.31049193 | 0.061589005 | -5.041353265 | 4.62E-07 | 2.62E-05 | | Rgs22 | 51.25571681 | -0.18011032428151 | 0.0926402353934048 | -1.9441911337623 | 0.0518724065708327 | 0.405765418511323 | NM_001195748 |
| Saysd1 | 197.8495034 | -0.311639463 | 0.096236384 | -3.238270692 | 0.001202566 | 0.013112291 | | Bdnf | 930.5590876 | 0.181393303988336 | 0.0933807601200214 | 1.94251260918408 | 0.0520750749786385 | 0.406272097828096 | NM_001048141,NM_001048142,NM_001048139,NM_007540 |
| Mrpl54 | 467.531179 | -0.311910518 | 0.095653665 | -3.260831851 | 0.001110859 | 0.012373579 | | Emb | 1139.043993 | 0.151047434527481 | 0.0777688713167416 | 1.94226085540429 | 0.0521055292753506 | 0.406272097828096 | NM_010330 |
| Dclk3 | 2929.17992 | -0.313456316 | 0.095372552 | -3.286651232 | 0.001013863 | 0.011471264 | | Mtmr2 | 2525.118719 | 0.133154192973489 | 0.0685514566078556 | 1.94239771935394 | 0.0520889711906621 | 0.406272097828096 | NM_023858,NR_024025 |
| Gfra1 | 1802.792151 | -0.313592962 | 0.09727164 | -3.223888925 | 0.001264624 | 0.013635415 | | Reep4 | 113.2158005 | -0.184763728860115 | 0.0951000044106283 | -1.94283617550985 | 0.0520359554858096 | 0.406272097828096 | NM_180588 |
| Bmpr1b | 995.1696079 | -0.314259492 | 0.067679555 | -4.643344534 | 3.43E-06 | 0.000136544 | | Srrt | 4831.873867 | -0.121879322941211 | 0.0627496986931549 | -1.94230929358242 | 0.0520996686207768 | 0.406272097828096 | NM_001109909,NM_001109910,NM_031405 |
| Hspa12b | 317.0013868 | -0.314429983 | 0.117948008 | -2.665835459 | 0.007679727 | 0.048706382 | | Ttf1 | 634.5481669 | -0.142370838983211 | 0.0732968917648695 | -1.94238576227659 | 0.052090417607222 | 0.406272097828096 | NM_009442 |
| D3Bwg0562e | 9157.264238 | -0.314432927 | 0.072080833 | -4.362226601 | 1.29E-05 | 0.000393448 | | 2810008D09Rik | 347.0601091 | -0.160615413312763 | 0.082746343786002 | -1.94105752549195 | 0.0522513002215754 | 0.406355486979988 | NR_027059 |
| Gap43 | 8199.855136 | -0.314654271 | 0.088533242 | -3.554080516 | 0.000379303 | 0.005618472 | | Atp10d | 202.8343792 | -0.163884840127359 | 0.0844095582474791 | -1.9415436300101 | 0.0521923726925835 | 0.406355486979988 | NR_003966,NM_153389 |
| Rgs20 | 1447.0646 | -0.314922014 | 0.098659601 | -3.192005745 | 0.001412885 | 0.014761459 | | Chrac1 | 399.29143 | 0.135790176891136 | 0.0700080719376547 | 1.93963600386057 | 0.0524239420763113 | 0.406355486979988 | NM_053068 |
| Epha7 | 2519.553871 | -0.315456576 | 0.094767146 | -3.328754631 | 0.000872352 | 0.010270688 | | Fas | 72.79650208 | 0.184121556868778 | 0.0949197401215269 | 1.93976044006278 | 0.05240881044706 | 0.406355486979988 | NM_001146708,NM_007987 |
| Vps13d | 5553.789079 | -0.315506096 | 0.070727663 | -4.460858505 | 8.16E-06 | 0.000266837 | | Hmg20b | 469.5491222 | -0.156566203876824 | 0.0807053599108322 | -1.9399728103537 | 0.0523829943349585 | 0.406355486979988 | NM_010440,NM_001163165,NM_001163166 |
| Gm561 | 324.0118655 | -0.316575242 | 0.087693079 | -3.610036774 | 0.000306154 | 0.004743544 | | Mrps25 | 1349.032936 | 0.111878189616145 | 0.0576722375429395 | 1.93989680967115 | 0.0523922318917634 | 0.406355486979988 | NM_025578 |
| Sh3rf1 | 1589.979885 | -0.316720585 | 0.068587584 | -4.617753944 | 3.88E-06 | 0.000149512 | | Nmrk1 | 194.6561968 | -0.170985060700074 | 0.0880887720176909 | -1.94105397071189 | 0.0522517313510326 | 0.406355486979988 | NM_145497 |
| 1810037I17Rik | 1330.502723 | -0.316759182 | 0.062926593 | -5.033788823 | 4.81E-07 | 2.69E-05 | | Plag1 | 148.1946885 | -0.178708727701475 | 0.0920868197541316 | -1.94065478836842 | 0.0523001637562088 | 0.406355486979988 | NM_019969 |
| Snx7 | 565.7914671 | -0.316998941 | 0.079621006 | -3.981348086 | 6.85E-05 | 0.001468323 | | Sde2 | 982.9217198 | 0.139593750596055 | 0.0719664543214011 | 1.93970582422515 | 0.0524154513652986 | 0.406355486979988 | NM_145943 |
| Sqle | 2287.280488 | -0.317297592 | 0.111352659 | -2.849483754 | 0.004379024 | 0.03306462 | | Tmub1 | 699.3632772 | 0.126372803682204 | 0.0651384611555013 | 1.94006430978653 | 0.052371874778343 | 0.406355486979988 | NM_022418 |
| Cetn2 | 1116.672825 | -0.317396487 | 0.094241256 | -3.367914437 | 0.000757391 | 0.009189519 | | Trim12c | 149.7194165 | -0.177867633448124 | 0.0916312080547758 | -1.94112505143222 | 0.0522431111316028 | 0.406355486979988 | NM_175677,NM_001146007 |
| Abcb10 | 1224.746943 | -0.317413339 | 0.07792678 | -4.073225365 | 4.64E-05 | 0.001077066 | | 0610009D07Rik | 1146.335862 | 0.108551464002906 | 0.056022341214634 | 1.93764597568355 | 0.0526664292303211 | 0.40686417718513 | NM_025323 |
| Tmem121 | 383.1719074 | -0.317529577 | 0.101484475 | -3.128848775 | 0.001754926 | 0.01736241 | | Asah2 | 796.25584 | 0.139667761902913 | 0.0721197942122862 | 1.93660827894448 | 0.0527932451874948 | 0.40686417718513 | NM_018830 |
| Glra2 | 556.5823404 | -0.317769293 | 0.104558944 | -3.039140224 | 0.002372544 | 0.021674644 | | Bhlhe40 | 6666.085036 | 0.184054229575673 | 0.0949424806227909 | 1.93858669342047 | 0.0525516851316764 | 0.40686417718513 | NM_011498 |

| | | | | | | | | | | | | | | | |
|---|---|---|---|---|---|---|---|---|---|---|---|---|---|---|---|
| Clmp | 1014.097593 | -0.318827512 | 0.072016174 | -4.427165394 | 9.55E-06 | 0.000301295 | | Cd164 | 2461.93788 | 0.134158743271121 | 0.0693096620985104 | 1.93564272583583 | 0.0529114736505713 | 0.40686417718513 | NM_016898 |
| Ppp1r3c | 1711.517539 | -0.319668846 | 0.070525229 | -4.532687786 | 5.82E-06 | 0.000206485 | | Chfr | 1441.505566 | -0.139191250875674 | 0.0718578294177934 | -1.93703667371293 | 0.0527408605619494 | 0.40686417718513 | NM_172717 |
| Tgfa | 2331.659008 | -0.319697036 | 0.110520295 | -2.892654573 | 0.003820012 | 0.030182183 | | Chst14 | 75.04760106 | -0.183936518813907 | 0.0949731052004179 | -1.93672217440667 | 0.0527793136719637 | 0.40686417718513 | NM_028117 |
| Crip2 | 3895.34984 | -0.3197841 | 0.096295179 | -3.320873407 | 0.000897362 | 0.010498156 | | Emc10 | 7147.984889 | 0.10972018026096 | 0.0566970681693543 | 1.93520024586149 | 0.0529657275958337 | 0.40686417718513 | NM_197991 |
| Cd59a | 534.7187711 | -0.320208716 | 0.109050668 | -2.936329706 | 0.003321211 | 0.027270189 | | Havcr2 | 222.6576837 | -0.173431045454205 | 0.0894553545679308 | -1.93874415111176 | 0.0525324996537291 | 0.40686417718513 | NM_134250 |
| Ptprt | 4164.382298 | -0.320666898 | 0.070842605 | -4.526469608 | 6.00E-06 | 0.000211202 | | Lmo2 | 689.6803646 | -0.137898338481642 | 0.0711951997039644 | -1.93690500279562 | 0.0527569568130557 | 0.40686417718513 | NM_008505,NM_001142335,NM_001142336,NM_001142337 |
| Kcns2 | 1728.273532 | -0.32077895 | 0.096319519 | -3.330362866 | 0.000867329 | 0.010225679 | | Mdh2 | 10944.66689 | 0.0873769020143274 | 0.0451451972025868 | 1.93546395693495 | 0.0529333875080322 | 0.40686417718513 | NM_008617 |
| Mnf1 | 1654.800376 | -0.321255486 | 0.06902765 | -4.654011609 | 3.26E-06 | 0.00013101 | | Morn1 | 172.5101294 | -0.170870786033185 | 0.0882948748980513 | -1.93522881402209 | 0.0529622233552129 | 0.40686417718513 | NM_001081100 |
| Sec61g | 782.4474239 | -0.321353053 | 0.080999317 | -3.967355084 | 7.27E-05 | 0.001534089 | | Rasl11a | 122.3092335 | 0.184195156882329 | 0.0950356605651127 | 1.93816884932503 | 0.0526026258647653 | 0.40686417718513 | NM_026864 |
| Diras2 | 15122.23639 | -0.321489069 | 0.096877479 | -3.318511924 | 0.000904985 | 0.010519003 | | Sf3a3 | 1242.746955 | 0.10999210020563 | 0.0567666859597724 | 1.93761707850109 | 0.0526699572784413 | 0.40686417718513 | NM_029157 |
| Bmpr2 | 2439.555218 | -0.322533917 | 0.089753414 | -3.593555976 | 0.000326196 | 0.004989214 | | Shpk | 190.9802717 | -0.178206893634944 | 0.0919974786997587 | -1.93708453920282 | 0.0527350102116819 | 0.40686417718513 | NM_029031 |
| Capn11 | 48.3427266 | -0.323204291 | 0.11165458 | -2.894680108 | 0.003795451 | 0.030064784 | | Slc8a1 | 8098.614869 | 0.139943512486599 | 0.0722659725384635 | 1.93650631923778 | 0.0528057193470007 | 0.40686417718513 | NM_011406,NM_001112798 |
| Gas1 | 235.7426758 | -0.323258158 | 0.109574237 | -2.950129214 | 0.003176411 | 0.026388904 | | Trim62 | 1125.869008 | 0.121521421246146 | 0.0627712787274248 | 1.93593987106484 | 0.0528750657540778 | 0.40686417718513 | NM_178110 |
| Elovl6 | 2488.470841 | -0.323325155 | 0.094816497 | -3.410009495 | 0.000649606 | 0.008229761 | | Zbtb8b | 266.9117646 | 0.160894084432061 | 0.0831103317072228 | 1.93590954490292 | 0.0528787805256179 | 0.40686417718513 | NM_153541 |
| Pts | 678.6571504 | -0.323508391 | 0.095355466 | -3.392657018 | 0.000692182 | 0.008641962 | | Pard3 | 993.9469024 | 0.119793072125025 | 0.0619243584348616 | 1.93450647132721 | 0.0530508871812664 | 0.40708779038303 | NM_033620,NM_001013580,NM_001122850,NM_001013581 |
| Sema5a | 1903.395182 | -0.323749049 | 0.093814785 | -3.450938464 | 0.000558641 | 0.007404262 | | Sfr1 | 3131.023416 | 0.105782515295952 | 0.0546786301808775 | 1.93462262946278 | 0.0530366210091366 | 0.40708779038303 | NM_026377 |
| Epha6 | 822.1901054 | -0.324265701 | 0.118628544 | -2.733454283 | 0.006267382 | 0.042079243 | | 2700081O15Rik | 1567.491693 | -0.120638830073845 | 0.0624056408027332 | -1.93313983354788 | 0.053218974179976 | 0.407317205569974 | NM_175381 |
| Shisa3 | 99.94989799 | -0.324471273 | 0.121878028 | -2.662262256 | 0.007761738 | 0.049146043 | | A730036I17Rik | 30.53031476 | -0.168104869391955 | 0.08696309742276 | -1.93305981932468 | 0.0532288291469307 | 0.407317205569974 | NR_045838 |
| Otud1 | 634.8731627 | -0.324611732 | 0.088772631 | -3.65666455 | 0.000255518 | 0.00410251 | | Gm14325 | 864.9481125 | -0.127973519765361 | 0.0662081812821785 | -1.93289586403135 | 0.0532490274939681 | 0.407317205569974 | NM_001024849 |
| Mettl23 | 436.3612169 | -0.324932491 | 0.103862482 | -3.128487649 | 0.001757084 | 0.01736241 | | Sox4 | 334.798716 | -0.172263152268486 | 0.0890857694930069 | -1.93367754747866 | 0.0531527860732813 | 0.407317205569974 | NM_009238 |
| Adamts19 | 27.20734008 | -0.325162619 | 0.11460095 | -2.837346626 | 0.00454902 | 0.033853271 | | Tm2d2 | 1934.821684 | 0.0916404037102986 | 0.0474079726977514 | 1.93301671629264 | 0.0532341385718015 | 0.407317205569974 | NM_027194 |
| Aldh1b1 | 370.2292102 | -0.3254819 | 0.119094786 | -2.732965154 | 0.006276697 | 0.042100064 | | Tor4a | 378.2787119 | -0.178085317478235 | 0.0920821543599067 | -1.93398296028329 | 0.0531152229332251 | 0.407317205569974 | NM_146115 |
| 2010107E04Rik | 2034.57687 | -0.325830826 | 0.079358878 | -4.105789238 | 4.03E-05 | 0.000969426 | | Pycrl | 609.7080917 | 0.133023466103941 | 0.0688443686583578 | 1.93223452689462 | 0.0533305654255457 | 0.407726207037619 | NM_025412 |
| Atp5j2 | 3066.282303 | -0.326085467 | 0.069561038 | -4.687760236 | 2.76E-06 | 0.000115354 | | 6330403K07Rik | 10108.15504 | 0.139517027212177 | 0.072224615484152 | 1.93171021094312 | 0.0533952837591736 | 0.407791741264856 | NM_134022 |
| Slc17a8 | 256.0028571 | -0.326233967 | 0.104302897 | -3.12775556 | 0.001761466 | 0.017372473 | | Nav1 | 4801.826613 | -0.131560627603027 | 0.0681017875522099 | -1.93182341215591 | 0.0533813053488289 | 0.407791741264856 | NM_173437 |
| Tma7 | 2265.681651 | -0.326271379 | 0.079397247 | -4.109353819 | 3.97E-05 | 0.000956068 | | Gpr161 | 137.9846312 | 0.174777616800529 | 0.0905051765316621 | 1.93113392513394 | 0.053466492588407 | 0.407906655115126 | NM_001081126 |
| Heatr3 | 1037.872929 | -0.327157151 | 0.064976203 | -5.035030323 | 4.78E-07 | 2.69E-05 | | Rasa4 | 300.9072449 | -0.183020959892575 | 0.0947693638977529 | -1.93122494828642 | 0.0534552400297729 | 0.407906655115126 | NM_133914,NM_001039103 |
| Kcnj4 | 5238.358052 | -0.32720964 | 0.073359252 | -4.460373212 | 8.18E-06 | 0.000266879 | | 4930402H24Rik | 4271.764783 | -0.112968074332145 | 0.0585089867472037 | -1.93078158779676 | 0.0535100682508558 | 0.407961168367773 | NM_029432 |
| Adcy2 | 6997.718518 | -0.327563177 | 0.072208261 | -4.536367064 | 5.72E-06 | 0.000204791 | | Mif4gd | 387.4049863 | -0.153741180286781 | 0.0796423356538087 | -1.93039517267634 | 0.0535578926116854 | 0.407961168367773 | NM_027162,NM_001243584,NM_001243587,NM_001243586 |
| Bcl2 | 1069.470493 | -0.327564366 | 0.082453532 | -3.972714782 | 7.11E-05 | 0.001510253 | | Wdhd1 | 289.806395 | -0.162522036071256 | 0.0841825582510646 | -1.93059036750288 | 0.0535337300196327 | 0.407961168367773 | NM_172598 |
| Fam101b | 582.9213059 | -0.328662294 | 0.082556683 | -3.981050146 | 6.86E-05 | 0.001468323 | | Cep250 | 1718.800348 | -0.137062594118594 | 0.0710403307278939 | -1.92936311971274 | 0.0536857988234614 | 0.408507026563437 | NM_177217,NM_001130000,NM_001129999,NM_008383 |
| Rem2 | 459.8983737 | -0.328914623 | 0.101348219 | -3.245391257 | 0.001172894 | 0.012845805 | | Crlf2 | 411.0540021 | -0.165344720179361 | 0.0856930658063225 | -1.92949941309209 | 0.0536688928687944 | 0.408507026563437 | NM_016715,NM_001164735 |
| Slc9a7 | 796.9766947 | -0.328946606 | 0.080878241 | -4.067182966 | 4.76E-05 | 0.001102063 | | Dcaf17 | 621.4321147 | -0.127753210657925 | 0.0662292840425939 | -1.92895352116088 | 0.0537366325626789 | 0.408679751102342 | NM_198005,NM_001165980,NM_001165981,NM_001165982 |
| Sik2 | 1392.284125 | -0.328960483 | 0.080615205 | -4.080625771 | 4.49E-05 | 0.001048053 | | Col2a1 | 86.5619747 | -0.182737948283315 | 0.0947554916496549 | -1.92852092371557 | 0.0537903642356037 | 0.408874322808152 | NM_031163,NM_001113515 |
| Zfp521 | 527.1197681 | -0.329066726 | 0.102453683 | -3.211858448 | 0.001318793 | 0.013995469 | | Isoc1 | 764.4282219 | 0.14184314230391 | 0.0735601949596912 | 1.928259466708 | 0.0538228607926146 | 0.408907361858535 | NM_025478 |
| 6330403M23Rik | 1107.133131 | -0.329682636 | 0.088084085 | -3.742817305 | 0.000181969 | 0.003175023 | | Arhgap24 | 286.4299536 | 0.181249385574188 | 0.0940714479445059 | 1.92672048251144 | 0.0540144738436641 | 0.409506391990117 | NM_146161,NM_029270 |
| Snca | 8355.118203 | -0.329724937 | 0.086191442 | -3.825495059 | 0.00013051 | 0.002399355 | | Csrnp3 | 4322.544056 | 0.12247604466453 | 0.0635633993449922 | 1.92683283031777 | 0.0540004666086894 | 0.409506391990117 | NM_178634,NM_153409 |
| Cntnap3 | 499.1323413 | -0.329807183 | 0.113604303 | -2.903122289 | 0.003694623 | 0.029386291 | | Ifnar1 | 2193.148729 | -0.104391443622634 | 0.0541646007139411 | -1.9273001600059 | 0.053942233678092 | 0.409506391990117 | NM_010508 |
| Slco1c1 | 2632.233386 | -0.329877494 | 0.061858678 | -5.332760131 | 9.67E-08 | 7.24E-06 | | Srprb | 1711.861855 | 0.103708586753203 | 0.0538139427472626 | 1.92716945569795 | 0.0539585151737563 | 0.409506391990117 | NM_009275 |
| 4833424O15Rik | 1496.574996 | -0.329946024 | 0.068975623 | -4.783516391 | 1.72E-06 | 7.87E-05 | | Pcdhb7 | 294.6513207 | 0.160652586870839 | 0.0834382626337597 | 1.92540666355914 | 0.0541785026079556 | 0.410426203096955 | NM_053132 |
| Penk | 13357.46749 | -0.33015191 | 0.109026064 | -3.028192515 | 0.002460213 | 0.022123355 | | Traf3ip3 | 30.89429908 | -0.16775612359031 | 0.0871326291791024 | -1.92529624287458 | 0.0541923074170426 | 0.410426203096955 | NM_153137 |
| Rn45s | 16977.48593 | -0.330302067 | 0.09646701 | -3.423989897 | 0.000617089 | 0.007974295 | | A330023F24Rik | 2444.630706 | -0.161319236112877 | 0.0838589924135615 | -1.92369633202019 | 0.0543926580800228 | 0.410949742048056 | NR_015566 |
| Cxxc4 | 871.6700059 | -0.331081165 | 0.096470744 | -3.431933353 | 0.000599295 | 0.007802919 | | Lhfpl3 | 1161.671131 | 0.135390091690672 | 0.0703675785810216 | 1.92404079294534 | 0.0543494704463344 | 0.410949742048056 | NM_029990,NM_001081231 |
| Epha5 | 3203.284868 | -0.331149666 | 0.093232823 | -3.551857121 | 0.000382523 | 0.005649956 | | Ppp2cb | 4528.981371 | 0.102251010939183 | 0.0531451407389117 | 1.92399548702892 | 0.0543551491522513 | 0.410949742048056 | NM_017374 |
| Gucy1a2 | 2335.100972 | -0.33122769 | 0.113187003 | -2.926375646 | 0.003429366 | 0.027877541 | | Tagln3 | 9297.751364 | 0.100819553869994 | 0.0524115095161132 | 1.923614770893 | 0.0544028881976051 | 0.410949742048056 | NM_019754 |
| Hist3h2ba | 274.7343234 | -0.331726121 | 0.094726918 | -3.501920322 | 0.000461918 | 0.006438851 | | Tm2d1 | 692.4928063 | 0.140969576562576 | 0.0732596162886418 | 1.92424672287605 | 0.0543236651431562 | 0.410949742048056 | NM_053157 |
| Gdpd4 | 10.93000515 | -0.331879283 | 0.121410321 | -2.733534355 | 0.006265859 | 0.042079243 | | Slit3 | 1343.43177 | 0.134678280416883 | 0.0700405385049007 | 1.92286168388273 | 0.0544974002746927 | 0.411449707063506 | NM_011412 |
| Tnrc6b | 3887.764246 | -0.332568813 | 0.08534825 | -3.896609611 | 9.75E-05 | 0.001922925 | | Msantd2 | 400.5723454 | -0.16021826861217 | 0.0833562975336715 | -1.92208955235146 | 0.0545944904604613 | 0.411968606975928 | NM_146222 |
| Egfr | 548.6533899 | -0.332614649 | 0.076792366 | -4.331350421 | 1.48E-05 | 0.000439549 | | Gpr135 | 206.8982888 | 0.158828945731546 | 0.0826462141106562 | 1.92179336271699 | 0.0546317638285842 | 0.412035826258574 | NM_181752 |
| Hopx | 775.0994898 | -0.332639432 | 0.107364508 | -3.098225278 | 0.001946834 | 0.018724864 | | Vcam1 | 1265.873557 | -0.122413476639905 | 0.063705681030963 | -1.92154725699279 | 0.0546627506339654 | 0.412055586771656 | NM_011693 |
| Tet1 | 504.1623979 | -0.332654926 | 0.113721779 | -2.925164645 | 0.00344274 | 0.027955792 | | 4933408B17Rik | 30.30399302 | -0.163723280828935 | 0.0852422555671836 | -1.92068217504987 | 0.0547717881589995 | 0.412110851836089 | NM_177773 |
| Id4 | 1572.418092 | -0.332726286 | 0.083622698 | -3.978899175 | 6.92E-05 | 0.001479626 | | Cobll1 | 563.4614645 | -0.171663503949989 | 0.0893773252753542 | -1.9206605637518 | 0.0547745144324211 | 0.412110851836089 | NM_027225,NM_177025 |
| Nrarp | 600.8433816 | -0.332884072 | 0.097955924 | -3.398304457 | 0.000678049 | 0.008486015 | | Eefsec | 692.6424699 | 0.131222664674452 | 0.068361593614282 | 1.91953782433528 | 0.0549163041668154 | 0.412110851836089 | NM_023060 |
| Deptor | 457.7023164 | -0.333184604 | 0.096653321 | -3.447213208 | 0.000566401 | 0.00748152 | | Nipal3 | 6381.032078 | 0.0880343987921669 | 0.0458639423226469 | 1.91946863557555 | 0.0549250519536989 | 0.412110851836089 | NM_028995 |
| Hras1 | 3978.317564 | -0.333284423 | 0.075184265 | -4.432901234 | 9.30E-06 | 0.000296317 | | Qpct | 417.6700302 | 0.156077728818668 | 0.0812735066765332 | 1.92040106550163 | 0.0548072590732351 | 0.412110851836089 | NM_027455 |
| 1810041L15Rik | 2878.681301 | -0.334149583 | 0.086290148 | -3.872395544 | 0.000107771 | 0.002067036 | | Rec8 | 225.4806122 | 0.182645382185329 | 0.0950730748121092 | 1.92110523979882 | 0.0547184416673101 | 0.412110851836089 | NM_020002 |
| Syt17 | 1341.397189 | -0.334364332 | 0.117097199 | -2.855442618 | 0.004297689 | 0.032721568 | | Shq1 | 154.9286031 | 0.170401204601693 | 0.0887647576299068 | 1.91969436014413 | 0.0548965170616035 | 0.412110851836089 | NM_181590 |
| 6430573F11Rik | 390.8860614 | -0.334494284 | 0.120093666 | -2.785278309 | 0.00534818 | 0.037563326 | | Tha1 | 154.1998129 | -0.175446616849817 | 0.0914028614865224 | -1.91946574753228 | 0.0549254171247879 | 0.412110851836089 | NM_027919 |
| Fgf10 | 207.8299588 | -0.33477406 | 0.105604932 | -3.17006084 | 0.00152407 | 0.015659114 | | Tsg101 | 2520.577723 | 0.0924821630953219 | 0.0481629098783262 | 1.92019467530013 | 0.0548333139577428 | 0.412110851836089 | NM_021884 |
| Lhfp | 947.0737585 | -0.335311787 | 0.086044609 | -3.896952867 | 9.74E-05 | 0.001922649 | | Pcdhgb4 | 196.5609926 | 0.168815591452081 | 0.0879785577256862 | 1.91882653928519 | 0.0550062899635868 | 0.41218506003754 | NM_033576 |
| Mpped1 | 8845.1666 | -0.335800884 | 0.090686895 | -3.702860089 | 0.000213182 | 0.003556118 | | Pcolce2 | 244.8938731 | 0.181005617154073 | 0.0943308522543071 | 1.91883792872028 | 0.0550048481002586 | 0.41218506003754 | NM_029620 |
| 4932438A13Rik | 5820.787091 | -0.336130938 | 0.078166329 | -4.300201164 | 1.71E-05 | 0.000493273 | | Tshz2 | 1588.398541 | 0.176403540222264 | 0.0919383840047109 | 1.91871482332368 | 0.0550204344907607 | 0.41218506003754 | NM_080455 |
| A730017L22Rik | 214.5836589 | -0.336162728 | 0.110801236 | -3.033925784 | 0.002413938 | 0.021882001 | | Cnot3 | 1771.112429 | -0.114034321808428 | 0.0594430924237284 | -1.91837801767708 | 0.055063096317111 | 0.412292029434203 | NM_146176 |
| Dnahc5 | 195.5043215 | -0.336390702 | 0.111281408 | -3.022883227 | 0.002503788 | 0.022398209 | | Il21r | 81.86647701 | -0.182375337879159 | 0.0950914114890167 | -1.91789494995795 | 0.0551243327142469 | 0.412537896448815 | NM_021887 |
| Itpr1 | 36498.7663 | -0.336504459 | 0.095099834 | -3.538433727 | 0.000402508 | 0.005876062 | | Dscam | 3166.689704 | 0.107331757780512 | 0.0559721941616336 | 1.91759067851735 | 0.0551629330164928 | 0.412614194128514 | NM_031174 |
| Fam210b | 1125.487404 | -0.336838743 | 0.095255394 | -3.536164503 | 0.000405982 | 0.005884268 | | Atp6v1d | 9754.728429 | 0.112453388721962 | 0.058692252256473 | 1.9159835310215 | 0.0553671919922067 | 0.413057568054036 | NM_023721 |
| Gnb5 | 5733.635549 | -0.337105168 | 0.054449604 | -6.191140879 | 5.97E-10 | 8.99E-08 | | C4a | 103.8722857 | -0.179770440649499 | 0.0938538333299054 | -1.9154299219467 | 0.0554376982414061 | 0.413057568054036 | NM_011413 |
| Mri1 | 247.1534631 | -0.337303676 | 0.111878734 | -3.014904307 | 0.002570603 | 0.022850792 | | Cblb | 1042.034924 | -0.121637556864578 | 0.0635897778023676 | -1.91284764734701 | 0.0557675593429186 | 0.413057568054036 | NM_001033238 |
| Rftn1 | 391.7976625 | -0.337371338 | 0.098655556 | -3.419689196 | 0.000626927 | 0.008054403 | | Ccdc91 | 1608.608868 | 0.110472947347645 | 0.0577508243671516 | 1.91292416269787 | 0.0557577617788474 | 0.413057568054036 | NM_025911 |
| Flrt3 | 1201.59772 | -0.337482795 | 0.094715042 | -3.563138323 | 0.000366447 | 0.00546773 | | Elf1 | 286.8425927 | -0.157700067922896 | 0.0823827443390395 | -1.91423664249268 | 0.0555899257540777 | 0.413057568054036 | NM_007920 |
| Gpr88 | 8447.602446 | -0.337957101 | 0.121851955 | -2.773505777 | 0.005545586 | 0.038692429 | | Fbxw10 | 33.6379734 | -0.172297767327169 | 0.0900635895483721 | -1.91306795777477 | 0.0557393531245667 | 0.413057568054036 | NM_001033669 |
| Scube1 | 3362.952801 | -0.338442086 | 0.114239744 | -2.962559906 | 0.003050924 | 0.025593407 | | Gm10516 | 205.9208208 | -0.16284703369973 | 0.0849876389931148 | -1.91612610526718 | 0.0553490462035676 | 0.413057568054036 | NR_033536 |
| Rbp1 | 312.982462 | -0.33850729 | 0.113336939 | -2.986734009 | 0.002819749 | 0.024320521 | | Gm10814 | 29.00067005 | -0.169218680812626 | 0.0883803615652802 | -1.91466382141508 | 0.055535390259311 | 0.413057568054036 | NR_045783 |
| Ms4a15 | 71.18807246 | -0.338766177 | 0.104307027 | -3.247779052 | 0.001163096 | 0.012799007 | | Il16 | 179.9539773 | -0.151816870091603 | 0.0793812494162812 | -1.91250290475355 | 0.0558117203999189 | 0.413057568054036 | NM_010551 |
| Myl9 | 462.6617639 | -0.338894808 | 0.114570415 | -2.957960914 | 0.003096814 | 0.025824561 | | Klf11 | 371.8072583 | -0.157917176925584 | 0.0825735210217613 | -1.91244329866916 | 0.0558193588111023 | 0.413057568054036 | NM_178357 |
| Pcsk1 | 1339.158831 | -0.339400625 | 0.110525785 | -3.07078232 | 0.002134987 | 0.020096898 | | Matk | 6568.965917 | -0.144901733944663 | 0.0757635580468532 | -1.91255186108147 | 0.0558054473868888 | 0.413057568054036 | NM_010768 |
| Xylt1 | 914.3304387 | -0.339876823 | 0.093645366 | -3.629403554 | 0.000284077 | 0.004450441 | | Mgp | 519.2752032 | 0.181623933207917 | 0.0949649387511702 | 1.91253672772656 | 0.0558073864347323 | 0.413057568054036 | NM_008597 |
| Slc4a4 | 7143.742797 | -0.34000955 | 0.092199481 | -3.687759935 | 0.000226237 | 0.003709328 | | Ndst1 | 4721.258535 | 0.0789363381005196 | 0.0412033088638671 | 1.91577667612375 | 0.0553935277581436 | 0.413057568054036 | NM_008306 |
| Galnt13 | 1256.303999 | -0.340688511 | 0.096620387 | -3.526052028 | 0.000421804 | 0.006034564 | | Ptma | 8748.120686 | 0.0845996691639598 | 0.0441698185621489 | 1.91532752268213 | 0.0554507477483915 | 0.413057568054036 | NM_008972 |
| Slc13a3 | 791.7545529 | -0.341391271 | 0.099487206 | -3.431509288 | 0.000600233 | 0.007807531 | | St14 | 40.55400058 | -0.168808072072523 | 0.0882599969761172 | -1.91262268135134 | 0.0557963738796931 | 0.413057568054036 | NM_011176 |
| Zfp704 | 3463.765205 | -0.341544049 | 0.065952517 | -5.178635519 | 2.24E-07 | 1.45E-05 | | Stat6 | 783.5728508 | -0.117325760693447 | 0.0612474792139687 | -1.91560146166697 | 0.0554158434070944 | 0.413057568054036 | NM_009284 |
| Kcnj2 | 964.3876273 | -0.341666527 | 0.110229966 | -3.099579366 | 0.001937956 | 0.01868494 | | Tmem66 | 9931.541202 | 0.0935397592658097 | 0.0488161642854421 | 1.91616364446121 | 0.0553442693187701 | 0.413057568054036 | NM_026432 |
| Ldha | 10348.72167 | -0.341789967 | 0.072453446 | -4.717373528 | 2.39E-06 | 0.00010311 | | Trpc4ap | 7430.420812 | 0.0693788113164479 | 0.0362642557569415 | 1.91314587514092 | 0.0557293802539508 | 0.413057568054036 | NM_001163452,NM_019828 |
| Eomes | 106.5929677 | -0.341877523 | 0.106565971 | -3.208130325 | 0.001336009 | 0.014158776 | | Ttll12 | 779.7917044 | 0.134947641124614 | 0.07053613076298 | 1.91317045129784 | 0.0557262349884364 | 0.413057568054036 | NM_183017 |
| Mll2 | 4384.654433 | -0.342139293 | 0.085793516 | -3.987938792 | 6.66E-05 | 0.001436263 | | Ubl3 | 6609.28289 | 0.103920246147981 | 0.0543094439989701 | 1.91348388965189 | 0.0556861340021149 | 0.413057568054036 | NM_011908 |
| Triqk | 384.2498901 | -0.342213695 | 0.115235638 | -2.969686302 | 0.00298104 | 0.025280915 | | Zbtb43 | 1426.348108 | -0.109313656254943 | 0.0570457051659098 | -1.91624691003501 | 0.055333674951186 | 0.413057568054036 | NM_001025594,NM_027947 |
| Luzp2 | 6138.995764 | -0.342319498 | 0.103493108 | -3.307655025 | 0.000940806 | 0.010837808 | | Tgoln1 | 4185.144269 | 0.10579635720154 | 0.0554068307630404 | 1.90944610519236 | 0.0562045679616752 | 0.415696310698215 | NM_009443 |
| Prkca | 9860.561643 | -0.342682493 | 0.07636685 | -4.487319986 | 7.21E-06 | 0.00024453 | | Cldn14 | 28.41128285 | -0.167995925809257 | 0.0880388459706055 | -1.90820227090831 | 0.0563650782399435 | 0.416282454007734 | NM_001165926,NM_019500,NM_001165925 |
| Npy5r | 160.6696558 | -0.34308874 | 0.105499772 | -3.252032989 | 0.001145827 | 0.012644905 | | Rasl10a | 1057.893851 | -0.174626539277938 | 0.0915153913080599 | -1.90816579355607 | 0.0563697912042691 | 0.416282454007734 | NM_145216 |
| D10Bwg1379e | 2815.14733 | -0.343116092 | 0.091070613 | -3.767582975 | 0.000164836 | 0.002915485 | | Ube2z | 4514.000159 | 0.0766240450383656 | 0.0401480109770591 | 1.90853900787636 | 0.0563215864785656 | 0.416282454007734 | NM_172300 |
| Synpr | 4159.978486 | -0.343251904 | 0.110036058 | -3.119449301 | 0.001811894 | 0.017756794 | | Mfsd7b | 465.8565763 | -0.131371298676673 | 0.0688743975124422 | -1.90740396172527 | 0.0564682967627979 | 0.416586327463892 | NM_001081259 |
| Ncan | 10604.43296 | -0.343589469 | 0.107443711 | -3.197855546 | 0.001384536 | 0.014524038 | | Ncoa3 | 2277.492465 | -0.12670009960811 | 0.0664236241758991 | -1.90745538473767 | 0.0564616432134238 | 0.416586327463892 | NM_008679 |
| Vipr2 | 77.4543463 | -0.344556224 | 0.128877676 | -2.673513635 | 0.007506122 | 0.047820667 | | Acrbp | 244.8180833 | -0.180935236284078 | 0.0948929177263104 | -1.90673066672826 | 0.0565554736631538 | 0.416806093572285 | NM_001127340,NM_016845 |
| Smim18 | 170.5820966 | -0.344591101 | 0.109425331 | -3.149098084 | 0.001637752 | 0.016466796 | | Cckar | 33.0881941 | 0.113360970826015 | 0.0594471571311886 | 1.90691996550564 | 0.0565309523207278 | 0.416806093572285 | NM_009827 |
| Asb2 | 130.4562002 | -0.345288696 | 0.129780489 | -2.660559376 | 0.007801097 | 0.04927652 | | Nup205 | 1429.457373 | -0.111771205976807 | 0.0586532269813017 | -1.90562756269897 | 0.0566985434744272 | 0.417225452132487 | NM_027513 |
| Nkain2 | 2179.655224 | -0.345359911 | 0.084382008 | -4.092814571 | 4.26E-05 | 0.001011188 | | St8sia6 | 75.50765712 | -0.180692983977813 | 0.0948143137483098 | -1.9057563867147 | 0.0566818198065915 | 0.417225452132487 | NM_145838 |
| Dynll1 | 7315.923822 | -0.345548646 | 0.07011187 | -4.928532769 | 8.28E-07 | 4.19E-05 | | Synrg | 4257.245829 | 0.0876116142507977 | 0.0459667270961211 | 1.90597895011304 | 0.0566529367581062 | 0.417225452132487 | NM_001115009,NM_194341 |
| Ptpn13 | 833.3590085 | -0.3456532 | 0.097127691 | -3.5587503 | 0.000372624 | 0.005535407 | | Isca1 | 7548.434374 | 0.0981233132313779 | 0.0515137921960585 | 1.9047969300712 | 0.0568064731090665 | 0.417808014370785 | NM_026921 |
| Mina | 427.2017802 | -0.345781538 | 0.115019988 | -3.006273462 | 0.00264471 | 0.02323841 | | Ipmk | 1542.368614 | 0.132027734716937 | 0.0693459378525887 | 1.90390005248172 | 0.0569232022870311 | 0.418112798290665 | NM_027184 |
| Lamc3 | 334.9395416 | -0.345974504 | 0.089536817 | -3.864047391 | 0.000111524 | 0.002126409 | | Nars | 8630.069556 | 0.0761493741565521 | 0.0399970240234711 | 1.90387600117114 | 0.0569263353256143 | 0.418112798290665 | NM_027350,NM_001142950 |
| Abcc9 | 473.2776881 | -0.346499658 | 0.088645285 | -3.908833482 | 9.27E-05 | 0.00185175 | | Tor3a | 360.6142234 | -0.170865134607967 | 0.089748805339725 | -1.90381514228733 | 0.0569342637352978 | 0.418112798290665 | NM_023141 |
| Zfp536 | 615.2204977 | -0.346516081 | 0.074493238 | -4.651644781 | 3.29E-06 | 0.00013218 | | Kcnj12 | 761.5273563 | 0.126809197127343 | 0.0666289459426171 | 1.90321481652366 | 0.0570125205954825 | 0.418421023954386 | NM_001267593,NM_010603 |
| Ccdc85c | 588.5678491 | -0.346848127 | 0.112194185 | -3.091498263 | 0.001991491 | 0.019082356 | | Sgpp2 | 1440.479995 | 0.142298513116468 | 0.0747738692562562 | 1.90305135379312 | 0.0570338446530142 | 0.418421023954386 | NM_001004173 |
| Tbx21 | 33.81222109 | -0.347168039 | 0.117782485 | -2.947535361 | 0.003203181 | 0.026528802 | | Cpsf2 | 2419.025691 | 0.114502623091723 | 0.0601809385654418 | 1.90263937089002 | 0.0570876181261169 | 0.418604109490143 | NM_016856 |
| Usmg5 | 3726.275804 | -0.347225825 | 0.087881505 | -3.951068253 | 7.78E-05 | 0.001620271 | | Coq7 | 726.3801508 | 0.133982569017211 | 0.0704589845765512 | 1.90156826446518 | 0.057227620130706 | 0.419207468491495 | NM_009940 |
| Rasl10b | 13699.88058 | -0.347642039 | 0.059982646 | -5.79571026 | 6.80E-09 | 7.64E-07 | | Lmcd1 | 204.7086064 | -0.166788718825511 | 0.0877107858659364 | -1.90157592568425 | 0.057226617736294 | 0.419207468491495 | NM_144799 |
| Kcnj12 | 735.6707858 | -0.348058417 | 0.076020035 | -4.578509062 | 4.68E-06 | 0.000175263 | | Hspb6 | 266.8516857 | 0.169185664178932 | 0.0889823407689877 | 1.90133977952058 | 0.0572575218288942 | 0.419215101858123 | NM_001012401 |
| Sema3d | 234.8302422 | -0.350798246 | 0.124180808 | -2.824899058 | 0.004729551 | 0.034663987 | | Zswim4 | 968.0245772 | 0.163548552736273 | 0.0860352991655835 | 1.90094710336867 | 0.0573089414947196 | 0.419380193527606 | NM_172503 |
| Uqcr11 | 2475.351203 | -0.351491905 | 0.08354877 | -4.20702666 | 2.59E-05 | 0.000684149 | | A230077H06Rik | 184.0051021 | -0.168958978905096 | 0.0889426463583511 | -1.89963966469311 | 0.0574804232115316 | 0.419820382139875 | NR_040329 |
| Prdm12 | 19.02797062 | -0.351805734 | 0.131179794 | -2.681859178 | 0.007321427 | 0.04706979 | | Arhgap18 | 384.8337566 | -0.134199090942683 | 0.0706573358292904 | -1.89929452289159 | 0.0575257626106232 | 0.419820382139875 | NM_176837 |
| Serpinf1 | 377.8529466 | -0.352462618 | 0.127519349 | -2.763993225 | 0.005709872 | 0.039425264 | | Ky | 184.789662 | 0.179852103368138 | 0.0946856436485885 | 1.89946539346167 | 0.0575033125591358 | 0.419820382139875 | NM_024291 |
| Sdk2 | 620.3097411 | -0.352630849 | 0.123021839 | -2.866408521 | 0.004151581 | 0.031907044 | | Mafk | 404.2418589 | 0.158993169246878 | 0.0837047300011721 | 1.89945262644837 | 0.0575049897178846 | 0.419820382139875 | NM_010757 |
| Pde10a | 13760.16363 | -0.353121874 | 0.109508378 | -3.224610566 | 0.001261441 | 0.013610566 | | Map4k2 | 2956.528494 | -0.139495929306931 | 0.073443859964658 | -1.89935454609899 | 0.0575178755534644 | 0.419820382139875 | NM_009006 |
| Mdn1 | 2486.672297 | -0.35327161 | 0.119410922 | -2.958453077 | 0.003091873 | 0.025824561 | | Pex14 | 1956.033731 | 0.0945789122566775 | 0.0498117337361887 | 1.89872757205327 | 0.0576003043924529 | 0.419820382139875 | NM_019781 |
| Gldc | 1325.11144 | -0.353320775 | 0.08649868 | -4.084695558 | 4.41E-05 | 0.001036094 | | Ppme1 | 10400.41949 | 0.095414488225363 | 0.0502418489433552 | 1.89910383936979 | 0.0575508244159316 | 0.419820382139875 | NM_028292 |
| 9430020K01Rik | 7366.799805 | -0.353481689 | 0.072474501 | -4.877324925 | 1.08E-06 | 5.16E-05 | | Prosc | 2950.538372 | 0.109485619101293 | 0.0576567940587522 | 1.89891964838917 | 0.0575750415162135 | 0.419820382139875 | NM_054057,NM_001039078,NM_001039077 |
| Agpat5 | 1575.95567 | -0.353563213 | 0.083486386 | -4.234980475 | 2.29E-05 | 0.000621314 | | Dmpk | 625.5666504 | -0.155776900320375 | 0.0820542214721218 | -1.89846296175391 | 0.057635122404359 | 0.419863484476288 | NM_001190490,NM_001190491,NM_032418 |
| Spryd4 | 278.1430381 | -0.354754786 | 0.083811892 | -4.232749998 | 2.31E-05 | 0.000625316 | | Nudt11 | 929.3578584 | 0.128972026249506 | 0.0679575298812794 | 1.89783275635265 | 0.0577181166970778 | 0.420257324883084 | NM_021431 |
| Efnb2 | 2587.22642 | -0.355468123 | 0.077254026 | -4.601289321 | 4.20E-06 | 0.000160129 | | Zfp467 | 956.3990433 | -0.124681330647933 | 0.0657112087158845 | -1.89741344109218 | 0.0577733930389435 | 0.420449051745337 | NM_001085417,NM_001085416,NM_001085415,NM_020589 |
| Tsku | 98.42037718 | -0.355821286 | 0.130009131 | -2.736894584 | 0.006202216 | 0.041781366 | | Zfp74 | 760.0134155 | 0.117066219825497 | 0.0617210672124531 | 1.89669792038005 | 0.0578678183460751 | 0.420925352676559 | NM_178384 |
| Abl2 | 3772.098611 | -0.357096436 | 0.095228881 | -3.749875369 | 0.000176922 | 0.003098526 | | Arhgef39 | 32.5049761 | -0.169145718503394 | 0.0892399876132224 | -1.89540275640213 | 0.0580390638077911 | 0.421959680116103 | NM_001013377 |
| Nptx2 | 2550.751919 | -0.357913128 | 0.124579499 | -2.87296971 | 0.004066331 | 0.031404095 | | Hexim1 | 1945.487051 | 0.125303935431956 | 0.0661406362927832 | 1.8945075592753 | 0.0581576719522922 | 0.422610476672812 | NM_138753 |
| Cacna1e | 7933.606378 | -0.358035221 | 0.085280561 | -4.198321569 | 2.69E-05 | 0.000704965 | | Snora34 | 23.80395049 | -0.143316630413882 | 0.0756698163510979 | -1.89397354618798 | 0.0582285212737591 | 0.422913750011312 | NR_034051 |
| 2810459M11Rik | 514.48768 | -0.358523092 | 0.133399484 | -2.687589801 | 0.007196975 | 0.04650122 | | Lpxn | 67.92121673 | -0.176552206129162 | 0.0932989636318051 | -1.89232762355118 | 0.0584473426470537 | 0.423355579337543 | NM_134152 |
| Bmp2 | 124.860465 | -0.358831968 | 0.12117024 | -2.961386958 | 0.003062569 | 0.025630477 | | Pip5kl1 | 238.3424525 | -0.17561245264951 | 0.0927897241343034 | -1.89258513577785 | 0.0584130620235815 | 0.423355579337543 | NM_198191 |
| Adam19 | 1045.091785 | -0.360017146 | 0.098732276 | -3.646397708 | 0.000265942 | 0.004243573 | | Plxnb2 | 2899.084593 | -0.116824774550696 | 0.0617401546922006 | -1.8922008720761 | 0.058464222232625 | 0.423355579337543 | NM_001159521,NM_138749 |
| Lrba | 1814.378767 | -0.36073029 | 0.090968057 | -3.965461076 | 7.33E-05 | 0.001540028 | | Rnasek | 4038.60441 | 0.0918437825653465 | 0.0485157921722178 | 1.89306983258825 | 0.0583485833651218 | 0.423355579337543 | NM_173742 |
| Gadd45g | 325.0803605 | -0.361428 | 0.122032982 | -2.961723904 | 0.00305922 | 0.025621379 | | Tmem164 | 1451.758121 | 0.118489308412414 | 0.0625881947575339 | 1.89315746957458 | 0.0583369314316734 | 0.423355579337543 | NM_001199357,NM_177592,NM_001199360 |
| N6amt2 | 650.4626173 | -0.36153623 | 0.077386298 | -4.671837757 | 2.99E-06 | 0.000122685 | | Tmem248 | 2261.480952 | 0.0996035323041404 | 0.0526216734693763 | 1.89282335085952 | 0.0583813651532846 | 0.423355579337543 | NM_027854,NM_001081394 |
| Frzb | 793.3125109 | -0.361718163 | 0.112287675 | -3.221352337 | 0.001275872 | 0.013689999 | | Ezh2 | 151.6984262 | -0.173458884142859 | 0.0917139770369043 | -1.89130260999435 | 0.0585839605035837 | 0.424011265707552 | NM_001146689,NM_007971 |
| Hivep3 | 2030.706399 | -0.36176914 | 0.104736569 | -3.454086225 | 0.000552161 | 0.00735613 | | Fam135a | 1285.901534 | -0.120943326428506 | 0.0639580643256327 | -1.89097852950555 | 0.0586272103740555 | 0.424112977038611 | NM_026604 |
| Mpp7 | 237.6069352 | -0.361769372 | 0.105895298 | -3.416293056 | 0.000634799 | 0.00811516 | | Tnik | 6917.43409 | -0.092922548660022 | 0.0491559003215285 | -1.89036408757069 | 0.0587092829659169 | 0.424495293361328 | NM_026910,NM_001163009,NM_001163008,NM_001163007 |
| Tmem64 | 2003.421858 | -0.362001073 | 0.067657687 | -5.350479566 | 8.77E-08 | 6.80E-06 | | Fam117b | 3423.313327 | 0.0961733211492962 | 0.0509015568780404 | 1.88939842016476 | 0.058838462431689 | 0.425190580844407 | NM_001037725 |
| Sorl1 | 8861.932392 | -0.362056233 | 0.082617379 | -4.38232533 | 1.17E-05 | 0.000362409 | | Scrn1 | 5482.763182 | 0.115685404737386 | 0.0612348746580263 | 1.88920783105126 | 0.058863985823909 | 0.425190580844407 | NM_027268 |
| Gm4980 | 214.2852052 | -0.362969315 | 0.120706946 | -3.007029228 | 0.002638144 | 0.023198299 | | Fan1 | 442.7553385 | -0.128983493377955 | 0.0682868262501584 | -1.88884885212622 | 0.0589120846757036 | 0.425326511928067 | NM_177893 |
| Hivep2 | 13497.52563 | -0.363061563 | 0.068329487 | -5.313395149 | 1.08E-07 | 7.90E-06 | | Abca7 | 1240.612167 | -0.157426085738885 | 0.0833698865217112 | -1.88828475492633 | 0.0589877328247486 | 0.425661106315101 | NM_013850 |
| Cntnap4 | 1105.743853 | -0.363064872 | 0.086915666 | -4.177208636 | 2.95E-05 | 0.000759536 | | Dusp15 | 786.4301022 | -0.147022987222432 | 0.0778990681814745 | -1.88735232210898 | 0.0591129533551711 | 0.42635290984966 | NM_001159376,NM_145744 |
| Sulf1 | 1077.691463 | -0.363179031 | 0.129208545 | -2.810797309 | 0.004941891 | 0.035738209 | | Pcdh11x | 741.1099835 | 0.161456909995531 | 0.0855829001073206 | 1.88655572308328 | 0.0592201068589378 | 0.426913782745871 | NM_001271809,NM_001271810,NR_073452,NR_073451 |
| Cdh8 | 1626.825762 | -0.36322873 | 0.081142534 | -4.476428218 | 7.59E-06 | 0.000254002 | | Vat1l | 4169.797223 | 0.179281569250561 | 0.0950795456418637 | 1.88559556148765 | 0.0593494759746355 | 0.427634170638668 | NM_173016 |
| Ttc9b | 2727.648613 | -0.363386937 | 0.100327329 | -3.622013476 | 0.000292319 | 0.004556528 | | Col27a1 | 407.6247667 | -0.176896375341796 | 0.0939430983384287 | -1.88301619246716 | 0.0596981735909428 | 0.42823489855903 | NM_025685 |
| Sez6 | 11127.77284 | -0.364118691 | 0.074159636 | -4.909930947 | 9.11E-07 | 4.50E-05 | | Gm2382 | 105.5620084 | -0.177183157635352 | 0.0940673545598612 | -1.8835775542364 | 0.0596221403453348 | 0.42823489855903 | NM_001128601 |
| Slc25a34 | 311.6324944 | -0.36492733 | 0.107799804 | -3.385231844 | 0.000711182 | 0.008815238 | | Gmps | 1644.285673 | 0.124442945220777 | 0.066081247205656 | 1.88318093987375 | 0.0596758511615131 | 0.42823489855903 | NM_001033300 |
| Dok6 | 618.8126888 | -0.366073005 | 0.088340616 | -4.143881076 | 3.41E-05 | 0.000847891 | | Map3k3 | 1235.001309 | -0.110778153656867 | 0.0588254641420457 | -1.88316667403373 | 0.0596777838358741 | 0.42823489855903 | NM_011947 |
| Smpdl3a | 1047.420008 | -0.366096783 | 0.098882415 | -3.702344684 | 0.000213616 | 0.003556118 | | Olfr613 | 97.49314752 | -0.178868851131472 | 0.0949241130330117 | -1.88433523807873 | 0.0595196438033682 | 0.42823489855903 | NM_147100 |
| Lmo1 | 218.6309662 | -0.366243549 | 0.11983009 | -3.056357112 | 0.002240442 | 0.020811399 | | Pcbp1 | 4410.849631 | 0.0760787445829107 | 0.040377239358563 | 1.88419876622339 | 0.0595380943867025 | 0.42823489855903 | NM_011865 |
| Sema3c | 1128.107834 | -0.366573899 | 0.099952804 | -3.667469904 | 0.000244962 | 0.00397429 | | Pdzd7 | 227.2644544 | -0.175949188778945 | 0.0934067042861378 | -1.88369582380242 | 0.0596061316579328 | 0.42823489855903 | NM_001195265 |
| Glyctk | 254.6023416 | -0.367477908 | 0.093092854 | -3.947434097 | 7.90E-05 | 0.001635816 | | Smg1 | 6914.010593 | -0.131832554007512 | 0.0700089295698661 | -1.88308198421957 | 0.0596892583150278 | 0.42823489855903 | NM_001031814 |
| Pkig | 2019.345202 | -0.367646146 | 0.077835234 | -4.723389713 | 2.32E-06 | 0.000100932 | | Vasp | 539.6331441 | -0.14296419080673 | 0.0758595023020544 | -1.88459173166574 | 0.0594849794803185 | 0.42823489855903 | NM_009499 |
| Rab40b | 1054.869768 | -0.368774995 | 0.081176314 | -4.542889153 | 5.55E-06 | 0.000199939 | | Hspa5 | 11784.71785 | 0.14135741744766 | 0.0750936225097375 | 1.88241574614848 | 0.0597795896238597 | 0.428607264993182 | NM_022310,NM_001163434 |
| Gabrg1 | 986.6643979 | -0.369344065 | 0.110323594 | -3.347824823 | 0.000814485 | 0.009714878 | | Gltpd1 | 929.1307266 | 0.120247631418298 | 0.0638880354301116 | 1.88216198242375 | 0.0598140257635868 | 0.428642594100573 | NM_024472 |
| Mas1 | 674.4205572 | -0.370121437 | 0.104321857 | -3.547880059 | 0.000388345 | 0.005703334 | | Psmd11 | 2300.146664 | 0.100204231888446 | 0.0532525940089941 | 1.88167794927552 | 0.0598797554509375 | 0.428722425083892 | NM_178616 |
| Dhrs13 | 214.7413904 | -0.3701961 | 0.121884313 | -3.037274377 | 0.00238728 | 0.021757955 | | Rac2 | 106.7236335 | -0.178945464970153 | 0.0951005308097455 | -1.88164528048896 | 0.0598841938933786 | 0.428722425083892 | NM_009008 |
| Fkbp1b | 856.709925 | -0.370981242 | 0.096098518 | -3.860426248 | 0.000113189 | 0.002143957 | | Calb2 | 843.1563154 | 0.146860096073126 | 0.0780948761843139 | 1.88053433526826 | 0.0600352914694655 | 0.429169608211346 | NM_007586 |
| Slc8a1 | 8127.002437 | -0.37164547 | 0.092166064 | -4.032346106 | 5.52E-05 | 0.001243636 | | Notch1 | 1327.081381 | -0.122298045764008 | 0.0650317062253033 | -1.88059106466079 | 0.0600275679140502 | 0.429169608211346 | NM_008714 |
| Pcbd1 | 137.197122 | -0.372247651 | 0.128860721 | -2.888759649 | 0.003867646 | 0.030449849 | | Plch1 | 555.6515981 | 0.145228190217252 | 0.0772128781857113 | 1.88088041308279 | 0.0599881882103813 | 0.429169608211346 | NM_183191,NM_001177733,NM_001177732 |
| BC056474 | 989.7080538 | -0.372481266 | 0.069760232 | -5.339449967 | 9.32E-08 | 7.12E-06 | | Fgf12 | 7081.309505 | 0.122911879645836 | 0.0653730464875704 | 1.88016141590105 | 0.060086082349827 | 0.429321412795665 | NM_001276419,NM_183064,NM_010199,NM_001276420 |
| Pcp4 | 8281.138623 | -0.373278771 | 0.112722367 | -3.311488063 | 0.000928012 | 0.010720099 | | E130311K13Rik | 284.5516993 | -0.162685914949265 | 0.0865433676181393 | -1.879819556677 | 0.0601326741969869 | 0.429443080327155 | NM_177856 |
| Lix1 | 1952.682821 | -0.373446176 | 0.109862585 | -3.399211637 | 0.000675804 | 0.008464759 | | Iars2 | 2397.818177 | 0.0999076543123741 | 0.0531786914723321 | 1.87871592207264 | 0.0602832927615035 | 0.430307179682358 | NM_198653 |
| Cx3cl1 | 32935.2499 | -0.373655608 | 0.073195004 | -5.104933211 | 3.31E-07 | 2.01E-05 | | Atp6v0e2 | 16826.7016 | 0.0813902980845178 | 0.0433367129681856 | 1.87809117281849 | 0.0603686936892631 | 0.43049368901828 | NM_133764 |
| Tesc | 1480.463594 | -0.373697521 | 0.092954967 | -4.020199594 | 5.81E-05 | 0.001299865 | | Nsf | 42381.32647 | 0.0720314577053258 | 0.0383517118470997 | 1.87818103120143 | 0.0603564041805585 | 0.43049368901828 | NM_008740 |
| 9830001H06Rik | 2647.79046 | -0.373976182 | 0.08940989 | -4.182716048 | 2.88E-05 | 0.000747571 | | Adamts18 | 86.40353207 | -0.177599041587948 | 0.0945777854244704 | -1.87780926346365 | 0.0604072625690173 | 0.430557358232358 | NM_172466 |
| Ctxn2 | 275.479193 | -0.37397704 | 0.101350102 | -3.689952266 | 0.000224296 | 0.003685307 | | 4931406C07Rik | 1114.244132 | -0.118662178017256 | 0.0632364760662208 | -1.87648309012339 | 0.0605889745359455 | 0.431188114253562 | NM_133732,NM_001199484,NM_001199485 |
| Syt9 | 581.687642 | -0.374321283 | 0.100781407 | -3.714189902 | 0.000203856 | 0.003448207 | | Hsbp1 | 7209.135297 | 0.0805732648396411 | 0.0429427004317502 | 1.87629711288646 | 0.0606144932745266 | 0.431188114253562 | NM_024219 |
| Gabrb1 | 808.6128594 | -0.374957743 | 0.08598263 | -4.360854547 | 1.30E-05 | 0.000395144 | | Ngb | 366.2476873 | 0.178191944593445 | 0.0949537312510643 | 1.87661866727799 | 0.0605703770216086 | 0.431188114253562 | NM_022414 |
| Tbc1d16 | 2909.047956 | -0.374965198 | 0.075411919 | -4.972227239 | 6.62E-07 | 3.51E-05 | | Ssc5d | 126.3808835 | -0.178155835388402 | 0.0949313356604656 | -1.87668101527192 | 0.0605618261533227 | 0.431188114253562 | NM_173008 |
| Kcna3 | 349.8183196 | -0.375096828 | 0.115586656 | -3.245156829 | 0.00117386 | 0.012845805 | | Hspd1 | 7221.753944 | 0.0995426808072316 | 0.0530724927805099 | 1.87559836729183 | 0.0607104507813908 | 0.431659328463281 | NM_010477 |
| Akap5 | 4962.845284 | -0.375216383 | 0.08368297 | -4.483784241 | 7.33E-06 | 0.000246462 | | Bod1 | 2741.268763 | 0.0864515749099352 | 0.0461354346028252 | 1.8738649728606 | 0.060949038350458 | 0.432354503034509 | NM_001024919 |
| Adamtsl1 | 344.74579 | -0.375271588 | 0.120521011 | -3.1137441 | 0.001847296 | 0.018012592 | | Hmox2 | 2795.38223 | 0.0934889789076789 | 0.0498925361842265 | 1.87380690695847 | 0.0609570440737074 | 0.432354503034509 | NM_010443,NM_001136066 |
| Slc35f1 | 3619.591104 | -0.375296682 | 0.069374304 | -5.409736203 | 6.31E-08 | 5.12E-06 | | Ptprr | 811.9118465 | 0.142398025476935 | 0.0759762900666795 | 1.87424294279126 | 0.0608969477672526 | 0.432354503034509 | NM_001161838,NM_001161839,NM_011217,NM_001161840,NM_001161837 |
| 1700020I14Rik | 2926.67797 | -0.375547293 | 0.058990056 | -6.366281362 | 1.94E-10 | 3.49E-08 | | Slc16a6 | 659.2616309 | -0.159305952488892 | 0.0850169980801419 | -1.87381295607169 | 0.0609562100233482 | 0.432354503034509 | NM_001029842,NM_134038 |
| Cep112 | 105.3829029 | -0.376242297 | 0.124610622 | -3.019343695 | 0.00253323 | 0.022583349 | | Ube2j2 | 67.59203655 | -0.178139910043451 | 0.0950675270395114 | -1.8738250125031 | 0.060954547713519 | 0.432354503034509 | NM_001039159,NM_001039158,NM_021402,NM_001039157 |
| Chst11 | 3137.898792 | -0.377129483 | 0.0509906 | -7.39605901 | 1.40E-13 | 5.30E-11 | | 4930538K18Rik | 86.58954393 | -0.177977476633452 | 0.0950020423589148 | -1.87340684699239 | 0.0610122252425227 | 0.432534691982862 | NM_029198 |
| Ankrd6 | 1816.454802 | -0.377901926 | 0.091874181 | -4.113254925 | 3.90E-05 | 0.000945932 | | Cyyr1 | 123.9333307 | -0.177611536648187 | 0.0948405756418348 | -1.87273785978415 | 0.0611045925972907 | 0.4329782010089 | NM_144853 |
| Fat4 | 647.3182522 | -0.378370004 | 0.120168372 | -3.148665473 | 0.001640178 | 0.016480495 | | Dbpht2 | 1752.426363 | 0.143359723631399 | 0.0765733150311664 | 1.87218907230345 | 0.0611804503739226 | 0.433304350137299 | NM_198866 |
| Klhdc8a | 716.0204553 | -0.378634207 | 0.107015985 | -3.538108883 | 0.000403004 | 0.005876062 | | Chchd4 | 1237.459649 | 0.0965131638019305 | 0.0515788694702754 | 1.87117640989689 | 0.0613206333863148 | 0.434085536339965 | NM_133928 |
| Kcnmb4 | 1936.313485 | -0.37869042 | 0.088499472 | -4.279013348 | 1.88E-05 | 0.000529798 | | Col3a1 | 103.163387 | -0.157576845594406 | 0.0842361502491646 | -1.87065583040422 | 0.0613928007351447 | 0.434384716745597 | NM_009930 |
| Fam181b | 491.066758 | -0.379117375 | 0.095949726 | -3.951208537 | 7.78E-05 | 0.001620271 | | Fbrsl1 | 1719.305775 | -0.126832319068215 | 0.0678385757404916 | -1.8696194264661 | 0.0615368856291831 | 0.435077841664156 | NM_001142642,NM_028596 |
| Herc2 | 11846.88981 | -0.379328855 | 0.070680203 | -5.366833145 | 8.01E-08 | 6.25E-06 | | Ndufb10 | 3378.546648 | 0.090870522915914 | 0.0486119714342127 | 1.86930338834109 | 0.0615806169944586 | 0.435077841664156 | NM_026684 |
| Dpysl3 | 636.4767929 | -0.380114796 | 0.104183706 | -3.648505209 | 0.000263771 | 0.004213259 | | Tmem177 | 474.034511 | 0.141254050745263 | 0.0755645031653036 | 1.86931753440181 | 0.0615786500446192 | 0.435077841664156 | NM_175106 |
| Igsf21 | 1819.117811 | -0.380179591 | 0.090194782 | -4.215095202 | 2.50E-05 | 0.000666971 | | Calcrl | 206.963729 | -0.165633810525435 | 0.0886302271818514 | -1.86881852604966 | 0.061648066493555 | 0.435129208410339 | NM_018782 |
| Atp5l | 3829.251598 | -0.380180633 | 0.064842099 | -5.863175872 | 4.54E-09 | 5.50E-07 | | Gm14378 | 132.3152922 | -0.174723029865977 | 0.0935045155446032 | -1.86860526305418 | 0.0616777530026772 | 0.435129208410339 | NM_001195258 |
| Mll3 | 3858.140982 | -0.380975137 | 0.099004723 | -3.848050104 | 0.000119062 | 0.002236051 | | Sbds | 2490.967058 | 0.100624280081657 | 0.0538426187354978 | 1.86885932454316 | 0.0616423886339038 | 0.435129208410339 | NM_023248 |
| Idi1 | 1620.958335 | -0.381210985 | 0.136776682 | -2.787105073 | 0.005318123 | 0.037437075 | | Ddit3 | 909.2465648 | -0.112149475227169 | 0.0600282321898919 | -1.86827882707587 | 0.0617232162689494 | 0.435133476753597 | NM_007837 |
| Notch2 | 1020.963249 | -0.38136072 | 0.108151464 | -3.526172506 | 0.000421612 | 0.006034564 | | Ogdh | 14131.17915 | 0.0913036347225593 | 0.0488732803539096 | 1.86817078905684 | 0.0617382690065513 | 0.435133476753597 | NM_001252288,NM_001252282,NM_001252283,NM_010956,NM_001252287 |
| Cldn5 | 924.0491 | -0.381964078 | 0.121045932 | -3.155530075 | 0.001602067 | 0.01619582 | | Tmem231 | 594.0926225 | 0.122692279278964 | 0.0656833724353184 | 1.86793513685348 | 0.0617711125349325 | 0.435153821863448 | NM_001033321 |
| Glt25d2 | 341.1285305 | -0.382371349 | 0.109785418 | -3.482897429 | 0.000496018 | 0.006795145 | | Pltp | 3082.505687 | -0.128302797396725 | 0.0687392573683387 | -1.86651416248528 | 0.0619696464893832 | 0.436339525532128 | NM_011125 |
| Frmd7 | 152.2623315 | -0.383141152 | 0.121595123 | -3.15095821 | 0.001627358 | 0.016383548 | | Atp13a4 | 558.1329455 | -0.141022248694702 | 0.0755935554037262 | -1.86553268914972 | 0.0621067745905958 | 0.436531911416255 | NM_001164613,NM_001164612,NM_172613 |
| Chsy3 | 503.0898584 | -0.38456634 | 0.079928622 | -4.811372056 | 1.50E-06 | 6.93E-05 | | Degs1 | 3403.016293 | 0.0923212965000519 | 0.0494938320442362 | 1.86530912412556 | 0.0621380869654019 | 0.436531911416255 | NM_007853 |
| Cxcr7 | 390.6885535 | -0.384626638 | 0.091040786 | -4.224772815 | 2.39E-05 | 0.000642268 | | Lepr | 164.811167 | -0.176619548755849 | 0.0946897229472992 | -1.86524517401058 | 0.0621470461798716 | 0.436531911416255 | NM_010704,NM_146146,NM_001122899 |
| Myo18b | 93.87978013 | -0.385053993 | 0.133524328 | -2.883774051 | 0.003929406 | 0.03076413 | | Plagl1 | 651.5089354 | 0.157603882100941 | 0.0844922881331889 | 1.86530493590732 | 0.062138673689196 | 0.436531911416255 | NM_009538 |
| Tekt5 | 278.2030778 | -0.385541228 | 0.112524633 | -3.426282924 | 0.000611903 | 0.007927107 | | Sntg2 | 116.0599844 | -0.169941239282569 | 0.0910715190753229 | -1.86601959655482 | 0.0620386238288215 | 0.436531911416255 | NM_172951 |
| Pcdhb21 | 92.0942294 | -0.385794202 | 0.124810754 | -3.091033338 | 0.001994612 | 0.019100448 | | Arf1 | 16005.64003 | 0.0710957888116577 | 0.0381291905957763 | 1.86460262336472 | 0.0622371248819686 | 0.436873272197415 | NM_007476,NM_001130408 |
| Kcna4 | 1156.017063 | -0.38607634 | 0.096110039 | -4.01702408 | 5.89E-05 | 0.001310159 | | Kctd21 | 890.6862418 | 0.109133377893947 | 0.0585601132392013 | 1.86361282206147 | 0.0623760957274844 | 0.436873272197415 | NM_001039039 |
| Huwe1 | 16699.80886 | -0.386175084 | 0.059640604 | -6.475036394 | 9.48E-11 | 1.93E-08 | | Nxpe4 | 514.7222274 | -0.119751479662191 | 0.064253680546183 | -1.86372949602659 | 0.0623597010461012 | 0.436873272197415 | NM_172921 |
| Ubr4 | 9260.184575 | -0.386378259 | 0.081334063 | -4.750509733 | 2.03E-06 | 8.98E-05 | | Slfn9 | 22.44223661 | -0.154697307802653 | 0.0829906955155539 | -1.8640319476978 | 0.0623172180253562 | 0.436873272197415 | NM_172796 |
| Hpcal4 | 50831.04813 | -0.387150414 | 0.0846597 | -4.573018979 | 4.81E-06 | 0.000179055 | | Sod1 | 6047.24217 | 0.10962454189885 | 0.0588100666164649 | 1.86404384563269 | 0.0623155473050522 | 0.436873272197415 | NM_011434 |
| Rab27b | 1357.156021 | -0.388126841 | 0.089318309 | -4.345434265 | 1.39E-05 | 0.000416938 | | Tusc3 | 6926.764975 | 0.098292523180762 | 0.0527297862308517 | 1.86407966259981 | 0.0623105152649983 | 0.436873272197415 | NM_030254 |
| Slc19a3 | 164.9094588 | -0.388430463 | 0.12763295 | -3.043340012 | 0.002339678 | 0.021437595 | | Txndc12 | 927.9028842 | 0.121550072613742 | 0.0652413878723238 | 1.86308226384781 | 0.0624506931778354 | 0.437184948964452 | NM_025334 |
| Cd93 | 427.9824535 | -0.388615555 | 0.121395492 | -3.20123547 | 0.001368397 | 0.014413281 | | Cyp2s1 | 271.8208914 | 0.157448363898823 | 0.0845255705179263 | 1.86273056702328 | 0.0625001830484349 | 0.437320644971852 | NM_028775 |
| C1galt1 | 341.715848 | -0.388718125 | 0.098468531 | -3.947638115 | 7.89E-05 | 0.001635816 | | Stmn1 | 24211.67332 | 0.0860158135477267 | 0.0461875700263419 | 1.86231519646238 | 0.0625586746957 | 0.437519166408155 | NM_019641 |
| Pcdha7 | 117.7062215 | -0.389436537 | 0.115319264 | -3.377029347 | 0.000732732 | 0.008981768 | | Pdzd2 | 3666.639281 | 0.130579800142646 | 0.0701272859854844 | 1.86202459843886 | 0.0625996230344727 | 0.437594862463306 | NM_001081064 |
| Foxo1 | 1649.49725 | -0.389994885 | 0.07347332 | -5.307979605 | 1.11E-07 | 8.06E-06 | | Syt11 | 39148.85559 | 0.0752138247270907 | 0.0404037764463194 | 1.8615543234435 | 0.0626659367198452 | 0.437847713704892 | NM_018804 |
| Aox3 | 105.9682648 | -0.390147998 | 0.13041688 | -2.991545242 | 0.002775694 | 0.024026036 | | Rap2a | 5366.838403 | 0.123521073184911 | 0.0664245040044495 | 1.85957087728704 | 0.0629462628146426 | 0.43959491040649 | NM_029519 |
| Nrp2 | 1666.46064 | -0.390314308 | 0.140470442 | -2.778622333 | 0.005458995 | 0.038236657 | | Trp53inp2 | 8184.98424 | 0.114930656065051 | 0.0618258040654755 | 1.85894316786136 | 0.0630351944303131 | 0.4400004437431393 | NM_178111 |
| Grm5 | 6632.334105 | -0.390362452 | 0.081951778 | -4.76331891 | 1.90E-06 | 8.55E-05 | | Chml | 509.0721088 | 0.148562767310778 | 0.0799536427923487 | 1.85811130202807 | 0.0631532101869472 | 0.44034222344404 | NM_021350 |
| Pcdh11x | 720.7165591 | -0.390751015 | 0.117009381 | -3.339484517 | 0.00083934 | 0.009942461 | | Kat6b | 2640.872198 | -0.0973112191748147 | 0.0523784347368904 | -1.85784893465474 | 0.0631904697748177 | 0.44034222344404 | NM_001205241,NM_017479 |
| Emx2 | 339.3077002 | -0.393800679 | 0.113404218 | -3.472539953 | 0.000515558 | 0.006970384 | | Lxn | 1349.490411 | -0.11665733363551 | 0.0628022453641771 | -1.85753443939844 | 0.063235156141007 | 0.44034222344404 | NM_016753 |
| Tmem158 | 1495.954543 | -0.39404104 | 0.119948969 | -3.285072337 | 0.001019562 | 0.011522314 | | Smyd3 | 1073.198134 | 0.113858666311098 | 0.0612777586910244 | 1.85807491565084 | 0.063158376442179 | 0.44034222344404 | NM_027188 |
| Mll1 | 6969.910578 | -0.395232328 | 0.09221934 | -4.285785712 | 1.82E-05 | 0.000518465 | | Ttc5 | 1429.725689 | -0.109957560342328 | 0.0591898699649394 | -1.85770910474141 | 0.063210334866123 | 0.44034222344404 | NM_177625,NM_001080949 |
| Zfp365 | 23643.60402 | -0.395234793 | 0.08759501 | -4.512069749 | 6.42E-06 | 0.000222978 | | C030046E11Rik | 1858.451244 | -0.112486030429271 | 0.0605792675761824 | -1.85684038335084 | 0.0633338663991426 | 0.440607156759552 | NM_001081319 |
| Kcnq5 | 4308.845105 | -0.39538801 | 0.068030102 | -5.81195679 | 6.17E-09 | 7.00E-07 | | Ergic1 | 6427.809363 | 0.0777169623675249 | 0.0418532602152625 | 1.85689148151914 | 0.0633265947598501 | 0.440607156759552 | NM_026170 |
| Grid2 | 425.8573006 | -0.395399531 | 0.086979373 | -4.545900005 | 5.47E-06 | 0.000198718 | | Bnip3 | 4047.164247 | 0.119252869487088 | 0.064247008413627 | 1.85616221566815 | 0.0634304399336381 | 0.440995265947177 | NM_009760 |
| Rpl41 | 7640.167928 | -0.395787114 | 0.06806047 | -5.815227438 | 6.06E-09 | 6.95E-07 | | Cdh4 | 1449.650056 | 0.140465197013518 | 0.075689452538123 | 1.85580939355809 | 0.0634807311782698 | 0.440995265947177 | NM_009867 |
| Gm13889 | 575.8387015 | -0.396463769 | 0.118752699 | -3.338566402 | 0.000842119 | 0.009967755 | | Pitpna | 15067.37498 | 0.0883581401047178 | 0.0476112645638781 | 1.85582426583464 | 0.063478610619892 | 0.440995265947177 | NM_008850 |
| Lmcd1 | 230.7347163 | -0.396926575 | 0.102309739 | -3.87965583 | 0.000104604 | 0.002033551 | | Fgf10 | 217.2360493 | 0.164840576728219 | 0.0888619097912718 | 1.8550195141587 | 0.0635934399880155 | 0.440996088261682 | NM_008002 |
| Lrrc7 | 2685.07043 | -0.397002339 | 0.08500096 | -4.670563019 | 3.00E-06 | 0.000122805 | | Gm13446 | 193.7996208 | -0.164552981524713 | 0.0886962453903905 | -1.85524179519036 | 0.0635617057465892 | 0.440996088261682 | NR_045894 |
| Ndst3 | 1154.764642 | -0.397976084 | 0.084522923 | -4.708498856 | 2.50E-06 | 0.000106515 | | Gmip | 404.1568533 | -0.152674213074696 | 0.0823137573806278 | -1.8547836708354 | 0.0636271247778184 | 0.440996088261682 | NM_198101 |
| Mlf1ip | 70.99086984 | -0.398573578 | 0.142581828 | -2.795402353 | 0.005183516 | 0.03674392 | | Mroh8 | 55.47332217 | -0.175575840050641 | 0.0946468914822996 | -1.85506187578782 | 0.0635873911654465 | 0.440996088261682 | NM_001039557 |
| Gpx8 | 124.3018002 | -0.401060086 | 0.136067522 | -2.947507833 | 0.003203466 | 0.026528802 | | Tmem115 | 1239.319519 | 0.101157049420138 | 0.0545395985564403 | 1.85474503108885 | 0.0636326449811706 | 0.440996088261682 | NM_019704 |
| Cdh11 | 2027.712629 | -0.401060434 | 0.079197557 | -5.06405058 | 4.10E-07 | 2.38E-05 | | 2210404J11Rik | 259.2478093 | -0.152038414725786 | 0.0820815662233636 | -1.85228452283736 | 0.0639849769982583 | 0.441368101351686 | NM_001034891,NM_001039552 |
| Tubb3 | 12282.5718 | -0.401124115 | 0.083111191 | -4.826355043 | 1.39E-06 | 6.48E-05 | | 5530601H04Rik | 427.3242011 | -0.129088770863681 | 0.069635727856229 | -1.8537721201134 | 0.0637717687407959 | 0.441368101351686 | NR_015467 |
| Tmeff1 | 2840.785162 | -0.401895606 | 0.066498549 | -6.043674779 | 1.51E-09 | 1.93E-07 | | Abcf2 | 4539.1199 | 0.0706892814373429 | 0.038140212058774 | 1.85340556912507 | 0.0638242497037012 | 0.441368101351686 | NM_001190443,NM_013853 |
| E030013I19Rik | 54.8386463 | -0.402257886 | 0.147105812 | -2.734479894 | 0.006247182 | 0.041997755 | | Ano1 | 177.2527774 | -0.175183474686824 | 0.094567052425533 | -1.8524789574548 | 0.0639570764721664 | 0.441368101351686 | NM_001242349,NM_178642 |
| Rcn3 | 152.2682099 | -0.402298342 | 0.12762043 | -3.152303613 | 0.001619837 | 0.016318843 | | Chl1 | 5342.755183 | 0.115561631981192 | 0.0623925892649222 | 1.85216919737873 | 0.0640015304505658 | 0.441368101351686 | NM_007697 |
| Myoc | 854.6466574 | -0.403915176 | 0.132811117 | -3.041275355 | 0.002355783 | 0.021559115 | | Haus7 | 207.7427599 | 0.162731379427392 | 0.0877997034614016 | 1.85343882737522 | 0.0638194864804812 | 0.441368101351686 | NM_028633,NM_207104 |
| Chl1 | 5226.664705 | -0.404302611 | 0.085256146 | -4.742210736 | 2.11E-06 | 9.29E-05 | | Metap2 | 2819.049468 | 0.0945829102196964 | 0.0510333435867837 | 1.85335515120336 | 0.0638314710829714 | 0.441368101351686 | NM_019648 |
| Angpt1 | 212.6973788 | -0.404486436 | 0.11377804 | -3.555048361 | 0.00037791 | 0.005603187 | | Mmaa | 794.8851144 | -0.130738712467155 | 0.0705918590452676 | -1.85203668291719 | 0.0640205555244391 | 0.441368101351686 | NM_133823 |
| Hivep1 | 3954.315007 | -0.405279431 | 0.089564869 | -4.524982145 | 6.04E-06 | 0.00021221 | | Opa3 | 1879.266386 | 0.0967169154673463 | 0.0521914890423038 | 1.85311661426162 | 0.0638656459750579 | 0.441368101351686 | NM_207525 |
| Ablim3 | 1535.617072 | -0.405721488 | 0.123020089 | -3.298010034 | 0.000973727 | 0.011167224 | | Pkd2 | 1510.992194 | -0.103349624735257 | 0.0558009335276743 | -1.85211282682218 | 0.0640096229891125 | 0.441368101351686 | NM_008861 |
| Gypc | 83.28495826 | -0.406910072 | 0.1300786 | -3.128186119 | 0.001758888 | 0.017366567 | | Yars | 4185.921649 | 0.0862684848715419 | 0.0465656592428055 | 1.85262028443999 | 0.0639368029673967 | 0.441368101351686 | NM_134151 |
| Lrp1b | 1665.482045 | -0.40891699 | 0.107015652 | -3.821095157 | 0.00013286 | 0.002433261 | | Faah | 4164.460774 | -0.113117575676743 | 0.0610848242635231 | -1.85181142846132 | 0.0640529059759102 | 0.44138164715658 | NM_010173 |
| Adora2a | 2534.098192 | -0.408970203 | 0.127281241 | -3.213122379 | 0.001313003 | 0.013943573 | | Rn45s | 16791.75629 | 0.107740209852339 | 0.0581984126457889 | 1.85125684626615 | 0.0641326111398816 | 0.441721341592186 | NR_046233 |
| Ptprf | 4144.378741 | -0.409466477 | 0.073773422 | -5.550325109 | 2.85E-08 | 2.58E-06 | | Tex264 | 2412.686386 | 0.103718725342212 | 0.0560424883508778 | 1.85071591919575 | 0.0642104326480037 | 0.442047746277205 | NM_001081654,NM_011573 |
| Flt1 | 2472.320054 | -0.409549538 | 0.07180206 | -5.70386893 | 1.17E-08 | 1.21E-06 | | Il3ra | 55.70510742 | -0.164814073206894 | 0.0890929102580746 | -1.84991233005498 | 0.0643261864631091 | 0.442634857680304 | NM_008369 |
| Vstm4 | 181.439375 | -0.410569589 | 0.129038195 | -3.181767923 | 0.001463791 | 0.015180704 | | 9830166K06Rik | 69.52807151 | -0.174057317300203 | 0.0941916320442468 | -1.84790637472381 | 0.0646158881874224 | 0.443022969298812 | NR_045314 |

| | | | | | | | | | | | | | | | |
|---|---|---|---|---|---|---|---|---|---|---|---|---|---|---|---|
| Trpc5 | 397.2779555 | -0.410827097 | 0.111469554 | -3.68555433 | 0.000228205 | 0.003729762 | | Colec12 | 600.8469558 | -0.155301384698933 | 0.0840420100816869 | -1.84790183561749 | 0.0646165449478333 | 0.443022969298812 | NM_130449 |
| Apcdd1 | 971.7775461 | -0.411438765 | 0.11797788 | -3.487422939 | 0.000487699 | 0.006728776 | | Dok4 | 673.9559663 | 0.145685479388637 | 0.0787847309176351 | 1.84915881150806 | 0.0644348842487846 | 0.443022969298812 | NM_053246 |
| Cyp51 | 2246.898954 | -0.412298282 | 0.078483852 | -5.253288076 | 1.49E-07 | 1.03E-05 | | Hlf | 10355.61112 | 0.139412653688514 | 0.0754485047659221 | 1.84778550775843 | 0.0646333782316642 | 0.443022969298812 | NM_172563 |
| Tmem91 | 538.0354061 | -0.412307367 | 0.124362831 | -3.315358492 | 0.000915257 | 0.010598645 | | Kctd20 | 1926.167099 | 0.107647426072472 | 0.058262693054494 | 1.84762178040393 | 0.0646570766152748 | 0.443022969298812 | NM_025888 |
| Grm1 | 2078.453675 | -0.412796979 | 0.085241855 | -4.842656 | 1.28E-06 | 6.02E-05 | | Mapk6 | 3899.669533 | 0.147112200244727 | 0.0796149834173928 | 1.84779540144436 | 0.0646319464198668 | 0.443022969298812 | NM_015806,NM_027418 |
| Nid1 | 433.6626812 | -0.413040271 | 0.129061037 | -3.200348309 | 0.001372616 | 0.014433037 | | Mei1 | 291.6338544 | -0.162555165333581 | 0.0879733458960205 | -1.84777745665957 | 0.0646345434040321 | 0.443022969298812 | NM_028897 |
| Rab3c | 16973.54471 | -0.413947202 | 0.083106995 | -4.980894816 | 6.33E-07 | 3.39E-05 | | Men1 | 2639.718043 | 0.0780149327308336 | 0.042197222527196 | 1.84881677178049 | 0.0644842747254287 | 0.443022969298812 | NM_001168488,NM_001168489,NM_001168490,NM_008583 |
| Ccdc138 | 107.5147142 | -0.414068605 | 0.129214595 | -3.204503359 | 0.001352957 | 0.014299262 | | Plcl2 | 3310.397728 | 0.135279814462293 | 0.073178952101216 | 1.84861644746133 | 0.0645132160285642 | 0.443022969298812 | NM_013880 |
| Mrpl33 | 498.3041142 | -0.414337965 | 0.098859114 | -4.19119643 | 2.77E-05 | 0.000725023 | | Klhdc7a | 848.7677886 | 0.144042292695294 | 0.0779948918657109 | 1.84681700621242 | 0.0647736662879626 | 0.443414795151935 | NM_173427 |
| Ccr2 | 10.19247887 | -0.414605138 | 0.144121417 | -2.876776728 | 0.004017598 | 0.031139898 | | Lsamp | 2368.805213 | 0.10234857402244 | 0.0554192446239145 | 1.84680564877773 | 0.0647753129087433 | 0.443414795151935 | NM_175548 |
| Shisa2 | 160.5973259 | -0.415446709 | 0.133586815 | -3.109937983 | 0.001871266 | 0.018189084 | | Map3k13 | 1161.403042 | 0.15651007067834 | 0.0847868249158339 | 1.84592442084846 | 0.0649031802223005 | 0.444080827088618 | NM_172821 |
| Neto2 | 2520.131009 | -0.415486635 | 0.082960973 | -5.008217963 | 5.49E-07 | 3.04E-05 | | Cd99l2 | 4666.777429 | 0.116985411601538 | 0.0633987632284915 | 1.8452317623282 | 0.0650038319755659 | 0.44435090036568 | NM_001199349,NM_138309 |
| Pcdhb18 | 168.648665 | -0.41682465 | 0.101759428 | -4.096177217 | 4.20E-05 | 0.001001219 | | Ms4a6b | 34.46474964 | -0.165131653960723 | 0.0894820924010794 | -1.84541565278296 | 0.0649770978974995 | 0.44435090036568 | NM_027209 |
| Gpr26 | 1957.299373 | -0.416980635 | 0.085134091 | -4.897927846 | 9.69E-07 | 4.76E-05 | | Wars | 4304.605108 | 0.082295084440196 | 0.0446083737384769 | 1.84483489406413 | 0.0650615597876449 | 0.444536320543429 | NM_001164314,NM_011710,NM_001164488 |
| Trpc7 | 196.435079 | -0.419613083 | 0.123285402 | -3.403590979 | 0.000665063 | 0.008364026 | | Dclre1b | 376.4343612 | 0.137556049867114 | 0.0745772940325226 | 1.84447627996702 | 0.0651137595746635 | 0.444683813625558 | NM_133865,NM_001025312 |
| Col1a2 | 891.7664996 | -0.42031474 | 0.125401943 | -3.351740251 | 0.000803053 | 0.009608114 | | Fam96b | 507.6215908 | 0.127559956615875 | 0.069195096643009 | 1.8434825992654 | 0.065258580008632 | 0.444691459208727 | NM_026753 |
| Qpct | 420.8748326 | -0.421673988 | 0.10303889 | -4.092377056 | 4.27E-05 | 0.001011549 | | Gpr115 | 107.5706986 | -0.174229942938039 | 0.0945032358231885 | -1.84363997084731 | 0.0652356267637283 | 0.444691459208727 | NM_030067 |
| Rxrg | 969.8600865 | -0.425065021 | 0.124636469 | -3.410438572 | 0.000648585 | 0.008223547 | | Postn | 84.73479375 | -0.168422423002818 | 0.0913311691654847 | -1.84408482385297 | 0.0651707792675748 | 0.444691459208727 | NM_001198765,NM_001198766,NM_015784 |
| Pbx1 | 7473.231177 | -0.426554828 | 0.074716752 | -5.708958421 | 1.14E-08 | 1.18E-06 | | Rlim | 2302.878944 | 0.124773413757675 | 0.0676858895232208 | 1.8434183939456 | 0.065267946511979 | 0.444691459208727 | NM_011276 |
| Osbpl8 | 4083.251491 | -0.427470449 | 0.084718741 | -5.045760143 | 4.52E-07 | 2.57E-05 | | Urgcp | 3426.48322 | 0.103902295859067 | 0.0563588460407301 | 1.84358451526807 | 0.0652437144114282 | 0.444691459208727 | NM_178623,NM_001077661 |
| Plcxd3 | 345.601027 | -0.427541785 | 0.135266563 | -3.160735196 | 0.001573715 | 0.016024696 | | Defb1 | 25.82907545 | 0.153632180651947 | 0.0833816019151841 | 1.84251893850903 | 0.0653992789082371 | 0.445248049719934 | NM_007843 |
| Mpped2 | 1304.708112 | -0.428323525 | 0.070755847 | -6.053542503 | 1.42E-09 | 1.84E-07 | | Pinx1 | 166.9244807 | 0.160750090280617 | 0.087248521337032 | 1.84243913612766 | 0.0654109416289646 | 0.445248049719934 | NM_028228 |
| Akap9 | 4475.988126 | -0.428959502 | 0.109911541 | -3.902770318 | 9.51E-05 | 0.001886616 | | Dnajb1 | 2422.213773 | 0.127505845598684 | 0.0692282295853124 | 1.84181866794028 | 0.0655016784803089 | 0.445445382863633 | NM_018808 |
| Grk5 | 614.130081 | -0.429136276 | 0.070058257 | -6.125420371 | 9.04E-10 | 1.24E-07 | | Fam98a | 1455.342868 | -0.114613361523958 | 0.0622246432525686 | -1.84192878468976 | 0.0654855675104354 | 0.445445382863633 | NM_133747 |
| Grm3 | 2977.855656 | -0.42957074 | 0.126783367 | -3.388226303 | 0.000703462 | 0.008740529 | | Polr1b | 883.297398 | -0.100803701412555 | 0.0547366669247262 | -1.84161197741873 | 0.0655319277970249 | 0.445445382863633 | NM_009086 |
| Lhx8 | 117.5787567 | -0.430135873 | 0.159779415 | -2.692060626 | 0.007101204 | 0.04597829 | | Boc | 1067.790138 | -0.133781296349726 | 0.0726646879442193 | -1.84107714674867 | 0.0656102541389852 | 0.445560800197709 | NM_172506 |
| B230216N24Rik | 134.2511896 | -0.430364302 | 0.134862419 | -3.191135871 | 0.001417146 | 0.014776085 | | Nhp2l1 | 4171.408381 | 0.100621218282734 | 0.0546474706545469 | 1.8412786004097 | 0.0655807420476483 | 0.445560800197709 | NM_011482 |
| Npffr2 | 9.440078024 | -0.430607862 | 0.147228368 | -2.924761496 | 0.003447203 | 0.027963856 | | Plekho2 | 755.2281215 | -0.146297082647292 | 0.079471938840243 | -1.8408639676123 | 0.0656414958840619 | 0.445564658510226 | NM_153119 |
| Epha3 | 224.8833475 | -0.431022572 | 0.124395574 | -3.464934954 | 0.00053036 | 0.007120792 | | Rexo4 | 1994.453235 | -0.105605613821185 | 0.0573943376799862 | -1.84000056608389 | 0.0657681542095428 | 0.445799443792729 | NM_207234 |
| Sv2c | 1345.056425 | -0.431200026 | 0.12502424 | -3.44893138 | 0.00056281 | 0.007440421 | | Tango6 | 279.844921 | 0.15163246349255 | 0.0823954459653406 | 1.84030150836653 | 0.065723984059135 | 0.445799443792729 | NM_173037 |
| Slc6a11 | 8376.849193 | -0.432399271 | 0.12511868 | -3.455912985 | 0.000548432 | 0.007319044 | | Ttbk2 | 4437.3676 | 0.135849748393549 | 0.0738292983424413 | 1.84005200433356 | 0.0657606027388717 | 0.445799443792729 | NM_001024856,NM_001024857,NM_080788 |
| Sla | 505.497341 | -0.432427188 | 0.08690375 | -4.975932423 | 6.49E-07 | 3.46E-05 | | Kdm4a | 2450.191678 | 0.103144197204733 | 0.056063153842018 | 1.83978585106692 | 0.0657996834943026 | 0.445805131734253 | NM_001161823,NM_172382 |
| Prkar2b | 2750.511227 | -0.432440352 | 0.086429695 | -5.003377067 | 5.63E-07 | 3.10E-05 | | Ngfrap1 | 5366.659253 | 0.110511844182461 | 0.0600769277858874 | 1.83950558484486 | 0.0658408572236547 | 0.44587612682089 | NM_009750,NM_001110234,NM_001110233 |
| Atp5e | 2029.569215 | -0.432453949 | 0.107616115 | -4.018486901 | 5.86E-05 | 0.001303924 | | 1700037C18Rik | 137.7539736 | 0.169889227546152 | 0.0925321944009249 | 1.83600128199763 | 0.0663574670491757 | 0.446279966747306 | NM_028484 |
| Chst15 | 2008.341138 | -0.432548801 | 0.067974309 | -6.363415923 | 1.97E-10 | 3.51E-08 | | Arl3 | 2890.875837 | 0.0952552203473376 | 0.0518347368749178 | 1.83767153245511 | 0.0661108211260511 | 0.446279966747306 | NM_019718 |
| Foxo6 | 392.2482541 | -0.432731942 | 0.112847204 | -3.834671354 | 0.000125732 | 0.002335847 | | Baz2a | 3446.236635 | -0.104798084005681 | 0.0570437369893769 | -1.83715320097624 | 0.0661872821812982 | 0.446279966747306 | NM_054078 |
| Gpr52 | 453.4012127 | -0.433044922 | 0.110652606 | -3.913553761 | 9.09E-05 | 0.001825314 | | Clcn4-2 | 9510.150246 | 0.109621432092465 | 0.0596388308715046 | 1.83808821351061 | 0.0660494076970242 | 0.446279966747306 | NM_011334 |
| Kctd12 | 3117.747668 | -0.43396618 | 0.10364205 | -4.187163237 | 2.82E-05 | 0.000736781 | | Ddx51 | 864.4577872 | -0.12739392427043 | 0.0693368448058113 | -1.83731931597431 | 0.0661627699942773 | 0.446279966747306 | NM_027156 |
| Shisa9 | 2795.824996 | -0.43396713 | 0.093018132 | -4.665403627 | 3.08E-06 | 0.000125258 | | Dusp27 | 26.55567117 | -0.157206404909308 | 0.0855597078142236 | -1.83738828613607 | 0.0661525948478142 | 0.446279966747306 | NM_001033344,NM_001160049 |
| Ddr2 | 446.4424151 | -0.436666849 | 0.100952102 | -4.325485468 | 1.52E-05 | 0.000446615 | | Eif2d | 1148.357056 | -0.105323786846456 | 0.0573665454484272 | -1.83597924579828 | 0.0663607261946895 | 0.446279966747306 | NM_001136070,NM_010709 |
| Myo16 | 1566.656143 | -0.436991914 | 0.137263231 | -3.183605043 | 0.001454533 | 0.015115049 | | Park7 | 3140.114478 | 0.0910596924168434 | 0.0495773603836748 | 1.83671925475944 | 0.0662513512638036 | 0.446279966747306 | NM_020569 |
| 4930426D05Rik | 40.55002337 | -0.437723138 | 0.163424648 | -2.678440138 | 0.007396595 | 0.047350844 | | Pld2 | 993.6330256 | -0.105834567695443 | 0.0575787588453736 | -1.83808351930022 | 0.0660500993013565 | 0.446279966747306 | NM_008876 |
| Alk | 360.9203877 | -0.438310853 | 0.115176759 | -3.80554945 | 0.00014149 | 0.002582146 | | Prpf38b | 1688.819877 | -0.11853839200029 | 0.0645503819446598 | -1.83637011012445 | 0.0663029370791007 | 0.446279966747306 | NM_025845 |
| AI848285 | 105.4950121 | -0.438589757 | 0.149987607 | -2.924173309 | 0.003453724 | 0.027983052 | | Rab11b | 10097.97235 | 0.0682021150013345 | 0.0370895062954839 | 1.83885205853061 | 0.0659369488444464 | 0.446279966747306 | NM_008997 |
| Lrrc9 | 107.9937792 | -0.438954142 | 0.138081648 | -3.178946287 | 0.001478115 | 0.015298537 | | Slc9a2 | 223.4621993 | -0.170855458559061 | 0.0930597886471841 | -1.83597514074335 | 0.0663613333453243 | 0.446279966747306 | NM_001033289 |
| Shc3 | 1198.963897 | -0.439128157 | 0.088359746 | -4.969776145 | 6.70E-07 | 3.53E-05 | | Strap | 4315.975215 | 0.0999164335462423 | 0.0543996274807699 | 1.83671172346837 | 0.0662524636564543 | 0.446279966747306 | NM_011499 |
| Pcsk9 | 238.3146009 | -0.439615476 | 0.130431555 | -3.370468726 | 0.000750404 | 0.009126188 | | Tacr3 | 128.6748117 | 0.174187235115788 | 0.0947362740141052 | 1.83865406285509 | 0.065966084055127 | 0.446279966747306 | NM_021382 |
| Alcam | 6704.339778 | -0.44062504 | 0.078174827 | -5.636405673 | 1.74E-08 | 1.70E-06 | | Tceal1 | 626.8341425 | 0.141182508758429 | 0.0768627036436468 | 1.83681424235327 | 0.0662373226526824 | 0.446279966747306 | NM_146236 |
| Dgkb | 6402.564201 | -0.440645555 | 0.110167197 | -3.999789118 | 6.34E-05 | 0.001377705 | | Bcas3 | 2919.233344 | 0.0919014737725794 | 0.0500738890994946 | 1.83531727663432 | 0.0664586926763118 | 0.446727889780706 | NM_138681,NM_001166642,NM_001166643 |
| Atp6v1c2 | 64.02860352 | -0.44097743 | 0.152744845 | -2.887020045 | 0.003889095 | 0.030510193 | | Dopey1 | 2566.895081 | -0.110069163470966 | 0.0599981310295611 | -1.83454320296636 | 0.0665734008475044 | 0.447108619213183 | NM_177208 |
| Gng13 | 609.5214455 | -0.44235934 | 0.114011452 | -3.87995532 | 0.000104476 | 0.002033551 | | Impa1 | 2621.57463 | 0.115925064276037 | 0.0631909632946055 | 1.8345196564828 | 0.0665768926998564 | 0.447108619213183 | NM_018864 |
| Prok2 | 11.95770798 | -0.442527696 | 0.152307242 | -2.905493458 | 0.003666745 | 0.029209537 | | 2900011O08Rik | 12361.1454 | 0.0921162303146599 | 0.0502281557448651 | 1.83395605410173 | 0.0666605177759615 | 0.447318172986166 | NM_144518 |
| Slc7a2 | 1535.146295 | -0.442693669 | 0.085380956 | -5.184922871 | 2.16E-07 | 1.42E-05 | | Lrrc42 | 693.7852614 | -0.122565896241762 | 0.0668336751257836 | -1.83389430569377 | 0.066669685014116 | 0.447318172986166 | NM_029985 |
| Tnfrsf8 | 59.45853693 | -0.442858344 | 0.154643175 | -2.863743216 | 0.004186673 | 0.032100014 | | Arl6ip1 | 10622.56379 | 0.111227080235061 | 0.0607570124600768 | 1.83068712122979 | 0.0671472564465922 | 0.447688158576222 | NM_019419 |
| Slc7a11 | 2016.721578 | -0.442901646 | 0.082067634 | -5.396788286 | 6.78E-08 | 5.39E-06 | | Atxn10 | 10556.49163 | 0.09342346289799 | 0.0510323787572315 | 1.83067035425527 | 0.0671497605418038 | 0.447688158576222 | NM_016843 |
| Cped1 | 221.5049818 | -0.443426851 | 0.12388598 | -3.579314243 | 0.000344497 | 0.005197309 | | Cyb5b | 8032.945354 | 0.0749320745803995 | 0.0409081965959113 | 1.83171297724449 | 0.0669941942090941 | 0.447688158576222 | NM_025558 |
| Efcab6 | 338.9362617 | -0.444110901 | 0.111687666 | -3.976364773 | 7.00E-05 | 0.001491362 | | Gatsl2 | 4772.356991 | 0.109120396744479 | 0.0595804602169267 | 1.83147958822712 | 0.0670289916151362 | 0.447688158576222 | NM_030719 |
| Trps1 | 1044.161014 | -0.444463009 | 0.099518756 | -4.46612303 | 7.96E-06 | 0.000262449 | | Gpr165 | 438.2791147 | 0.162244717041531 | 0.0886021700744258 | 1.83115963079962 | 0.0670767202175353 | 0.447688158576222 | NM_029536 |
| 9630013A20Rik | 48.92984847 | -0.444707705 | 0.158306451 | -2.809157183 | 0.004967139 | 0.035795744 | | Ints5 | 977.2759659 | 0.122317578730945 | 0.0668038733847374 | 1.83099530810875 | 0.0671012433884513 | 0.447688158576222 | NM_176843 |
| Mgp | 522.789843 | -0.449818358 | 0.114977346 | -3.912234649 | 9.14E-05 | 0.001832942 | | Morf4l1 | 12585.97659 | 0.106158923486167 | 0.0579381849487962 | 1.83227906742309 | 0.0669098541155413 | 0.447688158576222 | NM_001039147,NM_024431 |
| Hey2 | 181.2224759 | -0.450155663 | 0.141726805 | -3.176221064 | 0.001492072 | 0.015401841 | | Ncf4 | 36.28782518 | -0.160778110846909 | 0.0877372078947729 | -1.83249632287977 | 0.0668775090912596 | 0.447688158576222 | NM_008677 |
| Foxp2 | 946.2517522 | -0.450656073 | 0.123285046 | -3.655399317 | 0.000256782 | 0.004114354 | | Pcsk7 | 1115.953367 | -0.109460056846604 | 0.0597874403970007 | -1.83082025455124 | 0.067127376129343 | 0.447688158576222 | NM_008794 |
| Fam126a | 875.7450863 | -0.450834309 | 0.113663263 | -3.966403014 | 7.30E-05 | 0.001538131 | | Pola2 | 251.8079128 | -0.149321792633123 | 0.0815686815967519 | -1.83062653104189 | 0.0671563057646694 | 0.447688158576222 | NM_001164057,NM_008893 |
| Tmem45a | 27.50803503 | -0.450852648 | 0.166069686 | -2.714840125 | 0.006630777 | 0.043659664 | | Rabif | 1304.75897 | 0.109983182142863 | 0.0600185236119417 | 1.83248729765455 | 0.0668788525117805 | 0.447688158576222 | NM_145510 |
| Lum | 82.67209727 | -0.45117631 | 0.139031925 | -3.245127397 | 0.001173981 | 0.012845805 | | Slc39a8 | 138.9324356 | -0.172761025605677 | 0.0943607058124086 | -1.83085770838902 | 0.0671217841565394 | 0.447688158576222 | NM_001135149,NM_001135150,NM_026228 |
| Ntrk1 | 121.6366259 | -0.451497015 | 0.160994552 | -2.804424186 | 0.005040654 | 0.036129802 | | Srp68 | 3038.460287 | 0.0898133502381753 | 0.0490069124465054 | 1.83266698011659 | 0.066852110645607 | 0.447688158576222 | NM_146032 |
| Ccdc80 | 303.8183343 | -0.452397036 | 0.104375668 | -4.334315143 | 1.46E-05 | 0.000435663 | | Zscan2 | 302.1421328 | 0.146438023123724 | 0.0799581299105616 | 1.83143381776844 | 0.067035817559476 | 0.447688158576222 | NM_009553 |
| Aplnr | 10.97508254 | -0.452603225 | 0.164309783 | -2.754572602 | 0.005876885 | 0.040218743 | | Celf6 | 954.836482 | 0.157256112086287 | 0.0859218158596055 | 1.83022333167679 | 0.0672165503133957 | 0.447884224702929 | NM_175235 |
| Cdca3 | 23.50886033 | -0.453112299 | 0.16658306 | -2.720038265 | 0.006527436 | 0.043239925 | | Plod2 | 174.527182 | -0.166920451671614 | 0.0912334432801293 | -1.82959719232662 | 0.0673101938914533 | 0.448302556839638 | NM_001142916,NM_011961 |
| Dcbld1 | 435.4604426 | -0.453384754 | 0.097761674 | -4.637653323 | 3.52E-06 | 0.000139281 | | Banf1 | 1306.28899 | 0.0980294835459051 | 0.0536307909545782 | 1.8278582471202 | 0.0675708287481936 | 0.448803776129977 | NM_001038231,NM_011793 |
| Gng4 | 2627.038902 | -0.454575489 | 0.166667594 | -2.727437754 | 0.00638283 | 0.042645781 | | Lad1 | 84.69601444 | 0.173553480493524 | 0.0948973131988546 | 1.82885557707885 | 0.0674212465724776 | 0.448803776129977 | NM_133664 |
| Arhgap10 | 997.5865014 | -0.454906765 | 0.083015142 | -5.479804684 | 4.26E-08 | 3.62E-06 | | Mthfr | 1618.505898 | -0.130244555639242 | 0.0712549187676535 | -1.82786757590645 | 0.0675694283282502 | 0.448803776129977 | NR_027809,NM_001161798,NM_010840 |
| Zdbf2 | 1074.617536 | -0.454963452 | 0.130089915 | -3.497299928 | 0.000469993 | 0.006519313 | | Pptc7 | 2907.057664 | 0.124881320397408 | 0.0683192355067001 | 1.82790863321591 | 0.067563265165566 | 0.448803776129977 | NM_177242 |
| Adra2c | 780.1633966 | -0.455282537 | 0.09841723 | -4.626045005 | 3.73E-06 | 0.000145462 | | Scg5 | 7806.697272 | 0.110508130373412 | 0.0604327882748378 | 1.82861214132369 | 0.0674577325481207 | 0.448803776129977 | NM_009162 |
| Kcnn3 | 1048.614526 | -0.456094211 | 0.099884443 | -4.566218681 | 4.97E-06 | 0.000184075 | | Tmod2 | 22410.72088 | 0.106719066194526 | 0.0583793794655948 | 1.82802673086683 | 0.067545539961986 | 0.448803776129977 | NM_001038710,NM_016711 |
| AI504432 | 1115.839049 | -0.457149223 | 0.103519501 | -4.416068646 | 1.01E-05 | 0.000314613 | | Atp5c1 | 7568.670092 | 0.0963363968366425 | 0.0527207269188823 | 1.82729644424039 | 0.0676552095780666 | 0.449158854813069 | NM_020615,NM_001112738 |
| Fbln1 | 304.2890215 | -0.458543731 | 0.138040043 | -3.321816781 | 0.000894334 | 0.010481817 | | Notum | 98.52384054 | -0.172698948551561 | 0.0945435165002823 | -1.82666093820453 | 0.0677507648368368 | 0.449587761544034 | NM_175263 |
| Mei4 | 39.15728793 | -0.458646448 | 0.163790126 | -2.800208171 | 0.005106966 | 0.036330272 | | Ist1 | 3595.0028 | -0.0922868665844338 | 0.0505333619473356 | -1.82625622020978 | 0.0678116764414085 | 0.449662797420469 | NM_028018 |
| Kif27 | 203.4189338 | -0.459781272 | 0.098654765 | -4.660507489 | 3.15E-06 | 0.000127939 | | Trim16 | 90.07251888 | 0.173027855176663 | 0.0947488078986292 | 1.82617448191838 | 0.0678239838323177 | 0.449662797420469 | NM_053169 |
| Artn | 20.6351935 | -0.459889477 | 0.166615661 | -2.760181571 | 0.005776924 | 0.039763513 | | Aph1b | 1189.514552 | 0.113646265783073 | 0.0622724863185125 | 1.82498359230099 | 0.0680035053828293 | 0.450415406447951 | NM_177583 |
| Dmkn | 119.8857115 | -0.460134067 | 0.134827447 | -3.412762587 | 0.000643079 | 0.008180519 | | Fam69b | 1946.793305 | 0.114911977117816 | 0.0629681886361782 | 1.82492111662545 | 0.0680129341031961 | 0.450415406447951 | NM_019833 |
| Egln3 | 461.6554543 | -0.460395318 | 0.090483754 | -5.088154495 | 3.62E-07 | 2.15E-05 | | Syne4 | 109.7673514 | -0.167040893794751 | 0.0915390588977772 | -1.82480457857107 | 0.0680305246968749 | 0.450415406447951 | NM_153577 |
| Map2 | 26102.54727 | -0.460995134 | 0.059517995 | -7.745474847 | 9.52E-15 | 4.76E-12 | | Pcbd1 | 155.3952881 | 0.172139524613878 | 0.0943700251834965 | 1.82409111663543 | 0.0681382983053474 | 0.450923426507278 | NM_025273 |
| Tcfl5 | 24.02995482 | -0.461004371 | 0.168119905 | -2.742116528 | 0.006104468 | 0.041300088 | | Dennd4b | 2585.307737 | -0.116496620810029 | 0.0639013117827213 | -1.82307088164549 | 0.0682926560892728 | 0.451155090641708 | NM_201407 |
| Ccdc109b | 64.13640588 | -0.46163123 | 0.158658664 | -2.909587278 | 0.003619063 | 0.028933832 | | Fam92a | 1493.784776 | 0.111921764379216 | 0.0613819882115487 | 1.82336492577409 | 0.0682481388396138 | 0.451155090641708 | NM_026558 |
| Col3a1 | 152.2011881 | -0.46220536 | 0.152597807 | -3.028912202 | 0.00245436 | 0.02211392 | | Moxd1 | 415.1310954 | 0.137127704576042 | 0.0752193077064001 | 1.82303864203704 | 0.0682975385048269 | 0.451155090641708 | NM_021509 |
| Pcdhb22 | 225.5714205 | -0.463201053 | 0.09985626 | -4.638678149 | 3.51E-06 | 0.000138949 | | Myh9 | 3925.928972 | -0.12738654443765 | 0.0698562904840506 | -1.82355151633387 | 0.0682199020589988 | 0.451155090641708 | NM_022410 |
| Slc26a7 | 33.00970186 | -0.463455945 | 0.167154853 | -2.77261435 | 0.005560798 | 0.038738759 | | Ttc9b | 2568.618704 | -0.115100242515036 | 0.06314421094734 | -1.82281543768162 | 0.0683313487848762 | 0.451173260204142 | NM_028417 |
| Clec3b | 43.15523841 | -0.464923471 | 0.151398503 | -3.070859105 | 0.002134438 | 0.020096898 | | Dnajb12 | 1423.930332 | 0.100568815791269 | 0.055182335043096 | 1.82248206265152 | 0.0683818729883225 | 0.451301720594445 | NM_019965 |
| 2810011L19Rik | 576.5191333 | -0.464945404 | 0.132981853 | -3.496307144 | 0.000471745 | 0.006531924 | | Ociad1 | 11536.28788 | 0.0877920893498366 | 0.0481783219158831 | 1.82223219627601 | 0.0684197612814822 | 0.45134670861708 | NM_001159888,NM_001159889,NM_001159887,NM_023429 |
| Cdca7 | 149.4369424 | -0.465055668 | 0.139861983 | -3.325104207 | 0.000883855 | 0.010382448 | | Cct6a | 5553.718965 | 0.09500520953282 | 0.0521564203721215 | 1.82154390303216 | 0.0685242193396982 | 0.451830599241242 | NM_009838 |
| 2610034M16Rik | 150.2699268 | -0.466483812 | 0.130439618 | -3.576243311 | 0.000348567 | 0.005248495 | | Panx2 | 4005.646339 | 0.0882782340258585 | 0.0484754450938993 | 1.82109176831403 | 0.0685929083707627 | 0.452078306258484 | NM_001002005 |
| Pdgfc | 331.1595035 | -0.467951917 | 0.091213411 | -5.130297329 | 2.89E-07 | 1.80E-05 | | Gfod2 | 764.6225987 | 0.111020938205045 | 0.0609866853312466 | 1.82041272782804 | 0.0686961755981462 | 0.45255358128738 | NM_027469 |
| Gm10754 | 95.65462875 | -0.46856232 | 0.141257437 | -3.317080716 | 0.000909633 | 0.010547482 | | Pnpla2 | 951.4351372 | 0.149283358246259 | 0.0820262242262077 | 1.81994672623932 | 0.0687671181621744 | 0.452815574988099 | NM_025802,NR_028142,NM_001163689 |
| Dok4 | 686.064647 | -0.468600865 | 0.076086053 | -6.158827366 | 7.33E-10 | 1.05E-07 | | Alg9 | 840.4244159 | 0.112153875426174 | 0.0616320112658464 | 1.8197341466337 | 0.0687995005787616 | 0.452823536659307 | NM_133981,NR_045601 |
| Sox4 | 436.3485372 | -0.469625086 | 0.123897127 | -3.790443718 | 0.000150378 | 0.002721919 | | Acat3 | 168.0958657 | -0.168663194476341 | 0.0927287688691052 | -1.8188874556765 | 0.0689286020354626 | 0.453262504828941 | NM_153151 |
| Rgs2 | 2708.104062 | -0.46972086 | 0.083137382 | -5.649935656 | 1.61E-08 | 1.60E-06 | | Pank4 | 1580.248206 | -0.122699354224814 | 0.0674541223756881 | -1.8190045308341 | 0.0689107388398216 | 0.453262504828941 | NM_172990 |
| Csmd3 | 1312.055909 | -0.470074532 | 0.108601877 | -4.328419938 | 1.50E-05 | 0.000443226 | | Mrap2 | 433.3509991 | 0.170568847341609 | 0.0937925119042003 | 1.81857638609602 | 0.0689760831657597 | 0.453369495052409 | NM_001177731,NM_001101482 |
| Casq2 | 108.8846149 | -0.471048162 | 0.153932084 | -3.060103845 | 0.002212603 | 0.020626994 | | Vwa3a | 317.3566456 | -0.161345640890755 | 0.0887474727999962 | -1.81803082161414 | 0.0690594220871218 | 0.453711969804401 | NM_177697 |
| Gpc6 | 541.3085926 | -0.471154641 | 0.092917191 | -5.070694006 | 3.96E-07 | 2.32E-05 | | Arrdc1 | 342.0227425 | -0.155909671908905 | 0.0858286428401364 | -1.81652262636031 | 0.0692902403219714 | 0.454875446195876 | NM_001162485,NR_027876,NM_178408 |
| Fam102b | 2977.074862 | -0.471578725 | 0.106390595 | -4.432522658 | 9.31E-06 | 0.000296317 | | Ccnd2 | 3005.335572 | 0.151521510120396 | 0.0834156171967948 | 1.81646453280954 | 0.0692991437719588 | 0.454875446195876 | NM_009829 |
| Wwc1 | 3706.947063 | -0.472068145 | 0.07617525 | -6.197132839 | 5.75E-10 | 8.73E-08 | | Mapre2 | 15257.26421 | 0.0918848238574725 | 0.05059949536505 | 1.81592372008 | 0.0693820741291592 | 0.45498512424667 | NM_153058,NM_001162941,NM_001162942 |
| Tox | 707.9212671 | -0.472174658 | 0.0884043 | -5.341082457 | 9.24E-08 | 7.09E-06 | | Nxpe3 | 1212.16638 | 0.102268397273778 | 0.0563231705509771 | 1.81574290426738 | 0.0694098193123104 | 0.45498512424667 | NM_001134457 |
| Hmgcr | 4416.600257 | -0.47244834 | 0.095628936 | -4.940432896 | 7.79E-07 | 4.04E-05 | | Rab7 | 5146.013305 | 0.0993269045226862 | 0.0546909268792814 | 1.81614959172166 | 0.069347428174675 | 0.45498512424667 | NM_009005 |
| Ccdc3 | 807.4774521 | -0.472896806 | 0.113035946 | -4.183596657 | 2.87E-05 | 0.00074593 | | Ap2m1 | 22677.83199 | 0.0831546958842399 | 0.0458188655058328 | 1.8148571547162 | 0.0695458643323817 | 0.455055506888368 | NM_009679 |
| Fam19a4 | 22.10397271 | -0.473192585 | 0.168066973 | -2.815500131 | 0.004870138 | 0.035393018 | | Clcn5 | 290.7588735 | 0.16587498725423 | 0.0913858728486576 | 1.81510535582378 | 0.0695077202927675 | 0.455055506888368 | NM_016691,NM_001243762 |
| Stk32a | 356.039342 | -0.474526001 | 0.109599209 | -4.329648035 | 1.49E-05 | 0.000441602 | | Naglu | 402.8857193 | -0.158487023151243 | 0.0873188308403427 | -1.81503830990393 | 0.0695180223491506 | 0.455055506888368 | NM_013792 |
| Rspo2 | 614.9985749 | -0.474531863 | 0.129848705 | -3.65449825 | 0.000257685 | 0.004124564 | | Zcchc11 | 1449.433703 | -0.107482946829355 | 0.059207014296256 | -1.81537522381282 | 0.0694662659391367 | 0.455055506888368 | NM_175472 |
| Cdca5 | 8.941967987 | -0.474538934 | 0.160964518 | -2.948096503 | 0.003197372 | 0.026528802 | | Pvrl3 | 603.7438764 | -0.163683149598943 | 0.0902007936409461 | -1.81465309773772 | 0.0695772370869822 | 0.455055806359975 | NM_021495,NM_021497,NM_021496 |
| Ust | 1081.167932 | -0.475240707 | 0.097621669 | -4.868188722 | 1.13E-06 | 5.37E-05 | | Cep152 | 166.8543505 | -0.162803364829481 | 0.0897346306412435 | -1.81427575581566 | 0.0696352821591096 | 0.455184690642753 | NM_001081091 |
| Kcnf1 | 6721.608509 | -0.476051288 | 0.067337726 | -7.069607413 | 1.55E-12 | 4.38E-10 | | Dpp7 | 932.6501703 | -0.155240427987162 | 0.0855735172941243 | -1.81411764872988 | 0.0696596149868402 | 0.455184690642753 | NM_031843 |
| St3gal1 | 1253.639761 | -0.476123517 | 0.070666416 | -6.737620794 | 1.61E-11 | 3.78E-09 | | 1700001L05Rik | 276.5574763 | -0.17172939824233 | 0.0946853496806875 | -1.81368499795863 | 0.0697262360325625 | 0.455201065941378 | NR_027980 |
| Fjx1 | 2017.784391 | -0.476842571 | 0.123456393 | -3.862437249 | 0.000112261 | 0.002134206 | | Asns | 4199.982374 | 0.093377948756584 | 0.0515247677234027 | 1.81229247375016 | 0.0699410986820619 | 0.455201065941378 | NM_012055 |
| Dst | 15807.14299 | -0.477766466 | 0.052654442 | -9.073621226 | 1.15E-19 | 1.11E-16 | | Atxn7l3b | 8682.103927 | 0.0982650012330765 | 0.0541938847598047 | 1.81321198265453 | 0.069799132391994 | 0.455201065941378 | NM_001033474 |
| Steap3 | 225.1031783 | -0.477927192 | 0.153256027 | -3.118488718 | 0.001817811 | 0.017803517 | | Cd44 | 157.9527791 | -0.169316190264836 | 0.0933764047606853 | -1.81326525366635 | 0.0697909196748885 | 0.455201065941378 | NM_001039151,NM_001039150,NM_001177785,NM_001177787,NM_001177786,NM_009851 |
| Pde3a | 113.374785 | -0.478911227 | 0.15777332 | -3.035438618 | 0.002401861 | 0.021823847 | | Edf1 | 2775.077818 | 0.10008965182816 | 0.0552350025431578 | 1.81206928975797 | 0.0699754908610146 | 0.455201065941378 | NM_021519 |
| Aoc3 | 43.96556511 | -0.479526232 | 0.163358724 | -2.935418565 | 0.00333098 | 0.027292548 | | Gpr179 | 35.56088188 | 0.140586796791346 | 0.0775340687635137 | 1.81322609574571 | 0.0697969565192955 | 0.455201065941378 | NM_001081220 |
| Chat | 302.373814 | -0.480156554 | 0.140264511 | -3.423221949 | 0.000618835 | 0.007990195 | | Hivep3 | 1921.721076 | 0.147675798504957 | 0.0814902335134578 | 1.81219015013093 | 0.0699568204956946 | 0.455201065941378 | NM_010657 |
| Macf1 | 14882.16153 | -0.480417915 | 0.077097708 | -6.231286622 | 4.63E-10 | 7.10E-08 | | Plscr3 | 645.1219281 | -0.136082142554274 | 0.0750578261835199 | -1.81303069216877 | 0.0698270876358749 | 0.455201065941378 | NM_001168497,NM_023564,NR_033133 |
| Hes5 | 357.7953987 | -0.480614976 | 0.137200044 | -3.503023495 | 0.000460009 | 0.006432651 | | Tarbp2 | 686.8695132 | -0.150408732194517 | 0.0830037394021246 | -1.8120717605967 | 0.0699751091278701 | 0.455201065941378 | NM_001253795,NM_009319 |
| Acta2 | 546.0208198 | -0.480993306 | 0.13431229 | -3.581156325 | 0.000342077 | 0.005171348 | | Zswim3 | 383.3939188 | 0.150544316096109 | 0.0830649242449551 | 1.81236926975531 | 0.0699291578311921 | 0.455201065941378 | NM_178375 |
| Lhfpl5 | 123.7034971 | -0.481057187 | 0.140140875 | -3.432668647 | 0.000597672 | 0.007794891 | | Cisd3 | 677.1743188 | 0.110755361011338 | 0.0611549010536932 | 1.81106271293116 | 0.0701311455547906 | 0.456009402776584 | NM_001085500 |
| 2810417H13Rik | 14.22484206 | -0.481435825 | 0.155255465 | -3.100926759 | 0.00192916 | 0.01865818 | | Armcx2 | 4223.581534 | 0.0890331239450795 | 0.0492076073058764 | 1.80933658065604 | 0.0703987292288796 | 0.45601700931274 | NM_001166397,NM_001166398,NM_026139 |
| Gpx6 | 40.98740255 | -0.481960137 | 0.168030922 | -2.868282411 | 0.00412707 | 0.031734401 | | Atp5f1 | 7781.49038 | 0.097245485869673 | 0.0537519022182401 | 1.80915431559693 | 0.0704270327466867 | 0.45601700931274 | NM_009725 |
| Scgn | 60.29875569 | -0.482377882 | 0.130562632 | -3.694609063 | 0.000220225 | 0.003633836 | | Ctnna2 | 6788.090262 | 0.0893901706306234 | 0.0493905190482675 | 1.80986497718855 | 0.0703167284951896 | 0.45601700931274 | NM_009819,NM_001109764,NM_145732 |
| Mis18bp1 | 10.88304371 | -0.482902521 | 0.163821547 | -2.947735075 | 0.003201113 | 0.026528802 | | Fibin | 98.75554232 | 0.171114368419836 | 0.094573670107009 | 1.80932354878712 | 0.0704007526077424 | 0.45601700931274 | NM_026271 |
| Tlr5 | 20.9814671 | -0.485553471 | 0.167894193 | -2.89202064 | 0.003827728 | 0.030212337 | | Fubp3 | 1318.454189 | -0.128982712016873 | 0.0713073165848457 | -1.80882857740694 | 0.0704776391234408 | 0.45601700931274 | NM_001033389 |
| Ptprb | 2208.873738 | -0.485830172 | 0.112549699 | -4.316583484 | 1.58E-05 | 0.000464125 | | Nfkbia | 986.4424724 | 0.162138806807507 | 0.0895997324041894 | 1.80959030185592 | 0.0703593449900429 | 0.45601700931274 | NM_010907 |
| Zic4 | 115.8168944 | -0.486301673 | 0.165587854 | -2.936819704 | 0.003315968 | 0.027241576 | | Pigb | 310.1499431 | -0.137577971957064 | 0.0760384274207582 | -1.80932163675318 | 0.070401049481614 | 0.45601700931274 | NM_018889 |
| Pcdh10 | 7770.362955 | -0.486566448 | 0.063181285 | -7.701116683 | 1.35E-14 | 6.33E-12 | | Ppl | 96.16915691 | -0.172177868955964 | 0.0951016203551302 | -1.81046199121544 | 0.0702241734667075 | 0.45601700931274 | NM_008909 |
| Celsr1 | 381.849956 | -0.487017468 | 0.102295997 | -4.760865371 | 1.93E-06 | 8.63E-05 | | Ptpdc1 | 1359.416188 | 0.109556179681697 | 0.0605420940546869 | 1.80958688978839 | 0.070359874513204 | 0.45601700931274 | NM_207232 |
| Tyrp1 | 9.268890722 | -0.488700192 | 0.139250044 | -3.509515538 | 0.000448924 | 0.006323296 | | Ramp3 | 491.3450762 | 0.161019060186856 | 0.0889861199200708 | 1.80948512342696 | 0.0703756692691118 | 0.45601700931274 | NM_019511 |
| Grm4 | 833.9925681 | -0.488740893 | 0.091609711 | -5.335033688 | 9.55E-08 | 7.22E-06 | | Ssr1 | 6041.725923 | 0.0895030917065608 | 0.0494781461507506 | 1.80894190000292 | 0.0704600300496013 | 0.45601700931274 | NM_025965 |
| Lgr5 | 199.6824086 | -0.488976081 | 0.164818895 | -2.966747724 | 0.003009678 | 0.025407408 | | 1700037H04Rik | 2126.14505 | 0.10741971020325 | 0.0594197160146738 | 1.80781258154654 | 0.0706356749313403 | 0.456632761038117 | NM_026091 |
| Adamtsl3 | 71.63363357 | -0.489227947 | 0.164556529 | -2.973008418 | 0.002948964 | 0.025077521 | | Ppard | 1146.326812 | 0.117147026152545 | 0.0647950647350359 | 1.80796217476732 | 0.0706123878207896 | 0.456632761038117 | NM_011145 |
| Tln2 | 6148.558021 | -0.489390087 | 0.055067926 | -8.887026012 | 6.28E-19 | 5.40E-16 | | Grn | 3077.845787 | -0.139476643540246 | 0.0772049569386409 | -1.80657627529145 | 0.0708283714149285 | 0.457674787888457 | NM_008175 |
| Bcl11b | 4015.449145 | -0.48964874 | 0.083071657 | -5.894293619 | 3.76E-09 | 4.63E-07 | | AI480653 | 1258.920962 | 0.104397134709815 | 0.0578356951710962 | 1.80506405950469 | 0.0710646583485543 | 0.458300422397288 | NM_198626 |
| Sox1 | 622.5741073 | -0.489797483 | 0.134878784 | -3.631390126 | 0.000281899 | 0.004420787 | | Espl1 | 53.35582345 | -0.166821860283104 | 0.0924315390113816 | -1.80481534839058 | 0.071103581803521 | 0.458300422397288 | NM_001014976 |
| Map1b | 39800.8502 | -0.491352995 | 0.070977794 | -6.922629834 | 4.43E-12 | 1.21E-09 | | Fam5c | 1664.899203 | 0.11788904437508 | 0.0653202601781334 | 1.80478528489611 | 0.0711082879442011 | 0.458300422397288 | NM_001145807,NM_153539 |
| Isoc1 | 852.470721 | -0.491575813 | 0.141160478 | -3.482389819 | 0.00049696 | 0.006796205 | | Mir143hg | 33.55061287 | -0.161892757444072 | 0.0897039195658067 | -1.80474563684264 | 0.071114494842591 | 0.458300422397288 | NR_045402 |
| Gnal | 15366.84569 | -0.492662101 | 0.087409768 | -5.63623623 | 1.74E-08 | 1.70E-06 | | Prkacb | 24312.29171 | 0.114110538779478 | 0.0632047931321449 | 1.80540957615196 | 0.0710106137556914 | 0.458300422397288 | NM_001164198,NM_001164200,NM_011100,NM_001164199 |
| Fstl5 | 873.7978585 | -0.49283804 | 0.125740178 | -3.919495332 | 8.87E-05 | 0.001787846 | | Zfp40 | 675.4366362 | -0.133585509739419 | 0.0739904761220623 | -1.80544195335414 | 0.0710055511460488 | 0.458300422397288 | NM_009555 |
| Chst3 | 145.9086893 | -0.493034599 | 0.135474847 | -3.639307305 | 0.000273372 | 0.004326488 | | Crim1 | 2308.182542 | 0.1377755027964 | 0.0763975797032678 | 1.80340140789181 | 0.0713251966790902 | 0.45917408885104 | NM_015800 |
| Dnahc8 | 21.32577821 | -0.493411844 | 0.16812101 | -2.934861286 | 0.003336968 | 0.027312724 | | Decr1 | 819.737038 | -0.11318854475703 | 0.0627693429867559 | -1.80324565100055 | 0.0713496439154211 | 0.45917408885104 | NM_026172 |
| Khdrbs2 | 187.6412855 | -0.495382893 | 0.106001766 | -4.673345669 | 2.96E-06 | 0.000122111 | | Mif | 3916.093695 | 0.103459275555228 | 0.0573793658085677 | 1.80307457388767 | 0.071376503691701 | 0.45917408885104 | NM_010798 |
| Chrna5 | 101.8248127 | -0.495730856 | 0.150933333 | -3.28443589 | 0.001021867 | 0.011539952 | | Tsc2 | 5898.314783 | -0.102669637710769 | 0.056933490772826 | -1.8033258863476 | 0.0713370495059817 | 0.45917408885104 | NM_001039363,NM_011647 |
| Fzd5 | 126.8742834 | -0.497164192 | 0.134494163 | -3.696548455 | 0.000218551 | 0.003621628 | | Cebpb | 68.19415658 | -0.171189959789565 | 0.0950206810372752 | -1.80160737558185 | 0.0716072000319151 | 0.459800193456028 | NM_009883 |
| Gabrg3 | 247.3246116 | -0.497234577 | 0.099394789 | -5.002622211 | 5.66E-07 | 3.10E-05 | | Kcnk13 | 415.2137043 | 0.14316492215985 | 0.0794666797979269 | 1.80157170934912 | 0.0716128156439768 | 0.459800193456028 | NM_146037,NM_001164426,NM_001164427 |
| Prc1 | 48.80920691 | -0.49727722 | 0.163712073 | -3.037510981 | 0.002385407 | 0.021753678 | | Mid1 | 398.7893642 | 0.12132938904056 | 0.0673421201175976 | 1.80168650509794 | 0.0715947424583879 | 0.459800193456028 | NM_010797,NM_183151 |
| Cgref1 | 771.3410501 | -0.49743962 | 0.061802736 | -8.048828404 | 8.36E-16 | 5.18E-13 | | Spata1 | 34.03759494 | -0.152127292461512 | 0.0844471699698633 | -1.8014492672259 | 0.0716320967775706 | 0.459800193456028 | NM_027617 |
| E130309F12Rik | 754.8683557 | -0.498024044 | 0.123386833 | -4.036281931 | 5.43E-05 | 0.001230118 | | Taf12 | 515.9449681 | 0.133432980358866 | 0.0740622466943492 | 1.80163290089668 | 0.0716031813185843 | 0.459800193456028 | NM_025579 |
| Adcy5 | 13613.71904 | -0.498463838 | 0.082429008 | -6.047189559 | 1.47E-09 | 1.90E-07 | | Cep135 | 428.4065888 | -0.141515315091512 | 0.0785929066892518 | -1.80061179886181 | 0.0717640881699508 | 0.460037574914698 | NM_199032 |
| Dsel | 630.5473734 | -0.499066489 | 0.106674023 | -4.678425678 | 2.89E-06 | 0.000119762 | | Gorasp2 | 4209.114924 | 0.0891852706121533 | 0.0495243084857407 | 1.80083828202936 | 0.0717283730565691 | 0.460037574914698 | NR_027343,NM_027352 |
| Lmo4 | 6647.496987 | -0.501948986 | 0.085117901 | -5.897102487 | 3.70E-09 | 4.59E-07 | | Tsnax | 9032.506058 | 0.0963931642639089 | 0.053530079985662 | 1.80072894136769 | 0.0717456136413611 | 0.460037574914698 | NM_016909 |
| Gm7244 | 11.75286263 | -0.50199937 | 0.162214113 | -3.09467137 | 0.001970311 | 0.018926221 | | Thoc3 | 1164.185172 | 0.11110184343897 | 0.0617180292062945 | 1.80015215760712 | 0.0718366157814355 | 0.460196881222947 | NM_028597 |
| Fam64a | 10.3431049 | -0.503334519 | 0.159811692 | -3.149547516 | 0.001635235 | 0.016452164 | | Zfp661 | 400.365089 | -0.124083288780612 | 0.068933138315434 | -1.80005280207632 | 0.0718523011574862 | 0.460196881222947 | NM_001111029,NM_028141 |
| Hs3st5 | 247.2097574 | -0.503862373 | 0.128444998 | -3.922787033 | 8.75E-05 | 0.001768185 | | Hras1 | 4073.663734 | 0.0797574353094391 | 0.0443298935933169 | 1.79917948915319 | 0.0719902928549543 | 0.460419542077244 | NM_001130443,NM_008284,NM_001130444 |
| Abcc12 | 48.0976981 | -0.503892538 | 0.156422549 | -3.221354846 | 0.001275861 | 0.013689999 | | Ppp2r1a | 24334.62988 | 0.0780846867965765 | 0.0434037585056887 | 1.79903053295125 | 0.0720138509981755 | 0.460419542077244 | NM_016891 |
| Cdyl2 | 1144.80427 | -0.505191515 | 0.091465725 | -5.523287732 | 3.33E-08 | 2.96E-06 | | Rbmx | 2489.05226 | -0.106166500824499 | 0.059010529350624 | -1.79911114156743 | 0.0720011015721437 | 0.460419542077244 | NM_011252,NR_029425,NM_001166623 |
| Slc22a2 | 32.02283487 | -0.505432554 | 0.167308239 | -3.020966318 | 0.002519694 | 0.02250152 | | Zdhhc8 | 5578.868938 | -0.121838732495417 | 0.0677064180189806 | -1.79951526103864 | 0.0719372120599516 | 0.460419542077244 | NM_172151 |
| Des | 85.14853111 | -0.506622969 | 0.144284167 | -3.511285955 | 0.000445944 | 0.006298508 | | Smoc2 | 551.9565571 | -0.170333154195494 | 0.0947039309662106 | -1.79858589245115 | 0.072084210600192 | 0.460666627003251 | NM_022315 |
| Zic3 | 127.3222856 | -0.507331392 | 0.141520435 | -3.584863146 | 0.000337255 | 0.005132306 | | Armcx3 | 4059.873981 | 0.103067796973558 | 0.0573464176369928 | 1.79728396682047 | 0.0722905502303483 | 0.461579135229028 | NM_027870 |
| Kcnh8 | 27.42040681 | -0.508852299 | 0.168121452 | -3.026692844 | 0.002472451 | 0.022169068 | | Myo10 | 1681.414871 | -0.103554520487371 | 0.0576133857059692 | -1.79740383625193 | 0.0722715321679962 | 0.461579135229028 | NM_019472 |
| Mest | 1061.817526 | -0.510000942 | 0.093028309 | -5.482212309 | 4.20E-08 | 3.60E-06 | | Kif3c | 12909.76968 | 0.0694091213361969 | 0.0386362151621626 | 1.79647827937792 | 0.0724184841274026 | 0.462192838503801 | NM_008445 |
| Reln | 3254.117389 | -0.510583866 | 0.10675359 | -4.782826174 | 1.73E-06 | 7.88E-05 | | Sdhb | 4338.001759 | 0.087863415476675 | 0.0489144494293049 | 1.7962670847121 | 0.0724520500565841 | 0.462203987317497 | NM_023374 |
| 1700001L19Rik | 365.364897 | -0.511723503 | 0.106732134 | -4.794465192 | 1.63E-06 | 7.50E-05 | | Leo1 | 933.673296 | 0.109130001197708 | 0.0607823194400166 | 1.79542131548745 | 0.0725865988697618 | 0.462655961027714 | NM_001039522 |
| Fxyd2 | 90.91497288 | -0.513027388 | 0.142391264 | -3.602941464 | 0.000314636 | 0.004850723 | | Plekha8 | 829.4381955 | -0.128189767324588 | 0.0713943196080039 | -1.79551773906417 | 0.0725712490461941 | 0.462655961027714 | NM_001001335,NM_001164361 |
| AI606473 | 41.40187103 | -0.513742882 | 0.162320478 | -3.164991187 | 0.001550876 | 0.015892381 | | Rock1 | 1115.86303 | -0.131715823757142 | 0.0733877503834167 | -1.79479304201299 | 0.0726866798171946 | 0.463090662730074 | NM_009071 |
| Serpinh1 | 598.5418462 | -0.513822388 | 0.0920663 | -5.581004015 | 2.39E-08 | 2.23E-06 | | Tubb5 | 17841.21834 | 0.0898981363068189 | 0.0501015064357179 | 1.7943200255298 | 0.0727621035759069 | 0.46336795990514 | NM_011655 |
| Kif26b | 346.9705902 | -0.513883098 | 0.08442379 | -6.086946563 | 1.15E-09 | 1.54E-07 | | Gskip | 1316.288008 | 0.118489286735787 | 0.0660494399348386 | 1.79394839460687 | 0.0728214060541875 | 0.463542394541248 | NM_178613 |
| Alx4 | 94.55830759 | -0.513895215 | 0.162403315 | -3.164314823 | 0.001554485 | 0.01589782 | | Gad1 | 20625.89624 | 0.110438485852724 | 0.0615760728592711 | 1.79352921881078 | 0.0728883429018873 | 0.463573784267297 | NM_008077 |
| E130008D07Rik | 184.7770836 | -0.514051321 | 0.137886041 | -3.728088198 | 0.000192938 | 0.003311243 | | Tmem55a | 1522.057584 | 0.110922332794037 | 0.0618462396325094 | 1.79351781859557 | 0.0728901640690146 | 0.463573784267297 | NM_028264 |
| Agbl2 | 73.2121449 | -0.515412708 | 0.16243905 | -3.172960617 | 0.00150893 | 0.015534459 | | Faim2 | 29969.82156 | 0.0928490943621978 | 0.0517862328447645 | 1.79293007546087 | 0.0729841056043126 | 0.463821103580248 | NM_028224,NM_001038658 |
| Gm19522 | 49.98368768 | -0.51541897 | 0.156610782 | -3.291082293 | 0.000998027 | 0.011345145 | | Nomo1 | 6213.586004 | 0.0908212257554318 | 0.0506567525937954 | 1.79287500886024 | 0.0729929122115136 | 0.463821103580248 | NM_153057 |
| Olfml2a | 107.5034229 | -0.516083838 | 0.145312584 | -3.551542635 | 0.00038298 | 0.005651325 | | Sez6l2 | 19002.18718 | 0.0863625745728964 | 0.0481779752341331 | 1.79257376743617 | 0.0730411040801517 | 0.463924389098506 | NM_144926,NM_001252567,NM_001252566 |
| Cntn5 | 300.0182222 | -0.517378742 | 0.097436637 | -5.309899434 | 1.10E-07 | 8.02E-06 | | Extl3 | 9215.509741 | 0.0880575647720878 | 0.0491306307551247 | 1.79231496560631 | 0.0730825273535869 | 0.463984612035928 | NM_018788 |
| Gm5607 | 604.0169666 | -0.518529593 | 0.149627345 | -3.465473462 | 0.000529299 | 0.007115035 | | Hcrtr2 | 81.84969887 | 0.170142566094146 | 0.094971479032715 | 1.79151222900864 | 0.0732111339589248 | 0.464395166763031 | NM_198962 |
| Dcaf12l2 | 37.56002595 | -0.519668939 | 0.164844884 | -3.15247235 | 0.001618942 | 0.016318843 | | Itga1 | 292.2322211 | -0.152635367396693 | 0.0851941977357625 | -1.79161693464272 | 0.0731943485540551 | 0.464395166763031 | NM_001033228 |
| Tnnc1 | 238.9691526 | -0.520337808 | 0.150507367 | -3.457224837 | 0.000545769 | 0.007296074 | | Nin | 2215.790977 | -0.144416989570565 | 0.0806270081672821 | -1.79117386162877 | 0.0732653993106111 | 0.464536530068065 | NM_001081453,NM_008697 |
| Pcsk5 | 504.2211175 | -0.520831141 | 0.088730841 | -5.869787019 | 4.36E-09 | 5.32E-07 | | Alox5ap | 130.837655 | 0.167821028750101 | 0.0937156862056145 | 1.79074641124537 | 0.073333998298369 | 0.464565922059155 | NM_009663 |
| Klhl13 | 2168.725409 | -0.520965248 | 0.116114226 | -4.48666167 | 7.23E-06 | 0.00024475 | | Ccdc160 | 85.89560752 | 0.168844395828142 | 0.0942850576334722 | 1.79078636706694 | 0.0733275837993152 | 0.464565922059155 | NM_001034059 |
| Neb | 86.64729108 | -0.523143601 | 0.159120429 | -3.287721153 | 0.001010018 | 0.011447859 | | Nfu1 | 1223.426483 | 0.114321990842861 | 0.0638529296867423 | 1.79039538833561 | 0.0733903711718314 | 0.464720371247613 | NM_020045,NM_001170591 |
| Mrc1 | 208.9341952 | -0.524572285 | 0.147747236 | -3.550471063 | 0.000384542 | 0.005668982 | | Ninl | 318.5706394 | -0.151976554223361 | 0.0849281922496599 | -1.78947120146632 | 0.0735389614067685 | 0.465458367492252 | NM_207204 |
| P2ry1 | 208.3984836 | -0.526538526 | 0.150096264 | -3.508005537 | 0.00045148 | 0.006347755 | | Heca | 991.8241885 | 0.140894816507188 | 0.0787460160860234 | 1.78923104317141 | 0.0735776141674889 | 0.465500184406335 | NM_001033432 |
| Apc | 14761.67817 | -0.526857181 | 0.071061431 | -7.414108793 | 1.22E-13 | 4.86E-11 | | Galnt15 | 130.5742503 | 0.170051361397203 | 0.0950826352894768 | 1.78845864841131 | 0.0737020414572733 | 0.465836524871565 | NM_030166 |
| Cdhr1 | 1095.455868 | -0.52730108 | 0.149507297 | -3.526925383 | 0.000420415 | 0.006030802 | | Nfat5 | 3803.664495 | -0.118131354677991 | 0.0660577611227884 | -1.78830394294484 | 0.0737269840754322 | 0.465836524871565 | NM_018823,NM_133957 |
| Slc10a4 | 185.0474056 | -0.527329775 | 0.161383949 | -3.267547836 | 0.001084835 | 0.012171208 | | Supt7l | 765.0733134 | 0.117604666347571 | 0.0657589824348401 | 1.78841980202635 | 0.0737083038744284 | 0.465836524871565 | NM_028150 |
| Loxl1 | 85.82401008 | -0.527388471 | 0.152112538 | -3.467094027 | 0.000526118 | 0.007094577 | | Nalcn | 4703.648239 | 0.124709038034284 | 0.069774170217194 | 1.78732384270695 | 0.0738851626047557 | 0.466430192088953 | NM_177393 |
| Ranbp3l | 206.6090881 | -0.527575365 | 0.102062114 | -5.169159691 | 2.35E-07 | 1.50E-05 | | Ppm1a | 6948.59036 | 0.0887390173521611 | 0.0496479076372551 | 1.78736671040639 | 0.0738782383827901 | 0.466430192088953 | NM_008910 |
| Kcnb2 | 433.5559473 | -0.528183751 | 0.119116744 | -4.434168814 | 9.24E-06 | 0.000295885 | | Pcdha5 | 186.7102108 | 0.169865228069174 | 0.0950582536886693 | 1.78695927475599 | 0.073944071034227 | 0.466583524838712 | NM_009959 |
| Dcc | 1013.2957 | -0.5288814 | 0.123470889 | -4.283450157 | 1.84E-05 | 0.000521593 | | Srp72 | 4345.937837 | 0.104583889127357 | 0.0585321755397097 | 1.78677604519253 | 0.0739736925308794 | 0.466583524838712 | NM_025691 |
| Col12a1 | 633.761478 | -0.530149493 | 0.100484782 | -5.275918211 | 1.32E-07 | 9.39E-06 | | Gtf3c2 | 2837.630826 | -0.078527711059819 | 0.0439636980447694 | -1.78619439565462 | 0.0740677882031386 | 0.466974258436976 | NM_027901 |
| Igsf3 | 906.4658855 | -0.530934293 | 0.108739997 | -4.882603538 | 1.05E-06 | 5.05E-05 | | Psme3 | 4834.970964 | 0.0891102082747775 | 0.0498959820879433 | 1.78591951788258 | 0.0741122902622997 | 0.467052116420896 | NM_011192 |
| Bche | 207.2537614 | -0.531509482 | 0.15877572 | -3.34754888 | 0.000815296 | 0.009717075 | | Adc | 1198.999713 | 0.113636341058238 | 0.063645473953757 | 1.78545832089808 | 0.0741870060530227 | 0.467117663600437 | NM_172875 |
| Adcyap1 | 578.8085298 | -0.532217481 | 0.11357997 | -4.685839231 | 2.79E-06 | 0.000116128 | | Srsf2 | 3560.401533 | -0.118940643870984 | 0.066610634474408 | -1.78561043307104 | 0.0741623564604363 | 0.467117663600437 | NM_011358 |
| Sp8 | 119.9236596 | -0.534441798 | 0.151541326 | -3.526706627 | 0.000420763 | 0.006030802 | | Gm13845 | 70.93188522 | -0.169139234260456 | 0.0947616692634807 | -1.7848908274312 | 0.0742790267743737 | 0.467418831598541 | NR_040300 |
| Homer2 | 2865.701092 | -0.536601976 | 0.086029228 | -6.237437978 | 4.45E-10 | 7.00E-08 | | Trappc13 | 1961.07788 | 0.10394068272879 | 0.0582376913670429 | 1.78476653673829 | 0.0742991933196359 | 0.467418831598541 | NM_001093760,NR_003546,NM_025879,NM_001093759 |
| Hs3st2 | 1664.786729 | -0.537448585 | 0.116381354 | -4.617995626 | 3.87E-06 | 0.000149512 | | Zfp951 | 127.4613959 | -0.162794846468433 | 0.0912315594426494 | -1.78441372111775 | 0.0743564631115859 | 0.467576616086103 | NM_001039231 |
| Fam198b | 254.324318 | -0.539808915 | 0.129329578 | -4.173901478 | 2.99E-05 | 0.0007681 | | Cds1 | 5172.605435 | 0.10368215690421 | 0.0581155540450879 | 1.78406897443962 | 0.0744124579740959 | 0.467726250338259 | NM_173370 |
| Draxin | 33.65976406 | -0.539857461 | 0.165193998 | -3.268021039 | 0.001083023 | 0.012159682 | | Rgs17 | 3403.223187 | 0.123359486706709 | 0.0691693000822792 | 1.78344274931174 | 0.0745142596056737 | 0.468021138002773 | NM_001161822,NM_019958 |
| Tenm1 | 1124.117494 | -0.54090152 | 0.109304171 | -4.948589938 | 7.48E-07 | 3.91E-05 | | Sstr2 | 685.0597589 | 0.152728596835602 | 0.0856397695151809 | 1.78338402473781 | 0.0745238119372446 | 0.468021138002773 | NM_009217,NM_001042606 |
| Tenm4 | 3813.222659 | -0.541281378 | 0.092773732 | -5.834424958 | 5.40E-09 | 6.29E-07 | | Top3b | 1493.740954 | -0.109988256871455 | 0.0616916788098956 | -1.78287021837071 | 0.0746074320423017 | 0.468343802007984 | NM_011624 |
| Fbn2 | 109.6071113 | -0.541846825 | 0.160485255 | -3.376302851 | 0.00073467 | 0.008991293 | | Cd24a | 324.9068361 | 0.160546073109739 | 0.090071848047507 | 1.78242232828465 | 0.0746803870308948 | 0.468388672023642 | NM_009846 |
| Nusap1 | 21.25068205 | -0.542459079 | 0.167309341 | -3.242252202 | 0.00118589 | 0.012948685 | | Cdk9 | 3500.095966 | -0.111336000121194 | 0.0624865792692864 | -1.7817586170794 | 0.0747886058576606 | 0.468388672023642 | NM_130860 |
| Tfcp2l1 | 474.7655235 | -0.54322369 | 0.085687923 | -6.339559512 | 2.30E-10 | 4.06E-08 | | G3bp2 | 12083.91797 | 0.105259684992866 | 0.0590802396984122 | 1.78163943697904 | 0.0748080489532459 | 0.468388672023642 | NM_001080797,NM_001080796,NM_001080795,NM_001080794,NM_011816 |
| Pcdh12 | 16.6357907 | -0.54394977 | 0.167050933 | -3.256191151 | 0.001129177 | 0.012541555 | | Insrr | 22.40174239 | -0.13548915690103 | 0.0760218909752332 | -1.78223870996803 | 0.0747103127070939 | 0.468388672023642 | NM_011832 |
| Itm2a | 1626.313659 | -0.544095902 | 0.141268752 | -3.851495065 | 0.000117399 | 0.002212869 | | Shisa7 | 8120.467184 | -0.102026745833686 | 0.0572566616148488 | -1.78191922050913 | 0.0747624056881055 | 0.468388672023642 | NM_172737 |
| Gprin3 | 396.3241914 | -0.545681997 | 0.127894143 | -4.266669179 | 1.98E-05 | 0.000556922 | | Stk24 | 2074.355092 | 0.121615979283212 | 0.068238516223658 | 1.78221898736199 | 0.0747135276329821 | 0.468388672023642 | NM_145465 |
| Slc5a7 | 432.889718 | -0.546010666 | 0.150739207 | -3.622220644 | 0.000292085 | 0.004556528 | | Gm20743 | 46.4972765 | -0.159336749513851 | 0.0894868007054847 | -1.78056147116325 | 0.0749841187306474 | 0.46858965670874 | NR_040303 |
| Asic4 | 1140.803113 | -0.547545147 | 0.087860002 | -6.232018359 | 4.60E-10 | 7.10E-08 | | Gm4636 | 356.4736005 | 0.153525897389184 | 0.086198667846477 | 1.78107041819508 | 0.0749009476329265 | 0.46858965670874 | NM_001254951 |
| Zic1 | 538.5450475 | -0.549320527 | 0.166857702 | -3.292149659 | 0.000994247 | 0.011333586 | | Plcd1 | 508.4397483 | -0.142150025748639 | 0.0798391333395057 | -1.78045552103083 | 0.0750014423687058 | 0.46858965670874 | NM_019676 |
| Gng7 | 14281.16362 | -0.550281163 | 0.085891828 | -6.40667657 | 1.49E-10 | 2.74E-08 | | Slc25a27 | 2135.055923 | -0.0976684280011309 | 0.0548432952261081 | -1.78086359688023 | 0.07493473686309 | 0.46858965670874 | NM_028711 |
| Nes | 625.753946 | -0.550390045 | 0.097642289 | -5.636799898 | 1.73E-08 | 1.70E-06 | | Zhx3 | 1376.129605 | 0.128929271977238 | 0.0724135964768322 | 1.78045668562377 | 0.0750012519312971 | 0.46858965670874 | NM_177263 |
| Zfhx3 | 290.9259067 | -0.550950876 | 0.137251898 | -4.01415851 | 5.97E-05 | 0.001322382 | | Crlf1 | 44.16998074 | -0.12502956131152 | 0.0702610820749693 | -1.7794995126621 | 0.0751579047203707 | 0.468962080742313 | NM_018827 |
| Sfrp1 | 505.7902792 | -0.550998761 | 0.096104219 | -5.73334621 | 9.85E-09 | 1.04E-06 | | Dirc2 | 3140.053643 | 0.0900565143722637 | 0.0505998522298801 | 1.77977820889927 | 0.0751122652075755 | 0.468962080742313 | NM_153550 |
| Pclo | 10455.35487 | -0.552742744 | 0.065382255 | -8.454017728 | 2.81E-17 | 1.98E-14 | | Jag2 | 1517.115497 | -0.131227555609899 | 0.0737413369795781 | -1.77956572236059 | 0.0751470601184817 | 0.468962080742313 | NM_010588 |
| Mxd3 | 34.97445607 | -0.552872241 | 0.166518824 | -3.320178628 | 0.000899599 | 0.010498156 | | 5031414D18Rik | 72.34616054 | -0.169120375264654 | 0.0950704750733018 | -1.77889481602209 | 0.0752570081765455 | 0.469360462669917 | NM_198642 |
| Lama3 | 279.7090196 | -0.553979536 | 0.13261753 | -4.17727231 | 2.95E-05 | 0.000759536 | | C530005A16Rik | 289.1343129 | -0.144844434288412 | 0.0814320296914847 | -1.77871575640658 | 0.0752863746400184 | 0.469360462669917 | NR_029450 |
| Robo1 | 1844.399059 | -0.555201819 | 0.085857011 | -6.466586857 | 1.00E-10 | 2.02E-08 | | Zfhx3 | 280.731514 | 0.168559185466613 | 0.0947937499559921 | 1.77817228060781 | 0.0753755640479687 | 0.469714904916685 | NM_007496 |
| Ccbe1 | 294.2530545 | -0.557108616 | 0.13324599 | -4.181053524 | 2.90E-05 | 0.0007518 | | Stard4 | 931.0726841 | 0.113732141250555 | 0.0639924126053732 | 1.77727540844437 | 0.0755229376719977 | 0.470431471965454 | NM_133774 |
| Wnt7a | 484.6670115 | -0.557153429 | 0.113010566 | -4.930100346 | 8.22E-07 | 4.18E-05 | | Usp5 | 9597.998221 | 0.0843866997124551 | 0.0474905153160334 | 1.77691690963743 | 0.0755819117865827 | 0.470597021265281 | NM_013700 |
| Pcp4l1 | 6945.445593 | -0.557778212 | 0.120608778 | -4.624690016 | 3.75E-06 | 0.000146048 | | Zfp28 | 473.5198544 | -0.122131281991566 | 0.0687408389571458 | -1.77669175768577 | 0.075618969160084 | 0.470626026572142 | NM_175247 |
| Tead2 | 98.07639631 | -0.55836474 | 0.143939994 | -3.879149396 | 0.000104822 | 0.002035235 | | Etv4 | 344.1712377 | -0.158227325233433 | 0.0890828643852826 | -1.77618138264056 | 0.0757030258175462 | 0.470947388876093 | NM_008815 |
| Sebox | 96.58272711 | -0.559710047 | 0.158231969 | -3.537275369 | 0.000404278 | 0.005876062 | | Arl4d | 1029.957751 | 0.167092356813539 | 0.0941015205268087 | 1.77566054063813 | 0.0757888849345128 | 0.47125813670925 | NM_025404 |
| Rhcg | 37.07514021 | -0.559916956 | 0.162121536 | -3.453686484 | 0.00055298 | 0.007360713 | | Dusp14 | 2265.382841 | 0.141595664973405 | 0.0797504206520411 | 1.77548486660905 | 0.0758178621430206 | 0.47125813670925 | NM_019819 |
| Cyp2f2 | 15.7864392 | -0.560032805 | 0.159131162 | -3.519315753 | 0.000432661 | 0.006144506 | | Bpnt1 | 2012.712104 | 0.0813045346084234 | 0.0458152673316204 | 1.77461661458667 | 0.0759612119964625 | 0.471947205072974 | NM_011794 |
| Gjb2 | 394.7619326 | -0.56119568 | 0.122243893 | -4.590787043 | 4.42E-06 | 0.000166574 | | Abhd6 | 2259.205042 | 0.109694560181944 | 0.0618376255469839 | 1.77391287604663 | 0.0760775625786662 | 0.472435997154356 | NM_025341 |
| Phex | 44.42078854 | -0.565549626 | 0.168071315 | -3.364938434 | 0.000765607 | 0.009245769 | | Add3 | 4880.199115 | -0.0817695710553651 | 0.0461296693292277 | -1.77260258407177 | 0.0762945832448956 | 0.472435997154356 | NM_013758,NM_001164101,NM_001164100,NM_001164099 |
| Nlrp10 | 20.16829148 | -0.565975847 | 0.160674734 | -3.522494378 | 0.000427506 | 0.006088033 | | C330007P06Rik | 1993.078756 | 0.11156069003339 | 0.0629297811891435 | 1.77278051703501 | 0.0762650830210342 | 0.472435997154356 | NM_029951 |
| Adam12 | 210.3778322 | -0.566639059 | 0.121878369 | -4.649217627 | 3.33E-06 | 0.000133399 | | Fam5b | 5197.762044 | 0.109028635641397 | 0.0614840815619459 | 1.77328233376227 | 0.0761819349156376 | 0.472435997154356 | NM_207583 |
| Tmem40 | 91.26251538 | -0.569360844 | 0.163430244 | -3.483815662 | 0.00049432 | 0.006777868 | | Hexim2 | 356.6578104 | 0.144739773787042 | 0.0816643797958336 | 1.77237339154355 | 0.0763325597126032 | 0.472435997154356 | NM_001130515,NM_027658,NM_001130516 |
| Pde1c | 593.6563004 | -0.57011992 | 0.128332442 | -4.442523735 | 8.89E-06 | 0.000286993 | | Rnf14 | 15670.97569 | 0.088857410335441 | 0.0501297083951017 | 1.77254991461557 | 0.0763033173114604 | 0.472435997154356 | NM_020012,NM_001164622,NM_001164621 |
| Oprm1 | 65.46653988 | -0.570622301 | 0.161664629 | -3.529666979 | 0.000416083 | 0.005984147 | | Snx18 | 945.6355821 | -0.105022501633287 | 0.059215826582508 | -1.77355460008574 | 0.0761368529251726 | 0.472435997154356 | NM_130796 |
| Pcdh17 | 3009.395467 | -0.571025763 | 0.114511842 | -4.986608851 | 6.14E-07 | 3.32E-05 | | Tmco4 | 169.7375145 | -0.164732861526156 | 0.0929275326205846 | -1.77270241531911 | 0.0762780306738508 | 0.472435997154356 | NM_029857 |
| Inhba | 185.1593459 | -0.571767514 | 0.109933705 | -5.201021065 | 1.98E-07 | 1.32E-05 | | Vsnl1 | 23661.49269 | 0.115139901413104 | 0.0649447790130961 | 1.7728892631983 | 0.0762470581367419 | 0.472435997154356 | NM_012038 |

| | | | | | | | | | | | | | | | |
|---|---|---|---|---|---|---|---|---|---|---|---|---|---|---|---|
| Pdgfrl | 34.54616712 | -0.572529831 | 0.159532252 | -3.588803046 | 0.0003322 | 0.005071036 | | 0610009L18Rik | 76.91325961 | -0.168203041584531 | 0.0950602723772044 | -1.76943572091917 | 0.0768211888842952 | 0.472560914790984 | NR_038126 |
| C1ql1 | 215.1760049 | -0.572670363 | 0.128824901 | -4.445339043 | 8.78E-06 | 0.000284411 | | 1700025G04Rik | 4653.057972 | 0.0908116343526437 | 0.0512561533089597 | 1.77172160784781 | 0.0764407809875989 | 0.472560914790984 | NM_197990 |
| Ddah2 | 303.6091624 | -0.572737076 | 0.101963355 | -5.617087404 | 1.94E-08 | 1.89E-06 | | Aamdc | 649.9617597 | 0.109011685038306 | 0.061613476211875 | 1.76928316416496 | 0.0768466345879688 | 0.472560914790984 | NM_001177947,NM_001177946,NM_001177945,NM_183251 |
| Asxl3 | 467.935201 | -0.57285369 | 0.109521558 | -5.230510802 | 1.69E-07 | 1.15E-05 | | Calu | 3851.00062 | 0.0876913415677531 | 0.0495420890645124 | 1.77003721933413 | 0.076720940696125 | 0.472560914790984 | NM_184053,NM_007594 |
| Lrp4 | 1783.262611 | -0.572980762 | 0.076153661 | -7.524008139 | 5.31E-14 | 2.29E-11 | | Dcx | 1140.929455 | -0.122732692092056 | 0.0693627726266925 | -1.76943174911127 | 0.076821851196754 | 0.472560914790984 | NM_001110223,NM_001110222,NM_010025,NM_001110224 |
| Fn1 | 2353.942416 | -0.574702634 | 0.089283382 | -6.436837645 | 1.22E-10 | 2.37E-08 | | Eif3i | 2662.762687 | 0.0867815685704576 | 0.0490321775931588 | 1.76989015846944 | 0.0767454406066582 | 0.472560914790984 | NM_018799 |
| Ankrd63 | 2370.300801 | -0.576063712 | 0.09372285 | -6.146459599 | 7.92E-10 | 1.12E-07 | | Epb4.1l4a | 152.7379955 | 0.16737103158055 | 0.0945825355739791 | 1.76957649279383 | 0.0767977177018913 | 0.472560914790984 | NM_013512 |
| Rrm2 | 71.02413499 | -0.576089466 | 0.141869642 | -4.060695857 | 4.89E-05 | 0.001129761 | | Glt25d1 | 2440.226471 | -0.0985339961850945 | 0.0556393344491992 | -1.77094131625639 | 0.0765704608494769 | 0.472560914790984 | NM_146211 |
| Sall3 | 314.0693231 | -0.576720517 | 0.139375943 | -4.137877051 | 3.51E-05 | 0.000862091 | | Hdac9 | 1347.020357 | 0.115356844724875 | 0.0651151135259909 | 1.77158325430594 | 0.0764637614590401 | 0.472560914790984 | NM_024124,NM_001271386 |
| Hdac9 | 1424.869924 | -0.57841612 | 0.091322279 | -6.333789815 | 2.39E-10 | 4.13E-08 | | Itk | 26.07622598 | 0.149956319587582 | 0.0847188653776097 | 1.77004636357198 | 0.0767194175033934 | 0.472560914790984 | NM_010583 |
| Marcksl1 | 1937.714658 | -0.579730554 | 0.093865646 | -6.176173917 | 6.57E-10 | 9.78E-08 | | Kctd13 | 4852.530545 | 0.0731775029628895 | 0.0413553115770155 | 1.76948256880164 | 0.0768133771919926 | 0.472560914790984 | NM_172747 |
| Emp2 | 902.8232293 | -0.586712933 | 0.121568292 | -4.826200332 | 1.39E-06 | 6.48E-05 | | Lap3 | 2010.651727 | -0.0959648316197645 | 0.0542442852628423 | -1.76912334921115 | 0.0768732921417524 | 0.472560914790984 | NM_024434 |
| C730002L08Rik | 168.9063842 | -0.587160071 | 0.12655628 | -4.639517463 | 3.49E-06 | 0.00013874 | | Masp2 | 29.31607488 | -0.147773040563529 | 0.0835013103302587 | -1.76970924143665 | 0.0767755896051487 | 0.472560914790984 | NM_010767,NM_001003893 |
| Fabp7 | 865.4012846 | -0.58950708 | 0.167964267 | -3.509717219 | 0.000448583 | 0.006323296 | | Nip7 | 1008.380848 | 0.110848121622084 | 0.0625813004851027 | 1.77126586956229 | 0.0765165002316043 | 0.472560914790984 | NR_028367,NM_001164472,NM_025391 |
| Th | 283.198081 | -0.591860304 | 0.148074614 | -3.997040999 | 6.41E-05 | 0.001387951 | | Ryr3 | 2044.482804 | -0.151871148616226 | 0.0858315628786547 | -1.76940910223125 | 0.0768256277298587 | 0.472560914790984 | NM_177652 |
| Pnp2 | 38.38040982 | -0.592811953 | 0.164781111 | -3.597572244 | 0.000321201 | 0.004932302 | | Ubxn1 | 2033.47382 | 0.0906650204815626 | 0.0512396493349229 | 1.76943093206863 | 0.0768219874419657 | 0.472560914790984 | NM_146093 |
| Kntc1 | 23.58020688 | -0.595938928 | 0.166545025 | -3.578245152 | 0.000345909 | 0.005213534 | | Ptprg | 3065.153055 | 0.0895993650513512 | 0.0506524261962988 | 1.76890569277209 | 0.0769096140218 | 0.472576677289215 | NM_008981 |
| Wnt7b | 1369.551654 | -0.597448458 | 0.109164793 | -5.472904218 | 4.43E-08 | 3.73E-06 | | Slc9a7 | 814.2707811 | 0.131522535255602 | 0.0743603697390322 | 1.76871814539358 | 0.0769409226069801 | 0.472576677289215 | NM_177353 |
| Hspg2 | 463.3904859 | -0.60344803 | 0.094049578 | -6.416275816 | 1.40E-10 | 2.61E-08 | | Tgif1 | 66.66692358 | -0.165120953434225 | 0.0933865698142586 | -1.76814453901286 | 0.0770367431773237 | 0.472965228822402 | NM_001164077,NM_009372,NM_001164076,NM_001164075,NM_001164074 |
| Gpr6 | 378.4883978 | -0.604242328 | 0.118307558 | -5.107385693 | 3.27E-07 | 1.99E-05 | | Esrrg | 1486.445535 | 0.153997537962004 | 0.0871522773156189 | 1.76699384921758 | 0.0772292583943477 | 0.473555200376883 | NM_001243792,NM_011935 |
| Cenpf | 48.7826805 | -0.604886858 | 0.163210916 | -3.706166674 | 0.00021042 | 0.003536058 | | Pdss1 | 216.7109522 | -0.147897346569508 | 0.0836867598300398 | -1.76727294580258 | 0.0771825283952856 | 0.473555200376883 | NM_019501 |
| Prss35 | 131.5449012 | -0.605525326 | 0.156222717 | -3.876038888 | 0.000106171 | 0.002048579 | | Spata2 | 3374.979222 | 0.101705340184624 | 0.0575586694322495 | 1.76698560247885 | 0.0772306395217429 | 0.473555200376883 | NM_170756 |
| Tagln | 312.7328641 | -0.610695238 | 0.132151033 | -4.621191565 | 3.82E-06 | 0.000148161 | | Lekr1 | 139.8236272 | -0.157808000720626 | 0.0893431505122387 | -1.76631336387683 | 0.0773432907791774 | 0.473845905465344 | NM_001166659,NM_001037923 |
| Thbd | 558.9698971 | -0.613631151 | 0.083926667 | -7.311515829 | 2.64E-13 | 8.90E-11 | | Repin1 | 1734.687305 | -0.108550111520672 | 0.061453495887338 | -1.76637813607343 | 0.0773324306692143 | 0.473845905465344 | NM_001079904,NM_001079905,NM_001079902,NM_001079903,NM_175099,NM_001079901 |
| Stac | 55.24673797 | -0.614169807 | 0.167970987 | -3.65640412 | 0.000255778 | 0.00410251 | | Ranbp10 | 1886.47753 | -0.0994216114015304 | 0.056321755260841 | -1.7652434825776 | 0.0775228535804223 | 0.474745771968472 | NM_145824 |
| Tenm3 | 4115.8393 | -0.615308749 | 0.100536173 | -6.120272259 | 9.34E-10 | 1.27E-07 | | Dock4 | 4960.376975 | -0.101286259541138 | 0.0573939309959146 | -1.76475557229819 | 0.0776048543422412 | 0.475047667161987 | NM_172803 |
| Kif2c | 9.386927755 | -0.617577957 | 0.159496715 | -3.872041865 | 0.000107927 | 0.002067036 | | Bud31 | 1189.150247 | 0.111790840068898 | 0.0634056975895356 | 1.76310401618147 | 0.0778829479988737 | 0.47533760590767 | NM_001008705 |
| Gpr149 | 173.3419624 | -0.618746005 | 0.165008278 | -3.749787655 | 0.000176984 | 0.003098526 | | Edc4 | 3337.042521 | -0.0965912318318636 | 0.0548085675739829 | -1.76233819104067 | 0.0780121748921854 | 0.47533760590767 | NM_181594 |
| Drd1a | 2588.755088 | -0.624010223 | 0.115327163 | -5.41078273 | 6.27E-08 | 5.12E-06 | | Fxyd5 | 453.6522966 | -0.154882052635132 | 0.0878250990249454 | -1.76352835755004 | 0.0778114188586961 | 0.47533760590767 | NM_008761,NM_001111073 |
| Aldh1a7 | 35.50894822 | -0.62717093 | 0.167725019 | -3.739280728 | 0.000184548 | 0.003205583 | | Polm | 286.0286224 | -0.154229627357763 | 0.0874714602635197 | -1.76319941262127 | 0.0778668628257576 | 0.47533760590767 | NM_017401 |
| Il31ra | 12.33124961 | -0.627189031 | 0.163553465 | -3.834764553 | 0.000125685 | 0.002335847 | | Polr2c | 1819.836529 | 0.105496597229948 | 0.059857495936263 | 1.76246258851659 | 0.0779911719407616 | 0.47533760590767 | NM_009090 |
| Cpa6 | 24.12097752 | -0.628285644 | 0.155224756 | -4.047586605 | 5.17E-05 | 0.00118084 | | Ppp5c | 4830.762121 | 0.0929193555163895 | 0.0527019020213034 | 1.76311199316543 | 0.0778816028642242 | 0.47533760590767 | NM_011155 |
| Pf4 | 26.6843616 | -0.629235918 | 0.166919108 | -3.769705723 | 0.00016344 | 0.002904061 | | Prkra | 529.9367838 | 0.121275918161521 | 0.0688029557298977 | 1.76265564284213 | 0.0779585862596809 | 0.47533760590767 | NM_011871 |
| Klhl30 | 18.95397982 | -0.629489713 | 0.148865787 | -4.228572105 | 2.35E-05 | 0.00063482 | | Psme2 | 813.5823533 | -0.108299210807948 | 0.0614446870035845 | -1.76254801007661 | 0.0779767522496457 | 0.47533760590767 | NM_001029855,NM_011190 |
| Htr2a | 977.2380404 | -0.63005408 | 0.124338833 | -5.067234946 | 4.04E-07 | 2.35E-05 | | Rnf32 | 305.0838004 | -0.138088111542703 | 0.0782873871414521 | -1.76386154379122 | 0.0777552927929848 | 0.47533760590767 | NM_021470 |
| Slc6a3 | 22.44684923 | -0.631175305 | 0.157404684 | -4.00988897 | 6.07E-05 | 0.001336958 | | Sbk1 | 3940.383251 | 0.154488880359035 | 0.0875983287403703 | 1.76360534019912 | 0.0777984480066153 | 0.47533760590767 | NM_145587 |
| Jsrp1 | 140.1323547 | -0.632165231 | 0.126419935 | -5.000518574 | 5.72E-07 | 3.12E-05 | | Ythdf2 | 1556.631747 | 0.112149797084088 | 0.0636065782401922 | 1.76317922118977 | 0.077870267157611 | 0.47533760590767 | NM_145393 |
| Rarb | 1014.211528 | -0.633381103 | 0.102704361 | -6.167032229 | 6.96E-10 | 1.03E-07 | | Ankrd27 | 1839.694607 | 0.101439393320887 | 0.05758226409797 | 1.76164301473625 | 0.0781296314725205 | 0.475440314921092 | NM_178263,NM_145633 |
| Drd5 | 112.4693383 | -0.634705805 | 0.157372468 | -4.033143866 | 5.50E-05 | 0.001241224 | | Gm11696 | 94.98962498 | -0.165766929850926 | 0.0941281688324234 | -1.76107675212552 | 0.0782254132356746 | 0.475440314921092 | NR_038097 |
| Fat1 | 2472.149787 | -0.635631482 | 0.121384662 | -5.23650575 | 1.64E-07 | 1.12E-05 | | Pkm | 55423.20069 | 0.0895233058263372 | 0.0508085910575713 | 1.76197182332606 | 0.0780740582345711 | 0.475440314921092 | NM_011099,NM_001253883 |
| Zbtb20 | 153.9481137 | -0.636141247 | 0.164648558 | -3.863630854 | 0.000111714 | 0.002126409 | | Plec | 8355.872823 | -0.119665079863573 | 0.0679493474579779 | -1.76109240692234 | 0.0782227639856888 | 0.475440314921092 | 163549,NM_201387,NM_201394,NM_201385,NM_001163540,NM_201386,NM_201392,NM_201393,NM_201390,NM_201391,NM_011117,NM_001164203, |
| Slc38a5 | 115.841841 | -0.640446634 | 0.159959996 | -4.003792509 | 6.23E-05 | 0.001364151 | | Tcf25 | 29243.05939 | 0.0661526094429697 | 0.0375631587327851 | 1.76110347677528 | 0.0782208906865363 | 0.475440314921092 | NM_001037878,NM_025804,NM_001037877 |
| Htr1b | 308.4134959 | -0.641298213 | 0.12870037 | -4.982877767 | 6.26E-07 | 3.37E-05 | | Zic3 | 99.6178413 | -0.163198659088359 | 0.0926450295376281 | -1.76154792008647 | 0.0781457097970402 | 0.475440314921092 | NM_009575 |
| Kif26a | 392.4709979 | -0.642109763 | 0.104477397 | -6.145920392 | 7.95E-10 | 1.12E-07 | | Phlda3 | 686.3395049 | 0.133633385584224 | 0.075890858591506 | 1.76086274505768 | 0.0782616368221005 | 0.475461537631883 | NM_013750 |
| Hs6st2 | 1483.031801 | -0.64377485 | 0.08678249 | -7.418257443 | 1.19E-13 | 4.84E-11 | | Fam134b | 1545.574591 | 0.105792978369757 | 0.0601053960009745 | 1.76012447148742 | 0.0783867044069486 | 0.47574440492648 | NM_001034851,NM_025459 |
| Islr | 664.4305848 | -0.645081214 | 0.129633481 | -4.976192938 | 6.48E-07 | 3.46E-05 | | Gnas | 51957.59659 | 0.0907793003985932 | 0.0515722601991629 | 1.76023505752937 | 0.0783679601740769 | 0.47574440492648 | NM_019690,NM_022000,NM_001077507,NM_001077510,NM_010309,NR_003258,NM_201618,NM_201617,NM_201616 |
| Tshr | 28.92724504 | -0.650990687 | 0.164948729 | -3.946624442 | 7.93E-05 | 0.001635816 | | Zc3h8 | 260.7687989 | -0.137608523250496 | 0.0781863057318114 | -1.76000799580569 | 0.0784064508738808 | 0.47574440492648 | NM_020594 |
| Tll1 | 119.4395333 | -0.657391051 | 0.157577941 | -4.171846943 | 3.02E-05 | 0.00077378 | | Alkbh1 | 396.3504379 | -0.131032841367876 | 0.0745408070868452 | -1.75786722050398 | 0.0787701044226919 | 0.476674705543282 | NM_001102565 |
| Strip2 | 2925.504315 | -0.660054149 | 0.128948252 | -5.118752218 | 3.08E-07 | 1.88E-05 | | B3galnt1 | 1597.044895 | 0.0962095578598739 | 0.0547411865276029 | 1.75753512049581 | 0.0788266410165598 | 0.476674705543282 | NM_020026 |
| Wnt2 | 209.5323649 | -0.661132857 | 0.10901458 | -6.064627822 | 1.32E-09 | 1.75E-07 | | Cwh43 | 106.1098031 | -0.16484475617224 | 0.0937731402075395 | -1.75791016283985 | 0.0787627963432378 | 0.476674705543282 | NM_181323 |
| Hs3st3b1 | 71.39058243 | -0.66506915 | 0.149847078 | -4.438319113 | 9.07E-06 | 0.000292048 | | D8Ertd738e | 1018.093266 | 0.107662778215992 | 0.0612674450089164 | 1.75725914799162 | 0.0788736475945196 | 0.476674705543282 | NM_001007571 |
| Kdr | 638.963809 | -0.666399224 | 0.091302834 | -7.298779149 | 2.90E-13 | 9.37E-11 | | Fam180a | 74.91564944 | -0.161533599885518 | 0.0919197932121949 | -1.7573320635374 | 0.0788612256256367 | 0.476674705543282 | NM_173375 |
| Isl1 | 132.1635622 | -0.667520014 | 0.162895717 | -4.097836504 | 4.17E-05 | 0.000997139 | | Foxm1 | 66.7735972 | -0.166236935011432 | 0.0945361837021839 | -1.75844770225891 | 0.0786713626369529 | 0.476674705543282 | NM_008021 |
| Man1a | 898.6485704 | -0.669742817 | 0.091331252 | -7.333117668 | 2.25E-13 | 7.92E-11 | | Klhl8 | 1453.565078 | 0.100227799412317 | 0.057034721155489 | 1.75731199139335 | 0.078864644978387 | 0.476674705543282 | NM_178741 |
| Gbx2 | 15.59767381 | -0.670987313 | 0.154923248 | -4.331095049 | 1.48E-05 | 0.000439549 | | Mrpl43 | 1130.444247 | 0.102938063928293 | 0.0585674013519652 | 1.75759988825419 | 0.0788155953575083 | 0.476674705543282 | NM_053164 |
| Myh11 | 489.6863226 | -0.672791051 | 0.122388438 | -5.497178198 | 3.86E-08 | 3.36E-06 | | Odc1 | 2765.060226 | 0.0950432408482285 | 0.0540833224155738 | 1.75734841358156 | 0.0788584404334229 | 0.476674705543282 | NM_013614 |
| Dlx6as1 | 766.3631331 | -0.677375407 | 0.139112429 | -4.869265911 | 1.12E-06 | 5.36E-05 | | Snx10 | 3697.980186 | 0.107544104745751 | 0.0612095373586668 | 1.75698280670837 | 0.0789207398342002 | 0.476674705543282 | NM_001127348,NM_001127349,NM_028035 |
| Mgat4c | 187.1044456 | -0.680272279 | 0.123838817 | -5.493207205 | 3.95E-08 | 3.42E-06 | | Tex9 | 407.8750126 | -0.149368311463564 | 0.0850114960478109 | -1.75703661748946 | 0.0789115679666656 | 0.476674705543282 | NM_009359 |
| Fndc9 | 209.234282 | -0.680379426 | 0.134433053 | -5.06110225 | 4.17E-07 | 2.41E-05 | | 4931406P16Rik | 1826.150435 | -0.0969171922738262 | 0.0552521748539154 | -1.75408827851703 | 0.079415381828244 | 0.477013957208569 | NM_172741 |
| Crybg3 | 395.3215998 | -0.681990337 | 0.111283829 | -6.128386699 | 8.88E-10 | 1.23E-07 | | Abtb1 | 1327.6168 | -0.129042181493989 | 0.0735803249251947 | -1.75375933206574 | 0.0794717542055703 | 0.477013957208569 | NM_030251 |
| Zfhx4 | 667.4133547 | -0.682173035 | 0.109323866 | -6.239927848 | 4.38E-10 | 6.99E-08 | | Afap1l1 | 1308.383929 | -0.109549184000752 | 0.0623798658499524 | -1.75616254552807 | 0.0790606580004163 | 0.477013957208569 | NM_178928 |
| Kcnq1 | 30.48005247 | -0.682977703 | 0.166955539 | -4.090775955 | 4.30E-05 | 0.00101496 | | Bag5 | 1904.450722 | 0.107299453430246 | 0.0611691297265716 | 1.7541438616158 | 0.0794058596252392 | 0.477013957208569 | NM_027404 |
| Prss12 | 892.3969642 | -0.68309962 | 0.105701891 | -6.462510894 | 1.03E-10 | 2.05E-08 | | Cdh9 | 770.2065615 | -0.103981754180449 | 0.0592159801053373 | -1.75597455273862 | 0.0790927537588024 | 0.477013957208569 | NM_009869 |
| Fras1 | 1534.465963 | -0.685119703 | 0.11947191 | -5.734567244 | 9.78E-09 | 1.04E-06 | | Cmc2 | 444.5011765 | 0.124976111292832 | 0.0711881660883119 | 1.75557425004872 | 0.0791611322126415 | 0.477013957208569 | NM_026844 |
| Top2a | 76.86885423 | -0.688508204 | 0.16806215 | -4.09674757 | 4.19E-05 | 0.001000295 | | Cyb561d2 | 472.4059869 | 0.13645750329545 | 0.0777628168214055 | 1.75479115692074 | 0.0792950367536471 | 0.477013957208569 | NM_019720 |
| Pbk | 23.22985463 | -0.696667648 | 0.168056817 | -4.145429283 | 3.39E-05 | 0.000846251 | | Edil3 | 5534.306273 | 0.103050305720435 | 0.0587562136842854 | 1.75386225998419 | 0.0794541116940556 | 0.477013957208569 | NM_010103,NM_001037987 |
| Emid1 | 158.6438141 | -0.701681435 | 0.149157122 | -4.704310631 | 2.55E-06 | 0.000108426 | | Hps6 | 300.3633759 | 0.130347941288218 | 0.0743324317885668 | 1.75358101641264 | 0.0795023262014281 | 0.477013957208569 | NM_176785 |
| Ccnb1 | 19.32549349 | -0.706826457 | 0.168121338 | -4.204263805 | 2.62E-05 | 0.000691379 | | Letm2 | 465.6033568 | -0.125191439789103 | 0.0713846380315063 | -1.75375883721436 | 0.0794718390339922 | 0.477013957208569 | NM_173012 |
| Klhl14 | 159.6690267 | -0.732093069 | 0.133319796 | -5.491255566 | 3.99E-08 | 3.44E-06 | | Mmp11 | 73.33387872 | 0.166424468241223 | 0.0948859593633385 | 1.75394198844477 | 0.0794404479085985 | 0.477013957208569 | NM_008606 |
| Kcna5 | 349.5773752 | -0.73646154 | 0.144640631 | -5.091664306 | 3.55E-07 | 2.14E-05 | | Plod3 | 1221.492183 | -0.122546974407144 | 0.0698561577118508 | -1.75427590668008 | 0.0793832420843573 | 0.477013957208569 | NM_011962 |
| Slc47a1 | 86.07693662 | -0.740341337 | 0.165514367 | -4.47297326 | 7.71E-06 | 0.00025648 | | Setdb1 | 2097.389607 | -0.109627848971235 | 0.0625146750103349 | -1.7536338300265 | 0.079493270376892 | 0.477013957208569 | NM_001163641,NM_001163642,NM_018877 |
| Ntn1 | 632.0933595 | -0.741314527 | 0.12959358 | -5.720302872 | 1.06E-08 | 1.12E-06 | | Top2a | 47.2468573 | -0.161495530775893 | 0.0919854008674032 | -1.75566480390344 | 0.0791456598799732 | 0.477013957208569 | NM_011623 |
| D8Ertd738e | 1180.125411 | -0.747207375 | 0.06646913 | -11.24141952 | 2.55E-29 | 3.60E-26 | | Unc13a | 16404.80586 | -0.106704840165304 | 0.0607571842818199 | -1.75625058051338 | 0.0790456315471679 | 0.477013957208569 | NM_001029873 |
| Syt6 | 1942.15138 | -0.750614703 | 0.108671269 | -6.907204738 | 4.94E-12 | 1.32E-09 | | Zfp691 | 214.8860211 | -0.145707135922668 | 0.0830125033399369 | -1.75524324722501 | 0.079217709387094 | 0.477013957208569 | NM_183140,NM_001145936,NM_001145935 |
| Asgr1 | 40.00032472 | -0.753497358 | 0.155438588 | -4.847556642 | 1.25E-06 | 5.91E-05 | | Cox8a | 7784.657486 | -0.119133917926726 | 0.0680003825462469 | -1.75195952531154 | 0.0797807678454898 | 0.47848696685532 | NM_007750 |
| Prkch | 493.7340023 | -0.76355125 | 0.118629193 | -6.436453226 | 1.22E-10 | 2.37E-08 | | Adcy7 | 425.6547598 | -0.156104876035054 | 0.0891447890108544 | -1.7511385440157 | 0.0799220483096209 | 0.478875136013939 | NM_007406,NM_001037724,NM_001109756,NM_001037723 |
| Dkk2 | 28.60362561 | -0.766268069 | 0.162712269 | -4.709344115 | 2.49E-06 | 0.000106367 | | Glrb | 8548.445517 | 0.126250391896281 | 0.0721169157747027 | 1.75063493134806 | 0.0800088142316391 | 0.478875136013939 | NM_010298 |
| Il15 | 26.90211668 | -0.76659756 | 0.166631324 | -4.60056094 | 4.21E-06 | 0.000160129 | | Rnft1 | 595.2602143 | -0.133384529107565 | 0.0761789817152177 | -1.75093609949002 | 0.0799569176764173 | 0.478875136013939 | NM_029788 |
| Serpina9 | 394.2533876 | -0.771060176 | 0.134906068 | -5.715533682 | 1.09E-08 | 1.14E-06 | | Wisp1 | 189.2163406 | 0.166524243782803 | 0.0950850076954765 | 1.75131966456921 | 0.0798908622928673 | 0.478875136013939 | NM_018865 |
| Pbx3 | 825.1991979 | -0.780984077 | 0.150987552 | -5.172506404 | 2.31E-07 | 1.48E-05 | | Wnt4 | 983.8790969 | 0.148533679869478 | 0.0848494709808488 | 1.7505551673151 | 0.0800225635578855 | 0.478875136013939 | NM_009523 |
| Gpr126 | 192.649184 | -0.79448622 | 0.116826207 | -6.800582165 | 1.04E-11 | 2.69E-09 | | Zdhhc9 | 2916.866403 | 0.097569810559549 | 0.0557285780624361 | 1.75080387750492 | 0.0799796984748228 | 0.478875136013939 | NM_172465 |
| St8sia2 | 229.9925071 | -0.797983371 | 0.109064861 | -7.316594566 | 2.54E-13 | 8.76E-11 | | Gm11974 | 58.3790729 | -0.166364099188314 | 0.0950622132797382 | -1.7500549739859 | 0.0801088281677059 | 0.478984868730352 | NR_045893 |
| Tac1 | 1862.400821 | -0.810545373 | 0.140841918 | -5.755000951 | 8.66E-09 | 9.39E-07 | | Gm16532 | 406.282521 | 0.122084290261644 | 0.0697645923368262 | 1.74994630044158 | 0.0801275802708768 | 0.478984868730352 | NM_001134752 |
| Padi6 | 30.03708738 | -0.827113313 | 0.16778201 | -4.929690102 | 8.24E-07 | 4.18E-05 | | Adh5 | 1949.050324 | 0.0876724174983163 | 0.0501106775438712 | 1.74957557541624 | 0.080191577360354 | 0.479170239710614 | NM_007410 |
| Hmcn1 | 198.4489543 | -0.837015838 | 0.113932474 | -7.346595828 | 2.03E-13 | 7.33E-11 | | Cd53 | 281.9478423 | -0.150075401836767 | 0.0858019053016406 | -1.74909171660025 | 0.0802751668343094 | 0.479231996890504 | NM_007651 |
| Meis2 | 7112.184552 | -0.842296325 | 0.097514958 | -8.637611569 | 5.74E-18 | 4.24E-15 | | Eva1b | 50.42556125 | -0.165473128085914 | 0.0946132239506343 | -1.74894291914471 | 0.0803008867156468 | 0.479231996890504 | NM_172145 |
| Slc35d3 | 344.9785632 | -0.844357013 | 0.145888698 | -5.78767942 | 7.14E-09 | 7.95E-07 | | St13 | 4256.461169 | 0.0878465775221799 | 0.0502266019580623 | 1.74900499132968 | 0.0802901566284707 | 0.479231996890504 | NM_133726 |
| Clec12a | 40.07983665 | -0.853805357 | 0.158441556 | -5.388771599 | 7.09E-08 | 5.58E-06 | | 9530091C08Rik | 35.25207141 | -0.158218280652841 | 0.0905685624775944 | -1.74694481533791 | 0.0806469110215815 | 0.479417113666935 | NR_033299 |
| Fat3 | 3840.366726 | -0.866225878 | 0.117425621 | -7.376804733 | 1.62E-13 | 5.98E-11 | | Bre | 1475.790629 | 0.0963112589187885 | 0.0550975240939671 | 1.7480142801795 | 0.0804615547038185 | 0.479417113666935 | NM_181281,NM_181280,NM_181282,NM_144541,NM_181279 |
| 2610028E06Rik | 33.5463469 | -0.876846204 | 0.164692608 | -5.3241382 | 1.01E-07 | 7.56E-06 | | Gnl3l | 9949.833549 | 0.0937093651273905 | 0.0536090504761998 | 1.7480138949484 | 0.0804616214085096 | 0.479417113666935 | NM_001168600,NM_198110 |
| Cbln2 | 1327.489852 | -0.877116881 | 0.11218767 | -7.818300175 | 5.35E-15 | 2.96E-12 | | Ints12 | 499.0943925 | 0.125428463744303 | 0.071824221096301 | 1.74632542935801 | 0.0807544195509233 | 0.479417113666935 | NM_027927 |
| Aldh1a3 | 52.87269128 | -0.877649708 | 0.152503366 | -5.754953008 | 8.67E-09 | 9.39E-07 | | Klk8 | 74.87076071 | -0.164054436566586 | 0.0939134654099722 | -1.74686809660807 | 0.0806602209323949 | 0.479417113666935 | NM_008940 |
| Pou3f4 | 172.8316529 | -0.890303679 | 0.114616852 | -7.767650806 | 8.00E-15 | 4.13E-12 | | March3 | 52.63980417 | 0.162107103559396 | 0.0927335656680294 | 1.74809522744652 | 0.0804475392907416 | 0.479417113666935 | NM_177115 |
| Acsm5 | 16.26062706 | -0.898732704 | 0.166657553 | -5.392691101 | 6.94E-08 | 5.49E-06 | | Pifo | 31.1456515 | 0.139663163676309 | 0.0799525364997306 | 1.74682592686449 | 0.0806675378020491 | 0.479417113666935 | NM_029604,NM_001200028 |
| Ndnf | 509.431718 | -0.899595623 | 0.145982712 | -6.16234356 | 7.17E-10 | 1.04E-07 | | Plekhn1 | 250.2026367 | -0.15599490631444 | 0.0893122476368935 | -1.74662390032609 | 0.0807025986041074 | 0.479417113666935 | NM_001008233 |
| Pgm5 | 360.0563638 | -0.900829426 | 0.086790141 | -10.37939813 | 3.08E-25 | 3.41E-22 | | Prdm1 | 56.44363688 | -0.164703244613551 | 0.0942252546903606 | -1.74797346162441 | 0.0804686228915913 | 0.479417113666935 | NM_007548 |
| Drd3 | 30.83890732 | -0.91427374 | 0.159037359 | -5.748798552 | 8.99E-09 | 9.67E-07 | | Ssb | 3826.952959 | 0.108361527337662 | 0.0620525027640071 | 1.74628778068426 | 0.0807609580850192 | 0.479417113666935 | NM_001110145,NM_009278 |
| Mki67 | 78.65430023 | -0.937193313 | 0.160355298 | -5.844479863 | 5.08E-09 | 6.01E-07 | | Wdr83 | 781.056369 | -0.127786742457403 | 0.0731724064502719 | -1.74637884219712 | 0.0807451439545419 | 0.479417113666935 | NM_026399 |
| Ly6g6e | 76.49027072 | -0.940068672 | 0.166030917 | -5.662009743 | 1.50E-08 | 1.52E-06 | | Zbtb8a | 258.8554085 | 0.150283885112561 | 0.0860398638309364 | 1.74667739372369 | 0.0806933138599202 | 0.479417113666935 | NM_028603 |
| Crym | 2658.596979 | -0.997109884 | 0.111323971 | -8.956829986 | 3.34E-19 | 3.05E-16 | | Zfp956 | 905.1544437 | -0.124217832880463 | 0.0710957776534178 | -1.74719001578418 | 0.0806043830336267 | 0.479417113666935 | NM_178898 |
| Wdfy1 | 1568.823883 | -1.083603933 | 0.135574278 | -7.992695577 | 1.32E-15 | 7.87E-13 | | Pyroxd2 | 90.603692 | -0.160267896183311 | 0.0917897757853822 | -1.74603211318462 | 0.0808053718366446 | 0.479484816707148 | NM_029011 |
| Ndst4 | 629.6451472 | -1.135988357 | 0.167953715 | -6.763698916 | 1.35E-11 | 3.26E-09 | | Zbtb4 | 8131.29642 | 0.080887513821502 | 0.0463400662255908 | 1.74552003071659 | 0.0808943888315822 | 0.479817024159887 | NM_029348 |
| Tfap2d | 47.43301765 | -1.166659431 | 0.155288071 | -7.512872207 | 5.78E-14 | 2.42E-11 | | Cflar | 912.8302355 | -0.108650660101456 | 0.0622888958664111 | -1.74430223220644 | 0.0811064026286644 | 0.479898997115796 | NM_009805,NM_207653 |
| Abi3bp | 698.4032325 | -1.202316504 | 0.167972945 | -7.157798552 | 8.20E-13 | 2.54E-10 | | Hs3st5 | 241.1319307 | 0.161605112862759 | 0.0926159514498764 | 1.74489502437623 | 0.0810031436327667 | 0.479898997115796 | NM_001253355,NM_001081208,NM_001253356 |
| Dsg1c | 34.39440554 | -1.223430605 | 0.168116335 | -7.277285709 | 3.41E-13 | 1.08E-10 | | Psmb5 | 1846.434549 | 0.105795992153348 | 0.060672022279525 | 1.74373604469504 | 0.0812051270763374 | 0.479898997115796 | NM_011186 |
| Igfbpl1 | 59.42862806 | -1.277622235 | 0.165657338 | -7.712439749 | 1.23E-14 | 5.98E-12 | | Qtrt1 | 384.1259543 | -0.154122415622101 | 0.0883943078213679 | -1.74357851111363 | 0.0812326130808659 | 0.479898997115796 | NM_021888 |
| Prss56 | 24.32590147 | -1.293674925 | 0.168068177 | -7.697322308 | 1.39E-14 | 6.33E-12 | | Rnf26 | 1562.797909 | 0.109275464371249 | 0.062615818298597 | 1.74517346160271 | 0.0809546792556951 | 0.479898997115796 | NM_153762 |
| Uba1y | 19.48016777 | -1.404381475 | 0.16188274 | -8.675300844 | 4.12E-18 | 3.20E-15 | | Spred2 | 2662.482463 | 0.156007489564237 | 0.0894313558682322 | 1.74443838013703 | 0.0810826774521167 | 0.479898997115796 | NM_033523 |
| Nmbr | 121.7881212 | -1.663346432 | 0.154595971 | -10.75931297 | 5.36E-27 | 6.38E-24 | | Trmt1 | 1980.034791 | -0.120769959537997 | 0.0692669287856523 | -1.74354430974877 | 0.081238581433825 | 0.479898997115796 | NM_001164559,NM_198020,NM_001164560 |
| 1-Mar | 461.9146356 | -3.175418061 | 0.110532883 | -28.72826602 | 1.69E-181 | 4.37E-178 | | Ubr5 | 7058.405845 | -0.0991807486957737 | 0.0568693366745355 | -1.74401100258631 | 0.0811571713036469 | 0.479898997115796 | NM_001112721,NM_001081359 |
| Uty | 237.0766996 | -4.767794529 | 0.149628733 | -31.86416421 | 8.38E-223 | 2.60E-219 | | Wdfy4 | 137.6597233 | -0.153787608584322 | 0.0881808351885811 | -1.74400263112222 | 0.0811586310407535 | 0.479898997115796 | NM_001146022 |
| Eif2s3y | 429.3459162 | -4.920495119 | 0.153483317 | -32.05882711 | 1.65E-225 | 6.41E-222 | | Zfp182 | 456.125923 | -0.142193708221547 | 0.0815311631448701 | -1.74404120751825 | 0.0811519046228199 | 0.479898997115796 | NM_001111076,NM_001013387 |
| Kdm5d | 409.4792754 | -5.308457086 | 0.146524767 | -36.22907704 | 2.12E-287 | 1.10E-283 | | 2010107G23Rik | 919.6297053 | 0.109782936765794 | 0.0630008810001865 | 1.74256192765928 | 0.0814101649172982 | 0.480521761718274 | NM_027251 |
| Ddx3y | 757.8046647 | -6.167616207 | 0.138786601 | -44.43956533 | 0 | 0 | | Bves | 29.7111861 | -0.155551489024517 | 0.0892630547546895 | -1.74261893066509 | 0.0814002010339254 | 0.480521761718274 | NM_024285 |
| Raver2 | 814.6625797 | -0.26381753 | 0.099408373 | -2.653876356 | 0.007957298 | 0.050077324 | | Gnl3 | 1014.18192 | -0.131952671443874 | 0.0757585580140036 | -1.74175267986863 | 0.0815517301044039 | 0.481161832451897 | NM_153547 |
| 1700017B05Rik | 531.693281 | 0.270928801 | 0.102101613 | 2.653521263 | 0.007965675 | 0.050082274 | | Yipf5 | 1466.195301 | 0.111758525510002 | 0.0641863452025165 | 1.74115733116426 | 0.0816560042244114 | 0.481581452441657 | NM_023311 |
| Pak3 | 4269.614667 | -0.213193997 | 0.080344689 | -2.653492112 | 0.007966363 | 0.050082274 | | Fut9 | 3594.986023 | 0.142342120866121 | 0.0817992100166326 | 1.74014053237407 | 0.0818343445939892 | 0.482437374014727 | NM_010243 |
| Srsf5 | 7136.155534 | 0.28367845 | 0.106915528 | 2.653295146 | 0.007971014 | 0.050082274 | | 2610507I01Rik | 971.5975824 | -0.100059239071372 | 0.0575326576764431 | -1.73906383601868 | 0.0820235349281605 | 0.482582908262268 | NR_037964 |
| Tmem147 | 1766.800841 | -0.147999754 | 0.055777139 | -2.653412434 | 0.007968244 | 0.050082274 | | Grk1 | 31.27034136 | -0.155220193688248 | 0.0892748850253459 | -1.73867704947679 | 0.0820915851794069 | 0.482582908262268 | NM_011881 |
| Purb | 18895.6317 | -0.20428154 | 0.077012755 | -2.652567597 | 0.007988213 | 0.050169994 | | Nrbp1 | 3402.692718 | 0.0787250684902716 | 0.0452783094900519 | 1.73869275105265 | 0.0820888217923083 | 0.482582908262268 | NM_147201 |
| Fyb | 160.8124834 | 0.303036249 | 0.114259086 | 2.652185127 | 0.007997268 | 0.050206512 | | Nup93 | 1009.620214 | -0.112463335483256 | 0.0646700392544435 | -1.73903304803993 | 0.0820289500112204 | 0.482582908262268 | NM_172410 |
| Mcm10 | 28.48433067 | -0.435125881 | 0.164145914 | -2.650848077 | 0.008028995 | 0.050385275 | | Pdia5 | 156.6531153 | -0.161728083887681 | 0.0929726251100361 | -1.73952368986324 | 0.081942688938865 | 0.482582908262268 | NM_028295 |
| Nckap1l | 701.9189656 | 0.185009587 | 0.069800477 | 2.650549035 | 0.008036106 | 0.050409485 | | Stx5a | 2106.675557 | -0.0870961522466673 | 0.0500894011103705 | -1.73881400703421 | 0.0820674839772578 | 0.482582908262268 | NM_001167799,NM_019829 |
| St6galnac2 | 159.7571515 | 0.286484234 | 0.108094307 | 2.6503175 | 0.008041616 | 0.050423633 | | Tmem194b | 213.3653175 | -0.149974051160327 | 0.0862485814199373 | -1.7388581782014 | 0.0820597121491131 | 0.482582908262268 | NR_024590,NM_001142647 |
| Gas5 | 2884.296592 | 0.277857686 | 0.104866538 | 2.649631538 | 0.00805796 | 0.050505675 | | AI846148 | 884.4784894 | 0.104315701276713 | 0.0600079289793234 | 1.73836529690363 | 0.0821464674611312 | 0.482710188649026 | NM_001033139,NM_001167831 |
| Adamts9 | 130.577071 | 0.352779046 | 0.133160035 | 2.649286226 | 0.008066198 | 0.050516443 | | Mvb12b | 6136.781309 | 0.0792433930594907 | 0.0455965925982691 | 1.73792357156308 | 0.0822242816595986 | 0.482972064451003 | NM_175184 |
| Otogl | 29.5151961 | -0.44502907 | 0.167974698 | -2.649381576 | 0.008063923 | 0.050516443 | | Ccdc22 | 1487.104486 | -0.106617854891065 | 0.0613665197128025 | -1.73739451723904 | 0.0823175583565698 | 0.483019452842213 | NM_138603 |
| Kdelc1 | 321.9522424 | 0.348158275 | 0.131443852 | 2.648722399 | 0.008079667 | 0.050580347 | | Gm2011 | 27.39982895 | -0.144552308693145 | 0.0832089817418303 | -1.7372200172049 | 0.0823483429797173 | 0.483019452842213 | NR_038067 |
| Traip | 491.372109 | 0.379155153 | 0.143170718 | 2.648273043 | 0.008090415 | 0.050627178 | | Rer1 | 2362.472884 | 0.0915637683024229 | 0.0527153488853936 | 1.73694702280143 | 0.0823965223231999 | 0.483019452842213 | NM_026395 |
| Gga3 | 3563.319896 | 0.164927639 | 0.062293414 | 2.647593498 | 0.008106694 | 0.050688101 | | Rhof | 1902.266352 | 0.10943278838933 | 0.0630020012764137 | 1.73697320993356 | 0.0823918997035208 | 0.483019452842213 | NM_175092 |
| Sdf4 | 11115.16578 | 0.137190851 | 0.051816848 | 2.647610906 | 0.008106277 | 0.050688101 | | Trhr2 | 171.1160795 | -0.161025524915592 | 0.0927066968698215 | -1.73693519834607 | 0.0823986096752721 | 0.483019452842213 | NM_133202 |
| Ddx60 | 44.22567283 | 0.43813977 | 0.165593691 | 2.645872355 | 0.008148056 | 0.050926169 | | Gramd1c | 52.60213821 | -0.160958735919924 | 0.0927234855180043 | -1.73590040344924 | 0.082581446459676 | 0.4838959625951 | NM_153528,NM_001172107 |
| Smc5 | 1099.575572 | 0.242633942 | 0.091709728 | 2.645672889 | 0.008152861 | 0.050935658 | | Pmpcb | 1765.269315 | 0.100240321855086 | 0.0577551251482099 | 1.73560911863409 | 0.082632972529717 | 0.484002514603577 | NM_028431 |
| C130074G19Rik | 2663.999699 | 0.21285121 | 0.080472198 | 2.645027903 | 0.008168418 | 0.051012281 | | Vmp1 | 4214.215385 | 0.125916472152159 | 0.0725567311834438 | 1.7354209609279 | 0.0826662700489793 | 0.484002514603577 | NM_029478 |
| Acot9 | 684.6268536 | -0.193904785 | 0.073365175 | -2.64300857 | 0.008217296 | 0.051276189 | | Uchl1 | 19027.9955 | 0.0793814338269735 | 0.0457473372764773 | 1.73521429995425 | 0.0827028545489923 | 0.484021621748051 | NM_011670 |
| Akt2 | 4377.766625 | -0.303837945 | 0.114958666 | -2.643019067 | 0.008217041 | 0.051276189 | | Agt | 1137.312258 | 0.135154798263217 | 0.0779360285969874 | 1.73417610181437 | 0.0828868418887018 | 0.484423387256539 | NM_007428 |
| Lrrcc1 | 640.0532499 | 0.29699788 | 0.112403188 | 2.642254937 | 0.008235604 | 0.051369746 | | Atp2b3 | 4817.397347 | 0.0964268311260333 | 0.0556130758901518 | 1.73388775180315 | 0.0829380014986071 | 0.484423387256539 | NM_177236 |
| Slc36a2 | 19.51190585 | -0.444029795 | 0.168079158 | -2.64178973 | 0.008246924 | 0.051419653 | | Ctbp1 | 7754.502305 | 0.0788927032354396 | 0.0454862533432167 | 1.73442957897971 | 0.0828418906093114 | 0.484423387256539 | NM_013502,NM_001198859,NM_001198861,NM_001198860 |
| Zfp169 | 444.1690551 | 0.260368571 | 0.098570027 | 2.641457851 | 0.008255008 | 0.051449353 | | Hibadh | 2083.484178 | 0.101140734460372 | 0.058331812299952 | 1.7338863730187 | 0.0829382461866317 | 0.484423387256539 | NM_145567 |
| Pdyn | 686.5699239 | -0.359056341 | 0.135960645 | -2.640884357 | 0.008268994 | 0.051499554 | | Zfp114 | 85.77669782 | -0.163816694579374 | 0.0944752667169795 | -1.73396382219468 | 0.0829245024604167 | 0.484423387256539 | NM_001029933 |
| Trim72 | 35.96261804 | 0.41254655 | 0.156217039 | 2.640855008 | 0.00826971 | 0.051499554 | | Mfsd10 | 723.1081004 | -0.156640754947175 | 0.090352000145161 | -1.73367224516904 | 0.082976253802979 | 0.48445058791884 | NM_026660 |
| Phf7 | 154.8161466 | 0.289422653 | 0.109621966 | 2.640188488 | 0.008285993 | 0.051580222 | | Gm14322 | 618.2081494 | -0.117832258375649 | 0.0679982367188025 | -1.7328722634813 | 0.0831183751407491 | 0.485085382601254 | NM_001243903 |
| Rrad | 64.20442094 | 0.401675596 | 0.152156479 | 2.639884926 | 0.008293418 | 0.051605711 | | Inpp5f | 4599.175924 | 0.118129419997199 | 0.0681927324598383 | 1.73228752883265 | 0.0832238313165289 | 0.485168543369645 | NM_178641 |
| Atp10d | 182.1289409 | 0.268496422 | 0.101761134 | 2.638496747 | 0.00832745 | 0.051796672 | | Ntrk2 | 25271.18833 | 0.100735871102943 | 0.0581506113610849 | 1.73232694799073 | 0.0832153665571835 | 0.485168543369645 | NM_008745,NM_001025074 |
| Kank2 | 569.1855253 | -0.220487412 | 0.083593819 | -2.637604242 | 0.008349396 | 0.051912336 | | Ppm1l | 3064.522481 | 0.110580073049407 | 0.0638368730334362 | 1.73222884823146 | 0.0832328245957011 | 0.485168543369645 | NM_178726 |
| Zfand4 | 154.8127168 | 0.381100808 | 0.144506464 | 2.637257863 | 0.008357927 | 0.051944534 | | Apex1 | 1336.546402 | 0.0914705924808841 | 0.0528223589135066 | 1.73166428691043 | 0.0833333527895613 | 0.485559680152895 | NM_009687 |
| Hspe1 | 1480.166787 | -0.216229002 | 0.082033293 | -2.635868858 | 0.008392216 | 0.052136725 | | P2rx7 | 521.1487907 | -0.115524399355015 | 0.0667403332891038 | -1.73095328809028 | 0.0834600961681233 | 0.486103190432301 | NM_001038839,NM_001038845,NM_011027,NM_001038887 |
| Ccdc88c | 1462.97487 | -0.173846835 | 0.065961987 | -2.635560912 | 0.008399835 | 0.05214083 | | Zfp418 | 169.2569071 | 0.157881779346597 | 0.0912290146323453 | 1.73060928020393 | 0.0835214753863786 | 0.486265712009032 | NM_146179 |
| Chchd1 | 622.8809436 | -0.217752002 | 0.082628978 | -2.635298278 | 0.008406338 | 0.05214083 | | Ccl27a | 1774.101833 | -0.121234835712591 | 0.0700934776355071 | -1.72961650359288 | 0.0836988154397488 | 0.48658073359459 | NM_011336,NM_001164044,NM_001164045,NM_001048179 |
| Gstm6 | 203.9989223 | 0.30629529 | 0.116215053 | 2.635590508 | 0.008399102 | 0.05214083 | | Emc4 | 3446.759207 | 0.0738197255469663 | 0.0426759163464242 | 1.72977482071458 | 0.0836705147768474 | 0.48658073359459 | NM_026519 |
| Ndufb2 | 1098.622625 | -0.270415999 | 0.10260894 | -2.635403882 | 0.008403722 | 0.05214083 | | Nat8l | 22445.44556 | 0.0775495185235409 | 0.0448445210941926 | 1.72929750683821 | 0.0837558626296673 | 0.48658073359459 | NM_001001985 |
| AA387883 | 79.88772551 | -0.396262419 | 0.150410824 | -2.634533923 | 0.008425288 | 0.052216568 | | Snx30 | 2714.019246 | 0.121677425621096 | 0.0703569661509806 | 1.72942968234284 | 0.0837322214510639 | 0.48658073359459 | NM_172468 |
| Hbq1a | 10.44393273 | 0.418286261 | 0.158764935 | 2.634626225 | 0.008422998 | 0.052216568 | | Tspyl3 | 1588.911426 | 0.117947856209146 | 0.068185495662489 | 1.72980859144847 | 0.0836644789458552 | 0.48658073359459 | NM_198617 |
| Nsmce1 | 377.6811585 | -0.268384111 | 0.101878967 | -2.634342686 | 0.008430036 | 0.0522251 | | Vps25 | 1606.358034 | 0.0961235699441317 | 0.0555890487375592 | 1.72918177459628 | 0.0837765671705954 | 0.48658073359459 | NM_026776 |
| Pafah1b3 | 287.940247 | -0.233452186 | 0.088644119 | -2.633589097 | 0.008448767 | 0.052301612 | | Tpra1 | 1137.534381 | -0.106212687358084 | 0.061436587753073 | -1.72881813985138 | 0.0838416485285485 | 0.486764103327616 | NM_011906 |
| Ppp4r1 | 1371.808668 | 0.184171279 | 0.069932064 | 2.633574191 | 0.008449137 | 0.052301612 | | Crb1 | 70.57484767 | 0.147646677990082 | 0.0854360948674829 | 1.72815340189755 | 0.0839607254951851 | 0.487260686593312 | NM_133239 |
| Fuk | 711.9884873 | 0.202200007 | 0.076790511 | 2.633137919 | 0.008459999 | 0.052347931 | | Sepp1 | 9299.574653 | 0.133246529275556 | 0.0771528276446253 | 1.72704660792609 | 0.0841592933066481 | 0.488218009014525 | NM_001042613,NM_001042614,NM_009155 |
| Scml4 | 135.9778093 | 0.393618592 | 0.149496017 | 2.632970428 | 0.008464172 | 0.052352845 | | Recql4 | 120.6063395 | -0.164208937420189 | 0.0950992320436933 | -1.72671149799341 | 0.0842194896530866 | 0.488372178323647 | NM_058214 |
| Npnt | 664.1860647 | -0.285122725 | 0.108323606 | -2.632138409 | 0.008484929 | 0.05246029 | | Ak3 | 3584.321445 | 0.108218287585076 | 0.0627597369280307 | 1.72432666040609 | 0.0846488890036694 | 0.488499377116194 | NM_021299 |
| Milr1 | 13.49964511 | 0.414403684 | 0.157451209 | 2.631949844 | 0.008489639 | 0.052468478 | | C1qtnf4 | 2556.842732 | -0.137871073603975 | 0.0799072102392011 | -1.72538965121245 | 0.084457275059374 | 0.488499377116194 | NM_026161 |
| Fam196b | 185.8191022 | -0.430293652 | 0.163515443 | -2.63151691 | 0.008500464 | 0.052514428 | | Cept1 | 1330.661152 | 0.112910860027489 | 0.0655439815518364 | 1.72267319369653 | 0.0849476405476736 | 0.488499377116194 | NM_133869 |
| Ccl12 | 22.0148445 | 0.402526166 | 0.152994618 | 2.630982517 | 0.008513842 | 0.05257611 | | Ddx27 | 1226.073768 | -0.108301903918985 | 0.0627810572081038 | -1.72507295568456 | 0.0845143256234066 | 0.488499377116194 | NM_153065 |
| Wisp1 | 183.6152939 | -0.373963245 | 0.14217559 | -2.63029149 | 0.008531168 | 0.052662121 | | Etfdh | 1224.487096 | 0.0957478201871268 | 0.0554808279869965 | 1.72578210638039 | 0.0843866201456895 | 0.488499377116194 | NM_025794 |
| Pced1a | 716.7169775 | 0.206927018 | 0.078677092 | 2.63007965 | 0.008536487 | 0.052673963 | | Eya2 | 66.18497213 | 0.1589466034374 | 0.092218453309754 | 1.72358782578484 | 0.0847822778491491 | 0.488499377116194 | NM_001271962,NM_001271963,NM_010165 |
| Irgm1 | 512.1636084 | 0.442031927 | 0.16811371 | 2.629362752 | 0.008554506 | 0.052732358 | | Fcho2 | 1069.710424 | -0.118334280549076 | 0.0686106334421442 | -1.72472216932469 | 0.0845775538126294 | 0.488499377116194 | NM_172591 |
| Pacrgl | 229.4907987 | -0.255296154 | 0.097096736 | -2.62929698 | 0.00855616 | 0.052732358 | | Gm13826 | 65.35142094 | -0.152057592280858 | 0.0880932641287213 | -1.72609783261831 | 0.0843298137017565 | 0.488499377116194 | NM_001271591,NM_001271590 |
| Tcerg1l | 382.5813999 | -0.352364744 | 0.134003168 | -2.629525484 | 0.008550412 | 0.052732358 | | Kcnip4 | 2166.222137 | 0.117200786290972 | 0.0680341881842381 | 1.72267487007546 | 0.0849473372250254 | 0.488499377116194 | NM_030265,NM_001199242,NM_001199243,NM_001199244,NM_001199245 |
| Wnk4 | 258.0342013 | 0.344728973 | 0.131119628 | 2.629117995 | 0.008560665 | 0.052739143 | | Lyar | 351.7243037 | 0.137458646204882 | 0.0797218024872413 | 1.72422903040709 | 0.0846665053355058 | 0.488499377116194 | NM_025281 |
| Pdzrn4 | 280.4762648 | -0.277583845 | 0.105593137 | -2.628805732 | 0.008568529 | 0.052745648 | | Mcfd2 | 2670.903835 | 0.0848104677982789 | 0.0491535413769236 | 1.7254192764563 | 0.0844519398640268 | 0.488499377116194 | NM_176808,NM_139295 |
| Tmem59l | 9252.696358 | 0.197827516 | 0.075251367 | 2.628889333 | 0.008566423 | 0.052745648 | | Mrpl39 | 936.164238 | 0.119744649978773 | 0.069392716793052 | 1.72560832768494 | 0.0844179001571878 | 0.488499377116194 | NM_017404 |
| E130114P18Rik | 127.5588609 | -0.381051638 | 0.144980705 | -2.62829208 | 0.00858148 | 0.052804388 | | Ptges | 166.419141 | -0.156187180935834 | 0.090510980513633 | -1.72561583190792 | 0.0844165492098902 | 0.488499377116194 | NM_022415 |
| Ppp2r2b | 4975.855252 | -0.170584938 | 0.064912261 | -2.627930926 | 0.008590596 | 0.052839496 | | Rasl11b | 2450.804693 | -0.119677165556824 | 0.0694697573207406 | -1.72272324206167 | 0.0849385852142439 | 0.488499377116194 | NM_026878 |
| Gskip | 1211.490148 | -0.181869351 | 0.069216394 | -2.627547312 | 0.008600288 | 0.052851067 | | Renbp | 156.4308354 | -0.163768563021981 | 0.0950196499775227 | -1.72352311401611 | 0.0847939689740749 | 0.488499377116194 | NM_001164704,NM_023132 |
| H2-T23 | 775.1088678 | 0.348438446 | 0.132612019 | 2.627502757 | 0.008601414 | 0.052851067 | | Rimbp2 | 4806.382694 | -0.121222770168274 | 0.070296142085258 | -1.72445836389215 | 0.0846251291589545 | 0.488499377116194 | NM_001081388 |
| Incenp | 738.8310002 | 0.196877904 | 0.074931127 | 2.62745151 | 0.00860271 | 0.052851067 | | Rnd2 | 2243.273133 | 0.0977718747447316 | 0.0567246832678104 | 1.72362134281346 | 0.0847762230216193 | 0.488499377116194 | NM_009708 |
| Osbpl6 | 2933.617494 | 0.163703201 | 0.062341703 | 2.625901967 | 0.00864197 | 0.05307122 | | Trit1 | 533.7245096 | -0.122754685973753 | 0.0712384591012089 | -1.72315189747935 | 0.0848610598547798 | 0.488499377116194 | NM_025873 |
| Fcgbp | 12.15364838 | 0.424997902 | 0.161883486 | 2.62533204 | 0.008656451 | 0.053139083 | | Ube2v1 | 8981.914963 | 0.0737309487014546 | 0.0427991391567394 | 1.72272036667458 | 0.0849391054416441 | 0.488499377116194 | NM_023230 |
| Map4k1 | 71.572548 | 0.343200707 | 0.130757842 | 2.624704575 | 0.008672418 | 0.053216016 | | Zfp455 | 76.63077951 | 0.163941662202597 | 0.0951017942220992 | 1.72385456597938 | 0.0847341011057196 | 0.488499377116194 | NM_001048204 |
| 4933407K13Rik | 118.1492937 | 0.374167069 | 0.142587059 | 2.62413063 | 0.008687046 | 0.053284676 | | Zfp651 | 4113.652274 | 0.107738988174129 | 0.0624584157422943 | 1.72497151734786 | 0.0845326056474914 | 0.488499377116194 | NM_001166644 |
| Acss3 | 40.66408045 | 0.437113118 | 0.166613061 | 2.623522514 | 0.008702569 | 0.053358769 | | Tnfaip6 | 393.3999466 | 0.153332864263467 | 0.0890312568742387 | 1.72223632066722 | 0.0850267178646547 | 0.488760626712297 | NM_009398 |
| 1700001O22Rik | 145.43033 | 0.330566459 | 0.126030799 | 2.622902185 | 0.00871843 | 0.053413742 | | Fbxo25 | 2539.706752 | 0.0931427396584143 | 0.0540885978417745 | 1.72204019654725 | 0.0850622371672442 | 0.488771383343113 | NM_025785 |
| Ssfa2 | 2341.970013 | 0.23448622 | 0.089395045 | 2.623033735 | 0.008715064 | 0.053413742 | | D17H6S53E | 607.735037 | 0.111488352426873 | 0.0647714310966446 | 1.72125813092693 | 0.0852039934729601 | 0.489390873315253 | NM_033477 |
| Calm2 | 49721.6461 | -0.148800135 | 0.056734253 | -2.622756587 | 0.008722156 | 0.053415451 | | Rcor2 | 502.3597743 | -0.140118107354381 | 0.0814131887318306 | -1.72107381539765 | 0.0852374300900171 | 0.489390873315253 | NM_054048 |
| Plau | 140.7666596 | 0.407199065 | 0.155305391 | 2.621924856 | 0.008743471 | 0.053524826 | | 6430584L05Rik | 180.838553 | -0.155270754669749 | 0.0902398160378578 | -1.72064575585355 | 0.0853151287885748 | 0.489643445587846 | NR_046179 |
| Fam96a | 568.5432062 | -0.170081777 | 0.064881837 | -2.621408141 | 0.008756736 | 0.053584859 | | Sod2 | 5251.63261 | 0.0991965829456308 | 0.0576587437059633 | 1.72040832959341 | 0.0853582439568883 | 0.489697417790584 | NM_013671 |
| Pfdn1 | 1427.856507 | -0.151237023 | 0.057696744 | -2.621240169 | 0.008761052 | 0.053590105 | | Aldob | 66.26202399 | -0.163161070825436 | 0.0948605241313661 | -1.72001021836528 | 0.0854305841662797 | 0.489918936280845 | NM_144903 |
| Als2cl | 481.7774557 | 0.3241591 | 0.123672779 | 2.621103066 | 0.008764576 | 0.053590506 | | Mchr1 | 906.7964038 | 0.108480088211991 | 0.0630805090470714 | 1.71970850981945 | 0.0854854401904177 | 0.490040056908448 | NM_145132 |
| Usp6nl | 844.9485865 | -0.190985448 | 0.072880641 | -2.620523712 | 0.008779482 | 0.053660474 | | Ubald1 | 3725.507584 | 0.0852774435840642 | 0.0495954493895424 | 1.71946105204655 | 0.0855304537017177 | 0.490104682631618 | NM_145359 |
| Cdk11b | 2774.297479 | 0.163480168 | 0.062389417 | 2.620318907 | 0.008784757 | 0.053671542 | | Ntn5 | 250.090193 | -0.153896029878614 | 0.0895209474370939 | -1.71910635761263 | 0.085595007379932 | 0.490281181861551 | NM_001033356 |
| Plch2 | 3864.863156 | 0.258169869 | 0.09853474 | 2.620089832 | 0.008790661 | 0.053683867 | | Rnf103 | 1770.206256 | 0.113124628798651 | 0.0658142527490826 | 1.71884696814745 | 0.0856422406693298 | 0.490358371289982 | NM_009543 |
| Plvap | 322.912716 | 0.265729141 | 0.101424587 | 2.619967684 | 0.00879381 | 0.053683867 | | Tet2 | 1638.690964 | -0.114368851446668 | 0.0665507489512745 | -1.7185208769086 | 0.0857016498433504 | 0.490505187401303 | NM_001040400 |
| Tnik | 6364.252787 | 0.181047146 | 0.069106252 | 2.619837439 | 0.008797169 | 0.053683867 | | Slmo2 | 1601.198099 | 0.103762205823332 | 0.060389963081627 | 1.71820283584344 | 0.0857596244692892 | 0.490643680599013 | NM_025531 |
| Ihh | 10.92781065 | 0.424628891 | 0.162100461 | 2.619541536 | 0.008804805 | 0.05370931 | | Lgals12 | 37.76544752 | -0.156972822288266 | 0.0914077410821881 | -1.71728149530712 | 0.0859277514206426 | 0.491218621462516 | NM_019516 |
| Igf1 | 330.1659199 | -0.260893377 | 0.099615649 | -2.618999933 | 0.008818796 | 0.053773488 | | Pigt | 3689.18136 | 0.0841771948386085 | 0.049013264090836 | 1.71743703260822 | 0.0858993501805367 | 0.491218621462516 | NM_133779 |
| Enoph1 | 1422.581605 | -0.141499956 | 0.054070542 | -2.616950951 | 0.008871909 | 0.054017095 | | Scube1 | 3220.3861 | -0.132793000167775 | 0.0773375290974823 | -1.71705770429247 | 0.0859686291573559 | 0.491258972124214 | NM_022723,NM_001271472,NM_001271473 |
| Gpnmb | 184.0650342 | 0.435719212 | 0.166499562 | 2.616939091 | 0.008872217 | 0.054017095 | | Sec61a2 | 2412.738344 | -0.0885938588749086 | 0.0516111782975999 | -1.71656338408047 | 0.0860589775254201 | 0.491581874767696 | NM_021305 |
| Sema6d | 2951.992139 | -0.214548365 | 0.081975295 | -2.617232 | 0.008864607 | 0.054017095 | | Itfg1 | 11150.74297 | 0.114426572257933 | 0.0666689126079984 | 1.71634076185915 | 0.0860996918934123 | 0.49162111809841 | NM_028007 |
| Tyk2 | 736.6277638 | 0.27729153 | 0.105960995 | 2.616920778 | 0.008872693 | 0.054017095 | | Arrdc3 | 2271.91487 | 0.139646071788857 | 0.08138010060619 | 1.71597320166295 | 0.086166947364505 | 0.491660395221328 | NM_001042591 |
| Lig1 | 436.0703522 | 0.272480381 | 0.10414853 | 2.616267181 | 0.008889697 | 0.054099356 | | Dennd2a | 1162.726807 | 0.104936216846891 | 0.0611559783855421 | 1.71587831013589 | 0.0861843173277979 | 0.491660395221328 | NM_172477 |
| Dnajc21 | 1006.072871 | 0.266241814 | 0.101769293 | 2.616131118 | 0.00889324 | 0.05409967 | | H3f3b | 5847.095526 | 0.0990553395996972 | 0.0577502560026141 | 1.71523637220263 | 0.0863018988392007 | 0.491660395221328 | NM_008211 |
| Acad10 | 105.7457144 | 0.373817046 | 0.142924898 | 2.615478846 | 0.008910244 | 0.054181837 | | Pgr | 1112.645487 | 0.136840011850006 | 0.0797770738362747 | 1.71527990774443 | 0.0862939204936829 | 0.491660395221328 | NM_008829 |
| Vars2 | 962.9283214 | 0.233949595 | 0.089555844 | 2.612331968 | 0.008992882 | 0.054661717 | | Prph | 41.02283042 | 0.14589272140669 | 0.0850399219503642 | 1.71557920163642 | 0.0862390878676112 | 0.491660395221328 | NM_013639,NM_001163588,NM_001163589 |
| Mmrn2 | 241.7296841 | -0.302078912 | 0.115650479 | -2.611998809 | 0.009001457 | 0.054687291 | | Sf3b1 | 11806.16552 | -0.0962431761918095 | 0.0561121204983287 | -1.71519406746847 | 0.0863096521970527 | 0.491660395221328 | NM_031179 |
| Rad9b | 94.01030266 | 0.36932564 | 0.141400927 | 2.611903939 | 0.009003955 | 0.054687291 | | Gapdh | 53989.88843 | 0.0877513656684597 | 0.0511772790089366 | 1.71465477195723 | 0.0864085403523703 | 0.492030755452188 | NM_008084 |
| Pax6 | 334.7859712 | -0.3320233 | 0.127133991 | -2.611601329 | 0.009011928 | 0.054714267 | | Sirt2 | 5310.516425 | -0.0814545545158568 | 0.0475189154329268 | -1.71415011840559 | 0.0865011592272603 | 0.492365140648583 | NM_001122765,NM_001122766,NM_022432 |
| Ccdc60 | 28.91964419 | -0.438635755 | 0.167987908 | -2.611115051 | 0.009024753 | 0.054762618 | | Nat10 | 1616.7261 | -0.106670090475187 | 0.0622465189565409 | -1.71367157976595 | 0.0865890592818528 | 0.492672414856323 | NM_153126 |
| Raf1 | 2729.586524 | 0.123934005 | 0.047465536 | 2.6110314 | 0.00902696 | 0.054762618 | | Ccnd3 | 942.0067076 | 0.112642648470911 | 0.0657783627723655 | 1.71245746660989 | 0.0868123965210334 | 0.493466363632674 | NM_001081636,NM_001081635,NM_007632 |
| Glt25d1 | 2120.166764 | 0.185958894 | 0.071242961 | 2.61020726 | 0.009048738 | 0.054851779 | | Dio3 | 74.36877849 | 0.107178413351376 | 0.0625807521236594 | 1.71264182219466 | 0.0867784542243501 | 0.493466363632674 | NM_172119 |
| Mybpc2 | 9.082871711 | 0.407381604 | 0.156069127 | 2.610263872 | 0.009047241 | 0.054851779 | | Slc25a17 | 1233.66415 | 0.103040743291405 | 0.060174728325777 | 1.71235909422902 | 0.0868305125599004 | 0.493466363632674 | NM_011399 |
| Plk5 | 810.1352229 | 0.346710775 | 0.132842479 | 2.609939058 | 0.009055835 | 0.054873333 | | Olfr78 | 43.23795534 | -0.159643774873726 | 0.0932537506441369 | -1.71192872963296 | 0.0869098034324615 | 0.493723818795438 | NM_001168503,NM_130866 |
| Ankrd29 | 738.9602421 | -0.234865224 | 0.090027946 | -2.608803532 | 0.009085939 | 0.055034222 | | Pms2 | 703.3010103 | -0.0978196319348503 | 0.0571520503057751 | -1.71156820116681 | 0.0869762725983979 | 0.493908262613107 | NM_008886 |
| C1ql3 | 4383.28079 | -0.296416989 | 0.113632313 | -2.6085625 | 0.009092341 | 0.055051475 | | Scamp1 | 10363.30544 | 0.103788863987419 | 0.0606559674240575 | 1.71110722316588 | 0.0870613210066661 | 0.494198026159762 | NM_029153 |
| 4930432K21Rik | 109.0370228 | 0.32596401 | 0.124971087 | 2.608315396 | 0.009098907 | 0.055069716 | | Ptar1 | 225.1959051 | -0.152639884539616 | 0.0892723709678438 | -1.70982223150092 | 0.0872987506875187 | 0.495352208002694 | NM_028208 |
| Arl5a | 3846.892932 | -0.152465184 | 0.058460246 | -2.608014729 | 0.009106904 | 0.055092094 | | 0610037L13Rik | 1098.277923 | -0.124491303954932 | 0.0728529818353792 | -1.70880176512523 | 0.0874876756684166 | 0.495597905649833 | NM_028754 |
| Timm13 | 1429.025292 | -0.203204087 | 0.077918396 | -2.607909019 | 0.009109716 | 0.055092094 | | 2610015P09Rik | 258.4250176 | -0.152133399662986 | 0.0890646850890632 | -1.70812257979529 | 0.0876136000049169 | 0.495597905649833 | NM_027801 |
| Fam19a3 | 16.13808345 | -0.361628583 | 0.1386794 | -2.607658979 | 0.009116373 | 0.055110838 | | Ctps | 1852.536497 | 0.104828240518125 | 0.0613637042915323 | 1.70831017664933 | 0.0875788040090207 | 0.495597905649833 | NM_016748 |
| Fgf9 | 733.2931817 | 0.180801991 | 0.069343119 | 2.607353033 | 0.009124523 | 0.0551171 | | Dbn1 | 9725.131013 | -0.104735427679684 | 0.0612851225739357 | -1.70898618263068 | 0.0874535089651314 | 0.495597905649833 | NM_001177371,NM_001177372,NM_019813 |
| Ido1 | 138.6857691 | -0.323516708 | 0.124074147 | -2.607446555 | 0.009122031 | 0.0551171 | | Exoc3 | 5163.121147 | 0.0882686939876082 | 0.0516816456870868 | 1.70793117777329 | 0.0876491132875135 | 0.495597905649833 | NM_177333 |
| Zmynd15 | 15.0249667 | 0.433056287 | 0.166102801 | 2.607158247 | 0.009129716 | 0.055126973 | | Ifi30 | 126.8666609 | -0.160009453657141 | 0.0936800527256852 | -1.70804188300024 | 0.0876285713051915 | 0.495597905649833 | NM_023065 |
| Mcin | 9.571698715 | -0.417649728 | 0.160259325 | -2.606086901 | 0.009158322 | 0.055278163 | | Mtch1 | 14983.17587 | 0.0876855140411519 | 0.0513309971550329 | 1.70823710625217 | 0.0875923559875268 | 0.495597905649833 | NM_019880 |
| Chrna2 | 40.96448014 | -0.436581027 | 0.167547535 | -2.605714408 | 0.009168287 | 0.055316761 | | Sar1b | 1303.027518 | 0.114793438989015 | 0.0671757993915492 | 1.70885110454608 | 0.087478533586608 | 0.495597905649833 | NM_025535 |
| Edar | 13.27787921 | -0.432903459 | 0.166166133 | -2.605244837 | 0.009180863 | 0.055371074 | | Tmem121 | 414.6115833 | 0.136010011008024 | 0.0796067476446562 | 1.70853649466672 | 0.0875392220283116 | 0.495597905649833 | NM_153776 |
| Ccdc142 | 321.5963556 | 0.363215045 | 0.139434441 | 2.604916286 | 0.009189671 | 0.055394692 | | Arl5b | 423.9588839 | 0.158251848889716 | 0.0926847907714627 | 1.70741982122962 | 0.0877440488096162 | 0.495775125474115 | NM_029466 |
| Pank4 | 1389.069964 | 0.200540362 | 0.076987827 | 2.604832091 | 0.009191929 | 0.055394692 | | Mrpl49 | 1418.23281 | 0.0985481168645219 | 0.0577184157901976 | 1.70739469396973 | 0.0877487159296399 | 0.495775125474115 | NM_026246 |
| Mecp2 | 6988.573667 | -0.194302606 | 0.074623054 | -2.603787911 | 0.009219979 | 0.05554213 | | Bub3 | 2552.057001 | 0.0878672931427885 | 0.0514873770917412 | 1.70657932304893 | 0.0879002710673485 | 0.495973913612079 | NM_009774 |
| Cspp1 | 637.7998325 | 0.278385826 | 0.106944651 | 2.603083213 | 0.009238953 | 0.055586285 | | Ddx20 | 527.7872978 | -0.133699395686923 | 0.0783484855902512 | -1.70647070814039 | 0.08792047553016 | 0.495973913612079 | NM_017397 |
| Hbp1 | 973.3822187 | 0.20064778 | 0.077087471 | 2.602858517 | 0.00924501 | 0.055586285 | | Pcdhb2 | 136.315463 | 0.158812854316918 | 0.0930494584335971 | 1.7067574276131 | 0.0878671482962254 | 0.495973913612079 | NM_053127 |
| Pms2 | 631.075283 | 0.188045762 | 0.072238828 | 2.603112034 | 0.009238176 | 0.055586285 | | Pglyrp1 | 179.7573408 | 0.161905501580957 | 0.0948614178903634 | 1.70675818664317 | 0.0878670071581125 | 0.495973913612079 | NM_009402 |
| Strada | 967.8267689 | 0.185366986 | 0.071206353 | 2.603236616 | 0.009234819 | 0.055586285 | | Notch4 | 343.3685974 | 0.159936953965385 | 0.0937570457710148 | 1.70586597145992 | 0.0880330366767239 | 0.496223473327936 | NM_010929 |
| Vwa8 | 2146.162872 | 0.169843402 | 0.065252865 | 2.602849717 | 0.009245247 | 0.055586285 | | Tinagl1 | 346.5218157 | 0.159978566047787 | 0.0937800762239866 | 1.70589076581353 | 0.0880284193583732 | 0.496223473327936 | NM_001168333,NM_023476 |
| Gmppb | 389.0998828 | -0.295911639 | 0.113697228 | -2.602628429 | 0.009251216 | 0.055600597 | | Dgkk | 177.885269 | 0.0978097682080496 | 0.0573588299952559 | 1.70522599948673 | 0.0881522827867798 | 0.496559782097955 | NM_177914 |
| Galntl6 | 814.2043737 | -0.293602475 | 0.112835248 | -2.60204573 | 0.009266949 | 0.055673561 | | Inhbb | 238.5355568 | 0.152845363503046 | 0.0896359704976373 | 1.70517887690048 | 0.0881610682934479 | 0.496559782097955 | NM_008381 |
| Slc44a5 | 335.7946527 | 0.333842752 | 0.128351789 | 2.600998037 | 0.009295299 | 0.055822231 | | Chchd3 | 1680.613068 | 0.0969291439713408 | 0.056865777823383 | 1.7045250708148 | 0.0882830363717627 | 0.496728132978743 | NM_025336 |
| Ldb1 | 3544.473516 | 0.198727217 | 0.076435945 | 2.599918378 | 0.009324594 | 0.055976464 | | Speg | 6329.527422 | -0.120308207192325 | 0.0705858428667552 | -1.70442403612621 | 0.0883018966080532 | 0.496728132978743 | NM_001173477,NM_007463,NM_001085370,NM_001085371 |
| 2700069I18Rik | 15.74530004 | -0.436988679 | 0.168120626 | -2.5992568 | 0.009342585 | 0.056062749 | | Wdr25 | 321.9795314 | 0.12763676155514 | 0.0748916498784068 | 1.70428561478308 | 0.0883277411182771 | 0.496728132978743 | NM_177602 |

| | | | | | | | | | | | | | | | |
|---|---|---|---|---|---|---|---|---|---|---|---|---|---|---|---|
| Ctsb | 25983.30086 | 0.1769532 | 0.068092287 | 2.598726034 | 0.009357042 | 0.056127761 | | Zdhhc24 | 964.5183211 | -0.099379422294745 | 0.0583028902235651 | -1.70453680621441 | 0.088280845924684 | 0.496728132978743 | NM_027476,NM_001168516,NM_001168517 |
| Flt3l | 109.2265948 | 0.373314579 | 0.143662666 | 2.598549706 | 0.009361849 | 0.056134864 | | Limk1 | 2738.60956 | 0.0886975931140307 | 0.0520663923668977 | 1.70354789494542 | 0.0884655829203137 | 0.497310780766439 | NM_010717 |
| A730085K08Rik | 8.980426969 | -0.408040231 | 0.157036292 | -2.598381728 | 0.009366431 | 0.056140609 | | Serinc5 | 3721.50551 | -0.0869926982114742 | 0.0510820829920311 | -1.70299825527955 | 0.0885683949120798 | 0.497517106718603 | NM_172588 |
| Nudt21 | 834.2462704 | -0.199600703 | 0.076848224 | -2.597336558 | 0.009394982 | 0.056289966 | | Spock3 | 3966.969148 | 0.136213977205242 | 0.0799939968115175 | 1.70280249311946 | 0.088605036147668 | 0.497517106718603 | NM_001252620,NM_001252621,NM_023689 |
| Ccser2 | 4761.90317 | -0.154981226 | 0.059675538 | -2.597064586 | 0.009402424 | 0.05629968 | | Wnt10b | 45.20657458 | -0.152228868503196 | 0.0893986341165354 | -1.70280978012212 | 0.0886036720044096 | 0.497517106718603 | NM_011718 |
| Erp29 | 2311.972383 | -0.14123071 | 0.054382006 | -2.597011754 | 0.009403871 | 0.05629968 | | Galnt3 | 40.54175999 | -0.158288585402653 | 0.0929812853846 | -1.70237037214447 | 0.0886859604327598 | 0.497760972110522 | NM_015736 |
| Tjp3 | 16.8448153 | 0.430437402 | 0.165776449 | 2.596493077 | 0.00941808 | 0.056362973 | | Hspb2 | 23.59026371 | -0.141107796143527 | 0.0828970779470918 | -1.7022047053768 | 0.0887170010184593 | 0.497760972110522 | NM_024441,NM_001164708 |
| Ascc2 | 672.8611184 | 0.246690875 | 0.09503656 | 2.595747106 | 0.009438551 | 0.05646367 | | Cpne6 | 7355.165087 | -0.142396689351677 | 0.0836811225063029 | -1.70165845159345 | 0.0888194133155781 | 0.4978907096955 | NM_001136057,NM_001146183,NM_009947 |
| Rpa1 | 1820.384798 | 0.153781597 | 0.059252609 | 2.595355712 | 0.009449307 | 0.056506199 | | Prdm9 | 83.83594222 | -0.1618317006575 | 0.0951027169549125 | -1.70165170711393 | 0.0888206783735701 | 0.4978907096955 | NM_144809 |
| Bub1 | 9.631153939 | -0.399966074 | 0.15418346 | -2.594091964 | 0.009484111 | 0.056676538 | | Syngr1 | 29505.42441 | 0.0811069510569312 | 0.0476669876014921 | 1.70153297151911 | 0.0888429519159257 | 0.4978907096955 | NM_009303,NM_207708 |
| D430042O09Rik | 1191.97309 | 0.155866674 | 0.060089161 | 2.593923275 | 0.009488766 | 0.056676538 | | Mum1l1 | 154.8451774 | 0.159159855183421 | 0.0935755865492858 | 1.70086943670497 | 0.088967506897377 | 0.498204319657401 | NM_175541,NM_001164631,NM_001164630,NM_001164633,NM_001164632 |
| Hprt | 4053.716439 | -0.18862834 | 0.072715897 | -2.594045419 | 0.009485396 | 0.056676538 | | Rnf41 | 3563.521923 | 0.0781263924592285 | 0.0459283295762746 | 1.70105016185014 | 0.0889335682680878 | 0.498204319657401 | NM_001164237,NM_026259 |
| Hmgb3 | 1246.713194 | -0.24021185 | 0.092613929 | -2.593690318 | 0.009495197 | 0.056693095 | | Tmem59l | 10839.8556 | 0.0788621738951604 | 0.046395642210859 | 1.69977545599536 | 0.0891731698920153 | 0.499163570655651 | NM_182991 |
| Cd151 | 936.4358161 | -0.184831119 | 0.071271076 | -2.593353862 | 0.009504492 | 0.056704891 | | Cnih | 2298.233143 | 0.108695793975738 | 0.0639651640053521 | 1.69929672917971 | 0.089263288558006 | 0.49945978318924 | NM_009919 |
| Tnfsf13b | 35.04158362 | 0.416750175 | 0.160697549 | 2.593382269 | 0.009503707 | 0.056704891 | | Col5a2 | 201.7357889 | -0.161587285791325 | 0.0951012040167917 | -1.69910872803245 | 0.0892986991774324 | 0.49945978318924 | NM_007737 |
| Exoc3 | 4666.46096 | -0.12884067 | 0.049687981 | -2.592994685 | 0.009514424 | 0.056720464 | | Grb2 | 6910.897565 | 0.0783581321047011 | 0.0461220127131267 | 1.69893132357599 | 0.0893321242507435 | 0.49945978318924 | NM_008163 |
| Rps24 | 8193.711025 | -0.18831099 | 0.072621383 | -2.593051553 | 0.009512851 | 0.056720464 | | Srp54c | 631.7076961 | 0.128984021811147 | 0.07592814947711 | 1.69876419614351 | 0.0893636222297146 | 0.49945978318924 | NM_001100110 |
| Wsb1 | 2660.787774 | 0.351343111 | 0.135525846 | 2.592443584 | 0.009529681 | 0.056789568 | | Inpp5k | 1989.657461 | -0.0807518797294575 | 0.0475563361941734 | -1.69802567211541 | 0.0895029166259372 | 0.500045910349371 | NM_008916 |
| Gm15328 | 88.29767242 | 0.431562799 | 0.166534181 | 2.591436763 | 0.009557611 | 0.056934109 | | Tmco1 | 1203.332133 | 0.110451931112283 | 0.0650569443319037 | 1.69777311625313 | 0.0895505917587339 | 0.500119913835974 | NM_001039483 |
| Nip7 | 963.137193 | -0.171352023 | 0.066134946 | -2.590945246 | 0.009571272 | 0.056971681 | | Slc36a1 | 4282.048322 | 0.117570205757736 | 0.0692809840879066 | 1.69700542371854 | 0.0896956351335021 | 0.500737431187261 | NM_153139 |
| Rps9 | 5525.150169 | -0.156961625 | 0.060579089 | -2.591019888 | 0.009569196 | 0.056971681 | | 3110056K07Rik | 215.2790016 | -0.156852165691653 | 0.092461745555674 | -1.69640065466002 | 0.0898100298322319 | 0.500876679955115 | NR_045056,NR_045055 |
| Snx31 | 9.655172613 | -0.421946834 | 0.162894287 | -2.590310816 | 0.009588931 | 0.057023545 | | Hsf1 | 1654.797284 | -0.119388651159522 | 0.0703686505726423 | -1.69661703312438 | 0.0897690874180787 | 0.500876679955115 | NM_008296 |
| Tmem223 | 775.9271524 | -0.200153233 | 0.077272211 | -2.590235619 | 0.009591026 | 0.057023545 | | Tep1 | 312.0351233 | -0.152658431148765 | 0.089993530815978 | -1.69632672220547 | 0.0898240225308463 | 0.500876679955115 | NM_009351 |
| Tob2 | 1206.785894 | 0.316793369 | 0.122299999 | 2.590297401 | 0.009589305 | 0.057023545 | | Dnajc6 | 15129.80773 | 0.0882323898903885 | 0.0520302914981562 | 1.69578888278024 | 0.0899258686139852 | 0.500949827324692 | NM_198412,NM_001164585,NM_001164584,NM_001164583 |
| Pmm2 | 636.2002941 | 0.175984287 | 0.067957567 | 2.58962018 | 0.009608188 | 0.057103668 | | Fam20b | 5170.843302 | 0.0815690170862998 | 0.0480953766461285 | 1.69598457844421 | 0.0898888006361239 | 0.500949827324692 | NM_145413 |
| Dkk3 | 12875.11359 | -0.297464086 | 0.114898069 | -2.588938958 | 0.009627216 | 0.057150989 | | Mrpl15 | 983.0350172 | 0.112740032890993 | 0.0665068060647166 | 1.69516534565331 | 0.0900440588699416 | 0.500949827324692 | NR_033530,NM_025300,NM_001177658 |
| Sc4mol | 2804.140156 | -0.334658407 | 0.129260044 | -2.589032126 | 0.009624611 | 0.057150989 | | Nceh1 | 6254.402864 | 0.108003227954675 | 0.0637009471375967 | 1.69547287454586 | 0.0899857517475076 | 0.500949827324692 | NM_178772 |
| Susd5 | 301.7701421 | -0.276897413 | 0.106952588 | -2.58897347 | 0.009626251 | 0.057150989 | | Pmm1 | 4604.255351 | 0.0919563850679793 | 0.0542436019143452 | 1.69524850531101 | 0.0900282888962187 | 0.500949827324692 | NM_013872 |
| Mme | 633.4823142 | -0.307714059 | 0.118873424 | -2.588585822 | 0.009637093 | 0.057187713 | | St6galnac3 | 398.2283711 | 0.140021810040571 | 0.0825855734334357 | 1.69547542287672 | 0.0899852687139107 | 0.500949827324692 | NM_011372 |
| Dvl3 | 1577.463177 | 0.205365568 | 0.079356245 | 2.587894226 | 0.009656463 | 0.057280718 | | Sacm1l | 3507.399694 | 0.113169353120422 | 0.0668167286168221 | 1.69372783527642 | 0.0903170125677882 | 0.502091889581153 | NM_030692 |
| Fap | 93.35080752 | 0.388599408 | 0.150174951 | 2.587644645 | 0.009663461 | 0.057300295 | | Slc39a6 | 2041.152423 | 0.103219452904024 | 0.0609424479070391 | 1.69372016466214 | 0.0903184708436978 | 0.502091889581153 | NM_139143 |
| Cdk7 | 2304.743907 | 0.166007308 | 0.064169842 | 2.586998864 | 0.009681591 | 0.057385835 | | Cttn | 4965.778795 | 0.0853418161952408 | 0.0504055005997273 | 1.69310522026047 | 0.0904354408169767 | 0.502238743515435 | NM_007803,NM_001252572 |
| Rpl27a | 7433.819566 | -0.160243273 | 0.061947075 | -2.586777067 | 0.009687825 | 0.057400826 | | Fam83f | 28.82903363 | -0.150140442533131 | 0.0886811867118434 | -1.69303601023057 | 0.0904486130411937 | 0.502238743515435 | NM_145986 |
| Gpr45 | 387.6549953 | -0.191637781 | 0.074093314 | -2.586438233 | 0.009697355 | 0.057435328 | | Ptpro | 1674.014219 | 0.149951637951169 | 0.0885552606396219 | 1.69333146301293 | 0.0903961973379273 | 0.502238743515435 | NM_011216,NM_001164402,NM_001164403,NM_001164401 |
| Klhl17 | 1642.833048 | 0.330448527 | 0.127798292 | 2.585703784 | 0.009718041 | 0.057535851 | | Mesdc2 | 2260.582711 | 0.10426731245669 | 0.0616380179051124 | 1.6916071606521 | 0.090720900281748 | 0.502789697631694 | NM_023403 |
| Jazf1 | 1124.852943 | -0.177542804 | 0.068676883 | -2.585190167 | 0.00973253 | 0.057577633 | | Psme4 | 2665.152143 | -0.108215834846921 | 0.063962668037474 | -1.69185930117112 | 0.0906728035351134 | 0.502789697631694 | NM_134013 |
| Sestd1 | 2856.895091 | 0.136523196 | 0.052807495 | 2.585299627 | 0.009729441 | 0.057577633 | | Rad54l | 22.67863056 | -0.134174570110102 | 0.079311398970311 | -1.69174383319512 | 0.090694826936876 | 0.502789697631694 | NM_009015,NM_001122958,NM_001122959 |
| Armc5 | 600.5726449 | 0.179284029 | 0.069391078 | 2.583675501 | 0.009775372 | 0.057809012 | | Urod | 1494.053199 | 0.0920131248985846 | 0.0543825384571163 | 1.6919608298745 | 0.0906534423486611 | 0.502789697631694 | NM_009478 |
| Cfh | 1354.701137 | -0.219380876 | 0.08491732 | -2.583464433 | 0.009781355 | 0.057822326 | | Zan | 40.55497771 | -0.152462211503846 | 0.0901041407971669 | -1.69206664815831 | 0.0906332666928253 | 0.502789697631694 | NM_011741 |
| Rpap1 | 1001.342642 | 0.202987301 | 0.078641981 | 2.581157015 | 0.009846978 | 0.058188056 | | 1110051M20Rik | 3264.514111 | 0.0977179457562198 | 0.0578482531201148 | 1.68921169587127 | 0.0911788684261014 | 0.503727415234457 | NM_175123 |
| 6330403A02Rik | 15730.07279 | -0.271155728 | 0.105080012 | -2.580469133 | 0.009866618 | 0.058255183 | | Arf6 | 1448.021783 | 0.100139159936352 | 0.0592837913473393 | 1.68914904260499 | 0.0911908714583246 | 0.503727415234457 | NM_007481 |
| Man2b2 | 1129.911015 | 0.261686296 | 0.101414483 | 2.580364163 | 0.009869618 | 0.058255183 | | Arl6ip5 | 5402.766538 | 0.083476793187798 | 0.0494370493814021 | 1.68854723799923 | 0.0913062291279309 | 0.503727415234457 | NM_022992 |
| Stxbp3a | 725.6685839 | 0.300120832 | 0.11629909 | 2.58059485 | 0.009863026 | 0.058255183 | | Btk | 62.94161719 | -0.160588536930275 | 0.095093382702062 | -1.68874565576678 | 0.0912681822136675 | 0.503727415234457 | NM_013482 |
| Mt1 | 4270.630358 | 0.248165825 | 0.096189298 | 2.579973353 | 0.009880794 | 0.058298942 | | Cybb | 43.24797018 | -0.125071133758958 | 0.0740262237675132 | -1.68955172091118 | 0.0911137490175221 | 0.503727415234457 | NM_007807 |
| BC021891 | 150.8479476 | 0.351776438 | 0.136361582 | 2.579732746 | 0.00988768 | 0.058317366 | | Hnrnph3 | 1604.236155 | -0.0940615009499577 | 0.0556947801335074 | -1.68887462567373 | 0.0912434588696165 | 0.503727415234457 | NM_001079824 |
| Cxxc1 | 1519.476865 | 0.185084255 | 0.07176082 | 2.579182542 | 0.009903444 | 0.05838173 | | Lrrtm4 | 1101.401546 | 0.111360928072474 | 0.0658761564606093 | 1.69045879504313 | 0.0909402149378195 | 0.503727415234457 | NR_027323,NM_178731 |
| Oprl1 | 2084.905807 | 0.218369929 | 0.084669408 | 2.579088886 | 0.009906129 | 0.05838173 | | Pnmal2 | 16599.49626 | 0.110322231714442 | 0.065296169321078 | 1.68956667537354 | 0.0911108858937793 | 0.503727415234457 | NM_001099636 |
| Tmem128 | 678.8815323 | -0.180229013 | 0.069924117 | -2.57749431 | 0.009951952 | 0.058616351 | | Ppid | 2502.175388 | 0.0944494582628761 | 0.0559103585287998 | 1.68930160256841 | 0.0911616464358769 | 0.503727415234457 | NM_026352 |
| Unc13c | 2710.778719 | -0.33558413 | 0.130200541 | -2.577440354 | 0.009953506 | 0.058616351 | | Rpn1 | 5446.249637 | 0.0934957283631356 | 0.0553516632117353 | 1.68912229440132 | 0.0911959962317409 | 0.503727415234457 | NM_133933 |
| Tll2 | 33.70995509 | -0.432461503 | 0.167836353 | -2.576685532 | 0.009975266 | 0.058722175 | | Tigd2 | 534.3147574 | 0.122408838760567 | 0.0724693299813326 | 1.68911232920325 | 0.0911979055549365 | 0.503727415234457 | NM_001081145 |
| Recql4 | 102.2178267 | 0.380794852 | 0.1478128 | 2.576196731 | 0.009989379 | 0.058782925 | | Vamp1 | 9079.100309 | 0.121508420611361 | 0.0719600168773499 | 1.68855464303826 | 0.0913048089712186 | 0.503727415234457 | NM_009496,NM_001080557 |
| C030030A07Rik | 248.1172667 | 0.251824773 | 0.097796165 | 2.574996391 | 0.010024113 | 0.058942547 | | Eno2 | 46505.65641 | 0.0724430695857334 | 0.0429176336892016 | 1.68795582045244 | 0.0914197100049435 | 0.504161999822251 | NM_013509 |
| Tmem174 | 8.879555436 | 0.333825345 | 0.12963844 | 2.575049083 | 0.010022586 | 0.058942547 | | Gng10 | 1012.641527 | -0.11378217673132 | 0.0674425513375725 | -1.6870977517117 | 0.0915845573049426 | 0.504530343399053 | NM_025277 |
| Pqlc2 | 346.4230629 | 0.231023711 | 0.089726842 | 2.574744703 | 0.01003141 | 0.058963075 | | Knstrn | 39.21201519 | -0.153638271012172 | 0.0910608940840996 | -1.68720362958746 | 0.091564203736782 | 0.504530343399053 | NM_026412 |
| Pdlim7 | 1572.99402 | 0.25052761 | 0.097342551 | 2.573670061 | 0.010062618 | 0.059124082 | | Thbs2 | 379.766097 | -0.157472064968754 | 0.0933407962332902 | -1.68706579891582 | 0.0915907005055281 | 0.504530343399053 | NM_011581 |
| Nob1 | 906.9091006 | 0.186463878 | 0.072461454 | 2.573283695 | 0.010073859 | 0.059167695 | | Adamts6 | 152.558716 | -0.152949886791681 | 0.0906900859214935 | -1.68651165381058 | 0.091697292364883 | 0.504543510944049 | NM_001081020 |
| 2610017I09Rik | 867.3877344 | -0.268207579 | 0.104299096 | -2.571523523 | 0.010125214 | 0.059426857 | | Tmem192 | 613.1924042 | 0.125774600677693 | 0.0745665484170845 | 1.68674296112217 | 0.0916527874337987 | 0.504543510944049 | NM_028427,NM_001163747 |
| Ptchd4 | 180.3007202 | -0.338575576 | 0.131664191 | -2.571508422 | 0.010125655 | 0.059426857 | | Vgf | 9574.497451 | 0.129496583929287 | 0.0767785760843392 | 1.68662393252825 | 0.0916756870894378 | 0.504543510944049 | NM_001039385 |
| A730056A06Rik | 137.7093341 | -0.324882338 | 0.126369686 | -2.57088823 | 0.010143806 | 0.059471971 | | Itpk1 | 2774.904524 | 0.103525073154234 | 0.0614257979231314 | 1.68536798307099 | 0.0919175968312285 | 0.505527386898715 | NM_172584 |
| Acot7 | 8096.137784 | -0.169784467 | 0.066042092 | -2.570852334 | 0.010144857 | 0.059471971 | | Spry2 | 1994.832337 | 0.13320439338867 | 0.0790426288261375 | 1.68522220688872 | 0.0919457081224291 | 0.505527386898715 | NM_011897 |
| Mmaa | 689.8980468 | 0.21822006 | 0.084882156 | 2.570859069 | 0.01014466 | 0.059471971 | | Adam9 | 2716.25473 | 0.0913848386528477 | 0.0542388124647505 | 1.68486060995967 | 0.0920154678252198 | 0.505579537879826 | NM_007404,NM_001270996 |
| Baiap2l2 | 80.87529522 | -0.397249873 | 0.154531262 | -2.570676424 | 0.010150011 | 0.059479681 | | Arpc1a | 6032.413455 | 0.0677028482206112 | 0.0401894150303546 | 1.68459401983025 | 0.0920669259297934 | 0.505579537879826 | NM_019767 |
| Hhip | 429.5842646 | -0.25608849 | 0.099637449 | -2.570203204 | 0.010163888 | 0.059538479 | | Atp5l | 3807.233768 | 0.0950123610804529 | 0.0563940229876355 | 1.68479487801898 | 0.0920281534741788 | 0.505579537879826 | NM_013795 |
| Ftsj3 | 1372.985003 | 0.198778514 | 0.077354473 | 2.569709392 | 0.010178386 | 0.059578357 | | Erlin1 | 1188.737533 | -0.115145616371883 | 0.0683652364592813 | -1.68427145630461 | 0.0921292191083505 | 0.505579537879826 | NM_001164359,NM_145502,NM_001164360 |
| Zfp748 | 455.5767041 | -0.219001892 | 0.085222772 | -2.569757915 | 0.01017696 | 0.059578357 | | Ssr2 | 2046.677488 | 0.086798765743628 | 0.0515346738878635 | 1.68427893678918 | 0.0921277741005052 | 0.505579537879826 | NM_025448 |
| Alox12b | 252.6211388 | 0.431291711 | 0.167960582 | 2.567815059 | 0.010234174 | 0.059882286 | | Mir1249 | 47.23218661 | -0.151998604209908 | 0.0902680439857335 | -1.68385840102984 | 0.0922090374034165 | 0.505826464245479 | NR_037206 |
| Cep152 | 152.3413094 | 0.301226692 | 0.117318733 | 2.567592442 | 0.010240748 | 0.059898131 | | Fam84a | 2780.703902 | 0.118547530313665 | 0.0704146058295301 | 1.68356449513701 | 0.0922658652326717 | 0.505947134152431 | NM_029007 |
| Mif4gd | 360.4830886 | 0.242167759 | 0.094355308 | 2.566551525 | 0.010271536 | 0.060055541 | | Ankrd24 | 1477.511836 | -0.127954591534845 | 0.0760465977959508 | -1.68258140723368 | 0.0924561534738334 | 0.506608363976617 | NM_027480 |
| Aaas | 504.3766696 | 0.239070128 | 0.093153461 | 2.566411659 | 0.01027568 | 0.060057103 | | Cstf2 | 4481.077771 | -0.0806874668270235 | 0.0479545763413255 | -1.6825811629054 | 0.0924562008055907 | 0.506608363976617 | NM_133196 |
| Dlx5 | 239.4980981 | -0.393896048 | 0.153521872 | -2.565732439 | 0.010295821 | 0.060129459 | | Idh3a | 8301.072174 | 0.0774777344384828 | 0.0460930685781027 | 1.68089773210044 | 0.0927827805680963 | 0.50686824766159 | NM_029573 |
| Kcnv1 | 3139.357477 | -0.286843005 | 0.111793647 | -2.565825632 | 0.010293056 | 0.060129459 | | Mfsd5 | 1393.953558 | 0.0870006834887948 | 0.0517207881330693 | 1.68212215299109 | 0.0925451554763393 | 0.50686824766159 | NM_134100 |
| Plin4 | 421.809484 | 0.32650994 | 0.127321191 | 2.564458729 | 0.010333687 | 0.060327861 | | Ptpra | 7503.281894 | 0.0729431665499292 | 0.0433952923889905 | 1.68090045104605 | 0.0927822523563648 | 0.50686824766159 | NM_008980,NM_001163688 |
| Mycbp | 157.7033539 | -0.333435452 | 0.130042086 | -2.564058016 | 0.010345625 | 0.060352076 | | Rnf157 | 17133.33016 | 0.077961849248135 | 0.0463807342943452 | 1.68091019761281 | 0.0927803589029059 | 0.50686824766159 | NM_027258 |
| Slc2a4rg-ps | 304.7819679 | 0.430953188 | 0.168071699 | 2.564103246 | 0.010344277 | 0.060352076 | | Smn1 | 542.9803023 | 0.123942904741191 | 0.0737117942457885 | 1.68145282596037 | 0.0926749920946023 | 0.50686824766159 | NM_001252629,NM_011420 |
| Gm16702 | 149.8892959 | 0.349793136 | 0.136442073 | 2.563675037 | 0.010357046 | 0.060365743 | | Tceb2 | 3249.967871 | 0.0850152724150107 | 0.0505734298021316 | 1.68102643517817 | 0.0927577799658707 | 0.50686824766159 | NM_026305 |
| Lrrtm4 | 1013.18802 | -0.198608988 | 0.077473065 | -2.563587583 | 0.010359656 | 0.060365743 | | Ube2g2 | 1887.431077 | 0.10305347367553 | 0.0613032016688333 | 1.68104553873456 | 0.0927540695566633 | 0.50686824766159 | NM_019803 |
| Plekhm1 | 1364.389354 | 0.190249282 | 0.074209956 | 2.563662495 | 0.010357421 | 0.060365743 | | Zfp821 | 1293.281238 | -0.0993958052085322 | 0.0591322043692109 | -1.68090816618171 | 0.0927807535439523 | 0.50686824766159 | NR_030773,NR_030772,NM_001167946,NM_029468 |
| Ednra | 173.6189008 | -0.274665869 | 0.107191301 | -2.562389549 | 0.010395465 | 0.060460709 | | Atp8a2 | 1318.699026 | 0.100394954169224 | 0.0597411075100399 | 1.68050038497114 | 0.0928599994704679 | 0.507099380566924 | NM_015803 |
| Jrk | 431.8810855 | -0.256834836 | 0.100230364 | -2.562445414 | 0.010393792 | 0.060460709 | | Fosl2 | 1641.438958 | 0.143312447218703 | 0.0852892227400739 | 1.68031132908151 | 0.0928967579675316 | 0.50710947246763 | NM_008037 |
| Med10 | 658.123239 | -0.190768045 | 0.07444064 | -2.56268679 | 0.01038657 | 0.060460709 | | Tbccd1 | 594.5363998 | 0.105321423680576 | 0.0626893261625689 | 1.68005352948684 | 0.0929469012558242 | 0.507192594906876 | NM_001081368 |
| Sirt2 | 4875.095061 | 0.144992453 | 0.056581755 | 2.562530146 | 0.010391257 | 0.060460709 | | Tmem59 | 7226.004225 | 0.0911249158520678 | 0.0542510581320388 | 1.67968918929256 | 0.0930178042594026 | 0.507388893981255 | NM_029565 |
| Trim43a | 19.89857973 | 0.425084875 | 0.165889881 | 2.5624521 | 0.010393592 | 0.060460709 | | Fam211b | 924.8721535 | 0.124110332227132 | 0.0739074281309991 | 1.67926736683558 | 0.0930999479078586 | 0.507646337578661 | NM_198860 |
| Marcks | 3298.500684 | -0.308585171 | 0.120475793 | -2.561387328 | 0.010425505 | 0.060612674 | | 2010111I01Rik | 728.4696742 | -0.153911905199629 | 0.0916782325543772 | -1.67882714261904 | 0.0931857370972156 | 0.507923458564411 | NM_028079 |
| Chp2 | 53.17426593 | -0.422928585 | 0.165147491 | -2.560914378 | 0.010439708 | 0.060672482 | | Grik3 | 5968.400168 | 0.111040973561439 | 0.0661597978691033 | 1.67837534481488 | 0.0932738476527559 | 0.508213019881445 | NM_001081097 |
| Cish | 288.3801045 | 0.215417436 | 0.084131875 | 2.560473508 | 0.010452963 | 0.060726739 | | Gcnt4 | 691.4381233 | -0.139253782412262 | 0.083002592184215 | -1.67770401800468 | 0.0934048946692228 | 0.508545539717065 | NM_001166065 |
| 5330434G04Rik | 212.669487 | -0.288553416 | 0.112707455 | -2.560198139 | 0.01046125 | 0.060752102 | | Mmp15 | 1145.306193 | 0.125058961728521 | 0.0745349268810184 | 1.67785717329749 | 0.093374984839553 | 0.508545539717065 | NM_008609 |
| Mrpl28 | 1260.636699 | -0.18211918 | 0.071146573 | -2.559774472 | 0.010474011 | 0.06080342 | | 9530082P21Rik | 80.94987343 | -0.159483894769307 | 0.0950730396727055 | -1.67748812195696 | 0.0934470702498383 | 0.508585451944231 | NR_015472 |
| Fam76b | 522.1441007 | 0.265488141 | 0.103730059 | 2.559413766 | 0.010484886 | 0.060843757 | | Casp6 | 95.60868786 | -0.155851836936512 | 0.0929210866479855 | -1.67724940117121 | 0.0934937224588501 | 0.508647869826688 | NM_009811 |
| Cybrd1 | 143.0888403 | 0.416886143 | 0.162902812 | 2.559109555 | 0.010494066 | 0.060851447 | | Rpap3 | 1050.366857 | 0.105612274407883 | 0.0629798464969297 | 1.67692175008766 | 0.0935577843659332 | 0.508805831411286 | NM_028003 |
| Klhl9 | 3548.340711 | -0.159257735 | 0.062228971 | -2.559221742 | 0.01049068 | 0.060851447 | | B230206H07Rik | 185.4442793 | -0.159405303872356 | 0.0950753903272697 | -1.67662003094228 | 0.0936168072330505 | 0.508936280638956 | NR_033532 |
| Dcun1d4 | 4374.56235 | -0.14823998 | 0.057952616 | -2.557951467 | 0.010529079 | 0.061031629 | | 1700030J22Rik | 395.528747 | 0.14367278755351 | 0.0857314957656272 | 1.67584603850005 | 0.0937683536875283 | 0.509378872724396 | NM_027103 |
| Slc9a5 | 1700.740138 | 0.201862697 | 0.078950813 | 2.556815941 | 0.010563509 | 0.061208308 | | Tomm40 | 2343.95292 | 0.092324536510856 | 0.0550871590191465 | 1.67597200790055 | 0.09374367568784 | 0.509378872724396 | NM_001109748,NM_016871 |
| Chrdl1 | 767.7015845 | -0.286801204 | 0.112177523 | -2.556672651 | 0.010567861 | 0.061210633 | | Elp3 | 3903.957752 | 0.0814276117054879 | 0.0485989874894029 | 1.67550016804863 | 0.0938361381342281 | 0.509556539266466 | NM_001253812,NM_028811,NR_045599 |
| Fam160b1 | 1619.75783 | 0.191079641 | 0.074746301 | 2.556375869 | 0.01057688 | 0.061239978 | | Oxtr | 483.0012484 | 0.142316486832292 | 0.0849930915173081 | 1.67444770265017 | 0.0940426443635352 | 0.510487089695333 | NM_001081147 |
| Cmpk1 | 1722.468066 | -0.229079894 | 0.0896373 | -2.555631344 | 0.010599535 | 0.061325316 | | Akr1c14 | 64.30038214 | -0.158908283021777 | 0.0949382491930553 | -1.67380675726008 | 0.0941685839029158 | 0.51097977204847 | NM_134072 |
| Lman2 | 3320.594971 | -0.13155135 | 0.051472644 | -2.555752706 | 0.010595839 | 0.061325316 | | Tbc1d22b | 2224.341879 | 0.103846262878076 | 0.0620656216205584 | 1.67316881981696 | 0.0942940666424707 | 0.511469608681303 | NM_198647 |
| Trip10 | 252.6770645 | 0.374066109 | 0.146405936 | 2.554992775 | 0.010619 | 0.061415002 | | Arf2 | 2080.706031 | 0.100254550810731 | 0.0599365477702123 | 1.67267809943095 | 0.0943906828687981 | 0.511705397359582 | NM_007477 |
| Fam83f | 21.3748766 | 0.428149172 | 0.167595318 | 2.554660696 | 0.010629135 | 0.06145068 | | Hyou1 | 5364.034184 | 0.102059901938976 | 0.0610348843358219 | 1.67215688289715 | 0.0944933902619953 | 0.511705397359582 | NM_021395 |
| Pan3 | 1214.579562 | 0.197435271 | 0.077290167 | 2.554468167 | 0.010635015 | 0.061461741 | | Ift52 | 1321.571298 | 0.0831480097028301 | 0.0497197230182293 | 1.67233453155692 | 0.0944583739652206 | 0.511705397359582 | NM_172150 |
| Creg1 | 1042.278544 | 0.202608046 | 0.07935328 | 2.553240974 | 0.010672563 | 0.061609839 | | Mbd4 | 401.739327 | -0.149918904848436 | 0.0866711860408219 | -1.67205401781602 | 0.0945136707363182 | 0.511705397359582 | NM_010774 |
| Pdk3 | 1153.130711 | -0.1629687 | 0.063828177 | -2.553240726 | 0.01067257 | 0.061609839 | | Stat5a | 79.6012095 | -0.158967819944483 | 0.095046476204069 | -1.67252723397312 | 0.0944204021858631 | 0.511705397359582 | NM_001164062,NM_011488 |
| Prkd2 | 180.3744169 | 0.319078259 | 0.124966081 | 2.55331892 | 0.010670175 | 0.061609839 | | Atp6v1c1 | 10630.10081 | 0.084702522485838 | 0.0506705024554309 | 1.67163375891805 | 0.0945965635699241 | 0.511963369007719 | NM_025494 |
| Slc25a10 | 592.3006943 | 0.171000333 | 0.066985681 | 2.55278933 | 0.010686411 | 0.061666761 | | Agpat6 | 4066.440426 | 0.0653195705564282 | 0.0390975022276196 | 1.67068397812596 | 0.094784115158308 | 0.512787358208411 | NM_018743 |
| Mrpl52 | 435.2197962 | -0.238107697 | 0.093311752 | -2.551743915 | 0.010718527 | 0.06182906 | | Slc7a5 | 3439.387918 | 0.0981716975985679 | 0.0587693744905686 | 1.67045673787667 | 0.0948290320448102 | 0.512839359450079 | NM_011404 |
| Aldh6a1 | 2641.05394 | 0.216962061 | 0.085057343 | 2.550774019 | 0.010748399 | 0.061955245 | | Adcyap1 | 526.0617032 | 0.158012918386164 | 0.0946354875205264 | 1.66970047416822 | 0.0949786398968268 | 0.513457284384557 | NM_009625 |
| Wdr81 | 1394.502408 | 0.221594261 | 0.086869445 | 2.550888415 | 0.010744872 | 0.061955245 | | Alkbh2 | 115.5286466 | -0.152521816297288 | 0.0913827617791864 | -1.66904362844533 | 0.0951087337746565 | 0.513969295688489 | NM_175016 |
| Tst | 770.4894314 | -0.358146629 | 0.140417724 | -2.550579935 | 0.010754386 | 0.0619667 | | Shisa9 | 2812.682748 | 0.108691759912473 | 0.0651420535493332 | 1.66853444112195 | 0.0952096808342035 | 0.514323474822477 | NM_001174086,NM_028277 |
| 0610031J06Rik | 1276.506886 | 0.163391465 | 0.064069732 | 2.550213016 | 0.010765712 | 0.061981468 | | Atrip | 847.8544455 | -0.101144115390446 | 0.0606335835429157 | -1.66812036301396 | 0.0952918356218843 | 0.514330430400178 | NM_172774 |
| Dgat1 | 680.026395 | 0.264866495 | 0.103856614 | 2.550309366 | 0.010762737 | 0.061981468 | | BC065397 | 113.2797228 | -0.158555582574349 | 0.0950581725381485 | -1.66798475439571 | 0.0953187532639172 | 0.514330430400178 | NR_033324 |
| Scx | 43.14213191 | 0.406398603 | 0.159365241 | 2.550108171 | 0.01076895 | 0.061981468 | | Furin | 1583.895526 | 0.107139849658725 | 0.0642396761176726 | 1.66781428758247 | 0.0953525987241967 | 0.514330430400178 | NM_011046,NM_001081454 |
| Vat1l | 3673.592589 | -0.300640851 | 0.117900093 | -2.549962797 | 0.010773441 | 0.061984294 | | Tmem198b | 399.249995 | -0.127971554770267 | 0.0767165263652799 | -1.66810934792511 | 0.0952940218355073 | 0.514330430400178 | NM_178066 |
| Coa6 | 315.2250508 | -0.309658564 | 0.121450604 | -2.549666721 | 0.010782594 | 0.062013925 | | Inpp5e | 1573.982648 | -0.107267748390826 | 0.0643308541363532 | -1.66743858496679 | 0.0954272268611508 | 0.514541906972931 | NM_033134 |
| Slbp | 728.4719232 | 0.196212327 | 0.076966887 | 2.549308331 | 0.010793682 | 0.062054663 | | Gm10451 | 30.7481403 | -0.126289938382705 | 0.0757534733760645 | -1.66711746345624 | 0.0954910502831544 | 0.514694989392616 | NR_028520 |
| Plekhf1 | 125.9128497 | 0.416564314 | 0.163426973 | 2.548932449 | 0.010805323 | 0.062098543 | | Stx4a | 1821.855474 | 0.105658285299857 | 0.0633952171673412 | 1.66666019963234 | 0.0955819911967051 | 0.514803116100608 | NM_009294 |
| Gdpd1 | 1626.402637 | -0.182492948 | 0.07160358 | -2.548656764 | 0.010813867 | 0.062124605 | | Tufm | 2433.355961 | 0.0919414914100752 | 0.0551623341932553 | 1.66674403385412 | 0.0955653130051117 | 0.514803116100608 | NM_172745,NM_001163713 |
| Trafd1 | 2106.066564 | 0.203573624 | 0.07989025 | 2.54816606 | 0.010829091 | 0.062189005 | | Med26 | 385.6557246 | 0.121990549084328 | 0.0732088199649832 | 1.66633677667086 | 0.0956463556166094 | 0.514958844213072 | NM_027485 |
| Ryr3 | 2208.2611 | -0.319152816 | 0.125308133 | -2.546944141 | 0.010867083 | 0.062360955 | | Ammecr1l | 1705.929883 | -0.0996299885668624 | 0.0598131017442949 | -1.66568838032823 | 0.0957754975259113 | 0.515046225629311 | NM_153515,NM_001242430 |
| Snrpe | 600.7237106 | -0.190282579 | 0.07470868 | -2.546994271 | 0.010865522 | 0.062360955 | | Cul9 | 3585.09837 | -0.103958483633952 | 0.062405137009015 | -1.66586419991249 | 0.095740465533773 | 0.515046225629311 | NM_001081335 |
| Acad11 | 1080.329029 | 0.159731568 | 0.062719169 | 2.546774281 | 0.010872373 | 0.062368216 | | Fam111a | 108.8155152 | -0.155882019041846 | 0.0935804593613177 | -1.66575394164266 | 0.0957624332567668 | 0.515046225629311 | NM_026640 |
| Mpzl1 | 716.7581133 | -0.177858949 | 0.069857709 | -2.546017494 | 0.010895973 | 0.06248046 | | Mir3072 | 47.22242229 | -0.148648438683207 | 0.089259676122448 | -1.66534817445773 | 0.0958433125676976 | 0.515046225629311 | NR_037233 |
| Grin2b | 2290.504797 | -0.411914197 | 0.161816638 | -2.545561463 | 0.010910216 | 0.062538986 | | Pex11b | 1687.408521 | 0.0825121059969598 | 0.0495481583006202 | 1.66529107896079 | 0.095854697481962 | 0.515046225629311 | NR_027845,NR_027844,NM_011069,NM_001162388,NM_001162387 |
| Pnma1 | 161.9144658 | -0.323543101 | 0.127108994 | -2.545398952 | 0.010915295 | 0.062544964 | | Zfp784 | 198.2798639 | -0.146651295867699 | 0.0880689314306483 | -1.66518763751757 | 0.0958753265938081 | 0.515046225629311 | NM_001039532 |
| Rfx5 | 882.8432278 | 0.230846569 | 0.090699272 | 2.545186567 | 0.010921937 | 0.062559884 | | Klf8 | 218.2008194 | -0.140950074452182 | 0.0846854141106174 | -1.66439611747155 | 0.0960332954501025 | 0.51551354386851 | NM_173780 |
| Gm3435 | 25.27857174 | -0.42355297 | 0.166435507 | -2.54484741 | 0.01093255 | 0.062597535 | | Mex3b | 558.2294407 | -0.134721042909141 | 0.0809414130848736 | -1.66442662383315 | 0.0960272032378175 | 0.51551354386851 | NM_175366 |
| Slc25a14 | 1897.253094 | -0.162011966 | 0.063682641 | -2.54405226 | 0.010957469 | 0.062717039 | | Ccdc6 | 2635.932676 | 0.130959799866575 | 0.0786966724098559 | 1.66410847951144 | 0.0960907528271009 | 0.51563142798909 | NM_001111121 |
| Hif1an | 2500.047356 | 0.161885317 | 0.063637323 | 2.543873758 | 0.01096307 | 0.062725925 | | Gimap8 | 57.84724191 | -0.155927349562781 | 0.0937453492323767 | -1.66330757567788 | 0.0962508830497797 | 0.516109386187191 | NM_001077410,NM_212486 |
| ORF63 | 26.2326013 | -0.420781454 | 0.165422421 | -2.543678492 | 0.0109692 | 0.06273783 | | Tex2 | 5126.287909 | 0.0819951816814649 | 0.0492934107457137 | 1.66341059466238 | 0.0962302737954997 | 0.516109386187191 | NM_198292 |
| Arrb2 | 1392.625592 | 0.182312848 | 0.071682048 | 2.543354325 | 0.010979383 | 0.0627729 | | Akap12 | 1383.777628 | 0.150082476308217 | 0.0902902132858923 | 1.66222307874055 | 0.0964680542477813 | 0.516701679942209 | NM_031185 |
| Ccnk | 1079.140416 | 0.175585103 | 0.069050189 | 2.542862004 | 0.010994864 | 0.062791901 | | Chrnb1 | 31.52406175 | -0.13993580525144 | 0.0841853019102097 | -1.66223559310499 | 0.0964655460045394 | 0.516701679942209 | NM_009601 |
| Col28a1 | 43.53800897 | 0.410365977 | 0.16137734 | 2.542897146 | 0.010993759 | 0.062791901 | | Nrbp2 | 6763.313593 | -0.125042784262061 | 0.0752130223191362 | -1.66251508590484 | 0.096409541098215 | 0.516701679942209 | NM_144847 |
| Khk | 266.1523313 | 0.244798417 | 0.096268524 | 2.542870787 | 0.010994588 | 0.062791901 | | Apoo | 217.9382852 | 0.157978656191016 | 0.0950998794511434 | 1.66118671340878 | 0.0966759531974717 | 0.516802453707324 | NM_026673,NM_001199338,NM_001199339,NM_001199337 |
| Cdk5rap3 | 887.2964331 | 0.18031411 | 0.070933567 | 2.54201386 | 0.01102158 | 0.062813732 | | C030039L03Rik | 662.6159213 | -0.140622175897571 | 0.0846696360685281 | -1.66083359309301 | 0.0967468723214332 | 0.516802453707324 | NM_001112731,NM_198417 |
| Fam171b | 9868.905014 | -0.197599745 | 0.077724557 | -2.542307747 | 0.011012316 | 0.062813732 | | Carkd | 1344.265715 | 0.100290879159352 | 0.0603827540671067 | 1.66091925929535 | 0.0967296636781779 | 0.516802453707324 | NM_026995,NM_001190357 |
| Fbxl12 | 377.1574979 | 0.300081804 | 0.118054697 | 2.541887879 | 0.011025554 | 0.062813732 | | Ccdc65 | 292.4224862 | 0.149140826661838 | 0.0847907255636052 | 1.66190443265086 | 0.09653193778961 | 0.516802453707324 | NM_153518 |
| Kcnip4 | 2007.893234 | -0.173751897 | 0.06834652 | -2.542220098 | 0.011015079 | 0.062813732 | | Fezf2 | 913.969935 | 0.137332450857523 | 0.0826563002842289 | 1.66148799771197 | 0.0966154774692973 | 0.516802453707324 | NM_080433 |
| Rab32 | 76.80530418 | 0.363455411 | 0.142960533 | 2.542347904 | 0.011011051 | 0.062813732 | | Oxt | 130.0454043 | 0.0488213794217526 | 0.0293978769386231 | 1.66071106167575 | 0.096771490712097 | 0.516802453707324 | NM_011025 |
| Slc22a3 | 136.9519009 | -0.390945649 | 0.153786613 | -2.542130564 | 0.011017901 | 0.062813732 | | Rnf182 | 387.859798 | -0.13995723898108 | 0.0842297152533144 | -1.66161358328436 | 0.0965902780610556 | 0.516802453707324 | NM_183204 |
| Socs5 | 5009.815253 | -0.128816419 | 0.050678415 | -2.541839951 | 0.011027065 | 0.062813732 | | Smyd4 | 573.6050314 | -0.110551470464216 | 0.0665443503747638 | -1.66132015477817 | 0.0966491642508408 | 0.516802453707324 | NM_001102611 |
| Zkscan6 | 479.693555 | 0.217682586 | 0.085666722 | 2.541040237 | 0.01105232 | 0.062934453 | | Ccdc37 | 133.6101788 | -0.157559230109355 | 0.0949214818360481 | -1.65989012246455 | 0.0969365589139302 | 0.517493735679438 | NM_173775 |
| Flywch1 | 8319.120313 | 0.243925749 | 0.096003878 | 2.540790584 | 0.011060214 | 0.062956268 | | Scamp5 | 21340.20422 | 0.0536650508196507 | 0.0323360851834384 | 1.65960259305404 | 0.0969944263505897 | 0.517612430995101 | NM_020270 |
| Epn3 | 63.74274807 | 0.426404101 | 0.167877896 | 2.539965728 | 0.011086333 | 0.063081765 | | Fam155a | 3678.797636 | 0.0767668924882668 | 0.0462737357914083 | 1.65897330689519 | 0.0971211712653704 | 0.5179082723204 | NM_173446 |
| Tmem196 | 418.8113109 | -0.251940083 | 0.099196999 | -2.539795423 | 0.011091733 | 0.063089319 | | Slc22a18 | 28.66644486 | -0.134900213590233 | 0.0813120152730838 | -1.65904403103496 | 0.097106920070981 | 0.5179082723204 | NM_001042760,NM_008767 |
| Rpl37a | 4165.19202 | -0.184393539 | 0.072616005 | -2.539296091 | 0.011107577 | 0.063156258 | | Cpxm1 | 284.5811399 | -0.146047555204943 | 0.0880649828720648 | -1.65840667245813 | 0.0972354107279005 | 0.518327183938893 | NM_019696 |
| Ap3b2 | 6380.834835 | 0.168990645 | 0.066571804 | 2.538471761 | 0.011133779 | 0.063282013 | | 1810032O08Rik | 101.6062169 | -0.156723449986426 | 0.094631037054144 | -1.65615272605282 | 0.0976908939732376 | 0.518456434333801 | NR_027819,NR_027821,NR_027820 |
| Epha10 | 1995.285619 | 0.224738566 | 0.088574383 | 2.537286274 | 0.011171556 | 0.063439698 | | Abcc1 | 1127.263707 | -0.104333591165938 | 0.0629879173289175 | -1.6564064282538 | 0.0976395401814705 | 0.518456434333801 | NM_008576 |
| Hfe2 | 38.01671704 | 0.42547106 | 0.167692105 | 2.537215806 | 0.011173805 | 0.063439698 | | Atp8b2 | 4277.578005 | -0.0871042661627589 | 0.0525850937563057 | -1.6564440593455 | 0.0976319248236893 | 0.518456434333801 | NM_001081182 |
| Vstm2l | 1890.240467 | -0.241979262 | 0.095366246 | -2.537368026 | 0.011168947 | 0.063439698 | | B230216G23Rik | 70.26702126 | -0.156067999958068 | 0.0941696395447856 | -1.65730696976751 | 0.0974574289048787 | 0.518456434333801 | NM_001242345 |
| Camkk1 | 6415.11481 | 0.328870141 | 0.129636443 | 2.536864897 | 0.011185011 | 0.063480059 | | Bcap31 | 2315.805919 | 0.102646599627165 | 0.0619957366457712 | 1.6557041690409 | 0.0977817426237358 | 0.518456434333801 | NM_012060 |
| Pfdn5 | 3814.140752 | -0.167776788 | 0.066143121 | -2.536571966 | 0.011194373 | 0.06350993 | | Bpgm | 2395.308967 | 0.0954768271668194 | 0.0576198499390265 | 1.65701277021466 | 0.0975168932876594 | 0.518456434333801 | NM_007563 |
| Ptpmt1 | 834.8181617 | -0.173528558 | 0.068417407 | -2.536321736 | 0.011202376 | 0.06353207 | | D17Wsu104e | 1010.517933 | 0.119811651175532 | 0.0723242374344657 | 1.65659059017519 | 0.0976022760823719 | 0.518456434333801 | NM_080837 |
| Tmem200a | 575.710673 | -0.219054855 | 0.086372351 | -2.536168732 | 0.011207272 | 0.06353658 | | Dqx1 | 29.97147434 | -0.14475847400445 | 0.0873702938675198 | -1.65683858433564 | 0.0975521138580779 | 0.518456434333801 | NM_033606 |
| Tagap | 35.7175708 | 0.423186793 | 0.166926005 | 2.535175947 | 0.011239086 | 0.063685682 | | Fam134a | 9301.710451 | 0.0675455340943275 | 0.0407956886451752 | 1.65570275530367 | 0.0977820290621818 | 0.518456434333801 | NM_170755 |
| Usp2 | 3083.909682 | 0.198220353 | 0.078190608 | 2.53509159 | 0.011241793 | 0.063685682 | | Lrch3 | 1714.790976 | -0.114429176247295 | 0.0690902355623826 | -1.65622790711105 | 0.0976756737536577 | 0.518456434333801 | NM_001081255 |
| Tsc22d4 | 2292.721168 | 0.187720992 | 0.074068948 | 2.53440877 | 0.011263725 | 0.063786608 | | Mpc1 | 3181.559217 | 0.0857510871042708 | 0.0517933611596637 | 1.65563858348419 | 0.0977950316724917 | 0.518456434333801 | NM_018819 |
| Opn3 | 167.9629247 | -0.329088965 | 0.129862169 | -2.534140365 | 0.011272357 | 0.063812166 | | Phc1 | 2332.564676 | -0.0959742976714518 | 0.0578853526060787 | -1.65800661740071 | 0.0973161307138167 | 0.518456434333801 | NM_001271579,NM_001042623,NR_073364,NM_007905 |
| Atp5o | 5375.182098 | -0.174056831 | 0.068694918 | -2.53376578 | 0.011284413 | 0.063833939 | | Pvalb | 2358.121772 | 0.147813292285803 | 0.0892607290724755 | 1.65597227158861 | 0.0977274342680761 | 0.518456434333801 | NM_013645 |
| C1galt1c1 | 635.1293776 | -0.203575367 | 0.080345012 | -2.533764849 | 0.011284443 | 0.063833939 | | Sf3b3 | 4155.579579 | 0.074976221483193 | 0.0452514054968295 | 1.65688160754357 | 0.0975434135742575 | 0.518456434333801 | NM_133953 |
| Ccdc78 | 11.35947521 | 0.402335971 | 0.158822798 | 2.533238148 | 0.011301414 | 0.06390661 | | Slc50a1 | 1072.214212 | -0.121444293036409 | 0.0733236707242086 | -1.65627677715694 | 0.0976657811498537 | 0.518456434333801 | NM_009057 |
| Rfx3 | 1815.425875 | -0.237778996 | 0.09387261 | -2.532996536 | 0.011309207 | 0.063927345 | | Mrpl21 | 595.0490793 | 0.113432267918435 | 0.0685344039910905 | 1.65511423916638 | 0.0979013270393955 | 0.518695741252113 | NM_172252 |
| Ifitm7 | 17.85656859 | 0.424667829 | 0.167734358 | 2.531787969 | 0.011348259 | 0.064124699 | | Prokr2 | 126.60478 | 0.15739267045298 | 0.0951009389361814 | 1.65500648272884 | 0.097923182913461 | 0.518695741252113 | NM_144944 |
| Tchp | 403.4008906 | 0.221415473 | 0.087460698 | 2.531599658 | 0.011354355 | 0.064135753 | | Wls | 1761.751678 | -0.114507175368901 | 0.0691933240165101 | -1.65488762097307 | 0.0979472957630832 | 0.518695741252113 | NM_026582,NR_037590 |
| Jarid2 | 1600.541071 | -0.175556257 | 0.069362069 | -2.531012393 | 0.011373383 | 0.064196429 | | Kank3 | 762.9556119 | -0.121194313068577 | 0.0732664778029051 | -1.6541577635901 | 0.0980954620444064 | 0.519101887671055 | NM_030697 |
| Ywhaq | 12706.5606 | -0.216844961 | 0.085671146 | -2.531131795 | 0.011369512 | 0.064196429 | | Lrrtm2 | 2846.858699 | 0.13982493419376 | 0.0845208413166627 | 1.65432492170655 | 0.0980615119508705 | 0.519101887671055 | NM_178005 |
| Hes7 | 13.36109971 | 0.403484179 | 0.159435372 | 2.530706793 | 0.011383296 | 0.064228984 | | Pde4c | 145.1428472 | -0.150195799566839 | 0.0908186187774629 | -1.65379964580689 | 0.0981682279717388 | 0.519297771127996 | NM_201607 |
| Yaf2 | 2488.34511 | -0.143218749 | 0.056612429 | -2.529811036 | 0.011412397 | 0.064369742 | | 9030025P20Rik | 66.65325334 | -0.156725071008869 | 0.0948304949816345 | -1.65268641737261 | 0.0983947001373545 | 0.520050240757615 | NM_001123370 |
| Srrm3 | 1699.212928 | 0.159191173 | 0.062947262 | 2.528961047 | 0.011440072 | 0.064502358 | | Asb13 | 2384.192693 | 0.0905660729731318 | 0.0548339467899989 | 1.6516424272719 | 0.0986074655847531 | 0.520050240757615 | NM_178283,NM_001267724 |
| Kif18b | 9.903794219 | -0.383794341 | 0.151872233 | -2.527086972 | 0.011501302 | 0.064823997 | | Dhdds | 3227.124544 | 0.0885797369676877 | 0.0536350787234364 | 1.65152618539892 | 0.0986311784187399 | 0.520050240757615 | NM_026144 |
| Acvr1 | 1144.243057 | -0.138757411 | 0.054923354 | -2.526382714 | 0.011524386 | 0.064930487 | | Fam189a1 | 3618.714526 | 0.0767468601039555 | 0.0464531661278257 | 1.65213410626889 | 0.098507215561289 | 0.520050240757615 | NM_183087 |
| Arhgap9 | 33.10334903 | 0.411850165 | 0.163031636 | 2.526197825 | 0.011530453 | 0.064941056 | | Kcna6 | 5513.77789 | 0.0996802160596892 | 0.0603462604413915 | 1.65180435921956 | 0.0985744397628911 | 0.520050240757615 | NM_013568 |
| Ppp2r5a | 3271.737304 | -0.161666005 | 0.064014641 | -2.525453593 | 0.011554904 | 0.065055118 | | Pik3cg | 200.8150597 | -0.140670351231905 | 0.0851763552765738 | -1.65151879033963 | 0.0986326871325368 | 0.520050240757615 | NM_020272,NM_001146200,NM_001146201 |
| Cap1 | 10568.90027 | -0.1504396 | 0.059591789 | -2.52450216 | 0.011586229 | 0.065184109 | | Ppp1r1a | 2942.904391 | -0.0922508666905582 | 0.0558467143987588 | -1.65185844294912 | 0.098563411420369 | 0.520050240757615 | NM_021391 |
| Dapk2 | 53.41060275 | 0.380436077 | 0.150690744 | 2.52461477 | 0.011582518 | 0.065184109 | | Stxbp1 | 46683.02007 | 0.0674524102894451 | 0.0408402177023834 | 1.65161730480954 | 0.0986125900716428 | 0.520050240757615 | NM_009295,NM_001113569 |
| Slc22a8 | 768.4648216 | -0.358356485 | 0.141976198 | -2.524060303 | 0.011600803 | 0.065242409 | | Zdhhc5 | 2866.36717 | 0.100598338348919 | 0.0608712406559815 | 1.6526414981002 | 0.0984038471423368 | 0.520050240757615 | NM_144887 |
| Fgf3 | 24.28948821 | -0.424131483 | 0.168065943 | -2.523601606 | 0.011615949 | 0.065248465 | | Brpf3 | 1905.495867 | -0.0841553164636606 | 0.0509624529566907 | -1.65131997345322 | 0.098673255956318 | 0.52007536865801 | NM_001081315 |
| Mybl1 | 132.3014691 | 0.28996759 | 0.11490611 | 2.523517601 | 0.011618724 | 0.065248465 | | 1810037I17Rik | 1371.377611 | 0.0882269505194287 | 0.0534758483546081 | 1.64984667348108 | 0.098974299884135 | 0.520271327036625 | NM_024461 |
| Syce2 | 34.21120355 | -0.409118455 | 0.162112881 | -2.523664082 | 0.011613885 | 0.065248465 | | Galnt10 | 640.0203449 | 0.137392680378579 | 0.0833077383095346 | 1.64921870604731 | 0.099102837030122 | 0.520271327036625 | NM_134189 |
| Tia1 | 2159.189722 | 0.295062542 | 0.116923094 | 2.523560847 | 0.011617295 | 0.065248465 | | Gm527 | 151.1447623 | -0.151938553955678 | 0.0921475642850578 | -1.64886131428995 | 0.099176050155887 | 0.520271327036625 | NM_001025605 |
| Nop10 | 1007.761977 | -0.170212648 | 0.067481186 | -2.522371906 | 0.01165664 | 0.065437671 | | Gucy2g | 362.0691768 | 0.1271455564206 | 0.0771103466564554 | 1.64887802913224 | 0.0991726250919654 | 0.520271327036625 | NM_001081076 |
| Stxbp3b | 32.6130831 | 0.411312499 | 0.163102092 | 2.521810081 | 0.011675272 | 0.065518534 | | Mrto4 | 750.287953 | 0.10319816229683 | 0.0625286052953991 | 1.65041522690772 | 0.0988580388588409 | 0.520271327036625 | NM_023536 |
| Doc2g | 378.8774493 | -0.338442561 | 0.134223839 | -2.521478765 | 0.011686273 | 0.065556521 | | Nkain2 | 2263.081525 | 0.106936186151714 | 0.0647908700449706 | 1.6504823299562 | 0.0988443244429055 | 0.520271327036625 | NM_001013411,NM_001025286 |
| Cald1 | 998.5525057 | -0.186006667 | 0.073778024 | -2.521166297 | 0.011696656 | 0.06559102 | | P2ry1 | 179.8881599 | 0.141408640482161 | 0.085710653076447 | 1.64983739367889 | 0.0989761983754206 | 0.520271327036625 | NM_008772 |
| Tril | 2113.073739 | -0.314263014 | 0.124664126 | -2.520877695 | 0.011706253 | 0.065621088 | | Pcdh15 | 687.7594912 | -0.12048616397716 | 0.0730452926459704 | -1.64947198666342 | 0.0990509776075946 | 0.520271327036625 | NM_001142741,NM_001142740,NM_001142743,NM_001142742,NM_001142747,NM_001142746,NM_001142760,NM_001142738,NM_001142739,NM_0011 |
| Adamts2 | 248.2203904 | 0.28839671 | 0.114424805 | 2.52040377 | 0.011722028 | 0.065665269 | | Psmb1 | 4142.98066 | 0.0807025499027062 | 0.0489139537884301 | 1.6498880922972 | 0.0989658266461002 | 0.520271327036625 | NM_011185 |
| Rps10 | 4001.365644 | -0.185016026 | 0.073407806 | -2.520386283 | 0.011722611 | 0.065665269 | | Rab3b | 2577.998487 | 0.0921645060844319 | 0.0558460702812252 | 1.6503310908058 | 0.0988752366097595 | 0.520271327036625 | NM_023537 |
| Fam102a | 4165.251602 | 0.13011946 | 0.051637862 | 2.519846009 | 0.011740619 | 0.065742372 | | Rhebl1 | 239.1888389 | -0.150572392076156 | 0.0913108419869458 | -1.64900891065797 | 0.0991458092214568 | 0.520271327036625 | NM_026967 |
| Vamp3 | 976.72662 | -0.172823222 | 0.068625034 | -2.51836992 | 0.011789942 | 0.065994714 | | Tlr3 | 627.1140405 | -0.124752057630988 | 0.0756072441297453 | -1.6500013863342 | 0.0989426525208954 | 0.520271327036625 | NM_126166 |
| Dok5 | 737.8890983 | -0.272487185 | 0.108217286 | -2.517963581 | 0.011803553 | 0.066030994 | | Trpc7 | 197.9805688 | 0.154979846526788 | 0.093950104733933 | 1.64959737900975 | 0.0990253114344252 | 0.520271327036625 | NM_012035 |
| Mga | 2773.137326 | -0.202494937 | 0.080421451 | -2.51792196 | 0.011804947 | 0.066030994 | | Dnttip2 | 1033.816921 | 0.116365348885074 | 0.0705928620895256 | 1.64840106266692 | 0.0992703981122317 | 0.520390401652211 | NM_153806 |
| Scn3a | 1873.139203 | -0.262959427 | 0.104495511 | -2.516466252 | 0.011853825 | 0.066260306 | | Map2k5 | 1353.735923 | 0.0872345248172584 | 0.0529176823439666 | 1.6484948121921 | 0.0992511743919696 | 0.520390401652211 | NM_011840 |
| Tspan3 | 10279.70557 | -0.089707079 | 0.035648318 | -2.516446299 | 0.011854496 | 0.066260306 | | Dbndd2 | 5936.280301 | 0.102400906415529 | 0.0621398575916702 | 1.64791022033587 | 0.099371095827753 | 0.520411862568074 | NM_001048227,NM_026797,NM_001048229,NM_001048228 |
| Ddx5 | 29185.77943 | 0.173138709 | 0.068823408 | 2.515695078 | 0.011879791 | 0.066377745 | | Lama2 | 673.162143 | -0.131905234208505 | 0.0800494516808508 | -1.64779684855803 | 0.0993943956956276 | 0.520411862568074 | NM_008481 |
| Ttc25 | 61.15023481 | 0.361636371 | 0.143766143 | 2.515448792 | 0.011888095 | 0.066400194 | | Nap1l3 | 1884.967831 | 0.117440337693728 | 0.0712760634480488 | 1.64768271439861 | 0.099417796958998 | 0.520411862568074 | NM_138742 |
| BB031773 | 39.13247164 | -0.415664011 | 0.165260255 | -2.515208581 | 0.011896198 | 0.066421511 | | Psma7 | 2358.18614 | 0.0827247529190235 | 0.050194773927054 | 1.64807501751565 | 0.0993372781283102 | 0.520411862568074 | NM_011969 |
| Fgd3 | 267.2925308 | 0.264657791 | 0.105271191 | 2.514057146 | 0.011935109 | 0.066566805 | | Cdh18 | 998.1466103 | 0.12793949160313 | 0.0776760816453017 | 1.64708992643771 | 0.0995395633757829 | 0.520673999854744 | NM_001081299 |
| Tcof1 | 1188.179985 | 0.1757286 | 0.069892616 | 2.514265597 | 0.011928057 | 0.066566805 | | Mettl16 | 1006.342627 | 0.119916863504889 | 0.072800377696486 | 1.64720111762108 | 0.099516714188361 | 0.520673999854744 | NM_026197 |
| Zbtb12 | 216.6930755 | -0.235494192 | 0.093668995 | -2.514110365 | 0.011933308 | 0.066566805 | | Agpat5 | 1638.658267 | 0.0932415905651957 | 0.0566568369661279 | 1.64572530974399 | 0.0998203257351569 | 0.521115714131388 | NM_026792 |
| Abcg1 | 3352.703806 | 0.240963463 | 0.095892992 | 2.512837034 | 0.011976465 | 0.066761405 | | Atp6ap2 | 9301.666133 | 0.104304283063348 | 0.06340516223231 | 1.64504402151338 | 0.0999607332190369 | 0.521115714131388 | NM_027439 |
| Aff3 | 2436.383502 | -0.18925168 | 0.075325463 | -2.512452912 | 0.01198951 | 0.066761405 | | Card10 | 320.6189843 | -0.13700189938921 | 0.0832115343490858 | -1.64642919351138 | 0.0996754267619806 | 0.521115714131388 | NM_130859 |
| Atp9b | 2880.378434 | 0.116586739 | 0.046405772 | 2.512332701 | 0.011993596 | 0.066761405 | | Chrnb2 | 3214.461829 | 0.0791617488397172 | 0.048098943653609 | 1.6458105485603 | 0.0998027698447365 | 0.521115714131388 | NM_009602 |
| Cdk3-ps | 52.34865923 | 0.419423307 | 0.166950614 | 2.512259742 | 0.011996076 | 0.066761405 | | Crym | 1785.395939 | 0.137969766842571 | 0.0838441948931535 | 1.64554942674794 | 0.0998565585956128 | 0.521115714131388 | NM_016669 |
| Ddah1 | 4334.129333 | -0.262785255 | 0.104606153 | -2.512139557 | 0.012000162 | 0.066761405 | | Dcdc2b | 135.7061619 | -0.155897983694426 | 0.094713683230972 | -1.64599219855327 | 0.099765365206867 | 0.521115714131388 | NM_001195730 |
| Mtm1 | 199.0857281 | 0.303855477 | 0.120949852 | 2.512243481 | 0.011996629 | 0.066761405 | | Dnase1l2 | 164.6030468 | -0.153087076249774 | 0.0930163342823247 | -1.64580852847976 | 0.0998031858744945 | 0.521115714131388 | NM_025718 |
| Zfp503 | 213.5535354 | -0.368404696 | 0.146642708 | -2.512260597 | 0.011996047 | 0.066761405 | | Dzip1 | 2983.436817 | -0.115214549723645 | 0.0700419122982493 | -1.64493723747924 | 0.0999827547352456 | 0.521115714131388 | NM_025943 |
| Cntnap2 | 4282.51122 | -0.19729569 | 0.078582026 | -2.510697417 | 0.012049293 | 0.066986632 | | Sema6c | 800.0252612 | -0.129841427109096 | 0.0789144879030131 | -1.64534334010597 | 0.0998990269171524 | 0.521115714131388 | NR_073549,NM_001272024,NR_073551,NM_011351,NR_073550 |
| Rnpepl1 | 1144.618239 | 0.258594411 | 0.102994666 | 2.510755368 | 0.012047315 | 0.066986632 | | Ubqln2 | 7488.477001 | 0.0707019742744951 | 0.0429777302150747 | 1.64508395209982 | 0.0999524995354516 | 0.521115714131388 | NM_018798 |
| Htr6 | 122.96441 | -0.320303293 | 0.127588227 | -2.510445523 | 0.012057893 | 0.067010397 | | 4932418E24Rik | 111.4204912 | -0.156130365286051 | 0.0950254762990339 | -1.64303691354018 | 0.100375296051568 | 0.521168499961477 | NM_177841 |
| Pde6b | 10.45540635 | 0.355273494 | 0.141529801 | 2.510238066 | 0.012064979 | 0.06702574 | | Arhgef25 | 7295.418901 | -0.114823027785156 | 0.0698990765633383 | -1.64269734924339 | 0.100445567513228 | 0.521168499961477 | NM_001166413,NM_028027 |
| BC028528 | 60.91635151 | -0.355981096 | 0.141838512 | -2.509763326 | 0.01208121 | 0.067067815 | | Blvra | 614.1318862 | 0.123797297612949 | 0.0753358501098137 | 1.64327206014792 | 0.100326656379296 | 0.521168499961477 | NM_026678 |
| Gm17296 | 262.7183125 | 0.290662707 | 0.115809251 | 2.509840145 | 0.012078583 | 0.067067815 | | Dtx3l | 327.908652 | -0.140425799273294 | 0.0854503750877018 | -1.64336082936053 | 0.100308299506126 | 0.521168499961477 | NM_001013371 |
| Slc22a23 | 3858.830133 | -0.133431783 | 0.053178236 | -2.509142695 | 0.012102458 | 0.067149427 | | Dync1i2 | 5682.971103 | -0.0760626597432701 | 0.0462560113938053 | -1.64438431787174 | 0.100096842355054 | 0.521168499961477 | NM_001198874,NM_001198875,NM_001198876,NM_001198877,NM_001198872,NM_001198873,NM_001198878,NM_010064 |
| Traf3ip3 | 26.70997236 | 0.408718251 | 0.162895611 | 2.509080804 | 0.012104579 | 0.067149427 | | Egln1 | 3971.342359 | 0.112732676559937 | 0.0686132075757527 | 1.6430171470336 | 0.100379385575867 | 0.521168499961477 | NM_053207 |
| Otud3 | 370.8996506 | 0.237510325 | 0.094676272 | 2.508657339 | 0.012119098 | 0.067205909 | | Gosr1 | 2347.842871 | 0.0792427868349356 | 0.0482339130742708 | 1.64288530173609 | 0.10040666665889 | 0.521168499961477 | NM_016810 |
| Fezf1 | 10.38889332 | 0.379263999 | 0.151280671 | 2.507022185 | 0.012175307 | 0.067478396 | | Gpr68 | 621.9785986 | 0.146404180731284 | 0.0890880783524032 | 1.64336444829528 | 0.100307551191687 | 0.521168499961477 | NM_175493,NM_001177674,NM_001177673 |
| Pmepa1 | 2294.364198 | -0.187419108 | 0.074759077 | -2.506974615 | 0.012176946 | 0.067478396 | | Kpna2 | 1313.979789 | 0.102894894678293 | 0.0626215649752959 | 1.64312237675447 | 0.100357615958365 | 0.521168499961477 | NM_010655 |
| Mphosph9 | 967.7446748 | 0.193494165 | 0.077204778 | 2.506245982 | 0.01220207 | 0.067593447 | | Rnpepl1 | 1378.572454 | -0.116646749418136 | 0.0710121437186668 | -1.64263101083475 | 0.100459300557079 | 0.521168499961477 | NM_181405 |
| Arfgap3 | 751.124775 | 0.191290932 | 0.076338215 | 2.505834488 | 0.012216279 | 0.067647972 | | Syt4 | 8476.126474 | 0.115836158741883 | 0.0704992426655643 | 1.6430837319968 | 0.10036561023571 | 0.521168499961477 | NM_009308 |
| Chrna6 | 14.2017664 | 0.395650315 | 0.157906165 | 2.505596837 | 0.012224492 | 0.067669266 | | Tmem163 | 707.7050858 | 0.122310861054381 | 0.074384329549424 | 1.64430951781467 | 0.100112284321587 | 0.521168499961477 | NM_028135 |
| Gad2 | 11188.00529 | -0.323058189 | 0.128948649 | -2.505324346 | 0.012233915 | 0.06767942 | | Trex1 | 435.5875693 | -0.122256780531986 | 0.0743480299130714 | -1.64438493763628 | 0.100096714416786 | 0.521168499961477 | NM_011637,NM_001012236 |
| Nif3l1 | 740.932996 | -0.180847298 | 0.072202754 | -2.504714695 | 0.01225502 | 0.06778982 | | Zc3hc1 | 952.1529602 | 0.100747738563435 | 0.0613399228345588 | 1.64244971150622 | 0.100496839866861 | 0.521177115854347 | NM_172735 |
| Map3k5 | 1871.954592 | 0.172442273 | 0.068852441 | 2.50451939 | 0.012261788 | 0.067803051 | | Cacna1g | 4397.059948 | -0.110343600089143 | 0.0672022046085587 | -1.6419639910904 | 0.100597466822964 | 0.521512777683931 | NM_009783,NM_001177890,NM_001112813,NM_001177888 |
| Recql5 | 903.1199365 | 0.257056064 | 0.10267852 | 2.503503793 | 0.012297036 | 0.067973084 | | C3 | 46.09510738 | -0.122625716175677 | 0.074731935352004 | -1.64087435442576 | 0.100823499565158 | 0.52154254668122 | NM_009778 |
| Xab2 | 1423.771143 | 0.167553701 | 0.066930969 | 2.503380771 | 0.012301312 | 0.067973084 | | Glrx5 | 965.3516248 | 0.104285603334977 | 0.0635211323448383 | 1.64174660440308 | 0.100642528942783 | 0.52154254668122 | NM_028419 |
| Pnp | 490.9293828 | -0.209790879 | 0.08382003 | -2.50287288 | 0.012318978 | 0.067998647 | | Higd1b | 77.64062183 | -0.154783749861418 | 0.0943684561121804 | -1.6402064443803 | 0.100962249846316 | 0.52154254668122 | NM_080846 |
| Polr2k | 282.3981513 | -0.236112859 | 0.094336873 | -2.502869259 | 0.012319104 | 0.067998647 | | Loxl2 | 103.3420568 | -0.155871195764232 | 0.0950084937754641 | -1.64060274581982 | 0.100879904645817 | 0.52154254668122 | NM_033325 |
| Stat4 | 83.8514627 | 0.316676075 | 0.126519264 | 2.502987019 | 0.012315006 | 0.067998647 | | Palm2 | 2859.497511 | 0.119216027355198 | 0.0726344335752347 | 1.6413155783987 | 0.100731923957937 | 0.52154254668122 | NM_172868 |
| Nras | 2028.226768 | -0.179577793 | 0.071753357 | -2.502709277 | 0.012324674 | 0.068005162 | | Pfn2 | 16175.77163 | 0.0877167589211769 | 0.0534431041814239 | 1.64131107772864 | 0.100732857732922 | 0.52154254668122 | NM_019410 |
| Itgb2 | 293.5952065 | 0.311751747 | 0.124587808 | 2.502265281 | 0.012340142 | 0.068066273 | | Prmt5 | 2413.694046 | 0.0725328927723298 | 0.0442173480123773 | 1.64037184575296 | 0.1009278755409 | 0.52154254668122 | NM_013768 |
| Apbb2 | 2084.027992 | 0.237634991 | 0.094981039 | 2.501920325 | 0.012352172 | 0.068108382 | | Rspo3 | 375.1703646 | 0.142175749652085 | 0.086658606286801 | 1.64064200596012 | 0.100871749920769 | 0.52154254668122 | NM_028351 |
| Ipo9 | 8393.62714 | 0.158542409 | 0.063393143 | 2.500939392 | 0.012386437 | 0.068273021 | | Ssbp1 | 505.1876082 | -0.117260081053794 | 0.07147667587377 | -1.64053629551658 | 0.10089370823743 | 0.52154254668122 | NM_212468,NM_028358 |
| Tmem81 | 157.2749937 | 0.280364923 | 0.112133319 | 2.500282027 | 0.012409447 | 0.068375525 | | Sv2a | 21942.79796 | 0.0599428749554563 | 0.0365254612700294 | 1.64112574820901 | 0.10077131490229 | 0.52154254668122 | NM_022030 |
| Ccdc92 | 3187.824356 | 0.171530864 | 0.068611311 | 2.500037693 | 0.012418009 | 0.068398378 | | Csnk1d | 6775.670902 | 0.0828727033718588 | 0.0505456826552881 | 1.6395604731889 | 0.101096587268119 | 0.521924471882977 | NM_139059,NM_027874 |
| Opa3 | 1763.470155 | -0.144945427 | 0.057983727 | -2.49976044 | 0.012427731 | 0.068427601 | | Extl2 | 3686.921849 | 0.0942045150878455 | 0.0574591060104729 | 1.63950540878021 | 0.101108045152051 | 0.521924471882977 | NM_021388,NM_001163515,NM_001163514 |
| Ophn1 | 1698.860574 | 0.165645463 | 0.066268001 | 2.499269682 | 0.012432953 | 0.068428542 | | Mrrf | 714.6948683 | 0.100923846580685 | 0.0615779096938992 | 1.63896187906301 | 0.101221199136007 | 0.522322962220119 | NM_026422 |
| Lamtor5 | 1115.640312 | -0.178619744 | 0.071475739 | -2.499026201 | 0.01245351 | 0.068450002 | | Apba1 | 9401.40803 | 0.0855644103677557 | 0.0522169658291393 | 1.63863268026039 | 0.101289847032954 | 0.52249159019911 | NM_177034 |
| Parp9 | 165.8688331 | 0.33617349 | 0.134522284 | 2.499017118 | 0.01245383 | 0.068450002 | | Abca8b | 1072.347326 | -0.116749939547699 | 0.0712716467500892 | -1.63809796561987 | 0.101401259316084 | 0.522600575884305 | NM_013851 |

| | | | | | | | | | | | | | | | |
|---|---|---|---|---|---|---|---|---|---|---|---|---|---|---|---|
| Prepl | 18068.8158 | 0.16466208 | 0.065897302 | 2.498768142 | 0.012462582 | 0.068450002 | | Ascc1 | 611.910022 | 0.11741991236191 | 0.0717404027440569 | 1.63673338691477 | 0.101686189661507 | 0.522606075884305 | NM_026937,NM_001199187 |
| Solh | 1586.392046 | 0.212238545 | 0.084937408 | 2.498764086 | 0.012462725 | 0.068450002 | | Atp5o | 5567.383678 | 0.102648044322661 | 0.0626876158437503 | 1.63745331420025 | 0.101535786177086 | 0.522606075884305 | NM_138597 |
| Thap3 | 333.063253 | 0.219743868 | 0.087923999 | 2.499247875 | 0.012445722 | 0.068450002 | | Chsy1 | 591.3559668 | 0.113352125231662 | 0.0692695762148006 | 1.63639120412928 | 0.101757738883818 | 0.522606075884305 | NM_001081163 |
| Tmem63b | 5132.135135 | 0.116352341 | 0.046563764 | 2.498774383 | 0.012462362 | 0.068450002 | | Gm1673 | 244.4016644 | -0.155153505774322 | 0.0948331059418938 | -1.63606901021873 | 0.10182514513383 | 0.522606075884305 | NM_001033458 |
| Srbd1 | 409.9676013 | -0.2315317 | 0.092684742 | -2.49805625 | 0.012487638 | 0.068540534 | | Mgat3 | 8853.65715 | 0.0877155723146111 | 0.0535953857068807 | 1.63662545119719 | 0.101708754316193 | 0.522606075884305 | NM_010795 |
| Tshz1 | 1816.077142 | -0.341866006 | 0.136853455 | -2.498044397 | 0.012488055 | 0.068540534 | | Napg | 6789.598311 | 0.0866274879251752 | 0.0528965874060688 | 1.63767630717147 | 0.10148923558849 | 0.522606075884305 | NM_028017 |
| Nxph3 | 987.9824782 | -0.286587713 | 0.114756777 | -2.497348921 | 0.012512577 | 0.068626502 | | Nrg1 | 233.3843663 | 0.139641040022016 | 0.085336457111957 | 1.63635853593985 | 0.101764571783223 | 0.522606075884305 | NM_178591 |
| Zfp653 | 597.3371463 | 0.188896803 | 0.075637417 | 2.497398912 | 0.012510813 | 0.068626502 | | Pip5k1b | 842.7611792 | 0.098301834387382 | 0.0600032855290958 | 1.63827419649738 | 0.101364507899522 | 0.522606075884305 | NM_008846 |
| Pcdha12 | 300.6996188 | 0.270678478 | 0.108472907 | 2.495355622 | 0.012583095 | 0.068988845 | | Rad17 | 610.5109504 | 0.107393205755088 | 0.065647012269 | 1.63591916894902 | 0.101856505562045 | 0.522606075884305 | NM_001044371,NM_011233 |
| Pcdh18 | 305.0441844 | -0.288593816 | 0.115730733 | -2.493666191 | 0.012643138 | 0.069293521 | | Slc31a2 | 386.0321703 | -0.139934865198882 | 0.085546771646143 | -1.63577026351983 | 0.101887677743656 | 0.522606075884305 | NM_025286 |
| Hunk | 815.9018934 | 0.27625373 | 0.110815616 | 2.492913356 | 0.012669976 | 0.069378457 | | Smu1 | 2624.158638 | 0.0901974170372039 | 0.055124811137821 | 1.63623992854498 | 0.101789382852504 | 0.522606075884305 | NM_021535 |
| Itpr2 | 1022.74456 | -0.166243746 | 0.066684959 | -2.492972138 | 0.012667879 | 0.069378457 | | Sv2c | 1217.817121 | 0.126197978428071 | 0.0770461755516982 | 1.63795253332713 | 0.101431595947083 | 0.522606075884305 | NM_029210 |
| Wdr83 | 687.0332849 | 0.208141689 | 0.083495315 | 2.492854712 | 0.012672069 | 0.069378457 | | Sytl2 | 3757.830791 | -0.104618017136843 | 0.0639199751296094 | -1.63670303257645 | 0.101692535027924 | 0.522606075884305 | NM_001040087,NM_001040086,NM_001040085,NM_001040088,NM_031394 |
| Ttc18 | 53.54381262 | 0.401007819 | 0.160910933 | 2.492110464 | 0.012698654 | 0.069499449 | | Trpc4 | 617.3699563 | 0.119461928393368 | 0.0729925587342027 | 1.63663160279639 | 0.101707468177768 | 0.522606075884305 | NM_001253683,NM_016984,NR_046161,NM_001253682 |
| Matn2 | 1282.015897 | -0.282512739 | 0.113371097 | -2.491929128 | 0.012705139 | 0.069510388 | | Tubd1 | 195.6050602 | -0.149637511194746 | 0.0914460129691666 | -1.63634811771619 | 0.10176675094237 | 0.522606075884305 | NM_001199045,NM_001199046,NM_001199047,NM_019756 |
| Ggnbp2 | 3504.600686 | 0.122975054 | 0.049358214 | 2.49148105 | 0.012721175 | 0.069549009 | | Adck5 | 528.3807269 | -0.138634712191654 | 0.0847824544644366 | -1.6351816312396 | 0.102010977638953 | 0.522723164244731 | NM_172960,NR_073123 |
| Unc5a | 4732.578681 | 0.308737744 | 0.123915751 | 2.491513327 | 0.012720019 | 0.069549009 | | Bcl2l13 | 2701.947822 | 0.0956338191001055 | 0.0584779489617773 | 1.63538258092147 | 0.101968871674475 | 0.522723164244731 | NM_153516 |
| Mrp63 | 553.5465704 | -0.246135716 | 0.098814765 | -2.490879945 | 0.012742717 | 0.069620454 | | Clvs1 | 2726.460098 | 0.116127482664921 | 0.0710494707379821 | 1.63445950348003 | 0.102162402815007 | 0.522723164244731 | NM_028940 |
| Tuba8 | 541.234206 | 0.321543116 | 0.129088904 | 2.490865634 | 0.01274323 | 0.069620454 | | Crhbp | 766.7828428 | 0.115315598880446 | 0.0705473434351183 | 1.63458456782999 | 0.102136164889293 | 0.522723164244731 | NM_198408 |
| Cdc42se2 | 6584.412517 | -0.138622381 | 0.055658216 | -2.490600492 | 0.012752743 | 0.069647865 | | Obsl1 | 623.4127347 | -0.139906737804663 | 0.0855662049290151 | -1.63507003636223 | 0.10203436663247 | 0.522723164244731 | NM_178884 |
| Rnf170 | 1485.407178 | 0.169525524 | 0.068078644 | 2.490142475 | 0.01276919 | 0.069713116 | | Tle6 | 201.826859 | -0.148602939799952 | 0.090915628334958 | -1.63451479708701 | 0.102150801808978 | 0.522723164244731 | NM_053254 |
| Trib2 | 1610.208354 | -0.246468202 | 0.098998322 | -2.48962 | 0.012787974 | 0.069791078 | | Usp43 | 125.7584053 | -0.150346743648907 | 0.0919800547384549 | -1.6345581014973 | 0.10214171695681 | 0.522723164244731 | NM_173754 |
| Msi2 | 4042.303133 | -0.203190732 | 0.081646379 | -2.488668 | 0.012822264 | 0.069953577 | | Ncam2 | 2660.693913 | 0.127331205329146 | 0.0779282174047847 | 1.63395506235881 | 0.102268286699998 | 0.522856085706222 | NM_001113208,NM_010954 |
| Emb | 907.4011932 | 0.22033939 | 0.088563728 | 2.487919103 | 0.012849296 | 0.070002457 | | Ndufb11 | 2548.119861 | 0.0824879479427368 | 0.050487739788243 | 1.63382136512171 | 0.10229636483389 | 0.522856085706222 | NM_019435 |
| Irf7 | 110.9203906 | 0.415644269 | 0.167062091 | 2.487962804 | 0.012847717 | 0.070002457 | | Sema3a | 1086.555683 | 0.117424685108709 | 0.0718571904446181 | 1.63413966482883 | 0.10222952789488 | 0.522856085706222 | NM_009152,NM_001243073,NM_001243072 |
| Nfasc | 11960.15694 | -0.126155516 | 0.050699635 | -2.488292384 | 0.012835816 | 0.070002457 | | Nefm | 10432.9716 | 0.116564840005688 | 0.0713525290543171 | 1.63364692955666 | 0.102333007761407 | 0.522859398783747 | NM_008691 |
| Smap1 | 3609.743707 | -0.166875616 | 0.067073733 | -2.487942863 | 0.012848437 | 0.070002457 | | 1110004F10Rik | 3058.448699 | 0.0872477809199801 | 0.053504643374216 | 1.63065811521744 | 0.102962479139136 | 0.522975270877619 | NM_019772 |
| 1700024P16Rik | 47.98445793 | 0.417937256 | 0.167998174 | 2.487748804 | 0.01285545 | 0.070009434 | | 1700095B22Rik | 42.22440262 | -0.144901058311464 | 0.0889135812026809 | -1.62968419842586 | 0.103168258716473 | 0.522975270877619 | NR_040759 |
| Omp | 81.77302807 | -0.262601788 | 0.105562887 | -2.487633628 | 0.012859613 | 0.070009434 | | 2210013O21Rik | 811.3279508 | 0.106740225794079 | 0.0653783984598986 | 1.63265280747969 | 0.102542038116784 | 0.522975270877619 | NM_027327 |
| Adi1 | 1039.209929 | 0.164347411 | 0.066084653 | 2.486922502 | 0.012885346 | 0.07003183 | | Abcb7 | 1456.020763 | 0.110499131094287 | 0.067799307489808 | 1.62979734137989 | 0.103144335885859 | 0.522975270877619 | NM_009592 |
| Atox1 | 818.690892 | -0.213881832 | 0.085999427 | -2.487014629 | 0.01288201 | 0.07003183 | | Arhgef6 | 655.0655295 | -0.126934738243957 | 0.0777206849553236 | -1.63321692695996 | 0.102423381161341 | 0.522975270877619 | NM_152801 |
| Ctnnbip1 | 1379.052416 | -0.195603699 | 0.078653769 | -2.486895436 | 0.012886327 | 0.07003183 | | Asic2 | 2824.856602 | 0.0767314277277336 | 0.0470449906979936 | 1.63102227440776 | 0.102885619576547 | 0.522975270877619 | NM_001034013,NM_007384 |
| Polr2m | 6566.03823 | -0.098324381 | 0.039536789 | -2.486908617 | 0.012885849 | 0.07003183 | | B4galt6 | 6811.098659 | 0.0978341203336155 | 0.0600098244049041 | 1.63030172648898 | 0.103037742868649 | 0.522975270877619 | NM_019737 |
| Rps6ka1 | 603.1626318 | 0.212235267 | 0.085329401 | 2.487246658 | 0.012873611 | 0.07003183 | | Caprin1 | 8944.849851 | 0.0918307104903882 | 0.0562563403913047 | 1.6323619675869 | 0.102603256124332 | 0.522975270877619 | NM_016739,NM_001111292,NM_001111289,NM_001111291,NM_001111290 |
| Adam21 | 15.93885895 | 0.417787618 | 0.168025432 | 2.486454657 | 0.012902301 | 0.070094058 | | Eef1e1 | 602.3583082 | 0.118342847718805 | 0.07246297953398 | 1.63314907115171 | 0.102437648177067 | 0.522975270877619 | NM_025380 |
| Nop14 | 908.6755389 | 0.260668092 | 0.104887262 | 2.48522163 | 0.01294708 | 0.070288038 | | Evc | 470.111337 | -0.133365238833688 | 0.0816801873434577 | -1.63277341998371 | 0.10251665928193 | 0.522975270877619 | NM_021292 |
| Sidt2 | 3600.827955 | 0.141182687 | 0.056806689 | 2.48531801 | 0.012943575 | 0.070288038 | | Ezr | 2661.976918 | 0.118367562157271 | 0.072547767487053 | 1.63158104318504 | 0.102767774380038 | 0.522975270877619 | NM_009510 |
| Tbcc | 459.6201582 | -0.174540663 | 0.070238042 | -2.484987598 | 0.012955595 | 0.070309627 | | Gdpd1 | 1676.744932 | 0.094211350713364 | 0.0577343804138001 | 1.63180673349436 | 0.102720206396267 | 0.522975270877619 | NM_025638 |
| Fgfr1op | 589.25957 | 0.240970209 | 0.096992609 | 2.484418254 | 0.01297633 | 0.070380366 | | Gpr137c | 839.1832896 | 0.114316950967135 | 0.0701613971470828 | 1.62934256750171 | 0.103240519582317 | 0.522975270877619 | NM_027518 |
| Hrh2 | 210.7312372 | 0.248166321 | 0.099895645 | 2.484255655 | 0.012982257 | 0.070380366 | | Gtpbp2 | 2882.20412 | -0.117354657017968 | 0.07202895531096 | -1.62927056919443 | 0.103255753605742 | 0.522975270877619 | NM_001145979,NM_019581 |
| Mlxip | 1561.133755 | 0.188017591 | 0.075681143 | 2.484338697 | 0.012979229 | 0.070380366 | | Iqsec1 | 15142.34491 | 0.0603821936540919 | 0.0370394190447197 | 1.63021438271451 | 0.103056195182018 | 0.522975270877619 | NM_001134384,NM_001134383 |
| Flrt2 | 2910.060439 | -0.193894886 | 0.078069785 | -2.483609853 | 0.013005821 | 0.070483454 | | Lrrn1 | 2959.855618 | 0.0967850721497552 | 0.0593301580409642 | 1.63129638189958 | 0.102827796461445 | 0.522975270877619 | NM_008516 |
| Eif2d | 1019.429815 | 0.175129687 | 0.070527406 | 2.483143726 | 0.013022853 | 0.070551079 | | Mki67ip | 945.1054064 | 0.101978912854386 | 0.0625309838697926 | 1.63085412292146 | 0.10292110401414 | 0.522975270877619 | NM_026472 |
| Yes1 | 420.3026607 | -0.273738464 | 0.110261944 | -2.482619608 | 0.013042028 | 0.07063026 | | Nop16 | 684.8504176 | 0.101976385133084 | 0.0625036880910849 | 1.63152588667211 | 0.102779402195664 | 0.522975270877619 | NM_178605 |
| Skiv2l | 3115.62374 | 0.157956795 | 0.063631803 | 2.482356104 | 0.013051677 | 0.070657821 | | P4hb | 4890.533973 | 0.0764557190141187 | 0.0468452522920456 | 1.63209109297723 | 0.102660297845375 | 0.522975270877619 | NM_011032 |
| Gpr4 | 149.6672003 | -0.30580959 | 0.123237838 | -2.481458578 | 0.013084592 | 0.070737147 | | Pafah1b2 | 8430.692115 | 0.0930115585662837 | 0.05700200067079 | 1.63172445654081 | 0.102737545602749 | 0.522975270877619 | NM_008775 |
| Hrk | 2477.51507 | -0.222880156 | 0.089816716 | -2.481499714 | 0.013083082 | 0.070737147 | | Pnldc1 | 53.75440173 | -0.146649156387064 | 0.0899865365222359 | -1.62967886146865 | 0.103169387266203 | 0.522975270877619 | NM_001034866 |
| Tor3a | 313.396796 | 0.291020632 | 0.117275586 | 2.481510779 | 0.013082675 | 0.070737147 | | Ppp4r1l-ps | 318.4820442 | -0.128874852494216 | 0.0790945782400022 | -1.62937656868416 | 0.103233325939827 | 0.522975270877619 | NR_027957 |
| Zc3h7b | 9793.226436 | -0.149364354 | 0.060187444 | -2.481653031 | 0.013077454 | 0.070737147 | | Rab11fip4 | 6477.947085 | 0.0971144588997658 | 0.0594820718080294 | 1.63266772571726 | 0.102538898805823 | 0.522975270877619 | NM_175543 |
| 1110058L19Rik | 478.7329802 | -0.199866805 | 0.080561664 | -2.480917044 | 0.013104487 | 0.070795299 | | Sepsecs | 254.8427767 | -0.137145375323609 | 0.0841684284499545 | -1.62941587539744 | 0.10322501030091 | 0.522975270877619 | NM_172490 |
| Pcdha8 | 21.3861232 | 0.40507413 | 0.163272321 | 2.48097245 | 0.01310245 | 0.070795299 | | Slc2a5 | 238.6637701 | 0.139702816145148 | 0.0856945996946051 | 1.63024060609438 | 0.10305065493449 | 0.522975270877619 | NM_019741 |
| Astn2 | 1763.96251 | 0.163001582 | 0.065709132 | 2.480653411 | 0.013114182 | 0.070822981 | | Akr1a1 | 11959.34314 | 0.0753020821835004 | 0.046259708644079 | 1.62781142360564 | 0.103564877526848 | 0.523462117129643 | NM_021473 |
| Glp2r | 100.010374 | 0.416774735 | 0.168041158 | 2.480194374 | 0.013131078 | 0.07088952 | | Armcx4 | 3517.916265 | 0.096712817623855 | 0.0594116709148778 | 1.62784207436987 | 0.103558376525092 | 0.523462117129643 | NM_001202500 |
| Dhrs11 | 171.3207158 | 0.285428243 | 0.115177068 | 2.478169028 | 0.013205857 | 0.071268389 | | Ggnbp1 | 110.105889 | -0.154692215256121 | 0.0950415992769662 | -1.62762639131654 | 0.103604129612262 | 0.523462117129643 | NM_027544,NM_001251881 |
| Sult2b1 | 247.4431425 | 0.318202091 | 0.128426785 | 2.477692565 | 0.013223503 | 0.071338773 | | Prr7 | 574.3429618 | -0.1507237533226 | 0.0925947519375275 | -1.62777857458155 | 0.1035718451378 | 0.523462117129643 | NM_001030296 |
| Lin28b | 187.4496486 | 0.344265007 | 0.138970208 | 2.47725763 | 0.01323963 | 0.071400912 | | Puf60 | 5330.702898 | 0.0887876769815055 | 0.0545503989279658 | 1.62762653851075 | 0.103604098382312 | 0.523462117129643 | NM_028364,NM_133691,NM_001164600 |
| 2810032G03Rik | 311.6239633 | -0.259296296 | 0.104683894 | -2.476945456 | 0.013251215 | 0.071413679 | | Pygm | 1787.068689 | 0.111937023114121 | 0.0687683264290009 | 1.62774098086695 | 0.10357981960288 | 0.523462117129643 | NM_011224 |
| Mrpl20 | 1369.550086 | -0.188633913 | 0.076154788 | -2.476980358 | 0.013249919 | 0.071413679 | | Syp | 60443.48884 | 0.0736749028284978 | 0.0452640948151549 | 1.62766764980862 | 0.103595376159745 | 0.523462117129643 | NM_009305 |
| Vps39 | 3660.325859 | 0.126201465 | 0.050958627 | 2.476547594 | 0.013265994 | 0.071468465 | | Atp6v0a1 | 25402.80009 | 0.0710922926903662 | 0.043712397552781 | 1.62636452517904 | 0.103872132729471 | 0.523645898092662 | NM_001243049,NM_001243051,NM_016920,NM_001243050 |
| Peli2 | 1083.471299 | -0.238156198 | 0.096170516 | -2.476395126 | 0.013271661 | 0.071474145 | | Cela1 | 83.13800065 | -0.153281813910892 | 0.0943323583672974 | -1.62491234782942 | 0.104181236781481 | 0.523645898092662 | NM_033612 |
| Rwdd2a | 576.0691686 | -0.23918775 | 0.096593874 | -2.476220683 | 0.013278148 | 0.071484232 | | Ddx54 | 2012.712132 | 0.0821457837993477 | 0.0505518990134901 | 1.62497918777347 | 0.104166993502569 | 0.523645898092662 | NM_028041 |
| Abi2 | 9088.960439 | -0.15878794 | 0.064136178 | -2.475793622 | 0.01329404 | 0.07154493 | | Faim | 598.2527767 | 0.105901512439833 | 0.0651543251204079 | 1.62539497177083 | 0.104078426617919 | 0.523645898092662 | NM_011810,NM_001122851 |
| Ctnnd1 | 3646.11893 | -0.165739105 | 0.066948813 | -2.475609316 | 0.013300904 | 0.071557014 | | Gria4 | 4579.412275 | 0.0905502415885355 | 0.0557098997154706 | 1.62538870202614 | 0.10407976170293 | 0.523645898092662 | NM_001113181,NM_019691,NM_001113180 |
| C1qtnf2 | 30.89117522 | -0.39480758 | 0.159515363 | -2.475044239 | 0.013321967 | 0.071620597 | | Khdrbs2 | 182.033972 | 0.147119750863193 | 0.0904970694976522 | 1.6256852479296 | 0.104016629849617 | 0.523645898092662 | NM_133235 |
| Upf3b | 753.8853479 | 0.233964094 | 0.094529104 | 2.475048264 | 0.013321817 | 0.071620597 | | Mobp | 10624.55495 | 0.154309916980448 | 0.094962104572174 | 1.62496311213457 | 0.104170419004612 | 0.523645898092662 | NM_008614,NM_001039364,NM_001039365 |
| Zfp811 | 396.1164798 | -0.258104341 | 0.104291008 | -2.474847513 | 0.013329307 | 0.071635202 | | Pim2 | 1837.573036 | 0.081128025650019 | 0.0499231706229164 | 1.62505755619573 | 0.104150295529115 | 0.523645898092662 | NM_138606 |
| Bud31 | 1100.227061 | -0.1565717 | 0.063292745 | -2.473770088 | 0.01336957 | 0.071801776 | | Plcxd2 | 5910.421711 | 0.109842889799392 | 0.0675454565420438 | 1.62620693415582 | 0.103905641567586 | 0.523645898092662 | NM_001134480 |
| Ttc8 | 658.9703297 | 0.25338441 | 0.102426796 | 2.473809775 | 0.013368085 | 0.071801776 | | Reck | 542.3347409 | -0.1324699537058 | 0.081464362692922 | -1.62610924982181 | 0.103926416659731 | 0.523645898092662 | NM_016678 |
| 4930430F08Rik | 475.2644837 | 0.198677714 | 0.08033327 | 2.473168527 | 0.013392097 | 0.071872933 | | Terf1 | 322.3265696 | -0.127185676288132 | 0.078248776018174 | -1.62540147923326 | 0.104077040927466 | 0.523645898092662 | NM_009352 |
| Ap3s1 | 1172.46874 | -0.194856077 | 0.078785741 | -2.473240403 | 0.013389404 | 0.071872933 | | Tmem131 | 3882.273018 | -0.0747230073155702 | 0.0459814117895313 | -1.62506987948949 | 0.104147669996236 | 0.523645898092662 | NM_018872 |
| Ybx2 | 120.0278905 | 0.396590963 | 0.160368622 | 2.472996008 | 0.013398564 | 0.071882739 | | Tspan2 | 4472.108502 | 0.122512463790811 | 0.0753669073692954 | 1.6255471806811 | 0.104046019291347 | 0.523645898092662 | NM_027533,NM_001243132 |
| Zfp692 | 1056.399207 | 0.356536924 | 0.144182675 | 2.472813909 | 0.013405393 | 0.07189448 | | Uaca | 420.0117618 | -0.144049897485853 | 0.0885223069699601 | -1.62727229346538 | 0.103679279623489 | 0.523645898092662 | NM_028283 |
| Ank3 | 10268.14058 | -0.169225074 | 0.068438798 | -2.472648266 | 0.013411607 | 0.071902919 | | Vdac3 | 5628.43881 | 0.0866402378621298 | 0.0532923835815177 | 1.6257527256142 | 0.104002268730689 | 0.523645898092662 | NM_001198998,NM_011696 |
| Nanos3 | 18.31028373 | 0.412040216 | 0.166715198 | 2.471521617 | 0.013453942 | 0.072080004 | | Rbm39 | 6462.848051 | -0.0952894210964247 | 0.0586718088574281 | -1.62410914120574 | 0.104352517374647 | 0.524144075858963 | NM_133242 |
| Pdia3 | 7525.655605 | -0.19437627 | 0.078642814 | -2.471634215 | 0.013449705 | 0.072080004 | | Sh3bp5 | 2206.68011 | 0.0996319874122924 | 0.0613419441986574 | 1.62420654763781 | 0.104331733934094 | 0.524144075858963 | NM_011894 |
| 4930528A17Rik | 18.35362442 | 0.413291494 | 0.167244334 | 2.471183838 | 0.013466657 | 0.072123188 | | Eif2s1 | 1565.471226 | 0.115601920688625 | 0.0711967422546303 | 1.62369677358527 | 0.10444053996608 | 0.524404868138918 | NM_026114 |
| Cd47 | 2275.66031 | -0.138062371 | 0.055894144 | -2.470068631 | 0.013508713 | 0.072298447 | | Col17a1 | 35.55108943 | -0.130983976400702 | 0.0806833143364758 | -1.62343326470769 | 0.10449681854595 | 0.524427521558789 | NM_007732 |
| Sptssa | 543.9821358 | -0.186123272 | 0.075350302 | -2.470106511 | 0.013507283 | 0.072298447 | | Gm20257 | 29.28663572 | -0.136383198621156 | 0.0840228151625659 | -1.62316864005668 | 0.104553359660901 | 0.524427521558789 | NR_045007 |
| 6330416G13Rik | 578.9097369 | 0.185940245 | 0.07531168 | 2.46894301 | 0.01355128 | 0.072501219 | | Lonrf2 | 7786.819566 | 0.102396652515311 | 0.0630783326695577 | 1.62332528749177 | 0.104519886600771 | 0.524427521558789 | NM_001029878 |
| A330023F24Rik | 1931.706753 | 0.413220619 | 0.167398962 | 2.468477786 | 0.013568907 | 0.072520404 | | Txnrd1 | 2888.306394 | 0.0977586333283011 | 0.0602568516383916 | 1.62236543513694 | 0.104725125497858 | 0.525107757328919 | NM_001042513,NM_001042523,NM_015762,NM_001042514 |
| Atg9a | 6782.900769 | 0.178708864 | 0.072393574 | 2.468573596 | 0.013565275 | 0.072520404 | | Tmem191c | 2802.905502 | -0.117928382429014 | 0.0727192122473536 | -1.62169499344797 | 0.104868671334357 | 0.525646073085878 | NM_177473 |
| Kcng4 | 291.6605299 | 0.378820487 | 0.153462339 | 2.468491547 | 0.013568386 | 0.072520404 | | Timm22 | 1028.551786 | 0.0886822351413361 | 0.0546920016974731 | 1.62148453866945 | 0.104913763240549 | 0.52569069500939 | NM_019818,NM_023355 |
| Map3k15 | 88.538187 | 0.379608426 | 0.153880836 | 2.466898654 | 0.013628893 | 0.072815883 | | Drg2 | 2262.261811 | 0.0861745129025159 | 0.0531757685290671 | 1.62055980169635 | 0.105112079219293 | 0.526359793223222 | NM_021354 |
| Nefh | 2990.109655 | 0.26784837 | 0.108590642 | 2.466587965 | 0.013640722 | 0.072838195 | | Gnb1 | 26820.25464 | 0.066095787679808 | 0.0407909259496363 | 1.62035516824048 | 0.105156004401335 | 0.526359793223222 | NM_008142,NM_001160017,NM_001160016 |
| Pbrm1 | 2209.209884 | -0.189171265 | 0.076694928 | -2.466542055 | 0.013642471 | 0.072838195 | | Tmem167b | 2337.092422 | 0.0865290658579173 | 0.0533959476179881 | 1.6205174684224 | 0.105121164987977 | 0.526359793223222 | NM_026198 |
| Aurkaip1 | 1582.870454 | -0.164223904 | 0.066626798 | -2.464832599 | 0.013707728 | 0.073125764 | | Lzts2 | 1716.032477 | 0.11065130745141 | 0.0683074727460083 | 1.61990047359607 | 0.10525365811293 | 0.526667116000145 | NM_145503,NM_001130526,NM_001130525 |
| Birc2 | 1324.401187 | 0.137264127 | 0.055693094 | 2.464652563 | 0.013714616 | 0.073125764 | | Tulp4 | 8135.880014 | 0.0973957952528202 | 0.0601347309759613 | 1.61962635688441 | 0.105312564284311 | 0.526780409364293 | NM_001103181,NM_054040 |
| Ogfod2 | 870.7012968 | 0.161055929 | 0.065345278 | 2.464691149 | 0.01371314 | 0.073125764 | | Bzw1 | 6708.941635 | 0.108437262694729 | 0.0669628143739215 | 1.61936537029662 | 0.105368673173026 | 0.526879637353313 | NM_025824 |
| Pten | 7969.889839 | -0.174139849 | 0.070655373 | -2.464637042 | 0.01371521 | 0.073125764 | | Ppil6 | 96.63396439 | -0.152691386657888 | 0.0943021548198731 | -1.61917176706666 | 0.105410310796105 | 0.526906460641509 | NM_028430 |
| Atp13a4 | 487.8134072 | 0.256517149 | 0.104088902 | 2.464404409 | 0.013724117 | 0.073148078 | | Fam160b2 | 2731.568352 | -0.0990096120061696 | 0.0611766251261693 | -1.61842226180301 | 0.105571627596954 | 0.527531290840508 | NM_194345 |
| Inpp4b | 293.5930867 | -0.338546576 | 0.137405485 | -2.463850529 | 0.013745342 | 0.073236014 | | Fv1 | 216.0727659 | -0.137595696059041 | 0.0850668710893317 | -1.61750037702159 | 0.10577031436186 | 0.527979239319718 | NM_010244 |
| Mrto4 | 694.4030856 | -0.158323657 | 0.064279903 | -2.463035076 | 0.013776644 | 0.073357204 | | Mb21d2 | 1917.30955 | 0.0818660419813829 | 0.0506080761671062 | 1.61764777841117 | 0.105738526163269 | 0.527979239319718 | NM_177718 |
| Slc25a36 | 1231.374073 | 0.174070295 | 0.070675618 | 2.462946892 | 0.013780033 | 0.073357204 | | Ofd1 | 438.2630768 | -0.120202346626659 | 0.0742986266488595 | -1.61782730109863 | 0.105699821005447 | 0.527979239319718 | NM_177429 |
| Tcf19 | 228.0321951 | 0.280380618 | 0.113842206 | 2.462888127 | 0.013782291 | 0.073357204 | | M6pr | 4314.786775 | 0.0766825120711317 | 0.0474507978841922 | 1.61604262710781 | 0.106085097237119 | 0.528508890269676 | NM_010749 |
| Tex11 | 8.98652736 | 0.362451817 | 0.147175228 | 2.462722984 | 0.01378864 | 0.073365794 | | Prmt2 | 2018.358488 | 0.11968710893081 | 0.0740429501215898 | 1.61645516196026 | 0.105995949215127 | 0.528508890269676 | NM_133182,NM_001077638 |
| Mrrf | 645.4882133 | -0.205969073 | 0.083646494 | -2.462375436 | 0.013802011 | 0.073411724 | | Syt16 | 2937.660993 | 0.0894081178564087 | 0.055326870254924 | 1.61598111166712 | 0.106094721466775 | 0.528508890269676 | NM_172804 |
| Sdhb | 4067.832509 | -0.122950966 | 0.049939249 | -2.462070102 | 0.013816054 | 0.07346124 | | Tm7sf2 | 650.3721785 | -0.1331786709584 | 0.0823935174109188 | -1.61637314070614 | 0.106013655494577 | 0.528508890269676 | NM_028454 |
| Pfdn4 | 459.6045942 | -0.195004902 | 0.079238018 | -2.461001771 | 0.013854968 | 0.07364284 | | Ugcg | 3144.18393 | 0.103553819712811 | 0.0640574294546866 | 1.61657782078289 | 0.105969442657429 | 0.528508890269676 | NM_011673 |
| Trpv6 | 48.93898725 | 0.384465999 | 0.15623221 | 2.460862581 | 0.013860344 | 0.07364615 | | Zdhhc4 | 796.508067 | -0.11199733714296 | 0.0692967092770724 | -1.61620008357418 | 0.106051067374761 | 0.528508890269676 | NM_028379 |
| A130049A11Rik | 59.85166435 | 0.355505057 | 0.144492452 | 2.46037113 | 0.013879341 | 0.073687366 | | Rasgrf1 | 15912.5894 | 0.0986690899562268 | 0.0610733916173395 | 1.61558229113009 | 0.106184655230187 | 0.528775557721527 | NM_011245,NM_001039655 |
| Endog | 260.3943405 | -0.263422642 | 0.107072395 | -2.460229272 | 0.013884828 | 0.073687366 | | Trmt2a | 1349.754728 | -0.121248560917379 | 0.0750931215990182 | -1.61464270409241 | 0.106388091945178 | 0.529607067716128 | NM_001081000,NM_001080999,NM_001195205 |
| Tmed2 | 3821.910848 | -0.123388507 | 0.050154465 | -2.460169928 | 0.013887125 | 0.073687366 | | Prkar1b | 41686.5328 | 0.0870899285279985 | 0.053957964440714 | 1.61403287597493 | 0.106520295466704 | 0.530021850183169 | NM_001253890,NM_008923 |
| Zbtb48 | 345.5334953 | 0.260667431 | 0.105950705 | 2.460270843 | 0.01388322 | 0.073687366 | | Ssna1 | 898.1755689 | 0.0972959421015624 | 0.0602854129186506 | 1.61392179550486 | 0.106544389545288 | 0.530021850183169 | NM_023464 |
| Bet1 | 408.0194431 | -0.242510048 | 0.098591669 | -2.459741793 | 0.013903701 | 0.07373614 | | Ola1 | 3623.287017 | 0.103105545656585 | 0.0639138362193224 | 1.61319601131083 | 0.106701929757363 | 0.530442242181882 | NM_025942,NM_030091 |
| Zfp641 | 510.5076364 | 0.287424347 | 0.116854047 | 2.459686707 | 0.013905835 | 0.07373614 | | Zkscan1 | 2664.512498 | 0.106914835838569 | 0.0662719371988697 | 1.61327464319833 | 0.106684852399765 | 0.530442242181882 | NM_133906,NM_029869 |
| Atp6v0e | 524.2076979 | 0.22488637 | 0.091498695 | 2.457809602 | 0.013978726 | 0.074097292 | | Ppp2r2b | 5452.214812 | 0.0723104932334089 | 0.0448338602096291 | 1.6128545009354 | 0.106776122139418 | 0.530447999383445 | NM_027531,NM_028392,NR_073583 |
| Dleu7 | 149.5291586 | -0.258533973 | 0.105207965 | -2.457361215 | 0.013996188 | 0.074164478 | | Rheb | 3351.622749 | 0.112889598823209 | 0.0699934861559057 | 1.61285863904185 | 0.106775222900218 | 0.530447999383445 | NM_053075 |
| Sat2 | 210.9892027 | 0.27544826 | 0.112147826 | 2.456117688 | 0.014044715 | 0.074396178 | | Dlc1 | 3515.895399 | 0.110618581323943 | 0.068631149371773 | 1.61178389603714 | 0.107008973710839 | 0.531423026367058 | NM_001194940,NM_001194941,NM_015802 |
| Fance | 446.1045254 | 0.191122812 | 0.077828104 | 2.45570433 | 0.014060879 | 0.074456344 | | Hnrnpab | 4230.260515 | 0.0871505027097549 | 0.0540973803953038 | 1.61099302916646 | 0.107181241817814 | 0.532011007433082 | NM_001048061,NM_010448 |
| Ndufa4 | 6503.007162 | -0.179506724 | 0.073112178 | -2.455223297 | 0.01407971 | 0.074530586 | | Zfp26 | 881.5879932 | -0.114970273240543 | 0.0713700284689144 | -1.61090412470018 | 0.107200620869932 | 0.532011007433082 | NM_011753 |
| Cyld | 7809.164 | -0.186456332 | 0.075950789 | -2.454962395 | 0.014089933 | 0.074559227 | | Ankrd10 | 1426.40361 | -0.118360083549684 | 0.0734960482863132 | -1.6104278571359 | 0.107304483171807 | 0.532178903759375 | NR_030780,NM_133971,NR_030781,NR_030779,NM_001167967 |
| Xdh | 499.2138611 | 0.411058141 | 0.167470051 | 2.454517315 | 0.014107387 | 0.074626103 | | Rab3il1 | 418.9360023 | -0.129383647829389 | 0.0803502823889307 | -1.61024509164903 | 0.107344361008879 | 0.532178903759375 | NM_144538 |
| Dph1 | 490.8284242 | 0.241461236 | 0.098402522 | 2.45381146 | 0.014135107 | 0.074747218 | | Vkorc1l1 | 687.2155858 | 0.124219649369046 | 0.0771382189548714 | 1.61035153588028 | 0.107321134377695 | 0.532178903759375 | NM_027121,NM_001001327 |
| Ptpro | 1929.185085 | -0.410669089 | 0.167370121 | -2.453658315 | 0.014141128 | 0.074753542 | | 2510003E04Rik | 7058.514774 | 0.0803871106471732 | 0.04993463046329 | 1.60984691187953 | 0.107431281010968 | 0.532254442574493 | NM_028197 |
| Ints9 | 616.16819 | 0.175365434 | 0.071480145 | 2.453344691 | 0.014153464 | 0.074793239 | | Gabrq | 76.52522193 | 0.0756976207652702 | 0.0470218408262666 | 1.60983958592674 | 0.107432880739943 | 0.532254442574493 | NM_020488 |
| Rnf103 | 1638.098587 | -0.147157476 | 0.059987474 | -2.453136731 | 0.01416165 | 0.074810979 | | Itga8 | 231.2052956 | -0.138978482123676 | 0.0863757842962778 | -1.60899820772643 | 0.107616733526354 | 0.53280186475931 | NM_001001309 |
| Rassf5 | 1161.603384 | 0.195203027 | 0.079601494 | 2.452253306 | 0.014196469 | 0.074969354 | | Tmem9 | 2111.890346 | 0.11424112984 1175 | 0.0709982631436495 | 1.60906936004946 | 0.107601176120863 | 0.53280186475931 | NM_025439,NM_001160145,NM_001160146 |
| Zfp94 | 359.2682145 | 0.212097666 | 0.086511193 | 2.451678902 | 0.014219148 | 0.075063538 | | Nus1 | 3916.543561 | 0.118092951262543 | 0.0734044170432374 | 1.60879897994398 | 0.107660304017512 | 0.53283597143386 | NM_030250 |
| Edn1 | 113.0413102 | -0.384223666 | 0.156776199 | -2.450778041 | 0.014254782 | 0.075226021 | | 4921524J17Rik | 271.9149484 | 0.143350855775178 | 0.0891564963449127 | 1.60785654048818 | 0.107866601928373 | 0.533015564958536 | NM_025722,NR_033144 |
| Lpcat4 | 3971.975542 | 0.224941928 | 0.091804571 | 2.450225795 | 0.014276666 | 0.075315852 | | Atp6v1b2 | 33561.13915 | 0.0754739345247565 | 0.0469425079356317 | 1.60779510605286 | 0.107880060648361 | 0.533015564958536 | NM_007509 |
| Ccdc174 | 765.345761 | 0.20562648 | 0.083977854 | 2.448579835 | 0.014342064 | 0.075617409 | | Ccdc88b | 163.6501073 | -0.149009705227356 | 0.0926746038419496 | -1.60788068197715 | 0.107861313506883 | 0.533015564958536 | NM_001081291 |
| Edn3 | 212.6537943 | -0.3561097 | 0.145437475 | -2.448541541 | 0.014343589 | 0.075617409 | | Lrrc24 | 1143.920524 | 0.115166083366398 | 0.0716099074013266 | 1.60824231654103 | 0.107782118601307 | 0.533015564958536 | NM_198119 |
| Nckap5 | 153.2648546 | -0.326354035 | 0.133341381 | -2.447507532 | 0.014384812 | 0.075799835 | | Mapt | 24603.75781 | 0.0749487307315899 | 0.0465982307996011 | 1.60840292529371 | 0.107746961412249 | 0.533015564958536 | NM_001038609,NM_010838 |
| Nit2 | 310.1077565 | -0.265269897 | 0.108387201 | -2.44742825 | 0.014387977 | 0.075799835 | | D15Ertd621e | 3359.223735 | 0.103847244260087 | 0.0646149498357238 | 1.60717054681784 | 0.108016961141086 | 0.533425002070014 | NM_145959 |
| Capza1 | 1868.540963 | -0.1876753 | 0.07670898 | -2.446588393 | 0.014421543 | 0.075950845 | | Nlrp1b | 27.28018392 | -0.140394226397518 | 0.0873597135279021 | -1.60708203733609 | 0.108036373130248 | 0.533425002070014 | NM_001040696,NM_001162414 |
| Nfrkb | 1308.327002 | 0.204422341 | 0.083569642 | 2.446131587 | 0.014439829 | 0.076009456 | | Cd72 | 47.248882 | -0.140494521372335 | 0.0874779030982651 | -1.60605726013478 | 0.108261329439397 | 0.533447785426283 | NM_007654,NM_001110320,NM_001110321,NM_001110322 |
| Psmb8 | 138.1169236 | 0.409618625 | 0.167460215 | 2.446065316 | 0.014442483 | 0.076009456 | | Cstad | 60.85532511 | -0.152172716024563 | 0.0947353188774317 | -1.60629338485094 | 0.108209463142621 | 0.533447785426283 | NM_030137 |
| Mmd2 | 5521.812297 | -0.215051694 | 0.087926605 | -2.445809142 | 0.014452749 | 0.076037652 | | Ift172 | 3035.974669 | -0.0935790558265109 | 0.0582604459149933 | -1.60621935443217 | 0.108225722278495 | 0.533447785426283 | NM_026298 |
| Usp34 | 6631.766072 | -0.207727185 | 0.084956903 | -2.445088958 | 0.014481642 | 0.076161523 | | Pkmyt1 | 196.2881275 | -0.141645281504067 | 0.0881760248147669 | -1.60639223418865 | 0.108187756097088 | 0.533447785426283 | NM_023058 |
| Zkscan16 | 1879.419916 | 0.283218615 | 0.115836908 | 2.444977342 | 0.014486124 | 0.076161523 | | Poln | 35.07360966 | -0.14769218015515 | 0.0919410564666316 | -1.6063789761732 | 0.108190667321057 | 0.533447785426283 | NM_181857 |
| Mrpl21 | 537.6861446 | -0.18141174 | 0.074201945 | -2.444838068 | 0.014491719 | 0.076165094 | | Srsf1 | 5136.720653 | -0.0848541446136326 | 0.0528232054835213 | -1.6063800717301 | 0.108190426753931 | 0.533447785426283 | NM_173374,NM_001078167 |
| Ltbp1 | 359.2203421 | -0.219947925 | 0.0899924 | -2.444072218 | 0.014522519 | 0.076301089 | | Sugt1 | 2221.110583 | 0.114284731975704 | 0.0711914605169775 | 1.60531517608702 | 0.108424461013246 | 0.534070437666468 | NM_026474 |
| Oacyl | 63.12126134 | 0.386398094 | 0.158138969 | 2.443408447 | 0.014549261 | 0.076415676 | | 2210016L21Rik | 4256.002858 | 0.0811019278298304 | 0.0505946903462884 | 1.60297310399055 | 0.108940592788808 | 0.534621386980457 | NM_028211 |
| Tmtc2 | 594.9520114 | -0.202848873 | 0.083056307 | -2.442305475 | 0.014593792 | 0.07662359 | | 9430037G07Rik | 99.76322476 | -0.148982311574462 | 0.092862711318012 | -1.60432868543183 | 0.108641621326 | 0.534621386980457 | NR_040766 |
| Dcaf12l1 | 1517.828151 | -0.203775821 | 0.083456922 | -2.441688687 | 0.014618747 | 0.076728612 | | B3gnt2 | 1393.508236 | 0.133853024627816 | 0.0834285519860817 | 1.60440306635247 | 0.108625235557532 | 0.534621386980457 | NM_016888,NM_001169114 |
| Asic3 | 12.41795117 | 0.395718361 | 0.16209393 | 2.441290428 | 0.01463488 | 0.076784262 | | Capn12 | 31.51583232 | -0.140831415942086 | 0.0878247809364184 | -1.60354986873266 | 0.10881330845454 | 0.534621386980457 | NM_001110807 |
| Fkbp1a | 18582.69165 | -0.129539891 | 0.053064406 | -2.441182341 | 0.014639261 | 0.076784262 | | Elmsan1 | 1288.056934 | 0.0970570116206481 | 0.0605262863537334 | 1.60355140663047 | 0.108812969219557 | 0.534621386980457 | NM_001163502,NM_001163501 |
| Sh3rf2 | 573.6638858 | -0.31875945 | 0.130597306 | -2.440781206 | 0.014655531 | 0.076843586 | | Fxyd1 | 822.7900187 | -0.143922108400866 | 0.0897846254172643 | -1.60297052788274 | 0.108941161053432 | 0.534621386980457 | NM_194321,NM_019503,NM_052992,NM_052991 |
| Evc2 | 946.38831 | 0.279478095 | 0.114558123 | 2.439618306 | 0.014702788 | 0.077013181 | | Shc4 | 297.1892486 | 0.132595666667361 | 0.0826919134604071 | 1.60349012519643 | 0.108826487544266 | 0.534621386980457 | NM_199022 |
| Oaf | 413.275506 | -0.208282308 | 0.085372673 | -2.439683585 | 0.014700131 | 0.077013181 | | Sowahb | 1002.507321 | 0.143154821760432 | 0.0893025432374917 | 1.60303185744358 | 0.108927620837774 | 0.534621386980457 | NM_175270 |
| Sepn1 | 1139.232577 | 0.198731793 | 0.081455034 | 2.439772996 | 0.014696494 | 0.077013181 | | Tnmd | 26.7821999 | 0.13474680717869 | 0.0840179628550034 | 1.60378569772315 | 0.108761298226239 | 0.534621386980457 | NM_022322 |
| Cpne7 | 2053.134167 | 0.410051022 | 0.168109858 | 2.439184865 | 0.014720436 | 0.077053523 | | Zbtb7b | 703.2548198 | 0.112370822433778 | 0.0700835386178941 | 1.60338397075582 | 0.108849907734605 | 0.534621386980457 | NM_009565 |
| Gm12522 | 55.98720344 | 0.370994087 | 0.152093449 | 2.439250925 | 0.014717745 | 0.077053523 | | Zbtb7c | 309.7536658 | 0.131750238522671 | 0.0821588294236557 | 1.603604134174 | 0.108801338896067 | 0.534621386980457 | NM_145356 |
| 1700037H04Rik | 2019.460639 | -0.172623529 | 0.070779241 | -2.438900548 | 0.014732022 | 0.077088129 | | Abcf1 | 3821.328531 | 0.0632666662735388 | 0.039489355894226 | 1.60211947855015 | 0.109129191433906 | 0.535010276275204 | NM_013854 |
| Rhpn1 | 332.6034345 | 0.291871561 | 0.119689548 | 2.43857163 | 0.014745427 | 0.077132225 | | Gm5113 | 922.7197448 | 0.106658099955606 | 0.066587289592573 | 1.60177866692294 | 0.109204562106291 | 0.535010276275204 | NM_001033540 |
| Wbp7 | 2845.788741 | 0.157958422 | 0.064784311 | 2.438220301 | 0.014759776 | 0.077181224 | | Gpt2 | 2744.899661 | 0.125827956713571 | 0.0785533444835174 | 1.60181539743318 | 0.109196437149993 | 0.535010276275204 | NM_173866 |
| Odf3b | 38.94316064 | 0.408559974 | 0.167591173 | 2.437839073 | 0.014775432 | 0.077237027 | | Ptges3 | 3277.057921 | 0.0743008016291877 | 0.0463680752159154 | 1.60241289471694 | 0.1090643352572 | 0.535010276275204 | NM_019766 |
| 1700009P17Rik | 48.37430814 | 0.399258538 | 0.163790613 | 2.437615512 | 0.01478449 | 0.077258309 | | Tmem194 | 222.7605689 | -0.135175365540425 | 0.0843833706855834 | -1.60191948297604 | 0.109173415568803 | 0.535010276275204 | NM_173732,NM_001113211 |
| Efr3a | 5894.387988 | 0.235314772 | 0.096565769 | 2.436834244 | 0.014816469 | 0.077274849 | | Arhgap19 | 221.4718431 | -0.141877366708577 | 0.0886494193394208 | -1.60043199116011 | 0.109502782873786 | 0.535243338467863 | NM_001163495,NM_027667 |
| Grip1 | 635.2277811 | -0.221985929 | 0.091089124 | -2.437019034 | 0.0148089 | 0.077274849 | | Crip3 | 52.05594756 | -0.132982234644569 | 0.083068338921586 | -1.6008774988278 | 0.109404054210532 | 0.535243338467863 | NM_181664,NM_053250 |
| Il20ra | 33.82920702 | -0.408764873 | 0.167728096 | -2.437068588 | 0.014806871 | 0.077274849 | | Kif9 | 281.1062971 | -0.136393217392451 | 0.0852003919804229 | -1.60085199401185 | 0.109409704416361 | 0.535243338467863 | NM_001163569,NM_010628 |
| Lair1 | 365.8334691 | 0.250499848 | 0.102798388 | 2.436807151 | 0.014817579 | 0.077274849 | | Plxnd1 | 6209.809292 | -0.1219557235643 | 0.0761643525247555 | -1.60121788634205 | 0.109328668592516 | 0.535243338467863 | NM_026376 |
| Map3k12 | 5585.219302 | 0.121523388 | 0.049863868 | 2.437103125 | 0.014805456 | 0.077274849 | | Ptgs1 | 818.9620055 | -0.132155223300127 | 0.0825764146246577 | -1.60039914424505 | 0.109510064844175 | 0.535243338467863 | NM_008969 |
| Mex3a | 188.1628675 | -0.294263314 | 0.120728865 | -2.437389879 | 0.014793719 | 0.077274849 | | Scn1b | 17860.08847 | 0.0864813763162751 | 0.0540286912221514 | 1.6006565097179 | 0.109453018662921 | 0.535243338467863 | NM_011322 |
| Rgl3 | 141.7725437 | 0.285974135 | 0.117383098 | 2.436246267 | 0.014840577 | 0.077368742 | | Zfp667 | 748.1717169 | 0.130812224012375 | 0.0816989909472509 | 1.60113095006251 | 0.109347918462764 | 0.535243338467863 | NM_001024928 |
| Ppp2r2c | 28246.88256 | -0.172591299 | 0.070853347 | -2.435894804 | 0.01485500 4 | 0.077417905 | | Cog2 | 1136.517522 | -0.11193442637814 | 0.0699951559960694 | -1.59917389689689 | 0.109781968639114 | 0.535311472457794 | NM_029746 |
| Dbndd1 | 566.9889549 | -0.224054829 | 0.092032615 | -2.434515497 | 0.014911741 | 0.077621071 | | Hnrnpc | 7184.531359 | 0.0940312387760587 | 0.0587716508067761 | 1.59994210619004 | 0.109611427253411 | 0.535311472457794 | NM_016884,NM_001170984,NM_001170982,NM_001170983,NM_001170981 |
| Itgb1 | 2566.177853 | -0.142438265 | 0.058512103 | -2.434338487 | 0.014919036 | 0.077621071 | | Il17rc | 124.1346834 | -0.152081662646188 | 0.0950896894506969 | -1.59934966161648 | 0.109742930628204 | 0.535311472457794 | NM_134159,NM_178942 |
| Map3k4 | 2136.478402 | 0.196021586 | 0.080519747 | 2.434453581 | 0.014914293 | 0.077621071 | | Mospd2 | 367.9263573 | -0.116072696776632 | 0.0725670755554783 | -1.59952286747305 | 0.109704471684436 | 0.535311472457794 | NM_029730 |
| Rnf25 | 1295.622171 | 0.17347204 | 0.071251352 | 2.434649107 | 0.014906237 | 0.077621071 | | Oxld1 | 142.2193499 | 0.147941435727585 | 0.0925051353164383 | 1.59927808571396 | 0.109758826583209 | 0.535311472457794 | NM_025560 |
| Wdr8 | 705.5456928 | 0.173419964 | 0.071231183 | 2.434607393 | 0.014907955 | 0.077621071 | | She | 245.7796465 | -0.145619767818241 | 0.091050762458965 | -1.59932507851177 | 0.109748389968961 | 0.535311472457794 | NM_172530 |
| 4833439L19Rik | 5476.223068 | -0.123407144 | 0.050699939 | -2.434068889 | 0.014930153 | 0.077652835 | | Wbp11 | 3941.03823 | 0.0756969232203495 | 0.0473247633630606 | 1.59952037455796 | 0.109705025140275 | 0.535311472457794 | NM_021714 |
| Resp18 | 2126.149205 | 0.201774641 | 0.082927871 | 2.433134212 | 0.014968751 | 0.077827461 | | Adra2c | 758.8377062 | 0.118057346138243 | 0.073858025885154 | 1.59843625284294 | 0.109945921875873 | 0.535492106803922 | NM_007418 |
| Iqcd | 37.51066752 | 0.378204507 | 0.155502314 | 2.432147139 | 0.015009609 | 0.078013712 | | Bora | 66.56986819 | -0.151829945219417 | 0.0949739183257544 | -1.59864884903085 | 0.109898649124829 | 0.535492106803922 | NM_175265 |
| Mrfap1 | 9271.574401 | -0.140539604 | 0.057788431 | -2.431967823 | 0.015017041 | 0.07802617 | | Cdca7 | 128.187617 | -0.145448997363783 | 0.0909817799877478 | -1.5986607140833 | 0.109896011292737 | 0.535492106803922 | NM_025866 |
| Rln1 | 14.84427357 | -0.406432031 | 0.167138247 | -2.431711707 | 0.015027663 | 0.078055184 | | Ywhaq | 13941.4118 | 0.087516376096819 | 0.0547544105873973 | 1.59834386231093 | 0.109966470783154 | 0.535492106803922 | NM_011739 |
| Pnmal1 | 3362.872028 | -0.179500649 | 0.073820328 | -2.431588339 | 0.015032782 | 0.078055605 | | A230072C01Rik | 339.5441913 | -0.117831003569798 | 0.07375908481587 | -1.59751173518419 | 0.110151683927388 | 0.535496439333747 | NR_027445,NR_027446 |
| Mobp | 7684.129417 | 0.28533024 | 0.117361106 | 2.431216363 | 0.015048225 | 0.078109616 | | Ebf1 | 139.0909639 | 0.147584257246255 | 0.0923769721224975 | 1.59763038185046 | 0.110125260741183 | 0.535496439333747 | NM_007897 |
| Kif5b | 10623.88286 | 0.145622524 | 0.059915567 | 2.430462242 | 0.015079577 | 0.078219943 | | Kcnj10 | 10969.15985 | 0.121509344729083 | 0.0760432781337274 | 1.59789724629442 | 0.110065847038888 | 0.535496439333747 | NM_001039484 |
| Ntn4 | 289.4956289 | -0.27829617 | 0.114503163 | -2.430467088 | 0.015079376 | 0.078219943 | | Lrrc8c | 690.8206575 | 0.105140111999986 | 0.0657886425228049 | 1.59814989287157 | 0.110009622091007 | 0.535496439333747 | NM_133897 |
| Bag1 | 3521.01707 | -0.197817365 | 0.081400908 | -2.43016166 | 0.01509209 | 0.078232464 | | Mapk1ip1l | 1877.698815 | 0.0894036647941941 | 0.0559615977459615 | 1.59758956847593 | 0.110134349511913 | 0.535496439333747 | NM_178684 |
| Got1l1 | 20.09367141 | 0.4081008 | 0.16792333 | 2.430280529 | 0.01508714 | 0.078232464 | | Rfc5 | 353.7908623 | -0.117289137858388 | 0.0734767633506208 | -1.59627523736587 | 0.110427356178857 | 0.536657000954862 | NM_028128 |
| Tsc2 | 5349.248144 | 0.134022019 | 0.05515311 | 2.429999313 | 0.015098851 | 0.078241339 | | Trim41 | 2516.760384 | -0.0777147016331723 | 0.0487006988979327 | -1.59576152687351 | 0.110542045989881 | 0.537034702357531 | NM_145377 |
| Ints7 | 1312.884188 | 0.138506812 | 0.057007558 | 2.429621933 | 0.01511458 | 0.078296657 | | Arl4c | 2454.617196 | 0.0911440984766528 | 0.0571527785169176 | 1.59474483729034 | 0.110769307067422 | 0.537406303238926 | NM_177305 |
| Slx1b | 625.3364865 | 0.212525869 | 0.087511071 | 2.428559803 | 0.015158925 | 0.078500128 | | Cdh19 | 492.9611507 | 0.136757508538864 | 0.0857666600691658 | 1.59452995404715 | 0.110817387210411 | 0.537406303238926 | NM_001081386 |
| Acin1 | 4532.398558 | 0.192209252 | 0.079157166 | 2.42819776 | 0.015174067 | 0.078552285 | | Gpbp1 | 3672.725782 | 0.11115780251572 | 0.0696813559143766 | 1.59523018829181 | 0.110660770361078 | 0.537406303238926 | NM_001122963,NM_028487 |
| Zfp259 | 1358.522468 | 0.141211477 | 0.058161207 | 2.427932367 | 0.015185175 | 0.078583533 | | H2-Ob | 29.49696169 | -0.139454638528781 | 0.0874638168735873 | -1.59442662707411 | 0.1108405124951 | 0.537406303238926 | NM_010389 |
| Ctc1 | 1948.230254 | 0.144651782 | 0.059588366 | 2.427517172 | 0.015202567 | 0.07864267 | | Tspan11 | 359.1697115 | 0.116972320532633 | 0.0733407973693762 | 1.59491476406385 | 0.110731297602606 | 0.537406303238926 | NM_026743 |
| Gm6297 | 68.78318162 | 0.356088453 | 0.146694371 | 2.427417295 | 0.015206754 | 0.07864267 | | Usp29 | 2547.031806 | 0.136654111874856 | 0.0856875307830035 | 1.59479577280528 | 0.110757912657193 | 0.537406303238926 | NM_021323 |
| Ldhd | 562.3665884 | 0.22028979 | 0.090755228 | 2.427295863 | 0.015211845 | 0.07864267 | | Dusp4 | 411.700476 | 0.133256718431035 | 0.0835952811733897 | 1.59406986328139 | 0.110920387968516 | 0.537434808415834 | NM_176933 |
| Kctd14 | 98.58017415 | -0.377131029 | 0.155396757 | -2.426891248 | 0.01522882 | 0.078704249 | | Sgta | 4902.574164 | 0.0913312939635728 | 0.0572893061881123 | 1.5942119051623 | 0.11088858091675 | 0.537434808415834 | NM_024499 |
| Taf10 | 779.6938078 | -0.213746011 | 0.088089529 | -2.426463307 | 0.015246792 | 0.078770856 | | Ndufs7 | 2402.143527 | 0.103279061771355 | 0.0648006981577533 | 1.59379551004109 | 0.110981843535255 | 0.537553270821312 | NM_029272 |
| Mapk15 | 38.46176991 | 0.406271894 | 0.167450058 | 2.426227253 | 0.015256713 | 0.078795839 | | Ddx55 | 883.7208682 | -0.111831123274449 | 0.0701834016615494 | -1.59341269626315 | 0.111067639355433 | 0.537789509759008 | NM_001190795,NM_026409 |
| Pcf11 | 1185.623025 | 0.23151459 | 0.095443993 | 2.425659105 | 0.015280616 | 0.078892273 | | Laptm4b | 3837.876889 | 0.0754412888584181 | 0.0473595422389406 | 1.59294801621721 | 0.111171853298705 | 0.538114742091631 | NM_033521 |
| Tdrd7 | 1617.375225 | -0.132017837 | 0.054428192 | -2.42554148 | 0.015285569 | 0.078892273 | | Acvr2a | 2021.480382 | 0.12214269827095 3 | 0.0767260819025104 | 1.5919319120993 3 | 0.111400004208442 | 0.538859960416861 | NM_007396 |
| Slc26a2 | 209.8289778 | -0.317340961 | 0.130871528 | -2.424828112 | 0.015315636 | 0.079021136 | | Tac1 | 1346.900866 | 0.135466710542676 | 0.0850882921886241 | 1.59207227055836 | 0.111368466847925 | 0.538859960416861 | NM_009311 |
| Ccr1 | 12.79084354 | -0.400636525 | 0.165324645 | -2.423332134 | 0.015378859 | 0.079320919 | | 1300002E11Rik | 274.1472159 | -0.12300511696826 | 0.0774739555856034 | -1.58769636009984 | 0.112355016524074 | 0.538992394329159 | NR_037957,NR_037958 |
| Gltscr1l | 2607.37818 | 0.162670186 | 0.06714602 | 2.422633316 | 0.015408471 | 0.079447204 | | Agmat | 85.96677476 | -0.150758520686452 | 0.0949356971035815 | -1.58800667489663 | 0.112284829764133 | 0.538992394329159 | NM_001081408 |
| Sharpin | 1175.889754 | 0.143380368 | 0.059224009 | 2.420983834 | 0.015478566 | 0.079782068 | | Ccdc93 | 1488.835031 | -0.08478711168946 | 0.053343726536586 | -1.58944860425731 | 0.111959148771325 | 0.538992394329159 | NM_001025156,NM_029955 |
| Prkcd | 636.8251155 | 0.398575485 | 0.164658697 | 2.420616061 | 0.015494232 | 0.079836261 | | Cnksr2 | 9262.92383 | -0.127474202983748 | 0.0802068726632869 | -1.58931770745996 | 0.111988682957524 | 0.538992394329159 | NM_177751 |
| Sdr42e1 | 339.5243965 | -0.257939479 | 0.1065796 | -2.420158057 | 0.015513762 | 0.079903737 | | Galnt11 | 1623.099852 | 0.0964204569725368 | 0.0605836940530426 | 1.59152488932283 | 0.11149149864364 | 0.538992394329159 | NM_144908 |
| Ubxn11 | 416.7485753 | 0.285569623 | 0.118000703 | 2.420067135 | 0.015517642 | 0.079903737 | | Kcnc4 | 3245.25378 | 0.0896588217151579 | 0.0563478252724875 | 1.5911673836849 | 0.111571911054232 | 0.538992394329159 | NM_145922 |
| Ccdc177 | 540.8340145 | -0.207786405 | 0.085921041 | -2.418341336 | 0.015591443 | 0.080108712 | | Klhl25 | 449.2875211 | 0.114939995486026 | 0.0723660101452775 | 1.58831466948751 | 0.112215201977997 | 0.538992394329159 | NM_001122780,NM_182782,NM_029652 |
| Cd36 | 46.25815205 | 0.40286397 | 0.166566815 | 2.418632851 | 0.015578956 | 0.080108712 | | Mcf2 | 347.9225612 | 0.126974376171469 | 0.0799093385409503 | 1.58898044321064 | 0.112064808055148 | 0.538992394329159 | NM_133197 |
| Chst8 | 356.3214266 | 0.276724097 | 0.114429396 | 2.418295523 | 0.015593407 | 0.080108712 | | Ndrg3 | 25733.7713 | 0.0629973452116248 | 0.0395945209596717 | 1.59106218953349 | 0.111595580694363 | 0.538992394329159 | NM_013865,NM_180956 |
| Gucy1b3 | 7341.462739 | -0.179666739 | 0.074282354 | -2.418700124 | 0.015576075 | 0.080108712 | | Ndufv1 | 6185.171454 | 0.0854775358143664 | 0.0538316186304171 | 1.58786857963933 | 0.112316059771645 | 0.538992394329159 | NM_133666 |
| Ppfia4 | 3262.994381 | 0.207909628 | 0.08597265 | 2.418322879 | 0.015592234 | 0.080108712 | | Nrg2 | 253.7092912 | -0.125315291835826 | 0.0787658601391953 | -1.59098487103891 | 0.111612980584546 | 0.538992394329159 | NM_001167891 |
| Spats2l | 994.1113361 | 0.166492514 | 0.068828975 | 2.418930592 | 0.01556621 | 0.080108712 | | Pcdh20 | 938.0747184 | -0.138986357121109 | 0.087416846196078 | -1.58992646348004 | 0.111851381758278 | 0.538992394329159 | NM_178685 |
| Vnn1 | 8.709869866 | -0.385208634 | 0.159289674 | -2.418290055 | 0.015593641 | 0.080108712 | | Pcnxl4 | 1887.423036 | 0.0960271225859415 | 0.0604665068436434 | 1.58810435063252 | 0.112262744697389 | 0.538992394329159 | NM_026327 |
| Acadm | 1255.945702 | -0.135525956 | 0.056059465 | -2.417539242 | 0.015625849 | 0.080247565 | | Pknox2 | 3354.179219 | 0.0788215016929123 | 0.049632359688522 | 1.58810707747068 | 0.112262128192162 | 0.538992394329159 | NM_001029838,NM_148950 |
| Rassf6 | 22.13237468 | 0.404004639 | 0.167142443 | 2.417127765 | 0.015643525 | 0.080311722 | | Prdm8 | 997.1153801 | -0.128129384427759 | 0.0805944984105318 | -1.5898031125536 | 0.111879192072856 | 0.538992394329159 | NM_029947 |
| Acap2 | 3723.617134 | 0.17621286 | 0.072911864 | 2.416792696 | 0.015657932 | 0.080313518 | | Psmd13 | 2707.256983 | 0.0759202530391715 | 0.0477763791786562 | 1.58907506898489 | 0.112043445621613 | 0.538992394329159 | NM_011875 |
| Cep95 | 333.0930267 | 0.336130248 | 0.139083124 | 2.416757972 | 0.015659425 | 0.080313518 | | Ptdss1 | 3262.927915 | 0.0692611779228843 | 0.0435578388747204 | 1.59009674750143 | 0.111812999015436 | 0.538992394329159 | NM_008959 |
| Pex13 | 1161.566649 | -0.162748948 | 0.067339157 | -2.41685457 | 0.015655271 | 0.080313518 | | Pus7 | 719.80552 | -0.111891351722061 | 0.070424014913548 | -1.58882381044901 | 0.112100176061359 | 0.538992394329159 | NM_178403 |
| Ube2v2 | 2400.334127 | -0.240008877 | 0.099322888 | -2.416450837 | 0.015672643 | 0.080354707 | | Sdc3 | 10453.98971 | 0.0951140789015354 | 0.0598093993805158 | 1.5902864748132 | 0.111770245911065 | 0.538992394329159 | NM_011520 |
| Gmfg | 32.90891728 | 0.387747904 | 0.16052807 | 2.415452353 | 0.015715679 | 0.080512887 | | Sox13 | 398.9577887 | -0.128476739561344 | 0.0807794157715858 | -1.5904638370328 | 0.111730290817566 | 0.538992394329159 | NM_011439 |
| Gpr98 | 84.16520156 | -0.357341342 | 0.147941467 | -2.415423811 | 0.01571691 | 0.080512887 | | Spg21 | 1343.345702 | 0.0895968800318837 | 0.0564022543870067 | 1.58853366776992 | 0.112165714143701 | 0.538992394329159 | NM_138584 |
| Msrb2 | 666.4183593 | -0.177172882 | 0.073352169 | -2.415373453 | 0.015719084 | 0.080512887 | | Sppl2b | 1536.446298 | -0.123781245793032 | 0.0778870446916912 | -1.58924049927698 | 0.11200610628664 | 0.538992394329159 | NM_175195 |
| C87436 | 860.8515892 | 0.166001194 | 0.06874021 | 2.414906693 | 0.015739241 | 0.080589491 | | Tcerg1l | 343.1125089 | 0.133816385780983 | 0.0841127128728142 | 1.59091748691217 | 0.11162814657275 | 0.538992394329159 | NM_183289 |
| Atxn2 | 3684.107935 | 0.139564059 | 0.057821622 | 2.413700166 | 0.015791451 | 0.080824093 | | Tsen15 | 586.1426554 | -0.10264335801118 | 0.0645868999574841 | -1.58922874574794 | 0.112008758856277 | 0.538992394329159 | NM_025677 |
| Impad1 | 3170.301991 | -0.170567844 | 0.070669272 | -2.413606937 | 0.015795492 | 0.080824093 | | Zranb2 | 5116.067085 | -0.0931886990637303 | 0.0586898988330797 | -1.58781495481478 | 0.112328188780557 | 0.538992394329159 | NM_017381 |
| Il17rd | 239.2957254 | -0.254104508 | 0.105306643 | -2.412996006 | 0.015821993 | 0.080932968 | | Cab39l | 1458.728205 | 0.132755597070671 | 0.0836400133268794 | 1.58722591963059 | 0.11246148635321 | 0.539325041520875 | NM_026908 |
| Plekho2 | 787.8535139 | -0.243894579 | 0.101103919 | -2.412315779 | 0.015851546 | 0.081057377 | | Gps2 | 814.1890042 | -0.111176726578229 | 0.0700609417148537 | -1.58685743949483 | 0.112544936116547 | 0.539369099976563 | NM_019726 |
| Atp1a2 | 52691.32525 | -0.107353226 | 0.044513556 | -2.411697356 | 0.015878456 | 0.081061217 | | Jagn1 | 895.4174378 | 0.0975138806247659 | 0.0614473967104935 | 1.58694893266509 | 0.112524211087255 | 0.539369099976563 | NM_001205025,NM_026365,NM_001243739 |
| Cdh24 | 337.5611126 | 0.312915997 | 0.129731843 | 2.412021509 | 0.015864346 | 0.081061217 | | Ankrd28 | 2644.417499 | -0.0886577556707421 | 0.0559039901075901 | -1.58589316254735 | 0.112763547128972 | 0.540238550658129 | NM_001024604 |
| Dgcr8 | 1538.815447 | 0.19196399 | 0.07959411 | 2.411786371 | 0.01587458 | 0.081061217 | | Atp6v1g1 | 2705.23013 | 0.0774272129014157 | 0.0488295213070424 | 1.58566397599005 | 0.112815555159917 | 0.540309513436515 | NM_024173 |
| Dpf1 | 2061.231557 | -0.224251388 | 0.092978076 | -2.411873842 | 0.015870772 | 0.081061217 | | Bace1 | 4372.096654 | 0.0844801072435897 | 0.0533444126410493 | 1.5836730233033 | 0.113268146821779 | 0.540327971588701 | NM_001145947,NM_011792 |
| Slc18a3 | 98.06159218 | -0.401011073 | 0.166262273 | -2.411918631 | 0.015868823 | 0.081061217 | | Bcl2l2 | 6863.580407 | 0.0751724254252122 | 0.047470289869154 | 1.58356786175976 | 0.113292092299225 | 0.540327971588701 | NM_007537 |
| Sh3gl3 | 1173.60265 | -0.181831846 | 0.075415265 | -2.411074826 | 0.015905585 | 0.081172968 | | Cldn12 | 1969.163834 | 0.103435847994832 | 0.0653085865163048 | 1.58380166395131 | 0.113238860531311 | 0.540327971588701 | NM_022890,NM_001193659,NM_001193660,NM_001193661 |
| Fbxo7 | 963.58793 | 0.162027488 | 0.067212974 | 2.410657916 | 0.015923776 | 0.081184757 | | E2f3 | 1396.285099 | 0.0936346877230001 | 0.0591036802312201 | 1.58424462498259 | 0.113138061796988 | 0.540327971588701 | NM_010093 |
| Lhfpl2 | 773.9357478 | -0.20110315 | 0.08342874 | -2.410478112 | 0.015931627 | 0.081184757 | | Gmppa | 1349.816661 | -0.109174871729795 | 0.0689159541402106 | -1.58417413053118 | 0.113154098547168 | 0.540327971588701 | NM_133708 |
| Nbl1 | 1679.748589 | -0.212533337 | 0.08817268 | -2.410421646 | 0.015934094 | 0.081184757 | | Mgst3 | 2307.03499 | 0.0855064231989058 | 0.053997688575528 | 1.58352007795179 | 0.113302974078148 | 0.540327971588701 | NM_025569 |
| Psma5 | 2210.527955 | -0.125512509 | 0.052066677 | -2.410611503 | 0.015925821 | 0.081184757 | | Nuak1 | 4415.899705 | 0.124597208667082 | 0.0786571807649913 | 1.58405383278798 | 0.113181469161214 | 0.540327971588701 | NM_001004363 |
| Tyrobp | 580.6805622 | 0.328746552 | 0.136359634 | 2.410878815 | 0.015914135 | 0.081184757 | | Plxna3 | 1635.821575 | -0.127068507657117 | 0.0802058508400145 | -1.58427977917196 | 0.11313006525592 | 0.540327971588701 | NM_008883 |
| Adcy3 | 2639.524277 | -0.143760275 | 0.059655115 | -2.409856628 | 0.01595879 | 0.081256799 | | Pqbp1 | 2239.848351 | 0.084746693989775 | 0.0534569183193603 | 1.58532696335925 | 0.112892065899838 | 0.540327971588701 | NM_019478,NM_001252529,NM_001252528 |
| Crhr1 | 894.8175787 | 0.263526267 | 0.109356162 | 2.409798069 | 0.015961352 | 0.081256799 | | Rab8a | 1423.917367 | 0.0817748780771805 | 0.0516373919368634 | 1.5836368764938 | 0.113276377067203 | 0.540327971588701 | NM_023126 |
| E2f3 | 1301.380637 | -0.151646768 | 0.062930803 | -2.40973831 | 0.015963966 | 0.081256799 | | Rpl7l1 | 1589.410929 | 0.0814823889133021 | 0.0514518619737062 | 1.58366258843932 | 0.113270522681343 | 0.540327971588701 | NM_025433 |
| Lpcat2 | 380.1552825 | -0.352239162 | 0.14619062 | -2.409451182 | 0.015976533 | 0.081294058 | | Rtf1 | 4677.623887 | 0.0873627634094374 | 0.0551151709158121 | 1.58509466975432 | 0.112944827423362 | 0.540327971588701 | NM_030112 |
| Pld5 | 435.0255449 | 0.253304421 | 0.105152778 | 2.408918012 | 0.015999892 | 0.081386187 | | Wnt3 | 108.4961069 | 0.149123565304046 | 0.0941413464820461 | 1.58403901023962 | 0.113184842006457 | 0.540327971588701 | NM_009521 |
| Rbms3 | 826.5907613 | -0.214626477 | 0.089114251 | -2.408441691 | 0.016020785 | 0.081438992 | | Rpl8 | 8358.142175 | 0.0926478345790107 | 0.0585155834195393 | 1.58330190292649 | 0.113352669413719 | 0.540387553627724 | NM_012053 |
| Snx20 | 24.04377191 | 0.398792949 | 0.165573149 | 2.408560523 | 0.016015571 | 0.081438992 | | Spice1 | 595.9757425 | -0.110994010599971 | 0.0701177541915991 | -1.58296585336542 | 0.113429247493317 | 0.540575212955353 | NM_144550 |
| Skil | 2894.546778 | -0.192410958 | 0.079900694 | -2.408126247 | 0.016034635 | 0.081482662 | | Eid1 | 4955.580054 | 0.0908924364336365 | 0.057486971176021 | 1.58109628276171 | 0.113856024556177 | 0.542263927186338 | NM_025613 |
| Slc1a4 | 2773.012119 | -0.133413984 | 0.055407541 | -2.407866908 | 0.016046029 | 0.08151383 | | Gnptg | 2524.363774 | -0.0877277600933043 | 0.0554857422762521 | -1.58108653672733 | 0.113858252644798 | 0.542263927186338 | NM_172529 |
| Qars | 2054.775251 | 0.160040046 | 0.066487306 | 2.407076729 | 0.016080791 | 0.081663643 | | Scn3b | 5991.110544 | 0.109790126544095 | 0.0694626010994111 | 1.5805645744099 | 0.113977631171598 | 0.542483530329304 | NM_178227,NM_153522,NM_001083917 |
| Ranbp9 | 3503.501605 | 0.203832846 | 0.084695413 | 2.406657426 | 0.016099264 | 0.081730666 | | Slc18a2 | 55.4905087 | 0.0737271304927947 | 0.0466462564714954 | 1.58055835708591 | 0.113979053735718 | 0.542483530329304 | NM_172523 |

| | | | | | | |
|---|---|---|---|---|---|---|
| Sp5 | 9.472806124 | -0.335855633 | 0.139562085 | -2.40649624 | 0.01610637 | 0.081739959 |
| Lamtor1 | 2249.70844 | -0.14891505 | 0.061909457 | -2.405368379 | 0.016156171 | 0.081931735 |
| Pcdh19 | 1546.796578 | -0.232428509 | 0.096628375 | -2.405385661 | 0.016155407 | 0.081931735 |
| Rcor1 | 671.7970909 | 0.209192579 | 0.08697219 | 2.40528128 | 0.016160022 | 0.081931735 |
| Pfkfb2 | 2073.858066 | 0.204303754 | 0.084971259 | 2.40438656 | 0.016199633 | 0.082105697 |
| Acrbp | 203.1258016 | 0.361400614 | 0.150324828 | 2.404131237 | 0.016210952 | 0.082136199 |
| Nts | 202.9411025 | -0.367821888 | 0.15300851 | -2.403930914 | 0.016219838 | 0.082154356 |
| Dnaic2 | 91.06605114 | 0.37318443 | 0.155253479 | 2.403710575 | 0.016229617 | 0.082177021 |
| Lonrf2 | 7268.172454 | -0.150200575 | 0.062499791 | -2.403217246 | 0.01625153 | 0.082261091 |
| Mettl24 | 43.19378507 | 0.392309417 | 0.16327016 | 2.402823744 | 0.016269027 | 0.082322763 |
| Vmn2r1 | 18.13350838 | 0.380466106 | 0.158350002 | 2.402690882 | 0.016274938 | 0.08232579 |
| Nudt9 | 1537.403646 | 0.17800782 | 0.074110374 | 2.401928507 | 0.016308895 | 0.082470635 |
| Slc26a10 | 188.0081752 | 0.315051767 | 0.131204755 | 2.40122218 | 0.016340412 | 0.082603047 |
| Pfn4 | 202.1715907 | -0.272239119 | 0.113392527 | -2.400855922 | 0.016356775 | 0.082631847 |
| Snora34 | 19.81984439 | 0.40333706 | 0.167996416 | 2.400867056 | 0.016356278 | 0.082631847 |
| Psme4 | 2407.205705 | 0.158239066 | 0.065913981 | 2.400690453 | 0.016364173 | 0.082642272 |
| Phf21b | 414.1327973 | 0.275519302 | 0.114782946 | 2.400350504 | 0.01637938 | 0.082692118 |
| Dmrt2 | 41.99156631 | -0.39500297 | 0.16456982 | -2.400215121 | 0.016385439 | 0.082695764 |
| A230073K19Rik | 80.43695457 | 0.403028543 | 0.167957825 | 2.399581812 | 0.016413811 | 0.082811982 |
| 1110018G07Rik | 7783.527919 | -0.161827239 | 0.067443496 | -2.39944915 | 0.01641976 | 0.082815028 |
| Nfatc2ip | 220.1889419 | 0.251648219 | 0.104883965 | 2.399301168 | 0.016426398 | 0.082821547 |
| Map3k13 | 1153.666959 | -0.255266218 | 0.106401922 | -2.399075247 | 0.016436537 | 0.082842953 |
| Tspan4 | 401.8621278 | 0.243314355 | 0.10142458 | 2.398968316 | 0.016441337 | 0.082842953 |
| Lbp | 116.5046095 | 0.403104753 | 0.168057154 | 2.398617041 | 0.016457116 | 0.08286856 |
| Ldb2 | 1654.582872 | -0.264946849 | 0.110453564 | -2.398717055 | 0.016452622 | 0.08286856 |
| Sh3rf3 | 1447.751029 | -0.168427211 | 0.07022869 | -2.39826786 | 0.016472814 | 0.082920657 |
| C1qa | 1861.750467 | 0.296544238 | 0.123656096 | 2.398136834 | 0.016478708 | 0.082923385 |
| Ebf1 | 134.4177754 | -0.281703872 | 0.117526054 | -2.396948276 | 0.016532257 | 0.083145631 |
| Fig4 | 1480.784664 | -0.159717361 | 0.06663446 | -2.396918386 | 0.016533606 | 0.083145631 |
| Csda | 719.5115841 | 0.190862918 | 0.079640287 | 2.396562412 | 0.016549675 | 0.083172449 |
| Rbm47 | 32.37786064 | 0.402868712 | 0.168094529 | 2.396679499 | 0.016544388 | 0.083172449 |
| Mien1 | 1600.045223 | -0.199650502 | 0.083326339 | -2.396007122 | 0.016574768 | 0.08327155 |
| 3110047P20Rik | 1871.079331 | -0.217675064 | 0.090877389 | -2.395260983 | 0.016608539 | 0.083309297 |
| Armcx1 | 2475.894419 | 0.167935039 | 0.070115253 | 2.395128464 | 0.016614543 | 0.083309297 |
| Itih2 | 212.6515848 | -0.396906565 | 0.165692602 | -2.395439266 | 0.016600464 | 0.083309297 |
| Ms4a6c | 44.90943626 | 0.388101713 | 0.162037356 | 2.395137284 | 0.016614143 | 0.083309297 |
| Sart3 | 1745.175727 | 0.132209025 | 0.055189928 | 2.395528132 | 0.016596441 | 0.083309297 |
| Zfp933 | 538.8123092 | -0.245267113 | 0.102401918 | -2.395141779 | 0.01661394 | 0.083309297 |
| Elovl1 | 564.0734513 | 0.216782878 | 0.09051626 | 2.394960624 | 0.01662215 | 0.083320477 |
| Gxylt1 | 695.3580599 | 0.204777153 | 0.085523828 | 2.394387121 | 0.016648167 | 0.08337513 |
| Ppm1h | 4265.55814 | -0.153782593 | 0.06422684 | -2.39436649 | 0.016649104 | 0.08337513 |
| Ryr2 | 7766.159326 | -0.236834453 | 0.098913285 | -2.394364443 | 0.016649197 | 0.08337513 |
| Ralgps2 | 1778.011528 | -0.19264432 | 0.080463925 | -2.394170059 | 0.016658023 | 0.083392379 |
| Cript | 1371.747109 | -0.178507991 | 0.074563298 | -2.39404636 | 0.016663642 | 0.083393565 |
| Xkr6 | 25.48750705 | -0.390555182 | 0.163160681 | -2.393684426 | 0.016680093 | 0.083448939 |
| Heatr8 | 244.8854986 | 0.371077819 | 0.155057308 | 2.393165615 | 0.016703699 | 0.083513107 |
| Stat1 | 1256.354587 | 0.393510542 | 0.164430882 | 2.393166885 | 0.016703642 | 0.083513107 |
| Sub1 | 6694.016904 | -0.15812645 | 0.066085423 | -2.392758381 | 0.016722249 | 0.083578881 |
| Ralgapa1 | 6602.499185 | -0.136234445 | 0.056945068 | -2.39238358 | 0.016739338 | 0.08363731 |
| Adamts5 | 102.8277036 | -0.345467528 | 0.144438198 | -2.391801692 | 0.016765898 | 0.083743014 |
| Rtkn | 2199.586687 | 0.175367651 | 0.073331748 | 2.391428738 | 0.016782942 | 0.083801127 |
| Calb2 | 983.2274565 | -0.383300186 | 0.160322094 | -2.390813245 | 0.016811102 | 0.08386066 |
| E4f1 | 939.9410418 | 0.258207259 | 0.107996697 | 2.39088107 | 0.016807997 | 0.08386066 |
| Pcdhb17 | 511.158627 | -0.235700822 | 0.098584568 | -2.390849066 | 0.016809462 | 0.08386066 |
| Ctnnbl1 | 1049.036077 | -0.12531657 | 0.052436346 | -2.389879921 | 0.016853882 | 0.084028459 |
| Sppl2b | 1359.276204 | 0.194552752 | 0.08140818 | 2.38984279 | 0.016855586 | 0.084028459 |
| Bmp4 | 122.7559221 | -0.28323722 | 0.118536011 | -2.389461376 | 0.016873098 | 0.084062773 |
| Rassf10 | 38.68127571 | -0.382789235 | 0.160199288 | -2.389456537 | 0.01687332 | 0.084062773 |
| Canx | 12922.55125 | -0.169127917 | 0.070804606 | -2.388656977 | 0.016910083 | 0.084204199 |
| Plcxd1 | 780.7515543 | -0.189170859 | 0.079197287 | -2.388602783 | 0.016912577 | 0.084204199 |
| 6430411K18Rik | 87.11345067 | 0.370725609 | 0.155249352 | 2.387936597 | 0.016943265 | 0.084250743 |
| Dchs1 | 799.0475162 | -0.28273371 | 0.118407135 | -2.387809742 | 0.016949114 | 0.084250743 |
| Fbxl17 | 3843.373087 | -0.105197285 | 0.044053897 | -2.387922326 | 0.016943922 | 0.084250743 |
| Pnldc1 | 48.39150929 | 0.382192295 | 0.160030016 | 2.388253803 | 0.016928646 | 0.084250743 |
| Slc24a5 | 9.463707027 | 0.332381988 | 0.139198955 | 2.387819568 | 0.01694866 | 0.084250743 |
| Cpsf6 | 2306.861519 | 0.212878436 | 0.089166974 | 2.387413486 | 0.016967396 | 0.084304335 |
| Fam84b | 382.6851611 | -0.335619369 | 0.140582965 | -2.387340238 | 0.016970777 | 0.084304335 |
| Fam81a | 6024.713289 | -0.186458773 | 0.078108762 | -2.387168461 | 0.016978709 | 0.084316705 |
| Adam22 | 9401.099789 | -0.12601844 | 0.052792857 | -2.387035827 | 0.016984836 | 0.084320105 |
| Nov | 3583.438044 | -0.293309142 | 0.122918387 | -2.386210474 | 0.017023005 | 0.084482526 |
| Phf1 | 1700.216856 | 0.179616256 | 0.075278112 | 2.386035613 | 0.017031102 | 0.084495642 |
| Tmem184c | 2671.823804 | -0.129623878 | 0.054338208 | -2.385501543 | 0.017055851 | 0.084591342 |
| Nhs | 237.0490173 | -0.323038881 | 0.135428146 | -2.385315683 | 0.017064471 | 0.084607013 |
| Gm16039 | 1125.569957 | -0.147624871 | 0.061902768 | -2.384786269 | 0.017089046 | 0.084701755 |
| Cdc42ep4 | 1294.033511 | -0.157829581 | 0.066201732 | -2.384070275 | 0.017122332 | 0.084804452 |
| Qsox1 | 1016.655878 | 0.172560494 | 0.072383138 | 2.383987453 | 0.017126186 | 0.084804452 |
| Tmem50b | 3523.581493 | -0.175715821 | 0.07370171 | -2.384148509 | 0.017118692 | 0.084804452 |
| Tbc1d15 | 1146.362106 | 0.199495369 | 0.083690137 | 2.383738121 | 0.017137793 | 0.084834813 |
| Apaf1 | 631.7768199 | 0.329962863 | 0.13844804 | 2.383297461 | 0.017158323 | 0.084909314 |
| Cox5b | 5388.752058 | -0.2144712 | 0.090001726 | -2.382967629 | 0.017173704 | 0.084958294 |
| Vip | 1290.127885 | 0.28483478 | 0.11959138 | 2.381733362 | 0.017231369 | 0.085216353 |
| Arid4b | 1778.595775 | 0.16017041 | 0.067274306 | 2.380855629 | 0.01727248 | 0.085350894 |
| Cap2 | 9975.219708 | -0.173346537 | 0.072815455 | -2.380628359 | 0.017283139 | 0.085350894 |
| Rps8 | 7515.485747 | -0.142776088 | 0.05996433 | -2.381016993 | 0.017264915 | 0.085350894 |
| Sec11a | 632.4795626 | -0.163359317 | 0.068619099 | -2.380668345 | 0.017281263 | 0.085350894 |
| Shkbp1 | 489.418943 | 0.326661472 | 0.137220152 | 2.380564863 | 0.017286118 | 0.085350894 |
| Supt4a | 1111.186566 | -0.146423761 | 0.061522721 | -2.379994863 | 0.017312879 | 0.085455799 |
| Cd48 | 36.80707499 | 0.396416895 | 0.166595508 | 2.379517307 | 0.017335329 | 0.08551673 |
| Lrrc16b | 1711.862804 | 0.29925681 | 0.125764711 | 2.379497464 | 0.017336262 | 0.08551673 |
| Sgcd | 65.75656928 | -0.389417905 | 0.163699255 | -2.378861802 | 0.017366185 | 0.08563707 |
| Tro | 8701.391647 | 0.214940728 | 0.090361496 | 2.37867606 | 0.017374937 | 0.085652969 |
| Fam20b | 4776.139503 | -0.10263425 | 0.043157967 | -2.378106684 | 0.01740179 | 0.08575806 |
| Lmnb1 | 277.6137763 | -0.281157251 | 0.118272576 | -2.377197327 | 0.017444753 | 0.08594245 |
| Mfsd7b | 409.0526651 | 0.2394743 | 0.100810099 | 2.375499108 | 0.017525235 | 0.086311503 |
| Bhmt2 | 12.83070366 | 0.386528659 | 0.162847833 | 2.373557272 | 0.017617661 | 0.086684037 |
| Gm11517 | 13.85634754 | 0.396246462 | 0.166936829 | 2.373631184 | 0.017614135 | 0.086684037 |
| Rpl27 | 2818.026145 | -0.129140066 | 0.054403625 | -2.373740087 | 0.017608941 | 0.086684037 |
| Nfxl1 | 399.6758742 | 0.215797945 | 0.090923553 | 2.373399837 | 0.017625173 | 0.086693469 |
| Nsdhl | 937.791978 | -0.268150234 | 0.112993896 | -2.373139103 | 0.01763762 | 0.086727161 |
| Rps11 | 5109.247231 | -0.142522398 | 0.060061821 | -2.372928359 | 0.017647687 | 0.086749129 |
| Tbc1d5 | 1853.950946 | 0.135300785 | 0.057037671 | 2.372130237 | 0.017685855 | 0.08690918 |
| Tmem194 | 200.9220233 | 0.218518251 | 0.09216963 | 2.370827025 | 0.017748335 | 0.087188555 |
| Mrps16 | 597.0168219 | -0.20335199 | 0.085823464 | -2.36942185 | 0.01781592 | 0.087492825 |
| Arhgef33 | 47.38334487 | 0.397435742 | 0.167782099 | 2.368761297 | 0.017847768 | 0.087510545 |
| Dock1 | 1545.824465 | -0.17984882 | 0.075923406 | -2.368819184 | 0.017844975 | 0.087510545 |
| P4ha1 | 2028.397069 | -0.246445453 | 0.104026001 | -2.369075517 | 0.017832612 | 0.087510545 |
| Pde4a | 5161.492529 | -0.143069632 | 0.060388922 | -2.369137061 | 0.017829645 | 0.087510545 |
| Srebf2 | 8693.91765 | -0.214956159 | 0.090744975 | -2.368794064 | 0.017846187 | 0.087510545 |
| Parp10 | 110.4097956 | 0.366015921 | 0.154531118 | 2.368558029 | 0.017857579 | 0.087530948 |
| St8sia4 | 444.7314695 | -0.253243227 | 0.106933595 | -2.368228873 | 0.017873475 | 0.087581159 |
| Sgk1 | 3281.027366 | 0.397235072 | 0.167764677 | 2.367811148 | 0.017893667 | 0.087652379 |
| Dis3 | 550.9342643 | 0.210665726 | 0.088975059 | 2.367694136 | 0.017899327 | 0.087652391 |
| AW549542 | 49.97035661 | 0.388257674 | 0.164026386 | 2.367044005 | 0.017930801 | 0.087751051 |
| Akirin2 | 1963.988884 | -0.16145039 | 0.068206361 | -2.367086986 | 0.017928718 | 0.087751051 |
| Taok1 | 5420.766461 | -0.179412309 | 0.075827577 | -2.366056216 | 0.017978714 | 0.087957751 |
| Slco5a1 | 220.8090313 | -0.391912741 | 0.165677936 | -2.365509565 | 0.018005278 | 0.088059905 |
| Nol9 | 905.9099469 | 0.173542634 | 0.073376932 | 2.365084341 | 0.018025965 | 0.088077675 |
| Prr14l | 3069.974686 | -0.179816362 | 0.076029004 | -2.365102157 | 0.018025098 | 0.088077675 |
| Slfn5 | 452.7647933 | -0.265530051 | 0.112260463 | -2.365303372 | 0.018015307 | 0.088077675 |
| Crlf1 | 37.34904651 | 0.390460175 | 0.165117316 | 2.364743952 | 0.01804254 | 0.088130869 |
| Asb11 | 146.4962814 | 0.325906669 | 0.137870844 | 2.363854889 | 0.018085895 | 0.088276641 |
| Gipc3 | 23.77989283 | 0.39649886 | 0.167739214 | 2.363781549 | 0.018089476 | 0.088276641 |
| Moxd1 | 321.0978014 | 0.214130699 | 0.090586375 | 2.363828988 | 0.01808716 | 0.088276641 |
| Pik3ip1 | 1244.090556 | 0.17450964 | 0.073843736 | 2.36322875 | 0.018116483 | 0.088380602 |
| Tial1 | 3053.806012 | 0.198488621 | 0.083995043 | 2.363099227 | 0.018122816 | 0.08838367 |
| Efna1 | 139.3645602 | 0.278449517 | 0.117849208 | 2.362761034 | 0.018139362 | 0.088436524 |
| Chchd2 | 3885.550337 | -0.208896162 | 0.088439196 | -2.362031453 | 0.0181751 | 0.088582888 |
| Trmt12 | 356.3919084 | -0.201327592 | 0.085243686 | -2.361788901 | 0.018186995 | 0.088612988 |
| Cckbr | 2206.5492 | -0.257810182 | 0.10919737 | -2.360955974 | 0.018227894 | 0.088728557 |
| Grin2a | 1830.157059 | -0.287698731 | 0.121853738 | -2.361016868 | 0.018224902 | 0.088728557 |
| L3mbtl2 | 1855.018214 | 0.159174355 | 0.067417335 | 2.36103006 | 0.018224253 | 0.088728557 |
| Pde4d | 3059.079742 | -0.158648385 | 0.067202141 | -2.360763859 | 0.018237339 | 0.088746651 |
| Phf11b | 13.45256209 | 0.392874646 | 0.166435552 | 2.360521183 | 0.018249276 | 0.08874899 |
| Spg7 | 1905.747602 | 0.187363884 | 0.079373664 | 2.360529601 | 0.018248862 | 0.08874899 |
| Tmcc2 | 8279.87536 | -0.166629579 | 0.070613241 | -2.359749766 | 0.018287266 | 0.088905835 |
| 4930523C07Rik | 85.01359722 | -0.316609504 | 0.134191827 | -2.359379936 | 0.018305503 | 0.088966584 |
| Dagla | 5635.274089 | 0.225332447 | 0.095521164 | 2.358979269 | 0.018325279 | 0.08903477 |
| Larp1b | 484.5853246 | 0.238364649 | 0.101053756 | 2.358790577 | 0.018334599 | 0.089052126 |
| Wdr76 | 142.1958694 | 0.307757177 | 0.130486929 | 2.358528776 | 0.018347537 | 0.089087039 |
| Cep250 | 1589.249899 | 0.180283868 | 0.076463223 | 2.357785361 | 0.018384319 | 0.08923767 |
| Rasgrp1 | 23287.50574 | -0.168732706 | 0.071581867 | -2.357199009 | 0.018413376 | 0.089322744 |
| Tmem178b | 1927.931031 | -0.196679036 | 0.083445036 | -2.357204765 | 0.01841309 | 0.089322744 |
| Ehd3 | 5290.596227 | 0.220522838 | 0.093557643 | 2.357079874 | 0.018419284 | 0.089323441 |
| Clptm1l | 3937.739573 | -0.120814068 | 0.051262127 | -2.356790002 | 0.018433668 | 0.089365222 |
| Bmper | 378.8123935 | -0.193070621 | 0.081956666 | -2.355764689 | 0.018484623 | 0.089556206 |
| Mrpl43 | 1035.364622 | -0.142243595 | 0.060378146 | -2.355878804 | 0.018478946 | 0.089556206 |
| Ncf1 | 330.0553702 | 0.245922461 | 0.104418038 | 2.355172205 | 0.018514124 | 0.089659874 |
| Sft2d2 | 1819.452844 | -0.158601572 | 0.067343807 | -2.355102567 | 0.018517594 | 0.089659874 |
| Phactr4 | 514.3173277 | 0.247274775 | 0.105016147 | 2.35463576 | 0.01854087 | 0.089744528 |
| Akt3 | 6685.974434 | -0.148908785 | 0.063251394 | -2.354237204 | 0.018560763 | 0.089806328 |
| Sulf2 | 4035.195783 | -0.197211446 | 0.083771907 | -2.354147759 | 0.01856523 | 0.089806328 |
| Phf11d | 26.57630504 | 0.389959622 | 0.165661044 | 2.353960918 | 0.018574564 | 0.089823437 |
| Dlx1 | 1060.589347 | -0.39213317 | 0.166602584 | -2.353704016 | 0.018587405 | 0.089857488 |
| P4ha3 | 177.3235348 | 0.336193558 | 0.1428468 | 2.353525312 | 0.018596342 | 0.089872651 |
| Smoc2 | 629.4587014 | -0.318369396 | 0.135297912 | -2.353099114 | 0.018617671 | 0.089947675 |
| Ctsd | 15207.27433 | 0.212034274 | 0.090115035 | 2.352928937 | 0.018626193 | 0.089960798 |
| Phtf1 | 1949.7097 | 0.115509282 | 0.049098763 | 2.352590448 | 0.018643155 | 0.09001466 |
| Srpx2 | 68.3261335 | 0.368245104 | 0.156544772 | 2.352330896 | 0.01865617 | 0.090049441 |
| B3gnt2 | 1318.552831 | -0.199267049 | 0.084723758 | -2.35196187 | 0.018674689 | 0.090094901 |
| Dnahc1 | 455.383315 | 0.265751721 | 0.112993929 | 2.351911494 | 0.018677218 | 0.090094901 |
| Filip1 | 747.7268209 | -0.296640352 | 0.126149484 | -2.351498729 | 0.018697953 | 0.090108701 |
| Mbp | 60860.84191 | 0.24399672 | 0.103758657 | 2.351579401 | 0.018693899 | 0.090108701 |
| Trrap | 6414.980351 | -0.189725952 | 0.08068667 | -2.351391525 | 0.018703342 | 0.090108701 |
| Uba7 | 156.18167 | 0.384020309 | 0.163295871 | 2.351684135 | 0.018688637 | 0.090108701 |
| Abcg4 | 2641.15172 | 0.169940472 | 0.072316238 | 2.349962849 | 0.018775285 | 0.090427188 |
| Golga2 | 4426.497719 | 0.161026414 | 0.068526363 | 2.34984621 | 0.018781169 | 0.090427419 |
| Ngfrap1 | 5031.795109 | -0.135800006 | 0.057797487 | -2.349583234 | 0.018794442 | 0.090463211 |
| Atp5j | 4039.508351 | -0.161324763 | 0.068669692 | -2.349286254 | 0.01880944 | 0.090479188 |
| Pkdcc | 276.8951276 | 0.228382905 | 0.097210808 | 2.349357127 | 0.01880586 | 0.090479188 |
| Mob3c | 211.4845485 | 0.24700404 | 0.105207693 | 2.347775457 | 0.018885903 | 0.090762462 |
| Tfap2b | 65.40873001 | -0.371337572 | 0.158161416 | -2.347839195 | 0.018882671 | 0.090762462 |
| Wbscr17 | 2191.991231 | 0.169314681 | 0.072110622 | 2.347985305 | 0.018875266 | 0.090762462 |
| 6720483E21Rik | 17.09136666 | 0.394455097 | 0.168046637 | 2.347295387 | 0.018910256 | 0.090767572 |
| Rccd1 | 337.5748647 | 0.289164052 | 0.123188115 | 2.347337253 | 0.018908131 | 0.090767572 |
| Rnf128 | 471.3599408 | -0.242232611 | 0.10318986 | -2.347445867 | 0.01890262 | 0.090767572 |
| Tbkbp1 | 1793.174353 | 0.225606722 | 0.096113592 | 2.347292576 | 0.018910399 | 0.090767572 |
| Pdzd4 | 5768.544316 | -0.153401151 | 0.0653716 | -2.346602348 | 0.018945462 | 0.090907708 |
| Ctnna2 | 6187.254253 | -0.144311079 | 0.061509419 | -2.346162293 | 0.018967846 | 0.090958778 |
| Mreg | 176.4369828 | 0.263304315 | 0.112224253 | 2.346233618 | 0.018964217 | 0.090958778 |
| Itga10 | 156.0538537 | 0.354144757 | 0.150976209 | 2.345699094 | 0.018991433 | 0.091015545 |
| Tgif1 | 62.43609728 | 0.370820503 | 0.158078952 | 2.345793019 | 0.018986648 | 0.091015545 |
| Slc22a4 | 300.0352939 | 0.215163169 | 0.091750275 | 2.345095618 | 0.019022201 | 0.09113481 |
| Opa1 | 7295.160246 | -0.131847867 | 0.056230324 | -2.344782268 | 0.019038194 | 0.09113625 |
| Tatdn3 | 425.392812 | 0.341771954 | 0.145757558 | 2.344797469 | 0.019037418 | 0.09113625 |
| Tcea1 | 3073.037939 | -0.136132654 | 0.058058642 | -2.344744009 | 0.019040147 | 0.09113625 |
| Neu1 | 2044.273565 | 0.193882397 | 0.082734616 | 2.343425353 | 0.019107585 | 0.091430799 |
| Apoa1 | 11.25856687 | 0.373030793 | 0.159195694 | 2.343221635 | 0.019118022 | 0.091452498 |
| Cluap1 | 949.3418951 | 0.150693935 | 0.064316562 | 2.343003568 | 0.0191292 | 0.091469296 |
| Gprin1 | 5672.165665 | -0.160646118 | 0.068566543 | -2.342922808 | 0.019133341 | 0.091469296 |
| Hs3st3a1 | 86.32654093 | -0.353433256 | 0.150870872 | -2.342620887 | 0.019148829 | 0.091515103 |
| Rfng | 2281.401231 | 0.173372229 | 0.074022098 | 2.342168547 | 0.019172055 | 0.091597846 |
| Atxn7l1 | 914.5557991 | 0.15307298 | 0.065380346 | 2.341269046 | 0.019218312 | 0.091773148 |
| Cep350 | 2563.563985 | -0.199874439 | 0.085372085 | -2.34121538 | 0.019221075 | 0.091773148 |
| Ddrgk1 | 1813.813338 | 0.188886509 | 0.080682467 | 2.34110972 | 0.019226516 | 0.091773148 |
| Wdr1 | 5961.029606 | -0.108109727 | 0.046184409 | -2.340827349 | 0.019241063 | 0.091814297 |
| Fgf17 | 23.5495849 | -0.392848127 | 0.167851189 | -2.340454841 | 0.019260268 | 0.091850854 |
| Tmem144 | 364.3399557 | 0.235535402 | 0.100636854 | 2.340448785 | 0.01926058 | 0.091850854 |
| Srcrb4d | 36.00718693 | 0.371011918 | 0.158549812 | 2.340033794 | 0.019281995 | 0.091924687 |
| Adtrp | 8.362910432 | -0.350154788 | 0.149681546 | -2.339331714 | 0.019318273 | 0.092069307 |
| Trim67 | 192.0195953 | -0.345264256 | 0.147606554 | -2.339084866 | 0.019331042 | 0.092101833 |
| Lars2 | 55452.26981 | -0.20975622 | 0.089686313 | -2.338776262 | 0.019347016 | 0.092121443 |
| Stra6 | 286.7295106 | -0.278588165 | 0.119117098 | -2.338775624 | 0.019347049 | 0.092121443 |
| Slc5a6 | 1070.879787 | 0.148637508 | 0.06356205 | 2.338463084 | 0.019363239 | 0.092170205 |
| Cd209a | 27.40794034 | -0.392845576 | 0.168044844 | -2.337742513 | 0.019400609 | 0.092251522 |
| Cisd1 | 2162.321146 | -0.174341958 | 0.074573586 | -2.337851351 | 0.019394961 | 0.092251522 |
| Gcfc2 | 230.4716507 | 0.304663873 | 0.130311634 | 2.337963718 | 0.01938913 | 0.092251522 |
| Zfp574 | 1180.306197 | 0.176115369 | 0.075337848 | 2.337674542 | 0.019404138 | 0.092251522 |
| Hapln3 | 25.33553149 | -0.38670956 | 0.165480156 | -2.336893854 | 0.019444704 | 0.092416026 |
| Arpp21 | 24260.21947 | -0.173545691 | 0.074275486 | -2.336513701 | 0.019464484 | 0.092452392 |
| Ctdspl | 1551.811491 | 0.115394139 | 0.049389485 | 2.336411051 | 0.019469829 | 0.092452392 |
| Tpm2 | 380.6511522 | -0.191802801 | 0.082097233 | -2.336288247 | 0.019476224 | 0.092452392 |
| Zfp384 | 1531.1735 | 0.166615135 | 0.071315091 | 2.336323645 | 0.01947438 | 0.092452392 |
| Slc29a3 | 699.4996895 | 0.201262103 | 0.086154418 | 2.33606246 | 0.019487986 | 0.092479897 |
| Rps6kb2 | 948.0719135 | 0.2910023 | 0.124617222 | 2.335169207 | 0.019534583 | 0.092672636 |
| Ankrd50 | 1673.840755 | -0.199697847 | 0.085521909 | -2.335048985 | 0.019540862 | 0.092674048 |
| Ildr2 | 7442.478408 | -0.160886919 | 0.068918821 | -2.334440982 | 0.019572643 | 0.092796368 |
| Map3k9 | 4234.828424 | 0.149148133 | 0.063912065 | 2.333645966 | 0.019614268 | 0.092938739 |
| Zbed3 | 367.3401818 | 0.180911127 | 0.077523207 | 2.333638314 | 0.019614669 | 0.092938739 |
| Sema3f | 467.4219986 | 0.233406931 | 0.100025757 | 2.333468286 | 0.019623581 | 0.092952541 |
| Runx1 | 54.67550922 | 0.360086902 | 0.154347323 | 2.332964991 | 0.019649983 | 0.093049155 |
| Appl2 | 3152.496046 | 0.189032993 | 0.081034841 | 2.332737266 | 0.019661939 | 0.093077325 |
| Aldh5a1 | 4858.897513 | -0.166172626 | 0.071241434 | -2.332527813 | 0.019672942 | 0.093100964 |
| Fn3k | 1116.376025 | 0.147083721 | 0.063069432 | 2.332092049 | 0.019695849 | 0.093180912 |
| Hars2 | 972.309758 | 0.224481144 | 0.096290101 | 2.331300324 | 0.019737529 | 0.093349596 |
| Clca1 | 27.68452846 | -0.269466156 | 0.115609893 | -2.33082263 | 0.019762714 | 0.093440188 |
| Fbxo38 | 2060.759806 | 0.204880161 | 0.087936997 | 2.329851696 | 0.019813991 | 0.093654049 |
| Tnfaip6 | 311.5758205 | 0.281125414 | 0.120688801 | 2.329341351 | 0.019840989 | 0.093753062 |
| Ptdss1 | 2983.413897 | -0.098019523 | 0.042088286 | -2.328902726 | 0.019864219 | 0.093834212 |
| Chgb | 18358.45937 | 0.219727192 | 0.094354167 | 2.328749211 | 0.019872355 | 0.093844034 |
| Anapc4 | 1962.50138 | 0.142325252 | 0.061139315 | 2.32788432 | 0.019918247 | 0.093909252 |
| Ccdc159 | 387.8843533 | 0.349554159 | 0.150146839 | 2.328082036 | 0.019907748 | 0.093909252 |
| Gpr68 | 502.0228375 | 0.315392845 | 0.135486361 | 2.32785678 | 0.01991971 | 0.093909252 |
| H2afy2 | 649.8774578 | -0.174979958 | 0.0751704 | -2.32777741 | 0.019923926 | 0.093909252 |
| Rit2 | 2399.884391 | -0.174737199 | 0.075068944 | -2.327689575 | 0.019928593 | 0.093909252 |
| Sass6 | 185.8474938 | 0.25923209 | 0.111361407 | 2.327844951 | 0.019920338 | 0.093909252 |

| | | | | | | | |
|---|---|---|---|---|---|---|---|
| Cnpy2 | 1005.149383 | 0.108434329490193 | 0.0686163622100772 | 1.58029843025208 | 0.114038539193809 | 0.542588870071822 | NM_019953 |
| Copz1 | 3746.707802 | 0.0673834498201745 | 0.0426538128126591 | 1.57977553181778 | 0.114158280960601 | 0.542980743036574 | NM_019817 |
| Aldoa | 63778.62338 | 0.0715169664223763 | 0.045330627044834 | 1.57767432494686 | 0.114640447649192 | 0.543130934168015 | NM_001177307,NM_007438,NM_001177308 |
| Arap3 | 539.8077453 | -0.130938324506215 | 0.0830152696492054 | -1.57727999992672 | 0.114731122255375 | 0.543130934168015 | NM_139206,NM_001205336 |
| Foxf2 | 168.6593079 | -0.149283069284897 | 0.0946495951794946 | -1.57721825436014 | 0.114745314122159 | 0.543130934168015 | NM_010225 |
| Fubp1 | 3120.832801 | -0.0954670676966324 | 0.0605178325056231 | -1.57750308866006 | 0.114679811975485 | 0.543130934168015 | NM_057172 |
| Grid2 | 430.4439247 | 0.135341764107615 | 0.0857080331758796 | 1.57910243757296 | 0.11431256268018 | 0.543130934168015 | NM_008167 |
| Klhl18 | 1667.391587 | 0.0754056912009208 | 0.0477829794819984 | 1.57808684218465 | 0.114545660572002 | 0.543130934168015 | NM_177771 |
| Mfap3l | 2861.347111 | -0.10082438589105 | 0.0639139990537487 | -1.57750081959761 | 0.114680333665956 | 0.543130934168015 | NM_001177882,NM_027756,NM_001177881 |
| Mybbp1a | 2811.953097 | -0.0805714511352813 | 0.0510731336290571 | -1.57757015107923 | 0.114664394192704 | 0.543130934168015 | NM_016776 |
| Nans | 522.342846 | 0.111879915574565 | 0.0709511120361067 | 1.57685922551334 | 0.114827920409883 | 0.543130934168015 | NM_053179 |
| Ndufs2 | 7542.694531 | 0.064374117656215 | 0.0408170864853472 | 1.57713651804434 | 0.114764116113733 | 0.543130934168015 | NM_153064 |
| Nup50 | 1959.262861 | 0.0971831686287156 | 0.0615349370881902 | 1.57931694135699 | 0.114263377891231 | 0.543130934168015 | NM_016714 |
| Prepl | 21130.94236 | 0.0914029882588799 | 0.0579220488073043 | 1.57803444700239 | 0.11455769637263 | 0.543130934168015 | NM_001163624,NM_001163623,NM_001163622,NM_145984 |
| Ralbp1 | 3531.024216 | 0.0798532832282877 | 0.050645596412364 | 1.57670732993262001 | 0.114862882401497 | 0.543130934168015 | NM_001198949,NM_009067 |
| Slc25a16 | 1719.159798 | 0.101199366708635 | 0.0640817590634419 | 1.57922267097017 | 0.114284991630643 | 0.543130934168015 | NM_175194 |
| Soga3 | 2511.269222 | 0.0979299246250112 | 0.0620870525672021 | 1.57730026753989 | 0.114726450878989 | 0.543130934168015 | NM_026138 |
| Sp140 | 35.20064515 | -0.139860080373859 | 0.0887029295642625 | -1.57672447867164 | 0.114858935384727 | 0.543130934168015 | NM_001013817 |
| Tox2 | 789.2119309 | 0.140369659689927 | 0.0889835957640265 | 1.57747794393665 | 0.114685593217565 | 0.543130934168015 | NM_001098799 |
| Vcpip1 | 2477.210919 | 0.11332002998592 | 0.0718418915758247 | 1.57735309441731 | 0.114714301852753 | 0.543130934168015 | NM_173443 |
| Mdm1 | 243.7982616 | -0.125169246900802 | 0.0794044605152346 | -1.57635032199214 | 0.114945090402641 | 0.543342786589247 | NM_010785,NM_001162905,NM_001162904,NM_148922 |
| Cpt2 | 290.0582229 | 0.137729521296469 | 0.0874079315463287 | 1.57570965082807 | 0.115092732300752 | 0.54339411768961 | NM_009949 |
| Hpca | 28728.24186 | -0.0971088513387041 | 0.0616259981704883 | -1.57577733783804 | 0.115077126867283 | 0.54339411768961 | NM_001130419,NM_010471 |
| Slc25a33 | 1167.586742 | 0.102732092846512 | 0.065198157908982 | 1.57569011366745 | 0.115097236958482 | 0.54339411768961 | NM_027460 |
| Tmx4 | 11598.73543 | 0.098511190525404 | 0.0625208298923574 | 1.57565391718267 | 0.1151055831014 | 0.54339411768961 | NM_029148 |
| Gt(ROSA)26Sor | 106.5920242 | -0.149479828668135 | 0.0948991135519524 | -1.57514462541636 | 0.115223065473592 | 0.543478515937577 | NR_027009,NR_027008,NR_027010 |
| Proca1 | 127.22638 | 0.147454088685168 | 0.0936163022331314 | 1.57508986328006 | 0.115235703503218 | 0.543478515937577 | NM_001045516 |
| Usp39 | 1312.175654 | 0.0842383081723401 | 0.0534788287960111 | 1.57517114844937 | 0.115216944868342 | 0.543478515937577 | NM_138592 |
| Ppapdc3 | 1032.819432 | 0.0965967147628913 | 0.0614169834604502 | 1.57280135428819 | 0.115764821979373 | 0.54561966387812 | NM_145521 |
| Siglech | 701.2074 | -0.128998760789656 | 0.082017769870595 | -1.57281478139659 | 0.115761711982512 | 0.54561966387812 | NM_178706 |
| Ell | 973.6806343 | -0.0951364157112074 | 0.060511471575835 | -1.57220462886908 | 0.115903102297139 | 0.545881860313367 | NM_007924 |
| Plekhg5 | 3689.655429 | -0.11970689153542 | 0.0761684229302721 | -1.57160785178662 | 0.116041524395744 | 0.545881860313367 | NM_001004156 |
| Sik3 | 6774.248664 | 0.0748170333523613 | 0.0475919360166829 | 1.57205273864326 | 0.115938320828775 | 0.545881860313367 | NM_027498 |
| Tmem200a | 600.9512971 | 0.116235239623284 | 0.0739326314419816 | 1.57217776989988 | 0.115909329429162 | 0.545881860313367 | NM_029881 |
| Vps9d1 | 2375.520406 | -0.112068480314391 | 0.0713090317523736 | -1.57158886553891 | 0.116045930376406 | 0.545881860313367 | NM_028200 |
| Zmym5 | 1473.952487 | -0.0962672606391451 | 0.0612427895690294 | -1.5718954233892 | 0.115974806119038 | 0.545881860313367 | NM_001253753,NM_001253752,NM_144842 |
| B230209K01Rik | 66.34035792 | -0.149199405129263 | 0.0949585945594874 | -1.57120485924838 | 0.116135071746796 | 0.54612432897182 | NR_038008 |
| Leprel1 | 101.5267641 | -0.144742889844452 | 0.0921605209600773 | -1.57055199272531 | 0.116286748534697 | 0.546307021091528 | NM_173379 |
| Net1 | 1994.40915 | 0.11895435508281 | 0.0757393211197117 | 1.57057593498619 | 0.116281183416633 | 0.546307021091528 | NM_001047159,NM_019671 |
| Pcdhga11 | 944.8951425 | 0.0838444655466263 | 0.0533818098062051 | 1.57065610647169 | 0.116262549950569 | 0.546307021091528 | NM_033594 |
| Hsp90aa1 | 50276.28609 | 0.0975880409859255 | 0.0621477315507893 | 1.5702590995807 | 0.116354845299893 | 0.546450204599499 | NM_010480 |
| Cyp4f13 | 367.8737408 | -0.131341275955155 | 0.0836848071731208 | -1.56947575541933 | 0.116537124366643 | 0.546452316803884 | NM_130882 |
| Pole | 88.9540909 | -0.148866132787988 | 0.0948377533570201 | -1.56969273858245 | 0.116486611370059 | 0.546452316803884 | NM_011132 |
| Smox | 1600.132051 | 0.13197223130039 | 0.0840773749650818 | 1.5696521371558 | 0.116496061949646 | 0.546452316803884 | NM_145533,NM_001177833,NM_001177837,NM_001177836,NM_001177835,NM_001177834,NM_001177840,NM_001177838,NM_001177839 |
| Syt15 | 39.74547114 | -0.144732190800456 | 0.0922114855298182 | -1.56956793363506 | 0.116515663478198 | 0.546452316803884 | NM_176931,NM_181529 |
| Znhit2 | 570.150694 | -0.127105031438139 | 0.08098704944775809 | -1.56944884774951 | 0.116543389608869 | 0.546452316803884 | NM_013859 |
| Dynlt1a | 46.88893179 | 0.147946264438655 | 0.0942928483256433 | 1.56901214742942 | 0.116645108564678 | 0.546752774124078 | NM_001166629 |
| Ank1 | 3575.952339 | 0.0992021725163748 | 0.0632353680071392 | 1.56877670902738 | 0.116699977267184 | 0.546833506381648 | NM_031158,NM_001110783 |
| Tspyl1 | 7929.130684 | 0.0954475891941461 | 0.0608682551863726 | 1.56810128533987 | 0.116857496633322 | 0.547395032600981 | NM_009433 |
| Snrpd3 | 1291.323822 | 0.0894620722157599 | 0.0570850100237515 | 1.56717275127984 | 0.117074317790248 | 0.548233894333057 | NM_026095 |
| Brca2 | 201.6242448 | -0.133333153412211 | 0.0851622083208508 | -1.5656375761168 | 0.117433487852707 | 0.548468184404505 | NM_001081001,NM_009765 |
| Cplx1 | 33806.59349 | 0.0876513846688009 | 0.0559751926390218 | 1.56589697214721 | 0.117372738827154 | 0.548468184404505 | NM_007756 |
| Ermp1 | 3175.548393 | 0.0904244386973952 | 0.0577471816016036 | 1.5658675659919 | 0.117379624335916 | 0.548468184404505 | NM_001081213 |
| Grk4 | 87.88060051 | -0.148554814917305 | 0.0948686783562831 | -1.56589948854776 | 0.117372149621771 | 0.548468184404505 | NM_019497,NM_001080743 |
| Icosl | 203.3414158 | -0.145354936114328 | 0.0928172131417178 | -1.56603426448921 | 0.117340595752338 | 0.548468184404505 | NM_015790 |
| Phlda1 | 746.4759918 | 0.12680568912691 | 0.0809997789083193 | 1.56550660799256 | 0.11746416919196 | 0.548468184404505 | NM_009344 |
| Them6 | 1756.445251 | 0.0906537014113129 | 0.0578758860126443 | 1.56634667141869 | 0.117267480356243 | 0.548468184404505 | NM_198607 |
| Trim21 | 103.8013417 | -0.147920100006453 | 0.0944724335043164 | -1.56574880649914 | 0.117407435329325 | 0.548468184404505 | NM_009277,NM_001082552 |
| Zhx2 | 684.0154422 | 0.115276993109242 | 0.0736103506496361 | 1.56604325467661 | 0.117338491198181 | 0.548468184404505 | NM_199449 |
| Agxt2l2 | 113.9138907 | -0.145626229191938 | 0.093081837653527 | -1.56449671453623 | 0.11770096394016 | 0.548537626626739 | NM_028398 |
| Gm14169 | 395.8632721 | -0.136432162818012 | 0.0871866007934121 | -1.56482947581919 | 0.1176228984011 | 0.548537626626739 | NR_040372 |
| Kcnmb2 | 135.1212676 | 0.142497785914313 | 0.0910592577022946 | 1.56489070425095 | 0.117608538685622 | 0.548537626626739 | NM_028231 |
| Lca5l | 42.14975927 | -0.145413719028501 | 0.0928997101713395 | -1.56527634758287 | 0.11751812656225 | 0.548537626626739 | NM_001001492 |
| Mx2 | 42.49774316 | -0.140664449794343 | 0.0899053692379362 | -1.5645834170117 | 0.117680619691015 | 0.548537626626739 | NR_003508,NM_013606 |
| Psmd4 | 3415.509955 | 0.0745518265204943 | 0.0476528781259264 | 1.56447688896115 | 0.117705616287729 | 0.548537626626739 | NM_008951 |
| Tsga10 | 351.9140887 | -0.121094331281294 | 0.0774243796612777 | -1.564033600412 | 0.117809677815191 | 0.548846497736841 | NM_207228 |
| Shfm1 | 1147.914637 | 0.0970846290944457 | 0.062083796073085 | 1.56376760499886 | 0.117872154607758 | 0.548913245850142 | NM_009169 |
| Vcp | 15351.01958 | 0.0673430333317699 | 0.0430678194941757 | 1.56365086792651 | 0.117899581925681 | 0.548913245850142 | NM_009503 |
| Dimt1 | 505.5144749 | -0.10843794189784 | 0.0694053017046436 | -1.56238701128772 | 0.118196845080662 | 0.549241001804766 | NM_025447 |
| Gnb5 | 5789.706147 | 0.0987661840293805 | 0.0632055349792951 | 1.56261922411913 | 0.118142183834314 | 0.549241001804766 | NM_010313,NM_138719,NR_073186 |
| Htra1 | 5480.094645 | 0.121187790426434 | 0.0775542921728463 | 1.56261874141461 | 0.11814229743895 | 0.549241001804766 | NM_019564 |
| Klhdc4 | 829.8275319 | -0.120568197684927 | 0.077152577758387 | -1.56272416538694 | 0.118117487913648 | 0.549241001804766 | NM_145605 |
| Map7d2 | 7576.269299 | 0.0956669396546845 | 0.0612048681674992 | 1.56306095444684 | 0.11803825839999 | 0.549241001804766 | NM_001081124 |
| Zfp174 | 366.2062469 | -0.123091456121277 | 0.0787802792827023 | -1.56246534338327 | 0.118178404046636 | 0.549241001804766 | NM_001081217 |
| Als2cl | 589.9148434 | -0.133578314123585 | 0.0855218887326257 | -1.56191959863284 | 0.118306930838985 | 0.549576743641539 | NM_001146060,NM_001146059,NM_146228 |
| Ado | 2342.422587 | 0.107158914705581 | 0.0686426724790666 | 1.56111221832542 | 0.118497275710144 | 0.550284983045253 | NM_001005419 |
| Hnrnpa2b1 | 16295.16623 | -0.0602554649639214 | 0.0386020413773544 | -1.56093985742603 | 0.118537941940803 | 0.550297904963918 | NM_016806,NM_182650 |
| Galc | 1252.709598 | -0.0832397559213351 | 0.0533528453195817 | -1.56017463403745 | 0.118718618239684 | 0.550960590590944 | NM_008079 |
| Pcx | 3872.697909 | 0.0843215535955668 | 0.0540583699973113 | 1.55982419743253 | 0.118801431598427 | 0.550992846551327 | NM_001162946,NM_008797 |
| Pmm2 | 714.8707174 | -0.109347177368027 | 0.0700972593454123 | -1.55993512997714 | 0.118775211698128 | 0.550992846551327 | NM_016881 |
| Sprtn | 419.4713878 | -0.120340464610298 | 0.0771657674969512 | -1.55950583417774 | 0.11887670480237 | 0.551165979559278 | NM_001111141 |
| Dact2 | 2835.294794 | -0.135445312317796 | 0.0868692733051107 | -1.55918551133808 | 0.118952479054332 | 0.551341324423491 | NM_172826 |
| Mrpl22 | 444.2835168 | 0.115485212488598 | 0.074081487044054 | 1.55889436209511 | 0.11902138496003 | 0.551385848496957 | NM_175001 |
| Nt5dc3 | 7321.82067 | 0.0812264632802901 | 0.052107521118469 | 1.55882416850377 | 0.119038002263972 | 0.551385848496957 | NM_175331 |
| Ier3 | 360.4441047 | 0.134882756445597 | 0.0865507685937413 | 1.55842355460435 | 0.119132876540024 | 0.551649398986417 | NM_133662 |
| Tbk1 | 2062.750641 | -0.0909947512798069 | 0.0584096097366714 | -1.55787295429706 | 0.119263367613869 | 0.55207765390665 | NM_019786 |
| AW551984 | 819.1546187 | 0.093472700386269 | 0.0600271880651905 | 1.557172731209 | 0.119429480726076 | 0.552670480097794 | NM_178737,NM_001199556 |
| Gcn1l1 | 3771.260464 | -0.07768500779571 | 0.0499083714077155 | -1.5565526504778 | 0.119576733054078 | 0.55282355962557 | NM_172719 |
| Irak4 | 71.2646764 | -0.147917661708456 | 0.0950272575011759 | -1.55658140199011 | 0.119569902207869 | 0.55282355962557 | NM_029926 |
| Mmp24 | 2053.985057 | 0.111389536075058 | 0.0715582039733566 | 1.55662844915074 | 0.11955872530152 | 0.55282355962557 | NM_010808 |
| Rnf6 | 2783.786354 | 0.0926337523545936 | 0.059521168754493 | 1.55631608540283 | 0.119632948300477 | 0.55290747916409 | NM_001256086,NM_001256087,NM_001256085,NM_028774 |
| Rpa2 | 469.9877859 | -0.125443929297144 | 0.0806477785346964 | -1.55545424283665 | 0.119837924113209 | 0.55367865320244 | NM_011284 |
| 9430083A17Rik | 107.2306028 | -0.147758807727757 | 0.0950641193074621 | -1.55430680686019 | 0.120111250639726 | 0.553708672419124 | NR_029463 |
| Dlat | 3900.732607 | 0.0802057519036658 | 0.0515941897662657 | 1.55455008145331 | 0.120053260350172 | 0.553708672419124 | NM_145614 |
| Mtap | 448.4432552 | 0.107062860912407 | 0.0688558582808983 | 1.55488383393091 | 0.119973738178011 | 0.553708672419124 | NM_024433 |
| Nhsl2 | 2838.33764 | 0.120126618987124 | 0.0772775223845347 | 1.55447037224325 | 0.120072258522157 | 0.553708672419124 | NM_001163610 |
| Sec23b | 1864.130103 | 0.0734129252289261 | 0.0472080120134927 | 1.55509461419269 | 0.119923537472333 | 0.553708672419124 | NM_001252543,NM_019787,NR_045534,NM_001252545,NM_001252544 |
| Ttc39c | 882.0724182 | 0.112842558411695 | 0.0725833912428442 | 1.55466087323137 | 0.120026857759962 | 0.553708672419124 | NM_028341 |
| Wdr54 | 570.4358012 | 0.109883190947536 | 0.0706930608920638 | 1.55437025304802 | 0.120096124616913 | 0.553708672419124 | NM_023790 |
| Nfkbib | 793.8737636 | 0.111250624689899 | 0.0716032409418581 | 1.55370934648383 | 0.120253762622114 | 0.554014004392269 | NM_010908 |
| Scpep1 | 710.6691955 | -0.115036544803126 | 0.0740386765583667 | -1.55373583308771 | 0.120247441980048 | 0.554014004392269 | NM_029023 |
| Fastkd5 | 386.1333111 | 0.143114647387401 | 0.0921373297210605 | 1.5532753968524 | 0.12035735542475 | 0.554315454946074 | NM_001146084,NM_198176 |
| Fyttd1 | 2745.040308 | 0.107130297648928 | 0.0690128727835283 | 1.55232340472135 | 0.120584860484808 | 0.554583367538074 | NM_027226,NM_001159349 |
| Higd1a | 2821.966518 | 0.0892631913973696 | 0.0574930636790689 | 1.55259062024672 | 0.120520967932157 | 0.554583367538074 | NM_019814 |
| Lingo4 | 44.95769299 | 0.14602471626605 | 0.0940616964866253 | 1.55243549415264 | 0.120558056120624 | 0.554583367538074 | NM_177250 |
| Ndufa11 | 2132.943261 | 0.0912882693187481 | 0.0588109220207707 | 1.55223326181669 | 0.120606420077983 | 0.554583367538074 | NM_027244 |
| Zfp580 | 386.0110556 | -0.129404114476097 | 0.0833327513449876 | -1.55286021867176 | 0.120456532473962 | 0.554583367538074 | NM_026900 |
| Fancf | 79.06984216 | -0.147436664645067 | 0.0950242854998936 | -1.55156825299394 | 0.120765564324341 | 0.554612895155036 | NM_001115087 |
| Grlf1 | 11459.87932 | 0.0828847245930101 | 0.0534160468691198 | 1.55168211522831 | 0.12073830413044 | 0.554612895155036 | NM_172739 |
| Usp16 | 1083.14042 | -0.106815651618937 | 0.0688344136317352 | -1.55177687995428 | 0.120715619816218 | 0.554612895155036 | NM_024258 |
| Xylt1 | 860.2847853 | 0.103934874751625 | 0.0669817868586851 | 1.55168859515352 | 0.12073675289155 | 0.554612895155036 | NM_175645 |
| Clpp | 1333.019528 | 0.0957600935412043 | 0.061749866602502 | 1.55077409571803 | 0.120955830535018 | 0.555002663504315 | NM_017393 |
| Larp1 | 6702.731412 | 0.0664341857256896 | 0.0428359373814174 | 1.55089837614968 | 0.120926031964357 | 0.555002663504315 | NM_028451 |
| Swap70 | 596.138351 | -0.114401260505482 | 0.0737798350377814 | -1.55057625768477 | 0.121003265545791 | 0.555002663504315 | NM_009302 |
| Trim59 | 639.2742277 | -0.14243609697684 | 0.0918568810260698 | -1.5506306700792 | 0.120990217804166 | 0.555002663504315 | NM_025863 |
| Tmco6 | 359.8733908 | -0.117312953757823 | 0.0756682232515737 | -1.55035956596725 | 0.121055237745994 | 0.555065777619413 | NM_028036 |
| Gli1 | 137.8874235 | -0.139385868327147 | 0.0899374540593539 | -1.54980891759689 | 0.121187386012837 | 0.555321125937689 | NM_010296 |
| Rmdn3 | 1903.948619 | 0.0762903159639641 | 0.0492219394078129 | 1.54992503102904 | 0.12115951095218 | 0.555321125937689 | NM_001033136 |
| Ctu2 | 567.3627597 | -0.10691526308545 | 0.068997311322393 | -1.54955694702195 | 0.121247893213085 | 0.555423177802986 | NM_153775 |
| Lef1 | 244.7501972 | -0.130780139289263 | 0.0844198178640681 | -1.54916395934239 | 0.121342310861674 | 0.555673148028673 | NM_001276402,NM_001276403,NM_010703 |
| Thumpd1 | 2285.854121 | 0.0869819822817807 | 0.0561532211533374 | 1.54901144574163 | 0.121378968655857 | 0.555673148028673 | NM_145585 |
| Lin7a | 3041.350817 | 0.0957769069373746 | 0.061839039372761 | 1.54880974718962 | 0.121427461725697 | 0.55572000914539 | NM_001033223,NM_001039354 |
| Gaa | 16557.79508 | 0.129773687508231 | 0.0838102667002283 | 1.54842231885986 | 0.121520651101665 | 0.555796277677199 | NM_008064,NM_001159324 |
| Gnl2 | 2195.410266 | -0.0824551900967897 | 0.0532460718936676 | -1.54856850776622 | 0.121485481247827 | 0.555796277677199 | NM_145552 |
| Clec3b | 46.27576837 | 0.1293506108974 | 0.0835541643280912 | 1.54810489623809 | 0.121597043457119 | 0.555970617959747 | NM_011606 |
| Fabp3 | 2402.067766 | 0.0829536292226649 | 0.0535989293411669 | 1.54767325098324 | 0.121700985394379 | 0.555997647284637 | NM_010174 |
| Phip | 1901.960678 | -0.117054686602672 | 0.0756327978998351 | -1.54767098207439 | 0.121701531940375 | 0.555997647284637 | NM_001081216 |
| S100a4 | 38.77029213 | -0.134451994822077 | 0.0868775451831452 | -1.54760352100927 | 0.121717783172597 | 0.555997647284637 | NM_011311 |
| Pnpo | 2287.37089 | 0.0925746362855283 | 0.0598280947742802 | 1.54734387974069 | 0.12178034605483 | 0.556108552905519 | NM_134021 |
| Copb2 | 4402.403848 | 0.075457235010459 | 0.0487719423020991 | 1.54714435080457 | 0.121828441416551 | 0.556153343814212 | NM_015827 |
| 2310061J03Rik | 102.693532 | -0.143747034469134 | 0.0929444809813977 | -1.54659031877217 | 0.12196206568501 | 0.556510459908872 | NR_027965 |
| Cdc37l1 | 2757.542099 | 0.104324454015989 | 0.0674721715590114 | 1.54618491750701 | 0.1220599149986 | 0.556510459908872 | NM_025950 |
| Has1 | 37.24989401 | -0.139377451958938 | 0.0901274921174645 | -1.54644768964929 | 0.121996484241377 | 0.556510459908872 | NM_008215 |
| Serpinb9 | 412.0131733 | -0.10978852935091 | 0.070999977005459 | -1.54631781560252 | 0.122027831410045 | 0.556510459908872 | NM_009256 |
| Adamts15 | 294.0478007 | 0.127182916717077 | 0.0823195721610914 | 1.54499001122349 | 0.122348679494388 | 0.556819993252963 | NM_001024139 |
| Dcp2 | 3672.514735 | 0.103409098599692 | 0.0669313437271862 | 1.54500257788325 | 0.122345639823866 | 0.556819993252963 | NM_027490 |
| Dnajc5 | 16585.20147 | 0.0746816485850305 | 0.0483205782263452 | 1.54554542446085 | 0.122214390397724 | 0.556819993252963 | NR_073369,NM_001271584,NM_001271585,NM_016775 |
| Dus2l | 509.7713556 | -0.106335569085732 | 0.0688130472986084 | -1.54528208326973 | 0.122278047271474 | 0.556819993252963 | NM_025518 |
| Gpr101 | 462.2642137 | 0.102395063396341 | 0.0662596340040779 | 1.54536113782396 | 0.122258934870659 | 0.556819993252963 | NM_001033360 |
| Ptger4 | 45.308755 | 0.139924954582063 | 0.0905691097351115 | 1.54495230207411 | 0.122357801078305 | 0.556819993252963 | NM_001136079,NM_008965 |
| Lhfp | 919.6957953 | 0.0954526103907114 | 0.0617997230852425 | 1.54454754205048 | 0.122455743214099 | 0.557091176300657 | NM_175386 |
| Sap30bp | 1655.346027 | 0.0767313538820612 | 0.0496840899379393 | 1.54438481167526 | 0.122495137290292 | 0.557095918684651 | NM_020483 |
| Fam115a | 9072.443785 | 0.067449541338691 | 0.043681172273495 | 1.54413304012031 | 0.122556106134578 | 0.557198747327348 | NM_029930 |
| BC030867 | 43.18538234 | -0.139754674866626 | 0.0905372316767425 | -1.54361550798915 | 0.122681505853678 | 0.557245639159014 | NM_153544 |
| Kcnd1 | 297.2103252 | -0.12597565732446 | 0.081607834453574 | -1.54367112138145 | 0.122668025742582 | 0.557245639159014 | NM_008423 |
| Notch2 | 868.8237973 | -0.116527395270608 | 0.0754794718159167 | -1.54382897060082 | 0.122629771027754 | 0.557245639159014 | NM_010928 |
| Dnajb11 | 2030.517939 | 0.0947938358509757 | 0.0614413496736196 | 1.54283453007668 | 0.122870929083192 | 0.55748489235324 | NM_001190805,NM_001190804,NM_026400 |
| Enho | 1422.275599 | 0.102546919476853 | 0.0664882358731827 | 1.54233178441443 | 0.12299298895342 | 0.55748489235324 | NM_027147 |
| Erg | 174.8738988 | -0.140836061342662 | 0.0913414353917121 | -1.54186389494204 | 0.123106671272571 | 0.55748489235324 | NM_133659 |
| Marveld2 | 49.32369062 | 0.145509212208909 | 0.094361903350451 | 1.54203345886848 | 0.123065463133351 | 0.55748489235324 | NM_178410,NM_001038602 |
| Mlx | 1093.55356 | -0.089112528036436 | 0.0577450254669149 | -1.5432070119095 | 0.122780556512236 | 0.55748489235324 | NM_001159385,NM_001159384,NM_011550 |
| Slc25a11 | 4125.392369 | 0.0790432374784762 | 0.0512662754512867 | 1.54181743812427 | 0.1231179632844 | 0.55748489235324 | NM_024211 |
| Stk39 | 3283.030164 | 0.0947454313439786 | 0.0614200149887386 | 1.54258235465019 | 0.122932142045168 | 0.55748489235324 | NM_016866 |
| Themis2 | 55.50492914 | -0.141255726967469 | 0.0915505822816803 | -1.54292548935251 | 0.122848855508591 | 0.55748489235324 | NM_001033308 |
| Ttll7 | 10352.65285 | 0.0981479800326335 | 0.0636421609851993 | 1.54218490562347 | 0.123028667013724 | 0.55748489235324 | NM_027594 |
| Zfp948 | 406.5880409 | 0.136592083626416 | 0.0885731781902112 | 1.54213822307566 | 0.123040008269228 | 0.55748489235324 | NM_001002008 |
| Nfam1 | 147.1800003 | -0.138015064511862 | 0.0895529237725854 | -1.54115643239458 | 0.12327871807989 | 0.558038846627762 | NM_001271412,NM_001271413,NM_001271411,NM_028728,NM_001271414 |
| Ctrl | 25.50937916 | -0.126331573305597 | 0.0819853282664321 | -1.5409036301247 | 0.12334024221346 | 0.558143413829507 | NM_023182 |
| Nsun2 | 3189.606666 | 0.0671265721713103 | 0.0435708264332149 | 1.54063114396514 | 0.123406583631149 | 0.558269708447856 | NM_145354 |
| G3bp1 | 2436.892835 | 0.0878055400136071 | 0.0570008739510031 | 1.5404244518967 | 0.123456924930218 | 0.558323565235474 | NM_013716 |
| Amotl1 | 1873.057676 | 0.132251727740058 | 0.0858629599716016 | 1.54026518284251 | 0.123495726957993 | 0.558325219356303 | NM_001081395 |
| Fbxl12 | 458.8441898 | -0.133020591694152 | 0.0864016406536144 | -1.53956094685092 | 0.123667411042046 | 0.558428988904225 | NM_001002846,NM_013911 |
| Sar1a | 5713.453125 | 0.0706648561715309 | 0.045899972419796 | 1.53954027521494 | 0.123672453346062 | 0.558428988904225 | NM_009120 |
| Tcerg1 | 2868.606778 | -0.0864015698336334 | 0.0561125355198001 | -1.53979086906785 | 0.123611338358814 | 0.558428988904225 | NM_001039474 |
| Zfp503 | 202.8575333 | 0.145801153485371 | 0.0946989024228995 | 1.53962855748915 | 0.123650920318919 | 0.558428988904225 | NM_145459 |
| Heatr5b | 2357.337869 | 0.0952696978843307 | 0.06189615182079 | 1.53918612194452 | 0.123758864685671 | 0.558645515358625 | NM_001081179 |
| Rttn | 235.9238054 | -0.129419946228741 | 0.0841186761147121 | -1.53853997954332 | 0.12391664108577 | 0.559183947937838 | NM_175542 |
| Cbr1 | 2009.427586 | 0.0815597790168466 | 0.0530422225327151 | 1.53763879269091 | 0.124136956855275 | 0.559205950660562 | NM_007620 |
| Cx3cl1 | 32735.73569 | 0.0909994596453239 | 0.0591826360295747 | 1.53760402966589 | 0.12414546159113 | 0.559205950660562 | NM_009142 |
| Fendrr | 27.27575016 | -0.111546696114575 | 0.0725471451088586 | -1.53757526842989 | 0.124152498339551 | 0.559205950660562 | NR_045471 |
| Prss53 | 108.4733162 | -0.145991490670716 | 0.0949347509243592 | -1.53780875021242 | 0.124095383478431 | 0.559205950660562 | NM_001081268 |
| Serpini1 | 10679.45 | 0.118362615596327 | 0.0769473091033138 | 1.53822943226523 | 0.123992526997832 | 0.559205950660562 | NM_009250 |
| Wdr27 | 44.6007339 | -0.130831052998023 | 0.085061691580353 | -1.53807255143091 | 0.124030876475667 | 0.559205950660562 | NM_175173 |
| Cib2 | 693.6805191 | 0.105423688556307 | 0.0686397879191105 | 1.5358976440974 | 0.124563486054264 | 0.560776597509588 | NM_019686 |
| Ifi35 | 117.0458995 | -0.145984565933437 | 0.0951008275332457 | -1.5350504272153 | 0.124771441769998 | 0.560776597509588 | NM_027320 |
| Ltbp2 | 26.70398679 | 0.131539748541577 | 0.0856860967786198 | 1.53513526099135 | 0.124750606496455 | 0.560776597509588 | NM_013589 |
| N6amt2 | 666.0738179 | 0.0946975395532315 | 0.0616682748717724 | 1.53559572973522 | 0.124637562167183 | 0.560776597509588 | NM_026526 |
| Sema3g | 318.3997811 | 0.127957346416407 | 0.083343704124787 | 1.5352970900456 | 0.124710868617866 | 0.560776597509588 | NM_001025379 |
| Trim45 | 443.6801179 | 0.116716375127922 | 0.0760161258250887 | 1.53541601141012 | 0.124681675670061 | 0.560776597509588 | NM_001165953,NM_001165952,NM_194343 |
| Ube2d3 | 4695.295466 | 0.104300822798889 | 0.0679411872008827 | 1.53516338315521 | 0.12474370025928 | 0.560776597509588 | NM_025356 |
| Asna1 | 3497.021696 | 0.0797549212706352 | 0.0519779354214228 | 1.53439956058286 | 0.124931385653143 | 0.560889588876706 | NM_019652 |
| Ddx31 | 301.9873232 | -0.113983202733041 | 0.074282248457236 | -1.53446085841977 | 0.124916315515154 | 0.560889588876706 | NM_001033294 |
| Oprm1 | 51.94165578 | 0.137947902711802 | 0.0898942854713402 | 1.53455697421147 | 0.124892688199896 | 0.560889588876706 | NM_001039652 |
| Plcb3 | 814.5461466 | -0.10522571395744 | 0.0685813505632596 | -1.53431965240142 | 0.124951033292374 | 0.560889588876706 | NM_008874 |
| Cdc5l | 2626.899919 | 0.0963090105023759 | 0.0627928124821212 | 1.53375850985808 | 0.125089073592868 | 0.561335768616003 | NM_152810 |
| Ift140 | 1379.124975 | -0.0816804472922248 | 0.0532725089847752 | -1.53325699969507 | 0.125212544988752 | 0.561716315165723 | NM_134126 |
| Gm17066 | 522.2092127 | -0.122808678703792 | 0.0801309628915135 | -1.53259956291875 | 0.125374549262425 | 0.562122953479606 | NR_040628 |
| Stk11ip | 1907.26664 | -0.0943286120817044 | 0.0615490967168435 | -1.53257508417488 | 0.125380584419243 | 0.562122953479606 | NM_027886 |
| Htr4 | 155.9868428 | 0.143050476171061 | 0.0933583854711606 | 1.5322723872003 | 0.12545523211236 | 0.562279816022514 | NM_008313 |
| Med23 | 1719.670642 | -0.0811013698320346 | 0.0529341076740651 | -1.5321193346907 | 0.125492989366996 | 0.562279816022514 | NM_027347,NM_001166416 |
| Donson | 639.320635 | -0.12164416739608 | 0.0794149888106662 | -1.53175325235004 | 0.1255833358901 | 0.562511112284796 | NM_021720 |
| Nsfl1c | 3215.548756 | 0.0863805756965818 | 0.0564011331264385 | 1.53153972107149 | 0.125636057292747 | 0.562573787988422 | NM_198326 |
| Ap4m1 | 529.2497043 | -0.104675981639089 | 0.0684605173427189 | -1.52899781804267 | 0.12626498483723 | 0.562787717013074 | NM_021392 |
| Aspdh | 79.22646283 | -0.1452971877939 | 0.0949908169612623 | -1.52959193785177 | 0.12611776628198 | 0.562787717013074 | NM_026690 |
| BC029722 | 2401.273901 | 0.0850032642794712 | 0.055582545549274 | 1.52931584003736 | 0.126186164669862 | 0.562787717013074 | NR_015528 |
| Dld | 4292.888471 | 0.0875209080702221 | 0.0572003680211788 | 1.53007596101159 | 0.125997927727158 | 0.562787717013074 | NM_007861 |
| Dnaja1 | 11565.0829 | 0.0998422146751946 | 0.0652523845591632 | 1.53009296671249 | 0.125993718925148 | 0.562787717013074 | NM_001164671,NM_008298,NM_001164672 |
| Dus3l | 2510.923503 | -0.120075567797099 | 0.0784472306671065 | -1.53065400494051 | 0.125854926953155 | 0.562787717013074 | NM_144858 |
| Fgfr3 | 5085.562193 | 0.0823043464472444 | 0.0537611316867361 | 1.53092659817558 | 0.125787534712972 | 0.562787717013074 | NM_001163216,NM_001163217,NM_001163215,NM_008010,NM_001205270 |
| March7 | 1401.699691 | -0.0995916981436844 | 0.0650747384963818 | -1.53042025899531 | 0.125912737516485 | 0.562787717013074 | NM_020575 |
| Mrpl42 | 1150.204179 | 0.110244763886289 | 0.0720686312253446 | 1.52971913038802 | 0.126086266280475 | 0.562787717013074 | NM_026065 |
| Nup62 | 1183.074619 | 0.0895744000533589 | 0.0585722866362831 | 1.52929661978915 | 0.126190927224402 | 0.562787717013074 | NM_053074 |
| Rasd1 | 315.0932593 | 0.144711502527986 | 0.094640998813312 | 1.52905721983601 | 0.126250259487388 | 0.562787717013074 | NM_009026 |
| Rps4y2 | 457.7483298 | 0.138406859017676 | 0.0904865062010917 | 1.52958562362977 | 0.126119330196957 | 0.562787717013074 | NR_003634 |
| Sdr42e1 | 369.396636 | 0.125606237878928 | 0.0820492692204952 | 1.53086357833827 | 0.125803112380451 | 0.562787717013074 | NM_028725 |
| Tmem178 | 2505.288308 | 0.112292692042469 | 0.0734335903365703 | 1.52917338683557 | 0.126221466247029 | 0.562787717013074 | NM_026516 |
| Ubc | 5673.517106 | 0.102639231018713 | 0.0670977053504241 | 1.52969807957918 | 0.12609147921749 | 0.562787717013074 | NM_019639 |
| Sycp2 | 64.43173072 | -0.143265515865908 | 0.0937117560792564 | -1.52878914940769 | 0.126316723147735 | 0.562845619768096 | NM_177191 |
| Araf | 5262.505894 | 0.0716513546642901 | 0.0468771447416609 | 1.52849229745453 | 0.126390354503314 | 0.563001008744294 | NM_009703,NM_001159645 |
| Atp5g1 | 4529.742687 | 0.0879806160257236 | 0.0575796400600278 | 1.52798134781673 | 0.126517169055215 | 0.563050456015075 | NM_007506,NM_001161419 |
| Gmppb | 419.5956503 | 0.118425554020888 | 0.0775046975223821 | 1.52797904909814 | 0.126517739806774 | 0.563050456015075 | NM_177910 |
| Stat3 | 2544.860005 | -0.0817988988014379 | 0.0535271166467879 | -1.5281768181389 | 0.126468642830349 | 0.563050456015075 | NM_213660,NM_011486,NM_213659 |
| Tspan31 | 2375.177829 | 0.0848592754664303 | 0.0555618602473962 | 1.52729363431289 | 0.126688011687536 | 0.563635546025466 | NM_025982 |
| Fbxo21 | 6541.096973 | 0.0617987786884312 | 0.0404897296316488 | 1.52628281914054 | 0.126939445540541 | 0.563718246995381 | NM_145564 |
| Nfkbiz | 192.3359947 | -0.138005906354552 | 0.0904106734423185 | -1.52643378375672 | 0.126901869399738 | 0.563718246995381 | NM_001159394,NM_001159395,NM_030612 |
| Secisbp2 | 677.0310162 | -0.113179072210971 | 0.0741406243250477 | -1.52654598260153 | 0.126873947937092 | 0.563718246995381 | NM_029279 |
| Slc9a3r1 | 1633.505924 | 0.0952631813107289 | 0.0624103697776717 | 1.5263998859499 | 0.126910306045326 | 0.563718246995381 | NM_012030 |
| Spin2 | 207.8280499 | 0.135074208319877 | 0.0884755563487044 | 1.52668391015837 | 0.126839630257959 | 0.563718246995381 | NM_001005370 |
| Unc13d | 65.15363377 | -0.143108672645771 | 0.0937290861106018 | -1.52683311642344 | 0.126802514465655 | 0.563718246995381 | NM_001009573 |
| Kctd12 | 2781.719016 | -0.0914170591665934 | 0.0599114973524172 | -1.52586837596215 | 0.127042647843946 | 0.564004126705734 | NM_177715 |
| Myd88 | 260.7387984 | -0.129439174289308 | 0.0848462867569108 | -1.52557264715848 | 0.127116328473992 | 0.564158810697589 | NM_010851 |
| Lilrb3 | 24.10685186 | -0.12085037540406 | 0.0792721139312128 | -1.52450047577798 | 0.127383738097358 | 0.564894808028353 | NM_011095 |
| Nudcd3 | 7560.839367 | 0.0629051220430639 | 0.041258089293617 | 1.52467366085215 | 0.127340514499183 | 0.564894808028353 | NM_173748 |
| Trim37 | 18607.40739 | 0.0951984043065902 | 0.0624481140518856 | 1.52444002115891 | 0.12739882907207 | 0.564894808028353 | NM_197987 |
| Hprt | 4394.174498 | 0.0975008942180787 | 0.0639653353658023 | 1.52427707383217 | 0.127439511699541 | 0.56490276074078 | NM_013556 |
| Mtf1 | 1532.249891 | 0.113078386348823 | 0.0741957566957633 | 1.52405462771269 | 0.127495065547973 | 0.564976608343457 | NM_008636 |
| Calcoco1 | 3555.78984 | -0.0689066984447991 | 0.045228040578875 | -1.52353932566744 | 0.127623829805729 | 0.565235453070151 | NM_026192 |
| Ppp1r17 | 28.25592435 | 0.0924236346544732 | 0.0606649571131567 | 1.52350943695679 | 0.12763130153312 | 0.565235453070151 | NM_011153 |
| Gdap2 | 1380.232387 | 0.0924222577835294 | 0.0606894372078729 | 1.52287221690597 | 0.127790677911841 | 0.565596400776175 | NM_010269 |
| Sulf2 | 4225.877953 | 0.102750525720509 | 0.0674679316406935 | 1.52295354580775 | 0.127770327965513 | 0.565596400776175 | NM_001252578,NM_028072,NM_001252579 |
| Ate1 | 3285.588373 | 0.0833468785156463 | 0.0547480446096941 | 1.52237178715399 | 0.127915949866815 | 0.565771338542876 | NM_001136054,NM_001029895,NM_001271343,NM_013799 |
| Plxnc1 | 3006.303328 | 0.11715888486425 | 0.0769493065096194 | 1.52254633834294 | 0.12787224384484 | 0.565771338542876 | NM_018797 |
| Rsph3b | 120.3288833 | 0.141657637875048 | 0.0930582101115287 | 1.52224760937561 | 0.127947049918308 | 0.565771338542876 | NM_001083945 |
| Zfp354a | 323.4616906 | -0.128202618599394 | 0.0842284578835801 | -1.5220819897362 | 0.12798853864391 | 0.565782566141642 | NM_009329 |
| B3galt2 | 1512.850995 | 0.119483204529992 | 0.0785337925539265 | 1.5214240983962 | 0.128153445243399 | 0.566072312884823 | NM_020025 |

| | | | | | | | | | | | | | | |
|---|---|---|---|---|---|---|---|---|---|---|---|---|---|---|
| Zfp619 | 206.7300945 | -0.236074103 | 0.101403426 | -2.328068323 | 0.019908476 | 0.093909252 | | Cds2 | 31122.44181 | 0.0782700385215531 | 0.0514476129037359 | 1.52135413295083 | 0.128170992501596 | 0.566072312884823 | NM_138651 |
| Park7 | 2892.032928 | -0.129919656 | 0.055836587 | -2.326783624 | 0.019976784 | 0.094107722 | | Ptpn13 | 763.3909734 | -0.114553201933172 | 0.0752899008497929 | -1.52149492349195 | 0.128135684288136 | 0.566072312884823 | NM_011204 |
| Fam20c | 1566.98457 | -0.204420323 | 0.087927875 | -2.324863669 | 0.020079251 | 0.094532944 | | 1500009L16Rik | 413.8167108 | 0.116309946737461 | 0.0764812452841196 | 1.52076429071443 | 0.128318998560084 | 0.566288196453558 | NM_001145198 |
| Mysm1 | 1109.88114 | 0.1743695 | 0.075001166 | 2.324890528 | 0.020077815 | 0.094532944 | | Acyp2 | 366.4468367 | 0.124656627496984 | 0.0819608415930041 | 1.5209290811824 | 0.128277635173663 | 0.566288196453558 | NM_029344 |
| Mapkbp1 | 3164.165961 | 0.14445887 | 0.062144386 | 2.324568316 | 0.020095055 | 0.094578608 | | Aspn | 31.53221453 | -0.133998934747133 | 0.0881170034452834 | -1.52069327721 | 0.128336826567886 | 0.566288196453558 | NM_001172481,NM_025711 |
| Ank1 | 3077.521641 | 0.189734374 | 0.081627019 | 2.324406515 | 0.020103717 | 0.094590643 | | Esyt1 | 711.3280434 | -0.114974863778252 | 0.0756324893310097 | -1.52017823021874 | 0.12846618723379 | 0.566659163704225 | NM_011843 |
| Stard4 | 836.0553654 | -0.300823348 | 0.129432835 | -2.324165639 | 0.020116618 | 0.094622611 | | Tspyl5 | 2293.416301 | 0.0811768307997751 | 0.0534041241141979 | 1.52004797656055 | 0.128498918163412 | 0.566659163704225 | NM_001085421 |
| Gon4l | 2228.141765 | 0.141601194 | 0.060936012 | 2.323768655 | 0.020137897 | 0.094693951 | | Mal2 | 6535.906745 | 0.106459906281803 | 0.0700477158234834 | 1.51981981182764 | 0.128556268402338 | 0.56674001663501 | NM_178920 |
| Tomm40l | 2502.469417 | -0.216782943 | 0.093313704 | -2.323162986 | 0.020170398 | 0.094789249 | | Dnajc28 | 578.8346001 | 0.11191813851961 | 0.0736678260986548 | 1.51922683818213 | 0.128705408043233 | 0.567225351103157 | NM_138664,NM_001099738 |
| Ube2n | 6979.926633 | -0.141080851 | 0.060726818 | -2.323205073 | 0.020168138 | 0.094789249 | | Cxx1a | 1181.439099 | 0.0982990166840649 | 0.0647166913544049 | 1.51891288981006 | 0.128784424060261 | 0.567401438853307 | NM_024170 |
| Col16a1 | 895.8295544 | 0.388258667 | 0.167136259 | 2.323006806 | 0.020178787 | 0.094799916 | | Dnaaf3 | 52.94093693 | -0.141623414643931 | 0.093251041801436 | -1.51873278740946 | 0.128829770098946 | 0.567429120817857 | NM_001033548 |
| Cyb5 | 1543.927305 | -0.135032844 | 0.058157211 | -2.321858998 | 0.020240529 | 0.09495708 | | Pappa | 32.14503079 | 0.124579325928274 | 0.0820613536858274 | 1.51812418797314 | 0.128983094576285 | 0.567618221082904 | NM_021362 |
| Dnm2 | 2164.425059 | 0.186049992 | 0.080133958 | 2.321737197 | 0.020247091 | 0.09495708 | | Slc2a13 | 6957.325846 | 0.122089846789935 | 0.0804260522374201 | 1.51803853842685 | 0.129004683644007 | 0.567618221082904 | NM_001033633 |
| Frmd6 | 1549.798478 | 0.259564973 | 0.111803277 | 2.321622232 | 0.020253286 | 0.09495708 | | Trim68 | 222.8768647 | -0.139425762922307 | 0.091827220209045 | -1.51834894495231 | 0.128926455041256 | 0.567618221082904 | NM_198012 |
| Hsd17b7 | 897.2663332 | -0.236159908 | 0.101723444 | -2.321587815 | 0.020255141 | 0.09495708 | | Ubap1 | 2096.181693 | 0.0750705338564112 | 0.0494554675323305 | 1.51794205175263 | 0.129029007711397 | 0.567618221082904 | NM_023305 |
| Spata18 | 21.16729931 | 0.390023094 | 0.167989837 | 2.321706489 | 0.020248745 | 0.09495708 | | Tprn | 879.2536157 | 0.113440477672919 | 0.0747646691562433 | 1.51730060405739 | 0.129190805780042 | 0.568157930596696 | NM_175286 |
| Tmem167 | 2042.608962 | -0.173801376 | 0.074853363 | -2.32189136 | 0.020238786 | 0.09495708 | | Maf | 534.6474511 | -0.108909778148954 | 0.0717904538944373 | -1.51705097601274 | 0.129253814268224 | 0.568262986095708 | NM_001025577 |
| Zbtb7b | 645.1483283 | 0.343790826 | 0.14807031 | 2.321807967 | 0.020243278 | 0.09495708 | | Pdlim2 | 422.6319256 | 0.122993897543384 | 0.0810920019912413 | 1.51672044743289 | 0.129337279541147 | 0.568274875118074 | NM_001253736,NM_145978 |
| Lrrc36 | 30.73228488 | 0.388998769 | 0.16758813 | 2.321159438 | 0.02027824 | 0.095036617 | | Phactr3 | 5582.731063 | 0.0765658206951627 | 0.05047966922557 | 1.51676549925527 | 0.12932590056495 | 0.568274875118074 | NM_001177789,NM_001007154,NM_001177791,NM_001177790,NM_028806 |
| Dnali1 | 38.28962184 | 0.389283526 | 0.167736763 | 2.320800284 | 0.020297625 | 0.095098701 | | Tcea1 | 3289.484507 | 0.0902676111082271 | 0.0595206820169687 | 1.51657555070509 | 0.12937388214343 | 0.568274875118074 | NM_011541,NM_001159750,NM_001159751 |
| Iars | 2497.750222 | 0.148549795 | 0.064026202 | 2.320140678 | 0.020333268 | 0.095179354 | | Scg2 | 11339.01683 | 0.144085040220562 | 0.0950199990002357 | 1.51636541503442 | 0.12942697915567 | 0.568336245228315 | NM_009129 |
| Nudt5 | 440.0324851 | -0.206018146 | 0.088794644 | -2.320164113 | 0.020332001 | 0.095179354 | | Pef1 | 2749.388397 | 0.085303350511156 | 0.056262630084519 | 1.51616357754715 | 0.129477995318122 | 0.568388443635856 | NM_026441 |
| Zfp661 | 332.8856708 | 0.244570203 | 0.105403217 | 2.32032959 | 0.020323055 | 0.095179354 | | Dnmbp | 643.3614615 | -0.109384168801035 | 0.0721852234494557 | -1.51532631713228 | 0.129689786852722 | 0.56888561936693 | NM_028029 |
| Ndst1 | 4764.696882 | -0.187275845 | 0.080722557 | -2.319993972 | 0.020341203 | 0.09518774 | | Dync1li1 | 2857.555847 | 0.081179077569754 | 0.0535858012333239 | 1.51493633950313 | 0.129788526500028 | 0.56888561936693 | NM_146229 |
| Hist4h4 | 30.38192396 | 0.389378455 | 0.167853673 | 2.319749381 | 0.020354439 | 0.095220916 | | Rfk | 2251.865167 | 0.0868430771127067 | 0.0573232323450095 | 1.5149717411263 | 0.129779560645267 | 0.56888561936693 | NM_019437 |
| Ctif | 4353.583903 | 0.167721258 | 0.072305452 | 2.319621167 | 0.02036138 | 0.095224636 | | Rlbp1 | 836.9629526 | -0.130349758368043 | 0.086021240234106 | -1.51532061166867 | 0.129691231016266 | 0.56888561936693 | NM_020599,NM_001173483 |
| G3bp2 | 10947.83141 | -0.160406708 | 0.069159788 | -2.319363781 | 0.02037532 | 0.095261076 | | Slc29a1 | 925.1203206 | -0.108527658544548 | 0.0716454556209464 | -1.51478775037365 | 0.129826163594444 | 0.56888561936693 | NM_001199114,NM_001199115,NM_001199116,NM_001199113,NM_022880 |
| Ndufaf6 | 194.1908238 | -0.241920636 | 0.104313501 | -2.31916898 | 0.020385876 | 0.095281677 | | Snx11 | 1366.228973 | -0.097258957692449 | 0.0642071443137225 | -1.5147684690232 | 0.129831048112473 | 0.56888561936693 | NM_001163389,NM_028965 |
| Siah1b | 132.2060302 | -0.262408193 | 0.113182272 | -2.318456666 | 0.020424515 | 0.095433487 | | Synpo2 | 373.0600541 | 0.141316096066621 | 0.0933005584863813 | 1.51463290637481 | 0.129865394039704 | 0.56888561936693 | NM_080451 |
| BC068157 | 1419.481458 | 0.167027024 | 0.07205617 | 2.318011402 | 0.020448701 | 0.09551769 | | Dgcr8 | 1663.910489 | -0.085674683093487 | 0.0565829252177697 | -1.51414375915972 | 0.129989382208308 | 0.568914060246426 | NM_033324 |
| Dnajc15 | 693.4766325 | -0.208030818 | 0.089802526 | -2.316536374 | 0.020529001 | 0.095835515 | | Rtfdc1 | 3305.120197 | 0.0721426940242784 | 0.0476369721337713 | 1.51442652193955 | 0.129917696799715 | 0.568914060246426 | NM_025542 |
| Foxred1 | 1213.646557 | 0.165478151 | 0.071436956 | 2.316422189 | 0.020535228 | 0.095835515 | | Zcchc17 | 2162.619925 | 0.0820452972260551 | 0.0541809759213579 | 1.51428238105459 | 0.129954235249875 | 0.568914060246426 | NM_153160 |
| Zbtb49 | 300.237136 | 0.230282306 | 0.09941298 | 2.316420909 | 0.020535298 | 0.095835515 | | Scml2 | 37.62780447 | -0.135728574907247 | 0.0896766812601601 | -1.51353253710947 | 0.130144442808931 | 0.569421137422448 | NM_133194 |
| Mctp1 | 2187.580258 | -0.264173528 | 0.114086334 | -2.315558037 | 0.020582412 | 0.096026467 | | 2700060E02Rik | 2032.891568 | 0.0874040886301034 | 0.0578072897567356 | 1.51199077137013 | 0.130536210283341 | 0.569619261334004 | NM_026528 |
| Col14a1 | 87.76774834 | -0.31054446 | 0.134125525 | -2.315327074 | 0.020595039 | 0.096056453 | | Atp6v1g2 | 20292.21403 | 0.0740732973296716 | 0.0489895961394827 | 1.51202098336902 | 0.13052852454137 | 0.569619261334004 | NM_023179 |
| Ubr1 | 2645.337175 | -0.156985098 | 0.067809236 | -2.31509906 | 0.020607511 | 0.0960857 | | Auh | 1685.521634 | 0.0909134037893699 | 0.0601003230678289 | 1.51269409461852 | 0.130357380315248 | 0.569619261334004 | NM_016709 |
| Gpr63 | 176.0202225 | -0.279605251 | 0.120782353 | -2.314951186 | 0.020615604 | 0.096094514 | | E130008D07Rik | 156.9718806 | -0.13362048253529 1 | 0.0883335812458139 | -1.51268046252368 | 0.130360844662297 | 0.569619261334004 | NR_045155,NR_045154,NR_045153,NR_045152 |
| Slc35b3 | 325.1914811 | 0.209037459 | 0.090323883 | 2.314309914 | 0.020650728 | 0.096229289 | | Garem | 1414.446302 | 0.105839901223896 | 0.0699464816820684 | 1.51315546799016 | 0.130240172847413 | 0.569619261334004 | NM_001033445 |
| Tert | 46.79737606 | 0.346361431 | 0.14967512 | 2.314088218 | 0.020662884 | 0.096256981 | | Lrrc16b | 2062.692951 | -0.133789706070496 | 0.0884528238679201 | -1.51255437893395 | 0.130392889881807 | 0.569619261334004 | NM_001024645 |
| Alg9 | 767.8969419 | -0.165194971 | 0.071393437 | -2.313867726 | 0.020674979 | 0.096284377 | | Map1lc3b | 8770.993507 | 0.0659667490325087 | 0.0436297988893603 | 1.51196546194934 | 0.130542649110622 | 0.569619261334004 | NM_026160 |
| Acsl4 | 2880.736316 | -0.178455681 | 0.077148044 | -2.313158853 | 0.020713907 | 0.096374032 | | Map3k19 | 214.3781292 | 0.143693491763316 | 0.0950325781258956 | 1.5120445493224 | 0.130522529755191 | 0.569619261334004 | NM_011737 |
| Ccm2l | 137.4218963 | 0.294227869 | 0.127210273 | 2.312925381 | 0.020726742 | 0.096374032 | | Podxl2 | 5774.139812 | 0.0934779553077985 | 0.0617828349857329 | 1.51300851327695 | 0.130277496399824 | 0.569619261334004 | NM_176973 |
| Hmg20b | 419.5021256 | 0.205502803 | 0.088847791 | 2.312975937 | 0.020723962 | 0.096374032 | | Gsta3 | 69.05903904 | -0.143033790384779 | 0.0946111427398875 | -1.51180702655731 | 0.130582961366141 | 0.569624053094465 | NM_010356,NM_001077353 |
| Rpl10 | 8230.635221 | -0.092596678 | 0.040035419 | -2.312868956 | 0.020729845 | 0.096374032 | | 3110001D03Rik | 626.2366294 | -0.0952873019848795 | 0.063054575793262 | -1.51118774150982 | 0.130740624731386 | 0.570140592869443 | NM_025849 |
| S100a16 | 1735.708879 | -0.255047108 | 0.110259799 | -2.313146873 | 0.020714565 | 0.096374032 | | Dancr | 98.46820895 | -0.140488847857583 | 0.0929931572380032 | -1.51074393030899 | 0.130853705126609 | 0.570291305331268 | NR_015531 |
| Tpgs1 | 835.5395268 | -0.247129665 | 0.106851267 | -2.312837942 | 0.020731551 | 0.096374032 | | Slx1b | 707.9200385 | -0.105892787798265 | 0.0700910380481505 | -1.5107892641784 | 0.130842150854895 | 0.570291305331268 | NR_033446,NM_029420 |
| Tk2 | 1575.91435 | 0.116025761 | 0.050179674 | 2.312206355 | 0.020766314 | 0.096506681 | | Zbtb16 | 1879.614671 | 0.142553130622407 | 0.0944106742424052 | 1.50992598841517 | 0.131062310465875 | 0.571029130722048 | NM_001033324 |
| Ifi27l1 | 867.8324243 | 0.229996086 | 0.099477926 | 2.312031383 | 0.020775954 | 0.096522528 | | Phf19 | 42.09001923 | -0.141092188398187 | 0.0934741871918592 | -1.50942407349946 | 0.131190444893156 | 0.571416012749019 | NM_028716 |
| Zfp389 | 15.40260289 | 0.378352219 | 0.16366456 | 2.311754118 | 0.020791237 | 0.096564577 | | Rab9b | 1241.718515 | 0.113218102690907 | 0.0750171278963056 | 1.50923003673782 | 0.131240006783758 | 0.571460533135752 | NM_176971 |
| Fmn2 | 3592.397969 | -0.129413846 | 0.05598414 | -2.311616227 | 0.020798842 | 0.096570947 | | Cryzl1 | 1780.900909 | -0.0716343816866026 | 0.0474894851476548 | -1.50842615926191 | 0.131445492069982 | 0.572183763203042 | NM_133679,NM_026994 |
| Zkscan17 | 536.5637811 | 0.198175459 | 0.085747943 | 2.311139502 | 0.020825151 | 0.096664136 | | Prdm15 | 826.3184645 | -0.1097325339222 | 0.0727619362638936 | -1.50810354364707 | 0.131528028418915 | 0.57237152211299 | NM_144789 |
| Apba3 | 491.4972891 | 0.223875631 | 0.096939851 | 2.309428258 | 0.020919828 | 0.097003177 | | Inf2 | 9258.228609 | -0.111901544981361 | 0.0742313899467459 | -1.50746934769293 | 0.131690395007044 | 0.572734933494707 | NM_198411 |
| Arpc5l | 2403.559658 | -0.128639042 | 0.055708612 | -2.309141018 | 0.020935757 | 0.097003177 | | Phactr2 | 1636.561417 | 0.105443104388176 | 0.0699464659120703 | 1.50748294446682 | 0.131686912337769 | 0.572734933494707 | NM_001033257,NM_001195096,NM_001195065,NM_001195066 |
| Commd3 | 1411.706556 | -0.112348189 | 0.048646953 | -2.309459913 | 0.020918074 | 0.097003177 | | Atp6v1a | 22768.36613 | 0.0936620844733779 | 0.0621616276087715 | 1.50675083771715 | 0.13187453528522 | 0.572833272921038 | NM_007508 |
| Dclk1 | 39858.5784 | -0.167581142 | 0.072562389 | -2.309476638 | 0.020917147 | 0.097003177 | | Ephb2 | 878.6866315 | 0.105160076741039 | 0.069786678087008 | 1.50687895775649 | 0.131841685976143 | 0.572833272921038 | NM_010142 |
| Gba2 | 3788.971686 | 0.15620259 | 0.067641551 | 2.309269773 | 0.020928616 | 0.097003177 | | Gnb4 | 1402.306681 | 0.0938221169553503 | 0.0622651343586809 | 1.50681626116607 | 0.131857760262194 | 0.572833272921038 | NM_013531 |
| Ube2c | 11.96565997 | -0.354196721 | 0.15338789 | -2.309157005 | 0.02093487 | 0.097003177 | | Pdzd3 | 44.77838466 | -0.139379099735124 | 0.0924871198697422 | -1.50701092142802 | 0.131807857809376 | 0.572833272921038 | NM_133226 |
| Camsap3 | 1210.491617 | 0.15632296 | 0.067707543 | 2.30879681 | 0.020954859 | 0.097062657 | | Stc2 | 38.73799743 | -0.13852318517268 | 0.0919435131205271 | -1.50661183667294 | 0.13191018159995 | 0.572833272921038 | NM_011491 |
| Cxcl14 | 3211.43624 | -0.196702193 | 0.085239511 | -2.307641026 | 0.021019111 | 0.097310346 | | Sertad1 | 378.4443686 | 0.138663635041202 | 0.0920903671369396 | 1.50573441449101 | 0.132135365683266 | 0.573639665844328 | NM_018820 |
| Ifi27l2a | 24.83160593 | 0.38134825 | 0.165256875 | 2.307608996 | 0.021020894 | 0.097310346 | | Entpd1 | 741.7345441 | -0.111052353445183 | 0.0737648476221268 | -1.50549153187528 | 0.132197752357694 | 0.573705114144058 | NM_009848 |
| Ppard | 954.4322572 | 0.211687087 | 0.09173922 | 2.307487308 | 0.021027669 | 0.097312635 | | Plch2 | 4545.689559 | -0.121776057119595 | 0.0808945309429018 | -1.5053682331819 | 0.132229431512757 | 0.573705114144058 | NM_175556,NM_001113360 |
| 1500004A13Rik | 1024.808388 | -0.228858893 | 0.099195208 | -2.307156746 | 0.021046083 | 0.097333324 | | Cdc42bpg | 283.9781604 | -0.12708995900431 | 0.0844497013240091 | -1.50491898741835 | 0.132344906057029 | 0.573884731784883 | NM_001033342 |
| Ccdc71 | 1118.464219 | -0.123299462 | 0.053444024 | -2.307076682 | 0.021050546 | 0.097333324 | | Cyb561d1 | 1413.748332 | -0.106221271404029 | 0.0705836181935281 | -1.5048997787672 | 0.13234984520717 | 0.573884731784883 | NM_001081320,NM_028061 |
| Shmt2 | 737.9068257 | 0.195520617 | 0.084748499 | 2.30706879 | 0.021050986 | 0.097333324 | | Glo1 | 3551.456179 | 0.0703028169503056 | 0.0467284267335065 | 1.50449783707131 | 0.132453229851913 | 0.574161628418049 | NM_001113560,NM_025374 |
| Ccni | 7855.03282 | -0.145028156 | 0.062868778 | -2.306839103 | 0.021063791 | 0.09736348 | | Pdha1 | 9971.562582 | 0.0724084032692975 | 0.048136704205659 | 1.50422436401005 | 0.132523606437134 | 0.57429531834899 | NM_008810 |
| Mpeg1 | 1795.949944 | 0.323878968 | 0.140426767 | 2.306390549 | 0.02108882 | 0.097450096 | | Emx1 | 578.9228127 | -0.127068216791976 | 0.0845041271180384 | -1.50369243639997 | 0.132660577630494 | 0.574717432347315 | NM_010131 |
| Krtcap3 | 18.82308023 | 0.387596463 | 0.168072434 | 2.306127501 | 0.021103509 | 0.0974889 | | Ggct | 377.9822669 | -0.10482606734434 | 0.0697278918482348 | -1.50335919480396 | 0.132746443073485 | 0.574746596746778 | NM_026637 |
| Crebrf | 2340.210071 | 0.235486087 | 0.10212464 | 2.305869438 | 0.021117929 | 0.097526435 | | Slc25a35 | 217.7082853 | -0.131496138976979 | 0.0874634867456675 | -1.50344039403955 | 0.132725516726004 | 0.574746596746778 | NM_028048 |
| Cog3 | 1428.913701 | 0.123468493 | 0.053556404 | 2.305391767 | 0.021144642 | 0.097620705 | | B630019K06Rik | 1216.700635 | 0.122345576401073 | 0.0814094946693069 | 1.50284161445851 | 0.132879891891436 | 0.575107585198328 | NR_045448 |
| Grcc10 | 2318.265014 | -0.159861024 | 0.069348659 | -2.305178317 | 0.021156589 | 0.097646764 | | Zfp697 | 749.5312194 | -0.114027426853619 | 0.0758802450257377 | -1.50272876444907 | 0.132909002031584 | 0.575107585198328 | NM_172863 |
| Pdzd7 | 182.1496397 | 0.347004106 | 0.150615533 | 2.303906508 | 0.021227892 | 0.097946683 | | Calr | 13796.90374 | 0.0832887608165882 | 0.0554541980366319 | 1.50193788325222 | 0.133113151769709 | 0.575193212045581 | NM_007591 |
| Cacng5 | 775.2408815 | -0.38687089 | 0.167952724 | -2.303451124 | 0.021253474 | 0.098006348 | | Cul3 | 4967.403187 | 0.107574958990877 | 0.0716339333151356 | 1.50173184707906 | 0.133166375609892 | 0.575193212045581 | NM_016716 |
| Haus3 | 172.35982 | 0.261271121 | 0.113425639 | 2.303457336 | 0.021253125 | 0.098006348 | | Erap1 | 468.4370834 | -0.126569855181067 | 0.0842434514532959 | -1.50242960132321 | 0.132986196325992 | 0.575193212045581 | NM_030711 |
| Gprasp2 | 5002.273529 | 0.231178062 | 0.100368527 | 2.303292369 | 0.021262399 | 0.098018329 | | Fignl2 | 48.83703541 | -0.138171034285418 | 0.0920046021131124 | -1.50178394462863 | 0.133152916069558 | 0.575193212045581 | NM_001214911 |
| Trim54 | 21.05048235 | -0.377035161 | 0.163725307 | -2.30285206 | 0.021287168 | 0.098103327 | | Nxph1 | 1436.010892 | 0.096706804255435 | 0.0643886946712447 | 1.50192211147003 | 0.13311722539849 | 0.575193212045581 | NM_008751 |
| Ccdc62 | 94.66594518 | 0.357224991 | 0.15513585 | 2.302659188 | 0.021298026 | 0.09812418 | | Pdhb | 4512.572357 | 0.0720220923305268 | 0.0479582592973463 | 1.5017661897189 | 0.133157502979368 | 0.575193212045581 | NM_024221 |
| Cwf19l1 | 569.905139 | 0.163959084 | 0.071213299 | 2.302366091 | 0.021314536 | 0.098164262 | | Efhd2 | 10865.1192 | 0.0771421327706193 | 0.0513844503023187 | 1.50127387403691 | 0.133284739484244 | 0.57536229591326 | NM_025994 |
| Ftsjd1 | 254.8538347 | -0.224675702 | 0.097588356 | -2.302279818 | 0.021319397 | 0.098164262 | | Mrps22 | 574.5290449 | 0.104754064773055 | 0.0697755171147316 | 1.50130115984391 | 0.133277685142905 | 0.57536229591326 | NM_025485 |
| Gabbr1 | 31161.74343 | -0.105320341 | 0.045761985 | -2.301481059 | 0.021364455 | 0.098322779 | | Ankrd50 | 1681.66127 | 0.107148948899455 | 0.0714333515306344 | 1.49998490345933 | 0.133618313117294 | 0.575883579749171 | NM_001167883 |
| Pkp2 | 243.646644 | 0.358904411 | 0.15594745 | 2.301444557 | 0.021366516 | 0.098322779 | | Bicd2 | 3613.19503 | -0.084115913390095 | 0.0560814557883893 | -1.49988819312194 | 0.133643366882449 | 0.575883579749171 | NM_001039180,NM_001039179,NM_029791 |
| B9d2 | 413.8175876 | -0.237292133 | 0.103112314 | -2.301297715 | 0.021374809 | 0.098331736 | | Diap3 | 35.79586107 | -0.13118123811647 | 0.087426206610115 | -1.50047958390193 | 0.133490218107677 | 0.575883579749171 | NM_019670 |
| Cep85 | 634.410626 | 0.180991359 | 0.078674222 | 2.300516666 | 0.021418966 | 0.098505628 | | Nono | 5363.146507 | 0.0714036098934923 | 0.0476036834342032 | 1.49995976660472 | 0.133624824717445 | 0.575883579749171 | NM_023144,NM_001252518 |
| Sod3 | 512.2862201 | 0.212686547 | 0.092478603 | 2.299846014 | 0.021456946 | 0.098651013 | | Nr2f6 | 633.5407631 | -0.120146700505092 | 0.0800994747275987 | -1.49996864416009 | 0.133622524994718 | 0.575883579749171 | NM_010150 |
| Hyou1 | 5007.967909 | -0.146743993 | 0.063812629 | -2.29960737 | 0.021470474 | 0.09868393 | | Psmd14 | 2781.675319 | 0.090086327938503 | 0.0600528603754404 | 1.50011718634714 | 0.133584049817705 | 0.575883579749171 | NM_021526 |
| Sec24d | 580.0889352 | 0.191826566 | 0.083437764 | 2.299037704 | 0.021502798 | 0.098803191 | | Pml | 774.6111754 | -0.097407338787043 | 0.0649511661505738 | -1.49970115334384 | 0.133691831690607 | 0.575921573884269 | NM_178087,NM_008884 |
| Ube2o | 6890.490458 | 0.126023472 | 0.054828193 | 2.298515867 | 0.021532446 | 0.098910085 | | Ralyl | 2428.283489 | 0.0962348238602771 | 0.0641825846421031 | 1.49939153115919 | 0.133772089320262 | 0.576096462930958 | NM_001163330,NM_178631,NM_001163328,NM_001163329 |
| Zfp563 | 561.011329 | -0.183045923 | 0.079651452 | -2.298086452 | 0.021556869 | 0.098992926 | | Spsb4 | 147.1023634 | 0.141151808713718 | 0.0941539029509861 | 1.49916046270751 | 0.133832009203278 | 0.576183688703856 | NM_145134 |
| Gpc1 | 3432.502557 | -0.179242687 | 0.078003263 | -2.297887047 | 0.021568219 | 0.099015698 | | Pcdhb19 | 283.4569432 | 0.123334697796421 | 0.0822813544950456 | 1.49893859372292 | 0.133889563048832 | 0.576260679362174 | NM_053144 |
| Exoc8 | 1455.031752 | -0.143907311 | 0.062640976 | -2.297335079 | 0.021599662 | 0.099101322 | | Csrp2bp | 1538.387215 | 0.0735986000734221 | 0.049108050386533 | 1.49870743175757 | 0.133949547900126 | 0.576348084359371 | NM_001166640,NM_181417 |
| Lilrb3 | 21.61742934 | 0.386224109 | 0.168110547 | 2.297441266 | 0.02159361 | 0.099101322 | | Polg | 2988.642689 | -0.0902756795341287 | 0.060264848780529 | -1.49798234561066 | 0.134137837357007 | 0.576690607057762 | NM_017462 |
| Cyp2a5 | 36.56313288 | -0.379531379 | 0.165232693 | -2.296950874 | 0.021621572 | 0.099172481 | | Pomt2 | 1246.709531 | -0.0800538221608561 | 0.0534425290610723 | -1.49794224875423 | 0.134148255627714 | 0.576690607057762 | NM_153415 |
| Rttn | 212.7957745 | 0.227401694 | 0.099009805 | 2.29675933 | 0.021632503 | 0.099193252 | | Shc2 | 2111.498626 | -0.0919065075115309 | 0.0613540006055147 | -1.49797090009596 | 0.134140811154016 | 0.576690607057762 | NM_001024539 |
| Sbk1 | 3810.46787 | -0.247954091 | 0.107984491 | -2.296200948 | 0.021664394 | 0.099310097 | | Alg11 | 1517.288033 | 0.107532391774227 | 0.0718127403927915 | 1.497399920767 | 0.134289228895811 | 0.577125839923241 | NM_183142,NM_001243161,NR_040650 |
| Xrcc5 | 769.0484957 | 0.17878257 | 0.077865796 | 2.296034716 | 0.021673897 | 0.099324269 | | Fam171b | 9992.470243 | 0.0960386489267942 | 0.0641783069792997 | 1.4964347528089 | 0.134540398914461 | 0.57786288772156 | NM_175514 |
| Mtor | 5602.318909 | -0.161219701 | 0.070225631 | -2.295738735 | 0.021690824 | 0.099372452 | | Mkx | 556.2513477 | -0.129066444176277 | 0.086246809905797 | -1.49647789080257 | 0.134529165171443 | 0.57786288772156 | NM_177595 |
| Ift172 | 2648.092459 | 0.162624228 | 0.070842678 | 2.295568607 | 0.021700559 | 0.099387664 | | Wbp2 | 18407.93541 | 0.0610526231811505 | 0.0408028740102143 | 1.49628242279862 | 0.134580073603335 | 0.57786288772156 | NM_016852 |
| Lcorl | 548.1683062 | 0.25019637 | 0.108998777 | 2.295405296 | 0.021709908 | 0.09939954 | | Adprh | 3415.308842 | 0.0715791294221967 | 0.0478535177490308 | 1.49579660574998 | 0.134706666152587 | 0.577947814261831 | NM_007414 |
| Rps25 | 5419.827679 | -0.166363745 | 0.0724802 | -2.295299205 | 0.021715983 | 0.09939954 | | Cdk3-ps | 68.55457882 | -0.140113610740234 | 0.0936684080793362 | -1.49584703757919 | 0.134693520518509 | 0.577947814261831 | NR_004853 |
| Klhl5 | 1692.046546 | -0.144636657 | 0.063021699 | -2.295029463 | 0.021731436 | 0.099440894 | | Slc45a1 | 2395.776975 | 0.0771748906472032 | 0.0515961688116308 | 1.49574847173162 | 0.134719213760881 | 0.577947814261831 | NM_173774 |
| Sirt7 | 1041.731851 | 0.246087994 | 0.107238261 | 2.294777915 | 0.021745855 | 0.099477495 | | Emc3 | 3108.815265 | 0.0777160028701597 | 0.052013994258757 | 1.49520535729798 | 0.134860855811201 | 0.578384644674787 | NM_175101 |
| Zfp54 | 44.68072237 | 0.358887679 | 0.156418853 | 2.294401676 | 0.021767437 | 0.099546833 | | Nkiras1 | 3453.731083 | 0.0742228158711074 | 0.0496496702807477 | 1.49493069040356 | 0.134932531636322 | 0.578521238060572 | NM_023526 |
| Abtb1 | 1151.761487 | 0.217798402 | 0.094939088 | 2.294085689 | 0.021785577 | 0.099600041 | | Axl | 2448.928071 | -0.0947327150439953 | 0.0633832151798057 | -1.49460255014292 | 0.135018200219631 | 0.578547013684471 | NM_001190975,NM_001190974,NM_009465 |
| Ccdc116 | 98.70811567 | 0.320592707 | 0.139754222 | 2.29397511 | 0.021791928 | 0.099600041 | | Zfp593 | 335.9758142 | 0.113961029250828 | 0.0762422527301991 | 1.49472274454043 | 0.134986815823067 | 0.578547013684471 | NM_024215 |
| Jagn1 | 820.3731136 | -0.142870285 | 0.062288835 | -2.293674072 | 0.021809227 | 0.09964971 | | Atp9b | 3232.450001 | -0.0657048091733533 | 0.0439790187997171 | -1.49400352637645 | 0.135174697581844 | 0.578555714267335 | NM_015805,NM_001201569 |
| Znhit3 | 589.3560631 | -0.151001283 | 0.065847699 | -2.293189966 | 0.021837071 | 0.099747515 | | Gm10033 | 162.0509378 | 0.142043125958466 | 0.0950715079013303 | 1.49406619389992 | 0.135158318878532 | 0.578555714267335 | NR_038043,NR_038044 |
| Mfsd2b | 37.85891873 | 0.38097137 | 0.166143569 | 2.293025074 | 0.021846561 | 0.099761457 | | Mrpl20 | 1388.2695 | 0.0870264647468694 | 0.0582512316011292 | 1.49398497430537 | 0.13517954662146 | 0.578555714267335 | NM_025570 |
| Hebp1 | 727.5126955 | -0.296055439 | 0.129139937 | -2.292516515 | 0.021875856 | 0.099857587 | | Tmem229b | 1572.212926 | 0.0892936900872704 | 0.0597558165041824 | 1.49430959714224 | 0.135094717864237 | 0.578555714267335 | NM_001170401,NM_178745 |
| Irf5 | 171.2624384 | 0.250595372 | 0.109314015 | 2.292435896 | 0.021880503 | 0.099857587 | | Avl9 | 2741.159983 | 0.0967868627905833 | 0.0648105494082942 | 1.493381304035 | 0.135337404029506 | 0.578660429559486 | NM_030235 |
| Adat3 | 60.72315107 | 0.349030866 | 0.152292052 | 2.291852146 | 0.021914176 | 0.099952384 | | Eps8 | 1272.72727 | 0.0962387628564317 | 0.0644478110361835 | 1.49328210390884 | 0.135363358091913 | 0.578660429559486 | NM_001271595,NM_001271587,NM_001271588,NM_001271589,NM_007945 |
| Dusp14 | 2108.113259 | -0.192906287 | 0.084169756 | -2.29187175 | 0.021913045 | 0.099952384 | | Klhl13 | 1985.3741 | 0.120805688807502 | 0.0808905310547694 | 1.49344660286266 | 0.135320321776096 | 0.578660429559486 | NM_026167 |
| Poln | 34.04675409 | 0.368445979 | 0.160787641 | 2.291506836 | 0.021934117 | 0.100013893 | | Zcchc24 | 3614.830739 | 0.0950188462928787 | 0.0636224198852635 | 1.49348054450358 | 0.135311443259164 | 0.578660429559486 | NM_001101433 |
| Nav1 | 5110.355421 | -0.189417418 | 0.082665683 | -2.291367003 | 0.021942196 | 0.100021297 | | Ctsk | 130.4838285 | -0.13743424558154 | 0.0920909115157411 | -1.49237577649612 | 0.135600661657862 | 0.579179191366766 | NM_007802 |
| Gm13749 | 17.29913083 | -0.372685326 | 0.162772536 | -2.289608149 | 0.022044042 | 0.100455997 | | Luzp1 | 5599.610212 | 0.112587914923514 | 0.0754504677273196 | 1.49220963520611 | 0.135644197234596 | 0.579179191366766 | NM_024452 |
| Gstm5 | 4756.847382 | 0.170063798 | 0.074302431 | 2.288805289 | 0.022090668 | 0.100638874 | | Tacc3 | 249.398851 | -0.124654753140603 | 0.0835330244548142 | -1.4922810942639 | 0.135625470817598 | 0.579179191366766 | NM_001040435 |
| Ralgds | 4074.398618 | 0.171228875 | 0.074840813 | 2.287907751 | 0.022142894 | 0.100819843 | | Tmem86b | 203.0136207 | -0.141940839310423 | 0.095096533094351 | -1.49259741329997 | 0.135542600868431 | 0.579179191366766 | NM_023440 |
| Wif1 | 30.54631762 | -0.384285238 | 0.167964252 | -2.287898964 | 0.022143406 | 0.100819843 | | Ccser1 | 453.684784 | 0.111184435592436 | 0.0745648287311406 | 1.49111099005316 | 0.135932357408618 | 0.57943550743131 | NM_001164316,NM_183310 |
| Tnfaip8l1 | 271.4987942 | -0.30559385 | 0.133580011 | -2.287721407 | 0.02215375 | 0.100837311 | | Cgrrf1 | 671.7941936 | 0.0972334271012925 | 0.0652105789406977 | 1.49106830027864 | 0.135943563907883 | 0.57943550743131 | NM_026832 |
| Ddhd2 | 2908.00737 | 0.120799049 | 0.05280702 | 2.287556609 | 0.022163355 | 0.100848185 | | Mto1 | 891.9436817 | 0.085810242602984 | 0.0575488417403172 | 1.49108548509444 | 0.135939052633469 | 0.57943550743131 | NM_026658 |
| Rps15 | 8574.089299 | -0.169463042 | 0.074083593 | -2.287457094 | 0.022169157 | 0.100848185 | | Osbpl8 | 4014.019068 | 0.127062695131812 | 0.0852106584479435 | 1.49115964418273 | 0.135919586075378 | 0.57943550743131 | NM_001003717,NM_175489 |
| Wdfy4 | 147.8910775 | -0.255176733 | 0.111566703 | -2.287212279 | 0.022183436 | 0.10088352 | | St7l | 869.7450796 | 0.106897175380387 | 0.0716608769576797 | 1.49170900383366 | 0.135775447693953 | 0.57943550743131 | NM_001253702,NM_001253703,NR_045578,NM_153091 |
| 1700109H08Rik | 25.25710211 | 0.382473543 | 0.167275903 | 2.286483207 | 0.022226005 | 0.101047455 | | Zdhhc1 | 1561.675942 | -0.108240625760447 | 0.0725776193033024 | -1.49137746318336 | 0.135862421628575 | 0.57943550743131 | NM_175160 |
| Clcc1 | 950.504799 | 0.170172252 | 0.074448784 | 2.28576268 | 0.022268146 | 0.101179664 | | Fam110b | 1262.629062 | 0.0925451119083359 | 0.0620856334826483 | 1.49060429469888 | 0.136065416103983 | 0.579784756329261 | NM_173426 |
| Nptx1 | 17352.25587 | 0.305360794 | 0.133587659 | 2.285845843 | 0.022263279 | 0.101179664 | | Agap2 | 43160.91705 | -0.100454355584786 | 0.0674272100264514 | -1.48981925168457 | 0.13627176772788 | 0.579855847513867 | NM_001033263 |
| Bmp7 | 363.1587904 | -0.183043304 | 0.080157967 | -2.283532255 | 0.022399035 | 0.101714727 | | D830030K20Rik | 111.633145 | 0.136222972571029 | 0.0914432273725291 | 1.48969996450445 | 0.136303143959996 | 0.579855847513867 | NM_177135 |
| Svop | 8267.993609 | -0.194458064 | 0.085152909 | -2.283633846 | 0.022393059 | 0.101714727 | | Polrmt | 1343.863219 | -0.110983572223523 | 0.0745012663606108 | -1.48968705694646 | 0.136306539382831 | 0.579855847513867 | NM_172551 |
| Adra1b | 822.8302902 | -0.280916644 | 0.123032054 | -2.283280118 | 0.022413873 | 0.101752287 | | Rab36 | 1169.495628 | -0.10463023582857 | 0.0702390877185099 | -1.48962976637576 | 0.136321610853632 | 0.579855847513867 | NM_029781 |
| Csrp2 | 129.957248 | -0.292018185 | 0.127902764 | -2.28312646 | 0.02242292 | 0.101763541 | | Rps6kc1 | 2528.111936 | 0.0816532782133403 | 0.0548015676875757 | 1.48998070053117 | 0.136229310548143 | 0.579855847513867 | NM_178775 |
| 9330159M07Rik | 90.4850428 | 0.297178948 | 0.130200108 | 2.282478515 | 0.022461105 | 0.101906985 | | Uba3 | 2869.617247 | 0.0918512982277658 | 0.0616554683507951 | 1.4897510421163 | 0.136289708279565 | 0.579855847513867 | NM_001111106,NM_011666 |
| Tekt1 | 60.4284102 | 0.315131722 | 0.1380739 | 2.282340993 | 0.022469216 | 0.101913944 | | Pdyn | 627.3360027 | 0.125444517075807 | 0.0842487295032675 | 1.48897814620387 | 0.136493123562066 | 0.580415431165857 | NM_018863 |
| Atn1 | 7379.82213 | 0.163388492 | 0.071592029 | 2.282216247 | 0.022476576 | 0.101917493 | | Lgi1 | 6964.864269 | 0.0754825518024992 | 0.0507053196351977 | 1.48865153292717 | 0.136579153957618 | 0.580611293645995 | NM_020278 |
| Trmu | 448.0677832 | 0.228520784 | 0.10013765 | 2.282066587 | 0.022485409 | 0.101927716 | | Nr2f1 | 2411.914694 | -0.0896000556365047 | 0.0601980228787325 | -1.48842190078239 | 0.136639664434968 | 0.580698585600452 | NM_010151 |
| Slc35g1 | 89.00178292 | -0.293588764 | 0.128665947 | -2.281790722 | 0.022501699 | 0.101971723 | | Pdxp | 6369.921958 | 0.0782327929028096 | 0.0525788091009835 | 1.4879148889157 | 0.136773340638426 | 0.580996971744474 | NM_020271 |
| Abcb1a | 1676.413515 | -0.143784215 | 0.063036439 | -2.28096984 | 0.022550232 | 0.10216178 | | Reep5 | 18085.46623 | 0.074209176756436 | 0.0498767119040705 | 1.48785222448434 | 0.136789869431784 | 0.580996971744474 | NM_007874 |
| Bbip1 | 1246.773524 | -0.176061364 | 0.077212061 | -2.280231366 | 0.02259397 | 0.102330012 | | Nolc1 | 3019.987935 | 0.0675509740747751 | 0.0454088140982424 | 1.48761810710643 | 0.136851635446714 | 0.581089405582861 | NM_001039353,NM_001039352,NM_001039351,NM_053086 |
| Oas1c | 42.32408205 | 0.355131174 | 0.155755809 | 2.280050909 | 0.022604669 | 0.102348552 | | Nedd1 | 147.1069924 | -0.130937583975578 | 0.0880323783250327 | -1.48737983077239 | 0.13691452079038 | 0.581186536820882 | NM_008682 |
| Haus6 | 222.4744739 | 0.234055149 | 0.10266909 | 2.279704126 | 0.022625242 | 0.102411775 | | Tmem179b | 167.0778251 | -0.127417049160192 | 0.085684387908836 | -1.48705093506366 | 0.137001358811839 | 0.581385257990295 | NR_028418,NR_028419,NM_026325 |
| Lrrc71 | 28.53714169 | 0.381040799 | 0.1671777 | 2.27925614 | 0.022651844 | 0.102502239 | | Cox7c | 3208.653348 | 0.0766817853081559 | 0.051572505123206 | 1.48687338582767 | 0.137048254623693 | 0.581414411992921 | NM_007749 |
| Cdh4 | 1447.533733 | -0.243666862 | 0.106917477 | -2.279018054 | 0.022665992 | 0.102536316 | | 2900026A02Rik | 3081.60413 | 0.0939797096776952 | 0.0632166585956271 | 1.4866288691221 | 0.13711285872037 | 0.581518652780173 | NM_172884 |
| Tomm5 | 813.1185908 | -0.191169011 | 0.083886465 | -2.278901739 | 0.022672907 | 0.10253766 | | Eif3g | 2183.800845 | 0.0710492432314669 | 0.0478030364963128 | 1.4862914249589 | 0.137202053884382 | 0.581727097117491 | NM_016876 |
| Pstpip1 | 69.545495 | 0.35288232 | 0.154861784 | 2.278692082 | 0.022685376 | 0.102564112 | | Nuf2 | 85.55900615 | -0.141177778832672 | 0.0950275212293594 | -1.48565149344393 | 0.137371327129108 | 0.58201244734081 | NM_023284 |
| Neil2 | 71.58632919 | 0.320843557 | 0.140827823 | 2.278268241 | 0.022710601 | 0.102593826 | | Rab2a | 9564.63237 | 0.0968580314871232 | 0.065218614909874 | 1.48512861889158 | 0.137509756249047 | 0.58201244734081 | NM_021518 |
| Pgap1 | 1539.052029 | -0.243968089 | 0.107091067 | -2.278136703 | 0.022718434 | 0.102593826 | | Stx16 | 3800.761947 | -0.0806513691519923 | 0.0543028984018555 | -1.48521297252223 | 0.137487416661026 | 0.58201244734081 | NM_001102424,NM_001102425,NM_001102423,NM_172675 |
| Ptger1 | 43.11766507 | 0.379145039 | 0.166411614 | 2.278356837 | 0.022705326 | 0.102593826 | | Tecr | 12862.80772 | 0.0755944575840179 | 0.0508919838330435 | 1.48539026955627 | 0.137440471755996 | 0.58201244734081 | NM_134118,NM_027179 |
| Qprt | 14.58791849 | -0.380900307 | 0.167195668 | -2.278170908 | 0.022716397 | 0.102593826 | | Tirap | 115.9880912 | 0.137403723591296 | 0.0925074906506924 | 1.48532537878615 | 0.137457652171667 | 0.58201244734081 | NM_054096,NM_001177846,NM_001177847,NM_001177845 |
| Ist1 | 3331.807927 | 0.152871136 | 0.067125405 | 2.277396094 | 0.022762583 | 0.102763247 | | Tmem108 | 913.8428323 | 0.106290937641705 | 0.0715595837653083 | 1.48534874085216 | 0.137451466663567 | 0.58201244734081 | NM_178638 |
| Ptpn4 | 1678.787902 | -0.199800312 | 0.087739613 | -2.277196174 | 0.022774514 | 0.102787158 | | 1700003E16Rik | 998.5377028 | 0.107428986976414 | 0.072348953161902 | 1.48487272146163 | 0.137577543345251 | 0.582129739188208 | NM_027948 |
| 2610507I01Rik | 852.8499956 | 0.16932114 | 0.074372164 | 2.276673572 | 0.022805726 | 0.102898053 | | Clic4 | 4257.059909 | 0.122834716290163 | 0.0827466104677017 | 1.48446825309067 | 0.137684739348861 | 0.582244111726799 | NM_013885 |
| Inha | 854.581333 | 0.32704529 | 0.143662473 | 2.276483785 | 0.02281707 | 0.102919267 | | Retsat | 465.5941776 | 0.126357774941501 | 0.0851174663872603 | 1.48451052768193 | 0.137673532327395 | 0.582244111726799 | NM_026159 |
| Ppa1 | 2501.598363 | -0.130458082 | 0.057317309 | -2.276067789 | 0.022841953 | 0.103001517 | | Necap2 | 1062.723274 | -0.102762372407776 | 0.0692402232739127 | -1.48414270706856 | 0.137771065373626 | 0.582270147110065 | NM_025383 |
| Fem1a | 2681.070713 | -0.152428715 | 0.066984495 | -2.275582046 | 0.022871037 | 0.103100551 | | Traip | 611.0538986 | -0.140185700613416 | 0.0944509684481201 | -1.48421665671345 | 0.137751452260675 | 0.582270147110065 | NM_011634 |
| Sipa1l1 | 22549.3782 | -0.138845709 | 0.061018239 | -2.275478799 | 0.022877223 | 0.103100551 | | Ss18 | 1127.744518 | -0.0910582495457486 | 0.0613633152807333 | -1.48391997937469 | 0.137830150785595 | 0.582350427664794 | NM_001161369,NM_001161371,NM_001161370,NM_009280 |
| Eif4g2 | 24694.28079 | -0.135253628 | 0.059448915 | -2.275123583 | 0.022898517 | 0.10316651 | | Itpkc | 408.807499 | 0.118598766792005 | 0.0799657855282098 | 1.48311888651636 | 0.138042826937233 | 0.582865024557558 | NM_181593 |
| Dip2a | 3955.500496 | 0.108694583 | 0.047793404 | 2.274259092 | 0.022950413 | 0.103340221 | | Mus81 | 759.9069314 | -0.101435174993874 | 0.0683982719229396 | -1.48300786177983 | 0.138072322008988 | 0.582865024557558 | NM_027877 |
| Tox2 | 624.4545049 | 0.215283405 | 0.094658371 | 2.274319775 | 0.022946767 | 0.103340221 | | Rusc2 | 6804.061672 | 0.0787188997276669 | 0.0530719704284234 | 1.48324810803535 | 0.138008503784337 | 0.582865024557558 | NM_001037709,NM_199057 |
| Fam83h | 104.1186562 | 0.331585515 | 0.145851829 | 2.273440915 | 0.022999623 | 0.103506474 | | Dsel | 598.6474598 | 0.13076737405953 | 0.0882080199077179 | 1.4824884879667 | 0.138210364489307 | 0.582904025247445 | NM_001081316 |
| Lepr | 160.3336933 | 0.310068002 | 0.136388165 | 2.273423085 | 0.023000696 | 0.103506474 | | Fzd8 | 122.8274174 | -0.138094544520818 | 0.0931375336710415 | -1.4826948822648 | 0.138155494969213 | 0.582904025247445 | NM_008058 |
| Mib1 | 2945.310977 | -0.183312291 | 0.080637586 | -2.273285952 | 0.023008954 | 0.103513567 | | Lrfn5 | 2135.818812 | 0.090355417254125 | 0.0609594358430689 | 1.48222200557649 | 0.138281233147232 | 0.582904025247445 | NM_178714 |
| Phpt1 | 1172.757362 | -0.162187285 | 0.071363296 | -2.272698896 | 0.023044331 | 0.103642632 | | Scfd2 | 786.918445 | 0.0936285437125248 | 0.0631742768739427 | 1.4820675177549 | 0.138322330650416 | 0.582904025247445 | NM_178672,NM_001114660 |
| Abat | 15670.01688 | -0.181085071 | 0.079702343 | -2.272016915 | 0.023085489 | 0.103767497 | | Tex40 | 276.8690588 | -0.136476593798196 | 0.0920741272329123 | -1.48224694493125 | 0.138274599556834 | 0.582904025247445 | NM_001039494 |
| Kpnb1 | 8956.002034 | -0.112280722 | 0.049416879 | -2.272112775 | 0.0230797 | 0.103767497 | | Trhde | 1377.912931 | 0.101418733792258 | 0.0684271336089544 | 1.48214207498217 | 0.138302495446872 | 0.582904025247445 | NM_146241 |
| Maneal | 2588.990214 | 0.116968119 | 0.051491776 | 2.271588388 | 0.023111383 | 0.103853761 | | 1600020E01Rik | 174.1720981 | -0.132597004967526 | 0.089533101313137 | -1.48098304451418 | 0.1386110917734 | 0.583109099075629 | NR_037961,NR_037960 |
| Alx3 | 33.15203347 | -0.377147353 | 0.166053571 | -2.271239035 | 0.023132512 | 0.103918566 | | AA474331 | 98.41892866 | -0.139625462313983 | 0.0942720329898255 | -1.48109102865165 | 0.138582318192476 | 0.583109099075629 | NR_033628 |
| Pcdhb9 | 527.5164641 | 0.154816404 | 0.068173983 | 2.27090156 | 0.023152938 | 0.103980179 | | AI464131 | 1668.549762 | 0.107355554457827 | 0.0724945667754499 | 1.48087724684745 | 0.138639287207832 | 0.583109099075629 | NM_001085515 |
| Usp18 | 49.00204733 | 0.377768514 | 0.166429005 | 2.269847821 | 0.023216818 | 0.104236852 | | Csnk1e | 4529.457715 | 0.0689100970432911 | 0.0465217493480162 | 1.48124475130533 | 0.138541365010221 | 0.583109099075629 | NM_013767 |
| Csad | 667.1608305 | 0.169729977 | 0.074780458 | 2.269710323 | 0.023225165 | 0.10424412 | | Helb | 227.6619354 | -0.121973236857922 | 0.0823190902835537 | -1.48171264329789 | 0.138416771492893 | 0.583109099075629 | NM_080446 |
| 2700049A03Rik | 357.4740666 | 0.197214509 | 0.086908654 | 2.269216003 | 0.023255194 | 0.104288269 | | Mlf2 | 15778.97725 | 0.0681900605996424 | 0.0460328616479588 | 1.48133438066773 | 0.138517491195731 | 0.583109099075629 | NM_001170341,NM_145385 |
| Eef1g | 10085.7516 | -0.106346126 | 0.046862866 | -2.269304774 | 0.023249798 | 0.104288269 | | Sox2 | 907.4834152 | -0.126634750853112 | 0.0855160866498885 | -1.48082958205943 | 0.138651991477848 | 0.583109099075629 | NM_011443 |
| Fam3a | 677.9260827 | 0.159665837 | 0.07035587 | 2.269403198 | 0.023243818 | 0.104288269 | | Rnf38 | 2337.252657 | -0.0888711947272236 | 0.0600209270979928 | -1.48067014330066 | 0.138694493789302 | 0.583119020776672 | NM_175201,NM_001038993 |
| Ahnak | 1713.340244 | -0.300875311 | 0.132615532 | -2.268778831 | 0.023281779 | 0.104316912 | | Ccng1 | 3587.237864 | 0.100248150281758 | 0.0677371102456578 | 1.47995906406687 | 0.13888417159496 | 0.583135914437549 | NM_009831 |
| Nrxn1 | 11631.82436 | -0.17355436 | 0.076494595 | -2.268844735 | 0.023277769 | 0.104316912 | | Hsd3b7 | 279.9291625 | -0.125506694071486 | 0.0847887895825391 | -1.48022745329215 | 0.138812556284542 | 0.583135914437549 | NM_133943,NM_001040684 |
| Syngr3 | 6050.950194 | -0.144596391 | 0.063729781 | -2.268898266 | 0.023274513 | 0.104316912 | | Ltbp3 | 1739.230394 | -0.114012333942168 | 0.0770542489539724 | -1.47963721001645 | 0.138970090644843 | 0.583135914437549 | NM_008520 |
| Ppp1r7 | 4112.657302 | -0.125712953 | 0.055420579 | -2.268344269 | 0.023308232 | 0.104405245 | | Slc24a3 | 5470.550443 | 0.0901104452796447 | 0.0609006135212556 | 1.47963115754481 | 0.138971706746252 | 0.583135914437549 | NM_053195 |
| E230025N22Rik | 25.68347632 | -0.374904068 | 0.165311349 | -2.267866477 | 0.023337346 | 0.104505445 | | Snx17 | 3266.924324 | 0.0640433801213301 | 0.0432824997304619 | 1.47965992075677 | 0.138964026662944 | 0.583135914437549 | NM_153680 |
| Eps8l2 | 193.1425897 | 0.268429574 | 0.118381137 | 2.26750292 | 0.023359521 | 0.10457452 | | Tmem189 | 1653.73224 | 0.0829513050664381 | 0.0560360228601422 | 1.48032106549518 | 0.138787584078315 | 0.583135914437549 | NM_145538 |
| AI182371 | 9.735186324 | 0.366356213 | 0.161692282 | 2.2657619 | 0.023465965 | 0.104990374 | | Trim46 | 3712.71765 | -0.102678966868067 | 0.0693963474856253 | -1.47960188955673 | 0.138979521945669 | 0.583135914437549 | NM_183037,NM_001039466 |
| Zdhhc20 | 1687.488271 | 0.203318651 | 0.089733391 | 2.265808169 | 0.023463131 | 0.104990374 | | Slc35d2 | 27.07756562 | -0.122263764919676 | 0.0826496948420999 | -1.47930086315815 | 0.13905992228753 | 0.583304773649626 | NM_001001321 |
| Pdia4 | 1610.538625 | -0.20287468 | 0.08956296 | -2.265162747 | 0.023502694 | 0.105124349 | | Murc | 27.06517411 | -0.12916122544294 | 0.0873242363579103 | -1.47909939817343 | 0.13911375103851 | 0.583362109580082 | NM_026509 |
| Trpv3 | 16.13461334 | 0.379824153 | 0.167691262 | 2.265020541 | 0.023511419 | 0.105133024 | | Begain | 4073.300766 | -0.107079999781177 | 0.072419365671383 | -1.47861002079296 | 0.139244572952659 | 0.583573764198018 | NM_001163175 |
| Ano8 | 2026.554746 | 0.194877158 | 0.086054624 | 2.264575095 | 0.023538767 | 0.105224943 | | Zeb2 | 5261.271556 | -0.104177930192215 | 0.0704522523760598 | -1.47870262026732 | 0.139219811704526 | 0.583573764198018 | NM_015753 |
| Fbxo25 | 2335.179661 | -0.143639676 | 0.063439304 | -2.264206351 | 0.023561426 | 0.105295857 | | Kat7 | 4346.627133 | 0.0681089023155571 | 0.0460700111739 | 1.478378246067 | 0.139306564750903 | 0.58366517437889 | NM_001195003,NM_177619,NM_001195004 |
| Adgb | 36.98437031 | 0.378601515 | 0.167276907 | 2.263322062 | 0.023615843 | 0.1054782 | | Pithd1 | 2922.276534 | 0.0729612697024151 | 0.0493637921025569 | 1.47803210804455 | 0.139399184369661 | 0.583884818960121 | NM_025411 |
| Naip1 | 9.595191009 | 0.354984398 | 0.156834619 | 2.263431381 | 0.02360911 | 0.1054782 | | Myo6 | 5251.353255 | -0.0931028779510776 | 0.0630080960500164 | -1.47763357072671 | 0.139505883695949 | 0.584163293908145 | NM_001039546 |
| Polr3a | 1337.685989 | 0.129585721 | 0.05727145 | 2.262658279 | 0.023656763 | 0.105618495 | | Cdc7 | 620.492941 | -0.121787888492658 | 0.0824518566391722 | -1.47707878823917 | 0.139654518771516 | 0.584617158407793 | NM_001271566,NM_001271567,NM_001271568,NM_009863 |
| Prdx1 | 4737.77481 | -0.160236217 | 0.070819776 | -2.262591417 | 0.023660888 | 0.105618495 | | Lrrn2 | 5347.457364 | 0.0969877735628348 | 0.0656855680363689 | 1.47654616473948 | 0.139797331747063 | 0.585046396127294 | NM_010732 |
| Rgs14 | 2369.57764 | -0.207662135 | 0.091787951 | -2.262411707 | 0.023671978 | 0.105637567 | | Six5 | 88.28833281 | -0.139150465127186 | 0.0942648849959732 | -1.47616437587687 | 0.139899770383721 | 0.585306470217144 | NM_011383 |
| Atp5g2 | 3144.914635 | -0.133653854 | 0.059079112 | -2.262286087 | 0.023679733 | 0.105641748 | | Car11 | 6266.504902 | -0.0842416653368163 | 0.0570938314232567 | -1.47549504450495 | 0.140079499582052 | 0.585376896927921 | NM_009800 |
| Trappc2l | 994.8086607 | -0.158576275 | 0.070114682 | -2.261670014 | 0.023717799 | 0.105781109 | | Gfpt1 | 3360.481044 | 0.0901984334615474 | 0.061130583770189 | 1.47550420589201 | 0.140077038363269 | 0.585376896927921 | NM_013528 |
| Lta4h | 1574.970836 | 0.15633639 | 0.069129857 | 2.261488711 | 0.023729011 | 0.105800661 | | Ppap2a | 846.6607414 | 0.111965050757118 | 0.0758861281052356 | 1.47543501760756 | 0.140095626710813 | 0.585376896927921 | NM_008903,NM_008247 |
| Id1 | 333.7416681 | -0.190702254 | 0.084343133 | -2.261028813 | 0.023757473 | 0.10589709 | | Rps6ka6 | 173.8396539 | 0.137903096371669 | 0.0934713568479783 | 1.47535139129256 | 0.140118096559162 | 0.585376896927921 | NM_025949 |
| Fbxl7 | 58.89310291 | -0.324580322 | 0.143573168 | -2.260731066 | 0.023775916 | 0.105948816 | | Stk40 | 1722.65826 | 0.120741717397022 | 0.0818238745872154 | 1.47562942974944 | 0.140043400143075 | 0.585376896927921 | NM_028800,NM_001145827 |
| Fgf16 | 22.28780832 | -0.348704573 | 0.154279957 | -2.260206568 | 0.023808433 | 0.106063217 | | Frmd4b | 936.6902775 | -0.104346475298177 | 0.0707397823616767 | -1.47507487038449 | 0.140192415674827 | 0.58551898507548 3 | NM_145148 |
| Rcor3 | 316.9149482 | 0.239877984 | 0.106141597 | 2.259980926 | 0.023822435 | 0.106095086 | | Slc35b4 | 3406.90302 | 0.0694948783414355 | 0.0471226455814949 | 1.47476605958487 | 0.140275449018109 | 0.585529072539381 | NM_021435 |
| Tmem43 | 1551.953568 | 0.184408023 | 0.081606133 | 2.259732385 | 0.023837865 | 0.106133299 | | Zfp689 | 292.5122166 | 0.115726889174973 | 0.0784653702296388 | 1.47487852075742 | 0.140245205971817 | 0.585529072539381 | NM_175163 |
| Atp7b | 87.92969401 | 0.339661473 | 0.150331046 | 2.259423335 | 0.023857064 | 0.106157769 | | Spsb2 | 89.85255695 | 0.139513299531402 | 0.094613048028684 | 1.47456722342467 | 0.140328932292002 | 0.585584047823506 | NM_013539 |
| Tmem106a | 49.46717726 | 0.345014133 | 0.152697029 | 2.259468548 | 0.023854255 | 0.106157769 | | Rbbp4 | 5103.57892 | 0.0754662509719937 | 0.0511952096471044 | 1.47408813231146 | 0.140457863424153 | 0.585785507923988 | NM_009030 |
| Arhgdig | 1680.47015 | -0.213866749 | 0.094668132 | -2.259120839 | 0.023875869 | 0.106210944 | | Rpusd4 | 316.7488129 | -0.108388971220377 | 0.0735272011567565 | -1.47413432736678 | 0.140445427623613 | 0.585785507923988 | NM_028040 |
| Psma2 | 2660.183007 | -0.134226994 | 0.059421723 | -2.25888761 | 0.023890377 | 0.106244977 | | Esd | 2219.540545 | 0.0849334811072095 | 0.057647859493977 | 1.47331543361264 | 0.140666001023218 | 0.586485169593358 | NM_016903 |
| Dctn3 | 2242.12878 | -0.136697296 | 0.06052717 | -2.258445188 | 0.023917918 | 0.106273139 | | 1700008J07Rik | 67.80167708 | -0.137925052811222 | 0.0936538969544723 | -1.47271023733557 | 0.140829184972678 | 0.586773565221472 | NR_024331 |
| Edf1 | 2588.590742 | -0.146276707 | 0.06477479 | -2.258235125 | 0.023931004 | 0.106273139 | | Neb | 68.51955825 | 0.139839394086907 | 0.0949435047962276 | 1.47286951737286 | 0.140786222907077 | 0.586773565221472 | NM_010889 |
| Fez1 | 7849.285884 | -0.103619154 | 0.045882335 | -2.258367063 | 0.023922784 | 0.106273139 | | Tcf3 | 886.1030749 | -0.106355691037418 | 0.0722226017209631 | -1.47260952254712 | 0.140856355633159 | 0.586773565221472 | NM_001164149,NM_001164148,NM_001164147,NM_001164150,NM_001164151,NM_001164152,NM_001164153,NM_011548 |
| Prpf3 | 831.2739396 | 0.186887404 | 0.082753599 | 2.258359832 | 0.023923235 | 0.106273139 | | 2610316D01Rik | 180.5611967 | -0.135928449491006 | 0.0924184017116178 | -1.47079420303282 | 0.141346780648362 | 0.586989325492173 | NR_045172 |
| Snhg7 | 123.5945182 | 0.3058899 | 0.135453124 | 2.258271279 | 0.023928752 | 0.106273139 | | Aamp | 5143.013101 | 0.102720905883355 | 0.0697887016708127 | 1.47188446588218 | 0.141052079119131 | 0.586989325492173 | NM_146110,NM_001190444 |
| C130026L21Rik | 149.7780706 | -0.304425588 | 0.134814901 | -2.258100444 | 0.023939398 | 0.106279952 | | Cep57 | 563.4902292 | -0.0976579121478001 | 0.0663736732427722 | -1.47133505464736 | 0.141200527640615 | 0.586989325492173 | NM_026665,NR_037622 |
| Foxj1 | 252.5669499 | 0.295971783 | 0.131080366 | 2.257941382 | 0.023949314 | 0.106293519 | | Chst8 | 427.7406975 | 0.121617955580974 | 0.0826809882001508 | 1.47093011620236 | 0.141310017088827 | 0.586989325492173 | NM_175140 |
| Kctd8 | 263.4639795 | -0.271206975 | 0.120134991 | -2.257518584 | 0.02397569 | 0.106380107 | | Cux2 | 3935.01911 | -0.120526863951084 | 0.0819066088970243 | -1.47151573693661 | 0.141151694827727 | 0.586989325492173 | NM_007804 |
| Foxc2 | 41.5185798 | -0.379023445 | 0.167956397 | -2.256677633 | 0.024028225 | 0.106582687 | | Gpr19 | 505.7297864 | 0.111023726307281 | 0.0754326844661616 | 1.47182520538674 | 0.141068085272302 | 0.586989325492173 | NM_001167699,NM_001167696,NM_001167697,NM_001167694,NM_001167695,NR_074084,NM_001167700,NM_008157,NR_072990 |
| Plekhg5 | 3272.728174 | 0.209370395 | 0.092815837 | 2.255761521 | 0.02408557 | 0.106774598 | | Mdga2 | 1534.84931 | 0.0939980124412909 | 0.0638771400302616 | 1.47154384803014 | 0.141144098438991 | 0.586989325492173 | NM_001193266,NM_207010 |
| Syndig1l | 2159.093536 | -0.264516952 | 0.117266858 | -2.255683799 | 0.024090441 | 0.106774598 | | Psmc2 | 3938.541794 | 0.0666433443367837 | 0.045311796893246 | 1.47077248986163 | 0.141352654589813 | 0.586989325492173 | NM_011188 |
| Zim1 | 138.8114445 | 0.35781858 | 0.158631694 | 2.255656301 | 0.024092164 | 0.106774598 | | Tmsb4x | 16178.6461 | -0.0940178190168979 | 0.0639009360740036 | -1.47130581793068 | 0.14120843063777 | 0.586989325492173 | NM_021278 |
| Mkks | 1273.617601 | -0.155551291 | 0.068981708 | -2.254964316 | 0.024135569 | 0.10693327 | | Tonsl | 340.7670827 | -0.122751926763276 | 0.0834472742573945 | -1.47101181980667 | 0.141287920373481 | 0.586989325492173 | NM_183091 |
| Zfp235 | 701.6162628 | 0.161423955 | 0.071589171 | 2.254865546 | 0.02414177 | 0.10693327 | | Zfp326 | 902.1285307 | -0.0884352327011606 | 0.060112544725781 | -1.47116102145702 | 0.141247575684231 | 0.586989325492173 | NM_018759,NR_030777 |
| Agap1 | 5826.95639 | -0.140217458 | 0.062246391 | -2.252619889 | 0.024283126 | 0.107467225 | | Socs5 | 5398.655832 | 0.0904272554111439 | 0.0614893221406483 | 1.47061721064844 | 0.141394666865296 | 0.586995979104112 | NM_019654 |
| Hrasls | 1182.565824 | -0.157457203 | 0.069899548 | -2.252621196 | 0.024283044 | 0.107467225 | | Aar2 | 1954.484811 | 0.0845924225479743 | 0.0575419494902307 | 1.4701000452259 | 0.141534660114644 | 0.587141260969906 | NM_001164818,NM_026661 |
| Jmjd4 | 1610.186459 | -0.113463571 | 0.050366794 | -2.252745541 | 0.024275198 | 0.107467225 | | Igf2bp2 | 36.73827311 | -0.133091454896579 | 0.0905359414345681 | -1.47003999503077 | 0.141550922202713 | 0.587141260969906 | NM_183029 |
| Cd274 | 153.6026106 | 0.368607662 | 0.163725804 | 2.251371829 | 0.024361997 | 0.107754719 | | Itfg2 | 806.2369996 | 0.106902818399753 | 0.0727086637524941 | 1.47028996109265 | 0.141483238780734 | 0.587141260969906 | NM_133927 |
| Ddx27 | 1118.556946 | 0.151578927 | 0.067326029 | 2.251416434 | 0.024359174 | 0.107754719 | | Fam192a | 1743.137961 | 0.0716356061069466 | 0.0487450709708862 | 1.46959691882964 | 0.141670955271195 | 0.587317992753072 | NM_028221 |
| R3hdm1 | 10953.51452 | 0.229705108 | 0.102035276 | 2.251232282 | 0.024370829 | 0.107763022 | | Gm4532 | 39.1722044 | -0.133490408762999 | 0.0908354934991751 | -1.46958422991572 | 0.141674393956131 | 0.587317992753072 | NR_030674 |
| Snhg3 | 141.4303023 | 0.339445503 | 0.150795213 | 2.251036322 | 0.024383237 | 0.107787125 | | Cd302 | 504.167 | 0.111197784733548 | 0.0756923482029625 | 1.46907563807349 | 0.141812274715016 | 0.58755422205086 | NM_025422 |
| Gfpt1 | 3102.42671 | -0.12891471 | 0.057275311 | -2.250790223 | 0.024398827 | 0.107805069 | | Dpm2 | 1386.119261 | -0.0835092200153007 | 0.0568390965576518 | -1.46922145271252 | 0.141772733397288 | 0.58755422205086 | NM_010073 |
| Myh2 | 38.87337291 | 0.361700974 | 0.160702236 | 2.250752589 | 0.024401212 | 0.107805069 | | Atg4b | 2778.554339 | 0.0696445327925586 | 0.0474484843892877 | 1.46779256890831 | 0.142160576617873 | 0.587991040988389 | NM_174874 |
| Cacna1c | 2432.963704 | -0.236236264 | 0.104964957 | -2.250620304 | 0.024409596 | 0.10781137 | | Dlst | 5427.784385 | 0.0570839609867012 | 0.0388813519884106 | 1.46815782032776 | 0.142061358662008 | 0.587991040988389 | NM_030225 |
| Hmg20a | 3633.93658 | 0.100931846 | 0.04485336 | 2.250262763 | 0.02443227 | 0.107850027 | | Grin3a | 2264.129697 | 0.119649079281218 | 0.0814865789513961 | 1.46832866001878 | 0.14201496953129 | 0.587991040988389 | NM_001033351,NM_001276355 |
| Rtn2 | 3534.263178 | -0.142948031 | 0.063524293 | -2.250289194 | 0.024430593 | 0.107850027 | | Ppp1r18 | 373.3267814 | -0.107711930815725 | 0.0733768560163292 | -1.4679278544144 | 0.142123821067202 | 0.587991040988389 | NM_175242,NM_001146710,NM_001146711 |
| Best1 | 41.44429403 | 0.366170272 | 0.162735601 | 2.250093213 | 0.024443029 | 0.107866787 | | Sp9 | 170.9697843 | -0.134037426462791 | 0.0912758313847579 | -1.46848759884507 | 0.141971822361768 | 0.587991040988389 | NM_001005343 |
| Erp44 | 1161.354551 | -0.151181503 | 0.067195873 | -2.249862938 | 0.024457647 | 0.10788722 | | Tmem204 | 363.0165172 | 0.119848901123785 | 0.0816382122327072 | 1.46804906484428 | 0.142090895755509 | 0.587991040988389 | NM_001001183 |
| Snrpc | 608.3236628 | -0.156663078 | 0.069634196 | -2.249800924 | 0.024461585 | 0.10788722 | | Lin37 | 737.8399561 | -0.0986413189225787 | 0.0672130816704874 | -1.46759107707884 | 0.142215333213949 | 0.588050079779623 | NM_029377 |
| Heatr6 | 831.4745969 | 0.219784429 | 0.097697698 | 2.24963775 | 0.02447195 | 0.107902218 | | Gpr83 | 1189.988427 | 0.095553561137266 | 0.0651374406673911 | 1.46695295606082 | 0.142388853242382 | 0.588600023391817 | NM_010287 |
| Gdpd5 | 1410.55536 | -0.174961923 | 0.077802348 | -2.24880002 | 0.024525222 | 0.108104516 | | Cplx3 | 925.090099 | 0.120923653850207 | 0.0824482896951608 | 1.46666054926431 | 0.142468419778039 | 0.58869797892616 | NM_146223 |
| Gm7120 | 403.7331441 | 0.356298068 | 0.158454182 | 2.248587346 | 0.024538762 | 0.108104516 | | Cs | 13075.76874 | 0.056965272853946 | 0.0388440530672332 | 1.46651207471444 | 0.142508834111284 | 0.58869797892616 | NM_026444 |
| Trpv4 | 92.3273928 | 0.372154975 | 0.165501429 | 2.248651129 | 0.024534701 | 0.108104516 | | Fem1b | 6906.590098 | 0.101133313791556 | 0.0689661707693414 | 1.46641915396171 | 0.142534131342648 | 0.58869797892616 | NM_010193 |
| Gtf3c2 | 2578.218854 | 0.115607465 | 0.051417213 | 2.248419506 | 0.024549453 | 0.108120869 | | Ccdc141 | 939.3841763 | -0.107920946027639 | 0.0736213971735458 | -1.46589103400523 | 0.142677974961475 | 0.589124577683452 | NM_001025576 |
| Nktr | 4122.652561 | 0.247895388 | 0.110269269 | 2.248091329 | 0.024570367 | 0.108182229 | | C1ql1 | 205.7041733 | 0.133341743451774 | 0.0910065459113819 | 1.46518848564693 | 0.142869499393616 | 0.589284353257248 | NM_011795 |
| Hspb2 | 19.77604066 | 0.377777505 | 0.168072862 | 2.247700794 | 0.024595275 | 0.108261135 | | Dpysl3 | 602.1363725 | 0.120316353297982 | 0.0821352495035716 | 1.46485649005973 | 0.142960075786765 | 0.589284353257248 | NM_001136086,NM_009468 |
| Armc9 | 1077.86333 | 0.185465527 | 0.082529149 | 2.246949329 | 0.024634526 | 0.108421517 | | Idh3b | 8393.420495 | 0.0618228226514906 | 0.0421959889374792 | 1.46513505686742 | 0.142884073741027 | 0.589284353257248 | NM_130884 |
| Morc2a | 2229.523804 | 0.166532527 | 0.074116634 | 2.246898144 | 0.02464637 | 0.10842517 | | Mpzl2 | 62.84164535 | 0.127894967362304 | 0.0873001317978422 | 1.46500314178753 | 0.142920060580328 | 0.589284353257248 | NM_007962 |

| | | | | | | |
|---|---|---|---|---|---|---|
| Clcn1 | 81.47254248 | 0.353925718 | 0.157560637 | 2.246282605 | 0.024685912 | 0.10850482 |
| D230025D16Rik | 1548.202437 | 0.158703384 | 0.070654664 | 2.2461841 | 0.024692218 | 0.10850482 |
| Sema7a | 7349.776995 | 0.232202172 | 0.103362643 | 2.246480601 | 0.024673241 | 0.10850482 |
| Zfp709 | 233.359498 | -0.260119626 | 0.115805475 | -2.246177279 | 0.024692655 | 0.10850482 |
| Fam134b | 1414.722468 | -0.179680923 | 0.080009181 | -2.245753804 | 0.024719782 | 0.108593223 |
| 1110008L16Rik | 243.4329302 | 0.209904995 | 0.093493833 | 2.245121283 | 0.024760348 | 0.108654552 |
| Elof1 | 1079.697887 | -0.15136324 | 0.067418034 | -2.24514467 | 0.024758847 | 0.108654552 |
| Fabp5 | 2531.955827 | -0.165762648 | 0.073833121 | -2.245098761 | 0.024761793 | 0.108654552 |
| Fdxr | 384.3491888 | 0.281316987 | 0.125296716 | 2.24520638 | 0.024754887 | 0.108654552 |
| Mks1 | 187.5287445 | 0.28398343 | 0.126508756 | 2.244772926 | 0.024782714 | 0.108703579 |
| Sema4g | 1196.75279 | 0.21654403 | 0.096468761 | 2.244706229 | 0.024786998 | 0.108703579 |
| Bend4 | 440.3532014 | -0.205521131 | 0.091568169 | -2.244460429 | 0.024802792 | 0.108716533 |
| Pcyt1b | 2515.354689 | -0.120359949 | 0.053625781 | -2.24444187 | 0.024803985 | 0.108716533 |
| Pcdhb7 | 279.2284691 | -0.264177614 | 0.117728546 | -2.243955468 | 0.024835267 | 0.108822857 |
| Azi2 | 2281.498188 | -0.117453903 | 0.052353627 | -2.243472127 | 0.024866385 | 0.108893068 |
| Gfod2 | 709.9540246 | -0.143782371 | 0.064091872 | -2.243379166 | 0.024872374 | 0.108893068 |
| Palm3 | 93.21106216 | -0.309117475 | 0.137777732 | -2.243595323 | 0.02485845 | 0.108893068 |
| Slc19a2 | 586.5521686 | 0.155833552 | 0.069472838 | 2.243086019 | 0.024891268 | 0.108945002 |
| Nnt | 1298.625809 | -0.206297799 | 0.091975259 | -2.242970566 | 0.024898713 | 0.10894681 |
| Trp53inp1 | 428.6488205 | 0.294006576 | 0.131091579 | 2.242757147 | 0.024912479 | 0.108976271 |
| Ppp2ca | 13062.08922 | -0.132143154 | 0.05892439 | -2.242588397 | 0.024923369 | 0.108993136 |
| Fanca | 45.7358904 | 0.344544539 | 0.153655542 | 2.242317685 | 0.024940848 | 0.109038796 |
| Heatr5b | 2264.040468 | -0.168058219 | 0.074952316 | -2.2422018 | 0.024948333 | 0.109040753 |
| Prl | 250.8284412 | 0.181551963 | 0.080983034 | 2.241851831 | 0.02497095 | 0.109108827 |
| Mapre3 | 6850.543587 | 0.096960811 | 0.043252492 | 2.241739314 | 0.024978225 | 0.109109846 |
| Eif2ak1 | 3114.058678 | 0.165566051 | 0.073867728 | 2.241385452 | 0.025001118 | 0.109148302 |
| Hdac3 | 3300.136138 | 0.128777774 | 0.057453682 | 2.241419016 | 0.024998946 | 0.109148302 |
| Cxcl12 | 2722.267082 | -0.364751399 | 0.162787794 | -2.240655701 | 0.025048386 | 0.109318646 |
| Sdc2 | 1910.222066 | -0.171081886 | 0.076356573 | -2.240565282 | 0.025054248 | 0.109318646 |
| Tcirg1 | 552.6248563 | 0.218779556 | 0.097667239 | 2.240050597 | 0.025087638 | 0.109402721 |
| Tmem68 | 1112.736729 | -0.217179393 | 0.096950545 | -2.240104925 | 0.025084112 | 0.109402721 |
| Plat | 1229.226899 | 0.305953387 | 0.136619033 | 2.239463851 | 0.025125751 | 0.109538093 |
| Nupr1 | 291.2407848 | 0.329040364 | 0.147013847 | 2.238158991 | 0.025210689 | 0.109877472 |
| Mark1 | 4279.98538 | -0.102047444 | 0.045597269 | -2.238016581 | 0.025219974 | 0.109887029 |
| Crhbp | 655.8090608 | 0.214743182 | 0.095958698 | 2.237870948 | 0.025229472 | 0.109897333 |
| Pip5kl1 | 188.6627441 | 0.342559891 | 0.15308141 | 2.237762846 | 0.025236525 | 0.109897333 |
| 4933408B17Rik | 25.61998967 | 0.376146819 | 0.168112913 | 2.237465355 | 0.025255941 | 0.109920099 |
| Rap2a | 5088.675245 | -0.171266981 | 0.076544248 | -2.237489887 | 0.02525434 | 0.109920099 |
| 4933406I18Rik | 15.8133254 | 0.375035024 | 0.167638497 | 2.237165272 | 0.02527554 | 0.109974507 |
| Gpc2 | 93.59860485 | -0.310421669 | 0.138833598 | -2.235926125 | 0.025356611 | 0.110296274 |
| Hbs1l | 2076.925141 | 0.121395091 | 0.0543149 | 2.23502375 | 0.02541579 | 0.110305951 |
| Ino80 | 1455.924769 | 0.144627437 | 0.064688895 | 2.235738239 | 0.025368923 | 0.110305951 |
| Mesdc1 | 1079.453052 | -0.22203143 | 0.099341433 | -2.235033492 | 0.02541515 | 0.110305951 |
| Pola1 | 178.6940085 | -0.223570684 | 0.100017216 | -2.235322002 | 0.025396217 | 0.110305951 |
| Tbk1 | 1929.698046 | 0.149591805 | 0.06691217 | 2.235644206 | 0.025375086 | 0.110305951 |
| Ticrr | 10.09579708 | -0.332275649 | 0.148639111 | -2.23545234 | 0.025387667 | 0.110305951 |
| Tmbim1 | 1423.449509 | 0.188642871 | 0.084394358 | 2.23525452 | 0.025400644 | 0.110305951 |
| Zfp90 | 491.7518715 | 0.188836482 | 0.084483367 | 2.235191228 | 0.025404797 | 0.110305951 |
| Clmn | 4242.048565 | 0.272355866 | 0.121893879 | 2.23436868 | 0.025458825 | 0.110461784 |
| Ablim1 | 6036.168743 | 0.107970883 | 0.048328802 | 2.234089769 | 0.025477167 | 0.110510422 |
| Cacfd1 | 1335.107569 | 0.13518056 | 0.060536811 | 2.233030765 | 0.025546916 | 0.110781951 |
| Zdhhc8 | 4860.566398 | 0.181615281 | 0.081338834 | 2.23282376 | 0.025560569 | 0.110810144 |
| Nemf | 1714.650708 | 0.150074619 | 0.067219819 | 2.232594829 | 0.025575675 | 0.11084462 |
| Cct6a | 5100.296493 | -0.101664453 | 0.04554255 | -2.232296004 | 0.025595406 | 0.11089911 |
| Sap30 | 122.1509836 | 0.346447399 | 0.155207329 | 2.232158759 | 0.025604472 | 0.110907378 |
| Ndufaf5 | 990.5174572 | -0.151513775 | 0.067881146 | -2.232045046 | 0.025611986 | 0.110908918 |
| Pcdhgc3 | 2693.8457 | -0.130921093 | 0.058675131 | -2.231287612 | 0.025662084 | 0.111084152 |
| Pkd1l3 | 54.05133579 | 0.316619364 | 0.141904367 | 2.231216503 | 0.025666791 | 0.111084152 |
| Manf | 1419.335301 | -0.211247705 | 0.094685529 | -2.231045293 | 0.025678129 | 0.111102187 |
| Igfbp2 | 1184.645477 | -0.224949541 | 0.100844007 | -2.230668406 | 0.025703102 | 0.111179191 |
| Fnbp1 | 4949.954058 | 0.186608782 | 0.083666244 | 2.230395122 | 0.025721223 | 0.111226523 |
| Anapc16 | 1500.402122 | -0.136250073 | 0.061145443 | -2.22829481 | 0.025860862 | 0.111705659 |
| Nup35 | 550.8978395 | -0.175379807 | 0.078705622 | -2.228300873 | 0.025860458 | 0.111705659 |
| Stk11ip | 1685.380121 | 0.146529722 | 0.06575817 | 2.228312043 | 0.025859714 | 0.111705659 |
| Zmym6 | 1472.148053 | 0.27296641 | 0.122483781 | 2.228592288 | 0.025841045 | 0.111705659 |
| Aff4 | 9631.926052 | -0.173643615 | 0.07793381 | -2.228090941 | 0.025874451 | 0.111708246 |
| Emx2os | 132.7881613 | -0.335447561 | 0.15055525 | -2.228069503 | 0.025875881 | 0.111708246 |
| Cpxm1 | 269.5940423 | 0.248384144 | 0.111519134 | 2.227278278 | 0.025928681 | 0.111905011 |
| Tfr2 | 193.9693938 | 0.298066561 | 0.133848209 | 2.226899877 | 0.025953966 | 0.111982944 |
| Snrnp27 | 1025.554411 | -0.143989029 | 0.0646688 | -2.226561024 | 0.025976626 | 0.112049512 |
| Gcnt4 | 608.7428908 | 0.2266443 | 0.10183281 | 2.225651048 | 0.026037564 | 0.112249866 |
| Hcst | 11.72193882 | 0.365057011 | 0.164016208 | 2.225737417 | 0.026031775 | 0.112249866 |
| Gldn | 68.72771525 | 0.34601944 | 0.15548218 | 2.225460438 | 0.026050344 | 0.112273722 |
| Dlec1 | 101.2776592 | 0.353951356 | 0.159070337 | 2.225124827 | 0.02607286 | 0.112277067 |
| Trim7 | 49.80621069 | 0.340425498 | 0.152989635 | 2.225153993 | 0.026070902 | 0.112277067 |
| Tulp3 | 790.6567406 | 0.187642751 | 0.084322636 | 2.225295122 | 0.026061433 | 0.112277067 |
| Atg16l1 | 2597.244093 | 0.126692179 | 0.056973266 | 2.223712773 | 0.026167775 | 0.11265449 |
| Ccdc17 | 80.61054843 | 0.300390948 | 0.135111718 | 2.223278287 | 0.026197041 | 0.112749152 |
| Ptges | 198.4160614 | -0.268708864 | 0.120880402 | -2.222931588 | 0.026220413 | 0.112818407 |
| Cd44 | 141.6867841 | 0.292620573 | 0.131670513 | 2.22236981 | 0.026258324 | 0.112918809 |
| Pam16 | 631.2810988 | -0.155790479 | 0.070097818 | -2.22247258 | 0.026251385 | 0.112918809 |
| Ccdc157 | 1355.056535 | 0.161451596 | 0.072682685 | 2.221321284 | 0.026329208 | 0.113124374 |
| Fam50a | 804.5235151 | 0.197980397 | 0.089130942 | 2.221230832 | 0.026335331 | 0.113124374 |
| Rps27l | 242.6100206 | -0.203622587 | 0.091658639 | -2.221531865 | 0.026314959 | 0.113124374 |
| Scd3 | 191.5562099 | 0.318535404 | 0.143398567 | 2.22132906 | 0.026328682 | 0.113124374 |
| BC037034 | 2044.917709 | 0.175448672 | 0.078993158 | 2.221061626 | 0.026346788 | 0.11314222 |
| Mov10 | 344.0133928 | 0.248015679 | 0.111689924 | 2.220573434 | 0.026379867 | 0.113252883 |
| Heatr2 | 288.3010222 | 0.20929798 | 0.094268439 | 2.220233866 | 0.026402897 | 0.113320354 |
| Lmbr1l | 316.1864538 | 0.240556361 | 0.108368893 | 2.219791616 | 0.026432916 | 0.113328907 |
| Neurod4 | 13.00091171 | 0.356342214 | 0.16051976 | 2.219927409 | 0.026423696 | 0.113328907 |
| Rbm3 | 2276.092606 | 0.362637818 | 0.163355859 | 2.219925384 | 0.026423833 | 0.113328907 |
| Zfp458 | 161.4387853 | -0.285936566 | 0.128813397 | -2.219773497 | 0.026434147 | 0.113328907 |
| Egflam | 178.901555 | -0.314138209 | 0.141546881 | -2.219322716 | 0.026464777 | 0.11339998 |
| Nup133 | 1622.495315 | 0.137090064 | 0.061771366 | 2.219314106 | 0.026465363 | 0.11339998 |
| Ube2l3 | 6215.829755 | 0.089964861 | 0.040542226 | 2.219040963 | 0.026483938 | 0.113448198 |
| S100a5 | 54.18756256 | -0.1853579 | 0.083554757 | -2.218400318 | 0.026527549 | 0.113572216 |
| Smim12 | 1177.075915 | -0.131427795 | 0.059244238 | -2.218406368 | 0.026527137 | 0.113572216 |
| Col4a3 | 23.88964803 | 0.37118997 | 0.167432038 | 2.216959041 | 0.026625891 | 0.113961754 |
| Cass4 | 24.06769474 | -0.372553205 | 0.168060494 | -2.216780378 | 0.026638103 | 0.113978519 |
| Slc25a11 | 3768.288145 | -0.099276632 | 0.044786049 | -2.216686525 | 0.02664452 | 0.113978519 |
| Ptp4a3 | 2140.510833 | -0.19988915 | 0.090192397 | -2.216252781 | 0.026674195 | 0.114073965 |
| Abcd2 | 1977.864815 | 0.178048968 | 0.080422716 | 2.213913886 | 0.026834703 | 0.114697968 |
| Elf5 | 15.80981951 | -0.366330138 | 0.16546743 | -2.213910846 | 0.026834913 | 0.114697968 |
| Cdk9 | 3174.136287 | 0.155866472 | 0.070408154 | 2.213755993 | 0.026845569 | 0.114711871 |
| Banf1 | 1171.790933 | -0.163946046 | 0.074064013 | -2.213572269 | 0.026858217 | 0.114712817 |
| Sdc1 | 43.28793774 | -0.358804079 | 0.16209531 | -2.213537693 | 0.026860598 | 0.114712817 |
| Asnsd1 | 732.5874788 | -0.18276482 | 0.082627522 | -2.21191216 | 0.026972735 | 0.115128252 |
| Jph4 | 14605.33135 | -0.194550381 | 0.087955619 | -2.211915323 | 0.026972516 | 0.115128252 |
| Tmem159 | 396.377332 | -0.201170883 | 0.09096012 | -2.211638283 | 0.026991668 | 0.115177335 |
| Mrpl46 | 697.3331217 | -0.138092039 | 0.06244189 | -2.21152882 | 0.026999238 | 0.115177918 |
| Tmem120b | 253.2755958 | 0.243852919 | 0.110297793 | 2.210859457 | 0.027045571 | 0.115343814 |
| Tacc2 | 3163.571524 | 0.163623244 | 0.074025472 | 2.210364065 | 0.027079906 | 0.115458464 |
| Tm2d2 | 1815.132718 | -0.118269979 | 0.053537462 | -2.209106957 | 0.027167202 | 0.115798799 |
| Pvrl3 | 630.7527255 | -0.260984263 | 0.118176949 | -2.208419358 | 0.027215054 | 0.115970858 |
| Star | 55.14597684 | 0.349944297 | 0.158470418 | 2.208262597 | 0.027225973 | 0.11598549 |
| 1700030J22Rik | 363.8494526 | -0.235212454 | 0.106539019 | -2.207758774 | 0.027261093 | 0.11600752 |
| Acer2 | 316.3749469 | 0.354407881 | 0.160511608 | 2.207989096 | 0.027245033 | 0.11600752 |
| Arvcf | 1406.994098 | 0.174154163 | 0.078877689 | 2.207901433 | 0.027251145 | 0.11600752 |
| C920021L13Rik | 20.7258669 | 0.370816472 | 0.167959409 | 2.207774338 | 0.027260008 | 0.11600752 |
| Col4a6 | 78.52534686 | -0.306122605 | 0.138677029 | -2.207449975 | 0.027282638 | 0.116022932 |
| Lgr4 | 1184.208919 | -0.161221743 | 0.07303744 | -2.207384908 | 0.02728718 | 0.116022932 |
| Rasgrp2 | 1904.708393 | -0.211117764 | 0.095639846 | -2.207424758 | 0.027284398 | 0.116022932 |
| Cftr | 63.01583471 | -0.325965839 | 0.147686902 | -2.207141147 | 0.0273042 | 0.116063449 |
| Itgal | 39.88549724 | 0.35198668 | 0.159490515 | 2.206944278 | 0.027317952 | 0.116090059 |
| Pura | 635.6300238 | -0.240533321 | 0.109027408 | -2.206172977 | 0.027371891 | 0.116266709 |
| Spop | 8264.730782 | -0.094208364 | 0.042702895 | -2.206135281 | 0.027374529 | 0.116266709 |
| Arhgap39 | 2893.848842 | 0.243804874 | 0.110577979 | 2.204823035 | 0.027466515 | 0.116615879 |
| Atp5f1 | 7229.764798 | -0.134479104 | 0.060995231 | -2.204747858 | 0.027471793 | 0.116615879 |
| Tmem170 | 105.3249497 | 0.319074137 | 0.144737442 | 2.204503087 | 0.027488983 | 0.11665689 |
| Aldh3b2 | 65.68496793 | -0.339844636 | 0.154174978 | -2.204278798 | 0.027504743 | 0.11669181 |
| Zfp691 | 190.2602248 | 0.221099462 | 0.10031452 | 2.204062401 | 0.027519956 | 0.11672439 |
| Mcfd2 | 2533.199159 | -0.129749982 | 0.058890908 | -2.203226029 | 0.027578822 | 0.116942053 |
| Trim11 | 745.8333968 | 0.176780221 | 0.080242685 | 2.203069626 | 0.027589842 | 0.116956773 |
| Dbc1 | 9205.968381 | 0.179412697 | 0.081470124 | 2.202190065 | 0.027651886 | 0.117179083 |
| Fam163a | 331.0198716 | -0.299460911 | 0.135988058 | -2.202111835 | 0.02765741 | 0.117179083 |
| Arsg | 1055.389691 | 0.295773048 | 0.134324704 | 2.201925933 | 0.027670541 | 0.117202668 |
| Lpin1 | 1310.382718 | 0.202410454 | 0.091931818 | 2.201745348 | 0.027683302 | 0.117224672 |
| Snx8 | 474.7296435 | 0.196230035 | 0.089181441 | 2.200346088 | 0.02778235 | 0.117611947 |
| Sars2 | 386.9580589 | 0.209592189 | 0.095279563 | 2.199760182 | 0.027823914 | 0.117691436 |
| Tuba1b | 9316.397266 | -0.197378638 | 0.089726167 | -2.199789034 | 0.027821866 | 0.117691436 |
| Zfp955a | 430.4128816 | -0.225461702 | 0.102489017 | -2.19986209 | 0.027816681 | 0.117691436 |
| Kcnj8 | 146.1291788 | -0.275365645 | 0.125232901 | -2.198828281 | 0.027890135 | 0.117939342 |
| Cobl | 5837.614543 | -0.215118876 | 0.097839358 | -2.198694689 | 0.027899639 | 0.117947341 |
| Ankrd17 | 6708.384731 | -0.168581064 | 0.076689881 | -2.198217837 | 0.027933586 | 0.118058643 |
| Slc25a1 | 1224.820941 | -0.198390644 | 0.090271031 | -2.197722149 | 0.027968912 | 0.118143491 |
| Tpp1 | 2942.456635 | 0.169895307 | 0.077302525 | 2.197797636 | 0.02796353 | 0.118143491 |
| Hdhd2 | 2718.824976 | -0.120245941 | 0.054723259 | -2.197346112 | 0.027995737 | 0.118192356 |
| Ypel2 | 2401.412933 | -0.18490087 | 0.084145606 | -2.197391852 | 0.027992473 | 0.118192356 |
| Sumo1 | 2861.623442 | -0.133617317 | 0.060818573 | -2.196982105 | 0.028021724 | 0.118269843 |
| Adssl1 | 796.5715147 | 0.227999521 | 0.103786935 | 2.196803679 | 0.02803447 | 0.118291417 |
| Cradd | 174.6535522 | -0.218811317 | 0.09962127 | -2.19643171 | 0.028061058 | 0.118339149 |
| Etv6 | 940.3203609 | -0.180605839 | 0.082223145 | -2.196532845 | 0.028053827 | 0.118339149 |
| Bcl2a1b | 26.19112841 | 0.366925932 | 0.167095648 | 2.195903582 | 0.028098845 | 0.118434034 |
| Pcdhga4 | 489.5401906 | 0.175197408 | 0.079781162 | 2.195974624 | 0.02809376 | 0.118434034 |
| Fam107b | 308.5769769 | -0.196917437 | 0.089690717 | -2.195516355 | 0.028126579 | 0.118518688 |
| Fam136a | 755.3807347 | -0.246695492 | 0.112442512 | -2.193969942 | 0.028237571 | 0.118942553 |
| Gtf2a2 | 515.160977 | -0.216575347 | 0.098717007 | -2.193901063 | 0.028242523 | 0.118942553 |
| Rbm26 | 1372.176365 | 0.140448401 | 0.064021074 | 2.193783889 | 0.02825095 | 0.11894571 |
| Nomo1 | 5839.617597 | -0.134350523 | 0.061252065 | -2.193403975 | 0.028278286 | 0.119028461 |
| Srsf1 | 4473.441582 | 0.170876057 | 0.077924142 | 2.19285132 | 0.028318093 | 0.119163643 |
| 4930581F22Rik | 90.18582821 | 0.292799766 | 0.13355759 | 2.19231094 | 0.028357062 | 0.119232798 |
| Napa | 9502.803461 | 0.150302855 | 0.068556873 | 2.192381998 | 0.028351935 | 0.119232798 |
| Pcdhga11 | 851.8229789 | -0.163819488 | 0.074724829 | -2.192303303 | 0.028357613 | 0.119232798 |
| Angptl7 | 10.84652463 | -0.342442467 | 0.156287964 | -2.191099417 | 0.028444599 | 0.119533663 |
| Iqsec1 | 13203.63182 | 0.141324368 | 0.064496855 | 2.191182315 | 0.028438602 | 0.119533663 |
| Trem2 | 476.4180915 | 0.325817019 | 0.148724926 | 2.190735791 | 0.028470917 | 0.119579397 |
| Upf3a | 1063.602098 | 0.128938115 | 0.058854048 | 2.190811349 | 0.028465447 | 0.119579397 |
| Cct3 | 4289.575189 | -0.112598699 | 0.051400846 | -2.190600115 | 0.028480743 | 0.119588246 |
| Zfp623 | 456.5720743 | 0.188231346 | 0.085940735 | 2.190245933 | 0.028506406 | 0.119663573 |
| Csmd1 | 3013.876959 | -0.165226144 | 0.075464325 | -2.189460306 | 0.028563401 | 0.119870349 |
| 1810055G02Rik | 691.3240904 | 0.176695464 | 0.080720275 | 2.188984924 | 0.028597936 | 0.119878408 |
| Malt1 | 273.8656572 | 0.282831435 | 0.129189224 | 2.189280391 | 0.028576467 | 0.119878408 |
| Pld4 | 421.1596779 | 0.28663865 | 0.130932695 | 2.189206057 | 0.028581867 | 0.119878408 |
| Rpe | 632.136526 | -0.223396439 | 0.102058703 | -2.188901414 | 0.028604006 | 0.119878408 |
| Scrg1 | 705.6959461 | 0.261889422 | 0.119642233 | 2.188937941 | 0.028601351 | 0.119878408 |
| Zrsr1 | 3262.177632 | 0.179349552 | 0.081940475 | 2.188778518 | 0.028612942 | 0.119883429 |
| Ccdc171 | 128.6605904 | 0.324776658 | 0.148432207 | 2.188047091 | 0.028666172 | 0.119981547 |
| Liph | 52.36618567 | 0.3420732 | 0.156328277 | 2.188172268 | 0.028657056 | 0.119981547 |
| Pdss1 | 185.5036119 | 0.275411798 | 0.125865161 | 2.188149568 | 0.028658709 | 0.119981547 |
| Yeats4 | 1002.570626 | -0.14457933 | 0.066077364 | -2.18803113 | 0.028667335 | 0.119981547 |
| Gm694 | 67.13368306 | -0.350053441 | 0.160035401 | -2.187350044 | 0.02871698 | 0.120059603 |
| Gpx4 | 8168.969268 | -0.167100753 | 0.076390596 | -2.187451892 | 0.028709552 | 0.120059603 |
| Slc15a3 | 103.5657395 | 0.332718787 | 0.152095787 | 2.18756084 | 0.028701607 | 0.120059603 |
| Zfp7 | 301.603456 | 0.264722707 | 0.121023097 | 2.18737343 | 0.028715274 | 0.120059603 |
| Gm15446 | 158.1206719 | 0.298393316 | 0.13642733 | 2.187196032 | 0.028728216 | 0.12007418 |
| Hps4 | 709.2835555 | 0.246830498 | 0.112862834 | 2.186995404 | 0.02874286 | 0.120102984 |
| Arhgef17 | 10567.95701 | 0.199501673 | 0.091234489 | 2.186691403 | 0.02876506 | 0.120144436 |
| Dcp2 | 3426.32966 | -0.141981504 | 0.064937446 | -2.186434982 | 0.028783797 | 0.120144436 |
| Idua | 712.229041 | 0.285695428 | 0.13066562 | 2.186462118 | 0.028781813 | 0.120144436 |
| Srsf6 | 3354.075592 | 0.200198755 | 0.091559821 | 2.186535003 | 0.028776487 | 0.120144436 |
| Mndal | 32.59179133 | 0.364191339 | 0.166597164 | 2.18605966 | 0.028811241 | 0.1202266 |
| Gucy2g | 321.9000507 | -0.291342177 | 0.133296228 | -2.185674568 | 0.028839423 | 0.120311798 |
| Sybu | 3144.56938 | -0.120873286 | 0.055311866 | -2.185304803 | 0.028866505 | 0.120392365 |
| Slc9a3r2 | 1420.823702 | -0.161487933 | 0.073907991 | -2.184986086 | 0.028889867 | 0.120457373 |
| Mrpl1 | 514.1898107 | -0.205601091 | 0.094104298 | -2.184821476 | 0.028901939 | 0.120475286 |
| 4932418E24Rik | 126.9537062 | -0.339092265 | 0.155274562 | -2.183823678 | 0.028975207 | 0.120734838 |
| Glis3 | 114.4998254 | 0.282770506 | 0.129487825 | 2.183761339 | 0.02897979 | 0.120734838 |
| Baalc | 9374.951189 | -0.134428225 | 0.061576897 | -2.183095148 | 0.029028804 | 0.120809102 |
| Hid1 | 4494.510704 | 0.139152894 | 0.063738422 | 2.183186985 | 0.029022043 | 0.120809102 |
| Vax1 | 17.44627343 | -0.36567689 | 0.167503196 | -2.183103961 | 0.029028155 | 0.120809102 |
| Zfhx2as | 207.2041324 | 0.35307947 | 0.161730916 | 2.183129111 | 0.029026303 | 0.120809102 |
| Otud7b | 2653.255313 | 0.272499779 | 0.124851124 | 2.182597717 | 0.029065448 | 0.120929122 |
| Dip2c | 4231.51781 | -0.170467156 | 0.078116531 | -2.182216143 | 0.029093584 | 0.120956391 |
| Gm11696 | 81.12209881 | 0.335325913 | 0.153669967 | 2.182117434 | 0.029100867 | 0.120956391 |
| Gramd1a | 3519.014238 | 0.172498597 | 0.07904937 | 2.182162831 | 0.029097517 | 0.120956391 |
| Rpl31 | 6469.000867 | -0.156567842 | 0.071751472 | -2.182085424 | 0.029103229 | 0.120956391 |
| Lpar5 | 13.08798549 | 0.360832445 | 0.165375952 | 2.181891871 | 0.029117514 | 0.120983309 |
| Trhr2 | 139.3885761 | 0.314770827 | 0.144331328 | 2.180890531 | 0.029191514 | 0.121258263 |
| Armc2 | 301.0133433 | 0.259087673 | 0.118822774 | 2.180454671 | 0.029223775 | 0.121359736 |
| Ttpal | 3341.621795 | -0.097990887 | 0.044945792 | -2.180201564 | 0.029242524 | 0.121405054 |
| Bach2 | 436.240805 | -0.276821559 | 0.1269851 | -2.179953079 | 0.02926094 | 0.121448969 |
| Stmn3 | 12242.57949 | -0.165323721 | 0.075861906 | -2.179271922 | 0.029311474 | 0.121626131 |
| Mdga1 | 1165.826457 | -0.322710677 | 0.148100213 | -2.179002109 | 0.029331512 | 0.12164478 |
| Mtf2 | 1270.930077 | 0.189255971 | 0.086854508 | 2.178999975 | 0.02933167 | 0.12164478 |
| Eif3i | 2554.72046 | -0.141959719 | 0.065161779 | -2.178573389 | 0.029363375 | 0.121743679 |
| S100a8 | 14.66361853 | -0.329496199 | 0.151277155 | -2.17809622 | 0.029398874 | 0.121858254 |
| Cpsf7 | 3436.261118 | 0.129095883 | 0.059289211 | 2.177392482 | 0.029451296 | 0.121935044 |
| Diablo | 895.0813416 | 0.15706399 | 0.072131155 | 2.177477813 | 0.029444935 | 0.121935044 |
| Mier3 | 698.1506846 | 0.147371713 | 0.067684932 | 2.177319343 | 0.029456749 | 0.121935044 |
| Peg13 | 15878.86871 | 0.156370879 | 0.071812259 | 2.177495619 | 0.029443608 | 0.121935044 |
| Tnnt2 | 247.4143422 | 0.261053163 | 0.119874508 | 2.177720406 | 0.029426859 | 0.121935044 |
| Mapkap1 | 1939.106199 | -0.112149329 | 0.051515321 | -2.17700922 | 0.02947988 | 0.121998198 |
| Lmtk3 | 8240.308986 | 0.194396014 | 0.08930318 | 2.176809534 | 0.029494781 | 0.122027275 |
| Gfm2 | 972.5204655 | 0.143153271 | 0.065772347 | 2.176496298 | 0.02951817 | 0.122064687 |
| Tubgcp6 | 2147.810617 | 0.195804022 | 0.089963728 | 2.176477414 | 0.029519581 | 0.122064687 |
| Naip2 | 70.40954443 | 0.341650077 | 0.156998999 | 2.176129015 | 0.029545615 | 0.122139743 |
| Smad6 | 60.95485357 | -0.312260336 | 0.143523692 | -2.175671007 | 0.02957987 | 0.122248733 |
| Tifa | 245.0144317 | 0.272155357 | 0.125102467 | 2.175459562 | 0.029595695 | 0.122281521 |
| 1700013F07Rik | 35.26067892 | 0.341339414 | 0.15694523 | 2.174895116 | 0.029637977 | 0.122423571 |
| 3110001D03Rik | 611.0541901 | -0.164105893 | 0.075487308 | -2.173953443 | 0.029708632 | 0.122552061 |
| Ngp | 19.97648518 | -0.350371588 | 0.161166251 | -2.173976164 | 0.029706925 | 0.122552061 |
| Nlrp6 | 43.20504359 | 0.363457951 | 0.167162711 | 2.174276474 | 0.029684378 | 0.122552061 |
| Pip4k2b | 7983.327125 | -0.123897001 | 0.056989656 | -2.174026141 | 0.029703172 | 0.122552061 |
| Tgm7 | 9.266306776 | 0.330495245 | 0.1520202 | 2.174021912 | 0.02970349 | 0.122552061 |
| Sugp2 | 3683.208703 | 0.134519115 | 0.06188212 | 2.173796165 | 0.029720447 | 0.122568166 |

| | | | | | | | |
|---|---|---|---|---|---|---|---|
| Pik3cd | 1269.98839 | -0.0997518967498 | 0.0680680802196928 | -1.46547245680863 | 0.142792061640235 | 0.589284353257248 | NM_001164049,NM_008840,NM_001164051,NM_001164050,NM_001164052,NM_001029837 |
| Susd3 | 50.77266966 | -0.138404263603711 | 0.0944771030473375 | -1.46495033335605 | 0.142934468832239 | 0.589284353257248 | NM_025491,NM_028340 |
| Lemd1 | 40.03279641 | 0.132577125155598 | 0.0905215225771839 | 1.46459230226221 | 0.143032183432116 | 0.589285583787395 | NM_001033250 |
| Xpr1 | 6590.654136 | 0.0781808615975039 | 0.0533818763716538 | 1.46455814054184 | 0.14304150959895 | 0.589285583787395 | NM_011273 |
| Rnf139 | 1283.466406 | 0.107824567209202 | 0.0736335057971359 | 1.46434107736584 | 0.14310077885749 | 0.589362606658321 | NM_175226 |
| Mak | 161.9345077 | -0.127709227746253 | 0.0872278829347628 | -1.46408720984053 | 0.143170121475684 | 0.589481061382025 | NM_001145803,NM_001145802,NM_008547 |
| 4930563E22Rik | 116.1757101 | -0.134695473772399 | 0.0920406303824803 | -1.46343493316662 | 0.143348405736754 | 0.589907431747263 | NM_001163728 |
| Efr3a | 6897.875722 | 0.100298958337088 | 0.0685522522924632 | 1.46310230492769 | 0.143439387273552 | 0.589907431747263 | NM_133766 |
| Gga3 | 4089.798059 | -0.0898550589469984 | 0.0614325911356895 | -1.46266106126845 | 0.143560145991154 | 0.589907431747263 | NM_173048,NM_001252067 |
| Glul | 69300.96669 | 0.0907479918201273 | 0.0620572540438209 | 1.46232689825507 | 0.143651650959546 | 0.589907431747263 | NM_008131 |
| Immt | 5781.222406 | 0.0787825224464764 | 0.0538667168316578 | 1.462545465555 | 0.143591794890418 | 0.589907431747263 | NM_029673,NM_001253689,NM_001253688,NM_001253681,NM_001253687,NM_001253686 |
| Kctd18 | 414.5962677 | -0.102419939247748 | 0.070027565604196 | -1.46256603901723 | 0.143586161699866 | 0.589907431747263 | NR_027630,NM_001159864,NM_001159865 |
| Mrps21 | 985.4771378 | 0.0935053998461249 | 0.0639046419381882 | 1.46320199926272 | 0.14341211391379 | 0.589907431747263 | NM_078479 |
| Mta2 | 2255.724013 | -0.0782108205755982 | 0.0534816105930834 | -1.46238715903056 | 0.143635146245981 | 0.589907431747263 | NM_011842 |
| Naalad2 | 76.04894275 | -0.138748129409857 | 0.0948928560870397 | -1.46215568938711 | 0.14369855097989 | 0.589907431747263 | NM_028279 |
| Spata13 | 857.7231946 | -0.123180644305085 | 0.0842505083744471 | -1.46207597653435 | 0.143720391088638 | 0.589907431747263 | NM_001033272 |
| Txnl4b | 732.4136241 | 0.0956251120931609 | 0.0653721744785231 | 1.46278004144679 | 0.143527576048681 | 0.589907431747263 | NM_175646 |
| Acvr1 | 1165.714448 | -0.0812452175273441 | 0.0556697662600901 | -1.45941366356318 | 0.144451287564274 | 0.589967031603128 | NM_007394,NM_001110205,NM_001110204 |
| Arhgap36 | 219.7885606 | 0.0796412038105646 | 0.0544927095311478 | 1.46150199716243 | 0.143877727868123 | 0.589967031603128 | NM_001081123 |
| Cdh22 | 1770.680554 | 0.108560190560682 | 0.0743768517261375 | 1.45959647445701 | 0.144401008771309 | 0.589967031603128 | NM_174988 |
| Cerk | 3846.688005 | 0.077538545948696 | 0.0530447399025889 | 1.46175749171525 | 0.143807676487251 | 0.589967031603128 | NM_145475 |
| Cmpk1 | 1876.177159 | 0.100385954922805 | 0.0687424210942536 | 1.46032032804262 | 0.144202057774444 | 0.589967031603128 | NM_025647 |
| Ints6 | 898.4791176 | -0.104118112443297 | 0.0712715899969469 | -1.46086417389814 | 0.144052720153891 | 0.589967031603128 | NM_008715 |
| Lrrc59 | 4408.116712 | 0.0711529934896301 | 0.0487493591096739 | 1.45956777256402 | 0.144408901814306 | 0.589967031603128 | NM_133807 |
| March4 | 1122.249799 | 0.0832295219368205 | 0.057027298926065 | 1.45946807062923 | 0.14443632249364 | 0.589967031603128 | NM_001045533 |
| Myadm | 2770.564171 | 0.0803825007783447 | 0.055041901484155 | 1.4603874250508 | 0.144183626823268 | 0.589967031603128 | NM_001093766,NM_016969,NM_001093764,NM_001093765 |
| Pcdh10 | 7219.670581 | 0.0798918658419144 | 0.0546931185577447 | 1.46072975812423 | 0.144089619083651 | 0.589967031603128 | NM_001098172,NM_001098170,NM_001098171,NM_011043 |
| Pnma2 | 3109.840353 | 0.0847391704247973 | 0.0580568139393729 | 1.45959043693455 | 0.14440266906724 | 0.589967031603128 | NM_175498 |
| Ppfibp2 | 502.9475231 | -0.120901614043961 | 0.0827793197962605 | -1.46052920393075 | 0.144144687365906 | 0.589967031603128 | NM_001163557,NM_008905 |
| Prss22 | 46.28756175 | -0.133836030741287 | 0.0916077383602693 | -1.46096861615495 | 0.144024054357647 | 0.589967031603128 | NM_133731 |
| Rep15 | 22.65732805 | -0.122611130079347 | 0.0840252767590027 | -1.45921721187559 | 0.14450533296309 | 0.589967031603128 | NM_025620 |
| Sec1 | 75.34318251 | -0.132146210884027 | 0.0905098355425916 | -1.46002045072596 | 0.144284453479721 | 0.589967031603128 | NM_001271578 |
| Sf3b2 | 10948.94095 | 0.0633237068370219 | 0.0433588746432017 | 1.46045549747566 | 0.144164929779291 | 0.589967031603128 | NM_030109 |
| Tars2 | 1321.514692 | -0.0872629013670721 | 0.0597250284471476 | -1.46107760240405 | 0.143994146055543 | 0.589967031603128 | NM_001163618,NM_001163619,NM_027931,NM_001163617 |
| Tbx15 | 42.44829937 | -0.126922690942127 | 0.0868418990764443 | -1.46153748699577 | 0.14386799571713 | 0.589967031603128 | NM_009323 |
| Vgll3 | 84.78048203 | 0.119170505540017 | 0.0816676757274971 | 1.45921265027374 | 0.144506588079577 | 0.589967031603128 | NM_028572 |
| Dmrta2 | 35.9385545 | -0.125226070191906 | 0.0858556057329348 | -1.45856603215215 | 0.144684588427144 | 0.590269908454338 | NM_172296 |
| Gpr3 | 313.7306394 | 0.124855617781751 | 0.0856401133238209 | 1.45791046900707 | 0.144865222610472 | 0.590269908454338 | NM_008154 |
| Ildr2 | 7701.035073 | 0.0890576492909404 | 0.0610618849661571 | 1.45848182283105 | 0.144707781838412 | 0.590269908454338 | NM_001164528 |
| Parp2 | 663.4165181 | -0.107657780181622 | 0.073838423123777 | -1.45801840866987 | 0.144835468987991 | 0.590269908454338 | NM_009632 |
| Tbc1d24 | 5339.608572 | -0.0899964514205448 | 0.0617119169953068 | -1.45833180692457 | 0.144749107132544 | 0.590269908454338 | NM_173186,NM_001163849,NM_001163848,NM_001163847,NM_001163850,NM_001163851,NM_001163852,NM_001163853 |
| Tmppe | 1038.824429 | 0.0840498092648491 | 0.057647691904013 | 1.45799088374253 | 0.144843055803323 | 0.590269908454338 | NM_001200002 |
| Zfp560 | 277.9704453 | -0.132156720050015 | 0.090618636379534 | -1.45838345543525 | 0.144734878354772 | 0.590269908454338 | NM_001004190 |
| Narg2 | 355.3566288 | -0.104985982505547 | 0.07203218973217 | -1.45748703317095 | 0.144981988130941 | 0.590580022319137 | NM_145618 |
| Rnf219 | 678.3980867 | 0.120377277953248 | 0.0826008573248833 | 1.45733690728878 | 0.145023403745771 | 0.59058311264678 | NM_026047 |
| Wdr82 | 6918.831816 | 0.0661412451029184 | 0.0453910074871772 | 1.45714423989384 | 0.145076568675557 | 0.590634034916237 | NM_029896 |
| Aplf | 282.6614399 | -0.115727429573204 | 0.0794526373575215 | -1.45655869234966 | 0.145238237176566 | 0.591055906989352 | NM_001170489,NM_024251 |
| Ube2j1 | 4426.341397 | 0.0778716707944547 | 0.0534658765575731 | 1.45647421885248 | 0.145261571523612 | 0.591055906989352 | NM_019586 |
| Trmt10b | 466.451101 | 0.104879333909071 | 0.0720250223954989 | 1.45615135436077 | 0.145350783731585 | 0.591253286050126 | NM_027266 |
| Fhdc1 | 278.1016824 | 0.128897458715204 | 0.0885440184072716 | 1.45574439734958 | 0.145463291705989 | 0.591545289843559 | NM_001033301,NM_001205355 |
| Plk4 | 218.0248584 | -0.125559925250687 | 0.0862835785127193 | -1.45520071623105 | 0.145613702680122 | 0.591991224498027 | NR_045576,NM_173169,NM_011495 |
| Lsp1 | 267.1561058 | -0.123258706125857 | 0.0847340879011621 | -1.45465312932419 | 0.145765314535079 | 0.592441790413139 | NM_001271523,NM_001271510,NM_001271508,NM_001271509,NM_019391,NM_001136071 |
| Clptm1 | 7576.310765 | 0.0667226929656256 | 0.045876446736574 | 1.4543997565625 | 0.145835507411788 | 0.592514473997113 | NM_019649 |
| Pja2 | 26391.161 | 0.0963012929886207 | 0.0662325429852555 | 1.45398755125647 | 0.145949757591705 | 0.592514473997113 | NM_001025309,NM_144859 |
| Sept11 | 4395.747911 | -0.0659502994924509 | 0.0453624354337311 | -1.45385270569955 | 0.145987147352035 | 0.592514473997113 | NM_001009818 |
| Smyd2 | 2263.896485 | 0.0729076606460702 | 0.0501428834140787 | 1.45399816847389 | 0.145946813977943 | 0.592514473997113 | NM_026796 |
| Zfp346 | 1619.865653 | -0.0809793648936215 | 0.0556854692669101 | -1.45422793342143 | 0.145883122989332 | 0.592514473997113 | NM_012017 |
| Srsf9 | 1176.312039 | -0.100074037017711 | 0.0688529970571745 | -1.45344489412148 | 0.146100269307382 | 0.592807964234367 | NM_025573,NR_036616 |
| Jun | 1971.857171 | -0.106063259765405 | 0.0729827403508736 | -1.45326496724421 | 0.146150200150109 | 0.592844961569529 | NM_010591 |
| Agbl3 | 192.6135161 | -0.122257978575733 | 0.0841389451009511 | -1.45304862604405 | 0.146210253486275 | 0.592922987755899 | NM_178630 |
| Phf8 | 1409.554889 | 0.0967335666380467 | 0.0665810075260854 | 1.45287027385622 | 0.146259775794836 | 0.592958276080319 | NM_001113354,NM_177201 |
| Fat2 | 22.35304398 | 0.076205833328942 | 0.0524592774854822 | 1.45266646781461 | 0.146316381481132 | 0.593022253737425 | NM_001029988 |
| Dcaf12 | 2113.902775 | 0.0824909545223135 | 0.0568053326226142 | 1.45216920161953 | 0.146454563993119 | 0.59325125392193 | NM_026893 |
| Pparg | 202.6920262 | -0.135415148101746 | 0.0932501102870034 | -1.45217145250518 | 0.146453938282272 | 0.59325125392193 | NM_001127330,NM_011146 |
| Pes1 | 1902.388603 | 0.0782007848381782 | 0.0538639598520929 | 1.45182019764074 | 0.146551606361618 | 0.593478849737626 | NM_022889 |
| 1700123O20Rik | 578.1495442 | 0.101263346987005 | 0.0697720705039803 | 1.45134501893888 | 0.146683811338211 | 0.593534374886927 | NM_021437 |
| 9130011E15Rik | 867.8905073 | 0.0865605837573061 | 0.0596482672062863 | 1.45118354331982 | 0.146728758104017 | 0.593534374886927 | NM_198296 |
| Cct4 | 4022.609911 | 0.0841794545056513 | 0.0579939273185612 | 1.45152188164897 | 0.146634593658591 | 0.593534374886927 | NM_009837 |
| Rasa2 | 1288.727484 | -0.100846221568006 | 0.0694861678565544 | -1.45131361649113 | 0.146692551389841 | 0.593534374886927 | NM_053268 |
| 0610009B22Rik | 459.2832703 | 0.0971721388571762 | 0.0670294424921125 | 1.44969337718437 | 0.147144043414444 | 0.5940317226829 | NM_025319 |
| Acin1 | 4924.250809 | -0.0931620359055677 | 0.0642392741053124 | -1.45023487894399 | 0.146993031763795 | 0.5940317226829 | NM_001085473,NM_001085472,NM_001242605,NM_019567,NM_001242606,NM_023190 |
| Arcn1 | 5628.987344 | 0.0823299647423826 | 0.0567689062658975 | 1.45026512148677 | 0.146984601350901 | 0.5940317226829 | NM_145985 |
| Ccl25 | 112.4924001 | -0.137131120400712 | 0.0945878211889089 | -1.44977565480482 | 0.147121090550751 | 0.5940317226829 | NR_033527,NM_009138 |
| Grhpr | 1031.2325 | 0.102554263338771 | 0.0707273904152004 | 1.44999359847342 | 0.147060304364536 | 0.5940317226829 | NM_080289 |
| Itga10 | 180.6534112 | -0.137634763070098 | 0.0949314812587615 | -1.44983267136575 | 0.147105186332701 | 0.5940317226829 | NM_001081053 |
| Kcnj5 | 84.50146023 | 0.131575774625207 | 0.090768920170447 | 1.44956857895999 | 0.147178863412898 | 0.5940317226829 | NM_010605 |
| Ppp1r14b | 569.7083115 | -0.12025405126639 | 0.0829427104755885 | -1.44984472507422 | 0.147101824235686 | 0.5940317226829 | NM_008889 |
| Ino80e | 1020.68219 | 0.0969343443195269 | 0.0668965413157492 | 1.44901877455817 | 0.147332339560138 | 0.594485990125159 | NM_153580 |
| Nudt5 | 483.986821 | 0.105362368151267 | 0.0727306654903255 | 1.44866498114584 | 0.147431164541579 | 0.594719549050535 | NM_016918 |
| Cdkn2c | 62.15281764 | -0.137667831162263 | 0.095062031394681 | -1.44818945211354 | 0.147564073702363 | 0.594737170804894 | NM_007671 |
| Kpna1 | 5852.434835 | 0.105041621877585 | 0.0725393776565126 | 1.44806345561685 | 0.147599304746774 | 0.594737170804894 | NM_008465 |
| Lancl3 | 253.86312 | 0.135438387355173 | 0.0935172874826607 | 1.44827112719971 | 0.147541239210946 | 0.594737170804894 | NM_173414 |
| Tmed2 | 4128.813841 | 0.0842917672871588 | 0.0582082242554102 | 1.44810752029296 | 0.147586982684658 | 0.594737170804894 | NM_019770 |
| Sncb | 16574.42666 | 0.0559505589615383 | 0.0386444202386731 | 1.44783020720663 | 0.147664542455761 | 0.594835037080528 | NM_033610 |
| 2700049A03Rik | 397.040383 | -0.10779640184024 | 0.0744837137097647 | -1.44724794819284 | 0.147827491796861 | 0.594842534722965 | NM_029818,NM_001163378 |
| Ddx5 | 33424.01137 | -0.0746758673694405 | 0.0516030594108534 | -1.44712093085191 | 0.147863056756276 | 0.594842534722965 | NM_007840 |
| Ext1 | 1738.094518 | 0.0950528570744656 | 0.0656832435085019 | 1.44714012276452 | 0.147857682585762 | 0.594842534722965 | NM_010162 |
| Fbxw2 | 3047.023294 | 0.0749023474583948 | 0.0517605976560617 | 1.44709201304253 | 0.147871154680203 | 0.594842534722965 | NM_013890,NM_001164770,NM_001164769,NM_001164768,NM_001164772 |
| Trim67 | 205.4035967 | 0.13200266935524 | 0.0912081091831263 | 1.44726900423083 | 0.147821596720672 | 0.594842534722965 | NM_198632 |
| Fam207a | 1423.328824 | 0.0834610949358631 | 0.0576941103928501 | 1.44661377682333 | 0.148005125471505 | 0.595114347077491 | NM_133998 |
| Hagh | 2432.764688 | 0.0942730564490414 | 0.0651705873229578 | 1.44655833745773 | 0.148020661984784 | 0.595114347077491 | NM_024284,NM_001159626 |
| Adarb1 | 11079.32106 | 0.080395419499021 | 0.0555846256199124 | 1.44636072659308 | 0.148076051251149 | 0.595162562286933 | NR_021486,NM_001024837,NM_130895,NR_004429 |
| Antxr2 | 76.6736182 | -0.137222020251287 | 0.0948830145716506 | -1.44622323469354 | 0.148114598834315 | 0.595162562286933 | NM_133738 |
| Frem3 | 111.1131637 | 0.0914301439650547 | 0.0632420788971934 | 1.44571692707452 | 0.148256614638557 | 0.595568469092831 | NM_001167898 |
| Rab37 | 211.8571935 | 0.133036401122232 | 0.0920326672520927 | 1.44553455956919 | 0.148307792941171 | 0.595609344833687 | NM_001163753,NM_021411 |
| Yipf1 | 1188.421963 | 0.0811636243809523 | 0.0561562740261112 | 1.44531712241473 | 0.148368830572582 | 0.595689782448129 | NM_145550,NM_001205156 |
| Tmem222 | 2831.533113 | 0.0828046799100606 | 0.0573131444903082 | 1.44477642339207 | 0.148520695480432 | 0.596134739582413 | NM_025667 |
| Rangap1 | 11689.09376 | 0.0678460313351305 | 0.0469708955938045 | 1.44442703247241 | 0.14861889123537 | 0.596199396322836 | NM_001146174,NM_011241 |
| Srp9 | 2282.325396 | 0.0690668907728518 | 0.0478156075130726 | 1.44444239789214 | 0.148614571766983 | 0.596199396322836 | NM_012058 |
| Mrps31 | 638.6470024 | 0.104289271776508 | 0.0722176785160469 | 1.44409615373242 | 0.148711929870787 | 0.596407921950042 | NM_020560 |
| Akap10 | 1338.924471 | 0.110420221868759 | 0.0764810182948063 | 1.44375982865617 | 0.14880654551283 | 0.596622655291021 | NM_019921 |
| 1700047M11Rik | 186.2978794 | -0.136051109476378 | 0.0942494547113137 | -1.44352144946729 | 0.148873634649882 | 0.596726936237359 | NR_015458 |
| Cirh1a | 1279.991974 | 0.087461908077196 | 0.060597713850833 | 1.44332025944892 | 0.148930275281124 | 0.596789290685135 | NM_011574 |
| Rhbdd3 | 659.5876952 | -0.110686860800712 | 0.0767008017352946 | -1.44309913712125 | 0.148992546391491 | 0.5968741668182 | NM_177370 |
| Rangrf | 349.1368184 | 0.114924049834615 | 0.0796611024666233 | 1.44266205558435 | 0.149115693076918 | 0.597202800561157 | NM_021329 |
| Eif4g1 | 12867.15742 | 0.054893764289507 | 0.0380595775335391 | 1.44231144555226 | 0.149214532765093 | 0.597429753682506 | NM_001005331,NM_145941 |
| Uvrag | 1742.796682 | 0.0862620864491144 | 0.0598141190337925 | 1.44216930457497 | 0.149254617658944 | 0.597429753682506 | NM_178635 |
| Tbc1d13 | 2229.255033 | 0.0742067736794791 | 0.051468527300808 | 1.44178933362086 | 0.149361812831737 | 0.597694130356421 | NM_146252 |
| Negr1 | 10429.72397 | 0.105784806508097 | 0.0733823245356096 | 1.44155703948522 | 0.149427375224863 | 0.597791807357854 | NM_177274,NM_001039094 |
| Tmem41a | 567.7626192 | 0.09902166791552 | 0.0687079792583297 | 1.44119604425006 | 0.149529305626626 | 0.598034882580499 | NM_025693,NR_037773 |
| C330006A16Rik | 3211.590967 | 0.0741268832552145 | 0.0514497434809064 | 1.44076293174764 | 0.149651669017167 | 0.598098567494676 | NM_001256521,NM_001256522 |
| Hsd17b11 | 1663.063004 | 0.0872850484285862 | 0.0605859875095537 | 1.44068046121758 | 0.149674977332133 | 0.598098567494676 | NM_053262 |
| Hypk | 1083.625834 | 0.103581018597383 | 0.0719094561212477 | 1.44043668502697 | 0.149743891011756 | 0.598098567494676 | NM_026318 |
| Mks1 | 226.5437538 | -0.127579831635128 | 0.088573775686791 | -1.44037928434109 | 0.14976012127042 | 0.598098567494676 | NM_001039684 |
| Pde4d | 3241.869874 | 0.106948234182809 | 0.0742302252958547 | 1.44076397123345 | 0.149651375249135 | 0.598098567494676 | NM_011056 |
| Prtg | 168.2658957 | -0.131230111021269 | 0.0911152170163021 | -1.4402655815196 | 0.149792275130499 | 0.598098567494676 | NM_175485 |
| Aplp2 | 26177.45044 | 0.0648450827281867 | 0.0450495322917743 | 1.43941744629449 | 0.150032284262734 | 0.598234686028129 | NM_009691,NM_001102455,NM_001102456 |
| Ccdc163 | 132.9788488 | -0.135919620239769 | 0.0944239020827413 | -1.43946201376709 | 0.150019665061539 | 0.598234686028129 | NM_026714 |
| Eif1a | 1622.631862 | 0.0918028563290279 | 0.0637718614163579 | 1.43955114826678 | 0.14999442913857 | 0.598234686028129 | NM_010120 |
| Kcnk1 | 3794.988084 | 0.0868723218348928 | 0.0603372597730572 | 1.43977903805443 | 0.149929923553072 | 0.598234686028129 | NM_008430 |
| Ptpn9 | 2091.81978 | 0.102508740771702 | 0.0711899555111863 | 1.43993264268292 | 0.149886456720333 | 0.598234686028129 | NM_019651 |
| Tbx2 | 87.22981441 | -0.136659531846862 | 0.0949624072465341 | -1.43909085510098 | 0.15012478268733 | 0.598439240756353 | NM_009324 |
| 4933407L21Rik | 108.7815758 | -0.134226464740775 | 0.0933585183220543 | -1.43775273165476 | 0.150504225387394 | 0.598511432111577 | NR_037692 |
| Ccdc149 | 2581.669464 | 0.0785238380707542 | 0.0546229992212707 | 1.43755998737206 | 0.150558940781364 | 0.598511432111577 | NM_001256059 |
| Cfdp1 | 2870.268979 | 0.0890494866342202 | 0.0619370455305185 | 1.43774191796641 | 0.150507294727946 | 0.598511432111577 | NM_011801 |
| Exoc4 | 3026.058104 | 0.0762751422562586 | 0.0530251505242642 | 1.43847101803804 | 0.150300454932325 | 0.598511432111577 | NM_009148 |
| Itpr2 | 991.8711943 | -0.0944556821317883 | 0.0657168752985988 | -1.43731243615294 | 0.150629236769201 | 0.598511432111577 | NM_010586,NM_019923 |
| Kcna1 | 10999.59489 | 0.0987800721264075 | 0.068664092507375 | 1.43859866954184 | 0.150264263539476 | 0.598511432111577 | NM_010595 |
| Mapkapk2 | 1431.093553 | 0.0794794141838155 | 0.0552975676167037 | 1.43730398296592 | 0.150631637623749 | 0.598511432111577 | NM_008551 |
| Pam | 9043.512383 | 0.0750859807262479 | 0.0522267945041176 | 1.4376907761461 | 0.15052181138952 | 0.598511432111577 | NM_013626 |
| Phf16 | 303.4101277 | -0.114766884358947 | 0.0798462392435193 | -1.43734865218793 | 0.150618951105244 | 0.598511432111577 | NM_199317 |
| Rpf2 | 582.2218958 | 0.0959027911309339 | 0.0666868841762129 | 1.43810574321513 | 0.150404053336928 | 0.598511432111577 | NM_001042556,NM_023323 |
| Top1 | 3385.101512 | 0.105310250331453 | 0.073270316506288 | 1.43728395553492 | 0.150637325884616 | 0.598511432111577 | NM_009408 |
| Traf3ip2 | 29.39320858 | 0.121705414504849 | 0.0846634985973187 | 1.43751931494953 | 0.150570488625986 | 0.598511432111577 | NM_134000 |
| Adprm | 447.5005488 | -0.10536740856164 | 0.0733801549097407 | -1.43591150347454 | 0.151027524725155 | 0.598846773234123 | NM_025510 |
| Apbb3 | 1628.624627 | -0.0911339933343605 | 0.0634856678884395 | -1.43550499452107 | 0.151143246118541 | 0.598846773234123 | NM_146085 |
| Casp7 | 110.3607678 | -0.136254384279451 | 0.094943955691514 | -1.43510330159574 | 0.151257662880077 | 0.598846773234123 | NM_007611 |
| Cnrip1 | 2729.794564 | 0.0821308806792828 | 0.057166006401162 | 1.43670838405135 | 0.150800871688591 | 0.598846773234123 | NM_029861 |
| Cox6a2 | 186.6024851 | 0.125936644832499 | 0.0877195744461427 | 1.43567323060625 | 0.151095345957786 | 0.598846773234123 | NM_009943 |
| Lhfpl5 | 125.6131416 | 0.136543021932019 | 0.0950965128882109 | 1.43583626554772 | 0.15104893770325 | 0.598846773234123 | NM_026571 |
| Mybpc1 | 210.2743889 | 0.134767167689048 | 0.0938994939385822 | 1.4352278381521 | 0.151222183281626 | 0.598846773234123 | NM_175418,NM_001252372 |
| Ndufs3 | 3870.133386 | 0.0608460572486296 | 0.042395293933638 | 1.43520781678912 | 0.15122788680086 | 0.598846773234123 | NM_026688 |
| Nupl2 | 358.6505436 | -0.11148071546436 | 0.0776158050460767 | -1.43631461914464 | 0.150912835986841 | 0.598846773234123 | NM_153092 |
| Setbp1 | 1661.148975 | 0.0869703070158876 | 0.060556298314077 | 1.43618928892935 | 0.150948486045304 | 0.598846773234123 | NM_053099 |
| Timp4 | 1002.295067 | 0.109617011696746 | 0.0763778467721695 | 1.43519379413414 | 0.15123188155568 | 0.598846773234123 | NM_080639 |
| Txndc15 | 2139.404086 | 0.0766795760360963 | 0.0533878185474627 | 1.43627475559667 | 0.150924174438542 | 0.598846773234123 | NM_175150 |
| Zfand1 | 597.6578116 | -0.0965079793061323 | 0.0671914764590618 | -1.43631282406679 | 0.150913346549702 | 0.598846773234123 | NM_025512 |
| A230046K03Rik | 2221.019049 | -0.0920379351128475 | 0.064181514771228 | -1.43402559819463 | 0.151564958032293 | 0.598905574699364 | NM_001033375 |
| Cab39 | 7711.489134 | 0.0811457261056749 | 0.0565791950521631 | 1.43419725273332 | 0.151515980835441 | 0.598905574699364 | NM_133781 |
| Ewsr1 | 7833.508998 | -0.0870705445186385 | 0.0607231151406973 | -1.43389456085863 | 0.151602354272998 | 0.598905574699364 | NM_007968 |
| Pear1 | 282.6197591 | -0.123881206938778 | 0.0863916386004013 | -1.43394903657034 | 0.151586806806012 | 0.598905574699364 | NM_001032414,NM_152799,NM_001032413,NM_028460 |
| Ppap2b | 11016.5852 | -0.0771141837435525 | 0.0537744958567389 | -1.43402894838834 | 0.15156400202532 | 0.598905574699364 | NM_080555 |
| Rad51ap2 | 34.49530744 | -0.130979567848307 | 0.0913038006191383 | -1.4345467216055 | 0.151416306158933 | 0.598905574699364 | NM_001111118 |
| Rpusd2 | 342.5415737 | 0.106847258786158 | 0.0744865868914041 | 1.43444965389451 | 0.151443986567994 | 0.598905574699364 | NM_173450 |
| Trim39 | 1191.768146 | -0.116851263385492 | 0.0814352319682098 | -1.43489814618699 | 0.151316124057131 | 0.598905574699364 | NM_178281,NM_024468 |
| Anxa7 | 4177.441217 | 0.0689350951122193 | 0.0480948499286752 | 1.43331552576732 | 0.151767687010543 | 0.599395710036743 | NM_001110794,NM_009674 |
| Sclt1 | 346.5906517 | -0.109468033961501 | 0.0763875479056215 | -1.4330612378964 | 0.151840337574787 | 0.599519636752204 | NM_001081411 |
| Cyp2u1 | 218.6056096 | -0.120393822477091 | 0.0840547834551075 | -1.43232565153643 | 0.15205064520565 | 0.600023817510803 | NM_027816 |
| Pink1 | 16539.76894 | 0.0594576411929845 | 0.0415098041028299 | 1.43237585621203 | 0.152036284402839 | 0.600023817510803 | NM_026880 |
| Zbtb33 | 1033.829169 | 0.124251555805916 | 0.086759481857681 | 1.43213805736803 | 0.152104314742537 | 0.600072589883239 | NM_020256,NM_001079513 |
| Ddx24 | 8997.721972 | 0.0660340004231798 | 0.0461305314592879 | 1.4314597801991 | 0.152298486012354 | 0.600512434260439 | NM_020494,NM_001159502 |
| Manf | 1513.108 | 0.0922464376186789 | 0.0644402477529503 | 1.43150345995458 | 0.152285976071874 | 0.600512434260439 | NM_029103 |
| 2210408I21Rik | 425.569192 | 0.13015327000088 | 0.090953523278464 | 1.43098656664901 | 0.152434065153561 | 0.600557968652191 | NM_001145676,NM_001081353 |
| Kifap3 | 13367.37025 | 0.0908796749276725 | 0.063496184162399 | 1.43126199040933 | 0.152355143063934 | 0.600557968652191 | NM_010629 |
| Nav2 | 2474.555096 | -0.0990399543221843 | 0.0692097170802942 | -1.43101226966847 | 0.152426698692174 | 0.600557968652191 | NM_175272,NM_001111016 |
| Adssl1 | 950.4999389 | 0.109958822219152 | 0.0768868875262349 | 1.43013751443161 | 0.152677555060541 | 0.601351762007321 | NM_007421 |
| Arhgap44 | 6756.842509 | 0.0652298682234197 | 0.0456200406901602 | 1.42985116270379 | 0.152759741278192 | 0.601351762007321 | NM_175003,NM_001099288 |
| Arxes2 | 629.3982382 | 0.0974990706170397 | 0.0681851681709084 | 1.42991611273371 | 0.152741096926486 | 0.601351762007321 | NM_029823 |
| Ube2cbp | 70.60453136 | -0.135949818721224 | 0.0950958066271696 | -1.42960897586395 | 0.152829277939067 | 0.601462501041153 | NM_027394 |
| Cd34 | 3461.669379 | -0.104182637095253 | 0.0728923638292197 | -1.4292668205869 | 0.152927558569323 | 0.601523345729215 | NM_133654,NM_001111059 |
| Lamb2 | 1312.192823 | -0.107279382509654 | 0.0750564714392746 | -1.42931556003735 | 0.152913555720908 | 0.601523345729215 | NM_008483 |
| Caly | 5312.493231 | 0.0882713311893247 | 0.0617778479860632 | 1.42885085944137 | 0.153047103934769 | 0.601704475601213 | NM_026769,NM_001190386,NM_001190385 |
| Taf15 | 1805.090109 | -0.0989919334014455 | 0.0692823787873452 | -1.4288183392965 | 0.153056453073557 | 0.601704475601213 | NM_027427 |
| Plrg1 | 1435.288892 | 0.0856523871003355 | 0.0599553196440173 | 1.42860362698245 | 0.153118191107261 | 0.601784319270584 | NM_016784 |
| E030019B06Rik | 86.68092014 | 0.134762024297421 | 0.0943911388749655 | 1.42769783163579 | 0.15337885057234 | 0.602567676251561 | NM_001243018 |
| Zc3h11a | 4861.906056 | -0.0879592817847435 | 0.0616124126437186 | -1.42762274695164 | 0.153400472723274 | 0.602567676251561 | NM_144530 |
| Arhgap26 | 5313.781019 | 0.0869051986075249 | 0.0609763223141494 | 1.4252286020103 | 0.154091131036528 | 0.603402950266781 | NM_175164 |
| Bend5 | 291.137294 | 0.118828488711692 | 0.0833559975231815 | 1.42555415617989 | 0.153997077211268 | 0.603402950266781 | NM_026279,NR_033793 |
| Golim4 | 1016.226606 | -0.0861686939428879 | 0.0604515862487138 | -1.42541659020108 | 0.154036815215751 | 0.603402950266781 | NM_175193 |
| Hsdl1 | 2227.238256 | 0.0800489056834008 | 0.0561322889552716 | 1.4260759212436 | 0.153846428333873 | 0.603402950266781 | NM_175185 |
| Inpp4b | 301.5240164 | 0.123415792875715 | 0.0865398465796215 | 1.42611522614806 | 0.153835084394092 | 0.603402950266781 | NM_001024617 |
| Ndufaf1 | 751.5937688 | 0.0860923141787177 | 0.0603994582268295 | 1.42538222537359 | 0.154046743230127 | 0.603402950266781 | NM_027175 |
| Pcnxl3 | 3268.614451 | -0.0814824267304841 | 0.0571242108422924 | -1.42640791932233 | 0.153750629086717 | 0.603402950266781 | NM_144868 |
| Rasgef1c | 1674.1498 | -0.120238890818298 | 0.0843664400335714 | -1.42519810922983 | 0.154099942751144 | 0.603402950266781 | NM_029004 |
| Rpa1 | 2097.611207 | 0.0623016006436804 | 0.0436673409807178 | 1.4267321811784 | 0.153657105938058 | 0.603402950266781 | NM_026653,NM_001164223 |
| Tspan3 | 11131.18746 | 0.056314639190349 | 0.0395146690735031 | 1.42515780875187 | 0.154111589253047 | 0.603402950266781 | NM_019793 |
| Ypel1 | 1056.045679 | -0.0964383623804447 | 0.0676531192238271 | -1.42548286741049 | 0.154017669073016 | 0.603402950266781 | NM_023249 |
| Zfp830 | 383.6474421 | 0.105633668414908 | 0.0741050922649868 | 1.42545761952743 | 0.154024962461518 | 0.603402950266781 | NM_025884 |
| Abhd15 | 61.77201023 | 0.123303924663736 | 0.0865343616353944 | 1.42491285927857 | 0.154182391996988 | 0.603517495593706 | NM_026185 |
| 1810020O05Rik | 42.49780954 | -0.132662327648818 | 0.0931609822901397 | -1.42401168802252 | 0.154443088683399 | 0.604212309780515 | NR_045482 |
| Magee2 | 305.942482 | 0.126059383580998 | 0.0885162898914697 | 1.4241376783365 | 0.15440662126092 | 0.604212309780515 | NM_053206 |
| Eef1a2 | 40259.61988 | 0.0721736635455487 | 0.0506982638225667 | 1.42359240936024 | 0.154564494438296 | 0.604361735184571 | NM_007906 |
| Rac1 | 12289.47755 | 0.0578302547310826 | 0.0406199378287349 | 1.42369136493786 | 0.154535834462585 | 0.604361735184571 | NM_009007 |
| Cdipt | 5713.994163 | 0.0815615530971154 | 0.0573153491971524 | 1.42303160040361 | 0.154726994833849 | 0.604402956255828 | NM_026638 |
| Chek2 | 50.07923011 | -0.134029110536758 | 0.0941712354917322 | -1.42324893410287 | 0.154664004675582 | 0.604402956255828 | NM_016681 |
| Mrpl18 | 1067.727661 | 0.0766123509975863 | 0.0538267906697203 | 1.42331263009302 | 0.154645547260948 | 0.604402956255828 | NM_026310 |
| Wdr6 | 8302.65968 | 0.121826284502316 | 0.0856133901025477 | 1.4229816662603 | 0.154741470075411 | 0.604402956255828 | NM_031392 |
| Capzb | 8003.255689 | 0.0635996357911305 | 0.0447067058043056 | 1.42259722891516 | 0.154852947775504 | 0.604489542765733 | NM_001271405,NM_001037761,NM_001271406,NM_009798 |
| Cyp4v3 | 379.85215 | -0.125271902297753 | 0.0880645354056182 | -1.42250114328953 | 0.154880819850236 | 0.604489542765733 | NM_133969 |
| Dll4 | 130.1596886 | -0.134726885731774 | 0.0947416785320056 | -1.42204453013001 | 0.155013324163731 | 0.604489542765733 | NM_019454 |
| Slc25a46 | 3807.521163 | 0.0931325629583028 | 0.0654862760275478 | 1.42216917204339 | 0.154977145858601 | 0.604489542765733 | NM_026165 |
| Sox17 | 194.9755461 | -0.135049548127542 | 0.0949446911615586 | -1.42240231102275 | 0.154909492632738 | 0.604489542765733 | NM_011441 |
| Syngr3 | 6599.134168 | 0.0777641342093891 | 0.0546837496320632 | 1.42207026278595 | 0.155005854530698 | 0.604489542765733 | NM_011522 |
| A830018L16Rik | 2979.518549 | 0.102647290174845 | 0.0722194811531044 | 1.42132411554209 | 0.155222555889893 | 0.604578395570044 | NM_001160370,NM_001160371,NM_001160369,NM_177173 |
| Cfl1 | 27611.35003 | 0.0523801818451914 | 0.0368646769591816 | 1.42087727781229 | 0.155352439737212 | 0.604578395570044 | NM_007687 |
| Coa3 | 2016.328721 | 0.0880756614784281 | 0.0619873142443309 | 1.42086590703489 | 0.155355745996033 | 0.604578395570044 | NM_026618 |
| Hsp90b1 | 15752.54307 | 0.0859467841995512 | 0.0604716971823371 | 1.42127289631711 | 0.155237439771789 | 0.604578395570044 | NM_011631 |
| Slc41a1 | 3421.622285 | 0.0873840647052249 | 0.0614945058856754 | 1.42100604674637 | 0.155315001572394 | 0.604578395570044 | NM_173865 |
| Tle1 | 1452.305997 | -0.0865786117843622 | 0.0609284598004332 | -1.42098802543088 | 0.1553202406613 | 0.604578395570044 | NM_011599 |
| Ttyh3 | 11442.48905 | 0.0821505245888261 | 0.0578190905219924 | 1.42082007598474 | 0.155369072742873 | 0.604578395570044 | NM_175274 |
| Usp2 | 3594.314808 | -0.0936200261138449 | 0.0658758748600238 | -1.42115799316173 | 0.155270833617542 | 0.604578395570044 | NM_016808,NM_198092,NM_198091 |
| Cdr2l | 1234.485715 | 0.0890910894651641 | 0.0627349593265992 | 1.42011870927268 | 0.15557312434181 | 0.605210284999766 | NM_001080929 |
| Vstm2a | 3151.2393 | 0.094872917281302 | 0.0668222771221536 | 1.41977977056769 | 0.155671806090274 | 0.605432036216149 | NM_145967 |
| Crispld2 | 138.3228603 | 0.134439896789671 | 0.0947611566701811 | 1.41872367870723 | 0.155979591068062 | 0.606466686256602 | NM_030209 |
| Atp13a1 | 2347.602284 | -0.093216172392435 | 0.0657178138568087 | -1.4184308168793 | 0.156065023762616 | 0.606636482519604 | NM_133224 |
| Hipk2 | 1418.640082 | 0.11876248468355 | 0.0837490687277435 | 1.41807528713699 | 0.156168785439595 | 0.606877415006839 | NM_010433,NM_001136065 |
| Fars2 | 596.5474434 | 0.102211010259335 | 0.0721249024886928 | 1.41713897326043 | 0.156442299889816 | 0.607194109660642 | NM_024274,NM_001039189 |
| Fbln1 | 262.7160553 | 0.133243209453608 | 0.0940252405710607 | 1.41710043648235 | 0.156453564968723 | 0.607194109660642 | NM_010180 |
| Gm5089 | 91.72100402 | -0.133483890664523 | 0.0942153908628844 | -1.41679495711202 | 0.156542884529747 | 0.607194109660642 | NR_033325,NR_033326 |
| Mrps18a | 1397.951415 | 0.0878111012558455 | 0.0619765302177095 | 1.41684442396799 | 0.156528418219734 | 0.607194109660642 | NM_026768 |
| Nrarp | 632.5462193 | 0.106990175662834 | 0.0755137020866891 | 1.41683128632748 | 0.156532260151581 | 0.607194109660642 | NM_025980 |
| Prr11 | 28.0682471 | -0.124837431607602 | 0.0880711756582216 | -1.41746071486612 | 0.156348272324784 | 0.607194109660642 | NM_175563 |
| Tubg2 | 2184.491002 | 0.079307407830257 | 0.055956922160129 | 1.41729396058109 | 0.156397000176554 | 0.607194109660642 | NM_134028 |
| Irf3 | 736.2685615 | -0.118937138114748 | 0.0839618667288907 | -1.41656138373854 | 0.156611205475437 | 0.607279635638395 | NM_016849,NR_045611 |
| Lcat | 751.3325746 | -0.113821459643061 | 0.0803590120662048 | -1.41641188357676 | 0.156654946609032 | 0.607279635638395 | NM_008490 |
| Zfp703 | 1487.189313 | -0.0690757666901052 | 0.0487723017132833 | -1.41629089182995 | 0.156690353460877 | 0.607279635638395 | NM_001110508,NM_001101502 |
| Hdac6 | 2432.659667 | -0.0874275875243987 | 0.0617396596027715 | -1.41606850583404 | 0.156755448007215 | 0.607369868699068 | NM_001130416,NM_010413 |
| 5031434O11Rik | 94.5497811 | -0.134343275960792 | 0.0949162018523815 | -1.41538824077399 | 0.156954695507034 | 0.607833353330809 | NR_033624 |
| Csrnp1 | 539.1580003 | 0.132489065658773 | 0.0936441251832526 | 1.41481449476414 | 0.157122893115229 | 0.607833353330809 | NM_153287 |
| Mtmr9 | 2889.962594 | 0.079496360209891 | 0.0561828148757911 | 1.41495865569643 | 0.157080618495533 | 0.607833353330809 | NM_177594 |
| Plcl1 | 1943.281384 | 0.0977512136573054 | 0.0690813040786697 | 1.4150169132011 | 0.157063537160447 | 0.607833353330809 | NM_001114663 |
| Prdx6 | 9152.52253 | -0.0742320407755071 | 0.0524606699912098 | -1.41500367395127 | 0.157067418838201 | 0.607833353330809 | NM_007453 |
| Zmynd10 | 141.5679697 | -0.134254289249742 | 0.0948925380608819 | -1.41480343969308 | 0.157126135326806 | 0.607833353330809 | NM_053253 |
| Birc2 | 1439.072025 | -0.091520859482969 | 0.0647072338647295 | -1.4143837406849 | 0.157249261416426 | 0.60787851646781 | NM_007465 |
| Dis3 | 610.474263 | -0.106585888476216 | 0.0753566109062243 | -1.41441987895202 | 0.157238656743539 | 0.60787851646781 | NM_028315 |

| | | | | | | |
|---|---|---|---|---|---|---|
| Baz1b | 5566.544064 | 0.127988935 | 0.058905203 | 2.17279508 | 0.029795744 | 0.122845996 |
| 1700029I15Rik | 24.57306288 | 0.362550407 | 0.166994842 | 2.171027572 | 0.029929088 | 0.12336294 |
| Tacr2 | 11.90303852 | 0.32757969 | 0.15095477 | 2.170051932 | 0.030002912 | 0.123629389 |
| Yap1 | 992.1040176 | -0.13301709 | 0.061299259 | -2.169962465 | 0.03000969 | 0.123629389 |
| Gpam | 2286.281706 | -0.121895686 | 0.056186263 | -2.169492681 | 0.030045299 | 0.123743187 |
| Krt10 | 219.7425032 | -0.275184004 | 0.126857458 | -2.16923789 | 0.030064628 | 0.123756999 |
| Slc22a18 | 27.00296219 | 0.364452515 | 0.168002133 | 2.169332664 | 0.030057437 | 0.123756999 |
| Rxfp3 | 150.4545007 | 0.334139919 | 0.154055201 | 2.168962277 | 0.030085548 | 0.123810221 |
| Efcab7 | 76.63017477 | -0.303992738 | 0.140186215 | -2.168492369 | 0.030121245 | 0.123924207 |
| Trim39 | 1034.30367 | 0.237531354 | 0.109550277 | 2.168240567 | 0.030140388 | 0.123954281 |
| Zfp687 | 1571.2009 | 0.160230701 | 0.073900817 | 2.168185775 | 0.030144555 | 0.123954281 |
| Zdhhc2 | 1183.227332 | -0.183945346 | 0.084855925 | -2.167737202 | 0.030178687 | 0.124061709 |
| Gmps | 1498.636654 | -0.159203919 | 0.073448285 | -2.167564829 | 0.030191812 | 0.124082742 |
| Pram1 | 15.56077495 | 0.362507314 | 0.167341322 | 2.166274946 | 0.030290183 | 0.124421022 |
| Tmco1 | 1072.143361 | -0.175486315 | 0.081006394 | -2.166326728 | 0.030286229 | 0.124421022 |
| Psma3 | 2974.071512 | -0.104196435 | 0.048102314 | -2.166141839 | 0.03030035 | 0.124422122 |
| Tshz3 | 1065.65911 | 0.190918939 | 0.088141065 | 2.166061189 | 0.030306511 | 0.124422122 |
| Psph | 516.4974505 | -0.189436127 | 0.087511551 | -2.164698555 | 0.030410778 | 0.12481711 |
| Uggt1 | 2930.191023 | -0.100544724 | 0.046453976 | -2.164394382 | 0.030434095 | 0.12487973 |
| Ankrd10 | 1254.189588 | 0.221486218 | 0.102366386 | 2.163661601 | 0.03049033 | 0.125077356 |
| Gm16677 | 10.84447229 | 0.326240713 | 0.150795438 | 2.163465409 | 0.030505401 | 0.125106059 |
| Amh | 28.90897381 | 0.35625165 | 0.164675532 | 2.163355082 | 0.03051388 | 0.125107714 |
| Polr3c | 705.3069206 | 0.166878633 | 0.077165186 | 2.16261557 | 0.03057076 | 0.125307765 |
| Mrpl17 | 921.7257126 | -0.171142186 | 0.07914584 | -2.162364888 | 0.030590062 | 0.125320576 |
| Plek2 | 11.25747401 | 0.35157972 | 0.162582855 | 2.162464922 | 0.030582358 | 0.125320576 |
| Xkr4 | 789.4695445 | -0.223462837 | 0.103347007 | -2.16225745 | 0.030598338 | 0.125321343 |
| Dync1li2 | 10732.73287 | 0.102004893 | 0.04718036 | 2.162020238 | 0.030616616 | 0.125363069 |
| Cdon | 544.6461913 | -0.177771902 | 0.082233189 | -2.161802347 | 0.030633414 | 0.125372535 |
| Fam73b | 3269.264083 | 0.117966836 | 0.054569298 | 2.161780341 | 0.030635112 | 0.125372535 |
| Gpsm3 | 71.84833671 | 0.298388646 | 0.138090008 | 2.160827206 | 0.03070869 | 0.125640464 |
| Mthfsd | 859.9507372 | 0.227498736 | 0.105304651 | 2.160386398 | 0.03074277 | 0.125746693 |
| Atrnl1 | 5897.582313 | -0.188175153 | 0.087123057 | -2.159877751 | 0.030782135 | 0.125826058 |
| Bcl7b | 1595.923052 | 0.165337685 | 0.076551571 | 2.159820915 | 0.030786536 | 0.125826058 |
| Donson | 560.7720782 | 0.201083461 | 0.093093407 | 2.160018273 | 0.030771255 | 0.125826058 |
| Spata6 | 133.7433974 | 0.254896927 | 0.118027923 | 2.159632412 | 0.030801137 | 0.125851079 |
| Vcam1 | 1240.248454 | -0.200790008 | 0.092978474 | -2.159532187 | 0.030808903 | 0.125851079 |
| Mtr | 485.3805122 | -0.20434941 | 0.094638776 | -2.159256681 | 0.030830259 | 0.125905121 |
| Ccdc40 | 73.14121455 | 0.282765068 | 0.130976776 | 2.158894713 | 0.030858336 | 0.125986577 |
| Cpne4 | 2625.237474 | -0.245964323 | 0.113970216 | -2.158145632 | 0.030916511 | 0.126126654 |
| Tpt1 | 13957.01151 | -0.135086271 | 0.062593885 | -2.158138479 | 0.030917067 | 0.126126654 |
| Wdr52 | 157.8999337 | 0.301827207 | 0.139854776 | 2.158147303 | 0.030916381 | 0.126126654 |
| Rgp1 | 2087.672047 | 0.14488676 | 0.067141648 | 2.157926782 | 0.030933525 | 0.126160577 |
| Nrsn1 | 16341.20936 | -0.1716593 | 0.079557709 | -2.157670224 | 0.030953481 | 0.126208744 |
| Prss16 | 32.13904646 | 0.353231679 | 0.163735951 | 2.157325116 | 0.030980341 | 0.126256066 |
| Zfp951 | 104.2960381 | 0.310324764 | 0.143847901 | 2.157311727 | 0.030981384 | 0.126256066 |
| Rimklb | 837.0055581 | -0.223689156 | 0.103719395 | -2.15667624 | 0.0310309 | 0.126424601 |
| Itsn1 | 7000.462147 | -0.157981748 | 0.073276796 | -2.155958725 | 0.031086888 | 0.12661057 |
| Wdr91 | 1052.4502 | 0.153298334 | 0.071107019 | 2.155881893 | 0.031092889 | 0.12661057 |
| Afg3l1 | 1201.194055 | -0.171476552 | 0.079567202 | -2.155116013 | 0.031152757 | 0.126821024 |
| Gpr22 | 1745.345108 | 0.268318094 | 0.124517116 | 2.154869172 | 0.031172073 | 0.126833011 |
| Prph | 24.85868427 | 0.357155118 | 0.165737679 | 2.1549422 | 0.031166358 | 0.126833011 |
| Lpar3 | 15.18389111 | -0.355501921 | 0.165015973 | -2.154348543 | 0.031212849 | 0.126965576 |
| Ptgr2 | 1821.818859 | -0.16889695 | 0.078413163 | -2.153936207 | 0.031245175 | 0.127063712 |
| 1700007F19Rik | 9.385540833 | -0.331857463 | 0.154142748 | -2.152922969 | 0.031324733 | 0.127353821 |
| Snord19 | 11.2840475 | 0.309713979 | 0.143919721 | 2.151991245 | 0.031398044 | 0.127590955 |
| Syvn1 | 2158.563423 | 0.117850297 | 0.054763852 | 2.151972382 | 0.03139953 | 0.127590955 |
| Sh3bp2 | 166.1989452 | 0.230226356 | 0.106995197 | 2.15174477 | 0.031417462 | 0.12760348 |
| Urb1 | 646.6566295 | 0.165379619 | 0.076859116 | 2.151724197 | 0.031419083 | 0.12760348 |
| Cnpy2 | 895.8461855 | -0.175583918 | 0.081614213 | -2.151388979 | 0.03144551 | 0.127612295 |
| Emcn | 252.8276854 | -0.284150403 | 0.132079902 | -2.151352307 | 0.031448403 | 0.127612295 |
| Mir23b | 17.19551822 | 0.345642061 | 0.160650046 | 2.151521705 | 0.031435044 | 0.127612295 |
| Tmem246 | 2757.362831 | -0.126535627 | 0.058818794 | -2.151278824 | 0.031454199 | 0.127612295 |
| Mthfd1l | 561.3525795 | 0.270311079 | 0.125661834 | 2.151099264 | 0.031468366 | 0.12763635 |
| Ost4 | 907.8199939 | -0.21681163 | 0.100809282 | -2.150710986 | 0.031499019 | 0.127727243 |
| Gas8 | 579.5695564 | 0.204625127 | 0.0951529 | 2.150487549 | 0.03151667 | 0.12776538 |
| Msrb3 | 365.3495788 | 0.187945766 | 0.087407203 | 2.15023201 | 0.031536867 | 0.127809097 |
| Sfrp5 | 51.93560134 | 0.361439008 | 0.168108187 | 2.150038108 | 0.031552201 | 0.127809097 |
| Vangl2 | 256.5011432 | -0.214502586 | 0.099763719 | -2.150106151 | 0.031546819 | 0.127809097 |
| Optn | 714.5885709 | 0.188042198 | 0.087467358 | 2.149855705 | 0.03156663 | 0.127834127 |
| Rexo1 | 2200.499592 | 0.118562951 | 0.05515246 | 2.149730991 | 0.0315765 | 0.12784068 |
| Spock3 | 3533.792728 | -0.169960277 | 0.079080596 | -2.149203288 | 0.031618289 | 0.127976428 |
| Avpr1a | 16.05992101 | -0.322575475 | 0.150124128 | -2.148725058 | 0.031656201 | 0.128096417 |
| Gmfb | 6240.88514 | -0.211655877 | 0.098521695 | -2.148317452 | 0.031688545 | 0.128193818 |
| Bcl11a | 3115.532359 | -0.12476968 | 0.058104159 | -2.147345101 | 0.031765818 | 0.128439348 |
| Col8a1 | 132.0863238 | 0.348900557 | 0.162472147 | 2.147448433 | 0.031757598 | 0.128439348 |
| Dlx2 | 261.0675869 | -0.330615824 | 0.15400934 | -2.146725799 | 0.031815118 | 0.128605122 |
| Cfdp1 | 2588.755063 | -0.124974872 | 0.058225377 | -2.146398653 | 0.031841187 | 0.128676929 |
| 1500032L24Rik | 2159.602029 | -0.14373654 | 0.066983896 | -2.145837272 | 0.031885964 | 0.12881459 |
| Evl | 4412.839041 | -0.092590995 | 0.043152707 | -2.145659028 | 0.031900192 | 0.12881459 |
| Mir1249 | 35.2881246 | 0.36065409 | 0.168079106 | 2.14574017 | 0.031893714 | 0.12881459 |
| Ftsjd2 | 4483.906161 | 0.111759339 | 0.052094083 | 2.14533651 | 0.031925951 | 0.128885016 |
| Ube2b | 4780.210052 | -0.131668946 | 0.061380497 | -2.145126752 | 0.031942714 | 0.128919096 |
| Stk10 | 312.8997559 | 0.244600359 | 0.114054351 | 2.144594718 | 0.031985265 | 0.129057212 |
| Lipe | 378.8672569 | 0.262818147 | 0.122562588 | 2.144358662 | 0.03200416 | 0.129066229 |
| Trim41 | 2269.13392 | 0.114405839 | 0.053350139 | 2.144433751 | 0.031998148 | 0.129066229 |
| Mmp16 | 840.4209679 | -0.175923226 | 0.082047971 | -2.144150843 | 0.032020802 | 0.129099743 |
| Tcf7 | 220.7814781 | -0.248466971 | 0.115895428 | -2.143889336 | 0.032041755 | 0.129150611 |
| Fbxo21 | 5791.203927 | 0.153083436 | 0.071429652 | 2.143135683 | 0.032102205 | 0.129360615 |
| Slc39a8 | 150.486718 | -0.285833982 | 0.133389652 | -2.142849742 | 0.032125166 | 0.129419479 |
| Ypel1 | 1040.194633 | -0.229757057 | 0.107226049 | -2.142735459 | 0.032134346 | 0.129422813 |
| Crk | 5730.020793 | -0.136785917 | 0.063847258 | -2.14239299 | 0.032161871 | 0.12950001 |
| Dact1 | 272.5672186 | -0.269576542 | 0.125851595 | -2.142019265 | 0.032191932 | 0.129587371 |
| Gm2011 | 22.61763906 | 0.360078253 | 0.16811821 | 2.141815884 | 0.032208301 | 0.129592498 |
| Sptbn4 | 7911.858652 | 0.158553552 | 0.07402833 | 2.141795603 | 0.032209933 | 0.129592498 |
| Car9 | 29.91606227 | 0.359611238 | 0.167924643 | 2.141503668 | 0.032233443 | 0.129653418 |
| Ano3 | 5943.717289 | -0.219979756 | 0.102759521 | -2.140723842 | 0.032296315 | 0.129817618 |
| Cd101 | 11.69764137 | 0.336334746 | 0.157106166 | 2.14081187 | 0.032289212 | 0.129817618 |
| Stmn4 | 7949.148062 | 0.175676529 | 0.082069525 | 2.140581779 | 0.032307779 | 0.129817618 |
| Ulk4 | 221.0536387 | 0.267230666 | 0.124836327 | 2.140648262 | 0.032302414 | 0.129817618 |
| Lekr1 | 126.4758392 | 0.229553784 | 0.10725569 | 2.140248074 | 0.032334724 | 0.129858531 |
| Tpcn2 | 125.4452981 | 0.291424806 | 0.136163378 | 2.140258345 | 0.032333894 | 0.129858531 |
| Ccdc113 | 93.21470943 | 0.267401669 | 0.124953793 | 2.140004421 | 0.032354409 | 0.129903918 |
| Cacna1h | 3142.803043 | -0.206384923 | 0.096465576 | -2.139467056 | 0.032397861 | 0.130044679 |
| Dad1 | 1878.900998 | -0.136342985 | 0.063737411 | -2.139135892 | 0.032424664 | 0.130084865 |
| Sox6 | 980.9697261 | -0.158671739 | 0.074174645 | -2.13916412 | 0.032422379 | 0.130084865 |
| Cldn1 | 67.1888557 | 0.348308357 | 0.162897936 | 2.138199943 | 0.032500519 | 0.130355434 |
| Ninl | 274.9449821 | 0.236374537 | 0.110610234 | 2.137004219 | 0.032597648 | 0.130652867 |
| Prss23 | 807.4464755 | -0.276371626 | 0.129334667 | -2.136871979 | 0.032608406 | 0.130652867 |
| Slc40a1 | 463.2404825 | -0.317214027 | 0.148444877 | -2.136914607 | 0.032604938 | 0.130652867 |
| Spcs2 | 2695.72003 | -0.108284025 | 0.050669884 | -2.137049014 | 0.032594005 | 0.130652867 |
| Cutc | 179.1464652 | 0.217602821 | 0.101840251 | 2.136707432 | 0.032621795 | 0.130672723 |
| AI607873 | 16.91663762 | 0.334973702 | 0.156833065 | 2.135861486 | 0.032690706 | 0.130914912 |
| Klhdc4 | 733.686677 | 0.173227135 | 0.081109447 | 2.135720841 | 0.032702175 | 0.130927001 |
| Uqcrb | 2152.724217 | -0.165155774 | 0.077338333 | -2.135496957 | 0.032720438 | 0.130966281 |
| A530058N18Rik | 85.55421437 | -0.284378705 | 0.13321336 | -2.134761144 | 0.032780526 | 0.131105179 |
| Dnajc11 | 2472.726338 | 0.160488766 | 0.075174513 | 2.134882679 | 0.032770595 | 0.131105179 |
| Ppfibp2 | 493.6886026 | 0.278439172 | 0.130427023 | 2.134827312 | 0.032775119 | 0.131105179 |
| 5031425E22Rik | 336.4705233 | 0.229416101 | 0.107475847 | 2.134582862 | 0.032795099 | 0.131129615 |
| Tmco5 | 62.76067215 | -0.317871874 | 0.148968127 | -2.133824731 | 0.032857131 | 0.131343752 |
| Dbpht2 | 1590.15741 | -0.180788399 | 0.084755382 | -2.13306099 | 0.032919723 | 0.131560019 |
| Spsb1 | 725.7245026 | 0.28719305 | 0.134687542 | 2.132291126 | 0.032982921 | 0.131778592 |
| Spink8 | 48.28079395 | 0.288764373 | 0.135459893 | 2.131733359 | 0.033028773 | 0.131927766 |
| D17Wsu104e | 908.0261084 | -0.156197351 | 0.073280264 | -2.131506376 | 0.033047448 | 0.131968339 |
| Esr2 | 15.55691602 | 0.35598228 | 0.167077388 | 2.130643073 | 0.033118558 | 0.132184168 |
| Tnfrsf10b | 35.00027664 | 0.35699303 | 0.167546508 | 2.130710062 | 0.033113036 | 0.132184168 |
| Xpo5 | 2047.955227 | 0.10528917 | 0.049420804 | 2.130462515 | 0.033133447 | 0.132209537 |
| Zwilch | 121.533348 | 0.272662856 | 0.128065239 | 2.129093408 | 0.033246533 | 0.132626618 |
| Acbd4 | 1079.654013 | 0.153632732 | 0.072172236 | 2.128695766 | 0.033279439 | 0.132723715 |
| Kansl1l | 422.3079601 | 0.22806521 | 0.107149577 | 2.128475127 | 0.03329771 | 0.13273691 |
| Ldlr | 1005.89088 | -0.282710226 | 0.13282453 | -2.128448907 | 0.033299882 | 0.13273691 |
| 2900011O08Rik | 10355.8693 | 0.120134992 | 0.056452802 | 2.128060753 | 0.033332047 | 0.13283095 |
| Ttll8 | 51.63338781 | 0.352841496 | 0.165840956 | 2.127589615 | 0.033371124 | 0.13295248 |
| Cox14 | 1304.567424 | -0.162555134 | 0.076411324 | -2.127369689 | 0.033389379 | 0.132974238 |
| Ttll3 | 327.4116906 | 0.265522551 | 0.124815693 | 2.127317039 | 0.03339375 | 0.132974238 |
| Nek6 | 1075.976128 | 0.174827436 | 0.08219217 | 2.127057075 | 0.033415341 | 0.133026027 |
| Pitrm1 | 1510.592592 | 0.161486221 | 0.075940755 | 2.126476378 | 0.033463614 | 0.133183981 |
| Elfn1 | 1310.65856 | -0.18230414 | 0.085756048 | -2.125845871 | 0.033516095 | 0.133358597 |
| Vwa1 | 1101.852367 | -0.218734173 | 0.10292484 | -2.125183504 | 0.033571304 | 0.133543975 |
| Gnai1 | 5219.941118 | -0.131434769 | 0.061853726 | -2.124928878 | 0.033592548 | 0.133581176 |
| Tmc4 | 164.3382969 | 0.348565855 | 0.164049404 | 2.124761482 | 0.03360652 | 0.133581176 |
| Upf1 | 2918.368698 | 0.128247191 | 0.060357382 | 2.12479709 | 0.033603548 | 0.133581176 |
| Dtx3l | 286.2162849 | 0.238376962 | 0.112222907 | 2.124138187 | 0.03365859 | 0.133702991 |
| Elac1 | 1688.93885 | 0.181404783 | 0.085417852 | 2.123733846 | 0.033692405 | 0.133702991 |
| Golm1 | 1385.983488 | 0.272291981 | 0.128202677 | 2.123918054 | 0.033676996 | 0.133702991 |
| Gpank1 | 502.0097412 | 0.177390287 | 0.083526005 | 2.123773155 | 0.033689117 | 0.133702991 |
| Os9 | 5637.186447 | 0.110874366 | 0.052202793 | 2.12391636 | 0.033677138 | 0.133702991 |
| Tmem86b | 177.0758991 | 0.310099785 | 0.146020558 | 2.123672101 | 0.033697572 | 0.133702991 |
| Zbtb41 | 841.5755393 | -0.183872378 | 0.086569341 | -2.123989569 | 0.033671016 | 0.133702991 |
| Bex1 | 1063.092331 | -0.140067901 | 0.065983478 | -2.122772329 | 0.033772935 | 0.133967703 |
| Scd2 | 69037.32649 | -0.124472267 | 0.058640426 | -2.122635785 | 0.033784384 | 0.133978819 |
| Dhrs3 | 584.3765511 | 0.179884572 | 0.08476699 | 2.12210641 | 0.033828803 | 0.134120644 |
| Nup214 | 1897.38163 | 0.14690944 | 0.069234808 | 2.121901458 | 0.033846014 | 0.134154551 |
| Tmem132c | 347.3546684 | 0.181323759 | 0.085458332 | 2.121779761 | 0.033856237 | 0.134160751 |
| Ctsc | 315.5048445 | 0.228576538 | 0.107750807 | 2.121344096 | 0.033892856 | 0.134237196 |
| Serpine2 | 7362.563866 | -0.129019944 | 0.060819141 | -2.121370712 | 0.033890618 | 0.134237196 |
| Mcf2l | 6946.176185 | 0.145005634 | 0.068365758 | 2.121027221 | 0.033919512 | 0.134308437 |
| B3galt6 | 431.4777387 | -0.243382035 | 0.114776717 | -2.120482632 | 0.033965365 | 0.134455636 |
| Fam65a | 3955.520274 | 0.171814918 | 0.081051445 | 2.119825476 | 0.034020766 | 0.134573122 |
| Nfx1 | 3496.302576 | 0.152311891 | 0.071853617 | 2.119752581 | 0.034026916 | 0.134573122 |
| Sstr5 | 36.12959242 | 0.356362869 | 0.16811801 | 2.11971858 | 0.034029785 | 0.134573122 |
| St3gal6 | 434.5377962 | 0.167941318 | 0.079221096 | 2.119906512 | 0.03401393 | 0.134573122 |
| Vwa5a | 689.854735 | 0.19774887 | 0.093298451 | 2.119530046 | 0.034045697 | 0.134601693 |
| Grm8 | 689.702143 | -0.218380042 | 0.103052155 | -2.119121558 | 0.034080195 | 0.134650794 |
| Krt9 | 560.4347328 | -0.238687063 | 0.11263742 | -2.119074298 | 0.034084188 | 0.134650794 |
| Nucks1 | 4583.423902 | -0.114280163 | 0.053928729 | -2.11909618 | 0.034082339 | 0.134650794 |
| Atp6v0a2 | 1955.901211 | -0.114580418 | 0.054074652 | -2.118930285 | 0.034096359 | 0.134664539 |
| Ddx58 | 248.7023126 | 0.227925767 | 0.107572777 | 2.118805264 | 0.034106928 | 0.134669647 |
| Syngr2 | 319.9617977 | 0.229603547 | 0.108369534 | 2.118709372 | 0.034115036 | 0.134669647 |
| Tmem50a | 2850.485168 | -0.126500733 | 0.059721187 | -2.118188534 | 0.034159104 | 0.13480926 |
| Timm8a1 | 454.9576045 | -0.20390612 | 0.096270249 | -2.118059538 | 0.034170026 | 0.134818023 |
| Pld2 | 909.8046348 | 0.144597792 | 0.068285633 | 2.117543411 | 0.034213755 | 0.134921843 |
| Zkscan14 | 332.4571017 | 0.204141601 | 0.096402166 | 2.117603876 | 0.03420863 | 0.134921843 |
| Gins3 | 105.321586 | -0.264198333 | 0.124830979 | -2.116448469 | 0.034306683 | 0.135219474 |
| Med28 | 1866.959401 | -0.153260739 | 0.072411867 | -2.116514128 | 0.034301105 | 0.135219474 |
| Cep97 | 820.6691898 | 0.154162583 | 0.072844373 | 2.116327945 | 0.034316926 | 0.135225444 |
| Cntnap5b | 582.6756277 | -0.247419356 | 0.116926732 | -2.116020449 | 0.034343068 | 0.13525966 |
| Rasd1 | 258.9847984 | 0.355586793 | 0.16804389 | 2.116035246 | 0.03434181 | 0.13525966 |
| Mepce | 1608.981537 | 0.124422084 | 0.058812329 | 2.115578255 | 0.034380693 | 0.135373432 |
| Cenpb | 5206.01625 | -0.141645111 | 0.066959877 | -2.115372934 | 0.034398175 | 0.135407856 |
| Coa4 | 75.58419141 | -0.318092756 | 0.150423219 | -2.114651971 | 0.034459621 | 0.135580844 |
| Gsg2 | 8.813447853 | -0.326455306 | 0.154371431 | -2.11473913 | 0.034452187 | 0.135580844 |
| Mtch1 | 13618.66032 | -0.1249414 | 0.059090352 | -2.114412843 | 0.034480022 | 0.135626671 |
| Gm13845 | 77.19867594 | -0.297208446 | 0.140572827 | -2.1142667 | 0.034492495 | 0.135641298 |
| Slc39a10 | 10441.43147 | -0.190042271 | 0.089899384 | -2.113944079 | 0.034520044 | 0.135715189 |
| Unc79 | 3327.730961 | -0.175094186 | 0.082859111 | -2.113155489 | 0.034587462 | 0.135945746 |
| Crh | 156.4182257 | -0.298948569 | 0.141483793 | -2.112952744 | 0.034604813 | 0.135979449 |
| Arpc5 | 2849.237723 | -0.169123045 | 0.080058778 | -2.112485973 | 0.034644788 | 0.136074988 |
| Camkv | 21322.28554 | -0.143717619 | 0.068034471 | -2.112423559 | 0.034650136 | 0.136074988 |
| Zfp445 | 7620.513962 | 0.209869766 | 0.099353159 | 2.112361282 | 0.034655473 | 0.136074988 |
| Shc4 | 279.8511066 | -0.224452601 | 0.106278014 | -2.111938239 | 0.034691747 | 0.136182906 |
| Ccnh | 1231.281942 | -0.170079393 | 0.080563575 | -2.111120241 | 0.034761978 | 0.136424034 |
| Eif2ak3 | 1323.162797 | 0.169226157 | 0.080169849 | 2.110845393 | 0.034785603 | 0.136482181 |
| Plekhb2 | 11615.06826 | -0.091263217 | 0.043241388 | -2.110552441 | 0.034810799 | 0.136517045 |
| 9-Sep | 2993.252897 | 0.222840286 | 0.105584629 | 2.110537195 | 0.034812111 | 0.136517045 |
| 4930444P10Rik | 9.365659291 | 0.342426437 | 0.162264369 | 2.110299632 | 0.034832555 | 0.136562655 |
| Zfp608 | 907.356129 | -0.22061875 | 0.104552115 | -2.110131874 | 0.034846999 | 0.136584719 |
| Sptbn2 | 46970.45474 | -0.19170332 | 0.090856201 | -2.109964078 | 0.03486145 | 0.136606805 |
| Hcn3 | 659.0854656 | -0.264051746 | 0.125196686 | -2.109095332 | 0.034936354 | 0.136796529 |
| Ppp1r3b | 60.83814376 | -0.333822599 | 0.158277189 | -2.109101137 | 0.034935853 | 0.136796529 |
| Trappc12 | 3000.529438 | 0.103227884 | 0.048940575 | 2.109249507 | 0.034923051 | 0.136796529 |
| Faim2 | 25017.38475 | 0.132112843 | 0.062644419 | 2.108932373 | 0.034950419 | 0.136817028 |
| S1pr4 | 21.01762786 | -0.354352448 | 0.168057935 | -2.108513638 | 0.034986584 | 0.136916399 |
| Slc25a18 | 2144.488223 | 0.240378948 | 0.114013823 | 2.108331625 | 0.035002314 | 0.136916399 |
| Synj2 | 3778.113495 | 0.221607935 | 0.105106608 | 2.108411071 | 0.034995448 | 0.136916399 |
| 2610306M01Rik | 164.644197 | -0.220220185 | 0.104476238 | -2.10784949 | 0.03504401 | 0.137044901 |
| Tmem63a | 1369.15628 | 0.258778364 | 0.122776292 | 2.10772259 | 0.035054992 | 0.137053255 |
| Mrpl19 | 1162.108425 | -0.124755558 | 0.059201774 | -2.107294247 | 0.035092082 | 0.137163651 |
| Gm16532 | 363.1437107 | -0.238761384 | 0.113314492 | -2.10706839 | 0.035111652 | 0.137205532 |
| Pcdhb16 | 298.4048613 | -0.236387646 | 0.112224897 | -2.106374362 | 0.035171847 | 0.137406099 |
| Fam96b | 467.3104517 | -0.150167119 | 0.071299882 | -2.106134185 | 0.035192698 | 0.137452903 |
| Anln | 1763.645618 | 0.334644743 | 0.158902135 | 2.105980159 | 0.035206076 | 0.137470499 |
| Cdkn2aip | 561.3969644 | 0.168279137 | 0.079918806 | 2.105626259 | 0.03523683 | 0.137555919 |
| Col4a1 | 2146.192049 | 0.238851785 | 0.113465747 | 2.105056294 | 0.035286408 | 0.137714762 |
| Ankrd13c | 2029.176599 | -0.147584055 | 0.070116083 | -2.104853098 | 0.035304097 | 0.137749103 |
| Aph1b | 1152.370522 | -0.184948802 | 0.087881153 | -2.104533184 | 0.035331963 | 0.137823121 |
| Six2 | 15.82095811 | -0.347471306 | 0.165118456 | -2.104375943 | 0.035345666 | 0.137841871 |
| Pacrg | 389.5055084 | -0.204618257 | 0.097265478 | -2.103708956 | 0.035403843 | 0.138034006 |
| Gatsl3 | 106.3687495 | 0.251485107 | 0.119561674 | 2.103392323 | 0.035431489 | 0.138107041 |
| D330023K18Rik | 84.87307231 | -0.282915503 | 0.13452858 | -2.10301412 | 0.035464535 | 0.138201084 |
| Ap2b1 | 18164.42652 | -0.12870801 | 0.061205065 | -2.102898016 | 0.035474685 | 0.138205877 |
| Atp8b5 | 14.69977951 | 0.352580741 | 0.167683754 | 2.102652964 | 0.035496116 | 0.138254608 |
| Zc3hav1 | 176.081695 | 0.269888488 | 0.128370375 | 2.102420346 | 0.03551647 | 0.138299119 |
| Irf1 | 293.1465801 | 0.315925057 | 0.15029788 | 2.101992768 | 0.035553909 | 0.138398496 |
| Nme3 | 498.88601 | -0.169691223 | 0.080734604 | -2.101840024 | 0.035567292 | 0.138398496 |
| Plxna2 | 7399.574262 | -0.219102504 | 0.104244035 | -2.101822941 | 0.035568789 | 0.138398496 |
| Wnt10b | 49.21543297 | 0.352603055 | 0.167776231 | 2.101626986 | 0.035585964 | 0.138430561 |
| Calr3 | 41.82614921 | 0.337932808 | 0.160827449 | 2.101213508 | 0.035622228 | 0.138536848 |
| Klhl21 | 780.5739772 | 0.178947664 | 0.08517621 | 2.100911325 | 0.035648751 | 0.138605207 |
| Mrps28 | 258.853129 | -0.189892783 | 0.090394247 | -2.100717583 | 0.035665765 | 0.138636568 |
| Pdxk | 9376.750051 | -0.142052554 | 0.067630061 | -2.100435111 | 0.035690583 | 0.138698243 |
| Eif5b | 3558.308912 | 0.151722216 | 0.072246124 | 2.100074127 | 0.035722321 | 0.13878677 |
| Ap5z1 | 861.5989474 | 0.221172604 | 0.105380299 | 2.098804101 | 0.035834173 | 0.139186432 |
| AU040972 | 8.768139707 | 0.313071648 | 0.149203251 | 2.098289722 | 0.03587956 | 0.139327795 |
| Shroom2 | 4334.40958 | -0.158300015 | 0.075449051 | -2.098104792 | 0.035895889 | 0.139356279 |
| Lmo2 | 707.1828253 | -0.182322705 | 0.08690583 | -2.097934116 | 0.035910966 | 0.139356415 |

| | | | | | | | |
|---|---|---|---|---|---|---|---|
| Lrwd1 | 479.747393 | -0.114623294792422 | 0.0810457407015879 | -1.41430374749078 | 0.157272737122258 | 0.60787851646781 | NM_027891 |
| Prkar2b | 2690.899252 | 0.105763770890771 | 0.0747873593996822 | 1.4141931435972 | 0.157305200564677 | 0.60787851646781 | NM_011158 |
| Mfap2 | 116.0131626 | -0.132934899474835 | 0.0940234446584979 | -1.41384842852404 | 0.157406410756502 | 0.608107851768338 | NM_001161799,NM_008546 |
| Alcam | 6409.684531 | 0.0905248915367011 | 0.0640534322365482 | 1.41327151997717 | 0.15757590458517 | 0.60814117265812 | NM_009655 |
| Aurkaip1 | 1699.031199 | 0.0731870520470223 | 0.0517905118461723 | 1.41313629539715 | 0.157615653119082 | 0.60814117265812 | NM_025338 |
| C7 | 32.80899545 | -0.122208141403369 | 0.0864987318662598 | -1.41283159610156 | 0.15770524567273 | 0.60814117265812 | NM_001243837 |
| Fignl1 | 30.27093336 | -0.128171182359383 | 0.0907291257800027 | -1.4126795696254 | 0.157749961350392 | 0.60814117265812 | NM_021891,NM_001163360,NM_001163359 |
| Map6 | 5643.015192 | 0.0574811005107878 | 0.0406788792649344 | 1.41304533333928 | 0.15764239519751 | 0.60814117265812 | NM_010837,NM_001048167,NM_001043355 |
| Prkag1 | 1995.653115 | 0.0773996051019833 | 0.0547882159618267 | 1.41270533714606 | 0.157742381651513 | 0.60814117265812 | NM_016781 |
| Rad54b | 59.82860688 | -0.130006810801078 | 0.0920112263996379 | -1.4129450925522 | 0.157671869125465 | 0.60814117265812 | NM_001039556,NM_001256145 |
| Spsb1 | 903.7217685 | 0.113407034626348 | 0.0802253834792036 | 1.41360539156221 | 0.157477797365292 | 0.60814117265812 | NM_029035 |
| Gdap1l1 | 2891.989408 | 0.0777311340425882 | 0.0550348677174253 | 1.41239794454847 | 0.1578328213517 | 0.608299167671742 | NM_144891 |
| St8sia1 | 2494.507917 | 0.0918124628931074 | 0.0650249785234985 | 1.41195683532488 | 0.157962671188646 | 0.608476733409247 | NM_011374 |
| Yars2 | 547.6729949 | 0.108412221620464 | 0.0767780689392108 | 1.4120206866143 | 0.157943870203051 | 0.608476733409247 | NM_198246,NR_038046 |
| E130309F12Rik | 685.5043686 | 0.10920387089911 | 0.0773551524486607 | 1.41172071209589 | 0.158032212300398 | 0.608583222660545 | NM_178756 |
| Rfc2 | 1170.782329 | -0.0771847969256984 | 0.0546846921553406 | -1.41145161257272 | 0.158111493592613 | 0.6087271550295 | NM_020022 |
| 9630013A20Rik | 38.18985316 | -0.124609162091727 | 0.0883258968986699 | -1.41078852824651 | 0.158306978056942 | 0.609156864438447 | NR_015539 |
| Slc35e1 | 4289.669986 | 0.0626912913059381 | 0.0444335897469935 | 1.4108986391355 | 0.158274503504568 | 0.609156864438447 | NM_177766 |
| Snx32 | 3313.166463 | -0.0946401099836406 | 0.0670947240227438 | -1.41054473898066 | 0.158378895775072 | 0.609272203397432 | NM_001024560 |
| Cybasc3 | 596.6680083 | -0.100955134842225 | 0.0715815818251242 | -1.41035071128858 | 0.158436151529294 | 0.609331092696461 | NM_201351 |
| Tbp | 670.3561402 | -0.10043721274457 | 0.0712242350581028 | -1.41015502184946 | 0.158493913521404 | 0.609391897250374 | NM_013684 |
| Drap1 | 3784.359968 | 0.0783525093924473 | 0.0555778794628673 | 1.40977867723068 | 0.158605044629431 | 0.609496528647387 | NM_024176 |
| Kri1 | 1452.862903 | -0.0793509350884947 | 0.0562829911912558 | -1.40985639549347 | 0.158582090308911 | 0.609496528647387 | NM_145416 |
| Cenpe | 68.50988644 | -0.134045865500141 | 0.0950969518568217 | -1.40957057910711 | 0.158666519414195 | 0.609571505160169 | NM_173762 |
| Acad10 | 133.1651685 | -0.133984852105709 | 0.0950839267116197 | -1.40912199084996 | 0.158799099333109 | 0.609919544397868 | NM_028037 |
| D130043K22Rik | 1874.79223 | -0.0971309243316727 | 0.0689576481391107 | -1.40855912219812 | 0.158965573326931 | 0.61039754643061 | NM_001081051 |
| Mtmr7 | 3616.279528 | 0.0836274755080139 | 0.0593940357908469 | 1.40801133303189 | 0.159127714192551 | 0.610858661829015 | NM_001040699 |
| Lzts1 | 1702.330054 | 0.117243010179412 | 0.0832796887423383 | 1.40782238682656 | 0.15918366965457 | 0.610912017279337 | NM_199364 |
| Rgag1 | 109.057169 | 0.132633036802684 | 0.0942235542924552 | 1.40764204660559 | 0.159237090382551 | 0.610955619359994 | NM_001040434 |
| Elovl6 | 2653.400992 | 0.108064993599244 | 0.076783746627057 | 1.4073941211038 | 0.15931055349264 | 0.611076076058646 | NM_130450 |
| Scn1a | 7794.940686 | 0.0991656311419825 | 0.0704821552352631 | 1.40696082307609 | 0.159439006124536 | 0.611407339747892 | NM_018733 |
| Cdh24 | 405.6839497 | -0.126997202664552 | 0.0902937929628159 | -1.40648873524287 | 0.159579047288808 | 0.611570722944484 | NM_199470 |
| Cops5 | 2455.900228 | 0.0685730929744916 | 0.0487676257607674 | 1.40611916009365 | 0.15968874377732 | 0.611570722944484 | NM_013715 |
| Kcnip2 | 7927.653454 | -0.0870227159285258 | 0.0618890791231933 | -1.40610778446554 | 0.159692121170896 | 0.611570722944484 | NM_001276358,NM_145704,NM_030716,NM_145703 |
| Kcnj2 | 868.5950087 | -0.126487135513828 | 0.089921680701411 | -1.40663669236604 | 0.159535146958777 | 0.611570722944484 | NM_008425 |
| Mertk | 2079.525796 | 0.0955512877302779 | 0.0679524577923826 | 1.40614910533801 | 0.159679853372599 | 0.611570722944484 | NM_008587 |
| Slc45a4 | 2108.80798 | 0.0837323389496933 | 0.0595634758139999 | 1.40576650045014 | 0.159793472607571 | 0.611797570663567 | NM_001033219,NM_001168255 |
| Abcd1 | 635.2655377 | -0.106508684181913 | 0.0757834140214785 | -1.40543528629796 | 0.159891880092519 | 0.612013030362035 | NM_007435 |
| Efna3 | 1430.919074 | 0.0867965644130342 | 0.0617683966426448 | 1.40519374195817 | 0.159963674549035 | 0.612126537539325 | NM_010108 |
| Gprasp1 | 27616.76366 | 0.0871963750839792 | 0.0620689074535006 | 1.40483180164405 | 0.160071300040404 | 0.612215825273014 | NM_026081,NM_001004359,NM_001005385 |
| Mpv17l2 | 435.6982943 | 0.108664116330151 | 0.0773456547727786 | 1.40491559156591 | 0.160046379650701 | 0.612215825273014 | NM_183170 |
| Egln2 | 2929.720113 | 0.088977860534332 | 0.0633481364784675 | 1.40458528822827 | 0.160144633869554 | 0.612291376698903 | NM_053208 |
| Sae1 | 3776.683371 | 0.0644032198437483 | 0.0458601202734364 | 1.40434040424993 | 0.160217508111836 | 0.612291376698903 | NM_019748 |
| Vti1a | 2991.907455 | 0.0740447332555419 | 0.0527226410347393 | 1.40442003288024 | 0.160193808931705 | 0.612291376698903 | NM_016862 |
| Clk3 | 2222.913405 | -0.0940916240788232 | 0.0670211190633535 | -1.40391007183692 | 0.160345630091512 | 0.612373044071061 | NM_007713 |
| Map4k4 | 4434.556103 | -0.0776282505003142 | 0.0552959581733161 | -1.40386843929897 | 0.160358029368337 | 0.612373044071061 | NM_001252200,NM_008696,NM_001252202,NM_001252201 |
| Pllp | 1971.773029 | -0.108619453909361 | 0.0773728879155972 | -1.40384386360051 | 0.160365349004978 | 0.612373044071061 | NM_026385 |
| 4732491K20Rik | 167.2815483 | -0.129208945499948 | 0.0921213584834035 | -1.40259487731313 | 0.160737680264537 | 0.612387888326862 | NR_045290 |
| Cyc1 | 5281.680731 | 0.0736849741723901 | 0.0525093897086576 | 1.40327234007523 | 0.160535643052543 | 0.612387888326862 | NM_025567 |
| Gnai1 | 5740.643234 | 0.0758508009938609 | 0.054078550694351 | 1.40260417522218 | 0.160734906083801 | 0.612387888326862 | NM_010305 |
| Lpar1 | 1154.151294 | 0.0970328607496112 | 0.0691446071261408 | 1.40333230287351 | 0.160517769818552 | 0.612387888326862 | NM_172989,NM_010336 |
| Mmp17 | 9912.905129 | 0.0917082494104174 | 0.0653826428778989 | 1.40263907015324 | 0.160724494942454 | 0.612387888326862 | NM_011846 |
| Pcdh7 | 5410.637317 | 0.100493309415753 | 0.0716336629062359 | 1.40287827452426 | 0.160653140400544 | 0.612387888326862 | NM_018764,NM_001122758 |
| Rcn2 | 4249.444776 | 0.0702731547724948 | 0.050096845456442 | 1.40274610371616 | 0.160692563925409 | 0.612387888326862 | NM_011992 |
| Spats1 | 81.99153171 | -0.133225875522596 | 0.0949877777963105 | -1.40255808287548 | 0.160748658831772 | 0.612387888326862 | NM_027649 |
| Zfp59 | 355.6464229 | 0.103640601697359 | 0.0738876108361353 | 1.40267902188919 | 0.160712575694459 | 0.612387888326862 | NM_011762 |
| Abcb1a | 1676.852 | -0.0971786304598092 | 0.069312317476263 | -1.40203983935596 | 0.160903350317181 | 0.612415570556777 | NM_011076 |
| Arhgap31 | 1721.422996 | 0.110217915284472 | 0.0786236401904132 | 1.40184192715503 | 0.160962455169914 | 0.612415570556777 | NM_020260 |
| Cenpt | 572.5364358 | -0.102151463931479 | 0.0728620026562125 | -1.40198539989991 | 0.160919606578405 | 0.612415570556777 | NM_177150 |
| Cgref1 | 730.8393697 | 0.103368742647738 | 0.0737287401746886 | 1.4020142267835 | 0.160910998377704 | 0.612415570556777 | NM_026770,NM_001160149 |
| Zfp384 | 1736.961917 | -0.0718506503085345 | 0.0512549822044221 | -1.40182763154555 | 0.160966725071305 | 0.612415570556777 | NM_001252083,NM_175557 |
| Canx | 14191.647 | 0.0781531501881964 | 0.0557775510981328 | 1.40115778928151 | 0.161166893671935 | 0.6127243874567 | NM_001110499,NM_001110500,NM_007597 |
| Galr1 | 22.65099282 | 0.0958754481651185 | 0.0684271076079044 | 1.40113255574818 | 0.161174437868102 | 0.6127243874567 | NM_008082 |
| Micall2 | 87.66224489 | -0.133190882094686 | 0.0950467757784393 | -1.40131930834943 | 0.161118609822893 | 0.6127243874567 | NM_174850 |
| Lgalsl | 3164.649639 | 0.0890743006632633 | 0.0635834071349881 | 1.40090480640897 | 0.161242541300337 | 0.612822908144608 | NM_173752 |
| Cox4i1 | 13085.91229 | 0.0612096990624452 | 0.0437014362764787 | 1.40063357815518 | 0.161323674497745 | 0.61297088562758 | NM_009941 |
| Wdr35 | 2113.092058 | 0.0823941690568442 | 0.0588326019590792 | 1.40048487255677 | 0.161368170265507 | 0.612979613304591 | NM_001159527,NM_172470 |
| Pip4k2a | 4329.728961 | 0.0951818812522575 | 0.0679847987783825 | 1.40004652455518 | 0.161499386913791 | 0.613176432015876 | NM_008845 |
| Rpain | 355.9660153 | -0.106117973839678 | 0.0757969416983671 | -1.40002975663547 | 0.161504407881918 | 0.613176432015876 | NM_001252413,NM_027186,NM_001252415,NM_001252414 |
| Hes6 | 379.405116 | 0.118683449519854 | 0.0848128701790609 | 1.39935659846535 | 0.161706075017454 | 0.613781668592512 | NM_019479 |
| Ctdsp2 | 5139.616083 | -0.0918834103188485 | 0.0656880659962531 | -1.39878391798123 | 0.161877790306416 | 0.613952162399738 | NM_001113470,NM_146012 |
| Htr7 | 466.6932084 | 0.123787968573896 | 0.0884968210332412 | 1.39878435325263 | 0.161877659740285 | 0.613952162399738 | NM_008315 |
| Thsd1 | 192.0053297 | -0.119176689425182 | 0.0851945729424799 | -1.39887654000737 | 0.16185000874285 | 0.613952162399738 | NM_001205253,NM_019576 |
| 1500011B03Rik | 3262.675584 | 0.0714240246891991 | 0.0510970550044538 | 1.39781098309821 | 0.162169835324222 | 0.613957643347272 | NR_027818,NR_027817 |
| Adcy1 | 39493.82823 | 0.0982658806368412 | 0.0703097566295708 | 1.39761372172795 | 0.162229095549332 | 0.613957643347272 | NM_009622 |
| Appbp2 | 1861.088909 | 0.100950959526721 | 0.0722388031743427 | 1.397461683897 | 0.162274781106427 | 0.613957643347272 | NM_025825 |
| Fgf2 | 24.95742788 | 0.118072267513064 | 0.0844545558276558 | 1.3980568171362 | 0.162096006020578 | 0.613957643347272 | NM_008006 |
| Fli1 | 154.3597814 | -0.124846335614247 | 0.0893436993005415 | -1.39737146090491 | 0.162301896630423 | 0.613957643347272 | NM_008026 |
| Mid2 | 1281.701213 | 0.0972727139235238 | 0.0696084399878301 | 1.39742700656027 | 0.162285202592585 | 0.613957643347272 | NM_011845 |
| Ndufa12 | 2541.607643 | 0.0856014629397471 | 0.0612588446635047 | 1.39737312072986 | 0.162301397757495 | 0.613957643347272 | NM_025551 |
| Pfkm | 19955.37311 | 0.0702177187638573 | 0.0502308986864431 | 1.39789891481293 | 0.1621434246049 | 0.613957643347272 | NM_001163487,NM_021514,NM_001163488 |
| Rpusd3 | 268.4154605 | -0.105632664196236 | 0.075560169055366 | -1.3979940161176 | 0.162114864118278 | 0.613957643347272 | NM_001033204 |
| Sgcb | 2615.726059 | -0.0625590976543151 | 0.0447487936168222 | -1.39800635051751 | 0.162111160173193 | 0.613957643347272 | NM_011890 |
| Rsg1 | 43.84270447 | 0.130824187306735 | 0.093636628828194 | 1.39714755800076 | 0.162369202946057 | 0.614052341055563 | NM_001081174 |
| 6330409D20Rik | 71.98671376 | 0.132131511355011 | 0.0949607095219409 | 1.3914334888629 | 0.164094015367313 | 0.614351407822208 | NM_027529 |
| BC051142 | 89.41708228 | -0.132274599140657 | 0.0950011506690986 | -1.39234733694318 | 0.163817243505659 | 0.614351407822208 | NM_001163855,NM_001001177 |
| Cage1 | 147.550461 | -0.12798020306494 | 0.0916927101839726 | -1.39575112141587 | 0.162789454016602 | 0.614351407822208 | NM_027724 |
| Card9 | 64.37166191 | -0.132554426134323 | 0.0951005105439133 | -1.3938350633051 | 0.163367417213821 | 0.614351407822208 | NM_001037747 |
| Cbfb | 675.8876827 | 0.108244174901307 | 0.077571229847846 | 1.39541651091037 | 0.162890275225885 | 0.614351407822208 | NM_001161456,NM_001161457,NM_022309,NM_001161458 |
| Cdc23 | 2157.126353 | -0.0696506714260206 | 0.0500622339484709 | -1.39128172941128 | 0.164140011961861 | 0.614351407822208 | NM_178347 |
| Chpf | 3308.311216 | 0.0873555248646646 | 0.0626478229942468 | 1.39439043033765 | 0.163199736325449 | 0.614351407822208 | NM_001001566,NM_001001565 |
| Cops2 | 3156.898681 | 0.0928597974926208 | 0.0666688235809335 | 1.39285189845734 | 0.163664580704833 | 0.614351407822208 | NM_009939 |
| Ctcf | 2899.360203 | 0.0838069384455466 | 0.0602335493217249 | 1.39136642932844 | 0.16411433916549 | 0.614351407822208 | NM_181322 |
| Eif2c2 | 2829.732244 | 0.107958387186089 | 0.077325986262825 | 1.39614626859264 | 0.162670453174579 | 0.614351407822208 | NM_153178 |
| Fank1 | 169.4674352 | -0.132398502119962 | 0.0948417353658003 | -1.39599409067439 | 0.16271627465684 | 0.614351407822208 | NM_025850 |
| Fbf1 | 3256.886916 | -0.0823479676161803 | 0.0590422171574169 | -1.39473027235116 | 0.163097192506558 | 0.614351407822208 | NM_172571 |
| Hnrnph2 | 5113.969162 | 0.066954523735043 | 0.0480939768074079 | 1.39216026994736 | 0.163873870745641 | 0.614351407822208 | NM_019868 |
| Irak2 | 841.2951888 | 0.097321969195208 | 0.0698745347885844 | 1.39281026327645 | 0.163677174005026 | 0.614351407822208 | NM_172161,NM_001113553 |
| Itch | 3050.060612 | 0.0975490243903421 | 0.0700740181898262 | 1.39208549631174 | 0.163896509674596 | 0.614351407822208 | NM_001243712,NM_008395 |
| Krr1 | 1824.523853 | 0.0869671686358402 | 0.0623699388185894 | 1.39437636597328 | 0.16320398114948 | 0.614351407822208 | NM_178610 |
| Lrmp | 67.03359217 | -0.132497984117924 | 0.0951014477876875 | -1.39322783406751 | 0.163550905364052 | 0.614351407822208 | NM_008511 |
| Mcm7 | 512.0253477 | 0.0949721517671848 | 0.0682573454892371 | 1.39138360987332 | 0.164109132060398 | 0.614351407822208 | NM_008568 |
| Mllt11 | 10874.70604 | 0.0730351868027351 | 0.052474897609834 | 1.39181189729559 | 0.163979366295315 | 0.614351407822208 | NM_019914 |
| Morn2 | 289.4863291 | 0.123895495784663 | 0.08890872245939 | 1.39351339618285 | 0.163464596924429 | 0.614351407822208 | NM_194269 |
| Mrpl12 | 1621.573403 | 0.0717667890301111 | 0.0515430243192795 | 1.39236666800064 | 0.163811392622403 | 0.614351407822208 | NM_027204 |
| Ninj1 | 407.287414 | 0.114314141908209 | 0.0819486340948226 | 1.39494871599613 | 0.163031305048508 | 0.614351407822208 | NM_013610 |
| Nploc4 | 3402.602247 | 0.0590758880399725 | 0.0424280627627233 | 1.3923776904534 | 0.163808056554439 | 0.614351407822208 | NM_199469,NM_001195023 |
| Paics | 4063.189962 | 0.0768344441541906 | 0.0551260486607468 | 1.39379560155019 | 0.163379336765256 | 0.614351407822208 | NM_025939 |
| Pdhx | 1779.536542 | 0.0737042139377627 | 0.0528323403287103 | 1.39505866064597 | 0.162998150893164 | 0.614351407822208 | NM_175094 |
| Pla2g4b | 168.9457738 | -0.132556652071104 | 0.0950396937722908 | -1.39475041227196 | 0.163091117018704 | 0.614351407822208 | NM_145378 |
| Psmc4 | 3494.689964 | 0.0639514207107364 | 0.0458611874545337 | 1.39445627687109 | 0.163179864017811 | 0.614351407822208 | NM_011874 |
| Ptges2 | 1220.606399 | 0.0844035589379489 | 0.060473923260413 | 1.39570172377423 | 0.162804335016238 | 0.614351407822208 | NM_133783 |
| Rab5c | 5730.428263 | 0.0532600949008553 | 0.0381764459596929 | 1.39510353994418 | 0.162984618860925 | 0.614351407822208 | NM_024456 |
| Rassf9 | 44.66290788 | -0.125362544533741 | 0.0900483459720896 | -1.3921693194964 | 0.163871131007994 | 0.614351407822208 | NM_146240 |
| Rbx1 | 3532.351187 | 0.0580380910865191 | 0.0415932217694493 | 1.39537378008906 | 0.162903153803895 | 0.614351407822208 | NM_019712 |
| Rin3 | 156.7518132 | 0.132168765820051 | 0.0948902128675679 | 1.39285983059718 | 0.163662181571029 | 0.614351407822208 | NM_001161365,NM_177620 |
| Robo2 | 4895.781865 | 0.098572990105996 | 0.070728833783115 | 1.39367475516793 | 0.163415842886256 | 0.614351407822208 | NM_175549 |
| Sfrp5 | 101.0748739 | 0.0965783828799665 | 0.0693261027861394 | 1.39310272752381 | 0.163588728448731 | 0.614351407822208 | NM_018780 |
| Shcbp1l | 61.97514073 | -0.131263197542078 | 0.0940201342846597 | -1.39611795431667 | 0.162678977998253 | 0.614351407822208 | NM_001033162 |
| Spon1 | 3256.201236 | 0.0869903714727845 | 0.0624978098906593 | 1.39189471799052 | 0.16395428157768 | 0.614351407822208 | NM_145584 |
| Stac3 | 66.10831339 | -0.120494373358177 | 0.0865786566716317 | -1.39173299737345 | 0.164003266179572 | 0.614351407822208 | NM_177707 |
| Ube2l6 | 217.2373141 | -0.12662334016326 | 0.0908962041812032 | -1.39305421281217 | 0.163603397529361 | 0.614351407822208 | NM_019949 |
| Wnt2b | 147.142491 | 0.128082734638493 | 0.0920211664026259 | 1.39188340732484 | 0.163957707179646 | 0.614351407822208 | NM_009520 |
| Zcwpw1 | 291.4654278 | -0.106746354960035 | 0.0764766574339467 | -1.39580309262643 | 0.162773798838936 | 0.614351407822208 | NM_001005426 |
| 2810029C07Rik | 412.8439442 | -0.120281270693636 | 0.0864676760743473 | -1.3910548856758 | 0.164208783874144 | 0.614450488036016 | NR_045295 |
| Dlk1 | 647.2709613 | 0.0764158979550128 | 0.0549440737678773 | 1.39079417878346 | 0.164287848841114 | 0.614588022731398 | NR_033813,NM_001190704,NM_001190705,NM_001190703,NM_010052 |
| Bfsp2 | 148.9153738 | -0.131630761217304 | 0.0946665028115935 | -1.39046819421731 | 0.164386751007205 | 0.614641427318059 | NM_001002896 |
| Mettl7a1 | 1222.799213 | 0.118394174584681 | 0.085141508264549 | 1.39055763748994 | 0.164359609880664 | 0.614641427318059 | NM_027334 |
| 4930590J08Rik | 24.96602838 | -0.103986080544828 | 0.0747975431000773 | -1.39023390655622 | 0.164457860462163 | 0.614688762098491 | NM_198668 |
| Chtf8 | 3517.679086 | 0.0754512897003417 | 0.0542762454418881 | 1.39013465441572 | 0.164487991806408 | 0.614688762098491 | NM_145412 |
| Glyr1 | 10006.71581 | 0.0597476059959457 | 0.0429922515858467 | 1.38972963248138 | 0.164610993016875 | 0.614688762098491 | NM_001079814,NM_028720 |
| Gsto2 | 30.5830771 | -0.122859787007521 | 0.088401222077893 | -1.38979738197811 | 0.164590413332869 | 0.614688762098491 | NM_030051,NM_026619 |
| Nudcd2 | 405.8377132 | -0.105067378312954 | 0.0755973110354817 | -1.38982957030892 | 0.164580636438465 | 0.614688762098491 | NM_026023 |
| Exoc5 | 2021.174596 | 0.102120150908141 | 0.0735359670226104 | 1.38871024673874 | 0.164920877061868 | 0.615071432573389 | NM_207214 |
| Gopc | 2556.728171 | 0.0905655303026788 | 0.0652162349485402 | 1.38869608731845 | 0.164925184488046 | 0.615071432573389 | NM_001199272,NM_053187 |
| Kitl | 1895.328179 | 0.119601689349484 | 0.0861205861750407 | 1.38877003352476 | 0.164902690304244 | 0.615071432573389 | NM_013598 |
| Plekhg4 | 123.8804362 | -0.131731198256966 | 0.0948560999687588 | -1.38874778006214 | 0.164909459488711 | 0.615071432573389 | NM_001081333 |
| Rbm38 | 82.18855437 | -0.131280497230662 | 0.0945084475693078 | -1.38908743723028 | 0.164806163414676 | 0.615071432573389 | NM_019547 |
| Gpr146 | 1111.181766 | 0.0888054317256396 | 0.0639801331499683 | 1.38801573790854 | 0.165132252872733 | 0.615077498412757 | NM_001038703,NM_030258 |
| Nudt1 | 141.1066652 | 0.127136700457935 | 0.0915608358744153 | 1.38854892753835 | 0.164969956864402 | 0.615077498412757 | NM_008637 |
| Prss50 | 29.61765786 | 0.122216667781037 | 0.0880373652715941 | 1.38823631765899 | 0.165065096687845 | 0.615077498412757 | NM_146227 |
| Rpl22 | 3478.564206 | -0.0680299761798161 | 0.0490131229892997 | -1.38799513335781 | 0.165138527041839 | 0.615077498412757 | NM_009079 |
| Zfp536 | 610.5845141 | 0.110110512961065 | 0.07933050851245 | 1.38799706475832 | 0.165137938914946 | 0.615077498412757 | NM_172385 |
| Chic1 | 1526.332113 | 0.0981378180217304 | 0.0707370434639463 | 1.38736103766833 | 0.165331699489506 | 0.615639135294684 | NM_009767 |
| Eif2s3x | 8043.224867 | 0.073604096433881 | 0.0530705947620838 | 1.38690920582008 | 0.165469450368032 | 0.615678595298677 | NM_012010 |
| Plekho1 | 1431.114809 | -0.09543635613395 | 0.0688101075613532 | -1.38695257886141 | 0.165456223396035 | 0.615678595298677 | NM_023320 |
| Rfx5 | 1003.066723 | -0.0991204928604116 | 0.071458710076763 | -1.38710162489546 | 0.165410776625211 | 0.615678595298677 | NM_017395 |
| Dpp3 | 1559.908959 | 0.0826993869570228 | 0.0596501732586996 | 1.38640648365529 | 0.165622817764818 | 0.616091434277015 | NM_133803 |
| Aldh3a1 | 54.35531189 | 0.127340577028757 | 0.0918735695254181 | 1.38604146640375 | 0.165734241995559 | 0.616168889440416 | NM_007436,NM_001112725 |
| Bcl6b | 83.06419659 | -0.124116656030883 | 0.0895667095113694 | -1.38574540371083 | 0.165824658749041 | 0.616168889440416 | NM_007528 |
| Polr2h | 571.0716747 | 0.0915113760087272 | 0.0660402263873507 | 1.38569143406594 | 0.165841144932038 | 0.616168889440416 | NM_145632 |
| Ptgfrn | 2200.037166 | 0.0903223063928302 | 0.0651762071301649 | 1.38581716196595 | 0.165802740563819 | 0.616168889440416 | NM_011197 |
| Scd2 | 76254.07008 | 0.0894688352209905 | 0.0645684284683342 | 1.38564368598297 | 0.165855731633762 | 0.616168889440416 | NM_009128 |
| Srsf7 | 2788.49791 | -0.0812113387983113 | 0.05861718530925 | -1.38545271953711 | 0.165914080181765 | 0.616228056435777 | NM_001195486,NM_001195487,NM_001195485,NM_146083,NR_036615 |
| Azin1 | 6655.628201 | 0.100058375583423 | 0.072240653866632 | 1.38507018178638 | 0.166031008531613 | 0.616354840087141 | NM_001102458,NM_018745 |
| Cdk5rap2 | 625.3186453 | -0.110638938447652 | 0.0798960601391756 | -1.38478591128177 | 0.166117940172185 | 0.616354840087141 | NM_145990 |
| Hspa9 | 10110.92277 | 0.0744925421271577 | 0.0537883287310203 | 1.38492018407326 | 0.166076874475019 | 0.616354840087141 | NM_010481 |
| Rab6b | 48049.33502 | 0.0634379693943027 | 0.0458086827501474 | 1.38484596338013 | 0.166099573025751 | 0.616354840087141 | NM_173781 |
| Tuba1b | 10246.85473 | 0.0617377774043935 | 0.0445941592353784 | 1.3844364029497 | 0.166224868847374 | 0.61659408704723 | NM_011654 |
| Dnmt3b | 34.18673748 | 0.115914328192945 | 0.0837513863695937 | 1.38402876916469 | 0.166349645818558 | 0.616741948739248 | NM_001271746,NM_001271747,NM_001271744,NM_001271745,NM_001003961,NM_001003960,NM_001003963,NM_010068,NM_001122997 |
| Naif1 | 113.8994186 | 0.129633101810576 | 0.0936599396197593 | 1.38408269679503 | 0.166333134494033 | 0.616741948739248 | NM_194335 |
| Fbxo4 | 213.3445042 | -0.118600191081674 | 0.0857195085341668 | -1.38358459013331 | 0.166485689483035 | 0.616902656356161 | NM_134099 |
| Ndel1 | 4114.894361 | 0.0810876487567071 | 0.0586001911228383 | 1.38374375924355 | 0.166436929371228 | 0.616902656356161 | NM_023668 |
| Tenm1 | 1003.244866 | 0.111278087472992 | 0.0804339683889974 | 1.3834713082267 | 0.16652039879245 | 0.616902656356161 | NM_011855 |
| 4833439L19Rik | 5964.124729 | 0.056942915474344 | 0.0411996161689499 | 1.38212247514235 | 0.166934098622173 | 0.617356018956859 | NM_001252648,NM_133797,NM_001252645,NM_001252646,NM_001252647 |
| Card6 | 109.4846776 | -0.125524173603595 | 0.0908371298591521 | -1.3818597505032 | 0.167014768513859 | 0.617356018956859 | NM_001163138 |
| Dido1 | 3355.769063 | -0.0714161960062582 | 0.0516928498068503 | -1.38154882683203 | 0.16711027581842 | 0.617356018956859 | NM_175551,NM_011805,NM_177852 |
| Nudt4 | 6825.888024 | 0.0926393132777455 | 0.0669884172208054 | 1.38291539225939 | 0.166690810101198 | 0.617356018956859 | NM_027722 |
| Nutf2 | 543.9446335 | 0.105847491389413 | 0.0765566267955499 | 1.38260390798156 | 0.166786350137747 | 0.617356018956859 | NM_026532 |
| Pcif1 | 1932.132689 | 0.0748170877021379 | 0.0541218333358842 | 1.38238272967986 | 0.166854216053553 | 0.617356018956859 | NM_146129 |
| Pigs | 2605.706601 | 0.0801721915443771 | 0.0580275285630182 | 1.38162340409362 | 0.167087363969273 | 0.617356018956859 | NM_201406 |
| Ptgs2 | 485.1128347 | 0.12848282644592 | 0.0929332281978484 | 1.38252839094741 | 0.166809519303157 | 0.617356018956859 | NM_011198 |
| Rcor3 | 341.6986057 | -0.113183979940703 | 0.0819160737012803 | -1.38170660319299 | 0.167061806079167 | 0.617356018956859 | NM_144814 |
| Slc25a39 | 2361.069664 | 0.0919123162560068 | 0.0664893529595188 | 1.38236141825544 | 0.166860756305303 | 0.617356018956859 | NM_026542 |
| Znhit1 | 1114.694852 | 0.0879587472325806 | 0.0636443389014053 | 1.38203568064148 | 0.16696074573011 | 0.617356018956859 | NM_027318 |
| Lyz1 | 50.1612296 | -0.123363900699671 | 0.0893184595697393 | -1.38116914794471 | 0.167226958507212 | 0.617511665690607 | NM_013590 |
| Mapk9 | 12157.16594 | 0.0735132064248203 | 0.0532319883626531 | 1.38099681574916 | 0.167279939728248 | 0.617511665690607 | NM_001163671,NM_001163672,NM_016961,NM_207692 |
| Mfsd6 | 5086.404996 | 0.0742160631040825 | 0.0537367223729495 | 1.38110513307827 | 0.16724663754685 | 0.617511665690607 | NM_133829,NM_178081 |
| Acbd3 | 1498.978765 | 0.0770927528414054 | 0.0559168930545456 | 1.37870236756901 | 0.167986539242196 | 0.617953075861397 | NM_133225 |
| Cdca4 | 272.8290627 | -0.107918542841314 | 0.0782267447513052 | -1.37956070119502 | 0.167721944429839 | 0.617953075861397 | NM_028023 |
| Dhx33 | 1959.898251 | -0.0859179733085081 | 0.062273798211578 | -1.37968095372307 | 0.167684899712256 | 0.617953075861397 | NM_178367 |
| Fam78a | 405.6506848 | 0.117281511066868 | 0.0850327589074439 | 1.3792509213364 | 0.167817402812287 | 0.617953075861397 | NM_175511 |
| Hnmt | 690.5537955 | 0.100252916884654 | 0.0726769483150517 | 1.37943211993521 | 0.167761561676733 | 0.617953075861397 | NM_080462 |
| Irf2bp1 | 1667.780787 | 0.0669499410803109 | 0.0485529806813677 | 1.37890486105631 | 0.167924089215129 | 0.617953075861397 | NM_178757 |
| Nid2 | 167.5688479 | -0.129644438675215 | 0.0939180751857931 | -1.38039923006031 | 0.167463757120506 | 0.617953075861397 | NM_008695 |
| Nol4 | 3975.930586 | 0.0982660353143789 | 0.0711942089044357 | 1.38025320916596 | 0.167508696215811 | 0.617953075861397 | NM_199024,NM_001161483 |
| Pex2 | 878.8786157 | 0.0805331283492109 | 0.0583734733122096 | 1.37961857980389 | 0.167704113711377 | 0.617953075861397 | NM_001267714,NM_001267715,NM_008994,NM_001163301,NM_001163302,NM_001163305,NM_001163306 |
| Ppp1r2 | 5178.062344 | 0.0960704339460476 | 0.0696754638682725 | 1.37882733192386 | 0.167947997536188 | 0.617953075861397 | NM_025800 |
| Set | 6580.399084 | 0.0744517014759568 | 0.0539750551188797 | 1.3793724028994 | 0.167779963517195 | 0.617953075861397 | NM_023871,NM_001204875 |
| Ubr2 | 3919.731524 | -0.0867970433972302 | 0.0629432036550207 | -1.37897403304966 | 0.167902760213859 | 0.617953075861397 | NM_001177374,NM_146078 |
| Usp24 | 4109.229977 | -0.0780432259332707 | 0.0565668468149001 | -1.37966371342292 | 0.16769021034213 | 0.617953075861397 | NM_183225 |
| Usp4 | 4348.846473 | 0.0576700574468408 | 0.0418300704839081 | 1.37867464194272 | 0.167995091324291 | 0.617953075861397 | NM_011678 |
| Abl2 | 3482.953622 | 0.0898423399499547 | 0.0651841583301588 | 1.3782848816564 | 0.168115349070676 | 0.618082399544581 | NM_001136104,NM_009595 |
| Krt77 | 263.3887187 | -0.129322419578035 | 0.093826766852992 | -1.37831051751636 | 0.16810743732441 | 0.618082399544581 | NM_001003667 |
| Cpne3 | 678.7827198 | -0.0928438084806819 | 0.0673775048993252 | -1.37796447967921 | 0.168214255200788 | 0.618133132063408 | NM_027769 |
| Pin1 | 2659.625089 | 0.0695167128095011 | 0.0504441630742161 | 1.37809230192251 | 0.168174792020223 | 0.618133132063408 | NM_023371 |
| Tert | 53.62264683 | -0.129583874332031 | 0.0940700264244413 | -1.37752565038466 | 0.168349789997441 | 0.61847472167497 | NM_009354 |
| Trp53bp2 | 1457.055896 | 0.089637340452099 | 0.0650964430173169 | 1.3769929092478 | 0.168514440116231 | 0.618923073863053 | NM_173378 |
| Slc38a11 | 84.89212454 | -0.130442433006218 | 0.094756092694233 | -1.37661261980421 | 0.168632047095708 | 0.619198462111289 | NM_177074 |
| Cnpy3 | 2429.083737 | 0.0768733084672212 | 0.0558533668052779 | 1.37634153255659 | 0.168715920183396 | 0.619205089431689 | NM_028065 |
| Gpc4 | 333.9421758 | 0.128848391185462 | 0.0936266529520688 | 1.3761934996376 | 0.168761734067194 | 0.619205089431689 | NM_008150 |
| Lingo1 | 14657.53753 | 0.0988745655015291 | 0.071843470706746 | 1.37624984607327 | 0.168744294622339 | 0.619205089431689 | NM_181074 |
| Actr10 | 5211.666836 | 0.0788234534682745 | 0.0572865796284931 | 1.37594972469031 | 0.168837198960678 | 0.619267833995098 | NM_019785 |
| Gm11549 | 2066.04018 | 0.102704970619658 | 0.0746476805452417 | 1.37586285159137 | 0.168864098200095 | 0.619267833995098 | NR_040411 |
| Lyz2 | 401.8701488 | -0.116318096560416 | 0.0845650821364224 | -1.37548611816836 | 0.168980786508007 | 0.619378696669384 | NM_017372 |
| S100a13 | 454.5866654 | -0.105445143183641 | 0.0766677326633079 | -1.37535230951346 | 0.169022246564604 | 0.619378696669384 | NM_009113 |
| Txlnb | 117.3307999 | -0.128871046543499 | 0.0936940962930571 | -1.37544468266618 | 0.168993624310877 | 0.619378696669384 | NM_138628 |
| E530011L22Rik | 308.3559851 | -0.124643873354224 | 0.0906481493907344 | -1.37502943184149 | 0.169122320294448 | 0.619512712108871 | NR_033503 |
| Traf3 | 2535.95311 | 0.102169509255159 | 0.0743073062115754 | 1.37495913207043 | 0.169144115119357 | 0.619512712108871 | NM_011632,NM_001048206 |
| Dcun1d3 | 843.8940995 | 0.0949100854880864 | 0.0690754198391372 | 1.37400663954143 | 0.169439621178892 | 0.619945581247306 | NM_173408,NM_001163703 |
| Dpp6 | 12714.81085 | 0.0840703847774532 | 0.0611900827645584 | 1.37392173664696 | 0.169465980660354 | 0.619945581247306 | NM_207282,NM_001136060,NM_010075,NM_001198886 |
| Eftud1 | 643.6036202 | 0.0966692176283984 | 0.0703988212198152 | 1.37316528818794 | 0.169700968126768 | 0.619945581247306 | NM_001159672,NM_175317 |
| Emc6 | 974.5942158 | 0.102170604191159 | 0.0743620487913531 | 1.3739616625926 | 0.169453584619574 | 0.619945581247306 | NM_001168470,NM_025318 |
| Fam120aos | 500.6851342 | -0.100924070533855 | 0.0734949851376424 | -1.37321029924481 | 0.169686978802912 | 0.619945581247306 | NR_015601 |
| Fem1a | 2788.302274 | 0.0785663177975979 | 0.0571793195054382 | 1.37403380238069 | 0.169431188683857 | 0.619945581247306 | NM_010192 |
| Inadl | 519.6572493 | 0.130300337030572 | 0.0948804362614886 | 1.37331089700586 | 0.169655716396144 | 0.619945581247306 | NM_007704,NM_172696,NM_001005787,NM_001005784 |
| Josd1 | 4249.719417 | 0.0749263266007896 | 0.0545427551390462 | 1.37371730507158 | 0.169529462376879 | 0.619945581247306 | NM_028792 |
| Paf1 | 2280.481871 | -0.0843845096208348 | 0.0614569847769997 | -1.3730662174044 | 0.169731762124503 | 0.619945581247306 | NM_019458 |
| Timm8b | 1826.243628 | 0.0837788065056831 | 0.0609948672190658 | 1.37353863243587 | 0.169584959824503 | 0.619945581247306 | NM_013897 |
| Vash2 | 121.7421358 | 0.130354642849667 | 0.0949231815087674 | 1.37326457855426 | 0.169670110079382 | 0.619945581247306 | NM_144879,NR_027352 |
| Kpna6 | 5832.198914 | 0.0617840468797862 | 0.0450033701986115 | 1.37287600033325 | 0.169790898706378 | 0.620005679891617 | NM_008468 |
| Fam26e | 243.9636622 | 0.118488649713186 | 0.0863636250279268 | 1.37197401886352 | 0.170071526005167 | 0.620316914769489 | NM_178908 |
| Hdac2 | 2977.59833 | 0.0801279734616424 | 0.058419396050917 | 1.37159879899828 | 0.170188367916757 | 0.620316914769489 | NM_008229 |
| Micalcl | 115.7806354 | 0.130064390176314 | 0.0947764506434275 | 1.37232813946208 | 0.169961309474781 | 0.620316914769489 | NM_027587 |
| Rnf20 | 2781.386681 | -0.0776337480142087 | 0.0566097203131396 | -1.37138547204921 | 0.170254823863364 | 0.620316914769489 | NM_182999,NM_001163263 |
| Slu7 | 2860.774012 | 0.0790034301379363 | 0.0576067460611465 | 1.37142670849831 | 0.170241976303912 | 0.620316914769489 | NM_198936,NM_148673 |
| Smg8 | 774.6402723 | 0.0906562096284193 | 0.0660810079157256 | 1.37189507980924 | 0.170096102304423 | 0.620316914769489 | NM_024262 |
| Snhg1 | 270.7960553 | -0.119049615532723 | 0.0868108859120791 | -1.37136736115439 | 0.170260466693236 | 0.620316914769489 | NR_002896 |
| Tgfb1 | 227.2592222 | -0.128095808046304 | 0.093358763203698 | -1.37208124497979 | 0.170038147284961 | 0.620316914769489 | NM_011577 |
| Tspan9 | 2185.312416 | 0.0659046428853817 | 0.0480335490162808 | 1.372054412699 | 0.170046499520681 | 0.620316914769489 | NM_175414 |
| Haus8 | 288.3939228 | 0.112536772154556 | 0.0820786409899103 | 1.37108474016266 | 0.170348541361481 | 0.620482174477653 | NM_001163042,NM_029621 |
| 2810407C02Rik | 6146.307487 | 0.0804236071503055 | 0.0586729201545671 | 1.37071083113707 | 0.170465117056409 | 0.620595561494086 | NM_001040396 |
| Zfyve9 | 2725.618332 | 0.0919262874886576 | 0.0670609559532912 | 1.37078701282883 | 0.170441360619721 | 0.620595561494086 | NM_183300 |
| Ube2e1 | 2125.513654 | 0.0682822925348788 | 0.0498222933518582 | 1.37051684980961 | 0.170525619236753 | 0.620660271869974 | NM_009455 |
| Blcap | 3738.659455 | 0.0794115456243172 | 0.057961197739719 | 1.37008117018084 | 0.170661565018516 | 0.620885919172814 | NM_016916 |

| | | | | | | | | | | | | | | |
|---|---|---|---|---|---|---|---|---|---|---|---|---|---|---|
| Mau2 | 3754.47143 | 0.137005892 | 0.065306184 | 2.09790076 | 0.035913913 | 0.139356415 | | H2-DMb1 | 76.61832183 | -0.124418110235058 | 0.0908134469875901 | -1.37004060920691 | 0.170674225451254 | 0.620885919172814 | NM_010387 |
| Mapk1 | 26624.26341 | -0.14331351 | 0.06832622 | -2.097489219 | 0.035950291 | 0.139462645 | | Nmral1 | 443.9773331 | 0.109271811575135 | 0.0797788369584684 | 1.36968418870308 | 0.170785506434161 | 0.620885919172814 | NM_026393 |
| Gm608 | 3367.322114 | -0.187122987 | 0.089230745 | -2.097068532 | 0.035987509 | 0.139502243 | | Pbxip1 | 2896.771158 | -0.105087810572318 | 0.0767422897035251 | -1.36935985332596 | 0.170886817076477 | 0.620885919172814 | NM_146131 |
| Mpdu1 | 1268.129219 | -0.125709975 | 0.059944062 | -2.097121381 | 0.035982832 | 0.139502243 | | Pknox1 | 707.8692117 | -0.0956541492995348 | 0.0698505548632441 | -1.36941144543246 | 0.170870698558352 | 0.620885919172814 | NM_016670,NR_027493 |
| Rbm12b2 | 278.4539419 | 0.213172543 | 0.101645474 | 2.097216284 | 0.035974434 | 0.139502243 | | Ptpla | 204.9237983 | -0.113663140606408 | 0.0829870471489099 | -1.36964917431577 | 0.170796441495692 | 0.620885919172814 | NM_001012396,NM_013935 |
| Sema4f | 2283.982794 | 0.179410527 | 0.085560811 | 2.096877372 | 0.036004432 | 0.139532934 | | Xrcc6 | 636.2151619 | -0.0991999130725957 | 0.0724417768761881 | -1.36937437691707 | 0.170882279469389 | 0.620885919172814 | NM_010247 |
| Kif13b | 1303.216676 | 0.207962713 | 0.099195203 | 2.096499696 | 0.036037887 | 0.139595056 | | 9130023H24Rik | 154.5902409 | -0.123996170975113 | 0.0905877709348814 | -1.36879591688206 | 0.171063077589493 | 0.621020522044039 | NM_177001 |
| Rxfp1 | 234.9115249 | 0.334085991 | 0.15935469 | 2.096492992 | 0.036038481 | 0.139595056 | | Fam213a | 4319.33581 | 0.0805701119432309 | 0.0588521564321052 | 1.36902565390583 | 0.170991255968415 | 0.621020522044039 | NM_027464 |
| Cd34 | 2991.573717 | 0.220957282 | 0.105465292 | 2.095071076 | 0.036164675 | 0.140031964 | | Rgma | 2267.380066 | 0.0825892999970013 | 0.0603416730429424 | 1.36869423455041 | 0.1710948732769 | 0.621020522044039 | NM_177740 |
| Nrcam | 14856.3349 | -0.134758598 | 0.064323346 | -2.095018469 | 0.036169351 | 0.140031964 | | Zfp790 | 812.5874415 | 0.108153813352579 | 0.0790079418596822 | 1.36889799692112 | 0.171031161991542 | 0.621020522044039 | NM_146185,NM_001145880 |
| Hnmt | 623.5794288 | -0.159824358 | 0.076291791 | -2.094908971 | 0.036179085 | 0.140034661 | | Kif21a | 9545.683983 | 0.0657348530398868 | 0.0480403084213005 | 1.36832704035557 | 0.171209730385195 | 0.62128217426314 | NM_001109041,NM_001109042,NM_016705,NM_001109040 |
| AA474331 | 89.25430222 | 0.270157041 | 0.129005756 | 2.094147188 | 0.036246871 | 0.140261991 | | Aatf | 545.0507167 | 0.0964854732428362 | 0.0705720175812608 | 1.36719165116311 | 0.171565241549104 | 0.622106015662079 | NM_019816 |
| Lemd1 | 34.69114722 | -0.348274167 | 0.166350251 | -2.093619731 | 0.036293868 | 0.140373739 | | Ankrd44 | 462.7419247 | -0.101099316364342 | 0.0739320874945543 | -1.36746194772046 | 0.171480556668663 | 0.622106015662079 | NM_001081433 |
| Tmem37 | 43.53594592 | 0.341532582 | 0.163128477 | 2.093641704 | 0.03629191 | 0.140373739 | | Hjurp | 231.0132728 | -0.125343205307469 | 0.0916724056389703 | -1.36729481934949 | 0.171532914893218 | 0.622106015662079 | NM_198652 |
| Zfp459 | 88.20822079 | 0.30127737 | 0.143932948 | 2.093178621 | 0.036333212 | 0.14049084 | | Nif3l1 | 807.8588399 | 0.0924634570949354 | 0.0676493060803774 | 1.36680569916083 | 0.171686215952267 | 0.622234025180298 | NM_022988 |
| Cd33 | 258.4955267 | 0.263993573 | 0.126143108 | 2.092810121 | 0.036366108 | 0.140570291 | | Prcc | 1725.958996 | 0.0745881412326306 | 0.0545692220359736 | 1.36685366676952 | 0.171671177311793 | 0.622234025180298 | NM_033573 |
| Dlx6 | 312.3877761 | -0.30106151 | 0.143873574 | -2.092542096 | 0.03639005 | 0.140570291 | | Rest | 116.2152373 | -0.126268277291421 | 0.092394988418164 | -1.36661391979349 | 0.171746351814918 | 0.622296709020577 | NM_011263 |
| Ints10 | 1813.801952 | 0.192552608 | 0.092015182 | 2.092617806 | 0.036383286 | 0.140570291 | | Phyh | 3090.140028 | 0.100327225962827 | 0.0734338781284388 | 1.36622535156526 | 0.171868242682506 | 0.622583065637427 | NM_010726 |
| Plekhg4 | 112.0195022 | 0.325947559 | 0.155761514 | 2.092606519 | 0.036384294 | 0.140570291 | | Col28a1 | 52.06209842 | -0.126202637499896 | 0.0924555503592077 | -1.36500877459032 | 0.172250292172499 | 0.623232234675676 | NM_001037865 |
| Cnot6l | 674.8338572 | -0.201885377 | 0.09648741 | -2.092349423 | 0.03640727 | 0.140601753 | | E330009J07Rik | 740.4031864 | 0.117062259528696 | 0.085757286148285 | 1.36504155840801 | 0.17223998853668 | 0.623232234675676 | NM_175528 |
| Atrn | 10964.70067 | -0.147997934 | 0.070741493 | -2.092095154 | 0.036430005 | 0.140654496 | | Elk1 | 3440.205577 | -0.100376480891417 | 0.0735093950790886 | -1.36549186377363 | 0.172098508588035 | 0.623232234675676 | NM_007922 |
| Vps54 | 1169.758751 | 0.147177211 | 0.070367653 | 2.091546391 | 0.036479112 | 0.14080901 | | Fam214a | 2110.588997 | -0.0910232097710078 | 0.0666674138638918 | -1.36533284397143 | 0.172148460557688 | 0.623232234675676 | NM_001113283,NM_153584 |
| BC023829 | 1684.395711 | -0.148095199 | 0.070813887 | -2.091329905 | 0.036498501 | 0.14084876 | | Polr2f | 783.1488773 | 0.0802782519952165 | 0.0588131294459475 | 1.36497161010615 | 0.172261973166931 | 0.623232234675676 | NM_027231 |
| Ccrn4l | 1920.709929 | -0.151046686 | 0.072229716 | -2.091198668 | 0.036510259 | 0.140859051 | | Faf2 | 2483.602393 | 0.0643711654145351 | 0.0471648537316182 | 1.36481214975935 | 0.172312099125461 | 0.623258354555888 | NM_178397 |
| Ralgapb | 7204.384797 | -0.139651299 | 0.066816747 | -2.090064319 | 0.036612022 | 0.141216498 | | Clec4a3 | 33.6772112 | -0.120337866881772 | 0.0882122377663171 | -1.36418562694848 | 0.172509150649319 | 0.623815763587755 | NM_001204241,NM_153197 |
| Arhgap28 | 36.64853367 | -0.349432309 | 0.167249526 | -2.08928729 | 0.03668187 | 0.141256 | | Ccdc157 | 1516.276481 | -0.0847304361690406 | 0.0621211988423423 | -1.36395365427635 | 0.172582152531876 | 0.623924426998016 | NM_001164620,NM_177616 |
| Cdkl4 | 747.6635222 | 0.261075319 | 0.124950562 | 2.089428923 | 0.03666913 | 0.141256 | | Rab14 | 9290.168509 | 0.0812135663891533 | 0.0595655357274167 | 1.36343214910048 | 0.172746354747668 | 0.624056899186717 | NM_026697 |
| E130309D14Rik | 1810.362553 | 0.194954643 | 0.093294402 | 2.089671394 | 0.036647328 | 0.141256 | | Tmem126b | 566.2873938 | 0.0898453313944906 | 0.0658934376821792 | 1.3634943714401 | 0.172726757155509 | 0.624056899186717 | NM_026734 |
| Lin9 | 160.3153697 | 0.246873967 | 0.118154376 | 2.089418745 | 0.036670045 | 0.141256 | | Ugt1a7c | 22.51773913 | -0.107890503929082 | 0.0791317975059107 | -1.3634279433754 | 0.172747679445807 | 0.624056899186717 | NM_201642 |
| Lpar1 | 1030.004795 | -0.251821759 | 0.120511983 | -2.08959933 | 0.036653807 | 0.141256 | | Prickle1 | 3910.302872 | 0.0915737915236156 | 0.0671803427034114 | 1.36310396521638 | 0.172849747289161 | 0.624135563113996 | NM_001033217 |
| Paip2 | 4003.920561 | -0.13919817 | 0.066609336 | -2.089769666 | 0.036638495 | 0.141256 | | Rcl1 | 626.2873029 | 0.0982649269242444 | 0.072090038566786 | 1.36308606400882 | 0.172855388297371 | 0.624135563113996 | NM_021525 |
| Vcpip1 | 2361.674583 | -0.163547228 | 0.07828071 | -2.08924048 | 0.036686081 | 0.141256 | | Tm9sf2 | 5086.132671 | 0.0947257520024422 | 0.0695078257903799 | 1.36280700662561 | 0.172943342341857 | 0.624297959954725 | NM_080556 |
| Eya3 | 1534.958712 | 0.110832499 | 0.053069258 | 2.088450124 | 0.03675725 | 0.141494865 | | Erf | 2143.379264 | -0.0811624956162812 | 0.0595650754601298 | -1.36258528994239 | 0.173013247442558 | 0.624395138472197 | NM_010155 |
| Emp3 | 74.40874918 | -0.295604849 | 0.141556801 | -2.08824194 | 0.036776016 | 0.14153194 | | Gm3414 | 64.50861906 | -0.129562686717491 | 0.0951021646414436 | -1.36235265733405 | 0.173086616935234 | 0.624504768405666 | NR_027993 |
| Ttll7 | 9320.98452 | -0.160881823 | 0.0770463 | -2.088118752 | 0.036787124 | 0.141539532 | | Ism1 | 250.2263968 | 0.116536421771965 | 0.0855574671527206 | 1.36208358720983 | 0.173171507382889 | 0.624655901724321 | NM_001276489,NM_001126490 |
| Trim30a | 119.7318879 | 0.314386889 | 0.150608902 | 2.087438956 | 0.036848474 | 0.14174038 | | Sphkap | 7496.390236 | -0.0872926095962675 | 0.064095914016931 | -1.3619059956491 | 0.173227553790424 | 0.624702941002905 | NM_172430 |
| Ano2 | 275.5066339 | -0.277772484 | 0.133084854 | -2.087183304 | 0.036871568 | 0.141794012 | | Abi1 | 4462.462995 | 0.0880263757224516 | 0.0646642259681154 | 1.36128399288125 | 0.1734239595536 | 0.625247872406875 | NM_001077193,NM_001077192,NM_001077190,NM_145994,NM_007380 |
| Grin1 | 19513.86456 | 0.132746074 | 0.063612526 | 2.08679143 | 0.036906992 | 0.141895021 | | Pa2g4 | 2935.840477 | 0.0609409176985012 | 0.0447714792347103 | 1.36115488565888 | 0.173464747748844 | 0.625247872406875 | NM_011119 |
| AW495222 | 106.1211263 | 0.309749742 | 0.1484885 | 2.086018394 | 0.036976957 | 0.142128745 | | 2610524H06Rik | 376.6804476 | -0.105455620723049 | 0.0775544035767812 | -1.35976315798296 | 0.173904884522418 | 0.626678827232112 | NM_181075 |
| Lhfpl3 | 1084.297505 | -0.267995616 | 0.128481242 | -2.085873479 | 0.036990085 | 0.142143943 | | 4933407K13Rik | 140.8161676 | -0.12792587557444 | 0.0941204491141474 | -1.35917196293118 | 0.174092103443958 | 0.626964545679512 | NR_029443 |
| Il17ra | 923.6852512 | 0.262359945 | 0.125785646 | 2.08577015 | 0.036999449 | 0.142144671 | | Grm5 | 6530.84366 | 0.103658926835567 | 0.076277696233982 | 1.35896771865779 | 0.174156818244309 | 0.626964545679512 | NM_001143834,NM_001081414 |
| Orai3 | 653.6987739 | 0.171125454 | 0.082056993 | 2.085446325 | 0.037028806 | 0.142186943 | | Mrpl1 | 563.4331806 | 0.0872366055004754 | 0.0641913578599666 | 1.35900857076091 | 0.174143872817772 | 0.626964545679512 | NM_053158,NM_001039084 |
| Zfr | 13336.01006 | -0.119098728 | 0.057107496 | -2.085518307 | 0.037022278 | 0.142186943 | | Nlgn3 | 3094.930638 | 0.0792661102751665 | 0.0583251147477003 | 1.35903907978667 | 0.174134205428074 | 0.626964545679512 | NM_172932 |
| Gsdmd | 107.5053849 | 0.272777818 | 0.130850735 | 2.084648724 | 0.037101198 | 0.142429625 | | Car10 | 6162.508541 | 0.0779791955774403 | 0.0573914617459927 | 1.35872468142676 | 0.17423384800098 | 0.627072383832326 | NM_028296 |
| Slc17a9 | 15.0504936 | 0.344447085 | 0.165260549 | 2.084266854 | 0.037135901 | 0.142527531 | | Hlx | 25.60525893 | -0.116839890936971 | 0.0860001614406429 | -1.35860083259975 | 0.174273111216515 | 0.627072383832326 | NM_008250 |
| 9530082P21Rik | 77.08171998 | 0.311681198 | 0.149597658 | 2.08346309 | 0.037209033 | 0.142696449 | | Las1l | 2388.982548 | -0.0677936231350718 | 0.0499051304115959 | -1.3584499745004 | 0.174320945981099 | 0.627089168232154 | NM_152822 |
| Dok1 | 62.94948048 | 0.28780514 | 0.138133455 | 2.083529584 | 0.037202978 | 0.142696449 | | Tysnd1 | 558.5025793 | -0.103612828625894 | 0.0762835265062595 | -1.35825955316044 | 0.174381339633828 | 0.627151111542705 | NM_001272092,NM_001272091,NM_001272090,NM_027912,NR_073587 |
| Hsph1 | 12453.36614 | -0.191940088 | 0.092120782 | -2.083569895 | 0.037199308 | 0.142696449 | | Ndufs1 | 6494.930839 | 0.0699619374987508 | 0.0515173121127481 | 1.3580277120366 | 0.174454890991929 | 0.627260333304148 | NM_001160040,NM_001160038,NM_001160039,NM_145518 |
| Plxnc1 | 2773.045919 | -0.169109159 | 0.081173307 | -2.083309963 | 0.037222979 | 0.142696449 | | Rrs1 | 711.4447262 | 0.0916070532454055 | 0.0674728970017005 | 1.35768667592703 | 0.174563126440837 | 0.627494178341897 | NM_021511 |
| Vma21 | 1197.871174 | -0.165462762 | 0.079424265 | -2.083277225 | 0.037225961 | 0.142696449 | | Ncaph | 45.18572361 | -0.123799453348925 | 0.0911948887406078 | -1.35752622826326 | 0.174614065439687 | 0.627521997668702 | NM_144818 |
| Hcfc1 | 5627.77657 | -0.098920364 | 0.047498877 | -2.082583208 | 0.037289231 | 0.142903623 | | Prkab2 | 2316.790048 | -0.101663500651529 | 0.0749159979749659 | -1.35703325590752 | 0.174770643959103 | 0.62792935299281 | NM_182997 |
| Itgbl1 | 66.67599685 | -0.316598505 | 0.152035238 | -2.082402137 | 0.037305753 | 0.142931588 | | Khnyn | 807.3312474 | -0.0826818526620758 | 0.0609371731937907 | -1.35683767934448 | 0.174832792270425 | 0.627997314668692 | NM_027143 |
| A830010M20Rik | 6754.455905 | 0.159734705 | 0.07674677 | 2.081321532 | 0.037404486 | 0.143269134 | | Afg3l2 | 3627.167781 | 0.0530210282859727 | 0.0390889398064308 | 1.35642021882747 | 0.174965503766612 | 0.628026740791054 | NM_027130 |
| Iqcc | 468.1690868 | 0.199771906 | 0.095987166 | 2.081235586 | 0.037412348 | 0.143269134 | | Dip2c | 4275.984735 | 0.0829612935997564 | 0.0611964601567159 | 1.35565510467932 | 0.175208930165548 | 0.628026740791054 | NM_001081426 |
| Mprip | 11807.21537 | -0.137034402 | 0.065859125 | -2.080720055 | 0.037459537 | 0.143414399 | | Fcrls | 1297.364406 | -0.126111201184856 | 0.0930053076563497 | -1.35595703474076 | 0.175112838843837 | 0.628026740791054 | NM_030707 |
| Slc2a9 | 36.3942228 | 0.331073227 | 0.159122886 | 2.080613516 | 0.037469296 | 0.143416321 | | Fermt3 | 194.6057532 | -0.1274392501546 | 0.0939917709577063 | -1.35585539942581 | 0.175145180590944 | 0.628026740791054 | NM_153795 |
| Serf1 | 205.8413084 | -0.232513544 | 0.111767864 | -2.080325567 | 0.037495681 | 0.143481868 | | Heg1 | 913.3161586 | 0.0953314282467257 | 0.0703194064518531 | 1.3556915943538 | 0.175197314985565 | 0.628026740791054 | NM_175256 |
| Mid1 | 838.8949807 | -0.26148131 | 0.125702576 | -2.080158726 | 0.037510977 | 0.143504956 | | P2ry13 | 563.7015259 | -0.110253331755776 | 0.0813324415507992 | -1.35558861450032 | 0.175230096408243 | 0.628026740791054 | NM_028808 |
| Crim1 | 2342.157684 | -0.213936953 | 0.102854646 | -2.079993095 | 0.037526166 | 0.143527628 | | Paqr9 | 943.4207947 | 0.0957569644461814 | 0.0706201027400077 | 1.35594484758422 | 0.175116716728537 | 0.628026740791054 | NM_198414 |
| AI480653 | 1140.265587 | -0.140316368 | 0.067466658 | -2.07978834 | 0.037544951 | 0.14353686 | | Rtn1 | 41995.43105 | 0.0738272872111326 | 0.0544298026675998 | 1.35637616880576 | 0.174979511736741 | 0.628026740791054 | NM_153457,NM_001007596 |
| Cgnl1 | 898.7511999 | -0.213409 | 0.102614826 | -2.079709222 | 0.037552212 | 0.14353686 | | Zfp442 | 97.62701692 | -0.127504904296122 | 0.0940174673632326 | -1.35618314204862 | 0.175040904392451 | 0.628026740791054 | NM_001177550 |
| Nfu1 | 1124.871718 | -0.116124526 | 0.055838122 | -2.079663893 | 0.037556372 | 0.14353686 | | Pcdha7 | 122.486808 | 0.128868667211044 | 0.0950792411050993 | 1.35538173962278 | 0.175295964410289 | 0.628089033312773 | NM_009957 |
| Map3k10 | 6515.185642 | -0.160653786 | 0.077255056 | -2.07952455 | 0.037569164 | 0.143550339 | | Zfand4 | 165.314276 | -0.117015455759535 | 0.0863415466186386 | -1.35526244713197 | 0.175333954983016 | 0.628089033312773 | NM_001081317 |
| Gnrh1 | 14.28045599 | -0.30038036 | 0.144462069 | -2.07930263 | 0.037589544 | 0.143592798 | | Tsku | 98.6130307 | 0.127150883345328 | 0.0938342787395436 | 1.35505792822537 | 0.175399101543326 | 0.628167492361527 | NM_001024619,NM_001168540,NM_001168541,NM_001168539 |
| Lyrm2 | 659.133057 | -0.19992991 | 0.096169859 | -2.078924857 | 0.037624258 | 0.14368998 | | Coq10b | 876.6332329 | 0.115850153896951 | 0.0855109382475599 | 1.35479923704681 | 0.175481529767708 | 0.62830778935315 | NM_026424,NM_001039710 |
| Zfp518b | 707.9571535 | -0.153793675 | 0.073994057 | -2.07845982 | 0.037667028 | 0.143817874 | | Hyi | 151.1822505 | -0.125665831592838 | 0.0927930379629727 | -1.35425926719829 | 0.175653676500712 | 0.628459434691956 | NR_003946,NM_026601 |
| BC027072 | 11.5356439 | 0.339063625 | 0.163163936 | 2.078054952 | 0.037704298 | 0.143924709 | | Ifitm1 | 69.53064478 | 0.127520728240697 | 0.0941601743920807 | 1.35429579505347 | 0.175642027155692 | 0.628459434691956 | NM_001112715,NM_026820 |
| Thyn1 | 1036.639465 | 0.163830892 | 0.078848561 | 2.077791789 | 0.03772854 | 0.143981775 | | Serinc1 | 35054.02552 | 0.0892472554324462 | 0.0659001112024402 | 1.35428080171648 | 0.175646808712337 | 0.628459434691956 | NM_019760 |
| Ndufb4 | 1828.479802 | -0.184555102 | 0.088845212 | -2.077265584 | 0.037777054 | 0.144025017 | | Skil | 3017.793096 | 0.110603231539083 | 0.0816830886380193 | 1.35405300391155 | 0.175719468133753 | 0.628540013324524 | NM_001039090,NM_011386,NM_001271772 |
| Scamp1 | 9555.709007 | -0.135383924 | 0.065169459 | -2.07741367 | 0.037763396 | 0.144025017 | | Dgat1 | 823.8501774 | -0.107758143947551 | 0.0796168054260474 | -1.35345977989085 | 0.175908790760224 | 0.628658304763903 | NM_010046 |
| Trmt1 | 1826.348701 | 0.149392122 | 0.071908492 | 2.077530996 | 0.037752577 | 0.144025017 | | Lrrn3 | 2639.393133 | 0.0839995058107993 | 0.0620652231850658 | 1.35340697253807 | 0.175925651167924 | 0.628658304763903 | NM_001271709,NM_001271708,NM_010733 |
| Uba52 | 9253.365395 | -0.159656524 | 0.076856047 | -2.07734499 | 0.037769729 | 0.144025017 | | Tmem14a | 1139.379778 | 0.0865994517015074 | 0.0639718363447227 | 1.35371214349471 | 0.175828232384028 | 0.628658304763903 | NM_029398 |
| Blm | 167.0256856 | 0.246108471 | 0.118493294 | 2.076982279 | 0.037803195 | 0.144089225 | | Txnip | 1391.184153 | 0.123395222576152 | 0.0911599870502876 | 1.35361167293807 | 0.175860300846917 | 0.628658304763903 | NM_023719,NM_001009935 |
| Dhx58 | 58.61062843 | 0.335109513 | 0.16136934 | 2.076661613 | 0.037832802 | 0.144115079 | | Eif2ak1 | 3572.986522 | 0.0716431523122905 | 0.0529479139929595 | 1.35308734394743 | 0.176027728367204 | 0.628713740413574 | NM_013557 |
| Fmo5 | 179.1854441 | 0.232115975 | 0.111784269 | 2.076463688 | 0.037851086 | 0.144115079 | | Pgm1 | 330.7713981 | -0.10437898448721 | 0.0771362558725174 | -1.35317670408733 | 0.175999185695202 | 0.628713740413574 | NM_025700 |
| Kcnip2 | 6803.816169 | 0.157291989 | 0.075752066 | 2.076405267 | 0.037856485 | 0.144115079 | | Habp4 | 9677.312808 | 0.0555961905955244 | 0.0410969191398164 | 1.35280677382117 | 0.176117368172442 | 0.628879274845845 | NM_019986 |
| Nphp1 | 586.6773478 | -0.14887548 | 0.071692802 | -2.076575016 | 0.037840801 | 0.144115079 | | BC035044 | 29.66246364 | -0.119874310936233 | 0.0886282637626237 | -1.35255172387554 | 0.176198884021284 | 0.629015726048948 | NM_001254947,NM_001254946 |
| Pdhb | 4134.041323 | -0.105541905 | 0.050826487 | -2.076513864 | 0.03784645 | 0.144115079 | | Aars | 7940.137466 | 0.058236826236673 | 0.0431077079702096 | 1.35096086010694 | 0.176707970853896 | 0.629083075677498 | NM_146217 |
| Dclre1c | 575.403637 | 0.231559321 | 0.11155931 | 2.0756611 | 0.037925306 | 0.144306162 | | Baz2b | 1411.674929 | -0.0790994886677496 | 0.0585448768153141 | -1.35109155524022 | 0.176666106273821 | 0.629083075677498 | NM_001001182 |
| Tmbim6 | 12345.85279 | -0.09837705 | 0.047394854 | -2.075690534 | 0.037922582 | 0.144306162 | | Cacng8 | 1241.053453 | -0.0922660945179188 | 0.0683012954629679 | -1.35086888019488 | 0.176737438513002 | 0.629083075677498 | NM_133190 |
| Gal | 32.03576589 | -0.254908765 | 0.12282327 | -2.075410992 | 0.03794846 | 0.144358813 | | Chst11 | 3124.08881 | 0.0797726412192453 | 0.058988476042885 | 1.35234280609742 | 0.176265676642596 | 0.629083075677498 | NM_021439 |
| Rps29 | 7473.767094 | -0.180924724 | 0.087195942 | -2.074921379 | 0.037993822 | 0.144495895 | | Csnk1a1 | 7095.191343 | 0.0884745278384838 | 0.0654457721382079 | 1.35187537632903 | 0.176415185893073 | 0.629083075677498 | NM_146087 |
| Crat | 2102.288094 | 0.123808695 | 0.059672431 | 2.074805606 | 0.038004555 | 0.144501245 | | Fhl4 | 33.77905248 | -0.122344750990938 | 0.0905442096260984 | -1.35121562710813 | 0.176626370113514 | 0.629083075677498 | NM_010214 |
| Fut4 | 14.45237673 | -0.345527258 | 0.166546253 | -2.074662454 | 0.038017829 | 0.144516254 | | Ppm1d | 602.6116995 | 0.0931004774462997 | 0.0688646545172002 | 1.35193413949454 | 0.176396385070328 | 0.629083075677498 | NM_016910 |
| Psmd10 | 497.8241099 | -0.16111569 | 0.077665986 | -2.074469127 | 0.038035763 | 0.144548961 | | Psmb7 | 4476.414775 | 0.0692331472033059 | 0.0512416905604213 | 1.35110973986446 | 0.176660281916962 | 0.629083075677498 | NM_011187 |
| Snx27 | 6814.824847 | -0.1281985 | 0.061807087 | -2.074171537 | 0.038063383 | 0.144618453 | | Ptcd2 | 1142.282667 | 0.0859422369881023 | 0.0635834980283389 | 1.35164373859705 | 0.176489311151951 | 0.629083075677498 | NM_026873 |
| Rplp0 | 6285.950758 | -0.139631579 | 0.067332787 | -2.073753166 | 0.038102241 | 0.1447306 | | Samd14 | 1809.334686 | 0.100674844342538 | 0.0744712222956355 | 1.35186238709605 | 0.176419341899804 | 0.629083075677498 | NM_146025 |
| Qrich1 | 3312.669495 | 0.138448378 | 0.066767334 | 2.073594514 | 0.038116985 | 0.14475112 | | Sox2ot | 1372.800111 | -0.113695033096565 | 0.084115049116894 | -1.35166102011739 | 0.176483780175839 | 0.629083075677498 | NR_015580 |
| Luc7l | 4650.219192 | 0.107774756 | 0.051980552 | 2.073366916 | 0.038138145 | 0.144795988 | | Ubn2 | 1896.573159 | -0.108663311310037 | 0.0804079375237636 | -1.35140030519901 | 0.176567236099094 | 0.629083075677498 | NM_177185 |
| Rnf126 | 1931.667215 | 0.134453649 | 0.064885173 | 2.072178312 | 0.038248815 | 0.145180584 | | Amh | 31.78051295 | -0.111855508887641 | 0.0828483074157075 | -1.35012424968907 | 0.176976131060206 | 0.629241039295 | NM_007445 |
| Jph2 | 51.37896047 | -0.326585815 | 0.157641939 | -2.071693729 | 0.038294013 | 0.145316539 | | Kcns2 | 1598.376963 | -0.113600756728871 | 0.0841716463596428 | -1.34963211059797 | 0.177134018833818 | 0.629241039295 | NM_181317,NM_001271704 |
| Myo1a | 39.67449104 | 0.347060107 | 0.167547343 | 2.071415168 | 0.038320015 | 0.145344016 | | Lss | 979.3031997 | -0.093267123528448 | 0.0690909113851543 | -1.34991884834925 | 0.177042015044255 | 0.629241039295 | NM_146006 |
| Trim21 | 92.5275991 | 0.264612676 | 0.127741932 | 2.071462914 | 0.038315557 | 0.145344016 | | Prr22 | 35.59219749 | -0.118551528318441 | 0.0877943710151073 | -1.35033176897003 | 0.176909586282774 | 0.629241039295 | NM_001195673 |
| Ptges2 | 1125.619704 | -0.134810802 | 0.065092311 | -2.071071065 | 0.038352155 | 0.14540359 | | Setd2 | 3289.44041 | -0.0856054349222511 | 0.0634337070491362 | -1.34952596820395 | 0.177168085125731 | 0.629241039295 | NM_001081340 |
| Rexo2 | 1257.780076 | -0.139513211 | 0.06736451 | -2.071019461 | 0.038356977 | 0.14540359 | | Shroom4 | 198.2018885 | -0.121559260445131 | 0.09004504311859 | -1.34998281261412 | 0.177021496528119 | 0.629241039295 | NM_001040459 |
| Zfp142 | 1730.06038 | 0.153060614 | 0.073908561 | 2.070945656 | 0.038363875 | 0.14540359 | | Taf9b | 1296.709213 | 0.0943569368181339 | 0.0698642525699937 | 1.35057534214085 | 0.176831503963921 | 0.629241039295 | NM_001001176,NM_001167988 |
| Tusc1 | 74.60636128 | -0.314279261 | 0.151818838 | -2.07009397 | 0.038443545 | 0.145669918 | | Vps18 | 2504.571297 | 0.0725387169519483 | 0.0537381211441086 | 1.34985584548858 | 0.17706222733467 | 0.629241039295 | NM_172269 |
| Cdh5 | 726.7102373 | -0.171569334 | 0.08288477 | -2.069974185 | 0.038454762 | 0.145676793 | | Zbed3 | 416.4942774 | -0.0928149278300699 | 0.068776519092188 | -1.34951476252616 | 0.177171681861252 | 0.629241039295 | NM_028106 |
| Mapk8 | 4222.53239 | -0.153275211 | 0.074059681 | -2.069617485 | 0.038488179 | 0.145767747 | | Kcnn1 | 779.9608141 | 0.102435868368955 | 0.0759373615717089 | 1.34895216595355 | 0.177352330873831 | 0.62972866249652 | NM_032397 |
| Amot | 963.9198527 | -0.195909313 | 0.09467201 | -2.069347776 | 0.038513463 | 0.145792898 | | Nol11 | 1113.205864 | 0.087592722550105 | 0.0649456197045995 | 1.34870870350478 | 0.177430548857764 | 0.629852432235551 | NM_001161329,NM_133702 |
| Npat | 1035.616379 | 0.154550374 | 0.074685617 | 2.069345896 | 0.03851364 | 0.145792898 | | Steap2 | 699.9225502 | 0.109886791877639 | 0.0815159070282999 | 1.34804108650217 | 0.177645186186228 | 0.630460227968031 | NM_001103157,NM_001103156,NM_028734 |
| 2810029C07Rik | 395.1301443 | 0.19715963 | 0.095331328 | 2.068151508 | 0.038625779 | 0.146136809 | | Cic | 8212.125144 | -0.0787816390500669 | 0.0584618267574977 | -1.34757402256445 | 0.177795430453024 | 0.630839380254184 | NM_001110131,NM_001110132,NM_027882 |
| Shc2 | 1861.497472 | 0.149335781 | 0.07220524 | 2.068212507 | 0.038620045 | 0.146136809 | | Atoh8 | 59.00301956 | 0.126296727778069 | 0.0937427594127891 | 1.34726914984368 | 0.177893564126624 | 0.631033434066749 | NM_153778 |
| Tusc5 | 27.52011876 | 0.345693761 | 0.167157105 | 2.068076982 | 0.038632785 | 0.146136809 | | Enah | 3098.111227 | 0.0903639277825111 | 0.0670882741669689 | 1.34694071213717 | 0.177999328105093 | 0.631188774993418 | NM_010135,NM_001083121,NM_001083120,NM_008680 |
| Mfsd8 | 716.81208 | -0.23168742 | 0.11204428 | -2.067820155 | 0.038656938 | 0.146156806 | | Gm12522 | 69.08467253 | -0.127979569103345 | 0.0950204585311692 | -1.34686330798293 | 0.178024260715134 | 0.631188774993418 | NR_040560 |
| Rcn1 | 987.3937432 | -0.24803602 | 0.119948955 | -2.067846437 | 0.038654465 | 0.146156806 | | Ubfd1 | 2776.912894 | 0.0796961479264912 | 0.0591822831476367 | 1.34662172001172 | 0.178102095203379 | 0.631310647858535 | NM_138589 |
| Ctrl | 19.49000184 | 0.336853736 | 0.163004335 | 2.066532375 | 0.038778239 | 0.146579661 | | Tnfsf12 | 168.4604228 | 0.124960934982453 | 0.0928085248234271 | 1.34643811244924 | 0.178161266579185 | 0.631366323085934 | NM_011614 |
| Arl3 | 2709.021731 | -0.116939638 | 0.056617174 | -2.065444642 | 0.038880948 | 0.146932052 | | Wasl | 4956.996373 | 0.0854075853277661 | 0.063456464398077 | 1.34592410935448 | 0.178326992633673 | 0.63179948658457 | NM_001167745,NM_028459 |
| Klhl2 | 6899.263053 | -0.178274447 | 0.086317321 | -2.065338036 | 0.038891027 | 0.146934302 | | Kcp | 68.67068517 | -0.12424293469894 | 0.0923308936351294 | -1.34562690565868 | 0.178422870057472 | 0.631985030591279 | NM_001029985 |
| Casp3 | 265.9314145 | -0.192732773 | 0.093331764 | -2.065028708 | 0.038920284 | 0.147008991 | | Aim2 | 86.43046516 | -0.12695093000118 | 0.0943562421959509 | -1.34544283501179 | 0.178482270177038 | 0.632041310724441 | NM_001013779 |
| Inhbb | 212.5935797 | -0.243073745 | 0.117717988 | -2.064881915 | 0.038934175 | 0.147025616 | | Fam107b | 335.1305349 | 0.102007013312488 | 0.0758263291985254 | 1.34527168057177 | 0.178537515400293 | 0.632082853693556 | NM_025626 |
| Rell2 | 4389.244603 | 0.139530728 | 0.067585666 | 2.064501793 | 0.038970164 | 0.147125663 | | Hspe1 | 1547.537039 | 0.0750383254484136 | 0.0557872223923571 | 1.34508086673794 | 0.178599121279382 | 0.63209710823724 | NM_008303 |
| Rab11a | 3043.073394 | -0.164090131 | 0.079495678 | -2.064139033 | 0.039004536 | 0.147183702 | | Zfp133-ps | 100.8699413 | -0.125216867249361 | 0.0930987577982244 | -1.3449896670023 | 0.178628571479683 | 0.63209710823724 | NR_033459 |
| Ulk3 | 899.5216897 | 0.223949033 | 0.108494704 | 2.064147138 | 0.039003768 | 0.147183702 | | Fzd3 | 5143.528497 | 0.103658139204674 | 0.077085854469267 | 1.3447102573923 | 0.178718820864514 | 0.632262443224044 | NM_021458 |
| Sypl | 1365.43255 | 0.135849065 | 0.065820912 | 2.063919499 | 0.03902535 | 0.147226386 | | Magi2 | 5193.750052 | 0.0738272064072105 | 0.0549257133948142 | 1.34412831157111 | 0.178906898419666 | 0.632619670507318 | NM_015823,NM_001170745,NM_001170746 |
| Rora | 422.3171381 | 0.235699476 | 0.114210417 | 2.063730101 | 0.039043314 | 0.147258302 | | Srm | 1750.316817 | 0.0831483123538371 | 0.0618560498438515 | 1.34422279734538 | 0.178876351801738 | 0.632619670507318 | NM_009272 |
| 1700028P14Rik | 27.78884104 | 0.341124138 | 0.165460644 | 2.061663303 | 0.039239803 | 0.147923531 | | Abhd1 | 131.4923319 | -0.125142222156422 | 0.0931139618533446 | -1.34396839813907 | 0.178958606200876 | 0.632648506613271 | NM_021304,NR_003522 |
| Btd | 363.461182 | -0.198842706 | 0.096449797 | -2.06161871 | 0.039244051 | 0.147923531 | | Map3k1 | 441.4800993 | -0.0998255671406232 | 0.0743002026669962 | -1.34354367225657 | 0.179095994675019 | 0.632845581351673 | NM_011945 |
| H2-K1 | 1086.124872 | 0.262307857 | 0.127236705 | 2.061573799 | 0.039248331 | 0.147923531 | | Slc22a4 | 335.8526383 | -0.106677327807677 | 0.0794009733441279 | -1.34352670143389 | 0.179101485951861 | 0.632845581351673 | NM_019687 |
| Trim30d | 68.83175796 | 0.334053504 | 0.162052288 | 2.061393316 | 0.039265532 | 0.147952372 | | Grwd1 | 688.8024045 | 0.0844257340172651 | 0.0628589293121324 | 1.34309850551288 | 0.179240079443584 | 0.633029217835538 | NM_153419 |
| Crybb1 | 107.1500267 | -0.335723838 | 0.16289818 | -2.060942849 | 0.039308493 | 0.148006266 | | Kcnj16 | 1347.101363 | 0.0929923520624241 | 0.0692372645810691 | 1.34309685145837 | 0.179240614963346 | 0.633029217835538 | NM_010604,NM_001252209,NM_001252210,NM_001252208,NM_001252207 |
| D4Ertd617e | 12.44412095 | 0.330935737 | 0.160565539 | 2.061063284 | 0.039297003 | 0.148006266 | | Dcaf12l1 | 1647.653784 | 0.0926856094717005 | 0.0690249321398459 | 1.34278450696507 | 0.179341761521972 | 0.633232481251379 | NM_001190718,NM_178739 |
| Wfdc18 | 43.90132137 | -0.346057875 | 0.16790843 | -2.06099167 | 0.039303835 | 0.148006266 | | Lrpap1 | 7849.252989 | 0.0635472255433 | 0.0473299672396508 | 1.34264250007896 | 0.179387761661143 | 0.633240978345022 | NM_013587 |
| Galc | 1180.328706 | 0.170353831 | 0.082669759 | 2.060654735 | 0.039335991 | 0.14807382 | | Clcn7 | 2811.327118 | -0.0768494361080251 | 0.0572748338561875 | -1.34176619876345 | 0.179671815108241 | 0.633459544269818 | NM_011930 |
| Parn | 1017.867542 | 0.133238188 | 0.064679518 | 2.059974975 | 0.039400933 | 0.148282259 | | Depdc5 | 2900.832751 | 0.0705162697563994 | 0.0525544947782313 | 1.34177428693707 | 0.179669191795557 | 0.633459544269818 | NM_177786,NM_001025426,NM_001170567 |
| Me2 | 836.0819075 | -0.179221466 | 0.087028043 | -2.05935306 | 0.039460429 | 0.148470103 | | Nos2 | 29.41007555 | -0.105363845263067 | 0.0785017932827966 | -1.34218392799643 | 0.179536366326757 | 0.633459544269818 | NM_010927 |
| Zfp580 | 394.0301461 | -0.224126701 | 0.108855639 | -2.058935155 | 0.039500451 | 0.148584604 | | Phf21b | 482.1711151 | -0.121251479313439 | 0.0903821582014509 | -1.34154219954766 | 0.179744478172967 | 0.633459544269818 | NM_001081166,NR_030731 |
| Glb1l | 367.3573928 | 0.274506489 | 0.133348241 | 2.058568511 | 0.039535592 | 0.148618256 | | Porcn | 2363.459832 | -0.07885137586076 | 0.058756123929345 | -1.34201119113269 | 0.179592367079719 | 0.633459544269818 | NM_145908,NM_023638,NM_016913,NM_145907 |
| Lima1 | 625.3236354 | -0.18689693 | 0.090794696 | -2.058456469 | 0.039546336 | 0.148618256 | | Prdm5 | 112.232596 | -0.125451285546605 | 0.0935179994578984 | -1.34146673660489 | 0.179768962500985 | 0.633459544269818 | NM_027547 |
| Ttpa | 100.0505957 | -0.316447154 | 0.153731424 | -2.058441567 | 0.039547765 | 0.148618256 | | Sucla2 | 5968.439682 | 0.0842030469458192 | 0.0627736550907821 | 1.34137556310918 | 0.179798547502716 | 0.633459544269818 | NM_011506 |
| Uhmk1 | 2209.202703 | -0.157342064 | 0.076434866 | -2.058511665 | 0.039541043 | 0.148618256 | | Ttc32 | 48.95293652 | -0.126214002225523 | 0.0940761388287571 | -1.34161545952971 | 0.17972071097765 | 0.633459544269818 | NM_029321 |
| Slc45a1 | 2084.392877 | 0.174520166 | 0.084789659 | 2.058271825 | 0.039564047 | 0.148643392 | | Armc1 | 3211.537618 | 0.0720878620101514 | 0.0537541087659946 | 1.34106701171381 | 0.179898696575794 | 0.633658697007755 | NM_028840 |
| Med15 | 2489.456488 | 0.140354159 | 0.068200372 | 2.057967635 | 0.03959324 | 0.14871701 | | Zdhhc15 | 523.7139974 | 0.105423677364007 | 0.0786382037437528 | 1.34061654952771 | 0.180044981235822 | 0.634020217559162 | NM_175358 |
| Gsk3b | 8399.269487 | -0.152525168 | 0.074121588 | -2.05776985 | 0.039612232 | 0.148752282 | | Aldh1l2 | 162.6740741 | -0.122262115659799 | 0.0912234459008341 | -1.34024881928607 | 0.18016446474836 | 0.634219999609488 | NM_153543 |
| Angel2 | 2216.015571 | 0.136629356 | 0.066402439 | 2.057595449 | 0.039628984 | 0.148779132 | | Lamc2 | 696.5336057 | -0.109727573735375 | 0.081880940314961 | -1.34008687874492 | 0.180217101414635 | 0.634219999609488 | NM_008485 |
| 1110059E24Rik | 675.0913613 | -0.160649418 | 0.078110951 | -2.056682398 | 0.039716786 | 0.148989976 | | Tmem252 | 38.17854281 | 0.116419799043876 | 0.0868779247245189 | 1.34003890416387 | 0.180232697121573 | 0.634219999609488 | NM_183160 |
| Ammecr1l | 1534.285692 | 0.129763724 | 0.063108719 | 2.056193293 | 0.039763889 | 0.148989976 | | 2610002M06Rik | 2624.318357 | 0.0950002436914888 | 0.0710026671368893 | 1.33798133960705 | 0.18090251981613 | 0.634469715840679 | NM_025921 |
| Arf5 | 3231.440178 | -0.14812729 | 0.072039023 | -2.056209042 | 0.039762371 | 0.148989976 | | Abat | 16573.93197 | 0.0909921663892764 | 0.0679835126750105 | 1.33844461412662 | 0.180751543728847 | 0.634469715840679 | NM_172961,NM_001170978 |
| Cenpa | 71.02014864 | 0.313388339 | 0.152432699 | 2.055912818 | 0.03979092 | 0.148989976 | | Bad | 772.3944833 | 0.0898041361029951 | 0.0671029484102011 | 1.33830387830385 | 0.180797398092758 | 0.634469715840679 | NM_007522 |
| Emc6 | 927.6355634 | -0.16836515 | 0.081890454 | -2.055980186 | 0.039784426 | 0.148989976 | | Bmx | 61.72282659 | -0.124171428459356 | 0.0927063665169381 | -1.33940562147551 | 0.180438660359755 | 0.634469715840679 | NM_009759 |
| Foxs1 | 15.15181821 | -0.340001084 | 0.165303196 | -2.056833095 | 0.039702283 | 0.148989976 | | Cpsf4 | 695.2247647 | -0.101551195614027 | 0.0758587499175194 | -1.33868796584434 | 0.180672275584178 | 0.634469715840679 | NM_178576 |
| H2-Q5 | 15.99317857 | 0.342633203 | 0.166628699 | 2.056267642 | 0.039756725 | 0.148989976 | | D930048N14Rik | 240.2578163 | -0.125862136010264 | 0.0941461085527917 | -1.33688091780966 | 0.181261510526805 | 0.634469715840679 | NR_027958 |
| Nrip3 | 5403.021662 | -0.23497779 | 0.114285153 | -2.056065749 | 0.039776179 | 0.148989976 | | Dnm3 | 8411.969349 | 0.0896164542429439 | 0.0669864281755801 | 1.33783001547788 | 0.18095185497232 | 0.634469715840679 | NM_001038619,NM_172646 |
| Parva | 3232.852688 | 0.14645348 | 0.07120938 | 2.05665997 | 0.039718945 | 0.148989976 | | Dvl1 | 4406.221795 | -0.0902439710329857 | 0.0673870748083877 | -1.33918813495898 | 0.180509434058184 | 0.634469715840679 | NM_010091 |
| Rab27a | 236.2654994 | 0.277880511 | 0.135139861 | 2.056243872 | 0.039759015 | 0.148989976 | | Gabra3 | 7375.196708 | 0.0872425101413305 | 0.0652717260852747 | 1.33660491875689 | 0.181351632613396 | 0.634469715840679 | NM_008067 |
| Sorcs3 | 2260.869031 | -0.133706503 | 0.065032821 | -2.055984968 | 0.039783965 | 0.148989976 | | Glod4 | 3157.613902 | 0.0732672318332738 | 0.054692242703418 | 1.33962749032954 | 0.18036648181722 | 0.634469715840679 | NM_026029 |
| Snx3 | 4900.082904 | -0.123860628 | 0.060250666 | -2.055755328 | 0.039806106 | 0.149010825 | | Katna1 | 410.951674 | -0.104285324398426 | 0.0780293931126731 | -1.33648770339454 | 0.181389917054401 | 0.634469715840679 | NM_011835 |
| Ofd1 | 397.7542304 | 0.168656765 | 0.08204905 | 2.055560246 | 0.039824923 | 0.149045255 | | Large | 7447.765403 | 0.0784458119779505 | 0.058580431290706 | 1.33911291278588 | 0.180533917391843 | 0.634469715840679 | NM_010687 |
| Fam189a2 | 454.9321733 | 0.179894717 | 0.087601767 | 2.053551242 | 0.040019145 | 0.149735967 | | Lphn3 | 3854.946134 | 0.0856044011784198 | 0.064042282505768 | 1.33668566810856 | 0.181325262052758 | 0.634469715840679 | NM_198702 |
| Alpl | 382.8306343 | -0.310857659 | 0.151407185 | -2.053123554 | 0.040060596 | 0.149854871 | | Mbp | 75747.34624 | 0.114314726227881 | 0.0855433657182821 | 1.33633655009965 | 0.181439295029752 | 0.634469715840679 | NM_001025259,NM_001025258,NM_001025245,NM_001025255,NM_001025254,NM_001025251,NM_001025256,NM_010777 |
| Shroom3 | 189.2798877 | -0.229261134 | 0.111678822 | -2.052861311 | 0.04008603 | 0.149913819 | | Ncf1 | 365.2881546 | -0.121644781943186 | 0.0909675859480857 | -1.33723216545075 | 0.181146865565652 | 0.634469715840679 | NM_010876 |
| 5830417I10Rik | 387.1399517 | -0.190771539 | 0.092973683 | -2.051887518 | 0.040180595 | 0.150194968 | | Nmnat1 | 135.1929783 | 0.121484616621352 | 0.0908532574771427 | 1.33715201848339 | 0.181173020277075 | 0.634469715840679 | NM_133435 |
| Lrig2 | 1348.127048 | 0.128804825 | 0.062773343 | 2.051903212 | 0.04017907 | 0.150194968 | | Nr1d1 | 6665.987313 | -0.0827047691367368 | 0.0617958340316647 | -1.33835509193643 | 0.180780710733536 | 0.634469715840679 | NM_145434 |
| Rnf8 | 1177.6768 | 0.15525831 | 0.075681459 | 2.051470877 | 0.040221113 | 0.15031016 | | Prkcq | 571.2505346 | 0.116222184315851 | 0.0869153472749253 | 1.33718828676165 | 0.181161184343518 | 0.634469715840679 | NM_008859 |
| Nedd4 | 22168.6739 | -0.110910215 | 0.0540705 | -2.051214909 | 0.040246023 | 0.150366982 | | Pum1 | 4730.226026 | 0.0740351919424218 | 0.0553946088981008 | 1.33650536424241 | 0.181384148350897 | 0.634469715840679 | NM_001159604,NM_001159605,NM_001159606,NM_030722,NM_001159603 |
| Klhl1 | 213.3337146 | -0.30218832 | 0.147350892 | -2.050807544 | 0.040285693 | 0.150442642 | | Slc38a5 | 86.31757808 | -0.126607452727577 | 0.0946096220681227 | -1.33820905273689 | 0.180828298906878 | 0.634469715840679 | NM_172479 |
| Tfe3 | 1677.973936 | 0.166504046 | 0.08118834 | 2.050836943 | 0.040282829 | 0.150442642 | | St5 | 783.2227207 | -0.0906116815984061 | 0.0677890838994751 | -1.33667069070847 | 0.18133015305272 | 0.634469715840679 | NM_029811,NR_037617,NM_001001326 |
| Dlgap5 | 8.392274212 | -0.325903943 | 0.158924102 | -2.050689213 | 0.040297222 | 0.150449436 | | Tars | 1271.444276 | 0.0730078151937641 | 0.0546091327441776 | 1.33691585134261 | 0.181250106036095 | 0.634469715840679 | NM_033074 |
| Tpi1 | 22186.03448 | -0.120481373 | 0.05881293 | -2.048552457 | 0.040505898 | 0.151192093 | | Ubiad1 | 443.7472002 | 0.0998822173711054 | 0.0746069477678346 | 1.33877903277753 | 0.180642618519055 | 0.634469715840679 | NM_027873 |
| Esyt2 | 1953.413713 | -0.12015939 | 0.058661405 | -2.048355108 | 0.040525217 | 0.151227772 | | Vbp1 | 1499.768177 | 0.0891312908420723 | 0.0666973863477493 | 1.33635357729546 | 0.181433732175104 | 0.634469715840679 | NM_011692 |
| Ssr2 | 1915.146639 | -0.108873402 | 0.053172237 | -2.047561037 | 0.04060303 | 0.151481665 | | Wnt7a | 424.0786088 | 0.10728222701208 | 0.0802133130283646 | 1.33746161281411 | 0.181072004499238 | 0.634469715840679 | NM_009527 |
| Zcchc14 | 4021.892156 | -0.158933487 | 0.077639762 | -2.047063046 | 0.040651895 | 0.151627457 | | Zfp213 | 328.7713105 | 0.100689866479799 | 0.0752697410868431 | 1.3377203777495 | 0.180987605640586 | 0.634469715840679 | NM_001033496 |
| Spata13 | 936.1997267 | -0.225817002 | 0.110356098 | -2.046257591 | 0.040731034 | 0.151886074 | | Pfn4 | 219.8162655 | 0.116018748030846 | 0.086860766375131 | 1.33568644248185 | 0.181651782541223 | 0.635059878024985 | NM_028376 |
| Ywhae | 23309.57805 | -0.115628404 | 0.056512293 | -2.046075262 | 0.040748966 | 0.151916383 | | Arhgef10 | 2361.119044 | 0.0953913368172899 | 0.071430839613083 | 1.33543630921872 | 0.181733587706225 | 0.635192996876954 | NM_001037736,NM_172751 |
| Tyw3 | 494.2534413 | 0.164115723 | 0.080222406 | 2.045759179 | 0.04078007 | 0.151995768 | | Cfl2 | 2387.032931 | 0.0751236910222889 | 0.0562823919841634 | 1.33476365118645 | 0.18195371362917 | 0.635245711079828 | NM_007688 |
| Hspb1 | 228.5666377 | -0.314677189 | 0.153827395 | -2.045651155 | 0.040790705 | 0.15199884 | | D10Wsu102e | 1130.866456 | 0.0884568750361668 | 0.066332364889904 | 1.33354019840819 | 0.182354592896622 | 0.635245711079828 | NM_026579 |
| Atp11a | 3091.794114 | 0.167647855 | 0.082022494 | 2.043925368 | 0.040960921 | 0.152596418 | | Dzip1l | 919.2976628 | -0.0998047471687604 | 0.0748220724463963 | -1.33389444993337 | 0.182238450745105 | 0.635245711079828 | NM_028258 |
| Kcnj3 | 2077.376718 | 0.132006438 | 0.064624825 | 2.042658362 | 0.041086269 | 0.1530266 | | Gm872 | 37.45083222 | -0.122386995914264 | 0.0917341062802329 | -1.33414932435698 | 0.182154923537429 | 0.635245711079828 | NR_040653 |
| Aanat | 21.86623983 | 0.342445078 | 0.167665629 | 2.042428612 | 0.041109034 | 0.153074591 | | Gtf2h3 | 533.8571523 | -0.0948035134064921 | 0.0710933843448623 | -1.33350682739502 | 0.1823655364858 | 0.635245711079828 | NM_181410 |
| Rab23 | 1149.407251 | 0.124970323 | 0.06120776 | 2.041739847 | 0.041177344 | 0.153292111 | | Icmt | 3910.656538 | 0.0591258915981658 | 0.0443047228728381 | 1.33452796370867 | 0.182030888632553 | 0.635245711079828 | NR_037908,NM_133788 |
| 4921536K21Rik | 14.33233936 | 0.341254896 | 0.167152506 | 2.041578104 | 0.041193399 | 0.153300321 | | Mrpl41 | 1357.814261 | 0.0746275151381361 | 0.0559668625236293 | 1.33342323962913 | 0.182392950142751 | 0.635245711079828 | NM_001031808 |
| Cd200r1 | 18.71250043 | 0.343159838 | 0.168104434 | 2.041349114 | 0.041216139 | 0.153300321 | | Nars2 | 543.75718 | -0.0883568265958367 | 0.0661760419901195 | -1.33517847152341 | 0.181817941183723 | 0.635245711079828 | NM_153591 |
| Clip3 | 22367.47786 | 0.178267084 | 0.08733362 | 2.041219443 | 0.04122902 | 0.153300321 | | Pgs1 | 3158.641893 | 0.0715407083259964 | 0.0536232516013857 | 1.33413596135128 | 0.182159302143705 | 0.635245711079828 | NM_133757 |
| Fbxw10 | 30.80793728 | 0.336851377 | 0.165018611 | 2.041293249 | 0.041221688 | 0.153300321 | | Prss23 | 716.5379386 | 0.0874159922348342 | 0.0655282923658855 | 1.33401907906795 | 0.182197603853439 | 0.635245711079828 | NM_029614 |
| Fuom | 534.6117465 | -0.149344106 | 0.073160289 | -2.041327436 | 0.041218292 | 0.153300321 | | Raf1 | 3018.940928 | -0.0691629745391814 | 0.0518701203377133 | -1.33338758593346 | 0.182404644149385 | 0.635245711079828 | NM_029780 |
| Suv420h2 | 422.1991259 | 0.29922621 | 0.146621651 | 2.040805078 | 0.041270206 | 0.153416643 | | Rhobtb1 | 249.0277662 | -0.119588919443676 | 0.0896472229184643 | -1.33399469108423 | 0.182205596420587 | 0.635245711079828 | NM_001081347,NM_001252637,NM_001252636,NM_001252638 |
| Lrrtm3 | 2125.520524 | -0.198607117 | 0.097346624 | -2.040205495 | 0.041329863 | 0.153597903 | | Slc7a7 | 77.2852782 | -0.125718959932451 | 0.0942637736979941 | -1.33369326306874 | 0.182304403574068 | 0.635245711079828 | NM_011405,NM_001253679,NM_001253680 |
| Map3k19 | 227.3353093 | -0.292635506 | 0.143440635 | -2.040115806 | 0.041338793 | 0.153597903 | | Ston1 | 176.6388677 | 0.114265738360037 | 0.0856636454407626 | 1.3338883463587 | 0.18224045135109 | 0.635245711079828 | NM_029858 |
| Mmp17 | 9402.962004 | -0.195883 | 0.096034564 | -2.03971354 | 0.041378866 | 0.153709936 | | Tmed4 | 2554.19138 | 0.0613875130169729 | 0.0460022263333502 | 1.33444656726252 | 0.182057547241334 | 0.635245711079828 | NM_134020 |
| Gli1 | 129.1466443 | 0.32973034 | 0.161687569 | 2.039305448 | 0.041419552 | 0.153824195 | | Arxes1 | 477.7195351 | 0.108321219863017 | 0.0812715439984984 | 1.3328308351693 | 0.182587324197634 | 0.635424885312609 | NM_029541 |
| 0610040J01Rik | 155.6834325 | -0.260775388 | 0.127892818 | -2.0390151 | 0.041448521 | 0.153858111 | | Eif3c | 8001.520838 | 0.0643879591678834 | 0.048305557402534 | 1.3329306736145 | 0.182554555411286 | 0.635424885312609 | NM_146200 |
| Cenpw | 65.44475925 | -0.304885783 | 0.14953332 | -2.038915364 | 0.041458475 | 0.153858111 | | Spg11 | 1229.777915 | -0.0825105569779553 | 0.0618959046680978 | -1.33305357471385 | 0.182514223031785 | 0.635424885312609 | NM_145531 |
| Ptplad2 | 45.32862608 | -0.333715575 | 0.163671011 | -2.038941249 | 0.041455891 | 0.153858111 | | Entpd3 | 626.8333617 | 0.100864964226052 | 0.0757006241264387 | 1.33241918927357 | 0.182722479897289 | 0.63574293245222 | NM_178676 |
| Zfp59 | 321.3881677 | -0.190503546 | 0.09345569 | -2.038437115 | 0.041506238 | 0.153998479 | | Cenpn | 37.07319102 | 0.112318629261857 | 0.0843176226919704 | 1.33208961158903 | 0.182830743589758 | 0.635967284814373 | NM_028131 |
| Paqr5 | 186.9900009 | 0.311843371 | 0.153073504 | 2.037213258 | 0.041628676 | 0.154415777 | | Psmc3 | 5329.132943 | 0.0686295469845676 | 0.0515315688356579 | 1.3317961889233 | 0.182927170645172 | 0.636150366481153 | NM_008948 |
| Arid1a | 5937.290299 | -0.120325495 | 0.059075324 | -2.03681481 | 0.041668604 | 0.15452689 | | Per3 | 2381.507038 | -0.0882435201619325 | 0.0662691823072906 | -1.33159210796878 | 0.182994259688852 | 0.636231358602264 | NM_011067 |
| Mkrn3 | 22.03490136 | -0.342168667 | 0.168032471 | -2.036324671 | 0.041717764 | 0.154598191 | | Aass | 233.1911046 | 0.123645688269915 | 0.092931994805739 | 1.33049644020209 | 0.183354758458977 | 0.636722739989267 | NM_013930 |
| Oscp1 | 1192.673446 | 0.118042985 | 0.057968518 | 2.036329178 | 0.041717312 | 0.154598191 | | Agpat1 | 5249.82963 | 0.0709024426573452 | 0.0532808731973836 | 1.33072974226006 | 0.183277952897703 | 0.636722739989267 | NM_001163379,NM_018862 |
| Zfp599 | 198.290578 | -0.252770041 | 0.124123163 | -2.036445381 | 0.041705652 | 0.154598191 | | Pls3 | 7486.020904 | 0.0865313764854615 | 0.0650274007524932 | 1.330690993091 | 0.183290707892174 | 0.636722739989267 | NM_145629,NM_001166454,NM_001166453 |
| Stt3b | 3959.722342 | -0.130627974 | 0.064158709 | -2.036013125 | 0.041749037 | 0.154677089 | | Rpl15 | 7214.929103 | 0.0494431624269779 | 0.0371580067928688 | 1.33061933872263 | 0.183314295964701 | 0.636722739989267 | NM_025586 |
| Mdfic | 103.7097133 | -0.318846741 | 0.156612798 | -2.035891987 | 0.041761203 | 0.154685172 | | Zfp92 | 166.895916 | 0.122207418918249 | 0.0918352727705181 | 1.33072418942585 | 0.183279780673808 | 0.636722739989267 | NM_009566 |
| Cd209c | 67.30200929 | 0.301880452 | 0.14830189 | 2.035580608 | 0.041792487 | 0.15472707 | | Sec61b | 420.5226203 | 0.106381166695525 | 0.0799757344604648 | 1.33016804926141 | 0.183462908712989 | 0.636946035364456 | NM_024171 |
| Wdr77 | 2141.318358 | 0.12022915 | 0.05906093 | 2.035679934 | 0.041782505 | 0.15472707 | | Asic4 | 1061.735998 | 0.100562421860042 | 0.0756223448350906 | 1.32979772155093 | 0.183584926854139 | 0.637217359016303 | NM_183022 |
| Erich1 | 157.1363321 | 0.230219243 | 0.113104119 | 2.035462936 | 0.041804314 | 0.154733886 | | Casc1 | 34.02248517 | -0.108961289448451 | 0.0819785210149889 | -1.32914436732188 | 0.183800345102215 | 0.637355821354308 | NM_177222 |
| Ranbp10 | 1736.692805 | 0.12688284 | 0.062364441 | 2.034538227 | 0.041897358 | 0.155009396 | | Ccdc88a | 2761.361918 | -0.0963036840011181 | 0.0724725794900642 | -1.32882925761351 | 0.18390430721529 | 0.637355821354308 | NM_176841 |
| Syt15 | 45.21824293 | -0.328966779 | 0.161692231 | -2.034524335 | 0.041898758 | 0.155009396 | | Elavl3 | 10388.289 | 0.0668882086225088 | 0.0503109767137112 | 1.32949533067363 | 0.183684605272154 | 0.637355821354308 | NM_010487 |
| Heg1 | 839.6954979 | -0.12724698 | 0.062548184 | -2.034383278 | 0.041912967 | 0.155024947 | | Fam149a | 1401.334263 | 0.0780708781830719 | 0.0587489487178732 | 1.32888979099843 | 0.18388433244764 | 0.637355821354308 | NM_153535 |
| Art3 | 123.7680018 | -0.249032628 | 0.122423366 | -2.034191962 | 0.041932245 | 0.155059236 | | Glb1 | 1138.176301 | 0.098649195993164 | 0.0742312572575398 | 1.32894416230764 | 0.183866392407529 | 0.637355821354308 | NM_009752 |
| Slc5a3 | 1095.900571 | -0.227189116 | 0.111698213 | -2.033954795 | 0.041956153 | 0.155110628 | | Pdcd6 | 2014.775614 | 0.0713288494707234 | 0.0536707668250229 | 1.3290074595593 | 0.183845508848396 | 0.637355821354308 | NM_011051 |
| Atad3a | 1205.017581 | 0.158478003 | 0.077941827 | 2.033285726 | 0.042023664 | 0.15528611 | | Rhob | 9163.563858 | 0.0770470412365687 | 0.0579874304229457 | 1.32868521116744 | 0.183951846030696 | 0.637355821354308 | NM_007483 |
| Tspan31 | 2134.9313 | -0.132047695 | 0.06494222 | -2.033310438 | 0.042021169 | 0.15528611 | | Spin4 | 25.24535738 | 0.109704127044106 | 0.0825704439483178 | 1.32861253734777 | 0.183975833604476 | 0.637355821354308 | NM_178753 |
| Raph1 | 2476.631941 | -0.131714117 | 0.064804895 | -2.032471724 | 0.042105924 | 0.155552975 | | Aggf1 | 2812.029544 | 0.0751186146510962 | 0.056558482346029 | 1.32815824497402 | 0.184125835175269 | 0.637571366330384 | NM_025630 |
| Tmem51 | 169.4469357 | -0.237369465 | 0.11679908 | -2.03228882 | 0.042124426 | 0.155584231 | | Mc3r | 26.64443498 | 0.0899230407997781 | 0.0677001020898042 | 1.32825561592943 | 0.184093676900756 | 0.637571366330384 | NM_008561 |
| 4930453N24Rik | 538.6892909 | -0.151396161 | 0.074502576 | -2.032092979 | 0.042144244 | 0.155620333 | | C030018K13Rik | 77.24511084 | -0.126243639492385 | 0.0950681901703199 | -1.32792724113305 | 0.18420214442988 | 0.637683591512975 | NR_045412,NR_045413,NR_045411 |
| Aip | 2449.67361 | -0.169610819 | 0.083485661 | -2.031616171 | 0.042192528 | 0.155724401 | | Atp2c2 | 50.60192321 | -0.117333812184812 | 0.088515953868529 | -1.32556682786343 | 0.184983220053215 | 0.637780733057772 | NM_026922 |
| Rpl22l1 | 1299.492745 | -0.248052455 | 0.122090621 | -2.031707692 | 0.042183257 | 0.155724401 | | Chga | 6516.638227 | 0.0844182182310267 | 0.0637064790720038 | 1.32511197386397 | 0.185134015291285 | 0.637780733057772 | NM_007693 |
| Cyp2j6 | 913.1263962 | 0.155482873 | 0.076538436 | 2.031435207 | 0.042210866 | 0.155754979 | | Ctnnb1 | 19029.40035 | 0.0603611993492314 | 0.0455333807071798 | 1.32564721555396 | 0.184956579017552 | 0.637780733057772 | NM_001165902,NM_007614 |
| Apopt1 | 799.6641376 | -0.138486203 | 0.068177529 | -2.031258742 | 0.042228754 | 0.155783885 | | Dyrk1b | 1048.075109 | -0.0886111791325204 | 0.0668500218777054 | -1.32552206631472 | 0.18499805557002 | 0.637780733057772 | NM_010092,NM_001037957,NM_001271370 |
| Brdt | 484.4950791 | -0.171624159 | 0.084514985 | -2.030695007 | 0.042285943 | 0.155883901 | | E130003G02Rik | 23.37096452 | -0.0738726318305445 | 0.0557245863379992 | -1.32567393829481 | 0.184947723545448 | 0.637780733057772 | NM_001242940 |
| C1qtnf6 | 76.43563219 | -0.285823747 | 0.140748208 | -2.030745188 | 0.04228085 | 0.155883901 | | Eif4a1 | 10970.88632 | 0.059783350727655 | 0.0451212533241263 | 1.32494880623559 | 0.185188131510861 | 0.637780733057772 | NM_001159375,NM_144958 |
| Ncam1 | 10462.49979 | -0.134933477 | 0.066446978 | -2.030693969 | 0.042286049 | 0.155883901 | | Enpp5 | 10572.3316 | 0.0760089099274472 | 0.0572809393070781 | 1.32694943286405 | 0.184525410592023 | 0.637780733057772 | NM_032003,NM_001168620 |
| C030046E11Rik | 1741.86237 | 0.148057358 | 0.072913793 | 2.030580941 | 0.042297523 | 0.15588911 | | Fam203a | 327.2135359 | -0.101699724034252 | 0.0766027616707953 | -1.32762477247638 | 0.184302096597318 | 0.637780733057772 | NM_021555 |
| Pyroxd2 | 71.89463314 | 0.332430795 | 0.163754665 | 2.030053896 | 0.042351061 | 0.156049309 | | Fam98c | 740.0335082 | -0.110217736023883 | 0.0830257558421016 | -1.32751258818402 | 0.184339178621683 | 0.637780733057772 | NM_001146023 |
| Fry | 7183.985585 | -0.19676752 | 0.096940996 | -2.029765814 | 0.042380349 | 0.15610783 | | Homer3 | 360.0002061 | -0.126056648650031 | 0.0950965683443575 | -1.32556464281196 | 0.184983944233993 | 0.637780733057772 | NM_001146153,NM_011984 |
| Gpr27 | 243.9572567 | -0.25853388 | 0.127387643 | -2.029505171 | 0.042406863 | 0.15610783 | | Hsp90ab1 | 67936.94723 | 0.0449685585469354 | 0.0339284624584013 | 1.32539335085019 | 0.18504072120719 | 0.637780733057772 | NM_008302 |
| Itpkb | 1430.804117 | 0.169435783 | 0.083490786 | 2.029395003 | 0.042418074 | 0.15610783 | | Kif11 | 79.24705467 | -0.126059684555547 | 0.0950936488352788 | -1.32563726494403 | 0.18495987656443 | 0.637780733057772 | NM_010615 |
| Kdm4d | 66.92940261 | 0.308450388 | 0.152001941 | 2.029252953 | 0.042432533 | 0.15610783 | | Lrrc36 | 36.94045088 | -0.110767798356947 | 0.0834734618200686 | -1.326982204185 | 0.184514569504919 | 0.637780733057772 | NM_001170788,NM_001170789,NM_001033371 |
| Lrfn5 | 1950.505291 | -0.138796829 | 0.068392277 | -2.029422563 | 0.042415269 | 0.15610783 | | Lrrn4 | 25.47533856 | 0.110884930985252 | 0.0835432397968385 | 1.32727592627367 | 0.184417424290407 | 0.637780733057772 | NM_177303 |
| Pm20d2 | 433.3267568 | -0.167525885 | 0.082557421 | -2.029204442 | 0.042437471 | 0.15610783 | | Ormdl1 | 301.3183507 | 0.106516507633195 | 0.0803043836789114 | 1.32640962738834 | 0.184704051593431 | 0.637780733057772 | NM_145517 |
| Smim7 | 3264.276886 | -0.12635036 | 0.06226426 | -2.029259808 | 0.042431835 | 0.15610783 | | Rnf125 | 55.41134693 | 0.121941693567804 | 0.0920366999413278 | 1.32492466207003 | 0.185196140165061 | 0.637780733057772 | NM_026301 |
| Spef2 | 33.16150609 | 0.336043686 | 0.16564792 | 2.028662266 | 0.042492702 | 0.156273895 | | Rrbp1 | 2318.581809 | -0.0861353320167959 | 0.0649799358410497 | -1.32556812963767 | 0.184982788613631 | 0.637780733057772 | NM_133626,NM_024281 |
| Tsr1 | 1422.760314 | 0.111765865 | 0.055099628 | 2.028432274 | 0.042516149 | 0.156323021 | | S100a16 | 1773.125564 | -0.092758285582578 | 0.0699191904393077 | -1.32664987966495 | 0.184624527695493 | 0.637780733057772 | NM_026416 |
| Gm1943 | 322.2542003 | -0.173508753 | 0.085550723 | -2.028138942 | 0.042546069 | 0.156395919 | | Slco1c1 | 2473.91219 | -0.0824327042720959 | 0.0621328408781667 | -1.32671712909014 | 0.184602272565172 | 0.637780733057772 | NM_001177772,NM_021471 |
| Prmt10 | 715.1386915 | 0.14515952 | 0.071582348 | 2.027867534 | 0.042573769 | 0.156460622 | | Txndc16 | 3279.849748 | 0.0865449929949785 | 0.0652462006568726 | 1.32643728100147 | 0.184694896911355 | 0.637780733057772 | NM_172597 |
| Ctsl | 3243.483941 | 0.158072311 | 0.077964814 | 2.027482697 | 0.042613072 | 0.156567925 | | Vangl1 | 152.6091285 | 0.124923587435922 | 0.0941358949212244 | 1.32705582222872 | 0.184490217624486 | 0.637780733057772 | NM_001165951,NM_177545 |
| Rgs11 | 1257.449651 | 0.33439738 | 0.164950575 | 2.027258043 | 0.04263603 | 0.156615137 | | Wrn | 535.1215393 | -0.0896920547682825 | 0.0676810691144746 | -1.32521628192041 | 0.185099426583411 | 0.637780733057772 | NM_001122822,NM_011721 |
| Ncapd2 | 982.0402497 | 0.150763629 | 0.074402206 | 2.026332788 | 0.042730693 | 0.156888475 | | 4922501C03Rik | 467.1714974 | -0.089975258166426 | 0.0679166399629849 | -1.32478959818188 | 0.185240945774953 | 0.63778382989499 | NM_199316 |
| Vtn | 2747.298832 | -0.145165664 | 0.071637494 | -2.026392258 | 0.042724604 | 0.156888475 | | Tpst1 | 1361.203929 | 0.0753491642942569 | 0.0569182783705371 | 1.32381313088451 | 0.1855651140944 | 0.638748542022571 | NM_001130476,NM_013837 |
| Stmn1 | 21663.38994 | -0.196651353 | 0.097059708 | -2.02608638 | 0.042755934 | 0.156943955 | | Galnt2 | 3487.605994 | 0.0722080658371776 | 0.0545534524081142 | 1.32362046121278 | 0.18562912623406 | 0.638817504779899 | NM_139272 |
| Ptbp1 | 1249.047071 | 0.183584784 | 0.090628159 | 2.025692513 | 0.042796304 | 0.157054936 | | Meis3 | 1354.697952 | -0.077487096463358 | 0.0585791502713951 | -1.32277604069644 | 0.185909867217357 | 0.639632101184113 | NM_008627 |
| Mical1 | 463.3421518 | 0.193284563 | 0.095441851 | 2.025155211 | 0.042851429 | 0.157219996 | | Gpx4 | 8268.645602 | 0.0673096288538597 | 0.0509012513642048 | 1.32235705508006 | 0.186049282079388 | 0.639960187422493 | NM_008162,NM_001037741 |
| Ltf | 12.61114583 | -0.292428603 | 0.144405243 | -2.025055299 | 0.042861686 | 0.157220399 | | Nop2 | 1030.056657 | 0.0831470818219391 | 0.062920327708704 | 1.32146612787646 | 0.186345989419588 | 0.640677359126374 | NM_138747 |
| Intu | 567.0976319 | 0.190679101 | 0.094195267 | 2.02429599 | 0.042939706 | 0.157469303 | | Xbp1 | 3725.960326 | 0.100286469006357 | 0.0758837973300322 | 1.32157947460368 | 0.186308221921493 | 0.640677359126374 | NM_013842,NM_001271730 |
| Col4a2 | 2738.725012 | 0.249082591 | 0.123085881 | 2.023648759 | 0.043006304 | 0.15756436 | | 5430416O09Rik | 33.44224007 | -0.120458452282582 | 0.0911829989511402 | -1.32106262864999 | 0.186480482593268 | 0.640826268877189 | NR_033355 |
| Gm11549 | 1767.2102 | 0.26309713 | 0.130010194 | 2.023665384 | 0.043004593 | 0.15756436 | | Chd5 | 12701.51623 | 0.0765701506219878 | 0.0579695690454518 | 1.32086803270785 | 0.18654537036983 | 0.640826268877189 | NM_029216,NM_001081376 |

| | | | | | | |
|---|---|---|---|---|---|---|
| Golga7 | 2775.652516 | -0.111159614 | 0.054922306 | -2.023942959 | 0.042976021 | 0.15756436 |
| Upk1a | 16.28662789 | 0.326624922 | 0.16140266 | 2.023665047 | 0.043004627 | 0.15756436 |
| Zfp617 | 1263.994757 | -0.164717047 | 0.081409746 | -2.023308692 | 0.043041331 | 0.157655411 |
| Jam2 | 1956.495174 | -0.24755798 | 0.122394582 | -2.022622047 | 0.043112129 | 0.157877413 |
| Rasgrp4 | 51.39808938 | -0.339459666 | 0.167850791 | -2.022389435 | 0.043136136 | 0.157927998 |
| Postn | 120.7945585 | -0.312983343 | 0.154799778 | -2.02185912 | 0.043190908 | 0.158091172 |
| Fam120c | 3208.517417 | -0.145307309 | 0.071873827 | -2.021699899 | 0.043207364 | 0.158114054 |
| Nphp3 | 384.7314286 | 0.266943604 | 0.132048408 | 2.021558675 | 0.043221965 | 0.158130136 |
| Znrf3 | 588.5286384 | -0.163761878 | 0.081022303 | -2.021195052 | 0.043259578 | 0.158230382 |
| Dctpp1 | 253.6777673 | -0.206635155 | 0.102270197 | -2.020482612 | 0.043333352 | 0.158434995 |
| Gli3 | 513.1655299 | -0.173513374 | 0.085878266 | -2.020457354 | 0.043335969 | 0.158434995 |
| Il15ra | 142.4527134 | 0.234679826 | 0.116164043 | 2.020244992 | 0.043357982 | 0.158478076 |
| Pdrg1 | 1435.278895 | 0.129591425 | 0.064153633 | 2.020016937 | 0.043381631 | 0.158527119 |
| Mtss1l | 8229.557001 | -0.114818329 | 0.056848331 | -2.019730862 | 0.043411312 | 0.158598177 |
| Ap2s1 | 3728.417212 | -0.127101907 | 0.062937294 | -2.019500682 | 0.043435206 | 0.158648064 |
| BC026585 | 220.8586463 | 0.247962607 | 0.122824393 | 2.01883844 | 0.043504014 | 0.158861937 |
| Insl5 | 10.69493984 | 0.316806094 | 0.156933841 | 2.018723894 | 0.043515925 | 0.158867988 |
| Krt77 | 223.2176987 | 0.284855348 | 0.141156874 | 2.018005499 | 0.043590689 | 0.159103446 |
| Ptchd1 | 685.1208519 | -0.229514764 | 0.113776267 | -2.017246396 | 0.043669807 | 0.159354683 |
| Medag | 1007.076983 | 0.192068537 | 0.095229516 | 2.016901333 | 0.043705811 | 0.159448513 |
| 1700007K13Rik | 55.92737795 | 0.30922307 | 0.153340161 | 2.016582401 | 0.043739111 | 0.159532436 |
| Cacna1g | 3791.003629 | 0.18733001 | 0.092916827 | 2.016104254 | 0.043789075 | 0.159677085 |
| Plekha8 | 753.9403306 | 0.154218815 | 0.07650107 | 2.015904014 | 0.043810014 | 0.159715848 |
| Anapc11 | 1156.828199 | -0.162388687 | 0.080557924 | -2.015800302 | 0.043820862 | 0.159717816 |
| Ovca2 | 425.3348675 | 0.192104233 | 0.09531099 | 2.015551763 | 0.043846868 | 0.159775017 |
| Hsf1 | 1426.93034 | 0.189229035 | 0.093944272 | 2.014269011 | 0.043981298 | 0.160220654 |
| Lcmt2 | 264.054934 | -0.192312656 | 0.095479728 | -2.014172644 | 0.043991411 | 0.160220654 |
| Ttc12 | 96.85539026 | 0.31045393 | 0.154141115 | 2.014089039 | 0.044000186 | 0.160220654 |
| Agxt2l1 | 1045.036883 | 0.336873255 | 0.167333948 | 2.013179382 | 0.044095761 | 0.160530951 |
| Tcte2 | 77.01315574 | -0.279749535 | 0.138988175 | -2.012757808 | 0.044140115 | 0.160654671 |
| Hif3a | 336.8584827 | 0.326394034 | 0.16219839 | 2.012313643 | 0.044186885 | 0.160758077 |
| Ndufc2 | 1794.110212 | -0.190512449 | 0.094674405 | -2.012290945 | 0.044189276 | 0.160758077 |
| Pla2g4b | 135.6899402 | 0.3172061 | 0.157652247 | 2.012062029 | 0.044213399 | 0.160808075 |
| Cggbp1 | 2189.807455 | -0.15385137 | 0.076484087 | -2.011547461 | 0.044267663 | 0.160892133 |
| Hfm1 | 63.81814348 | 0.300837231 | 0.14954987 | 2.011618143 | 0.044260205 | 0.160892133 |
| Nr2e1 | 643.0924687 | -0.173989398 | 0.086489654 | -2.011678737 | 0.044253813 | 0.160892133 |
| Nlgn3 | 2936.442729 | -0.138825714 | 0.06902172 | -2.011333731 | 0.044290218 | 0.160936359 |
| Gbx1 | 22.84130351 | -0.313692828 | 0.155989785 | -2.010983144 | 0.044327237 | 0.160962837 |
| Mfge8 | 6197.098281 | -0.331623779 | 0.164914616 | -2.010881674 | 0.044337956 | 0.160962837 |
| Pak1 | 20968.67747 | 0.145174681 | 0.072198447 | 2.010772914 | 0.044349448 | 0.160962837 |
| Stard13 | 562.6615297 | 0.238439508 | 0.118576175 | 2.010855105 | 0.044340764 | 0.160962837 |
| Vps11 | 3324.454847 | 0.19387288 | 0.096405922 | 2.011005917 | 0.044324832 | 0.160962837 |
| 2700094K13Rik | 342.5801323 | -0.158498317 | 0.07884808 | -2.010173455 | 0.044412834 | 0.161089456 |
| Ccnt2 | 1861.946825 | 0.181050254 | 0.090068121 | 2.010148012 | 0.044415526 | 0.161089456 |
| Sncaip | 197.6563248 | -0.255667888 | 0.127178605 | -2.010305817 | 0.044398832 | 0.161089456 |
| Col4a4 | 24.0155392 | 0.330824276 | 0.164616927 | 2.009661354 | 0.044467043 | 0.161200833 |
| Lingo3 | 1859.619775 | -0.159675709 | 0.079451944 | -2.009714328 | 0.044461433 | 0.161200833 |
| Gpn3 | 475.815018 | 0.174698591 | 0.086934563 | 2.009541265 | 0.044479763 | 0.161209228 |
| Wdhd1 | 257.8627816 | 0.221780541 | 0.110423174 | 2.008460119 | 0.04459442 | 0.161586984 |
| Mgat2 | 886.3042739 | -0.137740176 | 0.068591565 | -2.008121212 | 0.044630413 | 0.161679592 |
| Ebpl | 293.036126 | -0.211397147 | 0.105288361 | -2.007792179 | 0.04466538 | 0.161768444 |
| Abhd13 | 1634.40341 | -0.138230229 | 0.068856129 | -2.007522517 | 0.044694056 | 0.161780242 |
| Snrk | 3852.552766 | -0.122770718 | 0.061154533 | -2.007548933 | 0.044691246 | 0.161780242 |
| Stard8 | 1650.700487 | -0.141928071 | 0.070700077 | -2.007466988 | 0.044699962 | 0.161780242 |
| Pknox1 | 704.1357927 | -0.181212017 | 0.090289989 | -2.00700009 | 0.044749653 | 0.161922263 |
| Son | 12725.10692 | 0.125524688 | 0.062547786 | 2.006860588 | 0.044764509 | 0.161938199 |
| Enpep | 153.8192603 | -0.215293271 | 0.107307461 | -2.006321557 | 0.044821951 | 0.16210815 |
| Fam83d | 61.14365969 | 0.328408788 | 0.163728936 | 2.005807865 | 0.044876751 | 0.16226813 |
| Rnf122 | 143.6863559 | -0.21293693 | 0.10616533 | -2.005710625 | 0.044887131 | 0.16226813 |
| Mcm4 | 498.0310746 | 0.155942427 | 0.077792157 | 2.004603454 | 0.045005457 | 0.162657931 |
| Cc2d1a | 2045.275316 | 0.161427888 | 0.08053816 | 2.004365239 | 0.04503095 | 0.162712113 |
| Cck | 6035.833451 | -0.238624809 | 0.11905924 | -2.004252741 | 0.045042994 | 0.162717683 |
| Neurod6 | 1988.816101 | -0.270906694 | 0.135173897 | -2.00413467 | 0.045055636 | 0.162725415 |
| Dnaja4 | 2447.624409 | -0.096046943 | 0.047940747 | -2.003451119 | 0.045128889 | 0.162927061 |
| Snrnp25 | 347.8560901 | -0.159980756 | 0.079853929 | -2.003417456 | 0.045132499 | 0.162927061 |
| Zfp358 | 1033.27465 | 0.17414886 | 0.086937125 | 2.003158706 | 0.045160256 | 0.162989288 |
| Nmt2 | 1892.492432 | -0.102410349 | 0.0511308 | -2.002909163 | 0.045187039 | 0.163047971 |
| Tmem33 | 3564.515493 | -0.178316354 | 0.089040133 | -2.002651485 | 0.04521471 | 0.163109828 |
| Aagab | 2213.748289 | 0.140096055 | 0.06996328 | 2.002422625 | 0.045239297 | 0.163122567 |
| Snora16a | 17.46999578 | 0.336011578 | 0.167801377 | 2.002436351 | 0.045237822 | 0.163122567 |
| Fnip2 | 841.8416717 | 0.167854208 | 0.083848977 | 2.001863523 | 0.045299412 | 0.163301323 |
| Ankrd39 | 278.2449259 | 0.194742428 | 0.097298468 | 2.001495317 | 0.045339038 | 0.163406153 |
| Adam18 | 22.40908354 | -0.33421128 | 0.166998151 | -2.001287308 | 0.045361437 | 0.163448861 |
| 2700081L22Rik | 8.448350352 | -0.314548567 | 0.15718244 | -2.001168615 | 0.045374222 | 0.163456916 |
| Cpne6 | 6542.291153 | 0.219116175 | 0.109524089 | 2.000620846 | 0.045433265 | 0.16363157 |
| Caln1 | 2811.706554 | -0.194167392 | 0.097115078 | -1.999353724 | 0.045570095 | 0.163895788 |
| Khsrp | 3689.281601 | 0.099440111 | 0.049731189 | 1.999552247 | 0.045548635 | 0.163895788 |
| Notch3 | 830.0802891 | -0.158251547 | 0.079146926 | -1.999465488 | 0.045558012 | 0.163895788 |
| Rdx | 2902.333307 | -0.121978458 | 0.061005865 | -1.999454603 | 0.045559189 | 0.163895788 |
| Tmem151b | 5857.611601 | -0.200889489 | 0.100472917 | -1.999439206 | 0.045560853 | 0.163895788 |
| Ush1g | 40.95785739 | -0.309837868 | 0.15493333 | -1.999814167 | 0.045520334 | 0.163895788 |
| Ppp1r2 | 5308.081901 | -0.213294204 | 0.106687012 | -1.999251836 | 0.045581113 | 0.163897368 |
| Cenpq | 126.1802555 | -0.249183453 | 0.12465119 | -1.999045914 | 0.045603386 | 0.163939412 |
| Phb | 2552.368448 | -0.098448631 | 0.049259345 | -1.998577731 | 0.045654062 | 0.164058455 |
| Rsad2 | 72.32131737 | 0.307005928 | 0.153618987 | 1.998489479 | 0.045663619 | 0.164058455 |
| Tnc | 173.9900211 | -0.239212446 | 0.119705049 | -1.998348834 | 0.045678855 | 0.164058455 |
| Ube2j2 | 56.51516385 | 0.304627048 | 0.152433849 | 1.998421287 | 0.045671006 | 0.164058455 |
| Fibcd1 | 593.5034961 | 0.297614188 | 0.148963591 | 1.997898857 | 0.045727627 | 0.164195561 |
| Slc7a5 | 2806.160557 | 0.159015666 | 0.079598908 | 1.997711639 | 0.045747932 | 0.164230411 |
| BC030307 | 54.72657536 | 0.297645319 | 0.149009445 | 1.997492979 | 0.045771656 | 0.164277518 |
| Elf2 | 1022.852604 | 0.168832015 | 0.084540472 | 1.997055502 | 0.045819154 | 0.164396772 |
| Hspa13 | 1504.671721 | -0.179867593 | 0.090069282 | -1.99699152 | 0.045826104 | 0.164396772 |
| Ift88 | 736.6688389 | 0.15206118 | 0.076152116 | 1.996808326 | 0.045846009 | 0.164406322 |
| Meis3 | 1271.050039 | 0.170654726 | 0.085465317 | 1.996771709 | 0.045849988 | 0.164406322 |
| Rbfox1 | 13232.77284 | 0.165871508 | 0.08308063 | 1.996512399 | 0.045878177 | 0.164469339 |
| Rpl19 | 9860.232707 | -0.13040246 | 0.065319733 | -1.996371599 | 0.04589349 | 0.164486174 |
| Samm50 | 2864.796395 | 0.097083249 | 0.048663974 | 1.994971675 | 0.046045969 | 0.164994507 |
| C030013G03Rik | 46.06686688 | 0.298595565 | 0.149756531 | 1.993873409 | 0.046165891 | 0.165337144 |
| Naga | 731.0207856 | 0.155423364 | 0.077957042 | 1.993705251 | 0.046184275 | 0.165337144 |
| Palm | 7941.578703 | -0.137511798 | 0.068970956 | -1.993763855 | 0.046177867 | 0.165337144 |
| Tmem177 | 431.8959443 | -0.175927009 | 0.08823633 | -1.993816026 | 0.046172164 | 0.165337144 |
| Cacna2d2 | 2876.383263 | -0.240180839 | 0.120496014 | -1.993267921 | 0.046232117 | 0.165470183 |
| Rps17 | 4226.715614 | -0.154005298 | 0.077268588 | -1.993116508 | 0.046248691 | 0.165491274 |
| Htr2c | 1970.653092 | -0.316648584 | 0.158879426 | -1.993011886 | 0.046260146 | 0.165494042 |
| Polr2d | 452.104452 | -0.148986958 | 0.074774744 | -1.992476989 | 0.046318748 | 0.165635037 |
| Sorcs1 | 2117.357958 | 0.160516821 | 0.080562251 | 1.992457004 | 0.046320938 | 0.165635037 |
| Large | 6934.451898 | -0.131842467 | 0.066175857 | -1.992304627 | 0.046337644 | 0.165656544 |
| Kcnc2 | 4303.805015 | 0.143732081 | 0.07215472 | 1.991998312 | 0.046371243 | 0.165700194 |
| Tmem67 | 416.2735996 | 0.225418979 | 0.113157496 | 1.992081719 | 0.046362092 | 0.165700194 |
| Psmc3ip | 74.25716914 | -0.281242974 | 0.141199098 | -1.991818488 | 0.046390977 | 0.165732487 |
| Gcnt1 | 181.2957898 | -0.329673308 | 0.165545103 | -1.991441019 | 0.046432423 | 0.165842316 |
| Rasal3 | 139.0768824 | 0.315412102 | 0.158420353 | 1.990982193 | 0.046482845 | 0.165984144 |
| A730090H04Rik | 9.780923368 | 0.319257936 | 0.160395594 | 1.990440807 | 0.046542398 | 0.166158507 |
| Reep4 | 102.4562888 | 0.270508231 | 0.13592741 | 1.990093314 | 0.046580657 | 0.166256785 |
| Arhgef28 | 1517.815833 | -0.189757151 | 0.09538292 | -1.989424854 | 0.046654329 | 0.166481384 |
| Ebna1bp2 | 1368.056494 | 0.123876422 | 0.062276592 | 1.989132979 | 0.046686527 | 0.166557921 |
| 2410004P03Rik | 82.92383588 | 0.287438338 | 0.144529236 | 1.988790266 | 0.046724358 | 0.166654511 |
| Dgkh | 1088.011768 | -0.173193913 | 0.087106264 | -1.988306057 | 0.046777851 | 0.166752575 |
| Pwwp2b | 623.7011022 | 0.182338228 | 0.091704238 | 1.98832936 | 0.046775276 | 0.166752575 |
| Traf7 | 1966.455543 | 0.163488604 | 0.082227422 | 1.988249178 | 0.046784138 | 0.166752575 |
| Setx | 3444.651695 | -0.164354068 | 0.08267597 | -1.987930321 | 0.046819397 | 0.166839865 |
| Araf | 4917.438419 | -0.11679196 | 0.058760325 | -1.987598968 | 0.046856061 | 0.166932123 |
| F13a1 | 131.5722786 | -0.329264237 | 0.165674592 | -1.987415412 | 0.046876381 | 0.166966127 |
| Wdr69 | 11.8005696 | 0.311543471 | 0.156781512 | 1.987118673 | 0.046909247 | 0.16704479 |
| Clic1 | 364.8363775 | -0.191635675 | 0.096452227 | -1.986845522 | 0.046939518 | 0.167114177 |
| Med25 | 2499.322148 | 0.14116631 | 0.071054483 | 1.986733328 | 0.046951956 | 0.167120058 |
| Pld6 | 36.86469182 | -0.308005386 | 0.155039215 | -1.986628906 | 0.046963536 | 0.16712288 |
| Cd248 | 165.7146057 | -0.222926467 | 0.112226413 | -1.986399299 | 0.046989004 | 0.167136739 |
| Tmem180 | 961.2820099 | 0.135053364 | 0.06798697 | 1.986459533 | 0.046982322 | 0.167136739 |
| C530008M17Rik | 1901.093845 | -0.184500995 | 0.092903508 | -1.985942174 | 0.047039745 | 0.167278818 |
| Dse | 263.7638344 | -0.20499017 | 0.10322687 | -1.985821812 | 0.047053113 | 0.16728796 |
| Akr1b10 | 569.4884522 | 0.169308433 | 0.085272613 | 1.98549602 | 0.047089313 | 0.167378254 |
| Ptpn5 | 11064.15916 | -0.122204564 | 0.061558276 | -1.985184953 | 0.047123898 | 0.167424924 |
| Smad9 | 704.6195877 | 0.190695394 | 0.096059326 | 1.985183547 | 0.047124054 | 0.167424924 |
| Slc39a4 | 16.2225224 | 0.306690028 | 0.154508269 | 1.984942492 | 0.04715087 | 0.167481793 |
| Anxa5 | 1897.922498 | 0.158452801 | 0.079850528 | 1.9843676 | 0.047214875 | 0.167632542 |
| Pex11a | 179.6917316 | -0.226782288 | 0.114284452 | -1.984366942 | 0.047214949 | 0.167632542 |
| 4930538K18Rik | 82.48910951 | 0.281672876 | 0.141974512 | 1.983967916 | 0.047259417 | 0.167751982 |
| Klhl32 | 370.0551405 | -0.165605898 | 0.083479051 | -1.983801886 | 0.04727793 | 0.167779259 |
| Zw10 | 802.3558829 | 0.165294638 | 0.083346857 | 1.983213815 | 0.047343551 | 0.167973663 |
| Hic1 | 113.8407213 | -0.25751591 | 0.129855756 | -1.98309199 | 0.047357155 | 0.167983462 |
| Atp5c1 | 7163.682559 | -0.123835687 | 0.062461893 | -1.982579804 | 0.047414385 | 0.168109493 |
| Cpm | 558.9512007 | 0.257887833 | 0.130073407 | 1.98263302 | 0.047408436 | 0.168109493 |
| Mif | 3642.633674 | -0.129419009 | 0.065304194 | -1.98178709 | 0.047503075 | 0.168385414 |
| Zc3hav1l | 266.6681085 | -0.21745024 | 0.109738165 | -1.981537045 | 0.047531079 | 0.168446144 |
| Alkbh5 | 2966.234666 | 0.128891541 | 0.065058234 | 1.981171826 | 0.047572008 | 0.168514103 |
| Papln | 219.0317001 | -0.289349593 | 0.14604702 | -1.981208472 | 0.0475679 | 0.168514103 |
| Fgfr1op2 | 3395.188119 | 0.150465693 | 0.0759575 | 1.980919507 | 0.047600301 | 0.168575786 |
| Cdk5r1 | 11967.49081 | -0.162551194 | 0.082068347 | -1.980680744 | 0.047627088 | 0.168596639 |
| Slit2 | 1244.625576 | -0.151362628 | 0.076419795 | -1.980673041 | 0.047627952 | 0.168596639 |
| Htr5a | 1082.182592 | 0.167303507 | 0.084483003 | 1.980321504 | 0.047667414 | 0.168697787 |
| Hgsnat | 2269.134365 | 0.16665925 | 0.084162435 | 1.980209453 | 0.047679998 | 0.168703789 |
| Mlxipl | 138.2012056 | 0.316318441 | 0.159750117 | 1.980082687 | 0.047694238 | 0.168715646 |
| Amfr | 7135.668803 | -0.088707573 | 0.044814399 | -1.979443548 | 0.04776609 | 0.168930373 |
| Sf3b2 | 9717.684122 | 0.130929835 | 0.066147934 | 1.979348829 | 0.047776746 | 0.168930373 |
| Klrg2 | 9.538053492 | 0.256422926 | 0.129569588 | 1.979036362 | 0.047811912 | 0.169016147 |
| Plek | 699.0642541 | 0.24916027 | 0.125928895 | 1.978579024 | 0.047863423 | 0.169159644 |
| Hspa4l | 5052.41623 | -0.139048345 | 0.070314058 | -1.97753263 | 0.047981456 | 0.169538125 |
| Smim15 | 1050.91118 | -0.167000402 | 0.084457456 | -1.97733167 | 0.048004152 | 0.169579647 |
| Bcl2l12 | 45.417328 | 0.308718979 | 0.156147238 | 1.977101754 | 0.04803013 | 0.169632739 |
| Nrip2 | 627.5544101 | 0.245791969 | 0.124328283 | 1.976959412 | 0.048046218 | 0.16965089 |
| Abcb4 | 269.0683441 | 0.192117333 | 0.097189697 | 1.976725306 | 0.048072689 | 0.169669432 |
| C230081A13Rik | 1956.393236 | -0.213684673 | 0.108105962 | -1.97662246 | 0.048084321 | 0.169669432 |
| Cln8 | 1155.060086 | -0.117999897 | 0.059697483 | -1.976631016 | 0.048083354 | 0.169669432 |
| Hsp90b1 | 14339.63203 | -0.132173292 | 0.066882266 | -1.976208346 | 0.048131185 | 0.169796125 |
| Pop7 | 438.6963896 | -0.16670934 | 0.084366339 | -1.976017244 | 0.048152824 | 0.169833794 |
| Rnf7 | 2090.483091 | -0.121468392 | 0.061478871 | -1.975774595 | 0.048180312 | 0.16989207 |
| Ctxn3 | 90.45444598 | -0.329805695 | 0.166973903 | -1.975193067 | 0.048246242 | 0.170085843 |
| Ifi35 | 101.166282 | 0.288747467 | 0.146234144 | 1.97455573 | 0.048318587 | 0.170302136 |
| Lgals12 | 30.87150873 | 0.328908982 | 0.16659678 | 1.974281744 | 0.048349716 | 0.170373094 |
| Atm | 1009.187116 | -0.122168279 | 0.061887023 | -1.974053242 | 0.048375689 | 0.170387117 |
| Myo9b | 2369.877461 | 0.212320076 | 0.107553621 | 1.974085807 | 0.048371987 | 0.170387117 |
| Yipf5 | 1333.845546 | -0.147197142 | 0.074592642 | -1.973346668 | 0.048456079 | 0.170631476 |
| Ppef2 | 23.68480295 | 0.331557416 | 0.168074293 | 1.972683688 | 0.048531612 | 0.17085862 |
| Insm1 | 157.8201947 | -0.272510667 | 0.138155226 | -1.972496256 | 0.048552983 | 0.17089503 |
| Lcp1 | 522.7735732 | 0.20998012 | 0.106459666 | 1.972391311 | 0.048564953 | 0.170898338 |
| Rhbdl3 | 999.0210003 | -0.253956059 | 0.128769713 | -1.972172284 | 0.048589942 | 0.17094745 |
| Abhd2 | 2519.656477 | -0.174119571 | 0.088298034 | -1.97195297 | 0.048614976 | 0.170996693 |
| Efcc1 | 227.1449971 | 0.201138806 | 0.102020206 | 1.971558516 | 0.048660027 | 0.171098935 |
| Sra1 | 916.6028501 | -0.133549026 | 0.06774295 | -1.971408476 | 0.048677172 | 0.171098935 |
| Zranb1 | 1454.379031 | 0.161525682 | 0.081932964 | 1.971437055 | 0.048673906 | 0.171098935 |
| Ccdc12 | 869.8145279 | -0.124794102 | 0.063318754 | -1.970886902 | 0.048736814 | 0.171242905 |
| Fam114a1 | 242.5322714 | -0.20404975 | 0.103548025 | -1.970580801 | 0.048771844 | 0.171242905 |
| G0s2 | 101.2016957 | -0.253709971 | 0.128736618 | -1.970767715 | 0.048750451 | 0.171242905 |
| Gm20063 | 142.0319636 | -0.258530634 | 0.131196045 | -1.97056728 | 0.048773392 | 0.171242905 |
| Matk | 5682.054505 | 0.196078418 | 0.09950268 | 1.970584291 | 0.048771445 | 0.171242905 |
| Zfp770 | 956.9522956 | -0.14659421 | 0.074407862 | -1.970144088 | 0.048821859 | 0.171374239 |
| 1700040N02Rik | 11.50819633 | 0.321821602 | 0.163387803 | 1.969679471 | 0.048875117 | 0.171405855 |
| Alkbh2 | 105.0922082 | 0.224946878 | 0.114199728 | 1.969767196 | 0.048865058 | 0.171405855 |
| Rnasel | 572.0764952 | 0.152402547 | 0.077366927 | 1.969866881 | 0.048853629 | 0.171405855 |
| Ubxn6 | 2593.555208 | -0.11185126 | 0.056785729 | -1.969707212 | 0.048871936 | 0.171405855 |
| Fes | 160.2739849 | 0.218028581 | 0.110708068 | 1.969401011 | 0.04890706 | 0.171440268 |
| Mrps27 | 1001.614593 | 0.118987386 | 0.060416389 | 1.969455432 | 0.048900816 | 0.171440268 |
| Nuak2 | 44.19769673 | 0.32460354 | 0.164849962 | 1.969084719 | 0.048943363 | 0.17152872 |
| Ppp2r2a | 4260.257295 | -0.130473113 | 0.066269238 | -1.96883375 | 0.048972185 | 0.171557863 |
| Ramp1 | 1929.127689 | -0.113369023 | 0.057582234 | -1.968819486 | 0.048973824 | 0.171557863 |
| Bbs5 | 524.899273 | 0.150943252 | 0.076703051 | 1.967891115 | 0.049080567 | 0.171892927 |
| Ftsj2 | 263.8125868 | -0.180145443 | 0.091554665 | -1.967627121 | 0.049110957 | 0.171950425 |
| Hsbp1l1 | 27.58993814 | 0.330358192 | 0.167911064 | 1.967459346 | 0.049130278 | 0.171950425 |
| Nars2 | 495.2251346 | 0.141739182 | 0.072040901 | 1.967482101 | 0.049127657 | 0.171950425 |
| Ecscr | 99.61098136 | -0.304544713 | 0.154810576 | -1.967208707 | 0.049159155 | 0.172012634 |
| Ctnnd2 | 8807.93021 | -0.113783568 | 0.057851182 | -1.966832219 | 0.049202557 | 0.17212563 |
| Laptm5 | 1847.709138 | 0.248314523 | 0.126262927 | 1.966646334 | 0.049223998 | 0.172129261 |
| Sfpq | 5744.535786 | 0.158354228 | 0.080520576 | 1.966630597 | 0.049225813 | 0.172129261 |
| Mxd1 | 1176.025157 | 0.163074897 | 0.082936443 | 1.96626345 | 0.049268186 | 0.172238556 |
| As3mt | 443.246386 | -0.259577785 | 0.132059511 | -1.965612194 | 0.049343424 | 0.17246267 |
| Bai3 | 4434.438047 | -0.112880655 | 0.057435617 | -1.965342431 | 0.049374618 | 0.172532776 |
| Mbd6 | 1771.428608 | 0.245499738 | 0.124931154 | 1.965080209 | 0.049404955 | 0.172599858 |
| Lrrfip2 | 1288.180849 | 0.156155486 | 0.079489945 | 1.964468407 | 0.049475797 | 0.172797243 |
| Ugdh | 828.1515649 | 0.171460508 | 0.087283921 | 1.964399686 | 0.049483759 | 0.172797243 |
| Chmp2b | 1486.703382 | -0.13423174 | 0.068361453 | -1.963558923 | 0.049581265 | 0.17307478 |
| E430025E21Rik | 2080.747214 | 0.140794663 | 0.071715712 | 1.963233147 | 0.049619089 | 0.17307478 |
| Man2b1 | 2189.857327 | 0.172735843 | 0.087982692 | 1.963293448 | 0.049612086 | 0.17307478 |
| Mblac1 | 96.29201085 | -0.243885639 | 0.12421453 | -1.963422794 | 0.049597067 | 0.17307478 |
| Nova2 | 4273.98137 | -0.10618815 | 0.054083893 | -1.963396942 | 0.049600069 | 0.17307478 |
| Ccdc9 | 1436.275166 | 0.171404608 | 0.087325271 | 1.962829374 | 0.049666003 | 0.173082559 |
| Il1r1 | 171.0612422 | 0.264798604 | 0.134890148 | 1.963068529 | 0.049638212 | 0.173082559 |
| Ptprv | 350.0405874 | 0.308234059 | 0.157023282 | 1.96298317 | 0.04964813 | 0.173082559 |
| Yars | 3649.680323 | 0.118662748 | 0.060452426 | 1.962911276 | 0.049656484 | 0.173082559 |
| Ugp2 | 3065.911484 | -0.191061611 | 0.097350778 | -1.962610002 | 0.049691508 | 0.173132498 |
| 0610012G03Rik | 891.0230695 | -0.127174795 | 0.064821846 | -1.961912558 | 0.049772665 | 0.173206003 |
| Clcn5 | 255.3842286 | -0.227508971 | 0.115966409 | -1.961852338 | 0.049779678 | 0.173206003 |
| Drg1 | 2567.631486 | -0.101245918 | 0.051599604 | -1.96214524 | 0.049745577 | 0.173206003 |
| Ept1 | 3299.788463 | -0.137760338 | 0.070216862 | -1.961926728 | 0.049771015 | 0.173206003 |
| Fam166b | 17.81977071 | 0.319080167 | 0.16260298 | 1.962326686 | 0.049724462 | 0.173206003 |
| Stau2 | 4178.629782 | -0.116969946 | 0.059622026 | -1.961857934 | 0.049779026 | 0.173206003 |
| Frmpd4 | 3631.074728 | -0.14079531 | 0.071804764 | -1.960807358 | 0.049901499 | 0.173512979 |
| N4bp3 | 494.4538448 | -0.204494235 | 0.104286936 | -1.960880645 | 0.049892948 | 0.173512979 |
| I7Rn6 | 850.4619408 | -0.149346161 | 0.07616521 | -1.960818605 | 0.049900187 | 0.173512979 |
| Cox6a1 | 7227.825755 | -0.156906215 | 0.080031155 | -1.960564161 | 0.049929887 | 0.173572731 |
| Mob4 | 1669.997492 | -0.150756313 | 0.076902212 | -1.960363791 | 0.049953285 | 0.173615118 |
| 1700029J07Rik | 142.7684202 | 0.308294051 | 0.157360927 | 1.959152481 | 0.050094932 | 0.174068374 |
| Ss18l1 | 2050.965675 | 0.109917807 | 0.056113772 | 1.958838329 | 0.050131723 | 0.174157157 |
| S100b | 4453.939724 | 0.159251788 | 0.081321438 | 1.958300202 | 0.050194797 | 0.17432844 |
| Tek | 683.292159 | -0.221948589 | 0.113341677 | -1.958225737 | 0.05020353 | 0.17432844 |

| | | | | | | | |
|---|---|---|---|---|---|---|---|
| Myo1g | 25.41201949 | -0.109463575103112 | 0.0828778514255853 | -1.3207820089472 | 0.186574060203268 | 0.640826268877189 | NM_178440 |
| Sfpq | 6225.734069 | -0.0963164637418361 | 0.0729085981904282 | -1.32105768225401 | 0.1864821317563 | 0.640826268877189 | NM_023603,NR_045010 |
| Slc41a2 | 1050.023712 | 0.0988570437587846 | 0.0748534440205415 | 1.32067462028408 | 0.186609880032391 | 0.640826268877189 | NM_177388 |
| Nfatc3 | 711.0180161 | -0.0947100873231063 | 0.071724704285286 | -1.32046675224196 | 0.186679229512181 | 0.640912901889374 | NM_010901 |
| Cyb5 | 1680.104763 | 0.0704657179475365 | 0.0533832614598178 | 1.31999649366828 | 0.186836188662942 | 0.641054259107977 | NM_025797 |
| Gnptab | 3873.801898 | 0.0642503639469311 | 0.0486765481178613 | 1.31994495154752 | 0.186853397906323 | 0.641054259107977 | NM_001004164 |
| Tkt | 6101.803659 | 0.0810589057678217 | 0.0614052684413527 | 1.32006434993831 | 0.186813534122736 | 0.641054259107977 | NM_009388 |
| Tmem185b | 570.3799743 | 0.097806303560612 | 0.0741060935498925 | 1.31981459115455 | 0.186896928770638 | 0.641054259107977 | NM_146103 |
| Sept9 | 3420.557151 | 0.0780119795823709 | 0.0591326603507279 | 1.31927058785561 | 0.187078667053299 | 0.641526137303168 | NM_017380,NM_001113486,NM_001113487,NM_001113488 |
| Dcps | 492.3686965 | 0.088834622070812 | 0.0673590953821368 | 1.31882148308023 | 0.18722880036514 | 0.641692692382393 | NM_027030 |
| Meis2 | 5451.105888 | 0.0951849214821498 | 0.0721792946990024 | 1.31872889419444 | 0.187259763390401 | 0.641692692382393 | NM_001159567,NM_001159569,NM_001159568,NM_010825,NM_001136072,NM_001159570 |
| Oxr1 | 11650.02434 | 0.0878816573893556 | 0.066631378518975 | 1.31892299608253 | 0.187194857334586 | 0.641692692382393 | NM_130885,NM_001130163,NM_001130164,NM_001130165,NM_001130166 |
| Dsp | 54.60585995 | -0.0565620000737161 | 0.0429013412802888 | -1.31842031940628 | 0.187362982436636 | 0.641710497933817 | NM_023842 |
| Ercc4 | 1161.076942 | -0.0721413942394834 | 0.0547223345721914 | -1.31831718810008 | 0.187397489483357 | 0.641710497933817 | NM_015769 |
| Zcchc12 | 3203.941783 | 0.124486030204969 | 0.0944146254757586 | 1.31850367014305 | 0.187335097264569 | 0.641710497933817 | NM_028325 |
| AU019823 | 690.9884575 | -0.103624273544606 | 0.0786453360353885 | -1.31761498861137 | 0.187632565489697 | 0.641776648960143 | NM_001134902,NM_212449 |
| Ing1 | 1326.788792 | 0.0778436955071163 | 0.0590671210136102 | 1.31788538481805 | 0.187542018930214 | 0.641776648960143 | NM_011919 |
| Itga11 | 98.90851397 | -0.12481100719406 | 0.0947100890017409 | -1.31782166514241 | 0.18756335359246 | 0.641776648960143 | NM_176922 |
| Mecp2 | 7058.127029 | 0.0884676665894672 | 0.0671338754431514 | 1.31777982435084 | 0.187577363727296 | 0.641776648960143 | NM_001081979,NM_010788 |
| Slc35b3 | 357.4900336 | -0.108675674413765 | 0.0824800439723963 | -1.31759961779525 | 0.187637713626169 | 0.641776648960143 | NM_134060,NM_001170430,NM_001170431 |
| Abcc10 | 488.1744212 | -0.118077024470768 | 0.0896590459488726 | -1.31695606640853 | 0.187853351411926 | 0.641781415887309 | NM_170680,NM_145140 |
| Alox12b | 298.4545643 | 0.119784496172809 | 0.0909731381139986 | 1.31670181611969 | 0.187938594632593 | 0.641781415887309 | NM_009659 |
| Bak1 | 486.581913 | -0.0876442123094292 | 0.066535227064401 | -1.31726028716482 | 0.187751391951712 | 0.641781415887309 | NM_007523 |
| Cmbl | 399.0173737 | 0.112224398934867 | 0.0852302909127261 | 1.31671965134769 | 0.18793261403394 | 0.641781415887309 | NM_181588 |
| Hnrnpu | 14763.34268 | -0.0758555759147422 | 0.0576115684262046 | -1.31667264035532 | 0.187948378299918 | 0.641781415887309 | NM_016805 |
| Mfge8 | 6569.319254 | 0.0878038148363668 | 0.066677052361754 | 1.31685207618343 | 0.187888213062461 | 0.641781415887309 | NM_008594,NM_001045489 |
| Tasp1 | 765.7294994 | 0.0937883392697146 | 0.0712259499601293 | 1.3167720377505 | 0.187915048373108 | 0.641781415887309 | NM_001159640,NM_001159641,NM_175225 |
| Scaper | 1705.806083 | 0.079853965681182 | 0.0606629106257118 | 1.31635565879584 | 0.188054697670848 | 0.641854954600828 | NM_001081341 |
| Zfp335 | 1721.091444 | -0.0928007780567799 | 0.0704988289770161 | -1.31634495782951 | 0.188058287675969 | 0.641854954600828 | NM_199027 |
| Dctn6 | 2037.150579 | 0.0696724873519213 | 0.0529490087478959 | 1.3158412026871 | 0.18822734679637 | 0.642130211264458 | NM_011722 |
| Psma5 | 2362.589153 | 0.0666976262582791 | 0.050683680933514 | 1.31595860896078 | 0.188187935491091 | 0.642130211264458 | NM_011967 |
| Chd8 | 4799.843963 | -0.0661968031456662 | 0.0503420568493344 | -1.31494037567401 | 0.18852994184932 | 0.64225748951764 | NM_201637 |
| Imp3 | 742.6083373 | 0.0909755784031674 | 0.0691850289644011 | 1.31496047287887 | 0.188523187125522 | 0.64225748951764 | NM_133976 |
| Ints1 | 4733.489996 | -0.0823639527526655 | 0.062617333020185 | -1.31535389292474 | 0.188390993553343 | 0.64225748951764 | NM_026748 |
| Olfr287 | 186.170199 | 0.120692742472289 | 0.0917636856493428 | 1.31525604729406 | 0.188423864398687 | 0.64225748951764 | NM_001011780 |
| Zdhhc3 | 3617.501925 | 0.0625916344110025 | 0.0475975345318711 | 1.3150184148528 | 0.188503713673778 | 0.64225748951764 | NM_026917 |
| Zswim6 | 1254.49152 | -0.0974597722289723 | 0.0740815153826171 | -1.31557476552161 | 0.188316807846709 | 0.64225748951764 | NM_145456 |
| Arhgef15 | 1087.043917 | -0.0830506343563993 | 0.0631742522167271 | -1.31462789731937 | 0.188634989618279 | 0.642464679764388 | NM_177566 |
| Gjd2 | 255.6622307 | 0.117560327227613 | 0.0894723569156792 | 1.31392903104592 | 0.188870088014572 | 0.643114603492656 | NM_010290 |
| Nup35 | 607.2336138 | 0.0883590847356135 | 0.0672724465174134 | 1.31345133572245 | 0.189030908836181 | 0.643290795561659 | NM_027091,NM_001190179 |
| Rundc1 | 2217.399086 | 0.1048537608738 | 0.0798350080308406 | 1.31338072682719 | 0.189054688575845 | 0.643290795561659 | NM_172566 |
| Xkr7 | 147.1795436 | 0.124876230119025 | 0.0950626327565282 | 1.31362057306844 | 0.188973921877857 | 0.643290795561659 | NM_001011732 |
| Gm11627 | 83.0364012 | 0.124513755704096 | 0.0948441486977805 | 1.31282485439199 | 0.189241972920603 | 0.643777259635755 | NR_040286 |
| Dusp5 | 197.3401857 | 0.0962712422687875 | 0.0733623987402832 | 1.31226955391148 | 0.189429201074715 | 0.644077614678967 | NM_001085390 |
| Foxp2 | 926.6374136 | 0.118335481752483 | 0.0901921776655957 | 1.3120370836508 | 0.189507622548389 | 0.644077614678967 | NM_212435,NM_053242 |
| Mob3a | 516.2944968 | 0.113909757915775 | 0.0868013372202121 | 1.31230418290434 | 0.18941752138385 | 0.644077614678967 | NM_172457 |
| Rrp15 | 278.3243043 | -0.109836365980462 | 0.0837064066476162 | -1.31216200024983 | 0.189465480233056 | 0.644077614678967 | NM_026041 |
| Psmd6 | 2218.497418 | 0.0758206310837964 | 0.0577958426081301 | 1.31186998341522 | 0.189564006888174 | 0.644118541300026 | NM_025550 |
| E530001F21Rik | 58.4658805 | -0.121675762707759 | 0.0927843676154856 | -1.31138214156942 | 0.189728689236618 | 0.644323531445567 | NR_002167 |
| Ndfip1 | 11585.41847 | 0.0776742065980845 | 0.0592346357972907 | 1.31129710772422 | 0.189757405171702 | 0.644323531445567 | NM_022996 |
| Tmem130 | 9335.829822 | 0.124616112714135 | 0.0950191721835708 | 1.31148388109912 | 0.189694335998146 | 0.644323531445567 | NM_177735 |
| Hpdl | 44.09153172 | 0.119781102585874 | 0.0913709733790833 | 1.3109316685171 | 0.189880850468783 | 0.644592015403023 | NM_146256 |
| Rac3 | 640.4748772 | 0.108455027270765 | 0.0827510266194548 | 1.31061851074688 | 0.189986682169607 | 0.644800594671897 | NM_133223 |
| Adcy6 | 2596.996269 | -0.0889667681972894 | 0.067894885673139 | -1.31036038009689 | 0.190073950114156 | 0.644946087212853 | NM_007405 |
| 8430419L09Rik | 14249.83385 | 0.0632392744465885 | 0.0482668855000575 | 1.31020002205266 | 0.190128178290768 | 0.644966497709401 | NM_028982 |
| Zfr2 | 2434.405785 | -0.094000110193981 | 0.0717514264877851 | -1.3100800192451 | 0.190168767017029 | 0.644966497709401 | NM_001034895 |
| Opn3 | 180.122375 | 0.117513635555372 | 0.0897328692292929 | 1.30959409372157 | 0.1903331875655 | 0.645373455316633 | NM_010098 |
| Gas2l3 | 106.7412231 | -0.124414185340916 | 0.0950205651449318 | -1.30933956403176 | 0.190419353460516 | 0.645514942442814 | NM_001079876,NM_001033331 |
| Dbf4 | 59.50202385 | -0.123354757135817 | 0.0942452019007608 | -1.30887042149593 | 0.190578247466276 | 0.645902851772079 | NM_013726,NM_001190717 |
| Tpx2 | 143.7183305 | -0.115815697138154 | 0.0885224092996107 | -1.30832066201641 | 0.190764569843251 | 0.646232775546425 | NM_001141977,NM_001141976,NM_001141975,NM_028109,NM_001141978 |
| Tspo | 103.8516472 | -0.124272268938738 | 0.0949851409744766 | -1.30833378425086 | 0.19076012094419 | 0.646232775546425 | NM_009775 |
| Deptor | 452.0526655 | 0.112215440098529 | 0.0857914995540444 | 1.30800184962193 | 0.190872681931107 | 0.64644825780631 | NM_001037937,NM_145470 |
| D2hgdh | 585.4577529 | -0.0898803625357036 | 0.0687529947218703 | -1.30729378260977 | 0.19111295498372 | 0.646547945193856 | NM_178882 |
| Ethe1 | 439.7883704 | 0.0878050479683757 | 0.067148011065221 | 1.30763438224686 | 0.190997349292147 | 0.646547945193856 | NM_023154 |
| Gars | 5594.50856 | 0.0695291432807954 | 0.0531869566299785 | 1.30725929224546 | 0.191124664509322 | 0.646547945193856 | NM_180678 |
| Pcdhgb7 | 363.3764578 | 0.117204922808912 | 0.0896429674900531 | 1.30746366492069 | 0.191055287412411 | 0.646547945193856 | NM_033579 |
| Slc38a10 | 3431.298764 | 0.0693935376522437 | 0.053068442316697 | 1.30762341276428 | 0.191001071731104 | 0.646547945193856 | NM_001164802,NM_001164800,NM_001164801,NM_001164798,NM_001164799,NM_024249 |
| 3110070M22Rik | 35.41209493 | 0.106466091431538 | 0.0814903712638315 | 1.30648676377784 | 0.191387077344037 | 0.646682643754242 | NR_027974 |
| Mkrn2 | 1275.752305 | 0.0800247235961859 | 0.0612506021036711 | 1.30651325615931 | 0.191378074012232 | 0.646682643754242 | NM_023290 |
| Pfkp | 12577.65326 | 0.0555456344285836 | 0.0425131233402129 | 1.30655266102369 | 0.191364683000287 | 0.646682643754242 | NM_019703 |
| Scrn3 | 1336.159214 | 0.0869717551895777 | 0.0665545858652302 | 1.30677329080961 | 0.191289718797571 | 0.646682643754242 | NM_029022 |
| Tmem106a | 58.76656118 | -0.121559144801971 | 0.0930071227156249 | -1.30698747851437 | 0.191216964117532 | 0.646682643754242 | NM_144830 |
| Eif4e | 3911.510224 | 0.0723440887545866 | 0.0554064460657951 | 1.30569805305105 | 0.191655260347685 | 0.646997680772897 | NM_007917 |
| Fgf5 | 121.1004977 | 0.124096809094937 | 0.0950821291876972 | 1.30515387229038 | 0.191840457631585 | 0.646997680772897 | NM_010203 |
| Gpr61 | 888.6900959 | -0.0788097526218667 | 0.0603790386609816 | -1.30525020552862 | 0.191807663621519 | 0.646997680772897 | NM_175470 |
| Plcd3 | 650.9711437 | -0.103667837890346 | 0.0794327958501175 | -1.30510120890064 | 0.191858387182147 | 0.646997680772897 | NM_152813 |
| Smarce1 | 2149.81454 | 0.0649830996969012 | 0.0497767766878236 | 1.30549031136436 | 0.191725944117421 | 0.646997680772897 | NM_020618 |
| Ssfa2 | 2525.46506 | -0.0749185248536549 | 0.0573796728131351 | -1.30566315875724 | 0.19166713173455 | 0.646997680772897 | NM_080558 |
| Susd4 | 4078.104016 | 0.0649815300192021 | 0.0497929683416063 | 1.30503426856166 | 0.191881179180066 | 0.646997680772897 | NM_144796 |
| Ttc23 | 130.3513877 | -0.121688665911592 | 0.0931784573503581 | -1.30597424954175 | 0.191561314628703 | 0.646997680772897 | NM_025905,NM_001168477,NM_001168476,NM_001168475 |
| Zfp169 | 519.6256822 | -0.0948598224732675 | 0.0726776921721168 | -1.30521236487007 | 0.191820544944625 | 0.646997680772897 | NM_001164576,NM_001164575,NM_026450 |
| Adam23 | 7538.113085 | 0.0741347092032084 | 0.0568166841529942 | 1.30480527521776 | 0.19195916240827 | 0.647018137855609 | NM_001177600,NM_011780 |
| Luc7l | 5198.976072 | -0.0703451094196058 | 0.053914425566853 | -1.30475487181773 | 0.191976330315137 | 0.647018137855609 | NR_037906,NM_028190,NR_037905,NM_025881 |
| Gm15417 | 71.88039497 | 0.123797313015436 | 0.0949059984881118 | 1.30442032102896 | 0.192090310310668 | 0.647252110316112 | NR_040404,NR_040403 |
| Dhps | 1139.219003 | 0.0741453953777665 | 0.0568499719206342 | 1.30422923482315 | 0.192155434871121 | 0.647321393074653 | NM_001039514 |
| 2610008E11Rik | 1272.190184 | -0.0893907610853105 | 0.0686573827851317 | -1.30198323121441 | 0.192922118017932 | 0.649602848013091 | NM_001004362 |
| Acta1 | 328.3779266 | 0.104224746917064 | 0.0800441847729705 | 1.30209017947621 | 0.192885559871733 | 0.649602848013091 | NM_009606,NM_001272041 |
| Aldh1a1 | 2479.789611 | 0.11548698323072 | 0.0888196169932445 | 1.30024185129625 | 0.193518090769227 | 0.649744561065956 | NM_013467 |
| Dusp26 | 3137.88749 | 0.0873151918609949 | 0.0671306590130136 | 1.30067539167647 | 0.19336958893931 | 0.649744561065956 | NM_025869 |
| Hnrnpf | 3467.150263 | 0.06735698501086 | 0.0517637806745097 | 1.30123774062023 | 0.193177090659705 | 0.649744561065956 | NM_001166427,NM_001166431,NM_001166430,NM_133834,NM_001166429,NM_001166428,NM_001166432 |
| Nlrp5-ps | 25.13263652 | 0.102071353133227 | 0.0784646721427509 | 1.30085744763616 | 0.193307253748942 | 0.649744561065956 | NR_045119,NR_045120 |
| Pex10 | 399.3846228 | 0.0993146280573831 | 0.0763483925462673 | 1.30080836966271 | 0.193324056283384 | 0.649744561065956 | NM_001042407 |
| Pomp | 3108.002956 | 0.061101164402351 | 0.0469950647026521 | 1.30016129968011 | 0.193545691569076 | 0.649744561065956 | NM_025624 |
| Rpl38 | 3409.281554 | -0.0992509495379072 | 0.0762607335363911 | -1.30146859249059 | 0.193098108273526 | 0.649744561065956 | NM_023372,NM_001048058,NM_001048057 |
| Sf3b5 | 821.3050851 | 0.0740165518417349 | 0.056913403524334 | 1.30051178208115 | 0.193425620767093 | 0.649744561065956 | NM_175102 |
| Specc1 | 4427.686361 | 0.078445435047925 | 0.0603295194174228 | 1.30028277707895 | 0.193504068764626 | 0.649744561065956 | NM_001029936 |
| Timm10 | 588.8626174 | 0.0994901569458489 | 0.0764796759844089 | 1.30087053410282 | 0.193302773566798 | 0.649744561065956 | NM_013899 |
| Tm2d3 | 1074.154362 | 0.085961300242115 | 0.0660614157651618 | 1.30123309115406 | 0.193178681646189 | 0.649744561065956 | NM_026795,NM_178056 |
| Ttc4 | 2070.909051 | -0.061702583299073 | 0.047448046263523 | -1.30042410927483 | 0.193455651220603 | 0.649744561065956 | NM_028209,NM_001172073 |
| Usp11 | 7947.496259 | 0.0766804368397474 | 0.0589739928412318 | 1.30024156658655 | 0.193518188319166 | 0.649744561065956 | NM_145628 |
| Gins3 | 92.42081169 | -0.12316193244969 | 0.0947746231390613 | -1.29952437024177 | 0.193764035577389 | 0.649958942357025 | NM_030198 |
| Hdac3 | 3708.585767 | -0.067559046479524 | 0.051995077518688 | -1.29933543142121 | 0.193828839936847 | 0.649958942357025 | NM_010411 |
| Mri1 | 225.6595815 | -0.119894347677414 | 0.0922745101189753 | -1.29932250545494 | 0.193833274011471 | 0.649958942357025 | NM_026423 |
| Spred1 | 4980.922216 | 0.12352931841073 | 0.0950635642953458 | 1.29943916290522 | 0.193793258977343 | 0.649958942357025 | NM_033524 |
| Tmem50a | 2992.071349 | 0.0584090056731579 | 0.0449494761300087 | 1.29943685003624 | 0.193794052262773 | 0.649958942357025 | NM_027935 |
| Ankrd13b | 3787.191639 | -0.090754120523369 | 0.0698569332612603 | -1.29914263748102 | 0.193894982969947 | 0.650015814129113 | NM_172945 |
| Angel2 | 2386.112988 | -0.0756858817675016 | 0.0582832838332517 | -1.29858643490368 | 0.194085895752926 | 0.650138480379698 | NM_001199020,NR_037578,NM_021421,NR_037579 |
| Fam3a | 774.6212779 | -0.0852593613329997 | 0.0656767515769337 | -1.29816653969444 | 0.194230113325413 | 0.650138480379698 | NM_025473 |
| Hspa4 | 11379.9623 | 0.076124943248291 | 0.0586337107256107 | 1.29831358626668 | 0.194179599636256 | 0.650138480379698 | NM_008300 |
| Ndst2 | 795.594213 | -0.0824108535910553 | 0.0634637666975897 | -1.29854967455288 | 0.194098518354608 | 0.650138480379698 | NM_010811 |
| Pdrg1 | 1565.039365 | -0.0824640540691907 | 0.0635138592907016 | -1.29836314451865 | 0.194162577473661 | 0.650138480379698 | NM_178939 |
| Ptplad1 | 7755.320922 | 0.0673912598673859 | 0.0518924131661757 | 1.29867269135429 | 0.19405627977178 | 0.650138480379698 | NM_021345 |
| Zbtb14 | 1212.036255 | 0.0791451645874371 | 0.0609688981893391 | 1.29812358330065 | 0.194244871599056 | 0.650138480379698 | NM_009547 |
| Col9a1 | 24.96099362 | -0.109821497613466 | 0.0846205644404586 | -1.29781097939543 | 0.194352295857758 | 0.650198399269873 | NM_007740 |
| Entpd6 | 2256.551258 | 0.0757491512143171 | 0.058366561846826 | 1.29781759996604 | 0.194350020290778 | 0.650198399269873 | NM_172117 |
| Gtdc1 | 2572.938064 | -0.0707246093760762 | 0.0545152810389388 | -1.29733549984929 | 0.194515774828587 | 0.650595474363356 | NM_172662 |
| Dclk1 | 41359.77701 | 0.0889447997227138 | 0.0685694297290147 | 1.29714947425146 | 0.194579761442267 | 0.650659671894652 | NM_001195538,NM_001195539,NM_001111051,NM_019978,NM_001111053,NM_001111052,NM_001195540 |
| Dvl3 | 1713.447173 | -0.0796060317395933 | 0.061406861185837 | -1.29637031110709 | 0.194847935437785 | 0.650822539981727 | NM_007889 |
| Imp4 | 777.9491588 | 0.0898200863157729 | 0.0692790404584495 | 1.29649726268428 | 0.194804222500864 | 0.650822539981727 | NM_178601 |
| Myh7 | 704.7068128 | 0.111640906467343 | 0.0861145049629304 | 1.29642394757306 | 0.194829466042356 | 0.650822539981727 | NM_080728 |
| Spag7 | 1756.931158 | 0.0751400693351341 | 0.0579599487471325 | 1.29641366080144 | 0.194833008130947 | 0.650822539981727 | NM_001167669,NM_172561 |
| Usp19 | 6148.282076 | -0.080633337528935 | 0.0621999722833822 | -1.29635709356887 | 0.194852487015044 | 0.650822539981727 | NM_027804,NM_145407,NM_001168372,NM_001168373,NM_001168371 |
| H2-K2 | 53.63784969 | -0.12119232185531 | 0.0935080618188735 | -1.2960628153117 | 0.194953844558462 | 0.650861766503384 | NR_004446 |
| Mrps2 | 1328.675318 | 0.073612764635856 | 0.0567930755511798 | 1.2961573910453 | 0.194921265852776 | 0.650861766503384 | NM_001166031,NM_080452 |
| Emc2 | 2158.64419 | 0.0704726222873927 | 0.0543891119134532 | 1.29571195057445 | 0.195074742578598 | 0.650938719800711 | NM_025736 |
| Foxred1 | 1394.365395 | -0.0765802857708807 | 0.0591078932396335 | -1.29560167980292 | 0.195112750107034 | 0.650938719800711 | NM_172291 |
| Gm12942 | 92.09976788 | -0.122910345097549 | 0.094864696127374 | -1.29563842098348 | 0.195100085753964 | 0.650938719800711 | NM_001099319 |
| Swsap1 | 542.8868675 | 0.0835763286665297 | 0.0645140019691794 | 1.29547580549192 | 0.195156142415813 | 0.650938719800711 | NM_025870 |
| 9130024F11Rik | 358.1781239 | -0.121420867021683 | 0.0937757262479986 | -1.29480060437575 | 0.195389023457305 | 0.651003085650191 | NR_024325,NR_024326 |
| Ftsj2 | 289.3321847 | 0.102081730568035 | 0.0788530063831453 | 1.29458260693348 | 0.195464255617508 | 0.651003085650191 | NM_026510 |
| Mapk12 | 240.2295634 | -0.117468117017588 | 0.0907432729132154 | -1.29451047164599 | 0.195489154592189 | 0.651003085650191 | NM_013871 |
| Mrpl50 | 1335.729573 | 0.0761325259040966 | 0.0588025903017846 | 1.29471381300334 | 0.19541897311075 | 0.651003085650191 | NM_178603 |
| P4ha1 | 2147.692088 | 0.114243870340708 | 0.0882193649745446 | 1.29499765016074 | 0.195321040102945 | 0.651003085650191 | NM_011030 |
| Slc27a1 | 2890.273291 | -0.102216848416469 | 0.0789329978103661 | -1.29487911916853 | 0.195361932755454 | 0.651003085650191 | NM_011977 |
| Tmem30a | 12720.46596 | 0.101175546819784 | 0.0781416319347996 | 1.29477135701752 | 0.195399115654116 | 0.651003085650191 | NM_133718 |
| Mcts2 | 221.0839935 | 0.103986415765607 | 0.0803480966975761 | 1.29419886767205 | 0.195596737814805 | 0.651212059018533 | NM_025543 |
| Ahdc1 | 3159.577334 | 0.0918210827636328 | 0.0709744945264248 | 1.29371943225952 | 0.195762350642458 | 0.651423790816696 | NM_146155 |
| Pitrm1 | 1766.573322 | -0.0671358943055076 | 0.0518985651738988 | -1.29359827348892 | 0.195804219147439 | 0.651423790816696 | NM_145131 |
| Ppp1r13l | 166.3876679 | -0.119404996975604 | 0.0923118714707845 | -1.29349557183871 | 0.195839714614933 | 0.651423790816696 | NM_001010836 |
| Rhbdd2 | 3657.143109 | 0.0746836540383244 | 0.0577207312904751 | 1.2938792071515 | 0.195707147702738 | 0.651423790816696 | NM_146002 |
| Rbm15b | 1568.469615 | 0.0805774781716931 | 0.0623030677954788 | 1.29331477602682 | 0.195902212233913 | 0.651482309093349 | NM_175402 |
| Setd7 | 9696.638615 | 0.082302898314104 | 0.0636435500407735 | 1.29318522083347 | 0.195947005949941 | 0.651482309093349 | NM_080793 |
| Dzank1 | 14206.39871 | 0.0815137167993858 | 0.0630399376188783 | 1.29304881759552 | 0.195994175488701 | 0.651490021315531 | NM_172859 |
| Ccnb2 | 35.40722651 | -0.115364112603971 | 0.0892420784902636 | -1.29270983549041 | 0.196111434761834 | 0.651730656909267 | NM_007630 |
| P4ha3 | 229.5619378 | -0.113122726320428 | 0.087517172219522 | -1.29257748452702 | 0.196157231004036 | 0.651733746012036 | NM_177161 |
| 1700040L02Rik | 73.09868875 | -0.122630294681329 | 0.0948882107490218 | -1.29236597163461 | 0.196230435220283 | 0.651827876059874 | NM_028491 |
| Mboat7 | 5438.495463 | 0.0694227822878355 | 0.0537235702475393 | 1.29222205389478 | 0.196280256322093 | 0.651844308032629 | NM_029934 |
| Ifrd1 | 1513.312934 | 0.104767760004376 | 0.08109389306625 | 1.2919315628217 | 0.196380846061316 | 0.652029295974098 | NM_013562 |
| Mtm1 | 233.977424 | -0.114964180120818 | 0.0890196878265953 | -1.29144667800634 | 0.196548833571185 | 0.652437924235611 | NM_001164190,NM_001164191,NM_001164192,NM_001164193,NM_019926 |
| Spty2d1 | 922.5526142 | 0.0860386950436342 | 0.0666330785810874 | 1.29123097530203 | 0.19662359721763 | 0.652536982678387 | NM_175318 |
| Ltbp1 | 344.0631741 | -0.106580426833155 | 0.0825677579650816 | -1.29082379684124 | 0.196764784092152 | 0.652856385043993 | NM_206958,NM_019919 |
| Mfn2 | 10744.93433 | 0.0604969515293512 | 0.046873920722548 | 1.29063134887818 | 0.19683154018756 | 0.652874708532728 | NM_133201 |
| Pcyt1b | 2767.012006 | 0.0677790856316292 | 0.052524857487258 | 1.29041922004399 | 0.196905142371051 | 0.652874708532728 | NM_177546,NM_211138 |
| Stim1 | 4242.418292 | 0.0615958745045325 | 0.0477321698874127 | 1.29044781852198 | 0.196895218403332 | 0.652874708532728 | NM_009287 |
| Chodl | 56.31277321 | 0.113980880748731 | 0.0883471619834414 | 1.29014761979671 | 0.196999408722709 | 0.653038204268843 | NM_139134 |
| Got2 | 12495.49033 | 0.0601525494998547 | 0.0466394322797517 | 1.28973588569966 | 0.197142375393942 | 0.653065027359726 | NM_010325 |
| Gpc3 | 206.7921396 | 0.121224816505105 | 0.0939826200172412 | 1.289864194921 | 0.197097814363591 | 0.653065027359726 | NM_016697 |
| Slc35a4 | 2298.56425 | 0.0648294004149642 | 0.0502609756221713 | 1.28985559099188 | 0.197100802226389 | 0.653065027359726 | NM_026404,NM_001083317 |
| 3010026O09Rik | 383.8398411 | -0.101710452393318 | 0.0788874644610388 | -1.28931070466274 | 0.197290090954344 | 0.653301922599689 | NM_026543 |
| Evi2a | 708.0564637 | -0.109285872629952 | 0.0847656232601198 | -1.28927115057702 | 0.197303836874903 | 0.653301922599689 | NM_010161,NM_001033711 |
| Cdc34 | 1278.906297 | 0.0810909836497619 | 0.0629045967006951 | 1.28911061993767 | 0.19735963202236 | 0.65333774265196 | NM_177613 |
| 4732471J01Rik | 160.2173552 | -0.117652935758875 | 0.0912900006939589 | -1.28878228573243 | 0.197473786109717 | 0.65341781709106 | NR_073028,NR_073027,NR_015569 |
| Ulk4 | 268.5801607 | -0.103254100504529 | 0.0801119061751712 | -1.2888733452273 | 0.197442122021931 | 0.65341781709106 | NM_177589 |
| Bub1b | 99.85335276 | -0.0814281461317337 | 0.0632000269731418 | -1.28841948384513 | 0.197599979989291 | 0.653498553372773 | NM_009773 |
| Mfhas1 | 2443.776761 | 0.0776842661755119 | 0.0602986960172087 | 1.28832414806021 | 0.197633150555321 | 0.653498553372773 | NM_001081279 |
| Rnf208 | 4277.136376 | 0.0755952186694228 | 0.058667258265836 | 1.28854186992823 | 0.19755740367552 | 0.653498553372773 | NM_176834 |
| 1700003D09Rik | 30.59079242 | -0.082766534548933 | 0.0642825663338105 | -1.28754247487784 | 0.197905274947126 | 0.65353528213834 | NR_045477 |
| Cacul1 | 2839.608649 | 0.0752828644886823 | 0.0584679005374629 | 1.28759308606344 | 0.197887647346975 | 0.65353528213834 | NM_001172096,NM_030197,NM_001172097 |
| Hectd3 | 2714.305273 | -0.0643742444089924 | 0.0499876710712598 | -1.28780243266833 | 0.197814745267494 | 0.65353528213834 | NM_175244 |
| Iws1 | 1501.797471 | -0.0831080300303389 | 0.064563740853944 | -1.28722451535678 | 0.198016044794401 | 0.65353528213834 | NM_173441 |
| Psenen | 1446.834647 | 0.0700233493583154 | 0.054396395848177 | 1.287279207868 | 0.19799698794161 | 0.65353528213834 | NM_025498 |
| Sez6l | 10063.62599 | 0.0798430453765184 | 0.0620038852249037 | 1.2877103601961 | 0.197846805820234 | 0.65353528213834 | NM_001253917,NM_001253916,NM_019982 |
| Tomm22 | 2136.873924 | 0.0711509783930094 | 0.0552788057116172 | 1.28712944277768 | 0.198049174719329 | 0.65353528213834 | NM_172609 |
| Uap1 | 2151.64394 | 0.0843480905293526 | 0.0655041366094034 | 1.2876757850014 | 0.1978588462295 | 0.65353528213834 | NM_133806 |
| Xpa | 208.8159764 | 0.120654882283574 | 0.0937280595466752 | 1.2872866766594 | 0.197994385648159 | 0.65353528213834 | NM_011728 |
| 9930104L06Rik | 791.8893989 | -0.0779596301250612 | 0.0605871270159106 | -1.28673587880456 | 0.19818636302063 | 0.653542590065304 | NM_177573 |
| Cdnf | 68.14441381 | -0.122376322949156 | 0.0950888506739504 | -1.28696815748433 | 0.198105387050368 | 0.653542590065304 | NM_177647 |
| Gas6 | 4190.759783 | 0.0919633267157808 | 0.0714696903418289 | 1.28674583975296 | 0.198182889981932 | 0.653542590065304 | NM_019521 |
| Ccdc77 | 176.9098655 | -0.111293360327708 | 0.0865124103876498 | -1.28644387353234 | 0.198288194955052 | 0.653581647360356 | NR_045558,NM_026028 |
| Cldn25 | 2475.4394 | 0.0631912054978093 | 0.0491155968592424 | 1.28657180962399 | 0.198243574682833 | 0.653581647360356 | NR_045517,NM_171826,NM_001252451,NM_001252450,NR_045518 |
| Tmtc2 | 572.3756769 | -0.103049718317371 | 0.0801198245401478 | -1.28619500740086 | 0.198375013037658 | 0.653679944123788 | NM_177368 |
| Zfp715 | 1200.454681 | -0.0896587326317169 | 0.0697136330727553 | -1.28610041795048 | 0.198408018287332 | 0.653679944123788 | NM_027264 |
| Ntrk3 | 3308.804061 | -0.0676651088614999 | 0.0526180003541385 | -1.28596883967632 | 0.198453936784753 | 0.653682967286921 | NM_008746,NM_182809 |
| Ajap1 | 2019.212675 | 0.0867581642732509 | 0.0675723549806572 | 1.28392986004537 | 0.199166498214127 | 0.65377550969071 | NM_001099299 |
| Bbs9 | 760.1745907 | 0.0851851216774353 | 0.0663214378299035 | 1.28442814970194 | 0.198992188733533 | 0.65377550969071 | NM_181316,NM_178415 |
| Ccdc110 | 30.46062274 | -0.107516788212636 | 0.0837973749273499 | -1.28305675811266 | 0.199472191859371 | 0.65377550969071 | NM_001033246 |
| Cherp | 2781.574521 | -0.0736428782553308 | 0.057356746777036 | -1.28394447721389 | 0.19916138331292 | 0.65377550969071 | NM_138585 |
| Flad1 | 1266.736573 | -0.0869690199770928 | 0.0676493582862691 | -1.28558529127607 | 0.198587832752697 | 0.65377550969071 | NM_177041 |
| Ghitm | 18209.57201 | 0.0773554376799856 | 0.0602585536539887 | 1.28372542965716 | 0.199238043416804 | 0.65377550969071 | NM_078478,NM_001199122 |
| Gm12216 | 25.78671874 | -0.109329936761706 | 0.0850581403046966 | -1.28535536246223 | 0.198668132094498 | 0.65377550969071 | NR_033332 |
| Gpr85 | 1424.532415 | 0.0857488820982056 | 0.0667377568385625 | 1.28486311437813 | 0.198840122484295 | 0.65377550969071 | NM_145066 |
| Kansl2 | 2056.595868 | -0.0667083622565205 | 0.0518853803680816 | -1.28568706219908 | 0.198552298294013 | 0.65377550969071 | NM_133714 |
| Lyl1 | 69.29809234 | -0.12142060209167 | 0.0946154142203241 | -1.28330677503484 | 0.19938462001541 | 0.65377550969071 | NM_008535 |
| Mettl18 | 165.0242488 | 0.11280892220338 | 0.08783752193738 | 1.28429081006864 | 0.19904022114127 | 0.65377550969071 | NM_027279 |
| Ms4a6c | 35.41701924 | -0.10992283698262 | 0.0856625896318633 | -1.28320702718673 | 0.19941955469405 | 0.65377550969071 | NM_001166376,NM_028595 |
| Ogdhl | 5555.216912 | 0.0773040801728137 | 0.0601661807522064 | 1.28484273401347 | 0.1988472456842 | 0.65377550969071 | NM_001081130 |
| Pmaip1 | 104.2672402 | 0.120765626849703 | 0.094002385286154 | 1.28470811120461 | 0.198894302775076 | 0.65377550969071 | NM_021451 |
| Rab4a | 1557.677251 | 0.0789504381206509 | 0.0614195615909264 | 1.28542822637657 | 0.198642682848158 | 0.65377550969071 | NM_009003 |
| Rnd1 | 952.8233762 | 0.0940508483819579 | 0.0732597023029727 | 1.28380058102067 | 0.19921174025426 | 0.65377550969071 | NM_172612 |
| Rps21 | 4041.956774 | -0.0772576759646533 | 0.0601908046434077 | -1.28354615663233 | 0.199300799664985 | 0.65377550969071 | NM_025587 |
| Scrib | 2246.425418 | -0.100985666845788 | 0.0785858438617422 | -1.28503636130006 | 0.198779577985916 | 0.65377550969071 | NM_134089 |
| Snord96a | 57.4566841 | -0.122033851058226 | 0.0950794913354897 | -1.28349288941426 | 0.199319449150527 | 0.65377550969071 | NR_028563 |
| Tsen54 | 277.0314471 | -0.116488518913002 | 0.0907827781621457 | -1.28315657739559 | 0.199437225422053 | 0.65377550969071 | NM_029557 |
| Ubac2 | 1307.471024 | 0.0685535623167634 | 0.0534245426340001 | 1.28318482361953 | 0.199427331655608 | 0.65377550969071 | NM_026861 |
| Zfp758 | 227.9333098 | 0.120819650262155 | 0.0941016945406843 | 1.28392640378989 | 0.199167707655641 | 0.65377550969071 | NM_145484 |
| Anp32e | 4576.2442 | 0.0676551250847793 | 0.052743740787036 | 1.28271381732197 | 0.199592357259741 | 0.654021786951274 | NM_001253758,NM_001253757,NM_023210 |
| Mob4 | 1850.649568 | 0.0774266930485596 | 0.060372012246362 | 1.28249316475658 | 0.199669701193898 | 0.65412766791668 | NM_025283 |
| Sfxn3 | 5856.743427 | -0.0580996484164045 | 0.0453128403646015 | -1.28218950630586 | 0.199776176442627 | 0.654328915221105 | NM_053197,NM_001178012,NM_001178013 |
| Baalc | 9988.794845 | 0.0620086410235367 | 0.0483848552384842 | 1.28157128336754 | 0.199993079197373 | 0.654380963092721 | NM_080640 |
| Cd63 | 1624.442451 | -0.107274543719496 | 0.0837175064217556 | -1.2813872307551 | 0.200057687036945 | 0.654380963092721 | NM_001042580,NM_007653 |
| Ddost | 3644.25303 | 0.0683332288294338 | 0.0533073810419076 | 1.28187180637731 | 0.19988761960033 | 0.654380963092721 | NM_007838 |
| Eri3 | 4970.993266 | 0.0621991118023624 | 0.0485282346221351 | 1.28170975694203 | 0.199944480975371 | 0.654380963092721 | NM_080469 |
| Rhbdf2 | 63.95651894 | -0.121841223842395 | 0.0950614606322975 | -1.28170999090455 | 0.199944398871965 | 0.654380963092721 | NM_001167680,NM_172572 |
| Snrnp35 | 362.8354171 | 0.0966512093180478 | 0.0754277951365926 | 1.28137391717498 | 0.200062361083214 | 0.654380963092721 | NM_029532 |
| Trappc9 | 3660.918189 | 0.0648949607084456 | 0.0506529934960228 | 1.28116733542197 | 0.200134896697447 | 0.654470848587823 | NM_029640,NM_001164641,NM_001164642,NM_001164643,NM_180662 |
| 1500012F01Rik | 532.9781181 | -0.103481162776571 | 0.0807983751600319 | -1.28073321488969 | 0.200287388992166 | 0.654475061639329 | NM_001081005 |
| Ccdc25 | 1917.416365 | 0.0777937912330833 | 0.0607759333441839 | 1.28000981560455 | 0.200541683832896 | 0.654475061639329 | NM_145944 |
| Chchd6 | 1417.396167 | 0.0717321264943711 | 0.0560057387853902 | 1.28079957607993 | 0.200264072993365 | 0.654475061639329 | NM_025351,NM_001167736 |
| Fbxo9 | 3219.305294 | 0.0614031228744815 | 0.0479567519828864 | 1.28038535421234 | 0.200409642235353 | 0.654475061639329 | NM_023605,NM_001081490 |
| Gpsm2 | 665.9245944 | -0.0914017208970552 | 0.0713996766117883 | -1.28014194509621 | 0.200495219083052 | 0.654475061639329 | NM_029522 |
| Lmbrd1 | 8667.37909 | 0.0747104232648696 | 0.0583641580758705 | 1.28007369124986 | 0.200519220297324 | 0.654475061639329 | NM_026719 |
| Med24 | 5183.182722 | -0.0731866478913644 | 0.0571466393315163 | -1.28068157196082 | 0.200305535102212 | 0.654475061639329 | NM_011869 |
| Rfx3 | 1781.699283 | -0.0966888028304755 | 0.0755254223858702 | -1.28021532056423 | 0.200469419208282 | 0.654475061639329 | NM_011265,NM_001166414 |
| Zfp711 | 336.8441745 | 0.105648225616936 | 0.0825330964310847 | 1.28007102829533 | 0.200520156758004 | 0.654475061639329 | NM_177747 |
| Atl1 | 4990.246012 | 0.080994323915974 | 0.0633780663860415 | 1.27795511183049 | 0.20126525409506 | 0.654716628936691 | NM_178628 |
| Cttnbp2nl | 1602.565155 | 0.0680823985125503 | 0.0532573882770333 | 1.27836532573472 | 0.201120644072353 | 0.654716628936691 | NM_001163333,NM_001163332,NM_030249 |
| D6Wsu163e | 1397.929567 | 0.0696729192634983 | 0.0544868600820303 | 1.27871048466741 | 0.200999026186248 | 0.654716628936691 | NM_138594 |
| Gas1 | 213.5148891 | -0.118897688530539 | 0.0930429305318229 | -1.27787987599846 | 0.201291784719211 | 0.654716628936691 | NM_008086 |
| Il1rl2 | 25.36087056 | -0.106444010950233 | 0.0832150264505308 | -1.27914411003055 | 0.200846312921097 | 0.654716628936691 | NM_133193 |
| Mapk10 | 20050.86103 | 0.0596400189593931 | 0.0466326254438386 | 1.27893333029726 | 0.200920534304513 | 0.654716628936691 | NM_009158,NM_001081567 |
| Nynrin | 640.9739895 | 0.121156178497654 | 0.0946804702089899 | 1.27963220113107 | 0.200674519331243 | 0.654716628936691 | NM_001040072 |
| Ppargc1a | 2273.64967 | 0.086831102080338 | 0.0679291455662698 | 1.27825988913151 | 0.20115780570845 | 0.654716628936691 | NR_027710,NM_008904 |
| Prdm2 | 3739.80308 | 0.0886150295066205 | 0.0693389177735284 | 1.27799845097166 | 0.201249972449064 | 0.654716628936691 | NM_001081355,NM_001256380 |
| Psmd8 | 3455.655223 | 0.0551124144831619 | 0.0430983225775194 | 1.27876008130092 | 0.200981555059049 | 0.654716628936691 | NM_026545 |
| Sepn1 | 1329.224137 | -0.0864579351935928 | 0.0676117808060021 | -1.27874068931368 | 0.200988386033428 | 0.654716628936691 | NM_029100 |

| | | | | | | | | | | | | | | | |
|---|---|---|---|---|---|---|---|---|---|---|---|---|---|---|---|
| Camk2d | 4208.383815 | 0.203724484 | 0.104046394 | 1.958015798 | 0.050228159 | 0.174374881 | | Sgce | 1011.990903 | 0.0773585493318854 | 0.0605318639413131 | 1.27798062532629 | 0.201256257778072 | 0.654716628936691 | NM_011360,NM_001130188,NM_001130189,NM_001130191,NM_001130190 |
| Cartpt | 282.2789716 | -0.329116406 | 0.168121218 | -1.957613742 | 0.050275354 | 0.174469598 | | Slc26a6 | 282.4218142 | -0.105780567377975 | 0.0827657151133121 | -1.27807229398253 | 0.201223936901823 | 0.654716628936691 | NM_134420 |
| Id3 | 856.2582333 | -0.181585752 | 0.09276432 | -1.957495646 | 0.050289223 | 0.174469598 | | Ubl7 | 3060.986888 | 0.0630788823913975 | 0.0493611562099426 | 1.27790528493925 | 0.201282824407323 | 0.654716628936691 | NM_027086,NM_001122873 |
| Zfp639 | 855.7434068 | 0.157068435 | 0.080236416 | 1.95757041 | 0.050280442 | 0.174469598 | | Wwp2 | 1891.50686 | 0.0709722303917445 | 0.0554684371333147 | 1.27950658175509 | 0.200718723373713 | 0.654716628936691 | NM_025830 |
| Nfs1 | 1413.552651 | -0.108994161 | 0.055701353 | -1.956759658 | 0.050375732 | 0.174730601 | | Smarcd3 | 2632.696689 | -0.070795394807454 | 0.0554154778453087 | -1.27753829002572 | 0.20141227122936 | 0.654961865206555 | NM_025891 |
| Abtb2 | 574.5916698 | 0.132612553 | 0.067777061 | 1.956599353 | 0.050394591 | 0.174738704 | | Tsc22d4 | 2688.805799 | 0.089287802851859 | 0.0699044747520138 | 1.27728308049817 | 0.201502324780323 | 0.655108050527971 | NM_023910,NM_029805 |
| Esf1 | 1048.481956 | 0.147034838 | 0.075152906 | 1.956475746 | 0.050409137 | 0.174738704 | | Agpat4 | 2055.838811 | 0.0739150972082045 | 0.0578953308148031 | 1.27670221705177 | 0.201707398442223 | 0.655168686616826 | NM_026644 |
| Yae1d1 | 1020.417012 | -0.124867651 | 0.06382351 | -1.956452253 | 0.050411902 | 0.174738704 | | Cfb | 25.2761475 | -0.0886532979006505 | 0.06944365424163 | -1.27662201634983 | 0.201735725227536 | 0.655168686616826 | NM_008198,NM_001142706 |
| Cnih | 2057.535605 | -0.105072906 | 0.053717947 | -1.956011185 | 0.050463837 | 0.174879598 | | Clip4 | 2567.078921 | 0.0736030874974362 | 0.057672628699348 | 1.27622218038881 | 0.201876990053981 | 0.655168686616826 | NM_001271484,NM_175378,NR_073185,NM_030179,NM_001271483 |
| 4930590J08Rik | 22.20173233 | 0.328639685 | 0.168060635 | 1.955482819 | 0.050526109 | 0.174998787 | | Mea1 | 2177.96862 | 0.0765689321107555 | 0.0599950554134412 | 1.27625404432248 | 0.201865729660672 | 0.655168686616826 | NM_010787 |
| Nudt8 | 162.3650279 | 0.23195965 | 0.118611746 | 1.955621245 | 0.050509788 | 0.174998787 | | Ndrg1 | 5432.108449 | 0.0953947947709011 | 0.074731201976684 | 1.27650555922631 | 0.201776862898182 | 0.655168686616826 | NM_008681 |
| Tut1 | 878.7096007 | 0.147284227 | 0.075320561 | 1.9554319 | 0.050532114 | 0.174998787 | | Nr2c1 | 482.3084072 | -0.084311656301638 | 0.066028152566165 | -1.27690466906751 | 0.20163590555027 | 0.655168686616826 | NM_011629 |
| 4932415G12Rik | 1363.432851 | 0.185961481 | 0.0951049 | 1.955330179 | 0.050544111 | 0.17500122 | | Pak1ip1 | 1562.194914 | 0.0713962751552451 | 0.0559349455733098 | 1.2764162800815 | 0.201808404277966 | 0.655168686616826 | NM_026550 |
| Irf2 | 1015.662364 | 0.151006688 | 0.077246862 | 1.954858545 | 0.050599769 | 0.175154785 | | Sema4b | 715.5615369 | 0.096728000726081 | 0.0757932545046307 | 1.27620856708523 | 0.201881800997998 | 0.655168686616826 | NM_013659 |
| 1110002L01Rik | 492.2336206 | -0.206830442 | 0.10585816 | -1.953845054 | 0.050719544 | 0.175451801 | | Arl1 | 3332.411309 | 0.080755777453456 | 0.0632930618857082 | 1.2759025246603 | 0.201989978510014 | 0.65537333657281 | NM_025859 |
| Ahrr | 32.7642281 | 0.317457945 | 0.162472501 | 1.953918004 | 0.050710915 | 0.175451801 | | Ctr9 | 2483.776996 | 0.0757027393154231 | 0.0593488684964175 | 1.27555488812719 | 0.202112909639201 | 0.655443358895353 | NM_009431 |
| Rassf7 | 38.15156428 | 0.322784518 | 0.165191754 | 1.953998976 | 0.050701338 | 0.175451801 | | Llgl1 | 2885.323746 | -0.0771152139664369 | 0.0604563302705529 | -1.27555234698058 | 0.202113808439444 | 0.655443358895353 | NM_001159404,NM_001159405,NM_008502 |
| Sh3d19 | 624.1746922 | 0.152592762 | 0.078104129 | 1.953709281 | 0.050735608 | 0.175468195 | | Shmt1 | 76.58329287 | -0.121136067805198 | 0.0949745106315326 | -1.27545872044724 | 0.202146926053365 | 0.655443358895353 | NM_009171 |
| Tango2 | 2402.549352 | -0.178558271 | 0.091399236 | -1.953607911 | 0.050747604 | 0.175470516 | | Rnf130 | 2071.30084 | 0.0748599876409421 | 0.0587225365152734 | 1.27480848211438 | 0.202377037736392 | 0.656043037303912 | NM_021540 |
| 9530036O11Rik | 34.24309716 | 0.310200124 | 0.158806874 | 1.95331673 | 0.050782076 | 0.175550533 | | Bend4 | 456.6978383 | 0.0991909996361432 | 0.0778260442224045 | 1.27452192421195 | 0.202478507775925 | 0.656065990220277 | NM_001164806 |
| 2010204K13Rik | 430.0839923 | -0.189618986 | 0.097095325 | -1.952915707 | 0.050829584 | 0.175582561 | | Fpgt | 982.5347785 | 0.0838531624658599 | 0.0658035496812805 | 1.27429542740479 | 0.202558736432149 | 0.656065990220277 | NM_029330 |
| AU021092 | 91.35431889 | -0.294418953 | 0.150748994 | -1.953040907 | 0.050814748 | 0.175582561 | | Hgsnat | 2559.376835 | 0.0656779571964985 | 0.0515435660867526 | 1.27422221981996 | 0.202584672638115 | 0.656065990220277 | NM_029884 |
| Amph | 9332.729008 | 0.134261476 | 0.068743149 | 1.953088823 | 0.050809071 | 0.175582561 | | Plekha3 | 1598.702769 | 0.0737635108169519 | 0.0578922880496296 | 1.27415089819418 | 0.202609943007584 | 0.656065990220277 | NM_031256 |
| Deb1 | 782.2850529 | -0.175856968 | 0.090055582 | -1.952760319 | 0.050848003 | 0.175582561 | | Rptor | 3791.592767 | 0.0605208221323162 | 0.0474810368295133 | 1.27463143548495 | 0.202439725501986 | 0.656065990220277 | NM_028898 |
| Ptgdr | 35.1114014 | -0.322775333 | 0.165286493 | -1.952823407 | 0.050840524 | 0.175582561 | | Eftud2 | 4504.08206 | 0.0685296373227523 | 0.0538155994197238 | 1.27341585082551 | 0.202870515619547 | 0.656606302071025 | NM_001109995,NM_011431 |
| Homer3 | 320.4432491 | 0.270769391 | 0.138676829 | 1.952520779 | 0.050876406 | 0.175611834 | | Fbxl7 | 57.6289795 | 0.121069853566451 | 0.0950948133006398 | 1.27314886442536 | 0.202965222096229 | 0.656606302071025 | NM_176959 |
| Mff | 6370.222147 | -0.114510096 | 0.058648007 | -1.952497662 | 0.050879148 | 0.175611834 | | Gm129 | 856.2818815 | 0.112817373264604 | 0.0886202440110902 | 1.273042909366 | 0.203002815819976 | 0.656606302071025 | NM_001033302 |
| Slc2a13 | 6321.850495 | -0.143679681 | 0.073621028 | -1.951611994 | 0.050984288 | 0.175926033 | | Hdhd3 | 141.8154252 | 0.116581559989718 | 0.0915653216967301 | 1.27320646975766 | 0.202944785379972 | 0.656606302071025 | NM_024257 |
| Zfp791 | 185.1506712 | -0.190745854 | 0.097741215 | -1.951539616 | 0.050992888 | 0.175926033 | | Nt5c3 | 1731.78508 | 0.0872555917692602 | 0.0685330982747703 | 1.27318907164281 | 0.202950957573096 | 0.656606302071025 | NM_001252374,NM_026004 |
| Pcdhb2 | 101.9541132 | 0.228107507 | 0.116902992 | 1.951254652 | 0.051026761 | 0.176003702 | | Smc4 | 280.6869833 | -0.102821875500111 | 0.0808014553343342 | -1.2725250439446 | 0.20318663168683 | 0.657054544052291 | NM_133786 |
| Slc25a53 | 346.0892947 | -0.150067152 | 0.076913006 | -1.95112843 | 0.05104177 | 0.176016289 | | Tmem150c | 1587.241756 | 0.0830445117040487 | 0.0652855970756895 | 1.27201887435862 | 0.20336641322703 | 0.657417489018977 | NM_182841 |
| Lxn | 1110.797032 | 0.211208684 | 0.108274387 | 1.950680035 | 0.05109512 | 0.176121866 | | Ugdh | 959.7120464 | 0.0721169832093811 | 0.0566977819417862 | 1.27195422359602 | 0.203389384252463 | 0.657417489018977 | NM_009466 |
| Nup88 | 2085.678942 | 0.128589284 | 0.065920222 | 1.950680377 | 0.051095079 | 0.176121866 | | Calm3 | 59475.43716 | 0.0590142323504623 | 0.0464339120870304 | 1.27092957922332 | 0.203753702319965 | 0.657622143623547 | NM_007590 |
| Rem1 | 17.46079429 | -0.327883235 | 0.168113534 | -1.950367869 | 0.051132289 | 0.176210783 | | Comt | 3679.304222 | 0.0594196415213877 | 0.0467592236125086 | 1.27075765871981 | 0.203814876125101 | 0.657622143623547 | NM_007744,NM_001111062,NM_001111063 |
| Pnrc2 | 1169.353648 | -0.133555708 | 0.068493792 | -1.949895074 | 0.051188626 | 0.176365705 | | Fgd5 | 388.7214099 | -0.11059201927052 | 0.08700662562932 | -1.27107583440464 | 0.203701671427876 | 0.657622143623547 | NM_172731 |
| Adarb1 | 9992.912615 | -0.11145373 | 0.057186738 | -1.948943651 | 0.051302154 | 0.17667828 | | Htr1d | 168.7756339 | -0.116467622441508 | 0.0916505112209558 | -1.27077984497786 | 0.203806980923937 | 0.657622143623547 | NM_008309 |
| Prss53 | 89.31206515 | 0.296956509 | 0.152362949 | 1.949007353 | 0.051294547 | 0.17667828 | | Ngfr | 202.8552945 | 0.10286215412678 9 | 0.0808948303036371 | 1.27155411218119 | 0.203531589634514 | 0.657622143623547 | NM_033217 |
| Yars2 | 488.5549125 | -0.141642457 | 0.072681945 | -1.948798385 | 0.051319507 | 0.176698764 | | Noxo1 | 68.95237976 | -0.120464875457586 | 0.0947661030797863 | -1.27118106097665 | 0.203664242617585 | 0.657622143623547 | NM_027988 |
| Map3k1 | 492.6648547 | -0.287225341 | 0.14739469 | -1.948681746 | 0.051333443 | 0.17670748 | | Pcyox1 | 6091.937461 | 0.0635837037879022 | 0.0500241721524571 | 1.27105959083381 | 0.203707449668646 | 0.657622143623547 | NM_025823 |
| Itpr3 | 282.3315393 | -0.204669152 | 0.105058167 | -1.948150802 | 0.051396922 | 0.176886696 | | Rps6ka1 | 711.7026847 | -0.0859134270767242 | 0.0675778768051182 | -1.27132474618109 | 0.203613142259649 | 0.657622143623547 | NM_009097 |
| Itga3 | 861.3854862 | 0.174372866 | 0.089557344 | 1.947052675 | 0.05152842 | 0.177299875 | | Ak5 | 8514.663961 | 0.0766870762908825 | 0.0604011061376226 | 1.26963032955312 | 0.20421634051357 | 0.657789520087591 | NM_001081277 |
| 4930427A07Rik | 71.19622977 | 0.30095723 | 0.154609623 | 1.946562088 | 0.051587258 | 0.177462915 | | Bmper | 413.1133067 | 0.109836515727193 | 0.0865133547942131 | 1.26959029606999 | 0.204230607805736 | 0.657789520087591 | NM_028472 |
| Cox16 | 287.4988112 | -0.189578446 | 0.097410692 | -1.946176982 | 0.051633484 | 0.177543098 | | Ciapin1 | 2439.991056 | 0.0701475312675813 | 0.0552471355817169 | 1.26970440239069 | 0.204189944052837 | 0.657789520087591 | NM_134141 |
| Gfra2 | 3100.363617 | -0.191536148 | 0.098415821 | -1.946192644 | 0.051631603 | 0.177543098 | | Dennd4c | 951.1500377 | -0.086202412909569 | 0.0679077963100407 | -1.26940377384656 | 0.204297090899075 | 0.657789520087591 | NM_184088,NM_001081014 |
| Ptpn6 | 215.9239118 | 0.271051454 | 0.139294015 | 1.945894479 | 0.051667416 | 0.177620357 | | Dok5 | 799.5301928 | 0.0886521635388724 | 0.0698411116664829 | 1.26934067089624 | 0.204319586578219 | 0.657789520087591 | NM_001163686,NM_029761 |
| Ankhd1 | 2892.331898 | 0.140780244 | 0.072364822 | 1.945423774 | 0.051723996 | 0.177775419 | | Flot2 | 5091.606352 | 0.0665779547932703 | 0.0524090476543849 | 1.2703523107751 | 0.20395916242045 | 0.657789520087591 | NM_001040403,NM_008028 |
| Fbxo10 | 1728.955282 | 0.191800337 | 0.098619529 | 1.944851469 | 0.051792857 | 0.177969403 | | Rora | 485.8677261 | 0.103725018135874 | 0.0816969329987874 | 1.26963172702473 | 0.204215842490171 | 0.657789520087591 | NM_013646 |
| Lrrc43 | 19.75802498 | 0.312608214 | 0.160751419 | 1.944668453 | 0.051814895 | 0.177969403 | | Rprd1b | 1381.262039 | 0.0778084859629513 | 0.0612955878162275 | 1.26939782674459 | 0.204299210914762 | 0.657789520087591 | NM_027434 |
| Rnf111 | 1595.775972 | 0.130464744 | 0.067085643 | 1.944749112 | 0.051805181 | 0.177969403 | | Sgsh | 402.8846555 | 0.11459505257183 | 0.0877592264763591 | 1.27006025158173 | 0.204063168894731 | 0.657789520087591 | NM_018822 |
| Aptx | 1258.035436 | 0.132327986 | 0.068055542 | 1.944411618 | 0.051845834 | 0.178036204 | | Zfp607 | 41.00724343 | -0.11757902235072 | 0.0925798485806758 | -1.27002824214234 | 0.204074570262103 | 0.657789520087591 | NM_001024726 |
| Bloc1s5 | 576.6016301 | -0.158223458 | 0.081389129 | -1.944036763 | 0.051891018 | 0.17815188 | | Plscr4 | 145.3188051 | -0.116857986189727 | 0.0920810186942606 | -1.26907790385914 | 0.204413280225706 | 0.65794533759331 | NM_178711 |
| Hnrnpa0 | 2771.090071 | -0.160155718 | 0.08239676 | -1.943713767 | 0.051929977 | 0.17820666 | | Nckap5l | 1074.527991 | -0.0921844640305655 | 0.0726590271281306 | -1.26872692457061 | 0.204538476061388 | 0.658202459740302 | NM_001001884 |
| Samd4b | 3285.990571 | 0.206448254 | 0.106208913 | 1.943794051 | 0.051920291 | 0.17820666 | | Ache | 2560.752976 | 0.098312401928094 | 0.0775511722008621 | 1.26771006985503 | 0.204901507386351 | 0.658461307576266 | NM_009599 |
| Zfp410 | 842.1599106 | 0.188130226 | 0.096803601 | 1.94342178 | 0.051965218 | 0.178288106 | | Ehmt1 | 1363.054681 | -0.0645602721157261 | 0.0509206912210474 | -1.26785930370562 | 0.20484819950518 | 0.658461307576266 | NM_172545,NM_001109687,NM_001109686,NM_001012518 |
| Nkd1 | 598.2562397 | -0.189394202 | 0.097478557 | -1.942931945 | 0.052024382 | 0.178451576 | | Gm561 | 325.8544624 | 0.0959601792800963 | 0.0756823643979315 | 1.26793315779018 | 0.204821821789839 | 0.658461307576266 | NM_001033297 |
| Trabd | 640.1246797 | 0.166361458 | 0.085630749 | 1.942777101 | 0.052043096 | 0.178476256 | | Maged2 | 1815.205831 | 0.0851570977258094 | 0.0671690528618483 | 1.2678025682595 | 0.204868464807409 | 0.658461307576266 | NM_001199246,NM_030700 |
| Rps23 | 6034.698607 | -0.134123714 | 0.069048024 | -1.942469984 | 0.05208023 | 0.178564082 | | Nr2e1 | 649.6091836 | -0.101334529707348 | 0.0799412262800633 | -1.26761290041181 | 0.204936222742138 | 0.658461307576266 | NM_152229 |
| 1700001C02Rik | 8.842111319 | 0.294694388 | 0.15172822 | 1.94225166 | 0.052106642 | 0.178615113 | | Rbbp7 | 5730.861776 | 0.076848445138805 | 0.0606060109674965 | 1.26800038332863 | 0.204797813672546 | 0.658461307576266 | NM_009031 |
| Simc1 | 759.4926414 | 0.12974995 | 0.066828451 | 1.941537597 | 0.052193104 | 0.17887192 | | Slc27a4 | 4262.290948 | 0.0674126415539552 | 0.0531730882734695 | 1.26779624322837 | 0.204870724132048 | 0.658461307576266 | NM_011989 |
| Camk2b | 36746.42362 | -0.087179452 | 0.044906992 | -1.94133357 | 0.05221783 | 0.178877528 | | Zmpste24 | 1353.600645 | 0.0664236468193997 | 0.0524077286353254 | 1.26743990913254 | 0.204998037259095 | 0.658514261217519 | NM_172700 |
| Cntn3 | 1043.372782 | 0.142504997 | 0.073402502 | 1.941418788 | 0.052207501 | 0.178877528 | | Eif2b5 | 2359.64404 | 0.0598964294712827 | 0.0472638648589361 | 1.26727743594498 | 0.205056105717632 | 0.658534107842166 | NM_172265 |
| Supt5 | 6515.283452 | 0.119931084 | 0.061800536 | 1.940615608 | 0.052304919 | 0.179136256 | | Elavl2 | 2761.412839 | 0.0772836017140845 | 0.0609952888538276 | 1.26704214647284 | 0.205140220155982 | 0.658534107842166 | NM_207686,NM_010486,NM_207685,NM_001177883 |
| Zc3h18 | 1661.076001 | 0.117374738 | 0.060486506 | 1.940511141 | 0.052317602 | 0.179140092 | | Gpr116 | 2036.928417 | -0.0867877754654166 | 0.0684917505051768 | -1.26712742520513 | 0.205109730672652 | 0.658534107842166 | NM_001081178 |
| Tiprl | 1499.16675 | -0.126014516 | 0.064944409 | -1.940344343 | 0.052337856 | 0.179169849 | | Ckap4 | 898.1424583 | 0.0754879774060398 | 0.0596148145600988 | 1.26626205185859 | 0.205419278284251 | 0.659018859003342 | NM_175451 |
| Usp30 | 1564.160514 | 0.114781631 | 0.059177243 | 1.939624524 | 0.052425338 | 0.179429687 | | Shmt2 | 845.928287 | -0.0942819728834948 | 0.0744582432873641 | -1.26623955546758 | 0.20542732986143 | 0.659018859003342 | NM_001252316,NM_028230 |
| Lzts1 | 1672.519523 | -0.201752275 | 0.104024065 | -1.93947694 | 0.05244329 | 0.179451487 | | Srpk1 | 2485.967827 | -0.0788939076583087 | 0.0623013104667202 | -1.26632822114475 | 0.205395597276619 | 0.659018859003342 | NM_016795 |
| Fam184b | 338.9119661 | -0.228070027 | 0.117616403 | -1.939100513 | 0.0524891 | 0.179568585 | | Pafah2 | 334.1567681 | -0.106764105939897 | 0.0843561632266641 | -1.26563492050988 | 0.205643817838233 | 0.659550986840317 | NM_133880 |
| Mrpl44 | 742.2556415 | -0.12330663 | 0.063604078 | -1.938659201 | 0.05254285 | 0.179712785 | | Spdya | 43.11098216 | 0.118094325743603 | 0.0933166389124039 | 1.26552270977589 | 0.205684012831243 | 0.659550986840317 | NM_001142631,NM_029254 |
| Lhx5 | 11.62742051 | -0.279132071 | 0.144008915 | -1.938297159 | 0.052586979 | 0.179744684 | | Bag3 | 395.8046842 | -0.104418173261478 | 0.0825527705421516 | -1.26486576496136 | 0.205919451499574 | 0.660160219042775 | NM_013863 |
| Sh3bgrl | 1592.247401 | -0.19222561 | 0.099171138 | -1.938322109 | 0.052583937 | 0.179744684 | | Adcy4 | 180.5294153 | -0.119779707983569 | 0.0947208962158628 | -1.26455420893188 | 0.20603117668631 | 0.66020374946784 | NM_080435 |
| Tmem74 | 248.116035 | -0.22695138 | 0.117083633 | -1.938369812 | 0.052578121 | 0.179744684 | | Fam189b | 2209.379766 | -0.0850608315948527 | 0.0672720864746846 | -1.26442981112028 | 0.206075798516907 | 0.66020374946784 | NM_001014995 |
| Ppfia1 | 1585.951337 | 0.168106661 | 0.086740762 | 1.938035332 | 0.052618912 | 0.179774515 | | Gabra5 | 2072.204624 | 0.0931271288508514 | 0.0736573754410287 | 1.26432863366698 | 0.206112096318498 | 0.66020374946784 | NM_176942 |
| Rtn4rl1 | 3001.220934 | 0.209983393 | 0.108346983 | 1.938064052 | 0.052615409 | 0.179774515 | | Ssu72 | 2100.945725 | 0.0631747001219384 | 0.0499672941042137 | 1.26432101746752 | 0.206114828847353 | 0.66020374946784 | NM_026899 |
| H2afz | 2827.780593 | -0.107414852 | 0.055430501 | -1.937829363 | 0.052644044 | 0.179820727 | | B3gat3 | 4202.738223 | 0.0826201470432341 | 0.0653593422337711 | 1.26409085862165 | 0.20619741731274 | 0.660267934329878 | NM_024256 |
| Kcnmb2 | 105.5058952 | 0.224632957 | 0.11594042 | 1.937486136 | 0.052685947 | 0.179844912 | | Ptcd1 | 1172.676065 | 0.080499426265378 | 0.0636856581851453 | 1.26401184441483 | 0.206225775716278 | 0.660267934329878 | NM_133735 |
| Nt5e | 376.6529847 | -0.229063299 | 0.118218648 | -1.93762408 | 0.052669102 | 0.179844912 | | Irf2bpl | 2773.590539 | 0.0913798167073479 | 0.0723094551741863 | 1.26373261266072 | 0.206326015408599 | 0.660297796833072 | NM_145836 |
| Phlpp2 | 2884.868924 | -0.103297362 | 0.053312975 | -1.937565168 | 0.052676295 | 0.179844912 | | Spryd3 | 7378.919815 | 0.0732430475261701 | 0.0579554520202886 | 1.2637818354092 | 0.206308342668045 | 0.660297796833072 | NM_001033277 |
| Adamts3 | 1327.525764 | -0.173194674 | 0.089424991 | -1.936759193 | 0.052774786 | 0.180028734 | | Dstn | 6049.520356 | 0.070811050540945 | 0.0560536611763284 | 1.26327253304997 | 0.206491253732355 | 0.660681046633521 | NM_019771 |
| Lrrc4 | 4343.673094 | 0.131126749 | 0.067707493 | 1.936665263 | 0.052786275 | 0.180028734 | | Cldn23 | 26.66720276 | -0.106312721974886 | 0.0841845418735489 | -1.26285324608139 | 0.206641924979925 | 0.661001099262553 | NM_027998 |
| Med20 | 737.7315826 | 0.119846274 | 0.061882708 | 1.93666823 | 0.052785912 | 0.180028734 | | Trpv6 | 56.584789 | -0.119700939491952 | 0.0947945338868599 | -1.26274094701303 | 0.206682293325796 | 0.661001099262553 | NM_022413 |
| Pgk1 | 17070.29627 | -0.101468003 | 0.052391754 | -1.936717042 | 0.052779941 | 0.180028734 | | Spire1 | 8944.91141 | 0.0695100999565238 | 0.0550598193738048 | 1.26244693039429 | 0.206788011118895 | 0.661193627451699 | NM_194355,NM_176832 |
| Klhl40 | 240.4674714 | 0.270899519 | 0.139912968 | 1.936200221 | 0.052843183 | 0.180143517 | | Fh1 | 2814.765895 | 0.0572260279000455 | 0.0453396480525236 | 1.26216303738732 | 0.206890126069382 | 0.661374553539579 | NM_010209 |
| Slc39a13 | 781.0906564 | 0.155554808 | 0.08033644 | 1.936292016 | 0.052831946 | 0.180143517 | | Gpr149 | 127.3449685 | 0.119066421442825 | 0.094351979370484 | 1.26193877687819 | 0.206970817369629 | 0.661486929177388 | NM_177346 |
| Prom1 | 656.1375165 | -0.181085359 | 0.093532007 | -1.936079048 | 0.05285802 | 0.180154458 | | Scfd1 | 1274.900107 | 0.0801999106599004 | 0.0635681637472487 | 1.26163642194827 | 0.207079644021723 | 0.661689157294225 | NM_029825 |
| Nfkbia | 855.7529168 | 0.32023547 | 0.165412556 | 1.935980424 | 0.052870099 | 0.180155995 | | Plcz1 | 62.93226129 | 0.118248747743376 | 0.0937678873427829 | 1.26107936410149 | 0.207280254654014 | 0.661893378567642 | NM_054066 |
| 2310061I04Rik | 1496.157731 | 0.136723546 | 0.070637586 | 1.935563683 | 0.052921162 | 0.180250711 | | Rabep2 | 439.4031515 | -0.0940776082185728 | 0.07459819813904 | -1.26112440468369 | 0.207264029162169 | 0.661893378567642 | NM_030566 |
| Zfp804a | 993.7938261 | -0.25964602 | 0.134143586 | -1.935582819 | 0.052918816 | 0.180250711 | | Usp35 | 1137.337273 | -0.0932497686011556 | 0.0739354649052407 | -1.2612319232816 | 0.20722530021523 | 0.661893378567642 | NM_001177412 |
| Lpl | 965.7956696 | 0.194636179 | 0.100564908 | 1.935428408 | 0.052937746 | 0.180267569 | | Zcchc9 | 409.8157982 | -0.10295017651689 | 0.0816813354017975 | -1.26038801900666 | 0.207529421578501 | 0.6625433797471 | NM_145453 |
| Eif2c3 | 775.9016469 | -0.166105969 | 0.085833733 | -1.935206162 | 0.052965002 | 0.180320752 | | Nedd4 | 23601.30461 | 0.0672191896513528 | 0.0533576176602915 | 1.25978618609461 | 0.207746504358463 | 0.662799411884697 | NM_010890 |
| Rars2 | 648.1360739 | 0.14452757 | 0.074707664 | 1.93457488 | 0.053042485 | 0.180544873 | | Slc7a3 | 409.7692653 | 0.115380747661428 | 0.0915813943031267 | 1.25987105284211 | 0.207715882719268 | 0.662799411884697 | NM_007515 |
| Chkb | 19.72388696 | 0.325014092 | 0.168080493 | 1.933681216 | 0.053152335 | 0.180810506 | | Smad1 | 854.750301 | 0.0820655623842565 | 0.0651395904832912 | 1.2598415460611 | 0.207726528991453 | 0.662799411884697 | NM_008539 |
| Foxd1 | 42.46373653 | -0.324907681 | 0.168026044 | -1.933674523 | 0.053153158 | 0.180810506 | | 6430548M08Rik | 17036.46239 | 0.0656867428974517 | 0.0521528169327763 | 1.25950517652997 | 0.20784792166319 | 0.66296573644721 | NM_172286,NM_001163762,NM_001163760,NM_001163761 |
| Wibg | 577.1176748 | -0.134563918 | 0.069590441 | -1.933655207 | 0.053155535 | 0.180810506 | | Adcy9 | 3015.763352 | 0.0901160950934602 | 0.0715554179012324 | 1.25938884485095 | 0.20788991666784 | 0.66296573644721 | NM_009624 |
| Cox5a | 4091.806955 | -0.176719548 | 0.091406271 | -1.933341625 | 0.053194127 | 0.180886756 | | Fkbp2 | 2292.30727 | 0.0841530436569232 | 0.0668417952741991 | 1.25898838162126 | 0.208034528425886 | 0.662990250090922 | NM_001166368,NM_008020 |
| Fas | 55.81484384 | 0.317099586 | 0.164029311 | 1.933188549 | 0.053212975 | 0.180886756 | | Lynx1 | 20758.29343 | 0.0731733937118046 | 0.0581145263754 | 1.25912397941832 | 0.20798555437694 | 0.662990250090922 | NM_011838 |
| Zfp46 | 1822.538192 | 0.119573355 | 0.061850261 | 1.933271639 | 0.053202744 | 0.180886756 | | Tango2 | 2667.494885 | 0.0764252883131351 | 0.0607013000981026 | 1.2590387387028 | 0.208016339911255 | 0.662990250090922 | NM_138583 |
| Dctn5 | 2976.175281 | -0.087172058 | 0.045106778 | -1.932571143 | 0.05328905 | 0.181105625 | | Atp6v0b | 7148.792276 | 0.0763868183645623 | 0.0607080244787386 | 1.25826559207695 | 0.208295720435828 | 0.663292927281844 | NM_033617 |
| 6720401G13Rik | 2989.036801 | 0.267554415 | 0.138516188 | 1.931575074 | 0.053411975 | 0.181483583 | | Ccdc12 | 954.6078983 | 0.0791801594039454 | 0.0629213656745838 | 1.25839861476384 | 0.20824763263623 | 0.663292927281844 | NM_028312 |
| Cdc45 | 68.87931556 | 0.307432915 | 0.159183391 | 1.931312767 | 0.053444385 | 0.181549961 | | Dhdh | 545.3377747 | -0.0928620164577882 | 0.0738042679921406 | -1.25822014070618 | 0.208312152984977 | 0.663292927281844 | NM_027903 |
| Kpna1 | 5393.446868 | -0.1336058 | 0.069185204 | -1.931132549 | 0.053466663 | 0.181549961 | | Plcd4 | 782.5803795 | -0.0963120943660852 | 0.0765325612084719 | -1.25844598488916 | 0.208230510244455 | 0.663292927281844 | NM_001081456,NM_148937 |
| Zyx | 1682.711905 | 0.173030328 | 0.089600385 | 1.931133753 | 0.053466514 | 0.181549961 | | 2610044O15Rik8 | 293.8725639 | -0.105685117984965 | 0.0840238082532274 | -1.25779966633333 | 0.208464216413399 | 0.66340509522158 | NM_153780 |
| Cbl | 1413.689457 | -0.20003168 | 0.103598273 | -1.930839909 | 0.053502853 | 0.181633043 | | Fads1 | 10507.90044 | 0.0788043791974409 | 0.0626652932535571 | 1.25754424987021 | 0.208556626364942 | 0.66340509522158 | NM_146094 |
| Zfp821 | 1157.101426 | 0.124839769 | 0.064664783 | 1.930568113 | 0.053536484 | 0.181707402 | | Grm1 | 2015.654262 | 0.0842114686165607 | 0.066967825699474 | 1.25749145559047 | 0.208575731094379 | 0.66340509522158 | NM_001114333,NM_016976 |
| Fkbp4 | 4208.822188 | -0.104501867 | 0.054133232 | -1.930456817 | 0.053550261 | 0.181714354 | | Hsph1 | 13809.52882 | 0.0787168565641209 | 0.0625779208483892 | 1.25790143707127 | 0.208427403920866 | 0.66340509522158 | NM_013559 |
| Zar1 | 8.165889161 | -0.242975907 | 0.125872553 | -1.93033272 | 0.053565625 | 0.181726692 | | Tcf7l2 | 240.0164466 | 0.105698559306394 | 0.0840542511867452 | 1.25750402643599 | 0.20857118195236 | 0.66340509522158 | NM_001142921,NM_001142919,NM_009333,NM_001142918,NM_001142920,NM_001142923,NM_001142922,NM_001142924 |
| 3110002H16Rik | 933.6123716 | 0.117692194 | 0.060981798 | 1.929956114 | 0.053612276 | 0.18184514 | | Sox21 | 331.4291973 | -0.11513772580086 3 | 0.0915786897902884 | -1.25725456505792 | 0.208661470570551 | 0.663532513464935 | NM_177753 |
| Nfe2l2 | 683.6596151 | 0.153081171 | 0.079329641 | 1.929684399 | 0.053645954 | 0.181889635 | | Mospd1 | 383.6920243 | 0.0974482104616147 | 0.0775348848027845 | 1.25683053130834 | 0.20881500789874 | 0.663584949625268 | NM_027409 |
| Tacr1 | 526.9605875 | -0.262203813 | 0.135880775 | -1.929660859 | 0.053648873 | 0.181889635 | | Pim3 | 1536.526028 | 0.0893881750756225 | 0.0711148761159233 | 1.2569546620585 | 0.208770053231167 | 0.663584949625268 | NM_145478 |
| Snhg6 | 278.3514881 | -0.226604268 | 0.117442211 | -1.929495921 | 0.053669326 | 0.181919173 | | Uba1 | 15890.43162 | 0.046673631693735 | 0.0371331677555819 | 1.25692566820451 | 0.20878055289493 | 0.663584949625268 | NM_001136085,NM_001276316,NM_001276317,NM_009457 |
| Kansl2 | 1930.837496 | 0.145748618 | 0.075548345 | 1.92920995 | 0.053704803 | 0.181999612 | | Igf2r | 1765.609842 | -0.0713646753645499 | 0.0567914611969208 | -1.25660924830051 | 0.208895164213966 | 0.663694478427838 | NM_010515 |
| P2ry13 | 554.6192313 | -0.288704034 | 0.149680247 | -1.928805163 | 0.053755055 | 0.182130071 | | 1700052K11Rik | 258.3134951 | -0.10646118599683 | 0.0847436922242864 | -1.25627268770713 | 0.209017120771972 | 0.663936736569794 | NR_027956 |
| Col7a1 | 159.1235025 | 0.316953638 | 0.164336542 | 1.928686306 | 0.053769817 | 0.18214026 | | Cog3 | 1568.831826 | -0.0703524872506733 | 0.0560067918744266 | -1.25614206591964 | 0.209064466954314 | 0.663941942933616 | NM_177381 |
| Stard5 | 265.7469477 | -0.210513111 | 0.109161738 | -1.928451449 | 0.053798998 | 0.182199272 | | Arl8a | 8534.214947 | 0.0527719647418299 | 0.0420299455256765 | 1.25558013653839 | 0.209268236828221 | 0.664196936803936 | NM_026823 |
| Pou2f1 | 991.4750727 | -0.184918205 | 0.09589965 | -1.928246915 | 0.053824421 | 0.182245538 | | Magoh | 453.656887 | 0.0906946913990632 | 0.0722354681443373 | 1.25554237729645 | 0.209281934445244 | 0.664196936803936 | NM_010760 |
| Gm10033 | 143.2451691 | -0.23362153 | 0.121181494 | -1.927864747 | 0.053871952 | 0.182326784 | | Slc14a2 | 28.96766282 | -0.0930357512792121 | 0.0740974641153985 | -1.25558617140148 | 0.209266047669579 | 0.664196936803936 | NM_207651,NM_001110273,NM_030683,NM_001110274 |
| Hck | 98.0211987 | 0.269538044 | 0.139807802 | 1.927918477 | 0.053865267 | 0.182326784 | | Chrm5 | 37.81740929 | 0.0958532429764851 | 0.0764146690277107 | 1.2543827539412 | 0.209702917968872 | 0.664226905018717 | NM_205783 |
| Scel | 98.31533492 | 0.257462711 | 0.13355852 | 1.927714615 | 0.053890633 | 0.182350179 | | Inmt | 61.00851825 | -0.115702190221246 | 0.0922259803409014 | -1.25455093883055 | 0.209641823030576 | 0.664226905018717 | NM_009349 |
| Pawr | 75.04228703 | -0.25842569 | 0.134120272 | -1.926820506 | 0.054002003 | 0.182650548 | | Minpp1 | 2565.450014 | 0.0653119474228603 | 0.0520475657580604 | 1.25485114378756 | 0.209532802444291 | 0.664226905018717 | NM_010799 |
| Snord32a | 9.975550076 | 0.313568986 | 0.162746928 | 1.926727523 | 0.054013596 | 0.182650548 | | Mpped2 | 1288.648609 | 0.0808498580568506 | 0.064422519165227 | 1.2549937367319 | 0.209481033651581 | 0.664226905018717 | NM_001143683,NM_029837 |
| Stk36 | 196.1488964 | 0.236499321 | 0.122747234 | 1.926718127 | 0.054014768 | 0.182650548 | | Nrp2 | 1511.405105 | 0.107625121355889 | 0.0857879459744151 | 1.25454829502489 | 0.209642783321203 | 0.664226905018717 | NM_001077405,NM_001077404,NM_001077407,NM_001077406,NM_001077403,NM_010939 |
| 1810020O05Rik | 44.58619223 | 0.322496803 | 0.167396242 | 1.926547456 | 0.054036052 | 0.182682652 | | Srp19 | 917.26169 | 0.0768193953475184 | 0.0612357223232678 | 1.25448663677034 | 0.20966517991144 | 0.664226905018717 | NM_025527 |
| Hemk1 | 377.2024423 | 0.231487924 | 0.120163041 | 1.926448617 | 0.054048382 | 0.182684474 | | Tfe3 | 1792.472289 | -0.0676389938890455 | 0.0538946312695882 | -1.25502285284607 | 0.209470464098578 | 0.664226905018717 | NM_001105197,NM_001105196,NM_001271490,NM_001271491,NM_001271489,NM_172472 |
| Hiat1 | 2059.639386 | -0.094476602 | 0.049044751 | -1.926334632 | 0.054062604 | 0.18269269 | | Tmem33 | 3971.856095 | 0.0815439929152535 | 0.0649965817251701 | 1.25458894530872 | 0.20962801856129 | 0.664226905018717 | NM_028975,NM_030108 |
| Fam161b | 675.7785654 | 0.169937168 | 0.08822425 | 1.926195655 | 0.054079948 | 0.182711451 | | Zfp146 | 1425.527078 | 0.0743761225730432 | 0.0592737411649861 | 1.25479042002799 | 0.209554851198411 | 0.664226905018717 | NM_011980 |
| Ube2a | 1086.427789 | -0.176804357 | 0.091836041 | -1.925217535 | 0.054202149 | 0.183084391 | | Apobr | 54.23075335 | -0.113239860193809 | 0.0903011425529667 | -1.25402466671324 | 0.209833039879771 | 0.66442122108336 | NM_138310 |
| Bnip3l | 5072.891126 | -0.114040868 | 0.059242818 | -1.924973715 | 0.054232647 | 0.183107568 | | Tceal3 | 3202.221898 | 0.0885742542281647 | 0.0706355058854085 | 1.25396219815935 | 0.20985574572012 | 0.66442122108336 | NM_001029978 |
| Gars | 4880.853975 | 0.119938573 | 0.062305067 | 1.925021166 | 0.05422671 | 0.183107568 | | Nadsyn1 | 180.5197248 | -0.111415870812807 | 0.0888688195340493 | -1.25371160995471 | 0.209946846478666 | 0.664564805393138 | NM_030221 |
| Fam207a | 1319.647085 | -0.116936272 | 0.060753169 | -1.924776511 | 0.054257324 | 0.183111081 | | Arhgdia | 12099.5019 | 0.070514445642989 | 0.0562628677682985 | 1.25330343866191 | 0.210095297490348 | 0.664608510640737 | NM_133796 |
| Pex26 | 727.3870079 | 0.153721993 | 0.079864129 | 1.924793965 | 0.054255139 | 0.183111081 | | Cacng5 | 648.5670185 | 0.10750584912819 | 0.0858036555973548 | 1.2529285422602 | 0.210231713421192 | 0.664608510640737 | NM_001199301,NM_080644 |
| 1500011K16Rik | 514.0058728 | -0.1718725 | 0.089312457 | -1.924395609 | 0.054305014 | 0.183192226 | | Fbln2 | 321.7883772 | -0.115588251528388 | 0.0922354957201712 | -1.25318621237821 | 0.210137946474995 | 0.664608510640737 | NM_001081437,NM_007992 |
| Isyna1 | 1202.531947 | -0.138505414 | 0.071970617 | -1.924471687 | 0.054295486 | 0.183192226 | | Gsto1 | 952.1495535 | 0.0840198492774928 | 0.0670592804378309 | 1.2529190401228 | 0.210235171856959 | 0.664608510640737 | NM_010362 |
| 3110039M20Rik | 123.326606 | -0.245175185 | 0.127442363 | -1.923812304 | 0.054378115 | 0.183328028 | | Sub1 | 7119.340532 | 0.0798806458026874 | 0.063747956826889 | 1.25306989868879 | 0.21018026963467 | 0.664608510640737 | NM_011294 |
| Cdc37 | 4111.477235 | -0.098702326 | 0.051306155 | -1.923791145 | 0.054380768 | 0.183328028 | | Tmem203 | 257.8051516 | -0.107316366485992 | 0.085631747059091 | -1.25323107575906 | 0.210121623647541 | 0.664608510640737 | NM_177344 |
| Foxk2 | 4801.766519 | -0.105842915 | 0.055017698 | -1.923797617 | 0.054379956 | 0.183328028 | | Med14 | 3767.419252 | 0.0810004486459835 | 0.0646718575290357 | 1.25248371920687 | 0.210393657146003 | 0.664820157429376 | NM_001048208,NM_012005 |
| A4galt | 97.69614179 | -0.259376347 | 0.134845829 | -1.923502926 | 0.054416919 | 0.183409996 | | Taf1b | 820.5573803 | -0.0723537035932213 | 0.05776811165976 | -1.25248517762476 | 0.2103931260422 | 0.664820157429376 | NR_028341,NM_020614 |
| Kctd21 | 781.806974 | -0.203300937 | 0.10571073 | -1.923181649 | 0.054457241 | 0.183466078 | | Tm9sf3 | 6955.598121 | 0.0747537648199223 | 0.0596960316208311 | 1.25224010357561 | 0.21048238689054 | 0.664955883421484 | NM_133352 |
| Smpd4 | 2441.290896 | 0.270340154 | 0.14056236 | 1.923275571 | 0.054445451 | 0.183466078 | | Acyp1 | 391.5395527 | 0.091862449237143 | 0.0733747894746611 | 1.25196201440368 | 0.210583705686629 | 0.665097900291184 | NM_025421 |
| Necap1 | 7860.732042 | -0.117872081 | 0.061298973 | -1.922904671 | 0.054492023 | 0.183543349 | | Eif3l | 2655.700377 | 0.0546432831926379 | 0.0436538681219775 | 1.25173977801815 | 0.21066470048463 | 0.665097900291184 | NM_145139 |
| Cldn26 | 35.29108699 | -0.309915378 | 0.161190041 | -1.922670762 | 0.054521411 | 0.183602422 | | Ppfia4 | 3742.786209 | -0.0807290017211188 | 0.0644933697701 | -1.25174110158756 | 0.210664218038607 | 0.665097900291184 | NM_001144855 |
| Crocc | 2268.532449 | 0.249401255 | 0.129727136 | 1.922506443 | 0.054542064 | 0.183632059 | | Coq5 | 1514.554985 | 0.0677745736381388 | 0.0541624875407258 | 1.2513194411017 | 0.210817955306969 | 0.665414392106873 | NM_026504 |
| Dnlz | 1381.016075 | -0.164606726 | 0.085661015 | -1.92160607 | 0.054655344 | 0.18395243 | | Lsm14a | 2668.717621 | 0.0759332584586438 | 0.0606876871306737 | 1.2512135829982 | 0.210856563876355 | 0.665414392106873 | NM_025948 |
| Hnrnpc | 6421.726388 | -0.140460454 | 0.073101315 | -1.921449082 | 0.054675116 | 0.18395243 | | Ecm1 | 285.5404961 | -0.112501127364725 | 0.0899387210315761 | -1.25086421148047 | 0.210984022955567 | 0.665621429296965 | NM_007899,NM_001252653 |
| Kcng1 | 686.3751542 | -0.223019908 | 0.116078725 | -1.921281508 | 0.054696227 | 0.18395243 | | Klhl1 | 236.4634064 | 0.117609518286566 | 0.094028747287507 | 1.25078257106794 | 0.21101381535953 | 0.665621429296965 | NM_053105 |
| Pde6d | 1136.34169 | -0.131790317 | 0.068594784 | -1.92128774 | 0.054695442 | 0.18395243 | | 1700017B05Rik | 657.650475 | 0.0923164749245 | 0.0739767993130734 | 1.24791117156282 | 0.212063589021564 | 0.666270994326506 | NM_028820 |
| Pex10 | 362.5420839 | -0.171126456 | 0.089069044 | -1.92127869 | 0.054696582 | 0.18395243 | | 2700097O09Rik | 157.470258 | -0.10901234327853 | 0.0873710750093547 | -1.247693739225 | 0.212143234946766 | 0.666270994326506 | NR_030764,NM_028314 |
| Doc2a | 1540.800862 | -0.164897964 | 0.085841764 | -1.920952657 | 0.054737676 | 0.184050685 | | 3110035E14Rik | 14883.03804 | 0.0979106674927408 | 0.0783511745913447 | 1.24963879614329 | 0.211431524230835 | 0.666270994326506 | NM_178399 |
| Gm14204 | 422.6636717 | -0.272268548 | 0.141792887 | -1.920184805 | 0.05483456 | 0.184315114 | | AI854703 | 1416.499807 | 0.0741089682335084 | 0.0593670440589339 | 1.24831831209147 | 0.211914510694634 | 0.666270994326506 | NR_027236 |
| Ptger3 | 66.47488622 | -0.259299325 | 0.135041825 | -1.92014085 | 0.054840111 | 0.184315114 | | Abca17 | 35.67291926 | -0.112899242554171 | 0.0903366046082879 | -1.24976185504999 | 0.211386554197019 | 0.666270994326506 | NM_001031621 |
| Rps6ka2 | 4589.566528 | -0.167064883 | 0.087013203 | -1.919994645 | 0.054858576 | 0.184337188 | | Akip1 | 96.5449448 | -0.11652577843623 | 0.0934771747191483 | -1.24656931698807 | 0.212555458071631 | 0.666270994326506 | NM_020616 |
| Dhtkd1 | 225.5715021 | 0.276196911 | 0.143871795 | 1.919743278 | 0.054890335 | 0.18437962 | | Alms1 | 526.7761204 | -0.102169912814611 | 0.0819501893523511 | -1.24673186995729 | 0.212495828971628 | 0.666270994326506 | NM_145223 |
| Ndufs7 | 2212.169063 | -0.113438455 | 0.059091567 | -1.919706335 | 0.054895004 | 0.18437962 | | Cd36 | 73.97923793 | 0.100198003729527 | 0.0802307265304516 | 1.24887319438017 | 0.211711457299262 | 0.666270994326506 | NM_001159558,NM_007643,NM_001159555,NM_001159556,NM_001159557 |
| Rbm43 | 620.6710339 | 0.12511251 | 0.065176047 | 1.919608769 | 0.054907336 | 0.18438107 | | Ctdnep1 | 2785.643481 | 0.060307667488322 | 0.0482588219825289 | 1.24967135563639 | 0.211419625179042 | 0.666270994326506 | NM_026017 |
| 2900052N01Rik | 1484.438359 | 0.163976915 | 0.085433045 | 1.919361709 | 0.054938573 | 0.184445993 | | Dcaf7 | 12022.97118 | 0.0623305459928375 | 0.049908140175809 | 1.24890540447447 | 0.211699674670625 | 0.666270994326506 | NM_027946 |
| Clec4a2 | 29.23073781 | -0.30901615 | 0.161012276 | -1.919208624 | 0.054957937 | 0.18447 1029 | | Ercc3 | 1365.771599 | 0.0715781312398886 | 0.0573780636500663 | 1.2474825166012 | 0.21222062693362 7 | 0.666270994326506 | NM_133658 |
| Gas2l3 | 96.30230857 | 0.280726229 | 0.146295329 | 1.918900837 | 0.054996885 | 0.184561778 | | Fam169a | 2637.963899 | 0.0787859334359305 | 0.0631018341134212 | 1.24855219412986 | 0.211828906846513 | 0.666270994326506 | NM_001100458,NM_001146045 |
| Dpagt1 | 500.9827251 | 0.139901283 | 0.072933301 | 1.918208579 | 0.055084569 | 0.184659741 | | Gabbr1 | 33901.78661 | 0.0539743797834322 | 0.043297901779486 | 1.2465818796098 | 0.212550849310146 | 0.666270994326506 | NM_019439 |
| Inca1 | 79.34815103 | 0.250837208 | 0.130741411 | 1.918575043 | 0.055038137 | 0.184659741 | | Gabrg2 | 7909.983652 | 0.0827616654342335 | 0.0662617035956513 | 1.24901203777177 | 0.2116606710092 | 0.666270994326506 | NM_008073,NM_177408 |
| Klhl29 | 2881.312996 | -0.162431267 | 0.084679009 | -1.918199915 | 0.055085667 | 0.184659741 | | Gm10432 | 120.1451003 | -0.112253131484086 | 0.0900779935471882 | -1.2461770856972 | 0.212699389539096 | 0.666270994326506 | NR_045741 |
| Lgr6 | 129.2390869 | -0.293804905 | 0.153157641 | -1.918316995 | 0.055070829 | 0.184659741 | | Hic2 | 470.2948362 | -0.096943289707247 | 0.0776064222569948 | -1.24916581499168 | 0.211604432485005 | 0.666270994326506 | NM_178922 |
| Tbp | 604.1238734 | 0.138932645 | 0.072424466 | 1.91831094 | 0.055071596 | 0.184659741 | | Hirip3 | 1187.130136 | -0.085218552094275 | 0.0683849977737382 | -1.24615858548732 | 0.212706180032376 | 0.666270994326506 | NM_172746 |
| Snhg12 | 510.4127891 | 0.207883298 | 0.108442691 | 1.916987633 | 0.055239503 | 0.185095341 | | Mb21d1 | 44.28449536 | -0.112497637927844 | 0.0902174910077891 | -1.2469603917286 | 0.212412021053067 | 0.666270994326506 | NM_173386 |
| Stk3 | 508.0199471 | 0.154728721 | 0.080711309 | 1.917063703 | 0.055229839 | 0.185095341 | | Mkrn1 | 5056.16282 | 0.0687451966037962 | 0.0551689674940993 | 1.24608452407867 | 0.2127333658052 | 0.666270994326506 | NM_018810 |
| Rc3h1 | 1819.729054 | -0.185674347 | 0.096865672 | -1.916822999 | 0.055260422 | 0.185125401 | | Ndufa8 | 3069.848201 | 0.058214268807485 | 0.0466837870547461 | 1.24699113932675 | 0.2124007465225 | 0.666270994326506 | NM_026703 |
| Sstr4 | 329.0088604 | -0.228038703 | 0.118987259 | -1.916496816 | 0.055301888 | 0.185220192 | | Nipa1 | 2195.766785 | 0.0763078151687419 | 0.0612117331594641 | 1.24662072498341 | 0.21253659883528 | 0.666270994326506 | NM_153578 |
| Tmem169 | 636.4382066 | -0.138416192 | 0.072226726 | -1.916412382 | 0.055312626 | 0.185220192 | | Nkiras2 | 369.8900942 | 0.0914094456612874 | 0.0732655272902928 | 1.24764604913174 | 0.212160706818405 | 0.666270994326506 | NM_028024 |
| Cadm2 | 6052.078997 | -0.187142657 | 0.097672415 | -1.916023647 | 0.055362086 | 0.185345756 | | Rab21 | 1693.249215 | 0.0768022824163586 | 0.0615160712813412 | 1.24849134245109 | 0.211851176941627 | 0.666270994326506 | NM_024454 |
| BC031361 | 243.8049379 | -0.183084504 | 0.095566112 | -1.915788982 | 0.055391961 | 0.185395636 | | Rpl13 | 6139.449868 | -0.0869839303367333 | 0.0696969714088627 | -1.24803027417734 | 0.212019970641896 | 0.666270994326506 | NM_016738 |
| Cd2bp2 | 3550.236217 | 0.099128891 | 0.051745015 | 1.915718659 | 0.055400916 | 0.185395636 | | Slc45a3 | 57.05666145 | 0.117632847532354 | 0.0943071436010356 | 1.24733761452894 | 0.212273730859681 | 0.666270994326506 | NM_145977,NM_001177628 |
| Nrxn2 | 13944.33774 | -0.09638522 | 0.05031907 | -1.915480945 | 0.055431197 | 0.185456913 | | Taf1c | 575.708282 | -0.10541797504605 | 0.0843358772796855 | -1.24997780833475 | 0.21130765402087 | 0.666270994326506 | NM_021441 |
| Arpc4 | 2862.254256 | -0.129759253 | 0.067757571 | -1.915051721 | 0.055485908 | 0.185599883 | | Tbx1 | 42.98428412 | -0.112493402588769 | 0.0900457755590943 | -1.24929128424179 | 0.211558554591747 | 0.666270994326506 | NM_011532 |
| Dhx37 | 669.6491214 | 0.137770273 | 0.07194479 | 1.914944405 | 0.055499594 | 0.185605592 | | Tob2 | 1394.767803 | 0.0852049542593922 | 0.0682516969710422 | 1.24839319812873 | 0.211887098717036 | 0.666270994326506 | NM_020507 |
| Slc14a2 | 22.251202 | 0.316605857 | 0.16536284 | 1.914613085 | 0.055541865 | 0.185706875 | | Tpgs2 | 2529.103205 | 0.0731362570774249 | 0.058678611296751 | 1.24638697919346 | 0.212622359193905 | 0.666270994326506 | NM_001142697,NM_001004361,NM_001142698 |
| Tdp1 | 546.7199545 | 0.180791819 | 0.094450863 | 1.91413623 | 0.055602751 | 0.185870341 | | Trp53inp1 | 486.8551964 | 0.0974230638377448 | 0.0781332746593279 | 1.24688315269676 | 0.212440344971371 | 0.666270994326506 | NM_001199105,NM_021897 |
| Fam163b | 7216.941356 | -0.132125997 | 0.069058757 | -1.9132403 | 0.055717297 | 0.18621307 | | Ube2b | 5183.109642 | 0.0726173628633742 | 0.058171638321777 | 1.24832933983552 | 0.211910473836325 | 0.666270994326506 | NM_009458 |
| Rpl35a | 4398.826847 | -0.147861957 | 0.077312211 | -1.912530447 | 0.055808191 | 0.186476626 | | Zbtb42 | 48.93665689 | 0.117879184608554 | 0.094455097584828 | 1.24799177199187 | 0.212034070403584 | 0.666270994326506 | NM_001100460 |
| Bcr | 8057.270284 | -0.170533062 | 0.089170883 | -1.91242989 | 0.055821077 | 0.186479468 | | Zbtb49 | 351.0730015 | -0.0996499959277877 | 0.079813505832958 | -1.2485355064636 | 0.211835013919916 | 0.666270994326506 | NM_029162 |
| Ero1lb | 839.4966842 | 0.119662704 | 0.06258112 | 1.912121494 | 0.055860612 | 0.186571315 | | Ccdc151 | 231.7858622 | -0.112520586215306 | 0.0903184286290325 | -1.24582090192761 | 0.212830154132317 | 0.666286814423716 | NM_001163787,NM_029939 |
| Kif20a | 32.45467724 | -0.305284402 | 0.159670823 | -1.911961103 | 0.055881183 | 0.186599796 | | Kdm8 | 225.0498728 | -0.105590225282668 | 0.0847521480892019 | -1.24587078514558 | 0.212811837173402 | 0.666286814423716 | NM_029842 |
| 1500017E21Rik | 11.82646337 | -0.301023447 | 0.157461505 | -1.911727236 | 0.055911189 | 0.186630603 | | 4931414P19Rik | 166.2483499 | 0.11008386843116 | 0.0883894124083316 | 1.24544179480012 | 0.212969398265458 | 0.66657907330404 | NM_028890 |
| Rabl2 | 879.8374207 | 0.216828932 | 0.113421964 | 1.911701439 | 0.0559145 | 0.186630603 | | AI314180 | 5055.791805 | -0.0717359233218958 | 0.0576396198166258 | -1.24455927277307 | 0.213293798890488 | 0.666732455924949 | NM_172381 |
| Slc36a1 | 4031.638546 | -0.153157541 | 0.080132322 | -1.911307915 | 0.055965023 | 0.186759005 | | Csrp2 | 130.316805 | 0.118159371782999 | 0.0949389894785963 | 1.24458215146305 | 0.213285384558614 | 0.666732455924949 | NM_007792 |
| Bcl2l13 | 2576.132641 | -0.13033495 | 0.068212755 | -1.91071231 | 0.056041562 | 0.186826112 | | Insig2 | 1804.509986 | 0.0796325779830132 | 0.063969769459814 | 1.24484703720933 | 0.213187982239051 | 0.666732455924949 | NM_178082,NM_133748,NM_001271531,NM_001271532 |
| Camk1 | 2452.02424 | -0.121009745 | 0.063333263 | -1.910682291 | 0.056045422 | 0.186826112 | | Lats1 | 1466.823874 | -0.0745163907546259 | 0.0598438327769721 | -1.2451807863366 | 0.213065303578072 | 0.666732455924949 | NM_010690 |
| Samd8 | 3560.075637 | -0.130104381 | 0.06808566 | -1.910892555 | 0.05601839 | 0.186826112 | | Ndufaf5 | 1041.485861 | 0.0732988590420038 | 0.0588855223650693 | 1.24476876654973 | 0.213216760149836 | 0.666732455924949 | NM_027093 |
| Spag6 | 241.9998119 | -0.184620348 | 0.096612823 | -1.910930054 | 0.05601357 | 0.186826112 | | Poli | 558.0106938 | -0.0861201718460543 | 0.0691867628660058 | -1.24474925952011 | 0.213223932770134 | 0.666732455924949 | NM_001136090,NM_011972 |
| Strn | 2962.699837 | -0.205291544 | 0.107440441 | -1.910747404 | 0.05603705 | 0.186826112 | | Dhx35 | 844.1011976 | -0.095559318259983 | 0.076809282273931 | -1.24411158952355 | 0.21345849619473 | 0.667103725414081 | NM_145742 |
| Acat2 | 1161.34528 | -0.148267617 | 0.077615464 | -1.910284493 | 0.056096593 | 0.186952759 | | Nostrin | 165.2980796 | 0.117795388194349 | 0.0947141509021407 | 1.2436936516071 | 0.213612333361266 | 0.667262578611936 | NM_181547 |
| Drp2 | 4164.108918 | 0.115476915 | 0.060452807 | 1.910199381 | 0.056107547 | 0.186952759 | | Rnf168 | 1513.098342 | 0.0816127845257581 | 0.0656262793327539 | 1.24359913978889 | 0.213647132942594 | 0.667262578611936 | NM_027355 |
| Rock1 | 1002.287694 | 0.160893504 | 0.084235043 | 1.910054277 | 0.056126225 | 0.186974787 | | Slc25a14 | 2019.543258 | 0.0766036266751984 | 0.0615980672664112 | 1.24360438686961 | 0.213645200841919 | 0.667262578611936 | NM_011398,NM_001166450 |
| Ttc4 | 1829.630066 | 0.132493186 | 0.069378311 | 1.909720547 | 0.056169204 | 0.18707774 | | Ar | 349.2862274 | 0.109156591109442 | 0.0877915892009161 | 1.24335664038771 | 0.213736440780823 | 0.667398009196525 | NM_013476 |
| Kansl1 | 4233.579039 | 0.119722708 | 0.062697792 | 1.90952031 | 0.056195004 | 0.187088776 | | Dcc | 987.4288771 | 0.0889898282791691 | 0.0715993480005963 | 1.24288601452667 | 0.213909840034991 | 0.667795902933222 | NM_007831 |
| Rhob | 8018.836568 | 0.135126611 | 0.070765167 | 1.909507406 | 0.056196667 | 0.187088776 | | Lbh | 2123.78035 | 0.0811616680988033 | 0.0653267442104355 | 1.24239573056571 | 0.214090590099721 | 0.668216568927491 | NM_029999 |
| Gabra5 | 2107.397166 | -0.202989678 | 0.106350302 | -1.90868925 | 0.056302191 | 0.187399816 | | Atpif1 | 3983.457429 | 0.067711739979 6407 | 0.0545078327807582 | 1.2422387118561 | 0.214148500533777 | 0.668253731203791 | NM_007512 |
| Gtf2f2 | 415.4198925 | -0.143983024 | 0.07545138 | -1.908288811 | 0.056353898 | 0.187451116 | | Crnkl1 | 795.1444645 | 0.0870244069950583 | 0.0700693575073174 | 1.24197523840532 | 0.214245698169463 | 0.668413447510656 | NM_025820 |
| Osbpl2 | 2536.114067 | 0.128350707 | 0.067255087 | 1.908416336 | 0.056337427 | 0.187451116 | | Rerg | 560.8866757 | 0.101583678288737 | 0.0818200100296183 | 1.24155054798898 | 0.214402437103585 | 0.668471732424699 | NM_001164212,NM_181988,NM_001164214 |
| Zfp451 | 719.5329128 | 0.158631688 | 0.083126501 | 1.908316678 | 0.056350299 | 0.187451116 | | Sirt5 | 395.1571157 | -0.105493848379456 | 0.0849648504556354 | -1.24161753729608 | 0.214377708114786 | 0.668471732424699 | NM_178848 |
| Zfp36 | 211.9632348 | 0.236575276 | 0.123988723 | 1.908038651 | 0.056386221 | 0.187518375 | | Tmem38b | 374.2449343 | 0.0983093251904819 | 0.079175777999244 | 1.24165909921871 | 0.214362366635609 | 0.668471732424699 | NM_028053 |
| Gpr17 | 2323.309889 | -0.13287439 | 0.069681571 | -1.906879939 | 0.056536137 | 0.187855651 | | Cinp | 1603.433467 | 0.0803620008152141 | 0.0647342220889651 | 1.24141448250929 | 0.214452671775714 | 0.668484873436484 | NM_026048,NM_027223 |
| Psmb1 | 3885.826125 | -0.104059121 | 0.054568735 | -1.906936644 | 0.056528792 | 0.187855651 | | Flrt3 | 1151.47031 | 0.100298945278047 | 0.0808046892891182 | 1.24125154320164 | 0.214512839291954 | 0.668528964504381 | NM_178382,NM_001172160 |
| Ptrh2 | 467.5424674 | -0.139794228 | 0.073307353 | -1.906960527 | 0.056525699 | 0.187855651 | | Carf | 448.7465277 | -0.0912523687443986 | 0.0735827032936162 | -1.24013340988949 | 0.214926053306187 | 0.668817995040068 | NM_139150 |
| Zfp160 | 483.885269 | 0.195104778 | 0.102315152 | 1.90690015 | 0.056533519 | 0.187855651 | | Cyp2j6 | 1024.468826 | -0.0803231216296495 | 0.0647520749812569 | -1.2404717787484 | 0.21480094622924 | 0.668817995040068 | NM_010008 |
| Ppic | 51.79372059 | -0.270184318 | 0.141705413 | -1.906661935 | 0.056564379 | 0.187909196 | | Jrk | 469.5244119 | 0.105465455100186 | 0.0850439755799658 | 1.24012846743058 | 0.214927881099204 | 0.668817995040068 | NM_008415 |
| 9530091C08Rik | 31.21152481 | 0.312162873 | 0.163777123 | 1.906022446 | 0.056647294 | 0.188063674 | | Klf10 | 1245.69717 | 0.106384982769108 | 0.0857784707672867 | 1.24022941674638 | 0.214890550799905 | 0.668817995040068 | NM_013692 |
| Nat9 | 454.2763008 | 0.188114915 | 0.098686737 | 1.906182335 | 0.056626553 | 0.188063674 | | Pank1 | 665.8333221 | 0.0804075495677271 | 0.0648232413675957 | 1.24041235629914 | 0.214822913034776 | 0.668817995040068 | NM_023792,NM_001114339 |
| Tmem132d | 1383.685315 | 0.198112965 | 0.103935563 | 1.906113359 | 0.0566355 | 0.188063674 | | Plk5 | 1052.280773 | -0.108169173704048 | 0.0872182329072303 | -1.2402128557122 | 0.21489667462558 | 0.668817995040068 | NM_183152 |
| Npsr1 | 176.4913471 | -0.233248693 | 0.122398695 | -1.905646888 | 0.056696034 | 0.188149434 | | Zfp783 | 303.1728064 | -0.108467756997519 | 0.0874596467342002 | -1.24020346580138 | 0.214900146818301 | 0.668817995040068 | NR_027963 |
| Ocm | 18.19864196 | 0.320262934 | 0.168069141 | 1.905542751 | 0.056709556 | 0.188149434 | | Xylb | 357.1806506 | -0.0953287604528321 | 0.0769064173633616 | -1.2395423388718 | 0.215144719394044 | 0.669349366870397 | NM_001033209,NM_001199568 |
| Zfp831 | 1421.431672 | -0.18033233 | 0.094632678 | -1.905603156 | 0.056701712 | 0.188149434 | | Nasp | 734.0167164 | -0.0827136733366206 | 0.066754011843487 | -1.23908168291897 | 0.215315249691746 | 0.66959308863676 | NM_001081475,NM_016777 |
| Cnga2 | 18.59482687 | 0.311509509 | 0.163516642 | 1.905063023 | 0.056771879 | 0.188315885 | | Stk4 | 2216.817596 | 0.08000149696935 | 0.064561011964862 | 1.23916101964862 | 0.215285873075835 | 0.66959308863676 | NM_021420 |
| Ralgapa2 | 1155.217317 | -0.130537993 | 0.068534936 | -1.90469272 | 0.056820026 | 0.188435248 | | Thap11 | 1181.388088 | 0.0778596346543337 | 0.0628471874596777 | 1.23887222008603 | 0.21539282297038 | 0.669690956007649 | NM_021513 |
| Fst | 125.0172191 | 0.239679681 | 0.125842828 | 1.904595474 | 0.056832675 | 0.188436866 | | 1110057K04Rik | 1771.079259 | 0.0651027350212585 | 0.0525779786769415 | 1.23821296785245 | 0.215637104407586 | 0.669812325609328 | NM_172401,NM_001167768,NM_001167767 |
| Herc6 | 651.1058136 | 0.218920852 | 0.114953139 | 1.904435623 | 0.056853473 | 0.188465494 | | Cdk14 | 8884.548188 | 0.0642094490824463 | 0.0518628819843722 | 1.23806172402437 | 0.215693174911322 | 0.669812325609328 | NM_011074 |
| Rnf38 | 2064.121112 | 0.122146061 | 0.064145432 | 1.904205133 | 0.056883473 | 0.188507836 | | Dcaf15 | 984.6894673 | -0.102388028794308 | 0.0826827864966217 | -1.23832339393268 | 0.215596172866412 | 0.669812325609328 | NM_172502 |
| Smyd2 | 1888.516259 | 0.111397738 | 0.058502589 | 1.90415055 | 0.05689058 | 0.188507836 | | Psg23 | 41.42622043 | -0.115277384409497 | 0.0930867063139773 | -1.23838718732481 | 0.215572529174614 | 0.669812325609328 | NM_020261 |
| Ccdc148 | 638.1659427 | -0.196952667 | 0.103438836 | -1.904049527 | 0.056903734 | 0.188511109 | | Rb1 | 1270.501365 | -0.0908797102805482 | 0.0734072848984096 | -1.23802031918657 | 0.215708526724524 | 0.669812325609328 | NM_009029 |
| Ipo4 | 3301.817512 | 0.10951018 | 0.057521633 | 1.903808611 | 0.056935115 | 0.188574747 | | Tbc1d14 | 4529.808611 | 0.0652749772109645 | 0.0527112661529579 | 1.23834963519087 | 0.215586446864604 | 0.669812325609328 | NM_001113364,NM_133910,NM_001113362 |
| Exoc6 | 1321.68762 | -0.11828129 | 0.062167265 | -1.902629791 | 0.057088869 | 0.189003192 | | Fbxo42 | 2005.240222 | 0.0631999615546552 | 0.0510546930439086 | 1.23788740636049 | 0.215757812585033 | 0.669822181151996 | NM_172518 |
| Ids | 17630.07972 | -0.108517145 | 0.057033093 | -1.902704888 | 0.057079064 | 0.189003192 | | 4833424O15Rik | 1509.422047 | 0.0950704981190162 | 0.0769113810919081 | 1.23610442003906 | 0.216419752122821 | 0.67004971252877 | NM_029425 |
| Dus1l | 983.1271972 | 0.145325867 | 0.076385574 | 1.902530272 | 0.057101865 | 0.189005832 | | 5830454E08Rik | 25.7830862 | -0.104893632271708 | 0.0848645918517261 | -1.23601174509831 | 0.216454197919077 | 0.67004971252877 | NR_073359 |
| Lrrc49 | 3070.245269 | 0.153221102 | 0.080540638 | 1.902407363 | 0.057117919 | 0.18901859 | | Adamts13 | 44.6965783 | 0.11421362831127 | 0.0924140632718703 | 1.23589012610849 | 0.216499407743319 | 0.67004971252877 | NM_001001322 |
| Mia | 20.75224388 | 0.319169748 | 0.167793882 | 1.902153662 | 0.057151069 | 0.189047536 | | Agpat3 | 9118.494202 | 0.0505924664808121 | 0.0408954636630649 | 1.2371168327529 | 0.216043711440195 | 0.67004971252877 | NM_053014 |
| Plk3 | 484.3415647 | 0.263795764 | 0.13867894 | 1.902204942 | 0.057144367 | 0.189047536 | | Arr3 | 52.72600707 | -0.110854317487158 | 0.0897223133701495 | -1.23552674160138 | 0.216634530342094 | 0.67004971252877 | NM_133205 |
| L1cam | 7016.747899 | -0.163857031 | 0.08617517 | -1.901441349 | 0.057244228 | 0.189315276 | | Car9 | 39.76128406 | -0.108566588758327 | 0.0877839114182132 | -1.2367481353286 | 0.216180602302744 | 0.67004971252877 | NM_139305 |
| Ccdc94 | 231.0846861 | -0.193315818 | 0.101682774 | -1.901165847 | 0.057280293 | 0.189364081 | | Ccdc47 | 4784.637751 | 0.0607102505260325 | 0.0490886585837292 | 1.23674698550747 | 0.216181029308742 | 0.67004971252877 | NM_026009 |
| Lrrk1 | 275.9001775 | -0.185025073 | 0.097327904 | -1.901048574 | 0.057295651 | 0.189364081 | | Cntnap5c | 227.1770104 | -0.112023520965688 | 0.0906580419893727 | -1.23567108341938 | 0.216580850339854 | 0.67004971252877 | NM_001081653 |
| Mansc1 | 437.1690142 | 0.204964203 | 0.107811204 | 1.901140106 | 0.057283664 | 0.189364081 | | Klhl9 | 3830.805735 | 0.0789847271934311 | 0.063930014042346 | 1.23548740566697 | 0.216649160839967 | 0.67004971252877 | NM_172871 |
| Tctn3 | 1017.692717 | 0.184004382 | 0.096802785 | 1.90081702 | 0.057325984 | 0.189423927 | | Map6d1 | 3223.150811 | -0.0818544719469993 | 0.0662545419093848 | -1.23545449999413 | 0.216661400230458 | 0.67004971252877 | NM_198599 |
| Cercam | 281.4003672 | 0.182222885 | 0.095876162 | 1.900606798 | 0.057353534 | 0.189474555 | | Pcdhb18 | 168.4858115 | 0.112917371733038 | 0.0912917980238984 | 1.23688408134407 | 0.216130120674818 | 0.67004971252877 | NM_053143 |
| Bgn | 776.7670467 | -0.2046629 | 0.107703648 | -1.900241121 | 0.057401484 | 0.189592538 | | Pld3 | 28867.81376 | 0.0708845367176453 | 0.0573247490289539 | 1.23654334154769 | 0.216256665598546 | 0.67004971252877 | NM_011116 |
| Pcbp3 | 2305.681298 | -0.232359288 | 0.122329151 | -1.899459662 | 0.057504065 | 0.189890876 | | Rnf128 | 465.328836 | 0.0943232117087824 | 0.0762844140549269 | 1.23646767006517 | 0.216284775927492 | 0.67004971252877 | NM_001254761,NM_023270 |
| D430019H16Rik | 8308.975802 | -0.137791496 | 0.070443134 | -1.899283669 | 0.057527185 | 0.189894005 | | Rnf215 | 881.0947206 | -0.0924898960443128 | 0.0748033561773738 | -1.23643998305371 | 0.216295061714268 | 0.67004971252877 | NM_027859 |
| Elf4 | 86.52765036 | 0.248006321 | 0.130580095 | 1.899265892 | 0.057529525 | 0.189894005 | | Sema3c | 1083.476865 | 0.0849916057009761 | 0.0687886285793596 | 1.23554732019295 | 0.216626876681353 | 0.67004971252877 | NM_013657 |
| Cfb | 17.33494544 | 0.276270862 | 0.145503403 | 1.898724405 | 0.057600721 | 0.190021638 | | Trpc3 | 811.4107849 | 0.0931549257065994 | 0.0753733877114636 | 1.23591268131937 | 0.216491022707827 | 0.67004971252877 | NM_019510 |
| Mt3 | 5441.060187 | -0.162084143 | 0.085366212 | -1.898691995 | 0.057604985 | 0.190021638 | | Ttll5 | 1615.190491 | 0.0784457243367351 | 0.063431145343019 | 1.23670672999078 | 0.216195979274988 | 0.67004971252877 | NM_001081423 |
| Palmd | 637.6644949 | -0.188837182 | 0.099455749 | -1.898705546 | 0.057603202 | 0.190021638 | | Vhl | 2014.393404 | -0.0728405709904209 | 0.0589026394908072 | -1.23662660315569 | 0.216225738740305 | 0.67004971252877 | NM_009507 |

| | | | | | | | | | | | | | | | |
|---|---|---|---|---|---|---|---|---|---|---|---|---|---|---|---|
| Obsl1 | 534.1506429 | 0.211319491 | 0.111303941 | 1.898580484 | 0.057619656 | 0.190029578 | | 1110008J03Rik | 538.0184866 | 0.0895355401213382 | 0.0725179007932397 | 1.23466811838112 | 0.216954045920859 | 0.670185967172541 | NM_133908,NM_029096 |
| Hmbox1 | 1038.481021 | -0.176439774 | 0.092942694 | -1.898371628 | 0.057647144 | 0.190071252 | | Atp2b4 | 14382.58423 | 0.0888030170677374 | 0.0719362419884154 | 1.23446839330359 | 0.217028417291727 | 0.670185967172541 | NM_213616,NM_001167949 |
| Phc1 | 2125.808898 | 0.118793078 | 0.062578728 | 1.898298076 | 0.057656827 | 0.190071252 | | Dnajc9 | 1023.235866 | 0.0921708994402418 | 0.0746495379806707 | 1.2347149350624 | 0.216936615506339 | 0.670185967172541 | NM_134081 |
| Chtf8 | 3198.402237 | -0.119842552 | 0.063137715 | -1.898113543 | 0.057681127 | 0.190110907 | | Hnrnpk | 13442.92726 | 0.0650362085207517 | 0.0526664973401685 | 1.23486868892549 | 0.216879378178475 | 0.670185967172541 | NM_025279 |
| Mgrn1 | 9530.207061 | 0.080326418 | 0.042323626 | 1.897909669 | 0.057707982 | 0.190158971 | | Lsmd1 | 726.4412722 | 0.0793761651506436 | 0.0642684666334333 | 1.23507171259243 | 0.216803816027601 | 0.670185967172541 | NM_030083 |
| Mgll | 10416.2096 | -0.179000451 | 0.094376225 | -1.896668906 | 0.05787165 | 0.190617208 | | Sdhaf2 | 1601.830204 | 0.0681803237169794 | 0.0552285484383118 | 1.23451232460209 | 0.217012057077263 | 0.670185967172541 | NM_025333 |
| Zkscan2 | 757.3201273 | 0.272440915 | 0.143640118 | 1.896690973 | 0.057868736 | 0.190617208 | | Zmynd8 | 6001.194261 | 0.0707656767951757 | 0.0573147763035537 | 1.23468468969298 | 0.216947876106592 | 0.670185967172541 | NM_001252587,NR_045545,NM_001252585,NM_001252584,NM_027230 |
| Llph | 871.8248981 | 0.145967383 | 0.076992615 | 1.89586212 | 0.057978279 | 0.190927833 | | Mrps18c | 610.0785649 | -0.0999285150324751 | 0.080962031492109 | -1.23426392829847 | 0.2171045726602 | 0.670278644519037 | NM_026826 |
| Dffb | 224.2666979 | 0.257508973 | 0.135834506 | 1.895755222 | 0.057992419 | 0.190933818 | | Amigo2 | 503.1895601 | 0.102127363461889 | 0.0827648320927489 | 1.23394636199397 | 0.217222892033832 | 0.670311137572657 | NM_001164563,NM_178114,NM_001164602 |
| A2m | 181.9421982 | 0.289126543 | 0.152552968 | 1.895253481 | 0.058058828 | 0.191086137 | | Csnk1g2 | 5800.137817 | 0.0750920275358852 | 0.0608958831164404 | 1.23312157888079 | 0.217530407718402 | 0.670311137572657 | NM_134002,NM_001159591 |
| Nfatc2 | 536.096439 | -0.200615958 | 0.105855567 | -1.89518571 | 0.058067803 | 0.191086137 | | Ebp | 624.9217604 | 0.0751189936697182 | 0.0609003455239937 | 1.23347401436536 | 0.217398965874055 | 0.670311137572657 | NM_007898 |
| Opcml | 7708.301497 | -0.174376588 | 0.092013179 | -1.895126217 | 0.058075682 | 0.191086137 | | Klhl11 | 1080.696485 | 0.0922613109162091 | 0.0747671516537105 | 1.23398188744068 | 0.217209653601159 | 0.670311137572657 | NM_172565 |
| Dync1i2 | 4919.938148 | 0.130294326 | 0.068755784 | 1.895030773 | 0.058088325 | 0.191087156 | | Klhl5 | 1754.876263 | -0.0593739046112049 | 0.0481289908981652 | -1.2336411693491 | 0.217336644907553 | 0.670311137572657 | NM_175174 |
| 2810008D09Rik | 302.6760673 | 0.244508595 | 0.129040273 | 1.894823905 | 0.058115735 | 0.191110028 | | Lpcat4 | 4758.637429 | -0.0794928554504895 | 0.0644436881753282 | -1.23352430162311 | 0.217380215751644 | 0.670311137572657 | NM_207206 |
| Acan | 206.1497196 | 0.245217216 | 0.129416421 | 1.894792128 | 0.058119947 | 0.191110028 | | Mir27b | 28.15194852 | -0.0950893075068793 | 0.0771095057376687 | -1.23317231250812 | 0.217511482943515 | 0.670311137572657 | NR_029531 |
| 0610010K14Rik | 598.1875366 | -0.196714219 | 0.103836012 | -1.894470096 | 0.05816264 | 0.191169271 | | Tnrc6a | 4046.26566 | -0.0752158305961304 | 0.0609776430497531 | -1.23349848951623 | 0.217389839916486 | 0.670311137572657 | NM_144925 |
| Gaa | 14079.33723 | 0.198588813 | 0.104824743 | 1.894484133 | 0.058160778 | 0.191169271 | | Vps29 | 2383.36857 | 0.0695818611880434 | 0.056425744673484 | 1.23315804852358 | 0.217516803608341 | 0.670311137572657 | NM_019780 |
| Traf2 | 386.3406078 | 0.152790484 | 0.0806681 | 1.894063251 | 0.058216615 | 0.191306093 | | 4932415G12Rik | 1625.615971 | -0.084980244090422 | 0.0689582646065892 | -1.23234313646434 | 0.217820932578303 | 0.670929685831227 | NR_015524 |
| Pvr | 274.7015454 | 0.190328726 | 0.100496364 | 1.89388669 | 0.058240052 | 0.191342527 | | Cdc25a | 300.1301301 | 0.105194786831435 | 0.0853711212362518 | 1.23220575421898 | 0.217872234368402 | 0.670929685831227 | NM_007658 |
| Ptplad1 | 6938.024722 | -0.127159362 | 0.067161704 | -1.893331382 | 0.058313814 | 0.191544253 | | Fosb | 840.5505186 | 0.113091681460238 | 0.0918015951715004 | 1.23191412141548 | 0.217981165764736 | 0.670929685831227 | NM_008036 |
| Hhatl | 370.9686276 | 0.19665292 | 0.103872035 | 1.893222949 | 0.058328227 | 0.191550985 | | Ido2 | 82.65260953 | -0.117063038685241 | 0.0950107175187499 | -1.23210351150268 | 0.217910419861666 | 0.670929685831227 | NM_145949 |
| Eif4b | 8814.46401 | 0.179647409 | 0.094905857 | 1.892901171 | 0.058371013 | 0.191610272 | | Tmem104 | 897.804177 | 0.0697889221894254 | 0.0566563594330477 | 1.23179326888973 | 0.218026318365969 | 0.670929685831227 | NM_001033393 |
| Gpr56 | 3082.723401 | -0.225314019 | 0.119029308 | -1.892928914 | 0.058367324 | 0.191610272 | | Tmem81 | 189.6696858 | -0.103964874272125 | 0.0844024543613762 | -1.23177548637378 | 0.218032962789893 | 0.670929685831227 | NM_029025 |
| Psma6 | 2735.384466 | -0.094700142 | 0.050064 | -1.891581597 | 0.05854675 | 0.192146439 | | Vwa7 | 44.55976635 | -0.098861536122654 | 0.0802631276656422 | -1.23171796312858 | 0.218054457304779 | 0.670929685831227 | NM_138582 |
| Npr2 | 2013.40115 | 0.197299483 | 0.104335219 | 1.891015183 | 0.058622317 | 0.192353703 | | Kif17 | 1588.607466 | -0.0824523084889742 | 0.0669514741253735 | -1.23152342149441 | 0.218127162299176 | 0.67101125784791 | NM_010623,NM_001190978 |
| Abca5 | 2050.874975 | 0.132493436 | 0.070091918 | 1.890281222 | 0.058720359 | 0.192512323 | | Radil | 671.7074251 | -0.094954686169052 | 0.0771191258369088 | -1.23127285402407 | 0.218220831195897 | 0.671157271639128 | NM_178702 |
| Fkbp11 | 19.06464606 | -0.317754455 | 0.168096292 | -1.890312097 | 0.058716232 | 0.192512323 | | Pop7 | 467.7312279 | 0.0799038719420436 | 0.0649104477156083 | 1.23098630119031 | 0.218327987822146 | 0.671344537974238 | NM_028753 |
| LOC100503496 | 19.6772291 | 0.317584767 | 0.167997942 | 1.890408678 | 0.058703324 | 0.192512323 | | Smim3 | 335.3800915 | 0.114812240535267 | 0.0932872110276238 | 1.23073934005026 | 0.218420369438679 | 0.671344537974238 | NM_134133 |
| Rps27a | 5929.417106 | -0.12540071 | 0.06633368 | -1.890453092 | 0.058697388 | 0.192512323 | | Trim56 | 161.6113384 | -0.115116628754951 | 0.0935263712077826 | -1.23084673625583 | 0.218380191914157 | 0.671344537974238 | NM_201373 |
| Sp7 | 84.27513707 | -0.292822935 | 0.154918418 | -1.8901751 | 0.058734546 | 0.192518098 | | 4930444A02Rik | 1576.567122 | 0.0791980522185501 | 0.0643816688976383 | 1.23013357023206 | 0.218647090803574 | 0.671604513701991 | NM_029037 |
| Acat1 | 3271.644261 | -0.098701248 | 0.052223027 | -1.889994784 | 0.058758658 | 0.192518619 | | Anxa4 | 187.2217748 | -0.107488152391756 | 0.0874224642975584 | -1.22952553734817 | 0.218874829124689 | 0.671604513701991 | NM_013471 |
| Lpin3 | 42.51150674 | 0.296036156 | 0.156633875 | 1.889988073 | 0.058759555 | 0.192518619 | | Atp6v1e1 | 13924.93199 | 0.0567256233802356 | 0.0461046666244037 | 1.23036619790262 | 0.218560005298329 | 0.671604513701991 | NM_007510 |
| Dio3os | 10.14829702 | -0.293686963 | 0.155432508 | -1.889482234 | 0.058827241 | 0.192658904 | | Clpx | 1250.72679 | 0.0867313076067199 | 0.0705321118295006 | 1.22967121438783 | 0.218820250371067 | 0.671604513701991 | NM_001044389,NM_011802 |
| Popdc2 | 14.50784079 | 0.31612689 | 0.167303401 | 1.889542518 | 0.058819171 | 0.192658904 | | Efnb3 | 5471.412496 | 0.0903377451541122 | 0.0734726544655938 | 1.22954241698748 | 0.218868504568476 | 0.671604513701991 | NM_007911 |
| Prdm5 | 93.43478864 | 0.27123949 | 0.143565721 | 1.889305391 | 0.05885092 | 0.19269572 | | Hspbp1 | 2474.504057 | 0.0719206227351453 | 0.0584612121056803 | 1.23022804599286 | 0.218611720254056 | 0.671604513701991 | NM_024172 |
| Acadvl | 1676.708446 | 0.119474593 | 0.063242951 | 1.889136923 | 0.058873484 | 0.192728874 | | Rnf123 | 2914.793028 | -0.075227397765847 | 0.0611731399998309 | -1.22974556751631 | 0.218792397305517 | 0.671604513701991 | NM_032543 |
| Gucy2e | 172.4471379 | 0.263571354 | 0.139527867 | 1.889023025 | 0.058888743 | 0.192738106 | | Tfr2 | 227.407892 | -0.111499047963934 | 0.0906800731646965 | -1.22958709750295 | 0.218851764060457 | 0.671604513701991 | NM_015799 |
| Ptrhd1 | 451.6157846 | -0.149268459 | 0.079032081 | -1.888707193 | 0.058931074 | 0.192835916 | | Bdh1 | 1944.60528 | 0.0756509632013262 | 0.0615402311909885 | 1.22929280142847 | 0.218962045282917 | 0.671730236489895 | NM_001122683,NM_175177 |
| AB124611 | 10.95766446 | 0.310409476 | 0.164361242 | 1.888580745 | 0.058948029 | 0.192839133 | | 1700063D05Rik | 65.10438178 | -0.113514335512787 | 0.0925012927002054 | -1.22716485574623 | 0.219760635776684 | 0.672421385190151 | NR_040392 |
| Mast1 | 3557.106992 | 0.125912761 | 0.066673989 | 1.888483987 | 0.058961006 | 0.192839133 | | 6430550D23Rik | 30.84052153 | -0.107461000297879 | 0.0875754286039901 | -1.22706793458939 | 0.21979705852979 | 0.672421385190151 | NM_001145352,NM_001145351 |
| Pou3f1 | 774.5875996 | -0.229262117 | 0.121404106 | -1.888421439 | 0.058969395 | 0.192839133 | | Ak1 | 3955.673936 | 0.0713231768929213 | 0.0581046209682064 | 1.22749577752082 | 0.219636307877692 | 0.672421385190151 | NM_001198791,NM_001198790,NM_001198792,NM_021515 |
| 1700063D05Rik | 51.31994931 | 0.302110997 | 0.16001509 | 1.888015676 | 0.059023846 | 0.192976465 | | Cyth1 | 3722.072819 | -0.0649171310050724 | 0.0528415451006366 | -1.22852446652417 | 0.219250150616819 | 0.672421385190151 | NM_011180,NM_001112699,NM_001112700 |
| Entpd7 | 454.0441139 | 0.185391939 | 0.09820686 | 1.887769738 | 0.059056869 | 0.193002981 | | Esyt2 | 1958.151436 | -0.0729168301571413 | 0.0594097073467456 | -1.22735548471163 | 0.219689009884807 | 0.672421385190151 | NM_028731 |
| Rcbtb2 | 871.1915896 | -0.1221368 | 0.064697367 | -1.887817176 | 0.059050499 | 0.193002981 | | Fam19a5 | 4645.213229 | 0.0590635552578605 | 0.0481281156078955 | 1.22721520491384 | 0.21974171607833 | 0.672421385190151 | NM_134096,NM_001252310,NR_045510 |
| Il10ra | 283.7247094 | 0.225499409 | 0.119460867 | 1.887642484 | 0.059073963 | 0.193018131 | | Ifnar2 | 667.0942986 | -0.0848552479061665 | 0.0691461448446772 | -1.22718696894493 | 0.219752326047554 | 0.672421385190151 | NM_001110498,NM_010509 |
| Sp9 | 200.6967147 | -0.287026507 | 0.152073108 | -1.887424486 | 0.059103254 | 0.193073123 | | Prkd3 | 1003.976072 | -0.0860742003299845 | 0.0700939447155254 | -1.22798339684426 | 0.219453200740271 | 0.672421385190151 | NM_001171004,NM_029239,NM_001171005 |
| Megf11 | 749.2686644 | -0.230170233 | 0.121978977 | -1.886966414 | 0.059164843 | 0.193233575 | | Ryr2 | 7444.547493 | -0.0920539560961466 | 0.0750101055515296 | -1.22722072471833 | 0.219739641995344 | 0.672421385190151 | NM_023868 |
| Anxa3 | 449.5001652 | 0.177199447 | 0.093913798 | 1.886830799 | 0.059183087 | 0.193252424 | | Ska3 | 29.27804344 | -0.100534731529614 | 0.0818538193122061 | -1.22822285355012 | 0.21936332189164 | 0.672421385190151 | NM_198605 |
| 1700016K19Rik | 63.72603215 | -0.309943663 | 0.164284188 | -1.88663113 | 0.059209957 | 0.193299424 | | Unk | 1033.390654 | -0.0728556379200902 | 0.0593239782397193 | -1.22809764418852 | 0.219410315287502 | 0.672421385190151 | NM_172569 |
| Dlx4 | 8.926601048 | 0.271744376 | 0.144063255 | 1.886285137 | 0.059256541 | 0.193410753 | | Usp30 | 1787.147395 | 0.0601868459787085 | 0.0490533976540135 | 1.22696589547623 | 0.219835409491121 | 0.672421385190151 | NM_001033202 |
| Fam211a | 636.9511286 | -0.177412549 | 0.094061441 | -1.886134706 | 0.059276805 | 0.193436144 | | Zfp112 | 464.3141376 | -0.0891036639222798 | 0.0725770062516806 | -1.22771203338546 | 0.219555087575807 | 0.672421385190151 | NM_021307 |
| 4930520O04Rik | 20.07905756 | 0.316859162 | 0.168069205 | 1.88528983 | 0.059390718 | 0.193522564 | | Zfp598 | 1738.806411 | -0.0943329711296875 | 0.0767914701384105 | -1.22843033164569 | 0.219285467422024 | 0.672421385190151 | NM_183149 |
| BC029214 | 944.8818816 | 0.1411845 | 0.074873057 | 1.885651601 | 0.059341919 | 0.193522564 | | 1600002H07Rik | 373.6520725 | -0.0945520548786051 | 0.0771996315447521 | -1.22477339576153 | 0.220660611444334 | 0.6727577950103 | NM_028056 |
| Cib2 | 642.7847484 | -0.166206681 | 0.088153235 | -1.885429174 | 0.059371918 | 0.193522564 | | Actrt3 | 34.85827009 | 0.112533281892229 | 0.0918067176537469 | 1.22576304619291 | 0.220287857199189 | 0.6727577950103 | NM_029690 |
| Eps8 | 1235.879272 | -0.156062706 | 0.082779021 | -1.885292967 | 0.059390295 | 0.193522564 | | Aff1 | 1100.134742 | 0.0975316039642565 | 0.0795908965977162 | 1.2254115499819 | 0.220420197344822 | 0.6727577950103 | NM_133919,NM_001080798 |
| Lmna | 1394.965905 | 0.154033715 | 0.081688699 | 1.885618423 | 0.059346393 | 0.193522564 | | Csnk2b | 3037.11541 | 0.0635480386894931 | 0.0519549655514082 | 1.22313696130957 | 0.221277970382839 | 0.6727577950103 | NM_009975 |
| Stmn2 | 13344.33827 | -0.1485685 | 0.078796231 | -1.885477237 | 0.059365435 | 0.193522564 | | Cxx1b | 2037.078947 | 0.077329979951666 | 0.0631880503824872 | 1.22380702496082 | 0.221025033620213 | 0.6727577950103 | NM_001018063 |
| Susd1 | 448.3676406 | -0.171615097 | 0.091005373 | -1.88576885 | 0.05932611 | 0.193522564 | | Ddx47 | 1771.279005 | 0.0630158469176939 | 0.0514091116784314 | 1.22577194704051 | 0.220284506724019 | 0.6727577950103 | NM_026360 |
| Gnai3 | 1200.225593 | -0.144171047 | 0.076475706 | -1.885187519 | 0.059404525 | 0.193526853 | | Def8 | 2557.571065 | 0.0576821689360743 | 0.0470906318692197 | 1.22491813438115 | 0.220606067070666 | 0.6727577950103 | NM_054046,NM_001253783,NR_045593,NM_001253784 |
| Taldo1 | 2618.861743 | -0.121129659 | 0.064275643 | -1.884534361 | 0.059492732 | 0.193773467 | | Dixdc1 | 4073.25443 | 0.0789236243389069 | 0.0644858825035119 | 1.22388996280866 | 0.220993740521475 | 0.6727577950103 | NM_178118 |
| Abce1 | 3437.329459 | -0.099196014 | 0.052646285 | -1.88419778 | 0.059538228 | 0.193827647 | | Dnajc10 | 3684.866483 | 0.075271483284699 | 0.0615123252782191 | 1.2236813182439 | 0.221072469803557 | 0.6727577950103 | NM_024181 |
| Fnip1 | 2124.379587 | 0.188303115 | 0.099941483 | 1.884133679 | 0.059546896 | 0.193827647 | | Enox2 | 417.7508755 | 0.0932924976122743 | 0.0761948787145493 | 1.22439328188681 | 0.220803902419129 | 0.6727577950103 | NM_001271447,NM_145951,NM_001271450,NM_001271451,NM_001271449,NM_001271448 |
| Slc16a7 | 392.9278288 | -0.195107571 | 0.103549546 | -1.884195334 | 0.059538558 | 0.193827647 | | Fa2h | 1629.162906 | 0.104515127143593 | 0.085326742132407 | 1.2248812568211 | 0.220619963365594 | 0.6727577950103 | NM_178086 |
| Hdhd3 | 126.959801 | -0.228985312 | 0.121544052 | -1.883969709 | 0.059569073 | 0.193859109 | | Fbxo45 | 3096.504317 | 0.0759549717798953 | 0.0621301192979161 | 1.22251450082831 | 0.221513123629127 | 0.6727577950103 | NM_173439 |
| Cd302 | 481.3590355 | -0.164447525 | 0.087309408 | -1.883502932 | 0.059632243 | 0.193869931 | | Fbxo46 | 473.968943 | 0.0870027444206557 | 0.0711299249336732 | 1.22315248472121 | 0.221272108232705 | 0.6727577950103 | NM_175530 |
| Cpq | 505.3974892 | -0.17057896 | 0.090554264 | -1.883720914 | 0.059602736 | 0.193869931 | | Fbxw17 | 310.215332 | -0.0976562622399818 | 0.0797410923347335 | -1.22466672302462 | 0.220700816978409 | 0.6727577950103 | NM_175401 |
| S1pr1 | 5966.649379 | -0.208131518 | 0.110503065 | -1.883490912 | 0.05963387 | 0.193869931 | | Fgd6 | 959.1096156 | -0.0891631438513368 | 0.072813463545153 | -1.22454199416081 | 0.220747834626479 | 0.6727577950103 | NM_053072 |
| Slc39a3 | 3107.628942 | -0.122884246 | 0.065243092 | -1.883482853 | 0.059634961 | 0.193869931 | | Fscn1 | 8062.839807 | 0.0755840152460442 | 0.0617492566371017 | 1.22404737097078 | 0.220934357925307 | 0.6727577950103 | NM_007984 |
| Vegfa | 2576.659863 | -0.181689829 | 0.096463503 | -1.883508503 | 0.059631489 | 0.193869931 | | Gatad1 | 2443.114777 | 0.0691888144162294 | 0.0565354426666763 | 1.22381308348739 | 0.221022747583209 | 0.6727577950103 | NM_026033 |
| Galns | 303.3851779 | 0.192011039 | 0.101976369 | 1.88289738 | 0.059714276 | 0.194087057 | | Gpr125 | 883.6500047 | 0.083229251631706 | 0.0679807340873923 | 1.22430645607197 | 0.220836642378472 | 0.6727577950103 | NM_133911 |
| Comt | 3370.854634 | -0.109400856 | 0.058106742 | -1.882756687 | 0.059733349 | 0.194108329 | | Gxylt1 | 758.2814787 | -0.0819097701363956 | 0.0669697986445591 | -1.22308520847031 | 0.221297514758987 | 0.6727577950103 | NM_001033275 |
| B3gnt8 | 16.81483042 | -0.312297759 | 0.165885245 | -1.882613243 | 0.0597528 | 0.194118889 | | Ifrd2 | 309.08056 | -0.0993783079380498 | 0.0812686678174803 | -1.22283667952 | 0.221391388388777 | 0.6727577950103 | NM_025903 |
| Dclk2 | 2962.966052 | -0.125748843 | 0.066797153 | -1.882547949 | 0.059761656 | 0.194118889 | | Ikbip | 352.222328 | -0.0860249981996 | 0.070311901443428 | -1.22347705628206 | 0.221149564842379 | 0.6727577950103 | NM_027078,NM_026166 |
| Rnf146 | 2108.828772 | 0.154422141 | 0.082043042 | 1.882208873 | 0.059807661 | 0.194227605 | | Ndufs4 | 2193.709452 | 0.0688940789193077 | 0.0562757986880096 | 1.22422214389623 | 0.220868437834819 | 0.6727577950103 | NM_010887 |
| Enah | 2999.104855 | -0.148470405 | 0.078886075 | -1.882086342 | 0.059824294 | 0.194240906 | | Parp6 | 6094.856895 | -0.0571203830785841 | 0.0466618692946446 | -1.22413405082209 | 0.220901662633879 | 0.6727577950103 | NM_001205239,NR_038074,NR_038075,NM_029922 |
| 2410137M14Rik | 12.35145875 | 0.296092344 | 0.157339932 | 1.881863941 | 0.059854491 | 0.194298238 | | Pcdha12 | 346.8053768 | -0.10386560336913 | 0.0848979721513673 | -1.22341677589124 | 0.221172320284397 | 0.6727577950103 | NM_138663 |
| Baat1 | 11.83307264 | 0.305663517 | 0.162462501 | 1.881440432 | 0.059912031 | 0.194316101 | | Pde3b | 451.3955057 | -0.083635230790709 | 0.0684124708400106 | -1.2225144007194 | 0.221513161462688 | 0.6727577950103 | NM_011055 |
| Foxd2 | 12.66986214 | -0.309538808 | 0.164525325 | -1.881405234 | 0.059916815 | 0.194316101 | | Pdzrn4 | 298.1133761 | 0.109622813398805 | 0.0896677529296504 | 1.22254444677353 | 0.221501806545373 | 0.6727577950103 | NM_001164594,NM_001164593 |
| Pcdhb11 | 130.2491745 | -0.231463859 | 0.123032097 | -1.881329061 | 0.05992717 | 0.194316101 | | Plk3 | 638.339206 | 0.114970636874871 | 0.0939144309042703 | 1.22420628829732 | 0.220874417598838 | 0.6727577950103 | NM_013807 |
| Rnf115 | 1852.633198 | 0.143977512 | 0.076534688 | 1.881205963 | 0.059943907 | 0.194316101 | | Rhbdf1 | 499.1221713 | -0.108276340792 | 0.0883580327914654 | -1.22542724607217 | 0.220414286469372 | 0.6727577950103 | NM_010117 |
| Sepp1 | 7957.313696 | 0.158243279 | 0.084114333 | 1.881287927 | 0.059932762 | 0.194316101 | | Rps3 | 8622.623395 | 0.0575980588359308 | 0.0470876901082934 | 1.22320841611609 | 0.221250987623814 | 0.6727577950103 | NM_012052 |
| Slc3a1 | 11.91776095 | 0.275348298 | 0.146362706 | 1.881273629 | 0.059934706 | 0.194316101 | | Runx2 | 335.0311196 | -0.10172937472066 | 0.0832144764704361 | -1.22249612129449 | 0.221520069773803 | 0.6727577950103 | NM_001271627,NM_001271631,NR_073392,NM_001271630,NM_001146038,NM_001145920,NM_009820,NR_073425 |
| Tmem119 | 945.1961778 | -0.245432916 | 0.130467711 | -1.881177452 | 0.059947784 | 0.194316101 | | Slitrk5 | 3009.392674 | 0.0687374057894056 | 0.0560473854610239 | 1.2264159197436 | 0.220042198194866 | 0.6727577950103 | NM_198865 |
| Fbxl6 | 854.3088637 | 0.166382667 | 0.088468901 | 1.880691019 | 0.060013962 | 0.194471862 | | Tbc1d9 | 5240.074349 | 0.059924677512998 | 0.0489127488669449 | 1.2251341194503 | 0.220524691637511 | 0.6727577950103 | NM_027758,NM_001111304 |
| Smg6 | 2145.106377 | 0.155527583 | 0.082699296 | 1.880639755 | 0.06002094 | 0.194471862 | | Tmem8 | 854.7835553 | -0.0930408813617169 | 0.0760954567975106 | -1.22268641621139 | 0.221448159509984 | 0.6727577950103 | NM_021793 |
| Ncmap | 26.711026 | 0.312396094 | 0.166162701 | 1.880061484 | 0.060099699 | 0.194686334 | | Vrk1 | 618.3192573 | -0.0776953622468814 | 0.0634580869013105 | -1.22435714722558 | 0.220817527521634 | 0.6727577950103 | NM_001029843,NM_011705,NM_001029844 |
| Eln | 306.2422749 | -0.21962433 | 0.116835939 | -1.879766895 | 0.060139854 | 0.194734984 | | Dph3 | 1272.873016 | 0.0733433699940838 | 0.0600059290983428 | 1.22226871737762 | 0.221606025045519 | 0.672878160495655 | NM_172254,NM_001047433 |
| Pdia2 | 12.85152103 | 0.300466568 | 0.159839879 | 1.879797261 | 0.060135714 | 0.194734984 | | Gde1 | 7272.4787 | 0.0616397581265982 | 0.0504541462665372 | 1.22176467358126 | 0.221796631225094 | 0.672987912193223 | NM_019580 |
| Bmp3 | 491.5306072 | -0.234464236 | 0.124770913 | -1.879157813 | 0.060222948 | 0.194963301 | | H2-M3 | 122.443436 | -0.11194296606885 | 0.0916482704833045 | -1.22144111916703 | 0.221919046497373 | 0.672987912193223 | NM_013819 |
| Rab11fip5 | 4713.725086 | 0.161916675 | 0.086187446 | 1.878657298 | 0.060291302 | 0.195143813 | | Kctd10 | 2756.050876 | 0.0588350348361382 | 0.048157117860336 | 1.22173081467977 | 0.221809439312574 | 0.672987912193223 | NM_026145,NM_001159941 |
| Lrrc56 | 140.5814483 | 0.247937519 | 0.131995178 | 1.878375477 | 0.060329818 | 0.195227694 | | Muc1 | 25.78984972 | -0.107398676442936 | 0.0879280475909756 | -1.22143820300132 | 0.221920150034802 | 0.672987912193223 | NM_013605 |
| Tbc1d17 | 1920.666227 | 0.130834234 | 0.069677688 | 1.877706317 | 0.060421352 | 0.195483072 | | Nfatc2 | 519.05603 | -0.0988648664812058 | 0.0809296622585067 | -1.22161471730118 | 0.221853360453912 | 0.672987912193223 | NM_010899,NM_001037178,NM_001136073,NM_001037177 |
| Gprin2 | 21.30903031 | -0.313286926 | 0.166864879 | -1.87748871 | 0.060451143 | 0.195497811 | | Taf10 | 839.0475278 | 0.0875987254761997 | 0.0716973779572463 | 1.22178422659243 | 0.221789234986724 | 0.672987912193223 | NM_020024 |
| Mtus2 | 2227.374698 | 0.163713903 | 0.087198015 | 1.877495747 | 0.06045018 | 0.195497811 | | Slc18a3 | 92.61240397 | 0.115948566403202 | 0.0949642585004352 | 1.22097058656574 | 0.222097156624525 | 0.673384115451441 | NM_021712 |
| Erlec1 | 1838.985182 | 0.115390291 | 0.061464274 | 1.877355463 | 0.060469391 | 0.195516016 | | Snx22 | 301.5839025 | 0.1132400202839 | 0.0927587350691654 | 1.22080168729622 | 0.22216111481614 | 0.673437469494834 | NM_001025612 |
| Eif3f | 3817.035764 | 0.112108131 | 0.059719971 | 1.877230145 | 0.060486558 | 0.195530717 | | Acpl2 | 636.7795203 | 0.0817876446449384 | 0.0670549743490791 | 1.21971032632364 | 0.222574705627052 | 0.673748940276311 | NM_153420 |
| Gas2 | 229.5652362 | 0.202196197 | 0.107716455 | 1.877115219 | 0.060502304 | 0.195540821 | | Agpat9 | 247.2578 | 0.111542782320032 | 0.0914779705091435 | 1.21934036904418 | 0.222715032590285 | 0.673748940276311 | NM_172715 |
| Mir1188 | 15.61899507 | 0.300274796 | 0.160009958 | 1.876600679 | 0.060572844 | 0.195687166 | | Cutc | 196.0940845 | -0.102966228666661 | 0.0844267197519582 | -1.21959290813585 | 0.222619236175286 | 0.673748940276311 | NM_001113562,NM_025530 |
| Uhrf1 | 47.76490722 | -0.305476104 | 0.162778862 | -1.876632504 | 0.060568479 | 0.195687166 | | Lrrc49 | 3466.944977 | 0.0679003829275977 | 0.0556777064202679 | 1.21952550299163 | 0.22264480228559 | 0.673748940276311 | NM_145616,NM_001146046,NM_001146047 |
| Nalcn | 4157.041132 | -0.151598637 | 0.080798807 | -1.876248458 | 0.060621171 | 0.195720863 | | Npc1l1 | 101.9992682 | -0.115920479728083 | 0.0950416822780083 | -1.21968042809893 | 0.222586043847343 | 0.673748940276311 | NM_207242 |
| Pappa | 35.03436232 | -0.303916027 | 0.16198067 | -1.876248734 | 0.060621133 | 0.195720863 | | Pax6 | 300.9743099 | -0.101156578110901 | 0.0829623526289296 | -1.21930700981142 | 0.222727689054581 | 0.673748940276311 | NM_013627,NM_001244202,NM_001244201,NM_001244198,NM_001244200 |
| Vps8 | 2478.922151 | 0.115517912 | 0.061564686 | 1.876366478 | 0.060604974 | 0.195720863 | | Tmem178b | 2051.458862 | 0.0973542727906028 | 0.0798185921545272 | 1.2196941860629 | 0.222580826401717 | 0.673748940276311 | NM_001004182 |
| Drd2 | 1323.643414 | -0.201698693 | 0.107540767 | -1.875555642 | 0.060716322 | 0.195987229 | | Ube2ql1 | 7668.109934 | 0.0690892708875469 | 0.0566431594040581 | 1.21972841229964 | 0.222567847133569 | 0.673748940276311 | NM_001145162 |
| Cdc42 | 10102.09644 | -0.101926372 | 0.054354927 | -1.875200249 | 0.06076518 | 0.196104082 | | Wwc1 | 3544.21486 | 0.0678906725896489 | 0.055657770297235 | 1.21978786119424 | 0.222545304221322 | 0.673748940276311 | NM_170779 |
| Wipf2 | 4199.261539 | 0.141215323 | 0.075310627 | 1.87510487 | 0.060778298 | 0.19610557 | | Zfp867 | 221.6743728 | -0.111719602974811 | 0.0915930679007181 | -1.21973862799212 | 0.222563973245256 | 0.673748940276311 | NM_178417 |
| 1110065P20Rik | 844.9550364 | -0.160536785 | 0.085626217 | -1.874855516 | 0.060812603 | 0.19612631 | | Uqcrfs1 | 3442.393437 | 0.0595274382740505 | 0.0488355107457726 | 1.21893755926784 | 0.222867892698108 | 0.674032689846496 | NM_025710 |
| Ptplb | 1411.394056 | 0.179649834 | 0.095824379 | 1.874782145 | 0.0608227 | 0.19612631 | | Atf3 | 58.51936373 | -0.103745412061228 | 0.0851376876427596 | -1.21856036889969 | 0.22301109869721 | 0.674048714143021 | NM_007498 |
| Twist1 | 14.89220231 | -0.315145412 | 0.168082165 | -1.874948551 | 0.060799802 | 0.19612631 | | Sec23a | 4500.418186 | 0.076886226903677 | 0.0630961297055686 | 1.21855694259694 | 0.223012399846576 | 0.674048714143021 | NM_009147 |
| Cox6c | 5532.557521 | -0.110883571 | 0.059157006 | -1.874394575 | 0.060876061 | 0.196257529 | | Ttc28 | 818.2303 | 0.0878753612016495 | 0.0721112396224991 | 1.21860838423629 | 0.222992865295363 | 0.674048714143021 | NM_001267622 |
| Ccdc89 | 17.93294122 | 0.315019443 | 0.168097351 | 1.874029793 | 0.060926319 | 0.196376894 | | Lysmd2 | 1189.765707 | 0.0680947117962264 | 0.0558907520427101 | 1.21835382970317 | 0.223089542305807 | 0.674141604230113 | NM_027309 |
| Fdft1 | 3616.44637 | -0.128260109 | 0.06845655 | -1.873598771 | 0.060985748 | 0.196529362 | | Mob3c | 241.1274825 | -0.100059318274347 | 0.0821586623346968 | -1.21787910648699 | 0.223269917071556 | 0.674214151927605 | NM_175308 |
| Aif1l | 360.3120694 | -0.224849613 | 0.120021468 | -1.873411632 | 0.061011565 | 0.196571676 | | Rpl18 | 3226.862672 | 0.0719623817873101 | 0.0590819643771914 | 1.21800929515227 | 0.223220440498599 | 0.674214151927605 | NM_009077 |
| Alox12 | 32.32659329 | -0.301456198 | 0.160927201 | -1.873245775 | 0.061034454 | 0.196604538 | | Slc8a2 | 13259.86438 | -0.0729491833411273 | 0.0599083393949692 | -1.21767994369167 | 0.223345621580313 | 0.674214151927605 | NM_148946 |
| Trak2 | 8844.268508 | -0.119625853 | 0.063874644 | -1.872822229 | 0.061092937 | 0.196752021 | | Tomm7 | 1414.1116 | 0.0618045606431852 | 0.0507462261190637 | 1.21791442181682 | 0.223256495147066 | 0.674214151927605 | NM_025394 |
| Rad50 | 756.7166771 | 0.156711187 | 0.083704664 | 1.87219183 | 0.061180069 | 0.196991685 | | Vps51 | 2107.419478 | -0.0720457807043765 | 0.0591608500465348 | -1.21779488712056 | 0.223301927766756 | 0.674214151927605 | NM_001081041 |
| Cyb5r4 | 1654.952277 | -0.125535886 | 0.067056729 | -1.872084843 | 0.061194867 | 0.196998392 | | Csnk1g3 | 2512.461737 | 0.0769099880403627 | 0.0632452917284809 | 1.21605871264767 | 0.223962556945873 | 0.675094439135868 | NM_152809 |
| Ankra2 | 1058.327639 | 0.139904036 | 0.074754908 | 1.871503027 | 0.061275391 | 0.197198933 | | Krt222 | 1759.015995 | 0.0798208694139588 | 0.0656256577560171 | 1.21630581914648 | 0.223868445619031 | 0.675094439135868 | NM_172946 |
| Sycp2 | 53.01852041 | 0.294368965 | 0.157294521 | 1.871450848 | 0.061282617 | 0.197198933 | | Ktn1 | 3754.44259 | -0.0790856564808858 | 0.0650283025120352 | -1.2161728574454 | 0.223919081007496 | 0.675094439135868 | NM_008477 |
| AI464131 | 1478.414859 | -0.178861285 | 0.095596558 | -1.871001308 | 0.0613449 | 0.197304828 | | Sap30 | 139.2768409 | 0.11143393476281 | 0.0916015918711863 | 1.21650653101654 | 0.223792024655207 | 0.675094439135868 | NM_021788 |
| Cdc42ep3 | 527.157063 | -0.188360737 | 0.100670429 | -1.871063229 | 0.061336318 | 0.197304828 | | Tmem40 | 67.11100441 | -0.105021298167818 | 0.0863376180364377 | -1.21640254356754 | 0.223831615581559 | 0.675094439135868 | NM_144805,NM_001168259,NM_001168258,NM_001168257,NM_001168256 |
| Nudt22 | 183.8568566 | 0.218509077 | 0.116791215 | 1.870937618 | 0.061353728 | 0.197304828 | | Tspan6 | 495.6796091 | 0.0971206773100739 | 0.0798231754365833 | 1.21669774196383 | 0.223719238517313 | 0.675094439135868 | NM_019656 |
| Stard3nl | 1181.515492 | -0.160391505 | 0.085745155 | -1.870560562 | 0.061406015 | 0.197431999 | | Zfp738 | 615.8333753 | -0.104660109458115 | 0.0860577315391111 | -1.21616161135446 | 0.223923364196699 | 0.675094439135868 | NM_001001187 |
| Hmmr | 36.56939789 | -0.30710375 | 0.164219593 | -1.870079838 | 0.061472732 | 0.197564511 | | Cdkal1 | 483.8980883 | 0.0950267173652377 | 0.0781597333854826 | 1.21580145234845 | 0.224060565432743 | 0.67514807635414 | NM_144536 |
| Id2 | 3466.397025 | 0.135528975 | 0.072472047 | 1.870086193 | 0.061471849 | 0.197564511 | | Galnt4 | 186.0708219 | -0.101878484682802 | 0.0837976356167014 | -1.21576800983746 | 0.22407330828193 | 0.67514807635414 | NM_015737 |
| Eno4 | 119.4745534 | 0.248726777 | 0.133013739 | 1.869932976 | 0.061493126 | 0.197589069 | | Il15ra | 166.5050854 | -0.10585310556856 | 0.0870885688268877 | -1.2154649800133 | 0.224188797612313 | 0.675355967257664 | NR_073192,NR_073191,NM_001271500,NM_001271501,NM_001271499,NM_133836,NM_001271498,NM_008358,NM_001271497 |
| Fam108a | 3298.831558 | -0.159243833 | 0.085170177 | -1.869713538 | 0.061523609 | 0.197596753 | | Aptx | 1455.9738 | -0.0613771365563052 | 0.0510173965134212 | -1.21405084928764 | 0.224728307215406 | 0.676420097515744 | NM_025545,NM_001025444 |
| Pcmtd2 | 2460.232626 | 0.101034274 | 0.054037084 | 1.869721048 | 0.061522565 | 0.197596753 | | BC037034 | 2377.968677 | -0.0835102592359113 | 0.0687748942377458 | -1.21425500048357 | 0.224650363586311 | 0.676420097515744 | NM_153161 |
| Slc12a2 | 2931.716607 | 0.140533033 | 0.07516581 | 1.86964036 | 0.061533777 | 0.197596753 | | Gtpbp4 | 1421.970142 | 0.0748842872206408 | 0.0616773570181589 | 1.21412931488931 | 0.224698347261586 | 0.676420097515744 | NM_027000 |
| Dhfr | 200.4083898 | 0.183910668 | 0.098386846 | 1.869260744 | 0.061586547 | 0.197725229 | | Tmem196 | 447.4272723 | 0.0883537744177537 | 0.0727710754804002 | 1.21413314059857 | 0.22469688659163 | 0.676420097515744 | NM_001160385 |
| Gm2694 | 42.53070387 | -0.299966686 | 0.160508271 | -1.868855014 | 0.061642988 | 0.197824454 | | Tdrd5 | 68.67454168 | -0.114536656452644 | 0.094354045770789 | -1.21390297063556 | 0.224784778411127 | 0.676449904951321 | NM_001134741 |
| Tti2 | 833.010732 | 0.11973991 | 0.064070076 | 1.868889776 | 0.061638151 | 0.197824454 | | Spns2 | 2586.070195 | 0.0937947870853887 | 0.0772773668811802 | 1.21374201620515 | 0.224846254445205 | 0.676494758092596 | NM_001276383,NM_153060 |
| Ssrp1 | 3885.961094 | 0.119589714 | 0.063996387 | 1.868694764 | 0.061665293 | 0.197855053 | | Golga4 | 4711.22674 | -0.0705300911511968 | 0.0581274621222872 | -1.21336952579862 | 0.224988572037921 | 0.676642649569948 | NM_018748 |
| Ttll6 | 12.52533153 | 0.280051347 | 0.149920671 | 1.867996889 | 0.061762504 | 0.198125931 | | Ttc1 | 1514.306948 | 0.0703012571820658 | 0.0579355850189364 | 1.21343828942243 | 0.224962294644407 | 0.676642649569948 | NM_133795 |
| Bfsp2 | 154.0966454 | -0.288637625 | 0.154526529 | -1.867883964 | 0.061778247 | 0.198135407 | | Cnn2 | 253.1925632 | -0.107387299700198 | 0.0885817038511385 | -1.21229661466732 | 0.225398859301967 | 0.677510814222137 | NM_007725 |
| AI854517 | 1112.35996 | 0.246495483 | 0.132017663 | 1.867140188 | 0.061882014 | 0.198427135 | | Gfm2 | 1099.750046 | -0.0753437559904063 | 0.0621520578889795 | -1.21224877259882 | 0.225417166813685 | 0.677510814222137 | NM_001271463,NM_001146043,NM_177266,NM_001271465,NM_001271464 |
| Xirp2 | 9.439446424 | 0.297108763 | 0.15916505 | 1.86667087 | 0.061947564 | 0.198596225 | | Zfp422 | 870.8523047 | 0.0753895634253086 | 0.0621881659131292 | 1.21228150594794 | 0.225404640773749 | 0.677510814222137 | NM_026057 |
| Prpsap2 | 1190.020333 | -0.152197835 | 0.081539915 | -1.866543954 | 0.061965301 | 0.198611991 | | Psd4 | 75.42475833 | -0.114747963577734 | 0.0946835289173002 | -1.21191050745435 | 0.225546639528402 | 0.677759719857173 | NM_177611 |
| BC037704 | 43.3263424 | 0.294091773 | 0.15758653 | 1.866224059 | 0.062010024 | 0.198714233 | | 4930479D17Rik | 64.00689176 | -0.115187110510248 | 0.0950679161499638 | -1.21162969774732 | 0.225654161208101 | 0.677942574086633 | NR_046277 |
| Klhl3 | 749.309244 | -0.191745726 | 0.102754171 | -1.866062706 | 0.062032593 | 0.19874545 | | Gm20605 | 38.62469015 | -0.0971717796445914 | 0.0802337395194608 | -1.21110869599967 | 0.225853749036737 | 0.67824895845055 | NR_033148 |
| Sclt1 | 308.1469929 | 0.173555084 | 0.093037225 | 1.865437023 | 0.062120172 | 0.198984896 | | Ift27 | 668.0443243 | 0.083206332127712 | 0.0687089000103832 | 1.21099787822448 | 0.225896217884054 | 0.67824895845055 | NM_025931 |
| Tctn1 | 414.3623719 | 0.21346218 | 0.114456849 | 1.865001373 | 0.062181212 | 0.199139251 | | Map3k8 | 55.64481751 | -0.114661733844038 | 0.0946834311619278 | -1.21100104249437 | 0.225895005157534 | 0.67824895845055 | NM_007746 |
| Lrrc48 | 448.9524157 | 0.186115266 | 0.099807084 | 1.864750066 | 0.062216445 | 0.199180439 | | Spata2l | 2172.480329 | 0.0874580055500317 | 0.0722372590644203 | 1.21070492821492 | 0.226008512966045 | 0.678445889511216 | NM_030176 |
| Ppp4r1l-ps | 276.4139006 | 0.19099051 | 0.102422813 | 1.864726263 | 0.062219784 | 0.199180439 | | Wdr12 | 1413.923785 | 0.0682099534453013 | 0.0563479356540001 | 1.21051379529037 | 0.226081800480588 | 0.678525668136575 | NM_021312,NM_001199060,NM_001199061 |
| Snx29 | 314.8396535 | -0.152474186 | 0.081788666 | -1.864245898 | 0.062287181 | 0.199355004 | | Lsm11 | 1448.385807 | 0.0912564483123184 | 0.0754040976697954 | 1.210231952008 | 0.226189900681417 | 0.678709873434881 | NM_028185 |
| BC035044 | 33.32726087 | -0.311608985 | 0.167196997 | -1.863723571 | 0.062360534 | 0.199469076 | | Pfdn2 | 1844.946409 | 0.0841676829035082 | 0.0696034687303939 | 1.20924552236798 | 0.226568533439809 | 0.679565252270632 | NM_011070 |
| Ern1 | 324.762199 | 0.18835061 | 0.101061803 | 1.863717095 | 0.062361443 | 0.199469076 | | Trappc6b | 2322.942246 | 0.0682100172340222 | 0.0564023927970686 | 1.20934616159702 | 0.226529883216096 | 0.679565252270632 | NM_030057 |
| Rps14 | 6562.15397 | -0.134561784 | 0.072197533 | -1.863800303 | 0.062349753 | 0.199469076 | | Coro7 | 3117.855941 | -0.0580766117761387 | 0.0480504688189208 | -1.20865858759883 | 0.226794037870306 | 0.679834423595694 | NM_030205 |
| Ube2s | 1938.556743 | -0.115161906 | 0.06179625 | -1.863574342 | 0.062381504 | 0.199492057 | | Cript | 1485.38317 | 0.0776589826855434 | 0.0642528497501557 | 1.2086465112056 | 0.226798679384878 | 0.679834423595694 | NM_019936 |
| Zfp113 | 381.5417402 | -0.173760584 | 0.093253499 | -1.863314374 | 0.062418049 | 0.199567736 | | Spry4 | 730.1003383 | 0.106710690897229 | 0.088288993530772 | 1.2086522544668 | 0.226796472752798 | 0.679834423595694 | NM_011898 |
| Ret | 241.7556416 | -0.278543101 | 0.149509734 | -1.863043247 | 0.062456182 | 0.199631622 | | C130071C03Rik | 84.37537473 | -0.114802355641954 | 0.0950065079715114 | -1.20836306999494 | 0.226907638363264 | 0.679880436234484 | NR_015561 |
| Trdn | 18.77577821 | -0.242071037 | 0.129936904 | -1.862989106 | 0.062463799 | 0.199631622 | | Myo7a | 1158.941619 | -0.0788536394487514 | 0.0652548352673065 | -1.20839534918354 | 0.226895227884253 | 0.679880436234484 | NM_008663,NM_001256082,NM_001256083,NM_001256081 |
| Klhdc1 | 482.8569889 | -0.165187689 | 0.08867643 | -1.862813915 | 0.062488452 | 0.199669224 | | Psmd9 | 912.3056086 | 0.0736912843965768 | 0.0609906297560601 | 1.2082394408996 | 0.226955174899637 | 0.679882629530237 | NM_026000 |
| Socs7 | 6538.312827 | -0.144533508 | 0.07760352 | -1.862460736 | 0.062538175 | 0.199786904 | | Tmem128 | 731.7569816 | 0.0808231527007996 | 0.066915171120033 | 1.20784496771021 | 0.227106900987799 | 0.680196874999745 | NM_025480 |
| Leprot | 985.4391701 | -0.103351429 | 0.05550781 | -1.861925876 | 0.062613539 | 0.199865303 | | Adi1 | 1210.517759 | 0.0644944243095564 | 0.0534179327097615 | 1.20735530257184 | 0.227295341312162 | 0.680203999115725 | NM_134052 |
| Orc4 | 1209.872764 | 0.160729129 | 0.08632358 | 1.861937712 | 0.062611871 | 0.199865303 | | Cd93 | 344.719405 | -0.112587945627235 | 0.0932279144965543 | -1.20766345825956 | 0.227176739195659 | 0.680203999115725 | NM_010740 |
| Serpind1 | 162.3737336 | 0.226070317 | 0.1214016 | 1.862169173 | 0.062579248 | 0.199865303 | | Exosc10 | 1855.397905 | 0.0618933907436251 | 0.0512809593213594 | 1.20694681930112 | 0.227452625261935 | 0.680203999115725 | NM_016699 |
| Tgfb1i1 | 500.8310573 | -0.199955784 | 0.107392231 | -1.861920379 | 0.062614314 | 0.199865303 | | Gda | 11549.53277 | 0.0897358626651844 | 0.0743416578645671 | 1.20707373554491 | 0.227403748644362 | 0.680203999115725 | NM_010266 |
| Nus1 | 3637.783549 | -0.139575458 | 0.074971647 | -1.861709902 | 0.062643992 | 0.19991849 | | Otud6b | 2715.267217 | 0.0767892612557069 | 0.0636160170765634 | 1.20707433103347 | 0.227403519333863 | 0.680203999115725 | NM_152812 |
| Mirlet7b | 14.65435737 | 0.310266133 | 0.166683322 | 1.861410782 | 0.062686189 | 0.200012318 | | Shc1 | 541.0776682 | -0.0819838123486738 | 0.0679035076035529 | -1.20735754664291 | 0.227294477460718 | 0.680203999115725 | NM_011368,NM_001113331 |
| Tex14 | 14.111638 | 0.306881355 | 0.164940217 | 1.860561121 | 0.06280618 | 0.200353911 | | Spryd7 | 1374.202366 | 0.080183174423257 | 0.066415537963155 | 1.2072954143413 | 0.227318396056034 | 0.680203999115725 | NM_025697 |
| Aatf | 462.1363089 | 0.142297026 | 0.076501609 | 1.86005273 | 0.062878066 | 0.200541942 | | Tax1bp1 | 6667.016245 | 0.0740533323019374 | 0.0613600470267195 | 1.20686563798901 | 0.227483892861365 | 0.680203999115725 | NM_025816 |
| Eftud2 | 3969.339645 | 0.141832944 | 0.076265243 | 1.859732421 | 0.062923393 | 0.200645204 | | 1700008O03Rik | 25.79014218 | 0.101567998985595 | 0.0842364609339316 | 1.20574864921328 | 0.227914420687732 | 0.680415543250314 | NM_027049 |
| Hspb11 | 183.7228903 | -0.199980017 | 0.107539864 | -1.859589651 | 0.062943605 | 0.200668356 | | 9430041J12Rik | 118.6500358 | -0.112559829712508 | 0.0933159690811027 | -1.20622258784754 | 0.227731676764681 | 0.680415543250314 | NR_033568 |
| Skp1a | 10753.18397 | -0.118571382 | 0.063767512 | -1.859432446 | 0.062965866 | 0.200698032 | | Fam199x | 412.9782982 | 0.101745361194197 | 0.08437136327326 | 1.20592292511224 | 0.22784721026439 | 0.680415543250314 | NM_146261 |
| Ppm1e | 8996.644957 | -0.158140131 | 0.085051738 | -1.859340386 | 0.062978906 | 0.200698307 | | St8sia3 | 7756.41902 | 0.0805402064306314 | 0.0667677922227425 | 1.20627332055466 | 0.227712121150854 | 0.680415543250314 | NM_009182 |
| 2310003H01Rik | 449.7535824 | 0.195024619 | 0.104912976 | 1.858917991 | 0.063038764 | 0.200845046 | | Stau2 | 4515.831692 | 0.0754894224974552 | 0.0626099388653304 | 1.20570988992383 | 0.227929370332922 | 0.680415543250314 | NM_001111272,NM_025303 |
| Cmtm3 | 299.2609895 | 0.186730654 | 0.100455877 | 1.858832544 | 0.063050878 | 0.200845046 | | Tmco3 | 1708.603178 | 0.0568421132562564 | 0.0471302836587039 | 1.20606346585735 | 0.227793020269164 | 0.680415543250314 | NM_172282 |
| Afmid | 121.571033 | 0.221595043 | 0.119218283 | 1.858733728 | 0.06306489 | 0.200848388 | | Trim32 | 8998.297681 | 0.0649987870821312 | 0.0539072500763609 | 1.20575223165824 | 0.227913038956673 | 0.680415543250314 | NM_001161782,NM_053084 |
| Zdhhc1 | 1347.672259 | 0.175090026 | 0.094219643 | 1.858317667 | 0.063123916 | 0.20099506 | | Zmynd11 | 8394.373164 | -0.0440180674779387 | 0.0364873984456628 | -1.20639095559226 | 0.227666781725821 | 0.680415543250314 | NM_001199141,NM_144516 |
| Map3k14 | 224.1868386 | 0.213200847 | 0.114743529 | 1.858064242 | 0.063159892 | 0.20106829 | | Clcc1 | 1042.476318 | -0.0622998478512946 | 0.0516809814997022 | -1.20546951786613 | 0.22802209859888 | 0.680552497277035 | NM_145543,NM_001177770,NM_001177771 |
| N4bp2l1 | 846.8445253 | 0.169234373 | 0.091100045 | 1.857676071 | 0.063215029 | 0.201202476 | | Tmem50b | 3839.535159 | 0.0640295668653523 | 0.0531625135495014 | 1.20441195478337 | 0.22843039367601 | 0.681631039140864 | NM_030018 |
| Carf | 403.2810094 | 0.147827258 | 0.079590189 | 1.857355274 | 0.063260625 | 0.201223594 | | Ctnnbl1 | 1130.314371 | 0.0682083698483966 | 0.0566393802589308 | 1.20425699463125 | 0.228490263097677 | 0.681669657374584 | NM_025680 |
| Crtac1 | 2539.1676 | -0.162124343 | 0.087280541 | -1.857508458 | 0.063238849 | 0.201223594 | | Syt2 | 2133.005195 | 0.100936533054159 | 0.0838281132148955 | 1.20408928679339 | 0.228555070224336 | 0.681722987695834 | NM_009307 |
| Tnfrsf23 | 45.52341289 | 0.291890417 | 0.157153621 | 1.857357246 | 0.063260345 | 0.201223594 | | Ipo5 | 7103.601652 | 0.0703587558096807 | 0.0584453208364144 | 1.20383898664209 | 0.228651817737728 | 0.681731589585023 | NM_023579 |
| Cbx6 | 18585.556 | -0.127970737 | 0.068907997 | -1.857124614 | 0.063293427 | 0.201286609 | | Slc22a15 | 346.7887761 | -0.0983431911721025 | 0.0816910324187779 | -1.20384316687246 | 0.228650201730654 | 0.681731589585023 | NM_001039371 |
| Zfp346 | 1483.856533 | 0.098768689 | 0.053191762 | 1.856841837 | 0.063333659 | 0.201373224 | | Cmtm4 | 8458.842364 | 0.0770614129984988 | 0.0640447911952028 | 1.20324247390584 | 0.228882502958878 | 0.681764163004386 | NM_153582 |
| Ecm1 | 303.8980988 | -0.215726306 | 0.116237515 | -1.85590948 | 0.063466462 | 0.201754074 | | Esrrb | 145.9412624 | 0.109055377074835 | 0.0906197108094425 | 1.20343991501098 | 0.228806129596839 | 0.681764163004386 | NM_011934,NM_001159500 |
| Ak5 | 7210.188462 | 0.148731125 | 0.080152164 | 1.855609606 | 0.063509224 | 0.20183165 | | Gm5083 | 22.41448252 | -0.092383900934264 | 0.0767815815443718 | -1.20320393349642 | 0.228897413119399 | 0.681764163004386 | NR_045285 |
| Chmp1b | 855.8538076 | 0.109283123 | 0.058897991 | 1.855464359 | 0.063529944 | 0.20183165 | | Neto2 | 2539.318839 | 0.104226397911004 | 0.0866157180564683 | 1.20331967741761 | 0.228852637251277 | 0.681764163004386 | NM_001081324 |
| Mrpl23 | 705.7515705 | -0.137250239 | 0.073968223 | -1.855529758 | 0.063520614 | 0.20183165 | | Rcan3 | 2166.267965 | 0.08556199799059 | 0.0710968317883245 | 1.2034572545417 | 0.228799423257111 | 0.681764163004386 | NM_022980 |
| Clgn | 320.7063197 | -0.208077488 | 0.112155123 | -1.855265126 | 0.063558376 | 0.20188058 | | Hebp1 | 800.0581332 | 0.0890411579055242 | 0.074054086096125 | 1.20238007920246 | 0.229216303701547 | 0.682442899114487 | NM_013546 |
| Hmgb2 | 196.0277998 | -0.185478397 | 0.099982295 | -1.855112405 | 0.063580177 | 0.201908436 | | Ptbp1 | 1345.265501 | -0.0899897009876362 | 0.0748434484278957 | -1.20237245714743 | 0.22921925545771 | 0.682442899114487 | NM_008956,NM_001077363 |
| Ncapg2 | 206.2178232 | -0.183202758 | 0.098772628 | -1.854792798 | 0.063625821 | 0.202011981 | | Amt | 838.1097188 | -0.093988523319681 | 0.0782078843003663 | -1.20177810920837 | 0.229449509001215 | 0.682503526431835 | NM_001013814 |
| Tusc2 | 3120.752287 | -0.122993171 | 0.066326371 | -1.854363045 | 0.063687238 | 0.202165554 | | Bai1 | 20664.8244 | -0.0803485156673813 | 0.0668616228342988 | -1.20171351309415 | 0.229474543789968 | 0.682503526431835 | NM_174991 |
| Ephx2 | 809.6284299 | -0.239168335 | 0.128991671 | -1.854137814 | 0.063719446 | 0.202226362 | | Fam43b | 964.9042267 | 0.083444858168359 | 0.0694321016923226 | 1.20181956378235 | 0.229433443942449 | 0.682503526431835 | NM_001081672 |
| Olfr856-ps1 | 145.9329317 | 0.234974569 | 0.126747902 | 1.853873438 | 0.063757269 | 0.202275644 | | Mpp6 | 1409.470282 | 0.0855403472484516 | 0.0711668960816757 | 1.20196821778317 | 0.229375842036945 | 0.682503526431835 | NM_019939,NM_001164734,NM_001164733 |
| Xkrx | 86.70127137 | -0.270238717 | 0.145771874 | -1.853846773 | 0.063761085 | 0.202275644 | | Orc4 | 1376.088701 | -0.0952306090946215 | 0.0792204095288711 | -1.20209690483758 | 0.229325985429674 | 0.682503526431835 | NM_001177313,NM_011958,NR_033442 |
| Nat8l | 18840.71131 | 0.127437016 | 0.068751495 | 1.853589027 | 0.063797979 | 0.202351255 | | Avpi1 | 697.2754858 | 0.0866539340219726 | 0.0721250551030715 | 1.20144010840807 | 0.229580525687001 | 0.682539237848827 | NM_027106 |
| 2-Mar | 1646.665163 | 0.162864162 | 0.087875576 | 1.853349591 | 0.063832268 | 0.202418574 | | Fam69c | 221.2046751 | -0.107123782148906 | 0.0891608064348858 | -1.20146717411241 | 0.229570032453456 | 0.682539237848827 | NM_173770 |
| Gm14322 | 510.2207469 | 0.214519882 | 0.115791939 | 1.852632257 | 0.063935086 | 0.202703135 | | Cnnm2 | 1253.078455 | 0.065995250220951 | 0.0549431910775943 | 1.20115430004326 | 0.229691352938425 | 0.682724729317983 | NM_001102471,NM_033569 |
| Bin3 | 376.9463805 | 0.167952039 | 0.090663598 | 1.852474883 | 0.063957661 | 0.202733224 | | Sox11 | 470.8615444 | 0.0900740817094593 | 0.0749969360707561 | 1.20103682135066 | 0.229736918415689 | 0.682724729317983 | NM_009234 |
| Gtf2h5 | 1631.198143 | -0.123859066 | 0.06688244 | -1.85189216 | 0.06404131 | 0.202956853 | | Smndc1 | 970.5266762 | 0.0776823715533636 | 0.0647025148156254 | 1.20060822635295 | 0.229903208489435 | 0.68307915862498 | NM_172429 |
| 1500012K07Rik | 10.17293774 | -0.295095903 | 0.159365173 | -1.851696307 | 0.064069445 | 0.202962988 | | 4632415L05Rik | 1007.194435 | 0.0727149868678661 | 0.0605980541009069 | 1.19995580628352 | 0.23015650450545 | 0.683133098579111 | NR_027985 |
| Spata7 | 688.8245567 | 0.184075699 | 0.099406 | 1.851756419 | 0.064060808 | 0.202962988 | | Ccdc146 | 37.03253974 | 0.101714554436201 | 0.0847578281171462 | 1.20006088753797 | 0.230115694285024 | 0.683133098579111 | NM_029195 |
| Rev1 | 1288.111845 | 0.143858078 | 0.077704025 | 1.851359415 | 0.064117864 | 0.203074853 | | Gm10635 | 89.77542047 | -0.105310503955703 | 0.0877417004036253 | -1.20023322401159 | 0.230048775416514 | 0.683133098579111 | NR_045336 |
| Dnahc11 | 70.51764961 | -0.276178785 | 0.149186393 | -1.851233079 | 0.064136029 | 0.203090871 | | Sltm | 2753.138603 | -0.0685895343292524 | 0.0571489518451371 | -1.20018884194828 | 0.23006600780723 | 0.683133098579111 | NM_026337,NM_025690 |
| Ctps | 1646.714087 | -0.106523461 | 0.057580096 | -1.85000491 | 0.064312842 | 0.203609149 | | Trdmt1 | 175.3824903 | -0.103270122668676 | 0.086027335280703 | -1.20043381945646 | 0.22997090091826 | 0.683133098579111 | NM_010067 |
| Grhpr | 865.732597 | 0.148658271 | 0.080378741 | 1.849472496 | 0.064389616 | 0.203768935 | | 1810041L15Rik | 2953.868386 | 0.0885835383943627 | 0.0739474613260152 | 1.19792534869827 | 0.230946080374063 | 0.68340205891701 | NM_001163145 |
| Gstz1 | 1249.876322 | 0.152141651 | 0.082260737 | 1.849505086 | 0.064384914 | 0.203768935 | | 5033411D12Rik | 150.5386032 | -0.107460130351405 | 0.0896209156312975 | -1.19905191321073 | 0.230507760391731 | 0.68340205891701 | NM_138654 |
| Naaa | 1279.892325 | 0.129878576 | 0.070245734 | 1.848912624 | 0.064469709 | 0.2039399 | | 9030624J02Rik | 2454.331406 | 0.0660968030545999 | 0.0552008431578266 | 1.19738756282434 | 0.231155529031564 | 0.68340205891701 | NM_027815 |
| Nck2 | 1144.520761 | -0.144503849 | 0.078154679 | -1.848946874 | 0.064465484 | 0.20393909 | | C2cd4b | 23.7708747 | -0.0852757184564786 | 0.0710873527168356 | -1.19959057690838 | 0.230298388016384 | 0.68340205891701 | NM_001081314 |
| Perp | 176.6429364 | -0.215854242 | 0.116779613 | -1.848389761 | 0.064545979 | 0.204138681 | | Cox20 | 381.8797593 | -0.0890498352464256 | 0.0743576591963931 | -1.1975879312073 | 0.231077476835521 | 0.68340205891701 | NM_025511 |
| Gm12216 | 19.82177914 | 0.309278009 | 0.167357985 | 1.848002704 | 0.064601952 | 0.204190665 | | Cradd | 191.3212028 | 0.111812921550348 | 0.0933102115864954 | 1.19829244462383 | 0.230803186905795 | 0.68340205891701 | NM_009950 |
| Rtn4 | 26325.62958 | -0.107214358 | 0.058014493 | -1.848061616 | 0.06459343 | 0.204190665 | | Crkl | 2231.219008 | 0.0823645909621531 | 0.0687917069308045 | 1.19730407394893 | 0.231188057105754 | 0.68340205891701 | NM_007764 |
| Sin3b | 2985.410322 | 0.107666575 | 0.058256184 | 1.848157002 | 0.064579634 | 0.204190665 | | Cryab | 3885.595258 | 0.0916067448736387 | 0.0765000389847067 | 1.19747317896076 | 0.231122175532857 | 0.68340205891701 | NM_009964 |
| Ybx1 | 3558.259041 | -0.098014092 | 0.053044161 | -1.847782853 | 0.064633762 | 0.204249544 | | Eif2b3 | 578.4021489 | 0.077602639569292 | 0.0647871112175038 | 1.19780984382374 | 0.230991054079551 | 0.68340205891701 | NM_175135,NM_001111277 |
| Fancl | 178.5188047 | -0.265246507 | 0.143577607 | -1.847408605 | 0.064687943 | 0.204337408 | | Eif3f | 4145.123622 | -0.0565354302857099 | 0.0471835868339471 | -1.19820119832506 | 0.230838699001235 | 0.68340205891701 | NM_025344 |
| St8sia5 | 1811.642984 | -0.177987347 | 0.096343622 | -1.847422201 | 0.064685974 | 0.204337408 | | Elfn1 | 1417.850479 | 0.0751631580319513 | 0.0627312782792123 | 1.19817673246519 | 0.230848221515525 | 0.68340205891701 | NM_175522 |
| Tm9sf3 | 6516.898438 | -0.122599143 | 0.066376491 | -1.847026573 | 0.064743289 | 0.204470551 | | G530011O06Rik | 168.3457632 | 0.0747917153608504 | 0.0624249145578935 | 1.19810681184814 | 0.230875437306373 | 0.68340205891701 | NR_029457 |
| Ddx42 | 3622.450174 | 0.098768192 | 0.053479716 | 1.846834631 | 0.064771111 | 0.20447506 | | Lin9 | 173.5007878 | -0.108815477431424 | 0.090734818010128 | -1.19926925316892 | 0.230423266575471 | 0.68340205891701 | NM_001103182,NM_175186 |
| Zmynd8 | 5386.935427 | 0.185803174 | 0.100604639 | 1.846864872 | 0.064766727 | 0.20447506 | | Mcm3 | 143.257507 | -0.110789669442848 | 0.0924559289835247 | -1.19829707689801 | 0.230801384177183 | 0.68340205891701 | NM_008563 |
| Tex9 | 376.305537 | 0.21130704 | 0.114422439 | 1.846727286 | 0.064786675 | 0.204482531 | | Ppp1r8 | 1361.276789 | 0.068323339014543 | 0.0570411307473018 | 1.19779075413534 | 0.230998487560023 | 0.68340205891701 | NM_146154 |
| Hsd17b10 | 819.480604 | -0.130592587 | 0.070739464 | -1.846106541 | 0.064876737 | 0.204600075 | | Prpf6 | 2918.388802 | 0.0500349624934452 | 0.0417518420395987 | 1.19838934162451 | 0.230765479867652 | 0.68340205891701 | NM_133701 |
| Kctd1 | 3439.56405 | -0.14883443 | 0.080612223 | -1.846301039 | 0.064848507 | 0.204600075 | | Rnf216 | 2433.823386 | 0.0638072545999126 | 0.0532754929424155 | 1.19768492182476 | 0.231039701500041 | 0.68340205891701 | NM_080561,NM_207110 |
| Map3k6 | 198.3903886 | 0.294120017 | 0.159310514 | 1.846205936 | 0.064862309 | 0.204600075 | | Tgm3 | 180.2894973 | -0.109219513392781 | 0.0911250601432073 | -1.19856725714241 | 0.230696256232693 | 0.68340205891701 | NM_009374 |
| Nkiras2 | 333.6770808 | -0.17130121 | 0.092787369 | -1.846169501 | 0.064867598 | 0.204600075 | | Ubxn2a | 2180.107869 | 0.0879473387946724 | 0.073395630575959 | 1.19826395801114 | 0.230814273177615 | 0.68340205891701 | NM_145441 |
| G6pdx | 1504.992627 | 0.169169513 | 0.091652446 | 1.845771937 | 0.064925327 | 0.204711643 | | Zfp9 | 1145.238766 | 0.0684692888896825 | 0.0571515578133819 | 1.19803014142251 | 0.23090528300289 | 0.68340205891701 | NM_011763 |
| Atp2a1 | 14.42976958 | 0.303495879 | 0.164479788 | 1.845186466 | 0.065010419 | 0.204938235 | | Kcnj6 | 1248.893749 | 0.0947761823149198 | 0.0791670380953392 | 1.19716721245505 | 0.231241386703182 | 0.683420627314431 | NM_010606,NM_001025584,NM_001025590,NM_001025585 |
| Psmg4 | 131.4781166 | -0.199272336 | 0.10802976 | -1.844605934 | 0.065094883 | 0.205162758 | | A830010M20Rik | 8095.231638 | 0.089836025330256 | 0.0750574726695519 | 1.19689648658659 | 0.231346903777077 | 0.683460324215196 | NM_001007574,NM_001168557 |
| 5730508B09Rik | 89.82824416 | -0.256028774 | 0.138808451 | -1.844475404 | 0.065113887 | 0.205180917 | | Dnaja3 | 2923.744921 | 0.0554411615914225 | 0.0463209657740785 | 1.19689131400727 | 0.231348920154628 | 0.683460324215196 | NM_001135112,NM_023646 |
| Rnf144b | 1281.670203 | 0.134215975 | 0.07277155 | 1.84434679 | 0.065132617 | 0.205198203 | | Yif1b | 969.1401045 | 0.0893930090761468 | 0.0746954156477861 | 1.19676700773264 | 0.231397381045769 | 0.683464489034331 | NM_001110201,NM_029887 |
| Nppa | 10.69718905 | -0.246384523 | 0.133597082 | -1.844235809 | 0.065148782 | 0.205207405 | | Cxxc1 | 1722.187743 | -0.0848272816080454 | 0.0709053573750144 | -1.19634516697234 | 0.231561889722191 | 0.683672359777346 | NM_028868 |
| Endod1 | 4501.532508 | 0.155651974 | 0.084403491 | 1.844141424 | 0.065162532 | 0.205208999 | | Tlr13 | 153.0650109 | -0.109805736088382 | 0.0917769305918574 | -1.19644158265328 | 0.231524282393344 | 0.683672359777346 | NM_205820 |
| Gm10635 | 103.7482685 | -0.309179806 | 0.167672898 | -1.843946224 | 0.065190978 | 0.20525686 | | Tcf7 | 207.9404799 | -0.111290568765277 | 0.0930399449779273 | -1.19615901311721 | 0.231634512064984 | 0.683747799686235 | NM_009331 |
| Cdh23 | 70.23838618 | -0.298660992 | 0.162048793 | -1.843031271 | 0.065324445 | 0.205426618 | | Calcr | 71.67556096 | 0.0430658915702746 | 0.0360109305491835 | 1.19591165553069 | 0.231731036230454 | 0.683754830851833 | NM_001042725,NM_007588 |
| Col1a1 | 435.5905914 | -0.309012049 | 0.167651477 | -1.843181196 | 0.065302559 | 0.205426618 | | S1pr3 | 508.6509617 | -0.0849740498396851 | 0.0710537232422965 | -1.19591269763471 | 0.231730629519506 | 0.683754830851833 | NM_010101 |
| Pkd2l2 | 14.90525563 | 0.308358599 | 0.167305662 | 1.843085254 | 0.065316564 | 0.205426618 | | Irak3 | 64.55234941 | -0.11155362446791 | 0.0932921614574486 | -1.19574488065421 | 0.23179613143492 | 0.68380800268555 | NM_028679 |
| Prtg | 145.3713886 | 0.24166175 | 0.131118127 | 1.843084213 | 0.065316716 | 0.205426618 | | Msln1 | 24.46440051 | -0.0941473419952265 | 0.0787452626723432 | -1.19559372589778 | 0.231855141043939 | 0.683843203818123 | NM_177822 |
| Sesn3 | 1899.004887 | -0.183346935 | 0.099473216 | -1.843178919 | 0.065302892 | 0.205426618 | | Notch3 | 812.5460388 | -0.0921060690198661 | 0.0770550716509873 | -1.19532779668353 | 0.231958983571571 | 0.684010595891321 | NM_008716 |
| Slc23a2 | 7989.154743 | 0.104660058 | 0.056781008 | 1.843222983 | 0.065296461 | 0.205426618 | | Mtfr1l | 2532.57332 | -0.0573041716729036 | 0.0479511473557227 | -1.19505319127811 | 0.232066248706716 | 0.684188010698955 | NM_029759,NM_001256112 |
| Slfn8 | 55.12547367 | 0.301916286 | 0.163867331 | 1.842443418 | 0.065410316 | 0.205654918 | | Doc2g | 157.3495688 | -0.113565018556169 | 0.0950540113202737 | -1.19474198909423 | 0.232187851669408 | 0.684268763106072 | NM_021791 |
| D130043K22Rik | 1687.534086 | 0.134314272 | 0.07290444 | 1.842333228 | 0.065426422 | 0.205663824 | | Kbtbd7 | 1390.901919 | 0.090459965757758 | 0.0757095173570155 | 1.19482951306089 | 0.232153646909513 | 0.684268763106072 | NM_001024135 |
| Dennd6a | 1039.359057 | 0.10373584 | 0.056351837 | 1.840859939 | 0.065642086 | 0.206258059 | | Tsfm | 468.9294897 | -0.0792562478212247 | 0.0663449204461063 | -1.19460913191699 | 0.232239779687706 | 0.684282969522031 | NM_025537 |
| Mfap3 | 1203.227814 | -0.139482226 | 0.07576852 | -1.840899443 | 0.065636296 | 0.206258059 | | 4933426M11Rik | 2473.335655 | 0.0643610417712264 | 0.0539004102273812 | 1.19407331965966 | 0.232449288751884 | 0.684327864506297 | NM_001242419,NM_178682 |
| Cbln1 | 249.5870599 | -0.243159523 | 0.132123754 | -1.840392179 | 0.065710681 | 0.206389882 | | Bcar1 | 2138.492097 | 0.0731539902358031 | 0.0612730647620362 | 1.19390127652188 | 0.232516588134315 | 0.684327864506297 | NM_001198839,NM_009954 |

| | | | | | | | | | | | | | | | |
|---|---|---|---|---|---|---|---|---|---|---|---|---|---|---|---|
| Dynlt3 | 3548.401282 | -0.123747553 | 0.067236475 | -1.840482471 | 0.065697435 | 0.206389882 | | Lrrc29 | 53.1732765 | -0.100564405840805 | 0.0842327237842461 | -1.19388761662737 | 0.232521932168346 | 0.684327864506297 | NM_177449 |
| Eny2 | 1756.549699 | 0.143589292 | 0.078036003 | 1.840039042 | 0.065762506 | 0.206448923 | | Mms22l | 50.12708291 | -0.111552052405153 | 0.0934020773743539 | -1.19432089243641 | 0.232352468143254 | 0.684327864506297 | NM_199467 |
| Ncor1 | 9715.332586 | -0.12921109 | 0.070218768 | -1.84012187 | 0.065750347 | 0.206448923 | | Orai3 | 752.1311172 | -0.0943294250057571 | 0.0790002721442852 | -1.19403924120002 | 0.232462618376188 | 0.684327864506297 | NM_198424 |
| Ulk1 | 4568.641924 | 0.162672493 | 0.088409361 | 1.839991729 | 0.065769452 | 0.206448923 | | Tpd52l2 | 1911.520936 | 0.0582561310780662 | 0.0487969675090225 | 1.19384736494731 | 0.232537679967168 | 0.684327864506297 | NM_025482 |
| Fam195a | 203.0140037 | -0.193970368 | 0.105449671 | -1.839459207 | 0.065847673 | 0.206568909 | | Crbn | 3272.868249 | 0.075731774139109 | 0.0634419250514291 | 1.19371809852424 | 0.232588258417969 | 0.684338068012846 | NM_021449,NM_175357 |
| Paox | 222.5881453 | 0.184660079 | 0.100386483 | 1.839491472 | 0.065842931 | 0.206568909 | | Zfp157 | 1070.404989 | -0.0677247193283515 | 0.0567439420938945 | -1.1935145291155 | 0.232667925439484 | 0.684433836560133 | NM_028130 |
| Pole4 | 885.1868671 | -0.205743416 | 0.111848285 | -1.83948656 | 0.065843653 | 0.206568909 | | Rsrc1 | 1739.978208 | 0.0644687337092879 | 0.0540331122495617 | 1.19313382156348 | 0.232816967545475 | 0.684628271664154 | NM_025822 |
| Gtf2e1 | 681.6185281 | -0.150024125 | 0.081563938 | -1.839343822 | 0.065864631 | 0.206580283 | | Tubb6 | 43.63800768 | -0.10892322182781 | 0.0912939171824061 | -1.19310492078219 | 0.232828284594584 | 0.684628271664154 | NM_026473 |
| Cecr6 | 3430.962028 | -0.189540693 | 0.103065352 | -1.839034058 | 0.065910177 | 0.206681295 | | Erbb2 | 136.9700901 | -0.109058975382538 | 0.0914247855834292 | -1.19288193772154 | 0.232915614048601 | 0.684746450044522 | NM_001003817 |
| Nasp | 643.147386 | 0.159691161 | 0.086842 | 1.838870145 | 0.065934288 | 0.206715066 | | 9330179D12Rik | 90.07962867 | -0.113171126268186 | 0.0949514403143099 | -1.19188425045018 | 0.233306634480916 | 0.685064073104533 | NR_040275,NR_040274,NR_040273 |
| Ucp3 | 124.0103508 | 0.249051149 | 0.135447425 | 1.8387293 | 0.065955012 | 0.206738206 | | Acsl4 | 3122.823219 | 0.0951040614176736 | 0.0797574979316953 | 1.1924153074501 | 0.233098441079007 | 0.685064073104533 | NM_207625,NM_019477,NM_001033600 |
| Caly | 4771.261857 | -0.125809417 | 0.068429204 | -1.838533983 | 0.065983759 | 0.206786481 | | Ank | 9859.875926 | 0.0570973225116591 | 0.0478935477600559 | 1.19217149662233 | 0.233194007307452 | 0.685064073104533 | NM_020332 |
| Nynrin | 615.166242 | -0.213663436 | 0.116230549 | -1.838272626 | 0.066022243 | 0.206865243 | | Axin2 | 1141.390776 | -0.0796663514319901 | 0.0668582451445749 | -1.19157108086996 | 0.233429469981687 | 0.685064073104533 | NM_015732 |
| Kcne1l | 147.3753399 | -0.23825585 | 0.129627903 | -1.837998176 | 0.066062674 | 0.206950075 | | Bag4 | 3059.825128 | 0.0807480432096223 | 0.0678004219858366 | 1.19096667608486 | 0.23366666722157 | 0.685064073104533 | NM_026121 |
| Pank3 | 2353.705901 | -0.122291843 | 0.06654915 | -1.837616913 | 0.066118875 | 0.207084262 | | Cisd2 | 1588.63032 | 0.0703039846102964 | 0.0590237222829082 | 1.19111404518543 | 0.233608816819406 | 0.685064073104533 | NM_025902 |
| BC005561 | 449.3696948 | 0.182940572 | 0.099559288 | 1.83750382 | 0.066135553 | 0.207094636 | | Fgf9 | 830.2465435 | -0.0787695217759012 | 0.0661224593922127 | -1.19126727136193 | 0.233548677962921 | 0.685064073104533 | NM_013518 |
| Sbno1 | 6932.784722 | -0.094466648 | 0.05141325 | -1.837398892 | 0.06615103 | 0.207101246 | | Lipa | 1010.983124 | 0.0682183601987311 | 0.0573029441182806 | 1.19048613030981 | 0.233855378167372 | 0.685064073104533 | NM_021460,NM_001111100 |
| Cinp | 1486.545104 | -0.121074594 | 0.065907875 | -1.837027744 | 0.0662058 | 0.207230841 | | Map2k7 | 4069.945911 | -0.0611526881863048 | 0.0513672863731422 | -1.19049871044539 | 0.233850436555466 | 0.685064073104533 | NM_001042557,NM_011944,NM_001164172 |
| Rfx2 | 107.8067867 | 0.231938287 | 0.126294275 | 1.836490901 | 0.066285087 | 0.20743711 | | Pcdhga1 | 289.0068331 | 0.110668794485743 | 0.0929059348990685 | 1.19119187171381 | 0.233578269763323 | 0.685064073104533 | NM_033584 |
| Xpc | 746.2894489 | 0.150057225 | 0.081714803 | 1.836353012 | 0.066305464 | 0.207458979 | | Pdia3 | 8246.25866 | 0.0781060663261239 | 0.0655828958677381 | 1.19095177626254 | 0.233672516778944 | 0.685064073104533 | NM_007952 |
| Herpud2 | 1524.596479 | 0.103876005 | 0.056597761 | 1.835337701 | 0.066455668 | 0.207886962 | | Peg10 | 2341.023213 | 0.0958788793579228 | 0.0805405445236015 | 1.19044240295429 | 0.233872555316355 | 0.685064073104533 | NM_130877,NM_001040611 |
| Mir1839 | 10.54534105 | 0.266920683 | 0.145460684 | 1.835002252 | 0.066505356 | 0.207916461 | | Psma1 | 2696.475169 | 0.0727219934697924 | 0.0610699189965896 | 1.19079891810326 | 0.233732533729369 | 0.685064073104533 | NM_011965 |
| Mrpl49 | 1302.075244 | -0.106045471 | 0.057788752 | -1.835053846 | 0.066497712 | 0.207916461 | | Snx16 | 991.1614126 | 0.0856057471561096 | 0.0718609424346227 | 1.19126947484709 | 0.233547813209827 | 0.685064073104533 | NM_001127191,NM_029068 |
| Nrbp1 | 2988.748294 | 0.108995408 | 0.059395942 | 1.835064896 | 0.066496074 | 0.207916461 | | Syt10 | 294.1553375 | 0.108196648985434 | 0.0908209948504476 | 1.19131759306973 | 0.233528929881005 | 0.685064073104533 | NM_018803 |
| Clic4 | 3794.803205 | -0.178113647 | 0.097103955 | -1.834257395 | 0.066615795 | 0.208219716 | | Ttc3 | 47349.97241 | 0.0661297421667346 | 0.0555405081129733 | 1.19065785340354 | 0.233787929887956 | 0.685064073104533 | NM_009441 |
| Morf4l2 | 6786.830027 | -0.11642313 | 0.063474937 | -1.834159047 | 0.066630389 | 0.208223324 | | Uhmk1 | 2383.265487 | 0.0818499293490807 | 0.0686838093990092 | 1.19169175480039 | 0.233382132232946 | 0.685064073104533 | NM_010633 |
| Pam | 8632.802494 | -0.115128475 | 0.062778118 | -1.833894964 | 0.066669587 | 0.208303808 | | Wee1 | 1090.670465 | 0.0935065004456747 | 0.0784759586918678 | 1.19153052736601 | 0.233445379764248 | 0.685064073104533 | NM_009516 |
| Fbxl5 | 3218.027649 | 0.087317537 | 0.047619426 | 1.833653717 | 0.066705413 | 0.208373724 | | Fzr1 | 2770.037971 | 0.0696929432377545 | 0.0585546011035621 | 1.19022146721643 | 0.233959357820484 | 0.685180167681522 | NM_019757 |
| Cir1 | 1159.914216 | 0.156487456 | 0.085375064 | 1.832941 | 0.066811346 | 0.20864524 | | Btbd11 | 1433.131587 | 0.0980992987367002 | 0.0824781187020949 | 1.18939787037369 | 0.23428313846933 | 0.685572979785801 | NM_001017525,NM_028709 |
| Fam211b | 803.2051039 | -0.200867393 | 0.10959608 | -1.832797236 | 0.06683273 | 0.20864524 | | Emx2os | 126.53938 | -0.111996005711548 | 0.094189039188757 | -1.18905561279912 | 0.234417783528342 | 0.685572979785801 | NR_002863 |
| Nrgn | 37700.61387 | 0.224139106 | 0.122289771 | 1.83285245 | 0.066824517 | 0.20864524 | | Erich1 | 172.4693353 | -0.0995674350608225 | 0.0837190362679372 | -1.18930460143095 | 0.234319825292302 | 0.685572979785801 | NM_001034862 |
| Zfp106 | 13024.46515 | -0.152737244 | 0.0833406 | -1.832687114 | 0.066849115 | 0.208654348 | | Foxr2 | 26.4338135 | 0.0974208914718354 | 0.0818985665041277 | 1.18953109474664 | 0.23423074245681 | 0.685572979785801 | NM_001034894 |
| Stt3a | 1343.634757 | -0.119831603 | 0.065392099 | -1.832508905 | 0.066875636 | 0.208695087 | | Lactb2 | 342.2572352 | -0.101631305260086 | 0.0854733993138877 | -1.18904017011024 | 0.234423860016252 | 0.685572979785801 | NM_145381 |
| Mtmr2 | 2142.206137 | 0.141810762 | 0.077392807 | 1.832350668 | 0.066899193 | 0.208726559 | | Trim28 | 5088.491881 | -0.0510598129243143 | 0.042935521526851 | -1.18922074563325 | 0.234352812981054 | 0.685572979785801 | NM_011588 |
| Fgd6 | 987.0892307 | -0.130065645 | 0.07099844 | -1.831950742 | 0.06695876 | 0.208828306 | | Zswim1 | 1193.107524 | 0.0675984916500898 | 0.0568324439241487 | 1.18943488934437 | 0.234268578390058 | 0.685572979785801 | NM_028028 |
| Nub1 | 4433.947982 | 0.093523485 | 0.051049279 | 1.832023614 | 0.066947903 | 0.208828306 | | Accs | 134.5138766 | -0.112529867637788 | 0.0946872130026139 | -1.18843784782937 | 0.2346609526893 | 0.685908078335004 | NM_183220 |
| Mars | 2027.289153 | 0.111587879 | 0.060917405 | 1.831789755 | 0.06698275 | 0.208861085 | | Fam181a | 75.04079135 | -0.110218263454208 | 0.0927546346826812 | -1.18827769449226 | 0.234724022565951 | 0.685908078335004 | NM_001195726 |
| Ggact | 498.145158 | 0.160277808 | 0.087518318 | 1.831362991 | 0.067046381 | 0.209017431 | | March6 | 12220.5601 | 0.073377241323507 | 0.0617492396949409 | 1.18831003727352 | 0.234711284710006 | 0.685908078335004 | NM_172606 |
| Bmpr1a | 2459.308416 | -0.155184909 | 0.084761963 | -1.830831928 | 0.067125633 | 0.209138259 | | Tpcn1 | 2245.824144 | -0.0652875603004937 | 0.0549434031083465 | -1.18826932091827 | 0.23472732048763 | 0.685908078335004 | NM_145853 |
| Cdc37l1 | 2382.826196 | 0.147611459 | 0.080621439 | 1.830920667 | 0.067112385 | 0.209138259 | | Vti1b | 4881.474513 | 0.0591238911322719 | 0.0497725941783796 | 1.18788044119979 | 0.234880516430938 | 0.686217695429567 | NM_016800 |
| Notum | 105.7742969 | -0.294432796 | 0.160813266 | -1.830898678 | 0.067115668 | 0.209138259 | | Cacnb2 | 2206.629121 | 0.0685660073493764 | 0.0577383729231475 | 1.18752926135693 | 0.235018921643801 | 0.686276190501263 | NM_001252533,NR_045533,NM_023116 |
| Ap4m1 | 470.3342553 | 0.167112541 | 0.091291102 | 1.830545779 | 0.067168368 | [illegible] | | Cd109 | 85.86945781 | -0.105626849189934 | 0.0890166051891144 | -1.18659714067428 | 0.235386564304127 | 0.686276190501263 | NM_153098 |
| Gpsm2 | 637.8916253 | 0.183511703 | 0.100258592 | 1.830383806 | 0.067192567 | [illegible] | | Chn2 | 2140.899479 | 0.0939802316944672 | 0.0791927977176877 | 1.18672700552258 | 0.2353353192304 | 0.686276190501263 | NM_001163640,NM_023543 |
| Rnd3 | 600.9682254 | -0.22134018 | 0.120922056 | -1.830436789 | 0.067184651 | [illegible] | | Egfr | 494.8709966 | -0.0897185482358083 | 0.0755810632159213 | -1.18705062377197 | 0.235207652819328 | 0.686276190501263 | NM_207655,NM_007912 |
| Pfn2 | 15152.19096 | -0.101909387 | 0.055691678 | -1.829885396 | 0.067267078 | [illegible] | | Gbas | 2806.405437 | 0.0616262312153507 | 0.0519261801504629 | 1.186804633747 | 0.235304690662349 | 0.686276190501263 | NM_008095 |
| Zfp949 | 874.7461313 | 0.124626456 | 0.068110456 | 1.829769789 | 0.06728437 | [illegible] | | Myzap | 27.77159268 | -0.101644537416781 | 0.0855875787640658 | -1.18760851614903 | 0.234987681104319 | 0.686276190501263 | NM_001033208 |
| Bnc2 | 22.69267497 | -0.303905976 | 0.166099486 | -1.829662354 | 0.067300444 | [illegible] | | Nfil3 | 359.0604024 | 0.103402344656655 | 0.0871481779132763 | 1.18651183688033 | 0.235420229744444 | 0.686276190501263 | NM_017373 |
| Ado | 2096.056265 | -0.139079588 | 0.07603844 | -1.829069453 | 0.067389204 | [illegible] | | Poldip2 | 3786.933966 | 0.058838631894926 | 0.0495769904896921 | 1.18681330419118 | 0.235301269874451 | 0.686276190501263 | NM_026389 |
| D430036J16Rik | 118.889761 | -0.269958568 | 0.147571144 | -1.829345221 | 0.067347908 | [illegible] | | Rab31 | 3938.760262 | 0.0728801065887865 | 0.0613900270896842 | 1.18716524562396 | 0.235162446608283 | 0.686276190501263 | NM_133685 |
| Ldlrad3 | 549.1791533 | -0.150068117 | 0.082045535 | -1.829083302 | 0.06738713 | [illegible] | | Rtn3 | 54052.64243 | 0.064493013629402 | 0.0543208940219362 | 1.18725979736928 | 0.235125160563726 | 0.686276190501263 | NM_001003933,NM_001271487,NM_001271486,NM_053076,NM_001003934 |
| Mir3060 | 12.52381966 | 0.29724734 | 0.162496035 | 1.829259035 | 0.067360812 | [illegible] | | Tjp1 | 4685.570654 | -0.0767137753792439 | 0.0646301684471996 | -1.18696542531707 | 0.235241258594122 | 0.686276190501263 | NM_009386,NM_001163574 |
| Pnkd | 7470.073027 | 0.144463812 | 0.078981473 | 1.829084811 | 0.067386904 | [illegible] | | Pth2r | 56.227155 | 0.104541213787625 | 0.0881207231558353 | 1.18634085200086 | 0.235487719785084 | 0.686335196147297 | NM_139270 |
| Ptprn2 | 16650.61737 | -0.104549355 | 0.057186739 | -1.82820976 | 0.067518077 | [illegible] | | Cops8 | 3852.302818 | 0.0522414756025524 | 0.0440577492901541 | 1.18574998596733 | 0.235721047997518 | 0.686877420905104 | NM_133805 |
| Agtpbp1 | 7828.426726 | -0.132185532 | 0.072313022 | -1.827963042 | 0.067555099 | [illegible] | | Athl1 | 297.5027925 | -0.108136372798233 | 0.0912832971881486 | -1.18462387018457 | 0.236166194867009 | 0.687039130954991 | NM_145387 |
| Gm4532 | 32.62562595 | 0.303912691 | 0.166256364 | 1.827976293 | 0.06755311 | [illegible] | | Cdkl3 | 368.2099831 | -0.1045327053429 | 0.0882375755144155 | -1.18467336317307 | 0.236146618100364 | 0.687039130954991 | NM_153785,NM_001166653,NM_001166655,NM_001166654,NM_001166657,NM_001166656 |
| Gm6277 | 128.5533372 | 0.23542559 | 0.1287997 | 1.827842693 | 0.067573164 | [illegible] | | Dad1 | 2049.219719 | 0.0580265927534587 | 0.0489812830785636 | 1.18466869600755 | 0.236148464131227 | 0.687039130954991 | NM_010015,NM_001113358 |
| Dleu2 | 126.1663542 | -0.23774803 | 0.130083839 | -1.827652314 | 0.067601749 | [illegible] | | Gpr98 | 79.6564586 | 0.112336858145523 | 0.094770486489161 | 1.18535698514507 | 0.235876331395343 | 0.687039130954991 | NM_054053 |
| Ebf2 | 32.69457032 | -0.307123054 | 0.168094373 | -1.827087057 | 0.067686681 | [illegible] | | Kcnip3 | 6584.170717 | 0.0706113299509119 | 0.0595761040817243 | 1.18522906187437 | 0.235926892343782 | 0.687039130954991 | NM_001111331,NM_019789 |
| Lrrc4c | 4524.979428 | -0.13881811 | 0.075998458 | -1.826591142 | 0.067761266 | [illegible] | | Mlip | 1133.432614 | -0.0808253963086273 | 0.06823398720673 | -1.18453280567863 | 0.236202218090956 | 0.687039130954991 | NM_027150 |
| Spag17 | 21.12215067 | 0.305896151 | 0.167464861 | 1.826628879 | 0.067755588 | [illegible] | | Slc25a20 | 1446.197791 | 0.0632396123873923 | 0.0533793483731482 | 1.18472057667913 | 0.236127944044209 | 0.687039130954991 | NM_020520 |
| Dnmt3a | 3630.299426 | -0.124700538 | 0.068285195 | -1.826172391 | 0.067824299 | [illegible] | | Stambp | 1484.918157 | 0.0637437837287036 | 0.053770539823698 | 1.18547784600463 | 0.235828568867964 | 0.687039130954991 | NM_024239 |
| Sssca1 | 496.8088328 | -0.146431418 | 0.080181928 | -1.826239676 | 0.067814167 | [illegible] | | Tmprss6 | 24.85148337 | 0.0953501285447747 | 0.0804921559806269 | 1.18458907434066 | 0.236179958920196 | 0.687039130954991 | NM_027902 |
| Il13ra1 | 284.5274408 | -0.203399808 | 0.111407287 | -1.825731627 | 0.067890697 | [illegible] | | Socs2 | 488.6314499 | 0.0882345643420646 | 0.0745061665472759 | 1.18425854437267 | 0.236310733633764 | 0.687217160513325 | NM_001168657,NM_001168656,NM_001168655,NM_007706 |
| Rasl10a | 1100.205473 | -0.228133168 | 0.124973421 | -1.825453496 | 0.067932623 | [illegible] | | Slx4ip | 323.9713199 | -0.0928055404918162 | 0.0783739773495589 | -1.1841371795882 | 0.236358764656901 | 0.687219258488019 | NM_028201,NM_001038641,NM_028834 |
| Emc2 | 1929.570916 | -0.177658102 | 0.097332885 | -1.825262868 | 0.067961371 | [illegible] | | Myo1b | 1935.035603 | -0.0775550156720177 | 0.0655038132722213 | -1.18397711213718 | 0.236422123106704 | 0.687227575729535 | NM_001161817,NM_010863 |
| Fcho1 | 3799.827286 | 0.168307397 | 0.092268192 | 1.824110711 | 0.068135337 | [illegible] | | Numa1 | 4603.519742 | -0.0697355883793069 | 0.0589037277819384 | -1.18389091837223 | 0.236456245593846 | 0.687227575729535 | NM_133947 |
| Ndufc1 | 1115.936163 | -0.222307805 | 0.12187734 | -1.824029024 | 0.068147685 | [illegible] | | Acer2 | 396.7680386 | 0.0888938765680391 | 0.0751134341933116 | 1.18346175384902 | 0.236626195575044 | 0.687308961592298 | NM_139306 |
| Atp5d | 5227.551254 | -0.138125865 | 0.07573787 | -1.82373581 | 0.068192022 | [illegible] | | BC052040 | 327.0796409 | 0.0919846845418117 | 0.0777192178829498 | 1.18355134093535 | 0.236590711785576 | 0.687308961592298 | NM_207264,NM_001145898 |
| Mettl13 | 443.3139483 | 0.1828547 | 0.100278692 | 1.823465143 | 0.068232972 | [illegible] | | Smim14 | 4212.661797 | 0.0584253115634739 | 0.049362949894189 | 1.18358630691055 | 0.236576863429668 | 0.687308961592298 | NM_133697 |
| Tfam | 1232.283988 | -0.104974753 | 0.057576242 | -1.82323039 | 0.068268504 | [illegible] | | Trafd1 | 2338.666339 | -0.0806988082698783 | 0.0682018065793643 | -1.1832356401875 | 0.236715771749877 | 0.687431687412777 | NM_172275,NM_001163470 |
| Pcdhga10 | 388.1040694 | -0.17971501 | 0.098583764 | -1.822967618 | 0.068308296 | [illegible] | | Nes | 525.7631733 | -0.106990092938253 | 0.0904315001563071 | -1.18310647012739 | 0.23676695393918 | 0.687442888850795 | NM_016701 |
| Il1b | 13.10329417 | 0.299700174 | 0.164421222 | 1.822758461 | 0.068339982 | [illegible] | | Ccdc14 | 153.8660788 | -0.104460801289935 | 0.0883040540997339 | -1.1829672188318 | 0.23682213946706 | 0.687465707014092 | NM_172824 |
| Ube3a | 1897.642477 | -0.160274013 | 0.087960477 | -1.822113949 | 0.068437698 | [illegible] | | Ubac1 | 1611.599458 | 0.0588815490993357 | 0.0497875002677582 | 1.18265726904684 | 0.236945005739391 | 0.68768494572835 | NM_133835 |
| Lifr | 1642.096512 | 0.164788054 | 0.090458205 | 1.821703788 | 0.068499943 | [illegible] | | Yeats4 | 1081.547413 | 0.0812752143813985 | 0.0687431570849768 | 1.18230261494872 | 0.237085648376679 | 0.687955678849307 | NM_026570 |
| Rab31 | 3468.811264 | 0.136574205 | 0.074972035 | 1.821668644 | 0.068505278 | [illegible] | | Erlin2 | 2559.467494 | 0.0640782335575146 | 0.0542325978095415 | 1.18154460869733 | 0.237386443258763 | 0.688415946252103 | NM_153592 |
| Adap2 | 198.4606307 | 0.187790893 | 0.10313186 | 1.820881461 | 0.068624878 | [illegible] | | Ermap | 46.8852731 | -0.11116784802412 | 0.0940862229435534 | -1.18155288358015 | 0.237383158133815 | 0.688415946252103 | NM_013848 |
| Sall1 | 1305.996014 | -0.217735005 | 0.119577007 | -1.820876869 | 0.068625576 | [illegible] | | Smarca2 | 12135.33654 | 0.0706062087968261 | 0.0597521904598848 | 1.18165055127524 | 0.237344386527277 | 0.688415946252103 | NM_011416,NM_026003 |
| Prep | 926.9609521 | -0.125408852 | 0.068881812 | -1.8206381 | 0.068661887 | [illegible] | | Sult2b1 | 302.7605581 | 0.106962003305395 | 0.090538528229562 | 1.18139763697275 | 0.237444796322245 | 0.688447726821743 | NM_017465 |
| Arhgap29 | 1000.002038 | -0.14808598 | 0.081343248 | -1.820507348 | 0.068681778 | [illegible] | | B9d1 | 309.1032205 | -0.0924544294458496 | 0.0782948836266745 | -1.18084892860548 | 0.237662742826846 | 0.688529916693811 | NM_013717 |
| Ephb1 | 1882.668525 | -0.195263892 | 0.107266795 | -1.820357292 | 0.068704612 | [illegible] | | Mrc1 | 149.5262205 | -0.112311446634378 | 0.0950947147513456 | -1.1810482520301 | 0.237583555393506 | 0.688529916693811 | NM_008625 |
| A230065H16Rik | 45.1085192 | -0.292660974 | 0.160838206 | -1.819598602 | 0.068820155 | [illegible] | | Slc6a9 | 2515.510375 | 0.0909248154710448 | 0.0769952309520799 | 1.18091490014017 | 0.237636531517922 | 0.688529916693811 | NM_008135 |
| C030023E24Rik | 89.62456163 | 0.271605076 | 0.149268528 | 1.819573618 | 0.068823962 | [illegible] | | Stab2 | 28.11291031 | 0.103406030396536 | 0.0875603042030584 | 1.18096929125247 | 0.237614922790199 | 0.688529916693811 | NM_138673 |
| Klhl33 | 111.2078045 | 0.245443127 | 0.134868183 | 1.819874202 | 0.068778164 | [illegible] | | Gadd45b | 426.8603743 | 0.110789900214562 | 0.0938367111960012 | 1.18066691386007 | 0.237735070131541 | 0.688602119387988 | NM_008655 |
| Rnf20 | 2546.767764 | 0.106864631 | 0.058723358 | 1.819797692 | 0.068789819 | [illegible] | | Frem1 | 172.8655989 | -0.109718560447585 | 0.0929688827798689 | -1.18016434280893 | 0.237934857800806 | 0.688758313982319 | NM_177863,NM_001198811 |
| Tcp11 | 15.6994748 | 0.301670042 | 0.165742421 | 1.820113648 | 0.0687417 | [illegible] | | Lrfn2 | 1095.753781 | 0.103959864516203 | 0.0881064343347811 | 1.1799349877352 | 0.238026072985766 | 0.688758313982319 | NM_027452 |
| Tcp11l1 | 474.8577324 | -0.17207038 | 0.094546939 | -1.819946601 | 0.068767137 | [illegible] | | Mall | 34.37002237 | 0.0975238981506637 | 0.0826481938233607 | 1.17998825671975 | 0.238004885553445 | 0.688758313982319 | NM_145532 |
| Tlr13 | 142.6334863 | 0.215177165 | 0.118250469 | 1.819672829 | 0.068808843 | [illegible] | | Reln | 2766.245626 | -0.0787061402865927 | 0.0666851448239534 | -1.18026496747326 | 0.237894846868228 | 0.688758313982319 | NM_011261 |
| Pou6f1 | 5530.442943 | 0.139318 | 0.076606063 | 1.818628886 | 0.068968068 | [illegible] | | Trim11 | 843.3912074 | -0.0863378430715135 | 0.073168494945819 | -1.17998659307461 | 0.238005547238591 | 0.688758313982319 | NM_053168 |
| Uck2 | 421.0909984 | -0.172919845 | 0.095100493 | -1.81828548 | 0.069020511 | [illegible] | | Abcd3 | 3023.997832 | 0.0641386840055072 | 0.0543642292053307 | 1.17979570285562 | 0.238081479039467 | 0.688781430895697 | NM_008991 |
| Basp1 | 12091.76112 | -0.159252378 | 0.087591188 | -1.818132406 | 0.069043898 | [illegible] | | Usp21 | 1660.797329 | -0.0719899563284925 | 0.0610319842457975 | -1.17954474556428 | 0.238181330181777 | 0.688933094826859 | NM_013919 |
| Pigz | 319.8683362 | 0.287220481 | 0.157997659 | 1.817878077 | 0.06908277 | [illegible] | | Prpf38a | 686.1645718 | 0.082804125794024 | 0.070211859054152 | 1.17934672161522 | 0.23826014101832 | 0.689023851967374 | NM_172697 |
| Ppapdc1a | 148.8631456 | 0.214553355 | 0.118027043 | 1.817832162 | 0.069089789 | [illegible] | | Mettl6 | 585.084674 | 0.0829970823555479 | 0.0704109524925952 | 1.17875244429162 | 0.23849676582173 | 0.689459925903681 | NM_025907 |
| Mrps12 | 940.9270518 | -0.136092749 | 0.074880723 | -1.817460396 | 0.069146647 | [illegible] | | Nrxn3 | 8655.758785 | 0.0699799342709327 | 0.0593749475265697 | 1.17861046091228 | 0.238553324218085 | 0.689459925903681 | NM_172544,NM_001198587,NM_001252074 |
| Qtrtd1 | 502.8900693 | 0.199208259 | 0.109631609 | 1.817069552 | 0.069206464 | [illegible] | | Snx25 | 2636.958146 | 0.0759543795735056 | 0.0644399744894136 | 1.17868419680991 | 0.238523950696479 | 0.689459925903681 | NM_207213 |
| Mmp9 | 54.49764597 | 0.295826007 | 0.16282162 | 1.816871782 | 0.069236748 | [illegible] | | Casp1 | 137.9201493 | -0.108209769630465 | 0.0919050891224718 | -1.17740780911779 | 0.239032774468581 | 0.689537017427783 | NM_009807 |
| Ptprd | 8954.94578 | 0.115464711 | 0.063555474 | 1.816754762 | 0.069254672 | [illegible] | | Fbll1 | 855.920131 | 0.0835685519798351 | 0.0709415522309705 | 1.17799159098963 | 0.238799958625729 | 0.689537017427783 | NM_001004147 |
| Cby1 | 1139.546193 | -0.104396726 | 0.057473331 | -1.816437712 | 0.069303255 | [illegible] | | Gdpgp1 | 396.8884283 | 0.0962999595499874 | 0.0817811202839921 | 1.17753289775901 | 0.238982874865585 | 0.689537017427783 | NM_178752 |
| Ryr1 | 957.7133464 | -0.261121914 | 0.143910946 | -1.814468753 | 0.069605589 | [illegible] | | Igsf1 | 408.7876096 | 0.0723517805876356 | 0.061429299503066 | 1.17780572418906 | 0.238874066094128 | 0.689537017427783 | NM_177591,NM_183336,NM_183335,NM_177915 |
| Tnks2 | 6422.241663 | -0.127708987 | 0.0704325 | -1.813211039 | 0.069799278 | [illegible] | | Nfe2l2 | 757.9884638 | -0.0887456545268746 | 0.0753123123761454 | -1.17836847292162 | 0.238649740784345 | 0.689537017427783 | NM_010902 |
| Sdf2l1 | 276.0709188 | -0.243203096 | 0.134137292 | -1.813090845 | 0.069817811 | [illegible] | | Pcbp3 | 2295.292946 | 0.070039119139438 | 0.0594642753011656 | 1.17783524283638 | 0.238862295548498 | 0.689537017427783 | NM_021568 |
| Asah1 | 2023.310133 | 0.131641504 | 0.072622415 | 1.812684209 | 0.069880541 | [illegible] | | Rab12 | 3414.613538 | 0.0619600696464903 | 0.0526125262619923 | 1.17766764017281 | 0.23892913246026 | 0.689537017427783 | NM_024448 |
| Retsat | 420.6220446 | 0.256372433 | 0.141432252 | 1.812687205 | 0.069880079 | [illegible] | | Snap25 | 116859.6071 | 0.0734461694993691 | 0.062382460569191 | 1.17735287818452 | 0.239054689506149 | 0.689537017427783 | NM_011428 |
| Acap3 | 6236.515341 | 0.200764211 | 0.110766488 | 1.81249956 | 0.069909042 | [illegible] | | Ube2l3 | 7198.042473 | 0.053504365981464 | 0.0454088127759115 | 1.17828154295738 | 0.238684383466878 | 0.689537017427783 | NM_009456 |
| Tjp1 | 4623.461295 | -0.085007727 | 0.046906179 | -1.812292753 | 0.069940974 | [illegible] | | Zfp2 | 951.7781392 | -0.0808918055498843 | 0.0686845046502141 | -1.17773005660938 | 0.238904240372158 | 0.689537017427783 | NM_001044698,NM_001044699,NM_001044700,NM_001044697 |
| Arid1b | 3767.703507 | -0.129924078 | 0.071698182 | -1.81209723 | 0.069971174 | [illegible] | | Caln1 | 2814.775874 | 0.0941511702804398 | 0.0799863638676133 | 1.17709026548914 | 0.239159480060699 | 0.689702324272725 | NM_181045,NM_021371 |
| Ms4a7 | 15.27995935 | -0.294099683 | 0.162295311 | -1.812126795 | 0.069966607 | [illegible] | | Ywhaz | 47863.67434 | 0.0667548924520477 | 0.0567194211419506 | 1.17693183583418 | 0.239222714040933 | 0.689747745962406 | NM_001253805,NM_001253807,NM_001253806,NM_011740 |
| Tspan5 | 8568.557634 | -0.134725605 | 0.074359785 | -1.811807339 | 0.070015971 | [illegible] | | Arv1 | 204.5755036 | -0.101046435678946 | 0.0859230539125284 | -1.17601075704099 | 0.239590577520402 | 0.690203878384272 | NM_026855 |
| Ccdc63 | 12.99156708 | 0.25812517 | 0.142475623 | 1.811714628 | 0.070030302 | [illegible] | | Kdr | 510.7818089 | -0.110325837862324 | 0.0938005545166454 | -1.17617468714161 | 0.239525077416855 | 0.690203878384272 | NM_010612 |
| Ebp | 562.7961153 | -0.136795433 | 0.075514733 | -1.811506533 | 0.070062479 | [illegible] | | Osbpl10 | 1283.052683 | 0.0727062391584882 | 0.0618225289857991 | 1.1760476375075 | 0.239575840416936 | 0.690203878384272 | NM_148958 |
| Zcchc9 | 374.5193095 | 0.163005773 | 0.089999675 | 1.81118179 | 0.070112716 | [illegible] | | Tceanc2 | 811.2170427 | 0.0739422037906641 | 0.0628791829112999 | 1.17594091346528 | 0.239618488137952 | 0.690203878384272 | NM_025617 |
| Lsm1 | 466.1104349 | -0.13064203 | 0.072147882 | -1.810753499 | 0.070179017 | [illegible] | | Unc13c | 2723.02985 | 0.111718814220265 | 0.0949983680067729 | 1.17600772059895 | 0.239591790883872 | 0.690203878384272 | NM_001081153 |
| Il17d | 208.1002727 | -0.184210522 | 0.101779137 | -1.809904536 | 0.070310593 | [illegible] | | Csde1 | 9247.926197 | 0.0681563015573892 | 0.0579940093505822 | 1.17523003359493 | 0.239902697600194 | 0.690751361057437 | NM_144901,NM_001161854 |
| Depdc7 | 43.41762778 | 0.279870471 | 0.154675101 | 1.809408689 | 0.070387534 | [illegible] | | Ndfip2 | 4230.045981 | 0.0885010997140296 | 0.0753117113578239 | 1.17513064194677 | 0.23994245326421 | 0.690751361057437 | NM_001190989,NM_029561 |
| Rp2h | 433.8713756 | -0.157173745 | 0.086864952 | -1.809403463 | 0.070388345 | [illegible] | | Safb | 4065.979174 | -0.0657836933201059 | 0.0559809414561357 | -1.17510873538365 | 0.239951216294632 | 0.690751361057437 | NM_001163300 |
| Akr1c18 | 56.15791418 | 0.270253969 | 0.149368467 | 1.80931072 | 0.070402745 | [illegible] | | Dyrk2 | 684.1796817 | 0.0912210313088467 | 0.0776442675709262 | 1.1748585460674 | 0.240051312649089 | 0.690902589169937 | NM_001014390 |
| Car12 | 939.8742721 | -0.203626327 | 0.11255609 | -1.809109814 | 0.070433945 | [illegible] | | Pdk3 | 1237.852272 | 0.0776185271854944 | 0.0661109265785638 | 1.17406503285437 | 0.240368978040734 | 0.691466899864115 | NM_145630 |
| Nudt13 | 390.4019595 | 0.172487548 | 0.095358954 | 1.808823817 | 0.070478379 | [illegible] | | Pik3c2b | 1106.362243 | -0.0860563556067038 | 0.0733010817407516 | -1.17401208226455 | 0.240390186170576 | 0.691466899864115 | NM_001099276 |
| Tnks1bp1 | 3512.590875 | 0.166285906 | 0.091955706 | 1.808326125 | 0.070555758 | [illegible] | | Zfp362 | 1156.497971 | -0.0710955785391691 | 0.0605493609589674 | -1.17417553898461 | 0.240324721621812 | 0.691466899864115 | NM_001081098 |
| Fam120a | 5814.374724 | -0.089348145 | 0.04941354 | -1.808171313 | 0.070579842 | [illegible] | | Trappc3 | 2255.91596 | 0.0596036908903559 | 0.0507863904898921 | 1.17361541774107 | 0.240549102873571 | 0.691787026003066 | NM_013718 |
| Fam210a | 1709.962413 | -0.109942583 | 0.060816158 | -1.807785718 | 0.070639857 | [illegible] | | Abcg3 | 23.78328439 | -0.0896480937223672 | 0.0764509963897483 | -1.17262165250718 | 0.240947562472274 | 0.691863634771044 | NM_030239 |
| Cst6 | 148.1844828 | -0.208011676 | 0.115079417 | -1.807548921 | 0.070676734 | [illegible] | | C230004F18Rik | 179.8341295 | -0.101302455876618 | 0.0863758148134623 | -1.17281042263267 | 0.24087183755965 | 0.691863634771044 | NR_030706 |
| Trmt61a | 696.1599846 | -0.212348806 | 0.117486298 | -1.807434644 | 0.070694536 | [illegible] | | Csk | 1182.876189 | 0.0643249670303145 | 0.0548343128453231 | 1.17307874745805 | 0.240764228221441 | 0.691863634771044 | NM_007783 |
| BC003965 | 943.7624337 | -0.158273294 | 0.087638484 | -1.80597937 | 0.070921562 | [illegible] | | Eif1b | 3107.627447 | 0.0760392727068619 | 0.0648130784321012 | 1.17320878048589 | 0.240712091789147 | 0.691863634771044 | NM_026892 |
| Dopey1 | 2445.400882 | 0.148205057 | 0.082042526 | 1.806441911 | 0.07084934 | [illegible] | | Kctd12b | 176.3092853 | -0.10208258452337 | 0.0870453217074597 | -1.1727521079932 | 0.24089522861892 | 0.691863634771044 | NM_175429 |
| Gm14169 | 344.1799889 | 0.195286282 | 0.108130094 | 1.806030826 | 0.070913525 | [illegible] | | Mbtps1 | 4206.309889 | 0.05176361223531 | 0.0441319964012298 | 1.17292704741242 | 0.240825062044594 | 0.691863634771044 | NM_001167910,NM_019709 |
| Gpr30 | 135.3860865 | -0.208085434 | 0.115217685 | -1.806019916 | 0.070915229 | [illegible] | | Mrps7 | 1489.562191 | 0.0674407141516864 | 0.0575138918981269 | 1.17259868748132 | 0.240956776008999 | 0.691863634771044 | NM_025305 |
| Hypk | 989.0446584 | -0.107361136 | 0.059439825 | -1.806215536 | 0.070884679 | [illegible] | | Scn7a | 215.7009389 | -0.110459492032454 | 0.0941904131703871 | -1.17272542198787 | 0.240905933393079 | 0.691863634771044 | NM_009135 |
| Insig1 | 2788.342048 | -0.206925112 | 0.114533735 | -1.806673927 | 0.070813135 | [illegible] | | Adamts17 | 237.0262788 | -0.10212757305004 | 0.0872159023833205 | -1.1709742175364 | 0.241609140732271 | 0.692393570570603 | NM_001033877 |
| Mecr | 430.2632265 | 0.178319266 | 0.098714091 | 1.806421593 | 0.070852511 | [illegible] | | Cmss1 | 339.4917291 | 0.0864757932253803 | 0.0738562685545104 | 1.17086599306809 | 0.24165264631122 | 0.692393570570603 | NM_025599 |
| Ppp1r9a | 16974.30638 | -0.156462478 | 0.086618937 | -1.806331081 | 0.07086664 | [illegible] | | Ddit4l | 1349.198628 | -0.097872341539401 | 0.0835357588223186 | -1.17162210434428 | 0.241348809454608 | 0.692393570570603 | NM_030143 |
| Samhd1 | 728.2284059 | 0.148788861 | 0.082370271 | 1.80634176 | 0.070864973 | [illegible] | | Hdlbp | 9996.648839 | 0.0498895534662741 | 0.0425739840508327 | 1.1718319198576 | 0.241264544573062 | 0.692393570570603 | NM_133808 |
| Slc7a4 | 2729.214994 | -0.149560265 | 0.082812141 | -1.806018573 | 0.070915438 | [illegible] | | Pgap1 | 1532.732071 | 0.0922097916541315 | 0.0787292122892158 | 1.17122715918196 | 0.241507481226608 | 0.692393570570603 | NM_001163314 |
| Gripap1 | 4069.472909 | 0.105299179 | 0.058314888 | 1.805699769 | 0.070965249 | [illegible] | | Rftn2 | 1222.204383 | -0.0740189141753073 | 0.0632007581404734 | -1.17117130162883 | 0.241529928285187 | 0.692393570570603 | NM_028713 |
| C630043F03Rik | 218.6019754 | -0.169619995 | 0.093961455 | -1.805208259 | 0.071042099 | [illegible] | | Sik1 | 806.0338662 | 0.0977940306364603 | 0.08345690635165 | 1.17179074700421 | 0.241281078540338 | 0.692393570570603 | NM_010831 |
| Ankrd13a | 1759.749148 | 0.165266704 | 0.091597802 | 1.804264954 | 0.071189781 | [illegible] | | Smad9 | 818.0579377 | 0.080954760782661 | 0.0691090180741522 | 1.17140661289383 | 0.241435375438051 | 0.692393570570603 | NM_019483 |
| St3gal4 | 811.2725644 | -0.213668586 | 0.11845671 | -1.803769378 | 0.071267468 | [illegible] | | Ssbp2 | 3899.289479 | 0.0737026271040739 | 0.0629488476756151 | 1.17083361849409 | 0.241665661764626 | 0.692393570570603 | NM_024186,NM_024272 |
| Ticam2 | 32.74025237 | 0.287790444 | 0.159536005 | 1.803921588 | 0.0712436 | [illegible] | | Stam2 | 1229.189055 | 0.0758757670487249 | 0.0647861447217983 | 1.17117274649614 | 0.241529347628629 | 0.692393570570603 | NM_019667 |
| Txnrd3 | 205.7562633 | -0.201720817 | 0.111829378 | -1.803826697 | 0.071258479 | [illegible] | | Yipf4 | 1230.732479 | 0.086832289641833 | 0.0741601454943619 | 1.17087539490378 | 0.241648866611705 | 0.692393570570603 | NM_026417 |
| Tha1 | 127.5549731 | 0.263038811 | 0.145835559 | 1.803667173 | 0.071283499 | [illegible] | | Aftph | 3788.783783 | 0.0826565495190776 | 0.0706065359621427 | 1.17066427906045 | 0.241733748831472 | 0.692421614793766 | NM_181411,NM_001252503 |
| Tbc1d8 | 1332.128332 | -0.136452016 | 0.0756564 | -1.803575324 | 0.071297907 | [illegible] | | Scd1 | 8228.779937 | 0.0961914484399007 | 0.0821747262999134 | 1.17057219919814 | 0.241770785504198 | 0.692421614793766 | NM_009127 |
| Zcchc8 | 1010.072386 | 0.143142743 | 0.07939945 | 1.802817818 | 0.071416831 | [illegible] | | 2610318N02Rik | 34.96250787 | 0.105750638060573 | 0.090476797277207 | 1.16881500277436 | 0.242478174443835 | 0.692772033309095 | NM_183287 |
| Dhcr7 | 1275.772649 | -0.189293831 | 0.105008725 | -1.802648605 | 0.071443419 | [illegible] | | Ahnak | 1507.113792 | 0.10755864524331 | 0.0921687790528026 | 1.16697482974892 | 0.243220534452765 | 0.692772033309095 | NM_001039959,NM_009643 |
| Pcdha5 | 157.4569188 | -0.208283486 | 0.115552977 | -1.802493471 | 0.071467801 | [illegible] | | Arl2 | 1753.651004 | 0.0661978127850897 | 0.0565983406172381 | 1.16960695425279 | 0.24215917746687 | 0.692772033309095 | NM_019722 |
| 2210404J11Rik | 207.4924549 | 0.252900032 | 0.140364909 | 1.801732601 | 0.071587486 | [illegible] | | Cdca8 | 45.5225276 | -0.106948394949938 | 0.0916557187554452 | -1.1668491219331 | 0.243271305545905 | 0.692772033309095 | NM_026560 |
| Cebpd | 66.40139963 | 0.294210054 | 0.163287342 | 1.801793399 | 0.071577917 | [illegible] | | Cenpq | 132.8411834 | 0.10961615324875 | 0.0939231405289156 | 1.16708356036075 | 0.243176626149243 | 0.692772033309095 | NM_031863 |
| Cept1 | 1186.752667 | -0.135487793 | 0.075193828 | -1.801847255 | 0.071569441 | [illegible] | | Chrdl1 | 756.5385378 | -0.0959950779515826 | 0.0822004515511323 | -1.16781691754904 | 0.242880622582247 | 0.692772033309095 | NM_031258,NM_001114385 |
| Eng | 1088.065793 | -0.152174425 | 0.084451817 | -1.801908251 | 0.071559842 | [illegible] | | Crybb1 | 120.631174 | 0.110495191005968 | 0.094584868369523 | 1.16821213488701 | 0.242721206840252 | 0.692772033309095 | NM_023695 |
| Mettl4 | 287.6794777 | -0.202374631 | 0.112307719 | -1.801965464 | 0.071550839 | [illegible] | | Dok6 | 550.4432998 | -0.0918639336212148 | 0.0785373696247367 | -1.1696843688572 | 0.24212801081693 | 0.692772033309095 | NM_001039173 |
| Mitf | 354.9048011 | -0.200906085 | 0.111514429 | -1.801615151 | 0.071605976 | [illegible] | | Drg1 | 2757.967359 | 0.0565765396556755 | 0.0484606600841529 | 1.16747356634081 | 0.243019177227399 | 0.692772033309095 | NM_007879 |
| Rab3b | 2491.016844 | -0.151044999 | 0.083870021 | -1.800941484 | 0.071712104 | [illegible] | | Dus4l | 198.0370166 | -0.105124012617805 | 0.089981583782217 | -1.16828364426478 | 0.242692370520925 | 0.692772033309095 | NM_028002 |
| Tmem176a | 435.6672337 | 0.203299998 | 0.11288579 | 1.800935242 | 0.071713087 | [illegible] | | Eif2b2 | 1169.748277 | 0.0627528100615982 | 0.0537790728054925 | 1.16686299684938 | 0.243265701354731 | 0.692772033309095 | NM_145445 |
| Acaca | 4652.113352 | -0.144226353 | 0.080124318 | -1.800032204 | 0.071855553 | [illegible] | | Faxc | 5801.365171 | 0.0830047844499946 | 0.0710554254144369 | 1.16816955166846 | 0.242738379764453 | 0.692772033309095 | NM_175234 |
| Anapc7 | 1392.99074 | 0.124698562 | 0.06927925 | 1.799940992 | 0.071869956 | [illegible] | | Gramd2 | 184.8347472 | -0.109587732382878 | 0.0939462350143291 | -1.16649413748367 | 0.243414717500141 | 0.692772033309095 | NM_001033498 |
| Dag1 | 5137.596882 | -0.108576958 | 0.060324173 | -1.799891357 | 0.071877795 | [illegible] | | Hdgf | 6494.240487 | 0.0450381595116327 | 0.038614776654902 | 1.16634520287754 | 0.243474904028673 | 0.692772033309095 | NM_008231 |
| Icam1 | 124.6123467 | 0.226280155 | 0.125728911 | 1.7997464 | 0.071900691 | [illegible] | | Hs6st3 | 542.3178099 | 0.0915563905411953 | 0.0784748082968305 | 1.16669785537906 | 0.243332409220185 | 0.692772033309095 | NM_015820 |
| Trp53inp2 | 7270.061936 | -0.116579594 | 0.064796686 | -1.799159803 | 0.071993406 | [illegible] | | Isl1 | 104.568127 | 0.104110595728412 | 0.0892681481610318 | 1.16626812444464 | 0.243506056592361 | 0.692772033309095 | NM_021459 |
| Irx6 | 10.11734141 | -0.294792748 | 0.163866147 | -1.798985042 | 0.072021047 | [illegible] | | Maz | 5859.916412 | -0.0669691292840715 | 0.0573439713032483 | -1.16784951865163 | 0.242867469744359 | 0.692772033309095 | NM_010772 |
| Rin2 | 1353.983476 | 0.15508627 | 0.086223724 | 1.798649643 | 0.072074119 | [illegible] | | Mff | 6757.570028 | 0.0599909591581112 | 0.0513181437628265 | 1.16900095676428 | 0.242403245893862 | 0.692772033309095 | NM_029409 |
| Efna2 | 223.3869736 | -0.176778395 | 0.098300295 | -1.798350608 | 0.072121465 | [illegible] | | Otud7b | 2952.429699 | 0.0851990627209185 | 0.0729544129840002 | 1.1678397404089 | 0.242871414700735 | 0.692772033309095 | NM_001025614,NM_001025613 |
| Plcd4 | 623.9555561 | 0.230022035 | 0.127942576 | 1.797853705 | 0.072200194 | [illegible] | | Pcdhgb6 | 652.2091989 | 0.0843849721651527 | 0.0721969917471059 | 1.16881562684411 | 0.242477922953094 | 0.692772033309095 | NM_033578 |
| 2810433D01Rik | 9.490779877 | -0.284828909 | 0.158459124 | -1.797491374 | 0.072257646 | [illegible] | | Pim1 | 123.7989194 | 0.111023487665517 | 0.0950812151063937 | 1.16767005492394 | 0.242939880170856 | 0.692772033309095 | NM_008842 |
| Sprn | 7659.373044 | -0.164744405 | 0.091675066 | -1.797047022 | 0.072328155 | [illegible] | | Ppef2 | 26.65041848 | -0.0975979846382859 | 0.0835118678608407 | -1.16867203594245 | 0.242535792765366 | 0.692772033309095 | NM_011148 |
| Txlna | 2141.476462 | 0.122646427 | 0.068252895 | 1.79694103 | 0.072344982 | [illegible] | | Riok3 | 2489.775814 | -0.053455971073811 | 0.0457957277677929 | -1.16726982361454 | 0.243101421056694 | 0.692772033309095 | NM_024182 |
| Creb5 | 82.11932191 | 0.276062391 | 0.15364232 | 1.796786139 | 0.072369578 | [illegible] | | Ripk2 | 196.4209525 | -0.101161928378716 | 0.0864873134356885 | -1.16967361292751 | 0.242132340920142 | 0.692772033309095 | NM_138952 |
| A330048O09Rik | 35.85012277 | -0.286253595 | 0.159379273 | -1.796052833 | 0.072486115 | [illegible] | | Sesn1 | 2731.704333 | 0.0683342624075574 | 0.0584420735991118 | 1.16926484977761 | 0.242296940444916 | 0.692772033309095 | NM_001162908,NM_001013370 |
| Lmln | 482.4323135 | 0.175087779 | 0.097480622 | 1.796129059 | 0.072473994 | [illegible] | | Slc35d3 | 258.2641998 | 0.106844247655853 | 0.0914699409096046 | 1.16808042722409 | 0.24277432455971 | 0.692772033309095 | NM_029529 |
| Ctsf | 2235.774225 | 0.12834959 | 0.071470109 | 1.795849938 | 0.072518386 | [illegible] | | Slc5a3 | 1060.568687 | 0.0860671727909585 | 0.0736886607043461 | 1.16798394716763 | 0.242813240166405 | 0.692772033309095 | NM_017391 |
| Rhoa | 5673.035807 | -0.066981468 | 0.037296701 | -1.795908616 | 0.072509052 | [illegible] | | Stk17b | 297.7872634 | -0.104605441128847 | 0.0894444331907426 | -1.16950197342939 | 0.24220144661768 | 0.692772033309095 | NM_133810 |
| Spint2 | 584.0856254 | 0.196822467 | 0.109611019 | 1.795644908 | 0.072551009 | [illegible] | | Tmem159 | 419.5400146 | 0.0935883120367323 | 0.0802458608302994 | 1.16626965015241 | 0.243505439924366 | 0.692772033309095 | NM_145586 |
| Fam193b | 3189.060558 | 0.250319872 | 0.139413497 | 1.795521073 | 0.072570718 | [illegible] | | Tmem17 | 191.8316393 | 0.0996361781416039 | 0.0853937387843317 | 1.16678552268618 | 0.243296994958479 | 0.692772033309095 | NM_153596 |
| Mier2 | 1251.130113 | 0.123846682 | 0.068983394 | 1.795311527 | 0.07260408 | [illegible] | | Trak1 | 8494.42346 | 0.076783014840251 | 0.0658377050636337 | 1.16624683023259 | 0.24351466350518 | 0.692772033309095 | NM_175114 |
| Cdk5rap1 | 346.3580878 | -0.233335046 | 0.129978187 | -1.795186184 | 0.072624041 | [illegible] | | Tug1 | 4490.365497 | 0.068588721328359 | 0.0586584332330983 | 1.1692900329574 | 0.242286797482743 | 0.692772033309095 | NR_002321,NR_002322 |
| Tmem52 | 16.76190222 | 0.289077849 | 0.161055541 | 1.794895397 | 0.072670367 | [illegible] | | Ube2k | 4452.124036 | 0.0717378852209789 | 0.0613960740579607 | 1.16844417695592 | 0.242627644121991 | 0.692772033309095 | NM_016786 |
| Zxdc | 1113.026313 | 0.132666312 | 0.073926414 | 1.794572532 | 0.072721833 | [illegible] | | Zfp385a | 3383.174848 | 0.0640103299215392 | 0.054777931106312 | 1.16854230579299 | 0.242588084786897 | 0.692772033309095 | NM_013866 |
| Arid5a | 402.4964374 | 0.245193778 | 0.136665037 | 1.794122213 | 0.072793664 | [illegible] | | Gm973 | 297.4700787 | -0.10430560368011 | 0.0894586126040351 | -1.16596491543851 | 0.243628630862928 | 0.692824880954363 | NM_001013771 |
| Slc29a2 | 482.166667 | 0.136083072 | 0.075861634 | 1.793832589 | 0.072839894 | [illegible] | | Med27 | 775.3672512 | 0.0841606248191271 | 0.072178956808432 | 1.16599946217698 | 0.243614662924926 | 0.692824880954363 | NM_026896 |
| Spns2 | 2413.318249 | 0.294936588 | 0.164424085 | 1.793755388 | 0.07285222 | [illegible] | | Ecsit | 1564.704508 | 0.0637502274265364 | 0.0547273561980481 | 1.16486948859426 | 0.244071825669027 | 0.693677820322499 | NM_001253898,NM_012029,NM_001253897 |
| Tcp11l2 | 1042.101507 | 0.156114009 | 0.087051908 | 1.793343914 | 0.07291795 | [illegible] | | Luzp2 | 6396.981403 | 0.0654689126278003 | 0.0561999053278541 | 1.16492923334788 | 0.244047639165597 | 0.693677820322499 | NM_178705 |
| D330045A20Rik | 12.48394589 | -0.292375158 | 0.163074466 | -1.79289355 | 0.072989947 | [illegible] | | Trim47 | 118.268768 | -0.107525362636571 | 0.0922895968303987 | -1.16508649218796 | 0.243983984024167 | 0.693677820322499 | NM_001205081,NM_172570 |
| Metrn | 1159.415383 | -0.243680497 | 0.135934137 | -1.792636518 | 0.073031063 | [illegible] | | Arid5a | 453.440432 | 0.100409561070899 | 0.0862151965891833 | 1.16463877649496 | 0.244165221784063 | 0.693692714478533 | NR_033310,NM_001172206,NM_001172205,NM_145996 |
| 1700010I14Rik | 39.20362389 | 0.294295457 | 0.164228593 | 1.791986719 | 0.073135093 | [illegible] | | Dysf | 223.9631151 | -0.1044156681347 | 0.0896563762006199 | -1.16462066123501 | 0.244172576698936 | 0.693692714478533 | NM_001077694,NM_021469 |
| Nudt4 | 5753.555696 | 0.134523403 | 0.075075622 | 1.791838677 | 0.073158811 | [illegible] | | F730043M19Rik | 304.5455292 | -0.0939881134626361 | 0.0807278414580366 | -1.16425896896416 | 0.244319079182 | 0.693973199882232 | NR_015602 |
| Dcaf11 | 3429.380127 | 0.110007429 | 0.061406978 | 1.791448343 | 0.073221377 | [illegible] | | BC005624 | 2975.489165 | 0.0580306613492426 | 0.0498629485541685 | 1.16380324533358 | 0.244503756669645 | 0.694361988149222 | NM_144885 |
| 1700020D05Rik | 33.30895522 | 0.296543604 | 0.165569081 | 1.791056651 | 0.073284205 | [illegible] | | Pgam5 | 1989.944492 | 0.0638749977355358 | 0.0548919370218525 | 1.16364991292086 | 0.244565915138227 | 0.694402752794739 | NM_028273,NM_001163538 |
| Pcdha9 | 131.5197065 | -0.211547008 | 0.118110339 | -1.791096443 | 0.07327782 | [illegible] | | Cep97 | 903.1019127 | -0.0906641436025997 | 0.0779415440878989 | -1.16323258236138 | 0.244735150344076 | 0.694476029282682 | NM_001159366,NM_001159365,NM_001159364,NM_028815 |
| Cd2ap | 926.1391301 | 0.177029037 | 0.098885809 | 1.790237035 | 0.073415814 | [illegible] | | Insc | 180.1991168 | -0.110563209240889 | 0.0950392646691583 | -1.16334243142317 | 0.244690596564303 | 0.694476029282682 | NM_173767 |
| Dzip1l | 891.7527709 | 0.188417453 | 0.10524079 | 1.790346234 | 0.073398268 | [illegible] | | Polr2e | 1712.271554 | 0.0660540143930456 | 0.0567836732315981 | 1.16325715886039 | 0.244725181845339 | 0.694476029282682 | NM_025554 |
| Gm973 | 244.1198557 | 0.226242566 | 0.126375623 | 1.790238972 | 0.073415502 | [illegible] | | BC005561 | 496.1957447 | -0.103552969574296 | 0.0890359420873942 | -1.16304682296338 | 0.244810505619966 | 0.694524329407536 | NM_001166581 |
| Gnb2 | 9511.257982 | -0.135920683 | 0.075935172 | -1.789956872 | 0.073460845 | [illegible] | | Ssr4 | 1073.851778 | -0.0655961869024232 | 0.0564047550279502 | -1.16295491168995 | 0.24484779642682 | 0.694524329407536 | NM_001166480,NM_009279 |
| Gpaa1 | 1297.802725 | 0.101538642 | 0.056708289 | 1.790543205 | 0.073366628 | [illegible] | | Fuca1 | 1187.416877 | 0.0734735320266937 | 0.0631876848541959 | 1.16278246617568 | 0.244917772836539 | 0.69456858400912 | NM_024243 |
| Gpld1 | 1918.831183 | 0.141416037 | 0.078989325 | 1.790318348 | 0.073402748 | [illegible] | | Tead3 | 83.96183618 | -0.110381342355451 | 0.0949369260311431 | -1.16268081314579 | 0.244959029043007 | 0.69456858400912 | NM_001098226,NM_011566,NM_001204156 |
| Gpr111 | 29.67823194 | 0.293877491 | 0.16412221 | 1.790601591 | 0.073357252 | [illegible] | | Tmbim6 | 13537.32555 | 0.0455913098250195 | 0.0392175643358718 | 1.1625227266681 | 0.245023198632878 | 0.694614946007257 | NM_001171035,NM_001171034,NM_001171036,NM_026669 |
| Hnrpdl | 5253.598948 | 0.18635293 | 0.104113273 | 1.789905604 | 0.073469088 | [illegible] | | Ap1g1 | 5333.316232 | 0.0683915205989752 | 0.0588470217454861 | 1.16219170606066 | 0.24515760289386 | 0.694718387691939 | NM_009677 |
| Hspd1 | 6431.176792 | -0.111463777 | 0.062242023 | -1.79081226 | 0.073323427 | [illegible] | | Cmtm8 | 39.76824599 | 0.10065101446175 | 0.0865962526785281 | 1.16230219378427 | 0.245112735825684 | 0.694718387691939 | NM_027294 |
| Iffo2 | 1155.182222 | 0.162503999 | 0.090768797 | 1.790306847 | 0.073404596 | [illegible] | | Srsf3 | 4027.577882 | 0.0628912601327248 | 0.0541195838760203 | 1.16207952146859 | 0.245203164924726 | 0.694718387691939 | NM_013663,NR_036613 |
| Mettl7a1 | 1016.330917 | 0.206442701 | 0.115321449 | 1.790150085 | 0.073429787 | [illegible] | | Chordc1 | 1628.71302 | 0.0821766475341508 | 0.0707803478651022 | 1.16100937637052 | 0.245638086372323 | 0.695046269862496 | NM_025844 |
| Pstk | 400.5562997 | 0.15686529 | 0.087636121 | 1.789961581 | 0.073460088 | [illegible] | | Gm166 | 222.8934967 | -0.10029292376484 | 0.0863871303113988 | -1.1609706608301 | 0.245653831025338 | 0.695046269862496 | NM_001033040 |
| Camta1 | 10685.64734 | 0.144657653 | 0.080836112 | 1.789517691 | 0.073531481 | [illegible] | | Hspa12a | 11063.82947 | 0.0640360160467257 | 0.0551528395682942 | 1.1610647166667 | 0.24561558207097 | 0.695046269862496 | NM_175199 |
| Myadml2 | 245.120023 | -0.240829747 | 0.134581435 | -1.78947227 | 0.073538789 | [illegible] | | Lrpprc | 3268.486324 | 0.0646206281276305 | 0.0556526097427898 | 1.16114281839232 | 0.245583824224763 | 0.695046269862496 | NM_028233 |
| Wls | 1725.69847 | -0.189765128 | 0.106048271 | -1.789422183 | 0.073546849 | [illegible] | | Nov | 3392.544286 | 0.103529678360845 | 0.0891345259050568 | 1.16149917565188 | 0.245438958223506 | 0.695046269862496 | NM_010930 |
| Engase | 400.3477346 | 0.177444989 | 0.099202984 | 1.788706166 | 0.073662149 | [illegible] | | Trmt61b | 308.3098543 | 0.0879594319135792 | 0.0757384310089647 | 1.16135798826826 | 0.245496346180646 | 0.695046269862496 | NR_027953,NR_027952,NR_015549 |
| Aldh3a1 | 36.85734319 | 0.299921761 | 0.167778961 | 1.787600542 | 0.073840478 | [illegible] | | Tstd3 | 791.1208429 | 0.0756676812901843 | 0.065161850060582 | 1.16122671808481 | 0.245549712008687 | 0.695046269862496 | NM_029840 |
| Cyp20a1 | 346.5384567 | -0.151260007 | 0.084617228 | -1.787579328 | 0.073843903 | [illegible] | | Aip | 2679.825726 | 0.0494596928896745 | 0.0426150266332114 | 1.16061626138065 | 0.245797989409443 | 0.695249198097981 | NM_001276284,NM_016666 |
| Iqce | 1185.993373 | 0.178945935 | 0.100102602 | 1.787625222 | 0.073836493 | [illegible] | | Sptssb | 111.0992747 | 0.109187698148745 | 0.0940819863578253 | 1.16055902278112 | 0.245821277807465 | 0.695249198097981 | NM_001164210,NM_133675 |
| Zfp229 | 96.24828073 | -0.245242905 | 0.137192439 | -1.787583244 | 0.073843271 | [illegible] | | Naa38 | 474.835151 | 0.0781768452453732 | 0.0673778373899952 | 1.16027537056245 | 0.245936708873273 | 0.695335424875491 | NM_133939 |
| Pet100 | 501.1683226 | -0.191611301 | 0.107216686 | -1.78714068 | 0.073914754 | [illegible] | | Tmem151b | 6275.11787 | 0.0890620247268857 | 0.076761140059229 | 1.16024885323701 | 0.245947501928285 | 0.695335424875491 | NM_001013749 |
| Glra1 | 23.17712663 | -0.300248562 | 0.168068071 | -1.786469975 | 0.074023194 | [illegible] | | 1600029O15Rik | 61.65260869 | -0.104290824982255 | 0.0899248389202929 | -1.15975548285047 | 0.246148373612869 | 0.695662109029072 | NR_033522 |
| H2-Oa | 21.22476706 | 0.285674903 | 0.159952825 | 1.785994736 | 0.074100111 | [illegible] | | Sec24d | 690.3225815 | 0.0862146675661209 | 0.0743403057442078 | 1.15972979533836 | 0.246158835220255 | 0.695662109029072 | NM_027135 |
| Casc4 | 3805.791949 | -0.102064369 | 0.057167964 | -1.785342021 | 0.074205857 | [illegible] | | Prdx3 | 2343.64249 | 0.0698846350397937 | 0.0602834427220379 | 1.15926748513727 | 0.246347170956036 | 0.696058938982178 | NM_007452 |
| Gh | 126.4188396 | 0.144199548 | 0.08076689 | 1.785379467 | 0.074199787 | [illegible] | | Atrnl1 | 6466.36453 | 0.0833987259420816 | 0.0720365014588287 | 1.15772871048918 | 0.246974763697607 | 0.696511264509423 | NM_181415 |
| Mirlet7c-2 | 17.75585441 | 0.28033651 | 0.157039523 | 1.785133484 | 0.074239668 | [illegible] | | Ftsjd1 | 264.299445 | 0.101044393638342 | 0.0872841999397302 | 1.15764816207416 | 0.247007646369279 | 0.696511264509423 | NM_146215 |
| H60b | [illegible] | [illegible] | [illegible] | [illegible] | [illegible] | [illegible] | | Mll5 | 5163.908404 | -0.0701060316986066 | 0.0605565293172165 | -1.1576956686432 | 0.246988252160351 | 0.696511264509423 | NM_026984 |
| Anxa2 | [illegible] | [illegible] | [illegible] | [illegible] | [illegible] | [illegible] | | Nol10 | 975.9402986 | 0.0744046870053577 | 0.0642508464882943 | 1.15803434619205 | 0.246850020436863 | 0.696511264509423 | NM_001008421 |
| Upb1 | [illegible] | [illegible] | [illegible] | [illegible] | [illegible] | [illegible] | | [illegible] | [illegible] | [illegible] | [illegible] | [illegible] | [illegible] | [illegible] | [illegible] |
| Smarcc1 | [illegible] | [illegible] | [illegible] | [illegible] | [illegible] | [illegible] | | [illegible] | [illegible] | [illegible] | [illegible] | [illegible] | [illegible] | [illegible] | [illegible] |
| Ache | [illegible] | [illegible] | [illegible] | [illegible] | [illegible] | [illegible] | | [illegible] | [illegible] | [illegible] | [illegible] | [illegible] | [illegible] | [illegible] | [illegible] |
| Brms1 | [illegible] | [illegible] | [illegible] | [illegible] | [illegible] | [illegible] | | [illegible] | [illegible] | [illegible] | [illegible] | [illegible] | [illegible] | [illegible] | [illegible] |
| Lrrc20 | [illegible] | [illegible] | [illegible] | [illegible] | [illegible] | [illegible] | | [illegible] | [illegible] | [illegible] | [illegible] | [illegible] | [illegible] | [illegible] | [illegible] |
| Trak2 | [illegible] | [illegible] | [illegible] | [illegible] | [illegible] | [illegible] | | [illegible] | [illegible] | [illegible] | [illegible] | [illegible] | [illegible] | [illegible] | [illegible] |
| [illegible] | [illegible] | [illegible] | [illegible] | [illegible] | [illegible] | [illegible] | | [illegible] | [illegible] | [illegible] | [illegible] | [illegible] | [illegible] | [illegible] | [illegible] |
| Prkaa2 | [illegible] | [illegible] | [illegible] | [illegible] | [illegible] | [illegible] | | [illegible] | [illegible] | [illegible] | [illegible] | [illegible] | [illegible] | [illegible] | [illegible] |
| Ap5b1 | [illegible] | [illegible] | [illegible] | [illegible] | [illegible] | [illegible] | | [illegible] | [illegible] | [illegible] | [illegible] | [illegible] | [illegible] | [illegible] | [illegible] |
| Dll1 | [illegible] | [illegible] | [illegible] | [illegible] | [illegible] | [illegible] | | [illegible] | [illegible] | [illegible] | [illegible] | [illegible] | [illegible] | [illegible] | [illegible] |
| Brpf1 | [illegible] | [illegible] | [illegible] | [illegible] | [illegible] | [illegible] | | [illegible] | [illegible] | [illegible] | [illegible] | [illegible] | [illegible] | [illegible] | [illegible] |
| Cadm4 | [illegible] | [illegible] | [illegible] | [illegible] | [illegible] | [illegible] | | [illegible] | [illegible] | [illegible] | [illegible] | [illegible] | [illegible] | [illegible] | [illegible] |
| Tomm40 | [illegible] | [illegible] | [illegible] | [illegible] | [illegible] | [illegible] | | [illegible] | [illegible] | [illegible] | [illegible] | [illegible] | [illegible] | [illegible] | [illegible] |
| Zfp768 | [illegible] | [illegible] | [illegible] | [illegible] | [illegible] | [illegible] | | [illegible] | [illegible] | [illegible] | [illegible] | [illegible] | [illegible] | [illegible] | [illegible] |
| Cep290 | [illegible] | [illegible] | [illegible] | [illegible] | [illegible] | [illegible] | | [illegible] | [illegible] | [illegible] | [illegible] | [illegible] | [illegible] | [illegible] | [illegible] |
| D330050I16Rik | [illegible] | [illegible] | [illegible] | [illegible] | [illegible] | [illegible] | | [illegible] | [illegible] | [illegible] | [illegible] | [illegible] | [illegible] | [illegible] | [illegible] |
| Mertk | [illegible] | [illegible] | [illegible] | [illegible] | [illegible] | [illegible] | | [illegible] | [illegible] | [illegible] | [illegible] | [illegible] | [illegible] | [illegible] | [illegible] |
| Pak7 | [illegible] | [illegible] | [illegible] | [illegible] | [illegible] | [illegible] | | [illegible] | [illegible] | [illegible] | [illegible] | [illegible] | [illegible] | [illegible] | [illegible] |
| Zfp369 | [illegible] | [illegible] | [illegible] | [illegible] | [illegible] | [illegible] | | [illegible] | [illegible] | [illegible] | [illegible] | [illegible] | [illegible] | [illegible] | [illegible] |
| [illegible] | [illegible] | [illegible] | [illegible] | [illegible] | [illegible] | [illegible] | | [illegible] | [illegible] | [illegible] | [illegible] | [illegible] | [illegible] | [illegible] | [illegible] |
| Pias1 | [illegible] | [illegible] | [illegible] | [illegible] | [illegible] | [illegible] | | Incenp | [illegible] | [illegible] | [illegible] | [illegible] | [illegible] | [illegible] | [illegible] |

| | | | | | | | | | | | | | | |
|---|---|---|---|---|---|---|---|---|---|---|---|---|---|---|
| Tmie | 183.1645321 | 0.216605652 | 0.121707867 | 1.779717455 | 0.075122164 | 0.225258455 | Map2k1 | 12014.39273 | 0.0681147963631587 | 0.058977021032338 | 1.15493789226509 | 0.248115860865948 | 0.697476022761195 | NM_008927 |
| Ctf1 | 52.01891411 | 0.254629911 | 0.143097707 | 1.779378201 | 0.075176161 | 0.225339415 | Sars | 7387.941725 | 0.0514404766195317 | 0.0445546751528667 | 1.15454732049966 | 0.248275849894146 | 0.697784357560906 | NM_001204979,NM_011319 |
| Dusp19 | 341.7103473 | -0.204155733 | 0.115099763 | -1.778593874 | 0.075306369 | 0.225566144 | Alkbh4 | 236.472494 | -0.0914080345139056 | 0.0792314504621922 | -1.15368372004699 | 0.248629860728123 | 0.697784357560906 | NM_028070 |
| Hinfp | 438.785681 | 0.160010512 | 0.089965359 | 1.778579168 | 0.075308784 | 0.225566144 | Bcan | 10817.40836 | -0.0818101053856441 | 0.0709116706004076 | -1.15369028388359 | 0.248627168720944 | 0.697784357560906 | NM_007529,NM_001109758 |
| Irf3 | 636.0657427 | 0.191265285 | 0.107529334 | 1.77872659 | 0.075284598 | 0.225566144 | Ctsa | 4537.358838 | -0.0789993277729249 | 0.0685066655846387 | -1.15316264561618 | 0.248843632513974 | 0.697784357560906 | NM_008906,NM_001038492 |
| Zc3h14 | 3252.519103 | 0.125990473 | 0.070838012 | 1.778571538 | 0.075310034 | 0.225566144 | Fndc4 | 3402.041871 | 0.0649551964489944 | 0.0562858456800696 | 1.15402363888772 | 0.248490477615654 | 0.697784357560906 | NM_022424 |
| Kcnj13 | 96.45954198 | 0.234910057 | 0.132100423 | 1.778268776 | 0.075359722 | 0.225627736 | Gpc5 | 1630.89767 | 0.0647426917354077 | 0.0561148680399993 | 1.15258996012187 | 0.249078726049266 | 0.697784357560906 | NM_175500 |
| Lca5l | 36.8289009 | 0.285357985 | 0.160468909 | 1.778275844 | 0.075358562 | 0.225627736 | Ing3 | 605.6728061 | 0.0740415921273917 | 0.0642375175701493 | 1.15262225142085 | 0.249065465995463 | 0.697784357560906 | NM_023626 |
| Commd8 | 1249.20763 | -0.113245066 | 0.063690615 | -1.778049492 | 0.075395727 | 0.225649903 | Kcnj11 | 2167.87526 | -0.0719332019077198 | 0.0623693033967116 | -1.15334303880509 | 0.248769611321263 | 0.697784357560906 | NM_010602,NM_001204411 |
| Srgap2 | 3636.92434 | 0.101019028 | 0.056817456 | 1.777957612 | 0.075410817 | 0.225649903 | Lpcat1 | 1031.936005 | -0.0727850812596662 | 0.0631343593397363 | -1.15286013649553 | 0.248967796410939 | 0.697784357560906 | NM_145376 |
| Zfp708 | 125.9846947 | -0.224784619 | 0.126426412 | -1.777987806 | 0.075405858 | 0.225649903 | Necab1 | 4790.977774 | 0.0942070944194674 | 0.0816319095787026 | 1.15404741731101 | 0.248480729356949 | 0.697784357560906 | NM_178617 |
| Map4k3 | 3000.786688 | 0.149373219 | 0.084023341 | 1.777758629 | 0.075443506 | 0.225660556 | Ptrhd1 | 499.59164 | 0.0786367464083988 | 0.0682062202565493 | 1.15292631834188 | 0.248940628583689 | 0.697784357560906 | NM_001204912 |
| Plekha1 | 3581.341946 | 0.169010225 | 0.09506708 | 1.777799679 | 0.075436761 | 0.225660556 | Ranbp17 | 268.2204097 | -0.0985664762844513 | 0.0854376291034725 | -1.15366586501457 | 0.248637183663686 | 0.697784357560906 | NM_023146 |
| Rilp | 49.5669599 | 0.288422388 | 0.16225641 | 1.77757161 | 0.07547424 | 0.225708912 | Rapgefl1 | 8032.985176 | -0.0729557943088902 | 0.0632772683398494 | -1.15295423179552 | 0.248929170656855 | 0.697784357560906 | NM_001080925 |
| Dnpep | 1314.939629 | -0.110771258 | 0.062336579 | -1.77698648 | 0.075570464 | 0.225909468 | Sec22c | 2309.318821 | 0.0711202730371887 | 0.0616479365849702 | 1.15365212490385 | 0.248642819037334 | 0.697784357560906 | NM_001164562,NM_178677 |
| Tm9sf2 | 4546.496132 | -0.132447828 | 0.074531538 | -1.777070913 | 0.075556573 | 0.225909468 | Srp14 | 2308.149641 | 0.070506084482399 | 0.0611774394443316 | 1.15249070359944 | 0.249119487698669 | 0.697784357560906 | NM_009273 |
| Slc10a7 | 178.2498962 | -0.180385441 | 0.101531308 | -1.776648445 | 0.0756261 | 0.226032174 | Trmt1l | 1206.196472 | -0.0679989866065058 | 0.0589992595349017 | -1.15253966138813 | 0.24909938163397 | 0.697784357560906 | NM_026876 |
| 0610010B08Rik | 10.01045317 | -0.289089807 | 0.162758166 | -1.776192336 | 0.075701221 | 0.22621306 | Zfp46 | 1980.934466 | -0.0695595051159208 | 0.0602837918630521 | -1.15386744871558 | 0.248554516325272 | 0.697784357560906 | NM_009557 |
| Fam78a | 326.6722316 | 0.194875555 | 0.109735052 | 1.775873349 | 0.075753795 | 0.226274597 | Zfp507 | 1314.407529 | 0.0641064234660673 | 0.0555745950587735 | 1.15352029822747 | 0.248696891014845 | 0.697784357560906 | NM_177739 |
| Sqstm1 | 16117.37628 | 0.125654935 | 0.070758565 | 1.775826507 | 0.075761517 | 0.226274597 | Dbc1 | 10570.58606 | 0.0657400693156217 | 0.0570508401250184 | 1.15230677009422 | 0.249195035949132 | 0.697861402004451 | NM_019967 |
| Srebf1 | 2762.316142 | 0.187581243 | 0.10563187 | 1.775801588 | 0.075765626 | 0.226274597 | Gem | 62.74031448 | -0.108677603528267 | 0.0943228277454941 | -1.15218771665228 | 0.249243944110263 | 0.697863826550825 | NM_010276 |
| Rab28 | 1946.928689 | -0.103866167 | 0.058498182 | -1.775545216 | 0.075807907 | 0.226357237 | Btbd1 | 5306.902157 | 0.0639907545514805 | 0.0555455678297407 | 1.1520406947252 | 0.249304351220406 | 0.697864808274404 | NM_146193 |
| Myo7a | 997.6894743 | 0.149078933 | 0.083975921 | 1.775258095 | 0.075855281 | 0.226455053 | Pde12 | 532.1722369 | 0.0818397283885064 | 0.0710557312183342 | 1.15176815416952 | 0.249416357414934 | 0.697864808274404 | NM_178668 |
| Shf | 1052.943489 | 0.127751401 | 0.071974595 | 1.7749513 | 0.075905929 | 0.226562601 | Prrg3 | 2507.966827 | 0.073350176049931 | 0.0636776734868359 | 1.1518978636224 | 0.24936304623955 | 0.697864808274404 | NM_001081135 |
| Pet112 | 653.5904501 | -0.137714294 | 0.077594611 | -1.774792006 | 0.075932237 | 0.226597472 | Smoc1 | 800.0629246 | 0.108149374819571 | 0.0939025513575262 | 1.15171923718878 | 0.249436464584931 | 0.697864808274404 | NM_001146217,NM_022316 |
| Rpl39 | 2422.037477 | -0.178053153 | 0.100336403 | -1.774561855 | 0.07597026 | 0.226667285 | Ostm1 | 2451.759829 | 0.0690680624120642 | 0.0599837207805721 | 1.1514467844488 | 0.24954847614723 | 0.698043744370241 | NM_172416 |
| Dgat2l6 | 24.73938272 | 0.294652895 | 0.166065327 | 1.774319175 | 0.076010371 | 0.226743296 | Galnt18 | 1412.165473 | 0.0828779488612144 | 0.0720066329766929 | 1.15097658972668 | 0.249741866633031 | 0.698181361568786 | NM_173739 |
| Mrpl50 | 1208.897597 | -0.117463597 | 0.066212628 | -1.774036185 | 0.076057165 | 0.226839214 | Pelo | 480.9243971 | 0.0781132747987931 | 0.0678619442079802 | 1.15106155166134 | 0.249706914149502 | 0.698181361568786 | NM_134058 |
| Dennd5b | 3530.395108 | -0.140820806 | 0.079382762 | -1.773946908 | 0.076071933 | 0.226839593 | Sox8 | 1947.678855 | 0.0942005128081764 | 0.0818319983685474 | 1.15114520830745 | 0.249672501988225 | 0.698181361568786 | NM_011447 |
| Porcn | 2063.71414 | 0.135296561 | 0.076292113 | 1.773401684 | 0.07616217 | 0.227064973 | Gabarap | 5130.223609 | 0.0652438669210139 | 0.0566933955503454 | 1.1508195317579 | 0.249806487697929 | 0.698227639080262 | NM_019749 |
| Bai1 | 18410.74252 | 0.12316402 | 0.069455657 | 1.773275577 | 0.076183054 | 0.227083539 | Sept7 | 14513.41068 | 0.0847104018569134 | 0.0736167186077747 | 1.1506951608132 | 0.24985766806496 | 0.698236338266949 | NM_001205367,NM_009859 |
| Sptan1 | 44149.52124 | -0.102459586 | 0.05779254 | -1.772886015 | 0.076247596 | 0.227147116 | BC028528 | 59.29283911 | 0.107090030985035 | 0.0931271146099419 | 1.14993395246461 | 0.250171075473404 | 0.698461385936824 | NM_153513 |
| Styk1 | 12.15494332 | -0.283381963 | 0.159842596 | -1.772881384 | 0.076248364 | 0.227147116 | Car13 | 43.47522351 | 0.0942124907426776 | 0.0819196422832385 | 1.15005984055614 | 0.250119225440646 | 0.698461385936824 | NM_024495 |
| Trp53bp1 | 4489.486908 | -0.13492119 | 0.076098767 | -1.772974727 | 0.076232895 | 0.227147116 | Fbxw8 | 2203.642126 | 0.0631164755111536 | 0.0548934900395736 | 1.14979891906403 | 0.250226700565554 | 0.698461385936824 | NM_172721 |
| Camk1d | 9260.571807 | 0.148961351 | 0.084032239 | 1.772669072 | 0.076283559 | 0.227208278 | Kdm6a | 1783.92851 | -0.0757254237841946 | 0.0658589124924562 | -1.1498128486842 | 0.25022096205677 | 0.698461385936824 | NM_009483 |
| Dbndd2 | 5316.625998 | 0.198133771 | 0.111797637 | 1.772253655 | 0.076352461 | 0.227282425 | Slc1a5 | 144.8432452 | -0.103153761270129 | 0.089671500707832 | -1.15035167757731 | 0.249999054227256 | 0.698461385936824 | NM_009201 |
| Glp1r | 23.35983173 | -0.280566485 | 0.158303151 | -1.772336707 | 0.076338681 | 0.227282425 | Tet3 | 2652.316234 | 0.0950373108975135 | 0.0826336603764882 | 1.15010409153501 | 0.250101001395654 | 0.698461385936824 | NM_183138 |
| Ltbp3 | 1450.22034 | 0.21445592 | 0.12099558 | 1.772427721 | 0.076323584 | 0.227282425 | Upf3b | 831.0781464 | -0.0756709329619625 | 0.0658247389631686 | -1.14958196802426 | 0.250316088552137 | 0.698576657504005 | NM_026573 |
| Cpne2 | 875.7064033 | 0.203209377 | 0.114692508 | 1.771775515 | 0.076431829 | 0.227474981 | Npsr1 | 195.7970387 | 0.101097710393008 | 0.0879593634507481 | 1.14936837224404 | 0.250404115889311 | 0.698688088245895 | NM_175678 |
| Mknk1 | 1053.224268 | 0.150876964 | 0.085160069 | 1.77168673 | 0.076446574 | 0.22747517 | Ccdc102a | 50.34937698 | -0.108283764596817 | 0.0942696821284709 | -1.14865948576391 | 0.250696417899897 | 0.699068333470275 | NM_001033533 |
| Atp8b1 | 170.9281834 | 0.252528541 | 0.142569116 | 1.77127101 | 0.076515646 | 0.227636985 | Ino80 | 1662.866576 | -0.0670382372622509 | 0.0583704082352372 | -1.14849697456426 | 0.250763461261129 | 0.699068333470275 | NM_026574 |
| Mcm2 | 199.4468085 | -0.215811905 | 0.121869952 | -1.770837695 | 0.076587696 | 0.227807594 | Pcdhb20 | 449.6009236 | 0.0856658438484843 | 0.0745917387250364 | 1.14846289029767 | 0.250777524179879 | 0.699068333470275 | NM_053145 |
| Ak3 | 3233.742515 | -0.086519 | 0.048867002 | -1.770499449 | 0.076643976 | 0.227841624 | Scd3 | 217.6522718 | 0.09603209872119 | 0.083618554721 | 1.14845441949587 | 0.250781019256065 | 0.699068333470275 | NM_024450 |
| Atp1a1 | 45155.12372 | 0.15842596 | 0.089484278 | 1.770433464 | 0.076654959 | 0.227841624 | Slc22a21 | 48.4837803 | -0.104820170934192 | 0.0912416919857459 | -1.14881879821526 | 0.250630706316572 | 0.699068333470275 | NM_019723 |
| Jup | 2085.422935 | -0.125929905 | 0.071121502 | -1.77063055 | 0.076622158 | 0.227841624 | Mamstr | 92.64262701 | -0.10919765347403 | 0.0951019481866267 | -1.14821678794363 | 0.25087908054623 | 0.699207506526197 | NM_172418 |
| Shcbp1l | 51.92287297 | 0.288357389 | 0.162875549 | 1.770415455 | 0.076657957 | 0.227841624 | 3000002C10Rik | 141.0389828 | -0.1050694872921 | 0.0916324012742854 | -1.14664120803288 | 0.25152993803578 | 0.699465245047928 | NR_033215 |
| Arhgef7 | 5540.58052 | -0.099684972 | 0.056342827 | -1.769257568 | 0.076850901 | 0.22832749 | AI413582 | 907.536474 | 0.0827574877182062 | 0.0721648336128308 | 1.14678415476166 | 0.251470839526782 | 0.699465245047928 | NM_001002895 |
| Slc25a24 | 112.6002428 | -0.228709636 | 0.129267655 | -1.769271955 | 0.076848501 | 0.22832749 | AW112010 | 34.69728357 | -0.102305753380556 | 0.0891647960229646 | -1.1473783145784 | 0.251225299664112 | 0.699465245047928 | NM_001177351 |
| Ccnb2 | 35.20569327 | 0.287771184 | 0.162681435 | 1.76892455 | 0.076906467 | 0.228357688 | Adamts2 | 300.1633795 | 0.100126121317816 | 0.0873393241022282 | 1.14640366578314 | 0.251628166624475 | 0.699465245047928 | NM_175643 |
| Cpt1c | 4011.529703 | 0.147851538 | 0.083576399 | 1.769058491 | 0.076884114 | 0.228357688 | Iglon5 | 1542.183123 | -0.074985634811238 | 0.0653825811693101 | -1.14687480167019 | 0.251433368367371 | 0.699465245047928 | NM_001164518 |
| Gjb6 | 2136.287412 | 0.241260456 | 0.136394472 | 1.768843358 | 0.076920019 | 0.228357688 | Itgb1bp1 | 826.6367653 | 0.0669688565193635 | 0.0584304975209012 | 1.14612846648119 | 0.25174200062719 | 0.699465245047928 | NM_008403 |
| Usp16 | 989.3432752 | 0.152441886 | 0.086181486 | 1.76884727 | 0.076919366 | 0.228357688 | Kcmf1 | 2307.880175 | 0.0604033469252616 | 0.0526560607056065 | 1.14713000015267 | 0.251327896632678 | 0.699465245047928 | NM_019715 |
| Nelfa | 732.0148985 | 0.127137232 | 0.071889813 | 1.768501369 | 0.076977124 | 0.228483439 | Mau2 | 4059.539204 | -0.0670088407112435 | 0.0583774335916121 | -1.14785520000783 | 0.251028345149515 | 0.699465245047928 | NM_001167939,NM_028993 |
| Mb21d2 | 1650.162609 | 0.116339475 | 0.06579615 | 1.768180589 | 0.077030718 | 0.228574612 | Nfkbie | 112.017 | 0.107198343243451 | 0.0935229691781997 | 1.14622476366415 | 0.251702163979032 | 0.699465245047928 | NM_008690 |
| Sdhaf2 | 1483.630257 | -0.096436536 | 0.054541204 | -1.768140938 | 0.077037345 | 0.228574612 | Pcdh19 | 1549.851102 | 0.0914663423650686 | 0.0797570773698415 | 1.14681161072302 | 0.251459489509445 | 0.699465245047928 | NM_001105245,NM_001105246 |
| Me1 | 2311.926913 | -0.133664183 | 0.075627099 | -1.76741123 | 0.077159384 | 0.22884906 | Sdk1 | 425.4508709 | -0.0941313780518092 | 0.0820382721422038 | -1.14740810104634 | 0.251212994644776 | 0.699465245047928 | NM_177879 |
| Zcchc17 | 1995.663777 | -0.100844692 | 0.057056029 | -1.767467748 | 0.077149926 | 0.22884906 | Srd5a3 | 737.2633937 | 0.0745436134153542 | 0.0650331116792215 | 1.14624091467503 | 0.251695482987379 | 0.699465245047928 | NM_020611 |
| Mycn | 226.7310954 | -0.246298758 | 0.139363113 | -1.767316709 | 0.077175203 | 0.228852172 | Stard3nl | 1314.109297 | 0.0609435031472465 | 0.0531644321141395 | 1.14632096542301 | 0.251662371196661 | 0.699465245047928 | NM_024270 |
| Bzw1 | 6140.216656 | -0.118852292 | 0.067261184 | -1.767026474 | 0.077223795 | 0.228920619 | Synj2 | 4495.854977 | 0.0902387950802193 | 0.0786409380703791 | 1.14747861984379 | 0.251183864462759 | 0.699465245047928 | NM_001113351,NM_011523,NM_001113352,NM_001113353 |
| Elavl3 | 9588.02679 | -0.084443616 | 0.047791579 | -1.766914132 | 0.07724261 | 0.228920619 | Vps35 | 7451.349308 | 0.0840531897614934 | 0.073336507360614 | 1.14613025533639 | 0.25174126138465 | 0.699465245047928 | NM_022997 |
| Tomm34 | 2639.895375 | 0.156128171 | 0.088359037 | 1.766974575 | 0.077232486 | 0.228920619 | Zfp184 | 125.6997692 | 0.10661264675449 | 0.0929748116008413 | 1.1466831168446 | 0.251512610659436 | 0.699465245047928 | NM_183014 |
| Tpm1 | 7433.48525 | -0.112442107 | 0.063641549 | -1.766803437 | 0.077261153 | 0.228931785 | Adam15 | 4670.192678 | -0.0768462718709562 | 0.0670938962828345 | -1.14535413992072 | 0.252062487329676 | 0.699923340323103 | NM_009614,NM_001037722 |
| Cd180 | 154.7772305 | 0.282967035 | 0.160203859 | 1.766293502 | 0.077346621 | 0.229102277 | Fkbp15 | 1848.142498 | -0.0601553600846348 | 0.0525466474403293 | -1.14479920332317 | 0.252292345441841 | 0.699923340323103 | NM_001045528 |
| Osbpl11 | 841.690223 | 0.120839024 | 0.068414278 | 1.766283702 | 0.077348265 | 0.229102277 | Hdac1 | 996.6535555 | -0.0740089313728752 | 0.0646257250976436 | -1.14519305216389 | 0.252129195813111 | 0.699923340323103 | NM_008228 |
| Znhit6 | 499.0651278 | 0.135743936 | 0.076858202 | 1.766160694 | 0.077368893 | 0.229119578 | Klhl32 | 396.8660416 | 0.0994646548354475 | 0.0868734075236304 | 1.14493787766288 | 0.252234892000019 | 0.699923340323103 | NM_001163020,NM_001033531 |
| Asb13 | 2061.758141 | 0.141351626 | 0.080052176 | 1.765743696 | 0.077438858 | 0.229195353 | Myom3 | 128.3771191 | -0.102477137584273 | 0.0895142533953564 | -1.14481363243531 | 0.25228636696922 | 0.699923340323103 | NM_001085509 |
| Edem2 | 742.886214 | 0.149447263 | 0.084631185 | 1.765865177 | 0.077418471 | 0.229195353 | Pias3 | 1116.777526 | -0.0709823608225594 | 0.0619997840517659 | -1.14488077512163 | 0.252258548752361 | 0.699923340323103 | NM_001165949,NM_018812,NM_146135 |
| Them4 | 691.2040602 | -0.129697352 | 0.073449288 | -1.765808153 | 0.07742804 | 0.229195353 | Ptges3l | 91.91072577 | 0.108603280271341 | 0.0948397158826822 | 1.14512447934454 | 0.252157596422004 | 0.699923340323103 | NM_026865 |
| Wdr70 | 474.513572 | 0.220964137 | 0.12516349 | 1.765404098 | 0.077495875 | 0.2293203 | Txndc5 | 2663.715615 | 0.0685751084925699 | 0.05988862105803 | 1.14504403810071 | 0.252190915385035 | 0.699923340323103 | NM_145367 |
| Cds2 | 29423.09203 | -0.101659891 | 0.057606048 | -1.764743359 | 0.077606908 | 0.229605009 | Ap1s2 | 378.9447483 | 0.100131009866455 | 0.0874822187908442 | 1.14458699436798 | 0.252380282293641 | 0.700033603322023 | NM_026887 |
| Gm13483 | 13.12149439 | 0.294333829 | 0.1668306 | 1.764267638 | 0.07768693 | 0.229752167 | Pigf | 238.4987365 | 0.101438306582184 | 0.0886498391429599 | 1.14425821369626 | 0.252516567252251 | 0.70027790299851 | NM_008838 |
| Gpr85 | 1315.320084 | -0.124291966 | 0.070456498 | -1.764095147 | 0.077715961 | 0.229752167 | Adm | 75.39743338 | 0.108607158621597 | 0.0949886335116901 | 1.1433720979303 | 0.252884132080148 | 0.700378198924256 | NM_009627 |
| Mutyh | 74.26758702 | 0.266479097 | 0.151050333 | 1.764174172 | 0.07770266 | 0.229752167 | Ap3s2 | 4330.097968 | 0.0496417527189472 | 0.0434100482175406 | 1.1435544247769 | 0.252808471637502 | 0.700378198924256 | NM_009682 |
| Vamp5 | 73.6921128 | -0.22991462 | 0.130326783 | -1.764139456 | 0.077708503 | 0.229752167 | B3gat1 | 10024.52128 | -0.0689306331971511 | 0.0602971210868368 | -1.14318282456439 | 0.25296269181757 | 0.700378198924256 | NM_029792 |
| Nudt1 | 126.2387949 | -0.200420406 | 0.113618962 | -1.763969695 | 0.077737082 | 0.229770763 | Cep78 | 387.3189306 | -0.089603616769685 | 0.0784049183337264 | -1.14283158090022 | 0.253108523937806 | 0.700378198924256 | NM_198019 |
| Gng10 | 982.6368485 | -0.16782708 | 0.095149605 | -1.763823196 | 0.077761751 | 0.229799841 | Dbr1 | 440.7555685 | 0.0793028605124661 | 0.0693948880953784 | 1.14277668988341 | 0.253131319313806 | 0.700378198924256 | NM_031403 |
| Fh1 | 2570.023455 | -0.085548831 | 0.048513216 | -1.76341291 | 0.077830874 | 0.229960252 | Flrt1 | 1402.872398 | 0.0783262713282845 | 0.0685287212353016 | 1.14296998275135 | 0.25305105417373 | 0.700378198924256 | NM_201411 |
| Bclaf1 | 4264.653653 | 0.151883325 | 0.086143827 | 1.763136508 | 0.077877469 | 0.229976969 | Ift122 | 1712.386468 | -0.0543253897958995 | 0.0475208277832211 | -1.14319115070384 | 0.252959235615563 | 0.700378198924256 | NM_031177,NM_001167763 |
| Cox4i2 | 46.29549677 | -0.279826982 | 0.158719607 | -1.763027191 | 0.077895904 | 0.229976969 | Mmp2 | 112.341225 | -0.106209254041532 | 0.0928858740029918 | -1.14343817272055 | 0.252856711103771 | 0.700378198924256 | NM_008610 |
| Itgb8 | 1216.768137 | -0.149410214 | 0.08473923 | -1.763176442 | 0.077870736 | 0.229976969 | Nsg2 | 27605.96569 | 0.0635993369926663 | 0.0556368694137076 | 1.14311494630927 | 0.252990869484037 | 0.700378198924256 | NM_008741 |
| Sertad4 | 843.9375329 | -0.179624895 | 0.101880514 | -1.763093727 | 0.077884683 | 0.229976969 | Scoc | 4350.104393 | 0.0790289788962869 | 0.0691071157052347 | 1.14357223695129 | 0.252801080938786 | 0.700378198924256 | NM_001039137,NM_019708 |
| Gja5 | 21.89368011 | -0.292537797 | 0.166010414 | -1.762165342 | 0.078041366 | 0.230362531 | Snx3 | 5302.910604 | 0.0594309806376031 | 0.0519488929482138 | 1.14402785631732 | 0.252612084700048 | 0.700378198924256 | NM_017472 |
| Cidea | 78.41526408 | -0.241258635 | 0.136940221 | -1.761780677 | 0.078106361 | 0.230510467 | Trappc2 | 984.1130138 | 0.0801256299824602 | 0.070069774521632 | 1.14351202825298 | 0.252826063578569 | 0.700378198924256 | NM_025432 |
| Rpl32 | 4341.525042 | -0.14986941 | 0.085106487 | -1.760963421 | 0.078244594 | 0.230874452 | Hopx | 831.9402275 | 0.0995867396285884 | 0.0872350561671708 | 1.14159083273659 | 0.253624140060127 | 0.701608123883717 | NM_175606,NM_001159901,NM_001159900 |
| Slc27a3 | 129.5515956 | 0.245658421 | 0.139537321 | 1.760521265 | 0.078319465 | 0.23105137 | 4933406C10Rik | 58.94514443 | -0.10805626150763 | 0.0949017083346597 | -1.13861167610648 | 0.254865156257036 | 0.701619623536836 | NR_044986 |
| Cdkn3 | 10.11112459 | -0.287411934 | 0.163270431 | -1.760342842 | 0.078349694 | 0.231096548 | Arhgap6 | 581.7381422 | 0.104153787288669 | 0.0912794659113316 | 1.1410429087068 | 0.25385206737333 | 0.701619623536836 | NM_178754,NM_009707 |
| Vamp4 | 2356.414829 | -0.107493764 | 0.061074314 | -1.760048658 | 0.078399557 | 0.231199606 | C530008M17Rik | 1973.451997 | 0.0858903094898307 | 0.0753372968465148 | 1.14007667676283 | 0.254254277804538 | 0.701619623536836 | NM_001163793 |
| Pfkm | 17444.5524 | 0.117216345 | 0.066603625 | 1.759909392 | 0.078423171 | 0.231225234 | Cep85l | 435.2991762 | -0.0992389199269415 | 0.0870627907032237 | -1.13985457076862 | 0.254346894231682 | 0.701619623536836 | NM_001204983 |
| Ifitm10 | 293.4467867 | 0.19639919 | 0.111623377 | 1.759480818 | 0.078495875 | 0.231393521 | Clip3 | 24785.60995 | 0.0482330369855017 | 0.042261774151447 | 1.14129228963877 | 0.253748309662442 | 0.701619623536836 | NM_001081114 |
| Tdrd6 | 41.72990122 | 0.281605759 | 0.160058118 | 1.759396911 | 0.078510116 | 0.231393521 | Cystm1 | 639.176107 | 0.0779355847337947 | 0.0684164365712265 | 1.13913539844564 | 0.254646685135141 | 0.701619623536836 | NM_001081365 |
| 4430402I18Rik | 127.6785205 | 0.238942977 | 0.135907003 | 1.758135876 | 0.078724393 | 0.231932319 | D8Ertd82e | 1726.467339 | 0.0830954707443011 | 0.0729463242395067 | 1.13913170554655 | 0.254648225174281 | 0.701619623536836 | NR_172911 |
| Arv1 | 171.6763115 | 0.220134535 | 0.125205143 | 1.758190833 | 0.078715044 | 0.231932319 | Dpp10 | 7596.747983 | 0.0824562333654524 | 0.0723809736326231 | 1.13919762649184 | 0.254620735325409 | 0.701619623536836 | NM_199021 |
| Surf4 | 2861.176253 | -0.077940001 | 0.044333037 | -1.758056868 | 0.078737834 | 0.231932319 | Fbxl6 | 983.8707619 | -0.0944418166202005 | 0.0827863710984802 | -1.14078942423814 | 0.253957562668062 | 0.701619623536836 | NM_013909 |
| Unc13d | 54.85576989 | 0.277869845 | 0.15806524 | 1.757944037 | 0.078757032 | 0.231944773 | Fxn | 144.8573165 | -0.104706486998539 | 0.0918787620714661 | -1.13961577885752 | 0.254446408697784 | 0.701619623536836 | NM_008044 |
| Csnk1e | 3999.008011 | 0.112287099 | 0.063885205 | 1.757638535 | 0.078809032 | 0.232053809 | Kremen1 | 1377.676778 | -0.0786450735572196 | 0.0689101635279315 | -1.14126958246649 | 0.253757756012165 | 0.701619623536836 | NM_032396 |
| Chst9 | 17.42401148 | 0.288968446 | 0.164441413 | 1.757272943 | 0.078871297 | 0.232193023 | Lst1 | 51.74836349 | -0.10773688637102 | 0.0944867737221682 | -1.14023245916312 | 0.254189467848519 | 0.701619623536836 | NM_010734 |
| Tspan6 | 448.4341502 | -0.158168607 | 0.090031815 | -1.756807935 | 0.078950552 | 0.232382191 | Map4k5 | 1956.737268 | -0.0701035754275097 | 0.0615519742611895 | -1.13893301179963 | 0.25473109541088 | 0.701619623536836 | NM_201519 |
| Mrps18c | 612.3440634 | -0.165895987 | 0.094473588 | -1.756003883 | 0.079087746 | 0.232741791 | Med25 | 2756.575204 | -0.0710506198817084 | 0.0623714345935851 | -1.13915320923236 | 0.25463925764468 | 0.701619623536836 | NM_029365 |
| Hpca | 26254.09681 | 0.157561668 | 0.089741502 | 1.755727995 | 0.079134864 | 0.23283623 | Med7 | 506.0736496 | 0.0911262159707831 | 0.0799514314775045 | 1.13976966124018 | 0.254382276441131 | 0.701619623536836 | NM_001104530,NM_001104556,NM_001104557,NM_025426 |
| Ptch1 | 1756.380356 | -0.189774142 | 0.108131908 | -1.755024457 | 0.079255124 | 0.233145794 | Mllt3 | 1382.972384 | 0.0774465365073324 | 0.0678974941273831 | 1.14063909872777 | 0.254020139627265 | 0.701619623536836 | NM_027326,NM_029931 |
| AI593442 | 6817.382807 | 0.18306877 | 0.10432836 | 1.7547364 | 0.079304407 | 0.233202217 | Nos3 | 513.7208743 | -0.0930760035754945 | 0.0817437399799553 | -1.13863157714974 | 0.254856852141845 | 0.701619623536836 | NM_008713 |
| D19Ertd737e | 450.9877335 | -0.140929139 | 0.080313392 | -1.754740227 | 0.079303752 | 0.233202217 | Nucb1 | 3357.893159 | 0.0643104351065177 | 0.0564168880317268 | 1.13991461333975 | 0.254321876269975 | 0.701619623536836 | NM_008749,NM_001163662 |
| Prtn3 | 37.87172653 | -0.283378666 | 0.16150502 | -1.754612128 | 0.079325676 | 0.233220496 | Ptrf | 1005.690595 | -0.0729089502898007 | 0.0640362783909357 | -1.13855695742807 | 0.254887989708848 | 0.701619623536836 | NM_008986 |
| Rhog | 931.1972982 | 0.189281544 | 0.107896615 | 1.75428621 | 0.079381478 | 0.233340279 | Rnaseh1 | 366.0368811 | -0.0844496019073316 | 0.0741555317407339 | -1.1388172928567 | 0.254779367555897 | 0.701619623536836 | NM_011275 |
| Pnmt | 10.95260652 | -0.280500405 | 0.159915388 | -1.754055115 | 0.079421064 | 0.23341236 | Sart3 | 1957.660363 | -0.0653771490195539 | 0.0574081370400545 | -1.13881328310548 | 0.25478104033724 | 0.701619623536836 | NM_016926 |
| Gm9079 | 65.59934935 | -0.247607113 | 0.141203312 | -1.753550315 | 0.079507591 | 0.233622343 | Shisa6 | 1175.071858 | -0.10446934739477 | 0.0917645067726 | -1.13845048667513 | 0.254932422761078 | 0.701619623536836 | NM_001034874 |
| Elk1 | 3070.005193 | 0.145642306 | 0.083061832 | 1.753420338 | 0.079529882 | 0.233643534 | Srbd1 | 446.9008292 | 0.0835811300911149 | 0.0732466368296994 | 1.14109171026437 | 0.253831760618624 | 0.701619623536836 | NM_030133 |
| Kcnb1 | 13905.89271 | -0.159775534 | 0.091137429 | -1.753127506 | 0.079580123 | 0.233746811 | Tmem220 | 27.97544786 | -0.0889293550046675 | 0.0781041935987841 | -1.13859897794338 | 0.254870454922277 | 0.701619623536836 | NM_177392 |
| Caskin1 | 6826.030657 | 0.109912428 | 0.062705288 | 1.752841443 | 0.079629227 | 0.233846711 | Tpm4 | 1270.514664 | 0.0818888938561106 | 0.0718114671833004 | 1.140331719544 | 0.254148127725371 | 0.701619623536836 | NM_001001491 |
| Commd1 | 565.1336904 | -0.124615271 | 0.07109706 | -1.752748578 | 0.079645173 | 0.233849216 | Trim26 | 869.5804525 | 0.0790931752863571 | 0.0693665812586716 | 1.14022017304579 | 0.254194585115862 | 0.701619623536836 | NM_001025599 |
| Chst12 | 732.352546 | 0.193701285 | 0.110565975 | 1.751906813 | 0.079789833 | 0.2341852 | Trpc6 | 381.867002 | -0.093889510610107 | 0.0823701864978868 | -1.13984822180158 | 0.254349539758375 | 0.701619623536836 | NM_013838 |
| Rab8b | 1795.947007 | 0.158758168 | 0.090619865 | 1.751913533 | 0.079788677 | 0.2341852 | Cox7a2l | 5273.18288 | 0.0537160049694487 | 0.047196907081082 | 1.13812553176773 | 0.255068068317232 | 0.701727037950021 | NM_009187,NM_001159529 |
| Trmt44 | 275.583264 | 0.198680217 | 0.113454405 | 1.751189975 | 0.079913192 | 0.234502839 | Mgat2 | 947.6314625 | 0.0744548335033859 | 0.0654147974556204 | 1.13819558261666 | 0.255038822818601 | 0.701727037950021 | NM_146035 |
| Fam216b | 34.59430406 | 0.293243179 | 0.167504079 | 1.750662914 | 0.080003991 | 0.234716536 | Lyst | 3053.330445 | 0.0839413577658805 | 0.073763116379368 | 1.13798551208391 | 0.255126532042063 | 0.701754971490816 | NM_010748 |
| Slitrk5 | 2985.110693 | -0.146142241 | 0.083481639 | -1.750591432 | 0.080016312 | 0.234716536 | Acsl5 | 3928.803513 | 0.07418418031097 | 0.0652329622422262 | 1.13721924869065 | 0.25544664215281 | 0.701952733578609 | NM_027976 |
| 1700037C18Rik | 117.4821038 | -0.259480027 | 0.148245984 | -1.750334279 | 0.080060649 | 0.234762853 | Aldoc | 34640.54346 | 0.0756202297462538 | 0.0665023002196787 | 1.1371069797053 | 0.255493566483559 | 0.701952733578609 | NM_009657 |
| St18 | 362.9649109 | 0.212716002 | 0.121529494 | 1.750324093 | 0.080062406 | 0.234762853 | Fah | 1013.739191 | -0.0757084826960088 | 0.0666188265295812 | -1.13644275409732 | 0.255771311078046 | 0.701952733578609 | NM_010176 |
| Terf1 | 287.4459959 | 0.192759645 | 0.110136703 | 1.750185355 | 0.080086335 | 0.234788586 | Fam53b | 2144.404432 | 0.0942949049779361 | 0.0829020574225274 | 1.13742538978668 | 0.255360498310376 | 0.701952733578609 | NM_212473,NM_175268 |
| Rnf139 | 1165.440606 | -0.118316236 | 0.0676097 | -1.749989066 | 0.0801202 | 0.234843433 | Golph3 | 3170.186263 | 0.0779473031409516 | 0.0685412207719178 | 1.13723248963327 | 0.255441108317886 | 0.701952733578609 | NM_025673 |
| Dlgap3 | 13078.72048 | -0.156333211 | 0.089362666 | -1.749424217 | 0.080217718 | 0.235084797 | Impad1 | 3350.812867 | 0.0805351629192818 | 0.0708916808365021 | 1.13603122353694 | 0.255943496974272 | 0.701952733578609 | NM_177730 |
| Cabin1 | 4050.570807 | 0.093188177 | 0.053280633 | 1.749006577 | 0.080289883 | 0.235162843 | Kcnj8 | 156.4024681 | 0.107800247603695 | 0.094835759415127 | 1.13670463829808 | 0.255661779686499 | 0.701952733578609 | NM_008428 |
| Med12l | 1782.38903 | -0.153352328 | 0.08767589 | -1.749082085 | 0.080276832 | 0.235162843 | Lphn2 | 4747.061589 | -0.0635498728907048 | 0.0558588858206522 | -1.13768600925458 | 0.255251617529583 | 0.701952733578609 | NM_001081298 |
| Ndufab1 | 1319.195636 | -0.17758346 | 0.101530667 | -1.749062279 | 0.080280255 | 0.235162843 | Pvt1 | 369.3005322 | 0.0854207087774366 | 0.0751508726360277 | 1.13665624604451 | 0.255682016981396 | 0.701952733578609 | NR_003368 |
| 1700034H15Rik | 102.541835 | 0.205900707 | 0.117731834 | 1.748895771 | 0.080309038 | 0.235174491 | Rabgef1 | 1621.840128 | 0.0627512888003117 | 0.0552101776011215 | 1.13658914944402 | 0.255710078142366 | 0.701952733578609 | NM_019983,NM_001199059 |
| Csnk1g2 | 5486.518084 | -0.142913907 | 0.081735492 | -1.748492662 | 0.080378755 | 0.235245265 | Rgs14 | 2353.830769 | -0.0867003611864921 | 0.0763168392551707 | -1.13605807096653 | 0.255932261456264 | 0.701952733578609 | NM_016758 |
| Dlgap2 | 2863.632138 | -0.144091853 | 0.082407279 | -1.748533065 | 0.080371765 | 0.235245265 | Sh3bgr | 37.20414397 | 0.0969829068437246 | 0.0853749791278243 | 1.13596404688313 | 0.255971611576889 | 0.701952733578609 | NM_015825 |
| Macrod2 | 1030.462823 | -0.141116515 | 0.080703595 | -1.748577807 | 0.080364025 | 0.235245265 | Snx29 | 329.1881835 | 0.0832399035414451 | 0.0732694034358959 | 1.13611818863771 | 0.255907103747629 | 0.701952733578609 | NM_028964 |
| BC089491 | 32.12494045 | -0.290797486 | 0.166353773 | -1.748066672 | 0.080452483 | 0.235416577 | Unc5a | 5471.624618 | 0.0885851069820639 | 0.0779211310685454 | 1.1368560205336 | 0.255598479904299 | 0.701952733578609 | NM_153131 |
| Clec16a | 4319.508769 | 0.140805275 | 0.080562823 | 1.747769874 | 0.080503884 | 0.235510228 | Zfp709 | 256.4531521 | 0.0994337270728426 | 0.0874636493802082 | 1.1368577435021 | 0.255597759515679 | 0.701952733578609 | NM_145624 |
| Hectd1 | 6626.104573 | -0.13574514 | 0.077674349 | -1.747618624 | 0.080530088 | 0.235510228 | Zfp940 | 695.0291367 | -0.074770268240327 | 0.0658154262903162 | -1.13605992477372 | 0.255931485659668 | 0.701952733578609 | NM_173738 |
| Lad1 | 71.51435038 | 0.27868292 | 0.159457202 | 1.747697292 | 0.080516458 | 0.235510228 | Mettl22 | 614.4825177 | 0.0785302357241377 | 0.0691661096945143 | 1.1353860448561 | 0.256213604170827 | 0.70221864418593 | NM_146247 |
| Ubfd1 | 2511.440972 | -0.094958721 | 0.054341983 | -1.747428334 | 0.080563066 | 0.23556221 | Socs3 | 144.3490188 | -0.107989661022708 | 0.0950939409205018 | -1.13561032361659 | 0.25611968633347 | 0.70221864418593 | NM_007707 |
| Gtf2h1 | 1673.021898 | 0.134257438 | 0.076841711 | 1.747194796 | 0.080603554 | 0.235636126 | Wac | 5085.862409 | 0.0733680735869929 | 0.064613099693077 | 1.13549843507746 | 0.256166537221473 | 0.70221864418593 | NR_027465,NM_153085,NM_001146298 |
| 2310042E22Rik | 9.810184355 | 0.277456616 | 0.158856577 | 1.746585631 | 0.080709241 | 0.235811614 | Fndc5 | 3823.986315 | 0.0701821419202595 | 0.0618303845367223 | 1.13507529422815 | 0.256343772058791 | 0.702442866049048 | NM_027402 |
| Cd55 | 156.9865048 | -0.231021268 | 0.132265496 | -1.746648025 | 0.080698411 | 0.235811614 | Ywhah | 36175.1318 | 0.0629564988067707 | 0.0554870011876716 | 1.13461707176669 | 0.256535797046953 | 0.702836474519811 | NM_011738 |
| Tmem179b | 143.5442264 | 0.205468242 | 0.117637869 | 1.746616493 | 0.080703884 | 0.235811614 | Capn6 | 94.09951572 | -0.107264149900044 | 0.0945977851772364 | -1.13389705371089 | 0.256837733145433 | 0.703000737077551 | NM_007603 |
| A330050F15Rik | 251.7232627 | 0.241221715 | 0.13813586 | 1.746264259 | 0.080765043 | 0.235930162 | Fras1 | 1254.848581 | -0.0857568125992271 | 0.0756001682786345 | -1.13434684805408 | 0.256649085151379 | 0.703000737077551 | NM_175473 |
| Emc3 | 2833.228246 | -0.086632608 | 0.049616628 | -1.746039804 | 0.080804036 | 0.235999572 | Gtpbp3 | 860.0821673 | -0.0808651939583171 | 0.0713069952146909 | -1.13404293246193 | 0.256776539675747 | 0.703000737077551 | NM_032544 |
| Ccdc151 | 202.892511 | 0.202473171 | 0.115975712 | 1.745823916 | 0.080841554 | 0.236064651 | Lpgat1 | 14608.22065 | 0.0734438999175049 | 0.0647687688045913 | 1.13394003426384 | 0.256819702537194 | 0.703000737077551 | NM_172266,NM_001134829 |
| BC002163 | 22.01179899 | -0.29340867 | 0.168121487 | -1.745218148 | 0.080946903 | 0.236238712 | Rapgef2 | 6139.684569 | 0.0697549404961997 | 0.061503420300609 | 1.13416359863012 | 0.256725930073238 | 0.703000737077551 | NM_001099624 |
| Hsd11b1 | 405.5762954 | 0.238585833 | 0.136695646 | 1.745379903 | 0.080918762 | 0.236238712 | Cdh5 | 688.4143168 | -0.0873399518458824 | 0.0770889107598656 | -1.13297685730635 | 0.25722397205774 | 0.703199912481652 | NM_009868 |
| Rdh9 | 23.28569845 | 0.293008104 | 0.167888759 | 1.745251476 | 0.080941104 | 0.236238712 | Hscb | 149.9744566 | -0.104035630189872 | 0.0918258226426316 | -1.13296703689504 | 0.257228096203613 | 0.703199912481652 | NM_153571 |
| Sgce | 913.0014181 | -0.102756393 | 0.058893205 | -1.744792018 | 0.081021079 | 0.236410658 | Jsrp1 | 116.6918845 | 0.105161447769833 | 0.0928236622574513 | 1.13291638373589 | 0.25724936905738 | 0.703199912481652 | NM_028001 |
| Ubiad1 | 402.0264575 | -0.152012554 | 0.087153799 | -1.744187359 | 0.081126425 | 0.236673475 | Lamtor5 | 1213.304515 | 0.0661861640881537 | 0.0584143990397423 | 1.13304536511835 | 0.257195203031206 | 0.703199912481652 | NM_026774 |
| Akr1c14 | 68.93427523 | -0.282257214 | 0.161879771 | -1.74362252 | 0.081224934 | 0.236871659 | Tsen34 | 1780.339706 | 0.0539382420590387 | 0.0476080409957569 | 1.13292297802618 | 0.257246599578111 | 0.703199912481652 | NM_024168,NM_001164204,NM_001164205 |
| Frk | 19.72830282 | -0.292873764 | 0.167961697 | -1.743693768 | 0.081212503 | 0.236871659 | Zfp532 | 2445.603619 | 0.0616205529640866 | 0.0543676798095026 | 1.13340413238154 | 0.257044579527604 | 0.703199912481652 | NM_207255 |
| Ap1s2 | 343.5443525 | -0.220914229 | 0.126721301 | -1.743307769 | 0.081279869 | 0.236948306 | Zfp874b | 516.1229943 | 0.0938663866118706 | 0.0828036457198359 | 1.13360209922961 | 0.256961492085404 | 0.703199912481652 | NM_001076791 |
| Ptpn7 | 78.13905147 | -0.272341447 | 0.156222087 | -1.743296694 | 0.081281802 | 0.236948306 | Acsl6 | 7157.849071 | 0.0537934773462818 | 0.0475010116376916 | 1.13247014098532 | 0.257436830788551 | 0.703271881179499 | NM_001033597,NM_144823,NM_001033599,NM_001033598 |
| Spo11 | 11.35404189 | 0.278583727 | 0.159826951 | 1.743033477 | 0.081327767 | 0.237037703 | Calm2 | 53347.05689 | 0.0698936116902546 | 0.0617221612147785 | 1.1323908676341 | 0.257470142573803 | 0.703271881179499 | NM_007589 |
| Coq6 | 573.516756 | 0.13237303 | 0.075991674 | 1.741941229 | 0.081518729 | 0.237524069 | Prrt1 | 1579.274773 | -0.0862430230663 | 0.076177417738188 | -1.13227753768444 | 0.257517770617772 | 0.703271881179499 | NM_030890 |
| Gmeb1 | 902.3367923 | -0.123320153 | 0.07079968 | -1.741817941 | 0.081540306 | 0.237524069 | Rbm3 | 2305.54484 | -0.103199297801467 | 0.0911094861668966 | -1.13269542111591 | 0.257342181210215 | 0.703271881179499 | NM_001166409,NM_001166411,NM_001166410,NM_016809 |
| Rnf165 | 2839.671923 | -0.165894178 | 0.095242074 | -1.741816092 | 0.08154063 | 0.237524069 | Slfn5 | 447.7216576 | -0.101338140234185 | 0.0894867957822699 | -1.13243679526476 | 0.25745084276835 | 0.703271881179499 | NM_183201 |
| Dmrtb1 | 18.99287888 | -0.291887645 | 0.167591286 | -1.741663616 | 0.081567323 | 0.237557161 | Cdc42 | 11081.24545 | 0.0624184642627428 | 0.0551343504488358 | 1.13211570925582 | 0.257585791227397 | 0.703325414167136 | NM_009861,NM_001243769 |
| Cyp3a13 | 18.75590997 | 0.273748167 | 0.157190544 | 1.741505306 | 0.081595044 | 0.237579737 | Epdr1 | 5575.033613 | 0.0596671911083136 | 0.0527126684898162 | 1.13193266092839 | 0.257662746122494 | 0.703403317078621 | NM_134065 |
| Mid1ip1 | 3131.064246 | -0.159414625 | 0.09154162 | -1.741444227 | 0.081605741 | 0.237579737 | A430078G23Rik | 232.1620452 | -0.0951137364327536 | 0.0841463573097615 | -1.13033967994282 | 0.258334272730236 | 0.703616071992063 | NM_001033378 |
| Col2a1 | 75.45778949 | 0.270525101 | 0.155354482 | 1.741340821 | 0.081623855 | 0.237587828 | Atad1 | 4481.84066 | 0.0675768522250796 | 0.0598023522736419 | 1.13000324662587 | 0.258474856448218 | 0.703616071992063 | NM_026487 |
| Gng12 | 2557.481828 | -0.120081468 | 0.068979681 | -1.740823754 | 0.081714477 | 0.237806931 | Brix1 | 775.9668915 | 0.0725538171514834 | 0.0642139784424366 | 1.12987575153162 | 0.25852858286962 | 0.703616071992063 | NM_026396 |
| Idh3g | 4234.613239 | 0.095159389 | 0.054666576 | 1.740723413 | 0.081732072 | 0.23781347 | Dennd3 | 366.2206977 | -0.0917922944196466 | 0.0811954251335901 | -1.13051066939574 | 0.258261105236534 | 0.703616071992063 | NM_001081066 |
| Pptc7 | 2924.356523 | -0.174469247 | 0.100234687 | -1.740607488 | 0.081752404 | 0.237827967 | Dgcr2 | 6337.521283 | 0.062560747658436 | 0.0552999859476545 | 1.13129771348684 | 0.257929806405603 | 0.703616071992063 | NM_001109750,NM_010048 |
| Ckap2l | 33.76202319 | -0.292047909 | 0.167867222 | -1.739755413 | 0.081901975 | 0.238218358 | Dnahc9 | 284.6487361 | -0.104312961037027 | 0.0922077100026896 | -1.13128241699077 | 0.25793624251244 | 0.703616071992063 | NM_001099633 |
| Tmsb10 | 3506.581788 | -0.222501212 | 0.127943281 | -1.739061327 | 0.082023976 | 0.238528432 | Dusp3 | 9168.745663 | 0.0615875792243026 | 0.054492331889811 | 1.13020634442363 | 0.258389287035132 | 0.703616071992063 | NM_028207 |
| Gpatch4 | 639.7236164 | 0.110290565 | 0.063428231 | 1.738824544 | 0.08206563 | 0.23860478 | Dusp6 | 2377.454242 | 0.105744549248157 | 0.0935089476733044 | 1.13084952701639 | 0.258118430149345 | 0.703616071992063 | NM_026268 |
| Cbr3 | 331.8053387 | -0.167865728 | 0.096548428 | -1.738668673 | 0.082093059 | 0.238639749 | Gstm6 | 273.1035258 | 0.104171822813379 | 0.0922310483548092 | 1.12946588672217 | 0.258701352301483 | 0.703616071992063 | NM_008184 |
| Gm4926 | 8.407138136 | -0.256981227 | 0.147815974 | -1.738521349 | 0.082118992 | 0.238670354 | Kif20a | 27.86900993 | -0.0991039240658382 | 0.0877576494078382 | -1.12929100465385 | 0.258775094312363 | 0.703616071992063 | NM_009004,NM_001166406,NM_001166407 |
| 1700001L05Rik | 227.8205524 | 0.269368012 | 0.154967109 | 1.738226998 | 0.082170824 | 0.238776208 | Loxl3 | 185.8043309 | -0.100234281451169 | 0.0886145901664957 | -1.13112616401928 | 0.258001993413822 | 0.703616071992063 | NM_013586 |
| Bend7 | 123.5091421 | -0.266467684 | 0.153360284 | -1.737527318 | 0.082294136 | 0.238932488 | Lrrc32 | 279.8456611 | 0.094123178609555 | 0.0833293332872362 | 1.12953236149283 | 0.258673325899359 | 0.703616071992063 | NM_001113379 |
| Dnaaf1 | 12.73161115 | 0.274401899 | 0.157930571 | 1.737484375 | 0.08230171 | 0.238932488 | Mmgt1 | 1676.177443 | 0.071744907800301 | 0.0635355652922101 | 1.12920861678549 | 0.258809839611193 | 0.703616071992063 | NM_146234 |
| Exoc3l | 24.56076315 | 0.291049235 | 0.167500976 | 1.737597247 | 0.082281805 | 0.238932488 | Pcdhga8 | 582.2285774 | 0.10080641256232 | 0.0890860614638397 | 1.13156211988604 | 0.257818573179802 | 0.703616071992063 | NM_033591 |
| Rabif | 1221.467124 | -0.114279482 | 0.065767776 | -1.737621187 | 0.082277584 | 0.238932488 | Rab18 | 3678.106911 | 0.0763697885850411 | 0.0675785155666845 | 1.13008976217718 | 0.258438403210601 | 0.703616071992063 | NM_181070 |
| Slit3 | 1367.408427 | -0.193461997 | 0.111340446 | -1.737571601 | 0.082286327 | 0.238932488 | Rbm12b1 | 270.4289666 | -0.106225559223036 | 0.0939927297968995 | -1.13014654912747 | 0.258414478025467 | 0.703616071992063 | NM_028226 |
| Mppe1 | 555.175926 | 0.127773723 | 0.073545466 | 1.737343297 | 0.082326593 | 0.238959955 | Serbp1 | 5437.705046 | 0.0635772046437606 | 0.0562567247904225 | 1.13012630722122 | 0.258423006064942 | 0.703616071992063 | NM_025814,NM_001113566,NM_001113565,NM_001113564 |
| Rab11fip3 | 5705.762383 | 0.10643118 | 0.061266444 | 1.737185535 | 0.082354427 | 0.238965437 | Skiv2l2 | 2990.258478 | 0.0651825144193922 | 0.05770847821431 | 1.12951365962773 | 0.258681210572677 | 0.703616071992063 | NM_028151 |
| Rbm15b | 1468.056825 | -0.123364246 | 0.071014993 | -1.73715776 | 0.082359328 | 0.238965437 | Slitrk3 | 2605.993439 | -0.0763272944106713 | 0.0675999929655436 | -1.12910210581791 | 0.258854763095524 | 0.703616071992063 | NM_198864 |
| Scube2 | 60.84118263 | -0.276252144 | 0.159067369 | -1.736699018 | 0.082440311 | 0.239155623 | Tfip11 | 2396.274955 | 0.0690033168095795 | 0.0611100159048682 | 1.12916542055887 | 0.258828057975817 | 0.703616071992063 | NM_018783 |
| 9430021M05Rik | 571.9625612 | 0.11622468 | 0.066943221 | 1.736168037 | 0.082534127 | 0.239382959 | Tmem240 | 791.9259352 | -0.091516356507246 | 0.0810472808128158 | -1.12917244834665 | 0.258825093887865 | 0.703616071992063 | NM_001101506 |
| 4930447N08Rik | 9.062319972 | -0.246309647 | 0.141906086 | -1.735722923 | 0.082612838 | 0.239411823 | Wbp1l | 1878.191305 | 0.0596592285169006 | 0.0527456245670054 | 1.13107445416847 | 0.258023755362997 | 0.703616071992063 | NM_001177813,NM_001177812,NM_146099 |
| Creb1 | 2043.444031 | -0.130065769 | 0.074934277 | -1.735731292 | 0.082611358 | 0.239411823 | Wdr89 | 90.58502795 | -0.107460544625068 | 0.0950585791500884 | -1.13046655636834 | 0.258279682929167 | 0.703616071992063 | NM_028203 |
| Cystm1 | 605.3853037 | -0.133335126 | 0.076814752 | -1.735801045 | 0.082599019 | 0.239411823 | Lgr4 | 1183.475228 | -0.0694416700823619 | 0.0615286332563276 | -1.12860738825562 | 0.259063492594141 | 0.703828283955767 | NM_172671 |
| Fbxl22 | 11.76143722 | -0.281036788 | 0.16192038 | -1.735648023 | 0.082626089 | 0.239411823 | Plcb4 | 2518.398446 | 0.0762279285123602 | 0.0675436666357594 | 1.12857255623139 | 0.25907819319231 | 0.703828283955767 | NM_013829 |
| Wdr13 | 9471.838789 | -0.098860436 | 0.056960789 | -1.735587542 | 0.08263679 | 0.239411823 | Tmem69 | 514.9401212 | -0.0798757383450148 | 0.0707683836649134 | -1.12869242179141 | 0.259027607246777 | 0.703828283955767 | NM_177670 |
| Zfp712 | 238.0018675 | -0.188751773 | 0.108749362 | -1.735656666 | 0.082624206 | 0.239411823 | AI837181 | 2137.540521 | 0.0591459598829054 | 0.0524561521799885 | 1.12753142243378 | 0.259517862821731 | 0.703893403935547 | NM_001256515,NM_134149 |
| 2500004C02Rik | 228.0279746 | -0.170396755 | 0.098198953 | -1.735219684 | 0.082701901 | 0.239496535 | Cnbp | 8508.111433 | 0.0520474730325162 | 0.0461708168464778 | 1.12728074977705 | 0.259623798736976 | 0.703893403935547 | NM_001109745,NM_001109746,NM_013493 |
| Ankrd13d | 1975.047324 | 0.147265751 | 0.084873429 | 1.73512196 | 0.082719205 | 0.239496535 | Commd6 | 665.6802608 | 0.0792604655473725 | 0.0703270467133508 | 1.12702678772271 | 0.25973115530464 | 0.703893403935547 | NM_001168592,NM_001033132 |
| Mrvi1 | 167.324469 | -0.214155939 | 0.123427618 | -1.735073092 | 0.08272786 | 0.239496535 | Efemp1 | 521.472493 | -0.10285230110333 | 0.0912648850390459 | -1.12696467057759 | 0.259757418564535 | 0.703893403935547 | NM_146015 |
| Nde1 | 796.1804172 | 0.141740845 | 0.081688234 | 1.735143945 | 0.082715312 | 0.239496535 | Galt | 1829.338374 | -0.0767720842623311 | 0.0680394759420289 | -1.12834620195697 | 0.259173738419701 | 0.703893403935547 | NM_016658 |
| Pigo | 1041.196464 | 0.118884433 | 0.068543919 | 1.734427151 | 0.082842321 | 0.239783098 | Hdc | 125.4646549 | 0.0719413195804174 | 0.0638042707413692 | 1.12753141356371 | 0.259517866569755 | 0.703893403935547 | NM_008230 |
| Mapk1ip1l | 1709.227097 | -0.105331999 | 0.06075374 | -1.733753325 | 0.08296186 | 0.240084249 | Hyls1 | 224.6006606 | -0.0937909682196042 | 0.083190571478318 | -1.12742305471539 | 0.259563656059333 | 0.703893403935547 | NM_029762 |
| Armcx5 | 938.1143283 | 0.177207274 | 0.102225371 | 1.733496016 | 0.083007545 | 0.240171597 | Nfkb1 | 785.1846131 | -0.0781280141678412 | 0.0693242246850939 | -1.12699441678199 | 0.259744841577198 | 0.703893403935547 | NM_008689 |
| Atp11c | 418.8469482 | 0.174097972 | 0.100453873 | 1.733113586 | 0.083075482 | 0.240323286 | Nipsnap1 | 2808.491747 | 0.0547999445393885 | 0.0485875144442943 | 1.1278606276972 | 0.259378783984971 | 0.703893403935547 | NM_008698 |
| Jakmip2 | 2276.774044 | 0.135418468 | 0.078161305 | 1.732551268 | 0.083175457 | 0.240567581 | Npas1 | 312.4928033 | -0.0975053065827891 | 0.0864488944651425 | -1.12789535581747 | 0.259364115460488 | 0.703893403935547 | NM_008718 |
| Chrna1 | 20.39074904 | 0.29117839 | 0.168091697 | 1.732259213 | 0.08322742 | 0.240672947 | Poll | 463.4766388 | 0.0884476970088892 | 0.0784159497499843 | 1.12792993378119 | 0.259349510930223 | 0.703893403935547 | NM_020032 |
| Cand1 | 6275.053374 | -0.102570919 | 0.059221644 | -1.731983644 | 0.083276475 | 0.240765129 | Pten | 8594.251715 | 0.0830234922498207 | 0.0736735836100474 | 1.12690992051184 | 0.259780568532181 | 0.703893403935547 | NM_008960 |
| Rpl13 | 6060.979544 | -0.145344054 | 0.083921466 | -1.731905573 | 0.083290376 | 0.240765129 | Sppl3 | 3528.074657 | 0.0604121598057413 | 0.0535931006729647 | 1.12723763035074 | 0.259642024308351 | 0.703893403935547 | NM_029012 |
| Rundc3b | 1526.247285 | -0.12809498 | 0.073967848 | -1.731765676 | 0.083315292 | 0.240792227 | Zfp788 | 1372.347446 | 0.0725984569169552 | 0.0643667819189968 | 1.1278870055725 | 0.259367642399172 | 0.703893403935547 | NM_023363 |
| Clpp | 1210.58312 | -0.106611847 | 0.061567876 | -1.731614828 | 0.083342164 | 0.240824971 | Heatr2 | 337.5494778 | -0.0908395822479595 | 0.0806549828256886 | -1.12627365434175 | 0.260049705683632 | 0.704052662567117 | NM_001081265 |
| Gbp11 | 24.98873814 | 0.290601079 | 0.167831924 | 1.731500612 | 0.083362516 | 0.240838862 | Hspa4l | 5388.261287 | 0.0740777038733088 | 0.0657767060074517 | 1.12619965896311 | 0.260081017851898 | 0.704052662567117 | NM_011020 |
| 1600002H07Rik | 318.8052919 | 0.179739272 | 0.103823405 | 1.731201853 | 0.083415769 | 0.240857981 | Lypd6b | 701.3095382 | 0.0731772258200578 | 0.0649564439419226 | 1.1265583732614 | 0.259929247270159 | 0.704052662567117 | NM_027990 |
| Aspg | 119.6052189 | 0.230326552 | 0.133032896 | 1.731350362 | 0.083389294 | 0.240857981 | Obscn | 366.1199401 | -0.0984924133824195 | 0.0874645697910597 | -1.12608355151925 | 0.260130155582935 | 0.704052662567117 | NM_001171512,NM_199152 |
| Smu1 | 2437.84386 | -0.099688795 | 0.05758353 | -1.73120326 | 0.083415518 | 0.240857981 | Sms | 4895.616278 | 0.0754185521092957 | 0.0669707793748517 | 1.12614117400605 | 0.26010576841699 | 0.704052662567117 | NM_009214 |
| Dnajc12 | 389.7266943 | -0.164868018 | 0.095252804 | -1.730846874 | 0.083479079 | 0.240950978 | Spin1 | 9660.627803 | 0.0700782810500194 | 0.0622143854236974 | 1.12639995674259 | 0.259996265103446 | 0.704052662567117 | NM_146043,NM_011462 |
| Ugcg | 2867.810196 | -0.115452817 | 0.066702757 | -1.730855248 | 0.083477585 | 0.240950978 | Actn1 | 4391.513086 | -0.0809431325466226 | 0.0719455597508066 | -1.12506084916123 | 0.260563250123815 | 0.704058139176516 | NM_134156 |
| C920006O11Rik | 101.3385917 | 0.247973202 | 0.143307184 | 1.730361275 | 0.083565748 | 0.241156212 | Aldh3a2 | 2429.358575 | 0.0531494815046394 | 0.0472127986086763 | 1.12574308388641 | 0.260274281639934 | 0.704058139176516 | NM_007437 |
| Tex10 | 775.4214835 | 0.127786691 | 0.073851599 | 1.729958675 | 0.083637659 | 0.241318787 | Baz1b | 6061.131448 | -0.0569099949441157 | 0.0505569409170921 | -1.12566136146255 | 0.260308884430482 | 0.704058139176516 | NM_011714 |
| Haus2 | 804.8281369 | -0.111951843 | 0.064721066 | -1.729758944 | 0.083673353 | 0.241376824 | Eif4e2 | 2345.62571 | 0.052296274732227 | 0.0464703366577034 | 1.12536896638897 | 0.260432716000469 | 0.704058139176516 | NM_023314,NM_001039169,NM_001039170 |
| Wdr86 | 27.86530411 | 0.274875208 | 0.158918415 | 1.729662413 | 0.083690608 | 0.24138166 | Gba | 1572.777947 | -0.0662152655413798 | 0.0588321075163915 | -1.12549538529423 | 0.260379171617735 | 0.704058139176516 | NM_008094,NM_001077411 |
| B130034C11Rik | 46.54259718 | 0.269918802 | 0.156166636 | 1.728402484 | 0.083916091 | 0.24166991 | Med4 | 864.6842326 | 0.0785516334141706 | 0.0698320316062051 | 1.12486536060037 | 0.260646092483384 | 0.704058139176516 | NM_026119 |
| Cacna1s | 51.44499538 | -0.277099445 | 0.160303836 | -1.728588983 | 0.083882682 | 0.24166991 | Rce1 | 571.8295019 | -0.0846705701834105 | 0.0752462882066203 | -1.12524580549292 | 0.260484887788501 | 0.704058139176516 | NM_023131 |
| Cdc7 | 527.7060332 | 0.245618134 | 0.142079761 | 1.728734139 | 0.083856688 | 0.24166991 | Slc16a4 | 94.85891166 | -0.104015652900413 | 0.0924630138713741 | -1.12494335351333 | 0.260613039173661 | 0.704058139176516 | NM_146136 |
| Chst2 | 5698.599544 | -0.167252404 | 0.096785849 | -1.728066711 | 0.083976265 | 0.24166991 | Smc2 | 446.9914081 | -0.0924848224822374 | 0.0821500484822374 | -1.12580368236838 | 0.260248625173692 | 0.704058139176516 | NM_008017 |
| Gm4951 | 19.11771252 | 0.21312278 | 0.123326583 | 1.728117125 | 0.083967228 | 0.24166991 | Strip1 | 3434.407204 | 0.0610444966748507 | 0.0542441111892909 | 1.12536633629832 | 0.26043383004909 | 0.704058139176516 | NM_153563 |
| Hist3h2a | 897.803703 | -0.164007138 | 0.094862687 | -1.728891275 | 0.083828807 | 0.24166991 | Tmem238 | 34.78583793 | -0.0980925015199906 | 0.0872072912403748 | -1.1248199562765 | 0.260665336120839 | 0.704058139176516 | NM_029384 |
| Metap1d | 468.1833227 | 0.20837278 | 0.120539245 | 1.728671682 | 0.083867872 | 0.24166991 | Ahsa2 | 1164.377702 | -0.0737595456970417 | 0.0656850064646817 | -1.12292819422499 | 0.261467992093193 | 0.704173027536904 | NM_172391 |
| Nol6 | 5653.385621 | 0.103159026 | 0.059682512 | 1.728463196 | 0.083905214 | 0.24166991 | Dner | 7463.302369 | 0.0715378236253302 | 0.0637282011550722 | 1.12254578551895 | 0.26163045170408 | 0.704173027536904 | NM_152915 |
| Nudt11 | 826.3418739 | -0.13450816 | 0.077828706 | -1.728258976 | 0.083941805 | 0.24166991 | Gadd45gip1 | 989.685134 | 0.06170455663395 | 0.0549652399586785 | 1.12261052040049 | 0.261602945324121 | 0.704173027536904 | NM_183358 |
| Ptk2b | 23606.20628 | 0.163603913 | 0.094628435 | 1.728908566 | 0.083825461 | 0.24166991 | Gm14403 | 291.6731994 | -0.100351302115569 | 0.089232873144353 | -1.12460014543328 | 0.260758512073673 | 0.704173027536904 | NR_036450 |
| Smarcc2 | 12777.15184 | 0.113452898 | 0.065634059 | 1.728567447 | 0.08388654 | 0.24166991 | Ip6k2 | 2616.516774 | 0.0653661825481786 | 0.058204702106141 | 1.12303955149497 | 0.261420697028992 | 0.704173027536904 | NM_029634 |
| Vav2 | 364.429964 | -0.177733745 | 0.102851689 | -1.728058593 | 0.08397772 | 0.24166991 | Lama3 | 237.7956647 | -0.0964442318584185 | 0.0859171741130493 | -1.12252565164117 | 0.261639007160743 | 0.704173027536904 | NM_010680 |
| Patl1 | 1056.21034 | 0.150743542 | 0.087238832 | 1.727940853 | 0.083998829 | 0.241685767 | Lancl2 | 4268.142315 | 0.0773201111585837 | 0.0688604325999001 | 1.12285253285928 | 0.261500129958283 | 0.704173027536904 | NM_133737 |
| 1700110C19Rik | 15.92851434 | 0.288982788 | 0.167259371 | 1.727752452 | 0.084032615 | 0.241738088 | Mtmr6 | 6627.085125 | 0.073165538523568 | 0.0651122197011893 | 1.12368367810738 | 0.261147243060331 | 0.704173027536904 | NM_144843 |
| Prrc2c | 7871.128241 | -0.150285035 | 0.086993979 | -1.727533743 | 0.084071851 | 0.241761183 | Pcsk4 | 196.4044227 | -0.101072638347388 | 0.0900447655933834 | -1.12247100296538 | 0.261662229910322 | 0.704173027536904 | NM_008793 |
| Zfp706 | 7503.403422 | -0.115734692 | 0.066992187 | -1.727584914 | 0.084062669 | 0.241761183 | Phc3 | 2926.023858 | -0.0683034193124651 | 0.0608214162848245 | -1.12301592900439 | 0.261430729352624 | 0.704173027536904 | NM_001165954,NM_001165956,NM_001165955,NM_153421 |
| Psmf1 | 1353.005288 | 0.126587136 | 0.073297256 | 1.727037854 | 0.084160865 | 0.241972249 | Prorsd1 | 66.8005408 | 0.103202742031268 | 0.0919453823574604 | 1.12243529131285 | 0.261677406212599 | 0.704173027536904 | NM_026465,NM_001163454 |
| Gm715 | 106.2922074 | -0.204186773 | 0.118258931 | -1.72660764 | 0.084238153 | 0.242149526 | Rad18 | 188.7378379 | 0.100500768585046 | 0.0894878908652699 | 1.12306556354487 | 0.261409650182383 | 0.704173027536904 | NM_001167730,NM_021385 |
| Sdc3 | 9385.650652 | -0.107325703 | 0.062164935 | -1.726466918 | 0.084263446 | 0.242177301 | Rpusd1 | 2529.715019 | 0.0572236311692859 | 0.050931112440221 | 1.12354960313208 | 0.261204146206389 | 0.704173027536904 | NM_028009 |
| Tmem55b | 2363.998148 | 0.078968194 | 0.045747731 | 1.726166362 | 0.084317488 | 0.242287677 | Rtbdn | 51.52634375 | -0.103849045409093 | 0.092502122653159 | -1.12266662029454 | 0.261579109637595 | 0.704173027536904 | NM_144929 |
| Ppil4 | 1502.633859 | 0.12394462 | 0.071808997 | 1.726031918 | 0.084341671 | 0.242312228 | Sik2 | 1375.924179 | 0.102535972701616 | 0.0912450738096304 | 1.12374255859053 | 0.261122256133999 | 0.704173027536904 | NM_178710 |
| Gm5111 | 14.32056056 | 0.284464322 | 0.164834671 | 1.725755389 | 0.084391429 | 0.242410233 | Stk32b | 180.6844995 | 0.106497669862797 | 0.0948589418743764 | 1.12269510663353 | 0.261567006955056 | 0.704173027536904 | NM_022416 |
| Ash1l | 7780.332323 | -0.184987166 | 0.107203665 | -1.725567561 | 0.084425239 | 0.242417469 | Tmem64 | 1994.492545 | 0.0753465843511296 | 0.0670706195254459 | 1.12339180529776 | 0.26127112862125 | 0.704173027536904 | NM_181401 |
| Gm16701 | 85.94746744 | -0.233393204 | 0.135251452 | -1.72562439 | 0.084415008 | 0.242417469 | Tmx3 | 2317.274502 | 0.0880451142138697 | 0.0783718647094688 | 1.12342757876517 | 0.261255942370477 | 0.704173027536904 | NM_198295 |
| Nlrp1a | 100.7171996 | 0.215419245 | 0.12489912 | 1.72474589 | 0.084573277 | 0.242797546 | Unc45a | 1138.512037 | -0.0824701648518602 | 0.0734453065988692 | -1.12287862452915 | 0.261489046971672 | 0.704173027536904 | NM_133952 |
| Smad4 | 1619.785957 | -0.1297286 | 0.075224023 | -1.724563444 | 0.084606176 | 0.242846998 | Zfp595 | 131.4045542 | -0.105776314357878 | 0.0941407648543825 | -1.12359735467939 | 0.261183878851932 | 0.704173027536904 | NM_177622 |
| Exosc10 | 1608.586723 | 0.116924178 | 0.067819146 | 1.724058538 | 0.084697276 | 0.242973448 | Kcng3 | 59.66639143 | -0.0990024112640458 | 0.0882190343079599 | -1.12223413054424 | 0.261762904425871 | 0.704272633763698 | NM_153512 |
| Ptprm | 1892.207669 | -0.113742451 | 0.065970605 | -1.724138365 | 0.084682868 | 0.242973448 | 1110001A16Rik | 392.4718473 | 0.0834711879021139 | 0.0744081261403485 | 1.12180204275902 | 0.261946617494365 | 0.704317238569845 | NM_001177402,NM_001177403 |
| U2af2 | 5550.764923 | -0.071339787 | 0.041377253 | -1.724130594 | 0.08468427 | 0.242973448 | Agk | 1736.406618 | 0.0596841112097376 | 0.0531997770136606 | 1.12188649201315 | 0.261910704751807 | 0.704317238569845 | NM_023538 |
| Ppp1r14c | 389.274034 | -0.146594941 | 0.085038445 | -1.723866681 | 0.084731913 | 0.243027817 | Cep76 | 329.5109579 | 0.0927879140526412 | 0.082717917589346 | 1.12173899871716 | 0.26197342971175 | 0.704317238569845 | NM_001081073 |

| | | | | | | |
|---|---|---|---|---|---|---|
| Gdf11 | 315.2618099 | -0.181563704 | 0.105331166 | -1.723741517 | 0.084754516 | 0.243047655 |
| Dr1 | 986.7148141 | -0.116842897 | 0.067796968 | -1.723423622 | 0.084811946 | 0.243122348 |
| Prosc | 2558.399287 | 0.134012246 | 0.077759074 | 1.723429041 | 0.084810967 | 0.243122348 |
| Jdp2 | 1410.357959 | 0.19699364 | 0.114327036 | 1.723071345 | 0.084875624 | 0.243259881 |
| Lpxn | 59.93703947 | 0.266518226 | 0.154709572 | 1.7227003 | 0.084942736 | 0.243407204 |
| Plscr1 | 74.9228647 | -0.240693305 | 0.139734687 | -1.722502196 | 0.084978585 | 0.243464904 |
| 2810021J22Rik | 496.7967671 | -0.127263983 | 0.07389278 | -1.722278991 | 0.085018992 | 0.243535636 |
| C630031E19Rik | 8.382087779 | 0.24778992 | 0.143893501 | 1.722036914 | 0.085062832 | 0.243586371 |
| Fam216a | 1427.014866 | -0.114429891 | 0.066451444 | -1.722007576 | 0.085068146 | 0.243586371 |
| Arl16 | 781.8319362 | -0.118182119 | 0.068648342 | -1.721558244 | 0.085149573 | 0.243774479 |
| Arr3 | 45.77845771 | 0.266524337 | 0.154872671 | 1.720925547 | 0.085264335 | 0.243893179 |
| Cyp11a1 | 102.8846507 | 0.254125712 | 0.147666278 | 1.720946145 | 0.085260597 | 0.243893179 |
| Endov | 1698.480515 | 0.12325798 | 0.071620656 | 1.720983675 | 0.085253786 | 0.243893179 |
| Smim3 | 266.2875888 | 0.235028424 | 0.136566497 | 1.720981561 | 0.08525417 | 0.243893179 |
| Ywhaz | 43501.72649 | -0.10130076 | 0.058865134 | -1.720895766 | 0.08526974 | 0.243893179 |
| Qser1 | 734.9057902 | -0.156089326 | 0.090712843 | -1.720697105 | 0.085305802 | 0.243951291 |
| Parp16 | 135.3823216 | 0.228478349 | 0.132794051 | 1.720546571 | 0.085333136 | 0.243984426 |
| Evi2b | 30.92163632 | 0.286383343 | 0.166468837 | 1.720342068 | 0.085370281 | 0.244006785 |
| Klhdc8b | 953.6710966 | -0.170224476 | 0.098948728 | -1.720330112 | 0.085372453 | 0.244006785 |
| Afap1l1 | 1149.790652 | 0.154661903 | 0.089959323 | 1.719242635 | 0.0855702 | 0.244481779 |
| Efemp1 | 528.4960136 | -0.19802695 | 0.115179445 | -1.719290707 | 0.085561451 | 0.244481779 |
| Hps1 | 447.4882487 | 0.288530188 | 0.167850587 | 1.718970379 | 0.085619766 | 0.244578291 |
| Abcf3 | 2781.846147 | 0.143925132 | 0.08374923 | 1.718524842 | 0.085700927 | 0.244765008 |
| Rfk | 2072.416895 | -0.108640251 | 0.063231761 | -1.718127875 | 0.085773293 | 0.244912142 |
| Rnf167 | 2517.952094 | 0.087490967 | 0.050924017 | 1.718068848 | 0.085784058 | 0.244912142 |
| BC026590 | 1112.873473 | 0.096096911 | 0.055941145 | 1.717821653 | 0.085829151 | 0.244983891 |
| Gmip | 348.0045897 | 0.207550487 | 0.120826402 | 1.717757743 | 0.085840812 | 0.244983891 |
| Snora70 | 15.62937499 | 0.28831741 | 0.167873992 | 1.717463235 | 0.085894566 | 0.245092156 |
| Vps33a | 5485.893068 | 0.141343311 | 0.082306721 | 1.717275445 | 0.085928856 | 0.245144854 |
| Zmiz2 | 13108.27426 | 0.180321843 | 0.105016855 | 1.71707525 | 0.085965424 | 0.245204027 |
| Cnr1 | 6686.17567 | -0.149898659 | 0.087337256 | -1.716319781 | 0.08610353 | 0.24546239 |
| Lacc1 | 213.2393604 | 0.173639152 | 0.1011673 | 1.716356494 | 0.086096814 | 0.24546239 |
| Prkce | 17906.87353 | -0.12173971 | 0.070926253 | -1.716426645 | 0.086083983 | 0.24546239 |
| Bub1b | 84.9209946 | 0.287069083 | 0.167284176 | 1.716056409 | 0.086151718 | 0.245554585 |
| Cav1 | 487.2728416 | -0.214382734 | 0.124954418 | -1.715687507 | 0.086219252 | 0.245701876 |
| Tdrd12 | 10.45086136 | -0.275340066 | 0.160500396 | -1.715510194 | 0.086251728 | 0.245749223 |
| Ankrd32 | 556.3327726 | 0.151652874 | 0.088417496 | 1.715190772 | 0.086310256 | 0.245799414 |
| Nat10 | 1415.120491 | 0.146184636 | 0.085231185 | 1.715154326 | 0.086316936 | 0.245799414 |
| Scarf1 | 81.19007347 | -0.222305284 | 0.129599506 | -1.715325079 | 0.086285643 | 0.245799414 |
| Gpatch8 | 3282.021016 | 0.166133577 | 0.096874362 | 1.714938534 | 0.086356497 | 0.245830535 |
| Phf11c | 10.97839985 | 0.278360647 | 0.162316833 | 1.714921625 | 0.086359597 | 0.245830535 |
| Per1 | 4117.479076 | 0.278894717 | 0.16264552 | 1.714739617 | 0.086392977 | 0.245880378 |
| Btbd17 | 479.3951368 | -0.263238497 | 0.153538218 | -1.714481912 | 0.086440256 | 0.24595758 |
| Slc6a13 | 430.6781508 | -0.244530653 | 0.142631814 | -1.714418729 | 0.086451851 | 0.24595758 |
| Gpd2 | 4965.184424 | -0.126180885 | 0.073613728 | -1.714094474 | 0.086511376 | 0.246081745 |
| Eif1a | 1454.699863 | -0.085229881 | 0.049728924 | -1.713889515 | 0.086549019 | 0.246143631 |
| Nfyc | 1406.76252 | 0.165767821 | 0.096762465 | 1.713141761 | 0.086686463 | 0.246489274 |
| Brpf3 | 1802.637395 | 0.12672318 | 0.073981456 | 1.712904647 | 0.086730083 | 0.246568057 |
| Kcnh6 | 112.8628633 | -0.228815708 | 0.133606253 | -1.712612263 | 0.086783896 | 0.246654237 |
| Spag1 | 417.2929861 | 0.195909759 | 0.114395386 | 1.712566962 | 0.086792236 | 0.246654237 |
| Bud13 | 416.7066335 | 0.150777184 | 0.088052082 | 1.71236364 | 0.086829675 | 0.246670149 |
| Lphn1 | 26793.65873 | -0.112266258 | 0.065560935 | -1.712395626 | 0.086823785 | 0.246670149 |
| Syn2 | 21412.88207 | -0.116800739 | 0.068230399 | -1.711857763 | 0.086922884 | 0.246889673 |
| Accsl | 12.88073023 | 0.267324761 | 0.156179386 | 1.71165202 | 0.086960816 | 0.24695214 |
| Uba6 | 811.0414721 | -0.144619229 | 0.084502152 | -1.711426575 | 0.087002395 | 0.247012669 |
| Zfp935 | 241.3951015 | -0.234630026 | 0.137101214 | -1.711363586 | 0.087014015 | 0.247012669 |
| Asna1 | 3079.53666 | 0.130863614 | 0.076473274 | 1.711233319 | 0.08703805 | 0.247035638 |
| Cnksr2 | 9288.689169 | -0.12679058 | 0.07411365 | -1.71075881 | 0.087125646 | 0.247103197 |
| 10-Mar | 65.14516492 | 0.270613255 | 0.158182043 | 1.71077102 | 0.087123391 | 0.247103197 |
| Stip1 | 6665.272649 | -0.1224701 | 0.071585376 | -1.7108257 | 0.087113294 | 0.247103197 |
| Tmem198 | 2238.030794 | -0.1770627 | 0.103497711 | -1.710788562 | 0.087120152 | 0.247103197 |
| Gm5069 | 199.2472907 | -0.206926675 | 0.120973724 | -1.710509266 | 0.087171741 | 0.247188683 |
| Snord37 | 8.138696652 | 0.259866645 | 0.151948813 | 1.710224905 | 0.087224292 | 0.247292439 |
| Med11 | 249.5512667 | -0.180696107 | 0.105675843 | -1.709909299 | 0.087282647 | 0.24741261 |
| Aspdh | 70.01622749 | 0.235485771 | 0.137741791 | 1.709617466 | 0.087336634 | 0.247497921 |
| Car5a | 10.59898722 | -0.273371719 | 0.159906345 | -1.709573928 | 0.08734469 | 0.247497921 |
| Aven | 293.4004143 | 0.174125292 | 0.101883496 | 1.70906279 | 0.087439319 | 0.247630198 |
| Cirbp | 1966.46323 | 0.187980087 | 0.10998787 | 1.70909835 | 0.087432733 | 0.247630198 |
| Mmadhc | 1357.367012 | -0.087956001 | 0.051460653 | -1.709189364 | 0.087415878 | 0.247630198 |
| Pdcd4 | 2624.658169 | 0.117849781 | 0.06897043 | 1.70870009 | 0.087506517 | 0.247729944 |
| Vps51 | 1899.31038 | 0.095968191 | 0.056163467 | 1.708729821 | 0.087501008 | 0.247729944 |
| Slc25a37 | 1326.688165 | 0.181151753 | 0.106046128 | 1.708235429 | 0.087592667 | 0.247928532 |
| Atp6v0b | 6473.483573 | -0.105527731 | 0.061782244 | -1.708059208 | 0.087625357 | 0.247946635 |
| Gm17066 | 425.019724 | 0.227499013 | 0.133193927 | 1.708028424 | 0.087631068 | 0.247946635 |
| Clip2 | 6694.133926 | 0.107741688 | 0.063100312 | 1.707466804 | 0.087735323 | 0.248196292 |
| Cldn25 | 2262.204554 | -0.099565007 | 0.058338558 | -1.706675832 | 0.087882322 | 0.248566607 |
| Zbtb24 | 498.7825236 | 0.145553783 | 0.085289258 | 1.706589857 | 0.087898312 | 0.248566607 |
| Tmem117 | 260.3696503 | -0.204823427 | 0.120036562 | -1.706341994 | 0.087944424 | 0.248651624 |
| Ptpn9 | 1957.263255 | -0.133314797 | 0.078142011 | -1.706057933 | 0.087997294 | 0.248755714 |
| Cox7a2l | 5000.333194 | -0.092935886 | 0.054477911 | -1.705937038 | 0.088019803 | 0.248773956 |
| E2f2 | 31.56679426 | -0.282757058 | 0.165762538 | -1.705795903 | 0.088046086 | 0.248802856 |
| Ttk | 16.13728722 | -0.285761127 | 0.167535658 | -1.705673472 | 0.088068891 | 0.248821919 |
| Mll5 | 4892.70503 | 0.143957177 | 0.084440236 | 1.704840997 | 0.088224083 | 0.24921494 |
| Btn2a2 | 54.53343545 | 0.282058515 | 0.165466014 | 1.704631109 | 0.088263246 | 0.249234687 |
| Mir425 | 20.00114247 | 0.286323532 | 0.167964694 | 1.704664982 | 0.088256924 | 0.249234687 |
| Mns1 | 51.0060736 | 0.261629624 | 0.153497178 | 1.704458858 | 0.088295396 | 0.249280041 |
| Ces5a | 12.64150111 | 0.281436161 | 0.165158654 | 1.704035203 | 0.088374511 | 0.249457947 |
| Ercc1 | 596.3682081 | 0.13629815 | 0.079998832 | 1.703751761 | 0.088427474 | 0.249561981 |
| Fam203a | 293.0175487 | 0.15441258 | 0.090645581 | 1.703476092 | 0.088479008 | 0.249660098 |
| Mapk7 | 706.7302424 | 0.180048277 | 0.10569976 | 1.703393431 | 0.088494466 | 0.249660098 |
| 2610035D17Rik | 508.3643898 | 0.149135239 | 0.087573995 | 1.702962613 | 0.088575065 | 0.249841992 |
| Lpin2 | 3929.11538 | 0.109076289 | 0.064057444 | 1.702788648 | 0.088607628 | 0.249888349 |
| Atpaf2 | 724.2690409 | 0.104382188 | 0.061308887 | 1.702562109 | 0.088650046 | 0.249962478 |
| Wdr27 | 39.61471571 | 0.2859686 | 0.16798501 | 1.702345939 | 0.088690538 | 0.250031149 |
| Itpripl2 | 394.3116563 | -0.148583038 | 0.087288735 | -1.702201751 | 0.088717555 | 0.250061814 |
| Deaf1 | 1531.56352 | 0.191353811 | 0.112461907 | 1.701498905 | 0.088849343 | 0.250382744 |
| Smim8 | 326.08302 | -0.177342641 | 0.104232001 | -1.701422203 | 0.088863735 | 0.250382744 |
| Mchr1 | 935.5354576 | -0.185497225 | 0.109037848 | -1.701218684 | 0.088901931 | 0.250444821 |
| Tbc1d24 | 4934.773546 | 0.095244628 | 0.055990611 | 1.701082132 | 0.088927566 | 0.250471497 |
| Oat | 3123.510016 | -0.081749545 | 0.048065735 | -1.700786337 | 0.088983116 | 0.25053687 |
| Pyurf | 790.8390421 | -0.163091188 | 0.095887967 | -1.700851447 | 0.088970886 | 0.25053687 |
| Tctex1d4 | 9.039008638 | 0.252861868 | 0.148815917 | 1.699158757 | 0.089289275 | 0.251353203 |
| Rgs6 | 1300.23808 | 0.186043315 | 0.109500796 | 1.699013352 | 0.089316668 | 0.251384642 |
| Zfp110 | 653.3694407 | 0.135341731 | 0.079664612 | 1.69889399 | 0.08933916 | 0.251402277 |
| Ube2ql1 | 7009.672199 | -0.122611389 | 0.072186557 | -1.698534949 | 0.089406842 | 0.251547052 |
| Fbxl15 | 443.9386806 | -0.159324198 | 0.093817997 | -1.698226387 | 0.089465042 | 0.251665099 |
| Laptm4a | 6182.679753 | -0.070823991 | 0.041708299 | -1.698079089 | 0.089492836 | 0.251697585 |
| Hcrtr2 | 71.45472533 | -0.228224846 | 0.134426194 | -1.697770644 | 0.089551059 | 0.251815626 |
| Myt1l | 10664.48447 | -0.114453512 | 0.067430982 | -1.697343091 | 0.089631815 | 0.251996977 |
| Fut7 | 8.200417491 | 0.262861688 | 0.154927185 | 1.696678915 | 0.089757381 | 0.25230422 |
| Poli | 486.3938117 | 0.137688196 | 0.081164658 | 1.696405779 | 0.08980906 | 0.252403696 |
| Nlgn1 | 2666.56176 | -0.139480346 | 0.082241428 | -1.695986435 | 0.089888449 | 0.2525352 |
| Procr | 23.99475095 | 0.284006224 | 0.167455729 | 1.696007808 | 0.089884401 | 0.2525352 |
| Khdrbs1 | 3962.977145 | -0.120122518 | 0.070834017 | -1.695830939 | 0.089917902 | 0.252572148 |
| Gulp1 | 84.21214276 | -0.238828431 | 0.140861721 | -1.695481419 | 0.089984132 | 0.25271237 |
| Fcgrt | 623.3309307 | -0.192368948 | 0.113488766 | -1.695048368 | 0.090066246 | 0.252897139 |
| Eif1 | 11852.91944 | -0.084580022 | 0.049904135 | -1.694849989 | 0.090103882 | 0.252956975 |
| Cdnf | 57.20405843 | 0.249764279 | 0.147402062 | 1.694442229 | 0.090181281 | 0.253004419 |
| Clptm1 | 7050.626363 | -0.091941924 | 0.054254783 | -1.694632594 | 0.09014514 | 0.253004419 |
| Lancl1 | 7738.949675 | 0.116943511 | 0.069023971 | 1.694244898 | 0.090218756 | 0.253004419 |
| Nipa1 | 1966.409344 | -0.115504272 | 0.068170053 | -1.694355037 | 0.090197838 | 0.253004419 |
| Spc24 | 64.44160373 | -0.235616644 | 0.139041758 | -1.69457469 | 0.090156132 | 0.253004419 |
| Tagap1 | 781.8964997 | 0.158589837 | 0.09360384 | 1.694266359 | 0.09021468 | 0.253004419 |
| Slmo1 | 1523.658433 | 0.141895766 | 0.083756934 | 1.694137539 | 0.09023915 | 0.253015816 |
| Ilf3 | 3439.728647 | 0.1349301 | 0.07965155 | 1.694004685 | 0.090264393 | 0.2530408 |
| Tsga14 | 300.2737847 | -0.230782144 | 0.136260613 | -1.693681966 | 0.090325733 | 0.253166952 |
| Rps15a-ps4 | 18.00614132 | -0.281231652 | 0.16609747 | -1.693172398 | 0.090422657 | 0.253392774 |
| Triobp | 359.9888177 | -0.146389837 | 0.086478298 | -1.692792748 | 0.090494924 | 0.25354943 |
| Stard6 | 68.74060725 | 0.239120501 | 0.141269276 | 1.692657504 | 0.090520679 | 0.253575736 |
| Acvr2a | 1655.222413 | 0.183139655 | 0.108294347 | 1.691128492 | 0.090812265 | 0.253890779 |
| Bbs1 | 2222.538176 | -0.096605983 | 0.057110658 | -1.691557859 | 0.090730307 | 0.253890779 |
| Bora | 62.18882426 | 0.257714923 | 0.152318692 | 1.691945483 | 0.090656369 | 0.253890779 |
| Clec2l | 1261.473911 | -0.104337716 | 0.06168593 | -1.691434584 | 0.090753832 | 0.253890779 |
| Glrx | 1262.084873 | 0.139164225 | 0.082299508 | 1.690948449 | 0.090846649 | 0.253890779 |
| Larp6 | 1330.007546 | 0.127269969 | 0.075247005 | 1.691362592 | 0.090767572 | 0.253890779 |
| Lyrm1 | 243.3445357 | -0.168175872 | 0.099456247 | -1.690953319 | 0.090845719 | 0.253890779 |
| Myo15 | 17.91348268 | 0.283144409 | 0.167359873 | 1.691829737 | 0.090678442 | 0.253890779 |
| Nmnat1 | 130.6143512 | -0.208403461 | 0.123252558 | -1.690865199 | 0.090862551 | 0.253890779 |
| Numb | 1976.209571 | -0.113268722 | 0.06698573 | -1.690938089 | 0.090848628 | 0.253890779 |
| Plekhn1 | 207.0382843 | 0.24882339 | 0.147084711 | 1.691701251 | 0.09070295 | 0.253890779 |
| Sema4a | 2258.716149 | -0.182332175 | 0.107809793 | -1.691239455 | 0.090791078 | 0.253890779 |
| Slc30a1 | 1495.440478 | -0.178173011 | 0.105356867 | -1.691138086 | 0.090810433 | 0.253890779 |
| Smap2 | 7926.175978 | -0.091111014 | 0.053880377 | -1.690986944 | 0.090839296 | 0.253890779 |
| Ttyh3 | 10576.59309 | -0.102344423 | 0.060543742 | -1.69042116 | 0.09094741 | 0.254082071 |
| Zfp292 | 1653.139393 | -0.144323409 | 0.085389287 | -1.690181681 | 0.090993202 | 0.254164174 |
| Tars | 1134.249308 | 0.137324071 | 0.081256751 | 1.690001992 | 0.091027573 | 0.254214352 |
| Exoc4 | 2710.992845 | -0.114294124 | 0.067638051 | -1.689790317 | 0.091068077 | 0.254235817 |
| Mlh3 | 1253.078643 | -0.097877938 | 0.057921284 | -1.689844075 | 0.091057789 | 0.254235817 |
| Ccdc64 | 1259.548578 | 0.164889955 | 0.097657463 | 1.688452159 | 0.091324465 | 0.254676254 |
| Ccnd1 | 2340.566587 | -0.165080195 | 0.097754083 | -1.68872941 | 0.091271297 | 0.254676254 |
| Cyp4b1 | 25.99196954 | 0.279023365 | 0.165253764 | 1.688453911 | 0.091324129 | 0.254676254 |
| Fbxo9 | 2856.870341 | -0.171881828 | 0.101784533 | -1.688683173 | 0.091280162 | 0.254676254 |
| Stap2 | 64.40375916 | 0.264974388 | 0.156915488 | 1.688643944 | 0.091287684 | 0.254676254 |
| Telo2 | 747.7107359 | 0.160999739 | 0.095350581 | 1.688502969 | 0.091314719 | 0.254676254 |
| Gltscr2 | 2054.907563 | 0.16010978 | 0.094832123 | 1.688349631 | 0.091344133 | 0.254685262 |
| Adamts20 | 375.589424 | 0.242882019 | 0.143886561 | 1.688010449 | 0.091409223 | 0.254729227 |
| Gpr50 | 20.09085074 | -0.271576234 | 0.160882254 | -1.688043445 | 0.091402889 | 0.254729227 |
| Snora81 | 17.20572739 | 0.283557819 | 0.167979443 | 1.688050714 | 0.091401494 | 0.254729227 |
| 4632415K11Rik | 375.4798491 | -0.146112754 | 0.086567871 | -1.687840443 | 0.091441862 | 0.254774358 |
| Dgkk | 156.548618 | -0.248187396 | 0.147064732 | -1.687606488 | 0.091486793 | 0.254853717 |
| 2810055G20Rik | 31.86903466 | 0.278631864 | 0.165117767 | 1.687473542 | 0.091512333 | 0.254879039 |
| Kansl3 | 3727.441082 | 0.11167621 | 0.066193858 | 1.687108342 | 0.091582521 | 0.255028682 |
| Eif4a2 | 22339.59389 | -0.095587061 | 0.056681885 | -1.686377582 | 0.091723097 | 0.255236649 |
| Smim13 | 10071.96034 | -0.154590836 | 0.091669678 | -1.686390072 | 0.091720692 | 0.255236649 |
| Syt14 | 226.0519141 | 0.160612625 | 0.095240412 | 1.686391547 | 0.091720409 | 0.255236649 |
| Usp19 | 5607.861882 | 0.116414079 | 0.069027883 | 1.686479052 | 0.091703566 | 0.255236649 |
| Chchd6 | 1272.479513 | -0.156476816 | 0.092827899 | -1.685665814 | 0.091860185 | 0.255572222 |
| Dars2 | 774.7369334 | 0.116000611 | 0.068848229 | 1.684874286 | 0.092012829 | 0.255766056 |
| Hps3 | 617.1261928 | 0.135431363 | 0.080389381 | 1.684692211 | 0.09204797 | 0.255766056 |
| Mrpl3 | 1772.663144 | -0.110929785 | 0.065846875 | -1.684662873 | 0.092053633 | 0.255766056 |
| Pdss2 | 349.9318552 | -0.149079277 | 0.08846996 | -1.685083583 | 0.091972447 | 0.255766056 |
| Rreb1 | 2192.52444 | 0.283213521 | 0.168117156 | 1.684619981 | 0.092061914 | 0.255766056 |
| Samd12 | 271.7780415 | -0.186682655 | 0.110811357 | -1.684688816 | 0.092048625 | 0.255766056 |
| Snord53 | 8.176687852 | 0.265309303 | 0.157476388 | 1.684756085 | 0.092035641 | 0.255766056 |
| Tfdp2 | 1685.002767 | 0.1838495 | 0.109123784 | 1.68477938 | 0.092031145 | 0.255766056 |
| Itfg3 | 693.6113591 | 0.134801563 | 0.080051588 | 1.683933656 | 0.092194491 | 0.256088463 |
| 1110032F04Rik | 439.0759845 | -0.200196716 | 0.118937634 | -1.683207487 | 0.092334932 | 0.256340698 |
| Lrrc55 | 904.3925211 | -0.248289592 | 0.147503336 | -1.683281191 | 0.09232067 | 0.256340698 |
| Npdc1 | 5687.219208 | 0.119685509 | 0.071100989 | 1.683317083 | 0.092313725 | 0.256340698 |
| Mgl2 | 16.76142413 | -0.264379531 | 0.157120404 | -1.682655623 | 0.092441777 | 0.256585309 |
| Vps37a | 2157.695645 | 0.088730677 | 0.052734851 | 1.682581361 | 0.092456163 | 0.256585309 |
| Gas6 | 3456.335157 | 0.126577536 | 0.07524262 | 1.682258483 | 0.092518728 | 0.256666996 |
| Lbr | 469.8781697 | 0.16947915 | 0.100740591 | 1.682332302 | 0.092504421 | 0.256666996 |
| Dvl1 | 4085.95285 | 0.113373245 | 0.067406494 | 1.681933568 | 0.092581723 | 0.256795777 |
| Gcn1l1 | 3511.489097 | 0.097722954 | 0.058105414 | 1.681821836 | 0.092603393 | 0.256809911 |
| Fam169b | 23.80812707 | -0.282461339 | 0.167970012 | -1.68161767 | 0.092643002 | 0.256873778 |
| Ccdc101 | 405.2147884 | 0.197104158 | 0.117230919 | 1.681332539 | 0.092698341 | 0.25698123 |
| Arf6 | 1314.212621 | -0.097676965 | 0.058105088 | -1.681039785 | 0.092755187 | 0.257092821 |
| Ahcyl1 | 16672.60622 | 0.064063629 | 0.038114061 | 1.680839758 | 0.092794044 | 0.257149547 |
| Hcls1 | 207.587284 | 0.235534442 | 0.14013538 | 1.680763574 | 0.092808847 | 0.257149547 |
| Sync | 56.5783377 | -0.250785839 | 0.149246423 | -1.680347405 | 0.092889743 | 0.257327673 |
| Fbxo22 | 1358.376155 | 0.126111582 | 0.075055755 | 1.680238677 | 0.092910887 | 0.257340236 |
| Slc19a1 | 591.0200935 | 0.153935568 | 0.091620845 | 1.680136961 | 0.092930671 | 0.257349029 |
| Tnfrsf12a | 138.216222 | 0.267571498 | 0.159278904 | 1.679892886 | 0.092978158 | 0.257388527 |
| Ywhag | 44860.09164 | -0.123350282 | 0.073424641 | -1.679957575 | 0.09296557 | 0.257388527 |
| Snord12 | 9.91430037 | 0.255197567 | 0.151924276 | 1.679768194 | 0.093002426 | 0.257409715 |
| Clec4a1 | 12.98577273 | -0.274277118 | 0.163331061 | -1.679271022 | 0.093099236 | 0.25758027 |
| Nfix | 3580.065298 | 0.164274252 | 0.097829132 | 1.679195642 | 0.093113921 | 0.25758027 |
| Wdr16 | 33.77385267 | 0.281232313 | 0.167471163 | 1.679287986 | 0.093095931 | 0.25758027 |
| Ccdc146 | 22.32748987 | 0.2815093 | 0.167677426 | 1.678874174 | 0.093176569 | 0.257707561 |
| 3110035E14Rik | 13419.38378 | -0.204437901 | 0.121786019 | -1.678664779 | 0.093217394 | 0.25777446 |
| Acadsb | 2051.519311 | 0.092259062 | 0.054972526 | 1.678275821 | 0.093293266 | 0.257938234 |
| Pard3b | 180.126834 | -0.202975981 | 0.120951318 | -1.678162625 | 0.093315356 | 0.257953278 |
| 0610009O20Rik | 1413.035145 | 0.127585189 | 0.076072933 | 1.677143021 | 0.093514518 | 0.258457713 |
| Il3ra | 50.05184338 | 0.259278391 | 0.154606426 | 1.677022085 | 0.093538163 | 0.258476958 |
| Smo | 614.0120718 | -0.156566513 | 0.093365655 | -1.676917636 | 0.093558589 | 0.2584873 |
| Ovol2 | 193.9946592 | 0.245832726 | 0.14662403 | 1.676619622 | 0.093616887 | 0.258602255 |
| Helz | 1130.598144 | -0.191131486 | 0.114020502 | -1.676290515 | 0.093681302 | 0.258661178 |
| Lpgat1 | 13165.73612 | -0.125768623 | 0.075024918 | -1.676358027 | 0.093668085 | 0.258661178 |
| Pinx1 | 152.7327591 | -0.180269092 | 0.107544482 | -1.676228183 | 0.093693506 | 0.258661178 |
| Rnf182 | 369.3571025 | 0.186645429 | 0.111352359 | 1.676169508 | 0.093704995 | 0.258661178 |
| Daam2 | 4173.112571 | 0.139629735 | 0.083356724 | 1.675086637 | 0.093917234 | 0.259077536 |
| Lrrc3b | 510.4011606 | -0.144039441 | 0.085983786 | -1.675193062 | 0.093896358 | 0.259077536 |
| Mapk6 | 3269.280817 | 0.14910326 | 0.089013752 | 1.67505871 | 0.093922713 | 0.259077536 |
| Plcg2 | 214.2150024 | 0.16588359 | 0.099024836 | 1.675171562 | 0.093900575 | 0.259077536 |
| Trim68 | 176.7693508 | 0.248730674 | 0.148538422 | 1.674520784 | 0.094028293 | 0.259322601 |
| Ica1 | 1696.847826 | 0.102245203 | 0.061064526 | 1.6743797 | 0.094056 | 0.25935285 |
| Med24 | 4834.310498 | 0.096890942 | 0.057871725 | 1.674236274 | 0.094084173 | 0.259384373 |
| Il6ra | 187.2570572 | 0.208198493 | 0.12438064 | 1.673881826 | 0.094153827 | 0.259530224 |
| Lcmt1 | 2147.991473 | -0.10364376 | 0.061921432 | -1.673794617 | 0.094170971 | 0.259531309 |
| 1810009A15Rik | 352.695977 | -0.156593695 | 0.093571565 | -1.673517966 | 0.094225373 | 0.259635057 |
| Tal1 | 92.897028 | 0.244637791 | 0.146192514 | 1.673394794 | 0.094249602 | 0.259655643 |
| 5830418K08Rik | 1081.683019 | 0.136893089 | 0.081829492 | 1.6729065 | 0.094345704 | 0.259862697 |
| Rbp4 | 320.082795 | -0.227421118 | 0.13594891 | -1.672842519 | 0.094358302 | 0.259862697 |
| Zfp595 | 113.6468897 | 0.21534906 | 0.128742975 | 1.672705321 | 0.094385321 | 0.259890913 |
| Plekhj1 | 677.2831688 | -0.118613368 | 0.070916705 | -1.672573015 | 0.094411383 | 0.259916483 |
| Mboat2 | 1234.118184 | -0.105202223 | 0.062907261 | -1.672338297 | 0.094457632 | 0.25999761 |
| Ssc5d | 108.8474851 | 0.25751548 | 0.153999744 | 1.672181217 | 0.094488593 | 0.260036636 |
| Smurf2 | 1431.563926 | -0.126691951 | 0.075770054 | -1.672058341 | 0.094512818 | 0.260057113 |
| Afap1l2 | 420.5962748 | -0.141768097 | 0.084791488 | -1.671961423 | 0.094531929 | 0.260063514 |
| Tmem141 | 460.5743786 | -0.161024697 | 0.096316831 | -1.671823047 | 0.094559221 | 0.260092414 |
| Psmb9 | 60.88580777 | 0.278691384 | 0.16671232 | 1.671690392 | 0.09458539 | 0.260118216 |
| Trmt10b | 401.3786115 | 0.147487503 | 0.088238264 | 1.671468775 | 0.094629121 | 0.260192299 |
| Nfkb1 | 716.9560893 | 0.134400113 | 0.080430958 | 1.670999785 | 0.09472172 | 0.260400698 |
| Fam98c | 739.5984597 | -0.158710717 | 0.094995146 | -1.670724499 | 0.094776108 | 0.260503993 |

| | | | | | | | |
|---|---|---|---|---|---|---|---|
| Ttc33 | 1492.726429 | 0.0682836084582754 | 0.0608663195880392 | 1.12186195781901 | 0.261921137769786 | 0.704317238569845 | NM_026213 |
| Zfp618 | 75.99833589 | -0.106623797387721 | 0.0950672437600528 | -1.12156188788681 | 0.262048763932166 | 0.704389405047861 | NM_028326 |
| Jazf1 | 1207.873628 | 0.0609989315347766 | 0.0543979467464673 | 1.12134621218471 | 0.262140521957713 | 0.70450568398481 | NM_173406,NM_001168277 |
| Nfkb2 | 122.3311822 | -0.104326335706368 | 0.0930777321197341 | -1.12085171534008 | 0.262350986664808 | 0.704810510873497 | NM_001177370,NM_001177369,NM_019408 |
| Ppib | 3046.794178 | -0.0610541851373525 | 0.0544693981450437 | -1.12088965945198 | 0.262334832992097 | 0.704810510873497 | NM_011149 |
| Nudt21 | 884.7805253 | 0.0727025467829034 | 0.0648729026655054 | 1.12069205778827 | 0.262418964048501 | 0.704862772146546 | NM_026623 |
| Epha10 | 2210.258782 | -0.0840865157073809 | 0.0750398331756054 | -1.12055840410259 | 0.262475879121691 | 0.704885305994026 | NM_177671,NM_001256432 |
| Uhrf1 | 37.03406341 | -0.0927804969197549 | 0.0828326526992103 | -1.12009568565513 | 0.262672988948556 | 0.705284258311779 | NM_001111078,NM_001111080,NM_001111079,NM_010931 |
| Far2 | 569.9657833 | 0.0842555091269871 | 0.0752520529959105 | 1.11964399338801 | 0.262865500381483 | 0.705670718636374 | NM_178797 |
| Cep170b | 13060.53013 | 0.0667832528797866 | 0.0596601415259207 | 1.11939481153881 | 0.262971743469289 | 0.705682020443724 | NM_001024602 |
| Kctd6 | 1241.738139 | -0.071618660413275 | 0.0639856407692165 | -1.11929269680348 | 0.263015290454054 | 0.705682020443724 | NM_027782 |
| Tnfrsf1a | 459.7421971 | -0.0948097141629959 | 0.0847050521811414 | -1.11929231753787 | 0.263015452201729 | 0.705682020443724 | NM_011609 |
| Pnpla6 | 2479.164265 | -0.0769019180447005 | 0.0687154399599558 | -1.11913593384912 | 0.263082152094886 | 0.705730626284453 | NM_015801,NM_001122818 |
| Pnoc | 343.0954923 | 0.105069642329946 | 0.0939080425809534 | 1.11885669685182 | 0.263201279298029 | 0.705919827009447 | NM_010932,NM_001205075 |
| Arrb2 | 1543.056135 | -0.0756699209349946 | 0.0676943659718313 | -1.11781711592486 | 0.263645109870747 | 0.706200556819541 | NM_001271360,NM_001271358,NM_145429,NM_001271359 |
| Glra3 | 204.1073634 | 0.103968416258079 | 0.0929821138782804 | 1.11815500768439 | 0.263500796396584 | 0.706200556819541 | NM_080438 |
| Golm1 | 1432.156782 | -0.0895180554227561 | 0.0800777600032955 | -1.1178891045288 | 0.263614358988424 | 0.706200556819541 | NM_027307,NM_001035122 |
| Magel2 | 203.5709119 | 0.0654584453388077 | 0.0585433873512581 | 1.1181185151802 | 0.263516379708827 | 0.706200556819541 | NM_013779 |
| Palb2 | 56.55460306 | -0.105286734417361 | 0.094135880743339 | -1.11845487168091 | 0.263372770225301 | 0.706200556819541 | NM_001081238 |
| Rpap1 | 1099.856327 | -0.0817992081088685 | 0.0731778074588356 | -1.1178144160015 | 0.263646263226791 | 0.706200556819541 | NM_177294,NM_001163701 |
| Rybp | 2625.415409 | 0.084731528620352 | 0.0757710097483085 | 1.11825787859774 | 0.263456871083731 | 0.706200556819541 | NM_019743 |
| Kdm4d | 77.18909466 | -0.104859783469338 | 0.0938823035627448 | -1.11692810561744 | 0.264025066346906 | 0.706954491014774 | NM_173433 |
| Rab11a | 3334.297483 | 0.0723677617646898 | 0.0647858156523383 | 1.11703095864438 | 0.263981088410791 | 0.706954491014774 | NM_017382 |
| Rassf5 | 1359.212837 | 0.0687583099336674 | 0.0615767594086573 | 1.11662761395658 | 0.2641535796284 | 0.707085906453636 | NM_018750 |
| Rcn1 | 1099.054125 | 0.0908575934668921 | 0.0813709076569784 | 1.11658571943065 | 0.264171500366507 | 0.707085906453636 | NM_009037 |
| Kif7 | 103.1764909 | -0.105945377457529 | 0.0949005633179393 | -1.11638301979923 | 0.264258218690345 | 0.707187709044942 | NM_010626 |
| Pex5 | 2346.030101 | 0.0584860626747569 | 0.0524025811838385 | 1.11609125645121 | 0.264383074430946 | 0.707261241102012 | NM_175933,NM_008995 |
| Zfp280b | 1047.607389 | 0.071711042561257 | 0.064247421589915 | 1.11616996895193 | 0.264349386593753 | 0.707261241102012 | NM_177475 |
| 4930453N24Rik | 575.0473276 | 0.0883964426247107 | 0.0793132489155165 | 1.11452303156651 | 0.265054869455899 | 0.707370514515615 | NM_026273 |
| Ahi1 | 10451.49133 | 0.105836512103642 | 0.0948648335975732 | 1.11565590841187 | 0.264569450689325 | 0.707370514515615 | NM_026203,NM_001177776 |
| Ergic3 | 3341.878652 | 0.0730628316833883 | 0.0654937538092938 | 1.11556946172507 | 0.264606470037598 | 0.707370514515615 | NM_025516 |
| Fbxo32 | 1494.481048 | -0.0822653693050011 | 0.0737933876330951 | -1.11480678613142 | 0.26493322781225 | 0.707370514515615 | NM_026346 |
| Gamt | 340.1396624 | 0.0866940981542243 | 0.0777383198447954 | 1.11520416607034 | 0.264762941213749 | 0.707370514515615 | NM_010255 |
| Gas2l1 | 2169.210666 | -0.0779794355474735 | 0.0699162722432441 | -1.11532598986652 | 0.264710751978225 | 0.707370514515615 | NM_030228,NM_144560,NM_001190406,NM_001190408 |
| Ift20 | 1350.604697 | -0.0612273658032942 | 0.0549417772582735 | -1.11440453619607 | 0.265105678164877 | 0.707370514515615 | NM_018854 |
| Mpv17l | 2804.856987 | 0.0636103749489388 | 0.0570786992698316 | 1.11443280527872 | 0.265093556275206 | 0.707370514515615 | NM_033564 |
| Nipal2 | 310.9845804 | 0.0972183884251852 | 0.0871818139961117 | 1.11512233995867 | 0.264797999447954 | 0.707370514515615 | NM_145469 |
| Rgsl1 | 125.5817812 | -0.10468418648202 | 0.093920842249421 | -1.1146001672772 | 0.265021798639458 | 0.707370514515615 | NM_001243223 |
| Siva1 | 190.2692243 | 0.103817192241122 | 0.0931576970171426 | 1.11442420288704 | 0.265097244972906 | 0.707370514515615 | NM_001161737,NM_013929 |
| Tmem183a | 2221.324488 | 0.0648219704102302 | 0.0580913653990184 | 1.11586239994499 | 0.264481038591594 | 0.707370514515615 | NM_020588,NM_001042485 |
| Wipi2 | 5439.478921 | 0.0431251959538023 | 0.038672085243031 | 1.1151505196264 | 0.26478592556442 | 0.707370514515615 | NM_178398 |
| Zscan29 | 368.3683436 | -0.0932288159379058 | 0.0836186940921135 | -1.11492791115831 | 0.264881314911058 | 0.707370514515615 | NM_178889 |
| Zfp800 | 439.6752478 | -0.101074460999079 | 0.0907085643828131 | -1.1142769339014 | 0.265160399264288 | 0.707386585805885 | NM_001081678 |
| Cdh13 | 2807.203556 | 0.0879823134137344 | 0.0789701076705969 | 1.11412173553985 | 0.265226965176611 | 0.707434244611724 | NM_019707 |
| Kctd16 | 689.3346116 | -0.0746831730511581 | 0.0670438319216934 | -1.11394547284211 | 0.265302579724346 | 0.707506016720368 | NM_026135 |
| Exosc3 | 226.2091564 | 0.0962796964158497 | 0.0864622086663562 | 1.11354657602349 | 0.265473756419667 | 0.707651445295817 | NM_025513 |
| Ncan | 10594.13281 | -0.0616849721048565 | 0.0553984731670486 | -1.11347783753627 | 0.265503261521561 | 0.707651445295817 | NM_007789 |
| Srgap2 | 3935.287052 | -0.0619300375487319 | 0.0556168160770509 | -1.11351281711155 | 0.265488246711477 | 0.707651445295817 | NM_001081011 |
| Igfbp3 | 460.7723442 | 0.094942596212422 | 0.0852783888724691 | 1.1133253977676 | 0.265568702367882 | 0.707696013684803 | NM_008343 |
| Cdv3 | 4001.5089 | 0.0604665866918025 | 0.0543321823465363 | 1.11290553922794 | 0.265749000806495 | 0.708046585787811 | NM_175833,NM_175565,NM_001134427,NM_001134426 |
| Nts | 153.6098757 | 0.0834989495168027 | 0.0750677384576264 | 1.11231470712196 | 0.266002862574931 | 0.708592991337511 | NM_024435 |
| Ythdc2 | 1359.310303 | -0.0754362258972963 | 0.0678334422142526 | -1.1120801692332 | 0.266103682320451 | 0.708731589546547 | NM_001163013 |
| Slc13a5 | 585.7188719 | -0.086578575933784 | 0.077873636562999 | -1.11178287999614 | 0.266231514501425 | 0.708942067763097 | NM_001004148 |
| Man1a | 749.4448074 | 0.0821537798049486 | 0.0739067280090469 | 1.11158729412148 | 0.266315638032021 | 0.709036099349916 | NM_008548 |
| Stmn2 | 14804.05942 | 0.053863867251028 | 0.0484743705545038 | 1.11118239669486 | 0.266489846779637 | 0.709239925672593 | NM_025285 |
| Trib3 | 26.9510519 | -0.0933310844366788 | 0.0839884039138445 | -1.11123774339632 | 0.266466029012974 | 0.709239925672593 | NM_175093 |
| Bend6 | 6011.222302 | 0.0695810705390026 | 0.06266725505688 | 1.11032580692815 | 0.266858656347972 | 0.709441588966076 | NM_177235 |
| Exog | 1188.821768 | 0.0698341426490949 | 0.0628876721143182 | 1.11045838240203 | 0.266801552275094 | 0.709441588966076 | NM_172456,NM_001172136 |
| Golga1 | 1670.501187 | -0.0590141844126465 | 0.0531393216289777 | -1.1105558483544 | 0.266759576245503 | 0.709441588966076 | NM_029793 |
| Pgap3 | 446.2472421 | 0.0799192113358591 | 0.0719420559980648 | 1.11088306036192 | 0.266618687844163 | 0.709441588966076 | NM_001033537 |
| Slc12a7 | 234.8767107 | -0.103290955383055 | 0.0929920209060205 | -1.11075073298432 | 0.266675658170636 | 0.709441588966076 | NM_011390 |
| Tcp1 | 7801.027021 | 0.0614960167240704 | 0.0553843376482046 | 1.11035031446411 | 0.266848099613412 | 0.709441588966076 | NM_013686 |
| Ebf3 | 58.43011339 | 0.0837351379044196 | 0.0754290886150809 | 1.1101173226648 | 0.266948473526007 | 0.709550508040033 | NM_001113415,NM_001113414,NM_010096 |
| 4930507D05Rik | 35.55420516 | -0.102877950983722 | 0.092692865298442 | -1.10987993145413 | 0.2670507694297 | 0.709692549713836 | NR_027926 |
| Gpx1 | 1942.661347 | 0.0598818472953114 | 0.0539718415456277 | 1.10950165086895 | 0.267213832658418 | 0.709971476777887 | NM_008160 |
| Plxnb3 | 1433.286335 | -0.0927547512935475 | 0.0836150990110081 | -1.10930624242084 | 0.267298093063408 | 0.709971476777887 | NM_019587 |
| Trhr | 199.8131296 | 0.081683945218026 | 0.0736358183084804 | 1.10929636003812 | 0.267302354846362 | 0.709971476777887 | NM_013696 |
| 1600012H06Rik | 606.1495293 | 0.0795532437139959 | 0.0717628705700826 | 1.10855715611746 | 0.267621269396643 | 0.710152313924697 | NM_001083882,NM_026451,NM_001083880,NM_001083881 |
| Ccnyl1 | 841.7177033 | -0.0733860192139135 | 0.066212718045908 | -1.10833721042885 | 0.267716210972808 | 0.710152313924697 | NM_001097644 |
| Mapk4 | 4369.782043 | 0.0729674487160943 | 0.0658020403830323 | 1.10889340651676 | 0.267476168564888 | 0.710152313924697 | NM_172632 |
| Mknk1 | 1205.156163 | -0.0879911750669539 | 0.0793977785561622 | -1.10823220330671 | 0.267761546424726 | 0.710152313924697 | NM_021461 |
| Npff | 41.96432999 | -0.100311834016655 | 0.0904544746478054 | -1.10897591752348 | 0.267440571175465 | 0.710152313924697 | NR_033409,NM_018787 |
| Rad21l | 24.28709945 | 0.0871558101136837 | 0.0786168290242504 | 1.10861518068605 | 0.267596226417405 | 0.710152313924697 | NM_001276400,NM_001114677 |
| Stat5b | 1599.268525 | 0.0693002583819897 | 0.0625068661024388 | 1.10868233688788 | 0.267567244303182 | 0.710152313924697 | NM_001113563,NM_011489 |
| Usp32 | 7933.86317 | 0.0782859075705965 | 0.070631004021468 | 1.10837880128112 | 0.267698256124871 | 0.710152313924697 | NM_001029934 |
| Prr13 | 1389.21344 | 0.0742176073790381 | 0.0669810434491984 | 1.10803898472153 | 0.267844979808481 | 0.710243916885359 | NM_025385,NM_001170911 |
| Lpar2 | 55.59011525 | 0.104499277360985 | 0.0943205170815459 | 1.10791671414013 | 0.26789778648644 | 0.710254288465418 | NM_020028 |
| Cmip | 15078.90582 | 0.0550295247028105 | 0.0496851437957677 | 1.10756496809209 | 0.268049739784462 | 0.710375070231693 | NM_001163262,NM_028941 |
| Gbp11 | 33.12723373 | -0.0976534906905291 | 0.088189654809526 | -1.10731231346175 | 0.268158922409045 | 0.710375070231693 | NM_001039647 |
| Lrrc33 | 315.7996311 | 0.104865205156252 | 0.094706048154163 | 1.10727041408751 | 0.268177031832892 | 0.710375070231693 | NM_146069 |
| Pigv | 266.2513255 | -0.0916067010162424 | 0.0827338768402437 | -1.10724535722086 | 0.268187862119689 | 0.710375070231693 | NM_001145955,NM_178698,NM_001145956 |
| Polr2g | 2037.095245 | 0.0602547657081785 | 0.0544154339760809 | 1.10731021155991 | 0.268159830856801 | 0.710375070231693 | NM_026329 |
| Hcrtr1 | 115.5685759 | 0.105049687871869 | 0.0949051271297061 | 1.10689159847284 | 0.268340798729297 | 0.710391551365366 | NM_198959,NM_001163027 |
| Per2 | 2341.741477 | 0.0931457558594508 | 0.0841458025574144 | 1.10695665176995 | 0.268312670461911 | 0.710391551365366 | NM_011066 |
| Rbak | 566.285696 | -0.0884506443714093 | 0.0799047635494223 | -1.10695083049337 | 0.268315187429746 | 0.710391551365366 | NM_021326,NM_001045482 |
| Elof1 | 1132.889576 | 0.0662160015139927 | 0.0598433548391702 | 1.10648879381761 | 0.268515011743792 | 0.71072322532622 6 | NM_170777 |
| Amacr | 367.7320397 | 0.0904730223635935 | 0.0818060096626654 | 1.10594591689128 | 0.268749928887192 | 0.710903000999521 | NM_008537 |
| Cadps | 12491.5789 | 0.0597625669875402 | 0.0540408258513252 | 1.10587812169926 | 0.268779275561504 | 0.710903000999521 | NM_001042617,NM_012061 |
| Celf1 | 10265.42286 | -0.0675614347962455 | 0.0610782417431447 | -1.10614570537844 | 0.268663458741118 | 0.710903000999521 | NM_198683,NM_001244903,NM_017368,NM_001244891 |
| Ebpl | 320.9371989 | 0.095475641772503 | 0.0863403329335218 | 1.10580580973686 | 0.268810579846573 | 0.710903000999521 | NM_026598 |
| Zfp180 | 2806.235038 | 0.0809051262402246 | 0.0731665608934305 | 1.10576642187769 | 0.268827632141702 | 0.710903000999521 | NM_001045486 |
| Fndc3b | 842.1388725 | 0.0706013709556691 | 0.0639128844619213 | 1.10465004904813 | 0.269311255394745 | 0.711922710803289 | NM_173182 |
| Serp2 | 1897.907762 | 0.0697434800808657 | 0.0631327769676149 | 1.10471110935991 | 0.269284788064548 | 0.711922710803289 | NM_001160326,NR_027699 |
| Anapc1 | 4050.637853 | -0.0481679715089269 | 0.0436160534109249 | -1.10436336490871 | 0.269435545969052 | 0.711972822484593 | NM_008569 |
| Fam222b | 1724.690129 | 0.0541581127391413 | 0.0490549788384837 | 1.10402886763971 | 0.269580615453664 | 0.711972822484593 | NM_145430 |
| Fgf1 | 2645.397517 | 0.090339193001137 | 0.0818253589781184 | 1.10404884414006 | 0.269571950259337 | 0.711972822484593 | NM_010197 |
| Jpx | 169.9052701 | -0.0965212186883047 | 0.0874175489749729 | -1.10414007050161 | 0.269532381485429 | 0.711972822484593 | NR_015508 |
| Paqr6 | 283.0685876 | 0.101741725252371 | 0.0921633555625329 | 1.10392817873628 | 0.269624294126927 | 0.711972822484593 | NM_198410 |
| Rnf7 | 2227.328874 | 0.066488498523155 | 0.0602092737144109 | 1.1042899942379 | 0.269467361807024 | 0.711972822484593 | NM_011279 |
| Nacc1 | 6470.869709 | 0.0532814358064915 | 0.0482742008194472 | 1.10372486549849 | 0.269712505860277 | 0.712076310455542 | NM_025788 |
| Ak2 | 764.8392738 | -0.0818273889351698 | 0.0741724254626588 | -1.10320497711599 | 0.269938160429005 | 0.712163729607985 | NM_001033966,NM_016895 |
| Entpd5 | 1925.334834 | 0.0569432101108403 | 0.0516042546593142 | 1.10345959818184 | 0.269827627441801 | 0.712163729607985 | NM_001026214,NM_007647 |
| Ing2 | 1118.753983 | 0.0809785363685649 | 0.0734439797136157 | 1.1025891663868 | 0.270205617035331 | 0.712163729607985 | NM_023503 |
| Ino80b | 330.7870061 | -0.0897392876124467 | 0.0813518359690192 | -1.10310095087002 | 0.269983327967271 | 0.712163729607985 | NM_023547 |
| Mien1 | 1744.956767 | 0.0583554014573731 | 0.0529396475642024 | 1.10230052790969 | 0.270331039943012 | 0.712163729607985 | NM_025559 |
| Mocos | 112.3910636 | -0.101361208407284 | 0.0919642085150807 | -1.10218105547728 | 0.270382966321633 | 0.712163729607985 | NM_026779 |
| Ndufb5 | 3366.529859 | 0.0616897977965417 | 0.055957146226726 | 1.10244717531856 | 0.270267311835105 | 0.712163729607985 | NM_025316 |
| Nsmce4a | 1113.572872 | 0.0717020836817752 | 0.0650476557171577 | 1.10230081147816 | 0.270330916703601 | 0.712163729607985 | NM_001162855 |
| Pdcd2 | 322.0599349 | -0.0854774424846086 | 0.0775428335320733 | -1.10232549664636 | 0.270320188629128 | 0.712163729607985 | NM_008799 |
| Pou2f1 | 974.2152843 | -0.0895139459097397 | 0.0812110262736946 | -1.10223882170954 | 0.270357858493859 | 0.712163729607985 | NM_198934,NM_011137,NM_198933,NM_198932 |
| Slc35c2 | 2070.874074 | 0.0647210001390629 | 0.0587157643708259 | 1.10227637896886 | 0.270341535260992 | 0.712163729607985 | NR_045544,NM_144893,NM_001252574,NM_001252575,NM_001252573 |
| Tmem25 | 2937.882917 | -0.0650750295888072 | 0.0590073146845259 | -1.10282987688427 | 0.270101050929885 | 0.712163729607985 | NM_027865 |
| Zfp708 | 142.9321592 | 0.101355901458378 | 0.0918657297537746 | 1.10330480942175 | 0.269894818747053 | 0.712163729607985 | NM_001012325,NM_001012448,NM_001012449 |
| 4933421O10Rik | 404.6307658 | -0.0823114257034828 | 0.0747160932950257 | -1.1016559093697 | 0.270611291995265 | 0.712294860704995 | NR_036602 |
| Ak4 | 2389.606713 | 0.0913111976023547 | 0.0828645336429266 | 1.10193340369024 | 0.270490625152573 | 0.712294860704995 | NM_009647,NM_001177602,NM_001177604,NM_001177605 |
| Gmfb | 6847.586738 | 0.0835177276006052 | 0.0758293879479859 | 1.10138997373806 | 0.270726967227886 | 0.712294860704995 | NM_022023 |
| Kif22 | 47.15587626 | -0.101994333164303 | 0.092568456962753 | -1.10182600543231 | 0.270537322304084 | 0.712294860704995 | NM_145588 |
| Mmrn2 | 225.8445753 | -0.0952722009546479 | 0.0864902612690215 | -1.10153674595005 | 0.270663120860516 | 0.712294860704995 | NM_153127 |
| Otx1 | 229.877529 | -0.0969424260185701 | 0.0880168438575107 | -1.10140765982826 | 0.270719273176856 | 0.712294860704995 | NM_011023 |
| Anapc2 | 3906.988553 | 0.0575314633691357 | 0.0522549866774646 | 1.10097556285373 | 0.270907292963932 | 0.712640227742497 | NM_175300 |
| 0610007N19Rik | 74.14745248 | -0.104534314125065 | 0.0949728317991272 | -1.10067597380019 | 0.271037706660123 | 0.712854196441235 | NR_038186 |
| Pcdhac2 | 1555.088788 | 0.0718384952456322 | 0.0652776441985943 | 1.10050686000674 | 0.271111342343479 | 0.712918783287722 | NM_001003672 |
| Cars2 | 406.9816056 | -0.0861940965753637 | 0.0783651058903016 | -1.09990404014794 | 0.27137393422479 | 0.713480139103944 | NM_024248 |
| Api5 | 4076.523725 | 0.0659884301790134 | 0.0600225605822895 | 1.09939378691692 | 0.271596339641325 | 0.713790098959437 | NM_007466 |
| Atp5g3 | 7057.107928 | 0.0520873281107085 | 0.0473921832123909 | 1.09907002758822 | 0.271737522184062 | 0.713790098959437 | NM_175015 |
| Clec12a | 29.47095516 | 0.0932971283925696 | 0.084881990710857 | 1.0991392592379 | 0.271707327941512 | 0.713790098959437 | NM_177686 |
| Lrrc47 | 2637.521065 | 0.0585017724966235 | 0.0532235005104215 | 1.09917183078119 | 0.271693123194672 | 0.713790098959437 | NM_201226 |
| Tmem57 | 2812.267771 | 0.060436189159824 | 0.0549858036879427 | 1.0991235029102 | 0.271714199602111 | 0.713790098959437 | NM_025382 |
| 1810055G02Rik | 824.4212943 | 0.072898560620067 | 0.0663568322230546 | 1.09858409718871 | 0.271949517376163 | 0.714078853635281 | NM_028077 |
| Sec11a | 686.2205541 | 0.0718069934478506 | 0.0653694108590284 | 1.09848004600661 | 0.271994926143743 | 0.714078853635281 | NM_019951 |
| Zfp763 | 779.0492561 | 0.091177354814341 | 0.0829971506456138 | 1.09856006025623 | 0.271960006824609 | 0.714078853635281 | NM_028543 |
| 9330020H09Rik | 60.76014836 | -0.102950552298903 | 0.0938448191664275 | -1.09702968382652 | 0.272628416246932 | 0.714142044060982 | NR_028442 |
| AI854517 | 1113.309525 | -0.0859401367204909 | 0.0783038086581157 | -1.09752179610722 | 0.272413358081611 | 0.714142044060982 | NR_040312,NR_040313,NR_040311,NR_040314 |
| Aarsd1 | 1900.21336 | 0.0660418377540188 | 0.0602033998243 | 1.09697854185574 | 0.272650772479439 | 0.714142044060982 | NM_144829 |
| Cabp7 | 322.1994472 | -0.0965944993693459 | 0.0879792372449284 | -1.09792381014208 | 0.272237759961944 | 0.714142044060982 | NM_138948 |
| Cd81 | 10156.31008 | 0.0704886080433056 | 0.0642659551489824 | 1.09682658384051 | 0.272717206899626 | 0.714142044060982 | NM_133655 |
| Clic1 | 348.5701239 | -0.0914273864754913 | 0.0833026587119979 | -1.09753263448149 | 0.272408622906714 | 0.714142044060982 | NM_033444 |
| Ctnna3 | 78.39929691 | 0.103130013012276 | 0.0939897468406864 | 1.09724748154799 | 0.272533222061567 | 0.714142044060982 | NM_177612,NM_001164376,NM_001164519 |
| Dok3 | 225.0450658 | -0.103450732402485 | 0.0943066093488682 | -1.09696163521043 | 0.272658163336605 | 0.714142044060982 | NM_013739 |
| Galnt1 | 2503.410677 | 0.064396827940646 | 0.0586950266721832 | 1.09714283461029 | 0.272578957901131 | 0.714142044060982 | NM_013814,NM_001160404 |
| H2-T10 | 57.64705417 | 0.104144481065392 | 0.0949200926123226 | 1.09718056735094 | 0.27256246623916 | 0.714142044060982 | NR_046286,NM_010395 |
| Htra2 | 1057.634354 | -0.0793031321177423 | 0.0723143991644489 | -1.09664372564861 | 0.272797165227115 | 0.714142044060982 | NM_019752 |
| Myl6 | 8971.997501 | 0.0626141197553882 | 0.0570923428648496 | 1.09671659303963 | 0.272765300610753 | 0.714142044060982 | NM_010860 |
| Poglut1 | 2210.157318 | 0.0632939075593482 | 0.0577033571603962 | 1.09688431789873 | 0.272691964882038 | 0.714142044060982 | NM_172380 |
| Rad50 | 810.4534724 | -0.0720219406528535 | 0.0656716225579672 | -1.09669805385547 | 0.272773407479742 | 0.714142044060982 | NM_009012 |
| Rbm45 | 897.5321314 | -0.0723989406988507 | 0.0660198138685148 | -1.09662442919092 | 0.272805603916728 | 0.714142044060982 | NM_153405 |
| Sc5d | 1801.222618 | 0.0714152306002318 | 0.0651062445246041 | 1.09690293337751 | 0.27268382631389 | 0.714142044060982 | NM_172769 |
| Nanp | 584.5423537 | 0.0707913359029934 | 0.0645630573424669 | 1.09646814783707 | 0.272873955156339 | 0.71419226533351 | NM_026086 |
| Nr3c2 | 1335.798233 | -0.0849740673314422 | 0.0775207699143537 | -1.09614581260381 | 0.273014968743962 | 0.714368645153821 | NM_001083906 |
| Pianp | 7731.401596 | 0.0517487224171518 | 0.0472121414153868 | 1.09608928690293 | 0.27303970245725 | 0.714368645153821 | NM_001145926,NM_175696,NM_001145927 |
| Dffa | 1828.956757 | 0.0577947358465108 | 0.052755325395053 | 1.09552420374096 | 0.27328704776023 | 0.714887026069711 | NM_010044,NM_001025296 |
| Thbs4 | 340.5523734 | 0.103991060897217 | 0.0949590300809641 | 1.09511502811846 | 0.273466245623539 | 0.71522698666322 | NM_011582 |
| Clint1 | 2227.856707 | 0.0536463909390888 | 0.0490353855385719 | 1.09403424383989 | 0.273939959773475 | 0.716079153440614 | NM_001045520 |
| Gcat | 394.0764756 | 0.091594951367041 | 0.083710254971791 | 1.09419032826871 | 0.273871512421083 | 0.716079153440614 | NM_001161712,NM_013847 |
| Sec14l3 | 112.1952406 | -0.10267447315236 | 0.0938431128375956 | -1.09410770857578 | 0.273907741992504 | 0.716079153440614 | NM_001029937 |
| Bcor | 1492.728439 | 0.0932040355066698 | 0.0852472357655221 | 1.09333792080993 | 0.274245459139726 | 0.716185716418173 | NM_029510,NM_175044,NM_175046,NM_001168321,NM_175045 |
| Dleu7 | 155.6975325 | 0.0958039743578193 | 0.0876700229096303 | 1.0927791641685 | 0.274490771955036 | 0.716185716418173 | NM_173419 |
| Eif4g2 | 26798.44199 | 0.0610094347463967 | 0.0558017676205439 | 1.09332441153594 | 0.274251388394528 | 0.716185716418173 | NM_001040131,NM_013507 |
| Gm16677 | 22.92660709 | 0.0846559175220147 | 0.0774620904480439 | 1.09286900253222 | 0.274451319806587 | 0.716185716418173 | NR_045091 |
| Hmgcll1 | 621.4183225 | 0.0772012492149996 | 0.0706268621083867 | 1.0930862126725 | 0.274355948868893 | 0.716185716418173 | NM_173731 |
| Mpped1 | 8852.494075 | 0.0637048778853321 | 0.0582777129977071 | 1.09312590711716 | 0.274338522587995 | 0.716185716418173 | NM_172610 |
| Naa50 | 2302.023933 | 0.0833595868539422 | 0.0762330607904435 | 1.09348340457021 | 0.274181611496155 | 0.716185716418173 | NM_028108 |
| Slc25a19 | 1616.561762 | 0.0656144497468697 | 0.0600415982491472 | 1.09281650822481 | 0.274474371995692 | 0.716185716418173 | NM_001252396,NM_001252394,NM_026071,NM_001252395,NM_001252384 |
| Spint2 | 707.3367213 | 0.101233272195356 | 0.0925680684816806 | 1.09360899342293 | 0.274126503187302 | 0.716185716418173 | NM_001082548,NM_011464 |
| Uprt | 146.9649442 | 0.0996415454012938 | 0.0911879124808906 | 1.09270563049872 | 0.274523066846784 | 0.716185716418173 | NM_001081189 |
| Vprbp | 1622.949327 | 0.0722273436649708 | 0.0660845328307888 | 1.09295383611035 | 0.274414069044809 | 0.716185716418173 | NM_001015507 |
| Tspan12 | 480.1323501 | -0.0783269747262989 | 0.0716978368559268 | -1.09245938456544 | 0.274631233286066 | 0.716339252058428 | NM_173007 |
| Gltscr2 | 2265.139876 | 0.0630940417578744 | 0.0577656296051925 | 1.0922419125196 | 0.274726784654803 | 0.716459833733513 | NM_133831 |
| Stk16 | 1692.271455 | -0.0657830952945905 | 0.0602371511928378 | -1.09206849912272 | 0.274802994104912 | 0.716529939394715 | NM_011494 |
| Gm20063 | 127.866124 | -0.102995440006265 | 0.0943316547402847 | -1.09184388093089 | 0.274901727824264 | 0.716658739837628 | NR_045049 |
| Sft2d1 | 806.0837213 | -0.0747888206588665 | 0.0685115236424069 | -1.09162395785508 | 0.274998420814432 | 0.716782174905874 | NM_134114 |
| Gipr | 56.4302575 | -0.100340436114286 | 0.09194182063458 | -1.09134706515206 | 0.27512019539853 | 0.716917117827604 | NM_001080815 |
| Kcnd2 | 4327.19521 | 0.0775586941061049 | 0.0710778389949358 | 1.09117968698557 | 0.275193824216084 | 0.716917117827604 | NM_019697 |
| Zc3h12a | 48.03006646 | -0.101796828659588 | 0.0932914802238384 | -1.09116961608223 | 0.275198254784987 | 0.716917117827604 | NM_153159 |
| Fahd2a | 631.3133607 | -0.0716361481490274 | 0.0656786422141704 | -1.09070689852921 | 0.275401874139106 | 0.717061771597895 | NM_029629 |
| Matr3 | 14816.56003 | 0.0766706192380224 | 0.0702943045065095 | 1.09070883873561 | 0.27540102013461 | 0.717061771597895 | NM_010771 |
| Pias2 | 2562.443234 | 0.0668370426309183 | 0.061278501916745 | 1.09070947461681 | 0.275400740244469 | 0.717061771597895 | NM_001164168,NM_001164167,NM_001164169,NM_001164170,NM_008602 |
| Esyt3 | 128.6670753 | 0.0751409551982659 | 0.0689120780334729 | 1.09038875829238 | 0.275541931722364 | 0.717137213491157 | NM_177775 |
| Slc3a2 | 4633.022832 | 0.0474852087503053 | 0.0435463891509612 | 1.09045111836229 | 0.275514474589272 | 0.717137213491157 | NM_008577,NM_001161413 |
| Trappc4 | 1632.86807 | 0.050387844689508 | 0.0462144626783251 | 1.09030467453947 | 0.275578956747049 | 0.717137213491157 | NM_021789 |
| Cog4 | 2217.900071 | 0.0513706802777765 | 0.047133487501106 | 1.0898977139464 | 0.275758203739292 | 0.717346645328967 | NM_133973 |
| Rsl1d1 | 1937.993221 | 0.0668744205307324 | 0.0613552410204454 | 1.08995449155595 | 0.275733191100653 | 0.717346645328967 | NM_025546 |
| Kbtbd2 | 2311.899907 | 0.0813485398212945 | 0.0746570465960321 | 1.08962976075748 | 0.275876267927369 | 0.717396825619937 | NM_145958 |
| Timeless | 226.1921685 | -0.0910324203061343 | 0.0835421877803559 | -1.0896580844336 | 0.27586378646198 | 0.717396825619937 | NM_001164080,NM_001164081,NM_011589,NM_001136082 |
| A030009H04Rik | 4598.088121 | 0.0654591586296326 | 0.0601385610824783 | 1.08847231213027 | 0.276386653252638 | 0.71747510541445 | NR_027827 |
| Aifm1 | 959.9196346 | 0.0618760985592784 | 0.0568748647661255 | 1.08793399006254 | 0.276624249458913 | 0.71747510541445 | NM_012019 |
| Atp5sl | 1701.822389 | 0.0597525653730073 | 0.0548668638682301 | 1.08904648744843 | 0.276133386088422 | 0.71747510541445 | NM_025504 |
| Ccdc55 | 715.9519455 | 0.0721739038584493 | 0.0663465531227331 | 1.087832004248 | 0.27666927805111 | 0.71747510541445 | NM_001012309 |
| Fbxo44 | 3909.458502 | 0.0569865957560119 | 0.0523602592928894 | 1.08835587381728 | 0.276438033184172 | 0.71747510541445 | NM_001161852,NM_173401,NM_001161851 |
| Ly6c1 | 1099.017821 | 0.090242591869492 | 0.0828935103671254 | 1.0886568981072 | 0.276305215614394 | 0.71747510541445 | NM_010741,NM_001252058,NM_001252055,NM_001252057,NM_001252056 |
| Mvp | 349.2382312 | -0.0839287868645776 | 0.0771374870085107 | -1.08804149732435 | 0.276576788456157 | 0.71747510541445 | NM_080638 |
| Nacad | 2168.965158 | -0.0738596053485766 | 0.0678301391881067 | -1.08889066471984 | 0.276202103414989 | 0.71747510541445 | NM_001081652 |
| Pnkp | 807.7235735 | -0.0877423480238624 | 0.0806193220423371 | -1.08835383132823 | 0.276438934517342 | 0.71747510541445 | NM_021549 |
| Prr14 | 1040.055817 | -0.0733055636887884 | 0.0673576519619971 | -1.08830343032366 | 0.276461176689865 | 0.71747510541445 | NM_145589 |
| Ptp4a1 | 421.5572657 | 0.0862429858437461 | 0.0792618915002267 | 1.08807630263907 | 0.27656142421726 | 0.71747510541445 | NM_011200 |
| Resp18 | 2680.184614 | 0.10296077687505 | 0.0946530751182229 | 1.08777001430171 | 0.276696650181175 | 0.71747510541445 | NM_009049 |
| Rrn3 | 2879.715981 | 0.0596237303912484 | 0.0547770499286166 | 1.08848012934154 | 0.276383204038976 | 0.71747510541445 | NM_001039521 |
| Sncg | 118.5122658 | 0.0751412933817849 | 0.0689866338231704 | 1.08921524674461 | 0.276058976929247 | 0.71747510541445 | NM_011430 |
| Usp51 | 46.00038944 | 0.0998199637196204 | 0.0917634210771515 | 1.0877968862527 | 0.276684784441149 | 0.71747510541445 | NM_001137547 |
| Vps13c | 2606.870453 | -0.0869100132883265 | 0.0798899231598761 | -1.08787203505506 | 0.276651603126028 | 0.71747510541445 | NM_177184 |
| Prr24 | 1075.278677 | -0.0612993537718463 | 0.0563847743260048 | -1.08716146343739 | 0.276965458426345 | 0.717792699930078 | NM_001136270 |
| Sema3e | 1228.362807 | 0.0869717448394266 | 0.0799884014569869 | 1.0873044498357 | 0.27690228268389 | 0.717792699930078 | NM_011348 |
| Ttc38 | 373.6582259 | -0.0859459168872787 | 0.0790556532792485 | -1.08715712693805 | 0.276967374577178 | 0.717792699930078 | NM_001033337 |
| Lamp5 | 8675.516963 | 0.0814317123106468 | 0.0749176312914253 | 1.08694990627616 | 0.277058948834507 | 0.717815689184511 | NM_029530 |
| Serf1 | 226.3717939 | 0.0973887724143972 | 0.0896012240406746 | 1.08691341504651 | 0.277075077052014 | 0.717815689184511 | NM_011353 |
| Zfpm1 | 1365.746766 | -0.0819357686869659 | 0.0753929053995685 | -1.08678354087459 | 0.277132483401412 | 0.717836386214144 | NM_009569 |
| Klf2 | 388.53213 | -0.0975181589441766 | 0.0897417828171345 | -1.08665279296811 | 0.27719028413986 | 0.717858097239366 | NM_008452 |
| Gcc1 | 1001.496428 | 0.0690792615701988 | 0.0635975967890057 | 1.08619295143348 | 0.277393633621309 | 0.71801550125743 | NM_028900 |
| Lipg | 293.5611412 | 0.0973145362292531 | 0.089583270180309 | 1.0863025655726 | 0.277345152337825 | 0.71801550125743 | NM_010720 |
| Tmem242 | 1299.104209 | 0.0581204568407784 | 0.053509045873447 | 1.08618002605088 | 0.277399352406492 | 0.71801550125743 | NM_027457 |
| Ctnnbip1 | 1387.161585 | 0.0682663457183129 | 0.0628644761856525 | 1.08592880845308 | 0.277510490224659 | 0.718077080380369 | NM_001141930,NM_023465 |
| Gpcpd1 | 3931.357867 | -0.0679063794215827 | 0.0625441967440793 | -1.08573429601222 | 0.277596562701221 | 0.718077080380369 | NM_001042672,NM_028802,NM_001042671 |
| Ppp1r11 | 2192.794445 | 0.0607075523742149 | 0.0559166500027203 | 1.08567935259465 | 0.277620878660068 | 0.718077080380369 | NM_029632 |
| Vapa | 4465.055123 | 0.0699559438351644 | 0.0644331614780027 | 1.08571335365947 | 0.277605830854002 | 0.718077080380369 | NM_013933 |
| Adk | 2149.417007 | 0.0588823003482093 | 0.0542524638937585 | 1.08533873159231 | 0.277771657501524 | 0.718083484048254 | NM_001243041,NM_134079 |
| Kif13a | 1721.15488 | 0.0792329761659895 | 0.0729946284371396 | 1.08546310683973 | 0.277716595257515 | 0.718083484048254 | NM_010617 |
| Pdss2 | 390.3417781 | 0.0825061409008366 | 0.0760167971659485 | 1.08536723430641 | 0.277759038390499 | 0.718083484048254 | NM_027772,NM_001168289 |
| Smcr8 | 2637.69212 | 0.0603418906078872 | 0.0556094516769347 | 1.0851013413771 | 0.277876773306696 | 0.718227403746097 | NM_175491,NM_001085440 |
| D17Wsu92e | 13171.73342 | 0.0557681519602558 | 0.051401183228526 | 1.08495852541593 | 0.277940024910628 | 0.718263085189786 | NM_001033279,NM_001271511,NM_001044719 |
| Ddb1 | 11658.2777 | 0.0428681307117293 | 0.0395251871569655 | 1.08457755156195 | 0.278108801949594 | 0.718319851514567 | NM_015735 |
| Fads2 | 3884.422893 | 0.0762209426990603 | 0.0702845364067467 | 1.08446248059401 | 0.278159793802109 | 0.718319851514567 | NM_019699 |
| Iqgap3 | 34.88402574 | -0.096658642182373 | 0.0891114616554658 | -1.08469371264593 | 0.278057333483919 | 0.718319851514567 | NM_001033484 |
| Lyrm9 | 1951.887335 | -0.0610576363828772 | 0.056297373014404 | -1.08455569262984 | 0.278118487894914 | 0.718319851514567 | NM_001076681 |
| Rbm10 | 2289.094833 | -0.0627603526405126 | 0.0578841062574502 | -1.08424154225297 | 0.278257716920848 | 0.718445004620022 | NM_001167775,NM_001167776,NM_145627 |
| Man2b2 | 1384.31428 | -0.0826230420857469 | 0.0762115124056441 | -1.08412809925588 | 0.278308005639323 | 0.718447145888894 | NM_008550 |
| Cops6 | 3369.72653 | 0.0519120232941064 | 0.0479417952812489 | 1.0828135031149 | 0.278891210754588 | 0.718495890058647 | NM_012002 |
| Dock11 | 386.3337423 | -0.0940712739298425 | 0.0868417185403694 | -1.08324979642258 | 0.278697562263737 | 0.718495890058647 | NM_001009947 |
| Erc1 | 2701.031583 | 0.0654255231012433 | 0.0603942909653016 | 1.08330641945664 | 0.278672436876409 | 0.718495890058647 | NM_053204,NM_178085 |
| Grina | 16946.74044 | 0.0542282432267334 | 0.0500303885477696 | 1.08390609789 | 0.27840643564046 | 0.718495890058647 | NM_023168 |
| H2afx | 830.2712995 | 0.0721982096162717 | 0.0666805498129169 | 1.08274766503128 | 0.278920440867459 | 0.718495890058647 | NM_010436 |
| Kif13b | 1485.332264 | -0.0931588670656264 | 0.0860216408780988 | -1.08297012373481 | 0.278821684269863 | 0.718495890058647 | NM_001081177 |
| Pgm2l1 | 14064.81734 | 0.0824611719678951 | 0.0761479671422914 | 1.08290706978174 | 0.27884967354492 | 0.718495890058647 | NM_027629 |
| Rcan1 | 2685.903935 | 0.0699594501627044 | 0.0645757406071527 | 1.08337046551744 | 0.278644019520518 | 0.718495890058647 | NM_001081549,NM_019466 |
| Slc9a6 | 6194.681242 | 0.0680994217522005 | 0.0628641115268963 | 1.08327979348065 | 0.278684251451547 | 0.718495890058647 | NM_172780 |
| Tsks | 36.56542411 | -0.100163985711523 | 0.0924967878219701 | -1.08289150434403 | 0.278856583244354 | 0.718495890058647 | NM_011651,NM_001077591 |

| | | | | | | |
|---|---|---|---|---|---|---|
| Snupn | 446.1974328 | -0.156512608 | 0.09368971 | -1.67054214 | 0.094812149 | 0.260556836 |
| A330021E22Rik | 410.8324754 | 0.232607212 | 0.13930208 | 1.6698043 | 0.094958089 | 0.260796651 |
| Alpk2 | 16.92066073 | 0.274301548 | 0.164260664 | 1.669916218 | 0.094935941 | 0.260796651 |
| Pou3f3 | 2247.655894 | -0.163345511 | 0.097811683 | -1.670000009 | 0.094919362 | 0.260796651 |
| Rps7 | 7261.134736 | -0.122287136 | 0.07323633 | -1.669760582 | 0.094966742 | 0.260796651 |
| Dstn | 5523.656187 | -0.105081943 | 0.062940057 | -1.669555901 | 0.095007262 | 0.260797001 |
| Man1c1 | 1950.140761 | 0.119225183 | 0.071411095 | 1.669561049 | 0.095006242 | 0.260797001 |
| Ppfibp1 | 1329.126568 | 0.142649563 | 0.085444233 | 1.669504869 | 0.095017366 | 0.260797001 |
| Lrrc32 | 254.0460927 | 0.224420371 | 0.134456031 | 1.669098585 | 0.095097844 | 0.260971659 |
| Asah2 | 633.1986194 | 0.148934045 | 0.089236293 | 1.668985122 | 0.095120328 | 0.260987138 |
| Ptgfr | 39.5446382 | -0.265555451 | 0.159124298 | -1.668855448 | 0.095146031 | 0.261011438 |
| Hspa1a | 204.7952997 | 0.191485969 | 0.114784316 | 1.668224155 | 0.095271238 | 0.261308648 |
| Abhd4 | 2784.680551 | 0.127218042 | 0.076288653 | 1.66758799 | 0.095397544 | 0.261469936 |
| Ier3ip1 | 1020.957 | -0.111607613 | 0.06692402 | -1.667676473 | 0.095379968 | 0.261469936 |
| Ldlrap1 | 141.3888867 | 0.183495992 | 0.110027514 | 1.667728235 | 0.095369688 | 0.261469936 |
| Mta1 | 2384.853693 | 0.109763848 | 0.065820406 | 1.667626417 | 0.095389911 | 0.261469936 |
| 4930506M07Rik | 1237.030035 | -0.197325837 | 0.118342856 | -1.667408111 | 0.095433282 | 0.261521626 |
| Ccna1 | 28.83746251 | 0.279397809 | 0.167574807 | 1.66730199 | 0.095454371 | 0.261533161 |
| Ctla2b | 33.41014683 | 0.279741394 | 0.167808803 | 1.667024549 | 0.095509524 | 0.261638005 |
| Tnfrsf11b | 90.12403785 | -0.242632981 | 0.145616931 | -1.666241551 | 0.095665313 | 0.262018448 |
| Cdk2ap2 | 546.7483056 | -0.122875382 | 0.073754778 | -1.665998945 | 0.095713624 | 0.262058119 |
| Stab2 | 18.85622956 | 0.279942041 | 0.168024782 | 1.666075909 | 0.095698296 | 0.262058119 |
| Cln6 | 474.0626384 | 0.150705707 | 0.090480879 | 1.665608349 | 0.095791447 | 0.262132229 |
| Nup85 | 1158.636761 | 0.130013632 | 0.078052784 | 1.665714219 | 0.095770349 | 0.262132229 |
| Scai | 1751.717248 | 0.161891181 | 0.097192618 | 1.66567363 | 0.095778437 | 0.262132229 |
| Gclc | 2495.130115 | 0.146402846 | 0.087907954 | 1.665410679 | 0.09583085 | 0.262170049 |
| Trip4 | 1613.510281 | -0.100649912 | 0.060436994 | -1.665369257 | 0.095839109 | 0.262170049 |
| Ptk7 | 462.7486134 | 0.145133658 | 0.087154885 | 1.665238366 | 0.09586521 | 0.262195156 |
| Gpr21 | 128.4859317 | -0.234514251 | 0.140856456 | -1.664916592 | 0.095929398 | 0.262324406 |
| Cfl1 | 25411.21969 | -0.101448016 | 0.060943755 | -1.664617093 | 0.095989173 | 0.262441547 |
| Klhl6 | 83.04347163 | 0.263439489 | 0.158267135 | 1.66452428 | 0.096007703 | 0.262445899 |
| 2610019F03Rik | 844.1471793 | -0.162051938 | 0.097374352 | -1.664215831 | 0.096069305 | 0.262567969 |
| 2810004N23Rik | 410.9821461 | -0.132501116 | 0.079645009 | -1.663646189 | 0.096183156 | 0.262816719 |
| Pcx | 3334.009733 | 0.100340958 | 0.060315891 | 1.66359074 | 0.096194244 | 0.262816719 |
| Max | 1876.150222 | 0.142999426 | 0.085967707 | 1.663408634 | 0.096230666 | 0.262869876 |
| Sec14l4 | 9.240876223 | 0.259855823 | 0.156238079 | 1.663204164 | 0.096271575 | 0.262935268 |
| Fkbp9 | 1805.118995 | -0.11539349 | 0.069397035 | -1.66280145 | 0.096352186 | 0.263062692 |
| Zfyve16 | 679.8277866 | 0.184099099 | 0.11071433 | 1.662829914 | 0.096346487 | 0.263062692 |
| Kif11 | 80.91806891 | -0.252754848 | 0.152086863 | -1.661911118 | 0.096530597 | 0.26350336 |
| Mbd3 | 3324.325022 | -0.101582375 | 0.061136017 | -1.661579869 | 0.096597043 | 0.263591859 |
| Ppan | 486.319828 | 0.157365243 | 0.09470476 | 1.661640268 | 0.096584924 | 0.263591859 |
| Pcdhb13 | 262.2417469 | 0.182978204 | 0.110139394 | 1.661332949 | 0.096646596 | 0.263680641 |
| Mycbpap | 73.12116443 | 0.265221296 | 0.159657956 | 1.661184341 | 0.09667643 | 0.263715599 |
| Nudt16 | 450.2908964 | -0.140003 | 0.084293297 | -1.660903118 | 0.096732906 | 0.263776777 |
| Sdk1 | 418.7774382 | -0.191225753 | 0.115127831 | -1.660986328 | 0.096716193 | 0.263776777 |
| Fryl | 3309.426139 | -0.146011677 | 0.087933611 | -1.660476305 | 0.096818671 | 0.263964189 |
| Nxpe4 | 440.0706167 | 0.162353464 | 0.097784949 | 1.660311378 | 0.096851828 | 0.264008132 |
| Mta2 | 2121.537522 | 0.11827454 | 0.07125716 | 1.659826747 | 0.096949311 | 0.264227375 |
| Unk | 917.6351036 | 0.134163005 | 0.080841102 | 1.659589021 | 0.096997159 | 0.264311287 |
| Sri | 1756.294573 | -0.116511864 | 0.070232058 | -1.658955584 | 0.097124743 | 0.264565887 |
| Tbc1d2b | 966.2958825 | 0.138335165 | 0.083385052 | 1.658992366 | 0.097117331 | 0.264565887 |
| Slc39a6 | 1916.927495 | -0.115552753 | 0.06967654 | -1.658416922 | 0.097233343 | 0.264815156 |
| Etfa | 1054.696266 | -0.130899609 | 0.078939437 | -1.658228306 | 0.097271394 | 0.264825685 |
| Tbx6 | 28.85650867 | 0.278755713 | 0.168101703 | 1.658256331 | 0.097265739 | 0.264825685 |
| Rmdn3 | 1731.020164 | -0.082753092 | 0.049924944 | -1.657550024 | 0.097408324 | 0.265151893 |
| Glrx3 | 298.516579 | -0.170866607 | 0.103150643 | -1.656476403 | 0.09762538 | 0.265692103 |
| Gm16119 | 10.93657159 | 0.271059605 | 0.163643924 | 1.656398838 | 0.097641076 | 0.265692103 |
| 1110006O24Rik | 14.94616826 | 0.277173437 | 0.167377325 | 1.655979612 | 0.097725948 | 0.265876353 |
| Gnpat | 2228.485269 | 0.086625459 | 0.052325344 | 1.655516294 | 0.097819814 | 0.266049876 |
| Trim65 | 121.9407076 | 0.190633447 | 0.115157595 | 1.655413583 | 0.097840633 | 0.266049876 |
| Tvp23b | 820.6722041 | -0.126988222 | 0.076711013 | -1.65541058 | 0.097841241 | 0.266049876 |
| Mmgt1 | 1526.887222 | -0.096164132 | 0.058094999 | -1.655291065 | 0.09786547 | 0.266069065 |
| Uchl5 | 1438.335401 | 0.157747969 | 0.09531047 | 1.655095912 | 0.097905044 | 0.266129957 |
| Dctn1 | 16711.8334 | 0.107091467 | 0.06471608 | 1.654789146 | 0.097967276 | 0.266252409 |
| Acads | 280.1342732 | -0.202933092 | 0.122640205 | -1.654702801 | 0.097984799 | 0.266253327 |
| Hspa4 | 10218.49202 | -0.11926238 | 0.072083004 | -1.654514568 | 0.098023006 | 0.266310442 |
| Mrpl27 | 1108.121268 | -0.099673201 | 0.060253965 | -1.654218113 | 0.098083204 | 0.266427272 |
| Chtop | 4697.498399 | -0.113266727 | 0.068488913 | -1.653796529 | 0.098168861 | 0.266566481 |
| Murc | 30.43337837 | 0.27791478 | 0.168039049 | 1.653870228 | 0.098153883 | 0.266566481 |
| Adrbk2 | 5162.258101 | 0.162571337 | 0.098338468 | 1.653181514 | 0.098293928 | 0.266812564 |
| Enkd1 | 207.8366467 | 0.183399203 | 0.110934706 | 1.653217546 | 0.098286597 | 0.266812564 |
| Sdc4 | 2016.916396 | 0.170193801 | 0.103076237 | 1.651144881 | 0.098708995 | 0.267892304 |
| Ilvbl | 833.5757193 | 0.154204126 | 0.093407674 | 1.650872129 | 0.098764688 | 0.267979485 |
| Shank3 | 10452.37904 | -0.137840967 | 0.083498577 | -1.650818167 | 0.098775709 | 0.267979485 |
| Gm16287 | 32.87559487 | 0.277425468 | 0.16808693 | 1.650488046 | 0.098843156 | 0.268056398 |
| Mtch2 | 2861.503088 | -0.083989874 | 0.050889834 | -1.650425394 | 0.098855961 | 0.268056398 |
| Tapbp | 1210.054025 | 0.123261947 | 0.074684142 | 1.650443373 | 0.098852286 | 0.268056398 |
| Ptpn22 | 30.47229796 | 0.267952296 | 0.162387308 | 1.650081517 | 0.098926265 | 0.268200095 |
| Gpx7 | 97.8630262 | -0.242367122 | 0.146918743 | -1.64966781 | 0.099010897 | 0.268382583 |
| Stambp | 1277.816659 | 0.091428978 | 0.055427489 | 1.649524088 | 0.099040312 | 0.268415358 |
| Zfp180 | 2697.99205 | -0.13231962 | 0.080231498 | -1.649222859 | 0.099101987 | 0.268535533 |
| Tmem258 | 679.886564 | -0.134641814 | 0.08164708 | -1.649070785 | 0.099133134 | 0.268572963 |
| Htr1d | 187.7952992 | -0.245978958 | 0.149196067 | -1.648695992 | 0.099209932 | 0.268690886 |
| Olfr1344 | 26.25325534 | -0.274760288 | 0.16665379 | -1.648689104 | 0.099211343 | 0.268690886 |
| Zkscan1 | 2427.397193 | -0.088917912 | 0.053955694 | -1.647980134 | 0.099356748 | 0.269037653 |
| Flna | 4240.357641 | 0.144791459 | 0.087876462 | 1.647670548 | 0.099420295 | 0.269162685 |
| Rbm17 | 1618.195647 | 0.094683426 | 0.057469407 | 1.647544841 | 0.099446107 | 0.269185532 |
| Ccp110 | 2983.730692 | 0.114993342 | 0.069821268 | 1.646967252 | 0.099564777 | 0.269443841 |
| Polr3g | 222.9594598 | 0.159454538 | 0.096820367 | 1.646911122 | 0.099576315 | 0.269443841 |
| Gtf2f1 | 2311.363784 | 0.134456263 | 0.0816542 | 1.646654588 | 0.099629063 | 0.269539498 |
| Alg14 | 270.3960585 | -0.160009401 | 0.097186503 | -1.646415883 | 0.099678165 | 0.26954004 |
| Surf1 | 1918.790619 | 0.116056145 | 0.07049086 | 1.646399904 | 0.099681453 | 0.26954004 |
| Zfp629 | 1427.430551 | 0.123633409 | 0.075087975 | 1.64651409 | 0.099657962 | 0.26954004 |
| Cep76 | 298.3568594 | 0.213710294 | 0.129824409 | 1.646148799 | 0.099733128 | 0.269632714 |
| Fbxw17 | 276.9335303 | 0.161529411 | 0.09813291 | 1.646026902 | 0.09975822 | 0.269653501 |
| Scg3 | 11225.70952 | 0.097127957 | 0.059018601 | 1.645717726 | 0.099821888 | 0.269778533 |
| Rnf208 | 3597.718028 | 0.131494715 | 0.07990859 | 1.64556419 | 0.099853517 | 0.26981695 |
| Ufd1l | 2795.565205 | 0.104695017 | 0.063627904 | 1.645426131 | 0.099881964 | 0.269846758 |
| Mettl21d | 384.4012589 | 0.155807361 | 0.094705373 | 1.645179734 | 0.099932752 | 0.269936899 |
| Ubap2l | 5235.096056 | 0.114252964 | 0.069465681 | 1.644739718 | 0.100023498 | 0.270134928 |
| Grp | 126.130614 | -0.214133673 | 0.130271278 | -1.643751991 | 0.100227442 | 0.270638547 |
| Alkbh7 | 328.1805171 | -0.170491382 | 0.103742255 | -1.643413114 | 0.100297489 | 0.270733325 |
| Rexo4 | 1828.670651 | 0.096415039 | 0.058664801 | 1.643490423 | 0.100281505 | 0.270733325 |
| Zfp780b | 538.6399964 | -0.140652744 | 0.085600399 | -1.643131875 | 0.100355651 | 0.270843139 |
| Abca6 | 130.33335 | 0.193659521 | 0.117867329 | 1.64302968 | 0.100376793 | 0.270853017 |
| Cers5 | 1343.337097 | 0.100336246 | 0.061080942 | 1.642676789 | 0.100449824 | 0.270934267 |
| Cpsf1 | 2493.481069 | 0.138837219 | 0.084521261 | 1.642630711 | 0.100459363 | 0.270934267 |
| Parp8 | 721.2038107 | 0.19070326 | 0.116089452 | 1.642726843 | 0.100439462 | 0.270934267 |
| Vldlr | 3675.032005 | -0.113434124 | 0.069066998 | -1.642378077 | 0.100511675 | 0.271028176 |
| Sult1a1 | 346.3899469 | 0.266950437 | 0.162594498 | 1.641817155 | 0.100627903 | 0.271294367 |
| Dcp1a | 1118.767705 | 0.111789935 | 0.068106215 | 1.641405785 | 0.10071321 | 0.271477118 |
| Rpl11 | 6142.279163 | -0.108505091 | 0.066118627 | -1.641066914 | 0.100783526 | 0.271619403 |
| Fmn1 | 922.1155731 | 0.166057032 | 0.101209058 | 1.640732911 | 0.10085287 | 0.271759021 |
| Zfp28 | 438.2880191 | 0.16645658 | 0.101477453 | 1.640330682 | 0.100936429 | 0.271936887 |
| Prlhr | 8.540878702 | -0.23887891 | 0.145665214 | -1.639917337 | 0.101022356 | 0.272073766 |
| Samd3 | 86.06833068 | 0.249731962 | 0.152281811 | 1.639932964 | 0.101019106 | 0.272073766 |
| Cyp2j12 | 118.0317607 | 0.269874905 | 0.16461162 | 1.639464487 | 0.101116561 | 0.272280152 |
| Nars | 7719.223909 | 0.128784099 | 0.078561565 | 1.63927615 | 0.101155761 | 0.272308699 |
| Oraov1 | 496.1631857 | 0.124784681 | 0.07612328 | 1.639244678 | 0.101162312 | 0.272308699 |
| Tmem208 | 566.5572256 | -0.154465583 | 0.09423794 | -1.63910186 | 0.101192048 | 0.272341426 |
| Hcn4 | 173.6593608 | -0.257635468 | 0.157206282 | -1.638836971 | 0.101247217 | 0.272442581 |
| Aifm2 | 322.1027906 | -0.191119396 | 0.116658596 | -1.638279586 | 0.101363384 | 0.272471248 |
| Gdf10 | 299.4413777 | 0.180914617 | 0.110428646 | 1.638294263 | 0.101360324 | 0.272471248 |
| Nbn | 687.5640095 | 0.151197438 | 0.092280615 | 1.638452861 | 0.10132726 | 0.272471248 |
| Snx11 | 1261.113865 | 0.100661921 | 0.061437226 | 1.63845159 | 0.101327525 | 0.272471248 |
| Synpo2 | 312.2082807 | 0.218128442 | 0.133112701 | 1.638674897 | 0.101280985 | 0.272471248 |
| Tspan7 | 44196.6614 | -0.142686644 | 0.087087 | -1.63843793 | 0.101330372 | 0.272471248 |
| Mtmr14 | 614.9868866 | 0.142755699 | 0.087147712 | 1.638088883 | 0.101403154 | 0.27253087 |
| Cct4 | 3636.356469 | -0.099049486 | 0.060477892 | -1.637780074 | 0.10146758 | 0.272656726 |
| Vipr1 | 1258.568698 | 0.166901229 | 0.101923714 | 1.637511258 | 0.101523688 | 0.272760192 |
| Nt5c3l | 1087.079698 | -0.173658306 | 0.106084901 | -1.636974758 | 0.101635744 | 0.273013907 |
| Zfp324 | 449.481359 | -0.13297048 | 0.081279985 | -1.635956013 | 0.101848794 | 0.273538778 |
| Pars2 | 338.0320474 | -0.147187755 | 0.089980349 | -1.635776655 | 0.101886339 | 0.273592191 |
| Dclre1a | 339.9308686 | 0.15528437 | 0.094939458 | 1.635614676 | 0.101920257 | 0.273635845 |
| Mab21l1 | 35.19397979 | 0.273704729 | 0.167458102 | 1.634466924 | 0.102160846 | 0.27423426 |
| Maats1 | 54.53201673 | 0.231707882 | 0.141806226 | 1.63397538 | 0.10226402 | 0.274407405 |
| Ppat | 611.5321949 | 0.122774278 | 0.075144969 | 1.63383231 | 0.102294066 | 0.274407405 |
| Rufy3 | 8947.127118 | 0.084619323 | 0.051792247 | 1.633822196 | 0.10229619 | 0.274407405 |
| Tbl1xr1 | 2726.036943 | -0.104113545 | 0.063723885 | -1.633822948 | 0.102296032 | 0.274407405 |
| Ska3 | 23.86533943 | 0.273642705 | 0.167526936 | 1.633425117 | 0.102379618 | 0.27458366 |
| Fam108b | 787.2235606 | 0.130812463 | 0.080101386 | 1.633086141 | 0.102450881 | 0.274727233 |
| Rnf169 | 2399.90417 | 0.09922354 | 0.060797838 | 1.632024151 | 0.102674399 | 0.275278964 |
| Fars2 | 539.8178072 | -0.119081426 | 0.072970849 | -1.631904084 | 0.102699694 | 0.27528099 |
| Limd1 | 524.0460605 | -0.120440024 | 0.073805732 | -1.631851903 | 0.102710688 | 0.27528099 |
| Mir341 | 8.448284105 | 0.244100317 | 0.149656406 | 1.631071625 | 0.102875207 | 0.275674241 |
| Gpr146 | 1009.536684 | -0.107575819 | 0.065965892 | -1.630779417 | 0.102936872 | 0.275791786 |
| Klhl20 | 839.4055186 | 0.104363365 | 0.064000947 | 1.630653443 | 0.102963466 | 0.275815342 |
| Nup54 | 583.0974803 | 0.128133089 | 0.078592877 | 1.630339729 | 0.103029715 | 0.275945101 |
| A630007B06Rik | 992.004553 | 0.131611308 | 0.080740132 | 1.6300606 | 0.10308869 | 0.276018401 |
| Arfgap1 | 4322.646027 | 0.10776752 | 0.066113357 | 1.630041566 | 0.103092712 | 0.276018401 |
| Ino80d | 1514.926483 | -0.122269419 | 0.075049053 | -1.629193377 | 0.103272089 | 0.276330832 |
| Prdm10 | 441.9958178 | -0.122675886 | 0.075290974 | -1.629357146 | 0.103237435 | 0.276330832 |
| Rasl11a | 98.72205403 | 0.266911162 | 0.163834365 | 1.629152479 | 0.103280744 | 0.276330832 |
| Zfp385b | 1428.745521 | -0.097456541 | 0.059818291 | -1.62920971 | 0.103268632 | 0.276330832 |
| Bahcc1 | 1312.242964 | -0.151152501 | 0.092785219 | -1.629057969 | 0.103300748 | 0.276336634 |
| Arid4a | 1556.25375 | 0.108063078 | 0.066351003 | 1.628657789 | 0.103385483 | 0.276515567 |
| Senp1 | 631.1351365 | 0.153322157 | 0.094147637 | 1.628529005 | 0.103412764 | 0.276540795 |
| Kat6a | 3335.655735 | -0.13386933 | 0.08221254 | -1.628332259 | 0.103454453 | 0.276556814 |
| Snord52 | 14.4949575 | 0.272168302 | 0.167144481 | 1.628341536 | 0.103452487 | 0.276556814 |
| Col15a1 | 262.4203989 | 0.264690014 | 0.162613258 | 1.627727144 | 0.103582755 | 0.276852028 |
| Rab9b | 1127.154169 | -0.153943389 | 0.09459196 | -1.627446861 | 0.103642226 | 0.276963202 |
| Rnf4 | 3554.416021 | -0.083085133 | 0.051055538 | -1.627348114 | 0.103663185 | 0.27697144 |
| 4932438H23Rik | 40.0754293 | 0.261261902 | 0.160585826 | 1.626930027 | 0.10375196 | 0.277160839 |
| Pitpnm1 | 8211.257429 | 0.125473512 | 0.077144411 | 1.626475721 | 0.103848494 | 0.277370896 |
| Tnfrsf25 | 128.2417777 | 0.250627097 | 0.15412376 | 1.62614185 | 0.103919483 | 0.277512663 |
| Dpy19l1 | 4659.300404 | -0.144977362 | 0.089169963 | -1.625854235 | 0.103980668 | 0.27758037 |
| S100a9 | 30.33289457 | -0.270831916 | 0.166571589 | -1.625919023 | 0.103966883 | 0.27758037 |
| Fpgs | 218.55434 | 0.166955835 | 0.102724857 | 1.625272019 | 0.104104611 | 0.277824139 |
| Parpbp | 17.34880898 | 0.27251942 | 0.167677757 | 1.625256834 | 0.104107845 | 0.277824139 |
| Actr1b | 9938.384078 | 0.105916773 | 0.065206301 | 1.624333421 | 0.104304668 | 0.278301451 |
| Gm960 | 33.73445343 | 0.262638297 | 0.16171207 | 1.624110662 | 0.104352193 | 0.278380316 |
| Nagpa | 360.2536231 | 0.161959403 | 0.099761933 | 1.623458948 | 0.104491332 | 0.278703512 |
| Ikbkb | 1582.845805 | 0.099804399 | 0.061483754 | 1.623264568 | 0.10453286 | 0.278766289 |
| Barhl2 | 24.57405904 | 0.253802879 | 0.156455026 | 1.622209817 | 0.104758431 | 0.279301801 |
| Kank1 | 766.814029 | -0.152027644 | 0.093724301 | -1.622072861 | 0.104787748 | 0.279301801 |
| Rrp36 | 453.1182602 | -0.124515159 | 0.076761488 | -1.622104547 | 0.104780965 | 0.279301801 |
| Fam217b | 1447.552844 | 0.112132806 | 0.069148096 | 1.621632585 | 0.104882041 | 0.279410144 |
| Gnaq | 9300.135729 | -0.095338787 | 0.058794983 | -1.621546295 | 0.10490053 | 0.279410144 |
| Pdp1 | 5947.328751 | -0.145849678 | 0.089940593 | -1.62162237 | 0.10488423 | 0.279410144 |
| Phactr2 | 1506.256242 | -0.177257012 | 0.109308578 | -1.621620326 | 0.104884668 | 0.279410144 |
| Fam92a | 1392.222016 | -0.108361745 | 0.066832737 | -1.621387208 | 0.104934622 | 0.279442249 |
| Pyhin1 | 11.48654586 | 0.258656511 | 0.15953435 | 1.621321744 | 0.104948654 | 0.279442249 |
| Aasdhppt | 688.7303287 | -0.124549731 | 0.076829969 | -1.621108687 | 0.104994331 | 0.279485598 |
| Cul4a | 2699.803422 | 0.105825932 | 0.065281228 | 1.621077536 | 0.105001011 | 0.279485598 |
| Ralgps1 | 1313.652456 | 0.11130089 | 0.068673167 | 1.620733317 | 0.105074845 | 0.279634086 |
| Secisbp2 | 614.2564613 | 0.150339178 | 0.092780748 | 1.620370394 | 0.105152736 | 0.279793317 |
| Cul2 | 2153.166555 | -0.108986352 | 0.06726401 | -1.620277364 | 0.105172709 | 0.279798413 |
| Gm996 | 4853.639863 | 0.132823927 | 0.081998841 | 1.619826885 | 0.105269469 | 0.280007753 |
| Fn3krp | 1287.518685 | 0.121115781 | 0.074791854 | 1.619371289 | 0.105367401 | 0.280220135 |
| Ggt5 | 104.1716105 | -0.207771206 | 0.128317794 | -1.619192477 | 0.105405856 | 0.280274298 |
| Hepacam | 3685.666239 | 0.114038877 | 0.07043509 | 1.619063394 | 0.105433624 | 0.280300029 |
| Fcho2 | 997.492919 | 0.11376431 | 0.070282633 | 1.618668878 | 0.105518526 | 0.280429511 |
| Hdac2 | 2718.900431 | -0.097569775 | 0.060276474 | -1.618704096 | 0.105510945 | 0.280429511 |
| Wdr78 | 403.0844226 | -0.195617035 | 0.120881205 | -1.618258479 | 0.105606905 | 0.280616255 |
| Cnst | 2285.990699 | -0.100642592 | 0.062200558 | -1.618033589 | 0.105655359 | 0.280696868 |
| Zzef1 | 2995.721911 | -0.111225503 | 0.06874491 | -1.617945289 | 0.105674389 | 0.280699294 |
| Eef1d | 2105.631821 | 0.103196851 | 0.06382977 | 1.616751094 | 0.10593202 | 0.281226693 |
| Jmjd6 | 742.4531876 | 0.159096313 | 0.098408763 | 1.616688485 | 0.105945541 | 0.281226693 |
| Mdm4 | 1509.62579 | 0.132349029 | 0.081852058 | 1.616929777 | 0.10589344 | 0.281226693 |
| Nr1d2 | 4649.563805 | 0.086801627 | 0.053688617 | 1.616760326 | 0.105930026 | 0.281226693 |
| Adam1a | 188.4424594 | 0.248655352 | 0.153830811 | 1.616420992 | 0.106003323 | 0.281283693 |
| Pgpep1 | 490.9257719 | -0.117532316 | 0.072708192 | -1.616493441 | 0.10598767 | 0.281283693 |
| Srsf3 | 3632.505017 | -0.099182794 | 0.061368076 | -1.616195276 | 0.1060521 | 0.281364937 |
| Cdc25a | 264.5112454 | 0.202523829 | 0.125356436 | 1.615583818 | 0.106184325 | 0.281667511 |
| Zbtb46 | 246.7347384 | -0.182507568 | 0.112982868 | -1.61535612 | 0.106233597 | 0.281749975 |
| 1600029O15Rik | 57.85542757 | 0.2657691 | 0.164589469 | 1.614739393 | 0.106367143 | 0.281828499 |
| Fut9 | 3221.643881 | -0.176435101 | 0.109248661 | -1.614986391 | 0.106313642 | 0.281828499 |
| Glt1d1 | 589.2324156 | 0.118366582 | 0.073303573 | 1.614745058 | 0.106365916 | 0.281828499 |
| Kcnk10 | 131.0118844 | -0.185661629 | 0.114978134 | -1.614755981 | 0.106363549 | 0.281828499 |
| Limch1 | 4941.186566 | -0.085310077 | 0.052832888 | -1.614715397 | 0.106372342 | 0.281828499 |
| Mir17hg | 14.55705698 | 0.268030947 | 0.165968934 | 1.614946484 | 0.106322285 | 0.281828499 |
| Gm9899 | 906.215188 | 0.167771914 | 0.103922348 | 1.614396887 | 0.106441366 | 0.281963162 |
| Rpa2 | 430.6330283 | 0.151017925 | 0.093570754 | 1.61394366 | 0.106539647 | 0.282175264 |
| Pdcd6 | 1886.986529 | -0.093517844 | 0.057958438 | -1.613532859 | 0.10662879 | 0.282363097 |
| 4921525O09Rik | 12.00856785 | -0.246502749 | 0.152810011 | -1.613132202 | 0.106715789 | 0.282545187 |
| Cnn1 | 20.07974851 | -0.269708087 | 0.167212491 | -1.612966143 | 0.106751864 | 0.28259241 |
| Cyba | 175.2882944 | 0.190403832 | 0.118089047 | 1.612375038 | 0.106880353 | 0.282884215 |
| Rgcc | 339.1961596 | -0.144263739 | 0.089493131 | -1.612009077 | 0.106959965 | 0.282977548 |
| Slit1 | 2332.836669 | 0.177013416 | 0.109810421 | 1.611991043 | 0.106963889 | 0.282977548 |
| Usp10 | 2357.068351 | -0.173119344 | 0.107396727 | -1.611961087 | 0.106970408 | 0.282977548 |
| Rbm10 | 1987.045772 | 0.121322071 | 0.075297426 | 1.611237966 | 0.107127866 | 0.28332306 |
| Zfp426 | 1048.022472 | -0.104493999 | 0.064855034 | -1.611193339 | 0.107137589 | 0.28332306 |
| Tsc22d3 | 2637.300384 | 0.190729054 | 0.118424203 | 1.610558057 | 0.107276082 | 0.283640889 |
| Ankrd34b | 911.0774415 | -0.163504125 | 0.101528259 | -1.61042972 | 0.107304077 | 0.283666502 |
| Fam135a | 1157.582834 | 0.139736327 | 0.086795917 | 1.609941244 | 0.107410684 | 0.283899887 |
| Ppp6r2 | 3635.610539 | 0.1120764 | 0.069633407 | 1.609520563 | 0.107502562 | 0.284094269 |
| Usp13 | 1088.398101 | 0.116160892 | 0.072177716 | 1.609373343 | 0.107534731 | 0.284130818 |
| Cyb561d2 | 412.6147067 | -0.147486041 | 0.091653837 | -1.609163859 | 0.107580517 | 0.28415488 |
| Tirap | 106.33414 | -0.183850693 | 0.114248167 | -1.609222256 | 0.107567751 | 0.28415488 |
| 4930412C18Rik | 74.90942367 | 0.248916461 | 0.154741011 | 1.608600461 | 0.107703733 | 0.284383375 |
| Alyref | 516.9348803 | -0.124872457 | 0.077624627 | -1.608670631 | 0.107688381 | 0.284383375 |
| Chrnb4 | 21.99394608 | -0.238360466 | 0.148195175 | -1.608422586 | 0.107742658 | 0.284437681 |
| Atg5 | 799.6865395 | -0.098898112 | 0.061498129 | -1.608148302 | 0.107802703 | 0.284480801 |
| Blmh | 2098.61921 | -0.083090393 | 0.051668105 | -1.608156374 | 0.107800935 | 0.284480801 |
| Tbxas1 | 95.92271577 | 0.259149782 | 0.161153142 | 1.60809637 | 0.107814074 | 0.284480801 |

| | | | | | | | |
|---|---|---|---|---|---|---|---|
| Unc80 | 12761.92811 | 0.0770344239011059 | 0.0711060833933612 | 1.08337318306436 | 0.278642813783516 | 0.718495890058647 | NM_175510 |
| Zfp334 | 986.7695268 | 0.0650353985846275 | 0.0600154000707892 | 1.08364517287092 | 0.278522153723766 | 0.718495890058647 | NM_178411 |
| Abhd4 | 3205.8619 | -0.0685095212401944 | 0.0632904751261222 | -1.08246179387454 | 0.279047383278987 | 0.718516358316667 | NM_001205181,NM_134076 |
| Otud4 | 2617.621183 | -0.0751280197798745 | 0.069406182323725 | -1.08243988164428 | 0.279057115125755 | 0.718516358316667 | NM_001256033,NM_001081164 |
| Pds5b | 4477.816906 | 0.0704771245302456 | 0.0651121678852201 | 1.08239560775311 | 0.279076779128641 | 0.718516358316667 | NM_175310 |
| Vamp2 | 52658.47258 | 0.0427461138633056 | 0.0394979630032818 | 1.0822359082101 | 0.279147716623927 | 0.718571634180252 | NM_009497 |
| Rfx2 | 121.3348945 | -0.102199273693815 | 0.0944511735980633 | -1.08203286206611 | 0.27923792610177 | 0.718676490884889 | NM_009056,NM_027787 |
| Enpep | 150.8106724 | -0.0962746070977095 | 0.0889886300033996 | -1.08187537097752 | 0.279307909998486 | 0.71872926494916 | NM_007934 |
| 1810062O18Rik | 35.17006265 | -0.0969367532253659 | 0.0896467593523029 | -1.0813191009439 | 0.27955519368537 | 0.718934998182959 | NR_033571 |
| A730046J19Rik | 29.30870069 | 0.0808205392684427 | 0.0747578298496825 | 1.08109798573542 | 0.27965352932897 | 0.718934998182959 | NR_040271 |
| Cpt1c | 4635.541332 | -0.0766228086464415 | 0.0708834001551254 | -1.08096971193193 | 0.27971058677728 | 0.718934998182959 | NM_153679,NM_001252470 |
| Mrgpre | 581.8105472 | 0.0944229539824521 | 0.0873355085958815 | 1.08115193350926 | 0.279629535192522 | 0.718934998182959 | NM_175534 |
| Nme3 | 542.7119345 | 0.0725506212624745 | 0.0671195502148603 | 1.08091638025327 | 0.279734311560656 | 0.718934998182959 | NM_019730 |
| Nxph4 | 238.3032636 | 0.102390866275888 | 0.0946809940429861 | 1.08142998825511 | 0.279505888085317 | 0.718934998182959 | NM_183297 |
| Wdr78 | 422.9098256 | -0.0782455225380008 | 0.0723838619632985 | -1.08098021320933 | 0.279705915408548 | 0.718934998182959 | NM_146254 |
| Aig1 | 918.7739701 | 0.0656310364794119 | 0.0607461279315115 | 1.08041514272988 | 0.279957355656783 | 0.719332627922656 | NM_025446 |
| Slc5a5 | 991.2797174 | 0.0698365894082961 | 0.0646427903943572 | 1.08034614505738 | 0.279988062172566 | 0.719332627922656 | NM_053248 |
| Slc4a11 | 157.559984 | -0.101851011147566 | 0.0942862790895701 | -1.08023152606128 | 0.280039092892196 | 0.719336492193111 | NM_001081162 |
| D10Wsu52e | 4741.037952 | 0.04903243036698 | 0.0454136261942589 | 1.07968542651145 | 0.280282285749249 | 0.719833890168598 | NM_145422 |
| 2010320M18Rik | 148.4111201 | 0.100124798384337 | 0.0929474158558659 | 1.07721981791943 | 0.281382073634713 | 0.720080691785073 | NR_029440 |
| Cercam | 312.8030461 | -0.0970519932710026 | 0.0899533489911239 | -1.07891473035183 | 0.28062574158483 | 0.720080691785073 | NM_207298 |
| Fam166b | 24.15292771 | -0.0737723324540534 | 0.0683909334819627 | -1.07868585349124 | 0.280727794121395 | 0.720080691785073 | NM_001162381,NM_177377 |
| Hspa1b | 183.8677698 | -0.0983977574957784 | 0.0912831256674741 | -1.07794027402415 | 0.281060410843893 | 0.720080691785073 | NM_010478 |
| Hydin | 87.07739964 | -0.102227848252024 | 0.0949364095190135 | -1.07680339682058 | 0.281568107485833 | 0.720080691785073 | NM_172916 |
| Igdcc4 | 2561.996501 | -0.0565267897713872 | 0.0524076178604652 | -1.07859872436655 | 0.280766650228591 | 0.720080691785073 | NM_020043 |
| Kdm4b | 3203.538991 | -0.0668786128793461 | 0.0620621539412963 | -1.07760702186723 | 0.281209167242808 | 0.720080691785073 | NM_172132 |
| Myg1 | 930.0516972 | 0.0659996616790923 | 0.0611957157157823 | 1.07850134453238 | 0.280810082063996 | 0.720080691785073 | NM_021713 |
| N4bp1 | 3181.350204 | 0.0685545517536374 | 0.0635488572341272 | 1.07876922949327 | 0.280690615176054 | 0.720080691785073 | NM_030563 |
| Nnat | 5669.7866 | 0.101389063250359 | 0.094124470899949 | 1.07718069786689 | 0.281399546753624 | 0.720080691785073 | NM_010923,NM_180960 |
| Pcdhb22 | 208.1356312 | 0.0876431759774927 | 0.0813598995597421 | 1.07722817323707 | 0.281378341796022 | 0.720080691785073 | NM_053147 |
| Pnpla8 | 4483.503475 | 0.0722294934665977 | 0.0669843300916741 | 1.07830433427856 | 0.280897963455614 | 0.720080691785073 | NM_026164 |
| Prox1 | 446.3904098 | -0.0990644047848055 | 0.0917966543292216 | -1.07917228039182 | 0.280510934265873 | 0.720080691785073 | NM_008937 |
| Rnf2 | 957.4969753 | 0.0749755642725361 | 0.069545662350211 | 1.07807678780859 | 0.280999489533034 | 0.720080691785073 | NM_011277 |
| Sh3tc2 | 157.3458656 | -0.0973976999316512 | 0.09035317669527 | -1.07796652529595 | 0.281048695126366 | 0.720080691785073 | NM_172628 |
| Slc35a5 | 1830.40936 | 0.0661013232467845 | 0.0613332896140549 | 1.0777397342085 | 0.281149920984276 | 0.720080691785073 | NM_028756 |
| Snw1 | 3377.999625 | 0.0644195733133818 | 0.0597762479456468 | 1.07767843461766 | 0.281177285671104 | 0.720080691785073 | NM_025507 |
| Srxn1 | 4972.62886 | 0.0562017859903008 | 0.052124369419696 | 1.0782247654216 | 0.280933462443991 | 0.720080691785073 | NM_029688 |
| Stx8 | 763.2681556 | 0.0664653837643751 | 0.0617137765288877 | 1.07699427101602 | 0.281482825124709 | 0.720080691785073 | NM_018768 |
| Tmem43 | 1770.52713 | 0.058595204657232 | 0.0543892363222733 | 1.07733089521679 | 0.281332464554685 | 0.720080691785073 | NM_028766 |
| Tubgcp2 | 1804.279564 | -0.0587284032195828 | 0.0544998839790615 | -1.07758767417094 | 0.281217805267932 | 0.720080691785073 | NM_133755 |
| Vta1 | 1445.067913 | 0.0541893530688288 | 0.0502092490590631 | 1.07927033533371 | 0.280467232990567 | 0.720080691785073 | NM_025418 |
| Xist | 21529.01429 | -0.082026289019893 | 0.0761736931454302 | -1.07683224526464 | 0.281555216911187 | 0.720080691785073 | NR_001463,NR_001570 |
| Zfp318 | 2633.932134 | -0.0679393288737206 | 0.063088562015077 | -1.0768882140234 | 0.281530209097436 | 0.720080691785073 | NM_021346,NM_207671 |
| Poc1b | 257.1290289 | -0.0824442758094743 | 0.0765752623399869 | -1.07664372657882 | 0.281639461423672 | 0.720136387368466 | NM_027740 |
| Atf6 | 4921.29566 | 0.0741231331390071 | 0.0688627057202014 | 1.07639007738353 | 0.281752838193548 | 0.720163706248182 | NM_001081304 |
| Atp1a3 | 106168.954 | 0.0507858754326971 | 0.0471861789137612 | 1.0762870951156 | 0.281798878308871 | 0.720163706248182 | NM_144921 |
| Gpr180 | 937.7739912 | 0.0646531293919516 | 0.060059071448143 | 1.07649232985178 | 0.281707132079656 | 0.720163706248182 | NM_021434 |
| Ntm | 5651.835564 | 0.0524836967723878 | 0.0487707410183014 | 1.07613080458821 | 0.281868760610804 | 0.72021558779816 | NM_172290 |
| Btrc | 5824.703144 | 0.0533307739952803 | 0.0495667944715082 | 1.07593752155863 | 0.281955199655226 | 0.720294901658224 | NM_001037758,NM_009771 |
| Tfb2m | 969.69367 | 0.0682523967622689 | 0.0634410495582097 | 1.07583965330909 | 0.28199897464755 | 0.720294901658224 | NM_008249 |
| Cacna1i | 5366.095026 | 0.065881892340466 | 0.0612587859425983 | 1.07546846263326 | 0.282165044534195 | 0.720537371494138 | NM_001044308 |
| Ccdc80 | 277.2201982 | -0.0856215310649618 | 0.0796178848801529 | -1.07540574826681 | 0.28219310935083 | 0.720537371494138 | NM_026439 |
| Rev1 | 1412.347901 | -0.064475768570525 | 0.0599726229948074 | -1.07508668707232 | 0.282335919247277 | 0.720775318626704 | NM_019570 |
| Pdp1 | 6703.975173 | 0.102187471340937 | 0.0950730831284838 | 1.07483072998526 | 0.282450519537531 | 0.720814519817671 | NM_001098231,NM_001098230,NM_001033453 |
| Slc38a2 | 5628.950296 | 0.081197943553668 | 0.0755417029084614 | 1.07487573654595 | 0.2824303663557 | 0.720814519817671 | NM_175121 |
| Terf2ip | 2556.706724 | 0.0584976174833131 | 0.0544310056191654 | 1.07471131238325 | 0.282503997458903 | 0.720824357471988 | NM_020584 |
| Ring1 | 1162.220857 | -0.0912166686579989 | 0.0848899364565359 | -1.07452864810074 | 0.282585811963266 | 0.720906481309871 | NM_009066 |
| A530054K11Rik | 474.9885696 | 0.0897171407623539 | 0.0835669555952618 | 1.07359589832229 | 0.283003836756252 | 0.720920006908497 | NM_183146 |
| Brd8 | 2621.370711 | -0.0523163548362937 | 0.0487070386266349 | -1.07410255912551 | 0.282776717708717 | 0.720920006908497 | NM_030147 |
| Gm14634 | 33.46517389 | 0.0966462426218944 | 0.0899860573787535 | 1.0740135242853 | 0.282816620092462 | 0.720920006908497 | NR_045852 |
| Gramd1b | 4160.001306 | 0.0750973110492009 | 0.0699606748042451 | 1.07342176528926 | 0.283081923289024 | 0.720920006908497 | NM_172768 |
| Gss | 511.7818451 | 0.0711776347404354 | 0.066301429094877 | 1.07354601118146 | 0.283026206171178 | 0.720920006908497 | NM_008180 |
| Lrrc55 | 907.5960819 | -0.0899802181526502 | 0.0837576526875603 | -1.07429250063038 | 0.282691605146939 | 0.720920006908497 | NM_001033346 |
| Ndufa13 | 3693.434647 | 0.064184449947945 | 0.0597810707536317 | 1.07365841961012 | 0.282975803876078 | 0.720920006908497 | NM_023312 |
| Prkce | 18609.37405 | 0.0522070575747792 | 0.0486366629633153 | 1.07340953087503 | 0.283087410120203 | 0.720920006908497 | NM_011104 |
| Psd | 16405.87659 | -0.079220294156044 | 0.0738023212940949 | -1.07341195733342 | 0.283086321908103 | 0.720920006908497 | NM_028627 |
| Zfp619 | 221.0969102 | 0.0921871805837336 | 0.0858621465830574 | 1.07366498803472 | 0.28297285887867 | 0.720920006908497 | NM_001004139 |
| Cish | 352.4925579 | 0.0779964099449272 | 0.0726751934630539 | 1.07321915812413 | 0.283172796858853 | 0.720995619419525 | NM_009895 |
| Wipf1 | 619.9860346 | -0.071527794407842 | 0.0666539245916477 | -1.07312202313748 | 0.283216370949181 | 0.720995619419525 | NM_153138 |
| 5430405H02Rik | 177.5997933 | -0.0970823324422778 | 0.0905003199703759 | -1.07272916243894 | 0.283392651907602 | 0.721034820635665 | NR_027914,NR_015591 |
| Cltb | 5343.39248 | 0.0463195890420222 | 0.0431885893362356 | 1.07249599382399 | 0.283497312392863 | 0.721034820635665 | NM_028870 |
| Mxi1 | 3532.067226 | 0.0639127386719913 | 0.0595948580451979 | 1.07245391244189 | 0.283516203906313 | 0.721034820635665 | NM_001008542,NM_010847,NM_001008543 |
| Smap2 | 8620.253198 | 0.0506332558820847 | 0.047213836951853 | 1.07242408478088 | 0.2835295948968 | 0.721034820635665 | NM_133716 |
| Sort1 | 29328.97443 | 0.0606782974809372 | 0.0565698701813459 | 1.0726257155341 | 0.283439082041588 | 0.721034820635665 | NM_001271599,NM_019972 |
| Spcs2 | 2927.906277 | 0.0569733965758296 | 0.053106668913361 | 1.07281058559287 | 0.283356110329989 | 0.721034820635665 | NM_025668 |
| Gnaz | 2626.301107 | 0.0595136082815445 | 0.055501023075911 | 1.07229749981627 | 0.283586429397768 | 0.721053119802552 | NM_010311 |
| Atp6v1f | 3287.146837 | 0.0535326712851625 | 0.0499352668110835 | 1.07204135881938 | 0.283701455958793 | 0.721155193971164 | NM_025381 |
| I7Rn6 | 937.9543132 | 0.0689485756450622 | 0.0643184804830392 | 1.0719870109998 | 0.283725866277379 | 0.721155193971164 | NM_026304 |
| Wdr19 | 1060.867365 | -0.0681195860225376 | 0.0635615604666267 | -1.07171040991519 | 0.283850123669759 | 0.72134480343368 | NM_153391 |
| Akap6 | 10170.44936 | 0.0738244499609457 | 0.0688978324045574 | 1.07150613284116 | 0.283941914626498 | 0.721415456458809 | NM_198111 |
| Nr2c2 | 7033.064735 | 0.0638436309511017 | 0.0595874484649913 | 1.07142750018254 | 0.283977253203323 | 0.721415456458809 | NM_011630 |
| Klf14 | 29.49413904 | 0.0932715827047396 | 0.0870685814588097 | 1.07124270479661 | 0.284060314461644 | 0.721500284642392 | NM_001135093 |
| F8a | 494.3545362 | 0.0864025074228351 | 0.0806655798182161 | 1.07111989546902 | 0.284115523513848 | 0.721514352196182 | NM_007978 |
| Ap3m2 | 7879.773524 | 0.0472328716642818 | 0.0441161401868954 | 1.07064832653497 | 0.284327585273653 | 0.721548306024648 | NM_001122820,NM_029505 |
| Clic5 | 580.7018636 | 0.0950834683983137 | 0.0887867823422823 | 1.07091918289996 | 0.284205769678634 | 0.721548306024648 | NM_172621 |
| Grpel1 | 1636.217454 | 0.0557039927051826 | 0.0520272688048582 | 1.07066916993308 | 0.284318209858722 | 0.721548306024648 | NM_024478 |
| Zfp866 | 1189.702765 | -0.066581337190321 | 0.0621854060095407 | -1.07069072090815 | 0.28430851639382 | 0.721548306024648 | NM_177899 |
| 2810001G20Rik | 247.3767091 | 0.086152590969598 | 0.0805463317801164 | 1.06960291133786 | 0.284798084181193 | 0.721641599170286 | NR_033780 |
| A430005L14Rik | 498.6868919 | 0.0848215464890692 | 0.079317351955278 | 1.06939458262417 | 0.284891907370785 | 0.721641599170286 | NM_001163019,NM_175287 |
| Armc2 | 376.6844156 | -0.0857911177805199 | 0.0801990785993356 | -1.0697269754073 | 0.284742220457296 | 0.721641599170286 | NM_001034858 |
| Col13a1 | 58.2443769 | -0.0979502445064935 | 0.0915217811326478 | -1.07023971009184 | 0.28451142427362 | 0.721641599170286 | NM_007731 |
| Fbxo33 | 2126.161525 | 0.080647173837332 | 0.0753986360296911 | 1.06961051398296 | 0.284794660639448 | 0.721641599170286 | NM_001033156 |
| Gusb | 613.0373484 | -0.0885006964201932 | 0.0827610059809592 | -1.06935259390823 | 0.284910819994602 | 0.721641599170286 | NM_010368 |
| Htr5b | 46.64434564 | 0.0935584748926802 | 0.0874812951570947 | 1.06946833291245 | 0.284858690705047 | 0.721641599170286 | NM_010483 |
| Igsf3 | 793.4124178 | 0.0677673973361114 | 0.06332404764037 | 1.07016844092115 | 0.28454349693866 | 0.721641599170286 | NM_207205 |
| Lsm10 | 224.2928109 | -0.0932779227250405 | 0.0872019914583491 | -1.06967651959638 | 0.284764938868341 | 0.721641599170286 | NM_001163266,NM_138721 |
| Mki67 | 47.92295484 | -0.0972473708209061 | 0.0908586621053706 | -1.07031480067499 | 0.284477634537008 | 0.721641599170286 | NM_001081117 |
| Ube4a | 3319.742511 | 0.0616750436094852 | 0.0576661904175248 | 1.06951825953708 | 0.284836205554576 | 0.721641599170286 | NM_145400 |
| Mppe1 | 611.9790515 | -0.0781682161667932 | 0.0731116378690423 | -1.06916242563715 | 0.284996486025003 | 0.721671536078837 | NM_172630 |
| Zfp142 | 1875.690942 | -0.065002029036271 | 0.0608003712327706 | -1.06910579192707 | 0.285022002098602 | 0.721671536078837 | NM_029888 |
| Ly96 | 26.78980905 | -0.0909479931532159 | 0.0851036643628342 | -1.06869420763667 | 0.285207482436205 | 0.722015317160738 | NM_016923,NM_001159711 |
| 9330133O14Rik | 304.0635877 | -0.0931288945524105 | 0.0873824216361756 | -1.06576234451548 | 0.286531086938995 | 0.722148664676319 | NR_045696 |
| Abracl | 217.3400706 | 0.0957387083217132 | 0.0898455241132498 | 1.06559240726377 | 0.286607932966267 | 0.722148664676319 | NM_028440 |
| Ahcyl2 | 7881.915158 | 0.055718934005421 | 0.0522532086067216 | 1.0663255997308 | 0.286276481101868 | 0.722148664676319 | NM_021414,NM_001171000,NM_001171001 |
| Aldh1a3 | 39.09578032 | 0.0985765361696862 | 0.0922885968682096 | 1.06813343701016 | 0.285460324883769 | 0.722148664676319 | NM_053080 |
| BC016423 | 1489.49667 | -0.0812134111823859 | 0.0761713823505211 | -1.06619321687852 | 0.286336307811175 | 0.722148664676319 | NM_134063 |
| C030034I22Rik | 90.82420222 | -0.10057416888289 | 0.0943803955244495 | -1.06562563468846 | 0.286592906351684 | 0.722148664676319 | NR_026848 |
| Cand1 | 6742.394693 | 0.0622944509039118 | 0.058397387394069 | 1.06673352496996 | 0.286092183857092 | 0.722148664676319 | NM_027994 |
| Ccdc58 | 209.7816637 | -0.0922591335745589 | 0.0865720093841367 | -1.06569241295056 | 0.286562708473 | 0.722148664676319 | NM_001159421,NM_198645,NM_001159422 |
| Clstn1 | 53421.70596 | 0.0541644237094199 | 0.0507728652284326 | 1.06679864265545 | 0.286062771647295 | 0.722148664676319 | NM_023051 |
| Clybl | 660.442379 | 0.080330986076522 | 0.0752680180143167 | 1.06726586132828 | 0.285851799286488 | 0.722148664676319 | NM_029556 |
| Ddx1 | 6232.239062 | 0.0593441978777713 | 0.05554825196409 | 1.06833601021568 | 0.285368970409121 | 0.722148664676319 | NM_134040 |
| Epas1 | 7309.736955 | 0.0697317540331043 | 0.0653792701898297 | 1.06657284228834 | 0.286164769390598 | 0.722148664676319 | NM_010137 |
| Fam160b1 | 1768.928673 | -0.0520802899907458 | 0.0487711108932709 | -1.06785121431244 | 0.285587863187369 | 0.722148664676319 | NM_145505 |
| Fndc1 | 238.0907312 | -0.0975294003150239 | 0.0913674051685879 | -1.06744194097519 | 0.285772317896638 | 0.722148664676319 | NM_001081416 |
| Glrx | 1422.542194 | 0.0644315521213752 | 0.0604759042538473 | 1.06540866013221 | 0.286691039513097 | 0.722148664676319 | NM_053108 |
| Lrrc57 | 1424.401461 | -0.0783232618847653 | 0.0733821053626603 | -1.06733462467022 | 0.285820758112222 | 0.722148664676319 | NM_001159609,NR_027510,NM_025657,NM_001159612,NM_001159610 |
| Ms4a6d | 48.68311452 | -0.0974314538201567 | 0.091238255846227 | -1.06787939901402 | 0.285574916384881 | 0.722148664676319 | NM_026835 |
| Pcsk9 | 217.4469274 | -0.096194966677505 | 0.0901215536202875 | -1.06739112679978 | 0.285795253604196 | 0.722148664676319 | NM_153565 |
| Pebp1 | 15242.63197 | 0.0395604607067967 | 0.0371325587863098 | 1.06538471895942 | 0.286701869006494 | 0.722148664676319 | NM_018858 |
| Psmd3 | 5201.791336 | 0.0556185625175161 | 0.0521983890297116 | 1.06552258702577 | 0.286639509876602 | 0.722148664676319 | NM_009439 |
| Pttg1ip | 2492.675082 | -0.0483639762287833 | 0.0453141972409386 | -1.06730294639512 | 0.285835058048343 | 0.722148664676319 | NM_145925 |
| Pxdc1 | 159.0585758 | 0.0965972163706487 | 0.0905109330117619 | 1.06724362633734 | 0.285861837102076 | 0.722148664676319 | NM_025831 |
| Rslcan18 | 39.63508228 | 0.0986340799135334 | 0.0925645774336263 | 1.06557046602691 | 0.286617855859905 | 0.722148664676319 | NM_001256052 |
| Snx21 | 1002.04595 | 0.0701951262988166 | 0.0657571512779916 | 1.06749037837821 | 0.285750456137523 | 0.722148664676319 | NM_133924 |
| Sult1a1 | 361.0962343 | 0.087354408125255 | 0.0818754097512932 | 1.06691872920826 | 0.286008536581168 | 0.722148664676319 | NM_133670 |
| Ttc7b | 8638.808872 | 0.0522254718221629 | 0.0488918607728503 | 1.0681833539697 | 0.285437811988143 | 0.722148664676319 | NM_001033213 |
| Usp46 | 4549.777705 | 0.062307356799087 | 0.0584749806700235 | 1.06553873272212 | 0.286632207612964 | 0.722148664676319 | NM_177561 |
| Xk | 947.6192772 | 0.0773741525599013 | 0.0725898872319664 | 1.06590815209075 | 0.28646516350128 | 0.722148664676319 | NM_023500 |
| Zfp672 | 1188.307363 | -0.0555899878798225 | 0.0521574741130062 | -1.06581058276287 | 0.286509276025248 | 0.722148664676319 | NM_001256518,NM_001256519,NM_178761,NM_001256520,NM_001256516,NM_001256517,NR_028331,NR_046280 |
| Mocs2 | 1566.440986 | -0.0552683384374022 | 0.0518970989951524 | -1.0649600749854 | 0.2868939973569 | 0.72225688138758 | NM_001113375,NM_001113374,NM_013826 |
| Samd12 | 274.2030091 | -0.0944658837650778 | 0.0886985339203965 | -1.06502192978587 | 0.28686600600952 | 0.72225688138758 | NM_177225 |
| Zfp655 | 1626.959197 | 0.0768996509609805 | 0.0722055904452504 | 1.06500965488662 | 0.286871560661628 | 0.72225688138758 | NM_001083958,NM_028298 |
| Ociad2 | 3690.307749 | 0.0648712679462075 | 0.0609277382816654 | 1.06472400234769 | 0.287000484998407 | 0.722400671364903 | NM_026950 |
| Hrasls | 1302.88489 | 0.0720859583724991 | 0.0677856003249998 | 1.06344058364728 | 0.287582196605484 | 0.723579188552091 | NM_013751 |
| Kcnb1 | 13815.29858 | 0.0739495834106967 | 0.0695337609718585 | 1.06350616415852 | 0.287552471279596 | 0.723579188552091 | NM_008420 |
| Lypla2 | 2763.008458 | 0.0707914041287459 | 0.06657328664665 | 1.06336051131867 | 0.287618493370492 | 0.723579188552091 | NM_011942 |
| Ano4 | 954.4957674 | 0.0644409926561254 | 0.0606170561432134 | 1.06308350745172 | 0.287744082988736 | 0.72376979212024 | NM_178773 |
| Dnajc3 | 2560.571912 | 0.0593453083548097 | 0.0558669845762837 | 1.06226081119121 | 0.288117299854666 | 0.724103150688742 | NM_008929 |
| Epc2 | 2140.43699 | 0.0637104589584374 | 0.0599708567097747 | 1.06235699227644 | 0.28807365037921 | 0.724103150688742 | NM_172663 |
| Kcnab1 | 5216.535938 | 0.0869858797536753 | 0.0818889513355609 | 1.06224195492782 | 0.288125857839799 | 0.724103150688742 | NM_010597 |
| Trmu | 505.7268358 | -0.0891696906545692 | 0.0839440747912578 | -1.06225115800378 | 0.288121680968199 | 0.724103150688742 | NM_028063 |
| Whsc1 | 4268.70127 | -0.0634877699946818 | 0.0597577561140576 | -1.06241890799087 | 0.288045553778872 | 0.724103150688742 | NM_001177884,NM_001081102,NM_175231 |
| Fcgrt | 656.6161306 | -0.0782101813079137 | 0.0736518305600746 | -1.06189052890031 | 0.288285385214857 | 0.724378741676356 | NM_010189 |
| 1110018G07Rik | 8007.702185 | 0.0586619277027168 | 0.0553515894220939 | 1.05980565897357 | 0.289233021798237 | 0.724444298191034 | NM_178065 |
| Asxl2 | 1259.222647 | -0.0751167053521934 | 0.0708816364035633 | -1.05974846467311 | 0.289259047880215 | 0.724444298191034 | NM_172421,NM_001270988 |
| B630005N14Rik | 791.0056192 | 0.0845982003753213 | 0.0798197223832102 | 1.05986588088054 | 0.289205619718672 | 0.724444298191034 | NM_175312 |
| Baiap3 | 1923.307258 | 0.0794831865333417 | 0.0749314901573703 | 1.060744773211 | 0.288805906491745 | 0.724444298191034 | NM_001163270 |
| Bms1 | 1749.2367 | -0.0630274943836259 | 0.0594141010768765 | -1.06081709966585 | 0.2887730295841 | 0.724444298191034 | NM_194339 |
| Eif3h | 4317.106618 | 0.0585225036854928 | 0.0551306053497766 | 1.06152477946136 | 0.288451477834518 | 0.724444298191034 | NM_080635 |
| Fam50a | 873.3156325 | -0.0765338087840479 | 0.072199161781955 | -1.06003735909267 | 0.28912760354619 | 0.724444298191034 | NM_138607 |
| Gadd45a | 238.1080149 | 0.0999172791554868 | 0.0942533558804743 | 1.06009253699354 | 0.289102502680709 | 0.724444298191034 | NM_007836 |
| Gpr157 | 145.3458164 | -0.0968232208833307 | 0.0912267632133497 | -1.06134666486953 | 0.28853238584764 | 0.724444298191034 | NM_177366 |
| LOC100504703 | 22.90848881 | -0.0835529852781667 | 0.078777048549293 | -1.06062599217442 | 0.28885990539176 | 0.724444298191034 | NR_040660 |
| Nanos1 | 1200.633553 | 0.0772566675163248 | 0.0728657249435977 | 1.06026074091935 | 0.289025994438062 | 0.724444298191034 | NM_178421 |
| Nipa2 | 1219.979795 | 0.0851843864607796 | 0.0803507365746608 | 1.06015688333644 | 0.28907323287251 | 0.724444298191034 | NM_001256132,NM_001256133,NM_001256130,NM_001256131,NM_023647 |
| Ogfod1 | 6590.582051 | 0.0601100511645112 | 0.0566268809101926 | 1.06151089726879 | 0.288457783228759 | 0.724444298191034 | NM_001093757,NM_177767 |
| Pdap1 | 3759.029649 | 0.0579968700494205 | 0.0546944363272829 | 1.06037970118892 | 0.288971893110209 | 0.724444298191034 | NM_001033313 |
| Rpl10 | 8775.641122 | 0.0458280099113533 | 0.0432108521528053 | 1.06056714061766 | 0.28888666234558 | 0.724444298191034 | NM_052835 |
| Serpinb1b | 69.18320478 | 0.0936602043631416 | 0.0882996603712475 | 1.06070854598258 | 0.288822374986163 | 0.724444298191034 | NM_173052 |
| Thrap3 | 5817.448631 | 0.0452173272443827 | 0.0426606995371211 | 1.0599293432832 | 0.289176745046374 | 0.724444298191034 | NM_146153 |
| Wdr37 | 3243.682756 | 0.0625147520871577 | 0.0589460649015397 | 1.06054156781422 | 0.288898289581841 | 0.724444298191034 | NM_001039388,NM_172445,NM_001039389 |
| Zfp451 | 798.8260081 | -0.0763697272068997 | 0.07195468097537 | -1.06135870761544 | 0.28852691498469 | 0.724444298191034 | NM_133817 |
| Angptl4 | 94.08156506 | 0.0973241462572297 | 0.0918796966016582 | 1.05925628682881 | 0.289483077019572 | 0.724785793515301 | NM_020581 |
| Tm9sf4 | 4630.078282 | 0.0510809970474884 | 0.0482246657201939 | 1.05922967603067 | 0.289495193031514 | 0.724785793515301 | NM_133847 |
| Zyg11b | 11260.90386 | 0.0738183613875746 | 0.0697053862055674 | 1.05900512723475 | 0.289597444673129 | 0.724916850133012 | NM_001033634 |
| Atg5 | 865.2977639 | 0.0745965446173145 | 0.0704657181310491 | 1.05862178937257 | 0.289772059476686 | 0.725139224105962 | NM_053069 |
| Cdk4 | 1374.9262 | -0.0600884602832952 | 0.0567626821698586 | -1.0585909260504 | 0.289786121157587 | 0.725139224105962 | NM_009870 |
| 2700046A07Rik | 29.63482662 | 0.091716849699157 | 0.0866952791560178 | 1.05792207594259 | 0.290090969745284 | 0.725277353962134 | NR_037693 |
| A930012L18Rik | 93.92366578 | -0.0999195499748374 | 0.0944450060377167 | -1.05796541465553 | 0.290071210283345 | 0.725277353962134 | NR_026853 |
| Kcnh2 | 1783.783108 | 0.0798969245912693 | 0.07551596154795 | 1.05801373581845 | 0.290049180236283 | 0.725277353962134 | NM_013569 |
| Larp4 | 1703.300649 | 0.0757018826946828 | 0.071545602579675 | 1.05809274036625 | 0.290013163787747 | 0.725277353962134 | NM_001024526,NM_001080948 |
| Vldlr | 3836.995625 | -0.0556437748705478 | 0.052588337127833 | -1.058101052621 | 0.29000937458732 | 0.725277353962134 | NM_013703,NM_001161420 |
| Fgfr2 | 2245.695055 | -0.0791224210818637 | 0.0748349395242431 | -1.05729251048879 | 0.290378110313767 | 0.725761284743831 | NM_201601,NM_010207 |
| Lrrc43 | 28.61993563 | -0.0853344350017323 | 0.0807113986590823 | -1.05727860524605 | 0.290384454559489 | 0.725761284743831 | NM_001033461 |
| Fam83h | 152.6974081 | 0.0999390199997683 | 0.0946015981730798 | 1.05641999638234 | 0.290776374234386 | 0.726615794276395 | NM_001168253,NM_134087 |
| Gria2 | 21687.52453 | -0.0695166516903348 | 0.0658288886700112 | -1.05602043562986 | 0.290958878564738 | 0.726946795671034 | NM_001083806,NM_001039195,NM_013540 |
| Slc38a3 | 3569.611547 | -0.0823543284033537 | 0.0780311388310641 | -1.05540338492956 | 0.291240873392098 | 0.727526212707415 | NM_001199217,NM_023805,NM_001199218 |
| Mtag2 | 33.62318314 | -0.0884193088847001 | 0.0838069406054067 | -1.0550356360222 | 0.291409026876985 | 0.727742893699919 | NR_015480,NR_027802 |
| Tcta | 1224.574759 | -0.0717321270035239 | 0.0679928880885823 | -1.05499455928494 | 0.291427813069836 | 0.727742893699919 | NM_133986 |
| Adat3 | 78.0716222 | -0.100203662609186 | 0.0950905023588817 | -1.05377151369973 | 0.291987538448812 | 0.727847831330164 | NM_001100606 |
| Ciao1 | 2182.309198 | 0.0597439265349022 | 0.0567110405327404 | 1.05347963947886 | 0.29212122102814 | 0.727847831330164 | NM_025296 |
| Clec10a | 25.99063784 | -0.0803712503685294 | 0.0762636197222745 | -1.05386094524773 | 0.291946585745196 | 0.727847831330164 | NM_010796,NM_001204252 |
| Gstm7 | 734.2944795 | 0.0935483668222834 | 0.0887490053154715 | 1.0540779188426 | 0.291847244729976 | 0.727847831330164 | NM_026672 |
| Nt5c | 930.9865909 | 0.0701276953329006 | 0.066561434168697 | 1.05357849043885 | 0.292075941266802 | 0.727847831330164 | NM_015807 |
| Nudt12 | 282.852855 | 0.0822864163346908 | 0.0780268124469714 | 1.0545915404479 | 0.291612174433022 | 0.727847831330164 | NM_026497 |
| Pex19 | 3583.470828 | 0.0440457044593879 | 0.0418087391768489 | 1.05350472955132 | 0.292109727800007 | 0.727847831330164 | NM_001159525,NM_023041 |
| Pgm5 | 249.7812861 | -0.0963390977195143 | 0.0914017967351807 | -1.05401754847 | 0.291874882933211 | 0.727847831330164 | NM_175013 |
| Rilp | 53.30880602 | -0.0928801654374545 | 0.0880929907021696 | -1.05434228872385 | 0.291726234103468 | 0.727847831330164 | NM_001029938 |
| St3gal3 | 2211.205826 | 0.0552521872830445 | 0.0523933290167955 | 1.05456531050609 | 0.291624176062077 | 0.727847831330164 | NM_001161774,NM_009176 |
| Zfp653 | 675.9426359 | -0.0724111971900533 | 0.0686539806440788 | -1.05472685648707 | 0.291550265238414 | 0.727847831330164 | NM_177318 |
| Zfp94 | 403.1129281 | -0.0791124166480957 | 0.0750808850754689 | -1.05369584506861 | 0.292022191819676 | 0.727847831330164 | NM_009568,NM_001199321 |
| Zmynd19 | 727.4106122 | 0.0653631143981257 | 0.0620079380897338 | 1.05410881915693 | 0.291833098918261 | 0.727847831330164 | NM_026021 |
| Abcb10 | 1258.086358 | 0.0568006844442334 | 0.0539365720567996 | 1.05310149084776 | 0.292294479660525 | 0.728154623829324 | NM_019552 |
| Gfm1 | 1797.379044 | 0.0634002407953209 | 0.0602231323389608 | 1.05275561620538 | 0.292453011552239 | 0.728310617042778 | NM_138591 |
| Tmem179 | 5593.783962 | 0.065300850402103 | 0.0620290595422113 | 1.05274609810367 | 0.292457374997282 | 0.728310617042778 | NM_178915 |
| Arntl | 1403.538022 | -0.0753600347069747 | 0.0715970116193983 | -1.05255838201153 | 0.29254343983606 | 0.728400069773502 | NM_001243048,NM_007489 |
| BC037032 | 35.72962518 | -0.0918492851273084 | 0.0872879959900004 | -1.05225562903094 | 0.292682283089337 | 0.728496032240525 | NR_028266 |
| Ttpal | 3629.86097 | 0.0635872921120577 | 0.0604292645336947 | 1.05225990424891 | 0.292680322156042 | 0.728496032240525 | NM_181734,NM_029512 |
| Camta1 | 11932.51425 | 0.0633649301232772 | 0.0602579476108267 | 1.05156137299127 | 0.293000837638069 | 0.728526370670165 | NM_001081557,NM_001195565 |
| Ccdc97 | 2432.423734 | 0.0558412319542049 | 0.0531068186456471 | 1.05148893076806 | 0.293034090651676 | 0.728526370670165 | NM_028771 |
| Hemk1 | 456.4835844 | -0.0933839486347294 | 0.0888132646487368 | -1.05146397899086 | 0.293045544804198 | 0.728526370670165 | NM_133984 |
| Reps2 | 8885.740192 | 0.0738837451139906 | 0.0702298587203722 | 1.05202753444467 | 0.292786917041412 | 0.728526370670165 | NM_178256 |
| Tbpl1 | 2501.372182 | 0.0473208341453649 | 0.0449916192475011 | 1.0517969987467 | 0.292905098689271 | 0.728526370670165 | NM_011603 |
| Ttc37 | 1020.380132 | -0.0690395164222493 | 0.0656292937245045 | -1.05196189847875 | 0.292817030915909 | 0.728526370670165 | NM_001081352 |
| Zfp808 | 51.7455227 | 0.0978400801714482 | 0.0930358218152871 | 1.05163880172628 | 0.292965298476172 | 0.728526370670165 | NM_001039239 |
| Rere | 6847.793228 | -0.0613485128951214 | 0.0583636707457105 | -1.0511421250801 | 0.293193319286513 | 0.728769020526334 | NM_001085492 |
| Srcin1 | 14489.33571 | -0.0552924947924285 | 0.0526121345004431 | -1.05094566714382 | 0.293283544629811 | 0.728786362299893 | NM_018873 |
| Stk3 | 619.0792355 | 0.0717540637492031 | 0.0682781289914977 | 1.05090845140646 | 0.293300638441772 | 0.728786362299893 | NM_019635 |
| Zfp72 | 169.8288826 | 0.0994027482996231 | 0.0946269477533283 | 1.05046977271997 | 0.29350218130243 | 0.729162422712349 | NM_001081680 |
| 1700045I19Rik | 22.50710758 | 0.0786266783954371 | 0.0748655770229318 | 1.0502380602951 | 0.293608674810633 | 0.729168132441703 | NR_003640 |
| Slc4a3 | 7575.561451 | -0.0663723544312825 | 0.0631995284735906 | -1.05020331692851 | 0.293624644868158 | 0.729168132441703 | NM_009208 |
| Snx6 | 1403.770931 | 0.0638570700530372 | 0.0608083158931604 | 1.05013712540952 | 0.293655071925097 | 0.729168132441703 | NM_026998 |
| Casp8 | 129.3604155 | -0.0992609936300773 | 0.0945787036084988 | -1.04950681118405 | 0.293944922065477 | 0.72928872143263 | NM_009812,NM_001080126 |
| Dnajc14 | 2753.266975 | 0.0544123645945778 | 0.0518424131905497 | 1.04957237223086 | 0.293914764865766 | 0.72928872143263 | NM_028873 |
| Eef1g | 11097.45697 | 0.0596868842600105 | 0.0568725121004091 | 1.04948563120674 | 0.293954665013634 | 0.72928872143263 | NM_026007 |
| Pigo | 1164.23472 | -0.0740814645380722 | 0.0705744392058229 | -1.04969257101741 | 0.293859480430363 | 0.72928872143263 | NM_020035 |
| Sin3b | 3336.217132 | -0.0612712416425745 | 0.0583770030801588 | -1.04957840261929 | 0.293911991072412 | 0.72928872143263 | NM_009188,NM_001113248 |
| A230056J06Rik | 39.20412072 | 0.0973575860207662 | 0.0927772326625253 | 1.04936936818419 | 0.294008150732766 | 0.729296857504124 | NR_045633 |
| Atad3a | 1404.870929 | -0.0633710895763299 | 0.0604088451005673 | -1.04903660169022 | 0.294161272920408 | 0.729447814716597 | NM_179203 |
| Msantd1 | 84.47720805 | 0.0997522720271058 | 0.0950910192189733 | 1.04901885421376 | 0.294169440906638 | 0.729447814716597 | NM_207277 |
| Ngf | 79.01724799 | 0.0997187752152879 | 0.0950784827277972 | 1.04880486472187 | 0.294267938046771 | 0.729567514604437 | NM_013609,NM_001112698 |
| Brms1l | 2701.797083 | 0.062221353802744 | 0.0593403821500868 | 1.04854993426518 | 0.294385308745655 | 0.729576943800697 | NM_001037756 |
| Ccdc134 | 350.5550437 | -0.0946569161281719 | 0.0902810525005651 | -1.04846934662818 | 0.294422418047617 | 0.729576943800697 | NM_172428 |
| Tomm6 | 2467.727167 | 0.0563294917400711 | 0.053714479677344 | 1.04868355941331 | 0.294323783438755 | 0.729576943800697 | NM_025365,NM_001164729 |
| Btbd2 | 3229.529778 | 0.064297314705926 | 0.0613418928450321 | 1.04817950219372 | 0.294555912636521 | 0.729621729617702 | NM_145361 |
| Tmem251 | 316.279217 | 0.0860942543662168 | 0.0821292472044564 | 1.04827765134496 | 0.29451070322223 | 0.729621729617702 | NM_177140 |
| Tmem62 | 1349.026158 | -0.0627109473121888 | 0.059832812546549 | -1.04810294958808 | 0.294591177489179 | 0.729621729617702 | NM_175285 |
| Ncoa4 | 4332.564655 | 0.0537418044960287 | 0.0512810060674885 | 1.04798654740318 | 0.294644804938745 | 0.729630146017764 | NM_001033988,NM_019744 |
| Stxbp3a | 773.6545433 | -0.0695743999385662 | 0.0663980113875989 | -1.04783860818917 | 0.29471297119511 | 0.729674555919579 | NM_011504 |
| Arhgap1 | 3609.576663 | 0.0503973224159428 | 0.0481202282725722 | 1.04732093394221 | 0.294951584234309 | 0.7300089803703 | NR_027373,NR_027372,NR_027371,NM_001145902,NM_146124 |
| Frem2 | 137.7366006 | -0.0975968188293687 | 0.0931775034867235 | -1.04742899495343 | 0.294901764684391 | 0.7300089803703 | NM_172862 |
| Stk35 | 1688.099703 | 0.0686274289575398 | 0.0655398763196825 | 1.04710952798869 | 0.295049065383039 | 0.7300089803703 | NM_001038635,NM_183262 |
| Vstm2b | 1029.586552 | 0.0625069958132308 | 0.0596933168501355 | 1.04713557750794 | 0.295037052554303 | 0.7300089803703 | NM_021387 |
| Btbd7 | 661.7401532 | -0.072365061349992 | 0.0691278160816389 | -1.04682985015077 | 0.295178060460529 | 0.730203764688292 | NM_172806 |

| | | | | | | | | | | | | | | | |
|---|---|---|---|---|---|---|---|---|---|---|---|---|---|---|---|
| Ppp1cb | 7532.461685 | -0.135239393 | 0.084159318 | -1.606944968 | 0.108066441 | 0.285049614 | | Ghr | 391.8857752 | -0.0828725754983967 | 0.0792046412365606 | -1.04630958747583 | 0.295418120339615 | 0.730610821876573 | NM_010284,NM_001048147,NM_001048178 |
| Ucma | 14.40894042 | 0.264126222 | 0.164358892 | 1.607009025 | 0.108052388 | 0.285049614 | | Kansl1l | 452.742509 | -0.0776395106377572 | 0.0742070456336695 | -1.04625524402403 | 0.295443203063678 | 0.730610821876573 | NM_177645,NM_001122738 |
| Eml2 | 5288.958745 | 0.149140858 | 0.092819726 | 1.606779771 | 0.108102687 | 0.285096687 | | Prrc2c | 7530.016035 | -0.0699075053583574 | 0.0668385117569587 | -1.04591654602601 | 0.295599564409321 | 0.730624344497157 | NM_001081290 |
| Car8 | 462.128477 | -0.197300427 | 0.122830559 | -1.60628128 | 0.108212121 | 0.285336727 | | Ptpn18 | 33.24612234 | -0.0888584181241859 | 0.0849551069195407 | -1.04594557462381 | 0.295586161065285 | 0.730624344497157 | NM_011206 |
| Zbtb43 | 1305.362414 | 0.103506203 | 0.064468169 | 1.605539664 | 0.108375092 | 0.285717827 | | Slc27a3 | 167.297147 | 0.0971629386239573 | 0.0928956513197525 | 1.04593635163304 | 0.295590419543035 | 0.730624344497157 | NM_011988 |
| Coq10b | 800.5042631 | -0.13848206 | 0.08626639 | -1.605284057 | 0.108431306 | 0.285817397 | | Abcc9 | 481.7000829 | 0.0865318293096711 | 0.0827521419983619 | 1.04567479729267 | 0.295711202605265 | 0.730775932127266 | NM_001044720,NM_021042,NM_011511,NM_021041 |
| Iqgap3 | 37.62195037 | -0.269801793 | 0.168115361 | -1.60486104 | 0.108524389 | 0.286014099 | | Fam214b | 1821.241821 | -0.0752280616511725 | 0.0719580915779394 | -1.04544270146035 | 0.295818409702828 | 0.730916519704589 | NM_001253354,NM_172691,NM_001253353,NR_045567 |
| Nav2 | 2449.636448 | -0.119941298 | 0.074745086 | -1.604671353 | 0.10856615 | 0.286055805 | | Disp2 | 16945.79879 | 0.0695968504518586 | 0.0665862135928446 | 1.04521411710573 | 0.295924020246851 | 0.73092880770375 | NM_170593 |
| Rhbdd1 | 323.3373147 | 0.168454286 | 0.104980702 | 1.604621446 | 0.108577139 | 0.286055805 | | Plekhg2 | 316.7349957 | -0.0822357761659904 | 0.0786705039221368 | -1.04531904673424 | 0.295875537526425 | 0.73092880770375 | NM_138752,NM_001083912 |
| Zxdb | 159.9154465 | -0.246801335 | 0.153920583 | -1.603432949 | 0.108839101 | 0.286697218 | | Ttc30a1 | 148.3520825 | 0.0954261116304352 | 0.0913532397875641 | 1.04458377012509 | 0.296215383907055 | 0.731524084772845 | NM_030188 |
| Abl1 | 2637.536238 | 0.114638911 | 0.071500094 | 1.603339304 | 0.108859763 | 0.286702902 | | Arhgap15 | 497.2280867 | -0.0750036399106594 | 0.0718203099920552 | -1.04432353935378 | 0.2963357256735 | 0.731596910588832 | NM_153820,NM_001025377 |
| Aars | 7112.702703 | 0.118203671 | 0.073730469 | 1.60318621 | 0.108893549 | 0.286743143 | | Max | 2073.222817 | 0.0520613844030809 | 0.0498579951710717 | 1.04419329787427 | 0.296395967149613 | 0.731596910588832 | NM_001146176,NM_008558 |
| Nucb1 | 2952.363781 | 0.09829437 | 0.061318522 | 1.603012691 | 0.108931852 | 0.286793683 | | Pcdhb16 | 294.7106433 | -0.0804081480967099 | 0.0769977904291247 | -1.04429163029976 | 0.296350484027216 | 0.731596910588832 | NM_053141 |
| Spcs3 | 3606.289511 | -0.140460454 | 0.087636395 | -1.602763934 | 0.108986782 | 0.286793683 | | Hcn4 | 175.8588439 | 0.098614711716242 | 0.0944699662961125 | 1.04387368369687 | 0.296543835189779 | 0.731837538220648 | NM_001081192 |
| Srp14 | 2178.498353 | -0.099238903 | 0.061916805 | -1.602778171 | 0.108983638 | 0.286793683 | | Rap1gds1 | 18058.04604 | 0.0651394466013269 | 0.0624149171185291 | 1.04365189619052 | 0.296646473166896 | 0.7319664802484 | NM_001040690,NM_145544 |
| Zbtb7c | 271.8409204 | 0.219896549 | 0.137187569 | 1.602889756 | 0.108958996 | 0.286793683 | | Hiat1 | 2235.002827 | 0.0679988060319437 | 0.0651634344358797 | 1.04351169671473 | 0.296711366403135 | 0.732002260253387 | NM_008246 |
| Dmxl1 | 2575.373531 | -0.16583825 | 0.103484062 | -1.602548709 | 0.109034325 | 0.286870069 | | Adra1a | 404.5498637 | 0.0828831979665237 | 0.0795709513668915 | 1.04162632898983 | 0.297584956917933 | 0.732415974955929 | NM_013461,NM_001271759,NM_001271760,NM_001271761 |
| Card6 | 97.69139589 | 0.193199999 | 0.120587213 | 1.60215992 | 0.109120251 | 0.287047396 | | Arhgef16 | 25.15265204 | -0.0777359863558909 | 0.0745900054140586 | -1.04217697698731 | 0.297329635073813 | 0.732415974955929 | NM_001112744 |
| Daxx | 921.2361264 | 0.117905857 | 0.073630905 | 1.601309362 | 0.109308417 | 0.287493567 | | Cacng7 | 3114.178893 | 0.0586575805793248 | 0.0562921347043231 | 1.04202089488037 | 0.29740199160685 | 0.732415974955929 | NM_133189 |
| Bbs4 | 2443.151434 | 0.110659374 | 0.069121161 | 1.600947851 | 0.10938847 | 0.287606475 | | Ccdc115 | 650.314827 | 0.0701318234565899 | 0.0673260622761412 | 1.04167422073403 | 0.297562744885706 | 0.732415974955929 | NM_027159 |
| Galnt15 | 109.1327259 | 0.262745018 | 0.164111275 | 1.601017468 | 0.10937305 | 0.287606475 | | Col1a1 | 352.8872234 | -0.0878892415337454 | 0.0842796244163725 | -1.04282906031403 | 0.297027469618561 | 0.732415974955929 | NM_007742 |
| Polr1a | 1509.001665 | 0.17106375 | 0.106860287 | 1.600816864 | 0.109417487 | 0.287633958 | | Efha2 | 3504.806222 | 0.0730135514138177 | 0.0700423324146597 | 1.04242033205816 | 0.297216844025307 | 0.732415974955929 | NM_030110 |
| Cngb1 | 17.68476354 | -0.255775901 | 0.159900626 | -1.599592867 | 0.109688932 | 0.288151968 | | Fbxw9 | 567.7334341 | -0.0883703294778482 | 0.0848260032006383 | -1.04178348788669 | 0.297512071290522 | 0.732415974955929 | NM_026791 |
| Kalrn | 22849.18386 | 0.115375824 | 0.072123691 | 1.599693835 | 0.10966652 | 0.288151968 | | Gm16576 | 54.30924684 | -0.0939445643828112 | 0.0900939749998224 | -1.04273969910858 | 0.297068866090592 | 0.732415974955929 | NR_045069 |
| Mall | 35.88474245 | -0.266561964 | 0.166634022 | -1.599685105 | 0.109668458 | 0.288151968 | | Grhl3 | 52.8985134 | 0.0982517145575205 | 0.0943145873508351 | 1.04174462633272 | 0.29753009301425 | 0.732415974955929 | NM_001013756 |
| Ppara | 294.1924763 | -0.169215628 | 0.105783854 | -1.599635687 | 0.109679427 | 0.288151968 | | Lamc3 | 309.8579945 | -0.0945009825424872 | 0.0906800545169139 | -1.04213636665669 | 0.297348460075553 | 0.732415974955929 | NM_011836 |
| Mast3 | 13134.78879 | 0.146073519 | 0.091327988 | 1.599438712 | 0.109723156 | 0.288193013 | | Rabgap1l | 4336.257362 | 0.0782139224245032 | 0.0750440589077663 | 1.04224003289365 | 0.29730040695852 | 0.732415974955929 | NM_001038621,NM_013862 |
| Rusc2 | 6042.144061 | 0.130188654 | 0.0814055 | 1.59926115 | 0.109762588 | 0.288247719 | | Rbm8a | 1803.305449 | 0.0609231329447918 | 0.0584622842688379 | 1.04209292720479 | 0.297368597404151 | 0.732415974955929 | NM_025875,NM_001102407 |
| Polr2c | 1701.920615 | -0.11940682 | 0.074669419 | -1.59913953 | 0.109789603 | 0.288269803 | | Tbkbp1 | 1935.898345 | -0.0739465162876174 | 0.0709418529356664 | -1.04235388882041 | 0.297247636527311 | 0.732415974955929 | NM_198100 |
| 1810013D10Rik | 578.2923584 | -0.199145017 | 0.124550994 | -1.598903472 | 0.109842052 | 0.288358652 | | Usp14 | 6045.750312 | 0.0714452617622302 | 0.0685055943837933 | 1.04291134767721 | 0.296989353507081 | 0.732415974955929 | NM_021522,NM_001038589 |
| Ccdc120 | 456.0041186 | 0.121493063 | 0.076005343 | 1.598480551 | 0.10993607 | 0.288556577 | | Txnl1 | 2604.432853 | 0.0614336018402635 | 0.0589957700944822 | 1.04132214465337 | 0.297726062475043 | 0.732639129851342 | NM_016792 |
| Gtf2ird2 | 647.7119966 | 0.118619442 | 0.074215449 | 1.598311995 | 0.109973559 | 0.288606085 | | 0610011F06Rik | 724.1337056 | 0.0704120823040256 | 0.067644424171777 | 1.04091480068217 | 0.297915091916431 | 0.732916828930711 | NM_026686 |
| Ifi203 | 44.14980388 | 0.265172474 | 0.165922136 | 1.598174177 | 0.110004219 | 0.288637658 | | Efs | 436.2932335 | -0.08802789576364 | 0.0845809816964464 | -1.04075282643992 | 0.297990278925016 | 0.732916828930711 | NM_010112 |
| Tsen54 | 256.6289705 | 0.150724056 | 0.094347588 | 1.597540111 | 0.110145364 | 0.28895907 | | H2afy2 | 683.0636943 | 0.0679671500973547 | 0.0653011076290446 | 1.04082691037119 | 0.297955888247369 | 0.732916828930711 | NM_207000 |
| Znhit1 | 988.1576962 | -0.136235619 | 0.085289485 | -1.597331953 | 0.110191732 | 0.289031775 | | Gbe1 | 504.3321159 | 0.0761650118413641 | 0.0732064157812941 | 1.04041443674703 | 0.29814739732631 | 0.732930968617697 | NM_028803 |
| Aldh1l1 | 2605.606247 | 0.151131812 | 0.09463417 | 1.597011006 | 0.110263254 | 0.289170422 | | Mlf1 | 40.1905445 | 0.0882438681892251 | 0.0848157151895314 | 1.04041884209822 | 0.298145351513366 | 0.732930968617697 | NM_001039543,NM_010801 |
| 2210039B01Rik | 11.43319355 | 0.259867179 | 0.162760292 | 1.596625176 | 0.110349284 | 0.289208519 | | Timm8a1 | 497.3095096 | 0.078867897756423 | 0.0758026280123513 | 1.04043751284681 | 0.298136681058067 | 0.732930968617697 | NM_013898 |
| Rad23a | 2065.252559 | -0.084574152 | 0.052966097 | -1.596760131 | 0.110319186 | 0.289208519 | | Arhgef4 | 9928.076527 | 0.0577128269513024 | 0.0555175260621647 | 1.03954248405593 | 0.298552510399015 | 0.732994422831217 | NM_183019 |
| Smad7 | 741.1600625 | -0.144924406 | 0.090774773 | -1.596527327 | 0.11037111 | 0.289208519 | | Atp6ap1l | 377.8667171 | -0.0795318977323927 | 0.0764833724750857 | -1.03985866677493 | 0.298405568057449 | 0.732994422831217 | NM_001145879 |
| Wnt5b | 72.411179 | -0.231121844 | 0.144751634 | -1.596678658 | 0.110337355 | 0.289208519 | | Ccdc23 | 282.9085974 | -0.0808144903955195 | 0.0776893906288477 | -1.04022556672637 | 0.298235116004205 | 0.732994422831217 | NM_024462,NM_001038998 |
| Zfp474 | 11.13058761 | 0.252684338 | 0.15826779 | 1.596561988 | 0.110363378 | 0.289208519 | | Fam129b | 1678.739922 | 0.0781095888035052 | 0.0750970951618687 | 1.04011464937683 | 0.298286638423849 | 0.732994422831217 | NM_146119 |
| Ankrd23 | 97.24709786 | 0.244381866 | 0.153079791 | 1.596434539 | 0.11039181 | 0.289213849 | | Gm4285 | 144.1901653 | -0.0926211026701978 | 0.0890868015615934 | -1.03967255582928 | 0.298492055165426 | 0.732994422831217 | NR_045294 |
| AI197445 | 38.80726191 | -0.260256344 | 0.163075251 | -1.595927912 | 0.110504889 | 0.289412693 | | Lrtm2 | 3971.714544 | 0.0578942574587784 | 0.0556973759183433 | 1.03944317850191 | 0.298598671500094 | 0.732994422831217 | NM_001172207,NM_172492 |
| Ccdc18 | 54.73161788 | -0.248872317 | 0.155963705 | -1.595706617 | 0.110554311 | 0.289412693 | | Naga | 830.5125108 | -0.0841212948020792 | 0.0809065498240978 | -1.03973405101281 | 0.298463476055665 | 0.732994422831217 | NM_008669 |
| Esd | 2094.382511 | -0.113093612 | 0.070871032 | -1.59576642 | 0.110540953 | 0.289412693 | | Ncl | 11344.7469 | 0.05726291898678 | 0.0550932651045202 | 1.03938147209361 | 0.298627357449755 | 0.732994422831217 | NM_010880 |
| Scrt2 | 973.3405392 | 0.208517466 | 0.130676552 | 1.595676209 | 0.110561103 | 0.289412693 | | Slc39a14 | 1658.432656 | 0.0599252545360553 | 0.0576215753605458 | 1.03997945493672 | 0.298349445877263 | 0.732994422831217 | NM_001135152,NM_144808,NM_001135151 |
| Slc35d1 | 338.501674 | 0.147017706 | 0.092122712 | 1.595889901 | 0.110513377 | 0.289412693 | | BC020402 | 148.1229454 | -0.0934165823220594 | 0.0899326782218459 | -1.03873902311259 | 0.298926127124001 | 0.733276458391256 | NR_033219 |
| Uqcrc2 | 7766.102323 | -0.091580059 | 0.057405059 | -1.595330804 | 0.110638281 | 0.289565798 | | Dll3 | 43.08684187 | 0.0970829805318517 | 0.0934658328599296 | 1.03870021334259 | 0.29894418192159 | 0.733276458391256 | NM_007866 |
| Slc7a8 | 3396.663295 | 0.11507678 | 0.072140829 | 1.595168532 | 0.110674554 | 0.289611811 | | Reep1 | 5729.254141 | 0.0562894737652608 | 0.0541886535262632 | 1.03876863701696 | 0.298912350849399 | 0.733276458391256 | NM_178608 |
| Gpr165 | 363.8398862 | -0.227844482 | 0.142853687 | -1.594949957 | 0.110723427 | 0.289690776 | | Rpsa | 13392.24864 | 0.0544877331293993 | 0.0524520261809604 | 1.03881083528433 | 0.298892721109617 | 0.733276458391256 | NM_011029 |
| 1700028K03Rik | 13.99096087 | 0.267176986 | 0.167584107 | 1.594285941 | 0.110872005 | 0.289955312 | | Suclg2 | 1088.225728 | -0.0624349496609218 | 0.0601167659441624 | -1.03856135106989 | 0.299008788374405 | 0.733311102469458 | NM_011507 |
| Rabepk | 723.4305857 | 0.123670379 | 0.077575771 | 1.594188202 | 0.110893888 | 0.289955312 | | Abca6 | 147.6934762 | -0.0955553689295206 | 0.0921261654618504 | -1.03722290459479 | 0.299631985880051 | 0.733704077735471 | NM_147218,NM_001166556,NM_001166557 |
| Slc43a3 | 31.067413 | 0.268033989 | 0.168119864 | 1.594302913 | 0.110868206 | 0.289955312 | | Cacng2 | 2206.437561 | 0.0573070370194081 | 0.0552733919491487 | 1.03679247823492 | 0.299832581777501 | 0.733704077735471 | NM_007583 |
| Smarce1 | 1991.443973 | -0.092049414 | 0.05774151 | -1.594163622 | 0.110899392 | 0.289955312 | | Fam3c | 2579.360557 | 0.0640374011368611 | 0.0617219029229212 | 1.03751501661949 | 0.299495900995473 | 0.733704077735471 | NM_138587 |
| Lmo3 | 3527.69097 | -0.154016004 | 0.096620537 | -1.594029694 | 0.110929384 | 0.289984795 | | Frmd7 | 31.29076485 | -0.0864901629166651 | 0.08343714320359 | -1.03659065490324 | 0.299926670356136 | 0.733704077735471 | NM_001190332 |
| Sec31a | 4436.886512 | 0.106588293 | 0.066873077 | 1.593889474 | 0.110960793 | 0.290017969 | | Gon4l | 2428.843195 | -0.0585490745969202 | 0.0564112312575283 | -1.03789747700475 | 0.299317788276871 | 0.733704077735471 | NM_001242372,NM_027389 |
| Nr2c1 | 430.9400013 | 0.138415155 | 0.086847146 | 1.593778972 | 0.110985549 | 0.290033749 | | Inpp4a | 7425.757087 | 0.0638816153972039 | 0.0615428187053399 | 1.03800275549714 | 0.299268772237779 | 0.733704077735471 | NM_172971,NM_030266 |
| Bag6 | 8447.948463 | 0.081649769 | 0.051234763 | 1.593640021 | 0.111016685 | 0.290066193 | | Mypop | 1573.842797 | 0.050970457159592 | 0.0491484508839589 | 1.03707148939312 | 0.299702541210888 | 0.733704077735471 | NM_145579 |
| Ckmt1 | 7384.853641 | -0.097900492 | 0.061442461 | -1.593368674 | 0.111077509 | 0.29017618 | | Nhlrc2 | 455.9609038 | -0.0805635476130811 | 0.0776725826427199 | -1.03721988985044 | 0.299633390560119 | 0.733704077735471 | NM_025811 |
| Rnf14 | 14053.44576 | -0.091591823 | 0.05750755 | -1.592692149 | 0.11122927 | 0.290523652 | | Nudt16l1 | 1535.975348 | 0.0632090257984616 | 0.060974901194724 | 1.0366400692738 | 0.299903631913798 | 0.733704077735471 | NM_025839 |
| Csnk2b | 2714.220313 | 0.125851187 | 0.079025593 | 1.592537079 | 0.111264079 | 0.290535356 | | Ppp1r35 | 211.0405517 | -0.0901907587697494 | 0.0869887133905121 | -1.03680989469131 | 0.299824463274109 | 0.733704077735471 | NM_027242 |
| Psap | 77343.38931 | 0.12094028 | 0.075943417 | 1.59250512 | 0.111271254 | 0.290535356 | | Simc1 | 850.6603714 | -0.0639350520974439 | 0.0616028462983541 | -1.03785873444539 | 0.299335827562002 | 0.733704077735471 | NM_176987 |
| Agbl4 | 328.9752428 | 0.135408223 | 0.085039479 | 1.592298357 | 0.111317682 | 0.29060761 | | Slc30a9 | 6540.362179 | 0.0495116844944447 | 0.047746551276079 | 1.03696881075576 | 0.299750392943991 | 0.733704077735471 | NM_178651 |
| Sap130 | 1802.951407 | 0.108596305 | 0.068222115 | 1.591805026 | 0.111428521 | 0.290847961 | | Thumpd2 | 156.6460917 | 0.091895952448669 | 0.0885803611148663 | 1.03743032080783 | 0.299535353598559 | 0.733704077735471 | NM_028138 |
| Fzd9 | 164.5389623 | -0.203302617 | 0.127786736 | -1.590952415 | 0.111620285 | 0.291299427 | | Wdr44 | 760.0425194 | -0.0859576119087812 | 0.0828871953057665 | -1.03704331642165 | 0.299715670265438 | 0.733704077735471 | NM_175180 |
| Nucb2 | 596.3961076 | -0.111188098 | 0.06990953 | -1.590456957 | 0.11173184 | 0.291541451 | | Zfand2a | 2944.117406 | 0.0555044376709907 | 0.0535429343844762 | 1.0366342134413 | 0.299906362014617 | 0.733704077735471 | NM_001159908,NM_133349 |
| Cand2 | 189.6004404 | -0.235667324 | 0.148286274 | -1.589272676 | 0.111998845 | 0.29211375 | | Hepacam | 4321.383488 | 0.0742090860649706 | 0.0716036321903631 | 1.03638717471316 | 0.300021551279427 | 0.733812603785984 | NM_175189 |
| Ddx24 | 8024.439951 | 0.112092565 | 0.070544774 | 1.58895632 | 0.112070255 | 0.29211375 | | Tamm41 | 283.0233369 | 0.0804646205646552 | 0.0776488590075225 | 1.03626275508903 | 0.300079576853696 | 0.733830965214946 | NM_026894 |
| Fmnl1 | 4952.338391 | 0.187283911 | 0.117844466 | 1.589246556 | 0.112004739 | 0.29211375 | | Fstl1 | 1630.596736 | 0.0734062081221117 | 0.0708513573363601 | 1.03605930615588 | 0.300174475442715 | 0.733939476566384 | NM_008047 |
| Rapgef2 | 6507.899594 | -0.175756685 | 0.110597198 | -1.589160383 | 0.112024188 | 0.29211375 | | Dusp12 | 474.7995372 | -0.074972146537914 | 0.0723786441042429 | -1.03583242634299 | 0.300280326940468 | 0.733951207998861 | NM_023173 |
| Tgfbrap1 | 2723.659081 | 0.089594043 | 0.056387492 | 1.588899231 | 0.112083145 | 0.29211375 | | Elac2 | 1294.900925 | -0.0623303889397528 | 0.0601730881547456 | -1.03585158832898 | 0.300271385893122 | 0.733951207998861 | NM_023479 |
| Trmt6 | 1044.242803 | 0.100409858 | 0.063191838 | 1.588968783 | 0.112067441 | 0.29211375 | | Ppfia1 | 1808.420609 | -0.0684828224270535 | 0.0661612544666234 | -1.03508953962778 | 0.300627097201608 | 0.734641346533552 | NM_001033319,NM_001195086 |
| Tspan17 | 1160.232996 | 0.214448934 | 0.134951203 | 1.589085016 | 0.1120412 | 0.29211375 | | Snapc2 | 1090.295093 | -0.0695673870950826 | 0.0672211260278792 | -1.03490362637231 | 0.300713920923716 | 0.734641346533552 | NM_133968 |
| Rab18 | 3410.992659 | -0.122121631 | 0.076864841 | -1.588784034 | 0.112109159 | 0.29213241 | | Zfp747 | 239.061103 | -0.0871816079521095 | 0.0842413660893472 | -1.03490259001313 | 0.300714404962722 | 0.734641346533552 | NM_175560 |
| Syt16 | 2748.663421 | -0.12009751 | 0.075609385 | -1.588394218 | 0.112197224 | 0.292312727 | | Ube2q1 | 4232.671677 | 0.0526647005544077 | 0.0509086278654542 | 1.03449459870721 | 0.300905000565858 | 0.734983359377779 | NM_027315 |
| Tmem150a | 333.2495482 | 0.201053746 | 0.126591936 | 1.588203419 | 0.112240348 | 0.292375918 | | Hps3 | 694.6830433 | -0.0706834721493165 | 0.0683462982280589 | -1.03419605716551 | 0.301044517029081 | 0.735062578155623 | NM_080634,NM_001146324,NM_001146323 |
| Csf2rb2 | 50.44082239 | 0.256465118 | 0.161513195 | 1.587889569 | 0.112311313 | 0.292511595 | | Nr1d2 | 5403.613271 | 0.0843403166323216 | 0.0815591203496073 | 1.03410037124962 | 0.301089242738948 | 0.735062578155623 | NM_011584 |
| Lrpap1 | 7340.852705 | -0.086299383 | 0.054370204 | -1.587255091 | 0.112454882 | 0.292787085 | | Pfas | 1278.675842 | -0.0708679573800076 | 0.0685223777381214 | -1.03423085595264 | 0.301028252405053 | 0.735062578155623 | NM_001159519 |
| Ubl5 | 2615.886908 | -0.109453569 | 0.068955204 | -1.587314129 | 0.112441517 | 0.292787085 | | Psmb10 | 1144.557962 | -0.0697184740701385 | 0.0674287686113392 | -1.03395739690869 | 0.301156080349947 | 0.735102205203045 | NM_013640 |
| Zfp788 | 1151.886933 | 0.111331736 | 0.070164821 | 1.586717305 | 0.112576685 | 0.293054967 | | Gm16793 | 24.44454894 | -0.079008128038622 | 0.0764375319237719 | -1.03363002506954 | 0.301309157275566 | 0.735352288068695 | NR_040376 |
| Kcnn4 | 10.6378526 | -0.261776997 | 0.164989843 | -1.586624922 | 0.11259762 | 0.293060225 | | Ddx3x | 12272.80136 | 0.0653678240681959 | 0.0632754928901716 | 1.03306700718485 | 0.301572541913327 | 0.735871450333108 | NM_010028 |
| Dgcr14 | 1093.894549 | 0.136336481 | 0.085934634 | 1.586513787 | 0.112622807 | 0.293076549 | | 4930441O14Rik | 161.7037849 | 0.0882805883642945 | 0.0854952149601843 | 1.03257929002702 | 0.301800824085973 | 0.735974432322909 | NR_027969 |
| Fbxo17 | 87.52423607 | 0.22350409 | 0.140921154 | 1.586022284 | 0.112734255 | 0.293317303 | | B230217O12Rik | 405.4880799 | 0.0769313702699503 | 0.0745535018010641 | 1.03189479248381 | 0.302121405751171 | 0.735974432322909 | NR_040316 |
| C330007P06Rik | 1739.499865 | -0.128283458 | 0.080894264 | -1.585816496 | 0.112780943 | 0.293389511 | | Bcap29 | 1485.330117 | 0.0795552369268226 | 0.077066516874932 | 1.03229314302513 | 0.301934812327779 | 0.735974432322909 | NM_007530,NM_001164090 |
| Dyrk1b | 945.5265789 | 0.106179739 | 0.06696748 | 1.585541809 | 0.112843286 | 0.293502412 | | Cdca3 | 22.69534928 | 0.0862071748075402 | 0.0835365977674591 | 1.03196894668269 | 0.302086664990751 | 0.735974432322909 | NM_013538 |
| Styx | 359.6190854 | 0.155269769 | 0.097960487 | 1.585024461 | 0.112960777 | 0.29375869 | | Ddx23 | 3015.502977 | -0.0568624185409474 | 0.0550549637244741 | -1.03283000649167 | 0.301683458717082 | 0.735974432322909 | NM_001080981 |
| Dynlrb1 | 4134.598283 | -0.095786912 | 0.060448432 | -1.584605393 | 0.113056019 | 0.293957032 | | Mtx3 | 2514.645902 | -0.0506201188422424 | 0.0490508110007975 | -1.03199351467236 | 0.302075155633487 | 0.735974432322909 | NM_001162945 |
| A330070K13Rik | 8.953398796 | 0.24534387 | 0.154843783 | 1.584460581 | 0.113088945 | 0.293993308 | | Nptx2 | 2494.699731 | 0.093333394155697 | 0.0904110428130202 | 1.03232294697364 | 0.301920854791707 | 0.735974432322909 | NM_016789 |
| Atp4a | 18.86154861 | 0.265457379 | 0.167559246 | 1.584259803 | 0.113134609 | 0.294050242 | | Sumf2 | 528.9120216 | -0.0818924139121454 | 0.0793440211758286 | -1.03211827052059 | 0.302016715809334 | 0.735974432322909 | NM_026445 |
| Ogn | 272.7007834 | -0.225049527 | 0.142066497 | -1.584113993 | 0.113167781 | 0.294050242 | | Svep1 | 70.16405231 | -0.0898158568796822 | 0.0869805637867311 | -1.03259685807398 | 0.301792599143351 | 0.735974432322909 | NM_022814 |
| Pih1d2 | 56.93986104 | 0.233160877 | 0.147185899 | 1.584125102 | 0.113165253 | 0.294050242 | | Wdtc1 | 3367.580455 | 0.0586991922232165 | 0.0568520429966925 | 1.03249046347607 | 0.301842412853275 | 0.735974432322909 | NM_199306 |
| Wfdc3 | 15.57352621 | 0.265996488 | 0.16793169 | 1.583956474 | 0.113203624 | 0.294094057 | | Dcaf11 | 3879.148454 | -0.0567699521268207 | 0.0550729400870457 | -1.03081389947754 | 0.302628098900937 | 0.735974512746528 | NM_133734,NR_037572,NM_001199009 |
| Etv1 | 2258.183266 | -0.118927222 | 0.075093216 | -1.58372791 | 0.113255651 | 0.294179891 | | Fnbp1l | 4984.004963 | -0.0713432548485909 | 0.0692069818442927 | -1.03086788279692 | 0.302602779592745 | 0.735974512746528 | NM_001114665,NM_153118 |
| Tbc1d1 | 971.3181185 | 0.119246107 | 0.075308443 | 1.583436106 | 0.113322099 | 0.294303151 | | Ilf2 | 3425.057396 | 0.0523624730923812 | 0.0507774253685453 | 1.03121559851315 | 0.302439727399953 | 0.735974512746528 | NM_026374 |
| Nckap1 | 22128.26872 | -0.106124461 | 0.067055258 | -1.582641889 | 0.11350311 | 0.294689071 | | Klf7 | 368.6588933 | 0.07760445313635 | 0.0752320589434331 | 1.03153435152826 | 0.302290307862987 | 0.735974512746528 | NM_033563 |
| Tuba1c | 34.48617399 | 0.263289301 | 0.166363223 | 1.582617215 | 0.113508737 | 0.294689071 | | Mctp1 | 2037.603057 | 0.0866365779485946 | 0.0840197962350306 | 1.03114482337286 | 0.302472910813091 | 0.735974512746528 | NM_030174 |
| Trmt2a | 1179.037258 | 0.144732149 | 0.091471504 | 1.582264887 | 0.113589113 | 0.294848336 | | Nek3 | 70.51091647 | 0.0980309714033764 | 0.095100282381269 | 1.03081682776038 | 0.302626725438772 | 0.735974512746528 | NM_001162947,NM_011848 |
| Ndufaf3 | 973.3157326 | -0.15393503 | 0.097339479 | -1.581424426 | 0.113781026 | 0.295297022 | | Pdlim3 | 73.72600403 | -0.0977422229037619 | 0.0947766177895948 | -1.03129047209461 | 0.302404625040581 | 0.735974512746528 | NM_016798 |
| Abcc6 | 57.87193829 | -0.264677566 | 0.167385734 | -1.581243274 | 0.113822424 | 0.295331486 | | Siah2 | 662.4420464 | 0.0784057104500011 | 0.0759967337123535 | 1.03169842465553 | 0.302213415644006 | 0.735974512746528 | NM_009174 |
| Krt222 | 1562.047825 | 0.143072473 | 0.090488277 | 1.581116124 | 0.113851489 | 0.295331486 | | Skap1 | 60.81650945 | 0.0980438897176976 | 0.095082023444917 | 1.03115064410147 | 0.3024701816329 | 0.735974512746528 | NM_001177899,NM_001177898,NM_001033186 |
| Pik3c3 | 2214.485819 | 0.094484951 | 0.059758343 | 1.581117316 | 0.113851216 | 0.295331486 | | Slc22a5 | 540.4700274 | -0.0829190599905511 | 0.0804242093754428 | -1.0310211394614 | 0.302530906683035 | 0.735974512746528 | NM_011396 |
| Mctp2 | 78.67363428 | -0.254341321 | 0.160884778 | -1.580891152 | 0.113902928 | 0.295415461 | | P2rx6 | 120.1929254 | -0.0976814238649572 | 0.0947941960682307 | -1.03045785413538 | 0.302795126920605 | 0.736257451230114 | NM_001159561,NM_011028 |
| E530001F21Rik | 51.21143859 | 0.2651921 | 0.167780598 | 1.580588597 | 0.113972135 | 0.295545482 | | Mlycd | 962.1934721 | 0.0694263420569263 | 0.0673831749967939 | 1.03032162049695 | 0.302859053068851 | 0.736289640983788 | NM_019966 |
| Trmt11 | 226.374449 | 0.135918447 | 0.086031371 | 1.579870743 | 0.114136471 | 0.295922101 | | Ttll1 | 1749.389657 | 0.0591892736746456 | 0.0574651735814294 | 1.03000252127966 | 0.303008822020147 | 0.736530480030899 | NM_178869 |
| 2810410L24Rik | 128.1307582 | 0.222185193 | 0.140655522 | 1.579640736 | 0.114189165 | 0.296009189 | | A330048O09Rik | 40.84209247 | 0.0905101778991467 | 0.0878942847487923 | 1.029761811679 | 0.303121831447086 | 0.736550420189535 | NR_045162 |
| Ttll12 | 698.2303168 | -0.118929229 | 0.075294976 | -1.579510815 | 0.114218938 | 0.296036839 | | Mbd1 | 1214.070183 | -0.0575031949348269 | 0.0558502068985547 | -1.02959681132918 | 0.303199312737964 | 0.736550420189535 | NM_013594 |
| Itga9 | 458.5966153 | -0.139134629 | 0.088148128 | -1.578418414 | 0.114469518 | 0.296434962 | | Plekhb1 | 11293.95654 | 0.0824009132314313 | 0.080023229971933 | 1.02971241301222 | 0.303145026820168 | 0.736550420189535 | NM_013746,NM_001163182,NM_001163183,NM_001163186,NM_001163187,NM_001163184,NM_001163185 |
| Med27 | 710.5066759 | -0.113920357 | 0.072176087 | -1.578367049 | 0.114481311 | 0.296434962 | | Slc43a3 | 37.65433235 | -0.0920160056685329 | 0.0893747073418043 | -1.02955308504259 | 0.303219848047186 | 0.736550420189535 | NM_021398 |
| Mfsd4 | 2043.72753 | 0.116119753 | 0.073580506 | 1.578132026 | 0.114535282 | 0.296434962 | | Egr3 | 2608.748014 | 0.0921270851125931 | 0.0894965477608274 | 1.02939261253736 | 0.303295219160954 | 0.736610324951014 | NM_018781 |
| Mrc2 | 361.1488894 | -0.197956235 | 0.125433896 | -1.578171781 | 0.114526151 | 0.296434962 | | Mlec | 7117.103988 | 0.0584176395648791 | 0.0567747388877566 | 1.02893717715498 | 0.303509196967791 | 0.736936169201399 | NM_175403 |
| Npc1 | 3406.395122 | 0.115573966 | 0.073236572 | 1.578090873 | 0.114544735 | 0.296434962 | | Tppp | 29335.93543 | 0.0488005774206632 | 0.0474302646112914 | 1.02889110614503 | 0.303530848157233 | 0.736936169201399 | NM_182839 |
| Ppt1 | 4526.349989 | 0.091360471 | 0.057876196 | 1.578550042 | 0.114439301 | 0.296434962 | | Camk4 | 11348.35482 | 0.0763620953961615 | 0.0742441515206998 | 1.02852674361658 | 0.303702117419447 | 0.737228769658236 | NM_009793 |
| Rab4b | 1880.342455 | 0.108504434 | 0.068732833 | 1.578640497 | 0.11441854 | 0.296434962 | | Fbxo34 | 3169.287603 | -0.0738079414427313 | 0.0717860810358246 | -1.02816507570455 | 0.303872183558293 | 0.737396793883169 | NM_001146086,NM_030236,NR_027414,NM_001146085 |
| Rnf43 | 128.3420004 | 0.190669077 | 0.120815239 | 1.57818731 | 0.114522585 | 0.296434962 | | Jkamp | 1681.531817 | 0.0525278725930286 | 0.0510890202769584 | 1.02816363101641 | 0.30387286301698 | 0.737396793883169 | NM_024205,NM_001205067 |
| Slc2a8 | 779.3806528 | 0.145726729 | 0.092327669 | 1.578364646 | 0.114481862 | 0.296434962 | | 1110034G24Rik | 269.6397315 | -0.0801944698388753 | 0.0780139035432366 | -1.02795099586871 | 0.303972879550049 | 0.737453291187883 | NM_028637 |
| Tmem127 | 5716.557241 | -0.06075356 | 0.038507852 | -1.577692798 | 0.114636202 | 0.296622129 | | Adnp | 6370.437369 | 0.0534365744842224 | 0.052002491133624 | 1.027577204848 | 0.304148751469156 | 0.737453291187883 | NM_009628 |
| P2ry12 | 1133.73008 | -0.264256064 | 0.167505466 | -1.577596667 | 0.114658299 | 0.296629768 | | Ap1m2 | 49.37254147 | -0.096420786692493 | 0.093926839552922 | -1.02655201805407 | 0.304631457532869 | 0.737453291187883 | NM_001110300,NM_009678 |
| Mir421 | 14.68276627 | 0.24082588 | 0.152666204 | 1.57746688 | 0.114688137 | 0.296657428 | | Arpc1b | 557.8208511 | -0.0700756110198209 | 0.0682866927703254 | -1.02619717220033 | 0.30479865402548 | 0.737453291187883 | NM_023142 |
| Agrn | 8392.062022 | 0.156854379 | 0.099443255 | 1.577325476 | 0.114720653 | 0.296692005 | | Cox6b1 | 6048.30261 | 0.059236074224738 | 0.0576977804341891 | 1.026661229929 | 0.30458001126764 | 0.737453291187883 | NM_025628 |
| Cda | 46.72362203 | 0.25572174 | 0.16214809 | 1.577087585 | 0.114775373 | 0.296753581 | | Creb3 | 1609.118852 | 0.048132336949239 | 0.0469151263992648 | 1.02594494874884 | 0.304917533854632 | 0.737453291187883 | NM_013497 |
| Ngb | 268.9008512 | 0.242539804 | 0.15379282 | 1.577055444 | 0.114782768 | 0.296753581 | | Ddr2 | 371.6227285 | -0.0934421426259016 | 0.0909851555090484 | -1.02700426353186 | 0.304418456482402 | 0.737453291187883 | NM_022563 |
| Leprel2 | 740.1303685 | 0.210226847 | 0.133337071 | 1.576657155 | 0.114874434 | 0.296941021 | | Dtx1 | 3033.08434 | 0.054500758050712 | 0.0530990203236239 | 1.02639856100065 | 0.304703756051011 | 0.737453291187883 | NM_008052 |
| Fxyd1 | 707.4762706 | 0.18430582 | 0.11690467 | 1.57654796 | 0.114899575 | 0.296956466 | | Erh | 1616.933183 | 0.0503157519176446 | 0.0490318576587073 | 1.02618489937449 | 0.304804437832756 | 0.737453291187883 | NM_007951 |
| Gtl3 | 1165.137151 | 0.103882366 | 0.065921201 | 1.575856693 | 0.115058833 | 0.29712047 | | Etohi1 | 564.6064396 | -0.0710081979320429 | 0.0691031111725778 | -1.02756875525774 | 0.30415272785562 | 0.737453291187883 | NM_001177400,NM_001177399 |
| Pkn1 | 1638.706657 | 0.101884524 | 0.064652082 | 1.575889283 | 0.115051321 | 0.29712047 | | Gabrg1 | 996.1057436 | 0.0822712281734799 | 0.080204520355169 | 1.02576797179459 | 0.305000966313689 | 0.737453291187883 | NM_010252 |
| Rab35 | 3391.477458 | 0.101148139 | 0.064186142 | 1.575856342 | 0.115058914 | 0.29712047 | | Man2a2 | 11686.05274 | -0.0401017241715457 | 0.0390496935460588 | -1.02694081643038 | 0.304448333180822 | 0.737453291187883 | NM_172903 |
| Timp4 | 863.0914008 | 0.14334641 | 0.090955526 | 1.576005497 | 0.115024536 | 0.29712047 | | Ppp1r13b | 3925.401745 | 0.0525519443039153 | 0.051233083296721 | 1.02574237040461 | 0.305013036861958 | 0.737453291187883 | NM_011625 |
| Zfp871 | 3792.312549 | -0.135722454 | 0.086124083 | -1.575894337 | 0.115050156 | 0.29712047 | | Ptprd | 10174.14231 | -0.0633022279606535 | 0.0616405531489098 | -1.02695749351453 | 0.304440479895674 | 0.737453291187883 | NM_001014288,NM_011211 |
| Lmod1 | 63.73070514 | -0.259564969 | 0.164733561 | -1.575665379 | 0.11510294 | 0.297184629 | | Rgag4 | 1451.827447 | 0.0766222128598127 | 0.0746387661329453 | 1.02657394849393 | 0.304621126332406 | 0.737453291187883 | NR_028404,NM_183318 |
| Dlg4 | 28797.77379 | -0.125702362 | 0.079788519 | -1.575444225 | 0.115153943 | 0.297227378 | | Snx4 | 1563.230313 | 0.0616904831681997 | 0.0601170484826811 | 1.02617285321271 | 0.304810114891027 | 0.737453291187883 | NM_080557 |
| Dysf | 236.8655917 | -0.173144072 | 0.109908734 | -1.575344065 | 0.115177048 | 0.297227378 | | Stt3b | 4295.42964 | 0.0640839527400023 | 0.0624126893613901 | 1.02677762159767 | 0.304525189177608 | 0.737453291187883 | NM_024222 |
| Taf13 | 737.2712828 | -0.112078377 | 0.071143346 | -1.575388047 | 0.115166902 | 0.297227378 | | Tor1b | 1299.475602 | -0.0654459941872175 | 0.0637372017217945 | -1.02680996999024 | 0.304509953799527 | 0.737453291187883 | NM_133673 |
| C230091D08Rik | 57.52774818 | 0.261137287 | 0.165799113 | 1.575022221 | 0.115251316 | 0.297319994 | | Trappc10 | 5613.073022 | 0.0447568005999407 | 0.0436204589737667 | 1.02605065725827 | 0.304867706790313 | 0.737453291187883 | NM_001081055 |
| St6galnac4 | 1125.17515 | -0.11761531 | 0.074673132 | -1.575068655 | 0.115240598 | 0.297319994 | | Xpot | 4214.112191 | 0.0570567333647637 | 0.0555116225996266 | 1.02783400471061 | 0.304027917641175 | 0.737453291187883 | NM_001081056 |
| Cacng7 | 2910.802726 | -0.091413999 | 0.058046497 | -1.574840937 | 0.115293165 | 0.29737844 | | Zfp942 | 547.3519317 | 0.0887216107576972 | 0.0864364540956839 | 1.02643741793808 | 0.304685448230237 | 0.737453291187883 | NM_001199048 |
| Mcam | 812.8395054 | -0.213092246 | 0.135335896 | -1.57454343 | 0.115361869 | 0.297506126 | | Zfyve1 | 2203.541697 | 0.0543201425981272 | 0.0529167086010858 | 1.02652156632834 | 0.304645803407534 | 0.737453291187883 | NM_183154 |
| Txlng | 457.8718545 | 0.176306458 | 0.111985731 | 1.574365378 | 0.115403003 | 0.297562678 | | 1600002K03Rik | 80.36374405 | -0.0972318999543152 | 0.0948027193773564 | -1.02562353266777 | 0.305069070651616 | 0.737466021016037 | NM_027207 |
| Dusp3 | 8412.359065 | -0.116230133 | 0.073842351 | -1.574030777 | 0.115480334 | 0.297712529 | | Plp1 | 64986.24614 | 0.0796943090187745 | 0.0777199347516776 | 1.02540370464032 | 0.305172740892786 | 0.737593882563829 | NM_011123 |
| B4galt6 | 6221.931128 | -0.099889696 | 0.063473177 | -1.573730832 | 0.115549691 | 0.297792233 | | Dcaf6 | 4088.089962 | 0.0594631675538761 | 0.0580059819713259 | 1.02512129840799 | 0.305305957126253 | 0.737793101516546 | NM_028759 |
| Vps4a | 4182.12895 | 0.089445533 | 0.056833743 | 1.573810337 | 0.115531304 | 0.297792233 | | Zfp106 | 12919.22506 | 0.060516645893955 | 0.0590438768380356 | 1.02494363742339 | 0.305389782857905 | 0.737872918462065 | NM_011743 |
| Kctd2 | 2405.076453 | 0.098221808 | 0.062423929 | 1.57346405 | 0.115611406 | 0.297901734 | | 4930578C19Rik | 31.89452137 | -0.0822613453843427 | 0.0802771327697341 | -1.02471703393169 | 0.305496723271448 | 0.737909530427187 | NM_175228 |
| Celsr2 | 10627.2883 | -0.12187196 | 0.077466787 | -1.573215632 | 0.115668897 | 0.297950771 | | Map3k15 | 95.18094176 | -0.095501591021731 | 0.0931999031486804 | -1.02469624747763 | 0.305506534213762 | 0.737909530427187 | NM_001163085 |
| Morn4 | 2251.879861 | -0.112421792 | 0.071457975 | -1.573257461 | 0.115659215 | 0.297950771 | | Cep128 | 282.0334221 | -0.0883914987537279 | 0.0862793404824926 | -1.02448046379844 | 0.305608393714377 | 0.738032839084795 | NM_181815 |
| Pold1 | 281.9869283 | 0.163744233 | 0.104089711 | 1.573106806 | 0.115694089 | 0.297966127 | | Ggcx | 677.4294235 | -0.0718012515867232 | 0.0701037447306367 | -1.02421421084721 | 0.305734107994655 | 0.73809101757194 | NM_019802 |
| Wnt6 | 53.02103312 | -0.264330773 | 0.168053893 | -1.572892886 | 0.115743623 | 0.298044156 | | Rdx | 3146.41282 | 0.0596453586393977 | 0.0582295341778748 | 1.02431454212218 | 0.305686731446029 | 0.73809101757194 | NM_009041,NM_001104617,NM_001104616 |
| Pnoc | 286.833138 | 0.171149561 | 0.108848495 | 1.57236497 | 0.115865933 | 0.298309534 | | 9430023L20Rik | 882.1102561 | 0.0746663943029319 | 0.0729224313107785 | 1.02391531605304 | 0.305875275332535 | 0.738179891025902 | NM_026566 |
| Zfp318 | 2481.8788 | 0.118364233 | 0.075290806 | 1.572094126 | 0.115928724 | 0.298421606 | | Mt1 | 5018.895583 | 0.0866958528797354 | 0.0846651864556036 | 1.02398466842327 | 0.305842516514527 | 0.738179891025902 | NM_013602 |
| Tmem214 | 2208.099155 | 0.103150636 | 0.065619695 | 1.571946299 | 0.115963006 | 0.298460268 | | Ube2s | 2058.72085 | 0.0536440281105529 | 0.0523962893211159 | 1.02381349529907 | 0.30592337491229 | 0.738179891025902 | NM_133777 |
| Gbp3 | 222.9413765 | 0.263317923 | 0.167556431 | 1.571517859 | 0.116062409 | 0.2986169 | | Anapc16 | 1611.637967 | 0.0500940603217039 | 0.0489716099389084 | 1.02292043051465 | 0.306345468813384 | 0.738584942735804 | NM_025514 |
| Taf4a | 888.4713477 | 0.11362369 | 0.072300046 | 1.571557637 | 0.116053178 | 0.2986169 | | B3gat2 | 565.0558174 | 0.074678885920909 | 0.0729822074388849 | 1.02324783726835 | 0.30619068005039 | 0.738584942735804 | NM_172124 |
| Txnip | 1117.997253 | 0.143118035 | 0.091082161 | 1.571306974 | 0.116111362 | 0.29869325 | | Gart | 1523.357011 | 0.0517443853841377 | 0.050570761363901 | 1.02320756082337 | 0.30620971883115 | 0.738584942735804 | NM_010256 |
| 9030617O03Rik | 336.7652492 | -0.135832636 | 0.086452388 | -1.571184311 | 0.116139843 | 0.298716921 | | Has3 | 104.9971422 | 0.0958856727792257 | 0.093731110956966 | 1.02298662418766 | 0.306314170113098 | 0.738584942735804 | NM_008217 |
| Smg9 | 1223.485021 | 0.136496286 | 0.086889367 | 1.570920488 | 0.116201119 | 0.298824919 | | Nfkbid | 136.4785951 | -0.097253074783062 | 0.0950694379701459 | -1.02296886212372 | 0.306322568434556 | 0.738584942735804 | NM_172142 |
| Mir374 | 17.90163355 | 0.253348668 | 0.161295332 | 1.570712957 | 0.116249338 | 0.298849717 | | Aars2 | 1255.119799 | -0.0790688430898518 | 0.0773430675339726 | -1.02231325457994 | 0.306632661863123 | 0.738806191753709 | NM_198608 |
| Yipf1 | 1065.456223 | -0.100717831 | 0.064121602 | -1.570731671 | 0.116244989 | 0.298849717 | | Gm3604 | 48.09479956 | 0.0923849092506054 | 0.090369992210331 | 1.02229630645076 | 0.306640680853856 | 0.738806191753709 | NM_001162910 |
| Dym | 2259.376154 | 0.109843648 | 0.069939332 | 1.57055615 | 0.116285782 | 0.2988829 | | Nhp2 | 854.9009451 | 0.0635021978944565 | 0.0621030681667352 | 1.02252915627236 | 0.306530520347155 | 0.738806191753709 | NM_026631 |
| Epc2 | 1962.074664 | 0.164583883 | 0.104797696 | 1.570491431 | 0.116300826 | 0.2988829 | | Nxt1 | 231.5933156 | -0.0905682790100323 | 0.0885918207373597 | -1.0223097150078 | 0.306634336596739 | 0.738806191753709 | NM_019761,NM_001110159 |
| Mgat5 | 1704.955291 | -0.124155965 | 0.079082218 | -1.569960576 | 0.116424283 | 0.299150555 | | Kctd14 | 92.35004801 | -0.0967872914660105 | 0.0946985547266005 | -1.0220566907852 | 0.306754069639745 | 0.738956818505297 | NM_001010826,NM_001012434,NM_001136235 |
| 6820431F20Rik | 2456.605678 | 0.173129933 | 0.110301363 | 1.569608279 | 0.116506271 | 0.299311585 | | Hcfc1r1 | 2489.989044 | 0.0688279839276272 | 0.0673677675728092 | 1.02167529676918 | 0.306934606793855 | 0.739163118548453 | NM_181821 |
| Fam109b | 25.30650372 | -0.25982024 | 0.16554518 | -1.569482358 | 0.116535587 | 0.299337266 | | Scyl3 | 1198.371902 | -0.0654918610256745 | 0.0641168211126588 | -1.02144585288469 | 0.307043250538439 | 0.739163118548453 | NM_028776 |
| Klf9 | 6826.07298 | 0.18393396 | 0.117213659 | 1.569219504 | 0.116596801 | 0.299444859 | | Zfp317 | 1171.778629 | -0.0524921968482631 | 0.051379995389077 | -1.02164658542228 | 0.306948200481144 | 0.739163118548453 | NM_172918 |
| Enthd2 | 613.7564151 | 0.124023421 | 0.079042039 | 1.56908176 | 0.116628889 | 0.299477629 | | Zfyve27 | 3661.631997 | -0.0620009960484473 | 0.0606967416821796 | -1.02148804581796 | 0.307023269893894 | 0.739163118548453 | NM_001164531,NM_177319 |
| Mllt11 | 10210.30031 | -0.100075246 | 0.063786391 | -1.568912192 | 0.116668401 | 0.299529444 | | Tnxb | 175.217121 | 0.0971023189017206 | 0.0950763066064558 | 1.02130932897563 | 0.307107907922391 | 0.739196266856778 | NM_031176 |
| Cep128 | 277.827191 | -0.159379782 | 0.101621663 | -1.568364229 | 0.116796154 | 0.299758094 | | Gdf10 | 382.3402594 | 0.0888767678722792 | 0.0870533188151937 | 1.0209463473869 | 0.307279858952888 | 0.739487612847856 | NM_145741 |
| Ntrk3 | 3249.607935 | -0.102884896 | 0.065598882 | -1.568394041 | 0.116789201 | 0.299758094 | | Dpagt1 | 564.6142597 | -0.0690568109691544 | 0.0676829250291898 | -1.02029885586907 | 0.307586745797163 | 0.739991032507178 | NM_007875 |
| Slc35c2 | 1829.521077 | 0.113195907 | 0.072198513 | 1.567842647 | 0.116917859 | 0.300020754 | | Zfp523 | 3933.452508 | -0.0698030536901603 | 0.0684149130546925 | -1.02029002995821 | 0.30759093035098 | 0.739991032507178 | NM_172617 |
| Slc44a2 | 1917.026584 | 0.104708628 | 0.066807996 | 1.567306818 | 0.117042993 | 0.300292122 | | Dtna | 5584.762442 | -0.0536407262389934 | 0.0525930026019983 | -1.01992135046793 | 0.307765762875777 | 0.740098991201351 | NM_207650,NM_010087 |
| Uprt | 132.9470319 | -0.196073262 | 0.125109547 | -1.56721263 | 0.117064999 | 0.300298857 | | Gad2 | 10636.33808 | 0.078941998857169 | 0.0773907059754032 | 1.02004495064638 | 0.307707142765148 | 0.740098991201351 | NM_008078 |
| Cwc22 | 788.6281913 | 0.105751507 | 0.067488547 | 1.566954852 | 0.117125245 | 0.300403666 | | Wrb | 2166.33031 | 0.0567734667522708 | 0.0556671879226255 | 1.01987308629965 | 0.307788655228374 | 0.740098991201351 | NM_207301 |
| Chd1 | 1049.629912 | 0.120765995 | 0.077101695 | 1.56632088 | 0.117273515 | 0.300734168 | | Ccdc153 | 105.6966644 | 0.0952516217733109 | 0.0934376138245075 | 1.0194141082435 | 0.308006411085974 | 0.740254984684789 | NM_001081369 |
| Fam178a | 2358.483085 | 0.164673061 | 0.105144297 | 1.566162544 | 0.117310569 | 0.300737957 | | Skiv2l | 3474.396982 | -0.0681207102880587 | 0.0668211269045949 | -1.01944869001266 | 0.307990000688951 | 0.740254984684789 | NM_021337 |
| Rgs19 | 571.321884 | -0.122350848 | 0.078122115 | -1.566148686 | 0.117313813 | 0.300737957 | | Snord104 | 33.49594113 | -0.0794278496224612 | 0.0779127491063473 | -1.01944611855559 | 0.307991220925623 | 0.740254984684789 | NR_030703 |
| Dnajc24 | 164.7118181 | -0.188778502 | 0.120544228 | -1.566051772 | 0.117336497 | 0.300746351 | | Ap3m1 | 2452.288812 | -0.05527953004733 | 0.0542588028451665 | -1.01881219541604 | 0.308292134845925 | 0.740628329834004 | NM_018829 |
| Arhgef25 | 6533.416218 | 0.137763466 | 0.087985022 | 1.565760419 | 0.117404716 | 0.300771935 | | Lmf1 | 619.919614 | 0.0696627988753498 | 0.0683796782166884 | 1.01876464897356 | 0.308314712274971 | 0.740628329834004 | NR_045520,NM_029624 |
| Cebpg | 1279.096436 | 0.102691778 | 0.065582622 | 1.565838243 | 0.117386491 | 0.300771935 | | Smad6 | 62.47383324 | 0.0958933441864697 | 0.0941226505348581 | 1.01881261993314 | 0.3082919332689 | 0.740628329834004 | NM_008542 |
| P2rx1 | 17.20879178 | -0.260036466 | 0.166071222 | -1.565812925 | 0.117392419 | 0.300771935 | | Brwd1 | 5062.06958 | -0.0597787906648802 | 0.0586990737779686 | -1.018394104326 | 0.308490702880866 | 0.740806185741832 | NM_176928,NM_001103179,NM_145125 |
| Fat2 | 14.99877462 | -0.256300803 | 0.163700588 | -1.565668185 | 0.117426318 | 0.300777546 | | Hint2 | 416.6419243 | 0.0794923252923489 | 0.0780647814003131 | 1.0182866571382 | 0.308541747469233 | 0.740806185741832 | NM_026871 |
| Zrsr2 | 1371.938477 | 0.09570301 | 0.061130634 | 1.565549118 | 0.11745421 | 0.300799261 | | Pcnx | 5937.974144 | 0.0622506038248646 | 0.0611267349330765 | 1.01838588128449 | 0.308494609177266 | 0.740806185741832 | NM_018814 |
| Bmp6 | 352.8333659 | -0.159304695 | 0.101772533 | -1.56530147 | 0.117512239 | 0.300871791 | | Prkcz | 12219.44863 | 0.0471034498521017 | 0.046267058527817 | 1.01807746917349 | 0.308641141741707 | 0.740922364062144 | NM_008860,NM_001039079 |
| Card9 | 53.13130101 | 0.251201328 | 0.16048511 | 1.565262516 | 0.117521368 | 0.300871791 | | Bin3 | 435.021654 | -0.072825188780461 | 0.0715599795688823 | -1.0176804020795 | 0.308829863764342 | 0.741252908301526 | NM_021328 |
| Exph5 | 1296.325006 | 0.16169477 | 0.103315269 | 1.565061695 | 0.117568444 | 0.300942586 | | Acot13 | 1503.917596 | 0.0640715464235275 | 0.0630202150746315 | 1.01668244622223 | 0.309304518974977 | 0.742145027028141 | NM_025790 |
| Trpm3 | 1676.607019 | 0.201965484 | 0.129062226 | 1.564869059 | 0.117613615 | 0.301008482 | | Csrp1 | 11311.99089 | 0.0759239852038 | 0.0746844139423274 | 1.01659745582833 | 0.309344965004844 | 0.742145027028141 | NM_007791 |
| Alkbh1 | 359.029531 | 0.123735856 | 0.079084076 | 1.564611508 | 0.117674029 | 0.301113361 | | Gm10767 | 54.45749165 | -0.0916515522979901 | 0.0901760853211464 | -1.01636206508177 | 0.309457003215576 | 0.742145027028141 | NM_001177750 |
| Reep5 | 16094.18526 | -0.088752353 | 0.056734078 | -1.564357018 | 0.117733749 | 0.301216431 | | Psen2 | 584.871434 | 0.0707255839937831 | 0.0695763908187357 | 1.01651699896365 | 0.309383256796167 | 0.742145027028141 | NM_001128605,NM_011183 |
| Bri3bp | 5313.813318 | -0.097565894 | 0.062381253 | -1.564025869 | 0.117811493 | 0.301365573 | | Ptger1 | 59.81588585 | -0.0957606639340308 | 0.0942140020945438 | -1.01641647531261 | 0.309431103363012 | 0.742145027028141 | NM_013641 |
| Ascc3 | 1061.176633 | -0.133019535 | 0.085063624 | -1.563765198 | 0.11787272 | 0.301421803 | | Als2 | 2161.274413 | 0.0629661069683495 | 0.0619666866880391 | 1.01612834788702 | 0.309568271393258 | 0.742289321600935 | NM_028717,NM_146109,NM_001159948 |
| Ccdc173 | 70.22491207 | 0.251090976 | 0.160567696 | 1.563770191 | 0.117871547 | 0.301421803 | | Bco2 | 177.3347568 | -0.0894730969145104 | 0.0880841314118052 | -1.01576862347897 | 0.309739580778097 | 0.742454975310665 | NM_133217 |
| Dtnb | 2502.897206 | -0.099672224 | 0.063742468 | -1.563670619 | 0.117894941 | 0.301421803 | | Zfp108 | 280.6798664 | -0.0795227605508756 | 0.0782845974955544 | -1.01581617706333 | 0.309716931030367 | 0.742454975310665 | NM_018791 |
| F830016B08Rik | 10.62410873 | 0.18692409 | 0.119553507 | 1.563518254 | 0.117930746 | 0.301421803 | | Glud1 | 10774.20738 | 0.0511384218939791 | 0.0503538178747776 | 1.01558181787035 | 0.30982856678402 | 0.742545745108839 | NM_008133 |
| Mtss1 | 1270.404357 | -0.128332416 | 0.082075319 | -1.563593269 | 0.117913116 | 0.301421803 | | Aldh5a1 | 5273.408651 | 0.0622981157649156 | 0.0613760046875243 | 1.01502396713644 | 0.310094402920486 | 0.743030104379923 | NM_172532 |
| Rhpn2 | 319.571945 | 0.215902796 | 0.138102452 | 1.56335237 | 0.117969737 | 0.301422015 | | Col4a6 | 69.39427782 | -0.0961108912085811 | 0.0947009554479794 | -1.01488829499061 | 0.310159078387975 | 0.743030104379923 | NM_053185 |
| Zc3h11a | 4325.937266 | 0.123912424 | 0.079258634 | 1.563393394 | 0.117960093 | 0.301422015 | | Sla | 467.2570232 | 0.0797993293315162 | 0.0786327522544892 | 1.01483576555036 | 0.310184121778869 | 0.743030104379923 | NM_001029841,NM_009192 |
| Nek8 | 121.8495658 | -0.235995359 | 0.150972184 | -1.563171133 | 0.118012348 | 0.301481174 | | Inpp5a | 2738.169681 | 0.056418074985216 | 0.0556046065611681 | 1.01462951496929 | 0.310282464577896 | 0.743039092797119 | NM_001127363,NM_183144 |
| Brms1l | 2485.106741 | -0.080862154 | 0.05173763 | -1.562927295 | 0.118069697 | 0.301577956 | | Svil | 484.8306531 | -0.0853678171563804 | 0.0841382760806939 | -1.014613338102 | 0.310290178776492 | 0.743039092797119 | NM_153153 |
| Lonp2 | 2878.654587 | -0.082723495 | 0.052932897 | -1.562799299 | 0.118099809 | 0.30160515 | | Lhfpl4 | 6876.209318 | 0.0521719458999717 | 0.0514371185095604 | 1.01428593614307 | 0.310446332864369 | 0.74329049467411 | NM_177763 |
| Adcy7 | 371.4909536 | 0.182363454 | 0.116706471 | 1.562582194 | 0.118150899 | 0.301607001 | | Nrip3 | 5387.897395 | 0.0700821001860268 | 0.0691079939672293 | 1.01409542026729 | 0.310537223121218 | 0.743385580596376 | NM_020610 |
| Arnt | 1349.111187 | 0.143855015 | 0.092055907 | 1.562691842 | 0.118125094 | 0.301607001 | | Parp16 | 153.4539701 | -0.0890337997894722 | 0.087821055687884 | -1.01380926353127 | 0.310673774188014 | 0.74358992319247 | NM_177460 |
| Cldn19 | 22.123372 | -0.245925229 | 0.157387305 | -1.562548064 | 0.118158932 | 0.301607001 | | Igf1 | 334.7252548 | 0.0878318046290417 | 0.0866518487266267 | 1.01361720401532 | 0.31076544524389 | 0.743686796970799 | NM_184052,NM_010512,NM_001111275,NM_001111274,NM_001111276 |
| Col6a5 | 68.37221718 | 0.254835273 | 0.163103368 | 1.562415765 | 0.118190076 | 0.301636803 | | Rrp1b | 700.3472306 | -0.0812699780618654 | 0.0801896998192996 | -1.01347153368824 | 0.310834986388651 | 0.743730688894999 | NM_001163734,NM_028244 |
| Cers4 | 1819.425559 | 0.152456894 | 0.097604678 | 1.561983477 | 0.118291881 | 0.301846905 | | Filip1 | 729.1666579 | 0.0777353942180801 | 0.0767345774114172 | 1.01304257924426 | 0.31103982336766 | 0.743730777652449 | NM_001081243 |
| Vsx2 | 13.97260011 | 0.255495115 | 0.163597924 | 1.561725902 | 0.118352574 | 0.301952047 | | Lin7b | 1427.492799 | -0.0753316020118201 | 0.0743406508175799 | -1.01332986977302 | 0.310902624771417 | 0.743730777652449 | NM_011698 |
| 1500009C09Rik | 3244.64739 | -0.097660925 | 0.062539 | -1.561600361 | 0.118382165 | 0.301977816 | | Mrps34 | 994.6701151 | 0.0599750143858631 | 0.0591954282200173 | 1.01316970227748 | 0.310979109507894 | 0.743730777652449 | NM_023260 |
| 2610034B18Rik | 703.1394747 | -0.136979119 | 0.087739515 | -1.561202138 | 0.118476065 | 0.302167596 | | Pik3ca | 3428.74435 | 0.0703091591980875 | 0.0693988457724884 | 1.0131171263076 | 0.311004218801166 | 0.743730777652449 | NM_008839 |
| Abi3 | 192.6927102 | 0.214108558 | 0.137216645 | 1.560368706 | 0.118672776 | 0.302391407 | | 8430410A17Rik | 585.3945338 | -0.0694146019708499 | 0.0685409022775674 | -1.01274712856484 | 0.311180960582532 | 0.743823372292556 | NM_173737 |
| Batf2 | 8.488925228 | 0.220847881 | 0.141526786 | 1.56046701 | 0.11864956 | 0.302391407 | | Tead2 | 78.14707801 | 0.0961494888841385 | 0.0949346774311763 | 1.01279628778265 | 0.311157474224243 | 0.743823372292556 | NM_011565 |
| Mpp3 | 2159.33731 | 0.103671353 | 0.06644176 | 1.56033423 | 0.118680918 | 0.302391407 | | Crk | 5978.367736 | 0.0698952656387561 | 0.0690320974339448 | 1.01250386757605 | 0.311297198411586 | 0.743856407982678 | NM_133656 |
| Pycr2 | 687.1900332 | -0.115885856 | 0.074266396 | -1.56040769 | 0.118663569 | 0.302391407 | | Rapsn | 31.53851093 | -0.0912281389999406 | 0.090098581125378 | -1.01253691079764 | 0.311281407629973 | 0.743856407982678 | NM_009023 |
| Susd4 | 3747.038484 | -0.085126346 | 0.054547722 | -1.56058481 | 0.118621745 | 0.302391407 | | Myo16 | 1426.063034 | 0.0661622281015181 | 0.065361195635353 | 1.01225547449643 | 0.311415918062396 | 0.744017701607626 | NM_001081397 |
| Usp7 | 7421.263275 | 0.110988449 | 0.071127665 | 1.560411826 | 0.118662592 | 0.302391407 | | Afap1 | 846.6710691 | -0.0646565064734282 | 0.0638996136790069 | -1.01184502927081 | 0.311612156076078 | 0.74424257972001 | NM_027373 |
| Htr3a | 283.118125 | 0.198532355 | 0.127279475 | 1.559814382 | 0.118803752 | 0.302604855 | | Cul2 | 2352.356581 | 0.0638869922715211 | 0.0631579835790285 | 1.0115426213945 | 0.311756792495222 | 0.74424257972001 | NM_029402 |
| Ppp1r9b | 37537.85477 | -0.134917029 | 0.086491455 | -1.559888539 | 0.118786223 | 0.302604855 | | Elfn2 | 4658.009703 | 0.076833605633704 | 0.0759583440553347 | 1.01152291547762 | 0.311766219027692 | 0.74424257972001 | NM_183141 |
| Gsto2 | 24.44548743 | 0.261434582 | 0.167665235 | 1.559265291 | 0.118933603 | 0.302885807 | | Fbxo5 | 43.11722563 | -0.0934299981375948 | 0.0923569450387058 | -1.01161854258431 | 0.311720476552563 | 0.74424257972001 | NM_025995 |
| Cxcr4 | 55.35712124 | -0.259527703 | 0.166460239 | -1.559097257 | 0.118973363 | 0.302937269 | | Vps33a | 6071.359379 | 0.0488099148431412 | 0.0482404059897295 | 1.01180563972726 | 0.311630992902056 | 0.74424257972001 | NM_029929 |
| P2ry6 | 93.35628654 | 0.223480542 | 0.143382489 | 1.55863204 | 0.119083495 | 0.303167873 | | Gipc1 | 2620.622055 | 0.074280674321371 | 0.0734641794186676 | 1.01111419074118 | 0.311961779167575 | 0.744587052939236 | NM_018771 |
| Ddx59 | 291.7719699 | 0.157890154 | 0.101325005 | 1.558254581 | 0.119172911 | 0.303345669 | | Itsn1 | 7098.836112 | 0.0633994741569693 | 0.0627248712329932 | 1.01075495111772 | 0.312133729201602 | 0.744800424038485 | NM_010587,NM_001110276,NM_001110275 |
| Rapgef5 | 3251.446077 | 0.118201077 | 0.075881472 | 1.557706682 | 0.119302796 | 0.303576534 | | Sncaip | 176.0570471 | -0.0959682772749945 | 0.0949611105598369 | -1.01060609663493 | 0.312204996693538 | 0.744800424038485 | NM_001199151,NM_001199153,NM_001199154,NM_026408 |
| Tuba4a | 22537.3309 | -0.152656231 | 0.097999661 | -1.557722032 | 0.119299155 | 0.303576534 | | Snrnp48 | 432.7975437 | -0.0762114633138627 | 0.0754116205230612 | -1.01060635993835 | 0.312204870621532 | 0.744800424038485 | NM_026382 |
| Myo1b | 1689.92804 | 0.117191249 | 0.075242773 | 1.557508372 | 0.119349834 | 0.30364636 | | Fam71e1 | 93.25075236 | -0.0960835904225939 | 0.0950885448908123 | -1.01046441012347 | 0.312272842314861 | 0.744839951964806 | NM_028169 |
| Icam2 | 96.25355347 | -0.189548943 | 0.121720764 | -1.557244113 | 0.119412539 | 0.303657683 | | D10Bwg1379e | 2606.834587 | 0.072786657439318 | 0.072048144829687 | 1.01025026544926 | 0.312375402470089 | 0.744839969842501 | NM_001033258 |
| Tln1 | 2213.03291 | 0.096867361 | 0.062204241 | 1.557246884 | 0.119411881 | 0.303657683 | | Noxred1 | 48.89121935 | -0.085563084767294 | 0.0846885409006561 | -1.01032659031951 | 0.312338845709686 | 0.744839969842501 | NM_027744 |
| Tmem220 | 22.64841627 | 0.261319863 | 0.167809429 | 1.557241832 | 0.11941308 | 0.303657683 | | Fam195b | 2091.122114 | 0.0655899563927515 | 0.0649332808242731 | 1.01011308161458 | 0.312441115486576 | 0.744874387585426 | NM_001033231 |
| 4933407L21Rik | 112.8923369 | -0.220969009 | 0.141950251 | -1.556665154 | 0.119550006 | 0.303955988 | | Gstt3 | 130.0093962 | -0.0922513286272719 | 0.0913515270417035 | -1.00984988006996 | 0.312567218283086 | 0.74493050250699 | NM_133994 |
| Ppp6c | 1257.078436 | -0.109697327 | 0.07048904 | -1.556232384 | 0.119652843 | 0.30416754 | | Lmo7 | 4062.487031 | -0.0651421660391391 | 0.0645053228598032 | -1.00987272291808 | 0.312556272691296 | 0.74493050250699 | NM_201529 |
| Sepw1 | 15798.26678 | -0.122107733 | 0.078485957 | -1.555790833 | 0.119757839 | 0.304384507 | | Dpysl4 | 2161.92881 | -0.0707757826140648 | 0.0701185898540505 | -1.00937258951417 | 0.312795979024925 | 0.745030948621872 | NM_011993 |
| Ggct | 368.0400453 | -0.12267539 | 0.078880359 | -1.55520832 | 0.119896463 | 0.304636898 | | Krt73 | 56.13520293 | 0.095467276129256 | 0.0945609012127254 | 1.00958509177585 | 0.312694115134555 | 0.745030948621872 | NM_212485 |
| Zfp420 | 499.0216027 | 0.134614754 | 0.086555879 | 1.555235249 | 0.119890052 | 0.304636898 | | Pkd1l3 | 64.09867124 | -0.0959058086867089 | 0.0950013356025762 | -1.00952063545629 | 0.312725010244999 | 0.745030948621872 | NM_001039700,NM_181544 |
| Gprc5b | 7573.804917 | -0.126227628 | 0.081170194 | -1.555098262 | 0.119922669 | 0.304653522 | | Slc39a11 | 682.2385319 | 0.070671377408876 | 0.0700178372257561 | 1.0093339098866 | 0.31281452262459 | 0.745030948621872 | NM_001166503,NM_027216 |
| Tmx2 | 8197.922598 | -0.064727991 | 0.041628482 | -1.554896747 | 0.119970662 | 0.304725482 | | Zfp521 | 539.2766053 | 0.0864778941910649 | 0.0857110513706238 | 1.00894683717185 | 0.313000131023024 | 0.74535080380991 | NM_145492 |
| Mamld1 | 1249.894525 | 0.176340991 | 0.113427233 | 1.554661844 | 0.120026626 | 0.304817269 | | Dnm2 | 2348.150913 | -0.0640202322051392 | 0.0634786016198609 | -1.00853249081512 | 0.313198897971877 | 0.745612332510311 | NM_001039520,NM_001253893,NM_007871,NM_001253894 |
| Rev3l | 1798.237886 | -0.162848394 | 0.104753954 | -1.554579937 | 0.120046145 | 0.304817269 | | Sdcbp2 | 34.95598718 | 0.0927181293953764 | 0.0919363120230302 | 1.00850390183316 | 0.313212615515484 | 0.745612332510311 | NM_145535 |
| Cd81 | 8603.855889 | 0.083752732 | 0.053885039 | 1.554285445 | 0.120116344 | 0.30494554 | | Cuedc2 | 3873.381626 | 0.0453182580258377 | 0.0449673378254754 | 1.00780389094245 | 0.313548617612601 | 0.745844950800566 | NM_001164293,NM_001164292,NM_001164291,NM_001164290,NM_001164295,NM_001164294,NM_024192 |
| Fbxl3 | 3064.172815 | 0.094700665 | 0.060936722 | 1.554082033 | 0.12016485 | 0.305018707 | | Dcakd | 1062.5341 | 0.0757246729830226 | 0.0751142942626802 | 1.00812599953622 | 0.313393977475969 | 0.745844950800566 | NM_026551 |
| 2310011J03Rik | 1020.517059 | -0.129375842 | 0.083261892 | -1.553842212 | 0.120222059 | 0.305063966 | | Fhl1 | 3777.8281 | -0.0540022022869819 | 0.0536067541356954 | -1.00737683446167 | 0.313753719272042 | 0.745844950800566 | NM_001077361,NM_001077362,NM_010211 |
| Fgf1 | 2459.369368 | -0.130785784 | 0.084166956 | -1.553885166 | 0.120211811 | 0.305063966 | | Kiss1r | 234.7777751 | -0.09047861999299 | 0.0897761220128388 | -1.00782499805517 | 0.313538482825741 | 0.745844950800566 | NM_053244 |
| Racgap1 | 579.7161744 | -0.145432995 | 0.093603371 | -1.553715358 | 0.120252328 | 0.30507634 | | Prkch | 366.601411 | 0.0836466779378622 | 0.0830373614505635 | 1.0073378594485 | 0.313772442127376 | 0.745844950800566 | NM_008856 |

| | | | | | | | | | | | | | | | |
|---|---|---|---|---|---|---|---|---|---|---|---|---|---|---|---|
| Sesn1 | 2575.33878 | 0.189647852 | 0.122065477 | 1.553656749 | 0.120266315 | 0.30507634 | | Rbm41 | 373.2292104 | -0.0860511093889889 | 0.0854124446426387 | -1.0074774202872 | 0.313705403150355 | 0.745844950800566 | NM_001172148,NM_153586,NM_001172147 |
| Ephx3 | 15.32974799 | -0.249478553 | 0.160592341 | -1.553489732 | 0.120306181 | 0.30512751 | | Sept2 | 2407.146423 | -0.0627457282662534 | 0.0622856021440562 | -1.00738735929907 | 0.313748663466307 | 0.745844950800566 | NM_010891,NM_001159718,NM_001159719,NM_001159717 |
| Lman2l | 1374.339392 | -0.170288391 | 0.109658798 | -1.552893107 | 0.120448674 | 0.305438912 | | Sf1 | 6189.680764 | -0.0492674086866391 | 0.0488912797202126 | -1.00769317081858 | 0.313601784495853 | 0.745844950800566 | NM_001110791,NM_011750 |
| Rab39 | 44.03358225 | -0.239382986 | 0.154194087 | -1.552478372 | 0.120547804 | 0.305640267 | | Tusc5 | 39.83922593 | -0.0915049898266493 | 0.0908034122223746 | -1.0077263352456 | 0.313585858591303 | 0.745844950800566 | NM_177709 |
| Mpp6 | 1329.244605 | -0.122056881 | 0.078640069 | -1.552095303 | 0.120639422 | 0.305822513 | | Fjx1 | 1902.522624 | -0.0650957391877975 | 0.0646602633852686 | -1.00673482877628 | 0.314062220271476 | 0.745923448604229 | NM_010218 |
| Zfp488 | 509.1308458 | 0.14181463 | 0.091423332 | 1.551186401 | 0.12085702 | 0.306324009 | | Gm13293 | 76.13686526 | -0.0932675880915134 | 0.0926420247158771 | -1.00675247953134 | 0.314053735942457 | 0.745923448604229 | NR_040369 |
| Ccdc41 | 1213.0545 | 0.106259065 | 0.068515376 | 1.550879104 | 0.120930659 | 0.306460522 | | Gria1 | 17289.37909 | 0.0526036804489963 | 0.0522387678373354 | 1.00698547509385 | 0.313941754240006 | 0.745923448604229 | NM_008165,NM_001113325,NM_001252403 |
| Zfp804b | 54.93675083 | -0.238858323 | 0.15407806 | -1.550242278 | 0.121083376 | 0.306797355 | | Prickle3 | 66.22636594 | -0.0951027958807127 | 0.0944498221423032 | -1.00691344592927 | 0.313976369898592 | 0.745923448604229 | NM_175097 |
| Pgp | 1016.332078 | -0.155094311 | 0.100077095 | -1.549748325 | 0.121201934 | 0.307047543 | | St3gal2 | 5290.971887 | 0.0445966658249416 | 0.044280525487952 | 1.007139489279 | 0.313867746777308 | 0.745923448604229 | NM_009179 |
| Aoc2 | 162.4974398 | 0.224714249 | 0.145019243 | 1.549547807 | 0.121250089 | 0.307084755 | | 2310045N01Rik | 535.2173316 | -0.0786462099656954 | 0.0783607230483432 | -1.00364323995806 | 0.315550604756507 | 0.745993051875022 | NM_001145552 |
| Atp1a3 | 93513.56428 | 0.080364203 | 0.051865865 | 1.549462321 | 0.121270623 | 0.307084755 | | Amn | 99.21245803 | -0.0951172461231778 | 0.0946433056389761 | -1.00500764931024 | 0.31489316656547 | 0.745993051875022 | NM_033603 |
| Cacng8 | 1197.939638 | -0.131248171 | 0.08470722 | -1.549433101 | 0.121277642 | 0.307084755 | | Ap3d1 | 8656.017006 | -0.0381541565492857 | 0.0380429975662945 | -1.00292193018696 | 0.315898530598964 | 0.745993051875022 | NM_007460 |
| Cdkl2 | 1519.309092 | -0.099399196 | 0.064155124 | -1.549357095 | 0.121295902 | 0.307084755 | | Arhgdib | 413.8444906 | -0.0881611232777768 | 0.0876571374182366 | -1.00574951309596 | 0.314536078922221 | 0.745993051875022 | NM_007486 |
| Tnpo3 | 3697.988227 | 0.086040326 | 0.055541382 | 1.549121096 | 0.121352612 | 0.307178137 | | Armc6 | 946.9346066 | 0.0667957450048792 | 0.0666098828162425 | 1.00279030949731 | 0.3159620453813 | 0.745993051875022 | NM_133972 |
| Pkp4 | 8915.032567 | -0.068740314 | 0.044379946 | -1.548904849 | 0.121404595 | 0.307259522 | | Asf1b | 26.25214772 | 0.0825013142363185 | 0.0823311923542246 | 1.00206631140919 | 0.316311567253222 | 0.745993051875022 | NM_024184 |
| Mfsd12 | 560.829234 | 0.12022505 | 0.077640879 | 1.548476163 | 0.121507696 | 0.307420027 | | Birc5 | 34.10042573 | 0.0890403621583287 | 0.0887312408015399 | 1.00348379391516 | 0.315627492336665 | 0.745993051875022 | NM_009689,NM_001012273 |
| Txn2 | 3823.552154 | -0.093641409 | 0.060470449 | -1.548548262 | 0.121490351 | 0.307420027 | | Bop1 | 2068.641063 | 0.0696554102196563 | 0.069417980230106 | 1.00342029527168 | 0.315658115882587 | 0.745993051875022 | NM_013481 |
| Epb4.9 | 14779.41777 | 0.122222227 | 0.078942987 | 1.54823413 | 0.121565937 | 0.307517164 | | Cep290 | 706.0802783 | -0.0816251878009722 | 0.0812269651928625 | -1.00490259124127 | 0.31494375659366 | 0.745993051875022 | NM_146009 |
| Acvrl1 | 788.8210393 | -0.133243794 | 0.086068957 | -1.548105135 | 0.121596986 | 0.307545495 | | Dio2 | 3249.051958 | 0.0950016281425637 | 0.094825654495666 | 1.00185575989782 | 0.316413262038038 | 0.745993051875022 | NM_010050 |
| Arhgap21 | 7951.638197 | 0.165602363 | 0.106978505 | 1.547996605 | 0.121623114 | 0.307561372 | | Dnaaf2 | 197.7179573 | -0.083221284360493 | 0.0827887490002945 | -1.00522456692995 | 0.31478872819922 | 0.745993051875022 | NM_027269 |
| Glo1 | 3207.684991 | -0.087063549 | 0.056265627 | -1.547366549 | 0.121774883 | 0.307800027 | | Emx2 | 311.0382275 | 0.0837164842123868 | 0.0835861679726743 | 1.00155906464997 | 0.316556600010852 | 0.745993051875022 | NM_010132 |
| Gm10421 | 33.71722981 | -0.250205852 | 0.16169881 | -1.547357409 | 0.121777086 | 0.307800027 | | Fam212b | 6679.721286 | 0.0697077065318689 | 0.0695730431461095 | 1.00193556842809 | 0.316374712597753 | 0.745993051875022 | NM_001163356,NM_175398 |
| Insig2 | 1755.715745 | -0.124858269 | 0.080686571 | -1.547447944 | 0.121755268 | 0.307800027 | | Fbxl19 | 2625.894953 | -0.0681296657177482 | 0.0678421533870231 | -1.00423795997578 | 0.315263929226225 | 0.745993051875022 | NM_172748 |
| Eif4e | 3552.586716 | -0.082661218 | 0.053424768 | -1.547245243 | 0.12180412 | 0.307818143 | | Fbxo3 | 3029.43806 | 0.058992419738294 | 0.0587480729970431 | 1.00415922989105 | 0.315301870008831 | 0.745993051875022 | NM_020593,NM_212433 |
| Pls1 | 500.9237097 | 0.15319471 | 0.099022882 | 1.547063742 | 0.121847876 | 0.307878504 | | Gm5595 | 159.1372985 | 0.0950192294976679 | 0.0947309397939693 | 1.00304324758443 | 0.315839995190654 | 0.745993051875022 | NM_001008427 |
| 1700012D14Rik | 13.98948182 | 0.246669586 | 0.159453457 | 1.546969192 | 0.121870675 | 0.307885901 | | Gm9079 | 69.45632179 | 0.0955748093689959 | 0.0950680692505144 | 1.00533028726129 | 0.314737835752475 | 0.745993051875022 | NR_004052 |
| Tapt1 | 2127.679978 | 0.135447573 | 0.087579673 | 1.546564043 | 0.121968406 | 0.308082569 | | Gpr107 | 2260.452708 | 0.0555397652356439 | 0.0554450845658223 | 1.00170764767631 | 0.316484811972944 | 0.745993051875022 | NM_178760 |
| Gm5089 | 86.76168879 | 0.205433783 | 0.132884535 | 1.54595704 | 0.122114943 | 0.308402434 | | Gpsm3 | 84.68939132 | -0.0956207800847305 | 0.0950621247681223 | -1.00587637921628 | 0.314474866839304 | 0.745993051875022 | NM_134116 |
| Dnajc16 | 2820.594457 | 0.117174119 | 0.075806998 | 1.545689993 | 0.122179455 | 0.30848017 | | Kndc1 | 6326.594644 | 0.0679479133116829 | 0.0675864923885115 | 1.0053475318869 | 0.314729534918224 | 0.745993051875022 | NM_177261 |
| Kcnk3 | 1013.639463 | 0.145276712 | 0.093994802 | 1.545582407 | 0.122205453 | 0.30848017 | | Mgat4b | 2108.838128 | -0.0659476733441439 | 0.0657068876559255 | -1.00366454258919 | 0.315540333198952 | 0.745993051875022 | NM_145926 |
| Smc1a | 3706.677148 | 0.094678476 | 0.061255582 | 1.54563017 | 0.122193911 | 0.30848017 | | Mrps36 | 668.5259063 | 0.0759364189637324 | 0.0757569922978508 | 1.00236845023066 | 0.316165673973496 | 0.745993051875022 | NM_001190264,NM_025369 |
| Dpp3 | 1385.176054 | -0.103101555 | 0.06672438 | -1.545185655 | 0.122301363 | 0.308671986 | | Nadkd1 | 1394.695373 | 0.053211899232007 | 0.0529708633872575 | 1.0045503476692 | 0.315113416165479 | 0.745993051875022 | NM_001040395,NM_001085410 |
| 4933432I09Rik | 40.61617095 | 0.25490658 | 0.164987583 | 1.545004634 | 0.122345142 | 0.308694072 | | Ncapg2 | 225.4243477 | 0.0867298481509226 | 0.0865065825336821 | 1.00258090899792 | 0.316063110809913 | 0.745993051875022 | NM_133762 |
| Pcid2 | 1122.09288 | 0.103467358 | 0.066969826 | 1.544984713 | 0.122349961 | 0.308694072 | | Ncrna00086 | 950.6696736 | 0.0568937366103018 | 0.056533026367137 | 1.00638052243696 | 0.31423255940867 | 0.745993051875022 | NR_028086 |
| Batf3 | 26.4141412 | 0.258439732 | 0.1673178 | 1.54460393 | 0.122442095 | 0.308775687 | | Ostf1 | 391.6205917 | -0.0809058985500301 | 0.0804499799861437 | -1.00566710599511 | 0.314575731487889 | 0.745993051875022 | NM_017375 |
| Dnaaf2 | 184.3207733 | 0.155324067 | 0.100554839 | 1.544670232 | 0.122426049 | 0.308775687 | | Pacsin3 | 1255.716577 | -0.0641573667630762 | 0.0638992730237244 | -1.00403907160659 | 0.31535978122059 | 0.745993051875022 | NM_030880,NM_028733 |
| Leo1 | 837.1390371 | 0.161459566 | 0.104528223 | 1.544650437 | 0.122430839 | 0.308775687 | | Pcmt1 | 5409.20742 | 0.0580033241511437 | 0.0576788541159011 | 1.0056254591083 | 0.314595772344974 | 0.745993051875022 | NM_008786 |
| Slc6a19 | 8.716080356 | 0.226218736 | 0.14655807 | 1.543543355 | 0.122698997 | 0.30937319 | | Pex5l | 8945.396087 | 0.0561073956241344 | 0.0559017175874171 | 1.00367927937806 | 0.315533227643002 | 0.745993051875022 | NM_021483,NM_001163517,NM_001163516 |
| Fech | 2277.719581 | 0.090720696 | 0.058778461 | 1.543434344 | 0.122725426 | 0.309389481 | | Ppm1b | 4285.478556 | 0.0584650002106859 | 0.0583409247726935 | 1.00212673073791 | 0.316282389136695 | 0.745993051875022 | NM_001159498,NM_011151,NM_001159497,NM_001159496 |
| Entpd2 | 418.0586274 | -0.192747317 | 0.124898611 | -1.543230268 | 0.122774916 | 0.309463893 | | Prep | 989.3502471 | 0.0583500225444755 | 0.058227759344179 | 1.00209974077096 | 0.316295423098291 | 0.745993051875022 | NM_011156 |
| Cyb5r3 | 2756.10856 | 0.125015232 | 0.081021813 | 1.542982407 | 0.122835045 | 0.30946442 | | Ptch1 | 1692.131562 | 0.07517909799246 | 0.0750486990990611 | 1.0017375237008 | 0.316470378631905 | 0.745993051875022 | NM_008957 |
| Scn9a | 107.6080588 | -0.257127493 | 0.166643052 | -1.542983576 | 0.122834761 | 0.30946442 | | Rexo1 | 2422.990928 | -0.0568137929764338 | 0.0566308274181742 | -1.00323084734236 | 0.315749492695092 | 0.745993051875022 | NM_025852 |
| Wwc2 | 1731.746708 | -0.082245775 | 0.053301072 | -1.543041675 | 0.122820665 | 0.30946442 | | Rfc4 | 137.692496 | -0.0906329064693155 | 0.0901738378087567 | -1.00509092960568 | 0.314853067277325 | 0.745993051875022 | NM_145480 |
| 9330020H09Rik | 57.48230374 | 0.235052301 | 0.152351787 | 1.542826014 | 0.122872996 | 0.30949535 | | Rpl7a | 8086.812699 | 0.0473345737190929 | 0.0472167301971272 | 1.00249580014274 | 0.316104193963846 | 0.745993051875022 | NM_013721 |
| Cep78 | 346.1385283 | 0.16220652 | 0.105139985 | 1.542767193 | 0.122887272 | 0.30949535 | | Rps9 | 5663.688088 | 0.0529822564771002 | 0.0526630936663471 | 1.00606046452142 | 0.314386485224681 | 0.745993051875022 | NM_029767 |
| 4922501L14Rik | 244.804466 | -0.164467501 | 0.106649421 | -1.54213216 | 0.123041481 | 0.309806965 | | Rqcd1 | 1361.020674 | 0.0600390646274555 | 0.0597220050356931 | 1.0053089240988 | 0.314748119275958 | 0.745993051875022 | NM_021383 |
| Il20rb | 14.31147771 | 0.254841724 | 0.165257038 | 1.542093017 | 0.123050992 | 0.309806965 | | Sfxn5 | 8006.291593 | -0.0510864353704403 | 0.0509183002588528 | -1.00330205664236 | 0.315715144128497 | 0.745993051875022 | NM_178639 |
| Asb8 | 1691.617119 | 0.092412689 | 0.059941274 | 1.541720468 | 0.123141536 | 0.309984558 | | Slbp | 847.7891587 | 0.0646794487364177 | 0.0644278941772398 | 1.00390443553045 | 0.31542467841624 | 0.745993051875022 | NM_009193 |
| Ngf | 72.43765657 | -0.219821194 | 0.142612906 | -1.541383595 | 0.123223454 | 0.310073074 | | Slc39a3 | 3381.566683 | 0.0581827012420408 | 0.0580174050912157 | 1.00284907866123 | 0.315933684741195 | 0.745993051875022 | NM_134135 |
| Pi4kb | 2173.92391 | 0.092247076 | 0.059842769 | 1.541490763 | 0.123197389 | 0.310073074 | | Smcr7l | 1296.828766 | 0.0628316178329101 | 0.0625635636581079 | 1.00428450937141 | 0.315241498037279 | 0.745993051875022 | NM_178719 |
| Srprb | 1526.322889 | -0.075985353 | 0.049298595 | -1.541328993 | 0.123236736 | 0.310073074 | | Stab1 | 786.0820661 | -0.088376124130399 | 0.0879293697529807 | -1.00508083224835 | 0.314857928954969 | 0.745993051875022 | NM_138672 |
| Lrp3 | 3362.099759 | 0.087898537 | 0.057080072 | 1.539916349 | 0.123580745 | 0.31078722 | | Syf2 | 1530.957202 | 0.0555829886787454 | 0.0554930108784955 | 1.00162142581247 | 0.316526468851719 | 0.745993051875022 | NM_026780 |
| Nkd2 | 127.792785 | 0.221538817 | 0.143863903 | 1.539919415 | 0.123579998 | 0.31078722 | | Syt13 | 11629.25394 | 0.0565141514961957 | 0.0564061457227156 | 1.0019147873356 | 0.316384750068594 | 0.745993051875022 | NM_030725 |
| Zscan18 | 940.5353363 | 0.127934049 | 0.083070255 | 1.54007051 | 0.123543167 | 0.31078722 | | Taf11 | 1237.001955 | 0.0620915638408246 | 0.0619617216414039 | 1.00209552278376 | 0.316297460075324 | 0.745993051875022 | NM_026836 |
| Zfp326 | 819.4951241 | 0.097594055 | 0.063391104 | 1.539554428 | 0.123669001 | 0.310958699 | | Thoc2 | 1621.198615 | -0.0686341190869642 | 0.0682778770371831 | -1.00521753260704 | 0.314792114626301 | 0.745993051875022 | NM_001033422 |
| Lfng | 616.7256274 | -0.166490022 | 0.10814989 | -1.539437742 | 0.123697466 | 0.310979805 | | Trak2 | 9143.460176 | 0.0534282634351181 | 0.0532604812907811 | 1.00315021832831 | 0.315788387923703 | 0.745993051875022 | NM_172406 |
| Utp20 | 1045.108786 | 0.113320828 | 0.073621071 | 1.539244502 | 0.123744617 | 0.311047874 | | Zfp606 | 507.0176629 | -0.0705902955107963 | 0.0702444845887923 | -1.0049229619098 | 0.314933946813614 | 0.745993051875022 | NM_001039951,NM_026112 |
| Serpinb8 | 419.5140329 | 0.230648982 | 0.149907876 | 1.538604833 | 0.123900798 | 0.311389937 | | Mpp2 | 7998.30909 | 0.0571876132853785 | 0.057104994574471 | 1.00144678607402 | 0.316610854604314 | 0.745999882235567 | NM_016695 |
| Ecd | 1678.802068 | -0.084937738 | 0.055215224 | -1.538302887 | 0.123974574 | 0.311524822 | | Tox4 | 3499.018986 | 0.0505751808225647 | 0.0505077013957531 | 1.00133602252621 | 0.316664383092648 | 0.746004999806001 | NM_023434 |
| Kctd17 | 5578.965209 | 0.109748318 | 0.071349199 | 1.538185702 | 0.124003216 | 0.311545238 | | Tut1 | 1004.334645 | 0.0575477201162439 | 0.05748198477553 | 1.00114358161032 | 0.316757397765463 | 0.746103123064879 | NM_197993 |
| Slc44a1 | 3219.778995 | 0.1299815 | 0.084507553 | 1.538105121 | 0.124022914 | 0.311545238 | | Mpp7 | 230.502208 | 0.0879443907412625 | 0.087861917261515 | 1.00093867152366 | 0.316856459006981 | 0.746215454529086 | NM_001081287,NM_001161620 |
| Mtg1 | 577.7272388 | 0.112033781 | 0.072844418 | 1.537987169 | 0.124051752 | 0.311567166 | | Bmp3 | 492.500929 | -0.0864676111529212 | 0.0864536925994063 | -1.00016099432073 | 0.317232602309732 | 0.746980188223564 | NM_173404 |
| Cyp46a1 | 8551.30337 | 0.120612562 | 0.07843031 | 1.537830998 | 0.124089942 | 0.311612571 | | Wdr36 | 1247.389238 | 0.0622381516810805 | 0.0622506198961803 | 0.999799709382485 | 0.317407446501561 | 0.747270756544842 | NM_001110016,NM_144863,NM_001110015 |
| Pter | 75.1868393 | -0.236713303 | 0.153944381 | -1.537654708 | 0.124133063 | 0.311670343 | | Dhcr24 | 5718.990761 | 0.0612344036449649 | 0.0612533511600996 | 0.999690669738457 | 0.317460228750936 | 0.747273907443866 | NM_053272 |
| Arl13b | 355.1801469 | 0.141426344 | 0.092025564 | 1.536815845 | 0.124338412 | 0.311791886 | | Zfp524 | 177.1816601 | -0.0869459513092139 | 0.0869855489577887 | -0.999544779000084 | 0.317530858315908 | 0.747319061551666 | NM_025324 |
| Casc3 | 2670.207624 | 0.095995838 | 0.062457073 | 1.536989069 | 0.124295986 | 0.311791886 | | Dock1 | 1565.847599 | -0.0710388637928934 | 0.0711519790103045 | -0.998410230903139 | 0.318080474571557 | 0.748272127339361 | NM_001033420 |
| Cd200 | 3439.861047 | -0.122262203 | 0.079543553 | -1.537047292 | 0.124281729 | 0.311791886 | | Fam131c | 110.7094655 | 0.094944356758266 | 0.0950974767604112 | 0.998389862619268 | 0.318090347399184 | 0.748272127339361 | NM_001085513 |
| Gabpa | 848.6126413 | -0.128708299 | 0.083729289 | -1.537195648 | 0.124245406 | 0.311791886 | | Srebf1 | 2843.644072 | -0.0830895596038413 | 0.0832074933141256 | -0.998582655172184 | 0.31799690585936 | 0.748272127339361 | NM_011480 |
| Ldlrad4 | 1397.217027 | -0.13889031 | 0.090376348 | -1.536799322 | 0.124342459 | 0.311791886 | | 3830408C21Rik | 109.9481873 | -0.0941639460409292 | 0.0943381970786995 | -0.998152911088338 | 0.318205216287439 | 0.748361825040592 | NR_015471 |
| Mgea5 | 9858.000217 | -0.153170218 | 0.099666538 | -1.536826916 | 0.1243357 | 0.311791886 | | Slc2a4 | 90.62613659 | -0.0949112338518504 | 0.0950920352784149 | -0.998098669083745 | 0.318231515439607 | 0.748361825040592 | NM_009204 |
| Pafah1b2 | 7673.676699 | -0.083864261 | 0.054555366 | -1.537232124 | 0.124236476 | 0.311791886 | | Cramp1l | 1956.627087 | -0.0573289156715296 | 0.0574706034477239 | -0.997534604342145 | 0.318505085749128 | 0.748510369766394 | NM_020608 |
| Wscd2 | 2302.40745 | -0.123894535 | 0.080610742 | -1.536948204 | 0.124305994 | 0.311791886 | | Gm10509 | 173.4146698 | 0.0919309364348262 | 0.0921808126874733 | 0.997289281300934 | 0.31862411499578 | 0.748510369766394 | NR_045766 |
| 1110032A03Rik | 1002.234043 | 0.091830314 | 0.059768463 | 1.536434262 | 0.124431909 | 0.311793329 | | H2afv | 991.4426575 | 0.0554087769780359 | 0.0555282803437767 | 0.997847882826536 | 0.318353127279098 | 0.748510369766394 | NM_029938 |
| A230070E04Rik | 556.1036033 | 0.128453368 | 0.083604159 | 1.536447108 | 0.124428761 | 0.311793329 | | Litaf | 877.2649918 | -0.0864709336215652 | 0.0866760127609533 | -0.997633957390799 | 0.31845688854069 | 0.748510369766394 | NM_019980 |
| BC017643 | 741.5838149 | 0.118854939 | 0.077346125 | 1.536663142 | 0.124375821 | 0.311793329 | | Pgrmc2 | 3387.956748 | 0.0499228194153793 | 0.0500611287663737 | 0.99723719072257 | 0.318649392776091 | 0.748510369766394 | NM_027558 |
| Myo1d | 825.1966961 | 0.20190028 | 0.131416296 | 1.536341271 | 0.124454702 | 0.311793329 | | Ppp6r1 | 8269.656714 | -0.0576146511385758 | 0.057746906476006 | -0.997709741603473 | 0.318420128033946 | 0.748510369766394 | NM_172894 |
| Pspc1 | 1437.575555 | -0.077416294 | 0.050383875 | -1.53652919 | 0.124408644 | 0.311793329 | | Ralgapb | 7376.947074 | 0.0591087650624694 | 0.0592764820968267 | 0.997170597369742 | 0.318681710170419 | 0.748510369766394 | NM_177658 |
| Scrn1 | 4910.807867 | -0.096246541 | 0.062648099 | -1.536304258 | 0.124463776 | 0.311793329 | | Wsb2 | 12913.47007 | 0.0552792745260283 | 0.055439012909563 | 0.99711866472451 | 0.318706914291969 | 0.748510369766394 | NM_021539 |
| Zmat3 | 4391.916504 | -0.10303965 | 0.067088879 | -1.535867821 | 0.124570802 | 0.312010993 | | Crip1 | 102.8186905 | -0.0940625479843059 | 0.0943729433756743 | -0.996710970535986 | 0.318904823113228 | 0.748792455885777 | NM_007763 |
| Tbx4 | 12.68325473 | -0.254396703 | 0.165672293 | -1.535541635 | 0.124650838 | 0.312160997 | | Tcap | 167.7699443 | 0.0944512529956377 | 0.0947678847780217 | 0.996658870427195 | 0.318930120099498 | 0.748792455885777 | NM_011540 |
| Jph1 | 777.9091553 | 0.147878178 | 0.096311941 | 1.535408559 | 0.124683503 | 0.312192338 | | Clu | 32161.34513 | -0.0690959063158567 | 0.0693399831934219 | -0.996479998028204 | 0.319016980828991 | 0.748875349631856 | NM_013492 |
| B3galt2 | 1415.88066 | -0.160744538 | 0.104725652 | -1.534910837 | 0.124805731 | 0.312304373 | | Cbx4 | 1767.639275 | 0.056816981541389 | 0.0570265694724648 | 0.9963247319098 | 0.319092390863404 | 0.748931341037617 | NM_007625 |
| Llgl1 | 2549.183849 | 0.106352478 | 0.069288177 | 1.534929647 | 0.12480111 | 0.312304373 | | Creb5 | 102.8163048 | -0.0946257053784815 | 0.0950466890838903 | -0.995570769382222 | 0.319458743132638 | 0.749427923569881 | NM_172728 |
| Rps3a1 | 10357.69369 | -0.094115367 | 0.061317015 | -1.534898049 | 0.124808873 | 0.312304373 | | Gak | 5662.56018 | 0.0382856844052962 | 0.0384558828542901 | 0.995574189529369 | 0.319457080653505 | 0.749427923569881 | NM_153569 |
| Zdhhc6 | 946.0737395 | 0.107580214 | 0.070086733 | 1.534958311 | 0.124794069 | 0.312304373 | | Pcdhga6 | 945.7823497 | 0.0747434248000518 | 0.0750693105056719 | 0.995658869071464 | 0.319415921079984 | 0.749427923569881 | NM_033589 |
| Uaca | 459.1879067 | -0.146249924 | 0.095319779 | -1.53430826 | 0.124953835 | 0.312616618 | | Lamp1 | 13801.0205 | 0.0410703779408472 | 0.0412719762008469 | 0.995115371771427 | 0.319680155225871 | 0.749632194119936 | NM_010684 |
| D630037F22Rik | 539.8813358 | -0.1095411 | 0.071410401 | -1.533965614 | 0.125038112 | 0.312625547 | | Mutyh | 92.79023509 | -0.0946222056625884 | 0.0950906930663098 | -0.99507325702848 | 0.319700636277916 | 0.749632194119936 | NM_133250,NM_001159581 |
| Mb21d1 | 47.71202055 | 0.257909291 | 0.168121508 | 1.534064821 | 0.125013707 | 0.312625547 | | Otub1 | 7709.163798 | 0.0450019947434752 | 0.0452163818492635 | 0.995258640850499 | 0.319610487679361 | 0.749632194119936 | NM_134150 |
| Med14 | 3442.869361 | -0.093624169 | 0.061032924 | -1.533994499 | 0.125031006 | 0.312625547 | | Aox3 | 92.43776219 | -0.0944236974327682 | 0.0949060328060073 | -0.994917758555718 | 0.319776265029828 | 0.749688512883035 | NM_023617 |
| Slc9a3 | 16.98757312 | 0.244308391 | 0.159240354 | 1.534211556 | 0.124977616 | 0.312625547 | | Ccnf | 246.8331458 | -0.0884086210091161 | 0.0889145380235583 | -0.994310075430992 | 0.320071932022107 | 0.750154747972766 | NM_007634 |
| Dedd2 | 448.3808762 | 0.150908013 | 0.098385787 | 1.533839567 | 0.125069126 | 0.312636269 | | H2afz | 3022.699343 | 0.0496047202270282 | 0.0498993551784197 | 0.994095415655414 | 0.320176417013001 | 0.750154747972766 | NM_016750 |
| Fam110b | 1221.767599 | -0.123869039 | 0.080764727 | -1.533702191 | 0.125102934 | 0.312636269 | | Phf17 | 1699.710454 | 0.0577238614167984 | 0.0580638161045119 | 0.99414515423682 | 0.320152204925435 | 0.750154747972766 | NM_001130186,NM_172303,NM_001130184,NM_001130185 |
| Nadsyn1 | 151.4422747 | 0.209686345 | 0.13671489 | 1.533749142 | 0.125091379 | 0.312636269 | | Wwtr1 | 594.5628202 | -0.0714114833418048 | 0.0718374190950452 | -0.994070837195907 | 0.320188381925949 | 0.750154747972766 | NM_133784,NM_001168281 |
| Dynll2 | 28312.97122 | -0.102867753 | 0.067084072 | -1.533415476 | 0.125173518 | 0.312711785 | | Zfp947 | 69.90886284 | 0.0935929469697073 | 0.0941599332963618 | 0.99397847569762 | 0.320233346563343 | 0.750154747972766 | NM_177596 |
| Mrpl36 | 659.9605518 | -0.177549201 | 0.115786299 | -1.533421503 | 0.125172034 | 0.312711785 | | Il33 | 3007.669493 | 0.0879031803194287 | 0.0884627484958383 | 0.993674533225294 | 0.320381344973948 | 0.750259457857741 | NM_001164724,NM_133775 |
| Prdm6 | 22.54000388 | -0.257689028 | 0.168071788 | -1.533208109 | 0.125224587 | 0.312788933 | | Ptpru | 1997.330321 | 0.0800475034466908 | 0.0805527667868386 | 0.993727548285401 | 0.320355527183816 | 0.750259457857741 | NM_011214,NM_001083119 |
| Ikzf4 | 477.0708976 | 0.130970968 | 0.08543456 | 1.532997507 | 0.125276469 | 0.312868087 | | Hipk3 | 4041.295141 | 0.0590040962382661 | 0.0593892802515511 | 0.993514250187011 | 0.320459409440642 | 0.750321305856668 | NM_001145824,NM_010434 |
| C78339 | 902.6663009 | -0.112901925 | 0.073678058 | -1.532368368 | 0.125431559 | 0.313154459 | | Atp5j2 | 3134.284665 | 0.051544176928609 | 0.0519240465905704 | 0.99268412832003 | 0.320863912040904 | 0.750352873820335 | NM_020582 |
| Tsnax | 8337.412509 | -0.090925277 | 0.059335431 | -1.532394315 | 0.125425159 | 0.313154459 | | Bach2 | 470.4892537 | 0.0882613281029977 | 0.0889395808438704 | 0.992374005651507 | 0.321015114480273 | 0.750352873820335 | NM_001109661 |
| Zscan29 | 344.3875504 | 0.170935063 | 0.11158209 | 1.531922036 | 0.125541675 | 0.313378881 | | Car8 | 533.890716 | 0.0893482146620937 | 0.0900262090663376 | 0.992468921980884 | 0.320968832428903 | 0.750352873820335 | NM_007592 |
| Kin | 303.8894087 | 0.162203401 | 0.105894822 | 1.531740623 | 0.125586454 | 0.313426464 | | Ccni | 8370.714697 | 0.055831391930265 | 0.0562106318292708 | 0.993253235434933 | 0.320586561023963 | 0.750352873820335 | NM_017367 |
| Tmeff2 | 2594.284649 | -0.086801265 | 0.056670593 | -1.531680911 | 0.125601195 | 0.313426464 | | Cox10 | 1120.894586 | 0.0600539346084575 | 0.0605133027915172 | 0.992408806628149 | 0.320998144706526 | 0.750352873820335 | NM_178379 |
| Mrps17 | 1171.016773 | -0.101954791 | 0.066587177 | -1.531147521 | 0.125732937 | 0.313704691 | | Fam179b | 2708.567775 | 0.0758739639201845 | 0.0764692938942731 | 0.992214783950906 | 0.321092762196558 | 0.750352873820335 | NM_177805 |
| Sft2d1 | 763.1272525 | -0.140007445 | 0.091451484 | -1.530947765 | 0.125782303 | 0.313777331 | | Frmpd3 | 169.1838599 | 0.0901294180596996 | 0.0908343605487816 | 0.992239253022501 | 0.321080828555929 | 0.750352873820335 | NM_177750 |
| Fam228b | 82.57758252 | 0.234938455 | 0.153473951 | 1.530803465 | 0.125817973 | 0.313815788 | | Npm1 | 4909.19626 | 0.0592033823835286 | 0.0596119899606185 | 0.993145547106214 | 0.32063903027701 | 0.750352873820335 | NM_001252260,NM_001252261,NM_008722 |
| Abhd8 | 7917.9002 | -0.120941129 | 0.079018438 | -1.530543153 | 0.125882341 | 0.313817527 | | Slc35e4 | 1044.884672 | 0.0734281310976027 | 0.0739776656842556 | 0.992571614938508 | 0.320918763334078 | 0.750352873820335 | NM_153142 |
| Gnptg | 2314.955989 | 0.095953097 | 0.06269506 | 1.53047302 | 0.125899687 | 0.313817527 | | Stx12 | 4962.126657 | 0.0598208430816495 | 0.0602730485277147 | 0.9924973855295 | 0.320954954197661 | 0.750352873820335 | NM_133887 |
| Prpf4 | 929.2660522 | 0.098546855 | 0.064386548 | 1.530550376 | 0.125880554 | 0.313817527 | | Tmem86a | 641.9822601 | -0.0851220388643642 | 0.0857818949683626 | -0.992307746241304 | 0.32104742574543 | 0.750352873820335 | NM_026436 |
| Txlnb | 93.76386259 | 0.234075461 | 0.152935189 | 1.530553317 | 0.125879827 | 0.313817527 | | Wdr46 | 381.8981641 | -0.0735768809748432 | 0.0740800459665531 | -0.993207820201176 | 0.320608688117304 | 0.750352873820335 | NM_020603 |
| Dusp23 | 126.8272405 | 0.182292233 | 0.119117398 | 1.530357746 | 0.125928202 | 0.313838115 | | Mcf2l | 7751.723824 | -0.0706316707421296 | 0.071202803502802 | -0.991978788297993 | 0.32120787286683 | 0.750501135799191 | NM_001159486,NM_001159485,NM_178076 |
| AI661453 | 25.76717833 | -0.254481273 | 0.166321647 | -1.530055034 | 0.126003107 | 0.313873335 | | Zfp933 | 585.3369672 | 0.0642374566224249 | 0.0647643566419192 | 0.991864351831555 | 0.321263700794357 | 0.750510858433392 | NM_198619 |
| Bbs7 | 707.1554011 | 0.095635938 | 0.062498452 | 1.530212913 | 0.125964036 | 0.313873335 | | Helz | 1077.913573 | 0.0805903456773254 | 0.0812831414066797 | 0.991476759911529 | 0.321452834923333 | 0.750831947271557 | NM_198298 |
| Setd4 | 226.5659771 | 0.194679247 | 0.127235392 | 1.530071506 | 0.12599903 | 0.313873335 | | Vipas39 | 2416.619174 | 0.053336362985441 | 0.0538094869255337 | 0.991207425174906 | 0.321584305627861 | 0.751018267451819 | NM_134044,NM_001142581,NM_001142580 |
| Masp2 | 26.44467142 | 0.256575984 | 0.167703897 | 1.529934539 | 0.126032933 | 0.313897165 | | Arhgap22 | 232.8388054 | -0.0847865841860496 | 0.0855589902906467 | -0.990972239130299 | 0.321699135998296 | 0.751165672642863 | NM_153800 |
| Magi3 | 2953.385851 | -0.095281396 | 0.062290191 | -1.529637249 | 0.126106544 | 0.313979558 | | Clec16a | 4805.594631 | 0.04989629901378 | 0.0503633511865826 | 0.990726348390276 | 0.321819221592898 | 0.751204565781526 | NM_001204229,NM_177562 |
| Pabpc4 | 1187.027774 | -0.128348213 | 0.083906512 | -1.529657365 | 0.126101562 | 0.313979558 | | Nr2f2 | 1013.33636 | 0.0933610380566391 | 0.094229241240012 | 0.990786265792362 | 0.321789957051287 | 0.751204565781526 | NM_183261,NM_009697 |
| Tmem165 | 787.9616893 | 0.151522882 | 0.099068689 | 1.529472961 | 0.126147237 | 0.314030413 | | Carm1 | 3486.052182 | 0.0445064716105298 | 0.0449282715130529 | 0.990611704204989 | 0.321875220343424 | 0.751214564702535 | NM_021531,NM_153141 |
| Gabra2 | 3495.688616 | -0.181129476 | 0.118433795 | -1.529373232 | 0.126171944 | 0.314041463 | | Wbscr17 | 2494.242365 | -0.0519018850741573 | 0.0524161036040246 | -0.99018968419034 | 0.322081413755251 | 0.751575038748398 | NM_145218 |
| Btbd9 | 3734.743756 | 0.142172099 | 0.092998591 | 1.528755414 | 0.126325089 | 0.314372138 | | Frrs1l | 8881.919733 | 0.0779369942505634 | 0.0787464010792415 | 0.989721348308177 | 0.322310337377476 | 0.7519884292877 | NM_001142965 |
| Vgll3 | 73.1959661 | 0.254259908 | 0.166337744 | 1.528576143 | 0.126369554 | 0.31443229 | | Fam120b | 3615.372369 | 0.0548985025623979 | 0.055492385802766 | 0.989297932828141 | 0.322517395161069 | 0.752229878309197 | NM_024203,NR_033586 |
| 2610301G19Rik | 3788.230515 | 0.081520342 | 0.053351053 | 1.527998741 | 0.12651285 | 0.314473278 | | Fsd1l | 1140.175678 | -0.0735529452315061 | 0.0743439947698056 | -0.989359604084381 | 0.322487231410219 | 0.752229878309197 | NM_176966,NM_001195284 |
| Gm16675 | 8.892258581 | 0.238735605 | 0.156244148 | 1.527965098 | 0.126521204 | 0.314473278 | | Pla2r1 | 32.16614687 | -0.0878417314024787 | 0.088813321645937 | -0.989060309585857 | 0.322633635203484 | 0.752380187022926 | NM_008867 |
| Gng5 | 394.9041953 | -0.142666495 | 0.093356796 | -1.528185425 | 0.126466507 | 0.314473278 | | 2610034B18Rik | 787.1820655 | 0.0697926118316832 | 0.0706239787628152 | 0.988228262614259 | 0.323040869537886 | 0.752792381840132 | NM_027420 |
| Mrpl38 | 1275.735991 | 0.115894706 | 0.07585595 | 1.527826181 | 0.1265557 | 0.314473278 | | Ccdc113 | 120.2000192 | 0.0931488701184244 | 0.0942564781339639 | 0.988248998504216 | 0.323030716561879 | 0.752792381840132 | NM_172914 |
| Myef2 | 2068.851141 | 0.108391306 | 0.070944305 | 1.52783659 | 0.126553115 | 0.314473278 | | Eif1ax | 1825.554996 | 0.0710935877860235 | 0.0719447102331511 | 0.988169770308763 | 0.323069510422097 | 0.752792381840132 | NM_025437 |
| Nr2c2 | 6569.09945 | -0.095226195 | 0.062312868 | -1.528194703 | 0.126464203 | 0.314473278 | | Rrp36 | 500.706703 | 0.0665017110596156 | 0.0672921760209866 | 0.988253241192195 | 0.323028639227476 | 0.752792381840132 | NM_144857 |
| Odc1 | 2486.604891 | -0.088383656 | 0.057844959 | -1.527940521 | 0.126527306 | 0.314473278 | | Twf2 | 1636.708815 | 0.0577775608178072 | 0.0584604885849737 | 0.988318131036935 | 0.322996868495603 | 0.752792381840132 | NM_011876 |
| Plekha7 | 690.5490425 | -0.17127636 | 0.112110803 | -1.527741793 | 0.126576659 | 0.314473278 | | AI118078 | 288.6246048 | -0.0832596468753628 | 0.0844303160724539 | -0.986134492306217 | 0.324067119682406 | 0.752925263857917 | NM_172923 |
| Prr7 | 580.9911503 | -0.220938615 | 0.144622473 | -1.527692144 | 0.126588992 | 0.314473278 | | Ankrd46 | 8837.500308 | 0.0443248417264863 | 0.0449412123058455 | 0.986284958777604 | 0.323993298618192 | 0.752925263857917 | NM_175134 |
| Zfp931 | 124.8478526 | -0.221603171 | 0.144993508 | -1.528366157 | 0.126421653 | 0.314473278 | | Asl | 551.5271567 | -0.0649405689966683 | 0.0657691949754888 | -0.98740100165238 | 0.323446093525733 | 0.752925263857917 | NM_133768 |
| Rnaseh2b | 272.0695826 | -0.151580484 | 0.099237444 | -1.527452519 | 0.126648525 | 0.314520316 | | BC053749 | 123.9181796 | -0.0920853744939924 | 0.0933944422754257 | -0.985983450946976 | 0.324141233812314 | 0.752925263857917 | NM_183321 |
| Zc3h15 | 3673.107715 | -0.095440584 | 0.062481449 | -1.527502734 | 0.126636048 | 0.314520316 | | Ccdc111 | 182.7030728 | -0.0869083038458685 | 0.0881317420301837 | -0.986118075552209 | 0.324075174646207 | 0.752925263857917 | NM_001001184 |
| Ttll5 | 1373.173005 | 0.100745602 | 0.065969067 | 1.527164264 | 0.12672017 | 0.314647807 | | Cwc25 | 730.0717149 | 0.0630464672837117 | 0.0638676142308501 | 0.987142983857667 | 0.323572548234139 | 0.752925263857917 | NM_026186 |
| Krit1 | 1476.235996 | 0.109521455 | 0.07172104 | 1.52704779 | 0.126749128 | 0.314669282 | | Flt1 | 2150.027807 | -0.0801622201408147 | 0.0812414322277234 | -0.986715988907191 | 0.323781889533048 | 0.752925263857917 | NM_010228 |
| 9130401M01Rik | 828.9907657 | -0.103335317 | 0.067682645 | -1.526762401 | 0.126820104 | 0.314691614 | | Gkap1 | 518.2502469 | 0.0819572238954156 | 0.0830950336765894 | 0.986307126541375 | 0.323982423713025 | 0.752925263857917 | NM_019832 |
| Dhcr24 | 5045.67271 | -0.167451222 | 0.109683771 | -1.526672726 | 0.126842413 | 0.314691614 | | Gzmm | 62.95828639 | 0.0929739514316943 | 0.0941305281139636 | 0.987713054357148 | 0.323293199306525 | 0.752925263857917 | NM_008504 |
| Elf1 | 281.7239634 | 0.173203506 | 0.113456784 | 1.52660334 | 0.126859676 | 0.314691614 | | Il17ra | 1136.140752 | -0.0829688901875821 | 0.084025529082373 | -0.987424787367241 | 0.323434437751263 | 0.752925263857917 | NM_008359 |
| Rps4y2 | 372.0673401 | 0.154649536 | 0.101296038 | 1.526708632 | 0.12683348 | 0.314691614 | | Kcna2 | 14557.04114 | 0.0672931738999031 | 0.0682589239378925 | 0.98585166623974 | 0.324205907957856 | 0.752925263857917 | NM_008417 |
| Selenbp1 | 339.79918 | 0.136134152 | 0.089156498 | 1.526912287 | 0.126782824 | 0.314691614 | | Klhdc3 | 7473.747242 | 0.0474695739706594 | 0.0481288748958014 | 0.986301343495576 | 0.323985260695913 | 0.752925263857917 | NM_001163729,NM_027910 |
| Ftl1 | 9881.961791 | 0.121371189 | 0.079512841 | 1.526435068 | 0.12690155 | 0.314745096 | | Matn2 | 1162.965927 | 0.0684675470782271 | 0.069395163891635 | 0.986632832010362 | 0.323822668832971 | 0.752925263857917 | NM_016762 |
| Szrd1 | 1303.567361 | -0.104345628 | 0.068370839 | -1.526171529 | 0.126967152 | 0.314857403 | | Mfsd11 | 1339.419073 | -0.0575245988779798 | 0.0582660290519257 | -0.987275086598313 | 0.323507800560875 | 0.752925263857917 | NM_178620 |
| Ankfy1 | 2567.334077 | -0.069960587 | 0.045846851 | -1.525962757 | 0.12701914 | 0.314885764 | | Nmt1 | 4675.311893 | 0.0531641736004901 | 0.0538469690637886 | 0.987319704801034 | 0.323485933695772 | 0.752925263857917 | NM_008707 |
| Pth1r | 202.2175711 | -0.201463055 | 0.132023606 | -1.525962375 | 0.127019235 | 0.314885764 | | Pfdn4 | 472.5566547 | 0.076441919955881 | 0.0774346771286813 | 0.987179423875552 | 0.323554687001508 | 0.752925263857917 | NM_001110152,NM_001199902,NM_001013369 |
| Msto1 | 658.1647394 | 0.111022208 | 0.072764263 | 1.525779326 | 0.127064831 | 0.314948407 | | Pi16 | 38.9080401 | 0.0719984009389345 | 0.0729792445476414 | 0.986559964894312 | 0.323858404879101 | 0.752925263857917 | NM_023734 |
| A530054K11Rik | 418.5197414 | -0.135319135 | 0.088708669 | -1.525433043 | 0.127151122 | 0.315111883 | | Pik3c2a | 1107.504017 | 0.0676153348289557 | 0.0685003476437235 | 0.987080170463208 | 0.323603337999853 | 0.752925263857917 | NM_011083 |
| Gpm6b | 32369.09858 | -0.106754575 | 0.06999386 | -1.52519914 | 0.127209435 | 0.315205979 | | Tceal7 | 24.2213352 | 0.0790382193432535 | 0.0801740385385263 | 0.985833079934884 | 0.324215029975993 | 0.752925263857917 | NM_001127169 |
| Dgki | 1495.198678 | -0.139036244 | 0.091182716 | -1.524809199 | 0.127306695 | 0.315245314 | | Uso1 | 4057.965747 | 0.0581716614729221 | 0.0589287302782225 | 0.987152806420127 | 0.323567733599343 | 0.752925263857917 | NM_019490 |
| Dhrs7 | 839.4037071 | -0.120004039 | 0.078701004 | -1.524809495 | 0.127306621 | 0.315245314 | | Zfp592 | 1673.355306 | 0.0599404210624036 | 0.0607568200089972 | 0.986562842715061 | 0.323856993467598 | 0.752925263857917 | NM_178707 |
| Epm2aip1 | 5274.824409 | -0.122051651 | 0.080043882 | -1.524809245 | 0.127306683 | 0.315245314 | | 2010002M12Rik | 35.30513387 | -0.0889098419693927 | 0.0904482006783772 | -0.982991826289008 | 0.32561146150868 | 0.752934080280062 | NM_053217 |
| Pi4k2a | 3613.738128 | 0.106515953 | 0.069845588 | 1.525020487 | 0.127253988 | 0.315245314 | | AW549877 | 2731.863108 | -0.0622813637592887 | 0.0632359760594955 | -0.984903968283676 | 0.324671243976161 | 0.752934080280062 | NM_145930 |
| Gareml | 584.3417319 | -0.151523935 | 0.099391866 | -1.524510412 | 0.127381258 | 0.315379548 | | Asb1 | 2696.664444 | 0.0599021270505996 | 0.0608459130280116 | 0.984488917489371 | 0.324875177894871 | 0.752934080280062 | NM_001039126,NM_023046 |
| Kcnab2 | 17102.65301 | 0.123800552 | 0.081231042 | 1.5240547 | 0.127495048 | 0.315610843 | | Ccnd1 | 2590.730164 | 0.0777446480523952 | 0.0790072475387383 | 0.984019194116033 | 0.325106075553924 | 0.752934080280062 | NM_007631 |
| Acox1 | 9186.537025 | 0.11379284 | 0.074669215 | 1.523959242 | 0.127518893 | 0.315619445 | | Chpf2 | 2494.170674 | -0.0819189035857678 | 0.0832358268924765 | -0.984178407833806 | 0.32502780035106 | 0.752934080280062 | NM_133913 |
| Zcchc11 | 1331.295381 | 0.118621235 | 0.077853425 | 1.52364824 | 0.127596606 | 0.31576135 | | Cryz | 631.4912646 | -0.0625023525724605 | 0.0634146003580576 | -0.985614546485409 | 0.324322297079569 | 0.752934080280062 | NM_009968 |
| Spsb4 | 138.2235986 | -0.201508258 | 0.132275992 | -1.523392527 | 0.127660531 | 0.315869093 | | Dnajc1 | 989.1359312 | -0.0763573505681839 | 0.0775030682677615 | -0.985217131073839 | 0.324517427580423 | 0.752934080280062 | NM_001190817,NM_007869 |
| Elavl4 | 2765.658371 | -0.122990911 | 0.080754398 | -1.523024299 | 0.127752626 | 0.316046494 | | Erc2 | 7315.038933 | 0.0626302051125272 | 0.0636752283016132 | 0.983588230196271 | 0.325318014480045 | 0.752934080280062 | NM_177814 |
| Gtpbp1 | 2413.407322 | -0.078252911 | 0.051389613 | -1.522737885 | 0.127824296 | 0.316173314 | | Fam131a | 11379.59752 | 0.0701063793237151 | 0.0712896566914131 | 0.98340183663922 | 0.325409706917266 | 0.752934080280062 | NM_133778 |
| A230056J06Rik | 34.02046686 | -0.242785555 | 0.159463805 | -1.522511992 | 0.127880843 | 0.316212222 | | Jtb | 909.6731735 | -0.0683848912278149 | 0.0695765597543053 | -0.982872557500708 | 0.325670165661547 | 0.752934080280062 | NM_206924 |
| C130071C03Rik | 83.45617654 | 0.217152485 | 0.142620159 | 1.522593202 | 0.127860511 | 0.316212222 | | Kcnq3 | 2021.236095 | 0.0721123025799521 | 0.073199225730044 | 0.985151166023252 | 0.324549823737573 | 0.752934080280062 | NM_152923 |
| Gm527 | 134.3110618 | 0.179355663 | 0.11782758 | 1.522187452 | 0.127962118 | 0.316362703 | | Kdelr2 | 848.2293423 | 0.0560665068154357 | 0.0568865739996488 | 0.985584169927016 | 0.324337209236853 | 0.752934080280062 | NM_025841 |
| Prkcz | 11507.05755 | -0.098739205 | 0.06488021 | -1.521869384 | 0.128041812 | 0.316509227 | | Mettl3 | 904.9520238 | -0.0673026709493681 | 0.0684028230209756 | -0.983916569185015 | 0.325156536169725 | 0.752934080280062 | NM_019721 |
| Pank1 | 605.9884951 | -0.119285948 | 0.078386132 | -1.521773625 | 0.128065812 | 0.316518057 | | Ndufa1 | 1428.3963 | 0.0632362600676815 | 0.0642695686850469 | 0.983922272414349 | 0.325153731761632 | 0.752934080280062 | NM_019443 |
| Accs | 109.8823617 | 0.225687114 | 0.148330583 | 1.521514373 | 0.128130807 | 0.316541113 | | Pld5 | 552.8031989 | 0.0782487181920512 | 0.0795018797262473 | 0.984237334532075 | 0.324998832971626 | 0.752934080280062 | NM_176916,NM_001195816 |
| Lipc | 9.028996654 | -0.242985268 | 0.159702207 | -1.521489734 | 0.128136986 | 0.316541113 | | Pmfbp1 | 23.20620155 | 0.069579690975586 | 0.0707451088751231 | 0.983526523344613 | 0.32534836795671 | 0.752934080280062 | NM_019938 |
| Rraga | 3250.114841 | -0.077485113 | 0.050929788 | -1.521410479 | 0.128156861 | 0.316541113 | | Ppp2r2a | 4567.930386 | 0.0690851563014067 | 0.0702357835493657 | 0.983617649155288 | 0.325303544001568 | 0.752934080280062 | NM_028032,NM_001205188 |
| Vwa3a | 281.1508617 | 0.18092853 | 0.11891681 | 1.521471435 | 0.128141574 | 0.316541113 | | Prrx1 | 244.7396249 | -0.0894798387020674 | 0.0909927883579581 | -0.983372861924629 | 0.325423961706787 | 0.752934080280062 | NM_175686,NM_011127,NM_001025570 |
| Bbs12 | 148.5834345 | -0.182136044 | 0.119755909 | -1.520894013 | 0.128286437 | 0.316760167 | | Rbm12 | 832.1186188 | 0.0749307912078331 | 0.0761671635308234 | 0.983767646507015 | 0.325229770421532 | 0.752934080280062 | NM_170598,NM_029397 |
| Src | 5026.393582 | 0.127580317 | 0.083883724 | 1.520918614 | 0.128280262 | 0.316760167 | | Rufy3 | 10205.35894 | 0.0499735336317066 | 0.050828094002046 | 0.983187243450343 | 0.325515291967002 | 0.752934080280062 | NM_027530 |
| 4930525G20Rik | 78.29518982 | -0.203919042 | 0.134090107 | -1.520761268 | 0.128319757 | 0.316791957 | | Scarf1 | 77.38514423 | -0.093323081508901 | 0.0947885767416292 | -0.984539420020326 | 0.324850359179366 | 0.752934080280062 | NM_001004157 |
| Ramp2 | 468.3035209 | 0.148535597 | 0.097707406 | 1.52020817 | 0.128458665 | 0.317033857 | | Scn2b | 15756.20764 | 0.0545182688264009 | 0.0554652724931572 | 0.982926187428843 | 0.325643768134531 | 0.752934080280062 | NM_001014761 |
| Zfp367 | 759.1093129 | 0.131086963 | 0.086225268 | 1.520284782 | 0.128439417 | 0.317033857 | | Sfxn1 | 2422.935702 | 0.0533426614489893 | 0.0541746018501008 | 0.984643349970279 | 0.324799288238493 | 0.752934080280062 | NM_027324 |
| Gm14378 | 106.4429904 | 0.25291999 | 0.166389607 | 1.520046804 | 0.128499213 | 0.317061653 | | Sytl1 | 80.41864933 | -0.0933703423028584 | 0.0949895100103827 | -0.982954247186375 | 0.325629957217819 | 0.752934080280062 | NM_031393 |
| Tmem9 | 1972.796631 | -0.10652992 | 0.070085451 | -1.520000482 | 0.128510855 | 0.317061653 | | Tmem218 | 569.4828748 | 0.0700628970094967 | 0.0711635575219758 | 0.984533368611607 | 0.324853332988841 | 0.752934080280062 | NM_025464 |
| Lancl3 | 265.7053155 | -0.238291614 | 0.15678147 | -1.519896542 | 0.12853698 | 0.317075619 | | Tmem44 | 3349.109905 | -0.0581493981682182 | 0.0591103961485379 | -0.983742318730114 | 0.325242226682795 | 0.752934080280062 | NM_172614 |
| Arih2 | 2070.082701 | 0.127742952 | 0.084075107 | 1.519390889 | 0.128664134 | 0.317241369 | | Tnip1 | 1238.210685 | 0.0587188021255391 | 0.0595984810338458 | 0.985239910597602 | 0.324506240796439 | 0.752934080280062 | NM_001271456,NM_021327,NM_001271455,NM_001199275,NM_001199276 |
| Caskin2 | 1261.914063 | -0.11045505 | 0.072697206 | -1.519385072 | 0.128665597 | 0.317241369 | | Ube2q2 | 2864.024775 | 0.057819161990883 | 0.0587663012913127 | 0.983882951970472 | 0.325173066841509 | 0.752934080280062 | NM_180600 |
| Oas2 | 35.49565973 | 0.248957196 | 0.16384446 | 1.519472775 | 0.128643536 | 0.317241369 | | Bcl2l11 | 283.4081948 | -0.0817085456287427 | 0.0831570461084306 | -0.982581145585677 | 0.325813627690948 | 0.753033837474394 | NM_009754,NM_207680,NM_207681 |
| Ccar1 | 1945.599696 | 0.16438698 | 0.108218864 | 1.51902334 | 0.128756621 | 0.317415289 | | Cwc27 | 684.7848398 | -0.0744945105048276 | 0.0758319004769913 | -0.982363755045682 | 0.325920675767693 | 0.753033837474394 | NM_026072 |
| Lcor | 166.1143048 | -0.19860799 | 0.13077495 | -1.518700568 | 0.128837884 | 0.317565092 | | Hfe2 | 55.13705992 | -0.0892445819231312 | 0.0908405857007424 | -0.982430718986456 | 0.325887698754171 | 0.753033837474394 | NM_027126 |
| Cct5 | 5433.127598 | -0.097712414 | 0.064346773 | -1.518528579 | 0.128881201 | 0.317621333 | | Trib1 | 979.817492 | 0.0841136883932034 | 0.0856078077051271 | 0.982546927062189 | 0.325830476158808 | 0.753033837474394 | NM_144549 |
| Mboat7 | 4885.682807 | -0.088584645 | 0.058366353 | -1.517734798 | 0.129081268 | 0.317941173 | | Abcb6 | 889.9380279 | -0.0648650793996795 | 0.066059570015727 | -0.98191797773187 | 0.326140258228427 | 0.753049595558931 | NM_023732 |
| Pdf | 989.3260155 | -0.105832165 | 0.06973249 | -1.51768803 | 0.129093063 | 0.317941173 | | Acat2 | 1274.60324 | 0.0593333860457831 | 0.0604176259918155 | 0.982054244465361 | 0.32607312530896 | 0.753049595558931 | NM_009338 |
| Rab33b | 1142.061642 | -0.124029826 | 0.081711226 | -1.517904349 | 0.129038513 | 0.317941173 | | LOC106740 | 505.4102921 | 0.0690999689490721 | 0.070354953270968 | 0.982162104250678 | 0.326019993673948 | 0.753049595558931 | NR_027905 |
| Ttc30a1 | 130.4108405 | -0.196261325 | 0.129311248 | -1.51774365 | 0.129079036 | 0.317941173 | | Rbms2 | 648.6625673 | 0.061620036450257 | 0.0627607946326881 | 0.981823713528366 | 0.326186703514856 | 0.753049595558931 | NM_001039080,NM_019711 |
| Col20a1 | 127.0899322 | 0.193234701 | 0.127357871 | 1.517257619 | 0.129201654 | 0.318158046 | | Zscan21 | 711.8900308 | 0.0642784352890694 | 0.0654641302043675 | 0.98188786879782 | 0.326155092852487 | 0.753049595558931 | NM_001044704,NM_001044705,NM_011757,NM_001044703 |
| Rnf187 | 10570.70324 | -0.096314512 | 0.063496432 | -1.5168492 | 0.129304762 | 0.318361351 | | Cenpf | 36.75370689 | -0.0869762910038819 | 0.0886232592728816 | -0.981416072005111 | 0.326387603662932 | 0.753373721590492 | NM_001081363 |
| Slc25a44 | 4663.980415 | 0.156252599 | 0.103019078 | 1.516734584 | 0.129333709 | 0.318382028 | | Lgals3 | 42.42317168 | 0.0743519200135507 | 0.075790976868016 | 0.981012820866905 | 0.326586419169443 | 0.753373721590492 | NM_010705,NM_001145953 |
| Lamc1 | 1559.00265 | -0.092684264 | 0.061120119 | -1.516428068 | 0.129411146 | 0.318468986 | | Nabp1 | 359.0363227 | -0.0750406181054225 | 0.076488811400548 | -0.981066599563931 | 0.326559900032702 | 0.753373721590492 | NM_028696 |
| Pou3f2 | 900.8554901 | -0.115701213 | 0.07630001 | -1.516398408 | 0.129418641 | 0.318468986 | | Rgmb | 1890.494421 | 0.0532310867103727 | 0.0542530733564158 | 0.981162603649582 | 0.326512562358841 | 0.753373721590492 | NM_178615 |
| Sdcbp2 | 31.37300917 | 0.254088054 | 0.167565493 | 1.516350707 | 0.129430696 | 0.318468986 | | Ubr3 | 8656.689115 | -0.0599005159435914 | [illegible] | [illegible] | [illegible] | [illegible] | [illegible] |
| Cd276 | 454.4088653 | -0.142511813 | 0.093995962 | -1.516148253 | 0.129481869 | 0.318544313 | | Yipf2 | 870.0851972 | 0.0640313272912415 | [illegible] | [illegible] | [illegible] | [illegible] | [illegible] |
| Cdc42bpa | 13614.45285 | -0.085738886 | 0.056560744 | -1.515872656 | 0.129551556 | 0.318665154 | | Cbwd1 | 441.3245274 | 0.076395554559744 | [illegible] | [illegible] | [illegible] | [illegible] | [illegible] |
| 4933427I22Rik | 14.89939805 | 0.25098492 | 0.165616121 | 1.515461896 | 0.129655473 | 0.318819536 | | Mthfd2 | 240.8989066 | 0.0838914864372429 | [illegible] | [illegible] | [illegible] | [illegible] | [illegible] |
| Rpl17 | 8876.804727 | -0.138518006 | 0.091402391 | -1.515474639 | 0.129652248 | 0.318819536 | | Efcc1 | 270.7722985 | 0.0820230992984304 | [illegible] | [illegible] | [illegible] | [illegible] | [illegible] |
| Tnfaip8 | 212.8371065 | -0.193765031 | 0.127873111 | -1.515291442 | 0.129698615 | 0.318875013 | | Eif4a2 | 24646.79695 | 0.0583074009171298 | [illegible] | [illegible] | [illegible] | [illegible] | [illegible] |
| C630020P19Rik | 134.9841611 | 0.183193383 | 0.120903562 | 1.51520253 | 0.129721122 | 0.318879751 | | Kcnip1 | 1203.638963 | 0.0837755911850598 | [illegible] | [illegible] | [illegible] | [illegible] | [illegible] |
| Agk | 1558.865458 | -0.094754138 | 0.06254512 | -1.514972509 | 0.129779366 | 0.318939271 | | Rps27l | 235.9756613 | -0.0806208747777376 | [illegible] | [illegible] | [illegible] | [illegible] | [illegible] |
| Rpl23a | 7023.904388 | -0.088383053 | 0.058340793 | -1.514944322 | 0.129786505 | 0.318939271 | | Eif4enif1 | 2523.070068 | 0.0477102783102491 | [illegible] | [illegible] | [illegible] | [illegible] | [illegible] |
| Thrsp | 494.995604 | 0.174464957 | 0.115182467 | 1.514683282 | 0.12985263 | 0.319051166 | | Rpia | 377.7627157 | -0.0736817883461582 | [illegible] | [illegible] | [illegible] | [illegible] | [illegible] |
| Gjd2 | 226.422726 | -0.170060557 | 0.112294426 | -1.514416726 | 0.12992018 | 0.319115926 | | Chd1 | 1127.176703 | -0.0737938757764013 | [illegible] | [illegible] | [illegible] | [illegible] | [illegible] |
| Pigt | 3224.80853 | 0.094457248 | 0.062370961 | 1.514442718 | 0.129913592 | 0.319115926 | | Abhd13 | 1780.513079 | 0.072929070332125 | [illegible] | [illegible] | [illegible] | [illegible] | [illegible] |
| Fam69c | 220.0243479 | -0.163187318 | 0.10776752 | -1.514253257 | 0.129961619 | 0.319167114 | | Mipep | 1225.524303 | 0.0506050432196633 | [illegible] | [illegible] | [illegible] | [illegible] | [illegible] |
| Eif4g3 | 11481.24289 | 0.073164337 | 0.048324587 | 1.51401887 | 0.130021054 | 0.319211885 | | Nexn | 435.134289 | -0.0930034332426272 | [illegible] | [illegible] | [illegible] | [illegible] | [illegible] |
| Rhbdl2 | 10.38032024 | -0.247437363 | 0.163429039 | -1.51403548 | 0.130016841 | 0.319211885 | | Nrip1 | 1017.411928 | 0.0709397345146866 | [illegible] | [illegible] | [illegible] | [illegible] | [illegible] |
| Bptf | 4846.532655 | -0.156658752 | 0.103479456 | -1.513911629 | 0.130048254 | 0.319228082 | | Plin5 | 29.82022386 | 0.0753329765137785 | [illegible] | [illegible] | [illegible] | [illegible] | [illegible] |
| Lyrm4 | 677.1186309 | -0.105418864 | 0.069653629 | -1.513472665 | 0.130159639 | 0.319450887 | | Thrb | 2834.719377 | 0.0643187982286281 | [illegible] | [illegible] | [illegible] | [illegible] | [illegible] |
| Gadd45gip1 | 873.3169231 | 0.108437587 | 0.071652131 | 1.51338957 | 0.130180733 | 0.319452055 | | Wipf2 | 4780.804919 | 0.0650200417068796 | [illegible] | [illegible] | [illegible] | [illegible] | [illegible] |
| Ccdc163 | 105.0446581 | 0.243481942 | 0.160955149 | 1.512731605 | 0.130347848 | 0.319760855 | | Wnk3 | 1698.643716 | 0.0719488324123271 | [illegible] | [illegible] | [illegible] | [illegible] | [illegible] |
| Ptma | 7991.508743 | -0.081564553 | 0.053918241 | -1.512745081 | 0.130344424 | 0.319760855 | | Ccdc61 | 254.5449087 | -0.0811123975150198 | [illegible] | [illegible] | [illegible] | [illegible] | [illegible] |
| Otud4 | 2386.775338 | 0.094461244 | 0.062461442 | 1.512312896 | 0.130454282 | 0.319971291 | | Ccnjl | 131.3319175 | -0.0904000474334512 | [illegible] | [illegible] | [illegible] | [illegible] | [illegible] |
| Kctd9 | 533.0305741 | 0.144373564 | 0.095472383 | 1.512202371 | 0.130482388 | 0.319989573 | | Lrrc61 | 1260.197272 | 0.0699201151446987 | [illegible] | [illegible] | [illegible] | [illegible] | [illegible] |
| Erbb2ip | 3637.366522 | -0.125720489 | 0.083160344 | -1.511784136 | 0.130588787 | 0.320149155 | | Msi1 | 668.948802 | -0.0711540341321274 | [illegible] | [illegible] | [illegible] | [illegible] | [illegible] |
| Gucy1a3 | 5356.131008 | -0.137173133 | 0.090731479 | -1.511858216 | 0.130569936 | 0.320149155 | | Npas4 | 1134.251097 | -0.0609180902616668 | [illegible] | [illegible] | [illegible] | [illegible] | [illegible] |
| Acot2 | 179.2618735 | 0.151733076 | 0.100390054 | 1.511435343 | 0.13067757 | 0.320316133 | | 1700034H15Rik | 119.9015299 | -0.0878333316244939 | [illegible] | [illegible] | [illegible] | [illegible] | [illegible] |
| Dcun1d5 | 997.3221253 | -0.104488932 | 0.069138869 | -1.511290739 | 0.130714393 | 0.32035571 | | 1700066M21Rik | 893.9267816 | 0.0790432434055341 | [illegible] | [illegible] | [illegible] | [illegible] | [illegible] |
| 1700028J19Rik | 32.1732815 | 0.253343476 | 0.167696048 | 1.510730153 | 0.130857217 | 0.320515687 | | 1700086L19Rik | 843.2468835 | 0.0826662401951309 | [illegible] | [illegible] | [illegible] | [illegible] | [illegible] |
| Arhgap4 | 176.6525619 | 0.211898162 | 0.140251096 | 1.510848527 | 0.130827048 | 0.320515687 | | 2810428I15Rik | 789.6459049 | -0.067342029402034 | [illegible] | [illegible] | [illegible] | [illegible] | [illegible] |
| Fam126b | 3376.297778 | -0.141260664 | 0.093499607 | -1.510815596 | 0.13083544 | 0.320515687 | | 4930513N10Rik | 29.95671132 | 0.0860119346040951 | [illegible] | [illegible] | [illegible] | [illegible] | [illegible] |
| Ppil1 | 545.5437127 | -0.123035981 | 0.081442501 | -1.510709765 | 0.130862413 | 0.320515687 | | Abce1 | 3781.612312 | 0.0532018609293147 | [illegible] | [illegible] | [illegible] | [illegible] | [illegible] |
| Adamtsl4 | 303.812651 | 0.239553883 | 0.158585626 | 1.510564916 | 0.130899338 | 0.320555452 | | B230219D22Rik | 4365.181058 | 0.065601653529154 | [illegible] | [illegible] | [illegible] | [illegible] | [illegible] |
| Ppm1j | 45.70529242 | -0.201916204 | 0.133715639 | -1.510041801 | 0.131032758 | 0.32078078 | | Bach1 | 648.7419822 | -0.0582446398872577 | [illegible] | [illegible] | [illegible] | [illegible] | [illegible] |
| Supt20 | 764.9853289 | 0.178301541 | 0.118074315 | 1.510078975 | 0.131023274 | 0.32078078 | | Bag2 | 339.2077925 | 0.0740679625732808 | [illegible] | [illegible] | [illegible] | [illegible] | [illegible] |
| Nolc1 | 2673.95883 | 0.084646996 | 0.056074795 | 1.509537319 | 0.131161526 | 0.321045289 | | Camk2a | 142857.9117 | -0.0606873639701999 | [illegible] | [illegible] | [illegible] | [illegible] | [illegible] |
| Ly6d | 12.79072422 | 0.230876688 | 0.152996738 | 1.509030133 | 0.131291083 | 0.321311646 | | Ccdc71 | 1207.031178 | 0.0519939249254419 | [illegible] | [illegible] | [illegible] | [illegible] | [illegible] |
| Phf15 | 3898.301202 | 0.191199265 | 0.126720172 | 1.508830532 | 0.131342096 | 0.321334982 | | Ctc1 | 2181.564333 | -0.0521195521706189 | [illegible] | [illegible] | [illegible] | [illegible] | [illegible] |
| Pkd1l2 | 21.00920327 | 0.248783768 | 0.164884912 | 1.508832824 | 0.13134151 | 0.321334982 | | Gabra2 | 3710.165519 | 0.09254785652188 | [illegible] | [illegible] | [illegible] | [illegible] | [illegible] |
| Nfkb2 | 113.9475407 | 0.192008867 | 0.127275974 | 1.508602616 | 0.131400365 | 0.321426786 | | Gpm6a | 38343.45946 | 0.0710093669573101 | [illegible] | [illegible] | [illegible] | [illegible] | [illegible] |
| Sv2b | 23288.16128 | -0.144848035 | 0.096021068 | -1.508502648 | 0.13142593 | 0.321438572 | | Gramd3 | 676.0670053 | -0.0718333736131296 | [illegible] | [illegible] | [illegible] | [illegible] | [illegible] |
| D630041G03Rik | 177.9648893 | 0.169805667 | 0.112590795 | 1.508166514 | 0.131511915 | 0.321598108 | | Gtf2f2 | 440.5862172 | 0.0711506450386127 | [illegible] | [illegible] | [illegible] | [illegible] | [illegible] |
| Ndufa12 | 2343.892655 | -0.090609356 | 0.060083786 | -1.508050045 | 0.131541719 | 0.32162023 | | Haus6 | 251.6933792 | -0.0822779138270524 | [illegible] | [illegible] | [illegible] | [illegible] | [illegible] |
| Mrpl47 | 399.6173265 | -0.122881382 | 0.08149133 | -1.507907423 | 0.131578223 | 0.321633594 | | Midn | 2442.124344 | 0.0916875607911683 | [illegible] | [illegible] | [illegible] | [illegible] | [illegible] |
| Ndp | 334.4153072 | -0.159681777 | 0.105899149 | -1.507866483 | 0.131588702 | 0.321633594 | | Napb | 24865.46334 | 0.0593147145175331 | [illegible] | [illegible] | [illegible] | [illegible] | [illegible] |
| Ppp1r16a | 2947.241788 | -0.123483577 | 0.081903221 | -1.507676691 | 0.131637294 | 0.321650881 | | Ogfrl1 | 4750.133711 | 0.0577917949397849 | [illegible] | [illegible] | [illegible] | [illegible] | [illegible] |
| Vipas39 | 2135.991985 | 0.089118736 | 0.059108882 | 1.50770464 | 0.131630137 | 0.321650881 | | Pkia | 6587.188655 | 0.0730081747680794 | [illegible] | [illegible] | [illegible] | [illegible] | [illegible] |
| Cdh19 | 421.8203825 | 0.189888339 | 0.125958039 | 1.507552356 | 0.131669135 | 0.321677952 | | Praf2 | 3131.873801 | 0.0665837340577966 | [illegible] | [illegible] | [illegible] | [illegible] | [illegible] |

| | | | | | | |
|---|---|---|---|---|---|---|
| Glcci1 | 2301.963577 | -0.100668595 | 0.066790735 | -1.507223947 | 0.131753264 | 0.321731296 |
| Med23 | 1542.533348 | 0.099308897 | 0.065887402 | 1.507251663 | 0.131746162 | 0.321731296 |
| Pgls | 414.3518463 | -0.14465356 | 0.095965359 | -1.507351838 | 0.131720497 | 0.321731296 |
| 1110015O18Rik | 10.08385814 | -0.225518337 | 0.149654078 | -1.506930779 | 0.131828401 | 0.321812686 |
| 1810021B22Rik | 27.90805086 | 0.245973203 | 0.163236429 | 1.506852391 | 0.131848497 | 0.321812686 |
| Rad9a | 396.4081893 | 0.109718297 | 0.07281298 | 1.506850801 | 0.131848905 | 0.321812686 |
| Tmem55a | 1400.683986 | -0.105926826 | 0.070320663 | -1.50633998 | 0.13197992 | 0.322048446 |
| Vps41 | 7906.513489 | 0.073674093 | 0.048910243 | 1.506312121 | 0.131987068 | 0.322048446 |
| Ap4e1 | 621.255981 | 0.16447983 | 0.10922014 | 1.505947804 | 0.132080573 | 0.322199735 |
| Mt2 | 2130.99088 | 0.170167945 | 0.113000188 | 1.505908509 | 0.132090662 | 0.322199735 |
| Gnl3l | 9004.24465 | -0.083307793 | 0.055330907 | -1.505628529 | 0.132162561 | 0.322324368 |
| Nipa2 | 1148.482193 | -0.129508783 | 0.086033871 | -1.505323213 | 0.132241 | 0.322464913 |
| Fam65b | 2769.490572 | -0.135133989 | 0.089788822 | -1.505020175 | 0.13231889 | 0.322604072 |
| Plekha3 | 1432.571969 | 0.12508276 | 0.083123676 | 1.504778973 | 0.132380912 | 0.322704506 |
| Grik4 | 970.7513924 | 0.158983934 | 0.105670082 | 1.504531191 | 0.132444648 | 0.322768524 |
| Mtfr1l | 2321.331311 | 0.072763078 | 0.048363148 | 1.504514911 | 0.132448837 | 0.322768524 |
| Lmnb2 | 839.6030421 | 0.120079388 | 0.079842137 | 1.50396009 | 0.132591643 | 0.323014924 |
| Rara | 437.8947372 | 0.195431332 | 0.129943204 | 1.503975008 | 0.132587802 | 0.323014924 |
| Tyw5 | 339.5131366 | 0.141452358 | 0.094062671 | 1.503809703 | 0.132630372 | 0.323058479 |
| Stard3 | 1229.24049 | 0.103924769 | 0.069126475 | 1.503400382 | 0.132735828 | 0.323264527 |
| Etnk2 | 288.9004254 | 0.221530414 | 0.147403446 | 1.502884904 | 0.132868726 | 0.323537333 |
| Eps8l1 | 84.87109809 | 0.181419469 | 0.120766791 | 1.502229775 | 0.133037778 | 0.323847184 |
| Frem2 | 142.4628563 | -0.191297497 | 0.127336278 | -1.502301616 | 0.133019231 | 0.323847184 |
| Akap3 | 9.703246949 | 0.240559855 | 0.160163117 | 1.501967867 | 0.133105408 | 0.323960915 |
| Eed | 450.5232675 | 0.134574595 | 0.089615694 | 1.501685567 | 0.133178333 | 0.324087497 |
| Slc16a1 | 1289.815611 | -0.145098572 | 0.096631711 | -1.501562681 | 0.133210087 | 0.324113865 |
| Emilin1 | 105.9379437 | -0.222389495 | 0.148155603 | -1.501053557 | 0.13334171 | 0.324383177 |
| Hist1h1c | 465.5860309 | -0.210482565 | 0.140269323 | -1.500560209 | 0.13346935 | 0.324642717 |
| Dexi | 1148.134135 | 0.126328838 | 0.084216932 | 1.500040845 | 0.133603822 | 0.324783353 |
| Ppp2r4 | 6962.000776 | -0.082813386 | 0.055207775 | -1.500031204 | 0.13360632 | 0.324783353 |
| Slc35f4 | 456.5015815 | 0.144828864 | 0.096551734 | 1.50001307 | 0.133611017 | 0.324783353 |
| Slc9a9 | 761.2705402 | -0.147131349 | 0.09808233 | -1.50008007 | 0.133593663 | 0.324783353 |
| Ube2m | 4277.199925 | -0.092006486 | 0.061342435 | -1.499883181 | 0.133644665 | 0.324814187 |
| Ldhb | 19038.34808 | -0.088784363 | 0.059210279 | -1.499475502 | 0.133750319 | 0.325019989 |
| Cul1 | 5092.786971 | 0.086041578 | 0.057385602 | 1.499358302 | 0.133780705 | 0.325042848 |
| Mpnd | 2254.745705 | -0.116910852 | 0.077979482 | -1.499251456 | 0.133808411 | 0.32505919 |
| Slc8a3 | 1369.26922 | -0.130635457 | 0.087154654 | -1.49889249 | 0.133901525 | 0.325234398 |
| Adck2 | 813.8792359 | 0.097136582 | 0.064816945 | 1.498629452 | 0.133969788 | 0.3253492 |
| Ivd | 2185.100833 | 0.124718805 | 0.083233722 | 1.49841677 | 0.134025002 | 0.32543228 |
| Abcd1 | 551.4438371 | 0.143331549 | 0.095678136 | 1.498059585 | 0.13411777 | 0.325535831 |
| Il1rap | 1271.653903 | 0.150209744 | 0.100270482 | 1.498045495 | 0.134121431 | 0.325535831 |
| Prr12 | 2933.101879 | 0.087954193 | 0.058715362 | 1.497975815 | 0.134139534 | 0.325535831 |
| Zfp51 | 189.8590095 | 0.175468017 | 0.117140407 | 1.497929033 | 0.13415169 | 0.325535831 |
| B3gat1 | 9724.129637 | -0.102930004 | 0.068734699 | -1.497496968 | 0.134263994 | 0.325757332 |
| Gm11744 | 82.37011331 | 0.199433944 | 0.133202514 | 1.497223575 | 0.134335093 | 0.32583618 |
| Ints3 | 3110.376072 | 0.083320882 | 0.055650755 | 1.49721028 | 0.134338551 | 0.32583618 |
| Cd74 | 315.5222307 | 0.200915333 | 0.134220005 | 1.49691049 | 0.13441655 | 0.325903062 |
| Fdx1 | 205.9021818 | -0.150699634 | 0.100677054 | -1.496861773 | 0.134429229 | 0.325903062 |
| Inpp4a | 6774.292024 | -0.083387642 | 0.05570275 | -1.497011225 | 0.134390337 | 0.325903062 |
| Pkia | 6377.68549 | -0.146340017 | 0.097776828 | -1.496673801 | 0.134478156 | 0.325970675 |
| Vcan | 874.5373205 | -0.153176089 | 0.102364696 | -1.496376152 | 0.134555661 | 0.326107525 |
| Hspb8 | 578.6441868 | -0.130451875 | 0.087187414 | -1.496223693 | 0.134595372 | 0.32613938 |
| Ift46 | 1283.569406 | 0.099752067 | 0.066671877 | 1.496164077 | 0.134610903 | 0.32613938 |
| Nrf1 | 632.5846544 | -0.115212085 | 0.077029513 | -1.495687568 | 0.134735091 | 0.326389229 |
| Fam195b | 1900.213392 | -0.095655413 | 0.063984213 | -1.4949846 | 0.134918461 | 0.326782342 |
| Cacng2 | 1890.182188 | 0.12557837 | 0.084020071 | 1.49462347 | 0.135012737 | 0.326959574 |
| Cryga | 11.60957235 | 0.248513498 | 0.166326278 | 1.494132502 | 0.13514099 | 0.327189831 |
| Irf2bp1 | 1543.190937 | -0.085977655 | 0.057544863 | -1.494097823 | 0.135150053 | 0.327189831 |
| 3425401B19Rik | 13.06232049 | 0.225545922 | 0.151035033 | 1.493335139 | 0.135349482 | 0.327438548 |
| Bcas1 | 3758.257839 | 0.172213289 | 0.115305108 | 1.493544319 | 0.135294762 | 0.327438548 |
| Hmbs | 559.2749129 | -0.126716131 | 0.084852204 | -1.49337465 | 0.135339145 | 0.327438548 |
| Lgals9 | 319.9079403 | 0.190961341 | 0.127881849 | 1.493263842 | 0.135368136 | 0.327438548 |
| Rybp | 2433.994646 | -0.107352797 | 0.071893486 | -1.493220078 | 0.135379588 | 0.327438548 |
| Ydjc | 418.2412156 | -0.198994206 | 0.13325569 | -1.493326151 | 0.135351833 | 0.327438548 |
| Tnrc6c | 3895.408319 | 0.10319807 | 0.069126956 | 1.492877401 | 0.135469282 | 0.327604347 |
| Ablim2 | 6340.608985 | 0.116557933 | 0.078094013 | 1.492533525 | 0.135559335 | 0.327770965 |
| Trim12a | 86.87936277 | 0.199133204 | 0.13345685 | 1.492116772 | 0.135668536 | 0.327932651 |
| Vps53 | 2842.578568 | 0.11310721 | 0.075803166 | 1.492117225 | 0.135668417 | 0.327932651 |
| Mrpl14 | 331.7658772 | -0.153104853 | 0.10262124 | -1.491941172 | 0.135714568 | 0.327984607 |
| Mrpl35 | 1025.362443 | -0.103626539 | 0.069460684 | -1.491873285 | 0.135732367 | 0.327984607 |
| Rph3al | 27.10653739 | 0.248390953 | 0.166532494 | 1.491546465 | 0.135818081 | 0.328140551 |
| Dusp10 | 244.6904732 | 0.189185488 | 0.126870012 | 1.491175768 | 0.135915354 | 0.328305122 |
| Nanp | 548.3087432 | -0.110251591 | 0.073938511 | -1.491125396 | 0.135928576 | 0.328305122 |
| Klhdc3 | 6645.267689 | 0.1260457 | 0.084536955 | 1.491013005 | 0.13595808 | 0.328325202 |
| Prkd3 | 980.7356597 | -0.112365257 | 0.075379359 | -1.490663466 | 0.136049873 | 0.328429143 |
| Rrp8 | 431.2569352 | 0.170286193 | 0.114237231 | 1.490636567 | 0.136056938 | 0.328429143 |
| Snx14 | 2195.322087 | -0.082880377 | 0.05560451 | -1.490533362 | 0.136084051 | 0.328429143 |
| Ttc9 | 1342.420241 | -0.119021212 | 0.079851803 | -1.490526284 | 0.136085911 | 0.328429143 |
| Apln | 1056.745517 | -0.245863329 | 0.16496298 | -1.49041518 | 0.136115104 | 0.328448437 |
| 1700019D03Rik | 124.2965629 | -0.167290506 | 0.112297598 | -1.489706894 | 0.136301321 | 0.328796121 |
| Synpo2l | 10.68006528 | -0.239987276 | 0.161097104 | -1.489705713 | 0.136301632 | 0.328796121 |
| Rpe65 | 128.0334479 | 0.214861855 | 0.144257171 | 1.489436224 | 0.136372536 | 0.328915951 |
| Svil | 492.4578633 | -0.143345548 | 0.096266423 | -1.489050317 | 0.136474119 | 0.329109728 |
| Mesdc2 | 2017.515739 | -0.086421271 | 0.058044682 | -1.488874912 | 0.136520311 | 0.329169888 |
| A230056P14Rik | 91.80491389 | -0.211846777 | 0.142390137 | -1.487791092 | 0.136805996 | 0.329602254 |
| Crip3 | 44.58264111 | 0.250006029 | 0.168011482 | 1.488029424 | 0.136743134 | 0.329602254 |
| Gtf3c4 | 2148.770041 | -0.079960077 | 0.053741497 | -1.487864712 | 0.136786576 | 0.329602254 |
| Maf | 490.6469589 | 0.134913201 | 0.09067481 | 1.487879613 | 0.136782645 | 0.329602254 |
| Ttc37 | 924.1078246 | 0.110218869 | 0.074071817 | 1.488000064 | 0.136750877 | 0.329602254 |
| Rfc1 | 1665.777719 | 0.072801802 | 0.04895558 | 1.487099153 | 0.136988625 | 0.329990945 |
| Cdc6 | 21.14970576 | -0.247812277 | 0.166675102 | -1.486798408 | 0.137068062 | 0.330130974 |
| Atraid | 1650.536124 | 0.088651342 | 0.059633796 | 1.486595637 | 0.137121641 | 0.330157374 |
| Gmeb2 | 1016.374487 | -0.111572871 | 0.075052035 | -1.486606876 | 0.137118671 | 0.330157374 |
| Il1rapl1 | 285.9203408 | -0.154272204 | 0.103799936 | -1.48624565 | 0.137214157 | 0.330296028 |
| Mocs1 | 327.7152867 | 0.135201017 | 0.090969933 | 1.486216509 | 0.137221862 | 0.330296028 |
| Glyr1 | 8968.868449 | 0.103972099 | 0.069978273 | 1.485776865 | 0.137338151 | 0.330443271 |
| H2-DMb1 | 66.37869106 | 0.240214581 | 0.161679727 | 1.485743363 | 0.137347016 | 0.330443271 |
| 11-Sep | 4113.730641 | -0.103462864 | 0.069636047 | -1.485765894 | 0.137341054 | 0.330443271 |
| Dnahc10 | 63.29001885 | 0.245303977 | 0.165140206 | 1.485428553 | 0.137430337 | 0.330592398 |
| Gsx1 | 13.95927557 | -0.226820968 | 0.152745941 | -1.48495578 | 0.137555538 | 0.330842209 |
| Smox | 1454.352962 | 0.1967776 | 0.132523305 | 1.4848528 | 0.137582822 | 0.33085647 |
| Eci1 | 414.9117712 | -0.162024724 | 0.109165183 | -1.484216126 | 0.137751593 | 0.331210922 |
| 9230110C19Rik | 305.2241294 | 0.167892279 | 0.113156708 | 1.483714775 | 0.137884605 | 0.331476498 |
| Utp14b | 273.3511431 | -0.220647913 | 0.1487208 | -1.483638557 | 0.137904835 | 0.331476498 |
| Bloc1s3 | 421.7080836 | -0.131865217 | 0.088891885 | -1.483433688 | 0.137959222 | 0.331518804 |
| Pigf | 234.4148012 | -0.175835968 | 0.11853489 | -1.483411069 | 0.137965228 | 0.331518804 |
| AF357359 | 192.7130186 | 0.24618781 | 0.166057054 | 1.482549541 | 0.138194132 | 0.332017349 |
| Avl9 | 2671.838888 | -0.132198631 | 0.089193508 | -1.482155298 | 0.138298978 | 0.332166232 |
| Eefsec | 616.4198431 | -0.115265898 | 0.077765102 | -1.482231679 | 0.13827866 | 0.332166232 |
| 4933426M11Rik | 2269.306056 | 0.155110707 | 0.104693745 | 1.481566135 | 0.138455775 | 0.332182376 |
| Actr6 | 459.6841273 | -0.108721481 | 0.073374578 | -1.481732282 | 0.138411544 | 0.332182376 |
| Ccdc23 | 273.3756038 | -0.140625497 | 0.094893463 | -1.481930288 | 0.138358845 | 0.332182376 |
| Fndc3b | 854.3663118 | -0.155083744 | 0.104660398 | -1.481780574 | 0.13839869 | 0.332182376 |
| Kptn | 689.1783022 | -0.183688551 | 0.123952645 | -1.481925231 | 0.138360191 | 0.332182376 |
| Pcdhgc4 | 516.3070115 | -0.132662486 | 0.089535769 | -1.481670257 | 0.138428055 | 0.332182376 |
| Plekhf2 | 286.1137699 | -0.132665315 | 0.089542399 | -1.48159214 | 0.138448852 | 0.332182376 |
| Elp4 | 495.6248542 | -0.158529584 | 0.107010623 | -1.481437822 | 0.138489942 | 0.332190947 |
| Narg2 | 300.9766093 | 0.174282357 | 0.117654115 | 1.481311191 | 0.138523668 | 0.332190947 |
| Tmco3 | 1536.837785 | -0.093056448 | 0.06281826 | -1.481359858 | 0.138510706 | 0.332190947 |
| Yipf2 | 695.9186715 | 0.148374185 | 0.100187933 | 1.480958637 | 0.138617596 | 0.332364753 |
| Snord88c | 9.83997425 | 0.225962295 | 0.152601018 | 1.480739107 | 0.138676109 | 0.332453601 |
| Gatc | 1161.18818 | 0.094318978 | 0.063710032 | 1.480441539 | 0.138755451 | 0.332592352 |
| Trpc4ap | 6582.544063 | 0.081173858 | 0.054855083 | 1.479787356 | 0.138930004 | 0.33295924 |
| B3gat3 | 3981.527165 | -0.119340014 | 0.080687923 | -1.479031923 | 0.139131783 | 0.333288164 |
| Rap1a | 1040.281206 | -0.113493791 | 0.07673337 | -1.479066941 | 0.139122425 | 0.333288164 |
| Rnf149 | 826.8663355 | -0.0962524 | 0.065073302 | -1.479138096 | 0.13910341 | 0.333288164 |
| Extl1 | 2960.89417 | 0.146080358 | 0.098804281 | 1.478482072 | 0.139278792 | 0.333588748 |
| 5730577I03Rik | 60.58208864 | -0.237438038 | 0.16063704 | -1.478102674 | 0.139380299 | 0.333742489 |
| C1qtnf4 | 2514.866642 | -0.199008237 | 0.134646921 | -1.478000657 | 0.139407602 | 0.333742489 |
| Rars | 1621.483231 | 0.142120403 | 0.096152826 | 1.478067882 | 0.13938961 | 0.333742489 |
| Clcn4-2 | 8399.115107 | -0.078001525 | 0.052794622 | -1.477452091 | 0.139554492 | 0.33393935 |
| Prss22 | 50.31606294 | -0.246671059 | 0.166954919 | -1.477471054 | 0.139549412 | 0.33393935 |
| Sncg | 104.430024 | -0.222219285 | 0.150393224 | -1.477588418 | 0.139517976 | 0.33393935 |
| Pard6g | 279.7804455 | -0.150404214 | 0.101812684 | -1.477264006 | 0.139604883 | 0.334008346 |
| 2310061J03Rik | 97.70773224 | 0.182050249 | 0.123263755 | 1.476916309 | 0.139698073 | 0.334076546 |
| Acot5 | 83.87108093 | 0.223598385 | 0.151380946 | 1.477057653 | 0.139660184 | 0.334076546 |
| Wdr17 | 1682.487955 | -0.107327642 | 0.072668652 | -1.476945559 | 0.139690231 | 0.334076546 |
| Med8 | 777.4886025 | 0.091217072 | 0.061772943 | 1.476650912 | 0.139769237 | 0.334195147 |
| Gm9159 | 8.588911443 | 0.218707177 | 0.148156089 | 1.476194317 | 0.139891735 | 0.334334817 |
| Mical3 | 4324.217106 | 0.103347557 | 0.070017195 | 1.476031106 | 0.139935542 | 0.334334817 |
| Nrtn | 18.37771453 | -0.242658189 | 0.164391041 | -1.476103489 | 0.139916113 | 0.334334817 |
| Slc9b2 | 109.740784 | 0.190901907 | 0.129317393 | 1.476227612 | 0.139882799 | 0.334334817 |
| Zfp1 | 667.1702377 | 0.122645909 | 0.083085796 | 1.476135698 | 0.139907467 | 0.334334817 |
| Fermt3 | 173.6451502 | 0.176687656 | 0.119728251 | 1.475739053 | 0.140013958 | 0.334412122 |
| Mcm3ap | 2557.832358 | 0.093178265 | 0.063143369 | 1.475661919 | 0.140034674 | 0.334412122 |
| Ninj1 | 402.0048245 | -0.161575471 | 0.109493927 | -1.475656912 | 0.140036019 | 0.334412122 |
| Siah2 | 583.0987224 | 0.130488291 | 0.088431319 | 1.475589106 | 0.140054231 | 0.334412122 |
| Leprel4 | 353.9038954 | -0.139929693 | 0.094865549 | -1.475031707 | 0.140204019 | 0.334666626 |
| Srsf10 | 1858.310009 | 0.119610515 | 0.081087826 | 1.475073631 | 0.140192749 | 0.334666626 |
| Nkx2-2as | 35.44538583 | -0.246611424 | 0.167262761 | -1.474395276 | 0.140375196 | 0.335023611 |
| Grlf1 | 10859.23531 | -0.094332221 | 0.063986128 | -1.474260496 | 0.140411467 | 0.335058566 |
| Ccbp2 | 172.4472642 | 0.224328543 | 0.152198361 | 1.473922201 | 0.140502539 | 0.335121051 |
| Hyls1 | 198.7642157 | 0.145699319 | 0.098850149 | 1.473941321 | 0.140497391 | 0.335121051 |
| Tmem134 | 723.3512697 | -0.125536595 | 0.0851679 | -1.473989551 | 0.140484404 | 0.335121051 |
| Mta3 | 2074.317999 | 0.111886047 | 0.075914742 | 1.473838205 | 0.140525159 | 0.335123413 |
| Cnnm3 | 1326.818294 | 0.10089703 | 0.068471206 | 1.473568754 | 0.140597739 | 0.335147509 |
| Dhx36 | 3148.423824 | -0.094506428 | 0.064134777 | -1.473559786 | 0.140600156 | 0.335147509 |
| Pelp1 | 1467.080903 | 0.085267668 | 0.057862297 | 1.473630889 | 0.140581 | 0.335147509 |
| Ptgis | 52.57602856 | -0.2307964 | 0.156642977 | -1.473391306 | 0.140645553 | 0.335204153 |
| Dok7 | 407.7664095 | 0.139904721 | 0.094964667 | 1.47322921 | 0.140689241 | 0.335256706 |
| Rapgef3 | 1414.817631 | 0.121258742 | 0.082374569 | 1.47204098 | 0.141009812 | 0.335925279 |
| Rnf26 | 1333.693709 | 0.10980276 | 0.07459282 | 1.472028553 | 0.141013167 | 0.335925279 |
| Ltbp2 | 30.89237574 | -0.232079361 | 0.157672446 | -1.471908167 | 0.141045678 | 0.335951073 |
| Snord69 | 11.00245022 | 0.203691037 | 0.138417999 | 1.471564675 | 0.141138471 | 0.336120422 |
| Esrrg | 1189.610342 | 0.155170217 | 0.105513146 | 1.47062449 | 0.141392697 | 0.336674112 |
| Fam19a5 | 4321.314485 | -0.078070435 | 0.053102979 | -1.470170522 | 0.141515576 | 0.336863164 |
| Fut8 | 2311.389292 | -0.088859967 | 0.060440768 | -1.470199184 | 0.141507816 | 0.336863164 |
| Zfp882 | 366.3064222 | -0.188521408 | 0.128265234 | -1.469777905 | 0.141621915 | 0.337064509 |
| A930024E05Rik | 97.08384028 | 0.206780295 | 0.140742288 | 1.469212267 | 0.141775224 | 0.337323079 |
| Fanci | 24.887643 | 0.24581976 | 0.167317513 | 1.469181292 | 0.141783623 | 0.337323079 |
| Gbp5 | 135.3492143 | 0.241442967 | 0.164352476 | 1.46905585 | 0.141817641 | 0.337323079 |
| Msl3 | 780.5531065 | 0.109167641 | 0.074311219 | 1.469060023 | 0.14181651 | 0.337323079 |
| Snx24 | 504.7597046 | 0.105920389 | 0.072110141 | 1.468869543 | 0.141868177 | 0.337391487 |
| Myocd | 10.85512103 | -0.231769465 | 0.157887627 | -1.467939376 | 0.142120691 | 0.33788829 |
| Prox2 | 34.96773978 | 0.233303477 | 0.158924603 | 1.468013591 | 0.142100531 | 0.33788829 |
| Srgn | 180.6997159 | -0.157762885 | 0.107494378 | -1.467638478 | 0.14220245 | 0.338030801 |
| 9330175M20Rik | 8.600210793 | 0.222748957 | 0.151796501 | 1.467418259 | 0.142262311 | 0.338118002 |
| Zbtb8os | 337.9869594 | 0.130888671 | 0.089201143 | 1.467342976 | 0.142282779 | 0.338118002 |
| Gchfr | 8.854347792 | -0.232579006 | 0.158540453 | -1.467001019 | 0.142375778 | 0.33828712 |
| 4930479D17Rik | 55.94142799 | 0.218033758 | 0.148650466 | 1.466754629 | 0.142442816 | 0.33839451 |
| Dgcr2 | 5620.361633 | 0.100082575 | 0.06824226 | 1.466577677 | 0.142490976 | 0.338457027 |
| Tpgs2 | 2281.973613 | -0.088335296 | 0.060259327 | -1.465919042 | 0.142670344 | 0.338831132 |
| Xlr3b | 38.53565111 | 0.224356493 | 0.153076058 | 1.465653716 | 0.142742649 | 0.338950898 |
| Emd | 1174.406291 | -0.127322958 | 0.086893099 | -1.465282745 | 0.142843792 | 0.339139093 |
| Nhlrc2 | 414.3914617 | 0.12197177 | 0.083258078 | 1.464984205 | 0.142925227 | 0.339280445 |
| Tmx3 | 2054.857352 | -0.115862564 | 0.079094244 | -1.464867205 | 0.142957152 | 0.339304245 |
| Commd6 | 604.0371885 | -0.121875864 | 0.083250145 | -1.463971785 | 0.143201658 | 0.339776502 |
| Tmem116 | 69.07292796 | 0.209097446 | 0.142826764 | 1.463993444 | 0.14319574 | 0.339776502 |
| Ythdc1 | 2309.92404 | 0.099543014 | 0.067998616 | 1.463897656 | 0.143221914 | 0.339776502 |
| Mthfd1 | 1478.002002 | 0.068227751 | 0.046622939 | 1.463394471 | 0.143359471 | 0.33999872 |
| Slc39a5 | 12.56485186 | -0.241095826 | 0.164743224 | -1.463464299 | 0.143340376 | 0.33999872 |
| Hlf | 8878.864707 | 0.137686221 | 0.094111109 | 1.463017727 | 0.143462528 | 0.340112608 |
| Mybbp1a | 2521.115236 | 0.089868607 | 0.061428279 | 1.462984271 | 0.143471683 | 0.340112608 |
| Prickle2 | 9523.924345 | -0.108750874 | 0.074335266 | -1.462978198 | 0.143473345 | 0.340112608 |
| Gyg | 1046.347057 | -0.117680742 | 0.080510424 | -1.461683295 | 0.143828017 | 0.340901223 |
| Spin1 | 9041.748063 | -0.089418595 | 0.061222337 | -1.460555088 | 0.144137579 | 0.341582694 |
| Atrx | 6226.999987 | -0.10433746 | 0.071443273 | -1.460423842 | 0.144173624 | 0.341588749 |
| Nrg2 | 238.634586 | 0.12804285 | 0.087677448 | 1.460385238 | 0.144184228 | 0.341588749 |
| Stoml1 | 2060.60092 | 0.135855039 | 0.093042842 | 1.460134232 | 0.144253186 | 0.341699873 |
| Dcun1d1 | 1650.657132 | -0.118147406 | 0.080921933 | -1.460017102 | 0.144285374 | 0.341723874 |
| Tnks | 3521.364202 | -0.113758683 | 0.077926413 | -1.459821892 | 0.14433903 | 0.341798706 |
| Megf6 | 260.4576749 | 0.17966712 | 0.123097742 | 1.459548457 | 0.144414214 | 0.341924485 |
| Taf8 | 655.0113974 | 0.136326653 | 0.093429783 | 1.459134858 | 0.144527994 | 0.342141594 |
| Dmbx1 | 9.447200537 | 0.193581456 | 0.132717876 | 1.458593686 | 0.144676972 | 0.342441951 |
| Itpkc | 350.3820626 | 0.151513032 | 0.103917796 | 1.45800852 | 0.144838195 | 0.342771191 |
| Chmp1a | 3707.007369 | -0.097240685 | 0.066703354 | -1.457808016 | 0.144893468 | 0.342806638 |
| Ntng1 | 805.7123801 | -0.209944347 | 0.144023059 | -1.457713431 | 0.144919548 | 0.342806638 |
| Prlr | 140.8214869 | 0.191816148 | 0.131579888 | 1.457792303 | 0.1448978 | 0.342806638 |
| Ntf3 | 38.74912685 | 0.23307668 | 0.159940283 | 1.457273156 | 0.145040994 | 0.343041545 |
| Larp7 | 651.984239 | -0.09758254 | 0.066966535 | -1.457183653 | 0.145065692 | 0.343047593 |
| Nos2 | 33.83073835 | -0.244945827 | 0.16812107 | -1.456960906 | 0.145127172 | 0.343140607 |
| G2e3 | 403.0692082 | 0.138689858 | 0.095210553 | 1.456664769 | 0.145208939 | 0.343281554 |
| Ssna1 | 813.9602516 | -0.089558858 | 0.061505343 | -1.456115086 | 0.145360808 | 0.343588155 |
| Smg7 | 5229.999556 | 0.087490206 | 0.060099121 | 1.455765154 | 0.145457552 | 0.343764385 |
| Dpcd | 1331.07876 | -0.113416508 | 0.07791404 | -1.455662013 | 0.145486076 | 0.34377936 |
| Tcf25 | 25576.61258 | 0.074442525 | 0.051168799 | 1.454842153 | 0.145712965 | 0.34426299 |
| Necap2 | 935.8443304 | 0.127938652 | 0.087983381 | 1.454122935 | 0.145912226 | 0.344681206 |
| Smoc1 | 695.0822383 | 0.216390918 | 0.148853881 | 1.453713645 | 0.146025714 | 0.344896708 |
| B4galnt1 | 5117.175979 | 0.113999828 | 0.078443606 | 1.453271138 | 0.146148488 | 0.345028899 |
| Ccdc33 | 35.87404782 | 0.244255818 | 0.168066274 | 1.453330354 | 0.146132053 | 0.345028899 |
| Scn8a | 15399.91458 | -0.101711784 | 0.06998797 | -1.45327524 | 0.146147349 | 0.345028899 |
| Mark4 | 2096.709039 | 0.108026824 | 0.074338385 | 1.453176904 | 0.146174643 | 0.345038074 |
| 5930403L14Rik | 1943.825698 | -0.165093299 | 0.113617386 | -1.4530637 | 0.146206068 | 0.345059684 |
| Trim8 | 2636.278165 | -0.115624603 | 0.079580696 | -1.452922745 | 0.146245205 | 0.345099483 |
| Frrs1 | 34.09456501 | 0.235973393 | 0.162442516 | 1.452657826 | 0.146318782 | 0.345180316 |
| Slc30a3 | 4284.380181 | 0.168160943 | 0.115762378 | 1.452638982 | 0.146324017 | 0.345180316 |
| Zfp810 | 600.5183561 | 0.142164028 | 0.097874521 | 1.452513134 | 0.14635898 | 0.345210235 |
| Hk2 | 215.2843771 | 0.143885874 | 0.09908935 | 1.452082127 | 0.146478771 | 0.345440194 |
| Fam159b | 12.77840498 | 0.238489955 | 0.16426088 | 1.451897465 | 0.146530118 | 0.345508696 |
| Matn4 | 266.4036146 | -0.160629407 | 0.110649812 | -1.451691632 | 0.146587367 | 0.345591094 |
| Dennd1a | 2590.607086 | 0.068984363 | 0.047535222 | 1.451226274 | 0.146716863 | 0.345620169 |
| Gabpb1 | 612.6494031 | -0.107061668 | 0.07376688 | -1.451351449 | 0.146682022 | 0.345620169 |
| Gtpbp10 | 350.838254 | -0.135457575 | 0.093328422 | -1.451407526 | 0.146666415 | 0.345620169 |
| H2-Ob | 23.2129287 | 0.243523642 | 0.16778348 | 1.451416087 | 0.146664033 | 0.345620169 |
| Ptges3l | 83.67605396 | -0.228968031 | 0.157773589 | -1.451244356 | 0.14671183 | 0.345620169 |

| | | | | | | | |
|---|---|---|---|---|---|---|---|
| Ripk1 | 288.431728 | -0.0751206139544915 | 0.0771156642336113 | -0.974129117619009 | 0.32999243363644 | 0.756141870819943 | NM_009068 |
| Rpl22l1 | 1316.492292 | -0.0817019081807382 | 0.0838382588595551 | -0.974518188857003 | 0.329799312466709 | 0.756141870819943 | NM_026517 |
| Rpl3 | 18791.30444 | 0.0463963706631865 | 0.0476855859106485 | 0.972964256958534 | 0.330571066866277 | 0.756141870819943 | NM_013762 |
| Rtn4 | 28672.34743 | 0.0631859298679267 | 0.0649271966284645 | 0.973181242207301 | 0.330463231817605 | 0.756141870819943 | NM_194054,NM_024226,NM_194052,NM_194053,NM_194051 |
| Terf2 | 1752.22177 | 0.0506495360738085 | 0.0520653669364608 | 0.972806666965775 | 0.33064939855764 | 0.756141870819943 | NM_001083118,NM_009353 |
| Tnfaip8l2 | 80.88242622 | -0.0925744167447287 | 0.0950014951993109 | -0.97445220783641 | 0.329832057951393 | 0.756141870819943 | NM_027206 |
| Trmt11 | 270.6850803 | 0.0811414042765748 | 0.0832792642600623 | 0.974329024127645 | 0.329893197988255 | 0.756141870819943 | NM_028604 |
| Vac14 | 1293.269055 | 0.061112679357374 | 0.0628145673141093 | 0.972906158085515 | 0.330599944097993 | 0.756141870819943 | NM_146216 |
| Wdr33 | 2555.275219 | 0.0480265288192347 | 0.0492809333221585 | 0.97454584525168 | 0.32978558759974 | 0.756141870819943 | NM_001170966,NM_001170967,NM_028866,NM_001170970 |
| Zfat | 300.6720454 | -0.0754290229756865 | 0.0773776231101433 | -0.974817007086363 | 0.329651039396326 | 0.756141870819943 | NM_001145888,NM_198644 |
| Erp29 | 2474.579857 | 0.0583203864063477 | 0.0599953920011549 | 0.97208109591525 | 0.331010205884629 | 0.75621299821795 | NM_026129 |
| Hp1bp3 | 10974.56032 | 0.0454489435578417 | 0.0467676966465822 | 0.971802051772912 | 0.33114903494867 | 0.75621299821795 | NM_001122897,NM_001122896,NM_010470 |
| Ier3ip1 | 1098.531755 | 0.0609869596618094 | 0.0627317735285664 | 0.972186122460535 | 0.330957963198601 | 0.75621299821795 | NM_025409 |
| Nop14 | 1040.10871 | 0.064927587909928 | 0.0667628327711419 | 0.972510979761674 | 0.330796405263892 | 0.75621299821795 | NM_029278 |
| Peli2 | 1058.558474 | 0.061467201289799 | 0.0632473318315328 | 0.971854456303779 | 0.331122959958492 | 0.75621299821795 | NM_033602 |
| Sec11c | 1470.986941 | -0.0641090171448356 | 0.0659343427262235 | -0.972316011566732 | 0.330893360664614 | 0.75621299821795 | NM_025468 |
| Slc30a4 | 3427.624056 | 0.0815204873492494 | 0.0838317097714185 | 0.972430212523745 | 0.330836567635567 | 0.75621299821795 | NM_011774 |
| Tmx1 | 1197.548909 | 0.0615214807019343 | 0.0633029420308315 | 0.971858159008952 | 0.331121117648921 | 0.75621299821795 | NM_028339 |
| Txlna | 2348.472007 | -0.0561711003525083 | 0.0577961563929045 | -0.971882973854716 | 0.331108771001751 | 0.75621299821795 | NM_001199695,NM_001005506 |
| Actl6a | 461.1349567 | -0.0637387658972878 | 0.065610600482282 | -0.971470546356306 | 0.331314013297726 | 0.756398469821747 | NM_019673 |
| Setd3 | 7327.255034 | 0.0436911525382154 | 0.0449761391265842 | 0.971429593261613 | 0.331334397871112 | 0.756398469821747 | NM_028262 |
| Kcnh8 | 24.73074827 | 0.0734379538065766 | 0.0756105052570656 | 0.971266539707642 | 0.331415566498718 | 0.756464883557569 | NM_001031811 |
| Zmat3 | 4692.980104 | 0.0656076229582771 | 0.0675644907196645 | 0.97103703823497 | 0.331529834913325 | 0.756606815703215 | NM_009517 |
| Zfp811 | 382.3456414 | -0.0765848072944703 | 0.0788802323638374 | -0.970899869326203 | 0.331598143252495 | 0.756643831116203 | NM_001267583,NM_183177 |
| Arl2bp | 5161.581138 | 0.041446496368778 | 0.042714232886087 | 0.970320513054983 | 0.331886755529381 | 0.75695848284981 | NM_024191,NM_024269 |
| Lrriq1 | 104.5457241 | -0.0899735348719797 | 0.0927160729077806 | -0.970420036679846 | 0.331837165268366 | 0.75695848284981 | NM_029134,NM_001163559 |
| Utp15 | 1315.891573 | -0.0573819627997713 | 0.0591378090343598 | -0.970309244402876 | 0.331892370733198 | 0.75695848284981 | NM_178918 |
| Acvrl1 | 858.0658386 | 0.0790309277239199 | 0.0814717470887168 | 0.970040910474903 | 0.332026100488426 | 0.757144605289619 | NM_009612 |
| Cklf | 49.38225594 | -0.0891639482843784 | 0.091953372777561 | -0.969664794135063 | 0.332213604432348 | 0.757166891883869 | NM_001037841,NM_029295 |
| Ly6e | 9473.272843 | -0.0436944214823134 | 0.0450690937464244 | -0.969498559881293 | 0.332296498400087 | 0.757166891883869 | NM_001164039,NM_001164038,NM_001164037,NM_001164036,NM_008529,NM_001164040 |
| Msh3 | 1010.487693 | 0.0614443315221599 | 0.063372923237442 | 0.969567575286748 | 0.33226208172424 | 0.757166891883869 | NM_010829 |
| Rpl11 | 6379.020324 | 0.0583775038338131 | 0.060199059621036 | 0.969741125547643 | 0.33217554567224 | 0.757166891883869 | NM_025919 |
| Trpc5 | 382.4622469 | 0.0847200994477849 | 0.0873854200343155 | 0.969499253016304 | 0.332296152735334 | 0.757166891883869 | NM_009428 |
| Pcgf5 | 339.9232043 | 0.0765074403035379 | 0.0789403029568395 | 0.969180981549669 | 0.332454898380106 | 0.757290238963371 | NM_029508 |
| Pnck | 2894.289033 | 0.0775321715300561 | 0.0799931630888477 | 0.969234976293433 | 0.332427963737204 | 0.757290238963371 | NM_012040,NM_001199352,NM_001199351 |
| 2410016O06Rik | 529.9190121 | 0.0674188618686097 | 0.0695921496559772 | 0.968771078374344 | 0.332659419725332 | 0.757637304943584 | NM_023633 |
| Eln | 317.1344407 | 0.0884510340823635 | 0.0913481838555877 | 0.968284538882522 | 0.33290228422236 | 0.757641998582448 | NM_007925 |
| Gm17644 | 54.84319258 | -0.0876947497880198 | 0.0905708719165216 | -0.968244513190149 | 0.332922268824987 | 0.757641998582448 | NR_045297 |
| Phkg2 | 1959.0334 | -0.0536521980979006 | 0.055404568059164 | -0.968371381229935 | 0.332858926990929 | 0.757641998582448 | NM_026888 |
| Prpsap2 | 1301.212155 | 0.056670647849198 | 0.0585215242208309 | 0.968372724458635 | 0.332858256394236 | 0.757641998582448 | NM_001164244,NM_144806,NM_001164242,NM_001164243 |
| Vps28 | 3066.578329 | 0.0566465481449065 | 0.0584978690315194 | 0.968352336294243 | 0.332868435123817 | 0.757641998582448 | NM_025842 |
| Cby1 | 1215.407912 | 0.0586877372911931 | 0.0606198566487666 | 0.96812728593589 | 0.332980804188392 | 0.757656510282047 | NM_028634 |
| Ctxn2 | 275.2576196 | 0.082255496794406 | 0.0849965264795311 | 0.96775127409724 | 0.333168603903683 | 0.757730861621216 | NM_001162934 |
| Dnajc18 | 5229.414324 | 0.0477070173327646 | 0.0493084520522855 | 0.967522105179398 | 0.333283096210319 | 0.757730861621216 | NM_029669 |
| Inca1 | 87.29347905 | -0.0919477882608199 | 0.0950428280681915 | -0.967435314475796 | 0.333326463290622 | 0.757730861621216 | NM_213729,NM_001252484,NM_001252485,NM_001252482,NM_001252483 |
| Nhs | 229.3773049 | 0.0906926331876322 | 0.0937231773186114 | 0.967664944598742 | 0.333211730957889 | 0.757730861621216 | NM_001081052 |
| Nmi | 121.5871984 | -0.0890372387987968 | 0.0920173010861736 | -0.967614108953424 | 0.333237128269047 | 0.757730861621216 | NM_001141948,NM_001141949,NM_019401 |
| Nr1h3 | 116.0648808 | -0.0910737516008407 | 0.0940948562009648 | -0.967892988818946 | 0.333097816232974 | 0.757730861621216 | NM_001177730,NM_013839 |
| Gprasp2 | 6018.148391 | 0.0736926621497324 | 0.0762047571901025 | 0.967034931505609 | 0.333526571500552 | 0.757852267925996 | NM_001163015,NM_001163016,NM_001163017 |
| Ppp4c | 1127.648505 | 0.0570816880291611 | 0.059028733232285 | 0.967015297525325 | 0.333536386400309 | 0.757852267925996 | NM_019674 |
| Utp14b | 316.6823049 | 0.0919795439101535 | 0.0950993740399516 | 0.967193999316048 | 0.333447061401577 | 0.757852267925996 | NM_001001981,NM_001136226 |
| Otud3 | 427.143216 | -0.0740004575101063 | 0.0765430084701966 | -0.966782714569153 | 0.333652667294471 | 0.757997911341803 | NM_028453 |
| Abcb1b | 153.8916806 | -0.0864868753442564 | 0.0895232792704074 | -0.966082521206808 | 0.334002889971536 | 0.758212398552346 | NM_011075 |
| Btf3l4 | 3095.723011 | 0.0585354008834084 | 0.06059940413863 | 0.965940205443277 | 0.33407410244171 | 0.758212398552346 | NM_027453 |
| Cbx5 | 9224.183557 | 0.0554677685470662 | 0.0574227705290268 | 0.965954237945165 | 0.334067080373214 | 0.758212398552346 | NM_007626,NM_001076789,NM_001110216 |
| Ltbr | 324.3509927 | -0.0773584136316247 | 0.0800589167321903 | -0.96626855307574 | 0.33390981744352 | 0.758212398552346 | NM_010736 |
| Napepld | 872.409783 | 0.0625806848889969 | 0.0647924690244077 | 0.965863561479999 | 0.334112457877845 | 0.758212398552346 | NM_178728 |
| Pitpnc1 | 3875.589455 | 0.0639917808610652 | 0.0662449797271426 | 0.965986873641472 | 0.334050749363154 | 0.758212398552346 | NM_145823 |
| Ppfia3 | 9435.835716 | -0.0544945062357665 | 0.0564075912911926 | -0.966084617129808 | 0.334001841279251 | 0.758212398552346 | NM_029741 |
| 9530036O11Rik | 39.84419328 | -0.0897988076619125 | 0.0930004211523392 | -0.965574204387932 | 0.334257288321596 | 0.758238351847355 | NR_015562 |
| Dazap2 | 5582.617312 | 0.048143788255914 | 0.0499107162176179 | 0.964598224677845 | 0.334746088973355 | 0.758238351847355 | NM_011873 |
| Ddx56 | 1579.473256 | 0.0517497300607295 | 0.0536121429712251 | 0.965261360444122 | 0.334413920052024 | 0.758238351847355 | NM_026538 |
| Eef2 | 45690.34342 | 0.0517751171444614 | 0.0536657917006397 | 0.964769464937273 | 0.334660293282394 | 0.758238351847355 | NM_007907 |
| Kcnd3 | 4289.439797 | 0.0733637117511962 | 0.0760149759471491 | 0.965121817603464 | 0.334483800290726 | 0.758238351847355 | NM_019931,NM_001039347 |
| March2 | 2445.142289 | -0.0476496662368144 | 0.0493579724338686 | -0.965389457613093 | 0.334349779860316 | 0.758238351847355 | NR_045522,NR_045523,NM_001252480,NM_145486 |
| Nat9 | 539.8454868 | -0.0691194339617108 | 0.0716180183083709 | -0.965112350136501 | 0.334488541748075 | 0.758238351847355 | NM_025400 |
| Pcdhb3 | 291.9673249 | -0.0768788959236569 | 0.0797130757176105 | -0.964445233502294 | 0.334822753400443 | 0.758238351847355 | NM_053128 |
| Prkg2 | 570.205071 | 0.0824976499260779 | 0.0855446178975174 | 0.964381535082782 | 0.334854676242653 | 0.758238351847355 | NM_008926 |
| Slc48a1 | 6386.780429 | 0.0482220525816552 | 0.0499970036462321 | 0.964498851228444 | 0.334795884077838 | 0.758238351847355 | NM_026353 |
| Smpdl3b | 176.0870777 | -0.0917599204330276 | 0.095030557744002 | -0.965583309320513 | 0.334252730458985 | 0.758238351847355 | NM_133888 |
| Vat1 | 2089.559691 | 0.0913694086704835 | 0.0947148528786814 | 0.964678779446736 | 0.334705727236062 | 0.758238351847355 | NM_012037 |
| Wasf3 | 2974.261158 | 0.0490462622064901 | 0.0508551650979667 | 0.964430301465113 | 0.334830236503811 | 0.758238351847355 | NM_145155 |
| Zc2hc1a | 2173.234254 | 0.0638887589813287 | 0.0662444476521356 | 0.964439454863038 | 0.33482564931903 | 0.758238351847355 | NM_173181 |
| Tmem41b | 1576.260596 | 0.0600894975943155 | 0.0623169970906192 | 0.964255346048454 | 0.334917922419142 | 0.758263363631619 | NM_153525 |
| Armcx1 | 2873.811515 | 0.0516626313922511 | 0.0536024746249189 | 0.963810565720301 | 0.335140908509055 | 0.758531760206066 | NM_030066,NM_001166377,NM_001166380,NM_001166378,NM_001166379 |
| Smarca5 | 3038.980075 | 0.0748015922417975 | 0.077607917087701 | 0.96383970925631 | 0.335126294766769 | 0.758531760206066 | NM_053124 |
| Stac | 40.35693625 | 0.0758304730731258 | 0.0787013862241529 | 0.963521441123663 | 0.33528590945838 | 0.758741723133265 | NM_016853 |
| Cited4 | 233.5489674 | 0.0888736183087839 | 0.0922964103925181 | 0.96291521989666 | 0.335590070970639 | 0.759126438682198 | NM_019563 |
| Npepps | 7205.206329 | 0.0492660792833828 | 0.0511671466753614 | 0.962845936982959 | 0.335624843839887 | 0.759126438682198 | NM_008942 |
| Tssc1 | 1624.957373 | 0.0652464887715552 | 0.0677698287866871 | 0.962766026411628 | 0.335664953576742 | 0.759126438682198 | NM_201357 |
| Vamp5 | 70.30868075 | -0.0915821128836389 | 0.0950961237672078 | -0.963047801063153 | 0.335523535380955 | 0.759126438682198 | NM_001080742,NM_016872 |
| Fam210b | 1147.989043 | 0.0736586776408752 | 0.0765398968322766 | 0.962356636072882 | 0.335870488417922 | 0.759354819417703 | NM_025912 |
| Hectd2 | 667.443337 | 0.0842492429727149 | 0.0875428696474801 | 0.962376985264156 | 0.33586027017261 | 0.759354819417703 | NM_172637,NM_001163471 |
| Cstf2t | 3550.870806 | 0.0552582409105047 | 0.0574381518581318 | 0.962047682992808 | 0.336025652254694 | 0.759375713552499 | NM_031249 |
| Pdcd2l | 418.2382283 | -0.0669218044719349 | 0.0695634077255983 | -0.962025965374159 | 0.336036561112176 | 0.759375713552499 | NM_026549 |
| Suv39h1 | 1288.646807 | 0.0643655151673273 | 0.0669059661072348 | 0.962029530582733 | 0.336034770276244 | 0.759375713552499 | NM_011514 |
| Rilpl1 | 2326.249729 | 0.0791529952633853 | 0.0823007612349963 | 0.961752893601761 | 0.336173745740839 | 0.759567558038796 | NM_021430 |
| Aldh3b1 | 198.6197626 | -0.084750687858096 | 0.0881397629548882 | -0.961548851696744 | 0.336276274961368 | 0.759681052890954 | NM_026316 |
| D330050G23Rik | 77.46233582 | -0.0913594126834543 | 0.0950248520281076 | -0.961426518785117 | 0.336337755794821 | 0.759701794538263 | NR_040335,NR_040336 |
| Degs2 | 120.4103886 | -0.087555556702814 | 0.0911250651283386 | -0.960828467771218 | 0.336638422294164 | 0.759707980266187 | NM_027299,NM_001171002 |
| Dnaja2 | 7142.598067 | 0.0595174081319133 | 0.0619266612312332 | 0.961095059035659 | 0.336504373813243 | 0.759707980266187 | NM_019794 |
| Fam219a | 7299.216877 | 0.0430666289289812 | 0.0448085623280225 | 0.961124987981329 | 0.336489326965704 | 0.759707980266187 | NM_001159583,NM_027993 |
| Mxd3 | 29.13045532 | 0.0778627331466074 | 0.0810380591189258 | 0.960816855600423 | 0.336644261952601 | 0.759707980266187 | NM_016662 |
| Shd | 847.1482601 | 0.0712164154325637 | 0.0741222362856832 | 0.9607969079357 | 0.336654293609627 | 0.759707980266187 | NM_001159523,NM_009168 |
| Six3 | 469.0312369 | 0.0869572792258727 | 0.0904817089783074 | 0.961048152248321 | 0.336527957181203 | 0.759707980266187 | NM_011381 |
| Gtse1 | 31.31342911 | -0.0792950460744162 | 0.0825670612906567 | -0.96037142214954 | 0.336868315696661 | 0.760072872601692 | NM_013882,NM_001168672 |
| 1110004E09Rik | 892.4639649 | 0.0573616365158875 | 0.0597824881590432 | 0.959505672686032 | 0.337304063307316 | 0.760113439446824 | NM_026502 |
| Aaed1 | 214.405573 | 0.082041967367791 | 0.0854848156888627 | 0.95972561567422 | 0.337193327646944 | 0.760113439446824 | NM_025370 |
| Dynlt3 | 3879.281735 | 0.0605216491889306 | 0.0630536671770081 | 0.959843446044629 | 0.337134012688425 | 0.760113439446824 | NM_025975 |
| Fam76a | 2283.434185 | 0.0455303376189169 | 0.0474174483764933 | 0.960202186701554 | 0.336953466530749 | 0.760113439446824 | NM_145553,NM_001163792 |
| Pccb | 2104.179748 | 0.052528500609963 | 0.0547454746432418 | 0.959503976397573 | 0.337304917435924 | 0.760113439446824 | NM_025835 |
| Tceal6 | 2325.855021 | 0.0573050911035527 | 0.0596953566183972 | 0.959958937340399 | 0.337075881712849 | 0.760113439446824 | NM_025355 |
| Tmem60 | 541.4205692 | 0.0757732803571259 | 0.0789294418563499 | 0.960012874473785 | 0.337048735396773 | 0.760113439446824 | NM_177601 |
| Wdr65 | 63.5095083 | 0.089017177956917 | 0.0927546168981855 | 0.95970616809974 | 0.337203118060591 | 0.760113439446824 | NM_026789 |
| Mll2 | 4004.258196 | -0.0732145129572991 | 0.0763228470072457 | -0.959273871842362 | 0.337420794398936 | 0.760256624699836 | NM_001033276 |
| 1190007I07Rik | 161.9971968 | -0.0855183607051405 | 0.0891758787802826 | -0.958985343063972 | 0.337566128991044 | 0.760312043979752 | NM_001135567,NM_001135569,NM_001135568 |
| Dlx1as | 618.7221959 | 0.0766308448033831 | 0.0799115370123593 | 0.9589459503392 | 0.337585974589462 | 0.760312043979752 | NR_002854 |
| Wdr1 | 6441.028026 | 0.0497708104712094 | 0.0519033468050699 | 0.958913317442332 | 0.337602415232645 | 0.760312043979752 | NM_011715 |
| Casr | 42.59387738 | 0.0763704216595752 | 0.0797088925233989 | 0.958116707457156 | 0.338003911539941 | 0.760581218590385 | NM_013803 |
| Cenpk | 24.15009258 | 0.0813561711221023 | 0.0849035795694035 | 0.958218387666431 | 0.33795264702441 | 0.760581218590385 | NM_181061,NM_021790,NR_075088 |
| Fbxo31 | 6093.084723 | 0.0547902273205173 | 0.057195346683662 | 0.957949037769679 | 0.338088457141548 | 0.760581218590385 | NM_133765 |
| Gm13152 | 77.82015888 | 0.0910753079578472 | 0.095029273207826 | 0.958392134165526 | 0.337865060119815 | 0.760581218590385 | NM_001039209 |
| Meaf6 | 1468.924919 | 0.0563018823040937 | 0.0587705027965524 | 0.957995586646512 | 0.338064984018956 | 0.760581218590385 | NM_027310 |
| Shc3 | 1132.804364 | 0.0710515804546815 | 0.0741239055031172 | 0.958551495262127 | 0.337784737826115 | 0.760581218590385 | NM_009167 |
| Slc2a6 | 1010.363617 | -0.0797582928315825 | 0.0832413408434317 | -0.958157233214198 | 0.337983478909207 | 0.760581218590385 | NM_172659,NM_001177627 |
| Mbtd1 | 918.9583902 | -0.0621095451628526 | 0.0648619163245992 | -0.957565682333953 | 0.338281810938567 | 0.760898356409667 | NM_134012 |
| BC006965 | 57.13497006 | -0.0909618562596562 | 0.0950516485267758 | -0.95697294754475 | 0.338580909584558 | 0.760991092254312 | NR_024085 |
| Cacnb1 | 4443.635518 | 0.0640704995629891 | 0.0669590389974174 | 0.95686109780429 | 0.338637368878691 | 0.760991092254312 | NM_145121,NM_001159320,NM_031173,NM_001159319 |
| Camk2d | 5114.463441 | 0.0738426027169563 | 0.0771609192809593 | 0.956994854455786 | 0.338569852164364 | 0.760991092254312 | NM_001025439,NM_001025438,NM_023813 |
| Gsk3b | 8614.97969 | 0.0576325248695733 | 0.0602063360817192 | 0.957250160370954 | 0.338441004682533 | 0.760991092254312 | NM_019827 |
| Hsf2 | 879.9272426 | -0.0701047583023422 | 0.0732646055605702 | -0.956870753154937 | 0.33863249483129 | 0.760991092254312 | NM_008297 |
| Upp1 | 55.44182546 | -0.0907443109626306 | 0.0948147054228556 | -0.957070008897124 | 0.338531920041435 | 0.760991092254312 | NM_009477,NM_001159402,NM_001159401 |
| Pex7 | 845.7996189 | 0.058676040398421 | 0.0613336290297675 | 0.956669959475955 | 0.338733865298465 | 0.761006690634724 | NM_001161825,NM_008822 |
| Tmem70 | 1165.906088 | 0.0657560788164814 | 0.0687365069162828 | 0.956639808545533 | 0.338749088644095 | 0.761006690634724 | NM_027415,NM_026392 |
| Alyref2 | 88.93795727 | 0.0905127445248856 | 0.0946481808407089 | 0.956307281565368 | 0.33891701219571 | 0.761116258869698 | NM_019484 |
| Atp5d | 5763.485063 | 0.0550487095608606 | 0.0575703770370434 | 0.956198524206985 | 0.338971945410516 | 0.761116258869698 | NM_025313 |
| Dap3 | 2718.75506 | 0.049523254926647 | 0.0517956198733879 | 0.956128241108117 | 0.33900744836066 | 0.761116258869698 | NM_001164533,NM_022994 |
| Heph | 360.6813782 | 0.0844679114432814 | 0.088322147274237 | 0.956361615405609 | 0.338889570376446 | 0.761116258869698 | NM_001159628,NM_010417,NM_001159627,NM_181273 |
| Tbc1d2b | 1090.465474 | 0.0526623915743251 | 0.0550877871050832 | 0.955972173539454 | 0.339086293186719 | 0.761175628933747 | NM_194334 |
| 9530077C05Rik | 268.9377397 | 0.075594223103433 | 0.0791119802455161 | 0.955534457219171 | 0.339307488808561 | 0.761467070181342 | NM_026739 |
| Lypla1 | 797.6182795 | 0.072016557775993 | 0.0753699322903561 | 0.955507794521501 | 0.339320965529659 | 0.761467070181342 | NM_008866 |
| Otud1 | 657.7845662 | 0.0781575024690662 | 0.0818086854747329 | 0.955369249799346 | 0.339390998801394 | 0.761506587672079 | NM_027715 |
| Desi1 | 5944.351073 | 0.0531367288050436 | 0.055630846626599 | 0.955166639143635 | 0.339493433592274 | 0.761618782449632 | NM_134095 |
| Arf5 | 3499.716555 | 0.0557042430975966 | 0.0583448442733573 | 0.954741482154123 | 0.339708446595307 | 0.761679979053987 | NM_007480 |
| Gtf2e2 | 568.4405945 | 0.073610224790495 | 0.0770834218658325 | 0.954942359961877 | 0.339606846558067 | 0.761679979053987 | NM_001167921,NM_026584,NM_001167922 |
| Lrrc48 | 535.1049744 | -0.0663604082390736 | 0.0695093213097346 | -0.95469797415185 | 0.33973045465309 | 0.761679979053987 | NM_029044 |
| Pdzd8 | 3081.970805 | 0.0648758779960111 | 0.0679520781187992 | 0.954729859513493 | 0.33971432569432 | 0.761679979053987 | NM_001033222 |
| Adck4 | 979.8631154 | 0.0745338859663906 | 0.0781140824383723 | 0.954167080246941 | 0.339999075292188 | 0.761914236978731 | NM_133770 |
| Ercc1 | 661.1613551 | -0.0694115402092786 | 0.0727342989597306 | -0.954316480697894 | 0.339923468187918 | 0.761914236978731 | NM_001127324,NM_007948 |
| Ptbp2 | 2955.577599 | 0.0605059373527287 | 0.0634183062298596 | 0.954076842314661 | 0.340044747234828 | 0.761914236978731 | NM_019550 |
| Ubtd2 | 1557.637216 | 0.0715247086512521 | 0.0749598093274752 | 0.954174100667516 | 0.339995522225747 | 0.761914236978731 | NM_173784 |
| Angptl1 | 35.88989857 | 0.0834930208391456 | 0.0875477226341996 | 0.953685810743522 | 0.340242704694122 | 0.762240211040532 | NM_028333 |
| Atp2a2 | 43170.81825 | 0.0555749082307642 | 0.0582825059948578 | 0.953543559634645 | 0.340314736806988 | 0.762284019561806 | NR_027838,NM_001110140,NM_009722 |
| Erbb4 | 1075.581145 | 0.0824490718281034 | 0.0865018151534738 | 0.953148459160309 | 0.340514856256917 | 0.762546123415894 | NM_010154 |
| Ltv1 | 639.4773597 | 0.0625129124911447 | 0.065594098523068 | 0.953026474922287 | 0.340576656829359 | 0.762546123415894 | NM_181470 |
| Nxt2 | 1089.310603 | 0.0590636881738653 | 0.0619773775561729 | 0.952987856259862 | 0.340596223604082 | 0.762546123415894 | NM_172782,NM_001161430 |
| Sdc2 | 2040.638072 | 0.0667926133093572 | 0.0700941866719167 | 0.952898043057224 | 0.340641731711809 | 0.762546123415894 | NM_008304 |
| Adam12 | 171.3464405 | -0.0883308459678812 | 0.0927556865298632 | -0.952295748891284 | 0.340947013179686 | 0.762632919878766 | NM_007400 |
| Apod | 8109.46498 | 0.0903625611623531 | 0.0948700865363635 | 0.952487390508675 | 0.340849857868147 | 0.762632919878766 | NM_007470 |
| Bahcc1 | 1277.197701 | -0.0676769304507025 | 0.0710742739446948 | -0.952200095682498 | 0.340995512502587 | 0.762632919878766 | NM_198423 |
| Polr2b | 3279.652756 | 0.0534155816806708 | 0.0560850629003236 | 0.952402991427556 | 0.340892642939465 | 0.762632919878766 | NM_153798 |
| Rabl3 | 347.8712514 | 0.0744421063085472 | 0.0781721988076822 | 0.952283643596726 | 0.340953150718015 | 0.762632919878766 | NM_001042499 |
| Vps45 | 1589.686145 | 0.0616318336642327 | 0.0647060607368659 | 0.952489348947777 | 0.340848865102473 | 0.762632919878766 | NM_013841 |
| Lipe | 424.3891044 | 0.0717341700935943 | 0.0753746463962363 | 0.951701580349651 | 0.341248347677606 | 0.762819666191009 | NM_001039507,NM_010719 |
| Ppp1ca | 8337.210691 | 0.0489953748416294 | 0.0514733090591366 | 0.951859822832445 | 0.341168077838069 | 0.762819666191009 | NM_031868 |
| Spag16 | 38.08820618 | 0.0742724778122815 | 0.0780376761882354 | 0.951751531313262 | 0.341223008322493 | 0.762819666191009 | NM_029160,NM_001271533,NM_025728 |
| Usp53 | 839.3599631 | 0.0694451080637481 | 0.0729755703420384 | 0.951621312971685 | 0.341289068606317 | 0.762819666191009 | NM_133857 |
| Gm9897 | 40.97565767 | -0.0868098851287704 | 0.0912911071232317 | -0.950912831099614 | 0.341648627719983 | 0.76303628017535 | NM_001042670 |
| Mustn1 | 41.43149693 | -0.080866565402365 | 0.0850387254541356 | -0.950938116375923 | 0.341635791109513 | 0.76303628017535 | NM_181390 |
| Tmem151a | 9943.82236 | 0.0506747564739111 | 0.0532831464123454 | 0.951046623293441 | 0.341580708759939 | 0.76303628017535 | NM_001001885 |
| Wdr81 | 1517.337103 | -0.0635217935062622 | 0.0667774567163537 | -0.951246073597557 | 0.341479474821344 | 0.76303628017535 | NM_138950 |
| Ypel3 | 4972.71571 | -0.0491002096673167 | 0.0516276052427615 | -0.951045655448077 | 0.341581200050926 | 0.76303628017535 | NM_025347,NM_026875 |
| Oprl1 | 2390.430432 | 0.057961499998342 | 0.0609711178911504 | 0.950638630274397 | 0.341787851445971 | 0.763229873959135 | NM_011012,NM_001252565,NR_045540,NR_045541,NR_045542 |
| Clpb | 2254.761775 | 0.0430196953609778 | 0.0452830492293493 | 0.950017635585712 | 0.342103291731496 | 0.763752913487252 | NM_009191 |
| Cnppd1 | 1292.686382 | -0.0543898665409511 | 0.0572542680626269 | -0.949970515411312 | 0.342127234480349 | 0.763752913487252 | NM_026977 |
| Actr6 | 501.0616266 | 0.073724826662342 | 0.0777554101218604 | 0.948163305251666 | 0.343046324637059 | 0.763762832218036 | NM_025914 |
| Adrb1 | 955.4151893 | -0.0822970722770086 | 0.0868185035465186 | -0.947920879941367 | 0.343169734422467 | 0.763762832218036 | NM_007419 |
| Aff2 | 564.5936772 | 0.0811611094032398 | 0.08549003210852 | 0.949363421693594 | 0.342435807387817 | 0.763762832218036 | NM_008032 |
| Atp1b2 | 24295.2928 | 0.0653369840225143 | 0.0689097080467903 | 0.948153545769631 | 0.34305129228152 | 0.763762832218036 | NM_013415 |
| Bckdhb | 790.4720255 | 0.0572024676585521 | 0.0602245208547704 | 0.949820220180649 | 0.342203609800287 | 0.763762832218036 | NM_199195 |
| Chrna2 | 26.99694746 | -0.082432195894977 | 0.086882806233317 | -0.94877455584954 | 0.342735285418603 | 0.763762832218036 | NM_144803 |
| Dnajc16 | 3140.019683 | 0.0494738505260913 | 0.0522688037216315 | 0.946527316553383 | 0.343879696266011 | 0.763762832218036 | NM_172338 |
| Efhd1 | 1008.073996 | 0.0806985105475735 | 0.085201727789662 | 0.947146409363837 | 0.343564178869256 | 0.763762832218036 | NM_028889 |
| Esrra | 1930.98202 | 0.0694674278485929 | 0.0731841741626336 | 0.949213797155364 | 0.342511885704545 | 0.763762832218036 | NM_007953 |
| Fis1 | 3826.085012 | 0.04692749285762 | 0.049573618669459 | 0.946622298656821 | 0.343831277125381 | 0.763762832218036 | NM_025562,NM_001163243 |
| Gcc2 | 3650.888607 | 0.0618563610442256 | 0.0652094521410686 | 0.948579676921849 | 0.342834431367293 | 0.763762832218036 | NM_027375 |
| Golga7 | 3026.161313 | 0.0526486574930351 | 0.0554922488225209 | 0.948756963543136 | 0.342744234868661 | 0.763762832218036 | NM_020585,NM_001042484 |
| Hs3st1 | 706.8753255 | -0.0722195934501214 | 0.0761649076197872 | -0.948200368214707 | 0.343027459748997 | 0.763762832218036 | NM_010474 |
| Il6ra | 193.0856311 | -0.0846959008552236 | 0.0894286347770014 | -0.947078092676028 | 0.343598987031519 | 0.763762832218036 | NM_010559 |
| Lonrf1 | 1468.410044 | 0.0723082914560963 | 0.0763402312835967 | 0.947184600312225 | 0.343544721109787 | 0.763762832218036 | NM_001081150 |
| Lrrc16a | 1015.362008 | -0.0583008906937637 | 0.0613906027431317 | -0.94967125404688 | 0.342279320474301 | 0.763762832218036 | NM_026825 |
| Manea | 1578.385944 | 0.0619278349284743 | 0.0653202477287818 | 0.948064912209255 | 0.343096409486705 | 0.763762832218036 | NM_172865 |
| Map4 | 19264.97133 | 0.0476208302419453 | 0.0502771082717556 | 0.947167247259872 | 0.343553562162468 | 0.763762832218036 | NM_001205330,NM_001205331,NM_001205332,NM_008633 |
| March1 | 71.76353904 | -0.0902681864376902 | 0.0950814417138124 | -0.94937755266049 | 0.342428622893697 | 0.763762832218036 | NM_175188,NM_001166375,NM_001166372 |
| Mul1 | 2047.812084 | 0.0450912700159182 | 0.0475289757577761 | 0.948711166125665 | 0.342767533350164 | 0.763762832218036 | NM_026689 |
| Oxsm | 466.5000625 | -0.0735309820318472 | 0.077497364695686 | -0.948819128503093 | 0.342712611360743 | 0.763762832218036 | NM_027695 |
| Pex16 | 739.1143716 | 0.0672194467185856 | 0.0708269341720953 | 0.949066163943448 | 0.342586962099594 | 0.763762832218036 | NR_028114,NM_145122 |
| Rps6kl1 | 1510.057494 | -0.0631795039481033 | 0.0666764913448254 | -0.947552918184657 | 0.343357104379565 | 0.763762832218036 | NM_146244 |
| S100a9 | 22.39428828 | -0.0535326312290327 | 0.0563721198503714 | -0.949629557503328 | 0.342300514279634 | 0.763762832218036 | NM_009114 |
| Scn9a | 127.1963583 | 0.0584983066026899 | 0.0617847903658091 | 0.94680755985962 | 0.343736848826466 | 0.763762832218036 | NM_018852 |
| Sertm1 | 1336.616456 | 0.0739003191879657 | 0.0780673844309753 | 0.946622199867679 | 0.343831327482979 | 0.763762832218036 | NM_177854 |
| Sidt2 | 4030.671608 | -0.0517682819705955 | 0.0546830624493162 | -0.946696831739768 | 0.343793285354734 | 0.763762832218036 | NM_172257 |
| Srpk2 | 10303.57613 | 0.0532964619351082 | 0.0562701530849771 | 0.947153313313754 | 0.343560661350236 | 0.763762832218036 | NM_009274 |
| Sumo3 | 5379.947736 | 0.0467475133168637 | 0.0493634856037777 | 0.947005924421314 | 0.343635760059022 | 0.763762832218036 | NM_019929 |
| Tnfrsf25 | 174.5804649 | -0.057684085835637 | 0.0608694028116822 | -0.947669652914125 | 0.343297654763548 | 0.763762832218036 | NM_033042 |
| Ube2h | 5839.538812 | 0.0461744624330055 | 0.0486529141840669 | 0.949058513911731 | 0.342590852681667 | 0.763762832218036 | NM_009459,NM_001169577,NM_001169576 |
| Ufl1 | 1485.623945 | -0.0588063616105699 | 0.062036713363796 | -0.94792838662676 | 0.343165912620229 | 0.763762832218036 | NM_026194 |
| Wibg | 603.8396559 | 0.0650821801073869 | 0.0687297726326031 | 0.946928494224555 | 0.343675217081581 | 0.763762832218036 | NM_001170869,NM_030100,NM_001253704,NM_001253705 |
| Wnk4 | 295.6890543 | -0.0883735923187596 | 0.0933738055865246 | -0.946449507585599 | 0.343919364280474 | 0.763762832218036 | NM_175638 |
| Alox12 | 27.82107848 | -0.0806157240363031 | 0.085209644244821 | -0.94608685144467 | 0.344104289593124 | 0.763939922150346 | NM_007440 |
| Klhl35 | 23.47374864 | -0.0812784379264 | 0.085905455739179 | -0.94613825428239 | 0.344078074441614 | 0.763939922150346 | NM_028145 |
| Kdm5c | 3416.711453 | 0.0532633658583116 | 0.0563196385641013 | 0.945733446028579 | 0.344284558836079 | 0.764106570153229 | NM_013668 |
| Taf6l | 360.5524529 | -0.0708771240417727 | 0.0749372245446508 | -0.945819977620617 | 0.344240414200018 | 0.764106570153229 | NM_001177798,NM_146092 |
| Fam150b | 39.22457132 | 0.0845413124294551 | 0.0894146005900477 | 0.945497847908129 | 0.344404769008211 | 0.76413986170968 | NM_001159743 |
| Vsig2 | 84.53584868 | -0.0898621826193443 | 0.0950349472702316 | -0.945569868774919 | 0.34436801867097 | 0.76413986170968 | NM_020518 |
| Hnrnpm | 4184.749016 | -0.0446464694060371 | 0.0472404285967363 | -0.945090269759356 | 0.344612792477865 | 0.764478237375078 | NM_029804,NM_001109913 |
| Popdc3 | 31.52475347 | 0.0828118172417389 | 0.0876322171625337 | 0.944992833949935 | 0.344662534529078 | 0.764478237375078 | NM_024286 |
| D14Abb1e | 1332.542042 | -0.0481990225122066 | 0.0510189664489873 | -0.944727536971956 | 0.344797994764554 | 0.764561710651453 | NM_028945,NM_001114879 |
| Fam132b | 29.80364986 | -0.0768093256058387 | 0.0813132759893247 | -0.944609901289953 | 0.344858070231882 | 0.764561710651453 | NM_173395 |
| Paqr4 | 2819.103881 | 0.0547915101324904 | 0.0580000804014578 | 0.944679899635333 | 0.344822321912502 | 0.764561710651453 | NM_023824 |
| 9130401M01Rik | 909.9953759 | 0.0549739533877569 | 0.0582067327006145 | 0.944460388637766 | 0.344934434611072 | 0.764614313621309 | NM_029418 |
| Lysmd1 | 883.2985224 | 0.0525477516544392 | 0.0556449151541764 | 0.944340583660595 | 0.344995633420691 | 0.764633288231455 | NM_153121 |
| Cd38 | 118.2301469 | 0.0877826153568307 | 0.0929748212087789 | 0.944154710012414 | 0.345090595150844 | 0.764727076302237 | NM_007646 |
| Dhx36 | 3413.794205 | 0.0588222336097292 | 0.0623207123894887 | 0.94386330570333 | 0.345239505408173 | 0.764823708335996 | NM_028136 |
| Uqcr11 | 2486.795524 | 0.0677948905571147 | 0.0718250997266306 | 0.943888568413339 | 0.345226594311474 | 0.764823708335996 | NM_025650 |
| Atg2b | 3011.223506 | 0.0569145455207516 | 0.060341414798124 | 0.943208668725498 | 0.345574180242112 | 0.765097831953754 | NM_029654 |
| C78339 | 972.2169703 | 0.0671149922173321 | 0.0712066717338784 | 0.942537975488614 | 0.345917278013351 | 0.765097831953754 | NM_001033192 |
| E130307A14Rik | 594.7339284 | -0.0617243097751585 | 0.0654979391288787 | -0.942385525347676 | 0.345995295201997 | 0.765097831953754 | NR_038037 |
| Epha2 | 61.88170521 | -0.0895453106257351 | 0.0949411899828992 | -0.943166086730786 | 0.34555959569061 | 0.765097831953754 | NM_010139 |
| Mmadhc | 1460.132399 | 0.0549544319951975 | 0.0583038919197654 | 0.942551692275067 | 0.345910258922952 | 0.765097831953754 | NM_133839 |
| Sec16b | 72.90354911 | -0.0891860447560085 | 0.0945741982136205 | -0.943027236187175 | 0.345666971905912 | 0.765097831953754 | NM_033354,NR_027641,NM_001159986 |
| Serpind1 | 176.0020591 | -0.0895229655524986 | 0.0949642894121542 | -0.942701368131764 | 0.345833673384947 | 0.765097831953754 | NM_008223 |
| Ssbp4 | 2722.811622 | -0.0621667194625049 | 0.0659303009018209 | -0.942915755156033 | 0.345723995520816 | 0.765097831953754 | NM_133772 |
| Thsd4 | 379.6541867 | -0.0894301072044457 | 0.0948361355250821 | -0.942996113341136 | 0.345682890936515 | 0.765097831953754 | NM_172444,NM_001040426 |
| Zbtb1 | 474.2136807 | -0.080599558261487 | 0.0855229962273516 | -0.942431413969919 | 0.345971810270785 | 0.765097831953754 | NM_178744 |
| Zc3h3 | 1023.463527 | -0.068617883839449 | 0.0727700968669018 | -0.942940669227812 | 0.345711251214401 | 0.765097831953754 | NM_172121 |
| Zscan18 | 1072.096508 | -0.0624118256328446 | 0.0661993683638245 | -0.942785817680859 | 0.345790467348837 | 0.765097831953754 | NM_001017955 |
| Krit1 | 1658.779825 | -0.0647766821539694 | 0.0687490478791921 | -0.942219334699688 | 0.346080356940113 | 0.765103440436257 | NM_030675,NM_001170552,NR_033173 |
| Slc39a7 | 3757.58495 | 0.0502257990631487 | 0.0533083682660062 | 0.942174759739866 | 0.346103174108953 | 0.765103440436257 | NM_008202,NM_001077709 |
| Myl6b | 650.6952316 | 0.0590591369395788 | 0.0626908167599418 | 0.942069987151873 | 0.346156809199955 | 0.765105570669286 | NM_172259 |
| 2410002F23Rik | 2425.453917 | 0.0510886832462324 | 0.0542707861641639 | 0.94136619085808 | 0.346517233202885 | 0.765286747186386 | NM_025880 |
| Cml3 | 202.9223107 | 0.0877836951480161 | 0.0932241347033273 | 0.941641297367634 | 0.346376318843294 | 0.765286747186386 | NM_001037842,NM_053097 |
| Nabp2 | 1730.64675 | 0.0549849945813665 | 0.0584207249409521 | 0.941189871179136 | 0.346607566414652 | 0.765286747186386 | NM_027257 |
| Pappa2 | 172.7907333 | 0.0863367920408615 | 0.0916855714049686 | 0.941661710974326 | 0.346365864092015 | 0.765286747186386 | NM_001085376 |
| Pcnt | 1609.595764 | 0.0512313044114763 | 0.0544298711182043 | 0.941235085789901 | 0.346584400350305 | 0.765286747186386 | NM_008787 |

| | | | | | | | | | | | | | | | |
|---|---|---|---|---|---|---|---|---|---|---|---|---|---|---|---|
| Sirt1 | 503.3344067 | 0.160635589 | 0.110694124 | 1.451166366 | 0.14673354 | 0.345620169 | | Plekhg1 | 890.4909819 | 0.0882750844561554 | 0.093786506228526 | 0.941234384411963 | 0.346584759699325 | 0.765286747186386 | NM_001159942,NM_001033253 |
| Dyrk2 | 664.1870862 | -0.151923421 | 0.104698429 | -1.451057307 | 0.146763903 | 0.345639142 | | Zfp426 | 1130.635745 | 0.0713201615090947 | 0.0757746570538191 | 0.941213913491413 | 0.346595248022581 | 0.765286747186386 | NM_001110309,NM_146221 |
| Gcdh | 968.8635853 | -0.132204118 | 0.091125713 | -1.450788292 | 0.146838821 | 0.345763023 | | Slc46a1 | 389.7501173 | 0.075541298542524 | 0.0803108282058517 | 0.940611623987958 | 0.346903922873009 | 0.765791249635359 | NM_026740 |
| Fosl2 | 1709.855283 | -0.242013956 | 0.166892044 | -1.450122785 | 0.147024282 | 0.346147125 | | Sugp2 | 4230.234792 | -0.0553394655234151 | 0.0588380758653618 | -0.940538328446489 | 0.346941498957063 | 0.765791249635359 | NM_172755,NM_001168290 |
| Stac2 | 4617.538443 | 0.140794788 | 0.097098875 | 1.450014616 | 0.147054444 | 0.346165535 | | Itpa | 1876.580369 | 0.052863699632084 | 0.0562205144132478 | 0.940291994546874 | 0.347067804793627 | 0.765953651235525 | NM_025922 |
| Gspt1 | 4663.855509 | -0.085997981 | 0.059328604 | -1.449519703 | 0.147192502 | 0.346377881 | | Bud13 | 456.559903 | -0.0647766633674767 | 0.0689063947505288 | -0.940067516258781 | 0.347182929811647 | 0.765974975462331 | NM_146000 |
| Lrrc73 | 1082.439978 | -0.129301114 | 0.089197751 | -1.4496006 | 0.147169929 | 0.346377881 | | Kcnn3 | 970.8322394 | 0.0751426677306702 | 0.0799308844836768 | 0.940095536488347 | 0.347168558144013 | 0.765974975462331 | NM_080466 |
| Mtx3 | 2105.61415 | 0.134088473 | 0.092509845 | 1.449450849 | 0.147211717 | 0.346377881 | | BC055324 | 56.86938224 | -0.0880897040245665 | 0.0937612836163903 | -0.939510431458808 | 0.347468739013158 | 0.766038371990845 | NM_201364 |
| Cyfip2 | 54647.31487 | 0.118318457 | 0.081647875 | 1.449130872 | 0.147301038 | 0.346535421 | | Cacng3 | 4551.092932 | 0.0559502670674565 | 0.059579454460107 | 0.939086595781428 | 0.347686285731491 | 0.766038371990845 | NM_019430 |
| Greb1l | 431.8401566 | 0.114688336 | 0.079162973 | 1.448762361 | 0.147403958 | 0.346662471 | | Galnt13 | 1293.149979 | 0.0833139585441156 | 0.0886606578458483 | 0.939694793252843 | 0.34737413667315 | 0.766038371990845 | NM_173030 |
| Htatsf1 | 2875.440198 | 0.069943214 | 0.048278811 | 1.448735206 | 0.147411545 | 0.346662471 | | Ndnl2 | 662.5974457 | 0.0591530892466664 | 0.0629461197124002 | 0.939741631683349 | 0.347350104883598 | 0.766038371990845 | NM_023239 |
| Ms4a6b | 43.70438488 | 0.242901223 | 0.167668733 | 1.448697193 | 0.147422165 | 0.346662471 | | Piga | 371.2521065 | -0.0742236556095801 | 0.0790114479646572 | -0.939403814530538 | 0.347523455283128 | 0.766038371990845 | NM_011081 |
| 4921531C22Rik | 57.27701201 | -0.206447803 | 0.142589981 | -1.447842277 | 0.147661166 | 0.347157599 | | Poldip3 | 3470.759358 | 0.0396673928732356 | 0.0422370608129565 | 0.939160824871302 | 0.347648179108613 | 0.766038371990845 | NM_178627 |
| Eif3g | 1941.83215 | 0.100211182 | 0.069224612 | 1.447623599 | 0.147722347 | 0.347157599 | | Syt1 | 72163.96653 | 0.0556607623817127 | 0.0592532094641185 | 0.939371265879171 | 0.347540160484112 | 0.766038371990845 | NM_009306,NM_001252341,NM_001252342 |
| Hist2h2be | 290.7999626 | -0.166032633 | 0.114692421 | -1.447633869 | 0.147719474 | 0.347157599 | | Tmem39a | 342.7355843 | -0.0827888633774802 | 0.0880986753417238 | -0.939728810408925 | 0.347356683098126 | 0.766038371990845 | NM_026407,NM_001205286,NM_001205287 |
| Hlcs | 674.2645561 | 0.106140198 | 0.073319248 | 1.447644387 | 0.147716531 | 0.347157599 | | Trim17 | 147.6025751 | -0.0892568238009112 | 0.0950355509396198 | -0.939194048105428 | 0.347631124320825 | 0.766038371990845 | NM_031172 |
| Rpn1 | 5038.823333 | -0.085150767 | 0.058825246 | -1.447520803 | 0.147751114 | 0.347172546 | | Rnf185 | 1554.603821 | 0.0512327466247719 | 0.0545628944962765 | 0.938966803314809 | 0.347747788603729 | 0.766057685359079 | NM_145355 |
| Gpd1l | 5929.346827 | -0.087578666 | 0.060508477 | -1.447378457 | 0.147790956 | 0.347213506 | | Wdr18 | 2451.001299 | 0.0508516319356548 | 0.0541726013527178 | 0.938696511998008 | 0.347886584784932 | 0.766247237389829 | NM_175450 |
| Rgs13 | 15.78194508 | -0.241833556 | 0.16710352 | -1.447208025 | 0.14783867 | 0.347272946 | | Rorb | 1769.736373 | -0.0636830445226962 | 0.0678592705366389 | -0.938457546317302 | 0.348009324401557 | 0.766401371476201 | NM_146095,NM_001043354 |
| Slc7a3 | 325.8718432 | 0.150012315 | 0.103687426 | 1.446774416 | 0.147960114 | 0.347505535 | | Spen | 3518.302242 | 0.0656467407641073 | 0.0700275485187898 | 0.937441651930639 | 0.348531424113943 | 0.767434813806145 | NM_019763 |
| Spp1 | 470.6882354 | 0.243183104 | 0.168119289 | 1.446491389 | 0.148039426 | 0.347586431 | | Sap18 | 1249.697788 | 0.0589996379233382 | 0.0629446271242959 | 0.937326037484223 | 0.348590873501934 | 0.767449382917412 | NM_009119 |
| Zfp961 | 432.8365317 | -0.13168431 | 0.09103623 | -1.446504432 | 0.14803577 | 0.347586431 | | H2-Ke2 | 1354.50902 | 0.0477437191608606 | 0.0509523443582427 | 0.937026936880028 | 0.348744702063929 | 0.767555385178884 | NM_010385,NM_001185182 |
| Cox20 | 361.2016516 | -0.187986171 | 0.130005917 | -1.445981648 | 0.148182349 | 0.347869291 | | Rnf121 | 924.7430275 | 0.0508257647387202 | 0.0542374854061265 | 0.937096628985295 | 0.348708855299899 | 0.767555385178884 | NM_029211 |
| Dcun1d3 | 797.825229 | -0.120103775 | 0.083090052 | -1.445465146 | 0.148327276 | 0.348065936 | | Rnf223 | 36.36356256 | 0.0855036176382521 | 0.0912688715087429 | 0.936832199465307 | 0.348844879422347 | 0.767659554384035 | NM_001220499 |
| Fbxw4 | 780.6484257 | -0.103151471 | 0.071363226 | -1.445442944 | 0.148333508 | 0.348065936 | | Cap2 | 10577.22333 | 0.0534399283038992 | 0.0570767260311134 | 0.936282299632398 | 0.349127859088673 | 0.768165901412006 | NM_026056 |
| Pbxip1 | 2533.337237 | 0.133857129 | 0.092600149 | 1.445539017 | 0.148306542 | 0.348065936 | | BC022687 | 533.1758843 | 0.0748483741512219 | 0.0799535561947669 | 0.936148155422772 | 0.349196912106486 | 0.768201475883509 | NM_145450 |
| Sash1 | 3563.382899 | -0.123791592 | 0.085648192 | -1.44534973 | 0.148359676 | 0.348074624 | | Gpr139 | 52.35536328 | 0.0828951007243138 | 0.0885598237276509 | 0.936035069121661 | 0.349255131944026 | 0.768213211177911 | NM_001024138 |
| Lats1 | 1494.675448 | -0.118916336 | 0.082300062 | -1.444911865 | 0.148482643 | 0.348265855 | | Rbfa | 966.5041841 | 0.0561130162923407 | 0.0599652807177298 | 0.935758419217237 | 0.349397584658011 | 0.768268270563435 | NM_199197 |
| Lsg1 | 599.9024546 | 0.137036087 | 0.094851767 | 1.444739415 | 0.148531094 | 0.348265855 | | Rnmt | 2775.54388 | 0.0532473692683604 | 0.0569018855785764 | 0.935775128134033 | 0.349388979847258 | 0.768268270563435 | NM_026440,NM_001170953 |
| Rfesd | 380.6642763 | -0.153913388 | 0.106530231 | -1.444786018 | 0.148518 | 0.348265855 | | Sun1 | 4737.759035 | -0.0516584830516422 | 0.0552154204156796 | -0.93558072478196 | 0.349489102623085 | 0.768268270563435 | NM_001256115,NM_001256116,NM_001256117,NM_001256118,NM_024451 |
| Zfp189 | 369.1714255 | 0.228644193 | 0.158249785 | 1.444830991 | 0.148505364 | 0.348265855 | | Zfp511 | 493.0675159 | -0.0705957679009487 | 0.0754570391867749 | -0.935575642269856 | 0.349491720493128 | 0.768268270563435 | NM_027201 |
| 1810024B03Rik | 13.25266823 | 0.236900952 | 0.164011081 | 1.44442041 | 0.148620753 | 0.348423354 | | Zfp935 | 269.2457117 | 0.0874678598774783 | 0.0935045936350533 | 0.935439174452367 | 0.349562016177572 | 0.768306528521019 | NM_001136496,NM_178875 |
| Zfp92 | 146.0444978 | -0.218500309 | 0.151351565 | -1.443660717 | 0.148834437 | 0.348871522 | | Ptprf | 3917.438714 | 0.0656977204024623 | 0.0702523133079441 | 0.935168072181236 | 0.349701689788046 | 0.76849723840562 | NM_011213 |
| Ceacam1 | 129.8818171 | -0.166012966 | 0.115001521 | -1.443571913 | 0.14885943 | 0.348877328 | | Arhgdig | 1809.750878 | 0.0565660598036258 | 0.0604955910662279 | 0.935044336399652 | 0.349765451018925 | 0.768521092346227 | NM_008113 |
| Fzd4 | 290.9696813 | 0.164973725 | 0.114323595 | 1.443041784 | 0.149008701 | 0.349174352 | | Rgs19 | 610.7127286 | 0.0625470618948847 | 0.066899696052627 | 0.934937908322958 | 0.349820299465701 | 0.768525358445065 | NM_026446 |
| Ddx31 | 273.2381838 | 0.144423656 | 0.100106296 | 1.442703014 | 0.14910415 | 0.349239561 | | 2700054A10Rik | 206.1290538 | 0.0759144868946248 | 0.081216012180405 | 0.934723151956736 | 0.349930992265187 | 0.768536074031464 | NR_045436,NR_045437 |
| Gsx2 | 10.77964239 | -0.226065627 | 0.156686147 | -1.442792684 | 0.149078881 | 0.349239561 | | Jund | 2348.946496 | -0.0818383887894283 | 0.0875505877952264 | -0.934755446540708 | 0.34991434510894 | 0.768536074031464 | NM_010592 |
| Tmem60 | 504.1477285 | -0.135518278 | 0.093931413 | -1.442736508 | 0.149094711 | 0.349239561 | | Edn3 | 210.3126546 | -0.080415283486789 | 0.0861304772665894 | -0.933644930793651 | 0.350487080301604 | 0.768667813196619 | NM_007903 |
| Nfat5 | 3984.97482 | -0.113576619 | 0.078730469 | -1.442600565 | 0.149133024 | 0.349254395 | | Fam210a | 1871.731944 | 0.0592336946696521 | 0.0634480913671221 | 0.933577250210967 | 0.350522004960482 | 0.768667813196619 | NM_153794 |
| Mapk8ip1 | 15560.35184 | -0.089832389 | 0.062276033 | -1.442487339 | 0.149164941 | 0.349276348 | | Hecw1 | 2968.508811 | 0.0704742147579347 | 0.0754951130318305 | 0.933493731285906 | 0.350565105591798 | 0.768667813196619 | NM_001081348 |
| Trim62 | 1058.313193 | -0.13624459 | 0.09448407 | -1.441984777 | 0.149306668 | 0.349555381 | | Hgs | 3591.652971 | 0.048798295605953 | 0.0522815099340138 | 0.933375789405144 | 0.350625976197218 | 0.768667813196619 | NM_008244,NM_001159328 |
| Sema3b | 139.7602293 | 0.23917753 | 0.165876814 | 1.441898505 | 0.149331008 | 0.349559546 | | Myh3 | 144.2511269 | -0.0880259570933765 | 0.0942672969083795 | -0.933791038677293 | 0.350411692966789 | 0.768667813196619 | NM_001099635 |
| Mapre2 | 13858.70105 | -0.068772258 | 0.047698897 | -1.441799749 | 0.149358874 | 0.349571962 | | Myo18b | 96.79263086 | 0.0880577898055765 | 0.0943290443821896 | 0.933517246806782 | 0.35055296987308 | 0.768667813196619 | NM_028901 |
| Spag8 | 34.14225429 | 0.240806585 | 0.167069472 | 1.441355995 | 0.149484136 | 0.349812294 | | Rab6a | 21026.24767 | 0.0588098353446781 | 0.0629461153773785 | 0.93428855763533 | 0.350155065076942 | 0.768667813196619 | NM_001163663,NM_024287 |
| Ppfia3 | 8313.470176 | 0.122683488 | 0.085128575 | 1.441155201 | 0.149540842 | 0.349892147 | | Slc2a8 | 895.9724486 | -0.0777165482842421 | 0.0832101316470643 | -0.933979393445461 | 0.350314522667874 | 0.768667813196619 | NM_019488 |
| Arhgap33 | 9924.79032 | -0.194264854 | 0.134853197 | -1.440565454 | 0.149707492 | 0.350068662 | | Sox5 | 1692.413849 | 0.0630804506492934 | 0.067583040985563 | 0.933376920147298 | 0.3506253925817 | 0.768667813196619 | NM_001243163,NM_011444,NM_001113559 |
| Ccdc170 | 12.84278924 | 0.23294575 | 0.161698697 | 1.440616118 | 0.149693164 | 0.350068662 | | Tm6sf2 | 51.37008414 | -0.0688685435141178 | 0.0737281767720865 | -0.934087163541409 | 0.350258932865075 | 0.768667813196619 | NM_181540 |
| Gramd1b | 3737.145377 | 0.124554444 | 0.086467483 | 1.440477274 | 0.149732415 | 0.350068662 | | Wnt10a | 339.8964078 | 0.081708046352111 | 0.0874821338712796 | 0.933996951564252 | 0.350305465484858 | 0.768667813196619 | NM_009518 |
| Ripk2 | 185.3302983 | 0.161142428 | 0.111867478 | 1.440476105 | 0.149732746 | 0.350068662 | | Zfp367 | 918.4298852 | 0.0697604038745387 | 0.074699454475553 | 0.933881035200455 | 0.350365262548389 | 0.768667813196619 | NM_175494 |
| Slc35c1 | 681.2339309 | 0.122829095 | 0.085257512 | 1.440683552 | 0.149674104 | 0.350068662 | | 1810021B22Rik | 30.36539337 | -0.0837521504871246 | 0.0897553131677664 | -0.933116353018333 | 0.350759896660657 | 0.768845368778136 | NR_040417 |
| Tnfsf10 | 141.2003222 | -0.240964294 | 0.167288853 | -1.440408553 | 0.149751845 | 0.350068662 | | Ccdc24 | 49.37526723 | -0.083003796133579 | 0.0889713216294026 | -0.932927539048139 | 0.350857382381748 | 0.768943019987518 | NM_001034876 |
| Tcaim | 693.161967 | -0.131831403 | 0.091533351 | -1.440255399 | 0.149795155 | 0.350117081 | | Mrpl35 | 1132.95293 | 0.0546494546332535 | 0.0585923382453852 | 0.932706498320329 | 0.350971528780624 | 0.769077149957361 | NM_025430 |
| Bcam | 725.60698 | -0.156634008 | 0.108761598 | -1.440159119 | 0.149822386 | 0.350127912 | | Amigo3 | 52.96610658 | 0.0876999090867309 | 0.0941418967122215 | 0.93157151225471 | 0.351558011222536 | 0.769244461938841 | NM_177275 |
| Kif9 | 224.0303608 | 0.211480811 | 0.146854091 | 1.440074356 | 0.149846364 | 0.350131135 | | BC037704 | 51.69833796 | -0.0817780967057773 | 0.0877678966068816 | -0.931754090815997 | 0.351463625349445 | 0.769244461938841 | NR_045645 |
| 1700047M11Rik | 137.4973179 | 0.240445908 | 0.166974812 | 1.439982002 | 0.149872491 | 0.350139382 | | Cep19 | 2411.332976 | 0.0581499242004526 | 0.0624013852122172 | 0.931869124422381 | 0.351404165782981 | 0.769244461938841 | NM_025892 |
| Chrnb1 | 33.31580632 | 0.238639109 | 0.165783043 | 1.439466332 | 0.150018442 | 0.350370787 | | Gemin2 | 197.4827138 | -0.0811531894392195 | 0.0871083154167129 | -0.931635390387187 | 0.351524986936208 | 0.769244461938841 | NM_025656 |
| Cops4 | 2312.059205 | -0.073026959 | 0.050748647 | -1.438993206 | 0.150152448 | 0.350370787 | | Ppp6c | 1330.257402 | 0.0651914257436118 | 0.0699574361347212 | 0.931872712116962 | 0.351402311446831 | 0.769244461938841 | NM_024209 |
| Cybb | 45.33866328 | 0.241927302 | 0.168118492 | 1.439028501 | 0.150142448 | 0.350370787 | | Prss12 | 770.3389609 | 0.070349257657714 | 0.0755023570719772 | 0.931749158382557 | 0.351466175010848 | 0.769244461938841 | NM_008939 |
| Egfl7 | 907.9797413 | -0.137597532 | 0.095599777 | -1.439308083 | 0.150063254 | 0.350370787 | | Sephs2 | 1207.621649 | 0.0564578873550849 | 0.0605561954819187 | 0.932322232362542 | 0.351170021423735 | 0.769244461938841 | NM_009266 |
| Ptprh | 94.63756118 | 0.207294364 | 0.144045992 | 1.439084567 | 0.150126564 | 0.350370787 | | Serinc3 | 13280.33102 | 0.0538544219321656 | 0.0577912330872471 | 0.9318787479558 | 0.351399191776604 | 0.769244461938841 | NM_012032 |
| Sf3b4 | 1633.095674 | 0.109744666 | 0.076252944 | 1.439218743 | 0.150088556 | 0.350370787 | | Slc35g1 | 79.7921708 | -0.0885728982947077 | 0.0950828480336614 | -0.931533921484461 | 0.351577446152552 | 0.769244461938841 | NM_175507 |
| Tubg2 | 1955.438157 | 0.13333591 | 0.092624341 | 1.439534233 | 0.149999218 | 0.350370787 | | Spop | 8966.12308 | 0.0401422974136202 | 0.0430603580468809 | 0.932233247338915 | 0.351215996794591 | 0.769244461938841 | NM_025287 |
| Zcchc12 | 3004.18388 | -0.225782076 | 0.15689959 | -1.439022729 | 0.150144083 | 0.350370787 | | Nbl1 | 1548.662864 | -0.0762741171314304 | 0.0819318301113563 | -0.930946092962451 | 0.351881449874946 | 0.769793665795703 | NM_008675 |
| Col11a1 | 740.0341048 | 0.23554942 | 0.16372862 | 1.438657574 | 0.150247565 | 0.350539946 | | Dalrd3 | 2531.951886 | -0.0734313954341132 | 0.0789062079802412 | -0.930616200090379 | 0.352052131477423 | 0.770051085957092 | NM_026378 |
| Ptpn14 | 626.8537632 | 0.198110501 | 0.137729439 | 1.43840346 | 0.150319612 | 0.350655234 | | Rgs7bp | 13059.16152 | 0.0764665863395604 | 0.0821878427697871 | 0.930388044783554 | 0.352170206252927 | 0.770193377902744 | NM_029879 |
| Cilp2 | 19.06767113 | -0.241646164 | 0.168054717 | -1.4379017 | 0.150461947 | 0.350776018 | | Btn2a2 | 71.13273365 | -0.0882457468330958 | 0.0948886963409177 | -0.929992193338235 | 0.352375126490921 | 0.770257961583715 | NM_175938 |
| Nenf | 1089.415352 | -0.11842915 | 0.082354944 | -1.438033271 | 0.150424614 | 0.350776018 | | Mknk2 | 1884.511799 | 0.066106707171929 | 0.0710884977693561 | 0.929921284683912 | 0.352411841710407 | 0.770257961583715 | NM_021462 |
| Slc16a9 | 123.8605851 | -0.212416251 | 0.14772177 | -1.437948186 | 0.150448756 | 0.350776018 | | Rnf8 | 1275.051034 | -0.0491555884975018 | 0.0528456919578814 | -0.930172104410692 | 0.352281982653967 | 0.770257961583715 | NM_021419 |
| Ubl3 | 6292.624756 | -0.104921536 | 0.072964442 | -1.437981752 | 0.150439232 | 0.350776018 | | Smc1a | 4187.087663 | 0.0527178813166218 | 0.0566803035594969 | 0.930091725096078 | 0.352323594830105 | 0.770257961583715 | NM_019710 |
| 1700003E16Rik | 895.7964678 | -0.13872458 | 0.096503376 | -1.43751013 | 0.150573097 | 0.350929536 | | March5 | 3008.401717 | 0.0562151866099897 | 0.0604701736332251 | 0.929634946162987 | 0.352560127248341 | 0.770466136363682 | NM_027314,NM_001164336,NM_001164337 |
| Vash2 | 113.3097375 | -0.232149435 | 0.161490255 | -1.437544543 | 0.150563326 | 0.350929536 | | Acot3 | 95.90063736 | 0.0879639537621661 | 0.0947254283182086 | 0.928620279938679 | 0.353085908142628 | 0.770519167563791 | NM_134246 |
| Errfi1 | 1257.646495 | 0.187593628 | 0.130609613 | 1.43629265 | 0.150919085 | 0.351663492 | | Actr8 | 1991.324311 | 0.0423408557956047 | 0.0455974638841451 | 0.928579183771823 | 0.353107213845336 | 0.770519167563791 | NM_027493 |
| Slc30a10 | 921.632983 | -0.160788357 | 0.111950719 | -1.436242295 | 0.150933408 | 0.351663492 | | Arih1 | 4352.880523 | 0.0675012988631154 | 0.0726942501978106 | 0.92856448315287 | 0.353114835362258 | 0.770519167563791 | NM_019927 |
| Orc2 | 1186.304747 | 0.094284747 | 0.065659263 | 1.435970225 | 0.151010814 | 0.351790942 | | Casz1 | 84.53915357 | -0.0877967313764413 | 0.0944999222191251 | -0.929066705185847 | 0.352854517934954 | 0.770519167563791 | NM_001159344,NM_027195 |
| Nudt15 | 242.2040111 | -0.132176537 | 0.092066527 | -1.435663343 | 0.151098161 | 0.351941507 | | Fam169b | 23.75927967 | 0.0708672386767986 | 0.0762836068488467 | 0.928996957592992 | 0.352890663038259 | 0.770519167563791 | NM_001013811 |
| Anks1 | 1390.870326 | 0.103593776 | 0.072189185 | 1.435031807 | 0.151278034 | 0.352254563 | | Lsm6 | 1284.17746 | 0.0581098781015855 | 0.0625287231963987 | 0.929330955936299 | 0.352717597241505 | 0.770519167563791 | NM_030145,NM_001191004 |
| Dancr | 88.79515554 | 0.187737546 | 0.130819108 | 1.435092688 | 0.151260687 | 0.352254563 | | Pmepa1 | 2491.679021 | 0.0648668682497251 | 0.0698543524064477 | 0.928601669260307 | 0.353095556474123 | 0.770519167563791 | NM_022995 |
| Gm514 | 53.02135781 | 0.241114965 | 0.168035962 | 1.434900972 | 0.151315319 | 0.352288437 | | Rhobtb3 | 1108.576675 | 0.060794007624826 | 0.0654489661982222 | 0.928876514881871 | 0.352953085247589 | 0.770519167563791 | NM_028493 |
| Celf6 | 885.9140165 | -0.135732105 | 0.094630247 | -1.43434166 | 0.151474787 | 0.352496758 | | Twistnb | 361.8585693 | 0.0731228561272786 | 0.0787179254830136 | 0.928922550722678 | 0.352929225288134 | 0.770519167563791 | NM_172253 |
| Eif4a1 | 10203.51352 | -0.076146183 | 0.053090502 | -1.434271286 | 0.151494861 | 0.352496758 | | Zfp704 | 3341.806908 | 0.0674734504520796 | 0.0726353297438304 | 0.9289343173638 | 0.352923126908306 | 0.770519167563791 | NM_133218 |
| Ralbp1 | 3194.750823 | 0.1247201 | 0.086957319 | 1.434267998 | 0.151495799 | 0.352496758 | | Fbn1 | 304.2643525 | -0.0874012959820094 | 0.0941723362204078 | -0.92809947687237 | 0.353355970960965 | 0.770624605620052 | NM_007993 |
| Ubxn10 | 70.54590193 | 0.214341644 | 0.149438465 | 1.434313744 | 0.15148275 | 0.352496758 | | Idi1 | 1713.710739 | 0.0686023020401227 | 0.0739199483942113 | 0.928062093256209 | 0.353375361285637 | 0.770624605620052 | NM_145360 |
| Adam32 | 8.848845749 | 0.221225018 | 0.154270577 | 1.43400655 | 0.151570394 | 0.35261737 | | Mtpn | 13536.04364 | 0.0684480069584654 | 0.073746983334284 | 0.928146533780248 | 0.353331564200406 | 0.770624605620052 | NM_008098 |
| Cd9 | 1314.85278 | 0.158436203 | 0.110519762 | 1.433555418 | 0.151699174 | 0.352863984 | | Slc39a10 | 10931.11749 | 0.0735039003178338 | 0.0791950495790392 | 0.928137562998486 | 0.353336216946679 | 0.770624605620052 | NM_172653 |
| Btg1 | 1044.495043 | -0.167947795 | 0.117182623 | -1.433214162 | 0.151796644 | 0.353037706 | | BC021891 | 185.8166326 | -0.0878695948803907 | 0.0947083458323327 | -0.927791464502516 | 0.353515752450681 | 0.77081504354527 | NM_145608 |
| 1810030O07Rik | 874.0631097 | 0.101554153 | 0.070862073 | 1.433124211 | 0.151822344 | 0.353044483 | | Surf2 | 936.6297192 | -0.0650730727799871 | 0.0701670586779506 | -0.927402031751914 | 0.353717836152618 | 0.771139920149022 | NM_013678 |
| Adck5 | 489.1439116 | 0.118437759 | 0.082649705 | 1.433008854 | 0.151855307 | 0.353068146 | | Gm5124 | 809.7497056 | 0.0661343889759704 | 0.0713275693953797 | 0.927192522282336 | 0.353826584607981 | 0.771261249702211 | NR_045668 |
| Tmtc1 | 4978.041604 | -0.104850221 | 0.073182979 | -1.432713226 | 0.151939808 | 0.353158624 | | Mta3 | 2303.530504 | 0.0575637262045085 | 0.0620940842298571 | 0.927040424518086 | 0.353905546050942 | 0.771317623696322 | NM_054082,NM_001171052,NM_001171053,NM_001171054 |
| Trpv2 | 741.5159218 | 0.139758326 | 0.097542685 | 1.432791461 | 0.151917442 | 0.353158624 | | Kif3a | 8945.720867 | 0.0470682313271146 | 0.0507865241328714 | 0.926785837990631 | 0.354037739376507 | 0.771423784546133 | NM_008443 |
| Fbxo36 | 85.7448077 | 0.186416612 | 0.130150843 | 1.432311981 | 0.152054556 | 0.353372324 | | Ubxn6 | 2782.362895 | 0.0485654489595167 | 0.0524102765277607 | 0.926639815262042 | 0.354113575337575 | 0.771423784546133 | NM_024432 |
| Xbp1 | 3317.519262 | -0.125886384 | 0.087902989 | -1.432105838 | 0.152113534 | 0.353456373 | | Zxdc | 1253.162486 | 0.0517130072070765 | 0.0558021145245614 | 0.926721283730474 | 0.354071263949059 | 0.771423784546133 | NM_030260,NM_173002 |
| E2f8 | 20.41412082 | -0.240016853 | 0.167613213 | -1.431968565 | 0.152152818 | 0.353494641 | | Rph3al | 36.11499381 | 0.0799600233104236 | 0.0862998694535015 | 0.926537013517803 | 0.354166970914993 | 0.771424414381644 | NM_029548 |
| 1810014B01Rik | 116.553151 | -0.206643687 | 0.144347865 | -1.431567326 | 0.152267686 | 0.353708475 | | Islr2 | 3048.861764 | 0.0783891669006875 | 0.0846396139115643 | 0.926152226811814 | 0.354366875592241 | 0.771744113171348 | NM_001161540,NM_001161535,NM_001161537,NM_001161536,NM_001161541,NM_001161539,NM_001161538,NM_177193 |
| Adm | 67.78721592 | 0.235468475 | 0.164510115 | 1.431331287 | 0.152335291 | 0.353735457 | | 2610100L16Rik | 198.2218503 | -0.0852125691329346 | 0.0920574563731009 | -0.925645488047981 | 0.354630245598355 | 0.77200441038975 | NR_033490 |
| Cops5 | 2306.220243 | -0.085536922 | 0.059765544 | -1.431207958 | 0.152370623 | 0.353735457 | | E2f5 | 290.4172252 | -0.0824985493644902 | 0.0891381592328446 | -0.92551327147097 | 0.354698983542695 | 0.77200441038975 | NM_007892 |
| Mapk10 | 17337.19708 | 0.101869732 | 0.071168756 | 1.431382774 | 0.152320543 | 0.353735457 | | Msl2 | 2151.242394 | 0.0534921675281849 | 0.0577818039365384 | 0.92576146613448 | 0.354569956792399 | 0.77200441038975 | NM_001100451 |
| Rbm15 | 312.457539 | 0.143164203 | 0.10002607 | 1.431268894 | 0.152353165 | 0.353735457 | | Zfyve26 | 1234.320293 | -0.0621330189962008 | 0.06713186264319 | -0.925536944005884 | 0.35468667583391 | 0.77200441038975 | NM_001008550 |
| Zdhhc12 | 162.1212063 | -0.183670499 | 0.128358489 | -1.430918203 | 0.152453659 | 0.353875206 | | Arhgef26 | 1222.559198 | -0.0676863718329503 | 0.0731525864301018 | -0.925276536840231 | 0.354822080288195 | 0.772141518860555 | NM_001081295 |
| Lemd3 | 1441.201453 | -0.115755275 | 0.080919845 | -1.430493034 | 0.152575564 | 0.353946068 | | Mpp1 | 1713.405663 | 0.0542990062171775 | 0.058689720250706 | 0.925187681679643 | 0.35486828995684 | 0.772141518860555 | NM_008621 |
| Mgat3 | 8345.692197 | -0.096636428 | 0.067550496 | -1.430580572 | 0.152550459 | 0.353946068 | | Ccdc60 | 24.45548615 | 0.0720462443965199 | 0.077882029443742 | 0.925068913985639 | 0.35493006175362 | 0.772160263147084 | NM_177759 |
| Psmd14 | 2563.940223 | -0.081487629 | 0.056961884 | -1.430564165 | 0.152555164 | 0.353946068 | | Bdp1 | 1651.38374 | -0.0728569274364376 | 0.0787874657913393 | -0.924727387848618 | 0.355107729384732 | 0.7722068672323 | NM_001081061 |
| Setd8 | 1535.172038 | 0.123200978 | 0.086116558 | 1.430630534 | 0.152536132 | 0.353946068 | | Rom1 | 193.8409752 | 0.0863361582348372 | 0.0933817458383001 | 0.92455069735296 | 0.355199668774905 | 0.7722068672323 | NM_009073 |
| Sfrp2 | 58.36713029 | -0.221246506 | 0.154680019 | -1.430349625 | 0.152616699 | 0.35397807 | | Slc4a4 | 7230.561922 | 0.0631379400719641 | 0.0682846544501598 | 0.924628535947966 | 0.355159164275853 | 0.7722068672323 | NM_001136260,NM_001197147,NM_018760 |
| Tbrg1 | 1728.696609 | 0.086098284 | 0.060196566 | 1.430285651 | 0.152635051 | 0.35397807 | | Uri1 | 1518.548269 | 0.0630929956792981 | 0.0682442900087637 | 0.924516844870038 | 0.35521728533982 | 0.7722068672323 | NM_011274 |
| Dis3l2 | 1045.927317 | 0.086570294 | 0.060534104 | 1.430107804 | 0.152686081 | 0.35404342 | | Wfdc18 | 40.43246144 | -0.0847065037727173 | 0.0916224529230608 | -0.924516874088155 | 0.355217270134705 | 0.7722068672323 | NM_007969 |
| Gria1 | 17469.31063 | -0.131456836 | 0.091941576 | -1.429786637 | 0.152778266 | 0.354204167 | | 1500015A07Rik | 149.6894128 | -0.0806362562866751 | 0.0872609183473632 | -0.924082141396713 | 0.355443550593036 | 0.772370993941384 | NR_029432 |
| Tank | 932.2985497 | -0.097084446 | 0.067907005 | -1.429667614 | 0.15281244 | 0.354230392 | | D5Ertd579e | 6869.052419 | 0.0667979277985781 | 0.0723109911375938 | 0.923758985290004 | 0.355611813815226 | 0.772370993941384 | NM_001081232 |
| Mrgbp | 291.6283852 | 0.173511612 | 0.121420616 | 1.429012774 | 0.153000562 | 0.354495595 | | Fam115c | 33.68736915 | 0.0789779609286966 | 0.0854495352060028 | 0.924264371225608 | 0.355348688021402 | 0.772370993941384 | NM_146174 |
| Prdm9 | 75.28310612 | 0.209599648 | 0.146670084 | 1.429055211 | 0.152988365 | 0.354495595 | | Ip6k1 | 11946.93314 | 0.0451459685035729 | 0.0488660539298224 | 0.923871785685991 | 0.355553074394526 | 0.772370993941384 | NM_013785 |
| Prss57 | 33.65445095 | -0.235352326 | 0.164702885 | -1.428950836 | 0.153018365 | 0.354495595 | | Kcnk6 | 249.3126616 | -0.086123412159206 | 0.093228176231043 | -0.92379166514821 | 0.355594795550675 | 0.772370993941384 | NM_001033525 |
| Slc52a3 | 90.45263649 | -0.195684671 | 0.136940824 | -1.42897249 | 0.15301214 | 0.354495595 | | Slc35e3 | 1507.801721 | 0.0495025147719577 | 0.053583039724639 | 0.92384670646438 | 0.355566133561968 | 0.772370993941384 | NM_029875 |
| Atp5b | 41779.31123 | -0.072270017 | 0.0505784 | -1.428871143 | 0.153041273 | 0.354495662 | | AI597468 | 1175.883627 | 0.0713134139473449 | 0.0772364448051514 | 0.923313004984256 | 0.355844112521954 | 0.772643676590014 | NM_001013028 |
| Jund | 2336.870245 | -0.181910866 | 0.127331192 | -1.42864339 | 0.153106756 | 0.354594332 | | Dnmt3a | 3698.615396 | -0.0517344984469226 | 0.0560297930572031 | -0.92333909557911 | 0.355830520023602 | 0.772643676590014 | NM_153743,NM_007872,NM_001271753 |
| 2410004B18Rik | 517.4208917 | -0.122705553 | 0.085904559 | -1.428393966 | 0.153178495 | 0.354648861 | | Vegfa | 2639.231711 | -0.0755058071168682 | 0.0817860230035353 | -0.923211623991111 | 0.355896932401472 | 0.772643676590014 | NM_001025257,NM_001025250,NM_001110268,NM_001110267,NM_001110266,NM_009505 |
| Mrpl34 | 449.7831171 | -0.133962439 | 0.093795255 | -1.428243241 | 0.153221858 | 0.354648861 | | Grhl1 | 592.7663582 | -0.0777511553274253 | 0.084244711736408 | -0.922920308288307 | 0.35604873649427 | 0.772857732563622 | NM_001161406,NM_145890 |
| Olfm4 | 26.32873218 | 0.237413377 | 0.166221325 | 1.428296739 | 0.153206466 | 0.354648861 | | Gm19757 | 176.5459794 | -0.0802968180312322 | 0.0870211052196741 | -0.922728087956741 | 0.35614892451933 | 0.772920905709106 | NR_040297 |
| Zfp473 | 20.98258571 | 0.237900756 | 0.16656559 | 1.428270727 | 0.15321395 | 0.354648861 | | Zfp71-rs1 | 536.2460382 | -0.0720457133649457 | 0.0780847653859389 | -0.922660303951163 | 0.35618425876475 | 0.772920905709106 | NM_145622 |
| Bbs9 | 699.0515345 | -0.107331834 | 0.075170994 | -1.427835769 | 0.153339135 | 0.354708354 | | Six4 | 90.61461893 | 0.0865152642669803 | 0.0937816009428873 | 0.922518525991766 | 0.356258171504179 | 0.772965825133638 | NM_011382 |
| Glipr1 | 41.71736313 | 0.228243156 | 0.159849318 | 1.42786443 | 0.153330884 | 0.354708354 | | Abcf3 | 3078.36581 | 0.0410452086752796 | 0.0445213813363838 | 0.921921275648666 | 0.356569640634452 | 0.773064268635232 | NM_013852 |
| Kcnma1 | 4028.386 | -0.101429813 | 0.071032635 | -1.427932558 | 0.153311271 | 0.354708354 | | Atg12 | 2611.715565 | 0.0582228411272376 | 0.0631368229849142 | 0.922169320131126 | 0.35644026332999 | 0.773064268635232 | NM_026217 |
| Pycrl | 524.9837824 | -0.133753861 | 0.093670607 | -1.427917094 | 0.153315723 | 0.354708354 | | Ly6h | 4944.000726 | 0.0683909997471218 | 0.0741682490944707 | 0.922106165132868 | 0.356473201482405 | 0.773064268635232 | NM_001135688,NM_001135689,NM_011837 |
| Emc1 | 3557.474732 | 0.070708584 | 0.049526699 | 1.427686178 | 0.153382206 | 0.354719364 | | Pcdhb21 | 86.36691118 | 0.0872476188733489 | 0.0946219169698028 | 0.922065644698285 | 0.356494335709085 | 0.773064268635232 | NM_053146 |
| Ppargc1b | 349.2025862 | -0.132000942 | 0.092459635 | -1.427660217 | 0.153389682 | 0.354719364 | | Ptprh | 98.52009466 | -0.0873706331450384 | 0.0947634091856526 | -0.921987018996637 | 0.35653534673716 | 0.773064268635232 | NM_207270 |
| Tbx3 | 124.6664972 | -0.189561433 | 0.132802981 | -1.427388381 | 0.153467978 | 0.354847464 | | AI504432 | 1077.13128 | 0.0767257193995245 | 0.0832483346594987 | 0.921648699800988 | 0.356711847355438 | 0.773088648244301 | NR_033498 |
| Afg3l2 | 3261.522888 | 0.090954493 | 0.063729872 | 1.427187755 | 0.153525783 | 0.35484834 | | Arfgef2 | 3991.424462 | 0.0509455709406575 | 0.0553104597359645 | 0.92108384533154 | 0.357006653753461 | 0.773088648244301 | NM_001085495 |
| Hsf2 | 813.9028862 | 0.108538789 | 0.076052899 | 1.427148608 | 0.153537064 | 0.35484834 | | Brip1 | 34.35903367 | -0.081077051726366 | 0.0879790422515787 | -0.921549606035989 | 0.356763554846771 | 0.773088648244301 | NM_178309 |
| Iyd | 8.898279509 | 0.194469607 | 0.136253709 | 1.427261022 | 0.153504671 | 0.35484834 | | Dph2 | 480.4501538 | 0.0746402440211233 | 0.0810286791345836 | 0.92115834564143 | 0.356967762090961 | 0.773088648244301 | NM_026344 |
| Hdac5 | 6247.872518 | -0.111143889 | 0.07788429 | -1.427038607 | 0.153568767 | 0.354868676 | | Fxr2 | 5451.267112 | 0.0412917461025802 | 0.0448162435424734 | 0.921356696561306 | 0.356864229264986 | 0.773088648244301 | NM_011814 |
| Arhgap23 | 5121.246177 | 0.10596346 | 0.074263036 | 1.426866786 | 0.153618296 | 0.35488508 | | Pdcd10 | 1216.994334 | 0.0756432254198878 | 0.0821217171145933 | 0.921111102856442 | 0.35699242410091 | 0.773088648244301 | NM_019745 |
| Hipk2 | 1520.108415 | -0.180208917 | 0.126305015 | -1.426775624 | 0.15364458 | 0.35488508 | | Slc35f2 | 173.2446045 | -0.0821399070453783 | 0.089153606708138 | -0.921330163604929 | 0.356878077522126 | 0.773088648244301 | NM_028060 |
| Rps27 | 24.71073765 | 0.239877794 | 0.168121362 | 1.426813287 | 0.15363372 | 0.35488508 | | Snx19 | 4306.221232 | 0.0501685817340263 | 0.0544554440321697 | 0.921277617429564 | 0.356905503770695 | 0.773088648244301 | NM_028874 |
| AU022754 | 77.12945137 | -0.222875108 | 0.156239527 | -1.426496305 | 0.153725133 | 0.354965306 | | Rhoa | 6160.156382 | 0.0442904922640501 | 0.048114045131209 | 0.920531461099728 | 0.357295100117243 | 0.773597946684017 | NM_016802 |
| Vbp1 | 1351.727283 | -0.105861679 | 0.074209034 | -1.426533574 | 0.153714383 | 0.354965306 | | Lbr | 496.4391211 | -0.0719474541351228 | 0.0781882284131673 | -0.920182687282969 | 0.35747729988263 | 0.773877087644574 | NM_133815 |
| Ifngr2 | 1606.269655 | 0.069614594 | 0.048821288 | 1.425906549 | 0.153895319 | 0.35530533 | | AI414108 | 2139.00495 | -0.0819647248854014 | 0.089112246158613 | -0.919791930050898 | 0.35768150129037 | 0.773905615389595 | NR_027907 |
| Pcdhb8 | 127.7875134 | 0.194120905 | 0.136171632 | 1.425560535 | 0.153995235 | 0.35548304 | | Cdyl2 | 1095.354892 | 0.0870669973299391 | 0.0946526592506531 | 0.919857910165776 | 0.35764701633639 | 0.773905615389595 | NM_029441 |
| Pmaip1 | 88.78510312 | 0.216012613 | 0.151545205 | 1.425400511 | 0.154041461 | 0.355536778 | | Golga3 | 2417.536845 | 0.0490334093980068 | 0.0533116897793981 | 0.919749675932338 | 0.357703586790978 | 0.773905615389595 | NM_008146 |
| Cntn2 | 7031.799008 | 0.103871886 | 0.072876432 | 1.425315196 | 0.154066109 | 0.355540706 | | Zbtb26 | 371.8672875 | 0.0826606933423808 | 0.089870286457981 | 0.919777788635724 | 0.357688892669156 | 0.773905615389595 | NM_199025 |
| Shd | 745.5049687 | 0.135173764 | 0.094847275 | 1.425172877 | 0.154107235 | 0.35558265 | | Ehmt2 | 8345.968623 | -0.0570760471559707 | 0.0620658600042005 | -0.91960454833153 | 0.357779449027112 | 0.773954471566318 | NM_145830,NM_147151 |
| Dach1 | 508.1055138 | -0.174540137 | 0.122502549 | -1.424787793 | 0.154218552 | 0.355680596 | | Edem3 | 2318.533988 | 0.064474788478666 | 0.0701240550949339 | 0.919438962726558 | 0.357866017598215 | 0.774026469867729 | NM_001039644 |
| Pom121 | 3552.348237 | 0.106968153 | 0.075071505 | 1.424883551 | 0.154190865 | 0.355680596 | | C130060K24Rik | 39.33274837 | -0.0818232824784669 | 0.089023430815438 | -0.919120749773188 | 0.358032417134725 | 0.77408780815794 | NM_175524 |
| Spats1 | 67.66641361 | 0.214413403 | 0.150487053 | 1.424796345 | 0.154216079 | 0.355680596 | | D430041D05Rik | 9268.905441 | 0.0655411890086603 | 0.0712976615051299 | 0.919261412296737 | 0.357958856053587 | 0.77408780815794 | NM_001033347 |
| Ift57 | 1459.063714 | -0.092687881 | 0.065072563 | -1.4243773 | 0.154337282 | 0.35590145 | | Paip1 | 2662.877352 | 0.0664460720190188 | 0.0722963653566306 | 0.919079011666037 | 0.3580542463867 | 0.77408780815794 | NM_001079849,NM_145457 |
| Ngfr | 192.2685964 | -0.222118924 | 0.155956617 | -1.424235333 | 0.15437836 | 0.355943201 | | Edc3 | 1305.808459 | 0.0533371538120996 | 0.0580430588888493 | 0.918923896037916 | 0.358135380023951 | 0.774147995569631 | NM_153799 |
| Aim1l | 49.24972859 | 0.213891793 | 0.150210931 | 1.423942927 | 0.154462994 | 0.356085349 | | Lemd2 | 1176.47004 | -0.0625217136745748 | 0.0680716582686433 | -0.918469084855172 | 0.358373336878796 | 0.774201530562373 | NM_146075 |
| Dync1i1 | 5145.170262 | -0.102702121 | 0.072138818 | -1.423673464 | 0.154541019 | 0.356106268 | | Myt1l | 11277.89934 | 0.057925896679523 | 0.063047987555115 | 0.91875885219787 | 0.3582217193382 | 0.774201530562373 | NM_001093775,NM_008666,NM_001093776,NM_001093778 |
| Snhg5 | 241.0829528 | 0.174127334 | 0.122303285 | 1.423732741 | 0.154523852 | 0.356106268 | | Nkx2-2 | 143.7980216 | -0.0854587795601871 | 0.0930255677982366 | -0.918659048075244 | 0.358273936179395 | 0.774201530562373 | NM_001077632,NM_010919 |
| Spsb3 | 1989.036105 | 0.157587452 | 0.11067909 | 1.423823169 | 0.154497667 | 0.356106268 | | Spag4 | 99.28848097 | -0.0873504334796775 | 0.0951010239753785 | -0.918501503225585 | 0.358356372321889 | 0.774201530562373 | NM_139151 |
| Dlst | 4916.537688 | -0.057721991 | 0.040547844 | -1.423552653 | 0.15457601 | 0.356133933 | | Aldh2 | 2824.1754 | 0.0614383209576724 | 0.0669282009160306 | 0.917973591352831 | 0.358632691851212 | 0.774627233750546 | NM_009656 |
| Gtf2ird1 | 1126.404695 | 0.13396454 | 0.094117888 | 1.423369593 | 0.154629042 | 0.356203149 | | C1qtnf1 | 410.6121988 | 0.0677562940967894 | 0.0738175348945063 | 0.917888875503915 | 0.358677046276069 | 0.774627233750546 | NM_019959,NM_001204129,NM_001204130 |
| Radil | 567.1796086 | 0.155900001 | 0.109565428 | 1.422894096 | 0.154766858 | 0.356467623 | | Hdgfrp3 | 4927.369758 | 0.0635398005268654 | 0.0692323571131415 | 0.917776068537241 | 0.358736113638124 | 0.774639629360396 | NM_013886 |
| Pik3c2b | 1102.846641 | -0.119117517 | 0.083724736 | -1.422727887 | 0.154815053 | 0.356525629 | | Sall2 | 3175.644506 | 0.0525716789224096 | 0.0572944746565682 | 0.917569787270627 | 0.358844141314302 | 0.774757728408375 | NM_015772,NM_001244916 |
| Smyd3 | 1021.534103 | -0.118353719 | 0.083197298 | -1.422566855 | 0.154861758 | 0.356580187 | | Alkbh8 | 632.4334323 | 0.0732261904132824 | 0.0798194875120026 | 0.917397401258324 | 0.358934434030426 | 0.774837507612715 | NM_026303 |
| Ccnc | 742.0571057 | -0.102364722 | 0.07199035 | -1.42192283 | 0.155048655 | 0.356862624 | | Arf4 | 3287.815003 | 0.0571369599302276 | 0.0622932328616598 | 0.917225793323599 | 0.359024333387469 | 0.774860308807848 | NM_007479 |
| Npm3 | 142.6160312 | -0.169277743 | 0.11904664 | -1.421944732 | 0.155042296 | 0.356862624 | | Stac2 | 5334.020699 | 0.0605438064891857 | 0.0660112840018287 | 0.917173592434719 | 0.359051682402976 | 0.774860308807848 | NM_146028 |
| Ubn1 | 2809.11202 | 0.14539162 | 0.102251212 | 1.421906085 | 0.155053516 | 0.356862624 | | Baiap2l2 | 72.44005485 | -0.0864407072007025 | 0.0943337893382704 | -0.916328155659429 | 0.359494804766581 | 0.775701356809173 | NM_177580 |
| Calca | 16.71029913 | -0.238994131 | 0.168109598 | -1.421656671 | 0.155125944 | 0.356976293 | | Eea1 | 2019.918117 | 0.0551715255657016 | 0.0602399865659326 | 0.915862182427888 | 0.35973918409822 | 0.776027411324571 | NM_001001932 |
| Was | 67.52605409 | 0.220237748 | 0.154974374 | 1.421123653 | 0.155280815 | 0.357279618 | | Sntg1 | 981.6187438 | 0.0639634096020524 | 0.0698415310788155 | 0.915836302756167 | 0.359752759731493 | 0.776027411324571 | NM_027671 |
| Apobr | 50.89052441 | 0.22740574 | 0.160028993 | 1.421028375 | 0.155308511 | 0.357290284 | | Zfp949 | 976.044994 | -0.0648658909098162 | 0.0708415563579999 | -0.915647456727439 | 0.359851831950804 | 0.776125866505921 | NM_001163061,NM_001142943 |
| Scamp2 | 913.8273747 | 0.103469888 | 0.072826477 | 1.420772949 | 0.155382777 | 0.357408069 | | Mtbp | 94.67729466 | -0.0866936888623689 | 0.0947164093109404 | -0.915297459997306 | 0.360035492194208 | 0.776328279376891 | NM_001168250,NM_134092 |
| Agfg1 | 1768.209748 | -0.098075735 | 0.069047085 | -1.420418195 | 0.155485969 | 0.357592341 | | Nat2 | 25.54230195 | -0.0768890399426037 | 0.0840087526937881 | -0.915250345673673 | 0.360060219856668 | 0.776328279376891 | NM_001168577,NM_010874 |
| Pttg1 | 970.4192634 | 0.106642203 | 0.075085556 | 1.420275865 | 0.155527385 | 0.357634506 | | St3gal1 | 1177.778037 | 0.0626268114627724 | 0.0684324045339167 | 0.915163099840122 | 0.360106013110388 | 0.776328279376891 | NM_009177 |
| Ubtd2 | 1496.755276 | -0.134246159 | 0.09452707 | -1.420187454 | 0.155553115 | 0.357640595 | | 4930414L22Rik | 400.521918 | -0.071998637358746 | 0.0787169865854189 | -0.914651849389806 | 0.360374429747704 | 0.776337124632091 | NR_046011 |
| Snord22 | 15.79444523 | 0.235612996 | 0.165913248 | 1.420097542 | 0.155579286 | 0.357647694 | | Apool | 133.6373054 | -0.0850037055269196 | 0.0929771506050062 | -0.91424296156418 | 0.360589194331896 | 0.776337124632091 | NM_026565 |
| Narf | 2217.128149 | 0.116637657 | 0.082192767 | 1.419074471 | 0.155877306 | 0.358209939 | | Chmp5 | 2487.650144 | 0.0470569081289522 | 0.0514650492840496 | 0.914346897235682 | 0.360534595461008 | 0.776337124632091 | NM_029814 |
| Rnaset2b | 10.89254366 | 0.229350487 | 0.161625987 | 1.419019866 | 0.155893224 | 0.358209939 | | Fam216a | 1516.370768 | 0.055582975540056 | 0.0608146865859237 | 0.91397290129127 | 0.360731085048339 | 0.776337124632091 | NM_026883 |
| Tsr3 | 558.8368277 | 0.123100011 | 0.08674683 | 1.419072155 | 0.155877981 | 0.358209939 | | Fam46a | 311.9802498 | 0.0858909992150452 | 0.0938934765196968 | 0.914770678419039 | 0.360312030948604 | 0.776337124632091 | NM_001160378,NM_001160379 |
| Tinf2 | 407.9743233 | 0.149023437 | 0.105032698 | 1.418828993 | 0.155948878 | 0.358284684 | | Mcm6 | 372.9984355 | -0.0775116017125675 | 0.0847986212364603 | -0.91406676880308 | 0.360681762721425 | 0.776337124632091 | NM_008567 |
| Cecr2 | 180.9749866 | 0.176848234 | 0.124687922 | 1.418326908 | 0.156095344 | 0.358438532 | | Mthfd2l | 149.0474252 | 0.0867286111550123 | 0.0948549607803849 | 0.914328680772034 | 0.360544164450726 | 0.776337124632091 | NM_026788 |
| Schip1 | 154.5538226 | -0.19375933 | 0.136612992 | -1.418308221 | 0.156100798 | 0.358438532 | | Ntng2 | 735.8011549 | -0.0810944849131394 | 0.0887097152336597 | -0.914155622070684 | 0.360635079008741 | 0.776337124632091 | NM_133501,NM_133500 |
| Slc1a6 | 357.0271818 | 0.170575641 | 0.120269179 | 1.418282244 | 0.156108378 | 0.358438532 | | Scn10a | 31.61648218 | 0.0792116404925812 | 0.0866111908141842 | 0.914565886324343 | 0.360419574393661 | 0.776337124632091 | NM_009134,NM_001205321 |
| Tada1 | 1166.557699 | -0.083836383 | 0.059105813 | -1.418411795 | 0.156070574 | 0.358438532 | | Sh2d3c | 955.425228 | -0.0794817948720274 | 0.0868712246393561 | -0.914938119060646 | 0.360224117018249 | 0.776337124632091 | NM_013781,NM_001252547,NR_045535 |
| Ly6g5b | 8.942715601 | 0.224501103 | 0.158318499 | 1.418034561 | 0.156180675 | 0.358551396 | | Tnfsf13b | 41.06284961 | -0.0822665069630912 | 0.0900136050747023 | -0.913934142453446 | 0.360751451966585 | 0.776337124632091 | NM_033622 |
| Gltp | 1383.841908 | 0.140166491 | 0.098852132 | 1.417940996 | 0.156207992 | 0.358560982 | | Vps26a | 2209.289948 | 0.0544508696495472 | 0.0595783838997265 | 0.913936667721482 | 0.360750124971708 | 0.776337124632091 | NM_001113355,NM_133672 |
| Pcdhgb4 | 166.9155259 | -0.182473915 | 0.128707324 | -1.417743055 | 0.156265795 | 0.358640531 | | 1500032L24Rik | 2263.057393 | -0.0470469213648425 | 0.0515848841086476 | -0.912029215104036 | 0.361753338464138 | 0.776449324221643 | NM_026914 |
| Sstr1 | 1258.008508 | -0.130769916 | 0.092271078 | -1.417236243 | 0.156413869 | 0.358927204 | | 1700011J10Rik | 303.4725756 | -0.0713717867936243 | 0.0782346563440679 | -0.912278395903455 | 0.361622184084544 | 0.776449324221643 | NR_045269 |
| Caprin2 | 262.8574708 | 0.200117283 | 0.141236777 | 1.416892171 | 0.156514456 | 0.359104839 | | A330069E16Rik | 25.64240387 | -0.0775008963836677 | 0.0850202046117724 | -0.911558572901111 | 0.362001138656253 | 0.776449324221643 | NR_015464 |
| 2810454H06Rik | 16.34501357 | 0.236585838 | 0.167012079 | 1.416579208 | 0.156605991 | 0.359261656 | | Acadl | 1427.329663 | 0.0616759160714288 | 0.0675148779416955 | 0.913515923478243 | 0.3609712627471 | 0.776449324221643 | NM_007381 |
| Jmy | 2409.68851 | 0.118163003 | 0.083426924 | 1.416365341 | 0.156668566 | 0.359352 | | Agbl5 | 599.6993829 | -0.077204935374784 | 0.0847906324679634 | -0.910536142113983 | 0.362539830232252 | 0.776449324221643 | NM_174849,NM_001048192 |
| Arhgap18 | 352.6553974 | 0.146746503 | 0.103627578 | 1.416095082 | 0.156747668 | 0.359427019 | | Aldh4a1 | 1107.694667 | -0.0676667661237367 | 0.0742627991392297 | -0.911179849238828 | 0.362200619566303 | 0.776449324221643 | NM_175438 |
| Azin1 | 5825.330562 | -0.121568448 | 0.085846733 | -1.416110358 | 0.156743196 | 0.359427019 | | Asrgl1 | 6631.199879 | -0.045874390874305 | 0.050347579251948 | -0.911153853986537 | 0.362214314282081 | 0.776449324221643 | NM_025610 |
| AI316807 | 737.9161315 | 0.110845092 | 0.078288219 | 1.415859156 | 0.156816745 | 0.359508355 | | BC003965 | 1048.920872 | 0.0542537212760551 | 0.059473568533916 | 0.912232486017983 | 0.361646346156187 | 0.776449324221643 | NM_183150 |
| Chmp3 | 4393.253408 | -0.085436907 | 0.060344664 | -1.415815449 | 0.156829545 | 0.359508355 | | BC061194 | 150.6377188 | -0.0836292540872241 | 0.0917807659395208 | -0.911184965947352 | 0.362197924040406 | 0.776449324221643 | NM_001001334 |
| AW209491 | 779.6424673 | -0.117685921 | 0.083141775 | -1.415484824 | 0.156926395 | 0.359528273 | | Eml4 | 1599.566124 | -0.0526333871945842 | 0.0577757837613806 | -0.910993910735422 | 0.362298582089371 | 0.776449324221643 | NM_199466,NM_001114361,NM_001114362 |
| Dcx | 1183.550688 | -0.231098195 | 0.163265092 | -1.415478301 | 0.156928306 | 0.359528273 | | Fam105a | 890.7173033 | -0.0611668272842317 | 0.0670927222335586 | -0.9116760394861 | 0.361939280775937 | 0.776449324221643 | NM_001242423,NM_001242424,NM_198301 |
| Dusp8 | 7035.265199 | -0.063386108 | 0.044776523 | -1.415610325 | 0.156889627 | 0.359528273 | | Gabre | 34.58322732 | 0.0374357337074942 | 0.0411208971611949 | 0.910382221495443 | 0.362620970364837 | 0.776449324221643 | NM_017369 |
| Mtus1 | 2083.952227 | 0.098869869 | 0.06984955 | 1.415468931 | 0.156931051 | 0.359528273 | | Gltscr1 | 847.7242948 | -0.0584013189683743 | 0.0640317783045462 | -0.912067734408492 | 0.361733062179327 | 0.776449324221643 | NM_001081418 |
| Ric8b | 2758.372174 | -0.119486561 | 0.08443528 | -1.415126017 | 0.157031551 | 0.359652159 | | Ik | 4905.7423 | 0.0554194785276454 | 0.0606695507464571 | 0.913464461921728 | 0.360998316115831 | 0.776449324221643 | NM_011879 |
| Tnni3 | 17.08130532 | -0.228857072 | 0.161717781 | -1.415163321 | 0.157020616 | 0.359652159 | | Kifc5b | 86.91991258 | -0.0864804334691322 | 0.0949870722358776 | -0.910444247132692 | 0.362588271828312 | 0.776449324221643 | NM_053173 |
| Smg5 | 4241.972481 | 0.124602918 | 0.088061193 | 1.414958319 | 0.157080717 | 0.359711592 | | Mocs1 | 364.3747968 | -0.0742100135769074 | 0.0813147819571816 | -0.91262636129289 | 0.361439085083827 | 0.776449324221643 | NM_028464,NM_020042 |
| Foxp1 | 4573.472792 | -0.115210126 | 0.081447426 | -1.414533663 | 0.157205271 | 0.35990366 | | Mrpl47 | 433.8482711 | 0.0702319832725387 | 0.077121156230145 | 0.910670777068648 | 0.362468865970599 | 0.776449324221643 | NM_029017 |
| Pex7 | 763.4412908 | -0.096575062 | 0.068274378 | -1.414513973 | 0.157211048 | 0.35990366 | | Phax | 1450.365537 | 0.0569481187394263 | 0.0623800866583622 | 0.912921443205341 | 0.361283858864016 | 0.776449324221643 | NM_001162989,NM_019996 |
| Cml3 | 173.098191 | -0.219454751 | 0.155177499 | -1.414217611 | 0.157298019 | 0.360049564 | | Phf21a | 1992.057811 | -0.0489336511800604 | 0.0536755947073773 | -0.911655500918649 | 0.361950095903634 | 0.776449324221643 | NM_001109690,NM_001109691,NM_138755 |
| Ms4a6d | 49.96779679 | 0.230590862 | 0.163093418 | 1.413857558 | 0.15740373 | 0.360238314 | | Pom121 | 3919.997651 | 0.045854256311672 | 0.0503047839513503 | 0.911528739612869 | 0.3620168499147 | 0.776449324221643 | NM_148932 |
| Ppp1r18 | 346.2834634 | 0.119506387 | 0.084533079 | 1.413723336 | 0.157443151 | 0.360275318 | | Prune2 | 4721.558093 | 0.0617004444467897 | 0.0675824021763314 | 0.912966134080365 | 0.361260353120191 | 0.776449324221643 | NM_181348 |
| Tbx2 | 83.4772758 | 0.193779568 | 0.137080537 | 1.413618393 | 0.157473978 | 0.360292648 | | Pycr2 | 749.0113552 | 0.065672621722829 | 0.0720679366678844 | 0.911259913343608 | 0.36215844255002 | 0.776449324221643 | NM_133705 |
| Atp2b1 | 27834.39602 | -0.137207053 | 0.097092836 | -1.413153222 | 0.157610677 | 0.360498942 | | Rhog | 1190.624217 | 0.0698425811593419 | 0.0766633157044433 | 0.911030008519367 | 0.362279562514048 | 0.776449324221643 | NM_019566 |
| Npy | 2427.223343 | 0.111458192 | 0.078869323 | 1.413200814 | 0.157596687 | 0.360498942 | | Rnf111 | 6596.111337 | 0.0596206227532283 | 0.0654785919246366 | 0.910536115710144 | 0.362539844150211 | 0.776449324221643 | NM_013876 |
| Pwp2 | 1220.571941 | 0.10931674 | 0.077362405 | 1.413047325 | 0.15764181 | 0.36051693 | | Rpf1 | 387.3146451 | 0.075170883827166 | 0.0823496924759842 | 0.912825313210347 | 0.361334422928452 | 0.776449324221643 | NM_027332,NM_027371 |
| Lhpp | 476.225887 | -0.191757496 | 0.135726912 | -1.412818532 | 0.157709088 | 0.360617563 | | Sdf2 | 1634.040642 | 0.0530335321772257 | 0.058137550297099 | 0.912207891564225 | 0.361659290472169 | 0.776449324221643 | NM_009143 |
| Hexdc | 1014.032459 | 0.156968226 | 0.11111271 | 1.4126937 | 0.157745805 | 0.360648295 | | Socs7 | 6749.84427 | 0.0631084022110676 | 0.0692385654566882 | 0.911463167886467 | 0.36205138379271 | 0.776449324221643 | NM_138657 |
| App | 63331.49167 | 0.099149455 | 0.070190208 | 1.412582444 | 0.157778534 | 0.360669903 | | Tes | 58.90274936 | -0.0855156881183826 | 0.093620133999502 | -0.913432660957711 | 0.36101503453598 | 0.776449324221643 | NM_207176 |
| Mir3101 | 14.14223009 | 0.236800373 | 0.167687111 | 1.412156075 | 0.157904011 | 0.360746278 | | Tmem110 | 1019.793154 | -0.0556309251231905 | 0.061100266119284 | -0.910485807289025 | 0.362566363271286 | 0.776449324221643 | NM_028839 |
| Nsmaf | 1254.379289 | 0.137897823 | 0.097648214 | 1.412189912 | 0.15789405 | 0.360746278 | | Tpp1 | 3505.648535 | -0.0519987522040154 | 0.0571119027071078 | -0.910471368301025 | 0.362573974731739 | 0.776449324221643 | NM_009906 |
| Rpf2 | 514.9392298 | -0.115015853 | 0.081447193 | -1.412152454 | 0.157905077 | 0.360746278 | | Twsg1 | 1285.260338 | 0.0692660268010915 | 0.0759538383258064 | 0.911948998600605 | 0.361795566140919 | 0.776449324221643 | NM_023053 |
| Tecta | 12.96739013 | 0.226707092 | 0.160515687 | 1.412367203 | 0.157841868 | 0.360746278 | | Ywhab | 27181.18443 | 0.0471301856757934 | 0.051663854760668 | 0.912246790219646 | 0.361638817840131 | 0.776449324221643 | NM_018753 |
| Klhl23 | 3329.147418 | -0.09481816 | 0.067153722 | -1.411956888 | 0.157962656 | 0.360824619 | | Zbtb37 | 152.6082528 | -0.086649258992793 | 0.0950371014642323 | -0.911741390023388 | 0.361904870063678 | 0.776449324221643 | NM_173424 |
| Ltbr | 323.9983843 | -0.123376612 | 0.087397173 | -1.411677382 | 0.158044976 | 0.360959443 | | Zfp316 | 1341.524016 | -0.0675012295489608 | 0.0741432942336085 | -0.910415840659572 | 0.362603246860013 | 0.776449324221643 | NM_017467 |
| Stk30 | 285.980693 | -0.119130434 | 0.084401178 | -1.411478333 | 0.15810362 | 0.36104016 | | Sertad4 | 835.1048019 | 0.0708817986618568 | 0.0778696260395963 | 0.910262476743035 | 0.362684102358472 | 0.776470047289486 | NM_001177794,NM_198247 |
| Iars2 | 2138.386259 | 0.122715897 | 0.086985622 | 1.410760693 | 0.158315188 | 0.361470015 | | Scyl1 | 3104.49187 | 0.0479830321894592 | 0.0527259228354862 | 0.910046322739089 | 0.362798080793742 | 0.776599605306498 | NM_023912 |
| Cd163 | 71.23336289 | -0.233354748 | 0.165425621 | -1.410632443 | 0.15835302 | 0.361503123 | | Nkain3 | 28.97645787 | 0.0808741285176799 | 0.0888896891348052 | 0.909825754875018 | 0.362914409780618 | 0.776734155955983 | NM_172987 |
| Csk | 1112.344283 | -0.103553549 | 0.07343015 | -1.41023203 | 0.158471181 | 0.361719576 | | Adam19 | 1067.919936 | 0.0674553568584851 | 0.0741624217542269 | 0.909562488156483 | 0.363053288996752 | 0.776821522420875 | NM_009616 |
| Efr3b | 10398.89226 | 0.094345797 | 0.066908431 | 1.410073378 | 0.158518017 | 0.361723971 | | Gjb2 | 315.8603766 | -0.0864505902444073 | 0.0951021887132181 | -0.90902839791732 | 0.363335135992609 | 0.776821522420875 | NM_008125 |
| Tma16 | 273.6743456 | 0.208686657 | 0.147997653 | 1.410067344 | 0.158519799 | 0.361723971 | | Immp2l | 102.2109603 | 0.0844681824890235 | 0.092888968033547 | 0.909345687407332 | 0.363167681323542 | 0.776821522420875 | NM_053122 |
| A230046K03Rik | 2006.60272 | 0.099979061 | 0.07091329 | 1.40987763 | 0.158575819 | 0.36174525 | | Mcm4 | 573.0215155 | -0.0581676508804455 | 0.0639920196348669 | -0.908982888996867 | 0.363359158023861 | 0.776821522420875 | NM_008565 |
| Mmgt2 | 496.9204622 | -0.12606898 | 0.089416025 | -1.40991484 | 0.15856483 | 0.36174525 | | Pdia4 | 1715.794411 | 0.0590461118119586 | 0.0649632849074541 | 0.90891511868704 | 0.363394932643466 | 0.776821522420875 | NM_009787 |
| 2310014L17Rik | 37.63967669 | 0.221264323 | 0.157062254 | 1.408653682 | 0.158937597 | 0.362517169 | | Rftn1 | 411.651816 | 0.0788756442447858 | 0.0867484543087484 | 0.909245529194765 | 0.363220536225329 | 0.776821522420875 | NM_181397 |
| 1810043H04Rik | 220.3387365 | -0.193863359 | 0.137638625 | -1.408495319 | 0.158984452 | 0.362540932 | | Tacc2 | 3579.037098 | 0.0423236878396328 | 0.0465547079032536 | 0.909117246049242 | 0.363288240081956 | 0.776821522420875 | NM_206856,NM_001004468,NM_021314 |
| Bard1 | 25.13931704 | -0.234291336 | 0.166345715 | -1.408460305 | 0.158994813 | 0.362540932 | | Tnpo3 | 4183.72099 | 0.0451593740875569 | 0.0496892269142775 | 0.908836319097188 | 0.363436532174877 | 0.776821522420875 | NM_177296 |
| Cep68 | 1356.355669 | -0.08176954 | 0.058069819 | -1.40812459 | 0.159094181 | 0.36271311 | | Xpo7 | 5228.282512 | 0.053475459426663 | 0.0588242341681015 | 0.90907191879195 | 0.363312164287369 | 0.776821522420875 | NM_023045 |
| Pink1 | 15505.55604 | 0.158786266 | 0.112770569 | 1.408047035 | 0.159117143 | 0.36271311 | | Rbpms2 | 100.9020771 | -0.0857766133454995 | 0.0943920111370631 | -0.908727468693791 | 0.363494000899234 | 0.776830051061096 | NM_028030 |
| Brd9 | 2376.18334 | 0.155971609 | 0.110831958 | 1.407280098 | 0.159344348 | 0.363177601 | | Ddx25 | 1819.778613 | 0.0525313208105576 | 0.0578287197175516 | 0.908395016647997 | 0.363669557676405 | 0.776984985235225 | NM_013932 |
| Pkmyt1 | 165.1550193 | 0.198108269 | 0.140863404 | 1.406385637 | 0.159609643 | 0.363728755 | | Trim65 | 133.7682302 | -0.0825204631370599 | 0.0908427893292746 | -0.908387597368361 | 0.363673476154089 | 0.776984985235225 | NM_178802 |
| Polr2i | 438.1155141 | -0.133113728 | 0.094665223 | -1.406152374 | 0.159678883 | 0.363833031 | | Tctn2 | 965.4424457 | 0.0545687581972775 | 0.0600857594807058 | 0.908181217461354 | 0.363782485873129 | 0.777103586734276 | NM_026486 |
| Cd320 | 385.3335548 | -0.122186781 | 0.086935839 | -1.405482285 | 0.159877914 | 0.364165372 | | 4632428N05Rik | 1447.858484 | -0.0629034721220756 | 0.0693178695851813 | -0.907464013226441 | 0.364161471522391 | 0.777191868882161 | NM_028732,NM_001159572 |
| Nebl | 1307.061628 | -0.096571991 | 0.068713779 | -1.405423941 | 0.159895252 | 0.364165372 | | Brcc3 | 1016.768983 | 0.0514620672766041 | 0.0567080254728194 | 0.907491785290079 | 0.364146791593359 | 0.777191868882161 | NM_001166457,NM_145956,NM_001166459 |
| Rnf5 | 1994.740429 | -0.079465369 | 0.056539879 | -1.405474703 | 0.159880167 | 0.364165372 | | F12 | 25.86089227 | -0.0779950514043503 | 0.0859549744116019 | -0.907394271690025 | 0.364198337565804 | 0.777191868882161 | NM_021489 |
| Hist1h2bc | 422.8362005 | -0.188747764 | 0.134325048 | -1.405156872 | 0.159974636 | 0.364292622 | | Fip1l1 | 2293.245223 | 0.0548977687988173 | 0.0604714134568092 | 0.90783009128813 | 0.363967997422156 | 0.777191868882161 | NM_001159574,NM_001159573,NM_024183 |
| Epb4.1l1 | 24999.08826 | -0.082391116 | 0.058639913 | -1.40503477 | 0.160010939 | 0.364321747 | | Lztr1 | 7065.51499 | -0.0606038772779037 | 0.0667725318456819 | -0.907616883810328 | 0.364080670831119 | 0.777191868882161 | NM_025808 |
| Arfgef1 | 5351.578562 | -0.106825742 | 0.076050509 | -1.404668344 | 0.160119923 | 0.364516322 | | Nupl1 | 1425.800427 | 0.0622485825913022 | 0.068601307689428 | 0.907396443127792 | 0.364197189687835 | 0.777191868882161 | NM_170591 |
| Zfp454 | 169.2438682 | 0.163172916 | 0.116183254 | 1.404444357 | 0.16018657 | 0.364614473 | | Rpl39 | 2378.199662 | -0.0716434163935944 | 0.078946594676765 | -0.907492168432693 | 0.364146589072055 | 0.777191868882161 | NM_026055 |
| 4732471J01Rik | 152.192627 | 0.163039621 | 0.116120598 | 1.404054264 | 0.160302692 | 0.364683468 | | Myh7b | 393.5028347 | -0.0673996858816388 | 0.074297333562998 | -0.907161571612654 | 0.364321361932159 | 0.777237063935001 | NM_001085378 |
| Asb5 | 19.78645864 | -0.236037344 | 0.168114622 | -1.40402626 | 0.16031103 | 0.364683468 | | Ncoa2 | 4926.514833 | 0.0541650529442044 | 0.0597132704342355 | 0.907085687156566 | 0.364361486302008 | 0.777237063935001 | NM_008678,NM_001077695 |
| Mynn | 560.4372921 | 0.114578841 | 0.081595181 | 1.404235393 | 0.160248766 | 0.364683468 | | Pcca | 1110.982566 | -0.0596473492503427 | 0.0657816826759919 | -0.906747088610306 | 0.364540556009167 | 0.777237063935001 | NM_144844 |
| Timm17a | 1668.377509 | -0.084906435 | 0.060473078 | -1.404036938 | 0.16030785 | 0.364683468 | | Ppdpf | 1267.634505 | 0.0578190980682485 | 0.0637529210815553 | 0.90692468811404 | 0.364446624675806 | 0.777237063935001 | NM_025598 |
| Haus5 | 221.398265 | 0.173607339 | 0.123692657 | 1.403537958 | 0.160456481 | 0.364960763 | | Snora61 | 25.19290436 | -0.0765057286424292 | 0.0843616734398243 | -0.906877797973047 | 0.3644714231259 | 0.777237063935001 | NR_034046 |

| | | | | | | | | | | | | | | |
|---|---|---|---|---|---|---|---|---|---|---|---|---|---|---|
| Nudt18 | 1324.545848 | -0.123193357 | 0.087783729 | -1.403373476 | 0.160505498 | 0.365018668 | | Zc3h10 | 629.1851297 | 0.0594331474481121 | 0.0655416938607442 | 0.906799076239763 | 0.36451305849158 | 0.777237063935001 | NM_134003 |
| Sp4 | 779.3606622 | -0.120323806 | 0.085764523 | -1.402955466 | 0.160630119 | 0.365248469 | | Rad23a | 2264.909881 | 0.0435218239275922 | 0.0480108555324193 | 0.906499653983548 | 0.364671447948345 | 0.777402033005234 | NM_009010 |
| Polr3k | 1265.501501 | -0.095838885 | 0.068329028 | -1.402608636 | 0.160733575 | 0.365430083 | | Derl3 | 56.16644826 | -0.0842169043770628 | 0.0929308050608227 | -0.906232377110511 | 0.364812869345915 | 0.777589396935988 | NM_024440 |
| Ppp2r5c | 7965.336575 | -0.059699519 | 0.042565835 | -1.402521968 | 0.160759435 | 0.365435254 | | Dlg1 | 8160.710735 | 0.060725095307733 | 0.0670395566873886 | 0.90580991743277 | 0.365036470855227 | 0.777634018770177 | NR_045516,NM_001252433,NM_001252436,NM_001252435,NM_001252434,NM_007862 |
| Rasgef1a | 12532.56497 | -0.127286908 | 0.090765019 | -1.402378458 | 0.160802263 | 0.365478987 | | Dnahc2 | 147.5302644 | -0.08569309313991 | 0.0946060990065369 | -0.90578825297499 | 0.365047939831601 | 0.777634018770177 | NM_001081330 |
| Tmem135 | 2305.754448 | -0.078522756 | 0.056000469 | -1.402180323 | 0.160861406 | 0.365559786 | | Kremen2 | 41.82555899 | 0.0843180647407769 | 0.0930713997268626 | 0.905950324033223 | 0.364962146272634 | 0.777634018770177 | NM_028416 |
| Mir541 | 9.145460242 | 0.222626521 | 0.158791248 | 1.402007504 | 0.160913006 | 0.365623421 | | Pld6 | 31.34537357 | -0.0793377139918071 | 0.0875809587033942 | -0.905878574137284 | 0.365000126077103 | 0.777634018770177 | NM_183139 |
| Hs6st3 | 431.527433 | 0.1350973 | 0.096382781 | 1.401674639 | 0.161012427 | 0.365745564 | | 2410015M20Rik | 1267.844495 | 0.0581864497187956 | 0.064263885935623 | 0.905429991847746 | 0.365237632804177 | 0.77773573326162 | NM_153152 |
| Mob2 | 885.9952601 | 0.120699643 | 0.086111345 | 1.401669465 | 0.161013973 | 0.365745564 | | Dnajb4 | 2817.04606 | 0.0690217665287555 | 0.0762218353034742 | 0.905537976800848 | 0.365180450189177 | 0.77773573326162 | NM_027287,NM_025926 |
| Ephb2 | 704.2007425 | 0.155176506 | 0.110738238 | 1.401291094 | 0.161127043 | 0.365948755 | | Drosha | 5441.449828 | 0.0366469910080047 | 0.0405079114360681 | 0.904687250189215 | 0.365631097510919 | 0.77773573326162 | NM_026799,NM_001130149 |
| Cox17 | 956.0233468 | -0.125414939 | 0.089505332 | -1.401200756 | 0.161154048 | 0.365956445 | | Ikbkb | 1775.762648 | -0.0549993799080981 | 0.0607921633637701 | -0.904711674414201 | 0.365618154660352 | 0.77773573326162 | NM_001159774,NM_010546 |
| Izumo4 | 834.1270696 | 0.196122768 | 0.139975755 | 1.401119556 | 0.161178325 | 0.365957937 | | Impg2 | 37.02668661 | -0.0795857753743168 | 0.0879297031203048 | -0.905106835916733 | 0.365408790984708 | 0.77773573326162 | NM_174876 |
| Zfp648 | 10.46353473 | 0.222518667 | 0.158826 | 1.401021661 | 0.161207596 | 0.365970767 | | Myt1 | 446.6396829 | -0.0708209463136809 | 0.0782224404813732 | -0.905378889713179 | 0.365264695501993 | 0.77773573326162 | NM_001171615,NM_008665,NM_001171616,NM_001171680 |
| Fhod1 | 175.437892 | 0.143669254 | 0.102565229 | 1.400759838 | 0.161285902 | 0.366094897 | | Nme7 | 667.6579572 | 0.0582700031482321 | 0.0644021995885702 | 0.904782810532663 | 0.365580459939131 | 0.77773573326162 | NM_178071,NM_138314 |
| Mkx | 506.7013427 | 0.139831736 | 0.099847639 | 1.400451101 | 0.161378277 | 0.366197278 | | Nr4a3 | 393.1229689 | 0.063048523451811 | 0.0696479064110826 | 0.905246499150741 | 0.365334812796121 | 0.77773573326162 | NM_015743 |
| Polq | 79.96117344 | -0.206492021 | 0.147440937 | -1.400506707 | 0.161361636 | 0.366197278 | | Smim12 | 1259.862754 | 0.055129935081691 | 0.0609351738873655 | 0.904730906054275 | 0.365607963658251 | 0.77773573326162 | NM_030252 |
| Srrm4 | 1968.051841 | 0.103446415 | 0.073891493 | 1.399977341 | 0.161520104 | 0.36646544 | | Tnfrsf10b | 36.82755066 | -0.0784854502450827 | 0.0867523246838456 | -0.904707171030977 | 0.365620541065281 | 0.77773573326162 | NM_020275 |
| Arl10 | 367.9783038 | -0.132141159 | 0.094399518 | -1.399807558 | 0.161570954 | 0.366527139 | | Zfp354b | 124.1056714 | 0.0827394643338511 | 0.0915032111827148 | 0.904224707137719 | 0.365876261657929 | 0.778143276258138 | NM_013744 |
| Comtd1 | 246.4065079 | -0.179501759 | 0.128282521 | -1.399269035 | 0.161732322 | 0.366839495 | | Bai3 | 4759.591146 | 0.0507084721335436 | 0.0561487408318022 | 0.903109693687427 | 0.36646767793893 | 0.778210102463893 | NM_175642 |
| 1110004E09Rik | 782.3391334 | 0.099118544 | 0.070845624 | 1.399077855 | 0.161789638 | 0.366915785 | | Bnip3l | 5460.576874 | 0.0559554152388486 | 0.0619094270239455 | 0.90382705718155 | 0.366087112159611 | 0.778210102463893 | NM_009761 |
| Car3 | 33.92960202 | 0.232086846 | 0.165955016 | 1.398492506 | 0.161965222 | 0.366938024 | | Creld1 | 3008.440181 | 0.0544183124519472 | 0.0602214277318658 | 0.903637037877716 | 0.366187896101112 | 0.778210102463893 | NM_133930 |
| E230008N13Rik | 22.53036371 | 0.221755695 | 0.158553958 | 1.398613431 | 0.161928936 | 0.366938024 | | Foxg1 | 2732.495687 | -0.0737167247033929 | 0.0816049146486673 | -0.903336827454139 | 0.366347156614038 | 0.778210102463893 | NM_008241,NM_001160112 |
| Foxj2 | 1738.892885 | 0.098123249 | 0.070150844 | 1.39874651 | 0.161889012 | 0.366938024 | | Nkain1 | 2879.051524 | 0.0538433416289817 | 0.0596042962285713 | 0.903346655122016 | 0.366341942354158 | 0.778210102463893 | NM_025998 |
| Hnrnpd | 5094.106278 | -0.104498594 | 0.074701067 | -1.398890232 | 0.161845902 | 0.366938024 | | Osbpl7 | 786.3433384 | -0.0623436925498047 | 0.0690504052250915 | -0.902872218440657 | 0.366593716763234 | 0.778210102463893 | NM_001081434 |
| Stag3 | 35.47783023 | 0.231903759 | 0.16582158 | 1.398513752 | 0.161958846 | 0.366938024 | | Pdpn | 347.1562361 | -0.0749682655765648 | 0.0830274136591267 | -0.902933889815608 | 0.366560982852411 | 0.778210102463893 | NM_010329 |
| Tet2 | 1530.998853 | 0.134516705 | 0.096171996 | 1.398709714 | 0.16190005 | 0.366938024 | | Phex | 27.54317274 | 0.0723701607223656 | 0.0801572284697758 | 0.902852582405012 | 0.366604139553932 | 0.778210102463893 | NM_011077 |
| Usp39 | 1208.387749 | -0.081742877 | 0.05844729 | -1.39857428 | 0.161940684 | 0.366938024 | | Picalm | 6284.066073 | 0.0536707731486886 | 0.0594149893579809 | 0.903320420126934 | 0.366355861942995 | 0.778210102463893 | NM_001252524,NM_001252521,NM_001252520,NM_001252523,NM_001252522,NM_146194 |
| 9330158H04Rik | 26.52249887 | -0.234941157 | 0.168045641 | -1.398079448 | 0.162089211 | 0.367061759 | | Ranbp6 | 1952.751867 | 0.0606525779542468 | 0.0671366879049518 | 0.903419275614478 | 0.366303413577661 | 0.778210102463893 | NM_177721 |
| Mrs2 | 882.881563 | 0.086908284 | 0.062162873 | 1.39807379 | 0.16209091 | 0.367061759 | | Sept3 | 25264.01924 | 0.0323636887345145 | 0.0358003366613552 | 0.904005150584229 | 0.36599267027832 | 0.778210102463893 | NM_011889 |
| Snw1 | 3082.408186 | 0.131491818 | 0.094043975 | 1.39819503 | 0.162054509 | 0.367061759 | | Sp8 | 45.52650057 | -0.0825053689780716 | 0.0913320980118529 | -0.903355674227085 | 0.366337157133698 | 0.778210102463893 | NM_177082 |
| Grasp | 1802.454415 | -0.143154617 | 0.102417439 | -1.397756266 | 0.162186272 | 0.367224038 | | Stub1 | 4233.447462 | 0.0405978214219437 | 0.044950563211284 | 0.903166023329211 | 0.366437787450196 | 0.778210102463893 | NM_019719 |
| Acpp | 15.58084136 | -0.226050936 | 0.161749778 | -1.397534751 | 0.162252824 | 0.367321049 | | Calhm2 | 196.0583231 | 0.0839704361918431 | 0.0930192929505397 | 0.902720645667474 | 0.366674176251707 | 0.778245044452411 | NM_133746 |
| B3galnt1 | 1454.943431 | -0.078826843 | 0.056413803 | -1.397297088 | 0.162324251 | 0.367321739 | | Pgap2 | 362.6601981 | 0.0641102876635345 | 0.071031925033201 | 0.902555965272921 | 0.366761606147795 | 0.778316886910572 | NM_145583 |
| Gpc4 | 333.8838493 | -0.171828741 | 0.122965997 | -1.397367927 | 0.162302959 | 0.367321739 | | Aqp4 | 5081.744615 | -0.0585438959394531 | 0.0648720575952859 | -0.902451658072695 | 0.366816990243735 | 0.77832071286598 | NM_009700 |
| Lpar4 | 37.38889357 | -0.223084359 | 0.159653923 | -1.39729958 | 0.162323502 | 0.367321739 | | Arid3b | 366.0580446 | -0.0802410842667733 | 0.0889322631806461 | -0.90227192468701 | 0.366912435686858 | 0.778335590524204 | NM_019689 |
| Prune2 | 4033.555022 | 0.118349033 | 0.084712822 | 1.397061626 | 0.16239504 | 0.367428264 | | Tubb4a | 33893.0329 | 0.0494323736099758 | 0.0547887001387248 | 0.902236656186645 | 0.366931166453927 | 0.778335590524204 | NM_009451 |
| Leng1 | 254.1338035 | 0.130700576 | 0.093572679 | 1.396781385 | 0.162479322 | 0.367565281 | | Ipo7 | 4807.592728 | 0.0655595501916797 | 0.0726895901787358 | 0.901911126895555 | 0.367104080121612 | 0.778490942774878 | NM_181517 |
| Dusp11 | 2401.220771 | 0.163882709 | 0.117351442 | 1.396512103 | 0.162560339 | 0.367607043 | | N4bp3 | 524.6189784 | 0.0785386708337784 | 0.0870816419692029 | 0.901896990660261 | 0.367111590114823 | 0.778490942774878 | NM_145974 |
| Kcnq1ot1 | 182.7966094 | -0.21344022 | 0.152841209 | -1.396483455 | 0.162568959 | 0.367607043 | | Efcab4a | 83.4619567 | -0.085718480465847 | 0.0950617125048376 | -0.901714036147674 | 0.367208794868064 | 0.778583411801708 | NM_001025103 |
| Mast4 | 2757.671133 | 0.10128957 | 0.072530955 | 1.396501267 | 0.162563599 | 0.367607043 | | Snrpc | 637.6955396 | 0.0563966067871911 | 0.0625586690007358 | 0.901499467428372 | 0.36732281685941 | 0.778711505793897 | NM_011432 |
| Kdm6b | 1164.698992 | 0.196562113 | 0.140788367 | 1.396153081 | 0.162668402 | 0.367778232 | | Ythdf1 | 2522.534626 | 0.0484632183737362 | 0.0537692446922507 | 0.901318563262629 | 0.367418966617939 | 0.778801679423929 | NM_173761 |
| Cdhr2 | 25.73751161 | 0.234604721 | 0.168076865 | 1.395894291 | 0.16274633 | 0.367900735 | | Mzt2 | 509.867807 | -0.0666950341490047 | 0.0740209046068707 | -0.901029709150758 | 0.367572523795859 | 0.778899851299585 | NM_029354 |
| Cdk2 | 98.35195754 | 0.196692369 | 0.140963301 | 1.395344512 | 0.162911975 | 0.368167758 | | Tac2 | 354.1943015 | 0.0847105858944792 | 0.0940149312218148 | 0.901033323043301 | 0.367570602374489 | 0.778899851299585 | NM_009312,NM_001199971 |
| Kcnip3 | 6085.073312 | -0.100640169 | 0.072122571 | -1.395404627 | 0.162893857 | 0.368167758 | | Dgkz | 21724.65729 | -0.0654850881659914 | 0.0727004942139141 | -0.90075162313626 | 0.367720394329921 | 0.778985919211964 | NM_001166597,NM_138306 |
| Cmss1 | 302.5169231 | -0.132819225 | 0.09519833 | -1.395184402 | 0.162960239 | 0.368223122 | | Foxk1 | 3315.925796 | -0.0499536365645447 | 0.0554524619772652 | -0.900837127574696 | 0.36767492392468 | 0.778985919211964 | NM_199068 |
| Kcnn2 | 1059.721471 | 0.151992936 | 0.108951941 | 1.395045702 | 0.163002058 | 0.368263909 | | Hist2h3c1 | 69.52019692 | 0.0847855990921959 | 0.0941487576781975 | 0.900549331481064 | 0.367827994527521 | 0.77910024037719 | NM_178216 |
| Tmem106c | 957.9491531 | 0.107431398 | 0.077015698 | 1.394928574 | 0.163037379 | 0.368290007 | | 2610035D17Rik | 607.6685015 | 0.066415783953438 | 0.0738201473746375 | 0.899697254956397 | 0.36828138462335 | 0.779243710606327 | NR_015556 |
| Rnf180 | 399.624126 | -0.117957858 | 0.084579116 | -1.394478628 | 0.163173119 | 0.368542901 | | A430033K04Rik | 739.1486408 | 0.0610309985050263 | 0.0678434017019095 | 0.899707035729805 | 0.368276178189011 | 0.779243710606327 | NM_183025 |
| Cyb5r2 | 12.54018756 | 0.229069399 | 0.164288266 | 1.394313816 | 0.163222861 | 0.368601516 | | Fiz1 | 898.0147067 | 0.0496084240627607 | 0.0551377975653686 | 0.899717185909492 | 0.368270775163294 | 0.779243710606327 | NM_011813,NM_001110328,NM_001110329,NM_001110330 |
| Snx1 | 2660.74592 | 0.122212276 | 0.087663329 | 1.394109454 | 0.163284555 | 0.368687102 | | Lyrm7 | 471.4010191 | 0.0667428917782721 | 0.0741719025812629 | 0.899840633117756 | 0.368205067129009 | 0.779243710606327 | NM_029327 |
| Cdk19 | 4218.700239 | 0.095913999 | 0.068812422 | 1.393847156 | 0.163363765 | 0.3686973 | | Npnt | 698.5263599 | 0.0738936366027757 | 0.0820922424869542 | 0.900129346746967 | 0.36805142017716 | 0.779243710606327 | NM_033525,NM_001029836 |
| Fank1 | 134.6905574 | 0.224096815 | 0.160782787 | 1.393786107 | 0.163382205 | 0.3686973 | | Rnf113a1 | 142.5391582 | 0.0839711623760735 | 0.0932943419346159 | 0.900067041953343 | 0.368084574020064 | 0.779243710606327 | NM_153503 |
| Prrg3 | 2127.449149 | 0.088679498 | 0.063625208 | 1.393779316 | 0.163384256 | 0.3686973 | | Tmem132e | 914.7755479 | 0.0807189554704821 | 0.0897096843690445 | 0.899779728779597 | 0.368237484158061 | 0.779243710606327 | NM_023438 |
| Zdhhc24 | 828.7894492 | 0.128101346 | 0.091902123 | 1.393888864 | 0.163351168 | 0.3686973 | | Txndc9 | 1655.2583 | 0.0485500695425925 | 0.0539675758260995 | 0.89961553394646 | 0.368324887582476 | 0.779243710606327 | NM_172054 |
| Aars2 | 1073.679402 | 0.136499551 | 0.097966497 | 1.393328902 | 0.163520354 | 0.368744952 | | B130034C11Rik | 51.49372361 | -0.0834049953374821 | 0.0927531170410816 | -0.899215012909387 | 0.368538145224388 | 0.779452462740169 | NR_040375 |
| Kcnd3 | 4122.20825 | -0.132737705 | 0.095267389 | -1.393317336 | 0.16352385 | 0.368744952 | | Sorl1 | 8356.353472 | 0.056718165464546 | 0.0630813092507874 | 0.899127905526755 | 0.368584535767742 | 0.779452462740169 | NM_011436 |
| Lyplal1 | 144.9522689 | -0.207083883 | 0.148593697 | -1.393624947 | 0.163430891 | 0.368744952 | | Tmod1 | 2863.068775 | -0.0496212798878761 | 0.0551768863463736 | -0.899312795150816 | 0.368486073929098 | 0.779452462740169 | NM_021883 |
| Ptdss2 | 1620.171482 | -0.102560619 | 0.073602719 | -1.393435197 | 0.163488229 | 0.368744952 | | 9530051G07Rik | 26.44202792 | -0.0753909699698228 | 0.0838872250917454 | -0.898718128861333 | 0.368802818169071 | 0.779573593818964 | NR_040272 |
| Ube2g1 | 1505.342069 | 0.101097312 | 0.072558802 | 1.393315623 | 0.163524368 | 0.368744952 | | Fbxw7 | 8953.072079 | -0.0540350054817723 | 0.0601214054725255 | -0.89876484185762 | 0.368777930737057 | 0.779573593818964 | NM_001177774,NM_080428,NM_001177773 |
| AI118078 | 300.9837202 | -0.136111217 | 0.097699208 | -1.393166018 | 0.163569593 | 0.36875934 | | Tiprl | 1613.516915 | 0.0611654134391085 | 0.0680494610632837 | 0.89883758788665 | 0.368739175683713 | 0.779573593818964 | NM_145513 |
| Slc22a7 | 8.699720755 | 0.197091147 | 0.141480683 | 1.393060468 | 0.163601506 | 0.36875934 | | Cacnb4 | 9153.097186 | 0.0711880511942005 | 0.0792379113187196 | 0.898408981375845 | 0.368967549998848 | 0.77969488380612 | NM_146123,NM_001037099 |
| Slc3a2 | 4225.439469 | 0.112148057 | 0.080504925 | 1.393058341 | 0.163602149 | 0.36875934 | | Ugt1a6a | 59.35965329 | -0.0830256758958164 | 0.0924097545346576 | -0.898451427708081 | 0.36894492940467 | 0.77969488380612 | NM_145079 |
| Nbr1 | 7676.570057 | 0.053720885 | 0.03857135 | 1.392766547 | 0.163690398 | 0.368860597 | | Tnfaip8 | 233.9844604 | 0.0832116540599264 | 0.0926515363748469 | 0.898114130814529 | 0.369124706226513 | 0.779913524748558 | NM_001177759,NM_134131,NM_001177760 |
| S1pr5 | 428.3868681 | 0.218075477 | 0.156578787 | 1.39275237 | 0.163694686 | 0.368860597 | | Alg10b | 1247.728985 | 0.0682032901134375 | 0.0760358154270059 | 0.896989000912503 | 0.369724786242829 | 0.780397654597347 | NM_001033441 |
| Gm1673 | 253.5963533 | -0.225490559 | 0.161956523 | -1.392290688 | 0.16383439 | 0.369048539 | | C030030A07Rik | 311.091108 | 0.0790222711016013 | 0.0881245320138506 | 0.896711384398402 | 0.369872944236385 | 0.780397654597347 | NM_001039558 |
| Nedd4l | 7230.270747 | -0.095936832 | 0.068900673 | -1.392393247 | 0.163803348 | 0.369048539 | | C920025E04Rik | 53.02889323 | 0.079523683634646 | 0.0886852131905255 | 0.896696086909128 | 0.369881109251419 | 0.780397654597347 | NM_001271005 |
| Traf6 | 866.9067289 | 0.157399416 | 0.113054751 | 1.392240613 | 0.163849548 | 0.369048539 | | Chmp4b | 6587.470511 | 0.037055126470904 | 0.0413185742176299 | 0.896815225901315 | 0.369817521929684 | 0.780397654597347 | NM_029362 |
| Mcmdc2 | 85.95875596 | 0.213788858 | 0.153580657 | 1.392029841 | 0.163913362 | 0.369138609 | | Cpt1a | 1483.018394 | -0.0560254836277694 | 0.0625120631726226 | -0.896234755091334 | 0.370127397128176 | 0.780397654597347 | NM_013495 |
| Coro1b | 2467.884385 | 0.069552689 | 0.049983781 | 1.39150516 | 0.164072296 | 0.369298607 | | Dkc1 | 1071.570197 | 0.0571682902275571 | 0.0637824529659455 | 0.896301217171439 | 0.370091909218823 | 0.780397654597347 | NM_001030307 |
| Filip1l | 186.7167389 | -0.228137115 | 0.163952785 | -1.391480569 | 0.164079748 | 0.369298607 | | Flnc | 117.9468674 | -0.0845568202332405 | 0.094353101046525 | -0.896174256524474 | 0.370159702614328 | 0.780397654597347 | NM_001081185 |
| Slc8a2 | 12625.56724 | -0.133388359 | 0.095856843 | -1.391537156 | 0.164062601 | 0.369298607 | | Gm16894 | 32.87984751 | -0.0762796934934067 | 0.0851028815758481 | -0.896323274616997 | 0.370080131953115 | 0.780397654597347 | NR_037980 |
| Sugt1 | 1997.242317 | -0.096936579 | 0.069661183 | -1.391543692 | 0.16406062 | 0.369298607 | | Gpx6 | 39.78474463 | 0.0727528060415399 | 0.0810599572246419 | 0.897518436136349 | 0.369442340251728 | 0.780397654597347 | NM_145451 |
| Ctbp2 | 701.0141151 | -0.138173664 | 0.099313377 | -1.391289562 | 0.164137638 | 0.369375245 | | Ly75 | 69.65168538 | -0.0853020928696432 | 0.0950624268405908 | -0.897327111295879 | 0.36954439378877 | 0.780397654597347 | NM_013825 |
| Hus1 | 641.202604 | -0.140188815 | 0.100768039 | -1.391203166 | 0.164163828 | 0.369380532 | | Oprk1 | 580.0382967 | -0.0703369777409331 | 0.0784683713231335 | -0.896373615953934 | 0.370053253769833 | 0.780397654597347 | NM_011011,NM_001204371 |
| Impg2 | 32.8384876 | 0.233153122 | 0.167628223 | 1.390894196 | 0.164257513 | 0.369537666 | | Rab5b | 7717.139106 | 0.0373474215897665 | 0.0416365347281674 | 0.896986788972634 | 0.369725966562022 | 0.780397654597347 | NM_177411 |
| 4632428C04Rik | 36.35364196 | -0.228562098 | 0.164420962 | -1.390103152 | 0.164497556 | 0.369577538 | | Siglec1 | 29.15343215 | -0.0620026561074998 | 0.0691710592945822 | -0.896367017359761 | 0.370056776813863 | 0.780397654597347 | NM_011426 |
| Dpysl2 | 5681.660567 | -0.096695993 | 0.069563572 | -1.390037764 | 0.16451741 | 0.369577538 | | Stard5 | 258.9675069 | -0.074094863716583 | 0.0826308816472354 | -0.896697000437511 | 0.369880621653687 | 0.780397654597347 | NM_023377 |
| Elovl2 | 661.7143353 | -0.122351221 | 0.087980197 | -1.390667731 | 0.164326207 | 0.369577538 | | Trim35 | 7009.517827 | 0.047407169508321 | 0.0528275884441561 | 0.897394162870693 | 0.36950862618048 | 0.780397654597347 | NM_029979 |
| Emc9 | 515.1193963 | -0.105578575 | 0.075923052 | -1.390599717 | 0.164346842 | 0.369577538 | | Srebf2 | 9405.293803 | 0.0442550140354359 | 0.0493909103886197 | 0.896015353578759 | 0.370244563163297 | 0.780463289001604 | NM_033218 |
| Gpatch2l | 1548.932771 | -0.08547773 | 0.06148641 | -1.390188994 | 0.164471495 | 0.369577538 | | Mfng | 144.3723482 | -0.0822362044741863 | 0.0917950544510327 | -0.895867483994517 | 0.37032354230843 | 0.780516508353491 | NM_008595 |
| Hint1 | 3231.433454 | -0.111751659 | 0.080384925 | -1.390206672 | 0.164466128 | 0.369577538 | | Fam181b | 452.6976127 | -0.0732202574069013 | 0.0817549945730643 | -0.895605923396698 | 0.370463270984468 | 0.780632010857947 | NM_021427 |
| Mak | 141.5644726 | 0.163369212 | 0.117484212 | 1.390563114 | 0.164357948 | 0.369577538 | | Trappc8 | 2843.956723 | 0.0597281760056221 | 0.0666933856872382 | 0.895563711305949 | 0.370485824236176 | 0.780632010857947 | NM_029491,NM_177038 |
| Mmp24 | 1916.687596 | -0.117524462 | 0.084551724 | -1.389971216 | 0.164537618 | 0.369577538 | | 0610007P14Rik | 1201.098142 | 0.0479528863471377 | 0.0535744783321194 | 0.895069589849625 | 0.370749888908193 | 0.78076125737553 | NM_021446 |
| Nek4 | 665.784405 | 0.096865281 | 0.069664837 | 1.390447249 | 0.164393107 | 0.369577538 | | 1810043H04Rik | 233.9736903 | -0.0759530155718505 | 0.08493165793172 | -0.89428391510869 | 0.371170003826004 | 0.78076125737553 | NM_001110242 |
| Nrde2 | 720.4543202 | -0.093501569 | 0.067265054 | -1.390046722 | 0.16451469 | 0.369577538 | | 4930451C15Rik | 59.47458308 | -0.0813887116794826 | 0.0910582798813544 | -0.893809017538318 | 0.371424083588413 | 0.78076125737553 | NM_029053,NM_001145435 |
| Zdhhc17 | 5080.160321 | 0.069936259 | 0.050314457 | 1.38998339 | 0.164533921 | 0.369577538 | | 6530402F18Rik | 687.8853714 | -0.070480256627457 | 0.0788263513427438 | -0.894120499387353 | 0.371257422353478 | 0.78076125737553 | NR_029460 |
| Rph3a | 20642.39654 | -0.132955792 | 0.095735167 | -1.38878738 | 0.164897414 | 0.370332009 | | A630007B06Rik | 1137.152647 | 0.0712855375229616 | 0.079765961628555 | 0.893683672428045 | 0.371491163740077 | 0.78076125737553 | NM_170757 |
| Ccdc91 | 1405.914582 | -0.097874138 | 0.07049673 | -1.388350041 | 0.165030481 | 0.370577142 | | Adamtsl4 | 406.1536915 | -0.0817054154118459 | 0.0913932557213663 | -0.893998301810628 | 0.371322799761616 | 0.78076125737553 | NM_144899 |
| Wdr20a | 576.673366 | 0.117950773 | 0.084971136 | 1.388127533 | 0.165098214 | 0.370675515 | | Ap5z1 | 1038.534672 | -0.0714090446227007 | 0.0798940630246329 | -0.893796634183993 | 0.371430710375466 | 0.78076125737553 | NM_172725 |
| Mecom | 96.69695863 | -0.198714663 | 0.143166187 | -1.38799997 | 0.165137054 | 0.370698637 | | B3gntl1 | 243.39908 | -0.0789069177050858 | 0.0882891885341897 | -0.893732505815583 | 0.371465028990693 | 0.78076125737553 | NM_178664 |
| Trim2 | 11261.98334 | -0.111038645 | 0.08000268 | -1.387936563 | 0.165156363 | 0.370698637 | | Frrs1 | 37.55165805 | -0.0793772025294684 | 0.0887461040042632 | -0.894430278603051 | 0.371091718157444 | 0.78076125737553 | NM_001113478,NM_009146 |
| Gipc1 | 2230.610072 | 0.120777915 | 0.087027541 | 1.387812568 | 0.165194127 | 0.370729693 | | Igsf8 | 7891.669454 | 0.0514721847225345 | 0.0575864403300545 | 0.893824734217355 | 0.371415673122679 | 0.78076125737553 | NM_080419 |
| Dnajc9 | 938.4275376 | -0.105057774 | 0.075710361 | -1.38762744 | 0.16525021 | 0.370802545 | | Mill2 | 37.6960303 | -0.0753178971386356 | 0.0842821661124726 | -0.89363978896942 | 0.371514650349695 | 0.78076125737553 | NM_153761,NM_153760 |
| Klhl11 | 1025.841382 | -0.108924759 | 0.078515662 | -1.387299758 | 0.165350377 | 0.370972884 | | Pbld2 | 49.91111262 | -0.0840082645227977 | 0.0939678049737531 | -0.894011140797241 | 0.371315930388879 | 0.78076125737553 | NM_026085 |
| Hgf | 262.5481898 | 0.16018627 | 0.115483065 | 1.387097489 | 0.165412038 | 0.37100378 | | Pfkfb3 | 2022.751178 | 0.0813713528022401 | 0.0910476738472818 | 0.893722479266498 | 0.371470396926105 | 0.78076125737553 | NM_133232,NM_001177758,NM_001177752,NM_001177753,NM_001177756,NM_001177757,NM_001177754,NM_001177755 |
| Pcdhga7 | 690.94397 | 0.128500994 | 0.0926378 | 1.387133474 | 0.165401067 | 0.37100378 | | Prpf18 | 2104.458144 | 0.055927621728129 | 0.0624872518595738 | 0.895024506019466 | 0.370773988085119 | 0.78076125737553 | NM_026045 |
| Zhx2 | 667.7196952 | -0.144526892 | 0.104204271 | -1.386957477 | 0.16545473 | 0.371045822 | | Prpf19 | 8673.016183 | 0.0361673566881793 | 0.040410577907286 | 0.894997264606264 | 0.370788550221545 | 0.78076125737553 | NM_134129,NM_001253843,NM_001253844 |
| Agpat2 | 48.81788497 | 0.211587975 | 0.152582236 | 1.386714342 | 0.165528885 | 0.371158401 | | Rnf4 | 3804.186659 | 0.047987179663123 | 0.0536371063582382 | 0.894663842277774 | 0.37096681281849 | 0.78076125737553 | NM_011278 |
| Xpot | 4002.586384 | -0.090459767 | 0.065237061 | -1.386631554 | 0.165554141 | 0.371161318 | | Syne3 | 105.8273005 | 0.0850724283340473 | 0.0950964263404641 | 0.894591222907485 | 0.371005645458521 | 0.78076125737553 | NM_172500,NM_001042699 |
| Rnf6 | 2495.987197 | 0.127909492 | 0.092258258 | 1.386428641 | 0.165616056 | 0.371246408 | | Tmem138 | 297.5613943 | -0.0682274484595355 | 0.0762040790101094 | -0.895325412311385 | 0.370613159608158 | 0.78076125737553 | NM_028411 |
| Sla2 | 41.12153579 | -0.228573039 | 0.164924845 | -1.385922417 | 0.165770595 | 0.371539071 | | Msi2 | 4026.501098 | 0.0498783226050634 | 0.0558412417546998 | 0.893216573230403 | 0.371741204447498 | 0.781124365080914 | NM_001201341,NM_054043 |
| Slc7a6os | 650.8039841 | 0.092988338 | 0.067113782 | 1.385532692 | 0.165889643 | 0.371752117 | | Pold2 | 499.4649789 | 0.0650072014787383 | 0.0728127983541145 | 0.892799108785589 | 0.371964763707063 | 0.781368063283991 | NM_008894 |
| Emc8 | 2625.726958 | 0.124404203 | 0.089802052 | 1.385315819 | 0.165955919 | 0.371806281 | | Yme1l1 | 2871.005648 | 0.0605769482492197 | 0.0678483041557261 | 0.892829216632783 | 0.371948637657904 | 0.781368063283991 | NM_013771 |
| Mafk | 355.311133 | 0.186057611 | 0.134316164 | 1.38521821 | 0.165985754 | 0.371806281 | | Dok7 | 491.2037876 | 0.0758096108246044 | 0.0849412501243274 | 0.892494644400016 | 0.372127862081577 | 0.781371686122721 | NM_172708 |
| Rnf144a | 1383.926541 | -0.090642256 | 0.065435371 | -1.385218047 | 0.165985804 | 0.371806281 | | Necab3 | 2929.808753 | -0.0518575391500922 | 0.0580989052912092 | -0.892573429570947 | 0.372085653441691 | 0.781371686122721 | NM_021546 |
| Gas2l2 | 10.40986047 | 0.20892815 | 0.150839005 | 1.385106919 | 0.166019777 | 0.371823021 | | Pgm3 | 598.7955767 | 0.0647823733784465 | 0.072575495146143 | 0.892620480893671 | 0.372060474418943 | 0.781371686122721 | NM_001163746,NM_028352 |
| Hrct1 | 29.17826643 | -0.222688112 | 0.160803935 | -1.38484243 | 0.166100654 | 0.371823021 | | 6330403A02Rik | 14847.70288 | 0.0660661979191485 | 0.0740787980183775 | 0.891836796579215 | 0.37248041560015 | 0.78144602643646 | NM_001081227 |
| Lpcat1 | 948.3618389 | 0.083014858 | 0.059949018 | 1.384757611 | 0.166126596 | 0.371823021 | | Dcun1d2 | 1903.148625 | 0.0488474937167365 | 0.0547724303976615 | 0.891826295858911 | 0.372486034819362 | 0.78144602643646 | NM_001024504,NM_001042650,NM_001042651,NM_001042649 |
| Rbm42 | 1548.172434 | -0.073385741 | 0.052994099 | -1.384790807 | 0.166116443 | 0.371823021 | | Syncrip | 2643.303813 | 0.0701368041922666 | 0.0786014296170406 | 0.89230952329983 | 0.372227051180063 | 0.78144602643646 | NM_019666,NM_019796 |
| Tacr3 | 129.9070936 | -0.232397046 | 0.167815691 | -1.384835023 | 0.166102919 | 0.371823021 | | Tenm2 | 2612.347159 | 0.0687374784636551 | 0.0770713091675979 | 0.891868572183973 | 0.372463381676041 | 0.78144602643646 | NM_011856 |
| Tmed3 | 834.059861 | -0.106948719 | 0.077234753 | -1.384722735 | 0.166137265 | 0.371823021 | | Xiap | 3706.010268 | 0.0595202406758439 | 0.0667191529637008 | 0.892101263758945 | 0.372338657592726 | 0.78144602643646 | NM_009688 |
| BC031181 | 2854.241449 | -0.07421025 | 0.053596587 | -1.384607769 | 0.166172435 | 0.371848022 | | Zfp1 | 729.4559288 | -0.0537464907191839 | 0.0602470056549411 | -0.8921022735472 | 0.372338116396565 | 0.78144602643646 | NM_001037665,NM_011742 |
| Arih1 | 4049.527506 | -0.094562557 | 0.068303544 | -1.384445827 | 0.166221985 | 0.371853802 | | 2700050L05Rik | 1137.809086 | -0.0680690570782849 | 0.0764299621273942 | -0.890606971187906 | 0.373140058014158 | 0.78222667069605 | NM_178115 |
| Ppp1r32 | 37.9829718 | 0.226323415 | 0.163476217 | 1.384442453 | 0.166223017 | 0.371853802 | | 5730577I03Rik | 67.81872095 | 0.0846233775063534 | 0.0950233246621128 | 0.890553743591482 | 0.373168624064235 | 0.78222667069605 | NR_033729 |
| Casp9 | 1314.619231 | 0.081238744 | 0.05868998 | 1.384201242 | 0.166296843 | 0.371915182 | | Ccdc28b | 532.5706997 | -0.0710941789658901 | 0.0798377432245214 | -0.890483323983213 | 0.373206418759986 | 0.78222667069605 | NM_025455 |
| Cd38 | 102.9231255 | -0.212982149 | 0.153875768 | -1.384117542 | 0.166322466 | 0.371915182 | | Hmox1 | 273.4490644 | -0.0719430633995771 | 0.0808023085771415 | -0.890359009122783 | 0.373273145199214 | 0.78222667069605 | NM_010442 |
| Cuta | 1148.770803 | 0.108676465 | 0.078512359 | 1.384195639 | 0.166298558 | 0.371915182 | | Lamtor3 | 1485.262467 | 0.0533488032652885 | 0.0598733914693384 | 0.891026914562019 | 0.37291473136766 | 0.78222667069605 | NM_019920 |
| Adss | 2971.132898 | 0.096359592 | 0.06965065 | 1.383470101 | 0.166520769 | 0.372076569 | | Pak1 | 24175.81546 | 0.0516281081920076 | 0.057957371314143 | 0.890794510195619 | 0.373039420850729 | 0.78222667069605 | NM_011035 |
| Atp10a | 681.2093407 | 0.170523665 | 0.123244979 | 1.383615519 | 0.166476214 | 0.372076569 | | Robo1 | 1626.330009 | 0.061490660293421 | 0.0690422496154367 | 0.89062364908331 | 0.373131107642601 | 0.78222667069605 | NM_019413 |
| Clybl | 587.4424075 | -0.11553569 | 0.083507466 | -1.383537254 | 0.166500193 | 0.372076569 | | Rtl1 | 34.31035068 | 0.0637756953667911 | 0.0716315595322193 | 0.890329566789696 | 0.373288949557003 | 0.78222667069605 | NM_184109 |
| Ddit4 | 1407.724289 | 0.231269922 | 0.167151693 | 1.383593052 | 0.166483097 | 0.372076569 | | Clec11a | 174.8118133 | -0.0787586137115594 | 0.0884787903209193 | -0.890141167465586 | 0.373390090287394 | 0.782325753860477 | NM_009131 |
| Mrpl15 | 930.9247488 | -0.11188929 | 0.080871307 | -1.383547452 | 0.166497068 | 0.372076569 | | Actl6b | 3352.566048 | -0.0510416723237851 | 0.0573862944587193 | -0.889440114668875 | 0.373766594091517 | 0.782348608562381 | NM_031404 |
| Scap | 4005.705034 | 0.081665954 | 0.059034814 | 1.383352448 | 0.166556824 | 0.372076569 | | Lamtor1 | 2409.271538 | 0.0462489809019359 | 0.0519650089738406 | 0.890000979798114 | 0.373465359967271 | 0.782348608562381 | NM_025605 |
| Snora41 | 8.144679043 | 0.205510392 | 0.14856175 | 1.383333146 | 0.166562739 | 0.372076569 | | Mrpl30 | 1430.491034 | 0.0534555152026963 | 0.0600944220717689 | 0.889525406182558 | 0.373720775322163 | 0.782348608562381 | NM_027098 |
| Smarca4 | 6525.420915 | -0.077188855 | 0.055825631 | -1.382677719 | 0.166763707 | 0.372471799 | | Ptp4a2 | 4967.961582 | 0.0549567777818453 | 0.0617852869091036 | 0.889480012655696 | 0.373745160381611 | 0.782348608562381 | NM_001164745,NM_008974 |
| Ap3m1 | 2073.641946 | 0.123489365 | 0.089376984 | 1.381668529 | 0.167073502 | 0.373002426 | | Sumo2 | 4830.276763 | 0.0536573104720404 | 0.0603285047723621 | 0.889418868816754 | 0.373778007945954 | 0.782348608562381 | NM_133354 |
| Lemd2 | 1048.888766 | 0.091916142 | 0.066522563 | 1.381728818 | 0.167054982 | 0.373002426 | | Tatdn2 | 1749.025391 | 0.0475097512632028 | 0.053403475719963 | 0.889637811447598 | 0.373660396301401 | 0.782348608562381 | NM_001033463 |
| Zscan25 | 504.5870473 | 0.11261997 | 0.081506498 | 1.38172996 | 0.167054632 | 0.373002426 | | Tmem28 | 289.7583628 | 0.076709749765532 | 0.0862353749718374 | 0.889539238283405 | 0.373713345017886 | 0.782348608562381 | NM_001081283 |
| Gm10767 | 48.58135333 | 0.215128415 | 0.155733795 | 1.381385554 | 0.167160445 | 0.373111272 | | Prdx4 | 444.2162913 | 0.0671070779805369 | 0.0754684718482841 | 0.88920679506329 | 0.373891951603971 | 0.782462265356963 | NM_016764 |
| Tmem170b | 3198.897884 | -0.093781587 | 0.067894955 | -1.38127475 | 0.167194499 | 0.373111272 | | Zfp292 | 1714.158627 | 0.0653358353386322 | 0.0734839320615267 | 0.889117300962171 | 0.373940041725736 | 0.782462265356963 | NM_013889 |
| Tmem19 | 1423.586912 | 0.079487051 | 0.057545001 | 1.38130246 | 0.167185982 | 0.373111272 | | Igfbp7 | 711.7604195 | -0.0716664837944819 | 0.0806346096773727 | -0.888780687117192 | 0.374120957195105 | 0.782728074926702 | NM_001159518,NM_008048 |
| Gps1 | 4715.618787 | -0.065009768 | 0.047068042 | -1.381186998 | 0.167221472 | 0.373117725 | | Arl14ep | 1176.477717 | -0.054267815679706 | 0.0610990113045381 | -0.888194661763296 | 0.374436049782917 | 0.782936239980806 | NM_001025102,NM_173750 |
| Ap1ar | 3579.98083 | -0.07798429 | 0.056532031 | -1.379470881 | 0.167749618 | 0.374188394 | | Fgf18 | 53.70592475 | 0.0830915520094613 | 0.0935467542719378 | 0.88823554228206 | 0.374414063928856 | 0.782936239980806 | NM_008005 |
| Tsks | 31.46846595 | 0.222123795 | 0.161013955 | 1.379531328 | 0.167730994 | 0.374188394 | | Kif3b | 4378.858949 | 0.0489337369185049 | 0.0550842041981902 | 0.888344265489319 | 0.374355595645629 | 0.782936239980806 | NM_008444 |
| St6gal2 | 497.1264103 | -0.145860059 | 0.105756379 | -1.379208141 | 0.167830589 | 0.374315121 | | Serpinb8 | 474.0693128 | 0.0799811690795055 | 0.0900351102380976 | 0.888331058474026 | 0.374361596927937 | 0.782936239980806 | NM_001159748,NM_011459 |
| Naglu | 351.5502816 | 0.162015318 | 0.11751364 | 1.378693724 | 0.167989205 | 0.374561052 | | Raver2 | 775.1529841 | -0.0716571698538729 | 0.0807142429935029 | -0.887788414984465 | 0.374654575775002 | 0.783149984077341 | NM_183024 |
| Rnft2 | 4379.39951 | -0.079391978 | 0.057584771 | -1.378697474 | 0.167988049 | 0.374561052 | | Rwdd4a | 1048.936627 | 0.0616859296198899 | 0.0694892998198899 | 0.887703964070901 | 0.374700013034388 | 0.783149984077341 | NM_203507 |
| Hpx | 18.96479061 | 0.231016709 | 0.167577379 | 1.378567381 | 0.168028179 | 0.374580427 | | Znrd1as | 118.9089995 | -0.0809431028015075 | 0.0911694962481905 | -0.887830975627597 | 0.374631678095449 | 0.783149984077341 | NR_029602 |
| Slc22a6 | 374.2911531 | -0.186981169 | 0.135640169 | -1.378508819 | 0.168046247 | 0.374580427 | | BC003331 | 2164.400692 | -0.0541087900078518 | 0.0609625060447632 | -0.887574896742615 | 0.374769461915676 | 0.78318237744 | NM_001077237,NM_145511 |
| B3gnt5 | 28.33625497 | -0.221153686 | 0.160450192 | -1.378332326 | 0.168100707 | 0.374647922 | | Recql | 770.9479037 | -0.0575694737283681 | 0.0648688610929553 | -0.887474710645414 | 0.374823375794519 | 0.783182444877903 | NM_001204906,NM_001204907,NM_023042 |
| Tfb1m | 163.3486628 | -0.170963891 | 0.124099046 | -1.377640652 | 0.168314263 | 0.375069926 | | 1700013F07Rik | 41.45600076 | -0.0812324051128466 | 0.0915733046207165 | -0.887075173861162 | 0.375038429157281 | 0.783489747868359 | NM_029314 |
| Pigg | 749.5945454 | -0.120599042 | 0.087563106 | -1.377281455 | 0.168425247 | 0.375263269 | | Entpd2 | 443.8454563 | -0.0727482497497282 | 0.0820530292052833 | -0.88660041505261 | 0.375294070333749 | 0.783489747868359 | NM_009849 |
| Uap1l1 | 527.777398 | 0.132658943 | 0.096348718 | 1.376862558 | 0.168554745 | 0.375497803 | | Snrpa | 1736.399821 | 0.0539766286559266 | 0.060857112221176 | 0.886940353984538 | 0.375111014029613 | 0.783489747868359 | NM_015782,NM_001046637 |
| Noc4l | 791.8126584 | 0.119901501 | 0.087092578 | 1.376713203 | 0.168600935 | 0.375546706 | | Taf5l | 877.2848025 | 0.0561109088208772 | 0.0632748468812493 | 0.886780633798814 | 0.375197010696051 | 0.783489747868359 | NM_133966 |
| Stxbp1 | 41661.2787 | 0.091586124 | 0.066545034 | 1.3763029 | 0.168727875 | 0.375775435 | | Tmem200c | 418.6899815 | -0.0655598952922718 | 0.0739549265922718 | -0.886889684678417 | 0.375138295830532 | 0.783489747868359 | NM_001206661 |
| Dhrs7b | 571.7599068 | -0.111686063 | 0.081170038 | -1.375951846 | 0.168836542 | 0.37587621 | | Usf1 | 1960.666402 | 0.0526921449360737 | 0.0594274438436501 | 0.886663499259642 | 0.375260098861588 | 0.783489747868359 | NM_009480 |
| Eps15 | 8095.577494 | -0.090764922 | 0.065966481 | -1.37592487 | 0.168844895 | 0.37587621 | | 2210016F16Rik | 343.1687005 | -0.0735686485883303 | 0.0831720425981156 | -0.884535792284046 | 0.376407049893342 | 0.783947144371458 | NM_027335 |
| Minos1 | 2198.352063 | -0.0850644 | 0.06182358 | -1.375921613 | 0.168845903 | 0.37587621 | | Camsap2 | 8275.832725 | 0.0572653789355776 | 0.0647265806429003 | 0.884727392777219 | 0.376303677849211 | 0.783947144371458 | NM_001081360 |
| Cdc42bpg | 275.7424976 | 0.148280217 | 0.107783872 | 1.375718041 | 0.168908944 | 0.375908529 | | Dnahc10 | 79.35187217 | -0.0774918620598149 | 0.0875356656152782 | -0.885260442302385 | 0.376016179860232 | 0.783947144371458 | NM_019536 |
| Slc39a1 | 1543.001741 | -0.070130135 | 0.050976717 | -1.375728757 | 0.168905625 | 0.375908529 | | Dock9 | 6538.722432 | -0.0529749855206286 | 0.0598652372505744 | -0.884903960188019 | 0.37620843194709 | 0.783947144371458 | NM_001081039,NM_001128308,NM_134074,NM_001128307 |
| Ano7 | 16.97265511 | -0.231076957 | 0.168006896 | -1.375401624 | 0.169006966 | 0.376018657 | | Hs1bp3 | 787.2071842 | -0.0631684659667059 | 0.0713113160991744 | -0.885812651092513 | 0.375718491438581 | 0.783947144371458 | NM_021429 |
| Srf | 2337.869843 | -0.113856634 | 0.082779233 | -1.375425094 | 0.168999694 | 0.376018657 | | Kdm6b | 1297.977244 | 0.0799822025916085 | 0.0904272671417698 | 0.884492090933305 | 0.376430630042057 | 0.783947144371458 | NM_001017426 |
| B3gnt3 | 34.03202724 | 0.228801256 | 0.166460379 | 1.374508801 | 0.16928378 | 0.376567454 | | Myom1 | 105.4540384 | -0.0839492606392682 | 0.0947885577313964 | -0.885647620856901 | 0.375807441794325 | 0.783947144371458 | NM_010867,NM_001083934 |
| Irf2bp2 | 1280.408169 | 0.129911239 | 0.094524144 | 1.374370973 | 0.169326543 | 0.376567454 | | Pip4k2b | 8379.547198 | 0.0431837729397073 | 0.0488136228761368 | 0.884666418825026 | 0.376336572534374 | 0.783947144371458 | NM_054051 |
| Tmem205 | 534.7581224 | -0.139785305 | 0.101708393 | -1.374373355 | 0.169325804 | 0.376567454 | | Pold3 | 1106.778103 | -0.0503390048138279 | 0.0568524459578889 | -0.885432525649195 | 0.375923396401019 | 0.783947144371458 | NM_133692 |
| Slc38a1 | 8583.691473 | -0.110439417 | 0.080369337 | -1.37414866 | 0.169395535 | 0.376666823 | | Prdx1 | 5178.568046 | 0.0490384050419488 | 0.0553585098137054 | 0.885833184581282 | 0.375707424914774 | 0.783947144371458 | NM_011034 |
| Vegfc | 127.3999181 | -0.153731329 | 0.11188721 | -1.373984826 | 0.169446393 | 0.376725845 | | Rabac1 | 3178.400989 | 0.0472470266120512 | 0.0533788575871985 | 0.885126223146862 | 0.376088557613873 | 0.783947144371458 | NM_010261 |
| Hdac11 | 7948.267176 | -0.075074663 | 0.054652412 | -1.373675184 | 0.169542544 | 0.376885536 | | Rwdd3 | 161.0728419 | -0.0809984764314285 | 0.0915657907084788 | -0.884593206750173 | 0.376376071878163 | 0.783947144371458 | NM_028456,NM_025637 |
| P2rx5 | 33.14490881 | 0.22988255 | 0.167362146 | 1.373563585 | 0.169577209 | 0.376908517 | | Smim15 | 1152.427635 | 0.0593080379417723 | 0.0670381499546815 | 0.884690851132753 | 0.376323391396649 | 0.783947144371458 | NM_001048250 |
| Ddx50 | 2405.674646 | 0.112862055 | 0.082173782 | 1.373455771 | 0.169610702 | 0.376928889 | | Snap29 | 1293.699706 | 0.0598356452922699 | 0.0675758855073942 | 0.885458545500269 | 0.375909368314517 | 0.783947144371458 | NM_023348 |
| Apobec1 | 73.30956365 | 0.197077977 | 0.143525002 | 1.373126461 | 0.169713036 | 0.377102221 | | Tapbpl | 430.704791 | -0.0815986133050758 | 0.0921270557544742 | -0.885718235938663 | 0.375769379076554 | 0.783947144371458 | NM_145391 |
| Pnkp | 701.4580796 | 0.138203856 | 0.100655391 | 1.37303978 | 0.16973998 | 0.377108009 | | Tmed8 | 2974.9104 | 0.0514355133713813 | 0.0580850840970847 | 0.8855201670262 | 0.375876147580362 | 0.783947144371458 | NM_001033475 |
| L3mbtl3 | 837.2152563 | 0.104824823 | 0.076353182 | 1.372893979 | 0.169785309 | 0.377154634 | | Usp10 | 2397.539498 | 0.0402763369963683 | 0.0455360817871933 | 0.884492811318161 | 0.376430241333135 | 0.783947144371458 | NM_009462 |
| Chrm5 | 31.3881219 | 0.209013675 | 0.152276953 | 1.372589031 | 0.169880144 | 0.377311202 | | Grik4 | 1044.036517 | -0.0677780796098938 | 0.0766472113658066 | -0.884286308687945 | 0.376541677212162 | 0.783953619490306 | NM_175481 |
| Cdk4 | 1293.096878 | -0.10558869 | 0.07695372 | -1.372106385 | 0.170030322 | 0.377590628 | | Rint1 | 732.0645556 | 0.0517266978794718 | 0.0584935220488475 | 0.884314981687634 | 0.376526203064566 | 0.783953619490306 | NM_177323 |
| Atp6v1f | 3001.833846 | -0.085126405 | 0.062049819 | -1.371904161 | 0.170093275 | 0.377591627 | | Katnal1 | 4484.73615 | 0.0474599301571562 | 0.053701557284393 | 0.883771952940166 | 0.376819329629824 | 0.783969862819081 | NM_153572 |
| E030019B13Rik | 20.85045332 | -0.223218191 | 0.162710881 | -1.371870091 | 0.170103883 | 0.377591627 | | Mrps14 | 958.6216779 | 0.055551083618643 | 0.0628400011902161 | 0.884008315825622 | 0.376691723802786 | 0.783969862819081 | NM_025474 |
| Tmem243 | 192.4514313 | 0.17749012 | 0.129376483 | 1.371888583 | 0.170098125 | 0.377591627 | | Rps6 | 8791.384766 | 0.0573560578229174 | 0.0648757790899626 | 0.884090466850229 | 0.376647378965173 | 0.783969862819081 | NM_009096 |
| Bcor | 1410.767925 | -0.116337148 | 0.084817524 | -1.371616875 | 0.170182738 | 0.377712554 | | Sipa1l2 | 2724.398469 | 0.0583867341131831 | 0.0660600709292116 | 0.883843043035013 | 0.37678094723775 | 0.783969862819081 | NM_001081337 |
| Rbm48 | 305.5668213 | 0.113267718 | 0.082585217 | 1.37152534 | 0.17021125 | 0.377721728 | | Zap70 | 53.7361053 | 0.0800068997321736 | 0.0905239836927465 | 0.883819916760738 | 0.376793433125051 | 0.783969862819081 | NM_009539 |
| Rfwd2 | 2973.851651 | -0.098098026 | 0.071536892 | -1.371292818 | 0.170283694 | 0.377828376 | | Eif2a | 1716.966961 | 0.056090507675048 | 0.0635372460051195 | 0.882797275640944 | 0.377345812725873 | 0.783997208767348 | NM_001005509 |
| A830082K12Rik | 269.4009045 | 0.165733523 | 0.120888835 | 1.370958059 | 0.170388031 | 0.377840546 | | Gabrb3 | 12095.47701 | 0.0557227685382706 | 0.0630720141299052 | 0.883478501629933 | 0.376977793022775 | 0.783997208767348 | NM_008071,NM_001038701 |
| Camkk2 | 12136.47269 | -0.141647393 | 0.103324689 | -1.370895903 | 0.170407409 | 0.377840546 | | Plxna2 | 6935.368802 | 0.054314730552999 | 0.0614965492221258 | 0.883215907884748 | 0.377119628198265 | 0.783997208767348 | NM_008882 |
| Cldn23 | 23.97528631 | 0.230175204 | 0.167866409 | 1.37118084 | 0.17031859 | 0.377840546 | | Prrc2b | 26688.02249 | 0.0492888709537581 | 0.0558071939022682 | 0.883199234852673 | 0.377128634940271 | 0.783997208767348 | NM_172661,NM_001159634 |
| Lbh | 1887.180088 | -0.095842544 | 0.069920926 | -1.370727617 | 0.170459882 | 0.377840546 | | Rwdd1 | 1412.193811 | 0.0565146114254779 | 0.0640202821475787 | 0.882761048993835 | 0.377365389699853 | 0.783997208767348 | NM_025614 |
| Ncaph | 49.54932276 | -0.210297314 | 0.153393095 | -1.370969889 | 0.170384343 | 0.377840546 | | Scx | 51.44310128 | -0.0803010201568115 | 0.0909649389927316 | -0.882768911253024 | 0.377361140861418 | 0.783997208767348 | NM_198885 |
| Sema4b | 661.954986 | 0.17598509 | 0.128374582 | 1.370871768 | 0.170414934 | 0.377840546 | | Sgip1 | 5922.293603 | 0.0458494537389165 | 0.051939432209053 | 0.882748458904504 | 0.377372193563355 | 0.783997208767348 | NM_144906 |
| Srp54b | 10.47395693 | -0.223428424 | 0.162996895 | -1.370752641 | 0.170452079 | 0.377840546 | | Tyw3 | 541.7239489 | -0.05833094119196 | 0.066056136292592 | -0.883051060291906 | 0.377208684398971 | 0.783997208767348 | NM_172474,NM_001168358 |
| Chac1 | 203.7055503 | -0.154680633 | 0.112871053 | -1.370418982 | 0.17055615 | 0.377867998 | | Wrnip1 | 2107.647197 | 0.0460697821898044 | 0.052176246000683 | 0.882964676725451 | 0.377255356876936 | 0.783997208767348 | NM_030215 |
| Lpp | 383.3123454 | -0.146847869 | 0.107156072 | -1.370411079 | 0.170558616 | 0.377867998 | | Zfp35 | 975.9423863 | 0.0682498195400371 | 0.0772984921780124 | 0.882938562150256 | 0.37726946711251 | 0.783997208767348 | NM_011755 |
| Mfn1 | 2932.826717 | 0.107384882 | 0.078361667 | 1.370375169 | 0.170569819 | 0.377867998 | | Gdi2 | 9130.207938 | 0.0496288184327429 | 0.0562373351040734 | 0.882488801094491 | 0.377512533201663 | 0.784176613368705 | NM_008112 |
| Ttc16 | 29.41492216 | 0.230378649 | 0.168111205 | 1.370394368 | 0.170563829 | 0.377867998 | | Cd97 | 486.4019319 | -0.0715576332737089 | 0.0810956339688075 | -0.882385768156553 | 0.377568229271754 | 0.784178571454463 | NM_001163029,NM_001163031,NM_001163030,NM_011925 |
| Lsp1 | 274.13026 | -0.145816798 | 0.10641808 | -1.370225793 | 0.170616429 | 0.37791722 | | Osbp2 | 5120.326757 | -0.0580942902482276 | 0.065849663605558 | -0.882226074778675 | 0.37765456403762 | 0.784178571454463 | NM_152818 |
| Gpr158 | 8615.76531 | -0.095816336 | 0.069948457 | -1.369813431 | 0.170745149 | 0.378069092 | | Slc25a1 | 1350.115354 | 0.0598018024321273 | 0.0677880876082385 | 0.882187483702659 | 0.377675429291989 | 0.784178571454463 | NM_153150 |
| Magohb | 89.40205552 | -0.183085333 | 0.133661209 | -1.369771641 | 0.170758197 | 0.378069092 | | Tnfrsf9 | 23.47195483 | -0.075750508329412 | 0.085879772458505 | -0.882052969644428 | 0.37774816331855 | 0.784217496121946 | NM_011612,NM_001077508,NM_001077509 |
| Oxtr | 474.6347513 | -0.180326277 | 0.131637179 | -1.369873453 | 0.170726408 | 0.378069092 | | Chrm4 | 2155.539563 | 0.0656965967915009 | 0.0745207255598563 | 0.881588626150959 | 0.377999506009878 | 0.78462715408681 | NM_007699 |
| Sipa1l2 | 2613.878189 | -0.096324992 | 0.070328715 | -1.369639577 | 0.170799439 | 0.378106373 | | Tgfa | 2346.638061 | -0.0619206139443276 | 0.0702863176787909 | -0.880976781673289 | 0.378330387065292 | 0.785201772040352 | NM_031199 |
| Noxred1 | 46.60955701 | 0.224708959 | 0.164076645 | 1.369536529 | 0.170831625 | 0.378123599 | | Rassf6 | 34.57196571 | 0.0784422450951597 | 0.0890564986935559 | 0.880814384642272 | 0.378418292732726 | 0.785272017476511 | NM_028478 |
| Fam108c | 915.7159766 | -0.08784519 | 0.064149906 | -1.369373643 | 0.170882509 | 0.37812819 | | Gm10406 | 66.60979705 | 0.0832408867283688 | 0.0945214003972154 | 0.880656511420255 | 0.378503761712551 | 0.785337186492861 | NM_001164727 |
| Zranb3 | 347.7690402 | 0.119756723 | 0.087452356 | 1.369393905 | 0.170876178 | 0.37812819 | | BC005764 | 2626.506919 | -0.0616448911331193 | 0.0700421405890296 | -0.880111467392453 | 0.378798927513783 | 0.785380516173885 | NM_181681,NR_033210,NM_001170935 |
| Sh3bp4 | 875.1662784 | -0.107251703 | 0.078326563 | -1.369289023 | 0.170908948 | 0.378132692 | | Cdc42ep5 | 35.338777 | -0.0770428451383333 | 0.0875961935371499 | -0.879522751244426 | 0.379117902828609 | 0.785380516173885 | NM_021454 |
| Spg20 | 1581.478148 | 0.084408152 | 0.061655729 | 1.369023663 | 0.170991878 | 0.378262159 | | Cetn4 | 241.5900601 | 0.076433500394181 | 0.0868606481506639 | 0.879955446125655 | 0.378883446105307 | 0.785380516173885 | NM_145825 |
| Mvd | 609.7741835 | -0.172704459 | 0.126168001 | -1.368845176 | 0.171047676 | 0.378277575 | | Eef1d | 2395.418703 | 0.0474180899495753 | 0.053907524295033 | 0.879619136097934 | 0.379065668746658 | 0.785380516173885 | NM_029663,NM_023240 |
| Vangl1 | 134.3235673 | -0.173816725 | 0.126978811 | -1.368864017 | 0.171041786 | 0.378277575 | | Fbxo11 | 5615.694671 | 0.0588258568222577 | 0.0668345245422461 | 0.880171697253406 | 0.378766303380367 | 0.785380516173885 | NM_001081034 |
| Prpf19 | 7750.999931 | 0.071758262 | 0.052440212 | 1.368382394 | 0.171192412 | 0.378543633 | | Foxred2 | 1028.714544 | 0.0640043892629596 | 0.0727304698861776 | 0.880021665790497 | 0.37884757270118 | 0.785380516173885 | NM_001017983,NM_001168260 |
| Adar | 8679.581518 | 0.083966096 | 0.061379097 | 1.367991712 | 0.17131467 | 0.378651855 | | Mvk | 709.1090954 | 0.0699037146658451 | 0.0794551842541443 | 0.879787962510438 | 0.37897418690307 | 0.785380516173885 | NM_023556 |
| F420014N23Rik | 13.53092543 | 0.215801919 | 0.157747792 | 1.368018633 | 0.171306244 | 0.378651855 | | Ptprj | 7011.397171 | 0.0631005008697653 | 0.0717257305162573 | 0.879747064485637 | 0.37899634703858 | 0.785380516173885 | NM_008982,NM_001135657 |
| Gca | 458.128898 | 0.160159214 | 0.117076125 | 1.367992096 | 0.17131455 | 0.378651855 | | Rela | 948.9665034 | -0.0577964032410615 | 0.0657135734318778 | -0.879520017291044 | 0.37911938451131 | 0.785380516173885 | NM_009045 |
| Espnl | 26.28826548 | -0.229198159 | 0.167566659 | -1.367802883 | 0.171373785 | 0.378728488 | | Uap1l1 | 602.9527432 | 0.0651840368607429 | 0.0740866281732679 | 0.879835382820982 | 0.378948493738485 | 0.785380516173885 | NM_001033293 |
| Cntnap5c | 228.6506983 | -0.185973181 | 0.135998381 | -1.367466139 | 0.171479244 | 0.378907502 | | Upp2 | 326.4756794 | 0.0753420539532678 | 0.0855755111445311 | 0.88041605531307 | 0.378633962026367 | 0.785380516173885 | NM_029692 |
| Gtpbp3 | 741.5823206 | 0.127157333 | 0.093016334 | 1.367043057 | 0.17161181 | 0.379146355 | | Rnaseh2a | 707.274422 | 0.0522627587831535 | 0.059439000496996 | 0.879267120008097 | 0.379256459142743 | 0.785552449095607 | NM_027187 |
| BC053749 | 111.9585432 | 0.169409237 | 0.123959297 | 1.36665212 | 0.171734372 | 0.37936304 | | 1700012B15Rik | 479.4558291 | -0.0744307199075287 | 0.0848612339309313 | -0.877087410349324 | 0.380439162301892 | 0.786275719104857 | NR_015551 |
| Kif6 | 78.27570001 | 0.197734926 | 0.144700514 | 1.366511568 | 0.171778452 | 0.379406321 | | Ablim2 | 6835.209315 | -0.0587157378824075 | 0.0668769767340364 | -0.877966390674546 | 0.379961958134264 | 0.786275719104857 | NM_001177700,NM_177678,NM_001177699,NM_001177698,NM_001177697,NM_001177696 |
| Rxra | 1716.936304 | 0.140151991 | 0.102575526 | 1.366329735 | 0.171835492 | 0.379478209 | | Aga | 212.3878963 | 0.0786474502671126 | 0.0895777596684903 | 0.877979652071802 | 0.379954761253627 | 0.786275719104857 | NM_001205054,NM_001005847 |
| Gm15645 | 13.69571187 | 0.225813767 | 0.165280985 | 1.366241659 | 0.171863126 | 0.379485147 | | Asb3 | 574.6769187 | 0.0616776107361198 | 0.0703027802986778 | 0.877313962180238 | 0.38031613062257 | 0.786275719104857 | NM_023906 |
| Scyl2 | 1251.100999 | 0.100677447 | 0.073711666 | 1.365827868 | 0.171992997 | 0.379717797 | | Atg7 | 1149.829656 | 0.0512857753987084 | 0.0584773674300127 | 0.877019223891852 | 0.380476196559334 | 0.786275719104857 | NM_001253717,NM_001253718,NM_028835 |
| Ccnyl1 | 810.2592657 | -0.095670884 | 0.070051662 | -1.365718964 | 0.17202719 | 0.379739176 | | Dgkh | 1021.258487 | -0.0735454762655011 | 0.0838826149302462 | -0.876766613995748 | 0.38061341641662 | 0.786275719104857 | NM_001081336,NR_045588 |
| Zfp763 | 678.794139 | -0.123934897 | 0.090762773 | -1.365481601 | 0.172101732 | 0.379849606 | | Fam163a | 361.6763684 | 0.076084948703815 | 0.0867827887048643 | 0.876728552277444 | 0.38063409450264 | 0.786275719104857 | NM_177838 |
| AW549877 | 2398.956076 | -0.105631639 | 0.077385802 | -1.365000239 | 0.172252975 | 0.38012088 | | Gnai3 | 1273.06518 | 0.0608402425225296 | 0.0693186399993128 | 0.877689500589347 | 0.380112243859734 | 0.786275719104857 | NM_010306 |
| Dap | 213.9628448 | -0.160372106 | 0.117501098 | -1.364856233 | 0.17229824 | 0.38012088 | | Kcnj14 | 94.14024467 | 0.0835240265746747 | 0.0950990929669442 | 0.878284155703852 | 0.379789532013934 | 0.786275719104857 | NM_145963 |
| Zfp873 | 157.6616045 | -0.155239689 | 0.113736462 | -1.364906961 | 0.172282294 | 0.38012088 | | Kif27 | 177.2641073 | -0.0784217652875181 | 0.0893409436774109 | -0.877780803062486 | 0.380062684208267 | 0.786275719104857 | NM_175214 |
| Armc6 | 850.4440859 | -0.105454244 | 0.077272274 | -1.364709992 | 0.172344218 | 0.38016523 | | Lrrc4 | 5048.97064 | 0.0468960837647511 | 0.053422996244959 | 0.87782578778846 | 0.380038267629873 | 0.786275719104857 | NM_138682 |
| Chuk | 1179.600658 | 0.174758284 | 0.128062176 | 1.364636217 | 0.172367416 | 0.38016523 | | Msantd3 | 366.0076895 | 0.0675254832132754 | 0.0770025215752328 | 0.876925610121765 | 0.380527044815395 | 0.786275719104857 | NM_001145924,NM_001145925,NM_028137 |
| Mzt2 | 459.3766003 | 0.1047469 | 0.076769907 | 1.364426561 | 0.172433353 | 0.380256529 | | Paxip1 | 1990.035042 | -0.0388182106583017 | 0.0442114616689332 | -0.878012379436412 | 0.379937000667998 | 0.786275719104857 | NM_018878 |
| Golga7b | 4394.643091 | -0.125494353 | 0.09200596 | -1.363980698 | 0.172573641 | 0.380349358 | | Rhot1 | 3456.474528 | 0.0520580042549626 | 0.0593537849256914 | 0.877097737769056 | 0.380443309864041 | 0.786275719104857 | NM_001163355,NM_001163354,NM_021536 |
| Grk4 | 75.81620961 | 0.21447059 | 0.157238583 | 1.363981951 | 0.172573246 | 0.380349358 | | Rnf19a | 1603.205691 | 0.0606146805422303 | 0.0690954482561796 | 0.877260110065343 | 0.380345373441945 | 0.786275719104857 | NM_013923 |
| Kremen2 | 36.69721408 | -0.225373703 | 0.165224448 | -1.36404573 | 0.172553173 | 0.380349358 | | Rps19-ps3 | 47.77891455 | -0.0829102004892248 | 0.0945295513900458 | -0.877082343775466 | 0.380441914044889 | 0.786275719104857 | NR_033639 |
| Slc16a6 | 547.4101983 | 0.185623218 | 0.136087453 | 1.363999501 | 0.172567723 | 0.380349358 | | Sdhc | 3109.837334 | 0.044340111851582 | 0.0505406043397755 | 0.87731661365763 | 0.380314690850838 | 0.786275719104857 | NM_025321 |
| Nudt3 | 8353.918462 | -0.099964681 | 0.073305733 | -1.363667979 | 0.172672086 | 0.380512204 | | Snap91 | 13767.80365 | 0.0497713380095887 | 0.056681888193928 | 0.878081863457054 | 0.379899294560614 | 0.786275719104857 | NM_013669 |
| D930016D06Rik | 436.7948959 | 0.136494909 | 0.100100938 | 1.363572728 | 0.17270208 | 0.38052418 | | Tfam | 1321.716206 | 0.0530896975374887 | 0.0605044102464272 | 0.877451698500335 | 0.380241343219312 | 0.786275719104857 | NM_009360 |
| Foxl2os | 18.91747006 | 0.229185069 | 0.168093196 | 1.363440487 | 0.172743729 | 0.380561827 | | 2310067B10Rik | 3732.224609 | -0.0504169540225704 | 0.0575258658971168 | -0.876422340391008 | 0.380800477724237 | 0.786507569944869 | NM_028014 |
| Dact3 | 4563.828074 | -0.11960407 | 0.087737931 | -1.363196828 | 0.172820487 | 0.380676801 | | Hs2st1 | 2451.787046 | 0.0537841309105342 | 0.0613861104546722 | 0.87616124416693 | 0.380942382183363 | 0.786576978478398 | NM_011828 |
| Emg1 | 832.3568103 | -0.102178072 | 0.074962769 | -1.363050933 | 0.172866459 | 0.380693484 | | Nup210 | 1505.744048 | -0.0552298213953611 | 0.063029518268795 | -0.876253268505537 | 0.380892363722157 | 0.786576978478398 | NM_018815 |
| Uox | 20.06799221 | 0.229152507 | 0.168121552 | 1.363016847 | 0.172877201 | 0.380693484 | | Acp1 | 2252.984393 | 0.0471678649858948 | 0.0538902963367417 | 0.875257498198918 | 0.381433814145063 | 0.786663047961363 | NM_021330,NM_001110239 |
| Cage1 | 136.907383 | 0.171363167 | 0.12573564 | 1.362884594 | 0.172918885 | 0.380731164 | | Ankrd37 | 143.7170454 | 0.0808985778021004 | 0.0923707175401565 | 0.875803284378853 | 0.381136984259558 | 0.786663047961363 | NM_001039562 |
| Pla2g6 | 799.279865 | 0.113012479 | 0.082938064 | 1.362612936 | 0.17300453 | 0.380811505 | | Ccndbp1 | 1928.43091 | 0.0446958423874481 | 0.0511292765254667 | 0.874173182661518 | 0.382023948008451 | 0.786663047961363 | NM_010761 |
| Tex26 | 11.558884 | 0.220034229 | 0.161475458 | 1.362648116 | 0.172993437 | 0.380811505 | | Dusp16 | 716.3156339 | -0.0693982766495596 | 0.079381406581983 | -0.874238438889496 | 0.381988418948913 | 0.786663047961363 | NM_130447,NM_001048054 |
| Ikzf2 | 112.2914945 | 0.159344968 | 0.116977838 | 1.362180822 | 0.173140827 | 0.381057382 | | Eltd1 | 937.0786802 | 0.0772112507513222 | 0.088215384013739 | 0.875258342006389 | 0.381433355124602 | 0.786663047961363 | NM_133222 |
| Setd7 | 8764.265114 | -0.072880207 | 0.053512057 | -1.361939924 | 0.173216845 | 0.381170544 | | Evi5l | 1581.017097 | 0.042687143564282 | 0.048799716206174 | 0.874741635462242 | 0.3817145003227 | 0.786663047961363 | NM_001039578 |
| Rnf123 | 2673.948487 | 0.074048245 | 0.054383903 | 1.361583859 | 0.173329252 | 0.381363736 | | Gal3st3 | 1972.708896 | 0.0626763305578676 | 0.0716057415331693 | 0.875297555976494 | 0.381412023600852 | 0.786663047961363 | NM_001024717 |
| Tyms | 144.0790501 | 0.179437363 | 0.131837841 | 1.361045975 | 0.173499161 | 0.381683373 | | Galns | 347.5656745 | -0.0704178755441917 | 0.0805376121975434 | -0.874347694484287 | 0.381928932941221 | 0.786663047961363 | NM_016722,NM_001193645 |
| Ireb2 | 2439.486669 | -0.103261426 | 0.075876745 | -1.360910065 | 0.173542112 | 0.381723664 | | Gar1 | 425.2484859 | -0.0632069015550975 | 0.0722286509154871 | -0.875094588559521 | 0.38152244125404 | 0.786663047961363 | NM_026578 |
| Ubr2 | 3684.569965 | 0.09565825 | 0.070303302 | 1.360650885 | 0.173624043 | 0.381795475 | | Greb1 | 66.914309 | -0.0804664908422395 | 0.0920213564664837 | -0.874432783128416 | 0.381882610659704 | 0.786663047961363 | NM_015764,NM_001252071 |
| Zbtb42 | 45.96939942 | 0.228160431 | 0.167679402 | 1.360694448 | 0.17361027 | 0.381795475 | | Id3 | 801.7745695 | -0.0742945808415721 | 0.0849135749443875 | -0.874943504501253 | 0.381604646230446 | 0.786663047961363 | NM_008321 |
| Bbox1 | 53.76858209 | 0.228016928 | 0.167609568 | 1.360405201 | 0.173701733 | 0.381912113 | | Llph | 997.4243182 | 0.0597862576868275 | 0.0683447372910245 | 0.874774855483701 | 0.381696421153484 | 0.786663047961363 | NM_025431 |

| | | | | | | | | | | | | | | |
|---|---|---|---|---|---|---|---|---|---|---|---|---|---|---|
| Cd24a | 309.6917897 | -0.177075851 | 0.130194034 | -1.360091902 | 0.173800843 | 0.381967414 | | Mllt10 | 1413.895637 | -0.0544541263930194 | 0.0622331759975209 | -0.875001565004309 | 0.381573054171115 | 0.786663047961363 | NR_045539,NR_045538,NM_001252561,NM_010804,NM_001252560 |
| Lrrd1 | 12.06337056 | 0.219379057 | 0.161294499 | 1.360114938 | 0.173793555 | 0.381967414 | | Pcdhga7 | 844.6752429 | 0.0620692598824277 | 0.0709816427925193 | 0.874441016585898 | 0.38187812854604 | 0.786663047961363 | NM_033590 |
| Tmem107 | 234.058013 | -0.160382631 | 0.117917526 | -1.360125481 | 0.173790219 | 0.381967414 | | Polr3d | 770.755334 | 0.0513533075938149 | 0.058657737688374 | 0.87547371612992 | 0.381316205445023 | 0.786663047961363 | NM_001164082,NM_025945 |
| Chic2 | 192.6727476 | -0.134574688 | 0.099016352 | -1.359115798 | 0.174109898 | 0.38253811 | | Rnf24 | 2614.386453 | 0.0531692674519246 | 0.0607252188146701 | 0.875571442800299 | 0.381263055702636 | 0.786663047961363 | NM_178607 |
| Sstr3 | 1169.601754 | -0.120391355 | 0.088578509 | -1.359148595 | 0.174099507 | 0.38253811 | | Slc36a4 | 1159.198057 | 0.0568458013645266 | 0.0650067294091124 | 0.874460257903056 | 0.381867654120837 | 0.786663047961363 | NM_172289 |
| Dpy30 | 564.7521342 | -0.1102446 | 0.081129917 | -1.358864929 | 0.174189394 | 0.38265851 | | Sox7 | 82.36668089 | -0.0831421703690985 | 0.0950765785021183 | -0.874475834942315 | 0.38185917455328 | 0.786663047961363 | NM_011446 |
| D030056L22Rik | 741.7312544 | -0.09137547 | 0.067256814 | -1.358605387 | 0.174271667 | 0.382784975 | | Spef1 | 317.3489419 | -0.0726716112949402 | 0.0831393744053995 | -0.87409379508418 | 0.382067176329851 | 0.786663047961363 | NM_027641 |
| Arhgap5 | 4139.438643 | -0.108665516 | 0.080001135 | -1.358299684 | 0.17436861 | 0.382943622 | | Stat4 | 96.98144758 | -0.0828251343502553 | 0.0945781653365936 | -0.875732089489043 | 0.381175696086926 | 0.786663047961363 | NM_011487 |
| Ccdc166 | 201.1955164 | 0.148985668 | 0.109729686 | 1.357751708 | 0.174542483 | 0.383216839 | | 1110019D14Rik | 71.87682369 | -0.0829336425834193 | 0.0949618196328655 | -0.873336704204398 | 0.382479580049557 | 0.786954444730859 | NR_045995 |
| Lepre1 | 309.5186252 | 0.136692973 | 0.100671144 | 1.357816823 | 0.174521816 | 0.383216839 | | Crh | 148.5363309 | 0.0801886681008041 | 0.0918027059350295 | 0.873489155728755 | 0.382396514483665 | 0.786954444730859 | NM_205769 |
| Bbc3 | 151.9406988 | 0.183656659 | 0.135297922 | 1.357424091 | 0.174646498 | 0.383336568 | | Emp2 | 807.2634246 | -0.0587360228203785 | 0.0672491886001725 | -0.873408646899687 | 0.382440379583519 | 0.786954444730859 | NM_007929 |
| Galnt11 | 1491.336784 | -0.075585634 | 0.055681825 | -1.357456126 | 0.174636325 | 0.383336568 | | Gpr151 | 28.60642181 | -0.0764867088419456 | 0.0875499294438317 | -0.873635299626554 | 0.382316896109135 | 0.786954444730859 | NM_181543 |
| Cdk1 | 15.43189986 | -0.220141621 | 0.162198593 | -1.357235081 | 0.174706527 | 0.383414014 | | Sigmar1 | 1383.255958 | 0.0482340599829442 | 0.0552229553754127 | 0.873442206326027 | 0.382422094411717 | 0.786954444730859 | NM_011014 |
| Icam5 | 7458.385829 | -0.103776131 | 0.076469162 | -1.35709778 | 0.174750144 | 0.383446291 | | Ip6k3 | 49.89378915 | -0.0804840286636239 | 0.09217816415294 | -0.873135513201224 | 0.382589219002077 | 0.787068544855427 | NM_173027 |
| Serinc4 | 14.02335306 | 0.223999096 | 0.165065329 | 1.357032982 | 0.174770731 | 0.383446291 | | Bbs12 | 147.5205878 | -0.0791130116533766 | 0.0906722019285142 | -0.872516713730512 | 0.382926554235801 | 0.787600928143281 | NM_001008502,NM_001255992 |
| Rcan3 | 2007.475451 | -0.125496669 | 0.092490259 | -1.356863631 | 0.174824545 | 0.383510052 | | Uqcrh | 4477.943128 | 0.0544329736160808 | 0.0623900871313994 | 0.872461894490383 | 0.382956447437422 | 0.787600928143281 | NM_025641 |
| Cdk10 | 2071.963481 | 0.130276903 | 0.096030656 | 1.356617863 | 0.174902663 | 0.383627104 | | Mcam | 840.6263845 | -0.0814395086970249 | 0.0933858665872271 | -0.872075311535032 | 0.383167293617906 | 0.787602461491642 | NM_023061 |
| Csnk1d | 6090.134862 | 0.112177904 | 0.082705268 | 1.356357418 | 0.174985475 | 0.383754415 | | Rab26 | 1273.04184 | -0.0686563500666037 | 0.0787417843434233 | -0.871917631014899 | 0.383253314551941 | 0.787602461491642 | NM_177375 |
| Map6d1 | 2694.846636 | 0.143352753 | 0.105703899 | 1.356172802 | 0.175044194 | 0.383828862 | | Sh3bgrl2 | 1297.067861 | 0.0600483715447978 | 0.0688735443272362 | 0.871864111704377 | 0.383282514132206 | 0.787602461491642 | NM_172507 |
| Snx13 | 3649.808047 | -0.096444185 | 0.071123555 | -1.356009064 | 0.175096284 | 0.383888755 | | Sparc | 12201.39713 | 0.0776648176946615 | 0.0890660103415307 | 0.871991654244414 | 0.383212930491878 | 0.787602461491642 | NM_009242 |
| E130112N10Rik | 18.53387761 | 0.227907771 | 0.168098154 | 1.355801747 | 0.175162255 | 0.383979058 | | Ubtd1 | 322.6724989 | -0.0686076402388571 | 0.0786718452470307 | -0.872073611892896 | 0.383168220776574 | 0.787602461491642 | NM_145500 |
| Gm13375 | 356.0449078 | 0.131658119 | 0.097129962 | 1.355484092 | 0.175263374 | 0.384146373 | | Zdhhc21 | 2053.886243 | -0.0613694342721847 | 0.070384436090473 | -0.87191768068298 | 0.383253287454185 | 0.787602461491642 | NM_026647 |
| Sec14l2 | 1913.44112 | 0.133470488 | 0.098483397 | 1.355258776 | 0.175335124 | 0.384249281 | | B830017H08Rik | 94.32034792 | -0.082089521118632 | 0.0941824670693077 | -0.871600879367742 | 0.383426150793602 | 0.787702474891803 | NR_027959 |
| 9230112J17Rik | 28.60085269 | 0.226479897 | 0.167126628 | 1.355139512 | 0.175373112 | 0.384278178 | | Rap1gap | 11284.05259 | 0.0559545460460132 | 0.0641992615359401 | 0.871576163141513 | 0.383439639264762 | 0.787702474891803 | NM_001256218,NM_029563,NM_001081155 |
| Rac3 | 550.7597499 | -0.152224285 | 0.112392611 | -1.354397622 | 0.175609556 | 0.384741863 | | Gm6682 | 30.22478552 | 0.0792296424641909 | 0.0909150419525831 | 0.871469019455694 | 0.383498114516963 | 0.787711200717393 | NR_033599 |
| Dtwd2 | 117.6136887 | -0.174414785 | 0.128800331 | -1.354148572 | 0.175688983 | 0.384861458 | | Chchd7 | 433.6110298 | -0.0682675759042726 | 0.0783708335709082 | -0.871083958071029 | 0.383708312538939 | 0.78787653971568 | NM_001190324,NM_181391,NM_001190322,NM_001190323 |
| Agbl3 | 164.4570717 | 0.166453871 | 0.122983502 | 1.353465047 | 0.175907109 | 0.385037626 | | Dpy30 | 598.2482697 | 0.0677298906186557 | 0.0777678402218416 | 0.870924156122228 | 0.383795566228015 | 0.78787653971568 | NM_001146222,NM_001146223,NM_001146224,NM_024428 |
| Ccna2 | 66.73768245 | -0.192048347 | 0.141886266 | -1.353537255 | 0.175884056 | 0.385037626 | | Rcor1 | 742.1190116 | -0.0642948764602426 | 0.0738172757417445 | -0.871000396779518 | 0.383753936443524 | 0.78787653971568 | NM_198023 |
| D930015E06Rik | 719.5345221 | -0.107954308 | 0.07974628 | -1.353722181 | 0.175825029 | 0.385037626 | | Rnf150 | 3982.987324 | 0.0689293482560368 | 0.0791430580309352 | 0.870946232948111 | 0.383783511305951 | 0.78787653971568 | NM_177378 |
| Prr24 | 976.3999939 | 0.074226323 | 0.054843146 | 1.353429346 | 0.175918507 | 0.385037626 | | Csnk2a2 | 1472.416881 | 0.0452175481273029 | 0.0519698032848562 | 0.870073490166147 | 0.384260243934048 | 0.788718991576371 | NM_009974 |
| Suco | 1543.067705 | -0.10598483 | 0.078308284 | -1.353430627 | 0.175918098 | 0.385037626 | | Tsn | 3269.542276 | 0.0420387963880662 | 0.0483324541311518 | 0.869784022842964 | 0.384418444406723 | 0.788932229931063 | NM_011650 |
| Ubqln4 | 6413.965212 | -0.095234619 | 0.07036055 | -1.353522934 | 0.175888628 | 0.385037626 | | Ccnj | 362.7271889 | -0.0688788653429151 | 0.0792004174639273 | -0.86967805913759 | 0.384476365941084 | 0.788939637188894 | NM_172839 |
| Diras1 | 4931.947634 | -0.07747065 | 0.05725702 | -1.353033217 | 0.176045019 | 0.385096926 | | Pisd | 3507.028886 | -0.0637250689536235 | 0.0732851443840446 | -0.869549613215984 | 0.384546583787935 | 0.788972270636094 | NM_177298 |
| E2f4 | 1058.617321 | 0.118910218 | 0.087880022 | 1.353097265 | 0.176024559 | 0.385096926 | | Ang | 52.85326398 | -0.0821864004941967 | 0.0946427081639686 | -0.86838597593603 | 0.38518306957257 | 0.789016184424233 | NM_001161731,NM_007447 |
| Sco2 | 106.4204618 | 0.157869998 | 0.116664553 | 1.35319592 | 0.175993048 | 0.385096926 | | Ccdc94 | 254.9011773 | 0.0759282207186096 | 0.0874677287453553 | 0.868071251051452 | 0.385355328210903 | 0.789016184424233 | NM_028381 |
| Slco4a1 | 92.25342619 | 0.215743764 | 0.15943787 | 1.353152577 | 0.176006892 | 0.385096926 | | Cnst | 2455.102067 | 0.0542661127722048 | 0.0624187612526795 | 0.869387852035838 | 0.384635025327114 | 0.789016184424233 | NM_146105 |
| Tmem39a | 285.0909019 | 0.172107576 | 0.127210349 | 1.352936909 | 0.176075787 | 0.385109859 | | Hes1 | 268.4224395 | 0.0779644522513387 | 0.0897977513803787 | 0.86822276786292 | 0.385272392510163 | 0.789016184424233 | NM_008235 |
| Rtn4r | 2835.298067 | 0.16630805 | 0.122933866 | 1.352825349 | 0.176111432 | 0.385133456 | | Ifngr2 | 1842.751798 | 0.0440318899348938 | 0.0507007406732999 | 0.868466404043719 | 0.385139056339655 | 0.789016184424233 | NM_008338 |
| Strn4 | 9488.093494 | 0.121808582 | 0.090055023 | 1.352601747 | 0.176182894 | 0.38523536 | | Lpin2 | 4291.813414 | -0.0450212072470271 | 0.0518664860213019 | -0.868021157796127 | 0.385382750136991 | 0.789016184424233 | NM_022882,NM_001164885 |
| Polr2a | 3517.095035 | 0.144973271 | 0.107187714 | 1.352517615 | 0.176209788 | 0.385239798 | | Nefh | 3638.723414 | 0.0757344820127081 | 0.0872365640538461 | 0.868150675512181 | 0.385311852312908 | 0.789016184424233 | NM_010904 |
| Vrk3 | 911.1796389 | -0.107930815 | 0.079843475 | -1.351780031 | 0.176445694 | 0.385701126 | | Nrgn | 46478.92447 | -0.0443483191842997 | 0.0510632958126778 | -0.868496999233822 | 0.385122314328061 | 0.789016184424233 | NM_022029 |
| Lrrc24 | 1002.111593 | -0.116837961 | 0.08645026 | -1.351505034 | 0.176533709 | 0.385784666 | | Pfkfb4 | 691.094326 | -0.0638361932467767 | 0.0734872817641637 | -0.868669948245475 | 0.385027683164878 | 0.789016184424233 | NM_173019 |
| Osgepl1 | 440.428318 | 0.127172156 | 0.094095274 | 1.351525433 | 0.176527178 | 0.385784666 | | Ppp2r4 | 7480.345278 | 0.0429046309464396 | 0.0493817816269611 | 0.868835216812324 | 0.384937267733783 | 0.789016184424233 | NM_138748 |
| C330018D20Rik | 333.3785293 | -0.161444008 | 0.119491277 | -1.35109451 | 0.17666516 | 0.385960345 | | Rab13 | 170.738375 | -0.0800024497776755 | 0.0920566257795052 | -0.869056942943989 | 0.384815985786101 | 0.789016184424233 | NM_026677 |
| Grpel1 | 1513.643788 | -0.079988668 | 0.059206105 | -1.351020613 | 0.17668883 | 0.385960345 | | Snca | 8558.941242 | 0.0506469687311424 | 0.0583073272744574 | 0.868620996684394 | 0.385054466166775 | 0.789016184424233 | NM_001042451,NM_009221 |
| Tm6sf2 | 43.56461501 | 0.226962542 | 0.167978105 | 1.351143599 | 0.176649438 | 0.385960345 | | Wdr13 | 9911.79914 | -0.0496967361313851 | 0.0572387628793335 | -0.868235678610876 | 0.385265326063254 | 0.789016184424233 | NM_026137,NR_029428 |
| Kcnj5 | 56.7339992 | 0.224246153 | 0.165994229 | 1.350927411 | 0.176718687 | 0.385971149 | | Ybx1 | 3674.499372 | -0.0422924621171101 | 0.048684047406474 | -0.868712943359059 | 0.385004160069867 | 0.789016184424233 | NM_011732 |
| Atxn1 | 3258.477935 | -0.079362919 | 0.058774436 | -1.350296568 | 0.176920873 | 0.386303833 | | Zbtb10 | 326.7619706 | 0.0765325561991757 | 0.0881534839060673 | 0.868173925839682 | 0.385299125958734 | 0.789016184424233 | NM_177660 |
| Tbcb | 1891.596574 | -0.101041664 | 0.074825962 | -1.350355687 | 0.176901918 | 0.386303833 | | Tmem134 | 781.9775544 | 0.0606777673829964 | 0.0699285674400579 | 0.86771071686845 | 0.385552717534282 | 0.789252927692078 | NM_001078649,NM_025889 |
| Flnc | 115.7884395 | 0.204079312 | 0.151164083 | 1.350051597 | 0.176999433 | 0.386420912 | | Enkd1 | 242.2400945 | -0.0768665755601197 | 0.0886212319568748 | -0.867360720030667 | 0.385744396832603 | 0.789534043735436 | NM_198299 |
| AI450353 | 53.87297458 | 0.216509014 | 0.160398901 | 1.349816065 | 0.177074991 | 0.386476955 | | Slc29a4 | 530.6355972 | 0.077119330265555 | 0.08893302077533 | 0.867161933702672 | 0.385853290084461 | 0.789645659589585 | NM_146257 |
| Egr4 | 824.0449137 | -0.221430232 | 0.164041265 | -1.349844697 | 0.177065804 | 0.386476955 | | D130040H23Rik | 111.4770081 | -0.0807820713734983 | 0.093223555989653 | -0.866541407007307 | 0.38619332943923 | 0.790230216007078 | NM_172491 |
| A730020M07Rik | 17.56249834 | -0.22273685 | 0.165022462 | -1.34973656 | 0.177100501 | 0.386478191 | | Mamld1 | 1354.721044 | 0.0644013763788448 | 0.0743372689264346 | 0.866340360749296 | 0.386303539012557 | 0.790344395450198 | NM_001081354,NM_001256048 |
| Pid1 | 1648.06395 | -0.124333604 | 0.092144028 | -1.349339795 | 0.177227849 | 0.386701633 | | Lrsam1 | 1770.587972 | 0.0544747843925959 | 0.0629125503988716 | 0.865881037205144 | 0.386555403104585 | 0.790748315096072 | NM_199302 |
| Cyp2d22 | 1298.653052 | 0.144444394 | 0.107059726 | 1.349194502 | 0.1772745 | 0.386748959 | | Rnft2 | 4777.991895 | 0.0432943715574972 | 0.0500203040474182 | 0.865535953489108 | 0.386744691184969 | 0.790912767021379 | NM_001109902,NM_172998 |
| Sike1 | 999.1619133 | -0.118689482 | 0.087991536 | -1.348873853 | 0.177377488 | 0.386919161 | | Sidt1 | 2841.12004 | 0.0565981311240472 | 0.0653898881706539 | 0.865548675910531 | 0.386737711576644 | 0.790912767021379 | NM_001159419,NM_198034 |
| Tie1 | 497.5000591 | -0.175389708 | 0.130040805 | -1.348728255 | 0.177424266 | 0.386966721 | | Plxnb1 | 4977.123788 | -0.0503268616073661 | 0.0581913197386508 | -0.864851696668755 | 0.387120192427603 | 0.791569244820293 | NM_172775 |
| Cacna2d4 | 90.67894028 | 0.18492413 | 0.137152015 | 1.34831508 | 0.177557064 | 0.387201851 | | Ap2s1 | 3990.713078 | 0.0419565517216902 | 0.0485598918564693 | 0.864016580714452 | 0.387578782183583 | 0.791583702425494 | NM_198613 |
| Olfr613 | 87.26211484 | 0.206169008 | 0.15295031 | 1.34794763 | 0.177675227 | 0.387405006 | | Dnajb3 | 28.74850084 | 0.0735332952931275 | 0.0850696765792835 | 0.864389030850444 | 0.387374216612132 | 0.791583702425494 | NM_008299 |
| 9430038I01Rik | 44.23061993 | -0.207523968 | 0.154031408 | -1.34728346 | 0.177888957 | 0.387414269 | | Iscu | 2207.896892 | 0.0617540900048348 | 0.0714334006865105 | 0.864498811639195 | 0.387313932864111 | 0.791583702425494 | NM_025526 |
| Bex4 | 368.9564351 | -0.145236703 | 0.10778515 | -1.347464873 | 0.177830559 | 0.387414269 | | Pvrl4 | 289.3050346 | -0.0779055297599281 | 0.0901331202939282 | -0.864338541768827 | 0.387401943520893 | 0.791583702425494 | NM_027893,NM_001122680 |
| Bicc1 | 172.4875477 | -0.18384948 | 0.136448355 | -1.347392422 | 0.17785388 | 0.387414269 | | Rims4 | 843.1632335 | 0.0619866091658738 | 0.0717188109734975 | 0.864305069466773 | 0.387422797431732 | 0.791583702425494 | NM_183023 |
| Ctgf | 1357.163445 | 0.132085927 | 0.098022581 | 1.347505102 | 0.177817612 | 0.387414269 | | Rtn4ip1 | 604.0147871 | 0.06044831546765 | 0.0699046978495917 | 0.864724651234626 | 0.387189935920588 | 0.791583702425494 | NM_130892 |
| Kif15 | 14.96683085 | -0.224300927 | 0.166489813 | -1.347235143 | 0.177904513 | 0.387414269 | | Sh3glb2 | 6230.015403 | -0.051399536596336 | 0.0594935044355139 | -0.863952074836152 | 0.387614218264785 | 0.791583702425494 | NM_139302 |
| Nat6 | 476.8784799 | 0.106609435 | 0.079128066 | 1.347302423 | 0.177882852 | 0.387414269 | | Tmed9 | 3910.371599 | 0.0413475508305254 | 0.0478493342100776 | 0.864119668812803 | 0.387522155193305 | 0.791583702425494 | NM_026211 |
| Sgip1 | 5698.137183 | -0.091304618 | 0.067766883 | -1.347333879 | 0.177872726 | 0.387414269 | | Tmem132b | 7390.676506 | 0.0446107093775264 | 0.0516360096842895 | 0.863945716376675 | 0.387617711369443 | 0.791583702425494 | NM_001190352 |
| Tmem140 | 73.32942144 | -0.208387297 | 0.154664876 | -1.34734726 | 0.177868418 | 0.387414269 | | Ankrd6 | 1781.811736 | 0.0636768936461524 | 0.0737692613966461 | 0.863190066439346 | 0.388032974393697 | 0.79188911531153 | NM_080471,NM_001012451,NM_001012450 |
| Vdr | 34.58514092 | 0.223353805 | 0.16577571 | 1.347325278 | 0.177875494 | 0.387414269 | | Ccdc66 | 597.8970399 | -0.0637542591521176 | 0.0738488204547656 | -0.863307751694814 | 0.387968283332001 | 0.79188911531153 | NM_177111 |
| Brap | 2872.796086 | 0.137302087 | 0.101954195 | 1.346703654 | 0.178075695 | 0.387578219 | | Cntln | 599.8497336 | -0.056829891224773 | 0.0658494010567909 | -0.863028217610679 | 0.388121952709265 | 0.79188911531153 | NM_177385,NM_175275 |
| Gpr39 | 10.16146523 | 0.19567215 | 0.145289428 | 1.34677487 | 0.178052751 | 0.387578219 | | Cxcr4 | 39.95362208 | -0.079286597798304 | 0.0918453380044985 | -0.863262082985862 | 0.387993386437934 | 0.79188911531153 | NM_009911 |
| Pgpep1l | 23.31495532 | -0.226220414 | 0.167970721 | -1.3467848 | 0.178049551 | 0.387578219 | | H2afy | 2482.03584 | 0.0517203272058345 | 0.0599333100360469 | 0.862964638107378 | 0.388156909696631 | 0.79188911531153 | NM_001159514,NM_001159515,NM_012015,NM_001159513 |
| Psmc6 | 2439.432814 | -0.089000215 | 0.066088088 | -1.346690729 | 0.178079859 | 0.387578219 | | Phtf2 | 731.7276285 | -0.0677790099381216 | 0.0785364112855855 | -0.863026573644341 | 0.38812285656305 | 0.79188911531153 | NM_172992 |
| Zfp879 | 106.5962502 | 0.175495497 | 0.130330671 | 1.34654027 | 0.178128342 | 0.387629289 | | Prtn3 | 39.77644207 | 0.0768726708251535 | 0.0890884721354993 | 0.86288011212308 | 0.388203386350916 | 0.79188911531153 | NM_011178 |
| Rbbp4 | 4690.467943 | -0.06586425 | 0.048919548 | -1.346378948 | 0.178180337 | 0.387687984 | | Zfp87 | 657.5178677 | 0.0578705249402065 | 0.0670154706597175 | 0.863539782240044 | 0.387840756426028 | 0.79188911531153 | NM_133228 |
| Cnr2 | 17.93677144 | 0.226260641 | 0.168116087 | 1.345859543 | 0.178347818 | 0.387849001 | | Sh3bp5l | 2155.749485 | 0.0565860974869328 | 0.0656045555567837 | 0.862533051351213 | 0.388394253421965 | 0.792167217805036 | NM_024480,NM_001161338 |
| Ctbp1 | 7139.353195 | -0.072209501 | 0.053653801 | -1.345841287 | 0.178353707 | 0.387849001 | | Gm16702 | 184.1232918 | 0.0812817495379458 | 0.094253006434519 | 0.862378322058248 | 0.38847936566012 | 0.792229575400661 | NR_045795 |
| Dazl | 116.084771 | -0.20606381 | 0.153111792 | -1.345838928 | 0.178354468 | 0.387849001 | | Clcn3 | 8891.236638 | 0.0500149894608216 | 0.0580487734303221 | 0.861602864371567 | 0.388906094240516 | 0.792357410251181 | NM_173876,NM_173874,NM_173873,NM_007711 |
| Ighmbp2 | 489.2183083 | 0.124361818 | 0.092404376 | 1.345843386 | 0.17835303 | 0.387849001 | | Commd5 | 471.4291037 | 0.0649779742422579 | 0.0753876298134021 | 0.861918253738578 | 0.388732503410739 | 0.792357410251181 | NM_025536 |
| Csdc2 | 4816.610841 | -0.136751135 | 0.101626101 | -1.345630044 | 0.178421857 | 0.387941097 | | Dcp1b | 547.2436678 | -0.0623130394677878 | 0.0723131722420853 | -0.861710771851915 | 0.388846696475025 | 0.792357410251181 | NM_001033379 |
| Rho | 13.72284992 | 0.222725178 | 0.165530808 | 1.345520999 | 0.178457045 | 0.387963161 | | Dohh | 1820.862141 | 0.0641958540142765 | 0.0744662035209459 | 0.86208039323798 | 0.38864327992461 | 0.792357410251181 | NM_133964 |
| Cdh2 | 4638.305819 | 0.098573414 | 0.073266161 | 1.345415302 | 0.178491156 | 0.387982881 | | Ralgps2 | 1816.028591 | 0.0555219621949342 | 0.0644250321184864 | 0.861807287776308 | 0.388793573880384 | 0.792357410251181 | NR_027640,NM_001159965,NM_001159967,NM_001159966,NM_023884,NM_001159968 |
| Cenpo | 128.4151259 | 0.187101823 | 0.139077903 | 1.3453023 | 0.178527631 | 0.388007731 | | Rbp1 | 300.3835604 | 0.0806060363436512 | 0.0935372703531133 | 0.861753139035966 | 0.388823376931956 | 0.792357410251181 | NM_011254 |
| Adra1d | 1013.454481 | 0.170745827 | 0.126944551 | 1.345042585 | 0.178611483 | 0.388110364 | | Zfp330 | 841.5815714 | 0.0560375063711783 | 0.0650411118191366 | 0.861570548286519 | 0.388923883731992 | 0.792357410251181 | NM_145600 |
| Xpo1 | 2634.424829 | -0.128843851 | 0.095794622 | -1.345000876 | 0.178624952 | 0.388110364 | | Yae1d1 | 1096.020737 | 0.0493407123077403 | 0.0572810006372753 | 0.861380069461149 | 0.389028749363272 | 0.79245990930457 | NM_025904 |
| Cacna2d1 | 7371.178065 | -0.119239768 | 0.088676852 | -1.344654954 | 0.178736688 | 0.388276877 | | Slc30a10 | 994.569136 | -0.0582917126542799 | 0.0676848734360898 | -0.861222156370297 | 0.389115699400358 | 0.792525890281773 | NM_001033286 |
| Dpp4 | 155.537685 | -0.171906064 | 0.127848417 | -1.344608466 | 0.178751708 | 0.388276877 | | Ppfia2 | 3852.872739 | 0.0471131396226059 | 0.0547135364892826 | 0.861087450120035 | 0.389189880643715 | 0.792565849744933 | NM_001205341,NM_177373 |
| 2310008H09Rik | 1116.031243 | -0.117656257 | 0.087533281 | -1.344131694 | 0.178905805 | 0.388502669 | | Vcan | 823.1120245 | -0.0656334571259897 | 0.0762399330117118 | -0.860880309481741 | 0.389303967493764 | 0.792687052398993 | NM_172955,NM_019389,NM_001134475,NM_001134474,NM_001081249 |
| En2 | 20.38769105 | 0.224546312 | 0.167051595 | 1.344173405 | 0.178892319 | 0.388502669 | | Camk2n2 | 3595.91626 | 0.0570100473812958 | 0.0662534441513164 | 0.86048428291653 | 0.3895221436802 | 0.792715562028886 | NM_028420 |
| Apba2 | 7754.8332 | 0.110598623 | 0.082349645 | 1.343037039 | 0.179259981 | 0.389108174 | | Gm1976 | 963.0075212 | 0.0644454155660897 | 0.0748890299713147 | 0.860545470955823 | 0.389488429536131 | 0.792715562028886 | NR_045963,NR_045964 |
| Il7r | 11.16661731 | -0.221007069 | 0.164542145 | -1.343163905 | 0.179218907 | 0.389108174 | | Nme2 | 2265.878812 | 0.0482416980388412 | 0.0560690849938597 | 0.860397455106042 | 0.389569988189848 | 0.792715562028886 | NM_008705,NM_001077529 |
| Tppp3 | 1088.163581 | 0.114943019 | 0.085584086 | 1.343041971 | 0.179258384 | 0.389108174 | | Pmvk | 1225.543727 | 0.0624318750615089 | 0.0725367534313217 | 0.860692898150068 | 0.389407155983469 | 0.792715562028886 | NM_026784,NM_027348 |
| Ppp1r3f | 1244.198598 | 0.107002623 | 0.079701502 | 1.342542122 | 0.17942082 | 0.389401577 | | Rcsd1 | 264.3387952 | 0.0709743096702969 | 0.0824987555869501 | 0.860307639373943 | 0.38961948288746 | 0.792715562028886 | NM_001038846,NM_178593 |
| Zc4h2 | 664.8062963 | -0.105607931 | 0.078677658 | -1.342286162 | 0.179503229 | 0.389527034 | | Sec23ip | 1910.857062 | 0.0526764117057482 | 0.0612353696596389 | 0.860228524764961 | 0.389663083692244 | 0.792715562028886 | NM_001029982 |
| Mdfi | 11.8349771 | -0.206073475 | 0.1535892 | -1.341718526 | 0.179687278 | 0.389871822 | | Zbtb38 | 1510.055168 | 0.0592772723920182 | 0.0689141122846969 | 0.860161589938688 | 0.389699974421608 | 0.792715562028886 | NM_175537 |
| Lst1 | 51.58919593 | -0.225045561 | 0.167752602 | -1.341532461 | 0.179747638 | 0.38994818 | | Zfp300 | 121.0078867 | 0.0816627102644015 | 0.0949696785106055 | 0.859881928054353 | 0.389854131373427 | 0.792918105898964 | NM_183185 |
| Wipf3 | 7852.592047 | 0.176668651 | 0.13171437 | 1.341301266 | 0.179822659 | 0.390056318 | | Fech | 2563.40109 | 0.0407936025758194 | 0.0474502283178021 | 0.859713515024641 | 0.389946982930597 | 0.792995922448529 | NM_007998 |
| Palb2 | 50.4393882 | 0.21378133 | 0.159413037 | 1.341052989 | 0.179903249 | 0.390176504 | | Camkmt | 295.8258925 | 0.0701721500382652 | 0.0816823587679679 | 0.859085745033401 | 0.390293211313813 | 0.793521684947212 | NM_028576 |
| 1700030L20Rik | 35.83407705 | -0.219105968 | 0.163398908 | -1.340926762 | 0.179944232 | 0.390210768 | | Ppp1r26 | 1500.51084 | 0.0556513691549466 | 0.0647827085972565 | 0.859046655503739 | 0.390314776190818 | 0.793521684947212 | NM_001005420 |
| Ckap4 | 808.0486773 | -0.08678407 | 0.064738126 | -1.340540345 | 0.180069737 | 0.390321947 | | Hcfc1 | 5767.180005 | -0.0416840120559958 | 0.0485613508982474 | -0.858378345844168 | 0.390683580690108 | 0.794106634999289 | NM_008224 |
| Drosha | 4820.658573 | 0.055293143 | 0.041247034 | 1.340536218 | 0.180071078 | 0.390321947 | | Zkscan16 | 2018.188199 | -0.0631971431426243 | 0.0736282690104899 | -0.858327161455073 | 0.390711835353154 | 0.794106634999289 | NM_001099323 |
| Pdpn | 328.1016569 | -0.159511444 | 0.118977976 | -1.340680436 | 0.180024229 | 0.390321947 | | Abca2 | 14392.90946 | 0.0505487580338426 | 0.0590673717316242 | 0.855781399306434 | 0.392118705706841 | 0.79418442318082 | NM_007379 |
| 3110045C21Rik | 21.71625581 | -0.22426974 | 0.167346274 | -1.340153771 | 0.180195357 | 0.390529473 | | Adam11 | 7243.000313 | -0.0587275417091475 | 0.0686953777666904 | -0.854898009420712 | 0.39260761270498 | 0.79418442318082 | NM_009613,NM_001110778 |
| Fam219b | 763.8556073 | -0.089389102 | 0.066726386 | -1.339636499 | 0.180363551 | 0.390529473 | | B430319G15Rik | 26.13548559 | -0.0729497700509918 | 0.0852902498227098 | -0.855311951865895 | 0.392378472669721 | 0.79418442318082 | NR_029474 |
| Kifap3 | 11935.4299 | -0.080254815 | 0.059908579 | -1.33962141 | 0.18036846 | 0.390529473 | | Barx2 | 191.7679776 | -0.0752474057869744 | 0.0878454787052132 | -0.856588260387144 | 0.391672475972298 | 0.79418442318082 | NM_013800 |
| Lrrc14 | 749.0038916 | -0.13591727 | 0.101449753 | -1.339749642 | 0.180326753 | 0.390529473 | | Bcl7b | 1801.100569 | 0.0463198244902256 | 0.0539793317632225 | 0.858102962322784 | 0.390835611756734 | 0.79418442318082 | NM_009745 |
| Pcnxl3 | 3024.164392 | 0.088050392 | 0.065719239 | 1.339796289 | 0.180311583 | 0.390529473 | | Cap1 | 11124.4634 | 0.0439783696869479 | 0.0514126864753408 | 0.855399176777922 | 0.392330199203551 | 0.79418442318082 | NM_007598 |
| Prpsap1 | 1435.487483 | 0.09663917 | 0.072134362 | 1.339710615 | 0.180339445 | 0.390529473 | | Casd1 | 2650.557358 | 0.0562801773299406 | 0.0657452922302849 | 0.856033571693756 | 0.391979210147916 | 0.79418442318082 | NM_145398 |
| Serf2 | 4269.68939 | -0.113814525 | 0.084934639 | -1.340024829 | 0.180237273 | 0.390529473 | | Cdc26 | 630.5878143 | -0.055038151963121 | 0.0643093411767168 | -0.855834486188883 | 0.392089336847915 | 0.79418442318082 | NM_139291 |
| She | 254.2414285 | -0.142873381 | 0.106635208 | -1.339833098 | 0.180299613 | 0.390529473 | | Cdk10 | 2421.355817 | -0.0612350668999796 | 0.0715271721059586 | -0.856109155402754 | 0.391937404964874 | 0.79418442318082 | NM_194444,NM_194446 |
| Pip5k1c | 20366.55346 | 0.119673638 | 0.089387303 | 1.338821437 | 0.18062881 | 0.391038534 | | Cdkl5 | 2591.540022 | 0.065120724456593 | 0.0761690407529576 | 0.854950040237502 | 0.392578806313848 | 0.79418442318082 | NM_001024624 |
| Tsc22d1 | 15341.75225 | -0.075361984 | 0.056296044 | -1.338672839 | 0.180677202 | 0.391088652 | | Cggbp1 | 2268.672101 | 0.0531849173032753 | 0.0621019011311183 | 0.856413673890334 | 0.391769003863658 | 0.79418442318082 | NM_178647 |
| Gusb | 548.0881935 | 0.135696257 | 0.101391709 | 1.338336815 | 0.180786666 | 0.391270932 | | Crybg3 | 318.6719045 | 0.0728102373353385 | 0.0850970336323915 | 0.855614282042658 | 0.392211167452088 | 0.79418442318082 | NM_174848 |
| Scaf1 | 6558.796832 | 0.091350133 | 0.068262305 | 1.338222219 | 0.180824008 | 0.391297092 | | Cwc22 | 883.0338666 | -0.0519481855134859 | 0.060645991979596 | -0.856580687656383 | 0.391676662593043 | 0.79418442318082 | NM_172667,NM_030560 |
| Ccsap | 2296.26145 | 0.122726303 | 0.091741538 | 1.337739758 | 0.180981286 | 0.391473411 | | Dmwd | 5259.762114 | 0.0512697798917839 | 0.0599719845702975 | 0.854895502610672 | 0.392609000610042 | 0.79418442318082 | NM_010058 |
| Stk38 | 1085.154668 | 0.155080863 | 0.115924793 | 1.337771318 | 0.180970994 | 0.391473411 | | Dync2li1 | 702.0541146 | 0.055591166528144 | 0.06483284552549 | 0.857453750141006 | 0.391194164721214 | 0.79418442318082 | NM_172256 |
| Theg | 13.58280173 | -0.220309945 | 0.164670957 | -1.337879791 | 0.180935626 | 0.391473411 | | Exoc2 | 2398.884347 | 0.0502501316682552 | 0.0586680560891355 | 0.85651605009563 | 0.391712398885607 | 0.79418442318082 | NM_025588 |
| Aspn | 37.84927768 | -0.221005457 | 0.165246545 | -1.337428612 | 0.18108277 | 0.391638252 | | Fam114a2 | 1469.806461 | -0.0452601485693406 | 0.0529173029847262 | -0.855299609324464 | 0.39238530377581 | 0.79418442318082 | NM_001168668,NM_001168667,NM_026342 |
| Atoh8 | 44.44842071 | 0.207104189 | 0.154895359 | 1.337058717 | 0.181203471 | 0.391771527 | | Fam161b | 807.9761638 | -0.0511043600230546 | 0.0597127943626503 | -0.855836015857598 | 0.39208849062051 | 0.79418442318082 | NM_172581 |
| Mad1l1 | 561.4880272 | 0.117642246 | 0.087998537 | 1.336865937 | 0.181266401 | 0.391771527 | | Fbxo27 | 576.8166377 | -0.0593038087781957 | 0.069282886437581 | -0.855966196379884 | 0.392016477565972 | 0.79418442318082 | NM_001163702,NM_207238 |
| Nosip | 1563.795502 | 0.095095645 | 0.071128821 | 1.33694955 | 0.181239105 | 0.391771527 | | Fkbp14 | 440.07528 | 0.0666901267656059 | 0.0779907014800502 | 0.855103563630146 | 0.392493816944551 | 0.79418442318082 | NM_153573 |
| Pgs1 | 2970.032842 | -0.093886322 | 0.070217222 | -1.337083971 | 0.181195229 | 0.391771527 | | Gpr12 | 728.3296246 | -0.0630663885649355 | 0.0737345035066064 | -0.855317192978521 | 0.39237557194402 | 0.79418442318082 | NM_008151,NM_001010941 |
| Tmem123 | 489.6015526 | -0.124389356 | 0.093046439 | -1.336852405 | 0.181270819 | 0.391771527 | | Grb14 | 832.5957558 | 0.0603482182550454 | 0.0705635187325004 | 0.855232552727702 | 0.392422418196919 | 0.79418442318082 | NM_016719 |
| C230035I16Rik | 22.01888649 | 0.224682847 | 0.168116861 | 1.336468249 | 0.181396272 | 0.391933319 | | Ifitm2 | 239.2989946 | -0.075856733445904 | 0.0884408417779087 | -0.857711572176057 | 0.391051748458551 | 0.79418442318082 | NM_030694 |
| Pcdha1 | 45.92473869 | -0.206383544 | 0.154424011 | -1.336473144 | 0.181394673 | 0.391933319 | | Katnal2 | 153.5936616 | 0.0802165328463141 | 0.0938056831603225 | 0.855135106358287 | 0.392476356514437 | 0.79418442318082 | NM_027721 |
| Gm3219 | 22.69986946 | -0.224512413 | 0.168010752 | -1.336297894 | 0.181451925 | 0.391944252 | | Map2k2 | 2731.698192 | -0.0560032692780132 | 0.0654287338309626 | -0.855943039073621 | 0.392029287101153 | 0.79418442318082 | NM_023138 |
| Prdm15 | 772.8845985 | 0.127231357 | 0.095211216 | 1.336306397 | 0.181449146 | 0.391944252 | | Map9 | 5566.631945 | 0.0562417608085056 | 0.0655903767326119 | 0.85746970226719 | 0.391185352139286 | 0.79418442318082 | NM_001081230 |
| Gm10451 | 27.12940873 | 0.224398593 | 0.167940205 | 1.336181484 | 0.181489962 | 0.391971768 | | Pnpla7 | 852.3864018 | -0.061041267175733 | 0.0712310225803116 | -0.856947787137411 | 0.391473741125781 | 0.79418442318082 | NM_146251 |
| Def8 | 2359.153676 | -0.072795148 | 0.054495022 | -1.335812798 | 0.181610469 | 0.392177366 | | Pogz | 2912.371053 | -0.0387640391689273 | 0.0451876500402832 | -0.857845874577913 | 0.390977574697303 | 0.79418442318082 | NM_001165948,NM_172683 |
| Mroh5 | 82.68337481 | 0.181598092 | 0.135956691 | 1.335705442 | 0.18164557 | 0.392198504 | | Rbm26 | 1520.764486 | -0.0558489717945748 | 0.0651190521289254 | -0.857644114413747 | 0.391089007873543 | 0.79418442318082 | NM_134077 |
| 9930014A18Rik | 91.23071809 | -0.199675205 | 0.149511735 | -1.335515266 | 0.181707762 | 0.39227812 | | Samd1 | 882.4757971 | -0.0543878177670814 | 0.0634790437650153 | -0.856783822522793 | 0.391564367908407 | 0.79418442318082 | NM_001081415 |
| Rbpjl | 14.43624082 | 0.219034158 | 0.1640175 | 1.335431635 | 0.181735117 | 0.392282516 | | Senp7 | 1268.732417 | 0.0507189881798 | 0.0592028781811363 | 0.856698014319858 | 0.391611801030065 | 0.79418442318082 | NM_025483,NM_001003972,NM_001003973,NM_001003971 |
| Tceb1 | 1815.195052 | -0.091137036 | 0.068250504 | -1.335331329 | 0.181767929 | 0.39229869 | | Snhg6 | 272.7049537 | -0.0778027700910459 | 0.090742530640401 | -0.857401370029745 | 0.391223102402603 | 0.79418442318082 | NR_024067 |
| Impg1 | 58.84893574 | -0.219461318 | 0.164361149 | -1.335238408 | 0.18179833 | 0.392309655 | | Sox18 | 207.7209836 | -0.080856414231669 | 0.0943490500181778 | -0.856993251444882 | 0.3914491116715 | 0.79418442318082 | NM_009236 |
| Amer3 | 1517.423964 | 0.105473052 | 0.078997433 | 1.335145306 | 0.181828794 | 0.392320753 | | Trim24 | 1215.57427 | -0.0457774718626143 | 0.0534289272877953 | -0.856791895072751 | 0.391559905737749 | 0.79418442318082 | NM_001272076,NM_145076,NM_001272064 |
| Ddx47 | 1581.43699 | 0.088057925 | 0.065964167 | 1.334935759 | 0.181897373 | 0.392412937 | | Ntmt1 | 661.0942672 | 0.065445014570028 | 0.0765822408926873 | 0.854571684076657 | 0.39278830899905 | 0.794436504667252 | NM_170592 |
| Orai2 | 3580.196339 | 0.137029286 | 0.102654425 | 1.334859999 | 0.181922172 | 0.392412937 | | Ppm1k | 3342.615244 | -0.0616980294612365 | 0.0722193514073051 | -0.854314366703045 | 0.39293082901385 | 0.79461411976266 | NM_175523 |
| Rpl36al | 1612.902668 | -0.095241728 | 0.071355903 | -1.334742095 | 0.181960771 | 0.392441563 | | Enkur | 103.0592239 | 0.0809153410717264 | 0.0947257165994977 | 0.854206692506092 | 0.392990475670281 | 0.794620350967734 | NM_027728 |
| Etfdh | 1128.961133 | -0.078895326 | 0.059148799 | -1.333844931 | 0.182254682 | 0.392903171 | | Exoc8 | 1571.457524 | 0.0484153641334455 | 0.0566850754353234 | 0.854111311692379 | 0.393043316928484 | 0.794620350967734 | NM_198103 |
| Rad51 | 42.90176671 | 0.198911849 | 0.149123663 | 1.333871802 | 0.182245874 | 0.392903171 | | 5031425E22Rik | 355.1181025 | -0.0761053091689717 | 0.0891986527992028 | -0.853211419462738 | 0.393542071825121 | 0.7947541882506 | NR_040469 |
| Spire2 | 598.3976047 | -0.096325466 | 0.072211083 | -1.33394295 | 0.182222554 | 0.392903171 | | AU022252 | 592.4238723 | 0.0571817726705433 | 0.0669760140880984 | 0.853764940316215 | 0.393235243915824 | 0.7947541882506 | NM_001012400 |
| Tmem5 | 759.4405109 | -0.096149382 | 0.072087931 | -1.333779182 | 0.182276236 | 0.392903171 | | Coq3 | 654.689873 | 0.0565382300811264 | 0.0662492390451075 | 0.853417048951021 | 0.393428070279711 | 0.7947541882506 | NM_172687 |
| Ppp4c | 1027.909594 | -0.077130189 | 0.057844715 | -1.333400791 | 0.182400313 | 0.393115934 | | E430025E21Rik | 2347.891125 | 0.0486541830125157 | 0.0570764127619492 | 0.852439399361686 | 0.39397026037618 | 0.7947541882506 | NM_153548 |
| Fam105b | 547.5202546 | 0.116969845 | 0.087736652 | 1.333192493 | 0.182468642 | 0.393208504 | | Fanca | 56.77712571 | -0.080377607558208 | 0.0942618179446228 | -0.85270589206574 | 0.393822422616402 | 0.7947541882506 | NM_016925 |
| 1110038B12Rik | 209.7950699 | 0.17088097 | 0.128213882 | 1.332780564 | 0.182603826 | 0.393364924 | | Gins4 | 540.988037 | 0.0656991284001074 | 0.0770173055597776 | 0.853043714300216 | 0.393635062830951 | 0.7947541882506 | NM_024240 |
| Gpr137c | 802.2638139 | -0.136980974 | 0.102769861 | -1.332890521 | 0.182567734 | 0.393364924 | | Gpkow | 1792.081372 | 0.042886789449618 | 0.0502831478152813 | 0.852905820597503 | 0.393711533617165 | 0.7947541882506 | NM_173747 |
| Igsf5 | 12.88947234 | -0.210519652 | 0.157969266 | -1.33266209 | 0.182642719 | 0.393364924 | | Gpr137 | 3289.924922 | 0.0515289056580365 | 0.0603708803078251 | 0.853539080352907 | 0.393360425198143 | 0.7947541882506 | NM_207220,NM_001177362,NM_001177361,NM_001177360 |
| Ivns1abp | 6966.512743 | 0.071344646 | 0.053535444 | 1.332661899 | 0.182642782 | 0.393364924 | | Guk1 | 2623.222087 | -0.0461887663925532 | 0.0541438284740323 | -0.853075367854818 | 0.393617510200727 | 0.7947541882506 | NM_001159410,NM_008193 |
| Rnf166 | 1070.203555 | 0.080183375 | 0.060172376 | 1.332561239 | 0.182675832 | 0.393381424 | | Mcm3ap | 2749.995317 | -0.0493882652592806 | 0.0579364013123566 | -0.852456558235472 | 0.393960740419192 | 0.7947541882506 | NM_019434 |
| Sytl4 | 44.75014824 | -0.214658849 | 0.161142389 | -1.332106656 | 0.182825144 | 0.393648245 | | Mdn1 | 2362.690612 | -0.0729835499573482 | 0.0855449908720423 | -0.853159831024081 | 0.393570675733986 | 0.7947541882506 | NM_001081392 |
| Sppl2a | 1492.445383 | -0.099935304 | 0.075027914 | -1.331974965 | 0.182868415 | 0.393686706 | | Pole3 | 391.5684666 | 0.0640074433008614 | 0.0750771930957671 | 0.852555092452839 | 0.393906075103362 | 0.7947541882506 | NM_021498 |
| Rnpep | 1496.059625 | -0.09373483 | 0.070383444 | -1.331773842 | 0.182934516 | 0.393774297 | | Sgk3 | 545.6037489 | 0.0762845838732709 | 0.0894925176988569 | 0.852412981943017 | 0.39398491736146 | 0.7947541882506 | NM_001037759,NM_133220,NM_177547 |
| Capn6 | 75.64997627 | 0.218579537 | 0.164138367 | 1.331678516 | 0.182965852 | 0.393787041 | | Slc25a47 | 85.76886298 | 0.080579059093878 | 0.09444626302775 | 0.853173609105131 | 0.393563036167026 | 0.7947541882506 | NM_001012310 |
| Rgs10 | 939.629422 | -0.130877379 | 0.098291964 | -1.331516571 | 0.183019096 | 0.393846927 | | Sra1 | 1002.011613 | 0.046612607052073 | 0.0546777168842396 | 0.852497318985692 | 0.393938126405344 | 0.7947541882506 | NM_001164406,NR_028355,NM_025291 |
| Gbp7 | 254.4399252 | 0.213098749 | 0.160052517 | 1.331430165 | 0.183047509 | 0.393853369 | | Stam | 2398.54688 | 0.0517462196479603 | 0.0606963516042904 | 0.852542505113315 | 0.39391305811597 | 0.7947541882506 | NM_011484 |
| Apbb1ip | 237.252086 | 0.179488293 | 0.1348604 | 1.330919184 | 0.183215604 | 0.394060885 | | Celf2 | 14886.33288 | -0.0455872277312368 | 0.0534930057055704 | -0.852209127715731 | 0.394098031427785 | 0.794871980633159 | NM_001110231,NM_001110232,NM_001160292,NM_001110230,NM_010160,NM_001110229,NM_001110228,NM_001160293 |
| Epb4.1l5 | 370.4985634 | -0.157486859 | 0.118329531 | -1.330917623 | 0.183216118 | 0.394060885 | | Ccny | 5140.962596 | 0.0578953640332691 | 0.0680232060004631 | 0.851111957776218 | 0.394707163526334 | 0.795437882544884 | NM_026484 |
| Zik1 | 680.9335632 | -0.098188408 | 0.073775667 | -1.330905054 | 0.183220254 | 0.394060885 | | Clmn | 4733.764666 | 0.0641951506592919 | 0.0754024329757671 | 0.851367099519494 | 0.394565461913912 | 0.795437882544884 | NM_053155,NM_001040682 |
| Ap2m1 | 20663.73222 | -0.078708536 | 0.059146917 | -1.33072931 | 0.183278095 | 0.394130577 | | Emcn | 254.3468512 | -0.0745200908198942 | 0.0875508654363076 | -0.851163383120488 | 0.394678600242518 | 0.795437882544884 | NM_016885,NM_001163522 |
| Prrt1 | 1410.196401 | 0.107085247 | 0.080496114 | 1.330315729 | 0.183414267 | 0.394368672 | | Ppp2r5e | 3900.520329 | 0.0538964548090536 | 0.0633008331229794 | 0.851433577569901 | 0.394528546130174 | 0.795437882544884 | NM_012024 |
| 0610007N19Rik | 77.7980029 | -0.188936775 | 0.142081187 | -1.329780383 | 0.183590641 | 0.394528903 | | Ufc1 | 2000.184659 | 0.0385037484784814 | 0.0452370182518247 | 0.851155756202572 | 0.394682836398125 | 0.795437882544884 | NM_025388 |
| Iqgap2 | 1292.417988 | 0.17548709 | 0.131951913 | 1.329932145 | 0.183540629 | 0.394528903 | | Zmat5 | 516.2055326 | 0.061483464565672 | 0.072201142442008 | 0.851558056924868 | 0.394459427385528 | 0.795437882544884 | NM_026015 |
| Ndufv2 | 4053.525859 | -0.102235739 | 0.076878238 | -1.329839772 | 0.183571069 | 0.394528903 | | Rb1cc1 | 4343.638024 | 0.0642121558403828 | 0.0754683541559071 | 0.850848763810715 | 0.394853369459803 | 0.795622145203649 | NM_009826 |
| Rabgap1l | 4028.004855 | -0.121230487 | 0.091151624 | -1.329987139 | 0.183522509 | 0.394528903 | | Emc7 | 2508.717385 | 0.0483360262263718 | 0.0568323816962513 | 0.850501506072902 | 0.395046323459676 | 0.795790167046908 | NM_133749 |
| Ang | 50.78240133 | 0.196562689 | 0.14788693 | 1.329141723 | 0.183801217 | 0.394698659 | | Gstm4 | 403.889741 | 0.0663258979296571 | 0.0779806738338859 | 0.850542764877157 | 0.395023394993057 | 0.795790167046908 | NM_001160411,NM_026764 |
| Chchd5 | 248.2440744 | -0.129615613 | 0.097528675 | -1.329000043 | 0.183847956 | 0.394698659 | | Lta4h | 1758.938399 | -0.0473791222022205 | 0.0557245636862685 | -0.850237652267084 | 0.395192972114089 | 0.795857763691923 | NM_008517 |
| Chrna3 | 33.68938477 | -0.216218444 | 0.162678288 | -1.329116788 | 0.183809442 | 0.394698659 | | Mesdc1 | 1161.295749 | 0.0624175217881043 | 0.0734058736513517 | 0.85030691255801 | 0.39515447439829 | 0.795857763691923 | NM_030705 |
| Cwh43 | 113.4586499 | -0.172133652 | 0.12948668 | -1.329354117 | 0.183731168 | 0.394698659 | | Zfx | 846.4792592 | -0.0554561409843152 | 0.065231359310362 | -0.850145414270189 | 0.395244245303148 | 0.795857763691923 | NM_011768,NM_001044386 |
| Kif1b | 39493.97554 | -0.120191802 | 0.090432898 | -1.329071656 | 0.18382433 | 0.394698659 | | Pcdha3 | 104.9119319 | 0.0791453192869993 | 0.0931580367046609 | 0.849581228701866 | 0.395557951864285 | 0.796239418040598 | NM_138662 |
| Pmp22 | 872.1830916 | 0.142176861 | 0.106964869 | 1.329192116 | 0.183784595 | 0.394698659 | | Rmdn2 | 232.0974923 | -0.0723434805078786 | 0.0851519999892557 | -0.849580520915618 | 0.395558345512224 | 0.796239418040598 | NM_201361 |
| Uckl1 | 689.9562338 | 0.139210228 | 0.104732425 | 1.329198937 | 0.183782346 | 0.394698659 | | Tst | 774.9081238 | -0.0630165472193374 | 0.0741799807009846 | -0.849508811189283 | 0.395598229381729 | 0.796239418040598 | NM_009437 |
| Map1s | 1873.206608 | 0.083711899 | 0.062995323 | 1.328858956 | 0.183894507 | 0.394743903 | | 2900079G21Rik | 225.9410858 | -0.0752744751610934 | 0.0886236845156832 | -0.849371988678405 | 0.395674334741687 | 0.796282266479654 | NR_033431,NR_033433,NR_033432,NR_015468 |
| Dis3l | 1213.094454 | 0.097767446 | 0.073609094 | 1.328197924 | 0.18411273 | 0.395156762 | | Sos1 | 2752.380641 | 0.0611957052802452 | 0.0720579593852281 | 0.849256706716987 | 0.395738465377177 | 0.796301004026718 | NM_009231 |
| Slc35f3 | 1609.412935 | 0.104049156 | 0.078343078 | 1.328121875 | 0.184137848 | 0.395156762 | | AA465934 | 36.43230754 | -0.0728665679724503 | 0.0858145810698885 | -0.849116397982609 | 0.395816526721789 | 0.796347765534723 | NR_028363 |
| Tubgcp4 | 872.2312489 | 0.093710664 | 0.070566033 | 1.327985442 | 0.184182916 | 0.395198741 | | Gas7 | 26669.47141 | 0.0453550540572735 | 0.0534240333401924 | 0.848963494921144 | 0.395901605562736 | 0.796351698323078 | NM_001109657,NM_008088 |
| Aoah | 12.6493359 | -0.217203121 | 0.163610264 | -1.327564151 | 0.184322134 | 0.395333215 | | Surf6 | 741.6188692 | 0.0538933183475005 | 0.0634848780952425 | 0.84891583577821 | 0.395928126482808 | 0.796351698323078 | NM_009298 |
| Mtmr12 | 2122.165134 | 0.121606481 | 0.091596959 | 1.327625749 | 0.184301774 | 0.395333215 | | Fam89b | 807.1961456 | 0.056070378353428 | 0.0660813866814374 | 0.848504869059875 | 0.396156861967711 | 0.796565438980175 | NM_023166,NM_181452 |
| Zmat5 | 458.8952039 | -0.129606617 | 0.097617738 | -1.327695356 | 0.184278768 | 0.395333215 | | Fblim1 | 80.38485613 | -0.0806556622189709 | 0.0950646775661084 | -0.848429346040569 | 0.396198905179111 | 0.796565438980175 | NM_001163256,NM_133754 |
| Atp2c1 | 5547.687789 | -0.087195973 | 0.065705137 | -1.327079998 | 0.184482221 | 0.395621804 | | Rnf10 | 10109.82369 | 0.0380016383255389 | 0.0447891939527787 | 0.848455508389011 | 0.396184340451829 | 0.796565438980175 | NM_016698 |
| Neu4 | 343.6359573 | 0.132082998 | 0.099552039 | 1.326773408 | 0.18458365 | 0.395751 | | Pstpip1 | 90.82614258 | -0.0800272801318156 | 0.0943498997941546 | -0.848196768691996 | 0.396328396515289 | 0.796715511732784 | NM_011193 |
| Rnf181 | 2419.354298 | 0.082715308 | 0.062344612 | 1.326743485 | 0.184593551 | 0.395751 | | Wfdc1 | 210.6951663 | 0.068759647904847 | 0.0810751234045642 | 0.848097974044849 | 0.396383409821665 | 0.796715844896846 | NM_023395 |
| Arl4a | 1342.237679 | -0.118568096 | 0.089433628 | -1.325766369 | 0.184917096 | 0.396245313 | | 2310057M21Rik | 631.1492045 | 0.0576093520989385 | 0.0679736588301394 | 0.847524660146654 | 0.396702747792254 | 0.796916624869352 | NM_026655 |
| Dcakd | 962.3464338 | -0.118113922 | 0.089084135 | -1.32586933 | 0.184882983 | 0.396245313 | | 2610034M16Rik | 148.4834789 | 0.0801719078122322 | 0.0945937085899295 | 0.847539535211409 | 0.396694460364618 | 0.796916624869352 | NM_027001 |
| Prelp | 915.86248 | 0.181563687 | 0.136950762 | 1.32575886 | 0.184919584 | 0.396245313 | | Fcf1 | 454.5302668 | -0.0663419301100445 | 0.0782617701614118 | -0.847692685371375 | 0.39660914103898 | 0.796916624869352 | NM_028632 |
| Zfp97 | 54.7323301 | -0.213877605 | 0.161327174 | -1.325738247 | 0.184926414 | 0.396245313 | | Lysmd4 | 1104.958597 | -0.0531996246178808 | 0.0627545753150295 | -0.847740971089637 | 0.396582243559956 | 0.796916624869352 | NM_001191051,NM_175215 |
| Gm9833 | 16.82237226 | 0.206212781 | 0.155569105 | 1.325538137 | 0.184992729 | 0.396332597 | | Zfp931 | 142.9720142 | 0.0779197333293206 | 0.0919517269017387 | 0.847398259443099 | 0.396773174335866 | 0.79694788860658 | NM_001162922 |
| 6430571L13Rik | 256.821184 | -0.212414272 | 0.160285676 | -1.325223049 | 0.185097183 | 0.396391949 | | 2310009B15Rik | 132.6620972 | -0.07748007382226 | 0.0914528651481669 | -0.847213192246422 | 0.396876301634567 | 0.79704481647224 | NM_001081226 |
| Commd9 | 788.7430815 | 0.094793024 | 0.071523943 | 1.325332753 | 0.18506081 | 0.396391949 | | Pcnp | 5536.58297 | 0.0452807042544984 | 0.0534665225105574 | 0.846898248255389 | 0.397051838971503 | 0.797284167685333 | NM_001024622 |
| Sec24c | 4083.199817 | 0.074141207 | 0.055945725 | 1.325234539 | 0.185093373 | 0.396391949 | | Pvr | 319.5300256 | 0.0690755724529501 | 0.0815813893672493 | 0.84670747812344 | 0.397158189478404 | 0.797284167685333 | NM_027514 |
| Clasp1 | 5144.145952 | 0.117042384 | 0.088349388 | 1.324767352 | 0.185248326 | 0.396605993 | | Sbf2 | 2876.940126 | -0.0425881560220374 | 0.0502987553149265 | -0.846703974191566 | 0.397160143010538 | 0.797284167685333 | NM_177324 |
| Trim28 | 4772.818636 | 0.077349696 | 0.058385304 | 1.324814481 | 0.185232691 | 0.396605993 | | Bahd1 | 2288.078718 | -0.0578314885274948 | 0.0683209968818184 | -0.846467281903565 | 0.397292118466887 | 0.797340698892446 | NM_001045523 |
| Dpp10 | 6783.735366 | -0.092696971 | 0.070002523 | -1.324194705 | 0.185438389 | 0.396849532 | | Pcna | 1566.822678 | 0.0467541335392892 | 0.05524154677472 | 0.846358153763449 | 0.397352975305137 | 0.797340698892446 | NM_011045 |
| Glce | 1795.413439 | -0.089308678 | 0.067443848 | -1.324193108 | 0.185438919 | 0.396849532 | | Slc25a51 | 3829.582885 | 0.0426698355502722 | 0.0504132924913088 | 0.846400491648674 | 0.39732936432213 | 0.797340698892446 | NM_001009949 |
| Tor1b | 1150.444994 | 0.099728257 | 0.075305772 | 1.324310936 | 0.1853998 | 0.396849532 | | Slitrk6 | 64.60520279 | 0.0658295783154476 | 0.0777957786265777 | 0.846184452133729 | 0.397449854049991 | 0.797424941979305 | NM_175499 |
| Ift81 | 1175.685554 | 0.101090735 | 0.076361656 | 1.323841583 | 0.185555663 | 0.397044529 | | C1d | 922.9927176 | 0.0591532012758673 | 0.0699305308722608 | 0.845885202615151 | 0.397616788115544 | 0.797649698130975 | NM_020558 |
| Ptprj | 6858.683197 | -0.11356251 | 0.08580147 | -1.323549691 | 0.185652643 | 0.39719719 | | Copg2 | 3890.38889 | 0.0497813656025504 | 0.0588675469514515 | 0.845650416580215 | 0.397747791304042 | 0.797651232461208 | NM_017478 |
| Mbtd1 | 856.3540546 | 0.111341611 | 0.084140436 | 1.323283027 | 0.185741274 | 0.397331948 | | Lgals8 | 1193.10997 | -0.0516019851313915 | 0.0610204349688683 | -0.845650889865305 | 0.39774752719975 | 0.797651232461208 | NM_001199043,NM_018886 |
| Pacsin3 | 1072.615313 | 0.12181277 | 0.092058916 | 1.323204481 | 0.185767386 | 0.397332949 | | Trap1 | 2960.171148 | 0.0393428886580539 | 0.0465272223224766 | 0.845588597259714 | 0.397782288857841 | 0.797651232461208 | NM_026508 |
| Mirlet7d | 12.29551379 | 0.184081662 | 0.139163115 | 1.322776241 | 0.185909801 | 0.397582671 | | 5830416P10Rik | 65.06033746 | -0.0713968414331505 | 0.084642803600897 | -0.843507520967723 | 0.398944661260721 | 0.797789837692759 | NR_028427 |
| Coro1c | 3806.55163 | -0.082203249 | 0.062156245 | -1.32252598 | 0.185993064 | 0.397705843 | | Aes | 15605.57331 | 0.0389418839450463 | 0.0461596029114716 | 0.843635592354033 | 0.398873068785371 | 0.797789837692759 | NR_074087,NM_010347,NM_001276288 |
| Cldn14 | 24.27679009 | 0.221554503 | 0.167554878 | 1.322283455 | 0.18607378 | 0.39777119 | | Arl16 | 820.1881203 | 0.0525886275469409 | 0.0622980125293824 | 0.844146151887877 | 0.398587740615317 | 0.797789837692759 | NM_197995 |
| Pcdhb19 | 250.5854634 | -0.12992888 | 0.098261254 | -1.32227988 | 0.18607497 | 0.39777119 | | Asic1 | 3833.652809 | 0.0531489306831267 | 0.0629943923183799 | 0.84370891958933 | 0.398832082016313 | 0.797789837692759 | NM_009597 |
| F8a | 451.8620611 | -0.139710665 | 0.105725915 | -1.321442002 | 0.186354029 | 0.398312777 | | Atf1 | 402.4124264 | -0.0615737957537458 | 0.0728386327742091 | -0.845345298347611 | 0.397918076504673 | 0.797789837692759 | NM_007497 |
| Brd4 | 3795.098712 | 0.093128873 | 0.070507963 | 1.320827736 | 0.186558809 | 0.398530566 | | Cntnap2 | 4295.383354 | 0.0530429826647378 | 0.0627890070102319 | 0.84478135887854 | 0.398232924368532 | 0.797789837692759 | NM_001004357,NM_025771 |
| Galk1 | 308.8816817 | 0.11948931 | 0.090457523 | 1.320943871 | 0.18652008 | 0.398530566 | | Dfna5 | 443.6536987 | 0.0610953899100376 | 0.0723758965764438 | 0.844140007930793 | 0.398591173458686 | 0.797789837692759 | NM_018769 |
| Map2k4 | 8593.68337 | 0.109823127 | 0.083144404 | 1.320872153 | 0.186543996 | 0.398530566 | | Gm5617 | 53.13242562 | -0.078828237789757 | 0.0927437922771645 | -0.84413453307789 | 0.398594232465186 | 0.797789837692759 | NM_001004191 |
| Syde2 | 137.069172 | -0.157867574 | 0.119515777 | -1.320893167 | 0.186536989 | 0.398530566 | | Itga9 | 506.9769934 | 0.0718575967434293 | 0.085168747797943 | 0.843708503427884 | 0.398832314625498 | 0.797789837692759 | NM_133721,NM_001113514 |
| Mum1 | 1249.928111 | 0.108082499 | 0.081846751 | 1.320547213 | 0.186652384 | 0.398675494 | | Klf6 | 2197.030923 | -0.0604010409195662 | 0.0715572515988566 | -0.844093918785044 | 0.398616925641567 | 0.797789837692759 | NM_011803 |
| Tead3 | 82.44750613 | 0.197368793 | 0.149569636 | 1.31957794 | 0.186975972 | 0.399311607 | | Mettl21a | 419.097279 | -0.0572960922762371 | 0.0678892998901104 | -0.843963516621617 | 0.398689792917545 | 0.797789837692759 | NM_025964 |
| BC055324 | 60.0320784 | -0.190652576 | 0.144510545 | -1.319298718 | 0.187069266 | 0.39941383 | | Parp8 | 833.1669233 | -0.0633122746713065 | 0.0749357914900309 | -0.844886981406333 | 0.398173943806083 | 0.797789837692759 | NM_001081009 |
| Ppp3ca | 31805.9687 | 0.104160973 | 0.07895287 | 1.319280381 | 0.187075394 | 0.39941383 | | Plcg2 | 233.2673947 | -0.0759698263710859 | 0.0898928021041902 | -0.845115788948631 | 0.398046193682073 | 0.797789837692759 | NM_172285 |
| Mcm6 | 325.8838534 | 0.146185996 | 0.110829777 | 1.319013718 | 0.187164526 | 0.399549073 | | Stra13 | 275.1533547 | -0.0713667746217018 | 0.0845466795111435 | -0.844110910497621 | 0.398607431452504 | 0.797789837692759 | NM_016665 |
| Tgoln1 | 3730.802909 | -0.075923824 | 0.05757173 | -1.318769201 | 0.187246284 | 0.399668538 | | Syngr2 | 377.2560454 | -0.0749474608616808 | 0.0886747063354477 | -0.845195478608769 | 0.398001706350396 | 0.797789837692759 | NM_009304 |
| Eif1ax | 1648.085098 | -0.085270625 | 0.064678507 | -1.318376523 | 0.187377636 | 0.399838005 | | Tcp11l1 | 478.8138155 | 0.0686348765251956 | 0.0813556711831402 | 0.843639730667223 | 0.398870755578869 | 0.797789837692759 | NM_177190 |
| Polr2h | 510.4137418 | -0.112481351 | 0.085319622 | -1.318352674 | 0.187385616 | 0.399838005 | | Timm13 | 1534.012172 | 0.0564652395336716 | 0.0668948073802508 | 0.844090023500713 | 0.398619102166843 | 0.797789837692759 | NM_013895 |
| Psg23 | 36.48122404 | 0.219168309 | 0.166250655 | 1.318300428 | 0.187403098 | 0.399838005 | | Ube2m | 4519.868387 | 0.0439759985949736 | 0.0521016722863268 | 0.844041979176824 | 0.398645947954898 | 0.797789837692759 | NM_001168469,NM_001243968,NM_145578 |
| Nkain3 | 31.52659367 | -0.221507462 | 0.168077162 | -1.317891493 | 0.187539974 | 0.400074949 | | Ucp2 | 1064.290598 | 0.0795735953936365 | 0.0941874039544532 | 0.844843281083704 | 0.398198345818067 | 0.797789837692759 | NM_011671 |
| Ppp1r35 | 206.0481219 | -0.1457966 | 0.110667251 | -1.317432201 | 0.187693793 | 0.400347967 | | Zscan22 | 434.4183922 | 0.0573324967810743 | 0.0679699054847281 | 0.843498256650601 | 0.398949840355239 | 0.797789837692759 | NM_001001447 |
| B3gnt1 | 4041.51153 | -0.072835989 | 0.055311524 | -1.31683209 | 0.187894914 | 0.400721788 | | 2310042D19Rik | 39.93868593 | -0.0779152089256668 | 0.0923824483009858 | -0.843398398273835 | 0.39900566743525 | 0.797791648336469 | NM_172417 |
| Kxd1 | 894.1259985 | 0.075306267 | 0.057206426 | 1.316395238 | 0.18804142 | 0.400979048 | | Serpine2 | 7949.444971 | 0.0441103559795422 | 0.0523073672783521 | 0.843291457297216 | 0.399065459345727 | 0.797801384868707 | NM_009255 |
| Zfp11 | 289.8121812 | -0.128867531 | 0.097912324 | -1.316152297 | 0.188122931 | 0.40109766 | | Pdpk1 | 7617.298576 | 0.0554282647086283 | 0.0657578665102523 | 0.842914584219159 | 0.399276216335311 | 0.798004075940947 | NM_011062,NM_001080773 |
| Htatip2 | 324.7246756 | 0.146147526 | 0.111107898 | 1.315365767 | 0.188387005 | 0.401605428 | | Psme1 | 1175.884263 | -0.0579051483332658 | 0.0686964150194653 | -0.842913685042491 | 0.3992767192578 | 0.798004075940947 | NM_011189 |
| Sema3a | 943.2010615 | -0.126413608 | 0.096112824 | -1.315262656 | 0.188421644 | 0.401624014 | | Zfp423 | 806.5306003 | 0.0716154197747548 | 0.0850040959088898 | 0.84249375290709 | 0.399511635017867 | 0.798363737827699 | NM_033327 |
| Oxld1 | 120.2555498 | -0.16486331 | 0.125376371 | -1.314947216 | 0.188527643 | 0.401684172 | | Eri2 | 456.7502944 | -0.0608139529295138 | 0.072201034670915 | -0.842286446540519 | 0.399627635672865 | 0.798421909005876 | NM_027698 |
| Tmc7 | 710.6562234 | 0.11188011 | 0.085081438 | 1.314976722 | 0.188517714 | 0.401684172 | | Jakmip3 | 1593.000602 | -0.0553338059287225 | 0.0656979711317983 | -0.842245277425873 | 0.399650674678626 | 0.798421909005876 | NM_028708 |
| Tmem189 | 1457.05935 | 0.10433157 | 0.079338654 | 1.315015626 | 0.188504651 | 0.401684172 | | Ctsd | 17702.75921 | -0.0712747869754141 | 0.0846569041029798 | -0.841925271549179 | 0.399829783670015 | 0.798662339247373 | NM_009983 |
| Gpn2 | 241.0598863 | 0.130629136 | 0.099357321 | 1.314740923 | 0.188596988 | 0.401747886 | | Pnrc2 | 1271.789725 | 0.058027043839992 | 0.0689371585292653 | 0.841738259568072 | 0.399934477582221 | 0.798662339247373 | NM_026383 |
| Swt1 | 291.0552203 | 0.137377087 | 0.104493676 | 1.31469283 | 0.188613157 | 0.401747886 | | Ptprs | 19412.04024 | -0.0481953391105156 | 0.0572570996705213 | -0.841735599390287 | 0.399935966934145 | 0.798662339247373 | NM_001252455,NM_001252456,NM_011218,NM_001252453 |
| Umps | 671.4531499 | -0.102194163 | 0.07773625 | -1.314626873 | 0.188635334 | 0.401747886 | | Alg8 | 304.9448542 | -0.065017366884292 | 0.0772783879479342 | -0.841339585500891 | 0.400157720160494 | 0.798904607890772 | NM_199035 |
| Tmem179 | 5003.319942 | -0.083623322 | 0.063622059 | -1.314376423 | 0.188719621 | 0.401872157 | | Kcnh7 | 1246.991139 | 0.0754278456895772 | 0.0896539013661415 | 0.841322514025732 | 0.400167281220794 | 0.798904607890772 | NM_133207 |
| Gcc2 | 3401.31645 | -0.090278049 | 0.068692938 | -1.314226053 | 0.188770143 | 0.401924502 | | Ankfn1 | 103.794273 | 0.0796878819842608 | 0.0947589286509665 | 0.840953808983867 | 0.400373811890342 | 0.799132593983304 | NM_001080933 |
| Mib2 | 3720.793943 | 0.101961977 | 0.077596897 | 1.313995541 | 0.188847705 | 0.402034396 | | Cse1l | 3062.585649 | 0.0620225334495531 | 0.0737553742088809 | 0.84092222586927 | 0.400391506196509 | 0.799132593983304 | NM_023565 |
| Abcc2 | 9.30674974 | 0.207817603 | 0.158173495 | 1.313858575 | 0.188893801 | 0.402059668 | | AI593442 | 8159.17285 | -0.0640685892671337 | 0.0762563047534044 | -0.840174323609268 | 0.400810652587365 | 0.799169897076002 | NM_178906 |
| Aurkb | 11.5885159 | -0.216811631 | 0.165031491 | -1.313759151 | 0.188927268 | 0.402059668 | | Abi3bp | 275.294842 | -0.073467767256532 | 0.0873878613612218 | -0.840709065448456 | 0.400510940708388 | 0.799169897076002 | NM_001014424,NM_178790,NM_001014399,NM_001014422,NM_001014423 |
| Sdf2 | 1512.917033 | -0.083802442 | 0.063779749 | -1.313728983 | 0.188937424 | 0.402059668 | | Gpr56 | 2823.990853 | 0.0691183615410357 | 0.0822422121781526 | 0.84042349655334 | 0.400670475043321 | 0.799169897076002 | NM_018882,NM_001198894 |
| Cacna1b | 3865.04407 | -0.085557818 | 0.065129789 | -1.313651078 | 0.188963551 | 0.40206026 | | Klf12 | 1574.795042 | 0.0572180839656363 | 0.0680732065393775 | 0.840537516512287 | 0.400607075708647 | 0.799169897076002 | NM_010636 |

| | | | | | | |
|---|---|---|---|---|---|---|
| Atp9a | 16676.76021 | -0.097948922 | 0.074569274 | -1.313529254 | 0.18900467 | 0.402092318 |
| Tnrc18 | 3087.901567 | -0.111751949 | 0.085101992 | -1.313153143 | 0.18913135 | 0.40230658 |
| Phf16 | 278.2449715 | 0.124862116 | 0.09510458 | 1.312892771 | 0.189219083 | 0.402437952 |
| Tjap1 | 661.8839185 | 0.159650177 | 0.121630242 | 1.312586194 | 0.189322424 | 0.402602476 |
| Emc7 | 2345.047201 | -0.077721244 | 0.059219195 | -1.312433309 | 0.189373974 | 0.402656835 |
| Zbtb16 | 1489.543646 | 0.217845786 | 0.16601577 | 1.31219935 | 0.189452881 | 0.402769339 |
| Cyp19a1 | 9.177297304 | 0.190243437 | 0.14500349 | 1.311992122 | 0.189522793 | 0.40286269 |
| Tmem130 | 8035.237347 | -0.126646244 | 0.096539694 | -1.311856688 | 0.189568494 | 0.40290456 |
| 2310050B05Rik | 8.43026902 | 0.157214462 | 0.119849176 | 1.311769232 | 0.189598009 | 0.402912022 |
| 1700007G11Rik | 14.94204235 | 0.214910619 | 0.163882654 | 1.311368922 | 0.189733153 | 0.403091979 |
| C030006K11Rik | 394.6467051 | 0.12303956 | 0.093825615 | 1.311364272 | 0.189734723 | 0.403091979 |
| Antxr1 | 782.4282343 | 0.097535626 | 0.074390122 | 1.311136791 | 0.189811553 | 0.403199918 |
| Tubgcp3 | 1577.693832 | 0.076413214 | 0.058292768 | 1.310852387 | 0.189907639 | 0.403348727 |
| Ncs1 | 14756.34968 | -0.080910095 | 0.06173254 | -1.310655531 | 0.189974169 | 0.403434727 |
| Oxsr1 | 2054.519657 | 0.087559878 | 0.066825734 | 1.310271845 | 0.190103888 | 0.403654878 |
| Pcdh20 | 877.5525293 | 0.165013334 | 0.125950193 | 1.310147534 | 0.190145931 | 0.403688825 |
| Osbpl1a | 11853.92705 | 0.13323612 | 0.101709386 | 1.309968777 | 0.190206398 | 0.403706566 |
| Slc9a4 | 33.10696575 | 0.196837665 | 0.150253307 | 1.310038816 | 0.190182705 | 0.403706566 |
| Bmp5 | 50.88284066 | -0.214228035 | 0.163618766 | -1.30931213 | 0.190428643 | 0.4040697 |
| Rfx1 | 1000.477133 | 0.15097262 | 0.115307082 | 1.309309163 | 0.190429647 | 0.4040697 |
| Ccdc51 | 206.1003606 | -0.123672822 | 0.094474575 | -1.309059313 | 0.19051426 | 0.404152589 |
| Gart | 1385.748606 | 0.110114012 | 0.084122107 | 1.308978301 | 0.190541701 | 0.404152589 |
| Tbc1d8b | 257.385404 | -0.164354109 | 0.125560569 | -1.308962764 | 0.190546964 | 0.404152589 |
| Lgmn | 4190.948676 | 0.080528371 | 0.061547783 | 1.308387842 | 0.190741794 | 0.404510452 |
| Ntm | 5099.340452 | -0.083027659 | 0.063465949 | -1.308223698 | 0.190797447 | 0.404573099 |
| Rngtt | 858.4208902 | -0.112664908 | 0.086161479 | -1.307601832 | 0.191008395 | 0.404854183 |
| Socs3 | 134.8086578 | 0.179299625 | 0.137114558 | 1.307662931 | 0.190987662 | 0.404854183 |
| Zfp191 | 1291.402695 | 0.119689087 | 0.091526526 | 1.30769835 | 0.190975643 | 0.404854183 |
| Cdc73 | 676.3146958 | -0.09884915 | 0.075609285 | -1.307367878 | 0.191087801 | 0.40491802 |
| Slc25a23 | 17586.02334 | 0.082417937 | 0.063041545 | 1.3073591 | 0.191090781 | 0.40491802 |
| Fstl1 | 1562.563221 | -0.112876819 | 0.08634555 | -1.307268512 | 0.191121534 | 0.404927807 |
| Slc22a21 | 39.18043603 | 0.215809952 | 0.165114155 | 1.30703483 | 0.191200883 | 0.405040535 |
| Msl1 | 6471.004428 | -0.072188088 | 0.055237356 | -1.30687081 | 0.191256591 | 0.405103161 |
| Grb14 | 749.9004811 | -0.103092093 | 0.078908183 | -1.306481644 | 0.191388817 | 0.40532782 |
| Ky | 146.1822601 | -0.195279351 | 0.149506235 | -1.306161918 | 0.1914975 | 0.405408899 |
| Lzts2 | 1495.878527 | 0.117368463 | 0.089859153 | 1.306138102 | 0.191505598 | 0.405408899 |
| Tspan12 | 450.6292005 | -0.135085611 | 0.103423379 | -1.306141914 | 0.191504302 | 0.405408899 |
| Zfp37 | 729.4334311 | -0.104673873 | 0.080155513 | -1.305884895 | 0.191591704 | 0.405535773 |
| Phf20 | 3747.939206 | 0.102331872 | 0.078389824 | 1.305422906 | 0.191748883 | 0.405813029 |
| Mospd1 | 332.9368397 | 0.123758518 | 0.094827101 | 1.30509651 | 0.191859987 | 0.405907402 |
| Nap1l1 | 9465.601092 | -0.069659588 | 0.053376498 | -1.30506103 | 0.191872067 | 0.405907402 |
| Tnfaip8l2 | 72.56482728 | 0.189134507 | 0.144912852 | 1.305160334 | 0.191838258 | 0.405907402 |
| Cacna1i | 5033.471203 | -0.083145549 | 0.063728303 | -1.304687946 | 0.191999128 | 0.406065313 |
| Dph3 | 1157.944532 | -0.074819093 | 0.057344577 | -1.304728309 | 0.191985378 | 0.406065313 |
| Atp5s | 451.7756526 | -0.12810893 | 0.09819807 | -1.304597236 | 0.19203003 | 0.40607524 |
| Blcap | 3417.506164 | -0.080511059 | 0.061728327 | -1.304280596 | 0.192137929 | 0.406161583 |
| Phyhip | 23217.32159 | 0.113835271 | 0.087275141 | 1.304326395 | 0.192122319 | 0.406161583 |
| Prpf38a | 639.8827561 | -0.097414343 | 0.074690124 | -1.304246637 | 0.192149503 | 0.406161583 |
| Casp8 | 110.4107026 | 0.18086627 | 0.138692274 | 1.304083233 | 0.192205205 | 0.406185958 |
| Ccdc55 | 620.2060811 | 0.083291594 | 0.063871032 | 1.304059001 | 0.192213466 | 0.406185958 |
| Setdb2 | 196.7613371 | 0.162319206 | 0.124490807 | 1.303865 | 0.192279616 | 0.406270335 |
| Sucla2 | 5340.981539 | -0.075243867 | 0.057719154 | -1.303620396 | 0.192363043 | 0.40639119 |
| Adam10 | 3317.692324 | -0.08437057 | 0.064746482 | -1.303091178 | 0.192543635 | 0.406446041 |
| Afap1 | 792.0776995 | -0.109039208 | 0.083694407 | -1.302825502 | 0.192634342 | 0.406446041 |
| B3galt1 | 1751.761484 | -0.115393646 | 0.088575217 | -1.302775753 | 0.192651331 | 0.406446041 |
| Cr2 | 12.8264472 | -0.213480543 | 0.163797304 | -1.303321468 | 0.192465035 | 0.406446041 |
| Dnmbp | 592.9266138 | 0.098756201 | 0.075783548 | 1.303135096 | 0.192528644 | 0.406446041 |
| Ext2 | 1448.120252 | 0.104562501 | 0.080242763 | 1.303077037 | 0.192548462 | 0.406446041 |
| Fam131c | 94.12592257 | -0.188968487 | 0.144990423 | -1.303317023 | 0.192466552 | 0.406446041 |
| Fkbp15 | 1623.275172 | 0.091037932 | 0.069873566 | 1.302895182 | 0.192610549 | 0.406446041 |
| Gatm | 3360.566133 | 0.159336252 | 0.12228484 | 1.302992686 | 0.192577259 | 0.406446041 |
| Limk2 | 4605.693885 | 0.085382953 | 0.065532954 | 1.302901034 | 0.192608551 | 0.406446041 |
| 9530077C05Rik | 247.7295275 | 0.165960679 | 0.127406149 | 1.302611227 | 0.192707523 | 0.406509239 |
| Gp9 | 8.848061451 | -0.180180833 | 0.138408278 | -1.301806766 | 0.19298245 | 0.407033772 |
| Iqcg | 154.975466 | -0.188553202 | 0.144856992 | -1.301650684 | 0.193035825 | 0.407090932 |
| Taf2 | 1495.806924 | 0.085717065 | 0.065867778 | 1.301350491 | 0.193138512 | 0.407252056 |
| Clspn | 132.096537 | -0.192607051 | 0.148087084 | -1.300633693 | 0.193383868 | 0.407700176 |
| Nxn | 397.5199592 | -0.135967565 | 0.104544121 | -1.300575905 | 0.193403659 | 0.407700176 |
| Usp12 | 3336.914348 | 0.098842553 | 0.076026026 | 1.300114697 | 0.193561661 | 0.407977742 |
| Kdm5b | 3030.021834 | 0.078681034 | 0.060529325 | 1.299882898 | 0.193641107 | 0.407978694 |
| Rcbtb1 | 1814.298909 | 0.092387666 | 0.071072977 | 1.299898646 | 0.193635709 | 0.407978694 |
| Slc7a7 | 70.89198641 | 0.193664147 | 0.148972457 | 1.299999681 | 0.193601078 | 0.407978694 |
| B430319G15Rik | 28.02414583 | -0.217945626 | 0.167729748 | -1.299385644 | 0.193811616 | 0.408226914 |
| Iqub | 65.85057378 | 0.20233166 | 0.155705348 | 1.299452225 | 0.193788779 | 0.408226914 |
| Wwox | 683.3330216 | 0.132084398 | 0.101666792 | 1.299189196 | 0.193879008 | 0.408313355 |
| Polr2l | 934.1371711 | -0.104719563 | 0.080626167 | -1.298828495 | 0.194002793 | 0.408518521 |
| Iqck | 59.4381145 | 0.204147772 | 0.15719264 | 1.298710756 | 0.194043211 | 0.408548107 |
| St7l | 796.9060952 | -0.108187131 | 0.083323744 | -1.298394987 | 0.194151641 | 0.408720859 |
| Ttc17 | 1792.053849 | 0.093986649 | 0.072398836 | 1.298179007 | 0.19422583 | 0.408821493 |
| Arap2 | 4196.060432 | -0.110667063 | 0.085286404 | -1.29759326 | 0.19442714 | 0.408911918 |
| Asxl2 | 1273.822881 | -0.108473499 | 0.083581406 | -1.297818555 | 0.194349692 | 0.408911918 |
| Cdh22 | 1699.455756 | -0.123988721 | 0.095544641 | -1.297704614 | 0.194388858 | 0.408911918 |
| Hvcn1 | 180.4419663 | 0.175687217 | 0.135376389 | 1.297768524 | 0.194366889 | 0.408911918 |
| Ifitm1 | 67.59696309 | -0.203179558 | 0.156545126 | -1.297897695 | 0.194322492 | 0.408911918 |
| Ube2d2a | 5396.318419 | -0.082513862 | 0.06358908 | -1.297610568 | 0.194421189 | 0.408911918 |
| Dnajc7 | 6010.056568 | 0.098723095 | 0.07611547 | 1.297017471 | 0.194625175 | 0.409106291 |
| Pcdhga8 | 458.6026719 | 0.136381813 | 0.105149724 | 1.297024928 | 0.19462261 | 0.409106291 |
| Pggt1b | 280.8867488 | -0.12251653 | 0.094456791 | -1.297064286 | 0.194609069 | 0.409106291 |
| Synm | 1851.600765 | 0.089034732 | 0.068641814 | 1.297091768 | 0.194599614 | 0.409106291 |
| Mocs3 | 219.2515407 | -0.144037339 | 0.111059376 | -1.296939931 | 0.194651856 | 0.409106871 |
| Ikbip | 352.9410744 | -0.10841349 | 0.083599654 | -1.296817447 | 0.194694006 | 0.409126645 |
| Psmb7 | 4245.350658 | -0.087118326 | 0.067181579 | -1.296759136 | 0.194714075 | 0.409126645 |
| Catsperd | 41.3197543 | 0.217994027 | 0.168121363 | 1.296646799 | 0.194752742 | 0.409152405 |
| Adck3 | 1583.360229 | 0.107443513 | 0.082910091 | 1.295903928 | 0.195008586 | 0.409594319 |
| Higd2a | 1985.032624 | -0.089682602 | 0.069205811 | -1.295882526 | 0.19501596 | 0.409594319 |
| Wdr4 | 839.2100333 | -0.096508108 | 0.074482941 | -1.29570754 | 0.195076263 | 0.409628147 |
| Zfp87 | 597.1264293 | -0.09490578 | 0.073247721 | -1.295682357 | 0.195084942 | 0.409628147 |
| Fads2 | 3501.915819 | -0.07563383 | 0.058378265 | -1.295582015 | 0.195119529 | 0.409645254 |
| B630019K06Rik | 1047.945508 | 0.11058091 | 0.085387249 | 1.295051786 | 0.195302366 | 0.409894454 |
| Klhdc7a | 705.7717208 | -0.155929196 | 0.120415065 | -1.294930963 | 0.195344046 | 0.409894454 |
| Prr5 | 225.4614639 | 0.174033376 | 0.134389745 | 1.294989998 | 0.19532368 | 0.409894454 |
| Tubgcp5 | 1217.154202 | 0.10336607 | 0.079822963 | 1.294941531 | 0.1953404 | 0.409894454 |
| Gopc | 2257.562715 | 0.106438685 | 0.082204767 | 1.294799436 | 0.195389427 | 0.409914495 |
| Pcdhgc5 | 4531.126476 | -0.122934663 | 0.094948576 | -1.29474993 | 0.19540651 | 0.409914495 |
| Usp25 | 1434.09782 | 0.104753182 | 0.080923967 | 1.29446425 | 0.19550511 | 0.410065815 |
| Gxylt2 | 25.41262791 | -0.217490247 | 0.168109592 | -1.29374085 | 0.19575495 | 0.410534272 |
| Rpl35 | 3466.266528 | -0.096046598 | 0.074277722 | -1.293074088 | 0.195985436 | 0.410962017 |
| Tfip11 | 2126.658619 | 0.098477116 | 0.076163679 | 1.292966895 | 0.196022509 | 0.410984135 |
| Aldh1a2 | 355.7754064 | -0.202655323 | 0.156764741 | -1.29273535 | 0.196102607 | 0.411040827 |
| Myrip | 3075.0621 | 0.125095231 | 0.096766529 | 1.292753103 | 0.196096465 | 0.411040827 |
| Prdx5 | 5059.420878 | -0.080202781 | 0.062052736 | -1.292493882 | 0.196186163 | 0.411160343 |
| Gm19345 | 43.84532391 | -0.202971539 | 0.157056264 | -1.292349209 | 0.196236237 | 0.411209665 |
| Mapre1 | 3769.6599 | -0.084571326 | 0.065444733 | -1.29225564 | 0.196268629 | 0.411221925 |
| Fkbp8 | 11013.73812 | -0.089289457 | 0.069102951 | -1.292122188 | 0.196314833 | 0.41124651 |
| Tcn2 | 868.1682202 | 0.132807141 | 0.102786461 | 1.292068431 | 0.196333447 | 0.41124651 |
| Dapl1 | 11.42358066 | -0.198415594 | 0.153607502 | -1.291705106 | 0.196459289 | 0.411454477 |
| Plin3 | 554.9483673 | 0.14907686 | 0.115448044 | 1.291289612 | 0.196603271 | 0.411700377 |
| Bcar1 | 1856.665221 | 0.100055665 | 0.077506127 | 1.290938775 | 0.196724909 | 0.411732505 |
| Cabyr | 18.83830968 | 0.216344707 | 0.167568908 | 1.291079053 | 0.196676267 | 0.411732505 |
| Gm20605 | 34.31490836 | 0.216601937 | 0.167769635 | 1.291067582 | 0.196680244 | 0.411732505 |
| Wdr46 | 333.1147059 | 0.11630262 | 0.090086718 | 1.29100741 | 0.196701108 | 0.411732505 |
| Ppp1r8 | 1308.095862 | -0.097877789 | 0.075827436 | -1.290796492 | 0.196774255 | 0.411780159 |
| E130311K13Rik | 282.5187032 | -0.148282433 | 0.11490703 | -1.290455713 | 0.196892479 | 0.411971921 |
| Lrch1 | 634.4789373 | 0.120086498 | 0.093081725 | 1.290118957 | 0.197009359 | 0.412160816 |
| Fyn | 4726.147293 | -0.059523954 | 0.046146034 | -1.289904011 | 0.197083988 | 0.412261281 |
| Ssh2 | 1948.613432 | -0.113956487 | 0.088368658 | -1.289557751 | 0.197204253 | 0.412457167 |
| Crip1 | 112.0889711 | -0.171616633 | 0.133103052 | -1.289351599 | 0.19727588 | 0.412495612 |
| Hap1 | 8067.595112 | 0.183209336 | 0.142093627 | 1.289356459 | 0.197274191 | 0.412495612 |
| 2900055J20Rik | 164.6071537 | -0.15861663 | 0.123060462 | -1.288932513 | 0.197421549 | 0.4127445 |
| Setd6 | 410.2096246 | 0.134518999 | 0.104379318 | 1.288751459 | 0.197484506 | 0.412820418 |
| Cntn4 | 2070.344764 | -0.10217319 | 0.079286561 | -1.288657103 | 0.197517322 | 0.412833318 |
| Plekhh1 | 1763.704274 | 0.154315936 | 0.119860537 | 1.287462411 | 0.197933163 | 0.413646672 |
| Ipo11 | 2272.361253 | -0.107871645 | 0.083813544 | -1.287043106 | 0.198079264 | 0.41386462 |
| Tmed8 | 2812.514556 | -0.079366542 | 0.061667396 | -1.287009792 | 0.198090875 | 0.41386462 |
| Qk | 9090.366197 | -0.112469478 | 0.087440123 | -1.286245644 | 0.198357346 | 0.414365474 |
| Ebf4 | 311.3494351 | 0.18448077 | 0.143436695 | 1.286147658 | 0.198391534 | 0.414381023 |
| 0610030E20Rik | 785.6958151 | 0.129796666 | 0.100930644 | 1.285998599 | 0.198443551 | 0.414386757 |
| Iscu | 2074.212315 | -0.089436219 | 0.069546778 | -1.285986512 | 0.198447769 | 0.414386757 |
| Gm13293 | 58.79498809 | 0.214951081 | 0.167182423 | 1.285727749 | 0.198538093 | 0.41450741 |
| Ttc7 | 467.9756467 | -0.112536031 | 0.087531195 | -1.28566771 | 0.198559055 | 0.41450741 |
| I830077J02Rik | 32.14324228 | 0.207225557 | 0.161206346 | 1.285467741 | 0.198628883 | 0.41459732 |
| Cox6a2 | 146.8647848 | 0.174131808 | 0.135511334 | 1.284998106 | 0.198792946 | 0.414883877 |
| Morc3 | 872.4706819 | 0.084124134 | 0.065477374 | 1.28478174 | 0.198868565 | 0.414985797 |
| Trpt1 | 325.6503713 | 0.123070271 | 0.095801459 | 1.28463879 | 0.198918537 | 0.415034179 |
| Rgs5 | 1547.971665 | -0.140044318 | 0.109021951 | -1.284551567 | 0.198949033 | 0.415041916 |
| Mettl16 | 879.8547229 | -0.109620721 | 0.085352507 | -1.284329244 | 0.199026779 | 0.415148211 |
| Cog7 | 2199.239511 | 0.069281782 | 0.053968473 | 1.283745454 | 0.199231035 | 0.415518327 |
| Stk33 | 43.39287773 | -0.215726171 | 0.168087477 | -1.283416081 | 0.199346343 | 0.415702858 |
| E130006D01Rik | 19.09981269 | -0.215445406 | 0.167902931 | -1.283154525 | 0.199437944 | 0.41583791 |
| Ctu2 | 504.8393859 | 0.108150054 | 0.084317656 | 1.282650149 | 0.199614672 | 0.416113389 |
| Pls3 | 6501.460095 | 0.101576466 | 0.079194255 | 1.282624169 | 0.199623778 | 0.416113389 |
| Sh3pxd2b | 1077.548197 | -0.109214287 | 0.085156744 | -1.282508951 | 0.199664167 | 0.416141593 |
| 1810032O08Rik | 92.29489524 | 0.171184334 | 0.133505778 | 1.282224167 | 0.199764021 | 0.416244936 |
| Glb1 | 971.8747349 | 0.099501057 | 0.077600957 | 1.282214301 | 0.199767481 | 0.416244936 |
| Aifm1 | 876.0443065 | -0.077196622 | 0.060219316 | -1.281924583 | 0.199869104 | 0.416400684 |
| Enpp5 | 9272.519054 | 0.067606666 | 0.052751562 | 1.281605014 | 0.19998124 | 0.416578292 |
| Foxg1 | 2800.768316 | -0.107087226 | 0.083583241 | -1.281204526 | 0.200121837 | 0.416815128 |
| Phf10 | 905.2166847 | 0.103371344 | 0.080696597 | 1.280987649 | 0.200198004 | 0.416917726 |
| Atad1 | 4031.429452 | -0.074869021 | 0.058454089 | -1.280817501 | 0.200257775 | 0.416986154 |
| Glg1 | 5521.770853 | -0.102845885 | 0.080316805 | -1.280502694 | 0.200368398 | 0.417104388 |
| Otop2 | 58.20368632 | -0.204551563 | 0.159735184 | -1.280566736 | 0.20034589 | 0.417104388 |
| Kras | 3234.429241 | -0.11733977 | 0.091654766 | -1.280236419 | 0.200462001 | 0.417243182 |
| Prickle3 | 59.98461639 | 0.187323594 | 0.146349025 | 1.279978422 | 0.200552725 | 0.417375946 |
| Fbxl20 | 2819.183261 | 0.101348656 | 0.079185778 | 1.279884582 | 0.200585731 | 0.417388573 |
| Myo1c | 920.1079587 | 0.107456264 | 0.083975456 | 1.279615122 | 0.200680529 | 0.4174737 |
| Naip6 | 9.324757374 | 0.194804635 | 0.152232483 | 1.279652226 | 0.200667473 | 0.4174737 |
| Cdc23 | 1918.103671 | 0.079867868 | 0.062437342 | 1.27916829 | 0.2008378 | 0.417688708 |
| Wdyhv1 | 574.3404132 | 0.111624668 | 0.087260041 | 1.279218608 | 0.200820085 | 0.417688708 |
| Cers2 | 2319.593313 | 0.108687398 | 0.084977949 | 1.279007079 | 0.200894563 | 0.417750686 |
| Eef1a2 | 34140.58874 | 0.089059679 | 0.06964435 | 1.278778238 | 0.200975159 | 0.417806134 |
| Stoml3 | 20.31063833 | -0.178587542 | 0.139652688 | -1.27879774 | 0.20096829 | 0.417806134 |
| BC039966 | 10.276408 | 0.196772817 | 0.153885058 | 1.278699956 | 0.201002735 | 0.417807403 |
| Agfg2 | 1700.58031 | 0.100002654 | 0.078237611 | 1.27819157 | 0.201181888 | 0.418011556 |
| Ccdc25 | 1653.855243 | 0.072847449 | 0.056991177 | 1.27822328 | 0.20117071 | 0.418011556 |
| Cpsf3 | 1763.54589 | 0.085101742 | 0.066575803 | 1.278268359 | 0.20115482 | 0.418011556 |
| Sptb | 9426.575295 | -0.091293915 | 0.071444313 | -1.277833195 | 0.201308247 | 0.418218019 |
| Ccdc77 | 158.9555165 | 0.14660309 | 0.114757387 | 1.277504602 | 0.201424157 | 0.418319339 |
| Gm10941 | 10.77644016 | 0.195979173 | 0.153412518 | 1.277465327 | 0.201438014 | 0.418319339 |
| Rab39b | 1598.446589 | -0.111965255 | 0.087637013 | -1.277602354 | 0.20138967 | 0.418319339 |
| Arf1 | 15095.00588 | -0.076736423 | 0.060073147 | -1.277383105 | 0.201467026 | 0.41832352 |
| Bhlhe40 | 5454.696346 | 0.200985706 | 0.157360355 | 1.277232164 | 0.201520295 | 0.418378058 |
| Arhgap20 | 4233.195755 | -0.097718924 | 0.076549329 | -1.27654841 | 0.201761725 | 0.41876707 |
| Arpc1a | 5809.927702 | -0.092114265 | 0.072155494 | -1.276607779 | 0.201740754 | 0.41876707 |
| Npffr1 | 18.09553131 | 0.206151964 | 0.16153275 | 1.276223954 | 0.201876363 | 0.418948885 |
| Ulk2 | 4997.595689 | -0.076275723 | 0.059795105 | -1.27561817 | 0.202090528 | 0.419337169 |
| C77370 | 1487.71064 | -0.122235719 | 0.095833808 | -1.275496842 | 0.202133441 | 0.41937005 |
| Arrdc3 | 1899.947523 | 0.075327392 | 0.059097159 | 1.274636415 | 0.202437962 | 0.419384114 |
| C030037D09Rik | 151.5551914 | -0.181272058 | 0.142190757 | -1.274851205 | 0.202361913 | 0.419384114 |
| Cpt2 | 228.4015257 | 0.136843735 | 0.107352711 | 1.274711491 | 0.202411378 | 0.419384114 |
| Fxr1 | 2301.335885 | 0.092615726 | 0.072660363 | 1.27463892 | 0.202437075 | 0.419384114 |
| Gne | 1566.792001 | -0.07248684 | 0.056867861 | -1.274653887 | 0.202431775 | 0.419384114 |
| Mre11a | 368.2836338 | -0.123552668 | 0.09690961 | -1.274926888 | 0.202335121 | 0.419384114 |
| Pglyrp1 | 164.9321696 | -0.178398158 | 0.139956176 | -1.274671564 | 0.202425516 | 0.419384114 |
| Timp2 | 15280.10831 | 0.130035084 | 0.10195732 | 1.275387431 | 0.202172145 | 0.419384114 |
| Tmem198b | 380.6174795 | -0.164902179 | 0.129310361 | -1.275243358 | 0.202223119 | 0.419384114 |
| Usp27x | 661.2433355 | -0.118277192 | 0.092790968 | -1.27466276 | 0.202428633 | 0.419384114 |
| Zfp456 | 74.49593103 | 0.167493131 | 0.13134475 | 1.275217557 | 0.202232249 | 0.419384114 |
| Gpr135 | 192.5791984 | -0.137371498 | 0.107811635 | -1.274180637 | 0.202599406 | 0.419662459 |
| Actr3 | 7471.781012 | -0.076258311 | 0.059859073 | -1.273964107 | 0.202676137 | 0.419709177 |
| Bfsp1 | 25.10416442 | 0.211963658 | 0.166373062 | 1.27402631 | 0.202654092 | 0.419709177 |
| Acn9 | 164.4981979 | -0.145937114 | 0.114564768 | -1.273839387 | 0.202720343 | 0.419744621 |
| A930016O22Rik | 25.27742302 | 0.212857275 | 0.167150813 | 1.273444448 | 0.202860373 | 0.419922328 |
| Plch1 | 496.244255 | -0.098423661 | 0.07728804 | -1.273465615 | 0.202852867 | 0.419922328 |
| Prkd1 | 451.9781371 | -0.112244895 | 0.088148391 | -1.273362952 | 0.202889278 | 0.419926058 |
| Ehbp1 | 2293.058515 | 0.098047567 | 0.077019895 | 1.273016109 | 0.203012326 | 0.420066199 |
| Gdap1l1 | 2605.936575 | 0.103509692 | 0.081312106 | 1.272992397 | 0.20302074 | 0.420066199 |
| H13 | 2476.754898 | 0.070909506 | 0.055705176 | 1.27294285 | 0.203038322 | 0.420066199 |
| F2rl2 | 76.6447412 | 0.194101749 | 0.152534722 | 1.272508623 | 0.203192462 | 0.420270158 |
| Irgm2 | 273.4950514 | 0.200435169 | 0.157530292 | 1.272359536 | 0.203245404 | 0.420270158 |
| Lgi2 | 1298.765978 | -0.123069176 | 0.096723787 | -1.272377567 | 0.203239001 | 0.420270158 |
| Zfp383 | 472.3349313 | 0.132145655 | 0.103850768 | 1.272457177 | 0.20321073 | 0.420270158 |
| Ambra1 | 3315.641175 | 0.069992812 | 0.055017953 | 1.272181305 | 0.203308708 | 0.420344959 |
| Slc16a4 | 95.11597242 | -0.21380562 | 0.168087687 | -1.271988591 | 0.203377173 | 0.420430409 |
| Actn1 | 4256.179937 | 0.110701743 | 0.087053284 | 1.27165499 | 0.203495729 | 0.420609802 |
| Ncdn | 59910.32366 | 0.100755164 | 0.079235471 | 1.271591651 | 0.203518245 | 0.420609802 |
| Dnajb3 | 24.78591691 | -0.213673033 | 0.168121366 | -1.270945142 | 0.203748165 | 0.420956615 |
| Mcu | 1099.339641 | -0.095702856 | 0.075295136 | -1.271036363 | 0.203715712 | 0.420956615 |
| Nbas | 2118.092973 | -0.085298373 | 0.067117006 | -1.270890624 | 0.203767563 | 0.420956615 |
| Supt16 | 2902.551955 | 0.081918121 | 0.064476081 | 1.270519553 | 0.203899623 | 0.421173277 |
| Abhd15 | 50.80253228 | 0.213443052 | 0.168022643 | 1.270323142 | 0.203969548 | 0.421222205 |
| Gm11602 | 88.43500679 | 0.17263427 | 0.135906138 | 1.270246304 | 0.203996908 | 0.421222205 |
| Kifc3 | 1983.704375 | -0.078573337 | 0.061857861 | -1.270223952 | 0.204004868 | 0.421222205 |
| Cml1 | 288.073289 | -0.154952733 | 0.122013177 | -1.269967207 | 0.204096311 | 0.421255955 |
| Kirrel3 | 2011.439999 | -0.106681158 | 0.084004282 | -1.269949053 | 0.204102779 | 0.421255955 |
| Ptpdc1 | 1189.789808 | 0.114566277 | 0.090205814 | 1.270054244 | 0.204065309 | 0.421255955 |
| Nr1h3 | 104.3251176 | 0.149233153 | 0.117530282 | 1.269742151 | 0.204176493 | 0.421351969 |
| Cplx1 | 29668.84106 | 0.088273913 | 0.069535855 | 1.269473322 | 0.2042723 | 0.421381309 |
| Dcps | 448.422341 | -0.109212699 | 0.086024141 | -1.269558725 | 0.20424186 | 0.421381309 |
| Garnl3 | 3559.918321 | 0.130368086 | 0.102691254 | 1.269514994 | 0.204257446 | 0.421381309 |
| Adcyap1r1 | 9518.920507 | -0.099753813 | 0.078594244 | -1.269225438 | 0.204360671 | 0.421507486 |
| Ccdc130 | 425.550829 | 0.129787316 | 0.102268198 | 1.269087741 | 0.204409772 | 0.421552643 |
| Itgb3 | 216.678942 | 0.146269701 | 0.11527878 | 1.268834565 | 0.204500074 | 0.421576043 |
| Msantd1 | 73.83391776 | -0.176489791 | 0.139096804 | -1.268827078 | 0.204502745 | 0.421576043 |
| Pkp1 | 31.71495909 | 0.212302449 | 0.167313576 | 1.26888956 | 0.204480457 | 0.421576043 |
| Dzip3 | 2387.637588 | -0.128883847 | 0.101583897 | -1.268742884 | 0.204532782 | 0.421581871 |

| | | | | | | | |
|---|---|---|---|---|---|---|---|
| Oaf | 429.4222265 | 0.0656570271289857 | 0.0781396748578843 | 0.840252115822068 | 0.400767043268908 | 0.799169897076002 | NM_178644 |
| Ppp2r5c | 8689.531824 | 0.0403757750953292 | 0.0480604741763293 | 0.840103552603213 | 0.400850328383296 | 0.799169897076002 | NM_012023,NM_001081458,NM_001081457,NM_001135001 |
| Tdp2 | 594.3210224 | -0.0577361030104672 | 0.0686933405564626 | -0.840490541626971 | 0.40063340257191 | 0.799169897076002 | NM_019551 |
| Tnpo1 | 1964.34257 | -0.0480700212436354 | 0.0572152300416273 | -0.840161285878982 | 0.40081796165092 | 0.799169897076002 | NM_001048267,NM_178716 |
| Akt3 | 7124.061253 | 0.0560724932646839 | 0.0668795247681072 | 0.838410462082456 | 0.401800215637935 | 0.799277435820584 | NM_011785 |
| Apba2 | 8985.215446 | 0.051001321173598 | 0.0607863655248852 | 0.839025671846077 | 0.401454903783905 | 0.799277435820584 | NM_007461 |
| Atp5g2 | 3372.638699 | 0.0468996061087915 | 0.0558637053161867 | 0.839536257814287 | 0.401168451664805 | 0.799277435820584 | NM_026468 |
| Bap1 | 6634.385386 | 0.0374824404265005 | 0.0447207539081387 | 0.838144198183544 | 0.401949722474829 | 0.799277435820584 | NM_027088 |
| Btg2 | 880.436352 | -0.067024769544615 | 0.0798718362508446 | -0.839153983315443 | 0.401382906136734 | 0.799277435820584 | NM_007570 |
| Cd3eap | 901.7751579 | 0.0533392259790257 | 0.063595272538856 | 0.83872941886421 | 0.401621165747096 | 0.799277435820584 | NM_145822 |
| Coch | 1916.920801 | 0.0698761832712977 | 0.0832437429903151 | 0.839416642755088 | 0.401235547839847 | 0.799277435820584 | NM_007728,NM_001198835 |
| Creb1 | 2236.596178 | 0.0639624156915342 | 0.0762798615189454 | 0.838522965535904 | 0.40173705512247 | 0.799277435820584 | NM_133828,NM_009952,NM_001037726 |
| Dcun1d1 | 1792.300614 | 0.0613873046779962 | 0.0732029735745783 | 0.838590315125048 | 0.401699247269548 | 0.799277435820584 | NM_033623,NM_001205362,NM_001205361 |
| Gyg | 1172.456874 | 0.0542373476571961 | 0.0647071258744002 | 0.838197446174221 | 0.401919821127102 | 0.799277435820584 | NM_013755 |
| Klhl15 | 234.4710552 | 0.0773794599487766 | 0.0923053556621493 | 0.838298703186805 | 0.401862964050352 | 0.799277435820584 | NM_001039061,NM_001039059,NM_153165,NM_001039060 |
| Odf3b | 56.49885954 | -0.0677318367472922 | 0.0807823735680412 | -0.838448212842411 | 0.401779021335826 | 0.799277435820584 | NM_001013022 |
| Psma3 | 3206.687555 | 0.04656438861025 | 0.0555043926963074 | 0.838931593487156 | 0.401507697602657 | 0.799277435820584 | NM_011184 |
| Sybu | 3415.872903 | 0.0444762234945336 | 0.0529578969734778 | 0.83984119529535 | 0.400997432291727 | 0.799277435820584 | NM_176998,NM_001032727,NM_178765 |
| Tank | 957.9198326 | -0.0579495516114547 | 0.0691130377736115 | -0.838474960415946 | 0.401764004927502 | 0.799277435820584 | NM_001164072,NM_001164071,NM_011529 |
| Tomm20 | 16551.82095 | 0.0536117735779591 | 0.0638642865459975 | 0.839464064776577 | 0.401208946401095 | 0.799277435820584 | NM_024214 |
| Uhrf1bp1 | 1300.239847 | -0.045944759081134 | 0.054719984592672 | -0.839633991550627 | 0.401113634473138 | 0.799277435820584 | NM_001080769 |
| Uqcr10 | 2198.325799 | 0.0485602974164014 | 0.0579168553722903 | 0.838448446557659 | 0.401778890123826 | 0.799277435820584 | NM_197979 |
| Zfp952 | 713.1664427 | -0.0579234854401545 | 0.0690295716512141 | -0.839111181695067 | 0.401406921958729 | 0.799277435820584 | NM_001045559 |
| 2310011J03Rik | 1111.564964 | 0.0486229064481436 | 0.0580589820135967 | 0.837474319421528 | 0.402326006110241 | 0.799551549968885 | NM_025521 |
| Atp5h | 6144.417114 | 0.0481562363099139 | 0.057512968943604 | 0.837310909077478 | 0.40241782884638 | 0.799551549968885 | NM_027862 |
| Btaf1 | 1780.846278 | -0.0658260319543977 | 0.0785916945240365 | -0.837569826596187 | 0.402272345006828 | 0.799551549968885 | NM_001080706 |
| Casc4 | 4107.250857 | 0.0518313695559953 | 0.061874325936517 | 0.837687825628585 | 0.402206052687288 | 0.799551549968885 | NM_177054,NM_199038,NM_001205370,NM_001205371,NM_001205369 |
| Gspt2 | 1328.582331 | 0.0505498613105453 | 0.0603709533997119 | 0.837320904572405 | 0.402412211866297 | 0.799551549968885 | NM_008179 |
| Neurod2 | 2832.278244 | -0.0558306371657921 | 0.0666624786285071 | -0.837512170480817 | 0.402304738821643 | 0.799551549968885 | NM_010895 |
| Fgf17 | 24.35879781 | 0.0692298054504435 | 0.0827072539651134 | 0.83704635484144 | 0.402566512496759 | 0.799737576658632 | NM_008004 |
| Olfm2 | 4335.832509 | 0.0586980788487726 | 0.0701342726476456 | 0.836938584130922 | 0.402627090774137 | 0.799748546504187 | NM_173777 |
| A530058N18Rik | 85.40079712 | 0.0793063352140311 | 0.0948440164155219 | 0.836176473870333 | 0.403055631439236 | 0.800009209064826 | NR_028423 |
| Arhgef18 | 2528.739716 | 0.0404821814853101 | 0.0483969207357232 | 0.836461925054438 | 0.402895087964704 | 0.800009209064826 | NM_133962 |
| Leprotl1 | 3127.497582 | 0.0469156793859666 | 0.0561003153620883 | 0.836281918972446 | 0.402996322533777 | 0.800009209064826 | NM_026609 |
| Morc2b | 58.45681588 | 0.079195178847079 | 0.094680591987758 | 0.836445750754482 | 0.402904183691278 | 0.800009209064826 | NM_177719 |
| Mtif3 | 668.8937555 | 0.0571101743847564 | 0.0683039998786038 | 0.83611756978007 | 0.403088765052008 | 0.800009209064826 | NM_001256102,NM_029581,NM_001256101,NM_001256100 |
| Zw10 | 899.2569113 | -0.0564198800251196 | 0.0674661043655719 | -0.836270013745018 | 0.403003018514703 | 0.800009209064826 | NM_012039 |
| Adam18 | 23.43330585 | 0.0701440231234731 | 0.0840102246020219 | 0.834946263454995 | 0.403747964908148 | 0.800224169498672 | NM_010084 |
| Clip2 | 7633.273073 | 0.043623083903988 | 0.0522270125220777 | 0.835259031627501 | 0.403571878877986 | 0.800224169498672 | NM_001039162,NM_009990 |
| Fbxl3 | 3282.596061 | -0.0517313103381269 | 0.0619220347173754 | -0.835426525860124 | 0.403477599848263 | 0.800224169498672 | NM_015822 |
| Morc3 | 961.643923 | -0.0552479574062829 | 0.0661385531823336 | -0.835336649321205 | 0.403528187847938 | 0.800224169498672 | NM_001045529 |
| Mrpl27 | 1210.001676 | 0.0535313153178041 | 0.0640613616825413 | 0.835625623805511 | 0.403365548903984 | 0.800224169498672 | NM_053161 |
| Nup37 | 122.9410103 | -0.0755386343413976 | 0.0904652996984418 | -0.83500120591209 | 0.403717029405375 | 0.800224169498672 | NM_028334,NM_027191 |
| Pde4b | 5750.642139 | 0.0533162706971083 | 0.0638168351228695 | 0.835457769011202 | 0.403460015186293 | 0.800224169498672 | NM_001177982,NM_001177983,NM_019840,NM_001177981,NM_001177980 |
| Ppp1r9a | 17055.89033 | 0.0608012378659526 | 0.0728078076457549 | 0.835092276940682 | 0.403665754736142 | 0.800224169498672 | NM_181595 |
| Tbc1d25 | 606.2077326 | 0.0533191803553829 | 0.0638534264307991 | 0.835024576373633 | 0.403703871032443 | 0.800224169498672 | NM_172478,NM_001166437 |
| Tnni3k | 29.04161803 | 0.073438453550481 | 0.0879400413603647 | 0.835096872988058 | 0.403663167180038 | 0.800224169498672 | NM_177066 |
| 1110037F02Rik | 2146.560702 | -0.0530169307239714 | 0.0635172584659502 | -0.83468543832685 | 0.403894842579067 | 0.80026024324482 | NM_001081183 |
| A530072M11Rik | 25.47318442 | -0.0686727803410237 | 0.0822802506009265 | -0.834620456786144 | 0.403931440415188 | 0.80026024324482 | NR_045765 |
| Acad11 | 1249.1382 | -0.0466536751824074 | 0.0558908287154582 | -0.834728635353085 | 0.403870514957753 | 0.80026024324482 | NM_175324 |
| 1810009A15Rik | 372.5924978 | 0.0659098326503094 | 0.0790927969238805 | 0.833322820961073 | 0.404662689078405 | 0.800290023356422 | NM_025463 |
| A930017M01Rik | 584.331785 | 0.0636935648348114 | 0.0764253748274337 | 0.833408602556804 | 0.404614324682366 | 0.800290023356422 | NR_033609 |
| Anapc5 | 8454.438082 | -0.0337951621221257 | 0.0405489472327624 | -0.833441172421369 | 0.404595962420245 | 0.800290023356422 | NM_021505,NM_001042491 |
| C230052I12Rik | 190.1832651 | 0.0693555155194727 | 0.0831620582944829 | 0.83398026626373 | 0.404292104105693 | 0.800290023356422 | NM_178643 |
| Casp8ap2 | 638.3047575 | 0.0720729364814495 | 0.0864498094166296 | 0.833696881089773 | 0.404451816118559 | 0.800290023356422 | NM_001122978,NM_011997 |
| Cxcr7 | 361.5060714 | -0.0626738332203619 | 0.0751344535758799 | -0.8341557067034 | 0.404193247185504 | 0.800290023356422 | NM_001271607,NM_007722 |
| Foxo1 | 1562.688142 | 0.0518797205131362 | 0.062187150400206 | 0.834251451936033 | 0.404139302904616 | 0.800290023356422 | NM_019739 |
| Gadd45g | 334.4906166 | 0.0722466240266464 | 0.0866496526448873 | 0.833778576386587 | 0.404405769893671 | 0.800290023356422 | NM_011817 |
| Mycbp | 167.0846487 | 0.077633770977679 | 0.0931235653737452 | 0.833664074889112 | 0.404470307682404 | 0.800290023356422 | NM_019660 |
| Qrich1 | 3703.13536 | 0.0380790974533117 | 0.04566593741216 | 0.833862165351541 | 0.404358659579808 | 0.800290023356422 | NM_001114119,NM_175143 |
| Rnf214 | 2692.522217 | 0.0471379001943385 | 0.0565245664885585 | 0.833936518626464 | 0.404316757213087 | 0.800290023356422 | NM_178709 |
| Rragc | 3379.268249 | 0.0483787858346676 | 0.058043538305043 | 0.833491328189142 | 0.404567686539512 | 0.800290023356422 | NM_017475 |
| Trmt12 | 377.8811517 | 0.0609116754837627 | 0.0730241460223579 | 0.834130610238335 | 0.404207387617977 | 0.800290023356422 | NM_026642 |
| Ctdspl2 | 886.5322965 | 0.0687128482984993 | 0.0824671258712344 | 0.833215024442451 | 0.404723470578117 | 0.80030127057143 | NM_212450 |
| Ebf2 | 36.50106641 | 0.0704235513920439 | 0.0845952016036364 | 0.832476902437178 | 0.405139810301911 | 0.800502700395235 | NM_001276387,NM_010095 |
| Eif2ak3 | 1498.699783 | 0.0473881620043903 | 0.0569462983648558 | 0.832155264961625 | 0.40532131084999 | 0.800502700395235 | NM_010121 |
| Exosc2 | 727.9633689 | 0.0542478843829646 | 0.065186583695116 | 0.832194008458662 | 0.405299445258035 | 0.800502700395235 | NM_144886 |
| Grin2b | 1839.507671 | 0.077003993422687 | 0.0924775536345313 | 0.832677664971595 | 0.405026544309358 | 0.800502700395235 | NM_008171 |
| Heatr1 | 1226.754739 | -0.0541671359247554 | 0.0650479298454717 | -0.83272651494729 | 0.404998987046018 | 0.800502700395235 | NM_144835 |
| Pacs1 | 4929.724261 | 0.0401541679860429 | 0.0482316305684644 | 0.832527690081811 | 0.40511115519383 | 0.800502700395235 | NM_153129 |
| Rpl4 | 17930.86843 | 0.0400972822179334 | 0.0481766493624167 | 0.832297030793799 | 0.405241306173132 | 0.800502700395235 | NM_024212 |
| Snx8 | 586.4766663 | 0.0708282168755851 | 0.0850956496285485 | 0.832336519960277 | 0.405219022385368 | 0.800502700395235 | NM_172277 |
| Zfp938 | 615.7400012 | 0.0653679974897899 | 0.0785287202242597 | 0.83240879646471 | 0.405178238557409 | 0.800502700395235 | NM_001105557 |
| Tmem123 | 521.6902769 | -0.0580171269543625 | 0.069736255722391 | -0.831950702735167 | 0.40543677092253 | 0.800621877979972 | NM_133739 |
| Adgb | 51.86077785 | -0.0733815701397345 | 0.0883128619512535 | -0.830927324948877 | 0.406014686158506 | 0.800733644061662 | NM_001127353 |
| Akr1b10 | 664.2299068 | -0.0597877972573375 | 0.0719229916531257 | -0.831275172001821 | 0.405818197073383 | 0.800733644061662 | NM_172398 |
| Copa | 11806.98066 | 0.0347736502016939 | 0.0418280548110253 | 0.831347533582366 | 0.405777329175266 | 0.800733644061662 | NM_009938 |
| Ext2 | 1642.06644 | 0.0410684300446588 | 0.0494091348560903 | 0.831191037128564 | 0.405865717305297 | 0.800733644061662 | NM_010163 |
| H2afj | 634.5104347 | -0.0701764932568546 | 0.0843953009703758 | -0.831521334126028 | 0.405679181289565 | 0.800733644061662 | NM_177688 |
| Mrpl38 | 1438.179278 | 0.0553094528463247 | 0.0665635456707003 | 0.830927083120672 | 0.406014822780263 | 0.800733644061662 | NM_024177 |
| Ppm1h | 4513.184505 | 0.0550671429573917 | 0.0662676488722502 | 0.830982792535711 | 0.405983350261122 | 0.800733644061662 | NM_001110218,NM_176919 |
| Rph3a | 21853.62779 | 0.047014144208352 | 0.056545511381341 | 0.831439013634348 | 0.405725667191005 | 0.800733644061662 | NM_011286 |
| Tbc1d8 | 1340.760951 | -0.0664824713851601 | 0.0799728000314481 | -0.831313538590831 | 0.405796528337092 | 0.800733644061662 | NM_018775 |
| Zfp513 | 873.6875304 | -0.0558436183422675 | 0.0672106653237605 | -0.830874357116734 | 0.406044611190844 | 0.800733644061662 | NM_175311,NM_001177901 |
| 1700084C01Rik | 485.6556735 | -0.0720483208444866 | 0.0867722544517602 | -0.830315188878039 | 0.406360602653528 | 0.800759396647402 | NM_001033185 |
| Atg3 | 1578.929811 | 0.0469945883624185 | 0.0565901349294605 | 0.83043782137995 | 0.406291289259004 | 0.800759396647402 | NM_026402 |
| Ccdc136 | 3238.855686 | 0.0364382938360093 | 0.0438892786821382 | 0.830232232794445 | 0.406407494451848 | 0.800759396647402 | NM_001201378,NM_145574 |
| Eps8l2 | 243.3265256 | 0.0721995830355087 | 0.0869421990041813 | 0.830431986566574 | 0.406294587007178 | 0.800759396647402 | NM_133191 |
| Mettl21d | 424.1845371 | -0.0644824855219146 | 0.0776739777041774 | -0.830168448016106 | 0.406443551664691 | 0.800759396647402 | NM_001033236 |
| Mrps35 | 568.8587691 | 0.0600968841503423 | 0.0723723017921268 | 0.830385142688389 | 0.406321063036956 | 0.800759396647402 | NM_145573 |
| Zswim5 | 887.4070209 | -0.0501700678282456 | 0.060398311825536 | -0.830653478745644 | 0.406169414191082 | 0.800759396647402 | NM_001029912 |
| Dtymk | 604.4486659 | -0.062168994401422 | 0.0749351956775216 | -0.82963677934414 | 0.406744175560496 | 0.8009171606604 | NM_001105667,NM_023136 |
| Ttpa | 109.7038733 | 0.0771648814902836 | 0.0929901625697439 | 0.82981768563324 | 0.406641869985893 | 0.8009171606604 | NM_015767 |
| Vil1 | 27.57050787 | 0.0697661288810612 | 0.0840853417806794 | 0.829706193774331 | 0.406704918707281 | 0.8009171606604 | NM_009509 |
| Xrn2 | 2046.313461 | 0.0495652151840873 | 0.0597234839600618 | 0.829911651122571 | 0.406588736967866 | 0.8009171606604 | NM_011917 |
| Atg16l1 | 2835.26567 | -0.0408113762174468 | 0.0492162787756053 | -0.829225151367509 | 0.406977015381603 | 0.80097579798674 | NM_029846,NM_001205392,NM_001205391 |
| Dpf3 | 114.3367124 | -0.0788233829861921 | 0.095032873699827 | -0.829432804854092 | 0.406859545011737 | 0.80097579798674 | NM_001267626,NM_058212,NM_001267625 |
| Fgfr1op | 648.231962 | -0.0627825188964762 | 0.0757328967285122 | -0.828999306886932 | 0.407104799431096 | 0.80097579798674 | NM_201230,NM_001197046 |
| Mt2 | 2668.855975 | 0.0788104924700348 | 0.0950381039610248 | 0.82925152318227 | 0.406962095623906 | 0.80097579798674 | NM_008630 |
| Pkp1 | 39.04563477 | -0.0724740791107665 | 0.0874143486603506 | -0.829086760027983 | 0.407055315124014 | 0.80097579798674 | NM_019645 |
| Ptpn1 | 2173.654056 | 0.0653820728322233 | 0.0788650622672581 | 0.829037230841921 | 0.407083340177134 | 0.80097579798674 | NM_011201 |
| A330102I10Rik | 93.60835047 | -0.0788244491329241 | 0.0951002615722927 | -0.828856281041917 | 0.407185736615415 | 0.801026545243163 | NR_045073,NR_045074,NR_045075,NR_045077 |
| 4930519F09Rik | 28.68851355 | 0.0718674772041741 | 0.0871160212485746 | 0.824962804477827 | 0.409392708147879 | 0.801055189437303 | NR_033601 |
| Alg5 | 542.5297613 | -0.059236688759506 | 0.0716480172574093 | -0.826773594399505 | 0.408365399280038 | 0.801055189437303 | NM_025442 |
| Arid3a | 290.3720964 | -0.064289570097204 | 0.0777502135407423 | -0.826873228631267 | 0.408308918750532 | 0.801055189437303 | NM_007880 |
| Atl2 | 2476.163667 | 0.0525909531956641 | 0.0636613340592225 | 0.826105107171335 | 0.408744470814419 | 0.801055189437303 | NM_178050,NM_019717 |
| Bmpr1a | 2471.165121 | 0.052917569432658 | 0.0640541542046376 | 0.826137977930348 | 0.408725826267283 | 0.801055189437303 | NM_009758 |
| Cyp2j9 | 1236.884403 | 0.0706063234087655 | 0.0854688618040254 | 0.826105811150981 | 0.408744071506448 | 0.801055189437303 | NM_028979 |
| Dbndd1 | 613.2349542 | 0.0575084459686741 | 0.0695210720390532 | 0.827208848798663 | 0.408118697043243 | 0.801055189437303 | NM_001170976,NM_001170975,NM_028146 |
| Dgat2 | 2718.270851 | 0.0531212090714777 | 0.0642462562590168 | 0.826837424694646 | 0.408329214706049 | 0.801055189437303 | NM_026384 |
| Dnajb9 | 1408.333355 | 0.0624836281760606 | 0.0757462107284099 | 0.824907643236403 | 0.409424026684422 | 0.801055189437303 | NM_013760 |
| Enox1 | 1157.572031 | 0.051799467759487 | 0.0626767426515039 | 0.826454368369191 | 0.408546393069761 | 0.801055189437303 | NM_001253759,NM_172813 |
| Ephx1 | 1207.60371 | 0.0583362774597685 | 0.0704700605341947 | 0.827816480042068 | 0.407774440127047 | 0.801055189437303 | NM_010145 |
| Eri1 | 610.877321 | 0.0709960658035526 | 0.0859631845675877 | 0.825889200832626 | 0.408866947133061 | 0.801055189437303 | NM_026067 |
| F13a1 | 88.92591791 | -0.076117499946331 | 0.0922587573464429 | -0.825043628763615 | 0.409346821646757 | 0.801055189437303 | NM_028784,NM_001166391 |
| Fmnl2 | 2158.147751 | 0.0466694644011146 | 0.056382147633991 | 0.82773477704455 | 0.407820719349079 | 0.801055189437303 | NM_172409 |
| Gab2 | 1188.732872 | 0.0492695606834239 | 0.059667679606438 | 0.825732808924377 | 0.408955676580943 | 0.801055189437303 | NM_010248,NM_001162477 |
| Gemin4 | 494.8568323 | 0.0615204959902662 | 0.0744673969087519 | 0.826140009508456 | 0.40872467395718 | 0.801055189437303 | NM_177367 |
| Hmgxb4 | 738.0119501 | -0.0510013316739913 | 0.061662769327386 | -0.827100894596705 | 0.408179877214476 | 0.801055189437303 | NM_178017 |
| Ift43 | 254.7814741 | 0.0656018030749301 | 0.0792165842513693 | 0.828132185891317 | 0.407595643531627 | 0.801055189437303 | NM_001199843,NM_029601 |
| Kcnq1ot1 | 177.0989626 | -0.0779991398985179 | 0.0944264274562627 | -0.826030826323979 | 0.408786605346691 | 0.801055189437303 | NR_001461 |
| Klhl14 | 119.7464575 | 0.0746067743762713 | 0.0900844092250334 | 0.828187419089373 | 0.407564367610617 | 0.801055189437303 | NM_001081403 |
| Magi1 | 4750.490204 | 0.0365419135990478 | 0.044107745452097 | 0.828469313597856 | 0.40740476650337 | 0.801055189437303 | NM_010367,NM_001029850,NM_001083321,NM_001083320 |
| Mphosph10 | 724.6335733 | 0.068145917177943 | 0.0825202870335882 | 0.82580804827067 | 0.40891298786750 | 0.801055189437303 | NM_026483 |
| Nat14 | 2264.713393 | 0.0481052767147751 | 0.0581795548459188 | 0.826841608571531 | 0.408326842986701 | 0.801055189437303 | NM_201355 |
| Nol12 | 660.6952756 | 0.0545949501716124 | 0.0660183662099402 | 0.826966089981672 | 0.408256281812848 | 0.801055189437303 | NM_133800 |
| Nudt10 | 500.7109192 | 0.0669955880807948 | 0.0811936577089108 | 0.825133267440446 | 0.409295934515806 | 0.801055189437303 | NM_001031664 |
| Pbx3 | 586.9501735 | 0.0732804094070728 | 0.0887723492160281 | 0.82548687800009 | 0.409095229453006 | 0.801055189437303 | NM_016768 |
| Prom1 | 681.4578636 | -0.0539815595502771 | 0.0654095245588976 | -0.825285918439443 | 0.409209284476097 | 0.801055189437303 | NM_001163583,NM_001163577,NM_001163581,NM_008935,NM_001163585,NM_001163582,NM_001163584,NM_001163578 |
| S100a3 | 22.71995486 | -0.0679235851985319 | 0.0821356526946934 | -0.826968350163489 | 0.408255000716138 | 0.801055189437303 | NM_011310 |
| S100pbp | 719.0841065 | 0.0619858222101723 | 0.0751487216445287 | 0.824842004676805 | 0.409461295716093 | 0.801055189437303 | NM_029036 |
| Slc16a13 | 200.9155014 | 0.0747003648770325 | 0.090226417625143 | 0.827921210253349 | 0.407715122123444 | 0.801055189437303 | NM_172371 |
| Slc16a3 | 105.5333292 | -0.0776565303975317 | 0.0937866006982736 | -0.828013061773772 | 0.40766310269759 | 0.801055189437303 | NM_001038654,NM_030696,NM_001038653 |
| Smim7 | 3574.660152 | 0.0445946457436861 | 0.0540479958588729 | 0.825093419932335 | 0.409318555149926 | 0.801055189437303 | NM_172396 |
| Smtnl2 | 49.70303081 | 0.0750252684463305 | 0.0909005393970988 | 0.825355591330242 | 0.409169739337293 | 0.801055189437303 | NM_177776 |
| Snapc1 | 1113.25254 | 0.0556770821565832 | 0.0673441969383989 | 0.826753969722323 | 0.408376524641137 | 0.801055189437303 | NM_178392 |
| Srgap3 | 17147.66138 | -0.0493849370093139 | 0.0598493378970087 | -0.82515427479411 | 0.409284009360303 | 0.801055189437303 | NM_080448 |
| Zdhhc6 | 1091.707145 | 0.0492609727609618 | 0.0595434806562426 | 0.827310936781745 | 0.4080608464214 | 0.801055189437303 | NM_001033573,NM_025883 |
| Zfp12 | 1588.944215 | 0.0442797272299502 | 0.053593952532463 | 0.82620753158911 | 0.408686376566595 | 0.801055189437303 | NM_177681 |
| Zfp120 | 557.4392678 | -0.0651398681832145 | 0.0788620528942842 | -0.825997622335999 | 0.408805440573567 | 0.801055189437303 | NM_023266,NM_181266 |
| Zfp13 | 639.4795662 | 0.0583163550840215 | 0.070505503157818 | 0.827117777650454 | 0.408170308832654 | 0.801055189437303 | NM_011747 |
| Zfp687 | 1696.216077 | -0.058216805759302 | 0.0703595800685264 | -0.827418323170804 | 0.407999998602526 | 0.801055189437303 | NM_030074 |
| Zmiz1 | 6664.247321 | 0.0508196446981535 | 0.0615811039358807 | 0.825247380285156 | 0.409231159048769 | 0.801055189437303 | NM_183208 |
| Fbxo47 | 32.37521958 | 0.0688702523872603 | 0.0835194966968161 | 0.824600902915711 | 0.409598208621674 | 0.801196796154758 | NM_001081435 |
| Gm20748 | 27.91853773 | 0.0702010497628606 | 0.0851517171955519 | 0.824423183406191 | 0.409699146484755 | 0.801196796154758 | NR_045420 |
| Plxdc2 | 1343.016489 | -0.0525275058002068 | 0.0637095078206729 | -0.824484564345705 | 0.40966428279401 | 0.801196796154758 | NM_026162 |
| Nfe2l3 | 162.2934901 | -0.0780533824071539 | 0.0947143193042556 | -0.824092734662634 | 0.409886868009638 | 0.801456002787454 | NM_010903 |
| Bri3 | 985.2649874 | 0.05045175052182 | 0.0612694122463716 | 0.823441072340429 | 0.410257214564878 | 0.80184318968123 | NM_018772,NM_001163709 |
| Ikzf4 | 523.2407709 | -0.0608388639725175 | 0.0738785102686117 | -0.823498792156421 | 0.410224403761625 | 0.80184318968123 | NM_011772 |
| Rpl19 | 10130.44555 | 0.0440132364558556 | 0.0534379119968111 | 0.823633162509831 | 0.410148027036361 | 0.80184318968123 | NM_009078,NM_001159483 |
| Rpp25l | 396.6345551 | 0.0646134660291195 | 0.0784757522078141 | 0.823355803688438 | 0.410305688345077 | 0.80184318968123 | NM_027278 |
| Ccnc | 816.7597675 | 0.0567585056821373 | 0.0689775875338553 | 0.822854317053048 | 0.410590843841095 | 0.80218461299741 | NM_016746,NM_001122982 |
| Strn | 2837.023923 | 0.0676350904724203 | 0.0821883396802704 | 0.822928054460459 | 0.410548907871181 | 0.80218461299741 | NM_011500 |
| Gm5148 | 99.90802547 | 0.0777196196078084 | 0.0944754334221417 | 0.822643694689774 | 0.410710643095964 | 0.802310758689077 | NM_198657 |
| Cdsn | 111.2833396 | 0.0761337564286724 | 0.0926339584431945 | 0.821877394728409 | 0.411146679670027 | 0.802622855649351 | NM_001008424 |
| Exoc1 | 2421.699006 | -0.0486566338867285 | 0.0592003112735423 | -0.821898277897637 | 0.411134793181062 | 0.802622855649351 | NM_027270 |
| Fam198b | 223.8975497 | -0.0724109282188519 | 0.0880803278420479 | -0.822101029740766 | 0.411019399486243 | 0.802622855649351 | NM_133187 |
| Pradc1 | 507.825454 | -0.057829334826184 | 0.0703539934625925 | -0.821976578443014 | 0.411090227118175 | 0.802622855649351 | NM_028505,NM_001163427,NR_028081 |
| Siglece | 40.75404433 | -0.0754991241981563 | 0.0918275798522988 | -0.822183534833368 | 0.410972448242268 | 0.802622855649351 | NM_031181 |
| AA387883 | 82.15072352 | 0.0780047982415043 | 0.0949547274611585 | 0.821494625145572 | 0.411364584387767 | 0.802795661018829 | NR_030678 |
| AW495222 | 132.0647537 | -0.0777406165228101 | 0.0947033735240053 | -0.820885398587248 | 0.411711548842929 | 0.802795661018829 | NR_045086 |
| Amer2 | 891.4527147 | 0.061271030226679 | 0.0746732117074322 | 0.82052223047185 | 0.411918461575965 | 0.802795661018829 | NM_001164705,NM_028113 |
| Bet1l | 1074.042338 | 0.0472911963990349 | 0.0576389444073086 | 0.820472978562015 | 0.411946527288394 | 0.802795661018829 | NM_018742 |
| Ccp110 | 3366.564975 | -0.0568972361848767 | 0.0693164248307057 | -0.820833393006624 | 0.411741174909642 | 0.802795661018829 | NM_182995 |
| Lmln | 543.5146396 | 0.0560989585385423 | 0.0683307622877436 | 0.820991258699958 | 0.411651247323055 | 0.802795661018829 | NM_172823 |
| Mmp16 | 862.3429657 | 0.058156869028902 | 0.0708605797859527 | 0.820722455342243 | 0.411804377096845 | 0.802795661018829 | NM_019724 |
| Pbrm1 | 2244.85273 | 0.0568849444456214 | 0.0693416179411874 | 0.820357905318403 | 0.412012105055675 | 0.802795661018829 | NM_001081251 |
| Rad21 | 5399.474624 | 0.0468413160199999 | 0.0570865806738431 | 0.820531120748356 | 0.411913395660895 | 0.802795661018829 | NM_009009 |
| Sec24c | 4396.030741 | -0.0358603793812527 | 0.0437144496161697 | -0.820332400296038 | 0.412026640657679 | 0.802795661018829 | NM_172596,NM_001168273 |
| Tcaim | 744.5200474 | 0.0614403284089033 | 0.0747868989841921 | 0.821538655077678 | 0.411339515349987 | 0.802795661018829 | NM_001013405 |
| Tubgcp4 | 960.196399 | -0.0542492755625012 | 0.0660974357509863 | -0.820747052380043 | 0.411790363446204 | 0.802795661018829 | NM_153387 |
| Ube3a | 1981.932883 | 0.0599915533504182 | 0.0731366602958092 | 0.820266513507395 | 0.412064191694643 | 0.802795661018829 | NM_001033962,NM_173010,NM_011668 |
| Zbtb48 | 414.094628 | -0.0677675844454589 | 0.0826050391386707 | -0.820380755848273 | 0.411999082537396 | 0.802795661018829 | NM_133879 |
| Znhit6 | 570.2062352 | -0.0528128306826486 | 0.0643827788460119 | -0.820294364879219 | 0.412048318038788 | 0.802795661018829 | NM_001081094 |
| Cftr | 60.39311418 | 0.0779371857511611 | 0.0950325087558053 | 0.820110789155612 | 0.412152952165572 | 0.80286090695415 | NM_021050 |
| Mpi | 2125.942331 | 0.048585384915026 | 0.0593010104794481 | 0.819301130321619 | 0.4126146278554 | 0.803544722379347 | NM_025837 |
| Tmem184c | 2909.542566 | 0.0538606910058424 | 0.0657324332199886 | 0.819392929295426 | 0.412562267748191 | 0.803544722379347 | NM_145599 |
| Hkdc1 | 630.9474858 | -0.07189447779582 | 0.0877762709240764 | -0.819065073498126 | 0.412749287537577 | 0.803699215921025 | NM_145419 |
| Tenm4 | 3283.136251 | 0.059942384245374 | 0.073205709589043 | 0.81882116274638 | 0.41288845488093 | 0.803781966244235 | NM_011858 |
| Zfp385b | 1546.082354 | 0.0559973304000181 | 0.0683897908573619 | 0.818796631748878 | 0.412902452988743 | 0.803781966244235 | NM_001113399,NM_001113400,NM_178723 |
| Znfx1 | 4274.098568 | -0.0452517942740777 | 0.0552743663740306 | -0.818675947687357 | 0.412971322953658 | 0.803808312638997 | NM_001033196 |
| Pphln1 | 1632.405479 | 0.0543482895705059 | 0.0663964635421445 | 0.818541932372781 | 0.413047808549241 | 0.8038494730689 | NM_001083114,NM_146062,NM_175363 |
| 4933411K20Rik | 1410.738907 | 0.0562686124337007 | 0.0687594681962533 | 0.818339843366718 | 0.413163161218832 | 0.803854924132843 | NM_025747 |
| Gucy1b3 | 7564.613198 | 0.0577536266328114 | 0.0705822129352829 | 0.8182461874037 | 0.413216626634995 | 0.803854924132843 | NM_017469,NM_001161796 |
| Jph1 | 883.4550962 | -0.0552857893248978 | 0.0675637165308751 | -0.818276319466051 | 0.413199424410561 | 0.803854924132843 | NM_020604 |
| Myh2 | 57.6902851 | 0.075630046329108 | 0.092471843436446 | 0.817871078574172 | 0.413430806226958 | 0.804056217867558 | NM_001039545 |
| Phf1 | 1917.197075 | -0.0532212144262402 | 0.0650716364052918 | -0.817886522704876 | 0.413421986641906 | 0.804056217867558 | NM_009343 |
| Cnot2 | 2460.178025 | 0.0390553005209432 | 0.0477598668635035 | 0.817743077733567 | 0.41350390717017 | 0.804090730325499 | NM_001037848,NR_024327,NM_028082,NM_001037847,NM_001037846 |
| Ndufb4 | 1792.126137 | 0.0543313294345178 | 0.0664515065266859 | 0.817608693532015 | 0.413580661867941 | 0.804132337611258 | NM_026610 |
| Hira | 2847.834971 | 0.0409987247273761 | 0.0501569646750663 | 0.817408409639213 | 0.413695071395108 | 0.804247136922557 | NM_010435 |
| Ccnh | 1317.370645 | 0.058190959901871 | 0.0712152526116164 | 0.817113718871779 | 0.413863443658769 | 0.804285711027944 | NM_023243 |
| Laptm4a | 6802.187437 | 0.0449489542677597 | 0.0550222565480336 | 0.816923134159719 | 0.413972356274556 | 0.804285711027944 | NM_008640 |
| Mmp19 | 26.94178065 | -0.0709775744532679 | 0.0868746003885233 | -0.817011809387781 | 0.413921679318337 | 0.804285711027944 | NM_021412,NM_001164197 |
| Pcgf3 | 1464.532127 | 0.0612197024986547 | 0.0749308496904366 | 0.817015992099022 | 0.413919289033712 | 0.804285711027944 | NM_172716 |
| Vwa8 | 2461.60373 | -0.0543513619471668 | 0.0665345590628756 | -0.81688918830595 | 0.413991756943132 | 0.804285711027944 | NM_027906,NM_173758 |
| Aasdhppt | 749.0952049 | 0.060028023097317 | 0.0735240649332809 | 0.816440483150505 | 0.414248250596284 | 0.804541293013405 | NM_026276 |
| Dpyd | 258.7254624 | 0.069120294347489 | 0.084668019367721 | 0.816368386359591 | 0.414289472101079 | 0.804541293013405 | NM_170778 |
| Lamtor2 | 1010.401644 | 0.0541222956992038 | 0.0662892002137203 | 0.816457216614138 | 0.414238685262357 | 0.804541293013405 | NM_031248 |
| Tbc1d12 | 623.5509863 | 0.0663470743733977 | 0.0813132984249066 | 0.815943709806209 | 0.414532331124813 | 0.804797733482896 | NM_145952 |
| Tmod3 | 934.4154551 | -0.0552401743549664 | 0.0676949795091625 | -0.816015822081601 | 0.414491086473346 | 0.804797733482896 | NM_016963 |
| Ppip5k1 | 6460.580396 | -0.0353437791133332 | 0.043332226065152 | -0.815646513525343 | 0.414702338294853 | 0.805020201265674 | NM_178795 |
| Glg1 | 5700.177223 | 0.0520421659344277 | 0.0638171399186671 | 0.815488848305546 | 0.41479254531466 | 0.805087722239545 | NM_009149 |
| Hist2h3c2 | 43.44213869 | -0.0735749163463708 | 0.090234216820489 | -0.815377125650018 | 0.414856473648959 | 0.805104226616536 | NM_054045 |
| Amn1 | 324.1682098 | 0.06387521805977 | 0.0783701556023659 | 0.815045186127481 | 0.415046445673632 | 0.805365304549183 | NM_001113424 |
| C77080 | 1226.815923 | 0.061679018672745 | 0.0757224517014992 | 0.814540698125914 | 0.415335267197021 | 0.805633085437867 | NM_001033189 |
| Commd3 | 1522.911871 | 0.0435106045446021 | 0.0534618891537883 | 0.813862084436253 | 0.415723963650454 | 0.805633085437867 | NM_147778 |
| Cyp4f15 | 612.5017202 | 0.0698236159701676 | 0.0857957117646027 | 0.813835732976286 | 0.415739061575261 | 0.805633085437867 | NM_134127 |
| Gca | 549.105828 | -0.0693027577686158 | 0.0851435004212047 | -0.813952414755973 | 0.415672211845369 | 0.805633085437867 | NM_145523 |
| Gm5820 | 38.52581905 | 0.0708638525381905 | 0.0870311442451252 | 0.814235560762028 | 0.41551001724959 | 0.805633085437867 | NM_001033789 |
| Hpcal4 | 50641.00071 | 0.0517061867539856 | 0.0634792318697703 | 0.814537057727203 | 0.415337351772166 | 0.805633085437867 | NM_174998 |
| Ntn1 | 527.119091 | 0.0686189134412986 | 0.0842715829308698 | 0.814259220662658 | 0.415496465835553 | 0.805633085437867 | NM_008744 |
| Pomc | 190.1549055 | -0.0468903082181316 | 0.0575993582429761 | -0.814076921140864 | 0.41560088621706 | 0.805633085437867 | NM_008895 |
| Scmh1 | 1895.125065 | -0.0385128911031767 | 0.0473054578382004 | -0.814132086722486 | 0.415569285973975 | 0.805633085437867 | NM_013883,NM_001159630 |
| T2 | 35.52360724 | -0.0694512140117593 | 0.0852744740076201 | -0.81444057385365 | 0.415392602056325 | 0.805633085437867 | NM_001161832 |
| Arhgef40 | 984.7553031 | -0.057807627118699 | 0.0710513936764378 | -0.81360300097631 | 0.415872418158745 | 0.805784013095094 | NM_198249,NM_001145921,NM_001145922 |
| 4922501L14Rik | 267.713592 | 0.0696300837912876 | 0.0856059422372443 | 0.813379094623106 | 0.416000741438453 | 0.805925149391166 | NM_175176 |
| 2810408M09Rik | 503.2475503 | 0.0578732456878598 | 0.0711647767277945 | 0.813228798415614 | 0.416086891015704 | 0.805984555126566 | NM_001007581 |
| Orai2 | 3712.413504 | -0.0569218738876321 | 0.0700059649770085 | -0.813100339468594 | 0.416160531852076 | 0.806019718091101 | NM_178751 |
| Aprt | 728.6856793 | 0.0574165590004876 | 0.0706798933311774 | 0.812346429188801 | 0.416592875889832 | 0.806153083919673 | NM_009698 |
| Arhgef7 | 5862.863316 | 0.045494322205904 | 0.0560005842608067 | 0.812390134967651 | 0.416567805041751 | 0.806153083919673 | NM_001113518,NM_001113517,NM_017402 |
| Btbd19 | 269.2447229 | -0.0630969152933103 | 0.0776950514068326 | -0.812109833905863 | 0.416728610913806 | 0.806153083919673 | NR_024078 |
| Ddx39b | 5870.487878 | -0.0472022482923464 | 0.0582030518749404 | -0.81099266742522 | 0.41736988165698 | 0.806153083919673 | NM_001252457,NM_019693 |
| Fam49b | 6191.594442 | 0.0576179913647714 | 0.0710385937978373 | 0.811080122570296 | 0.41731966008453 | 0.806153083919673 | NM_144846 |
| Gm20300 | 1708.167474 | 0.0591154013747034 | 0.0727450454143532 | 0.812638180895815 | 0.416425534148203 | 0.806153083919673 | NR_045008 |
| Gpr6 | 321.6066478 | 0.0749952839061204 | 0.092462355477475 | 0.811089913498798 | 0.41731403781458 | 0.806153083919673 | NM_199058 |
| Hrh3 | 4619.972979 | 0.0610238709918142 | 0.0752164647550076 | 0.811310012915111 | 0.417187661346627 | 0.806153083919673 | NM_133849 |
| Hunk | 969.1343016 | -0.0755562827936453 | 0.0931923860162804 | -0.810755964338609 | 0.417505827504316 | 0.806153083919673 | NM_015755 |
| Mplkip | 217.5241651 | -0.0723637290415973 | 0.0892473488457919 | -0.810822169817421 | 0.417467801037491 | 0.806153083919673 | NM_025479 |
| Mterfd1 | 552.4408775 | -0.0577316567971453 | 0.0711668538147152 | -0.811215526647437 | 0.417241910615467 | 0.806153083919673 | NM_025547 |
| Nrbf2 | 813.0591172 | 0.0496991097471058 | 0.0611747185743191 | 0.812412560373745 | 0.416554941395804 | 0.806153083919673 | NM_001036293 |
| Poc5 | 1381.643238 | 0.0467183940082443 | 0.0575100272743545 | 0.812352144876785 | 0.416589597426206 | 0.806153083919673 | NM_026173 |
| Ppil1 | 565.2194004 | 0.0569274726768306 | 0.0701146503635651 | 0.811919796813432 | 0.416837654087538 | 0.806153083919673 | NM_026845 |
| Ppp1r12b | 2043.516944 | 0.0545315510905044 | 0.0672428207676898 | 0.810964657162432 | 0.417385967449386 | 0.806153083919673 | NR_015606,NM_001081307 |
| Prex2 | 2661.798707 | -0.0669157069137269 | 0.0824413172227608 | -0.811676828657615 | 0.41697709358329 | 0.806153083919673 | NM_001033636,NM_029525 |
| Pvrl2 | 231.174141 | 0.0740867225499523 | 0.0913745316004016 | 0.810802761374391 | 0.41747894840412 | 0.806153083919673 | NM_008990,NM_001159724 |

| | | | | | | | | | | | | | | |
|---|---|---|---|---|---|---|---|---|---|---|---|---|---|---|
| Atf6 | 4761.193318 | -0.115320782 | 0.090913231 | -1.268470837 | 0.204629859 | 0.42171341 | | Rspry1 | 1018.228248 | 0.0488492834575847 | 0.0602003381527375 | 0.811445333307706 | 0.417109974416948 | 0.806153083919673 | NM_026274 |
| Fth1 | 49965.45387 | 0.136759225 | 0.10782577 | 1.268335254 | 0.204678253 | 0.42171341 | | Slc9a4 | 31.5009191 | -0.0644505089942483 | 0.0793726331180286 | -0.811999129453211 | 0.416792131018693 | 0.806153083919673 | NM_177084 |
| Syn3 | 2071.169055 | -0.137879317 | 0.108707557 | -1.268350803 | 0.204672702 | 0.42171341 | | Sspo | 136.8729808 | 0.0770973507002739 | 0.0950816118295729 | 0.810854477714003 | 0.417449245079366 | 0.806153083919673 | NM_173428 |
| Rmdn1 | 234.5719626 | 0.172887675 | 0.136324579 | 1.268206196 | 0.204724325 | 0.421752252 | | Tti1 | 1159.076545 | 0.0424866299486417 | 0.0523572881140187 | 0.811474991907875 | 0.417092948665495 | 0.806153083919673 | NM_029282 |
| Pip4k2c | 6537.314096 | 0.11331618 | 0.08937559 | 1.26786497 | 0.204846176 | 0.421947174 | | Vmac | 485.7351243 | -0.067878170449208 | 0.0836011988333216 | -0.811928194768342 | 0.416832834990505 | 0.806153083919673 | NM_001166474,NM_178926 |
| Faf2 | 2264.661499 | -0.055683753 | 0.043926347 | -1.267661829 | 0.204918742 | 0.42204054 | | Zfp612 | 2738.078057 | 0.0562080305679962 | 0.0691955076590207 | 0.812307510553666 | 0.416615202023382 | 0.806153083919673 | NM_175480 |
| Slitrk2 | 1109.666162 | -0.073120682 | 0.057733676 | -1.266517003 | 0.205328046 | 0.422827319 | | Ppp4r4 | 1706.312515 | -0.0523268452546028 | 0.0645490714406471 | -0.810652176502916 | 0.417565444272288 | 0.806161037147695 | NM_028980 |
| Cpsf3l | 967.1027995 | -0.075646284 | 0.05973531 | -1.266357936 | 0.205384964 | 0.422843318 | | Otof | 1692.062104 | -0.0746439072401024 | 0.0920939232015825 | -0.81051935507966 | 0.417641745539694 | 0.806201195443136 | NM_031875,NM_001100395 |
| Fam120b | 3165.000336 | 0.066631076 | 0.052616936 | 1.266342752 | 0.205390397 | 0.422843318 | | 1110008P14Rik | 3264.994854 | 0.0538124486981724 | 0.0664104414576856 | 0.81030102370965 | 0.417761871133688 | 0.806218970293285 | NM_198001 |
| Fen1 | 542.8136946 | 0.137131071 | 0.108365535 | 1.265449129 | 0.205710373 | 0.423436651 | | Bmp2k | 759.1263795 | -0.0537802344303618 | 0.0663797829012927 | -0.810189971701677 | 0.417831000227138 | 0.806218970293285 | NM_080708 |
| Gm16596 | 39.19155249 | -0.203875027 | 0.161116961 | -1.265385255 | 0.205733258 | 0.423436651 | | Ctsl | 3528.124934 | -0.0561226735570601 | 0.0692772484766864 | -0.810116954571988 | 0.417872960714453 | 0.806218970293285 | NM_009984 |
| Irs1 | 757.552852 | -0.098829405 | 0.078125649 | -1.265005869 | 0.205869224 | 0.423660215 | | Mad1l1 | 629.725253 | 0.0516738205138121 | 0.0637711289831757 | 0.810301171356788 | 0.417767102295995 | 0.806218970293285 | NM_010752 |
| Hmga2-ps1 | 38.21044611 | -0.212635318 | 0.168116438 | -1.264809798 | 0.205939518 | 0.423692324 | | Hbegf | 546.505629 | 0.0611286262451567 | 0.0754658737898089 | 0.81001680859636 | 0.417930515273376 | 0.806222930260432 | NM_010415 |
| Ipo7 | 4321.870839 | -0.087652807 | 0.069297535 | -1.264876261 | 0.205915688 | 0.423692324 | | Map3k10 | 6897.374397 | 0.0534501012157888 | 0.0660118914123302 | 0.809704131668086 | 0.418110242830877 | 0.806462539816932 | NM_001081292 |
| 4931428F04Rik | 1185.169472 | -0.107303625 | 0.084863466 | -1.264426611 | 0.206076946 | 0.423761947 | | Ccdc137 | 1012.09381 | -0.0582326170304157 | 0.0719496619293764 | -0.809352198034997 | 0.418312589687368 | 0.806745708682781 | NM_152807 |
| 6430550D23Rik | 26.0838998 | 0.21192096 | 0.16759192 | 1.264505833 | 0.206048528 | 0.423761947 | | 1700023F06Rik | 65.74930753 | -0.0758228589004099 | 0.0936986784084363 | -0.809220153243731 | 0.418388524692811 | 0.80678504044707 | NM_001254724,NM_027082 |
| Gpi1 | 23538.58571 | -0.055072837 | 0.043555426 | -1.264431141 | 0.206075322 | 0.423761947 | | Angpt2 | 58.78972417 | -0.0698324998192511 | 0.0864168185286656 | -0.808089223929098 | 0.419039220590664 | 0.806975047512235 | NM_007426 |
| Tpm4 | 1212.215722 | -0.108274213 | 0.085632175 | -1.264410408 | 0.206082759 | 0.423761947 | | Arpc2 | 6863.342616 | 0.0468721349675293 | 0.0579813806724232 | 0.808399772891616 | 0.41886048266339 | 0.806975047512235 | NM_029711 |
| Lsr | 288.1094879 | -0.167886114 | 0.132789541 | -1.264302237 | 0.206121567 | 0.423777155 | | Atrx | 6321.334407 | -0.057147780646107 | 0.0707001040406404 | -0.808312539586319 | 0.418910685674595 | 0.806975047512235 | NM_009530 |
| Slc35b4 | 3194.562976 | -0.07940213 | 0.062806349 | -1.264237326 | 0.206144857 | 0.423777155 | | Copg1 | 15157.77424 | 0.034835083077999 | 0.0431070898496727 | 0.80810564997341 | 0.419029762487429 | 0.806975047512235 | NM_017477,NM_201244 |
| Xlr | 26.76001555 | 0.212467996 | 0.168077244 | 1.26410923 | 0.206190824 | 0.423815419 | | Dicer1 | 2764.587117 | -0.0495971095797514 | 0.0613408086529984 | -0.808549979513957 | 0.4187740466282 | 0.806975047512235 | NM_148948 |
| Agpat1 | 4616.445789 | -0.108097188 | 0.085522482 | -1.263962245 | 0.206243578 | 0.423831546 | | Gtf3c4 | 2317.169582 | 0.0472340491086758 | 0.0584539867688425 | 0.808055219491937 | 0.419058794730725 | 0.806975047512235 | NM_172977,NM_001166033 |
| Xrcc6bp1 | 142.3074847 | -0.155929783 | 0.12336852 | -1.263934941 | 0.206253379 | 0.423831546 | | Idh1 | 1534.436858 | 0.0475874097866304 | 0.0588832149295502 | 0.808165957710794 | 0.418995051925865 | 0.806975047512235 | NM_010497,NM_001111320 |
| Zfr2 | 2026.180466 | 0.144306212 | 0.114179904 | 1.263849485 | 0.206284056 | 0.423838371 | | Mtif2 | 1010.563425 | -0.0465729333928032 | 0.0576171629468449 | -0.808317018937039 | 0.418908107709503 | 0.806975047512235 | NM_133767 |
| Darc | 1851.653767 | 0.102161572 | 0.080858896 | 1.263454942 | 0.20642573 | 0.424073224 | | Plscr2 | 70.57171207 | -0.0763373301052625 | 0.0944784312971489 | -0.807986850090367 | 0.419098152170749 | 0.806975047512235 | NM_008880,NM_001195084 |
| Ints2 | 802.8297114 | 0.135553645 | 0.107299276 | 1.26332302 | 0.206473117 | 0.42411434 | | Rogdi | 4500.089056 | -0.0479329353445227 | 0.0592892128334303 | -0.808459634625064 | 0.418826034116555 | 0.806975047512235 | NM_133185 |
| Bod1 | 2584.826831 | -0.086825154 | 0.068742407 | -1.263050832 | 0.206570912 | 0.424258976 | | Serpinb6a | 1191.053835 | 0.0462613128350929 | 0.0572183561599486 | 0.808504751618059 | 0.418800071806335 | 0.806975047512235 | NM_001243192,NM_001164118,NM_009254,NM_001164117 |
| Lrp2 | 21.29169247 | 0.211593368 | 0.167536841 | 1.262966204 | 0.206601326 | 0.424265201 | | Gstp1 | 1778.936799 | 0.0512678076213311 | 0.0634717965420944 | 0.807725799715318 | 0.41924844808645 | 0.80711450168009 | NM_013541 |
| Thoc5 | 1164.862807 | 0.066659386 | 0.052829585 | 1.261781374 | 0.207027466 | 0.42508396 | | Lmf2 | 746.427105 | -0.0565476856197149 | 0.0700135234673931 | -0.807668044960634 | 0.419281703819218 | 0.80711450168009 | NM_178919 |
| Gnpda2 | 634.7257135 | 0.09406153 | 0.074561371 | 1.261531653 | 0.207117363 | 0.425155859 | | D030056L22Rik | 791.7127024 | 0.0562961346290835 | 0.0697421019740814 | 0.807204443737659 | 0.419548705979511 | 0.807483874702979 | NM_177640 |
| Nfkbid | 118.0953421 | 0.206978927 | 0.164062301 | 1.261587373 | 0.207097302 | 0.425155859 | | Glt25d2 | 320.2818892 | -0.0687552463200297 | 0.0851835974924443 | -0.807141848242887 | 0.419584764302498 | 0.807483874702979 | NM_177756 |
| Eapp | 736.0195562 | -0.084250125 | 0.066797729 | -1.261272304 | 0.207210756 | 0.425234895 | | Aph1c | 433.2641787 | -0.0675364690334824 | 0.0837404692809173 | -0.80649737950385 | 0.419956118387195 | 0.807657282110808 | NM_026674 |
| Hnrnph3 | 1421.159744 | 0.101986743 | 0.080857996 | 1.261306833 | 0.20719832 | 0.425234895 | | Dctn2 | 9914.24023 | 0.0434977410509712 | 0.05395185439408 | 0.806232548250355 | 0.420108774728951 | 0.807657282110808 | NM_001190454,NM_027151,NM_001190453 |
| Nos3 | 517.8592264 | -0.110573619 | 0.087673846 | -1.26119275 | 0.20723941 | 0.425237375 | | Dhrs4 | 354.0008292 | -0.0609488941795086 | 0.0755598765079187 | -0.8066304101638 | 0.419879447994728 | 0.807657282110808 | NM_030686,NM_001037938 |
| Cryab | 3015.513412 | 0.144635962 | 0.114747925 | 1.260466903 | 0.20750098 | 0.425492348 | | Ggps1 | 1710.688767 | 0.0522370067017232 | 0.0647580465072118 | 0.806648895684428 | 0.419868794769982 | 0.807657282110808 | NM_010282 |
| Ocrl | 3556.615506 | 0.123412132 | 0.097905623 | 1.260521389 | 0.207481337 | 0.425492348 | | Nupr1 | 297.8289533 | -0.0686254048660663 | 0.0850687817140919 | -0.80670492139772 | 0.419836508063509 | 0.807657282110808 | NM_019738 |
| Pi16 | 32.79571635 | -0.205552426 | 0.163050861 | -1.260664458 | 0.207429764 | 0.425492348 | | Riok1 | 569.1897864 | -0.0510749821389718 | 0.0633744136520435 | -0.805924334375674 | 0.42028647910475 | 0.807657282110808 | NM_024242 |
| Rgl1 | 5243.726395 | -0.104392695 | 0.082818236 | -1.260503733 | 0.207487702 | 0.425492348 | | Tbcd | 2080.912493 | -0.041632431069203 | 0.0516571915270061 | -0.805936789022644 | 0.420279297373924 | 0.807657282110808 | NM_029878 |
| Ube4b | 5068.093557 | 0.097362036 | 0.077232749 | 1.260631491 | 0.207441647 | 0.425492348 | | Tph2 | 40.52028626 | 0.072137935213969 | 0.0894915564325424 | 0.806086496756212 | 0.420192976951614 | 0.807657282110808 | NM_173391 |
| Alg13 | 236.2906821 | 0.127746007 | 0.101381144 | 1.260056868 | 0.207648848 | 0.425682886 | | Zfp341 | 497.6247067 | -0.0666673689090071 | 0.0827111476987234 | -0.806026403500578 | 0.420227625047214 | 0.807657282110808 | NM_199304 |
| Ociad1 | 10450.6343 | -0.058569014 | 0.046478447 | -1.260132755 | 0.207621476 | 0.425682886 | | Zfp770 | 1035.512961 | 0.0661523843749734 | 0.0820274344069908 | 0.806466578568689 | 0.419973871255686 | 0.807657282110808 | NM_175466 |
| Gpcpd1 | 3351.000397 | 0.113677014 | 0.090247283 | 1.259617021 | 0.207807552 | 0.425908943 | | Zfp941 | 1454.252698 | 0.0562072082945263 | 0.0697412242433995 | 0.805939512882095 | 0.420277726722763 | 0.807657282110808 | NM_001001180 |
| Tdrkh | 1042.085744 | -0.067035608 | 0.053219805 | -1.25959889 | 0.207814096 | 0.425908943 | | Zfand2b | 567.8586234 | -0.0629123045736232 | 0.0780799986962322 | -0.805741619161414 | 0.420391846578271 | 0.807752905211107 | NM_026846,NM_001159906,NM_001159905 |
| Flot1 | 3744.698931 | -0.102507331 | 0.081400877 | -1.259290257 | 0.207925511 | 0.426024579 | | Wdr24 | 1128.922232 | -0.0561106688686237 | 0.0696506259918739 | -0.805601788491752 | 0.420472494044569 | 0.807801011571407 | NM_173741 |
| Tspan14 | 1617.400829 | 0.102041859 | 0.081026414 | 1.25936535 | 0.207898399 | 0.426024579 | | Akt1 | 7194.594152 | 0.0374991567873587 | 0.0465575722674241 | 0.805436257972508 | 0.420567975665949 | 0.80787760043951 | NM_001165894,NM_009652 |
| Syt11 | 35077.79624 | -0.06758898 | 0.053690513 | -1.258862638 | 0.208079951 | 0.426284644 | | Aebp1 | 1003.636885 | -0.0730221926697892 | 0.090690205902848 | -0.805182786198703 | 0.420714208397478 | 0.807944816391033 | NM_009636 |
| Zfyve26 | 1151.840258 | 0.085220255 | 0.067703345 | 1.258730336 | 0.20812775 | 0.426326198 | | Tmem255a | 1274.473093 | 0.0738987964750431 | 0.0917686931009906 | 0.805272408028282 | 0.420662500432576 | 0.807944816391033 | NM_172930 |
| Gjc1 | 147.5705827 | 0.158549696 | 0.125985061 | 1.258480137 | 0.208218166 | 0.426398661 | | AI450353 | 65.82770912 | -0.075331469136907 | 0.0936326951631463 | -0.804542355697963 | 0.421083818018343 | 0.808470524691774 | NR_028364 |
| Slc38a4 | 44.41413628 | -0.197880728 | 0.157228619 | -1.258554128 | 0.208191425 | 0.426398661 | | Ano3 | 6424.921394 | 0.0700535635352277 | 0.0870763386939387 | 0.804507453872816 | 0.421103966271858 | 0.808470524691774 | NM_001128103,NM_001081556 |
| 2010109I03Rik | 8.233823068 | 0.187583667 | 0.149105098 | 1.258058337 | 0.20837066 | 0.426414656 | | Cyp4f17 | 83.40454236 | 0.0765003154973909 | 0.0951000636887755 | 0.804419182596405 | 0.421154926362568 | 0.808470524691774 | NM_001101445 |
| Cd86 | 109.4505734 | 0.184677377 | 0.14680053 | 1.258015738 | 0.208386065 | 0.426414656 | | Adhfe1 | 767.5047039 | -0.0553051885698871 | 0.0687994944076146 | -0.803860392377623 | 0.421477606764963 | 0.808555694118839 | NR_027664,NM_175236 |
| Cltb | 4910.369744 | -0.058133552 | 0.046199614 | -1.258312505 | 0.20827876 | 0.426414656 | | Atg2a | 2667.081051 | -0.0538246580041151 | 0.0669449095321487 | -0.804014201830643 | 0.421388773109921 | 0.808555694118839 | NM_194348 |
| Gna12 | 1950.243551 | -0.12803034 | 0.10177278 | -1.258001801 | 0.208391105 | 0.426414656 | | Ccdc127 | 3596.312014 | 0.0435628741505306 | 0.0541917187404736 | 0.803865888792991 | 0.421474432085085 | 0.808555694118839 | NM_024201,NM_001168659,NM_001168658 |
| Imp3 | 709.8767354 | -0.107503157 | 0.085439298 | -1.258240176 | 0.208304909 | 0.426414656 | | Gm2115 | 410.5281722 | -0.0751109757918221 | 0.093435982402996 | -0.803876342497938 | 0.421468394158224 | 0.808555694118839 | NR_045098 |
| Plekhs1 | 24.33032145 | 0.209899798 | 0.166837569 | 1.258108708 | 0.208352444 | 0.426414656 | | Mrpl14 | 366.3344351 | 0.0643896788572631 | 0.0800909933749991 | 0.803956551715873 | 0.421422068023561 | 0.808555694118839 | NM_026732 |
| Rrp1b | 614.049503 | 0.146168428 | 0.116222404 | 1.257661372 | 0.208514248 | 0.426610294 | | Mrpl13 | 907.8816668 | 0.0564426112120055 | 0.0702326684548756 | 0.803651811240375 | 0.421598091711982 | 0.808573261184085 | NM_026759 |
| Zfp395 | 423.8942693 | -0.132157407 | 0.105088921 | -1.257576966 | 0.208544788 | 0.426616444 | | Npbwr1 | 68.85813751 | -0.0718917748943122 | 0.0894490968438933 | -0.803717169104321 | 0.421560336178923 | 0.808573261184085 | NM_010342 |
| Mrap | 10.06648932 | -0.205790046 | 0.163675422 | -1.257305732 | 0.208642949 | 0.426760905 | | Tpbg | 581.4412408 | 0.0669944394122285 | 0.0833832447971753 | 0.803452055328244 | 0.421713497784468 | 0.808687824266296 | NM_001164792,NM_011627 |
| Olfr558 | 15.58488801 | -0.197917308 | 0.157433288 | -1.257150315 | 0.208699211 | 0.426819637 | | Amph | 10572.47176 | 0.0342859538501639 | 0.0427200178592883 | 0.802573490561155 | 0.422221295671744 | 0.808695226068607 | NM_175007 |
| Spink10 | 36.6025272 | -0.207644648 | 0.165201445 | -1.256917867 | 0.208783378 | 0.426935418 | | Cdc16 | 1988.251122 | 0.0425516776114607 | 0.0530513391102477 | 0.802084892202866 | 0.422503853541111 | 0.808695226068607 | NM_027276 |
| Bank1 | 64.63687392 | -0.185609946 | 0.147724058 | -1.256463898 | 0.208947827 | 0.427134075 | | Cope | 2789.474421 | 0.0502333011775783 | 0.0626349691137028 | 0.802000893245251 | 0.42255244154349 | 0.808695226068607 | NM_021538 |
| Snapc3 | 568.6621205 | -0.09223849 | 0.073414525 | -1.256406547 | 0.208968609 | 0.427134075 | | Cry2 | 7734.337601 | 0.0486428773201525 | 0.0606227655440184 | 0.802386312858538 | 0.422329527999277 | 0.808695226068607 | NM_009963,NM_001113333 |
| Spin4 | 22.28352849 | -0.209865459 | 0.167045761 | -1.256335136 | 0.208994488 | 0.427134075 | | Cybrd1 | 144.2919682 | -0.0762962595830136 | 0.0950528160485977 | -0.802672269530716 | 0.42216418496391 | 0.808695226068607 | NM_028593 |
| Tmem231 | 522.1618862 | -0.093820457 | 0.074681807 | -1.256269253 | 0.209018365 | 0.427134075 | | Kat2b | 1183.077278 | 0.0583914665965447 | 0.072696364144774 | 0.803224030300315 | 0.42184525856841 | 0.808695226068607 | NM_001190846,NM_020005 |
| Zmynd10 | 122.6674478 | 0.17073535 | 0.135904402 | 1.256290061 | 0.209010824 | 0.427134075 | | Pcdha2 | 109.5392377 | 0.0756553450332397 | 0.0943171912771704 | 0.80213738353288 | 0.42247349233976 | 0.808695226068607 | NM_198117 |
| Trappc11 | 3099.56442 | 0.061281344 | 0.048783749 | 1.256183553 | 0.209049428 | 0.427141217 | | Ppp2r2d | 2211.522503 | 0.0507231536986595 | 0.0631715533060871 | 0.802942955259504 | 0.422007706714846 | 0.808695226068607 | NM_026391 |
| C030039L03Rik | 633.0159649 | 0.086995986 | 0.069286524 | 1.255597504 | 0.209261937 | 0.427406334 | | Ppp4r2 | 2868.505732 | 0.062614241221353 | 0.0779975330933382 | 0.802772071604158 | 0.422106487330069 | 0.808695226068607 | NM_182939 |
| Rab3d | 511.2456451 | 0.129293818 | 0.102970457 | 1.255639939 | 0.209246544 | 0.427406334 | | Rundc3b | 1692.040264 | 0.0515917526544935 | 0.0643134673879102 | 0.802192056343581 | 0.422441870720134 | 0.808695226068607 | NM_198620 |
| Top1mt | 474.0558515 | 0.123480843 | 0.098336196 | 1.255700829 | 0.209224459 | 0.427406334 | | Snhg4 | 238.8027137 | -0.0660859610251676 | 0.0823387380719046 | -0.802610807168993 | 0.422199719920715 | 0.808695226068607 | NR_038073 |
| Gm15708 | 11.17322191 | 0.203852718 | 0.162396039 | 1.255281347 | 0.209376644 | 0.427553024 | | Tmpo | 1088.801251 | -0.0465762167399745 | 0.0580408206968324 | -0.802473434744461 | 0.422279149225968 | 0.808695226068607 | NM_001080129,NM_011605,NM_001080134,NM_001080132,NM_001080131,NM_001080130 |
| Nt5c2 | 2825.381946 | 0.094679547 | 0.075426998 | 1.255247445 | 0.209388947 | 0.427553024 | | Toe1 | 484.9837271 | 0.0523600886518483 | 0.0652466611045488 | 0.802494530225056 | 0.422266951162536 | 0.808695226068607 | NM_026654 |
| Zfp597 | 840.7139711 | -0.102618347 | 0.0817669 | -1.255010844 | 0.209474824 | 0.427672014 | | Zbtb44 | 942.5713906 | -0.0524149499840105 | 0.0653315606661488 | -0.802292082870279 | 0.422384021041649 | 0.808695226068607 | NM_172765,NM_001115130 |
| Fktn | 1284.588412 | -0.126957351 | 0.101172434 | -1.254861088 | 0.209529192 | 0.427726653 | | Zfp780b | 557.9550371 | -0.0694414867644989 | 0.0865137411567249 | -0.802664245425491 | 0.422168824065134 | 0.808695226068607 | NM_001081021 |
| 5430427O19Rik | 12.51327503 | 0.200583704 | 0.159901245 | 1.2544224 | 0.209688515 | 0.427871411 | | Vps11 | 3655.071193 | 0.0391944369008492 | 0.0488905047847484 | 0.801677893763045 | 0.422739306492747 | 0.80894627402365 | NM_027889 |
| Foxn2 | 583.394082 | 0.106398223 | 0.084827741 | 1.254285698 | 0.20973818 | 0.427871411 | | 5730507C01Rik | 40.43037769 | -0.0731876738073804 | 0.0913562717535607 | -0.801123693015944 | 0.423060041098388 | 0.808999423486937 | NM_001033157,NM_001201330 |
| Neo1 | 4708.386084 | -0.063468694 | 0.05059726 | -1.254389936 | 0.209700309 | 0.427871411 | | Arpc5l | 2522.980218 | 0.0470361141320394 | 0.0588308696874982 | 0.79951417311845 | 0.423992331750382 | 0.808999423486937 | NM_028809 |
| Prr3 | 946.8654901 | 0.091850157 | 0.073228389 | 1.254297116 | 0.209734032 | 0.427871411 | | Asprv1 | 49.08449787 | -0.0751824257746343 | 0.0939297863072641 | -0.800410910429379 | 0.423472761739467 | 0.808999423486937 | NM_026414 |
| Slc38a3 | 3287.266443 | -0.164296544 | 0.130987973 | -1.254287242 | 0.209737619 | 0.427871411 | | Capza1 | 2008.85684 | 0.0521699846587032 | 0.0651634618834823 | 0.800601796632406 | 0.423362210304153 | 0.808999423486937 | NM_009797 |
| Gja4 | 108.1403361 | 0.154899605 | 0.123530132 | 1.253941872 | 0.209863134 | 0.428013611 | | Ccdc103 | 106.8740931 | -0.0758648834802745 | 0.0948490985013975 | -0.799848229228627 | 0.423798735925807 | 0.808999423486937 | NM_028492 |
| Nup107 | 615.7182374 | 0.097946962 | 0.078108889 | 1.253979708 | 0.209849381 | 0.428013611 | | Cdk7 | 2677.038501 | 0.0526806909739465 | 0.0658329555661291 | 0.800217619289914 | 0.423584723202546 | 0.808999423486937 | NM_009874 |
| Kif4 | 68.30568263 | -0.180228937 | 0.143777624 | -1.253525631 | 0.210014477 | 0.4282659 | | Cebpa | 385.6366623 | 0.0680959242200527 | 0.0851668444835295 | 0.799559084812592 | 0.423966301028632 | 0.808999423486937 | NM_007678 |
| Ajap1 | 1980.420023 | -0.117282236 | 0.093573885 | -1.253365043 | 0.210072887 | 0.428328637 | | Cntn4 | 2288.37415 | 0.0620458064609045 | 0.0775149665340619 | 0.800436473563336 | 0.423457955912196 | 0.808999423486937 | NM_001109749,NM_173004,NM_001109751 |
| Clrn1 | 12.84448777 | 0.199239629 | 0.158978796 | 1.253246554 | 0.210115992 | 0.428360156 | | D7Ertd715e | 154.4433967 | -0.0748157868991844 | 0.0935735621696783 | -0.799539796973683 | 0.423977480104245 | 0.808999423486937 | NR_015456 |
| Itga8 | 220.6725581 | 0.12558126 | 0.100240055 | 1.252805182 | 0.210276615 | 0.428462109 | | Kdm3b | 2794.420598 | -0.0359148339821579 | 0.0448160593526443 | -0.80138313142516 | 0.422909877634282 | 0.808999423486937 | NM_001081256 |
| Kirrel | 390.3754495 | -0.143370963 | 0.114428008 | -1.252935938 | 0.210229022 | 0.428462109 | | Lrif1 | 420.3430366 | 0.0676345198887699 | 0.0844301668131671 | 0.801070546721011 | 0.423090806131062 | 0.808999423486937 | NM_028081,NM_001039478,NM_001039488 |
| Naa40 | 1360.918461 | 0.146853873 | 0.117219154 | 1.252814643 | 0.210273171 | 0.428462109 | | Mapk8ip2 | 12672.88845 | 0.0369214472826629 | 0.0461206287962026 | 0.800540847910186 | 0.423397506822109 | 0.808999423486937 | NM_021921 |
| Trappc13 | 1792.133527 | -0.089112334 | 0.071124937 | -1.252898604 | 0.21024261 | 0.428462109 | | Mars2 | 373.0186572 | -0.0595844186531965 | 0.0744543248813166 | -0.800282397942319 | 0.423547199052758 | 0.808999423486937 | NM_175439 |
| Helb | 207.5760586 | 0.134792746 | 0.107634378 | 1.252320571 | 0.210453076 | 0.428708909 | | Pdik1l | 608.5190103 | 0.061226705810089 | 0.0764730995829249 | 0.800630628861811 | 0.423345513633703 | 0.808999423486937 | NM_146156,NM_001163794 |
| Phkg2 | 1809.566854 | 0.076746989 | 0.061283207 | 1.252333104 | 0.210448511 | 0.428708909 | | Rsph9 | 357.5396842 | 0.0641529567030931 | 0.0801389623992974 | 0.800521429057778 | 0.423408752995899 | 0.808999423486937 | NM_029338 |
| 9430037G07Rik | 83.24602483 | 0.184295946 | 0.14718523 | 1.252136144 | 0.210520259 | 0.428789391 | | Slc25a29 | 474.9895771 | 0.0582768183199307 | 0.07275025266859 | 0.801053139779565 | 0.423100882849615 | 0.808999423486937 | NM_181328 |
| Pcdha6 | 123.2704618 | -0.193646882 | 0.154666365 | -1.252029691 | 0.210559045 | 0.428812019 | | Slc37a2 | 185.4429621 | -0.0744355518144306 | 0.0929287679254933 | -0.800995789313705 | 0.423134083501724 | 0.808999423486937 | NM_020258,NM_001145960 |
| Doc2b | 2215.449168 | -0.148250556 | 0.118416551 | -1.251941178 | 0.210591299 | 0.428821341 | | Stox2 | 5930.490023 | 0.0538426861907914 | 0.0673441014348079 | 0.799515993882754 | 0.423991276421111 | 0.808999423486937 | NM_001114311,NM_175162 |
| Zfp30 | 421.0433743 | 0.115404778 | 0.092199979 | 1.251679011 | 0.210686851 | 0.428959537 | | Thg1l | 211.3284299 | 0.0663009007163857 | 0.0829040936777794 | 0.799730116369808 | 0.423867180062151 | 0.808999423486937 | NM_001080969 |
| Fam212b | 6107.305511 | -0.112119505 | 0.089597422 | -1.251369764 | 0.210799603 | 0.42913271 | | Trmt44 | 301.1491449 | -0.0657554279899202 | 0.082216413041985 | -0.799784684796955 | 0.423835557901681 | 0.808999423486937 | NM_030208 |
| Lypd6 | 1257.910282 | -0.085003515 | 0.067932838 | -1.251287554 | 0.210829585 | 0.42913736 | | Xrcc2 | 293.566991 | 0.0666509734929975 | 0.0833221157920588 | 0.799919359457141 | 0.423757520441995 | 0.808999423486937 | NM_020570 |
| Etohd2 | 27.88126795 | -0.204181158 | 0.16326906 | -1.250580843 | 0.211087444 | 0.429605786 | | Zfp266 | 1577.661144 | -0.0583488548727172 | 0.0728295487158695 | -0.801170073157449 | 0.423033193889431 | 0.808999423486937 | NM_001082485,NM_001135019 |
| Slc25a19 | 1428.756418 | 0.080687932 | 0.064524735 | 1.250496152 | 0.21111836 | 0.429612276 | | Cln5 | 565.5562391 | -0.0601948332191118 | 0.0752988203024364 | -0.799412699659043 | 0.424051148991885 | 0.809005383537709 | NM_001033242 |
| Fgf23 | 13.71045425 | 0.206268011 | 0.164965097 | 1.250373653 | 0.211163084 | 0.429646859 | | Cox6b2 | 73.49383735 | 0.0732242444380111 | 0.0916165522929227 | 0.799246889403714 | 0.424147268152786 | 0.80908249733255 | NM_183405,NM_183406 |
| 2900026A02Rik | 2590.265197 | 0.117818439 | 0.09424521 | 1.250126551 | 0.211253322 | 0.429728004 | | Ccdc124 | 2165.5407 | 0.0425588310105937 | 0.0532731614346216 | 0.798879395955492 | 0.424360347194099 | 0.809162684238523 | NM_026964 |
| Dapp1 | 235.504319 | -0.17849274 | 0.142781336 | -1.25011255 | 0.211258436 | 0.429728004 | | Scamp2 | 1047.989014 | -0.0496189790275597 | 0.062112707844389 | -0.798853902036765 | 0.424375131328833 | 0.809162684238523 | NM_022813 |
| Umodl1 | 13.08750624 | -0.203356792 | 0.162683275 | -1.250016584 | 0.211293489 | 0.429742888 | | Scara5 | 28.79602406 | 0.0541507598670507 | 0.0677981804575474 | 0.798705286295317 | 0.424461365867172 | 0.809162684238523 | NM_001168318,NM_028903 |
| Phf14 | 1462.85246 | -0.080866535 | 0.064781102 | -1.2483044 | 0.211919604 | 0.430959751 | | Sun2 | 3537.066185 | -0.0508484951350645 | 0.0636722022087918 | -0.798598028199556 | 0.424523531363196 | 0.809162684238523 | NM_001205345,NM_001205346,NM_194342 |
| Ap3s2 | 3882.244403 | 0.072734056 | 0.058291712 | 1.247759808 | 0.212119032 | 0.431308698 | | Susd5 | 321.2174868 | 0.0660994640952723 | 0.0827620704010565 | 0.798668566107169 | 0.424482618220957 | 0.809162684238523 | NM_001101510 |
| Trib3 | 35.81015496 | -0.204669034 | 0.164054018 | -1.247570996 | 0.212188206 | 0.431392739 | | Zbtb11 | 2332.297816 | 0.0646973122647453 | 0.0809674932027525 | 0.79905294890923 | 0.42425972432645 | 0.809162684238523 | NM_173026 |
| C030018K13Rik | 76.49972424 | -0.198822402 | 0.159388231 | -1.24740955 | 0.212247367 | 0.431456403 | | Cers5 | 1468.316719 | -0.0408109595917268 | 0.0511202448857688 | -0.798332630896454 | 0.424677486796324 | 0.809188912286515 | NM_028015 |
| Osgin1 | 29.00154326 | 0.205467253 | 0.164744461 | 1.247187622 | 0.212328711 | 0.431565137 | | Hmgb1 | 7109.140258 | 0.0495387097763671 | 0.0620562908777616 | 0.798286669661717 | 0.424704151955581 | 0.809188912286515 | NM_010439 |
| Serpinb1b | 59.42686677 | -0.205956575 | 0.165179681 | -1.246863864 | 0.212447419 | 0.431749778 | | Lrrc8b | 3012.355475 | 0.0479042143771029 | 0.0600304785365316 | 0.797998209325465 | 0.424871529258519 | 0.809188912286515 | NM_001033550 |
| Elp6 | 312.8409599 | -0.145801768 | 0.117002258 | -1.246144909 | 0.2127112 | 0.432229158 | | Nubpl | 180.0093613 | -0.0678466999955459 | 0.0850110842916085 | -0.798092396549318 | 0.424816873474914 | 0.809188912286515 | NM_029760 |
| Rhoc | 430.3330208 | 0.119532542 | 0.095936035 | 1.245960828 | 0.212778777 | 0.432309778 | | Sh2b1 | 3527.037226 | 0.0491208569330669 | 0.0615507891781399 | 0.798054055666154 | 0.424839121761152 | 0.809188912286515 | NM_001081459,NM_011363 |
| Clstn2 | 2801.826474 | 0.080000989 | 0.064214323 | 1.245843384 | 0.212821899 | 0.432340697 | | Slc44a5 | 366.5880458 | -0.0679419238250429 | 0.0851227888852934 | -0.798163734396584 | 0.424775474438924 | 0.809188912286515 | NM_001081263 |
| Cdc40 | 1903.615357 | 0.078558197 | 0.063068053 | 1.245609989 | 0.212907613 | 0.432401437 | | Pik3c3 | 2522.269866 | 0.0468894211249991 | 0.0587840285811237 | 0.797655796255786 | 0.42507026231445 | 0.809421961270407 | NM_181414 |
| Usp49 | 642.2855886 | 0.114959993 | 0.092288603 | 1.245657525 | 0.212890153 | 0.432401437 | | Sh3gl3 | 1189.490956 | 0.0536377385092622 | 0.0672493101937389 | 0.797595370937441 | 0.425105338188898 | 0.809421961270407 | NM_017400 |
| Ten1 | 565.9142909 | -0.103845946 | 0.083381426 | -1.245432602 | 0.212972776 | 0.432477088 | | 6030419C18Rik | 6202.466997 | 0.0509050919386669 | 0.0638414196607066 | 0.797367793655099 | 0.425237458121821 | 0.809519334216707 | NM_176921 |
| Ankrd54 | 470.1476557 | 0.128640573 | 0.103299468 | 1.245316897 | 0.213015287 | 0.43250673 | | Apoa1bp | 1593.065319 | 0.0414412662714708 | 0.0520401789175301 | 0.796332125167062 | 0.425839018001774 | 0.809519334216707 | NM_144897 |
| Ier2 | 255.8904783 | -0.202737524 | 0.162824149 | -1.24513179 | 0.21308331 | 0.432588157 | | Apoa2 | 25.11865789 | 0.0674721872526739 | 0.0846751178278621 | 0.796830659795506 | 0.425546249532471 | 0.809519334216707 | NM_013474 |
| Cox8a | 7639.345278 | -0.123370387 | 0.099095184 | -1.244968536 | 0.213143316 | 0.432653287 | | Gm19461 | 34.73284799 | 0.0573714354353631 | 0.0720945436449929 | 0.795780547802214 | 0.426159599941839 | 0.809519334216707 | NR_037984 |
| Tpd52 | 2445.70238 | 0.081903683 | 0.065793131 | 1.244866773 | 0.21318072 | 0.43267254 | | Hddc3 | 409.9594481 | 0.064149183019944 | 0.0805674575345607 | 0.796216865863978 | 0.425905996156046 | 0.809519334216707 | NM_026812 |
| Neat1 | 5031.528199 | 0.20872049 | 0.167709764 | 1.244533924 | 0.213303122 | 0.432864253 | | Mapk11 | 2234.308038 | -0.0708840434542079 | 0.0890496510876624 | -0.796005852784624 | 0.426028633566272 | 0.809519334216707 | NM_011161 |
| Fxyd5 | 428.8217881 | 0.129575573 | 0.104135066 | 1.244302983 | 0.213388074 | 0.432979939 | | Myof | 227.6737549 | 0.0746280107381076 | 0.0937690484172777 | 0.795870407109269 | 0.426107363271552 | 0.809519334216707 | NM_001099634 |
| Osgep | 675.7741031 | -0.119036735 | 0.095679157 | -1.244123991 | 0.213453933 | 0.433023546 | | Parn | 1158.35197 | -0.0496808452232609 | 0.0624221023775925 | -0.795885484963972 | 0.4260985963645 | 0.809519334216707 | NM_028761 |
| Stat3 | 2451.369591 | 0.106677692 | 0.085747384 | 1.244092665 | 0.21346546 | 0.433023546 | | Pcyt1a | 2007.202353 | 0.0494711914447769 | 0.0621525036804677 | 0.795964579305016 | 0.426052623458122 | 0.809519334216707 | NM_001163159,NM_001163160,NM_009981 |
| C230052I12Rik | 179.7640373 | -0.141211494 | 0.113539763 | -1.243718413 | 0.213603217 | 0.433246268 | | Pnp | 503.1314493 | -0.0544372802799926 | 0.0683104160257621 | -0.796910390056218 | 0.425503076185992 | 0.809519334216707 | NM_013632 |
| Slc25a45 | 138.513087 | 0.164022748 | 0.131906567 | 1.243476745 | 0.213692205 | 0.43337003 | | Ppp1r15b | 2765.588894 | 0.0561145536816453 | 0.0704932131908971 | 0.796027747092275 | 0.426015907988665 | 0.809519334216707 | NM_133819 |
| Ltc4s | 78.68254962 | 0.176001766 | 0.141581966 | 1.243108647 | 0.2138278 | 0.433588265 | | Qdpr | 6348.612752 | 0.0524988310798147 | 0.0659167450666904 | 0.796441496416421 | 0.42577546712949 | 0.809519334216707 | NM_024236 |
| Atp13a5 | 564.2392137 | -0.132571034 | 0.106667065 | -1.242848802 | 0.213923555 | 0.433676986 | | Rrad | 83.36854814 | 0.0740999951823634 | 0.0930637310992934 | 0.796228501770503 | 0.425899234153548 | 0.809519334216707 | NM_019662 |
| Smim14 | 3888.006172 | -0.061701528 | 0.049645672 | -1.242838008 | 0.213927533 | 0.433676986 | | Slc43a2 | 3460.132762 | 0.0488754638480823 | 0.0613378358759171 | 0.79682406707264 | 0.425553215521593 | 0.809519334216707 | NM_001199284,NM_173388,NM_001199283 |
| Adrm1 | 2276.747012 | 0.072019286 | 0.057951738 | 1.242745925 | 0.213961475 | 0.433689049 | | Taf3 | 819.8214554 | 0.0620376931891 39 | 0.0779954446058366 | 0.795403559731746 | 0.426111634280849 | 0.809519334216707 | NM_027748 |
| Pttg1ip | 2370.702691 | -0.081941526 | 0.065942333 | -1.242624009 | 0.214006418 | 0.433723407 | | Tef | 19815.82884 | 0.0493023376785729 | 0.0619494005938404 | 0.795848502026972 | 0.426120096520255 | 0.809519334216707 | NM_153484,NM_017376 |
| Camk4 | 11080.12719 | -0.121622918 | 0.097905609 | -1.242246693 | 0.214145557 | 0.433891786 | | Tmem129 | 1151.329135 | 0.0447067827794723 | 0.0561202311039899 | 0.796625065506793 | 0.425668815563963 | 0.809519334216707 | NM_026698 |
| Ddi2 | 494.9362502 | 0.11559289 | 0.093047499 | 1.242299799 | 0.214125969 | 0.433891786 | | Zranb3 | 391.5682378 | -0.0590394737645543 | 0.0741784044020051 | -0.795911886222218 | 0.426083252051686 | 0.809519334216707 | NM_027678 |
| Epsti1 | 34.54792512 | 0.205084006 | 0.165105882 | 1.242136276 | 0.214186286 | 0.433917667 | | G6pdx | 1680.788967 | 0.0474331912933218 | 0.0596293143905669 | 0.795467661805375 | 0.426341514785322 | 0.809717745604856 | NM_008062 |
| Pcbp2 | 7795.673722 | 0.075324138 | 0.060662337 | 1.241695285 | 0.21434901 | 0.434190556 | | Rsph4a | 227.3965194 | 0.0709386459688414 | 0.0891851011420818 | 0.79540915534567 | 0.426375536013462 | 0.809717745604856 | NM_001162957 |
| Tmed7 | 2889.924406 | -0.091853232 | 0.07401962 | -1.241540176 | 0.214406266 | 0.434249763 | | Etv6 | 945.4848535 | 0.0590732493402118 | 0.0742891819681906 | 0.795179698781795 | 0.426508979195043 | 0.809764423623784 | NM_007961 |
| Exosc4 | 623.9155933 | -0.084361682 | 0.067971964 | -1.241124682 | 0.214559693 | 0.434503709 | | Gm13375 | 412.7650055 | -0.0672356363331957 | 0.0845544956459304 | -0.795175180450997 | 0.426511607128292 | 0.809764423623784 | NR_033225 |
| 9630033F20Rik | 1406.572891 | 0.074935295 | 0.06041053 | 1.240434313 | 0.214814796 | 0.43465253 | | Pafah1b1 | 26402.36539 | 0.0510927056148561 | 0.0642688142170392 | 0.794984414094111 | 0.426622568498174 | 0.809869240721965 | NM_013625,NR_037610 |
| Aspscr1 | 1086.399461 | 0.086500306 | 0.06971483 | 1.24077339 | 0.214689474 | 0.43465253 | | Vapb | 7256.492685 | 0.0327471348552389 | 0.0412148434545865 | 0.794547112409201 | 0.42687699338356 | 0.810246335540258 | NM_019806 |
| Mrpl45 | 1242.918877 | 0.084511176 | 0.068112608 | 1.24075672 | 0.214695634 | 0.43465253 | | Coro2b | 9845.907753 | 0.0321548691598716 | 0.0404829286338995 | 0.794282188689333 | 0.427031170687476 | 0.810328132440584 | NM_175484 |
| Mtfr1 | 278.6830154 | 0.134542907 | 0.108467845 | 1.240394399 | 0.214829552 | 0.43465253 | | Cyp4b1 | 32.28568738 | -0.0714788361263791 | 0.0899918346171975 | -0.794281352640848 | 0.427031657292625 | 0.810328132440584 | NM_007823 |
| Phyhipl | 6731.696796 | -0.102745193 | 0.082819921 | -1.240585493 | 0.214758914 | 0.43465253 | | Evl | 4792.552457 | 0.0364612879388097 | 0.045927735311852 | 0.793883863230692 | 0.427263044688303 | 0.810334558001624 | NM_007965,NM_001163396,NM_001163395,NM_001163394 |
| Tmprss9 | 22.33669349 | 0.207977069 | 0.16765563 | 1.240501546 | 0.214789943 | 0.43465253 | | Fkrp | 2446.954951 | 0.0452497993530269 | 0.0570345534482858 | 0.793375184291671 | 0.427559264498952 | 0.810334558001624 | NM_173430 |
| Zfp449 | 211.9592111 | -0.134791601 | 0.108658302 | -1.240509001 | 0.214787187 | 0.43465253 | | Frmpd4 | 3872.182689 | 0.0567276110229707 | 0.0715069746630832 | 0.793315775003097 | 0.427593868202306 | 0.810334558001624 | NM_001033330 |
| Sumo2 | 4477.961626 | -0.062426532 | 0.050331084 | -1.240317651 | 0.214857926 | 0.434653181 | | Hbp1 | 1074.835405 | -0.0460071530414318 | 0.0579643224774833 | -0.793715014253874 | 0.427361357520475 | 0.810334558001624 | NM_153198,NM_177993 |
| Abca2 | 13134.97386 | 0.100405064 | 0.080965264 | 1.240100496 | 0.214938226 | 0.43470211 | | Ikbkap | 3529.200529 | -0.035634188605479 | 0.0448841005990453 | -0.793915621119451 | 0.427244555032936 | 0.810334558001624 | NM_026079 |
| Mthfd2 | 208.7466335 | 0.132143255 | 0.106556524 | 1.240123555 | 0.214929698 | 0.43470211 | | Mbnl2 | 8145.314078 | 0.0551468526852512 | 0.0695442651300544 | 0.792974842456404 | 0.427792480277752 | 0.810334558001624 | NM_207515,NM_175341 |
| Cela1 | 73.30835476 | 0.175558891 | 0.141584095 | 1.23996195 | 0.214989474 | 0.434748998 | | Pign | 767.9057362 | -0.0571779812623686 | 0.0720131036850886 | -0.793994125185972 | 0.427198851440532 | 0.810334558001624 | NM_013784 |
| 1700096K18Rik | 41.53409874 | 0.192993 | 0.155661157 | 1.239827605 | 0.215039166 | 0.434792748 | | Pitpnm3 | 5903.629934 | -0.0522314963535092 | 0.0658711431419338 | -0.792934415013339 | 0.427816035062265 | 0.810334558001624 | NM_001081641,NM_001024927 |
| Ankrd9 | 327.5863409 | 0.133416553 | 0.107636007 | 1.239516003 | 0.215154406 | 0.434969115 | | Plod1 | 910.0275586 | 0.0591557822126572 | 0.0745672230803428 | 0.793321512709672 | 0.427590526130249 | 0.810334558001624 | NM_011122 |
| Sp100 | 132.5544189 | 0.179542905 | 0.144868156 | 1.239353837 | 0.215214489 | 0.435030698 | | Psd3 | 22118.62539 | 0.05511410915002 | 0.0694853619448318 | 0.793175834052071 | 0.42767538505245 | 0.810334558001624 | NM_027626,NM_030263 |
| Mex3b | 548.8003879 | -0.131001924 | 0.105732947 | -1.23898867 | 0.215349694 | 0.435250216 | | Rorc | 112.9396857 | -0.0754218647964984 | 0.0949990787330579 | -0.793922065375283 | 0.427240803200959 | 0.810334558001624 | NM_011281 |
| Gm7854 | 14.17049051 | 0.194243483 | 0.156822812 | 1.238617529 | 0.215487173 | 0.435414484 | | Tmem9b | 3772.70643 | 0.0494707398193842 | 0.0623611855947108 | 0.793293766749362 | 0.427606687606432 | 0.810334558001624 | NM_020050 |
| Ppap2b | 10110.45147 | -0.137295345 | 0.110844773 | -1.23862715 | 0.215483609 | 0.435414484 | | Ube2e2 | 2404.979575 | 0.039543041078101 | 0.0498460169547173 | 0.793303927950183 | 0.427600768865518 | 0.810334558001624 | NM_144839 |
| Msx1 | 95.77028498 | 0.191404743 | 0.154580504 | 1.238220459 | 0.215634328 | 0.435631624 | | Wrap53 | 468.8541712 | 0.064699174460401 | 0.0815801857442462 | 0.793074615731237 | 0.427734351265069 | 0.810334558001624 | NM_144824 |
| Stim2 | 2855.945161 | 0.109504737 | 0.088440376 | 1.238175839 | 0.215650868 | 0.435631624 | | Fahd1 | 485.3630308 | 0.0568149679035391 | 0.0716753625026635 | 0.792670817053904 | 0.427969639252126 | 0.810464137520907 | NM_023480 |
| Zfp553 | 890.5760157 | 0.085422109 | 0.06900057 | 1.237991349 | 0.215719268 | 0.43571299 | | Rmnd5a | 6754.819649 | 0.0573988549084314 | 0.0724161073969628 | 0.792625521747373 | 0.427996034622255 | 0.810464137520907 | NM_024288 |
| Lipg | 256.3727519 | -0.160221485 | 0.129505703 | -1.237177057 | 0.216021357 | 0.436266281 | | Apln | 1177.523283 | 0.0668021817256593 | 0.0843441097597017 | 0.792019524730063 | 0.428349291755789 | 0.810486243852795 | NM_013912 |
| Uchl3 | 737.7666646 | -0.090484332 | 0.07314251 | -1.237096338 | 0.216051319 | 0.436269926 | | Arsk | 270.2615336 | -0.0630411842267068 | 0.0796153171600411 | -0.791822308513608 | 0.42846429265528 | 0.810486243852795 | NM_029847 |
| Fgfr3 | 4680.997222 | 0.121244705 | 0.098027728 | 1.236840911 | 0.216146151 | 0.436353563 | | Brk1 | 5586.905557 | 0.0346476939755832 | 0.0437246093121945 | 0.792407171170207 | 0.428123299363076 | 0.810486243852795 | NM_133937 |
| Icmt | 3484.665416 | 0.084248124 | 0.068116002 | 1.236833065 | 0.216149064 | 0.436353563 | | Gpr160 | 37.17106739 | -0.0727727222245316 | 0.0919041189412521 | -0.79183036693981 | 0.428459593266516 | 0.810486243852795 | NM_001134385,NM_001134386,NM_027965 |
| Zfp442 | 98.51912176 | -0.163704021 | 0.132370367 | -1.236712001 | 0.216194022 | 0.436387464 | | Mtrf1 | 296.6505976 | 0.0593346541292831 | 0.0749459839391619 | 0.791698917091737 | 0.428536233907271 | 0.810486243852795 | NM_145960 |
| Myb | 24.71277541 | 0.204879839 | 0.165679679 | 1.236602095 | 0.216234842 | 0.436413005 | | Ogfod3 | 210.80085 | 0.0720414128550012 | 0.0909433083639504 | 0.792157379701816 | 0.428268916304912 | 0.810486243852795 | NM_025402 |
| Abhd10 | 642.2947172 | 0.087592032 | 0.070862927 | 1.236076965 | 0.216429956 | 0.436443452 | | Prpsap1 | 1557.567479 | -0.0509604552602271 | 0.0643332066738427 | -0.792132988467169 | 0.428283136817674 | 0.810486243852795 | NM_026364 |
| Ccdc42 | 55.37902446 | 0.192187296 | 0.155495842 | 1.235964215 | 0.216471866 | 0.436443452 | | Rhoh | 33.26382315 | -0.0722820552551804 | 0.0913167794340486 | -0.791552830741084 | 0.428621459815308 | 0.810486243852795 | NM_001081105 |
| Emp1 | 90.47709606 | 0.191461842 | 0.154907238 | 1.235977376 | 0.216466973 | 0.436443452 | | Sel1l3 | 3410.75924 | -0.0514061205685723 | 0.0649361197393006 | -0.791641397344847 | 0.428569801534631 | 0.810486243852795 | NM_172710 |
| Klk8 | 56.52797813 | 0.206321403 | 0.166942328 | 1.235884306 | 0.216501571 | 0.436443452 | | Stt3a | 1460.620194 | 0.0478363206519171 | 0.0604314838680202 | 0.791579448162974 | 0.428605934279646 | 0.810486243852795 | NM_008408 |
| Map7d2 | 6884.898809 | 0.119973454 | 0.097047 | 1.236240734 | 0.216369094 | 0.436443452 | | Zfp939 | 252.3858476 | -0.0666506011497995 | 0.0841556821495405 | -0.792065572381589 | 0.428327690499484 | 0.810486243852795 | NM_001243021 |
| Pcdhga12 | 572.1809851 | 0.109887872 | 0.0889085 | 1.235965874 | 0.216471249 | 0.436443452 | | Emc1 | 3991.314413 | 0.0373281636668702 | 0.047179360698833 | 0.791196894446125 | 0.428829103452351 | 0.810705510743762 | NM_001039200,NM_146157 |
| Ppia | 29828.72814 | -0.091297336 | 0.073871607 | -1.235892114 | 0.216498669 | 0.436443452 | | P2rx3 | 118.1726567 | 0.073570891749362 | 0.0930020903109011 | 0.791067077131475 | 0.428904849928804 | 0.810705510743762 | NM_145526 |
| Sf1 | 5565.280103 | 0.06920738 | 0.055998496 | 1.235879268 | 0.216503445 | 0.436443452 | | Zbtb22 | 1521.383231 | 0.059229531683601 | 0.0748708663357264 | 0.791089172135117 | 0.42889195727275 | 0.810705510743762 | NM_020625 |
| Zdhhc16 | 966.5522336 | 0.077503447 | 0.06268591 | 1.236377477 | 0.216318284 | 0.436443452 | | Rnf115 | 2116.303904 | 0.0480116764648243 | 0.0607085388270134 | 0.790855411651921 | 0.429028370276775 | 0.810833477314655 | NM_026406 |
| Arpc3 | 3858.624131 | -0.088391259 | 0.071548099 | -1.235410308 | 0.216677838 | 0.436738185 | | Gm5141 | 122.5258018 | 0.075052928536952 | 0.0949327461615379 | 0.790595013511973 | 0.429182984420674 | 0.811020168036257 | NM_001256065 |
| 9930012K11Rik | 63.85689864 | 0.185303988 | 0.150043506 | 1.235001721 | 0.216829864 | 0.436760518 | | Fbxo36 | 106.9655308 | 0.0745916138911487 | 0.0944141187573439 | 0.790047239469114 | 0.429500180635014 | 0.811442205987932 | NM_025386 |
| Baz2a | 3263.369709 | 0.085138761 | 0.068930565 | 1.235138016 | 0.216779143 | 0.436760518 | | Hdac4 | 1353.182725 | 0.0593610058170953 | 0.0751662297921927 | 0.789729727791829 | 0.429685629105603 | 0.811442205987932 | NM_207225 |
| Dopey2 | 4222.116807 | 0.055606204 | 0.045024237 | 1.235028219 | 0.216820002 | 0.436760518 | | Klhdc5 | 1461.714013 | 0.0554025460512769 | 0.0701419834840498 | 0.789862836768444 | 0.429607877031246 | 0.811442205987932 | NM_001081237 |
| Mdm2 | 1693.121607 | 0.118123318 | 0.095625827 | 1.23526585 | 0.216731579 | 0.436760518 | | Mitd1 | 229.1314368 | -0.067261462833423 | 0.0851520266280296 | -0.789898555524015 | 0.42958701504199 | 0.811442205987932 | NM_026913 |
| Usp32 | 7624.303373 | -0.10061915 | 0.08146394 | -1.235137282 | 0.216779416 | 0.436760518 | | Pdxdc1 | 4410.610855 | 0.0388289421960243 | 0.0491632644814402 | 0.789795848700867 | 0.429647003843196 | 0.811442205987932 | NM_001039533,NM_053181 |
| Ccnjl | 118.9656999 | 0.140604968 | 0.11389019 | 1.234566104 | 0.216992031 | 0.436916767 | | Anxa8 | 23.55428055 | 0.0654478042195104 | 0.082889179405425 | 0.789582002967281 | 0.429771922618655 | 0.811499668264472 | NM_013473 |
| Mef2d | 4954.697462 | 0.110365112 | 0.089393119 | 1.234604115 | 0.216977877 | 0.436916767 | | Nln | 1367.088097 | -0.0445766164600484 | 0.0564662342050533 | -0.789438450918676 | 0.429855789972032 | 0.811552535109662 | NM_029447 |
| Tm4sf1 | 318.1300214 | -0.150932427 | 0.122241213 | -1.234709826 | 0.216938518 | 0.436916767 | | Bbip1 | 1293.897137 | -0.0474870411522377 | 0.0601951684378922 | -0.788884596298349 | 0.430179459468162 | 0.81186074378066 | NM_001195338,NM_001195348 |
| Atp6v1g1 | 2501.726398 | -0.069089332 | 0.055982643 | -1.234120567 | 0.217157981 | 0.437194095 | | Ift80 | 1121.762585 | 0.046040179111175 | 0.0583690618214949 | 0.788777096537473 | 0.430242298111135 | 0.81186074378066 | NM_026641 |
| Tmem17 | 164.4543393 | -0.158231722 | 0.128253417 | -1.233742739 | 0.217298783 | 0.437420727 | | Pigp | 713.0271422 | -0.0571521550506482 | 0.0724559360083366 | -0.787849944014464 | 0.430237681484749 | 0.81186074378066 | NM_001159617,NM_001159616,NM_001159620,NM_019543,NM_001159619,NM_001159618 |
| Stxbp5l | 5824.395622 | -0.087711069 | 0.071112267 | -1.233416866 | 0.217420276 | 0.437608437 | | Rnaseh2c | 343.4393081 | -0.0661944384302312 | 0.0838934904739541 | -0.789029495092736 | 0.430094767771373 | 0.81186074378066 | NM_026616 |
| 2610203C20Rik | 668.0500346 | 0.154430251 | 0.125229994 | 1.233173027 | 0.217511217 | 0.437645554 | | Cdhr1 | 458.8840209 | -0.0748822960660497 | 0.0949810260071482 | -0.788392158033901 | 0.430467356388319 | 0.812136658431946 | NM_130878 |
| Gprc5c | 188.5419426 | -0.139526353 | 0.113147192 | -1.233140218 | 0.217523455 | 0.437645554 | | Sept10 | 203.6225349 | -0.0731108049358431 | 0.0927407541233092 | -0.788335242978876 | 0.430500638160952 | 0.812136658431946 | NM_001024911,NM_001024910 |
| Shank2 | 6175.420076 | -0.102569106 | 0.083175533 | -1.233164403 | 0.217514433 | 0.437645554 | | Adarb2 | 512.1500155 | -0.0610901128015929 | 0.0776278552630305 | -0.786961234309964 | 0.431304559526434 | 0.812181605806493 | NM_052977 |
| Cyp2j9 | 1061.31183 | 0.110886798 | 0.089944053 | 1.232841912 | 0.217634751 | 0.437812624 | | Ccdc126 | 365.5588396 | 0.0686591469249991 | 0.0871656303273687 | 0.787686002695504 | 0.430880395001402 | 0.812181605806493 | NM_175098 |
| Prss30 | 21.39475252 | -0.205389491 | 0.166643416 | -1.232508888 | 0.217759049 | 0.438005803 | | Dguok | 385.4466469 | -0.0564254125546852 | 0.0716088380338196 | -0.787967157462276 | 0.430715916173163 | 0.812181605806493 | NM_013764,NM_001162521 |
| Dnajc8 | 2814.115513 | 0.070215213 | 0.056979998 | 1.232278262 | 0.217845157 | 0.438122127 | | Diablo | 1009.065092 | -0.0501688706770043 | 0.063661644792292 | -0.788054893031582 | 0.430664598005588 | 0.812181605806493 | NM_023232 |
| Lyrm5 | 565.1137524 | -0.142865011 | 0.11594493 | -1.232179887 | 0.217881895 | 0.438139141 | | Epor | 329.2007869 | -0.0610679385093931 | 0.0775549311788247 | -0.787415288508009 | 0.431038799874299 | 0.812181605806493 | NM_010149 |
| Zfp955b | 727.2684129 | -0.089808124 | 0.07290575 | -1.231838694 | 0.218009346 | 0.438338542 | | Gm10069 | 40.33905058 | -0.0715315801269052 | 0.0908257104787318 | -0.787566725188719 | 0.430948429128957 | 0.812181605806493 | NR_028592,NR_028593 |
| Gm711 | 9.963430847 | 0.192873315 | 0.156599637 | 1.231633217 | 0.218086127 | 0.438436026 | | Gm16701 | 92.31056591 | 0.0748290395875925 | 0.0950586337045727 | 0.787188257871611 | 0.431171674073184 | 0.812181605806493 | NR_037988 |
| Pappa2 | 173.1223308 | -0.19923811 | 0.161852373 | -1.230986647 | 0.218327859 | 0.438694312 | | Grip2 | 1598.212922 | -0.0526211088571079 | 0.0668669754397393 | -0.786952131527621 | 0.431309888444403 | 0.812181605806493 | NM_001159507 |
| Pdcd10 | 1146.373835 | -0.111034291 | 0.090185337 | -1.231178972 | 0.218255934 | 0.438694312 | | Gucy2e | 224.0846723 | -0.0677603171946545 | 0.0860077048690391 | -0.787840081279122 | 0.430790253011084 | 0.812181605806493 | NM_008192 |
| Stk19 | 633.4944735 | 0.103478736 | 0.084059226 | 1.231021746 | 0.218314731 | 0.438694312 | | L3hypdh | 54.47056413 | 0.0742103924966334 | 0.0943220737615542 | 0.786776515596602 | 0.431412703063481 | 0.812181605806493 | NM_026038 |
| Wasf1 | 8418.016047 | -0.071531343 | 0.05810213 | -1.231131164 | 0.218273812 | 0.438694312 | | Mir3064 | 47.05966274 | -0.0689758380143377 | 0.0876177410696971 | -0.787235977237414 | 0.431143740115662 | 0.812181605806493 | NR_037224 |
| Colec11 | 20.29710053 | 0.206367808 | 0.167685015 | 1.23068724 | 0.218439862 | 0.438862452 | | Nab1 | 1137.920649 | 0.0561592264272923 | 0.0713122886993933 | 0.787511205313725 | 0.430982671544464 | 0.812181605806493 | NM_008667 |
| Zbtb38 | 1411.78459 | 0.133209557 | 0.108250924 | 1.230562771 | 0.218486436 | 0.438899111 | | Ncs1 | 15706.94283 | 0.0415144326810167 | 0.0527356223381866 | 0.78721802911114 | 0.431154244904087 | 0.812181605806493 | NM_019681 |
| Myo1e | 864.1682578 | 0.13113825 | 0.106606044 | 1.230120218 | 0.21865209 | 0.439118015 | | Rdh9 | 23.50898887 | -0.0655289208035656 | 0.0832455021730728 | -0.787176713371574 | 0.431178426995316 | 0.812181605806493 | NM_153133 |
| Zc3hc1 | 860.113706 | -0.073450271 | 0.059709139 | -1.230134486 | 0.218646748 | 0.439118015 | | Tmem106b | 7221.466313 | 0.0513129364073106 | 0.0652201019134208 | 0.786765658162083 | 0.431419060333395 | 0.812181605806493 | NM_027992 |
| 0610038B21Rik | 13.94850875 | -0.203016803 | 0.16505963 | -1.229960366 | 0.218711947 | 0.439129313 | | Wdr45b | 2808.480853 | 0.0435940945 78898 | 0.0554086875452896 | 0.786773636232502 | 0.431414389198835 | 0.812181605806493 | NM_025793 |
| 5033411D12Rik | 156.1623094 | -0.132587892 | 0.107799082 | -1.229953817 | 0.2187144 | 0.439129313 | | Gm2694 | 45.60092276 | 0.0728355704567332 | 0.0926268570100503 | 0.786333173852917 | 0.431672322377155 | 0.812553084728594 | NR_033430 |
| Morn2 | 245.3325895 | -0.141555895 | 0.115110234 | -1.229742051 | 0.218793715 | 0.439231642 | | Bsg | 14125.52614 | 0.0480075076136506 | 0.061124177587194 | 0.785409464939919 | 0.43221353312816 | 0.81325466451425 | NM_009768,NM_001077184 |
| Sowahc | 386.5069707 | 0.129421395 | 0.105275324 | 1.229361163 | 0.218936425 | 0.439461195 | | Ehd3 | 5963.304865 | 0.0452426116888628 | 0.0575989267576869 | 0.785476644021408 | 0.43217415895243 | 0.81325466451425 | NM_020578 |
| Gm13704 | 23.72697792 | 0.172736499 | 0.140525394 | 1.229219107 | 0.218989667 | 0.439511127 | | Khk | 305.7951116 | -0.0673881492734232 | 0.0858103577841558 | -0.785314861906345 | 0.432268984215512 | 0.81325466451425 | NM_008439 |
| Sigmar1 | 1249.228201 | -0.083793938 | 0.068181177 | -1.228989309 | 0.219075815 | 0.439627078 | | Vps33b | 950.6436222 | -0.0441375907927587 | 0.0561974310618153 | -0.785402285456943 | 0.432217741201166 | 0.81325466451425 | NM_178070 |
| Gm11992 | 44.02785316 | 0.190593899 | 0.155185352 | 1.228169388 | 0.219383388 | 0.440187284 | | Acox1 | 10123.43949 | 0.0398532680222204 | 0.0507672382041266 | 0.785019422604339 | 0.432442180983311 | 0.813262564901102 | NM_015729,NM_001271898 |
| Ccdc155 | 147.7129948 | 0.143392039 | 0.11676308 | 1.22805975 | 0.219424539 | 0.440212846 | | Dbt | 869.4177858 | 0.0462792438589516 | 0.0589424543003662 | 0.785159769953185 | 0.432359899489241 | 0.813262564901102 | NM_010022 |
| Psme1 | 1074.220716 | 0.081853715 | 0.066660053 | 1.227927543 | 0.219474169 | 0.440255408 | | Fam168b | 17363.14694 | 0.0397477369897684 | 0.0506388525270615 | 0.78492570439915 | 0.43249713023964 | 0.813262564901102 | NM_001160235,NM_174997,NM_001160236 |
| Shc1 | 526.6805184 | 0.125435722 | 0.10216022 | 1.227833325 | 0.219509543 | 0.440269367 | | Tacr1 | 504.2905649 | 0.0719690280478284 | 0.0916718263765295 | 0.785072479653949 | 0.432411074140626 | 0.813262564901102 | NM_009313 |
| Rpap3 | 919.3708231 | 0.100317204 | 0.08171714 | 1.227615203 | 0.219591452 | 0.440376644 | | 4930506M07Rik | 1390.457924 | 0.0574354731642381 | 0.0732059613852806 | 0.78457371609338 | 0.432703545619222 | 0.81346077646298 | NM_001114312,NM_175172 |
| Dock11 | 422.6409196 | -0.166404045 | 0.135571978 | -1.227422119 | 0.219663977 | 0.440386617 | | Zdhhc7 | 1422.281772 | 0.048068061911071 | 0.0612679337818832 | 0.784554969361226 | 0.432714540804726 | 0.81346077646298 | NM_133967 |
| Gbp2 | 230.6271282 | 0.193073121 | 0.157300165 | 1.227418425 | 0.219665365 | 0.440386617 | | Cccd9 | 1583.548336 | -0.0536631671838593 | 0.0684781729894166 | -0.783653605830767 | 0.433243392222199 | 0.813783127125213 | NM_001136471,NM_172297 |
| Ppp2r3a | 1818.965405 | 0.093344963 | 0.076052524 | 1.227374959 | 0.219681694 | 0.440386617 | | Commd4 | 1347.17706 | -0.058363684629666 | 0.0744728267042567 | -0.783691008016083 | 0.433221440038085 | 0.813783127125213 | NM_025417 |
| Abca3 | 4752.108171 | 0.068026998 | 0.055431317 | 1.227230408 | 0.219736003 | 0.440438504 | | Fam122b | 243.288841 | 0.0642245736294326 | 0.0819061523317273 | 0.784123924773286 | 0.432967398281557 | 0.813783127125213 | NM_030167,NM_001166583,NM_001166365 |
| Zswim5 | 889.6096737 | -0.112497115 | 0.09171662 | -1.226572838 | 0.219983183 | 0.440876917 | | Gdf9 | 37.00989605 | 0.0723625654391562 | 0.092309018856752 | 0.78391652663377 | 0.4330890917239 | 0.813783127125213 | NM_008110 |
| Zfp703 | 1364.58744 | 0.063844223 | 0.052073152 | 1.226048757 | 0.220180328 | 0.44121495 | | Sdf4 | 12663.99184 | 0.0337126438942132 | 0.0430230817489049 | 0.783594352700487 | 0.433278170534655 | 0.813783127125213 | NM_011341 |
| Pcdhb15 | 126.4936235 | -0.145722863 | 0.118880652 | -1.225791249 | 0.220277241 | 0.441352072 | | Tuba4a | 24712.82242 | 0.0409929108247165 | 0.0523072374458333 | 0.783694815983479 | 0.433219205092431 | 0.813783127125213 | NM_009447 |
| Chsy1 | 564.985191 | -0.118547924 | 0.096741335 | -1.225411289 | 0.220420296 | 0.441581596 | | Zbtb5 | 508.0232383 | 0.0526687322294698 | 0.0672018643907668 | 0.783739152283196 | 0.433193184031655 | 0.813783127125213 | NM_001163283,NM_001163284,NM_173399 |
| Mcph1 | 529.1154347 | -0.081137281 | 0.066227824 | -1.225123769 | 0.220528591 | 0.441741433 | | Arl10 | 405.0621861 | 0.0677416555699204 | 0.0864713314783535 | 0.783400167567424 | 0.43339215779224 | 0.813891982429228 | NM_019968 |
| Fam71f2 | 17.1322985 | 0.205849466 | 0.168065478 | 1.224817066 | 0.220644153 | 0.441801565 | | Itga3 | 980.4745997 | 0.0534624531133931 | 0.0682559786873365 | 0.783264032566161 | 0.433472079788103 | 0.813936844752067 | NM_013565 |
| Gm15421 | 16.51977924 | 0.202191821 | 0.165075476 | 1.22484469 | 0.220633743 | 0.441801565 | | Nudc | 4642.277746 | 0.0353528634899583 | 0.0451624804424653 | 0.782792777181405 | 0.433748809723349 | 0.814351196851669 | NM_010948 |
| Ppm1a | 6535.341505 | -0.079083409 | 0.064561494 | -1.224931525 | 0.220601021 | 0.441801565 | | Sacs | 277.8933766 | -0.0739977516832326 | 0.0945719410642633 | -0.782449327469654 | 0.433950554111532 | 0.814624676792984 | NM_172809 |
| Mir22hg | 720.719731 | 0.123541393 | 0.100899286 | 1.224403055 | 0.220800217 | 0.44191594 | | Med18 | 114.5724284 | 0.0718260786165571 | 0.0918277413832524 | 0.782182786319264 | 0.434107159326143 | 0.814708087386506 | NM_026039 |
| Nkiras1 | 3047.50552 | 0.070592546 | 0.057656554 | 1.224362893 | 0.220815361 | 0.44191594 | | Vav3 | 319.3728672 | 0.072173549334461 | 0.0922669573021058 | 0.782225310607634 | 0.434082172156862 | 0.814708087386506 | NM_020505,NM_146139 |
| Rac2 | 99.62284882 | 0.167697062 | 0.13696095 | 1.224415149 | 0.220795657 | 0.44191594 | | Scarb2 | 3376.609283 | 0.0441036607474496 | 0.0564036858733807 | 0.781928699596988 | 0.434256477390184 | 0.814883037148922 | NM_007644 |
| Snhg1 | 246.8679815 | 0.127594805 | 0.104200454 | 1.224512948 | 0.220758785 | 0.44191594 | | Hist1h2bc | 456.3911049 | -0.0579712553461165 | 0.0741596018259531 | -0.781709366268857 | 0.434385395934484 | 0.815019666926416 | NM_023422 |
| Angptl6 | 239.6731691 | -0.167431002 | 0.136840253 | -1.223550807 | 0.221121727 | 0.442129311 | | Ankrd32 | 629.8214639 | -0.0537779204133968 | 0.06887162564325 | -0.780842907527151 | 0.434894894443997 | 0.815326687685887 | NM_134071 |
| Dhx16 | 1012.827522 | 0.088513761 | 0.07232198 | 1.223884652 | 0.220995744 | 0.442129311 | | Dnajc21 | 1218.6789 | 0.0611094433715875 | 0.078213943357815 | 0.781311371708017 | 0.43461938344857 | 0.815326687685887 | NM_030046 |
| Ercc6l2 | 869.0277835 | 0.105301385 | 0.086048953 | 1.223738129 | 0.221051031 | 0.442129311 | | Dr1 | 1070.264524 | 0.0559352944669139 | 0.0716219302329832 | 0.780979991532742 | 0.434814262796729 | 0.815326687685887 | NM_026106 |
| Nmnat2 | 7365.105149 | 0.079196775 | 0.064720231 | 1.223678808 | 0.221073417 | 0.442129311 | | Lrp12 | 2548.397071 | 0.0454163774936767 | 0.0581534678239845 | 0.780974535020685 | 0.434817472105804 | 0.815326687685887 | NM_172814 |
| Nrm | 85.00248411 | -0.177921108 | 0.145408767 | -1.223592719 | 0.221105908 | 0.442129311 | | Ncln | 2329.108165 | -0.0542135630445353 | 0.0694536118522135 | -0.780572263972294 | 0.435054110024854 | 0.815326687685887 | NM_134009 |
| Sctr | 16.60404906 | 0.193946337 | 0.158495881 | 1.223667996 | 0.221077497 | 0.442129311 | | Ncoa7 | 3389.027683 | 0.0525744775798291 | 0.0673456684007213 | 0.780660659574903 | 0.434998927656755 | 0.815326687685887 | NM_001111267,NM_172495 |
| Tmprss4 | 8.581361245 | 0.178545787 | 0.145897873 | 1.223772379 | 0.221038107 | 0.442129311 | | Ntrk1 | 102.6440613 | 0.0738247762241604 | 0.0945389604723903 | 0.780892616708225 | 0.434865654928576 | 0.815326687685887 | NM_001033124 |
| Mpzl2 | 77.66480809 | -0.2053997 | 0.167893936 | -1.223389632 | 0.221182567 | 0.442136847 | | Slc24a2 | 29718.16885 | 0.0605039160745532 | 0.0774796352106666 | 0.7809008897634 | 0.434860788731972 | 0.815326687685887 | NM_172426,NM_001110240 |
| Sox12 | 795.4184923 | 0.125155363 | 0.102301455 | 1.223397675 | 0.221179531 | 0.442136847 | | Tnfrsf11b | 78.06196496 | -0.0725613572571391 | 0.0929591956522307 | -0.780572128939846 | 0.435054189470833 | 0.815326687685887 | NM_008764 |
| Trhr | 156.7015947 | 0.204829705 | 0.167503173 | 1.222840745 | 0.221389853 | 0.442484962 | | 4932416H05Rik | 274.9223674 | -0.0694954872374306 | 0.0890481084967687 | -0.780426315736424 | 0.435139983164455 | 0.815382274954448 | NR_029452 |

| | | | | | | | | | | | | | | | |
|---|---|---|---|---|---|---|---|---|---|---|---|---|---|---|---|
| Unc119b | 1244.430031 | -0.091942969 | 0.075191918 | -1.222777273 | 0.221413832 | 0.442484962 | | Dcbld1 | 392.7101598 | 0.0582806132882764 | 0.0746974631190292 | 0.780222123412775 | 0.435260142422775 | 0.815502235113276 | NM_025705 |
| Lrp6 | 2236.124704 | -0.092163079 | 0.075406546 | -1.222215891 | 0.221625996 | 0.442851842 | | Prrt2 | 13263.8674 | -0.0329457881310485 | 0.0422407960163135 | -0.779951876814176 | 0.435419201494356 | 0.815695037517026 | NM_001102563 |
| Agbl5 | 502.7610006 | 0.157967749 | 0.129271279 | 1.221986433 | 0.221712758 | 0.442910968 | | Emg1 | 882.1332981 | 0.0489765770411152 | 0.0628258459116103 | 0.779560964607152 | 0.435649340064721 | 0.815960392627735 | NM_013536 |
| Rspo3 | 366.9279618 | -0.155057063 | 0.126887156 | -1.222007557 | 0.221704769 | 0.442910968 | | Fbxo2 | 2461.630796 | 0.0484931293506524 | 0.0622241534963153 | 0.779329675469575 | 0.435785538070079 | 0.815960392627735 | NM_176848 |
| 2610008E11Rik | 1137.069822 | 0.105900283 | 0.086670217 | 1.221876297 | 0.22175441 | 0.442929929 | | Idnk | 483.8567001 | 0.0557803520107465 | 0.0715670405239292 | 0.779413981665146 | 0.435735890284926 | 0.815960392627735 | NM_198004,NM_001048060 |
| 3300002I08Rik | 24.77765357 | -0.201265728 | 0.164732052 | -1.221776365 | 0.221792209 | 0.442929929 | | Ttc39b | 3331.717801 | 0.0536117200987457 | 0.068787754685085 | 0.779378834854892 | 0.435756587789359 | 0.815960392627735 | NM_027238 |
| Capza2 | 6975.491148 | -0.065476787 | 0.053593299 | -1.221734592 | 0.221808011 | 0.442929929 | | Car3 | 41.24078025 | -0.0718308681736258 | 0.0922153829922725 | -0.778946698943333 | 0.436011113405692 | 0.816020977050171 | NM_007606 |
| Socs6 | 417.0393565 | -0.120474573 | 0.098629754 | -1.221483049 | 0.22190318 | 0.44306287 | | Espn | 39.6417337 | -0.0685370518711852 | 0.0879971664774422 | -0.77885521335229 | 0.436065008894693 | 0.816020977050171 | NM_019585,NM_207688,NM_207689,NM_207687,NM_207691,NM_207690 |
| Nup98 | 1105.824405 | 0.093701023 | 0.076720268 | 1.221333372 | 0.221959823 | 0.443118863 | | Polq | 81.00769293 | -0.0740650531162417 | 0.0950816175740299 | -0.778962907527461 | 0.436001566095558 | 0.816020977050171 | NM_029977,NM_001159369 |
| Agl | 1917.02454 | -0.079990613 | 0.065502916 | -1.22117637 | 0.222019249 | 0.443158192 | | Rnf181 | 2668.684219 | 0.0344214709979936 | 0.044180957546383 | 0.779101968576767 | 0.435919650602528 | 0.816020977050171 | NM_025607 |
| Retn | 19.45650277 | 0.205185317 | 0.168029025 | 1.2211302 | 0.222036727 | 0.443158192 | | Tnnc1 | 214.1709953 | -0.0661679692433431 | 0.0849616693053331 | -0.778797895384451 | 0.436098777697954 | 0.816020977050171 | NM_009393 |
| Adprh | 3094.872523 | -0.0533568 | 0.043707137 | -1.22078001 | 0.222169324 | 0.443251547 | | Stk32c | 7229.015864 | 0.0532896158882224 | 0.0684384141112708 | 0.778650653373462 | 0.436185531827494 | 0.816078185899817 | NM_001162540,NM_021302 |
| Map3k8 | 52.985293 | 0.176807026 | 0.144828555 | 1.220802251 | 0.222160901 | 0.443251547 | | Lingo2 | 943.0089269 | 0.0651698292725955 | 0.0837101977260686 | 0.778517206300907 | 0.436264167057324 | 0.816120191973559 | NM_175516,NM_001166000,NM_001166001,NM_001165999 |
| Nfib | 2903.103431 | -0.083600642 | 0.068480574 | -1.22079355 | 0.222164197 | 0.443251547 | | Glipr2 | 127.1244699 | 0.0735028547231077 | 0.0944409658342117 | 0.778294186996559 | 0.436395601641164 | 0.816260946361002 | NM_027450 |
| Slc5a11 | 33.77533449 | -0.205024479 | 0.167974594 | -1.220568383 | 0.222249483 | 0.443297309 | | Mrps9 | 957.0270195 | 0.0469044816821742 | 0.0602843255057554 | 0.778054349761218 | 0.436536973194933 | 0.816420248825749 | NM_023514 |
| Ssr1 | 5415.770971 | -0.065470127 | 0.05363633 | -1.220630249 | 0.222226048 | 0.443297309 | | Cox16 | 304.7187999 | 0.0640718564321791 | 0.08236160148264 | 0.777933591367623 | 0.436608163962449 | 0.816448273650687 | NM_025461 |
| Tmem8b | 4246.408514 | -0.08740621 | 0.071638475 | -1.220101495 | 0.222426402 | 0.443593084 | | Nbas | 2370.687572 | 0.044837927190616 | 0.0576682954937688 | 0.777514348338262 | 0.436855372467845 | 0.816490110713834 | NM_027706 |
| Aamdc | 575.091094 | -0.087465392 | 0.071702283 | -1.219841097 | 0.222525119 | 0.443624247 | | Nmbr | 43.07922016 | 0.0697218873831422 | 0.0896659713707682 | 0.777573546767733 | 0.436820460965171 | 0.816490110713834 | NM_008703 |
| Actn3 | 39.49687424 | 0.189579087 | 0.155419359 | 1.21979069 | 0.222544232 | 0.443624247 | | Plaa | 1796.127145 | 0.0548912335316737 | 0.0705822133294304 | 0.777692154190153 | 0.436750518618038 | 0.816490110713834 | NM_172695 |
| Slc1a5 | 130.9747615 | 0.156570865 | 0.128359591 | 1.219783145 | 0.222547092 | 0.443624247 | | Slc1a4 | 2977.968852 | 0.047846943560698 | 0.0615364791237273 | 0.777537880652797 | 0.436841494399445 | 0.816490110713834 | NM_018861 |
| Wbp11 | 3524.039969 | 0.08552911 | 0.070119726 | 1.219758189 | 0.222556556 | 0.443624247 | | Crat | 2341.42568 | -0.0412216594445248 | 0.0530315790850231 | -0.777304016884657 | 0.436979425699987 | 0.816564673280286 | NM_007760 |
| A430107P09Rik | 20.59531828 | -0.204439548 | 0.167655397 | -1.219403318 | 0.222691151 | 0.443835438 | | Fam136a | 784.2721074 | -0.0552829496544741 | 0.0711432293504067 | -0.777065507979475 | 0.437120122499484 | 0.816564673280286 | NM_025591 |
| Scg5 | 7026.97994 | 0.121394655 | 0.099567463 | 1.219220123 | 0.222760656 | 0.443916863 | | Prmt8 | 4218.65397 | 0.0424654135636035 | 0.0546456080626777 | 0.777105701063776 | 0.437096410705105 | 0.816564673280286 | NM_201371 |
| Ubtf | 5537.519114 | 0.050944081 | 0.041801766 | 1.218706455 | 0.222955627 | 0.444248262 | | Zfp185 | 83.60433733 | -0.0635221463916287 | 0.0817412406788417 | -0.777112579453068 | 0.43709235289336 | 0.816564673280286 | NM_009549,NM_001109043 |
| Rbpj | 1607.941894 | 0.081825204 | 0.067152466 | 1.218498864 | 0.223034456 | 0.44429106 | | Gng11 | 349.3889454 | -0.0718085008678808 | 0.0924338221631858 | -0.776863913958979 | 0.43723906348394 | 0.816681835690846 | NM_025331 |
| Samd10 | 844.1962258 | 0.076043774 | 0.062407258 | 1.218508493 | 0.223030799 | 0.44429106 | | Nudt17 | 284.8727331 | 0.0679185428795154 | 0.0874729452471188 | 0.77645199538714 | 0.437482154407387 | 0.816715817365596 | NM_030094,NM_001162925 |
| Trpc1 | 1313.426421 | 0.087967359 | 0.072208966 | 1.218233193 | 0.223135369 | 0.444434942 | | Pfdn1 | 1520.077688 | 0.0424892314466949 | 0.0547087412614778 | 0.776644288773153 | 0.4373686641045 | 0.816715817365596 | NM_026027 |
| Lyz1 | 41.94389754 | 0.204237495 | 0.167680103 | 1.218018666 | 0.223216879 | 0.444540145 | | Prodh | 1220.572305 | 0.0610492565618994 | 0.0786082900908823 | 0.776626186516941 | 0.437379347215511 | 0.816715817365596 | NM_011172 |
| Tmem79 | 11.73558908 | 0.20150271 | 0.16544773 | 1.217923693 | 0.223252972 | 0.444554883 | | Suds3 | 3755.615839 | 0.0354722889322369 | 0.0456831887561157 | 0.776484520850969 | 0.437462956899858 | 0.816715817365596 | NM_178622,NM_001122666 |
| Btla | 14.16077839 | -0.202362112 | 0.166180286 | -1.217726345 | 0.223327982 | 0.444647103 | | B230216N24Rik | 133.1087361 | 0.0707234579109818 | 0.0912138726070788 | 0.775358570901123 | 0.438127808502515 | 0.816984131628323 | NR_037993 |
| AW554918 | 596.6801396 | 0.098631829 | 0.081002051 | 1.217646065 | 0.223358501 | 0.444650728 | | BC064078 | 82.99375902 | -0.0735390371389669 | 0.0948490489192118 | -0.775327090539454 | 0.438146405391486 | 0.816984131628323 | NR_015455 |
| Cd99l2 | 3997.885305 | -0.100250235 | 0.08234684 | -1.217414467 | 0.223446561 | 0.444711756 | | Chn1 | 46967.97579 | 0.0465394246984884 | 0.0600262072912455 | 0.775318428377131 | 0.438151522606451 | 0.816984131628323 | NM_001113246,NM_029716,NM_001166604,NM_175752,NM_001166603 |
| Ism1 | 229.6560945 | 0.16534679 | 0.13581646 | 1.217428213 | 0.223441334 | 0.444711756 | | Gata2 | 212.8275769 | -0.0707890964201905 | 0.0912515641681665 | -0.775757621970557 | 0.437892110139077 | 0.816984131628323 | NM_008090 |
| Dock5 | 616.5031507 | -0.123316932 | 0.101310274 | -1.217220388 | 0.223520375 | 0.444744406 | | Nfia | 1168.07113 | -0.0534492345533275 | 0.0688897306900784 | -0.775866504599725 | 0.437827811595933 | 0.816984131628323 | NM_001122952,NM_001122953,NM_010905 |
| Grk1 | 29.27377036 | 0.200595765 | 0.164797791 | 1.217223625 | 0.223519144 | 0.444744406 | | Rimbp3 | 126.5856325 | -0.0720653855659973 | 0.0929568692431352 | -0.775256160763173 | 0.438188308516884 | 0.816984131628323 | NM_001033338 |
| Foxm1 | 76.86541793 | -0.184648117 | 0.151731069 | -1.216943362 | 0.223625766 | 0.444896972 | | Sdk2 | 554.5047146 | 0.0682265913051366 | 0.0879413211308585 | 0.77581949449695 | 0.43785557184197 | 0.816984131628323 | NM_172800 |
| Smc6 | 1622.652002 | 0.095936059 | 0.07886773 | 1.216417136 | 0.223826059 | 0.44523828 | | Slc15a2 | 1109.383752 | -0.0491407119029143 | 0.0633625338210749 | -0.775548402860892 | 0.438015675736416 | 0.816984131628323 | NM_021301,NM_001145899 |
| Gltpd1 | 798.2591467 | 0.091876733 | 0.075541058 | 1.216248946 | 0.223890104 | 0.445308506 | | Slc26a1 | 40.89580576 | -0.0709053659405932 | 0.0914305549567345 | -0.775510615397074 | 0.438037995257649 | 0.816984131628323 | NM_174870 |
| Lrtm2 | 3457.775125 | 0.061726027 | 0.050760056 | 1.216035437 | 0.223971423 | 0.44541307 | | Txndc11 | 1214.490965 | 0.0450312847608488 | 0.0580331777106377 | 0.775957590766124 | 0.437774026584181 | 0.816984131628323 | NM_029582,NM_134105 |
| Gm4371 | 10.32010343 | 0.191327953 | 0.157354583 | 1.215903276 | 0.22402177 | 0.44545602 | | Crtac1 | 2643.917884 | 0.0543045280068562 | 0.0700705334369953 | 0.774998067564089 | 0.438340801469274 | 0.817094138947092 | NM_145123 |
| Depdc5 | 2723.774465 | -0.071128335 | 0.058531911 | -1.215206096 | 0.224287496 | 0.445927174 | | Rfwd2 | 3150.224455 | 0.0562392568014287 | 0.0725699763762321 | 0.774965896500525 | 0.438359811704164 | 0.817094138947092 | NM_011931 |
| Gpr75 | 446.1384369 | -0.108720181 | 0.089477831 | -1.215051594 | 0.224346414 | 0.445987085 | | Agtrap | 522.3748529 | 0.0686213822339537 | 0.0886067069224073 | 0.774449075215551 | 0.438665272039913 | 0.817139233348521 | NM_009642 |
| Slc7a1 | 2120.111597 | -0.102840137 | 0.084658529 | -1.214764052 | 0.224456095 | 0.446147882 | | B930003M22Rik | 25.89232793 | -0.0652968177798932 | 0.0842980862012689 | -0.774594308392619 | 0.438579421550342 | 0.817139233348521 | NR_037588 |
| Sox11 | 473.1803678 | -0.154346177 | 0.127097274 | -1.21439408 | 0.224597275 | 0.446371239 | | Lemd3 | 1495.260578 | 0.0544900531259042 | 0.0703521296725657 | 0.774533100554496 | 0.438615601657305 | 0.817139233348521 | NM_001081193 |
| 1500012F01Rik | 584.0385071 | 0.183924738 | 0.151529684 | 1.213786849 | 0.22482913 | 0.446526045 | | Plekha6 | 5832.610833 | -0.0417671322394699 | 0.0539085101425044 | -0.774778084741363 | 0.438470801195702 | 0.817139233348521 | NM_182930,NM_001160268 |
| 4930452G13Rik | 12.46373813 | -0.194670667 | 0.16037503 | -1.213846488 | 0.224806351 | 0.446526045 | | Sh3bgrl | 1643.953609 | 0.0606190350867377 | 0.0782587588638672 | 0.774597450391282 | 0.43857756435332 | 0.817139233348521 | NM_019989 |
| Aldh2 | 2521.000786 | -0.087702473 | 0.072258205 | -1.213737229 | 0.224848083 | 0.446526045 | | 2500004C02Rik | 233.8569072 | -0.0653789927399343 | 0.0844398474912087 | -0.774267063269402 | 0.438772876894125 | 0.81723487751815 | NR_040318 |
| Aup1 | 1851.796753 | 0.092134572 | 0.075897273 | 1.213937846 | 0.22477146 | 0.446526045 | | A830009L08Rik | 165.2782873 | -0.0689190455171788 | 0.0890732962349877 | -0.773734086761097 | 0.439088058042036 | 0.817545616940309 | NR_045161 |
| Dcaf13 | 734.6630133 | 0.07902049 | 0.065087868 | 1.214058648 | 0.224725329 | 0.446526045 | | Adamtsl1 | 352.0561341 | 0.0679508597324642 | 0.087871675534476 | 0.773296506742995 | 0.439346922675561 | 0.817545616940309 | NM_029967 |
| Elfn2 | 4314.869525 | -0.091738796 | 0.075576235 | -1.213857717 | 0.224802062 | 0.446526045 | | Cisd1 | 2350.291021 | 0.0360372586009013 | 0.0466089473248551 | 0.773183276372427 | 0.439413922037981 | 0.817545616940309 | NM_134007 |
| Bcl7a | 4015.134442 | 0.078181447 | 0.064438207 | 1.213277816 | 0.225023621 | 0.446817377 | | Dhx15 | 3629.886969 | 0.0527055624039648 | 0.0681521019267435 | 0.773351971750158 | 0.439314105704954 | 0.817545616940309 | NM_001042620,NM_007839 |
| Eif4h | 12957.06104 | -0.060242298 | 0.049660783 | -1.213075877 | 0.225100812 | 0.446913375 | | Gmeb1 | 960.2058012 | 0.0439473586855508 | 0.0567890747549132 | 0.773869954303995 | 0.43900769902359 | 0.817545616940309 | NM_001122992,NM_020273 |
| Stat5b | 1518.699839 | -0.094645898 | 0.078028556 | -1.21296488 | 0.225143248 | 0.446940357 | | Hip1 | 3005.812957 | 0.0455448113628629 | 0.0588802108440226 | 0.773516444812231 | 0.439216800234512 | 0.817545616940309 | NM_146001 |
| Cdk5r2 | 9618.630456 | -0.114806707 | 0.094666765 | -1.212745643 | 0.225227083 | 0.447049504 | | Pcdha11 | 151.4618078 | 0.069778228489765 | 0.0902341658159977 | 0.773301640889043 | 0.439343884897194 | 0.817545616940309 | NM_009960 |
| Cep192 | 381.4003644 | 0.134796892 | 0.11115772 | 1.212663341 | 0.225258561 | 0.447054713 | | Prlr | 276.2364173 | 0.0361924629805722 | 0.0468129889400849 | 0.7731286508301 | 0.439446246528289 | 0.817545616940309 | NM_001253782,NM_001253781,NM_011169 |
| Chrna4 | 1802.433821 | -0.098084619 | 0.080902752 | -1.212376788 | 0.225368182 | 0.447194849 | | Slc19a2 | 678.0016153 | 0.0474268663073855 | 0.0613301645904206 | 0.773304076780406 | 0.439342443629969 | 0.817545616940309 | NM_001276455,NM_054087 |
| Igj | 13.35680682 | 0.200064434 | 0.165025021 | 1.212327878 | 0.225386896 | 0.447194849 | | Aard | 58.11741459 | 0.0713941996052971 | 0.0923596210685528 | 0.773002300989026 | 0.439521018868763 | 0.817580121862962 | NM_175503 |
| Ublcp1 | 1739.040398 | 0.065800303 | 0.054291124 | 1.21199007 | 0.225516182 | 0.447394074 | | Map3k11 | 1127.767866 | -0.0481392621027571 | 0.0622852823252186 | -0.772883421341835 | 0.439591377115613 | 0.817606190010422 | NM_022012 |
| Scn4b | 15817.88756 | -0.131104229 | 0.108200814 | -1.211675076 | 0.225636783 | 0.447576024 | | Camk1d | 9941.852593 | -0.0538148872174019 | 0.0696490583467223 | -0.77265778597471 | 0.43972493590089 | 0.817695543515655 | NM_177343 |
| Ube2w | 1905.665933 | -0.095060138 | 0.078465643 | -1.211487393 | 0.225708663 | 0.447661294 | | Ern1 | 344.8220218 | -0.064904838421111 | 0.0840070216394154 | -0.772612064497454 | 0.439752002336796 | 0.817695543515655 | NM_023913 |
| Lims2 | 410.3786252 | -0.135358477 | 0.111757704 | -1.211178048 | 0.225827174 | 0.447839016 | | Ppp1r12c | 4314.09069 | -0.0494618722564414 | 0.0640330109258604 | -0.772443331045451 | 0.439851898311111 | 0.817776612679791 | NM_029834 |
| Hspa8 | 79645.73242 | -0.061707348 | 0.050991474 | -1.210150298 | 0.226221226 | 0.44856305 | | Zcchc2 | 1292.460647 | -0.0491342018880219 | 0.0636304973222248 | -0.772180070182484 | 0.440007783751304 | 0.817961743891917 | NM_001122676,NM_001122675 |
| Marveld3 | 19.68125727 | -0.203164071 | 0.167921583 | -1.209874679 | 0.226326985 | 0.448715329 | | Ube2d2a | 5972.446498 | 0.041113234782285 | 0.0532815413986379 | 0.771622473807337 | 0.44033805960114 | 0.818470972970717 | NM_019912 |
| Jak3 | 226.7371253 | 0.181152076 | 0.149740015 | 1.209777336 | 0.226364345 | 0.448731981 | | Med21 | 531.5227941 | 0.0615621642581735 | 0.0798006586285882 | 0.771449325308189 | 0.440440647956171 | 0.81855691558487 | NM_025315 |
| Ubxn8 | 610.2856804 | -0.109426809 | 0.090465819 | -1.209592865 | 0.226435157 | 0.448757525 | | Mcm10 | 24.72572504 | 0.0663810826288624 | 0.0860763940796371 | 0.771188005011539 | 0.440595502964619 | 0.818563571794836 | NM_027290 |
| Xk | 853.179271 | -0.080561556 | 0.066601265 | -1.209609987 | 0.226428584 | 0.448757525 | | Slc12a8 | 59.07772276 | 0.072970101815706 | 0.0946084855026125 | 0.77128495851133 | 0.4405380459399 | 0.818563571794836 | NM_001083902,NM_134251 |
| Gorab | 335.2005745 | -0.098535439 | 0.081493187 | -1.209124866 | 0.226614877 | 0.449017805 | | Svop | 8344.723636 | 0.0489628218692701 | 0.0634925939134872 | 0.77115800208108 | 0.4406132843084 | 0.818563571794836 | NM_026805 |
| Tgfbr1 | 1218.935316 | -0.109708693 | 0.090735835 | -1.209099945 | 0.226624451 | 0.449017805 | | Smad7 | 808.5485568 | 0.0597346501640683 | 0.0774794136171802 | 0.770974474061106 | 0.440722061799341 | 0.81866095520425 | NM_001042660 |
| Zfp52 | 162.640089 | 0.142788035 | 0.11811528 | 1.208887073 | 0.226706233 | 0.449122411 | | Ndst4 | 275.1989516 | 0.0727322711479735 | 0.0943576748228939 | 0.770814576392323 | 0.440816846072725 | 0.81873232400619 | NM_022565 |
| Gucy2f | 49.22275876 | -0.192234545 | 0.159039874 | -1.208719174 | 0.226770753 | 0.449192795 | | Osbpl1a | 14364.4813 | 0.0523209116327111 | 0.0678901599217191 | 0.770670030723743 | 0.440902540028972 | 0.81878679320645 | NR_045524,NM_207530,NM_001252489,NM_001252491,NM_001252490,NM_001252493,NM_001252492 |
| Mvk | 620.8295266 | -0.137366392 | 0.113681344 | -1.208345961 | 0.226914216 | 0.449416926 | | Cdpf1 | 322.4609421 | 0.0649425208040477 | 0.0843417717777912 | 0.769992370745385 | 0.441304418292399 | 0.818962635635223 | NM_197998,NM_001164626,NM_001164625 |
| Rps6 | 8462.512855 | -0.087572861 | 0.072477655 | -1.208273925 | 0.226941915 | 0.449416926 | | Ermn | 2834.863886 | 0.0689388431730175 | 0.0895181058786973 | 0.770110610544351 | 0.441234282464885 | 0.818962635635223 | NM_029972 |
| Fbxl18 | 613.091389 | 0.134106923 | 0.111000221 | 1.208168073 | 0.22698262 | 0.449440091 | | Gripap1 | 4444.940047 | -0.036454223123994 | 0.0473468381222694 | -0.769939970011385 | 0.441335502667805 | 0.818962635635223 | NM_207670 |
| Ccnd3 | 821.2100373 | 0.10805621 | 0.089448568 | 1.20802615 | 0.227037204 | 0.449490728 | | Lpar4 | 36.14557277 | -0.069689023127997 | 0.0904971138337887 | -0.770069012985222 | 0.441258955994142 | 0.818962635635223 | NM_175271 |
| Ncald | 20174.25527 | -0.111054383 | 0.09194722 | -1.207805769 | 0.227121982 | 0.449601123 | | Tiam2 | 2792.513929 | 0.0594089785612454 | 0.0771469468223839 | 0.770075563690462 | 0.441255070400807 | 0.818962635635223 | NM_001122998,NM_011878 |
| Ggh | 369.8772891 | -0.139542036 | 0.115597017 | -1.207142194 | 0.227377388 | 0.449991729 | | Ubl5 | 2733.87901 | 0.0481140964348562 | 0.0624528357298077 | 0.770406913835173 | 0.441058553484081 | 0.818962635635223 | NM_025401 |
| Msl2 | 2028.619259 | -0.072591261 | 0.060131089 | -1.207216811 | 0.227348658 | 0.449991729 | | Tbl1x | 1628.922442 | 0.0443199984984468 | 0.0575727062230196 | 0.769809192688707 | 0.441413085895162 | 0.819001977993758 | NM_020601 |
| Cops3 | 2205.103575 | -0.063366566 | 0.052521768 | -1.206481982 | 0.227631702 | 0.450437496 | | Gm11992 | 58.02173373 | 0.0720682620828338 | 0.0936587629530288 | 0.769476980162306 | 0.441610205033507 | 0.819100917722126 | NM_001037928 |
| Rpgrip1 | 26.68339034 | -0.200379139 | 0.166111191 | -1.20629524 | 0.227703672 | 0.450522372 | | Idh3g | 4824.478471 | -0.038790864171331 | 0.0504163547012154 | -0.769410331254985 | 0.441649757400077 | 0.819100917722126 | NM_008323 |
| Slc25a3 | 11865.39342 | -0.065791962 | 0.054547713 | -1.206136029 | 0.227765045 | 0.450586262 | | Nhlrc3 | 205.0077852 | -0.0686452626786685 | 0.0892196660091244 | -0.769396095605741 | 0.4416582057177 | 0.819100917722126 | NM_172501 |
| Bcas3 | 2529.635635 | 0.075234738 | 0.062400248 | 1.205680119 | 0.227940854 | 0.450876495 | | Slmo1 | 1622.759108 | -0.0428851961318173 | 0.0557428966363578 | -0.769339211264558 | 0.441691965339213 | 0.819100917722126 | NM_144867 |
| Clns1a | 2222.744848 | -0.057411706 | 0.047630614 | -1.205353049 | 0.228067039 | 0.450971516 | | Chic2 | 201.6589089 | 0.0653289996507947 | 0.0849456685753204 | 0.769068049571806 | 0.441852914223996 | 0.819294796019629 | NM_028850 |
| Lrch3 | 1585.957394 | 0.099208661 | 0.082308347 | 1.205329281 | 0.22807621 | 0.450971516 | | 1600014C10Rik | 1392.087473 | 0.0419930217675218 | 0.0546177179234652 | 0.768853466678448 | 0.441980304375415 | 0.819321835293169 | NM_028166,NM_001085385 |
| Micall2 | 90.45157461 | -0.179288958 | 0.148743301 | -1.205358205 | 0.228065049 | 0.450971516 | | Dnajb6 | 5594.444838 | 0.041912674197735 | 0.054510653699622 | 0.768889590440292 | 0.441958857524904 | 0.819321835293169 | NM_001037940,NM_001037941,NM_011847,NM_001127367 |
| 2310035C23Rik | 2608.006354 | 0.062100991 | 0.051525885 | 1.205238711 | 0.228111162 | 0.450983073 | | Mamdc4 | 66.0465452 | -0.072420564145067 | 0.094223686947077 | -0.768602529698756 | 0.442129303303128 | 0.819493461755932 | NM_001081199 |
| Lars | 1772.813595 | 0.07815956 | 0.064865644 | 1.204945414 | 0.228224375 | 0.451149333 | | Phf20 | 4099.1054 | 0.0437688208315069 | 0.0569594777428502 | 0.768420332593392 | 0.442237504543977 | 0.819589434933122 | NM_172674 |
| Elac2 | 1140.562648 | 0.09398 | 0.078015858 | 1.204626885 | 0.228347373 | 0.451219768 | | Dnalc1 | 5765.663337 | 0.0481347923039871 | 0.0626597044276369 | 0.768193733814624 | 0.442372095706409 | 0.819629727325421 | NM_028821 |
| Qtrt1 | 345.2620872 | 0.139174809 | 0.115529866 | 1.204665199 | 0.228332576 | 0.451219768 | | Evc2 | 1035.903142 | -0.0673504000988457 | 0.0876721645624173 | -0.768207337357301 | 0.442364015051958 | 0.819629727325421 | NM_145920 |
| Strap | 3966.3969 | -0.072851921 | 0.060475017 | -1.204661438 | 0.228334028 | 0.451219768 | | Cnnm4 | 629.9917767 | 0.0554618090106662 | 0.0722233033266826 | 0.767921245028071 | 0.442533974838013 | 0.819825088444966 | NM_033570 |
| Shb | 443.6533571 | -0.127203599 | 0.105613249 | -1.204428426 | 0.228424031 | 0.451313686 | | Hrk | 2473.255116 | 0.0620467754658102 | 0.0808444793081874 | 0.767483147850845 | 0.442794309005266 | 0.819995322824399 | NM_007545 |
| Dnajc25 | 91.29773315 | 0.165001571 | 0.13700588 | 1.20433934 | 0.228458447 | 0.451324134 | | Iqub | 79.34538647 | -0.072976175683201 | 0.095068736838443 | -0.767614865938679 | 0.442716027854069 | 0.819995322824399 | NM_172535 |
| Pgam2 | 164.7044779 | -0.17089323 | 0.141914944 | -1.204194742 | 0.228514318 | 0.451376956 | | Wdr3 | 936.1831544 | -0.0431998629277161 | 0.0562878142369845 | -0.767481621258818 | 0.442795216317953 | 0.819995322824399 | NM_175552 |
| Enpp1 | 162.6575017 | -0.164654593 | 0.136784207 | -1.203754417 | 0.228684513 | 0.451625538 | | Ankrd13d | 2302.250179 | -0.0573362961044707 | 0.0747176938497884 | -0.767372400702556 | 0.442860133077595 | 0.820010999755914 | NM_026720 |
| Slc25a32 | 283.5621273 | 0.111358399 | 0.092512007 | 1.203718338 | 0.228698462 | 0.451625538 | | Atp8a1 | 11217.54984 | 0.0465782432535167 | 0.0607180450469703 | 0.767123566272344 | 0.443008051607502 | 0.820159899181452 | NM_009727,NM_001038999 |
| Dcp1b | 482.0873202 | 0.10932046 | 0.09082758 | 1.203604229 | 0.228742584 | 0.451655103 | | Fdx1 | 215.459652 | 0.0653865798517333 | 0.0852445358540136 | 0.76704716843918 | 0.443053471628587 | 0.820159899181452 | NM_007996 |
| Cybasc3 | 504.6778091 | 0.133931874 | 0.111293367 | 1.203412906 | 0.228816576 | 0.451686079 | | Spag9 | 19197.79926 | 0.0486350485258306 | 0.0634313683141112 | 0.766734973853797 | 0.443239105138413 | 0.820398985886926 | NM_001025428,NM_001025429,NM_001199204,NM_001199205,NM_001199203,NM_027569,NM_001025430 |
| Gabrb2 | 11914.94012 | -0.093715257 | 0.077870368 | -1.20347777 | 0.228791489 | 0.451686079 | | Arl13b | 394.8629945 | -0.0572533908646654 | 0.0747348209542583 | -0.766087215218024 | 0.443624409591314 | 0.820717991751493 | NM_026577 |
| Med18 | 103.857058 | -0.143978704 | 0.119668454 | -1.203146681 | 0.228919564 | 0.451831812 | | Dclk2 | 3038.180846 | -0.038503006604062 | 0.0502570315847337 | -0.766121782165858 | 0.443603843399439 | 0.820717991751493 | NM_001195500,NM_027539,NM_001195497,NM_001195496,NM_001195499,NM_001195498 |
| Csrp1 | 9394.908719 | 0.104675533 | 0.087032138 | 1.202722745 | 0.229083629 | 0.452040467 | | Dnajc13 | 3378.115027 | -0.0492995599850279 | 0.0643618001734163 | -0.765975467625133 | 0.443690899443163 | 0.820717991751493 | NM_001163026 |
| Thoc2 | 1498.405905 | 0.098999443 | 0.082309917 | 1.202764458 | 0.229067482 | 0.452040467 | | Lasp1 | 7423.273949 | 0.0330600362745139 | 0.0431584959089366 | 0.766014560476568 | 0.443667638537355 | 0.820717991751493 | NM_010688 |
| Fam107a | 17423.09026 | 0.152999632 | 0.127235493 | 1.202491761 | 0.229173056 | 0.452159344 | | Rictor | 2611.119886 | -0.0558282355555819 | 0.0728856369232695 | -0.765970332569045 | 0.443693954923893 | 0.820717991751493 | NM_030168 |
| Lamtor2 | 921.8769241 | -0.075354664 | 0.062679885 | -1.202214451 | 0.229280452 | 0.452292337 | | Kin | 328.5047284 | -0.0573926925254142 | 0.0749483982437317 | -0.765762763051633 | 0.443817474456061 | 0.820841944735006 | NM_025280 |
| Tmem185b | 497.831753 | -0.095026376 | 0.079045905 | -1.202166967 | 0.229298845 | 0.452292337 | | Oard1 | 454.3637353 | 0.0511623697093032 | 0.0668279759244651 | 0.765583110988301 | 0.443924396833033 | 0.820935173570546 | NM_207219 |
| 8430419L09Rik | 12703.0853 | 0.073092282 | 0.060810598 | 1.201966169 | 0.229376636 | 0.452388187 | | Hyal3 | 28.9094863 | 0.0673568162161723 | 0.0880376776625944 | 0.765090788449897 | 0.444217484681276 | 0.821372604694528 | NM_178020 |
| L3mbtl1 | 367.5758206 | 0.184303225 | 0.153373002 | 1.201666672 | 0.229492699 | 0.452510398 | | Serpinb1a | 513.9535706 | -0.0708133891672466 | 0.092577107766561 | -0.76491252400979 | 0.4443236357171 | 0.821464316205498 | NM_025429 |
| Lsm5 | 105.4282647 | -0.142255748 | 0.118383133 | -1.201655543 | 0.229497012 | 0.452510398 | | Apbb1 | 17908.69019 | 0.0400117924658759 | 0.052344530074103 | 0.76439300169167 | 0.444633077964418 | 0.821493053967672 | NM_001253885,NM_001253887,NM_001253886,NM_009685 |
| Cul7 | 1397.301564 | -0.076178277 | 0.063401464 | -1.201522368 | 0.229548635 | 0.452554594 | | Cbl | 1362.001873 | 0.0654306227815998 | 0.0856843692601525 | 0.763623789806296 | 0.445091468123706 | 0.821493053967672 | NM_007619 |
| Map1lc3a | 4712.989746 | -0.1186927 | 0.098796922 | -1.201380546 | 0.229603619 | 0.452605404 | | Fpgs | 249.4047334 | -0.0691779664285363 | 0.0904627567201209 | -0.764712119514147 | 0.444442987764259 | 0.821493053967672 | NM_010236 |
| Gm10814 | 30.73187582 | 0.200253042 | 0.166700476 | 1.201274571 | 0.229644711 | 0.45262882 | | Mettl14 | 855.3611547 | 0.0482940636739995 | 0.063206605906124 | 0.764066751394106 | 0.444827487063371 | 0.821493053967672 | NM_201638 |
| Kcnk6 | 219.6721598 | 0.1424867 | 0.118641824 | 1.200982041 | 0.229758168 | 0.452794843 | | Mrpl55 | 720.534564 | 0.050049355000639 | 0.0655451337331475 | 0.763586129893394 | 0.445113917404785 | 0.821493053967672 | NM_026035 |
| Zfp781 | 95.51744913 | -0.16668869 | 0.138813608 | -1.200809441 | 0.229825129 | 0.452869203 | | Ndrg4 | 81956.22306 | 0.0447380876410698 | 0.0585335057012576 | 0.764315875242521 | 0.444679027151731 | 0.821493053967672 | NM_001195006,NM_145602 |
| 6430584L05Rik | 182.651583 | -0.143646055 | 0.119669254 | -1.200358909 | 0.22999998 | 0.452972079 | | Pmp22 | 1082.570996 | 0.0630339807414135 | 0.0825530135453274 | 0.763557598134239 | 0.445130925773065 | 0.821493053967672 | NM_008885 |
| Catsperg1 | 33.73933621 | 0.197809486 | 0.164800273 | 1.20029829 | 0.230023514 | 0.452972079 | | Ppat | 715.0373372 | 0.0529068289658299 | 0.0692609399283896 | 0.763876854985385 | 0.444940631324983 | 0.821493053967672 | NM_172146 |
| Map3k2 | 1202.154553 | -0.112070549 | 0.093367082 | -1.200321862 | 0.230014363 | 0.452972079 | | Pthlh | 106.2116703 | 0.0727087099565236 | 0.0950940033573789 | 0.764598264764102 | 0.444510802762217 | 0.821493053967672 | NM_008970 |
| Mlec | 6412.875781 | -0.078078137 | 0.065040992 | -1.20044505 | 0.229966541 | 0.452972079 | | Rassf8 | 330.3216505 | 0.0666756127191286 | 0.087252829868703 | 0.764165618690662 | 0.444768552415854 | 0.821493053967672 | NM_027760 |
| Ppp2r5e | 3608.369154 | -0.073089693 | 0.060881857 | -1.200516811 | 0.229938688 | 0.452972079 | | Solh | 1773.604461 | -0.0591315025299529 | 0.0774245252522775 | -0.763730902285559 | 0.445027621315447 | 0.821493053967672 | NM_015830 |
| Rpl3 | 17829.27268 | -0.070670019 | 0.058882734 | -1.200182372 | 0.23006852 | 0.453003132 | | Spata5 | 521.82967 | 0.0593412080431421 | 0.0777128863932868 | 0.76359547041954 | 0.445108349405606 | 0.821493053967672 | NM_001163511,NM_021343 |
| Lyz2 | 362.5027814 | 0.187782031 | 0.156472787 | 1.200093858 | 0.230102891 | 0.453013238 | | Tead4 | 27.3794714 | -0.0658400709616538 | 0.086154378130984 | -0.764210390579971 | 0.444741875530702 | 0.821493053967672 | NM_001080979,NM_011567 |
| Csde1 | 8449.548095 | -0.064218265 | 0.053526218 | -1.199753451 | 0.230235108 | 0.453215952 | | Thnsl1 | 911.4655345 | 0.0475467348654861 | 0.0622590176207603 | 0.763692340202163 | 0.44505060651303 | 0.821493053967672 | NM_001001297,NM_177588 |
| Chaf1b | 19.67297757 | -0.201608843 | 0.16806522 | -1.199586942 | 0.2302998 | 0.45328571 | | Gdnf | 46.41318351 | 0.070937895690208 | 0.0929391290083095 | 0.76327265433987 | 0.44530080695562 | 0.821629343314143 | NM_010275 |
| 3110009E18Rik | 32.94612746 | -0.198195807 | 0.165361456 | -1.198561088 | 0.230698656 | 0.453668953 | | Rbm18 | 1782.862473 | 0.0539252608922762 | 0.0706527163258957 | 0.76324398687417 | 0.445317900310632 | 0.821629343314143 | NM_001159635,NM_026434,NR_027515 |
| Crebl2 | 1367.477755 | 0.110686369 | 0.092361411 | 1.198404911 | 0.230759422 | 0.453668953 | | 2610306M01Rik | 176.9390866 | 0.0664918592598684 | 0.0871310559933835 | 0.76312468042299 | 0.445389042384026 | 0.821656239480614 | NR_028298 |
| Fis1 | 3738.827597 | -0.093193917 | 0.077745271 | -1.198708491 | 0.230641315 | 0.453668953 | | Dynlt1b | 37.47015881 | -0.0645812161930193 | 0.0846932233980388 | -0.762531092830206 | 0.445743093124983 | 0.821773577915295 | NM_009342 |
| Gdap1 | 4542.465689 | -0.076140457 | 0.063527034 | -1.19855204 | 0.230702177 | 0.453668953 | | Emp3 | 65.45624216 | -0.0692339441895692 | 0.0907776515592254 | -0.7626760882264 | 0.445656594499766 | 0.821773577915295 | NM_010129,NM_001146346 |
| Gyk | 824.5423227 | -0.086866861 | 0.072494289 | -1.198257987 | 0.230816597 | 0.453668953 | | Nr1h2 | 1799.39717 | -0.0432694415352336 | 0.0567295188627258 | -0.762732390520307 | 0.445623009314749 | 0.821773577915295 | NM_009473 |
| Klf3 | 652.7116837 | -0.099625875 | 0.08310645 | -1.198774279 | 0.230615726 | 0.453668953 | | Podxl | 2094.447959 | -0.0575159864334863 | 0.0754397802063941 | -0.762409252467724 | 0.445815785750968 | 0.821773577915295 | NM_013723 |
| Phf13 | 673.1897173 | 0.116667244 | 0.097354899 | 1.198370545 | 0.230772794 | 0.453668953 | | Sdf2l1 | 298.2674253 | 0.0687025043270496 | 0.0901063713332997 | 0.762460004886025 | 0.445785504927182 | 0.821773577915295 | NM_022324 |
| Rhobtb2 | 5472.627938 | -0.084579865 | 0.070584066 | -1.19828553 | 0.230805878 | 0.453668953 | | Tada3 | 1346.111409 | 0.0399860895648016 | 0.0524508050031688 | 0.762354163341932 | 0.445848655345617 | 0.821773577915295 | NM_133932 |
| Slc37a1 | 246.2057962 | 0.130362402 | 0.108755509 | 1.198674012 | 0.230654727 | 0.453668953 | | Timp2 | 17050.32347 | 0.0606876733165607 | 0.0795957419462546 | 0.762448722468424 | 0.44579222468424 | 0.821773577915295 | NM_011594 |
| Sparc | 10824.85432 | 0.199823053 | 0.166662501 | 1.198968283 | 0.230540278 | 0.453668953 | | 2810429I04Rik | 28.44363196 | 0.0648650323499567 | 0.0852512034780135 | 0.760869403640565 | 0.446735075076475 | 0.82221060703319 | NR_015522 |
| Zfp518a | 412.8097296 | -0.151926464 | 0.126765421 | -1.198485066 | 0.230728233 | 0.453668953 | | Cyp20a1 | 377.3691754 | 0.0543414219610551 | 0.0713956501880863 | 0.76113071059507 | 0.446578998948215 | 0.82221060703319 | NM_030013 |
| Bace2 | 146.2030808 | 0.162711894 | 0.135846295 | 1.197764686 | 0.231008639 | 0.453988819 | | Dlg5 | 1841.303824 | 0.056135837681159 | 0.0737567516568131 | 0.761094224191931 | 0.44660079006125 | 0.82221060703319 | NM_001163513,NM_027726 |
| 1110038F14Rik | 460.1336451 | -0.115895004 | 0.09677964 | -1.197514317 | 0.23110615 | 0.45406327 | | Matn4 | 290.632908 | 0.0658490283000463 | 0.0865609330285868 | 0.76072456703187 | 0.446821597956767 | 0.82221060703319 | NM_001252563,NM_013592 |
| Sod1 | 5637.486818 | -0.080963646 | 0.067611636 | -1.197480957 | 0.231119146 | 0.45406327 | | Mtmr3 | 5023.602389 | -0.0320043647269596 | 0.0420510258128231 | -0.761084042739336 | 0.44660687093263 | 0.82221060703319 | NM_028860 |
| Tmco6 | 316.6737769 | 0.120555092 | 0.100677213 | 1.197441697 | 0.23113444 | 0.45406327 | | Otop2 | 53.48251751 | -0.0617001648806662 | 0.081015526074842 | -0.761584450166598 | 0.446308058414611 | 0.82221060703319 | NM_172801 |
| Fignl2 | 54.35269759 | -0.18281599 | 0.152686377 | -1.197330068 | 0.231177929 | 0.45409113 | | Oxa1l | 1818.086703 | 0.0394576813535196 | 0.051830420704314 | 0.761284219138808 | 0.446487324245634 | 0.82221060703319 | NM_026936 |
| Diap2 | 1206.214312 | -0.118666661 | 0.099119184 | -1.197211842 | 0.231223995 | 0.454124044 | | Scn2a1 | 13451.62779 | 0.0540229461903324 | 0.0710066667152043 | 0.760815127500763 | 0.446767497584923 | 0.82221060703319 | NM_001099298 |
| Aldh16a1 | 230.2489788 | 0.11708456 | 0.097810565 | 1.197054323 | 0.231285382 | 0.454187035 | | Sesn3 | 1860.815458 | 0.0565678398633882 | 0.0742797520598693 | 0.761551274370313 | 0.4463278652556 | 0.82221060703319 | NM_030261 |
| Rab33a | 727.5078049 | 0.092328226 | 0.077139639 | 1.196897303 | 0.231346586 | 0.454249651 | | Thap7 | 654.3745925 | 0.0577568364707839 | 0.0759215741059933 | 0.760743400685426 | 0.446810346519219 | 0.82221060703319 | NM_026909 |
| Ctdsp2 | 4886.588314 | 0.090730267 | 0.075810439 | 1.196804403 | 0.231382802 | 0.454263194 | | Tmsb10 | 3647.695146 | -0.0637228171988502 | 0.0836838681550568 | -0.761470742255593 | 0.446375947785622 | 0.82221060703319 | NM_025284,NM_001039392,NM_001190327 |
| Gm1661 | 13.62924176 | 0.200626828 | 0.167655422 | 1.19666173 | 0.231438429 | 0.454314839 | | Tnks1bp1 | 3876.027794 | -0.0556817828871825 | 0.073144475812254 | -0.761257528594581 | 0.446503262965005 | 0.82221060703319 | NM_001081260 |
| Zufsp | 274.5067301 | 0.114339985 | 0.095580928 | 1.196263601 | 0.231593708 | 0.454562062 | | Vamp3 | 1049.342801 | 0.047866271929507 | 0.0628331785619506 | 0.761799307706725 | 0.446179379651195 | 0.82221060703319 | NM_009498 |
| Fscn1 | 7521.526885 | -0.091902387 | 0.076856102 | -1.19577216 | 0.231785483 | 0.454880845 | | Atp5s | 511.292725 | 0.0523344844913666 | 0.0688808298121682 | 0.759783014143094 | 0.447384297739458 | 0.822458097494885 | NM_026536 |
| Fendrr | 24.17456761 | 0.200854888 | 0.16808166 | 1.194983963 | 0.232093296 | 0.455245894 | | Fance | 509.8691662 | -0.0653986967355521 | 0.0860932745546104 | -0.759626080827819 | 0.447478124876236 | 0.822458097494885 | NM_001163819,NR_028297,NR_028296,NM_001163820 |
| Plcd1 | 438.6798023 | 0.145707908 | 0.121947172 | 1.194844509 | 0.232147787 | 0.455245894 | | Gareml | 636.2626547 | 0.0580786897672193 | 0.0764356961007948 | 0.759837258375087 | 0.447351868852779 | 0.822458097494885 | NM_001167879 |
| Six1 | 11.93115248 | -0.188837153 | 0.158037491 | -1.194888323 | 0.232130666 | 0.455245894 | | Jmjd1c | 3512.48661 | -0.0530348331743434 | 0.0698020078539368 | -0.759789507564319 | 0.447380415700146 | 0.822458097494885 | NM_207221,NM_001242396 |
| Slc44a4 | 16.98421301 | 0.200921069 | 0.168119469 | 1.195108875 | 0.232044495 | 0.455245894 | | Ldoc1 | 55.82609096 | 0.0707214180098719 | 0.0931064126404186 | 0.759576231156079 | 0.447507931290099 | 0.822458097494885 | NM_001018087 |
| Snord42b | 19.07482168 | 0.19967661 | 0.167111129 | 1.194873202 | 0.232136574 | 0.455245894 | | Nell2 | 23909.52563 | 0.0484076633113505 | 0.0637236118440267 | 0.759650338556384 | 0.447463620958337 | 0.822458097494885 | NM_016743 |
| Tax1bp1 | 6135.004462 | -0.073115983 | 0.061192157 | -1.194858722 | 0.232142233 | 0.455245894 | | Polr3g | 250.5924392 | -0.0621569414223388 | 0.0817663123667626 | -0.760177873052829 | 0.44714826936753 | 0.822458097494885 | NM_001081176 |
| Fbxl13 | 10.55143406 | 0.194401531 | 0.162788885 | 1.19419413 | 0.232402039 | 0.455686812 | | Ppp2r3c | 1141.669163 | 0.0532217223690539 | 0.0700461427250078 | 0.759809466996571 | 0.447368483235071 | 0.822458097494885 | NM_021529 |
| Rbm46 | 8.99877052 | 0.189380659 | 0.158635131 | 1.193812854 | 0.232551183 | 0.455921552 | | Rgs2 | 2590.691165 | 0.0617871813630197 | 0.0813468516021206 | 0.759552215557522 | 0.447522291243259 | 0.822458097494885 | NM_009061 |
| Olig1 | 2955.154384 | -0.129746246 | 0.10875886 | -1.192971743 | 0.23288044 | 0.456378333 | | Zfp706 | 8317.970152 | 0.0430600878726442 | 0.0566584893841027 | 0.759993574497346 | 0.44725842569545 | 0.822458097494885 | NM_026521 |
| Phrf1 | 2266.401425 | 0.080018319 | 0.067072739 | 1.193008081 | 0.232866208 | 0.456378333 | | Cntn6 | 889.7674486 | 0.05351508483573 | 0.0704983792544346 | 0.759096668628218 | 0.447794731835174 | 0.822750603925845 | NM_017383 |
| Shq1 | 133.1838679 | -0.141146944 | 0.118327804 | -1.192846806 | 0.232929375 | 0.456378333 | | Tmprss9 | 24.5538342 | -0.0625603286523354 | 0.0824055790262178 | -0.759175888229988 | 0.447747347652795 | 0.822750603925845 | NM_001081688 |
| Trappc6b | 2116.113181 | -0.065113913 | 0.054587229 | -1.192841514 | 0.232931448 | 0.456378333 | | Eps8l1 | 178.6142132 | 0.0318384522505953 | 0.04194881197345 | 0.758983407462081 | 0.447862482489265 | 0.822771015636659 | NM_026146 |
| Ube2z | 4246.787922 | -0.073573614 | 0.061671275 | -1.192996487 | 0.232870749 | 0.456378333 | | Eya1 | 469.4696465 | -0.0633175742362358 | 0.0834446315399195 | -0.75879745728094 | 0.447973726946487 | 0.822871314823554 | NM_001252192,NM_010164 |
| Slu7 | 2534.832178 | 0.076301914 | 0.06397157 | 1.192747253 | 0.232968374 | 0.456392967 | | Pcdhb12 | 326.9293572 | 0.0614372934045764 | 0.080987559732785 | 0.758601612485747 | 0.44809090781652 | 0.822982491711059 | NM_053137 |
| Pik3r4 | 2085.168807 | 0.087262973 | 0.073177564 | 1.192482616 | 0.233072063 | 0.456500657 | | Abhd8 | 8562.653295 | 0.0492513260935167 | 0.0649400911068172 | 0.758411718463796 | 0.448204544753541 | 0.823087132375466 | NM_022419 |
| Slc46a1 | 333.2542117 | -0.15136577 | 0.126936087 | -1.192456569 | 0.233082271 | 0.456500657 | | Aff4 | 9748.687109 | 0.0519635864465003 | 0.0685355253920034 | 0.758199286418008 | 0.448331688356768 | 0.823216547221643 | NM_033565 |
| Plk4 | 195.1377744 | 0.121641335 | 0.102031055 | 1.192199133 | 0.233183173 | 0.456594481 | | A430105I19Rik | 376.9978254 | -0.0630777912030386 | 0.0833955487388634 | -0.756368800936296 | 0.44942810827106 | 0.823263966309774 | NM_001001982 |
| Sh3kbp1 | 1657.543605 | -0.094327066 | 0.079121233 | -1.192183979 | 0.233189114 | 0.456594481 | | Adam22 | 10178.21649 | 0.0454156830716225 | 0.0600731984316508 | 0.756005743947443 | 0.449645751871707 | 0.823263966309774 | NM_001098225,NM_001007220,NM_001007221 |
| Tom1 | 1595.942335 | 0.082552377 | 0.069258935 | 1.191938293 | 0.233285442 | 0.456725377 | | Apoe | 42530.03042 | -0.0521636978998772 | 0.069070475870641 | -0.755224243677895 | 0.4501144444742 | 0.823263966309774 | NM_009696 |
| Mlst8 | 1685.50605 | 0.074354405 | 0.062393869 | 1.191694102 | 0.233381212 | 0.456855148 | | Asphd1 | 610.1073789 | 0.0552445897379656 | 0.0729390564591798 | 0.757407518273603 | 0.448805753260451 | 0.823263966309774 | NM_001039645 |
| Sar1a | 5238.054964 | -0.062128179 | 0.052139267 | -1.191581371 | 0.233425433 | 0.456883989 | | Camta2 | 15209.67056 | -0.0474917781091918 | 0.0628577887012621 | -0.755543252323132 | 0.449923090583294 | 0.823263966309774 | NM_001190379,NM_001190378,NM_001190376,NM_178116 |
| Bysl | 1094.647966 | -0.08658585 | 0.072702485 | -1.190961357 | 0.233668755 | 0.457143705 | | Cnot6l | 675.0700039 | 0.0649034808747171 | 0.085766836798768 | 0.756743320579703 | 0.449203655712982 | 0.823263966309774 | NM_178854,NM_144910 |
| Mtmr1 | 1823.012354 | 0.111903554 | 0.093962177 | 1.190942542 | 0.233676142 | 0.457143705 | | Cpeb3 | 2875.080441 | 0.057470405717842 | 0.0758466392716361 | 0.757718552459764 | 0.448619490084365 | 0.823263966309774 | NM_198300 |
| Pou6f2 | 32.79163728 | 0.197098775 | 0.165474358 | 1.1911137 | 0.233608952 | 0.457143705 | | Crem | 505.3920905 | 0.0634401956976457 | 0.0838242783012232 | 0.75682364326088 | 0.449155525983412 | 0.823263966309774 | NM_001110851,NM_001110856,NM_001110857,NM_001110854,NM_001271503,NM_001110858,NM_001110859,NR_073193,NM_001110850,NM_001110 |
| Wdr43 | 985.7000102 | 0.074050944 | 0.062176786 | 1.190974129 | 0.233663741 | 0.457143705 | | Dctn5 | 3213.869024 | 0.0344664842860265 | 0.0455250779050127 | 0.757087870512607 | 0.448997220410339 | 0.823263966309774 | NM_021608 |
| Adk | 1916.002114 | -0.079358351 | 0.066655862 | -1.190568221 | 0.233823133 | 0.457315783 | | Grin1 | 21475.63651 | -0.032485381066686 | 0.0430016590858251 | -0.755444830671531 | 0.449982122829488 | 0.823263966309774 | NM_001177657,NM_001177656,NM_008169 |
| Wnt16 | 19.30065258 | -0.198401661 | 0.166640175 | -1.190599211 | 0.233810961 | 0.457315783 | | Hipk1 | 6506.15531 | 0.0440947140039831 | 0.0582703010286586 | 0.756727067229263 | 0.449213395153156 | 0.823263966309774 | NM_010432 |
| Smad5 | 1586.461262 | -0.088665355 | 0.074484747 | -1.190382704 | 0.233896008 | 0.457400574 | | Ica1l | 1608.044001 | 0.0501164117946283 | 0.0661210665624024 | 0.757949234641133 | 0.448481374145898 | 0.823263966309774 | NM_027407 |
| Cib1 | 343.5618478 | 0.090044217 | 0.075659069 | 1.190131181 | 0.233994836 | 0.457478359 | | Kcnn2 | 1195.943831 | 0.0599810079560286 | 0.0794021133596672 | 0.755408205375253 | 0.4500040914082 | 0.823263966309774 | NM_080465 |
| Xpo6 | 4180.380516 | 0.105595049 | 0.088724359 | 1.190147224 | 0.233988532 | 0.457478359 | | Kti12 | 397.710618 | 0.0566954350073383 | 0.0748746072219903 | 0.757205107457139 | 0.448926990780597 | 0.823263966309774 | NM_029571 |
| Pcif1 | 1743.081628 | 0.097296818 | 0.081774898 | 1.189812761 | 0.234119993 | 0.4576653 | | Lamb3 | 90.20870048 | 0.0717685011762155 | 0.0950217432700221 | 0.755285040101526 | 0.450077972839549 | 0.823263966309774 | NM_008484 |
| D630003M21Rik | 285.4808509 | -0.157184957 | 0.132135502 | -1.189573995 | 0.234213872 | 0.45779106 | | Mettl24 | 52.73349745 | -0.0712779812878392 | 0.0943031230541924 | -0.755839032466385 | 0.44974571125919 | 0.823263966309774 | NM_177793 |
| Coq2 | 2160.851229 | -0.096675322 | 0.081285115 | -1.189336102 | 0.234307434 | 0.457916169 | | Mrpl37 | 1530.754725 | 0.038851194929886 | 0.0513068879882469 | 0.757231561945168 | 0.448911144343792 | 0.823263966309774 | NM_025500 |
| Atf7ip | 2403.281245 | 0.103620935 | 0.087132578 | 1.189232975 | 0.234348002 | 0.45793769 | | Myo1e | 941.8448037 | 0.0502291383837691 | 0.0663041164095959 | 0.757556862284039 | 0.448716312952518 | 0.823263966309774 | NM_181072 |
| Socs4 | 204.4773262 | -0.12864508 | 0.108217463 | -1.188764521 | 0.234532343 | 0.458182338 | | Nog | 205.7963875 | 0.0670707544737274 | 0.0887990696015411 | 0.755309202840605 | 0.450063478133022 | 0.823263966309774 | NM_008711 |
| Ubald1 | 3177.275875 | 0.077270078 | 0.064997052 | 1.188824342 | 0.234508797 | 0.458182338 | | Phldb1 | 5284.516827 | -0.0595242818876226 | 0.0785807212434189 | -0.757492180597766 | 0.448755048784101 | 0.823263966309774 | NM_153537 |
| Acvr2b | 133.268294 | -0.140943413 | 0.118579926 | -1.188594206 | 0.234599389 | 0.458255539 | | Ppa2 | 1298.709846 | 0.049290058162734 | 0.065120646869116 | 0.75690369387455 | 0.44910756218885 | 0.823263966309774 | NM_146141 |
| Gria2 | 20032.8255 | 0.061784645 | 0.052001328 | 1.188135912 | 0.234779868 | 0.458404859 | | Rasa3 | 3989.570226 | -0.036927006018171 | 0.0487591915682594 | -0.757334254947107 | 0.448849633473048 | 0.823263966309774 | NM_009025 |
| Insl6 | 14.63293602 | 0.171109139 | 0.144025691 | 1.188045946 | 0.234815309 | 0.458404859 | | Sec61g | 767.8473452 | 0.0603998979376775 | 0.0797741732059026 | 0.757135994149149 | 0.448968391668532 | 0.823263966309774 | NM_001109972,NM_001109971,NM_011343 |
| Llgl2 | 50.60767207 | 0.193687073 | 0.162996022 | 1.188293254 | 0.234717895 | 0.458404859 | | Smyd5 | 2323.517956 | -0.0346441580000272 | 0.0458362202704877 | -0.755824930493524 | 0.449754167310885 | 0.823263966309774 | NM_144918 |
| Mbd2 | 1497.050441 | 0.109147411 | 0.09186569 | 1.188119425 | 0.234786362 | 0.458404859 | | Snrpn | 52.49886374 | -0.0660022901165738 | 0.0871823768938235 | -0.757059998455373 | 0.449013917807194 | 0.823263966309774 | NM_001082962,NM_001082961,NM_013670 |
| Zfp287 | 714.1340679 | 0.095930248 | 0.080747702 | 1.188024488 | 0.234823762 | 0.458404859 | | Tbl2 | 1182.389318 | 0.0380370713177194 | 0.0503264222116415 | 0.755807181320365 | 0.449764810483573 | 0.823263966309774 | NM_013763 |
| Fam133b | 1070.402896 | 0.126122316 | 0.106177381 | 1.187845421 | 0.234894316 | 0.458484824 | | Tmem198 | 2333.325568 | 0.0603196048834236 | 0.0798044554840541 | 0.755842571916003 | 0.449743588891653 | 0.823263966309774 | NM_177056 |
| Kif16b | 696.4921236 | 0.094634023 | 0.079704785 | 1.187306663 | 0.235106681 | 0.458829708 | | Ubxn2b | 1710.044016 | 0.056291074477113 | 0.0744194761176945 | 0.756402455562688 | 0.449407936189039 | 0.823263966309774 | NM_026534 |
| Steap2 | 629.7891626 | -0.145559034 | 0.122602157 | -1.187246926 | 0.235130236 | 0.458829708 | | Ywhae | 25384.54213 | 0.0350597412937624 | 0.0464127061138673 | 0.755391017445697 | 0.450014401278607 | 0.823263966309774 | NM_009536 |
| Gm6225 | 16.97815084 | 0.192524317 | 0.162210906 | 1.186876524 | 0.235276328 | 0.459056974 | | Zfp219 | 1189.360411 | -0.0526169124072049 | 0.0696041156291593 | -0.755945419772881 | 0.449681920496889 | 0.823263966309774 | NM_027248,NM_001253694,NM_001253695,NM_001253696 |
| Alox5 | 32.70542633 | 0.199405685 | 0.168099884 | 1.18623333 | 0.235530167 | 0.459475465 | | Zfp934 | 73.70059542 | 0.0710748859085668 | 0.0939983287953207 | 0.756129250588389 | 0.449571706018123 | 0.823263966309774 | NM_001162911,NM_175252 |
| Cpeb2 | 2477.554734 | 0.096610582 | 0.081446623 | 1.186182783 | 0.235550124 | 0.459475465 | | Arhgap21 | 8222.876087 | -0.0398770932724906 | 0.0528339969602927 | -0.754762001111901 | 0.450391797491095 | 0.823365451229261 | NM_001081364,NM_001128084 |
| Psmc1 | 5287.499535 | 0.101010002 | 0.085173785 | 1.185928304 | 0.235650614 | 0.459613623 | | Myeov2 | 1353.482549 | -0.0501696318854283 | 0.0664715106747619 | -0.754753899469849 | 0.450396659470447 | 0.823365451229261 | NM_001163425,NR_028108 |
| 7-Mar | 1263.614658 | 0.093157831 | 0.078579933 | 1.185516799 | 0.235813177 | 0.459699215 | | Phf2 | 2613.648702 | 0.0342976495684072 | 0.0454264953614024 | 0.755014211321901 | 0.450240455286819 | 0.823365451229261 | NM_011078 |
| Phf17 | 1525.406485 | 0.091268276 | 0.076984626 | 1.185538993 | 0.235804407 | 0.459699215 | | Ppp1cb | 8133.580152 | 0.0580095512964487 | 0.0768552794761309 | 0.754789413191385 | 0.450375347100247 | 0.823365451229261 | NM_172707 |
| Slc30a5 | 1679.665966 | 0.075529165 | 0.063705157 | 1.185605197 | 0.235778249 | 0.459699215 | | Ssr3 | 5123.979975 | 0.0495540378035372 | 0.0656740591358666 | 0.754545073893175 | 0.450521990699468 | 0.823490932037306 | NM_026155 |
| Tmem138 | 256.0541739 | 0.121394932 | 0.102385467 | 1.185665657 | 0.235754362 | 0.459699215 | | AI317395 | 35.9728698 | -0.0646503662922718 | 0.0857761976404191 | -0.75370986439958 | 0.451023457346551 | 0.82399279856823 | NM_144821 |
| Ankrd13b | 3414.500451 | 0.09081033 | 0.076619735 | 1.185208091 | 0.235935182 | 0.459821346 | | Fam171a1 | 4095.646063 | 0.0480736572126601 | 0.0637775792583981 | 0.75377049069058 | 0.450987046192286 | 0.82399279856823 | NM_001081161 |
| Tspan8 | 14.84534253 | -0.189344769 | 0.159747411 | -1.185275974 | 0.23590835 | 0.459821346 | | Sema4c | 699.4879073 | -0.0532041865190441 | 0.0705744105769194 | -0.753873621956169 | 0.450925111071333 | 0.82399279856823 | NM_001126047 |
| Dlk1 | 581.2297118 | -0.172693054 | 0.14575703 | -1.184800855 | 0.236096195 | 0.460019421 | | Zfp366 | 149.960106 | -0.0712974505041204 | 0.0945682616318812 | -0.75392578095233 | 0.450893789002005 | 0.82399279856823 | NM_001004149 |
| Lamp2 | 2349.474284 | 0.080911833 | 0.068290644 | 1.184815786 | 0.23609029 | 0.460019421 | | 9330159F19Rik | 3337.18 | 0.0560471826503172 | 0.0743873624031384 | 0.753450328653574 | 0.451179349057849 | 0.824173946732181 | NM_001162537 |
| Cacng1 | 10.73293909 | 0.173121234 | 0.146134885 | 1.184667398 | 0.236148978 | 0.460064411 | | Irak1bp1 | 463.6792055 | 0.0558601002943155 | 0.0741552046752656 | 0.75328630726506 | 0.4512778852164 | 0.824250290538592 | NM_022986,NM_001168240 |
| Asb3 | 516.8455345 | -0.105046194 | 0.088703376 | -1.18424122 | 0.236317589 | 0.46027715 | | 2810032G03Rik | 317.5005057 | -0.0590341479042219 | 0.0784239354975701 | -0.752756763986309 | 0.451596092441743 | 0.824520467421915 | NR_015579 |
| Lipt1 | 150.8861907 | -0.154135068 | 0.130153688 | -1.184254325 | 0.236312404 | 0.46027715 | | Mybpc3 | 32.5271713 | -0.0663872177348811 | 0.0881816801748623 | -0.752845915424119 | 0.451542511674111 | 0.824520467421915 | NM_008653 |
| Sox21 | 384.6218319 | -0.158048435 | 0.133481651 | -1.184046152 | 0.236394794 | 0.460369651 | | Rnf13 | 4476.977457 | 0.0502606576508766 | 0.0667635315834878 | 0.752816042905608 | 0.451560464909625 | 0.824520467421915 | NM_011883,NM_001113413 |
| Slc7a10 | 1016.099328 | -0.171482637 | 0.144889057 | -1.183544434 | 0.236593447 | 0.460698614 | | Gdi1 | 28373.95432 | 0.0284861075182777 | 0.0378489598903744 | 0.752625900441777 | 0.451674749051158 | 0.824558474870953 | NM_010273 |
| Slc12a8 | 47.85542696 | 0.185708931 | 0.15696335 | 1.183135622 | 0.236755402 | 0.460956044 | | Gpatch4 | 739.5136368 | 0.0441185550236129 | 0.0586340752609665 | 0.752438830616011 | 0.45178720236114 | 0.824558474870953 | NM_001110809,NM_025663 |
| Pdlim4 | 267.9419049 | 0.122820664 | 0.103816382 | 1.183056674 | 0.236786687 | 0.460959031 | | Mtrr | 533.70605 | 0.0615475244006361 | 0.0817972369678914 | 0.752440139570922 | 0.451786415453898 | 0.824558474870953 | NM_172480 |
| Fam173a | 1457.52927 | 0.10274437 | 0.086866217 | 1.182788576 | 0.236892949 | 0.461107959 | | D430042O09Rik | 1337.313168 | -0.0414518518010926 | 0.0551140543157033 | -0.752110370317684 | 0.451984688042312 | 0.824608085719999 | NM_001081022 |
| Popdc3 | 27.07799168 | 0.198772558 | 0.168111292 | 1.182386708 | 0.237052295 | 0.461360163 | | Figf | 91.05785371 | 0.0715042764744713 | 0.0950565511632561 | 0.752228811159637 | 0.451913470247639 | 0.824608085719999 | NM_010216 |
| Htr5b | 38.5811311 | -0.198705342 | 0.168092112 | -1.18212175 | 0.237157395 | 0.461506742 | | Lrrc4c | 4815.938085 | 0.0512357120026788 | 0.0681191267209599 | 0.752148691109304 | 0.451961645277184 | 0.824608085719999 | NM_178725 |
| Gm13446 | 178.2292629 | 0.166637081 | 0.140975219 | 1.182031019 | 0.237193393 | 0.461518829 | | 2610020H08Rik | 30.95755699 | -0.06278775179359 | 0.0835039837923964 | -0.751913249428793 | 0.452103229761027 | 0.824720773013774 | NM_001004187,NM_028129 |
| Ednrb | 1909.673665 | -0.114989725 | 0.097290606 | -1.181920122 | 0.237237397 | 0.461546488 | | Mgme1 | 270.7837938 | -0.0591881086282877 | 0.0787334238746738 | -0.751753267106763 | 0.452199450541212 | 0.824792719558219 | NM_028984 |
| Fntb | 1190.957545 | 0.066074075 | 0.055919264 | 1.181597733 | 0.237365354 | 0.46173745 | | Tmprss5 | 67.62718429 | -0.0691539741586894 | 0.09203852386513 | -0.751359009842717 | 0.452436624524983 | 0.825121708455731 | NM_030709 |
| Dock6 | 797.1480952 | 0.139895853 | 0.118414946 | 1.181403684 | 0.237442395 | 0.461829333 | | Mast3 | 14541.41295 | -0.058628534785619 | 0.078081755054707 | -0.75086087325447 | 0.452736389839542 | 0.825461127000023 | NM_199308 |
| Pag1 | 1448.479325 | -0.08484273 | 0.071827087 | -1.181208007 | 0.237520101 | 0.461922486 | | Mmgt2 | 532.439205 | -0.0520811749759854 | 0.0693616673001386 | -0.75086394262442 | 0.452734542431161 | 0.825461127000023 | NM_175002 |
| Gm9897 | 32.39055432 | 0.197634405 | 0.167348816 | 1.180972826 | 0.237613518 | 0.462046166 | | D430020J02Rik | 131.8205057 | 0.0708468921867769 | 0.0943735349864088 | 0.75070719982016 | 0.452828889049787 | 0.825470560369691 | NR_028421 |
| Ctdnep1 | 2512.328089 | -0.065135092 | 0.055175636 | -1.180504599 | 0.237799582 | 0.462349947 | | Gtf2h2 | 637.5769268 | -0.0504840576900736 | 0.0672600613682244 | -0.75058001231506 | 0.45290545390415 | 0.825470560369691 | NM_022011 |
| Ccdc167 | 504.1807522 | -0.117868865 | 0.099859232 | -1.180350211 | 0.237860956 | 0.462353236 | | Ifngr1 | 698.3689365 | -0.0463666363776805 | 0.0617830795057422 | -0.750474672816708 | 0.45296887214008 | 0.825470560369691 | NM_010511 |
| Col6a3 | 150.8114551 | -0.17553962 | 0.14871421 | -1.180382292 | 0.237848202 | 0.462353236 | | Snhg3 | 147.7173678 | -0.069294668078547 | 0.0923307600054582 | -0.750504686352096 | 0.452950802381807 | 0.825470560369691 | NR_003270 |
| Spice1 | 510.8293749 | 0.130110711 | 0.110239126 | 1.180258911 | 0.237897255 | 0.462365789 | | Dnajc30 | 1293.297223 | 0.042900034289971 | 0.0571992653674268 | 0.750010232026532 | 0.453248542284991 | 0.825626289596525 | NM_025362 |
| Gm1821 | 1452.26643 | -0.095591784 | 0.081008187 | -1.180026204 | 0.237989793 | 0.462487627 | | Glt8d2 | 658.066425 | 0.0543597297185717 | 0.0724869734402688 | 0.749924119309044 | 0.453300407087001 | 0.825626289596525 | NM_029102 |
| Parm1 | 1084.843639 | -0.076407754 | 0.064766358 | -1.179744497 | 0.238101851 | 0.462589351 | | Nova2 | 4478.027945 | -0.0406758644636775 | 0.054218401911166 | -0.750222489595373 | 0.45312071604345 | 0.825626289596525 | NM_001029877 |
| Rcn2 | 3859.375806 | -0.070334184 | 0.059614369 | -1.179819303 | 0.238072091 | 0.462589351 | | Rab10 | 5913.93494 | 0.0436759112100368 | 0.0582416900635022 | 0.749908032586555 | 0.453310096325422 | 0.825626289596525 | NM_016676 |
| Fam181a | 75.399425 | -0.187033798 | 0.158614571 | -1.179171605 | 0.23832985 | 0.462671816 | | Tbxa2r | 33.72619568 | -0.06078563344364 | 0.0810625511517066 | -0.749860849183998 | 0.453338516165626 | 0.825626289596525 | NM_009325 |
| G630025P09Rik | 32.73196021 | -0.192935826 | 0.163599291 | -1.179319447 | 0.238270997 | 0.462671816 | | Efna5 | 1481.917923 | 0.0570889296601269 | 0.0761978651579502 | 0.749219542329532 | 0.453724892340732 | 0.825985605778632 | NM_010109,NM_207654 |
| Gimap9 | 12.78829912 | 0.192388452 | 0.16316376 | 1.179112639 | 0.238353326 | 0.462671816 | | Eme2 | 640.3440417 | -0.0626935438453632 | 0.0837331861961348 | -0.748729944403539 | 0.454019991535991 | 0.825985605778632 | NM_001163102 |
| Nek9 | 3048.216903 | 0.066993418 | 0.056813346 | 1.179184528 | 0.238324705 | 0.462671816 | | Id1 | 325.1614954 | -0.0604294684611948 | 0.0807110016935798 | -0.748714142969207 | 0.45402951746085 | 0.825985605778632 | NM_010495 |
| Ppp2r1b | 598.8823197 | 0.101666537 | 0.086202461 | 1.179392507 | 0.238241917 | 0.462671816 | | Itpr3 | 272.5163924 | -0.0651084956645787 | 0.0869593483173654 | -0.748723362403313 | 0.4540239594937 | 0.825985605778632 | NM_080553 |
| Rsl1 | 125.6456871 | 0.166518542 | 0.141211125 | 1.179216874 | 0.238311829 | 0.462671816 | | Olfml2a | 79.53306282 | -0.0709034407700346 | 0.0947037942744088 | -0.748686378547711 | 0.454046255568346 | 0.825985605778632 | NM_172854 |
| Snord100 | 12.15908257 | 0.190221045 | 0.161273774 | 1.179491498 | 0.23820252 | 0.462671816 | | Pde11a | 96.82414906 | 0.0711769664620098 | 0.0950694128219372 | 0.748684191363658 | 0.454047574152718 | 0.825985605778632 | NM_001081033 |
| Adamts1 | 560.1777684 | 0.198186585 | 0.168113011 | 1.178889038 | 0.238442363 | 0.462759891 | | Phka1 | 1276.536408 | 0.0552789786383054 | 0.0738142167686702 | 0.748893384764578 | 0.453921467824834 | 0.825985605778632 | NM_008832,NM_173021 |
| Farp2 | 372.6309053 | 0.14575656 | 0.123644295 | 1.178837732 | 0.238462796 | 0.462759891 | | Pitpnm2 | 12606.59897 | -0.0558046674595721 | 0.0744841885434847 | -0.749214948176454 | 0.45372766090766 | 0.825985605778632 | NM_011256 |
| Tnfrsf14 | 22.37885663 | 0.1972734 | 0.167354769 | 1.178773698 | 0.2384883 | 0.462759891 | | Plin3 | 704.4615415 | 0.0583308946477126 | 0.0778942623772364 | 0.748847127728359 | 0.453949350861922 | 0.825985605778632 | NM_025836 |
| Usp20 | 3754.427122 | 0.08707076 | 0.073873707 | 1.178643433 | 0.238540189 | 0.462802616 | | Pcdhga12 | 652.5348964 | -0.055225898041822 | 0.0737766760426519 | -0.748554977048501 | 0.454125477224911 | 0.826023877055981 | NM_033595 |
| Cryba4 | 18.35147458 | -0.197779779 | 0.167820757 | -1.178517975 | 0.238590171 | 0.462841631 | | Atp2a3 | 493.009536 | -0.0654179884650332 | 0.0874079009967434 | -0.748421912882573 | 0.454205709240604 | 0.826066374412047 | NM_016745,NM_001163337,NM_001163336 |

| | | | | | | | | | | | | | | |
|---|---|---|---|---|---|---|---|---|---|---|---|---|---|---|
| Mettl22 | 541.5173748 | 0.094214976 | 0.079954347 | 1.178359642 | 0.23865326 | 0.46290606 | | Aqr | 2160.438812 | 0.0425675843662779 | 0.0569634737447209 | 0.747278590435732 | 0.454895413125829 | 0.826168452034221 | NM_009702 |
| Tmem132e | 769.3993453 | 0.141009457 | 0.119681057 | 1.178210319 | 0.23871277 | 0.462963531 | | Bcl2l12 | 55.44640986 | -0.0693319098434426 | 0.0926795586771901 | -0.748082002471881 | 0.454410696993232 | 0.826168452034221 | NM_029410 |
| 2610203C22Rik | 12.82333829 | 0.1952206 | 0.16572931 | 1.177948549 | 0.238817119 | 0.463059624 | | Cdh2 | 5159.925315 | 0.0425226038806591 | 0.0568542864816749 | 0.74792256682994 | 0.454506864860324 | 0.826168452034221 | NM_007664 |
| Polr2e | 1580.044169 | -0.080243287 | 0.068121937 | -1.177936081 | 0.238822089 | 0.463059624 | | Cep85 | 702.7393547 | -0.0465777877697746 | 0.0623179121836346 | -0.747422147785087 | 0.454808780458761 | 0.826168452034221 | NM_144527 |
| Snph | 15311.15964 | 0.118994422 | 0.10103474 | 1.17775749 | 0.2388933 | 0.463139739 | | Exosc9 | 1289.148943 | 0.0479636522214949 | 0.0641508999882907 | 0.747669202306587 | 0.454659712034289 | 0.826168452034221 | NM_019393 |
| Rasl12 | 136.2705087 | -0.151465126 | 0.128618399 | -1.177631866 | 0.2389434 | 0.463178912 | | Lrrc28 | 941.815737 | 0.0481007839987383 | 0.064338624152075 | 0.747619095569159 | 0.45468994334722 | 0.826168452034221 | NR_028143,NM_175124,NM_027413,NR_028144 |
| Crabp1 | 69.84479758 | -0.193283719 | 0.164165422 | -1.177371681 | 0.239047188 | 0.463206244 | | Nrn1l | 48.15236825 | -0.0691396242720298 | 0.0924726651846985 | -0.747676344506077 | 0.454655402964131 | 0.826168452034221 | NM_175024 |
| Dgcr6 | 1403.777269 | -0.080740776 | 0.068574328 | -1.177419855 | 0.239027969 | 0.463206244 | | Prkg1 | 552.6847202 | -0.0607573730588118 | 0.0812239972577505 | -0.748022445460405 | 0.45444661905398 | 0.826168452034221 | NM_001013833,NM_011160 |
| Slmap | 5048.549344 | -0.091023048 | 0.077309809 | -1.177380336 | 0.239043735 | 0.463206244 | | Ralgds | 4700.522137 | 0.0476291973201897 | 0.0637171451601536 | 0.747509908054941 | 0.454755824277304 | 0.826168452034221 | NM_001145836,NM_001145834,NM_001145835,NM_009058 |
| Ethe1 | 384.7066135 | 0.110691863 | 0.094029477 | 1.177203856 | 0.23911415 | 0.463246349 | | Tsr3 | 641.8944857 | -0.0548389356142889 | 0.0733060400764927 | -0.748082089239388 | 0.454410644660215 | 0.826168452034221 | NM_026676,NM_001163718 |
| Rbm28 | 1861.026815 | 0.118623264 | 0.100776285 | 1.177095026 | 0.23915758 | 0.463246349 | | Zdhhc22 | 678.7872955 | 0.0478078441886972 | 0.0639885488940077 | 0.747131244808931 | 0.454984341573021 | 0.826168452034221 | NM_001080943 |
| Spata1 | 28.8230893 | 0.196293101 | 0.166751284 | 1.177160962 | 0.239131267 | 0.463246349 | | Zfand3 | 3205.45094 | 0.0365330153106854 | 0.0488836297510574 | 0.74734661678627 | 0.454854360051658 | 0.826168452034221 | NM_148926 |
| Atxn7l3 | 11988.02855 | -0.089381357 | 0.075944689 | -1.176927029 | 0.239224633 | 0.463318308 | | Zfp449 | 236.934497 | 0.0616442561580179 | 0.0825110219072321 | 0.747103292785842 | 0.455001212740862 | 0.826168452034221 | NM_030139 |
| Myom3 | 120.814855 | -0.178628916 | 0.151804406 | -1.176704425 | 0.239313501 | 0.463432494 | | Tom1 | 1860.478475 | 0.0522182376191597 | 0.0699115156045065 | 0.74691897561474 | 0.455112470965321 | 0.826267185757063 | NM_001136259,NM_011622 |
| Nap1l2 | 3288.145398 | 0.113018474 | 0.096069222 | 1.1764275 | 0.239424088 | 0.463588705 | | Tmie | 229.4721811 | 0.070915271968537 | 0.0949881373010568 | 0.746569771589237 | 0.455323300818242 | 0.826546646799024 | NM_146260 |
| Cmtm4 | 7870.075732 | -0.078249953 | 0.066547039 | -1.175859273 | 0.239651116 | 0.46397031 | | Cdc123 | 2383.029745 | 0.0350454529058375 | 0.046998625905742 | 0.74566973460294 | 0.455866946191982 | 0.826603827285574 | NM_133837 |
| 3230401D17Rik | 649.7001163 | 0.101794543 | 0.086584944 | 1.175661015 | 0.239730363 | 0.464065748 | | Dsn1 | 134.4841723 | -0.0695556009229823 | 0.0932426943892091 | -0.745963009527017 | 0.455689760535088 | 0.826603827285574 | NM_025853 |
| Casz1 | 95.01282322 | -0.173323462 | 0.147452426 | -1.17545344 | 0.239813354 | 0.464110431 | | Hsf4 | 811.0129347 | -0.0565021067966896 | 0.075750891800872 | -0.745893618588913 | 0.455731680424693 | 0.826603827285574 | NM_011939,NM_001256042,NR_045689,NM_001256044 |
| Wnt4 | 796.9972816 | 0.143473726 | 0.122053206 | 1.175501494 | 0.23979414 | 0.464110431 | | Pacsin2 | 3117.484608 | -0.0429675048130156 | 0.0576153110314161 | -0.745765388467426 | 0.455809151473034 | 0.826603827285574 | NM_011862,NM_001159510,NM_001159509 |
| Ppp1r1b | 17095.2017 | -0.139073304 | 0.11833323 | -1.175268382 | 0.23988736 | 0.464195673 | | Pcdhgb1 | 317.7048591 | 0.0703862283044808 | 0.0943738213920412 | 0.74582365314439 | 0.455773949111991 | 0.826603827285574 | NM_033574 |
| 4933413G19Rik | 42.05392704 | 0.19672884 | 0.167419207 | 1.175067327 | 0.239967781 | 0.464293306 | | Pdgfd | 90.63040677 | 0.0709435479176088 | 0.0950980305901655 | 0.746004385972483 | 0.455664765565766 | 0.826603827285574 | NM_027924 |
| Snhg4 | 221.1965992 | 0.124302035 | 0.105825998 | 1.174588823 | 0.240159257 | 0.464547756 | | Smarcad1 | 1069.59225 | -0.0555004014042713 | 0.0744291585769454 | -0.74568089261542 | 0.455860204232152 | 0.826603827285574 | NM_007958,NM_001253392 |
| Wdr74 | 836.342893 | 0.100002114 | 0.085137574 | 1.174594364 | 0.240157039 | 0.464547756 | | Topors | 1030.917881 | 0.0511970335269258 | 0.0686185577573935 | 0.746110603314298 | 0.455600604591789 | 0.826603827285574 | NM_134097 |
| 4933412O06Rik | 76.45580352 | 0.182330983 | 0.155296579 | 1.174082419 | 0.240362015 | 0.464881919 | | Tuba8 | 601.5741162 | -0.0583178006444867 | 0.0781731965912305 | -0.746007623935758 | 0.455662809587069 | 0.826603827285574 | NM_017379 |
| Clstn1 | 46959.18858 | 0.100653816 | 0.08575787 | 1.173697705 | 0.24051613 | 0.465121931 | | Efnb2 | 2446.109008 | 0.0543883096245127 | 0.0729614211204124 | 0.745439285437609 | 0.456006202082536 | 0.826649955254077 | NM_010111 |
| 2700089E24Rik | 3410.690197 | -0.117271687 | 0.099927115 | -1.173572234 | 0.240566408 | 0.465161104 | | Prrc1 | 849.3640376 | -0.054229384871236 | 0.0727436605816987 | -0.745486059370503 | 0.45597793558113 | 0.826649955254077 | NM_028447 |
| 2610019E17Rik | 67.25870507 | 0.179632898 | 0.15308875 | 1.173390586 | 0.240639211 | 0.465243815 | | Adrbk1 | 10986.45263 | 0.0441946190957992 | 0.0593310143271557 | 0.744882244084126 | 0.456342909999516 | 0.826749850168438 | NM_130863 |
| 6330549D23Rik | 53.85239455 | 0.165994379 | 0.141481009 | 1.173262622 | 0.240690507 | 0.465284929 | | Art3 | 123.2049317 | -0.0666600969865936 | 0.0895152326371268 | -0.744678810776458 | 0.456465911659894 | 0.826749850168438 | NM_181728 |
| Plcb3 | 815.8222209 | -0.078992239 | 0.067339014 | -1.173053092 | 0.240774516 | 0.465389265 | | Cdc42ep4 | 1389.80762 | 0.0502564872343277 | 0.0674478009195062 | 0.745116765101133 | 0.456201134938531 | 0.826749850168438 | NM_001163346,NM_020006 |
| A930003A15Rik | 9.119688138 | -0.184541146 | 0.157328307 | -1.172968485 | 0.240808444 | 0.465396786 | | Eml6 | 1272.270171 | -0.0514584658641681 | 0.0690877412427507 | -0.744827735550952 | 0.456375865607089 | 0.826749850168438 | NM_146016 |
| 9530026P05Rik | 9.714312093 | 0.184464751 | 0.157293651 | 1.172741238 | 0.240899589 | 0.465428677 | | Mrpl10 | 2045.047854 | 0.0407143848120615 | 0.0546799107891377 | 0.744594938515327 | 0.456516628679681 | 0.826749850168438 | NM_026154 |
| Fcgr1 | 152.8228721 | 0.163406527 | 0.139335542 | 1.172755528 | 0.240893857 | 0.465428677 | | Prpf40a | 1404.403085 | -0.0474916879474744 | 0.0637695144375047 | -0.744739682689871 | 0.456429104786946 | 0.826749850168438 | NM_018785 |
| Gm17821 | 356.3756072 | 0.117752483 | 0.100420059 | 1.172599223 | 0.240956561 | 0.465428677 | | Slc30a6 | 390.1327636 | -0.0564723288393674 | 0.0758285486784047 | -0.744737039328965 | 0.456430703089579 | 0.826749850168438 | NM_144798,NM_001252478 |
| Gm5431 | 8.869499416 | 0.178721378 | 0.152430469 | 1.172478047 | 0.241005181 | 0.465428677 | | Stag3 | 39.63246908 | -0.0637174089213317 | 0.0855491584541378 | -0.744804625465604 | 0.456389838261922 | 0.826749850168438 | NM_016964 |
| Sympk | 5660.307043 | 0.09579863 | 0.081705839 | 1.172482059 | 0.241003571 | 0.465428677 | | Slc5a6 | 1284.201278 | 0.0445727854747269 | 0.0598955362165651 | 0.744175414233949 | 0.456770359870974 | 0.827003146888417 | NM_177870,NM_001177622,NM_001177621 |
| Tmem30a | 11302.24965 | -0.101654496 | 0.086688873 | -1.172636029 | 0.240941795 | 0.465428677 | | Slc9b2 | 127.0227326 | -0.0706449420767421 | 0.0949253469571363 | -0.744215790000135 | 0.456745936882036 | 0.827003146888417 | NM_178877 |
| Tspan15 | 539.3444342 | -0.137528043 | 0.117340253 | -1.172044881 | 0.241179038 | 0.465706382 | | Lin52 | 1128.827138 | 0.0456371959069981 | 0.0613439347729726 | 0.743956123386876 | 0.456903020025622 | 0.827140237897829 | NM_173756 |
| 2810006K23Rik | 428.7555294 | -0.132474073 | 0.113139789 | -1.170888457 | 0.241643616 | 0.466262791 | | 6030458C11Rik | 1843.896651 | 0.0418108894529836 | 0.0562216111267325 | 0.743680030064153 | 0.45707007365831 | 0.82723646772497 | NM_001166360,NM_029998 |
| 4930441O14Rik | 138.8910197 | 0.128658148 | 0.109860547 | 1.171104202 | 0.241556895 | 0.466262791 | | Rhov | 336.224113 | 0.0611972109583179 | 0.0822807121619725 | 0.743761318422343 | 0.457020885582271 | 0.82723646772497 | NM_145530 |
| Bad | 708.7982409 | -0.098966411 | 0.084514178 | -1.171003647 | 0.241597311 | 0.466262791 | | Mpp5 | 1646.886733 | 0.0460235452739497 | 0.0619053407607201 | 0.743450317991825 | 0.45720908989369 | 0.827384980664725 | NM_019579 |
| Gm16938 | 53.0004067 | 0.185756986 | 0.158624865 | 1.171045825 | 0.241580358 | 0.466262791 | | Pusl1 | 670.1031865 | -0.0530011320265613 | 0.0713003758296173 | -0.743349967091551 | 0.457269827303071 | 0.827391817564076 | NM_001033490 |
| Phb2 | 2192.673428 | -0.067962653 | 0.0580464 | -1.170833216 | 0.241665823 | 0.466262791 | | Pex1 | 973.5915458 | -0.0397112108747984 | 0.0534451140604083 | -0.743027900172753 | 0.457464788996671 | 0.827641490218663 | NM_027777 |
| Prokr2 | 126.0538647 | -0.194174876 | 0.165793958 | -1.171153607 | 0.241537039 | 0.466262791 | | 1110065P20Rik | 853.2581114 | -0.0490670760205576 | 0.0660801147552408 | -0.742539207177543 | 0.457760706115082 | 0.827642075203399 | NM_001142727 |
| Rpp25 | 467.210933 | 0.132260126 | 0.11296528 | 1.170803325 | 0.241677841 | 0.466262791 | | B4galt7 | 684.1379096 | -0.048732163774179 | 0.0656497345801511 | -0.742305571924017 | 0.457902216668634 | 0.827642075203399 | NM_146045 |
| AI837181 | 1954.47698 | 0.111522637 | 0.095300282 | 1.170232583 | 0.241911005 | 0.46665452 | | C130036L24Rik | 25.72832874 | -0.0619478414681391 | 0.0834673377516647 | -0.742180631811317 | 0.457977901726898 | 0.827642075203399 | NR_015507 |
| Tmem57 | 2468.477217 | 0.058424917 | 0.04994407 | 1.169806875 | 0.242078696 | 0.466919871 | | Dhx9 | 5632.271574 | -0.0464261555462627 | 0.0625199385657548 | -0.742581592549621 | 0.45773503634654 | 0.827642075203399 | NM_007842 |
| C130060K24Rik | 44.99221469 | -0.195763888 | 0.167375701 | -1.169607578 | 0.242158926 | 0.467016481 | | Fbxl18 | 662.5350711 | 0.0550048102606148 | 0.0740631211673293 | 0.742674753557084 | 0.457678618274074 | 0.827642075203399 | NM_001033312 |
| Pdcl | 794.1859622 | -0.093180061 | 0.079692334 | -1.169247491 | 0.242303932 | 0.467063589 | | Fundc2 | 1653.157992 | 0.0461689619794351 | 0.0621881315975138 | 0.742407928867264 | 0.457840217063092 | 0.827642075203399 | NM_026126 |
| Sec24b | 2498.778609 | 0.066155039 | 0.056573647 | 1.169361397 | 0.242258056 | 0.467063589 | | Mterfd2 | 490.7996349 | 0.0518979051805926 | 0.0699238486509889 | 0.742206073919512 | 0.457962489074365 | 0.827642075203399 | NM_178051 |
| Zbtb10 | 289.8792558 | -0.121747407 | 0.104116308 | -1.169340412 | 0.242266507 | 0.467063589 | | R3hdm2 | 12523.14988 | 0.0429955005076056 | 0.0578814272070114 | 0.742820323932811 | 0.457590469025334 | 0.827642075203399 | NM_001168293,NM_001168292,NM_027900 |
| Zfp166 | 1258.687465 | 0.070317535 | 0.060136579 | 1.16929722 | 0.242283903 | 0.467063589 | | Vamp4 | 2584.076745 | 0.0420882858226572 | 0.0566823042611026 | 0.742529549059628 | 0.457766555454836 | 0.827642075203399 | NM_016796 |
| Cdca7l | 48.72390096 | -0.196087381 | 0.16776164 | -1.168845157 | 0.242466023 | 0.467201662 | | Pcdhga2 | 564.3070954 | -0.0602286323295871 | 0.0811788083282464 | -0.741925553847166 | 0.458132442272045 | 0.827715405030314 | NM_033585 |
| Pqlc1 | 599.4602038 | 0.110514817 | 0.094548533 | 1.16886853 | 0.242456555 | 0.467201662 | | Tmem180 | 1064.595744 | -0.0490323683630255 | 0.0660798218855008 | -0.74201726013453 | 0.45807687808898 | 0.827715405030314 | NM_029186 |
| Sirt5 | 342.3834422 | 0.120818487 | 0.103362244 | 1.1688493 | 0.242464353 | 0.467201662 | | Tomm40l | 2714.700796 | 0.0432947476320406 | 0.0583801199017361 | 0.741600868667505 | 0.458329197085039 | 0.827967904098654 | NM_001037170 |
| A930003O13Rik | 12.99113037 | -0.193225481 | 0.165385458 | -1.168334164 | 0.242672 | 0.467209584 | | Ccbe1 | 229.633856 | -0.0701520288062126 | 0.0946110663914165 | -0.741478047779167 | 0.458403637228929 | 0.827999407409528 | NM_178793 |
| Ddc | 344.5498891 | 0.196266797 | 0.167991214 | 1.168315845 | 0.242679386 | 0.467209584 | | Atad5 | 223.6187343 | -0.062659275781179 | 0.0845456786443961 | -0.741129254455777 | 0.458615073290556 | 0.828043599048601 | NM_001029856 |
| Gucy2c | 15.44328467 | -0.195584567 | 0.167350618 | -1.168711349 | 0.242519948 | 0.467209584 | | Bcl6 | 1205.100555 | 0.0619320809683849 | 0.0835634914581965 | 0.741138024365187 | 0.458609756363111 | 0.828043599048601 | NM_009744 |
| Hddc2 | 311.4537738 | -0.105868288 | 0.090601982 | -1.168498591 | 0.242605707 | 0.467209584 | | Flt4 | 199.4383297 | 0.0687998743457964 | 0.0928396218771853 | 0.741061552758258 | 0.458656119919113 | 0.828043599048601 | NM_008029 |
| Klk6 | 164.0258607 | 0.182093511 | 0.155835553 | 1.168497871 | 0.242605998 | 0.467209584 | | Thumpd3 | 913.4852124 | 0.0438105351715116 | 0.0591072952240487 | 0.741203518202718 | 0.458570050548295 | 0.828043599048601 | NM_008188 |
| Mrpl40 | 672.5870013 | -0.087740996 | 0.075099342 | -1.168332423 | 0.242672702 | 0.467209584 | | Gosr2 | 4441.795297 | 0.0317242889146516 | 0.0428235416197957 | 0.740814227751462 | 0.458806087247001 | 0.828211410879822 | NM_019650 |
| Rpusd3 | 254.1102956 | 0.113628644 | 0.097258875 | 1.168311312 | 0.242681214 | 0.467209584 | | Exosc1 | 504.1819983 | 0.048259266374991 | 0.0651565844466287 | 0.740665993849346 | 0.458895983129271 | 0.828270756826018 | NM_025644,NM_001164561 |
| Gab2 | 1169.921858 | -0.101827838 | 0.087170095 | -1.168151055 | 0.242745839 | 0.467217893 | | Ddhd2 | 3352.099267 | -0.0316868614782415 | 0.042790871286667 | -0.740505171440964 | 0.458993524426232 | 0.828343885677158 | NM_028102 |
| Rint1 | 676.9439882 | -0.076850861 | 0.065784926 | -1.168213842 | 0.242720518 | 0.467217893 | | Fitm2 | 645.3870066 | 0.0435237514322968 | 0.0587966010177928 | 0.740242644623722 | 0.459152775991268 | 0.828381304839699 | NM_173397 |
| Arfip2 | 3655.586091 | 0.097591109 | 0.083556206 | 1.167969604 | 0.242819026 | 0.467300706 | | Prr14l | 3091.086824 | 0.056265208118919 | 0.0760243079111769 | 0.740094973105925 | 0.459242368709493 | 0.828381304839699 | NM_194340 |
| Rbak | 515.5859663 | 0.106485555 | 0.091179276 | 1.167870155 | 0.242859144 | 0.467319869 | | Usp36 | 1899.463342 | -0.0445068942500952 | 0.0601292201328801 | -0.740187452152864 | 0.459186260269925 | 0.828381304839699 | NM_001033528 |
| Cry2 | 7162.290936 | 0.11048175 | 0.09461877 | 1.167651525 | 0.242947358 | 0.46741904 | | Wdsub1 | 380.6397965 | -0.0571505711872319 | 0.077210574567392 | -0.740190984297739 | 0.459184117340233 | 0.828381304839699 | NM_001159636,NM_028118 |
| Tnfsf9 | 13.65695359 | -0.194468548 | 0.166555097 | -1.167592895 | 0.242971018 | 0.46741904 | | Ilvbl | 985.300961 | -0.0542575513602746 | 0.0733457021569323 | -0.739750929702517 | 0.459451138774776 | 0.828552109477641 | NM_173751 |
| Cenpt | 505.7952431 | 0.103357442 | 0.088541899 | 1.167328057 | 0.243077912 | 0.467508587 | | Sgk2 | 91.95481426 | -0.0702807213051178 | 0.0949999676745415 | -0.739797318083429 | 0.45942298659498 | 0.828552109477641 | NM_013731 |
| St8sia6 | 72.38200638 | 0.18264235 | 0.15645479 | 1.167381004 | 0.243056539 | 0.467508587 | | Nsun4 | 540.4878928 | 0.0524788362424465 | 0.0709605289647039 | 0.739549676532847 | 0.459573286481984 | 0.828566657495011 | NM_028142 |
| Sf3b5 | 757.3952892 | -0.087072743 | 0.074617446 | -1.166922055 | 0.243241848 | 0.467765818 | | Vstm4 | 157.1341039 | -0.0700640181433373 | 0.0947318821571105 | -0.739603357897375 | 0.459540703574294 | 0.828566657495011 | NM_178791 |
| Grb2 | 6526.952233 | -0.078692652 | 0.067441833 | -1.166822547 | 0.24328204 | 0.467785049 | | Zfp597 | 939.1581275 | 0.0589373912794308 | 0.079731866930132 | 0.739194923543893 | 0.459788642958942 | 0.828852051082353 | NM_001033159 |
| 1190003J15Rik | 23.2997865 | 0.193342671 | 0.165761014 | 1.166394116 | 0.243455137 | 0.468001723 | | Ephb6 | 5517.078892 | -0.0512556721157803 | 0.0693584882199169 | -0.738996385752663 | 0.459909192032539 | 0.828966487587127 | NM_007680,NM_001146351 |
| Madd | 11031.96419 | 0.074169884 | 0.063585217 | 1.166464278 | 0.243426783 | 0.468001723 | | Apol8 | 265.1206459 | -0.0601158961413258 | 0.0814054057120875 | -0.738475481026596 | 0.460225561398851 | 0.829022384037663 | NM_001081970 |
| Plekhg3 | 678.2524021 | 0.148645176 | 0.127478741 | 1.166038937 | 0.243598703 | 0.4681697 | | Gm16386 | 26.48522602 | -0.0617592952399392 | 0.0835906234663109 | -0.738830417562679 | 0.460009978921973 | 0.829022384037663 | NR_030709 |
| Zfyve21 | 471.801476 | -0.086166219 | 0.073897185 | -1.16602843 | 0.243602951 | 0.4681697 | | Mink1 | 16581.19653 | -0.0471327196584538 | 0.0638177055559555 | -0.73855240090272 | 0.460178836763521 | 0.829022384037663 | NM_001045964,NM_176893,NM_001045959,NM_016713 |
| 9630028B13Rik | 16.62289437 | -0.189331641 | 0.16246711 | -1.165353658 | 0.243875868 | 0.468636077 | | Ppp1r32 | 54.73790139 | 0.0665505925700343 | 0.0901070101946681 | 0.738572839233476 | 0.460166422037232 | 0.829022384037663 | NM_133689 |
| Syt1 | 65383.76035 | -0.059910657 | 0.051424523 | -1.165021141 | 0.244010435 | 0.468836518 | | Ykt6 | 3987.132807 | 0.0380034413007302 | 0.0514542601957882 | 0.738586876131341 | 0.460157895812416 | 0.829022384037663 | NM_019661 |
| Akap1 | 1959.286914 | 0.071614173 | 0.061498376 | 1.164488848 | 0.24422596 | 0.469017976 | | Fbxo38 | 2236.84109 | -0.0453998542843122 | 0.0615177954067872 | -0.737995469182672 | 0.460517202403954 | 0.829239231699496 | NM_134136 |
| Apeh | 727.0379699 | -0.102376649 | 0.087901441 | -1.164675431 | 0.244150398 | 0.469017976 | | Fmod | 1033.732929 | -0.0595194428778153 | 0.0806471725334038 | -0.738022685831454 | 0.460500663609111 | 0.829239231699496 | NM_021355 |
| Gimap1 | 50.30328664 | 0.176623084 | 0.151663605 | 1.164571318 | 0.24419256 | 0.469017976 | | Sema4g | 1399.705581 | -0.0622079301123633 | 0.0842756565411625 | -0.738148270396199 | 0.460424353691253 | 0.829239231699496 | NM_011976 |
| Maz | 5522.533108 | -0.094324219 | 0.080995387 | -1.164562855 | 0.244195987 | 0.469017976 | | Wnt5b | 75.13131147 | 0.070144337364809 | 0.0950625367436959 | 0.737875716002925 | 0.460589977002 | 0.829267477185306 | NM_001271757,NM_009525,NM_001271758 |
| Cbr2 | 42.24324239 | -0.193941001 | 0.166559258 | -1.164396402 | 0.244263405 | 0.469022123 | | Mpzl3 | 58.39818457 | -0.0701048429567574 | 0.0950291805039792 | -0.737719115170332 | 0.460685153803834 | 0.82933604463434 | NM_176993,NM_001093749 |
| Gm10336 | 491.6472984 | -0.103847051 | 0.089190084 | -1.164334051 | 0.244288662 | 0.469022123 | | Nrip2 | 744.9196021 | 0.0662556347974274 | 0.0898356273320524 | 0.737520700471672 | 0.460805759481358 | 0.829450367066444 | NM_021717,NM_001162858 |
| Efhd1 | 787.2098693 | 0.127554614 | 0.109621389 | 1.163592391 | 0.244589236 | 0.469482858 | | 1200014J11Rik | 897.2719695 | 0.0524938575831678 | 0.0711877672904091 | 0.737399771669573 | 0.460879125296793 | 0.829479689266958 | NM_025818 |
| Timm23 | 2172.598336 | -0.083190362 | 0.071493403 | -1.163608926 | 0.244582532 | 0.469482858 | | Srp54a | 136.7049572 | 0.0662295256073199 | 0.0898431136500837 | 0.73716863671118 | 0.461019803644125 | 0.829630038123707 | NM_011899 |
| Fam117a | 197.7174087 | 0.158667461 | 0.136405892 | 1.163200936 | 0.244747987 | 0.469671205 | | Chmp3 | 4818.785208 | 0.0394467248514553 | 0.0535732448150464 | 0.736313900485945 | 0.461539688676599 | 0.830270578153304 | NM_025783 |
| Rrp7a | 3716.488022 | -0.071703732 | 0.061640871 | -1.163249825 | 0.244728157 | 0.469671205 | | Csgalnact2 | 496.0452852 | 0.0528210003814282 | 0.0717474276786443 | 0.736207583887366 | 0.461604377610222 | 0.830270578153304 | NM_030165 |
| Tbr1 | 4183.424161 | 0.117330753 | 0.100897903 | 1.162866125 | 0.244883823 | 0.469873679 | | Kif5a | 154107.5859 | 0.0434265100391117 | 0.0589830878925774 | 0.73625358709944 | 0.461576386075543 | 0.830270578153304 | NM_001039000,NM_008447 |
| Klf2 | 336.5266341 | 0.181856655 | 0.156422681 | 1.162597734 | 0.24499275 | 0.470024477 | | St6galnac4 | 1223.657706 | 0.0396339596676116 | 0.0538168175644176 | 0.736460486913232 | 0.46145050564552 | 0.830270578153304 | NM_011373,NM_001276425 |
| Suclg1 | 3147.773873 | -0.058896127 | 0.050664015 | -1.162484404 | 0.245038756 | 0.470054536 | | Wdr62 | 81.4855094 | -0.0695860120557744 | 0.0945434749368354 | -0.736021307681624 | 0.461717730623153 | 0.830371642321644 | NM_146186 |
| Cox19 | 688.9984811 | -0.073636046 | 0.063350785 | -1.16235412 | 0.245091651 | 0.470097802 | | Fnip1 | 2243.74396 | -0.0573186918580497 | 0.0779345974750172 | -0.735471712372978 | 0.462052261566983 | 0.830604783684898 | NM_173753 |
| Bdnf | 853.5535749 | -0.153944457 | 0.132472256 | -1.162088288 | 0.245199464 | 0.470246648 | | Gm694 | 72.2540243 | 0.062684064504831 | 0.0852307902932685 | 0.735462768938571 | 0.462057767309643 | 0.830604783684898 | NM_001033374 |
| Rasl11b | 2141.188047 | 0.124771132 | 0.107382438 | 1.161932391 | 0.245262929 | 0.470309878 | | Igfbp4 | 3271.215952 | 0.0578556834705356 | 0.0786622336981807 | 0.735495049537014 | 0.462038053813176 | 0.830604783684898 | NM_010517 |
| Sergef | 686.8141075 | 0.106543246 | 0.091712879 | 1.161704302 | 0.245355598 | 0.47037115 | | Penk | 12888.26142 | 0.0608657541184733 | 0.0827618412124004 | 0.735432576497025 | 0.462076088183785 | 0.830604783684898 | NM_001002927 |
| Zfp36l1 | 911.5156877 | -0.119133462 | 0.10254412 | -1.1617776 | 0.245325815 | 0.47037115 | | Peo1 | 700.8135561 | 0.0475767517292524 | 0.0647247897676051 | 0.735062282937916 | 0.462301563422628 | 0.830835164221796 | NM_153796 |
| Syt4 | 7771.551305 | -0.08661102 | 0.074560249 | -1.161624606 | 0.245387983 | 0.470375034 | | Tex261 | 886.5577403 | 0.0421761080390477 | 0.0573797872793177 | 0.735034234855902 | 0.462318644665068 | 0.830835164221796 | NM_009357 |
| Card10 | 316.0706401 | -0.14464655 | 0.124613179 | -1.160764466 | 0.245737697 | 0.470819915 | | Wdr7 | 10078.98971 | 0.0415575154345677 | 0.0565465236686505 | 0.734926088084301 | 0.462384509193239 | 0.830850739799726 | NM_001014981 |
| Hipk3 | 3908.148318 | -0.094460439 | 0.081368003 | -1.160903987 | 0.245680947 | 0.470819915 | | Itih2 | 149.4433111 | 0.0677822691548814 | 0.0922643159891503 | 0.734653136786406 | 0.462550767749083 | 0.830943909513132 | NM_010582 |
| Rpa3 | 143.4034611 | -0.141182894 | 0.121630246 | -1.160754824 | 0.245741619 | 0.470819915 | | Nptn | 23427.94185 | 0.048418855140992 | 0.0659032841814458 | 0.734695633797014 | 0.462524880017927 | 0.830943909513132 | NM_009145 |
| Tk1 | 13.75165579 | -0.192608955 | 0.165910083 | -1.160923744 | 0.245672912 | 0.470819915 | | 2810025M15Rik | 453.7925597 | 0.055348315998011 | 0.0753962285517686 | 0.734099265456066 | 0.462888241199959 | 0.831014287519936 | NR_027984 |
| Cdkl3 | 370.6494894 | 0.148509524 | 0.127960997 | 1.160584298 | 0.245810994 | 0.470874197 | | Chtop | 5182.952182 | 0.0354824115939934 | 0.0483494610775606 | 0.733873983353686 | 0.463025544708902 | 0.831014287519936 | NM_023215 |
| Ttll9 | 44.75876019 | 0.181120724 | 0.156066471 | 1.160535784 | 0.245830733 | 0.470874197 | | Cml5 | 73.57518549 | 0.0690588674978337 | 0.0940649835556564 | 0.734161266896654 | 0.462850456938837 | 0.831014287519936 | NM_023493 |
| Actn4 | 6690.69616 | -0.080022951 | 0.068979684 | -1.160094481 | 0.246010341 | 0.471159978 | | Dsc3 | 25.10332378 | 0.0614585693581462 | 0.083717743444034 | 0.734116411047699 | 0.462877792343098 | 0.831014287519936 | NM_007882 |
| Erv3 | 17.86889303 | 0.194552991 | 0.167738635 | 1.159857959 | 0.246106642 | 0.471286158 | | Gpr108 | 1100.030648 | -0.0472900886751373 | 0.0644115243692263 | -0.734186764530773 | 0.462834918939826 | 0.831014287519936 | NM_030084 |
| Sp110 | 73.17541967 | 0.167399242 | 0.144361376 | 1.159584687 | 0.246217938 | 0.471382768 | | Rdh12 | 89.66463097 | 0.0689691404695822 | 0.0939841783967144 | 0.733837776167582 | 0.463047614152992 | 0.831014287519936 | NM_030017 |
| Vps37d | 546.3852641 | 0.09037688 | 0.077938288 | 1.159595407 | 0.246213572 | 0.471382768 | | Tsacc | 29.05707932 | -0.0632524604747 | 0.0861731876424342 | -0.734015558727604 | 0.462939255595949 | 0.831014287519936 | NM_029801 |
| Cops8 | 3551.534459 | -0.049235661 | 0.042465529 | -1.159426539 | 0.246282364 | 0.471389617 | | Wnt9a | 745.567634 | -0.068813764443288 | 0.0937494749736451 | -0.734017598099967 | 0.462938012679523 | 0.831014287519936 | NM_139298 |
| Tep1 | 268.2153584 | 0.149907832 | 0.129293417 | 1.159439022 | 0.246277279 | 0.471389617 | | Manba | 553.2931419 | -0.0536811208896268 | 0.0731673033671908 | -0.733676361150384 | 0.463146008930035 | 0.831088193417874 | NM_027288 |
| 4931406P16Rik | 1701.835815 | 0.068318355 | 0.0589329 | 1.159256628 | 0.246351595 | 0.471424129 | | Shprh | 1754.95998 | -0.0539768240275255 | 0.0735905355660012 | -0.733475076910199 | 0.463268723368561 | 0.831205715866072 | NM_001077707,NM_172937 |
| Ascl5 | 8.90921616 | 0.176095055 | 0.151945671 | 1.158934334 | 0.246482953 | 0.471424129 | | Fhad1 | 281.7001767 | 0.0672329433481056 | 0.0916798187221402 | 0.733345072942092 | 0.463347990887851 | 0.831245265609105 | NM_177868 |
| Kcnh1 | 5097.029546 | -0.095831035 | 0.082678484 | -1.159080698 | 0.246423293 | 0.471424129 | | Zhx1 | 3481.895013 | 0.0433947372671065 | 0.0591925213313812 | 0.733111823775288 | 0.463490229204224 | 0.831397761470044 | NM_001042438,NM_009572 |
| Lnpep | 1836.85406 | -0.13570367 | 0.117064221 | -1.159224126 | 0.24636484 | 0.471424129 | | 8030462N17Rik | 1127.476326 | 0.0563377032154902 | 0.0769596860955207 | 0.732041749047224 | 0.464143086138657 | 0.831598959715103 | NM_178670 |
| Thoc7 | 1126.694672 | -0.080013991 | 0.069038166 | -1.158974597 | 0.24646654 | 0.471424129 | | Cln6 | 563.1927362 | 0.0547236662530071 | 0.0748170658527866 | 0.73143293777848 | 0.464514752727026 | 0.831598959715103 | NM_001033175 |
| Tppp | 25867.86801 | -0.107620489 | 0.092859908 | -1.158955366 | 0.246474379 | 0.471424129 | | Cpsf3l | 1046.438122 | 0.0455827046568761 | 0.0622894253789372 | 0.731788812941749 | 0.464297487253427 | 0.831598959715103 | NM_028020 |
| Pcmtd1 | 3956.216254 | 0.094319669 | 0.081402641 | 1.158680701 | 0.24658636 | 0.4715055 | | Ddhd1 | 3016.050097 | -0.0387824625274269 | 0.0529829534113745 | -0.73198000036557 | 0.464180774876259 | 0.831598959715103 | NM_176845,NM_001042719,NM_001039106 |
| Slc25a16 | 1602.049887 | -0.084887435 | 0.073259387 | -1.158724339 | 0.246568567 | 0.4715055 | | Diexf | 830.48708 | -0.0486442534360676 | 0.0665204870717361 | -0.73126724678984 | 0.464615932282394 | 0.831598959715103 | NM_145415 |
| Gucy1b2 | 9.715393783 | -0.169977947 | 0.146724758 | -1.158481699 | 0.246667516 | 0.471602479 | | Ecm2 | 167.2268394 | -0.0688235767315085 | 0.0941289739876089 | -0.731162508374609 | 0.464679897343033 | 0.831598959715103 | NM_001012324 |
| Atp5sl | 1481.649108 | 0.06204501 | 0.053565229 | 1.158307574 | 0.246738542 | 0.471680069 | | Endog | 280.7444189 | 0.058874523987691 | 0.0804931498283363 | 0.731422787072561 | 0.464520950925328 | 0.831598959715103 | NM_007931 |
| Nsg1 | 11377.81936 | -0.086285513 | 0.074503254 | -1.158144224 | 0.246805185 | 0.471749265 | | Fam175b | 1335.367393 | 0.0435327685781838 | 0.0594602156939535 | 0.73213270537408 | 0.464087573413991 | 0.831598959715103 | NM_198017 |
| Lck | 55.52871928 | -0.165752077 | 0.143174321 | -1.157694175 | 0.246988862 | 0.472042115 | | Gm16938 | 65.86533898 | -0.0694730587224921 | 0.0948036331846395 | -0.732810087427624 | 0.463674267962728 | 0.831598959715103 | NR_045971,NR_045970,NR_045972,NR_045969 |
| AI414108 | 1894.537704 | 0.115877987 | 0.100101058 | 1.157610007 | 0.247023224 | 0.472049559 | | Mrpl2 | 921.5959112 | 0.0492986620561441 | 0.067418829364864 | 0.731229873519559 | 0.464638756044011 | 0.831598959715103 | NM_025302 |
| Ch25h | 14.22578733 | 0.178042774 | 0.153831023 | 1.157391859 | 0.247112299 | 0.472061549 | | Nckap1 | 23486.7415 | 0.0502135729314337 | 0.0686418735846756 | 0.731529754494405 | 0.464455637069624 | 0.831598959715103 | NM_016965 |
| Haus1 | 227.4770436 | 0.120435151 | 0.104057336 | 1.157392223 | 0.24711215 | 0.472061549 | | Ndufb7 | 1664.546242 | 0.0414330030988541 | 0.0565818536731023 | 0.73226662629737 | 0.464005845143725 | 0.831598959715103 | NM_025843 |
| Nudt6 | 92.13492721 | 0.156262426 | 0.135015007 | 1.157370798 | 0.2471209 | 0.472061549 | | Prdm11 | 122.5796996 | 0.0658691859880331 | 0.0900226686497059 | 0.731695549310382 | 0.464354413555173 | 0.831598959715103 | NM_001177536 |
| Has1 | 33.57585029 | 0.188839618 | 0.163188562 | 1.157186604 | 0.24719613 | 0.472120974 | | Procr | 24.55659061 | -0.06313238724201 | 0.0862076321867129 | -0.732329442772243 | 0.463967512731158 | 0.831598959715103 | NM_011171 |
| Pus3 | 350.3693 | 0.112894565 | 0.097569278 | 1.157070833 | 0.247243422 | 0.472120974 | | Tnfaip8l3 | 321.7505918 | -0.0565935537944376 | 0.0774138523366375 | -0.731052028625808 | 0.464747374016743 | 0.831598959715103 | NM_001033535 |
| Zfat | 281.5159272 | 0.113376674 | 0.098291778 | 1.157133142 | 0.247217968 | 0.472120974 | | Vmn2r87 | 31.99456273 | -0.0618348923190529 | 0.0845739683774071 | -0.731133864301103 | 0.464697391490167 | 0.831598959715103 | NM_001103366 |
| Tsn | 3023.995351 | -0.056324541 | 0.048694958 | -1.156681165 | 0.247402648 | 0.472366805 | | Wdfy3 | 8573.562693 | 0.0538057551351328 | 0.0734339881604186 | 0.732709151212006 | 0.463735841310207 | 0.831598959715103 | NM_172882 |
| 11-Mar | 120.2965932 | -0.153835644 | 0.133016632 | -1.156514349 | 0.247470834 | 0.472380571 | | Wdr92 | 592.640039 | 0.0507564144420824 | 0.0693203538495603 | 0.732200740813218 | 0.464046052248271 | 0.831598959715103 | NM_178909 |
| Smek2 | 1686.972935 | 0.121858325 | 0.105364257 | 1.156543295 | 0.247459001 | 0.472380571 | | Zdhhc2 | 1286.90106 | 0.0557535188996362 | 0.0762509711793387 | 0.731184377553783 | 0.464666541157045 | 0.831598959715103 | NM_178395 |
| Numbl | 3628.403801 | 0.09705439 | 0.083936631 | 1.156281694 | 0.247565954 | 0.47244573 | | Zfp647 | 99.28900146 | 0.0691107791990988 | 0.0943342724704244 | 0.732615807481489 | 0.463792787125086 | 0.831598959715103 | NM_172817,NM_001168276 |
| Rapgefl1 | 7116.634556 | 0.088491813 | 0.076530262 | 1.156298317 | 0.247559156 | 0.47244573 | | Acss1 | 1243.746607 | -0.052996730183486 | 0.0725710923153983 | -0.73027328778676 | 0.465223152788571 | 0.831682622118967 | NM_080575 |
| Naa35 | 1716.203603 | 0.069132924 | 0.059800895 | 1.156051658 | 0.247660028 | 0.472567053 | | Ccdc132 | 3086.77643 | 0.0530663333961547 | 0.0726179219014599 | 0.730760837085975 | 0.464925248437874 | 0.831682622118967 | NM_024260,NM_001167751,NM_001167750 |
| Gm2115 | 330.6411161 | 0.171887328 | 0.148714327 | 1.155822246 | 0.247753872 | 0.472687906 | | Ccdc167 | 545.3180825 | -0.0504863080379704 | 0.0691321889349105 | -0.730286554147799 | 0.465215045318994 | 0.831682622118967 | NM_001163741,NM_026782 |
| Xrcc6 | 574.7283053 | 0.093835643 | 0.081222484 | 1.155291462 | 0.247971091 | 0.473044088 | | Fzd9 | 179.3480099 | 0.067812620591281 | 0.0928041827544427 | 0.730706511049306 | 0.464958437700368 | 0.831682622118967 | NM_010246 |
| Ankle1 | 8.668018369 | -0.176978086 | 0.153234712 | -1.154947757 | 0.248111821 | 0.473111538 | | Gp1bb | 314.5766309 | -0.0627017636158304 | 0.0858840801587274 | -0.730074345559126 | 0.465344741753625 | 0.831682622118967 | NM_010327,NM_001001999 |
| Glt8d1 | 1025.747975 | 0.15976582 | 0.138344538 | 1.154840096 | 0.248155914 | 0.473111538 | | Hcfc2 | 969.1486022 | 0.0460330596889486 | 0.063040143701922 | 0.730218190913564 | 0.465256824976244 | 0.831682622118967 | NM_001081218 |
| Gpc3 | 191.8428026 | -0.159597052 | 0.138179396 | -1.154998907 | 0.248090874 | 0.473111538 | | Heatr3 | 996.0667476 | 0.0526664445903165 | 0.0721512711995863 | 0.729944791196119 | 0.465423931929305 | 0.831682622118967 | NM_172757 |
| Ifnar1 | 2105.389605 | -0.068349525 | 0.059185674 | -1.154832259 | 0.248159124 | 0.473111538 | | Limd1 | 564.306984 | 0.0486134616396071 | 0.066590384037123 | 0.730037262024287 | 0.465367408319623 | 0.831682622118967 | NM_013860 |
| Sertm1 | 1156.884099 | 0.103529779 | 0.089635098 | 1.155013844 | 0.248084757 | 0.473111538 | | Nrxn1 | 12651.71961 | 0.0489120651264274 | 0.0669846672580899 | 0.730197926310072 | 0.465269209931333 | 0.831682622118967 | NM_020252,NM_177284 |
| Ick | 1013.036525 | 0.079945519 | 0.069248372 | 1.15447507 | 0.248305454 | 0.473241848 | | Papd4 | 814.6432859 | -0.0487775036663617 | 0.0668007150053679 | -0.730167389113159 | 0.465287873451063 | 0.831682622118967 | NM_133905 |
| Pygo1 | 452.7040918 | -0.113799418 | 0.098575279 | -1.154441753 | 0.248319016 | 0.473241848 | | Zfp60 | 1555.128762 | -0.0500758841709774 | 0.0685995941490089 | -0.729973475676939 | 0.465406397879097 | 0.831682622118967 | NM_009560,NM_029531 |
| Slc18a2 | 34.82142139 | -0.183146735 | 0.15864251 | -1.154461907 | 0.248310847 | 0.473241848 | | Sema4d | 2216.270637 | -0.0378167281979373 | 0.0518332509887468 | -0.72958433971559 | 0.465644297497875 | 0.831871733544967 | NM_013660 |
| Gpr137b | 358.3258256 | 0.117982374 | 0.102216349 | 1.154241721 | 0.248401082 | 0.473339856 | | Tanc1 | 1777.196687 | -0.0604137810417309 | 0.0828228251246566 | -0.729651288169495 | 0.465603363518128 | 0.831871733544967 | NM_198294 |
| Zfp799 | 854.1699229 | -0.126407821 | 0.109538427 | -1.154004349 | 0.248498386 | 0.473467043 | | Dhx8 | 2027.217469 | 0.0372058123649089 | 0.05101767697193 | 0.729272961318478 | 0.465834708413547 | 0.831904963660583 | NM_144831 |
| Mcts1 | 947.6168792 | -0.124216104 | 0.10768249 | -1.15354042 | 0.248688637 | 0.47377127 | | Gabrg3 | 233.9012017 | 0.0688578033804027 | 0.0944085599964162 | 0.729359746436304 | 0.465781634123635 | 0.831904963660583 | NM_008074 |
| Frem1 | 147.1729021 | 0.174053444 | 0.150897817 | 1.153452362 | 0.24872476 | 0.473781833 | | Lsm14b | 4152.216808 | -0.0401248471935059 | 0.0550149826138297 | -0.729343994801504 | 0.465791266940848 | 0.831904963660583 | NM_177727 |
| Ggn | 39.02249532 | -0.192168034 | 0.166640768 | -1.153187405 | 0.248833472 | 0.473930647 | | Cd82 | 842.8715933 | 0.0633961214936173 | 0.087016558401893 | 0.728552388854741 | 0.466275511287591 | 0.832050282035795 | NM_001271430,NM_001271431,NM_001271432,NM_001136055,NM_007656,NM_001271462,NM_001271461 |
| Bloc1s4 | 237.9935582 | -0.146209342 | 0.126866168 | -1.152469129 | 0.249128348 | 0.474379245 | | Dnm1l | 9942.944337 | 0.0420614022551526 | 0.0577260683374085 | 0.728637918129189 | 0.466223177515168 | 0.832050282035795 | NM_001025947,NM_001276341,NR_075074,NM_001276340,NM_152816 |
| Trim46 | 3440.888662 | 0.081833145 | 0.071007084 | 1.152464526 | 0.249130239 | 0.474379245 | | Gcdh | 1025.196964 | -0.0510194744531052 | 0.0700233281057055 | -0.728606820516835 | 0.466242205188522 | 0.832050282035795 | NM_008097,NM_001044744 |
| Smek1 | 1522.498237 | 0.107267806 | 0.09308837 | 1.152322314 | 0.249188651 | 0.474432165 | | Mcl1 | 2770.663348 | 0.0484362859377942 | 0.0664789170989316 | 0.728596193372298 | 0.466248707710529 | 0.832050282035795 | NM_008562 |
| Brca1 | 92.39825393 | -0.16600758 | 0.144109758 | -1.151952389 | 0.249340638 | 0.474623391 | | Ptpn3 | 3381.431787 | 0.0442292712483691 | 0.0606707006590034 | 0.729005446911804 | 0.465998330608032 | 0.832050282035795 | NM_011207 |
| Ewsr1 | 6927.860543 | 0.090705271 | 0.07874718 | 1.151854209 | 0.249380988 | 0.474623391 | | Zfp791 | 189.6368672 | 0.0624183936390202 | 0.085682525798999 | 0.728484519532563 | 0.466317041584291 | 0.832050282035795 | NM_001037745 |
| Minpp1 | 2334.537241 | -0.06057523 | 0.052587959 | -1.151884019 | 0.249368736 | 0.474623391 | | Zfpl1 | 808.9075741 | -0.0495084513862725 | 0.067947872858324 | -0.728624006957525 | 0.466231689280318 | 0.832050282035795 | NM_024231 |
| Ss18 | 1029.461351 | 0.07040148 | 0.061143965 | 1.151405202 | 0.249565575 | 0.474916361 | | Ascl1 | 155.1298305 | -0.0646737080384046 | 0.0888608432650895 | -0.727808848780223 | 0.466730607018306 | 0.832095317448889 | NM_008553 |
| Smarcal1 | 3495.249201 | -0.060823952 | 0.052835071 | -1.15120414 | 0.249648263 | 0.475015373 | | Elavl1 | 3163.139415 | 0.0402178360819347 | 0.0552461154795489 | 0.727975185536394 | 0.466628389637634 | 0.832095317448889 | NM_010485 |
| Magee2 | 270.467799 | -0.131115157 | 0.113903428 | -1.151108084 | 0.249687773 | 0.475032214 | | Mras | 10814.17554 | 0.0384254818414779 | 0.0527976176332819 | 0.727788176132699 | 0.466743263566952 | 0.832095317448889 | NM_008624 |
| Zdhhc21 | 1928.335284 | -0.070014963 | 0.060840926 | -1.150787273 | 0.249819762 | 0.475224971 | | Pigw | 143.4846665 | -0.0671296811740199 | 0.0922315767146765 | -0.727838377757428 | 0.466712528631951 | 0.832095317448889 | NM_001077636,NM_027388 |
| Plin5 | 21.71819375 | 0.192304778 | 0.167121885 | 1.150685793 | 0.249861523 | 0.475246064 | | Prim2 | 299.9606647 | 0.0646723893087749 | 0.0888589954429042 | 0.727809142451304 | 0.466730427223533 | 0.832095317448889 | NM_008922 |
| Zadh2 | 1225.306559 | -0.098995801 | 0.086039531 | -1.150585078 | 0.249902975 | 0.475266563 | | Tchh | 272.0762785 | 0.0608817044143221 | 0.0836470207353331 | 0.727840679549812 | 0.466711119432786 | 0.832095317448889 | NM_001163098 |
| Rasgrf1 | 14672.72925 | 0.121215387 | 0.105387552 | 1.15018695 | 0.25006668 | 0.475519912 | | Ube2w | 2093.497837 | 0.0521859454089583 | 0.0716973156676748 | 0.727864703482708 | 0.46669641168501 | 0.832095317448889 | NM_001271016,NM_001271017,NR_073122,NM_025773 |
| 1700052N19Rik | 905.2895477 | 0.076584929 | 0.066592683 | 1.150050205 | 0.250123194 | 0.47556863 | | Foxc1 | 149.3345406 | -0.0689655348980977 | 0.0947863023774085 | -0.727589674544949 | 0.466864803169843 | 0.832107746115968 | NM_008592 |
| Zfp800 | 391.3196483 | 0.118538292 | 0.10308418 | 1.1499174 | 0.250177894 | 0.475614268 | | Gm2a | 2675.517015 | 0.0427561084509532 | 0.0587622950160614 | 0.727611275891568 | 0.466851576131713 | 0.832107746115968 | NM_010299 |
| Trappc5 | 1176.839821 | -0.080309941 | 0.069882246 | -1.149218094 | 0.250466062 | 0.476103688 | | Arhgap27 | 794.1639527 | -0.0506540613432795 | 0.0696333446692477 | -0.727439728536406 | 0.466956624544754 | 0.832169295563379 | NM_183288,NM_133715,NM_001205236 |
| 3010001F23Rik | 35.96528432 | -0.184299034 | 0.1603938 | -1.149040887 | 0.250539122 | 0.476184145 | | Bcam | 825.1390905 | 0.0616428721567473 | 0.0847571840350275 | 0.72728787368953 | 0.467049625029656 | 0.832211278317856 | NM_020486 |
| Sepsecs | 234.3843779 | 0.11914594 | 0.103718805 | 1.148739997 | 0.250663208 | 0.476361553 | | Palm3 | 90.48982105 | -0.0690919977235094 | 0.0950199014294095 | -0.727129488779125 | 0.467146635666054 | 0.832211278317856 | NM_028877 |
| Sowaha | 3377.491107 | -0.117706522 | 0.102485035 | -1.148523992 | 0.250752315 | 0.476448049 | | Slirp | 620.2293177 | -0.0594261223599925 | 0.0817286048942906 | -0.727115340300442 | 0.467155302142055 | 0.832211278317856 | NM_026958 |
| Wnk2 | 5566.95994 | 0.110896016 | 0.096558895 | 1.148480585 | 0.250770223 | 0.476448049 | | Srsf4 | 2239.573156 | 0.0374398643026622 | 0.0514972050691876 | 0.727027112488162 | 0.467209347011023 | 0.832211278317856 | NM_020587 |
| Anapc13 | 507.7503024 | -0.1121543 | 0.097661479 | -1.148398535 | 0.250804078 | 0.476453946 | | Mpp3 | 2382.843616 | -0.0418426126228615 | 0.0575651794585784 | -0.726873652030735 | 0.467303359077918 | 0.83227667900513 | NM_007863 |
| Zfp710 | 916.9824609 | -0.07848273 | 0.068345727 | -1.148319494 | 0.250836694 | 0.47645749 | | BC017158 | 998.2834206 | 0.053601427491401 | 0.0737718709303639 | 0.72658354485814 | 0.467481115524575 | 0.832353543860922 | NM_145590 |
| Kat8 | 708.1168675 | 0.101092728 | 0.088054156 | 1.148074469 | 0.250937823 | 0.476591153 | | Ccna2 | 67.30770954 | -0.0689142447900078 | 0.0949154399290382 | -0.726059372864027 | 0.467802373551789 | 0.832353543860922 | NM_009828 |
| Apoc3 | 9.787055126 | 0.181510644 | 0.158112699 | 1.147982702 | 0.250975705 | 0.476604678 | | Ccs | 563.7553511 | 0.0535216322096208 | 0.0737446242845689 | 0.725769949048615 | 0.467979812163931 | 0.832353543860922 | NM_016892 |
| 2310045N01Rik | 490.8040076 | 0.085105901 | 0.074146475 | 1.147807779 | 0.251047925 | 0.476683401 | | Dpp9 | 3240.247971 | 0.0386839897148467 | 0.053295369365991 | 0.725841478819545 | 0.467935955555176 | 0.832353543860922 | NM_172624 |
| Ecsit | 1479.201988 | -0.082695151 | 0.072083289 | -1.147216676 | 0.251292081 | 0.476971641 | | Gja4 | 124.2609827 | -0.0680217235296099 | 0.0937550008070824 | -0.725526349997871 | 0.468129185631963 | 0.832353543860922 | NM_008120 |
| Gng11 | 324.7404262 | -0.174077344 | 0.151729929 | -1.147284158 | 0.251264199 | 0.476971641 | | Lars2 | 56155.43219 | -0.0617864282471161 | 0.0851052314211303 | -0.726000355270469 | 0.467838552758875 | 0.832353543860922 | NM_153168 |
| Ikbke | 44.81917586 | 0.184562125 | 0.160859772 | 1.147347923 | 0.251237855 | 0.476971641 | | Lrdd | 68.58760997 | -0.0690187889504256 | 0.0951009509786694 | -0.725742365668962 | 0.467996726026529 | 0.832353543860922 | NM_022654 |
| Immp2l | 89.72395888 | -0.156208972 | 0.136181776 | -1.147062233 | 0.251355901 | 0.477034338 | | Npat | 1155.997173 | -0.0501029121503604 | 0.0689930707242112 | -0.726202089926376 | 0.467714890960974 | 0.832353543860922 | NM_001081152 |
| Atf4 | 3968.653618 | 0.101730351 | 0.088710115 | 1.146772838 | 0.251475518 | 0.477149975 | | Nup54 | 657.4756469 | -0.0457457744891953 | 0.0629935143600339 | -0.726189164270361 | 0.467717297185105 | 0.832353543860922 | NM_183392 |
| Gstt3 | 129.3197066 | -0.154900131 | 0.13508442 | -1.146691319 | 0.251509219 | 0.477149975 | | Serac1 | 1333.203496 | -0.0475736251051255 | 0.0655740690226773 | -0.72549447997002 | 0.468148730093882 | 0.832353543860922 | NM_177311,NM_001111017 |
| Kif19a | 111.5012093 | -0.14367928 | 0.125294149 | -1.14673575 | 0.25149085 | 0.477149975 | | Sgtb | 6196.746426 | 0.0463276325689763 | 0.0637958854445338 | 0.726185274272196 | 0.467725198136074 | 0.832353543860922 | NM_144838 |
| Cep104 | 1444.81921 | 0.075935405 | 0.066226425 | 1.146602815 | 0.251545813 | 0.477160972 | | Tbca | 1126.067798 | 0.0609073434819982 | 0.0838736533101957 | 0.726179689070421 | 0.467728621619946 | 0.832353543860922 | NM_009321 |
| Bccip | 1333.87916 | 0.07227216 | 0.06305434 | 1.146188519 | 0.251717157 | 0.477427548 | | Unkl | 2161.799324 | -0.0452173212842817 | 0.0623207383642045 | -0.725558176477791 | 0.468109668327027 | 0.832353543860922 | NM_001197024,NM_028789 |
| Phospho1 | 1071.2511 | 0.142724744 | 0.124544657 | 1.145956376 | 0.251813203 | 0.477481256 | | Upf1 | 3214.246418 | -0.0414528255425295 | 0.0570767086534126 | -0.726265170895397 | 0.467676226528185 | 0.832353543860922 | NM_001122829,NM_030680 |
| Taf5l | 824.6266211 | -0.073183665 | 0.063868401 | -1.145850903 | 0.251856849 | 0.477575574 | | Phf23 | 1779.092683 | 0.0371120715476149 | 0.0511619788684588 | 0.725383817600815 | 0.468216597881636 | 0.832372329069715 | NM_030064 |
| Pecam1 | 954.9595997 | -0.120227574 | 0.104942717 | -1.145649526 | 0.251940196 | 0.477675159 | | Zbtb46 | 257.56855 | 0.0625838089401682 | 0.0862902152831318 | 0.725271211050069 | 0.468285663603133 | 0.83239323904786 | NM_027656,NM_028125 |
| Allc | 17.23619925 | 0.192585042 | 0.168121422 | 1.145511619 | 0.25199785 | 0.477720814 | | D10Jhu81e | 2408.169184 | 0.0346877782298376 | 0.0478737107577463 | 0.7245684046797 | 0.468716847939914 | 0.832446623447266 | NM_138601 |
| Sfxn1 | 2279.90105 | -0.070289246 | 0.061364278 | -1.145442404 | 0.252025941 | 0.477720814 | | Eci1 | 464.9017671 | 0.0600977091307963 | 0.0829075841395119 | 0.724875893496879 | 0.468528170994606 | 0.832446623447266 | NM_010023 |
| Ndfip1 | 10755.5344 | -0.068880355 | 0.060139514 | -1.14534274 | 0.252067208 | 0.477740589 | | Eif3b | 3850.415667 | 0.0310332637024667 | 0.0428143501364535 | 0.724833230063301 | 0.468554347015962 | 0.832446623447266 | NM_133916 |
| Actg1 | 43908.84406 | -0.099907634 | 0.087249905 | -1.145074411 | 0.252178334 | 0.477892749 | | Rpl6 | 9806.062646 | 0.0396549598407775 | 0.0546880806145538 | 0.725111567185344 | 0.468383588699174 | 0.832446623447266 | NM_011290 |
| A830052D11Rik | 38.60046104 | 0.189050091 | 0.165178702 | 1.144518567 | 0.252408642 | 0.478270699 | | Taf2 | 1626.599421 | -0.0422428931111255 | 0.0582994507507078 | -0.724584752809401 | 0.4687068155732 | 0.832446623447266 | NM_001081288 |
| AU041133 | 154.4654543 | -0.153002028 | 0.133711161 | -1.144272675 | 0.252510572 | 0.478384902 | | Tbc1d8b | 246.1292695 | 0.0656805733817648 | 0.0906458450545626 | 0.724584489694255 | 0.468706977082392 | 0.832446623447266 | NM_001081499 |
| Cpt1a | 1382.985871 | -0.090422331 | 0.079030154 | -1.144149747 | 0.25256154 | 0.478384902 | | Timmdc1 | 687.6647289 | 0.0473437787158133 | 0.0653119332154168 | 0.7248871130435 | 0.468521287409951 | 0.832446623447266 | NM_024273 |
| Sarm1 | 1331.449138 | 0.073969485 | 0.064647085 | 1.144204494 | 0.25253884 | 0.478384902 | | Haus1 | 254.1824458 | -0.0612036684781339 | 0.0844949896603789 | -0.724346718357353 | 0.46885290205171 | 0.832586461516276 | NM_146089 |
| Gsg1l | 4597.972467 | -0.114300142 | 0.099944061 | -1.143641158 | 0.252772485 | 0.478665461 | | Apip | 356.9861328 | 0.0595505516386635 | 0.0822551557507286 | 0.724237626489821 | 0.468919862319154 | 0.832603583919817 | NM_019735 |
| Ptges3 | 2927.3967 | -0.08819692 | 0.077121483 | -1.143610263 | 0.252785303 | 0.478665461 | | 2410018M08Rik | 246.9079346 | 0.06287976579585 | 0.0872470639549137 | 0.720709247343287 | 0.471088421697403 | 0.833245343374287 | NM_183088 |
| Ubqln1 | 6977.531444 | 0.06713441 | 0.058706032 | 1.1435692 | 0.252802341 | 0.478665461 | | Add1 | 19769.02436 | 0.0316376434980306 | 0.0438078301278127 | 0.722191521600713 | 0.470176735250613 | 0.833245343374287 | NM_001102444,NM_001024458,NM_013457 |
| Dmgdh | 12.8007829 | 0.189483914 | 0.165717444 | 1.143415618 | 0.252866071 | 0.478727628 | | Akirin1 | 2148.36703 | 0.0443902964610245 | 0.0614966558115198 | 0.721832689479524 | 0.470397348806417 | 0.833245343374287 | NM_023423 |
| Mmp15 | 852.9693779 | 0.140286041 | 0.122715838 | 1.143177954 | 0.252964713 | 0.478855867 | | Atg9a | 7701.874643 | 0.032811939111388 | 0.0455656107254786 | 0.720103134556271 | 0.471461497709928 | 0.833245343374287 | NM_001003917 |
| Fv1 | 203.345902 | 0.129291939 | 0.113110379 | 1.143059901 | 0.253013722 | 0.47889013 | | Col6a3 | 154.5346183 | -0.0647309678744271 | 0.0896014641634536 | -0.72242198806422 | 0.470035073323996 | 0.833245343374287 | NM_001243008,NM_001243009 |
| Zfp82 | 192.3624234 | 0.12942952 | 0.113245427 | 1.142911667 | 0.253075268 | 0.478948113 | | Cyp1b1 | 137.2443396 | -0.0680639372395752 | 0.0943717338908238 | -0.721232242255044 | 0.47076663755045 | 0.833245343374287 | NM_009994 |
| 4933427G17Rik | 16.66359955 | -0.190279604 | 0.166525582 | -1.142644877 | 0.253186065 | 0.478982282 | | Dnahc6 | 121.4181679 | 0.0653441119054129 | 0.0905227819100797 | 0.721852670969857 | 0.47038506248478 | 0.833245343374287 | NM_001164669 |
| Abcg2 | 736.6336593 | 0.086621723 | 0.075803142 | 1.142719436 | 0.253155098 | 0.478982282 | | Dnajb14 | 804.8074676 | 0.0462362164610138 | 0.0642236474192685 | 0.719925110434767 | 0.471571106494054 | 0.833245343374287 | NM_001033155 |
| Phldb2 | 328.8468264 | -0.113021291 | 0.09890888 | -1.142680931 | 0.25317109 | 0.478982282 | | Dpy19l3 | 2743.488024 | 0.0512867684647553 | 0.0711486655186983 | 0.720839499811515 | 0.471008269649529 | 0.833245343374287 | NM_178704 |
| Dennd2a | 1110.962134 | -0.102234355 | 0.089486118 | -1.142460499 | 0.253262656 | 0.479068685 | | Ecd | 1783.282302 | 0.0366020231778511 | 0.0507429177538992 | 0.721322793367326 | 0.470710936306138 | 0.833245343374287 | NM_027475 |
| Dnajc5 | 15041.48259 | -0.052234202 | 0.045725942 | -1.142331901 | 0.253316086 | 0.479111259 | | Elmo3 | 73.3383848 | -0.068648601870582 | 0.0951026087975526 | -0.721837209409099 | 0.470394569553342 | 0.833245343374287 | NM_172760 |
| Fgfrl1 | 654.4530236 | -0.133754374 | 0.117126867 | -1.141961506 | 0.253470021 | 0.479343891 | | Epb4.1l4b | 458.1813264 | -0.0571486937096006 | 0.0793497808169101 | -0.720212370106783 | 0.471394248745638 | 0.833245343374287 | NM_019427 |
| Hip1r | 3465.386484 | 0.064761806 | 0.056732106 | 1.14153715 | 0.253646462 | 0.479619023 | | Fam211a | 694.4062806 | 0.0474937181173548 | 0.065963370624068 | 0.720001383616771 | 0.471524143662592 | 0.833245343374287 | NM_198861 |
| Nodal | 22.12132244 | -0.191523343 | 0.167796873 | -1.141399953 | 0.253703524 | 0.479668384 | | Fktn | 1420.590095 | 0.0576468884475856 | 0.0799676444001587 | 0.720877660958976 | 0.470984788263657 | 0.833245343374287 | NM_139309 |
| Bach1 | 649.5594566 | -0.094071212 | 0.082431554 | -1.141203909 | 0.253785078 | 0.479754037 | | Frmpd1 | 650.2935885 | 0.0611733194663986 | 0.0847360296340866 | 0.721928083373291 | 0.470338694114481 | 0.833245343374287 | NM_001081172 |
| Ypel3 | 4549.843231 | 0.07965225 | 0.06980046 | 1.141142193 | 0.253810755 | 0.479754037 | | Homer2 | 2656.123097 | 0.0541096200793783 | 0.075017114028396 | 0.7212970104248 | 0.470726795950478 | 0.833245343374287 | NM_011983,NM_001164086,NM_001164087 |
| 2610001J05Rik | 947.0249733 | -0.079296447 | 0.069499555 | -1.140963383 | 0.253885161 | 0.479783814 | | Kif23 | 55.24900184 | -0.0681396600996547 | 0.0946130229427277 | -0.720193245922412 | 0.471406021834672 | 0.833245343374287 | NM_024245 |
| Gm8234 | 17.58122535 | 0.179671455 | 0.157474551 | 1.140955504 | 0.25388844 | 0.479783814 | | Lsm2 | 212.5278799 | -0.0641925699188478 | 0.0888582761967606 | -0.722415206172861 | 0.470039240674342 | 0.833245343374287 | NM_001110101,NM_001204274,NM_001204273,NM_030597 |
| Galnt12 | 55.70476269 | 0.179416384 | 0.157278959 | 1.140752616 | 0.253972884 | 0.479797818 | | Mrpl3 | 1898.789156 | 0.0462440368694719 | 0.0640532105387474 | 0.721962825602593 | 0.470317333218552 | 0.833245343374287 | NM_053159 |
| Kdm4b | 2988.166303 | 0.07199206 | 0.063103848 | 1.140850553 | 0.253932119 | 0.479797818 | | Mtss1 | 1398.216377 | 0.0549588748809344 | 0.0762521423609663 | 0.720751878954 | 0.471062187117331 | 0.833245343374287 | NM_144800,NM_001146180 |
| Nedd9 | 420.164579 | 0.147797831 | 0.129566014 | 1.1407145 | 0.253988751 | 0.479797818 | | Ncoa5 | 2222.023306 | 0.0396816578729216 | 0.0550996552827264 | 0.720179784597702 | 0.471414308893134 | 0.833245343374287 | NM_144892 |
| Rinl | 73.11171151 | 0.15574205 | 0.136581579 | 1.14028591 | 0.254167206 | 0.480076398 | | Ndrg2 | 33606.63464 | 0.0496111695872911 | 0.068587934656365 | 0.723322109578745 | 0.469482012375225 | 0.833245343374287 | NM_001145959,NM_013864 |
| Lrp8 | 3608.491016 | -0.089172568 | 0.078243372 | -1.139682063 | 0.254418783 | 0.480493006 | | Oaz2 | 7289.869475 | 0.0371593215279622 | 0.0515382085965071 | 0.721005299561005 | 0.470906254143114 | 0.833245343374287 | NM_010952 |
| Ercc4 | 1094.972193 | 0.076494507 | 0.067168899 | 1.138838176 | 0.254770656 | 0.481098908 | | Pex13 | 1241.952532 | 0.0482208333998046 | 0.0668784941817596 | 0.721021518049615 | 0.470896275667414 | 0.833245343374287 | NM_023651 |
| Tceal3 | 2795.574988 | 0.074586157 | 0.06550911 | 1.138561592 | 0.254886056 | 0.481258171 | | Pla2g12a | 559.9562487 | 0.0514202168741905 | 0.0712270549311386 | 0.721919738558655 | 0.470343824916188 | 0.833245343374287 | NM_183423,NM_023196 |
| Alg10b | 1176.518509 | -0.112026424 | 0.098423957 | -1.138202809 | 0.255035806 | 0.481364938 | | Polr1d | 1423.71042 | 0.0442327786543691 | 0.0612256603924615 | 0.722454904868864 | 0.470014840975651 | 0.833245343374287 | NM_009087,NM_181730 |
| Fkrp | 2062.800204 | 0.08064009 | 0.07084207 | 1.138307933 | 0.254991922 | 0.481364938 | | Ppp3cb | 11080.48906 | 0.0539250210026905 | 0.0749044453706113 | 0.719917499367106 | 0.471575792914775 | 0.833245343374287 | NM_008914 |
| Noxo1 | 58.59938679 | 0.179640091 | 0.157822823 | 1.138238992 | 0.255020701 | 0.481364938 | | Ptpre | 1445.596094 | -0.0424204290310713 | 0.058807323880263 | -0.72134602005429 | 0.470696649322296 | 0.833245343374287 | NM_011212 |
| Znfx1 | 4094.91522 | 0.098846913 | 0.086854668 | 1.138072534 | 0.255090196 | 0.481408952 | | Rbm12b2 | 314.8561596 | -0.0635216283762804 | 0.087952579276341 | -0.722225873293604 | 0.470155618486199 | 0.833245343374287 | NM_198957 |
| Cbll1 | 672.6888457 | 0.08797691 | 0.077314181 | 1.137914283 | 0.255156277 | 0.481475015 | | Rnf17 | 33.31415361 | 0.0591261253197827 | 0.0818199694461172 | 0.722636854059454 | 0.469903019923124 | 0.833245343374287 | NM_001033043 |
| Eef2 | 42445.04113 | -0.06412787 | 0.056384322 | -1.137335125 | 0.255398216 | 0.481872864 | | Sag | 76.26016113 | -0.0684843100241132 | 0.0950958503960085 | -0.720160866524916 | 0.471425955369725 | 0.833245343374287 | NM_009118 |
| Aff1 | 1053.116401 | 0.151000021 | 0.132810803 | 1.136955868 | 0.255556735 | 0.48192595 | | Siah3 | 83.9924798 | 0.0685170086204766 | 0.0949472621427445 | 0.721632273266264 | 0.470520591561591 | 0.833245343374287 | NM_001128093 |
| Emc10 | 6161.475952 | -0.099550422 | 0.087565207 | -1.136871875 | 0.255591851 | 0.48192595 | | Skida1 | 328.3612429 | -0.058053773145811 | 0.0803041639075549 | -0.722923573585073 | 0.469726839693637 | 0.833245343374287 | NM_028317 |
| Gm10516 | 212.0865773 | -0.108475642 | 0.095420131 | -1.136821349 | 0.255612977 | 0.48192595 | | Tcf12 | 2061.737527 | -0.0388659198903258 | 0.0537557904375468 | -0.723008992593644 | 0.469674359429234 | 0.833245343374287 | NM_001253863,NM_001253862,NM_011544,NM_001253865,NM_001253864 |
| Nanos2 | 23.6705391 | 0.189955656 | 0.167065623 | 1.137012224 | 0.255533176 | 0.48192595 | | Tinf2 | 433.4806691 | -0.0583355980195594 | 0.0808247598457071 | -0.721754053224791 | 0.470445702792512 | 0.833245343374287 | NM_145705 |
| Piezo2 | 37.05556738 | -0.189285302 | 0.166455599 | -1.137151911 | 0.255474786 | 0.48192595 | | Tmem101 | 698.9130905 | 0.0454079577923627 | 0.0627962487999769 | 0.72309984529489 | 0.469618544335589 | 0.833245343374287 | NM_029649 |
| Slc25a33 | 1043.06481 | -0.072826313 | 0.064058181 | -1.136877636 | 0.255589442 | 0.48192595 | | Tor2a | 794.266343 | -0.0535227221729239 | 0.0742394396022882 | -0.720947281655857 | 0.470941950795505 | 0.833245343374287 | NR_045485,NM_152800 |
| Ttf1 | 557.4432912 | 0.125439729 | 0.110366211 | 1.136577288 | 0.255715039 | 0.482059717 | | Tox | 667.1458423 | 0.0592275422175787 | 0.082160646123308 | 0.720874834999338 | 0.470986527116077 | 0.833245343374287 | NM_145711 |
| Gmnc | 17.54119322 | 0.190514796 | 0.167685908 | 1.136140762 | 0.255897658 | 0.482345293 | | Wscd1 | 3358.000689 | 0.0497674810110637 | 0.0689808900527478 | 0.721467655360896 | 0.47062183405203 | 0.833245343374287 | NM_177618 |
| Ganab | 5082.637687 | -0.078504461 | 0.069108657 | -1.135956975 | 0.255974571 | 0.482431579 | | Zfyve20 | 3069.757984 | 0.0368206557432156 | 0.0511451866089051 | 0.719924164609551 | 0.471571688874284 | 0.833245343374287 | NM_030081 |
| Ifrd2 | 270.9328065 | 0.131450574 | 0.115731709 | 1.135821593 | 0.256031238 | 0.482479689 | | Ric3 | 4145.216632 | 0.0383674240947621 | 0.0533052321966579 | 0.719768445116494 | 0.47166757636791 | 0.833304495485816 | NM_178780,NM_001038624 |

| | | | | | | | | | | | | | | |
|---|---|---|---|---|---|---|---|---|---|---|---|---|---|---|
| 9930021J03Rik | 2807.385656 | -0.092825997 | 0.081754717 | -1.135420684 | 0.256199097 | 0.482569738 | | Tpm3 | 6702.791957 | 0.0281062103982153 | 0.0390574310754303 | 0.719612366310899 | 0.471763695900742 | 0.833304495485816 | NM_001271764,NM_001253738,NM_001253740,NM_022314 |
| Idh2 | 1264.942591 | 0.121204499 | 0.106741638 | 1.135494085 | 0.256168359 | 0.482569738 | | Zfp710 | 936.9323652 | -0.0465313069963488 | 0.0646642064409478 | -0.719583670122694 | 0.471781369329158 | 0.833304495485816 | NM_001145999,NM_175433,NM_001146000 |
| Ipw | 377.3501248 | 0.190515584 | 0.167796409 | 1.135397269 | 0.256208903 | 0.482569738 | | Maats1 | 69.76084729 | 0.0682350791449675 | 0.094845002339363 | 0.719437792840333 | 0.471871217974575 | 0.83336186167765 | NM_001081025 |
| Sntb2 | 422.3698309 | 0.127446 | 0.112243188 | 1.135445296 | 0.25618879 | 0.482569738 | | Cacfd1 | 1535.679999 | 0.0373547399858614 | 0.0519329372449506 | 0.719288027358656 | 0.471963471247746 | 0.83342346016834 | NM_001243240,NM_001243239,NM_029862 |
| Tmem38a | 4020.161649 | -0.083054319 | 0.073153984 | -1.135335553 | 0.256234751 | 0.482569738 | | Calb1 | 5032.35623 | 0.0583452843559433 | 0.081146176527113 | 0.719014583964393 | 0.472131933883363 | 0.833441696707998 | NM_009788 |
| Sh3tc1 | 109.1683346 | 0.147275554 | 0.129773829 | 1.13486328 | 0.256432607 | 0.482883668 | | Psmg2 | 589.135085 | 0.0456915011035189 | 0.0635454751614474 | 0.719036264776246 | 0.472118575589035 | 0.833441696707998 | NM_134138 |
| Ogfod1 | 5916.947022 | -0.061404644 | 0.054114132 | -1.134724738 | 0.256490669 | 0.482934309 | | Sgcd | 52.74460964 | -0.0672728816814793 | 0.0935751105485749 | -0.718918538135821 | 0.472191113528328 | 0.833441696707998 | NM_011891 |
| 5330426P16Rik | 172.8342734 | -0.150525249 | 0.132688959 | -1.134421809 | 0.256617655 | 0.483114696 | | Thop1 | 1659.378584 | 0.0463160055238044 | 0.0644263257172445 | 0.718898757738831 | 0.472203301934932 | 0.833441696707998 | NM_022653 |
| Cox8b | 9.345137366 | 0.159576647 | 0.14070756 | 1.134101447 | 0.256751997 | 0.483132796 | | Nme4 | 69.47063149 | -0.0679915880336484 | 0.094614133145147 | -0.718619785157709 | 0.472375219433472 | 0.833643838839827 | NM_019731 |
| Pphln1 | 1538.654301 | -0.080552032 | 0.071027104 | -1.134102725 | 0.256751461 | 0.483132796 | | Acot9 | 715.5236344 | 0.0490697500984704 | 0.0683022716249542 | 0.71842047022844 | 0.472498068830717 | 0.833671042611583 | NM_019736 |
| Rbm18 | 1655.317741 | -0.077305264 | 0.068160059 | -1.13417249 | 0.256722201 | 0.483132796 | | Ipo8 | 2372.290738 | 0.046972514662637 | 0.0653841254547309 | 0.718408548496359 | 0.472505417446039 | 0.833671042611583 | NM_001081113 |
| Timp3 | 5575.209021 | -0.081477336 | 0.071829614 | -1.134313997 | 0.25666286 | 0.483132796 | | 1700018L02Rik | 23.30451592 | -0.0594372580339348 | 0.082974853753562 | -0.716328566368618 | 0.473788491800446 | 0.833983941564835 | NR_028360 |
| Cuedc2 | 3532.513372 | 0.086588125 | 0.076407451 | 1.133241906 | 0.25711268 | 0.483752747 | | 1700021F05Rik | 527.7904998 | 0.0571875051335533 | 0.0797048518233427 | 0.71749089077166 | 0.473071255097722 | 0.833983941564835 | NM_026411 |
| Tmprss6 | 23.56131969 | 0.180896363 | 0.159637955 | 1.133166375 | 0.257144392 | 0.483753668 | | 1700123M08Rik | 71.61222827 | -0.0680533723324539 | 0.0949369216145859 | -0.71682724882032 | 0.473480696110868 | 0.833983941564835 | NR_040577 |
| Csrnp1 | 484.7719378 | 0.190496116 | 0.168120817 | 1.133090592 | 0.257176212 | 0.483754792 | | 2310003H01Rik | 522.7574211 | -0.0635442314710159 | 0.0886961771690447 | -0.716425820133238 | 0.473728456411858 | 0.833983941564835 | NM_027980 |
| 4933430I17Rik | 15.77635331 | -0.189906023 | 0.167626914 | -1.132908898 | 0.257252513 | 0.483780852 | | 8430427H17Rik | 2891.697204 | -0.047478874929538 | 0.0661339258494189 | -0.717920104087624 | 0.472806551132688 | 0.833983941564835 | NM_001134300,NM_001001986 |
| Snx5 | 3450.071886 | 0.060665061 | 0.053545577 | 1.132961206 | 0.257230545 | 0.483780852 | | Alg6 | 332.0361137 | -0.0578852693797427 | 0.0807729465887982 | -0.716641794367452 | 0.473595149055457 | 0.833983941564835 | NM_001081264 |
| Med30 | 385.2275989 | -0.118197179 | 0.104350426 | -1.132694742 | 0.257342467 | 0.483891284 | | Asf1a | 377.6786308 | -0.0581687662723344 | 0.0811466687766751 | -0.716834925564492 | 0.473475958747332 | 0.833983941564835 | NM_025541 |
| A330009N23Rik | 450.3140024 | -0.137357242 | 0.121329597 | -1.132100047 | 0.257592375 | 0.484107923 | | Atp10b | 110.6800002 | 0.0681210314948376 | 0.0950660797543276 | 0.716565063699669 | 0.473642507724119 | 0.833983941564835 | NM_176999 |
| Fam65c | 17.1559238 | 0.186301137 | 0.164556354 | 1.132141861 | 0.257574798 | 0.484107923 | | Coq10a | 1329.075156 | 0.0366441590270332 | 0.0511604388820694 | 0.716259669145961 | 0.473831025040244 | 0.833983941564835 | NM_001081040 |
| Pcdhb12 | 315.8682183 | -0.112264333 | 0.099146818 | -1.132303941 | 0.257506674 | 0.484107923 | | Dctd | 315.5842074 | -0.0554909946816277 | 0.0773901233623394 | -0.717029412420234 | 0.473355948472806 | 0.833983941564835 | NR_027759,NM_001161516,NM_001161515,NM_178788 |
| Rnf40 | 2641.962262 | 0.072885979 | 0.06438413 | 1.132048835 | 0.257613904 | 0.484107923 | | Fxyd7 | 2095.80625 | -0.0501962299715482 | 0.0699512543621512 | -0.717588704152074 | 0.473010924563438 | 0.833983941564835 | NM_022007 |
| Vmn2r29 | 188.2571239 | -0.154452637 | 0.136413425 | -1.132239269 | 0.257533855 | 0.484107923 | | Gm5468 | 122.6306544 | -0.0682523733031017 | 0.095102727623028 | -0.717669987065389 | 0.472960793114498 | 0.833983941564835 | NR_027376 |
| Bnip3 | 3723.073268 | -0.088530915 | 0.078238104 | -1.131557521 | 0.257820508 | 0.484143855 | | Klhdc1 | 528.0980603 | 0.0534711400667833 | 0.0746000704774677 | 0.716770637407558 | 0.473515632148336 | 0.833983941564835 | NM_178253 |
| Efha2 | 3098.935447 | 0.096587954 | 0.085348994 | 1.131682397 | 0.257767985 | 0.484143855 | | Krt12 | 571.1490252 | -0.059871979781504 | 0.0835513924450036 | -0.716588653156365 | 0.473627947880352 | 0.833983941564835 | NM_010661 |
| Mob1a | 349.0685411 | -0.098130281 | 0.086720036 | -1.131575653 | 0.257812881 | 0.484143855 | | Mblac2 | 908.3606643 | 0.0449748997179447 | 0.0627855769326064 | 0.716325339595436 | 0.473790483780445 | 0.833983941564835 | NM_028372 |
| Tgtp2 | 27.72436825 | 0.166154913 | 0.146821277 | 1.131681429 | 0.257768392 | 0.484143855 | | Nup155 | 934.0509391 | -0.0421829405344259 | 0.0588542614895452 | -0.71673553395822 | 0.47353729590256 | 0.833983941564835 | NM_133227 |
| Tmem104 | 784.650637 | 0.070070818 | 0.061916861 | 1.131692016 | 0.257763939 | 0.484143855 | | Sil1 | 782.3109718 | 0.0527631289212808 | 0.0735489034529435 | 0.717388383023801 | 0.473134485625275 | 0.833983941564835 | NM_030749 |
| Trit1 | 493.5479999 | 0.103775716 | 0.091701479 | 1.131668946 | 0.257773642 | 0.484143855 | | Smcr7 | 616.8742522 | -0.0556642301734745 | 0.0776322817601958 | -0.717024270205273 | 0.473359121318292 | 0.833983941564835 | NM_001009927 |
| Msh3 | 933.6717271 | 0.116664109 | 0.103143382 | 1.131086713 | 0.258018596 | 0.48433973 | | Zfp644 | 2359.999673 | 0.05090837536638 | 0.0710493708551981 | 0.716521128246633 | 0.473669626144708 | 0.833983941564835 | NM_026856 |
| Pdcl3 | 689.5905013 | -0.093360019 | 0.082536205 | -1.131140186 | 0.257996092 | 0.48433973 | | Zzef1 | 2992.431442 | -0.0436329034544136 | 0.0608293234852507 | -0.717300488554558 | 0.473188705851104 | 0.833983941564835 | NM_001045536 |
| Phtf2 | 669.6101778 | 0.088740637 | 0.07845277 | 1.131134537 | 0.25799847 | 0.48433973 | | 9430020K01Rik | 7191.252919 | 0.0518630395003623 | 0.0724488077627921 | 0.715857735991591 | 0.474079197623271 | 0.833994865114109 | NM_001081963 |
| Cdkn1c | 212.2369083 | 0.145007707 | 0.128239723 | 1.130754989 | 0.25815823 | 0.484543139 | | Nphs1 | 88.27058411 | -0.0680767524822389 | 0.0950906346902806 | -0.715914376888655 | 0.47404422052502 | 0.833994865114109 | NM_019459 |
| Mpv17l2 | 380.9715886 | -0.099885252 | 0.088350273 | -1.130559631 | 0.258240487 | 0.48463882 | | Ssbp3 | 5175.466065 | 0.031834233767749 | 0.0444745937627493 | 0.715784700307089 | 0.474124300985289 | 0.833994865114109 | NM_198438,NM_023672 |
| Uvrag | 1592.244874 | 0.106659805 | 0.094358677 | 1.130365621 | 0.258322194 | 0.484733447 | | Supt6 | 8116.412477 | 0.0340618640397869 | 0.0475773205942712 | 0.715926487963854 | 0.474036741832692 | 0.833994865114109 | NM_009297 |
| Mblac2 | 834.3715977 | -0.084865749 | 0.075085179 | -1.130259662 | 0.258366826 | 0.48475849 | | Znf41-ps | 47.70766417 | -0.0635584175948301 | 0.0887881296846761 | -0.715843636086859 | 0.47408790486956 | 0.833994865114109 | NR_040355 |
| Lif | 24.43363156 | 0.189484926 | 0.167675669 | 1.130068106 | 0.258447527 | 0.484851192 | | Ankrd26 | 809.1464496 | -0.0505097098636575 | 0.0705818643006562 | -0.7156188117744 | 0.474226754606273 | 0.834074081294434 | NM_001081112 |
| Snx22 | 257.2681705 | 0.124685319 | 0.110387925 | 1.129519547 | 0.258678728 | 0.485226177 | | 2900056M20Rik | 885.1726572 | 0.0549301325081467 | 0.0768007839434349 | 0.71522879959928 | 0.47446767606676 | 0.834396787233143 | NR_040270,NR_040269 |
| Gkn3 | 100.8579886 | -0.171493567 | 0.151848205 | -1.129375001 | 0.258739674 | 0.485281746 | | Gm10336 | 480.9820674 | -0.0537000283356263 | 0.0751222425478384 | -0.714835267350143 | 0.474710840102272 | 0.834491965381406 | NR_045170 |
| Nudt2 | 442.3119106 | -0.114785199 | 0.101655614 | -1.129157501 | 0.258831398 | 0.485395024 | | Ndufaf7 | 1377.741036 | 0.034643034063075 | 0.0484674323220539 | 0.714769328667563 | 0.474751590383584 | 0.834491965381406 | NM_028611 |
| Bop1 | 1883.774401 | -0.086229302 | 0.07637473 | -1.129029232 | 0.258885503 | 0.485437732 | | Nt5e | 361.2258381 | 0.0652475236791125 | 0.0912830159501133 | 0.714782733677102 | 0.474743305896472 | 0.834491965381406 | NM_011851 |
| Zfp652 | 571.2856494 | -0.10969761 | 0.097170997 | -1.128913086 | 0.2589345 | 0.485470855 | | Rabgap1 | 4751.349746 | 0.040234117335969 | 0.0562848683848993 | 0.714830086495563 | 0.474714041814663 | 0.834491965381406 | NM_146121,NM_001033960 |
| Pdhx | 1677.161823 | -0.076196458 | 0.067518278 | -1.128530817 | 0.25909581 | 0.485714516 | | Cyb5d1 | 454.687513 | 0.0470924557697789 | 0.0659267157399476 | 0.714315209566001 | 0.475032289354675 | 0.834699977238266 | NM_001045525 |
| Pabpc1l | 14.42429854 | 0.174079906 | 0.154281651 | 1.128325402 | 0.259182519 | 0.485818287 | | Psap | 85413.27367 | 0.0335281455882832 | 0.0469379616960079 | 0.714307660086031 | 0.475036956590132 | 0.834699977238266 | NM_001146124,NM_001146121,NM_001146120,NM_001146123,NM_001146122,NM_011179 |
| Zswim1 | 1017.338473 | 0.066693435 | 0.059115374 | 1.128191037 | 0.259239249 | 0.485865842 | | Tmed1 | 408.6220989 | -0.0594647541238791 | 0.0832599425365012 | -0.714206043293986 | 0.475099780518104 | 0.834699977238266 | NM_010744 |
| Srfbp1 | 382.6947095 | 0.097060207 | 0.086059732 | 1.12782372 | 0.259394374 | 0.486097777 | | Tra2a | 1954.24342 | -0.0529609886563909 | 0.0741486959070844 | -0.714253811324696 | 0.475070247672481 | 0.834699977238266 | NM_198102 |
| Atg2a | 2466.860051 | 0.077779849 | 0.068979488 | 1.127579399 | 0.259497591 | 0.486232395 | | Mfsd12 | 655.1790922 | 0.0503444421915577 | 0.0705074550195693 | 0.714030057922026 | 0.475208593128202 | 0.834790183066908 | NM_028657 |
| Gm13157 | 76.53630962 | 0.160198076 | 0.142109827 | 1.127283587 | 0.259622599 | 0.486370038 | | Ccdc18 | 53.91926091 | -0.065297975440781 | 0.0914811239834556 | -0.713786326593344 | 0.475359315975103 | 0.834927401268865 | NM_028481 |
| Ngdn | 540.4091065 | 0.108032057 | 0.095836225 | 1.127257013 | 0.259633831 | 0.486370038 | | Golph3l | 919.1420795 | 0.044424257610688 | 0.0622434441273744 | 0.713717857896466 | 0.475401661565109 | 0.834927401268865 | NM_146133,NM_001177669,NM_001177670 |
| Enkur | 81.54132192 | 0.152097855 | 0.13497363 | 1.12687089 | 0.259797073 | 0.486583408 | | Selrc1 | 509.8833535 | 0.0537758886991232 | 0.0753603896963514 | 0.713582943450818 | 0.475485107675703 | 0.834973002187773 | NM_027250 |
| Epb4.1 | 1302.421956 | 0.116025431 | 0.102969113 | 1.126798394 | 0.25982773 | 0.486583408 | | Lrrc25 | 23.65945545 | -0.0603814641999524 | 0.0846331598370126 | -0.713449247508136 | 0.475567808053885 | 0.835017282701647 | NM_153074 |
| Epm2a | 457.3143976 | 0.106464173 | 0.0944866 | 1.126764778 | 0.259841946 | 0.486583408 | | 4921531C22Rik | 56.20515678 | -0.0674773031360242 | 0.0947028931769433 | -0.712515751867784 | 0.476145460798247 | 0.835195112086295 | NR_033782 |
| Gm17762 | 68.32139159 | 0.155802121 | 0.138294379 | 1.126597641 | 0.259912637 | 0.486656967 | | Adamts9 | 153.1298214 | -0.067052155880019 | 0.0940557465757229 | -0.712898024003629 | 0.475908862006978 | 0.835195112086295 | NM_175314 |
| Dmrta2 | 40.61036895 | -0.185180845 | 0.164400288 | -1.126402192 | 0.259995319 | 0.486752958 | | Adrb3 | 50.61204653 | -0.0640963989498476 | 0.0900685169506182 | -0.711640439078059 | 0.476687458803475 | 0.835195112086295 | NM_013462 |
| Sbspon | 13.20048409 | -0.187951554 | 0.166932654 | -1.125912458 | 0.260202576 | 0.487051764 | | Cbr4 | 375.8882964 | -0.051538246808246 | 0.0723412750450413 | -0.712432104302241 | 0.476197241183901 | 0.835195112086295 | NM_145595 |
| Spint1 | 165.7080776 | 0.137172023 | 0.121835763 | 1.125876508 | 0.260217794 | 0.487051764 | | Cep120 | 2315.040653 | 0.042499603223275 | 0.05973958389638 | 0.711414450040224 | 0.476827447192981 | 0.835195112086295 | NM_178686 |
| Ccdc43 | 670.2885991 | 0.07074529 | 0.062843459 | 1.125738325 | 0.260276297 | 0.48709443 | | Cgn | 226.6762952 | -0.0667630441761117 | 0.0938574705094648 | -0.71132371044859 | 0.476883661957533 | 0.835195112086295 | NM_001037711 |
| Hdgfrp3 | 4426.881292 | -0.081755507 | 0.072628039 | -1.125674161 | 0.260303465 | 0.48709443 | | Cyfip2 | 60715.76747 | 0.0382117456669565 | 0.0536215689628674 | 0.712618940587471 | 0.476081588095862 | 0.835195112086295 | NM_001252459,NM_001252460,NM_133769 |
| Clip1 | 4673.497504 | 0.102541485 | 0.091106108 | 1.125517128 | 0.260369963 | 0.487160031 | | Dus1l | 1097.469803 | -0.0493826326084469 | 0.0693155772612417 | -0.712431960601439 | 0.47619733014172 | 0.835195112086295 | NM_026824 |
| Amn1 | 307.1942675 | -0.110212315 | 0.09795196 | -1.125167021 | 0.260518265 | 0.487378653 | | Hhatl | 433.9208484 | -0.0625206864704002 | 0.0877836707640463 | -0.712213170470502 | 0.476332782488179 | 0.835195112086295 | NR_027967,NM_029095 |
| Scn2b | 13903.76542 | -0.092365518 | 0.082113698 | -1.124849082 | 0.260652992 | 0.487571828 | | Jakmip1 | 3250.050842 | 0.0311134113354583 | 0.043745155606971 | 0.711242442820348 | 0.476934011755185 | 0.835195112086295 | NM_178394 |
| Slc9a6 | 5570.718228 | -0.084104167 | 0.074810655 | -1.124227121 | 0.260916687 | 0.488006174 | | Lman2l | 1538.335193 | 0.050904956651662 | 0.0715341599590382 | 0.711617452148892 | 0.476701696975469 | 0.835195112086295 | NM_001013374 |
| Auts2 | 2725.837832 | 0.125829202 | 0.111936431 | 1.124113039 | 0.260965075 | 0.488037763 | | Mrpl54 | 488.5930813 | 0.060094994245381 | 0.0843932054080595 | 0.712083324182422 | 0.476413179936261 | 0.835195112086295 | NM_025317 |
| Wnt5a | 316.6419987 | -0.139372185 | 0.124002517 | -1.12394642 | 0.261035757 | 0.488111033 | | Nudt9 | 1738.601295 | 0.0403112793289181 | 0.0565482045866362 | 0.712865768658635 | 0.475928823241406 | 0.835195112086295 | NM_028794 |
| Brd7 | 1621.080479 | 0.068170222 | 0.060663046 | 1.123752043 | 0.261118231 | 0.488147427 | | P2ry2 | 32.81743136 | -0.0620298608714404 | 0.0870188160744552 | -0.712832737443432 | 0.475949265099135 | 0.835195112086295 | NM_008773 |
| Tsta3 | 671.0610766 | 0.078678205 | 0.070013705 | 1.123754351 | 0.261117252 | 0.488147427 | | Pipox | 83.58929742 | -0.0667132430610927 | 0.0936810438512665 | -0.712131721834895 | 0.476383212498175 | 0.835195112086295 | NM_008952 |
| Bpifa1 | 10.11102975 | -0.112744252 | 0.100358001 | -1.123420657 | 0.261258881 | 0.488351441 | | Rpp14 | 1341.261577 | 0.0457084979062341 | 0.064203156991907 | 0.711935363427624 | 0.476504802453457 | 0.835195112086295 | NM_025938 |
| Preb | 2677.564496 | 0.078397715 | 0.069812122 | 1.122981407 | 0.261445391 | 0.48864112 | | Sdr39u1 | 2173.377465 | 0.0438265205427732 | 0.0615884181527448 | 0.711603282845803 | 0.476710473599791 | 0.835195112086295 | NM_001082975 |
| Haus4 | 74.64047351 | -0.150406572 | 0.133947861 | -1.122874015 | 0.261491005 | 0.488667426 | | Ska2 | 291.5998259 | -0.0554438422350946 | 0.0777444449670498 | -0.713155033245053 | 0.475749828124981 | 0.835195112086295 | NM_025377 |
| Rnf135 | 141.6339528 | 0.121583571 | 0.108293637 | 1.122721281 | 0.261555887 | 0.488729729 | | Tab2 | 3129.76991 | -0.0435499466030384 | 0.0612240729334503 | -0.711320637723278 | 0.476885565628141 | 0.835195112086295 | NM_138667 |
| 1190002N15Rik | 1195.240707 | -0.136789066 | 0.12184543 | -1.122644204 | 0.261588634 | 0.488731978 | | Tmem255b | 53.58792924 | 0.0653728809580873 | 0.0918844216078586 | 0.711468601686188 | 0.476793901029845 | 0.835195112086295 | NM_001143672,NM_001143671 |
| Sacm1l | 2973.871318 | 0.067751503 | 0.060360038 | 1.122456254 | 0.261668498 | 0.488784616 | | Wdr52 | 191.477926 | -0.0673373303598575 | 0.0945187510311407 | -0.712422980892672 | 0.476202889038387 | 0.835195112086295 | NM_001033247 |
| Slc28a1 | 18.79909953 | 0.188668503 | 0.168089414 | 1.122429421 | 0.261679901 | 0.488784616 | | Zfp617 | 1363.988524 | 0.0488491338124718 | 0.0686409099933376 | 0.711662095056916 | 0.476674045239119 | 0.835195112086295 | NM_133358 |
| Cd37 | 131.2718526 | 0.174691872 | 0.15566509 | 1.122228956 | 0.261765104 | 0.488863021 | | Lgmn | 4612.288785 | -0.0430307764044396 | 0.0605125258105796 | -0.711105276602359 | 0.477019000427268 | 0.835243249783812 | NM_011175 |
| Khdc3 | 8.317956657 | 0.165802656 | 0.147750211 | 1.122182198 | 0.26178498 | 0.488863021 | | Fen1 | 586.1066246 | 0.0479184340016313 | 0.0674014845734313 | 0.710940334695833 | 0.47712120999202 | 0.835247565637132 | NM_001271614,NM_001271615,NM_007999 |
| Vps18 | 2248.408116 | 0.093538536 | 0.083380591 | 1.121826255 | 0.261936321 | 0.48908669 | | Fkbp9 | 1987.664237 | 0.0418102839828073 | 0.0588123629866117 | 0.710909779161996 | 0.477140145658113 | 0.835247565637132 | NM_012056 |
| Mipep | 1108.213878 | -0.078357325 | 0.069859288 | -1.12164505 | 0.262013389 | 0.489171641 | | Gpsm1 | 2152.435043 | 0.0409858567618552 | 0.0576597274659048 | 0.71082293592336 | 0.47719396580081 | 0.835247565637132 | NM_153410,NM_001199147,NM_001199146 |
| Mllt3 | 1287.262065 | -0.087853443 | 0.078339818 | -1.121440484 | 0.262100412 | 0.489275155 | | Bscl2 | 3235.318365 | 0.0358992279159592 | 0.0505328885743569 | 0.7104131374389 | 0.477447978734158 | 0.835289574743151 | NM_008144,NM_001136064 |
| Amdhd1 | 9.052811219 | -0.170854008 | 0.152403742 | -1.121061763 | 0.262261573 | 0.489476425 | | Kdm1a | 2591.94119 | -0.0322229610838997 | 0.0453527799162311 | -0.710495831642893 | 0.477396714907261 | 0.835289574743151 | NM_133872 |
| Cacng3 | 4089.641714 | -0.103018956 | 0.091908159 | -1.120890215 | 0.262334596 | 0.489476425 | | Oplah | 1347.169872 | -0.0469372010860721 | 0.0660636020743588 | -0.710485041872849 | 0.477403403535349 | 0.835289574743151 | NM_153122 |
| Efcab1 | 121.4431669 | -0.147215089 | 0.131327401 | -1.120977708 | 0.262297351 | 0.489476425 | | Snhg7 | 136.8122798 | -0.0641603186839748 | 0.0902868909047371 | -0.710627180103823 | 0.47731529550359 | 0.835289574743151 | NR_024068 |
| Snx4 | 1383.410419 | 0.064555749 | 0.057592864 | 1.120898412 | 0.262331107 | 0.489476425 | | Igdcc3 | 68.32232789 | 0.0655521724473911 | 0.0923465087677697 | 0.709850034636823 | 0.47779713773717 | 0.835457924704683 | NM_008988 |
| Cadm3 | 21401.14211 | 0.083737941 | 0.074714884 | 1.120766518 | 0.262387259 | 0.489515737 | | Lmbr1 | 1499.319357 | 0.0488905024839538 | 0.0688748438818137 | 0.709845565208798 | 0.47779990962091 | 0.835457924704683 | NM_020295 |
| Mrpl18 | 965.0895375 | -0.061743776 | 0.055105089 | -1.120473195 | 0.262512169 | 0.489689808 | | Map2k3 | 576.7383564 | -0.0593581241334474 | 0.0835869584404391 | -0.710136189196833 | 0.477619686540879 | 0.835457924704683 | NM_008928 |
| Gm410 | 61.60330962 | 0.179169572 | 0.159916445 | 1.120394914 | 0.262545511 | 0.489693048 | | Mrps6 | 614.6598125 | 0.0444888866717745 | 0.0626618729230062 | 0.709983353457034 | 0.477714459104683 | 0.835457924704683 | NM_080456 |
| Chia | 78.98573644 | -0.148750307 | 0.132788987 | -1.120200632 | 0.262628275 | 0.489788455 | | Sav1 | 756.9826623 | -0.0513700238330742 | 0.0723731265304898 | -0.709794177697058 | 0.477831780149147 | 0.835457924704683 | NM_022028 |
| Gsk3a | 9488.231549 | 0.084298352 | 0.075258199 | 1.120121849 | 0.262661841 | 0.489792101 | | Rnf111 | 1758.90836 | -0.0409237171709221 | 0.0576988055429209 | -0.70926454691475 | 0.478160324819073 | 0.835931746097227 | NM_033604 |
| Lphn2 | 4421.044804 | 0.072231882 | 0.064499489 | 1.119883012 | 0.262763618 | 0.489907889 | | Dst | 14511.95524 | 0.0574501388352701 | 0.0811171650178751 | 0.708236522104912 | 0.478798389427385 | 0.83593645948681 | NM_134448,NM_133833 |
| Tnmd | 18.02933645 | 0.186647311 | 0.16667502 | 1.119827745 | 0.262787173 | 0.489907889 | | Fam217a | 35.91140464 | -0.0641728344350588 | 0.090620702392188 | -0.708147616836293 | 0.478853592137529 | 0.83593645948681 | NM_027967 |
| Mst1r | 53.19347207 | 0.170693755 | 0.152455325 | 1.119631305 | 0.262870909 | 0.490005037 | | Fam43a | 578.6182751 | 0.0615315440272465 | 0.086864880186217 | 0.708359280474893 | 0.478722172495584 | 0.83593645948681 | NM_177632 |
| Msr1 | 10.90050293 | 0.159640154 | 0.142636345 | 1.119210917 | 0.263050169 | 0.490280202 | | Fbxl4 | 798.6534185 | 0.0450146016284931 | 0.0635587603395531 | 0.708235991199472 | 0.47879871906479 | 0.83593645948681 | NM_172988 |
| Zfp248 | 460.5362595 | 0.097612204 | 0.08722164 | 1.119128288 | 0.263085414 | 0.490286913 | | Gm16287 | 29.8090622 | -0.0543972959269244 | 0.0767362142254089 | -0.708886885755597 | 0.478394673910626 | 0.83593645948681 | NR_033543 |
| St6galnac6 | 4817.27972 | -0.089157997 | 0.079709098 | -1.118542282 | 0.263335459 | 0.490693878 | | Kdm4c | 1253.125264 | -0.0428567203352098 | 0.0604515070437935 | -0.708943787028546 | 0.478359361097266 | 0.83593645948681 | NM_144787,NM_001172095 |
| Hcfc1r1 | 2217.179695 | -0.08278541 | 0.074017499 | -1.118457279 | 0.263371742 | 0.490702474 | | Myh13 | 25.55260901 | -0.0580833990194977 | 0.0820032680151018 | -0.708305905671967 | 0.478755310470241 | 0.83593645948681 | NM_001081250 |
| 2410018M08Rik | 202.234848 | 0.116202357 | 0.10391667 | 1.118226334 | 0.26347034 | 0.490790797 | | N4bp2l2 | 1344.319234 | -0.0493477327251415 | 0.0696191648075333 | -0.708823969112046 | 0.478433721508911 | 0.83593645948681 | NM_201369 |
| Snora31 | 17.82126674 | 0.187180454 | 0.167394753 | 1.118197856 | 0.2634825 | 0.490790797 | | P4ha2 | 450.0449871 | 0.0599167700932821 | 0.0845363568126549 | 0.70876924855026 | 0.478467683835568 | 0.83593645948681 | NM_011031,NM_001136076 |
| Ppp1r3d | 312.6578132 | -0.104775384 | 0.093742794 | -1.117690011 | 0.26369941 | 0.491076763 | | Psmb3 | 2002.353287 | 0.0407360948382733 | 0.0575223414504975 | 0.708178662604162 | 0.478834314926362 | 0.83593645948681 | NM_011971 |
| Prpf40a | 1280.301304 | 0.089220735 | 0.07982561 | 1.117695632 | 0.263697009 | 0.491076763 | | Rcc2 | 4445.071524 | 0.0377262513684187 | 0.0532046129683248 | 0.709078579161527 | 0.478275715059943 | 0.83593645948681 | NM_173867 |
| Podxl2 | 5107.318845 | 0.072570579 | 0.06493731 | 1.117548281 | 0.263759968 | 0.491130507 | | Slc39a12 | 2374.079635 | 0.0447489579819883 | 0.0631895281484526 | 0.708170472120927 | 0.478839400592073 | 0.83593645948681 | NM_001012305 |
| Mir377 | 12.02549698 | 0.17105077 | 0.153086088 | 1.117350196 | 0.263844621 | 0.491170078 | | Gng13 | 556.7487848 | -0.0590529458749628 | 0.0834150258126315 | -0.707941348691886 | 0.478981680729068 | 0.836059588352612 | NM_022422 |
| Stox1 | 154.9708997 | -0.127097962 | 0.113745088 | -1.117392979 | 0.263826336 | 0.491170078 | | Trim43a | 24.27252925 | -0.0553295953887834 | 0.0781773013406579 | -0.707745016007708 | 0.479103616964747 | 0.836171950021617 | NM_001034906 |
| Pdk1 | 2690.528541 | 0.106245568 | 0.095099883 | 1.117199785 | 0.263908913 | 0.491230742 | | Il17rd | 233.8025272 | 0.0618605052405335 | 0.0874262478431236 | 0.707573603656594 | 0.479210089797501 | 0.836180652257999 | NM_134437 |
| Zfp3 | 467.284277 | -0.094603159 | 0.084700484 | -1.116914025 | 0.264031087 | 0.491399119 | | Orc6 | 607.8473079 | -0.0468645875068682 | 0.0662348649826874 | -0.707551642463796 | 0.479223731932249 | 0.836180652257999 | NR_028268,NM_019716,NM_001163791 |
| Il2ra | 11.75439319 | 0.177050731 | 0.158531544 | 1.116817048 | 0.264072558 | 0.491417273 | | Serping1 | 272.8311532 | -0.0572067411018319 | 0.0808643272199226 | -0.707441007284332 | 0.479292460932506 | 0.836200130615612 | NM_009776 |
| Eva1c | 63.83711738 | 0.177202167 | 0.158713713 | 1.116489326 | 0.264212737 | 0.49160707 | | Pou6f2 | 44.83223655 | 0.0609608378634109 | 0.0861986935211036 | 0.707213014179689 | 0.479434112215245 | 0.836346813262717 | NM_175006 |
| Mfsd7c | 113.0162254 | -0.161083228 | 0.14428716 | -1.116407229 | 0.26424786 | 0.49160707 | | 2900055J20Rik | 174.5013905 | -0.0628599674243192 | 0.0889179828262941 | -0.706943246194861 | 0.479601747581037 | 0.836438346183472 | NR_045177 |
| Npepps | 6607.557112 | -0.058274172 | 0.052200343 | -1.116356103 | 0.264269736 | 0.49160707 | | Fhl3 | 36.32210048 | -0.0639136658296747 | 0.0903990804258766 | -0.707016769734524 | 0.47955605646398 | 0.836438346183472 | NM_010213 |
| Fem1c | 788.8668946 | -0.107202481 | 0.09605959 | -1.11599977 | 0.264422233 | 0.491723661 | | 2310022B05Rik | 3287.896357 | 0.0561713638817562 | 0.0796248733020444 | 0.705449962459331 | 0.480530260785682 | 0.836914993047623 | NM_175149 |
| Slitrk6 | 56.31074322 | -0.187617554 | 0.168118019 | -1.115987185 | 0.26442762 | 0.491723661 | | Efcab4b | 42.4783988 | -0.0651621679590569 | 0.0924167274716162 | -0.705090623113332 | 0.480753841558233 | 0.836914993047623 | NM_001033464 |
| Swsap1 | 475.8911343 | -0.09404897 | 0.084268479 | -1.116063465 | 0.264394969 | 0.491723661 | | Ept1 | 3398.314274 | 0.0456860680327467 | 0.0647527070129884 | 0.705546843371086 | 0.480469991226146 | 0.836914993047623 | NM_027652 |
| Arhgef18 | 2285.955562 | 0.067397499 | 0.060414393 | 1.11558679 | 0.264599049 | 0.491924366 | | Gpr17 | 2329.894966 | -0.054618089959447 | 0.0774174653922745 | -0.705500879971942 | 0.480498584516767 | 0.836914993047623 | NM_001025381 |
| Podxl | 1895.966885 | -0.135633836 | 0.121577389 | -1.115617279 | 0.264585993 | 0.491924366 | | Gprin1 | 5979.255856 | 0.0457640929355689 | 0.0648405217740367 | 0.705794643279593 | 0.480315853789981 | 0.836914993047623 | NM_012014 |
| Adcy9 | 2872.887658 | 0.147459496 | 0.132195289 | 1.115467103 | 0.264650308 | 0.491960633 | | Lipt1 | 167.2489064 | 0.0634146745452981 | 0.0899005325922733 | 0.705387084111095 | 0.480569379574344 | 0.836914993047623 | NM_001037918 |
| Camkmt | 261.4541303 | -0.109159626 | 0.097887642 | -1.115152274 | 0.264785174 | 0.492152289 | | Lyrm4 | 738.8070238 | 0.053074354646795 | 0.0751511300435799 | 0.706234951038226 | 0.480042038397226 | 0.836914993047623 | NM_201358 |
| Lax1 | 12.95677503 | 0.185686561 | 0.166579495 | 1.114702389 | 0.264977977 | 0.492392514 | | Mier3 | 771.8650071 | -0.0487464792763954 | 0.0690435250542803 | -0.706025354847861 | 0.480172369939168 | 0.836914993047623 | NM_172593 |
| Tmem229a | 2038.721082 | -0.113266667 | 0.101608526 | -1.114735852 | 0.264963633 | 0.492392514 | | Mlxip | 1731.936039 | -0.0428894237907803 | 0.0607416033935045 | -0.706096339158651 | 0.480128228166275 | 0.836914993047623 | NM_133917,NM_177582 |
| Gm16897 | 14.19940257 | -0.1867849 | 0.167619212 | -1.114340636 | 0.26513308 | 0.49262165 | | Mpnd | 2481.101488 | 0.0471130907553748 | 0.0668210598013421 | 0.705063506856091 | 0.4807707155799 | 0.836914993047623 | NM_026530 |
| Amacr | 326.3761898 | -0.096915872 | 0.086989553 | -1.114109326 | 0.265232288 | 0.492706257 | | Mxra8 | 217.4505862 | -0.0670558129681726 | 0.0950938465088858 | -0.705154070741126 | 0.480714360354979 | 0.836914993047623 | NM_024263 |
| Sytl5 | 163.2693188 | -0.16108249 | 0.1445871 | -1.114086183 | 0.265242216 | 0.492706257 | | Naa20 | 1579.392049 | 0.0424837255586441 | 0.060151870672652 | 0.706274386541387 | 0.480017518686225 | 0.836914993047623 | NM_001141965,NM_026425 |
| Exd2 | 3314.991285 | 0.134143404 | 0.120428171 | 1.113887242 | 0.265327563 | 0.492805714 | | Slc1a6 | 389.7458701 | 0.0571930387290328 | 0.081022355792047 | 0.705892073489262 | 0.480255257265036 | 0.836914993047623 | NM_009200 |
| Cnrip1 | 2428.538817 | -0.075154155 | 0.067508455 | -1.113255445 | 0.265598736 | 0.493225838 | | Tmem29 | 519.2715152 | 0.0494081706335901 | 0.0700400322507675 | 0.705427582567235 | 0.480544183889394 | 0.836914993047623 | NM_001164684,NM_001164686,NM_001164683 |
| Dock2 | 238.472003 | 0.14060824 | 0.126308601 | 1.113211917 | 0.265617426 | 0.493225838 | | Zfp777 | 1290.086799 | 0.0433490000421837 | 0.0614860728514568 | 0.705021446839025 | 0.480796889507257 | 0.836914993047623 | NM_001081382 |
| Wdr41 | 1325.15404 | -0.078507369 | 0.070534112 | -1.113041143 | 0.26569076 | 0.493302892 | | Zglp1 | 33.7786923 | -0.0534155012024096 | 0.0757541315400953 | -0.705116673064066 | 0.480737631386223 | 0.836914993047623 | NM_001103168 |
| S100a1 | 1377.735024 | -0.073964897 | 0.066461404 | -1.112900005 | 0.265751378 | 0.49335632 | | Ptpn14 | 755.9632099 | 0.0666151797564809 | 0.0945090655357069 | 0.704854919249125 | 0.480900527162006 | 0.836995094363204 | NM_008976 |
| Lat2 | 86.05372282 | 0.174079649 | 0.156467072 | 1.112564113 | 0.26589568 | 0.493565074 | | Met | 362.4192209 | -0.0512743928045464 | 0.0728000369882495 | -0.704318224629783 | 0.481234619286345 | 0.837109199764217 | NM_008591 |
| Zcchc18 | 11679.72647 | -0.048454404 | 0.043556842 | -1.112440717 | 0.265948706 | 0.493604367 | | Ppil3 | 557.8932343 | 0.0498555152961234 | 0.0707750509542115 | 0.70442217453687 | 0.481169900658017 | 0.837109199764217 | NM_027374,NM_027351 |
| Ankmy2 | 1846.131722 | -0.080118702 | 0.072039054 | -1.112156491 | 0.266070871 | 0.493660492 | | Stard8 | 1635.835967 | -0.0569309771001803 | 0.0808455664566242 | -0.704194176569042 | 0.481311857113503 | 0.837109199764217 | NM_199018 |
| Ap2a1 | 11902.26648 | 0.072868458 | 0.065520469 | 1.11214798 | 0.26607453 | 0.493660492 | | Tubgcp6 | 2321.103811 | -0.0435583594621051 | 0.0618485897641128 | -0.704274093042936 | 0.481262096797357 | 0.837109199764217 | NM_001163319 |
| Samd1 | 829.0063508 | -0.085603399 | 0.076963339 | -1.112262024 | 0.266025507 | 0.493660492 | | Utp6 | 1859.445799 | 0.0446452809506519 | 0.0633720131605643 | 0.704495229424622 | 0.481124419930667 | 0.837109199764217 | NM_144826 |
| Sox3 | 28.574023 | -0.185351813 | 0.166709713 | -1.111823719 | 0.266213951 | 0.493860029 | | Yod1 | 382.2712303 | -0.0607483700466769 | 0.0862429461002761 | -0.704386535868612 | 0.481192088558897 | 0.837109199764217 | NM_178691 |
| 1110046J04Rik | 90.59358953 | -0.165135582 | 0.148623257 | -1.111101895 | 0.266524492 | 0.494085203 | | Pus1 | 525.1064458 | 0.0499221837081643 | 0.0709047169856108 | 0.704074225672395 | 0.481386550281212 | 0.837138875779346 | NM_001025561,NM_001025562,NM_019700 |
| Adamts8 | 135.329917 | -0.164131523 | 0.14771232 | -1.111156623 | 0.266500939 | 0.494085203 | | Acot2 | 207.4409855 | 0.0567616588227084 | 0.0806822285991796 | 0.70352120669217 | 0.48173099559998 | 0.837383764621454 | NM_134188 |
| Katnb1 | 1820.322883 | 0.066123759 | 0.059507466 | 1.111184257 | 0.266489046 | 0.494085203 | | Bloc1s3 | 431.5161675 | -0.0505323583457548 | 0.0718531272445057 | -0.70327291634504 | 0.481885685698155 | 0.837383764621454 | NM_177692 |
| Nkpd1 | 21.6648737 | 0.184141926 | 0.165740912 | 1.111022761 | 0.266558552 | 0.494085203 | | C230081A13Rik | 1816.429615 | 0.0478730834135838 | 0.0680511956830756 | 0.703486293415559 | 0.481752745671802 | 0.837383764621454 | NM_172924 |
| Rad54b | 60.60801379 | 0.166460019 | 0.149822599 | 1.111047469 | 0.266547918 | 0.494085203 | | Grm3 | 3024.754287 | 0.045215607329533 | 0.064299750908913 | 0.703200349774066 | 0.481930901296665 | 0.837383764621454 | NM_181850 |
| Tamm41 | 264.6218114 | 0.145091927 | 0.130591188 | 1.111039183 | 0.266551484 | 0.494085203 | | Ltn1 | 2885.553642 | -0.0448901980094655 | 0.0638220985784955 | -0.703364493009494 | 0.481828628369323 | 0.837383764621454 | NM_001081068 |
| Tnfrsf1a | 444.604296 | 0.108865091 | 0.097967965 | 1.11123153 | 0.266468703 | 0.494085203 | | Rala | 2296.559456 | 0.0421143066398613 | 0.0598720685273815 | 0.703404904418844 | 0.481803450994044 | 0.837383764621454 | NM_019491 |
| Pcbd2 | 157.8378927 | -0.123167001 | 0.11094656 | -1.110147088 | 0.266935649 | 0.494724993 | | Vps13a | 2475.684744 | 0.0529466220840572 | 0.0752880122922431 | 0.703254349159013 | 0.481897254530432 | 0.837383764621454 | NM_173028 |
| Acsl1 | 2334.86195 | 0.068754708 | 0.061946597 | 1.109902909 | 0.267040867 | 0.494860805 | | Arid1b | 3797.719548 | 0.0318195724502078 | 0.0452667256124312 | 0.702935147610267 | 0.482096166079751 | 0.837477953579289 | NM_001085355 |
| C1qtnf9 | 11.54995185 | 0.181990174 | 0.163994829 | 1.10973117 | 0.267114887 | 0.494879596 | | Gdap1 | 5040.736906 | 0.0463064346110726 | 0.0658764667444041 | 0.702928327816035 | 0.482100416345175 | 0.837477953579289 | NM_010267 |
| Dusp16 | 655.4525216 | 0.092147429 | 0.083032394 | 1.109776842 | 0.267095201 | 0.494879596 | | Scly | 606.9125546 | 0.0463403955417214 | 0.0659344014249473 | 0.702825756209684 | 0.482164343979648 | 0.837488850968356 | NM_016717 |
| Avpi1 | 647.841116 | -0.092833328 | 0.083676194 | -1.109435354 | 0.267242418 | 0.495056674 | | Fam108c | 976.5965541 | 0.0464832765993238 | 0.0661660043776112 | 0.702525066105585 | 0.482351775294709 | 0.837629227089782 | NM_133722 |
| 1110035M17Rik | 76.58679424 | -0.155990279 | 0.140623842 | -1.109273339 | 0.267312283 | 0.495119458 | | Mtmr10 | 974.8946923 | 0.0503935236216819 | 0.0717334211714259 | 0.702511086167957 | 0.482360490472672 | 0.837629227089782 | NM_172742 |
| Ap1g1 | 4965.994085 | -0.065520009 | 0.059069152 | -1.109208559 | 0.267340221 | 0.495119458 | | Cryl1 | 545.0190727 | 0.0603671164332406 | 0.0860248742938958 | 0.701740245815438 | 0.482841169476696 | 0.837880118991517 | NM_030004 |
| Suds3 | 3546.117545 | -0.07562864 | 0.068232454 | -1.108396886 | 0.267690449 | 0.495708835 | | Gspt1 | 5034.92348 | 0.0470147481033134 | 0.0669971782647805 | 0.701742212448197 | 0.482839942796992 | 0.837880118991517 | NM_001130008,NM_146066 |
| Lefty1 | 30.38633122 | 0.185931189 | 0.167764966 | 1.108283771 | 0.267739282 | 0.495740014 | | Scnn1a | 275.2606305 | 0.0564194027861694 | 0.0804010854666694 | 0.701724391638447 | 0.482851058521134 | 0.837880118991517 | NM_011324 |
| Ndnl2 | 586.6742595 | 0.086054037 | 0.077657709 | 1.108119685 | 0.267810131 | 0.495811945 | | Slitrk2 | 1168.210587 | -0.0386646363390418 | 0.055079543655594 | -0.701978153283319 | 0.48269278788382 | 0.837880118991517 | NM_198863,NM_001161431 |
| Grem2 | 1303.29523 | -0.132216185 | 0.119329631 | -1.107991235 | 0.267865601 | 0.495855392 | | Tbcb | 2023.565176 | 0.0381380579963187 | 0.0543315416815151 | 0.70195059473702 | 0.482709974730341 | 0.837880118991517 | NM_025548 |
| Sfmbt1 | 1615.190641 | 0.100071764 | 0.090332051 | 1.107821236 | 0.267939027 | 0.495872824 | | Tmem135 | 2537.888349 | 0.039698681185086 | 0.0565420862800054 | 0.702108531837539 | 0.482611481999317 | 0.837880118991517 | NM_028343 |
| Sh3tc2 | 141.6138503 | 0.137190373 | 0.123836112 | 1.107838186 | 0.267931705 | 0.495872824 | | Aimp2 | 769.3677838 | 0.0421011080671788 | 0.0600092944458619 | 0.701576455046656 | 0.482943339287376 | 0.837940151276687 | NM_146165,NM_001172146 |
| Syk | 185.1038212 | -0.142219626 | 0.128399755 | -1.107631599 | 0.268020951 | 0.495905972 | | 1700026D08Rik | 40.77998813 | 0.0609411560548863 | 0.0868948870703172 | 0.701320389605563 | 0.483103091947108 | 0.838104565866552 | NM_029335 |
| Traf5 | 87.39833263 | 0.164693593 | 0.148686884 | 1.107653803 | 0.268011358 | 0.495905972 | | D3Ertd254e | 1336.325613 | 0.0521986312664065 | 0.0744572886018638 | 0.70105468848754 | 0.4832688863898 | 0.838104565866552 | NM_001101478 |
| Fxr2 | 5047.86653 | 0.091854274 | 0.082940933 | 1.107466131 | 0.268092448 | 0.495979031 | | Hace1 | 2148.674895 | -0.0495812579044724 | 0.0707188456723011 | -0.701103891517452 | 0.483238181934525 | 0.838104565866552 | NM_172473 |
| 1110059G10Rik | 375.1338284 | -0.0971328 | 0.087716709 | -1.107346601 | 0.268144103 | 0.49601537 | | Tlk2 | 2443.836372 | 0.0482376622296137 | 0.0687923512943983 | 0.701206766769457 | 0.483173987510229 | 0.838104565866552 | NM_001112705,NM_011903 |
| Aldoa | 59874.47854 | -0.079585955 | 0.07187845 | -1.107229706 | 0.268194627 | 0.496049606 | | Btbd8 | 73.78480812 | -0.0665989801102838 | 0.0950600858302338 | -0.700598779483766 | 0.483553440327669 | 0.838397860372371 | NM_001255991 |
| Rrn3 | 2580.314146 | 0.068634152 | 0.062002175 | 1.106963606 | 0.268309663 | 0.496203142 | | Htr1a | 653.0955023 | -0.0574934043892429 | 0.0820535426036977 | -0.700681561878743 | 0.483501765243181 | 0.838397860372371 | NM_008308 |
| Zkscan5 | 610.4804891 | 0.115587839 | 0.1044567 | 1.106562233 | 0.268483244 | 0.496464898 | | A830052D11Rik | 43.59084611 | -0.0657202658016342 | 0.0938564465795919 | -0.700221116360955 | 0.483789226226641 | 0.838706564049192 | NR_045403 |
| Adamtsl2 | 238.6595451 | -0.160683087 | 0.145342161 | -1.105550417 | 0.268921161 | 0.497156003 | | Amfr | 7522.171861 | 0.0296369774789895 | 0.0423883502594144 | 0.699177422513801 | 0.484441158424788 | 0.838713365223021 | NM_011787 |
| Pin4 | 507.7052617 | 0.098275665 | 0.088891388 | 1.105570034 | 0.268912666 | 0.497156003 | | C1galt1 | 326.160726 | 0.0618944855455166 | 0.0885518887862058 | 0.698962915784963 | 0.484575206727557 | 0.838713365223021 | NM_052993 |
| Eif3m | 2033.687958 | -0.065618295 | 0.059363232 | -1.105369317 | 0.268999593 | 0.497241672 | | Fam72a | 54.50845018 | 0.0650827467606558 | 0.0931388365357986 | 0.698771143579815 | 0.484695064920692 | 0.838713365223021 | NM_175382 |
| Pard6a | 707.8057943 | -0.104503968 | 0.094582962 | -1.104892113 | 0.26920634 | 0.49756448 | | Itga4 | 484.2055957 | -0.0582854761295418 | 0.0833747807355583 | -0.699078013942934 | 0.484503277753997 | 0.838713365223021 | NM_010576 |
| Aqp11 | 405.1963468 | 0.09917172 | 0.089777223 | 1.104642328 | 0.269314602 | 0.497705206 | | Mllt1 | 3605.110729 | 0.0476910035538849 | 0.0681244139372055 | 0.700057450737772 | 0.483891426729903 | 0.838713365223021 | NM_022328 |
| Fbxo31 | 5644.01074 | 0.118411356 | 0.107204289 | 1.104539353 | 0.269359243 | 0.497728337 | | Myl12a | 827.6842992 | 0.0511986535469763 | 0.0731419042121077 | 0.699997011186612 | 0.483933105402039 | 0.838713365223021 | NM_026064 |
| Eif3c | 7152.185581 | 0.068612571 | 0.062140883 | 1.104145419 | 0.269530062 | 0.497865854 | | Myo5a | 24015.03842 | 0.0445446447434952 | 0.0637500349167048 | 0.698739183947936 | 0.48471506714145 | 0.838713365223021 | NM_010864 |
| Twf1 | 1965.123656 | -0.064450038 | 0.058368987 | -1.104182914 | 0.2695138 | 0.497865854 | | Neu3 | 135.1721684 | -0.0623070409560374 | 0.0890325646456261 | -0.699823350448103 | 0.484037630301401 | 0.838713365223021 | NM_016720 |
| Zfp300 | 105.8851236 | -0.152714818 | 0.138300398 | -1.104225445 | 0.269495354 | 0.497865854 | | Phyhd1 | 579.8196451 | -0.0616897603408713 | 0.0882876388270479 | -0.698736098965329 | 0.484716969643898 | 0.838713365223021 | NM_001252568,NM_172267,NM_001252571,NM_001252570,NR_045543 |
| Cebpa | 315.3716242 | 0.12156839 | 0.11011169 | 1.104046178 | 0.269573107 | 0.497867046 | | Pih1d2 | 66.16859713 | -0.0663332084313726 | 0.0949206968183046 | -0.698827659876396 | 0.484659740399399 | 0.838713365223021 | NM_028300 |
| Tbc1d23 | 1364.600189 | 0.085488382 | 0.077436823 | 1.10397584 | 0.269603618 | 0.497867046 | | Pkd1 | 6010.513083 | -0.0464010243689463 | 0.0663783759943948 | -0.699038258677262 | 0.484528121593399 | 0.838713365223021 | NM_013630 |
| Zc3h10 | 568.368523 | 0.10176061 | 0.092181003 | 1.103921697 | 0.269627106 | 0.497867046 | | Plk2 | 9175.85058 | 0.0469804436514903 | 0.0671585717879692 | 0.699545008190695 | 0.484211495654784 | 0.838713365223021 | NM_152804 |
| Anxa1 | 77.00415017 | -0.18361089 | 0.16639775 | -1.103445746 | 0.26983364 | 0.498070338 | | Rnaseh2b | 296.9996688 | 0.0593587220521152 | 0.0848997178785007 | 0.699162612683042 | 0.48445028767769 | 0.838713365223021 | NM_026001 |
| Gm20187 | 10.5949802 | -0.180475033 | 0.163547985 | -1.10349897 | 0.269810539 | 0.498070338 | | Rnf19b | 1793.699392 | 0.0383546813316891 | 0.0548392859825614 | 0.699401544795563 | 0.484301122701144 | 0.838713365223021 | NM_029219 |
| Smurf1 | 2478.401444 | -0.065454268 | 0.059310108 | -1.103593802 | 0.269769381 | 0.498070338 | | Tiam1 | 5361.423897 | 0.0469219281270667 | 0.0670921293341021 | 0.699364417384174 | 0.484323069503443 | 0.838713365223021 | NM_009384,NM_001145887,NM_001145886 |
| BC025920 | 150.3095689 | -0.14055945 | 0.127423645 | -1.103087653 | 0.269989102 | 0.498297933 | | Ube2f | 1189.314176 | 0.0406655661021105 | 0.0581986626939542 | 0.698737122465443 | 0.484716329895009 | 0.838713365223021 | NM_026454 |
| Stx18 | 831.1552124 | 0.096305984 | 0.087319215 | 1.102918581 | 0.270062524 | 0.498374077 | | Srpx2 | 79.37463046 | 0.0663032051027536 | 0.0949038926406833 | 0.698635253601081 | 0.484780006281497 | 0.838722531115416 | NM_001083895,NM_026838 |
| Ddx18 | 805.7829153 | 0.075858226 | 0.068789473 | 1.102759227 | 0.270131739 | 0.498442439 | | A630089N07Rik | 74.41043591 | -0.0661103242718041 | 0.0946463345619165 | -0.698497555256183 | 0.484866086663441 | 0.838771558279522 | NR_015491 |
| Pcgf5 | 318.7060356 | -0.098864155 | 0.089659499 | -1.102662355 | 0.27017382 | 0.498460217 | | 2310061I04Rik | 1706.823394 | 0.0397505190178927 | 0.0569480031669727 | 0.698014272753502 | 0.485168268193137 | 0.83898200351664 | NM_001033630 |
| 5730408K05Rik | 12.83389096 | -0.182556238 | 0.165573449 | -1.102569524 | 0.270214151 | 0.498475778 | | Caskin1 | 7679.504247 | -0.0406532612441487 | 0.0582394831593334 | -0.698035748925317 | 0.48515483762622 | 0.83898200351664 | NM_027937 |
| Cerk | 3508.060198 | 0.095362659 | 0.086532261 | 1.102047468 | 0.270441036 | 0.498730506 | | Cbx1 | 726.8946924 | 0.0466276960198762 | 0.0667976746058547 | 0.698043706939658 | 0.485149860968971 | 0.83898200351664 | NM_007622 |
| Crb3 | 11.58058855 | 0.170278904 | 0.154515456 | 1.102018581 | 0.270453594 | 0.498730506 | | Fam229b | 295.43065 | -0.0537773611997911 | 0.0770522682438216 | -0.697935235196427 | 0.485218767144657 | 0.83898200351664 | NM_183254 |
| Fgd1 | 632.3751565 | -0.070030327 | 0.063544369 | -1.102069754 | 0.270431347 | 0.498730506 | | Dnaic1 | 143.7616719 | -0.0656634432498203 | 0.0941003615048064 | -0.697801063615588 | 0.485294894563187 | 0.839013763202196 | NM_175138 |
| Fmo1 | 297.3660851 | -0.13978359 | 0.126872158 | -1.101767263 | 0.270562866 | 0.498730506 | | Mettl2 | 594.8594167 | -0.0442996747680353 | 0.0635194683550572 | -0.697418853073702 | 0.485540706326763 | 0.839338843282466 | NM_172567 |
| Klhl22 | 3293.423932 | 0.063908167 | 0.058005917 | 1.101752553 | 0.270569263 | 0.498730506 | | Mon1b | 1924.403994 | 0.0359885723943412 | 0.0516167159332976 | 0.697227085133582 | 0.485660691116285 | 0.839446358776197 | NM_173015,NM_001048143 |
| Mrpl30 | 1306.778361 | -0.069730832 | 0.063299493 | -1.101601746 | 0.270634849 | 0.498730506 | | Lamp2 | 2677.796491 | 0.0460426010096814 | 0.0660737487168053 | 0.696836524143376 | 0.485905105753948 | 0.839716418408557 | NM_001017959,NM_010685 |
| Ogfr | 1346.09774 | -0.068912409 | 0.062557479 | -1.101585461 | 0.270641932 | 0.498730506 | | Metrn | 1228.805674 | -0.0575384861256022 | 0.0825761923166323 | -0.696792677300682 | 0.485932549438409 | 0.839716418408557 | NM_133719 |
| S100a6 | 436.3462323 | 0.127637418 | 0.115849218 | 1.101754677 | 0.270568339 | 0.498730506 | | Ogfod2 | 984.8879972 | -0.0468914739440841 | 0.0673104786429416 | -0.696648499453232 | 0.486022796085669 | 0.839772467698397 | NM_025671 |
| Wdr11 | 2164.423184 | 0.065952152 | 0.059863866 | 1.101702191 | 0.270591164 | 0.498730506 | | 2210408F21Rik | 369.5793372 | -0.055088521595348 | 0.079241627911988 | -0.695196742556775 | 0.486932013498852 | 0.839801621369315 | NR_040262,NR_040260,NR_040261,NR_040259,NR_040258,NR_040257 |
| Wdr48 | 3446.581606 | 0.05341848 | 0.048502999 | 1.101339078 | 0.270749109 | 0.498866677 | | 9930014A18Rik | 92.62978358 | 0.0659857807878927 | 0.0947872013303137 | 0.696146524655222 | 0.486337072786219 | 0.839801621369315 | NR_030696 |
| Prx | 24.93524271 | -0.182984224 | 0.166169565 | -1.101189764 | 0.270814076 | 0.498929048 | | Abcb4 | 307.0692636 | -0.0578533962023279 | 0.0831403875758021 | -0.695851894478852 | 0.486521586192957 | 0.839801621369315 | NM_008830 |
| Cttnbp2 | 4671.253769 | 0.069106949 | 0.062764996 | 1.101042846 | 0.27087801 | 0.498987503 | | Acot6 | 144.7968604 | -0.0608168074357044 | 0.0875590835886886 | -0.694580222747257 | 0.487318409968947 | 0.839801621369315 | NM_172580 |
| Rab11fip2 | 2653.533019 | -0.073010552 | 0.066319134 | -1.10089724 | 0.270941384 | 0.499044912 | | Acox3 | 1843.456649 | -0.0397227623002746 | 0.0572483446510908 | -0.693867461637193 | 0.487765359090624 | 0.839801621369315 | NM_030721 |
| Lin54 | 515.7277537 | 0.130181905 | 0.118288548 | 1.100545289 | 0.271094608 | 0.499267783 | | Asb11 | 167.17861 | 0.059488109518734 | 0.0854502218533718 | 0.696172676490383 | 0.486320697911696 | 0.839801621369315 | NM_026853 |
| Dnajc28 | 502.7541736 | 0.086385741 | 0.078502738 | 1.10041692 | 0.27115051 | 0.499311386 | | Bod1l | 3380.943892 | -0.053150911607229 | 0.076586197408353 | -0.694001183056935 | 0.487681466996156 | 0.839801621369315 | NM_001081422 |
| Sgcz | 85.58715352 | -0.149697745 | 0.136088897 | -1.0999997 | 0.271332252 | 0.499586681 | | Cdc14b | 497.8220295 | 0.0547520782374678 | 0.0787777984985292 | 0.695021695015993 | 0.487043303029639 | 0.839801621369315 | NM_001122989,NM_172587 |
| Aktip | 4162.402379 | -0.060104083 | 0.054669387 | -1.09941024 | 0.271589166 | 0.499940898 | | Cdh12 | 493.1495629 | 0.0629566928120838 | 0.090690838068462 | 0.694189875343458 | 0.487563141186387 | 0.839801621369315 | NM_001008420 |
| Polr2g | 1924.032436 | -0.070315137 | 0.063955187 | -1.099443844 | 0.271574515 | 0.499940898 | | Clic6 | 517.7450935 | 0.0316904409862448 | 0.0456278456944626 | 0.694541688390315 | 0.487342566435706 | 0.839801621369315 | NM_172469 |
| Galnt1 | 2247.339256 | 0.078670354 | 0.071593412 | 1.098849066 | 0.271833906 | 0.500331972 | | Dpysl2 | 6304.401706 | 0.0394204752967485 | 0.056612136111846 | 0.696325523185829 | 0.486224992669539 | 0.839801621369315 | NM_009955 |
| Sgsm1 | 5754.829528 | -0.090003293 | 0.081913342 | -1.09876231 | 0.27187175 | 0.500342201 | | Elf4 | 98.92336941 | -0.0650455525938211 | 0.0936622038854047 | -0.694469308213204 | 0.487387941929215 | 0.839801621369315 | NM_019680 |
| 2900092D14Rik | 588.4398588 | -0.106264159 | 0.096749808 | -1.098339737 | 0.272056166 | 0.500535371 | | Esco1 | 876.4277949 | 0.0561804768173088 | 0.0808989863479018 | 0.694452170145756 | 0.487398682603027 | 0.839801621369315 | NM_001081222 |
| Mpi | 1944.212642 | 0.096604429 | 0.087957613 | 1.098306623 | 0.272070621 | 0.500535371 | | Fam65b | 2826.288801 | 0.051285070852996 | 0.0737471339601657 | 0.695417816277458 | 0.486793498448057 | 0.839801621369315 | NM_029679,NM_001080381,NM_178658 |
| Smyd5 | 2002.07832 | 0.085350483 | 0.077714468 | 1.098299719 | 0.272073634 | 0.500535371 | | Gm11744 | 91.79887892 | -0.0660022754207501 | 0.0950954861181881 | -0.694063179178874 | 0.487642588544825 | 0.839801621369315 | NM_001163318 |
| Htr4 | 145.1613738 | -0.183959957 | 0.167580892 | -1.097738258 | 0.272318799 | 0.500748572 | | Hmgn3 | 1160.918793 | -0.0447110566795621 | 0.0644387552587616 | -0.693853512533249 | 0.48774079546531 | 0.839801621369315 | NM_175074,NM_026122 |
| Syt13 | 10224.19455 | 0.089716882 | 0.08172365 | 1.097808058 | 0.272288312 | 0.500748572 | | Kcnh5 | 906.9526816 | 0.058133602076105 | 0.0837366555350261 | 0.694243180655674 | 0.487529717114821 | 0.839801621369315 | NM_172805 |
| Wdr5b | 164.8584227 | -0.136647072 | 0.12447684 | -1.097771056 | 0.272304473 | 0.500748572 | | Kdelr1 | 2246.175163 | 0.0353743862218048 | 0.0509599369692536 | 0.694143052868123 | 0.487592501340042 | 0.839801621369315 | NM_133950 |
| Xrcc4 | 270.6936457 | -0.128763548 | 0.117276423 | -1.097949146 | 0.272226696 | 0.500748572 | | Mtor | 5625.338735 | 0.0339693181661135 | 0.0488229612292202 | 0.695765215116651 | 0.486575876716606 | 0.839801621369315 | NM_020009 |
| Sardh | 530.9826734 | -0.116092465 | 0.105767201 | -1.09762255 | 0.272369342 | 0.500782079 | | Mxd4 | 1095.986639 | 0.0483327635381525 | 0.0696038975772045 | 0.69439737170669 | 0.487433041543381 | 0.839801621369315 | NM_010753 |
| Gck | 51.23499475 | 0.182242581 | 0.166053602 | 1.097492491 | 0.272426161 | 0.500827117 | | Nicn1 | 4695.002813 | 0.0325199483864109 | 0.0468086812447063 | 0.694741835715091 | 0.48721711498482 | 0.839801621369315 | NM_025449 |
| Heyl | 281.9341015 | -0.150458294 | 0.137106475 | -1.097382849 | 0.27247406 | 0.50085576 | | Nt5c3l | 1211.298709 | 0.04491428044335 | 0.0645235685816443 | 0.69609108066956 | 0.486371791928907 | 0.839801621369315 | NM_026561,NM_001102650 |
| Fam3c | 2355.629652 | -0.061344814 | 0.055907156 | -1.097262275 | 0.272526757 | 0.500893187 | | Pnma1 | 171.4126621 | 0.0638030420031421 | 0.0916827260474424 | 0.695911277077959 | 0.48648439704731 | 0.839801621369315 | NM_027438 |
| Hspb3 | 64.42158644 | 0.152883552 | 0.139375843 | 1.096915711 | 0.27267824 | 0.501112164 | | Ptprb | 2065.623283 | -0.0522450403939077 | 0.0752206790323875 | -0.694556883372626 | 0.487333040091531 | 0.839801621369315 | NM_029928 |
| Asb1 | 2493.072167 | -0.07374827 | 0.067250011 | -1.096628373 | 0.272803879 | 0.501283599 | | Rrm2b | 1630.948967 | 0.0480981242887903 | 0.069243350838553 | 0.694624447059694 | 0.487290687408787 | 0.839801621369315 | NM_199476 |
| Nol10 | 875.3819833 | -0.075808387 | 0.069138312 | -1.096474358 | 0.272871239 | 0.501347916 | | Scarb1 | 1134.969821 | -0.0428376923872204 | 0.0615825557358402 | -0.69561407244892 | 0.486670550849672 | 0.839801621369315 | NM_001205082,NM_001205083,NM_016741 |
| B430306N03Rik | 12.64067679 | 0.174293585 | 0.158970597 | 1.096388817 | 0.272908656 | 0.50135721 | | Thoc5 | 1334.102972 | 0.0366316790227731 | 0.0527280973295413 | 0.694727857231902 | 0.48722585545822 | 0.839801621369315 | NM_172438 |
| Mtap | 392.9806395 | -0.09166859 | 0.083627837 | -1.096149249 | 0.27301346 | 0.501430848 | | Usp15 | 2710.041592 | 0.047379203754507 | 0.0681411520062809 | 0.695309696996901 | 0.486861238548986 | 0.839801621369315 | NM_027604 |
| Trmt5 | 374.2889622 | 0.114788924 | 0.104717842 | 1.096173501 | 0.273002854 | 0.501430848 | | Usp54 | 4136.89888 | 0.055095439011089 | 0.0792443674009295 | 0.695260001664708 | 0.486892375932685 | 0.839801621369315 | NM_030180 |
| Mdc1 | 1059.907789 | 0.102434736 | 0.093466529 | 1.095951002 | 0.273100218 | 0.501471293 | | Zfp563 | 595.3001898 | 0.053992592895702 | 0.0775648192213592 | 0.696100712123284 | 0.486365760595766 | 0.839801621369315 | NM_001024950 |
| Nphs2 | 17.67623178 | -0.181882943 | 0.165958746 | -1.095952747 | 0.273099454 | 0.501471293 | | Hsd17b7 | 950.5711047 | 0.0502312771836547 | 0.072437998467043 | 0.693443956988471 | 0.488030984682161 | 0.839945270556051 | NM_010476 |
| Cyp2u1 | 218.4817147 | -0.099519639 | 0.090841596 | -1.095529399 | 0.273284773 | 0.501649393 | | Ppan | 568.9016234 | -0.0564168964588963 | 0.0813499533043043 | -0.693508652031574 | 0.487990398058745 | 0.839945270556051 | NM_145610 |
| Klhdc5 | 1314.3435 | 0.07608696 | 0.069447037 | 1.095611328 | 0.273248902 | 0.501649393 | | Utp23 | 409.4833124 | -0.0551502319513309 | 0.0795291942837844 | -0.693458954890068 | 0.488021575542464 | 0.839945270556051 | NM_030132 |
| Tmem101 | 625.9193563 | -0.098090398 | 0.08953877 | -1.095507546 | 0.273294341 | 0.501649393 | | Zbtb18 | 11090.63823 | 0.0497024736751875 | 0.0717200266446396 | 0.693006904772285 | 0.488305218356778 | 0.840031768769702 | NM_013915,NM_001012330 |
| Lct | 81.40493008 | 0.16550512 | 0.151117999 | 1.095204549 | 0.273427033 | 0.501851505 | | Ccdc90b | 434.0238802 | 0.0506816017354835 | 0.0731690997063227 | 0.692666739633316 | 0.48851871655989 | 0.840318276329969 | NM_025515,NM_001162918 |
| Mir568 | 19.02698099 | -0.182741848 | 0.166879308 | -1.095053963 | 0.273492996 | 0.501895118 | | Copz2 | 158.5035938 | 0.0642079210391287 | 0.0926943943281758 | 0.692683969774985 | 0.488507901178252 | 0.840318276329969 | NM_019877 |
| Zfp2 | 844.7152888 | 0.090957446 | 0.083072166 | 1.094920842 | 0.273551317 | 0.501942694 | | Entpd7 | 510.8477122 | 0.0501761606819195 | 0.0724421470396064 | 0.692637608036282 | 0.488536957428514 | 0.840318276329969 | NM_053103 |
| Nono | 4933.752965 | -0.056561904 | 0.051665443 | -1.09477247 | 0.27361633 | 0.502002537 | | Rdh11 | 903.3951386 | 0.0453816425856669 | 0.0655189206813439 | 0.692649422697053 | 0.48852985655266 | 0.840318276329969 | NM_021557 |
| Cenpc1 | 849.4423517 | 0.087356127 | 0.079811698 | 1.094527564 | 0.273723538 | 0.50213977 | | Fam73a | 4677.19906 | 0.0395620809615157 | 0.0571285001210642 | 0.692510408459516 | 0.488616851645178 | 0.840356190740926 | NM_174868,NM_001162375 |
| Gm14827 | 186.0112022 | 0.180362093 | 0.164830266 | 1.094229221 | 0.273854459 | 0.502142128 | | Gipc3 | 30.12514509 | -0.0615929442015562 | 0.0889705105019538 | -0.69228943636893 | 0.488758408067683 | 0.840500134437216 | NM_148951 |
| Lig3 | 1254.993708 | 0.096351953 | 0.088047335 | 1.094319923 | 0.273814691 | 0.502142128 | | Tnfrsf12a | 148.4324896 | 0.0647131775089711 | 0.0935068645383822 | 0.692068735578317 | 0.488894164429546 | 0.840634071082337 | NM_001161746,NM_013749 |
| Ranbp17 | 253.1732783 | 0.1169966 | 0.106920192 | 1.094242334 | 0.273848709 | 0.502142128 | | 1190002N15Rik | 1254.789725 | -0.0587411366812988 | 0.0849219401461255 | -0.691707426611106 | 0.489121082162813 | 0.840836702688179 | NM_001033145 |
| Sorbs2 | 6514.191753 | 0.078590386 | 0.071814735 | 1.094349042 | 0.273801924 | 0.502142128 | | Chmp2b | 1570.988367 | 0.0447661604220434 | 0.0647365921886478 | 0.691512464721515 | 0.48924355026301 | 0.840836702688179 | NM_026879 |
| Aes | 14483.80129 | -0.063478937 | 0.058017151 | -1.094140884 | 0.273893194 | 0.502153727 | | Gprc5c | 186.2160479 | -0.0646012111308796 | 0.0934010531669989 | -0.691653990403878 | 0.489154647238093 | 0.840836702688179 | NM_001110337,NM_147217,NM_001110338 |
| Hepacam2 | 13.35046926 | 0.18071809 | 0.165168649 | 1.094022256 | 0.273945217 | 0.502189682 | | Ndufb6 | 1618.016413 | 0.040964807241419 | 0.059238854791814 | 0.691519229826161 | 0.489239300389186 | 0.840836702688179 | NM_001033305 |
| Spc25 | 12.98815244 | -0.18065646 | 0.165142775 | -1.093941046 | 0.273980835 | 0.502195559 | | Cdc42ep3 | 512.6752049 | 0.0512737915317397 | 0.0742353342535913 | 0.690692539439319 | 0.489758778775264 | 0.841119426334601 | NM_026514 |
| Chchd10 | 3756.904024 | -0.099135315 | 0.090639121 | -1.0937365 | 0.274070561 | 0.5023006 | | Gm3230 | 135.7193894 | -0.0640107372745835 | 0.0926878250109476 | -0.690605667648616 | 0.48981338478345 | 0.841119426334601 | NR_033642 |
| Nxph2 | 68.82851891 | -0.180334122 | 0.165004149 | -1.092906593 | 0.274434813 | 0.502902752 | | Mrps26 | 1301.807157 | 0.0412034338069304 | 0.0596331697208573 | 0.690948242392675 | 0.489598067622418 | 0.841119426334601 | NM_207207 |
| Trmt2b | 605.6187536 | -0.110109235 | 0.100755124 | -1.09284006 | 0.274464029 | 0.502902752 | | Nab2 | 1946.717768 | 0.0621749957363191 | 0.0900018044297589 | 0.690847212789439 | 0.489661562065235 | 0.841119426334601 | NM_001122895,NM_008668 |
| Mocos | 98.88467813 | 0.147606879 | 0.135097164 | 1.09259791 | 0.274570381 | 0.503038132 | | Nxph2 | 72.06861153 | 0.0624520443954756 | 0.0903623813584651 | 0.691128802236077 | 0.489484601558919 | 0.841119426334601 | NM_008752 |

| | | | | | | | | | | | | | | | |
|---|---|---|---|---|---|---|---|---|---|---|---|---|---|---|---|
| Dnajc19 | 484.5306663 | -0.083942636 | 0.076848557 | -1.09231245 | 0.27469579 | 0.503089429 | | Sh3glb1 | 2315.150088 | -0.044653665087954 | 0.0646549956911533 | -0.690645241108011 | 0.489788509229194 | 0.841119426334601 | NM_019464 |
| Pcdhga6 | 774.3867739 | 0.115518954 | 0.105755336 | 1.092322694 | 0.274691289 | 0.503089429 | | Stk32a | 332.320069 | 0.0617834726881591 | 0.0894621961811161 | 0.690609836618347 | 0.489810764170994 | 0.841119426334601 | NM_178749 |
| Ung | 76.78285397 | -0.158001594 | 0.144646958 | -1.09232573 | 0.274689955 | 0.503089429 | | Rnls | 30.14557899 | 0.0591295744977825 | 0.0856545424090907 | 0.69032619677514 | 0.489989077250731 | 0.841321670939021 | NM_001146342,NM_001167818 |
| Gpr62 | 312.5098788 | 0.143680124 | 0.131568513 | 1.092055548 | 0.274808686 | 0.503236708 | | Nek2 | 69.47983999 | 0.0654030852968194 | 0.0947645927897766 | 0.690163734907909 | 0.490091226288494 | 0.841397607028326 | NM_010892 |
| Tmem175 | 1279.101553 | 0.088743909 | 0.081268697 | 1.091981444 | 0.274841257 | 0.503236876 | | Syne2 | 949.7837351 | -0.0424303340277606 | 0.0614869053794105 | -0.690071060918425 | 0.49014950083954 | 0.841398209547998 | NM_001005510 |
| Gm13476 | 145.465822 | -0.16251424 | 0.148875361 | -1.091612736 | 0.275003356 | 0.503474181 | | Ak7 | 118.0907536 | 0.0606640871395036 | 0.0879653120887488 | 0.689636468046623 | 0.490422827927114 | 0.841767930812863 | NM_030187 |
| Utp18 | 648.7310528 | -0.087174542 | 0.079899517 | -1.091052176 | 0.275249925 | 0.50386606 | | Vars2 | 1166.00833 | -0.0503676955215804 | 0.0730478478984377 | -0.689516487763051 | 0.490498301171414 | 0.841798006003775 | NM_175137 |
| Itga5 | 333.6946138 | 0.137994352 | 0.126553083 | 1.090406879 | 0.275533953 | 0.504326411 | | Arl6 | 801.9515331 | 0.0455816140257675 | 0.0662011678017683 | 0.688531872462678 | 0.491117906353827 | 0.841866734446035 | NM_019665 |
| 2410004I01Rik | 13.18370108 | -0.180510185 | 0.165568702 | -1.090243401 | 0.27560594 | 0.504398586 | | Draxin | 26.49340856 | 0.0601406256806166 | 0.0873144537624231 | 0.688782018201194 | 0.490960453172308 | 0.841866734446035 | NM_027426 |
| Dram2 | 569.6489118 | -0.086758711 | 0.07962072 | -1.089649923 | 0.275867383 | 0.504817436 | | Efcab6 | 299.9288608 | -0.0604449000374747 | 0.0877382544076595 | -0.688922983999985 | 0.490871734794189 | 0.841866734446035 | NM_029946,NM_001161629,NM_001161628 |
| Acad8 | 858.0057154 | 0.069773288 | 0.064060046 | 1.089185722 | 0.276071994 | 0.504893705 | | Fam108a | 3532.431138 | 0.0448659576210234 | 0.0651567587446628 | 0.688584860349557 | 0.491084551087416 | 0.841866734446035 | NM_145421 |
| Arhgap19 | 223.9301013 | 0.118099783 | 0.108419901 | 1.089281412 | 0.276029807 | 0.504893705 | | Grk5 | 572.4293992 | 0.0431489194120438 | 0.0626327390716205 | 0.68891956589513 | 0.490873885914212 | 0.841866734446035 | NM_018869 |
| Lman1 | 1450.340784 | 0.060465312 | 0.055507316 | 1.089321498 | 0.276012136 | 0.504893705 | | Lrrc71 | 39.11062007 | -0.0558465754288559 | 0.0810640590763084 | -0.688919060619523 | 0.490874203900345 | 0.841866734446035 | NM_028971 |
| Mnt | 1910.083417 | 0.092700821 | 0.085105458 | 1.089246485 | 0.276045205 | 0.504893705 | | Rae1 | 1604.014402 | -0.0355440578741857 | 0.0516214473996133 | -0.688552136073038 | 0.491105150500053 | 0.841866734446035 | NM_175112 |
| Ppp6r3 | 3447.019568 | 0.068649341 | 0.063020237 | 1.089322173 | 0.276011838 | 0.504893705 | | Trim7 | 60.20531669 | -0.0652153615469959 | 0.0946726682797925 | -0.688850992920792 | 0.490917042096439 | 0.841866734446035 | NM_053166 |
| Opn1mw | 11.9270485 | 0.178874493 | 0.164240407 | 1.089101618 | 0.276109077 | 0.504901927 | | Zfp664 | 4921.272471 | -0.0431420750807534 | 0.0626101551880461 | -0.689058746958518 | 0.490786299025206 | 0.841866734446035 | NM_001081750 |
| Atg14 | 932.5302857 | 0.064215073 | 0.058982259 | 1.088718426 | 0.276278074 | 0.505132102 | | Zfp799 | 933.4219438 | -0.0395522181949739 | 0.0574105914883377 | -0.688935911817046 | 0.490863598961403 | 0.841866734446035 | NM_177359 |
| Erap1 | 437.660611 | 0.097180674 | 0.089275927 | 1.088542881 | 0.276355517 | 0.505132102 | | Fn3krp | 1508.062582 | -0.0393005868330474 | 0.0571013106027124 | -0.688260679452436 | 0.491288638291688 | 0.841892493483271 | NM_181420 |
| Gtf2e2 | 518.3262086 | -0.073648188 | 0.067658808 | -1.088523289 | 0.276364161 | 0.505132102 | | Kif26a | 336.0786843 | 0.0557210970980268 | 0.0810109777638059 | 0.687821559943224 | 0.49156515740888 | 0.841892493483271 | NM_001097621 |
| Wdr59 | 1439.094584 | -0.063329483 | 0.058179408 | -1.08852058 | 0.276365356 | 0.505132102 | | Ptprz1 | 8914.845706 | -0.0452129827547756 | 0.0657647586440998 | -0.687495608391956 | 0.491770467245323 | 0.841892493483271 | NM_001081306 |
| Brf2 | 490.5940917 | -0.083204466 | 0.076492368 | -1.087748595 | 0.276706109 | 0.505605783 | | Rai1 | 3707.322845 | -0.0373435275930388 | 0.054275047911048 | -0.688042277811372 | 0.491426158128635 | 0.841892493483271 | NM_001037764,NM_009021 |
| Flt4 | 180.1858191 | 0.14325256 | 0.131719539 | 1.087557405 | 0.276790544 | 0.505605783 | | Rps6ka5 | 1014.088615 | 0.0551745869873863 | 0.0801491267375802 | 0.688399103436721 | 0.491201488275978 | 0.841892493483271 | NM_153587 |
| Lamp5 | 7298.585867 | 0.118890329 | 0.109316653 | 1.087577466 | 0.276781683 | 0.505605783 | | Setx | 3426.636317 | 0.0430064795198514 | 0.0625039525457095 | 0.68806015888996 | 0.491414898265142 | 0.841892493483271 | NM_198033 |
| Pik3ap1 | 124.1428116 | 0.155363944 | 0.142819354 | 1.087835367 | 0.276667793 | 0.505605783 | | Slc25a15 | 538.7781309 | -0.0470951853686472 | 0.0684835255199438 | -0.687686345162416 | 0.491650320691047 | 0.841892493483271 | NM_181325 |
| Pnmal2 | 15463.46883 | -0.091234812 | 0.083900024 | -1.087422963 | 0.276849927 | 0.505605783 | | Stradb | 1586.523883 | -0.0413819849841032 | 0.0601324414515632 | -0.688180688912099 | 0.491339003124711 | 0.841892493483271 | NM_172656 |
| Ptrh1 | 88.98114447 | -0.153960877 | 0.141571317 | -1.08751462 | 0.276809441 | 0.505605783 | | Tbc1d4 | 700.836372 | 0.0524528197782611 | 0.076250270672631 | 0.687903391234627 | 0.491513620874556 | 0.841892493483271 | NM_001081278 |
| Tpp2 | 2240.997529 | 0.078796419 | 0.072462065 | 1.087416142 | 0.276852941 | 0.505605783 | | Tmem53 | 78.30153419 | -0.0653770074829908 | 0.0950904217027657 | -0.687524635103067 | 0.491752182080924 | 0.841892493483271 | NM_026837 |
| Snap91 | 12515.20191 | -0.065517121 | 0.060262355 | -1.087198141 | 0.276949252 | 0.505722064 | | Vps16 | 2398.202141 | -0.0422078579375458 | 0.0613934413632703 | -0.687497833649661 | 0.491769065447589 | 0.841892493483271 | NM_030559 |
| Scyl3 | 1096.174775 | 0.076139637 | 0.070044793 | 1.087013526 | 0.277030832 | 0.50581142 | | Angptl2 | 241.7269622 | 0.0534011918451893 | 0.0777843852812989 | 0.686528429222261 | 0.492379943282217 | 0.842340249218877 | NM_011923 |
| Atp10b | 92.45460193 | -0.182558434 | 0.167984281 | -1.086759031 | 0.277143318 | 0.505957178 | | Atf2 | 5146.090263 | 0.0463443079659582 | 0.0674607839339862 | 0.686981461871366 | 0.49209441055288 | 0.842340249218877 | NM_009715,NM_001025093 |
| Eltd1 | 898.5104858 | -0.112099706 | 0.103184864 | -1.086396794 | 0.277303479 | 0.506189928 | | C2cd2 | 722.4431312 | 0.0498354052152304 | 0.0725752993383816 | 0.686671714337317 | 0.492289625417843 | 0.842340249218877 | NM_174847 |
| Tsnaxip1 | 10.97165806 | 0.158608089 | 0.146040863 | 1.086052802 | 0.277455632 | 0.506408006 | | Ero1l | 2386.528937 | 0.0383369155545959 | 0.0558250421498078 | 0.686733302443693 | 0.492250806905567 | 0.842340249218877 | NM_015774 |
| Arrdc4 | 309.222921 | 0.106063639 | 0.097673791 | 1.085896616 | 0.277524734 | 0.50641482 | | Pcdhga3 | 635.4462423 | 0.053934490290372 | 0.0785204025816369 | 0.686885045377816 | 0.492155171500977 | 0.842340249218877 | NM_033586 |
| Pigu | 816.8536786 | 0.063377251 | 0.058362133 | 1.08593102 | 0.277509512 | 0.50641482 | | Stard7 | 4933.793819 | 0.0313015785654291 | 0.0455901912645548 | 0.686585813684999 | 0.492343770683545 | 0.842340249218877 | NM_139308 |
| 2610018G03Rik | 206.9364362 | -0.114659825 | 0.105636213 | -1.085421572 | 0.277734982 | 0.506699118 | | Ak8 | 93.4145262 | 0.0648803854740666 | 0.0945355215461302 | 0.686306950159546 | 0.492519567141004 | 0.842479890754855 | NM_001033874 |
| Agap3 | 9651.161794 | 0.087592008 | 0.080700447 | 1.085396819 | 0.277745941 | 0.506699118 | | Dgki | 1512.431845 | 0.0546996019605383 | 0.0797405939804059 | 0.685969331680415 | 0.492732447937591 | 0.842744794388491 | NM_001081206 |
| 4930577N17Rik | 13.61345547 | 0.180981535 | 0.166799489 | 1.085024517 | 0.277910797 | 0.50694018 | | 5730409E04Rik | 4323.740723 | -0.0266120685957464 | 0.0388057745421277 | -0.685775993643838 | 0.492854376843148 | 0.842781258864066 | NM_001145950,NM_001013755 |
| Nipsnap3b | 315.3507926 | -0.122460509 | 0.1128939 | -1.084739829 | 0.278036902 | 0.507083953 | | Dennd5b | 3603.414314 | 0.0531674294256594 | 0.0775316233788649 | 0.685751530905681 | 0.492869805455751 | 0.842781258864066 | NM_177192 |
| Plekhh2 | 516.391218 | -0.113074927 | 0.10424546 | -1.084698819 | 0.278055071 | 0.507083953 | | Atg4d | 986.9685738 | 0.0507933409233385 | 0.0740981442337915 | 0.685487355298364 | 0.49303643710725 | 0.842867751608793 | NM_153583 |
| Lrdd | 54.02416605 | 0.176713671 | 0.162946616 | 1.084488131 | 0.278148427 | 0.507194507 | | Itgal | 53.02055003 | -0.0646543927861122 | 0.0943170388675772 | -0.685500664168307 | 0.493028041670161 | 0.842867751608793 | NM_001253872,NM_001253873,NM_001253874,NM_008400 |
| Col11a2 | 906.29531 | 0.158559349 | 0.146251358 | 1.08415642 | 0.27829545 | 0.507283495 | | Ndufa5 | 2222.605013 | 0.0475024144339119 | 0.0693071558381237 | 0.685389753185953 | 0.493098008341073 | 0.842873813739989 | NM_026614 |
| Leprel1 | 93.75609248 | 0.133951951 | 0.12353972 | 1.084282457 | 0.278239581 | 0.507283495 | | Eml2 | 5820.892064 | 0.0430790425284236 | 0.0628839868535819 | 0.685055841461328 | 0.493308684079018 | 0.843035522933155 | NM_001162996,NM_028153 |
| Pmpcb | 1645.025308 | -0.07844346 | 0.072351353 | -1.08420171 | 0.278275373 | 0.507283495 | | Fam49a | 10779.65902 | 0.052595169587545 | 0.076772128829591 | 0.685081557452824 | 0.493292457312687 | 0.843035522933155 | NM_029758,NM_001146119 |
| Mxi1 | 3276.322784 | 0.106781367 | 0.098510289 | 1.083961558 | 0.278381844 | 0.507381283 | | Tnfaip1 | 2538.141721 | 0.0314496125075114 | 0.0459207073410829 | 0.684867771611502 | 0.493427364666295 | 0.843139148234632 | NM_001159392,NM_009395 |
| C330013E15Rik | 34.49888404 | 0.174033226 | 0.160585567 | 1.083741391 | 0.278479478 | 0.507499533 | | 9930013L23Rik | 891.1469304 | -0.0476647224483073 | 0.0696810459882247 | -0.684041431530206 | 0.493949003777485 | 0.843404385363906 | NM_030728 |
| Grap | 152.2924145 | -0.128699355 | 0.118789294 | -1.083425542 | 0.278619584 | 0.50765721 | | C330027C09Rik | 119.8976515 | -0.0650407139805727 | 0.0950197281450759 | -0.684496948689105 | 0.493661415499505 | 0.843404385363906 | NM_172616 |
| Tfb2m | 872.882933 | -0.083069551 | 0.076674965 | -1.083398617 | 0.278631529 | 0.50765721 | | Capn2 | 3597.517025 | -0.0331285132237465 | 0.048415792445408 | -0.684250149599455 | 0.493817219634273 | 0.843404385363906 | NM_009794 |
| Acsl5 | 3392.512396 | 0.086671357 | 0.080038967 | 1.082864512 | 0.278868566 | 0.507917251 | | Dhx40 | 1249.71941 | 0.0442184468989441 | 0.0646279627338259 | 0.684199919484703 | 0.493848933102017 | 0.843404385363906 | NM_026191 |
| Klf5 | 218.619607 | -0.133305647 | 0.123105674 | -1.082855426 | 0.278872599 | 0.507917251 | | Fibcd1 | 687.0053464 | -0.0587082530316671 | 0.0858177435082574 | -0.684103900098681 | 0.493909559283542 | 0.843404385363906 | NM_178887 |
| Tmem203 | 238.0716132 | 0.105369233 | 0.097295534 | 1.082981188 | 0.278816773 | 0.507917251 | | Samhd1 | 829.7792567 | -0.0526475836920845 | 0.0769726206209299 | -0.683978059566923 | 0.493989020423799 | 0.843404385363906 | NM_018851,NM_001139520 |
| Emc4 | 3093.165791 | -0.05790194 | 0.053561252 | -1.081041575 | 0.27967862 | 0.509325405 | | Tor1a | 997.8321903 | 0.03792962100719 | 0.0554510004332741 | 0.684020499374612 | 0.493962221334045 | 0.843404385363906 | NM_144884 |
| Eif1ad | 968.5425612 | 0.084505396 | 0.07820587 | 1.080550561 | 0.279897084 | 0.50948372 | | Insr | 2698.427314 | -0.038139659125591 | 0.0557739020190185 | -0.683826265420442 | 0.494084787887018 | 0.843468909490901 | NM_010568 |
| Npr1 | 195.307958 | -0.174366882 | 0.161368347 | -1.080551948 | 0.279896467 | 0.50948372 | | 1110001J03Rik | 377.5177111 | -0.0509949483583454 | 0.0746266464502957 | -0.683334315341506 | 0.494395615271134 | 0.843503724151808 | NM_025363 |
| Rbm6 | 2030.424896 | 0.06928201 | 0.064116775 | 1.080559805 | 0.279892971 | 0.50948372 | | Dlx1 | 871.9934522 | 0.0560740543039484 | 0.0820399141423256 | 0.683497232903843 | 0.494292698109677 | 0.843503724151808 | NM_010053 |
| Twsg1 | 1225.273733 | -0.091452686 | 0.084622128 | -1.080718348 | 0.279822419 | 0.50948372 | | Fastk | 2975.529613 | -0.0483520843768604 | 0.0707490402610234 | -0.683430958046747 | 0.494334563423776 | 0.843503724151808 | NM_023229 |
| Mmp25 | 15.38811468 | -0.181530644 | 0.168026388 | -1.080369853 | 0.279977515 | 0.509510408 | | Rplp0 | 6413.227458 | 0.0347470131920079 | 0.0508486911374913 | 0.683341348906198 | 0.494391178839374 | 0.843503724151808 | NM_007475 |
| Tmx4 | 10364.73909 | -0.074817805 | 0.069251771 | -1.080373884 | 0.279975721 | 0.509510408 | | Telo2 | 856.9360634 | -0.0562191100063957 | 0.0822368285445555 | -0.683624490406223 | 0.494212315847456 | 0.843503724151808 | NM_001163661,NM_027880 |
| Rft1 | 413.7652269 | 0.08890548 | 0.082297779 | 1.080290148 | 0.280012996 | 0.509515133 | | Coro1b | 2753.538314 | -0.0368265895667674 | 0.0539137510474354 | -0.683064874012679 | 0.494565850037928 | 0.843609829416698 | NM_011778 |
| Eif5 | 8667.705849 | 0.07843978 | 0.072624612 | 1.08007159 | 0.280110302 | 0.509632341 | | Tgif2 | 46.33734617 | -0.0597127948531509 | 0.0874205640206617 | -0.683052042984278 | 0.494573957545959 | 0.843609829416698 | NM_173396 |
| Slc37a2 | 159.9636739 | 0.155326312 | 0.143869258 | 1.079635181 | 0.280304669 | 0.509926094 | | A730056A06Rik | 140.7574483 | 0.0630581420548631 | 0.0924071117797576 | 0.682394907164638 | 0.494989275212999 | 0.844219116090646 | NR_040325,NR_040324 |
| Gspt2 | 1181.382337 | 0.078553371 | 0.072777137 | 1.079368802 | 0.280423353 | 0.510082112 | | Fgf22 | 37.18324962 | 0.0583243240098349 | 0.0854947412175372 | 0.682197795785258 | 0.495113888275634 | 0.84433251245502 | NM_023304 |
| Vti1a | 2643.52934 | -0.078428135 | 0.072698809 | -1.078809077 | 0.280672848 | 0.510416091 | | 3110002H16Rik | 1039.863742 | -0.0372168697622517 | 0.0546059214199437 | -0.681553736197171 | 0.495521177113013 | 0.844369474264346 | NM_029623 |
| Wdr6 | 6999.013636 | 0.113026846 | 0.1047678 | 1.078831912 | 0.280662666 | 0.510416091 | | A230001M10Rik | 173.113857 | 0.0647536198001562 | 0.0950391448291161 | 0.681336305335928 | 0.495658715892336 | 0.844369474264346 | NR_040391 |
| Fgf13 | 3923.910022 | -0.102961604 | 0.095451917 | -1.078675073 | 0.280732601 | 0.510439319 | | Capn10 | 956.1157049 | 0.0529439194560887 | 0.0776451471512919 | 0.681870295807763 | 0.495320969563567 | 0.844369474264346 | NM_011796 |
| Igf2r | 1793.093347 | -0.086967674 | 0.080627702 | -1.078632676 | 0.280751509 | 0.510439319 | | Ccbp2 | 177.8455325 | -0.0646890997951251 | 0.0949415160875796 | -0.681357349881079 | 0.495645402995046 | 0.844369474264346 | NM_021609 |
| Inf2 | 8709.124272 | -0.107652313 | 0.099824856 | -1.078411911 | 0.280849974 | 0.51055843 | | Ddx19a | 728.4905667 | -0.0484424082030267 | 0.0710608913810066 | -0.681702794062819 | 0.495426900354918 | 0.844369474264346 | NM_007916 |
| Echdc3 | 23.31948962 | 0.180587445 | 0.167500472 | 1.078130961 | 0.280975317 | 0.510715963 | | Gja1 | 7234.807969 | 0.0474816032268187 | 0.0696377242732898 | 0.681837376541478 | 0.495341787280674 | 0.844369474264346 | NM_010288 |
| Zfx | 816.6267901 | -0.076564884 | 0.071020332 | -1.078069919 | 0.281002555 | 0.510715963 | | Hlcs | 763.7866106 | -0.0445653790242377 | 0.0653776318332469 | -0.681660956118857 | 0.495453361230157 | 0.844369474264346 | NM_139145 |
| Armc8 | 3112.98283 | -0.058833522 | 0.054593944 | -1.07765656 | 0.281187051 | 0.510991341 | | Klhdc2 | 4890.571383 | 0.0380845085810249 | 0.0558933696321466 | 0.681377931437524 | 0.495632383171645 | 0.844369474264346 | NM_027117 |
| Glra3 | 158.2189142 | 0.169521803 | 0.157373135 | 1.077196579 | 0.281392453 | 0.511269957 | | Ldlr | 1116.443046 | 0.0645330303944266 | 0.0947096821214347 | 0.681377330690264 | 0.495632763199755 | 0.844369474264346 | NM_001252659,NM_001252658,NM_010700 |
| Limd2 | 2551.599673 | -0.094195325 | 0.087447408 | -1.077165437 | 0.281406363 | 0.511269957 | | Cdk18 | 1753.158895 | 0.0524126141452047 | 0.0769583888404781 | 0.681051343913233 | 0.495839002877693 | 0.844379525885271 | NM_008795 |
| Psmb4 | 3436.261866 | -0.067113778 | 0.062311002 | -1.07707749 | 0.281445649 | 0.511281379 | | D3Bwg0562e | 8979.751846 | 0.0496929593538967 | 0.0729553528322436 | 0.681142060516967 | 0.495781605266203 | 0.844379525885271 | NM_177664 |
| Gpr19 | 471.8088751 | 0.137898167 | 0.128066439 | 1.076770527 | 0.281582796 | 0.511426532 | | Snupn | 488.7591427 | 0.0524084680539639 | 0.0769478940779406 | 0.681090349280765 | 0.495814323224076 | 0.844379525885271 | NM_178374 |
| Rbm22 | 1253.948939 | -0.061812634 | 0.057408118 | -1.076722872 | 0.281604091 | 0.511426532 | | Spg7 | 2078.474008 | -0.0488404430247913 | 0.071737113100442 | -0.680825320589747 | 0.495982026289424 | 0.844524078523055 | NM_153176 |
| Rerg | 507.1543271 | 0.140277186 | 0.130287153 | 1.076677035 | 0.281624575 | 0.511426532 | | Sdcbp | 6893.521911 | 0.0347806852605099 | 0.0511057195502658 | 0.680563458778833 | 0.496147755160305 | 0.844707254038746 | NM_001098227,NM_016807 |
| Ndrg1 | 4481.54286 | 0.09146947 | 0.084991009 | 1.076225259 | 0.281826526 | 0.511733293 | | Plekha4 | 113.3162512 | -0.0644042783560808 | 0.0946619169971743 | -0.680361019500622 | 0.496275896541875 | 0.84482640023055 | NM_148927 |
| Copz2 | 145.1021692 | -0.122908114 | 0.114236767 | -1.07590679 | 0.281968945 | 0.511811954 | | 2810047C21Rik1 | 57.0214405 | 0.0553911411460372 | 0.0814370114957423 | 0.68017158450041 | 0.496395822373825 | 0.844826505915688 | NR_015598 |
| Dhx9 | 5403.381649 | -0.074143485 | 0.068911978 | -1.075915793 | 0.281964918 | 0.511811954 | | Ahr | 418.2719965 | -0.0545532980748595 | 0.0802153733764829 | -0.680085322533114 | 0.49645043745672 | 0.844826505915688 | NM_013464 |
| Psma1 | 2499.518693 | -0.067229873 | 0.062482731 | -1.075975264 | 0.281938319 | 0.511811954 | | Nudt6 | 95.5998246 | -0.064359385954643 | 0.0946338591233892 | -0.680088358974428 | 0.496448514389689 | 0.844826505915688 | NM_153561 |
| Fam43a | 494.6485382 | 0.11692086 | 0.108693652 | 1.075691707 | 0.282065157 | 0.511926619 | | Ppp1cc | 8080.254641 | 0.0474448103156814 | 0.0697781992312856 | 0.67993744232954 | 0.496544072379211 | 0.844829674173311 | NM_013636 |
| Appl1 | 4111.574042 | -0.076600982 | 0.071227972 | -1.075433992 | 0.28218047 | 0.511995964 | | Sspn | 578.0131677 | 0.0538502350571139 | 0.0792033236942052 | 0.679898677800737 | 0.49656861889658 | 0.844829674173311 | NM_010656 |
| Card11 | 8.194410806 | 0.16204799 | 0.150679384 | 1.07544898 | 0.282173763 | 0.511995964 | | Eif6 | 1384.007953 | 0.0438580597131303 | 0.0645236668387618 | 0.679720509727496 | 0.496681446998805 | 0.844908800459849 | NM_010579 |
| Smg1 | 6906.342286 | -0.090164405 | 0.083843856 | -1.075384765 | 0.2822025 | 0.511995964 | | Hspb3 | 68.7556155 | -0.0635530779120541 | 0.093509701502553 | -0.679641544041491 | 0.496731457794789 | 0.844908800459849 | NM_019960 |
| Bbs10 | 188.2486504 | -0.112686953 | 0.104818364 | -1.075068804 | 0.282343925 | 0.512132612 | | Dym | 2518.918777 | 0.0361184505554629 | 0.0531796717437358 | 0.679177764193654 | 0.497025234471961 | 0.845038521354618 | NM_027727 |
| Foxk1 | 3050.404742 | 0.061187283 | 0.056919059 | 1.075140607 | 0.282311782 | 0.512132612 | | Fkbp3 | 3872.051503 | 0.0464250497240427 | 0.0683939324702765 | 0.678789010183304 | 0.497271558092832 | 0.845038521354618 | NM_013902 |
| 5730409E04Rik | 3866.729646 | 0.074627373 | 0.069428595 | 1.074879499 | 0.282428681 | 0.512226383 | | Frs3 | 713.8735248 | 0.0555817596945333 | 0.0818950776731613 | 0.678694755212969 | 0.497331290035841 | 0.845038521354618 | NM_144939 |
| Bcl2l1 | 4176.566229 | 0.120729868 | 0.112387303 | 1.074230495 | 0.282719388 | 0.512513653 | | Pde8b | 4126.022225 | -0.033114125841834 | 0.0487759805953861 | -0.678902308833673 | 0.497199762708545 | 0.845038521354618 | NM_172263,NM_001170669 |
| Def6 | 74.48293941 | -0.15722533 | 0.146349723 | -1.074312458 | 0.282682663 | 0.512513653 | | Rab11fip5 | 5190.927647 | 0.0263523420007001 | 0.0388100287700507 | 0.679008566492895 | 0.497132434095974 | 0.845038521354618 | NM_001003955,NM_177466 |
| Prkcq | 449.4749496 | 0.116619672 | 0.10856105 | 1.074231246 | 0.282719051 | 0.512513653 | | Sdhaf1 | 416.0814894 | 0.0572451585592886 | 0.084328663616587 | 0.678833935037348 | 0.497243089333467 | 0.845038521354618 | NM_001033140 |
| Tfpt | 602.2887793 | 0.089195786 | 0.083024959 | 1.074324964 | 0.28267706 | 0.512513653 | | Snrnp27 | 1097.872609 | 0.0454652627128753 | 0.0669237302076428 | 0.679359362842018 | 0.496910191609137 | 0.845038521354618 | NM_025665 |
| Cd63 | 1499.283891 | 0.087795291 | 0.081746637 | 1.073992698 | 0.282825954 | 0.51252694 | | Spata24 | 121.0507671 | -0.0635205284704262 | 0.0935911494767301 | -0.678702300650978 | 0.497326508145463 | 0.845038521354618 | NM_029485,NM_027733 |
| Folh1 | 262.0021354 | 0.115378132 | 0.107423125 | 1.074053015 | 0.282798921 | 0.51252694 | | Trappc6a | 205.4926891 | -0.0595887266010631 | 0.0877631520862674 | -0.678972042190212 | 0.497155776638617 | 0.845038521354618 | NM_025960 |
| Pcnp | 5172.49143 | 0.099841482 | 0.092950561 | 1.07413533 | 0.282762032 | 0.51252694 | | Irf4 | 23.08404237 | -0.0476714026512217 | 0.0702778834169237 | -0.678327239430519 | 0.4975642312963 | 0.845236583301374 | NM_013674 |
| Upk1b | 52.28559402 | 0.178717541 | 0.166424071 | 1.073868345 | 0.282881693 | 0.512567997 | | Kars | 3008.559926 | 0.0373550622290635 | 0.0550672337562907 | 0.678353708384638 | 0.497547452629409 | 0.845236583301374 | NM_001130868,NM_053092 |
| Cds1 | 4328.36351 | 0.080602494 | 0.075080916 | 1.073541696 | 0.283028141 | 0.512599701 | | Il13ra1 | 281.3380115 | -0.0609536933559242 | 0.0899508069996016 | -0.677634397792956 | 0.498000405676126 | 0.845489269103718 | NM_133990 |
| Ceacam9 | 11.68099402 | 0.170083307 | 0.158454844 | 1.073386605 | 0.283097692 | 0.512599701 | | Mecom | 96.23276985 | -0.0641676293599198 | 0.0946755832967158 | -0.677763232351226 | 0.497921826908323 | 0.845489269103718 | NM_021442,NM_007963 |
| Nup43 | 121.2885924 | 0.148127595 | 0.137989845 | 1.073467366 | 0.283061473 | 0.512599701 | | Pex3 | 1068.64928 | -0.0470567137586061 | 0.0694089556864707 | -0.677963143130512 | 0.497795062468548 | 0.845489269103718 | NM_001164195,NM_019961 |
| Sfn | 63.38660011 | -0.146903496 | 0.136824116 | -1.07366669 | 0.282972096 | 0.512599701 | | Shank3 | 10887.63873 | -0.044977041390319 | 0.0663541944943703 | -0.677832678597809 | 0.497877788743694 | 0.845489269103718 | NM_021423 |
| Usp29 | 2474.751376 | -0.136975551 | 0.127597801 | -1.073494603 | 0.283049259 | 0.512599701 | | Zfr | 13881.19244 | 0.0394683361734038 | 0.0582409269838606 | 0.67767355736527 | 0.497978695852899 | 0.845489269103718 | NM_011767 |
| Vps4b | 1655.086822 | 0.068669568 | 0.063971128 | 1.073446259 | 0.283070939 | 0.512599701 | | Arpp21 | 24241.97259 | -0.0531425056878089 | 0.0784566522761267 | -0.677348626866907 | 0.49818478509999 | 0.845625663417497 | NM_001177623,NM_001177620,NM_001177617,NM_001177616,NM_001177615,NM_001177619,NM_001177618,NM_028755,NM_033264 |
| AI317395 | 30.83641502 | 0.177339942 | 0.165256812 | 1.07311729 | 0.283218494 | 0.512698604 | | Dok1 | 72.71964384 | -0.0642169148335098 | 0.0948226718843815 | -0.677231653119945 | 0.498258987552689 | 0.845625663417497 | NM_010070 |
| Fam57a | 180.3291564 | 0.152207956 | 0.141828685 | 1.073181753 | 0.283189576 | 0.512698604 | | Snrpf | 479.975422 | 0.0496608567225039 | 0.0733258444461651 | 0.67726626362966 | 0.498239315421756 | 0.845625663417497 | NM_027246 |
| Klc4 | 828.0229109 | 0.087899926 | 0.081959518 | 1.072479781 | 0.283504591 | 0.513156557 | | Aoc2 | 172.6556978 | -0.0642928788695757 | 0.0950741994219317 | -0.676306382388986 | 0.498846141386392 | 0.84605234220734 | NM_178932 |
| 2410089E03Rik | 2325.516928 | 0.097997917 | 0.091396274 | 1.072230995 | 0.283616292 | 0.513238826 | | Galk1 | 370.2363397 | 0.0571502288427571 | 0.0844738225765133 | 0.676543657448583 | 0.498695537473133 | 0.84605234220734 | NM_016905 |
| Trim37 | 16383.24485 | -0.075344184 | 0.070268336 | -1.072235206 | 0.283614401 | 0.513238826 | | Kctd5 | 928.9931265 | 0.037643771038233 | 0.0556535076476445 | 0.676395300662172 | 0.498789700090581 | 0.84605234220734 | NM_027008 |
| Trp53bp2 | 1246.973607 | 0.07091751 | 0.066175887 | 1.071651822 | 0.283876446 | 0.513649618 | | Pcdh18 | 296.6320954 | 0.0611390107700057 | 0.0904042398487785 | 0.676284772568467 | 0.49885985894299 | 0.84605234220734 | NM_130448 |
| Pear1 | 268.8977448 | 0.092255535 | 0.086110539 | 1.071361721 | 0.284006818 | 0.513825506 | | Tnnt2 | 272.8664888 | -0.058055455350295 | 0.0858297351288878 | -0.676402592447885 | 0.498785071745596 | 0.84605234220734 | NM_001130179,NM_001130178,NM_011619,NM_001130180,NM_001130181,NM_001130177,NM_001130176,NM_001130175,NM_001130174 |
| Ddx52 | 918.7518901 | 0.074924604 | 0.069943116 | 1.071221996 | 0.284069314 | 0.51387913 | | Wdr5b | 187.8663093 | 0.0580048582615312 | 0.0857416617960459 | 0.676724324733548 | 0.498580880021393 | 0.84605234220734 | NM_027113 |
| Cisd2 | 1464.34161 | -0.06102846 | 0.056981551 | -1.071021395 | 0.28415981 | 0.513960079 | | Gins1 | 28.36905246 | -0.0588605239009754 | 0.0870511439791631 | -0.676160257182427 | 0.498938902426837 | 0.846087613431267 | NM_001163476,NM_027014 |
| Trim32 | 7948.22561 | 0.070500183 | 0.065828044 | 1.070974902 | 0.284180715 | 0.513960079 | | Adra1d | 1108.215432 | -0.0593512227896283 | 0.0877977270936758 | -0.67599953614179 | 0.499040939653175 | 0.846161863907398 | NM_013460 |
| Wasl | 4406.545273 | 0.09722989 | 0.090792994 | 1.070896393 | 0.284216018 | 0.513963933 | | AA986860 | 173.6189217 | 0.0620565656934873 | 0.0920086597141753 | 0.67446440244066 | 0.500016109983399 | 0.846241311321458 | NM_177604 |
| Ufsp1 | 292.5084045 | -0.111328534 | 0.10398497 | -1.070806081 | 0.284256632 | 0.513977391 | | Cdkn2b | 22.59345146 | -0.0539936664144951 | 0.0800034346585473 | -0.674891854892715 | 0.499744475754808 | 0.846241311321458 | NM_007670 |
| Rdh12 | 73.72730301 | 0.139705925 | 0.130480182 | 1.070706089 | 0.284301604 | 0.513998723 | | Dctn3 | 2403.652134 | 0.0317613420436781 | 0.0470896983627368 | 0.674485994771451 | 0.500002386774959 | 0.846241311321458 | NM_016890,NM_001159565 |
| Eml4 | 1497.282309 | 0.05923744 | 0.05534727 | 1.070286582 | 0.284490332 | 0.514279922 | | Dnaic2 | 124.140422 | -0.0642635374978847 | 0.0950995594175458 | -0.675750107481866 | 0.499199316552704 | 0.846241311321458 | NM_001034878 |
| Megf8 | 9102.421395 | -0.077508602 | 0.072429779 | -1.070120643 | 0.284565008 | 0.514354904 | | Fads3 | 1467.440238 | 0.0458529178494067 | 0.067984794875386 | 0.674458427556539 | 0.500019907412369 | 0.846241311321458 | NM_021890 |
| Xrra1 | 54.1838493 | 0.158590595 | 0.148221634 | 1.069955787 | 0.28463921 | 0.514429012 | | Itsn2 | 2384.277964 | -0.043012037346294 | 0.0637455035112193 | -0.674746217020998 | 0.499837015801784 | 0.846241311321458 | NM_011365,NM_001198968,NM_001198969 |
| Add1 | 18180.0682 | -0.055951844 | 0.052314653 | -1.069525293 | 0.284833038 | 0.514480389 | | Kcnma1 | 4212.108782 | 0.0449952259425438 | 0.0666061319796197 | 0.675541794805191 | 0.499331607034563 | 0.846241311321458 | [illegible]M_001253374,NM_001253359,NM_001253372,NM_001253371,NM_001253370,NM_010610,NM_001253368,NM_001253369,NM_001253364,NM_0012533[illegible] |
| Gdpd3 | 18.75690575 | 0.179845756 | 0.168120948 | 1.069740316 | 0.284736214 | 0.514480389 | | Kctd9 | 580.7946788 | -0.0495532324828836 | 0.0733883777181192 | -0.675219080033985 | 0.499536586160486 | 0.846241311321458 | NM_001111028,NM_134073 |
| Hgs | 3150.125042 | 0.063695475 | 0.059548653 | 1.069637539 | 0.284782491 | 0.514480389 | | Limch1 | 5200.64355 | -0.0347118178262633 | 0.0513750290454587 | -0.675655439446056 | 0.499259433869933 | 0.846241311321458 | NM_001001980,NM_001256122 |
| Lamtor4 | 845.064462 | -0.100972698 | 0.094409015 | -1.069523905 | 0.284833663 | 0.514480389 | | Qsox2 | 902.656472 | 0.0435540233080652 | 0.0645580164091899 | 0.674649342259921 | 0.49989857622117 | 0.846241311321458 | NM_153559 |
| R3hcc1l | 552.8045638 | 0.08412809 | 0.07863886 | 1.069803026 | 0.28470798 | 0.514480389 | | Snapc5 | 674.8120294 | 0.0504515177409045 | 0.0747509112621036 | 0.674928464269865 | 0.499721215148822 | 0.846241311321458 | NM_183316 |
| 0610007P14Rik | 1072.632876 | -0.072347914 | 0.06765099 | -1.069428755 | 0.284876516 | 0.514547813 | | Sumo1 | 3084.116667 | 0.0392449740058392 | 0.0581082138770876 | 0.675377393097152 | 0.499436024680893 | 0.846241311321458 | NM_009460 |
| Cpa2 | 42.80398667 | 0.172116622 | 0.161017526 | 1.068930977 | 0.285100772 | 0.514782819 | | Ttyh2 | 1928.785964 | 0.056872673799772 | 0.0842556567640529 | 0.67500125195198 | 0.499674969535768 | 0.846241311321458 | NM_053273 |
| Rgs7bp | 11765.7414 | -0.089389445 | 0.083619493 | -1.069002473 | 0.285068555 | 0.514782819 | | Wash | 1275.420718 | -0.0427769219717613 | 0.0634205544055446 | -0.674496184600075 | 0.499995910603178 | 0.846241311321458 | NM_026833,NM_001037757 |
| Elp5 | 2080.900377 | -0.063857343 | 0.059752801 | -1.068692045 | 0.285208457 | 0.51491725 | | Zfp553 | 1013.253898 | 0.0415234848565949 | 0.0615094024761153 | 0.675075386361998 | 0.499627870615417 | 0.846241311321458 | NM_146201 |
| Creb3l2 | 334.3391402 | 0.083752094 | 0.078377192 | 1.068577144 | 0.285260145 | 0.514950564 | | Zfp787 | 508.9485262 | -0.0532091132427902 | 0.0787985011766586 | -0.675255397605857 | 0.499513516054167 | 0.846241311321458 | NM_001013012 |
| Arf3 | 58150.79269 | -0.069143882 | 0.064709414 | -1.068055042 | 0.285495684 | 0.515019946 | | Nacc2 | 6757.798505 | 0.0321008818058738 | 0.0475602662042977 | 0.67435005597702 | 0.500088466935193 | 0.846258745421786 | NM_001037098,NM_026495 |
| Cldn10 | 1020.082093 | -0.093145657 | 0.087199015 | -1.068196214 | 0.285432012 | 0.515019946 | | Ido1 | 129.1848271 | -0.0641150248068245 | 0.0950983854710739 | -0.674196775115268 | 0.500186219606382 | 0.846268337560688 | NM_008324 |
| Dohh | 1667.43535 | -0.095377636 | 0.089304841 | -1.068000727 | 0.285520184 | 0.515019946 | | Lonrf3 | 1057.417065 | 0.0556892288687401 | 0.0826392617360346 | 0.673883426721528 | 0.500385429664653 | 0.846268337560688 | NM_028894 |
| Gins4 | 488.1354333 | -0.0843749 | 0.078985556 | -1.06823202 | 0.285415865 | 0.515019946 | | Mir22hg | 760.4971644 | -0.0506945983278604 | 0.0752104455767014 | -0.674036670560086 | 0.500288000181743 | 0.846268337560688 | NR_030711 |
| Gm5113 | 839.0118194 | -0.07231284 | 0.067687511 | -1.068333557 | 0.285370077 | 0.515019946 | | Tmem47 | 3358.238133 | 0.0401824323217571 | 0.059627936967331 | 0.673886006550457 | 0.500383789287482 | 0.846268337560688 | NM_138751 |
| Ly75 | 67.43123078 | -0.158075087 | 0.148013683 | -1.067976174 | 0.285531259 | 0.515019946 | | Zscan12 | 307.6519267 | 0.0499384951213793 | 0.0740764449381292 | 0.674148106906175 | 0.50021715748264 | 0.846268337560688 | NM_016684 |
| Prkaa1 | 1167.651232 | 0.074210211 | 0.069468757 | 1.068253042 | 0.285406384 | 0.515019946 | | Cnksr3 | 215.7862117 | 0.0620256193285309 | 0.0920846417673584 | 0.673571815430761 | 0.500583576883952 | 0.846307848907854 | NM_172546 |
| Gdf5 | 11.03954688 | 0.170574913 | 0.159739921 | 1.067828956 | 0.285597674 | 0.515055081 | | Gldc | 1296.12851 | -0.0418292272902898 | 0.0620987476779471 | -0.673592123100808 | 0.500570662380709 | 0.846307848907854 | NM_138595 |
| Parp14 | 215.4161879 | 0.140406133 | 0.131492811 | 1.067785627 | 0.285617223 | 0.515055081 | | Ythdf3 | 2216.958067 | 0.0499785591212111 | 0.0741833966443098 | 0.673716246857039 | 0.500491734508094 | 0.846307848907854 | NM_001145919,NM_172677,NR_027375 |
| Zfp963 | 148.2951088 | 0.122142118 | 0.114401099 | 1.067665596 | 0.285671383 | 0.515092797 | | 0610040J01Rik | 156.8869441 | -0.0632837554451583 | 0.0941367180432693 | -0.672253683372203 | 0.501422210340625 | 0.846601651291981 | NM_029554 |
| Sirt3 | 2202.687747 | -0.078015538 | 0.073106792 | -1.067144871 | 0.285906212 | 0.51545661 | | BC024139 | 62.59192886 | -0.0611476571390867 | 0.0909645506999321 | -0.672214139118837 | 0.501447381020365 | 0.846601651291981 | NM_001142968 |
| 9030612E09Rik | 43.50178527 | 0.170240496 | 0.159605634 | 1.066632123 | 0.286137989 | 0.515760851 | | Dll1 | 221.9250356 | -0.0576721434360309 | 0.0857645511144778 | -0.672447330355063 | 0.501298959963849 | 0.846601651291981 | NM_007865 |
| Npl | 156.6260435 | 0.131391557 | 0.123189201 | 1.066583398 | 0.28616 | 0.515760851 | | Erbb2ip | 3729.267965 | -0.0494045387287538 | 0.0734933298969966 | -0.672231599765529 | 0.501436266890462 | 0.846601651291981 | NM_001005868,NM_021563 |
| Susd2 | 614.9995738 | -0.135482097 | 0.127028346 | -1.066550113 | 0.286175038 | 0.515760851 | | Frat2 | 215.7218289 | 0.0587480081836858 | 0.0874032842933348 | 0.672148748856178 | 0.501489004651363 | 0.846601651291981 | NM_177603 |
| Lss | 908.2744817 | 0.074338016 | 0.06973654 | 1.065983715 | 0.286431004 | 0.516162127 | | Grtp1 | 103.7713524 | -0.0623717543512675 | 0.0927434811622307 | -0.67251901232998 | 0.501253340650759 | 0.846601651291981 | NM_025768 |
| B330016D10Rik | 65.98139691 | 0.157783602 | 0.148072018 | 1.065586895 | 0.286610426 | 0.516185277 | | Helz2 | 337.0376756 | -0.0582478653334284 | 0.086664458630067 | -0.672107877371776 | 0.501515021985922 | 0.846601651291981 | NM_183162 |
| Fahd1 | 435.695269 | -0.098830746 | 0.092728105 | -1.065812213 | 0.286508539 | 0.516185277 | | Rabl6 | 6754.425239 | 0.0301582043403436 | 0.0448580339283136 | 0.672303302203093 | 0.501390627944582 | 0.846601651291981 | NM_001024616 |
| Gramd3 | 644.2934578 | -0.091051577 | 0.085438945 | -1.065691724 | 0.28656302 | 0.516185277 | | Sema5a | 1891.247813 | 0.0440137510973697 | 0.065409885454444 | 0.67289142599132 | 0.501016367293648 | 0.846601651291981 | NM_009154 |
| Wtip | 178.795008 | 0.121683882 | 0.114189306 | 1.065632905 | 0.286589619 | 0.516185277 | | Slc35f5 | 745.3700354 | -0.0404960513827011 | 0.0602138445032525 | -0.672537216594992 | 0.50124175557702 | 0.846601651291981 | NM_028787 |
| Zic2 | 428.9986134 | -0.132173997 | 0.124020694 | -1.065741472 | 0.286540525 | 0.516185277 | | Trmt6 | 1162.970233 | -0.0440428303505594 | 0.0654893615896639 | -0.672518853161498 | 0.501253441945154 | 0.846601651291981 | NM_175113 |
| Six3 | 432.6716726 | -0.162211465 | 0.152250053 | -1.065427974 | 0.286682303 | 0.516254719 | | Ttll13 | 41.10665815 | -0.0587033313160603 | 0.087269057595526 | -0.672670622709575 | 0.501156861137266 | 0.846601651291981 | NM_177765 |
| Pkn3 | 61.75583106 | 0.153954202 | 0.144559765 | 1.064986523 | 0.286882029 | 0.516554346 | | Zcchc16 | 90.69428237 | -0.0638707051724926 | 0.0949349628793531 | -0.672783800986596 | 0.501084844875993 | 0.846601651291981 | NM_001033795 |
| Ormdl2 | 193.0846536 | -0.139136802 | 0.130760902 | -1.064055075 | 0.287303751 | 0.517169742 | | Trmt10a | 245.7641711 | -0.0549900396092016 | 0.0818295191308895 | -0.67200736596341 | 0.501579007016631 | 0.846611277704344 | NM_175389 |
| Rab20 | 11.744965 | 0.17216067 | 0.161803544 | 1.064010504 | 0.287323941 | 0.517169742 | | Cnot11 | 1286.641683 | 0.0426768333276083 | 0.0635220739807663 | 0.671842568309882 | 0.501683925683115 | 0.8466899854157 | NM_028043 |
| Tacc1 | 9975.023693 | -0.068847593 | 0.064705356 | -1.064016921 | 0.287321034 | 0.517169742 | | Trappc12 | 3437.1841 | 0.0288115586455355 | 0.0429077940977977 | 0.671476109442184 | 0.501917273891728 | 0.846985399692292 | NM_001161410,NM_001161411,NM_178811 |
| Rps13 | 3462.102145 | -0.090835222 | 0.085384112 | -1.063842205 | 0.287400188 | 0.517246895 | | Mcm9 | 222.9135382 | -0.0561605955985328 | 0.0836587568823193 | -0.671305643204005 | 0.502025840409577 | 0.847070200695726 | NM_027830 |
| Arrb1 | 17019.29764 | -0.063594399 | 0.05980773 | -1.063314041 | 0.28763956 | 0.517531236 | | Zfp583 | 215.8113838 | 0.0568555591098784 | 0.0847401141110171 | 0.67094070507614 | 0.502258303930482 | 0.847364009627663 | NM_001033249 |
| G3bp1 | 2210.560185 | -0.062266945 | 0.058559585 | -1.06330918 | 0.287641763 | 0.517531236 | | Asb8 | 1880.713549 | 0.0320088528851326 | 0.0477389228096108 | 0.670498012969169 | 0.502540372705392 | 0.847394259983065 | NM_001170711,NM_001170710,NM_030121 |
| Pes1 | 1671.896537 | 0.060019125 | 0.056447546 | 1.063272518 | 0.287658384 | 0.517531236 | | Ephb3 | 1082.27588 | 0.0548237565706179 | 0.0817988492859095 | 0.670226501341037 | 0.502713421368619 | 0.847394259983065 | NM_010143 |
| AU019823 | 610.2404072 | 0.111273904 | 0.104674243 | 1.063049527 | 0.287759492 | 0.517653032 | | Golgb1 | 4970.274831 | -0.0441871622588608 | 0.0659422606014728 | -0.670088678425955 | 0.502801261654553 | 0.847394259983065 | NM_030035 |
| Degs2 | 101.5245377 | 0.144082072 | 0.135588743 | 1.062640368 | 0.287945073 | 0.517926743 | | Pi4k2a | 3997.779369 | 0.0296429906841364 | 0.0442177947116263 | 0.670386003586523 | 0.502611754656033 | 0.847394259983065 | NM_145501 |
| Mpdz | 2635.595138 | -0.060536848 | 0.056972507 | -1.062562481 | 0.287980409 | 0.517930175 | | Ppapdc1a | 182.6902633 | 0.0608588925624455 | 0.0907903723610559 | 0.67032319594881 | 0.502651783395719 | 0.847394259983065 | NM_001080963 |
| Lmo7 | 4185.847381 | -0.102067498 | 0.096093763 | -1.062165693 | 0.288160471 | 0.518193865 | | Rab4b | 2186.126537 | 0.0373882891481215 | 0.0557710090638808 | 0.670389325488024 | 0.502609637579386 | 0.847394259983065 | NM_029391 |
| Wdr45b | 2729.671626 | -0.081151493 | 0.076411538 | -1.062031918 | 0.288221195 | 0.518242915 | | Rsbn1 | 1095.541202 | 0.0501745503280298 | 0.0748544589513615 | 0.670294743037711 | 0.502669917639432 | 0.847394259983065 | NM_172684 |
| Cbx1 | 675.7151607 | -0.091285221 | 0.08598073 | -1.061693947 | 0.288374648 | 0.518458668 | | Smim1 | 490.1394891 | 0.0557918564285811 | 0.0832504677623686 | 0.670168684070751 | 0.502750264511396 | 0.847394259983065 | NM_001163722,NM_001163721 |
| Dbp | 3109.427239 | 0.144451221 | 0.136101782 | 1.061347022 | 0.288532224 | 0.518626383 | | Wbscr27 | 368.8033623 | -0.0482471662412719 | 0.0719538502389704 | -0.670529319571297 | 0.502520422421669 | 0.847394259983065 | NM_024479 |
| Ect2 | 27.01609526 | -0.175421427 | 0.165282786 | -1.061341176 | 0.288534879 | 0.518626383 | | Psip1 | 6134.342547 | 0.0401810126508659 | 0.0599739488283912 | 0.66997443783199 | 0.50287408554473 | 0.84741867363898 | NM_133948 |
| Ppp2r5b | 5571.73657 | 0.063500555 | 0.059836661 | 1.061231589 | 0.288584667 | 0.518655704 | | Gpr155 | 3323.475644 | 0.0555924222910272 | 0.0830190371590057 | 0.669634630723932 | 0.503090732211158 | 0.847567617625473 | NM_001276444,NM_001080707,NM_001276443,NM_001190297 |
| Gm266 | 69.11805724 | -0.16209379 | 0.152888495 | -1.060209213 | 0.289049431 | 0.519370507 | | Rab5a | 4012.876439 | 0.0490569567240716 | 0.0732633170319976 | 0.669597811175353 | 0.503114209760589 | 0.847567617625473 | NM_025887 |
| Wbscr22 | 794.8289508 | 0.083165553 | 0.078438473 | 1.060264806 | 0.289024146 | 0.519370507 | | Top1mt | 552.9500387 | -0.0484198944532706 | 0.0723158541766634 | -0.669561260176554 | 0.503137516644944 | 0.847567617625473 | NM_028404 |
| Atp2b3 | 4270.213891 | 0.076759111 | 0.072407092 | 1.06010487 | 0.289096892 | 0.519395553 | | 1700007K13Rik | 77.13125273 | 0.0634502027159469 | 0.0950539593877996 | 0.667517724928046 | 0.504441491652754 | 0.847807552547121 | NM_027040 |
| Chi3l1 | 377.5241923 | -0.136415359 | 0.128703923 | -1.059916093 | 0.289182774 | 0.519489611 | | Acaca | 4608.817875 | 0.0368320322329787 | 0.0551037445322493 | 0.668412510721987 | 0.503870311717636 | 0.847807552547121 | NM_133360 |
| Cntln | 573.6743929 | -0.086474006 | 0.081634893 | -1.059277507 | 0.289473415 | 0.519937033 | | Acad9 | 1818.043312 | 0.0298014756405534 | 0.0446436456382669 | 0.667541264036213 | 0.504426461264287 | 0.847807552547121 | NM_172678 |
| Mkl2 | 12851.93544 | -0.09397182 | 0.088717825 | -1.059221417 | 0.289498953 | 0.519937033 | | Arhgap42 | 693.0421104 | -0.0524798523360166 | 0.078571632366651 | -0.667923660591637 | 0.504182323079521 | 0.847807552547121 | NM_027823 |
| Alox5ap | 113.7988001 | -0.136805406 | 0.129224252 | -1.058666651 | 0.289751621 | 0.520046504 | | B330016D10Rik | 65.98635061 | -0.0636090847999226 | 0.0950708380073424 | -0.669070412263936 | 0.503450562952496 | 0.847807552547121 | NR_030695 |
| Brip1 | 36.23683618 | -0.171314781 | 0.161810941 | -1.058734229 | 0.289720835 | 0.520046504 | | Ctla2b | 30.58074054 | -0.0566590918044933 | 0.0847913171140278 | -0.668156590890819 | 0.504033693728279 | 0.847807552547121 | NM_007797,NM_001145801 |
| Dennd1c | 63.65636058 | 0.168123752 | 0.158762198 | 1.058965889 | 0.289615315 | 0.520046504 | | Dach1 | 560.4255589 | 0.06265911156759 | 0.0937415671239869 | 0.668423981910973 | 0.503862991385084 | 0.847807552547121 | NM_001038610,NM_007826 |
| Fam13c | 2549.43466 | -0.067773954 | 0.064028414 | -1.058498089 | 0.289828422 | 0.520046504 | | Eif2s2 | 2672.910255 | 0.0417805789920129 | 0.0625255556321153 | 0.668216036940789 | 0.503995700059408 | 0.847807552547121 | NM_026030 |
| Fam196a | 472.6069251 | -0.12340581 | 0.116585074 | -1.058504367 | 0.289825561 | 0.520046504 | | Epb4.1l1 | 26497.45911 | 0.0293290877759838 | 0.0439070534505699 | 0.667981234700757 | 0.504145570977951 | 0.847807552547121 | NM_013510,NM_001003815,NM_001006664 |
| Fam69b | 1769.539417 | -0.070297 | 0.066407562 | -1.058569206 | 0.289796017 | 0.520046504 | | Hmbox1 | 1026.298303 | -0.0438751449701981 | 0.0655769217184679 | -0.669063808126905 | 0.503454775550346 | 0.847807552547121 | NM_177338 |
| Tet3 | 2418.117423 | -0.081618318 | 0.077089285 | -1.058750478 | 0.289713432 | 0.520046504 | | Mkrn3 | 22.78616889 | 0.0533539130522891 | 0.0799222865860282 | 0.667572404786731 | 0.504406577374383 | 0.847807552547121 | NM_011746 |
| Zfp266 | 1438.904755 | 0.07433462 | 0.070205908 | 1.058808617 | 0.289686948 | 0.520046504 | | Mob1b | 426.929653 | -0.0582152056323171 | 0.0870734187606617 | -0.668576087906617 | 0.503765930339538 | 0.847807552547121 | NM_026735 |
| U2surp | 2567.306888 | -0.067178764 | 0.063487241 | -1.058145909 | 0.289988927 | 0.520274251 | | Mrpl32 | 442.5518272 | 0.049330230966712 | 0.0738151961789431 | 0.668293705365565 | 0.503946130587325 | 0.847807552547121 | NM_029271 |
| Chn2 | 1879.531631 | -0.122778395 | 0.116039912 | -1.058070389 | 0.290023353 | 0.520275771 | | Nthl1 | 93.14107384 | 0.0631127227678407 | 0.0944767598690922 | 0.668023785482168 | 0.504118409677301 | 0.847807552547121 | NM_008743 |
| Ywhah | 32788.2713 | -0.068562843 | 0.064818046 | -1.057773988 | 0.290158494 | 0.520457943 | | Rhoc | 460.8635305 | -0.0530583398173589 | 0.0794750763692623 | -0.667609799716779 | 0.504382700632493 | 0.847807552547121 | NM_007484 |
| Rnd1 | 861.147822 | 0.118046492 | 0.111614517 | 1.057626695 | 0.290225667 | 0.520518172 | | Sass6 | 212.4065278 | -0.0588862898591372 | 0.0882222392707572 | -0.667476708207475 | 0.504467682556957 | 0.847807552547121 | NM_028349 |
| Fgf11 | 234.6615145 | -0.118671515 | 0.112260806 | -1.057105499 | 0.290463442 | 0.520884325 | | Slc20a2 | 2429.07958 | 0.0377883237488654 | 0.0566019440955031 | 0.667613769327538 | 0.504380166062628 | 0.847807552547121 | NM_011394 |
| BC018242 | 6584.968432 | 0.079277584 | 0.075007528 | 1.056928365 | 0.290544282 | 0.520968997 | | Slit2 | 1351.024103 | 0.0495641803838701 | 0.0741714385378622 | 0.668238089498144 | 0.503981625434582 | 0.847807552547121 | NM_178804 |
| Zfp12 | 1427.247254 | 0.059241508 | 0.05605576 | 1.056831766 | 0.290588374 | 0.520987765 | | Ubap2 | 2236.191356 | -0.0335504829377845 | 0.0501462936292511 | -0.669052097565472 | 0.503462245442012 | 0.847807552547121 | NM_026872 |
| Amdhd2 | 388.4523223 | 0.12473543 | 0.118045834 | 1.056669481 | 0.290662458 | 0.521060295 | | Vps37a | 2369.623989 | -0.0418986796906236 | 0.0627207507801401 | -0.668019422112696 | 0.504121194897091 | 0.847807552547121 | NM_033560 |
| 1700024F13Rik | 14.55627125 | -0.176283658 | 0.166882212 | -1.056335819 | 0.290814817 | 0.52115253 | | Zdhhc23 | 313.2306631 | -0.0506300856496687 | 0.0758597126399512 | -0.667417313982871 | 0.504505609542704 | 0.847807552547121 | NM_001007460 |
| Asb4 | 43.67168318 | -0.170024741 | 0.160955337 | -1.056347336 | 0.290809557 | 0.52115253 | | Acbd5 | 2832.858341 | 0.0450230368977055 | 0.067564817985228 | 0.666368063147141 | 0.505175870878634 | 0.84818327275139 | NM_001102437,NM_001102436,NM_028793,NM_001102438 |
| Stxbp5 | 5076.964502 | -0.080214205 | 0.075930928 | -1.05641017 | 0.290780861 | 0.52115253 | | Alg14 | 298.5990896 | 0.0509455649424662 | 0.0764597300519403 | 0.666305843714829 | 0.505215631162327 | 0.84818327275139 | NM_024178 |
| D14Abb1e | 1266.402198 | -0.085404898 | 0.080868753 | -1.056091378 | 0.290926942 | 0.521292322 | | Cyth3 | 1738.37911 | 0.0331040728956721 | 0.0496577890623427 | 0.666644116074353 | 0.50499948268562 | 0.84818327275139 | NM_011182,NM_001163548 |
| Erc2 | 6930.720012 | -0.079955428 | 0.075717528 | -1.055969876 | 0.290981978 | 0.521331495 | | Eci2 | 1204.114626 | 0.0467818999858805 | 0.0701821154811151 | 0.666578652769006 | 0.505041308471662 | 0.84818327275139 | NR_073427,NM_011868,NM_001110331 |
| Zfp334 | 897.2841765 | -0.060316864 | 0.057131245 | -1.055759665 | 0.29107803 | 0.521443288 | | Hook3 | 3095.875678 | -0.0454706440488533 | 0.0682486949133061 | -0.666249289399791 | 0.505251772859818 | 0.84818327275139 | NM_207659 |
| Itsn2 | 2250.135015 | 0.080215847 | 0.075986424 | 1.055660247 | 0.291123465 | 0.52146439 | | Mettl8 | 459.1805188 | -0.0475855619228849 | 0.0713574863987076 | -0.666861521116399 | 0.504860591449538 | 0.84818327275139 | NM_145524,NM_001110512 |
| Ccdc67 | 10.1339444 | 0.167614756 | 0.158813612 | 1.055420729 | 0.291232946 | 0.521539907 | | Pfdn5 | 3856.703368 | 0.0382361235231696 | 0.0573905106292481 | 0.666244699758486 | 0.5052547059836 | 0.84818327275139 | NM_027044 |
| Plekhd1 | 153.0634034 | 0.133579546 | 0.126563186 | 1.055437609 | 0.29122523 | 0.521539907 | | Rtn2 | 3668.26897 | 0.0343880208816896 | 0.0515727919836894 | 0.666786100945733 | 0.504908772047323 | 0.84818327275139 | NM_001025364,NM_013648 |
| Col10a1 | 14.92005335 | -0.174585196 | 0.16546099 | -1.055156239 | 0.291353876 | 0.521616167 | | Trove2 | 1824.928792 | 0.0510137258829595 | 0.0765085503938907 | 0.666771565012334 | 0.504918058327011 | 0.84818327275139 | NM_013835 |
| Arl6ip4 | 1037.535121 | 0.064236963 | 0.060887737 | 1.055006578 | 0.291422316 | 0.521743749 | | Gng5 | 434.9680834 | 0.0501238576478327 | 0.075267274504342 | 0.665944900727623 | 0.505446319394409 | 0.848406891093505 | NM_010318 |
| Clec4a3 | 31.03558248 | 0.171241203 | 0.162321496 | 1.054950869 | 0.291447796 | 0.521743749 | | Dmrtc1a | 196.1539943 | 0.0623136518809416 | 0.0936111876027149 | 0.665664580274665 | 0.505625181887776 | 0.848413569693639 | NM_027591,NM_029378,NM_001038616 |
| 0610040F04Rik | 44.09892936 | 0.162148361 | 0.153739248 | 1.054697243 | 0.291563813 | 0.521749609 | | Fbxl20 | 2979.165324 | -0.0346993138322705 | 0.0521273213487341 | -0.665664625276456 | 0.505625489117103 | 0.848413569693639 | NM_028149 |
| 1700123O20Rik | 548.3879099 | -0.090792832 | 0.086081968 | -1.054725324 | 0.291550966 | 0.521749609 | | Zik1 | 758.14953 | 0.0523397431422459 | 0.0786263259440473 | 0.66567700809753 | 0.505617525939937 | 0.848413569693639 | NM_009577 |
| Dalrd3 | 2156.714379 | 0.11693443 | 0.110868521 | 1.054712633 | 0.291556772 | 0.521749609 | | C1qbp | 1888.3787 | 0.0449323644128956 | 0.0676262407489086 | 0.664422033744656 | 0.506420234444348 | 0.848908639017717 | NM_007573 |
| Pafah1b1 | 24073.3691 | -0.062463313 | 0.059226622 | -1.054649259 | 0.291585766 | 0.521749609 | | Heatr5a | 647.5831416 | -0.0516600634374892 | 0.0777413015770057 | -0.664512458494384 | 0.506362377799398 | 0.848908639017717 | NM_177171 |
| H2-T24 | 76.18856033 | 0.174491835 | 0.165514419 | 1.054239475 | 0.291773291 | 0.521904338 | | Irgq | 618.786199 | 0.0366671600888878 | 0.0551596512968587 | 0.66474624838289 | 0.506212937249398 | 0.848908639017717 | NM_153134 |
| Pcdhga5 | 446.0402153 | 0.098277424 | 0.093211924 | 1.054343908 | 0.291725493 | 0.521904338 | | Lancl1 | 8647.956352 | 0.0342808062095808 | 0.0515823806149113 | 0.664585475502529 | 0.506315661721551 | 0.848908639017717 | NM_021295,NM_001190984,NM_001190985 |
| Tspyl3 | 1421.838961 | -0.070037599 | 0.066429928 | -1.05430792 | 0.291741964 | 0.521904338 | | Mib2 | 4235.614599 | -0.0487029874597354 | 0.0732373898437531 | -0.665001682387076 | 0.506049417011804 | 0.848908639017717 | NM_001256107,NM_145124,NM_001256108 |
| Rpl4 | 16421.58382 | 0.070496549 | 0.066916596 | 1.053498723 | 0.292112479 | 0.522450739 | | Rnf217 | 434.1666649 | 0.0554369590684002 | 0.0834153105203872 | 0.664589734457094 | 0.506312936924284 | 0.848908639017717 | NM_001146349 |
| Gsr | 943.8385654 | -0.082127873 | 0.07797622 | -1.053242545 | 0.292229844 | 0.522600324 | | Tlk1 | 5023.081272 | 0.045802673670955 | 0.0689042939395324 | 0.664728873227401 | 0.506223922855412 | 0.848908639017717 | NM_172664 |
| Unc13a | 15275.00453 | -0.10925496 | 0.103758155 | -1.052977091 | 0.292351492 | 0.522757532 | | Ttc17 | 1965.370352 | 0.0353976089724518 | 0.053230099782168 | 0.664992346760732 | 0.506055388140709 | 0.848908639017717 | NM_183106 |
| Fam219a | 6435.63558 | 0.053096704 | 0.050434657 | 1.052782112 | 0.292440865 | 0.522857001 | | Zcchc3 | 381.3339381 | 0.0469433067396216 | 0.0706572146157371 | 0.66438094106764 | 0.506446527999968 | 0.848908639017717 | NM_175126 |
| Ift122 | 1520.209681 | 0.066857082 | 0.063534825 | 1.052290336 | 0.292666364 | 0.523199798 | | Cav1 | 518.6253231 | -0.0527721262997154 | 0.0794579294632484 | -0.664151792731071 | 0.506593163995474 | 0.848943696869019 | NM_001243064,NM_007616 |
| Cndp2 | 1841.144209 | 0.053166481 | 0.05053673 | 1.052036422 | 0.292782839 | 0.523347637 | | Mis12 | 1069.869418 | 0.0458410971990238 | 0.069026295738465 | 0.66411063651325 | 0.506619502937496 | 0.848943696869019 | NM_025993 |
| Mybpc3 | 26.50999208 | 0.175763283 | 0.167115523 | 1.051747197 | 0.292915551 | 0.52352446 | | Rprd1a | 5534.724745 | 0.0412573248904159 | 0.0621275755807981 | 0.664074277882612 | 0.506642772143572 | 0.848943696869019 | NM_144861 |
| Gab3 | 53.76478252 | 0.159325663 | 0.15155395 | 1.051280179 | 0.293129928 | 0.523847186 | | Enpp4 | 1538.339634 | 0.0490578475795356 | 0.0738846118331293 | 0.663979174585559 | 0.506703640079892 | 0.848947759607901 | NM_199016 |
| Ermn | 2191.097111 | 0.135725789 | 0.129128193 | 1.051093383 | 0.293215703 | 0.523879624 | | Ccdc74a | 589.9788752 | -0.0558385343385339 | 0.08414411603312 | -0.663605929576291 | 0.506942561194565 | 0.848958234808791 | NM_001166164 |
| Hspa2 | 1295.38218 | 0.094765814 | 0.090154215 | 1.051152337 | 0.29318863 | 0.523879624 | | Fbxw11 | 6398.42148 | 0.0318928847857519 | 0.0480580290472839 | 0.663630322852291 | 0.50692694497433 | 0.848958234808791 | NM_134015,NM_001271347,NM_001271348,NM_001271349 |
| Nop2 | 907.9924419 | 0.082503027 | 0.078498328 | 1.051016353 | 0.293251079 | 0.523882419 | | Kcnk9 | 664.0534304 | -0.047461507435642 | 0.0715736936107283 | -0.66311384869661 | 0.507257641821005 | 0.848958234808791 | NM_001033876 |
| Prdm1 | 59.88455048 | -0.176371884 | 0.167951493 | -1.050135852 | 0.293655657 | 0.524423785 | | Mapk8ip3 | 23485.13487 | 0.0307642199697465 | 0.0463983513384448 | 0.663045541108611 | 0.507301387470373 | 0.848958234808791 | NM_001163450,NM_001163451,NM_013931,NM_001163453,NM_001163447,NM_001163449,NM_001163448 |
| Rab8a | 1261.907444 | 0.064183041 | 0.061118371 | 1.05014319 | 0.293652284 | 0.524423785 | | Mtmr4 | 4922.569922 | 0.0369135383725607 | 0.0556641168311831 | 0.663147831564656 | 0.507235879199063 | 0.848958234808791 | NM_133215 |
| Timm10b | 508.8226847 | -0.10536525 | 0.100325681 | -1.050232091 | 0.293611419 | 0.524423785 | | Myh1 | 29.11644539 | 0.059800305802647 | 0.0901132751668741 | 0.663612610815746 | 0.506938283887877 | 0.848958234808791 | NM_030679 |
| Lurap1 | 524.0651879 | -0.101821913 | 0.097028869 | -1.04939812 | 0.293994923 | 0.524969153 | | Ppcdc | 488.4705131 | -0.0517796532911004 | 0.0780774290397084 | -0.663183379985098 | 0.507213114522518 | 0.848958234808791 | NM_176831 |
| Ccne1 | 617.4708187 | -0.072640062 | 0.069250758 | -1.048942496 | 0.294204585 | 0.525282996 | | Rnf152 | 939.9366662 | 0.0495813749234591 | 0.074769871258681 | 0.663119704351538 | 0.507253891825781 | 0.848958234808791 | NM_178779,NM_001160368 |
| Ufc1 | 1879.275163 | -0.061967013 | 0.059089303 | -1.048701033 | 0.294315739 | 0.525420908 | | Syde2 | 145.6925192 | 0.0602235291350243 | 0.0908031519983529 | 0.663231702971243 | 0.507182170065857 | 0.848958234808791 | NM_001166064 |
| Ece2 | 1158.70352 | 0.134852348 | 0.12864854 | 1.048222912 | 0.294535916 | 0.525753397 | | Tet1 | 494.1505663 | -0.0499385758927659 | 0.0753260777608782 | -0.662965302020575 | 0.507352776839812 | 0.848958234808791 | NM_001253857,NM_027384 |
| Nprl2 | 1148.888695 | 0.079113261 | 0.075522796 | 1.047541472 | 0.294849915 | 0.526253264 | | Zfp953 | 53.13583901 | -0.0629465888364123 | 0.0948759827863917 | -0.663461784402626 | 0.507034847060628 | 0.848958234808791 | NM_001038651 |
| Asb10 | 9.120888748 | 0.162434301 | 0.155095043 | 1.047321033 | 0.294951538 | 0.526313387 | | Caap1 | 180.4764777 | -0.0581712876496734 | 0.0878027756211548 | -0.662522093247447 | 0.507636680503597 | 0.849139845807836 | NM_026368 |
| Hmgcs2 | 424.8222214 | 0.16168046 | 0.154369815 | 1.047357994 | 0.294934498 | 0.526313387 | | Lor | 122.7544798 | -0.0630034149193333 | 0.0950872543089677 | -0.662585278933556 | 0.507596200919783 | 0.849139845807836 | NM_008508 |
| Tmem80 | 811.6868597 | 0.115297272 | 0.110164902 | 1.046588073 | 0.295289605 | 0.526855958 | | Ppp1r10 | 2212.752825 | 0.0313220807549247 | 0.0472733099048693 | 0.662574311338805 | 0.507603227131228 | 0.849139845807836 | NM_001163818 |
| Scn7a | 192.6546179 | 0.128783902 | 0.123089241 | 1.046264486 | 0.295438937 | 0.527008738 | | Aktip | 4485.472246 | 0.0385083189393108 | 0.0581528704328248 | 0.662191198004466 | 0.507848694505799 | 0.849201120800189 | NM_010241 |
| Snx15 | 1058.0902 | 0.069191646 | 0.066132672 | 1.046255117 | 0.295443262 | 0.527008738 | | Dap | 223.1199938 | -0.0581052241783947 | 0.0877380831562882 | -0.662257734476505 | 0.507806058961677 | 0.849201120800189 | NM_146057 |
| Igdcc4 | 2304.655976 | -0.120627086 | 0.115306538 | -1.046142638 | 0.295495182 | 0.527040675 | | Ndufb9 | 4590.030676 | 0.0350970665866566 | 0.0529909175757003 | 0.662322303374322 | 0.507764686004895 | 0.849201120800189 | NM_023172 |
| Narfl | 921.6653927 | 0.074019404 | 0.070770677 | 1.045904981 | 0.295604905 | 0.52717569 | | Alk | 328.6680858 | 0.0575827370156747 | 0.0871308958647569 | 0.660876218982686 | 0.508691698060371 | 0.85005033310116 | NM_007439 |
| Pcdh9 | 4733.884826 | 0.085954252 | 0.082215981 | 1.045468917 | 0.295806299 | 0.52747414 | | Htatip2 | 366.6569648 | -0.0526653593818441 | 0.0797042082532233 | -0.66076008451805 | 0.50876618449549 | 0.85005033310116 | NM_001146052,NM_001146049,NM_001146050,NM_001146053,NM_016865 |
| Gucd1 | 553.0228871 | -0.079256255 | 0.07581715 | -1.045360517 | 0.295856377 | 0.527502729 | | Npr3 | 491.947377 | 0.0603037154835828 | 0.0912203573103596 | 0.661077387346895 | 0.508562686028598 | 0.85005033310116 | NM_001039181,NM_008728 |
| Brwd3 | 358.2704106 | -0.10505866 | 0.100551059 | -1.044828979 | 0.296102019 | 0.527758505 | | Prss35 | 107.196387 | 0.0627167743690448 | 0.0948626456918008 | 0.661132460640042 | 0.508527369761298 | 0.85005033310116 | NM_178738 |
| Gtpbp4 | 1212.11697 | 0.070424227 | 0.067393473 | 1.044971033 | 0.296036358 | 0.527758505 | | Synm | 2114.063102 | 0.0308349604709077 | 0.0466414342786233 | 0.661106609344559 | 0.50854394698758 | 0.85005033310116 | NM_201639,NM_207663,NM_183312 |

| | | | | | | | | | | | | | | | |
|---|---|---|---|---|---|---|---|---|---|---|---|---|---|---|---|
| Ndc80 | 12.07436441 | -0.170522977 | 0.163199897 | -1.04487184 | 0.296082206 | 0.527758505 | | Zfp748 | 487.7329107 | 0.0524019336351489 | 0.0792800686569545 | 0.66097235437437 | 0.508630042974708 | 0.85005033310116 | NM_001035231,NM_183145 |
| Wrap53 | 425.7405504 | 0.111778163 | 0.107010039 | 1.044557722 | 0.296227428 | 0.527921299 | | Zim1 | 166.340216 | 0.0621217946336502 | 0.0940044883014029 | 0.66083860203006 | 0.508715824231862 | 0.85005033310116 | NM_011769 |
| Mon1b | 1763.245246 | -0.050022077 | 0.047917145 | -1.04392858 | 0.296518434 | 0.528336622 | | Haus4 | 80.47024945 | 0.0628025557838813 | 0.0950592061471187 | 0.660667791467612 | 0.508825383577708 | 0.850051468872891 | NM_145462 |
| Plin2 | 237.0035293 | -0.139031432 | 0.133183804 | -1.043906451 | 0.296528673 | 0.528336622 | | 4932438H23Rik | 51.30655632 | 0.0586070259301746 | 0.088724715350342 | 0.660549044296806 | 0.508901556307659 | 0.850080957558078 | NM_028905,NM_001163695 |
| Cdk17 | 8567.236864 | -0.080647638 | 0.077297899 | -1.043335453 | 0.296792957 | 0.528685913 | | Dbp | 3952.261596 | -0.0441078947101504 | 0.0668036543045357 | -0.660261705281527 | 0.50908590034362 | 0.850095618851756 | NM_016974 |
| Lamp1 | 12406.18079 | 0.050288243 | 0.048197183 | 1.043385508 | 0.296769783 | 0.528685913 | | Prox2 | 41.86354855 | -0.059004944837298 | 0.0893571547658656 | -0.660327032478857 | 0.509043986223402 | 0.850095618851756 | NM_175198 |
| D830030K20Rik | 97.66554383 | -0.129967623 | 0.12461573 | -1.042947174 | 0.29697276 | 0.528945389 | | Tmcc3 | 2446.992838 | 0.0515089626926932 | 0.0779927762405602 | 0.660432480744363 | 0.508976334123561 | 0.850095618851756 | NM_172051,NM_001168684,NM_177026 |
| AI462493 | 511.8579732 | -0.071408784 | 0.068503842 | -1.042405539 | 0.2972237 | 0.529027456 | | Ankrd17 | 6670.933469 | 0.0442366312350386 | 0.067030748010878 | 0.659945361610164 | 0.509288892938061 | 0.850336834346928 | NM_198010,NM_030886 |
| Gm6402 | 10.93467629 | -0.171182337 | 0.164196714 | -1.042544229 | 0.297159431 | 0.529027456 | | E130309D14Rik | 2013.405399 | 0.0368818699892263 | 0.0559326138100555 | 0.65939829156698 | 0.509640038914688 | 0.850457636652046 | NM_001013784 |
| Mthfd2l | 145.3220107 | -0.148201758 | 0.142147397 | -1.042592141 | 0.297137231 | 0.529027456 | | Fam132a | 466.3786199 | -0.0506811518475287 | 0.0768599327165879 | -0.659396255711146 | 0.509641345899158 | 0.850457636652046 | NM_026125 |
| Pramef8 | 712.8073667 | -0.072916584 | 0.06994759 | -1.042445979 | 0.297204959 | 0.529027456 | | Gclc | 2728.399142 | 0.034184737449619 | 0.0518503216328228 | 0.659296536127542 | 0.50970536630637 | 0.850457636652046 | NM_010295 |
| Ttyh2 | 1507.013485 | 0.128866215 | 0.123593992 | 1.042657596 | 0.297106904 | 0.529027456 | | Sh3gl1 | 1345.788856 | 0.0475298738337742 | 0.0720930195072306 | 0.659285381007063 | 0.509712528204097 | 0.850457636652046 | NM_001252471,NM_013664 |
| Vstm2a | 2910.134783 | -0.089237156 | 0.08558649 | -1.042654689 | 0.297108251 | 0.529027456 | | Tmem18 | 1517.22293 | -0.0330495237107043 | 0.0501189219148176 | -0.659422079486774 | 0.509624767608866 | 0.850457636652046 | NM_172049 |
| Cpe | 76847.68497 | 0.085522537 | 0.082083873 | 1.04189208 | 0.297461716 | 0.529268699 | | Zfp846 | 450.2924265 | -0.0506908323159266 | 0.0768615854051273 | -0.659508024050531 | 0.509569595141411 | 0.850457636652046 | NM_172919 |
| Dnajb4 | 2550.780622 | 0.103682318 | 0.099508785 | 1.041941344 | 0.297438874 | 0.529268699 | | Cacna2d3 | 5442.984278 | 0.0451642887669274 | 0.0685442458675784 | 0.658907078125579 | 0.509955440358291 | 0.850575621900297 | NM_009785 |
| Gpsm1 | 2230.170285 | -0.138518816 | 0.13293744 | -1.041984981 | 0.297418642 | 0.529268699 | | Mro | 453.7825564 | -0.0517967360269436 | 0.078603689810862 | -0.658960618153907 | 0.509921058078601 | 0.850575621900297 | NM_027741 |
| Gm14405 | 13.1899039 | -0.173267246 | 0.16631882 | -1.041777746 | 0.297514734 | 0.529302249 | | Smug1 | 728.2450196 | -0.0479244249141604 | 0.0727338029829217 | -0.658901679119039 | 0.50995890755402 | 0.850575621900297 | NM_027885 |
| Klc2 | 10359.76807 | 0.070510219 | 0.06769405 | 1.041601424 | 0.297596508 | 0.529386946 | | Snx13 | 3805.381011 | 0.0430330424877651 | 0.0653443661234547 | 0.658557807515694 | 0.510179764358683 | 0.850846298171552 | NM_001014973 |
| D6Ertd474e | 19.21335687 | -0.174936944 | 0.167980232 | -1.041413873 | 0.297683507 | 0.529480915 | | Atxn3 | 1324.99662 | 0.0448555610370531 | 0.0681338457122647 | 0.658344770769027 | 0.510316615567225 | 0.850976829035645 | NM_029705,NM_001167914 |
| Bdh2 | 43.91755434 | 0.169639674 | 0.162931808 | 1.041169777 | 0.29779676 | 0.529540878 | | 1700010I14Rik | 44.44877067 | -0.0605448789207219 | 0.0920627812227415 | -0.657647728176237 | 0.510764518088072 | 0.851624320316363 | NM_025851 |
| Gm16973 | 162.0726494 | 0.122511424 | 0.117659486 | 1.041237124 | 0.29776551 | 0.529540878 | | Paqr7 | 2840.539603 | 0.0305429590122058 | 0.0464490689302212 | 0.65755804617073 | 0.510822160465129 | 0.851624320316363 | NM_027995 |
| Has2 | 12.76859917 | -0.170023531 | 0.163308257 | -1.041120233 | 0.29781975 | 0.529540878 | | Senp5 | 3310.27789 | 0.042851635984088 | 0.0652112152742085 | 0.657120647175488 | 0.511103343814554 | 0.851995314694768 | NM_177103 |
| Dzip1 | 2590.845607 | 0.113283595 | 0.108864986 | 1.040587976 | 0.298066814 | 0.529836392 | | Gyk | 914.9462957 | 0.0418021930300967 | 0.0636333573117265 | 0.656922639258471 | 0.511230660420123 | 0.852011997850242 | NM_212444,NM_008194 |
| Galt | 1601.633693 | 0.09190827 | 0.088333384 | 1.040470379 | 0.298121419 | 0.529836392 | | Iqcg | 179.2352012 | 0.0624580430348299 | 0.09506503734017 | 0.65700329776695 | 0.511178796012151 | 0.852011997850242 | NM_178378 |
| Pex19 | 3357.952168 | -0.059436821 | 0.057117371 | -1.040608485 | 0.298057292 | 0.529836392 | | 8430408G22Rik | 65.36938146 | 0.0488171378704163 | 0.0743873380977587 | 0.656256012364115 | 0.511659414869535 | 0.852417907766886 | NM_001166580,NM_145980 |
| Tmem167b | 2140.070685 | -0.058784323 | 0.056501432 | -1.040404129 | 0.298152184 | 0.529836392 | | Egr1 | 12389.93262 | 0.0515361799009884 | 0.0785259403614094 | 0.656294972894272 | 0.511634351460563 | 0.852417907766886 | NM_007913 |
| Zfp947 | 60.76940665 | -0.156419527 | 0.150346447 | -1.040393906 | 0.298156932 | 0.529836392 | | Pla1a | 63.81061276 | 0.0624029862286309 | 0.0951005508686231 | 0.65617901956044 | 0.511708946422088 | 0.852417907766886 | NM_134102 |
| Ccdc127 | 3301.711676 | -0.054274336 | 0.052185848 | -1.040020192 | 0.29833052 | 0.529906191 | | Zfp85-rs1 | 152.3697686 | 0.0604233327757948 | 0.0920686966118676 | 0.656285306508903 | 0.511640569811288 | 0.852417907766886 | NM_001001130 |
| Lrfn3 | 951.3312787 | -0.081109977 | 0.077984177 | -1.04008248 | 0.298301583 | 0.529906191 | | Marc2 | 1882.385599 | 0.0438261350238 | 0.0668501703231694 | 0.655587484847596 | 0.512089580156826 | 0.852888435374525 | NM_133684 |
| Pmf1 | 90.89429907 | -0.133170712 | 0.128046936 | -1.040014825 | 0.298333013 | 0.529906191 | | Ptgfr | 38.25095048 | 0.0572615163713435 | 0.0873501269515207 | 0.655540161986526 | 0.512120037288726 | 0.852888435374525 | NM_008966 |
| Zbtb8b | 220.0219526 | 0.105224029 | 0.101159197 | 1.040182517 | 0.298255112 | 0.529906191 | | Tdrkh | 1128.905826 | 0.0375284793499765 | 0.0572546245211201 | 0.655466342917557 | 0.512167549344071 | 0.852888435374525 | NM_028307 |
| 1500015O10Rik | 66.23948984 | 0.144417364 | 0.138910222 | 1.039645332 | 0.298504708 | 0.530150383 | | Cdk5r1 | 13027.1673 | 0.0404454461656938 | 0.0617346470983494 | 0.655149872344135 | 0.512371264871913 | 0.852934333432204 | NM_009871 |
| Brd1 | 2186.468735 | 0.07476211 | 0.071923259 | 1.039470559 | 0.298585944 | 0.530212546 | | Eif3e | 2830.372619 | 0.0412030374255068 | 0.0628818874308546 | 0.655244922010555 | 0.512310075935726 | 0.852934333432204 | NM_008388 |
| Slc2a3 | 7099.238062 | -0.087717485 | 0.084390573 | -1.039422788 | 0.29860815 | 0.530212546 | | Epha5 | 3231.079127 | 0.0392440089579433 | 0.0598950717568765 | 0.655212654510891 | 0.512330847953594 | 0.852934333432204 | NM_007937 |
| E2f7 | 18.63773243 | -0.174131358 | 0.167631201 | -1.038776531 | 0.298908679 | 0.530536947 | | Aldh3b2 | 68.08524532 | 0.0602995360149055 | 0.0921010454345026 | 0.654710657522203 | 0.512654062190975 | 0.852934797674339 | NM_001177438 |
| Eri1 | 551.4673448 | 0.09985909 | 0.096142793 | 1.038653929 | 0.298965715 | 0.530536947 | | Col25a1 | 2176.426035 | 0.0552634279202846 | 0.0844346667783657 | 0.654511115266751 | 0.512782568366378 | 0.852934797674339 | NM_198711,NM_001244952,NM_029838 |
| Fpgt | 885.2394797 | -0.079644988 | 0.076675203 | -1.038732022 | 0.298929384 | 0.530536947 | | Krt9 | 499.7172269 | -0.0597632145894386 | 0.091264887179251 | -0.654832503896698 | 0.512575600792761 | 0.852934797674339 | NM_201255 |
| Sh3bgrl2 | 1284.155747 | -0.124403938 | 0.119761229 | -1.038766379 | 0.298913401 | 0.530536947 | | Lif | 26.42078716 | -0.0550681308056287 | 0.0840666396791468 | -0.655053312655348 | 0.512433429798078 | 0.852934797674339 | NM_001039537,NM_008501 |
| Tbl3 | 1042.503587 | 0.081852281 | 0.078804693 | 1.038672675 | 0.298956993 | 0.530536947 | | Ppih | 367.6000354 | 0.0476737084703032 | 0.0728323918928507 | 0.654567387275152 | 0.512746327222067 | 0.852934797674339 | NM_028677,NM_001110130,NM_001110129 |
| Vps36 | 523.2545972 | -0.082839167 | 0.079761321 | -1.038588198 | 0.298996297 | 0.530536947 | | Riiad1 | 53.78677097 | -0.0603199007869197 | 0.092136111326248 | -0.654682511760571 | 0.512672187180411 | 0.852934797674339 | NM_025506 |
| Kif1a | 64998.7361 | -0.067076298 | 0.064636638 | -1.03774423 | 0.299389147 | 0.531173188 | | Trerf1 | 1588.513038 | -0.0459513620129331 | 0.0701714604600855 | -0.654844030773322 | 0.512568178533633 | 0.852934797674339 | NM_001097623,NM_172622 |
| 2310022B05Rik | 2743.952165 | 0.087462104 | 0.084313831 | 1.037339938 | 0.299577459 | 0.531244904 | | Tmem119 | 1052.632164 | 0.0490899661758818 | 0.0750160665922108 | 0.654392697536863 | 0.512858837891893 | 0.852963988918896 | NM_146162 |
| Cnot6 | 1373.409647 | -0.093430199 | 0.090053105 | -1.037501142 | 0.299502364 | 0.531244904 | | Pik3cb | 1659.310372 | 0.043794505790455 | 0.0669402411873446 | 0.654232865219114 | 0.512961790759806 | 0.853023560191314 | NM_029094 |
| Dnahc7a | 36.481052 | 0.173842719 | 0.167600984 | 1.037241632 | 0.29962326 | 0.531244904 | | Ush1g | 33.00694938 | 0.0542332888337953 | 0.0829058994315007 | 0.654154761068656 | 0.513012103939923 | 0.853023560191314 | NM_176847 |
| Naa50 | 2059.631266 | -0.08100948 | 0.078089274 | -1.037395734 | 0.299551466 | 0.531244904 | | Kctd7 | 683.2730962 | 0.0486129993259104 | 0.0743261888404683 | 0.654049401486896 | 0.513079978613463 | 0.853038774103143 | NM_172509 |
| Phf5a | 689.710471 | -0.070722291 | 0.068188749 | -1.037154828 | 0.299663706 | 0.531244904 | | Hs3st2 | 1577.557515 | 0.0456592883540861 | 0.0698303320497278 | 0.65386039295318 | 0.513201753276024 | 0.853143587558654 | NM_001081327 |
| Soga3 | 2329.752946 | -0.077486309 | 0.07469334 | -1.037392473 | 0.299552985 | 0.531244904 | | 4930432K21Rik | 135.5021809 | 0.0600535823151282 | 0.0919826041598189 | 0.652879779428572 | 0.513833785772264 | 0.853195313198702 | NM_001163752,NM_029045 |
| Zcchc3 | 344.5151295 | -0.082685514 | 0.079724374 | -1.037142225 | 0.299669579 | 0.531244904 | | Arl8b | 9487.114692 | 0.0453629208902615 | 0.0694000965759207 | 0.653643483631704 | 0.51334152244106 | 0.853195313198702 | NM_026011 |
| 4833418N02Rik | 26.13202795 | 0.173643961 | 0.167482199 | 1.036790553 | 0.299833479 | 0.531324702 | | Bmp2 | 124.8062992 | 0.0597141288171031 | 0.0915118883120799 | 0.652528648665429 | 0.514060197681492 | 0.853195313198702 | NM_007553 |
| Gm17644 | 45.26630935 | 0.170073511 | 0.16404083 | 1.036775487 | 0.299840502 | 0.531324702 | | Cacnb3 | 12386.01636 | -0.0400105868373299 | 0.0613288283781477 | -0.652394443125971 | 0.514146748186269 | 0.853195313198702 | NM_001044741,NM_007581 |
| Mrpl41 | 1250.811805 | 0.097612001 | 0.094150194 | 1.036768985 | 0.299843533 | 0.531324702 | | Camk2b | 38256.90206 | -0.0268769267088356 | 0.0412517074539565 | -0.651534890739603 | 0.514701261844983 | 0.853195313198702 | NM_001174054,NM_007595,NM_001174053 |
| Nnmt | 19.29785546 | -0.174289747 | 0.168111428 | -1.036751334 | 0.299851761 | 0.531324702 | | Cd300a | 62.86571438 | -0.06198302319475 | 0.0948470685024124 | -0.65350489133118 | 0.513430837094891 | 0.853195313198702 | NM_170758 |
| Rragc | 3104.88049 | 0.084810199 | 0.08182099 | 1.036533512 | 0.299953314 | 0.53144387 | | Dact3 | 4727.598341 | -0.0426477151680661 | 0.0653109588433122 | -0.652994779488423 | 0.51375964410019 | 0.853195313198702 | NM_001081655 |
| Rpl12 | 5007.443237 | -0.063586093 | 0.061356658 | -1.036335659 | 0.300045576 | 0.531546553 | | Ebag9 | 667.1245661 | 0.0506027212050633 | 0.0775473396430092 | 0.652539744600059 | 0.514053042140646 | 0.853195313198702 | NM_019480 |
| Spa17 | 64.94272403 | -0.160552586 | 0.154945048 | -1.036190491 | 0.300113282 | 0.531605715 | | Ggn | 36.34540413 | -0.0549624319969636 | 0.0841341787997745 | -0.653271153068068 | 0.513581486097942 | 0.853195313198702 | NM_182696,NM_182694 |
| Aim1 | 58.24524131 | -0.172799704 | 0.166797706 | -1.035983697 | 0.300209748 | 0.531655028 | | Hnrnpul2 | 11648.1059 | 0.0385758076137755 | 0.0591236756584727 | 0.652459563519161 | 0.514104750441976 | 0.853195313198702 | NM_001081196 |
| Zfp964 | 72.86458967 | -0.158597476 | 0.153088317 | -1.035986802 | 0.3002083 | 0.531655028 | | Homer1 | 6575.036986 | 0.0582168019187114 | 0.0893177416074044 | 0.651794378933169 | 0.514533828312367 | 0.853195313198702 | NM_011982,NM_152134,NM_147176 |
| Dhx35 | 735.5665357 | 0.105318057 | 0.101715399 | 1.035418996 | 0.300473278 | 0.532060911 | | Mapre1 | 3938.325166 | 0.0377080703597409 | 0.0578099107963778 | 0.652276916540469 | 0.514222548489649 | 0.853195313198702 | NM_007896 |
| Pim3 | 1371.05235 | 0.106568519 | 0.102935776 | 1.035291351 | 0.300532868 | 0.532105617 | | Mapre3 | 7756.615354 | 0.0230561818598352 | 0.0353793483030862 | 0.651684752989753 | 0.514604560481095 | 0.853195313198702 | NM_133350 |
| Ccdc61 | 233.1276546 | 0.100886208 | 0.097459323 | 1.035162211 | 0.300593163 | 0.532151562 | | Mbtps2 | 1247.552409 | 0.0490659253517571 | 0.0751300855003513 | 0.653079588888897 | 0.51370497021063 | 0.853195313198702 | NM_172307 |
| Ano6 | 1219.189938 | 0.095973522 | 0.092740257 | 1.034863668 | 0.300732584 | 0.53233756 | | Pmpca | 4443.360325 | 0.0251111978203759 | 0.0384604372605831 | 0.652909836938115 | 0.513814406862224 | 0.853195313198702 | NM_173180 |
| Kbtbd12 | 12.05431378 | -0.168943012 | 0.163295806 | -1.034582675 | 0.300863848 | 0.532509078 | | Rnf126 | 2154.196257 | -0.0438781911663528 | 0.0672655750582448 | -0.652312733940934 | 0.514199446974653 | 0.853195313198702 | NR_027505,NM_144528 |
| Nedd1 | 137.7732087 | 0.117502826 | 0.113594164 | 1.034408999 | 0.300944999 | 0.53259187 | | Rnf187 | 11503.69444 | 0.0264320144364598 | 0.0405495575381252 | 0.651844706606431 | 0.514501357896973 | 0.853195313198702 | NM_022423 |
| Peli3 | 555.9250354 | 0.072759787 | 0.070396504 | 1.033571028 | 0.301336749 | 0.533163366 | | Sec22b | 2301.213228 | 0.0369936493650782 | 0.0567669323222043 | 0.651676034827907 | 0.514610185776132 | 0.853195313198702 | NM_011342 |
| Sdha | 16296.74015 | -0.038967777 | 0.037700589 | -1.033611879 | 0.301317644 | 0.533163366 | | Slco2a1 | 87.21819133 | -0.0616886353250907 | 0.0943840676227956 | -0.653591616454044 | 0.513374946864436 | 0.853195313198702 | NM_033314 |
| Ticam1 | 247.8635776 | 0.096963592 | 0.093826039 | 1.033440109 | 0.301397985 | 0.533210821 | | Sox3 | 28.58704366 | -0.0545104078792559 | 0.0834329303931265 | -0.6533440408051 | 0.513534506019557 | 0.853195313198702 | NM_009237 |
| Fam129a | 121.1580265 | 0.153595739 | 0.148716511 | 1.03280892 | 0.301693329 | 0.533658642 | | Ttc22 | 54.06649903 | -0.0565224763722354 | 0.0866798495303336 | -0.652083231321893 | 0.514347480975554 | 0.853195313198702 | NM_177667 |
| Kat2b | 1097.528834 | 0.113679373 | 0.110074597 | 1.032748483 | 0.301721618 | 0.533658642 | | Vps36 | 547.2749956 | 0.0455473859462388 | 0.0698755135396929 | 0.651836153166381 | 0.514506876331508 | 0.853195313198702 | NM_027338 |
| Zfp651 | 3709.737553 | 0.104963691 | 0.101642191 | 1.032678359 | 0.301754444 | 0.533658642 | | Zfp296 | 82.59661152 | 0.0619494828905025 | 0.0950680474932647 | 0.65163306204318 | 0.514637913945026 | 0.853195313198702 | NM_022409 |
| Rbfa | 854.4184493 | 0.076721498 | 0.074318928 | 1.032327834 | 0.301918566 | 0.53382703 | | Zfp369 | 665.8517258 | -0.0443565691446639 | 0.068077737480357 | -0.651557627887713 | 0.514686589675736 | 0.853195313198702 | NM_178364 |
| Zfp568 | 174.7873816 | 0.117591698 | 0.113904544 | 1.032370561 | 0.301898558 | 0.53382703 | | Zfp628 | 527.2431936 | -0.0458187058685147 | 0.0702277881888263 | -0.652429857897827 | 0.514123908106954 | 0.853195313198702 | NM_170759 |
| Cdc42bpb | 7875.405969 | 0.077269899 | 0.074867645 | 1.03208667 | 0.302031518 | 0.533904888 | | 4930581F22Rik | 111.6509545 | -0.0612750578582783 | 0.0950803026352799 | -0.644455856365164 | 0.519279874070244 | 0.853297834676958 | NR_029475 |
| Cox15 | 1121.951279 | -0.060251962 | 0.058375778 | -1.032139754 | 0.302006653 | 0.533904888 | | A230072E10Rik | 23.54851746 | -0.0516010808053072 | 0.0800417838585546 | -0.644676796515353 | 0.519136656116383 | 0.853297834676958 | NR_015600 |
| Efcab2 | 416.6612204 | 0.101926046 | 0.098777455 | 1.031875612 | 0.302130392 | 0.534018742 | | A830082K12Rik | 321.639868 | -0.0569913025621201 | 0.0879505338753594 | -0.647992684648013 | 0.516989681742076 | 0.853297834676958 | NR_045195 |
| 2610204G22Rik | 21.89452361 | -0.173355629 | 0.168020589 | -1.031752297 | 0.302188172 | 0.534025881 | | Aaas | 560.5213386 | -0.0420925337802156 | 0.0646377439808885 | -0.651206728264854 | 0.514913047672109 | 0.853297834676958 | NM_153416 |
| B230217C12Rik | 1065.703605 | 0.078573397 | 0.076157685 | 1.031719874 | 0.302203364 | 0.534025881 | | Acadsb | 2329.275907 | 0.0370625696038973 | 0.0570715000848644 | 0.649405912737284 | 0.516076043222208 | 0.853297834676958 | NM_025826 |
| Vsig10 | 87.83310894 | 0.133484428 | 0.129441718 | 1.031231901 | 0.302432084 | 0.534369109 | | Actn2 | 1065.725798 | -0.0541611663893322 | 0.0842252923149862 | -0.643051094281513 | 0.520190946648315 | 0.853297834676958 | NM_033268 |
| Tnfsf12 | 146.4199015 | -0.12575955 | 0.122040992 | -1.030469751 | 0.302789545 | 0.534939705 | | Alad | 1146.885414 | 0.0349381134977588 | 0.0541419830573705 | 0.645305390102489 | 0.518729300397861 | 0.853297834676958 | NM_008525,NM_001276446 |
| Pthlh | 98.13622443 | -0.145384087 | 0.141098824 | -1.030370655 | 0.302836043 | 0.534960854 | | Alx3 | 25.26391443 | 0.0469972698484364 | 0.072334574918998 | 0.649720688910734 | 0.515872657511319 | 0.853297834676958 | NM_007441 |
| Mpp2 | 7689.336452 | -0.079835437 | 0.077493839 | -1.03021657 | 0.302908353 | 0.53502759 | | Arhgef10l | 2123.39725 | 0.0415797588459806 | 0.064575805306344 | 0.643890674668763 | 0.519646329281637 | 0.853297834676958 | NM_001112722,NM_001112723,NM_172415 |
| Traf3ip2 | 27.69905862 | 0.169099557 | 0.16416393 | 1.030065233 | 0.302979385 | 0.535092054 | | Asph | 3724.573828 | 0.0318898732425995 | 0.0491466426849599 | 0.648871855744453 | 0.516421207529395 | 0.853297834676958 | NM_023066,NM_001177856,NM_001177855,NM_001177854,NM_001177849,NM_001177851,NM_001177850,NM_001177853,NM_001177852,NM_133723 |
| Adcy8 | 929.920884 | 0.106548636 | 0.103454933 | 1.029903873 | 0.303055133 | 0.535164831 | | BC029214 | 1075.448506 | 0.0480079942292988 | 0.0744275143661045 | 0.645030196670956 | 0.518907617268983 | 0.853297834676958 | NM_153557 |
| Gramd1c | 48.885542 | 0.170026361 | 0.165178708 | 1.029347928 | 0.303316209 | 0.535503799 | | Bend3 | 279.9162906 | -0.0569725573973745 | 0.0885865470804039 | -0.643128773781694 | 0.520140545311178 | 0.853297834676958 | NM_199028 |
| Rusc1 | 6695.074574 | 0.077030951 | 0.074833847 | 1.029359766 | 0.303310648 | 0.535503799 | | Brd7 | 1809.589395 | 0.0340237093677044 | 0.0528491945425133 | 0.643788607607537 | 0.519712522231889 | 0.853297834676958 | NM_012047 |
| A930011G23Rik | 22.28760303 | -0.1724977 | 0.167626678 | -1.029058749 | 0.303452069 | 0.535633715 | | Btbd3 | 7228.764526 | 0.0436423659558681 | 0.0677676950148238 | 0.643999562716743 | 0.519575717552853 | 0.853297834676958 | NM_001025431,NM_145534 |
| Jmjd1c | 3415.37995 | -0.065656916 | 0.06380379 | -1.029044135 | 0.303458936 | 0.535633715 | | Cad | 1015.473458 | -0.0501640560677689 | 0.0775327361803223 | -0.647004846457366 | 0.517628806840714 | 0.853297834676958 | NM_023525 |
| 2810428I15Rik | 750.6016863 | -0.094722807 | 0.092056787 | -1.028960605 | 0.303498187 | 0.535641977 | | Camkk1 | 7183.229903 | 0.049912613226744 | 0.0774528070291247 | 0.644426136911672 | 0.51929914038804 | 0.853297834676958 | NM_018883 |
| Zmpste24 | 1253.062325 | -0.069347227 | 0.067401166 | -1.028872803 | 0.30353945 | 0.535653786 | | Cask | 3734.499423 | 0.0332115045118408 | 0.0513578064203301 | 0.646669062148533 | 0.517846150250827 | 0.853297834676958 | NM_009806 |
| Pck2 | 826.6243246 | 0.0852042 | 0.082832129 | 1.028637085 | 0.303650245 | 0.53578828 | | Caskin2 | 1331.201844 | -0.0377498457104869 | 0.0580969157045935 | -0.649773662726507 | 0.515838433732126 | 0.853297834676958 | NM_080643 |
| Ptger4 | 38.430263 | 0.171444027 | 0.166722614 | 1.028318971 | 0.30379981 | 0.535930122 | | Cc2d1a | 2305.969251 | 0.0388718995461733 | 0.0604114223556666 | 0.643452811246831 | 0.51993032496142 | 0.853297834676958 | NM_145970 |
| Txndc16 | 2946.69625 | -0.05471531 | 0.053205285 | -1.028381108 | 0.303770591 | 0.535930122 | | Cenpb | 5443.275758 | 0.0398028175080543 | 0.0613818933892381 | 0.64844558077827 | 0.516696797507663 | 0.853297834676958 | NM_007682 |
| 1810011H11Rik | 23.24774067 | -0.171906427 | 0.167345425 | -1.02725501 | 0.304300401 | 0.536752096 | | Cnp | 13254.7067 | 0.0478093447647207 | 0.074312077858783 | 0.643359008956443 | 0.519991174992978 | 0.853297834676958 | NM_009923,NM_001146318 |
| Slc35f6 | 1440.670803 | 0.065485305 | 0.063757818 | 1.027094504 | 0.304375966 | 0.536824271 | | Cspg4 | 857.3446284 | 0.05255028504367 4 | 0.0815858575679413 | 0.644110224617206 | 0.519503960589134 | 0.853297834676958 | NM_139001 |
| Nod1 | 246.3475725 | 0.126030284 | 0.122759496 | 1.026643866 | 0.30458819 | 0.537031958 | | Ctf1 | 58.91079099 | -0.0611021758868893 | 0.0945187999132692 | -0.6464552654388 | 0.517984559232759 | 0.853297834676958 | NM_007795 |
| Pycr1 | 35.7755793 | -0.172566614 | 0.168091405 | -1.026623667 | 0.304597705 | 0.537031958 | | Dcxr | 172.772978 | 0.0570526832352146 | 0.0883943105715169 | 0.645433884447293 | 0.518646050861746 | 0.853297834676958 | NM_026428 |
| Trmt61b | 278.5082268 | -0.086996514 | 0.084736018 | -1.026676915 | 0.304572623 | 0.537031958 | | Dkk3 | 12643.19957 | 0.0389938697613035 | 0.0603841898910072 | 0.645762903032847 | 0.518432916186951 | 0.853297834676958 | NM_015814 |
| Ccdc164 | 665.6362659 | 0.074260724 | 0.07236679 | 1.026171315 | 0.30481084 | 0.537102144 | | Dlx6as1 | 619.8218595 | 0.0515011934434936 | [illegible] | [illegible] | [illegible] | [illegible] | [illegible] |
| Cdk5rap2 | 535.7926005 | 0.117344401 | 0.114345344 | 1.026228065 | 0.304784096 | 0.537102144 | | Ehd2 | 320.6655307 | -0.055862612183295 | [illegible] | [illegible] | [illegible] | [illegible] | [illegible] |
| Kcnj11 | 1972.834453 | 0.084349306 | 0.082175516 | 1.026453015 | 0.3046781 | 0.537102144 | | Eif2b4 | 1193.599382 | 0.0355795514877824 | [illegible] | [illegible] | [illegible] | [illegible] | [illegible] |
| Loh12cr1 | 1183.854724 | 0.082881373 | 0.080753826 | 1.026346088 | 0.304728481 | 0.537102144 | | Eif5b | 3892.350032 | -0.0402466509058894 | [illegible] | [illegible] | [illegible] | [illegible] | [illegible] |
| Tspyl5 | 2036.013803 | -0.067637533 | 0.065911633 | -1.026185048 | 0.304804368 | 0.537102144 | | Epn3 | 107.8147269 | 0.0364876450288666 | [illegible] | [illegible] | [illegible] | [illegible] | [illegible] |
| Jam3 | 1129.693405 | 0.071629019 | 0.069846394 | 1.025522074 | 0.305116916 | 0.537580338 | | Exoc7 | 3151.697393 | 0.0307784849821165 | [illegible] | [illegible] | [illegible] | [illegible] | [illegible] |
| Igsf10 | 178.454035 | -0.14291475 | 0.139432401 | -1.02497518 | 0.305374899 | 0.537895292 | | Fam171a2 | 2852.710484 | 0.038883727280011 | [illegible] | [illegible] | [illegible] | [illegible] | [illegible] |
| Prmt8 | 3775.69855 | -0.078096094 | 0.076202553 | -1.02484879 | 0.305434541 | 0.537895292 | | Fam63a | 1009.562378 | 0.0385020536161805 | [illegible] | [illegible] | [illegible] | [illegible] | [illegible] |
| Rab11b | 8795.318171 | 0.052832795 | 0.051551451 | 1.024855621 | 0.305431317 | 0.537895292 | | Fbxl17 | 4113.752539 | 0.0375100410239719 | [illegible] | [illegible] | [illegible] | [illegible] | [illegible] |
| Zeb2 | 5077.629658 | -0.061970956 | 0.060455842 | -1.025061498 | 0.305334171 | 0.537895292 | | Gm9776 | 129.0754226 | -0.0610559949887339 | [illegible] | [illegible] | [illegible] | [illegible] | [illegible] |
| Cdc20 | 43.03935707 | -0.160043097 | 0.156196562 | -1.024626247 | 0.305539575 | 0.538019113 | | Gpr137b-ps | 500.4556499 | -0.0486768429157442 | [illegible] | [illegible] | [illegible] | [illegible] | [illegible] |
| Lasp1 | 6573.675095 | -0.090442132 | 0.088299833 | -1.024261642 | 0.30571171 | 0.53826105 | | Hat1 | 605.623402 | 0.0492923909293856 | [illegible] | [illegible] | [illegible] | [illegible] | [illegible] |
| Kdelr1 | 2049.578652 | -0.052421481 | 0.051186536 | -1.024126359 | 0.305775596 | 0.53831236 | | Htr5a | 1224.273156 | 0.0409646794998739 | [illegible] | [illegible] | [illegible] | [illegible] | [illegible] |
| Mr1 | 243.285977 | 0.117763123 | 0.115010956 | 1.023929609 | 0.305868524 | 0.538414782 | | Hyal2 | 353.5065767 | 0.0482096577816341 | [illegible] | [illegible] | [illegible] | [illegible] | [illegible] |
| Cacna2d3 | 5020.281322 | -0.069244524 | 0.067661122 | -1.023401946 | 0.30611784 | 0.538792436 | | Icam5 | 7735.155972 | -0.0491926657026903 | [illegible] | [illegible] | [illegible] | [illegible] | [illegible] |
| Coro7 | 2969.545098 | -0.053459663 | 0.052270785 | -1.022744594 | 0.306428621 | 0.539175836 | | Isoc2a | 705.626877 | 0.0443989185727169 | [illegible] | [illegible] | [illegible] | [illegible] | [illegible] |
| Cstf2t | 3262.027784 | -0.055004182 | 0.053776915 | -1.022821441 | 0.306392278 | 0.539175836 | | Kdm5b | 3260.792503 | -0.0376918558744329 | 0.0581603094061897 | [illegible] | [illegible] | [illegible] | [illegible] |
| Smc4 | 260.7716712 | 0.093878977 | 0.091793395 | 1.022720389 | 0.306440068 | 0.539175836 | | Klhl38 | 52.76888205 | -0.0582610056647507 | 0.0899705268396032 | [illegible] | [illegible] | [illegible] | [illegible] |
| Sntg1 | 851.5365817 | 0.087164691 | 0.085254996 | 1.022399804 | 0.306591713 | 0.539381402 | | Lcmt1 | 2291.03139 | 0.0321739018012383 | 0.0497848000698686 | [illegible] | [illegible] | [illegible] | [illegible] |
| Cdo1 | 1087.267559 | -0.129061249 | 0.12627095 | -1.022097713 | 0.306734656 | 0.539510359 | | Lrrc73 | 1195.646956 | 0.0424539429680305 | 0.0656000808366278 | [illegible] | [illegible] | [illegible] | [illegible] |
| Ttc19 | 3180.604789 | -0.068336332 | 0.066858899 | -1.022097781 | 0.306734623 | 0.539510359 | | Lsr | 263.782766 | -0.060560927506383 | 0.0939083858356319 | [illegible] | [illegible] | [illegible] | [illegible] |
| Ctage5 | 1853.612535 | 0.081111865 | 0.079400244 | 1.021556865 | 0.306990682 | 0.539605236 | | Meis1 | 266.601734 | 0.0598293356799976 | 0.0926728235771665 | [illegible] | [illegible] | [illegible] | [illegible] |
| Focad | 1570.081249 | 0.061097212 | 0.05981159 | 1.021494532 | 0.307020198 | 0.539605236 | | Mnat1 | 446.239477 | 0.0510039256897227 | 0.0788384679941404 | [illegible] | [illegible] | [illegible] | [illegible] |
| Rock2 | 10323.25194 | -0.093841596 | 0.091835502 | -1.021844423 | 0.30685454 | 0.539605236 | | Mrvi1 | 170.3799913 | -0.0601869442486082 | 0.0935253008519206 | [illegible] | [illegible] | [illegible] | [illegible] |
| Sbds | 2180.472258 | 0.063441463 | 0.062100151 | 1.021599174 | 0.306970649 | 0.539605236 | | Msl3 | 848.3414427 | -0.0462495289521107 | 0.0717307845323629 | [illegible] | [illegible] | [illegible] | [illegible] |
| Sec13 | 1589.475292 | -0.057353256 | 0.056139614 | -1.02161829 | 0.306961597 | 0.539605236 | | Ndufaf2 | 460.8916777 | 0.0476853657125505 | 0.0737100350348703 | [illegible] | [illegible] | [illegible] | [illegible] |
| Sntg2 | 123.3358075 | -0.129321219 | 0.1266032 | -1.021468799 | 0.307032384 | 0.539605236 | | Neurl2 | 62.99390615 | -0.0603912266884054 | 0.0931200795402877 | [illegible] | [illegible] | [illegible] | [illegible] |
| Srxn1 | 4706.465379 | 0.106359472 | 0.10410044 | 1.021700496 | 0.306922676 | 0.539605236 | | Nfatc4 | 68.46158207 | -0.060571123601453 | 0.0940241538397305 | [illegible] | [illegible] | [illegible] | [illegible] |
| Senp8 | 246.4874988 | -0.133538199 | 0.130760775 | -1.021240496 | 0.30714051 | 0.539734044 | | Nmd3 | 790.7195706 | 0.0435647022901305 | 0.0673847325926171 | [illegible] | [illegible] | [illegible] | [illegible] |
| Myadm | 2615.153929 | -0.08102286 | 0.079348378 | -1.02110292 | 0.30720568 | 0.539787344 | | Npas3 | 467.527508 | -0.046752810613434 | 0.0720071803861509 | [illegible] | [illegible] | [illegible] | [illegible] |
| Kdm8 | 197.4295471 | 0.116074232 | 0.113698756 | 1.020892717 | 0.30730527 | 0.539901106 | | Numb | 2059.618356 | -0.04233938792614 | 0.0652977626064988 | [illegible] | [illegible] | [illegible] | [illegible] |
| Tfcp2 | 800.3342327 | 0.071489512 | 0.070041223 | 1.020677668 | 0.307407178 | 0.54000629 | | P2rx5 | 39.36389795 | 0.0552195496149553 | 0.0853802974863666 | [illegible] | [illegible] | [illegible] | [illegible] |
| Tnfrsf11a | 230.7027707 | 0.128924245 | 0.126319623 | 1.020619294 | 0.307434845 | 0.54000629 | | Paqr3 | 361.9867078 | 0.0491195988938976 | 0.0758004695082658 | [illegible] | [illegible] | [illegible] | [illegible] |
| Dydc2 | 13.71324319 | 0.169039449 | 0.165675292 | 1.020305724 | 0.30758349 | 0.540141653 | | Parg | 1568.447523 | 0.0432390293863411 | 0.0670200591599455 | 0.645165491172573 | 0.518819946640953 | 0.853297834676958 | NM_011960 |
| Eid2b | 880.905984 | -0.090801327 | 0.089007419 | -1.020154591 | 0.307655149 | 0.540141653 | | Pcdhac1 | 170.611437 | 0.057657723917641 | 0.089117240820059 | 0.64698731005441 | 0.517640156475591 | 0.853297834676958 | NM_001003671 |
| Frs3 | 661.4379201 | -0.08015175 | 0.078576006 | -1.020053761 | 0.307702965 | 0.540141653 | | Pcdhb13 | 301.4471925 | 0.0534641087246108 | 0.0831136960033323 | 0.643264724053028 | 0.520052341798818 | 0.853297834676958 | NM_053138 |
| Gak | 5243.514958 | 0.082640413 | 0.081020392 | 1.019995223 | 0.307730726 | 0.540141653 | | Pcdhga9 | 500.020102 | 0.0535368983982931 | 0.0829801272729316 | 0.645177347369013 | 0.51881226420868 | 0.853297834676958 | NM_033592 |
| Olfm3 | 950.2529625 | -0.098492834 | 0.096573435 | -1.019875017 | 0.30778774 | 0.540141653 | | Pelp1 | 1653.068789 | -0.0335481008606227 | 0.0521351982534625 | -0.643482752238209 | 0.519910902858469 | 0.853297834676958 | NM_029231 |
| Tmem222 | 2504.715506 | -0.072026698 | 0.070607211 | -1.020103991 | 0.307679144 | 0.540141653 | | Phactr4 | 593.0931151 | 0.0513189983313613 | 0.0788454788976589 | 0.650880672536381 | 0.515123518699779 | 0.853297834676958 | NM_175306,NM_001161797 |
| Uqcc | 2140.230045 | -0.051676576 | 0.0506635 | -1.019996175 | 0.307730275 | 0.540141653 | | Pitpnb | 3674.827026 | 0.0335062568895515 | 0.051688007015237 | 0.648240449272388 | 0.516829443727028 | 0.853297834676958 | NM_019640 |
| Uxs1 | 1469.772436 | -0.092311 | 0.090512643 | -1.019868564 | 0.3077908 | 0.540141653 | | Pomt1 | 686.8474727 | -0.0469479630825266 | 0.072341234673883 | -0.648979289532033 | 0.516351762799979 | 0.853297834676958 | NM_145145 |
| Mxd4 | 1037.146477 | -0.081546294 | 0.079998291 | -1.01935046 | 0.308036616 | 0.540511815 | | Prkca | 8923.908718 | -0.0459876142779887 | 0.0714936131147685 | -0.6432408752956 | 0.520067814135229 | 0.853297834676958 | NM_011101 |
| Cbx3 | 3375.00603 | -0.070724001 | 0.069406183 | -1.018987043 | 0.308209118 | 0.540692037 | | Rab15 | 12818.25421 | 0.040328443797168 | 0.0625274453492244 | 0.644971877100177 | 0.51894541062069 | 0.853297834676958 | NM_134050 |
| Tbc1d14 | 4057.6187 | 0.06526454 | 0.064047359 | 1.019004383 | 0.308200886 | 0.540692037 | | Rab3ip | 1377.595051 | 0.0384800873315379 | 0.0592103072347653 | 0.649883239866979 | 0.515764360760898 | 0.853297834676958 | NM_001003950 |
| Braf | 5522.824927 | -0.099331544 | 0.097512324 | -1.018656302 | 0.308366165 | 0.540783855 | | Rbks | 56.17545918 | 0.0611816842155649 | 0.0946456032685765 | 0.646429227588619 | 0.518001417077537 | 0.853297834676958 | NM_153196 |
| Plcb1 | 11559.21202 | -0.080304932 | 0.078830956 | -1.018697942 | 0.30834639 | 0.540783855 | | Rmi1 | 717.6004462 | -0.0414683483631365 | 0.0644660838048822 | -0.643258375809638 | 0.520056460319659 | 0.853297834676958 | NM_001168248,NR_031761,NM_028904 |
| Sec22c | 2109.11264 | -0.06611504 | 0.064901112 | -1.018704265 | 0.308343387 | 0.540783855 | | Rnf167 | 2904.971766 | 0.0329478154274476 | 0.0507842302231759 | 0.648780443902673 | 0.516480299555359 | 0.853297834676958 | NM_027445 |
| 9930111J21Rik1 | 101.6560598 | -0.146997172 | 0.144329255 | -1.018484939 | 0.308447554 | 0.54086537 | | Rpl27 | 2911.385572 | 0.031343219221585 | 0.0482240246360383 | 0.649950298801106 | 0.515724326357268 | 0.853297834676958 | NM_011289 |
| Bnipl | 8.470775996 | -0.157050817 | 0.154311912 | -1.017749145 | 0.308797186 | 0.541417177 | | Sfmbt2 | 123.4476883 | 0.0613730571622776 | 0.0949946336723545 | 0.646068675562865 | 0.518234880648598 | 0.853297834676958 | NM_001198809,NM_001198808,NM_177386 |
| Tmem212 | 28.36674606 | 0.170598393 | 0.167657163 | 1.017543119 | 0.308895131 | 0.541527626 | | Slc29a2 | 548.3372384 | -0.0480351726337064 | 0.0745818746866799 | -0.64405960342916 | 0.519536784465344 | 0.853297834676958 | NM_007854 |
| Tefm | 149.5189205 | -0.117759211 | 0.115749721 | -1.01736064 | 0.308981899 | 0.541618457 | | Slc52a3 | 99.73116728 | 0.0608160675951802 | 0.0943532254501281 | 0.64455737792796 | 0.519214063170476 | 0.853297834676958 | NM_001164819,NM_027172,NM_001164820 |
| 4921508A21Rik | 15.44481682 | 0.170694098 | 0.167834178 | 1.017040157 | 0.309134326 | 0.54172842 | | Slc9a8 | 1833.656651 | -0.0390290532206203 | 0.0600599115354123 | -0.649835343124143 | 0.515798586542567 | 0.853297834676958 | NM_178371,NM_148929 |
| Mapk8ip3 | 21408.68188 | 0.067402232 | 0.066278643 | 1.016952492 | 0.30917603 | 0.54172842 | | Slfn2 | 94.42620863 | -0.0603099649312709 | 0.0934819995068653 | -0.645150566412967 | 0.518829617486663 | 0.853297834676958 | NM_011408 |
| Mkl1 | 2989.255276 | 0.10324546 | 0.101530474 | 1.016891343 | 0.309205122 | 0.54172842 | | Snx20 | 30.53145945 | -0.0585533389630371 | 0.0909683654415104 | -0.643667044899086 | 0.519791364254645 | 0.853297834676958 | NM_027840 |
| Nkain4 | 756.8624741 | -0.078117337 | 0.076822023 | -1.01686123 | 0.309219449 | 0.54172842 | | Sox10 | 1784.436896 | -0.0527697295417454 | 0.0817361934598207 | -0.645610314207813 | 0.518531755945101 | 0.853297834676958 | NM_011437 |
| Prdx6 | 7950.481737 | 0.082979582 | 0.081590947 | 1.017019466 | 0.309144169 | 0.54172842 | | Sptan1 | 44838.78761 | -0.0331434910687395 | 0.0512689070762544 | -0.646463772271248 | 0.517979051663635 | 0.853297834676958 | NM_001076554,NM_001177668,NM_001177667 |
| Ssu72 | 1939.082096 | -0.063124309 | 0.062112097 | -1.016296531 | 0.3094882 | 0.542076664 | | Sptssa | 556.7791981 | -0.0475424851462889 | 0.0735560998969918 | -0.64634320216634 | 0.51805711506075 | 0.853297834676958 | NM_134054 |
| Ubxn7 | 1711.513803 | -0.06587598 | 0.064817321 | -1.016332952 | 0.309470862 | 0.542076664 | | Steap3 | 223.414919 | 0.0584527797988133 | 0.0908844728648426 | 0.643154743118117 | 0.520123696005326 | 0.853297834676958 | NM_001085409,NM_133186 |
| Mal2 | 5768.464621 | -0.105939136 | 0.104271622 | -1.015992018 | 0.309633188 | 0.542269313 | | Tbc1d7 | 576.9956443 | -0.0404908173770802 | 0.0629491805273476 | -0.643230253958419 | 0.520074705006773 | 0.853297834676958 | NM_001252640,NM_001252639,NM_025935 |
| 2210408I21Rik | 382.71195 | -0.108460451 | 0.106765543 | -1.015875045 | 0.309688894 | 0.542305575 | | Tchp | 464.5350482 | -0.0467710151921411 | 0.0720528920192363 | -0.649120581858872 | 0.51626043943022 | 0.853297834676958 | NM_029992 |
| Mmp2 | 96.40096998 | 0.163622301 | 0.161162527 | 1.015262691 | 0.309980624 | 0.542755089 | | Tmem200b | 199.0587817 | -0.0584146442957705 | 0.0901968925638568 | -0.647634775825727 | 0.5172211992728 | 0.853297834676958 | NM_001201367 |
| 3110021N24Rik | 14.03335348 | 0.163931952 | 0.161537922 | 1.014820232 | 0.310191528 | 0.543062998 | | Tspan18 | 144.0682901 | -0.0604199518470247 | 0.0930081775576528 | -0.649619780041087 | 0.515937853032247 | 0.853297834676958 | NM_183180 |
| Rai2 | 379.299967 | 0.105614415 | 0.104098542 | 1.014561902 | 0.310314708 | 0.543217273 | | Vps72 | 972.708412 | 0.0377585420900336 | 0.0587090358403353 | 0.643147030939521 | 0.520128699756324 | 0.853297834676958 | NM_009336 |
| Smarca1 | 1076.946111 | -0.102600523 | 0.101201852 | -1.013820609 | 0.310668359 | 0.543774916 | | Wif1 | 29.11675392 | 0.0532550994610135 | 0.0821274882912492 | 0.648444273276136 | 0.516697642934864 | 0.853297834676958 | NM_011915 |
| Abcc10 | 438.592471 | 0.133228317 | 0.131428281 | 1.013695957 | 0.310727854 | 0.543817617 | | Zfhx2 | 2695.866948 | -0.0529308059685006 | 0.082273159139577 | -0.64335448549755 | 0.519994109477006 | 0.853297834676958 | NM_001039198 |
| Zbtb5 | 468.1316776 | -0.083617425 | 0.082501528 | -1.013525777 | 0.31080909 | 0.543898356 | | Zfp53 | 179.6023293 | -0.057374719840384 | 0.0881295626947603 | -0.651026943581954 | 0.515029094292272 | 0.853297834676958 | NM_013843 |
| Ccdc93 | 1386.479814 | 0.063966861 | 0.063127539 | 1.013295651 | 0.310918964 | 0.543987214 | | Zfp654 | 624.5531815 | 0.0561457257201458 | 0.0864864803768732 | 0.649184999499174 | 0.516218806290094 | 0.853297834676958 | NM_028059 |
| Zfp114 | 81.24335085 | 0.136211409 | 0.134427242 | 1.013272358 | 0.310930087 | 0.543987214 | | Zfp773 | 89.22095658 | -0.0607426605568875 | 0.0935124418309954 | -0.649567687117694 | 0.515971511065259 | 0.853297834676958 | NM_029584 |
| 3110070M22Rik | 24.49791855 | 0.169404272 | 0.167217731 | 1.01307601 | 0.311023856 | 0.54402222 | | Apc | 12874.77053 | 0.0422515143235975 | 0.0657481606781581 | 0.64262656600101 | 0.520466444684436 | 0.853555599970098 | NM_007462 |
| Cflar | 972.3233144 | -0.098995561 | 0.097724179 | -1.013009102 | 0.311055431 | 0.54402222 | | Bloc1s1 | 1003.903174 | -0.0397414003704198 | 0.06184869165717 | -0.642558464950369 | 0.520510641411817 | 0.853555599970098 | NM_015740 |
| Pfkl | 6657.9081 | -0.098207655 | 0.096945497 | -1.013019251 | 0.311050966 | 0.54402222 | | Zfp804b | 57.20184686 | 0.0582405584093176 | 0.0906491814928248 | 0.642483003709499 | 0.520559621377881 | 0.853555599970098 | NM_001163223 |
| Pm20d1 | 103.7336959 | -0.153249659 | 0.151300735 | -1.012881128 | 0.311116944 | 0.544068389 | | 1810026J23Rik | 3110.388477 | 0.0318967668953311 | 0.0496582027981671 | 0.64232642416501 | 0.520661377588844 | 0.853626091518685 | NM_178619 |
| Ank | 8933.881206 | 0.076336318 | 0.07537437 | 1.012762268 | 0.311173727 | 0.544106278 | | Glrx3 | 329.5228097 | 0.0550419023422225 | 0.0857046516340572 | 0.642227712181144 | 0.520725342200883 | 0.853634614694733 | NM_023140 |
| Nrbf2 | 722.0221238 | 0.069228841 | 0.068383119 | 1.012367398 | 0.31136242 | 0.544374784 | | 3110047P20Rik | 1917.873608 | -0.0497261764120591 | 0.0774525751629265 | -0.64202095679139 | 0.520859576022714 | 0.853725740110665 | NM_177006 |
| Tmed4 | 2310.752443 | 0.065589141 | 0.064835846 | 1.011618493 | 0.3117205 | 0.544939347 | | Bcorl1 | 1182.22536 | -0.0433162956084617 | 0.0674821963342257 | -0.641892202113944 | 0.520943177683901 | 0.853725740110665 | NM_178782 |
| Eif1b | 2800.356998 | 0.082030262 | 0.081127931 | 1.01112233 | 0.311957884 | 0.545231298 | | Ids | 18601.13721 | 0.0346124131477724 | 0.0539365268268489 | 0.641724916008918 | 0.52105180849551 | 0.853725740110665 | NM_010498 |
| Fam63a | 909.364996 | 0.067744407 | 0.066996064 | 1.011169952 | 0.311935094 | 0.545231298 | | Keap1 | 2927.65566 | 0.0308442036004046 | 0.0480671762912456 | 0.641689526622395 | 0.52107479084 6 | 0.853725740110665 | NM_016679,NM_001110306,NM_001110307,NM_001110305 |
| 1700007L15Rik | 11.58169702 | 0.168237654 | 0.166434663 | 1.010833025 | 0.312096354 | 0.545255252 | | Topbp1 | 1012.20112 | 0.0457145832107109 | 0.0712354719242264 | 0.641739037811637 | 0.52104263775004 | 0.853725740110665 | NM_176979 |
| 9930013L23Rik | 907.141211 | -0.098904596 | 0.097846185 | -1.010817096 | 0.312103979 | 0.545255252 | | Acy3 | 104.3460704 | 0.0603012808169639 | 0.094024841786484 | 0.641333499437296 | 0.521306028785568 | 0.853911972675263 | NM_027857 |
| Copb1 | 2535.3313 | -0.057278923 | 0.056675186 | -1.010652586 | 0.312182738 | 0.545255252 | | Crtc1 | 10452.1949 | -0.0410206387150055 | 0.0639561241402552 | -0.641387189521485 | 0.52127115395318 | 0.853911972675263 | NM_001004062 |
| Lrrn4cl | 63.12654398 | 0.161951712 | 0.160233403 | 1.010723787 | 0.312148649 | 0.545255252 | | S100a10 | 782.2015871 | 0.0419154473296497 | 0.0653759289680943 | 0.641155113798724 | 0.521424492119903 | 0.854009716149928 | NM_009112 |
| Scmh1 | 1788.742641 | -0.04470194 | 0.044229338 | -1.010685262 | 0.312167093 | 0.545255252 | | 0610040B10Rik | 40.58498063 | -0.0593312102525137 | 0.0927190098831885 | -0.639903406294586 | 0.522235399263905 | 0.854113377026378 | NR_027874 |
| Tab1 | 933.2343497 | -0.066752835 | 0.066038861 | -1.010811434 | 0.31210669 | 0.545255252 | | BC068281 | 329.5600589 | -0.0493978554212206 | 0.0771896888891801 | -0.639954068115759 | 0.522202461269196 | 0.854113377026378 | NM_001170858,NM_173416 |
| 1700021F05Rik | 476.2414204 | -0.079031827 | 0.078209086 | -1.01051977 | 0.312246332 | 0.545304855 | | Fadd | 124.0480374 | -0.0608626487811161 | 0.0949900080331768 | -0.640726851605896 | 0.521700165263155 | 0.854113377026378 | NM_010175 |
| Fam115a | 8457.355551 | -0.056667904 | 0.056087778 | -1.010343186 | 0.31233098 | 0.545391066 | | Fam35a | 239.4821845 | 0.0574765107649204 | 0.0897875091356979 | 0.640139272357526 | 0.522082059045478 | 0.854113377026378 | NM_029389 |
| Arf4 | 3042.880835 | -0.069512884 | 0.068830013 | -1.009921116 | 0.312533085 | 0.545436767 | | Gstt2 | 88.25032805 | 0.0606503271282611 | 0.0947976593606055 | 0.639787179739853 | 0.522310968478956 | 0.854113377026378 | NM_010361 |
| Cab39l | 1263.50508 | 0.10678985 | 0.105735694 | 1.00996973 | 0.312509793 | 0.545436767 | | Kif16b | 793.6921808 | 0.0427762671427016 | 0.0667663126257279 | 0.640686380008622 | 0.521726464938367 | 0.854113377026378 | NM_001081133 |
| Mylip | 589.5415802 | -0.074689961 | 0.073937799 | -1.0101729 | 0.31241246 | 0.545436767 | | Pak2 | 2368.191581 | 0.0386388232122112 | 0.0603636241839132 | 0.640101116104099 | 0.522106863453418 | 0.854113377026378 | NM_177326 |
| Orc5 | 834.8111864 | 0.0814945 | 0.080689935 | 1.00997108 | 0.312509146 | 0.545436767 | | Pcdhga4 | 595.6335306 | 0.0482248923836272 | 0.0752573750582895 | 0.640799554147023 | 0.521652922652962 | 0.854113377026378 | NM_033587 |
| Tubb4a | 28586.72965 | 0.079976394 | 0.079186134 | 1.009979774 | 0.31250498 | 0.545436767 | | Smad3 | 4334.874365 | 0.0541159807352389 | 0.0845794234016383 | 0.639824422522496 | 0.522286753023434 | 0.854113377026378 | NM_016769 |
| Aldoc | 30197.82806 | 0.055072745 | 0.054570676 | 1.009200343 | 0.312878562 | 0.545944885 | | Spef2 | 45.1903929 | -0.059680855871795 | 0.0931481667874166 | -0.640708861270443 | 0.521711855846273 | 0.854113377026378 | NM_177123 |
| Rtbdn | 43.71684162 | 0.168882901 | 0.167348869 | 1.009166674 | 0.312894706 | 0.545944885 | | Sv2b | 24553.25358 | 0.0479559124978419 | 0.0749239898351641 | 0.640060848379096 | 0.522133041132261 | 0.854113377026378 | NM_001109753,NM_153579 |
| Pdk4 | 511.773732 | 0.131811772 | 0.13066764 | 1.008756047 | 0.313091645 | 0.546226996 | | Tmem147 | 1875.510172 | 0.0377350122526485 | 0.0589261132950575 | 0.640378435860969 | 0.521926598864786 | 0.854113377026378 | NM_027215 |
| Stard7 | 4586.21541 | -0.046003591 | 0.04560974 | -1.008635233 | 0.313149604 | 0.546266602 | | Tspan17 | 1499.313098 | -0.0433895393558625 | 0.0677425581838498 | -0.6405063599474 | 0.521843455713639 | 0.854113377026378 | NM_028841 |
| Gm9962 | 65.13246285 | -0.155949688 | 0.154667504 | -1.008251143 | 0.313333911 | 0.546413192 | | Utrn | 3483.326188 | -0.040217655171645 | 0.0628013594979759 | -0.640394658541448 | 0.521916054594931 | 0.854113377026378 | NM_011682 |
| Slc2a1 | 4373.888318 | 0.157931061 | 0.156637657 | 1.008257296 | 0.313330958 | 0.546413192 | | Rps27a | 6003.498312 | -0.0387985684934987 | 0.0606635704966047 | -0.639569484220686 | 0.522452526807416 | 0.854248694777637 | NM_024277,NM_001033865 |
| Slc6a15 | 2374.12819 | 0.058244586 | 0.057769948 | 1.008216005 | 0.313350776 | 0.546413192 | | Aagab | 2475.645321 | 0.031463909047999 | 0.0492345898922875 | 0.639061056806483 | 0.522783212744305 | 0.854337224801527 | NM_025857 |
| Zmat1 | 497.4053865 | 0.105533172 | 0.104678352 | 1.008166158 | 0.313374701 | 0.546413192 | | Ap5b1 | 150.934432 | 0.0604971926842316 | 0.0946923892018574 | 0.638881257450044 | 0.522900181648043 | 0.854337224801527 | NM_001033448 |
| Crot | 1317.053435 | -0.065255041 | 0.064762357 | -1.007607565 | 0.313642896 | 0.54681929 | | Atp13a3 | 2927.098222 | 0.0526571752838326 | 0.0824330820500977 | 0.638786928430395 | 0.522961552992761 | 0.854337224801527 | NM_001128094,NM_001128096 |
| Csrnp2 | 1194.565587 | 0.081863065 | 0.081270598 | 1.00729006 | 0.313795405 | 0.546931265 | | Cd151 | 1006.814956 | 0.0436465738559248 | 0.0683224469193003 | 0.638832128133464 | 0.52293214821378 | 0.854337224801527 | NM_009842,NM_001111049,NM_001111050 |
| Nkain1 | 2343.763054 | 0.094310489 | 0.093625649 | 1.007314662 | 0.313783586 | 0.546931265 | | Cyhr1 | 5852.625988 | 0.0300566669399989 | 0.0470278612406994 | 0.639124683688293 | 0.52274182334263 | 0.854337224801527 | NM_180962,NM_019396,NM_001276321,NM_001276322,NM_001276333,NM_001276334,NM_001276335 |
| Psmg3 | 298.9814814 | -0.103600751 | 0.102854706 | -1.007253393 | 0.313813021 | 0.546931265 | | Fgf11 | 240.764764 | -0.0597758908708401 | 0.093580717916448 | -0.638762901180241 | 0.522977185937986 | 0.854337224801527 | NM_010198 |
| Hspa14 | 966.3662692 | -0.055907426 | 0.055509579 | -1.007167173 | 0.313854445 | 0.546941938 | | Ghdc | 309.0362297 | 0.0518956698947262 | 0.0812124163700349 | 0.639011523290597 | 0.522815435551067 | 0.854337224801527 | NM_031871 |
| Smad1 | 768.6985072 | 0.091320141 | 0.090718922 | 1.006627281 | 0.314113919 | 0.547332554 | | Smurf2 | 1426.264771 | -0.038812354048119 | 0.060702318474334 | -0.639388330202001 | 0.522570338745913 | 0.854337224801527 | NM_025481 |
| Hlx | 25.96144508 | 0.165883455 | 0.16483163 | 1.006381208 | 0.31423223 | 0.547477136 | | Amer1 | 511.1855869 | -0.0419931767681998 | 0.0657955588295123 | -0.638237253626972 | 0.523319250125912 | 0.854628731286121 | NM_175179 |
| Clta | 4951.280947 | -0.046246537 | 0.045963563 | -1.006156484 | 0.314340301 | 0.547542285 | | Sfrp2 | 60.84432214 | 0.0601877346509286 | 0.0943059982315514 | 0.638217459966316 | 0.52333213305728 | 0.854628731286121 | NM_009144 |
| Ppt2 | 968.5547507 | 0.076943899 | 0.076472667 | 1.006162093 | 0.314337603 | 0.547542285 | | Tmem19 | 1602.191049 | 0.0328858911043981 | 0.0515201869088409 | 0.638310788013754 | 0.523271390851172 | 0.854628731286121 | NM_133683 |
| Dram1 | 47.83103794 | 0.159312357 | 0.158372551 | 1.005934143 | 0.314447251 | 0.547605452 | | Dlgap4 | 11716.3133 | 0.0341747056707479 | 0.0535627952057368 | 0.638030661758438 | 0.52345372083106 | 0.854731187004122 | NM_146128,NM_001042487,NM_001042488 |
| Zfp946 | 470.3382727 | 0.100323317 | 0.099727133 | 1.005978144 | 0.314426084 | 0.547605452 | | Rgs12 | 2815.904419 | 0.0477999978612291 | 0.0749286402336718 | 0.637940281742207 | 0.523512554782849 | 0.854731187004122 | NM_173402,NM_001163512 |
| Kifc2 | 23131.26605 | 0.086171913 | 0.085688188 | 1.005645173 | 0.314586286 | 0.547727297 | | Eaf1 | 1064.410219 | 0.0398377812330998 | 0.0625101512262451 | 0.6373009895451 | 0.523928806523104 | 0.855218546303474 | NM_028932 |
| Zar1l | 15.67516203 | -0.167345046 | 0.166406219 | -1.005641779 | 0.314587919 | 0.547727297 | | Taf5 | 259.3027581 | 0.0580800005749671 | 0.0911242408739123 | 0.637371571142434 | 0.523882841564826 | 0.855218546303474 | NM_177342 |
| Mettl5 | 197.4552752 | -0.110134903 | 0.109571325 | -1.005143482 | 0.314827765 | 0.548083303 | | 1500009C09Rik | 3577.375291 | 0.0368293184945355 | 0.0578168444019101 | 0.63699980300756 | 0.524124971913159 | 0.855442622697814 | NR_037697,NR_037698 |
| Tfeb | 283.0313521 | 0.105804094 | 0.105282874 | 1.004950666 | 0.314920606 | 0.548183335 | | 2610001J05Rik | 1015.115364 | -0.0401418805128302 | 0.0630520631175513 | -0.636646582650144 | 0.52435507533291 | 0.855487854117416 | NR_024619 |
| Ly6h | 4382.958666 | -0.075728958 | 0.075393505 | -1.004449359 | 0.315162069 | 0.548542023 | | AI467606 | 75.53129156 | 0.0597720055818912 | 0.0939639690980803 | 0.636116227907537 | 0.524700669134726 | 0.855487854117416 | NM_178901 |
| Slc22a5 | 462.5372488 | 0.11428417 | 0.113788457 | 1.004356448 | 0.315206835 | 0.548558317 | | Chgb | 20595.60617 | 0.0390783369097029 | 0.0614066437471862 | 0.636386138779871 | 0.524524773178324 | 0.855487854117416 | NM_007694 |
| Bax | 962.4061326 | -0.065063283 | 0.064794499 | -1.004148268 | 0.315307153 | 0.548643115 | | Fig4 | 1541.341694 | 0.0392533286490956 | 0.0616774747788549 | 0.636428919793471 | 0.524496896351265 | 0.855487854117416 | NM_133999 |
| Krcc1 | 413.4035114 | -0.105323343 | 0.104892406 | -1.004108372 | 0.315326381 | 0.548643115 | | Grm2 | 1760.554063 | -0.0500360017856557 | 0.0785766036367467 | -0.636779899739216 | 0.524268220582299 | 0.855487854117416 | NM_001160353 |
| 5430416N02Rik | 131.9862758 | 0.129475274 | 0.129003086 | 1.003660282 | 0.315542388 | 0.54876192 | | Rock2 | 10632.62065 | 0.0509263575418676 | 0.0800651524013626 | 0.63606145763111 | 0.524736365604041 | 0.855487854117416 | NM_009072 |
| 6230400D17Rik | 13.95585025 | 0.166073984 | 0.165489678 | 1.003530773 | 0.315604837 | 0.54876192 | | Sp6 | 24.33279068 | -0.0502652881010241 | 0.0790049817059177 | -0.636229349284934 | 0.524626946334473 | 0.855487854117416 | NM_031183 |
| Dact2 | 2524.153756 | 0.117386355 | 0.116939399 | 1.003856458 | 0.315447807 | 0.54876192 | | Sulf1 | 1106.627434 | 0.0591675807098994 | 0.0930107347471377 | 0.636137117621471 | 0.524687054613545 | 0.855487854117416 | NM_001198566,NM_001198565,NM_172294 |
| Fam131b | 7869.792651 | -0.081399406 | 0.081116533 | -1.003487239 | 0.315625831 | 0.54876192 | | Tmem201 | 3618.514701 | -0.0387529150455199 | 0.0609271412642835 | -0.636053394946293 | 0.524741620555292 | 0.855487854117416 | NM_177672,NM_001025106 |
| Gngt2 | 64.09989949 | -0.157316928 | 0.156774004 | -1.003463095 | 0.315637475 | 0.54876192 | | Tom1l2 | 17058.29249 | 0.0320039351807968 | 0.0502962049748112 | 0.636309144931008 | 0.524574945575595 | 0.855487854117416 | NM_153080,NM_001039092,NM_001039093 |
| Mob3a | 455.0014859 | -0.076877388 | 0.07658975 | -1.003755569 | 0.315496445 | 0.54876192 | | Dcaf5 | 4420.56428 | 0.0352582407271905 | 0.0554459378055606 | 0.635903045789126 | 0.524839617353032 | 0.855551597090129 | NM_177267 |
| Ppp1r14b | 556.3094291 | -0.092210354 | 0.091899819 | -1.00337906 | 0.315678004 | 0.54876192 | | Ndufa6 | 2063.605394 | 0.0411007298587803 | 0.0646536247185268 | 0.635706505825692 | 0.524967735207305 | 0.855664421187311 | NM_025987 |
| Tor4a | 386.1379459 | -0.141134381 | 0.140561526 | -1.003432987 | 0.315651995 | 0.54876192 | | Cog5 | 1060.625363 | 0.0391056344374519 | 0.0615360388687373 | 0.635491577884437 | 0.525107857896496 | 0.855700778977395 | NM_001163126 |
| Fam46c | 58.19601801 | 0.164145641 | 0.163663813 | 1.002944008 | 0.315887877 | 0.549010123 | | Slc44a1 | 3778.883667 | 0.0482202381774695 | 0.0758769504538988 | 0.635505748307151 | 0.525098618872455 | 0.855700778977395 | NM_133891,NM_001159633 |
| Nup37 | 116.8913676 | -0.122477188 | 0.122118625 | -1.002936187 | 0.315891651 | 0.549010123 | | Ptgds | 21239.52221 | 0.0575204318258828 | 0.0905290038317903 | 0.635381252319534 | 0.525179792297298 | 0.855722003691592 | NM_008963 |
| Col9a3 | 458.4563731 | 0.168281886 | 0.167849686 | 1.002574922 | 0.316066001 | 0.54917695 | | Olig2 | 1007.657254 | 0.0520824359350703 | 0.081991595704435 | 0.635216762005934 | 0.525287052525064 | 0.855800776691238 | NM_016967 |
| E530011L22Rik | 279.2476365 | 0.103979005 | 0.103712678 | 1.002567927 | 0.316069377 | 0.54917695 | | Htr1f | 104.9981316 | -0.0595418212772434 | 0.093764056920063 | -0.635017545454595 | 0.525416971914596 | 0.855879197314464 | NM_008310 |
| Il1rl2 | 23.97345068 | 0.16805198 | 0.16763006 | 1.002516971 | 0.316093974 | 0.54917695 | | Snrpd2 | 1394.798995 | 0.0395667548263856 | 0.0623135533748685 | 0.634962262356606 | 0.525453027788131 | 0.855879197314464 | NM_026943 |
| Chordc1 | 1450.491861 | -0.088049555 | 0.087860721 | -1.002149235 | 0.316271521 | 0.549267267 | | 2900008C10Rik | 174.412632 | 0.0553957373136438 | 0.0872952617479676 | 0.63457897031775 | 0.525703047303972 | 0.855892498712622 | NR_045434 |
| Cited4 | 208.874593 | -0.113351887 | 0.113091824 | -1.002299579 | 0.316198926 | 0.549267267 | | B3gnt4 | 23.48582515 | 0.0479511878754198 | 0.0755775198051018 | 0.634463634147769 | 0.525778292646666 | 0.855892498712622 | NM_198611 |
| Eif4ebp2 | 861.2109892 | 0.074729236 | 0.074576965 | 1.002042203 | 0.31632321 | 0.549267267 | | Bcas2 | 1611.087504 | 0.0311998036335859 | 0.049184224026103 | 0.634345753163201 | 0.525855203472064 | 0.855892498712622 | NM_026602 |
| Sox8 | 1588.200315 | 0.101264828 | 0.101043554 | 1.002189884 | 0.316251893 | 0.549267267 | | Cdc42ep1 | 982.8570644 | -0.0539464156217993 | 0.0850120978838254 | -0.634573395606829 | 0.52570668410932 | 0.855892498712622 | NM_027219 |
| Stk16 | 1555.575337 | -0.090116962 | 0.089932484 | -1.00205129 | 0.316318822 | 0.549267267 | | Dcun1d4 | 4603.67483 | 0.0339215303262158 | 0.0535031409278007 | 0.634010073763537 | 0.526074247854405 | 0.855892498712622 | NM_001190733,NM_178896,NM_001190734 |
| Nek10 | 53.75182265 | 0.155585334 | 0.155289427 | 1.001905517 | 0.316389228 | 0.549320338 | | Dynlrb2 | 48.78579972 | 0.0539917902654318 | 0.0851326338815443 | 0.634207919111974 | 0.52594514012162 | 0.855892498712622 | NM_029297 |
| Col25a1 | 2026.797708 | -0.113904606 | 0.113724126 | -1.001586998 | 0.316543103 | 0.549376037 | | Fkbp1a | 19507.56803 | 0.0296556443991691 | 0.0467760636646391 | 0.633991876951964 | 0.526086123342116 | 0.855892498712622 | NM_008019 |
| Igsf9 | 214.7619593 | 0.153796077 | 0.153555974 | 1.00156362 | 0.316554399 | 0.549376037 | | Mios | 963.7661168 | -0.0488351337447218 | 0.0770089806588896 | -0.634148554193133 | 0.525983878123902 | 0.855892498712622 | NM_145374 |
| Rasa1 | 1842.721249 | -0.086068588 | 0.085922159 | -1.001704204 | 0.316486476 | 0.549376037 | | Rbpj | 1748.451627 | -0.0390814426989706 | 0.0615994945210181 | -0.634444210993254 | 0.525790964601648 | 0.855892498712622 | NM_009035,NM_001080927,NM_001080928 |

| | | | | | | | | | | | | | | |
|---|---|---|---|---|---|---|---|---|---|---|---|---|---|---|
| Slc25a13 | 130.6656785 | 0.138834643 | 0.1386204 | 1.001545534 | 0.316563138 | 0.549376037 | | Tead1 | 768.502256 | 0.0499971614839969 | 0.0788653098175816 | 0.633956318686152 | 0.526109329547363 | 0.855892498712622 | NM_001166585,NM_009346,NM_001166584 |
| Gm5860 | 9.605743896 | 0.157774278 | 0.157559207 | 1.001365015 | 0.316650371 | 0.549465881 | | Wdr70 | 512.7918887 | 0.0426880216605053 | 0.0672573841166908 | 0.634696431048255 | 0.52562642178166 | 0.855892498712622 | NM_001081402 |
| Rab22a | 1161.229521 | -0.066705367 | 0.066619767 | -1.001284911 | 0.316689086 | 0.549471522 | | Dhfr | 242.5307081 | 0.0534436088616203 | 0.0843167368319393 | 0.633843420294413 | 0.526183013302108 | 0.855916511895457 | NM_010049 |
| Nt5c3 | 1563.310431 | -0.062223811 | 0.062179799 | -1.000707824 | 0.316968084 | 0.549877862 | | Col23a1 | 886.3005708 | -0.0581262518081958 | 0.0917367318643672 | -0.633620259048856 | 0.526328676225451 | 0.855977055876102 | NM_153393 |
| Pitpna | 13582.28152 | 0.074948357 | 0.074899395 | 1.000653697 | 0.31699426 | 0.549877862 | | Copb1 | 2748.780706 | 0.0402006332574166 | 0.0634473842238165 | 0.633605841268494 | 0.526338087779523 | 0.855977055876102 | NM_033370 |
| Galk2 | 826.3542025 | 0.074700651 | 0.074684486 | 1.000216448 | 0.317205771 | 0.550183165 | | Banp | 1087.501475 | -0.0568441166481227 | 0.0897297875602596 | -0.633503301341799 | 0.526405025671305 | 0.85599008204426 | NM_001110100,NM_016812 |
| H2-Q1 | 29.07651382 | 0.167735517 | 0.167737863 | 0.999986016 | 0.317317275 | 0.550314962 | | Anks1b | 12133.90948 | 0.0379859469723983 | 0.0600073929051984 | 0.633021118454683 | 0.526719852143225 | 0.855999809537752 | NM_001177396,NM_001177398,NM_001128086,NM_001177397,NM_181398 |
| Pde8b | 3689.303714 | 0.056620484 | 0.056656368 | 0.999366636 | 0.317617116 | 0.550773318 | | Mrgprf | 40.48612488 | 0.0561667516679884 | 0.088737686584984 | 0.632952625085595 | 0.526764580576756 | 0.855999809537752 | NM_145379 |
| Ddx19a | 704.1560766 | 0.095096287 | 0.095181652 | 0.999103141 | 0.31774473 | 0.550877097 | | Opalin | 685.7542955 | -0.0589836460067413 | 0.0931683998906273 | -0.633086390621538 | 0.526677229065825 | 0.855999809537752 | NM_153520 |
| Spata5 | 478.7337254 | -0.07163996 | 0.071704764 | -0.999096243 | 0.317748071 | 0.550877097 | | Ranbp9 | 3938.882557 | 0.0386474431449493 | 0.0610412839305653 | 0.633136144202192 | 0.526644740891936 | 0.855999809537752 | NM_019930 |
| 1110007C09Rik | 155.6381591 | 0.114037742 | 0.114176945 | 0.998780809 | 0.317900885 | 0.55103906 | | Rnf138 | 290.6421065 | -0.0565023026135684 | 0.0892105649379075 | -0.633358870139374 | 0.52649931749334 | 0.855999809537752 | NM_019706,NM_207623 |
| Arhgef15 | 1007.840861 | -0.098504274 | 0.098630497 | -0.99872024 | 0.317930233 | 0.55103906 | | Spa17 | 75.88814891 | 0.0601434275542794 | 0.0950172835785418 | 0.632973552696489 | 0.526750913949506 | 0.855999809537752 | NM_011449 |
| Zfp202 | 185.6069599 | -0.123680448 | 0.123843527 | -0.998683191 | 0.317948186 | 0.55103906 | | Mdp1 | 1535.814053 | -0.0380750867822308 | 0.0601817015585783 | -0.632668831159088 | 0.526949928050605 | 0.856196077858577 | NM_023397,NR_028316 |
| Pifo | 19.54342346 | 0.165223906 | 0.165468737 | 0.998520377 | 0.318027088 | 0.55111416 | | Mroh1 | 3836.331428 | -0.0411156366027076 | 0.0650029020152443 | -0.632520015691995 | 0.527047133605313 | 0.856196077858577 | NM_175457,NM_001162489 |
| Ppp1r13b | 3713.408321 | 0.103247166 | 0.103424328 | 0.998287034 | 0.318140193 | 0.551248507 | | Nfx1 | 3823.565079 | -0.0268324480837126 | 0.0424290986863013 | -0.632406742412743 | 0.527121129305332 | 0.856196077858577 | NM_023739 |
| Supt7l | 692.2647744 | -0.069431107 | 0.069588232 | -0.997742075 | 0.318404445 | 0.551644691 | | Taf7 | 293.2422514 | 0.0549745119146186 | 0.0869265873671325 | 0.632424596199032 | 0.527109465978496 | 0.856196077858577 | NM_175770 |
| 2810013P06Rik | 590.3418468 | -0.076898951 | 0.077084131 | -0.997597685 | 0.318474484 | 0.551704345 | | Lrg1 | 46.51723309 | 0.0588189415294509 | 0.0930246514944573 | 0.632294134775183 | 0.527194695429548 | 0.856219828467086 | NM_029796 |
| Pim1 | 119.8554952 | -0.150106651 | 0.150487835 | -0.997467018 | 0.318537875 | 0.55175247 | | Jak2 | 2287.642963 | 0.0498865642781086 | 0.0789508122874878 | 0.631868917275405 | 0.527472535570921 | 0.856575299240156 | NM_001048177,NM_008413 |
| 1810058I24Rik | 421.7928355 | 0.082251594 | 0.082497137 | 0.99702362 | 0.318753045 | 0.552030005 | | Fbxo28 | 2423.43761 | 0.0385914399862407 | 0.0611448533243909 | 0.631147805384406 | 0.527943885865582 | 0.856965326619609 | NM_175127 |
| Gng3 | 6920.720767 | 0.065998038 | 0.066199203 | 0.996961219 | 0.318783335 | 0.552030005 | | Gfra4 | 1867.594379 | -0.0354585540094155 | 0.0561694569141568 | -0.631278206296466 | 0.527858634224913 | 0.856965326619609 | NM_001271001,NM_001136063,NM_001271002,NM_020014 |
| Mag | 6216.197743 | 0.159866272 | 0.160360726 | 0.996916611 | 0.318804988 | 0.552030005 | | Grik1 | 616.9820497 | 0.0534419651506263 | 0.0847300280233105 | 0.630732296417081 | 0.528215578162656 | 0.856965326619609 | NM_146072,NM_010348 |
| Pla2g12a | 515.1491366 | -0.074536233 | 0.074773849 | -0.996822213 | 0.318850814 | 0.552047661 | | Ldhd | 661.500099 | -0.0476344072715515 | 0.0755254098564318 | -0.630707034388837 | 0.528232098751431 | 0.856965326619609 | NM_027570 |
| Hspbp1 | 2226.094474 | -0.077901295 | 0.078159696 | -0.996693938 | 0.318913093 | 0.552068788 | | Mterf | 67.36165212 | -0.0598427440878266 | 0.0948846604220331 | -0.63068934242537 | 0.528243668907612 | 0.856965326619609 | NM_172135,NM_001013023 |
| Ormdl3 | 2134.65925 | 0.074199137 | 0.074448517 | 0.996650305 | 0.318934279 | 0.552068788 | | Pcsk6 | 957.9676874 | 0.0502639851208477 | 0.0796953819339379 | 0.630701351836335 | 0.528235815002093 | 0.856965326619609 | NM_011048 |
| Prkx | 455.7821389 | 0.093961629 | 0.094294745 | 0.99646729 | 0.319023152 | 0.552160939 | | Plekhd1 | 193.7042958 | 0.0592613509550629 | 0.0939291748875157 | 0.630915272342496 | 0.528095925398976 | 0.856965326619609 | NM_001177503 |
| Bag2 | 309.4004141 | -0.081618744 | 0.081926334 | -0.996245522 | 0.319130866 | 0.552285674 | | Rbfox2 | 12535.67322 | 0.0343791710415237 | 0.0544927720381855 | 0.630894148997095 | 0.528109737804182 | 0.856965326619609 | NM_175387,NM_001110829,NM_001110828,NM_001110830,NM_001110827,NM_053104 |
| AW146154 | 141.5832176 | -0.109802699 | 0.110238199 | -0.996049462 | 0.319226113 | 0.552388809 | | Zfp622 | 1188.747025 | 0.0409731707012576 | 0.0648941540081499 | 0.631384618961392 | 0.527789070476754 | 0.856965326619609 | NM_144523 |
| Snrpf | 443.1899895 | -0.085946329 | 0.086311917 | -0.99576434 | 0.31936466 | 0.55256684 | | Riok2 | 935.5331215 | 0.0426049830205302 | 0.0675989195299952 | 0.630261301759791 | 0.528523637459164 | 0.857228043516282 | NM_025934 |
| Degs1 | 2986.684062 | 0.052246758 | 0.052498355 | 0.995207539 | 0.319635336 | 0.552973413 | | Zfp7 | 346.1904972 | -0.0492404960938776 | 0.0781253158972925 | -0.630275801490667 | 0.528514152384144 | 0.857228043516282 | NM_145916 |
| Pla2g5 | 94.28904912 | -0.147990685 | 0.148733189 | -0.995007815 | 0.319732463 | 0.55307969 | | 1810014B01Rik | 123.7914981 | -0.0572318222615018 | 0.0908375181872303 | -0.630046080117889 | 0.528664436016684 | 0.857360678528341 | NR_015572 |
| Sgol1 | 8.790862732 | -0.158526088 | 0.159366072 | -0.994729219 | 0.319867979 | 0.553252341 | | Adra1b | 829.6518513 | 0.0489010921719828 | 0.0777186165148367 | 0.62920693091143 | 0.529213591926829 | 0.857868166089624 | NM_007416 |
| Arhgef6 | 605.9003969 | 0.117585814 | 0.118266086 | 0.994247954 | 0.320102167 | 0.553376867 | | Dusp19 | 379.4090556 | 0.0513365726987874 | 0.0815767088922531 | 0.629304287901501 | 0.529149864701602 | 0.857868166089624 | NM_024438 |
| Fbxo48 | 28.08582094 | 0.166413655 | 0.167389604 | 0.994169591 | 0.32014031 | 0.553376867 | | Naa16 | 213.272083 | 0.0553174425282574 | 0.0878810422844156 | 0.629458198154156 | 0.529049127221131 | 0.857868166089624 | NM_025832 |
| Hdx | 87.05584079 | 0.145602817 | 0.146476581 | 0.994034789 | 0.320205931 | 0.553376867 | | Zfp541 | 34.0941933 | -0.0532442352977265 | 0.0846123661754188 | -0.629272501224469 | 0.529170670962503 | 0.857868166089624 | NM_001099277 |
| Msc | 41.99876292 | -0.160239097 | 0.161187728 | -0.99411475 | 0.320167005 | 0.553376867 | | Alkbh7 | 346.6968999 | -0.0503912499769442 | 0.0802179416562588 | -0.62817929426406 | 0.529886492803829 | 0.857915133398583 | NM_027372,NM_025538 |
| Rps6ka4 | 1611.330204 | 0.067187067 | 0.067597787 | 0.993924055 | 0.320259842 | 0.553376867 | | BC048546 | 337.2759692 | 0.0560522951985946 | 0.0892563217944181 | 0.627992438761912 | 0.530008893287421 | 0.857915133398583 | NM_001001179 |
| Snx19 | 3923.275107 | -0.05514018 | 0.055475553 | -0.993954592 | 0.320244975 | 0.553376867 | | Chek1 | 43.12398007 | -0.0586130359035468 | 0.0932690828500913 | -0.628429422831935 | 0.529722667519668 | 0.857915133398583 | NM_007691 |
| Tarsl2 | 1813.202553 | -0.062830788 | 0.063202139 | -0.994124394 | 0.32016231 | 0.553376867 | | Creb3l1 | 110.1748762 | 0.0596939895371187 | 0.0950862947840437 | 0.627787523666721 | 0.530143140296659 | 0.857915133398583 | NM_011957 |
| Unc45a | 1106.358599 | -0.085413042 | 0.085934394 | -0.993933134 | 0.320255422 | 0.553376867 | | Elovl5 | 2882.541524 | 0.0366221329192302 | 0.0582782069171171 | 0.628401847903693 | 0.52974072685068 | 0.857915133398583 | NM_134255 |
| Zfp26 | 809.9906615 | 0.074063048 | 0.074516044 | 0.993920825 | 0.320261415 | 0.553376867 | | Fndc3a | 2498.566194 | 0.0434420530695061 | 0.0691787885163169 | 0.627967820790323 | 0.530025020464308 | 0.857915133398583 | NM_207636 |
| Tsr2 | 3326.378483 | -0.046446497 | 0.046742711 | -0.993662877 | 0.320387022 | 0.553532172 | | Gatad2b | 1173.615804 | 0.0460345620937602 | 0.0732627374046319 | 0.628348922311082 | 0.529775389677833 | 0.857915133398583 | NM_139304 |
| Pomp | 2825.773596 | -0.04697457 | 0.047280567 | -0.993528044 | 0.320452691 | 0.553583899 | | Ginm1 | 456.5552701 | 0.0446311484432521 | 0.0709841736118879 | 0.628747876599956 | 0.529514128929542 | 0.857915133398583 | NM_145418 |
| A830082N09Rik | 943.8772894 | -0.084824373 | 0.08538533 | -0.993430292 | 0.320500306 | 0.553584696 | | Hnrnpa0 | 2886.35905 | -0.0445935494805322 | 0.0710606373315989 | -0.627542211202524 | 0.530303875741702 | 0.857915133398583 | NM_029872 |
| Ppap2a | 743.5537633 | 0.09742333 | 0.098072531 | 0.993380399 | 0.32052461 | 0.553584696 | | Kansl1 | 4712.068806 | 0.0280362685389236 | 0.0446702058360376 | 0.62762792367313 | 0.530247711770562 | 0.857915133398583 | NM_001081045 |
| Ndfip2 | 3669.301209 | -0.076149415 | 0.076667635 | -0.993240691 | 0.320592673 | 0.553640534 | | Map3k2 | 1214.875371 | -0.0492128216721949 | 0.0782327869765828 | -0.62905622532565 | 0.5293122473866 | 0.857915133398583 | NM_011946 |
| Ccdc134 | 312.4128003 | 0.122674435 | 0.123531581 | 0.99306132 | 0.320680072 | 0.553706097 | | Ppp1r3e | 283.8356564 | -0.0517300951238472 | 0.0824450515054654 | -0.627449363900488 | 0.530364718292667 | 0.857915133398583 | NM_001167908 |
| Xylt2 | 965.3961535 | 0.076848562 | 0.07738904 | 0.993016094 | 0.320702111 | 0.553706097 | | Rbm15 | 329.4823395 | -0.050613879534085 | 0.0806285275859453 | -0.627741582904805 | 0.530173240058585 | 0.857915133398583 | NM_001045807 |
| Alkbh8 | 584.0131489 | -0.070418267 | 0.070924838 | -0.992857648 | 0.320779332 | 0.55377712 | | Serpinb6b | 54.99398414 | 0.058676518733564 | 0.0935056541006473 | 0.627518403009148 | 0.530319476839526 | 0.857915133398583 | NM_011454 |
| Grsf1 | 3935.872799 | 0.076262765 | 0.07682419 | 0.992692085 | 0.320860033 | 0.55385532 | | Snx2 | 2214.221924 | 0.0410175435696828 | 0.0653307450803483 | 0.627844417191884 | 0.530105865636545 | 0.857915133398583 | NM_026386 |
| Dbf4 | 56.78960586 | 0.146320648 | 0.147410479 | 0.992606825 | 0.320901597 | 0.553865361 | | Spp1 | 503.9173172 | 0.0563967196851558 | 0.0897640582934051 | 0.628277294469197 | 0.529822303099823 | 0.857915133398583 | NM_001204233,NM_009263,NM_001204203,NM_001204202,NM_001204201 |
| Cntn6 | 765.6208954 | -0.122141397 | 0.123061623 | -0.992522232 | 0.32094284 | 0.553874845 | | Stx18 | 900.6309211 | 0.0344330363666526 | 0.0548606130671854 | 0.627645854494625 | 0.530235962798902 | 0.857915133398583 | NM_026959 |
| Adra2b | 46.40447594 | -0.166374692 | 0.167716848 | -0.991997488 | 0.321198751 | 0.554073138 | | Tnfaip3 | 299.167344 | -0.0584086481937698 | 0.0929603934048386 | -0.628317566809368 | 0.52979592603959 | 0.857915133398583 | NM_001166402,NM_009397 |
| Fip1l1 | 2102.694995 | -0.054873898 | 0.055309061 | -0.992132156 | 0.321133062 | 0.554073138 | | Yipf6 | 1798.166531 | 0.0395086260657126 | 0.0628498925566683 | 0.628618832245255 | 0.529598628246165 | 0.857915133398583 | NM_207633 |
| Nudt10 | 440.4827162 | -0.090983846 | 0.091718204 | -0.991993323 | 0.321200783 | 0.554073138 | | Plekhf2 | 306.6307872 | 0.0510726250744728 | 0.0814136042042752 | 0.627322983347185 | 0.530447540775882 | 0.857953566118524 | NM_175175 |
| Themis2 | 47.28465368 | 0.166413051 | 0.167745806 | 0.992054912 | 0.32117074 | 0.554073138 | | Ankrd54 | 527.228082 | 0.0480027371206409 | 0.076563947065819 | 0.626962675779696 | 0.530683701498334 | 0.858116380871637 | NM_144849 |
| Gpr123 | 2680.995365 | 0.083363915 | 0.084104034 | 0.991199953 | 0.321587954 | 0.554679256 | | Capn1 | 1104.069116 | -0.0370570761137395 | 0.0591180762908684 | -0.62683156216745 | 0.530769652761383 | 0.858116380871637 | NM_001110504,NM_007600 |
| Chadl | 351.7282444 | -0.107505992 | 0.108514956 | -0.990702077 | 0.321831077 | 0.554975037 | | Ckb | 49969.98867 | 0.0319408386667038 | 0.0509578567990183 | 0.626808909815044 | 0.530784502418536 | 0.858116380871637 | NM_021273 |
| Pogk | 3354.731617 | -0.056328934 | 0.056854623 | -0.990753798 | 0.321805814 | 0.554975037 | | Rbbp8 | 189.3168293 | -0.0532148183497471 | 0.0848685549242537 | -0.627026327916647 | 0.530641977331869 | 0.858116380871637 | NR_045526,NM_001252495,NR_045527,NM_001081223 |
| Nsmf | 18516.86405 | -0.083431518 | 0.084246676 | -0.990324155 | 0.322015704 | 0.555231617 | | Cyp27a1 | 296.6152398 | -0.0580965114180417 | 0.0927340752044804 | -0.626485046515402 | 0.530996842031137 | 0.858310785186028 | NM_024264 |
| Elk3 | 358.178704 | -0.089342733 | 0.090247608 | -0.989973417 | 0.322187113 | 0.555465352 | | Rbm34 | 769.6456439 | 0.0420444692719011 | 0.0671159500797427 | 0.626445267063144 | 0.531022926233432 | 0.858310785186028 | NM_172762 |
| Dnajc4 | 337.6439497 | 0.106998797 | 0.108104196 | 0.989774685 | 0.322284261 | 0.555571021 | | 4732415M23Rik | 26.35568566 | 0.0538277467616166 | 0.0861236955917778 | 0.625005074291721 | 0.531967727733711 | 0.858328942120579 | NM_177279 |
| Bcap29 | 1298.491283 | -0.091975779 | 0.092960745 | -0.989404493 | 0.322465277 | 0.555799581 | | A330050F15Rik | 292.3073731 | 0.0531609737276205 | 0.0849633298110227 | 0.625693153103372 | 0.531516225074966 | 0.858328942120579 | NM_001145192 |
| Il1rl1 | 11.62216241 | -0.159309111 | 0.161034844 | -0.989283482 | 0.322524463 | 0.555799581 | | Acad12 | 89.93531835 | -0.0581724826030151 | 0.0932306827360291 | -0.623962851025378 | 0.532651982269278 | 0.858328942120579 | NM_178799 |
| Limk1 | 2455.64453 | -0.072090263 | 0.072866682 | -0.98934467 | 0.322494536 | 0.555799581 | | Ap4b1 | 686.7151916 | -0.0411497566136272 | 0.0659086784741466 | -0.624345041749982 | 0.532401009528107 | 0.858328942120579 | NM_026193,NM_001163553,NM_001163552 |
| Lpar6 | 140.1215552 | -0.141376609 | 0.142960461 | -0.988921045 | 0.322701773 | 0.556002754 | | Apba3 | 579.6376837 | -0.0497437906459181 | 0.0795986477843303 | -0.624932609165638 | 0.532015289111802 | 0.858328942120579 | NM_018758 |
| Zfp622 | 1083.438995 | 0.083250714 | 0.084185528 | 0.988895786 | 0.322714132 | 0.556002754 | | Blzf1 | 878.427211 | -0.0382097983522485 | 0.0611048146852994 | -0.625315673552661 | 0.531763895060156 | 0.858328942120579 | NM_001160208,NM_001160209,NM_025505 |
| Fgf22 | 28.47812483 | 0.164339062 | 0.16623539 | 0.988592512 | 0.322862551 | 0.556196616 | | Cep72 | 68.102927 | -0.0595520582334176 | 0.0951027430830683 | -0.626186546285015 | 0.531192590611945 | 0.858328942120579 | NM_028959 |
| Shfm1 | 1038.680663 | -0.062438497 | 0.063179023 | -0.988278927 | 0.323016063 | 0.556399208 | | Efna1 | 161.7267738 | -0.0564082168061385 | 0.090145557953308 | -0.625745938977446 | 0.531481596133514 | 0.858328942120579 | NM_010107,NM_001162425 |
| Ankrd26 | 765.9500307 | 0.085813637 | 0.086919995 | 0.987271532 | 0.323509543 | 0.557187289 | | Fam122a | 394.1339247 | 0.0557342793118039 | 0.0893014732799225 | 0.624113771752684 | 0.532552870177088 | 0.858328942120579 | NM_026520 |
| Lyl1 | 71.17327148 | -0.148380005 | 0.150323365 | -0.987072134 | 0.323607277 | 0.557293671 | | Gpd2 | 5447.839175 | 0.0293686234238392 | 0.0469055569699246 | 0.62612247505493 | 0.531234611605762 | 0.858328942120579 | NM_001145820,NM_010274 |
| Lrig3 | 108.3421125 | 0.124008179 | 0.12565297 | 0.986910053 | 0.323686736 | 0.557368558 | | Ing4 | 1483.557911 | -0.0358934633753519 | 0.0575156354598902 | -0.624064449403206 | 0.532585259938042 | 0.858328942120579 | NM_133345 |
| Exosc5 | 317.1288025 | -0.091673044 | 0.092930867 | -0.986464962 | 0.323905001 | 0.557682418 | | Josd2 | 1105.178138 | -0.0508877302354472 | 0.0813010060006698 | -0.625917596087654 | 0.531368992398281 | 0.858328942120579 | NM_001205073,NM_001205072,NM_001205071,NM_001205070,NM_025368 |
| Gmpr | 1499.908432 | -0.058413494 | 0.059221719 | -0.986352548 | 0.323960142 | 0.557715382 | | Kcnab3 | 4350.224734 | 0.0463916260896419 | 0.0743488318531704 | 0.623972494702534 | 0.532645648830734 | 0.858328942120579 | NM_010599 |
| Rabgef1 | 1485.687142 | 0.083891134 | 0.085081662 | 0.986007227 | 0.324129566 | 0.557945062 | | Ksr2 | 965.5400077 | 0.0452067597152131 | 0.0723183964226257 | 0.625107330243147 | 0.531900617267106 | 0.858328942120579 | NM_001114545 |
| Sertad1 | 380.1337679 | -0.142143477 | 0.144231772 | -0.985521258 | 0.324368095 | 0.558293631 | | Lnp | 2714.706288 | 0.0483996822056558 | 0.077441968576754 | 0.6249800088396 | 0.531984178816487 | 0.858328942120579 | NM_001110209,NM_027133 |
| Cdc14b | 436.2300373 | 0.11357372 | 0.115271443 | 0.985271956 | 0.324490504 | 0.558442282 | | Map3k12 | 6354.295899 | -0.026927922339572 | 0.0431207607692896 | -0.624476977195401 | 0.532314385533979 | 0.858328942120579 | NM_001163643,NM_009582 |
| Parp4 | 390.0265235 | 0.092112743 | 0.093504183 | 0.985119855 | 0.324565644 | 0.558509561 | | Mgat5 | 1827.879643 | 0.0473199754423709 | 0.0757387429927402 | 0.624778991208062 | 0.532116121033238 | 0.858328942120579 | NM_145128 |
| Klhl28 | 484.1084474 | 0.088199877 | 0.089549443 | 0.984929377 | 0.324658762 | 0.558546456 | | Myc | 140.1656095 | 0.0578567399622111 | 0.0926500497709823 | 0.624465287446955 | 0.532322060307669 | 0.858328942120579 | NM_001177354,NM_001177352,NM_001177353,NM_010849 |
| Slc2a6 | 854.0496946 | 0.112991328 | 0.114720333 | 0.98492852 | 0.324659183 | 0.558546456 | | Pkn2 | 1177.920086 | -0.0410782562998072 | 0.0656816944449026 | -0.625414076889656 | 0.531699325522287 | 0.858328942120579 | NM_178654 |
| Nefm | 8992.632089 | 0.09130271 | 0.092777131 | 0.984107929 | 0.325062449 | 0.559178148 | | Pomgnt1 | 3328.139462 | 0.0354039113144044 | 0.056618060918222 | 0.625311265349428 | 0.531766787693724 | 0.858328942120579 | NM_029786,NM_026651 |
| Mtrr | 475.9707847 | 0.097027502 | 0.098631512 | 0.983737353 | 0.325244669 | 0.559429496 | | Prrg4 | 37.51370285 | 0.0566151893824074 | 0.0907429927714554 | 0.623907010924777 | 0.532688655735717 | 0.858328942120579 | NM_178695 |
| Vegfb | 1866.206416 | -0.096105436 | 0.097704639 | -0.983632269 | 0.325296353 | 0.559456287 | | Sipa1 | 594.1322035 | -0.0447326451322029 | 0.0716970648265201 | -0.623911805043052 | 0.532685507107158 | 0.858328942120579 | NM_001164568,NM_011379,NM_001164482,NM_001164480,NM_001164481 |
| Aak1 | 20381.28435 | 0.062103128 | 0.063197021 | 0.982690741 | 0.325759669 | 0.559970267 | | Sp3 | 1855.246975 | 0.0439259209119343 | 0.0702514846882019 | 0.625266798372893 | 0.531795967073097 | 0.858328942120579 | NM_001018042,NM_001098425 |
| Cmc1 | 306.8782707 | -0.100486729 | 0.102265053 | -0.982610643 | 0.325799104 | 0.559970267 | | Thap1 | 305.8990433 | -0.0516440510982565 | 0.0827205021745708 | -0.62431984502788 | 0.532417553587001 | 0.858328942120579 | NM_199042 |
| Hecw2 | 2452.726486 | -0.086744005 | 0.088277071 | -0.982633468 | 0.325787866 | 0.559970267 | | Thnsl2 | 194.162328 | 0.0568767209775377 | 0.0908446185279116 | 0.626087949943481 | 0.531257255533073 | 0.858328942120579 | NM_178413,NM_001033929 |
| Pdia5 | 133.9333575 | 0.147946442 | 0.150529521 | 0.982840051 | 0.325686166 | 0.559970267 | | Trip4 | 1733.957382 | 0.035581317071514 | 0.056954490578451 | 0.624732426014823 | 0.532146687456743 | 0.858328942120579 | NM_001170907,NM_019797 |
| Smpd2 | 462.145808 | 0.082124492 | 0.083580094 | 0.982584343 | 0.325812053 | 0.559970267 | | Zfp275 | 1480.222797 | 0.0453698873323489 | 0.0727012397728109 | 0.624059333432668 | 0.532588619629528 | 0.858328942120579 | NM_031494,NM_001160229 |
| Tbc1d22a | 1737.311906 | 0.084584179 | 0.086052867 | 0.982932727 | 0.325640549 | 0.559970267 | | Zswim7 | 271.9425458 | 0.0496286042361958 | 0.079351004714892 | 0.62543132773821 | 0.53168800640374 | 0.858328942120579 | NM_027198 |
| Sall2 | 2990.642919 | -0.072206844 | 0.073533545 | -0.981957885 | 0.326120597 | 0.56043839 | | Fbxl2 | 2273.018148 | -0.0293481006689133 | 0.0470582776102206 | -0.623654374093352 | 0.532854593114034 | 0.858501089127601 | NM_178624 |
| St7 | 660.3346173 | -0.076847013 | 0.078277946 | -0.981719854 | 0.326237881 | 0.560577768 | | Chm | 2007.146936 | 0.0514255429356268 | 0.082548795058125 | 0.62297145463379 | 0.533303280352704 | 0.859128695841563 | NM_018818 |
| Nup188 | 944.641777 | 0.059704531 | 0.060827825 | 0.981533214 | 0.326329864 | 0.560673643 | | Ganab | 5678.249337 | 0.0280868587882524 | 0.0450983423774091 | 0.622791377856091 | 0.533421625025173 | 0.859128786463651 | NM_008060 |
| Nelfe | 688.2859197 | -0.097270607 | 0.099130794 | -0.981235025 | 0.326476856 | 0.560801819 | | Lysmd3 | 275.1516301 | 0.0540091032484471 | 0.08671202634689 | 0.622855969625073 | 0.533379174428418 | 0.859128786463651 | NM_030257 |
| Sptssb | 83.84425516 | 0.138018383 | 0.140649494 | 0.981293137 | 0.326448206 | 0.560801819 | | Dctpp1 | 264.955618 | 0.0491753592133481 | 0.0790601027574056 | 0.621977336207573 | 0.533956770709216 | 0.859470724637387 | NM_023203 |
| Cdc14a | 138.9501469 | 0.113355112 | 0.115580503 | 0.980745966 | 0.32671803 | 0.561007261 | | Emc9 | 565.8434124 | 0.0447288222336243 | 0.0719011131469892 | 0.622088035579978 | 0.533883981735321 | 0.859470724637387 | NM_033146 |
| Sgsm2 | 5583.705884 | -0.085509939 | 0.087192864 | -0.980698815 | 0.326741288 | 0.561007261 | | Lmbrd2 | 1637.994182 | 0.0481933510346125 | 0.0774901893303118 | 0.621928420244042 | 0.533988936379762 | 0.859470724637387 | NM_177178 |
| Tbc1d7 | 512.1636642 | 0.083292329 | 0.084920268 | 0.98082979 | 0.326676685 | 0.561007261 | | Nom1 | 1217.881048 | 0.0385296122675656 | 0.0619426447140747 | 0.622020781408626 | 0.533928203268559 | 0.859470724637387 | NM_001033457 |
| Vapa | 4059.633244 | -0.069371227 | 0.070721514 | -0.980906987 | 0.326638612 | 0.561007261 | | Shb | 464.1924432 | 0.0501910933918537 | 0.0806755981392256 | 0.622134753872374 | 0.533853264216371 | 0.859470724637387 | NM_001033306 |
| Smad3 | 3867.658412 | 0.091008676 | 0.092840304 | 0.980271205 | 0.326952264 | 0.5613073 | | Specc1l | 2746.471139 | -0.0372206436306351 | 0.0598413518078098 | -0.621988683153956 | 0.5339493092007 | 0.859470724637387 | NM_153406,NM_001145826 |
| Ncl | 10173.18772 | 0.059429508 | 0.060631904 | 0.980168912 | 0.327002747 | 0.561331771 | | Map3k4 | 2406.184743 | 0.0344769377095605 | 0.0554583248376225 | 0.621672901417526 | 0.534156973802116 | 0.859551797656774 | NM_011948 |
| Plk1 | 39.20613737 | -0.160407184 | 0.163686481 | -0.979965988 | 0.327102908 | 0.561441504 | | Rad51d | 1095.186318 | 0.0374325508667086 | 0.0601982377091666 | 0.621671867663494 | 0.534157653686334 | 0.859551797656774 | NM_011235 |
| Dars | 1779.59311 | -0.080032708 | 0.081716107 | -0.979399423 | 0.327382662 | 0.561859434 | | Chchd1 | 624.0586705 | 0.0435155785881137 | 0.0700275311614272 | 0.621406721990553 | 0.534332050367663 | 0.859648542365729 | NM_025366 |
| Pomgnt1 | 2992.520243 | -0.066354142 | 0.067757572 | -0.97928748 | 0.327437955 | 0.561892089 | | Peg3 | 11309.99947 | 0.0515211564500952 | 0.0829113516199719 | 0.621400513240272 | 0.534336134449964 | 0.859648542365729 | NM_008817 |
| Dio2 | 3857.314582 | -0.163490064 | 0.167035254 | -0.978775794 | 0.327690772 | 0.562263658 | | Ubr7 | 1946.984824 | -0.0373407862598875 | 0.0601359081924158 | -0.620939923953737 | 0.534639151522524 | 0.860040787930917 | NM_025666 |
| Acp5 | 9.529002625 | 0.149507356 | 0.152771755 | 0.978632184 | 0.327761751 | 0.562323173 | | Slc29a3 | 772.4344231 | -0.0386181713842291 | 0.0622043483493379 | -0.620827521049662 | 0.534713113424516 | 0.8600645206073 | NM_023596 |
| 1110012L19Rik | 303.9756137 | -0.087166783 | 0.089105219 | -0.978245541 | 0.327952897 | 0.562588816 | | Thrsp | 618.7899953 | 0.0461168522547333 | 0.0743241317296862 | 0.620482897722626 | 0.534939913459197 | 0.860334054794984 | NM_009381 |
| Herc3 | 7350.090016 | 0.082855261 | 0.084715982 | 0.978035775 | 0.32805663 | 0.562704465 | | Ampd3 | 1149.829412 | -0.0343065756090262 | 0.0553088557336316 | -0.620272742112896 | 0.535078237463554 | 0.860420882847261 | NM_009667,NM_001276301 |
| Brd2 | 8207.414125 | 0.073183656 | 0.074839012 | 0.977881102 | 0.328133133 | 0.562773385 | | Elk4 | 383.8420086 | 0.0510672043890815 | 0.0823371236681437 | 0.620220893249875 | 0.535112367867421 | 0.860420882847261 | NM_007923 |
| Rcc1 | 452.0523291 | 0.093637627 | 0.095789261 | 0.977537845 | 0.328302951 | 0.563002317 | | Xrcc6bp1 | 141.7020967 | -0.0576401078013251 | 0.0929515004020394 | -0.620109493144453 | 0.535185702610951 | 0.860443554634939 | NM_026858,NM_001159559 |
| Bri3 | 929.1652977 | -0.078051438 | 0.079888679 | -0.977002486 | 0.328567922 | 0.563394354 | | Ebf4 | 369.5028124 | -0.055553455828949 | 0.0896041629718597 | -0.619987442395904 | 0.535266054493859 | 0.860477501945306 | NM_001110513 |
| Rab4a | 1447.861186 | 0.099692449 | 0.102055185 | 0.976848447 | 0.328644187 | 0.563434496 | | 4930452B06Rik | 566.8694402 | 0.0469056491429923 | 0.0757516043634529 | 0.619203375785166 | 0.535782388252526 | 0.860926434928783 | NM_028934 |
| Trf | 11240.14151 | 0.163752615 | 0.167640482 | 0.976808309 | 0.328664062 | 0.563434496 | | Mrfap1 | 9980.236079 | 0.0322919370927195 | 0.0521334747473638 | 0.619408877869825 | 0.53564703408787 | 0.860926434928783 | NM_026242 |
| Ager | 18.68590809 | 0.164020095 | 0.167942052 | 0.97664696 | 0.328743962 | 0.563509121 | | Nfs1 | 1520.615002 | 0.0337965933610329 | 0.0545675378795959 | 0.619353459479986 | 0.53568353377236 | 0.860926434928783 | NM_010911 |
| Gm13031 | 52.40378175 | -0.147575768 | 0.151130701 | -0.976477752 | 0.328827768 | 0.563590424 | | Slc19a3 | 156.7515836 | 0.0584103207795118 | 0.0943298839543834 | 0.619213321705752 | 0.535775836964775 | 0.860926434928783 | NM_030556 |
| Gm15997 | 18.51521607 | -0.163773338 | 0.167720295 | -0.976176016 | 0.328977246 | 0.563784255 | | Arl4a | 1364.464105 | -0.0475774830720311 | 0.0768675305170794 | -0.618954228183943 | 0.535946512267718 | 0.860987999916764 | NM_007487,NM_001039515 |
| Mospd3 | 454.8439959 | 0.086188777 | 0.088299353 | 0.97609749 | 0.329016155 | 0.563788576 | | F2rl2 | 88.43269118 | 0.0585376203763057 | 0.0945643667275347 | 0.619024082770715 | 0.535900493855107 | 0.860987999916764 | NM_010170 |
| Cysltr2 | 11.12522589 | -0.154290943 | 0.158088402 | -0.975978888 | 0.329074927 | 0.563826929 | | Folh1 | 314.6547868 | 0.0538748635562009 | 0.0870528594771083 | 0.618875288873975 | 0.535998518742069 | 0.860987999916764 | NM_016770,NM_001159706 |
| Zfp61 | 526.4502326 | 0.088913977 | 0.09113482 | 0.975631229 | 0.329247243 | 0.564059794 | | Grsf1 | 4335.633866 | 0.0411106074242304 | 0.0664462021312734 | 0.61870514951345 | 0.536110617006535 | 0.86107284637737 | NM_001098476,NM_178700 |
| Tbca | 1058.238075 | -0.084332333 | 0.086461752 | -0.975371549 | 0.329375991 | 0.564217977 | | Brd3 | 2337.284655 | 0.0328840467580485 | 0.0531719206806414 | 0.618447600483629 | 0.536280328583288 | 0.861250199336743 | NM_023336,NM_001113573,NM_001113574 |
| Cps1 | 18.15517449 | -0.163413052 | 0.167609236 | -0.974964483 | 0.329577878 | 0.564501398 | | Tctex1d2 | 290.3038354 | 0.0500304007762486 | 0.0809192599529528 | 0.61827556010408 | 0.536393709414343 | 0.861337057589293 | NM_025329 |
| Alg1 | 893.8590091 | -0.081416518 | 0.083595768 | -0.9739311 | 0.330090751 | 0.565317353 | | Uqcrb | 2312.6422 | 0.0447521724114372 | 0.0724200874384023 | 0.617952476921567 | 0.536606665504677 | 0.861583776182263 | NM_026219 |
| Irf6 | 58.86127556 | -0.148229066 | 0.152211167 | -0.973838314 | 0.330136826 | 0.565333774 | | E2f6 | 1429.334547 | 0.0418622069473103 | 0.0677721497271494 | 0.617690409937526 | 0.536779434793346 | 0.861670689557758 | NR_027677,NM_033270 |
| Gm15910 | 8.59686301 | 0.153783761 | 0.157944595 | 0.973656371 | 0.330227187 | 0.565426018 | | Marcks | 3502.891669 | 0.0445241951132353 | 0.072073766546846 | 0.617758683172133 | 0.536734422544494 | 0.861670689557758 | NM_008538 |
| Acsl3 | 6925.88872 | 0.054268077 | 0.055759997 | 0.973243906 | 0.330432094 | 0.56571435 | | Tex15 | 41.8687796 | 0.057295511949692 | 0.0927852221655976 | 0.617506868145817 | 0.536900452515181 | 0.861769720799505 | NM_031374 |
| Slc25a12 | 6321.78113 | 0.081454692 | 0.083707409 | 0.973088191 | 0.330509473 | 0.565744467 | | 9930021J03Rik | 2943.656426 | 0.0471970538226293 | 0.0764897287608064 | 0.617037798241132 | 0.53720979464569 | 0.861996453104079 | NM_172836 |
| Zfp260 | 2441.661549 | 0.063790847 | 0.065561797 | 0.972988079 | 0.330559227 | 0.565744467 | | Cog6 | 2107.264933 | 0.0317371438520217 | 0.0514726486443931 | 0.616582683966444 | 0.537510018910822 | 0.861996453104079 | NM_026225 |
| Zscan21 | 641.3482048 | 0.075605983 | 0.077703557 | 0.973005427 | 0.330550605 | 0.565744467 | | Eif3m | 2165.208712 | 0.0348001363435819 | 0.0563914796602968 | 0.617116921797734 | 0.537157607978138 | 0.861996453104079 | NM_145380 |
| Ncoa2 | 4805.521027 | -0.079476351 | 0.081719388 | -0.972551966 | 0.330776026 | 0.566052987 | | Gtf2f1 | 2556.536917 | 0.029312423225838 | 0.0475280238461965 | 0.61673978536736 | 0.537406374639271 | 0.861996453104079 | NM_133801 |
| Prr13 | 1190.916385 | 0.073538236 | 0.075625809 | 0.972396028 | 0.330853567 | 0.566123156 | | Lrp11 | 8240.367894 | 0.0360981448101478 | 0.0585451950219397 | 0.616585952043035 | 0.537507862765293 | 0.861996453104079 | NM_172784 |
| Dnase2a | 153.3761961 | 0.133574201 | 0.137382443 | 0.972279996 | 0.330911273 | 0.566143084 | | Rbm42 | 1653.921883 | 0.0364001419464403 | 0.0590253213667636 | 0.616686891381108 | 0.537441269188252 | 0.861996453104079 | NM_133693 |
| Unc45b | 54.48967995 | 0.148276526 | 0.152512458 | 0.97222567 | 0.330938293 | 0.566143084 | | Smg9 | 1394.519165 | -0.0417140787260451 | 0.0676238415595246 | -0.616854614645445 | 0.537330624844052 | 0.861996453104079 | NM_028047 |
| 4930507D05Rik | 31.82166681 | 0.154840563 | 0.159298146 | 0.972017359 | 0.331041913 | 0.566257827 | | Trmt61a | 731.0937212 | -0.0510403891250359 | 0.0827807797228971 | -0.616572944781266 | 0.537516444459366 | 0.861996453104079 | NM_177374,NM_001099793,NM_001099792 |
| Ankrd44 | 412.8962872 | 0.100296663 | 0.103192648 | 0.971936129 | 0.331082325 | 0.566264437 | | Spc24 | 67.91946935 | 0.0585844369852433 | 0.0950345616258862 | 0.616454014023521 | 0.537594913618311 | 0.86202712387897 | NM_026282 |
| LOC100504703 | 25.03549314 | 0.162639557 | 0.167363491 | 0.971774402 | 0.331162793 | 0.56633955 | | Kirrel | 380.0182984 | -0.0476893950234449 | 0.0773858831404998 | -0.616254452208825 | 0.537726595155749 | 0.862037665157456 | NM_130867,NM_001170985 |
| Dscam | 2964.854063 | 0.11010571 | 0.113316144 | 0.971668344 | 0.33121557 | 0.566339732 | | Ptprm | 2015.425282 | 0.0430677301929352 | 0.0698953748001889 | 0.61617425067186 | 0.537779520973211 | 0.862037665157456 | NM_008984 |
| Paqr6 | 220.7617775 | 0.128704093 | 0.132472425 | 0.971553839 | 0.331272557 | 0.566339732 | | Usp42 | 1091.588729 | 0.0347168856680571 | 0.0563267146446187 | 0.616348492666328 | 0.537664540224337 | 0.862037665157456 | NM_029749 |
| Zfp870 | 450.9737637 | -0.079162468 | 0.081478728 | -0.971572217 | 0.33126341 | 0.566339732 | | Adar | 9583.344352 | 0.0311529886843385 | 0.0506187679960658 | 0.615443439610378 | 0.538261911171796 | 0.862082843901637 | NM_001146296,NM_001038587,NM_019655 |
| Riok2 | 806.2805603 | 0.066655769 | 0.068622996 | 0.971332834 | 0.331382563 | 0.566465295 | | Cbx3 | 3589.586358 | 0.0358186493955826 | 0.0582341531583793 | 0.615079767678026 | 0.53850204284229 | 0.862082843901637 | NM_007624 |
| 8-Mar | 2597.930665 | -0.068114055 | 0.070130099 | -0.971252789 | 0.331422412 | 0.566470916 | | Cnot4 | 1447.677367 | 0.0440895342497128 | 0.0716504506319189 | 0.615342037082343 | 0.538328861606028 | 0.862082843901637 | NM_001164411,NM_001164413,NM_001164412,NM_001164414,NM_016877 |
| Ak4 | 2110.117776 | -0.086760008 | 0.089375159 | -0.97073963 | 0.331677952 | 0.566732816 | | Ctxn3 | 73.23637755 | 0.0565254963298255 | 0.0917569042110022 | 0.616035347049642 | 0.537871191086937 | 0.862082843901637 | NM_001134697 |
| Cdk16 | 11345.36541 | 0.068118343 | 0.070176592 | 0.970670442 | 0.331712415 | 0.566732816 | | Gngt2 | 68.70333517 | -0.0584557269946834 | 0.0950418087083175 | -0.615052762454085 | 0.538519876467262 | 0.862082843901637 | NM_001038664,NM_023121 |
| Cenpk | 20.61194501 | -0.162615038 | 0.167520993 | -0.970714389 | 0.331690525 | 0.566732816 | | Hk1 | 18313.52239 | 0.0295254458474566 | 0.0479936198633776 | 0.615195226605245 | 0.538425799779233 | 0.862082843901637 | NM_010438,NM_001146100 |
| Habp4 | 8761.111421 | 0.059828579 | 0.061637561 | 0.9706513 | 0.331721951 | 0.566732816 | | Iqcd | 45.21330901 | -0.0558812689478162 | 0.0908333462309869 | -0.615206543263434 | 0.538418327138869 | 0.862082843901637 | NM_029408 |
| 2-Mar | 2226.807536 | 0.053725044 | 0.055353777 | 0.970575937 | 0.331759493 | 0.566734464 | | Lmo3 | 3682.135072 | 0.0466885925497346 | 0.0758727713204635 | 0.615353726207471 | 0.538321143715548 | 0.862082843901637 | NM_207222 |
| Pebp1 | 14166.94793 | -0.053667235 | 0.055320877 | -0.970108169 | 0.331992577 | 0.567070113 | | Ogfr | 1421.363754 | 0.0338365673975193 | 0.0550031396985695 | 0.615175198778686 | 0.538439024724761 | 0.862082843901637 | NM_031373 |
| 8430429K09Rik | 207.4729391 | -0.088628472 | 0.091390969 | -0.969772751 | 0.332159778 | 0.567168129 | | Ogg1 | 226.6285956 | 0.0538208955497207 | 0.0874365831192976 | 0.615542072089985 | 0.538196793654494 | 0.862082843901637 | NM_010957 |
| Mapkapk5 | 1177.370908 | 0.06873533 | 0.07086761 | 0.969911773 | 0.332090471 | 0.567168129 | | Rpp25 | 522.0898787 | 0.0441506837964711 | 0.0717724744606663 | 0.615147856169667 | 0.538457080093007 | 0.862082843901637 | NM_133982 |
| Trappc2 | 871.6540559 | -0.07851808 | 0.080962647 | -0.969806232 | 0.332143086 | 0.567168129 | | Rtn4r | 3476.872765 | 0.0426473706479693 | 0.0693293285443609 | 0.615141838863781 | 0.538461053590501 | 0.862082843901637 | NM_022982 |
| Athl1 | 246.9337955 | 0.138415607 | 0.142742931 | 0.969684495 | 0.332203781 | 0.567180759 | | 9330159M07Rik | 108.1508232 | -0.0576912774376629 | 0.0938711857633121 | -0.614579191351927 | 0.538832660000168 | 0.862393479414107 | NR_037983,NR_037982 |
| Tgm2 | 711.1601974 | -0.08493551 | 0.087611348 | -0.969457857 | 0.332316797 | 0.567311201 | | Skp1a | 11697.12948 | 0.0401985009199989 | 0.0654003444622052 | 0.614652739990225 | 0.538784076732377 | 0.862393479414107 | NM_011543 |
| 1700013G23Rik | 16.04302459 | -0.162031731 | 0.167167941 | -0.969275145 | 0.332407927 | 0.567404255 | | Grin2a | 1618.02739 | 0.0538454163906726 | 0.0876341897794379 | 0.614433893052397 | 0.538928644658432 | 0.862452075829942 | NM_008170 |
| Serp1 | 1327.144052 | -0.063963584 | 0.06599801 | -0.969174434 | 0.332458165 | 0.567427495 | | 9030617O03Rik | 368.5386635 | 0.0524237827103634 | 0.0854057066820386 | 0.61382060692425 | 0.53933387842095 | 0.862815409465058 | NM_145448 |
| 5730522E02Rik | 50.74946407 | 0.149145737 | 0.153903829 | 0.969083992 | 0.332503284 | 0.567441996 | | Mrpl45 | 1371.286693 | 0.0341684964445604 | 0.0556497190571219 | 0.613992254111616 | 0.539220445775595 | 0.862815409465058 | NM_025927 |
| 4930402H24Rik | 3950.366378 | -0.105681531 | 0.109062515 | -0.968999582 | 0.332545398 | 0.567451365 | | Tmed5 | 756.2455296 | 0.0489942332662925 | 0.0798183616015284 | 0.613821585450262 | 0.539333231730431 | 0.862815409465058 | NM_028876 |
| Thtpa | 1248.771011 | 0.053658528 | 0.055379604 | 0.968922207 | 0.332584004 | 0.567454748 | | Phf6 | 1002.65708 | 0.0484793716891042 | 0.0789959684644836 | 0.613694250876871 | 0.539417388159215 | 0.862853978680846 | NM_027642 |
| Pcdhac2 | 1395.671182 | -0.076511209 | 0.078977835 | -0.968768128 | 0.332660892 | 0.567523438 | | 1110058L19Rik | 493.4279497 | -0.0418537753330112 | 0.0683425434296379 | -0.612411731151062 | 0.540265382173683 | 0.863544777888966 | NM_026503 |
| A230077H06Rik | 179.8350525 | -0.130696424 | 0.135025926 | -0.967935772 | 0.333076448 | 0.567976249 | | 1700003M07Rik | 740.3104653 | -0.0417202298211725 | 0.0681191993490954 | -0.612459192413078 | 0.540233989206535 | 0.863544777888966 | NR_040648,NR_040647 |
| Cenpj | 138.8635373 | -0.10538385 | 0.108882305 | -0.967869392 | 0.333109602 | 0.567976249 | | Jph4 | 15315.35718 | -0.0343696465322325 | 0.0561118992113452 | -0.612519750985069 | 0.540193934427241 | 0.863544777888966 | NM_177049 |
| Mrpl42 | 1060.984099 | -0.075646349 | 0.07815639 | -0.967884391 | 0.333102111 | 0.567976249 | | March10 | 84.42266805 | -0.0580963202710148 | 0.0947910646912856 | -0.61288814995614 | 0.539950299211357 | 0.863544777888966 | NM_172568,NM_001039242 |
| Rimbp2 | 4717.511023 | -0.066892202 | 0.069098032 | -0.968076804 | 0.333006014 | 0.567976249 | | Pak6 | 3992.494436 | 0.0361889762277447 | 0.0590864397811191 | 0.612475152705153 | 0.540223432572076 | 0.863544777888966 | NM_001033254,NM_001145854 |
| Zfp637 | 943.4473277 | -0.067274999 | 0.069491538 | -0.968103471 | 0.332992697 | 0.567976249 | | Rps6ka4 | 1758.470363 | -0.0387643913242263 | 0.0632799707798882 | -0.612585480784491 | 0.54015046096518 | 0.863544777888966 | NM_019924 |
| Ergic1 | 5651.866556 | 0.051060161 | 0.052762678 | 0.967732558 | 0.333177954 | 0.568030283 | | Tcf20 | 5061.403951 | 0.0404003870099829 | 0.0659421719460235 | 0.612663881363392 | 0.540098609382494 | 0.863544777888966 | NM_001114140,NM_013836 |
| Tmem126b | 486.4551292 | 0.083845527 | 0.086652729 | 0.967603994 | 0.333242182 | 0.568077277 | | Spata6 | 169.6138203 | 0.0577738926975019 | 0.0943798532575758 | 0.612142217892932 | 0.540443667403612 | 0.8637347026851 | NM_026470 |
| Dock7 | 1723.705442 | 0.082603308 | 0.085385162 | 0.967419936 | 0.333334148 | 0.5681402 | | Angel1 | 582.0436381 | -0.0454403438542351 | 0.0743319245156482 | -0.611316660376109 | 0.540989963490423 | 0.863847445274474 | NM_144524 |
| Slc37a3 | 2308.314995 | -0.04881401 | 0.050459841 | -0.967383349 | 0.333352431 | 0.5681402 | | Atp2b2 | 34053.61818 | 0.0398068145534763 | 0.0650950120731069 | 0.611518659966912 | 0.540856268830794 | 0.863847445274474 | NM_001036684,NM_009723 |
| 4933436C20Rik | 11.90254202 | 0.146792999 | 0.151779216 | 0.967148221 | 0.333469943 | 0.56815299 | | Dsc2 | 24.77883008 | -0.0500262539806844 | 0.0818238820349913 | -0.611389398015742 | 0.54094181973818 | 0.863847445274474 | NM_013505 |
| Lrrc16a | 943.3036367 | 0.089885163 | 0.092936892 | 0.967163424 | 0.333462344 | 0.56815299 | | Epha3 | 201.2201566 | -0.0559116898187262 | 0.0914446770780006 | -0.611426499664213 | 0.540917263640309 | 0.863847445274474 | NM_010140 |
| Mrpl10 | 1835.139608 | -0.05203485 | 0.053799042 | -0.967207745 | 0.333440191 | 0.56815299 | | Fam172a | 1842.565285 | 0.0397510503662654 | 0.0649643221688893 | 0.61189048140799 | 0.540610219722115 | 0.863847445274474 | NM_138312,NR_028109,NM_001163419,NM_001163420 |
| Cisd3 | 616.9010725 | -0.074531986 | 0.077076045 | -0.966992881 | 0.333547592 | 0.568222803 | | Me3 | 1976.287777 | -0.0386633641872891 | 0.0632269500444236 | -0.611505602597054 | 0.540864910431377 | 0.863847445274474 | NM_181407 |
| Snx12 | 1715.34453 | -0.051862648 | 0.053643024 | -0.966810665 | 0.333638692 | 0.568284318 | | Ptprk | 3031.743985 | -0.0347401874159854 | 0.0568061649741 | -0.611556640583372 | 0.540831132972727 | 0.863847445274474 | NM_008983 |
| Tagln3 | 8194.455318 | -0.070310652 | 0.072727087 | -0.966773936 | 0.333657057 | 0.568284318 | | Tnks2 | 6593.270637 | 0.0436083857125843 | 0.0713310352531205 | 0.61135220535968 | 0.540966436629413 | 0.863847445274474 | NM_001163635 |
| Gga2 | 1352.494945 | 0.049248976 | 0.050948575 | 0.966640891 | 0.333723585 | 0.568335154 | | Polr3h | 1551.535296 | 0.0364858805816504 | 0.0597054798201947 | 0.611097686368638 | 0.541134911420809 | 0.863862965709127 | NM_030229 |
| Alg11 | 1359.126618 | -0.064948477 | 0.067197353 | -0.966533256 | 0.333777414 | 0.568364354 | | Tbc1d17 | 2175.508884 | 0.0349824412948794 | 0.0572513635441773 | 0.611032456334173 | 0.541178093621992 | 0.863862965709127 | NM_001042655 |
| Exoc7 | 2813.596921 | 0.047492834 | 0.049142827 | 0.966424534 | 0.333831792 | 0.568394483 | | Zfp865 | 956.7671921 | -0.0413888393937965 | 0.0677250207829197 | -0.611130700519988 | 0.541113056749973 | 0.863862965709127 | NM_001033383 |
| AA414768 | 269.4824003 | 0.093797578 | 0.097067412 | 0.966313783 | 0.333887191 | 0.568426343 | | Lpcat3 | 677.1899507 | 0.0489345871756886 | 0.0801094929869725 | 0.610846297375099 | 0.541301340084749 | 0.863964758386008 | NM_145130 |
| BC029722 | 2080.928881 | -0.078525213 | 0.081280564 | -0.966100739 | 0.333993775 | 0.568482868 | | Sept5 | 27566.69083 | 0.0364668368776586 | 0.0597237824998039 | 0.610591549148587 | 0.541470018756494 | 0.864139034348102 | NM_213614 |
| Slc45a4 | 1874.210921 | 0.076614925 | 0.079301858 | 0.966117656 | 0.333985311 | 0.568482868 | | Slc39a1 | 1617.050862 | -0.0316649275095948 | 0.0518937800987929 | -0.610187337467277 | 0.541737716818536 | 0.864471281391416 | NM_013901 |
| 6720456H20Rik | 1048.617676 | -0.072813577 | 0.07537793 | -0.965980053 | 0.334054163 | 0.568523198 | | Acsf2 | 767.1893369 | -0.0428027864337854 | 0.070159251492092 | -0.610080431639296 | 0.541808528593728 | 0.86448931089109 | NM_153807 |
| Mllt10 | 1337.155133 | 0.075851891 | 0.078548311 | 0.965671826 | 0.334208422 | 0.56872326 | | Cdc42bpa | 14426.72585 | 0.0374300596893401 | 0.0613829343602916 | 0.609779576024203 | 0.542007832695078 | 0.864560610941176 | NM_001033285 |
| Lhfpl4 | 6285.666257 | -0.052984995 | 0.054889947 | -0.965295068 | 0.334397042 | 0.568981744 | | Med13 | 2694.629995 | 0.0477884131609634 | 0.078382988440991 | 0.609678376793964 | 0.54207488111778 | 0.864560610941176 | NM_001080931 |
| Ccdc47 | 4314.204223 | 0.056049966 | 0.058072549 | 0.965171435 | 0.334458952 | 0.569024596 | | Rc3h1 | 1765.830521 | 0.0448846309183439 | 0.0736231687015547 | 0.609653614615423 | 0.542091287653327 | 0.864560610941176 | NM_001024952 |
| Nxt2 | 996.5639337 | -0.075297978 | 0.078040552 | -0.964857061 | 0.334616411 | 0.569229982 | | Senp6 | 3112.176232 | 0.0416431623794138 | 0.068277811492821 | 0.609907691370458 | 0.541922957233949 | 0.864560610941176 | NM_146003 |
| Ccdc132 | 2729.82367 | 0.06632758 | 0.068753778 | 0.964711783 | 0.334689192 | 0.569238517 | | Mre11a | 402.6208153 | 0.0457458189075736 | 0.0750514936224795 | 0.609525767907292 | 0.542176001571784 | 0.864600790298797 | NM_018736 |
| Ensa | 11811.77905 | -0.050964785 | 0.05283105 | -0.964674837 | 0.334707703 | 0.569238517 | | Hap1 | 9885.436271 | 0.0577643970774137 | 0.094830964610801 | 0.60913012236548 | 0.542438192887186 | 0.864833083907413 | NM_177981,NM_010404 |
| Mef2c | 18167.07783 | 0.099942273 | 0.103607164 | 0.964627049 | 0.334731646 | 0.569238517 | | Pla2g4a | 153.5100082 | -0.0572921419498358 | 0.0940562643090858 | -0.60912627532775 | 0.542440742632551 | 0.864833083907413 | NM_008869 |
| Ccdc65 | 243.6260114 | -0.101425111 | 0.105162211 | -0.964463468 | 0.334813616 | 0.569315426 | | Ilf3 | 3784.795342 | 0.0279765354403233 | 0.0459582385446176 | 0.60873820072897 | 0.542697981987632 | 0.865148253550521 | NM_001042708,NM_001042709,NM_010561,NM_001042707 |
| Corin | 8.551559676 | -0.147118153 | 0.152558428 | -0.964339729 | 0.334875629 | 0.569354094 | | Armc5 | 685.3417738 | -0.0434117028401978 | 0.0713642145367324 | -0.608311926670937 | 0.542980612302036 | 0.865503826873628 | NM_146205 |
| Erlin2 | 2205.048776 | 0.050799989 | 0.052685657 | 0.964209085 | 0.33494111 | 0.569354094 | | Ecscr | 100.5309614 | -0.0562871596320744 | 0.0925510966527186 | -0.608173880891783 | 0.543072155804402 | 0.86555476576857 | NM_001033141 |
| Serinc1 | 32227.4425 | -0.065388027 | 0.067815964 | -0.964198141 | 0.334946596 | 0.569354094 | | Adora1 | 11969.92825 | 0.0351753469761679 | 0.0578520696092416 | 0.608022274980959 | 0.543172700410682 | 0.865558162856155 | NM_001008533,NM_001039510 |
| Nr6a1 | 472.9886945 | 0.080398316 | 0.083405429 | 0.96394584 | 0.33507308 | 0.569506615 | | Ddx41 | 1530.30833 | -0.0436920419305375 | 0.0718653719064388 | -0.607970720410659 | 0.543206893366097 | 0.865558162856155 | NM_134059 |
| Gramd4 | 2327.811112 | -0.071029662 | 0.0736987 | -0.963784461 | 0.335153999 | 0.569519199 | | Slc25a38 | 698.8869466 | 0.0398233365589042 | 0.0655095612721913 | 0.607901133598481 | 0.543253047691007 | 0.865558162856155 | NM_144793 |
| Scg2 | 11742.93462 | -0.153365967 | 0.15912377 | -0.963815567 | 0.335138401 | 0.569519199 | | Rabl5 | 1422.308888 | 0.0321251041415503 | 0.0528939718179166 | 0.607349061479795 | 0.543619285618496 | 0.866046692574498 | NM_026073 |
| Pex2 | 783.2432322 | -0.060464712 | 0.062745263 | -0.963653811 | 0.335219518 | 0.569568069 | | 2310015A10Rik | 431.3356608 | -0.0472353655674418 | 0.0778507668066281 | -0.60674245746055 | 0.544021840959956 | 0.866049979903839 | NR_033514 |
| Igfbp5 | 7577.156347 | -0.09547549 | 0.099086393 | -0.963558035 | 0.335267555 | 0.569587226 | | Mboat1 | 150.3644128 | -0.0575060989307339 | 0.0947463860199681 | -0.606947677335306 | 0.543885636090888 | 0.866049979903839 | NM_153546 |
| Slc29a4 | 380.7054243 | 0.140603717 | 0.145941266 | 0.963426728 | 0.335333418 | 0.569636661 | | Plbd2 | 3808.60445 | -0.0360401796034963 | 0.0593976049960156 | -0.606761494944349 | 0.544009205027542 | 0.866049979903839 | NM_023625 |
| Pcyt1a | 1869.743387 | 0.092989451 | 0.096544757 | 0.963174529 | 0.335459745 | 0.569789124 | | Plk1 | 39.92907796 | -0.0549401807343975 | 0.090553214540146 | -0.606717067013013 | 0.544038693833301 | 0.866049979903839 | NM_011121 |
| 2310042D19Rik | 37.40346073 | 0.15052373 | 0.15629227 | 0.963091329 | 0.335501693 | 0.56979757 | | Sec14l1 | 11248.84812 | 0.0290183099077883 | 0.0477860162143028 | 0.607255264336156 | 0.543681521710734 | 0.866049979903839 | NM_028777,NR_029459,NM_001166507,NM_001166506 |
| Trim27 | 896.0382314 | 0.067958681 | 0.070582218 | 0.962830065 | 0.33563281 | 0.569957778 | | Surf1 | 2135.032187 | 0.0338556123093165 | 0.0557820273574528 | 0.606926888697837 | 0.543899432781828 | 0.866049979903839 | NM_013677,NM_001271724,NR_073420 |
| Gm15663 | 295.8298831 | -0.139180781 | 0.144610731 | -0.962451268 | 0.335822971 | 0.569968355 | | Vmn1r58 | 68.0715051 | -0.0574054674280508 | 0.0945567826705782 | -0.607100472401255 | 0.543784236696533 | 0.866049979903839 | NM_030739 |
| L2hgdh | 802.0516853 | 0.069257158 | 0.071956589 | 0.962485287 | 0.33580589 | 0.569968355 | | Clec2l | 1332.169927 | 0.035832077235785 | 0.0590907193827597 | 0.606390946159971 | 0.544255179164447 | 0.866248040495723 | NM_001101507 |
| Malsu1 | 205.3364872 | -0.098952201 | 0.102810773 | -0.962469187 | 0.335813974 | 0.569968355 | | Dctn4 | 4263.605287 | 0.0308929592856182 | 0.0509779002450201 | 0.606006899796468 | 0.544510171467334 | 0.866248040495723 | NM_026302 |
| Pld1 | 432.5780714 | 0.080363347 | 0.083475132 | 0.962722005 | 0.335687051 | 0.569968355 | | Fam184b | 340.7193542 | -0.0477468545437195 | 0.078810508604885 | -0.605843755977994 | 0.544618510773814 | 0.866248040495723 | NM_021416 |
| Slc15a4 | 847.300736 | 0.101729605 | 0.10569365 | 0.962494953 | 0.335801037 | 0.569968355 | | Gid4 | 3702.575253 | 0.03330074237731 | 0.0549291754483842 | 0.606248721293878 | 0.544349604190599 | 0.866248040495723 | NM_025757 |
| Dld | 3960.59639 | -0.062111774 | 0.064544536 | -0.962308784 | 0.335894518 | 0.570014191 | | Mycl1 | 519.5995909 | 0.0482287887339087 | 0.0795876409458321 | 0.605983393410712 | 0.544525780748861 | 0.866248040495723 | NM_008506 |
| Impdh2 | 1686.970016 | 0.09571097 | 0.099473278 | 0.962177706 | 0.335960346 | 0.570014191 | | Polg2 | 73.91133661 | -0.0562596674334855 | 0.0928515956006343 | -0.605909538436636 | 0.544574825170324 | 0.866248040495723 | NR_027785,NM_015810 |
| Rest | 112.5168288 | 0.11260827 | 0.117031691 | 0.962203219 | 0.335947532 | 0.570014191 | | Srcrb4d | 43.76959127 | -0.0515975536129887 | 0.0851271092598639 | -0.606123643356302 | 0.544432651866118 | 0.866248040495723 | NM_001160366 |
| Smtn | 320.1119419 | 0.103250563 | 0.107326162 | 0.962026045 | 0.336036521 | 0.57008101 | | Xpnpep1 | 1543.474683 | 0.0309078500000224 | 0.0510189548229842 | 0.605811116814535 | 0.544640186826893 | 0.866248040495723 | NM_133216 |
| Dimt1 | 452.2432509 | 0.100711638 | 0.104720061 | 0.961722493 | 0.336189021 | 0.570214854 | | Amer3 | 1676.415074 | 0.0390477713811871 | 0.0644900638387376 | 0.605485078737542 | 0.54485673607825 | 0.866307897140178 | NM_213727 |
| Rpl38 | 3323.726093 | -0.094302322 | 0.098055556 | -0.961723384 | 0.336188573 | 0.570214854 | | Dhrs3 | 665.0815163 | -0.0485963194043222 | 0.0802581626555935 | -0.605500023877451 | 0.544846808821414 | 0.866307897140178 | NM_011303,NM_001172424 |
| B130024G19Rik | 10.9025305 | 0.158352981 | 0.164678543 | 0.961588427 | 0.336256387 | 0.570266688 | | Msl1 | 6812.601661 | -0.0371546108917362 | 0.0613626572662016 | -0.605492208894298 | 0.544851999885394 | 0.866307897140178 | NM_028722 |
| Ift27 | 585.8502549 | 0.071208877 | 0.074064354 | 0.961446 | 0.336327965 | 0.570325652 | | Cul1 | 5708.404643 | 0.0352611483850984 | 0.058254928947347 | 0.605290385247381 | 0.544986068754438 | 0.866418697026045 | NM_012042 |
| Adamts14 | 16.96499327 | 0.152922706 | 0.159080643 | 0.961290469 | 0.336406139 | 0.570346257 | | Zic1 | 434.1994382 | 0.05152306676341 | 0.0851555745823281 | 0.60504631688672 | 0.545148222127057 | 0.866581645285362 | NM_009573 |
| Mvb12b | 5594.942764 | -0.051352611 | 0.053421333 | -0.961275354 | 0.336413737 | 0.570346257 | | Papss2 | 839.2769651 | 0.045064114940584 | 0.0744949342386373 | 0.604928582072783 | 0.545226450976315 | 0.866611163899984 | NM_001201470,NM_011864 |
| Slc25a30 | 211.4833031 | 0.10695524 | 0.111272528 | 0.961200767 | 0.336451231 | 0.570347414 | | Jarid2 | 1618.567028 | -0.0401243343748117 | 0.0663559714884699 | -0.604683097462958 | 0.545389581045005 | 0.86668078484408 | NM_001205044,NM_021878,NM_001205043 |
| Sox13 | 392.3321883 | -0.100144755 | 0.104209122 | -0.960997975 | 0.336553186 | 0.570457834 | | Ndufv3 | 1733.566461 | 0.0379770323097361 | 0.0628042221520602 | 0.604689159556613 | 0.545385552355177 | 0.86668078484408 | NM_001083891,NM_030087 |
| Ogdh | 12661.32192 | -0.049389638 | 0.051401346 | -0.960862736 | 0.336621189 | 0.570503566 | | Itm2c | 32996.07567 | 0.0287926411120954 | 0.0476256883734145 | 0.604561153769443 | 0.545470624387104 | 0.8667147549603 | NM_022417 |
| Stag1 | 1436.256735 | 0.081873755 | 0.085214339 | 0.960797874 | 0.336653808 | 0.570503566 | | H1f0 | 3829.460374 | 0.0345311591943653 | 0.0571488375606653 | 0.604232048599578 | 0.545689376324891 | 0.866967503061945 | NM_008197 |
| Psenen | 1262.275784 | 0.062207624 | 0.064755004 | 0.960661274 | 0.336722509 | 0.570557585 | | Hebp2 | 113.550386 | -0.0569615166321301 | 0.0943175768083047 | -0.603933206934496 | 0.545888050191551 | 0.866989658236774 | NM_019487 |
| 3110057O12Rik | 290.8721778 | 0.104434009 | 0.108770248 | 0.960133955 | 0.336987801 | 0.570594737 | | Iqca | 46.96585163 | -0.0567477286296369 | 0.0939737022896392 | -0.603868180639867 | 0.545931285277662 | 0.866989658236774 | NM_029122 |
| Cyp2r1 | 10.86477611 | -0.151934272 | 0.158254866 | -0.960060669 | 0.337024682 | 0.570594737 | | Med19 | 1345.32423 | 0.03487825320756 | 0.0577514880028485 | 0.603936875286048 | 0.545885611205809 | 0.866989658236774 | NM_025885 |
| Efcab4a | 84.15184665 | -0.13566493 | 0.141312907 | -0.96003212 | 0.33703905 | 0.570594737 | | Vps4a | 4687.123892 | 0.0279265834961094 | 0.0462473997175361 | 0.603851971498414 | 0.545942062776523 | 0.866989658236774 | NM_126165 |
| Il11 | 12.61849689 | 0.154623603 | 0.161054766 | 0.960068471 | 0.337020755 | 0.570594737 | | Atxn7l1 | 1047.350242 | 0.035247898265672 | 0.058439992949013 | 0.603146860343133 | 0.546410995034684 | 0.867128614080498 | NM_028139,NM_001033436 |
| Nol3 | 900.6758856 | 0.08656701 | 0.090128553 | 0.960483746 | 0.336811808 | 0.570594737 | | Fbl | 1049.739853 | -0.0382634380151608 | 0.0634723021965821 | -0.60283677590036 | 0.546617279008849 | 0.867128614080498 | NM_007991 |
| Pla2g4a | 145.9192873 | -0.137989811 | 0.143729098 | -0.960068719 | 0.337020631 | 0.570594737 | | Gabrr2 | 39.86393692 | 0.051780017653807 | 0.0859199476629367 | 0.602654203851934 | 0.546738753278107 | 0.867128614080498 | NM_008076 |
| Rnf19a | 1455.798487 | 0.071992704 | 0.074979427 | 0.960166096 | 0.336971627 | 0.570594737 | | Gna12 | 2189.563876 | 0.0408319286224249 | 0.0677489072193967 | 0.60269501455124 | 0.546711598728918 | 0.867128614080498 | NM_010302 |
| Rpp25l | 367.1585145 | -0.082163591 | 0.085565671 | -0.960240133 | 0.336934372 | 0.570594737 | | Htatsf1 | 3244.153662 | 0.0311388790381327 | 0.0516253219832393 | 0.603170650407609 | 0.546395170264592 | 0.867128614080498 | NM_028242,NM_029371 |
| Chpf | 3037.308032 | -0.070365586 | 0.073317779 | -0.959734286 | 0.337188963 | 0.570700002 | | Irf2bp2 | 1449.896811 | 0.0464072262750273 | 0.0769721555967121 | 0.602909271739424 | 0.546569047627536 | 0.867128614080498 | NM_001164598 |
| Ddx20 | 492.3717722 | 0.078589549 | 0.081887575 | 0.959724949 | 0.337193663 | 0.570700002 | | Mrps11 | 607.1378782 | 0.0403207267066864 | 0.0668111602021875 | 0.603502866656793 | 0.546174209062011 | 0.867128614080498 | NM_026498 |
| Lmf2 | 663.5639468 | 0.078025354 | 0.081302744 | 0.959689065 | 0.337211728 | 0.570700002 | | Nkx2-1 | 63.39790507 | 0.0535103053882417 | 0.088792642682911 | 0.602643459766518 | 0.546745902269261 | 0.867128614080498 | NM_009385,NM_001146198 |
| Hmga1 | 12.2203665 | 0.159939799 | 0.166754199 | 0.959135062 | 0.337490709 | 0.571052559 | | Rhpn1 | 394.2323382 | -0.056588349780795 | 0.0938990920812462 | -0.602650659623332 | 0.546741111562035 | 0.867128614080498 | NM_008164,NM_001163465 |
| Tigd3 | 126.1530219 | -0.112466886 | 0.117259394 | -0.959129009 | 0.337493758 | 0.571052559 | | Trrap | 6070.238127 | -0.0428089536799897 | 0.071014255435591 | -0.602821974509284 | 0.546627126612953 | 0.867128614080498 | NM_001081362 |
| Cog5 | 947.7284861 | -0.066155161 | 0.069015923 | -0.958549253 | 0.337785868 | 0.571422015 | | Wasf2 | 1017.781954 | -0.0386540365577094 | 0.0641090203280868 | -0.602942243695068 | 0.546547112128526 | 0.867128614080498 | NM_153423 |
| Rnf215 | 781.404933 | 0.087210461 | 0.090981664 | 0.958549859 | 0.337785562 | 0.571422015 | | Zfp964 | 81.87597017 | 0.0571314856538968 | 0.0947910495119562 | 0.602709706750221 | 0.546701823023674 | 0.867128614080498 | NM_001177527 |
| Psd | 16358.7105 | -0.089139071 | 0.093021492 | -0.958263178 | 0.337930067 | 0.571603543 | | Hsd17b4 | 2495.058912 | 0.0270364678558142 | 0.044884952258006 | 0.60235037569839 | 0.546940934897123 | 0.86734323365891 | NM_008292 |
| Ccdc114 | 94.50101332 | 0.143312188 | 0.149574278 | 0.958133914 | 0.337995236 | 0.571651369 | | Nap1l1 | 10159.15565 | 0.0419909230263594 | 0.0697446569008183 | 0.602066522258091 | 0.547129857833312 | 0.867487125252749 | NM_015781,NM_001146707 |
| Oxr1 | 10782.37388 | -0.076594708 | 0.079955689 | -0.957964449 | 0.338080685 | 0.57173348 | | Pkig | 2033.509431 | 0.0400162538759814 | 0.0664683630771586 | 0.602034592450085 | 0.547151111218896 | 0.867487125252749 | NM_001039391,NM_001039390,NM_001164055,NM_001164053,NM_011106 |

| | | | | | | | | | | | | | | | |
|---|---|---|---|---|---|---|---|---|---|---|---|---|---|---|---|
| Epb4.1l3 | 7817.170012 | 0.055234037 | 0.057676194 | 0.957657445 | 0.338235522 | 0.571901923 | | Desi2 | 994.540767 | -0.0460427712129554 | 0.0765622197593238 | -0.601377172157399 | 0.54758879959204 | 0.867553753435222 | NM_024282 |
| Tes | 66.70952328 | -0.143071681 | 0.149403305 | -0.95762059 | 0.338254112 | 0.571901923 | | Enthd2 | 706.8027797 | -0.043688953882785 | 0.0725995035460033 | -0.601780339380711 | 0.547320363849435 | 0.867553753435222 | NM_183137 |
| 3000002C10Rik | 119.6177392 | 0.136646771 | 0.142789826 | 0.956978342 | 0.338578187 | 0.57220009 | | Ints3 | 3454.674953 | -0.0298190997477193 | 0.0495926387236765 | -0.601280764951171 | 0.547652998829633 | 0.867553753435222 | NM_145540,NM_178876 |
| Hnrnpm | 3771.98518 | 0.059706888 | 0.06237964 | 0.957153462 | 0.338489803 | 0.57220009 | | Itgb8 | 1244.87451 | -0.0412685181403364 | 0.0686190825095638 | -0.601414601172853 | 0.547563875961803 | 0.867553753435222 | NM_177290 |
| Ndufaf1 | 692.6295926 | -0.068762983 | 0.071852196 | -0.957006002 | 0.338564225 | 0.57220009 | | Nol7 | 996.2078482 | -0.0458943779734007 | 0.0762773080927508 | -0.60167799730943 | 0.547388498812794 | 0.867553753435222 | NM_023554 |
| Tmem59 | 6563.297528 | -0.049101735 | 0.05130597 | -0.957037457 | 0.338548349 | 0.57220009 | | Scel | 121.6691407 | -0.056092183513727 | 0.0932838096840153 | -0.601306740191366 | 0.547635701098526 | 0.867553753435222 | NM_022886 |
| Lclat1 | 1691.523974 | -0.06346229 | 0.066323991 | -0.956852701 | 0.338641608 | 0.572226272 | | Synj1 | 18192.74367 | 0.0336244423573439 | 0.0559238712858079 | 0.601253839983659 | 0.547670929299255 | 0.867553753435222 | NM_001164483,NM_001045515 |
| Nudc | 4225.465817 | 0.060714794 | 0.06345601 | 0.956801326 | 0.338667543 | 0.572226272 | | Ttll8 | 54.20827939 | -0.0557091046871377 | 0.0926068543006988 | -0.601565673597417 | 0.547463283954044 | 0.867553753435222 | NM_172818 |
| Cox18 | 301.3237464 | 0.089520702 | 0.093584584 | 0.956575302 | 0.33878166 | 0.57223188 | | Rcc1 | 479.1378801 | -0.043802480898456 | 0.0728716686772606 | -0.601090680281408 | 0.547779590424099 | 0.867631264911183 | NM_133878,NM_001197082 |
| Zdhhc14 | 1697.470456 | -0.07787312 | 0.081396314 | -0.956715563 | 0.338710841 | 0.57223188 | | Rnf5 | 2138.50311 | 0.029614681310801 | 0.0492763500200315 | 0.600991780007291 | 0.547845461232629 | 0.867640991045047 | NM_019403 |
| Zfp9 | 1044.595998 | 0.08677288 | 0.090708368 | 0.956613831 | 0.338762205 | 0.57223188 | | Sharpin | 1333.895627 | -0.0413026062245403 | 0.0687449901149458 | -0.600808963031049 | 0.547967233609755 | 0.867726281486481 | NM_025340 |
| C7 | 29.62906944 | 0.158802575 | 0.166080605 | 0.956177723 | 0.338982453 | 0.572508625 | | Zfp386 | 636.6647695 | -0.042839844652709 | 0.0713127904400215 | -0.600731571270373 | 0.548018787440243 | 0.867726281486481 | NM_019565,NM_001004066 |
| Mfhas1 | 2211.643343 | 0.075916797 | 0.079432194 | 0.955743428 | 0.339201876 | 0.572754344 | | Cox6a1 | 7265.205531 | -0.0329869728029569 | 0.0549367810387722 | -0.60045332433431 | 0.548204158951388 | 0.86792518941993 | NM_007748 |
| Prkci | 2568.762703 | 0.06427433 | 0.067246366 | 0.95580376 | 0.339171388 | 0.572754344 | | C87436 | 977.7366355 | 0.0332938771130781 | 0.055461150093748 | 0.60030989362464 | 0.54829972634279 | 0.867981890241431 | NM_001243741,NM_001243742,NM_146170 |
| Sirpa | 21604.14535 | -0.05384758 | 0.056365447 | -0.955329608 | 0.339411039 | 0.573045074 | | 1200011I18Rik | 2053.083574 | 0.0359059314491077 | 0.0598325361543025 | 0.600107128277325 | 0.548434842239439 | 0.868006593851612 | NM_026177 |
| Wdr3 | 815.922808 | 0.076336509 | 0.079914479 | 0.955227517 | 0.339462653 | 0.57306977 | | Rnd3 | 598.3951675 | 0.0542373020475048 | 0.0903793548652914 | 0.600107205106126 | 0.548434791040238 | 0.868006593851612 | NM_028810 |
| Ipmk | 1410.877791 | -0.071711367 | 0.075082653 | -0.955099003 | 0.339527633 | 0.573117022 | | Gcsh | 1659.720252 | 0.0383631345898087 | 0.0639484567585811 | 0.599907121052783 | 0.548568136325434 | 0.868028403950246 | NM_026572 |
| Igfbp4 | 2956.996421 | 0.114333663 | 0.119798985 | 0.954379229 | 0.339891716 | 0.57366909 | | Shisa3 | 37.41123878 | 0.05441966645833 | 0.0907132011873752 | 0.599909007134716 | 0.548566879278295 | 0.868028403950246 | NM_001033415 |
| A730043L09Rik | 10.87153933 | -0.14310545 | 0.149999483 | -0.954039624 | 0.340063585 | 0.573834153 | | Pard3b | 178.5138623 | 0.0538110713117051 | 0.0897144029352451 | 0.599804151297148 | 0.548636766380923 | 0.868042442920084 | NM_001081050 |
| Egr2 | 457.2895258 | -0.143054743 | 0.14994123 | -0.954072093 | 0.340047151 | 0.573834153 | | Actg1 | 49361.89195 | 0.0256914082328448 | 0.0428586312891851 | 0.599445373313352 | 0.548875927504922 | 0.868267066631208 | NM_009609 |
| Xpo4 | 326.9004302 | -0.094752395 | 0.09932931 | -0.953921804 | 0.340123225 | 0.573872292 | | Glmn | 472.2990274 | -0.0484967877998977 | 0.0809213025346265 | -0.599308047212237 | 0.548967482590925 | 0.868267066631208 | NM_001161739,NM_001161738,NM_133248 |
| Anapc15 | 363.5676395 | -0.078851116 | 0.082678491 | -0.953707736 | 0.340231603 | 0.573900218 | | Leprel4 | 386.6313519 | 0.0533572826274993 | 0.0890426998411849 | 0.599232533634609 | 0.549017830580176 | 0.868267066631208 | NM_176830 |
| Mfap2 | 108.5860249 | 0.112047521 | 0.117490919 | 0.953669631 | 0.340250897 | 0.573900218 | | Rhoj | 152.6372243 | -0.0517815979209646 | 0.0864013092906452 | -0.599314968095872 | 0.548962868268855 | 0.868267066631208 | NM_023275 |
| Nfkbib | 706.8351407 | 0.094851478 | 0.099446043 | 0.953798408 | 0.340185695 | 0.573900218 | | Sep15 | 4827.64494 | 0.0334835348073227 | 0.0558971075542674 | 0.599020884485219 | 0.549158957861473 | 0.868365901699098 | NM_053102 |
| Rdh13 | 816.7595936 | 0.062689058 | 0.065752123 | 0.953414974 | 0.340379858 | 0.574055243 | | Uba52 | 9305.035077 | 0.0344225667977824 | 0.0574706073223588 | 0.59895950994049 | 0.549199885647089 | 0.868365901699098 | NM_019883 |
| Anxa6 | 4378.250549 | 0.079779906 | 0.083715165 | 0.95299228 | 0.340593982 | 0.574189552 | | Rai2 | 432.3785076 | 0.051040191056534 | 0.0852975797234986 | 0.59837795189484 | 0.549587773904256 | 0.868884632535178 | NM_198409,NM_001103367 |
| Fam78b | 1655.30227 | -0.059016924 | 0.061929041 | -0.952976546 | 0.340601954 | 0.574189552 | | Lck | 59.2696806 | 0.0564073460346557 | 0.0942822514330515 | 0.59828170389747 | 0.549651982527624 | 0.868891576689113 | NM_010693,NM_001162433,NM_001162432 |
| Rab37 | 151.6813864 | 0.159722572 | 0.167605891 | 0.952965146 | 0.34060773 | 0.574189552 | | Ccz1 | 1297.994928 | 0.0370074462454559 | 0.061889362969201 | 0.597961337295918 | 0.549865730995041 | 0.869134886663109 | NM_177682 |
| Wscd1 | 2875.445517 | -0.112550338 | 0.118078885 | -0.953179206 | 0.34049928 | 0.574189552 | | Klhl12 | 1969.221069 | 0.0319810706405462 | 0.0534989131221674 | 0.597789165688578 | 0.54998062074068 | 0.869221901520958 | NM_153128 |
| Zfp551 | 215.5689118 | -0.089116822 | 0.093541884 | -0.95269432 | 0.340744972 | 0.57435842 | | Fkbp1b | 837.0261835 | -0.040262336094834 | 0.0673699534203273 | -0.597630457655709 | 0.550086536746295 | 0.869294716359517 | NM_016863 |
| Iqcb1 | 888.7408228 | 0.064793681 | 0.068018705 | 0.952586221 | 0.340799761 | 0.574388284 | | Gm3219 | 23.59290988 | 0.0484430975168561 | 0.0810954698872086 | 0.597358860910887 | 0.550267813904678 | 0.869486594667612 | NR_027380 |
| Rab30 | 418.3964392 | -0.091184417 | 0.095739223 | -0.952424864 | 0.340881555 | 0.574463651 | | 6330549D23Rik | 60.94252516 | -0.0562963209864243 | 0.0942962676717777 | -0.597015368438314 | 0.550497119964441 | 0.869565154915558 | NR_003619 |
| Arpp19 | 12780.46044 | -0.091533975 | 0.096117456 | -0.952313752 | 0.340937885 | 0.574496095 | | Chmp1a | 4010.33592 | 0.03349907442858 | 0.0561085245231141 | 0.597040729787503 | 0.550480187822067 | 0.869565154915558 | NM_145606 |
| Zmiz1 | 6541.841961 | -0.081869958 | 0.085979187 | -0.952206694 | 0.340992167 | 0.574525079 | | Thap6 | 135.0167403 | -0.0555789223634979 | 0.0930702875983789 | -0.597171490468951 | 0.550392891390472 | 0.869565154915558 | NR_028429 |
| 2410002F23Rik | 2062.920712 | 0.058637391 | 0.061617055 | 0.951642216 | 0.341278464 | 0.574801598 | | Agps | 1835.596191 | 0.038276296169151 | 0.0641856305098334 | 0.596337464711623 | 0.550949807691622 | 0.869642173800581 | NM_172666 |
| Lrrtm1 | 1675.315308 | -0.091196779 | 0.09582835 | -0.951668054 | 0.341265356 | 0.574801598 | | Ccdc64 | 1343.910024 | -0.0398468429420959 | 0.0667939672114607 | -0.596563501250737 | 0.550798845605446 | 0.869642173800581 | NM_001080808 |
| Slc16a11 | 468.338856 | 0.125892858 | 0.132297298 | 0.951590543 | 0.341304679 | 0.574801598 | | Cxcl12 | 2637.23013 | -0.0511158547246783 | 0.0856740298863782 | -0.596631847392595 | 0.550753203558698 | 0.869642173800581 | NM_021704,NM_001012477,NM_013655 |
| Zc3h4 | 2795.186763 | -0.049352145 | 0.051860112 | -0.951639768 | 0.341279706 | 0.574801598 | | Edn1 | 109.5330764 | -0.0565513107049581 | 0.09483869423344 | -0.596289427665044 | 0.550981892616863 | 0.869642173800581 | NM_010104 |
| AA987161 | 397.7206078 | -0.101460087 | 0.106632199 | -0.951495776 | 0.341352762 | 0.574820095 | | Galm | 111.5457009 | 0.0560351580941597 | 0.0939816167940322 | 0.596235306495791 | 0.551018042353019 | 0.869642173800581 | NM_176963 |
| Dmwd | 4789.244944 | -0.056659352 | 0.059572099 | -0.951105501 | 0.341550822 | 0.575028625 | | Jam3 | 1245.852579 | -0.0414838441340912 | 0.0695428903336215 | -0.596521713939107 | 0.550826752390465 | 0.869642173800581 | NM_023277 |
| Mapk11 | 1900.398859 | 0.119783395 | 0.125934426 | 0.951156874 | 0.341524747 | 0.575028625 | | Mut | 1791.671159 | 0.0400740965205562 | 0.0671903875917342 | 0.596426035879665 | 0.550890651607838 | 0.869642173800581 | NM_008650 |
| Fto | 8641.253163 | 0.052531707 | 0.055241559 | 0.950945415 | 0.341632086 | 0.575055208 | | Rgcc | 374.2452108 | 0.0505221862502406 | 0.084736757109529 | 0.596225156279425 | 0.55102482222639 | 0.869642173800581 | NM_025427 |
| Rsph9 | 316.0894234 | -0.096273299 | 0.1012414 | -0.950928169 | 0.341640841 | 0.575055208 | | BC089491 | 32.0808767 | 0.0487542663310602 | 0.0818516154111797 | 0.595642078487325 | 0.551414359988815 | 0.869755927298638 | NM_175033 |
| Mast2 | 6044.896164 | -0.064680904 | 0.068032641 | -0.950733408 | 0.341739725 | 0.575159167 | | Bcl7c | 316.0941806 | 0.0464151323745167 | 0.0779607590867652 | 0.595365321197802 | 0.551599301057747 | 0.869755927298638 | NM_009746 |
| Mir331 | 20.14501712 | 0.15804545 | 0.166305628 | 0.950331336 | 0.341943921 | 0.575421758 | | Ier5l | 356.1154228 | -0.0552166707036649 | 0.0927526876666293 | -0.595310735384015 | 0.551635781233812 | 0.869755927298638 | NM_030244 |
| Ring1 | 1000.153844 | 0.105992612 | 0.111538302 | 0.950279951 | 0.341970023 | 0.575421758 | | Pigz | 473.9264018 | 0.0543137183222183 | 0.0911708483769826 | 0.595735580935217 | 0.551351884539238 | 0.869755927298638 | NM_172822 |
| Gpkow | 1637.78344 | 0.070801316 | 0.07453651 | 0.949887728 | 0.342169303 | 0.575602106 | | Sec14l5 | 489.4635947 | -0.0555209674170798 | 0.0932407394892067 | -0.595458248414115 | 0.55153719971464 | 0.869755927298638 | NM_001127725 |
| Rtn4ip1 | 523.6757993 | -0.102606799 | 0.10801362 | -0.949943161 | 0.342141134 | 0.575602106 | | Spink8 | 51.52842064 | -0.0313076430620555 | 0.052559052174838 | -0.5956660511668 | 0.551398341852635 | 0.869755927298638 | NM_183136 |
| Ube2l6 | 203.3456761 | 0.122410855 | 0.128873927 | 0.94984965 | 0.342188654 | 0.575602106 | | Srr | 4600.389344 | 0.0306235131124399 | 0.0514202580116541 | 0.595553470492102 | 0.551473568327358 | 0.869755927298638 | NM_001163311,NM_013761 |
| Commd7 | 884.1599229 | -0.067432206 | 0.070998159 | -0.949773996 | 0.342227102 | 0.575604289 | | Sypl | 1575.14594 | -0.0350492704081073 | 0.0588639221912511 | -0.595428729574475 | 0.551556926174298 | 0.869755927298638 | NM_198710,NM_013635 |
| Ergic2 | 1695.388389 | -0.064762091 | 0.068198448 | -0.949612386 | 0.342309243 | 0.575617474 | | Zfp551 | 232.9542769 | 0.050233979975067 | 0.0843437632164714 | 0.595586182776071 | 0.551451709439583 | 0.869755927298638 | NM_001033820 |
| Vamp8 | 237.828524 | 0.106224776 | 0.111857594 | 0.94964296 | 0.342293702 | 0.575617474 | | 4933431E20Rik | 5657.442047 | 0.0279784992570673 | 0.0470656146703051 | 0.594457322039813 | 0.552206278930015 | 0.870265824568551 | NR_015459 |
| Acvr1b | 4792.300795 | 0.082927805 | 0.087399488 | 0.948836275 | 0.342703889 | 0.575828939 | | Adra2b | 47.8449036 | 0.0544640934111301 | 0.0916482286146427 | 0.594273286146427 | 0.552329342912191 | 0.870265824568551 | NM_009633 |
| Alg2 | 5429.982202 | -0.042606362 | 0.044920771 | -0.948477964 | 0.342886186 | 0.575828939 | | Cacna2d2 | 2902.459956 | 0.0491133316462963 | 0.0826721816030599 | 0.594073250444845 | 0.552463121160853 | 0.870265824568551 | NM_001174047,NM_020263,NM_001174050,NM_001174049,NM_001174048 |
| Alg5 | 514.8841768 | 0.090130678 | 0.095091499 | 0.947831082 | 0.343215454 | 0.575828939 | | Dcn | 537.0362102 | 0.0503435995008414 | 0.0846623948493204 | 0.594639445180371 | 0.552084507250615 | 0.870265824568551 | NM_007833,NM_001190451 |
| Apoe | 40943.05504 | 0.07785142 | 0.082112978 | 0.948101282 | 0.343077896 | 0.575828939 | | H2-Q1 | 31.01731263 | -0.0417360901144415 | 0.070281543978078 | -0.59384139493947 | 0.552618199491967 | 0.870265824568551 | NM_010390 |
| Arhgdia | 10904.11064 | -0.064519901 | 0.068058187 | -0.948010866 | 0.343123923 | 0.575828939 | | Map3k7 | 2824.394722 | 0.032378069439906 | 0.0545025965099369 | 0.594064714586631 | 0.552468830056356 | 0.870265824568551 | NM_172688,NM_009316 |
| Bcl3 | 21.62825498 | 0.159389503 | 0.167939048 | 0.94909138 | 0.342574138 | 0.575828939 | | Mrpl23 | 732.1607989 | -0.0389569819717455 | 0.0655876772990192 | -0.593968007040982 | 0.552533511370983 | 0.870265824568551 | NM_011288 |
| Cdh6 | 408.2508943 | 0.083402727 | 0.087950648 | 0.94829008 | 0.342981799 | 0.575828939 | | Pcdhb15 | 129.2942163 | -0.0550732039790451 | 0.0927318506865153 | -0.593897388775544 | 0.552580745629752 | 0.870265824568551 | NM_053140 |
| Derl3 | 45.61461867 | 0.157745631 | 0.166421684 | 0.947867053 | 0.34319714 | 0.575828939 | | Plagl2 | 448.9712372 | -0.0455322985322975 | 0.0766089486838406 | -0.594346996200221 | 0.552280051711495 | 0.870265824568551 | NM_018807 |
| Eif2b4 | 1096.711984 | 0.069478071 | 0.073287425 | 0.94802172 | 0.343118397 | 0.575828939 | | Rtkn | 2633.854717 | 0.034566770964874 | 0.0581688814169375 | 0.594248507498528 | 0.552345913311859 | 0.870265824568551 | NM_133641,NM_001136227,NM_009106 |
| Grin2c | 1938.439969 | -0.096050451 | 0.101195371 | -0.949158545 | 0.342539982 | 0.575828939 | | Sh2d5 | 4827.366357 | -0.0410483633693147 | 0.0690979793753559 | -0.594060256759908 | 0.552471811521666 | 0.870265824568551 | NM_001099631 |
| Invs | 538.7378875 | 0.132359116 | 0.139603295 | 0.948108826 | 0.343074055 | 0.575828939 | | Gm4944 | 364.9740824 | 0.0544981008675862 | 0.0917895393506047 | 0.593728885155661 | 0.552693460218277 | 0.870289994919317 | NM_001205095 |
| Kcnj10 | 9747.172351 | -0.071039328 | 0.074887283 | -0.948616713 | 0.342815588 | 0.575828939 | | Fhit | 100.8615289 | -0.0549707963102433 | 0.0926491274396559 | -0.593322331568064 | 0.552965456410253 | 0.870600462219512 | NM_010210 |
| Lgalsl | 2861.428381 | -0.051697191 | 0.054511094 | -0.948379266 | 0.342936411 | 0.575828939 | | Gm3558 | 35.07799301 | 0.0494187507035416 | 0.0833707321370714 | 0.592758986718399 | 0.553342458974566 | 0.870600462219512 | NM_001270842 |
| Lypla1 | 708.5421395 | -0.082748583 | 0.087248435 | -0.948424847 | 0.342913215 | 0.575828939 | | Gm410 | 75.96427275 | -0.0560424716739017 | 0.0945289589691148 | -0.592860349728514 | 0.553274615362867 | 0.870600462219512 | NM_001033349 |
| Npr3 | 471.1122436 | 0.139552352 | 0.147070521 | 0.948880517 | 0.342681385 | 0.575828939 | | Gstm1 | 8579.775442 | 0.0441248741725372 | 0.0744562571537555 | 0.592628153217766 | 0.553430033604225 | 0.870600462219512 | NM_010358 |
| Pmvk | 1107.991716 | -0.073743898 | 0.077726974 | -0.948755543 | 0.342744957 | 0.575828939 | | Ndufs6 | 1352.204156 | 0.0369682775133902 | 0.0623426469330459 | 0.592985369278481 | 0.553190943729381 | 0.870600462219512 | NR_033306,NM_010888 |
| Ppwd1 | 249.1012572 | 0.086297592 | 0.091043398 | 0.947873042 | 0.343194039 | 0.575828939 | | Snrpa1 | 702.8857397 | 0.0360422958143146 | 0.060812782768296 | 0.592676312012231 | 0.553397797274087 | 0.870600462219512 | NM_021336 |
| Rabggta | 1571.602124 | 0.059837741 | 0.063106648 | 0.94820027 | 0.34302751 | 0.575828939 | | Tmem63c | 2152.68242 | -0.0374006861594456 | 0.0630742973422603 | -0.592962560166531 | 0.553206208684545 | 0.870600462219512 | NM_172583 |
| Tbc1d4 | 603.7358776 | 0.095120212 | 0.100325572 | 0.948115325 | 0.343070747 | 0.575828939 | | Tspan14 | 1793.641982 | -0.0361932827263571 | 0.0610638516059055 | -0.592712083737228 | 0.553373853143653 | 0.870600462219512 | NM_145928 |
| Wdr33 | 2321.336549 | 0.063525239 | 0.067013131 | 0.947952115 | 0.343153832 | 0.575828939 | | Zfp775 | 338.0714683 | 0.0506829614338156 | 0.0854933058006511 | 0.592829590096745 | 0.553295202762939 | 0.870600462219512 | NM_173429 |
| Zfpl1 | 713.7963941 | 0.070635119 | 0.074510916 | 0.947983501 | 0.343137854 | 0.575828939 | | Tmem161a | 1059.273809 | 0.03738917590056 | 0.0631090474561956 | 0.592453497678159 | 0.553546951509091 | 0.870690093949221 | NM_145597 |
| AU023762 | 83.87971847 | -0.135232141 | 0.142714597 | -0.947570491 | 0.343348154 | 0.575863522 | | Ap5m1 | 765.1691755 | -0.040853199968775 | 0.0689757635080048 | -0.592283403489023 | 0.553660827583818 | 0.870774922204692 | NM_144535 |
| Ankrd28 | 2538.424663 | -0.057586953 | 0.060783718 | -0.947407545 | 0.343431148 | 0.575863522 | | Hmgb2 | 201.1767463 | -0.0515783905715738 | 0.0871253231448659 | -0.592002287162973 | 0.553849056832146 | 0.870976659039055 | NM_008252 |
| D1Ertd622e | 2134.236505 | 0.075898622 | 0.08011081 | 0.947420477 | 0.343424561 | 0.575863522 | | Sec13 | 1716.225172 | 0.0350254416634387 | 0.05917823025101 | 0.591863618004036 | 0.553941918159924 | 0.871028393937113 | NM_024206 |
| Stk35 | 1475.626354 | 0.062217685 | 0.065655056 | 0.947644989 | 0.343310215 | 0.575863522 | | Acsl1 | 2597.754848 | 0.03060114001545 | 0.0519161684742516 | 0.589433714289354 | 0.555570365806118 | 0.871360988694929 | NM_007981 |
| Trpm1 | 8.455845851 | 0.142682192 | 0.150611474 | 0.947352733 | 0.343459068 | 0.575863522 | | Axin1 | 1195.636397 | 0.0343115528466626 | 0.0581789139480518 | 0.589759253280312 | 0.555352063925142 | 0.871360988694929 | NM_001159598,NM_009733 |
| Utp11l | 452.4697294 | 0.075470387 | 0.07965204 | 0.947501003 | 0.343385546 | 0.575863522 | | Cyb5r3 | 3033.814293 | 0.0372001094824966 | 0.062925273664537 | 0.591179144977824 | 0.554400394688984 | 0.871360988694929 | NM_029787 |
| Ezh2 | 161.820538 | 0.126836654 | 0.133932214 | 0.94702126 | 0.343627946 | 0.576022005 | | Flywch2 | 240.1638379 | -0.0489458685856324 | 0.0829041900261647 | -0.590390769998295 | 0.55492869718909 | 0.871360988694929 | NM_029798 |
| Man1a2 | 5643.189334 | -0.054662834 | 0.057719054 | -0.947050074 | 0.343613263 | 0.576022005 | | Fnta | 1886.784149 | 0.0339012381057728 | 0.0574600670696729 | 0.589996493820064 | 0.55519300014495 | 0.871360988694929 | NM_008033 |
| Taf6 | 1280.89925 | 0.060821124 | 0.064231066 | 0.946911327 | 0.343683965 | 0.576053587 | | Gipc2 | 56.72068496 | 0.056026493232982 | 0.0947596336778795 | 0.591248520687989 | 0.554353916713366 | 0.871360988694929 | NM_016867 |
| Igbp1 | 1072.545944 | 0.113278166 | 0.1196852 | 0.946467616 | 0.343910132 | 0.576370318 | | Gm4980 | 191.8590147 | -0.0510008430306507 | 0.0864159063195521 | -0.590178882601286 | 0.555070728217752 | 0.871360988694929 | NM_001195529 |
| Rfxap | 783.1271041 | 0.071302838 | 0.07534299 | 0.946376538 | 0.343956568 | 0.576385795 | | Hhat | 30.44325075 | 0.0530363929268242 | 0.0897502411387516 | 0.590933152422747 | 0.554565211765513 | 0.871360988694929 | NM_144881 |
| Arxes2 | 544.5424958 | -0.096986269 | 0.102491033 | -0.946290281 | 0.344000549 | 0.576397157 | | Ift46 | 1446.78044 | 0.0355001812457627 | 0.0601619767351009 | 0.590076709780556 | 0.555139222408808 | 0.871360988694929 | NM_023831 |
| Ap3b1 | 2026.488278 | 0.051859681 | 0.05481674 | 0.946055547 | 0.344120255 | 0.576410728 | | Irs1 | 795.9643901 | 0.0437220622078746 | 0.0740107751700411 | 0.59075265875033 | 0.554686159281367 | 0.871360988694929 | NM_010570 |
| Cited1 | 63.63005366 | 0.145856988 | 0.154170425 | 0.946076318 | 0.344109662 | 0.576410728 | | Lcmt2 | 280.5618335 | 0.0542260370746034 | 0.092016092667244 | 0.589310364119675 | 0.555653093644577 | 0.871360988694929 | NM_177846 |
| Syn1 | 25587.49247 | 0.071229605 | 0.07528933 | 0.946078348 | 0.344108626 | 0.576410728 | | Megf11 | 829.0870972 | 0.0502523819889743 | 0.0852477033100924 | 0.5894866376186 | 0.555534873311819 | 0.871360988694929 | NM_172522,NM_001134399 |
| Nup153 | 2207.825796 | 0.067535235 | 0.071403533 | 0.945824841 | 0.344237933 | 0.576510979 | | Msh6 | 860.6782147 | -0.0376124676932804 | 0.0635945278383354 | -0.591441889291098 | 0.554224380264661 | 0.871360988694929 | NM_010830 |
| Tmem255a | 1070.200471 | 0.158741358 | 0.16783955 | 0.945792322 | 0.344254523 | 0.576510979 | | Olfm4 | 31.16336309 | -0.0524270201560341 | 0.0887478373281272 | -0.590741382938671 | 0.554693715549963 | 0.871360988694929 | NM_001030294 |
| Lrp10 | 1160.211294 | 0.07431359 | 0.078580213 | 0.945703595 | 0.344299788 | 0.57652447 | | Pin4 | 547.7474896 | -0.0419115120200433 | 0.071111650693927 | -0.589376165664265 | 0.555608961464763 | 0.871360988694929 | NM_027181 |
| 9530052E02Rik | 12.81494132 | -0.15757016 | 0.16672746 | -0.945076236 | 0.344619957 | 0.576722182 | | Rp2h | 447.9411686 | -0.0452979649032624 | 0.0768316564663945 | -0.589574232635155 | 0.555476131034889 | 0.871360988694929 | NM_133669 |
| Arx | 599.5358166 | -0.089944953 | 0.09518283 | -0.944970358 | 0.344674009 | 0.576722182 | | Sbf1 | 9060.890295 | -0.0344061603324347 | 0.0583122972023468 | -0.590032668633229 | 0.555168747803871 | 0.871360988694929 | NM_001170561,NM_001081030 |
| Cst3 | 29140.80583 | -0.078609023 | 0.083193939 | -0.94488882 | 0.34471564 | 0.576722182 | | Sel1l | 9475.377267 | 0.0325273716268701 | 0.0551476678540692 | 0.589823158305507 | 0.555309215031994 | 0.871360988694929 | NM_001039089,NM_011344 |
| Map2k6 | 407.4579139 | 0.113521286 | 0.120093927 | 0.945270825 | 0.344520629 | 0.576722182 | | Selm | 1961.786054 | 0.0373842686337345 | 0.0632415094604306 | 0.59113498322056 | 0.554429981672082 | 0.871360988694929 | NM_053267 |
| Pgd | 1322.551735 | 0.070111783 | 0.074191763 | 0.945007643 | 0.344654974 | 0.576722182 | | Senp8 | 284.4279286 | 0.0491992622931836 | 0.0834858331709834 | 0.589312706413565 | 0.555651522671301 | 0.871360988694929 | NM_001172070,NM_001172071,NM_001172069,NM_001172068,NM_027838 |
| Ppm1b | 4134.029688 | -0.072974648 | 0.077225078 | -0.944960497 | 0.344679044 | 0.576722182 | | Shoc2 | 2545.503623 | 0.0391192417163529 | 0.0663524453913188 | 0.589567445263609 | 0.555480682619873 | 0.871360988694929 | NM_019658,NM_001168505 |
| Slc27a4 | 3908.810715 | -0.061761106 | 0.065333258 | -0.945324131 | 0.344493422 | 0.576722182 | | Sys1 | 985.8547656 | 0.0380657701119949 | 0.0644341377465066 | 0.590770225897199 | 0.55467438709562 | 0.871360988694929 | NM_025575 |
| Zfp457 | 8.839519458 | 0.14682617 | 0.155377233 | 0.944965795 | 0.344676339 | 0.576722182 | | Tmem206 | 414.4254114 | -0.0424312294786516 | 0.0719076767421466 | -0.590079271102104 | 0.555137505310245 | 0.871360988694929 | NM_025864 |
| Ginm1 | 402.0289567 | 0.077387696 | 0.081908439 | 0.944807369 | 0.344757229 | 0.576729487 | | Yes1 | 424.6763001 | 0.0444681190782986 | 0.0753750554168198 | 0.589958028321073 | 0.555218788776893 | 0.871360988694929 | NM_001205132,NM_001205133,NM_009535 |
| AI467606 | 66.90449025 | -0.141277021 | 0.149673796 | -0.943899498 | 0.345221009 | 0.576948985 | | Zfp358 | 1129.338888 | -0.0325444306403484 | 0.0551670665829373 | -0.589924979814209 | 0.555240946132997 | 0.871360988694929 | NM_080461 |
| Akip1 | 85.8667189 | 0.126675906 | 0.134298615 | 0.943240599 | 0.345557852 | 0.576948985 | | Rab39b | 1724.127224 | 0.0453646819626002 | 0.0769965478885167 | 0.589178128197172 | 0.555741787611916 | 0.871401030114683 | NM_175122 |
| Cort | 96.80326922 | 0.151737693 | 0.160827547 | 0.943480738 | 0.345435063 | 0.576948985 | | Xrra1 | 61.52171339 | -0.0556122202851575 | 0.0944030575924808 | -0.589093422431553 | 0.555798605535766 | 0.871401030114683 | NM_001164258 |
| Dbi | 3219.290115 | -0.081454249 | 0.086313455 | -0.943702798 | 0.345321544 | 0.576948985 | | Ccdc13 | 200.4838213 | -0.0525193015910474 | 0.0892218130453395 | -0.588637462056043 | 0.556104497902412 | 0.871546692028314 | NM_028384 |
| Denr | 2073.786632 | 0.054314256 | 0.057594773 | 0.943041408 | 0.345659723 | 0.576948985 | | Fam173a | 1698.059809 | -0.044061904545159 | 0.0748383758287996 | -0.58876083369253 | 0.556021722864748 | 0.871546692028314 | NM_145410 |
| Eif6 | 1247.762082 | 0.066326854 | 0.070310189 | 0.943346266 | 0.345503818 | 0.576948985 | | Fam183b | 77.89235173 | 0.053507509299368 | 0.0908794243955931 | 0.588774738123921 | 0.55601239419442 | 0.871546692028314 | NM_029283,NM_001162878 |
| Fignl1 | 31.85959655 | -0.154849762 | 0.16420584 | -0.943022257 | 0.345669519 | 0.576948985 | | Fam65a | 4345.699577 | -0.0403986043839111 | 0.0686523906961754 | -0.588451530591223 | 0.556229258224256 | 0.871546692028314 | NM_001081241 |
| Galnt6 | 655.1739531 | 0.130607894 | 0.138293314 | 0.94442667 | 0.344951658 | 0.576948985 | | Ston2 | 1043.639745 | -0.0409310745833341 | 0.0695521797315889 | -0.588494490629805 | 0.556200430754213 | 0.871546692028314 | NM_175367 |
| Hectd2 | 599.4413907 | -0.096675158 | 0.102511497 | -0.943066492 | 0.345646893 | 0.576948985 | | Ylpm1 | 4580.971564 | -0.041034952584231 | 0.0697377142728821 | -0.588418376083594 | 0.556251506388166 | 0.871546692028314 | NM_178363 |
| Htra1 | 5332.229608 | 0.151162668 | 0.160268474 | 0.943184046 | 0.345586773 | 0.576948985 | | Gcnt2 | 1899.370583 | 0.0486250311948176 | 0.0826582873485744 | 0.588265650723722 | 0.556353997569511 | 0.871613262370008 | NM_008105,NM_133219,NM_023887 |
| Il2rg | 45.1150865 | 0.145869968 | 0.154662416 | 0.943150715 | 0.345603818 | 0.576948985 | | Mmd2 | 5951.510708 | 0.0299059796966977 | 0.0508495812957301 | 0.588126370653302 | 0.556447473892522 | 0.871665696728435 | NM_175217 |
| Lrguk | 44.74658474 | -0.137989968 | 0.146252031 | -0.943508042 | 0.345421104 | 0.576948985 | | B3gnt9 | 145.8148949 | -0.054144839418966 | 0.0920913390308532 | -0.587947140184659 | 0.556567773762915 | 0.871666143577369 | NM_178879,NM_001271915 |
| Mapk9 | 10646.0154 | 0.048227689 | 0.051136339 | 0.943119709 | 0.345619676 | 0.576948985 | | Rell2 | 5067.287949 | 0.0366494710607052 | 0.062329916724355 | 0.587991657726453 | 0.556537892311038 | 0.871666143577369 | NM_153793 |
| Med17 | 1016.252218 | 0.07463759 | 0.07908583 | 0.943754277 | 0.345295231 | 0.576948985 | | Tcf4 | 9927.453659 | -0.037998863725398 | 0.0646447153116772 | -0.587810829426516 | 0.556659274320119 | 0.871715461273615 | NM_013685,NM_001083967 |
| Mybph | 11.78802157 | 0.13783049 | 0.146132091 | 0.943191118 | 0.345583156 | 0.576948985 | | Peli3 | 627.8951574 | 0.0393044258400174 | 0.0668880567821848 | 0.587615005291735 | 0.556790736919343 | 0.871733374055871 | NM_172835 |
| Nudt14 | 159.7638798 | -0.097003265 | 0.102827598 | -0.943358272 | 0.345497679 | 0.576948985 | | Sgsm2 | 5861.383871 | -0.0405169680274146 | 0.0689471957980898 | -0.587652152613539 | 0.556765797648262 | 0.871733374055871 | NM_197943 |
| Rnaseh2c | 324.2460017 | -0.112256357 | 0.118918903 | -0.94397404 | 0.345182914 | 0.576948985 | | Smarcb1 | 2339.505658 | 0.0303132967125543 | 0.0516047443035327 | 0.587412981532382 | 0.556926377366115 | 0.871851768253064 | NM_001161853,NM_011418 |
| Rom1 | 170.1285862 | 0.116924707 | 0.123978269 | 0.943106467 | 0.345626448 | 0.576948985 | | Txndc17 | 919.9945139 | -0.0389302180563364 | 0.0663457309993022 | -0.586778040877202 | 0.557352786632728 | 0.87242527786929 | NM_026559 |
| Setd5 | 4738.151442 | 0.057926541 | 0.061360644 | 0.944034122 | 0.345152212 | 0.576948985 | | Ppp3r1 | 31725.43799 | 0.0463766418691777 | 0.0790735673797137 | 0.586499931721502 | 0.557539607397142 | 0.872623676010223 | NM_024459 |
| Syngr1 | 26888.13004 | -0.065002566 | 0.068930173 | -0.943020502 | 0.345670416 | 0.576948985 | | 1110054M08Rik | 134.4103889 | 0.0554239189647361 | 0.0946123036824649 | 0.585800332594673 | 0.558009700205733 | 0.872673742619629 | NR_037954 |
| Ugt1a7c | 21.65682924 | 0.156759933 | 0.166201109 | 0.943194267 | 0.345581545 | 0.576948985 | | Arhgef17 | 11788.18575 | 0.0336768654675766 | 0.0574582239283342 | 0.586110449734414 | 0.557801294459885 | 0.872673742619629 | NM_001081116 |
| Mapk4 | 4008.792027 | 0.086076525 | 0.091286516 | 0.942927044 | 0.345718221 | 0.576966622 | | Fam32a | 3125.426146 | -0.0281487522609064 | 0.0480169281318286 | -0.586225594932376 | 0.557723923919364 | 0.872673742619629 | NM_026455 |
| Cdc123 | 2186.89042 | -0.041046658 | 0.043539859 | -0.942737499 | 0.345815188 | 0.577066292 | | Klf4 | 304.0337988 | 0.0485527029718725 | 0.0828053747953903 | 0.586347240041421 | 0.557642191508535 | 0.872673742619629 | NM_010637 |
| Gm14295 | 1694.566296 | -0.1345732 | 0.142844308 | -0.942097043 | 0.346142958 | 0.577551044 | | Lrba | 1736.386791 | 0.0398661630805092 | 0.0680719193975785 | 0.58564770074527 | 0.558112286172129 | 0.872673742619629 | NM_030695,NM_001077687,NM_001077688 |
| Fchsd2 | 1900.741819 | -0.08244709 | 0.087534574 | -0.941880287 | 0.346253934 | 0.577674001 | | Mex3c | 1629.261196 | 0.045493934826525 | 0.0776374391806235 | 0.585979332995301 | 0.557889403258981 | 0.872673742619629 | NM_001039214 |
| Rab3gap2 | 4312.17338 | 0.06972247 | 0.074034526 | 0.941756149 | 0.346317501 | 0.577717846 | | Ncbp2 | 2101.159569 | 0.0325288335027724 | 0.0555392956005484 | 0.585690422448412 | 0.558083571403297 | 0.872673742619629 | NM_026554 |
| D17Wsu92e | 12150.4624 | -0.073340457 | 0.077893463 | -0.941548294 | 0.346423953 | 0.577815952 | | Pkp2 | 253.3057477 | -0.0552453859122369 | 0.094287866398236 | -0.585922537253112 | 0.557927571384604 | 0.872673742619629 | NM_026163 |
| Fam115c | 30.53282782 | -0.156690759 | 0.166427981 | -0.941492879 | 0.346452337 | 0.577815952 | | Slc5a11 | 25.12181864 | 0.0460637602078121 | 0.0786237087804612 | 0.585876206074616 | 0.557958708008592 | 0.872673742619629 | NM_146198 |
| Fam131a | 10788.97297 | -0.100897889 | 0.107175947 | -0.941422886 | 0.34648819 | 0.577815952 | | Egr2 | 417.2830477 | 0.0302804590350026 | 0.0517234227493736 | 0.585430302664364 | 0.558258418246531 | 0.872714354654445 | NM_010118 |
| Ccdc160 | 74.65780813 | -0.142444032 | 0.151346401 | -0.941178852 | 0.346613212 | 0.577962237 | | Pds5a | 1871.246883 | -0.0379701731061396 | 0.064853345007697 | -0.585477481564493 | 0.558226703642211 | 0.872714354654445 | NM_001081321 |
| Cyp4v3 | 352.2845021 | 0.094479369 | 0.100392941 | 0.941095737 | 0.3466558 | 0.57797105 | | Myrf | 3175.776699 | 0.0493814249423469 | 0.0843644330692662 | 0.585334638612494 | 0.558322728232565 | 0.872720967427767 | NM_001033481 |
| Cyp1b1 | 176.1955793 | -0.145863784 | 0.155028118 | -0.940885989 | 0.346763289 | 0.578088056 | | Osbpl5 | 1328.486502 | -0.0380188864725779 | 0.0649760261606125 | -0.585121755193215 | 0.558465851649485 | 0.872850144597668 | NM_001199227,NM_024289 |
| Haus7 | 184.251154 | -0.10380642 | 0.110353837 | -0.940668873 | 0.346874575 | 0.578166078 | | Osbpl9 | 1901.94764 | 0.0361923869305586 | 0.0618638412952663 | 0.585032971650337 | 0.558525546883886 | 0.872850144597668 | NM_173350,NM_001134791,NM_133885 |
| Lancl2 | 3771.136896 | -0.069516636 | 0.073908557 | -0.940576292 | 0.346922036 | 0.578166078 | | Zfp91 | 3881.186594 | 0.0383210128459634 | 0.0655189145064683 | 0.58488473343343 | 0.558625224463247 | 0.872912006298745 | NM_053009 |
| Vps26a | 2073.219914 | -0.061399443 | 0.06527626 | -0.940609077 | 0.346905229 | 0.578166078 | | Atg4c | 1802.674312 | 0.0360449280396247 | 0.061673309051932 | 0.58444939299873 | 0.558918003794107 | 0.873275564495343 | NM_175029,NM_001145967 |
| Gpm6a | 34413.24414 | -0.081525031 | 0.086735917 | -0.939922398 | 0.347257367 | 0.578662386 | | Sema5b | 883.0659627 | -0.0418817381802963 | 0.0717201765028843 | -0.583960333374417 | 0.559246999728381 | 0.87369562465632 | NM_013661 |
| Mfsd1 | 1041.927889 | 0.060548989 | 0.064427664 | 0.939798001 | 0.347321184 | 0.578686732 | | Coq4 | 687.3960142 | -0.0412924075979606 | 0.0709861012586477 | -0.581697076833477 | 0.56077074085759 | 0.873713575125364 | NM_178693 |
| Mms22l | 41.23666157 | 0.156890907 | 0.166949858 | 0.939748666 | 0.347346496 | 0.578686732 | | Cyr61 | 302.8813045 | -0.0421377709150449 | 0.0723907868747809 | -0.582087482871727 | 0.560507755948216 | 0.873713575125364 | NM_010516 |
| Rlbp1 | 768.8120905 | -0.139955276 | 0.148952384 | -0.939597424 | 0.347424098 | 0.578753787 | | D030028A08Rik | 275.0976525 | -0.0464494510883288 | 0.0796990682129573 | -0.582810466042276 | 0.560020898593679 | 0.873713575125364 | NR_003293 |
| Nos1 | 1237.92031 | -0.118587493 | 0.126230879 | -0.939449158 | 0.347500184 | 0.578818303 | | Darc | 2039.61281 | -0.038467128077533 | 0.0659320018208821 | -0.583436374069711 | 0.559599577340512 | 0.873713575125364 | NM_010045 |
| 4930578M01Rik | 19.03879293 | -0.151617445 | 0.161457616 | -0.939054153 | 0.347702941 | 0.579093774 | | Ddx4 | 110.3017683 | -0.0553804637264743 | 0.0950818917778219 | -0.58245016680865 | 0.560263498713808 | 0.873713575125364 | NM_010029,NM_001145885 |
| Foxp4 | 1421.150567 | 0.06521685 | 0.0694586 | 0.938931248 | 0.347766044 | 0.579099155 | | Igfbp2 | 1285.98226 | 0.0497017971014129 | 0.0852866190572968 | 0.582761957863783 | 0.560053557616761 | 0.873713575125364 | NM_008342 |
| Psmd8 | 3259.088566 | -0.057892524 | 0.061659798 | -0.938902269 | 0.347780924 | 0.579099155 | | Kank2 | 545.4459165 | -0.0423599506597232 | 0.0727926043933719 | -0.581926570875383 | 0.560616142082244 | 0.873713575125364 | NM_145611 |
| Asb18 | 12.39904003 | -0.155193671 | 0.165319444 | -0.938750264 | 0.34785898 | 0.579166886 | | Lrrfip2 | 1462.78372 | 0.0373055307866417 | 0.0640455829050813 | 0.582484054238469 | 0.560240679136447 | 0.873713575125364 | NM_027742,NM_001164838 |
| Cabp1 | 3864.496703 | 0.112253423 | 0.119603718 | 0.938544595 | 0.347964611 | 0.579228723 | | Mybl1 | 152.2716196 | -0.0548888949535839 | 0.0943113471902622 | -0.581996722439475 | 0.560568888565916 | 0.873713575125364 | NM_008651 |
| Zbtb7a | 5091.879057 | 0.066954358 | 0.071339423 | 0.938532372 | 0.347970889 | 0.579228723 | | Nos1ap | 3601.636947 | 0.0364134338565283 | 0.0625155325423375 | 0.582470185819308 | 0.560250017986295 | 0.873713575125364 | NM_027528,NM_001109985 |
| Zxda | 40.77376756 | 0.147185747 | 0.156868626 | 0.938273959 | 0.348103639 | 0.57938745 | | Nrn1 | 6797.787196 | 0.0343055275614063 | 0.0588774171381675 | 0.582660198576674 | 0.560122071927592 | 0.873713575125364 | NM_153529 |
| Gal3st1 | 351.8062454 | -0.115409653 | 0.123016758 | -0.93816204 | 0.348161143 | 0.579420918 | | Olfr1344 | 27.38216268 | 0.0500347398842019 | 0.0859812913826815 | 0.581925894337985 | 0.56061659780166 | 0.873713575125364 | NM_177061 |
| Cpa4 | 9.689432344 | 0.152774147 | 0.162868192 | 0.938023225 | 0.348232475 | 0.579477388 | | Pex26 | 823.2083883 | 0.0386497690307324 | 0.0663638515605596 | 0.582391891396824 | 0.56030274205472 | 0.873713575125364 | NM_028730 |
| Clec11a | 147.4760546 | 0.126518184 | 0.134909562 | 0.937799973 | 0.348347215 | 0.579556372 | | Pms1 | 147.438202 | -0.051886248719634 | 0.0891703809014978 | -0.581877616704926 | 0.560649118351442 | 0.873713575125364 | NM_153556 |
| Efhc2 | 71.84877043 | -0.139692285 | 0.14895977 | -0.937785315 | 0.34835475 | 0.579556372 | | Prkag2 | 3506.670689 | -0.0314761809349737 | 0.0539547485644907 | -0.583381106805661 | 0.559636773537745 | 0.873713575125364 | NM_001170555,NM_145401,NM_001170556 |
| Zbtbd6 | 24.99524599 | -0.157418688 | 0.167895635 | -0.937598454 | 0.348450806 | 0.579653939 | | Qpctl | 726.6513806 | 0.0369161330904315 | 0.0633285339621112 | 0.582930486161547 | 0.559940096810016 | 0.873713575125364 | NM_026111 |
| Armc3 | 27.73129786 | 0.156524308 | 0.166960247 | 0.937494472 | 0.348504266 | 0.579680633 | | Rnf180 | 417.7981334 | 0.044251435932586 | 0.0760904187205553 | 0.581563837821697 | 0.560860506854396 | 0.873713575125364 | NM_027934 |
| Pcnxl4 | 1700.789845 | -0.054882366 | 0.058555237 | -0.937275104 | 0.348617066 | 0.579806013 | | Sept8 | 9478.225228 | 0.0243994925633988 | 0.0418551931099502 | 0.582950184922174 | 0.559926835448587 | 0.873713575125364 | NM_033144,NM_001252332,NM_001252333 |
| Zfp105 | 169.2982824 | 0.097163648 | 0.103687264 | 0.937083728 | 0.348715491 | 0.579907462 | | Setd1b | 2073.4321 | 0.0384568602607468 | 0.0659248903401252 | 0.583343560563194 | 0.559662043745604 | 0.873713575125364 | NM_001040398 |
| Cpxm2 | 95.17164078 | -0.14541902 | 0.155210673 | -0.93691379 | 0.348802905 | 0.579990579 | | Sgpp1 | 1269.901095 | 0.0383455836865624 | 0.0659215043437662 | 0.581685507154063 | 0.560778535324412 | 0.873713575125364 | NM_030750 |
| Rai14 | 557.9948775 | 0.08854415 | 0.094524977 | 0.936727548 | 0.348898722 | 0.580087649 | | Slc25a26 | 622.5259969 | -0.0382283217956333 | 0.0657242036203507 | -0.581647546715901 | 0.560804109554044 | 0.873713575125364 | NM_026255 |
| Cdh20 | 859.2066795 | 0.066956818 | 0.07150201 | 0.936432669 | 0.349050464 | 0.58015801 | | Smyd1 | 119.4640895 | 0.0553119753522729 | 0.0950762901287967 | 0.581764131492127 | 0.56072556731896 | 0.873713575125364 | NM_001160127,NM_009762 |
| Ldoc1 | 49.41917001 | -0.147985206 | 0.158042266 | -0.936364741 | 0.349085425 | 0.58015801 | | Tceanc | 80.06431823 | -0.0553340189545797 | 0.0950393369293057 | -0.582222274927481 | 0.560416971339972 | 0.873713575125364 | NM_001007577 |
| Mpp5 | 1461.62597 | 0.064194039 | 0.068555212 | 0.936384516 | 0.349075247 | 0.58015801 | | Tmem56 | 2851.762777 | 0.0479113158392538 | 0.0823882112258237 | 0.581531206059691 | 0.560882492636928 | 0.873713575125364 | NM_178936 |
| Ppp1r15a | 626.2953724 | 0.118407297 | 0.12645566 | 0.936354265 | 0.349090817 | 0.58015801 | | Trmt112 | 762.4916843 | 0.0406593581988546 | 0.0698194141676949 | 0.582350320230379 | 0.560330737376236 | 0.873713575125364 | NM_001166370,NM_026306 |
| C5ar1 | 43.91905704 | 0.154961783 | 0.165626487 | 0.935609914 | 0.349474068 | 0.580421397 | | Wdr41 | 1475.84898 | 0.0311477505703838 | 0.0535030146313691 | 0.582168141084926 | 0.560453430493881 | 0.873713575125364 | NM_172590 |
| Chrnb2 | 2884.325367 | 0.05968216 | 0.063776433 | 0.935802722 | 0.34937477 | 0.580421397 | | Zfp652 | 560.4461167 | -0.0416084399069669 | 0.0713052824703989 | -0.583525349952017 | 0.559539696952662 | 0.873713575125364 | NM_201609 |
| Dscaml1 | 2670.205175 | -0.070897893 | 0.075773579 | -0.935654539 | 0.349451084 | 0.580421397 | | Bai2 | 19031.54906 | -0.0400363103749845 | 0.0688812408362972 | -0.581236776354459 | 0.561080884713027 | 0.87387211030248 | NM_001199696,NM_173071 |
| Dspp | 9.467365553 | -0.152568173 | 0.163035384 | -0.935797915 | 0.349377245 | 0.580421397 | | Gm6277 | 165.1035631 | 0.0534272023207337 | 0.0919308774975752 | 0.581167109191827 | 0.561127832678954 | 0.87387211030248 | NR_045421 |
| Gm7008 | 14.04952023 | 0.153019619 | 0.163550541 | 0.935610598 | 0.349473716 | 0.580421397 | | Stk19 | 698.5386231 | -0.0459278561030394 | 0.0790343844322513 | -0.58111234031827 | 0.561164742179646 | 0.87387211030248 | NM_019442 |
| Rapgef1 | 7243.845863 | 0.114884867 | 0.122768703 | 0.935783017 | 0.349384917 | 0.580421397 | | 2410006H16Rik | 382.6052973 | -0.0437828411859925 | 0.0753865093890415 | -0.580778199452713 | 0.561389949769449 | 0.873934985333935 | NR_030738 |
| Rasgef1b | 2657.181794 | -0.099782422 | 0.106668371 | -0.935445264 | 0.349558879 | 0.580500029 | | Apitd1 | 74.17766495 | 0.0552146638660531 | 0.095088753563636 | 0.580664503390529 | 0.561466589741561 | 0.873934985333935 | NM_027263 |
| Itm2c | 31448.61079 | -0.061524295 | 0.065784924 | -0.935233953 | 0.349667744 | 0.580618587 | | BC056474 | 945.8924814 | -0.033645550930844 | 0.0579338243468348 | -0.580758327456803 | 0.561403344672421 | 0.873934985333935 | NM_001001493 |
| Ddx39b | 5243.881011 | 0.062976922 | 0.067344147 | 0.935150634 | 0.349710675 | 0.580627647 | | Larp1b | 559.2875416 | 0.0440787681910244 | 0.0759302077346605 | 0.580516891841762 | 0.561566098933426 | 0.873934985333935 | NM_001040399 |
| Nfatc3 | 658.7756281 | 0.071883518 | 0.076878143 | 0.935031917 | 0.349771851 | 0.580666995 | | Rprml | 1158.558784 | 0.0438412783087933 | 0.0755112988226116 | 0.580592295356801 | 0.561515266189033 | 0.873934985333935 | NM_001033212 |
| Fbxo30 | 296.0676914 | -0.10254503 | 0.109705753 | -0.934727911 | 0.349928539 | 0.580864879 | | Tmem48 | 488.6741707 | 0.0455654588338893 | 0.07847452108933 | 0.580640164493909 | 0.561482996704849 | 0.873934985333935 | NM_028355 |
| Mrpl48 | 1717.277959 | -0.061875357 | 0.066203354 | -0.934625704 | 0.349981228 | 0.580890106 | | Dars | 2006.621147 | 0.0294504217444856 | 0.0507535315743156 | 0.58026349755313 | 0.561736939196391 | 0.874107207152306 | NM_145507,NM_177445 |
| Mras | 9924.456202 | 0.083695954 | 0.089561407 | 0.93450914 | 0.350041323 | 0.58092762 | | Pofut1 | 1031.483556 | -0.0345925602393004 | 0.0596356987139105 | -0.580064642241399 | 0.561871026479691 | 0.874222207652527 | NM_080463 |
| Btg2 | 1063.098737 | -0.153347691 | 0.164138438 | -0.934258257 | 0.350170691 | 0.581080078 | | B4galt5 | 2724.848681 | 0.0290182411652763 | 0.0500404899833441 | 0.579895224346024 | 0.56198527644218 | 0.874306321687812 | NM_019835 |
| Enox1 | 1149.524669 | -0.09404502 | 0.100700678 | -0.933906519 | 0.350352116 | 0.581256632 | | Maml1 | 566.0718395 | 0.0402970529142209 | 0.0695385287574374 | 0.579492457408527 | 0.562256934529615 | 0.874635278536859 | NM_175334 |
| Pdzd11 | 1014.442888 | -0.075956426 | 0.081327208 | -0.933960814 | 0.350324107 | 0.581256632 | | Cstf3 | 975.7086267 | -0.0380769725202946 | 0.0657179460429734 | -0.579399917571919 | 0.562319359722302 | 0.87463872141837 | NM_177253,NM_145529,NM_001037326 |
| Rfc3 | 185.327006 | 0.104096933 | 0.111499729 | 0.933607048 | 0.350506628 | 0.581450717 | | Sdad1 | 1005.894066 | -0.0356591728820117 | 0.0615555940871768 | -0.579300279865874 | 0.562386576712529 | 0.874649615987816 | NM_172713 |
| 2210016L21Rik | 3756.668437 | 0.047652239 | 0.051055673 | 0.93333878 | 0.350645078 | 0.581493615 | | Ccdc107 | 605.8396544 | -0.0388817509209655 | 0.0672028437287293 | -0.5785730002426 | 0.562877327209546 | 0.875103820297693 | NM_001037913 |
| Fubp1 | 2712.921755 | 0.089737273 | 0.096142189 | 0.933380802 | 0.350623389 | 0.581493615 | | Ccdc164 | 751.6754194 | -0.0370142382191485 | 0.0639777004925207 | -0.578549055908561 | 0.56289348777328 | 0.875103820297693 | NM_001033460 |
| Vwf | 898.2523373 | -0.088861773 | 0.095199262 | -0.933429225 | 0.350598397 | 0.581493615 | | Cpeb4 | 2136.11034 | -0.0363304757891632 | 0.0628278153600238 | -0.578254640575639 | 0.563092213529212 | 0.875103820297693 | NM_026252 |
| Rab34 | 525.7814607 | 0.119759827 | 0.128377575 | 0.932871854 | 0.350886136 | 0.581831099 | | Nr4a1 | 4042.832383 | 0.0403589916613021 | 0.0697957420946794 | 0.578244323365088 | 0.563099178099042 | 0.875103820297693 | NM_010444 |
| Psmb6 | 2584.905107 | -0.070262929 | 0.075330075 | -0.932734086 | 0.350957281 | 0.5818475 | | Sssca1 | 534.7077253 | 0.0391174523450683 | 0.0676423882069723 | 0.57829791912989 | 0.563062999058793 | 0.875103820297693 | NM_020491 |
| Spata2 | 2941.008464 | 0.068397382 | 0.0733321 | 0.932707265 | 0.350971133 | 0.5818475 | | Syne1 | 18771.97789 | 0.0357843194589622 | 0.0618846112381135 | 0.578242615458548 | 0.563100331014907 | 0.875103820297693 | NM_153399,NM_022027,NM_001079686 |
| Rundc3a | 11918.53311 | -0.071072622 | 0.076228514 | -0.932362683 | 0.351149124 | 0.582080295 | | Zfp958 | 269.7278944 | -0.05002132962041 | 0.0864935223502036 | -0.57832457577434 | 0.563045005296832 | 0.875103820297693 | NM_145591 |
| Fam53c | 2991.643276 | 0.071990715 | 0.077229664 | 0.932164036 | 0.351251759 | 0.582188141 | | Cntnap3 | 499.5988788 | 0.0441416731899789 | 0.0763986422641464 | 0.577780859473394 | 0.563412079410472 | 0.875108780622359 | NM_001081129 |
| 2810025M15Rik | 400.8737746 | -0.092732004 | 0.099542953 | -0.931577791 | 0.351554765 | 0.582452457 | | Dnajc24 | 180.7641425 | 0.0539990428329453 | 0.0934160647302967 | 0.578048786242998 | 0.563231182091906 | 0.875108780622359 | NM_026992,NR_033993 |
| A630019I02Rik | 24.69201386 | -0.155532108 | 0.166953218 | -0.931590956 | 0.351547959 | 0.582452457 | | Gpr156 | 50.06294547 | -0.0532297198002968 | 0.0921545968807224 | -0.577613288995156 | 0.563525232957562 | 0.875108780622359 | NM_153394 |
| Siah3 | 75.07763029 | 0.136417096 | 0.14643869 | 0.931564578 | 0.351561596 | 0.582452457 | | Isg20l2 | 607.9341382 | 0.0421900422826672 | 0.0730415402774151 | 0.577617094634471 | 0.563522663042138 | 0.875108780622359 | NM_177663 |
| St8sia1 | 2322.547516 | -0.082102702 | 0.088118668 | -0.931728821 | 0.351476688 | 0.582452457 | | Mpdz | 2847.237261 | 0.0295130279238409 | 0.0510911097988105 | 0.57765486089574 | 0.563497160117661 | 0.875108780622359 | NM_010820 |
| Grm7 | 2091.015162 | -0.076380724 | 0.082016532 | -0.931284486 | 0.351706425 | 0.582567816 | | Slc25a53 | 357.6231718 | -0.0415183307146303 | 0.0718594355761601 | -0.577771455922822 | 0.563418428956644 | 0.875108780622359 | NM_001114179,NM_001082412 |
| Srp9 | 2172.704684 | -0.066741084 | 0.0716644 | -0.9313004 | 0.351698195 | 0.582567816 | | Slc4a1ap | 1207.091011 | 0.0287699195279594 | 0.049789807646351 | 0.57782748895733 | 0.563380594359559 | 0.875108780622359 | NM_009206 |
| H2-Q2 | 14.8448339 | -0.144805222 | 0.155527595 | -0.931058069 | 0.351823527 | 0.582637209 | | Isca2 | 873.2575052 | -0.0382923356606547 | 0.06634656264532 | -0.577156285629452 | 0.56383388445515 | 0.875120312597021 | NM_028863 |
| Thop1 | 1521.106972 | -0.072733641 | 0.07811403 | -0.93112136 | 0.351790791 | 0.582637209 | | Ispd | 298.1157838 | 0.0470448914622806 | 0.0814942936176006 | 0.577278351328886 | 0.563751435601091 | 0.875120312597021 | NM_178629 |
| AI413582 | 826.7788496 | -0.076297709 | 0.081987257 | -0.930604474 | 0.352058199 | 0.582963529 | | Ivd | 2538.754856 | -0.0360940063311459 | 0.0625081955905821 | -0.577428383432397 | 0.563650104894974 | 0.875120312597021 | NM_019826 |
| 1700026D08Rik | 29.59193764 | 0.155263786 | 0.166871025 | 0.930441854 | 0.352142357 | 0.583040573 | | Krt1 | 111.3443878 | 0.0548822123676097 | 0.0950791435179102 | 0.577226617078975 | 0.563786378611776 | 0.875120312597021 | NM_008473 |
| Chtf18 | 57.01780333 | 0.141373421 | 0.151982159 | 0.930197477 | 0.352268848 | 0.583187684 | | Zfp946 | 550.3286806 | 0.0474676537175314 | 0.0822306713129489 | 0.577249984216737 | 0.56377059555063 | 0.875120312597021 | NM_198003 |
| Mmp11 | 63.58565418 | -0.139471409 | 0.149954218 | -0.930093267 | 0.352322796 | 0.58321468 | | Mn1 | 1883.530982 | 0.0436224966740324 | 0.0756259505052257 | 0.576819152455057 | 0.564061630064772 | 0.875251365912015 | NM_001081235 |
| Ebag9 | 628.66907 | -0.08193653 | 0.088120579 | -0.929822873 | 0.352462801 | 0.583347711 | | Mtcp1 | 284.4361514 | -0.0479000782723864 | 0.083049742723849 | -0.576763716114814 | 0.564099083551445 | 0.875251365912015 | NM_001039373,NM_010839 |
| Psmd4 | 3069.981823 | 0.06427493 | 0.069128242 | 0.929792634 | 0.352478461 | 0.583347711 | | Slc25a10 | 674.1428173 | -0.0473984892121233 | 0.0821680512241499 | -0.576848160641207 | 0.564042032247726 | 0.875251365912015 | NM_013770 |
| Aatk | 11601.78781 | 0.06194257 | 0.066641413 | 0.92949064 | 0.352634874 | 0.583544242 | | Kpna3 | 3892.249974 | 0.0392436142729597 | 0.0680624675630487 | 0.576582302670806 | 0.564221657120569 | 0.875323715765288 | NM_008466 |
| F8 | 110.2658355 | 0.117167003 | 0.126079358 | 0.929311545 | 0.352727654 | 0.583635441 | | Ntf3 | 50.06571224 | 0.0493838198347456 | 0.0856702498148395 | 0.576440712399925 | 0.564317332725883 | 0.875323715765288 | NM_008742,NM_001164034,NM_001164035 |
| Caprin1 | 8137.635469 | -0.059269218 | 0.06380535 | -0.928906719 | 0.352937431 | 0.583857842 | | Tsga14 | 334.2112915 | 0.0452197331385002 | 0.0784483040017853 | 0.57642716071301 | 0.564326490303038 | 0.875323715765288 | NM_031998 |
| Rab40c | 1603.503978 | 0.062076016 | 0.066826882 | 0.928907851 | 0.352936844 | 0.583857842 | | Arg2 | 96.20349935 | -0.0545631308610755 | 0.0947636119631579 | -0.575781460106106 | 0.564762906551866 | 0.875364911761399 | NM_009705 |
| Fam92b | 10.78926911 | -0.149749933 | 0.161241592 | -0.928730184 | 0.353028934 | 0.583928053 | | Atad2b | 755.7661509 | -0.042765214234046 | 0.0743868478401111 | -0.574902895817909 | 0.565356971348957 | 0.875364911761399 | NM_001099628 |
| Ltv1 | 582.8716848 | -0.070607947 | 0.076030485 | -0.928679424 | 0.353055247 | 0.583928053 | | Cog8 | 875.6406049 | 0.0356263319909179 | 0.061916632991653 | 0.575391946711971 | 0.565026249421332 | 0.875364911761399 | NM_139229 |
| Bhlha9 | 38.28866341 | -0.155742895 | 0.167737907 | -0.928489558 | 0.353153682 | 0.583966184 | | Eef1a1 | 88995.03367 | 0.0305957870810857 | 0.0531733739479133 | 0.57539675987189 | 0.565022994971846 | 0.875364911761399 | NM_010106 |
| Spag4 | 80.56865009 | 0.146682589 | 0.157973889 | 0.928524265 | 0.353135687 | 0.583966184 | | Esr1 | 133.6615975 | 0.0423239003557499 | 0.0734466166949195 | 0.576253914207564 | 0.564443568247648 | 0.875364911761399 | NM_007956 |
| Mtfr2 | 14.19835155 | 0.155272669 | 0.167249112 | 0.928391588 | 0.35320448 | 0.583987858 | | Fgl2 | 77.25385921 | -0.0472667477335307 | 0.082046636989555 | -0.576096101776213 | 0.56455022620044 | 0.875364911761399 | NM_008013 |
| E330020D12Rik | 19.0215704 | 0.155986143 | 0.168101072 | 0.927930687 | 0.353443525 | 0.584133757 | | Hs3st3a1 | 82.98658041 | 0.0541532241876679 | 0.0942113828625009 | 0.574805533495917 | 0.565422823971999 | 0.875364911761399 | NM_178870 |
| Gdi1 | 25701.98993 | 0.042550766 | 0.045855452 | 0.927932528 | 0.35344257 | 0.584133757 | | Kif24 | 35.59236734 | -0.0512813273609708 | 0.0891593276547133 | -0.575165029951411 | 0.565179690897721 | 0.875364911761399 | NM_024241 |
| Mylk4 | 14.31164527 | 0.155372527 | 0.16742547 | 0.928010095 | 0.353402333 | 0.584133757 | | Mier2 | 1367.757666 | -0.0385936281899459 | 0.0670323948312173 | -0.575746413200934 | 0.56478659870505 | 0.875364911761399 | NM_027422 |
| Uevld | 639.2699104 | 0.071819674 | 0.077381435 | 0.928125387 | 0.353342532 | 0.584133757 | | Mtfmt | 433.2959033 | -0.0438559687753339 | 0.0762157234219917 | -0.575418913660531 | 0.565008015659314 | 0.875364911761399 | NM_027134 |
| Ggt1 | 75.74652659 | -0.134274831 | 0.144753131 | -0.927612624 | 0.353608547 | 0.584344158 | | Ntsr2 | 3431.572111 | -0.038331012435574 | 0.0666630293021002 | -0.574996558615232 | 0.565293624439347 | 0.875364911761399 | NM_008747 |
| Klhl4 | 532.9334684 | 0.092632412 | 0.099869897 | 0.927530871 | 0.353650971 | 0.58435194 | | Paip2b | 3715.459485 | 0.0351503134202968 | 0.061141596162665 | 0.574900160060929 | 0.565358821673286 | 0.875364911761399 | NM_146169 |
| Zc3h8 | 246.4634479 | 0.082308285 | 0.088761907 | 0.92729289 | 0.353774485 | 0.584493695 | | Slc2a3 | 7196.587038 | 0.0379378375493839 | 0.0660037060707254 | 0.57478344486796 | 0.565437764495195 | 0.875364911761399 | NM_011401 |
| Lrrc14b | 8.642557725 | 0.135688498 | 0.14634983 | 0.927151729 | 0.353847762 | 0.584552427 | | Slc35f6 | 1597.487692 | 0.030541777814474 | 0.0531356154172164 | 0.574789198820086 | 0.565433872562254 | 0.875364911761399 | NM_175675 |
| 4933431E20Rik | 5094.994154 | 0.048455625 | 0.052329659 | 0.925968665 | 0.354462265 | 0.585380817 | | Usp31 | 5864.895531 | 0.0418666002416157 | 0.0727030807411123 | 0.575857306387031 | 0.564711635139207 | 0.875364911761399 | NM_001033173 |
| Heatr5a | 605.1863091 | -0.077715469 | 0.08393108 | -0.925943867 | 0.354475152 | 0.585380817 | | Xpnpep3 | 434.2429119 | -0.0450100187540622 | 0.0781813379366572 | -0.57571307861896 | 0.564809133749598 | 0.875364911761399 | NM_177310 |
| Tctn2 | 877.0459494 | -0.062434727 | 0.067431728 | -0.925895412 | 0.354500336 | 0.585380817 | | Zc4h2 | 723.2434763 | 0.0450425756292739 | 0.0783592367196572 | 0.574821520919366 | 0.565412010359787 | 0.875364911761399 | NM_001003916 |
| Zp2 | 8.626441853 | 0.142252359 | 0.153601453 | 0.926113366 | 0.354387069 | 0.585380817 | | Zfp143 | 490.4812056 | 0.0453224087331657 | 0.0787225676683413 | 0.575723202069695 | 0.564802289987832 | 0.875364911761399 | NM_009281 |
| Cdk20 | 332.7509843 | 0.080240359 | 0.086678301 | 0.925726023 | 0.354588381 | 0.585463807 | | Nebl | 1433.551425 | 0.0360651174513986 | 0.0627616817385123 | 0.574635934833306 | 0.565537543644628 | 0.87542608258545 | NM_028757 |
| Pcdhgb2 | 290.8558481 | -0.105199565 | 0.113697419 | -0.925259044 | 0.354831177 | 0.585802265 | | Yap1 | 1000.379734 | -0.038948821257884 | 0.0677935713670137 | -0.57452086492135 | 0.565615385428274 | 0.875453285959628 | NM_009534,NM_001171147 |
| Gm3279 | 12.06548888 | -0.150350391 | 0.162510563 | -0.925173036 | 0.354875907 | 0.58581369 | | Elmod2 | 662.7499074 | 0.0428018885943268 | 0.074530543687415 | 0.574286547188494 | 0.565773911150282 | 0.875605351946409 | NM_178736,NM_001170691 |
| Klc3 | 78.96828778 | -0.12615745 | 0.136375254 | -0.925075819 | 0.35492647 | 0.585834742 | | Tcf19 | 265.9168357 | -0.0481516077960575 | 0.0838706211302644 | -0.574117696365577 | 0.56588815900228 | 0.875688867334306 | NM_025674,NM_001163764,NM_001163763 |
| Vash1 | 2451.662182 | -0.0795175 | 0.085977914 | -0.924859605 | 0.355038941 | 0.585957963 | | Capns1 | 5410.627837 | 0.0347695748581961 | 0.0605945649269659 | 0.573806824095587 | 0.566098530378486 | 0.875752430301446 | NM_009795 |
| Stub1 | 3850.149428 | -0.049531576 | 0.053568833 | -0.924634218 | 0.355156208 | 0.58608907 | | Dph5 | 340.4255467 | 0.0423250846554695 | 0.0737638924911397 | 0.573791366291489 | 0.566108991858506 | 0.875752430301446 | NM_027193 |
| Galnt4 | 180.4409276 | -0.100999958 | 0.109285024 | -0.924188454 | 0.355388206 | 0.586284589 | | Rnf166 | 1172.231463 | -0.0384170733884525 | 0.0669532260797128 | -0.573789728111295 | 0.566110100545957 | 0.875752430301446 | NM_001033142 |
| Gm20753 | 10.65851557 | 0.149760445 | 0.162033675 | 0.924255068 | 0.35535353 | 0.586284589 | | Mmp28 | 116.8465574 | -0.0521922516106451 | 0.0909943593153092 | -0.573576779960515 | 0.566254228448956 | 0.875844211697171 | NM_172797,NM_080453 |
| Tmem42 | 362.3844539 | -0.11214618 | 0.121326656 | -0.924332573 | 0.355313188 | 0.586284589 | | Zfp473 | 24.81086895 | -0.0497653730613696 | 0.0867712277078748 | -0.573523901596857 | 0.566290020395142 | 0.875844211697171 | NM_178734 |
| S100pbp | 680.6379026 | -0.078409147 | 0.084872822 | -0.923842816 | 0.355568159 | 0.58651901 | | Abca8a | 612.5531924 | 0.0473306077951611 | 0.0826508636461607 | 0.572657147271802 | 0.566876857822431 | 0.875959314031032 | NM_153145 |
| Capn3 | 225.8964674 | -0.131089671 | 0.141949226 | -0.923496906 | 0.355748312 | 0.586748225 | | Akap7 | 1255.625868 | -0.0357166424280654 | 0.0623287941106295 | -0.573035993038317 | 0.566620323819563 | 0.875959314031032 | NM_018747 |
| Gm20748 | 26.3654268 | -0.154096999 | 0.166874479 | -0.923430591 | 0.355782856 | 0.586748225 | | Drd1a | 2137.950667 | 0.0541233011027123 | 0.0944973663770211 | 0.572749306967707 | 0.566814447102697 | 0.875959314031032 | NM_010076 |
| Ccdc32 | 1552.422371 | -0.069814802 | 0.075648837 | -0.922880035 | 0.356069726 | 0.586971416 | | Gm11602 | 96.77181737 | -0.0541962507426659 | 0.0946473591581128 | -0.572612392197109 | 0.566907167231566 | 0.875959314031032 | NR_045915,NR_045914 |
| Cirh1a | 1136.86539 | 0.068885336 | 0.074634826 | 0.922965044 | 0.356025422 | 0.586971416 | | Nsd1 | 6768.020628 | -0.0364247789040325 | 0.0636072610637304 | -0.572651271173856 | 0.566880837238741 | 0.875959314031032 | NM_008739 |

| | | | | | | | | | | | | | | |
|---|---|---|---|---|---|---|---|---|---|---|---|---|---|---|
| Cnksr3 | 186.5974524 | 0.096782176 | 0.104864924 | 0.922922291 | 0.356047703 | 0.586971416 | | Pcdha6 | 123.6958951 | 0.0544141436748167 | 0.0949339236994234 | 0.573179128749602 | 0.566523414497345 | 0.875959314031032 | NM_007767 |
| Ncoa3 | 2191.942367 | 0.062385889 | 0.067588219 | 0.92302903 | 0.355992077 | 0.586971416 | | Rp9 | 732.2698952 | 0.0404662667980821 | 0.070608346957363 | 0.573108825540382 | 0.566571011942476 | 0.875959314031032 | NM_018739 |
| Ccbl2 | 217.0903522 | 0.08856334 | 0.095997015 | 0.922563475 | 0.356234737 | 0.58718096 | | Stoml2 | 1631.313808 | 0.0301152244360293 | 0.0525752942842408 | 0.572801823480351 | 0.566778884294794 | 0.875959314031032 | NM_023231 |
| Luzp1 | 5035.997611 | 0.088388029 | 0.095823335 | 0.922406101 | 0.356316789 | 0.587253731 | | Tnfrsf23 | 58.70531957 | -0.0541279544643316 | 0.0944694831031955 | -0.572967615427766 | 0.566666213466848 | 0.875959314031032 | NM_024290 |
| Nmi | 110.0686612 | 0.111154731 | 0.120540156 | 0.9221386 | 0.356456285 | 0.587394165 | | Ddah2 | 283.3190953 | 0.0490090566458607 | 0.085637614228929 | 0.572284236163425 | 0.56712942755342 | 0.876209536762495 | NM_001190449,NM_016765 |
| Zfp366 | 144.0507112 | -0.131325731 | 0.142420685 | -0.922097311 | 0.356477819 | 0.587394165 | | Supt4a | 1163.845714 | 0.0326137701470087 | 0.05701355891914 | 0.572035332740119 | 0.567298037864232 | 0.876376826333706 | NM_009296 |
| Traf4 | 320.2069641 | -0.091164213 | 0.098902747 | -0.921756126 | 0.356655797 | 0.587624939 | | Sugp1 | 2063.69791 | -0.0311666482361044 | 0.0545064922024498 | -0.571796991087671 | 0.567459516000835 | 0.87653306352915 | NM_027481 |
| Iqsec3 | 9183.362736 | 0.074415191 | 0.080759698 | 0.921439679 | 0.35682092 | 0.587709508 | | Ripk4 | 32.07509919 | 0.0504092161388495 | 0.0881775080597701 | 0.571678846998945 | 0.567539567606168 | 0.876563504417564 | NM_023663 |
| Nrg1 | 208.6563094 | -0.097912105 | 0.106244268 | -0.921575413 | 0.356750088 | 0.587709508 | | Tdrd3 | 828.9038367 | 0.0430605359583422 | 0.075345873908895 | 0.571504897672421 | 0.567657441345056 | 0.876652348817593 | NM_001253755,NM_172605 |
| Sh3glb1 | 2151.9357 | -0.053545071 | 0.058109941 | -0.921444264 | 0.356818528 | 0.587709508 | | Serpinf1 | 346.2986498 | -0.0531256761044271 | 0.0930286676559599 | -0.571067794939269 | 0.567953688133598 | 0.877016612504374 | NM_011340 |
| Acsl6 | 6394.044766 | -0.045500494 | 0.049398309 | -0.92109415 | 0.357001274 | 0.587944063 | | Tyro3 | 11049.08311 | 0.0401647244201176 | 0.0703542315581595 | 0.570892802473647 | 0.568072310174915 | 0.877106545238183 | NM_019392 |
| Galnt3 | 37.57596139 | 0.146370274 | 0.158958793 | 0.920806402 | 0.357151512 | 0.588003988 | | L3mbtl3 | 937.4724868 | 0.0368964172003466 | 0.0646427025850658 | 0.570774669450016 | 0.568152395640028 | 0.877136964509198 | NM_172787 |
| Sec61a2 | 2158.715885 | 0.066429803 | 0.072141486 | 0.920826657 | 0.357140935 | 0.588003988 | | Crebbp | 2204.735788 | -0.0418694165269493 | 0.0733718836217412 | -0.570646608213051 | 0.568239217800457 | 0.877177776596114 | NM_001025432 |
| Yipf6 | 1598.590955 | 0.058888556 | 0.063943784 | 0.920942625 | 0.357080383 | 0.588003988 | | Trps1 | 968.7568448 | -0.0407088576466584 | 0.0713673698964965 | -0.570412748931312 | 0.568397784633358 | 0.877329318837973 | NM_032000 |
| Zscan20 | 270.9988051 | 0.102881159 | 0.111759597 | 0.920557712 | 0.357281389 | 0.588155316 | | Bcl10 | 493.3104382 | 0.0435135117864822 | 0.0763672814816597 | 0.569792598901565 | 0.568818375916975 | 0.877419106782413 | NM_009740 |
| Copg1 | 13598.53206 | 0.042672584 | 0.046362187 | 0.92041785 | 0.357354444 | 0.588213084 | | Bdh2 | 57.46826962 | 0.0525252613977688 | 0.0921439278524934 | 0.570034972698936 | 0.568653978144061 | 0.877419106782413 | NM_001172055,NM_027208 |
| Adal | 570.5848755 | 0.075274805 | 0.081791672 | 0.920323595 | 0.357403682 | 0.58823164 | | Fam196b | 141.2264603 | 0.0428825802834242 | 0.0752480689074064 | 0.569882801061536 | 0.568757190766657 | 0.877419106782413 | NM_001025382 |
| Gm13238 | 12.61156714 | 0.15086826 | 0.163967168 | 0.920112619 | 0.357513911 | 0.588350561 | | Fermt2 | 3651.786226 | -0.0306902524047418 | 0.0538377120614732 | -0.570051200721512 | 0.568642971780363 | 0.877419106782413 | NM_146054 |
| Haghl | 2481.200639 | 0.096041249 | 0.104403726 | 0.9199025 | 0.357623712 | 0.588468755 | | Ggact | 535.331065 | -0.0450664806717892 | 0.0790838590487452 | -0.5698568736259 | 0.56877477731886 | 0.877419106782413 | NM_145466 |
| 2810429I04Rik | 23.29190464 | -0.153982532 | 0.167420514 | -0.91973515 | 0.357711179 | 0.588549826 | | Hectd1 | 6625.172399 | 0.0322863306379171 | 0.0566279640246669 | 0.57014818021452 | 0.568577199310317 | 0.877419106782413 | NM_144788 |
| Nqo2 | 358.4824109 | -0.110164933 | 0.119795151 | -0.919610954 | 0.3577761 | 0.588549826 | | Arih2 | 2199.04863 | -0.0296301483388831 | 0.0520414494616677 | -0.569356707881625 | 0.569114090060672 | 0.87745335001908 | NM_011790 |
| Optc | 28.97770028 | 0.153871157 | 0.167325786 | 0.919590224 | 0.357786937 | 0.588549826 | | Bnip1 | 648.7408361 | -0.0352516169445393 | 0.0619387586504188 | -0.569136639361774 | 0.56926341529532 | 0.87745335001908 | NM_172149 |
| Rfx7 | 1605.954663 | -0.061505566 | 0.066896671 | -0.919411473 | 0.357880391 | 0.588578577 | | Hapln3 | 23.67634687 | 0.0441007701248556 | 0.0774860564777828 | 0.569144593614729 | 0.56925801769281 | 0.87745335001908 | NM_178255 |
| Tubb2b | 3384.666494 | -0.118855402 | 0.129266822 | -0.919457912 | 0.35785611 | 0.588578577 | | Mbd2 | 1569.64353 | -0.0333091696192609 | 0.0584715896668596 | -0.569664170395214 | 0.568905495858676 | 0.87745335001908 | NM_010773 |
| Chrnb3 | 63.76631455 | 0.153734267 | 0.167261609 | 0.91912465 | 0.358030377 | 0.588749683 | | Piezo2 | 39.419641 | 0.0502145786366691 | 0.0881989436264483 | 0.569333107314124 | 0.569130103087918 | 0.87745335001908 | NM_001039485 |
| Gas7 | 24337.08852 | 0.068422491 | 0.074447757 | 0.919067194 | 0.358060427 | 0.588749683 | | Samd4b | 3655.579713 | 0.0410259021812432 | 0.0720656621306188 | 0.56928502380071 | 0.569162728503481 | 0.87745335001908 | NM_175021 |
| E030030I06Rik | 19.33197641 | 0.154467083 | 0.168119263 | 0.918794672 | 0.35820298 | 0.588859095 | | Slc37a3 | 2494.096341 | 0.0320611309171577 | 0.0562986128859931 | 0.569483496548712 | 0.569028067406495 | 0.87745335001908 | NM_028123 |
| Gpr81 | 53.61712825 | -0.154126373 | 0.167735598 | -0.918865016 | 0.358166181 | 0.588859095 | | Znrf2 | 377.4218758 | -0.0443864939484319 | 0.0780149268962939 | -0.568948734739384 | 0.569390930851052 | 0.877556781065504 | NM_199143 |
| A830009L08Rik | 138.8637268 | 0.136399395 | 0.148494535 | 0.918548251 | 0.35833191 | 0.588946072 | | Ccdc17 | 99.41722785 | -0.0527511377331208 | 0.0927558975590293 | -0.568709258616686 | 0.569553463540509 | 0.877633217243605 | NM_001037916 |
| Fbxo28 | 2208.429998 | 0.064980338 | 0.070738633 | 0.918597592 | 0.358306092 | 0.588946072 | | Lix1l | 851.3488499 | 0.0341375311548 | 0.0600275608880029 | 0.568697622388697 | 0.569561361624361 | 0.877633217243605 | NM_001163170 |
| Zfp775 | 307.9812816 | -0.092527817 | 0.100776443 | -0.918149261 | 0.358540728 | 0.589226776 | | Ctnna1 | 3628.537208 | 0.0257174946468315 | 0.0452451114167424 | 0.568403830636054 | 0.569760789944182 | 0.877708859955916 | NM_009818 |
| Crbn | 2892.93264 | 0.073460316 | 0.080064453 | 0.917514735 | 0.358872975 | 0.589710243 | | Fam73b | 3675.49661 | -0.0320542887260696 | 0.0563980337222845 | -0.568358267309664 | 0.569791721697941 | 0.877708859955916 | NM_001242407,NM_175392 |
| Zfp69 | 38.9184144 | -0.152338069 | 0.166133303 | -0.916962861 | 0.359162102 | 0.590122757 | | Katnbl1 | 411.9931978 | -0.0455936601369927 | 0.0801959510127141 | -0.568528205741514 | 0.56967635900745 | 0.877708859955916 | NM_024254 |
| B4galt2 | 3045.527199 | -0.095734772 | 0.104466084 | -0.916419648 | 0.359446834 | 0.590476142 | | Arvcf | 1554.134042 | -0.036356488152294 | 0.0640021945512013 | -0.568050649000935 | 0.570000576701565 | 0.877844399614816 | NM_001272031,NM_001272030,NM_001272032,NM_001272028,NM_001272029,NM_033474 |
| D730039F16Rik | 15.19634192 | 0.15294994 | 0.166901735 | 0.916407125 | 0.3594534 | 0.590476142 | | Elac1 | 1910.310858 | 0.0348562658262672 | 0.0613587595589317 | 0.568073182652751 | 0.569985276418865 | 0.877844399614816 | NM_053255 |
| Arhgap22 | 216.9443347 | -0.118740956 | 0.129622745 | -0.916050309 | 0.359640509 | 0.590660962 | | AK010878 | 140.4038519 | 0.0504388827327293 | 0.0889204956679477 | 0.567235735179449 | 0.570554032338564 | 0.878246677165849 | NM_001142938 |
| Rsf1 | 2426.264364 | -0.080291427 | 0.087649882 | -0.916047174 | 0.359642153 | 0.590660962 | | Col20a1 | 152.7110427 | -0.0523849326886589 | 0.0924383807524978 | -0.566701106858618 | 0.570917268724082 | 0.878246677165849 | NM_028518 |
| Caap1 | 173.9304366 | -0.097581525 | 0.106535495 | -0.91595318 | 0.359691452 | 0.590679317 | | Cyp39a1 | 191.5228943 | -0.0494858848032265 | 0.0872321625707076 | -0.567289441702364 | 0.570517549218529 | 0.878246677165849 | NM_018887 |
| Zmym4 | 2469.541407 | -0.057525416 | 0.062811719 | -0.915838912 | 0.359751391 | 0.590715139 | | Ednra | 178.5656625 | 0.0505618497031351 | 0.0892516150935251 | 0.566509072694677 | 0.571047767164494 | 0.878246677165849 | NM_010332 |
| Zfp385c | 25.43576974 | -0.153876458 | 0.168038004 | -0.915724148 | 0.359811596 | 0.59075139 | | Grin2c | 2122.892377 | -0.0453257530191122 | 0.0799404777047125 | -0.56699377237322 | 0.570718412712378 | 0.878246677165849 | NM_010350 |
| Klf12 | 1512.472478 | -0.077771916 | 0.084944807 | -0.915558216 | 0.359898655 | 0.590831719 | | Gstk1 | 349.1316588 | 0.0505388379445669 | 0.0890940039875284 | 0.567252965212355 | 0.570542327766604 | 0.878246677165849 | NM_029555 |
| Ier3 | 339.5199949 | 0.142514121 | 0.155687725 | 0.91538444 | 0.359989844 | 0.59091881 | | Krt2 | 60.6929632 | 0.0253964199222492 | 0.0448273006053742 | 0.566539130826106 | 0.571027339969737 | 0.878246677165849 | NM_010668 |
| Smug1 | 682.6406575 | -0.061793375 | 0.067511608 | -0.915300002 | 0.360034158 | 0.590928946 | | Lrig2 | 1503.097145 | -0.0316312818043286 | 0.0558010463277461 | -0.566858220158509 | 0.570810511614135 | 0.878246677165849 | NM_001025067 |
| Stab1 | 809.6821873 | -0.100242142 | 0.109570368 | -0.914865436 | 0.360262277 | 0.591240729 | | Lrrc3b | 551.619615 | 0.0421827407738254 | 0.0743581481766012 | 0.567291437565673 | 0.570516193439634 | 0.878246677165849 | NM_146052 |
| Mmp14 | 463.3730223 | -0.131912334 | 0.144208782 | -0.914731625 | 0.360332538 | 0.591293406 | | Mlst8 | 1899.537259 | 0.0353665749403657 | 0.0623826593375914 | 0.566929581327644 | 0.570762025465034 | 0.878246677165849 | NM_001252464,NM_001252465,NM_019988,NM_001252463 |
| 2310010J17Rik | 40.57674791 | -0.145823676 | 0.159490498 | -0.914309493 | 0.360554244 | 0.591344072 | | Tpmt | 402.9915102 | 0.0425992925184203 | 0.0751759307411299 | 0.56666132495402 | 0.570944301687742 | 0.878246677165849 | NM_016785 |
| Aspm | 29.66348298 | -0.153139769 | 0.167488881 | -0.914327977 | 0.360544534 | 0.591344072 | | Unc45b | 59.65655461 | -0.0535375673795333 | 0.0943932091729332 | -0.567176048453335 | 0.570594579145676 | 0.878246677165849 | NM_178680 |
| Gpr35 | 14.15077631 | 0.152111434 | 0.166363963 | 0.914329231 | 0.360543875 | 0.591344072 | | Vgll4 | 457.1810566 | -0.0383928395198318 | 0.0677403426548164 | -0.566764766978959 | 0.570874011084801 | 0.878246677165849 | NM_177683 |
| Ide | 1990.478111 | -0.053357272 | 0.058353284 | -0.91438335 | 0.360515447 | 0.591344072 | | Tcf23 | 22.82379261 | 0.0435677694519383 | 0.0769464310425838 | 0.56620909978043 | 0.571251644705062 | 0.87837423425275 | NM_053085 |
| Tmem238 | 30.32097445 | 0.152757839 | 0.167037014 | 0.914514908 | 0.360446348 | 0.591344072 | | Trp53bp1 | 4594.611014 | 0.0386584600401309 | 0.0682672400880692 | 0.566281279135629 | 0.571202584613466 | 0.87837423425275 | NM_013735 |
| Baiap2l1 | 27.14052769 | 0.152800143 | 0.167151059 | 0.914144034 | 0.360641167 | 0.591424031 | | Agpat2 | 52.07817461 | -0.0524619896870189 | 0.0927155197341787 | -0.565838274297882 | 0.571503725283173 | 0.878389917835506 | NM_026212 |
| Grpr | 27.76504019 | 0.153658733 | 0.168119026 | 0.913987766 | 0.360723274 | 0.591496075 | | Dexi | 1272.578744 | 0.0345714351637976 | 0.0610895578673913 | 0.565913985477563 | 0.571452253869277 | 0.878389917835506 | NM_021428 |
| Zfp783 | 247.6483314 | 0.134817471 | 0.147533049 | 0.913812006 | 0.360815637 | 0.591584918 | | Enc1 | 39062.24961 | 0.0440552618934088 | 0.0778247918687653 | 0.566082617576395 | 0.571337619115502 | 0.878389917835506 | NM_007930 |
| 6530411M01Rik | 14.86124717 | -0.151176225 | 0.165505709 | -0.913420005 | 0.361021688 | 0.591756282 | | Pus3 | 378.7075977 | -0.0462746995592606 | 0.0817558202833985 | -0.566011072959136 | 0.571386253238527 | 0.878389917835506 | NM_023292 |
| Foxq1 | 140.5985014 | 0.115071749 | 0.125978491 | 0.913423777 | 0.361019705 | 0.591756282 | | Tmem170b | 3350.451087 | 0.0372738966560498 | 0.0659584123080801 | 0.565112096421455 | 0.571997521311326 | 0.87905586061874 | NM_001163572 |
| Tns1 | 2632.262513 | -0.069266211 | 0.075839812 | -0.913322553 | 0.361072924 | 0.591756282 | | AI115009 | 102.1412007 | 0.0529239934926211 | 0.0937193259903244 | 0.564707363538712 | 0.572272824898446 | 0.879277717687715 | NR_040386 |
| Tti1 | 1024.096521 | 0.047527835 | 0.052034688 | 0.913387528 | 0.361038763 | 0.591756282 | | Abi2 | 9618.214371 | 0.0405498223783837 | 0.0718615488726351 | 0.564277044045527 | 0.572565601790452 | 0.879277717687715 | NM_001198571,NM_001198570,NM_198127 |
| Tstd1 | 15.74263358 | 0.151469001 | 0.165884591 | 0.913098681 | 0.361190644 | 0.591886604 | | Cstf1 | 902.3240849 | -0.0318377939896455 | 0.0564215594012558 | -0.564284190786419 | 0.572560738774687 | 0.879277717687715 | NM_024199 |
| Airn | 21.33529681 | -0.151527921 | 0.166012011 | -0.912752762 | 0.361372588 | 0.592122131 | | Pip4k2c | 7301.830824 | 0.0345740940113881 | 0.0612668324925056 | 0.564319920009205 | 0.572536427039535 | 0.879277717687715 | NM_054097 |
| Fmr1 | 1838.848056 | -0.056009467 | 0.061392221 | -0.91232189 | 0.361599294 | 0.592430947 | | Pls1 | 608.9812672 | 0.0444094134589254 | 0.0786764171774762 | 0.564456479490516 | 0.572443510477591 | 0.879277717687715 | NM_001033210 |
| A730020E08Rik | 53.90710479 | -0.140813726 | 0.154364607 | -0.912215103 | 0.361655495 | 0.592460376 | | Zdhhc20 | 1927.717238 | 0.0351913651318661 | 0.0623213958739238 | 0.564675496085778 | 0.572294504149849 | 0.879277717687715 | NM_029492 |
| 4931406C07Rik | 1016.011891 | 0.064761047 | 0.071021666 | 0.911849166 | 0.361848124 | 0.592600427 | | Zfp287 | 798.9211102 | -0.0387420465639846 | 0.0686165244118365 | -0.564616860094149 | 0.572334394908875 | 0.879277717687715 | NM_133208 |
| A930004D18Rik | 103.8126059 | -0.123724394 | 0.135694267 | -0.911787925 | 0.361880368 | 0.592600427 | | Cdkn2aip | 631.2838396 | -0.0390910359432404 | 0.0693808750093205 | -0.563426678288346 | 0.573144375010501 | 0.87931064707661 | NM_172407 |
| Blvrb | 220.9626668 | -0.091489807 | 0.100345579 | -0.911747267 | 0.361901776 | 0.592600427 | | Dlgap3 | 13361.05398 | -0.0417933824982 | 0.0741398128349848 | -0.563710385824965 | 0.572951248048502 | 0.87931064707661 | NM_198618 |
| Gle1 | 2315.111185 | 0.074962899 | 0.082223923 | 0.911692074 | 0.361930837 | 0.592600427 | | Dnajc25 | 104.9663613 | 0.0534820333792184 | 0.0948809712071197 | 0.563675020383911 | 0.572975320522198 | 0.87931064707661 | NM_001033165 |
| Tbpl1 | 2235.077881 | 0.044395098 | 0.048695417 | 0.911689446 | 0.361932221 | 0.592600427 | | Etfb | 969.2403766 | 0.035304435507245 | 0.0625947784384822 | 0.564015663733709 | 0.572743472079466 | 0.87931064707661 | NR_075104,NM_026695 |
| Gbp9 | 157.8052779 | 0.140137899 | 0.153794212 | 0.911203985 | 0.362187905 | 0.592893756 | | Mrpl24 | 1246.338287 | -0.0346899400380572 | 0.0615769016876621 | -0.563359621664886 | 0.573190026677131 | 0.87931064707661 | NM_026591 |
| Gclm | 1129.860608 | 0.074474674 | 0.081725717 | 0.911275865 | 0.36215004 | 0.592893756 | | Scrg1 | 800.3154471 | 0.0484463528241318 | 0.085973322346157 | 0.563504486066861 | 0.573091406196614 | 0.87931064707661 | NM_009136 |
| Dguok | 352.7933601 | -0.101127529 | 0.111032355 | -0.910793329 | 0.362404278 | 0.593183458 | | Slc26a2 | 188.7104057 | 0.048909589608162 | 0.0867502428775746 | 0.563797725352598 | 0.572891800019469 | 0.87931064707661 | NM_007885 |
| Gatad2b | 1097.479438 | 0.092663101 | 0.101752089 | 0.910675165 | 0.362466553 | 0.593183458 | | Spg20 | 1759.757005 | -0.0367604377240917 | 0.0652525702977372 | -0.56335616446003 | 0.573192380364066 | 0.87931064707661 | NM_001144988,NM_001144987,NM_144895 |
| Rad1 | 326.3528614 | -0.086148899 | 0.094601531 | -0.910650159 | 0.362479733 | 0.593183458 | | Tbc1d22a | 1908.940176 | 0.0331594442960712 | 0.0588309258318848 | 0.563639681463242 | 0.572999375423395 | 0.87931064707661 | NM_145476 |
| Birc3 | 92.61354309 | -0.133320801 | 0.146440463 | -0.910409585 | 0.362606545 | 0.593328314 | | Vwa1 | 1164.474292 | -0.0338687300822366 | 0.0601189939446475 | -0.563361557803513 | 0.573188708543709 | 0.87931064707661 | NM_147776 |
| 1700007P06Rik | 12.41491263 | 0.150587525 | 0.165450342 | 0.910167509 | 0.362734176 | 0.593467136 | | Fam126b | 3396.38772 | 0.0476571448404932 | 0.0846212282761902 | 0.563181908503477 | 0.573311020859737 | 0.879352236276764 | NM_172513 |
| Pithd1 | 2665.824443 | 0.071728979 | 0.0788141 | 0.910103383 | 0.36276799 | 0.593467136 | | Myo5b | 2000.474351 | 0.0324164262307342 | 0.0575638588383414 | 0.563138519288125 | 0.573340563801269 | 0.879352236276764 | NM_201600 |
| Echs1 | 1664.194968 | -0.060295196 | 0.066262589 | -0.909943254 | 0.362852437 | 0.593542616 | | Prl | 547.63629 | -0.0188365546560417 | 0.0334621210980099 | -0.56292171679344 | 0.573488191560544 | 0.879485797150387 | NM_011164,NM_001163530 |
| Acacb | 231.3087356 | 0.117915214 | 0.12968594 | 0.909236682 | 0.363225205 | 0.593603784 | | Fbxl14 | 1140.831171 | -0.0403236206377747 | 0.0717413668536316 | -0.562069311002172 | 0.574068796593071 | 0.879633263640396 | NM_133940 |
| Atg7 | 1010.549692 | 0.061669283 | 0.067830907 | 0.909162001 | 0.363264619 | 0.593603784 | | Foxj2 | 1922.068892 | 0.0287622261577369 | 0.0511530655638788 | 0.562277662945132 | 0.573926854693677 | 0.879633263640396 | NM_021899 |
| Cnnm2 | 1152.05273 | -0.050804349 | 0.055872316 | -0.909293772 | 0.363195077 | 0.593603784 | | Lrrc2 | 73.68806584 | -0.0531535494539204 | 0.0945369027228113 | -0.562251860628122 | 0.573944431884941 | 0.879633263640396 | NM_028838 |
| Ddx41 | 1342.247922 | 0.076534291 | 0.084175072 | 0.909227514 | 0.363230044 | 0.593603784 | | Nrap | 59.02367751 | -0.0525937744892736 | 0.0935643351891703 | -0.562113484619309 | 0.574038701474314 | 0.879633263640396 | NM_198059,NM_008733 |
| Fbxl19 | 2495.799893 | -0.065705216 | 0.072250742 | -0.909405409 | 0.363136168 | 0.593603784 | | Otx2 | 72.66662828 | 0.0226848113719358 | 0.0403277032459392 | 0.562511860236426 | 0.573767325235303 | 0.879633263640396 | NM_144841 |
| Gm10509 | 153.3196779 | -0.116114288 | 0.127717947 | -0.909146216 | 0.36327295 | 0.593603784 | | Pigq | 4841.783727 | -0.0315096176661914 | 0.0560094374270012 | -0.562576935489824 | 0.573723001297303 | 0.879633263640396 | NM_011822 |
| Hebp2 | 109.604699 | 0.117870089 | 0.129641778 | 0.909198336 | 0.363245442 | 0.593603784 | | Tril | 2251.460726 | 0.0293176496744446 | 0.0521258430914906 | 0.562444767886491 | 0.57381302475317 | 0.879633263640396 | NM_025817 |
| Mustn1 | 43.87914477 | 0.147318982 | 0.161959102 | 0.909606066 | 0.363030298 | 0.593603784 | | Wfs1 | 7041.56952 | -0.0534586051045369 | 0.095080811919748 | -0.562243885229526 | 0.57394986497987 | 0.879633263640396 | NM_011716 |
| Myom1 | 95.16259355 | 0.132270392 | 0.145438372 | 0.909460072 | 0.363107325 | 0.593603784 | | Gtf3c5 | 857.5217751 | 0.0384787342010945 | 0.0685100318793915 | 0.561651091753021 | 0.574353762917656 | 0.879791496376871 | NM_148928 |
| Olfr1393 | 20.7944011 | -0.152406229 | 0.167552542 | -0.909602605 | 0.363032216 | 0.593603784 | | Pdcd6ip | 3843.519286 | 0.032020846233739 | 0.0570050315750202 | 0.561719647354264 | 0.574307045898553 | 0.879791496376871 | NM_001164678,NM_011052,NM_001164677 |
| Ak1 | 3584.317141 | -0.055009774 | 0.060535171 | -0.908724188 | 0.363495733 | 0.593901493 | | Taf13 | 784.6500024 | 0.039849587604215 | 0.0709283754892176 | 0.561828567612872 | 0.574232826198434 | 0.879791496376871 | NM_025444 |
| Pth2r | 48.31474261 | -0.138261536 | 0.152160504 | -0.908655874 | 0.363531803 | 0.593901493 | | Braf | 5472.075405 | 0.0393768818486104 | 0.0701781588183079 | 0.561098861379006 | 0.574730175000014 | 0.880182448292062 | NM_139294 |
| Tecr | 10991.7111 | 0.05347133 | 0.058857614 | 0.908486191 | 0.363621406 | 0.593985247 | | Pih1d1 | 729.8706901 | 0.0381976634085281 | 0.0680690106646443 | 0.561160843026154 | 0.574687893933328 | 0.880182448292062 | NM_029406 |
| Npc2 | 2178.494441 | 0.068632222 | 0.075577532 | 0.90810302 | 0.363823541 | 0.594252787 | | Ano10 | 1059.778182 | 0.0380994009253382 | 0.0679265404940924 | 0.560892878894971 | 0.574870564839801 | 0.880304641035521 | NM_133979,NM_001271873 |
| Pex12 | 281.2537421 | -0.087386963 | 0.096251337 | -0.907903887 | 0.363929004 | 0.594362389 | | Bmf | 164.2064518 | -0.0529020852022261 | 0.0943626051495397 | -0.560625526588529 | 0.575052584029291 | 0.880490949870505 | NM_138313 |
| 3110052M02Rik | 451.5443453 | 0.109638583 | 0.120813853 | 0.907500091 | 0.364142402 | 0.594378872 | | Apaf1 | 740.1200539 | 0.0516696852695418 | 0.0922192355150455 | 0.560291841294996 | 0.575280391477283 | 0.880746518402088 | NM_009684,NM_001042558 |
| Capn10 | 871.8828527 | -0.07436083 | 0.081927857 | -0.907637922 | 0.364069552 | 0.594378872 | | Cd59b | 23.37160762 | 0.0459126579984082 | 0.0820756053645466 | 0.559385541777783 | 0.575862455353426 | 0.880811222300151 | NM_181858 |
| Mcm9 | 195.634284 | 0.092140776 | 0.101537224 | 0.907458091 | 0.364164602 | 0.594378872 | | Fancl | 196.0795827 | 0.0485737840161794 | 0.0868107362524674 | 0.559536597811066 | 0.575795561432803 | 0.880811222300151 | NM_025923 |
| Mtmr7 | 3213.639047 | 0.066922746 | 0.073748196 | 0.907449264 | 0.364169268 | 0.594378872 | | Igf1r | 3143.198715 | 0.0404669987999804 | 0.072388468705858 | 0.559439735644774 | 0.575861649231723 | 0.880811222300151 | NM_010513 |
| Prorsd1 | 55.42666473 | -0.149665218 | 0.164867284 | -0.9077921 | 0.363988073 | 0.594378872 | | Lrrc8d | 2088.047168 | 0.0373712662642438 | 0.0667938746818006 | 0.559501398029038 | 0.575819577372958 | 0.880811222300151 | NM_001122768,NM_178701 |
| Txndc12 | 852.2420588 | -0.067591535 | 0.074484617 | -0.907456295 | 0.364165552 | 0.594378872 | | Pard6b | 225.482892 | 0.0491037911128168 | 0.087772979890677 | 0.55972799809677 | 0.575664982033755 | 0.880811222300151 | NM_021409 |
| Bivm | 855.8328053 | 0.059796794 | 0.065902725 | 0.907349336 | 0.364222092 | 0.594402475 | | Prkd2 | 212.221959 | -0.052995191293561 | 0.0946297935551024 | -0.5600246491685223 | 0.575461368076733 | 0.880811222300151 | NM_178900,NM_001252458 |
| Pygo2 | 2096.871229 | -0.059134358 | 0.065181957 | -0.907219745 | 0.364290604 | 0.594451672 | | Pyurf | 845.8525161 | -0.0400377578720325 | 0.0715688607360404 | -0.559429861817968 | 0.575868386216751 | 0.880811222300151 | NM_025574 |
| Runx2 | 326.2508076 | 0.105241605 | 0.116038296 | 0.906955795 | 0.364430174 | 0.594616798 | | Rbm43 | 732.7565843 | 0.0384645584792828 | 0.0687089062980715 | 0.559819105727235 | 0.575602830406993 | 0.880811222300151 | NM_001141982,NM_030243,NM_001141981 |
| Ankrd2 | 13.10159108 | 0.151048264 | 0.166652118 | 0.906368707 | 0.36474073 | 0.594935559 | | Ucp3 | 153.4182366 | -0.0531826336441286 | 0.095065050614541 | -0.559434127477273 | 0.575865475721305 | 0.880811222300151 | NM_009464 |
| Slc38a9 | 880.5416938 | 0.070559978 | 0.077848658 | 0.906373721 | 0.364738077 | 0.594935559 | | Htr6 | 117.8693707 | -0.0512822996840987 | 0.0916985932029076 | -0.559248488912179 | 0.575992144910896 | 0.880907759209905 | NM_021358 |
| Sptlc1 | 1147.111639 | -0.051133321 | 0.056413362 | -0.906404432 | 0.364721828 | 0.594935559 | | Bmpr2 | 2291.268515 | 0.0420884643042778 | 0.0753289854150981 | 0.558773182878137 | 0.576316526628255 | 0.880966011161667 | NM_007561 |
| Car11 | 5670.378617 | -0.106740073 | 0.117798015 | -0.906127939 | 0.364868139 | 0.595018097 | | Cd47 | 2415.201853 | 0.0403088521929964 | 0.0721247007613674 | 0.558879214142479 | 0.576245517717102 | 0.880966011161667 | NM_010581 |
| Tgfbr2 | 1000.487006 | 0.08021665 | 0.08852127 | 0.906185034 | 0.364837923 | 0.595018097 | | Msrb3 | 419.2345191 | -0.0457592366522059 | 0.0819172757475277 | -0.558604018746344 | 0.576431996726623 | 0.880966011161667 | NM_177092 |
| Bsdc1 | 2566.512623 | 0.068556669 | 0.075709406 | 0.905523798 | 0.365187958 | 0.595372142 | | Mylk3 | 190.0963092 | 0.0520595035088708 | 0.0931615098549789 | 0.558809143281489 | 0.576291981747935 | 0.880966011161667 | NM_175441 |
| 3-Mar | 47.59453514 | -0.138299334 | 0.15271968 | -0.905576376 | 0.365160118 | 0.595372142 | | Pcdhb9 | 595.4219 | -0.0411494140672144 | 0.0736691287066489 | -0.55857066303949 | 0.576454766356302 | 0.880966011161667 | NM_053134 |
| Snora64 | 17.77571683 | 0.148909793 | 0.164450337 | 0.905500078 | 0.365200519 | 0.595372142 | | Pygl | 215.0604082 | -0.0494639778320616 | 0.088471909024029 | -0.559092466498345 | 0.576098615896293 | 0.880966011161667 | NM_133198 |
| Epb4.1l2 | 3876.282409 | -0.06819043 | 0.075340447 | -0.905097225 | 0.365413882 | 0.595469362 | | Ttc27 | 805.9985363 | 0.0395360336671564 | 0.0707713537248876 | 0.558644586916429 | 0.576404302419603 | 0.880966011161667 | NM_152817 |
| Med19 | 1206.742111 | 0.057873618 | 0.063939368 | 0.905132781 | 0.365395048 | 0.595469362 | | Epc1 | 1758.733766 | 0.0272997636052347 | 0.0488823087237576 | 0.558479433983813 | 0.576517044277713 | 0.880968502543452 | NM_027497,NM_001276350,NM_007935 |
| Papd7 | 1928.753028 | 0.068268442 | 0.075421503 | 0.905158859 | 0.365381234 | 0.595469362 | | Adipor1 | 3183.892681 | 0.0269672995664227 | 0.048321116939886 | 0.558085186647722 | 0.57678621546006 | 0.881151497931004 | NM_028320 |
| Srpk1 | 2292.899991 | 0.063970652 | 0.070661278 | 0.905314101 | 0.365299008 | 0.595469362 | | Ss18l1 | 2321.18701 | 0.0276679751810277 | 0.0495805783165097 | 0.558037496627879 | 0.576818779693372 | 0.881151497931004 | NM_178750 |
| 2310015A10Rik | 384.8017204 | 0.096937008 | 0.107141887 | 0.904753595 | 0.365595941 | 0.595703387 | | Ttc18 | 67.77618991 | -0.0520372821238401 | 0.0932442675020612 | -0.558074866347037 | 0.576793262409962 | 0.881151497931004 | NM_029698,NM_001163638,NM_001163639 |
| Micalcl | 98.89648915 | 0.130908284 | 0.144717128 | 0.904580444 | 0.3656877 | 0.595790254 | | Beta-s | 118.1808697 | 0.0211741425184769 | 0.0379557347274722 | 0.557864119098481 | 0.576937174583854 | 0.881236189389762 | NM_001201391 |
| 2410127L17Rik | 267.3160764 | 0.088952929 | 0.098375373 | 0.904219481 | 0.365879033 | 0.596039294 | | Dhx34 | 539.6156268 | -0.0430677283034348 | 0.0772129398448166 | -0.557778636456438 | 0.576995552615037 | 0.881236189389762 | NM_027883 |
| Ccdc13 | 182.055429 | -0.134674973 | 0.148975801 | -0.904005701 | 0.365992379 | 0.596098593 | | Slc39a4 | 22.5590391 | -0.0292295360414734 | 0.0524196275486239 | -0.557606709707358 | 0.577112973712417 | 0.881322861243332 | NM_028064 |
| Prkcb | 52543.29474 | -0.069594897 | 0.076984314 | -0.904013985 | 0.365987986 | 0.596098593 | | Dazap1 | 1337.641959 | -0.0400216724339584 | 0.0718164335625306 | -0.557277359075643 | 0.577337942296586 | 0.881375634638492 | NM_133188,NM_001122604,NM_001122605 |
| Bmp2k | 671.2787017 | 0.069716519 | 0.077146446 | 0.903690606 | 0.366159482 | 0.596153796 | | Fnip2 | 947.1811742 | 0.0333978214591791 | 0.0599218491413628 | 0.557356322238817 | 0.577284001392878 | 0.881375634638492 | NM_001162999 |
| Med29 | 601.0314982 | -0.060564036 | 0.067010778 | -0.903795442 | 0.366103879 | 0.596153796 | | Mfsd1 | 1181.134448 | -0.0319141351814459 | 0.0572595669982109 | -0.557359003124195 | 0.577282170082122 | 0.881375634638492 | NM_025813 |
| Tdp2 | 538.5785242 | 0.069825332 | 0.07726536 | 0.903708104 | 0.366150201 | 0.596153796 | | Tmem256 | 867.6918605 | -0.0352852470201212 | 0.0633259072282095 | -0.557200813451636 | 0.577390234009355 | 0.881375634638492 | NM_026982 |
| Xpr1 | 6145.866821 | -0.067020007 | 0.07416576 | -0.903651586 | 0.366180178 | 0.596153796 | | Hhip | 436.432172 | 0.048345920304 0336 | 0.0867884883593666 | 0.55705452667 6934 | 0.57749017520 1237 | 0.881435566352125 | NM_020259 |
| Dlg1 | 7741.701007 | -0.065809517 | 0.072848302 | -0.903377496 | 0.366325579 | 0.596239482 | | Srf | 2467.390456 | 0.0390869539102596 | 0.0701853255451988 | 0.556910630628733 | 0.577588491026308 | 0.881493004901046 | NM_020493 |
| Lingo2 | 821.3108854 | -0.092988944 | 0.102931094 | -0.90340965 | 0.36630852 | 0.596239482 | | 5730480H06Rik | 28.15024117 | -0.0496582456269029 | 0.0892371788919042 | -0.556474848751723 | 0.577886283567856 | 0.881703921295932 | NR_045500 |
| Tmem97 | 544.2609843 | -0.103154892 | 0.114202576 | -0.90326223 | 0.366386737 | 0.596239482 | | Best1 | 54.25564751 | -0.0503846845722836 | 0.0905299532190604 | -0.556552641205557 | 0.577833118619959 | 0.881703921295932 | NM_011913 |
| Ttc21b | 502.6656256 | 0.083581849 | 0.092529023 | 0.903304136 | 0.366364592 | 0.596239482 | | Layn | 157.3448347 | -0.0509074348671566 | 0.0915020221669681 | -0.556353112877259 | 0.577969484963504 | 0.881703921295932 | NM_001033534 |
| 2310039H08Rik | 70.39366605 | -0.12901247 | 0.142933085 | -0.902607464 | 0.366734263 | 0.596573703 | | Vps41 | 8954.391805 | 0.0297852039086936 | 0.053530195085647 | 0.556418743870408 | 0.577924628215359 | 0.881703921295932 | NM_172120 |
| Ap5s1 | 556.7820422 | -0.080052954 | 0.08868468 | -0.902669482 | 0.366701338 | 0.596573703 | | Vwa5b1 | 219.5862542 | 0.0525316082027691 | 0.0944592142847256 | 0.556130056771642 | 0.578121949135405 | 0.881843897211056 | NM_029401 |
| Rcl1 | 557.3551244 | -0.059187019 | 0.065575 | -0.902585116 | 0.366746129 | 0.596573703 | | Fmo2 | 183.6569665 | 0.0424905647122467 | 0.0764170997804945 | 0.556034772770746 | 0.578187083789997 | 0.881850648796041 | NM_018881 |
| Zfp64 | 455.2570908 | -0.080909675 | 0.089628167 | -0.902725984 | 0.366671342 | 0.596573703 | | Rpl9 | 9497.936054 | -0.0293892651285606 | 0.0528643501612964 | -0.555937319552588 | 0.578253704857368 | 0.881859665801378 | NM_011292 |
| Tor1aip1 | 1753.537879 | 0.067464274 | 0.074764475 | 0.902357352 | 0.366867069 | 0.596707785 | | Acad8 | 989.6005796 | 0.0425405619275057 | 0.0765547959341248 | 0.555687745077549 | 0.578424335669276 | 0.881901419080596 | NM_025862 |
| Mrpl32 | 430.5742038 | -0.083868737 | 0.092971166 | -0.902094061 | 0.367006904 | 0.596872569 | | Ctbs | 361.2927296 | 0.0427161878860365 | 0.0768556721858054 | 0.555797466487136 | 0.578349317657765 | 0.881901419080596 | NM_028836 |
| Decr1 | 749.6465442 | -0.112999003 | 0.125314254 | -0.901725058 | 0.367202939 | 0.596940754 | | Prpf4 | 1011.135452 | -0.0326804466214814 | 0.0588168296433709 | -0.555630876734355 | 0.578463219124323 | 0.881901419080596 | NM_027297 |
| Fahd2a | 554.0878826 | 0.083683649 | 0.092799163 | 0.901771594 | 0.367178309 | 0.596940754 | | Slc47a1 | 55.17437671 | -0.0464200407564931 | 0.0835817670832984 | -0.555384773215317 | 0.578631505379676 | 0.882065405304353 | NM_026183 |
| Rilpl1 | 2138.008236 | 0.107907165 | 0.119665344 | 0.901741147 | 0.36719439 | 0.596940754 | | C2 | 107.610072 | -0.051290203132497 | 0.092423859082586 | -0.554945483142683 | 0.578931950297253 | 0.882430798532832 | NM_013484 |
| Ubr7 | 1801.222714 | 0.050009306 | 0.055458305 | 0.901746028 | 0.367191797 | 0.596940754 | | Hsd11b1 | 483.2881546 | -0.0513832341368088 | 0.0926185255968387 | -0.554783546873507 | 0.579042722314365 | 0.88245268230009 | NM_008288,NM_001044751 |
| Gm12070 | 155.6847635 | -0.136545648 | 0.151443889 | -0.90162534 | 0.367255925 | 0.596964258 | | St6gal2 | 515.920625 | -0.0491031305638997 | 0.0885596281424328 | -0.554464055392439 | 0.579261298684635 | 0.88245268230009 | NM_172829 |
| Cyp4x1 | 316.0346238 | 0.095073216 | 0.105487699 | 0.901273013 | 0.367443179 | 0.597143341 | | Thbd | 459.9607729 | -0.0461994012055153 | 0.0833259959037934 | -0.55444163258314 | 0.57927664044249 | 0.88245268230009 | NM_009378 |
| Oxt | 42.17968518 | -0.089592175 | 0.099399926 | -0.901330404 | 0.367412673 | 0.597143341 | | Ubash3b | 1389.855187 | 0.0354286921907052 | 0.0639055244447503 | 0.554391697721457 | 0.579310806719927 | 0.88245268230009 | NM_176860 |
| Nadk | 2655.002254 | -0.056516961 | 0.062718214 | -0.901125154 | 0.36752178 | 0.597208439 | | Usp12 | 3559.360058 | -0.0334415852754856 | 0.0602861888456459 | -0.554713872543975 | 0.579090385891888 | 0.88245268230009 | NM_011669 |
| A630001G21Rik | 21.30169277 | 0.151383517 | 0.168062746 | 0.900755942 | 0.367718097 | 0.597464401 | | Zfp382 | 402.6788821 | -0.0433134766794121 | 0.0781101673468929 | -0.554517781111053 | 0.579224540152109 | 0.88245268230009 | NM_001081007 |
| Ano1 | 176.7209534 | -0.137324404 | 0.152463291 | -0.900704714 | 0.367745342 | 0.597446401 | | Sf3a1 | 3634.542239 | 0.023336904756732 | 0.0420981251459335 | 0.554269112811716 | 0.579394685400706 | 0.882487910258012 | NM_026175 |
| Ptprs | 17729.73001 | 0.053152758 | 0.059053808 | 0.900073329 | 0.368081228 | 0.597837201 | | Zc3h13 | 2902.623559 | -0.0271420924851753 | 0.0489778307826377 | -0.554170980042607 | 0.579461836820459 | 0.882497655866427 | NM_026083 |
| Sntb1 | 73.64341992 | -0.146285423 | 0.162526967 | -0.900066822 | 0.368083733 | 0.597837201 | | Chp2 | 48.74438397 | -0.0522432137604275 | 0.0942952734341813 | -0.554038520254079 | 0.579552483719298 | 0.882543178373678 | NM_027363 |
| Tmco4 | 151.9737644 | 0.119691211 | 0.132993857 | 0.900034961 | 0.368101646 | 0.597837201 | | Gucy2f | 46.96239142 | -0.0518300308836214 | 0.0935688182794598 | -0.55392417940795 | 0.579630736537561 | 0.882569819595871 | NM_001007576 |
| Abhd16a | 2655.929107 | 0.058327757 | 0.064814695 | 0.899915634 | 0.368165149 | 0.597863573 | | A330041J22Rik | 44.01756463 | -0.0514751054853875 | 0.0930317140569379 | -0.553307073906897 | 0.580053157984438 | 0.883120445748029 | NR_045835 |
| Kif1c | 4476.555011 | 0.07240266 | 0.080459966 | 0.89985944 | 0.368195057 | 0.597863573 | | Adck2 | 941.0342725 | 0.0368721431941457 | 0.0666855016106267 | 0.55292593297776 | 0.580314128885115 | 0.883332603603183 | NM_178873 |
| Cbwd1 | 396.4672691 | -0.085496613 | 0.095021807 | -0.899757811 | 0.368249151 | 0.597863573 | | Dhcr7 | 1410.856342 | 0.0403401073595724 | 0.0729523765447853 | 0.552964951522966 | 0.58028740998011 | 0.883332603603183 | NM_007856 |
| Rarg | 416.4014242 | 0.084606254 | 0.094039675 | 0.899686798 | 0.368286951 | 0.597887471 | | Csad | 766.724909 | -0.0401262414885718 | 0.0726041391039786 | -0.552671541647313 | 0.580488343712174 | 0.883489182239975 | NM_144942 |
| Syt10 | 286.742948 | -0.142103008 | 0.15799951 | -0.899388917 | 0.368445541 | 0.598082264 | | Rpe | 681.6518674 | 0.0441968125185396 | 0.07998002853876 3 | 0.552598108878133 | 0.580538637235341 | 0.883489182239975 | NM_025683 |
| Tnxb | 131.5199224 | 0.129261628 | 0.143746332 | 0.899234273 | 0.368527888 | 0.598153269 | | Ppp1r7 | 4320.688302 | 0.0342827869981517 | 0.0620753561707006 | 0.552276927801714 | 0.580758635597944 | 0.883731399611957 | NM_023200 |
| Tcf23 | 19.27040081 | 0.150452214 | 0.167345333 | 0.8988955 | 0.368708325 | 0.598383449 | | Ndufb2 | 1120.611042 | -0.0375758555418806 | 0.0680518767742625 | -0.552164867789391 | 0.580835402166423 | 0.883755635473915 | NM_026612 |
| Bpifc | 9.204371208 | 0.133297007 | 0.15058815 | 0.898457196 | 0.368941855 | 0.598699739 | | Gpx8 | 117.2531456 | 0.049956622368319 | 0.0905095817983537 | 0.55194843878096 | 0.580983680028469 | 0.883796097611639 | NM_027127 |
| Scn5a | 391.0487005 | 0.148446042 | 0.165276723 | 0.898166655 | 0.369096707 | 0.598888301 | | Pdcd11 | 1211.880023 | -0.0410651585164754 | 0.0743966760636761 | -0.55197571570708 | 0.580964991334738 | 0.883796097611639 | NM_011053 |
| Adarb2 | 496.8110099 | 0.116396069 | 0.129603438 | 0.898093991 | 0.369135442 | 0.598888434 | | Col9a2 | 265.0029341 | 0.0516323286336905 | 0.0936147473312503 | 0.551540543617477 | 0.581263181578942 | 0.883958169006879 | NM_007741 |
| Rasa4 | 261.0113403 | 0.140240175 | 0.15623989 | 0.897595198 | 0.3694014 | 0.599257177 | | Kif26b | 315.9457154 | 0.0479178426345836 | 0.0868822071284427 | 0.551526534814477 | 0.581272781932652 | 0.883958169006879 | NM_001161665 |
| 4930550C14Rik | 8.489261048 | 0.125781799 | 0.140159774 | 0.89741725 | 0.369496311 | 0.599285653 | | Mccc2 | 703.2649048 | -0.0436923169853979 | 0.0792065609270083 | -0.551624972401743 | 0.58120532337446 | 0.883958169006879 | NM_030026 |
| Atp6v0a4 | 12.77901486 | -0.147225179 | 0.164044251 | -0.897472347 | 0.369466923 | 0.599285653 | | A930004D18Rik | 119.3633853 | 0.0515359672091783 | 0.0934910767255466 | 0.551239423206857 | 0.581469558334296 | 0.884020488736626 | NR_028376,NR_028377 |
| F9 | 11.29878812 | -0.134282892 | 0.149774014 | -0.896570031 | 0.369948257 | 0.599956086 | | Acsbg1 | 6981.680987 | 0.033984614596038 | 0.0617750203445068 | 0.550135223048291 | 0.582226629198905 | 0.884020488736626 | NM_053178 |
| Cpox | 1274.346763 | 0.062806156 | 0.070073576 | 0.896288718 | 0.370098583 | 0.600106315 | | Adrm1 | 2533.798306 | 0.0322408909082183 | 0.0584745591712606 | 0.551366121697318 | 0.58138271971492 | 0.884020488736626 | NM_019822 |
| Poc5 | 1261.143326 | 0.077195975 | 0.086132053 | 0.896251428 | 0.370118494 | 0.600106315 | | Cmya5 | 414.7496218 | 0.0448015907021303 | 0.0814044896317943 | 0.550357737082747 | 0.582074030250548 | 0.884020488736626 | NM_023821 |
| Slc13a5 | 555.2860016 | -0.087283054 | 0.097404004 | -0.896093072 | 0.370203057 | 0.600180618 | | Dis3l2 | 1205.049517 | 0.0305450476345259 | 0.0554692076448285 | 0.5506667380235 | 0.58186215003285 | 0.884020488736626 | NM_153530,NM_001172157 |
| Snx30 | 2416.01707 | 0.074818313 | 0.083511867 | 0.895900379 | 0.370305972 | 0.600284654 | | Gm10790 | 71.50012643 | -0.0523469764689673 | 0.0951025672467094 | -0.550426534051093 | 0.582026853435042 | 0.884020488736626 | NR_033545 |
| Gpr12 | 653.1292314 | 0.080412412 | 0.089780054 | 0.895660096 | 0.370434329 | 0.60040892 | | Gstz1 | 1496.802548 | -0.0324091245284478 | 0.0588960985261818 | -0.550276254954996 | 0.582129908094712 | 0.884020488736626 | NM_010363,NM_001252555,NM_001252556 |
| Nid2 | 161.3037206 | 0.119703353 | 0.133666224 | 0.895539271 | 0.370498882 | 0.60040892 | | Hiatl1 | 1583.161187 | 0.0351636922734545 | 0.0638728333976159 | 0.550526576057092 | 0.581958253843123 | 0.884020488736626 | NM_133680,NM_001083901 |
| Reck | 502.4898129 | 0.08066125 | 0.09006616 | 0.895577764 | 0.370478316 | 0.60040892 | | Map3k14 | 272.2316413 | 0.0495324658105525 | 0.0900152671833202 | 0.550267386416544 | 0.582135990006696 | 0.884020488736626 | NM_016896 |
| Arnt2 | 17196.76819 | 0.058151351 | 0.064947903 | 0.895353788 | 0.370597996 | 0.600443935 | | Pdk4 | 538.2805523 | 0.0466099352546414 | 0.0846660708552381 | 0.550514920366802 | 0.581966246047374 | 0.884020488736626 | NM_013743 |
| Atp6v1c1 | 9366.141147 | 0.052128815 | 0.058217608 | 0.895413202 | 0.370566246 | 0.600443935 | | Sh3bp2 | 187.5032248 | -0.0507020620495169 | 0.0921251848862928 | -0.550360491673335 | 0.582072141283972 | 0.884020488736626 | NM_011893,NM_001145858,NM_001145859,NM_001136088 |
| Zfp40 | 552.4256265 | 0.129559105 | 0.144726263 | 0.895201068 | 0.370679614 | 0.600513377 | | Ttc16 | 37.56412862 | -0.0459491913909576 | 0.0833790685581742 | -0.551087847172322 | 0.581573455887416 | 0.884020488736626 | NM_177384 |
| Actr10 | 4676.209296 | -0.04056411 | 0.045323177 | -0.894997061 | 0.370788659 | 0.60051934 | | Twist2 | 26.28527379 | -0.0474919782851484 | 0.0863270573358641 | -0.550140126986792 | 0.582223265902471 | 0.884020488736626 | NM_007855 |
| Ccdc117 | 712.7152667 | 0.064237742 | 0.071776764 | 0.894965694 | 0.370805427 | 0.60051934 | | Xpo1 | 2931.18204 | 0.0416156932269556 | 0.075546834419579 | 0.550859523720432 | 0.581729976219197 | 0.884020488736626 | NM_134014,NM_001035226 |
| Chd6 | 3971.662892 | -0.070233742 | 0.078481859 | -0.894904151 | 0.370838327 | 0.60051934 | | Zfp719 | 751.0793206 | 0.0387128398944926 | 0.0703261240077671 | 0.550475949594735 | 0.581992968335176 | 0.884020488736626 | NM_172482 |
| Rbm4 | 439.9062325 | -0.078136384 | 0.087298221 | -0.895051276 | 0.370759679 | 0.60051934 | | 9830001H06Rik | 2546.660144 | 0.0438937569552675 | 0.0799177132005661 | 0.549236898772481 | 0.58284288513206 | 0.884065160480968 | NM_001164663 |
| Cygb | 2361.323845 | 0.057958251 | 0.064783555 | 0.8946445 | 0.370977156 | 0.600555851 | | Akirin2 | 2119.745569 | 0.0330183239836828 | 0.060104229293756 | 0.549351091789359 | 0.582764531145612 | 0.884065160480968 | NM_001007589 |
| Fblim1 | 68.90810906 | 0.139212243 | 0.155602641 | 0.89466504 | 0.370966172 | 0.600555851 | | Brf1 | 1495.667284 | 0.0266846312576591 | 0.0485727083587967 | 0.549374991827615 | 0.582748132662603 | 0.884065160480968 | NM_028193 |
| Pde4dip | 10132.57593 | -0.062375303 | 0.069715383 | -0.894713623 | 0.370940194 | 0.600555851 | | Cir1 | 1291.543461 | 0.0354644428353487 | 0.0644925144566856 | 0.549900141653917 | 0.582387867069788 | 0.884065160480968 | NM_025854 |
| Rabl6 | 6200.363873 | -0.045985589 | 0.05141323 | -0.89443104 | 0.371091311 | 0.600677891 | | Pde9a | 672.8891056 | 0.0363580532678907 | 0.0661428862584336 | 0.549689548258183 | 0.582532326787744 | 0.884065160480968 | NM_001163748,NM_008804 |
| C2 | 102.5879124 | 0.143228444 | 0.16017618 | 0.89419316 | 0.371218551 | 0.600821083 | | Snx14 | 2383.831237 | 0.0366963050907668 | 0.0667840900302658 | 0.549476755229224 | 0.582678312363987 | 0.884065160480968 | NM_172926 |
| F2rl1 | 15.39855513 | 0.14658492 | 0.164082769 | 0.893359623 | 0.371664618 | 0.601354594 | | Srfbp1 | 423.9854066 | -0.0384473236020953 | 0.0700053931776246 | -0.549205166301158 | 0.582864659364328 | 0.884065160480968 | NM_026040 |
| Ppp1r42 | 21.6721438 | 0.148473235 | 0.166190864 | 0.893389879 | 0.371648421 | 0.601354594 | | Tmem169 | 661.7897955 | 0.0348374208782296 | 0.0633961841933801 | 0.549519207212276 | 0.582649187033521 | 0.884065160480968 | NM_175564 |
| Tax1bp3 | 285.4937813 | -0.092249312 | 0.103255132 | -0.893411401 | 0.3716369 | 0.601354594 | | Trpm3 | 1983.67992 | 0.0520565811683274 | 0.0946968730901479 | 0.54971805794233 | 0.582512769160788 | 0.884065160480968 | NM_177341,NM_001035239,NM_001035241,NM_001035240,NM_001035243,NM_001035242,NM_001035245,NM_001035244,NM_001035246 |
| Dusp26 | 2819.367714 | -0.054465463 | 0.060981226 | -0.893151327 | 0.371776139 | 0.601409428 | | Zfp235 | 795.6725572 | -0.03645182212213 | 0.0663655403641578 | -0.549258273527396 | 0.58282821838626 1 | 0.884065160480968 | NM_019941 |
| Nyx | 19.80501746 | 0.150071207 | 0.168019179 | 0.893179025 | 0.371761309 | 0.601409428 | | A930005H10Rik | 224.4698388 | -0.0441003078613859 | 0.0807401441750214 | -0.546200509201335 | 0.58492811616784 5 | 0.884313345041393 | NR_027895,NR_027894,NR_015487 |
| A930015D03Rik | 45.55080053 | 0.149636905 | 0.167637716 | 0.892620756 | 0.3720603 | 0.601694982 | | Adal | 659.9342684 | 0.0379243408131086 | 0.0693057503965198 | 0.54720337917318 | 0.584239014891601 | 0.884313345041393 | NM_029475 |
| Nlrc4 | 13.23053909 | 0.147484398 | 0.165242752 | 0.892531721 | 0.372107998 | 0.601694982 | | Arhgef9 | 20715.92114 | 0.0331236018530708 | 0.0605082071766559 | 0.54742329013922 4 | 0.584087958160723 | 0.884313345041393 | NM_001033329 |
| 5-Sep | 24702.30582 | -0.083668077 | 0.093735768 | -0.892594985 | 0.372074106 | 0.601694982 | | Atxn1 | 3469.274185 | 0.0391654858699839 | 0.0714509312141317 | 0.548145212448088 | 0.583592197948453 | 0.884313345041393 | NM_001199305,NM_001199304,NM_009124 |
| Usp22 | 11181.6756 | -0.048126999 | 0.053919492 | -0.892571443 | 0.372086718 | 0.601694982 | | Bambi | 147.8322686 | 0.0512701862709974 | 0.0937429936484745 | 0.54692286085139 | 0.584431729167761 | 0.884313345041393 | NM_026505 |
| Rab7 | 4673.944339 | 0.078383223 | 0.087848204 | 0.89225755 | 0.372254902 | 0.601869712 | | Birc3 | 82.12341621 | -0.0519094686236619 | 0.0946208772565719 | -0.54860481247606 | 0.583276682492073 | 0.884313345041393 | NM_007464 |
| Acss2 | 1948.905593 | -0.102484707 | 0.11495243 | -0.891540151 | 0.372639461 | 0.602302923 | | Cdkl1 | 586.1453496 | 0.0408847360107062 | 0.0748216242407401 | 0.546429410288645 | 0.584770798266835 | 0.884313345041393 | NM_183294 |
| Ddr1 | 2983.888881 | 0.071921656 | 0.080661945 | 0.891642958 | 0.372584337 | 0.602302923 | | Creb3l2 | 366.4912308 | -0.0423647016913645 | 0.0773431011750731 | -0.547750233022958 | 0.583863414874188 | 0.884313345041393 | NM_178661 |
| Rgs9bp | 13.98943723 | 0.140177943 | 0.157224007 | 0.891581038 | 0.372617538 | 0.602302923 | | Ddx49 | 982.4630439 | 0.0358784100157917 | 0.0655622676928935 | 0.547241748620612 | 0.584212657628323 | 0.884313345041393 | NM_001024922 |
| Ikzf1 | 206.7245943 | 0.09423357 | 0.105720341 | 0.891347575 | 0.372742733 | 0.602407 | | Dnajc11 | 2756.87791 | 0.0278128115722458 | 0.0508754282660164 | 0.54668456895967 | 0.584595457386837 | 0.884313345041393 | NM_172704 |
| Secisbp2l | 6270.940956 | -0.064474644 | 0.072361495 | -0.891007631 | 0.372925077 | 0.602638834 | | Dynll1 | 7325.422381 | 0.0273371392879344 | 0.0500049111106986 | 0.546689088746037 | 0.584592351683734 | 0.884313345041393 | NM_019682 |
| Akap10 | 1212.387809 | -0.056643615 | 0.063607538 | -0.890517328 | 0.373188168 | 0.602699272 | | Efcab7 | 72.69426466 | -0.0520261376609923 | 0.0949382697259503 | -0.547999640304921 | 0.583692149816193 | 0.884313345041393 | NM_145549 |
| Gm16576 | 46.51514197 | 0.14178154 | 0.159188837 | 0.890650015 | 0.373116958 | 0.602699272 | | Enoph1 | 1489.5508 | -0.0317042084847641 | 0.0580204720747942 | -0.54643141206942 | 0.584769422580551 | 0.884313345041393 | NM_026421,NM_001163035,NR_027990 |
| Golga5 | 1130.748679 | 0.051055695 | 0.057338214 | 0.890430504 | 0.373234769 | 0.602699272 | | Eva1c | 67.47001664 | 0.0520305728333031 | 0.0950533903505322 | 0.54738260930439 | 0.5841159004349 | 0.884313345041393 | NM_001199210,NM_027627 |
| Mbnl1 | 2822.82628 | 0.068525618 | 0.076935126 | 0.890693512 | 0.373093616 | 0.602699272 | | Fbxw5 | 4034.172031 | -0.0338918507661771 | 0.0620581784527903 | -0.546130286308028 | 0.584976382521186 | 0.884313345041393 | NM_013908 |
| Mpp1 | 1515.943759 | 0.056942792 | 0.063938867 | 0.890581808 | 0.373153562 | 0.602699272 | | Khsrp | 4078.584727 | -0.0287144467109233 | 0.0524350843728237 | -0.547618966468289 | 0.583953563477877 | 0.884313345041393 | NM_010613 |
| Snrnp35 | 318.0252707 | -0.08381657 | 0.094122713 | -0.890503123 | 0.373195792 | 0.602699272 | | Mdh1b | 53.08518482 | -0.050669433380892 | 0.0924062040330182 | -0.548333674249697 | 0.583462809288184 | 0.884313345041393 | NM_029696 |
| Strbp | 6754.389725 | 0.06667563 | 0.074877361 | 0.89049029 | 0.37320268 | 0.602699272 | | Mmel1 | 40.06437495 | 0.0505018030145094 | 0.0920477670771079 | 0.548647779496968 | 0.583247189693813 | 0.884313345041393 | NM_013783 |
| H1fx | 474.0105955 | -0.112432984 | 0.126278849 | -0.89035484 | 0.373275383 | 0.602702041 | | Nob1 | 1018.121201 | -0.0313175447995201 | 0.0571741520431981 | -0.547757048952087 | 0.583858734144503 | 0.884313345041393 | NM_026277 |
| Uhrf1bp1 | 1194.434209 | 0.05728906 | 0.064362468 | 0.890100421 | 0.373411967 | 0.602859749 | | Nras | 2116.090972 | -0.035142967168651 | 0.0641070914696745 | -0.548191570744949 | 0.583560369368238 | 0.884313345041393 | NM_010937 |
| B4galt1 | 267.8086848 | 0.103779404 | 0.116617621 | 0.889911863 | 0.373513214 | 0.602960381 | | Pigm | 612.8545717 | -0.0373009144775069 | 0.0681425114217606 | -0.54739565213024 | 0.584106941696116 | 0.884313345041393 | NM_026234 |
| Nkx3-1 | 58.5219506 | -0.139556198 | 0.156841524 | -0.889791135 | 0.373578047 | 0.603002215 | | Ppp2r2c | 29407.02663 | 0.031324520113716 | 0.0572668180057254 | 0.54699250289381 | 0.584383882774597 | 0.884313345041393 | NM_172994 |
| Inpp5f | 4186.828688 | -0.071817665 | 0.080726659 | -0.88964 | 0.373659221 | 0.603070413 | | Pygo2 | 2275.277368 | 0.0378023894415599 | 0.0689434716388498 | 0.548309920329834 | 0.583479116832046 | 0.884313345041393 | NM_026869 |
| Cnbp | 7709.209246 | -0.039814388 | 0.044764279 | -0.889423185 | 0.373775689 | 0.603120383 | | Rpn2 | 5699.977562 | 0.0282972107119014 | 0.0517377362100373 | 0.546935617689659 | 0.584422964660381 | 0.884313345041393 | NM_019642 |
| Ikzf3 | 9.384554634 | -0.136430991 | 0.15340326 | -0.889361748 | 0.373808696 | 0.603120383 | | Slc37a4 | 926.4906572 | -0.0393173212896921 | 0.0716337508556037 | -0.548865874257322 | 0.583097498978534 | 0.884313345041393 | NM_008063 |
| Mink1 | 15871.55482 | -0.060108929 | 0.067591852 | -0.889292528 | 0.373845886 | 0.603120383 | | Smarca4 | 7084.989487 | 0.0269621069476106 | 0.0491748689066956 | 0.548290367560888 | 0.583492540360659 | 0.884313345041393 | NM_011417,NM_001174078,NM_001174079 |
| Txndc17 | 847.6222636 | -0.069317453 | 0.077942788 | -0.88933762 | 0.373821659 | 0.603120383 | | Stap2 | 73.36613954 | -0.0511760766385224 | 0.0936340206047793 | -0.54655430054138 | 0.584684972665451 | 0.884313345041393 | NM_145934 |
| Igfbp3 | 441.5004479 | 0.120660584 | 0.135731142 | 0.888967577 | 0.374020505 | 0.603339272 | | Tbrg1 | 1967.670236 | 0.0272388777099698 | 0.0497369099401559 | 0.547659228181726 | 0.583925912659219 | 0.884313345041393 | NM_025289 |
| Apc2 | 6697.520261 | -0.050476854 | 0.056794634 | -0.888760972 | 0.374131555 | 0.603455581 | | Tomm70a | 8423.242691 | 0.0426265718454105 | 0.078028296694491 | 0.546296326476393 | 0.584862260995465 | 0.884313345041393 | NM_138599 |
| A630089N07Rik | 75.64373217 | -0.131501867 | 0.14804331 | -0.888266192 | 0.374397581 | 0.603821809 | | Ttyh1 | 20577.17498 | -0.022904501953998 | 0.0417766242722877 | -0.548261195177314 | 0.583512568294174 | 0.884313345041393 | NM_021324,NM_001109765,NM_001001454 |
| Fam71e1 | 90.28755069 | 0.116949653 | 0.131725119 | 0.887831064 | 0.374631631 | 0.604136395 | | Vps26b | 6106.798283 | 0.0321305641858299 | 0.0588122192752898 | 0.546324634264052 | 0.58484280572466 | 0.884313345041393 | NM_178027 |
| Pot1b | 256.6620085 | 0.115011607 | 0.129566037 | 0.887667865 | 0.374719436 | 0.604215105 | | Yy2 | 42.8618466 | 0.050855041450765 | 0.0930968530883112 | 0.546259511076321 | 0.584887563796511 | 0.884313345041393 | NM_001098723 |
| Fundc2 | 1532.45724 | -0.054619256 | 0.061544791 | -0.887471618 | 0.37482504 | 0.60431583 | | Zfp454 | 200.3169338 | -0.0461198283355601 | 0.0844663469852795 | -0.546014240956789 | 0.585056148115463 | 0.884341894643622 | NM_172794 |
| St13 | 3932.308687 | -0.054693001 | 0.061632387 | -0.887406826 | 0.37485991 | 0.60431583 | | AI429214 | 124.5127771 | -0.0504900827789396 | 0.0925546823644529 | -0.5455162449818 | 0.585398510843915 | 0.884512516052311 | NM_001039220 |
| Srd5a3 | 648.8377223 | 0.074867009 | 0.084377246 | 0.887289072 | 0.374923288 | 0.60431583 | | Evpl | 245.8334212 | 0.0483131139033734 | 0.0885603246395943 | 0.545539033421443 | 0.585382842195197 | 0.884512516052311 | NM_025276 |
| Ift74 | 745.7418799 | 0.067470071 | 0.076072884 | 0.886913544 | 0.375125449 | 0.604492326 | | Hmmr | 38.39553436 | 0.0487318348829731 | 0.0892979755404616 | 0.54572160889462 | 0.585257315799355 | 0.884512516052311 | NM_013552 |
| Ushbp1 | 189.5071984 | 0.1150115 | 0.129669009 | 0.88695597 | 0.375102606 | 0.604492326 | | Impdh1 | 2636.700416 | 0.0331659773977621 | 0.0608095694059101 | 0.545407206822594 | 0.585473484912775 | 0.884512516052311 | NM_011829 |
| Zfp820 | 21.99147199 | 0.148809429 | 0.167767829 | 0.886991454 | 0.375083502 | 0.604492326 | | Nfya | 1069.335458 | -0.0310775930336849 | 0.0569785050870058 | -0.54542661282934 | 0.585460141117695 | 0.884512516052311 | NM_001110832,NM_010913 |
| Ppp6r2 | 7548.466109 | 0.057916824 | 0.06530892 | 0.886813376 | 0.375179385 | 0.604516367 | | Ankrd45 | 3385.170126 | 0.0375631195422617 | 0.0688865298934493 | 0.545289762749882 | 0.585554243831561 | 0.884542527651545 | NM_028664 |
| Tmem38b | 326.8767344 | 0.094572367 | 0.10667503 | 0.88654624 | 0.37532315 | 0.604685129 | | Omd | 26.13690973 | -0.0431233430776746 | 0.0790970713063505 | -0.545195193266433 | 0.58561927708992 | 0.884548780181779 | NM_012050 |
| Cltc | 27104.65603 | -0.061640891 | 0.069572857 | -0.885990507 | 0.375622642 | 0.60487861 | | 2210015D19Rik | 524.7517609 | 0.0342014913556251 | 0.0628070871672608 | 0.544548281064899 | 0.586064233629945 | 0.884588668914926 | NR_015577 |
| Fbxo4 | 201.5458163 | 0.09099916 | 0.102693332 | 0.886125306 | 0.37555008 | 0.60487861 | | Actr5 | 256.7364122 | -0.0425379515610566 | 0.0781605334483585 | -0.544238245113333 | 0.586277536887095 | 0.884588668914926 | NM_175419 |
| Golga4 | 4496.747278 | 0.06180334 | 0.069751229 | 0.886053774 | 0.375588551 | 0.60487861 | | Fam118b | 582.6463534 | 0.0369365159929255 | 0.0678749634880453 | 0.544184690418744 | 0.586314385907939 | 0.884588668914926 | NM_175411,NM_194257 |
| Nr1h2 | 1600.857567 | 0.064447198 | 0.072742688 | 0.885961193 | 0.37563844 | 0.60487861 | | G0s2 | 105.9997458 | 0.0510917962925999 | 0.0938847660544736 | 0.544196874953659 | 0.586306002082044 | 0.884588668914926 | NM_008059 |
| Zfp646 | 1325.193226 | 0.070336248 | 0.079382428 | 0.886043046 | 0.375594332 | 0.60487861 | | Mctp2 | 69.19939814 | -0.049564857175361 | 0.0909385271435506 | -0.545036946740084 | 0.585728107085611 | 0.884588668914926 | NM_001024703 |
| Zdhhc23 | 298.45051 | -0.079088798 | 0.089278689 | -0.885864237 | 0.37569069 | 0.60489988 | | Mtch2 | 3082.504962 | 0.0296935817693855 | 0.0544889918509868 | 0.544946433411536 | 0.585790359529425 | 0.884588668914926 | NM_019758 |
| Gsto1 | 846.2183455 | 0.075825399 | 0.085615582 | 0.885649519 | 0.375806419 | 0.605023343 | | Nav3 | 2945.86555 | 0.0324291055357823 | 0.0595163779645841 | 0.544877001000962 | 0.58583811521011 | 0.884588668914926 | NM_001081035 |
| Lrrc39 | 10.27489823 | -0.137085745 | 0.154835176 | -0.88536564 | 0.375959458 | 0.60520684 | | Osbpl11 | 947.7527536 | 0.0336020770720687 | 0.0616870583148526 | 0.544718422145577 | 0.58594719267897 | 0.884588668914926 | NM_176840 |
| Bod1l | 3375.267634 | -0.07048511 | 0.079624759 | -0.885215997 | 0.376040146 | 0.605210971 | | Ranbp3 | 3290.02563 | 0.0321286455849789 | 0.0590298910500274 | 0.544277568761732 | 0.586250480410936 | 0.884588668914926 | NM_001252466,NM_001252467,NR_045519,NM_027933 |
| Zbtb34 | 912.7039682 | 0.082867169 | 0.09361005 | 0.885228924 | 0.376033176 | 0.605210971 | | Syn1 | 29550.45756 | -0.0236330017068046 | 0.0434284089654451 | -0.544182996103054 | 0.58631555172194 | 0.884588668914926 | NM_013680,NM_001110780 |
| Ngef | 15915.5029 | -0.075964828 | 0.085822763 | -0.885136131 | 0.376083215 | 0.605217421 | | Wdr11 | 2466.112379 | 0.0265690254409313 | 0.048823539703818 | 0.544184743713975 | 0.586314349236912 | 0.884588668914926 | NM_172255 |
| Cpne8 | 1122.290982 | -0.078305448 | 0.088487018 | -0.88493713 | 0.376190541 | 0.605327265 | | Nkap | 651.0486679 | 0.0398578957048049 | 0.073270227253739 | 0.543984878971028 | 0.586451878313069 | 0.884593011538357 | NM_025937 |
| Dapk3 | 943.9414897 | -0.061916606 | 0.07000619 | -0.884444735 | 0.376456183 | 0.605377488 | | Osbpl3 | 1679.310054 | 0.0450561653382633 | 0.0828370322888027 | 0.54391332105162 4 | 0.58650112172146 | 0.884593011538357 | NM_001163645,NM_027881 |
| Gabrd | 2705.939155 | -0.076987306 | 0.08704121 | -0.884492597 | 0.376430357 | 0.605377488 | | Rbbp5 | 2570.970094 | -0.0257557145781653 | 0.0473457629666861 | -0.543991963890998 | 0.58644700284788 | 0.884593011538357 | NM_172517 |
| Gamt | 277.0015097 | -0.141197049 | 0.159604474 | -0.884668493 | 0.376335453 | 0.605377488 | | Epha7 | 2474.650684 | -0.0405451803820886 | 0.0745603125159614 | -0.543790376058429 | 0.586585732213294 | 0.884628773476983 | NM_001122889,NM_010141 |

| | | | | | | | | | | | | | | |
|---|---|---|---|---|---|---|---|---|---|---|---|---|---|---|
| Morf4l1 | 11811.04303 | -0.067005255 | 0.07575947 | -0.884447249 | 0.376454826 | 0.605377488 | Churc1 | 676.9848993 | 0.0361636862115567 | 0.0665365911729914 | 0.543515764393958 | 0.58677473982451 | 0.884638284451565 | NM_206534 |
| Rab6a | 18501.34492 | 0.056206814 | 0.063536573 | 0.88463717 | 0.376352353 | 0.605377488 | Cops4 | 2498.06974 | 0.029384935410729 | 0.0540514626750837 | 0.543647367831077 | 0.58668415729859 | 0.884638284451565 | NM_012001 |
| Zfp689 | 262.1371146 | -0.078221841 | 0.08842833 | -0.88457897 | 0.376383753 | 0.605377488 | Ngly1 | 954.6577561 | 0.0379352989403623 | 0.0697874165130493 | 0.543583654988416 | 0.586728009980984 | 0.884638284451565 | NM_021504 |
| D10Wsu102e | 1076.971978 | -0.075043932 | 0.084857146 | -0.884356075 | 0.376504027 | 0.605391593 | Fuom | 571.0488298 | 0.0379243913353803 | 0.0698214379883578 | 0.543162564794268 | 0.587017878814514 | 0.884663352599169 | NM_026928 |
| Nfe2l1 | 16143.58044 | -0.055122777 | 0.062349903 | -0.884087617 | 0.376648917 | 0.605561723 | Mcat | 598.5064316 | -0.0386275870730392 | 0.0711213140113155 | -0.543122516927816 | 0.587045450300919 | 0.884663352599169 | NM_001030014 |
| Rspry1 | 913.1829943 | -0.062414152 | 0.070660873 | -0.883291545 | 0.377078771 | 0.606189923 | Snrpb | 2195.882687 | 0.0350352508267448 | 0.0645157862730879 | 0.543049272908875 | 0.587095877671485 | 0.884663352599169 | NM_009225 |
| Polh | 209.9739348 | 0.085744076 | 0.097086162 | 0.883175049 | 0.3771417 | 0.606228188 | Tfap2b | 52.86375893 | -0.0460798879428731 | 0.084821823500547 | -0.543255096874638 | 0.586954176166655 | 0.884663352599169 | NM_001025305,NM_009334 |
| Aarsd1 | 1730.625939 | -0.058259771 | 0.06599927 | -0.882733562 | 0.377380244 | 0.60642289 | Ttc21b | 568.6819447 | 0.0418076137004924 | 0.0769616892765268 | 0.543226299909761 | 0.586974000763161 | 0.884663352599169 | NM_001047604 |
| Cep57 | 532.5306237 | 0.076288111 | 0.086410227 | 0.882859746 | 0.377312055 | 0.60642289 | Strip2 | 2330.248677 | -0.0480701792165532 | 0.0885449528775818 | -0.542890110100491 | 0.587205465710348 | 0.884736707282286 | NM_177204,NM_001037740 |
| Lrrtm2 | 2554.357053 | -0.070070412 | 0.079374903 | -0.882777927 | 0.377356269 | 0.60642289 | Igf2 | 3280.12683 | 0.0374885569072902 | 0.0690796774457246 | 0.54268575496382 | 0.587346183813762 | 0.884773742666262 | NM_001122736,NM_001122737,NM_010514 |
| C130036L24Rik | 30.66970351 | -0.145672941 | 0.165131492 | -0.88216329 | 0.37768851 | 0.606792387 | Inip | 805.6205584 | 0.0374427398309853 | 0.0690075470055711 | 0.542589056642782 | 0.587412775318286 | 0.884773742666262 | NM_001013577 |
| Dusp12 | 399.4322907 | 0.102665589 | 0.116375743 | 0.882190622 | 0.377673733 | 0.606792387 | Tle4 | 2549.456615 | 0.0383481915770009 | 0.0706692640468607 | 0.542643143298793 | 0.587375527996921 | 0.884773742666262 | NM_011600 |
| Bace1 | 3957.113346 | -0.044797396 | 0.050789824 | -0.882015193 | 0.377768591 | 0.606858119 | Get4 | 1852.306889 | 0.0314957602995931 | 0.058059462204938 | 0.542474199785377 | 0.587491876277239 | 0.884801139948489 | NM_026269,NM_001163316 |
| Brix1 | 681.9424542 | 0.054364027 | 0.061652989 | 0.881774396 | 0.37789882 | 0.607004387 | Tigd3 | 133.7414706 | 0.0498601029777955 | 0.0919832599333356 | 0.542056272129639 | 0.587779741154627 | 0.885142911052469 | NM_198634 |
| Mrpl51 | 1156.618859 | -0.071771688 | 0.081411834 | -0.881587906 | 0.377999698 | 0.607065669 | 2510039O18Rik | 1040.104313 | 0.031177659465692 | 0.0578040048124553 | 0.539368501660873 | 0.589632610656717 | 0.88518841122125 | NM_029841 |
| Zfp516 | 810.4667674 | -0.10131241 | 0.11492414 | -0.881559004 | 0.378015333 | 0.607065669 | 5430417L22Rik | 1538.067701 | 0.0343734913666958 | 0.0635758430877067 | 0.540669060717937 | 0.588735707447088 | 0.88518841122125 | NR_030716 |
| Maml3 | 481.9819355 | -0.084458181 | 0.095815576 | -0.881466082 | 0.378065605 | 0.607083478 | Bcl11a | 3219.854303 | 0.032600747387656 | 0.0603528790865493 | 0.540168884750382 | 0.589080568738444 | 0.88518841122125 | NM_001159289,NM_001159290,NM_016707,NM_001242934 |
| Kcnt2 | 806.4586586 | 0.091755723 | 0.104114543 | 0.881295931 | 0.378157668 | 0.607105471 | Car12 | 1000.483699 | 0.0461732145000396 | 0.085407625056484 | 0.540621688865638 | 0.588768365393037 | 0.88518841122125 | NM_178396 |
| Tsga10 | 324.3073191 | -0.122063375 | 0.138500777 | -0.881319065 | 0.37814515 | 0.607105471 | Cetn2 | 1159.612994 | 0.0344067330441738 | 0.0637697619539174 | 0.539546204814713 | 0.589510024266026 | 0.88518841122125 | NM_019405 |
| Snn | 10928.64642 | 0.058813638 | 0.066770081 | 0.880838202 | 0.3784054 | 0.607440247 | Cks1b | 50.97863319 | 0.0489364542407799 | 0.0906733482403389 | 0.539700531528502 | 0.589403573367705 | 0.88518841122125 | NM_016904 |
| Bola1 | 204.0374117 | -0.098879515 | 0.112306913 | -0.880440143 | 0.378620918 | 0.607597359 | Cyb5r4 | 1799.251807 | 0.0396795002859374 | 0.0733376279691451 | 0.541052409039348 | 0.588471459593432 | 0.88518841122125 | NM_024195 |
| Gm10406 | 53.61151871 | 0.13589837 | 0.154331107 | 0.880563692 | 0.378554017 | 0.607597359 | D130020L05Rik | 95.17930375 | -0.0507750081057612 | 0.0937295780680869 | -0.541718091047816 | 0.588012725038272 | 0.88518841122125 | NR_038049,NR_038048,NR_038050,NR_038047 |
| Tcap | 119.6426229 | 0.135511762 | 0.153902293 | 0.880505154 | 0.378585714 | 0.607597359 | D630003M21Rik | 305.2512108 | -0.0473658422359724 | 0.0877907222533685 | -0.539531296932177 | 0.589520307838513 | 0.88518841122125 | NM_177657,NM_001131021 |
| Tmod2 | 22212.14452 | -0.090804394 | 0.103151978 | -0.880297164 | 0.378698349 | 0.60765868 | Egf | 58.18632245 | -0.0494999198702925 | 0.0915239385616951 | -0.540841233978642 | 0.588617019011561 | 0.88518841122125 | NM_010113 |
| Adra2a | 656.2493234 | 0.119401664 | 0.135703407 | 0.879872261 | 0.378928513 | 0.607752012 | Ergic2 | 1887.519071 | 0.0354943165567535 | 0.0657095400376976 | 0.540169913476649 | 0.589079859356678 | 0.88518841122125 | NM_026168,NM_026355 |
| Atxn10 | 9654.259973 | -0.048584118 | 0.055221832 | -0.879799088 | 0.378968159 | 0.607752012 | Gnal | 14501.02437 | 0.0399153157183603 | 0.0739633346563331 | 0.539663549565806 | 0.589429081838728 | 0.88518841122125 | NM_177137,NM_010307 |
| Eml5 | 2808.250243 | 0.118050904 | 0.134175169 | 0.879826757 | 0.378953167 | 0.607752012 | Ikzf2 | 127.4279702 | -0.0506327421585338 | 0.0935129116641602 | -0.541451883568494 | 0.588196154006462 | 0.88518841122125 | NM_011770 |
| Eral1 | 636.2589448 | 0.058320565 | 0.066297255 | 0.879682949 | 0.379031089 | 0.607752012 | Lrig1 | 1781.006751 | 0.0434570890946932 | 0.0803341659169646 | 0.540954008778924 | 0.588539283169335 | 0.88518841122125 | NM_008377 |
| Sp2 | 665.6809862 | 0.056653326 | 0.064397857 | 0.879739294 | 0.379000558 | 0.607752012 | Mef2a | 3344.526183 | -0.0370268667362284 | 0.0685730375605535 | -0.539962469994592 | 0.589222914736575 | 0.88518841122125 | NM_001033713 |
| Ube2cbp | 73.61089701 | 0.133374218 | 0.151551325 | 0.880059731 | 0.378826952 | 0.607752012 | Ndufv2 | 4270.231008 | 0.0387074067860068 | 0.0716876448226598 | 0.539945298548457 | 0.589234757078939 | 0.88518841122125 | NM_028388 |
| Utp3 | 914.6769507 | 0.070332181 | 0.079933458 | 0.879884123 | 0.378922086 | 0.607752012 | Nrde2 | 752.438431 | -0.0355279717623474 | 0.0655955343506968 | -0.541621805722358 | 0.588079066920408 | 0.88518841122125 | NM_183155 |
| AA415398 | 228.7321846 | 0.086903696 | 0.098810876 | 0.879495243 | 0.379132812 | 0.607852213 | Nup160 | 596.9576961 | -0.0387065868729652 | 0.0717233133748453 | -0.539665348011381 | 0.589427841341225 | 0.88518841122125 | NM_021512 |
| Gabbr2 | 10647.79382 | 0.076475222 | 0.086975368 | 0.879274484 | 0.379252467 | 0.607968717 | Pcdhb8 | 156.514974 | 0.0488021935327102 | 0.0903778753609281 | 0.539979429011984 | 0.58921121900414 | 0.88518841122125 | NM_053133 |
| Kiss1r | 215.6941594 | 0.098593473 | 0.112137892 | 0.879216396 | 0.379283956 | 0.607968717 | Polr3c | 795.4062611 | 0.0309032356460496 | 0.0571084679427872 | 0.541132283166996 | 0.588416408034853 | 0.88518841122125 | NM_028925 |
| Gmcl1 | 986.1843835 | 0.053697237 | 0.061089424 | 0.878994009 | 0.379404524 | 0.60803617 | Pou3f1 | 759.3548512 | 0.0477691905720247 | 0.0885663426468976 | 0.539360541989118 | 0.589638101815835 | 0.88518841122125 | NM_011141 |
| Lrrc33 | 276.0205107 | 0.099017076 | 0.112644801 | 0.879020385 | 0.379390223 | 0.60803617 | Rab22a | 1220.573556 | 0.0343728596862221 | 0.0637224964333386 | 0.539414831654944 | 0.589600649333136 | 0.88518841122125 | NM_024436 |
| Zfp592 | 1554.329327 | 0.085111292 | 0.096869444 | 0.87861857 | 0.379608124 | 0.608299541 | Sh2b2 | 233.7494355 | -0.0473857575532494 | 0.0876407147648522 | -0.540682006991722 | 0.588726782490175 | 0.88518841122125 | NM_018825 |
| Bcl2l15 | 10.4603284 | 0.140700492 | 0.160181273 | 0.878382907 | 0.379735957 | 0.608347533 | Slc2a10 | 80.56127861 | -0.0513048145432845 | 0.0949849759242518 | -0.540136100936624 | 0.589103175774781 | 0.88518841122125 | NM_130451 |
| Ly6g6d | 8.26898452 | -0.129573537 | 0.14751989 | -0.878346215 | 0.379755863 | 0.608347533 | Soat1 | 882.862521 | -0.0364836844402016 | 0.0676107122064326 | -0.539613964260688 | 0.589463284319388 | 0.88518841122125 | NM_009230 |
| Tceb3 | 1485.396308 | 0.070418846 | 0.080166352 | 0.878409017 | 0.379721793 | 0.608347533 | Tnfrsf8 | 55.29894472 | 0.050018829606259 | 0.0924899174967096 | 0.540803051403289 | 0.588643339393593 | 0.88518841122125 | NM_009401 |
| Pgf | 80.08302212 | -0.117978236 | 0.134402292 | -0.877799281 | 0.380052655 | 0.608760037 | Tnk2 | 10948.64348 | -0.0335187398221047 | 0.0618774555219118 | -0.541695509929867 | 0.588028283421171 | 0.88518841122125 | NM_001110147,NM_016788 |
| Fam110a | 174.4860926 | -0.088387736 | 0.100739871 | -0.877385834 | 0.380277105 | 0.609056591 | Trp53i11 | 4444.985621 | 0.0436367272349942 | 0.080891865063753 | 0.539445186491905 | 0.589579709103799 | 0.88518841122125 | NM_001025246 |
| Ptrf | 1017.436632 | -0.074373572 | 0.084787463 | -0.877176529 | 0.380390762 | 0.609175656 | Wdr91 | 1162.438254 | -0.0320505726385496 | 0.0592874774425205 | -0.54059599128033 | 0.588786081544206 | 0.88518841122125 | NM_001013366 |
| AW046200 | 15.71622264 | -0.145930205 | 0.166447391 | -0.876734714 | 0.380630747 | 0.609245124 | Zfp740 | 3031.988753 | -0.0256207082734714 | 0.0473889610882014 | -0.540647182068111 | 0.588750790388464 | 0.88518841122125 | NM_153194 |
| Cdpf1 | 278.2672707 | -0.092724063 | 0.105750186 | -0.876821745 | 0.380583466 | 0.609245124 | Mthfd1 | 1675.742466 | 0.0248597801318844 | 0.0461034127865881 | 0.539217785176987 | 0.589736589825737 | 0.885244776667216 | NM_138745 |
| Etf1 | 3280.720858 | -0.066075972 | 0.075356396 | -0.876846232 | 0.380570164 | 0.609245124 | Med9 | 1976.026139 | 0.0260973339502 | 0.0484155587185874 | 0.539027838176756 | 0.589867646129469 | 0.885258541964735 | NM_138675 |
| Rpgrip1l | 1074.141273 | -0.071885334 | 0.08199131 | -0.876743318 | 0.380626073 | 0.609245124 | Tspan4 | 447.7299201 | -0.040285480076568 | 0.0747370099030827 | -0.539029861226844 | 0.589866250230103 | 0.885258541964735 | NM_053082,NM_001252588 |
| Taok2 | 6324.153893 | 0.059262427 | 0.067580996 | 0.876909636 | 0.380535722 | 0.609245124 | Lyrm1 | 266.1242574 | 0.0427768770432611 | 0.0794413310524334 | 0.538471302992484 | 0.590251711771673 | 0.885570623285077 | NM_029610 |
| Slx4ip | 303.5110228 | -0.090421441 | 0.103144397 | -0.876649081 | 0.380677272 | 0.609256652 | Trim2 | 11434.13753 | 0.0314081439090015 | 0.0583179785517636 | 0.538567088382929 | 0.59018560194776 | 0.885570623285077 | NM_001271728,NM_030706,NM_001271725,NM_001271726,NM_001271727 |
| Qrsl1 | 480.0168613 | -0.073711066 | 0.084127017 | -0.876187794 | 0.380927951 | 0.609594885 | Ttll11 | 1961.386907 | 0.0300650238394975 | 0.0558350495568005 | 0.538461487509071 | 0.590258486482591 | 0.885570623285077 | NM_028921,NM_029774 |
| Magi1 | 4281.700778 | -0.044734357 | 0.05107209 | -0.875906132 | 0.381081066 | 0.609769958 | Gpr153 | 1400.0198 | -0.0318247998318092 | 0.0591837486671757 | -0.537728693239395 | 0.590764366975358 | 0.885693150538513 | NM_178406 |
| Tkt | 5370.194231 | -0.058845324 | 0.067187164 | -0.875841763 | 0.381116063 | 0.609769958 | Gstm2 | 90.11721129 | 0.0485184031166635 | 0.0901998976273026 | 0.537898649476709 | 0.590647020856693 | 0.885693150538513 | NM_008183 |
| Dbh | 16.74623483 | -0.145841352 | 0.16660807 | -0.875355871 | 0.381380303 | 0.610129728 | Hspbap1 | 218.3290219 | -0.0445386998487841 | 0.0828120994159696 | -0.537828411100458 | 0.59069551557497 | 0.885693150538513 | NM_175111 |
| 1700084E18Rik | 8.614635622 | 0.131192908 | 0.149892756 | 0.875245152 | 0.38144053 | 0.610149688 | Ick | 1139.948388 | -0.0359395211720603 | 0.0668031805997917 | -0.537991168225489 | 0.590583145885463 | 0.885693150538513 | NM_001163780,NM_019987 |
| Casp4 | 14.13253228 | 0.143676469 | 0.164166378 | 0.875188151 | 0.381471539 | 0.610149688 | Scai | 1962.294568 | -0.0393140220392586 | 0.0730832158332836 | -0.537935031881208 | 0.590621902054717 | 0.885693150538513 | NM_178778 |
| Ufm1 | 2214.470129 | -0.047873268 | 0.054727896 | -0.874750758 | 0.381709535 | 0.610467335 | Sec22a | 595.3917864 | 0.0358636288103251 | 0.066697716454555 | 0.537703998228502 | 0.590781418514496 | 0.885693150538513 | NM_133704 |
| Dctd | 252.8615026 | 0.123822886 | 0.141581649 | 0.874568747 | 0.381808599 | 0.610501468 | Slc25a32 | 325.0354166 | 0.0461587206384853 | 0.0858460031569847 | 0.537692134065646 | 0.590789610623495 | 0.885693150538513 | NM_172402 |
| Etv3 | 1301.764456 | 0.078333424 | 0.089568262 | 0.874566757 | 0.381809682 | 0.610501468 | Vps53 | 3132.143895 | 0.0278875407168349 | 0.0518706116467012 | 0.537636627591387 | 0.590827938091573 | 0.885693150538513 | NM_026664 |
| Atcay | 10536.10617 | -0.04597697 | 0.052590166 | -0.874250335 | 0.38198194 | 0.610650864 | Ehbp1 | 2522.165322 | 0.0315766826770728 | 0.058760398027645 | 0.537380340109624 | 0.591004920572587 | 0.885827500260987 | NM_001252515,NM_153078 |
| Nmd3 | 704.6826246 | 0.063522349 | 0.072654728 | 0.874304409 | 0.381952499 | 0.610650864 | Pgls | 436.991174 | -0.0433598733598679 | 0.0807348174450506 | -0.537065354602174 | 0.591222471088412 | 0.885827500260987 | NM_025396 |
| Cd209f | 19.35221151 | -0.130029265 | 0.148773703 | -0.874007045 | 0.382114417 | 0.610799626 | Plekhh2 | 513.0118713 | -0.0456993916941267 | 0.0850801994736594 | -0.537133104727559 | 0.591175675114746 | 0.885827500260987 | NM_177606 |
| D2hgdh | 548.5688286 | -0.077502461 | 0.088688564 | -0.873872099 | 0.382187911 | 0.610799825 | Taok3 | 2523.449173 | 0.027348434825876 | 0.0509208863653509 | 0.537076959533942 | 0.59121445527639 | 0.885827500260987 | NM_183306,NM_001081308,NM_001199685 |
| Stau1 | 2670.191467 | 0.056858007 | 0.065065198 | 0.873862048 | 0.382193385 | 0.610799825 | Zbtb21 | 854.5866132 | 0.0342347804912265 | 0.063741089958185 | 0.537091231318526 | 0.591204597471987 | 0.885827500260987 | NM_001081684,NM_001081685,NM_175428 |
| Asphd1 | 533.4901059 | -0.089285269 | 0.102186892 | -0.873744836 | 0.382257228 | 0.610838644 | Rnf165 | 2977.436261 | 0.0485204216545804 | 0.0903876627783895 | 0.536803587604004 | 0.591403293268934 | 0.886007037750055 | NM_001164504 |
| Lrrc59 | 4083.792501 | -0.052956511 | 0.060613672 | -0.873672698 | 0.382296523 | 0.610838644 | Pla2g16 | 1937.548566 | 0.0319098034851916 | 0.0594729640728795 | 0.536543015513547 | 0.591583315273452 | 0.886185339554725 | NM_139269 |
| Igdcc3 | 54.00682994 | 0.139854926 | 0.160131499 | 0.873375491 | 0.382458445 | 0.611034353 | 0610031J06Rik | 1494.254466 | 0.0318065345698422 | 0.0593466824794473 | 0.535944609555174 | 0.591996832555139 | 0.88622671269197 | NM_020003 |
| Pyroxd1 | 449.4111143 | 0.088397327 | 0.101243752 | 0.873113901 | 0.382600998 | 0.611199078 | Ankrd23 | 113.3906172 | -0.0505039435210272 | 0.0944307692557934 | -0.534825077875014 | 0.592770820175263 | 0.88622671269197 | NM_153502 |
| Pdlim2 | 309.9829665 | 0.121168021 | 0.138812731 | 0.87288839 | 0.382723915 | 0.611332406 | B3gnt5 | 28.36080055 | 0.0452891361212521 | 0.0846436838312865 | 0.535056298016558 | 0.592610928248966 | 0.88622671269197 | NM_001159407,NM_001159408,NM_054052 |
| 0610009L18Rik | 66.79084824 | 0.13958861 | 0.160011987 | 0.872363458 | 0.383010129 | 0.611369871 | Bptf | 4632.690115 | 0.0383624617469021 | 0.0715901039356099 | 0.535862635168212 | 0.592053489759087 | 0.88622671269197 | NM_176850 |
| Gch1 | 55.1187699 | 0.132064223 | 0.151347778 | 0.872587789 | 0.382887799 | 0.611369871 | Ccdc155 | 172.194081 | -0.0486872961391799 | 0.0910489017593904 | -0.534737873805913 | 0.592831128130791 | 0.88622671269197 | NM_201374 |
| Ino80c | 938.1483269 | 0.061354546 | 0.070322682 | 0.872471654 | 0.382951126 | 0.611369871 | Ccser2 | 4922.800074 | 0.0301775016805388 | 0.056405746287894 | 0.53500757753498 | 0.592644617491307 | 0.88622671269197 | NM_028407,NM_027045 |
| Jmjd7 | 99.31871252 | 0.122120622 | 0.140009805 | 0.872229071 | 0.383083423 | 0.611369871 | Csmd1 | 2938.989296 | -0.0410328295123695 | 0.0766492869007029 | -0.535332175569049 | 0.592420180985015 | 0.88622671269197 | NM_053171 |
| Rian | 20955.46053 | 0.083914877 | 0.096211241 | 0.872194103 | 0.383102496 | 0.611369871 | Dmrt2 | 35.66620313 | 0.0492662630384063 | 0.0920530664439311 | 0.535194154215536 | 0.592515608192135 | 0.88622671269197 | NM_145831 |
| Sh3pxd2a | 2468.916073 | -0.06881927 | 0.078890837 | -0.872335398 | 0.383025432 | 0.611369871 | Ece1 | 2822.164523 | 0.031758714539451 | 0.0592507208247691 | 0.536005538791261 | 0.591954722475482 | 0.88622671269197 | NM_199307 |
| Snord104 | 31.13770234 | 0.146642835 | 0.168074926 | 0.872484896 | 0.382943904 | 0.611369871 | Fam134c | 3206.731703 | 0.0296390438419673 | 0.0552715973131911 | 0.536243663703449 | 0.591790160174356 | 0.88622671269197 | NM_028933,NM_026501 |
| Tmub2 | 2377.634671 | 0.055590164 | 0.063726138 | 0.872329087 | 0.383028874 | 0.611369871 | Gldn | 79.88109502 | -0.045889493203454 | 0.0855802037769369 | -0.536216217982655 | 0.59180912616933 | 0.88622671269197 | NM_177350 |
| Zfp329 | 1179.10925 | 0.093573542 | 0.107259802 | 0.872400853 | 0.382989736 | 0.611369871 | Grpr | 29.1294315 | 0.0432222983874047 | 0.0806256216602339 | 0.536086389132585 | 0.591898846427023 | 0.88622671269197 | NM_008177 |
| Kiss1 | 15.11924878 | -0.140421768 | 0.161038633 | -0.87197576 | 0.383221656 | 0.611434079 | Lcorl | 617.7680595 | -0.0397161873186114 | 0.0741295292737022 | -0.535767429089838 | 0.592119295268267 | 0.88622671269197 | NM_001163073,NM_172153,NM_178142 |
| Zfp697 | 697.7158205 | 0.076718047 | 0.087976003 | 0.872033786 | 0.383189947 | 0.611434079 | Lipt2 | 126.9123227 | -0.0483809797790666 | 0.0903350306042319 | -0.535572739118551 | 0.592253873520283 | 0.88622671269197 | NM_026010 |
| 2210404O07Rik | 16.92486679 | -0.145834294 | 0.167363484 | -0.871362681 | 0.383556156 | 0.611876648 | Nol9 | 1003.416548 | -0.0319035014146083 | 0.0596153642079125 | -0.535155690793782 | 0.592542202847084 | 0.88622671269197 | NM_028727,NM_001159599 |
| Lenep | 202.3775702 | 0.105221215 | 0.120760339 | 0.871322621 | 0.383578023 | 0.611876648 | Pbx4 | 247.1347867 | -0.0479561846891724 | 0.0895560184152244 | -0.535488128411702 | 0.59231236452878 | 0.88622671269197 | NM_001024954 |
| Abcb6 | 808.9765916 | 0.058087327 | 0.066672864 | 0.871228922 | 0.383629171 | 0.611880995 | Ptpmt1 | 868.16637 | -0.0321906912167189 | 0.0601797685116781 | -0.534908857458835 | 0.592712883149465 | 0.88622671269197 | NM_025576 |
| Pde8a | 707.938274 | 0.0926361 | 0.106334914 | 0.871172944 | 0.38365973 | 0.611880995 | Snora65 | 48.06668916 | -0.0499596055589359 | 0.0933458097281962 | -0.535209943589412 | 0.592504691148491 | 0.88622671269197 | NR_002898 |
| Prrx2 | 24.40641086 | -0.146042356 | 0.167655623 | -0.871085345 | 0.383707555 | 0.611894284 | Vsig10 | 99.82133378 | -0.049120830380891 | 0.0917571989393873 | -0.535334894140993 | 0.592418301450078 | 0.88622671269197 | NM_001033311 |
| Med22 | 1276.131055 | 0.051127493 | 0.058740023 | 0.870403012 | 0.384080202 | 0.612425506 | Zfp61 | 567.339226 | -0.0397388097014959 | 0.0743005597360611 | -0.534838631669271 | 0.592761446993404 | 0.88622671269197 | NM_009561 |
| Commd4 | 1210.491808 | 0.064859904 | 0.074530122 | 0.870250876 | 0.384163318 | 0.612431984 | 2410127L17Rik | 307.9129223 | -0.0446780688658655 | 0.083585390994195 | -0.53452006785451 | 0.592981769069046 | 0.886253111213807 | NM_026120 |
| Tmcc3 | 2080.475811 | 0.069556178 | 0.079924338 | 0.870275314 | 0.384149966 | 0.612431984 | 2700089E24Rik | 3890.883384 | 0.0362877990655609 | 0.0679060640615079 | 0.534382305425509 | 0.593077058631746 | 0.886253111213807 | NR_073368,NM_001163445,NM_001271582,NM_001271583 |
| Fbxo16 | 251.9236949 | 0.089498209 | 0.102851977 | 0.870165181 | 0.384210141 | 0.612443614 | Hmbs | 595.4495541 | -0.0394546171091846 | 0.073820402009457 | -0.534467654404403 | 0.593018022355714 | 0.886253111213807 | NM_013551,NM_001110251 |
| Mpv17 | 1718.988999 | -0.068905707 | 0.079209807 | -0.869913835 | 0.384347494 | 0.612599535 | Ocm | 24.283821 | 0.0443685842888373 | 0.0830450450387583 | 0.534271301413948 | 0.593153844638622 | 0.886253111213807 | NM_033039 |
| Abca9 | 496.966352 | -0.102663233 | 0.118025564 | -0.869838952 | 0.384388421 | 0.612601749 | Tob1 | 1105.484094 | -0.0397125198251342 | 0.074328545942853 | -0.534283555818069 | 0.593145367543334 | 0.886253111213807 | NM_009427 |
| Snai2 | 18.28562654 | 0.145309436 | 0.167077263 | 0.869714007 | 0.384456716 | 0.612647573 | Egfem1 | 200.1171509 | 0.0479050715158955 | 0.0896893116350887 | 0.534122412610355 | 0.593256844253421 | 0.886291956220826 | NM_001167748,NM_029412 |
| Ahsa2 | 1026.268876 | 0.076520764 | 0.088014664 | 0.869409254 | 0.384623323 | 0.61266645 | G6pc3 | 1902.431932 | 0.0393385610239625 | 0.0736598093955726 | 0.534057328504668 | 0.593301871285372 | 0.886291956220826 | NM_175935 |
| Cgrrf1 | 611.1345087 | 0.084452569 | 0.097135434 | 0.869431114 | 0.384611371 | 0.61266645 | Ankrd11 | 8942.456333 | -0.030071637907508 | 0.0563883678005713 | -0.533768355057015 | 0.59350181021955 | 0.886313552411855 | NR_037865,NM_001081379 |
| Depdc1b | 12.48791962 | 0.130122024 | 0.149678264 | 0.869344826 | 0.384658551 | 0.61266645 | Klhdc8b | 1044.528724 | 0.0406912185900192 | 0.0762131817901299 | 0.533913132010045 | 0.593401636088704 | 0.886313552411855 | NM_030075 |
| Prr5l | 477.5146725 | 0.127081614 | 0.146135895 | 0.869612589 | 0.384512156 | 0.61266645 | Myo1d | 1104.629728 | 0.0425258915712706 | 0.0796655143838679 | 0.533805523006602 | 0.593476092217415 | 0.886313552411855 | NM_177390 |
| Spr | 689.7928637 | -0.100698521 | 0.115834538 | -0.869330706 | 0.384666272 | 0.61266645 | Naa60 | 4791.452585 | 0.0212968226873103 | 0.0399053275036351 | 0.53368369637789 | 0.593560390875845 | 0.886313552411855 | NM_029090 |
| Copa | 10607.43666 | 0.037655959 | 0.043332944 | 0.868991457 | 0.384851803 | 0.61273055 | Fam13b | 3325.002583 | 0.039910625711161 | 0.0748173569934477 | 0.533440732404597 | 0.593728527603178 | 0.886473490796974 | NM_146084 |
| Ppp2r3c | 1019.619137 | -0.062624013 | 0.072067121 | -0.868967873 | 0.384864703 | 0.61273055 | Rps23 | 6104.436091 | 0.0313378171818076 | 0.0587727108161381 | 0.533203535223064 | 0.593892694599712 | 0.886549756375718 | NM_024175 |
| Rhobtb1 | 234.4168945 | -0.118532138 | 0.136392604 | -0.869051063 | 0.384819202 | 0.61273055 | Tma16 | 265.1160707 | -0.0456252562395366 | 0.0855702616651104 | -0.533190565877858 | 0.593901671436879 | 0.886549756375718 | NM_025465 |
| Rmst | 59.05506884 | 0.136783083 | 0.157378188 | 0.869136217 | 0.38477263 | 0.61273055 | Shisa5 | 2460.162017 | 0.0305915969249624 | 0.0574528787047222 | 0.532464127379713 | 0.594404580802073 | 0.887209303404328 | NM_025858,NM_026381 |
| Rab5b | 7255.267845 | -0.046255411 | 0.053242829 | -0.868763215 | 0.384976657 | 0.612814857 | Rgp1 | 2385.926871 | -0.0360971890398971 | 0.0678045346002556 | -0.53237131192935 | 0.594468850447981 | 0.887214067770202 | NM_172866 |
| Zfp467 | 861.7523536 | 0.08513891 | 0.098004278 | 0.868726467 | 0.384996761 | 0.612814857 | Nqo2 | 415.8770042 | 0.0434756716885298 | 0.081708207375698 | 0.532084512497339 | 0.594667463505626 | 0.887269096170567 | NM_020282,NM_001163242,NM_001163241,NM_001163239 |
| Slc35e4 | 878.0769059 | 0.071013162 | 0.081755883 | 0.868600023 | 0.385065942 | 0.612862014 | Plekhj1 | 725.218458 | 0.0344186531472776 | 0.0646796634643925 | 0.53214026331825 | 0.594628852823173 | 0.887269096170567 | NM_023900 |
| Sema6b | 4179.248385 | -0.061644257 | 0.070976594 | -0.868515285 | 0.385112308 | 0.612872854 | Slc46a3 | 309.3021528 | 0.0441766993076977 | 0.0830305640160319 | 0.532053465265729 | 0.594688966013813 | 0.887269096170567 | NM_027872 |
| 1810026J23Rik | 2909.346808 | 0.075320861 | 0.086731862 | 0.86843357 | 0.385157024 | 0.612881065 | Lypd6 | 1328.079387 | -0.0348281550997751 | 0.0654972577832209 | -0.531749820962694 | 0.594899280912722 | 0.88731416582475 | NR_033304,NM_177139 |
| Btk | 53.61841418 | 0.139379883 | 0.160587761 | 0.8679359 | 0.385429425 | 0.613251541 | Trp53rk | 128.4257079 | 0.0503503940142985 | 0.09468893481429 | 0.531745278506395 | 0.594902427438196 | 0.88731416582475 | NM_023815 |
| Mrap2 | 343.041319 | 0.103259108 | 0.118985573 | 0.867828808 | 0.385488057 | 0.613272331 | Zdhhc16 | 1066.128811 | -0.0348323189301873 | 0.0654952107599204 | -0.531830015142158 | 0.594843732256483 | 0.88731416582475 | NM_023740 |
| Zfp58 | 214.3843029 | 0.095143361 | 0.109641542 | 0.867767447 | 0.385521654 | 0.613272331 | Ddx6 | 7483.665534 | 0.0343472121931617 | 0.0646052724315776 | 0.53164719999499 | 0.594970367542391 | 0.887324390032933 | NM_001110826,NM_181324,NM_007841 |
| Arcn1 | 5231.268893 | -0.059444408 | 0.068516604 | -0.867591276 | 0.385618124 | 0.613299857 | 5730455P16Rik | 3618.322984 | 0.0323431220660743 | 0.060894579543372 | 0.531133022160667 | 0.595326602355629 | 0.887381665349327 | NM_027472 |
| Trappc3 | 2099.628001 | 0.079071342 | 0.091134366 | 0.86763475 | 0.385594317 | 0.613299857 | Cat | 2082.831661 | -0.0286083579647877 | 0.0538700159493928 | -0.531062734261363 | 0.595375307069706 | 0.887381665349327 | NM_009804 |
| Gprasp1 | 24211.94543 | 0.062994794 | 0.07262107 | 0.867445137 | 0.38569816 | 0.613364188 | Ccdc86 | 364.2331175 | 0.0433171577561366 | 0.0815381366335162 | 0.531250278024272 | 0.595245356096563 | 0.887381665349327 | NM_023731 |
| Slfn9 | 22.35631945 | 0.145115473 | 0.167414662 | 0.866802649 | 0.386050151 | 0.613704655 | Eng | 1161.075779 | 0.0420958272171953 | 0.0792027258940979 | 0.531494677007477 | 0.595076028994811 | 0.887381665349327 | NM_001146348,NM_007932,NM_001146350 |
| Txnl4b | 642.5932217 | -0.066037365 | 0.076178746 | -0.866873875 | 0.386011119 | 0.613704655 | Gigyf2 | 3040.131504 | -0.0291492175004171 | 0.054858342262938 | -0.531354326397686 | 0.595173265508275 | 0.887381665349327 | NM_146112,NM_001110212 |
| Upp2 | 259.1219636 | 0.095849956 | 0.110573779 | 0.866841647 | 0.38602878 | 0.613704655 | Itgb4 | 629.0901408 | 0.0470147954197254 | 0.0885258642427597 | 0.531085415792149 | 0.595359590119163 | 0.887381665349327 | NM_133663,NM_001005608 |
| Vps25 | 1406.679059 | -0.049372304 | 0.056961568 | -0.866765169 | 0.38607069 | 0.613704655 | Btd | 381.7954214 | 0.0432212225760945 | 0.0814178015058267 | 0.530857156257178 | 0.595517769021439 | 0.88749770826135 | NM_025295 |
| Dll4 | 135.2778033 | 0.121448688 | 0.140147787 | 0.866575854 | 0.386174448 | 0.613806617 | Gm10653 | 28.22248124 | -0.0466663918663678 | 0.0879407753387345 | -0.530657043750364 | 0.595656458412916 | 0.88749770826135 | NR_003965 |
| Grpel2 | 843.9911768 | 0.062438892 | 0.072076825 | 0.866282499 | 0.386335261 | 0.61393626 | N4bp2 | 249.8630082 | -0.0500580599948177 | 0.0943469375053352 | -0.530574296510547 | 0.595713811278589 | 0.88749770826135 | NM_001024917 |
| Pla2r1 | 30.86322092 | -0.144740997 | 0.167074593 | -0.866325596 | 0.386311633 | 0.61393626 | Rars2 | 723.8722016 | -0.034174190473498 | 0.0643932608870583 | -0.530710667587364 | 0.595619292587802 | 0.88749770826135 | NM_181406 |
| Abca8a | 447.9011592 | 0.134078213 | 0.154811242 | 0.866075431 | 0.386448798 | 0.613972588 | Zfp438 | 264.8196908 | 0.0423602065746372 | 0.0798481423359257 | 0.530509606553217 | 0.595758650231063 | 0.88749770826135 | NM_178722 |
| Rbl2 | 2333.894507 | 0.089041429 | 0.102816353 | 0.866024 | 0.386477001 | 0.613972588 | Kcnq4 | 280.9423353 | 0.0460932305433795 | 0.0869479761106436 | 0.530124249065034 | 0.596025787345858 | 0.887804613103561 | NM_001081142 |
| Slc25a35 | 191.5654539 | 0.107422148 | 0.124039686 | 0.866030471 | 0.386473452 | 0.613972588 | Cmc1 | 346.3058325 | 0.0405955042787616 | 0.0765931183688819 | 0.530015034552433 | 0.596101506843217 | 0.887826359930747 | NM_026442 |
| 5830403L16Rik | 34.29917644 | 0.137508677 | 0.158883116 | 0.865470672 | 0.386780506 | 0.614237786 | Chsy3 | 492.8939178 | 0.0389480413287625 | 0.0735978771985636 | 0.529200607562125 | 0.596666295268221 | 0.887922450933845 | NM_001081328 |
| Cd7 | 45.25950252 | 0.144649621 | 0.167113255 | 0.865578382 | 0.386721415 | 0.614237786 | Fstl4 | 868.3593811 | -0.043817780548545 | 0.0827676363944606 | -0.529407174800965 | 0.596523022065989 | 0.887922450933845 | NM_177059 |
| Hells | 60.93598857 | -0.12655782 | 0.146236826 | -0.865430568 | 0.38680251 | 0.614237786 | Gemin8 | 385.8397393 | -0.0391920240680449 | 0.0740863969509511 | -0.529004320374656 | 0.596802452831209 | 0.887922450933845 | NM_146238 |
| Rnf157 | 15890.60018 | -0.06125963 | 0.070784461 | -0.865438952 | 0.386797909 | 0.614237786 | Hexa | 2478.786815 | -0.0357978347754915 | 0.0676123150898958 | -0.52945731451283 | 0.596488247969963 | 0.887922450933845 | NM_010421 |
| Nelfb | 2027.964669 | 0.059670656 | 0.068968536 | 0.86518664 | 0.386936357 | 0.614387366 | Mboat2 | 1302.608727 | -0.0308999017926406 | 0.0583612026398401 | -0.529459647761729 | 0.596486629781681 | 0.887922450933845 | NM_001083341,NM_026037 |
| Copb2 | 3983.521931 | 0.060407696 | 0.069842753 | 0.86491001 | 0.387088183 | 0.614565459 | Mrps5 | 1319.754416 | 0.0297246385992177 | 0.056201130162138 | 0.528897524186138 | 0.596876539554677 | 0.887922450933845 | NM_029963 |
| Itfg1 | 9927.09633 | -0.053174404 | 0.061496088 | -0.86467946 | 0.387214746 | 0.61470341 | Pmel | 24.36044251 | -0.0427655575268063 | 0.0808351514935859 | -0.529046543943196 | 0.596773162623428 | 0.887922450933845 | NM_021882 |
| Ahsa1 | 4011.031162 | -0.058227673 | 0.067349411 | -0.864560981 | 0.387279796 | 0.614743691 | Rpl10a | 4451.543823 | 0.0281257334396484 | 0.0531823482121227 | 0.528854674251432 | 0.596906266616528 | 0.887922450933845 | NM_011287 |
| 2810002D19Rik | 122.2878211 | -0.118402661 | 0.137006591 | -0.864211421 | 0.387471759 | 0.614790257 | Sec63 | 4151.258461 | 0.0304698126566485 | 0.057586080124566 | 0.529117672026615 | 0.59672382302598 | 0.887922450933845 | NM_153055 |
| Cmc2 | 412.1291523 | -0.077400188 | 0.089553308 | -0.864291777 | 0.387427626 | 0.614790257 | Tenc1 | 1277.263668 | -0.0395971186883989 | 0.0748674782244358 | -0.528896119216085 | 0.596877514239333 | 0.887922450933845 | NM_153533 |
| Fam180a | 90.37512287 | -0.134907198 | 0.156076651 | -0.864365026 | 0.387387399 | 0.614790257 | Trim13 | 300.4118027 | -0.0411088484317501 | 0.0777081403753734 | -0.529015985110074 | 0.596794361016669 | 0.887922450933845 | NM_023233,NM_001164220 |
| Ifitm3 | 342.900116 | 0.090888873 | 0.105177645 | 0.864146298 | 0.387507529 | 0.614790257 | Yy1 | 1059.946247 | 0.0384641577286257 | 0.0727418391732993 | 0.528776260894218 | 0.596960667480375 | 0.887922450933845 | NM_009537 |
| Rad18 | 178.6788782 | 0.12069146 | 0.139647286 | 0.86425926 | 0.387445485 | 0.614790257 | Zfp879 | 122.6001804 | -0.0494312970589038 | 0.0933084674478296 | -0.529762179263545 | 0.596276830725972 | 0.887922450933845 | NM_173387 |
| Slc25a29 | 425.0970198 | 0.074833869 | 0.086607514 | 0.864057474 | 0.387556319 | 0.61480471 | Dnlz | 1463.803995 | -0.0307574494578063 | 0.0582022100527799 | -0.528458445648616 | 0.597181181377079 | 0.888159500428325 | NM_001139504,NM_026828,NM_001139503 |
| Dhrs1 | 2171.286006 | 0.053096087 | 0.061465024 | 0.8638423 | 0.387674527 | 0.614866324 | 2900009J06Rik | 23.55765944 | -0.0439835415715116 | 0.0832818029401563 | -0.528129075244887 | 0.597409751823009 | 0.888165440564526 | NR_045298 |
| Prss45 | 11.33269832 | 0.141207951 | 0.163460679 | 0.863864949 | 0.387662083 | 0.614866324 | Iffo2 | 1273.298386 | 0.0284721732211706 | 0.0539117249442638 | 0.528125806595992 | 0.597412020339451 | 0.888165440564526 | NM_183148,NM_001205173 |
| Awat2 | 8.462637929 | -0.129504818 | 0.149995647 | -0.86339051 | 0.387922795 | 0.615197113 | Klhl3 | 769.4199014 | 0.0482543329837245 | 0.0913373518626478 | 0.528308868164789 | 0.597284977492596 | 0.888165440564526 | NM_001195075 |
| 1600020E01Rik | 169.7082311 | 0.101997638 | 0.118159353 | 0.863221027 | 0.388015955 | 0.615281875 | Snx9 | 434.3776852 | 0.0360047339736088 | 0.0681778375583365 | 0.528100263414798 | 0.597429748021202 | 0.888165440564526 | NM_025664 |
| 2700060E02Rik | 1891.665054 | -0.060312545 | 0.069916485 | -0.862636982 | 0.388337091 | 0.615635694 | Jak1 | 8130.569314 | 0.0299100082271012 | 0.0566566227025079 | 0.527917246041164 | 0.597556774188892 | 0.888198349104731 | NM_146145 |
| Gm11190 | 9.155661789 | -0.13920346 | 0.161376897 | -0.862598445 | 0.388358286 | 0.615635694 | Kif20b | 27.2371596 | -0.0421850996366915 | 0.0799123410142313 | -0.527892176618614 | 0.597574174982827 | 0.888198349104731 | NM_183046 |
| Zfp236 | 2156.341849 | -0.077706654 | 0.090079313 | -0.862647055 | 0.388331551 | 0.615635694 | Snx5 | 3813.032458 | -0.0295810694058468 | 0.056073017720563 | -0.52754552204168 | 0.597814813019126 | 0.88846510885163 | NM_024225,NM_001199188 |
| Spef1 | 310.2043035 | -0.084012702 | 0.097469334 | -0.861939845 | 0.388720621 | 0.616047658 | D2Wsu81e | 1270.826053 | 0.0335968998917682 | 0.0637033374722226 | 0.527396227967145 | 0.597918462403578 | 0.888528243976918 | NM_172660 |
| Taf1c | 523.8634455 | 0.080157641 | 0.093007887 | 0.861837031 | 0.388777204 | 0.616047658 | Slc16a9 | 138.0628451 | 0.047325988816831 | 0.0897602914952294 | 0.527248608805446 | 0.59802095698221 | 0.888589650278598 | NM_025807 |
| Txndc5 | 2372.095655 | -0.044666186 | 0.051822527 | -0.86190676 | 0.388738829 | 0.616047658 | Casc3 | 2902.936162 | -0.0266156743775234 | 0.0505365011234874 | -0.526662388290143 | 0.598428058919012 | 0.88871702396247 | NM_138660 |
| Vps35 | 6580.227258 | -0.061725001 | 0.071604252 | -0.862029824 | 0.388671107 | 0.616047658 | Enpp6 | 261.8927145 | -0.0476803960329992 | 0.0905270820697749 | -0.526697590851862 | 0.598403608885228 | 0.88871702396247 | NM_177304 |
| Pa2g4 | 2612.572635 | 0.046666893 | 0.054154072 | 0.861743006 | 0.388828954 | 0.616066655 | Mef2d | 5409.4268 | 0.0339445901761377 | 0.0644603314038976 | 0.526596581755787 | 0.598473766239906 | 0.88871702396247 | NM_133665 |
| 9430016H08Rik | 378.1038584 | -0.082347612 | 0.095585238 | -0.861509724 | 0.388957368 | 0.616109828 | Rin1 | 3428.076075 | -0.0422165577787745 | 0.0801315402379342 | -0.526840712825699 | 0.598304207802359 | 0.88871702396247 | NM_145495 |
| Adc | 1058.489422 | 0.076524202 | 0.088829092 | 0.861476795 | 0.388975496 | 0.616109828 | Slc16a14 | 238.2213052 | 0.0490177539574611 | 0.093038620531234 | 0.526853834220439 | 0.59829509510544 | 0.88871702396247 | NM_027921 |
| Pard6b | 208.2692062 | -0.086792102 | 0.100734482 | -0.861592777 | 0.388911647 | 0.616109828 | Uchl3 | 773.9448398 | 0.0404612459950759 | 0.0768166964010848 | 0.526724630070196 | 0.598384829031343 | 0.88871702396247 | NM_016723 |
| 2310008H04Rik | 113.8882889 | -0.111470888 | 0.12942089 | -0.861305219 | 0.389069962 | 0.616133482 | 2010001M06Rik | 77.64222168 | 0.0498145468763356 | 0.0947928982123844 | 0.525509271430077 | 0.599229209824172 | 0.889150486911646 | NR_026688 |
| Rcc2 | 4119.707011 | -0.048891691 | 0.056760775 | -0.861364041 | 0.389037574 | 0.616133482 | B230120H23Rik | 439.9706128 | -0.0414470896303106 | 0.0788550347920276 | -0.525611202120742 | 0.599158371869157 | 0.889150486911646 | NM_001164791,NM_023057,NM_178084 |
| Eea1 | 1946.249702 | -0.067612702 | 0.078510764 | -0.861190222 | 0.389133284 | 0.616170783 | Gabrd | 2951.430796 | -0.0362422772739975 | 0.0689464547391468 | -0.525658315733812 | 0.599125630981578 | 0.889150486911646 | NM_008072 |
| Hes6 | 343.663794 | -0.07639193 | 0.088721425 | -0.861031368 | 0.389220767 | 0.616246328 | H13 | 2822.159125 | 0.0285841540748585 | 0.0543572192666837 | 0.525857548647235 | 0.598987186068908 | 0.889150486911646 | NM_001159551,NM_010376,NM_001159553,NM_001159552 |
| Rbbp6 | 3230.57207 | 0.062822498 | 0.072972368 | 0.860908037 | 0.389288695 | 0.6162909 | Il17rb | 58.79160167 | -0.0499653450051835 | 0.0951026007298861 | -0.525383581749745 | 0.599316564598095 | 0.889150486911646 | NM_019583 |
| Lmbrd1 | 7694.251998 | 0.052068133 | 0.060505463 | 0.860552588 | 0.389484508 | 0.616537901 | Nfasc | 12948.48597 | 0.0309512323522454 | 0.0588391931318847 | 0.52603087678089 | 0.59886675392017 | 0.889150486911646 | NM_182716,NM_001160318,NM_001160316,NM_001160317 |
| Ube2e1 | 1937.880494 | -0.043270408 | 0.050298037 | -0.86028025 | 0.389634577 | 0.616712446 | Ppp2r5b | 6227.352538 | 0.0292021808344697 | 0.0555482866895429 | 0.525708038443733 | 0.599091077821334 | 0.889150486911646 | NM_198168 |
| Ddx39 | 452.8116589 | 0.07669283 | 0.089169079 | 0.860083238 | 0.38974316 | 0.616770739 | Rassf3 | 2645.947553 | -0.0318312623628119 | 0.060530195489182 | -0.525874104743323 | 0.598975682059817 | 0.889150486911646 | NM_138956 |
| Wdr45 | 1383.571032 | 0.057390541 | 0.066727835 | 0.860068979 | 0.38975102 | 0.616770739 | Rfwd3 | 1566.267095 | 0.0297452603874481 | 0.0566141915246294 | 0.525402899633526 | 0.599303138225105 | 0.889150486911646 | NM_146218 |
| BC048943 | 9.662921329 | -0.110708632 | 0.128757081 | -0.85982558 | 0.389885196 | 0.616888543 | Gm14204 | 424.8587609 | 0.0441648149536973 | 0.084082397626819 | 0.525256369944551 | 0.599404983128097 | 0.889190848133041 | NR_040358,NR_040359 |
| Gm11110 | 21.1354512 | -0.14367875 | 0.167109218 | -0.859789493 | 0.389905092 | 0.616888543 | Slc35b2 | 915.3244772 | 0.0311400040679532 | 0.0592971205252927 | 0.525152044350463 | 0.599477499012891 | 0.889207612647938 | NM_028662 |
| Aph1a | 2104.661621 | 0.042365146 | 0.049283879 | 0.859614684 | 0.390001478 | 0.616915049 | Rab42 | 32.59435577 | -0.0453730772171585 | 0.0864375997872956 | -0.524922919294519 | 0.599636775948004 | 0.889353053647203 | NM_001081651 |
| Mmab | 1334.902532 | 0.063382972 | 0.073732672 | 0.85963211 | 0.389991869 | 0.616915049 | Kifc2 | 24984.93137 | -0.0311765179911875 | 0.0594434108420993 | -0.524473908033344 | 0.599948962957173 | 0.889725230823471 | NM_010630 |
| Gm11974 | 58.68638944 | 0.124187915 | 0.144537052 | 0.859211622 | 0.390223773 | 0.617203668 | Nrg3 | 1863.906284 | 0.0365038181117794 | 0.0696154624446211 | 0.524363651836948 | 0.600025632752996 | 0.889748095280729 | NM_001190188,NM_008734,NM_001190187 |
| C330005M16Rik | 69.68397893 | 0.134363448 | 0.156454484 | 0.858802153 | 0.390449679 | 0.617448986 | Mex3a | 176.6476821 | 0.0469880396648448 | 0.0896256834732443 | 0.524269805751295 | 0.600090894801768 | 0.889754040817646 | NM_001029890 |
| Siglec1 | 31.19796171 | 0.14194772 | 0.165288808 | 0.858786032 | 0.390458575 | 0.617448986 | 2810474O19Rik | 529.8974601 | -0.0422499734107411 | 0.0806554572760206 | -0.523832792443944 | 0.600394843036304 | 0.889841392709453 | NM_026054 |
| Pros1 | 576.2951601 | 0.067880141 | 0.079064285 | 0.858543668 | 0.390592329 | 0.617597463 | Dlg2 | 19247.23475 | 0.0314881606078251 | 0.060103489498263 | 0.523899042646021 | 0.600348760713992 | 0.889841392709453 | NM_001243046,NM_011807,NM_001243047 |
| 1700021K19Rik | 3764.971208 | -0.047498268 | 0.055363641 | -0.85793253 | 0.390929721 | 0.617802551 | Glt1d1 | 682.1278789 | 0.0324450094721791 | 0.0619359019810391 | 0.523848179075712 | 0.600384140258823 | 0.889841392709453 | NM_177005 |
| Ap5m1 | 665.0013434 | 0.074198356 | 0.086472295 | 0.858059282 | 0.39085973 | 0.617802551 | Mllt4 | 4181.841489 | -0.0251551278870931 | 0.0479994185349875 | -0.524071512840455 | 0.600228801380658 | 0.889841392709453 | NM_010806 |
| Foxred2 | 883.3243643 | 0.064174369 | 0.07479815 | 0.857967331 | 0.390910503 | 0.617802551 | Mapk3 | 7574.559889 | 0.0282125068779906 | 0.0538870573684964 | 0.523548849310228 | 0.600592366344132 | 0.890043329271053 | NM_011952 |
| Itprip | 51.84479614 | 0.140808702 | 0.16415036 | 0.857803187 | 0.39100115 | 0.617802551 | BC030500 | 278.3131111 | 0.0433045566935073 | 0.0827373863392619 | 0.52339768766611 | 0.600697532997867 | 0.890108371348262 | NM_173411 |
| Mrps11 | 571.4761288 | -0.067440192 | 0.078611285 | -0.857894534 | 0.390950703 | 0.617802551 | 2310069G16Rik | 93.24951268 | 0.0497125005719476 | 0.0950128833072767 | 0.523218524072938 | 0.600822192031129 | 0.89020228084906 | NR_040309 |
| Piga | 338.5768355 | -0.110443131 | 0.128746547 | -0.857833737 | 0.390984278 | 0.617802551 | Aox4 | 41.60249158 | 0.0479222488993323 | 0.0916928373675919 | 0.522638957143561 | 0.601225525043627 | 0.890297392009909 | NM_023631 |
| Pik3cb | 1515.474815 | 0.064791901 | 0.07553095 | 0.857819226 | 0.390992292 | 0.617802551 | Csrnp2 | 1371.523593 | 0.0355566506433695 | 0.0680301813025328 | 0.522659942434193 | 0.601210918798731 | 0.890297392009909 | NM_153407 |
| Dusp5 | 165.9994193 | 0.135039658 | 0.157448441 | 0.85767542 | 0.391071716 | 0.617851042 | Ensa | 12617.45953 | 0.0214526283200293 | 0.0410349202566894 | 0.522789570098705 | 0.601120698514416 | 0.890297392009909 | NM_001026212,NM_019561 |
| Klhdc9 | 159.1412293 | 0.110494336 | 0.128871002 | 0.857402626 | 0.391222408 | 0.617995175 | Pde4a | 5372.665172 | 0.029132046500685 | 0.055732730376797 | 0.522709838612418 | 0.601176190554486 | 0.890297392009909 | NM_183408,NM_019798 |
| Rpl8 | 7851.347703 | -0.064669852 | 0.075428536 | -0.857365865 | 0.391242718 | 0.617995175 | Rexo2 | 1344.878489 | 0.0313735476532475 | 0.0600338225492775 | 0.522597867685258 | 0.60125412471549 | 0.890297392009909 | NM_024233 |
| Etv5 | 5349.696101 | -0.099395333 | 0.115941991 | -0.857285028 | 0.391287381 | 0.618002719 | Slc16a10 | 118.8375983 | -0.0490721726424748 | 0.0938254436018967 | -0.523015620908643 | 0.600963382772572 | 0.890297392009909 | NM_028247,NM_001114332 |
| Bola3 | 365.3355468 | -0.093106366 | 0.108616554 | -0.857202353 | 0.391333063 | 0.618011871 | Olfml2b | 304.6100365 | -0.0495468963490977 | 0.0948317689714233 | -0.522471497542434 | 0.601342086516567 | 0.890336881942682 | NM_177068 |
| Utp6 | 1740.195392 | -0.053579324 | 0.06251254 | -0.857097208 | 0.391391164 | 0.618040633 | Cand2 | 196.4728651 | -0.0489605678783158 | 0.0937471257521638 | -0.522262069215341 | 0.601487874979137 | 0.890461306738273 | NM_025958 |
| Abhd12 | 11366.5171 | 0.060819173 | 0.070981994 | 0.856825364 | 0.391541406 | 0.618204152 | Uros | 879.607677 | -0.031726093678445 | 0.0607576277292438 | -0.522174661259439 | 0.601548726629676 | 0.890461306738273 | NM_009479 |
| Ppp4r4 | 1618.613171 | -0.047565304 | 0.055517298 | -0.856765464 | 0.391574516 | 0.618204152 | Ppic | 52.69714413 | 0.0489385411432572 | 0.0938020899864115 | 0.521721223379421 | 0.601864445477362 | 0.890837878031807 | NM_008908 |
| Cnksr1 | 10.07086803 | 0.132673402 | 0.154867723 | 0.856688526 | 0.391617046 | 0.618208305 | Dapk3 | 1042.095434 | 0.0336488819770317 | 0.064579720528275 | 0.521044094055799 | 0.602336054817775 | 0.891122970143255 | NM_007828,NM_001190473,NM_001190474 |
| Bbx | 1503.858441 | 0.085222315 | 0.099520875 | 0.856326019 | 0.391817473 | 0.618286373 | Denr | 2322.584796 | 0.0336710793924113 | 0.0646124281182907 | 0.521123882401807 | 0.602280474905855 | 0.891122970143255 | NM_026603 |
| Mettl18 | 149.908694 | -0.10139854 | 0.118402483 | -0.856388631 | 0.391782851 | 0.618286373 | Insig1 | 3059.414677 | 0.0447936048259824 | 0.0859755031273529 | 0.521004276760452 | 0.602363792085682 | 0.891122970143255 | NM_153526 |
| Prelid1 | 3542.928675 | -0.05069804 | 0.059205215 | -0.856310383 | 0.39182612 | 0.618286373 | Nucks1 | 4895.875906 | 0.0333419983364253 | 0.0639741710996941 | 0.521179059662482 | 0.602242040227143 | 0.891122970143255 | NM_175294,NM_001145804 |
| Ptpn20 | 18.26636562 | -0.142736729 | 0.166668255 | -0.85641221 | 0.391769813 | 0.618286373 | Preb | 3055.602201 | -0.0325682846646678 | 0.0624864816199746 | -0.521205288253211 | 0.602223770634545 | 0.891122970143255 | NM_016703 |
| Ahctf1 | 2607.237059 | 0.08826043 | 0.103149688 | 0.855653878 | 0.392189259 | 0.618521669 | A830082N09Rik | 1046.350898 | 0.0401375890888909 | 0.0771504369277662 | 0.520250962758262 | 0.602888669224209 | 0.891281204889515 | NR_015526 |
| Fam26f | 8.347921985 | -0.129605396 | 0.151415843 | -0.855956642 | 0.392021762 | 0.618521669 | C030013G03Rik | 56.19889518 | -0.0494016724882646 | 0.0948944405948889 | -0.520596066308709 | 0.602648190205675 | 0.891281204889515 | NR_077217,NR_077216 |
| Higd1b | 77.09183923 | -0.121247743 | 0.141709488 | -0.8556078 | 0.392214754 | 0.618521669 | Eps15 | 8837.963978 | 0.0340899413170998 | 0.0655335573593961 | 0.520190612118693 | 0.602930727886229 | 0.891281204889515 | NM_001159964,NM_007943 |
| Pcdhb14 | 289.7421541 | 0.095575863 | 0.111694448 | 0.855690365 | 0.392169071 | 0.618521669 | Nudt18 | 1459.118101 | 0.0375133185807579 | 0.0720911543359902 | 0.520359521584607 | 0.602813017360276 | 0.891281204889515 | NM_153136 |
| Slc12a4 | 841.7785675 | 0.073880706 | 0.086343887 | 0.855656474 | 0.392187823 | 0.618521669 | Ptchd2 | 1934.662846 | -0.0331585012469636 | 0.063687582687528 | -0.520643112012117 | 0.602615410624016 | 0.891281204889515 | NM_001083342 |
| Slc25a42 | 1429.537997 | 0.067754017 | 0.079183773 | 0.855655318 | 0.392188462 | 0.618521669 | Rplp2 | 3432.831938 | 0.0285592863811579 | 0.0548817539466973 | 0.520378528880391 | 0.602799772102559 | 0.891281204889515 | NM_026020 |
| Pgbd1 | 79.64814995 | 0.121570066 | 0.142138141 | 0.855295173 | 0.392387759 | 0.618634473 | Suox | 552.0955786 | 0.0361263725765864 | 0.0694497086264374 | 0.520180333238061 | 0.60293789142106 | 0.891281204889515 | NM_173733 |
| Rab36 | 1043.391537 | 0.082919509 | 0.096952162 | 0.855262095 | 0.392406067 | 0.618634473 | Tbx3 | 123.9436085 | -0.0489876333272772 | 0.094180490791182 | -0.52014629479786 | 0.602961613689196 | 0.891281204889515 | NM_011535,NM_198052 |
| Sowahb | 845.8800677 | 0.109789711 | 0.128357529 | 0.855342975 | 0.392361303 | 0.618634473 | BC026585 | 276.0836269 | -0.0429959799526892 | 0.0827127249116349 | -0.519823038095087 | 0.60318692056615 | 0.891523525452167 | NM_001033284 |
| Dom3z | 547.7972238 | 0.081507961 | 0.095318417 | 0.855112406 | 0.392488922 | 0.618702142 | Aen | 1011.007635 | 0.0305223885688349 | 0.0587459461160453 | 0.519565869422577 | 0.603366191752263 | 0.891540502556126 | NM_001162939,NM_026531 |
| Gla | 537.0982149 | 0.085558961 | 0.100075507 | 0.854944071 | 0.392582111 | 0.618743087 | Dab2 | 451.4014109 | -0.0492535592932497 | 0.0948364803631165 | -0.519352459145091 | 0.603514977335769 | 0.891540502556126 | NM_001037905,NM_001102400,NM_023118,NM_001008702 |
| Myc | 126.9688261 | 0.112205255 | 0.13124631 | 0.854921216 | 0.392594765 | 0.618743087 | Dctn1 | 18642.13113 | 0.0245394739333265 | 0.0472553275366516 | 0.519295394033476 | 0.603554764840114 | 0.891540502556126 | NM_001198867,NM_001198866,NM_007835 |
| Frrs1l | 8030.202146 | -0.068674304 | 0.080357634 | -0.854608344 | 0.392768007 | 0.618824677 | Il1rapl1 | 311.0167253 | 0.0479101958110586 | 0.0922630363845493 | 0.519278333864609 | 0.603566659929571 | 0.891540502556126 | NM_001160403 |
| Gm16516 | 25.61308369 | 0.14329964 | 0.167651181 | 0.854748767 | 0.392690247 | 0.618824677 | Sash1 | 3501.780517 | -0.0332484606376345 | 0.0639829115677723 | -0.519645946440197 | 0.603310367835734 | 0.891540502556126 | NM_175155 |
| Pxdn | 3137.70331 | 0.09477035 | 0.110897595 | 0.854575341 | 0.392786284 | 0.618824677 | Snrnp200 | 6441.840876 | -0.0223812045076031 | 0.0430849052492117 | -0.519467418534305 | 0.603434827788267 | 0.891540502556126 | NM_177214 |
| Ripply3 | 8.639553801 | -0.128456112 | 0.15032208 | -0.854539212 | 0.392806293 | 0.618824677 | Snx33 | 493.2610494 | -0.0447644118092295 | 0.0862476057480015 | -0.519022080914597 | 0.603745343274954 | 0.89171376272618 | NM_175483 |
| Aebp2 | 2516.979197 | 0.06253734 | 0.073196471 | 0.854376441 | 0.392896445 | 0.618903774 | Cnn3 | 2731.482496 | 0.0320025990906576 | 0.0617010221217648 | 0.518672106071461 | 0.603989416617799 | 0.891718992982693 | NM_028044 |
| Fert2 | 589.2462759 | -0.066331956 | 0.077646125 | -0.854285461 | 0.392946841 | 0.618920235 | F630042J09Rik | 31.03482154 | 0.0469561352309708 | 0.0905795709188814 | 0.518396529754181 | 0.604181635422228 | 0.891718992982693 | NR_033540 |
| Uap1 | 2014.437567 | -0.073329089 | 0.085866515 | -0.853989344 | 0.393110893 | 0.619115693 | Foxd1 | 29.37742409 | -0.039962602693049 | 0.0770348529775192 | -0.518760030667042 | 0.603928093613531 | 0.891718992982693 | NM_008242 |
| Fzd10 | 48.87105983 | -0.141747422 | 0.165999552 | -0.853902436 | 0.39315905 | 0.619128602 | Kcnq5 | 4116.736189 | 0.0378524577144247 | 0.072968684039481 | 0.518749354092009 | 0.603935539841 | 0.891718992982693 | NM_001160139,NM_023872 |
| Bloc1s6 | 1166.614612 | -0.060155429 | 0.070480656 | -0.853502676 | 0.393380604 | 0.61923566 | Kctd1 | 3554.850212 | 0.0388676671663142 | 0.0749888118711002 | 0.518312881568581 | 0.604239986777409 | 0.891718992982693 | NM_134112,NM_001142731 |
| Nom1 | 1118.685903 | 0.062730093 | 0.073481179 | 0.85368925 | 0.393277192 | 0.61923566 | Myh14 | 1766.220153 | -0.0379083569066277 | 0.0731031788580554 | -0.518572392187516 | 0.604058965444815 | 0.891718992982693 | NM_001271540,NM_001271538,NM_028021 |
| Ralb | 642.3797652 | -0.067539251 | 0.079132909 | -0.853491322 | 0.393386898 | 0.61923566 | Prob1 | 269.0963437 | 0.0475092809513686 | 0.091657150607443 | 0.518336962653852 | 0.604223188019097 | 0.891718992982693 | NM_001270646 |
| Wdr44 | 740.0976696 | -0.063655628 | 0.074574883 | -0.85357999 | 0.39333775 | 0.61923566 | Zfp260 | 2647.713686 | -0.0373603086464348 | 0.072065993254537 | -0.518418007706883 | 0.604166653226382 | 0.891718992982693 | NM_011981 |
| Med1 | 1988.570684 | -0.054005313 | 0.063316165 | -0.852946684 | 0.393688871 | 0.619648047 | Fancb | 101.6228171 | -0.0488380214074827 | 0.0943104273069438 | -0.517843284163415 | 0.604567615798012 | 0.891930650729426 | NM_175027,NM_001146081 |
| Lrwd1 | 446.5857992 | 0.071174853 | 0.083459639 | 0.852805673 | 0.393767077 | 0.619708186 | Orc3 | 1644.42117 | 0.0316233995826791 | 0.0610599361518955 | 0.51790751146564 | 0.604522800947371 | 0.891930650729426 | NM_001159563,NM_015824 |
| Frmd3 | 198.1408387 | -0.117557271 | 0.137870222 | -0.852666145 | 0.39384447 | 0.619762733 | Tssc4 | 1052.323024 | -0.0354081625092824 | 0.0683584831438925 | -0.517977592257997 | 0.604473903501829 | 0.891930650729426 | NM_138631,NM_020285,NM_001115085 |
| Slco4c1 | 68.47457103 | -0.118970093 | 0.139538163 | -0.852598964 | 0.393881737 | 0.619762733 | Saysd1 | 201.6698073 | 0.04321639984411 | 0.0835055392765266 | 0.517527342719145 | 0.604788086870535 | 0.891988592901812 | NM_026209 |
| Topors | 956.07908 | 0.082413743 | 0.096681063 | 0.852429011 | 0.393976024 | 0.619848143 | Setd8 | 1700.175687 | 0.0304761481572959 | 0.0588713414259064 | 0.517673751253864 | 0.604685915206369 | 0.891988592901812 | NM_030241 |
| Zfp74 | 680.9881335 | 0.079248055 | 0.092976038 | 0.852349242 | 0.394020283 | 0.619854835 | St7 | 696.9975574 | 0.0371996039666629 | 0.071880096342272 | 0.517523012066223 | 0.60479110914842 | 0.891988592901812 | NM_001083315,NM_022332 |
| 2810408M09Rik | 471.2845247 | -0.074984786 | 0.087990609 | -0.852190782 | 0.394108212 | 0.619921228 | Anapc15 | 403.2932147 | 0.0388544460168894 | 0.0751085838943242 | 0.517310325961632 | 0.604939546951004 | 0.892113813439142 | NM_027532 |
| Col27a1 | 360.1577219 | 0.118233397 | 0.138768386 | 0.852019687 | 0.394203165 | 0.619921228 | Apopt1 | 833.1010329 | 0.0335952303043335 | 0.0649528131799934 | 0.517225177164173 | 0.604998978544863 | 0.892113813439142 | NM_001163388,NM_026511 |
| Itih5 | 2035.547038 | -0.087442503 | 0.10263119 | -0.852007098 | 0.394210153 | 0.619921228 | Ebi3 | 37.06821898 | -0.0476717256310392 | 0.0921838700291472 | -0.517137386572793 | 0.605060256781112 | 0.892113813439142 | NM_015766 |
| Pilra | 11.09924532 | 0.136306715 | 0.159990486 | 0.851967625 | 0.394232061 | 0.619921228 | Ddx10 | 796.650767 | 0.0338861312104899 | 0.0655195530576504 | 0.516935475860347 | 0.605201201943091 | 0.892231062562199 | NM_029936 |
| Prrc1 | 757.2402699 | 0.072219284 | 0.084773437 | 0.851909349 | 0.394264407 | 0.619921228 | Rpl37a | 4144.273669 | -0.031671668118576 | 0.0612855242255631 | -0.516788728150672 | 0.605303649423607 | 0.892291537601717 | NM_009084 |
| Shh | 104.7925646 | 0.126097751 | 0.148029744 | 0.851840635 | 0.394302549 | 0.619921228 | 1700020I14Rik | 2869.59612 | 0.0417780355685435 | 0.0808576181265047 | 0.516686448804122 | 0.605375057254588 | 0.892306248775255 | NR_027832,NR_015473 |
| Apoh | 14.97679845 | 0.135816569 | 0.159468663 | 0.851681872 | 0.394390685 | 0.619996882 | Nt5c2 | 3069.938243 | 0.0255887778044349 | 0.0495384391696211 | 0.516543884574525 | 0.605474596865412 | 0.892362418229198 | NM_029810,NM_001164365,NM_001164363,NR_028353 |
| Irf4 | 20.36222075 | 0.141962996 | 0.166743425 | 0.851385873 | 0.394555037 | 0.620018297 | Usp9x | 11182.577 | 0.0372430256046214 | 0.0721421538999687 | 0.516244991191447 | 0.605683310672265 | 0.892498451170969 | NM_009481 |
| Rnf216 | [illegible] | [illegible] | [illegible] | [illegible] | [illegible] | [illegible] | Zfp772 | [illegible] | [illegible] | [illegible] | [illegible] | [illegible] | [illegible] | [illegible] |
| Timm21 | [illegible] | [illegible] | [illegible] | [illegible] | [illegible] | [illegible] | Adsl | [illegible] | [illegible] | [illegible] | [illegible] | [illegible] | [illegible] | [illegible] |
| Tmem200b | [illegible] | [illegible] | [illegible] | [illegible] | [illegible] | [illegible] | [illegible] | [illegible] | [illegible] | [illegible] | [illegible] | [illegible] | [illegible] | [illegible] |
| Aox1 | [illegible] | [illegible] | [illegible] | [illegible] | [illegible] | [illegible] | [illegible] | [illegible] | [illegible] | [illegible] | [illegible] | [illegible] | [illegible] | [illegible] |
| Zfp945 | [illegible] | [illegible] | [illegible] | [illegible] | [illegible] | [illegible] | [illegible] | [illegible] | [illegible] | [illegible] | [illegible] | [illegible] | [illegible] | [illegible] |
| Gbp3 | [illegible] | [illegible] | [illegible] | [illegible] | [illegible] | [illegible] | [illegible] | [illegible] | [illegible] | [illegible] | [illegible] | [illegible] | [illegible] | [illegible] |
| Scd4 | [illegible] | [illegible] | [illegible] | [illegible] | [illegible] | [illegible] | [illegible] | [illegible] | [illegible] | [illegible] | [illegible] | [illegible] | [illegible] | [illegible] |
| Fam150b | [illegible] | [illegible] | [illegible] | [illegible] | [illegible] | [illegible] | [illegible] | [illegible] | [illegible] | [illegible] | [illegible] | [illegible] | [illegible] | [illegible] |
| Raver1 | [illegible] | [illegible] | [illegible] | [illegible] | [illegible] | [illegible] | [illegible] | [illegible] | [illegible] | [illegible] | [illegible] | [illegible] | [illegible] | [illegible] |
| Sap18 | [illegible] | [illegible] | [illegible] | [illegible] | [illegible] | [illegible] | Gpr4 | [illegible] | [illegible] | [illegible] | [illegible] | [illegible] | [illegible] | [illegible] |

| | | | | | | | | | | | | | | |
|---|---|---|---|---|---|---|---|---|---|---|---|---|---|---|
| Wfikkn2 | 74.01261014 | 0.137370412 | 0.161563604 | 0.850255929 | 0.395182813 | 0.620486675 | | Kif18a | 35.22322763 | -0.0464063349772938 | 0.0901787484060047 | -0.514603892797028 | 0.606829844304508 | 0.893150687578348 | NM_139303 |
| Zfand1 | 534.8712866 | 0.070431921 | 0.082846162 | 0.850153096 | 0.395239975 | 0.620513545 | | Mtr | 450.3690184 | -0.0408680196618011 | 0.0794389643382201 | -0.514458112618399 | 0.606931738750232 | 0.893150687578348 | NM_001081128 |
| 4932443I19Rik | 15.12654626 | 0.127288671 | 0.149758044 | 0.849962167 | 0.39534612 | 0.620554431 | | Tfb1m | 172.4304156 | -0.0456326737121634 | 0.0886609664722488 | -0.514687302968286 | 0.606771547407879 | 0.893150687578348 | NM_146074 |
| E130317F20Rik | 148.7940329 | -0.100128734 | 0.11779587 | -0.850019054 | 0.395314493 | 0.620554431 | | Tmed7 | 3154.944175 | 0.0364859082440158 | 0.0709158711520952 | 0.514495664387503 | 0.606905490852399 | 0.893150687578348 | NM_025698 |
| Ccdc69 | 11.51478732 | 0.134188344 | 0.157926657 | 0.849687739 | 0.395498717 | 0.620711462 | | Tmem107 | 250.3442531 | -0.0429228630494184 | 0.0834718741213705 | -0.514219472142285 | 0.607098555279986 | 0.893305674027256 | NM_028336,NM_025838 |
| Tmem62 | 1254.595658 | 0.044556896 | 0.052442202 | 0.849638167 | 0.395526285 | 0.620711462 | | Parva | 3776.591033 | 0.031954026106145 | 0.0621962396060616 | 0.513761383462013 | 0.607418829751424 | 0.893686409497538 | NM_020606 |
| 4933434E20Rik | 1666.659053 | -0.050859555 | 0.059880895 | -0.849345271 | 0.395689197 | 0.620841359 | | Vopp1 | 8373.689951 | 0.0487126070679815 | 0.0948644045291797 | 0.513497210146909 | 0.607603561820916 | 0.89386766649895 | NM_146168 |
| Klhl35 | 25.30494885 | -0.140898127 | 0.165882397 | -0.849385648 | 0.395666737 | 0.620841359 | | Ivns1abp | 7768.071221 | -0.0369358739234599 | 0.0719613010642649 | -0.513274126192833 | 0.607759580291275 | 0.893916126297191 | NM_001039511,NM_054102,NM_001039512 |
| Tyw1 | 583.3612702 | 0.056095471 | 0.066053221 | 0.849246556 | 0.395744112 | 0.620864649 | | Neurl4 | 5589.489115 | -0.0261339412684066 | 0.0509146430236765 | -0.513289295895754 | 0.607748970473265 | 0.893916126297191 | NM_001013414 |
| Efnb3 | 4529.005573 | 0.065755681 | 0.077455606 | 0.848946689 | 0.395910957 | 0.620922658 | | Btbd17 | 534.9062839 | 0.0406575408215166 | 0.0792756814533333 | 0.512862709927637 | 0.608047359789002 | 0.894000455572205 | NM_028055 |
| Krt73 | 47.42333646 | 0.130639722 | 0.153894396 | 0.848892006 | 0.395941388 | 0.620922658 | | Ski | 7297.235713 | -0.0303498953578881 | 0.0591804706562742 | -0.512836329642648 | 0.60806581447428 | 0.894000455572205 | NM_011385 |
| Rps28 | 2071.510394 | -0.086876321 | 0.102339671 | -0.848901702 | 0.395935992 | 0.620922658 | | Sobp | 3600.309384 | 0.027481220712222 | 0.0535787427814442 | 0.512912757664397 | 0.608012348903807 | 0.894000455572205 | NM_175407 |
| Zfp747 | 217.5707374 | 0.089366503 | 0.105259309 | 0.849012822 | 0.395874157 | 0.620922658 | | Zdhhc14 | 1823.234888 | 0.0399733679101899 | 0.07792038439657 | 0.513002704231391 | 0.607949429058213 | 0.894000455572205 | NM_146073 |
| Ikzf5 | 949.026983 | 0.076638024 | 0.090303191 | 0.848674593 | 0.396062388 | 0.620986707 | | 1700021K19Rik | 3982.560209 | 0.029064503504331 | 0.0567039615712316 | 0.512565660299062 | 0.608255179296643 | 0.894011406907116 | NM_001200038,NM_172615 |
| Rsph3b | 103.4851568 | -0.116320305 | 0.137054727 | -0.848714288 | 0.396040294 | 0.620986707 | | Cd209a | 24.39922291 | -0.0435446327310349 | 0.084947527077754 | -0.512606243277426 | 0.60822678508824 | 0.894011406907116 | NM_133238 |
| Sh3gl1 | 1180.694777 | 0.072512235 | 0.085490766 | 0.848187913 | 0.396333328 | 0.621348637 | | Ercc6l2 | 1018.063026 | -0.028300735343024 | 0.055209382226612 | -0.512607354070021 | 0.608226007921539 | 0.894011406907116 | NM_001013608,NM_023507 |
| Pdzd3 | 39.55240884 | 0.140668742 | 0.165955898 | 0.84762725 | 0.396645593 | 0.621775276 | | Pde6d | 1196.248058 | 0.0325356072201292 | 0.0634913182920357 | 0.512441828195753 | 0.608341823077937 | 0.894048292394791 | NM_008801 |
| Asphd2 | 3393.145856 | 0.087989758 | 0.103820386 | 0.847519072 | 0.396705861 | 0.62180684 | | Ube2i | 7606.254705 | 0.0202113396054687 | 0.0394724411185509 | 0.512036728226823 | 0.608625304893011 | 0.894374423760838 | NM_011665,NM_001177609,NM_001177610 |
| Orc6 | 562.4572421 | -0.066079333 | 0.078028652 | -0.846859858 | 0.397073239 | 0.622256779 | | Camsap1 | 5603.648446 | 0.0273218576013505 | 0.0535067363922301 | 0.510624632402697 | 0.609613924203196 | 0.894400210292768 | NM_001276361,NM_001276360,NM_001115076,NM_001276359 |
| Snx33 | 427.2596767 | 0.091359905 | 0.107876303 | 0.846895026 | 0.397053635 | 0.622256779 | | Cdh20 | 964.4008414 | 0.0370992120475535 | 0.0726581125337601 | 0.510599721817927 | 0.60963137070317 | 0.894400210292768 | NM_011800 |
| Col18a1 | 256.657546 | 0.102625264 | 0.121273815 | 0.84622772 | 0.397425721 | 0.622746169 | | Creg2 | 7177.531454 | -0.036031663324798 | 0.070545555452195 | -0.510757383563515 | 0.609520953688327 | 0.894400210292768 | NM_170597 |
| Oaz2 | 6821.680643 | -0.047594173 | 0.056258329 | -0.845993364 | 0.397556446 | 0.622825033 | | Ddo | 303.6005966 | 0.0447264202525234 | 0.0874974361526041 | 0.511174066569404 | 0.609229176221765 | 0.894400210292768 | NM_027442 |
| Ubac1 | 1496.873644 | -0.050837793 | 0.060087591 | -0.846061438 | 0.397518471 | 0.622825033 | | Fn1 | 2002.061496 | 0.0394911869410937 | 0.0771647564645044 | 0.511777510232401 | 0.60880673193147 | 0.894400210292768 | NM_001276413,NM_001276412,NM_001276411,NM_001276410,NM_001276408,NM_001276409,NM_010233 |
| Defb1 | 17.68320313 | 0.139318926 | 0.164699955 | 0.845895347 | 0.397611129 | 0.622847723 | | Gemin7 | 688.1867338 | 0.0305950611053156 | 0.05980092404268 | 0.511615189816796 | 0.60892035247435 | 0.894400210292768 | NM_027189 |
| H2-K2 | 50.65935898 | 0.136458513 | 0.16134673 | 0.845746999 | 0.397693898 | 0.622914402 | | Gorasp1 | 1512.549264 | 0.0290607738060256 | 0.0569242477221442 | 0.510516607050752 | 0.609689582976661 | 0.894400210292768 | NM_028976 |
| Mphosph8 | 2039.5236 | 0.058566974 | 0.069267007 | 0.845524824 | 0.397817879 | 0.622982638 | | Gpr150 | 154.9789695 | 0.0457224662833588 | 0.0895549835014584 | 0.510551892208369 | 0.609664869506425 | 0.894400210292768 | NM_175495 |
| Satb2 | 2145.809708 | 0.105521973 | 0.124791485 | 0.845586329 | 0.397783555 | 0.622982638 | | Hmgn2 | 2752.527149 | 0.0305289965796102 | 0.0596370787949333 | 0.511913011108183 | 0.608711891675042 | 0.894400210292768 | NM_016957 |
| Camk2n1 | 59769.82772 | -0.08461625 | 0.100116797 | -0.845175362 | 0.398012936 | 0.623225107 | | Krcc1 | 452.3601433 | -0.041161962461466 | 0.0805362929674822 | -0.511098300465428 | 0.609282225940526 | 0.894400210292768 | NM_145568 |
| Pcsk6 | 822.8987079 | -0.096593811 | 0.114344959 | -0.844757937 | 0.398246004 | 0.62352704 | | Pan3 | 1383.827862 | -0.0365330686561047 | 0.0714256535953939 | -0.511483855129338 | 0.609012290629192 | 0.894400210292768 | NM_028291 |
| Baz1a | 149.1198714 | -0.116228252 | 0.137677505 | -0.844206554 | 0.398553993 | 0.623934497 | | Rps11 | 5199.604328 | -0.0313351941141469 | 0.0613162934104921 | -0.511041884159048 | 0.609321728701382 | 0.894400210292768 | NM_013725 |
| Dmrtc1a | 174.2173123 | -0.113584734 | 0.134566984 | -0.844075792 | 0.398627054 | 0.623934497 | | Sept6 | 3990.18888 | 0.0270033681001293 | 0.0528429895904404 | 0.511011362328644 | 0.609343100595953 | 0.894400210292768 | NM_001253706,NM_001177324,NM_001177323,NM_019942 |
| Mrpl9 | 1326.883885 | -0.046736221 | 0.055365494 | -0.84413987 | 0.398591251 | 0.623934497 | | Slx4 | 1330.91441 | 0.0335803719870942 | 0.0657582843623017 | 0.510663748495625 | 0.609586529111317 | 0.894400210292768 | NM_177472 |
| Dstyk | 3294.429537 | -0.058456905 | 0.06926609 | -0.843946948 | 0.398699052 | 0.623984153 | | Sos2 | 1722.276081 | 0.0353713666638545 | 0.0691824762395079 | 0.511276389434223 | 0.609157535318488 | 0.894400210292768 | NM_001135559 |
| Syndig1 | 1617.480047 | -0.067242728 | 0.07968393 | -0.843868118 | 0.398743106 | 0.62399007 | | Sparcl1 | 95552.39038 | 0.0280339213500312 | 0.0548321310659736 | 0.511268134304337 | 0.609163314972353 | 0.894400210292768 | NM_010097 |
| Efcab3 | 11.08370414 | 0.129832478 | 0.153883846 | 0.843704402 | 0.398834607 | 0.624070229 | | Sync | 56.60589655 | 0.0477093024772977 | 0.093373475488092 | 0.510951340601883 | 0.609385129782127 | 0.894400210292768 | NM_023485 |
| 6330418K02Rik | 93.2653855 | -0.120863674 | 0.143315191 | -0.84334168 | 0.399037378 | 0.624324461 | | Emp1 | 95.23179129 | 0.0485081691010703 | 0.0950375570231898 | 0.510410522118471 | 0.609763886780259 | 0.894418885122695 | NM_010128 |
| Gabra4 | 4613.988801 | -0.075777038 | 0.089862689 | -0.84325362 | 0.399086616 | 0.624338452 | | Akap1 | 2144.653175 | -0.0291476333595107 | 0.057248135031077 | -0.509145552840943 | 0.610650203970076 | 0.894800514414105 | NM_001042541,NM_009648 |
| Pitpnb | 3472.50823 | 0.078174353 | 0.092718012 | 0.84314095 | 0.399149619 | 0.624370251 | | Clock | 3366.772998 | 0.0374863149277803 | 0.0735970635950225 | 0.509345252333077 | 0.610510243937063 | 0.894800514414105 | NM_007715 |
| Rps6ka6 | 151.7500459 | -0.110047466 | 0.130531339 | -0.843073144 | 0.399187538 | 0.624370251 | | Fbxo16 | 292.1191571 | -0.044007862360595 | 0.0863541931929275 | -0.509620444976832 | 0.610317397611016 | 0.894800514414105 | NM_015795 |
| Cfl2 | 2136.518648 | -0.060204536 | 0.071426804 | -0.842884351 | 0.399293126 | 0.624385379 | | Fkbp8 | 11878.36172 | 0.0296033157262404 | 0.058140799961112 | 0.509165951380801 | 0.610635906934976 | 0.894800514414105 | NM_001199631,NM_010223,NM_001111066 |
| Gdnf | 44.73546183 | -0.134004782 | 0.158974385 | -0.84293317 | 0.399265821 | 0.624385379 | | Galnt12 | 59.06607515 | -0.0474316729936903 | 0.0931985693216903 | -0.508931342389732 | 0.61080034987548 | 0.894800514414105 | NM_172693 |
| Plekho1 | 1263.699703 | 0.075412156 | 0.089473902 | 0.842839694 | 0.399318105 | 0.624385379 | | Hrsp12 | 1293.283937 | -0.0314109593098143 | 0.0615974207745131 | -0.50993952205887 | 0.610093832311666 | 0.894800514414105 | NM_008287 |
| Begain | 4014.077453 | -0.074929023 | 0.088930364 | -0.842558384 | 0.399475474 | 0.624521618 | | Ipo11 | 2539.709815 | 0.0353942391774213 | 0.0694515424494712 | 0.509624954739803 | 0.610314237536542 | 0.894800514414105 | NM_029665 |
| Hist2h3c1 | 57.38393019 | -0.1351642 | 0.160424699 | -0.842539839 | 0.39948585 | 0.624521618 | | Map4k3 | 3338.460901 | 0.0301301567728612 | 0.0591975100683967 | 0.508976758280017 | 0.610768515280698 | 0.894800514414105 | NM_001081357 |
| Ribc1 | 11.80555578 | -0.139888957 | 0.166097453 | -0.842210128 | 0.399670346 | 0.624747007 | | Nlk | 5254.564151 | 0.0386002717698649 | 0.0757853476170534 | 0.509336870299965 | 0.610516118225813 | 0.894800514414105 | NM_008702 |
| Trim6 | 8.56547513 | 0.134421179 | 0.159627654 | 0.84209205 | 0.399736431 | 0.624787275 | | Sgms2 | 168.7468205 | 0.048346922153872 | 0.0949112602129291 | 0.509390793520262 | 0.610478328242376 | 0.894800514414105 | NM_028943 |
| Taf6l | 344.2571011 | 0.082435939 | 0.097909393 | 0.841961495 | 0.399809507 | 0.624838461 | | Syvn1 | 2446.585617 | -0.0304440838655822 | 0.0597989466234127 | -0.509107360323678 | 0.610676972940219 | 0.894800514414105 | NM_028769,NM_001164709 |
| Dnahc6 | 87.7134062 | 0.137655266 | 0.163599466 | 0.841416355 | 0.400114726 | 0.625252402 | | Tmem186 | 921.8479003 | -0.0325219336026394 | 0.0639067893002831 | -0.508896377970523 | 0.610824858937785 | 0.894800514414105 | NM_025708 |
| Gm14393 | 23.2362761 | 0.140867971 | 0.167441477 | 0.841296756 | 0.400181707 | 0.625294007 | | Zbtb2 | 430.9781667 | 0.0404914926019019 | 0.0794432238022373 | 0.509690954922723 | 0.610267990816413 | 0.894800514414105 | NM_001033466 |
| Mtpn | 12001.84505 | -0.072605269 | 0.086313068 | -0.841185125 | 0.400244232 | 0.625328641 | | Zfp408 | 519.1081987 | -0.0413748411270735 | 0.0813197048430652 | -0.508792318011 | 0.610897804586717 | 0.894817133147792 | NM_001033451 |
| 3110056K07Rik | 195.460663 | 0.103851405 | 0.123514592 | 0.840802718 | 0.400458464 | 0.625348059 | | Rimkla | 731.3550249 | 0.0384098138712513 | 0.0755293789685983 | 0.508541370202717 | 0.611073733963925 | 0.894894350192557 | NM_177572 |
| Gmnn | 119.5645859 | -0.101598785 | 0.120803952 | -0.841022034 | 0.400335591 | 0.625348059 | | Sirt6 | 598.9806409 | -0.0430240417936245 | 0.0845891291255247 | -0.508623770434847 | 0.61101596401153 | 0.894894350192557 | NM_181586,NM_001163430 |
| Hnrnph1 | 6858.105057 | 0.070253469 | 0.083554821 | 0.840806882 | 0.400456131 | 0.625348059 | | Chmp7 | 4021.862568 | 0.0230818602363523 | 0.0454128235058703 | 0.508267455190683 | 0.611265790349701 | 0.894995148736997 | NM_134078 |
| Nmt1 | 4366.552331 | 0.079519104 | 0.094553606 | 0.840994942 | 0.400350768 | 0.625348059 | | Mfn1 | 3182.393317 | -0.0295116775882412 | 0.0580602947270655 | -0.508293623499021 | 0.611247441201255 | 0.894995148736997 | NM_024200 |
| Ppp2cb | 4182.77292 | -0.054445237 | 0.064750159 | -0.840851029 | 0.400431395 | 0.625348059 | | 4930579G24Rik | 63.23709868 | 0.0482297525203042 | 0.0949470823063518 | 0.507964556137579 | 0.611478200122337 | 0.895166660839403 | NM_029482 |
| Slco2a1 | 81.00811144 | 0.125315225 | 0.149102428 | 0.840464016 | 0.400648269 | 0.625581405 | | Ercc5 | 1071.955071 | -0.0302604132719058 | 0.0596000433307968 | -0.507724685768299 | 0.611646433844425 | 0.895166660839403 | NM_011729 |
| Nsmce2 | 859.5167393 | -0.077299928 | 0.092049401 | -0.839765684 | 0.401039777 | 0.626129616 | | Nlgn1 | 2887.615208 | 0.0381272114278142 | 0.0751036741364344 | 0.507661068066414 | 0.611691055727104 | 0.895166660839403 | NM_138666,NM_001163387 |
| P4htm | 2748.021378 | 0.0622595 | 0.074164906 | 0.839473865 | 0.401203449 | 0.626195854 | | Pecam1 | 940.1205918 | 0.0402273925722873 | 0.079230354665947 | 0.507727028888045 | 0.611644790392043 | 0.895166660839403 | NM_001032378,NM_008816 |
| Tgif2 | 50.43011266 | -0.134506822 | 0.160217595 | -0.839525908 | 0.401174257 | 0.626195854 | | Zfhx4 | 547.5503279 | 0.0478730401798017 | 0.0942706126547366 | 0.507825703383676 | 0.61157558237299 | 0.895166660839403 | NM_030708 |
| Ttc32 | 49.19572702 | -0.138188 | 0.164604902 | -0.839513276 | 0.401181342 | 0.626195854 | | Adamtsl3 | 50.51023068 | 0.045476034922859 | 0.0896470519940437 | 0.507278643428012 | 0.611959321297898 | 0.895166660839403 | NM_001190374 |
| Atf1 | 359.4638159 | 0.070719189 | 0.084265513 | 0.839242372 | 0.401333314 | 0.626335452 | | Hrct1 | 27.93839499 | 0.0441062877772381 | 0.0869177735620917 | 0.507448430506907 | 0.611840211594934 | 0.895245243652933 | NM_027511 |
| Far2 | 492.2033448 | -0.087224227 | 0.103945577 | -0.83913361 | 0.401394338 | 0.626367597 | | Phf10 | 972.3977261 | -0.0344028127069272 | 0.067823879356325 | -0.507237466116996 | 0.611988209713814 | 0.895245243652933 | NM_024250 |
| Fam58b | 389.2620919 | -0.069551039 | 0.082896428 | -0.839011289 | 0.401462975 | 0.626411615 | | Prickle2 | 9784.658351 | 0.0339208637699484 | 0.0668743109093378 | 0.507233096067865 | 0.611991275607436 | 0.895245243652933 | NM_001081146,NM_001134461,NM_001134460,NM_001134459 |
| Alg3 | 403.5236729 | 0.068596976 | 0.081785218 | 0.838745398 | 0.401612197 | 0.626453747 | | Adamts8 | 153.9663283 | 0.0457524210578865 | 0.0902817497373073 | 0.506773752070738 | 0.612313575283343 | 0.895344806682315 | NM_013906 |
| Appbp2 | 1715.530223 | -0.061445164 | 0.07324987 | -0.838843322 | 0.401557236 | 0.626453747 | | Ap1b1 | 9858.007086 | 0.0249378379852368 | 0.0492250154024094 | 0.506609043824011 | 0.612429161448126 | 0.895344806682315 | NM_001243043,NM_001243044,NM_007454 |
| Foxf2 | 165.5578192 | 0.091897445 | 0.109574559 | 0.83867501 | 0.401651705 | 0.626453747 | | Gck | 70.06369567 | 0.0313411654787329 | 0.0618515228635287 | 0.506716149057237 | 0.61235399785401 | 0.895344806682315 | NM_010292 |
| Prcp | 359.6245277 | 0.100797836 | 0.120186228 | 0.838680419 | 0.401648669 | 0.626453747 | | Pfkl | 7480.034835 | 0.0386904965485995 | 0.0763185340751215 | 0.50696068808811 | 0.612182402178477 | 0.895344806682315 | NM_008826 |
| Nuak1 | 3820.564498 | 0.088958331 | 0.106084138 | 0.838563921 | 0.401714064 | 0.626466868 | | Tfeb | 325.0689475 | -0.0429750740004989 | 0.0848220669828817 | -0.506649690689239 | 0.612400636082704 | 0.895344806682315 | NM_011549,NM_001161722,NM_001161723 |
| Twistnb | 317.1388932 | -0.089103327 | 0.106263125 | -0.838515968 | 0.401740984 | 0.626466868 | | Trnt1 | 1275.858953 | 0.0361668055343239 | 0.0713624851668539 | 0.506804176588884 | 0.61229222553566 | 0.895344806682315 | NM_001242358,NM_027296,NM_001242360 |
| Plscr3 | 556.5837572 | 0.100077574 | 0.119369532 | 0.838384569 | 0.401814753 | 0.62651847 | | Ahctf1 | 2659.381714 | -0.0344742774464634 | 0.0680623924930474 | -0.506509926902511 | 0.612498722691869 | 0.89535638983819 | NM_026375 |
| 2310069G16Rik | 78.5563848 | -0.127001029 | 0.151529161 | -0.838129292 | 0.401958093 | 0.626679281 | | 2010300C02Rik | 11894.21745 | 0.0319565946187671 | 0.0631451503545436 | 0.506081534992618 | 0.612799412565069 | 0.895428201148793 | NM_028096 |
| Zfp932 | 1375.390122 | 0.112269897 | 0.134011239 | 0.837764786 | 0.402162819 | 0.626935378 | | 4930455F23Rik | 972.9030029 | 0.0331854406051878 | 0.0656092582382936 | 0.505804233979568 | 0.61299408593919 | 0.895428201148793 | NM_029115 |
| Frs2 | 2386.398179 | 0.057258711 | 0.068357235 | 0.837639367 | 0.402233276 | 0.626982131 | | Fam69a | 1070.903349 | 0.0404632009937085 | 0.0799354003155228 | 0.506198765928378 | 0.612717121254453 | 0.895428201148793 | NM_026062 |
| Rbx1 | 3255.55558 | -0.04023311 | 0.04805151 | -0.837291289 | 0.402428855 | 0.627223888 | | Isy1 | 677.3220545 | 0.0327926659223752 | 0.0648476646683259 | 0.505687692688684 | 0.613075909464327 | 0.895428201148793 | NM_133934 |
| Prr11 | 32.43841625 | -0.137223594 | 0.163922488 | -0.837124884 | 0.402522374 | 0.627306545 | | Mdga1 | 1152.557439 | 0.0424567993346949 | 0.0839648582834735 | 0.505649627744938 | 0.613102635868504 | 0.895428201148793 | NM_001081160 |
| Rnasek | 3555.760641 | 0.052844543 | 0.063153225 | 0.83676713 | 0.402723477 | 0.627556829 | | Nr2c2ap | 536.9398693 | 0.0375173889196713 | 0.0741618437710759 | 0.505885331485023 | 0.612937150306607 | 0.895428201148793 | NM_001025587,NM_001025586 |
| Pald1 | 544.845052 | 0.075867341 | 0.090701416 | 0.836451564 | 0.402900915 | 0.62777019 | | Nyap2 | 608.9440898 | -0.0420119725021183 | 0.0830602066436535 | -0.50580144451553 | 0.612996044362797 | 0.895428201148793 | NM_172849 |
| Jak1 | 7445.739883 | -0.045586735 | 0.054521185 | -0.836067509 | 0.403116129 | 0.628043604 | | Srpx | 23.12686749 | 0.0431206852510018 | 0.0851783460591433 | 0.50623999227528 | 0.612688183209686 | 0.895428201148793 | NM_016911 |
| Sh3yl1 | 271.9032342 | 0.086129093 | 0.103029299 | 0.835966995 | 0.403173471 | 0.628068546 | | Zc3h14 | 3640.903968 | 0.0328615538277682 | 0.0649524130698562 | 0.505932763305124 | 0.612903851219621 | 0.895428201148793 | NM_001008506,NM_001160107,NR_027648,NM_029334,NM_001160108 |
| 1810033B17Rik | 10.64209597 | 0.136616931 | 0.163451521 | 0.835825387 | 0.403253142 | 0.628129505 | | C030037D09Rik | 172.7147809 | 0.0460619993300892 | 0.091141201478816 | 0.505391618529358 | 0.613283804524532 | 0.895602749022153 | NR_038058 |
| Lrig1 | 1592.887045 | 0.092431273 | 0.110600655 | 0.835720852 | 0.403311962 | 0.628157975 | | BC024978 | 315.0953449 | -0.0388467131163159 | 0.0769508285560358 | -0.5048251441247 | 0.613681653827117 | 0.895819717368722 | NM_001243888 |
| Arhgap42 | 617.572874 | 0.078470543 | 0.093936623 | 0.835356222 | 0.403517171 | 0.628351261 | | Coasy | 988.5220244 | 0.0385754409091229 | 0.0764275095849607 | 0.504732407461747 | 0.613746795972635 | 0.895819717368722 | NM_027896 |
| Suv39h2 | 159.0044346 | -0.092478045 | 0.110703183 | -0.835369343 | 0.403509785 | 0.628351261 | | Gprc5b | 8119.836315 | 0.0326267262223431 | 0.0646517701952789 | 0.504653254254214 | 0.613802398937828 | 0.895819717368722 | NM_001195774,NM_022420 |
| Fbxo8 | 669.6041852 | -0.05545176 | 0.066413896 | -0.834942138 | 0.403750288 | 0.628651086 | | Rasgef1a | 12153.44476 | -0.0367361509988231 | 0.072766980928735 | -0.504846436253842 | 0.613666697767543 | 0.895819717368722 | NM_027526 |
| Ccnf | 236.3197071 | 0.081991426 | 0.098298192 | 0.834109199 | 0.404219452 | 0.629203086 | | Satb1 | 5160.266937 | 0.0358366535308289 | 0.0709602173986133 | 0.505024573551112 | 0.613541576513401 | 0.895819717368722 | NM_009122,NM_001163630,NM_001163631,NM_001163632 |
| Fundc1 | 1211.453258 | -0.056799443 | 0.068096972 | -0.834096447 | 0.404226637 | 0.629203086 | | Slc22a8 | 777.7109672 | -0.0445836099478352 | 0.0883164475688719 | -0.504816613157674 | 0.613687646211001 | 0.895819717368722 | NM_001164635,NM_001164634,NM_031194 |
| Nlrp1c-ps | 12.858969 | -0.129481885 | 0.155222695 | -0.834168516 | 0.40418603 | 0.629203086 | | Kif1b | 38937.04614 | 0.0311043470928675 | 0.0616542002586016 | 0.504496805771607 | 0.613912306254762 | 0.89589011057431 | NM_008441,NM_207682 |
| Ciapin1 | 2196.596086 | 0.058225964 | 0.069826236 | 0.833869439 | 0.40435456 | 0.629275769 | | Scn5a | 470.6619929 | 0.0477300338704136 | 0.0946442384889728 | 0.504309978424885 | 0.614043566527396 | 0.895991647149871 | NM_001253860,NM_021544 |
| Terf2 | 1601.221744 | 0.062927132 | 0.075462416 | 0.833887048 | 0.404344637 | 0.629275769 | | Gfod1 | 6975.219711 | 0.0436532390295678 | 0.0865925347214993 | 0.504122430068206 | 0.614175345803256 | 0.896093920564292 | NM_001033399 |
| Supt3 | 326.066491 | -0.073586553 | 0.088258128 | -0.833765161 | 0.404413331 | 0.629304022 | | Ccdc135 | 119.0861456 | 0.0266837299671802 | 0.0529745736854354 | 0.503708253050396 | 0.614466407938625 | 0.89611763962382 | NM_001042715 |
| Rnf32 | 289.0511987 | -0.093731207 | 0.112436647 | -0.833635733 | 0.404486283 | 0.629354336 | | Cygb | 2590.194946 | 0.0261713934619451 | 0.0519647769157268 | 0.50363717531951 | 0.614516363789111 | 0.89611763962382 | NM_030206 |
| Gm1653 | 16.33567347 | 0.139972077 | 0.167936053 | 0.833484382 | 0.404571603 | 0.629423879 | | Esam | 680.2306681 | -0.0374564423473432 | 0.0743351281083671 | -0.503886161233738 | 0.614341375848333 | 0.89611763962382 | NM_027102 |
| Prkar2a | 2727.27992 | 0.083240697 | 0.099901491 | 0.83322777 | 0.404716283 | 0.629585753 | | Ier5 | 3092.861158 | 0.0477387420814204 | 0.0948143459130406 | 0.503497035408589 | 0.614614864131012 | 0.89611763962382 | NM_010500 |
| Acat3 | 148.3868612 | 0.120445676 | 0.144591962 | 0.833003955 | 0.404842499 | 0.629676416 | | Plekhm1 | 1531.579563 | -0.0340203233032711 | 0.0675049783247081 | -0.503967620575015 | 0.614284130779363 | 0.89611763962382 | NM_183034 |
| Adhfe1 | 683.4762905 | -0.102481473 | 0.123030262 | -0.832977766 | 0.404857269 | 0.629676416 | | Sardh | 580.0463207 | -0.0387519463940603 | 0.0769150422971905 | -0.503827928018649 | 0.614382300335186 | 0.89611763962382 | NM_138665 |
| Pcyox1l | 549.8555373 | -0.088794779 | 0.106608117 | -0.832908235 | 0.404896485 | 0.629676416 | | Tmem214 | 2525.032949 | -0.0305340637240224 | 0.0606454481805273 | -0.503484839177537 | 0.61462343684236 | 0.89611763962382 | NM_144525 |
| Wdr25 | 276.6885318 | -0.074841217 | 0.089868059 | -0.832789953 | 0.404963202 | 0.629716966 | | Cachd1 | 622.7228952 | -0.0447446410098351 | 0.0889086829191925 | -0.503265142848902 | 0.614777870055197 | 0.896163393343238 | NM_198037 |
| Fgf14 | 1626.235692 | -0.071056444 | 0.085356198 | -0.832469646 | 0.405143904 | 0.62986703 | | Rpl13a | 8693.027279 | 0.0311407285313794 | 0.0618774387812508 | 0.503264665518367 | 0.614778205608245 | 0.896163393343238 | NM_009438 |
| Gm16907 | 9.2819162 | -0.130223704 | 0.156455542 | -0.832336792 | 0.405218869 | 0.62986703 | | Sh3pxd2a | 2480.92748 | -0.0338794581558444 | 0.0673380471475069 | -0.50312504729503 | 0.614876357661146 | 0.896216533351978 | NM_008018,NM_001164717 |
| Gm6682 | 25.81745905 | 0.138428811 | 0.166283737 | 0.832485571 | 0.405134919 | 0.62986703 | | Zfp282 | 722.8025257 | -0.0329285050747748 | 0.0654619898342884 | -0.503017173143232 | 0.614952198253051 | 0.896237145763402 | NM_146175 |
| Gpr155 | 2826.171408 | 0.066526168 | 0.079927569 | 0.832330684 | 0.405222316 | 0.62986703 | | Nudcd1 | 592.4015767 | -0.0350696956401199 | 0.0697849853646321 | -0.502539270544773 | 0.615288235703318 | 0.896636929356581 | NM_001113554,NM_026149 |
| Tra2b | 1924.557411 | -0.042586383 | 0.051171671 | -0.832225762 | 0.405281525 | 0.629895872 | | Itgbl1 | 58.09859479 | 0.0460586279339104 | 0.0916847682844736 | 0.502358557432383 | 0.615415325252561 | 0.896719391231212 | NM_145467 |
| Wdsub1 | 350.394992 | 0.071328449 | 0.085714873 | 0.832052984 | 0.405379038 | 0.629984235 | | Pdk2 | 5268.65198 | 0.0216739717038588 | 0.0431508944405762 | 0.50228325472387 | 0.615468286560318 | 0.896719391231212 | NM_133667 |
| Kdm2a | 4287.285582 | 0.05439588 | 0.065389213 | 0.831878496 | 0.405477531 | 0.630019591 | | Actb | 39092.0296 | 0.0260088460037848 | 0.0530228635596045 | 0.501837211675169 | 0.615782035111093 | 0.896906632109068 | NM_007393 |
| Lonrf3 | 1019.693794 | 0.110772883 | 0.133161514 | 0.831868607 | 0.405483113 | 0.630019591 | | Camk2n1 | 64376.91164 | -0.0398792269823055 | 0.0794657423178595 | -0.501841747388331 | 0.615778844317872 | 0.896906632109068 | NM_025451 |
| Anxa7 | 3630.638085 | 0.04954536 | 0.059572917 | 0.831675904 | 0.405591905 | 0.63006226 | | Gm3086 | 36.31725815 | -0.0440455736121055 | 0.0877643852698297 | -0.501861586299367 | 0.615764880883558 | 0.896906632109068 | NR_036607 |
| Ptcra | 8.433840029 | -0.133940647 | 0.161047886 | -0.831682119 | 0.405588396 | 0.63006226 | | Farp2 | 383.1558713 | -0.0381228423066706 | 0.0760154061557437 | -0.501514682807362 | 0.616008947027992 | 0.89714717911857 | NM_145519 |
| Has3 | 91.8190172 | -0.113709155 | 0.136746673 | -0.831531421 | 0.405673486 | 0.630125813 | | Cnbd2 | 193.1065437 | -0.0429879965748393 | 0.0857558930956385 | -0.501283294046012 | 0.616171760860856 | 0.897204390363351 | NM_027585 |
| A030009H04Rik | 4192.115389 | -0.048413281 | 0.058262174 | -0.830955626 | 0.405998697 | 0.630504542 | | Gmpr | 1633.466118 | 0.0320497097829964 | 0.0639346275002233 | 0.501288754405968 | 0.616167918528671 | 0.897204390363351 | NM_025508 |
| Zcchc4 | 198.293236 | 0.085829196 | 0.103287244 | 0.830975751 | 0.405987328 | 0.630504542 | | 4932441J04Rik | 29.20089624 | 0.0449798021276432 | 0.0898712398543018 | 0.500491616679195 | 0.616728957633622 | 0.897296157710936 | NR_015588 |
| Crispld1 | 205.1665269 | -0.10369086 | 0.124798171 | -0.830868433 | 0.406047958 | 0.630517846 | | Apold1 | 234.8755958 | 0.0373660237232423 | 0.0746219893399543 | 0.500737437500017 | 0.616555920867474 | 0.897296157710936 | NM_001109914 |
| Pygb | 8233.664118 | -0.060086542 | 0.072340785 | -0.830603964 | 0.406197395 | 0.630686686 | | Elovl2 | 701.7164468 | 0.036836498147479 | 0.0736005865865096 | 0.500491909859735 | 0.616728751246994 | 0.897296157710936 | NM_019423 |
| Ccny | 4798.18846 | -0.058392179 | 0.070352312 | -0.829996595 | 0.406540709 | 0.631050988 | | Nkx6-2 | 320.7178134 | 0.047222036598789 | 0.0942883833144405 | 0.500825604797031 | 0.616493863844158 | 0.897296157710936 | NR_027857,NM_183248 |
| Ibtk | 1130.100054 | -0.062412295 | 0.075198008 | -0.829972712 | 0.406554212 | 0.631050988 | | Slc35b1 | 1551.572583 | 0.02755682651319347 | 0.0550437698232257 | 0.500842606174519 | 0.616481897648856 | 0.897296157710936 | NM_016752 |
| Pkhd1l1 | 22.88857945 | -0.136599592 | 0.164557592 | -0.830102039 | 0.406481094 | 0.631050988 | | Slc7a4 | 2869.482937 | 0.0380856481481648 | 0.0760787458073123 | 0.500608254565945 | 0.616646851914186 | 0.897296157710936 | NM_144852 |
| Ciao1 | 1976.322657 | 0.054739074 | 0.065996602 | 0.829422606 | 0.406865314 | 0.631280911 | | Stk11 | 2285.012166 | 0.02782739264169 | 0.055582558438273 | 0.500649725805507 | 0.616617659931127 | 0.897296157710936 | NM_011492 |
| Mtx1 | 837.147316 | -0.055769515 | 0.067236752 | -0.829449865 | 0.406849895 | 0.631280911 | | Zfp414 | 965.478051 | -0.0429100541399153 | 0.0856687368116772 | -0.500883469710112 | 0.616453136805079 | 0.897296157710936 | NR_034153,NM_026712 |
| Sec23ip | 1721.098748 | 0.055115871 | 0.066447725 | 0.829462126 | 0.40684296 | 0.631280911 | | Bivm | 938.9893374 | -0.0327886358961642 | 0.0655308339814433 | -0.500354320310483 | 0.616825611763758 | 0.89734690400404 | NM_144558 |
| Serinc5 | 3207.766111 | 0.081253667 | 0.097950585 | 0.829537335 | 0.40680042 | 0.631280911 | | Ptov1 | 5068.583792 | 0.0291013983112483 | 0.0581819679627465 | 0.500178995833927 | 0.616949046638956 | 0.897436596382684 | NM_133949 |
| Cdk2ap1 | 2855.375536 | 0.069799049 | 0.084194564 | 0.829020855 | 0.407092606 | 0.631443872 | | Ddn | 42259.69258 | -0.037096112320821 | 0.0742077153485689 | -0.499895626035283 | 0.617148572277548 | 0.89762105097879 | NM_001013741 |
| Fbxw9 | 509.4931913 | 0.091271662 | 0.110091067 | 0.829056018 | 0.40707271 | 0.631443872 | | Zer1 | 8544.117056 | 0.0260981157267153 | 0.052214674717014 | 0.499823390036581 | 0.6171994394311 | 0.89762105097879 | NM_178694 |
| Slirp | 588.2824161 | -0.078975431 | 0.095248099 | -0.829154927 | 0.407016746 | 0.631443872 | | Arhgap20 | 4311.892568 | -0.0389004916163944 | 0.0778919882498102 | -0.499415825561356 | 0.617486472636547 | 0.897948593604813 | NM_175535 |
| Chid1 | 1987.555034 | -0.088688777 | 0.107017514 | -0.828731428 | 0.407256398 | 0.631561428 | | AK129341 | 779.6873915 | 0.0426008877943169 | 0.0853402942186729 | 0.499188433603919 | 0.617646642123919 | 0.898001713891708 | NM_001045524 |
| Hnrnph2 | 4573.004573 | 0.048013597 | 0.057942722 | 0.828638963 | 0.407308734 | 0.631561428 | | Wdr8 | 817.9945723 | 0.0364488909390226 | 0.0730038109709855 | 0.499273811246769 | 0.617586502010498 | 0.898001713891708 | NM_021499 |
| Tmem63c | 1999.589695 | 0.058400106 | 0.070468729 | 0.8287379 | 0.407252735 | 0.631561428 | | Evi5 | 3050.968603 | -0.0298599038881976 | 0.0598487830603463 | -0.498922490338517 | 0.61783398930262 | 0.898138096331922 | NM_007964 |
| Urod | 1326.414156 | 0.06802593 | 0.082097544 | 0.828598846 | 0.407331441 | 0.631561428 | | Gbf1 | 4642.931356 | 0.0246692462882761 | 0.0494492843865847 | 0.498879743039693 | 0.617864105510457 | 0.898138096331922 | NM_178930 |
| Fgl2 | 88.05299186 | -0.137535334 | 0.166032383 | -0.828364513 | 0.407464098 | 0.631703895 | | Cox11 | 547.5801596 | -0.0396059707764808 | 0.079404020520441 | -0.498790459588388 | 0.617926983685606 | 0.898139620273876 | NM_199008 |
| Zscan22 | 383.1638721 | -0.067642695 | 0.081696674 | -0.827973675 | 0.407685409 | 0.631983764 | | Zfp516 | 793.040433 | 0.0322116341803832 | 0.0645923771096004 | 0.498690954285928 | 0.617997118118259 | 0.898151689623395 | NM_001177464,NM_183033 |
| Arhgap6 | 512.0272363 | 0.138490896 | 0.167326705 | 0.827667619 | 0.407858762 | 0.631999566 | | Atrn | 11077.82109 | 0.0335605371348318 | 0.0673267119473023 | 0.498472837365058 | 0.618150809710358 | 0.898285180257368 | NM_009730 |
| Phf8 | 1338.729295 | -0.068989141 | 0.083349794 | -0.827706198 | 0.407836908 | 0.631999566 | | Rab1b | 5925.596634 | 0.0248352626282236 | 0.0498343180066005 | 0.498356626952018 | 0.61823270180632 | 0.898314316938942 | NM_029576 |
| Taf1 | 3466.955789 | -0.060961701 | 0.073653559 | -0.827681669 | 0.407850803 | 0.631999566 | | Cebpd | 72.12625372 | 0.0463693910644176 | 0.0930885775514489 | 0.49812116893493 | 0.618398640809827 | 0.898465558752106 | NM_007679 |
| Tph2 | 37.55128812 | -0.138969695 | 0.167881074 | -0.827786553 | 0.407791391 | 0.631999566 | | Asah1 | 2196.0434 | -0.0245460336890316 | 0.04928876644207 | -0.498004625818359 | 0.618480781755871 | 0.898495033081886 | NM_019734 |
| Gm17769 | 50.18222238 | -0.123454455 | 0.149194745 | -0.827471871 | 0.407969659 | 0.63206613 | | Dapk1 | 5507.924766 | 0.0328479897236284 | 0.0660161942676344 | 0.497574725232725 | 0.618783821918541 | 0.89857580697678 | NM_134062,NM_029653 |
| Prkra | 457.9782439 | -0.075604401 | 0.0913706 | -0.827447782 | 0.407983307 | 0.63206613 | | Krtcap3 | 25.75287548 | -0.042665352972492 | 0.0857409304705499 | -0.497607767239552 | 0.618760528058477 | 0.89857580697678 | NM_027221 |
| 4933404O12Rik | 655.7883706 | 0.063711143 | 0.077029018 | 0.827105733 | 0.408177135 | 0.632239981 | | Zfp319 | 1013.948839 | 0.030546250968755 | 0.0613700449592315 | 0.497738774495719 | 0.618668174705173 | 0.89857580697678 | NM_024467 |
| St3gal3 | 1930.834518 | 0.050444638 | 0.060984619 | 0.827169836 | 0.408140806 | 0.632239981 | | Zfp365 | 23209.16665 | 0.0313218167223023 | 0.0629178891692898 | 0.497820526655406 | 0.618610546703801 | 0.89857580697678 | NM_178679 |
| Tmed10 | 5675.602002 | -0.054685228 | 0.066124616 | -0.827002575 | 0.408235602 | 0.632267335 | | Fam174a | 862.0156519 | 0.0350510118836983 | 0.0704967536461593 | 0.497200368397501 | 0.619047761680977 | 0.898695176608308 | NM_026321 |
| 2410016O06Rik | 483.3300907 | -0.064512124 | 0.078022959 | -0.826835142 | 0.408330509 | 0.632351115 | | Polb | 1348.761451 | -0.0338596728036062 | 0.0681014100111223 | -0.497194886245031 | 0.619051627229983 | 0.898695176608308 | NM_011130 |
| Naa16 | 187.9179767 | 0.091631645 | 0.110848847 | 0.826635982 | 0.408443416 | 0.632395817 | | Xrcc4 | 307.0890215 | 0.0389294436074446 | 0.0782733143925951 | 0.497352691776745 | 0.61894036036566 | 0.898695176608308 | NM_028012 |
| Snora61 | 22.18609337 | 0.138771248 | 0.167888433 | 0.826568247 | 0.408481821 | 0.632395817 | | Trp73 | 50.25457952 | -0.0445901764309979 | 0.0897392603426092 | -0.496885936665404 | 0.619269489298628 | 0.89892161502467 | NM_011642,NM_001126330,NM_001126331 |
| Tars2 | 1138.734632 | 0.081558311 | 0.09865561 | 0.826697146 | 0.40840874 | 0.632395817 | | Slk | 4382.972673 | -0.0340527383085846 | 0.0685845234405148 | -0.496507617175754 | 0.619536314521509 | 0.899129234163197 | NM_001164639,NM_009289 |
| Cox10 | 1018.529404 | -0.055061133 | 0.066623721 | -0.826449379 | 0.408549223 | 0.632436972 | | Zfp760 | 380.6097328 | -0.0412172833514879 | 0.0830109024385177 | -0.496525853969349 | 0.619521546684913 | 0.899129234163197 | NM_001008501 |
| 2510003E04Rik | 6279.516197 | 0.054624626 | 0.066103435 | 0.826350795 | 0.408605127 | 0.632460323 | | Anxa2 | 434.1638064 | -0.0424897041414083 | 0.0857014236056786 | -0.495787611847709 | 0.620044266044287 | 0.899216176925702 | NM_007585 |
| 2610020C07Rik | 21.68067674 | -0.13764763 | 0.166595625 | -0.826237962 | 0.408669117 | 0.632496185 | | Capn5 | 2665.903296 | 0.0292395587089329 | 0.0589560474969672 | 0.495955206468633 | 0.619926014686214 | 0.899216176925702 | NM_007602 |
| Srl | 333.4644312 | 0.077880545 | 0.094311529 | 0.825779685 | 0.40892908 | 0.632835314 | | Elp4 | 544.531901 | 0.0334445143864307 | 0.0674689313152286 | 0.495702447548355 | 0.620104360008309 | 0.899216176925702 | NM_023876 |
| Gm16793 | 21.22041269 | 0.138792206 | 0.168119772 | 0.825555522 | 0.409056275 | 0.632908847 | | Fap | 125.5701645 | -0.0471394883011578 | 0.0950770424011199 | -0.495803057296222 | 0.62003336763866 | 0.899216176925702 | NM_007986 |
| Plgrkt | 338.2347216 | 0.09266329 | 0.11224466 | 0.825547423 | 0.409060871 | 0.632908847 | | Manbal | 1492.350986 | 0.029760426714746 | 0.0600348324619046 | 0.495719326503171 | 0.620092449612998 | 0.899216176925702 | NM_026968 |
| Rwdd2b | 340.5223163 | -0.077898276 | 0.094367723 | -0.825479984 | 0.409099142 | 0.632908847 | | Rgs8 | 6027.649285 | 0.0306446133584975 | 0.0618292417433337 | 0.495633012704729 | 0.620153356770046 | 0.899216176925702 | NM_026380 |
| Arhgef2 | 6543.23415 | 0.052414169 | 0.063518036 | 0.825185602 | 0.409266226 | 0.633104124 | | Rpl18a | 7142.635575 | 0.0272820531253729 | 0.0549792894140682 | 0.496224185800261 | 0.619736428883546 | 0.899216176925702 | NM_029751 |
| Eepd1 | 809.3980271 | -0.05504483 | 0.066767997 | -0.824419368 | 0.409701314 | 0.633656044 | | Rprm | 2076.397932 | 0.0298711402800853 | 0.0602526595497527 | 0.49576467666825 | 0.620060449405155 | 0.899216176925702 | NM_023396 |
| H2-Eb1 | 98.66044544 | 0.105210809 | 0.127616839 | 0.824427324 | 0.409696795 | 0.633656044 | | Slco5a1 | 163.4932876 | -0.044902472375438 | 0.0905091860306474 | -0.496109559091975 | 0.619817115306817 | 0.899216176925702 | NM_172841 |
| H2-Aa | 119.954637 | 0.095744281 | 0.116245891 | 0.823635833 | 0.410146509 | 0.63423556 | | Hist3h2ba | 263.7124902 | -0.042730771184349 | 0.0863043061090124 | -0.495117487305617 | 0.620517190469247 | 0.899412757063786 | NM_030082 |
| Rapsn | 26.77104769 | 0.137175592 | 0.166554128 | 0.823609681 | 0.410161373 | 0.63423556 | | Inhba | 148.7563255 | -0.0470514858157526 | 0.0950084993176513 | -0.495234491163161 | 0.620434606472193 | 0.899412757063786 | NM_008380 |
| Hira | 2749.59249 | 0.090775602 | 0.110247198 | 0.823382393 | 0.410290572 | 0.634372031 | | Lrig3 | 116.7363399 | -0.0468865450597 | 0.0947278224229689 | -0.494960655279787 | 0.620627893602016 | 0.899412757063786 | NM_177152 |
| Cmtm6 | 1148.603129 | -0.076090166 | 0.092424224 | -0.823270811 | 0.410354008 | 0.634393404 | | Nup88 | 2331.248676 | -0.0279248666860977 | 0.0564398793868453 | -0.494771905777784 | 0.620761137741159 | 0.899412757063786 | NR_075087,NM_001083331,NM_001276406,NM_172394 |
| G6pc3 | 1571.324411 | 0.079214362 | 0.096225718 | 0.823214044 | 0.410386284 | 0.634393404 | | Syn3 | 1987.677941 | 0.0332409202062963 | 0.0671677395166749 | 0.49489413289015 | 0.620674852403906 | 0.899412757063786 | NM_013722,NM_001164495 |
| D3Ertd751e | 525.1926616 | 0.102338956 | 0.124337611 | 0.823073201 | 0.410466367 | 0.6344539 | | Synpo | 11437.95751 | 0.0391126178321079 | 0.0790190768324294 | 0.494976901780964 | 0.620616425273719 | 0.899412757063786 | NM_177340,NM_001109975 |
| Ints12 | 465.0077612 | -0.075047122 | 0.091204132 | -0.822847899 | 0.410594494 | 0.634502692 | | Tmem132a | 10450.60301 | 0.0394071032740501 | 0.0796522856156135 | 0.494739139496094 | 0.620784269745388 | 0.899412757063786 | NM_133804 |
| Prkdc | 1137.202951 | -0.084613713 | 0.102836098 | -0.822801668 | 0.410620788 | 0.634502692 | | Tnr | 3379.038304 | 0.0282296673945216 | 0.0570360504626115 | 0.494944288139777 | 0.620639447181947 | 0.899412757063786 | NM_022312 |
| Soga2 | 3993.213776 | 0.060975996 | 0.074099546 | 0.82289297 | 0.410568861 | 0.634502692 | | Bzw2 | 1307.094933 | 0.0288576128994422 | 0.0583405335757749 | 0.494640719975785 | 0.620853759559938 | 0.899423727067667 | NM_025840 |
| Palm2 | 2545.371537 | 0.072461144 | 0.08808389 | 0.822637871 | 0.410713956 | 0.63458337 | | 3110052M02Rik | 539.4399635 | -0.0388648742588673 | 0.078681205630932 | -0.493953720551384 | 0.621338863956096 | 0.899558901531836 | NM_001166497 |
| Dynlrb2 | 31.81926436 | 0.13701994 | 0.166564717 | 0.822453982 | 0.410818567 | 0.634618433 | | Azi2 | 2415.718208 | 0.0302704188399093 | 0.0613038646794485 | 0.493776681098168 | 0.621463903474595 | 0.899558901531836 | NM_001048146,NM_013727 |
| Zdhhc13 | 801.9356219 | -0.070044432 | 0.085160933 | -0.822494886 | 0.410795296 | 0.634618433 | | BC031361 | 259.2318633 | 0.0406671727215211 | 0.082334997655654 | 0.493923287538085 | 0.621360357422643 | 0.899558901531836 | NR_033221 |
| Parp2 | 605.9660255 | 0.076559146 | 0.093112825 | 0.822219137 | 0.410952189 | 0.634761561 | | Ccdc36 | 29.77106284 | 0.0441085250888468 | 0.0893026345385066 | 0.493921879424818 | 0.62136135191744 | 0.899558901531836 | NM_001135198 |
| Cbfb | 631.8414927 | -0.080026391 | 0.097342158 | -0.822114409 | 0.411011785 | 0.634790331 | | Elmod3 | 945.246657 | 0.0294480191024939 | 0.0595818870577959 | 0.494244485306895 | 0.621133526198191 | 0.899558901531836 | NM_144917,NM_001253692 |
| Nae1 | 1356.568928 | 0.05116393 | 0.062259943 | 0.821779271 | 0.411202534 | 0.635021634 | | Krt18 | 42.28911412 | 0.0218063265395842 | 0.0441500493385408 | 0.493913987286589 | 0.621366925835831 | 0.899558901531836 | NM_010664 |
| Wdr92 | 561.8195381 | 0.095752398 | 0.116530929 | 0.821690848 | 0.411252869 | 0.635036073 | | L3mbtl4 | 30.56816909 | -0.0418381064447724 | 0.084686083496838 | -0.494037564582079 | 0.621279653064982 | 0.899558901531836 | NM_177278 |
| Kctd13 | 4420.710679 | 0.066249705 | 0.080633958 | 0.821610479 | 0.411298623 | 0.635043435 | | Ranbp3l | 167.6799781 | -0.04554864877937 | 0.0922330586653195 | -0.49384298253243 | 0.621417074782122 | 0.899558901531836 | NM_198024 |
| Kank3 | 734.7644958 | -0.093258285 | 0.113529925 | -0.821442322 | 0.411394365 | 0.635152969 | | Mr1 | 277.5835903 | -0.0461186519982923 | 0.0934831349626812 | -0.493336600411433 | 0.621774771246635 | 0.899950211949842 | NM_008209 |
| Brwd1 | 4632.877631 | -0.06796798 | 0.082766808 | -0.821198522 | 0.411533199 | 0.635152443 | | Htra4 | 46.59015206 | -0.0464239773748498 | 0.09414616452013405 | -0.493129021439587 | 0.621921425998489 | 0.900001855385281 | NM_001081187 |
| Eif2b2 | 1077.268774 | 0.070060379 | 0.085313779 | 0.821208248 | 0.411527659 | 0.635152443 | | Mapk8 | 4551.62203 | 0.0329988695725707 | 0.0669330300256501 | 0.493013233674388 | 0.62200323669371 | 0.900001855385281 | NM_016700 |
| Il1f9 | 50.38023512 | -0.121062945 | 0.147398691 | -0.82132985 | 0.411458409 | 0.635152443 | | Ubxn10 | 94.48964215 | 0.0430883765501515 | 0.0873869259076683 | 0.493075778814763 | 0.621959044385714 | 0.900001855385281 | NM_178671 |
| Gadl1 | 27.09782797 | 0.137294218 | 0.167249951 | 0.820892429 | 0.411707544 | 0.635358236 | | 4833422C13Rik | 135.6032684 | -0.046368089125352 | 0.0941224041940312 | -0.492636046883857 | 0.622269773171149 | 0.900114632385487 | NR_015501,NR_028313,NR_028314 |
| Gm7102 | 28.89618421 | -0.134903998 | 0.164490078 | -0.820089693 | 0.412164978 | 0.635937479 | | Cog1 | 2203.287363 | -0.027893605588942 | 0.0566470274670536 | -0.492410755448115 | 0.62242899475992 | 0.900114632385487 | NM_013581 |
| Uba1 | 14571.01628 | 0.050917348 | 0.062159994 | 0.820114145 | 0.412151039 | 0.635937479 | | F8 | 120.30421 | -0.0458488680253639 | 0.0930703138193606 | -0.492626124742113 | 0.622276785253796 | 0.900114632385487 | NM_001161373,NM_007977,NM_001161374 |
| Med16 | 2238.272062 | -0.064206655 | 0.078299744 | -0.820011051 | 0.412209807 | 0.635943319 | | Nudt16 | 465.4821431 | -0.0368154036157194 | 0.0747693024767642 | -0.492386613171366 | 0.622446061015573 | 0.900114632385487 | NM_029385 |
| Ergic3 | 3016.973837 | -0.046143681 | 0.05629643 | -0.819655547 | 0.412412498 | 0.636114585 | | Orai1 | 205.9631901 | 0.0429318081230589 | 0.0871911383566649 | 0.49238728765579 | 0.622445584292971 | 0.900114632385487 | NM_175423 |
| Gm10532 | 9.209810764 | 0.122255733 | 0.149165061 | 0.819600329 | 0.412443986 | 0.636114585 | | Ube2t | 35.32875421 | -0.0412142168923538 | 0.0836769949464965 | -0.492539400679486 | 0.622338075526218 | 0.900114632385487 | NM_026024 |
| Htra4 | 49.21213292 | -0.122366267 | 0.150878911 | -0.819639185 | 0.412421828 | 0.636114585 | | Ccdc85c | 548.6124088 | 0.0389262912763988 | 0.0790749523703871 | 0.492270815340717 | 0.622527908755974 | 0.900143380707673 | NM_001159910 |
| Impa2 | 124.6967514 | 0.107657748 | 0.131404435 | 0.819285501 | 0.412623543 | 0.636328175 | | Hivep2 | 12754.31193 | 0.0382500312926048 | 0.0777414367444893 | 0.492016007091818 | 0.622708027689389 | 0.900224602927554 | NM_010437 |
| Lpcat3 | 610.468335 | -0.060381652 | 0.073733801 | -0.818914136 | 0.412835404 | 0.636528187 | | Ttbk1 | 5535.607757 | 0.0327387960675976 | 0.0665386395397086 | 0.492026832740694 | 0.622700374791976 | 0.900224602927554 | NM_001162864 |
| Myg1 | 842.258642 | -0.052160826 | 0.063691047 | -0.818966384 | 0.412805593 | 0.636528187 | | L3mbtl2 | 2120.13473 | 0.0296754209110261 | 0.0603319983355399 | 0.491868688750943 | 0.622812174426586 | 0.900285565302079 | NM_145993 |
| Nup93 | 1033.578815 | 0.090402395 | 0.110434653 | 0.818605326 | 0.413011628 | 0.636706509 | | Dzip3 | 2684.340463 | 0.0289033661956152 | 0.0587855594465092 | 0.491674596070065 | 0.622949399816176 | 0.900360726037068 | NM_001110017,NM_027341 |
| Paics | 3663.105181 | -0.042766144 | 0.052245107 | -0.818567446 | 0.413033247 | 0.636706509 | | Sptbn2 | 47246.85343 | -0.0326539467563035 | 0.0664211368394915 | -0.491619811253949 | 0.622988135577487 | 0.900360726037068 | NM_021287 |
| Bcl2l11 | 256.2857164 | 0.083086951 | 0.101556976 | 0.818131399 | 0.413282161 | 0.637026841 | | Sec14l2 | 2075.243697 | 0.0300852770937641 | 0.0612194854996833 | 0.491433027380143 | 0.623120209492649 | 0.900462013837069 | NM_144520 |
| Gm15760 | 10.72703723 | 0.126436118 | 0.154631334 | 0.817661691 | 0.413550391 | 0.637376878 | | Def6 | 72.95029853 | -0.0467151878524783 | 0.0950947255819289 | -0.491248989537604 | 0.623250353561066 | 0.900470920611502 | NM_027185 |
| Cops7a | 4926.393974 | -0.044115589 | 0.053976321 | -0.817313737 | 0.413749158 | 0.637388731 | | Fam84b | 389.9520169 | 0.0460979306862455 | 0.0938279201308759 | 0.49130291518714 | 0.623212218318589 | 0.900470920611502 | NM_001162926 |
| Derl1 | 2554.152324 | -0.042074598 | 0.051475243 | -0.817375411 | 0.413713923 | 0.637388731 | | Abcg2 | 790.7512602 | -0.0334164789795093 | 0.0680578182192683 | -0.491001325841042 | 0.623425509813882 | 0.900570704204234 | NM_011920 |
| Gm10778 | 11.63371234 | -0.136058959 | 0.166496877 | -0.817186254 | 0.413821997 | 0.637388731 | | Emid1 | 135.0211432 | 0.0464245634592067 | 0.0945556647750072 | 0.490976014707028 | 0.623443411915034 | 0.900570704204234 | NM_080595 |
| Gm3604 | 36.40572569 | 0.134794924 | 0.164888338 | 0.817492163 | 0.413647226 | 0.637388731 | | Kcnb2 | 401.9151345 | 0.0417130597994501 | 0.0849956761434309 | 0.490766844763479 | 0.623591362497284 | 0.90069485250428 | NM_001098528 |
| Mettl14 | 764.7438829 | -0.056641088 | 0.069315905 | -0.817144166 | 0.413846046 | 0.637388731 | | Adora2b | 146.2352356 | 0.0440898023923968 | 0.0898863679440159 | 0.490505995523785 | 0.623775888282605 | 0.90073387321099 | NM_007413 |
| Mrps10 | 622.5145387 | -0.067654191 | 0.082762877 | -0.817446107 | 0.413673536 | 0.637388731 | | Armcx5 | 998.3691025 | -0.0338173619165567 | 0.0689494957168273 | -0.490465688907185 | 0.623804403449164 | 0.90073387321099 | NM_001009575 |
| Zbtb9 | 677.2038759 | -0.054749862 | 0.066989883 | -0.817285527 | 0.413765276 | 0.637388731 | | Flot1 | 4075.040378 | 0.0298149726013376 | 0.0607750616457114 | 0.490579059798313 | 0.623724199944464 | 0.90073387321099 | NM_008027 |
| Tmem39b | 269.517906 | 0.082622632 | 0.101141043 | 0.816905077 | 0.413982676 | 0.63753579 | | 1810026B05Rik | 275.8374221 | -0.0425437942968122 | 0.0868041174584381 | -0.490112629935812 | 0.624054201305013 | 0.900825929460063 | NR_037569 |
| Eya2 | 101.8802245 | -0.123682022 | 0.151554111 | -0.816091496 | 0.414447807 | 0.63764026 | | 2700029M09Rik | 1097.028397 | -0.0373202573132087 | 0.0761455275098547 | -0.490117522770839 | 0.624050739209141 | 0.900825929460063 | NM_028299 |
| Fhad1 | 283.7044634 | 0.126426184 | 0.154869599 | 0.816339579 | 0.414305943 | 0.63764026 | | Rnf141 | 923.0134817 | -0.031050250593958 | 0.0633515352488179 | -0.490126253010377 | 0.62404456184465 | 0.900825929460063 | NM_025999 |
| Mboat1 | 120.3048774 | 0.12726768 | 0.155951279 | 0.816073329 | 0.414458197 | 0.63764026 | | Dync1h1 | 24765.90902 | -0.0330204863499245 | 0.0673930566111419 | -0.489968670518282 | 0.624156068521332 | 0.900883451047383 | NM_030238 |
| Mcl1 | 2514.764332 | 0.065630005 | 0.080406092 | 0.816231754 | 0.414367599 | 0.63764026 | | Tenm3 | 3557.641161 | 0.0336783485343524 | 0.0687655364666101 | 0.489756210230474 | 0.624306420809863 | 0.901010935785799 | NM_001145937,NM_011857 |
| Nceh1 | 5554.875065 | -0.050848198 | 0.062308884 | -0.816066586 | 0.414462054 | 0.63764026 | | St8sia5 | 1957.099554 | 0.0351015965680691 | 0.0717145834151923 | 0.489462462116638 | 0.624514324020524 | 0.901133756499869 | NM_153124,NM_013666 |
| Nploc4 | 3088.217232 | 0.053360686 | 0.065380278 | 0.816158751 | 0.414409345 | 0.63764026 | | Upf2 | 2669.223139 | 0.0360576899001494 | 0.0736682073254237 | 0.489460667080811 | 0.624515594567271 | 0.901133756499869 | NM_001081132 |
| Slc4a11 | 171.4043077 | 0.128383246 | 0.1572166 | 0.816601085 | 0.414156435 | 0.63764026 | | Shank1 | 17622.55666 | -0.033779312916902 | 0.0690424319300721 | -0.489254389983171 | 0.624661607273914 | 0.901254917288525 | NM_001034115 |
| Tbata | 96.78027954 | 0.122142266 | 0.149608558 | 0.816412296 | 0.414264367 | 0.63764026 | | Naa30 | 1064.978752 | 0.0401175496712416 | 0.082020640094668 | 0.489115296161284 | 0.624760072792762 | 0.901307460262952 | NM_001081430 |
| Tbx18 | 125.2194596 | -0.113794024 | 0.139347283 | -0.81662176 | 0.414144616 | 0.63764026 | | Pstk | 453.0986912 | -0.0390610607064281 | 0.0798992806049193 | -0.488878753484336 | 0.624927538440869 | 0.901459525659589 | NM_001039534 |
| Znrf2 | 369.8387647 | -0.071461407 | 0.087525121 | -0.816467388 | 0.414232868 | 0.63764026 | | Dennd1b | 505.6277922 | 0.0380705599289059 | 0.0779062538101226 | 0.48867142324278 | 0.625074338422982 | 0.901503145317256 | NM_001166501,NM_181347 |
| Hdlbp | 9242.996845 | -0.03802918 | 0.046613774 | -0.815835688 | 0.414594119 | 0.637716795 | | Lbp | 180.1178385 | 0.0211241138624891 | 0.0432285877418379 | 0.488660744335271 | 0.625081900016206 | 0.901503145317256 | NM_008489 |
| Svip | 1902.590772 | 0.045412122 | 0.055662273 | 0.815851016 | 0.414585351 | 0.637716795 | | Glrx2 | 2182.590388 | -0.0236133110863996 | 0.0484049066763433 | -0.487828873305937 | 0.62567105810208 | 0.902263257221366 | NM_001038592,NM_001038593,NM_023505,NM_001038594 |
| Rsbn1l | 845.8081726 | 0.06603205 | 0.08095079 | 0.815706064 | 0.41466827 | 0.637767538 | | Hspb8 | 590.054229 | 0.0425930603858622 | 0.0873364340855957 | 0.487689482995356 | 0.625769802248641 | 0.902316075785563 | NM_030704 |
| Scfd1 | 1128.86844 | -0.05175325 | 0.063462438 | -0.815494203 | 0.414789481 | 0.637890642 | | Ddx39 | 505.1748351 | -0.035330638020478 | 0.0724617974399085 | -0.48757606447421 | 0.625850152923384 | 0.902342364403481 | NM_197982 |
| Dhx57 | 1935.355047 | 0.043256257 | 0.053070945 | 0.81506475 | 0.415035248 | 0.638205253 | | Ap3s1 | 1213.010322 | -0.03623648096881 | 0.0743878652450712 | -0.487128926867691 | 0.626166968176196 | 0.902402690506287 | NM_009681 |
| Banp | 997.344282 | 0.080640342 | 0.098977226 | 0.814736329 | 0.415223254 | 0.63843099 | | Fam198a | 349.225277 | 0.0378207482149091 | 0.0776313576519075 | 0.487183908137923 | 0.626128007978789 | 0.902402690506287 | NM_177743,NM_001199927 |
| Apip | 317.582073 | -0.08141154 | 0.099955383 | -0.814478789 | 0.415370718 | 0.638594355 | | Rfng | 2652.543718 | -0.0341441002538432 | 0.0700997707832384 | -0.487078629107407 | 0.626202610512417 | 0.902402690506287 | NM_009053 |
| Usp42 | 995.7528294 | 0.069133817 | 0.084889449 | 0.814398229 | 0.415416853 | 0.638601917 | | Ric8b | 2849.531979 | 0.031371523225029 | 0.0643826978122845 | 0.487266366446719 | 0.626069579273522 | 0.902402690506287 | NM_183172,NM_001013441 |
| Zcchc16 | 86.20426533 | -0.121439751 | 0.149150197 | -0.814211138 | 0.415524006 | 0.638703269 | | Trappc2l | 1065.283706 | 0.02823669871822 | 0.0579503989823956 | 0.487256331502739 | 0.626076689756963 | 0.902402690506287 | NM_021502 |
| Zfp532 | 2241.153255 | -0.046819757 | 0.057524664 | -0.813907536 | 0.415697923 | 0.638907214 | | Man1a2 | 6144.086133 | 0.0328384230468696 | 0.0675003967378142 | 0.486492296844136 | 0.626618165615404 | 0.9028224036629 | NM_010763 |
| Gtf2h3 | 481.9449639 | 0.070706097 | 0.08688252 | 0.813812689 | 0.415752265 | 0.638927355 | | Morn4 | 2450.819665 | 0.0302087632975608 | 0.0620853896450103 | 0.486567990799244 | 0.626564511876316 | 0.9028224036629 | NM_198108 |
| Stk32b | 143.7311265 | 0.13274166 | 0.163132014 | 0.813706993 | 0.415812827 | 0.638957051 | | AW554918 | 669.0164213 | 0.0314647441460438 | 0.0647125873401406 | 0.486222934970282 | 0.626809111967839 | 0.902918401472118 | NM_001033532 |
| 2210408F21Rik | 354.4407285 | -0.079536041 | 0.097766334 | -0.813531998 | 0.415913108 | 0.638959676 | | Ptgr1 | 36.02164019 | -0.0438318786958343 | 0.0901347528728761 | -0.486292770532735 | 0.626759604245301 | 0.902918401472118 | NM_025968 |
| Fkbp10 | 265.8208026 | -0.073662097 | 0.090558929 | -0.813416169 | 0.415979492 | 0.638959676 | | Cpox | 1471.049512 | 0.0289638274621072 | 0.0595914710806882 | 0.486039813027765 | 0.626938938484677 | 0.902926335557051 | NM_007757 |
| Pnrc1 | 1025.794994 | 0.070751225 | 0.086960708 | 0.813599927 | 0.41587418 | 0.638959676 | | Tfpi | 174.7264153 | -0.0455308718297488 | 0.0936637092672888 | -0.486110065316942 | 0.62688913090784 | 0.902926335557051 | NM_001177319,NM_001177320,NM_011576 |
| Tssc1 | 1490.908257 | -0.069686911 | 0.085670156 | -0.813432751 | 0.415969988 | 0.638959676 | | Dhx57 | 2173.04168 | -0.0222637187851506 | 0.0458411076567227 | -0.485671484028498 | 0.627200104778614 | 0.903033870751724 | NM_001163759,NM_198942 |
| Cmya5 | 385.2793425 | 0.089467106 | 0.110013354 | 0.813238599 | 0.416081273 | 0.63898932 | | Mapkapk5 | 1292.864931 | -0.0285207283002264 | 0.0587169133335423 | -0.485732758774512 | 0.627156654243977 | 0.903033870751724 | NM_010765 |
| Zfp213 | 283.6166565 | -0.073856891 | 0.090816385 | -0.813255124 | 0.4160718 | 0.63898932 | | Zfp281 | 1769.42323 | 0.0354948146199803 | 0.0730833580557876 | 0.485675748408901 | 0.627197080821996 | 0.903033870751724 | NM_001160251,NM_177643 |
| Tecpr2 | 4500.166658 | 0.055940123 | 0.068813611 | 0.81292236 | 0.416262574 | 0.639204393 | | E030013I19Rik | 51.28353327 | 0.0453705196678456 | 0.0934415427686768 | 0.48554977072847 | 0.627286416730909 | 0.90306863126196 | NR_040353 |
| Srsf7 | 2473.840235 | 0.072041725 | 0.088636545 | 0.81277677 | 0.416346057 | 0.63926923 | | Pcdhb4 | 99.139789 | -0.045571311103428 | 0.0939034832300009 | -0.485299474906683 | 0.627463927915484 | 0.903234666227363 | NM_053129 |
| Egfl8 | 54.13830042 | -0.132163884 | 0.162650436 | -0.812563967 | 0.416468098 | 0.639292284 | | Gnpnat1 | 357.6021379 | -0.0342892941363004 | 0.070712917356405 | -0.484908492227475 | 0.627741258146362 | 0.903544343622081 | NM_019425 |
| Sp140 | 33.34216167 | 0.131497798 | 0.161836522 | 0.812534753 | 0.416484853 | 0.639292284 | | Fam151b | 201.978998 | -0.0406076175378926 | 0.083763742776805 | -0.484785724909133 | 0.627827072841509 | 0.903578327563238 | NM_001163627 |
| Zfp382 | 378.0466522 | 0.06456215 | 0.079451344 | 0.812599854 | 0.416447515 | 0.639292284 | | Naf1 | 456.1947769 | 0.0409814367433461 | 0.0845723865461872 | 0.484572310383663 | 0.6279797593199 | 0.903708538129668 | NM_001163564 |
| Gnl1 | 7600.382524 | -0.056992666 | 0.070154369 | -0.812389407 | 0.416568223 | 0.639356913 | | Fam108b | 881.8077151 | 0.0321340086736773 | 0.0663606943839926 | 0.484232556213707 | 0.628220834325479 | 0.903965907816929 | NM_146096 |
| Magt1 | 620.8881539 | -0.088780679 | 0.109296549 | -0.812291696 | 0.416624274 | 0.639379607 | | Pcbd2 | 174.9820814 | 0.0452816319813526 | 0.0935579473364199 | 0.483995569275658 | 0.628389013589555 | 0.904028801763086 | NM_028281 |
| Phf21a | 1924.937784 | 0.067762939 | 0.083440021 | 0.812115558 | 0.416725327 | 0.639471349 | | Rpl36a | 2593.246186 | -0.034534401871729 | 0.0713513531891068 | -0.484004862251181 | 0.628382418407852 | 0.904028801763086 | NM_019865 |
| Entpd5 | 1731.106209 | 0.048740507 | 0.060025355 | 0.811998648 | 0.416792407 | 0.639510949 | | Gm16861 | 31.75664318 | 0.0406943938274154 | 0.0841074332741706 | 0.48383825594536 | 0.628500662632722 | 0.904099883680225 | NR_037666 |
| Pla1a | 57.50642296 | 0.12631771 | 0.155580023 | 0.811914712 | 0.416840572 | 0.639521519 | | Cdk16 | 12558.35849 | 0.0219861533440446 | 0.0454798370810363 | 0.483426387497157 | 0.628793015794181 | 0.904346422581565 | NM_011049 |
| Prmt6 | 482.223881 | -0.075456355 | 0.093076967 | -0.810664217 | 0.417558528 | 0.640559587 | | Maoa | 1364.673694 | -0.0322104613208326 | 0.0666301944898505 | -0.483421391269375 | 0.628796562582528 | 0.904346422581565 | NM_173740 |
| Chpt1 | 857.727491 | -0.067225171 | 0.082964885 | -0.810284629 | 0.417776607 | 0.640787605 | | Chst15 | 1816.326528 | -0.031364159616803 | 0.0648961581755861 | -0.483297632687942 | 0.628884420697368 | 0.904383238793186 | NM_029935 |
| Copg2 | 3456.629089 | -0.047989215 | 0.059226821 | -0.810261533 | 0.417789879 | 0.640787605 | | Aldh6a1 | 2964.156377 | 0.0292185529257643 | 0.0604824521230007 | 0.483091407510127 | 0.62903083478763 | 0.904428764037583 | NM_134042 |

| | | | | | | |
|---|---|---|---|---|---|---|
| Mettl10 | 452.7251927 | -0.076607204 | 0.094567489 | -0.810079704 | 0.417894368 | 0.640820996 |
| Thoc6 | 160.2669248 | -0.085218915 | 0.105189004 | -0.810150418 | 0.41785373 | 0.640820996 |
| Sp1 | 1282.835051 | -0.066554821 | 0.082185005 | -0.80981709 | 0.418045309 | 0.640989017 |
| Elmo2 | 7295.211283 | 0.047224127 | 0.058338907 | 0.809479112 | 0.418239613 | 0.641099272 |
| F11r | 391.0655531 | 0.092592149 | 0.114385279 | 0.809476095 | 0.418241348 | 0.641099272 |
| Mex3d | 465.4396971 | -0.061132512 | 0.075511521 | -0.809578598 | 0.418182412 | 0.641099272 |
| Pdlim3 | 78.02237828 | -0.123899631 | 0.153112846 | -0.809204676 | 0.418397426 | 0.641275073 |
| Gm5820 | 31.40843979 | -0.135961341 | 0.168115768 | -0.808736401 | 0.418666785 | 0.641528526 |
| Magee1 | 6631.988667 | -0.046235952 | 0.057168679 | -0.8087637 | 0.41865108 | 0.641528526 |
| Tbxa2r | 34.20073439 | -0.133964905 | 0.165654379 | -0.808701257 | 0.418687005 | 0.641528526 |
| Fbxo46 | 409.3302056 | 0.067358406 | 0.083309328 | 0.808533785 | 0.418783365 | 0.641612722 |
| Rps6ka5 | 1005.819026 | 0.108152893 | 0.13380709 | 0.808274758 | 0.41893243 | 0.641714194 |
| Trerf1 | 1518.214119 | 0.068124435 | 0.084280453 | 0.808306462 | 0.418914184 | 0.641714194 |
| Plcl1 | 1722.303482 | 0.073141667 | 0.090515872 | 0.808053505 | 0.419059782 | 0.641845815 |
| Dus2l | 446.2408935 | 0.0823519 | 0.101973594 | 0.807580642 | 0.419332034 | 0.642189209 |
| Mrpl22 | 431.9054088 | -0.087065873 | 0.107828437 | -0.807448166 | 0.419408326 | 0.642189209 |
| P2rx4 | 513.6189763 | 0.063700618 | 0.07888877 | 0.807473836 | 0.419393542 | 0.642189209 |
| Htr7 | 386.6767317 | 0.08832782 | 0.109467193 | 0.806888507 | 0.419730721 | 0.642619347 |
| Lmf1 | 543.9081029 | -0.067134682 | 0.08322977 | -0.806618609 | 0.419886249 | 0.642730443 |
| Rtl1 | 37.96039997 | 0.123306471 | 0.15286794 | 0.806620875 | 0.419884943 | 0.642730443 |
| AI314180 | 4877.04855 | 0.056697489 | 0.070362072 | 0.805796181 | 0.420360381 | 0.643196272 |
| Clock | 3229.71826 | -0.061895595 | 0.076829729 | -0.805620372 | 0.420461775 | 0.643196272 |
| Gm10658 | 12.95065956 | 0.1323364 | 0.164225065 | 0.8058234 | 0.420344683 | 0.643196272 |
| Mbtps1 | 3831.798326 | 0.052606958 | 0.065271794 | 0.805967715 | 0.420261465 | 0.643196272 |
| Mrps34 | 931.8995168 | -0.058027684 | 0.072010255 | -0.805825282 | 0.420343598 | 0.643196272 |
| Trim44 | 14729.38232 | -0.038948926 | 0.048348518 | -0.805586769 | 0.420481157 | 0.643196272 |
| Uba3 | 2563.463054 | -0.048192957 | 0.059816968 | -0.805673693 | 0.420431022 | 0.643196272 |
| Vmn2r87 | 27.01138101 | 0.135315572 | 0.167989675 | 0.805499338 | 0.420531588 | 0.643209914 |
| Hps6 | 257.6418368 | 0.080173908 | 0.099567886 | 0.805218549 | 0.420693574 | 0.64339416 |
| Nxph4 | 182.2140989 | 0.106233213 | 0.131958544 | 0.805049902 | 0.420790884 | 0.643479466 |
| Gabra1 | 12773.10968 | 0.062913023 | 0.078175479 | 0.804766701 | 0.420954321 | 0.64353885 |
| Ndufaf4 | 922.3848322 | -0.070473345 | 0.087563488 | -0.804825689 | 0.420920275 | 0.64353885 |
| Txnl1 | 2334.867916 | -0.043560453 | 0.054126248 | -0.8047935 | 0.420938853 | 0.64353885 |
| B4galt4 | 734.0594434 | -0.066234702 | 0.082319884 | -0.804601493 | 0.421049681 | 0.643621128 |
| 2610305D13Rik | 9.24982342 | 0.124452936 | 0.154722605 | 0.804361687 | 0.421188121 | 0.64363258 |
| Ankrd33b | 4640.540299 | -0.097355853 | 0.121043369 | -0.804305552 | 0.421220532 | 0.64363258 |
| BC005537 | 6027.313899 | 0.046033743 | 0.057221606 | 0.804481846 | 0.42111875 | 0.64363258 |
| Lrrc3 | 645.2624931 | -0.078646845 | 0.097791636 | -0.804228753 | 0.421264876 | 0.64363258 |
| Zfp85-rs1 | 139.5306057 | -0.097269325 | 0.120945321 | -0.804242151 | 0.42125714 | 0.64363258 |
| Prnp | 34417.04835 | 0.047120746 | 0.058601764 | 0.804084083 | 0.421348417 | 0.643696743 |
| Bap1 | 6285.644644 | -0.051718513 | 0.064333318 | -0.80391489 | 0.42144613 | 0.643782543 |
| Morc2b | 47.51827889 | 0.120970678 | 0.150542705 | 0.803563866 | 0.421648898 | 0.644028789 |
| Rbpms | 128.2056268 | 0.097072423 | 0.120815602 | 0.803475884 | 0.42169973 | 0.644042939 |
| Zfyve28 | 2122.073797 | 0.053829886 | 0.067029907 | 0.803072666 | 0.421932736 | 0.644335285 |
| Fbxw5 | 3493.271695 | 0.063704111 | 0.079349226 | 0.802832163 | 0.422071749 | 0.644484053 |
| 5730455P16Rik | 3327.241694 | 0.057846722 | 0.072079829 | 0.80253689 | 0.422242458 | 0.644554152 |
| Rpl18 | 3073.162152 | -0.062750073 | 0.078186534 | -0.802568799 | 0.422224008 | 0.644554152 |
| Thg1l | 179.2911929 | 0.081577827 | 0.101641623 | 0.802602564 | 0.422204486 | 0.644554152 |
| Ube2k | 4131.808243 | -0.054454788 | 0.067863847 | -0.802412329 | 0.422314483 | 0.644600592 |
| Iba57 | 329.8193672 | -0.079272346 | 0.098806316 | -0.8023004 | 0.422379211 | 0.644635884 |
| Sh3bp5 | 1959.931409 | 0.067448681 | 0.084092219 | 0.802079933 | 0.422506722 | 0.64476698 |
| Kif5a | 139939.9325 | -0.067553511 | 0.084250127 | -0.801820885 | 0.422656576 | 0.644932144 |
| Dpm3 | 256.2929338 | -0.106714645 | 0.133132575 | -0.801566748 | 0.422803619 | 0.645029467 |
| Tex15 | 46.90644742 | -0.132946039 | 0.165853088 | -0.801589166 | 0.422790647 | 0.645029467 |
| Eif4enif1 | 2399.66444 | 0.080625066 | 0.100622826 | 0.801260207 | 0.422981023 | 0.645236582 |
| Cdc42ep2 | 426.3963463 | 0.095793329 | 0.119581952 | 0.801068454 | 0.423092017 | 0.645342362 |
| Il18 | 553.4087036 | -0.082596047 | 0.103159599 | -0.800662741 | 0.423326918 | 0.645637097 |
| Mum1l1 | 133.1896097 | -0.104859296 | 0.130986338 | -0.800536125 | 0.423400242 | 0.645685369 |
| Hsdl2 | 1290.016573 | -0.073503449 | 0.091868666 | -0.800092699 | 0.423657091 | 0.64601348 |
| Zfp182 | 402.7108213 | 0.100081773 | 0.125100801 | 0.80000905 | 0.423705554 | 0.646023799 |
| Tmem11 | 750.4501946 | -0.07513653 | 0.093973825 | -0.799547431 | 0.423973055 | 0.646368053 |
| Apol10b | 17.58885436 | -0.13170982 | 0.164808377 | -0.799169448 | 0.424192165 | 0.646517445 |
| H2-DMb2 | 15.99011988 | -0.12976181 | 0.162372258 | -0.799162441 | 0.424196227 | 0.646517445 |
| Ilkap | 1198.325067 | 0.053771961 | 0.067276972 | 0.799262494 | 0.424138222 | 0.646517445 |
| Eogt | 1142.148878 | -0.068373353 | 0.085572009 | -0.799015404 | 0.42428148 | 0.646583776 |
| Maged2 | 1599.491205 | -0.059231082 | 0.074184333 | -0.798431138 | 0.42462034 | 0.64703654 |
| Ccdc124 | 1960.003261 | -0.058966952 | 0.073871254 | -0.798239484 | 0.424731528 | 0.647078692 |
| Fam135b | 1055.031826 | -0.083288967 | 0.104337544 | -0.798264592 | 0.424716961 | 0.647078692 |
| Sypl2 | 106.7184176 | 0.112012242 | 0.140343435 | 0.79812955 | 0.424795315 | 0.647112242 |
| Cdkn2c | 66.11176658 | -0.112761109 | 0.141360132 | -0.797686785 | 0.425052274 | 0.647312752 |
| Gm19461 | 26.14918571 | 0.132504846 | 0.166086295 | 0.797807226 | 0.424982367 | 0.647312752 |
| Gm5141 | 103.6764198 | -0.124672156 | 0.156292105 | -0.797686843 | 0.425052241 | 0.647312752 |
| Dtx2 | 265.1188358 | 0.094533246 | 0.118566889 | 0.797298865 | 0.425277479 | 0.647528426 |
| Rpusd2 | 306.1194442 | -0.073517021 | 0.092206941 | -0.797304627 | 0.425274134 | 0.647528426 |
| Trappc9 | 3359.285744 | 0.063489554 | 0.079652655 | 0.797080193 | 0.425404459 | 0.64765812 |
| 1110057K04Rik | 1560.198489 | -0.054022346 | 0.067810312 | -0.796668594 | 0.425643528 | 0.647831123 |
| Sptlc2 | 943.6327482 | 0.051833211 | 0.065050784 | 0.796811474 | 0.42556053 | 0.647831123 |
| Zfp277 | 733.694385 | 0.06246178 | 0.07839822 | 0.796724463 | 0.425611073 | 0.647831123 |
| Hsf2bp | 12.81656627 | 0.132442267 | 0.166284782 | 0.796478577 | 0.425753923 | 0.647900885 |
| Tceal6 | 2052.673695 | 0.055291191 | 0.069422419 | 0.796445748 | 0.425772997 | 0.647900885 |
| Csnk2a1 | 5618.717578 | -0.048768771 | 0.061245492 | -0.796283452 | 0.425867302 | 0.647980749 |
| Gimap5 | 17.9763272 | -0.133553809 | 0.167742046 | -0.796185644 | 0.42592414 | 0.648003597 |
| Nufip2 | 1794.484977 | 0.052566288 | 0.066034326 | 0.796044891 | 0.426005943 | 0.648064417 |
| Ipo8 | 2268.817707 | -0.061660104 | 0.077472856 | -0.795892999 | 0.426094231 | 0.648135089 |
| Ccdc24 | 40.61955369 | 0.133742099 | 0.168121551 | 0.795508359 | 0.42631785 | 0.648329765 |
| Cep63 | 933.4702578 | 0.079610361 | 0.100081295 | 0.795456946 | 0.426347746 | 0.648329765 |
| Itga6 | 963.6195681 | -0.071565686 | 0.089962746 | -0.79550357 | 0.426320635 | 0.648329765 |
| Kif17 | 1533.067433 | 0.064231363 | 0.080757917 | 0.795356859 | 0.426405948 | 0.648354637 |
| Cdc42se1 | 2556.89788 | -0.044148109 | 0.0555128 | -0.795278009 | 0.426451803 | 0.648360733 |
| Dnaja1 | 10538.99716 | -0.048988992 | 0.061610565 | -0.795139479 | 0.426532372 | 0.648419601 |
| Dennd4a | 3518.47007 | -0.054108514 | 0.068065839 | -0.794943761 | 0.426646217 | 0.648529038 |
| Adpgk | 554.7744291 | 0.055645117 | 0.070025884 | 0.794636411 | 0.426825032 | 0.648609949 |
| Fkbp7 | 73.7795458 | 0.11358606 | 0.142915887 | 0.794775597 | 0.426744048 | 0.648609949 |
| Ttc9c | 1293.742898 | 0.044050928 | 0.055430888 | 0.794700018 | 0.426788022 | 0.648609949 |
| Lingo4 | 33.75730959 | 0.12399894 | 0.156141324 | 0.794145568 | 0.427110692 | 0.648980393 |
| Tmem47 | 2920.286654 | -0.086118744 | 0.108472167 | -0.793924804 | 0.427239209 | 0.649112013 |
| Fam179b | 2538.359323 | -0.060327629 | 0.076003387 | -0.79374921 | 0.427341446 | 0.649140036 |
| Rpp14 | 1191.101055 | -0.052073907 | 0.065600625 | -0.793801997 | 0.42731071 | 0.649140036 |
| Dirc2 | 2869.404216 | -0.041533648 | 0.052353233 | -0.793334933 | 0.427582709 | 0.649203686 |
| Kif2a | 3069.963507 | -0.060785011 | 0.076618908 | -0.793342173 | 0.427578492 | 0.649203686 |
| Phka1 | 1142.013112 | 0.069813433 | 0.08800188 | 0.793317523 | 0.42759285 | 0.649203686 |
| Prkrip1 | 521.9163775 | 0.056387579 | 0.071067721 | 0.79343447 | 0.427524734 | 0.649203686 |
| Vps9d1 | 2161.700972 | 0.062708275 | 0.079034942 | 0.793424688 | 0.427530432 | 0.649203686 |
| Nit1 | 1045.479204 | 0.071456721 | 0.090111623 | 0.792980076 | 0.427789431 | 0.649438511 |
| Tubd1 | 195.0895305 | 0.104811518 | 0.132228502 | 0.792654509 | 0.427979141 | 0.64966286 |
| Dpm2 | 1251.352067 | 0.056625348 | 0.071448467 | 0.792534114 | 0.428049309 | 0.649685981 |
| Mrps18a | 1242.767098 | -0.049163002 | 0.062041407 | -0.79242242 | 0.428114411 | 0.649685981 |
| Ube2e2 | 2183.232836 | 0.06496676 | 0.081986032 | 0.792412545 | 0.428120167 | 0.649685981 |
| Itk | 17.23646412 | 0.12899577 | 0.162821014 | 0.792255048 | 0.428211976 | 0.649761665 |
| Nppc | 83.93647574 | -0.10949529 | 0.138286584 | -0.791799802 | 0.428477418 | 0.650037121 |
| Unc93b1 | 637.3990027 | 0.108366776 | 0.136854019 | 0.791842119 | 0.42845274 | 0.650037121 |
| Farp1 | 3038.489655 | 0.058853598 | 0.074361428 | 0.791453299 | 0.428679518 | 0.650280052 |
| Zfyve1 | 2075.49585 | 0.076057328 | 0.096110057 | 0.791356606 | 0.428735925 | 0.65030195 |
| Pja1 | 7300.473163 | -0.06706629 | 0.084777432 | -0.791086588 | 0.428893465 | 0.650413561 |
| Zc2hc1a | 1988.105021 | -0.059568786 | 0.075298831 | -0.791098423 | 0.428886559 | 0.650413561 |
| 4930506C21Rik | 46.48612643 | -0.131335392 | 0.166116048 | -0.790624347 | 0.429163235 | 0.650758971 |
| Slc17a7 | 51375.26665 | -0.07474471 | 0.094620719 | -0.789940199 | 0.429562693 | 0.651267729 |
| Smcr7l | 1167.255954 | -0.039675623 | 0.050228301 | -0.78990574 | 0.429582819 | 0.651267729 |
| 2700029M09Rik | 1019.983567 | -0.070194314 | 0.088920537 | -0.789404969 | 0.429875353 | 0.651519976 |
| Evi2a | 597.570977 | 0.104166726 | 0.131952949 | 0.789423249 | 0.429864672 | 0.651519976 |
| Paxip1 | 1873.197795 | 0.045914075 | 0.058156277 | 0.789494747 | 0.429822899 | 0.651519976 |
| Cysltr1 | 21.95226128 | -0.130321431 | 0.165119077 | -0.789257263 | 0.42996166 | 0.651548009 |
| Gsta4 | 1193.945956 | -0.058510922 | 0.074143536 | -0.789157422 | 0.430020004 | 0.651548009 |
| Tbc1d10b | 3139.957104 | 0.043708827 | 0.055382308 | 0.789220024 | 0.42998342 | 0.651548009 |
| Yme1l1 | 2602.31824 | -0.043091299 | 0.05462341 | -0.788879695 | 0.430182324 | 0.651730217 |
| Rpl29 | 5053.820781 | -0.056816545 | 0.072055588 | -0.788509901 | 0.43039851 | 0.651993988 |
| Micall1 | 2490.690102 | 0.054575173 | 0.069279211 | 0.78775685 | 0.430838945 | 0.652275301 |
| Nmb | 100.1704317 | -0.132460987 | 0.168089738 | -0.78803732 | 0.430674877 | 0.652275301 |
| Prim1 | 205.4980727 | 0.091218618 | 0.115805437 | 0.787688555 | 0.430878902 | 0.652275301 |
| Prkg1 | 510.5347522 | -0.089047253 | 0.113010037 | -0.787958796 | 0.430720808 | 0.652275301 |
| Slc39a11 | 599.8407931 | -0.067915647 | 0.08618405 | -0.788030344 | 0.430678957 | 0.652275301 |
| Spsb2 | 81.55843434 | -0.116622515 | 0.148032108 | -0.787819053 | 0.430802555 | 0.652275301 |
| Tigd5 | 212.1538662 | -0.081588426 | 0.103568937 | -0.787769273 | 0.430831677 | 0.652275301 |
| Cep19 | 2233.142856 | -0.049470528 | 0.06286045 | -0.786989717 | 0.431287886 | 0.652830647 |
| Bhlhe22 | 906.2403092 | 0.087320296 | 0.110980089 | 0.786810473 | 0.431392822 | 0.652925699 |
| Ptpra | 6865.32804 | -0.03645477 | 0.04634141 | -0.786656473 | 0.431482991 | 0.652998385 |
| Helz2 | 301.8274867 | 0.075529993 | 0.09603831 | 0.786456918 | 0.431599849 | 0.653111443 |
| 4931408D14Rik | 14.52117776 | 0.129607582 | 0.164883063 | 0.786057583 | 0.431833753 | 0.653295873 |
| Pcdhb6 | 103.2351802 | -0.11320932 | 0.144006785 | -0.786138796 | 0.431786178 | 0.653295873 |
| Slc6a6 | 7755.263573 | 0.055035665 | 0.070017001 | 0.786032987 | 0.43184822 | 0.653295873 |
| Pum2 | 8430.12264 | -0.060077191 | 0.076452931 | -0.785806246 | 0.431981007 | 0.653320983 |
| Spin2 | 174.1517619 | 0.099657057 | 0.126824252 | 0.78578865 | 0.431991317 | 0.653320983 |
| Tram2 | 19.64504898 | -0.131018596 | 0.166708862 | -0.785912605 | 0.431918689 | 0.653320983 |
| Zmym1 | 444.954953 | 0.068228287 | 0.086849302 | 0.785593951 | 0.432105409 | 0.653429749 |
| AI836003 | 893.8624824 | -0.080851139 | 0.102933748 | -0.785467749 | 0.432179372 | 0.653477818 |
| Sspo | 103.1973181 | 0.110180227 | 0.140297654 | 0.78533193 | 0.43225898 | 0.65353441 |
| Susd3 | 48.65137964 | -0.129526879 | 0.165001749 | -0.785003067 | 0.432451771 | 0.653762097 |
| Epc1 | 1619.454179 | 0.06630153 | 0.084487951 | 0.784745386 | 0.432602867 | 0.653926714 |
| Ccrl2 | 34.80790341 | -0.126734792 | 0.161526355 | -0.784607516 | 0.432683722 | 0.653973357 |
| D10Wsu52e | 4349.760596 | 0.055816602 | 0.071144841 | 0.784548834 | 0.43271814 | 0.653973357 |
| Nrap | 46.11569408 | 0.124094617 | 0.158206571 | 0.784383454 | 0.432815144 | 0.654004155 |
| Wasf2 | 897.983756 | 0.058546246 | 0.074641092 | 0.784370167 | 0.432822938 | 0.654004155 |
| Mrpl55 | 633.8563917 | -0.065369722 | 0.083363764 | -0.784150322 | 0.432951911 | 0.654110921 |
| Sox9 | 828.6672561 | -0.081481609 | 0.1039166 | -0.784105803 | 0.432978031 | 0.654110921 |
| Vstm5 | 495.9183251 | 0.057746874 | 0.073658684 | 0.783979175 | 0.43305233 | 0.654159384 |
| Ramp3 | 393.2424967 | -0.114803431 | 0.146459159 | -0.783859692 | 0.433122444 | 0.654201515 |
| Ube2q1 | 3872.088485 | 0.057671907 | 0.073599467 | 0.783591371 | 0.433279921 | 0.654375581 |
| Hnrnpab | 3930.071135 | -0.052326905 | 0.066812671 | -0.783188335 | 0.433516524 | 0.654541515 |
| Ror1 | 47.42496023 | 0.116256765 | 0.148436816 | 0.783207081 | 0.433505517 | 0.654541515 |
| Scnm1 | 736.2852655 | 0.058016064 | 0.074072753 | 0.783230838 | 0.433491569 | 0.654541515 |
| Ubac2 | 1229.491544 | -0.058301729 | 0.07445232 | -0.783074715 | 0.433583238 | 0.654578456 |
| Zfp541 | 28.74802507 | 0.130324505 | 0.166460514 | 0.782915432 | 0.433676775 | 0.65465588 |
| St6gal1 | 1107.905847 | -0.042227757 | 0.053973572 | -0.782378403 | 0.433992222 | 0.655068241 |
| Srpr | 2922.297067 | -0.035239211 | 0.045053436 | -0.782164784 | 0.434117738 | 0.655094028 |
| U2af1 | 1358.081605 | 0.057173761 | 0.073094946 | 0.782184872 | 0.434105934 | 0.655094028 |
| Zfp764 | 175.0975746 | 0.087250291 | 0.111554224 | 0.782133453 | 0.434136148 | 0.655094028 |
| Spdya | 34.34806141 | -0.126264595 | 0.161565335 | -0.781507959 | 0.434503797 | 0.655584948 |
| Dnahc2 | 149.726623 | -0.110772701 | 0.141762964 | -0.781393803 | 0.434570915 | 0.655622371 |
| Slc38a8 | 15.38911121 | 0.13007309 | 0.166568723 | 0.780897447 | 0.434862813 | 0.655998874 |
| Ano10 | 890.6945432 | 0.066152424 | 0.084763946 | 0.780431148 | 0.43513714 | 0.656157163 |
| Arhgdib | 394.2675028 | -0.079195653 | 0.101473033 | -0.780460101 | 0.435120104 | 0.656157163 |
| BC003331 | 1954.609424 | 0.047425587 | 0.060753944 | 0.780617429 | 0.435027537 | 0.656157163 |
| Srms | 14.25514479 | -0.130523576 | 0.167231492 | -0.780496391 | 0.435098751 | 0.656157163 |
| Slc25a4 | 18279.35085 | -0.047108355 | 0.060374689 | -0.780266618 | 0.435233957 | 0.656239289 |
| Cdip1 | 10104.43992 | 0.058256377 | 0.074682953 | 0.780049185 | 0.435361925 | 0.65630935 |
| Tufm | 2094.984566 | -0.05967158 | 0.076497738 | -0.780043721 | 0.435365141 | 0.65630935 |
| D16Ertd472e | 400.9617831 | 0.086368931 | 0.110747438 | 0.779872953 | 0.43546566 | 0.656397017 |
| A930007I19Rik | 13.25595126 | 0.127599396 | 0.163631984 | 0.779794957 | 0.435511575 | 0.656402368 |
| Ccdc110 | 30.61906568 | -0.130671212 | 0.167590718 | -0.779704348 | 0.435564919 | 0.656418914 |
| Rap1gds1 | 16174.92772 | -0.053171994 | 0.068202448 | -0.779620016 | 0.435614571 | 0.656429893 |
| 6330408A02Rik | 572.5937471 | 0.053489501 | 0.068656029 | 0.779094012 | 0.435924337 | 0.656577397 |
| Ankrd66 | 12.44894353 | -0.129830402 | 0.166626417 | -0.779170581 | 0.435879238 | 0.656577397 |
| Frmpd3 | 152.5595989 | -0.085306761 | 0.109486309 | -0.779154596 | 0.435888653 | 0.656577397 |
| Nhlrc1 | 621.0874187 | 0.068127157 | 0.087420688 | 0.779302461 | 0.435801565 | 0.656577397 |
| Zfp707 | 321.7717877 | 0.070815065 | 0.090890419 | 0.779125739 | 0.435905649 | 0.656577397 |
| A730017C20Rik | 1430.689383 | -0.065004238 | 0.083457209 | -0.778893032 | 0.436042729 | 0.656626826 |
| Adcy1 | 36973.19334 | -0.087305181 | 0.112109302 | -0.77875055 | 0.436126673 | 0.656626826 |
| Tmem53 | 74.84516962 | -0.11475881 | 0.147327917 | -0.778934586 | 0.436018249 | 0.656626826 |
| Tom1l1 | 695.8877736 | 0.057251463 | 0.073513892 | 0.778784276 | 0.436106802 | 0.656626826 |
| Nsd1 | 6955.261315 | -0.071992636 | 0.092491807 | -0.778367703 | 0.436352273 | 0.656902654 |
| Mdk | 350.3763455 | -0.083545203 | 0.107350508 | -0.778246927 | 0.436423457 | 0.656945986 |
| B3galt4 | 144.5814249 | -0.081656723 | 0.104940662 | -0.778122815 | 0.436496614 | 0.65699228 |
| Tmem201 | 3131.757534 | 0.067500583 | 0.086763551 | 0.777983173 | 0.436578933 | 0.657052355 |
| Wnt8b | 11.34611189 | 0.119943681 | 0.154192859 | 0.77788091 | 0.436639223 | 0.657079267 |
| Smpdl3b | 174.321638 | 0.118836436 | 0.152863605 | 0.777401763 | 0.436921772 | 0.657440608 |
| Zbtb14 | 1126.30684 | -0.056167151 | 0.072264611 | -0.777242841 | 0.437015511 | 0.657517803 |
| Creb3l3 | 22.83512179 | 0.130541618 | 0.168121354 | 0.77647256 | 0.437470016 | 0.658137725 |
| Hnrnpul2 | 10937.83103 | -0.043864272 | 0.056500588 | -0.776350719 | 0.437541934 | 0.658182012 |
| Aco2 | 20870.41951 | -0.034722332 | 0.044752525 | -0.775874251 | 0.437823237 | 0.658477309 |
| Serping1 | 332.0367562 | -0.116937051 | 0.150708535 | -0.775915248 | 0.437799029 | 0.658477309 |
| Dos | 10310.52196 | -0.049510041 | 0.063841568 | -0.775514179 | 0.43803589 | 0.658642811 |
| Fkbp3 | 3518.309412 | -0.052677295 | 0.067932272 | -0.775438451 | 0.438080622 | 0.658642811 |
| Paqr3 | 335.4104287 | -0.064266667 | 0.08286089 | -0.775597113 | 0.437986905 | 0.658642811 |
| Sez6l | 9236.349388 | 0.071095884 | 0.091689298 | 0.775400029 | 0.438103318 | 0.658642811 |
| AI846148 | 793.8136019 | -0.049554184 | 0.063955196 | -0.774826559 | 0.438442153 | 0.658832579 |
| Cacnb4 | 8155.658781 | 0.073065968 | 0.094286952 | 0.774931907 | 0.438379897 | 0.658832579 |
| Cyp2s1 | 225.8167933 | -0.095843307 | 0.123695624 | -0.774831829 | 0.438439039 | 0.658832579 |
| Cyp7b1 | 381.1451626 | -0.060815763 | 0.078479716 | -0.774923323 | 0.438384969 | 0.658832579 |
| Itpripl1 | 79.86457257 | 0.101002903 | 0.130320645 | 0.775033788 | 0.438319694 | 0.658832579 |
| Gadd45a | 184.921943 | 0.115939255 | 0.149673258 | 0.774615698 | 0.438566779 | 0.658955942 |
| Zfp438 | 223.6393599 | 0.074643294 | 0.09637907 | 0.774476184 | 0.438649247 | 0.659015944 |
| Brcc3 | 925.9086605 | -0.050050252 | 0.064643106 | -0.774255063 | 0.438779972 | 0.659148428 |
| Fiz1 | 802.9746057 | 0.04558231 | 0.058918852 | 0.773645595 | 0.439140401 | 0.659561979 |
| Zfp958 | 246.6996955 | -0.088303562 | 0.11413254 | -0.773693126 | 0.439112286 | 0.659561979 |
| Nsun7 | 160.3040673 | 0.113009568 | 0.14611338 | 0.7734375 | 0.439263504 | 0.659682924 |
| Ndrg2 | 28502.24942 | 0.051030819 | 0.065995233 | 0.773250082 | 0.439374392 | 0.659781311 |
| Zfp715 | 1083.492043 | 0.069127197 | 0.089406016 | 0.773182836 | 0.439414183 | 0.659781311 |
| Pik3r2 | 5246.572417 | 0.060691791 | 0.078522438 | 0.772922913 | 0.439568003 | 0.659948318 |
| Exoc6b | 4653.121809 | 0.058608959 | 0.075904386 | 0.772141922 | 0.440030375 | 0.660415671 |
| Jhdm1d | 2455.537424 | -0.065413001 | 0.084698574 | -0.772303443 | 0.439934727 | 0.660415671 |
| Ptp4a1 | 371.9907172 | -0.072159339 | 0.093457433 | -0.772109145 | 0.440049786 | 0.660415671 |
| Rps4x | 8955.883965 | -0.03460931 | 0.044819992 | -0.772184651 | 0.440005071 | 0.660415671 |
| 9430091E24Rik | 46.92200696 | -0.123507922 | 0.15999401 | -0.771953413 | 0.440142021 | 0.660426154 |
| Ankrd12 | 2918.008475 | 0.065542548 | 0.084901195 | 0.771986161 | 0.440122624 | 0.660426154 |
| Lrrc38 | 89.10596748 | 0.106804936 | 0.138379928 | 0.7718239 | 0.440218734 | 0.660477299 |
| Gpr137 | 2816.511147 | 0.054472775 | 0.070602939 | 0.771536927 | 0.440388743 | 0.660668396 |
| Iqgap1 | 1226.191355 | 0.055852667 | 0.072409819 | 0.771341063 | 0.440504799 | 0.660778522 |
| Katnal2 | 124.0388856 | 0.090947834 | 0.117945272 | 0.771101988 | 0.440646482 | 0.660927067 |
| Cachd1 | 654.7657027 | -0.092980389 | 0.120648432 | -0.77067217 | 0.440901272 | 0.661093969 |
| Dnase1l1 | 65.63983326 | 0.108225055 | 0.140409763 | 0.77078013 | 0.440837267 | 0.661093969 |
| Ears2 | 194.2392617 | -0.09875068 | 0.128148274 | -0.770597033 | 0.44094582 | 0.661093969 |
| Tcea3 | 14.20852046 | -0.129183733 | 0.167650373 | -0.770554403 | 0.440971097 | 0.661093969 |
| Xkr8 | 250.7770511 | 0.100633531 | 0.130577934 | 0.770677927 | 0.440897859 | 0.661093969 |
| Angel1 | 503.8195606 | 0.08044321 | 0.104421005 | 0.770373835 | 0.44107817 | 0.661140563 |
| Tlr4 | 62.25508253 | -0.11202485 | 0.145419194 | -0.770358071 | 0.441087518 | 0.661140563 |

| | | | | | | | |
|---|---|---|---|---|---|---|---|
| Rhpn2 | 336.8227755 | 0.0398089218274992 | 0.0824131950217397 | 0.483040632231259 | 0.629066886049166 | 0.904428764037583 | NM_027897 |
| Ubn1 | 2992.444611 | 0.0281772390268578 | 0.058339181850962 | 0.482989958598351 | 0.629102866022011 | 0.904428764037583 | NM_026666 |
| Rsf1 | 2432.786515 | -0.0333546788962261 | 0.0690772546130632 | -0.482860517301427 | 0.629194777666249 | 0.904471384500735 | NM_001081267 |
| 2900052N01Rik | 1712.447503 | 0.0362218118201488 | 0.0750632926615859 | 0.482550265726426 | 0.629415099618827 | 0.904624801904692 | NR_015605 |
| Tada1 | 1252.211826 | 0.0291898765371502 | 0.0604927854193135 | 0.482534840060956 | 0.629426054856858 | 0.904624801904692 | NM_030245 |
| Tdrd7 | 1713.364556 | 0.0296135274220964 | 0.0613832822990135 | 0.482436362360707 | 0.629495995197247 | 0.904635815812743 | NM_146142 |
| Lrrk2 | 3319.544875 | 0.0423473025220012 | 0.0878253423474459 | 0.482176344436792 | 0.629680679796772 | 0.904807492467134 | NM_025730 |
| Lsg1 | 642.3273361 | -0.0314838926589445 | 0.0653185883455873 | -0.48200509925857 | 0.629802323856202 | 0.904807492467134 | NM_178069 |
| Polr1a | 1576.26449 | -0.0276810350765773 | 0.0574287024028853 | -0.482006974184855 | 0.629800991947428 | 0.904807492467134 | NM_009088 |
| Mrpl28 | 1316.683761 | 0.0306013701489431 | 0.0635384587155649 | 0.48161964843895 | 0.630076165684332 | 0.905068772049975 | NM_024227 |
| Zfp81 | 381.5889177 | -0.0435454703060522 | 0.0904232710601107 | -0.481573712115596 | 0.630108804333017 | 0.905068772049975 | NM_207541 |
| Crebrf | 2604.462717 | 0.0330657806400818 | 0.0687672841902164 | 0.480835924080103 | 0.630633115930889 | 0.905249211412947 | NM_029870 |
| Lnx2 | 262.238318 | -0.0419547866243752 | 0.0871836929305198 | -0.48122286650338 | 0.630358110758028 | 0.905249211412947 | NM_080795 |
| Nop56 | 3106.47539 | -0.0292554305999724 | 0.0607972121237407 | -0.481196909825878 | 0.630376556915238 | 0.905249211412947 | NM_024193 |
| Sephs1 | 1058.792201 | 0.0301767811910365 | 0.0627658938699147 | 0.480783102580828 | 0.630670660849444 | 0.905249211412947 | NM_175400 |
| Slc9a3r2 | 1504.747261 | 0.035565676953671 | 0.0739448907855043 | 0.480975447740374 | 0.630533948692465 | 0.905249211412947 | NM_023449,NM_023055 |
| Ttc39a | 150.325608 | -0.0445472685744476 | 0.092638276946935 | -0.480873242061326 | 0.630606591310762 | 0.905249211412947 | NM_001145948,NM_153392 |
| Znrf3 | 589.4041584 | 0.0313786177785067 | 0.0652333461205801 | 0.481021128680186 | 0.630501482149573 | 0.905249211412947 | NM_001080924 |
| Pdk1 | 3108.047109 | -0.0375430256128514 | 0.0781543245718128 | -0.48037041863697 | 0.630964024703515 | 0.905472959095347 | NM_172665 |
| Unc5d | 2605.060235 | 0.0394221712620577 | 0.0820780940139366 | 0.48030076399391 | 0.631013545705782 | 0.905472959095347 | NM_153135 |
| Zfp37 | 793.7462444 | 0.0380061911667379 | 0.0791183803711719 | 0.480371198050789 | 0.630963470588257 | 0.905472959095347 | NM_009554 |
| Mfap4 | 66.18306093 | -0.0418792819418975 | 0.0872250665655082 | -0.480128976576304 | 0.631135685134907 | 0.905558767509843 | NM_029568 |
| Oas1c | 52.29759082 | -0.0455374682872418 | 0.094885640957223 | -0.479919488637604 | 0.631284642955093 | 0.90568303442624 | NM_033541 |
| Pkhd1l1 | 22.79108359 | -0.0386800306916494 | 0.0806710274989764 | -0.479478592139417 | 0.631598194325111 | 0.906043390358144 | NM_138674 |
| Plcb1 | 12848.0359 | 0.0348093049150546 | 0.0726661531630481 | 0.479030516958144 | 0.631916918862494 | 0.906377073924305 | NM_019677,NM_001145830 |
| Trpt1 | 381.0615654 | -0.0388408662629992 | 0.0810914414473962 | -0.478976148034009 | 0.631955597184728 | 0.906377073924305 | NM_153597,NR_027351 |
| Ibtk | 1243.175737 | 0.0322785846038005 | 0.0674140819639239 | 0.478810712294118 | 0.632073295185528 | 0.906456381268139 | NM_001081282 |
| Cep170 | 3434.43801 | -0.0333112018227214 | 0.0695999866489868 | -0.478609313400009 | 0.632216591483029 | 0.906498455986076 | NM_001099637 |
| Eno4 | 141.1017829 | 0.0446354622937868 | 0.0932900949211442 | 0.478458750969393 | 0.632323726411756 | 0.906498455986076 | NM_178689 |
| Nhej1 | 67.37252186 | -0.0453238103675581 | 0.0947051037228582 | -0.478578329845794 | 0.632238637659382 | 0.906498455986076 | NM_029342 |
| Slc22a3 | 129.4828251 | -0.0454758629331037 | 0.0950545342719621 | -0.478418660208171 | 0.632352254888263 | 0.906498455986076 | NM_011395 |
| Crmp1 | 7886.429112 | 0.0243673446186583 | 0.0509661519232073 | 0.478108385647274 | 0.632573063922918 | 0.906637357927673 | NM_007765,NM_001136058 |
| Gm2061 | 144.5439885 | -0.0435035265760632 | 0.0910106837679216 | -0.478004611931031 | 0.632646922516109 | 0.906637357927673 | NR_040305 |
| Nop10 | 1051.846636 | 0.0360061783623973 | 0.0753651362465384 | 0.477756428975487 | 0.632823575973855 | 0.906637357927673 | NM_025403 |
| Ppp1r15a | 643.6401668 | 0.0371868776232359 | 0.0778337654596791 | 0.477773077065121 | 0.632811725420763 | 0.906637357927673 | NM_008654 |
| Shisa4 | 3874.67045 | 0.0312574362596757 | 0.0654130947708393 | 0.477846773175606 | 0.632759267700977 | 0.906637357927673 | NM_175259 |
| U2af2 | 5928.770422 | 0.0195190480075128 | 0.0408631539152849 | 0.477668660817971 | 0.632886053241539 | 0.906637357927673 | NM_001205231,NM_133671 |
| Zfp873 | 165.4441022 | -0.0411064788739137 | 0.0860465180458057 | -0.477723908038106 | 0.632846725514136 | 0.906637357927673 | NM_001024626 |
| Arfgap3 | 846.3549613 | -0.0350775488986035 | 0.073585139011044 | -0.476693383610227 | 0.633580475001351 | 0.906932171011455 | NM_025445 |
| Atp10a | 781.7540844 | 0.0404083382089626 | 0.0847968032171603 | 0.476531386513226 | 0.633695852272671 | 0.906932171011455 | NM_009728 |
| D130017N08Rik | 511.5963103 | 0.0337583026655654 | 0.0708382435509493 | 0.476554767218152 | 0.633679199560411 | 0.906932171011455 | NR_015486 |
| Gpr21 | 126.2389897 | -0.0453357942370707 | 0.0951022594684308 | -0.476705753264673 | 0.633571665475147 | 0.906932171011455 | NM_177383 |
| Gpr25 | 275.6256105 | -0.0407791555241017 | 0.0855254916569453 | -0.476807028338084 | 0.633499540481777 | 0.906932171011455 | NM_001101516 |
| Phb2 | 2321.231459 | 0.0248604585624962 | 0.0521288616381404 | 0.476903921959173 | 0.633430539084012 | 0.906932171011455 | NM_007531 |
| Ptcd3 | 2087.717201 | -0.0241611439322532 | 0.0506928745553965 | -0.476618146912347 | 0.633634058834571 | 0.906932171011455 | NM_027275 |
| Tmed3 | 924.044474 | 0.030769668763334 | 0.0644866061205934 | 0.477148211301321 | 0.63325658609884 | 0.906932171011455 | NM_025360 |
| Ubtf | 6101.358076 | -0.0247091411310733 | 0.051808027924973 | -0.476936531281531 | 0.633407317541475 | 0.906932171011455 | NM_001044383,NM_011551 |
| Vash1 | 2692.459352 | -0.0306274852151572 | 0.0642755596184043 | -0.476502816886988 | 0.633716201002772 | 0.906932171011455 | NM_177354 |
| Pyroxd1 | 521.2595987 | -0.0356750107613637 | 0.0748902919877663 | -0.476363622232792 | 0.633815346425005 | 0.906984703198662 | NM_183165 |
| Cgnl1 | 854.2880322 | -0.04275134288281 | 0.0898839909648529 | -0.475628000313503 | 0.634339423578965 | 0.907220905508528 | NM_026599 |
| Dda1 | 2175.793023 | 0.0295071378879687 | 0.0619994037882481 | 0.475926155495736 | 0.634126987566061 | 0.907220905508528 | NM_025600 |
| Ica1 | 1917.557601 | 0.0269781551576336 | 0.0567120411060084 | 0.475704182595102 | 0.634285140723348 | 0.907220905508528 | NM_001252266,NM_010492 |
| Med30 | 395.5684241 | -0.0410674136924113 | 0.0863634769376252 | -0.47551829950144 | 0.634417593154043 | 0.907220905508528 | NM_027212 |
| Naca | 6008.750296 | 0.0272158100342624 | 0.0572336647731146 | 0.475521009219857 | 0.634415662239203 | 0.907220905508528 | NM_013608,NM_001113199 |
| Nck2 | 1147.882809 | -0.0314171258773429 | 0.0660118945135148 | -0.47593128645794 | 0.634123332011606 | 0.907220905508528 | NM_010879 |
| Nsun6 | 300.8896949 | -0.0367727509013091 | 0.0773083173093526 | -0.475663579044947 | 0.634314072096286 | 0.907220905508528 | NM_001165941,NM_001165942,NM_028950,NM_001165943 |
| 1700003M02Rik | 70.19258166 | 0.0448457048688808 | 0.0944981119940597 | 0.474567204810397 | 0.635095486139919 | 0.907297111690447 | NM_027041 |
| Dgkd | 5251.983661 | -0.0283972704756631 | 0.0597806400502811 | -0.475024530546651 | 0.634769488875359 | 0.907297111690447 | NM_177646 |
| Exph5 | 1484.129813 | -0.0401169503069589 | 0.0845291626231683 | -0.474593016918913 | 0.635077084510599 | 0.907297111690447 | NM_176846 |
| Flcn | 2347.389585 | -0.0287073009936762 | 0.0604893505251726 | -0.47458438129088 | 0.635083240884002 | 0.907297111690447 | NR_073164,NM_146018,NM_001271357,NM_001271356 |
| Gabra4 | 5052.292896 | 0.0399131255003921 | 0.0840122418565028 | 0.475087018491492 | 0.634724950850818 | 0.907297111690447 | NM_010251 |
| Gas8 | 653.1457828 | -0.0372876638770282 | 0.0784454651938905 | -0.475332306142436 | 0.634550135873672 | 0.907297111690447 | NM_018855 |
| Gpn1 | 1018.430583 | 0.0284759942108566 | 0.0599860782670496 | 0.474710049956683 | 0.634993653682975 | 0.907297111690447 | NM_133756 |
| Kcnh1 | 5240.879835 | -0.0340450983562651 | 0.0716761377313389 | -0.474985112672716 | 0.634797584482191 | 0.907297111690447 | NM_001038607,NM_010600 |
| Mfap1a | 1219.592671 | 0.0348356751062324 | 0.0733979303083249 | 0.474613861179697 | 0.635062224658544 | 0.907297111690447 | NM_026220 |
| Rarres2 | 213.6741156 | 0.04516319177903 | 0.0950567736306561 | 0.475118080006712 | 0.634702812376121 | 0.907297111690447 | NM_027852 |
| Arc | 11827.65662 | 0.0259673179233813 | 0.0547563677635196 | 0.474233755524623 | 0.635333224685804 | 0.907547489604286 | NM_001276684,NM_018790 |
| Btbd9 | 4136.766328 | 0.0305283122086269 | 0.0644268967572252 | 0.473844213289743 | 0.635611003303187 | 0.907676507469731 | NM_027060,NM_172618 |
| Mecr | 485.2086906 | -0.0388480344204848 | 0.0819782481392728 | -0.473882222446202 | 0.635583897101051 | 0.907676507469731 | NM_025297 |
| Ptprt | 4017.176099 | 0.0335430362888319 | 0.0707717157885722 | 0.473961043830566 | 0.635527687244488 | 0.907676507469731 | NM_021464 |
| C1qtnf6 | 80.07360459 | 0.0450329827789542 | 0.0951021925609623 | 0.473522024742881 | 0.635840791435106 | 0.907889736447229 | NM_001204153,NM_001204152,NM_028331 |
| Elp6 | 345.0339147 | 0.0417442310806544 | 0.0881878129366078 | 0.473356007940252 | 0.635959209987093 | 0.907889736447229 | NM_001081381 |
| Fam184a | 1109.383772 | -0.0263028642828768 | 0.0555831114774705 | -0.473216838419313 | 0.636058485743143 | 0.907889736447229 | NM_001081428 |
| Pold4 | 311.8608596 | -0.0408911239764205 | 0.0863685580012207 | -0.473449191728343 | 0.635892741537336 | 0.907889736447229 | NM_027196 |
| Zfp930 | 448.155642 | -0.0413832463857486 | 0.0874546314852419 | -0.473196738502432 | 0.636072824440551 | 0.907889736447229 | NM_001013379 |
| Jmjd8 | 1479.815022 | 0.0319839273690042 | 0.0677286595775959 | 0.472236237487626 | 0.636758176880788 | 0.90877866745631 | NM_028101 |
| 1700029I15Rik | 31.27177332 | -0.0415080391125757 | 0.0882387996276527 | -0.470405754472296 | 0.638065153666478 | 0.908932006246768 | NM_183112 |
| 2410004B18Rik | 534.2046695 | 0.0364134971020524 | 0.0773383845169105 | 0.470833433223452 | 0.637759687332527 | 0.908932006246768 | NM_025555 |
| 6720456H20Rik | 1179.963032 | 0.0303400808683216 | 0.0644663842525693 | 0.470634133123612 | 0.637902028281234 | 0.908932006246768 | NM_172600 |
| Auts2 | 2886.007818 | 0.0357698373588893 | 0.0759831542909142 | 0.470760100613071 | 0.637812060230774 | 0.908932006246768 | NM_177047 |
| Bbs7 | 801.190529 | -0.0298855629944962 | 0.0635407423740054 | -0.470337013354167 | 0.638114257227848 | 0.908932006246768 | NM_027810 |
| Cacna2d1 | 8105.616857 | 0.0354879323545913 | 0.0753221686943092 | 0.471148573783331 | 0.637534639910945 | 0.908932006246768 | NM_001110845,NM_001110844,NM_001110846,NM_001110843,NM_009784 |
| Ctnnd1 | 3668.825215 | -0.0306487711295196 | 0.0651826764996414 | -0.470198107463235 | 0.638213486149203 | 0.908932006246768 | NM_001085453,NM_001085448,NM_001085449,NM_007615,NM_001085450 |
| Eif1 | 12720.74894 | 0.0246676957802429 | 0.0523464575228521 | 0.471239066549519 | 0.637470023607459 | 0.908932006246768 | NM_011508 |
| Eif2b1 | 961.0461242 | 0.0252588173137824 | 0.0535375249457597 | 0.471796507951624 | 0.637072043702117 | 0.908932006246768 | NM_145371 |
| Epn1 | 14982.43468 | 0.0297153184490479 | 0.0630161642117941 | 0.471550733382886 | 0.637247499188473 | 0.908932006246768 | NM_010147,NM_001252454 |
| Fam98b | 1720.417333 | -0.0267448136970568 | 0.0568218061181928 | -0.470678697565965 | 0.637870199014597 | 0.908932006246768 | NM_026620 |
| Gid8 | 1858.276807 | 0.0269636962763412 | 0.0571817190998742 | 0.471543995192696 | 0.637252309787239 | 0.908932006246768 | NM_029607 |
| Gtpbp6 | 925.8955132 | -0.0364838864665827 | 0.0774341494144659 | -0.471160163086481 | 0.637526364422399 | 0.908932006246768 | NM_145147 |
| Lrrc7 | 2448.769855 | 0.0353902141735268 | 0.0752693071180414 | 0.47018121367885 | 0.638225554848371 | 0.908932006246768 | NM_001081358 |
| Maob | 1186.620419 | 0.0314005739211271 | 0.0667551165531816 | 0.470384526946505 | 0.63808031686859 | 0.908932006246768 | NM_172778 |
| Nat1 | 57.60594234 | -0.0448112217289911 | 0.0950283924971407 | -0.471556137607394 | 0.637243640959395 | 0.908932006246768 | NM_008673 |
| Pip5k1a | 2277.641769 | 0.0319257543627949 | 0.0678679701126316 | 0.470409742766903 | 0.638062304773059 | 0.908932006246768 | NM_008847 |
| Ppie | 409.9375836 | -0.0335091581289637 | 0.0710234910534499 | -0.471803872661593 | 0.637066786438813 | 0.908932006246768 | NM_019489 |
| Psmd5 | 1809.073795 | 0.0261665412565598 | 0.0556547957768142 | 0.47015788830656 | 0.638242218347586 | 0.908932006246768 | NM_080554 |
| Trpm6 | 41.67163737 | -0.042179155433154 | 0.089521278435238 | -0.471163461586035 | 0.63752400909532 | 0.908932006246768 | NM_153417 |
| Zc3h7b | 10364.64474 | 0.0245977884163128 | 0.0523001303417406 | 0.470319830095745 | 0.638126531922496 | 0.908932006246768 | NM_001081016 |
| Zfp605 | 603.455958 | -0.036276332756122 | 0.0769630201742734 | -0.471347572821059 | 0.637392548404456 | 0.908932006246768 | NM_001163996 |
| Snhg12 | 543.5105242 | -0.0330259190711031 | 0.0702927318905918 | -0.46983405229586 | 0.638473583633477 | 0.909172363087921 | NR_029468 |
| Cdadc1 | 2388.935397 | 0.0256564094275203 | 0.0546481016149948 | 0.469484001626884 | 0.638723717628678 | 0.909380944976886 | NM_001168536,NM_001168537,NM_001168535,NM_027986,NM_001168538 |
| Pemt | 65.72776903 | -0.0445491224891846 | 0.0949191914863439 | -0.469337357299276 | 0.63882851676897 | 0.909380944976886 | NM_008819 |
| Pln | 68.66275445 | -0.043088368211331 | 0.0917965139043017 | -0.469390027776554 | 0.638790875064226 | 0.909380944976886 | NM_023129,NM_001141927 |
| Spag17 | 28.27072378 | -0.0368268221350472 | 0.0784753843749633 | -0.469278646143165 | 0.638870476627366 | 0.909380944976886 | NM_028892 |
| Dusp1 | 1507.804976 | 0.0361657564956516 | 0.0771111191234596 | 0.46900832080712 | 0.63906368843246 | 0.909490331847324 | NM_013642 |
| Pbx1 | 6943.50812 | 0.0252567986373434 | 0.0538629764964214 | 0.468908335190526 | 0.639135158164497 | 0.909490331847324 | NM_183355,NM_008783 |
| Tox3 | 567.1624606 | 0.0397842222244219 | 0.0848330837368341 | 0.46897060052466 | 0.639090650501725 | 0.909490331847324 | NM_172913 |
| 3300002I08Rik | 24.08072593 | 0.0400618248640447 | 0.0855340803611005 | 0.468372661457458 | 0.639518114872348 | 0.909767910746815 | NM_027017 |
| Abhd12 | 12355.33395 | 0.0204391471411525 | 0.0436347293279194 | 0.468414665473235 | 0.639488082449407 | 0.909767910746815 | NM_024465 |
| Upk1b | 68.50374693 | -0.0442927645069372 | 0.0945434067273561 | -0.468491310395324 | 0.63943328367103 | 0.909767910746815 | NM_178924 |
| Adamts1 | 511.678778 | -0.0430221993453712 | 0.0919625960704931 | -0.467822801700737 | 0.639911313206765 | 0.910092094811638 | NM_009621 |
| Calml4 | 53.76396739 | 0.0170309593648894 | 0.0364452331544761 | 0.467302796299924 | 0.640283256107836 | 0.910092094811638 | NM_001102468,NM_138304 |
| Coa4 | 80.24238244 | 0.044411908733895 | 0.0950481003692666 | 0.467257194634638 | 0.640315877803738 | 0.910092094811638 | NM_183270 |
| Epha8 | 514.1184521 | -0.042097378505825 | 0.0900003807259357 | -0.467746671361509 | 0.639965761103899 | 0.910092094811638 | NM_007939 |
| Fastkd2 | 772.7308968 | -0.0311141285789679 | 0.0666001565134564 | -0.467178009899741 | 0.640372525201002 | 0.910092094811638 | NM_172422 |
| Hist3h2a | 986.5866838 | 0.0327615118941744 | 0.0701256215911125 | 0.467183194256726 | 0.640368816337079 | 0.910092094811638 | NM_178218 |
| Kcnu1 | 145.7448643 | -0.0431504682531713 | 0.0923603449678623 | -0.467196915171613 | 0.640359000504253 | 0.910092094811638 | NM_008432 |
| Plin2 | 264.1238248 | -0.0391841364382313 | 0.0838207715438189 | -0.467475253645775 | 0.640159892971775 | 0.910092094811638 | NM_007408 |
| Tmem205 | 599.0355709 | 0.0300252810798768 | 0.0642589328012723 | 0.467254586576051 | 0.640317743530612 | 0.910092094811638 | NM_178577,NM_001253869,NM_001253868,NR_045606,NM_001253870,NM_001253867 |
| Zc3h6 | 845.0874689 | -0.0366041421244767 | 0.0783064682706757 | -0.467447235622351 | 0.640179934301679 | 0.910092094811638 | NM_178404 |
| 2810004N23Rik | 442.6856408 | 0.0362695480333424 | 0.0776634581327196 | 0.467009181735909 | 0.64049330896209 | 0.910174702209286 | NM_025615 |
| Mtx1 | 915.3971421 | -0.0258935868644627 | 0.0555000147825807 | -0.466550990407838 | 0.640821158133676 | 0.910551515509125 | NM_013604,NM_001161824 |
| Gpank1 | 555.0422649 | -0.0300630316792482 | 0.0644801293606743 | -0.466237148984123 | 0.641045761216567 | 0.910692491680376 | NM_032460,NM_001128597 |
| Morc4 | 163.4447336 | -0.0417800800758058 | 0.0895985772446459 | -0.46630294096888 | 0.640998673941857 | 0.910692491680376 | NM_001193309,NM_029413 |
| 1190005I06Rik | 155.7561113 | -0.0421019482514453 | 0.0903802200394972 | -0.465831442245287 | 0.641336157138555 | 0.911015941579762 | NM_197988 |
| C330018D20Rik | 375.0079947 | 0.0347803298106636 | 0.0747176157800752 | 0.465490359235184 | 0.641580339293971 | 0.91109559180768 | NM_029909 |
| Carns1 | 644.5477731 | -0.0387417507649498 | 0.0832486681935258 | -0.465373820454256 | 0.641663778578243 | 0.91109559180768 | NM_134148 |
| Ccdc106 | 1239.868657 | -0.0323919440622995 | 0.0695777283064182 | -0.465550469248528 | 0.641537303574414 | 0.91109559180768 | NM_146178 |
| Mtdh | 2534.115481 | -0.0340924428420202 | 0.0732674361251378 | -0.465315079181857 | 0.641705837793224 | 0.91109559180768 | NM_026002 |
| St6galnac2 | 202.76118 | 0.0433870097406638 | 0.0932361137700377 | 0.465345540330818 | 0.641684027226471 | 0.91109559180768 | NM_009180 |
| Fert2 | 655.3576907 | 0.0346058209911996 | 0.0743980700837818 | 0.465144068283344 | 0.641828289492025 | 0.911180388307385 | NM_001037997,NM_008000 |
| Anapc7 | 1562.940931 | 0.0223513783881834 | 0.0482963185855072 | 0.462796731568908 | 0.643510074230162 | 0.91125847294421 | NM_019805 |
| Ankrd34a | 2880.020847 | 0.0251880633386726 | 0.0543719596212151 | 0.463254653945646 | 0.64318184510033 | 0.91125847294421 | NM_001024851 |
| Atn1 | 8130.227989 | -0.0253020385150925 | 0.0546346776966889 | -0.463113165150528 | 0.643283253863293 | 0.91125847294421 | NM_007881 |
| Atp2b1 | 27462.82502 | 0.0367691716597333 | 0.079193587837967 | 0.464294807995874 | 0.642436543314644 | 0.91125847294421 | NM_026482 |
| C630020P19Rik | 161.2830284 | 0.0426878572913934 | 0.0920767129240835 | 0.463611872489293 | 0.642925846696349 | 0.91125847294421 | NR_040354 |
| Ddx59 | 326.562234 | 0.0384632429005435 | 0.0827640008869777 | 0.464733972359176 | 0.642121976790088 | 0.91125847294421 | NM_026500 |
| Dhx38 | 1673.409398 | 0.0274631291778462 | 0.0593337714326361 | 0.462858310111403 | 0.643465931976178 | 0.91125847294421 | NM_178380 |
| Dmxl2 | 10188.49897 | -0.0282263773893792 | 0.0608781039463188 | -0.463654016134744 | 0.642895647515297 | 0.91125847294421 | NM_172771 |
| Dolpp1 | 713.8268362 | -0.0355871098959085 | 0.0765755889213456 | -0.464731781984226 | 0.642123545561956 | 0.91125847294421 | NM_020329 |
| Fgf13 | 4360.349527 | 0.0310517515845086 | 0.0670914660267451 | 0.462827143650881 | 0.643488273332081 | 0.91125847294421 | NM_010200 |
| Gabpa | 892.6517351 | -0.0354586559626243 | 0.0765575646736995 | -0.463163321792624 | 0.643247304510493 | 0.91125847294421 | NM_008065 |
| Kcnmb1 | 58.44556278 | 0.0420968998997821 | 0.0908619631038353 | 0.463306079483167 | 0.64314498870381 | 0.91125847294421 | NM_031169 |
| Kdm1b | 385.9473133 | -0.0350137249260746 | 0.0755737524592027 | -0.463305364451451 | 0.643145501157042 | 0.91125847294421 | NM_172262 |
| Mark4 | 2265.909417 | -0.028733039322717 | 0.061994292884679 | -0.463478781444703 | 0.64302122059591 | 0.91125847294421 | NM_172279 |
| Ncaph2 | 4746.006042 | 0.0256492457769992 | 0.055398238157715 | 0.462997500100591 | 0.643366159001422 | 0.91125847294421 | NM_001115132,NR_073376,NM_001271601,NM_001271600 |
| Neo1 | 5003.096134 | 0.0252910512459224 | 0.0544656738730806 | 0.464348450087246 | 0.6423981168884 | 0.91125847294421 | NM_008684,NM_001042752 |
| Nrp1 | 1194.038201 | 0.038723820913975 | 0.0834595882838984 | 0.46398288932664 | 0.642660004713493 | 0.91125847294421 | NM_008737 |
| Nucb2 | 636.1071033 | 0.0358089654438139 | 0.077104466071112 | 0.464421417700862 | 0.642345848191754 | 0.91125847294421 | NM_001130479 |
| Pitpnm1 | 8973.384169 | -0.0301781769990715 | 0.0652091159755178 | -0.462790770088061 | 0.643514347753112 | 0.91125847294421 | NM_001136078,NM_008851,NR_075078 |
| Ppargc1b | 362.424251 | 0.0386152131774737 | 0.0834183356725826 | 0.46291037655125 | 0.643428609404517 | 0.91125847294421 | NM_133249 |
| Prkd1 | 497.8617416 | 0.0344200858179707 | 0.0743439294600518 | 0.462984483978159 | 0.643375488834223 | 0.91125847294421 | NM_008858 |
| Ptplb | 1555.183367 | 0.0251140417446886 | 0.0541475438679883 | 0.463807588501458 | 0.64278560603169 | 0.91125847294421 | NM_023587 |
| S100a11 | 155.8520556 | -0.0440218608790539 | 0.0950643158553255 | -0.463074503644974 | 0.643310964743242 | 0.91125847294421 | NM_016740 |
| Serp1 | 1428.499196 | 0.0332500654903251 | 0.0716157890524711 | 0.464284006784644 | 0.64244428086077 | 0.91125847294421 | NM_030685 |
| Slc6a13 | 380.4688131 | 0.0391139740502072 | 0.084450701706085 | 0.463157478387049 | 0.64325149267948 | 0.91125847294421 | NM_144512 |
| Tnfaip8l1 | 281.9734565 | 0.0395822350443129 | 0.0853112267572914 | 0.463974514830546 | 0.642666004725303 | 0.91125847294421 | NM_025566 |
| Alg13 | 270.8259585 | -0.0412487861392972 | 0.0891681903837008 | -0.46259530401816 | 0.643654475296447 | 0.911368058110556 | NR_037145,NM_026247 |
| Ankrd40 | 5968.972367 | 0.0233036391926482 | 0.0504766869254401 | 0.461671330114639 | 0.644317033790126 | 0.912152041147281 | NM_146024,NM_027799 |
| Dhx32 | 1455.253134 | 0.0268943991634072 | 0.0582573693045562 | 0.461648019545982 | 0.644333752871558 | 0.912152041147281 | NM_133941 |
| Ap1m1 | 2714.080932 | -0.0291742530109673 | 0.0632182599114105 | -0.461484594037387 | 0.644450971897375 | 0.912229079885136 | NM_007456 |
| Gm13251 | 30.20971057 | -0.0383869300726826 | 0.0832451577940977 | -0.461131086658885 | 0.644704559805187 | 0.91235393164436 | NM_001085522 |
| Lmo4 | 5907.638661 | -0.0257610655690227 | 0.0558727191530533 | -0.461066974357466 | 0.644750555083441 | 0.91235393164436 | NM_010723,NM_001161769,NM_001161770 |
| Mlph | 132.4334585 | 0.0437556429352925 | 0.094906165606449 | 0.461041099444852 | 0.644769118580138 | 0.91235393164436 | NM_053015 |
| Pbx2 | 2341.268921 | -0.0231939005356084 | 0.0503109017236239 | -0.461011425774496 | 0.644790407700744 | 0.91235393164436 | NM_017463 |
| C1ql2 | 148.9309008 | 0.0332083627900049 | 0.0721209126952168 | 0.460454000774277 | 0.645190381597211 | 0.912363974014534 | NM_207233 |
| Cetn3 | 1335.523653 | 0.027645313505322 | 0.0600651731931547 | 0.460255286643752 | 0.645332991465613 | 0.912363974014534 | NM_007684 |
| Chd6 | 4016.021013 | -0.0307003263754898 | 0.0666514749046554 | -0.460609857762434 | 0.645078537863556 | 0.912363974014534 | NM_173368 |
| Dclk3 | 2784.412726 | -0.0377692520806171 | 0.0821631815591369 | -0.459685851544498 | 0.645741726451579 | 0.912363974014534 | NM_172928 |
| Dusp7 | 2864.873115 | 0.0279364903901553 | 0.0607208381909518 | 0.460080776591095 | 0.645458241709358 | 0.912363974014534 | NM_153459 |
| Faf1 | 1463.780132 | 0.0259546484824679 | 0.0564198087000333 | 0.460027232996495 | 0.645496673307058 | 0.912363974014534 | NM_007983 |
| Fbxl16 | 44591.63305 | -0.0311888501057129 | 0.0676697536194356 | -0.460897940919281 | 0.644871829124597 | 0.912363974014534 | NM_001164225 |
| Hif1a | 2910.067772 | 0.0284093137776622 | 0.0618227093155315 | 0.459528773361685 | 0.645854494480441 | 0.912363974014534 | NM_010431 |
| Mcm5 | 262.3850434 | 0.0392162140695962 | 0.0851109194934499 | 0.460765954627177 | 0.644966530040733 | 0.912363974014534 | NM_008566 |
| Paip2 | 4320.151563 | 0.0294364458930029 | 0.0639232168887864 | 0.460496941889148 | 0.645159566042887 | 0.912363974014534 | NM_026420 |
| Pawr | 70.889627 | 0.0437239701758879 | 0.0950187608442118 | 0.460161443776094 | 0.645400343596426 | 0.912363974014534 | NM_054056 |
| Pkdrej | 99.70849658 | 0.043723199004037 | 0.0950898706917877 | 0.459809217174728 | 0.645653166716356 | 0.912363974014534 | NM_011105 |
| Plxna4 | 4677.338901 | 0.0329200323033463 | 0.071490052493563 | 0.460484097508677 | 0.645168783408636 | 0.912363974014534 | NM_175750 |
| Pou3f4 | 130.4732084 | 0.0433954218930662 | 0.0943903910928122 | 0.459744062829407 | 0.645699938044393 | 0.912363974014534 | NM_008901 |
| Serinc2 | 250.6216452 | 0.0430551405643112 | 0.0936582939610976 | 0.459704514607054 | 0.645728328589837 | 0.912363974014534 | NM_001253386,NM_172702 |
| Uxt | 183.5071012 | -0.0405850151382928 | 0.0883216496808755 | -0.459513780425694 | 0.64586525848764 | 0.912363974014534 | NM_013840 |
| Vstm2l | 1982.502138 | 0.0314275483687851 | 0.0682914760506163 | 0.460197233773241 | 0.645374656351034 | 0.912363974014534 | NM_198627 |
| Arfgef1 | 5610.477091 | 0.0348249705378546 | 0.0758310228894751 | 0.459244372697076 | 0.646058689326913 | 0.912548475414502 | NM_001102430 |
| Map1s | 2081.981465 | -0.024716455421135 | 0.0538500668821868 | -0.458986531534114 | 0.646243837954313 | 0.912721243570671 | NM_173013 |
| AW146154 | 154.9836677 | 0.0409745405843347 | 0.0895150815700208 | 0.457738962705222 | 0.647139992060519 | 0.912750424296538 | NM_001033530 |
| Cd6 | 150.4396791 | -0.0432422359844302 | 0.0945266607281262 | -0.457460738074752 | 0.647339916330343 | 0.912750424296538 | NM_001037801,NM_009852 |
| Dtnbp1 | 1937.890772 | 0.0221652693450166 | 0.0484045192922773 | 0.457917352947516 | 0.647011819323996 | 0.912750424296538 | NM_025772 |
| Esf1 | 1206.613117 | 0.0326122340795478 | 0.0712948662616547 | 0.457427523040154 | 0.647363785405718 | 0.912750424296538 | NM_001081090 |
| Hrh1 | 777.7074855 | 0.0357489579282143 | 0.0781029158717586 | 0.457716047207667 | 0.647156457520359 | 0.912750424296538 | NM_001252642,NM_001252643,NM_008285 |
| Ipo4 | 3777.456341 | -0.0265369241691328 | 0.0579570827270457 | -0.457871979066146 | 0.647044419299077 | 0.912750424296538 | NM_024267 |
| Katnb1 | 2060.248964 | 0.0288362012799266 | 0.062974095831823 | 0.457905761075725 | 0.647020147723449 | 0.912750424296538 | NM_028805 |
| Klc2 | 11693.53747 | 0.0193115376934123 | 0.0421471689375993 | 0.458192998016162 | 0.646813789939164 | 0.912750424296538 | NR_045528,NM_008451 |
| Mrps17 | 1281.809489 | 0.0234353575446828 | 0.0511271605900814 | 0.45837393029859 | 0.646683817865352 | 0.912750424296538 | NM_025450 |
| Myo1h | 83.67972504 | 0.0429359118494211 | 0.0938543693035373 | 0.457473766730676 | 0.647330553742443 | 0.912750424296538 | NM_001164573 |
| Parl | 802.1282467 | -0.0294049546724529 | 0.064154448439337 | -0.458346309379583 | 0.646703658559098 | 0.912750424296538 | NM_001005767 |
| Pxmp4 | 446.1113049 | -0.036128904720691 | 0.0789032116074299 | -0.457888899382759 | 0.64703226240448 | 0.912750424296538 | NM_021534 |
| Rbm27 | 1667.073722 | 0.0314743877432126 | 0.068799167412882 | 0.45748210228078 | 0.647324563720171 | 0.912750424296538 | NM_172626 |
| Shh | 120.1114167 | -0.0424728031784045 | 0.0928434135712953 | -0.457467057108895 | 0.647335375370496 | 0.912750424296538 | NM_009170 |
| Sri | 1928.940421 | 0.0246146425346915 | 0.0537141293696616 | 0.458252657606961 | 0.64677093248937 | 0.912750424296538 | NM_001080974,NM_025618 |
| Tmem109 | 1728.013501 | 0.0278424108216439 | 0.0607608563218376 | 0.458229401412128 | 0.646787638820676 | 0.912750424296538 | NM_134142 |
| Wdr59 | 1504.457292 | -0.0253838508916301 | 0.0553873790770111 | -0.458296660983655 | 0.646739322689669 | 0.912750424296538 | NM_176923,NM_001170742,NM_001170743 |
| Zfp119b | 91.30390689 | 0.0433777395706454 | 0.0948389157501002 | 0.457383334969217 | 0.647395540515437 | 0.912750424296538 | NM_146249 |
| Vezt | 2002.630666 | -0.025710080112529 | 0.0562402333166229 | -0.457147465370309 | 0.647565055601764 | 0.912900814991385 | NM_172538 |
| Crtap | 222.9143748 | -0.0407214052289468 | 0.0890976811324712 | -0.457042256446628 | 0.647640673190671 | 0.912918817929907 | NM_019922 |
| Commd8 | 1324.560879 | -0.0261290441258869 | 0.0571919739531482 | -0.456865576056037 | 0.64776766817637 | 0.912960942585905 | NM_178599 |
| Fnbp1 | 5979.701557 | -0.0293862168646419 | 0.0643269636610631 | -0.45682580355381 | 0.647796257416558 | 0.912960942585905 | NM_019406,NM_001038700,NM_001177648,NM_001177649,NM_001177650 |
| Gsk3a | 10343.82716 | 0.0189157370651219 | 0.0414322577104653 | 0.456546133626312 | 0.647997304210221 | 0.913155688878315 | NM_001031667 |
| 2310036O22Rik | 1344.550205 | -0.0332355875383586 | 0.0728282337747291 | -0.456355808945776 | 0.648134137915744 | 0.913259917292085 | NM_026760 |
| Lmnb1 | 284.5565653 | -0.0356950687790752 | 0.0782622570126567 | -0.456095570733445 | 0.64832125510815 | 0.913290059990352 | NM_010721 |
| Ramp1 | 2053.921042 | 0.0237556376727786 | 0.0520884164380016 | 0.456063733499248 | 0.64834414832855 | 0.913290059990352 | NM_001168392,NM_178401,NM_016894 |
| Stag2 | 2683.478246 | 0.0302795404629697 | 0.0663697185881441 | 0.456225235048363 | 0.648228020740239 | 0.913290059990352 | NM_001077712,NM_021465 |
| Ccdc81 | 43.09806391 | -0.0406360409590776 | 0.0894429315267433 | -0.454323670584605 | 0.649595881545614 | 0.913724196313701 | NM_001162979 |
| Dtwd2 | 114.0674526 | 0.0423140373824871 | 0.0930459782305791 | 0.454764818288308 | 0.649278443345446 | 0.913724196313701 | NM_001170960,NM_026854 |
| Fut11 | 773.763657 | -0.0285156775963543 | 0.0626767296123216 | -0.454964350768366 | 0.649134885970414 | 0.913724196313701 | NM_028428 |
| Gdpd2 | 363.143587 | 0.0414318180745081 | 0.0911159448721187 | 0.454715342442617 | 0.649314041683809 | 0.913724196313701 | NM_023608 |
| Gucd1 | 612.0142677 | 0.0313412990818842 | 0.0689757812429988 | 0.454381212029626 | 0.649554472636738 | 0.913724196313701 | NM_175133 |
| Lrrc18 | 24.4443545 | 0.0341200873262746 | 0.0750049637577111 | 0.454904390547976 | 0.64917802410311 | 0.913724196313701 | NM_001146021,NM_026253 |
| Mkl2 | 12853.78058 | 0.0330782656541614 | 0.0727024754373493 | 0.454981284408484 | 0.649122703346125 | 0.913724196313701 | NM_181860,NM_153588,NM_001122667 |
| Mrc2 | 359.6322112 | -0.0430085015380699 | 0.0946407300159718 | -0.454439663882682 | 0.649512409673734 | 0.913724196313701 | NM_008626 |
| Phf12 | 2617.917342 | -0.0218016053862127 | 0.0479759929942488 | -0.454427392233991 | 0.649521240471423 | 0.913724196313701 | NM_174852 |
| Pqlc2 | 404.6375803 | -0.0372620289904141 | 0.0818318345921978 | -0.455348815972492 | 0.648858312037635 | 0.913724196313701 | NM_145384 |
| St3gal5 | 6622.68357 | 0.0250857514312569 | 0.0550824767807929 | 0.455421631294622 | 0.648805936104311 | 0.913724196313701 | NM_011375,NM_001035228 |
| Tmem229a | 2307.726268 | 0.0292799582659284 | 0.0643126378363136 | 0.455275343245145 | 0.64891116260159 | 0.913724196313701 | NM_177013 |
| Tmem97 | 624.066566 | 0.031655888621117 | 0.0695502140674217 | 0.455151562731783 | 0.649000204670874 | 0.913724196313701 | NM_133706 |
| Zbtb20 | 92.19674938 | 0.0392865180951493 | 0.086352800004817 | 0.454953609992471 | 0.649142613290716 | 0.913724196313701 | NM_181058,NM_019778 |
| Zfp111 | 481.1942115 | 0.0342182919951793 | 0.0752647039253444 | 0.454639295852683 | 0.649368759485175 | 0.913724196313701 | NM_019940 |
| Atxn7l3 | 12622.98767 | 0.0244792461847524 | 0.0539106250739961 | 0.454070902556833 | 0.649777795398556 | 0.913891581715669 | NM_001098837,NM_001098836 |
| 2610002J02Rik | 736.6321611 | 0.0343424026995982 | 0.0757066596281696 | 0.45362459350696 | 0.650099049177874 | 0.91400066498865 | NM_001190445,NR_033802 |
| Emc8 | 2824.10095 | 0.0224214389879909 | 0.0494609371452343 | 0.453316097148622 | 0.650321143242357 | 0.91400066498865 | NM_010926 |
| Fam213b | 2031.652552 | 0.0299921216674582 | 0.0661738560723253 | 0.45323218938127 | 0.650381555867645 | 0.91400066498865 | NM_025582 |
| Iqsec3 | 9870.081603 | -0.0276740754478604 | 0.0610019241086979 | -0.453659058336399 | 0.650074239034868 | 0.91400066498865 | NM_001033354 |
| Nae1 | 1497.610285 | -0.0301614914948922 | 0.0664948499135551 | -0.453591391425094 | 0.650122950675584 | 0.91400066498865 | NM_144931 |
| Ndufa4l2 | 85.760714 | -0.0427206615882475 | 0.0942503844764795 | -0.45326776994643 | 0.650355937990267 | 0.91400066498865 | NM_001098789 |
| Tmbim4 | 1090.074995 | 0.0259300250686312 | 0.0572048480284758 | 0.45328369818802 | 0.6503444698482 | 0.91400066498865 | NM_026617 |
| Utp11l | 493.4128912 | -0.0319115088246951 | 0.0703092535276125 | -0.453873526223152 | 0.649919859090235 | 0.91400066498865 | NM_026031 |
| Zfp128 | 391.7335756 | -0.0357814897606451 | 0.0789570715675618 | -0.453176505286517 | 0.650421649042247 | 0.91400066498865 | NM_153802 |

| | | | | | | | | | | | | | |
|---|---|---|---|---|---|---|---|---|---|---|---|---|---|
| Kat6b | 2646.855897 | -0.059763926 | 0.077594231 | -0.770210944 | 0.441174773 | 0.661207384 | Ier2 | 240.7875535 | -0.0374546739163356 | 0.08268282984536 | -0.452992162778975 | 0.650554384939705 | 0.914098761427177 | NM_010499 |
| Efna3 | 1252.100074 | 0.065556317 | 0.08512484 | 0.770119711 | 0.441228885 | 0.661224523 | Rpl31 | 6542.748419 | -0.0285919084764999 | 0.063140579845633 | -0.452829361820905 | 0.650671619053212 | 0.91417505932556 | NM_053257,NM_001252219,NM_001252218 |
| Cnga4 | 53.81050859 | 0.123995865 | 0.161048584 | 0.769928315 | 0.441342417 | 0.661314189 | Pi4k2b | 84.54333072 | -0.0429826295402159 | 0.0949415864265235 | -0.452727104718022 | 0.65074525953075 | 0.914190100574824 | NM_025951,NM_028744 |
| Klf10 | 1024.191863 | -0.127876066 | 0.166099791 | -0.769874937 | 0.441374082 | 0.661314189 | Sec61a1 | 2824.68651 | -0.0246691046726713 | 0.0545113186980399 | -0.452550135675921 | 0.650872711884664 | 0.914280728443732 | NM_016906 |
| Kctd3 | 3248.118637 | 0.055856145 | 0.072569907 | 0.769687429 | 0.441485329 | 0.66141691 | Abcc4 | 680.8562222 | 0.0332226816382289 | 0.0735390174778216 | 0.451769452158487 | 0.651435078640717 | 0.914343204175973 | NM_001033336,NM_001163675,NM_001163676 |
| Sec16b | 61.88444107 | 0.119578043 | 0.155389342 | 0.769538253 | 0.441573845 | 0.661485561 | Arhgap32 | 20119.87912 | 0.0355555393792159 | 0.0786849024738033 | 0.451872446446172 | 0.651360875157759 | 0.914343204175973 | NM_177379,NM_001195632 |
| Lurap1l | 305.4903035 | 0.099801273 | 0.129716091 | 0.769382365 | 0.441666354 | 0.661560179 | Cd4 | 642.3354204 | -0.0428250011075506 | 0.094744943496429 | -0.45200302546135 | 0.651266802888195 | 0.914343204175973 | NM_013488 |
| 2510002D24Rik | 404.5670678 | 0.075080422 | 0.097635212 | 0.768989185 | 0.441899731 | 0.661742575 | Hyal1 | 186.4544206 | 0.0419352665362044 | 0.0927853542369916 | 0.45195997666931 | 0.651297815662056 | 0.914343204175973 | NM_008317 |
| Ccdc112 | 315.5923076 | 0.066937756 | 0.08704957 | 0.768961353 | 0.441916253 | 0.661742575 | Klhl23 | 3474.773143 | -0.0325811595491332 | 0.0721034074726739 | -0.451867126549893 | 0.651364707856941 | 0.914343204175973 | NM_177784 |
| Cpeb3 | 2802.3721 | -0.069076952 | 0.089830784 | -0.768967485 | 0.441912613 | 0.661742575 | Lrrc27 | 363.5664686 | -0.0324299894025743 | 0.0717951021664594 | -0.451701974424164 | 0.651483695664417 | 0.914343204175973 | NM_027164,NM_001143755 |
| Rps26 | 2652.39605 | -0.042582526 | 0.055394184 | -0.768718359 | 0.442060523 | 0.661830684 | Snx27 | 7077.109587 | 0.027645849193456 | 0.0611332062617293 | 0.452223118726932 | 0.651108254961894 | 0.914343204175973 | NM_001082484,NM_029721 |
| Scaf11 | 2813.169685 | 0.061890751 | 0.080504731 | 0.768784022 | 0.442021536 | 0.661830684 | Ttc9c | 1433.001892 | 0.0289852628123062 | 0.0641555165933445 | 0.451796889050585 | 0.651415311061836 | 0.914343204175973 | NM_027412 |
| Rap2c | 994.1624724 | -0.070376406 | 0.091571882 | -0.768537291 | 0.442168045 | 0.661927699 | Usp8 | 3229.341828 | -0.0228954126360867 | 0.0506703863802587 | -0.451849971386972 | 0.651377067292032 | 0.914343204175973 | NM_019729,NM_001252580 |
| Ccne2 | 125.365141 | 0.117554557 | 0.152987356 | 0.768393938 | 0.44225318 | 0.661991187 | Gna11 | 5997.842192 | 0.0249274592954855 | 0.0552435963914573 | 0.451228032274529 | 0.651825207925625 | 0.914447302961913 | NM_010301 |
| 4930502E18Rik | 8.639294525 | -0.121133309 | 0.157733343 | -0.767962601 | 0.442509404 | 0.66224676 | Gtl3 | 1283.418068 | -0.0258204292461704 | 0.0572420022581796 | -0.451074879067159 | 0.651935582367725 | 0.914447302961913 | NM_008187 |
| Scara3 | 1302.814859 | 0.076207022 | 0.099229964 | 0.767983975 | 0.442496705 | 0.66224676 | Maf1 | 2504.679052 | -0.0237640861278846 | 0.0526717333827728 | -0.451173420764183 | 0.651864564469968 | 0.914447302961913 | NM_026859,NM_001164607,NM_001164608 |
| Mlph | 115.2625895 | 0.092306639 | 0.120259713 | 0.767560781 | 0.44274817 | 0.662540095 | Snx24 | 570.3194726 | 0.0305164539709436 | 0.0676433352310903 | 0.451137630436023 | 0.651890357793664 | 0.914447302961913 | NM_029394 |
| Akap7 | 1098.413956 | 0.071226414 | 0.092820322 | 0.767357976 | 0.442868707 | 0.66265647 | Trim27 | 989.2562488 | 0.0270214017543113 | 0.0598490145517652 | 0.451492844902873 | 0.651634380777375 | 0.914447302961913 | NM_009054 |
| Zcchc2 | 1202.706098 | 0.04760825 | 0.062070649 | 0.767000992 | 0.443080925 | 0.66290999 | Zfand5 | 6485.344159 | 0.0325951862365728 | 0.0722580996083298 | 0.451093876164096 | 0.651921891127015 | 0.914447302961913 | NM_009551 |
| Naalad2 | 64.46796623 | 0.121366929 | 0.158355062 | 0.766422795 | 0.443424774 | 0.663360379 | Nek5 | 24.64126282 | -0.0302415319384249 | 0.067065273802291 | -0.450926839239891 | 0.652042278947145 | 0.914508655400814 | NM_177898 |
| Nlk | 4774.215292 | -0.059811614 | 0.07806211 | -0.76620544 | 0.443554072 | 0.663489747 | Pir | 210.8202146 | -0.0377939552913593 | 0.0838395627665323 | -0.450789031386101 | 0.652141607449127 | 0.914559663043639 | NM_027153 |
| Ankdd1b | 11.78846257 | -0.12746447 | 0.166414656 | -0.765944981 | 0.44370904 | 0.663529376 | Gls | 15759.56626 | -0.0303461870687585 | 0.067351716681437 | -0.45056293386437 | 0.652304586332618 | 0.914611623655175 | NM_001113383,NM_001081081 |
| Msh6 | 792.8022391 | 0.052456082 | 0.068477865 | 0.766029754 | 0.443658598 | 0.663529376 | Stil | 32.77807842 | 0.0407935007993821 | 0.0905366597879002 | 0.450574395995489 | 0.652296323634012 | 0.914611623655175 | NM_009185 |
| Tollip | 6820.188828 | 0.068751756 | 0.089759586 | 0.765954464 | 0.443703397 | 0.663529376 | AU023762 | 91.30270151 | -0.0424986005596267 | 0.094605433175378 | -0.449219448959591 | 0.653273358766122 | 0.914683957693943 | NR_040760 |
| Ttll11 | 1774.884556 | 0.060693172 | 0.079247897 | 0.765864764 | 0.443756773 | 0.663536716 | Apc2 | 7163.901947 | 0.0244650102059627 | 0.0544019589984325 | 0.449708257871148 | 0.652920816189099 | 0.914683957693943 | NM_011789 |
| 1110037F02Rik | 2068.876311 | -0.046992772 | 0.061366415 | -0.765773451 | 0.443811114 | 0.663553932 | Arid4b | 2003.799241 | -0.0316829986740321 | 0.0705830505602341 | -0.448875451295413 | 0.65352150586336 | 0.914683957693943 | NM_194262,NM_198122 |
| Cdk8 | 829.1003027 | 0.054992681 | 0.071828868 | 0.765606962 | 0.4439102 | 0.66363804 | Asgr1 | 30.54472829 | 0.035879056447726 | 0.0799318506196899 | 0.448870583748099 | 0.653525017405627 | 0.914683957693943 | NM_009714 |
| Kif22 | 46.70694851 | -0.117198916 | 0.153152451 | -0.765243493 | 0.444126565 | 0.663645311 | Atm | 1041.058208 | -0.0299659318398233 | 0.0666732823418888 | -0.449444376926926 | 0.653111124842376 | 0.914683957693943 | NM_007499 |
| Ppm1k | 3113.409394 | -0.059636659 | 0.07792741 | -0.765284761 | 0.444101996 | 0.663645311 | Cldn1 | 89.18507606 | 0.0279016171194471 | 0.0620613101699023 | 0.449581503243522 | 0.653012227711777 | 0.914683957693943 | NM_016674 |
| Rab14 | 8204.765543 | -0.053056971 | 0.069321286 | -0.765377767 | 0.444046628 | 0.663645311 | Daam1 | 3033.21054 | -0.0298006467228826 | 0.0664036297907293 | -0.448780387710719 | 0.653590087950257 | 0.914683957693943 | NM_026102,NM_172464 |
| Rprd1b | 1217.576775 | 0.04117444 | 0.053805986 | 0.765239024 | 0.444129226 | 0.663645311 | Foxf1 | 36.20602261 | -0.0400552570585132 | 0.0890888514058486 | -0.449610208532598 | 0.652991525887521 | 0.914683957693943 | NM_010426 |
| Zfp655 | 1418.393397 | 0.048859164 | 0.063825626 | 0.765510135 | 0.443967834 | 0.663645311 | Grap | 169.3459996 | 0.0421573277060719 | 0.0937387630097073 | 0.449732067636802 | 0.652903645902733 | 0.914683957693943 | NM_027817 |
| Dyrk1a | 3588.291365 | -0.054429787 | 0.0712147 | -0.764305511 | 0.444685202 | 0.664220593 | Gsn | 2603.920419 | 0.0347806334194971 | 0.0774526684817064 | 0.449056618723369 | 0.653390813653453 | 0.914683957693943 | NM_001206367,NM_146120,NM_001206368,NM_001206369 |
| Itgb3bp | 108.7609379 | -0.114894686 | 0.15038239 | -0.764016892 | 0.444857176 | 0.664220593 | Kcnab2 | 18802.99989 | -0.0207661770568703 | 0.0462835145660493 | -0.448673296562986 | 0.65366735063278 | 0.914683957693943 | NM_010598,NM_001252656,NM_001252655,NM_001252654 |
| L3hypdh | 48.98996932 | -0.11802213 | 0.154424861 | -0.764268973 | 0.444706971 | 0.664220593 | Mark1 | 4567.746752 | 0.0210723790551921 | 0.0469675959088509 | 0.44865781710622 | 0.653678518850394 | 0.914683957693943 | NM_145515 |
| Nhlrc4 | 14.10132117 | 0.122518592 | 0.160332545 | 0.76415298 | 0.444776083 | 0.664220593 | Megf10 | 1329.46255 | 0.0333802353302247 | 0.0742687760087467 | 0.449451803625975 | 0.653105768462551 | 0.914683957693943 | NM_001001979 |
| Pdcd2 | 300.8540736 | 0.066163023 | 0.086594747 | 0.764053547 | 0.444835333 | 0.664220593 | Parp1 | 3327.02438 | 0.0265027609610285 | 0.0589791454704393 | 0.449358171428777 | 0.653173300393501 | 0.914683957693943 | NM_007415 |
| Rtp3 | 8.874265687 | -0.11230421 | 0.146943157 | -0.76426975 | 0.444706508 | 0.664220593 | Peli1 | 1413.263538 | 0.0306430142371905 | 0.0682205664745176 | 0.449175605257347 | 0.653304983847687 | 0.914683957693943 | NM_023324 |
| Tm7sf3 | 1722.170353 | 0.045383658 | 0.05939317 | 0.764122502 | 0.444794244 | 0.664220593 | Pigc | 481.2161915 | 0.028836548271891 | 0.0641041383543854 | 0.449839105745008 | 0.652826458209327 | 0.914683957693943 | NM_001039045,NM_026078 |
| Tmem186 | 808.1796464 | 0.063198211 | 0.082713958 | 0.764057393 | 0.444833041 | 0.664220593 | Rps26 | 2871.064861 | 0.0310415167293497 | 0.0691302442880553 | 0.449029466755598 | 0.653410400110134 | 0.914683957693943 | NM_013765 |
| Txndc15 | 1971.664308 | -0.042780911 | 0.056013904 | -0.763755219 | 0.445013128 | 0.66438942 | Slc30a5 | 1867.518602 | 0.0283438239163012 | 0.0629860475900924 | 0.45000162735659 | 0.652709267164413 | 0.914683957693943 | NM_022885 |
| Gm17455 | 8.208940609 | -0.116341437 | 0.152407732 | -0.76335653 | 0.445250797 | 0.66461617 | Tmem8b | 4454.966356 | 0.025881372194204 | 0.0575104064466535 | 0.450029373696245 | 0.652689260696856 | 0.914683957693943 | NM_001085508 |
| Lrrc25 | 25.75659007 | -0.125630192 | 0.164564862 | -0.76340836 | 0.445219896 | 0.66461617 | Traf6 | 920.3990901 | 0.029986811645273 | 0.0667409713719331 | 0.449301396561385 | 0.65321425047384 | 0.914683957693943 | NM_009424 |
| Klhl36 | 207.8852861 | -0.078726075 | 0.103178651 | -0.763007411 | 0.445458976 | 0.664862863 | Zc3hav1l | 287.5552538 | 0.0358524515189884 | 0.0797963149964799 | 0.44929958884154 | 0.653215554347378 | 0.914683957693943 | NM_172467 |
| Atp6v1d | 8725.989307 | 0.059161482 | 0.077569036 | 0.762694558 | 0.445645577 | 0.66501325 | Bcl9 | 2544.904686 | 0.0270755504652982 | 0.0603896960127709 | 0.448347189222022 | 0.653902649038163 | 0.914733231888324 | NM_029933 |
| Brsk1 | 15401.49642 | -0.063853145 | 0.083718374 | -0.762713627 | 0.445634202 | 0.66501325 | Lima1 | 614.317978 | -0.0325579847625015 | 0.0726152397864725 | -0.448362972541843 | 0.653891260001654 | 0.914733231888324 | NM_001113545,NM_023063 |
| Zfp60 | 1434.194338 | -0.063850623 | 0.083776573 | -0.762153673 | 0.445968292 | 0.665430732 | Map1b | 35301.17457 | 0.0319340050075993 | 0.0712254859199039 | 0.448350819866787 | 0.653900029205396 | 0.914733231888324 | NM_008634 |
| Adora3 | 42.69240062 | 0.125459053 | 0.164643884 | 0.762002513 | 0.446058505 | 0.665501249 | Armc9 | 1231.224553 | 0.023351181711916 | 0.0520967083234361 | 0.448227584110363 | 0.653988957136805 | 0.914765873025444 | NM_030184,NM_027456 |
| Cdh1 | 80.88761612 | -0.127928053 | 0.168080335 | -0.761112556 | 0.446589841 | 0.666037444 | Mospd3 | 527.8526164 | 0.0316592842811428 | 0.0706507446955934 | 0.448109703833277 | 0.654074025107198 | 0.914796773416826 | NM_001254762,NM_030037 |
| Oprk1 | 535.5925476 | -0.08456735 | 0.111088689 | -0.761259772 | 0.446501923 | 0.666037444 | Rfesd | 417.2865474 | 0.0319649148766795 | 0.0713621418522429 | 0.447925385183416 | 0.654207047148509 | 0.914894730613194 | NM_001131068,NM_001131069,NM_178916 |
| Pdxdc1 | 3953.247432 | 0.032008219 | 0.042049 | 0.761212376 | 0.446530228 | 0.666037444 | BC027231 | 210.5270215 | -0.0391900504553747 | 0.0875155764697021 | -0.4478065738268 | 0.654292798656848 | 0.914926568472216 | NM_145972 |
| Thada | 582.6089293 | -0.061694294 | 0.081052339 | -0.761166123 | 0.44655785 | 0.666037444 | Rnf122 | 150.2536487 | 0.0416398707132443 | 0.0930263039363777 | 0.447613943059831 | 0.654431838653911 | 0.915032908680982 | NM_175136 |
| Gtf3c3 | 1175.367613 | 0.04796521 | 0.063025958 | 0.761038971 | 0.44663379 | 0.666038878 | Vps13b | 3019.33549 | 0.0292413982182348 | 0.0653484736870925 | 0.447468725256708 | 0.654536664118982 | 0.915091393935739 | NM_177151 |
| Fndc5 | 3195.470871 | 0.058381555 | 0.076756133 | 0.760610946 | 0.44688948 | 0.666356039 | 0610040F04Rik | 51.03505515 | -0.0420346842743075 | 0.0949510699350825 | -0.442698374047248 | 0.657983918680365 | 0.915208181488406 | NR_040758,NR_040757 |
| Carkd | 1176.044982 | -0.045815806 | 0.060253987 | -0.760377998 | 0.447028671 | 0.666499445 | 1110032F04Rik | 452.4438441 | -0.0387264866358967 | 0.0868958142435285 | -0.445665731692949 | 0.655838721700986 | 0.915208181488406 | NM_001167996 |
| Yif1b | 844.0457741 | -0.070199465 | 0.09235439 | -0.760109675 | 0.44718903 | 0.66667438 | 2610019E17Rik | 71.51897576 | -0.041638954310943 | 0.0944325922829392 | -0.440938380534808 | 0.659257607059871 | 0.915208181488406 | NR_045289 |
| Cetn4 | 216.8574237 | -0.076117077 | 0.100168132 | -0.759893147 | 0.447318458 | 0.666726681 | 2610019F03Rik | 823.8171039 | 0.0315392004785 | 0.0715712203477069 | 0.440668753799034 | 0.65945282045514 | 0.915208181488406 | NM_173744 |
| Grrp1 | 58.2147173 | -0.111688185 | 0.146990046 | -0.759835021 | 0.447353206 | 0.666726681 | 2900097C17Rik | 69371.02063 | 0.0261803893939933 | 0.0593045046873436 | 0.441457011267824 | 0.658882176672416 | 0.915208181488406 | NR_024329 |
| Zer1 | 7648.197387 | 0.0398468 | 0.052433906 | 0.759943374 | 0.447288433 | 0.666726681 | Abcg4 | 3172.312505 | 0.029627795040633 | 0.0670858585927561 | 0.441639947108499 | 0.658749772175115 | 0.915208181488406 | NM_138955 |
| Chrm4 | 1915.472243 | 0.059641921 | 0.078506447 | 0.759707295 | 0.447429567 | 0.666776351 | Abtb2 | 639.0764212 | -0.032082418722739 | 0.072640994574013 | -0.441657206249436 | 0.658737280982224 | 0.915208181488406 | NM_178890 |
| Cd28 | 11.68156491 | -0.121501462 | 0.159992915 | -0.759417764 | 0.44760269 | 0.666970195 | Aif1l | 371.367335 | -0.0375127259434224 | 0.0850449021601332 | -0.441093175376799 | 0.659145543990916 | 0.915208181488406 | NM_145144 |
| Ifltd1 | 19.59386864 | 0.122988064 | 0.161979713 | 0.759280661 | 0.447684684 | 0.667028223 | Atp7b | 107.4730051 | -0.0421450987626514 | 0.094591294130738 | -0.445549446700678 | 0.655922734536683 | 0.915208181488406 | NM_007511 |
| 3110062M04Rik | 145.0410871 | -0.090153871 | 0.118761517 | -0.759116871 | 0.447782647 | 0.667103964 | B3gnt7 | 49.60234899 | -0.0393643471729658 | 0.0888961813526068 | -0.442812577256013 | 0.657901305285244 | 0.915208181488406 | NM_145222 |
| Trim71 | 11.68374045 | 0.122390898 | 0.161241846 | 0.759051702 | 0.44782163 | 0.667103964 | B4galt1 | 283.7322375 | 0.0400799703211376 | 0.089759241481899 | 0.446527506911032 | 0.655216247617533 | 0.915208181488406 | NM_022305 |
| Zbtb18 | 9549.740872 | 0.078456807 | 0.10337229 | 0.758973293 | 0.447868533 | 0.667109696 | B4galt3 | 1717.055847 | 0.0246086587128438 | 0.0555337920294134 | 0.443129449899799 | 0.657672104840997 | 0.915208181488406 | NM_020579 |
| Ampd3 | 1129.76831 | 0.083751489 | 0.110392502 | 0.758670086 | 0.448049935 | 0.667315745 | Ccdc109b | 52.41279572 | -0.0421795233852206 | 0.0944705000006589 | -0.446483541263425 | 0.655247998916871 | 0.915208181488406 | NM_025779 |
| Stx7 | 3523.779352 | -0.052156992 | 0.068759657 | -0.758540604 | 0.448127415 | 0.66736699 | Cd164l2 | 127.5729421 | 0.0404278724699526 | 0.0913526547032008 | 0.442547319520164 | 0.658093196345805 | 0.915208181488406 | NM_027152 |
| Clasp2 | 9975.679647 | -0.05800507 | 0.076482189 | -0.758412781 | 0.448203909 | 0.667416757 | Clmp | 974.1308965 | -0.0285958512390168 | 0.0645127325918975 | -0.443259029499059 | 0.657578386544388 | 0.915208181488406 | NM_133733 |
| Cramp1l | 1824.153236 | 0.047939105 | 0.063228798 | 0.758184667 | 0.448340439 | 0.667491761 | Cotl1 | 2299.315704 | -0.0307605946206163 | 0.0693848974917481 | -0.443332709478179 | 0.657525099789123 | 0.915208181488406 | NM_028071 |
| Gpr173 | 499.4226631 | -0.065867235 | 0.08687481 | -0.758185656 | 0.448339847 | 0.667491761 | Cox6c | 5898.109783 | 0.0206707131419639 | 0.0469059077188393 | 0.440684641812436 | 0.65944131667577 | 0.915208181488406 | NM_053071 |
| Ptpru | 1744.754612 | 0.070821972 | 0.093457677 | 0.757797268 | 0.448572358 | 0.667719362 | Dcaf4 | 880.2003357 | 0.0291822248609214 | 0.066259425041596 | 0.440423756206782 | 0.659630222132822 | 0.915208181488406 | NM_030246,NM_001165256 |
| Rhoj | 142.3456911 | 0.09351735 | 0.123408762 | 0.757785333 | 0.448579504 | 0.667719362 | Diras2 | 15233.72614 | 0.0260269519441922 | 0.0585392720665562 | 0.44460668924275 | 0.656604012519816 | 0.915208181488406 | NM_001024474 |
| Epha1 | 11.23721149 | 0.123632624 | 0.163166482 | 0.757708461 | 0.448625533 | 0.667723727 | Dmp1 | 341.0126752 | 0.0394231176801637 | 0.0894329585585479 | 0.440811959210263 | 0.659349134915902 | 0.915208181488406 | NM_016779 |
| Kpna2 | 1186.229407 | -0.043412785 | 0.057311756 | -0.757484814 | 0.448759461 | 0.667794764 | Dmxl1 | 2477.941964 | 0.035181232483499 | 0.0788329752105192 | 0.446275589492206 | 0.655398186861111 | 0.915208181488406 | NM_001081371 |
| Tmppe | 929.6836041 | 0.052540025 | 0.069357552 | 0.757524206 | 0.448735869 | 0.667794764 | Dtwd1 | 245.0863596 | 0.0358796517121695 | 0.080900735132463 | 0.443502171561505 | 0.657402548853523 | 0.915208181488406 | NM_026981 |
| Spred3 | 2075.53102 | -0.08130203 | 0.107362776 | -0.757264608 | 0.44889135 | 0.667926877 | E130114P18Rik | 129.3936959 | 0.0417439476788307 | 0.0938120545680145 | 0.444974239942331 | 0.656338370229286 | 0.915208181488406 | NR_015513 |
| Acy3 | 99.41741866 | 0.111594653 | 0.147385308 | 0.757162669 | 0.448952412 | 0.667953589 | E430018J23Rik | 221.1699086 | -0.0383021498913185 | 0.0866030354741442 | -0.442272602589708 | 0.65829195406306 | 0.915208181488406 | NM_198011 |
| Dpm1 | 1262.101188 | 0.048755103 | 0.064409736 | 0.756952385 | 0.449078389 | 0.668076867 | Epb4.1 | 1351.238105 | -0.0276471694361633 | 0.0623988275826626 | -0.443071937522188 | 0.657713702326563 | 0.915208181488406 | NM_001128606,NM_001128607,NM_183428 |
| Actr2 | 11301.45998 | -0.054340828 | 0.071839025 | -0.756424914 | 0.449394475 | 0.668226295 | Etnk1 | 16398.34267 | 0.0267525240277717 | 0.0607614127188827 | 0.440288051753245 | 0.659728493368926 | 0.915208181488406 | NM_029250 |
| Dph5 | 296.1303938 | -0.076515831 | 0.101150342 | -0.75645647 | 0.449375562 | 0.668226295 | Exoc3l | 26.52875741 | -0.0360570522111533 | 0.0816501050062924 | -0.441604480586701 | 0.658775441142332 | 0.915208181488406 | NM_177788 |
| Pip5k1a | 2200.123925 | 0.083740687 | 0.110677713 | 0.756617426 | 0.449279098 | 0.668226295 | Fbxl8 | 29.131312 | 0.0382219301908477 | 0.0866276463401248 | 0.441220924331448 | 0.659053066440047 | 0.915208181488406 | NM_015821 |
| Samd5 | 242.1208187 | -0.090258587 | 0.119313603 | -0.756481948 | 0.449360292 | 0.668226295 | Fmn1 | 1075.232518 | 0.0354182000564623 | 0.0800518826350685 | 0.442440563427131 | 0.658170431516399 | 0.915208181488406 | NM_010230,NM_001043322 |
| Wdr85 | 775.5408446 | 0.061094838 | 0.080765117 | 0.756450809 | 0.449378955 | 0.668226295 | Fth1 | 55905.41649 | 0.036000069253018 | 0.0816633690183661 | 0.440834975163976 | 0.659332471200955 | 0.915208181488406 | NR_073181,NM_010239 |
| BC018473 | 10.49417964 | 0.124179718 | 0.164214747 | 0.756203209 | 0.449527369 | 0.668359759 | Gatsl3 | 134.6397733 | 0.0417498488080906 | 0.0938042081408755 | 0.445074369642251 | 0.656266010372893 | 0.915208181488406 | NM_028022 |
| Hspa9 | 9117.42648 | 0.051317977 | 0.067869481 | 0.756127448 | 0.449572787 | 0.668363151 | Gdf11 | 307.5632462 | -0.0358304772497022 | 0.0811462632207976 | -0.441554248187721 | 0.658811797631106 | 0.915208181488406 | NM_010272 |
| Bmyc | 1153.972216 | -0.066156315 | 0.087522135 | -0.755880955 | 0.449720574 | 0.668454582 | Gga2 | 1478.598732 | -0.0241645711023292 | 0.0546195813515754 | -0.442415897456019 | 0.658188277203923 | 0.915208181488406 | NM_028758 |
| Vimp | 1148.735055 | 0.052088414 | 0.068904326 | 0.755952749 | 0.449677526 | 0.668454582 | Git1 | 17368.26165 | 0.0284687283240616 | 0.0639913491626228 | 0.444884014739451 | 0.656403575250733 | 0.915208181488406 | NM_001004144 |
| 4921507P07Rik | 37.4614982 | 0.122283928 | 0.161807994 | 0.755734777 | 0.449808229 | 0.668456617 | Gpc1 | 3529.527674 | 0.0290625682802837 | 0.0651899057370297 | 0.445813933180385 | 0.655731656350846 | 0.915208181488406 | NM_016696 |
| Lrrc41 | 1858.501887 | 0.054462768 | 0.072061048 | 0.755786506 | 0.449777208 | 0.668456617 | Itpr1 | 33699.14409 | 0.0293937010119841 | 0.0658522736810484 | 0.446358179739561 | 0.655338536460225 | 0.915208181488406 | NM_010585 |
| Sumf2 | 482.3769159 | -0.089844066 | 0.118897915 | -0.755640382 | 0.449864837 | 0.668476627 | Klhl36 | 225.3818599 | -0.0362847492499338 | 0.0821506391372442 | -0.441685538067635 | 0.658716776216409 | 0.915208181488406 | NM_146219 |
| Omg | 4826.191376 | 0.057510484 | 0.076124164 | 0.755482634 | 0.449959448 | 0.668553096 | Mapk13 | 35.60025696 | -0.0395120806209609 | 0.0888930404097352 | -0.444490147247048 | 0.656688250735394 | 0.915208181488406 | NM_011950 |
| 5430417L22Rik | 1495.597843 | -0.065741334 | 0.087069608 | -0.755043417 | 0.450222932 | 0.668713036 | Mapkap1 | 2053.297051 | 0.0244720883360735 | 0.0547652651328888 | 0.446854192647322 | 0.654980339842746 | 0.915208181488406 | NM_177345 |
| AW112010 | 30.2035809 | 0.124817586 | 0.16530918 | 0.755055383 | 0.450215752 | 0.668713036 | Mgat1 | 2407.477624 | 0.0265303520517661 | 0.0596625713370171 | 0.444673292773515 | 0.65655587249841 | 0.915208181488406 | NM_010794,NM_001110148,NM_001110149,NM_001110150 |
| Dusp1 | 1384.910223 | 0.124014716 | 0.164254548 | 0.755015419 | 0.450239731 | 0.668713036 | Mon2 | 3992.45919 | -0.0251751065870666 | 0.0568138877943032 | -0.443115364296384 | 0.657682292561827 | 0.915208181488406 | NM_153395,NM_001163025,NM_001163024 |
| Rsbn1 | 979.6591244 | 0.056258342 | 0.074504275 | 0.7551022 | 0.450187663 | 0.668713036 | Mphosph6 | 384.8102637 | 0.0344685871195579 | 0.0771017481317274 | 0.447053250474539 | 0.654836612190786 | 0.915208181488406 | NM_026758 |
| Rbm4b | 376.333024 | -0.062033528 | 0.082177635 | -0.754871178 | 0.450326281 | 0.668777474 | Mpst | 796.8083172 | -0.0293796060228148 | 0.0661414877149319 | -0.444193304956189 | 0.656902832284878 | 0.915208181488406 | NM_138670,NM_001162493,NM_001162492 |
| Wbp1 | 773.3275554 | -0.087834444 | 0.116453158 | -0.754246996 | 0.450700923 | 0.669269704 | Mtmr12 | 2358.657323 | 0.0295326915988507 | 0.0663616534013845 | 0.445026458581795 | 0.656300633438525 | 0.915208181488406 | NM_172958 |
| Igbp1b | 9.97958867 | -0.115465599 | 0.153127055 | -0.754050932 | 0.45081864 | 0.669316213 | Myoc | 820.4377983 | -0.0413394409663883 | 0.0939089195536541 | -0.440207822248124 | 0.659786594833167 | 0.915208181488406 | NM_010865 |
| Nkx2-1 | 60.93497913 | -0.126681231 | 0.167991989 | -0.754090909 | 0.450794636 | 0.669316213 | Npepl1 | 517.3958374 | -0.0359363807014892 | 0.0806201224277054 | -0.445749517854112 | 0.655778191111032 | 0.915208181488406 | NM_213733 |
| Kcna1 | 10162.81788 | 0.076712958 | 0.101755747 | 0.753893124 | 0.4509134 | 0.669392758 | Nprl2 | 1259.52434 | 0.027790570983228 | 0.0626102369601755 | 0.443866248276696 | 0.657139287993107 | 0.915208181488406 | NM_018879 |
| Ndufa9 | 3540.181492 | -0.045633947 | 0.060539072 | -0.753793305 | 0.450973345 | 0.669417609 | Ocrl | 3778.350686 | 0.0242954732640248 | 0.0548364881777389 | 0.443053048643031 | 0.657727364484347 | 0.915208181488406 | NM_177215 |
| Saal1 | 363.4020811 | 0.056375369 | 0.074858639 | 0.753091026 | 0.451395217 | 0.669979643 | Otud5 | 2320.464022 | -0.0214236311601233 | 0.0483840760624424 | -0.442782686032382 | 0.657922927877142 | 0.915208181488406 | NM_138604 |
| Lect1 | 22.94122483 | 0.125817592 | 0.167171214 | 0.752627135 | 0.451674007 | 0.670329221 | Pcbp2 | 8530.959251 | 0.0185497309281044 | 0.04210613580665 | 0.440546979026622 | 0.659540994638249 | 0.915208181488406 | NM_001174073,NM_001103165,NM_001103166,NM_011042 |
| Smim5 | 9.752686901 | 0.118202927 | 0.157166069 | 0.752089354 | 0.451997326 | 0.670702205 | Pde1b | 16336.72887 | -0.0348449220789255 | 0.0782016846225537 | -0.445577640010023 | 0.655902365215522 | 0.915208181488406 | NM_008800 |
| Syne3 | 92.54021392 | 0.115653041 | 0.153780622 | 0.752065112 | 0.452011903 | 0.670702205 | Polr2j | 796.5881912 | 0.0286895768492491 | 0.0641767980883347 | 0.447039704438977 | 0.654846392560602 | 0.915208181488406 | NM_011293 |
| 2310009A05Rik | 116.6805004 | -0.095010814 | 0.12646792 | -0.751264145 | 0.452493703 | 0.671352814 | Pqlc1 | 662.5895276 | 0.0301178994611421 | 0.0674771573567563 | 0.446342149564877 | 0.655350114003259 | 0.915208181488406 | NM_025861,NM_001164420,NM_001164421,NM_001164422,NM_001164423,NM_001164424 |
| Fam19a1 | 1263.120234 | -0.070788238 | 0.094239633 | -0.751151454 | 0.452561513 | 0.67138913 | Purb | 18992.73068 | 0.032983578138829 | 0.0746255599343402 | 0.441987680465645 | 0.658498120751265 | 0.915208181488406 | NM_011221 |
| Mbnl2 | 7482.261466 | 0.060247556 | 0.080239273 | 0.750848728 | 0.4527437 | 0.671466535 | Pygb | 9084.037252 | -0.022129622245035 | 0.0501368902067918 | -0.441384021900051 | 0.658935007580221 | 0.915208181488406 | NM_153781 |
| Mfsd2a | 910.5516322 | 0.126152676 | 0.167983027 | 0.75098466 | 0.452661888 | 0.671466535 | Rab2b | 1865.848115 | 0.0242975639299886 | 0.0548874691572604 | 0.442679618919441 | 0.657997486341068 | 0.915208181488406 | NM_172601 |
| Thumpd1 | 2025.798939 | 0.04503672 | 0.059976705 | 0.750903534 | 0.452710713 | 0.671466535 | Rassf4 | 567.4108254 | 0.038592219999599 | 0.0868559820367646 | 0.444324260627939 | 0.656808163474309 | 0.915208181488406 | NM_178045 |
| Zfp808 | 37.07228873 | 0.12399859 | 0.165161424 | 0.750772108 | 0.452789818 | 0.671470659 | Rpgr | 733.3382894 | 0.0341141174881428 | 0.0774651912066119 | 0.440379955910198 | 0.659661939752066 | 0.915208181488406 | NM_011285,NM_001177950,NM_001177951,NM_001177952,NM_001177953,NM_001177954 |
| Cenpi | 18.30035922 | -0.125941809 | 0.167801896 | -0.750538653 | 0.452930353 | 0.671518227 | Rpl37 | 4554.883024 | 0.0237217562290518 | 0.0532455206035458 | 0.445516467116548 | 0.655946562207838 | 0.915208181488406 | NM_026069 |
| Inpp5a | 2412.675289 | 0.040350143 | 0.05377446 | 0.750358867 | 0.453038598 | 0.671518227 | Rps6kb1 | 2011.866647 | -0.0306222509032443 | 0.0689945920282969 | -0.443835523959402 | 0.657161502849636 | 0.915208181488406 | NM_028259,NM_001114334 |
| Khdrbs3 | 4830.722772 | -0.064503003 | 0.085937966 | -0.750576329 | 0.452907671 | 0.671518227 | Sh3d21 | 56.77660645 | 0.0414340055302901 | 0.0941171730135509 | 0.440238526122373 | 0.659764359129518 | 0.915208181488406 | NM_025856,NM_001162533 |
| Rif1 | 1092.440875 | -0.070684316 | 0.094198679 | -0.750374817 | 0.453028994 | 0.671518227 | Siah1b | 138.2267667 | 0.03927054624651 | 0.088123864960209 | 0.445628959467926 | 0.655865288191709 | 0.915208181488406 | NM_009173 |
| Trpa1 | 11.4919667 | 0.114936191 | 0.153160181 | 0.750431285 | 0.452994995 | 0.671518227 | Sigirr | 61.49286018 | -0.0416436438073421 | 0.0940935450355035 | -0.4425770523539 | 0.658071686087511 | 0.915208181488406 | NM_023059 |
| Zfp825 | 218.0749704 | 0.076267176 | 0.101663014 | 0.7501959 | 0.453136728 | 0.671599432 | Slc35a1 | 666.0283846 | -0.0333030279640376 | 0.0755630762561548 | -0.440731500278551 | 0.659407389085877 | 0.915208181488406 | NM_011895 |
| A330050B17Rik | 73.29345392 | 0.109410183 | 0.145874763 | 0.750028176 | 0.453237735 | 0.671684884 | Snora7a | 23.66617941 | -0.0372251195195051 | 0.0842690290306094 | -0.441741407818804 | 0.658676342000261 | 0.915208181488406 | NR_028546 |
| 2900056M20Rik | 777.2718262 | 0.065001817 | 0.08667997 | 0.749905856 | 0.453311407 | 0.67169764 | Snph | 16326.58469 | 0.0241137691000322 | 0.0541849991646049 | 0.445026658149032 | 0.656300489219155 | 0.915208181488406 | NM_198214 |
| Gpatch2 | 698.3224981 | 0.050207165 | 0.066954499 | 0.74986993 | 0.453333047 | 0.67169764 | Stx17 | 1290.193638 | -0.0268600965623365 | 0.06063439610917 | -0.442984482173714 | 0.657776958960703 | 0.915208181488406 | NM_026343 |
| Cd40 | 19.0224353 | -0.125966282 | 0.168112277 | -0.749298526 | 0.453677296 | 0.672143433 | Tbc1d30 | 5785.382973 | 0.0286083381055933 | 0.0646300113240449 | 0.442647889417124 | 0.658020440059997 | 0.915208181488406 | NM_029057 |
| Shisa5 | 2081.008554 | 0.064666672 | 0.08632419 | 0.749114142 | 0.453788412 | 0.672243777 | Tbl1xr1 | 2818.32923 | -0.0246853725420156 | 0.0552716299496373 | -0.446619225170464 | 0.655150012139437 | 0.915208181488406 | NM_030732 |
| Cd164 | 2162.98476 | -0.05063605 | 0.067606563 | -0.748981279 | 0.453868489 | 0.672298123 | Thtpa | 1362.259897 | -0.0302084916043581 | 0.0683605477690444 | -0.441899495984399 | 0.658561935387841 | 0.915208181488406 | NM_153083 |
| Apoo | 189.4165721 | -0.118959363 | 0.158872461 | -0.748772708 | 0.453994212 | 0.672303015 | Tmem202 | 70.58762089 | 0.041891670034794 | 0.0951009914567524 | 0.440496669835924 | 0.659577423693829 | 0.915208181488406 | NM_178388 |
| Phyh | 2629.588045 | -0.059667464 | 0.079685842 | -0.748783757 | 0.453987551 | 0.672303015 | Tmem74 | 252.6354662 | -0.039304313331423 | 0.0879244937940849 | -0.44702348157354 | 0.654858105705639 | 0.915208181488406 | NM_175502 |
| Pla2g7 | 5060.012468 | 0.083706747 | 0.111804599 | 0.748687871 | 0.454045356 | 0.672303015 | Tns3 | 3399.968016 | -0.0268172846091124 | 0.0601861617377287 | -0.44557226835586 | 0.655906246150221 | 0.915208181488406 | NM_001083587 |
| Synj2bp | 1954.494792 | -0.044544469 | 0.059491433 | -0.748754343 | 0.454005283 | 0.672303015 | Trf | 16025.29359 | 0.0386074458796397 | 0.0872009549516312 | 0.442741090404969 | 0.657953017629301 | 0.915208181488406 | NM_133977 |
| Smchd1 | 1717.056779 | -0.043576185 | 0.058233061 | -0.748306631 | 0.454275226 | 0.672579106 | Tsen2 | 560.9298955 | 0.0296348162806895 | 0.0666658545203552 | 0.444527659532828 | 0.656661135844273 | 0.915208181488406 | NM_199033 |
| Phf23 | 1702.402482 | 0.078905177 | 0.105482405 | 0.748041125 | 0.454435352 | 0.672687623 | Ube4b | 5507.336933 | 0.0213520307207843 | 0.0483860730964823 | 0.441284637383326 | 0.659006946468589 | 0.915208181488406 | NM_022022 |
| Ssb | 3407.762385 | 0.058324016 | 0.077964434 | 0.748084898 | 0.454408951 | 0.672687623 | Vip | 1440.1732 | -0.033883760276427 | 0.0761986908832897 | -0.444676409576712 | 0.656553619755272 | 0.915208181488406 | NM_011702 |
| Bcl7c | 288.9295897 | -0.072079842 | 0.096387633 | -0.747812145 | 0.454573475 | 0.672827802 | Wars2 | 338.9216324 | 0.036147832447976 | 0.081658078846877 | 0.442673069932975 | 0.658002223974685 | 0.915208181488406 | NM_027462 |
| Armcx6 | 272.0169892 | 0.083674559 | 0.111906206 | 0.747720458 | 0.454628789 | 0.672845396 | Wdfy2 | 334.8950578 | -0.0355311758500699 | 0.0804949433699628 | -0.44140879367745 | 0.658917077168501 | 0.915208181488406 | NM_175546 |
| Arhgef3 | 3335.436297 | 0.067389954 | 0.090161329 | 0.747437452 | 0.454799545 | 0.67296955 | Zfp280d | 1558.559409 | -0.0325165935766565 | 0.0736604453212732 | -0.441439003454758 | 0.658895210866243 | 0.915208181488406 | NM_146224 |
| Snap25 | 101385.2305 | 0.06834872 | 0.091442009 | 0.747454265 | 0.4547894 | 0.67296955 | Zfp58 | 250.2950668 | -0.0376897320148184 | 0.0848855641498355 | -0.444005976797543 | 0.657038262750346 | 0.915208181488406 | NM_001007575 |
| Pmel | 21.29431325 | 0.125550465 | 0.168014935 | 0.747257771 | 0.454907978 | 0.673065719 | Zfp608 | 838.0494475 | -0.0355953041432378 | 0.0807352159243619 | -0.440889440075144 | 0.659293038889546 | 0.915208181488406 | NM_175751 |
| B830017H08Rik | 84.04199776 | 0.102704983 | 0.137460085 | 0.747162225 | 0.454965643 | 0.673086764 | Rfx4 | 697.6099249 | 0.0317688911205913 | 0.0721980961473982 | 0.440023945447711 | 0.659919764448769 | 0.915305499702361 | NM_027689,NM_001024918 |
| Rpain | 344.4601638 | 0.077947033 | 0.104338965 | 0.747055838 | 0.455029856 | 0.67311749 | 1700048O20Rik | 344.7045669 | -0.0414148114829753 | 0.0949843873201961 | -0.436017040814974 | 0.66282436582714 | 0.915428982038356 | NR_033553 |
| Ubox5 | 714.7321449 | -0.070323952 | 0.094226718 | -0.746327086 | 0.455469853 | 0.673704048 | 2610507B11Rik | 14264.75371 | 0.0176110302419819 | 0.0405625759231331 | 0.434169424430913 | 0.664165414418479 | 0.915428982038356 | NM_001002004 |
| Abca13 | 8.498165381 | 0.10410297 | 0.139622664 | 0.745602232 | 0.455907734 | 0.674115317 | 4833418N02Rik | 28.90428122 | 0.037887651760116 | 0.0870129586315777 | 0.435425393596102 | 0.663253681588722 | 0.915428982038356 | NR_015506 |
| Loxhd1 | 10.00185357 | -0.114773383 | 0.153938649 | -0.745578737 | 0.455921932 | 0.674115317 | 9530080O11Rik | 62.95875076 | 0.0414094354205454 | 0.0949267846317803 | 0.436224987301235 | 0.66267350033709 | 0.915428982038356 | NR_045777,NR_045776 |
| Mpp4 | 18.80791091 | 0.123630176 | 0.165813797 | 0.745596433 | 0.455911238 | 0.674115317 | A330050B17Rik | 87.75817268 | -0.0412251363362577 | 0.0950982475021971 | -0.433500484173542 | 0.664651214411978 | 0.915428982038356 | NR_029456 |
| Zfp628 | 505.0823896 | 0.063133329 | 0.084673597 | 0.745608216 | 0.455904118 | 0.674115317 | AI661453 | 25.3059165 | 0.0326433049212988 | 0.0754523220881331 | 0.432634861564225 | 0.665280058809217 | 0.915428982038356 | NM_145489 |
| Zfp217 | 187.4399117 | 0.078960336 | 0.105937077 | 0.745351284 | 0.456059386 | 0.67425421 | Aak1 | 21706.13326 | -0.0229854191494353 | 0.0527340566913741 | -0.435874282988646 | 0.662927944765249 | 0.915428982038356 | NM_001040106,NM_177762 |
| Zfp937 | 450.9879097 | -0.072335934 | 0.097091689 | -0.745027043 | 0.456255372 | 0.674479604 | Abca3 | 5311.281086 | -0.0206377100757713 | 0.0475769051473281 | -0.433775799662966 | 0.664451256820772 | 0.915428982038356 | NM_013855,NM_001039581 |
| Skap1 | 61.13403916 | -0.114820048 | 0.154130273 | -0.744954549 | 0.456299197 | 0.674480037 | Ankrd13c | 2100.107927 | 0.0315984785556193 | 0.072961245955931 | 0.433085786044621 | 0.664952448609305 | 0.915428982038356 | NM_001013806 |
| Wnt11 | 10.45994348 | 0.119767023 | 0.160798235 | 0.74482797 | 0.456375724 | 0.674528805 | Asnsd1 | 766.9620794 | -0.030216206080967 | 0.0697221101368546 | -0.433380544875319 | 0.664738332024494 | 0.915428982038356 | NM_133728 |
| Atp1b1 | 54268.86191 | -0.044498021 | 0.059821785 | -0.743843089 | 0.456971409 | 0.675299957 | Bcas1 | 4716.833287 | 0.0389882297548358 | 0.0894486928387895 | 0.435872549027665 | 0.662929202892373 | 0.915428982038356 | NM_029815,NM_001164369 |
| Dsn1 | 129.2233824 | 0.101847916 | 0.136925263 | 0.743821217 | 0.456984642 | 0.675299957 | Car14 | 563.5479475 | 0.0413132516362541 | 0.0946705313872299 | 0.43638977230698 | 0.66255395828493 | 0.915428982038356 | NM_011797 |
| Ccdc97 | 2153.188899 | 0.04180549 | 0.056239593 | 0.743346234 | 0.457272087 | 0.675531438 | Ccdc117 | 806.0503039 | 0.0338507039391317 | 0.0775811034173357 | 0.436326662654396 | 0.662599739695725 | 0.915428982038356 | NM_134033 |
| Gphn | 2820.944486 | -0.039081406 | 0.052573252 | -0.743370537 | 0.457257377 | 0.675531438 | Cd248 | 162.9279379 | -0.0414412717354828 | 0.0949500642400043 | -0.436453330149752 | 0.662507853019406 | 0.915428982038356 | NM_054042 |
| Kat7 | 3929.506192 | 0.049144558 | 0.06610273 | 0.743457318 | 0.457204853 | 0.675531438 | Cd276 | 491.8485862 | 0.0315993156954533 | 0.0725027819355304 | 0.435835906593922 | 0.662955790125796 | 0.915428982038356 | NM_133983 |
| Bcl2l2 | 6352.74434 | -0.046105544 | 0.062051534 | -0.743020203 | 0.457469449 | 0.675565349 | Cdc42se2 | 6939.169026 | 0.0219247105473964 | 0.0500828166848332 | 0.437769119204431 | 0.661553658753377 | 0.915428982038356 | NM_178626 |
| Dcaf7 | 10961.57824 | -0.0435692 | 0.0586285 | -0.743140269 | 0.457396762 | 0.675565349 | Cnot7 | 3385.306766 | 0.0307381396499697 | 0.0709130859424123 | 0.433462163456429 | 0.664679048077693 | 0.915428982038356 | NM_001271542,NM_001271543,NM_011135 |
| Fcf1 | 422.8307612 | 0.06619056 | 0.089079235 | 0.743052633 | 0.457449815 | 0.675565349 | Cox18 | 343.936979 | -0.0343962689334892 | 0.0785852066115235 | -0.437693942875572 | 0.661608160928214 | 0.915428982038356 | NR_028088,NM_001033310,NM_001163456 |
| Zfp352 | 21.927997 | -0.124639382 | 0.167721434 | -0.743133295 | 0.457400984 | 0.675565349 | Cubn | 33.33847989 | -0.0338288417505893 | 0.0781176542012713 | -0.433049892453616 | 0.664978524040218 | 0.915428982038356 | NM_001081084 |
| Slx4 | 1209.719743 | 0.057207138 | 0.077013504 | 0.742819563 | 0.45759093 | 0.675680345 | Cyp11a1 | 133.8465451 | -0.0414542157189237 | 0.0946244189437191 | -0.438092156143964 | 0.661319480239883 | 0.915428982038356 | NM_019779 |
| Cav2 | 910.798954 | 0.080290919 | 0.108103857 | 0.742720208 | 0.457651093 | 0.675704787 | Dedd | 1124.857376 | 0.0230517917779146 | 0.0532123550029054 | 0.433203750832978 | 0.664866754196571 | 0.915428982038356 | NM_001128609,NM_011615 |
| Flad1 | 1082.015187 | 0.079695057 | 0.107361303 | 0.742307091 | 0.457901297 | 0.675945378 | Ears2 | 214.6948018 | -0.0391093176416649 | 0.0891201667363224 | -0.43883802144779 | 0.660778908240199 | 0.915428982038356 | NM_026140 |
| Prkg2 | 505.8075079 | -0.076940357 | 0.103649913 | -0.742309901 | 0.457899595 | 0.675945378 | Exosc4 | 655.0515078 | -0.028472665447066 | 0.0651416601817589 | -0.437088421873518 | 0.662047223686248 | 0.915428982038356 | NM_175399 |
| Atg10 | 361.1437581 | -0.079848031 | 0.107640304 | -0.741804212 | 0.458205968 | 0.6759631 | Fads6 | 433.448382 | -0.0354715508702857 | 0.0808546996707623 | -0.438707348054284 | 0.660873602079489 | 0.915428982038356 | NM_178035 |
| Decr2 | 1270.197979 | 0.057210203 | 0.07712867 | 0.741750161 | 0.458238722 | 0.6759631 | Fam129a | 120.6896662 | -0.041593959747289 | 0.0951023547712065 | -0.437359935485868 | 0.661850335018757 | 0.915428982038356 | NM_022018 |
| Gcat | 340.9674043 | 0.08025262 | 0.108195089 | 0.741739953 | 0.458244908 | 0.6759631 | Fam221b | 54.04660429 | 0.0412360488220161 | 0.0947026331745337 | 0.435426634294523 | 0.663252781187484 | 0.915428982038356 | NM_175517 |
| Gm5086 | 35.38221242 | 0.123315359 | 0.166216615 | 0.7418955 | 0.458150652 | 0.6759631 | Fntb | 1331.650984 | 0.0232124128908189 | 0.0530026423801013 | 0.437948220097297 | 0.661423819414245 | 0.915428982038356 | NM_145927 |
| Lactb | 531.7833558 | -0.056547828 | 0.076209459 | -0.742005376 | 0.458084079 | 0.6759631 | Fos | 1219.200699 | 0.0263064879669978 | 0.0600832834595104 | 0.437833727657802 | 0.661506819628122 | 0.915428982038356 | NM_010234 |
| Metrnl | 219.0008296 | 0.095062555 | 0.128166532 | 0.741711219 | 0.458262321 | 0.6759631 | Frat1 | 418.3901945 | -0.0370328226837054 | 0.0855214065127313 | -0.433024013446183 | 0.66499732448 | 0.915428982038356 | NM_008043 |
| Mrps7 | 1350.563149 | -0.041824081 | 0.056387685 | -0.74172368 | 0.458254769 | 0.6759631 | Frmd8 | 1030.395003 | -0.033354576054857 | 0.0759259381275464 | -0.439304101831779 | 0.660441202454959 | 0.915428982038356 | NM_026169 |
| Pikfyve | 2244.818641 | -0.060045446 | 0.080906107 | -0.742162098 | 0.45798913 | 0.6759631 | Fut8 | 2524.530269 | 0.0293023154374068 | 0.0670386143508362 | 0.437096078449022 | 0.662041671184168 | 0.915428982038356 | NM_016893,NM_001252616,NM_001252615,NM_001252614,NR_045554 |
| Efs | 402.1326853 | -0.098791713 | 0.133216138 | -0.741589674 | 0.458335982 | 0.676007397 | Gm12070 | 163.2437605 | -0.0414412657715292 | 0.0947926996167171 | -0.437177820012427 | 0.661982393856778 | 0.915428982038356 | NR_002890 |
| Arpc1b | 525.8277498 | 0.052373333 | 0.070631158 | 0.741504665 | 0.458387505 | 0.676019036 | Gm5126 | 85.66316328 | -0.0413313191293336 | 0.0943279936958511 | -0.438165994101404 | 0.661265957691416 | 0.915428982038356 | NR_026596 |
| Pitpnc1 | 3427.416826 | -0.058160377 | 0.078452961 | -0.741340753 | 0.458486858 | 0.676101207 | Hmgn5 | 358.7031296 | 0.0361286571033686 | 0.0832236366236645 | 0.434115337530149 | 0.664204688345741 | 0.915428982038356 | NM_016710 |
| Rasgef1c | 1581.274398 | -0.083748599 | 0.112983195 | -0.741248281 | 0.458542914 | 0.67611952 | Hspb11 | 196.1545947 | 0.0368731946026908 | 0.0848914422027943 | 0.434357028763933 | 0.664029197104876 | 0.915428982038356 | NM_028394 |
| Pias2 | 2380.599884 | 0.066396883 | 0.089606764 | 0.740980707 | 0.458705139 | 0.676294359 | Ints8 | 1112.859722 | 0.0289988560163031 | 0.0670259683095528 | 0.432651056712449 | 0.665268291439918 | 0.915428982038356 | NM_001159595,NM_178112 |
| Fkbp2 | 2010.592874 | -0.05400009 | 0.072931761 | -0.74041939 | 0.459045557 | 0.676731861 | Iqgap2 | 1317.455564 | -0.0401788389534217 | 0.0928450984465413 | -0.432751320486304 | 0.665195441774526 | 0.915428982038356 | NM_027711 |
| Cebpb | 70.74134214 | 0.116455164 | 0.157362168 | 0.740045499 | 0.459272387 | 0.676917731 | Kalrn | 25732.04975 | -0.0200641140623514 | 0.0458678246132723 | -0.437433303879548 | 0.661797135785446 | 0.915428982038356 | NM_177357,NM_001164268 |
| Etnk1 | 14726.91397 | -0.047221265 | 0.063812906 | -0.739995533 | 0.459302705 | 0.676917731 | Kbtbd11 | 6505.58947 | 0.0233804033581703 | 0.0539149108976221 | 0.433653751233436 | 0.664539895854597 | 0.915428982038356 | NM_029116 |
| Shoc2 | 2364.938354 | -0.05475215 | 0.073981004 | -0.740083898 | 0.459249088 | 0.676917731 | Kctd17 | 6277.326907 | -0.0235953548316492 | 0.0544691840802753 | -0.433187227421563 | 0.664878757209915 | 0.915428982038356 | NM_001081367 |
| 2610318N02Rik | 28.38692282 | -0.122458467 | 0.165578887 | -0.739577791 | 0.459556222 | 0.676957488 | Kdm3a | 2218.788272 | -0.031896232878246 | 0.0733974619958054 | -0.434568607836452 | 0.663875585331163 | 0.915428982038356 | NM_001038695,NM_173001 |
| Ddx21 | 1211.273294 | 0.076382116 | 0.103377935 | 0.738862857 | 0.459990279 | 0.676957488 | Klhdc9 | 178.3624245 | 0.0414977012420197 | 0.0948360812316386 | 0.437572922700812 | 0.661695903253849 | 0.915428982038356 | NM_001033039 |
| Fbxo47 | 24.33760891 | 0.124159702 | 0.167961339 | 0.739215953 | 0.459775875 | 0.676957488 | Lurap1 | 552.3042711 | 0.0312633279076674 | 0.0722309082362577 | 0.432824793029172 | 0.665142060091846 | 0.915428982038356 | NM_026547 |
| Lsm3 | 452.4564055 | -0.062188959 | 0.084157764 | -0.738956881 | 0.459933181 | 0.676957488 | Mettl23 | 435.3405592 | -0.0342902666857776 | 0.0780913186149246 | -0.439104721164539 | 0.660585658357268 | 0.915428982038356 | NM_028865 |
| Mrps5 | 1195.807599 | 0.051608911 | 0.069804248 | 0.739337681 | 0.459701973 | 0.676957488 | Mgat5b | 4626.939773 | -0.0334852125619901 | 0.0766171809582816 | -0.437045740174426 | 0.662078176535858 | 0.915428982038356 | NM_172948 |
| Nab2 | 1701.092071 | 0.082601356 | 0.111752768 | 0.73914371 | 0.459819738 | 0.676957488 | Mical3 | 4731.10071 | -0.026238927092128 | 0.0600141727045679 | -0.437212176885192 | 0.661957479580265 | 0.915428982038356 | NM_153396,NM_001270475 |
| Nek2 | 66.84815621 | -0.112700102 | 0.152361038 | -0.73969109 | 0.459487456 | 0.676957488 | Mns1 | 60.56858392 | -0.0410917983591318 | 0.0948681191931136 | -0.433146548162142 | 0.664908307992798 | 0.915428982038356 | NM_008613 |
| Pim2 | 1687.687445 | -0.045660911 | 0.061790684 | -0.738961095 | 0.459930622 | 0.676957488 | Myo3b | 135.7653998 | -0.0402818505526317 | 0.0918662081181273 | -0.438483871031607 | 0.661035559618057 | 0.915428982038356 | NM_177376 |
| Plekhg2 | 284.984652 | 0.075911792 | 0.102613046 | 0.739786945 | 0.459429282 | 0.676957488 | Napa | 11036.88616 | 0.0180356950626962 | 0.04164269681295 | 0.433105837110134 | 0.664937882392188 | 0.915428982038356 | NM_025898 |
| Psmd2 | 9006.134837 | -0.037239039 | 0.050374004 | -0.739251122 | 0.459754524 | 0.676957488 | Neu4 | 388.5525933 | 0.0346755985790899 | 0.0789062783582191 | 0.439452972571707 | 0.660333350412642 | 0.915428982038356 | NM_173772 |
| Ptpn1 | 2025.146636 | -0.065223546 | 0.088213893 | -0.739379525 | 0.459676572 | 0.676957488 | Nprl3 | 1099.887991 | -0.0307257476688276 | 0.0710251106422042 | -0.432604009919978 | 0.665302475793675 | 0.915428982038356 | NM_181569 |
| Sbf1 | 8349.287794 | 0.050025657 | 0.067644835 | 0.73953402 | 0.45958279 | 0.676957488 | Ocln | 373.1343401 | -0.0325434450178508 | 0.0751768928043338 | -0.432891594795665 | 0.665093526553858 | 0.915428982038356 | NM_008756 |
| Sf3a3 | 1109.444489 | 0.057067045 | 0.077242938 | 0.738799515 | 0.460028746 | 0.676957488 | Pcdhgb5 | 223.2947254 | 0.0371017481069757 | 0.0844914796505795 | 0.43911821949873 | 0.660575878102924 | 0.915428982038356 | NM_033577 |
| Slitrk4 | 1112.109099 | -0.081867402 | 0.110708613 | -0.739485394 | 0.459612306 | 0.676957488 | Pet100 | 529.6934827 | -0.0314287385228715 | 0.0724394449583222 | -0.433862221624612 | 0.664388494744545 | 0.915428982038356 | NM_001195244 |
| Tbl2 | 1084.360477 | 0.050285862 | 0.067992254 | 0.739582214 | 0.459553537 | 0.676957488 | Ppm1j | 32.70240461 | -0.037115534399698 | 0.0849493552208289 | -0.436913668187532 | 0.662173959050127 | 0.915428982038356 | NM_027982 |
| Trpc3 | 671.9431308 | 0.059761631 | 0.080876905 | 0.738920847 | 0.459955063 | 0.676957488 | Ppp1r42 | 24.34914671 | -0.037769072712792 | 0.0862276930222757 | -0.438015577003027 | 0.6613749916052 | 0.915428982038356 | NM_145692,NM_027033 |
| Bcorl1 | 1127.038631 | 0.055551297 | 0.075208399 | 0.738631561 | 0.460130754 | 0.677038414 | Prcp | 431.1652159 | -0.0379096387125891 | 0.0868809402868512 | -0.436340106212299 | 0.662589987277658 | 0.915428982038356 | NM_028243 |
| R3hdm4 | 10168.74753 | -0.059185012 | 0.080135134 | -0.738565083 | 0.460171133 | 0.677038414 | Ptp4a3 | 2307.548644 | 0.0284569691423593 | 0.0652092724804251 | 0.436394519673589 | 0.662550514470645 | 0.915428982038356 | NM_001166390,NM_008975,NM_001166389,NM_001166388 |
| Tceal1 | 559.1655831 | -0.065258208 | 0.088380471 | -0.738378135 | 0.460284698 | 0.677076911 | Rab35 | 3760.079226 | 0.0183615332541737 | 0.0418580711613047 | 0.438661714330205 | 0.660906672313216 | 0.915428982038356 | NM_198163 |
| Zfp777 | 1191.012896 | 0.058237068 | 0.078870105 | 0.738392173 | 0.46027617 | 0.677076911 | Rab39 | 42.32181912 | -0.0414204051799172 | 0.0942192655461156 | -0.439617151968182 | 0.660214415924478 | 0.915428982038356 | NM_175562 |
| Vsnl1 | 19691.90344 | 0.064898598 | 0.087905375 | 0.738277925 | 0.460345578 | 0.677102183 | Rgl1 | 5323.274968 | 0.0266823351608407 | 0.0613073640127606 | 0.43522235200468 | 0.663401039743528 | 0.915428982038356 | NM_016846 |
| Fchsd1 | 331.6514419 | -0.073932132 | 0.100188007 | -0.737933957 | 0.460554583 | 0.677345297 | Ripply2 | 58.78617703 | 0.0410048026518352 | 0.0944936187265358 | 0.433942558285368 | 0.664330154096683 | 0.915428982038356 | NM_001037907 |
| Rbm19 | 578.012566 | 0.051828692 | 0.070244664 | 0.737831024 | 0.460617138 | 0.677373001 | Rnase4 | 632.3964794 | 0.0313348652407729 | 0.0716676636249749 | 0.437224595526695 | 0.66194847415341 | 0.915428982038356 | NM_201239,NM_021472 |
| Unkl | 1988.315132 | 0.040756215 | 0.055268593 | 0.737420887 | 0.460866437 | 0.677675295 | Rpl29 | 5317.366329 | 0.0228738747029045 | 0.0525660402252945 | 0.435145478047588 | 0.663456834689258 | 0.915428982038356 | NM_009082 |
| Hadhb | 2779.176812 | -0.057138833 | 0.077507098 | -0.73720774 | 0.460996027 | 0.677767216 | Rpl35 | 3527.452421 | 0.0279973418911222 | 0.0643902379189937 | 0.434807244016466 | 0.663702346359274 | 0.915428982038356 | NM_025592 |
| Prg4 | 262.5092609 | -0.110654087 | 0.150105756 | -0.737174172 | 0.461016438 | 0.677767216 | Rps2 | 10390.10425 | -0.0216616211510042 | 0.0498343362319987 | -0.434672613078677 | 0.663800080064319 | 0.915428982038356 | NM_008503 |
| Smpd1 | 4579.140266 | -0.048544782 | 0.065868456 | -0.736995902 | 0.461124841 | 0.677862267 | Scara3 | 1499.410589 | -0.0337844090727971 | 0.0776613392914531 | -0.435022230894171 | 0.66354629109172 | 0.915428982038356 | NM_172604 |
| 2700099C18Rik | 10.48386147 | -0.121922001 | 0.165465166 | -0.736843915 | 0.461217274 | 0.677933825 | Slain2 | 1011.342311 | 0.0279305495469128 | 0.0644970562548291 | 0.43305154015958 | 0.664977327030935 | 0.915428982038356 | NM_153567,NM_001113423 |
| 1500011B03Rik | 3003.95319 | -0.039778395 | 0.054002921 | -0.736597101 | 0.461367399 | 0.677935394 | Slc37a1 | 279.9556221 | 0.0370428794725938 | 0.0850422983830075 | 0.435581824303039 | 0.663140160332987 | 0.915428982038356 | NM_153062,NM_001242427 |
| Gm5148 | 76.73333438 | 0.104289789 | 0.141582277 | 0.736602008 | 0.461364413 | 0.677935394 | Smad4 | 1649.336654 | 0.0331789427242137 | 0.0757695195932348 | 0.437893006347847 | 0.661463845581517 | 0.915428982038356 | NM_008540 |
| Nefl | 10522.66853 | -0.060512368 | 0.082156004 | -0.736554423 | 0.46139336 | 0.677935394 | St3gal4 | 897.0512347 | 0.0315653734011524 | 0.0721286964092254 | 0.437625729738201 | 0.661657616402599 | 0.915428982038356 | NM_009178 |
| Ythdf1 | 2327.926066 | 0.059228292 | 0.080407091 | 0.736605329 | 0.461362394 | 0.677935394 | Stk36 | 236.7802186 | -0.0388527410224308 | 0.0894861906572733 | -0.434175829109035 | 0.664160763873595 | 0.915428982038356 | NM_175031 |
| Pik3ca | 3204.848271 | -0.051476876 | 0.069904372 | -0.736389925 | 0.461493435 | 0.678018137 | Taf9 | 2574.138369 | 0.0268992608139684 | 0.061540802461631 | 0.437096361080755 | 0.662041466221706 | 0.915428982038356 | NM_027592,NM_001015889,NM_027139 |
| Akr7a5 | 586.1360266 | 0.057297725 | 0.07788128 | 0.735705999 | 0.461909638 | 0.678212469 | Tigd5 | 233.5608633 | 0.0368364466497875 | 0.0840945179204938 | 0.438036242559998 | 0.661360011189942 | 0.915428982038356 | NM_178646 |
| Ankfn1 | 93.16936004 | 0.120924567 | 0.164285981 | 0.736061385 | 0.461693341 | 0.678212469 | Traf5 | 110.0529589 | -0.0400886769614787 | 0.092217514603859 | -0.434718688024597 | 0.663766631869358 | 0.915428982038356 | NM_011633 |
| Cars2 | 371.1748432 | 0.063476957 | 0.086255909 | 0.735914309 | 0.461782849 | 0.678212469 | Trp63 | 27.38522602 | 0.0373986566464684 | 0.0853016557053913 | 0.438428261880792 | 0.661075862966639 | 0.915428982038356 | NM_001127259,NM_001127265,NM_001127264,NM_001127263,NM_001127262,NM_001127261,NM_001127260,NM_011641 |
| Dusp2 | 36.02515702 | 0.11949356 | 0.162428413 | 0.735669074 | 0.461932115 | 0.678212469 | Ttc21a | 95.80847237 | 0.0403131020260419 | 0.0931603123091586 | 0.432728283394545 | 0.665212179787879 | 0.915428982038356 | NM_028735 |
| Eri2 | 403.9451679 | 0.069333564 | 0.09424411 | 0.735680605 | 0.461925096 | 0.678212469 | Tusc1 | 74.93072418 | 0.0411606836917953 | 0.0946525926627489 | 0.434860604806173 | 0.663663611320796 | 0.915428982038356 | NM_026954 |
| Slc30a4 | 3010.914904 | 0.065745828 | 0.089360781 | 0.735734709 | 0.461892163 | 0.678212469 | Zfp202 | 206.2649099 | 0.0366042696295324 | 0.0834070640188532 | 0.438862943566248 | 0.660760848783309 | 0.915428982038356 | NM_030713 |
| Tnfrsf9 | 20.46195897 | 0.12358276 | 0.167949146 | 0.735834406 | 0.46183148 | 0.678212469 | Zfp239 | 2066.376569 | -0.0241904344331174 | 0.0554483719476417 | -0.436269516730982 | 0.662641195947292 | 0.915428982038356 | NM_008616,NM_001001792 |
| Esam | 641.119318 | 0.067162091 | 0.091327059 | 0.73540188 | 0.462094777 | 0.678387008 | Zfp609 | 2600.118983 | 0.0280051245976867 | 0.0641137946851702 | 0.436803417036933 | 0.662253920680906 | 0.915428982038356 | NM_172536 |
| Hs3st4 | 2485.035822 | 0.073739161 | 0.100297383 | 0.73520524 | 0.462214509 | 0.678498493 | Zfp839 | 554.1431413 | 0.0291280140910853 | 0.0666966506204667 | 0.436723790776789 | 0.662311673453818 | 0.915428982038356 | NM_028365,NM_001199785 |
| Arl14ep | 1061.27528 | 0.056590178 | 0.076993712 | 0.734997398 | 0.462341079 | 0.678564691 | Zfp93 | 450.2142992 | -0.0305310938408164 | 0.0703355674473345 | -0.434077593298401 | 0.664232095975673 | 0.915428982038356 | NM_009567 |
| Zbtb33 | 935.0810253 | -0.077115075 | 0.104920274 | -0.734987355 | 0.462347195 | 0.678564691 | Zyx | 1847.115968 | -0.026810682893807 | 0.0618086533863964 | -0.433769082885533 | 0.664456134833323 | 0.915428982038356 | NM_011777 |
| Srsf12 | 594.249664 | -0.050376903 | 0.068567932 | -0.734700629 | 0.462521837 | 0.67875671 | D19Ertd737e | 470.6863504 | 0.0312609874845675 | 0.0723666709414404 | 0.431980455299633 | 0.665755618323367 | 0.915591677044461 | NM_029648 |
| Irak1 | 2316.033811 | 0.058814162 | 0.080068061 | 0.73455209 | 0.462612325 | 0.678825208 | Gm765 | 118.2116051 | -0.0409454787285219 | 0.0948044347526883 | -0.43189412853243 | 0.665818362354343 | 0.915591677044461 | NM_001128092 |
| Myh9 | 3764.109509 | -0.053397277 | 0.072770944 | -0.733771942 | 0.463087743 | 0.679458475 | Nrm | 87.35412634 | -0.0410900200259113 | 0.0950965496712428 | -0.432087390846073 | 0.66567789866142 | 0.915591677044461 | NM_134122 |
| Cadps | 11494.73447 | -0.038891329 | 0.053030915 | -0.733370881 | 0.463332254 | 0.679675919 | Nrsn1 | 17735.90897 | 0.0259949771620777 | 0.0601892388753246 | 0.431887454432237 | 0.66582321332205 | 0.915591677044461 | NM_009513 |
| Evi5l | 1447.812306 | 0.053133093 | 0.072467563 | 0.733198284 | 0.463437502 | 0.679675919 | Scn3a | 1753.089045 | -0.0288664008257595 | 0.0667946185963248 | -0.432166564199048 | 0.665620358591421 | 0.915591677044461 | NM_018732 |
| Il10rb | 381.9501679 | -0.081815771 | 0.111557029 | -0.733398613 | 0.463315345 | 0.679675919 | Spink10 | 39.24904164 | 0.0394014696698573 | 0.0912475758288279 | 0.431808399422807 | 0.665880674310924 | 0.915591677044461 | NM_177829 |
| Prosapip1 | 12939.83964 | 0.057616422 | 0.078585444 | 0.733169134 | 0.463455278 | 0.679675919 | Srgn | 172.976683 | -0.0384118598953372 | 0.0889683510455177 | -0.431747463496149 | 0.66592496681638 | 0.915591677044461 | NM_011157 |
| Zfp78 | 129.718445 | -0.098165885 | 0.133881337 | -0.73323054 | 0.463417832 | 0.679675919 | Wipf3 | 8038.122761 | 0.0317535728846933 | 0.0734989752360296 | 0.432027423276611 | 0.665721482035661 | 0.915591677044461 | NM_001167861,NM_001167860 |
| Hadha | 2204.849255 | 0.046136431 | 0.06296267 | 0.732758489 | 0.463705744 | 0.679978875 | Rpl36al | 1720.17208 | 0.0311287238157982 | 0.0721620235088753 | 0.431372657003855 | 0.666197428085417 | 0.915879598747753 | NM_025589 |
| Megf9 | 3180.721121 | -0.05774376 | 0.078828923 | -0.732519966 | 0.463851261 | 0.680013495 | Taok1 | 5593.101119 | 0.0310334560379935 | 0.0719610267031657 | 0.431253658539426 | 0.666283941866135 | 0.915911851949226 | NM_144825 |
| Nrg3 | 1589.835353 | 0.065452548 | 0.089363305 | 0.732432042 | 0.463904908 | 0.680013495 | 1700028P14Rik | 36.30497859 | -0.0381051908754178 | 0.0884077900426229 | -0.431018650804096 | 0.666454809277472 | 0.91601740256939 | NM_026188 |
| Tiam1 | 4978.981321 | 0.089128124 | 0.121675568 | 0.732506333 | 0.463859579 | 0.680013495 | Kank4 | 318.2608084 | -0.0396644230846964 | 0.0920689209471409 | -0.430812294492609 | 0.666604859329058 | 0.91601740256939 | NM_172872 |
| Wasf3 | 2699.427375 | -0.034991156 | 0.047770299 | -0.732487689 | 0.463870954 | 0.680013495 | Pdlim5 | 1264.795765 | -0.0254262824190095 | 0.0590174100582483 | -0.430826808460666 | 0.666594305197455 | 0.91601740256939 | NM_019809,NM_019808,NM_001190852,NM_001190853,NM_001190856,NM_001190857,NM_001190854,NM_001190855,NM_022554 |
| Pknox2 | 2901.230965 | 0.051108112 | 0.069791442 | 0.732297695 | 0.463986886 | 0.680069322 | Tle3 | 1762.80627 | 0.0288799375200386 | 0.067037745567766 | 0.430801144570784 | 0.666612967269794 | 0.91601740256939 | NM_001083927,NM_001083928,NM_009389 |
| Csf2ra | 690.9675447 | 0.121148525 | 0.165455095 | 0.732213932 | 0.464038002 | 0.68007991 | Adamtsl5 | 248.1315761 | -0.0388766677245862 | 0.0903326705937039 | -0.430372172870264 | 0.66692493418974 | 0.916077198994267 | NM_001113548,NM_025629 |

| | | | | | | | | | | | | | | | |
|---|---|---|---|---|---|---|---|---|---|---|---|---|---|---|---|
| Snf8 | 1733.778739 | -0.042455545 | 0.057996896 | -0.73203133 | 0.464149445 | 0.680178899 | | Ccdc96 | 81.95383742 | -0.0407242889415698 | 0.0946704292823584 | -0.430169053317672 | 0.667072671679411 | 0.916077198994267 | NM_025725 |
| Slc41a1 | 3068.903233 | 0.051733333 | 0.070690564 | 0.731827979 | 0.464273569 | 0.680296452 | | Mcm2 | 213.3250939 | 0.0401767602003265 | 0.0933491246752522 | 0.430392468489613 | 0.666910173032151 | 0.916077198994267 | NM_008564 |
| Cstad | 48.16732965 | 0.122803126 | 0.167857338 | 0.731592239 | 0.464417486 | 0.680386485 | | Msrb1 | 672.8602655 | 0.0315163995949382 | 0.0732065375448492 | 0.430513457566956 | 0.666822179435914 | 0.916077198994267 | NM_013759 |
| Gm10433 | 12.8841283 | -0.111841828 | 0.152876373 | -0.731583473 | 0.464422839 | 0.680386485 | | Prx | 23.9803642 | 0.0361913632565075 | 0.0841291002888005 | 0.43018840249413 | 0.667058597644042 | 0.916077198994267 | NM_198048,NM_019412 |
| D330041H03Rik | 22.82113479 | 0.122943872 | 0.168107561 | 0.731340525 | 0.464571183 | 0.680539464 | | Slc6a8 | 5268.51837 | -0.024308623926607 | 0.0565140300542469 | -0.430134320685917 | 0.667097935492315 | 0.916077198994267 | NM_001142809,NM_001142810 |
| Ankib1 | 2277.824565 | -0.052207134 | 0.071398724 | -0.73120541 | 0.464653696 | 0.680542649 | | Tubb2b | 3947.512118 | 0.0304488236734046 | 0.0707415715928972 | 0.430423342142173 | 0.666887718638192 | 0.916077198994267 | NM_023716 |
| Cse1l | 2648.822125 | 0.048719927 | 0.066632248 | 0.731176393 | 0.464671418 | 0.680542649 | | Dnajc7 | 6508.945282 | 0.0210449317531699 | 0.0489712283800037 | 0.42974073653752 | 0.667384246996573 | 0.916320386773434 | NM_019795 |
| Epha8 | 413.0871082 | 0.106259388 | 0.145337583 | 0.7311212 | 0.464705126 | 0.680542649 | | Oat | 3335.263901 | 0.0213859759980977 | 0.0497675301480206 | 0.429717446987839 | 0.667401190421516 | 0.916320386773434 | NM_016978 |
| Pabpc1 | 5359.08856 | 0.044606864 | 0.061021802 | 0.730998793 | 0.46477989 | 0.68058781 | | Stag1 | 1550.300345 | -0.0268357093542809 | 0.0624802316365279 | -0.429507200139634 | 0.667554155183282 | 0.916443782080367 | NM_009282 |
| 5830415F09Rik | 100.8563872 | -0.091222234 | 0.124819481 | -0.730833306 | 0.464880977 | 0.680671505 | | Fam135b | 1024.59786 | -0.0293714347896402 | 0.0684171884555639 | -0.429299061429812 | 0.667705599778981 | 0.916478460019793 | NM_177819 |
| Grip2 | 1430.857763 | 0.089209857 | 0.122095481 | 0.730656504 | 0.464988989 | 0.680694055 | | Sco2 | 121.7233621 | -0.0386844392011647 | 0.0900976048536575 | -0.429361460429481 | 0.667660195985652 | 0.916478460019793 | NM_001111288 |
| Immt | 5287.129796 | -0.041106375 | 0.056264447 | -0.730592365 | 0.465028177 | 0.680694055 | | Endov | 1893.808257 | -0.0254156634281418 | 0.0592195988142366 | -0.429176555347278 | 0.667794743221041 | 0.91651421391051 | NM_001164636,NM_177394 |
| Vps45 | 1448.058916 | -0.047946261 | 0.065623976 | -0.730621095 | 0.465010623 | 0.680694055 | | Ccdc162 | 29.82213276 | 0.0371902795768276 | 0.0867698037253218 | 0.428608547906331 | 0.668208123832047 | 0.91699491797679 | NM_001177571 |
| Nicn1 | 4159.589224 | 0.040519226 | 0.055487332 | 0.730242818 | 0.465241774 | 0.680878063 | | Fam57a | 227.94405 | -0.0371805569001238 | 0.0867720067033575 | -0.428485617801036 | 0.668297602324621 | 0.917031076078541 | NM_027773 |
| Ppap2c | 103.2276227 | 0.103494484 | 0.141714307 | 0.730303708 | 0.465204562 | 0.680878063 | | Zfp110 | 751.2346559 | 0.0300779018501469 | 0.0702315774993752 | 0.42826749620446 | 0.668456380498961 | 0.917162310676103 | NM_022981 |
| Dhrs4 | 322.937256 | 0.057516431 | 0.078778691 | 0.730101383 | 0.465328216 | 0.680940251 | | Lats2 | 1090.915133 | 0.0329698539344963 | 0.077027847067492 | 0.428025125843229 | 0.668632827576617 | 0.917270892731121 | NM_153382,NM_015771 |
| 2410004N09Rik | 101.5600217 | 0.103079469 | 0.141208688 | 0.729979652 | 0.465402622 | 0.680984817 | | Stxbp6 | 2842.721118 | 0.02680463978718 | 0.0626298119858907 | 0.427985314616921 | 0.668661812138913 | 0.917270892731121 | NM_144552 |
| Elp3 | 3419.329212 | 0.042784055 | 0.058642446 | 0.729574873 | 0.465650086 | 0.68114179 | | Cox7a2 | 4093.687619 | 0.0247992782880482 | 0.0579771803110577 | 0.427742055667346 | 0.668838927537734 | 0.917427220152325 | NM_009945 |
| Rnf13 | 3773.583413 | 0.057811096 | 0.079243706 | 0.729535493 | 0.465674165 | 0.68114179 | | Ash1l | 7256.629629 | 0.0318937585809182 | 0.074618464172215 | 0.427424484472225 | 0.669070176976005 | 0.917484507764887 | NM_138679 |
| Rnf44 | 6588.656931 | -0.040808442 | 0.055939022 | -0.72951655 | 0.465685748 | 0.68114179 | | Lsm1 | 489.6887922 | 0.0303129588508945 | 0.0709155940546204 | 0.427451243340738 | 0.669050690454238 | 0.917484507764887 | NM_026032 |
| Zdhhc7 | 1305.510387 | -0.053237024 | 0.072955615 | -0.729717979 | 0.465562589 | 0.68114179 | | Sptb | 9619.602592 | 0.0233756005222941 | 0.0546802164292095 | 0.427496488653383 | 0.669017742117838 | 0.917484507764887 | NM_013675 |
| Rhbdl1 | 1714.675452 | -0.061271049 | 0.084014226 | -0.72929373 | 0.465822007 | 0.681148186 | | Plekhm3 | 2669.989955 | 0.0271091251214223 | 0.0634394703396438 | 0.427322690058488 | 0.669144308419665 | 0.917499549188603 | NM_001039493 |
| Zfp341 | 462.1009314 | 0.056351689 | 0.077262937 | 0.729349562 | 0.465787862 | 0.681148186 | | Fam57b | 2522.977672 | -0.0255438099333082 | 0.0598614592672162 | -0.426715456756291 | 0.669586591026775 | 0.917701189578706 | NM_029978,NM_001146347,NM_026884 |
| Zfpm1 | 1283.79896 | 0.072961232 | 0.100043325 | 0.729296358 | 0.465820399 | 0.681148186 | | Gjc3 | 1986.131971 | -0.034331304723444 | 0.0804601426703388 | -0.42668709728873 | 0.669607249645099 | 0.917701189578706 | NM_080450 |
| Cdh15 | 78.20387337 | -0.108845964 | 0.149288768 | -0.729096808 | 0.465942447 | 0.681260005 | | N4bp2l1 | 946.3497619 | 0.0282969998353359 | 0.0662726965985415 | 0.426978247267498 | 0.669395171650776 | 0.917701189578706 | NM_133898 |
| Dlk2 | 925.6786684 | 0.090260243 | 0.123851126 | 0.728780155 | 0.466136153 | 0.681389166 | | Nat6 | 524.8691185 | -0.0327782222748309 | 0.0768181155685442 | -0.426699119501091 | 0.66959849196438 | 0.917701189578706 | NM_019750 |
| Stac3 | 60.22126559 | 0.121871188 | 0.167220999 | 0.72880313 | 0.466122097 | 0.681389166 | | Zfp825 | 251.8944842 | -0.0383722779692551 | 0.0899201798615743 | -0.426737113163324 | 0.669570815460043 | 0.917701189578706 | NM_146231 |
| Tmem29 | 477.7873258 | -0.056812756 | 0.077960604 | -0.728736733 | 0.466162719 | 0.681389166 | | Adss | 3455.360806 | -0.0244608840804879 | 0.0573531143470061 | -0.426496178263156 | 0.669746332220125 | 0.91780520960656 | NM_007422 |
| 1110051M20Rik | 2904.480626 | -0.0388002 | 0.053267995 | -0.728396108 | 0.466371145 | 0.681565225 | | Cyld | 8073.553005 | 0.0326076920444364 | 0.0765253568945901 | 0.426103103176009 | 0.670032719094779 | 0.918093365654835 | NM_001128170,NM_001128171,NM_001128169,NM_001276279,NM_173369 |
| Myo6 | 4803.10504 | -0.069764313 | 0.095773355 | -0.728431337 | 0.466349586 | 0.681565225 | | Mast4 | 3053.898699 | 0.0326328767725319 | 0.0765968696398244 | 0.426034078494056 | 0.670083014090084 | 0.918093365654835 | NM_175171 |
| Aig1 | 835.0992092 | -0.045030032 | 0.061866126 | -0.727862486 | 0.466697769 | 0.681613104 | | Prpf3 | 926.5945504 | -0.0286841710873399 | 0.0673429251110587 | -0.425941864569074 | 0.670150208289154 | 0.918098832934854 | NM_027541 |
| Cdc16 | 1792.995564 | -0.035800389 | 0.04917973 | -0.727950101 | 0.466644132 | 0.681613104 | | Jam2 | 2125.005234 | 0.0261144070967264 | 0.0613235590502042 | 0.425846240844358 | 0.670219889919559 | 0.918099319576618 | NM_023844 |
| Dhx34 | 454.1688839 | 0.08413967 | 0.115572933 | 0.728022282 | 0.466599946 | 0.681613104 | | Rcbtb1 | 2124.003821 | 0.0270995200987055 | 0.0636485732603077 | 0.425767911369746 | 0.670276971231587 | 0.918099319576618 | NM_027764 |
| Dyrk3 | 309.0748985 | 0.109438214 | 0.150353011 | 0.727875111 | 0.46669004 | 0.681613104 | | Il34 | 2156.430176 | 0.0307316510024773 | 0.0721957887186957 | 0.425670964302646 | 0.670347622443074 | 0.918109519480303 | NM_001135100,NM_029646 |
| Lactb2 | 307.9668814 | 0.076263622 | 0.104732147 | 0.728177776 | 0.466504769 | 0.681613104 | | AA388235 | 139.7688716 | -0.0389230283977789 | 0.0914703540079882 | -0.425526158938662 | 0.670453156329761 | 0.918167488342236 | NR_033305 |
| Nthl1 | 85.69078635 | 0.101410065 | 0.139324711 | 0.727868476 | 0.466694102 | 0.681613104 | | Wtap | 2227.672651 | -0.0224031814315343 | 0.0526700156879823 | -0.425349815049438 | 0.670581684218089 | 0.918256932970584 | NM_175394,NM_001113532,NM_001113533 |
| Tcf7l1 | 157.1007233 | 0.093602624 | 0.128603389 | 0.727839485 | 0.466711851 | 0.681613104 | | Rbm20 | 252.8014645 | 0.0392176732262228 | 0.0922365186969321 | 0.425185965171594 | 0.670701114512029 | 0.918333904175863 | NM_001170847 |
| Inpp5j | 2931.417406 | -0.091602945 | 0.125902789 | -0.727568828 | 0.466877568 | 0.681756583 | | Polr3a | 1521.186209 | -0.0216924387965966 | 0.0510291821656179 | -0.425098696000902 | 0.670764728469592 | 0.918334443520198 | NM_001081247 |
| Klc1 | 23672.97381 | 0.039734665 | 0.054619878 | 0.72747626 | 0.46693425 | 0.681756583 | | Igsf11 | 1595.12903 | -0.0298870191522835 | 0.0703446721875394 | -0.42486543163645 | 0.670934775694148 | 0.918480685301403 | NM_170599 |
| Psmd12 | 2452.817284 | 0.060606443 | 0.083312011 | 0.72746345 | 0.466942097 | 0.681756583 | | Sall1 | 414.255196 | 0.029193474333911 | 0.0687366907734041 | 0.424714573911473 | 0.671044758335483 | 0.918544681453188 | NM_030233 |
| Fam118b | 550.5490566 | -0.061677035 | 0.084798606 | -0.727335478 | 0.46702047 | 0.681806761 | | Atp6v1c2 | 59.15687189 | 0.0400023859726621 | 0.0942363482359272 | 0.424489984188621 | 0.671208508254776 | 0.91868225675199 | NM_133699,NM_001159632 |
| 1500009L16Rik | 358.4435041 | -0.067344787 | 0.09263563 | -0.726985791 | 0.46723466 | 0.681990939 | | Tmem106c | 1131.354572 | 0.0258123514527547 | 0.0608549697179327 | 0.424161766448933 | 0.671447842154782 | 0.918836679711763 | NM_201359,NM_001252153 |
| Bambi | 123.8550893 | 0.108877305 | 0.149757988 | 0.727021681 | 0.467212674 | 0.681990939 | | Ypel2 | 2471.946979 | -0.0284990385901686 | 0.0671764489683171 | -0.424241516600703 | 0.671389685880258 | 0.918836679711763 | NM_001005341 |
| Polb | 1241.988059 | -0.04979657 | 0.068522612 | -0.726717333 | 0.467399133 | 0.682158898 | | Amz1 | 726.7206732 | 0.0365350538223881 | 0.0862578212671471 | 0.423556418254946 | 0.671889344878595 | 0.919094512073309 | NM_173405 |
| Zc3h13 | 2705.157401 | 0.037294999 | 0.05132427 | 0.726654254 | 0.467437784 | 0.682158898 | | Hist1h2be | 73.53797018 | 0.0402586931317094 | 0.095038418532303 | 0.423604417569128 | 0.671854333057005 | 0.919094512073309 | NM_001177653,NM_178194 |
| 3632454L22Rik | 35.34004446 | 0.119364743 | 0.164287142 | 0.726561688 | 0.467494505 | 0.682177421 | | Ino80d | 1560.808507 | -0.0326787085490121 | 0.0771293089067495 | -0.423687298800008 | 0.671793879281313 | 0.919094512073309 | NM_001114609,NM_001081436 |
| Dock10 | 2100.99105 | -0.08423246 | 0.116015112 | -0.726047316 | 0.467809765 | 0.682372648 | | Tcf7l1 | 159.4527196 | -0.0399736045660289 | 0.0943391987281562 | -0.423722112387398 | 0.671768486788765 | 0.919094512073309 | NM_009332,NM_001079822 |
| Rbm8a | 1615.786224 | 0.044516832 | 0.061307572 | 0.726122902 | 0.46776343 | 0.682372648 | | Aimp1 | 1475.149809 | 0.0247754115562989 | 0.0585064079815142 | 0.423464923092304 | 0.67195608552152 | 0.919099255959096 | NM_007926 |
| Thap1 | 288.0708085 | -0.081668496 | 0.112485799 | -0.726033838 | 0.467818027 | 0.682372648 | | Abcd4 | 536.0649424 | -0.0337567555611097 | 0.0797461170926323 | -0.423302811369463 | 0.672074335975269 | 0.919174457383406 | NM_008992 |
| Usp5 | 8651.93891 | -0.04236851 | 0.058360105 | -0.725984131 | 0.467848499 | 0.682372648 | | Ankrd29 | 777.7050138 | 0.0282363538816768 | 0.0667465380833117 | 0.423038477987169 | 0.672267187663376 | 0.91919030578735 | NM_001190371 |
| Zfp711 | 300.478785 | -0.071162502 | 0.09802167 | -0.725987442 | 0.467846469 | 0.682372648 | | Mark2 | 5192.638462 | 0.0250246551666415 | 0.0591364070436084 | 0.423168339398566 | 0.672172444693446 | 0.91919030578735 | NM_001080388,NM_001080389,NM_001080390,NM_007928 |
| Tymp | 11.21297542 | -0.119265847 | 0.164492406 | -0.725053821 | 0.468419013 | 0.683140454 | | Usp22 | 12406.03501 | 0.0186397281975676 | 0.044062768228171 | 0.423026717273983 | 0.672275768187031 | 0.91919030578735 | NM_001004143 |
| Park2 | 228.0408165 | 0.088954348 | 0.122716919 | 0.724874355 | 0.468529115 | 0.683236716 | | Adcyap1r1 | 10004.48219 | 0.0271062563246715 | 0.0640991083241782 | 0.42288039620743 | 0.672382526453235 | 0.919249748636207 | NM_007407,NM_001025372 |
| Nek7 | 1648.913311 | 0.053136986 | 0.073321231 | 0.724714862 | 0.468626976 | 0.683315111 | | 9030612E09Rik | 49.51962606 | 0.0397526387209024 | 0.0941428158756294 | 0.422258866501497 | 0.672836078393239 | 0.919353962749294 | NM_172419 |
| Lhx2 | 1566.516755 | -0.063930465 | 0.088229184 | -0.724595445 | 0.468700254 | 0.68335765 | | Add2 | 17912.42 | 0.0234094737152737 | 0.0555526128207191 | 0.421392847728358 | 0.673468241019525 | 0.919353962749294 | NM_001271857,NM_001271861,NM_001271860,NM_013458,NM_001271858,NM_001271859 |
| BC021785 | 26.81849828 | 0.121501822 | 0.167709338 | 0.724478576 | 0.468771975 | 0.68339791 | | Apeh | 789.4639851 | 0.0306174839077009 | 0.0725683113179317 | 0.421912586246654 | 0.673088822842262 | 0.919353962749294 | NM_146226 |
| Dlat | 3606.037603 | -0.047748792 | 0.065931103 | -0.724222564 | 0.468929108 | 0.683562668 | | Arid1a | 5989.866793 | -0.0239717827663312 | 0.0569079578705775 | -0.4212377963175 | 0.673581447350452 | 0.919353962749294 | NM_001080819 |
| 1700012B09Rik | 9.700104813 | 0.101539892 | 0.140347463 | 0.723489328 | 0.469379308 | 0.684154563 | | Ctdp1 | 766.9145408 | 0.0255372624748019 | 0.0605128814046825 | 0.422013658612555 | 0.673015047912633 | 0.919353962749294 | NM_026295 |
| Lins | 420.0686119 | -0.05970722 | 0.082579558 | -0.723026634 | 0.469663511 | 0.68450443 | | Efcab10 | 49.80047629 | 0.0379633552529596 | 0.0900761033466891 | 0.421458675969191 | 0.673420180658171 | 0.919353962749294 | NM_029152 |
| Egln1 | 3715.397893 | -0.059583472 | 0.082465758 | -0.722523792 | 0.469972503 | 0.684890328 | | Fastkd3 | 471.0320477 | 0.0294356321349845 | 0.0697679924506998 | 0.42190739766211 | 0.673092610188448 | 0.919353962749294 | NM_027123 |
| Ccdc14 | 134.7202439 | 0.103188345 | 0.142848518 | 0.722362028 | 0.470071926 | 0.684906378 | | Gbgt1 | 33.60790895 | 0.0373886708063605 | 0.0886978209690753 | 0.421528628300757 | 0.673369110811913 | 0.919353962749294 | NM_139197 |
| Krr1 | 1590.232229 | 0.043731286 | 0.060535178 | 0.722411117 | 0.470041754 | 0.684906378 | | Kifc3 | 2161.349083 | 0.0277328727014819 | 0.0656477165412494 | 0.422449921530114 | 0.672696646184606 | 0.919353962749294 | NM_001145832,NM_010631,NM_001145831 |
| Dnajc30 | 1132.198269 | -0.060017836 | 0.083095121 | -0.722278693 | 0.47012315 | 0.684916604 | | Lnpep | 1765.988565 | 0.0366931487028517 | 0.0869961169791984 | 0.421779154943495 | 0.673186222085384 | 0.919353962749294 | NM_172827 |
| H2afx | 768.7425469 | -0.064265939 | 0.088992513 | -0.722149953 | 0.470202289 | 0.684967495 | | Rpl27a | 7672.724678 | -0.0261407805752944 | 0.0620029176970288 | -0.421605652544108 | 0.673312879742469 | 0.919353962749294 | NM_011975 |
| Alpk3 | 20.60317969 | 0.120113303 | 0.166452617 | 0.721606577 | 0.470536394 | 0.685260917 | | Runx1t1 | 815.9607604 | 0.0356884105496596 | 0.0844482908498537 | 0.422606664865633 | 0.672582263097364 | 0.919353962749294 | NM_009822,NM_001111026,NM_001111027 |
| Tab2 | 3032.939389 | 0.059756374 | 0.082801707 | 0.721680462 | 0.470490957 | 0.685260917 | | Slc9a1 | 2934.392608 | -0.0233887300072744 | 0.0555268006098001 | -0.421215156472503 | 0.673597977801234 | 0.919353962749294 | NM_016981 |
| Tmcc1 | 3685.313977 | 0.055469553 | 0.076865526 | 0.721644103 | 0.470513317 | 0.685260917 | | Smad2 | 1785.587608 | -0.0235462696907174 | 0.0558889907882779 | -0.421304256144413 | 0.673532922712548 | 0.919353962749294 | NM_010754,NM_001252481 |
| Ppil2 | 2415.929794 | -0.035149189 | 0.048718387 | -0.721476854 | 0.470616176 | 0.685312691 | | Snx15 | 1189.626193 | -0.0247535667870129 | 0.0586499169234596 | -0.422056297527536 | 0.672983925780506 | 0.919353962749294 | NM_026912 |
| Btbd8 | 61.39246431 | 0.112776229 | 0.156380276 | 0.721166578 | 0.470807032 | 0.685526187 | | Stk38l | 1516.03285 | 0.0213565275628738 | 0.0506363197877446 | 0.421763028047758 | 0.67319799441278 | 0.919353962749294 | NM_172734 |
| Mterfd3 | 472.2392003 | -0.075013815 | 0.104044329 | -0.720979369 | 0.470922208 | 0.685615821 | | Ugp2 | 3276.739289 | -0.0275236772370026 | 0.065169278969398 | -0.422341288292097 | 0.672775925483232 | 0.919353962749294 | NM_139297 |
| Zmynd12 | 12.40994413 | 0.118485914 | 0.164353153 | 0.720922673 | 0.470957092 | 0.685615821 | | Zfp707 | 354.0981903 | 0.031882619700328 | 0.0755000990707626 | 0.42228579952519 | 0.672816421956267 | 0.919353962749294 | NM_001081065 |
| Apitd1 | 73.80665607 | -0.105737795 | 0.146697666 | -0.720787169 | 0.471040471 | 0.685672779 | | Cdkn2d | 103.2958927 | 0.039282049514139 | 0.0933367943255268 | 0.420863495452143 | 0.673854762834227 | 0.919618027520667 | NM_009878 |
| Arid2 | 1986.007884 | -0.06297668 | 0.08744718 | -0.720168224 | 0.471421426 | 0.686115494 | | Gadl1 | 34.54013076 | -0.0376734581442684 | 0.0895530045025 | -0.420683352318087 | 0.673986319178474 | 0.91971115757506 | NM_028638 |
| Trmt10c | 416.7274191 | 0.057323339 | 0.07959926 | 0.720149146 | 0.471433171 | 0.686115494 | | Mrs2 | 977.1558393 | -0.0240230205907243 | 0.0571175966183817 | -0.420588785470591 | 0.674055384190284 | 0.919719003451819 | NM_001013389 |
| Stx16 | 3368.826147 | 0.058306989 | 0.080995416 | 0.719880111 | 0.471598815 | 0.686292105 | | 2810021J22Rik | 519.9824151 | -0.0288274515422978 | 0.0686388151232813 | -0.419987604543015 | 0.674494508915563 | 0.919867577130338 | NM_172403 |
| Tfrc | 4034.068564 | 0.079625012 | 0.110636826 | 0.719697185 | 0.47171146 | 0.686391563 | | B930041F14Rik | 1229.641557 | -0.0320847419117589 | 0.0763806493190019 | -0.420063749101658 | 0.674438883985479 | 0.919867577130338 | NM_178699 |
| Crygn | 8.818391556 | 0.103731865 | 0.144156683 | 0.719577222 | 0.47178534 | 0.686434601 | | Echdc3 | 35.16945818 | 0.0348116109715881 | 0.0828729380620807 | 0.420060055618041 | 0.674441582098719 | 0.919867577130338 | NM_024208 |
| Cd59b | 19.22995161 | -0.120901374 | 0.16812038 | -0.719135746 | 0.472057285 | 0.686650062 | | Lcor | 156.7969155 | 0.0392511766652727 | 0.093462256671823 | 0.419968210302225 | 0.674508677031472 | 0.919867577130338 | NM_172154 |
| Mapt | 21616.63965 | -0.040552319 | 0.05638272 | -0.7192331 | 0.471997308 | 0.686650062 | | Lhx8 | 76.99132947 | 0.036195115224519 | 0.0861953565327505 | 0.419919548807324 | 0.674544226324684 | 0.919867577130338 | NM_010713 |
| Snx16 | 904.1328057 | -0.058741135 | 0.081684634 | -0.719120986 | 0.472066378 | 0.686650062 | | Mesp2 | 23.09763834 | -0.0330769187637461 | 0.0787075027088603 | -0.420251126606509 | 0.674302007561027 | 0.919867577130338 | NM_008589 |
| Gemin2 | 182.5589399 | 0.0689541 | 0.095912781 | 0.718925039 | 0.472187108 | 0.686761199 | | G2e3 | 464.2004146 | -0.0356025999410872 | 0.0848547786914684 | -0.419570948037448 | 0.674798915262278 | 0.919907397428193 | NM_001167964,NM_001167963,NM_001015099 |
| Cdc26 | 589.3188005 | -0.05185574 | 0.072141746 | -0.7188035 | 0.472262001 | 0.686805655 | | Gm14420 | 1125.63376 | -0.0241140216417044 | 0.0575495935177663 | -0.419012892493499 | 0.675206710132135 | 0.919907397428193 | NM_001177568 |
| Klhdc7b | 13.44270856 | 0.114654004 | 0.15954511 | 0.718630633 | 0.472368534 | 0.68686651 | | Neu1 | 2315.841425 | 0.0260487621122543 | 0.0621516677078949 | 0.419116060323277 | 0.675131313831788 | 0.919907397428193 | NM_010893 |
| Ostm1 | 2232.774663 | 0.053825168 | 0.074903684 | 0.718591733 | 0.472392509 | 0.68686651 | | Ppapdc1b | 683.0630844 | -0.0305587683392933 | 0.0728047255326235 | -0.419736055808631 | 0.674678282303746 | 0.919907397428193 | NM_028000 |
| 2300009A05Rik | 146.2069163 | 0.094152297 | 0.131111084 | 0.718110887 | 0.472688918 | 0.687213221 | | Rit1 | 1070.317851 | 0.0243711531303049 | 0.05815542202531 | 0.419069319447089 | 0.675165472226714 | 0.919907397428193 | NR_028056,NM_009069,NM_001163310 |
| Gale | 388.8984167 | 0.080758509 | 0.112467474 | 0.718061015 | 0.472719666 | 0.687213221 | | Tagln | 232.4140614 | -0.0398625931442235 | 0.095020036345 | -0.419517763595563 | 0.674837775276962 | 0.919907397428193 | NM_011526 |
| Ip6k2 | 2476.663663 | 0.07219634 | 0.100632098 | 0.717428551 | 0.473109708 | 0.687592909 | | Tmem168 | 631.8457754 | 0.0298413039269161 | 0.071174608337861 | 0.419268958745533 | 0.675019579807866 | 0.919907397428193 | NM_028990 |
| Ptgs2 | 560.6470203 | -0.12051165 | 0.168000174 | -0.717330506 | 0.473170188 | 0.687592909 | | Trim36 | 323.929746 | -0.0371308000642483 | 0.0885285306064376 | -0.419421849768601 | 0.674907858353092 | 0.919907397428193 | NM_178872,NM_001170855 |
| S100a3 | 22.16049593 | 0.120343571 | 0.167778202 | 0.717277746 | 0.473202736 | 0.687592909 | | Zbed4 | 632.0216474 | -0.0309914492314636 | 0.0739276856637578 | -0.419213031670175 | 0.675060448968404 | 0.919907397428193 | NM_181412 |
| Spag16 | 23.95808064 | 0.1181322 | 0.164656134 | 0.717447915 | 0.473097664 | 0.687592909 | | Zfp39 | 527.7564839 | -0.0331738681757836 | 0.0790992005811774 | -0.419395745241928 | 0.674926933103888 | 0.919907397428193 | NM_011758 |
| Ythdf3 | 2010.662846 | -0.045490972 | 0.063421518 | -0.71727978 | 0.473201481 | 0.687592909 | | Pou6f1 | 6116.836699 | -0.0229948455662112 | 0.0549392817129737 | -0.418550167553083 | 0.675544915210234 | 0.920229308631337 | NM_010127 |
| Kcnc4 | 2745.638006 | 0.051485883 | 0.071803341 | 0.717040208 | 0.473349288 | 0.68774136 | | Rps12 | 4592.81765 | -0.0250851516500571 | 0.059938314991094 | -0.418516130354755 | 0.675569795543034 | 0.920229308631337 | NM_011295 |
| Peg3 | 10592.90237 | 0.081924489 | 0.114303101 | 0.716730238 | 0.473540564 | 0.687954759 | | Tbc1d10b | 3497.007504 | 0.0181610211330943 | 0.0434028108269666 | 0.418429608291882 | 0.675633042582487 | 0.920229308631337 | NM_144522 |
| AI987944 | 282.1878581 | -0.073946081 | 0.103200717 | -0.716526813 | 0.473666117 | 0.687973655 | | Ammecr1 | 74.55235788 | -0.0396939750773899 | 0.0950965723618383 | -0.417407001025821 | 0.676380734893779 | 0.9206434178862 | NM_019496 |
| Rims2 | 4866.566337 | 0.048810077 | 0.068129504 | 0.716430817 | 0.473725372 | 0.687973655 | | Asb7 | 853.7700626 | 0.0289086269485196 | 0.0692132564614174 | 0.417674711847067 | 0.676184963849647 | 0.9206434178862 | NM_080443,NR_003961 |
| Slc15a2 | 1044.977058 | -0.049503713 | 0.069098595 | -0.716421405 | 0.473731182 | 0.687973655 | | B3galtl | 916.7758885 | 0.0251251257010384 | 0.0601339834632102 | 0.417819080893084 | 0.676079399021116 | 0.9206434178862 | NM_001081204 |
| Uba5 | 1692.407454 | 0.037968933 | 0.052984196 | 0.716608647 | 0.473615607 | 0.687973655 | | Bmp4 | 111.0548631 | -0.0395182603057863 | 0.0946600420160787 | -0.417475626083853 | 0.676330548810259 | 0.9206434178862 | NM_007554 |
| Lrrc4b | 5068.034591 | -0.046636981 | 0.065132406 | -0.716033446 | 0.473970697 | 0.68825698 | | Cep68 | 1434.321039 | -0.0270963480821921 | 0.0648642837197952 | -0.417739108925409 | 0.676137874946069 | 0.9206434178862 | NM_172260 |
| Fgf18 | 45.00914696 | 0.113317653 | 0.158322067 | 0.71574137 | 0.474151061 | 0.688454365 | | Psmf1 | 1575.218612 | 0.0283121103998536 | 0.067799264339921 | 0.417587280267629 | 0.676248898237063 | 0.9206434178862 | NM_212446 |
| BC030336 | 1961.022127 | 0.051140469 | 0.071461876 | 0.715632893 | 0.474218057 | 0.688487123 | | Zfp945 | 566.6313989 | -0.030830273404498 | 0.0738492649469785 | -0.417475697647542 | 0.676330496475881 | 0.9206434178862 | NM_001110254,NM_177358 |
| Rab5c | 5226.872411 | -0.035298514 | 0.049340645 | -0.715404395 | 0.474359198 | 0.688627509 | | Neto1 | 2810.44728 | 0.0285563866946498 | 0.0684676316469888 | 0.417078640048238 | 0.676620887937587 | 0.920884008074711 | NM_144946 |
| Nabp2 | 1609.101212 | 0.060531375 | 0.084641233 | 0.715152335 | 0.474514918 | 0.688789033 | | Zfp959 | 180.3042008 | -0.0374883550325419 | 0.0899291203220995 | -0.416865581452034 | 0.676776729988884 | 0.92100981635924 | NM_145490 |
| Crb2 | 272.1248721 | 0.076689554 | 0.107255774 | 0.715015625 | 0.474599388 | 0.688847112 | | Med20 | 846.9367139 | 0.0252697922505903 | 0.0606443387766069 | 0.416688395988216 | 0.676906343122613 | 0.921056061614246 | NM_020048 |
| Acot4 | 42.29219627 | 0.11581185 | 0.162013436 | 0.714828679 | 0.474714911 | 0.688950247 | | Slc7a8 | 3897.894192 | 0.0233684539432668 | 0.0560777072018949 | 0.416716457668846055 | 0.676937526765365 | 0.921056061614246 | NM_016972 |
| Plk1s1 | 541.5822641 | -0.07519067 | 0.105219509 | -0.714607693 | 0.47485149 | 0.688983917 | | Ccdc8 | 113.7481559 | -0.0388920035845806 | 0.0934896926173511 | -0.416003117517604 | 0.677407721980508 | 0.921064065908822 | NM_001101535 |
| Dll3 | 34.18248516 | 0.114463377 | 0.160212944 | 0.714445252 | 0.474951899 | 0.68916508 | | Grcc10 | 2489.403652 | 0.0244543029346922 | 0.0587923685487105 | 0.415944421286854 | 0.677450673287199 | 0.921064065908822 | NM_013535 |
| Rbm34 | 692.3277105 | -0.053044479 | 0.074301038 | -0.713913033 | 0.475280958 | 0.689577971 | | Jmy | 2634.920724 | 0.0273557930554344 | 0.0657300377705574 | 0.41618404588363 | 0.677275333200473 | 0.921064065908822 | NM_021310 |
| Galnt10 | 565.4176672 | -0.065374953 | 0.091587708 | -0.713796145 | 0.475353244 | 0.689618273 | | Ocel1 | 195.9780444 | -0.0368373956501613 | 0.088527712652342 | -0.416111458734123 | 0.677328445422588 | 0.921064065908822 | NM_029865 |
| Cdca2 | 13.31195699 | -0.117315753 | 0.164404742 | -0.713578891 | 0.475487615 | 0.689748628 | | Pth1r | 212.4402689 | -0.0386310563861225 | 0.0928649945086589 | -0.415991683308832 | 0.677416088946731 | 0.921064065908822 | NM_001083936,NM_011199,NM_001083935 |
| Ccdc158 | 12.74728167 | -0.116014375 | 0.162599527 | -0.713497614 | 0.475537889 | 0.689756979 | | Shf | 1201.896599 | 0.026049718662026 | 0.0625805119266091 | 0.416259277210383 | 0.677220287919619 | 0.921064065908822 | NM_001013829 |
| C130030K03Rik | 134.254635 | -0.107602224 | 0.15092261 | -0.712962916 | 0.475868705 | 0.689955989 | | Sufu | 405.9004075 | -0.0316901801332481 | 0.0761793668137248 | -0.415994270610539 | 0.677414195688854 | 0.921064065908822 | NM_001025391,NM_015752 |
| Lama4 | 411.0630674 | -0.076162468 | 0.10682618 | -0.712956952 | 0.475872395 | 0.689955989 | | Zfp629 | 1611.589342 | -0.0231206810916599 | 0.0555641861417214 | -0.416107617102292 | 0.677331256400139 | 0.921064065908822 | NM_177226 |
| Lsm14b | 3829.813136 | -0.042172921 | 0.059189626 | -0.712505282 | 0.476151942 | 0.689955989 | | Slc12a6 | 6686.683671 | -0.0225212600971641 | 0.0541574460715563 | -0.415847897764743 | 0.677521307213414 | 0.921073889432106 | NM_133648,NM_133649 |
| Ncf4 | 33.15740409 | 0.119514289 | 0.167644714 | 0.712902222 | 0.475906264 | 0.689955989 | | Mdk | 399.6866949 | 0.0318699738418515 | 0.076674062489608 | 0.415655213862849 | 0.677662317804296 | 0.921093181288033 | NM_010784,NM_001012336,NM_001012335 |
| Papd5 | 1534.384359 | 0.054410077 | 0.076374413 | 0.712412369 | 0.476209458 | 0.689955989 | | Polr3k | 1380.428261 | 0.0255506444704817 | 0.0614587140312765 | 0.41573672461612 | 0.677602664948026 | 0.921093181288033 | NM_025901 |
| Psat1 | 4389.474557 | -0.047235787 | 0.066265022 | -0.712831387 | 0.475950101 | 0.689955989 | | Agphd1 | 164.819238 | 0.0374565431092609 | 0.0901789118223138 | 0.415358118127044 | 0.67787976155056 | 0.921155235310422 | NM_177351 |
| Sapcd2 | 45.01868081 | -0.115406939 | 0.161854864 | -0.713027316 | 0.475828854 | 0.689955989 | | Fam118a | 765.8215258 | 0.0247240695365396 | 0.0595283183878033 | 0.415332907196742 | 0.677898214612998 | 0.921155235310422 | NM_133750,NM_177067 |
| Slc43a2 | 3017.572934 | 0.046669979 | 0.065489065 | 0.712637735 | 0.476069955 | 0.689955989 | | Zfp433 | 47.23895548 | -0.0383424608124674 | 0.0923120107084068 | -0.415357227279804 | 0.677880413600148 | 0.921155235310422 | NM_001243067 |
| Tmem2 | 634.9983919 | -0.079793489 | 0.111959827 | -0.712697504 | 0.476032961 | 0.689955989 | | Agbl4 | 389.620156 | 0.0304076732432481 | 0.0733511374512679 | 0.41454944394625 | 0.67847176436997 | 0.921316908708726 | NM_030231,NM_001048189 |
| Tmem65 | 3643.430786 | 0.050415881 | 0.070733921 | 0.712753938 | 0.475998034 | 0.689955989 | | Cirbp | 2120.298096 | -0.0355910770066657 | 0.0857893010109398 | -0.414866149825923 | 0.678239891120402 | 0.921316908708726 | NM_007705 |
| Trim34a | 63.19343896 | 0.114493511 | 0.16069376 | 0.712495065 | 0.476158266 | 0.689955989 | | Dynlrb1 | 4460.797246 | 0.0246372766032258 | 0.0594847185643575 | 0.414178249436791 | 0.678743569711336 | 0.921316908708726 | NM_025947 |
| Wipi1 | 641.2028401 | 0.052365154 | 0.073502445 | 0.712427375 | 0.476200169 | 0.689955989 | | Ggt5 | 112.2145457 | 0.0382418067365653 | 0.0923107677378124 | 0.4142724372652 | 0.678674597182491 | 0.921316908708726 | NM_011820 |
| Gm872 | 34.22522344 | 0.113574674 | 0.159504313 | 0.71204767 | 0.476435257 | 0.690211698 | | Hmgcs2 | 438.1283012 | 0.0392796821462755 | 0.0948848866223311 | 0.41413082638503 | 0.678778298017402 | 0.921316908708726 | NM_008256 |
| Kcnj14 | 82.77219175 | -0.103599745 | 0.145508648 | -0.711983419 | 0.476475044 | 0.690211698 | | Mthfs | 55.11688784 | 0.0384395060041005 | 0.0927158827631698 | 0.414594618079505 | 0.678438688688527 | 0.921316908708726 | NM_026829 |
| Twist2 | 23.73766272 | -0.119608093 | 0.168015856 | -0.711885746 | 0.47653553 | 0.690234785 | | Nbn | 775.8664975 | 0.0311805270042114 | 0.0752311383265682 | 0.414463049447171 | 0.678535022581044 | 0.921316908708726 | NM_013752 |
| Ube2g2 | 1821.928755 | -0.068630167 | 0.096429599 | -0.711712664 | 0.476642724 | 0.690325515 | | Ndst3 | 1080.902064 | -0.0303481852064421 | 0.0732392598679658 | -0.414370451874488 | 0.678602825208947 | 0.921316908708726 | NM_031186 |
| Sac3d1 | 370.7179773 | 0.064471001 | 0.090693604 | 0.710866018 | 0.477167266 | 0.69102062 | | Phgdh | 2934.791273 | 0.0275977433409389 | 0.0665102395364265 | 0.41493976767072 | 0.67818599686673 | 0.921316908708726 | NM_016966 |
| Car13 | 64.19912326 | -0.115349012 | 0.162449018 | -0.710062848 | 0.477665164 | 0.691328116 | | Slc4a10 | 9665.719981 | 0.028024233977229 | 0.0675963611158315 | 0.414581991021785 | 0.678447933928909 | 0.921316908708726 | NM_001242379,NM_001242378,NM_001242380,NM_001242381,NM_001242382,NM_001242383,NM_033552 |
| Csnk1g3 | 2281.688109 | -0.056135588 | 0.079047251 | -0.710152313 | 0.477609689 | 0.691328116 | | Trim23 | 2165.148236 | 0.0317381470880628 | 0.0765329893177127 | 0.414698907895885 | 0.678362331941504 | 0.921316908708726 | NM_030731 |
| Glmn | 424.3374238 | 0.066670834 | 0.093862552 | 0.710302805 | 0.477516381 | 0.691328116 | | Trim8 | 2799.340608 | 0.0255669919357049 | 0.0616861048772211 | 0.414469222632762 | 0.678530502488055 | 0.921316908708726 | NM_053100 |
| Il1r2 | 12.56616994 | 0.118245786 | 0.166554253 | 0.70995357 | 0.477732929 | 0.691328116 | | Dbi | 3334.027272 | 0.0325127032714574 | 0.0785492247935253 | 0.413915011343783 | 0.678936349830049 | 0.921359250526092 | NM_001037999,NM_007830 |
| Ktn1 | 3390.570224 | 0.049522787 | 0.069751938 | 0.70998438 | 0.47771382 | 0.691328116 | | Unc119b | 1267.817895 | -0.027823725025588 | 0.0672202156636798 | -0.413919008602968 | 0.678933422315096 | 0.921359250526092 | NM_175352 |
| Slc27a1 | 2519.537115 | 0.07720077 | 0.108693638 | 0.710260249 | 0.477542765 | 0.691328116 | | 1700113A16Rik | 40.78105288 | -0.0350203743470982 | 0.0847016337540326 | -0.413455712658327 | 0.679272763505616 | 0.921471438427585 | NR_045997 |
| Tmem132a | 9397.38563 | -0.075713894 | 0.106642327 | -0.709979761 | 0.477716687 | 0.691328116 | | Ank2 | 20077.30876 | 0.0237213007476291 | 0.0573568927839389 | 0.413573671729154 | 0.679186358193785 | 0.921471438427585 | NM_001034168,NM_178655 |
| Ttll13 | 40.55526167 | 0.11930872 | 0.168052823 | 0.709947727 | 0.477736552 | 0.691328116 | | Ncbp1 | 2495.615462 | 0.0253918791086685 | 0.0614009074433364 | 0.413542407856127 | 0.679209258647786 | 0.921471438427585 | NM_001033201 |
| Fuca1 | 1083.766197 | 0.062386907 | 0.087893339 | 0.709802448 | 0.477826651 | 0.691393923 | | Txnrd2 | 522.17354 | -0.0276120276400876 | 0.0667771010405331 | -0.4134954529297 | 0.679243653187542 | 0.921471438427585 | NM_013711 |
| Kcnh3 | 6891.977575 | -0.054160363 | 0.076332228 | -0.709534688 | 0.477992734 | 0.691569653 | | Msra | 1158.349555 | 0.0233656442721061 | 0.0565581102230038 | 0.413126325826257 | 0.679514062487121 | 0.921712696954704 | NM_001253715,NM_026322,NM_001253714,NM_001253716,NM_001253712 |
| 1700018L02Rik | 23.15032039 | 0.11739939 | 0.165487116 | 0.709417105 | 0.478065676 | 0.691616606 | | Fbxo6 | 651.997247 | -0.029389065594294 | 0.0711543860469286 | -0.413032382500089 | 0.679582888588808 | 0.921719985027174 | NM_001163704,NM_001163705,NM_001163706,NM_001163707,NM_015797 |
| Creld2 | 515.6167596 | -0.07910887 | 0.111533086 | -0.709286129 | 0.478146935 | 0.69166358 | | Cdh6 | 451.8518178 | 0.0319457876737007 | 0.0773744651017684 | 0.41287248489128 | 0.679700041200194 | 0.921792810986278 | NM_007666 |
| Ecel1 | 987.3731519 | 0.118540051 | 0.167159457 | 0.709143555 | 0.478235397 | 0.691726964 | | Astn1 | 13889.22965 | 0.0251828895427259 | 0.0610091961994701 | 0.412772026374355 | 0.679773648368393 | 0.921806573580964 | NM_007495,NM_001205204 |
| Hist1h2ac | 32.25540862 | -0.119030996 | 0.16795128 | -0.708723361 | 0.478496165 | 0.692032764 | | AI197445 | 37.85354342 | -0.0368675619960841 | 0.0893582480435735 | -0.41258152216801 | 0.679913241482184 | 0.921823758238772 | NR_045084,NR_045083 |
| Tada3 | 1206.407539 | 0.041124718 | 0.058031749 | 0.708658949 | 0.478536145 | 0.692032764 | | Dlg4 | 31099.60296 | 0.0248186047546677 | 0.060146395664646 | 0.4126366090671 | 0.679872875094584 | 0.921823758238772 | NM_007864,NM_001109752 |
| Xpnpep3 | 416.3586283 | -0.064177772 | 0.090572281 | -0.708580724 | 0.478584702 | 0.692038392 | | Cd209c | 82.108452 | -0.0391374968576791 | 0.0949181578846872 | -0.412328870785987 | 0.68009839024077 | 0.921988727637651 | NM_130903 |
| Pcsk2 | 18273.46689 | -0.067614199 | 0.095473807 | -0.708196322 | 0.47882335 | 0.692318867 | | Cbll1 | 760.4269545 | 0.0331783736019834 | 0.0805605645789373 | 0.411843856549361 | 0.68045387390549 | 0.922298495134006 | NM_001253848,NM_001253847,NM_134048 |
| Kit | 1756.975653 | -0.037005736 | 0.052263377 | -0.708062478 | 0.478906459 | 0.692349819 | | Gnb1l | 171.7942033 | -0.0360866809112667 | 0.0876153872752648 | -0.411876064621979 | 0.680430265295932 | 0.922298495134006 | NM_001081682,NM_023120 |
| Tradd | 157.1876433 | 0.077675068 | 0.109707771 | 0.708017924 | 0.478934126 | 0.692349819 | | 2610017I09Rik | 876.3191787 | -0.0353316753420475 | 0.085813288545301 | -0.411727320336825 | 0.680539292789649 | 0.922327641743822 | NR_027826 |
| Plekhb1 | 9509.201287 | 0.063923067 | 0.090312997 | 0.707794771 | 0.479072714 | 0.692473415 | | Rai14 | 620.4672859 | -0.0319178073814179 | 0.0775379162632783 | -0.411641283640402 | 0.680602367606501 | 0.922327641743822 | NM_001166408,NM_030690 |
| Tac2 | 322.0488343 | 0.091156004 | 0.128799385 | 0.707736333 | 0.47910901 | 0.692473415 | | Adora2a | 2395.651432 | -0.0391151452162239 | 0.0950889111195596 | -0.411335382383806 | 0.68081343155885 | 0.92235552199439 | NM_009630 |
| Zfp664 | 4553.089956 | 0.044048798 | 0.062254606 | 0.707558861 | 0.479219248 | 0.69256814 | | Enpp1 | 173.8683887 | -0.0390376577505181 | 0.0948977950418437 | -0.411365277067871 | 0.680804710924504 | 0.92235552199439 | NM_008813 |
| Catsper3 | 8.428020913 | 0.099828967 | 0.141133844 | 0.707335423 | 0.479358507 | 0.692574948 | | Rad23b | 8418.823109 | 0.021752333033574 | 0.0528645955525086 | 0.411472608581072 | 0.680726023363819 | 0.92235552199439 | NM_009011 |
| Cnnm4 | 597.8059973 | -0.059675592 | 0.084362673 | -0.707369628 | 0.479336806 | 0.692574948 | | Rab3gap2 | 4680.873836 | 0.0218153450886539 | 0.0530477705438314 | 0.411239621665697 | 0.680896837831581 | 0.922382492431367 | NM_001163754 |
| Mtmr6 | 6202.629554 | -0.049642609 | 0.070178453 | -0.707376791 | 0.479332356 | 0.692574948 | | Tmem176b | 1082.688639 | -0.0354594980708163 | 0.0862948767318532 | -0.410910814334908 | 0.681137932182667 | 0.922451016762131 | NM_001164208,NM_001164209,NM_023056,NM_001164207 |
| Bpnt1 | 1892.304297 | -0.050712623 | 0.071761695 | -0.706680954 | 0.479764788 | 0.692981569 | | Usp33 | 6258.813554 | 0.02443556170745 9 | 0.0594543733589784 | 0.410996875871855 | 0.681074825508657 | 0.922451016762131 | NM_001252486,NM_001076676,NM_133247 |
| Clpx | 1179.969015 | -0.060961967 | 0.08626694 | -0.706666627 | 0.479773673 | 0.692981569 | | Xkr4 | 782.0496379 | 0.0342856312111004 | 0.0834338811638363 | 0.410931754975834 | 0.681122576904273 | 0.922451016762131 | NM_001011874 |
| Dhdds | 2893.595931 | 0.045489735 | 0.064366396 | 0.706731119 | 0.479733587 | 0.692981569 | | Arl15 | 1643.109601 | -0.0310947390282422 | 0.0757922321435666 | -0.410262874556091 | 0.681613122804849 | 0.922538400546282 | NM_172595 |
| Cdh9 | 719.6300131 | -0.063904928 | 0.090460398 | -0.706440934 | 0.479913973 | 0.693055 | | Cpsf6 | 2564.17476 | -0.026979238058671 | 0.065768265251151 | -0.410216657251151 | 0.681647022791081 | 0.922538400546282 | NM_001013391 |
| Rchy1 | 1179.818977 | -0.044490008 | 0.062975287 | -0.706467719 | 0.479897321 | 0.693055 | | Kcnc1 | 10143.72355 | -0.0171495993717286 | 0.0417857692874696 | -0.410417222517697 | 0.681499914592163 | 0.922538400546282 | NM_008421,NM_001112739 |
| Eif3h | 4053.105411 | -0.044186516 | 0.062604796 | -0.705800818 | 0.480312013 | 0.693397062 | | Patl1 | 1169.223439 | 0.0282566541319489 | 0.0688174131980112 | 0.410603258954893 | 0.681363473648928 | 0.922538400546282 | NM_172635 |
| Kcnip1 | 1054.569209 | 0.076238188 | 0.108019298 | 0.705783031 | 0.480323077 | 0.693397062 | | Ppp3ca | 36009.71848 | 0.0295461026791804 | 0.0720071392081073 | 0.410321851473497 | 0.681569864688691 | 0.922538400546282 | NM_008913 |
| Map6 | 5059.765005 | -0.037078283 | 0.052538592 | -0.705734229 | 0.48035343 | 0.693397062 | | Smarcd1 | 8470.112969 | 0.0214392128058557 | 0.0522556988916084 | 0.410275113736974 | 0.681604145581083 | 0.922538400546282 | NM_031842 |
| Tmem150b | 12.82447305 | 0.084361304 | 0.119498655 | 0.705960282 | 0.480212838 | 0.693397062 | | Suco | 1617.833245 | 0.0280457513111408 | 0.0683465582989186 | 0.410346212145469 | 0.681551997041356 | 0.922538400546282 | NM_172645 |
| Zfand6 | 1080.919527 | -0.061682762 | 0.087406468 | -0.705700193 | 0.480374601 | 0.693397062 | | 2310022A10Rik | 986.4513492 | 0.0231073995809093 | 0.0563658849119191 | 0.409953638083654 | 0.68183995730238 | 0.922544285411829 | NM_001122767,NM_175107 |
| 1300002E11Rik | 259.1585931 | -0.07089978 | 0.100491036 | -0.705533378 | 0.480478368 | 0.693475978 | | Ctu1 | 325.250607 | 0.032280524097949 | 0.0788338964324226 | 0.409475182107994 | 0.682190976107624 | 0.922544285411829 | NM_145582 |
| 1700120C14Rik | 16.09144385 | 0.114354192 | 0.16209683 | 0.705468404 | 0.480518788 | 0.693475978 | | Dusp8 | 7565.240847 | -0.0161621110647809 | 0.0394548956218857 | -0.409635124109054 | 0.682073627132251 | 0.922544285411829 | NM_008748 |
| Lgi1 | 6475.521042 | -0.04512034 7 | 0.06398726 | -0.705145783 | 0.480719551 | 0.693670584 | | Fam58b | 404.789951 | -0.0327084746306692 | 0.079886985475798 | -0.409434332128065 | 0.682220940853957 | 0.922544285411829 | NM_197989 |
| Zfp930 | 401.8907281 | -0.078235825 | 0.11095584 | -0.705107767 | 0.480743173 | 0.693670584 | | Ggh | 403.983674 | -0.0322188078754899 | 0.0786245487297987 | -0.409778543016333 | 0.681966940380706 | 0.922544285411829 | NM_010281 |
| Otud7a | 1776.685688 | 0.060068642 | 0.08522696 | 0.704807989 | 0.480929736 | 0.693875159 | | Oma1 | 262.6430264 | -0.0340763342110449 | 0.08316183834434 | -0.409759270471491 | 0.68198254612 1 | 0.922544285411829 | NM_025909 |
| Antxr2 | 73.72692293 | 0.097955184 | 0.139066271 | 0.70437773 | 0.481197571 | 0.694196943 | | Prrg2 | 163.6529354 | 0.0371520829028913 | 0.0907790171811495 | 0.409258483474813 | 0.682349975917051 | 0.922544285411829 | NM_022999 |
| 1190005I06Rik | 151.472035 | 0.092081462 | 0.130751293 | 0.704248955 | 0.481277749 | 0.69424797 | | Rbfox1 | 14778.77875 | 0.0254698564538516 | 0.0622241818639104 | 0.409324086564433 | 0.682301841520816 | 0.922544285411829 | NM_021477,NM_183188 |
| Aplf | 273.6885648 | 0.062610636 | 0.08893046 | 0.704040388 | 0.481407622 | 0.694370666 | | Smarca1 | 1191.288121 | -0.022037580299833 | 0.0538118160519397 | -0.409530506804716 | 0.682150383670903 | 0.922544285411829 | NM_053123 |
| Mcmbp | 1005.053302 | 0.054622661 | 0.077605196 | 0.703853143 | 0.481524235 | 0.694413582 | | Tbc1d5 | 2166.671719 | 0.0239094707520917 | 0.0583995707607223 | 0.409411754926331 | 0.682237514572519 | 0.922544285411829 | NM_028162 |
| Reep2 | 4828.411119 | 0.051287785 | 0.07288527 | 0.703678327 | 0.48163312 | 0.694413582 | | Tdrd6 | 49.85002799 | -0.0386781731522193 | 0.0943706908845968 | -0.409853661021913 | 0.681913299672063 | 0.922544285411829 | NM_198418,NM_001161366,NM_001161367 |
| Stim1 | 3813.326368 | 0.041985817 | 0.059652206 | 0.703843503 | 0.481530239 | 0.694413582 | | Dnalc4 | 1844.547929 | 0.0302119994773424 | 0.0738939522880716 | 0.408856185680291 | 0.682645204372254 | 0.922857537339355 | NM_017470 |
| Vapb | 6504.891629 | -0.034653772 | 0.049242104 | -0.703742722 | 0.481593009 | 0.694413582 | | Dhx30 | 5114.721567 | 0.0203578252393286 | 0.0498143302965886 | 0.408674072663841 | 0.682778863354112 | 0.922882553072545 | NM_133347,NM_001252682,NM_001252683 |
| Wdr95 | 15.3164765 | 0.118252638 | 0.168060151 | 0.703632821 | 0.481661467 | 0.694413582 | | Dpp4 | 165.9206796 | -0.0379882480063877 | 0.0929782623786876 | -0.408571283593867 | 0.682854308166303 | 0.922882553072545 | NM_001159543,NM_010074 |
| Gemin4 | 437.4848065 | 0.068324306 | 0.097173784 | 0.703114602 | 0.481984333 | 0.694749768 | | Rcbtb2 | 917.4886961 | -0.0248072644947226 | 0.0607078278742351 | -0.408633702825183 | 0.6828084935089 | 0.922882553072545 | NM_134083,NR_033185,NM_001170694 |
| Pak1ip1 | 1443.290302 | -0.042326044 | 0.060196169 | -0.703135184 | 0.481971507 | 0.694749768 | | 4933403G14Rik | 180.3540019 | 0.0385563723966627 | 0.0943992616236079 | 0.408439342993974 | 0.682951154174412 | 0.922927571451996 | NM_028908 |
| Aacs | 1640.434787 | -0.057814811 | 0.082261171 | -0.702820172 | 0.482167824 | 0.694820338 | | 4930451G09Rik | 142.0340653 | -0.038518657971519 | 0.0943910946578614 | -0.408075127332057 | 0.683218519992992 | 0.922993771592081 | NM_001271586,NR_073370 |
| Foxc1 | 167.2301909 | -0.094577921 | 0.134543966 | -0.702951785 | 0.482085797 | 0.694820338 | | Map2 | 24471.4282 | 0.0249028879342168 | 0.0610325431784595 | 0.40802638456996 | 0.683254304414818 | 0.922993771592081 | NM_001039934,NM_008632 |
| Rtca | 1769.877754 | 0.072777431 | 0.103541631 | 0.702880862 | 0.482129999 | 0.694820338 | | Noa1 | 705.9277347 | 0.0259083859366666 | 0.0634694426718309 | 0.408202512469284 | 0.683125003756579 | 0.922993771592081 | NM_019836 |
| Zmym3 | 6315.213497 | 0.06587453 | 0.093748289 | 0.702674479 | 0.482258636 | 0.694886572 | | Sntb1 | 81.58741308 | 0.038818534893722 | 0.0951027446357144 | 0.408174706654515 | 0.683145416203133 | 0.922993771592081 | NM_016667 |
| Akirin1 | 2001.685598 | -0.044917674 | 0.06395737 | -0.702306456 | 0.482488068 | 0.694990786 | | Gucy1a2 | 2357.988973 | 0.0352882971994501 | 0.086518650820936 | 0.407869249746911 | 0.683369669542058 | 0.923036731521311 | NM_001033322 |
| Creb3 | 1435.465168 | 0.033927909 | 0.048317568 | 0.702185787 | 0.482563308 | 0.694990786 | | Rhoq | 1655.193187 | 0.0280455958313518 | 0.0687712363440664 | 0.407809970014761 | 0.683413193412619 | 0.923036731521311 | NM_145491 |
| Csnk1a1 | 6427.778591 | -0.040858372 | 0.058198261 | -0.702054862 | 0.48264495 | 0.694990786 | | Pde10a | 12955.69318 | 0.0386953938828415 | 0.0949578421921122 | 0.407500770758414 | 0.683640228166182 | 0.923110100965354 | NM_011866 |
| Ppm1d | 547.7392764 | 0.069359584 | 0.098741798 | 0.702433879 | 0.482408624 | 0.694990786 | | Sat2 | 241.6503722 | -0.0341614306112566 | 0.0838279116489745 | -0.407518569164719 | 0.683627158610893 | 0.923110100965354 | NM_026991 |
| Sh2b3 | 653.7765032 | 0.058749826 | 0.083674983 | 0.702119361 | 0.482604729 | 0.694990786 | | Zfp526 | 448.5382381 | 0.032537265815107 | 0.0798506857872294 | 0.407476347815046 | 0.683658162342371 | 0.923110100965354 | NM_175436 |
| Thumpd3 | 806.4924676 | -0.054572024 | 0.077717467 | -0.702184797 | 0.482563926 | 0.694990786 | | Tomm5 | 844.1061028 | -0.0280520907702975 | 0.0689343503179608 | -0.406939220300282 | 0.684052629178672 | 0.923556881815168 | NM_001134646,NM_001099675 |
| Zc3h12b | 484.9744567 | -0.079882287 | 0.113756612 | -0.70222104 | 0.482541326 | 0.694990786 | | Pm20d2 | 440.1140099 | 0.0285791911686084 | 0.0702530578635833 | 0.406803519130843 | 0.684152301864478 | 0.923605607517045 | NM_001034867 |
| 4931440F15Rik | 24.0031026 | -0.117951197 | 0.168059145 | -0.701843373 | 0.482776847 | 0.695105921 | | Etohd2 | 24.62952025 | -0.0341989232449713 | 0.0841199914475558 | -0.406549295315757 | 0.684339044396827 | 0.923655037989443 | NR_015349 |
| Ercc3 | 1259.498238 | 0.063714091 | 0.09079821 | 0.701710865 | 0.482859496 | 0.695105921 | | Hist1h1c | 521.3730531 | 0.0385399253225485 | 0.0948105674115978 | 0.406493984528502 | 0.684379676020954 | 0.923655037989443 | NM_015786 |
| Nbeal2 | 670.8633228 | 0.084120824 | 0.119867452 | 0.701782027 | 0.482815109 | 0.695105921 | | Tmem126a | 556.1418991 | 0.0255557434768487 | 0.0628455164948164 | 0.406643861045467 | 0.684269577981466 | 0.923655037989443 | NM_025460 |
| 4930444A02Rik | 1420.942895 | -0.046749129 | 0.066678531 | -0.701112161 | 0.483233022 | 0.69544979 | | Arsj | 118.8941233 | 0.0379470085846748 | 0.0934542021166281 | 0.406049248992764 | 0.68470641451252 | 0.923762012051692 | NM_173451 |
| Pgr15l | 17.05242208 | -0.102982754 | 0.146865479 | -0.701204633 | 0.483175319 | 0.69544979 | | Immp1l | 489.2569448 | 0.033592133209088 | 0.0827487604322237 | 0.405953310159878 | 0.684776906634491 | 0.923762012051692 | NM_028260 |
| Pik3c2a | 1028.897237 | -0.05259154 | 0.075005773 | -0.701166556 | 0.483199079 | 0.69544979 | | Rel | 63.67588292 | 0.0383498647868803 | 0.0944498925545997 | 0.406033957830278 | 0.684717649679878 | 0.923762012051692 | NM_009044 |
| Daglb | 1608.36277 | 0.054273313 | 0.077434744 | 0.700890974 | 0.483371058 | 0.695583837 | | Rpl23 | 6028.662333 | 0.0280755583844581 | 0.0691447692954791 | 0.406040235154762 | 0.684713037413135 | 0.923762012051692 | NM_022891 |
| Fastk | 2643.718488 | 0.051343948 | 0.073319793 | 0.700274047 | 0.483756176 | 0.69596223 | | Ten1 | 626.456009 | 0.0295760227084247 | 0.0728005122646562 | 0.406261189494178 | 0.684550698612018 | 0.923762012051692 | NM_027107 |
| Fgf7 | 18.96677089 | 0.11768871 | 0.168056051 | 0.700294391 | 0.483743474 | 0.69596223 | | Brwd3 | 357.4567916 | -0.0355955609516738 | 0.0877107733643703 | -0.405828834774968 | 0.684868370396503 | 0.923799605198217 | NM_001081477 |
| Hnrnpk | 12322.00153 | -0.038748741 | 0.055335274 | -0.700253889 | 0.483768763 | 0.69596223 | | 5830403L16Rik | 41.56615473 | -0.0379868313533081 | 0.0936913048426489 | -0.405446710525652 | 0.685149181849033 | 0.923835237681152 | NM_178243 |
| Vps37c | 843.2561517 | 0.061424213 | 0.087745904 | 0.700023708 | 0.483912499 | 0.696039756 | | Adap1 | 8298.297197 | 0.0232317331104121 | 0.0572820973821089 | 0.405567082424396 | 0.685060719517252 | 0.923835237681152 | NM_172723 |
| Zswim4 | 902.2578683 | -0.084714351 | 0.121005288 | -0.700088006 | 0.483872346 | 0.696039756 | | Hdgfrp2 | 3262.442751 | -0.0234981653529224 | 0.0579444839707106 | -0.405528943269217 | 0.68508874783932 | 0.923835237681152 | NM_008233 |
| Gadd45b | 404.817298 | -0.09219148 | 0.131802381 | -0.69946749 | 0.484259923 | 0.696474821 | | Sox1 | 530.3994289 | 0.0361490288133357 | 0.089119427834517 | 0.405624561239999 | 0.685018479370337 | 0.923835237681152 | NM_009233 |
| Bcl10 | 458.7087436 | -0.057500963 | 0.082254637 | -0.699060443 | 0.484514258 | 0.696517343 | | 1700030C10Rik | 47.57827614 | -0.0365883755832279 | 0.0918768265581535 | -0.398232905444 | 0.690458512517707 | 0.923860705630581 | NR_015521 |
| Dab2ip | 6414.175407 | 0.062690208 | 0.08967392 | 0.699090748 | 0.48449532 | 0.696517343 | | 2310044G17Rik | 4455.780238 | 0.0248652771533557 | 0.062024391046973 | 0.400895145566131 | 0.688497325595203 | 0.923860705630581 | NR_033623,NM_173735 |
| Naf1 | 391.4548438 | -0.089807015 | 0.128427459 | -0.699282032 | 0.484375794 | 0.696517343 | | 4930528A17Rik | 22.41968482 | -0.0331805333050421 | 0.0828688574685027 | -0.400398102720959 | 0.688863321442283 | 0.923860705630581 | NR_028384 |
| Pef1 | 2440.609711 | -0.049438666 | 0.070714605 | -0.699129485 | 0.48447111 | 0.696517343 | | 5330417C22Rik | 1357.815352 | -0.0228085737177252 | 0.0572941847789525 | -0.398095789402071 | 0.690559578016707 | 0.923860705630581 | NM_001033304 |
| Upf2 | 2512.143887 | 0.063583264 | 0.090941814 | 0.699164232 | 0.484449401 | 0.696517343 | | 5330426P16Rik | 199.4362824 | 0.0383172651239542 | 0.0948629952635272 | 0.403922151282592 | 0.686269966666065 | 0.923860705630581 | NR_028300 |
| Epha2 | 74.17867061 | -0.108005771 | 0.154520012 | -0.698975939 | 0.484567068 | 0.696528634 | | 5930412G12Rik | 97.831882 | 0.0375275940335169 | 0.0946760390842436 | 0.396379025803134 | 0.691825453095965 | 0.923860705630581 | NR_015517 |
| D17H6S53E | 556.4153014 | -0.055632623 | 0.079602002 | -0.698884726 | 0.484624073 | 0.696545955 | | A230056P14Rik | 100.7843497 | 0.0375738682078299 | 0.0947641708501811 | 0.396498675298208 | 0.691737184916287 | 0.923860705630581 | NR_015495 |
| N4bp1 | 2970.323635 | 0.065103133 | 0.093189346 | 0.698611329 | 0.484794961 | 0.69666232 | | Abl1 | 2854.731808 | -0.0196514649312799 | 0.0496425007510343 | -0.395838160265078 | 0.692224423451405 | 0.923860705630581 | NM_009594,NM_001112703 |
| Sik3 | 6740.653434 | -0.068955878 | 0.098701557 | -0.698630092 | 0.484783233 | 0.69666232 | | Adcy2 | 7070.330304 | 0.020270716345201 | 0.0505547308758727 | 0.400965765102614 | 0.688445329367079 | 0.923860705630581 | NM_153534 |
| A530072M11Rik | 26.08005159 | 0.116062129 | 0.166381569 | 0.697566021 | 0.485448638 | 0.697149058 | | Afg3l1 | 1268.676015 | -0.0217655189185575 | 0.0539326525348798 | -0.403568485798008 | 0.686530063762583 | 0.923860705630581 | NR_028260,NM_054070 |
| Ccnt1 | 703.144811 | 0.068234163 | 0.097769573 | 0.697907957 | 0.485234758 | 0.697149058 | | Akap17b | 881.485694 | -0.0243396177359282 | 0.0605833341886491 | -0.401754344852292 | 0.687864825059809 | 0.923860705630581 | NM_001081956 |
| Gm2762 | 14.25800011 | 0.116720984 | 0.167226569 | 0.6979811 | 0.485189014 | 0.697149058 | | Alox8 | 320.4272668 | 0.0349646341171668 | 0.0882691168541416 | 0.39611405849 8437 | 0.692020887595982 | 0.923860705630581 | NM_009661 |
| Npas1 | 284.2592587 | 0.07976764 | 0.114349625 | 0.697576755 | 0.485441923 | 0.697149058 | | Anxa9 | 37.50422041 | -0.0342234729733878 | 0.0864008636598836 | -0.396101051814809 | 0.692030482397087 | 0.923860705630581 | NM_001085383,NM_023628 |
| Ppif | 931.9832712 | 0.042582346 | 0.061038101 | 0.697635495 | 0.485405178 | 0.697149058 | | Atl3 | 1772.466941 | -0.02486707974425 | 0.061528041020054 | -0.404158483383943 | 0.686096181061581 | 0.923860705630581 | NM_001163505,NM_146091 |
| Ranbp3 | 3018.122364 | 0.050144317 | 0.071884526 | 0.697567611 | 0.485447643 | 0.697149058 | | Aven | 341.6808024 | 0.0325999636456255 | 0.0811688026296261 | 0.401631693329016 | 0.687955101561022 | 0.923860705630581 | NM_001165935,NM_028844 |
| Snai1 | 21.82421618 | -0.117208414 | 0.16797509 | -0.697772596 | 0.48531942 | 0.697149058 | | BC025920 | 169.8906753 | 0.0354397357112396 | 0.0890407224768275 | 0.398017162545628 | 0.690617534792873 | 0.923860705630581 | NR_030677 |
| Gna14 | 11.44649048 | 0.111340537 | 0.159662535 | 0.697349172 | 0.485584302 | 0.697252604 | | BC052688 | 31.22208005 | -0.034379039690431 | 0.0865093038844944 | -0.39740280116383 | 0.691070450188761 | 0.923860705630581 | NR_028430 |
| St6galnac3 | 318.4121481 | 0.080480323 | 0.115427831 | 0.697234991 | 0.485655744 | 0.697252604 | | Bank1 | 64.82785053 | -0.0379636822922024 | 0.0950377096532121 | -0.399459145540563 | 0.689554920949445 | 0.923860705630581 | NM_001033350 |
| Ube2d3 | 4246.032617 | -0.063907349 | 0.091654293 | -0.6972652 | 0.485636843 | 0.697252604 | | Bcdin3d | 103.5321561 | -0.0367240324601376 | 0.0923736834028463 | -0.397559468317208 | 0.690959494565163 | 0.923860705630581 | NM_029236 |
| Mis18a | 168.1543223 | -0.074169279 | 0.106417466 | -0.696965278 | 0.485824524 | 0.697365682 | | Bcl11b | 3719.918351 | 0.0336492699430938 | 0.0832433460780821 | 0.404227743458688 | 0.686045254163735 | 0.923860705630581 | NM_021399,NM_001079883 |
| Napg | 6044.882863 | -0.043624666 | 0.062591771 | -0.696971269 | 0.485820873 | 0.697365682 | | C130030K03Rik | 150.607495 | -0.0365094157378068 | 0.0914383960813842 | -0.399278829546249 | 0.689687764442017 | 0.923860705630581 | NR_046212 |
| Fads3 | 1236.608619 | 0.057728777 | 0.082854077 | 0.696752398 | 0.485957761 | 0.697492316 | | C1qtnf2 | 29.32233302 | 0.0344145026971837 | 0.08499560229897 | 0.404897450766118 | 0.685552892297516 | 0.923860705630581 | NM_026979 |
| Eci2 | 1086.257467 | 0.050724468 | 0.072812347 | 0.696646517 | 0.486024037 | 0.697522676 | | Cant1 | 2188.746886 | 0.02096970393438 | 0.0521618860773644 | 0.402011995948125 | 0.687675197880248 | 0.923860705630581 | NM_001025617,NM_001267591,NM_001267592,NM_029502,NM_001025618 |
| Enpp6 | 230.9769829 | -0.104901648 | 0.150628848 | -0.696424687 | 0.486162907 | 0.697592895 | | Ccdc43 | 763.6764426 | 0.0268590840235859 | 0.0677288002125181 | 0.396568135553974 | 0.691685953957871 | 0.923860705630581 | NM_025918 |
| Tnfaip3 | 307.4276896 | -0.082494378 | 0.118446115 | -0.696471789 | 0.486133419 | 0.697592895 | | Ccno | 76.73030648 | -0.0377639892710987 | 0.0945190491101862 | -0.399538395980636 | 0.689496538111305 | 0.923860705630581 | NM_001081062 |
| Atp13a3 | 2790.094682 | -0.064770141 | 0.09304768 | -0.696096245 | 0.486368558 | 0.697779778 | | Cda | 60.77324181 | -0.0373878916992601 | 0.0941022188771191 | -0.397311478362504 | 0.6911377840195 | 0.923860705630581 | NM_028176 |
| Rnf141 | 837.0160583 | -0.059977527 | 0.086165591 | -0.696072833 | 0.486383219 | 0.697779778 | | Cdc42ep2 | 507.5855451 | -0.0342065308371412 | 0.084419419386875 | -0.405197418859048 | 0.685332402142795 | 0.923860705630581 | NM_026772 |
| Gpbp1 | 3300.873221 | -0.05100778 | 0.073308753 | -0.695793853 | 0.486557939 | 0.697836592 | | Chmp2a | 2623.674373 | 0.0201903228758547 | 0.0502991948661014 | 0.401404494238967 | 0.68812234106449 | 0.923860705630581 | NM_026885 |
| Pde9a | 627.0343803 | 0.067054014 | 0.096357776 | 0.695885855 | 0.486500316 | 0.697836592 | | Col1a2 | 743.2950451 | -0.0372770405932652 | 0.0938802481905907 | -0.397070111125053 | 0.691315759793915 | 0.923860705630581 | NM_007743 |
| Tas1r3 | 12.49878659 | -0.109900186 | 0.157948378 | -0.695798133 | 0.486555259 | 0.697836592 | | Cpd | 5308.549896 | -0.0260494945733293 | 0.0649509095056799 | -0.401064354165068 | 0.688372744066135 | 0.923860705630581 | NM_007754 |
| Lyrm7 | 393.5479695 | -0.089515291 | 0.128672785 | -0.695681619 | 0.486628239 | 0.6978782 | | Dchs1 | 708.802189 | -0.0327359545528178 | 0.0817535171127151 | -0.400422583748711 | 0.688845293140111 | 0.923860705630581 | NM_001162943 |
| Exoc5 | 1869.578726 | -0.067715079 | 0.097463149 | -0.694782695 | 0.487191495 | 0.698562527 | | Dlgap1 | 21014.70375 | 0.0238475361631534 | 0.0595197916410958 | 0.400665652644653 | 0.688666302095313 | 0.923860705630581 | NM_001128181,NM_001128180,NM_177639,NM_027712 |
| Notch4 | 328.8568353 | -0.100929053 | 0.14528474 | -0.694698238 | 0.487244432 | 0.698562527 | | Dmd | 2562.630837 | -0.0272379403369521 | 0.0675512144304056 | -0.403219106667784 | 0.686787044988486 | 0.923860705630581 | NM_007868 |
| Rell1 | 513.9085803 | 0.080469465 | 0.115831296 | 0.694712641 | 0.487235405 | 0.698562527 | | Dnali1 | 60.54585601 | 0.0347303948786197 | 0.0877341362658501 | 0.395859540616897 | 0.692208649927187 | 0.923860705630581 | NM_175223 |

| | | | | | | | | | | | | | | |
|---|---|---|---|---|---|---|---|---|---|---|---|---|---|---|
| Fbxl2 | 1952.357578 | 0.060646918 | 0.087354716 | 0.694260376 | 0.487518936 | 0.698826754 | | Eif4ebp1 | 97.43826148 | -0.037620222352546 | 0.0935265907317067 | -0.40224092483457 | 0.68750672627576 | 0.923860705630581 | NM_007918 |
| Glrb | 7288.754177 | -0.058509877 | 0.084273369 | -0.694286675 | 0.487502446 | 0.698826754 | | Ell2 | 1279.667192 | 0.0312309030941165 | 0.0787195117358599 | 0.396736493982718 | 0.691561785729309 | 0.923860705630581 | NM_138953 |
| AK129341 | 687.0759947 | 0.064817492 | 0.093400547 | 0.693973372 | 0.487698908 | 0.698856463 | | Epb4.9 | 17497.56027 | 0.0234454907402916 | 0.0584839779167242 | 0.400887415245178 | 0.688503015783645 | 0.923860705630581 | NM_001252666,NM_001252664,NM_001252665,NM_001252662,NM_013514,NM_001252663 |
| Atp13a2 | 5925.12541 | 0.047645333 | 0.068671286 | 0.693817386 | 0.487796738 | 0.698856463 | | Esr2 | 22.54296179 | 0.0298727779767181 | 0.0747790137546698 | 0.399480769761458 | 0.689538990464768 | 0.923860705630581 | NM_207707,NM_010157 |
| BC024978 | 276.3213452 | 0.06492577 | 0.093561544 | 0.693936493 | 0.487722037 | 0.698856463 | | Fam149b | 1172.98435 | -0.0222150888661448 | 0.0557832361276434 | -0.39823951438228 | 0.690453641340542 | 0.923860705630581 | NM_001024512,NM_001177633,NM_172379 |
| Slitrk3 | 2429.371237 | 0.044089803 | 0.063539899 | 0.693891613 | 0.487750184 | 0.698856463 | | Fam212a | 60.47212867 | 0.0380379996064203 | 0.0941671900679589 | 0.403941113449057 | 0.686256022321467 | 0.923860705630581 | NM_026597 |
| Tsfm | 423.6609091 | 0.056486028 | 0.081415929 | 0.693795778 | 0.487810291 | 0.698856463 | | Fgfr1 | 2846.932721 | -0.0265579751272486 | 0.0665705459465842 | -0.398944829873538 | 0.689933855969632 | 0.923860705630581 | NM_001079908,NM_010206,NM_001079909 |
| Usp21 | 1565.290165 | -0.04647983 | 0.066992702 | -0.693804384 | 0.487804893 | 0.698856463 | | Fmo1 | 312.8572329 | -0.0352341331437117 | 0.0881414840738732 | -0.399745176904229 | 0.68934421381753 | 0.923860705630581 | NM_010231 |
| Dck | 935.9648853 | 0.050999497 | 0.073531545 | 0.693573034 | 0.48795001 | 0.698991998 | | Foxp4 | 1636.062095 | 0.0308248542322642 | 0.0769567358193126 | 0.400547839043462 | 0.68875305548472 | 0.923860705630581 | NM_028767,NM_001110825,NM_001110824 |
| Ppib | 2751.892343 | 0.04420312 | 0.063742938 | 0.69345909 | 0.488021491 | 0.699029766 | | Gal3st1 | 403.906242 | 0.0314703141255317 | 0.0786223357309858 | 0.40027193078072 | 0.688956239700781 | 0.923860705630581 | NM_016922,NM_001177703,NM_001177691 |
| Rragb | 2309.045396 | -0.049595229 | 0.071548161 | -0.693172656 | 0.488201206 | 0.699222544 | | Gemin6 | 171.3893674 | -0.0353691126490439 | 0.0883598717869147 | -0.40028478916695 | 0.688946770036194 | 0.923860705630581 | NM_026053 |
| Tmem199 | 1258.391153 | 0.053003911 | 0.076478879 | 0.693052928 | 0.488276336 | 0.69926551 | | Gm14295 | 1938.45606 | 0.0348390620958769 | 0.0867949345021908 | 0.40139510785618 | 0.688129250636835 | 0.923860705630581 | NM_001205057 |
| Rnf145 | 4630.530505 | 0.046978566 | 0.067806398 | 0.692833826 | 0.488413841 | 0.699364482 | | Gm9899 | 968.1097588 | -0.0282094444328928 | 0.0704286554037378 | -0.400539301384925 | 0.688759342446095 | 0.923860705630581 | NR_040427 |
| Tnfaip8l3 | 320.5580317 | -0.072750017 | 0.105008841 | -0.692798966 | 0.488435721 | 0.699364482 | | Gm9962 | 70.00460941 | 0.038288445566477 | 0.0950944853927343 | 0.402635814351884 | 0.687216158648863 | 0.923860705630581 | NR_033504 |
| 5730507C01Rik | 39.0862782 | -0.11520335 | 0.166377183 | -0.692422774 | 0.488671868 | 0.699637952 | | Gmpr2 | 793.5347113 | -0.0228141313700814 | 0.0571745158606215 | -0.399026227448906 | 0.689873879074494 | 0.923860705630581 | NM_177992 |
| Sh3d21 | 49.80100149 | 0.104703548 | 0.151238276 | 0.692308527 | 0.488743597 | 0.699675994 | | Gnao1 | 32710.19495 | 0.0168729001732125 | 0.0424731925555651 | 0.397259992903495 | 0.691175746187614 | 0.923860705630581 | NM_010308,NM_001113384 |
| Crem | 497.1539186 | -0.082040803 | 0.118564366 | -0.6919516 | 0.488967725 | 0.699932181 | | Grem1 | 34.47172527 | 0.0314941344837954 | 0.0782226062131986 | 0.402621901882904 | 0.6872263949377 | 0.923860705630581 | NM_011824 |
| Prdx2 | 6338.249413 | 0.035806443 | 0.051783127 | 0.691469305 | 0.489270664 | 0.700301124 | | Gstm5 | 5391.326145 | 0.0253466381670782 | 0.0636100814057157 | 0.398468884286016 | 0.690284590143188 | 0.923860705630581 | NM_010360 |
| D030028A08Rik | 252.3541778 | -0.074210178 | 0.107343975 | -0.691330631 | 0.489357786 | 0.700319482 | | Gstt1 | 354.3711448 | 0.033248909111777 | 0.0838802390412767 | 0.396385483539401 | 0.691820673412028 | 0.923860705630581 | NM_008185 |
| Gtf2h4 | 468.2559844 | 0.063502997 | 0.091859594 | 0.691305002 | 0.489373889 | 0.700319482 | | Herc1 | 9642.374489 | 0.0269386470807279 | 0.0680554687619096 | 0.395833686415005 | 0.692227724086689 | 0.923860705630581 | NM_145617 |
| Lctl | 9.927637418 | 0.111048717 | 0.160671414 | 0.691154166 | 0.489468664 | 0.700390421 | | Hey2 | 165.4119558 | 0.0359286974071152 | 0.0901612234379365 | 0.39849389834253 | 0.690266155109388 | 0.923860705630581 | NM_013904 |
| Nr2f6 | 635.2188479 | -0.076082784 | 0.110100853 | -0.691028107 | 0.489547878 | 0.700413803 | | Il12a | 96.52560462 | 0.03828363280158 | 0.0944556531897933 | 0.405308009720004 | 0.68525111960389 | 0.923860705630581 | NM_008351,NM_001159424 |
| Slc35f2 | 158.9181391 | 0.073327293 | 0.106120058 | 0.690984288 | 0.489575415 | 0.700413803 | | Kcne4 | 64.65174636 | 0.037685396221716 | 0.093209190588049 | 0.404309875281203 | 0.685984864524153 | 0.923860705630581 | NM_021342 |
| Psma4 | 2700.316828 | -0.038353297 | 0.0555307 | -0.690668343 | 0.489773988 | 0.700633198 | | Kcnj3 | 2310.849364 | -0.0235584853882678 | 0.0586979902964316 | -0.401350800415734 | 0.688161866903519 | 0.923860705630581 | NM_008426 |
| Tbcel | 2140.46758 | 0.043120133 | 0.062451705 | 0.690455646 | 0.489907693 | 0.700759767 | | Klhl40 | 257.7856711 | -0.0317273756200019 | 0.0791930609013808 | -0.400633278457465 | 0.688690140789431 | 0.923860705630581 | NM_028202 |
| Irak1bp1 | 419.1004121 | -0.061779613 | 0.089496193 | -0.690304367 | 0.490002802 | 0.700802864 | | Kpnb1 | 9373.113242 | 0.0226765180516768 | 0.056695977227095 | 0.399966966983325 | 0.689180847087459 | 0.923860705630581 | NM_008379 |
| Pcyox1 | 5603.39692 | 0.052684659 | 0.076333349 | 0.690191904 | 0.490073514 | 0.700802864 | | Lipo1 | 204.7496045 | -0.0348487464657696 | 0.0863187546364153 | -0.403721608502771 | 0.686417447787938 | 0.923860705630581 | NM_001013770 |
| Zfp101 | 281.3517477 | 0.077901472 | 0.112858109 | 0.690260295 | 0.490030512 | 0.700802864 | | Med15 | 2729.028774 | -0.0187162811925131 | 0.0463303130532952 | -0.403974848410438 | 0.686231214664568 | 0.923860705630581 | NM_001040683,NM_033609 |
| Acta1 | 269.2933722 | 0.08280491 | 0.120001403 | 0.690032844 | 0.490173533 | 0.700872964 | | Mme | 597.7960825 | 0.0372275616992807 | 0.09252127842834 | 0.402367567023143 | 0.687413535383823 | 0.923860705630581 | NM_008604 |
| Mrps9 | 875.5995501 | -0.039461226 | 0.057192663 | -0.689970076 | 0.490213006 | 0.700872964 | | Mreg | 199.8472193 | -0.0374782957103711 | 0.0946513169540796 | -0.395961693048115 | 0.692133288003228 | 0.923860705630581 | NM_001005423 |
| Mob3b | 205.0528134 | -0.098712707 | 0.143087605 | -0.689876015 | 0.490272161 | 0.700892864 | | Mvd | 685.1178201 | 0.0298765975800425 | 0.0748761063478755 | 0.399013771379021 | 0.689883057064366 | 0.923860705630581 | NM_138656,NR_028354 |
| Ptprn | 25116.23741 | 0.044602008 | 0.064702232 | 0.68934265 | 0.490607664 | 0.701243095 | | Mzt1 | 1943.707297 | 0.0237356093130319 | 0.0598847974883807 | 0.396354505793182 | 0.691843522772841 | 0.923860705630581 | NM_175245 |
| Rdm1 | 165.1227109 | 0.084698256 | 0.122856433 | 0.689408399 | 0.4905663 | 0.701243095 | | Neurod6 | 2024.986081 | 0.0367501258224188 | 0.0915542782190193 | 0.401402605506909 | 0.688123731409574 | 0.923860705630581 | NM_009717 |
| Pea15a | 21177.32046 | -0.046897261 | 0.06806366 | -0.689020569 | 0.490810324 | 0.701468052 | | Nme6 | 161.7417226 | 0.0354108712875627 | 0.0890104302410732 | 0.397828335304717 | 0.690756728970112 | 0.923860705630581 | NM_018757 |
| Sgk3 | 538.5891274 | 0.097071043 | 0.140918206 | 0.688846713 | 0.490919736 | 0.701559711 | | Npc1 | 3765.555935 | -0.0263865809739228 | 0.0661354539735994 | -0.398977846049958 | 0.689909528133644 | 0.923860705630581 | NM_008720 |
| Fnbp1l | 4561.594576 | -0.05922331 | 0.085988295 | -0.688736884 | 0.490988861 | 0.701593785 | | Orc2 | 1314.643951 | -0.0259620295781656 | 0.0649334273694223 | -0.399825338503407 | 0.689285166500388 | 0.923860705630581 | NM_001025378,NM_001271526,NM_008765 |
| Ppig | 2264.101357 | 0.05486297 | 0.079679897 | 0.688542171 | 0.491111423 | 0.701612191 | | Osgepl1 | 482.9409453 | -0.0293242057869648 | 0.0734563544519179 | -0.399205841424645 | 0.689741539413029 | 0.923860705630581 | NM_028091 |
| Stam | 2173.23903 | -0.041816449 | 0.060729529 | -0.688568646 | 0.491094757 | 0.701612191 | | Pam16 | 646.0095027 | -0.0276370216431885 | 0.069433820023629 | -0.398034007545363 | 0.690605117993731 | 0.923860705630581 | NM_025571 |
| Tm9sf1 | 1132.977918 | 0.040464371 | 0.05877173 | 0.688500604 | 0.49113759 | 0.701612191 | | Pank3 | 2538.385381 | 0.0238856943748664 | 0.0602012198515356 | 0.396764292048762 | 0.69154128481139 | 0.923860705630581 | NM_145962 |
| Ccdc90a | 997.2245169 | 0.040494006 | 0.058841747 | 0.68818498 | 0.491336302 | 0.70183135 | | Pcdhga5 | 533.179367 | 0.0334209333310021 | 0.0825109853355203 | 0.405048287753445 | 0.685442016921309 | 0.923860705630581 | NM_033588 |
| Mrpl2 | 834.0894578 | 0.0505041 | 0.073395579 | 0.688108208 | 0.491384642 | 0.701835697 | | Pcdhgb2 | 318.4397501 | -0.0317473091172861 | 0.0797120244394677 | -0.398275032412388 | 0.690427462677865 | 0.923860705630581 | NM_033575 |
| Tatdn2 | 1675.130667 | -0.052452271 | 0.076257602 | -0.687830062 | 0.491559803 | 0.702021162 | | Pdp2 | 314.8162547 | -0.0347606950516306 | 0.0860436202186392 | -0.403989220389644 | 0.686220646054138 | 0.923860705630581 | NM_001024606 |
| BC048403 | 460.5734932 | 0.059184455 | 0.086075918 | 0.687584363 | 0.491714558 | 0.702100612 | | Pi4kb | 2397.418482 | 0.016615417460606 | 0.0416646420569103 | 0.398789396484212 | 0.690048390738473 | 0.923860705630581 | NM_175356 |
| Fancf | 70.59874138 | 0.102181794 | 0.148607493 | 0.687595167 | 0.491707753 | 0.702100612 | | Pnpla1 | 27.1947314 | 0.03444319632516 | 0.0859596800413406 | 0.400690141106286 | 0.68864827025121 | 0.923860705630581 | NM_001034885 |
| Inadl | 461.9018188 | -0.104565899 | 0.152106035 | -0.68745398 | 0.491796692 | 0.702100612 | | Ppm1m | 766.2159165 | -0.0327917952205195 | 0.0816996267210144 | -0.401370196371839 | 0.688147588786046 | 0.923860705630581 | NM_026447,NM_198931 |
| Nol8 | 1052.822855 | 0.045714702 | 0.066494396 | 0.687497056 | 0.491769555 | 0.702100612 | | Pstpip2 | 432.1183724 | -0.034720361505134 | 0.0875213215830562 | -0.39670746370283 | 0.691583195639382 | 0.923860705630581 | NM_013831 |
| Tmem237 | 742.8737308 | 0.048734861 | 0.070922479 | 0.687156763 | 0.491983948 | 0.702303232 | | Ptpn5 | 11897.84777 | 0.0253223905730233 | 0.0626566083327795 | 0.404145568150321 | 0.686105677784023 | 0.923860705630581 | NM_001163565,NM_013643 |
| Tbrg4 | 1245.328305 | 0.041587837 | 0.060566366 | 0.686649037 | 0.492303919 | 0.702695248 | | Rab8b | 2052.392587 | 0.0295508751131586 | 0.0736419760926672 | 0.401277595755624 | 0.688215756701652 | 0.923860705630581 | NM_173413 |
| D7Ertd443e | 198.8571453 | 0.084820917 | 0.123550416 | 0.686528779 | 0.492379723 | 0.702738709 | | Racgap1 | 575.632949 | -0.0317719826639227 | 0.0793131387735576 | -0.400589147715274 | 0.688722636906801 | 0.923860705630581 | NM_001253809,NM_001253808,NM_012025 |
| Hyal1 | 154.1855313 | 0.080312305 | 0.117016871 | 0.68633099 | 0.492504411 | 0.702817162 | | Rasgrf2 | 8774.125166 | 0.0294935483710995 | 0.074114743449252 | 0.397944416974126 | 0.690671158020424 | 0.923860705630581 | NM_009027 |
| Nxpe2 | 10.76358979 | -0.10942783 | 0.159463315 | -0.686225732 | 0.492570774 | 0.702817162 | | Rasgrp2 | 1950.021513 | -0.0318523742560221 | 0.0787869028112919 | -0.404285142827787 | 0.686003049517164 | 0.923860705630581 | NM_011242 |
| Rtkn2 | 105.9123102 | 0.098882032 | 0.144093319 | 0.686236063 | 0.49256426 | 0.702817162 | | Rpl21 | 10665.76231 | 0.026389673544253 | 0.0657861600963382 | 0.401143242068051 | 0.688314665631753 | 0.923860705630581 | NM_019647 |
| Ncln | 2021.093847 | 0.059565151 | 0.086823689 | 0.686047221 | 0.492683331 | 0.702913033 | | Rps13 | 3524.942007 | -0.0272037200158622 | 0.0685665669739312 | -0.396749045729604 | 0.691552528858327 | 0.923860705630581 | NM_026533 |
| Ddx49 | 883.1983317 | 0.045061839 | 0.065736913 | 0.685487604 | 0.49303628 | 0.703173645 | | Slc22a23 | 4033.449442 | 0.0221660868669317 | 0.0552095753434385 | 0.401489899696649 | 0.688059472974152 | 0.923860705630581 | NM_001033167 |
| Fgfr4 | 8.546220118 | -0.109542516 | 0.15978629 | -0.685556414 | 0.492992875 | 0.703173645 | | Slc25a30 | 235.6215925 | 0.0361469046015594 | 0.0912260380712606 | 0.396234511174578 | 0.691932034130148 | 0.923860705630581 | NM_026232 |
| Hdac6 | 2138.40591 | 0.06155411 | 0.089785404 | 0.685569224 | 0.492984795 | 0.703173645 | | Smo | 665.4025983 | 0.033240104927557 | 0.0834366634031947 | 0.398387274511798 | 0.690344736761311 | 0.923860705630581 | NM_176996 |
| Lcn2 | 17.81525375 | 0.069090439 | 0.100792833 | 0.685469765 | 0.493047533 | 0.703173645 | | St8sia2 | 172.026629 | -0.0380316935850311 | 0.0950889805423288 | -0.399959000171437 | 0.689186715052202 | 0.923860705630581 | NM_009181 |
| Dus4l | 175.6141132 | 0.07865039 | 0.114777916 | 0.685239747 | 0.493192646 | 0.703315863 | | Taf6 | 1455.581009 | 0.0259581618838753 | 0.064474978787186 | 0.402608304371158 | 0.687236399547861 | 0.923860705630581 | NM_009315 |
| Cd83 | 638.0142853 | -0.067562283 | 0.098641547 | -0.684927237 | 0.493389838 | 0.703465993 | | Tbc1d10a | 901.1617876 | -0.0277009670164935 | 0.0690915462259525 | -0.400931351657728 | 0.688470666628655 | 0.923860705630581 | NM_134023 |
| Etaa1 | 367.5817337 | -0.058561911 | 0.08550453 | -0.684898342 | 0.493408072 | 0.703465993 | | Tbcel | 2379.610695 | 0.0230949345616068 | 0.0573545933965262 | 0.402669310231838 | 0.687191513833477 | 0.923860705630581 | NM_173038 |
| Rpl37 | 4322.367656 | -0.046435719 | 0.067810643 | -0.68478511 | 0.493479533 | 0.703465993 | | Tfcp2 | 878.7095978 | -0.0249569662285005 | 0.0624148037059616 | -0.39985652035491 | 0.689262198349647 | 0.923860705630581 | NM_033476 |
| Wdr54 | 517.7601756 | -0.053855166 | 0.078643876 | -0.684797964 | 0.493471421 | 0.703465993 | | Tgfbrap1 | 3014.852674 | 0.0158828405820641 | 0.0398529066814071 | 0.398536566204194 | 0.690234709875226 | 0.923860705630581 | NM_001013025 |
| Rassf4 | 506.7051678 | -0.088216266 | 0.128843645 | -0.684676887 | 0.493547837 | 0.703498637 | | Tlr4 | 64.97680533 | -0.0381429801402259 | 0.094873284582058 | -0.402041315511062 | 0.687653620384639 | 0.923860705630581 | NM_021297 |
| Ift52 | 1136.6903 | -0.052778744 | 0.077094813 | -0.684595271 | 0.493599352 | 0.703507346 | | Tmem199 | 1347.041587 | 0.0218845229685862 | 0.054912705232704 | 0.398532960192837 | 0.690237367402097 | 0.923860705630581 | NM_199199 |
| Gpr116 | 2065.947494 | -0.055398465 | 0.080957373 | -0.684291778 | 0.493790938 | 0.703710274 | | Tnnt1 | 123.0218306 | -0.0370529041518235 | 0.0928349408916477 | -0.399126705914208 | 0.68979984532505 | 0.923860705630581 | NM_011618 |
| Kdm1a | 2434.845811 | -0.030473402 | 0.044537052 | -0.684225839 | 0.493832568 | 0.703710274 | | Tpd52l1 | 672.686288 | 0.0273490680133896 | 0.0688098456610295 | 0.39745864491713 | 0.691029276858256 | 0.923860705630581 | NM_009413 |
| Nradd | 51.27723724 | -0.104907012 | 0.153342592 | -0.68413485 | 0.493890017 | 0.703727416 | | Trabd | 735.0852821 | -0.0281268775409017 | 0.0703702845572422 | -0.399698220887854 | 0.689378802663897 | 0.923860705630581 | NM_026485 |
| Meig1 | 26.73356607 | -0.111119066 | 0.162610033 | -0.683346925 | 0.494387649 | 0.7043717 | | Trim3 | 5302.547921 | 0.0244038098498761 | 0.0608908462279167 | 0.400779614041357 | 0.688582389216964 | 0.923860705630581 | NM_018880 |
| Copz1 | 3264.539672 | -0.045101272 | 0.066014221 | -0.683205391 | 0.494477067 | 0.70443432 | | Trim9 | 12978.39311 | 0.0294788118572468 | 0.0742191904330726 | 0.397185844863525 | 0.691230419694562 | 0.923860705630581 | NM_001110203,NM_001110202,NM_053167 |
| Msantd4 | 2134.200277 | -0.050370667 | 0.073753104 | -0.682963349 | 0.494630003 | 0.704483316 | | Ttc34 | 104.3626185 | -0.0375481765381344 | 0.0931460946590016 | -0.403110583171463 | 0.686866875390246 | 0.923860705630581 | NM_172878 |
| Pygl | 193.5745049 | 0.071474643 | 0.104647835 | 0.683001645 | 0.494605803 | 0.704483316 | | Ubox5 | 811.9144691 | 0.0273259761320056 | 0.0685306944584558 | 0.398740686169042 | 0.690084285547379 | 0.923860705630581 | NM_001255994,NM_080562,NM_001255993 |
| Tm4sf5 | 9.785699211 | -0.107873906 | 0.157956311 | -0.682935084 | 0.494647863 | 0.704483316 | | Vps37b | 1310.962151 | 0.0272930966858448 | 0.0677670682446994 | 0.402748671187788 | 0.687133124811625 | 0.923860705630581 | NM_177876 |
| Ripk4 | 25.50695218 | 0.114645106 | 0.16790055 | 0.682815546 | 0.494723405 | 0.704526143 | | Wdr77 | 2522.828476 | 0.020150700783238 | 0.0506154157413838 | 0.398113904392225 | 0.690546225502548 | 0.923860705630581 | NM_027432 |
| Cdr2 | 644.3913054 | 0.061940385 | 0.090774699 | 0.682352965 | 0.49501579 | 0.704812961 | | Zfp119a | 108.7530164 | -0.0372635233214333 | 0.0937961838904524 | -0.397281869856823 | 0.691159615391062 | 0.923860705630581 | NM_144546 |
| Numa1 | 4410.540145 | 0.043664244 | 0.063985739 | 0.682405872 | 0.494982344 | 0.704812961 | | Zfp276 | 1033.033181 | -0.0332165035535901 | 0.0830223431876999 | -0.40009113544884 | 0.68908939307386 | 0.923860705630581 | NM_020497 |
| Oxa1l | 1644.934595 | 0.03698911 | 0.054240441 | 0.681947068 | 0.495272422 | 0.705048778 | | Zfp3 | 487.1218953 | -0.02727387883025 | 0.0674731008694968 | -0.404218547521654 | 0.686052015849493 | 0.923860705630581 | NM_177565 |
| Tmem86a | 608.4299339 | -0.061299132 | 0.089883856 | -0.681981561 | 0.49525061 | 0.705048778 | | Zfp831 | 1404.375448 | 0.03483708745621 | 0.0877650270268965 | 0.396935871113374 | 0.691414751218265 | 0.923860705630581 | NM_001099328 |
| Dbnl | 2402.110858 | -0.056470416 | 0.082826629 | -0.681790596 | 0.495371371 | 0.705124853 | | Kctd2 | 2650.877016 | -0.0213589671341531 | 0.0540056300227233 | -0.395495194207829 | 0.692477467623639 | 0.924048414626801 | NM_183285 |
| Trp73 | 51.39660746 | -0.114338225 | 0.167753708 | -0.681583892 | 0.495502103 | 0.70524615 | | Lyplal1 | 162.8974637 | 0.0357843976185364 | 0.0904856022914022 | 0.395470624191629 | 0.692495596972832 | 0.924048414626801 | NM_146106 |
| Usp28 | 1193.144654 | 0.060929284 | 0.089411641 | 0.681446882 | 0.495588767 | 0.705304708 | | Tmem176a | 533.289147 | -0.0357238291974732 | 0.0903604918619581 | -0.395347883365308 | 0.692586165741512 | 0.924084379862332 | NM_001098271,NM_025326 |
| 2010002M12Rik | 33.77311532 | 0.114169956 | 0.167606065 | 0.681180335 | 0.495757389 | 0.705449141 | | Ipcef1 | 4819.100153 | 0.0297572259356212 | 0.0753433079371121 | 0.394955129398607 | 0.692876003021592 | 0.924216420559379 | NM_001033391,NM_001170802,NM_001170801,NM_001170800 |
| Smim1 | 457.2804021 | -0.058583579 | 0.086007814 | -0.681142519 | 0.495781315 | 0.705449141 | | Man2a1 | 1031.985349 | 0.030341662643701 | 0.0768140794478752 | 0.395001318271221 | 0.69284191508128 | 0.924216420559379 | NM_008549 |
| Cbs | 1067.763785 | 0.085121357 | 0.124996249 | 0.680991289 | 0.495877002 | 0.705520502 | | Plxdc1 | 2028.460571 | 0.0226498118075656 | 0.0573408689949747 | 0.395002939518595 | 0.692840718592629 | 0.924216420559379 | NM_001163608,NM_028199 |
| Ephb6 | 5020.532607 | -0.087830612 | 0.129049082 | -0.680598503 | 0.496125575 | 0.705679763 | | Cpne7 | 2449.14296 | 0.0368067725604716 | 0.093218502905228 | 0.394844064358046 | 0.692957972894962 | 0.924240888281353 | NM_170684 |
| Rnf223 | 34.54662009 | 0.114001415 | 0.16746958 | 0.680729089 | 0.496042926 | 0.705679763 | | Ccdc28a | 646.4057603 | 0.028002912859833 | 0.0709835697331258 | 0.394498515150964 | 0.693213023247234 | 0.92448250072754 | NM_144820 |
| Zfp575 | 295.2863084 | 0.065083914 | 0.095625086 | 0.680615477 | 0.496114831 | 0.705679763 | | Eif4g3 | 12517.89994 | -0.0177516158008685 | 0.0450061791792682 | -0.394426190460658 | 0.693266410603407 | 0.92448250072754 | NM_172703,NM_001256198,NM_001256195 |
| Ccdc149 | 2551.604648 | 0.087211145 | 0.12815581 | 0.68050871 | 0.496182409 | 0.705695818 | | Rps20 | 3833.126347 | -0.0252631168285431 | 0.0640714383505506 | -0.394296077611406 | 0.693362458831359 | 0.924525709288077 | NM_026147 |
| Zfp934 | 60.96080511 | -0.11133794 | 0.163630886 | -0.680421299 | 0.496237738 | 0.705709731 | | Eapp | 790.6372851 | 0.0259184502583383 | 0.065778618797488 | 0.394025455872419 | 0.693562245339778 | 0.92470722127633 | NM_025456 |
| Nf2 | 4600.747214 | -0.04833596 | 0.071065644 | -0.680159316 | 0.49640359 | 0.705880802 | | Micall1 | 2803.249276 | -0.0275242003683433 | 0.0699964389937552 | -0.393222866248937 | 0.694154882395367 | 0.925412428567832 | NM_177461 |
| Ar | 340.6826411 | -0.08499094 | 0.125002524 | -0.679913794 | 0.496559047 | 0.705999105 | | 9830147E19Rik | 107.3180719 | -0.0373513069270448 | 0.0950108143109165 | -0.393126900321211 | 0.694225756746829 | 0.925421982426763 | NM_001242388 |
| Entpd6 | 2035.345873 | -0.039947942 | 0.058756997 | -0.679883995 | 0.496577917 | 0.705999105 | | Galnt14 | 928.073522 | 0.0255524428120319 | 0.0650208951511734 | 0.392988173303713 | 0.694328216455207 | 0.925473634816328 | NM_027864 |
| Acbd5 | 2582.128235 | -0.039197761 | 0.057689726 | -0.679458262 | 0.496847545 | 0.706089882 | | 6430573F11Rik | 351.9590568 | 0.0321483855162548 | 0.0821918949449854 | 0.391138147353497 | 0.695695124697344 | 0.925527709389081 | NM_176952 |
| Cnot2 | 2212.564471 | 0.032589682 | 0.047972717 | 0.679337837 | 0.496923827 | 0.706089882 | | 9430021M05Rik | 632.8507329 | -0.0293033123290652 | 0.0747982666247693 | -0.391764590963949 | 0.695232160228803 | 0.925527709389081 | NR_033569 |
| Frat1 | 403.8305114 | -0.059811163 | 0.088045056 | -0.679324491 | 0.496932282 | 0.706089882 | | Akr1b8 | 43.63937458 | 0.035115374857521 | 0.0895651751739986 | 0.392065049717172 | 0.695010150653035 | 0.925527709389081 | NM_008012 |
| Hyal3 | 25.56481971 | -0.111406994 | 0.164045013 | -0.679124537 | 0.497058956 | 0.706089882 | | Cdh15 | 85.24311831 | -0.0372038300577997 | 0.0950781700406448 | -0.391297287714894 | 0.695577503462909 | 0.925527709389081 | NM_007662 |
| Nagk | 1130.060745 | -0.048418546 | 0.071339694 | -0.678704137 | 0.497325344 | 0.706089882 | | Dnajc19 | 515.7598156 | -0.0289697333186478 | 0.0738823314536491 | -0.39210637710888 | 0.694979615805865 | 0.925527709389081 | NM_001026211,NM_026332 |
| Pik3cg | 191.7182701 | -0.084855957 | 0.124937484 | -0.679187335 | 0.497019171 | 0.706089882 | | Fat4 | 570.4319975 | 0.0360594045123467 | 0.092130963048418 | 0.3913928967984 | 0.695506841954599 | 0.925527709389081 | NM_183221 |
| Reep3 | 1469.207403 | -0.046861229 | 0.068977802 | -0.679366799 | 0.496905481 | 0.706089882 | | Gabrb1 | 798.8713675 | 0.0278714142456871 | 0.0713598021556157 | 0.390575834065619 | 0.696110791169438 | 0.925527709389081 | NM_008069 |
| Rnf214 | 2482.895318 | 0.05063386 | 0.074559122 | 0.679110193 | 0.497068044 | 0.706089882 | | Grk6 | 3054.952951 | 0.0214188205100508 | 0.0547705537276506 | 0.391064523768683 | 0.695749542640672 | 0.925527709389081 | NM_011938,NM_001038018,NM_001112711 |
| Robo4 | 345.5569146 | 0.071926513 | 0.105839741 | 0.679579448 | 0.496770787 | 0.706089882 | | H2-Bl | 35.3620011 | -0.0341586767009094 | 0.0871338356191006 | -0.392025399297601 | 0.695039446930183 | 0.925527709389081 | NM_008199 |
| Rpp21 | 329.9904098 | 0.057542889 | 0.084765608 | 0.678847124 | 0.497234732 | 0.706089882 | | Ipo9 | 9290.927526 | -0.017847764445397 | 0.045482126097583 | -0.392412711910262 | 0.694753294998371 | 0.925527709389081 | NM_153774 |
| Sbf2 | 2792.065342 | 0.059826722 | 0.088076698 | 0.679257096 | 0.496974976 | 0.706089882 | | Lurap1l | 331.0264656 | 0.0335251521307068 | 0.0857999564793558 | 0.390736237013981 | 0.695992210716973 | 0.925527709389081 | NM_026821 |
| Srm | 1518.895173 | -0.047491712 | 0.069937121 | -0.679063022 | 0.497097931 | 0.706089882 | | Madd | 12085.89484 | -0.0209846351785605 | 0.0535774836119431 | -0.391668920671048 | 0.69530285668661 | 0.925527709389081 | NM_145527,NM_001177720,NM_001177719 |
| Tle3 | 1706.450691 | -0.058459421 | 0.086107871 | -0.67890914 | 0.497195434 | 0.706089882 | | March9 | 662.2350963 | -0.0269812055109881 | 0.0689534035764928 | -0.391296210361201 | 0.695578299714418 | 0.925527709389081 | NM_001033262 |
| Tmem256 | 824.4771174 | -0.052432146 | 0.077237034 | -0.678847222 | 0.49723467 | 0.706089882 | | Msl3l2 | 433.39366 | 0.0295383775178116 | 0.0752246951232315 | 0.392668623906318 | 0.694564247522795 | 0.925527709389081 | NM_001163833 |
| Ttc26 | 418.1469905 | 0.053874198 | 0.079370336 | 0.678769935 | 0.497283646 | 0.706089882 | | Nek7 | 1830.046854 | 0.0266446741244293 | 0.0681506972470956 | 0.390967006952593 | 0.695821623376265 | 0.925527709389081 | NM_021605 |
| Cd164l2 | 105.1785953 | 0.082834674 | 0.122101788 | 0.67840672 | 0.49751385 | 0.706229334 | | Pced1a | 856.4742029 | -0.0261665313841763 | 0.0669624388133486 | -0.390764312768133 | 0.695971456029551 | 0.925527709389081 | NM_178762,NM_001114541 |
| Lcp2 | 132.6095193 | 0.100106601 | 0.147561636 | 0.678405336 | 0.497514727 | 0.706229334 | | Pola1 | 183.4866778 | 0.0343990022087008 | 0.0875680200054212 | 0.392826082016828 | 0.6944479394044 | 0.925527709389081 | NM_008892 |
| Gm6981 | 33.44942253 | 0.108505841 | 0.160037032 | 0.678004578 | 0.497768791 | 0.706525251 | | Ppap2c | 130.2914855 | 0.03715976754106 | 0.0946911124822295 | 0.392431417975301 | 0.69473947579859 | 0.925527709389081 | NM_015817 |
| Acp2 | 3657.289588 | 0.036795179 | 0.054282895 | 0.677841132 | 0.497872428 | 0.706607622 | | Prim1 | 236.1473986 | -0.0358603831802667 | 0.0914132432969511 | -0.392288708800935 | 0.694844905484944 | 0.925527709389081 | NM_008921 |
| Myct1 | 31.35777722 | -0.113901694 | 0.168081502 | -0.677657521 | 0.497988866 | 0.706685488 | | Rabepk | 786.637916 | -0.0259167898773234 | 0.0663485440848684 | -0.390615803779695 | 0.696081242222114 | 0.925527709389081 | NM_145522 |
| Nat8 | 48.48660238 | 0.108489245 | 0.160105549 | 0.677610774 | 0.498018513 | 0.706685488 | | Relt | 829.6282319 | 0.0259099942114798 | 0.0659807214734126 | 0.392690374292443 | 0.694548180939182 | 0.925527709389081 | NM_177073 |
| 2900008C10Rik | 147.6573109 | 0.082562855 | 0.12191556 | 0.677213436 | 0.498270544 | 0.706913635 | | Rpl7 | 8751.792771 | 0.0209570471300174 | 0.0536381222750987 | 0.390711796780154 | 0.696010278072002 | 0.925527709389081 | NM_011291 |
| Hax1 | 1454.551465 | -0.058423862 | 0.086265361 | -0.677257497 | 0.498242593 | 0.706913635 | | Scrn2 | 129.3806998 | 0.0363670908744979 | 0.0928001798444407 | 0.391885995646338 | 0.695142450917878 | 0.925527709389081 | NM_146027 |
| Atr | 625.7483048 | 0.076173788 | 0.112508689 | 0.677048043 | 0.498375472 | 0.706925704 | | Ssx2ip | 3658.834567 | 0.024991581699607 | 0.0639150006007552 | 0.391012774226771 | 0.695787793599024 | 0.925527709389081 | NM_138744,NM_001253769,NM_001253770,NM_001253768 |
| Rhoq | 1524.833155 | -0.059773239 | 0.088302772 | -0.676912372 | 0.498461554 | 0.706925704 | | St3gal6 | 506.0283028 | -0.0297365530973199 | 0.076049910553506 | -0.391013649863511 | 0.695787146360876 | 0.925527709389081 | NM_018784 |
| Smcr7 | 549.8537263 | 0.056730942 | 0.083789488 | 0.677065148 | 0.49836462 | 0.706925704 | | Wwc2 | 1871.585389 | 0.0219527644578946 | 0.0561287611413664 | 0.39111435940309 | 0.695712707089249 | 0.925527709389081 | NM_133791 |
| Tom1l2 | 15169.7077 | 0.04228695 | 0.062469258 | 0.676924163 | 0.498454072 | 0.706925704 | | Zfp512 | 2100.601185 | 0.0214847857990166 | 0.0549234253229662 | 0.391177091972681 | 0.695666339907373 | 0.925527709389081 | NM_172993 |
| Ifngr1 | 649.8619592 | 0.047555694 | 0.070305793 | 0.67641218 | 0.498778986 | 0.707181699 | | Zfp955b | 783.3153974 | -0.0309854805762094 | 0.0793443622447877 | -0.390518994665496 | 0.696152812390548 | 0.925527709389081 | NM_001142957 |
| Pcolce | 269.2047136 | -0.100607071 | 0.148725856 | -0.676459854 | 0.498748727 | 0.707181699 | | 3010001F23Rik | 35.27948868 | -0.0346209558719712 | 0.0887576495153357 | -0.390061657344693 | 0.696490954621707 | 0.925553710258432 | NR_045452,NR_045451 |
| Thbs1 | 125.4159506 | 0.11352071 | 0.167809857 | 0.676484158 | 0.498733301 | 0.707181699 | | Git2 | 1528.896096 | 0.0214583152882654 | 0.0549672525315259 | 0.390383624794749 | 0.696252894740784 | 0.925553710258432 | NM_001077359,NM_001077360,NM_019834 |
| Ndufaf2 | 422.8828843 | -0.057665789 | 0.085278547 | -0.67620511 | 0.498910428 | 0.707303338 | | Slc39a13 | 917.0454348 | -0.0278278134615769 | 0.0713004222000797 | -0.390289602822938 | 0.696322410662003 | 0.925553710258432 | NM_026721 |
| Gabarapl1 | 23987.59184 | -0.037713735 | 0.055779365 | -0.676123429 | 0.498962283 | 0.707312133 | | Supv3l1 | 903.2659824 | 0.0249166098978966 | 0.0638744882699419 | 0.39008703745063 | 0.696472187710342 | 0.925553710258432 | NM_181423 |
| Slain1 | 1858.446771 | 0.06541958 | 0.096789479 | 0.675895566 | 0.499106953 | 0.707452487 | | Syt6 | 1559.464452 | 0.0331106605616613 | 0.0848573745781115 | 0.390191904077621 | 0.696394647738643 | 0.925553710258432 | NM_001276679,NM_018800,NM_001276680,NM_001276681,NM_001276677,NM_001276676 |
| BC022687 | 480.588446 | -0.061284354 | 0.090688275 | -0.675769328 | 0.499187111 | 0.707501382 | | Hrh2 | 248.5009117 | -0.0340056147182642 | 0.0872005986208208 | -0.389969968739925 | 0.696558753834555 | 0.925559134485981 | NM_001010973,NM_008286 |
| Aprt | 662.7637505 | -0.048744716 | 0.072191317 | -0.675215765 | 0.499538692 | 0.707924612 | | Clta | 5409.761339 | 0.0147593516563084 | 0.0378832011647139 | 0.389601490965234 | 0.696831249465802 | 0.925836525175179 | NM_016760,NM_001080384,NM_001080385,NM_001080386 |
| Birc5 | 29.62744576 | -0.109176504 | 0.161740225 | -0.675011453 | 0.499668488 | 0.707924612 | | Cckbr | 2222.692141 | -0.0291549122598948 | 0.0748972330031721 | -0.389265545479631 | 0.697079721005942 | 0.925997259015301 | NM_007627 |
| Mfap1b | 167.2659605 | 0.076534282 | 0.113379472 | 0.675027682 | 0.499658178 | 0.707924612 | | Tesk1 | 4154.805602 | 0.0225769086891827 | 0.0579886434383009 | 0.389333278906656 | 0.697029621485235 | 0.925997259015301 | NM_011571 |
| Pcdhb4 | 83.5469653 | 0.099891261 | 0.147983499 | 0.67501621 | 0.499665466 | 0.707924612 | | Fabp5 | 2735.544513 | 0.0230120477416709 | 0.0591832537451753 | 0.388827012464601 | 0.69740411708544 | 0.926115799890216 | NM_010634,NM_001272098,NM_001272097 |
| Fam184a | 1074.633142 | -0.085720288 | 0.127016958 | -0.674872781 | 0.499756595 | 0.707984701 | | Gle1 | 2481.239133 | 0.017058485944764 | 0.0438846405753917 | 0.387711989459235 | 0.697489212226699 | 0.926115799890216 | NM_028923 |
| Adamts16 | 161.0495322 | 0.100905745 | 0.14956402 | 0.674665905 | 0.499888051 | 0.70810314 | | Lonp1 | 2999.289013 | -0.0241409037215682 | 0.0620674923653929 | -0.3889460134695 | 0.697316082986756 | 0.926115799890216 | NM_028782 |
| Gm2061 | 137.8211397 | -0.078017227 | 0.115662389 | -0.674525461 | 0.499977304 | 0.70810314 | | Lrp1 | 16774.69883 | -0.0285342404375053 | 0.0734230534157182 | -0.388627809796273 | 0.697551491573651 | 0.926115799890216 | NM_008512 |
| Nsl1 | 56.10229366 | 0.103511294 | 0.153448585 | 0.674566624 | 0.499951144 | 0.70810314 | | Rps7 | 7373.48833 | 0.0218517537628372 | 0.0562098598499624 | 0.388753037655044 | 0.6974588439344 | 0.926115799890216 | NM_011300 |
| Aggf1 | 2507.456949 | -0.035177523 | 0.052159208 | -0.674425942 | 0.500040554 | 0.708127991 | | Trmt10c | 448.1149207 | -0.0323315154360567 | 0.0831695652341612 | -0.388742147984402 | 0.697466900287574 | 0.926115799890216 | NM_029092 |
| Fam134c | 3055.710573 | -0.047224347 | 0.070030126 | -0.674343311 | 0.500093075 | 0.708137644 | | Nradd | 50.61781557 | -0.0365241623056964 | 0.0940441401751803 | -0.388372547589474 | 0.697740356810737 | 0.9261972423497 | NM_026012 |
| Chrna10 | 25.70431038 | -0.10825703 | 0.160562564 | -0.674235806 | 0.500161409 | 0.708169685 | | Zfp637 | 993.9707072 | 0.0241338373547083 | 0.0621408874262716 | 0.38837291120669 | 0.697740087761697 | 0.9261972423497 | NM_177684 |
| Fyttd1 | 2470.862867 | -0.046237429 | 0.06862353 | -0.673783891 | 0.500448718 | 0.708447001 | | Bhlhb9 | 2827.598845 | 0.0273478282850102 | 0.0706861572068082 | 0.386890861884004 | 0.698837006538487 | 0.926409492782556 | NM_198161,NM_001098222 |
| Ksr1 | 1630.779762 | 0.08417143 | 0.124918287 | 0.673811911 | 0.500430901 | 0.708447001 | | Ccdc85b | 2000.741521 | -0.0276375598273153 | 0.0713933601260085 | -0.387116669933105 | 0.698669837085543 | 0.926409492782556 | NM_001243307,NM_198616 |
| B230378P21Rik | 10.94163064 | 0.098094216 | 0.145649491 | 0.673495084 | 0.500632376 | 0.708642246 | | Dock5 | 640.9229967 | 0.0346236642547666 | 0.0894242239352002 | 0.387184397371523 | 0.698619700185171 | 0.926409492782556 | NM_177780 |
| Lrfn2 | 1027.49568 | -0.078419834 | 0.116457544 | -0.673377017 | 0.500707467 | 0.708673794 | | Fam105b | 609.8190101 | 0.0239422852989286 | 0.0617198837170216 | 0.387918509514716 | 0.698076340032681 | 0.926409492782556 | NM_001013792 |
| Mavs | 351.3194906 | 0.060035676 | 0.089168381 | 0.673284357 | 0.500766403 | 0.708702471 | | Galnt7 | 530.5643584 | 0.0310613946756983 | 0.0800999321018438 | 0.387783033775922 | 0.698176602067893 | 0.926409492782556 | NM_001167981,NM_144731 |
| 1810026B05Rik | 257.3006401 | 0.062690012 | 0.093173516 | 0.672830802 | 0.501054939 | 0.708899185 | | Gm14446 | 162.5608299 | 0.0356434617714139 | 0.0919542615242293 | 0.387621641244132 | 0.698296051315926 | 0.926409492782556 | NM_001110517,NM_001101605 |
| Alad | 1011.09613 | 0.038651743 | 0.057448059 | 0.672811992 | 0.501066908 | 0.708899185 | | Gsta4 | 1327.874971 | 0.0217550892458893 | 0.0561565982857597 | 0.387400410815232 | 0.698459799710608 | 0.926409492782556 | NM_010357 |
| Chek1 | 36.6499643 | 0.107089065 | 0.159235192 | 0.672521344 | 0.501251857 | 0.708899185 | | Marf1 | 7072.053541 | 0.0270706965380512 | 0.069960640417986 | 0.386941805797016 | 0.698799290636636 | 0.926409492782556 | NM_001081154 |
| Cmpk2 | 1041.00115 | -0.042202774 | 0.062755932 | -0.672490595 | 0.501271426 | 0.708899185 | | Mitf | 334.7660166 | -0.0301208159779212 | 0.0778762670395614 | -0.386778606867672 | 0.698920668553217 | 0.926409492782556 | NM_001178049,NM_008601,NM_001113198 |
| Hic2 | 417.8648384 | 0.082403223 | 0.122502479 | 0.672665758 | 0.501159957 | 0.708899185 | | Mylk | 871.0037769 | -0.0309470072480329 | 0.0800032365560827 | -0.386821940964087 | 0.698888032750855 | 0.926409492782556 | NM_139300 |
| Pacsin2 | 2888.70102 | -0.073203915 | 0.108802882 | -0.672812273 | 0.501066729 | 0.708899185 | | Oip5 | 30.54585769 | -0.0328910222735683 | 0.0849597411042132 | -0.387136564284295 | 0.698655109666902 | 0.926409492782556 | NM_001042653 |
| Pigb | 283.8421601 | 0.066363003 | 0.098658107 | 0.672656364 | 0.501165934 | 0.708899185 | | Pik3r4 | 2273.824185 | 0.0186220094027074 | 0.0480232784699302 | 0.387770472904459 | 0.69818589830573 | 0.926409492782556 | NM_001081309 |
| Tpmt | 365.5804214 | -0.054787744 | 0.081458999 | -0.672580609 | 0.501214141 | 0.708899185 | | Sh3rf3 | 1476.488946 | 0.0308688233186388 | 0.0796832012796273 | 0.38739436697972 | 0.698464273380639 | 0.926409492782556 | NM_172788 |
| BC030867 | 41.06606576 | 0.109691075 | 0.163135234 | 0.672393528 | 0.501333202 | 0.708921843 | | Tsr2 | 3590.564462 | 0.0219085321686276 | 0.0564850019874322 | 0.387864590559849 | 0.698116243403553 | 0.926409492782556 | NR_028392,NM_175146,NM_001164578 |
| Elavl2 | 2462.61656 | 0.049074234 | 0.073038406 | 0.67189629 | 0.501649722 | 0.709201588 | | Uevld | 697.4532374 | -0.0277432145123704 | 0.0717042309446203 | -0.386911820221564 | 0.698821490117759 | 0.926409492782556 | NM_001040695 |
| Gfod1 | 6825.624607 | 0.089710547 | 0.133538547 | 0.671795142 | 0.501714122 | 0.709201588 | | Vps37c | 919.1336124 | 0.02416748674899 | 0.0624222415044402 | 0.387161469478326 | 0.698636672974263 | 0.926409492782556 | NM_181403 |
| Supv3l1 | 815.9103371 | 0.0495101 | 0.073683246 | 0.671931577 | 0.501627257 | 0.709201588 | | 6330418K02Rik | 103.1578767 | -0.0367433961794513 | 0.0950805234552744 | -0.386445034631464 | 0.699167102742752 | 0.926567040821204 | NR_045821 |
| Tdrd9 | 27.93414474 | 0.111086267 | 0.165351082 | 0.67182062 | 0.5016979 | 0.709201588 | | Mttp | 636.2254763 | 0.0320510465623776 | 0.08292325490177 | 0.386514574233066 | 0.699115610986346 | 0.926567040821204 | NM_008642,NM_001163457 |
| Cul3 | 4498.466325 | -0.048274243 | 0.071878405 | -0.671609822 | 0.501832124 | 0.709238979 | | Ftsjd2 | 5102.46938 | 0.0183397905306725 | 0.0474866377831536 | 0.386209497804849 | 0.699341520189401 | 0.926713640053935 | NM_028791 |
| Mgat4a | 1910.784035 | 0.045821082 | 0.068221692 | 0.671649749 | 0.501806699 | 0.709238979 | | Acss2 | 2236.448779 | 0.0220074823706472 | 0.0570484064241715 | 0.385768573569175 | 0.699668071775057 | 0.926781946381661 | NM_019811 |
| Tpo | 14.87062312 | -0.111443229 | 0.165993689 | -0.671370274 | 0.501984677 | 0.709389874 | | Aim1 | 61.89689853 | 0.0364766300862822 | 0.0946010060579736 | 0.385583955248092 | 0.699804817902214 | 0.926781946381661 | NM_172393 |
| Calm1 | 122655.312 | -0.043596818 | 0.065009068 | -0.670626717 | 0.502458358 | 0.709756076 | | Arpc5 | 2892.799947 | 0.0270072538645291 | 0.0700917618910449 | 0.385312811889518 | 0.700005670460092 | 0.926781946381661 | NM_026369 |
| Dlgap4 | 11175.95011 | -0.049099026 | 0.073222667 | -0.670544089 | 0.502511011 | 0.709756076 | | Cst6 | 149.307521 | -0.0349196498609774 | 0.0905573645366012 | -0.38560806224505 | 0.699786961383513 | 0.926781946381661 | NM_028623 |
| E2f5 | 288.1378865 | -0.076314653 | 0.113836521 | -0.670388135 | 0.502610396 | 0.709756076 | | Nr4a2 | 2329.717604 | 0.03330969985554 | 0.0864495175392939 | 0.385308105859581 | 0.700009156691142 | 0.926781946381661 | NM_013613,NM_001139509 |
| Figf | 87.47033351 | -0.090089064 | 0.134326303 | -0.670673296 | 0.502428678 | 0.709756076 | | Ppp2r5a | 3411.370689 | -0.024908183288996 | 0.0646351547990694 | -0.385365879704749 | 0.699966358214231 | 0.926781946381661 | NM_144880 |
| Impact | 9458.615061 | -0.049250785 | 0.073458779 | -0.670454715 | 0.502567965 | 0.709756076 | | Slc16a12 | 141.9235147 | 0.0367000735565285 | 0.0950968307333926 | 0.38592320347056 | 0.699553545456795 | 0.926781946381661 | NM_172838 |
| Kpna6 | 5357.157729 | -0.032088532 | 0.047870723 | -0.670316443 | 0.502656088 | 0.709756076 | | Sp2 | 749.7916449 | 0.0245732460366833 | 0.0637537671945914 | 0.38543990603222 | 0.699911521400116 | 0.926781946381661 | NM_030220,NM_001080964 |
| Mrps23 | 1172.19521 | -0.053830078 | 0.080300039 | -0.670361792 | 0.502627185 | 0.709756076 | | Tmem100 | 403.6363953 | 0.0310612556782378 | 0.0805913073863852 | 0.385416947380174 | 0.699928528416243 | 0.926781946381661 | NM_026433 |
| Mtmr3 | 4802.642707 | 0.047189878 | 0.070367437 | 0.670620955 | 0.50246203 | 0.709756076 | | Xpc | 821.4690657 | -0.0232553328634182 | 0.0603597979300475 | -0.385278507565076 | 0.700031083278231 | 0.926781946381661 | NM_009531 |
| Parp11 | 772.7999301 | -0.057095685 | 0.085143087 | -0.670585092 | 0.502484882 | 0.709756076 | | Bbox1 | 75.72484481 | 0.0354749691235543 | 0.0921011260705029 | 0.38517410847288 | 0.700108424723463 | 0.926799870366629 | NM_130452 |
| Msx2 | 9.676149083 | 0.10471414 | 0.156235091 | 0.67023445 | 0.502708346 | 0.709765183 | | Actr3b | 3699.222022 | 0.0204123215392808 | 0.0530544057706515 | 0.384743194137751 | 0.700427689689038 | 0.927053541724188 | NM_001004365 |
| Pop1 | 312.5982026 | 0.062983107 | 0.093992537 | 0.670086252 | 0.502802808 | 0.709833869 | | Tmem87b | 1729.103327 | -0.0194959036791732 | 0.0506647212490253 | -0.384802347640435 | 0.700383859649774 | 0.927053541724188 | NM_028248 |
| Dnajc2 | 1060.392244 | -0.044849501 | 0.066939773 | -0.669997799 | 0.502859193 | 0.709848792 | | Gab1 | 1477.560951 | -0.0322072801081776 | 0.083788779999291 | -0.384386550424175 | 0.700691967497489 | 0.927195030301845 | NM_021356 |
| 9530051G07Rik | 25.90577971 | -0.110119645 | 0.164416206 | -0.669761504 | 0.503009837 | 0.709867422 | | Slc38a7 | 1455.057658 | -0.0218987200941256 | 0.0569644332988938 | -0.384427946104941 | 0.700661290879754 | 0.927195030301845 | NM_172758 |
| Dpp9 | 2852.17505 | -0.04951136 | 0.073919533 | -0.669800773 | 0.502984801 | 0.709867422 | | Wdr74 | 915.4819834 | 0.0235678392121378 | 0.0613202045305561 | 0.384340518253972 | 0.70072608031486 | 0.927195030301845 | NM_134139 |
| Nyap2 | 594.4297271 | -0.068749988 | 0.102632721 | -0.669864226 | 0.502944346 | 0.709867422 | | Kdm5a | 2173.544979 | -0.029337441349342 | 0.0763758277991295 | -0.384119454999562 | 0.700889912544115 | 0.927327340342091 | NM_145997 |
| Ces2g | 10.95154869 | -0.10416784 | 0.155624706 | -0.669352848 | 0.503270423 | 0.70997253 | | 2900041M22Rik | 27.13538567 | -0.0324585430949429 | 0.0845257687557945 | -0.384007665031946 | 0.700972766381449 | 0.927352495852179 | NR_015489 |
| Gjb1 | 446.1865323 | 0.102863419 | 0.153643049 | 0.669496079 | 0.503179081 | 0.70997253 | | Cox15 | 1214.126381 | 0.0208403371057923 | 0.0543043789896624 | 0.383768997887989 | 0.701149667937603 | 0.927502055956799 | NM_144874 |
| Mrps18b | 757.3434769 | 0.04784847 | 0.071493438 | 0.669270797 | 0.503322752 | 0.70997253 | | Akap9 | 3850.443199 | 0.0309363071173434 | 0.0806903999969752 | 0.383395138932303 | 0.701426807064493 | 0.92751464986683 | NM_194462 |
| Nlgn2 | 11191.40095 | 0.031066762 | 0.046426362 | 0.669162088 | 0.503392088 | 0.70997253 | | Aldh1b1 | 384.6247408 | 0.0333946826118245 | 0.0871028896981513 | 0.383393509992049 | 0.701428014673884 | 0.92751464986683 | NM_028270 |
| Ogfrl1 | 4331.60472 | -0.043732319 | 0.065355203 | -0.669148245 | 0.503400917 | 0.70997253 | | Casq2 | 99.30415449 | -0.034702276918645 | 0.0905809573415055 | -0.383107862150447 | 0.701639790402179 | 0.92751464986683 | NM_009814 |
| Osbp | 3134.146657 | -0.044573331 | 0.066582804 | -0.669442075 | 0.50321352 | 0.70997253 | | Depdc7 | 58.01859656 | 0.0362765721329348 | 0.0945516857479051 | 0.383669226476362 | 0.701223623925203 | 0.92751464986683 | NM_144804 |
| Spg21 | 1194.089071 | 0.043940716 | 0.065667278 | 0.669141727 | 0.503405074 | 0.70997253 | | Gatc | 1337.303643 | -0.0228674805285476 | 0.0596764143575627 | -0.383191261987236 | 0.701577956412626 | 0.92751464986683 | NM_029645 |
| Fbxl16 | 43855.31174 | -0.06015995 | 0.089933785 | -0.668936041 | 0.503536279 | 0.710079244 | | Hacl1 | 240.2417037 | 0.0322921613526384 | 0.0843368849903226 | 0.382894878751377 | 0.701797708707581 | 0.92751464986683 | NM_019975 |
| Sox18 | 206.1547186 | -0.102445223 | 0.153159478 | -0.668879422 | 0.503572398 | 0.710079244 | | Mgat4a | 2058.318255 | 0.0215898863800887 | 0.0563780526033388 | 0.382948423777413 | 0.701758006092371 | 0.92751464986683 | NM_173870 |
| Klhl26 | 2033.278825 | 0.051605882 | 0.077165225 | 0.668771229 | 0.503641423 | 0.710111949 | | Phkb | 1910.919414 | -0.0234950712247177 | 0.061345624992394 | -0.382995058370189 | 0.701723428094848 | 0.92751464986683 | NM_199446 |
| Rtf1 | 4327.60451 | 0.062015958 | 0.092756708 | 0.668587304 | 0.503758774 | 0.710129757 | | Src | 5477.919689 | -0.0238371548740721 | 0.0621675045099168 | -0.383434320904253 | 0.701397759868315 | 0.92751464986683 | NM_001025395,NM_009271 |
| Slco3a1 | 1453.743363 | 0.059859682 | 0.089538465 | 0.668535936 | 0.503791551 | 0.710129757 | | Wapal | 1807.561045 | 0.0263371993025022 | 0.0687316866994614 | 0.383188607281896 | 0.701579924623707 | 0.92751464986683 | NM_001004436 |
| Zbtb44 | 887.5538306 | -0.04605729 | 0.068891649 | -0.668546782 | 0.50378463 | 0.710129757 | | E030003E18Rik | 338.8016284 | 0.0362991747615178 | 0.0948608858955749 | 0.382656923544619 | 0.701974157798695 | 0.927663447614979 | NR_015502 |
| Peli1 | 1350.465465 | -0.051217819 | 0.076647272 | -0.668227555 | 0.503988349 | 0.710277922 | | Ralgapa2 | 1230.757632 | 0.0240360901445841 | 0.0628443461136201 | 0.382470208236837 | 0.702112622612813 | 0.927762026387131 | NM_001033348 |
| Psmc4 | 3201.606307 | 0.047085127 | 0.070461701 | 0.668237152 | 0.503982224 | 0.710277922 | | 9430016H08Rik | 405.5448671 | 0.0288124178451805 | 0.0755713868902621 | 0.381260937913171 | 0.703009635613755 | 0.927855160570584 | NM_001081181 |
| A930005H10Rik | 206.330726 | 0.067641342 | 0.101313174 | 0.667646071 | 0.504359542 | 0.710709078 | | BC021785 | 31.81521089 | -0.0341471698312285 | 0.0895549613478182 | -0.381298470987062 | 0.702981788089813 | 0.927855160570584 | NM_001113388,NM_001001489,NM_001113385 |
| Zfp658 | 260.8800028 | 0.078024101 | 0.116871726 | 0.667604598 | 0.504386022 | 0.710709078 | | Bbs5 | 601.2924875 | -0.0285643421303243 | 0.0748376837211609 | -0.381683941966358 | 0.702695812407239 | 0.927855160570584 | NM_028284 |
| 9330117O12Rik | 76.88595692 | 0.11212559 | 0.168005001 | 0.667394356 | 0.50452027 | 0.710734511 | | Col6a5 | 86.77813917 | 0.0356697128498565 | 0.0934220322379063 | 0.381812640930576 | 0.702600341760687 | 0.927855160570584 | NM_001167923 |
| Cit | 3880.426694 | 0.046068181 | 0.069037182 | 0.667295216 | 0.504583581 | 0.710734511 | | Drd5 | 84.58536221 | -0.0340871942778579 | 0.0893289841108357 | -0.381591648188498 | 0.702764280077772 | 0.927855160570584 | NM_013503 |
| F2r | 194.0571893 | 0.069849632 | 0.104664324 | 0.667368112 | 0.504537029 | 0.710734511 | | Ep400 | 3993.024519 | -0.0224119163552719 | 0.0588074198354382 | -0.381106949054856 | 0.703123891242825 | 0.927855160570584 | NM_173066,NM_029337 |
| Flot2 | 4542.176595 | -0.055776444 | 0.083592173 | -0.667244809 | 0.504615773 | 0.710734511 | | Fmn2 | 3857.004185 | 0.0205746421310355 | 0.0539882629415005 | 0.381094723372178 | 0.703132962660904 | 0.927855160570584 | NM_019445 |
| Gnas | 45382.66248 | 0.026489301 | 0.039705443 | 0.667145341 | 0.504679301 | 0.710734511 | | Klhl28 | 552.3376202 | 0.0285669142191136 | 0.0749623345437541 | 0.381083572076315 | 0.703141236924204 | 0.927855160570584 | NM_025707 |
| Plag1 | 136.1812689 | 0.087822319 | 0.131632266 | 0.667179268 | 0.504657632 | 0.710734511 | | Myh15 | 76.44030467 | 0.0361356203131218 | 0.0948149432389147 | 0.381117354276817 | 0.703116170634163 | 0.927855160570584 | NM_001166210 |
| Serpina3g | 10.70017179 | 0.078740231 | 0.118094834 | 0.666754235 | 0.50492913 | 0.711021715 | | Plp2 | 261.2415118 | -0.0324962032085587 | 0.0851777573374232 | -0.381510434465047 | 0.702824530060726 | 0.927855160570584 | NM_019755 |
| Dynlt1b | 37.24422471 | -0.111925765 | 0.167931873 | -0.666495068 | 0.505094715 | 0.711125626 | | Pop5 | 1371.083626 | 0.0235539360086605 | 0.0618004138903837 | 0.381129098106664 | 0.703107456823725 | 0.927855160570584 | NM_026398 |
| Pex11b | 1510.497518 | 0.045639187 | 0.06847025 | 0.666555 | 0.505056421 | 0.711125626 | | Qk | 9641.975165 | -0.0252046035124482 | 0.0660036570286055 | -0.381866772950545 | 0.70256018729162 | 0.927855160570584 | NM_021881,NM_001159516,NM_001159517 |
| Dhps | 1024.364 | 0.047786738 | 0.071708958 | 0.66639844 | 0.505156459 | 0.711147936 | | Rad51c | 134.6666012 | 0.0340684210162644 | 0.0891692258818514 | 0.382064786461684 | 0.702413310367682 | 0.927855160570584 | NM_053269 |
| 4930452B06Rik | 518.6956291 | -0.053557077 | 0.080405419 | -0.666087902 | 0.505354917 | 0.711233452 | | Rps25 | 5319.221594 | -0.0244095201815184 | 0.0639048075309884 | -0.381966884704815 | 0.702485927806175 | 0.927855160570584 | NM_024266 |
| Ing4 | 1314.742555 | 0.047052411 | 0.070632683 | 0.666156365 | 0.505311161 | 0.711233452 | | Zmat4 | 2609.941324 | 0.0249570714224251 | 0.0652936902288868 | 0.382227917505324 | 0.702292315923223 | 0.927855160570584 | NM_177086 |
| Psmd5 | 1672.3725 | 0.050233529 | 0.075405133 | 0.666181821 | 0.505294891 | 0.711233452 | | Nufip2 | 1988.547595 | -0.025366827842159 | 0.0665905407435008 | -0.380937405807669 | 0.703249695569431 | 0.927913986542061 | NM_001024205 |
| 2310040G24Rik | 30.67612911 | 0.109945273 | 0.165112844 | 0.665879589 | 0.505488068 | 0.711356233 | | F930015N05Rik | 76.12422523 | 0.0361521168309138 | 0.0949380005141242 | 0.380797116382658 | 0.703353799144899 | 0.927967055983543 | NR_028445 |
| Ccl25 | 119.5070755 | 0.095292591 | 0.143154451 | 0.66566279 | 0.50562662 | 0.71142204 | | Hells | 65.96714403 | 0.0354415999137377 | 0.0931942286841798 | 0.380298226769423 | 0.703724051653531 | 0.928079891702509 | NM_008234 |
| Ndufa6 | 1943.327302 | -0.051208084 | 0.076921318 | -0.665720313 | 0.505589887 | 0.71142204 | | Mab21l1 | 47.28291418 | 0.0341567451930923 | 0.0897962920568058 | 0.380380352136194 | 0.703663097219474 | 0.928079891702509 | NM_010750 |
| Stradb | 1445.509465 | 0.0417672 | 0.062764797 | 0.665455822 | 0.505758991 | 0.711479009 | | Pcdhb10 | 174.3146878 | 0.0342954503792633 | 0.0901915067995661 | 0.380251440476303 | 0.70375877785372 | 0.928079891702509 | NM_053135 |
| Tnnt1 | 108.0620907 | 0.098918485 | 0.148642494 | 0.665479184 | 0.505744053 | 0.711479009 | | Spns1 | 2465.623301 | -0.0270560529334977 | 0.0711177918009505 | -0.380440003103922 | 0.703618824748223 | 0.928079891702509 | NM_023712,NR_045537 |
| Pdcd7 | 835.9637297 | -0.045995329 | 0.069149419 | -0.66515857 | 0.505949076 | 0.711681796 | | Yaf2 | 2636.947887 | 0.025376616237229 | 0.0667071522080263 | 0.380418221993527 | 0.703634990397743 | 0.928079891702509 | NR_028315,NM_024189 |
| Rarres2 | 168.1464343 | 0.095739281 | 0.143971481 | 0.664987816 | 0.506058286 | 0.711706189 | | Gjc2 | 569.559234 | -0.0338680605699111 | 0.0891293414567852 | -0.379987779740661 | 0.703954486366057 | 0.928127791674073 | NM_080454,NM_175452 |
| Slc2a10 | 80.26386372 | -0.098244808 | 0.147736752 | -0.664999109 | 0.506051063 | 0.711706189 | | Gnb2 | 10319.40907 | 0.024367338161983 | 0.064149074934474 | 0.379854864421246 | 0.704053153388408 | 0.928127791674073 | NM_010312 |
| 4931414P19Rik | 145.8254662 | 0.081946227 | 0.123267401 | 0.664784251 | 0.506188497 | 0.711824703 | | Polr3e | 1564.639121 | 0.0228160554438611 | 0.0600405245826683 | 0.380010927660139 | 0.703937303481611 | 0.928127791674073 | NM_025298,NM_001164096 |
| Col9a2 | 243.6309242 | -0.074043938 | 0.11146294 | -0.664291988 | 0.506503448 | 0.712202961 | | Rpp30 | 298.9269785 | -0.0317163254049355 | 0.0835052450667405 | -0.379812374415363 | 0.704084696043214 | 0.928127791674073 | NM_019428 |
| Gm889 | 16.39239648 | 0.11042917 | 0.166306464 | 0.664010089 | 0.506683854 | 0.712391981 | | Tmem88 | 111.0816766 | -0.0358221814346822 | 0.0943254612736004 | -0.379772130992038 | 0.704114571406326 | 0.928127791674073 | NM_025915 |
| Cend1 | 10170.52643 | -0.03501981 | 0.05279074 | -0.663370316 | 0.507093412 | 0.712838367 | | 1810058I24Rik | 481.4888708 | -0.0298276487089101 | 0.0785807761677385 | -0.379579461587908 | 0.70425760901295 | 0.928232104938038 | NR_027875,NR_015608 |
| Mon2 | 3861.898415 | 0.057654487 | 0.086909648 | 0.663384203 | 0.507084521 | 0.712838367 | | Pde2a | 24756.71641 | -0.0242016804509649 | 0.063814020854736 | -0.379253338479592 | 0.704499746376769 | 0.928382773824635 | NM_001143849,NM_001143848,NM_001008548,NM_001243757,NM_001243758 |
| Pofut2 | 972.0560114 | 0.050187487 | 0.075663496 | 0.66329855 | 0.507139365 | 0.712838367 | | Tmcc2 | 8369.536368 | 0.0214350099384222 | 0.0565130981288039 | 0.3792927772172 | 0.704470462615415 | 0.928382773824635 | NM_178874 |
| 9330182L06Rik | 1354.735189 | -0.051986092 | 0.078416796 | -0.662945889 | 0.50736521 | 0.713085955 | | Tmf1 | 1602.96625 | 0.0255850961108135 | 0.0675039843973792 | 0.379016088297846 | 0.704675916878062 | 0.928530693810842 | NM_001081111 |
| Pak4 | 331.9297388 | -0.059578832 | 0.08987879 | -0.662879776 | 0.507407555 | 0.713085955 | | Papola | 4359.404446 | 0.0284542790350919 | 0.075131093446574 | 0.378728402979065 | 0.704889559321972 | 0.928727958159724 | NM_011112 |
| Msh5 | 9.581838008 | 0.106791547 | 0.161154335 | 0.662666304 | 0.507544295 | 0.713213433 | | Abcb8 | 2049.627655 | 0.024534736066285 | 0.06483953620467 | 0.378391603370505 | 0.705139704866216 | 0.928783752712291 | NM_029020 |
| Cndp1 | 13.34192625 | 0.104297666 | 0.157441752 | 0.6624524 | 0.507681331 | 0.713276618 | | Ift57 | 1516.183053 | 0.0233957220037252 | 0.0618511621889359 | 0.378258405755717 | 0.705238641303187 | 0.928783752712291 | NM_028680 |
| Rxrb | 2497.312601 | 0.037240519 | 0.056215979 | 0.662454332 | 0.507680094 | 0.713276618 | | Kcne1l | 159.3121094 | 0.0359798498002266 | 0.09509646417653 | 0.378351078684022 | 0.705169805238582 | 0.928783752712291 | NM_021487 |
| 2700062C07Rik | 427.7599329 | -0.061498215 | 0.092919301 | -0.661845435 | 0.508070284 | 0.713389644 | | Mier1 | 1144.850262 | -0.0260986086383468 | 0.0690313730586198 | -0.378068803820322 | 0.705379482302216 | 0.928783752712291 | NM_001039081,NM_027696 |
| Aen | 879.2665494 | 0.049901505 | 0.07537438 | 0.662048632 | 0.507940055 | 0.713389644 | | Pxmp2 | 144.6131378 | -0.0355349278653263 | 0.0939865555859692 | -0.378085223400092 | 0.70536728503358 | 0.928783752712291 | NM_008993 |
| Car10 | 5023.263574 | 0.072979808 | 0.110258216 | 0.661899051 | 0.50803592 | 0.713389644 | | Rbp4 | 345.0147695 | 0.031086279182516 | 0.0822228280798417 | 0.378073582586212 | 0.705375932292877 | 0.928783752712291 | NM_001159487,NM_011255 |
| Celf2 | 14229.09489 | -0.037470604 | 0.056587338 | -0.662172939 | 0.507860395 | 0.713389644 | | Serf2 | 4756.04364 | 0.018043435176468 | 0.0476950921476355 | 0.378308005373305 | 0.705201799149168 | 0.928783752712291 | NM_011354 |
| Crkl | 2092.211695 | 0.068307821 | 0.103180086 | 0.662025237 | 0.507955048 | 0.713389644 | | D630037F22Rik | 568.9202018 | -0.0283680929697365 | 0.0751161881035881 | -0.37765618418517 | 0.705686021243078 | 0.929018954556548 | NM_001033385 |
| Gm5176 | 14.39025908 | -0.110402601 | 0.166746547 | -0.662098272 | 0.507908243 | 0.713389644 | | Palm | 8255.780365 | -0.0233175486836282 | 0.0617324965470919 | -0.377719191477064 | 0.705639209452037 | 0.929018954556548 | NM_001161747,NM_023128 |
| Papolg | 783.0237049 | 0.046287627 | 0.069946801 | 0.661754738 | 0.508128418 | 0.713389644 | | Ufd1l | 3117.908842 | 0.0212131869181665 | 0.0562085893015644 | 0.377401162024468 | 0.705875503453102 | 0.929184192402335 | NR_028403,NM_011672 |
| Tnfaip1 | 2356.170738 | -0.038717988 | 0.058508298 | -0.661752079 | 0.50813012 | 0.713389644 | | Klc4 | 940.4031696 | -0.0280635331703591 | 0.0743940318804366 | -0.377228286476821 | 0.706003960862315 | 0.929269077155308 | NM_029091 |
| Bambi-ps1 | 38.26029841 | -0.104440495 | 0.157896574 | -0.661448767 | 0.50832456 | 0.713450923 | | Plekha7 | 733.4470622 | -0.0309872444783409 | 0.0821930891215147 | -0.377005473447155 | 0.70616953731912 | 0.929402799592057 | NM_172743 |
| Fastkd5 | 353.1191091 | 0.089120076 | 0.134745273 | 0.661396675 | 0.508357957 | 0.713450923 | | Cluap1 | 1072.128166 | -0.0218079659802356 | 0.057877923701794 | -0.376792472594514 | 0.706327835170944 | 0.929442715254383 | NM_029738 |
| Maoa | 1284.047643 | -0.052655869 | 0.079606573 | -0.661451275 | 0.508322952 | 0.713450923 | | Sec31a | 4877.130781 | 0.0161139343546479 | 0.0427642736687611 | 0.376808325553742 | 0.706316053140089 | 0.929442715254383 | NM_026969 |
| 1-Sep | 21.21572057 | 0.11107061 | 0.167907928 | 0.661497117 | 0.508293563 | 0.713450923 | | Ddx19b | 1637.38085 | 0.0291139351849212 | 0.0773640286135585 | 0.376323928661321 | 0.706676092086929 | 0.929476901984127 | NM_001190786,NM_001190800,NM_172284 |
| Gls | 14595.87558 | -0.03950478 | 0.059746728 | -0.661204075 | 0.508481448 | 0.713494979 | | Fbn2 | 84.8601326 | 0.0357927924461438 | 0.095064581482414 | 0.376510282673102 | 0.706537572480081 | 0.929476901984127 | NM_010181 |
| Ipcef1 | 4322.602595 | -0.061627562 | 0.093203725 | -0.661213512 | 0.508475397 | 0.713494979 | | Itpkb | 1573.385111 | -0.0258707520149829 | 0.0687144424050021 | -0.376496572037957 | 0.706547763460451 | 0.929476901984127 | NM_001081175 |

| | | | | | | |
|---|---|---|---|---|---|---|
| Ints8 | 1020.09852 | 0.055765045 | 0.084351286 | 0.661104859 | 0.508545069 | 0.713519633 |
| Fbxl21 | 196.2633894 | -0.076841121 | 0.116252726 | -0.660983386 | 0.508622968 | 0.713564313 |
| Fam111a | 104.6795112 | 0.083906883 | 0.126958901 | 0.660897992 | 0.508677734 | 0.713576534 |
| D630023F18Rik | 135.7331276 | 0.081452907 | 0.123349451 | 0.660342681 | 0.509033947 | 0.713970788 |
| Pla2g16 | 1646.521191 | 0.052159074 | 0.078991058 | 0.660316191 | 0.509050942 | 0.713970788 |
| 5430402O13Rik | 17.68293405 | 0.110711856 | 0.167711237 | 0.660133795 | 0.509167973 | 0.714070292 |
| Pot1a | 673.4261343 | 0.039972889 | 0.060566312 | 0.659985522 | 0.50926312 | 0.714139088 |
| Arhgef39 | 30.90244382 | 0.108800081 | 0.164895704 | 0.659811497 | 0.509374804 | 0.714231061 |
| Acaa2 | 775.6814445 | -0.045611337 | 0.069142775 | -0.659668873 | 0.509466346 | 0.714294776 |
| Scand1 | 256.0297069 | -0.110544716 | 0.167669101 | -0.659302847 | 0.509701315 | 0.714559552 |
| Srd5a1 | 381.0906811 | 0.058189768 | 0.088290038 | 0.659075131 | 0.509847525 | 0.71469986 |
| Gm20300 | 1470.080242 | -0.057975235 | 0.087978549 | -0.658970122 | 0.509914955 | 0.714729719 |
| Sdr39u1 | 1880.914238 | 0.044169919 | 0.067043251 | 0.658827225 | 0.510006722 | 0.714793682 |
| Atp8a2 | 1186.789217 | -0.045066529 | 0.068412071 | -0.658751138 | 0.510055589 | 0.714797512 |
| Acot11 | 2217.641878 | -0.044911922 | 0.068212935 | -0.65840771 | 0.510276182 | 0.714904317 |
| Glrx2 | 2004.262127 | -0.036522736 | 0.055464243 | -0.65849156 | 0.510222318 | 0.714904317 |
| Lrp11 | 7320.484085 | 0.041665119 | 0.063274997 | 0.658476824 | 0.510231784 | 0.714904317 |
| Slamf1 | 10.01821087 | -0.102334892 | 0.155442613 | -0.658345162 | 0.510316364 | 0.714904317 |
| Asap2 | 2096.4364 | -0.055209727 | 0.083904847 | -0.658004019 | 0.510535548 | 0.715146713 |
| BC005624 | 2630.260416 | 0.035681332 | 0.054246569 | 0.657762002 | 0.510691074 | 0.715235244 |
| Cdc5l | 2395.855423 | 0.059046842 | 0.089768533 | 0.657767708 | 0.510687407 | 0.715235244 |
| Grin2d | 423.2078632 | -0.072146485 | 0.109738563 | -0.657439676 | 0.510898247 | 0.715460723 |
| Sgtb | 5588.182931 | -0.044001447 | 0.066966308 | -0.657068438 | 0.511136912 | 0.715730259 |
| Plcxd2 | 4866.534976 | 0.04618678 | 0.070314334 | 0.656861508 | 0.511269971 | 0.715851882 |
| Nupl2 | 324.9585881 | 0.062066563 | 0.094500389 | 0.656786324 | 0.511318319 | 0.715854887 |
| Stk40 | 1501.653346 | 0.068828464 | 0.104815264 | 0.656664514 | 0.511396657 | 0.715899874 |
| 9230114K14Rik | 74.72735619 | -0.097603388 | 0.148707796 | -0.656343451 | 0.511603166 | 0.715959495 |
| Efcab10 | 38.59291579 | 0.104703964 | 0.15951797 | 0.656377232 | 0.511581436 | 0.715959495 |
| Ficd | 569.7072464 | -0.047824657 | 0.0728484 | -0.656495647 | 0.511505267 | 0.715959495 |
| Frmd4a | 1934.257791 | -0.051296985 | 0.078159576 | -0.656310936 | 0.511624082 | 0.715959495 |
| Hn1l | 213.5734729 | -0.059117318 | 0.090159155 | -0.65569956 | 0.512017452 | 0.716445263 |
| Slc27a5 | 11.64357807 | 0.107210503 | 0.16352373 | 0.655626574 | 0.512064423 | 0.716446286 |
| Dnajc17 | 281.3286903 | -0.053439491 | 0.081521282 | -0.655528107 | 0.512127796 | 0.716470255 |
| Josd1 | 4038.679012 | -0.051677143 | 0.078875202 | -0.655176047 | 0.512354415 | 0.71672258 |
| Ankrd37 | 131.6074743 | -0.078180279 | 0.1195031 | -0.654211304 | 0.51297568 | 0.717275477 |
| Mkln1 | 2343.437338 | 0.062439143 | 0.095435254 | 0.654256582 | 0.512946513 | 0.717275477 |
| Plxnb2 | 2763.911546 | 0.043572347 | 0.066603725 | 0.65420285 | 0.512981126 | 0.717275477 |
| Rpusd1 | 2230.095845 | 0.052057969 | 0.079569104 | 0.654248525 | 0.512951703 | 0.717275477 |
| Zfp583 | 185.2716755 | 0.072050924 | 0.110092048 | 0.654460749 | 0.512815007 | 0.717275477 |
| Zglp1 | 29.73814788 | 0.10997057 | 0.16811999 | 0.654119535 | 0.513034796 | 0.717285791 |
| Dctn4 | 3796.728018 | 0.038959984 | 0.059587899 | 0.653823762 | 0.513225356 | 0.717391525 |
| Inmt | 57.25964956 | -0.108668331 | 0.166184529 | -0.653901613 | 0.513175195 | 0.717391525 |
| Slc6a1 | 19579.76945 | -0.042153669 | 0.064476194 | -0.653786562 | 0.513249326 | 0.717391525 |
| Myo18a | 8327.400638 | -0.05511082 | 0.084327908 | -0.653530034 | 0.513414634 | 0.717557851 |
| Rrp1 | 10899.49526 | 0.037032424 | 0.056672661 | 0.653444243 | 0.513469924 | 0.717570398 |
| Slc25a51 | 3600.783044 | -0.039178021 | 0.059998946 | -0.652978476 | 0.513770155 | 0.717925214 |
| 4930470H14Rik | 74.55849228 | 0.088963175 | 0.13630062 | 0.652698245 | 0.513950834 | 0.718048172 |
| Tlcd1 | 534.5844104 | -0.067796244 | 0.103861458 | -0.65275652 | 0.513913259 | 0.718048172 |
| 4930414L22Rik | 377.4400261 | -0.064185528 | 0.098506115 | -0.65158927 | 0.514666171 | 0.718669124 |
| Abhd5 | 737.1301244 | -0.043647516 | 0.067002206 | -0.651434018 | 0.514766357 | 0.718669124 |
| Crcp | 968.8392653 | 0.041772177 | 0.064100178 | 0.651670218 | 0.514613939 | 0.718669124 |
| Dsc2 | 23.8512984 | -0.105055756 | 0.161267211 | -0.651439034 | 0.51476312 | 0.718669124 |
| Nipbl | 2271.018137 | -0.056247006 | 0.086324626 | -0.651575435 | 0.514675099 | 0.718669124 |
| Sec23a | 4109.50894 | -0.042092547 | 0.064610338 | -0.651483154 | 0.514734648 | 0.718669124 |
| Stx4a | 1582.050392 | -0.052893746 | 0.081147853 | -0.651819418 | 0.514517673 | 0.718669124 |
| Tapbpl | 381.2379022 | 0.085170422 | 0.130660001 | 0.651847711 | 0.514499419 | 0.718669124 |
| Ammecr1 | 63.94162102 | 0.090486458 | 0.13900143 | 0.65097502 | 0.515062612 | 0.718950077 |
| Ccdc36 | 27.00021538 | -0.107688336 | 0.165413621 | -0.651024599 | 0.515030608 | 0.718950077 |
| Fhod3 | 2739.679919 | 0.060192422 | 0.092484962 | 0.650834689 | 0.515153205 | 0.718950077 |
| Pcgf1 | 377.5119576 | 0.054443897 | 0.083650978 | 0.650845907 | 0.515145963 | 0.718950077 |
| Ercc6l | 9.12442639 | -0.103193516 | 0.15862621 | -0.650545178 | 0.515340129 | 0.719146173 |
| Tube1 | 91.00867004 | 0.088141395 | 0.135519221 | 0.65039774 | 0.515435336 | 0.719214256 |
| Ccdc125 | 9.840496829 | 0.101039374 | 0.155404129 | 0.650171746 | 0.515581289 | 0.71929855 |
| Tmem200c | 400.2623587 | -0.055236186 | 0.084957778 | -0.650160435 | 0.515588595 | 0.71929855 |
| Sh3gl2 | 13216.14166 | 0.046136721 | 0.070986896 | 0.649932925 | 0.515735549 | 0.719438786 |
| Cnot7 | 3059.532693 | -0.040767707 | 0.062740284 | -0.649785182 | 0.515830992 | 0.719507147 |
| Acer3 | 338.6673427 | -0.069602883 | 0.107159734 | -0.649524604 | 0.515999349 | 0.719677191 |
| Mark2 | 4848.52171 | -0.04201597 | 0.064702656 | -0.649370098 | 0.516099187 | 0.719751647 |
| Dkc1 | 961.5832104 | -0.044313762 | 0.068256075 | -0.649228107 | 0.516190947 | 0.719814826 |
| Itpk1 | 2501.417543 | 0.057094067 | 0.087993865 | 0.64884145 | 0.516440863 | 0.720098518 |
| Fam129b | 1496.534332 | 0.054625308 | 0.084235839 | 0.648480606 | 0.516674151 | 0.720358642 |
| Tsen34 | 1580.711717 | -0.037735319 | 0.058196778 | -0.648409072 | 0.516720404 | 0.720358642 |
| Mcc | 653.3971365 | 0.063938071 | 0.098619825 | 0.648328779 | 0.516772324 | 0.720366207 |
| Dusp22 | 981.4855818 | -0.038997365 | 0.060158375 | -0.648244992 | 0.516826506 | 0.720376924 |
| Gpd1 | 4107.537257 | 0.086872363 | 0.134154185 | 0.647556114 | 0.51727209 | 0.720933144 |
| Ppid | 2146.842905 | -0.050151962 | 0.077474605 | -0.647334211 | 0.517415664 | 0.721003535 |
| Wdr93 | 12.57260587 | 0.101190881 | 0.156306952 | 0.647385671 | 0.517382368 | 0.721003535 |
| Tnip1 | 1096.68632 | 0.042473135 | 0.065673503 | 0.646731681 | 0.517805615 | 0.721482033 |
| Ogg1 | 187.7883143 | 0.069595619 | 0.107628123 | 0.646630428 | 0.51787116 | 0.721508475 |
| Sdad1 | 921.966244 | 0.041511823 | 0.064214799 | 0.646452582 | 0.517986296 | 0.721586379 |
| Synj1 | 17435.27485 | -0.045899033 | 0.071007147 | -0.646400185 | 0.51802022 | 0.721586379 |
| D15Ertd621e | 3147.90537 | -0.051168053 | 0.079185238 | -0.646181717 | 0.518161679 | 0.721718541 |
| Lnx2 | 238.7593756 | -0.078324106 | 0.121242407 | -0.646012465 | 0.518271283 | 0.721806315 |
| Dmap1 | 830.3518416 | 0.044788833 | 0.069347897 | 0.645857125 | 0.518371888 | 0.721881542 |
| Ncapg | 11.2407673 | -0.10423978 | 0.161429527 | -0.645729329 | 0.518454663 | 0.721931927 |
| Tssc4 | 951.2982165 | 0.041669878 | 0.064571401 | 0.645330247 | 0.518713195 | 0.722227017 |
| Lrrc2 | 62.76978077 | 0.092531546 | 0.143437642 | 0.645099465 | 0.518862731 | 0.722370307 |
| Mrps35 | 496.4656099 | 0.049927645 | 0.077407233 | 0.644999734 | 0.518927358 | 0.722395372 |
| Nek1 | 1385.700247 | 0.083251595 | 0.129122065 | 0.644751111 | 0.519088488 | 0.72255476 |
| 1700003D09Rik | 43.82252613 | 0.085030662 | 0.1319666 | 0.644334714 | 0.51935841 | 0.722636233 |
| 2410131K14Rik | 816.0703731 | 0.053641848 | 0.083255872 | 0.644301077 | 0.519380218 | 0.722636233 |
| 9330159F19Rik | 3103.338697 | 0.060516752 | 0.093896452 | 0.644505203 | 0.519247885 | 0.722636233 |
| Dbr1 | 401.35588 | 0.070488629 | 0.109393819 | 0.644356596 | 0.519344224 | 0.722636233 |
| Hist1h4i | 19.73407781 | -0.108068347 | 0.167665943 | -0.644545608 | 0.519221693 | 0.722636233 |
| Ythdc2 | 1294.903099 | 0.046270542 | 0.071824026 | 0.644220948 | 0.519432169 | 0.722643623 |
| Gm12942 | 78.55149213 | 0.101071267 | 0.156918825 | 0.644099052 | 0.519511205 | 0.722688688 |
| BC006965 | 54.26947882 | 0.093213468 | 0.144762801 | 0.643904835 | 0.519637146 | 0.722789818 |
| Brf1 | 1415.820743 | 0.058267616 | 0.090499717 | 0.643843069 | 0.519677202 | 0.722789818 |
| Pcdhgb8 | 28.37376782 | -0.107206568 | 0.166621689 | -0.643413045 | 0.519956121 | 0.723112839 |
| Gm2518 | 9.716638256 | -0.102856743 | 0.160009862 | -0.64281502 | 0.520344136 | 0.723587511 |
| Nipal3 | 5842.832626 | -0.037013593 | 0.057596829 | -0.642632487 | 0.520462598 | 0.723687292 |
| Prrt4 | 146.166614 | 0.085590584 | 0.133222897 | 0.642461514 | 0.52057357 | 0.723755825 |
| Sh3bgr | 32.17154788 | 0.106870918 | 0.166358685 | 0.642412617 | 0.52060531 | 0.723755825 |
| Cacnb2 | 2039.924815 | 0.055614241 | 0.086622801 | 0.642027743 | 0.52085517 | 0.723931462 |
| Gm16845 | 24.44215899 | 0.107867837 | 0.168017887 | 0.642002104 | 0.520871817 | 0.723931462 |
| Pcdhgb7 | 296.2653701 | -0.076019045 | 0.118403731 | -0.64203251 | 0.520852075 | 0.723931462 |
| Myof | 236.5276257 | -0.090006244 | 0.140356229 | -0.641270038 | 0.521347252 | 0.724527251 |
| Nkx2-2 | 131.7747054 | -0.094859034 | 0.147988809 | -0.640987889 | 0.521530551 | 0.724716983 |
| Tmf1 | 1538.944418 | 0.065097691 | 0.101581824 | 0.640839951 | 0.521626674 | 0.724785551 |
| Capn1 | 985.3789761 | 0.045722564 | 0.071415128 | 0.640236384 | 0.522018932 | 0.725200514 |
| Tlk2 | 2219.191218 | 0.05143114 | 0.08032691 | 0.640272854 | 0.521995226 | 0.725200514 |
| Ccdc106 | 1203.046213 | -0.047897149 | 0.074842159 | -0.639975511 | 0.52218852 | 0.725306045 |
| Ubxn2a | 1999.45366 | -0.049599362 | 0.077500924 | -0.639984135 | 0.522182914 | 0.725306045 |
| Rab9 | 906.0775379 | -0.040372734 | 0.063120025 | -0.639618478 | 0.522420666 | 0.725563446 |
| Mrpl39 | 836.5745324 | 0.052268282 | 0.081740295 | 0.639443275 | 0.522534605 | 0.725656643 |
| Gm10548 | 53.90553573 | -0.100998338 | 0.158018753 | -0.639154129 | 0.52272267 | 0.72574435 |
| Mtdh | 2391.944094 | 0.051508884 | 0.08057624 | 0.639256489 | 0.522656089 | 0.72574435 |
| Tnni1 | 96.48379536 | 0.103365278 | 0.161728064 | 0.639130127 | 0.522738282 | 0.72574435 |
| Rnf168 | 1398.027547 | -0.04373162 | 0.068449387 | -0.638889871 | 0.522894578 | 0.725896299 |
| Tecpr1 | 3991.744248 | 0.047856251 | 0.074920387 | 0.638761395 | 0.522978166 | 0.725947294 |
| Pddc1 | 997.235046 | 0.040880322 | 0.064018346 | 0.638571983 | 0.523101412 | 0.726053327 |
| Bub3 | 2277.784628 | -0.036113814 | 0.056578208 | -0.638299011 | 0.523279056 | 0.726072113 |
| Gm4890 | 15.07178263 | 0.103848043 | 0.162716186 | 0.638215814 | 0.523333205 | 0.726072113 |
| Lztfl1 | 1260.164141 | 0.05368292 | 0.084126796 | 0.638119158 | 0.523396117 | 0.726072113 |
| Mapkapk3 | 127.0741021 | 0.0883659 | 0.138446679 | 0.638266664 | 0.523300108 | 0.726072113 |
| Snx9 | 397.0628377 | 0.057217226 | 0.089661311 | 0.638148439 | 0.523377057 | 0.726072113 |
| Zfp455 | 59.92333224 | -0.097254234 | 0.152405338 | -0.638128791 | 0.523389846 | 0.726072113 |
| Layn | 166.8733655 | -0.088601945 | 0.138906352 | -0.637853808 | 0.523568849 | 0.72624671 |
| Pabpc4l | 12.60977279 | -0.093729619 | 0.146998562 | -0.637622693 | 0.52371932 | 0.726390398 |
| Ip6k3 | 43.50025501 | 0.100827371 | 0.158175094 | 0.63744151 | 0.523837297 | 0.726488998 |
| Cdh3 | 13.24534967 | -0.094464389 | 0.148239638 | -0.63724109 | 0.523967817 | 0.726543599 |
| Kars | 2775.272535 | 0.046829847 | 0.073488895 | 0.637237044 | 0.523970451 | 0.726543599 |
| Egr3 | 2768.242867 | -0.106301971 | 0.166948183 | -0.636736314 | 0.524296615 | 0.7267399 |
| Fbxo6 | 613.6449152 | -0.049924584 | 0.078384947 | -0.636915448 | 0.52417992 | 0.7267399 |
| Supt6 | 7694.061734 | 0.056849634 | 0.089273494 | 0.63680306 | 0.524253133 | 0.7267399 |
| Tef | 18092.39366 | 0.045345165 | 0.071215501 | 0.636731674 | 0.524299639 | 0.7267399 |
| Tgm4 | 13.83775446 | -0.105044406 | 0.165006746 | -0.636606735 | 0.524381037 | 0.726787708 |
| Golgb1 | 4833.234087 | -0.041135464 | 0.064624803 | -0.636527499 | 0.524432664 | 0.726794248 |
| Igf1r | 3133.508271 | -0.063507302 | 0.09978715 | -0.636427654 | 0.524497721 | 0.726819399 |
| 7-Sep | 12745.39314 | -0.05304618 | 0.083373158 | -0.636250101 | 0.524613423 | 0.726900682 |
| Tmem234 | 1187.011318 | -0.048532045 | 0.076285019 | -0.636193655 | 0.524650208 | 0.726900682 |
| Capns1 | 4823.071507 | -0.040283336 | 0.063337598 | -0.63600985 | 0.524770002 | 0.727001647 |
| Il33 | 2018.053977 | 0.105190685 | 0.165524752 | 0.635498216 | 0.52510353 | 0.727398667 |
| Golga3 | 2238.857407 | 0.050772148 | 0.079939138 | 0.635135044 | 0.525340343 | 0.727661655 |
| Exosc6 | 231.0221759 | 0.063950315 | 0.100700333 | 0.635055647 | 0.525392122 | 0.727668324 |
| 2810408A11Rik | 11.68559419 | 0.10448961 | 0.164794356 | 0.63406061 | 0.526041268 | 0.728341121 |
| Erg | 191.4667898 | -0.079169666 | 0.124868795 | -0.634022822 | 0.526065928 | 0.728341121 |
| Tbc1d22b | 2004.284419 | -0.039371482 | 0.062084872 | -0.634155807 | 0.525979145 | 0.728341121 |
| Zfp952 | 649.7016516 | -0.058069169 | 0.091560194 | -0.634218504 | 0.525938233 | 0.728341121 |
| Fbxo34 | 2960.242935 | 0.044421057 | 0.070095725 | 0.633719909 | 0.52626363 | 0.728549739 |
| Dsg2 | 94.47644739 | 0.080893946 | 0.127773901 | 0.63310227 | 0.52666686 | 0.728977696 |
| Fam173b | 187.0006998 | 0.066947971 | 0.105739806 | 0.633138777 | 0.526643021 | 0.728977696 |
| Aebp1 | 950.3378089 | 0.076083088 | 0.120233019 | 0.632796954 | 0.526866246 | 0.729188532 |
| Disc1 | 29.15871968 | 0.104730789 | 0.165628822 | 0.632322249 | 0.527176328 | 0.729452594 |
| Mdh1 | 26459.33545 | -0.034885547 | 0.055173452 | -0.632288648 | 0.52719828 | 0.729452594 |
| Zfp282 | 681.6733305 | -0.043995786 | 0.069570118 | -0.632394871 | 0.527128885 | 0.729452594 |
| Atpaf1 | 2076.735692 | 0.050074687 | 0.079266253 | 0.631727692 | 0.527564829 | 0.729883917 |
| Clic6 | 341.2041643 | 0.097109486 | 0.153735152 | 0.631667414 | 0.527604226 | 0.729883917 |
| Ankle2 | 1616.249255 | -0.030098133 | 0.04767808 | -0.631278217 | 0.527858627 | 0.729981578 |
| Cx3cr1 | 2564.477999 | -0.102867546 | 0.162997696 | -0.63109816 | 0.527976344 | 0.729981578 |
| Fam89b | 774.0120373 | -0.054735119 | 0.086741355 | -0.631015267 | 0.528030542 | 0.729981578 |
| Fbxw2 | 2766.81002 | 0.045963242 | 0.072843884 | 0.63098286 | 0.528051731 | 0.729981578 |
| Gtse1 | 27.99827502 | 0.105577266 | 0.167262851 | 0.631205704 | 0.527906033 | 0.729981578 |
| Ltb | 12.30251148 | -0.104778905 | 0.165967823 | -0.631320596 | 0.527830923 | 0.729981578 |
| Mx2 | 41.99859868 | 0.106071765 | 0.168065037 | 0.631135224 | 0.527952112 | 0.729981578 |
| Tgfb2 | 442.117993 | -0.061199233 | 0.096973648 | -0.631091373 | 0.527980782 | 0.729981578 |
| Eif2b3 | 522.7548729 | 0.055382848 | 0.087823562 | 0.630614908 | 0.528292349 | 0.730249054 |
| Vps28 | 2761.78234 | -0.033859525 | 0.053715083 | -0.63035414 | 0.528462908 | 0.730419652 |
| Scube3 | 199.9302187 | -0.086657406 | 0.137547331 | -0.630018814 | 0.528682275 | 0.730657672 |
| Fam183b | 61.43152185 | 0.095710287 | 0.151940714 | 0.629918635 | 0.52874782 | 0.730663399 |
| Med6 | 500.2857781 | -0.043768209 | 0.069495823 | -0.629796262 | 0.528827892 | 0.730663399 |
| Nelfcd | 1648.764371 | 0.033798656 | 0.053660373 | 0.629862483 | 0.528784561 | 0.730663399 |
| Slc12a7 | 226.2823927 | 0.060929185 | 0.09675972 | 0.629695759 | 0.528893658 | 0.730689107 |
| Agtrap | 449.8254462 | -0.072665158 | 0.115425684 | -0.629540632 | 0.528995176 | 0.730714258 |
| Iqca | 35.78274144 | 0.104248378 | 0.165635489 | 0.629384306 | 0.52909749 | 0.730714258 |
| Rnmt | 2482.068486 | 0.038223503 | 0.060732024 | 0.629379696 | 0.529100507 | 0.730714258 |
| Zfp512 | 1802.820692 | 0.047356844 | 0.075233379 | 0.629465861 | 0.529044112 | 0.730714258 |
| 2700097O09Rik | 142.7715771 | 0.082579155 | 0.131329005 | 0.628796016 | 0.529482609 | 0.731176786 |
| Recql | 678.4482361 | 0.05051108 | 0.080362617 | 0.628539506 | 0.529650575 | 0.731343554 |
| Cdyl | 295.4605896 | 0.0566197 | 0.090110153 | 0.628338737 | 0.529782061 | 0.731394747 |
| Mrps31 | 593.567952 | -0.053230227 | 0.084710477 | -0.628378314 | 0.52975614 | 0.731394747 |
| Mtmr4 | 4564.498698 | -0.032761556 | 0.052155775 | -0.628148198 | 0.529906861 | 0.731501863 |
| Sft2d3 | 171.5549895 | 0.073949223 | 0.11775251 | 0.628005495 | 0.53000034 | 0.731565726 |
| Myo3b | 126.3486871 | -0.099161342 | 0.157929823 | -0.627882309 | 0.530081041 | 0.731611941 |
| Ncbp1 | 2316.705374 | 0.048992197 | 0.07804169 | 0.627769554 | 0.530154914 | 0.731648725 |
| Carhsp1 | 2368.750631 | -0.052322053 | 0.083361223 | -0.627654573 | 0.53023025 | 0.731687522 |
| Bcl6b | 98.51193222 | 0.10437519 | 0.166538093 | 0.626734628 | 0.530833201 | 0.732282217 |
| Ddt | 708.1271216 | 0.043160492 | 0.068857649 | 0.626807524 | 0.530785411 | 0.732282217 |
| Mapkapk2 | 1275.648947 | 0.041659968 | 0.066474222 | 0.626708614 | 0.530850256 | 0.732282217 |
| Rpgr | 647.8147627 | 0.041930537 | 0.066899644 | 0.626767718 | 0.530811507 | 0.732282217 |
| Slc4a3 | 6871.914398 | -0.059824271 | 0.095496172 | -0.626457269 | 0.531015056 | 0.732379142 |
| Usp33 | 5577.351477 | 0.03282913 | 0.052402143 | 0.626484492 | 0.530997206 | 0.732379142 |
| Thrap3 | 5381.623998 | 0.043353046 | 0.069220579 | 0.626302856 | 0.531116313 | 0.732453596 |
| Bicd1 | 856.095204 | -0.05028039 | 0.080299082 | -0.626163944 | 0.531207414 | 0.732514033 |
| Trak1 | 7825.333289 | -0.04780542 | 0.076364062 | -0.626019864 | 0.531301912 | 0.732579143 |
| Anapc5 | 7395.660668 | 0.044228291 | 0.070669562 | 0.625846408 | 0.531415689 | 0.732628482 |
| Msrb1 | 619.2041207 | 0.06036402 | 0.096455707 | 0.62582114 | 0.531432264 | 0.732628482 |
| Dhx40 | 1122.204085 | -0.049268844 | 0.078778998 | -0.625405831 | 0.531704736 | 0.732808502 |
| Gm19705 | 15.87710644 | 0.10515029 | 0.168118011 | 0.625455238 | 0.531672318 | 0.732808502 |
| Slco1a4 | 1929.963197 | 0.055663767 | 0.088986499 | 0.625530481 | 0.531622949 | 0.732808502 |
| Kif21b | 4825.106593 | -0.043278303 | 0.069211968 | -0.62530086 | 0.531773616 | 0.732838246 |
| Rpl7l1 | 1511.464988 | -0.05115991 | 0.081843643 | -0.625093267 | 0.531909847 | 0.732960794 |
| Ccdc121 | 18.33301211 | 0.102064876 | 0.163376046 | 0.624723628 | 0.532152463 | 0.732989631 |
| Dpf3 | 109.7959633 | -0.09229753 | 0.147700148 | -0.624898016 | 0.532037995 | 0.732989631 |
| Krt1 | 97.58030842 | 0.077008609 | 0.123272748 | 0.624701004 | 0.532167314 | 0.732989631 |
| Lin37 | 699.3363979 | 0.052433496 | 0.083930314 | 0.624726558 | 0.53215054 | 0.732989631 |
| Ppp1r37 | 5928.409961 | -0.044169327 | 0.070680581 | -0.6249146 | 0.532027109 | 0.732989631 |
| Clec1a | 31.45513116 | -0.104845159 | 0.167860038 | -0.624598684 | 0.532234483 | 0.733016986 |
| 1700052K11Rik | 236.121215 | -0.079151835 | 0.12684526 | -0.624003096 | 0.532625552 | 0.733426462 |
| Klhl41 | 16.45577994 | 0.104322408 | 0.167182891 | 0.624001695 | 0.532626472 | 0.733426462 |
| Atp5a1 | 32801.1374 | -0.044386378 | 0.071151088 | -0.623832735 | 0.532737439 | 0.733514075 |
| Odf4 | 13.32431388 | 0.103264094 | 0.165675721 | 0.623290449 | 0.533093673 | 0.733939343 |
| Pfkfb1 | 25.14722184 | 0.104574802 | 0.167918419 | 0.622771479 | 0.533434703 | 0.734343607 |
| A330102I10Rik | 90.9404528 | -0.078080844 | 0.125446355 | -0.622424174 | 0.533662989 | 0.734388578 |
| Aplp2 | 23520.06434 | 0.03047712 | 0.048970144 | 0.622361238 | 0.533704362 | 0.734388578 |
| Gm20743 | 38.55297641 | 0.102459361 | 0.164586712 | 0.622525111 | 0.533596637 | 0.734388578 |
| Nol12 | 601.0562568 | -0.047457393 | 0.076232766 | -0.622532739 | 0.533591623 | 0.734388578 |
| Ttc33 | 1370.817082 | -0.039638386 | 0.063683826 | -0.622424692 | 0.533662648 | 0.734388578 |
| Ndufv3 | 1645.812697 | -0.047548695 | 0.07641216 | -0.622266074 | 0.533766926 | 0.734409444 |
| Apool | 125.0352762 | -0.081257525 | 0.130656141 | -0.621918907 | 0.533995192 | 0.734639672 |
| Rpl7a | 7655.240134 | -0.039408786 | 0.06337169 | -0.621867368 | 0.534029084 | 0.734639672 |
| Edc3 | 1210.040075 | 0.052171063 | 0.083924873 | 0.621640057 | 0.534178575 | 0.734780082 |
| Creld1 | 2625.371495 | -0.046709211 | 0.075166271 | -0.62141185 | 0.534328677 | 0.734914706 |
| Nup62 | 1045.564111 | 0.041574386 | 0.066917855 | 0.621274935 | 0.534418742 | 0.734914706 |
| Senp5 | 3054.03657 | 0.045383872 | 0.073047173 | 0.621295385 | 0.53440529 | 0.734914706 |
| Adamtsl5 | 207.6198469 | 0.083627526 | 0.134747096 | 0.620625813 | 0.534845852 | 0.735428175 |
| Tceal5 | 2163.737861 | 0.046243593 | 0.074518748 | 0.620563199 | 0.53488706 | 0.735428175 |
| Cyhr1 | 5301.302516 | -0.034284229 | 0.05526523 | -0.620358022 | 0.535022103 | 0.735548577 |
| Bahd1 | 2142.065436 | 0.029429057 | 0.047471668 | 0.619928861 | 0.535304624 | 0.735675877 |
| Kbtbd11 | 6051.746415 | -0.035748122 | 0.0576477 | -0.620113583 | 0.53518301 | 0.735675877 |
| Nipal4 | 192.2892279 | 0.101108096 | 0.163095055 | 0.619933545 | 0.53530154 | 0.735675877 |
| Pdp2 | 277.6550284 | 0.062889291 | 0.101441988 | 0.619953258 | 0.535288561 | 0.735675877 |
| Skiv2l2 | 2637.808367 | 0.036750094 | 0.059306006 | 0.619669009 | 0.535475723 | 0.735845752 |
| Ercc8 | 402.0540899 | -0.052930016 | 0.08545054 | -0.619422839 | 0.535637839 | 0.735872732 |
| Ghitm | 15912.52682 | -0.037622113 | 0.060728998 | -0.619508215 | 0.535581611 | 0.735872732 |

| | | | | | | | |
|---|---|---|---|---|---|---|---|
| Jup | 2165.810506 | 0.0245355765833665 | 0.0652123948441018 | 0.376240999000601 | 0.706737738015605 | 0.929476901984127 | NM_010593 |
| Nubp2 | 899.798738 | 0.0243709239651804 | 0.064772985185267 | 0.376251363056271 | 0.706730033768975 | 0.929476901984127 | NM_011956 |
| Slc19a1 | 661.9542411 | -0.0292076046041152 | 0.0775464773792984 | -0.376646439544298 | 0.706436371271291 | 0.929476901984127 | NM_001199271,NM_031196 |
| Hnrnph1 | 7883.664307 | -0.0194681089139744 | 0.0517998281101263 | -0.375833465558713 | 0.707040706757062 | 0.92970700700218 | NM_021510 |
| Mta1 | 2643.631082 | 0.0191749133429354 | 0.0510140136093103 | 0.375875411211242 | 0.707009521353187 | 0.92970700700218 | NM_054081 |
| 4930523C07Rik | 79.25576008 | 0.0355065453292879 | 0.0945416550479445 | 0.375565091506824 | 0.707240246892102 | 0.929868290049939 | NM_001162896,NR_028111 |
| Tbcc | 485.2069964 | 0.0306260714292513 | 0.0815989928202798 | 0.375324135393492 | 0.707419418514517 | 0.929868290049939 | NM_178385 |
| Tekt1 | 76.18079098 | 0.0356956299073431 | 0.095080645224045 | 0.375424775707307 | 0.707344581802481 | 0.929868290049939 | NM_011569 |
| Ttll3 | 404.6952139 | -0.0352671771865464 | 0.0939508838320826 | -0.375378876153832 | 0.707378712621874 | 0.929868290049939 | NM_133923,NM_001142732 |
| Atg14 | 1032.366883 | 0.0248168193240171 | 0.0661436621622111 | 0.375195725678995 | 0.70751490881584 | 0.929909660284011 | NM_172599 |
| 3110082J24Rik | 35.76163169 | -0.0326391212575973 | 0.0872276823987626 | -0.374183061615544 | 0.708268125060142 | 0.92995985388767 | NM_001256263 |
| Dab2ip | 7119.213559 | 0.0206022785733127 | 0.055066194053222 | 0.374136599188249 | 0.708302690517072 | 0.92995985388767 | NM_001001602,NM_001114124,NM_001114125 |
| Fzd5 | 100.5709769 | -0.0355072497923756 | 0.0947354250712248 | -0.374804353975086 | 0.707805975933287 | 0.92995985388767 | NM_022721,NM_001042659 |
| Gm9958 | 51.76059818 | 0.0353273592172807 | 0.0942015280283374 | 0.375018961546501 | 0.707646364809382 | 0.92995985388767 | NR_045618 |
| Gpr158 | 9090.89194 | 0.0260363561032071 | 0.0695677789745692 | 0.374258837740454 | 0.708211753134865 | 0.92995985388767 | NM_001004761 |
| Hdac8 | 496.8528209 | 0.0287016454515015 | 0.0767371405317694 | 0.374025475181981 | 0.708385363022654 | 0.92995985388767 | NM_027382 |
| Ints7 | 1506.187113 | -0.0250930702886105 | 0.0670864259819013 | -0.374040946753971 | 0.708373852489982 | 0.92995985388767 | NM_178632 |
| Mst1r | 59.42694795 | -0.0344555687749523 | 0.0920166460277148 | -0.374449300886 | 0.708070069467838 | 0.92995985388767 | NM_009074 |
| Mycbp2 | 14611.77692 | -0.0248663308954092 | 0.0664167169931917 | -0.374398675832745 | 0.70810772796297 | 0.92995985388767 | NM_207215 |
| Ptrh2 | 502.6035792 | 0.0258910240346501 | 0.0690981138187111 | 0.374699432499402 | 0.707884014361594 | 0.92995985388767 | NM_175004,NM_001098810 |
| Rbm4b | 420.2433066 | 0.0286095019476659 | 0.0764146872355277 | 0.374397946032087 | 0.708108270845534 | 0.92995985388767 | NM_025717 |
| Rps6ka3 | 1106.720843 | -0.033477315836235 | 0.0892935169639389 | -0.374913173705038 | 0.707725041305863 | 0.92995985388767 | NM_148945 |
| Smad5 | 1663.912966 | 0.0220451235746661 | 0.0589046308345439 | 0.374251111709507 | 0.708217500666743 | 0.92995985388767 | NM_008541,NM_001164042,NM_001164041 |
| Pla2g5 | 107.3205902 | 0.0343702895037384 | 0.0919383133990458 | 0.373840765977062 | 0.708522788031745 | 0.930056209917687 | NM_001122954,NM_011110 |
| Lpp | 372.5861268 | -0.033705698876182 | 0.0902259517860095 | -0.37356988991507 | 0.70872433897894 | 0.930152669679082 | NM_178665,NM_001145952,NM_001145954 |
| Xab2 | 1675.25958 | 0.0227010253161352 | 0.0607641490981304 | 0.373592416796201 | 0.708707576608468 | 0.930152669679082 | NM_026156 |
| Agtr1a | 29.92458302 | 0.0308044723065449 | 0.0825809255471641 | 0.373021640317553 | 0.709132337857438 | 0.930183884749607 | NM_177322 |
| Lrch2 | 640.3687954 | -0.0284447513480497 | 0.0762243168858484 | -0.373171613865006 | 0.709020721565076 | 0.930183884749607 | NM_001081173 |
| Pcdhgc5 | 4834.397716 | -0.0288046751551648 | 0.0771520348975381 | -0.37334951947047 | 0.708888325221827 | 0.930183884749607 | NM_033583 |
| Pik3r1 | 6482.194144 | 0.0211098592653607 | 0.0565280826369764 | 0.373440213794766 | 0.708820834403602 | 0.930183884749607 | NM_001024955,NM_001077495 |
| Rpl28 | 4515.451775 | 0.0242463354518993 | 0.0649621800005174 | 0.373237712338259 | 0.708971530429503 | 0.930183884749607 | NM_009081 |
| Zfp809 | 430.9711014 | -0.0299686846775102 | 0.0803303072682561 | -0.373068219164559 | 0.709097671354115 | 0.930183884749607 | NR_028420,NM_001164624,NM_172763 |
| Amotl2 | 890.2334175 | -0.0319897797577672 | 0.0859131439506572 | -0.372350239867139 | 0.709632097356894 | 0.93022482748166 | NM_019764 |
| Calm1 | 135982.678 | 0.023458413841748 | 0.0630400148448333 | 0.372119421283269 | 0.709803936927352 | 0.93022482748166 | NM_009790 |
| Cbr2 | 43.37557645 | -0.0334211075900094 | 0.0896945693819882 | -0.372610157117503 | 0.709438612121183 | 0.93022482748166 | NM_007621 |
| Cdk11b | 3081.523234 | -0.0148942281118308 | 0.0399918417717768 | -0.372431662358247 | 0.709571483551721 | 0.93022482748166 | NM_007661 |
| Clp1 | 279.6493112 | -0.0310036432544612 | 0.0833102390439014 | -0.372146852659052 | 0.709783514073281 | 0.93022482748166 | NM_133840 |
| Cox7b | 3745.964129 | -0.0248177369611478 | 0.0665657559066155 | -0.372830393392729 | 0.70927468050077 | 0.93022482748166 | NM_025379 |
| Cul4b | 2178.667822 | 0.0282297899681145 | 0.0758406457120138 | 0.372225074075848 | 0.709725278817918 | 0.93022482748166 | NM_028288,NM_001110142 |
| Emilin1 | 100.6967573 | -0.0351592470160038 | 0.0944730465491363 | -0.372161672564642 | 0.709772480635071 | 0.93022482748166 | NM_133918 |
| Mcc | 690.6135849 | 0.0267962184244627 | 0.0719840404207375 | 0.372252214071929 | 0.709705073691541 | 0.93022482748166 | NM_001085374,NM_001085373 |
| Rassf10 | 32.84561343 | 0.0332003563387898 | 0.0891008077272234 | 0.372615660684364 | 0.709434515408532 | 0.93022482748166 | NM_175279 |
| Ephx4 | 2125.282538 | 0.026961585958616 | 0.0724895433093257 | 0.371937588895631 | 0.709939317701523 | 0.930318315645677 | NM_001001804 |
| Cbr3 | 320.1270175 | -0.0325056130482999 | 0.0874689099073895 | -0.371624764533093 | 0.710172248151989 | 0.930478980863676 | NM_173047 |
| Eif3a | 6822.743775 | 0.0202295028685621 | 0.0544513966735159 | 0.371514857366391 | 0.710254091960377 | 0.930478980863676 | NM_010123 |
| Tmem158 | 1436.196577 | -0.0333341180554883 | 0.0897052021835383 | -0.371596264699189 | 0.71019347060487 | 0.930478980863676 | NM_001002267 |
| 2810013P06Rik | 639.4029957 | 0.0230641309330491 | 0.0627774103319741 | 0.367395386510583 | 0.713324112387242 | 0.930507332344087 | NR_045268 |
| 9230110C19Rik | 336.3578769 | 0.029927759171767 | 0.0814188549964985 | 0.367577745634645 | 0.713188111227934 | 0.930507332344087 | NR_199017 |
| A730017L22Rik | 214.4328625 | -0.0302145690258012 | 0.0820206237139552 | -0.368377703773307 | 0.712591620187625 | 0.930507332344087 | NR_015523 |
| Aloxe3 | 249.3677216 | -0.0329459415288893 | 0.0892063927880341 | -0.369322651653151 | 0.711887243584475 | 0.930507332344087 | NM_011786 |
| Ankrd63 | 2098.627005 | 0.0283198913096388 | 0.0770904343906219 | 0.367359342744655 | 0.713350994456758 | 0.930507332344087 | NM_001081971 |
| BC030307 | 69.63131569 | -0.0348981166399563 | 0.0949912646409714 | -0.367382377441305 | 0.713333814737037 | 0.930507332344087 | NM_153595,NM_001003939,NM_001003910 |
| Baz1a | 142.2263022 | -0.0350995749926977 | 0.0950163005500512 | -0.3694058260478 | 0.711825256055424 | 0.930507332344087 | NM_013815 |
| Casp9 | 1466.136495 | -0.0192985929058029 | 0.0523628800567271 | -0.368554840468972 | 0.712459561467146 | 0.930507332344087 | NM_015733 |
| Casq1 | 27.94500136 | 0.0286300778409408 | 0.0776362262173326 | 0.368772147177718 | 0.71229756702615 | 0.930507332344087 | NM_009813 |
| Cdk20 | 383.6576553 | 0.0285662547099557 | 0.0776414644136089 | 0.367925243627278 | 0.712928976800863 | 0.930507332344087 | NM_053180 |
| Col4a3bp | 3382.24449 | -0.0237245697166029 | 0.0643268007556425 | -0.368813145343963 | 0.712267005803267 | 0.930507332344087 | NM_001164222,NM_023420 |
| Col8a1 | 191.7011473 | 0.0228564335083469 | 0.0621978965686702 | 0.367479203788058 | 0.713261601375724 | 0.930507332344087 | NM_007739 |
| Coq6 | 672.0053583 | -0.0240560252269512 | 0.0647996284923302 | -0.371237085561355 | 0.710460953260282 | 0.930507332344087 | NM_172582 |
| Crabp2 | 82.22443329 | 0.0329104425202443 | 0.0892283733870663 | 0.368833827973992 | 0.712251588545995 | 0.930507332344087 | NM_007759 |
| Cyth2 | 3184.611371 | 0.0192684507596185 | 0.0520877550899422 | 0.369922848975672 | 0.711439976188236 | 0.930507332344087 | NM_011181,NM_001112701 |
| Dcaf8 | 6930.124131 | 0.0148968769771282 | 0.0404761300882096 | 0.368041039117708 | 0.712842633737274 | 0.930507332344087 | NM_153555 |
| Ddx42 | 4023.36249 | 0.0185288430419843 | 0.0502342737192667 | 0.368848630031607 | 0.712240554860398 | 0.930507332344087 | NM_028074 |
| Dgkg | 3999.322969 | 0.0245087399347529 | 0.0660689292980285 | 0.370957123040349 | 0.7106694676084 | 0.930507332344087 | NM_138650 |
| Ehbp1l1 | 968.1241115 | -0.0264366094157449 | 0.0716029308953274 | -0.369211275085809 | 0.711970252378985 | 0.930507332344087 | NM_001114597,NM_053252,NM_001114596,NM_001114595 |
| Grem2 | 1416.080935 | -0.029576197733349 | 0.0801241447822404 | -0.369129652662509 | 0.712031087620484 | 0.930507332344087 | NM_011825 |
| Gria3 | 10301.9165 | 0.0237094179822281 | 0.0645435964212672 | 0.367339585905316 | 0.713365729603279 | 0.930507332344087 | NM_016886 |
| Mxra7 | 605.976468 | -0.0288847443187046 | 0.0783237331671094 | -0.368786613593569 | 0.712286783287203 | 0.930507332344087 | NM_026280 |
| Ncor2 | 6690.949239 | 0.0235789346581433 | 0.0641654339038649 | 0.367470976561464 | 0.713267737164544 | 0.930507332344087 | NM_001253904,NM_001253905,NM_011424 |
| Nek10 | 68.8962501 | 0.0345693457567241 | 0.0936541457548111 | 0.369117090099006 | 0.712040450977264 | 0.930507332344087 | NM_001195229 |
| Nipsnap3b | 340.1446934 | -0.0309915155182118 | 0.0842734554074932 | -0.367749433891804 | 0.713060076599418 | 0.930507332344087 | NM_025623 |
| P2rx4 | 577.7130213 | -0.025847294013635 | 0.0700851883530063 | -0.36879823855858 | 0.712278117700224 | 0.930507332344087 | NM_011026 |
| Pea15a | 23765.23299 | 0.0193708052725183 | 0.0522863610888825 | 0.370475299277178 | 0.711028377656616 | 0.930507332344087 | NM_011063 |
| Pgam2 | 162.1047821 | -0.0345862296019083 | 0.0941468414967327 | -0.367364736321065 | 0.713346971810717 | 0.930507332344087 | NM_018870 |
| Pou3f3 | 2217.521314 | -0.0300460985641409 | 0.0811931206359972 | -0.370057196087372 | 0.711339874237417 | 0.930507332344087 | NM_008900 |
| Pts | 679.1509887 | -0.0265018649274236 | 0.0717883600895603 | -0.369166601582219 | 0.712003548435806 | 0.930507332344087 | NM_011220 |
| Rasa1 | 1942.2559 | 0.0260207717411112 | 0.0707365990523579 | 0.367854435888997 | 0.712981776498946 | 0.930507332344087 | NM_145452 |
| Rmst | 68.73692086 | -0.0343923660656841 | 0.0936422331356961 | -0.367274091123462 | 0.713414578019833 | 0.930507332344087 | NR_028262 |
| Rsl1 | 158.2020158 | -0.0344856485923982 | 0.0936817910949411 | -0.368114744491264 | 0.712787677142503 | 0.930507332344087 | NM_001013769 |
| Slain1 | 2183.122532 | -0.0241620505041315 | 0.0651709401495727 | -0.370748840643968 | 0.710824609094255 | 0.930507332344087 | NM_198014 |
| Snai3 | 30.63580156 | 0.030040701784868 | 0.08178404833122 | 0.367317372982139 | 0.713382296686464 | 0.930507332344087 | NM_013914 |
| Sstr5 | 43.44835017 | -0.0338728637607921 | 0.0913027875589311 | -0.370994847653791 | 0.710641369287663 | 0.930507332344087 | NM_001191008,NM_011425 |
| Susd2 | 709.458078 | 0.0309390040635284 | 0.0839391903479326 | 0.368588307026605 | 0.712434612490493 | 0.930507332344087 | NM_027890,NM_001162913 |
| Tab3 | 1142.917156 | 0.0269652200951182 | 0.0733924162409486 | 0.367411532093328 | 0.713312070845367 | 0.930507332344087 | NM_025729 |
| Tekt2 | 41.883037 | 0.0323606291700835 | 0.0879562551523455 | 0.367917314283481 | 0.712934889461346 | 0.930507332344087 | NM_011902 |
| Tfpt | 674.7398583 | 0.0262207419441926 | 0.071203069187007 | 0.368252973412238 | 0.712684614175858 | 0.930507332344087 | NM_023524 |
| Tmem117 | 263.5637299 | 0.0325002715914467 | 0.0878949567386274 | 0.369762643926119 | 0.711559351376075 | 0.930507332344087 | NM_178789 |
| Tmem140 | 73.59117716 | -0.0340164184678492 | 0.0916495959132689 | -0.371157320759385 | 0.710520359382838 | 0.930507332344087 | NM_197986 |
| Tor1aip2 | 2340.794684 | 0.0219903579323012 | 0.0597593056110667 | 0.367982152862045 | 0.712886541893459 | 0.930507332344087 | NM_172843,NM_022329,NM_001160182,NM_001160181,NM_001160180 |
| Txnrd3 | 213.6602322 | 0.0326352232243898 | 0.0883196486755182 | 0.369512602391455 | 0.711745681451792 | 0.930507332344087 | NM_001178058,NM_001178060,NM_001178059,NM_153162 |
| U2surp | 2823.684805 | 0.0222008272919663 | 0.0599378968169664 | 0.370397168919047 | 0.711086582922814 | 0.930507332344087 | NM_026476,NM_001114977 |
| Urb2 | 728.5705188 | -0.0235575125814535 | 0.0640419257089663 | -0.367845162690904 | 0.71298869141084 | 0.930507332344087 | NM_001029876 |
| Vwf | 877.8542035 | -0.0330639494004424 | 0.0898757818764412 | -0.367884970902371 | 0.712959007074733 | 0.930507332344087 | NM_011708 |
| Yrdc | 716.7426106 | 0.0227606356447692 | 0.0614082381145256 | 0.3706446617524 | 0.71090221240876 | 0.930507332344087 | NM_153566 |
| Zbtb41 | 861.0598178 | 0.0291700283831927 | 0.0791608237831631 | 0.368490712818945 | 0.712507368834546 | 0.930507332344087 | NM_172643 |
| Dtd2 | 321.3887439 | 0.0296064915950887 | 0.0806971425554436 | 0.366884014198487 | 0.713705536067189 | 0.930719689102432 | NM_029545 |
| Siah1a | 860.190694 | -0.0251525493034878 | 0.0685417086081043 | -0.366967060119564 | 0.713643588694749 | 0.930719689102432 | NM_009172 |
| Fam19a2 | 2490.017024 | -0.0308565738698678 | 0.0842096953118273 | -0.366425430653874 | 0.714047646396385 | 0.931082236225663 | NM_182807,NR_045514,NM_001252387 |
| Zkscan6 | 577.6449231 | 0.0251930539706191 | 0.0687893969981187 | 0.366234551689821 | 0.714190061968244 | 0.931184349712837 | NM_026107 |
| Mon1a | 819.6110487 | 0.0263348196737013 | 0.0719264567466483 | 0.366135367497142 | 0.714264067629097 | 0.931197257797546 | NM_028369 |
| Klf16 | 2097.18372 | -0.0263367789815225 | 0.0719898829356765 | -0.365840002893943 | 0.714484467984009 | 0.931210732902501 | NM_078477 |
| Mll3 | 3391.890313 | -0.0287612885050664 | 0.0786318234217194 | -0.365771608154289 | 0.714535507370537 | 0.931210732902501 | NM_001081383 |
| Nudt15 | 248.2148902 | -0.0295823652433786 | 0.0808294273106611 | -0.365985090178622 | 0.714376201224695 | 0.931210732902501 | NM_172527 |
| Relb | 196.3395466 | -0.033815570323477 | 0.0924700916431609 | -0.365691973724544 | 0.71459493595375 | 0.931210732902501 | NM_009046 |
| Tbc1d9b | 12654.23887 | 0.0217751680861222 | 0.0595069276158179 | 0.365926606507139 | 0.714419842107983 | 0.931210732902501 | NM_029745 |
| 1700088E04Rik | 143.1374331 | 0.0340169593409221 | 0.0942889383720786 | 0.360773595802786 | 0.718268702820473 | 0.931254890141504 | NM_138581 |
| 3110039I08Rik | 30.08179067 | -0.0302306978041796 | 0.0830847851267804 | -0.363853595553628 | 0.71596733822528 | 0.931254890141504 | NR_040725 |
| 9130221H12Rik | 304.159099 | 0.0293349886619891 | 0.0810639848563412 | 0.361874495979633 | 0.717445819401344 | 0.931254890141504 | NR_046001 |
| 9230114K14Rik | 84.3271121 | 0.0342768484652535 | 0.0949604756734455 | 0.360959106640602 | 0.718130017194278 | 0.931254890141504 | NR_015537 |
| Anxa11 | 2198.718769 | 0.0279394272219793 | 0.0764460384018545 | 0.365479072638269 | 0.714753825603176 | 0.931254890141504 | NM_013469 |
| Asap2 | 2252.773331 | 0.0271359920571976 | 0.0742817893625034 | 0.365311502187581 | 0.714878893360905 | 0.931254890141504 | NM_001135192,NM_001004364,NM_001098168 |
| Atp5j | 4228.157154 | 0.022609236484196 | 0.0621954067541653 | 0.363519392574466 | 0.71621692914107 | 0.931254890141504 | NM_016755 |
| Bhlha9 | 45.16825961 | 0.0318736983051576 | 0.0883135450869244 | 0.360915171888811 | 0.718162861436591 | 0.931254890141504 | NM_177182 |
| Bicc1 | 157.9998573 | -0.0345139468748448 | 0.0950993347985 | -0.362925218646106 | 0.716660747553663 | 0.931254890141504 | NM_031397 |
| Ccdc34 | 267.3767626 | -0.0304568864806923 | 0.083865961233452 | -0.363161478539685 | 0.71648426167741 | 0.931254890141504 | NM_026613 |
| Chka | 1860.220687 | -0.0203368849228666 | 0.0560775484173616 | -0.362656455155773 | 0.71686153197942 | 0.931254890141504 | NM_001271496,NR_073190,NM_013490 |
| Cldn10 | 1149.881478 | 0.0259251692612215 | 0.0710932835833774 | 0.364664113886606 | 0.715362149514972 | 0.931254890141504 | NM_001160097,NM_001160096,NM_001160099,NM_001160098,NM_021386,NM_023878 |
| Csda | 843.0578766 | -0.0229645926782714 | 0.0635441852943015 | -0.361395658344492 | 0.717803693144825 | 0.931254890141504 | NM_011733,NM_139117 |
| Ctdsp1 | 1224.629237 | -0.0250308589679776 | 0.0688790388442571 | -0.363403139590479 | 0.71630375680795 | 0.931254890141504 | NM_153088 |
| D830031N03Rik | 323.612467 | 0.0309745486201188 | 0.0852176975945356 | 0.363475539640782 | 0.716249681829974 | 0.931254890141504 | NM_001167918 |
| Daglb | 1819.046863 | -0.0237775452925859 | 0.0655158181336073 | -0.362928312122975 | 0.716658436632105 | 0.931254890141504 | NM_144915 |
| Dmap1 | 918.1405225 | -0.0230610831681171 | 0.0632816018495698 | -0.36442002879347 | 0.715544381422142 | 0.931254890141504 | NM_023178 |
| Dync2h1 | 1754.559538 | -0.024281887587029 | 0.0668908939852175 | -0.363007371263347 | 0.716599377923825 | 0.931254890141504 | NM_029851 |
| Eif3k | 2846.714229 | 0.0169516409310082 | 0.0467838698016871 | 0.362339434571462 | 0.717098392860758 | 0.931254890141504 | NM_028659 |
| Fgf14 | 1800.77957 | 0.0287688664691468 | 0.0796946395604721 | 0.360988726817907 | 0.718107874372786 | 0.931254890141504 | NM_010201,NM_207667 |
| Foxc2 | 27.5237018 | 0.028171310389658 | 0.077716535071915 | 0.362487987448098 | 0.716987398681138 | 0.931254890141504 | NM_013519 |
| Gm11837 | 27.84336635 | -0.0302493871167743 | 0.0837258866223433 | -0.361290735005509 | 0.717882119038837 | 0.931254890141504 | NM_001243100 |
| Gm5506 | 36.78957726 | -0.0320574022859017 | 0.0888601547907849 | -0.360762395264543 | 0.718277076502611 | 0.931254890141504 | NM_001025388 |
| Gpn3 | 547.8292056 | 0.0249213400357975 | 0.0690041200286348 | 0.361157276195332 | 0.717981878325202 | 0.931254890141504 | NM_024216 |
| Hapln2 | 1081.700083 | 0.0325708789903753 | 0.0901241323120276 | 0.361400194884496 | 0.717800302257929 | 0.931254890141504 | NM_022031 |
| Id4 | 1581.266745 | 0.0252761902297929 | 0.0698919949691949 | 0.361646426617718 | 0.717616266185464 | 0.931254890141504 | NM_031166 |
| Ldlrad4 | 1421.229055 | 0.02683949391998 | 0.0737199895968475 | 0.364073490334943 | 0.715803132004692 | 0.931254890141504 | NM_172631 |
| Marveld1 | 456.8062013 | -0.0293991996447957 | 0.0805874651240873 | -0.364811073279538 | 0.715252438672427 | 0.931254890141504 | NM_183195 |
| Mbd5 | 2671.562558 | 0.0211967726512493 | 0.0584978491672326 | 0.362351316381775 | 0.717089514914549 | 0.931254890141504 | NM_029924 |
| Ncmap | 28.29647334 | 0.0319352385103196 | 0.0881233009899568 | 0.362392671989888 | 0.717058614798055 | 0.931254890141504 | NM_181047,NM_145555,NM_001243306,NM_001243305,NM_001168500,NM_001168498 |
| Noc3l | 680.2204322 | -0.0306244094514882 | 0.0846278355217535 | -0.361871590625949 | 0.717447990618107 | 0.931254890141504 | NM_021315 |
| Nyap1 | 3618.903346 | 0.0213330632133923 | 0.0588844181089633 | 0.362287068438314 | 0.717137520663018 | 0.931254890141504 | NM_175521 |
| Opcml | 8410.674377 | 0.0246228235454202 | 0.0680809437423329 | 0.361669832877326 | 0.71759877295742 | 0.931254890141504 | NM_177906 |
| Pcmtd2 | 2713.829262 | -0.0247016396283794 | 0.0680947029440271 | -0.362754202021907 | 0.716788506222922 | 0.931254890141504 | NM_153594 |
| Phc2 | 3359.676217 | 0.0192767896464695 | 0.0530905258453239 | 0.363092836989998 | 0.716535535278074 | 0.931254890141504 | NM_001195083,NM_018774,NM_001195130 |
| Phyhip | 26751.42129 | 0.0205310479918833 | 0.0567331620897918 | 0.361887954691979 | 0.717435761522508 | 0.931254890141504 | NM_145981 |
| Plat | 1312.192342 | 0.0251917362815912 | 0.0693061109824608 | 0.363485065378526 | 0.716242567243897 | 0.931254890141504 | NM_008872 |
| Prmt7 | 1603.129662 | 0.0218802197229256 | 0.0601168051821614 | 0.363961785005471 | 0.715886546211806 | 0.931254890141504 | NM_145404 |
| Pwp1 | 876.6286458 | 0.0212246398655439 | 0.0586146375893475 | 0.362104770044697 | 0.71727373921204 | 0.931254890141504 | NM_133993 |
| Rbm14 | 1301.126615 | 0.0211264177603497 | 0.0585617765430139 | 0.360754386350152 | 0.718283064101983 | 0.931254890141504 | NM_019869 |
| Satb2 | 2447.434984 | 0.0277867791913207 | 0.0766270996617267 | 0.36262339712695 | 0.716886229905005 | 0.931254890141504 | NM_139146 |
| Scaf8 | 2806.283313 | 0.0201077220580001 | 0.0552619835836152 | 0.363861750050571 | 0.715961248627755 | 0.931254890141504 | NM_134123 |
| Slc5a7 | 351.9755979 | 0.034427172692716 | 0.0948537603920584 | 0.362950003778642 | 0.716642232376713 | 0.931254890141504 | NM_022025 |
| Smim13 | 11035.80032 | 0.0265534512262455 | 0.0726939392718419 | 0.365277373770429 | 0.714904366359663 | 0.931254890141504 | NM_001135577 |
| Sstr3 | 1180.740952 | -0.0273523028121315 | 0.0753291479232675 | -0.363103839167189 | 0.716527316828085 | 0.931254890141504 | NM_009218 |
| Swt1 | 329.5878548 | -0.0308258745648224 | 0.0844029763649633 | -0.365222601055318 | 0.7149452486529 | 0.931254890141504 | NM_025819 |
| Taco1 | 465.3766403 | 0.0240335157687644 | 0.0663553689811207 | 0.362193988787891 | 0.717207071303656 | 0.931254890141504 | NM_027346 |
| Tesk2 | 292.2783278 | 0.0291818900762953 | 0.0801433081606838 | 0.364121356430494 | 0.715767389784913 | 0.931254890141504 | NM_146151 |
| Tmem132d | 1668.172943 | -0.0296330739156265 | 0.0813547555806779 | -0.364245134830993 | 0.715674965774052 | 0.931254890141504 | NM_172885 |
| Tpgs1 | 863.4208907 | -0.0272087216563701 | 0.0749616547634561 | -0.36296853027362 | 0.716628392683254 | 0.931254890141504 | NM_148934 |
| Tpr | 4880.017398 | -0.0248689684261032 | 0.0684721809940349 | -0.363198134849389 | 0.716456880811749 | 0.931254890141504 | NM_133780 |
| Ube2e3 | 2049.9747 | 0.0233859978492591 | 0.0642732629010138 | 0.36385266273591 | 0.715968034834021 | 0.931254890141504 | NM_009454 |
| Vdr | 46.35660121 | -0.0329160001273368 | 0.0910995404278054 | -0.36131905795312 | 0.71786094850336 | 0.931254890141504 | NM_009504 |
| Vkorc1 | 410.1066254 | 0.0279250363599172 | 0.0773103017862851 | 0.361207183450306 | 0.717944572528276 | 0.931254890141504 | NM_178600 |
| Wfikkn2 | 82.59592107 | 0.0214990392948229 | 0.0590730878927045 | 0.363939656140406 | 0.715903070998072 | 0.931254890141504 | NM_181819 |
| Zfp658 | 288.1045133 | 0.0318062875081586 | 0.0874724587193263 | 0.363614879172606 | 0.716145614331058 | 0.931254890141504 | NM_001008549 |
| Zfp870 | 494.2907812 | 0.0280544716283409 | 0.0770888113354954 | 0.36392403958917 | 0.715914732777524 | 0.931254890141504 | NM_207245 |
| Zfp746 | 1114.524531 | 0.0185913607471172 | 0.0516212647942696 | 0.360149268353088 | 0.718735510523048 | 0.931758324485301 | NM_001163475 |
| Ndufa7 | 1762.67206 | 0.0211189512483796 | 0.0587051132880352 | 0.359746367318294 | 0.719036814120624 | 0.932065747092289 | NM_023202 |
| 1700102P08Rik | 26.36959013 | -0.0283574026082168 | 0.0788820197096598 | -0.359491335447438 | 0.719227558519675 | 0.932229813068332 | NM_053216 |
| 1700086O06Rik | 116.6824124 | -0.0336086183187654 | 0.0936741371974316 | -0.358782256493384 | 0.719757987401375 | 0.932251852429102 | NR_015475,NR_027903,NR_027904 |
| 2700046G09Rik | 108.110012 | -0.0341517239168329 | 0.095053699288482 | -0.359288740706288 | 0.719379096407873 | 0.932251852429102 | NR_033198 |
| Aspscr1 | 1235.737899 | -0.0225683517809044 | 0.0628914948478587 | -0.35884584768576 | 0.719710412284905 | 0.932251852429102 | NM_198223,NM_001164224,NM_026877 |
| Bax | 1044.168073 | 0.0244093709306111 | 0.0679698039535614 | 0.359120808223725 | 0.719504715797791 | 0.932251852429102 | NM_007527 |
| Fam114a1 | 259.8087014 | 0.0291605057142798 | 0.0811965601855648 | 0.359134742255547 | 0.719494292364754 | 0.932251852429102 | NM_026667 |
| Pde1a | 9866.673962 | 0.0256731055269334 | 0.0715127374196561 | 0.359000458565537 | 0.719594746225755 | 0.932251852429102 | NM_016744,NM_001009978,NM_001159582,NM_001009979 |
| Ror2 | 35.4003654 | -0.0300252160126963 | 0.0836442841865076 | -0.358963153366785 | 0.719622654057928 | 0.932251852429102 | NM_013846 |
| Usp1 | 1054.99503 | 0.0260033765618342 | 0.0724541084160143 | 0.358894438566948 | 0.719674060233366 | 0.932251852429102 | NM_146144 |
| Pcsk2 | 19534.81616 | 0.0211112725352692 | 0.0588760732902153 | 0.358571337053399 | 0.719915792447049 | 0.932373109939893 | NM_008792 |
| Eepd1 | 864.7383722 | 0.0245729367973807 | 0.0685535357501228 | 0.358448860857431 | 0.720007431781395 | 0.932408661322683 | NM_026189 |
| Cbx2 | 132.2336821 | -0.0329357911295173 | 0.0919329321367282 | -0.358258899874239 | 0.720149572649349 | 0.932426934628552 | NM_007623 |
| Pcdhb14 | 347.5135326 | 0.0302375644547012 | 0.0844015430392886 | 0.358258431846747 | 0.720149922869181 | 0.932426934628552 | NM_053139 |
| Commd10 | 842.6268179 | -0.0240919049499172 | 0.0672669550005834 | -0.358153642449078 | 0.720228337102939 | 0.932445349800115 | NM_178377 |
| Amz2 | 2246.322124 | -0.0189643422535415 | 0.0531075552146863 | -0.357093113717597 | 0.72102209955065 | 0.932513932898887 | NM_001252193,NM_025275 |
| Arfgap1 | 4764.928585 | -0.0199193478371947 | 0.0556781511579278 | -0.357758787296916 | 0.720523834878785 | 0.932513932898887 | NM_001177709,NM_001177708,NM_145760,NM_001177710,NM_001177706,NM_001177707 |
| Bean1 | 631.3278556 | -0.0267018520837485 | 0.0748594989246894 | -0.356692904271391 | 0.721321718158047 | 0.932513932898887 | NM_001141922,NM_001141924,NM_001141925 |
| Caml | 911.3940425 | -0.0207360569731944 | 0.0581534502897988 | -0.356574835540444 | 0.721410119018994 | 0.932513932898887 | NM_007596 |
| Cebpz | 1667.443777 | 0.0245235793228807 | 0.0688097846538791 | 0.356396687567576 | 0.721543509678228 | 0.932513932898887 | NM_001024806 |
| Clip1 | 4973.310359 | 0.0207974234233116 | 0.0583766163657891 | 0.356262913441137 | 0.721643680398936 | 0.932513932898887 | NM_019765 |
| Ephb4 | 275.9549459 | 0.0284042626847368 | 0.0796074879810835 | 0.356803906329594 | 0.721238611677971 | 0.932513932898887 | NM_010144,NM_001159571 |
| Fam126a | 788.9090188 | -0.0213849286208161 | 0.0600616600925678 | -0.356049576183165 | 0.721803438259043 | 0.932513932898887 | NM_053090 |
| Fam160a2 | 3795.860466 | 0.0226245650507513 | 0.063380868630391 | 0.356962054002252 | 0.721120213285023 | 0.932513932898887 | NM_001242364,NM_001242365,NM_001242363,NM_199009 |
| Gm1078 | 67.94103402 | -0.0338784714142288 | 0.0949406151303307 | -0.35683854973682 | 0.721212675070339 | 0.932513932898887 | NM_001200041 |
| Hdx | 102.7657981 | -0.0338323827211399 | 0.0946677393487705 | -0.357380274989944 | 0.720807141348858 | 0.932513932898887 | NM_001080549 |
| Ino80c | 1034.418129 | 0.021846983122208 | 0.0612544212715626 | 0.356659693597505 | 0.721346583401498 | 0.932513932898887 | NM_172625 |
| Lamtor4 | 945.7629724 | 0.0231071070392614 | 0.0649030893283785 | 0.356024763664955 | 0.721822019930819 | 0.932513932898887 | NM_001081108 |
| Mettl11b | 22.89154615 | 0.0292221605434858 | 0.081643602539801 | 0.357923457986069 | 0.720400595187576 | 0.932513932898887 | NM_001143956 |
| Mical2 | 15374.50107 | -0.0272841365324917 | 0.0766000083506349 | -0.35618973313422 | 0.721698480191248 | 0.932513932898887 | NM_177282,NM_001193305 |
| Mrp63 | 578.0353437 | -0.0228914693463121 | 0.064090598449871 | -0.357173593319103 | 0.720961853303293 | 0.932513932898887 | NM_026401 |
| Nrxn2 | 14980.64799 | 0.0170732358186472 | 0.047807756652341 | 0.357122714265891 | 0.720999940666623 | 0.932513932898887 | NM_020253,NM_001205235,NM_001205234 |
| Rnf149 | 865.2568957 | 0.0207689206044443 | 0.0582881809262715 | 0.356314441013605 | 0.721605095735059 | 0.932513932898887 | NM_001033135 |
| Slc9a5 | 1923.424505 | -0.024678552668228 | 0.0692626354708207 | -0.356303979778891 | 0.721612929216297 | 0.932513932898887 | NM_001081332 |
| Sprn | 8159.711286 | -0.0215242438781746 | 0.0603393952749026 | -0.356719582291329 | 0.721301744206971 | 0.932513932898887 | NM_183147 |
| Srsf10 | 2117.541498 | 0.0253895879974443 | 0.0710359197600348 | 0.357419008344122 | 0.720778148694808 | 0.932513932898887 | NM_010178,NM_001080387 |
| Tmem37 | 50.01888128 | 0.0338257090184355 | 0.0948197986044194 | 0.356736773503956 | 0.721288873171039 | 0.932513932898887 | NM_019432 |
| Trappc11 | 3444.219492 | 0.0156303089350389 | 0.0438028817358811 | 0.356832891253256 | 0.72121691140622 | 0.932513932898887 | NM_177240 |
| Ulk1 | 4935.197006 | -0.0171208232812376 | 0.048038994245323 | -0.356394290725694 | 0.72154530440315 | 0.932513932898887 | NM_009469 |
| Commd2 | 379.7697922 | 0.0277609695046294 | 0.0782667360796748 | 0.354696910784283 | 0.7228166655003 | 0.932559879833859 | NM_175095 |
| Gm7694 | 153.8197865 | -0.0337297676174028 | 0.0950244227595816 | -0.354958932007843 | 0.722620357918438 | 0.932559879833859 | NM_001198955 |
| Igbp1 | 1079.79087 | -0.0246043347695295 | 0.0693567725976341 | -0.35475028390189 | 0.722776676619494 | 0.932559879833859 | NM_008784 |
| Junb | 1824.061189 | 0.0280170033987031 | 0.0788594092871853 | 0.355277875550303 | 0.722381428495325 | 0.932559879833859 | NM_008416 |
| Myl9 | 442.0100652 | -0.0319193156898697 | 0.0899696581462492 | -0.354778670359996 | 0.722755408867504 | 0.932559879833859 | NM_172118 |
| Pgbd1 | 95.79576621 | -0.0332601900557599 | 0.0937939373519678 | -0.354609167658128 | 0.722882407157927 | 0.932559879833859 | NM_001162920,NM_001012311,NM_001162919 |
| Prkcd | 845.6229721 | -0.0284751607122121 | 0.0801914312502245 | -0.355089817805595 | 0.722522304475763 | 0.932559879833859 | NM_011103 |
| Rsu1 | 843.5821075 | -0.021994644604325 | 0.0620156150298984 | -0.354663008561975 | 0.722842066544348 | 0.932559879833859 | NM_009105 |
| Sh2b3 | 742.4839146 | 0.0242074728771254 | 0.0682658285615575 | 0.354606006946751 | 0.722884775363435 | 0.932559879833859 | NM_008507 |
| Smim8 | 349.2067987 | -0.0277476836898902 | 0.0781633290650521 | -0.354996185830276 | 0.722592448654658 | 0.932559879833859 | NM_025471 |
| Spry1 | 344.5534619 | 0.0290575377973692 | 0.0818043369004433 | 0.355207790911288 | 0.722433928515418 | 0.932559879833859 | NM_011896 |
| Supt5 | 7177.528473 | -0.018863799542004 | 0.0530643307310277 | -0.355489257701577 | 0.722223091198944 | 0.932559879833859 | NM_013676 |
| Unc50 | 1183.274345 | 0.0219317943906778 | 0.0618146347383995 | 0.35479938502417 | 0.722739889125612 | 0.932559879833859 | NM_026123 |
| Usp37 | 1158.450192 | 0.0282364813243908 | 0.0793985774329636 | 0.355629562106839 | 0.7221180017602 | 0.932559879833859 | NM_176972 |
| Zfp518a | 452.3210531 | -0.0268499453225148 | 0.0754592305071366 | -0.355820555577696 | 0.721974954118896 | 0.932559879833859 | NM_028319 |
| Zfp84 | 896.490999 | -0.0232862794418952 | 0.0655079350672639 | -0.355472652557048 | 0.722235528969449 | 0.932559879833859 | NM_023750 |
| Cltc | 29028.76708 | 0.0229589073305681 | 0.0648519349989264 | 0.354020390154097 | 0.72332360240459 | 0.932617157481839 | NM_001003908 |
| Dok2 | 24.02917382 | -0.0271020977967428 | 0.0764953016285858 | -0.354297547950513 | 0.723115905181077 | 0.932617157481839 | NM_010071 |
| Gan | 212.4499467 | 0.0300650039794714 | 0.0849421054274311 | 0.353947006942945 | 0.723378597917381 | 0.932617157481839 | NM_001081151 |
| Rnf44 | 7137.132753 | -0.0208291454412515 | 0.0588206104483692 | -0.354113044432523 | 0.72325416663429 | 0.932617157481839 | NM_001146025,NM_001146026,NM_001146027,NR_027395,NR_027396,NM_134064 |
| Ttc30b | 457.5661767 | 0.0314648252714459 | 0.0888933409691471 | 0.35396155581965 | 0.723367694452487 | 0.932617157481839 | NM_028235 |
| Usp7 | 8024.291347 | -0.0182539098814325 | 0.0515093740700612 | -0.354380347480135 | 0.723053860612013 | 0.932617157481839 | NM_001003918 |
| Zfp212 | 628.4210275 | -0.02439258900515 | 0.06890242068082 | -0.354016430252066 | 0.72332657003388 | 0.932617157481839 | NM_145576,NM_001145881 |
| 1810013D10Rik | 623.1290827 | -0.0294728113530194 | 0.0834069794437354 | -0.35336145187827 | 0.723817481067042 | 0.932714926679326 | NM_001145433 |
| 2310035C23Rik | 2928.944225 | 0.0230080794222037 | 0.0651946807846481 | 0.352913445472711 | 0.724153330566601 | 0.932714926679326 | NM_173187,NM_029349 |
| Abhd5 | 801.2237837 | -0.0214489662213279 | 0.0607365743307788 | -0.35314744793663 | 0.723977903208709 | 0.932714926679326 | NM_026179 |
| Ccl17 | 53.37243747 | -0.0324181397881164 | 0.0918573405487206 | -0.352918336133648 | 0.72414966397948 | 0.932714926679326 | NM_011332 |
| D1Ertd622e | 2417.002069 | -0.0257920788604017 | 0.0730853463418542 | -0.352903559350464 | 0.724160742330266 | 0.932714926679326 | NM_133825 |
| Filip1l | 159.3397364 | 0.033477987371844 | 0.0948376622802536 | 0.353003085134191 | 0.724086127643103 | 0.932714926679326 | NM_001040397,NM_001177871 |
| Igsf10 | 186.255584 | 0.0334879409623721 | 0.0947976606441768 | 0.353257039623257 | 0.723895749313916 | 0.932714926679326 | NM_001162884 |
| Iqsec2 | 9225.665809 | -0.0237756802611661 | 0.0672622407044472 | -0.35347737470771 | 0.723730587785888 | 0.932714926679326 | NM_001005475,NM_001114664 |
| Mlc1 | 7288.055738 | 0.0182347283519615 | 0.0516650901486251 | 0.352940995544681 | 0.724132676030034 | 0.932714926679326 | NM_133241 |
| Prdm16 | 575.7782272 | -0.0306340724468194 | 0.0866488548156266 | -0.353542727275546 | 0.723681602463898 | 0.932714926679326 | NM_001177995,NM_027504 |

| | | | | | | |
|---|---|---|---|---|---|---|
| Gm5595 | 130.8863771 | -0.082146449 | 0.132613413 | -0.619442988 | 0.535624569 | 0.735872732 |
| Plp2 | 252.1191319 | -0.074619661 | 0.120494753 | -0.619277263 | 0.53573372 | 0.735939202 |
| Card14 | 12.82305078 | -0.102694603 | 0.165891783 | -0.619045751 | 0.53588622 | 0.73608343 |
| Dpyd | 226.7685054 | 0.080379705 | 0.129957377 | 0.618508213 | 0.536240386 | 0.736158233 |
| H2-M5 | 17.88093326 | 0.100565126 | 0.162587423 | 0.618529554 | 0.536226323 | 0.736158233 |
| Lamb2 | 1323.052436 | -0.046695182 | 0.075474616 | -0.618687239 | 0.536122418 | 0.736158233 |
| Mro | 428.4483635 | -0.05254895 | 0.084949701 | -0.618588987 | 0.536187159 | 0.736158233 |
| Naa15 | 2153.295549 | -0.043399391 | 0.07017351 | -0.618458319 | 0.536273265 | 0.736158233 |
| Nrg4 | 15.28238527 | 0.097821348 | 0.158100612 | 0.618728457 | 0.53609526 | 0.736158233 |
| Trim17 | 126.1370945 | 0.094115192 | 0.152121909 | 0.618682693 | 0.536125413 | 0.736158233 |
| Agpat9 | 208.0184076 | 0.072858375 | 0.117843968 | 0.618261389 | 0.536403049 | 0.736259913 |
| Lamb3 | 83.49485168 | 0.087757245 | 0.141961031 | 0.618178415 | 0.536457737 | 0.736259913 |
| Tcf7l2 | 186.6188788 | -0.099109552 | 0.160337812 | -0.618129627 | 0.536489894 | 0.736259913 |
| Ehd1 | 1491.853617 | -0.052175095 | 0.084444918 | -0.617859503 | 0.536667956 | 0.73626355 |
| Hist1h2be | 58.81570444 | -0.103846947 | 0.168101004 | -0.617765179 | 0.53673014 | 0.73626355 |
| Vgf | 8548.986705 | 0.067707776 | 0.109574586 | 0.617914957 | 0.536631399 | 0.73626355 |
| Zfyve20 | 2757.05948 | 0.028945175 | 0.04684972 | 0.61783026 | 0.536687234 | 0.73626355 |
| Zkscan4 | 201.4303729 | -0.067147047 | 0.108669605 | -0.617900905 | 0.536640663 | 0.73626355 |
| Atl2 | 2255.425341 | -0.03755591 | 0.060810497 | -0.617589261 | 0.536846126 | 0.736292279 |
| Fam228a | 57.61225714 | 0.103365316 | 0.167353321 | 0.617647231 | 0.536807903 | 0.736292279 |
| Ubn2 | 1819.359699 | -0.070445448 | 0.114093678 | -0.617435157 | 0.536947739 | 0.73636646 |
| Nuf2 | 81.0874181 | -0.099348095 | 0.160925455 | -0.617354755 | 0.537000758 | 0.736373993 |
| Sgms2 | 149.0550072 | 0.084245542 | 0.136566766 | 0.616881723 | 0.537312743 | 0.736698119 |
| Slc25a21 | 19.9167664 | -0.103663506 | 0.168052424 | -0.616852194 | 0.537332222 | 0.736698119 |
| Slitrk1 | 3530.036347 | 0.040679851 | 0.065956403 | 0.616768794 | 0.537387238 | 0.736708358 |
| Scrn3 | 1159.564495 | -0.050132954 | 0.081332398 | -0.61639587 | 0.537633279 | 0.736980449 |
| Csnk2a2 | 1366.077064 | 0.047843406 | 0.077672166 | 0.615965911 | 0.537917019 | 0.737262992 |
| Strc | 17.33315745 | 0.103104234 | 0.16739349 | 0.615939329 | 0.537934563 | 0.737262992 |
| Actb | 35544.54328 | -0.031999904 | 0.051985318 | -0.615556571 | 0.538187221 | 0.737397186 |
| Dgkg | 3940.089015 | 0.072282542 | 0.117397913 | 0.61570551 | 0.5380889 | 0.737397186 |
| Dok3 | 195.4260009 | 0.096471435 | 0.156736028 | 0.61550261 | 0.538222846 | 0.737397186 |
| Lrrn4 | 18.82943768 | 0.103496811 | 0.168114409 | 0.615633195 | 0.538136638 | 0.737397186 |
| Trp53i11 | 3894.626471 | 0.061101017 | 0.099285958 | 0.61540441 | 0.53828768 | 0.737420805 |
| Zfp174 | 351.5371279 | -0.060609416 | 0.09850853 | -0.615270737 | 0.538375939 | 0.73747651 |
| Tbx15 | 55.83725186 | -0.100110884 | 0.162743339 | -0.615145818 | 0.538458426 | 0.737524298 |
| Rnase4 | 565.5582172 | -0.046372382 | 0.075431134 | -0.614764477 | 0.538710272 | 0.737738815 |
| Surf6 | 658.4674457 | 0.038865903 | 0.063220081 | 0.614771481 | 0.538705646 | 0.737738815 |
| Lyzl4 | 14.53070523 | 0.10292519 | 0.16746813 | 0.614595685 | 0.538821765 | 0.737826286 |
| Pde4b | 5451.238946 | -0.049282552 | 0.08020949 | -0.614422954 | 0.538935872 | 0.73791732 |
| Ppie | 367.9067388 | -0.058542812 | 0.095339549 | -0.614045401 | 0.539185326 | 0.738193642 |
| Nhsl1 | 1021.149801 | 0.047931214 | 0.078070729 | 0.613946035 | 0.539250988 | 0.738218308 |
| 9130024F11Rik | 331.9156161 | 0.085349611 | 0.1391079 | 0.613549704 | 0.539512928 | 0.738511645 |
| Dusp4 | 353.8899367 | -0.095902776 | 0.156397259 | -0.613199851 | 0.539744203 | 0.738762958 |
| Mtif3 | 610.4618012 | -0.044842761 | 0.073146366 | -0.613055218 | 0.539839829 | 0.738828576 |
| Fam172a | 1597.050508 | 0.042696262 | 0.069689995 | 0.612659845 | 0.540101279 | 0.73899057 |
| Gm4636 | 294.9924339 | -0.063781232 | 0.104089152 | -0.6127558 | 0.54003782 | 0.73899057 |
| Mlkl | 9.256713253 | -0.097027364 | 0.15835337 | -0.61272687 | 0.540056953 | 0.73899057 |
| Zmynd11 | 7853.757492 | 0.028440169 | 0.046432953 | 0.612499673 | 0.540207214 | 0.739070249 |
| Ppp3cc | 756.8253611 | -0.055801956 | 0.091184399 | -0.611968237 | 0.540558773 | 0.739420643 |
| Zfp251 | 2040.689552 | 0.056006586 | 0.091508369 | 0.612037854 | 0.540512713 | 0.739420643 |
| Adck1 | 1341.360054 | -0.04188061 | 0.068458864 | -0.611763142 | 0.540694479 | 0.739475705 |
| Smtnl2 | 45.80297574 | 0.101950226 | 0.16664359 | 0.611786063 | 0.540679312 | 0.739475705 |
| S100a7a | 13.59255941 | 0.098574394 | 0.161317442 | 0.6110585 | 0.541160853 | 0.740048213 |
| Gm16861 | 30.33648144 | -0.101760058 | 0.166561241 | -0.610946808 | 0.541234795 | 0.74008401 |
| Chrac1 | 344.6026352 | 0.050613122 | 0.082889082 | 0.610612644 | 0.54145605 | 0.740125279 |
| Entpd1 | 689.9804184 | -0.063964918 | 0.104739082 | -0.610707267 | 0.541393394 | 0.740125279 |
| Kdm2b | 1016.216389 | -0.038597526 | 0.06321073 | -0.610616676 | 0.54145338 | 0.740125279 |
| Pdzrn3 | 1294.359135 | 0.057466319 | 0.094090078 | 0.610758545 | 0.541359441 | 0.740125279 |
| Mex3c | 1538.122192 | -0.049977328 | 0.081866973 | -0.61046997 | 0.54155053 | 0.740189125 |
| Plk2 | 8538.54075 | 0.05065636 | 0.083068933 | 0.609811135 | 0.541986925 | 0.740720245 |
| Abhd1 | 112.777406 | 0.091925764 | 0.150822475 | 0.609496457 | 0.54219542 | 0.740777876 |
| Actl6b | 2984.150117 | -0.062738147 | 0.103029548 | -0.608933535 | 0.542568495 | 0.740777876 |
| Anxa9 | 27.88387546 | 0.101730878 | 0.166905576 | 0.609511561 | 0.542185412 | 0.740777876 |
| Cbx2 | 120.2436227 | 0.076487277 | 0.125549033 | 0.609222348 | 0.542377069 | 0.740777876 |
| Dnajc3 | 2453.633561 | -0.050748459 | 0.083272256 | -0.609428174 | 0.542240668 | 0.740777876 |
| Gosr1 | 2080.864821 | 0.031482748 | 0.051692449 | 0.609039595 | 0.542498194 | 0.740777876 |
| Hrh1 | 675.265137 | -0.06981984 | 0.114667816 | -0.608887854 | 0.542598775 | 0.740777876 |
| Polr2j | 723.8734926 | 0.047070878 | 0.077307094 | 0.60888175 | 0.542602822 | 0.740777876 |
| Tmed5 | 678.4442641 | -0.052612589 | 0.086313607 | -0.609551506 | 0.542158944 | 0.740777876 |
| Trim16 | 83.23713713 | -0.092699429 | 0.152170806 | -0.609180117 | 0.542405058 | 0.740777876 |
| Zfp472 | 168.3668712 | -0.080046087 | 0.131456198 | -0.608918317 | 0.542578582 | 0.740777876 |
| Zfp787 | 472.5939517 | 0.05261417 | 0.086355443 | 0.609274509 | 0.5423425 | 0.740777876 |
| Kif3a | 7972.371217 | -0.03141031 | 0.051606339 | -0.608652174 | 0.542755014 | 0.740883918 |
| Vmn1r58 | 66.96891914 | -0.088299379 | 0.145081219 | -0.608620325 | 0.54277613 | 0.740883918 |
| Stxbp4 | 923.3177638 | -0.042030996 | 0.069123392 | -0.60805748 | 0.543149352 | 0.741328053 |
| Cdc42ep5 | 38.70031211 | -0.099393421 | 0.163540657 | -0.6077597 | 0.543346861 | 0.741401697 |
| Fam122a | 363.3067163 | -0.060098042 | 0.098882124 | -0.607774584 | 0.543336988 | 0.741401697 |
| Snora3 | 12.30488376 | -0.100958196 | 0.166077369 | -0.607898575 | 0.543254745 | 0.741401697 |
| 2310036O22Rik | 1320.703145 | -0.058001615 | 0.095529829 | -0.607157109 | 0.543746653 | 0.741881881 |
| Arl8b | 8450.704777 | -0.039697046 | 0.065404586 | -0.606945915 | 0.543886806 | 0.741925327 |
| Ccdc28b | 479.6287133 | 0.060785236 | 0.100156768 | 0.606900933 | 0.543916659 | 0.741925327 |
| Dctn6 | 1851.989778 | 0.041590867 | 0.068530845 | 0.606892659 | 0.54392215 | 0.741925327 |
| Polr2f | 719.9464122 | -0.045252234 | 0.074574504 | -0.606805691 | 0.543979871 | 0.741938743 |
| Amhr2 | 28.53907677 | 0.100265793 | 0.16528411 | 0.606626936 | 0.54409852 | 0.742035249 |
| Pgbd5 | 6570.503865 | 0.045862223 | 0.075632281 | 0.606384238 | 0.544259632 | 0.742189645 |
| Aph1c | 379.025436 | 0.066473668 | 0.109699259 | 0.605962779 | 0.54453947 | 0.742315495 |
| Ddo | 249.0116207 | -0.094477976 | 0.155915412 | -0.60595662 | 0.54454356 | 0.742315495 |
| Myl2 | 9.41690481 | -0.0975707 | 0.161009831 | -0.605992191 | 0.544519939 | 0.742315495 |
| Slc30a7 | 823.8962026 | 0.040480681 | 0.066797378 | 0.606022005 | 0.544500141 | 0.742315495 |
| B4galnt3 | 32.25578941 | 0.101835785 | 0.168079941 | 0.605877089 | 0.544596374 | 0.74232218 |
| Gpr89 | 869.4841826 | -0.043166581 | 0.071268234 | -0.605691751 | 0.544719463 | 0.742359342 |
| Wdr89 | 84.88160923 | -0.087912768 | 0.145128984 | -0.605756106 | 0.544676721 | 0.742359342 |
| Zbtb17 | 1311.034451 | 0.051251892 | 0.084628607 | 0.60560954 | 0.544774066 | 0.742368459 |
| 6-Sep | 3378.162243 | 0.051355496 | 0.08481822 | 0.605477167 | 0.544861991 | 0.742422979 |
| Cyp2c44 | 8.441753471 | -0.09161044 | 0.151342894 | -0.605317091 | 0.544968328 | 0.742435638 |
| Fsd2 | 8.105248128 | -0.079721489 | 0.131695987 | -0.605344857 | 0.544949882 | 0.742435638 |
| Grm2 | 1633.962397 | -0.063800397 | 0.105419103 | -0.605207171 | 0.545041351 | 0.742435638 |
| Kif3b | 4106.107783 | -0.037845023 | 0.062548289 | -0.60505289 | 0.545143855 | 0.742435638 |
| Plod1 | 784.105156 | -0.056254742 | 0.09297837 | -0.605030416 | 0.545158787 | 0.742435638 |
| Ttf2 | 92.69814457 | 0.084375594 | 0.139443005 | 0.605090189 | 0.545119073 | 0.742435638 |
| Tasp1 | 703.8749663 | -0.05229266 | 0.0864533 | -0.604865979 | 0.54526805 | 0.742519174 |
| Usp38 | 1217.293996 | 0.041083323 | 0.067945779 | 0.604648646 | 0.545412477 | 0.742650577 |
| Ctso | 1352.317329 | 0.046407716 | 0.076794087 | 0.604313662 | 0.545635125 | 0.742888456 |
| Arntl | 1384.342415 | 0.058080458 | 0.096190312 | 0.603807775 | 0.54597145 | 0.743215748 |
| Tmem161b | 613.731862 | -0.049715447 | 0.082326754 | -0.603879594 | 0.545923696 | 0.743215748 |
| 2610528A11Rik | 12.6770639 | -0.10077751 | 0.167102881 | -0.603086612 | 0.546451072 | 0.743600379 |
| Bcap31 | 1976.59574 | -0.045356381 | 0.075196393 | -0.60317229 | 0.54639408 | 0.743600379 |
| Melk | 50.1864003 | 0.100809221 | 0.16713666 | 0.603154453 | 0.546405945 | 0.743600379 |
| Rai1 | 3606.17286 | -0.039825104 | 0.066042524 | -0.603022132 | 0.546493966 | 0.743600379 |
| Ssbp2 | 3609.698034 | -0.045588724 | 0.075598241 | -0.603039478 | 0.546482427 | 0.743600379 |
| Zfp606 | 451.3699727 | 0.057880202 | 0.096001543 | 0.60290908 | 0.546569175 | 0.743637408 |
| Slc16a12 | 127.8309968 | 0.076836147 | 0.12748981 | 0.602684612 | 0.54671852 | 0.743775288 |
| Nmral1 | 372.1136646 | -0.064388803 | 0.106898597 | -0.602335343 | 0.546950939 | 0.74402615 |
| Agtr2 | 20.24657334 | 0.100166497 | 0.166417765 | 0.601897865 | 0.547242126 | 0.744128707 |
| B4galt7 | 629.3408993 | 0.044530974 | 0.073989494 | 0.601855361 | 0.54727042 | 0.744128707 |
| B930059L03Rik | 35.36801585 | 0.099861392 | 0.165896823 | 0.601948789 | 0.547208227 | 0.744128707 |
| Chrna7 | 487.3587347 | 0.046675316 | 0.077562701 | 0.601775272 | 0.547323738 | 0.744128707 |
| Fos | 1299.876561 | -0.093032632 | 0.154567873 | -0.601888548 | 0.547248328 | 0.744128707 |
| Msra | 1102.766488 | 0.05869602 | 0.097559244 | 0.601644882 | 0.547410546 | 0.744128707 |
| Rab38 | 10.2809071 | 0.095869667 | 0.159341146 | 0.601662967 | 0.547398506 | 0.744128707 |
| Tnk2 | 10854.0892 | 0.06281706 | 0.104339423 | 0.602045307 | 0.547143979 | 0.744128707 |
| Ccdc122 | 55.59397627 | -0.093686299 | 0.155748372 | -0.601523454 | 0.547491395 | 0.74417332 |
| Sema4d | 1978.342213 | 0.037356054 | 0.062125614 | 0.601298752 | 0.547641021 | 0.744311401 |
| Tbc1d25 | 544.2520363 | 0.049692047 | 0.082659895 | 0.601162714 | 0.547731616 | 0.744369236 |
| Dda1 | 2029.744976 | -0.042381477 | 0.070512288 | -0.601050941 | 0.547806058 | 0.74440511 |
| Etohi1 | 519.2063723 | -0.062697066 | 0.104417053 | -0.60044853 | 0.548207354 | 0.744814712 |
| Folr2 | 15.07447431 | -0.098120864 | 0.16343073 | -0.600381976 | 0.548251697 | 0.744814712 |
| Lats2 | 1000.532933 | 0.060076734 | 0.100058014 | 0.60041901 | 0.548227022 | 0.744814712 |
| Usp14 | 5274.207639 | -0.053022814 | 0.088329773 | -0.600282464 | 0.548318003 | 0.744839483 |
| Pde3b | 412.9722291 | 0.046245963 | 0.077062522 | 0.600109648 | 0.548433163 | 0.744930607 |
| Ttc22 | 51.11747396 | -0.095444786 | 0.159067385 | -0.60002738 | 0.548487988 | 0.744939769 |
| Gtf2a1 | 1230.187129 | -0.044010712 | 0.073364303 | -0.599892727 | 0.54857773 | 0.744996349 |
| Lrrc46 | 31.38138717 | 0.100290693 | 0.167246033 | 0.599659622 | 0.548733103 | 0.745142043 |
| Fdxacb1 | 163.5125176 | -0.070184499 | 0.117100473 | -0.599352826 | 0.548937628 | 0.745354448 |
| Hsdl1 | 2041.973916 | -0.039696724 | 0.066252678 | -0.599171616 | 0.549058448 | 0.745387856 |
| Rab11fip4 | 5956.184506 | -0.042676082 | 0.071221942 | -0.59919852 | 0.54904051 | 0.745387856 |
| Ccdc71l | 520.1760712 | 0.052844221 | 0.088209005 | 0.599079662 | 0.549119763 | 0.745405783 |
| 3830408C21Rik | 106.2112079 | 0.079723742 | 0.133123966 | 0.598868443 | 0.549260617 | 0.745531669 |
| Relb | 197.8866786 | -0.07631247 | 0.127466896 | -0.598684618 | 0.549383217 | 0.745632757 |
| Col17a1 | 41.47010534 | -0.098689271 | 0.16487637 | -0.598565285 | 0.549462811 | 0.745675466 |
| Slc26a1 | 39.79662281 | -0.097293799 | 0.162580686 | -0.598433933 | 0.54955043 | 0.745729056 |
| Mul1 | 1909.589671 | -0.036172095 | 0.060453377 | -0.598346977 | 0.549608437 | 0.745742458 |
| Cntd1 | 20.69288458 | -0.099938 | 0.167131648 | -0.597959758 | 0.549866785 | 0.745962347 |
| Prkar1a | 25550.26677 | -0.035787418 | 0.059846173 | -0.597990088 | 0.549846547 | 0.745962347 |
| Atp6v1e1 | 12725.87037 | 0.044054776 | 0.073690262 | 0.597837149 | 0.5499486 | 0.746008021 |
| Ankrd55 | 134.7097154 | 0.085103344 | 0.142537748 | 0.597058291 | 0.550468464 | 0.746061754 |
| Arhgef26 | 1175.244826 | -0.050408777 | 0.084430311 | -0.597045968 | 0.55047669 | 0.746061754 |
| Ccr9 | 11.45335156 | -0.097094636 | 0.162572172 | -0.597240199 | 0.550347024 | 0.746061754 |
| Fam114a2 | 1393.450406 | -0.039003024 | 0.06533342 | -0.596984274 | 0.55051788 | 0.746061754 |
| Fbxl14 | 1060.2336 | 0.04885723 | 0.081839256 | 0.596990156 | 0.550513953 | 0.746061754 |
| Nrn1l | 44.28824163 | -0.100285174 | 0.167882692 | -0.597352668 | 0.550271948 | 0.746061754 |
| Rabggtb | 1352.468121 | 0.03658569 | 0.061232725 | 0.597485892 | 0.550183024 | 0.746061754 |
| Six5 | 87.77942271 | -0.086543129 | 0.144958192 | -0.597021303 | 0.550493158 | 0.746061754 |
| Speg | 6024.046489 | -0.051439 | 0.086140592 | -0.597151692 | 0.550406108 | 0.746061754 |
| Tipin | 378.0803007 | -0.06377995 | 0.106728494 | -0.597590651 | 0.550113104 | 0.746061754 |
| Tns3 | 3232.820089 | -0.03276958 | 0.05487283 | -0.59719137 | 0.55037962 | 0.746061754 |
| Hars | 3643.222555 | -0.040185484 | 0.067326228 | -0.596877112 | 0.55058943 | 0.74609346 |
| Junb | 1830.703675 | -0.100327165 | 0.168114651 | -0.596778237 | 0.55065545 | 0.746117669 |
| Tmpo | 1013.597682 | 0.039055684 | 0.06545632 | 0.596667886 | 0.550729137 | 0.746152261 |
| Tmem9b | 3357.727927 | -0.036735866 | 0.061577893 | -0.596575563 | 0.55079079 | 0.746170543 |
| Tmtc4 | 416.2612799 | -0.060452728 | 0.101355193 | -0.596444308 | 0.550878448 | 0.746224049 |
| Tbck | 739.5901732 | -0.038673106 | 0.0648698 | -0.596165029 | 0.551064986 | 0.746411477 |
| Lyrm9 | 1764.184818 | 0.046501224 | 0.078027172 | 0.595961921 | 0.551200666 | 0.746529992 |
| Erbb3 | 417.7410406 | -0.062136822 | 0.104313599 | -0.595673265 | 0.551393522 | 0.746628481 |
| Mkrn1 | 4765.783074 | -0.04183727 | 0.070233789 | -0.595685783 | 0.551385158 | 0.746628481 |
| Sphk2 | 1043.828299 | 0.039970846 | 0.067106083 | 0.595636707 | 0.551417949 | 0.746628481 |
| Zcwpw1 | 268.0337954 | -0.073548984 | 0.123519537 | -0.595444132 | 0.551546633 | 0.746737464 |
| Rnase1 | 9.796111194 | 0.077105906 | 0.129518354 | 0.595328026 | 0.551624226 | 0.746777261 |
| Cct7 | 6515.985863 | -0.036061263 | 0.060609536 | -0.594976712 | 0.551859038 | 0.747009187 |
| Zfp688 | 249.4758524 | -0.054709629 | 0.09196017 | -0.594927448 | 0.551891969 | 0.747009187 |
| Cdv3 | 3605.447501 | -0.028746537 | 0.048342498 | -0.594643178 | 0.552082012 | 0.74713588 |
| Iglon5 | 1361.487352 | 0.056889551 | 0.095660213 | 0.59470441 | 0.552041073 | 0.74713588 |
| B230206H07Rik | 138.9740127 | 0.086590284 | 0.145663526 | 0.594454122 | 0.552208419 | 0.747209042 |
| Serpinb6b | 50.78160011 | -0.090729714 | 0.152636198 | -0.594418071 | 0.552232525 | 0.747209042 |
| 2700046A07Rik | 25.95309645 | -0.099638578 | 0.167877997 | -0.593517793 | 0.552834679 | 0.74757312 |
| A730036I17Rik | 32.10432879 | -0.099794289 | 0.168096155 | -0.593673832 | 0.552730289 | 0.74757312 |
| Acsbg1 | 6036.166675 | -0.055681372 | 0.093811902 | -0.59354273 | 0.552817995 | 0.74757312 |
| Dtl | 367.2746342 | 0.07661291 | 0.129084268 | 0.593510818 | 0.552839345 | 0.74757312 |
| Gm101 | 9.413431964 | 0.086565102 | 0.145776355 | 0.593821281 | 0.552631654 | 0.74757312 |
| Hmgxb3 | 2440.229766 | 0.041451358 | 0.069831814 | 0.593588443 | 0.552787413 | 0.74757312 |
| Oas1a | 24.05345707 | 0.099548704 | 0.167659242 | 0.59375614 | 0.552675228 | 0.74757312 |
| Esco1 | 775.1239776 | 0.052547844 | 0.088552354 | 0.593409908 | 0.55290686 | 0.74759917 |
| Dnttip1 | 728.2437001 | 0.039966107 | 0.067424443 | 0.592753976 | 0.553345813 | 0.747996861 |
| Drg2 | 1969.198591 | 0.028360797 | 0.047841489 | 0.592807579 | 0.553309935 | 0.747996861 |
| Myh8 | 20.35493054 | -0.099456305 | 0.167757163 | -0.592858769 | 0.553275673 | 0.747996861 |
| Aco1 | 1314.751263 | -0.036587154 | 0.061748588 | -0.592518068 | 0.553503725 | 0.748145051 |
| Ppa2 | 1168.133258 | -0.04091596 | 0.069087834 | -0.592231049 | 0.553695881 | 0.748339496 |
| Chm | 1773.805331 | -0.052045066 | 0.087912514 | -0.592009753 | 0.553844058 | 0.748409195 |
| Hmgb1 | 6512.796655 | -0.034830853 | 0.058830453 | -0.592054819 | 0.55381388 | 0.748409195 |
| 2610002J02Rik | 666.0513271 | -0.047657641 | 0.080535374 | -0.591760349 | 0.554011079 | 0.748569604 |
| Ano4 | 825.0102154 | -0.05640346 | 0.095327501 | -0.591680888 | 0.554064297 | 0.748576231 |
| Epg5 | 2833.413321 | -0.037738014 | 0.063789735 | -0.591600102 | 0.554118406 | 0.748584059 |
| Gabrq | 51.31806127 | -0.09524499 | 0.161088787 | -0.591257728 | 0.554347748 | 0.748771246 |
| H2afy | 2227.298024 | 0.032958697 | 0.055757799 | 0.5911047 | 0.554450271 | 0.748771246 |
| Prdm2 | 3529.744522 | -0.048321468 | 0.081730676 | -0.591228045 | 0.554367634 | 0.748771246 |
| Rel | 58.45766292 | -0.087103075 | 0.147352973 | -0.591118546 | 0.554440994 | 0.748771246 |
| Bex2 | 5446.217425 | -0.034963757 | 0.059170432 | -0.590899128 | 0.55458801 | 0.748803416 |
| Cic | 7663.632863 | 0.038278682 | 0.064777095 | 0.590929279 | 0.554567807 | 0.748803416 |
| F630042J09Rik | 24.52632518 | 0.099316258 | 0.168089691 | 0.590852763 | 0.554619079 | 0.748803416 |
| Atf5 | 569.5793618 | -0.056606894 | 0.095954574 | -0.589934294 | 0.555234702 | 0.749154089 |
| Atp6ap2 | 8360.627703 | -0.044212537 | 0.074911027 | -0.590200653 | 0.555056134 | 0.749154089 |
| Ccdc107 | 562.3874758 | -0.045010429 | 0.076295817 | -0.589946219 | 0.555226706 | 0.749154089 |
| Cmtm8 | 33.45298972 | -0.097387998 | 0.165130103 | -0.589765257 | 0.555348038 | 0.749154089 |
| Dcn | 492.094384 | 0.084358289 | 0.142937005 | 0.590178096 | 0.555071256 | 0.749154089 |
| Dnajc13 | 3324.396462 | 0.055776136 | 0.094503867 | 0.590199515 | 0.555056897 | 0.749154089 |
| Fbxl8 | 26.49390958 | 0.096619156 | 0.163831194 | 0.589748223 | 0.55535946 | 0.749154089 |
| Itgb5 | 2081.987797 | 0.049714232 | 0.084259873 | 0.59001076 | 0.555183436 | 0.749154089 |
| Lat | 15.19571426 | 0.098919616 | 0.167753671 | 0.589671843 | 0.555410676 | 0.749154089 |
| Mki67ip | 842.9502212 | -0.04214926 | 0.071456085 | -0.589862428 | 0.555282885 | 0.749154089 |
| Scara5 | 20.67301429 | 0.098549616 | 0.167115306 | 0.589710294 | 0.555384892 | 0.749154089 |
| Akap17b | 812.6572692 | -0.052830968 | 0.089607865 | -0.58957959 | 0.555472538 | 0.749172311 |
| Plekha5 | 2291.425778 | 0.039020661 | 0.066204494 | 0.58939596 | 0.555595686 | 0.749273179 |
| B3gntl1 | 205.6849733 | 0.078149374 | 0.132624414 | 0.589253303 | 0.555691365 | 0.749276814 |
| Enc1 | 36674.0018 | -0.056774489 | 0.096362588 | -0.589175637 | 0.555743459 | 0.749276814 |
| Hilpda | 144.705607 | -0.068081984 | 0.115546649 | -0.589216428 | 0.555716098 | 0.749276814 |
| Lims1 | 993.3352613 | -0.043439393 | 0.073745253 | -0.589046635 | 0.555829991 | 0.749298616 |
| Zbtb4 | 7437.542528 | -0.030650191 | 0.052037027 | -0.589007339 | 0.555856351 | 0.749298616 |
| A930009A15Rik | 11.08351095 | 0.094855232 | 0.161188962 | 0.588472252 | 0.556215353 | 0.749717328 |
| Aldh4a1 | 1012.303573 | -0.054437956 | 0.09259229 | -0.587931851 | 0.556578037 | 0.750140927 |
| R3hcc1 | 656.1629 | 0.044806101 | 0.07622955 | 0.587778643 | 0.556680881 | 0.750184817 |
| Ube2v1 | 8181.743499 | -0.032435491 | 0.055186888 | -0.587739085 | 0.556707437 | 0.750184817 |
| Sos1 | 2607.209177 | -0.049906049 | 0.084932574 | -0.587596101 | 0.556803429 | 0.75024892 |
| Atl3 | 1635.074579 | 0.040952703 | 0.069713175 | 0.587445673 | 0.556904427 | 0.750319756 |
| Atat1 | 1838.465362 | 0.04205974 | 0.071654181 | 0.586982359 | 0.557215555 | 0.750608399 |
| Dnajc14 | 2559.125205 | -0.036750028 | 0.062607638 | -0.58698952 | 0.557210745 | 0.750608399 |
| Fam105a | 838.8125251 | 0.044857206 | 0.076473643 | 0.586570806 | 0.557491994 | 0.750915497 |
| E130309D02Rik | 733.5535901 | 0.045698277 | 0.077988885 | 0.58595885 | 0.557903168 | 0.751338695 |
| Thbs2 | 391.9664599 | -0.074719881 | 0.127512237 | -0.58598204 | 0.557887584 | 0.751338695 |
| Myl12b | 7018.32707 | 0.041512508 | 0.07088437 | 0.58563698 | 0.558119492 | 0.751456036 |
| Ostf1 | 357.7107995 | 0.051262081 | 0.087537493 | 0.585601425 | 0.558143391 | 0.751456036 |
| Pfas | 1182.207835 | -0.054379696 | 0.092882373 | -0.585468415 | 0.558232798 | 0.751456036 |
| Wtap | 2039.566126 | -0.036744336 | 0.06274556 | -0.585608536 | 0.558138611 | 0.751456036 |
| Zc3h12c | 621.7316892 | -0.069746806 | 0.119126938 | -0.585483078 | 0.558222942 | 0.751456036 |
| Ankrd45 | 3106.934529 | -0.033943296 | 0.057999316 | -0.585236144 | 0.558388945 | 0.751590158 |
| Rfx4 | 709.3999139 | -0.064414327 | 0.11008689 | -0.585122596 | 0.558465287 | 0.751590158 |

| | | | | | | | |
|---|---|---|---|---|---|---|---|
| Tmem63b | 5918.504264 | 0.0171974814134941 | 0.0486551244916457 | 0.353456734376395 | 0.723746059077625 | 0.932714926679326 | NM_198167 |
| Fam104a | 1365.994267 | 0.0214175150808468 | 0.0607367773731279 | 0.352628440413809 | 0.724367013202818 | 0.93288234633848 | NM_138598 |
| Ppp1r9b | 37517.9857 | -0.0235141056171047 | 0.0666955422876652 | -0.352558878908065 | 0.72441917022568 | 0.93288234633848 | NM_172261 |
| Alg12 | 458.3531367 | -0.0252042811456315 | 0.0715493243384465 | -0.352264418688411 | 0.724639969841713 | 0.932965302793691 | NM_145477,NM_001142357 |
| Hadhb | 3142.315585 | 0.0193110322712122 | 0.0548272415583153 | 0.352216010186699 | 0.724676270922568 | 0.932965302793691 | NM_145558 |
| Nova1 | 2529.555626 | 0.0269463673564493 | 0.0764923802249824 | 0.352275184498032 | 0.724631896746314 | 0.932965302793691 | NM_021361 |
| 4933412O06Rik | 78.36825715 | -0.0333986112929134 | 0.0949292136207958 | -0.351826482270542 | 0.724968396771069 | 0.933258678792676 | NR_045507 |
| Arhgap23 | 5795.798553 | -0.0215214827933492 | 0.0612103102922665 | -0.3515989821092 | 0.725139028682224 | 0.933334512171561 | NM_021493 |
| Mfap3 | 1234.658304 | 0.0286677916841483 | 0.0815715431254554 | 0.351443537607935 | 0.725255624547354 | 0.933334512171561 | NM_180599,NM_145426 |
| Phf13 | 765.4796978 | 0.0268328308956309 | 0.0763586429294265 | 0.351405287813075 | 0.725284315943314 | 0.933334512171561 | NM_172705 |
| Ttc19 | 3375.014419 | 0.0243715920645167 | 0.0693369236134355 | 0.351495145651287 | 0.725216913661935 | 0.933334512171561 | NM_029704,NM_028360 |
| Nudt8 | 191.0709493 | -0.030931167224159 | 0.0880621292847998 | -0.351242554266717 | 0.725406387643186 | 0.933408910169627 | NM_025529 |
| Aldh18a1 | 978.8416549 | 0.0206314504580139 | 0.0588830413329283 | 0.350380177229 | 0.726053400966534 | 0.933575663697777 | NM_019698,NM_153554 |
| Bloc1s4 | 256.1819043 | -0.0312861599203915 | 0.0893067764812756 | -0.350322351260221 | 0.726096792901982 | 0.933575663697777 | NM_133724 |
| Camsap3 | 1406.820521 | 0.01849532090995 | 0.0527640162238059 | 0.350529058127409 | 0.725941686515226 | 0.933575663697777 | NM_001163749,NM_027171 |
| Diap2 | 1244.134568 | -0.0280986649944793 | 0.0800982353175681 | -0.350802547435354 | 0.725736485939359 | 0.933575663697777 | NM_172493 |
| Fchsd1 | 366.9271334 | 0.0278782987146704 | 0.0794498766649675 | 0.350891655027112 | 0.725669632260164 | 0.933575663697777 | NM_175684 |
| Ireb2 | 2613.282499 | 0.0246587459473643 | 0.0703615208692953 | 0.350457830398107 | 0.725995132335235 | 0.933575663697777 | NM_022655 |
| Mmp14 | 411.7851467 | 0.0330761521187061 | 0.094279958604808 | 0.350829090383365 | 0.725716571663016 | 0.933575663697777 | NM_008608 |
| Rpgrip1 | 28.10911649 | 0.0291200807101525 | 0.0830695853291858 | 0.35055045230762 | 0.725925633631263 | 0.933575663697777 | NM_023879,NM_001168515 |
| Rpl35a | 4369.42981 | -0.0229239877648533 | 0.0654412246798093 | -0.350298880820396 | 0.726114405098271 | 0.933575663697777 | NM_021338,NM_001130485,NM_001130484 |
| B230209E15Rik | 1129.751398 | 0.0262364943052226 | 0.0752188239863472 | 0.348802240114585 | 0.727237781545654 | 0.933579400119738 | NR_045727 |
| Ccar1 | 2048.17983 | -0.02449989871615 | 0.0703217971720542 | -0.348396936673942 | 0.727542102777604 | 0.933579400119738 | NM_026201 |
| Cdk8 | 920.2883986 | 0.0228791257532755 | 0.0656850011946859 | 0.348315830663737 | 0.727603006215769 | 0.933579400119738 | NM_153599 |
| Chrnb3 | 91.35448139 | 0.0332524888474814 | 0.0950898139206264 | 0.349695592792284 | 0.726567161548033 | 0.933579400119738 | NM_173212,NM_027454 |
| Crcp | 1038.054103 | -0.0190647704454653 | 0.054618311766354 | -0.349054553846712 | 0.727048354036507 | 0.933579400119738 | NM_007761 |
| Dync1li2 | 11869.66118 | -0.0199184381580649 | 0.0571020585056372 | -0.348821718153971 | 0.727223157583971 | 0.933579400119738 | NM_001013380 |
| Fam217b | 1674.122277 | 0.0212655556848541 | 0.061070029121535 | 0.348215908699399 | 0.72767804113641 | 0.933579400119738 | NM_001081289 |
| Gnb2l1 | 5805.721191 | 0.0173628313295747 | 0.0496150422069444 | 0.349950953526439 | 0.726375506396189 | 0.933579400119738 | NM_008143 |
| Kat2a | 3550.559765 | -0.0182067065095883 | 0.0520480414676442 | -0.349805794727292 | 0.726484449909123 | 0.933579400119738 | NM_001038010,NM_020004 |
| Kcnt2 | 974.5968447 | 0.0289024446192647 | 0.0830161116338779 | 0.348154641917364 | 0.727724049811083 | 0.933579400119738 | NM_001081027 |
| Klhl7 | 3108.543804 | 0.0246108574803268 | 0.0706191262732394 | 0.3485013023965 | 0.727463735885305 | 0.933579400119738 | NM_001161800,NM_026448 |
| Mogs | 1111.601066 | 0.0219378419160939 | 0.0628530470531909 | 0.349033864619618 | 0.727063886089937 | 0.933579400119738 | NM_020619 |
| Ndufaf6 | 190.9813776 | 0.0303883944102785 | 0.0871129653148381 | 0.34883893919178 | 0.727210228244415 | 0.933579400119738 | NM_001085493 |
| Plaur | 40.02117959 | -0.0325217166573958 | 0.0931638026921221 | -0.349081034882938 | 0.727028474053479 | 0.933579400119738 | NM_011113 |
| Prnp | 38453.64813 | 0.0166392398833626 | 0.0477521758475184 | 0.348449878734213 | 0.727502348902435 | 0.933579400119738 | NM_011170 |
| Prr12 | 3253.785862 | -0.0215051066106201 | 0.0616062748656635 | -0.349073315299674 | 0.72703426932072 | 0.933579400119738 | NM_175022 |
| Sfn | 67.91907469 | 0.0331671228645999 | 0.0949964575724858 | 0.349140628104918 | 0.72698373658805 | 0.933579400119738 | NM_018754 |
| Smpdl3a | 1034.75466 | -0.0274906978079413 | 0.0788900059136713 | -0.34846870005339 | 0.727488216263572 | 0.933579400119738 | NM_020561 |
| Snrnp40 | 750.3586333 | 0.0240528566528052 | 0.0688086808173687 | 0.349561368814002 | 0.726667907144635 | 0.933579400119738 | NM_025645 |
| Srsf12 | 632.7183814 | 0.0220037990446178 | 0.0631639273068124 | 0.348360211006776 | 0.727569680291324 | 0.933579400119738 | NM_177774 |
| Tanc2 | 8520.550719 | 0.0219087948686734 | 0.062818166526242 | 0.348765270943097 | 0.727265537987603 | 0.933579400119738 | NM_181071 |
| Tapt1 | 2309.112017 | 0.0202436793849041 | 0.0579230060296863 | 0.349492900532975 | 0.726719299766855 | 0.933579400119738 | NM_173764 |
| Tmem208 | 605.2096735 | -0.0241001870253625 | 0.0690945439331475 | -0.348800146197949 | 0.727239353648022 | 0.933579400119738 | NM_025486 |
| Zbtb7a | 5580.955499 | 0.0220155409837671 | 0.0632053629449013 | 0.348317610373647 | 0.727601669792614 | 0.933579400119738 | NM_010731 |
| Zfp41 | 289.6158766 | -0.0276999932825924 | 0.0794984045307757 | -0.348434581122557 | 0.72751383571184 | 0.933579400119738 | NM_011759,NM_001044718 |
| Bsn | 20513.01519 | 0.0270491333436969 | 0.0778031977667163 | 0.347660946080912 | 0.728094829603051 | 0.933972579902324 | NM_007567 |
| Fam168a | 16128.76744 | 0.0183761621597746 | 0.0528707435458228 | 0.347567689186139 | 0.728164875365975 | 0.933979954045577 | NM_178764 |
| Acadm | 1317.635808 | -0.0182344208951178 | 0.052540327937014 | -0.347055711509404 | 0.72854946502805 | 0.933996867222924 | NM_007382 |
| Agfg2 | 1857.685967 | -0.0216240193502296 | 0.0624996939616959 | -0.34598600376319 | 0.729353233300092 | 0.933996867222924 | NM_145566,NM_178162 |
| Arsg | 1426.317249 | -0.02526156964315 | 0.07293252909121 | -0.346369033926655 | 0.729065393825009 | 0.933996867222924 | NM_028710,NM_001166177 |
| B3galt4 | 153.3745816 | -0.0307560906782652 | 0.0886315433134466 | -0.347010663793769 | 0.728583307443342 | 0.933996867222924 | NM_019420 |
| Cstb | 618.3963443 | 0.025862141491994 | 0.0747540704110209 | 0.345962987029281 | 0.72937053112878 | 0.933996867222924 | NM_007793 |
| Cyp4f16 | 150.9165522 | 0.0321364381128745 | 0.0925824152895312 | 0.34711168435577 | 0.728507415768084 | 0.933996867222924 | NM_024442 |
| Fam189a2 | 539.7591073 | 0.0277879721676756 | 0.0800460433484126 | 0.347149852825633 | 0.728478742406023 | 0.933996867222924 | NM_001114174 |
| Fopnl | 1258.917805 | 0.0172113259223083 | 0.0496512944674209 | 0.346644052424487 | 0.728858746496636 | 0.933996867222924 | NM_025345 |
| Frs2 | 2542.218086 | -0.0198623903515618 | 0.0573111123044245 | -0.346571363788177 | 0.728913362408069 | 0.933996867222924 | NM_177798 |
| Ift81 | 1359.555421 | -0.0233759025016214 | 0.0673972623358459 | -0.346837567157215 | 0.728713352438432 | 0.933996867222924 | NM_009879 |
| Il4ra | 364.9141222 | -0.0276862445879452 | 0.079788590869195 | -0.346995031323888 | 0.728595051570661 | 0.933996867222924 | NM_001008700 |
| Lrrc14 | 839.879937 | 0.0240182733971317 | 0.0694321987476133 | 0.345924136500968 | 0.729399728884541 | 0.933996867222924 | NM_145471 |
| Neurl1b | 2359.794983 | -0.0225262641910283 | 0.0650547835657258 | -0.346266069247155 | 0.729142765962545 | 0.933996867222924 | NM_001081656 |
| Ppp6r2 | 3995.113466 | 0.0175823175359271 | 0.0506348350955625 | 0.347237578689537 | 0.728412841401915 | 0.933996867222924 | NM_027805,NM_026813 |
| Reep6 | 464.2920379 | 0.0257156965897306 | 0.074307636471287 | 0.346070711045524 | 0.729289574192019 | 0.933996867222924 | NM_139292,NM_001204931 |
| Secisbp2l | 6589.039596 | 0.0224579016806383 | 0.0648617172343885 | 0.346242786010166 | 0.729160262381733 | 0.933996867222924 | NM_177608 |
| Slc4a2 | 1517.699465 | 0.0323401134454631 | 0.0933083087916012 | 0.346594144340274 | 0.728896245683365 | 0.933996867222924 | NM_001253892,NM_009207 |
| Usp6nl | 854.3966154 | 0.0250097498492233 | 0.0721821132168759 | 0.346481264327631 | 0.728981062164437 | 0.933996867222924 | NM_001080548,NM_181399 |
| Zfp518b | 726.4054561 | -0.0214602362951694 | 0.0618862390727798 | -0.346769114050244 | 0.728764782420429 | 0.933996867222924 | NM_001081144,NM_001177902 |
| Ahrr | 41.43317039 | -0.0313950768587494 | 0.0907936673508642 | -0.345784874372635 | 0.729504393274995 | 0.934048551495158 | NM_009644 |
| Chd2 | 2742.49414 | -0.0224432624076875 | 0.0649884171864831 | -0.345342499160843 | 0.729836899947239 | 0.934125507660513 | NM_001081345 |
| Fbxl5 | 3623.466486 | -0.0215321532902783 | 0.0623619433793334 | -0.345277137360892 | 0.729886032765167 | 0.934125507660513 | NM_001159963,NM_178729 |
| Hey1 | 1831.846278 | -0.0204375034296431 | 0.0591900439984216 | -0.34528616721738 | 0.729879244907274 | 0.934125507660513 | NM_010423 |
| Mmd | 7810.58178 | 0.0209696063353216 | 0.0607070560635214 | 0.345422883188074 | 0.729776476360623 | 0.934125507660513 | NM_026178 |
| Rif1 | 1124.871273 | 0.0280528210725088 | 0.0812083039849233 | 0.345442765036898 | 0.729761531702415 | 0.934125507660513 | NM_175238 |
| Atp2c1 | 6046.212058 | -0.0195621203416999 | 0.0567030680465603 | -0.34499227318065 | 0.730100179652316 | 0.93415266534216 | NM_001253834,NM_001253836,NM_001253831,NM_175025 |
| C1ql3 | 4464.274178 | 0.028553972328679 | 0.0827274626273839 | 0.345157114963021 | 0.729976257137067 | 0.93415266534216 | NM_153155 |
| Papd7 | 2082.607239 | 0.0216079715548524 | 0.0626322893423264 | 0.344997313394546 | 0.730096390484336 | 0.93415266534216 | NM_001169131,NM_198600 |
| Snn | 11743.64419 | -0.0179941442931131 | 0.0521757862747338 | -0.34487538335051 | 0.730188057771251 | 0.934182819022829 | NM_009223 |
| Elf2 | 1118.839471 | -0.0233352955511629 | 0.0677732784188294 | -0.344314102778885 | 0.730610079488543 | 0.934558120346123 | NM_023502 |
| Rdh10 | 329.5972594 | -0.0279691370010781 | 0.0812311210313465 | -0.344315536286703 | 0.73060900154321 | 0.934558120346123 | NM_133832 |
| Nup107 | 674.7274305 | -0.0224427841776464 | 0.0652276866979308 | -0.344068375160671 | 0.730794865510964 | 0.93471217895679 | NM_134010 |
| L2hgdh | 879.184282 | 0.0220792758313933 | 0.0643272786353199 | 0.343233481965927 | 0.731422817961299 | 0.93535063418486 | NM_145443 |
| Mrm1 | 584.6644151 | 0.0239186723811281 | 0.06967151828853 | 0.343306317541035 | 0.731368028599901 | 0.93535063418486 | NM_145433 |
| Aco1 | 1454.117593 | 0.0195989796617275 | 0.0571461907436382 | 0.342962136350433 | 0.731626945284722 | 0.935424324812553 | NM_007386 |
| Npl | 165.7533467 | -0.0304754278291829 | 0.0888335644799436 | -0.34306208478287 | 0.731551754072127 | 0.935424324812553 | NM_028749 |
| Rcn3 | 147.9949088 | 0.032581018795006 | 0.0950589186104132 | 0.342745523170084 | 0.731789912209684 | 0.935424324812553 | NM_026555 |
| Thap3 | 402.6732951 | 0.0259601748442859 | 0.0757455103914858 | 0.342728891918641 | 0.731802425111514 | 0.935424324812553 | NM_175152,NM_001145929 |
| Unc5c | 1563.304218 | 0.0188390554034927 | 0.0549453899464974 | 0.342868717863994 | 0.731697226348904 | 0.935424324812553 | NM_009472 |
| Smim18 | 173.7006034 | 0.0288564118721612 | 0.0842253364470248 | 0.342609636119542 | 0.731892151803375 | 0.935456700140416 | NM_001206849 |
| Fam216b | 48.56273082 | -0.0303338957415379 | 0.0885930662263849 | -0.342395822084143 | 0.732053032203042 | 0.935580005787559 | NM_177629 |
| Rab11fip3 | 6297.570274 | 0.014560775688594 | 0.0425557937505922 | 0.342157304688774 | 0.732232514084982 | 0.935722271504668 | NM_001162868,NM_001162869,NM_153140 |
| Uchl5 | 1581.706537 | 0.0254673008168107 | 0.0744490986595213 | 0.342076684276334 | 0.732293183427307 | 0.935722271504668 | NM_019562,NM_001159866 |
| Taf1 | 3574.728455 | 0.0194995020828838 | 0.0570791331745504 | 0.341622253149036 | 0.732635188126454 | 0.936076941043616 | NM_001081008 |
| Crtc3 | 787.9497277 | 0.0228366727450579 | 0.0668893360999075 | 0.341409768381443 | 0.732795122306641 | 0.936094952395418 | NM_173863 |
| Fam120c | 3417.49273 | 0.0211726847493829 | 0.0620176348593617 | 0.341397810435637 | 0.732804123222661 | 0.936094952395418 | NM_198105 |
| Sertad2 | 1442.249081 | 0.0226958413255101 | 0.0664891235572008 | 0.341346676136959 | 0.732842613151638 | 0.936094952395418 | NM_001038625,NM_021372 |
| 0610009O20Rik | 1660.489052 | -0.0195084661991941 | 0.0573903474052957 | -0.339925912304094 | 0.733912322172099 | 0.93645633651286 | NM_024179 |
| 1700009P17Rik | 72.46294828 | 0.0286897439366521 | 0.0844756386040215 | 0.339621509949572 | 0.734141577400512 | 0.93645633651286 | NM_001081275 |
| 1700101I11Rik | 66.6860692 | 0.0320447696897502 | 0.0944884580027407 | 0.339139513619967 | 0.734504632848542 | 0.93645633651286 | NR_045270 |
| B230312C02Rik | 39.55334325 | 0.0294310382873842 | 0.086935718961252 | 0.338537929351017 | 0.734957849113671 | 0.93645633651286 | NR_040745 |
| C130026L21Rik | 152.7157664 | -0.0314457881636267 | 0.0925206834359928 | -0.339878468206316 | 0.733948052291969 | 0.93645633651286 | NR_015546 |
| Ccdc148 | 693.7124892 | 0.0247998927324811 | 0.0731052419792823 | 0.339235492025445 | 0.734432334029911 | 0.93645633651286 | NM_001001178 |
| Chac1 | 218.7731004 | 0.0298911093900569 | 0.0880410418840552 | 0.33951335366319 | 0.734223039095024 | 0.93645633651286 | NM_026929 |
| Cth | 136.9878684 | -0.0314173243045603 | 0.0926708280120153 | -0.339020649524 | 0.734594174311006 | 0.93645633651286 | NM_145953 |
| Fam63b | 3658.079968 | 0.021990915039426 | 0.0648222355995156 | 0.339249562068341 | 0.734421735515715 | 0.93645633651286 | NM_172772 |
| Foxj3 | 3650.045617 | 0.0181528158727877 | 0.0535768497311243 | 0.338818276249681 | 0.734746632338634 | 0.93645633651286 | NM_172699 |
| Gabpb1 | 651.9286205 | 0.022780644450595 | 0.067225149382351 | 0.33887086344752 | 0.73470701435493 | 0.93645633651286 | NR_073183,NM_010249,NM_207669,NM_001271492,NM_001271469,NM_001271468,NM_001271470,NM_001271467 |
| Gga1 | 2976.624104 | -0.0216741513495581 | 0.0640374080331533 | -0.338460784333073 | 0.735015974629615 | 0.93645633651286 | NM_145929 |
| Gm16516 | 31.98343668 | -0.0304576461233869 | 0.0897767473352102 | -0.339259853218602 | 0.734413983552751 | 0.93645633651286 | NR_027800 |
| Hddc2 | 337.2468707 | 0.0270178046643702 | 0.0796635134032396 | 0.339149047163058 | 0.734497451295525 | 0.93645633651286 | NM_027168 |
| Hnrnpa1 | 117.8933061 | -0.0323117953126537 | 0.0948472837203563 | -0.340671804665703 | 0.733350666896703 | 0.93645633651286 | NM_010447,NM_001039129 |
| Kidins220 | 16056.68149 | 0.0162170580253026 | 0.0479004109543376 | 0.338557805709891 | 0.734942873361858 | 0.93645633651286 | NM_001081378 |
| Lhx9 | 67.62040579 | 0.020050961805368 | 0.059218058411065 | 0.338595393759507 | 0.734914553093042 | 0.93645633651286 | NM_010714,NM_001025565,NM_001042577 |
| Lyve1 | 34.99872084 | 0.027317502090404 | 0.0802423905199704 | 0.340437291478814 | 0.733527239433551 | 0.93645633651286 | NM_053247 |
| Oraov1 | 562.7162804 | -0.02304486671415 | 0.0679183540635808 | -0.339302490937528 | 0.73438186634117 | 0.93645633651286 | NM_028184 |
| Pcdhga10 | 409.5123834 | -0.0256204672572907 | 0.0754736426275638 | -0.339462444971932 | 0.734261383794007 | 0.93645633651286 | NM_033593 |
| Prima1 | 189.2849872 | -0.0319810520789061 | 0.0938889881474633 | -0.340626230082236 | 0.733384980365444 | 0.93645633651286 | NM_133364 |
| Rbfox3 | 12478.02584 | -0.0170541483252542 | 0.0502488235722454 | -0.339393982044864 | 0.734312951482234 | 0.93645633651286 | NM_001039168,NM_001024931,NM_001039167 |
| Rspo1 | 645.2136961 | 0.026670764973115 | 0.0784534246795205 | 0.339956669604471 | 0.733889159176608 | 0.93645633651286 | NM_138683 |
| Rxrb | 2738.495131 | -0.0214125407800087 | 0.0632248693483049 | -0.338672756475736 | 0.734856266209216 | 0.93645633651286 | NM_011306,NM_001205216,NM_001205215,NM_001205214 |
| Selenbp1 | 390.4675331 | 0.0290802160052747 | 0.0858727615121084 | 0.338643074861104 | 0.734878628852889 | 0.93645633651286 | NM_009150 |
| Sowahc | 431.6932025 | -0.0302466622826965 | 0.0887668013208041 | -0.340742956067379 | 0.733297097501911 | 0.93645633651286 | NM_172939 |
| Sptlc2 | 1073.555255 | 0.0218152342071237 | 0.0644815648436457 | 0.338317382030368 | 0.735124026246464 | 0.93645633651286 | NM_011479 |
| Stk30 | 308.8872255 | 0.0255136188262543 | 0.0748924949893296 | 0.340669901043484 | 0.733352100138449 | 0.93645633651286 | NM_011973 |
| Tbl3 | 1203.030503 | 0.0210984188521685 | 0.0621187639400997 | 0.339646469342393 | 0.734122778784683 | 0.93645633651286 | NM_145396 |
| Tfap4 | 123.1206976 | -0.0321983468216467 | 0.0950972736650635 | -0.338583279843022 | 0.734923680139943 | 0.93645633651286 | NM_031182 |
| Wdr75 | 1171.699408 | -0.0204714014670155 | 0.0605037512805898 | -0.338349292956699 | 0.735099981359036 | 0.93645633651286 | NM_028599 |
| Npy5r | 155.4980991 | 0.0303946618079005 | 0.0898734967031641 | 0.33819382713336 | 0.735217127326709 | 0.936492808799348 | NM_016708 |
| Gpr133 | 307.2523333 | 0.0316981961684901 | 0.0937640851817489 | 0.338063301178137 | 0.735315485464182 | 0.936535970350961 | NM_001081342 |
| Mtrf1l | 260.9656684 | 0.0275905183834294 | 0.0816845586819011 | 0.337769081802516 | 0.735537211075553 | 0.936736237776914 | NM_175374 |
| Dnahc11 | 48.48938375 | -0.0306724634303385 | 0.0908452273796283 | -0.337634285422204 | 0.735638801855143 | 0.936783486959569 | NM_010060 |
| E2f4 | 1165.986779 | -0.0226944943604688 | 0.0672772559620553 | -0.337327883486636 | 0.735869742252627 | 0.936995430922306 | NM_148952 |
| Cldn26 | 34.53931205 | 0.0299409657691607 | 0.0888386071368153 | 0.337026510591846 | 0.736096915462142 | 0.937120402629543 | NM_029070 |
| Psma4 | 2923.900567 | 0.0188418497002891 | 0.0559030451099324 | 0.337045140622249 | 0.736082871579666 | 0.937120402629543 | NM_011966 |
| Armc10 | 898.1849763 | 0.0221712127107206 | 0.0658496365733984 | 0.336694534160531 | 0.73634718415544 | 0.937175149961488 | NM_026034 |
| Cpne4 | 2682.254625 | -0.026234178974892 | 0.077901297485527 | -0.336761771904581 | 0.736296493048995 | 0.937175149961488 | NM_028719 |
| Pi4ka | 17009.24952 | 0.0144996749951509 | 0.0430592210662152 | 0.336737977049184 | 0.736314432061411 | 0.937175149961488 | NM_001001983 |
| Rps29 | 7567.206932 | -0.0234373590472639 | 0.0696240902785626 | -0.336627149503745 | 0.736397987171996 | 0.937175149961488 | NM_009093 |
| C5ar2 | 45.39900486 | -0.0310392523441293 | 0.0924071385000759 | -0.335896694215933 | 0.73694877001066 | 0.937206199382379 | NM_001146005,NM_176912 |
| Ccdc177 | 533.3644897 | -0.0247395276415753 | 0.0737929589273376 | -0.33525581520086 | 0.737432121261072 | 0.937206199382379 | NM_001008423 |
| Ccdc89 | 25.82032089 | 0.0286156859920009 | 0.0853981917617713 | 0.335085385318553 | 0.737560677047443 | 0.937206199382379 | NM_027298 |
| Cpq | 539.7141006 | 0.0268343764511998 | 0.0797593398597833 | 0.336446446251626 | 0.736534229664721 | 0.937206199382379 | NM_018755,NM_176073 |
| Eprs | 4179.033859 | 0.0188536197874234 | 0.056084677647882 | 0.336163468849595 | 0.736747599115767 | 0.937206199382379 | NM_029735 |
| Fam131b | 8051.693071 | 0.0187793364608381 | 0.0558906536294636 | 0.33600137485131 | 0.736869829706964 | 0.937206199382379 | NM_029528,NM_001113327 |
| Golga2 | 4877.686947 | -0.0159108471584591 | 0.0474657137784358 | -0.335207160956834 | 0.737468820558553 | 0.937206199382379 | NM_001080968,NM_133852 |
| Kdm2a | 4567.692233 | -0.020758925203034 | 0.0619568015221526 | -0.335054823571092 | 0.737583730644318 | 0.937206199382379 | NM_001001984 |
| Mrpl48 | 1878.717643 | 0.0195411747925591 | 0.0582698553455863 | 0.33535650082988 | 0.737356177231506 | 0.937206199382379 | NR_003559,NM_198831 |
| Nat8 | 59.47749196 | -0.0312253101630291 | 0.0931275846088072 | -0.335296038163069 | 0.737401782027704 | 0.937206199382379 | NM_023455 |
| Pcf11 | 1365.102876 | 0.02824243408951 | 0.0841328307244604 | 0.33568862293491 | 0.737105686070112 | 0.937206199382379 | NM_029078 |
| Pycr1 | 38.34277971 | -0.0290923898990595 | 0.0868246616339421 | -0.335070581924232 | 0.737571843640077 | 0.937206199382379 | NM_144795 |
| Slc17a7 | 54750.44893 | 0.0214659538989299 | 0.0638342514071112 | 0.336276425676679 | 0.736662425440963 | 0.937206199382379 | NM_182993 |
| St6gal1 | 1171.462898 | -0.0181635004412858 | 0.0540190824809192 | -0.33624229822307 | 0.736688158477111 | 0.937206199382379 | NM_001252506,NM_001252505,NM_145933 |
| Tatdn1 | 290.4920243 | -0.0286493920698874 | 0.0852373071984249 | -0.336113293715323 | 0.736785434082177 | 0.937206199382379 | NM_175151 |
| Tex30 | 105.3363976 | 0.031674356846114 | 0.0944155583960227 | 0.335478149832648 | 0.737264424603707 | 0.937206199382379 | NM_029368 |
| Vmn2r29 | 214.5813051 | 0.029079136352302 | 0.0866515067348742 | 0.335587198053864 | 0.73718217918075 | 0.937206199382379 | NR_003555,NM_001113468 |
| Yipf3 | 1809.574892 | 0.019457826954206 | 0.0579920103256067 | 0.335525994787153 | 0.737228339108084 | 0.937206199382379 | NM_145353 |
| Mycn | 236.4336439 | -0.0308881845355052 | 0.0922577486881009 | -0.334803146345246 | 0.737773586921348 | 0.937365444207076 | NM_008709 |
| Fam117a | 211.8257745 | 0.030203530275298 | 0.090237784725864 | 0.33471045822854 | 0.737843511545368 | 0.937372297420676 | NM_172543 |
| Abhd10 | 725.0112739 | -0.0216999166522381 | 0.0648738331250196 | -0.334494134336409 | 0.738006716382674 | 0.937497644265389 | NR_073572,NM_001272070,NM_172511 |
| Papd5 | 1664.056952 | 0.0194259155079424 | 0.0580980258723154 | 0.334364470919523 | 0.738104548439279 | 0.937539932723764 | NM_001164499,NM_001164498,NM_001164497 |
| Lix1 | 1962.722114 | -0.0204441905567547 | 0.0611918859671507 | -0.334099696939063 | 0.738304328752326 | 0.937711697075832 | NM_025681 |
| Macrod2 | 1075.7308 | 0.0259011464863852 | 0.0776084495390388 | 0.333741321212149 | 0.738574765744982 | 0.937973163770905 | NM_001013802,NM_028387 |
| Dgkb | 6243.640685 | 0.0286951768440869 | 0.0860642664548017 | 0.333415690693846 | 0.738820520643751 | 0.938076298642582 | NM_178681 |
| Pqlc3 | 38.60128659 | -0.0292509899030132 | 0.0877414723156737 | -0.333377012386741 | 0.73884971311922 | 0.938076298642582 | NM_172574,NM_001161111 |
| Ttc25 | 78.19065383 | -0.0314164691417534 | 0.0942194254357474 | -0.333439404840966 | 0.738802622563161 | 0.938076298642582 | NM_028918 |
| Ikbke | 56.07664125 | -0.0311497128108857 | 0.0934656107646594 | -0.333274586835138 | 0.738927020683849 | 0.938092457826743 | NM_019777 |
| Masp1 | 428.8092392 | 0.0237467575233988 | 0.0713314876663827 | 0.332907083537391 | 0.739204422257906 | 0.938362617995135 | NM_008555 |
| 1110035M17Rik | 80.62596021 | -0.0316098705464687 | 0.0950083002585361 | -0.332706410497316 | 0.739355910103971 | 0.938390908708631 | NR_045780 |
| Pigl | 272.1940361 | -0.0255326923302329 | 0.0767283612336892 | -0.332767335567988 | 0.739309916769031 | 0.938390908708631 | NM_001039536 |
| Mrps16 | 633.3032356 | 0.0233977050132843 | 0.0703652966916929 | 0.332517677226628 | 0.739498393861685 | 0.938418053976557 | NM_025440 |
| Tuba1c | 36.94862649 | -0.0252915741180703 | 0.076063297841492 | -0.332506936141203 | 0.73950650308892 | 0.938418053976557 | NM_009448 |
| Deaf1 | 1617.263987 | -0.022432027466594 | 0.0675086615616457 | -0.332283694383574 | 0.739675051096701 | 0.938470248011599 | NR_027769,NM_016874,NR_027770 |
| Rpl12 | 5248.686572 | -0.0171968264659605 | 0.0517538170246218 | -0.332281316714846 | 0.739676846309018 | 0.938470248011599 | NM_009076 |
| Smap1 | 3815.804078 | 0.0197094649162951 | 0.0593519514174003 | 0.332077790967406 | 0.739830519703884 | 0.938583242726518 | NM_028534 |
| Atp11c | 484.9555082 | -0.0244353061270396 | 0.0736202670567853 | -0.33191004466463 | 0.739957185412581 | 0.938661957497437 | NM_001037863,NM_001001798 |
| Cd55 | 157.4005395 | -0.0311027364907852 | 0.0937591262105111 | -0.331730229875994 | 0.740092971900938 | 0.938752227543925 | NM_010016 |
| Arhgef11 | 4328.282219 | 0.0161367098012543 | 0.0487432468206688 | 0.331055291836074 | 0.740602721153634 | 0.938975678096587 | NM_001003912 |
| Coil | 843.706678 | -0.0203979521563488 | 0.0616246272690611 | -0.331003254060891 | 0.740642027593045 | 0.938975678096587 | NM_016706 |
| Cpsf3 | 1961.611931 | 0.0169641624563614 | 0.0512580403753026 | 0.330956125754179 | 0.74067762627744 | 0.938975678096587 | NM_018813 |
| Rbm7 | 891.051306 | -0.0192019185039222 | 0.0580297416036563 | -0.330897880522568 | 0.740721622973206 | 0.938975678096587 | NM_144948,NR_037589 |
| Tmem98 | 426.8442742 | -0.031485610420877 | 0.0950990250943632 | -0.331082367980481 | 0.740582269608787 | 0.938975678096587 | NM_029537 |
| Vps37d | 628.7418342 | -0.0216135500323266 | 0.0652705373808865 | -0.331137920716059 | 0.740540309287369 | 0.938975678096587 | NM_177574,NM_001199677 |
| Zfp759 | 307.1316882 | -0.0256430252815825 | 0.0774633103196677 | -0.331034462324957 | 0.740618454525875 | 0.938975678096587 | NM_172392 |
| Nlrx1 | 398.7624383 | -0.0283133501102098 | 0.0856180553401608 | -0.330693683682966 | 0.740875873819567 | 0.939089262050876 | NM_178420,NM_001163742,NM_001163743 |
| Slc35e2 | 2203.66353 | 0.0240925974074053 | 0.0728990907794623 | 0.330492426583088 | 0.741027914171123 | 0.93920002453972 | NM_177186 |
| Eif2c3 | 786.2044031 | -0.0287206763023352 | 0.0869638149928698 | -0.330260077765563 | 0.741203455442126 | 0.939258605404547 | NM_153402 |
| Snhg8 | 339.3549297 | 0.0244794352241378 | 0.0741105893750282 | 0.330309547266753 | 0.741166079734649 | 0.939258605404547 | NR_028574 |
| Zdhhc12 | 168.0903733 | -0.0299493903677156 | 0.0907085529405297 | -0.330171625462387 | 0.741270285359978 | 0.939261354251486 | NM_025428,NM_001037762 |
| Sergef | 794.0302588 | -0.0255361404323515 | 0.0774250080318535 | -0.329817730491493 | 0.741537689347212 | 0.939436287760126 | NM_013789 |
| Ssh2 | 1957.216504 | 0.0245605400911079 | 0.0744560878586239 | 0.329866110313815 | 0.741501131581432 | 0.939436287760126 | NM_177710 |
| Foxk2 | 5019.084325 | 0.0143138773623402 | 0.0434127244267844 | 0.329716173111425 | 0.741614432140638 | 0.9394515776816 | NM_001080932 |
| Mag | 8433.319032 | 0.0267955382513339 | 0.0813027810719637 | 0.329577142356499 | 0.74171949621061 | 0.939502738224218 | NM_010758 |
| Tmem230 | 988.032073 | 0.022738600032069 | 0.0690341452201882 | 0.329381930630747 | 0.741867023770654 | 0.939560330483802 | NM_027478,NM_001141971 |
| Zfp955a | 457.8265945 | 0.0266062871908488 | 0.0807852623640884 | 0.329345804076712 | 0.741894326769187 | 0.939560330483802 | NM_029952 |
| Zfp148 | 4427.75554 | 0.0237465461747528 | 0.0721926184060241 | 0.328933161022059 | 0.742206208855021 | 0.939873366735946 | NM_011749 |
| 1700016K19Rik | 68.17025686 | 0.0309914258690945 | 0.094698105095854 | 0.327265533325347 | 0.743467058769364 | 0.939912915706461 | NM_198637 |
| 3110009E18Rik | 34.12500378 | 0.0279759831750057 | 0.0853634638291725 | 0.327727834837989 | 0.743117455532206 | 0.939912915706461 | NM_001172074,NM_028439 |
| 4932411E22Rik | 117.7389976 | 0.0312018316711645 | 0.0950008372074669 | 0.328437438956718 | 0.742580939333137 | 0.939912915706461 | NM_172534 |
| Ahcyl1 | 18950.55947 | 0.0145738680702423 | 0.0445227770871244 | 0.327335108538342 | 0.743414440979868 | 0.939912915706461 | NM_145542 |
| Ap1ar | 3803.455109 | 0.018068792623897 | 0.0551121241276975 | 0.32785513006232 | 0.743021201197453 | 0.939912915706461 | NM_145964 |
| Chdh | 254.3672679 | -0.0304694147378477 | 0.0927868444139058 | -0.328380762707362 | 0.742623786418314 | 0.939912915706461 | NM_172264,NM_001136240,NM_175343 |
| Dnahc5 | 199.361247 | 0.030464337405852 | 0.0931284742862052 | 0.327121620313763 | 0.743575899960639 | 0.939912915706461 | NM_133365 |
| Elmod1 | 10106.37323 | 0.0224803113943117 | 0.0686894160111127 | 0.327274749150216 | 0.743460089029684 | 0.939912915706461 | NM_177769 |
| Grb7 | 100.1777383 | -0.0308636197930844 | 0.0943815165878453 | -0.327009152945303 | 0.743660962426983 | 0.939912915706461 | NM_010346 |
| Iars | 2810.873002 | 0.0177841502159124 | 0.0541477953110634 | 0.328437198850805 | 0.742581120850851 | 0.939912915706461 | NM_172015 |
| Mtx2 | 1128.130582 | -0.0211842518703048 | 0.0644583913751195 | -0.328650023966961 | 0.742420232770375 | 0.939912915706461 | NM_016804 |
| Narf | 2416.933334 | 0.0190524537251827 | 0.0581882624204887 | 0.327427782385096 | 0.743344356200251 | 0.939912915706461 | NM_026272 |
| Ntn3 | 401.0350304 | 0.0264939657586896 | 0.0807541054883904 | 0.328081966835711 | 0.742849688438521 | 0.939912915706461 | NM_010947 |
| Piwil2 | 46.37866932 | -0.0307550143058858 | 0.0940474306452002 | -0.327015997086736 | 0.743655785906774 | 0.939912915706461 | NM_021308 |
| Pla2g15 | 1230.264543 | -0.0179347559397871 | 0.0546810038678593 | -0.327988783511139 | 0.742920143411747 | 0.939912915706461 | NM_133792 |
| Rims1 | 7019.241446 | 0.0190824175844486 | 0.0583139182327408 | 0.327236072669433 | 0.743489339401313 | 0.939912915706461 | NM_001012624,NM_001012625,NM_053270,NM_183018,NM_001012623 |
| Slc7a14 | 6587.15055 | 0.0240085759532407 | 0.0731663108262156 | 0.328137030309783 | 0.742808056512703 | 0.939912915706461 | NM_172861 |
| Soga2 | 4401.90274 | -0.021494961553727 | 0.0655083257657559 | -0.328125643610379 | 0.742816665610436 | 0.939912915706461 | NM_172963,NM_001114098 |
| Sord | 1036.389798 | 0.0218001702498287 | 0.0665411487428853 | 0.327619385322974 | 0.743199462844273 | 0.939912915706461 | NM_146126 |
| Syn2 | 23023.70974 | 0.0203524042780903 | 0.0619509179333892 | 0.328524660441248 | 0.74251500169426 | 0.939912915706461 | NM_013681,NM_001111015 |
| Tagap | 43.57632134 | -0.0276077618100485 | 0.0843286038850871 | -0.327383124327174 | 0.743378128679299 | 0.939912915706461 | NM_145968 |
| Zranb1 | 1620.476177 | 0.0181967505364671 | 0.0555850105822475 | 0.3273679422898 | 0.743389610147269 | 0.939912915706461 | NM_207302 |
| Osgin1 | 24.6887979 | -0.0254682002230371 | 0.0779255840851404 | -0.326827197024419 | 0.74379858780811 | 0.939923295911319 | NM_027950 |
| Zfp397 | 1535.507804 | -0.0245920737716025 | 0.0752282641503292 | -0.326899391463559 | 0.743743981365992 | 0.939923295911319 | NM_027007 |
| Pmf1 | 98.00850234 | 0.0309458867985073 | 0.0947433764209471 | 0.326628498661627 | 0.743948885961803 | 0.939962106929925 | NM_025928 |
| Zbtb3 | 168.3752935 | 0.0284328918675035 | 0.0870531001619877 | 0.326615500362374 | 0.743958718392768 | 0.939962106929925 | NM_133759,NM_001098237 |
| BC005537 | 6808.673588 | -0.0157175710033347 | 0.0481860884079224 | -0.326184828913204 | 0.744284519015387 | 0.940096833434738 | NM_024473 |
| Gins2 | 28.36377864 | 0.0289490705277174 | 0.0887291728795314 | 0.326263274954922 | 0.744225171598133 | 0.940096833434738 | NM_178856 |
| Kcnk2 | 2902.487803 | -0.0198828052468795 | 0.0609654498080084 | -0.326132347247403 | 0.744324224241562 | 0.940096833434738 | NM_001159850,NM_010607,NR_027627 |
| Rsbn1l | 916.3228262 | -0.027949330464132 | 0.0856853678059005 | -0.326185569132928 | 0.744283959003885 | 0.940096833434738 | NM_001080977 |
| Apex2 | 139.1705835 | 0.0305444746625354 | 0.0937935008046712 | 0.325656622265817 | 0.744684166976366 | 0.940439377003047 | NM_029943 |
| Ndufc1 | 1190.510756 | -0.0250615117073953 | 0.0769695904560368 | -0.325602768040059 | 0.744724917641887 | 0.940439377003047 | NM_025523 |
| BC017643 | 838.3609455 | -0.0213651142561088 | 0.0656506805784821 | -0.325436295067313 | 0.744850889701594 | 0.940516691916321 | NM_001252550,NM_001252549,NM_001252548,NM_001254735,NM_144832 |
| Ankib1 | 2367.992256 | 0.0205367932299751 | 0.063183833144541 | 0.325032404776623 | 0.745156546602333 | 0.94063202447114 | NM_001003909 |
| Dnph1 | 89.57710358 | 0.0303176555680024 | 0.093326913022496 | 0.324854370364679 | 0.745291292585624 | 0.94063202447114 | NM_207161 |
| Eif4ebp2 | 942.2518647 | 0.0229678606933786 | 0.0706218538381244 | 0.32522313483903 | 0.745012200521508 | 0.94063202447114 | NM_010124 |
| Fam163b | 7758.956276 | -0.0163459732686034 | 0.0503657252706633 | -0.32454557500683 | 0.745525023938884 | 0.94063202447114 | NM_175427 |
| Fbxl21 | 224.6001 | 0.0298988450725849 | 0.0920867742500994 | 0.3246812076551 | 0.745422358879869 | 0.94063202447114 | NM_178674 |
| Nfyc | 1525.24212 | 0.0183172976499967 | 0.0563798054173996 | 0.324891111531643 | 0.74526348426329 | 0.94063202447114 | NM_001048168,NM_008692 |
| Polr2a | 3679.467281 | -0.0184855768544879 | 0.0568576019850287 | -0.325120585622927 | 0.745089809499639 | 0.94063202447114 | NM_009089 |
| Sarm1 | 1501.799603 | -0.0193126071701822 | 0.059504684877813 | -0.324556078396831 | 0.745517073396982 | 0.94063202447114 | NM_001168521,NM_172795 |

| | | | | | | | | | | | | | | | |
|---|---|---|---|---|---|---|---|---|---|---|---|---|---|---|---|
| Synrg | 4110.187106 | -0.051789787 | 0.088513853 | -0.585103748 | 0.558477959 | 0.751590158 | | Tnrc18 | 3022.160248 | -0.0204805096611024 | 0.063077939941568 | -0.324685772554945 | 0.74541890362737 | 0.94063202447114 | NM_001122730,NM_178242 |
| Slc16a3 | 97.11809984 | -0.08073135 | 0.138036849 | -0.584853615 | 0.55864615 | 0.751751212 | | Slc23a2 | 8989.443049 | 0.0168668302817749 | 0.0520002859397968 | 0.324360337197039 | 0.745665244166861 | 0.940687916447383 | NM_018824 |
| Rab3a | 19382.83934 | 0.034549133 | 0.059092631 | 0.5846606 | 0.558775951 | 0.751795293 | | Ubxn7 | 1821.254951 | 0.021595024054264 | 0.0665863755081835 | 0.324315956371734 | 0.74569884055429 | 0.940687916447383 | NM_177633 |
| Slc38a11 | 91.18270251 | -0.081811439 | 0.139925795 | -0.58467732 | 0.558764707 | 0.751795293 | | Chd3 | 24837.14271 | 0.0168010415166904 | 0.0518184184295046 | 0.324229145270943 | 0.745764558156414 | 0.940689125719005 | NM_146019 |
| Lama5 | 633.9109276 | 0.056431028 | 0.096544123 | 0.584510234 | 0.558877082 | 0.751866068 | | Pum2 | 8756.970725 | 0.0219311822639181 | 0.0676612958774436 | 0.324131868589135 | 0.745838200593794 | 0.940700330105535 | NM_001160219,NM_001160222,NM_001160220,NM_001160221,NR_027670,NM_030723 |
| Leprotl1 | 2710.314875 | 0.032328651 | 0.055321311 | 0.584379695 | 0.558964885 | 0.751918903 | | Colq | 48.71367921 | -0.0282256549907453 | 0.0871754592529855 | -0.323779825568038 | 0.74610473100074 | 0.940755508444617 | NM_009937 |
| Abca1 | 1951.831964 | 0.096349053 | 0.164919817 | 0.584217563 | 0.559073946 | 0.752000323 | | Farsa | 1324.830006 | 0.0197475944224746 | 0.0609998374056189 | 0.323731919007633 | 0.74614100322115 | 0.940755508444617 | NM_025648 |
| Slc12a6 | 6120.359942 | -0.028780124 | 0.049275824 | -0.584061754 | 0.559178765 | 0.752076023 | | Gab3 | 61.37936639 | 0.030722410445875 | 0.0948633209953752 | 0.323859739713021 | 0.746044225644623 | 0.940755508444617 | NM_181584 |
| Fhdc1 | 232.9268354 | -0.068196278 | 0.116796872 | -0.583887878 | 0.559295749 | 0.75216807 | | Tubg1 | 1513.832931 | 0.0205417533848439 | 0.0634103494842316 | 0.323949537448174 | 0.7459762389993 | 0.940755508444617 | NM_134024 |
| Nlrc3 | 17.10053237 | -0.097851741 | 0.16762087 | -0.583768246 | 0.559376245 | 0.752211034 | | Myh11 | 371.4032305 | -0.0288133617349447 | 0.089071650620284 | -0.323485211448221 | 0.74632780556228 | 0.940909365005874 | NM_001161775,NM_013607 |
| Tifab | 151.6708886 | 0.087675122 | 0.150236193 | 0.583581893 | 0.559501645 | 0.75231437 | | Bambi-ps1 | 40.52470621 | -0.0297138684903823 | 0.091973101813973 | -0.323071288282549 | 0.746641253912365 | 0.940970690109281 | NR_027919 |
| Pus7 | 644.553115 | 0.058586493 | 0.100406888 | 0.583490779 | 0.559562963 | 0.75233153 | | Dynll2 | 31018.2382 | 0.0158530375520677 | 0.0491328481061792 | 0.322656596617569 | 0.746955326253894 | 0.940970690109281 | NM_001168472,NM_001168471,NM_026556 |
| Ccr5 | 200.4138042 | -0.093915922 | 0.161038603 | -0.583188879 | 0.559766157 | 0.752539422 | | Glb1l2 | 28.66460502 | 0.021591543535521 | 0.0668824530057738 | 0.322828224222832 | 0.746825336665436 | 0.940970690109281 | NM_153803 |
| Rictor | 2536.0841 | 0.049219016 | 0.084412376 | 0.583078193 | 0.559840663 | 0.752574288 | | Mdfic | 98.87685759 | -0.0288542990909846 | 0.0892680181747943 | -0.32323221329374 | 0.74651938650768 | 0.940970690109281 | NM_175088 |
| Manbal | 1336.123048 | -0.038859504 | 0.066689317 | -0.582694582 | 0.560098921 | 0.752835852 | | Mrpl53 | 662.4420921 | 0.0210607173676845 | 0.065318569299281 | 0.32243078183766 | 0.747126367808273 | 0.940970690109281 | NM_026744 |
| Rbbp9 | 1076.451062 | -0.039697608 | 0.068133459 | -0.582644831 | 0.560132419 | 0.752835852 | | Myadml2 | 265.3926673 | 0.0283440438515353 | 0.0878915114463917 | 0.322488979710212 | 0.747082285115123 | 0.940970690109281 | NM_026751,NM_001204820 |
| Lingo1 | 12552.16738 | -0.06746305 | 0.115869174 | -0.582234663 | 0.560408628 | 0.753141754 | | Nagpa | 424.1400848 | -0.0258573344693032 | 0.0802196763612696 | -0.322331573027727 | 0.7472015166464 | 0.940970690109281 | NM_013796,NR_027804 |
| Crabp2 | 96.09040249 | -0.094996009 | 0.163180928 | -0.582151419 | 0.560464693 | 0.753151774 | | Phf5a | 742.2565555 | -0.0191350231508334 | 0.0593272695330686 | -0.322533352730276 | 0.747048674782117 | 0.940970690109281 | NM_026737 |
| Greb1 | 70.58939524 | -0.087010581 | 0.149513922 | -0.581956382 | 0.560596062 | 0.753262976 | | Prpf31 | 1344.733885 | 0.0188093600381064 | 0.0582428015070917 | 0.322947378068965 | 0.746735094583053 | 0.940970690109281 | NM_027328,NM_001159714 |
| Bspry | 12.1063203 | 0.096968537 | 0.166675725 | 0.581779601 | 0.560715146 | 0.753357655 | | Ranbp2 | 6111.813253 | 0.0270294346575451 | 0.0838588675749947 | 0.322320530185705 | 0.747209881543581 | 0.940970690109281 | NM_011240 |
| Ikbkap | 3197.34942 | 0.02864111 | 0.049254961 | 0.581486812 | 0.560912404 | 0.753557339 | | Sap130 | 1986.377376 | 0.0178133870858585 | 0.0552680302707233 | 0.322309063641347 | 0.747218567424656 | 0.940970690109281 | NM_172965 |
| Npbwr1 | 59.09688859 | 0.096588096 | 0.166174801 | 0.58124394 | 0.561076058 | 0.753711846 | | Wipi1 | 728.1366486 | -0.0211189069500626 | 0.0654335290341862 | -0.322753598373532 | 0.746881856894663 | 0.940970690109281 | NM_145940 |
| Cpped1 | 1590.52956 | -0.033208067 | 0.057161546 | -0.58095117 | 0.561273364 | 0.75391153 | | Zfp410 | 934.1518762 | 0.0214769444487835 | 0.0665020680042865 | 0.322951527573536 | 0.746731951986394 | 0.940970690109281 | NR_045615,NR_045546,NM_001252583,NM_001252582,NM_144833 |
| Dnaic1 | 123.4087539 | 0.081263076 | 0.139936702 | 0.580713103 | 0.561433829 | 0.754061698 | | 0610012G03Rik | 911.0440581 | -0.0188465284867381 | 0.0589046426838112 | -0.319949797300404 | 0.749006387570318 | 0.941086400352566 | NR_027897 |
| Il17rc | 113.4405912 | 0.081734455 | 0.14081777 | 0.580427135 | 0.561626611 | 0.75425524 | | 2610018G03Rik | 218.2087193 | 0.0287475010464207 | 0.0895819349169588 | 0.320907346699636 | 0.748280606299324 | 0.941086400352566 | NM_133729 |
| 4933403G14Rik | 152.3025018 | -0.08478462 | 0.146094399 | -0.580341344 | 0.561684452 | 0.754267542 | | 4632415K11Rik | 409.7477116 | 0.02272867463042 | 0.0707109467836599 | 0.321430777895795 | 0.747883962183463 | 0.941086400352566 | NM_028883 |
| Nr3c1 | 4234.877718 | -0.038412334 | 0.066199567 | -0.580250535 | 0.561745679 | 0.754284388 | | AI316807 | 873.0553844 | 0.0215867859063616 | 0.0673820556747549 | 0.320364015169831 | 0.748692400907881 | 0.941086400352566 | NM_001146117,NM_001012667 |
| Tmem255b | 42.76691906 | -0.095978633 | 0.165465179 | -0.580053356 | 0.561878637 | 0.754332172 | | Acap3 | 6696.738566 | -0.0208345644110774 | 0.0655754044710496 | -0.317719190283844 | 0.750697953181196 | 0.941086400352566 | NM_207223 |
| Zfp867 | 222.1062431 | -0.072803195 | 0.125508187 | -0.580067296 | 0.561869237 | 0.754332172 | | Ace | 1335.84005 | 0.02343711207708 | 0.0739598133600283 | 0.316889821814324 | 0.751327205348252 | 0.941086400352566 | NM_207624,NM_009598 |
| Drd4 | 8.807071055 | 0.090545838 | 0.156173169 | 0.579778453 | 0.56206403 | 0.754515689 | | Amot | 963.6202368 | 0.0251787331211559 | 0.0789626129853939 | 0.318869046618472 | 0.749825817236125 | 0.941086400352566 | NM_153319 |
| Ppp1r3g | 276.4768181 | 0.078448295 | 0.135393772 | 0.579408442 | 0.562313609 | 0.75478533 | | Atp5e | 1967.148257 | -0.0220179349890702 | 0.069127788575807 | -0.318510622756649 | 0.75009763810929 | 0.941086400352566 | NM_025983 |
| Gm9767 | 11.64910974 | 0.095987575 | 0.165743099 | 0.579134669 | 0.562498308 | 0.754939222 | | Bend7 | 111.066064 | 0.0298083104178608 | 0.0939611761603678 | 0.317240711918991 | 0.751060960455858 | 0.941086400352566 | NM_178663,NM_001190400 |
| Vav1 | 186.4588382 | 0.074227743 | 0.128179079 | 0.57909406 | 0.562525708 | 0.754939222 | | Bhlhe22 | 997.3186921 | 0.0265432938278881 | 0.0827014125977826 | 0.320953330712512 | 0.748245758001397 | 0.941086400352566 | NM_021560 |
| Desi2 | 909.5545877 | -0.049879534 | 0.086177469 | -0.578800176 | 0.562724012 | 0.755139949 | | Bphl | 878.7874699 | -0.0198239705125099 | 0.0617860571962508 | -0.32084860908899 | 0.74832512046895 | 0.941086400352566 | NM_026512 |
| Tmem66 | 8924.509462 | -0.028586529 | 0.049396767 | -0.578712553 | 0.562783144 | 0.755153896 | | Ccdc129 | 67.64882905 | -0.030086140113314 | 0.0949418380927636 | -0.316890221610393 | 0.751326901978126 | 0.941086400352566 | NM_001081665 |
| Coq3 | 557.8929939 | 0.045680271 | 0.078966911 | 0.578473574 | 0.562944434 | 0.755201161 | | Ccdc59 | 614.2139827 | -0.022957061216576 | 0.0724360989781553 | -0.316928458882072 | 0.751297887248457 | 0.941086400352566 | NM_025602 |
| Fbn1 | 292.2813088 | 0.057767443 | 0.09986701 | 0.578443706 | 0.562964594 | 0.755201161 | | Chrna5 | 94.50771782 | 0.0300577786338344 | 0.0947761908832616 | 0.317144826709245 | 0.751133712337103 | 0.941086400352566 | NM_176844 |
| Hnrnpa2b1 | 14024.76091 | 0.055694257 | 0.096262925 | 0.578563942 | 0.562883441 | 0.755201161 | | Chst1 | 11021.95561 | 0.021285857377013 | 0.066822933405736 | 0.318541199737063 | 0.750074447980126 | 0.941086400352566 | NM_023850 |
| Nampt | 2108.800118 | -0.042527922 | 0.073562535 | -0.578119303 | 0.563183576 | 0.755298738 | | Cmklr1 | 180.1897721 | -0.0284509795373946 | 0.0897181397071911 | -0.317115130008812 | 0.751156244840875 | 0.941086400352566 | NM_008153 |
| Scaper | 1512.215008 | -0.033351944 | 0.05768812 | -0.578142322 | 0.563168036 | 0.755298738 | | Cntn2 | 8266.706136 | 0.021400250830656 | 0.0672348955312435 | 0.318290831889691 | 0.75026433808157 | 0.941086400352566 | NM_177129 |
| Sec23b | 1692.585276 | 0.041181269 | 0.071230234 | 0.57814311 | 0.563167503 | 0.755298738 | | Cox19 | 735.3874489 | -0.0205373937287468 | 0.0642757230297543 | -0.319520228799911 | 0.749332054320168 | 0.941086400352566 | NM_197980 |
| A730046J19Rik | 24.51861197 | 0.085028044 | 0.147133686 | 0.577896509 | 0.563333993 | 0.7553697 | | Cpped1 | 1702.520757 | 0.0172092601587374 | 0.0540022967967854 | 0.318676448586939 | 0.749971875541297 | 0.941086400352566 | NM_146067 |
| Rassf1 | 464.6100751 | 0.056587443 | 0.09791216 | 0.577940918 | 0.563304008 | 0.7553697 | | Cuedc1 | 1150.542418 | -0.0222330201158945 | 0.0699207476788797 | -0.317974576273163 | 0.750504222024334 | 0.941086400352566 | NM_001172099,NM_198013 |
| Gm20337 | 10.49401531 | 0.094580657 | 0.163767109 | 0.577531456 | 0.563580495 | 0.755589345 | | Ddx28 | 497.6684505 | 0.0227301587026983 | 0.0710675322145062 | 0.31983886304214 | 0.74909048535403 | 0.941086400352566 | NM_028038 |
| Snap29 | 1227.038063 | -0.047134949 | 0.081617618 | -0.577509487 | 0.563595331 | 0.755589345 | | Dennd5a | 15005.83809 | 0.0139328757272656 | 0.0434577771472279 | 0.320607187985322 | 0.74850808973242 | 0.941086400352566 | NM_021494 |
| Mesp2 | 21.38800553 | -0.09704882 | 0.168121542 | -0.577253924 | 0.563767934 | 0.755755353 | | Dennd6a | 1172.359513 | 0.0194430176246442 | 0.0609886308058683 | 0.318797411382015 | 0.749880141360917 | 0.941086400352566 | NM_001134465,NM_145969 |
| D8Ertd82e | 1533.56423 | -0.057099154 | 0.098954216 | -0.577025984 | 0.563921903 | 0.755826487 | | Dhrs7 | 919.8087509 | 0.0195768244651628 | 0.0614254082300214 | 0.318708902867245 | 0.749947262942067 | 0.941086400352566 | NM_025522 |
| Dnajc18 | 4726.297645 | 0.030132738 | 0.052214188 | 0.577098655 | 0.563872813 | 0.755826487 | | Ephx2 | 891.6867486 | 0.0217132063076016 | 0.0676200455060626 | 0.321106505878925 | 0.74813001899886 | 0.941086400352566 | NM_007940,NM_001271421,NM_001271403,NM_001271402 |
| Gar1 | 386.7581865 | 0.046771109 | 0.081064914 | 0.576958718 | 0.563967343 | 0.755826487 | | Epn2 | 6187.417822 | -0.0172012198847415 | 0.0539985310476675 | -0.318549774429179 | 0.75006794482072 | 0.941086400352566 | NM_001252189,NM_001252188,NM_010148 |
| 9230116N13Rik | 14.0430113 | 0.093568654 | 0.162203867 | 0.576858341 | 0.564035155 | 0.755839226 | | Exo1 | 23.71558815 | -0.027008552748186 | 0.0851266367657433 | -0.317274989055539 | 0.751034953580643 | 0.941086400352566 | NM_012012 |
| Usf2 | 3409.190482 | -0.032190213 | 0.055808253 | -0.57680023 | 0.564074414 | 0.755839226 | | Exosc6 | 259.940508 | 0.0270941337316209 | 0.085090673120357 | 0.318414848709436 | 0.750170277004827 | 0.941086400352566 | NM_028274 |
| 9430041J12Rik | 120.8946012 | 0.076742297 | 0.133138171 | 0.576410934 | 0.564337456 | 0.755931229 | | Exosc7 | 595.0792705 | 0.0207348873316907 | 0.0649288944765387 | 0.319347610934343 | 0.74946293288581 | 0.941086400352566 | NM_001081188 |
| Actr3b | 3441.665919 | 0.041823317 | 0.072536684 | 0.576581603 | 0.56422213 | 0.755931229 | | Gm11762 | 41.7222032 | -0.0293241426292532 | 0.0923494650983315 | -0.31753451520298 | 0.750838054126206 | 0.941086400352566 | NR_045099 |
| Naa25 | 1432.854971 | -0.03016889 | 0.052334246 | -0.576465559 | 0.564300543 | 0.755931229 | | Gm13157 | 86.8986811 | -0.0301751069650637 | 0.0950945722142439 | -0.3173168169586 | 0.751003218143215 | 0.941086400352566 | NM_001127189 |
| Rnf138 | 265.9028744 | -0.07478353 | 0.129745941 | -0.576384357 | 0.564355416 | 0.755931229 | | Hip1r | 3885.408244 | -0.0189614828355306 | 0.0596312125604791 | -0.31797915925824 | 0.750500745599036 | 0.941086400352566 | NM_145070 |
| Sstr2 | 570.4334335 | 0.071888166 | 0.124732738 | 0.576337592 | 0.564387018 | 0.755931229 | | Ift88 | 812.8360044 | -0.02201529846379 | 0.0687629419718717 | -0.320162253569598 | 0.748845335635236 | 0.941086400352566 | NM_009376 |
| Cdcp1 | 12.24717585 | 0.095193628 | 0.165307473 | 0.575857983 | 0.564711178 | 0.756103957 | | Lins | 454.6942513 | -0.0229892941535472 | 0.0717379333305882 | -0.320462175117399 | 0.748617999516912 | 0.941086400352566 | NR_034030,NM_152815,NM_001191001 |
| Fhit | 102.0758147 | -0.072298019 | 0.125519746 | -0.575989211 | 0.564622474 | 0.756103957 | | Map3k9 | 4708.941278 | -0.0163653467266851 | 0.0509612104754514 | -0.321133398794921 | 0.748109301043233 | 0.941086400352566 | NM_001174107,NM_177395 |
| Gstcd | 192.5794838 | 0.05660972 | 0.098281269 | 0.575997042 | 0.564617181 | 0.756103957 | | Mat2b | 4653.343362 | 0.0211449507601999 | 0.0660146612179736 | 0.320306888955795 | 0.748735701415038 | 0.941086400352566 | NM_134017,NM_001199274 |
| Plscr2 | 67.11826278 | 0.084112497 | 0.146046879 | 0.575928069 | 0.564663802 | 0.756103957 | | Myl4 | 639.5123345 | -0.030035868266754 | 0.0941416833467749 | -0.319049619370743 | 0.749688886547643 | 0.941086400352566 | NM_010858 |
| Rbm45 | 798.6069207 | 0.052797391 | 0.091700828 | 0.575756969 | 0.564779463 | 0.756130044 | | Naa15 | 2328.903842 | 0.0219476684215965 | 0.0682220190988825 | 0.32170945262973 | 0.747672816096652 | 0.941086400352566 | NM_053089 |
| Acot8 | 257.4529484 | -0.059792871 | 0.103980432 | -0.575039651 | 0.565264431 | 0.756568262 | | Narfl | 1035.28946 | -0.0220368032333935 | 0.0692655998448451 | -0.318149316294899 | 0.750371676492267 | 0.941086400352566 | NM_026238 |
| Cttn | 4453.828542 | 0.036685013 | 0.063789112 | 0.575098354 | 0.565224781 | 0.756568262 | | Nrd1 | 6773.270908 | -0.0167330346451093 | 0.052139298205376 | -0.320929418328534 | 0.748263879580562 | 0.941086400352566 | NM_146150 |
| Letmd1 | 2074.641289 | 0.03397266 | 0.059084529 | 0.574984023 | 0.565302102 | 0.756568262 | | Pacrgl | 226.9400187 | 0.0272368169861374 | 0.0851518730299842 | 0.31986163095375 | 0.749073225060079 | 0.941086400352566 | NM_025755 |
| Ppfia2 | 3551.241972 | -0.036899425 | 0.064172115 | -0.575007148 | 0.565286463 | 0.756568262 | | Patz1 | 1181.821587 | 0.0187633837343143 | 0.0585729742732302 | 0.32034200016525 | 0.748709087739288 | 0.941086400352566 | NM_001253690,NM_001253691,NR_045108,NR_045109,NM_019574 |
| Fam35a | 213.2553049 | 0.065684066 | 0.114265865 | 0.574835414 | 0.565402613 | 0.756637424 | | Pcdhb6 | 113.762484 | 0.0305886355106623 | 0.0950679820507758 | 0.321755388626267 | 0.747638013158251 | 0.941086400352566 | NM_053131 |
| Acsf3 | 629.9471688 | 0.061135083 | 0.106421963 | 0.57445927 | 0.565657055 | 0.756694044 | | Pcgf1 | 415.2666676 | 0.0234615029296746 | 0.07322208679744 | 0.320415654584728 | 0.748653259964688 | 0.941086400352566 | NM_197992 |
| Capn7 | 1771.112866 | 0.040199315 | 0.070001026 | 0.574267515 | 0.565786788 | 0.756694044 | | Pdgfb | 1267.80647 | 0.0254483548368332 | 0.0800293045096094 | 0.317987954447081 | 0.750494074018955 | 0.941086400352566 | NM_011057 |
| Hnrnpa1 | 104.0818431 | -0.085935867 | 0.149615764 | -0.574377091 | 0.565712652 | 0.756694044 | | Ppm1e | 8982.750348 | 0.0176439513298578 | 0.0554645453533023 | 0.318112250221616 | 0.750399791601702 | 0.941086400352566 | NM_177167 |
| Mir143hg | 32.34100031 | 0.09324339 | 0.162349678 | 0.574336771 | 0.565739931 | 0.756694044 | | Prdm10 | 460.8632286 | -0.0217493607478124 | 0.0679521315724093 | -0.320068852065906 | 0.74891613705885 | 0.941086400352566 | NM_001080817 |
| Ndufa5 | 2094.784946 | -0.046256126 | 0.080488749 | -0.574690587 | 0.565500575 | 0.756694044 | | Prps2 | 1619.561725 | 0.0222627900367784 | 0.0699052643209277 | 0.318470865578481 | 0.750127791002463 | 0.941086400352566 | NM_026662 |
| Nqo1 | 176.1937164 | 0.07684816 | 0.133779771 | 0.57443782 | 0.565671566 | 0.756694044 | | Prrt3 | 2076.388553 | 0.0225472184794008 | 0.0709541046879957 | 0.31777186927447 | 0.75065799057177 | 0.941086400352566 | NM_172487 |
| Rad51d | 1002.918748 | 0.042216804 | 0.073484898 | 0.574496332 | 0.565631982 | 0.756694044 | | Rab40c | 1730.927852 | -0.0222681600221018 | 0.069959147927185 | -0.318302333317138 | 0.750255614556185 | 0.941086400352566 | NM_139154 |
| Zfp790 | 700.9034284 | 0.041881542 | 0.072997137 | 0.573742259 | 0.566142227 | 0.757104062 | | Rps24 | 8186.68895 | 0.0234712481836201 | 0.0738742193035402 | 0.31771906904599 | 0.750698045153757 | 0.941086400352566 | NM_011297,NM_207635,NM_207634 |
| Adipor1 | 2909.099371 | -0.026157489 | 0.045618531 | -0.57339612 | 0.566376517 | 0.757352011 | | Slc13a3 | 769.6376132 | -0.029348321656694 | 0.0918527110009092 | -0.31951502940837 | 0.749335996384377 | 0.941086400352566 | NM_054055 |
| Cml2 | 10.05381525 | 0.088875638 | 0.155194272 | 0.572673442 | 0.566865823 | 0.757895294 | | Slc25a3 | 12847.5027 | 0.0160653603826544 | 0.0504434730170452 | 0.318482440279753 | 0.750119012404148 | 0.941086400352566 | NM_133668 |
| Fign | 148.8357352 | -0.087328793 | 0.15253438 | -0.572518753 | 0.566970585 | 0.757895294 | | Slc4a8 | 5422.833289 | 0.0189625083162456 | 0.0592742358956803 | 0.319911476372613 | 0.749035437817891 | 0.941086400352566 | NM_021530 |
| Hspbap1 | 199.383466 | 0.05535839 | 0.096702769 | 0.5724592 | 0.567010919 | 0.757895294 | | Sorbs1 | 6792.458409 | 0.019604950692221 | 0.0614961898426549 | 0.318799436881903 | 0.749878605318947 | 0.941086400352566 | NM_001034963,NM_001034962,NM_001034964,NM_178362,NM_009166 |
| Irak2 | 809.1641783 | -0.057672593 | 0.100731504 | -0.572537791 | 0.566957691 | 0.757895294 | | Spata7 | 773.5487497 | 0.0241234551468803 | 0.0759608637639729 | 0.317577420154636 | 0.750805504203518 | 0.941086400352566 | NM_178914 |
| Taf11 | 1117.580581 | -0.036889806 | 0.064443671 | -0.572434894 | 0.567027382 | 0.757895294 | | Stom | 630.8424942 | -0.025190881394828 | 0.0785283087466119 | -0.320787265088208 | 0.748371610781212 | 0.941086400352566 | NM_013515 |
| Colq | 36.44070418 | 0.096152554 | 0.168032009 | 0.572227602 | 0.56716779 | 0.758017574 | | Taf4b | 254.5287006 | -0.0268965666377102 | 0.0842401942567432 | -0.319284242813308 | 0.749510980295668 | 0.941086400352566 | NM_001100449 |
| Gpr37l1 | 7063.508031 | -0.064193749 | 0.112199939 | -0.572137111 | 0.567229089 | 0.758034113 | | Tgm2 | 717.8311895 | -0.0237868768665683 | 0.0739007277823769 | -0.321876084043664 | 0.747546571960037 | 0.941086400352566 | NM_009373 |
| Ghdc | 267.4533465 | 0.056710465 | 0.099202223 | 0.571665259 | 0.567548775 | 0.758395923 | | Traf3ip1 | 764.5769918 | -0.020108697981542 | 0.0626715231682558 | -0.320858612731587 | 0.748317539171035 | 0.941086400352566 | NM_028718 |
| Nt5m | 1145.553573 | -0.032159557 | 0.056314135 | -0.571074323 | 0.567949263 | 0.758865633 | | Wdr47 | 6805.344373 | 0.0190035499892396 | 0.0596871768704264 | 0.318385807231158 | 0.750192302779991 | 0.941086400352566 | NM_181400 |
| Wdr7 | 9534.380612 | -0.043259893 | 0.075766263 | -0.57096511 | 0.568023294 | 0.758899104 | | Zfand6 | 1221.727536 | 0.0201049323851597 | 0.0632762042781601 | 0.317732907883966 | 0.750687546860144 | 0.941086400352566 | NM_022985 |
| Pcdha3 | 94.09224325 | -0.074009865 | 0.129702684 | -0.570611675 | 0.568262903 | 0.759153769 | | AI597479 | 1429.046797 | 0.0188406300801359 | 0.0595784046706866 | 0.316232537347643 | 0.751826012745573 | 0.941374878843062 | NM_133818 |
| Mad2l2 | 269.0808264 | 0.05663899 | 0.09929174 | 0.570430025 | 0.56838607 | 0.759252846 | | Chrm1 | 9233.924903 | 0.0209812065899183 | 0.0663020327734574 | 0.316448918868106 | 0.751661791255559 | 0.941374878843062 | NM_007698,NM_001112697 |
| Sh3bp5l | 1956.098145 | -0.040165405 | 0.070434758 | -0.570249777 | 0.568508299 | 0.759350654 | | Coa6 | 323.8840699 | -0.0266766022007571 | 0.0843770733166227 | -0.316159368323358 | 0.751881546491615 | 0.941374878843062 | NM_174987 |
| Fcrlb | 12.36515853 | -0.094546337 | 0.165889165 | -0.569936782 | 0.568720577 | 0.759568711 | | Kat8 | 759.4679141 | -0.0195660967347243 | 0.0618435579156172 | -0.31638051551661 | 0.751713704364158 | 0.941374878843062 | NM_026370 |
| 1700088E04Rik | 121.0465836 | 0.071914765 | 0.126306046 | 0.569369142 | 0.569105654 | 0.759886504 | | Ptdss2 | 1784.164498 | 0.018807233104806 | 0.0594664848668081 | 0.316266097566219 | 0.751800541677402 | 0.941374878843062 | NM_013782 |
| Anp32a | 6641.390151 | -0.025714061 | 0.045160286 | -0.569395445 | 0.569087807 | 0.759886504 | | Pdpr | 837.334499 | -0.0186879723975537 | 0.0591756260697514 | -0.315805233315586 | 0.75215034564358 | 0.941549114169134 | NM_198308 |
| Prkcsh | 2831.081554 | 0.036551268 | 0.064195256 | 0.569376466 | 0.569100684 | 0.759886504 | | Slit1 | 2499.936468 | -0.0263982192345674 | 0.0835900544172623 | -0.315805742903259 | 0.75214995882956 | 0.941549114169134 | NM_015748 |
| Ccdc88b | 147.9719682 | 0.075765006 | 0.133100886 | 0.569229914 | 0.569200123 | 0.759947152 | | Nsmce1 | 395.5379807 | 0.0257706141326304 | 0.0816320831670172 | 0.315692227036573 | 0.752236127146556 | 0.941565790635865 | NM_026330 |
| 9830147E19Rik | 96.11558511 | -0.087596297 | 0.153973211 | -0.568906089 | 0.569419873 | 0.760138156 | | Tmem237 | 840.2946033 | 0.022916663387408 | 0.0726091130214819 | 0.31561690308251 | 0.752293306217806 | 0.941565790635865 | NM_001033449,NM_001037812 |
| Cacnb1 | 4101.574959 | 0.053322107 | 0.093745823 | 0.568794477 | 0.569495623 | 0.760138156 | | Pet112 | 690.4927614 | -0.0208059553463881 | 0.065979460118209 | -0.31533988470218 | 0.752503604979611 | 0.941691983419072 | NM_144896 |
| Eid1 | 4575.92487 | -0.047596371 | 0.083720798 | -0.568513111 | 0.569686606 | 0.760138156 | | Scrt2 | 1029.459953 | 0.0256003234769853 | 0.0811901162968172 | 0.315313299754306 | 0.752523787933952 | 0.941691983419072 | NM_001160410 |
| Lypla2 | 2466.515044 | 0.046223839 | 0.081287317 | 0.568647614 | 0.569595305 | 0.760138156 | | Rrp9 | 705.3567711 | -0.0226243319390801 | 0.0717961151913983 | -0.315119165971123 | 0.752671176961151 | 0.94179528956307 | NM_145620 |
| Siae | 1028.712101 | 0.054619302 | 0.096062436 | 0.56858127 | 0.569640338 | 0.760138156 | | AI848285 | 57.16966845 | -0.0294542139214341 | 0.093812766334533 | -0.313968077824306 | 0.753545283915156 | 0.941834640146532 | NM_001207021 |
| Ttc34 | 84.66276334 | 0.085025607 | 0.14954365 | 0.56856715 | 0.569649924 | 0.760138156 | | Birc6 | 4956.560049 | -0.0226996673024017 | 0.072258008712914 | -0.314147424025883 | 0.753409072182124 | 0.941834640146532 | NM_007566 |
| Zfp397 | 1336.586802 | 0.050606015 | 0.088993002 | 0.568651624 | 0.569592584 | 0.760138156 | | Farp1 | 3259.15115 | 0.0141811066132546 | 0.0450926442073945 | 0.314488246642435 | 0.75315024180099 | 0.941834640146532 | NM_134082 |
| Vmac | 445.9078806 | 0.050070105 | 0.088142392 | 0.568059295 | 0.569994706 | 0.760483766 | | Fkbp7 | 78.86721178 | -0.0295089431625392 | 0.0940057017433937 | -0.313905886720461 | 0.753592519257804 | 0.941834640146532 | NM_010222 |
| Tmem154 | 25.49268815 | 0.095338829 | 0.168040858 | 0.567355048 | 0.570472984 | 0.761056347 | | Klhl33 | 131.9150315 | 0.0286917939086505 | 0.0915089177471188 | 0.313540959886982 | 0.753869706800477 | 0.941834640146532 | NM_001166651 |
| Ermap | 51.51710048 | -0.083312079 | 0.146875805 | -0.567228064 | 0.570559243 | 0.76110589 | | Loh12cr1 | 1279.561591 | -0.0195799792631663 | 0.0624181100567465 | -0.31369067799979 | 0.753755981546841 | 0.941834640146532 | NM_001170479,NM_026371 |
| Capn5 | 2350.0524 | 0.04317872 | 0.076158635 | 0.56695764 | 0.570742961 | 0.761260281 | | Mas1 | 635.5610135 | -0.024157217695491 | 0.0769949245951333 | -0.313750780619869 | 0.753710329349741 | 0.941834640146532 | NM_008552 |
| Rslcan18 | 27.96306873 | 0.093307324 | 0.164588417 | 0.566913065 | 0.570773247 | 0.761260281 | | Nkx2-2as | 33.44167404 | -0.0271730108301846 | 0.086582710764801 | -0.313838762844919 | 0.753643502170431 | 0.941834640146532 | NR_030769 |
| Cks2 | 32.62427738 | -0.089742683 | 0.158386998 | -0.566603849 | 0.570983359 | 0.761290531 | | Pcdhgb8 | 30.14092795 | 0.0271892977172719 | 0.0866694285226173 | 0.313712668708512 | 0.753739277946078 | 0.941834640146532 | NM_033580 |
| Crlf2 | 362.7397428 | 0.07295355 | 0.128843157 | 0.566219828 | 0.571244353 | 0.761290531 | | Pibf1 | 441.4693469 | 0.0249051142225963 | 0.0793931044640517 | 0.313693668873637 | 0.753753709745955 | 0.941834640146532 | NM_029454,NM_029320 |
| D6Wsu163e | 1271.217306 | 0.03930445 | 0.069371637 | 0.566578096 | 0.57100086 | 0.761290531 | | Plk1s1 | 619.0908794 | 0.0257611788065801 | 0.0818045787604174 | 0.314911209090475 | 0.752829070675895 | 0.941834640146532 | NM_001033298 |
| Galr1 | 21.30730012 | -0.092683429 | 0.163560561 | -0.566661233 | 0.570944364 | 0.761290531 | | Ppp1r21 | 3123.027819 | 0.0136217975566967 | 0.0434424774954483 | 0.313559408717515 | 0.753855692857359 | 0.941834640146532 | NM_028658 |
| Gas2l1 | 2141.536354 | -0.045255056 | 0.079933801 | -0.566156684 | 0.571287273 | 0.761290531 | | Samd8 | 3659.628401 | 0.0218742080333808 | 0.0694661908524987 | 0.314889988423685 | 0.75284518329901 | 0.941834640146532 | NM_026283 |
| Lipt2 | 121.3747502 | -0.064072131 | 0.113078795 | -0.566614905 | 0.570975846 | 0.761290531 | | Sft2d2 | 1801.567993 | -0.0175281614360416 | 0.055813400909124 | -0.314049334936267 | 0.753483568961487 | 0.941834640146532 | NM_145512 |
| Pion | 258.504343 | 0.079091862 | 0.139669728 | 0.566277765 | 0.571204973 | 0.761290531 | | Tmem232 | 87.54835569 | 0.0298170632784023 | 0.0949292312433039 | 0.314097806206616 | 0.753446755675882 | 0.941834640146532 | NM_001008973 |
| Tlk1 | 4819.497231 | -0.046456874 | 0.082022307 | -0.566393173 | 0.571126534 | 0.761290531 | | Usp34 | 6414.367507 | 0.0224495122066873 | 0.0714554049904525 | 0.314175144758985 | 0.753388019233833 | 0.941834640146532 | NM_001190401 |
| Ugt8a | 1655.267426 | -0.093366209 | 0.164834559 | -0.56642375 | 0.571105754 | 0.761290531 | | Zfp354c | 1898.580178 | 0.0228930374318316 | 0.0728978030471837 | 0.314042899441207 | 0.753488456676594 | 0.941834640146532 | NM_013922 |
| Zbed4 | 596.1574726 | -0.046984692 | 0.082976179 | -0.566243138 | 0.571228509 | 0.761290531 | | Zfp459 | 120.211093 | 0.0299115984250117 | 0.0950849302464535 | 0.314577697511928 | 0.753082314863277 | 0.941834640146532 | NM_177811 |
| Lcat | 678.0830077 | -0.075892238 | 0.134071956 | -0.566056022 | 0.571355698 | 0.761316235 | | Lmna | 1618.032975 | 0.0195078939153843 | 0.0622613903808426 | 0.313322490809436 | 0.754035664578265 | 0.941879960758782 | NM_019390,NM_001111102,NM_001002011 |
| A330040F15Rik | 9.76531492 | -0.088919466 | 0.157163395 | -0.565777203 | 0.571545246 | 0.761503314 | | Zfp324 | 489.0004982 | 0.0217057227649015 | 0.069273526407206 | 0.31333359063187 | 0.75402723243883 | 0.941879960758782 | NM_178732 |
| Tdgf1 | 14.06972081 | 0.092015333 | 0.162699809 | 0.5655528 | 0.571697822 | 0.761616356 | | Tpt1 | 14978.53337 | 0.0189448479130913 | 0.0604897410482949 | 0.313191089675285 | 0.75413548756409 | 0.94192365368495 | NM_009429 |
| Tfap4 | 104.3016255 | 0.078262189 | 0.13839276 | 0.56550783 | 0.5717284 | 0.761616356 | | Far1 | 2472.868808 | -0.0208567865419954 | 0.0666245547386315 | -0.313049544928543 | 0.754243021058886 | 0.941966229553812 | NM_026143,NM_027379 |
| Foxl2 | 27.22668218 | -0.094949901 | 0.167942601 | -0.565371148 | 0.571821344 | 0.761674683 | | Limk2 | 5213.806105 | -0.0174758871201069 | 0.0558624658811774 | -0.312837731819413 | 0.754403947296805 | 0.941966229553812 | NM_173053,NM_001034030,NM_010718 |
| Necab1 | 4211.967138 | -0.054060909 | 0.095644574 | -0.56522714 | 0.571919279 | 0.761739646 | | Mad2l1 | 266.1245084 | -0.0256331043907406 | 0.0819683700082874 | -0.312719459812962 | 0.754493809779609 | 0.941966229553812 | NM_019499 |
| Mrgpre | 490.6500124 | -0.056769992 | 0.100461634 | -0.565091265 | 0.572011689 | 0.761797242 | | Robo4 | 385.2742879 | -0.0266020190330704 | 0.0850465400429871 | -0.312793665910739 | 0.754437427966696 | 0.941966229553812 | NM_028783 |
| Cthrc1 | 194.5271856 | -0.059476988 | 0.105271528 | -0.564986464 | 0.572082971 | 0.761826691 | | Tmem2 | 625.38734 | 0.0190990093752032 | 0.0610539211186107 | 0.312821994480897 | 0.754415904256665 | 0.941966229553812 | NM_031997,NM_001033759 |
| Clpb | 2039.223153 | -0.027308365 | 0.048364263 | -0.564639328 | 0.572319109 | 0.761944067 | | Rps15a | 5791.893228 | -0.0149360995383021 | 0.0477757593264901 | -0.312629244387969 | 0.754562357245846 | 0.941970849205324 | NM_170669 |
| Efnb1 | 601.8756222 | -0.05801971 | 0.102781538 | -0.564495449 | 0.572416996 | 0.761944067 | | 4930412C18Rik | 98.14085797 | -0.0296634045466484 | 0.0950298136804637 | -0.312148402672777 | 0.754927742903333 | 0.942099798455631 | NR_030693 |
| Mettl21a | 383.3669285 | -0.057191105 | 0.101262814 | -0.564778943 | 0.572224131 | 0.761944067 | | Serpine1 | 73.06677021 | 0.0271524264418431 | 0.0869877437164296 | 0.312140829061585 | 0.754933498435511 | 0.942099798455631 | NM_008871 |
| Mmp23 | 35.8829567 | 0.087738363 | 0.155423845 | 0.564510309 | 0.572406886 | 0.761944067 | | Tmem38a | 4352.647314 | 0.0149620825672108 | 0.0479431838793165 | 0.312079452313256 | 0.754980141924637 | 0.942099798455631 | NM_144534 |
| Sys1 | 874.9445353 | -0.052523263 | 0.093021647 | -0.564634843 | 0.572322161 | 0.761944067 | | Vipr1 | 1495.980132 | -0.0231959463499786 | 0.0743152868653455 | -0.312128867806272 | 0.754942588366061 | 0.942099798455631 | NM_011703 |
| Smc3 | 2271.429075 | 0.045807268 | 0.081159065 | 0.564413453 | 0.572472785 | 0.761952868 | | Zfp30 | 463.121794 | 0.0226768931382088 | 0.0726668452460646 | 0.312066569856173 | 0.754989932109459 | 0.942099798455631 | NM_013705 |
| Cln5 | 522.3420494 | 0.044750993 | 0.079312333 | 0.5642375 | 0.57259251 | 0.762018442 | | Eif5 | 9610.98575 | 0.0213152782551272 | 0.0683276937383057 | 0.311956647282206 | 0.755073470748673 | 0.942123109095665 | NM_173363,NM_178041 |
| Rilpl2 | 354.7589868 | 0.055963092 | 0.099190783 | 0.564196491 | 0.572620416 | 0.762018442 | | 2210018M11Rik | 2071.620838 | -0.0178708400137126 | 0.0576065240209202 | -0.310222502007285 | 0.756391759938329 | 0.942139347792531 | NM_172280 |
| Ddx10 | 707.8152486 | -0.03934421 | 0.06975832 | -0.564007415 | 0.572749086 | 0.762058766 | | Acn9 | 177.3891692 | 0.0271151426828187 | 0.0875543047714702 | 0.309695140102972 | 0.75679279884223 | 0.942139347792531 | NM_001077713 |
| Ptgfrn | 1921.118534 | -0.050798468 | 0.090062797 | -0.564033871 | 0.572731081 | 0.762058766 | | Aldh16a1 | 262.2091167 | 0.0256146094416784 | 0.0829436555471967 | 0.308819393993349 | 0.757458915495042 | 0.942139347792531 | NM_145954 |
| Asxl1 | 2618.141823 | 0.032641512 | 0.057958435 | 0.563188292 | 0.573306674 | 0.762342397 | | Bbs1 | 2348.631345 | -0.0183136827727939 | 0.0589549200519198 | -0.310638751721919 | 0.756075263999653 | 0.942139347792531 | NM_001033128 |
| Gab1 | 1391.512485 | 0.051137226 | 0.090787413 | 0.563263391 | 0.573255543 | 0.762342397 | | Cactin | 1112.606658 | 0.0197110763556181 | 0.0636343108102022 | 0.309755477896335 | 0.756746910898134 | 0.942139347792531 | NM_027381 |
| Glul | 66094.55951 | -0.051948118 | 0.09218573 | -0.56351583 | 0.573083684 | 0.762342397 | | Cd2bp2 | 4015.197645 | 0.0171446610691949 | 0.0551018235042731 | 0.311145076131016 | 0.75569033483279 | 0.942139347792531 | NM_027353 |
| Ranbp6 | 1750.719123 | -0.038645892 | 0.068581408 | -0.563503913 | 0.573091796 | 0.762342397 | | Cenpp | 23.99808206 | 0.0257744685662972 | 0.0829302874459536 | 0.310796807295461 | 0.755950969983112 | 0.942139347792531 | NM_025495 |
| Sppl3 | 3118.161838 | 0.031712853 | 0.05630317 | 0.563251636 | 0.573263546 | 0.762342397 | | Chrna3 | 33.24641455 | 0.0263682518758151 | 0.0850646426706168 | 0.30997898830795 | 0.75657693480615 | 0.942139347792531 | NM_145129 |
| Trim56 | 168.5985045 | -0.069081865 | 0.122622083 | -0.56337214 | 0.573181504 | 0.762342397 | | Chtf18 | 61.88512155 | -0.0290072021411539 | 0.0939944043902447 | -0.308605627423545 | 0.757621539621103 | 0.942139347792531 | NM_145409 |
| Zdhhc22 | 595.2748481 | -0.061170585 | 0.108613274 | -0.563196213 | 0.573301281 | 0.762342397 | | Cpeb2 | 2707.286743 | -0.0205086422127879 | 0.0664726159387696 | -0.308527683515256 | 0.757680838559297 | 0.942139347792531 | NM_001177379,NM_175937 |
| Ash2l | 1419.037037 | -0.034304855 | 0.06093622 | -0.562963288 | 0.573459883 | 0.762480685 | | Cpne8 | 1234.062279 | 0.0189965266488358 | 0.0609883630914616 | 0.311477889976281 | 0.755437348735983 | 0.942139347792531 | NM_025815,NM_001033851 |
| Rabl3 | 295.0727364 | -0.058000561 | 0.103098755 | -0.562572859 | 0.573725778 | 0.762659199 | | D430036J16Rik | 104.4409099 | 0.0289128823151223 | 0.0936677159542743 | 0.308675000992196 | 0.75756876211046 | 0.942139347792531 | NR_040393,NR_040396,NR_040397,NR_040394,NR_040395 |
| Tmem232 | 86.92880545 | -0.087757309 | 0.15599337 | -0.562570757 | 0.573727209 | 0.762659199 | | Dnase1l1 | 75.00606145 | 0.0294960728953225 | 0.0950303069992241 | 0.31038595819293 | 0.756267470967359 | 0.942139347792531 | NM_027109,NM_001172154 |
| Zfp354c | 1804.092385 | 0.050099332 | 0.08905767 | 0.562549319 | 0.573741811 | 0.762659199 | | Efhc1 | 59.87738477 | -0.0285089535578876 | 0.0922458799099426 | -0.309053950005357 | 0.757280488035738 | 0.942139347792531 | NM_027974 |
| Crmp1 | 7134.079123 | -0.030066659 | 0.053464533 | -0.56236644 | 0.57386638 | 0.762720771 | | Ercc8 | 429.8211814 | 0.0227059319537056 | 0.0734080079884157 | 0.309311375909952 | 0.757084678268056 | 0.942139347792531 | NM_028042 |
| Usp1 | 978.2084561 | 0.046926027 | 0.083448262 | 0.562336778 | 0.573886585 | 0.762720771 | | Fbln5 | 279.381418 | 0.0267323840888237 | 0.0860819155501311 | 0.310545878515629 | 0.756145876701673 | 0.942139347792531 | NM_011812 |
| Oard1 | 388.1861107 | 0.045811898 | 0.081482262 | 0.562231548 | 0.573958269 | 0.762750616 | | Fgd4 | 552.4815572 | -0.0252323636988926 | 0.0809510657508602 | -0.311698968566383 | 0.755269311899439 | 0.942139347792531 | NM_139234,NM_139233,NM_139232 |
| Psmd11 | 2044.35448 | 0.033792442 | 0.060115637 | 0.562123987 | 0.574031546 | 0.762779925 | | Fxyd2 | 86.41771495 | 0.0293566125911029 | 0.0950565859060248 | 0.308833020997888 | 0.757448549038242 | 0.942139347792531 | NM_007503,NM_052823,NR_028499 |
| Zfp760 | 346.1673176 | -0.067859959 | 0.120735518 | -0.56205465 | 0.574078785 | 0.762779925 | | Gm14326 | 855.107165 | -0.0231530050297428 | 0.0745468457647594 | -0.310583295540158 | 0.756117427808142 | 0.942139347792531 | NM_001190302 |
| Apmap | 2245.332474 | 0.042210736 | 0.075169045 | 0.561544133 | 0.574426653 | 0.762944234 | | Gpc2 | 89.06956176 | -0.0291854608262662 | 0.0945940606815228 | -0.3085337558827 | 0.757676218712252 | 0.942139347792531 | NM_172412 |
| C1qtnf7 | 28.82564837 | -0.089720617 | 0.159804576 | -0.5614396 | 0.574497895 | 0.762944234 | | Gpr88 | 8533.691 | 0.0288456112047951 | 0.0929611597877318 | 0.310297454019091 | 0.756334767073228 | 0.942139347792531 | NM_022427 |
| Glb1l2 | 22.6372275 | 0.094101933 | 0.16751737 | 0.561744333 | 0.574290224 | 0.762944234 | | Gramd4 | 2432.291924 | 0.0214603497375846 | 0.0690728234463941 | 0.310691653631903 | 0.756035042902735 | 0.942139347792531 | NM_001205353,NM_172611 |
| Magoh | 428.1393623 | -0.056083974 | 0.099885751 | -0.561481227 | 0.574469525 | 0.762944234 | | Gys1 | 896.1102894 | 0.0243979037672175 | 0.0785662759030147 | 0.310539140194646 | 0.756151000013668 | 0.942139347792531 | NM_030678 |
| Man1b1 | 3687.942776 | 0.0277558 | 0.049412646 | 0.561714512 | 0.574310546 | 0.762944234 | | Hnrnpul1 | 3696.431959 | 0.0149679280576921 | 0.0483585890380833 | 0.309519536351745 | 0.756926353436164 | 0.942139347792531 | NM_144922,NM_178089 |
| Spred1 | 4532.063589 | -0.064632023 | 0.115106533 | -0.561497434 | 0.574458479 | 0.762944234 | | Ldlrap1 | 159.7715177 | -0.0286640886854593 | 0.0925405403783829 | -0.309746285987272 | 0.756753901449906 | 0.942139347792531 | NM_145554 |
| Ptch2 | 74.83602779 | -0.087948297 | 0.156668534 | -0.56136542 | 0.574548453 | 0.762945983 | | Leng9 | 248.1971712 | 0.0276220019205813 | 0.0889606929067756 | 0.310496703859167 | 0.756183265654781 | 0.942139347792531 | NM_175529 |
| 5830454E08Rik | 25.96236218 | 0.093895773 | 0.167286423 | 0.561287466 | 0.574601584 | 0.762951148 | | Magt1 | 698.7700904 | -0.0201411648715095 | 0.0652174582255518 | -0.308830877797355 | 0.757450179430033 | 0.942139347792531 | NM_001190409,NM_025952 |
| Lrp12 | 2286.212438 | 0.031896463 | 0.056861927 | 0.560945874 | 0.574834436 | 0.763194923 | | Mprip | 11860.43447 | 0.0143445045815087 | 0.0463938941534367 | 0.309189492351465 | 0.757177386463937 | 0.942139347792531 | NM_201245,NM_012027 |
| Adprm | 419.3402751 | 0.047267874 | 0.084288369 | 0.560787617 | 0.574942329 | 0.763202562 | | Myo18a | 8373.099591 | -0.0138490107632623 | 0.0447742197610497 | -0.30930769619597 | 0.757087477114868 | 0.942139347792531 | NM_011586 |
| Lin7b | 1252.878162 | -0.07660111 | 0.136626815 | -0.560659416 | 0.575029739 | 0.763202562 | | Ost4 | 961.6343622 | -0.0209087761194446 | 0.0675395936380409 | -0.30957805626577 | 0.756881845588502 | 0.942139347792531 | NM_024460,NM_001134692 |
| Nfkbil1 | 275.7508445 | -0.060013639 | 0.107033864 | -0.560697679 | 0.575003649 | 0.763202562 | | Pacs2 | 6982.382603 | 0.0209329750646193 | 0.0672418688684838 | 0.311308644701137 | 0.755565996062541 | 0.942139347792531 | NM_001081170 |
| Pi4ka | 16689.23417 | -0.045810195 | 0.081711991 | -0.560630022 | 0.575049781 | 0.763202562 | | Pcyox1l | 626.0822316 | 0.0225213816945747 | 0.0727201057942198 | 0.309699517741418 | 0.756789469542998 | 0.942139347792531 | NM_172832 |
| Pkd2l1 | 47.52111772 | 0.091870842 | 0.163886448 | 0.560576201 | 0.575086479 | 0.763202562 | | Phpt1 | 1202.639008 | 0.0213335212055613 | 0.0689540786675936 | 0.309387372259785 | 0.757026874992757 | 0.942139347792531 | NM_029293 |
| Ap3d1 | 8086.209003 | -0.023396579 | 0.041763015 | -0.560222456 | 0.575327712 | 0.763230445 | | Pla2g7 | 6173.78166 | -0.0160324042999628 | 0.0514219327929308 | -0.311781440898441 | 0.755206629510026 | 0.942139347792531 | NM_013737 |
| Arhgap12 | 1533.532502 | -0.045165298 | 0.080592657 | -0.560414557 | 0.575196705 | 0.763230445 | | Plscr1 | 76.51569174 | -0.0295396602056688 | 0.095101062762888 | -0.310613355387195 | 0.75609457295964 | 0.942139347792531 | NM_011636 |
| Baiap2 | 12995.6329 | -0.049554778 | 0.088448175 | -0.560269087 | 0.57529591 | 0.763230445 | | Rab40b | 1007.969566 | 0.0256952798290909 | 0.0829832676657937 | 0.309644107202141 | 0.75683161092262 | 0.942139347792531 | NM_139147 |
| Ndrg3 | 23188.8046 | -0.032353624 | 0.057751553 | -0.560220855 | 0.575328804 | 0.763230445 | | Rhno1 | 503.2603625 | 0.0213383411230643 | 0.068964016044884 | 0.30941268137831 | 0.757007625028464 | 0.942139347792531 | NR_027360,NR_027359 |
| Psg16 | 94.31802111 | 0.069387578 | 0.123871175 | 0.560159198 | 0.575370855 | 0.763230445 | | Smchd1 | 1838.016417 | -0.021121228601853 | 0.0681336985224349 | -0.309996801287666 | 0.756563388826629 | 0.942139347792531 | NM_028887 |
| Spry1 | 312.6673071 | 0.053826642 | 0.096099784 | 0.560111997 | 0.575403048 | 0.763230445 | | Suz12 | 1576.199654 | -0.0236125036785648 | 0.076390398081339 | -0.309103032208611 | 0.757243152693156 | 0.942139347792531 | NM_001163018,NM_199196 |
| Rab7l1 | 114.9442478 | -0.0766331 | 0.136863597 | -0.559923176 | 0.57553184 | 0.763335929 | | Trub2 | 977.4974835 | -0.0170354788685242 | 0.0549753601891035 | -0.309874802273707 | 0.756656165153473 | 0.942139347792531 | NR_027344,NM_145520 |
| Dock4 | 4803.356088 | 0.034963136 | 0.062460396 | 0.559764878 | 0.575639823 | 0.763413799 | | Cideb | 192.9489446 | -0.0285240785110604 | 0.0925052175305948 | -0.308351023569308 | 0.757815244940228 | 0.942225819395853 | NM_009894 |
| Mettl20 | 75.15535246 | 0.082776521 | 0.14790545 | 0.559658356 | 0.575712493 | 0.763444827 | | 2610301B20Rik | 1554.303034 | 0.0179133449416993 | 0.0581815306283053 | 0.307887137864064 | 0.758168213205307 | 0.942510232563136 | NM_026005 |
| Cys1 | 262.3306415 | 0.064286837 | 0.114950688 | 0.559255783 | 0.575987167 | 0.76363536 | | C2cd2l | 8626.82246 | 0.0205154151103782 | 0.0666344854508475 | 0.307879845872168 | 0.75817376204738 | 0.942510232563136 | NM_027909 |
| Glis2 | 481.2954931 | 0.046764649 | 0.083623124 | 0.559231071 | 0.576004031 | 0.76363536 | | 6820408C15Rik | 92.84907989 | 0.029119918677607 | 0.0946316364729236 | 0.307718642125975 | 0.758296433237928 | 0.942526199240263 | NM_177656 |
| Wdr18 | 2175.271159 | 0.032421459 | 0.057972875 | 0.559252217 | 0.575989601 | 0.76363536 | | Ace2 | 71.91481858 | -0.0289686771599107 | 0.0942921069420802 | -0.307222715658533 | 0.7586738564716 | 0.942526199240263 | NM_027286,NM_001130513 |
| Ttll4 | 506.9156926 | 0.055167771 | 0.098672322 | 0.559100771 | 0.576092949 | 0.763687898 | | Adcy5 | 12511.62248 | 0.0227821904994714 | 0.0742403449639354 | 0.306870752156911 | 0.758941752051119 | 0.942526199240263 | NM_001012765 |
| Tmem18 | 1388.671869 | 0.030419555 | 0.054475334 | 0.558409706 | 0.576564646 | 0.764247808 | | Angptl6 | 264.4997176 | -0.0260162323438586 | 0.0848390893917647 | -0.306653837639893 | 0.759106870062366 | 0.942526199240263 | NM_145154 |
| Kif18a | 30.68812692 | 0.091822594 | 0.164519381 | 0.558126305 | 0.576758139 | 0.764373502 | | Anxa5 | 2299.22525 | 0.0196786862023623 | 0.0639699652452969 | 0.30762383763854 | 0.758368579436432 | 0.942526199240263 | NM_009673 |
| Zfp862 | 526.3511311 | 0.047363122 | 0.084855057 | 0.558164992 | 0.576731724 | 0.764373502 | | Coq9 | 2282.230968 | -0.0196229186923589 | 0.0640036950895833 | -0.306590403333645 | 0.75915515911628 | 0.942526199240263 | NM_026452 |
| Qsox2 | 809.0852883 | -0.041931095 | 0.075146308 | -0.557992753 | 0.576849333 | 0.764428974 | | Daam2 | 4682.238942 | 0.0232053090056835 | 0.0755184872330434 | 0.30727984439193 | 0.758630375901469 | 0.942526199240263 | NM_001008231 |
| 4930473A02Rik | 10.30533971 | 0.071634048 | 0.128406007 | 0.557871471 | 0.576932154 | 0.764473343 | | Gna13 | 3009.58833 | -0.0141086483694004 | 0.045989509101198 | -0.306779712267746 | 0.759011051401936 | 0.942526199240263 | NM_010303 |
| Nod2 | 47.73521017 | 0.086383119 | 0.154864579 | 0.557797786 | 0.576982475 | 0.764474642 | | Hbs1l | 2327.353797 | -0.0145158896353089 | 0.0473075903902744 | -0.306840604553241 | 0.758964700122181 | 0.942526199240263 | NM_001042593,NM_001145209,NM_019702 |
| 5330417C22Rik | 1266.269681 | 0.035132721 | 0.063014822 | 0.55753107 | 0.577164637 | 0.764560121 | | Mtmr1 | 1953.408088 | 0.0220091991374051 | 0.0716991262809437 | 0.306966071680774 | 0.758869197127058 | 0.942526199240263 | NM_016985 |
| Hscb | 141.5269484 | 0.066514615 | 0.119296817 | 0.557555658 | 0.577147843 | 0.764560121 | | Nudt22 | 206.937743 | -0.0268852603102595 | 0.0876976181400882 | -0.306567736735028 | 0.759172414180848 | 0.942526199240263 | NM_022675 |
| Sufu | 384.1833299 | 0.052978514 | 0.095031012 | 0.557486579 | 0.577195026 | 0.764560121 | | Pon2 | 2294.831359 | 0.0184984694131279 | 0.0603140078388928 | 0.306702705987305 | 0.759069669913584 | 0.942526199240263 | NM_183308 |
| Mfsd5 | 1241.565059 | 0.038387495 | 0.068921507 | 0.556974111 | 0.577545117 | 0.764827696 | | Ric8 | 2420.674927 | 0.0175135737921086 | 0.0569590371419427 | 0.307476647620719 | 0.75848059518358 | 0.942526199240263 | NM_053194 |
| Nat1 | 50.10764092 | -0.09064873 | 0.162720382 | -0.557082822 | 0.577470844 | 0.764827696 | | Sema7a | 9010.790392 | 0.0208460759469696 | 0.0677916733086466 | 0.307502012113791 | 0.758461291728708 | 0.942526199240263 | NM_011352 |
| Rsph1 | 116.4601944 | 0.088476213 | 0.158840237 | 0.557013857 | 0.577517962 | 0.764827696 | | Utp18 | 705.3550327 | -0.0226191346388288 | 0.0737984063963223 | -0.306498957678793 | 0.759224773324405 | 0.942526199240263 | NM_001013375 |
| 4930524L23Rik | 15.53073396 | -0.093567705 | 0.168044674 | -0.556802559 | 0.577662335 | 0.764852181 | | Zfp827 | 2269.638638 | -0.0200377497521738 | 0.065331342383822 | -0.306709597890273 | 0.759064423621512 | 0.942526199240263 | NM_178267 |
| Med26 | 352.0258037 | -0.049796488 | 0.08942817 | -0.55683224 | 0.577642054 | 0.764852181 | | Rwdd2b | 376.9818685 | 0.0231201982147922 | 0.0755368060628345 | 0.306078578376215 | 0.759544817693972 | 0.94272457014444 | NM_016924 |
| Ptbp3 | 879.5328262 | -0.045944812 | 0.08255374 | -0.556544276 | 0.577838835 | 0.7650205 | | Sarnp | 1141.920067 | -0.0236563954266856 | 0.0773431432800254 | -0.305862865451902 | 0.759709060864024 | 0.94272457014444 | NM_025364 |
| Ankrd42 | 513.0221254 | 0.048434334 | 0.087071724 | 0.556257901 | 0.578034562 | 0.765057436 | | Sart1 | 2400.448434 | -0.0188237426774806 | 0.0615366369816287 | -0.30589488800144 | 0.759684678306249 | 0.94272457014444 | NM_016882 |
| BC064078 | 74.75305515 | 0.084642994 | 0.152229031 | 0.556023994 | 0.578194452 | 0.765057436 | | Tifa | 271.1922725 | 0.023457389375789 | 0.0766825366569055 | 0.305902626575103 | 0.759678786049863 | 0.94272457014444 | NM_145133 |
| Coch | 1537.875167 | 0.072871161 | 0.13107666 | 0.555943071 | 0.578249773 | 0.765057436 | | Wdr26 | 6211.28101 | 0.0186399764032009 | 0.0609128674334935 | 0.306010489878064 | 0.759596658900257 | 0.94272457014444 | NM_145514 |
| Fam188b | 269.1670418 | -0.047395866 | 0.085190976 | -0.556348432 | 0.577972684 | 0.765057436 | | Lrp5 | 532.5823767 | 0.0221888602268318 | 0.07256544788006 | 0.305777210436388 | 0.759774281340798 | 0.942724968886686 | NM_008513 |
| Itga4 | 439.0183097 | 0.052010405 | 0.09352615 | 0.556105486 | 0.578138745 | 0.765057436 | | Fbxo30 | 310.8397207 | -0.0268020952407041 | 0.0877369029337252 | -0.305482577393344 | 0.759998637433277 | 0.94292805547983 | NM_001168297,NM_027968 |
| Pelo | 443.9514889 | -0.049211581 | 0.088521892 | -0.555925544 | 0.578261755 | 0.765057436 | | Arhgap10 | 901.297739 | -0.0180611253893945 | 0.0592363128923491 | -0.30489955413358 | 0.760442655353924 | 0.943070947807658 | NM_030113 |
| Ppme1 | 9416.025652 | -0.042093702 | 0.075656451 | -0.556379548 | 0.577951417 | 0.765057436 | | Gm17769 | 53.64765458 | -0.0283095873390227 | 0.0927936912804864 | -0.305080948374515 | 0.76030450095276 | 0.943070947807658 | NR_027377 |
| Pvt1 | 314.0901327 | -0.056931969 | 0.102406552 | -0.555940686 | 0.578251403 | 0.765057436 | | Prkrir | 3194.677027 | 0.0183959613608825 | 0.0603283632173386 | 0.304930556372122 | 0.760419042730384 | 0.943070947807658 | NM_028410 |
| Rps15a-ps6 | 19.82120477 | -0.089262934 | 0.160600384 | -0.555807726 | 0.578342303 | 0.765098672 | | Tmem125 | 368.4495179 | 0.0262367799356845 | 0.0860075909305211 | 0.305051910556117 | 0.760326616369319 | 0.943070947807658 | NM_172383 |
| Hapln1 | 576.4980755 | -0.072127569 | 0.12980944 | -0.55564194 | 0.578455655 | 0.76515473 | | Vtn | 2766.102544 | -0.0250029445246352 | 0.0819692976323156 | -0.305028165994411 | 0.760344700548151 | 0.943070947807658 | NM_011707 |
| Nrip1 | 1018.936234 | -0.061810276 | 0.111249337 | -0.555601295 | 0.578483446 | 0.76515473 | | Cacna1e | 7234.502493 | -0.0229079206423398 | 0.0752696491404229 | -0.3043447246526 | 0.760865274807365 | 0.943105712587767 | NM_009782 |
| Sf3a1 | 3348.285125 | 0.030487301 | 0.054971567 | 0.55460127 | 0.57916742 | 0.765970429 | | Nuak2 | 51.31285886 | -0.0278502925187363 | 0.0915326813606418 | -0.30426610588414 | 0.760925165326217 | 0.943105712587767 | NM_001195025,NM_028778 |
| Slc6a20a | 499.8538132 | 0.080291089 | 0.144784695 | 0.554555089 | 0.579199015 | 0.765970429 | | Osbp | 3279.439995 | 0.01593826602769 | 0.0523312997590614 | 0.304564688839593 | 0.760697717226592 | 0.943105712587767 | NM_001033174 |
| Trnt1 | 1140.047267 | 0.03383133 | 0.061023039 | 0.554402583 | 0.579303359 | 0.766043035 | | Pdgfa | 833.2626315 | 0.0214613366075812 | 0.070483760852508 | 0.304486258224649 | 0.76075746041083 | 0.943105712587767 | NM_008808 |
| Poglut1 | 1981.256679 | 0.034368849 | 0.062009936 | 0.554247457 | 0.579409504 | 0.766052637 | | Pik3r5 | 190.249903 | 0.0261500926625562 | 0.0859175775796836 | 0.304362548377292 | 0.760851697178567 | 0.943105712587767 | NM_177320 |
| Ylpm1 | 4415.793214 | -0.034087651 | 0.061498156 | -0.554287364 | 0.579382197 | 0.766052637 | | Slc25a22 | 9585.713366 | 0.0217774511583626 | 0.0714794791453053 | 0.304667177471913 | 0.760619650408529 | 0.943105712587767 | NM_001177576,NM_026646 |
| 1810044D09Rik | 13.82045169 | -0.089220727 | 0.161096983 | -0.553832385 | 0.579693562 | 0.766362804 | | Zfyve21 | 493.6370105 | -0.0204256321507834 | 0.0670392802365981 | -0.304681555033054 | 0.760608699043049 | 0.943105712587767 | NM_026752 |
| Alg12 | 387.0680473 | 0.059346128 | 0.107233796 | 0.553427463 | 0.579970738 | 0.766607924 | | Med12l | 1804.277381 | -0.0185153814542162 | 0.0608955544068444 | -0.304051447343999 | 0.761088696059718 | 0.943227915618416 | NM_177855 |
| Rab24 | 1704.021134 | 0.038338867 | 0.06927664 | 0.553416953 | 0.579977933 | 0.766607924 | | Polr3b | 809.7484371 | -0.018420007070809 | 0.0606627305990626 | -0.303646190814458 | 0.761397456858017 | 0.943530068104381 | NM_027423 |
| Dusp6 | 2135.10789 | -0.087438275 | 0.158105377 | -0.553037962 | 0.580237416 | 0.766885483 | | Elovl7 | 860.0128634 | -0.0284799730834436 | 0.0939138149007932 | -0.303256481631895 | 0.761694408130642 | 0.943656543497288 | NM_029001 |
| Plxna3 | 1390.807285 | 0.077597581 | 0.140349202 | 0.552889363 | 0.580339171 | 0.766954548 | | Rpp38 | 202.2329828 | 0.0272081250294729 | 0.089688587107535 | 0.303362176916121 | 0.76161386678609 | 0.943656543497288 | NM_001013376 |
| Use1 | 1638.54185 | -0.03784995 | 0.068476115 | -0.552746746 | 0.580436839 | 0.767018199 | | Umps | 719.8507417 | 0.0211483256094573 | 0.0697302975625944 | 0.303287471137968 | 0.761670793408552 | 0.943656543497288 | NM_009471 |
| Car5b | 153.5490642 | 0.067668483 | 0.122471997 | 0.552522089 | 0.580590705 | 0.767107029 | | Dlx2 | 228.8366357 | 0.0267725555374367 | 0.0883294524174736 | 0.303098851002732 | 0.761814529759558 | 0.94372487287117 | NM_010054 |
| Carm1 | 3215.166508 | 0.038379391 | 0.069464456 | 0.552504021 | 0.58060318 | 0.767107029 | | Ggt7 | 5862.850682 | -0.0187246052441812 | 0.0617980177118045 | -0.302996858758537 | 0.761892255495068 | 0.943740675647766 | NM_144786 |
| Atp6v1b2 | 31166.81289 | -0.036898066 | 0.066793327 | -0.552421444 | 0.580659642 | 0.767116345 | | Drd2 | 1353.429454 | 0.0275577881962431 | 0.0911169775741127 | 0.302444055212742 | 0.762313575008427 | 0.944182042170226 | NM_010077 |
| Dyx1c1 | 90.0565935 | 0.073325528 | 0.132765546 | 0.552293348 | 0.580747387 | 0.767166853 | | Cdc40 | 2072.64197 | -0.019199507383759 | 0.0635273883418574 | -0.302224093967809 | 0.762481238185852 | 0.944309187985991 | NM_027879 |
| AI597479 | 1296.250587 | -0.03802538 | 0.068890231 | -0.551970564 | 0.580968521 | 0.767185714 | | Trio | 7818.217123 | 0.0192427481464288 | 0.0636927676840305 | 0.302118260003656 | 0.762561912999416 | 0.944328588863873 | NM_001081302 |
| Arhgap24 | 244.9311633 | -0.067948597 | 0.123072058 | -0.552104177 | 0.58087698 | 0.767185714 | | BC017612 | 66.83134447 | 0.0286735053920188 | 0.0951011367643509 | 0.301505390656562 | 0.763029139999219 | 0.944431816409278 | NM_133214 |
| Mllt6 | 7928.291013 | 0.033608209 | 0.060879355 | 0.552046071 | 0.580916789 | 0.767185714 | | Fkbp10 | 277.7961793 | 0.0247160003930395 | 0.0819568608082024 | 0.301573292941033 | 0.762977369765861 | 0.944431816409278 | NM_010221,NM_001163481 |
| Rmi1 | 650.6745029 | 0.044246799 | 0.080170151 | 0.551911133 | 0.581009241 | 0.767185714 | | Gjc1 | 174.3377636 | 0.0277711819228176 | 0.0919952776988857 | 0.301876168184598 | 0.762746463761386 | 0.944431816409278 | NM_008122,NM_001159382,NM_001159383 |
| Snx17 | 2932.025867 | -0.025987725 | 0.04708678 | -0.551911278 | 0.581009141 | 0.767185714 | | Pdcd5 | 965.1622359 | -0.0227047253101863 | 0.0752949816605458 | -0.30154367297887 | 0.762999932562809 | 0.944431816409278 | NM_019746 |
| Dera | 65.37567884 | 0.082173201 | 0.148931672 | 0.551751012 | 0.581118955 | 0.767239587 | | Selplg | 1532.777003 | 0.019909224237435 | 0.0660345204884937 | 0.301497218275464 | 0.763035370878376 | 0.944431816409278 | NM_009151 |
| Dnmt1 | 2374.21126 | 0.040708004 | 0.073785542 | 0.551707054 | 0.581149076 | 0.767239587 | | Wdr48 | 3806.363821 | 0.0133975759330586 | 0.044432168667975 | 0.30158950382933 | 0.762965010375701 | 0.944431816409278 | NM_026236 |
| Zfp133-ps | 94.31778235 | -0.080237165 | 0.145478297 | -0.551540449 | 0.581263246 | 0.767324933 | | Cnot6 | 1425.858772 | 0.0228082609756553 | 0.075718989282149 | 0.301222469976002 | 0.763248856513699 | 0.944610614783845 | NM_212484 |
| 4930429B21Rik | 394.6208255 | -0.059014274 | 0.107063093 | -0.551210252 | 0.581489553 | 0.767558284 | | 4632434I11Rik | 25.0664515 | -0.0222973077546632 | 0.0740621592482012 | -0.301062080429534 | 0.763367155754266 | 0.944624337178718 | NM_001080995 |
| Gpr162 | 7192.702998 | -0.044067649 | 0.079997304 | -0.550864182 | 0.581726783 | 0.76767521 | | Basp1 | 12765.52319 | -0.0192624061609217 | 0.064000916886127 | -0.300970784452888 | 0.763436772832904 | 0.944624337178718 | NM_027395 |
| Slc1a1 | 5266.366623 | -0.036285107 | 0.065863416 | -0.550914438 | 0.58169233 | 0.76767521 | | Mrpl11 | 1340.784471 | 0.0170330478640974 | 0.0565972087899682 | 0.300952082766252 | 0.763451033903219 | 0.944624337178718 | NM_025553 |
| Ufl1 | 1370.167696 | 0.040723804 | 0.073916261 | 0.550945135 | 0.581671286 | 0.76767521 | | Cast | 557.0704538 | 0.0227406712661429 | 0.0756230829750842 | 0.300710713865439 | 0.763635098239824 | 0.944771606935669 | NM_009817 |
| Capzb | 7379.898525 | 0.041903888 | 0.076094683 | 0.55068089 | 0.581852447 | 0.767761516 | | Sp4 | 808.0456685 | -0.0221741638843584 | 0.0737636564085413 | -0.300610964314816 | 0.763711169666805 | 0.944785253839211 | NM_001166385,NM_009239 |
| Gpr151 | 27.81127556 | 0.0899005 | 0.163270141 | 0.550624254 | 0.581891279 | 0.767761516 | | Actr1b | 11157.37239 | 0.0153922209680634 | 0.0512323914073033 | 0.30043924449463 | 0.763842132710379 | 0.944786343643644 | NM_146107 |
| Cyr61 | 316.1565312 | -0.089105878 | 0.162037976 | -0.549907371 | 0.582382908 | 0.768194642 | | Zfp273 | 222.3728049 | -0.0261238445065055 | 0.0869421303849358 | -0.300473940434198 | 0.76381567112084 | 0.944786343643644 | NM_198322 |
| Dhx29 | 1111.169425 | -0.038505297 | 0.070027604 | -0.549858841 | 0.582416196 | 0.768194642 | | Melk | 46.62873738 | -0.0276616896788951 | 0.0921020530113912 | -0.30033738417833 | 0.763919820220842 | 0.944801984548996 | NM_010790 |
| Exd1 | 34.73727159 | 0.087742865 | 0.159537175 | 0.549983822 | 0.58233047 | 0.768194642 | | D3Ertd751e | 659.4676532 | 0.0229561129199894 | 0.0765702418029996 | 0.299804627874839 | 0.764326184387655 | 0.94487204460526 | NM_028667,NM_027271,NM_001099785 |
| Matn1 | 13.23215288 | 0.091795009 | 0.166987504 | 0.549711848 | 0.582517029 | 0.768194642 | | Ddr1 | 3414.39037 | -0.0215581354403699 | 0.0718427325228721 | -0.300073990552998 | 0.76412071805778 | 0.94487204460526 | NM_172962,NM_001198831,NM_001198833,NM_007584 |

| | | | | | | | | | | | | | | | |
|---|---|---|---|---|---|---|---|---|---|---|---|---|---|---|---|
| Slc20a2 | 2222.290289 | 0.034621587 | 0.062973108 | 0.549783673 | 0.582467758 | 0.768194642 | | Fam162a | 684.0135823 | -0.0213160755053081 | 0.0710284346717044 | -0.300106226525077 | 0.76409612990863 | 0.94487204460526 | NM_027342 |
| Stc1 | 158.6219822 | 0.08183276 | 0.148845355 | 0.549783769 | 0.582467693 | 0.768194642 | | Hilpda | 154.6785017 | -0.0270481681906286 | 0.0903089761702962 | -0.299506974142014 | 0.76455325152981 | 0.94487204460526 | NM_023516,NM_001190461 |
| Zfp93 | 415.5877512 | 0.045160486 | 0.08216654 | 0.54962136 | 0.582579105 | 0.768211119 | | Hmgn1 | 1651.298838 | -0.0191028848994837 | 0.0638091028769725 | -0.299375544212165 | 0.764653519867553 | 0.94487204460526 | NM_008251 |
| Pwwp2a | 831.1122896 | -0.033258183 | 0.060545393 | -0.549309894 | 0.582792798 | 0.768379032 | | P2ry14 | 104.6403911 | -0.0284367753983711 | 0.0949883572597517 | -0.299371167359058 | 0.764656859051473 | 0.94487204460526 | NM_133200,NM_001008497 |
| Spock2 | 29177.39221 | -0.028078876 | 0.05111838 | -0.549291198 | 0.582805627 | 0.768379032 | | Pikfyve | 2315.787756 | 0.0198156239948341 | 0.066096484841585 | 0.299798454370556 | 0.764330894363471 | 0.94487204460526 | NM_011086 |
| Akr1a1 | 10758.47722 | 0.035694782 | 0.065006293 | 0.549097337 | 0.582938652 | 0.768489022 | | Sema6d | 3027.15409 | 0.023206265703197 | 0.0775286253951767 | 0.299325127782296 | 0.764691983779392 | 0.94487204460526 | NM_199240,NM_199241,NM_199239,NM_199238,NM_172537 |
| L3mbtl4 | 32.50067235 | -0.089098696 | 0.162297464 | -0.548983907 | 0.583016494 | 0.768526251 | | Supt16 | 3192.354189 | 0.0164164691197027 | 0.0548299098963412 | 0.299407187623304 | 0.76462937864158 | 0.94487204460526 | NM_033618 |
| 1700023F06Rik | 60.33701765 | 0.087006132 | 0.158663745 | 0.548368072 | 0.583439195 | 0.769018025 | | Trnau1ap | 484.2052668 | -0.0214202607291581 | 0.0715125849770819 | -0.299531344532194 | 0.764534659710496 | 0.94487204460526 | NM_027925 |
| Wdr24 | 1070.551066 | 0.040267833 | 0.073465385 | 0.548119819 | 0.583609633 | 0.769177241 | | Zmat2 | 7864.083048 | 0.0171152138340128 | 0.0570987659325134 | 0.299747526141664 | 0.764369743820063 | 0.94487204460526 | NM_025594 |
| Fam5b | 4795.485344 | -0.046105215 | 0.084146621 | -0.547915226 | 0.583750114 | 0.76923649 | | Epha4 | 8052.688912 | -0.0199993053368127 | 0.0668553084028903 | -0.299143116897925 | 0.76483084910815 | 0.94492945980006 | NM_007936 |
| Tvp23a | 1624.162441 | -0.037747112 | 0.068892939 | -0.547909738 | 0.583753882 | 0.76923649 | | Papolg | 864.9832126 | -0.0218277969246068 | 0.07297979474434 | -0.299093701223374 | 0.764868552136108 | 0.94492945980006 | NM_172555 |
| Pgap3 | 404.8244089 | 0.051978025 | 0.094878736 | 0.547836401 | 0.583804242 | 0.769237428 | | Rabep1 | 6829.230607 | 0.0186577949554164 | 0.0624386078995298 | 0.298818240557809 | 0.765078732523986 | 0.945108739573384 | NM_019400 |
| 1110019D14Rik | 66.25876234 | -0.083052663 | 0.151832207 | -0.547002932 | 0.584376718 | 0.769926264 | | Senp2 | 4588.905566 | 0.0170260476050488 | 0.05699637705226 | 0.298721576451037 | 0.765152492707718 | 0.945119482063972 | NR_027488,NM_029457 |
| Psma7 | 2228.059734 | -0.043684699 | 0.079876391 | -0.546903771 | 0.584444845 | 0.769950551 | | Insm1 | 169.0599213 | 0.0280892065969527 | 0.094079217451194 | 0.298569730467036 | 0.765268364087077 | 0.945182234225736 | NM_016889 |
| 3110039I08Rik | 24.38128652 | 0.091646113 | 0.167685971 | 0.546534167 | 0.584698808 | 0.770154155 | | Ppef1 | 25.22957242 | -0.0252199926955449 | 0.0845251438706496 | -0.298372668068327 | 0.765418747262661 | 0.945287597580124 | NM_011147 |
| Gstt1 | 308.5353495 | -0.055963769 | 0.102388832 | -0.546580795 | 0.584666766 | 0.770154155 | | Rfc1 | 1856.858309 | -0.0166013118379045 | 0.0556711592665908 | -0.298203092168538 | 0.765548161891827 | 0.9453083329044 | NM_011258 |
| A830018L16Rik | 2572.598838 | -0.043413223 | 0.079495834 | -0.546106894 | 0.584992461 | 0.770330198 | | Tmem258 | 733.7281752 | 0.0202792551927857 | 0.0680100837441894 | 0.298180123833745 | 0.765565691056544 | 0.9453083329044 | NM_026919,NM_001163431 |
| Dlg5 | 1781.896664 | 0.064298479 | 0.117756654 | 0.546028417 | 0.585046403 | 0.770330198 | | Eya3 | 1773.960511 | 0.0137697832217356 | 0.0462233019989026 | 0.297897004892954 | 0.76578177398158 | 0.94545314186285 | NM_210071,NM_211357,NM_010166 |
| Morn5 | 18.0922029 | 0.09001505 | 0.164934911 | 0.545761046 | 0.585230203 | 0.770330198 | | Gm715 | 107.3477245 | -0.0275013429306336 | 0.0923574708161461 | -0.297770637151594 | 0.765878226648779 | 0.94545314186285 | NM_001271548 |
| Nfam1 | 153.1128724 | 0.075680768 | 0.138667365 | 0.545772022 | 0.585222657 | 0.770330198 | | Syce2 | 28.1511615 | 0.0256842541191031 | 0.0862513088144653 | 0.297783934784715 | 0.765868076797532 | 0.94545314186285 | NM_001168244,NM_027954,NM_001168246,NR_031759 |
| Sag | 79.64616371 | -0.074184569 | 0.135913318 | -0.545822663 | 0.585187843 | 0.770330198 | | Stbd1 | 51.4973384 | 0.0266878214476267 | 0.089731106450626 | 0.297419952826633 | 0.766145912373274 | 0.945703222564087 | NM_175096 |
| Scarb2 | 3127.367233 | -0.030378247 | 0.055658303 | -0.545799013 | 0.585204102 | 0.770330198 | | Rasgrp1 | 24575.28858 | 0.0219478959862202 | 0.0739023580510165 | 0.29698505656706 | 0.766477917859872 | 0.946032648044226 | NM_011246 |
| Tnn | 17.07874531 | 0.089641927 | 0.16422241 | 0.545856847 | 0.585164343 | 0.770330198 | | Pdcd4 | 2898.777094 | -0.0189877008017026 | 0.0639694077445703 | -0.296824708421883 | 0.766600340552537 | 0.946103359971636 | NM_001168491,NM_011050,NM_001168492 |
| Trim9 | 12174.33499 | -0.040158729 | 0.073527527 | -0.546172712 | 0.584947222 | 0.770330198 | | Etaa1 | 397.6356917 | 0.0257731514986583 | 0.0870595096984649 | 0.296040623108549 | 0.767199058307758 | 0.946118814923041 | NM_026576 |
| Oip5 | 25.49654365 | 0.091485345 | 0.167677769 | 0.545602112 | 0.585339472 | 0.770408578 | | Gmeb2 | 1103.799408 | 0.0200470831511139 | 0.0676932869101668 | 0.296145808043236 | 0.767118732325423 | 0.946118814923041 | NM_198169 |
| Arhgef5 | 176.5287083 | 0.070205621 | 0.128746005 | 0.545303294 | 0.585544939 | 0.770474879 | | Golga5 | 1261.465989 | -0.017301657391018 | 0.0583267201130702 | -0.296633470174178 | 0.766746334856704 | 0.946118814923041 | NM_001199004,NM_013747 |
| Cep170 | 3260.298136 | -0.036848226 | 0.067568943 | -0.545342654 | 0.585517873 | 0.770474879 | | Lace1 | 752.5503694 | 0.0176913274767708 | 0.0597217821869781 | 0.296229061306014 | 0.767055156552462 | 0.946118814923041 | NM_145743 |
| Evi5 | 2818.827892 | 0.031607306 | 0.057969574 | 0.545239576 | 0.585588755 | 0.770474879 | | Pag1 | 1520.402842 | -0.0193136531107809 | 0.0652002350748387 | -0.296220605471931 | 0.767061613719604 | 0.946118814923041 | NM_053182,NM_001195031 |
| Megf10 | 1315.816933 | -0.055617682 | 0.101977773 | -0.545390241 | 0.585485151 | 0.770474879 | | Pde7b | 3295.230313 | 0.0281627810653295 | 0.0950941443174582 | 0.296156837705086 | 0.76711030951106 | 0.946118814923041 | NM_013875 |
| Gm10863 | 13.13959842 | 0.086681568 | 0.159092479 | 0.544850196 | 0.585856552 | 0.770761775 | | S1pr5 | 597.2507353 | 0.0278523298148052 | 0.0940692017395034 | 0.296083407744162 | 0.76716638490597 | 0.946118814923041 | NM_053190 |
| Htra3 | 153.424567 | 0.068451039 | 0.1256957 | 0.544577414 | 0.586044192 | 0.770943175 | | Slc35f3 | 1729.264494 | -0.023588422263377 | 0.0795915567942364 | -0.296368399029546 | 0.766948755780748 | 0.946118814923041 | NM_175434 |
| 5430405H02Rik | 162.3550858 | 0.061463667 | 0.11296623 | 0.544088854 | 0.58638033 | 0.771031552 | | Ufsp1 | 292.1435533 | -0.0253678986472386 | 0.0855822458812716 | -0.296415435070862 | 0.76691283921224 | 0.946118814923041 | NM_027356 |
| Arhgap44 | 6359.016289 | 0.050628972 | 0.09300847 | 0.544347967 | 0.586202044 | 0.771031552 | | 5730508B09Rik | 91.63837487 | -0.0276568460161878 | 0.0943241915667566 | -0.293210528039501 | 0.769361237356626 | 0.946190740283965 | NM_027482 |
| C330006A16Rik | 2868.919883 | -0.035613193 | 0.06544319 | -0.544184851 | 0.586314276 | 0.771031552 | | AA414768 | 311.4804665 | -0.0234730083474326 | 0.0797655481306079 | -0.294275020952629 | 0.768547758429568 | 0.946190740283965 | NM_001272033 |
| Dtwd1 | 227.5989053 | -0.054349048 | 0.099884728 | -0.544117696 | 0.586360484 | 0.771031552 | | Ablim3 | 1428.205356 | 0.0182573504189156 | 0.0623233034888513 | 0.292945806734741 | 0.769563575207835 | 0.946190740283965 | NM_001164491,NM_198649 |
| Gpt | 472.2157925 | 0.068410455 | 0.125743924 | 0.544045805 | 0.586409953 | 0.771031552 | | Acot1 | 205.2443649 | 0.0250483580655947 | 0.0855157870825397 | 0.292909168238352 | 0.769591580819099 | 0.946190740283965 | NM_012006 |
| Katnbl1 | 392.328904 | -0.046582134 | 0.085597426 | -0.544200168 | 0.586303736 | 0.771031552 | | Ampd2 | 3185.007379 | -0.0197594777820107 | 0.0674371178853971 | -0.293005964691286 | 0.769517592522289 | 0.946190740283965 | NM_028779 |
| Wars2 | 316.8494004 | -0.044798139 | 0.082355991 | -0.543957249 | 0.586470892 | 0.771046245 | | Arhgef19 | 614.8035819 | -0.0238912303128773 | 0.0818571981267821 | -0.291864745674217 | 0.770390039308803 | 0.946190740283965 | NM_172520 |
| Eno2 | 41775.84434 | -0.026118498 | 0.048026571 | -0.543834328 | 0.586555484 | 0.771090368 | | B4galt2 | 3335.776508 | -0.0214857221091847 | 0.0729223140289534 | -0.294638512165891 | 0.768270038995882 | 0.946190740283965 | NM_017377,NR_045570,NM_001253381 |
| Wdfy2 | 312.6339017 | 0.049538485 | 0.09110294 | 0.543763852 | 0.586603987 | 0.771090368 | | Cald1 | 1005.736011 | -0.0212569400642727 | 0.0722157304772954 | -0.29435332058236 | 0.768487932384302 | 0.946190740283965 | NM_145575 |
| 4933417O13Rik | 9.174195133 | 0.085140678 | 0.156617588 | 0.543621437 | 0.586702005 | 0.771145473 | | Ccdc15 | 90.44954804 | -0.0276787675648785 | 0.0943994440098812 | -0.293209010446939 | 0.769362397272988 | 0.946190740283965 | NM_001081429 |
| Mamdc4 | 57.05258338 | 0.090454828 | 0.166412371 | 0.543558319 | 0.586745449 | 0.771145473 | | Cd163 | 58.30930589 | 0.0251515510733148 | 0.086152529686617 | 0.291942107385638 | 0.77033088811247 | 0.946190740283965 | NM_053094,NM_001170395 |
| Rbm7 | 838.1205129 | 0.037814397 | 0.069587693 | 0.543406386 | 0.58685003 | 0.771170181 | | Cdk12 | 1163.485224 | 0.0210122067294332 | 0.0711179571603924 | 0.295455712852443 | 0.767645779009948 | 0.946190740283965 | NM_001109626,NM_026952,NM_001109628 |
| Siva1 | 171.1826869 | -0.065988866 | 0.121440041 | -0.543386392 | 0.586863793 | 0.771170181 | | Cdk15 | 27.54721064 | 0.0247744908949864 | 0.0840144670811979 | 0.294883628447497 | 0.768082778753938 | 0.946190740283965 | NM_001033373 |
| Etfb | 839.5823339 | -0.050697988 | 0.093335066 | -0.543182643 | 0.587004056 | 0.771289081 | | Cep112 | 93.23294653 | -0.0276914598421216 | 0.0949028429799682 | -0.291787463605981 | 0.77044913094305 | 0.946190740283965 | NM_029586,NM_029606,NM_145688 |
| Thnsl2 | 163.7217201 | 0.066002719 | 0.121540918 | 0.543049375 | 0.587095808 | 0.771344225 | | Ckap5 | 8228.99655 | 0.018061055448831 | 0.0617587789616197 | 0.292445151159078 | 0.76994629063655 | 0.946190740283965 | NM_029437,NM_001165989 |
| Gm3230 | 126.6873184 | -0.076088414 | 0.140140668 | -0.542943137 | 0.587168954 | 0.771374917 | | Cpne5 | 5524.402414 | 0.0163645169760596 | 0.0555639353404507 | 0.294516881782961 | 0.768362965363635 | 0.946190740283965 | NM_153166 |
| Zcchc10 | 222.358831 | -0.064346037 | 0.118533635 | -0.542850453 | 0.587232772 | 0.771393351 | | Ddx18 | 900.3755974 | 0.0172192610676485 | 0.0588966232738953 | 0.292364147730021 | 0.770008217241668 | 0.946190740283965 | NM_025860 |
| Fam193a | 1954.879038 | 0.032421206 | 0.059774861 | 0.542388656 | 0.587550793 | 0.771745675 | | Ddx52 | 1008.516938 | -0.0202098038513572 | 0.0691491598089374 | -0.292263910468296 | 0.770084850019496 | 0.946190740283965 | NM_030096 |
| Exo1 | 22.50942625 | -0.090635404 | 0.16720242 | -0.542069927 | 0.587770335 | 0.771945244 | | Dnm1 | 56731.60448 | -0.0117554191821787 | 0.0397990514118973 | -0.295369330804316 | 0.767711759180187 | 0.946190740283965 | NM_010065 |
| Mss51 | 15.03948817 | 0.090493928 | 0.166992334 | 0.541904682 | 0.587884171 | 0.771945244 | | Eny2 | 1948.409617 | 0.0165050424905944 | 0.0563706422576553 | 0.292795005158079 | 0.769678846339489 | 0.946190740283965 | NM_175009 |
| Sec61a1 | 2555.351737 | -0.040791116 | 0.075264127 | -0.541972889 | 0.587837182 | 0.771945244 | | Fam86 | 163.9800653 | -0.0262355106225723 | 0.0896858199863524 | -0.292526852367125 | 0.769883832069817 | 0.946190740283965 | NM_027446 |
| Srp68 | 2779.255889 | 0.045355091 | 0.083699701 | 0.541878776 | 0.587902019 | 0.771945244 | | Fez2 | 1602.223408 | 0.0151496145238005 | 0.0518892219602723 | 0.291960718459017 | 0.770316658181229 | 0.946190740283965 | NM_199448 |
| Cbfa2t3 | 1293.621034 | 0.059280562 | 0.109460961 | 0.541568081 | 0.588116086 | 0.772160887 | | Hadh | 886.5788698 | 0.0191061576219562 | 0.0649005877760615 | 0.294391133835055 | 0.768459041076413 | 0.946190740283965 | NM_008212 |
| Nup210 | 1339.036602 | 0.045332907 | 0.083749101 | 0.541294254 | 0.58830478 | 0.772274883 | | Il1f9 | 55.55358002 | 0.0273645351336014 | 0.0936792706048714 | 0.292108755297871 | 0.770203472727519 | 0.946190740283965 | NM_153511 |
| Pex5 | 2129.756942 | 0.034057109 | 0.062929785 | 0.541192199 | 0.588375114 | 0.772274883 | | Itpripl1 | 95.76589268 | 0.0278166479175908 | 0.0940307718257399 | 0.295824945148183 | 0.767363771693959 | 0.946190740283965 | NM_001163528,NM_001163527 |
| Sned1 | 256.7051707 | 0.066000328 | 0.121948523 | 0.541214655 | 0.588359638 | 0.772274883 | | Jhdm1d | 2549.879036 | 0.0264113378011832 | 0.0895415473690455 | 0.29496182026349 | 0.768023045795978 | 0.946190740283965 | NM_001033430 |
| Stk32c | 6147.26102 | 0.046303171 | 0.085574114 | 0.541088522 | 0.58844657 | 0.772274883 | | Kcns1 | 919.9171472 | 0.023293135653688 | 0.079288563182086 | 0.293776740539431 | 0.768928509584812 | 0.946190740283965 | NM_008435 |
| Tceal7 | 17.06878168 | 0.090665906 | 0.167564556 | 0.541080457 | 0.588452128 | 0.772274883 | | Mettl1 | 211.697519 | 0.0245115914917358 | 0.0829670096189491 | 0.295437808404963 | 0.767659454612337 | 0.946190740283965 | NM_010792 |
| Klhl7 | 2902.61606 | -0.042744422 | 0.079017569 | -0.540948338 | 0.588543192 | 0.772328975 | | Mettl4 | 306.636589 | -0.0231093726801181 | 0.0791556481553305 | -0.291948499174306 | 0.770326000974923 | 0.946190740283965 | NM_176917 |
| Hint2 | 365.7362738 | -0.046319722 | 0.085678218 | -0.540624247 | 0.588766602 | 0.772556718 | | Mynn | 630.5088272 | 0.0193806759395807 | 0.0660830969279299 | 0.293277355943491 | 0.769310160404531 | 0.946190740283965 | NM_030557 |
| Zfp446 | 426.1447276 | 0.074587078 | 0.138009457 | 0.540449036 | 0.588887399 | 0.772649789 | | Nfatc1 | 295.3470599 | 0.0244118786871626 | 0.0832947399351566 | 0.293078274884666 | 0.769462322154972 | 0.946190740283965 | NM_001164110,NM_001164111,NM_001164112,NM_016791,NM_001164109,NM_198429 |
| Bcas2 | 1531.147994 | -0.03650223 | 0.067562929 | -0.540270094 | 0.589010779 | 0.772708749 | | Nlgn2 | 12292.3845 | -0.012541540769076 | 0.0425878332310491 | -0.294486427252185 | 0.768386198954185 | 0.946190740283965 | NM_198862 |
| Dnajb9 | 1266.195632 | 0.04245037 | 0.078595605 | 0.540111243 | 0.589120317 | 0.772708749 | | Nop58 | 1090.33897 | 0.0236076791166632 | 0.0802291704594954 | 0.294253062588773 | 0.76856453630521 | 0.946190740283965 | NM_018868 |
| Tcf3 | 820.5181599 | 0.04065741 | 0.075278317 | 0.540094559 | 0.589131822 | 0.772708749 | | Pacsin1 | 21579.59504 | 0.0157329302600265 | 0.0533108683244841 | 0.295116751133481 | 0.767904693755866 | 0.946190740283965 | NM_011861,NM_178365 |
| Ttc27 | 738.984848 | -0.03884873 | 0.071916861 | -0.540189462 | 0.58906638 | 0.772708749 | | Rabggta | 1792.489877 | -0.0141417956779306 | 0.0484488180855655 | -0.291891448269281 | 0.770369622204625 | 0.946190740283965 | NM_019519 |
| Gm13003 | 11.74540247 | 0.079124047 | 0.146622325 | 0.539645288 | 0.589441678 | 0.772918869 | | Ralb | 711.0196377 | 0.0211325881570578 | 0.0722625512760039 | 0.292441766639856 | 0.769948878050555 | 0.946190740283965 | NM_022327 |
| Med31 | 254.704196 | -0.053928676 | 0.099931124 | -0.539658456 | 0.589432595 | 0.772918869 | | Rps8 | 7734.12279 | 0.016578893764299 | 0.0562144042495971 | 0.294922520048192 | 0.768053068178952 | 0.946190740283965 | NM_009098 |
| Zfp35 | 888.5747094 | 0.04563404 | 0.084543301 | 0.53977121 | 0.589354824 | 0.772918869 | | Rtca | 1834.818347 | 0.017430497616588 | 0.0596833282474975 | 0.292049915695836 | 0.770248459504886 | 0.946190740283965 | NM_025517 |
| Clp1 | 258.3360283 | 0.060736835 | 0.112585026 | 0.539475252 | 0.589558969 | 0.772972905 | | Sars2 | 480.497672 | 0.0237004624509033 | 0.0804472659493352 | 0.294608675275921 | 0.768292834256865 | 0.946190740283965 | NM_023637 |
| Sfxn4 | 448.4389421 | 0.076798213 | 0.142366315 | 0.539440903 | 0.589582664 | 0.772972905 | | Sh3rf2 | 537.5569452 | 0.0276675147493143 | 0.0943372460672563 | 0.293283044637419 | 0.769305812549213 | 0.946190740283965 | NM_172966,NM_001146299 |
| Kazn | 4001.846865 | -0.050839996 | 0.094328086 | -0.538969863 | 0.589907649 | 0.773333541 | | Sirpa | 23465.59815 | 0.0139045940914775 | 0.0472350928880123 | 0.294369995724224 | 0.768475191661877 | 0.946190740283965 | NM_001177646,NM_001177647,NM_007547 |
| B9d1 | 283.9950284 | -0.059873947 | 0.111215744 | -0.538358554 | 0.590329534 | 0.773821134 | | Sirt1 | 564.2601079 | -0.0246791671454457 | 0.0841902863971176 | -0.293135564702042 | 0.76941853345841 | 0.946190740283965 | NM_019812,NM_001159589 |
| Ebi3 | 36.36160618 | -0.083975887 | 0.15612682 | -0.537869707 | 0.590667003 | 0.774197999 | | Slc17a8 | 257.5411261 | 0.0255131242581501 | 0.0873278520367295 | 0.29215334699197 | 0.770169379935559 | 0.946190740283965 | NM_182959 |
| Pign | 682.7833016 | 0.046930789 | 0.087271511 | 0.537756117 | 0.590745432 | 0.7742353 | | Snrpb2 | 744.4743191 | 0.0191163869743384 | 0.0654738845943215 | 0.291969647024675 | 0.77030983147317 | 0.946190740283965 | NM_021335 |
| Rab6b | 43874.09773 | -0.030336851 | 0.056443111 | -0.537476596 | 0.590938447 | 0.774422761 | | Tceb3 | 1572.42614 | -0.0168649459428623 | 0.0572869198862579 | -0.294394356972715 | 0.768456578445102 | 0.946190740283965 | NM_013736 |
| Skor1 | 14.50772559 | -0.087232606 | 0.162397116 | -0.537156126 | 0.591159774 | 0.774647289 | | Tomm34 | 3184.513982 | -0.0146091560426052 | 0.0499559325569017 | -0.292440863274143 | 0.769949568660479 | 0.946190740283965 | NM_025996 |
| Lefty2 | 13.17076397 | 0.089327036 | 0.1664186 | 0.536761131 | 0.591432624 | 0.774873759 | | Trip10 | 270.2350669 | -0.024952162829545 | 0.0845523350643733 | -0.295109092026233 | 0.767910544438002 | 0.946190740283965 | NM_134125,NM_001242389,NM_001242391,NM_001242390 |
| Tmem191c | 2468.554151 | 0.057219425 | 0.106594058 | 0.536797512 | 0.59140749 | 0.774873759 | | U2af1l4 | 994.6607711 | -0.0190929297403346 | 0.0652978653519192 | -0.292397456447226 | 0.769982752762144 | 0.946190740283965 | NM_170760 |
| Fgfr1 | 2835.623602 | -0.047563585 | 0.088686811 | -0.53630956 | 0.591744625 | 0.775206951 | | Wdyhv1 | 680.6220696 | -0.0179376675632244 | 0.0610927401969759 | -0.293613733896853 | 0.76905308000153 | 0.946190740283965 | NM_029734 |
| Serinc3 | 12117.60224 | 0.035948704 | 0.067037431 | 0.536248234 | 0.591787002 | 0.775206951 | | Wnk2 | 5935.243561 | 0.0171503727306633 | 0.058016715642231 | 0.295610886290492 | 0.767527258961987 | 0.946190740283965 | NM_029361 |
| Zfp362 | 1039.939701 | 0.037782068 | 0.070483892 | 0.536038328 | 0.591932061 | 0.775331419 | | Acbd6 | 1389.018181 | 0.0157354337005531 | 0.0540785014218308 | 0.290973922849882 | 0.771071263567763 | 0.946250219280113 | NM_028250,NM_001145781,NM_026683 |
| Gtpbp5 | 362.7761543 | 0.061191779 | 0.114211438 | 0.535776277 | 0.59211318 | 0.775372009 | | B4galnt4 | 3389.280536 | -0.0173865814176234 | 0.0597579838744125 | -0.290949933220021 | 0.77108961120625 | 0.946250219280113 | NM_177897 |
| Lsamp | 2075.803701 | 0.026264858 | 0.04901191 | 0.535887255 | 0.592036473 | 0.775372009 | | Dyrk1a | 3745.7956 | 0.0173174769049759 | 0.0594248951348228 | 0.291417879083944 | 0.770731742126944 | 0.946250219280113 | NM_007890,NM_001113389 |
| Slc24a3 | 4927.338196 | -0.030894416 | 0.057657563 | -0.535825916 | 0.592078869 | 0.775372009 | | Lrrc4b | 5580.999704 | 0.0145493192587277 | 0.049953218179606 | 0.291258897603271 | 0.770853320285932 | 0.946250219280113 | NM_198250 |
| A330069E16Rik | 25.34016021 | 0.087315723 | 0.163135436 | 0.535234555 | 0.592487674 | 0.7754411 | | Nemf | 1944.073279 | -0.0204631444648217 | 0.0703317279630461 | -0.2909518229891 | 0.771088165877099 | 0.946250219280113 | NM_025441 |
| Calb1 | 4230.795365 | 0.052287493 | 0.097771552 | 0.534792499 | 0.59279335 | 0.7754411 | | Phospho1 | 1110.957084 | -0.0223215154277702 | 0.076779231395544 | -0.29072335086003 | 0.771262911201328 | 0.946250219280113 | NM_153104 |
| Col6a6 | 22.64783505 | -0.088277921 | 0.164976005 | -0.53509552 | 0.592583808 | 0.7754411 | | Plxna1 | 5302.096037 | 0.0190461992958616 | 0.0653572889121703 | 0.291416605750839 | 0.77073271586261 | 0.946250219280113 | NM_008881 |
| Hk3 | 75.10203008 | 0.089639666 | 0.167502847 | 0.535153087 | 0.592544003 | 0.7754411 | | Prps1l3 | 593.845782 | 0.0229201961674152 | 0.0788122785970159 | 0.290820118075904 | 0.771188898049549 | 0.946250219280113 | NM_001037746 |
| Kcnk13 | 333.8588311 | -0.065918972 | 0.123208601 | -0.535019241 | 0.592636552 | 0.7754411 | | Pskh1 | 489.7598474 | -0.0230947936409831 | 0.0793547764653074 | -0.291032180666274 | 0.771026707625481 | 0.946250219280113 | NM_173432 |
| Mmel1 | 32.01723255 | 0.089821862 | 0.167989748 | 0.534686568 | 0.592866611 | 0.7754411 | | Scamp3 | 1740.9067 | 0.0176251397588514 | 0.0605928849027099 | 0.290878042647267 | 0.77114459499067 | 0.946250219280113 | NM_011886 |
| Nxph1 | 1242.408831 | 0.036095747 | 0.067490555 | 0.53482664 | 0.59276974 | 0.7754411 | | Slc23a3 | 68.69627719 | -0.0275051305027377 | 0.0945742597090833 | -0.290831042054628 | 0.771180542890951 | 0.946250219280113 | NM_194333 |
| Pold4 | 280.5835191 | -0.063054564 | 0.117916091 | -0.534740958 | 0.592828995 | 0.7754411 | | Zkscan8 | 960.9636284 | -0.0173484144428348 | 0.0596776637612323 | -0.290701970376137 | 0.771279264510294 | 0.946250219280113 | NM_001251833,NM_139141,NM_001251834 |
| Rlim | 2133.296295 | -0.045069941 | 0.084231059 | -0.535075088 | 0.592597936 | 0.7754411 | | Gtf2a2 | 544.0651869 | 0.0214780886831205 | 0.0739492593069438 | 0.290443594491875 | 0.77147689672744 | 0.946412752458643 | NM_001039519 |
| Tmub1 | 610.1568111 | 0.042684019 | 0.079816238 | 0.534778641 | 0.592802934 | 0.7754411 | | 2310047M10Rik | 847.0577223 | 0.0187821836688396 | 0.0647136140238367 | 0.290235431170964 | 0.771636132045773 | 0.946528158596258 | NM_028005 |
| Topbp1 | 949.0796511 | -0.038875688 | 0.072630162 | -0.535255425 | 0.592473245 | 0.7754411 | | Map7 | 4516.166447 | -0.0179374851173982 | 0.0618349770650881 | -0.290086387490968 | 0.771750149482127 | 0.946588083372235 | NM_008635,NM_001198635 |
| Trappc8 | 2578.254857 | -0.030042254 | 0.056117041 | -0.535349936 | 0.592407902 | 0.7754411 | | Adrbk2 | 5538.752118 | -0.0174645625249536 | 0.0607878382779129 | -0.287303563010552 | 0.773879894727128 | 0.947121017002802 | NM_177078,NM_001035531 |
| Zbtb39 | 538.7564335 | 0.038390297 | 0.071768007 | 0.534922154 | 0.592703688 | 0.7754411 | | Alg3 | 462.0286813 | 0.0221042037411387 | 0.0767902113414051 | 0.287851841465374 | 0.773460152215558 | 0.947121017002802 | NM_145939 |
| Zbtb8a | 221.2639498 | -0.059531116 | 0.111204198 | -0.535331559 | 0.592420607 | 0.7754411 | | Bmyc | 1269.620687 | -0.0175808942334984 | 0.0608234187712565 | -0.289048109900173 | 0.772544561679597 | 0.947121017002802 | NM_023326 |
| Fbxl4 | 704.5238708 | 0.038152817 | 0.071365891 | 0.534608574 | 0.592920553 | 0.775446193 | | Cables2 | 1746.602232 | -0.0178783550225726 | 0.062042865402235 | -0.28816133662854 | 0.773223242952832 | 0.947121017002802 | NM_145851 |
| D10Jhu81e | 2127.894553 | -0.038433928 | 0.07197716 | -0.533973948 | 0.593359558 | 0.775954845 | | Col4a4 | 34.80270465 | -0.0180071966786502 | 0.0626230644355865 | -0.28754895406264 | 0.773692023871796 | 0.947121017002802 | NM_007735 |
| Atpbd4 | 673.1346387 | -0.03762146 | 0.070480675 | -0.533784055 | 0.593490947 | 0.776046641 | | Eef2k | 2025.14673 | -0.0159133741016629 | 0.0553354716270074 | -0.287579985021689 | 0.773668237581015 | 0.947121017002802 | NM_007908,NM_001267710,NM_001267711 |
| Cdipt | 5101.79367 | -0.036329299 | 0.068067101 | -0.533727721 | 0.593529927 | 0.776046641 | | Epha6 | 810.9172101 | 0.0258034310825246 | 0.0893395514844835 | 0.288824273838068 | 0.772715855529511 | 0.947121017002802 | NM_007938 |
| 4732415M23Rik | 25.93944297 | -0.088772466 | 0.166451122 | -0.533324525 | 0.593808953 | 0.776345956 | | Fam133b | 1113.504475 | -0.0199438858834919 | 0.0689314505544462 | -0.289329264407964 | 0.772329419746564 | 0.947121017002802 | NM_001042501 |
| Papd4 | 742.9325855 | 0.039605544 | 0.074288652 | 0.533130466 | 0.593943271 | 0.776400816 | | Fbxo22 | 1494.902252 | 0.0150497186025605 | 0.0522288478822917 | 0.288149542116611 | 0.773232270910781 | 0.947121017002802 | NM_028049 |
| Sec63 | 3951.845278 | -0.037222037 | 0.069819365 | -0.533119106 | 0.593951134 | 0.776400816 | | Gm1943 | 342.0960388 | 0.0220202499387863 | 0.0766032510005247 | 0.287458425734902 | 0.773761330619893 | 0.947121017002802 | NR_002928 |
| Rnf121 | 854.5150217 | -0.0315951 | 0.059274079 | -0.533034018 | 0.594010032 | 0.776412303 | | Lamc1 | 1604.924255 | -0.0177240638921125 | 0.061379117531928 | -0.288763745795024 | 0.772762177410277 | 0.947121017002802 | NM_010683 |
| Ate1 | 2963.607927 | -0.030964589 | 0.058115128 | -0.532814606 | 0.594161922 | 0.776414339 | | Lrp1b | 1589.409674 | -0.0219360249574693 | 0.0761399175193672 | -0.288101506701653 | 0.773269039320276 | 0.947121017002802 | NM_053011 |
| Cxx1b | 1781.601242 | -0.028567546 | 0.053609856 | -0.532878618 | 0.594117607 | 0.776414339 | | Lrrc6 | 278.1191389 | -0.023776566866245 | 0.082554110891989 | -0.288011906485838 | 0.773337624607691 | 0.947121017002802 | NM_019457 |
| Rapgef6 | 1702.602406 | 0.04857852 | 0.091154832 | 0.532923147 | 0.594086781 | 0.776414339 | | Maml3 | 497.3678703 | 0.0268382123647899 | 0.0933763367520864 | 0.287419846379765 | 0.77379086678095 | 0.947121017002802 | NM_001004176 |
| Armcx3 | 3691.361337 | -0.034622052 | 0.065010731 | -0.532559033 | 0.594338867 | 0.776516842 | | Megf6 | 324.6525616 | 0.0272086317068941 | 0.0946353550969661 | 0.287510219399667 | 0.77372167816463 | 0.947121017002802 | NM_001162977 |
| Asap1 | 6369.518621 | -0.036852966 | 0.0692001 | -0.532556533 | 0.594340598 | 0.776516842 | | Mib1 | 2866.751943 | 0.0200290624907092 | 0.0693233919224196 | 0.288922136313294 | 0.772640963463155 | 0.947121017002802 | NM_144860 |
| Fxn | 137.9790933 | -0.067819046 | 0.127383135 | -0.532402078 | 0.594447546 | 0.776591086 | | Mir3061 | 41.01418997 | 0.0258113507529993 | 0.0898255060972308 | 0.287349906217729 | 0.773844413453731 | 0.947121017002802 | NR_037221 |
| Ripk1 | 283.2885721 | 0.05701631 | 0.107118324 | 0.532274111 | 0.59453616 | 0.776641368 | | Mlf1ip | 65.49633033 | -0.027284740455484 | 0.0946477836347367 | -0.288276591460196 | 0.773135024234587 | 0.947121017002802 | NM_027973 |
| 2700081O15Rik | 1492.068242 | -0.034457289 | 0.064774606 | -0.531956751 | 0.59475595 | 0.776753967 | | Mt3 | 5885.203173 | 0.0170624661228057 | 0.0592343209938583 | 0.288050336975666 | 0.773308207428373 | 0.947121017002802 | NM_013603 |
| 9130023H24Rik | 142.9165213 | -0.071457838 | 0.134336292 | -0.531932489 | 0.594772755 | 0.776753967 | | Pwwp2b | 702.4520872 | 0.0204665136439134 | 0.071116311735903 | 0.287789300996341 | 0.773508027632591 | 0.947121017002802 | NM_001098636,NM_001033206 |
| Mical2 | 13542.45555 | 0.056516875 | 0.106225916 | 0.532044124 | 0.594695435 | 0.776753967 | | Rbm48 | 342.686583 | -0.0205774437050225 | 0.0713905297438342 | -0.288237722550303 | 0.773164775071185 | 0.947121017002802 | NM_172991 |
| 6720468P15Rik | 14.28620424 | 0.089048715 | 0.167519094 | 0.531573525 | 0.595021405 | 0.777013203 | | Tm9sf1 | 1280.748852 | 0.0178386523933382 | 0.0620596133789949 | 0.287443820901662 | 0.773772511968542 | 0.947121017002802 | NM_028780 |
| Pomt1 | 581.9629841 | 0.060501188 | 0.113912629 | 0.531119227 | 0.595336161 | 0.777358712 | | Tmco5 | 59.86418556 | -0.0269499208997115 | 0.0935143348955653 | -0.288190264410355 | 0.773201100683013 | 0.947121017002802 | NM_026104 |
| Wapal | 1700.363611 | -0.03553505 | 0.066926028 | -0.53096009 | 0.595446436 | 0.777437185 | | Tmem116 | 78.15209665 | -0.027272440172074 | 0.0945118996401133 | -0.288560914296752 | 0.772917409484774 | 0.947121017002802 | NM_001161627,NM_001161626,NM_029912 |
| Utp23 | 378.6408259 | 0.046944919 | 0.088462092 | 0.530678369 | 0.595641678 | 0.777626572 | | Usmg5 | 3628.767415 | 0.0214994988270403 | 0.0743899853168854 | 0.289010660984231 | 0.772573219243451 | 0.947121017002802 | NM_023211 |
| Irs4 | 133.4081737 | -0.062918382 | 0.118659514 | -0.530243042 | 0.595943432 | 0.777954969 | | Utp14a | 688.4371045 | -0.0204424667706986 | 0.0706400956099154 | -0.289389002013599 | 0.772283710252429 | 0.947121017002802 | NM_028276 |
| Btf3 | 3333.605446 | -0.030492641 | 0.057554073 | -0.529808563 | 0.596244668 | 0.778217074 | | Taok2 | 6801.371059 | -0.0157013038881965 | 0.0546834019765843 | -0.287131073061618 | 0.774011960584817 | 0.947202842414074 | NM_001163775,NM_001163774 |
| Sphkap | 7731.660266 | -0.048917952 | 0.09232868 | -0.529824013 | 0.596233954 | 0.778217074 | | 2410066E13Rik | 1994.606962 | 0.0188777102440163 | 0.0659627272289796 | 0.28618753403699 | 0.774734491032097 | 0.94723891786813 | NM_026629 |
| Elk4 | 370.1737255 | -0.054002226 | 0.101971525 | -0.529581427 | 0.596402175 | 0.778291527 | | Ceacam1 | 129.3305535 | -0.0267138646922826 | 0.0931543828093548 | -0.286769810358294 | 0.774288580320615 | 0.94723891786813 | NM_011926,NM_001039186,NM_001039187,NM_001039185 |
| Fzd2 | 471.4328076 | -0.044463106 | 0.083952739 | -0.529620673 | 0.596374958 | 0.778291527 | | Crlf3 | 476.6597903 | -0.021278507503977 | 0.074417840768614 | -0.285932879591843 | 0.774929530304066 | 0.94723891786813 | NM_018776 |
| Fam103a1 | 2256.902235 | 0.046273345 | 0.087410981 | 0.529376791 | 0.596544095 | 0.778411168 | | Egfl7 | 995.6964821 | 0.0226636279193674 | 0.0791813186131227 | 0.286224431675622 | 0.774706232391568 | 0.94723891786813 | NM_001164564,NM_198724,NM_198725,NM_178444 |
| Glt8d2 | 570.2109351 | -0.061646914 | 0.116495469 | -0.529178641 | 0.596681532 | 0.778495955 | | Fabp7 | 625.7803171 | 0.0218394242223427 | 0.0763555207752028 | 0.286022857294626 | 0.774860614958935 | 0.94723891786813 | NM_021272 |
| Wdr35 | 1907.025135 | -0.033072314 | 0.06250222 | -0.529138231 | 0.596709562 | 0.778495955 | | Fhl2 | 1410.453471 | -0.0222322300121551 | 0.0775464482622714 | -0.28669565802631 | 0.774345362488705 | 0.94723891786813 | NM_010212 |
| Prmt5 | 2242.055961 | 0.043861185 | 0.082967139 | 0.528657325 | 0.597043186 | 0.778865633 | | Gm15663 | 337.4361147 | -0.0239643747390861 | 0.0836653973076936 | -0.286431135334876 | 0.774547930642538 | 0.94723891786813 | NR_038032 |
| Mocs2 | 1454.159588 | -0.029765777 | 0.056318893 | -0.528522057 | 0.597137042 | 0.778922489 | | Kdm2b | 1071.395185 | -0.0159697911489656 | 0.055874809152197 | -0.285813793215214 | 0.775020743168855 | 0.94723891786813 | NM_001005866,NM_001003953,NM_013910 |
| 2210417K05Rik | 12.93306885 | 0.087928029 | 0.166441081 | 0.528283213 | 0.597302781 | 0.778986324 | | Mcph1 | 567.6510999 | 0.0182466483233443 | 0.0638231883115376 | 0.285893713649618 | 0.774959528672333 | 0.94723891786813 | NM_173189 |
| A130077B15Rik | 365.3042928 | 0.050521924 | 0.095656161 | 0.528161735 | 0.597387085 | 0.778986324 | | Nagk | 1247.100166 | 0.0205852465885478 | 0.0719812102055687 | 0.285980834856195 | 0.774892800388214 | 0.94723891786813 | NM_019542,NM_001164187 |
| Gfer | 405.7905869 | -0.0418071 | 0.079141329 | -0.528258753 | 0.597319756 | 0.778986324 | | Nfkbil1 | 287.0920814 | 0.0225248455568972 | 0.0788560727831504 | 0.285645033564367 | 0.775150008033765 | 0.94723891786813 | NM_010909 |
| Trpm6 | 44.37186436 | -0.084973052 | 0.160863945 | -0.528229317 | 0.597340184 | 0.778986324 | | Pygo1 | 471.8702692 | 0.024579827565011 | 0.0860484571486173 | 0.285650996886072 | 0.775145440200769 | 0.94723891786813 | NM_028116 |
| Ccdc8 | 104.4766446 | -0.071152161 | 0.134735766 | -0.528086658 | 0.597439191 | 0.778988708 | | Rgs6 | 1591.991873 | -0.0185615857859775 | 0.0647484014473578 | -0.286672494935162 | 0.774363099879408 | 0.94723891786813 | NM_015812 |
| Pbx2 | 2157.282221 | 0.02903111 | 0.054998514 | 0.52785263 | 0.597601625 | 0.779069379 | | Smarcc1 | 3255.724166 | 0.0141869122721073 | 0.04958642586371 | 0.28610475598908 | 0.774797888995841 | 0.94723891786813 | NM_009211 |
| Steap4 | 21.08238514 | -0.088726547 | 0.168067567 | -0.52792189 | 0.597553551 | 0.779069379 | | Stk33 | 45.1613517 | -0.0239575420336649 | 0.0837809244362219 | -0.285954615503229 | 0.774912882263443 | 0.94723891786813 | NM_054103 |
| Imp4 | 679.1388081 | -0.04017686 | 0.076141348 | -0.52766153 | 0.597734279 | 0.779176755 | | Tnni1 | 128.838898 | 0.0267011716879114 | 0.0930669272035447 | 0.286902904073687 | 0.774186666801089 | 0.94723891786813 | NM_001112702,NM_021467 |
| Tpst1 | 1207.874192 | 0.030600785 | 0.058009549 | 0.527512896 | 0.597837463 | 0.779245702 | | Zrsr2 | 1585.658191 | 0.0184867725176916 | 0.0645624409893878 | 0.286339429463801 | 0.774618161430094 | 0.94723891786813 | NM_009453,NM_178794 |
| Ccdc88a | 2676.535623 | -0.039200295 | 0.074356931 | -0.527190871 | 0.598061047 | 0.779433947 | | 4933412E12Rik | 165.2499761 | -0.0258870457825506 | 0.0908483473247473 | -0.28494789993278 | 0.775684057628193 | 0.947241001013522 | NR_038026,NR_038025 |
| Tfdp1 | 1637.834743 | 0.039221455 | 0.074401428 | 0.527159982 | 0.598082496 | 0.779433947 | | Bloc1s2a | 909.6889597 | 0.016257337120201 | 0.0572389697611003 | 0.284025676703383 | 0.776390703432947 | 0.947241001013522 | NM_028607 |
| Mcts2 | 196.4677665 | -0.060324287 | 0.11454019 | -0.52666481 | 0.598426377 | 0.779816509 | | Egln3 | 420.4351376 | 0.0200470493685803 | 0.0703614158753995 | 0.284915377542727 | 0.775708974481266 | 0.947241001013522 | NM_028133 |
| Abhd11 | 772.606822 | 0.044935977 | 0.085418332 | 0.526069472 | 0.598839938 | 0.780027407 | | Fam186b | 46.56080866 | 0.0263677709603271 | 0.0924116701850706 | 0.285329449273246 | 0.775391752918337 | 0.947241001013522 | NM_001081254 |
| Atad5 | 214.4275116 | -0.054586302 | 0.103756875 | -0.526098169 | 0.598820001 | 0.780027407 | | Gen1 | 40.62602183 | 0.0253089804801077 | 0.0888273540237921 | 0.28492327344715 | 0.775702925055615 | 0.947241001013522 | NM_177331 |
| Nap1l3 | 1631.661027 | -0.040644714 | 0.077239806 | -0.526214597 | 0.598739113 | 0.780027407 | | Grik2 | 1950.134638 | 0.018918886895899 | 0.0664630102168785 | 0.284652874345653 | 0.775910098335158 | 0.947241001013522 | NM_010349,NM_001111268 |
| Ssx2ip | 3474.949982 | 0.048989644 | 0.093118218 | 0.526101608 | 0.598817612 | 0.780027407 | | Hr | 2295.637585 | 0.0210520559688737 | 0.07375861681906 | 0.285418258595023 | 0.775323720716586 | 0.947241001013522 | NM_021877 |
| Wdpcp | 348.7030177 | -0.054240656 | 0.10304961 | -0.526354792 | 0.59864172 | 0.780027407 | | Icam2 | 98.39521827 | 0.0270003392608279 | 0.0948558481168657 | 0.284646016000642 | 0.775915353241359 | 0.947241001013522 | NM_010494 |
| 4930405A21Rik | 9.256650748 | -0.0841704 | 0.160031708 | -0.525960766 | 0.598915467 | 0.780060209 | | Klhl21 | 895.21471 | 0.0200615745601783 | 0.0704012044591571 | 0.284960672396122 | 0.775674272137697 | 0.947241001013522 | NM_001033352 |
| Urb2 | 685.4355988 | -0.032119228 | 0.061091086 | -0.525759648 | 0.599055214 | 0.78017664 | | Ndufa2 | 1690.770932 | 0.0191793282089635 | 0.0674806425057623 | 0.284219703559072 | 0.776242016589074 | 0.947241001013522 | NM_010885 |
| Neu3 | 117.3537397 | 0.069355613 | 0.13193603 | 0.525676061 | 0.599113299 | 0.780186709 | | Ntpcr | 606.3977133 | 0.0192069859002079 | 0.0674960484173036 | 0.284564598233338 | 0.775977736819968 | 0.947241001013522 | NM_025636 |
| Gm10785 | 11.98684131 | 0.087050489 | 0.165702633 | 0.525341615 | 0.599345733 | 0.780423799 | | Polr2d | 477.3520165 | 0.0215628360926335 | 0.0755350620057036 | 0.28546790748653 | 0.775285688034898 | 0.947241001013522 | NM_027101,NM_027002 |
| Gm3086 | 32.84045887 | 0.083363092 | 0.158762537 | 0.525080377 | 0.599527317 | 0.780594643 | | Reep2 | 5541.734573 | -0.0158020878869574 | 0.0555965273920985 | -0.284227965813621 | 0.776235685232314 | 0.947241001013522 | NM_001204914,NM_144865 |
| Adrb3 | 43.32108901 | 0.084132134 | 0.160329151 | 0.524746332 | 0.599759544 | 0.78083139 | | Sebox | 72.45696277 | 0.0269387491578448 | 0.0948188369659505 | 0.284107567861421 | 0.776327947523457 | 0.947241001013522 | NM_008759 |
| Cobll1 | 552.2965673 | 0.061109291 | 0.116488241 | 0.524596219 | 0.599863916 | 0.780901656 | | Sez6 | 10720.64235 | -0.0187860575254139 | 0.066088916297372 | -0.284254283137048 | 0.776215518394828 | 0.947241001013522 | NM_021286 |
| Amigo3 | 51.85024728 | -0.085252003 | 0.162591809 | -0.524331475 | 0.600048008 | 0.780910185 | | Snd1 | 3344.794664 | 0.0131106471245135 | 0.0460591636326527 | 0.28464796341241 | 0.775913861121422 | 0.947241001013522 | NM_019776 |
| Rrp15 | 245.6297048 | 0.065456789 | 0.124833679 | 0.524352001 | 0.600033735 | 0.780910185 | | Srd5a1 | 441.5359827 | 0.0210816401778351 | 0.0742015503481907 | 0.284113203550459 | 0.77632362876179 | 0.947241001013522 | NM_175283 |
| Senp3 | 2653.709835 | -0.026994279 | 0.051472964 | -0.524436062 | 0.59997528 | 0.780910185 | | Use1 | 1731.002809 | 0.0183125389107984 | 0.0643008014140504 | 0.284794878260987 | 0.775801296365938 | 0.947241001013522 | NM_001145780,NM_025917 |
| Thsd1 | 184.5954078 | -0.057543246 | 0.109753174 | -0.524296874 | 0.600072071 | 0.780910185 | | Zfp90 | 549.8929987 | -0.0181625492309519 | 0.0638702879990963 | -0.284366170874459 | 0.776129781047824 | 0.947241001013522 | NM_011764 |
| Tm6sf1 | 519.364006 | 0.053925418 | 0.102929354 | 0.523907082 | 0.600343169 | 0.781197368 | | Rsrc2 | 2568.191703 | -0.0152285391563032 | 0.0536871630656511 | -0.28365326619477 | 0.776676112359783 | 0.947430064506064 | NM_001005523,NM_001005525,NM_025438 |
| Rgag1 | 104.1534605 | -0.08292019 | 0.158330717 | -0.523715119 | 0.600476698 | 0.781304013 | | Snf8 | 1869.125067 | 0.0140241195001545 | 0.0494326297581295 | 0.283701667679295 | 0.77663901659969 | 0.947430064506064 | NM_033568 |
| Ric3 | 3710.007343 | -0.026190142 | 0.05001514 | -0.52364428 | 0.600525977 | 0.781304013 | | Zfp69 | 39.55926206 | -0.0253632582408667 | 0.0894602881562527 | -0.283514157662525 | 0.776782730452407 | 0.947463203866492 | NM_001005788 |
| Cdkn2b | 23.094942 | -0.087617513 | 0.16742217 | -0.523332804 | 0.600742676 | 0.781520321 | | Znrd1 | 481.9252302 | -0.0215011422759386 | 0.0758557872299089 | -0.283447618976933 | 0.776833729729445 | 0.947463203866492 | NM_023162 |
| Gm5617 | 44.61026715 | 0.085582857 | 0.163584754 | 0.523171354 | 0.600855014 | 0.781537017 | | Cpne1 | 1669.331385 | 0.0131553120703945 | 0.046452957231672 | 0.2831964390294 | 0.777026257915407 | 0.947474492811931 | NM_170590,NM_170588 |
| Sult4a1 | 17284.15835 | -0.040430953 | 0.07728081 | -0.523169373 | 0.600856392 | 0.781537017 | | Dlk2 | 1118.542873 | -0.0230296649834177 | 0.0813408292761093 | -0.283125524885468 | 0.777080615735177 | 0.947474492811931 | NM_207666 |
| Mut | 1543.040115 | -0.050041275 | 0.09571147 | -0.522834671 | 0.60108931 | 0.781774344 | | Gm4349 | 25.20428387 | 0.0231448444847267 | 0.0817564112226865 | 0.2830951620624 | 0.77710389008408 | 0.947474492811931 | NR_033637 |
| Nufip1 | 496.2177907 | 0.038621918 | 0.073895179 | 0.522658159 | 0.60121216 | 0.781868492 | | Gpm6b | 34999.9339 | -0.0165891092565292 | 0.0585653702432564 | -0.28325799337774 | 0.776979075544899 | 0.947474492811931 | NM_001177958,NM_001177960,NM_023122,NM_001177961,NM_001177959,NM_001177962,NM_001177955,NM_001177956,NM_001177957 |
| Furin | 1472.765093 | 0.055827783 | 0.106862562 | 0.522426022 | 0.601373742 | 0.781953027 | | Crybb3 | 25.06219879 | -0.0233109170707436 | 0.0824534495337854 | -0.282716092565562 | 0.777394479230653 | 0.947506677101774 | NM_001159650,NM_021352 |
| Sspn | 492.1177506 | -0.060872372 | 0.116520041 | -0.522419762 | 0.601378099 | 0.781953027 | | Gpr111 | 34.30868479 | -0.0244943015762215 | 0.0866143002934514 | -0.282797430600192 | 0.777332124054163 | 0.947506677101774 | NM_001033493 |
| Aimp2 | 675.4891072 | -0.045706985 | 0.087513236 | -0.522286532 | 0.601470845 | 0.782007995 | | Itih5 | 2002.811949 | 0.0230067022702443 | 0.0813232824550012 | 0.282904250489085 | 0.777250236216078 | 0.947506677101774 | NM_172471 |
| B3gnt4 | 17.86325094 | -0.085253889 | 0.163372331 | -0.52183799 | 0.601783136 | 0.782282734 | | Lpo | 37.75473909 | 0.0260760113219132 | 0.0921784425905711 | 0.282886221431783 | 0.777264057069112 | 0.947506677101774 | NM_080420 |
| Stk11 | 2100.619902 | 0.040654048 | 0.07789851 | 0.521884798 | 0.601750544 | 0.782282734 | | Rxfp3 | 184.5066157 | -0.026241678801334 | 0.0928464354852843 | -0.282635285503159 | 0.777456428774339 | 0.947506677101774 | NM_178717 |
| Aurka | 42.59070592 | 0.085744342 | 0.164336306 | 0.521761404 | 0.601836466 | 0.782286426 | | 2310008H09Rik | 1135.296047 | 0.0161285011012685 | 0.057241922265096 | 0.281760298450058 | 0.778127315233245 | 0.947847159838836 | NM_001168220,NM_023197,NM_001168219,NM_001168218 |
| Ilf2 | 3089.11293 | 0.032060685 | 0.061476002 | 0.521515454 | 0.602007743 | 0.782377786 | | Atox1 | 861.5300554 | 0.0178106299991523 | 0.0631745690740332 | 0.281927209955012 | 0.777999324936176 | 0.947847159838836 | NM_009720 |
| Knstrn | 39.41195593 | -0.086390073 | 0.165643342 | -0.521542681 | 0.601988781 | 0.782377786 | | Cyb5d2 | 619.580471 | -0.0169256013850977 | 0.0600299857813943 | -0.281952447010967 | 0.77797997329942 | 0.947847159838836 | NM_001024926 |
| Mier1 | 1056.509272 | 0.042878802 | 0.082267926 | 0.521209229 | 0.602221026 | 0.782589329 | | Sac3d1 | 411.6348499 | 0.0199596761731256 | 0.0707971160917088 | 0.281927813941889 | 0.777998861800727 | 0.947847159838836 | NM_133678 |
| Tmem71 | 20.98049719 | -0.087535125 | 0.168114371 | -0.520687934 | 0.602584181 | 0.78292992 | | Spr | 779.7647022 | 0.0201624214429038 | 0.0714686947435931 | 0.282115428513703 | 0.777855003289293 | 0.947847159838836 | NM_011467 |
| Trim45 | 375.9160394 | -0.049450828 | 0.094967195 | -0.520714838 | 0.602565436 | 0.78292992 | | Whrn | 1504.147398 | -0.0161060546636992 | 0.0571452567997585 | -0.281844120853916 | 0.778063038178203 | 0.947847159838836 | [illegible]_028640,NM_001276371,NM_001008797,NM_001008796,NM_001008795,NM_001008794,NM_001008793,NM_001008792,NM_001008791,NM_00100879[illegible] |
| Wnt9a | 643.1987464 | 0.071522894 | 0.137392731 | 0.520572618 | 0.602664528 | 0.782968657 | | Erbb3 | 457.6064107 | 0.024839419682711 | 0.0882028944895575 | 0.281616831584272 | 0.778237332632775 | 0.947901684875373 | NM_010153 |
| Tmprss5 | 49.64822643 | 0.082641805 | 0.158781515 | 0.520474975 | 0.602732565 | 0.782991395 | | Rem2 | 419.141539 | -0.0249786466885261 | 0.0887494530246389 | -0.281451274765507 | 0.778364295211389 | 0.947976838453981 | NM_080726 |
| Rpf1 | 341.7337062 | 0.051209556 | 0.098431972 | 0.520253283 | 0.602887052 | 0.783126424 | | Rab33b | 1202.553819 | -0.0202481506003346 | 0.0719696510703485 | -0.281342903560037 | 0.778447406364629 | 0.947998576865576 | NM_016858 |
| Creg2 | 6770.976562 | -0.040110544 | 0.077120243 | -0.520103958 | 0.60299112 | 0.783168941 | | Rrm2 | 61.21839361 | -0.0263168602999082 | 0.0935866874185799 | -0.281203032459117 | 0.778554678904331 | 0.948049733067676 | NM_009104 |
| Napb | 23209.18757 | -0.037183007 | 0.071502906 | -0.52002092 | 0.603048994 | 0.783168941 | | Klhl24 | 2900.131161 | -0.0195051487310205 | 0.0693872818709793 | -0.281105531231054 | 0.778629458850921 | 0.948061317625187 | NM_029436 |
| Zfp771 | 388.78494 | -0.079758962 | 0.15338594 | -0.51998874 | 0.603071423 | 0.783168941 | | Anapc11 | 1225.968668 | 0.018335323000802 | 0.0653814186044521 | 0.280436298143484 | 0.779142791918101 | 0.948407833568281 | NM_025389,NM_001038230 |
| Hrh3 | 4025.517459 | 0.040434484 | 0.077798945 | 0.519730498 | 0.603251427 | 0.783302166 | | Crhr2 | 99.91973813 | -0.0263253227603188 | 0.0938930705546987 | -0.280375565574699 | 0.779189381409599 | 0.948407833568281 | NM_009953 |
| Rogdi | 4085.288766 | 0.031133932 | 0.059913725 | 0.519646067 | 0.603310284 | 0.783302166 | | Grid2ip | 155.894082 | 0.0266580859578291 | 0.0951025208443973 | 0.280308931047642 | 0.779240499355461 | 0.948407833568281 | NM_001159321,NM_133355 |
| Stk17b | 291.9778228 | 0.055418498 | 0.106651155 | 0.519623985 | 0.603325677 | 0.783302166 | | Ncstn | 3035.677189 | 0.0154296969179846 | 0.0550338444050031 | 0.280367419081883 | 0.779195630850887 | 0.948407833568281 | NM_021607 |
| A430078G23Rik | 208.6479762 | 0.053769645 | 0.103562485 | 0.519200025 | 0.603621262 | 0.783430843 | | Zcchc14 | 4062.821087 | 0.0160022951876637 | 0.0570321852987124 | 0.2805835880889 | 0.779029805367055 | 0.948407833568281 | NM_080855 |
| Dixdc1 | 3703.791155 | 0.038070504 | 0.073318918 | 0.519245304 | 0.60358969 | 0.783430843 | | Tmcc1 | 3964.630832 | 0.0190841887390566 | 0.0681413867044627 | 0.28006751347499 | 0.7794257081904 | 0.948553773221661 | NM_177412 |
| Dtx3 | 7443.176167 | 0.050161574 | 0.096593961 | 0.519303413 | 0.603549174 | 0.783430843 | | Tmem68 | 1179.90041 | 0.0211353734194098 | 0.0755386328475355 | 0.279795551265386 | 0.779634365019091 | 0.948728222021221 | NM_028097 |
| Mc4r | 83.67093784 | -0.087156133 | 0.167868876 | -0.519191734 | 0.603627043 | 0.783430843 | | Naa10 | 820.5820656 | 0.0181669890847048 | 0.0649687275528261 | 0.279626672231393 | 0.779763941614507 | 0.948806417816413 | NM_001177965,NM_019870 |
| Pias4 | 587.5839783 | 0.037531851 | 0.072306225 | 0.519068047 | 0.60371329 | 0.783477151 | | 4931428F04Rik | 1305.593642 | 0.0188860761005049 | 0.0679017757315473 | 0.278138176756553 | 0.780906290673351 | 0.948894514042564 | NM_001166394,NM_028888 |
| Anapc10 | 288.2411336 | -0.047797355 | 0.092175298 | -0.518548422 | 0.604075685 | 0.783867561 | | 5830415F09Rik | 108.5340196 | -0.0262206402167058 | 0.0943453560292327 | -0.277921895896831 | 0.781072315300647 | 0.948894514042564 | NM_029086 |
| Prdm11 | 108.9645971 | 0.066483355 | 0.128224551 | 0.51849162 | 0.604115306 | 0.783867561 | | Bag6 | 9398.78552 | -0.0137850847793922 | 0.0497075797000538 | -0.277323596573688 | 0.781531642407835 | 0.948894514042564 | NM_001252468,NM_001252469,NM_057171 |
| Itgb1bp2 | 11.37530105 | 0.079877374 | 0.154197566 | 0.518019682 | 0.604444537 | 0.784170038 | | Cdh10 | 1898.271721 | 0.0163365031654426 | 0.0586942658300606 | 0.278332183466477 | 0.780757372946615 | 0.948894514042564 | NM_009865 |
| Usp43 | 118.5062943 | 0.07683897 | 0.14833424 | 0.518012364 | 0.604449643 | 0.784170038 | | Eed | 522.3326084 | 0.0209708748681123 | 0.0752407565808674 | 0.278716959013738 | 0.780462046647598 | 0.948894514042564 | NM_021876 |
| Dap3 | 2463.786631 | -0.025541785 | 0.049316711 | -0.5179134 | 0.604518692 | 0.784193957 | | Eef1b2 | 4513.658928 | 0.0131544900902653 | 0.0474025465082686 | 0.277505979303679 | 0.781391615226678 | 0.948894514042564 | NM_018796 |
| 2310034G01Rik | 16.34958121 | -0.087034524 | 0.168117353 | -0.517701012 | 0.604666892 | 0.784264815 | | Efr3b | 11192.85693 | -0.0130768241451815 | 0.0470562346480584 | -0.27789780127937 | 0.781090811775751 | 0.948894514042564 | NM_001082483 |
| Zbtb6 | 532.0229142 | -0.04869831 | 0.094068471 | -0.517690038 | 0.60467455 | 0.784264815 | | Fam188a | 1277.61608 | 0.0196985782879476 | 0.0707658657499956 | 0.27836271172799 | 0.780733940472147 | 0.948894514042564 | NM_024185 |
| Lamc2 | 635.4194175 | 0.059462563 | 0.11508302 | 0.516692757 | 0.605370653 | 0.785101941 | | Fanci | 30.295939 | -0.0235068412062951 | 0.0843906045649453 | -0.278548083965966 | 0.78059165920651 | 0.948894514042564 | NM_145946 |
| Abcc4 | 634.1941226 | 0.045533751 | 0.088145393 | 0.516575505 | 0.605452518 | 0.785142394 | | Garnl3 | 3961.693254 | 0.0141004556917109 | 0.0505677719378802 | 0.278842732264981 | 0.780365518925955 | 0.948894514042564 | NM_178888 |
| Plec | 8670.419617 | -0.044566177 | 0.086341011 | -0.516164648 | 0.605739419 | 0.785251552 | | Gstcd | 215.2302535 | -0.0224631458089286 | 0.0809504413828666 | -0.277492567368298 | 0.781401912207566 | 0.948894514042564 | NM_026231 |
| Rpl24 | 5012.913876 | -0.039639601 | 0.076792423 | -0.516191568 | 0.605720619 | 0.785251552 | | Gtpbp10 | 373.8056523 | -0.0208790103871022 | 0.0752482340008338 | -0.277468443802559 | 0.781420433113158 | 0.948894514042564 | NM_153116 |
| Slc24a2 | 27643.09709 | 0.050717881 | 0.098242443 | 0.516252238 | 0.60567825 | 0.785251552 | | Icam4 | 28.88962123 | -0.022428967628082 | 0.0805385326583362 | -0.278487413263799 | 0.780638225804115 | 0.948894514042564 | NM_023892 |
| Zfp944 | 301.0423839 | 0.05280435 | 0.102280549 | 0.516269714 | 0.605666046 | 0.785251552 | | Ilk | 2853.277738 | 0.0140632969340345 | 0.0506042602732548 | 0.277907371001868 | 0.781083465467329 | 0.948894514042564 | NM_001161724,NM_010562 |
| Fbxo5 | 40.72755384 | -0.083500919 | 0.16186612 | -0.515864089 | 0.605949337 | 0.785457963 | | Lig1 | 496.7273337 | -0.0237156777033683 | 0.085245572568615 | -0.278204216228113 | 0.780855598493399 | 0.948894514042564 | NM_001199310,NM_010715,NM_001083188 |
| Sox2 | 952.4562778 | -0.056606599 | 0.109798114 | -0.515551651 | 0.606167586 | 0.785675136 | | Ormdl2 | 211.8579914 | -0.0237537274235788 | 0.0850861695464583 | -0.279172602905916 | 0.780112367815658 | 0.948894514042564 | NM_024180 |
| Slc36a4 | 1124.526106 | 0.061428518 | 0.119244774 | 0.515146421 | 0.606450706 | 0.785976348 | | Padi2 | 1620.712642 | -0.0223004260798173 | 0.0798129379540341 | -0.279408660443756 | 0.779931225572494 | 0.948894514042564 | NM_008812 |
| Nle1 | 304.3146394 | 0.055231098 | 0.107286068 | 0.514802146 | 0.606691286 | 0.786222381 | | Plekhh3 | 488.0551148 | -0.0230609069861758 | 0.0831493559341175 | -0.277343182362686 | 0.781516604773794 | 0.948894514042564 | NM_146030 |
| 2610316D01Rik | 168.1640833 | -0.07674214 | 0.149271994 | -0.514109435 | 0.607175481 | 0.786522522 | | Scyl2 | 1387.02377 | 0.0160902599381793 | 0.057902700445442 | 0.277884447778737 | 0.781101062777978 | 0.948894514042564 | NM_198021 |
| Shpk | 178.074681 | 0.065502483 | 0.127390848 | 0.514185155 | 0.607122545 | 0.786522522 | | Slc35a2 | 1076.28585 | 0.0186260330091871 | 0.0668454854127219 | 0.278643096002445 | 0.78051873617814 | 0.948894514042564 | NM_078484,NM_001083937 |
| Tgds | 158.1064716 | -0.070150925 | 0.136418302 | -0.514233969 | 0.607088421 | 0.786522522 | | Slc35c1 | 780.4085517 | 0.0185053256367819 | 0.0663319019963983 | 0.278980778174984 | 0.78025957615263 | 0.948894514042564 | NM_211358,NM_145832 |
| Zap70 | 47.65134935 | -0.079189652 | 0.1540332 | -0.514107683 | 0.607176706 | 0.786522522 | | Spred3 | 2143.257766 | -0.0235712929095789 | 0.0849444598669815 | -0.277490644433908 | 0.781403388538926 | 0.948894514042564 | NM_182927 |
| Zbtb11 | 2059.691094 | 0.029901383 | 0.058135869 | 0.514336221 | 0.607016942 | 0.786522522 | | Uhrf1bp1l | 4259.305814 | 0.0169859338720376 | 0.0610895581733829 | 0.278049708983466 | 0.780974200372319 | 0.948894514042564 | NM_029166 |
| Cwc15 | 1775.219599 | 0.047267965 | 0.091977772 | 0.513906395 | 0.607317436 | 0.786538225 | | Uox | 26.57085437 | 0.0237027028889341 | 0.0849253080431747 | 0.279100581853457 | 0.780167636612678 | 0.948894514042564 | NM_009474 |
| E130003G02Rik | 33.35411529 | 0.059060679 | 0.114944721 | 0.513818112 | 0.607379164 | 0.786538225 | | Wdr76 | 165.572237 | -0.0255784699387012 | 0.0922346497839511 | -0.277319532286573 | 0.78153476290825 | 0.948894514042564 | NM_030234 |
| Otud5 | 2167.729162 | 0.030670134 | 0.059692299 | 0.513803864 | 0.607389126 | 0.786538225 | | Zc3h12b | 474.8221754 | 0.0205972987425829 | 0.0739303209125979 | 0.278604211213062 | 0.780548580548753 | 0.948894514042564 | NM_001034907 |
| Pop5 | 1331.511493 | 0.055676971 | 0.108375896 | 0.513739426 | 0.607434184 | 0.786538225 | | Akap5 | 4841.566534 | 0.0219958609084241 | 0.0793768668350757 | 0.277106690972393 | 0.781698184281165 | 0.948979618993899 | NM_001101471 |
| Psmg1 | 324.5091036 | 0.042605189 | 0.082933471 | 0.513727319 | 0.607442649 | 0.786538225 | | Clec1a | 32.50005026 | -0.0236355300770837 | 0.0853876019961295 | -0.276802832314638 | 0.781931506246591 | 0.948979618993899 | NM_175526 |
| Gm3558 | 28.71803363 | 0.085765921 | 0.166988897 | 0.513602538 | 0.607529905 | 0.786585472 | | Dlx6 | 293.9240847 | 0.0256898895171463 | 0.0927539556087578 | 0.27696812872874 | 0.781804578731911 | 0.948979618993899 | NM_010057 |
| 1700102P08Rik | 26.9093646 | -0.086242428 | 0.168024785 | -0.513272065 | 0.607761022 | 0.786629774 | | Mog | 2643.008173 | 0.0237828224914891 | 0.0859135586097884 | 0.276822691043547 | 0.781916256854356 | 0.948979618993899 | NM_010814 |
| Hadh | 784.6530571 | -0.04178259 | 0.081405773 | -0.513263229 | 0.607767202 | 0.786629774 | | Nek6 | 1264.118413 | -0.0192419243015194 | 0.0694638790526501 | -0.277006187445061 | 0.781775355095793 | 0.948979618993899 | NM_021606,NM_001159631 |
| Nkx2-9 | 12.68229921 | 0.083809601 | 0.16328148 | 0.513282958 | 0.607753403 | 0.786629774 | | BC026590 | 1250.457145 | -0.0171806141679131 | 0.0622119818580153 | -0.276162463480491 | 0.782423286323295 | 0.949184473387406 | NM_001081420 |
| Pcdh15 | 634.1396526 | -0.063917332 | 0.124531094 | -0.513264034 | 0.607766639 | 0.786629774 | | Csmd3 | 1241.598349 | 0.0187612085211245 | 0.0679378212474059 | 0.276152637465407 | 0.782430833025093 | 0.949184473387406 | NM_001081391 |
| Msl3l2 | 386.9255868 | -0.048114702 | 0.093792369 | -0.512991645 | 0.607957165 | 0.786809916 | | Dmc1 | 24.18965986 | -0.0214745776715609 | 0.0777591432927101 | -0.27616787894285 | 0.782419127079373 | 0.949184473387406 | NM_010059 |
| Il21r | 77.53425147 | 0.073605775 | 0.143525142 | 0.512842379 | 0.608061583 | 0.786879326 | | Lrrc40 | 1039.431429 | 0.0200331003258193 | 0.0724983946518394 | 0.276324743768807 | 0.782298652702838 | 0.949184473387406 | NM_024194 |
| Per2 | 1888.81075 | -0.081215589 | 0.158435347 | -0.512610285 | 0.608223957 | 0.787023718 | | Tdrd9 | 31.32655883 | -0.0244498869139136 | 0.0884828001645089 | -0.276323611690135 | 0.78229952213636 | 0.949184473387406 | NM_029056 |
| Icam4 | 26.22460807 | -0.086024895 | 0.167868897 | -0.512452853 | 0.608334109 | 0.787100517 | | Vps39 | 4167.094667 | 0.0135466106649456 | 0.0490690288555008 | 0.276072524378623 | 0.782492363266844 | 0.949184473387406 | NR_027618,NM_178851,NM_147153 |
| Chrm3 | 1850.603948 | 0.047306267 | 0.092374584 | 0.512113447 | 0.608571614 | 0.787210601 | | Lrrc20 | 1122.299414 | -0.0177356966102327 | 0.0643104980951853 | -0.275782292713427 | 0.782715284863758 | 0.949296336973445 | NM_153542 |
| Rab43 | 963.3199815 | 0.044507317 | 0.086900483 | 0.512164202 | 0.608536094 | 0.787210601 | | Znhit3 | 620.8518741 | 0.0169933022627254 | 0.0616114403062388 | 0.275814072488169 | 0.782690874533801 | 0.949296336973445 | NM_001005223 |
| Zpbp | 51.09197502 | -0.078690559 | 0.153642353 | -0.512167105 | 0.608534063 | 0.787210601 | | Zfp68 | 1544.774309 | 0.021050658041524 | 0.0763577938071975 | 0.275684471642498 | 0.782790423448196 | 0.949308205961638 | NM_001163797,NM_001044747,NM_013844 |
| Zfp111 | 425.2946643 | -0.043600299 | 0.085150902 | -0.512035667 | 0.608626048 | 0.787215292 | | Rbm17 | 1805.559771 | 0.0144968188718959 | 0.0526392574425185 | 0.275399380162728 | 0.78300942023951 | 0.949438153421865 | NM_152824 |
| Nhlrc3 | 193.7832775 | 0.056429067 | 0.110262646 | 0.51176957 | 0.60881229 | 0.787324732 | | Smek2 | 1810.450033 | 0.0224710440680604 | 0.0816016706410658 | 0.275374804112796 | 0.783028299462615 | 0.949438153421865 | NM_134034 |
| Polr1e | 229.7882535 | 0.052726878 | 0.10301539 | 0.511834958 | 0.608766522 | 0.787324732 | | Aup1 | 2123.128696 | -0.0184549107507554 | 0.067090557644544 | -0.275074636412062 | 0.783258897394984 | 0.949478217058132 | NM_007517 |
| Fam69a | 992.2106911 | -0.043268008 | 0.084590552 | -0.511499298 | 0.60900148 | 0.787503666 | | Hist1h1e | 38.04432562 | 0.0239705625820742 | 0.0871542838435782 | 0.275035965244071 | 0.783288607144167 | 0.949478217058132 | NM_015787 |

| | | | | | | | | | | | | | | | |
|---|---|---|---|---|---|---|---|---|---|---|---|---|---|---|---|
| Wbscr25 | 9.26874711 | 0.078853994 | 0.154212657 | 0.51133283 | 0.60911802 | 0.787588634 | | Lrp8 | 3850.45885 | -0.0174804541301092 | 0.0635804226001841 | -0.27493453826239 | 0.783366531561621 | 0.949478217058132 | NR_033496,NM_001080926 |
| Dio3 | 57.08990426 | -0.085556973 | 0.167393794 | -0.511111978 | 0.609272649 | 0.787624958 | | Med22 | 1395.390669 | -0.0170731538698013 | 0.0620441716011007 | -0.275177400700413 | 0.78317994860715 | 0.949478217058132 | NM_001033908,NM_011513 |
| Hc | 8.886618883 | 0.080904288 | 0.158296519 | 0.511093288 | 0.609285735 | 0.787624958 | | Ttc26 | 475.2209501 | 0.0218171559455699 | 0.079362128752791 | 0.274906385305379 | 0.783388161327393 | 0.949478217058132 | NM_153600 |
| Rpl22 | 3239.791195 | 0.027045474 | 0.052918807 | 0.511074893 | 0.609298615 | 0.787624958 | | 4930529M08Rik | 40.80287016 | 0.0250570574557201 | 0.0916582395725789 | 0.273374849577805 | 0.784565083842297 | 0.949699053346043 | NM_175280 |
| Prkacb | 21325.71257 | 0.034400718 | 0.067330424 | 0.510923834 | 0.609404391 | 0.787695974 | | Amhr2 | 37.02119871 | 0.0242557176497231 | 0.0884700324684904 | 0.274168743617926 | 0.783954947083756 | 0.949699053346043 | NM_144547 |
| 4932411E22Rik | 106.238372 | 0.077424832 | 0.151626708 | 0.51062793 | 0.609611615 | 0.787891263 | | B3galt6 | 465.5454521 | 0.0214414456559601 | 0.0781540652782627 | 0.274348437021404 | 0.78381686451321 | 0.949699053346043 | NM_080445 |
| Cnot8 | 2030.352198 | -0.032390805 | 0.063450388 | -0.510490267 | 0.609708032 | 0.787891263 | | Gorab | 359.7611478 | 0.0199029942373591 | 0.0726994378589427 | 0.273770950966312 | 0.784260648785725 | 0.949699053346043 | NM_178883 |
| Cyp4f15 | 592.2127849 | -0.054744479 | 0.107235922 | -0.510505039 | 0.609697685 | 0.787891263 | | Ift74 | 829.4137554 | 0.0192762981161185 | 0.0704105798800411 | 0.273769909990227 | 0.784261448814438 | 0.949699053346043 | NM_026319 |
| Lias | 1053.786665 | 0.041999455 | 0.082367927 | 0.509900592 | 0.610121107 | 0.78826483 | | Ogn | 236.333343 | -0.0228407132836713 | 0.0834389613917847 | -0.273741581902291 | 0.784283220087735 | 0.949699053346043 | NM_008760 |
| Maml1 | 524.0835667 | 0.055819899 | 0.10945915 | 0.509961015 | 0.610078774 | 0.78826483 | | Papss1 | 2887.319488 | 0.0116473660146004 | 0.0425474295911455 | 0.27375016837737 | 0.784276621018953 | 0.949699053346043 | NM_011863 |
| Rlf | 1577.384865 | -0.0380831 | 0.074701671 | -0.509802514 | 0.610189824 | 0.78826483 | | Prr18 | 1469.842065 | 0.0226806920802619 | 0.0829858209014264 | 0.273308040263925 | 0.78461643530262 | 0.949699053346043 | NM_178774,NR_028280 |
| Slc39a9 | 2422.941793 | -0.023411018 | 0.045923126 | -0.509787109 | 0.610200618 | 0.78826483 | | Rap2c | 1047.083525 | 0.019942679558346 | 0.0729328767673792 | 0.273438817201103 | 0.78451591746172 | 0.949699053346043 | NM_172413 |
| Agphd1 | 135.995157 | 0.061732892 | 0.121127049 | 0.509654059 | 0.610293844 | 0.788319535 | | Rnf25 | 1492.510554 | -0.0186576911217912 | 0.0681939224060896 | -0.273597565054054 | 0.784393905430197 | 0.949699053346043 | NM_021313 |
| Itfg2 | 691.0574365 | 0.039473942 | 0.077470408 | 0.509535743 | 0.610376751 | 0.788360903 | | Rxrg | 867.438878 | -0.02571113947302 | 0.0940601151402795 | -0.273347948114616 | 0.784585760927641 | 0.949699053346043 | NM_009107,NM_001159731 |
| Rbbp7 | 5257.589115 | 0.041533546 | 0.081525071 | 0.50945735 | 0.610431686 | 0.788366137 | | Sall3 | 261.6470168 | -0.0229539434703035 | 0.0837773560395743 | -0.273987441898508 | 0.784094272433979 | 0.949699053346043 | NM_178280 |
| Ddx54 | 1823.593484 | -0.029239776 | 0.057403987 | -0.509368387 | 0.610494031 | 0.788366656 | | Ssrp1 | 4258.250854 | -0.011468836499959 | 0.0419209838264426 | -0.273582236224256 | 0.78440568678625 | 0.949699053346043 | NM_182990,NM_001136081 |
| Igsf11 | 1502.429436 | -0.040889313 | 0.080283496 | -0.509311566 | 0.610533852 | 0.788366656 | | Stra6 | 256.2085967 | -0.0259459461932284 | 0.0948667484145851 | -0.273498845768803 | 0.784469779474727 | 0.949699053346043 | NM_001162479,NM_009291,NM_001162475,NM_001162476 |
| Kdm1b | 349.8054666 | 0.041727701 | 0.081972736 | 0.509043651 | 0.610721628 | 0.788477702 | | Tmem241 | 273.9296371 | -0.0216466681194411 | 0.0790514346600201 | -0.273830174145958 | 0.784215133952102 | 0.949699053346043 | NM_178801 |
| Rbck1 | 3305.827814 | -0.02832629 | 0.055644539 | -0.509057857 | 0.610711671 | 0.788477702 | | Usb1 | 412.3402773 | -0.0226016417891446 | 0.0823968331307642 | -0.274302311513304 | 0.783852308285434 | 0.949699053346043 | NM_133954 |
| Agmo | 259.1224963 | -0.047498044 | 0.093342715 | -0.50885646 | 0.610852841 | 0.788581396 | | 9430091E24Rik | 54.23804104 | 0.0257092470286113 | 0.0941679109945755 | 0.273014944868983 | 0.784841727492084 | 0.949834245705751 | NR_040363,NR_040364 |
| Ehd4 | 642.7232822 | 0.046853917 | 0.092126313 | 0.508583435 | 0.611044242 | 0.788697059 | | Fggy | 487.5685305 | -0.0193456822196272 | 0.0708652260109603 | -0.272992598889547 | 0.784858904805599 | 0.949834245705751 | NM_029347,NM_001113412 |
| Rsrc1 | 1504.216196 | 0.030216163 | 0.059410722 | 0.508597801 | 0.611034171 | 0.788697059 | | Chd7 | 577.6250975 | -0.0245296937675597 | 0.0900309233389245 | -0.272458538220438 | 0.785269467425853 | 0.950251939672437 | NM_001081417 |
| 3830406C13Rik | 834.3781015 | -0.064507756 | 0.126908111 | -0.508302867 | 0.61124096 | 0.788819543 | | Coro1a | 3361.562305 | 0.018410590233354 | 0.0676098543987495 | 0.272306313880992 | 0.78538650182526 | 0.950314396127757 | NM_009898 |
| Nudt12 | 241.6882444 | 0.046178455 | 0.090837229 | 0.508364859 | 0.611197492 | 0.788819543 | | 2310033P09Rik | 452.8840533 | -0.0194236983435504 | 0.0716357745644126 | -0.271145226831954 | 0.78627933799608 | 0.95048482101323 | NM_024210 |
| Igfbp7 | 662.2002832 | 0.046327845 | 0.091172309 | 0.508135038 | 0.611358645 | 0.788905708 | | 9330182L06Rik | 1482.501909 | 0.0168917668983088 | 0.0620988398277044 | 0.27201421065475 | 0.785611092673424 | 0.95048482101323 | NM_172706 |
| Ahr | 392.3705427 | -0.044437943 | 0.087510603 | -0.507800669 | 0.61159314 | 0.789142581 | | Anapc13 | 556.168848 | -0.0205141888458412 | 0.0757580980198064 | -0.270785426007895 | 0.786556069614553 | 0.95048482101323 | NM_181394 |
| Jtb | 872.9558435 | -0.040657499 | 0.080109294 | -0.507525369 | 0.611786241 | 0.789194573 | | Ankfy1 | 2756.717284 | 0.0136355098639887 | 0.05030553044344 | 0.271053892957546 | 0.78634958255859 | 0.95048482101323 | NM_009671 |
| Mtmr10 | 894.9975315 | 0.045024736 | 0.088708566 | 0.50755793 | 0.611763401 | 0.789194573 | | Atxn2 | 4169.331033 | 0.0152976831792993 | 0.0564384461116338 | 0.27105075056533 | 0.786351999392371 | 0.95048482101323 | NM_009125 |
| Slc46a3 | 266.3226753 | 0.049830425 | 0.098178111 | 0.507551273 | 0.61176807 | 0.789194573 | | Lrrc56 | 176.6639672 | 0.0247767179960026 | 0.0911420483202395 | 0.271847280729816 | 0.785739448922542 | 0.95048482101323 | NM_153777,NM_001172065,NM_001172064 |
| AV039307 | 18.02152348 | 0.084830314 | 0.167171774 | 0.507443998 | 0.611843321 | 0.789202498 | | Mpp4 | 25.26352598 | -0.022931610745922 | 0.0844809834334348 | -0.271441096137222 | 0.786051797898763 | 0.95048482101323 | NM_145143,NM_001164682 |
| Gkap1 | 449.7771473 | 0.0559575 | 0.110308324 | 0.507282656 | 0.611956506 | 0.789217089 | | Ppwd1 | 285.8772863 | -0.0218402457769333 | 0.0806177759595875 | -0.270911043091556 | 0.786459451376332 | 0.95048482101323 | NM_172807 |
| Hs2st1 | 2226.048203 | 0.038530088 | 0.07594723 | 0.507327097 | 0.611925329 | 0.789217089 | | Rapgef5 | 3559.277601 | -0.0127499254828126 | 0.0470809945210928 | -0.270761746694905 | 0.786574282899631 | 0.95048482101323 | NM_175930 |
| Bicd2 | 3398.568428 | 0.031905894 | 0.062959104 | 0.506771729 | 0.612314995 | 0.789482277 | | Smg6 | 2334.176138 | -0.0134491850160306 | 0.0496665934663157 | -0.270789359152484 | 0.78655304439098 | 0.95048482101323 | NM_001002764 |
| Pmm1 | 4135.479462 | -0.030846082 | 0.060862778 | -0.506813574 | 0.612285631 | 0.789482277 | | Sstr1 | 1301.208042 | -0.0199032607431633 | 0.0732528487041349 | -0.271706303512532 | 0.785847854076324 | 0.95048482101323 | NM_009216 |
| Srrd | 514.4627484 | 0.039874869 | 0.078681282 | 0.506789766 | 0.612302338 | 0.789482277 | | St18 | 459.1804387 | 0.02478283467853 | 0.0913209396745687 | 0.271381730924431 | 0.786097451613639 | 0.95048482101323 | NM_173868,NR_045188,NR_045189,NM_001244692,NM_001244693 |
| Eaf1 | 958.6380153 | -0.028425419 | 0.056107822 | -0.506621318 | 0.612420547 | 0.789486976 | | Them4 | 750.1553198 | 0.0178317895072051 | 0.0656831018795203 | 0.271482146807153 | 0.786020229073322 | 0.95048482101323 | NM_029431 |
| Fosb | 880.3338224 | -0.084653084 | 0.167091056 | -0.506628457 | 0.612415538 | 0.789486976 | | Uggt1 | 3063.878217 | -0.015711496834974 | 0.0579066598041675 | -0.271324522742429 | 0.786141447199945 | 0.95048482101323 | NM_198899 |
| Cdh7 | 643.8437396 | -0.045129914 | 0.089102856 | -0.506492336 | 0.612511068 | 0.789511234 | | Wdr55 | 435.6905926 | 0.0200218211862252 | 0.0736968158846006 | 0.271678239363512 | 0.785869434643206 | 0.95048482101323 | NM_026464 |
| Hmgxb4 | 665.9315276 | -0.045462537 | 0.089767205 | -0.506449294 | 0.612541277 | 0.789511234 | | Zic4 | 88.98067463 | -0.02365658423793 | 0.0872189233123959 | -0.271232243411194 | 0.786212415468304 | 0.95048482101323 | NM_009576 |
| Tigd2 | 485.9670798 | 0.049546451 | 0.097848535 | 0.506358641 | 0.612604903 | 0.789527564 | | Camk1 | 2569.371688 | -0.0154368362212954 | 0.0570485112675588 | -0.270591394557104 | 0.786705315144478 | 0.950564083163258 | NM_133926 |
| Pnlip | 30.04926287 | 0.084538568 | 0.167023662 | 0.506147257 | 0.612753278 | 0.789587432 | | Commd9 | 863.0233957 | -0.017093327366074 | 0.063223998640315 | -0.270361377541446 | 0.786882250279202 | 0.950698791279688 | NM_029635 |
| Ttbk1 | 5301.237446 | -0.04602009 | 0.090913912 | -0.50619415 | 0.612720362 | 0.789587432 | | G630025P09Rik | 28.24946078 | -0.0240441784443537 | 0.0889640048482681 | -0.270268615777381 | 0.786953608180518 | 0.950705930840835 | NR_027913 |
| 1700084C01Rik | 465.8140345 | -0.045264016 | 0.089577582 | -0.505305177 | 0.613344507 | 0.790241343 | | Glyctk | 251.2299951 | 0.0234749444838761 | 0.0868908722912378 | 0.270165828295446 | 0.787032680553592 | 0.950722388168106 | NM_001039586,NM_174846 |
| Baiap3 | 1343.991877 | 0.079338609 | 0.157153156 | 0.504848973 | 0.613664916 | 0.790241343 | | 1110002L01Rik | 525.5766389 | -0.018358143808725 | 0.0680674134868356 | -0.269705324006112 | 0.787386964323418 | 0.950831248440814 | NR_030694 |
| Cacna1a | 6106.130465 | -0.027066457 | 0.053610892 | -0.504868624 | 0.613651113 | 0.790241343 | | Aoc3 | 35.31367698 | -0.0226184744527823 | 0.0838891728615408 | -0.2696232860719 | 0.787450083900798 | 0.950831248440814 | NM_009675 |
| Cdk6 | 23.68723309 | -0.083012259 | 0.164385073 | -0.504986603 | 0.613568246 | 0.790241343 | | Ccdc116 | 127.8115693 | -0.0250909087509856 | 0.093036578647539 | -0.269688644141143 | 0.78739979761523 | 0.950831248440814 | NM_029779,NM_001164606 |
| Gstp1 | 1583.387563 | -0.038249984 | 0.075765405 | -0.504847621 | 0.613665866 | 0.790241343 | | Nfib | 2996.006185 | -0.0190794495277941 | 0.0707483679480996 | -0.269680419225933 | 0.787406125789451 | 0.950831248440814 | NM_008687,NM_001113209,NM_001113210 |
| Ppcdc | 432.9987021 | 0.045868965 | 0.090842175 | 0.504930282 | 0.613607804 | 0.790241343 | | Slco4c1 | 66.13525866 | -0.025622793683786 | 0.0949512098035014 | -0.269852208695514 | 0.787273955465053 | 0.950831248440814 | NM_172658 |
| Prpf18 | 1951.620662 | -0.031370293 | 0.062115799 | -0.505029209 | 0.613538321 | 0.790241343 | | Iba57 | 357.1305783 | -0.0217662534578551 | 0.0807626052433099 | -0.269509055487758 | 0.787537974667865 | 0.950858334305162 | NM_173785,NM_001270791 |
| Speccl1 | 2616.77255 | 0.035188241 | 0.069701287 | 0.504843494 | 0.613668765 | 0.790241343 | | Gm960 | 38.4509825 | -0.0249700241951003 | 0.0927113970893138 | -0.269330686183549 | 0.787675220162854 | 0.950945000672009 | NM_001033447 |
| Nlrp5-ps | 18.20773253 | 0.080738788 | 0.159991356 | 0.504644688 | 0.613808416 | 0.790355489 | | Ppp1r16b | 9051.981931 | 0.0197605050383412 | 0.0733974602865726 | 0.269226005384769 | 0.787755769424364 | 0.950963210060526 | NM_153089,NM_001159662 |
| Kif20b | 28.38733913 | -0.084707091 | 0.167918519 | -0.50445354 | 0.613942702 | 0.790462708 | | 1700001O22Rik | 177.907996 | 0.0252571496346703 | 0.0939138062906272 | 0.268939686636799 | 0.78797609614727 | 0.95107110699088 | NM_198000,NR_028486,NR_028487 |
| 2310022A10Rik | 900.3087391 | 0.033423352 | 0.06627666 | 0.504300496 | 0.614050229 | 0.790490492 | | Hes5 | 343.0352855 | 0.0254506338629734 | 0.0946241453741882 | 0.268965534772649 | 0.787956204908377 | 0.95107110699088 | NM_010419 |
| Api5 | 3694.754164 | -0.033039719 | 0.065528346 | -0.504204983 | 0.614117339 | 0.790490492 | | Chadl | 386.8495163 | -0.0213168108482694 | 0.0792976411792995 | -0.268820238928293 | 0.788068018030357 | 0.951103026745511 | NM_001164320 |
| Itgb4 | 487.0040135 | 0.072639316 | 0.144053764 | 0.504251424 | 0.614084708 | 0.790490492 | | Arhgef12 | 11106.69345 | 0.0163358237167874 | 0.06080934867419 | 0.268640004751785 | 0.788206724183994 | 0.95114233061094 | NM_027144 |
| 6820408C15Rik | 78.01011039 | 0.075231982 | 0.149408924 | 0.503530713 | 0.614591192 | 0.791034718 | | Cox5b | 5438.992948 | -0.0193937616291637 | 0.0722191336883926 | -0.268540491123071 | 0.78828331160856 | 0.95114233061094 | NM_009942 |
| Hpgd | 187.983353 | -0.066146275 | 0.131559073 | -0.50278763 | 0.615113592 | 0.791541695 | | Hadha | 2494.464711 | -0.0156524410032456 | 0.0582909460949092 | -0.268522678938172 | 0.788297020392751 | 0.95114233061094 | NM_178878 |
| Sh2d4b | 12.79213699 | 0.083272381 | 0.165632978 | 0.502752423 | 0.615138347 | 0.791541695 | | Cabp1 | 4141.621742 | 0.0157394046210677 | 0.0586631224156524 | 0.26830151503951 | 0.788467240128923 | 0.951189696047897 | NM_013879 |
| Tmp1 | 4578.937256 | -0.041877256 | 0.083290896 | -0.502783117 | 0.615116765 | 0.791541695 | | Mrpl34 | 477.3698722 | -0.0228122630397347 | 0.0850110632269162 | -0.268344638613011 | 0.788434049093997 | 0.951189696047897 | NM_053162 |
| Pkdrej | 85.1338863 | 0.066608109 | 0.132558861 | 0.502479493 | 0.615330274 | 0.791722908 | | Ccdc41 | 1338.603656 | -0.0188034106439987 | 0.0701274976206147 | -0.268131778289367 | 0.788597885592542 | 0.951268301454681 | NM_029852 |
| Dcaf4 | 774.2871877 | 0.037820794 | 0.075300369 | 0.502265726 | 0.615480615 | 0.791809747 | | Timm9 | 869.9663582 | -0.0180233539480046 | 0.0672407841466603 | -0.268041995296983 | 0.788666993483509 | 0.951272668549485 | NM_001024854,NM_001024853,NM_013896 |
| Pwp1 | 814.6656963 | -0.033405376 | 0.066513012 | -0.502238202 | 0.615499974 | 0.791809747 | | Chaf1a | 225.9114935 | -0.0226795925690978 | 0.0846692569086071 | -0.267861008790693 | 0.788806307726155 | 0.9512827252827 | NM_013733 |
| AK010878 | 130.7434513 | -0.058204127 | 0.115984132 | -0.501828361 | 0.615788261 | 0.792073939 | | Hnrnpd | 5590.233005 | 0.0149308887017881 | 0.0557262905999554 | 0.267932577981426 | 0.7887512165817 | 0.9512827252827 | NM_001077265,NM_001077267,NM_001077266,NM_007516 |
| Mlx | 1004.767174 | 0.035331393 | 0.070409186 | 0.501800898 | 0.615807582 | 0.792073939 | | Alg1 | 983.4627927 | -0.018976422287606 | 0.0709914836221595 | -0.267305616383577 | 0.789233862807186 | 0.951324461964749 | NM_145362 |
| 4833417C18Rik | 10.52508099 | 0.079575372 | 0.158669616 | 0.501516132 | 0.616007927 | 0.79225276 | | Chid1 | 2279.898182 | 0.0187970500173866 | 0.0702478810853105 | 0.267581736658492 | 0.789021290513983 | 0.951324461964749 | NM_001142681,NM_026522 |
| Nans | 489.1273568 | -0.04719701 | 0.094119579 | -0.501457937 | 0.616048874 | 0.79225276 | | Ipo13 | 4453.326033 | 0.0152592036127538 | 0.0570759392114192 | 0.26734914613023 | 0.789200350216205 | 0.951324461964749 | NM_146152 |
| Matr3 | 13492.23547 | -0.035429334 | 0.070675888 | -0.501293091 | 0.616164867 | 0.792336165 | | Pcdha8 | 28.90709352 | -0.0187193674238907 | 0.0700099925216547 | -0.267381365854319 | 0.789175545213084 | 0.951324461964749 | NM_201243 |
| Mir343 | 8.169445386 | 0.073555497 | 0.146777519 | 0.501135992 | 0.616275418 | 0.792346803 | | Slc33a1 | 1237.409157 | 0.0190074076491949 | 0.0710381437354088 | 0.267566218227641 | 0.789033237023883 | 0.951324461964749 | NM_015728,NM_001272035 |
| Pkd1 | 5634.166979 | 0.042835207 | 0.085476088 | 0.501136732 | 0.616274897 | 0.792346803 | | Sorbs2 | 7309.487454 | -0.0183199147217403 | 0.0684985011339232 | -0.267449862675427 | 0.789122812266698 | 0.951324461964749 | NM_001205219,NM_172752 |
| Wdr53 | 302.3134645 | 0.043443621 | 0.086720082 | 0.500963792 | 0.616396606 | 0.792436857 | | Cnnm3 | 1439.650687 | -0.01605049578959 | 0.0600662046255282 | -0.267213417089592 | 0.789304846295618 | 0.95133108175348 | NM_053186,NM_001039551 |
| Serpinb1a | 424.8375972 | 0.074666584 | 0.149084195 | 0.50083501 | 0.616487244 | 0.792487626 | | Pde1c | 501.9408207 | -0.022674142078401 | 0.0848959590373225 | -0.267081523496693 | 0.789406393143379 | 0.951374534705112 | NM_001025568,NM_011054,NM_001159960,NM_001159955,NM_001159956,NM_001159957,NM_001159952,NM_001159953 |
| Zfp866 | 1114.712498 | -0.033918215 | 0.067709883 | -0.500631333 | 0.616630607 | 0.792606158 | | Klhl2 | 6725.650091 | 0.0178499996823891 | 0.0669026156695802 | 0.266805707127312 | 0.78961875984679 | 0.951472592744464 | NM_178633 |
| Sav1 | 710.2840649 | 0.037241568 | 0.074409893 | 0.500492158 | 0.616728577 | 0.792666327 | | Uba5 | 1886.300095 | -0.0137293665182064 | 0.0514457667402115 | -0.266870675434508 | 0.789568735645888 | 0.951472592744464 | NM_025692 |
| Thap6 | 117.911951 | 0.06433808 | 0.128596642 | 0.500309877 | 0.6168569 | 0.792765495 | | 2010204K13Rik | 462.0543593 | 0.0192544761257765 | 0.0723419417089042 | 0.266159238623347 | 0.790116573917971 | 0.951607109054254 | NR_027924,NR_027925 |
| Chdh | 274.1768713 | -0.070860563 | 0.141660984 | -0.50021227 | 0.616925619 | 0.792788052 | | 3110040N11Rik | 228.5261906 | 0.024049689731871 | 0.090338218868495 | 0.266218329662665 | 0.790071067211651 | 0.951607109054254 | NM_026077 |
| Idh3a | 7739.965482 | -0.033371084 | 0.066740963 | -0.500008314 | 0.617069223 | 0.792841077 | | Bcr | 7894.237924 | 0.0178868411407416 | 0.06727028927387 | 0.265895112594521 | 0.790319989235201 | 0.951607109054254 | NM_001081412 |
| Rims4 | 754.4587819 | -0.04480283 | 0.089597226 | -0.500047071 | 0.617041934 | 0.792841077 | | Chst12 | 837.0214946 | 0.0195712322526131 | 0.0735876317963842 | 0.265958174965685 | 0.790271420772124 | 0.951607109054254 | NM_021528 |
| 2810047C21Rik1 | 41.2318845 | 0.08392357 | 0.168057528 | 0.499374858 | 0.617515328 | 0.792926278 | | Cnih4 | 1462.772037 | -0.0158680839612069 | 0.0596353354072784 | -0.266085263926767 | 0.790173543713105 | 0.951607109054254 | NM_030131 |
| Alpk1 | 190.8934208 | 0.068747193 | 0.137708937 | 0.499221003 | 0.6176237 | 0.792926278 | | Map7d1 | 10819.92674 | -0.00978451671235331 | 0.0367850860688256 | -0.265991404615618 | 0.790245828769618 | 0.951607109054254 | NM_144941,NM_001145970 |
| Cdkn1b | 812.4614429 | -0.046973047 | 0.094058655 | -0.499401645 | 0.617496461 | 0.792926278 | | Mapk1 | 28002.99232 | 0.018916551084331 | 0.0711004740787183 | 0.266053796819802 | 0.790197777672414 | 0.951607109054254 | NM_011949,NM_001038663 |
| Gpr161 | 127.3023963 | 0.074031536 | 0.148223526 | 0.499458742 | 0.617456246 | 0.792926278 | | Zfp871 | 4066.687965 | -0.0187032161672163 | 0.0703062723624837 | -0.266024858646844 | 0.790220064187091 | 0.951607109054254 | NM_172458 |
| Plxdc2 | 1212.886961 | -0.046419868 | 0.092937544 | -0.499473805 | 0.617445636 | 0.792926278 | | Zic2 | 436.5149283 | 0.0251863275175601 | 0.0946785140276368 | 0.266019463615665 | 0.790224219148457 | 0.951607109054254 | NM_009574 |
| Psmd3 | 4656.737295 | 0.028526985 | 0.057146834 | 0.499187501 | 0.617647299 | 0.792926278 | | Clspn | 124.0718075 | -0.0243390600081924 | 0.091592741517906 | -0.265731318932453 | 0.790446141268244 | 0.951680120021758 | NM_175554 |
| Rps6kc1 | 2260.823025 | 0.033324981 | 0.066754295 | 0.499218523 | 0.617625447 | 0.792926278 | | Agmo | 281.8501621 | 0.0216840225198063 | 0.0829481773056517 | 0.261416503944432 | 0.793771331115046 | 0.951728597725432 | NM_178767 |
| Slc35e2 | 2068.476093 | -0.033272173 | 0.06658152 | -0.499720835 | 0.61727166 | 0.792926278 | | Arhgap5 | 4390.32342 | -0.0208068573446847 | 0.0790227241536133 | -0.263302202847854 | 0.792317661952038 | 0.951728597725432 | NM_009706 |
| Snora52 | 16.39225133 | 0.081809117 | 0.163840868 | 0.499320577 | 0.617553561 | 0.792926278 | | Baiap2 | 13106.60831 | -0.0204617811873166 | 0.0779575701184203 | -0.262473306392624 | 0.792956562547139 | 0.951728597725432 | NM_001037755,NM_001037754,NM_130862 |
| Zfp296 | 67.67082881 | 0.078205341 | 0.156497935 | 0.499721233 | 0.61727138 | 0.792926278 | | Chd1l | 393.6170937 | -0.0188648323933924 | 0.0717326451395149 | -0.262988104742292 | 0.792559747539965 | 0.951728597725432 | NM_026539 |
| Sdpr | 392.3578933 | 0.054297244 | 0.108817405 | 0.498975724 | 0.617796486 | 0.793052092 | | Chst2 | 6152.779704 | -0.0164205079634667 | 0.0628022429472485 | -0.26146371837801 | 0.793734925143181 | 0.951728597725432 | NM_018763 |
| Dnajc6 | 13878.45464 | -0.033656692 | 0.067465244 | -0.498874539 | 0.617867772 | 0.793071598 | | Cobl | 5964.665241 | -0.0209449495095869 | 0.0796964745252585 | -0.262808984140808 | 0.792697810547053 | 0.951728597725432 | NM_172496 |
| Gm5083 | 21.68296133 | -0.081342048 | 0.163072585 | -0.49880885 | 0.617914052 | 0.793071598 | | Ctsb | 29907.83768 | 0.0136696187088254 | 0.0523129207828261 | 0.261304826881564 | 0.793857444537003 | 0.951728597725432 | NM_007798 |
| Sgms1 | 1575.848696 | 0.03889691 | 0.078014549 | 0.498585341 | 0.618071534 | 0.793142318 | | Det1 | 146.1770879 | -0.0248438682779778 | 0.0936475272222512 | -0.26529123635071 | 0.79078511510126 | 0.951728597725432 | NM_029585 |
| Trpc6 | 366.7489013 | -0.056691472 | 0.113696148 | -0.498622626 | 0.618045263 | 0.793142318 | | Dnpep | 1366.595704 | -0.0167875838518448 | 0.0638145851801044 | -0.263068134102338 | 0.792498064419497 | 0.951728597725432 | NM_001110831,NM_016878 |
| Gpr157 | 139.8435857 | -0.062456867 | 0.125364511 | -0.498202135 | 0.618341578 | 0.793423139 | | Exo5 | 347.1928655 | 0.0216659665642363 | 0.0823138098653945 | 0.263211806131511 | 0.792387331579715 | 0.951728597725432 | NM_001160043,NM_028457 |
| Gbf1 | 4199.01657 | 0.024798568 | 0.049801673 | 0.497946487 | 0.61852176 | 0.793588618 | | Fam96a | 589.0532565 | 0.0161569403504823 | 0.0618428046606454 | 0.261258208438985 | 0.793893392443829 | 0.951728597725432 | NM_026635 |
| BC037032 | 34.27856213 | -0.083671215 | 0.168068474 | -0.497840034 | 0.618596796 | 0.793592753 | | Gk5 | 81.45858571 | -0.025158669582333 | 0.0951027335864821 | -0.264542023489313 | 0.791362287669668 | 0.951728597725432 | NM_177352 |
| Zbtb2 | 395.2091516 | 0.044160226 | 0.088711388 | 0.497796588 | 0.618627421 | 0.793592753 | | Gm1821 | 1591.852893 | 0.0169062510735873 | 0.0645157776764056 | 0.262048318079865 | 0.793284190391234 | 0.951728597725432 | NR_002875 |
| Amz2 | 2034.68262 | -0.031826994 | 0.064039231 | -0.496992139 | 0.619194595 | 0.794188088 | | Ifitm10 | 349.4580706 | -0.0196485623792824 | 0.0748199149795457 | -0.262611396773892 | 0.792850114950796 | 0.951728597725432 | NM_177265 |
| C2cd4b | 27.38339722 | -0.082529569 | 0.166106688 | -0.496846753 | 0.619297123 | 0.794188088 | | Iqck | 64.35724011 | 0.0247318811924701 | 0.0940432202238879 | 0.262984201663778 | 0.792562755894891 | 0.951728597725432 | NM_001081446 |
| Mtbp | 82.86856437 | 0.075192918 | 0.151362149 | 0.496774909 | 0.619347791 | 0.794188088 | | Irf2 | 1144.518555 | -0.0174868220648847 | 0.0660217979748006 | -0.26486437209055 | 0.791113945328169 | 0.951728597725432 | NM_008391 |
| Rfc4 | 129.2673028 | -0.062956715 | 0.126696492 | -0.496909693 | 0.619252736 | 0.794188088 | | Kat6a | 3308.261731 | 0.0167797017672925 | 0.0640339539514337 | 0.262043817878543 | 0.793287660367671 | 0.951728597725432 | NM_001081149 |
| Slc22a13b-ps | 17.59674693 | 0.081107373 | 0.163196266 | 0.496992821 | 0.619194114 | 0.794188088 | | Lmtk2 | 10525.96123 | 0.0155280616657191 | 0.0592583076319403 | 0.262040248637634 | 0.79329041209429 | 0.951728597725432 | NM_001081109 |
| Psmd9 | 842.9677412 | 0.044340706 | 0.089304603 | 0.49651087 | 0.61953402 | 0.794361148 | | Lsm7 | 707.4486536 | 0.0222469089115145 | 0.0843233741895989 | 0.263828495068199 | 0.791912076371585 | 0.951728597725432 | NM_025349 |
| Chrm2 | 316.0055412 | -0.071933242 | 0.144967127 | -0.496203814 | 0.61975062 | 0.794414831 | | Mfsd7a | 47.46066268 | -0.0236864279233882 | 0.0900300165531863 | -0.263094785830625 | 0.792477522724562 | 0.951728597725432 | NM_172883 |
| Fam86 | 145.1380965 | -0.064234813 | 0.129463776 | -0.496160781 | 0.619780979 | 0.794414831 | | Mga | 2703.808148 | -0.019615411603578 | 0.0746640542589439 | -0.262715597194191 | 0.792769794139756 | 0.951728597725432 | NM_001164274,NM_013720 |
| Hcfc2 | 871.4573886 | 0.038422176 | 0.07740906 | 0.496352446 | 0.61964577 | 0.794414831 | | Mlh3 | 1339.302655 | 0.0159642632129606 | 0.0610234094932443 | 0.261608837420464 | 0.793623029976381 | 0.951728597725432 | NM_175337 |
| Lrrc8c | 640.6475225 | 0.055251395 | 0.111327082 | 0.496297888 | 0.619684256 | 0.794414831 | | Mmachc | 658.7870081 | -0.0182997980131829 | 0.069502650249416 | -0.263296405928587 | 0.792322129644292 | 0.951728597725432 | NM_025962 |
| Wdr36 | 1102.116638 | -0.031199469 | 0.062913029 | -0.495914264 | 0.619954902 | 0.794572028 | | Mphosph8 | 2216.805036 | 0.0195991697999895 | 0.0743072888329316 | 0.263758375629277 | 0.79196611046556 | 0.951728597725432 | NM_023773 |
| Rtfdc1 | 2958.487745 | 0.032663688 | 0.065888868 | 0.495739103 | 0.620078495 | 0.794664698 | | Mvb12a | 575.5633187 | -0.0199862883184076 | 0.0760054947676612 | -0.262958466088578 | 0.792582592043687 | 0.951728597725432 | NM_028617 |
| C1qbp | 1738.514956 | 0.04082244 | 0.082362205 | 0.4956453 | 0.620144686 | 0.794683795 | | Myo1c | 1011.679864 | 0.0184899008282506 | 0.0700953105818833 | 0.263782279795325 | 0.791947689784394 | 0.951728597725432 | NM_008659,NM_001080775,NM_001080774 |
| 6720489N17Rik | 82.92300074 | 0.078261731 | 0.158108563 | 0.494987302 | 0.620609084 | 0.795015325 | | Ncoa1 | 6547.411819 | 0.0147357860961586 | 0.0557190256565777 | 0.264465968715643 | 0.791420884567901 | 0.951728597725432 | NM_010881 |
| Cckar | 20.82064125 | -0.067407057 | 0.136145758 | -0.495109489 | 0.620522836 | 0.795015325 | | Necab2 | 3085.613525 | -0.0194453826062808 | 0.0741737134395329 | -0.262160025493841 | 0.793198070850564 | 0.951728597725432 | NM_054095 |
| Ptgs1 | 763.4948245 | 0.046127001 | 0.093184869 | 0.495005266 | 0.620596403 | 0.795015325 | | Nod2 | 51.81415763 | -0.0248065350914635 | 0.0948304680343075 | -0.261588238523605 | 0.793638912657435 | 0.951728597725432 | NM_145857 |
| Sec14l3 | 101.4660051 | -0.078185962 | 0.157978191 | -0.494915234 | 0.620659957 | 0.795015325 | | Nol3 | 1079.207368 | 0.0164837353012895 | 0.0621052921925739 | 0.26541595280121 | 0.790689048170278 | 0.951728597725432 | NM_030152 |
| Slc25a22 | 8664.2695 | -0.038362429 | 0.077511862 | -0.494923331 | 0.620654241 | 0.795015325 | | Oscp1 | 1356.59232 | 0.0178002246378237 | 0.0672980869185659 | 0.264498226515159 | 0.791396031203872 | 0.951728597725432 | NM_172701 |
| Grik1 | 525.6646866 | 0.047450927 | 0.095965641 | 0.494457463 | 0.62098314 | 0.795363542 | | Pex6 | 2347.849023 | -0.01974628253405 | 0.0749289549313023 | -0.263533405670241 | 0.792139479235758 | 0.951728597725432 | NM_145488 |
| Mettl6 | 511.026129 | 0.037411678 | 0.075708701 | 0.494152947 | 0.621198167 | 0.795573186 | | Pigx | 316.4080413 | -0.0202118351149686 | 0.0770483850382184 | -0.262326525143167 | 0.793069713768721 | 0.951728597725432 | NM_001111025,NM_024464 |
| N28178 | 11524.72701 | 0.034449617 | 0.06972632 | 0.494069048 | 0.621257416 | 0.795583305 | | Polr3f | 1083.670432 | -0.0173979230501525 | 0.0664782233644257 | -0.261708604256451 | 0.793546106445171 | 0.951728597725432 | NM_029763 |
| Gk5 | 73.95411687 | -0.073299729 | 0.148435096 | -0.493816697 | 0.62143564 | 0.795745769 | | Pou3f2 | 955.504763 | -0.0210320747692373 | 0.0803229112503623 | -0.261844032106623 | 0.793441690305954 | 0.951728597725432 | NM_008899 |
| Ccndbp1 | 1787.864175 | -0.02925525 | 0.059273634 | -0.493562617 | 0.621615107 | 0.795909799 | | Ppp1r3b | 58.30968144 | 0.0237563765478959 | 0.0896287000030014 | 0.265053231242898 | 0.790968455150885 | 0.951728597725432 | NM_177741 |
| 9530080O11Rik | 54.61233321 | 0.068693607 | 0.139371135 | 0.492882594 | 0.622095547 | 0.796459131 | | Ppp1r3c | 1687.902341 | 0.0179263655544924 | 0.0678548127676727 | 0.264187090396524 | 0.791635758166496 | 0.951728597725432 | NM_016854 |
| Ina | 4755.291665 | 0.04501239 | 0.091355531 | 0.492716643 | 0.622212816 | 0.79654345 | | Ptpn12 | 1283.228061 | -0.0172742657577092 | 0.0653895242409544 | -0.264174819410753 | 0.791645213230339 | 0.951728597725432 | NM_011203 |
| Kbtbd4 | 1220.334743 | -0.043312808 | 0.087954439 | -0.492445963 | 0.622404114 | 0.796656699 | | Rilpl2 | 394.3579007 | 0.0213210079851545 | 0.0809227007513047 | 0.263473756245289 | 0.792185448641688 | 0.951728597725432 | NM_030259 |
| Mlycd | 890.1232525 | -0.037393334 | 0.07592338 | -0.492514083 | 0.622355969 | 0.796656699 | | Sept4 | 8225.011335 | 0.0211821611563916 | 0.080220802959735 | 0.26404823156686 | 0.791742753730993 | 0.951728597725432 | NM_011129 |
| Wars | 3766.71202 | -0.026732175 | 0.054384224 | -0.491542819 | 0.623042575 | 0.797408034 | | Slc7a2 | 1350.545931 | -0.0171999260639252 | 0.0658326532517568 | -0.261267398689665 | 0.793886305722946 | 0.951728597725432 | NM_001044740,NM_007514 |
| Ei24 | 4257.710656 | -0.030511106 | 0.062113674 | -0.491213999 | 0.623275098 | 0.797508 | | Snx12 | 1844.879756 | 0.0123278591822013 | 0.0466359733794857 | 0.264342272474667 | 0.791516189614867 | 0.951728597725432 | NM_001110311,NM_001110310,NM_018875 |
| Fastkd3 | 443.0948416 | -0.041714005 | 0.084913329 | -0.49125391 | 0.623246874 | 0.797508 | | Sox6 | 1006.161787 | 0.0204536935816145 | 0.0775671103924031 | 0.26369026674836 | 0.792018580766322 | 0.951728597725432 | NM_001025559,NM_011445,NM_001025560 |
| Trim47 | 130.614341 | -0.065780903 | 0.133901655 | -0.491262809 | 0.62324058 | 0.797508 | | Spast | 2121.908136 | 0.017520624273009 | 0.0663348352735051 | 0.264124033786618 | 0.791684344994215 | 0.951728597725432 | NM_016962,NM_001162870 |
| 1110020A21Rik | 23.16602162 | 0.082480142 | 0.168093298 | 0.490680729 | 0.623652278 | 0.797775617 | | Strn3 | 2907.962242 | 0.0187287819432023 | 0.0709974795222271 | 0.263795025810839 | 0.791937867682568 | 0.951728597725432 | NM_001172098,NM_052973 |
| Baz2b | 1323.226655 | 0.043906446 | 0.08949046 | 0.490627114 | 0.623690206 | 0.797775617 | | Thy1 | 56123.68496 | 0.0106885704322063 | 0.0407970549491989 | 0.261993676884664 | 0.793326317088517 | 0.951728597725432 | NM_009382 |
| Ctsh | 492.7938271 | 0.052286863 | 0.106555469 | 0.490717775 | 0.623626073 | 0.797775617 | | Tmem42 | 402.868127 | -0.0212409795331038 | 0.0805264003133433 | -0.263776593147728 | 0.791952071935622 | 0.951728597725432 | NM_025339,NM_001164823 |
| Rad51ap1 | 57.6017152 | 0.075242613 | 0.153349537 | 0.490660839 | 0.623666349 | 0.797775617 | | Tmtc3 | 602.0030987 | 0.0226521455858304 | 0.0863083561825262 | 0.262455999137851 | 0.792969904195058 | 0.951728597725432 | NM_001110013,NM_001033332 |
| Poc1b | 234.5608553 | 0.043416536 | 0.08853113 | 0.490409823 | 0.623843927 | 0.797906373 | | Traf2 | 427.2780109 | -0.0178596403824364 | 0.0683334747583978 | -0.261360050042553 | 0.793814862019833 | 0.951728597725432 | NM_009422 |
| Elavl1 | 2854.099808 | -0.025309829 | 0.051620689 | -0.490303976 | 0.623918814 | 0.797936286 | | Wnt5a | 312.8252163 | 0.0216826550644009 | 0.0816497729033098 | 0.265568324737713 | 0.790580534813018 | 0.951728597725432 | NM_009524,NM_001256224 |
| Pex11c | 12.99093749 | -0.080672044 | 0.164678016 | -0.489877434 | 0.624220633 | 0.79824887 | | Zdhhc18 | 1264.110425 | 0.0154917265956596 | 0.0584474697511128 | 0.265053845130133 | 0.790967982246505 | 0.951728597725432 | NM_001017968 |
| Zfp14 | 399.9540936 | 0.045222418 | 0.092325894 | 0.489812948 | 0.624266268 | 0.79824887 | | Zzz3 | 2501.565554 | -0.0161076928338634 | 0.0609617595834255 | -0.264226179558026 | 0.791605639314031 | 0.951728597725432 | NM_198416,NM_001080755 |
| Fam120aos | 479.6717005 | 0.039700411 | 0.081076687 | 0.48966494 | 0.624371015 | 0.798316926 | | Prosapip1 | 14260.20106 | -0.0184088926552284 | 0.0705298736697339 | -0.261008445037497 | 0.794085994738498 | 0.951878275713444 | NM_197945 |
| Rundc1 | 2024.472193 | -0.043011984 | 0.087971119 | -0.48893301 | 0.624889124 | 0.798913449 | | Vamp8 | 272.1153111 | 0.0247216644806879 | 0.094745757734143 | 0.260926347225561 | 0.794149306303953 | 0.951878275713444 | NM_016794 |
| Crtc1 | 9661.581225 | 0.03512708 | 0.071908554 | 0.48849654 | 0.625198176 | 0.799217394 | | Anxa6 | 5159.863783 | 0.0151629721359508 | 0.058219296857459 | 0.260445813577499 | 0.79451990777206 | 0.951979492661738 | NM_013472,NM_001110211 |
| Irak3 | 61.01279577 | 0.075890718 | 0.155369998 | 0.488451562 | 0.625230027 | 0.799217394 | | Dopey2 | 4705.202021 | -0.0109609848704283 | 0.0420536860779919 | -0.260642666378883 | 0.794368083576438 | 0.951979492661738 | NM_026700,NM_027293 |
| Kdm4c | 1158.809682 | 0.02814271 | 0.057645459 | 0.48820341 | 0.625405769 | 0.799244226 | | Klc3 | 78.06763474 | 0.0247451835854347 | 0.0950053202805618 | 0.260461030101885 | 0.794508171636131 | 0.951979492661738 | NM_146182 |
| Tmc3 | 18.55201597 | -0.07885378 | 0.161496857 | -0.4882682 | 0.625359883 | 0.799244226 | | Opa1 | 7734.422422 | 0.0151935880026016 | 0.0583489103412521 | 0.260391974995631 | 0.794561432536893 | 0.951979492661738 | NM_133752,NM_001199177 |
| Unc5c | 1442.031136 | -0.041471312 | 0.084936754 | -0.488261093 | 0.625364917 | 0.799244226 | | Rab33a | 789.3828867 | -0.0167965297131768 | 0.064480454285446 | -0.260490250872317 | 0.794485634495563 | 0.951979492661738 | NM_011228 |
| Zwint | 16037.34365 | -0.040401968 | 0.08277208 | -0.488111065 | 0.625471174 | 0.799261886 | | 4930525G20Rik | 78.35342472 | -0.0247410942496382 | 0.0950652264760217 | -0.260253882168773 | 0.794667943801494 | 0.952028581580248 | NR_045194 |
| Polr2b | 3018.961107 | 0.036225234 | 0.074301351 | 0.487544753 | 0.625872336 | 0.799642614 | | Mak16 | 1182.9102 | 0.01629908045458 | 0.0626665493997948 | 0.260092196086887 | 0.794792657454111 | 0.952099467440081 | NM_026594 |
| Tmem259 | 3360.760967 | 0.034607501 | 0.070979378 | 0.487571205 | 0.625853596 | 0.799642614 | | Mmp9 | 61.60077138 | 0.0245507271780716 | 0.0944825555429633 | 0.259844021332677 | 0.794984092791584 | 0.952171745703377 | NM_013599 |
| Ppp1r15b | 2559.183579 | 0.038022092 | 0.078014511 | 0.487372044 | 0.625994701 | 0.799667071 | | Trim33 | 2764.988841 | 0.01774365879476 | 0.068276324461659 | 0.259880111219579 | 0.794956253255118 | 0.952171745703377 | NM_001079830,NM_053170 |
| Rab12 | 3158.092854 | 0.037385648 | 0.07670024 | 0.48742544 | 0.625956869 | 0.799667071 | | Ccm2 | 1998.720623 | 0.0166146843368613 | 0.0639991412559369 | 0.259607926150416 | 0.795166221730689 | 0.95231136424935 | NM_146014,NM_001190344,NM_001190343 |
| Pbdc1 | 507.3006437 | 0.031533672 | 0.064727147 | 0.487178462 | 0.626131867 | 0.799776352 | | Cyp2j12 | 164.6802121 | -0.0244832075339365 | 0.0944491990112392 | -0.259220912302529 | 0.795464796742205 | 0.952590407046766 | NM_001100182 |
| Senp7 | 1172.903083 | 0.044169471 | 0.090700048 | 0.486983991 | 0.626269676 | 0.799820505 | | Lrfn4 | 1434.932946 | 0.0204169681463484 | 0.0788039735172217 | 0.259085516060767 | 0.795569259858602 | 0.95263696881593 | NM_153388 |
| Snrnp200 | 6163.730648 | 0.039245328 | 0.080587466 | 0.48699047 | 0.626265084 | 0.799820505 | | Islr | 521.8362234 | -0.024306679162088 | 0.0938685947213422 | -0.258943677960075 | 0.795678697033242 | 0.952689478495292 | NM_001195431,NM_012043 |
| Hcn1 | 2240.803533 | 0.046268868 | 0.095026946 | 0.486902606 | 0.626327352 | 0.799828236 | | Acads | 298.9708926 | -0.0225395093943082 | 0.0870770094516933 | -0.25884662939432 | 0.795753578530529 | 0.952700608401422 | NM_007383 |
| Rimbp3 | 117.7905104 | 0.061501661 | 0.126364253 | 0.48670142 | 0.626469939 | 0.799944391 | | 1110038B12Rik | 217.0000935 | -0.0225961122746582 | 0.0875413318983588 | -0.258119356704486 | 0.79631479314779 | 0.95273444749767 | NR_015536,NR_027943 |
| Gm11944 | 19.71670886 | 0.081734905 | 0.167990249 | 0.486545529 | 0.626580433 | 0.79996003 | | 4933434E20Rik | 1827.849909 | -0.0119481430934187 | 0.0462546904266133 | -0.25831203224729 | 0.796166100760015 | 0.95273444749767 | NM_027500,NM_025762 |
| Nubpl | 157.4451283 | 0.06135689 | 0.126127906 | 0.486465617 | 0.626637077 | 0.79996003 | | Ddah1 | 4560.723602 | -0.0141010920532511 | 0.0546179631809779 | -0.258176820078897 | 0.796270446540489 | 0.95273444749767 | NM_026993 |
| Snord49a | 10.00875279 | 0.077216398 | 0.158718681 | 0.486498489 | 0.626613776 | 0.79996003 | | Eif2c1 | 4271.622393 | -0.0153104521350262 | 0.0593495112276084 | -0.257970989455789 | 0.796429296671936 | 0.95273444749767 | NM_153403 |
| Ccbl1 | 394.5870832 | 0.048737932 | 0.100379363 | 0.485537365 | 0.627295214 | 0.799960418 | | Fasn | 13137.73045 | -0.0128069245718148 | 0.0496459889507653 | -0.257964940219353 | 0.796433965308006 | 0.95273444749767 | NM_007988 |
| Cdh26 | 9.193531413 | -0.075582973 | 0.155562262 | -0.485869592 | 0.627059629 | 0.799960418 | | Gm16845 | 27.63573501 | 0.021399647043432 | 0.0829572079451608 | 0.257960068491919 | 0.796437725179967 | 0.95273444749767 | NR_040406,NR_040407 |
| Ddx1 | 5637.070374 | 0.028103888 | 0.057870768 | 0.485631846 | 0.627228213 | 0.799960418 | | Kbtbd8 | 299.4242575 | 0.0222412024575042 | 0.0861287611985649 | 0.258232002271906 | 0.796227861024334 | 0.95273444749767 | NM_001008785,NM_001102670 |
| Enpp3 | 19.97968512 | -0.079736776 | 0.164059568 | -0.486022685 | 0.626951082 | 0.799960418 | | Pus7l | 117.127374 | -0.0241494952144182 | 0.0934572469310325 | -0.258401525910982 | 0.796097039072607 | 0.95273444749767 | NM_172437 |
| Gbp4 | 124.0646516 | 0.068543273 | 0.141106485 | 0.485755656 | 0.627140418 | 0.799960418 | | Wdr17 | 1776.008777 | -0.0188751712819029 | 0.073009729300816 | -0.258529533839703 | 0.795998258728686 | 0.95273444749767 | NM_001172152,NM_028220 |
| Hist1h2bg | 8.629779784 | 0.075029339 | 0.154463584 | 0.486077681 | 0.626912091 | 0.799960418 | | Zscan25 | 558.4562568 | -0.0197087353299167 | 0.0762899074167567 | -0.258340008492234 | 0.796144511605405 | 0.95273444749767 | NM_001081431 |
| Mgat1 | 2180.038983 | -0.031998641 | 0.065868076 | -0.485798926 | 0.627109736 | 0.799960418 | | Pde4dip | 10423.1031 | 0.014400842209088 | 0.0558689048599812 | 0.257761311863538 | 0.79659112438113 | 0.952839482276045 | NM_001110163,NM_177145,NM_001039376,NM_178080 |
| Mstn | 12.4107045 | 0.078706883 | 0.162104085 | 0.485533 | 0.62729881 | 0.799960418 | | Dennd4a | 3636.719824 | -0.0183157991031627 | 0.0711297618299861 | -0.25749838930912 | 0.79679405854157 | 0.952885371480875 | NM_001162917 |
| Myl12a | 780.6169583 | 0.048713657 | 0.100163714 | 0.486340364 | 0.626725866 | 0.799960418 | | Dhrs7b | 625.1057176 | 0.0162560129125263 | 0.0631456963719249 | 0.257435502387646 | 0.796842599223523 | 0.952885371480875 | NM_145428,NM_001172112 |
| Nol7 | 904.3571637 | -0.050142822 | 0.103276875 | -0.485518524 | 0.627308576 | 0.799960418 | | Dnaja4 | 2535.990096 | -0.0161903583281419 | 0.0628535004877702 | -0.257588808939619 | 0.796724267500053 | 0.952885371480875 | NM_021422 |
| Ocel1 | 172.3712685 | 0.055474294 | 0.114161472 | 0.485928335 | 0.627017978 | 0.799960418 | | Dock3 | 13962.45755 | 0.0152823925422631 | 0.0594572212723656 | 0.25703173164209 | 0.797154277449787 | 0.952885371480875 | NM_153413 |
| Olfm1 | 34448.53703 | -0.041748906 | 0.085977393 | -0.485580037 | 0.627264953 | 0.799960418 | | Lman1 | 1656.006792 | 0.0127758062854881 | 0.0496977466630808 | 0.257070131813016 | 0.797124634243546 | 0.952885371480875 | NM_027400,NM_001172062 |
| Tle1 | 1397.26853 | -0.031392714 | 0.064567189 | -0.486202273 | 0.62682376 | 0.799960418 | | Mycbpap | 99.74884166 | -0.024275536860441 | 0.0943837413400117 | -0.257200408839377 | 0.797024068414522 | 0.952885371480875 | NM_170671 |
| Gtdc2 | 1144.758456 | -0.033081127 | 0.068174674 | -0.485240707 | 0.627505609 | 0.800135255 | | Rapgef1 | 7209.017131 | -0.0139960493831265 | 0.0544286901792359 | -0.257144703226128 | 0.797067069299819 | 0.952885371480875 | NM_001039087,NM_054050,NM_001039086 |
| Rnf113a1 | 129.737977 | -0.058705129 | 0.120996701 | -0.485179585 | 0.627548962 | 0.800135255 | | Zfp943 | 514.265741 | 0.0205056176408554 | 0.0797202283667361 | 0.257219755399139 | 0.797009134350976 | 0.952885371480875 | NM_001025373 |
| A930006K02Rik | 18.86192921 | -0.081247997 | 0.167572541 | -0.484852689 | 0.627780844 | 0.800365046 | | 3830406C13Rik | 989.4279286 | 0.017490055095605 | 0.0681368440196605 | 0.256690126278205 | 0.797417994312642 | 0.952897915014139 | NM_146051,NM_178141 |
| Clic5 | 503.8828523 | 0.050238567 | 0.103636537 | 0.484757296 | 0.627848518 | 0.800370404 | | Cntnap5b | 598.0942971 | 0.0238270498389849 | 0.0927631216826159 | 0.256859077258179 | 0.797287562489566 | 0.952897915014139 | NM_172851 |
| Syt7 | 13591.17518 | -0.031452501 | 0.064890504 | -0.484701136 | 0.627888361 | 0.800370404 | | Itpka | 5081.016021 | -0.019439679920202 | 0.0757355969300818 | -0.256678242572623 | 0.797427168863546 | 0.952897915014139 | NM_146125 |
| Oxnad1 | 585.6124477 | 0.03282735 | 0.067861354 | 0.483741454 | 0.62856937 | 0.801172575 | | Qrfpr | 106.4185809 | -0.0244090057386184 | 0.0950854580074257 | -0.256705932544876 | 0.797405791478953 | 0.952897915014139 | NM_198192 |
| Atxn3 | 1219.881615 | 0.039749308 | 0.082326905 | 0.482822819 | 0.629221547 | 0.801937867 | | Acp2 | 4016.718641 | 0.0133026825625933 | 0.0520825508646537 | 0.255415371208734 | 0.798402299910514 | 0.952967259224917 | NM_007387 |
| Sh2d4a | 9.457385991 | -0.079018363 | 0.163706745 | -0.482682392 | 0.629321267 | 0.80199899 | | Ccdc40 | 88.92466881 | 0.0242080023574029 | 0.0944507704737553 | 0.256302857414271 | 0.797716991033712 | 0.952967259224917 | NM_175430 |
| Prrc2a | 20015.86181 | 0.025016776 | 0.051860087 | 0.482389787 | 0.629529075 | 0.802187836 | | Fzd7 | 338.1670163 | 0.0197666898482052 | 0.0773909021607677 | 0.255413611888681 | 0.79840365859612 | 0.952967259224917 | NM_008057 |
| Chst7 | 47.86388514 | 0.07112757 | 0.14757028 | 0.481991157 | 0.629812228 | 0.802480427 | | Gch1 | 65.3091783 | 0.0232536976256891 | 0.0909849894695933 | 0.25557729644471 | 0.798277251095595 | 0.952967259224917 | NM_008102 |
| Faxc | 5539.462891 | -0.043375767 | 0.090011472 | -0.481891539 | 0.629882997 | 0.802480427 | | Gpr182 | 68.95347525 | 0.0236331200015174 | 0.0921483584559333 | 0.256468160665272 | 0.797589362552425 | 0.952967259224917 | NM_007412 |
| Kctd5 | 866.1310624 | -0.035489372 | 0.07365097 | -0.481858852 | 0.629906219 | 0.802480427 | | Gpr26 | 1840.511067 | -0.0180028649630387 | 0.070235802789958 | -0.256320341590979 | 0.797703491473431 | 0.952967259224917 | NM_173410 |
| D330050G23Rik | 68.33213034 | 0.069044702 | 0.14334399 | 0.481671408 | 0.63003939 | 0.802584092 | | Lzic | 351.087845 | -0.018642310208069 | 0.0728606292827495 | -0.255862602225462 | 0.798056932685641 | 0.952967259224917 | NM_026963 |
| Gga1 | 2641.466426 | 0.03803169 | 0.079045771 | 0.481135035 | 0.630420529 | 0.802712142 | | Mtmr14 | 686.9601682 | -0.0184798202508857 | 0.0722923995010488 | -0.255626046146354 | 0.798239604534271 | 0.952967259224917 | NM_026849 |
| Parp6 | 5636.092162 | -0.025909948 | 0.053835404 | -0.481280837 | 0.630316856 | 0.802712142 | | Naa35 | 1880.207888 | 0.013390068478749 | 0.0523667943081644 | 0.255697692701067 | 0.798184276915474 | 0.952967259224917 | NM_030153 |
| Pdcd11 | 1096.934607 | -0.045792486 | 0.095163671 | -0.481197135 | 0.630376397 | 0.802712142 | | Nhlrc1 | 691.8977754 | -0.0188851478152817 | 0.0737445756872132 | -0.256088636205365 | 0.797882396475503 | 0.952967259224917 | NM_175340 |
| Phlpp1 | 4528.613778 | -0.033292588 | 0.06921154 | -0.481026546 | 0.630497632 | 0.802712142 | | Nsun3 | 198.5255624 | -0.0223035536125588 | 0.0871021578566765 | -0.256062009959139 | 0.797902955886482 | 0.952967259224917 | NM_178925 |
| Prim2 | 270.2410451 | 0.045044282 | 0.093643339 | 0.481019602 | 0.630502567 | 0.802712142 | | Onecut2 | 231.862696 | -0.0243050851723164 | 0.095100319651138 | -0.255573117540259 | 0.79828047824246 | 0.952967259224917 | NM_194268 |
| Serhl | 167.5050927 | 0.062252648 | 0.129302878 | 0.481448278 | 0.630197931 | 0.802712142 | | Slc30a7 | 925.8733624 | -0.0155515901141292 | 0.0607179455438376 | -0.256128397870464 | 0.797851694834049 | 0.952967259224917 | NM_023214 |
| Tmem194b | 185.7876316 | 0.069936275 | 0.145315758 | 0.481271104 | 0.630323831 | 0.802712142 | | Ugt8a | 1973.313662 | 0.0217803012167373 | 0.0851629755283131 | 0.255748476161408 | 0.798145061017819 | 0.952967259224917 | NM_011674 |
| Tagln2 | 439.2771569 | 0.05742314 | 0.119406052 | 0.480906446 | 0.630582992 | 0.802748572 | | Cited2 | 959.0137624 | 0.0191596207217149 | 0.0750668958494208 | 0.255233955059869 | 0.798542406972875 | 0.952978215470307 | NM_010828 |
| B4galt5 | 2576.209746 | -0.032869711 | 0.068362516 | -0.480814813 | 0.630648121 | 0.802765527 | | Zbtb8os | 396.7528987 | 0.0187477964012591 | 0.0734539903065975 | 0.255231830469735 | 0.798544047824906 | 0.952978215470307 | NM_025970 |
| Cep72 | 56.40899362 | 0.075539133 | 0.157148789 | 0.480685428 | 0.63074009 | 0.802816639 | | Ccpg1 | 4433.659618 | -0.0133633900670649 | 0.0524056106646714 | -0.254999224273397 | 0.798723698371891 | 0.95303601466651 | NM_001114328,NM_028181 |
| 1700095B22Rik | 42.66254303 | 0.078672339 | 0.163738344 | 0.480475965 | 0.630888989 | 0.802873394 | | Spata18 | 29.37841669 | -0.0194335985855627 | 0.0761934918624984 | -0.255055886146199 | 0.798679935286293 | 0.95303601466651 | NM_178387 |
| E2f6 | 1306.476256 | -0.030428982 | 0.063358824 | -0.480264316 | 0.631039459 | 0.802873394 | | Atpbd4 | 728.2421347 | -0.0190438032198802 | 0.0747357142500462 | -0.25481529695648 | 0.798865759927389 | 0.953053772399362 | NM_025675 |
| Gpn1 | 913.2205586 | -0.028208092 | 0.058728321 | -0.480314966 | 0.631003448 | 0.802873394 | | Nfic | 6148.513303 | 0.0138759453670462 | 0.0544711632289618 | 0.254739288542832 | 0.798924469157275 | 0.953053772399362 | NM_026756,NM_008688 |
| Lamtor3 | 1374.561926 | 0.037786124 | 0.078647099 | 0.480451596 | 0.630906314 | 0.802873394 | | Nipbl | 2325.133858 | -0.0189049379498548 | 0.0742170139106096 | -0.254725122190599 | 0.798935411435153 | 0.953053772399362 | NM_027707,NM_201232 |
| Ufsp2 | 1466.217874 | -0.03127833 | 0.065128146 | -0.480258251 | 0.631043771 | 0.802873394 | | Gpatch2 | 755.9572319 | -0.0161771581255805 | 0.0636054737667411 | -0.254335942609383 | 0.799236034327806 | 0.953180007018908 | NM_026367 |
| 1810013L24Rik | 1676.904736 | 0.032066051 | 0.066813671 | 0.479932487 | 0.6312754 | 0.802875181 | | Samd3 | 110.3620888 | -0.0241801586381262 | 0.0950185684963459 | -0.254478245892077 | 0.79912610829227 | 0.953180007018908 | NM_001115154,NM_001013766 |
| Bend6 | 5224.913692 | 0.033278877 | 0.069345259 | 0.479901264 | 0.631297602 | 0.802875181 | | Tram1 | 946.5027737 | -0.0153870336693102 | 0.0604994891919848 | -0.254333282393213 | 0.799238089321926 | 0.953180007018908 | NM_028173 |
| Klhl8 | 1348.671759 | 0.046262806 | 0.096402558 | 0.47989189 | 0.631304268 | 0.802875181 | | Syt17 | 1317.18047 | 0.0227949033919028 | 0.0896623320843501 | 0.254230543216949 | 0.799317455515875 | 0.953196400855726 | NM_138649 |
| Ptbp2 | 2706.003729 | 0.039697698 | 0.082702623 | 0.480005304 | 0.631223622 | 0.802875181 | | 1110059E24Rik | 717.1752098 | 0.0164254366958952 | 0.0651831226257857 | 0.251989101917358 | 0.801049487465365 | 0.953228630707512 | NM_025423 |
| Yy1 | 991.9881287 | 0.041597001 | 0.086643646 | 0.480092928 | 0.631161317 | 0.802875181 | | A330035P11Rik | 23.58671429 | -0.0216041698979657 | 0.0858848629446106 | -0.251548051161225 | 0.801390416686741 | 0.953228630707512 | NR_015586 |
| Lyar | 296.0254457 | 0.040210221 | 0.083808623 | 0.479786208 | 0.631379421 | 0.802904855 | | Aebp2 | 2673.855368 | -0.0162673372130791 | 0.0644142466516503 | -0.252542380494174 | 0.800621860489628 | 0.953228630707512 | NM_178803,NM_009637,NM_001005605 |
| B3galtl | 823.5198714 | 0.031448769 | 0.065587694 | 0.47949191 | 0.631588722 | 0.802907422 | | Aldh1a2 | 302.0078489 | 0.0187926318817204 | 0.0740466756323756 | 0.253794403614032 | 0.799654397194155 | 0.953228630707512 | NM_009022 |
| Bdkrb2 | 19.99913979 | 0.078976807 | 0.164686137 | 0.479559533 | 0.631540627 | 0.802907422 | | Coprs | 703.6916941 | 0.018002805740857 | 0.0713121831445856 | 0.252450632514731 | 0.800692768036261 | 0.953228630707512 | NM_025556 |
| Hoga1 | 27.29510821 | 0.080631299 | 0.168120509 | 0.479604182 | 0.631508872 | 0.802907422 | | Dusp9 | 43.9491792 | 0.0234072973899627 | 0.0929426657480753 | 0.251846632561725 | 0.801159611164222 | 0.953228630707512 | NM_029352 |
| Tbccd1 | 533.0767839 | 0.039146652 | 0.08163636 | 0.479524707 | 0.631565396 | 0.802907422 | | E130012A19Rik | 2455.219556 | -0.0209692891592935 | 0.0828452603160119 | -0.253113926847553 | 0.800180177156162 | 0.953228630707512 | NM_175332 |
| 6030407O03Rik | 10.30395131 | 0.072586031 | 0.151410387 | 0.479399285 | 0.631654602 | 0.802925294 | | E2f2 | 32.73276693 | 0.0216136874198825 | 0.0851071636950334 | 0.253958497516511 | 0.799527621306903 | 0.953228630707512 | NM_177733 |
| 3110007F17Rik | 53.34745158 | -0.069028704 | 0.144133387 | -0.478922377 | 0.631993851 | 0.803224734 | | Eda | 312.6311316 | 0.0218630369090081 | 0.0868706254966817 | 0.25167352927421 | 0.801293419109946 | 0.953228630707512 | NM_001177937,NM_001177938,NM_001177939,NM_010099,NM_001177943,NM_001177942,NM_001177941,NM_001177940,NM_001177944 |
| Pcdhb10 | 164.6923182 | -0.049382896 | 0.103110585 | -0.478931395 | 0.631987436 | 0.803224734 | | Fmo5 | 224.9472015 | 0.0219851248452373 | 0.0865702090616268 | 0.253957164751523 | 0.79952865095499 | 0.953228630707512 | NM_001161765,NM_010232,NM_001161763 |
| Aspa | 540.3752395 | 0.06824639 | 0.142625499 | 0.478500624 | 0.632293931 | 0.803503065 | | Fryl | 3175.323234 | -0.017333999347107 | 0.068344708776501 | -0.253626062023209 | 0.799784460252745 | 0.953228630707512 | NM_028194 |
| Ccser1 | 408.1849538 | 0.048068057 | 0.100462256 | 0.478468814 | 0.632316566 | 0.803503065 | | Gm13031 | 54.33205761 | -0.0239303495837165 | 0.094512045744213 | -0.253198937715109 | 0.800114487356352 | 0.953228630707512 | NR_045911 |
| Aard | 51.61656172 | -0.072927366 | 0.152457107 | -0.478346776 | 0.632403409 | 0.803547517 | | Gpatch3 | 220.0628412 | -0.0215387425569656 | 0.085470726070689 | -0.252001399159192 | 0.801039982324078 | 0.953228630707512 | NM_172876 |
| 0610011F06Rik | 622.9309509 | 0.035313831 | 0.073843706 | 0.478223976 | 0.6324908 | 0.803592657 | | Hdac5 | 6733.568788 | 0.0180572794249922 | 0.0717261486323601 | 0.251753088229325 | 0.801231919725066 | 0.953228630707512 | NM_001077696,NM_010412 |
| Igsf1 | 310.3915469 | -0.072092913 | 0.150848543 | -0.477915874 | 0.632710083 | 0.803750179 | | Il12rb1 | 43.07853109 | 0.021831397138594 | 0.0867978569802426 | 0.251520001738792 | 0.801412099978636 | 0.953228630707512 | NM_008353 |
| Sp3 | 1743.918816 | -0.037831906 | 0.079162145 | -0.477904003 | 0.632718532 | 0.803750179 | | Impa2 | 146.4191138 | -0.0237674721889288 | 0.0937530798788945 | -0.253511374982352 | 0.799873072239058 | 0.953228630707512 | NM_053261 |
| Acot1 | 191.2553657 | 0.049392266 | 0.103402015 | 0.477672184 | 0.632883545 | 0.80382799 | | Itprip | 48.97004031 | -0.0234154071757969 | 0.0923563234612254 | -0.253533340200875 | 0.799856100796525 | 0.953228630707512 | NM_001001738 |
| Zfp941 | 1268.396444 | 0.040564877 | 0.084920146 | 0.477682614 | 0.63287612 | 0.80382799 | | Letmd1 | 2315.809494 | -0.0139596710948503 | 0.0553696091072271 | -0.252117927504534 | 0.800949913313477 | 0.953228630707512 | NM_134093 |
| 2700070H01Rik | 9.773139921 | 0.072692835 | 0.152207849 | 0.477589269 | 0.63294257 | 0.803837064 | | Lrrc3 | 723.2899482 | 0.0200649270028998 | 0.079578937565442 | 0.252138664032796 | 0.800933885571346 | 0.953228630707512 | NM_145152 |
| Pigc | 432.2989833 | 0.035681176 | 0.074734576 | 0.477438656 | 0.633049793 | 0.803907343 | | Mettl5 | 214.3073317 | 0.021657231080157 | 0.0857383544565278 | 0.252596766259818 | 0.800579829154641 | 0.953228630707512 | NM_029280 |
| Add3 | 4566.887958 | -0.02369281 | 0.049658732 | -0.477112663 | 0.633281898 | 0.804136184 | | Mex3d | 495.7789623 | -0.0190185303368702 | 0.075133252817323 | -0.253130666166036 | 0.800167242198387 | 0.953228630707512 | NM_198615 |
| Usp24 | 4120.736361 | -0.037172854 | 0.077954153 | -0.476855339 | 0.633465136 | 0.804302944 | | Nnt | 1233.908979 | -0.0237209618413693 | 0.0938070591686583 | -0.252869688609689 | 0.800368913370957 | 0.953228630707512 | NM_008710,NR_003544 |
| Msi1 | 622.2950416 | 0.03817594 | 0.080075452 | 0.476749606 | 0.633540434 | 0.804332636 | | Osbpl2 | 2807.725604 | 0.0118274315009645 | 0.0470056823468676 | 0.251617057990706 | 0.801337072381218 | 0.953228630707512 | NM_144500 |
| Nyap1 | 3287.93037 | -0.029429713 | 0.061760871 | -0.476510652 | 0.633710621 | 0.804482782 | | Palld | 98.49167647 | 0.0240862980324045 | 0.0950677134023102 | 0.253359391641991 | 0.799990504869551 | 0.953228630707512 | NM_001081390 |
| Fam168b | 15674.11167 | 0.026558372 | 0.055747212 | 0.476407183 | 0.633784318 | 0.804510423 | | Ptn | 8097.002106 | -0.0130034993904717 | 0.0514889941292532 | -0.252549105112228 | 0.800616663424331 | 0.953228630707512 | NM_008973 |
| Ppp1r12a | 2709.059036 | -0.025807256 | 0.054189354 | -0.476242181 | 0.633901852 | 0.804593699 | | Raly | 2980.844934 | 0.0130173630488567 | 0.0513217290350985 | 0.25364233227517 | 0.799771889386905 | 0.953228630707512 | NM_001139512,NM_023130,NM_001139511,NM_001139513 |
| Gm14015 | 11.86561769 | 0.075352651 | 0.158292407 | 0.476034531 | 0.634049778 | 0.804715535 | | Sema6a | 1181.182194 | -0.0202396762163107 | 0.0804828174597045 | -0.251478226721426 | 0.801444393971557 | 0.953228630707512 | NM_018744 |
| Kcnj16 | 1135.384627 | -0.049040157 | 0.103049449 | -0.475889559 | 0.634153061 | 0.804729607 | | Slc41a3 | 237.5515732 | 0.0236379331787626 | 0.0934163777314922 | 0.253038425946095 | 0.800238519577508 | 0.953228630707512 | NM_027868,NM_001037493 |
| Pold3 | 1011.627632 | 0.028764296 | 0.060449941 | 0.47583662 | 0.634190778 | 0.804729607 | | Sptbn1 | 40931.29879 | 0.0131991260317931058 | 0.0522017319567054 | 0.252848445205874 | 0.800385329403612 | 0.953228630707512 | NM_009260,NM_175836 |
| Wbp4 | 2158.69212 | 0.033633764 | 0.07068883 | 0.475800267 | 0.63421668 | 0.804729607 | | Tmem143 | 829.4186436 | -0.0170208161566578 | 0.0676035315035732 | -0.25177406827864 | 0.801215702269743 | 0.953228630707512 | NM_144801 |
| 4930578C19Rik | 30.9467851 | -0.079363465 | 0.166828036 | -0.475720193 | 0.634273733 | 0.804736097 | | Usp20 | 4149.550936 | -0.0156302520712402 | 0.0617655670267938 | -0.253057695794484 | 0.800223628925985 | 0.953228630707512 | NM_028846 |
| Igsf9b | 686.5394148 | 0.055606091 | 0.11692159 | 0.475584454 | 0.634370453 | 0.804792909 | | Xpo6 | 4399.266636 | -0.0113262810199928 | 0.0449366495610499 | -0.252049966578063 | 0.801002442472128 | 0.953228630707512 | NM_028816 |
| Asic1 | 3309.80094 | 0.032802755 | 0.068985314 | 0.47550345 | 0.634428175 | 0.804800241 | | Nt5dc2 | 131.2597592 | 0.0239030665194089 | 0.0950985512463785 | 0.251350480171686 | 0.801543149983165 | 0.953268036405392 | NM_027289 |
| Mpv17l | 2447.938291 | 0.025568626 | 0.053785537 | 0.475381076 | 0.63451538 | 0.80484497 | | Etfa | 1151.581558 | 0.0142005981452991 | 0.0565321673313436 | 0.251195006589209 | 0.801663344989933 | 0.953332930767398 | NM_145615 |
| Cluh | 4325.21593 | 0.02371903 | 0.049914087 | 0.475197117 | 0.634646482 | 0.804945366 | | B9d2 | 430.515541 | -0.0194020572239418 | 0.07744987149143 | -0.250511160965433 | 0.802192074612211 | 0.953337330308904 | NM_172148 |
| Crls1 | 805.5518575 | -0.030234166 | 0.06367124 | -0.474848086 | 0.634895256 | 0.805134487 | | Chia | 84.09555846 | 0.0237497872811533 | 0.0945896305075899 | 0.25108235600146 | 0.801750436918639 | 0.953337330308904 | NM_023186 |

| | | | | | | | | | | | | | | | |
|---|---|---|---|---|---|---|---|---|---|---|---|---|---|---|---|
| Tmem229b | 1359.901048 | 0.029941176 | 0.063055014 | 0.474842104 | 0.63489952 | 0.805134487 | | E330033B04Rik | 48.43008222 | -0.0217447644238781 | 0.086642809348148 | -0.250970214233282 | 0.801837137917718 | 0.953337330308904 | NR_030690 |
| Esr1 | 113.5684499 | 0.071690651 | 0.151035662 | 0.474660422 | 0.635029032 | 0.805232819 | | Efcab14 | 2737.738613 | 0.0156985603284242 | 0.0626903850042748 | 0.250414163628979 | 0.802267077483246 | 0.953337330308904 | NM_172698 |
| Dnaja2 | 6513.125564 | -0.03016937 | 0.063595764 | -0.474392764 | 0.635219852 | 0.805306026 | | Egfl6 | 117.1479493 | 0.023349293130007 | 0.0930955787151779 | 0.250809903673764 | 0.801961084249838 | 0.953337330308904 | NM_019397 |
| Nipal2 | 255.5690212 | 0.055937573 | 0.117898888 | 0.474453782 | 0.635176349 | 0.805306026 | | Stard6 | 82.47310712 | -0.023821385568104 | 0.0950429770135452 | -0.250638040985491 | 0.802093967805719 | 0.953337330308904 | NM_029019 |
| Pomt2 | 1117.46432 | 0.035557092 | 0.074957916 | 0.474360731 | 0.635242691 | 0.805306026 | | Tbrg4 | 1354.86826 | -0.0143918855019014 | 0.0574450223963471 | -0.250533203775311 | 0.802175030336167 | 0.953337330308904 | NM_134011,NM_001130457 |
| Cd79b | 8.48564503 | -0.074215849 | 0.156505261 | -0.474206736 | 0.63535249 | 0.805379325 | | Tdrd1 | 34.96613852 | 0.0189016808896419 | 0.0755036153866897 | 0.250341401439354 | 0.802323341802723 | 0.953337330308904 | NM_031387,NM_001002241,NM_001002238,NM_001002240 |
| A430035B10Rik | 75.17145507 | -0.071311549 | 0.150429448 | -0.474053116 | 0.635462031 | 0.805452283 | | Thoc6 | 172.5996704 | -0.0211193886035482 | 0.0843333919153851 | -0.250427358889325 | 0.802256874184557 | 0.953337330308904 | NM_001008425 |
| Ppip5k1 | 5988.750691 | 0.020960222 | 0.044238303 | 0.473802571 | 0.635640701 | 0.805612845 | | Ubqln4 | 6796.688491 | 0.0141071508835588 | 0.0562388078299331 | 0.250843704337031 | 0.801934950383604 | 0.953337330308904 | NM_033526 |
| Cr1l | 589.6359846 | 0.038225272 | 0.080722558 | 0.473538902 | 0.635828754 | 0.805715213 | | Elovl1 | 695.5011703 | 0.0197580573343039 | 0.0790103850984189 | 0.25006911825189 | 0.802533897447413 | 0.953509520229112 | NM_019422,NM_001039176,NM_001039175 |
| Stam2 | 1073.711164 | 0.027878438 | 0.058881034 | 0.473470593 | 0.635877476 | 0.805715213 | | Zufsp | 314.2162158 | -0.0219925864239401 | 0.0879889381407917 | -0.249947173913494 | 0.802628201230668 | 0.953534571691886 | NM_028287 |
| Tbc1d20 | 1526.655238 | -0.043847879 | 0.092606286 | -0.473487072 | 0.635865722 | 0.805715213 | | Akap11 | 11900.41975 | 0.0180684368264423 | 0.0724777065480839 | 0.249296475937123 | 0.803131457086078 | 0.95375427199217 | NM_001164503 |
| Crlf3 | 419.024549 | 0.040256449 | 0.085074166 | 0.473192405 | 0.636075916 | 0.805900748 | | CK137956 | 94.87982515 | 0.0233523597580673 | 0.0938327173575052 | 0.248872252831538 | 0.803459599150972 | 0.95375427199217 | NM_001134733 |
| Maged1 | 19974.96192 | 0.031420306 | 0.066447588 | 0.472858483 | 0.636314147 | 0.806136663 | | Ccne1 | 647.5109777 | 0.0160687219517353 | 0.0645481408718212 | 0.248941669499736 | 0.803405902080077 | 0.95375427199217 | NM_007633 |
| Asns | 3716.161975 | 0.029218937 | 0.061814672 | 0.472686113 | 0.636437136 | 0.806226554 | | Ccrl2 | 35.01544031 | -0.0225387273095011 | 0.0905996944065448 | -0.248772663717428 | 0.803536637643633 | 0.95375427199217 | NM_017466 |
| Clcn3 | 8209.459066 | -0.031903345 | 0.067523717 | -0.472476142 | 0.636586967 | 0.806350431 | | Cyp46a1 | 9882.758265 | -0.0168535762454695 | 0.0677626953604693 | -0.248714667499802 | 0.803581502273766 | 0.95375427199217 | NM_010010 |
| Eif3a | 6309.261379 | 0.030162352 | 0.063869387 | 0.472250526 | 0.636747979 | 0.806488448 | | Efcab5 | 157.7713223 | -0.0226710442868433 | 0.0909833899537603 | -0.249177836727838 | 0.803223222557607 | 0.95375427199217 | NM_176965 |
| Gfpt2 | 318.0836347 | 0.065642091 | 0.139058743 | 0.47204577 | 0.636894119 | 0.806541682 | | Kirrel3 | 2169.599311 | 0.017519032430694 | 0.070424161701391 | 0.248764515010876 | 0.803542941269178 | 0.95375427199217 | NM_001190913,NM_001190912,NM_001190911,NM_026324,NM_001190914 |
| Tmem14a | 972.6597829 | -0.043375238 | 0.091885708 | -0.472056422 | 0.636886516 | 0.806541682 | | Pcdhgc3 | 2765.790565 | 0.0157814781085439 | 0.0634560831271075 | 0.248699215754183 | 0.803593455521973 | 0.95375427199217 | NM_033581 |
| BC027231 | 205.2101524 | 0.054256436 | 0.114974191 | 0.471901002 | 0.636997453 | 0.806606615 | | Tceal5 | 2332.687375 | 0.0167314168101131 | 0.0672050554199892 | 0.24896068763804 | 0.803391190822854 | 0.95375427199217 | NM_177919 |
| Cdkn2d | 96.0730874 | -0.060669831 | 0.128704038 | -0.471390266 | 0.637362066 | 0.806879358 | | Tpk1 | 94.61243359 | -0.0237064377160536 | 0.0951027075961112 | -0.249271953609689 | 0.803150424479398 | 0.95375427199217 | NM_013861 |
| Coasy | 858.2951255 | 0.03304097 | 0.070094062 | 0.471380448 | 0.637369076 | 0.806879358 | | Tvp23a | 1782.15453 | 0.0153810241405794 | 0.0617354294425055 | 0.249144199359686 | 0.803249241000724 | 0.95375427199217 | NM_001013778 |
| Ppcs | 165.8454288 | -0.057812227 | 0.122619701 | -0.47147584 | 0.637300969 | 0.806879358 | | Zfp944 | 347.5378677 | 0.0223574486234563 | 0.0898440134440937 | 0.248847394126804 | 0.803478828756072 | 0.95375427199217 | NM_176962 |
| Lrrc61 | 1055.723586 | 0.040841093 | 0.086700568 | 0.471059114 | 0.637598522 | 0.807097323 | | Ccdc53 | 447.1501063 | 0.0185204267040661 | 0.074540570095475 | 0.248461028408345 | 0.803777719637339 | 0.953895029040195 | NM_026070,NM_001122960,NM_027487 |
| Slc22a15 | 330.0559183 | 0.038261491 | 0.081235723 | 0.470993422 | 0.637645432 | 0.807097323 | | Zbtb39 | 588.903224 | 0.0171513294343437 | 0.0692337051561195 | 0.247730919436827 | 0.804342607150417 | 0.954487436603787 | NM_198035 |
| Pcnxl2 | 4929.598415 | -0.036151506 | 0.076798576 | -0.470731466 | 0.637832511 | 0.807202232 | | Tmem5 | 802.5904747 | 0.0168983908894656 | 0.0682715338395804 | 0.24751737567772 | 0.804507845927013 | 0.954605535854908 | NM_153059 |
| Pms1 | 136.2562566 | 0.052964061 | 0.112499873 | 0.47079218 | 0.63778915 | 0.807202232 | | Zcchc4 | 228.3440087 | 0.0210113528742409 | 0.0849668893338785 | 0.247288714921368 | 0.804684791821285 | 0.954735706003103 | NM_030185 |
| Ing5 | 728.3948221 | 0.032042498 | 0.068123889 | 0.47035626 | 0.638100509 | 0.807468732 | | Tgds | 180.535093 | -0.0219647795340552 | 0.0888879184435792 | -0.247106467545386 | 0.804825828543031 | 0.954826852778182 | NM_029578 |
| Thrb | 2558.303045 | -0.04054195 | 0.086219519 | -0.470217773 | 0.638199438 | 0.807468732 | | D430019H16Rik | 8571.480738 | -0.015104586112921 | 0.0612467908400794 | -0.246618408993222 | 0.805203556214044 | 0.955118965994219 | NM_001252508 |
| Zfp558 | 121.4400737 | -0.06567017 | 0.139644133 | -0.470268015 | 0.638163546 | 0.807468732 | | Ube2o | 7756.536398 | -0.0123552858862697 | 0.0500843342438908 | -0.246689630056864 | 0.805148432601906 | 0.955118965994219 | NM_173755 |
| Ppp1r14a | 238.0493898 | -0.075451213 | 0.160559174 | -0.46992776 | 0.63840663 | 0.807664925 | | Gpr126 | 154.8421911 | 0.0216818295573257 | 0.0879818981847666 | 0.246435119094529 | 0.805345423209736 | 0.955131255514748 | NM_001002268 |
| 6430531B16Rik | 20.30496894 | -0.078290249 | 0.166786641 | -0.469403593 | 0.638781181 | 0.808051631 | | Klc1 | 25785.81948 | 0.0102510122946044 | 0.0415950202137645 | 0.246448066184908 | 0.805335401908012 | 0.955131255514748 | NM_001081959,NM_008450,NM_001025360 |
| Insc | 163.0756786 | -0.075888158 | 0.161686389 | -0.469354029 | 0.638816602 | 0.808051631 | | Thyn1 | 1159.156575 | -0.0155287923317583 | 0.0630476710184223 | -0.246302394377437 | 0.805448156585934 | 0.955175110014473 | NM_144543 |
| E030019B06Rik | 63.2509931 | 0.069704359 | 0.148576651 | 0.469147466 | 0.638964233 | 0.808106426 | | Eaf2 | 42.00122174 | -0.0224147270183599 | 0.0912162410466757 | -0.245731755235236 | 0.805889888632521 | 0.955397949759603 | NM_134111,NM_001113405,NM_001113401 |
| Zfand2a | 2717.425749 | 0.038891908 | 0.082892948 | 0.469182345 | 0.638939304 | 0.808106426 | | Mxd1 | 1301.175897 | 0.0172753878807046 | 0.0703052362843856 | 0.245719789786715 | 0.805899151755777 | 0.955397949759603 | NM_010751 |
| Irs2 | 1557.312647 | 0.074600314 | 0.159069273 | 0.468980038 | 0.639083905 | 0.808191807 | | Sf3a2 | 1385.496062 | 0.0137836694513601 | 0.0560816469862994 | 0.245778613718806 | 0.805853613120501 | 0.955397949759603 | NM_013651 |
| Alg8 | 259.7344675 | 0.0578476 | 0.1233721 | 0.468887213 | 0.639150257 | 0.808209751 | | Zfp764 | 193.5119329 | -0.0221554201979923 | 0.090078735315129 | -0.245956164021224 | 0.805716166280753 | 0.955397949759603 | NM_001167832,NM_146203 |
| 2010001E11Rik | 12.37412882 | 0.078074646 | 0.166582229 | 0.468685326 | 0.639294578 | 0.808326276 | | 2810002D19Rik | 133.9927761 | -0.022332610339709 | 0.0912990213536005 | -0.244609526023449 | 0.80675878740431 | 0.955587482484189 | NR_027831 |
| Bcar3 | 581.860314 | -0.044159158 | 0.094300111 | -0.468283206 | 0.639582076 | 0.808434982 | | 4933404O12Rik | 765.9110929 | 0.0145087820059291 | 0.0593401617490086 | 0.244501895146444 | 0.806842134391886 | 0.955587482484189 | NR_015555 |
| Elmo3 | 65.58835888 | 0.067704816 | 0.144518226 | 0.468486352 | 0.639436829 | 0.808434982 | | Acaa2 | 857.0997139 | 0.0184813747392279 | 0.0754701912162258 | 0.244883104725121 | 0.806546943944896 | 0.955587482484189 | NM_177470 |
| Exosc7 | 540.6850564 | -0.032380062 | 0.069147808 | -0.468273159 | 0.639589261 | 0.808434982 | | Acbd4 | 1211.29055 | -0.0175567990724244 | 0.0718566803079465 | -0.244330784516953 | 0.806974643220735 | 0.955587482484189 | NM_025988 |
| Npm1 | 4408.65652 | 0.02708196 | 0.057823161 | 0.468358336 | 0.639528357 | 0.808434982 | | Dennd1a | 2903.319527 | 0.0141897721633646 | 0.0579950280539334 | 0.244672218283413 | 0.806710240906155 | 0.955587482484189 | NM_146122 |
| Hist1h1e | 29.58430736 | -0.071986683 | 0.153831611 | -0.467957676 | 0.639814857 | 0.808654164 | | Gng3 | 7853.265932 | 0.0120392481699208 | 0.0492531184143505 | 0.244436262261376 | 0.806892960133406 | 0.955587482484189 | NM_010316 |
| Nsfl1c | 2866.65993 | 0.030182816 | 0.064509754 | 0.467879884 | 0.639870489 | 0.808658513 | | LOC100503496 | 24.87964622 | -0.0207059970784216 | 0.0847316841242342 | -0.244371362288307 | 0.806943219113746 | 0.955587482484189 | NR_040680 |
| Col9a1 | 21.84694913 | 0.077559735 | 0.165872226 | 0.467587232 | 0.640079798 | 0.808726883 | | Mafg | 1079.825353 | 0.02015268109227 | 0.0823836020838643 | 0.244620137982218 | 0.806750569855303 | 0.955587482484189 | NM_010756 |
| Lage3 | 322.1109161 | 0.043188736 | 0.092365469 | 0.467585302 | 0.640081178 | 0.808726883 | | Medag | 1180.488223 | 0.0207158105380282 | 0.0846535074422923 | 0.24471296185985 | 0.806678691031714 | 0.955587482484189 | NM_027519 |
| Mttp | 568.8631284 | 0.037590598 | 0.080378834 | 0.467667872 | 0.64002212 | 0.808726883 | | Ncam1 | 11331.0157 | 0.0151126604651317 | 0.0617019987135649 | 0.244929836637679 | 0.80651075888436 | 0.955587482484189 | NM_010875,NM_001081445,NM_001113204 |
| Snora43 | 11.35438356 | 0.07516654 | 0.160794692 | 0.467469038 | 0.640164339 | 0.808766004 | | Ngdn | 585.8456838 | 0.0163098888106174 | 0.0666873669970267 | 0.244572391219833 | 0.806787543532861 | 0.955587482484189 | NM_026890 |
| Zfp828 | 1738.115065 | -0.027394807 | 0.058625849 | -0.467282053 | 0.640298095 | 0.808869032 | | Ninj2 | 89.44296393 | -0.0230286833145991 | 0.0941240890264183 | -0.24466301403603 | 0.806717368278103 | 0.955587482484189 | NM_016718 |
| Pgm3 | 533.2770265 | 0.039935569 | 0.085521257 | 0.466966581 | 0.640523788 | 0.80903468 | | Proser2 | 155.2120193 | 0.0221165108094884 | 0.090370604949583 | 0.244731246646263 | 0.806664532271007 | 0.955587482484189 | NM_001159657,NM_144883 |
| Smarcb1 | 2130.257326 | -0.019520604 | 0.041804235 | -0.466952793 | 0.640533653 | 0.80903468 | | Socs1 | 22.74104369 | -0.0186107815796674 | 0.0761725938381251 | -0.244323852476618 | 0.806980011540241 | 0.955587482484189 | NM_001271603,NM_009896 |
| Arg2 | 89.28678067 | 0.069168244 | 0.148211613 | 0.466685725 | 0.640724744 | 0.809078173 | | 1700028J19Rik | 45.18845501 | 0.0220057857142002 | 0.0907357116584347 | 0.242526181940786 | 0.808372472047237 | 0.955716697308331 | NR_029436 |
| Mfng | 136.8498088 | -0.052970887 | 0.113473793 | -0.466811642 | 0.640634646 | 0.809078173 | | 1810030O07Rik | 1013.083951 | -0.0147048329386393 | 0.0604359872168653 | -0.243312529765969 | 0.807763298659199 | 0.955716697308331 | NM_175141 |
| Txnrd1 | 2598.766296 | -0.025895551 | 0.055482052 | -0.466737438 | 0.640687741 | 0.809078173 | | 2810468N07Rik | 999.9604315 | -0.0200150243774063 | 0.0821593894252914 | -0.243612136329302 | 0.807531228009838 | 0.955716697308331 | NM_045176 |
| S100a10 | 748.0651074 | 0.047743427 | 0.102320977 | 0.466604486 | 0.640782877 | 0.80908564 | | 5031439G07Rik | 6193.871752 | 0.0159620691817365 | 0.065511971478889 | 0.243651180408762 | 0.807500986311694 | 0.955716697308331 | NM_001033273 |
| Orc3 | 1425.807233 | 0.032510318 | 0.069702206 | 0.466417347 | 0.640916797 | 0.809188791 | | Atf6b | 2405.003685 | 0.0160514258526917 | 0.0661532947804719 | 0.242639854990714 | 0.808284403828137 | 0.955716697308331 | NM_017406 |
| Kank4 | 269.9284852 | 0.061530062 | 0.132010896 | 0.466098357 | 0.6411451 | 0.809345134 | | Bccip | 1456.883299 | -0.0124682136157616 | 0.0511481435691351 | -0.24376668918411 | 0.807411520359678 | 0.955716697308331 | NM_025392 |
| Rbbp8 | 180.146347 | 0.051116358 | 0.109655671 | 0.46615334 | 0.641105745 | 0.809345134 | | Bicd1 | 908.8114037 | 0.0189718112151165 | 0.0782772888231492 | 0.242366738812062 | 0.808496004735359 | 0.955716697308331 | NM_001112796,NM_009753 |
| Gm11762 | 41.57431306 | -0.072042719 | 0.154622865 | -0.465925389 | 0.641268907 | 0.809435475 | | Ccnt1 | 740.4188523 | 0.0183278720906315 | 0.0752364753670596 | 0.243603544706401 | 0.807537882713382 | 0.955716697308331 | NM_009833 |
| 1700008J07Rik | 64.48689863 | 0.069664507 | 0.149982351 | 0.464484699 | 0.642300519 | 0.809467217 | | Cep104 | 1581.591695 | 0.0159511155143588 | 0.0658225328721011 | 0.242335182472439 | 0.808520454391718 | 0.955716697308331 | NM_177673 |
| 5430416O09Rik | 35.48401167 | -0.075149949 | 0.161694505 | -0.464765014 | 0.642099745 | 0.809467217 | | Dnajc2 | 1148.418212 | 0.016270544516274 | 0.0670040052097875 | 0.242829431842641 | 0.80813753452665 | 0.955716697308331 | NM_009584 |
| AI429214 | 106.2603508 | 0.063301095 | 0.136184494 | 0.464818666 | 0.642061319 | 0.809467217 | | Eif1ad | 1058.222509 | -0.0146791538204898 | 0.0603165026656002 | -0.243368782534893 | 0.807719724836425 | 0.955716697308331 | NM_027236 |
| Ap1s3 | 113.0627882 | -0.05548598 | 0.119177306 | -0.465575051 | 0.641519705 | 0.809467217 | | Fam185a | 708.0033206 | 0.0170629661207871 | 0.0703978035504938 | 0.24237923997939 | 0.808486318960105 | 0.955716697308331 | NM_177869 |
| Atg4d | 883.3401051 | 0.041653789 | 0.089686055 | 0.464439968 | 0.64233256 | 0.809467217 | | Kpna4 | 2943.215879 | 0.0180118323981035 | 0.0739035068004906 | 0.243720943401618 | 0.807446951911755 | 0.955716697308331 | NM_008467 |
| Cog6 | 1972.273183 | -0.027612334 | 0.059434837 | -0.464581632 | 0.642231088 | 0.809467217 | | Musk | 79.78569996 | -0.0224785646678213 | 0.0927252352096812 | -0.242421220253474 | 0.808453793292348 | 0.955716697308331 | NM_001165996,NM_001037128,NM_001037129,NM_001037130,NM_001037127,NM_010944 |
| Ctbs | 303.0671159 | 0.043223427 | 0.093044433 | 0.464546086 | 0.642256549 | 0.809467217 | | Nmt2 | 1994.73006 | -0.0151151936114553 | 0.0623428332525801 | -0.24245278603583 | 0.808429336829891 | 0.955716697308331 | NM_008708 |
| Exosc2 | 659.4288818 | 0.032920093 | 0.070725644 | 0.465461901 | 0.641600714 | 0.809467217 | | Pdf | 1072.684113 | -0.0147248173339575 | 0.0606118285029389 | -0.242936365683862 | 0.808054693541179 | 0.955716697308331 | NM_026513 |
| Fgf12 | 6128.982329 | -0.037239721 | 0.08019612 | -0.464358138 | 0.642391177 | 0.809467217 | | Phf14 | 1530.776321 | 0.0140469663016039 | 0.0576954595246132 | 0.243467448172614 | 0.807643299123617 | 0.955716697308331 | NM_029404,NM_001168382 |
| Kcnab1 | 4747.755296 | -0.044038515 | 0.094836166 | -0.464364147 | 0.642386873 | 0.809467217 | | Rps27 | 28.44165344 | 0.0211522189613089 | 0.0868858622506105 | 0.243448340309936 | 0.807658099797067 | 0.955716697308331 | NR_033727,NM_027015 |
| LOC106740 | 451.7221732 | -0.036432101 | 0.078422138 | -0.464563987 | 0.642243727 | 0.809467217 | | Slc25a36 | 1419.034993 | -0.01423807878659685 | 0.0584665664961015 | -0.243525295211544 | 0.807598492065569 | 0.955716697308331 | NM_138756 |
| Lrrc18 | 22.21560156 | -0.078098068 | 0.168101648 | -0.464588357 | 0.642226272 | 0.809467217 | | Timm50 | 870.6171966 | 0.0160868895121432 | 0.0663059191051544 | 0.24261619066966 | 0.808302737565636 | 0.955716697308331 | NM_025616 |
| Mmachc | 608.8961714 | -0.034772388 | 0.074882599 | -0.464358725 | 0.642390757 | 0.809467217 | | Ubr1 | 2769.435415 | 0.0150164053925589 | 0.0619707619100744 | 0.242314358089531 | 0.808536589096935 | 0.955716697308331 | NM_009461 |
| P2ry2 | 28.73660034 | 0.077240212 | 0.166331618 | 0.4643748 | 0.642379241 | 0.809467217 | | Usf2 | 3627.870295 | 0.0126881425959279 | 0.0522019264558551 | 0.243058895664659 | 0.807959772968602 | 0.955716697308331 | NM_011680 |
| Pla2g15 | 1118.920937 | -0.029002057 | 0.062279892 | -0.465672882 | 0.641449666 | 0.809467217 | | Cyb5rl | 117.9153398 | -0.0226951060462685 | 0.0937282820142629 | -0.242137224309893 | 0.80867383541361 | 0.95572338566578 | NM_175471 |
| Ppm1g | 2569.78848 | 0.029259172 | 0.062871196 | 0.465382777 | 0.641657365 | 0.809467217 | | Macf1 | 13108.53187 | -0.0198247864002685 | 0.0818600373747004 | -0.242179053859015 | 0.808641424624269 | 0.95572338566578 | NM_001199136,NM_001199137 |
| Rbm24 | 646.4649537 | -0.036240709 | 0.077871031 | -0.465393978 | 0.641649346 | 0.809467217 | | 1110059G10Rik | 410.793823 | 0.0196801383115124 | 0.0815224501113907 | 0.241407591217166 | 0.809239229775311 | 0.955725085142442 | NM_025419 |
| 3-Sep | 23623.84203 | 0.034826124 | 0.074864522 | 0.465188629 | 0.641796381 | 0.809467217 | | Bmp5 | 45.19765562 | -0.022415707292352 | 0.0928038762259651 | -0.241538480976514 | 0.809137795660727 | 0.955725085142442 | NM_007555 |
| Smn1 | 484.3892752 | 0.042986331 | 0.092375547 | 0.46534318 | 0.641685718 | 0.809467217 | | Ccbl1 | 462.604712 | 0.0192655325916894 | 0.0797230053285184 | 0.24165587476665 | 0.809046823088252 | 0.955725085142442 | NM_172404 |
| Spock1 | 12375.56968 | 0.036230737 | 0.07789592 | 0.465117261 | 0.641847485 | 0.809467217 | | Gjb1 | 586.3641472 | 0.0222957934989123 | 0.092263332140541 | 0.24165389414887 | 0.809048357917156 | 0.955725085142442 | NM_008124 |
| Txndc11 | 1126.17895 | 0.037741768 | 0.081223483 | 0.464665714 | 0.642170865 | 0.809467217 | | Herc4 | 1376.768933 | -0.0170090077940101 | 0.0705426869148271 | -0.241116528699094 | 0.80946480259586 | 0.955725085142442 | NM_026101,NM_030114 |
| Idh3b | 7487.130642 | -0.017931151 | 0.038642692 | -0.464024371 | 0.642630285 | 0.809702662 | | Itgb3bp | 124.6024209 | -0.0220099419804473 | 0.0912513949171559 | -0.241201156436341 | 0.809399214649817 | 0.955725085142442 | NM_026348 |
| Litaf | 749.9752729 | 0.061288984 | 0.13210472 | 0.463942422 | 0.642688998 | 0.809704323 | | Kansl3 | 4135.203094 | 0.0112481623729331 | 0.0466150191790329 | 0.241299104259351 | 0.809323305080615 | 0.955725085142442 | NM_172652 |
| Zfp874a | 506.9761365 | 0.039051574 | 0.084185255 | 0.463876652 | 0.642736122 | 0.809704323 | | Prr16 | 310.0110801 | 0.0226154271556237 | 0.0937577584292168 | 0.241211261174695 | 0.809391383393961 | 0.955725085142442 | NM_001081224 |
| Ppp1r12c | 4117.033617 | 0.034214404 | 0.073779011 | 0.463741693 | 0.642832822 | 0.809760305 | | Qars | 2318.228202 | 0.0124550268120411 | 0.0515003093441925 | 0.241843728137636 | 0.808901254280596 | 0.955725085142442 | NM_001168270,NM_133794 |
| Tdrd3 | 760.0894646 | 0.043109963 | 0.092996371 | 0.463566076 | 0.642958664 | 0.809852983 | | Slc31a1 | 1016.882345 | 0.0201082731980637 | 0.0833081568012395 | 0.241372201356452 | 0.809266655994619 | 0.955725085142442 | NM_175090 |
| Zfp280b | 939.3192414 | 0.032622734 | 0.070399926 | 0.463391597 | 0.643083701 | 0.809944632 | | Syndig1l | 2103.028786 | -0.0230018794480628 | 0.0950922352633701 | -0.241890196232701 | 0.808865246869429 | 0.955725085142442 | NM_001033334 |
| Kti12 | 353.5307854 | -0.04043999 | 0.087332835 | -0.463055957 | 0.643324259 | 0.810181749 | | Ypel5 | 3838.501272 | -0.0141375360831228 | 0.0584556720443545 | -0.241850543988198 | 0.808895972757648 | 0.955725085142442 | NM_027166 |
| Kifc1 | 43.49081701 | -0.071098126 | 0.15363888 | -0.462761286 | 0.643535484 | 0.810250186 | | Nhsl1 | 1087.797462 | -0.016522235915916 | 0.0685666303793615 | -0.240966135049991 | 0.809581363552432 | 0.955785020071735 | NM_001163592,NM_173390 |
| Ogfod3 | 184.4510648 | 0.050951897 | 0.110083739 | 0.462846716 | 0.643474243 | 0.810250186 | | 4930429B21Rik | 427.4943965 | -0.0176563070425829 | 0.0735636649882024 | -0.24001396675131 | 0.810319429132821 | 0.955876687722809 | NR_027966 |
| Shisa7 | 7628.324715 | 0.032565087 | 0.070367398 | 0.462786568 | 0.64351736 | 0.810250186 | | C030006K11Rik | 451.1963801 | -0.0189551139773829 | 0.0788418469435604 | -0.240419456319334 | 0.810005096503687 | 0.955876687722809 | NM_145472,NM_176828 |
| Tmem251 | 296.2046812 | 0.055662039 | 0.120338157 | 0.462546884 | 0.643689189 | 0.810377858 | | Cenpc1 | 933.9866073 | -0.0155477494723801 | 0.0647218656684878 | -0.24022406201974 | 0.810156560952796 | 0.955876687722809 | NM_007683 |
| Ascc1 | 527.2957859 | 0.034356631 | 0.074301184 | 0.462396822 | 0.643796778 | 0.810447455 | | Dpy19l1 | 5085.751615 | 0.0141609961101119 | 0.058877538320192 | 0.240516103664195 | 0.809930180689621 | 0.955876687722809 | NM_172920 |
| Kcnh5 | 839.7195182 | -0.041034811 | 0.088798372 | -0.462112201 | 0.644000861 | 0.810638504 | | Harbi1 | 235.111235 | -0.0199311527176014 | 0.0830474745621658 | -0.239997095910265 | 0.810332507952059 | 0.955876687722809 | NM_178724 |
| Atf6b | 2132.13656 | -0.038069855 | 0.082441046 | -0.461782779 | 0.644237101 | 0.810795483 | | Hfe | 170.5530703 | -0.0228329770385524 | 0.0950815652040015 | -0.240140946245083 | 0.810220992239962 | 0.955876687722809 | NM_010424 |
| Cacul1 | 2645.685604 | -0.03651697 | 0.079126606 | -0.461500519 | 0.644439549 | 0.810795483 | | Hist2h2be | 309.7576301 | -0.0194197933965255 | 0.0808786757060549 | -0.240110180180308 | 0.810244842396811 | 0.955876687722809 | NM_178214 |
| Ccng2 | 1612.423112 | -0.045487203 | 0.098526465 | -0.461674977 | 0.644314418 | 0.810795483 | | Pgbd5 | 7153.614185 | -0.0154477562954873 | 0.0643838538986245 | -0.239932146960487 | 0.810382858963678 | 0.955876687722809 | NM_171824 |
| Stambpl1 | 546.95164 | -0.031224655 | 0.067650664 | -0.461557259 | 0.644398851 | 0.810795483 | | Sh3gl2 | 14822.09849 | 0.0139558334859481 | 0.0581652468231951 | 0.239934226160334 | 0.810381247073022 | 0.955876687722809 | NM_019535 |
| Tprgl | 5487.78155 | 0.039783933 | 0.086202298 | 0.461518247 | 0.644426833 | 0.810795483 | | Tbc1d15 | 1280.815689 | 0.0159129647534413 | 0.0661609494794562 | 0.240518990108847 | 0.809927943300035 | 0.955876687722809 | NM_025706 |
| Trim35 | 6438.632577 | -0.025194698 | 0.054558144 | -0.461795362 | 0.644228077 | 0.810795483 | | Tnc | 181.5854343 | 0.0228082622465939 | 0.0947316783148057 | 0.240767002678862 | 0.809735705422722 | 0.955876687722809 | NM_011607 |
| Rbm41 | 352.0077338 | 0.038024619 | 0.082418553 | 0.46135995 | 0.64454038 | 0.8108565 | | Gm20754 | 31.73011875 | 0.0210636189084899 | 0.0879182403262617 | 0.23958189825369 | 0.81065439922501 | 0.956073262674805 | NR_040557 |
| Arl1 | 3060.159073 | -0.029576671 | 0.064158992 | -0.460990274 | 0.644805583 | 0.810992589 | | Man1b1 | 4020.703192 | 0.00936055681843211 | 0.039083804448001 | 0.239499632920481 | 0.810718181067677 | 0.956073262674805 | NM_001029983 |
| Mir453 | 8.11833049 | 0.071683246 | 0.155453174 | 0.461124367 | 0.644709381 | 0.810992589 | | Pm20d1 | 112.2183872 | -0.0226935755649898 | 0.0947688055347344 | -0.239462504955517 | 0.810746967480948 | 0.956073262674805 | NM_178079 |
| Prkrir | 3042.46139 | -0.031624395 | 0.068592867 | -0.46104495 | 0.644766356 | 0.810992589 | | Dpf2 | 2410.316016 | -0.0145010652876293 | 0.06059394738556 | -0.239315408771092 | 0.8108610180341 | 0.956130136209378 | NM_011262 |
| Rabgap1 | 4390.546154 | 0.031322135 | 0.067974765 | 0.460790625 | 0.644948828 | 0.811106911 | | 2700062C07Rik | 459.4380699 | 0.0179290596660128 | 0.074977375053439 | 0.239126265133104 | 0.811007675845162 | 0.956135608054323 | NM_026529 |
| 2010003O02Rik | 48.97811894 | -0.070286749 | 0.15259041 | -0.46062363 | 0.645068655 | 0.811191765 | | Acan | 235.8600573 | 0.0206139876946589 | 0.0862856322086504 | 0.238904058149698 | 0.811179978719721 | 0.956135608054323 | NM_007424 |
| Clec2d | 152.480538 | -0.062926963 | 0.136849732 | -0.459825251 | 0.645641657 | 0.811715225 | | Acvr1b | 5235.688607 | 0.0173664070671428 | 0.0726790823287034 | 0.238946427372326 | 0.811147124235923 | 0.956135608054323 | NM_007395 |
| Fbxo42 | 1831.312051 | 0.031972883 | 0.069519861 | 0.459910056 | 0.645580782 | 0.811715225 | | Cenpw | 64.5576932 | 0.0223131658581919 | 0.0934304100302218 | 0.238821234445768 | 0.811244203894539 | 0.956135608054323 | NM_001109747 |
| Fhl2 | 1283.653666 | -0.040536104 | 0.088169563 | -0.459751677 | 0.645694472 | 0.811715225 | | Dcdc2a | 446.3103645 | -0.0196247126445876 | 0.0821430923489832 | -0.23890886115186 | 0.811176254297721 | 0.956135608054323 | NM_001195617,NM_177577 |
| Scml2 | 30.08057921 | 0.073288376 | 0.159393222 | 0.459796062 | 0.64566261 | 0.811715225 | | E2f1 | 693.4371017 | 0.0173271870506827 | 0.0726449199725258 | 0.238518908923513 | 0.81147865129727 | 0.956135608054323 | NM_007891 |
| AA388235 | 141.9485314 | -0.059199505 | 0.128928199 | -0.459166463 | 0.646114631 | 0.812045758 | | Rab24 | 1944.884119 | 0.014419586734876 | 0.060434061651125 | 0.238600324732726 | 0.811415513299465 | 0.956135608054323 | NM_009000 |
| Hint3 | 355.0223802 | -0.045625599 | 0.099351294 | -0.459235078 | 0.646065363 | 0.812045758 | | Romo1 | 790.7697913 | -0.0166472359891807 | 0.0697876327991956 | -0.238541347821337 | 0.811461249799189 | 0.956135608054323 | NM_001164217,NM_001164216,NM_025946 |
| Zbtb3 | 148.6405204 | -0.05184315 | 0.112890686 | -0.459233192 | 0.646066718 | 0.812045758 | | Slc13a4 | 588.4630619 | -0.0156649379830943 | 0.0656586014576668 | -0.238581657777073 | 0.811429989423878 | 0.956135608054323 | NM_172892 |
| Hmgcll1 | 549.440076 | 0.036707509 | 0.07998431 | 0.458933864 | 0.64628166 | 0.812189799 | | Tbck | 794.6266304 | -0.015769735789838 | 0.0661314144485617 | -0.238460585204994 | 0.811523882121833 | 0.956135608054323 | NM_173032,NM_001163455 |
| S100a4 | 51.76095752 | -0.074293386 | 0.161955619 | -0.458726822 | 0.646430351 | 0.812310775 | | Gpr81 | 41.5017933 | 0.0206927781055347 | 0.0868175829076992 | 0.238347779476127 | 0.811611366259697 | 0.956161127841716 | NM_175520 |
| 1700049G17Rik | 235.2861814 | -0.043938461 | 0.095812596 | -0.458587526 | 0.646530396 | 0.812370607 | | Nek11 | 77.1371579 | -0.0226546649126631 | 0.0951017244040708 | -0.238215080269284 | 0.811714281381774 | 0.956204821292 | NM_172461 |
| Dmp1 | 287.5943455 | -0.061533929 | 0.13425517 | -0.458335642 | 0.646711321 | 0.812466165 | | 3110045C21Rik | 22.98764411 | 0.0204345202628337 | 0.0858228609335379 | 0.238101131103732 | 0.811802657470005 | 0.956219756241722 | NR_040438 |
| Rasgrp3 | 818.928538 | -0.053778524 | 0.11732108 | -0.458387562 | 0.646674026 | 0.812466165 | | Ankmy2 | 2044.826234 | 0.0137971883275477 | 0.0579977210137078 | 0.237891904826514 | 0.811964934316682 | 0.956219756241722 | NM_146033 |
| Cdc25c | 16.86389724 | -0.075574705 | 0.165025781 | -0.457956957 | 0.646983365 | 0.812742035 | | Ap3b2 | 7561.098576 | -0.0131564849252834 | 0.0554503394821294 | -0.237266084358662 | 0.812450371687478 | 0.956219756241722 | NM_021492 |
| Ccdc105 | 12.42612336 | -0.074603093 | 0.162967989 | -0.457777589 | 0.647112239 | 0.812772132 | | Dhrs1 | 2454.266828 | 0.0138091787504049 | 0.0581450299339307 | 0.237495427659011 | 0.812272465950449 | 0.956219756241722 | NM_026819 |
| Tec | 93.23658821 | -0.060406851 | 0.131940251 | -0.457834894 | 0.647071065 | 0.812772132 | | Eif4e3 | 620.1677231 | 0.0173221813751451 | 0.0729839047616152 | 0.237342485740163 | 0.812391104681251 | 0.956219756241722 | NM_025829 |
| Ahdc1 | 2918.780824 | 0.045619895 | 0.099764807 | 0.457274426 | 0.647473809 | 0.81311793 | | Gtf3c6 | 601.7845856 | 0.0150679562112793 | 0.0633945819684292 | 0.23768523655197 | 0.812125235094664 | 0.956219756241722 | NM_026113 |
| Cdkn2aipnl | 1485.392265 | 0.035777928 | 0.078258651 | 0.457175378 | 0.647544994 | 0.81311793 | | Itm2b | 20904.76061 | 0.0114369761268169 | 0.0481538905688111 | 0.237508869827946 | 0.812262038917944 | 0.956219756241722 | NM_008410 |
| Fam132a | 418.1455108 | -0.052599448 | 0.115052543 | -0.45717762 | 0.647543383 | 0.81311793 | | Mob1a | 373.9635971 | -0.0181671738045982 | 0.0763565078181876 | -0.237925676850702 | 0.811938740035172 | 0.956219756241722 | NM_145571 |
| Parg | 1424.952453 | -0.029365409 | 0.064262247 | -0.456962056 | 0.647698319 | 0.813244551 | | Nkd2 | 138.6003437 | 0.0222078880657388 | 0.0935040975644119 | 0.23750711085619 | 0.81226340334286 | 0.956219756241722 | NM_028186 |
| D630039A03Rik | 10.10624168 | 0.073131712 | 0.160078831 | 0.456848116 | 0.647780219 | 0.813278284 | | Pcdhb5 | 291.3369402 | 0.0214783436847448 | 0.0904716335396633 | 0.237404176805635 | 0.812343249723994 | 0.956219756241722 | NM_053130 |
| Eid2 | 1070.31375 | -0.028622927 | 0.06267827 | -0.456664276 | 0.647912372 | 0.813278284 | | Zcchc10 | 233.2471755 | 0.0205625967634874 | 0.086665040000387 | 0.237265185170347 | 0.81245106922314 | 0.956219756241722 | NM_026479 |
| Ntpcr | 535.700454 | -0.039067674 | 0.085538126 | -0.456728198 | 0.64786642 | 0.813278284 | | 1110008F13Rik | 1327.275663 | 0.0135381388035658 | 0.0579600027842022 | 0.233577262823317 | 0.815313181142326 | 0.956264994427604 | NM_026124 |
| S1pr3 | 456.3812282 | 0.040779623 | 0.089305108 | 0.456632597 | 0.647935145 | 0.813278284 | | 1700096K18Rik | 49.56198769 | -0.0223388553833948 | 0.0945442183546376 | -0.2362794443929 | 0.813215836413707 | 0.956264994427604 | NR_027388 |
| Pigl | 235.2228964 | 0.049615178 | 0.108711873 | 0.456391532 | 0.648108454 | 0.813429922 | | 4930427A07Rik | 79.52496722 | -0.0221262409998739 | 0.0951012970435129 | -0.232659718507837 | 0.816025650837288 | 0.956264994427604 | NM_134041 |
| Thra | 21059.7299 | -0.045967965 | 0.10074598 | -0.456275921 | 0.648191576 | 0.813468353 | | Ablim1 | 6891.982326 | 0.0105273735354199 | 0.0453456922159376 | 0.232158183522445 | 0.816415155191276 | 0.956264994427604 | NM_001103177,NM_178688,NM_001103178 |
| Gm5506 | 30.76186486 | 0.074593729 | 0.163577823 | 0.456013701 | 0.648380126 | 0.813573183 | | Acd | 1482.848626 | -0.0153192771269553 | 0.0646237381764859 | -0.23705340420139 | 0.812615360175439 | 0.956264994427604 | NM_001012638 |
| Mcat | 528.371208 | 0.038626753 | 0.084693925 | 0.456074666 | 0.648336287 | 0.813573183 | | Aff3 | 2412.274394 | 0.0145203281453947 | 0.0616037189290558 | 0.235705382691532 | 0.813661292784495 | 0.956264994427604 | NM_010678 |
| Zc3h6 | 789.3637171 | 0.043587911 | 0.095615814 | 0.455865083 | 0.648487 | 0.813641394 | | Alox5 | 33.5229515 | 0.0208827737617792 | 0.0891790991162404 | 0.234166682201617 | 0.8148555797276 | 0.956264994427604 | NM_009662 |
| Zfp865 | 883.9341585 | 0.034379732 | 0.075437343 | 0.455738912 | 0.648577737 | 0.813689349 | | Arsa | 1830.285942 | -0.0147389561882501 | 0.0626008505253947 | -0.235443385585809 | 0.813864615501618 | 0.956264994427604 | NM_009713 |
| Trim29 | 8.766087716 | -0.067707983 | 0.148618369 | -0.455582866 | 0.648689676 | 0.813764257 | | Atf5 | 621.403412 | -0.0173965276851521 | 0.0735193068969466 | -0.236625295033544 | 0.812947494815648 | 0.956264994427604 | NR_033136,NM_030693 |
| Foxn3 | 544.0779441 | 0.046529371 | 0.102177047 | 0.455379875 | 0.648835971 | 0.81388152 | | BC051226 | 109.1446425 | 0.0223755068255489 | 0.0949799385508641 | 0.235581399261132 | 0.813757508496787 | 0.956264994427604 | NR_045146 |
| Rassf9 | 39.91020945 | 0.074496909 | 0.163673543 | 0.455155472 | 0.648997392 | 0.814018101 | | Bmp6 | 358.350517 | 0.0221141009118737 | 0.0951016972684942 | 0.232531085638151 | 0.816125545944902 | 0.956264994427604 | NM_007556 |
| Pmfbp1 | 18.42114727 | -0.074498483 | 0.168101267 | -0.455073768 | 0.649056169 | 0.814025926 | | Ccdc104 | 3216.753499 | 0.0151224587152444 | 0.0646510722828226 | 0.233908861543544 | 0.815055733518363 | 0.956264994427604 | NM_025740 |
| Tmem176b | 965.4574841 | 0.038017261 | 0.083564418 | 0.454945561 | 0.649148404 | 0.814075708 | | Ccna1 | 36.88054035 | -0.0209530949422835 | 0.0897544770706064 | -0.233449022557398 | 0.815412750059041 | 0.956264994427604 | NM_007628 |
| Fgd2 | 205.1216026 | 0.069778304 | 0.153454823 | 0.454715615 | 0.649313845 | 0.814217281 | | Ctage5 | 2015.471967 | 0.0144895707185661 | 0.061804667164839 | 0.234441367994443 | 0.814642346329998 | 0.956264994427604 | NM_001165253,NM_001165254,NM_146034 |
| Nubp1 | 504.2843087 | -0.039714995 | 0.087378881 | -0.454514802 | 0.649458341 | 0.814332567 | | E130317F20Rik | 164.3759881 | 0.0215332692398942 | 0.0914579903113464 | 0.235443373603547 | 0.8138638487274 | 0.956264994427604 | NR_029447 |
| Amer1 | 482.1731231 | -0.032362914 | 0.071256227 | -0.454176649 | 0.649701689 | 0.814571773 | | Ep300 | 3234.413906 | 0.0163086391640709 | 0.0697831871976125 | 0.233704418198726 | 0.815214457506156 | 0.956264994427604 | NM_177821 |
| Poldip2 | 3466.790439 | -0.025661248 | 0.056548882 | -0.45378878 | 0.64998086 | 0.81485585 | | Fam102a | 4740.45706 | -0.0105452297707398 | 0.0450844338065394 | -0.233899571989529 | 0.815062945498857 | 0.956264994427604 | NM_153560 |
| Lrrc8d | 1805.216571 | 0.032432912 | 0.071524254 | 0.453453339 | 0.650222336 | 0.815026686 | | Fam53c | 3181.347342 | -0.0137544014712309 | 0.0590820845006888 | -0.232801560531781 | 0.815915501087449 | 0.956264994427604 | NM_175104 |
| Rdh11 | 833.1999514 | -0.032153673 | 0.070906748 | -0.453464216 | 0.650214505 | 0.815026686 | | Frmd3 | 212.3332619 | -0.0203746304137989 | 0.0873285578883868 | -0.233310052363848 | 0.81552065332256 | 0.956264994427604 | NM_172869,NM_001163732 |
| 1700101I11Rik | 59.12270159 | 0.06277025 | 0.138721497 | 0.452491153 | 0.650915193 | 0.815240171 | | Gclm | 1237.698375 | 0.0149793115483276 | 0.0645144141978206 | 0.232185500474306 | 0.816393939008648 | 0.956264994427604 | NM_008129 |
| Arhgef40 | 915.7118899 | 0.03356339 | 0.074153536 | 0.452620227 | 0.650822231 | 0.815240171 | | Golt1b | 1481.261687 | 0.0158814856263468 | 0.0677487758861991 | 0.234417307451041 | 0.814661023527329 | 0.956264994427604 | NM_025872 |
| Bcat2 | 390.3451884 | -0.042566175 | 0.094071781 | -0.452486122 | 0.650918817 | 0.815240171 | | Gtpbp5 | 430.0149818 | -0.019200784739257 | 0.0817761846116148 | -0.23479677892101 | 0.814366467913911 | 0.956264994427604 | NM_028422,NM_181424,NM_001083329,NM_001083328 |
| Cers3 | 9.894783385 | -0.069910092 | 0.154359736 | -0.452903679 | 0.650618102 | 0.815240171 | | Hint3 | 399.416157 | -0.0168667456541798 | 0.0717410267820771 | -0.23510599737323 | 0.814126463921458 | 0.956264994427604 | NM_025798 |
| Eif2a | 1564.040355 | 0.032534938 | 0.071846286 | 0.452840918 | 0.650663297 | 0.815240171 | | Hsdl2 | 1412.470834 | -0.0162094012225295 | 0.06938214732926 | -0.233624957521221 | 0.815276150559111 | 0.956264994427604 | NM_024255 |
| Ero1l | 2179.01182 | 0.032475295 | 0.071702753 | 0.452915592 | 0.650609523 | 0.815240171 | | Id2 | 3941.778486 | 0.0157566441616724 | 0.0678689953638986 | 0.232162625617025 | 0.816411705152612 | 0.956264994427604 | NM_010496 |
| Gpatch3 | 204.7593501 | 0.051966695 | 0.11483736 | 0.452524294 | 0.650891324 | 0.815240171 | | Ing5 | 814.2228683 | -0.0134399710942293 | 0.057522753945488 | -0.233646169078862 | 0.815259681851629 | 0.956264994427604 | NM_025454 |
| Irak4 | 67.12996675 | 0.062631244 | 0.138402896 | 0.452528426 | 0.650888348 | 0.815240171 | | Klf3 | 657.5651006 | -0.016763516753217 | 0.0714510076955204 | -0.23461553998864 | 0.814507147010312 | 0.956264994427604 | NM_008453 |
| Lpar2 | 47.08296158 | -0.072310639 | 0.159615308 | -0.453030725 | 0.650526617 | 0.815240171 | | Lhpp | 536.3084099 | 0.0172126641804296 | 0.0732047033369475 | 0.235130577624268 | 0.814107386382735 | 0.956264994427604 | NM_029609 |
| Rmdn2 | 206.9554802 | 0.047544689 | 0.105021284 | 0.452714792 | 0.650754127 | 0.815240171 | | Lmnb2 | 930.7598363 | -0.0142935196202505 | 0.0612467967944686 | -0.233375790544877 | 0.815469610543492 | 0.956264994427604 | NM_010722 |
| Slc39a12 | 2032.137974 | 0.033747344 | 0.074623246 | 0.452236339 | 0.651098732 | 0.815376338 | | Lrch1 | 707.4113842 | -0.016452288200666 | 0.0696067236076983 | -0.236360617882127 | 0.813152852851498 | 0.956264994427604 | NM_001252132,NM_001033439 |
| Tpcn1 | 2014.925785 | -0.043742618 | 0.096735242 | -0.45218906 | 0.651132789 | 0.815376338 | | Magi3 | 3046.552019 | 0.0141472844920304 | 0.0605581424088617 | 0.233614901799896 | 0.815283957875325 | 0.956264994427604 | NM_001159354,NM_133853 |
| Myl6b | 530.3553611 | 0.049067344 | 0.108626687 | 0.451706164 | 0.651480677 | 0.815691978 | | Mavs | 405.1361502 | 0.0210363200963226 | 0.090445995953318 | 0.232584315917978 | 0.816084207434975 | 0.956264994427604 | NM_001206385,NM_144888,NM_001206382,NM_001206383 |
| Stx6 | 3167.10966 | -0.024999767 | 0.055347107 | -0.451690593 | 0.651491896 | 0.815691978 | | Mcmbp | 1092.986833 | -0.0158516953907667 | 0.0674117391675219 | -0.23514740290817 | 0.814094327791829 | 0.956264994427604 | NM_145955 |
| Tshz2 | 1339.691874 | 0.063975908 | 0.141658725 | 0.451619964 | 0.651542785 | 0.815691978 | | Mcts1 | 1062.70383 | 0.0166857983898947 | 0.0711773191689679 | 0.234425777546978 | 0.814654448532754 | 0.956264994427604 | NM_026902 |
| Glipr2 | 136.9618425 | 0.074793951 | 0.165707345 | 0.451361713 | 0.651728873 | 0.815859025 | | Mettl20 | 80.3273201 | -0.0221029839828412 | 0.0949849910441763 | -0.232699753296406 | 0.815994560797373 | 0.956264994427604 | NM_001252095,NM_001252094,NM_001252097,NM_001252096,NM_177101 |
| Abr | 24718.65854 | 0.03105003 | 0.068817821 | 0.451191705 | 0.651851388 | 0.815946469 | | Mettl25 | 204.7093029 | 0.0203385492181991 | 0.0872611416125166 | 0.233076817955379 | 0.815701755800749 | 0.956264994427604 | NM_207522 |
| Lrrc47 | 2403.8609 | 0.032174108 | 0.071361493 | 0.450860912 | 0.652089797 | 0.816178957 | | Mfsd4 | 2358.489945 | 0.0147825196667639 | 0.0627849862965237 | 0.235446729206061 | 0.813862020607826 | 0.956264994427604 | NM_001114662,NM_172510 |
| 0610010F05Rik | 1557.816758 | 0.037724703 | 0.083720224 | 0.450604426 | 0.652274676 | 0.816214639 | | Mrps15 | 927.7097035 | -0.0165789186524426 | 0.0705300292211884 | -0.235061842955567 | 0.814160733889552 | 0.956264994427604 | NM_025544 |
| Gm2382 | 107.8433377 | 0.056205258 | 0.124728617 | 0.450620385 | 0.652263172 | 0.816214639 | | N6amt1 | 801.0526072 | 0.0151507154625442 | 0.0646934374403648 | 0.234192463130597 | 0.81483556589543 | 0.956264994427604 | NM_026366,NM_001159331 |
| Trip11 | 1137.316174 | -0.039455817 | 0.087562432 | -0.450602114 | 0.652276343 | 0.816214639 | | Ndp | 357.5170334 | -0.0194128387545864 | 0.0835477949796426 | -0.232356087426564 | 0.816261452749436 | 0.956264994427604 | NM_010883 |
| H2-Ke2 | 1238.780192 | 0.033041088 | 0.073340288 | 0.450517561 | 0.652337295 | 0.81622499 | | Psmd10 | 523.6470725 | 0.016537511379336 | 0.0697729789271723 | 0.237018852192014 | 0.812642164983892 | 0.956264994427604 | NM_001164177,NM_016883 |
| Gm12359 | 13.38375729 | 0.071538028 | 0.158849406 | 0.450351243 | 0.652457195 | 0.816309091 | | Rfxap | 864.8323688 | 0.0153592616379714 | 0.0651787909809123 | 0.235648151903729 | 0.813705705636174 | 0.956264994427604 | NM_133231 |
| 4930519F09Rik | 26.74828577 | 0.075445713 | 0.167621402 | 0.450095942 | 0.652641263 | 0.816472811 | | Rnpep | 1623.769919 | 0.013735907365554 | 0.058289013924383 | 0.235651736764208 | 0.813702923656592 | 0.956264994427604 | NM_001159624,NM_145417 |
| Pradc1 | 472.2996927 | 0.044539604 | 0.098971714 | 0.450023571 | 0.652693446 | 0.816472811 | | Sin3a | 1832.983482 | -0.0148085495581745 | 0.0637950352881459 | -0.23212699062377 | 0.816439381859503 | 0.956264994427604 | NM_001110351,NM_001110350,NM_011378 |
| Zfp408 | 477.5856246 | 0.043997839 | 0.097826035 | 0.449755927 | 0.65288644 | 0.8166483 | | Skap2 | 496.2721513 | -0.0156819803749235 | 0.0664218147329 | -0.236096837130768 | 0.81335752816963 | 0.956264994427604 | NM_018773 |
| Pcdhgb1 | 272.4836591 | 0.0589293 | 0.131071675 | 0.449595998 | 0.653001774 | 0.81672663 | | Slc26a11 | 210.6901297 | -0.0206320502470036 | 0.0886734535051052 | -0.232674486348 | 0.816014182459255 | 0.956264994427604 | NM_178743 |
| Sfrp4 | 50.09720636 | 0.074228866 | 0.165263391 | 0.449154925 | 0.653319901 | 0.817058564 | | Slc6a12 | 35.50689461 | -0.0179490459139091 | 0.0765254786801569 | -0.234549933218036 | 0.814558072980746 | 0.956264994427604 | NM_133661 |
| Tmem106b | 6553.444947 | 0.03241647 | 0.072221733 | 0.448846465 | 0.653542417 | 0.817270881 | | Snrpg | 523.4707825 | -0.0165445920062571 | 0.07009600298758 | -0.236027609294478 | 0.813411246215824 | 0.956264994427604 | NM_026506 |
| Ntsr2 | 2971.02922 | 0.042239034 | 0.094154842 | 0.44861245 | 0.653711251 | 0.817350075 | | Sorbs3 | 1544.346509 | 0.0185997181032641 | 0.0788283662940803 | 0.23595209412149 | 0.813469843986037 | 0.956264994427604 | NM_001271407,NM_011366,NM_001271409,NM_001271408 |
| Stx11 | 9.107791921 | 0.072160704 | 0.160849942 | 0.44862126 | 0.653704989 | 0.817350075 | | Stpg1 | 68.19018347 | 0.0222174920401145 | 0.0947224016359439 | 0.234553723901283 | 0.814555130515802 | 0.956264994427604 | NM_030189 |
| Cnpy3 | 2088.929784 | -0.036423744 | 0.081249169 | -0.448296817 | 0.653938998 | 0.817485766 | | Tgfbr3 | 583.1386451 | 0.0202704801890786 | 0.0856867150533361 | 0.23656502850484 | 0.812994253354206 | 0.956264994427604 | NM_011578 |
| Gm10653 | 27.69384023 | -0.072514605 | 0.161816506 | -0.448128601 | 0.654060387 | 0.817485766 | | Triobp | 375.6468225 | -0.0173682607334272 | 0.0744379833461655 | -0.233325245428239 | 0.815508856513003 | 0.956264994427604 | NM_138579,NM_001024716,NM_001039155,NM_001039156 |
| Hdgf | 5879.122691 | 0.021998027 | 0.049092169 | 0.448096459 | 0.654083583 | 0.817485766 | | Ubr4 | 8426.739225 | 0.0185950219922754 | 0.0794904777972753 | 0.23392766653336 | 0.81504113414802 | 0.956264994427604 | NM_001160319 |
| Lace1 | 668.8188396 | -0.033444079 | 0.074594206 | -0.44834687 | 0.653902879 | 0.817485766 | | Uck1 | 1602.709592 | -0.0135996947663327 | 0.0580132654336832 | -0.234423879860355 | 0.814655921629368 | 0.956264994427604 | NM_011675 |
| Zcrb1 | 1089.552523 | -0.039575854 | 0.088291846 | -0.44823906 | 0.653980676 | 0.817485766 | | Vps8 | 2868.748628 | -0.0116145393915542 | 0.0495930706082933 | -0.234196819214738 | 0.814832184262855 | 0.956264994427604 | NM_001081366 |
| Gm16291 | 9.455070472 | 0.07212736 | 0.16101337 | 0.447958827 | 0.654182912 | 0.817543961 | | Wdr63 | 24.662043 | -0.0150947558064506 | 0.0647768677562042 | -0.23302694818869 | 0.81574048008868 | 0.956264994427604 | NM_172864 |
| Sema3e | 1080.066587 | -0.044398145 | 0.099236718 | -0.447396349 | 0.654588911 | 0.817985369 | | Xkr8 | 283.5597063 | 0.020391384532388 | 0.086647611324892 | 0.235336949519923 | 0.8139472187384 | 0.956264994427604 | NM_201368 |
| Paqr7 | 2562.656074 | -0.032820397 | 0.073372063 | -0.447314621 | 0.654647856 | 0.817993125 | | Zbtb25 | 386.479464 | 0.0175789914190179 | 0.075726745642791 | 0.2321371566915 | 0.816431486134463 | 0.956264994427604 | NM_001172104,NM_028356 |
| Svep1 | 76.95649176 | -0.072267353 | 0.161631741 | -0.447111147 | 0.654794811 | 0.818110701 | | Zc3h4 | 2980.322011 | -0.0129829951036341 | 0.0555413253030966 | -0.233753786622557 | 0.815176128573734 | 0.956264994427604 | NM_198631 |
| Cul5 | 2491.067696 | -0.027085456 | 0.060595852 | -0.446985323 | 0.65488567 | 0.818111807 | | Zfp62 | 1387.853984 | -0.0171813046675918 | 0.0733524881317659 | -0.234229337070726 | 0.814806940726985 | 0.956264994427604 | NM_009562,NM_001024846 |
| Tob1 | 1055.776651 | -0.041060821 | 0.091866129 | -0.446963658 | 0.6549013 | 0.818111807 | | Ddrgk1 | 2009.506883 | -0.0151688671978344 | 0.0653755099376054 | -0.232026751490148 | 0.816517236030753 | 0.956279075270718 | NM_029832 |
| Ccdc74a | 536.1031166 | 0.041413375 | 0.0926869 | 0.446809364 | 0.655012709 | 0.818185015 | | As3mt | 477.9433224 | 0.020465186357783 | 0.0884158900907187 | 0.231465026668677 | 0.81695355242054 | 0.956302287868477 | NM_020577 |
| Scly | 513.5363888 | -0.048388588 | 0.108325395 | -0.44669662 | 0.655094123 | 0.818185942 | | Bmi1 | 1404.408899 | -0.0168687060323784 | 0.0728860049267636 | -0.231439575393496 | 0.816973322890545 | 0.956302287868477 | NM_007552 |
| Tcta | 1073.94419 | 0.041282172 | 0.092423723 | 0.44666208 | 0.655119065 | 0.818185942 | | Ccdc112 | 358.8377235 | 0.0174511371114971 | 0.0754727709676669 | 0.23122427979984 | 0.817140568484277 | 0.956302287868477 | NM_001160399 |
| Pigh | 384.060621 | -0.035081672 | 0.078600488 | -0.446328928 | 0.655359663 | 0.818420458 | | Fam53a | 655.547041 | 0.0157216728532612 | 0.0679994421068982 | 0.231202968232387 | 0.817157124156499 | 0.956302287868477 | NM_178390 |
| Adam9 | 2494.625411 | 0.038199862 | 0.085605948 | 0.446229067 | 0.655431788 | 0.818444562 | | Mtfr1 | 308.2393662 | 0.0185188428838755 | 0.0800864285999111 | 0.231235718805621 | 0.817131682243805 | 0.956302287868477 | NM_001253391,NM_001253390,NM_026182,NR_045575 |
| Brca2 | 180.8978839 | 0.060614413 | 0.136032602 | 0.445587393 | 0.655895319 | 0.818891384 | | Pabpc5 | 63.39373777 | -0.0218967368504757 | 0.0946239642927373 | -0.231407942101579 | 0.816997895693906 | 0.956302287868477 | NM_053114 |
| Zfp758 | 197.0190867 | -0.069051582 | 0.154948764 | -0.445641385 | 0.655856311 | 0.818891384 | | Phospho2 | 923.0710993 | 0.0189691187176412 | 0.0820677174243939 | 0.231139835649953 | 0.817206168533279 | 0.956302287868477 | NM_028521 |
| Gapdh | 48807.23821 | 0.033139709 | 0.07444133 | 0.445178889 | 0.656190482 | 0.819193886 | | Ptms | 15928.3821 | 0.0123145616620023 | 0.0532599255900708 | 0.231216276131976 | 0.817146786041862 | 0.956302287868477 | NM_026988 |
| Fads6 | 364.153409 | 0.054338956 | 0.122147139 | 0.444864748 | 0.6564175 | 0.819411275 | | Rbm19 | 648.9577913 | 0.0149331232038527 | 0.0645269759718824 | 0.231424500822103 | 0.816985032824808 | 0.956302287868477 | NM_028762 |
| 2900009J06Rik | 22.27651492 | -0.074648082 | 0.167850891 | -0.444728542 | 0.65651594 | 0.819468136 | | Rdm1 | 204.4538513 | 0.020003386100455 | 0.0865497256728124 | 0.231120156013834 | 0.817221456751967 | 0.956302287868477 | NM_025654 |
| Naa38 | 439.5377711 | -0.035635262 | 0.080149293 | -0.444611061 | 0.656600853 | 0.819508105 | | Schip1 | 154.7298963 | 0.0219357502605106 | 0.0949316177129381 | 0.231068960889739 | 0.817261228252738 | 0.956302287868477 | NM_001113420,NM_001113421,NM_013928,NR_036665 |
| Pigw | 134.5914783 | 0.049260062 | 0.110830745 | 0.444462066 | 0.656708549 | 0.819576501 | | Gm10125 | 56.78938564 | -0.0219227954680542 | 0.0949386387090038 | -0.230915418276111 | 0.817380512358648 | 0.956364826622772 | NR_033552 |
| Add2 | 16689.42055 | 0.03121146 | 0.070298841 | 0.443982568 | 0.657055187 | 0.819629267 | | Rab27a | 279.4633469 | 0.0202163454768867 | 0.0877209655993255 | 0.230461957854259 | 0.81773282113511 | 0.956663546249151 | NR_023635 |
| Alms1 | 509.4697043 | -0.041780623 | 0.094121387 | -0.443901477 | 0.657113817 | 0.819629267 | | Sapcd2 | 50.50678842 | -0.0214842554342325 | 0.0932406449007336 | -0.230417276254419 | 0.817767537778859 | 0.956663546249151 | NM_001081085 |
| Cyb561 | 3109.103695 | -0.028977994 | 0.065273367 | -0.443948198 | 0.657080037 | 0.819629267 | | Mpdu1 | 1355.891654 | 0.0144086592319667 | 0.0625664803033469 | 0.230293587910137 | 0.817863642830728 | 0.956698927021997 | NM_011900 |
| Phax | 1353.429561 | 0.038948649 | 0.08774123 | 0.443903616 | 0.65711227 | 0.819629267 | | Bclaf1 | 4679.09344 | 0.0129693946130658 | 0.0564296409194292 | 0.229833016864021 | 0.818221527650306 | 0.956955007593708 | NM_153787,NM_001025392,NM_001025393 |
| Poc1a | 326.96966 | -0.043654704 | 0.098335979 | -0.443934195 | 0.657090161 | 0.819629267 | | Ccdc30 | 426.5518498 | -0.0168602330554316 | 0.073382726361428 | -0.229757517762298 | 0.818280197530046 | 0.956955007593708 | NM_001270446,NM_028506,NM_029286,NM_028698,NM_001270435,NM_028857 |
| Sin3a | 1747.839872 | 0.039176265 | 0.088171367 | 0.444319585 | 0.656811544 | 0.819629267 | | Zfp51 | 208.1054058 | -0.0211813734166222 | 0.0921568829055514 | -0.2298403846659 | 0.818215802232065 | 0.956955007593708 | NM_009558 |
| Tjp2 | 1384.081889 | 0.029367771 | 0.066159816 | 0.443891365 | 0.657121128 | 0.819629267 | | Gm13629 | 232.4427224 | -0.020767777568694 | 0.0904785724146092 | -0.229532551348486 | 0.81845502355626 | 0.957030829757089 | NR_033495 |
| Siglech | 617.0011948 | -0.074283856 | 0.167396317 | -0.443760398 | 0.657215823 | 0.819681395 | | Trappc1 | 1173.285533 | 0.0152953459038241 | 0.0666450718774859 | 0.22950453008651 | 0.818476800087589 | 0.957030829757089 | NM_001024206 |
| 2310010M20Rik | 11.41739012 | -0.071824328 | 0.162022355 | -0.443298878 | 0.657549567 | 0.819772818 | | Uhrf2 | 1907.653354 | 0.0162916151725466 | 0.0710549636384723 | 0.229281873331726 | 0.818649841231853 | 0.957156116688176 | NM_144873 |
| Cdca4 | 252.2988388 | 0.041586973 | 0.093813695 | 0.443293203 | 0.657553671 | 0.819772818 | | Ptpn22 | 36.99774802 | -0.0197171975168022 | 0.0861648105461686 | -0.228831206055254 | 0.819000111310854 | 0.957334482731268 | NM_008979 |
| Chmp6 | 938.9167627 | 0.029873973 | 0.067374425 | 0.443402266 | 0.657474797 | 0.819772818 | | Rapgef6 | 1819.599547 | -0.0118875426124941 | 0.0519264636686249 | -0.228930332871423 | 0.818923064324708 | 0.957334482731268 | NM_175258,NM_001252498,NM_001252494,NM_001252497,NM_001252496 |
| Impa1 | 2415.723805 | 0.041544345 | 0.093658474 | 0.443572735 | 0.657351522 | 0.819772818 | | Stard13 | 605.8827332 | -0.0157154231332615 | 0.0686616886977253 | -0.228881978164661 | 0.818960648128152 | 0.957334482731268 | NM_146258,NM_001163493 |
| Rps19bp1 | 474.0858653 | -0.035015874 | 0.078968702 | -0.443414584 | 0.657465889 | 0.819772818 | | Bmp7 | 357.3878193 | -0.0214013862787811 | 0.0942543016877633 | -0.22706004814165 | 0.82037704965074 | 0.957571904906369 | NM_007557 |
| Dido1 | 3205.551805 | 0.02477542 | 0.055933145 | 0.442947013 | 0.657804061 | 0.819944081 | | Brd4 | 4110.533811 | 0.0117297561294372 | 0.0516734858646388 | 0.226997577832544 | 0.820425625622275 | 0.957571904906369 | NM_198094,NM_020508 |
| Ftsj1 | 776.2437647 | 0.030171656 | 0.068125449 | 0.442883774 | 0.657849804 | 0.819944081 | | Cpne2 | 969.2240376 | 0.0179045910721174 | 0.078608605862979 | 0.227768841280897 | 0.819825951082696 | 0.957571904906369 | NM_153507 |
| Nabp1 | 318.4965549 | 0.046069633 | 0.104016544 | 0.442906784 | 0.65783316 | 0.819944081 | | Fam109a | 452.4747573 | -0.0179758987299414 | 0.078820096235908 | -0.228062379880121 | 0.819597745928076 | 0.957571904906369 | NM_175474 |
| Exoc2 | 2049.909998 | 0.034194599 | 0.07724005 | 0.442705552 | 0.657978726 | 0.820038802 | | Gnaq | 9758.125776 | 0.0147385922251424 | 0.0645597148496997 | 0.228293948625007 | 0.819417728657936 | 0.957571904906369 | NM_008139 |
| Rgs4 | 19846.32822 | -0.056317083 | 0.127232733 | -0.442630464 | 0.658033046 | 0.820040539 | | Lactb | 576.9308083 | -0.0153007938584355 | 0.0672056991289306 | -0.227671076357357 | 0.819901959664584 | 0.957571904906369 | NM_030717 |
| Gria4 | 3805.706546 | 0.035597261 | 0.080464327 | 0.442398045 | 0.658201194 | 0.820133819 | | Lpin3 | 55.32720625 | -0.0213946517115889 | 0.0941500519062217 | -0.2272399353842 | 0.820237175889413 | 0.957571904906369 | NM_001199118,NM_022883 |
| Psd4 | 70.10342554 | 0.062411874 | 0.141081827 | 0.442380672 | 0.658213763 | 0.820133819 | | Mrps27 | 1139.056753 | 0.0138534960761675 | 0.0610773251569378 | 0.226818971534379 | 0.82056451098933 | 0.957571904906369 | NM_173757 |
| Pir | 195.0371162 | -0.045399639 | 0.102651384 | -0.442270114 | 0.658293755 | 0.82016429 | | Nf2 | 4734.377592 | 0.00910908798571073 | 0.040177457626853 | 0.226721363763509 | 0.820640413776272 | 0.957571904906369 | NM_010898,NM_001252253,NM_001252252,NM_001252251,NM_001252250 |
| Slc35a5 | 1722.931907 | -0.03219888 | 0.072827154 | -0.442127397 | 0.65839702 | 0.82016429 | | Nkd1 | 625.3136427 | -0.016805104682588 | 0.0738429974414876 | -0.227578853308389 | 0.819973661205454 | 0.957571904906369 | NM_001163660,NM_027280 |
| Srp19 | 856.6904535 | -0.03617017 | 0.081802648 | -0.442163805 | 0.658370676 | 0.82016429 | | Olfml3 | 1071.666135 | -0.0211527725266349 | 0.093047792786741 | -0.22733234065117 | 0.820165327145405 | 0.957571904906369 | NM_133859 |
| Srp72 | 3866.597356 | -0.025177673 | 0.056958238 | -0.442037425 | 0.658462124 | 0.820179448 | | P4htm | 2996.02623 | 0.0133679179920096 | 0.0586794622788877 | 0.227812550982071 | 0.819791968965584 | 0.957571904906369 | NM_028944 |
| 0610009B22Rik | 425.3823604 | -0.042064852 | 0.095201805 | -0.441849308 | 0.658598255 | 0.820224001 | | Prss16 | 43.87131591 | 0.0198328456442125 | 0.0872298757298987 | 0.22736299322062 | 0.820141493895577 | 0.957571904906369 | NM_019429 |
| Bpgm | 2144.097761 | -0.025306591 | 0.057347561 | -0.441284521 | 0.659007031 | 0.820224001 | | R3hdm4 | 10673.8584 | -0.0139971842084004 | 0.0615882483374011 | -0.227270373590091 | 0.820213508815714 | 0.957571904906369 | NM_177994 |
| Hspb7 | 16.85478504 | -0.074091056 | 0.167909247 | -0.441256555 | 0.659027275 | 0.820224001 | | Rpp40 | 136.9825899 | -0.0209204627785734 | 0.0921060544784999 | -0.227134501602789 | 0.82031915666062 | 0.957571904906369 | NM_145938 |
| Ipp | 389.1948952 | -0.041386966 | 0.093719792 | -0.441603261 | 0.658776324 | 0.820224001 | | Scamp4 | 2000.724114 | 0.0159209704603758 | 0.0697553402185161 | 0.228240166423124 | 0.819459537101358 | 0.957571904906369 | NM_019575 |
| Pdzd2 | 4433.401548 | -0.074207528 | 0.16812151 | -0.441392231 | 0.658929066 | 0.820224001 | | Serpina9 | 292.480607 | -0.0215680858946045 | 0.0950043391381459 | -0.227022113834636 | 0.820406546714415 | 0.957571904906369 | NM_027997 |
| Ssbp1 | 476.9496582 | 0.033570134 | 0.076008685 | 0.441661815 | 0.658733949 | 0.820224001 | | Smtn | 378.185092 | 0.0199625625159288 | 0.0880507768154889 | 0.226716483805254 | 0.820644208625109 | 0.957571904906369 | NM_001159284,NM_013870 |
| Suv420h1 | 2105.269485 | -0.027383208 | 0.062040035 | -0.441379639 | 0.65893818 | 0.820224001 | | Svip | 2125.12793 | 0.0122558080259535 | 0.0537495671263475 | 0.228016869366485 | 0.819633126079911 | 0.957571904906369 | NR_027704,NM_001160345 |
| Syp | 53687.97567 | -0.025026263 | 0.056687246 | -0.441479597 | 0.658865829 | 0.820224001 | | Trim66 | 2364.697612 | 0.0145366638689061 | 0.064121625434149 | 0.226704544223303 | 0.820653493334668 | 0.957571904906369 | NM_001170913,NM_001170912,NM_181853 |
| Yipf3 | 1622.891288 | 0.025383006 | 0.057460957 | 0.441743529 | 0.658674807 | 0.820224001 | | Zbtb6 | 570.8084497 | -0.0166967696197413 | 0.07345289326088 | -0.227312620082098 | 0.820180660539932 | 0.957571904906369 | NM_146253 |
| Zfp275 | 1303.542341 | -0.032993266 | 0.074683434 | -0.441774899 | 0.658652104 | 0.820224001 | | Zic5 | 48.2356047 | -0.0203773972627374 | 0.0894277208243841 | -0.2278644327054748 | 0.81975163056584 | 0.957571904906369 | NM_022987 |
| Trank1 | 17780.32987 | 0.054604414 | 0.123827865 | 0.440970325 | 0.65923448 | 0.820415987 | | Nwd1 | 3572.357827 | 0.0127567365471406 | 0.0562945156566155 | 0.22660709304178 | 0.82072927631755 | 0.957583411067368 | NM_176940 |
| 2210013O21Rik | 736.0575477 | -0.02988023 | 0.067844338 | -0.440423344 | 0.65963052 | 0.820537487 | | Ttc7 | 483.4525591 | 0.0190949326714588 | 0.0843044859181859 | 0.226499603947406 | 0.820812867231701 | 0.957604024528767 | NM_028639 |
| BC018507 | 5895.60377 | 0.048021587 | 0.109028701 | 0.440449044 | 0.659611911 | 0.820537487 | | Cenpl | 64.85413096 | 0.021450717217241 | 0.0948876580943196 | 0.226064354922941 | 0.821151367610052 | 0.957611693785944 | NM_027429,NM_001159930 |

| | | | | | | |
|---|---|---|---|---|---|---|
| Bdh1 | 1710.099705 | -0.026931231 | 0.061153039 | -0.440390724 | 0.659654142 | 0.820537487 |
| Car2 | 6400.215571 | -0.039021955 | 0.088575096 | -0.440552224 | 0.659537197 | 0.820537487 |
| Casp7 | 99.61756201 | 0.062270044 | 0.141418851 | 0.440323506 | 0.659702819 | 0.820537487 |
| Gramd2 | 159.7254439 | 0.064004135 | 0.145232215 | 0.440702047 | 0.659428714 | 0.820537487 |
| Sp6 | 23.35578097 | 0.073646472 | 0.167246017 | 0.440348136 | 0.659684982 | 0.820537487 |
| Atg3 | 1498.330904 | -0.033298519 | 0.075649098 | -0.440170736 | 0.659813453 | 0.820543354 |
| Atp6ap1l | 351.2087245 | -0.048241864 | 0.109590405 | -0.440201525 | 0.659791155 | 0.820543354 |
| Ccdc30 | 369.6223591 | 0.046228857 | 0.105058103 | 0.440031333 | 0.659914414 | 0.820603044 |
| Memo1 | 635.4228854 | 0.048737795 | 0.110897672 | 0.439484379 | 0.660310598 | 0.820963923 |
| Myl4 | 554.3424728 | -0.073618754 | 0.167484282 | -0.4395562 | 0.66025857 | 0.820963923 |
| Zfp644 | 2199.532098 | -0.031985698 | 0.072796012 | -0.43938805 | 0.660380384 | 0.820984808 |
| Mrpl11 | 1262.73555 | -0.027274992 | 0.062085362 | -0.43931437 | 0.660433763 | 0.820985296 |
| Kctd16 | 635.4272273 | -0.038013815 | 0.086599764 | -0.438959806 | 0.660690661 | 0.821172878 |
| Ttc3 | 44297.92609 | -0.030094623 | 0.068550872 | -0.439011529 | 0.660653183 | 0.821172878 |
| 1700012D01Rik | 13.51378355 | -0.072424001 | 0.165068673 | -0.438750732 | 0.660842163 | 0.821229425 |
| Gm5126 | 83.98629361 | 0.057808662 | 0.131757321 | 0.438751042 | 0.660841938 | 0.821229425 |
| Prkar1b | 36725.58424 | 0.033662713 | 0.076752218 | 0.438589451 | 0.660959042 | 0.821308797 |
| 4930451G09Rik | 121.2841846 | 0.060810001 | 0.13867221 | 0.438516128 | 0.661012181 | 0.821308961 |
| Traf1 | 29.56528989 | -0.0736312 | 0.168014578 | -0.438242926 | 0.661210195 | 0.821489115 |
| Fam89a | 24.49921398 | 0.073252606 | 0.167439637 | 0.437486651 | 0.661758455 | 0.822104354 |
| Capn12 | 29.29644343 | 0.073294317 | 0.167657976 | 0.437165704 | 0.66199118 | 0.822261612 |
| Zcchc24 | 3280.169977 | 0.035055057 | 0.080182189 | 0.437192561 | 0.661971704 | 0.822261612 |
| Zkscan7 | 118.9331238 | 0.069257742 | 0.158529021 | 0.436877371 | 0.662200284 | 0.822455407 |
| Kif5c | 40602.91736 | -0.028657895 | 0.065608428 | -0.436802046 | 0.662254915 | 0.822457331 |
| Epn2 | 5804.92083 | -0.023044414 | 0.052782502 | -0.436591929 | 0.662407317 | 0.822580666 |
| Eprs | 3836.950151 | -0.021387173 | 0.048996843 | -0.436501049 | 0.662473238 | 0.822596598 |
| Cebpz | 1534.359702 | -0.028665987 | 0.065683865 | -0.436423564 | 0.662529445 | 0.822600467 |
| Gm11127 | 44.66253358 | -0.072417545 | 0.166076139 | -0.436050269 | 0.662800258 | 0.822804839 |
| Kbtbd3 | 245.4250349 | -0.05177782 | 0.1187321 | -0.436089479 | 0.66277181 | 0.822804839 |
| Islr2 | 2860.805204 | 0.053766859 | 0.123370914 | 0.435814711 | 0.66297117 | 0.822951074 |
| Ppp1r26 | 1305.724365 | 0.031386915 | 0.07205673 | 0.435586162 | 0.663137013 | 0.823090994 |
| Psmd7 | 3401.350144 | -0.030246107 | 0.069500078 | -0.435195295 | 0.663420677 | 0.823377121 |
| Rpusd4 | 302.7999289 | 0.033730686 | 0.077541 | 0.435004528 | 0.663559141 | 0.823483005 |
| Fli1 | 161.2922819 | -0.048821016 | 0.11239652 | -0.434364123 | 0.664024046 | 0.823504463 |
| Mob1b | 410.517406 | -0.039304769 | 0.090503745 | -0.434288871 | 0.664078685 | 0.823504463 |
| Naprt1 | 174.5882294 | 0.063097146 | 0.145206834 | 0.434532894 | 0.663901514 | 0.823504463 |
| Nlrp1b | 26.08243885 | -0.071167018 | 0.163839653 | -0.434369926 | 0.664019833 | 0.823504463 |
| Peg10 | 1951.577894 | -0.073053613 | 0.168097894 | -0.434589701 | 0.663860272 | 0.823504463 |
| Serpinb6a | 1062.332578 | -0.027162265 | 0.062539976 | -0.43431844 | 0.664057215 | 0.823504463 |
| Slc25a39 | 2064.625561 | -0.03090553 | 0.071079727 | -0.434800904 | 0.663706949 | 0.823504463 |
| Spred2 | 2410.178305 | -0.051288458 | 0.118108518 | -0.434248593 | 0.66410793 | 0.823504463 |
| Ssr4 | 989.329928 | -0.031230269 | 0.071893319 | -0.434397376 | 0.663999903 | 0.823504463 |
| Tbc1d19 | 933.5075097 | 0.046031385 | 0.105898887 | 0.434672986 | 0.663799809 | 0.823504463 |
| Ccdc137 | 926.9431607 | 0.033287956 | 0.076698654 | 0.434009644 | 0.664281438 | 0.823653697 |
| Dicer1 | 2577.65294 | -0.030470405 | 0.070249101 | -0.433747965 | 0.664471472 | 0.823807252 |
| Faf1 | 1312.210893 | 0.029450667 | 0.067918228 | 0.433619477 | 0.664564789 | 0.823807252 |
| Kcnj6 | 1147.719299 | -0.036721062 | 0.084684705 | -0.43362095 | 0.664563719 | 0.823807252 |
| Ddx11 | 79.55756514 | -0.066983504 | 0.154544003 | -0.433426742 | 0.664704777 | 0.823829802 |
| Gmpr2 | 702.5060629 | -0.039427114 | 0.090964103 | -0.433435969 | 0.664698074 | 0.823829802 |
| Ripply2 | 55.23182112 | -0.067215765 | 0.155098457 | -0.433374817 | 0.664742493 | 0.823829802 |
| 2900041M22Rik | 26.91761232 | -0.070781222 | 0.16347465 | -0.432979806 | 0.66502944 | 0.82397451 |
| Elmsan1 | 1230.085004 | 0.048559769 | 0.112216694 | 0.432732128 | 0.665209386 | 0.82397451 |
| Ptms | 14660.56089 | 0.029352337 | 0.067835045 | 0.432701667 | 0.665231519 | 0.82397451 |
| Rhod | 17.59476613 | 0.072721638 | 0.168045086 | 0.432750756 | 0.665195852 | 0.82397451 |
| Spata5l1 | 64.7577502 | 0.060983039 | 0.140916036 | 0.432761523 | 0.665188029 | 0.82397451 |
| Tmod3 | 910.3499372 | -0.029455213 | 0.068007649 | -0.433116182 | 0.664930367 | 0.82397451 |
| Xkr7 | 116.7853846 | -0.062238588 | 0.143820361 | -0.432752272 | 0.665194751 | 0.82397451 |
| 2410006H16Rik | 385.6369425 | 0.047947586 | 0.110861425 | 0.432500178 | 0.665377923 | 0.824089971 |
| Caps2 | 8.66176849 | 0.06403012 | 0.148078978 | 0.432405198 | 0.665446941 | 0.824109576 |
| 2900002K06Rik | 9.300791402 | 0.069965871 | 0.162047258 | 0.431762139 | 0.665914299 | 0.824289729 |
| Dnttip2 | 943.6065321 | 0.039446226 | 0.091375719 | 0.431692645 | 0.665964814 | 0.824289729 |
| Nxt1 | 222.078285 | 0.045053516 | 0.10432217 | 0.431869045 | 0.665836594 | 0.824289729 |
| Slain2 | 938.3515146 | -0.030284584 | 0.070131568 | -0.431825279 | 0.665868405 | 0.824289729 |
| Tex261 | 795.783763 | -0.029464522 | 0.068210743 | -0.431963063 | 0.665768259 | 0.824289729 |
| Zfp607 | 38.94664459 | -0.072355068 | 0.167587331 | -0.431745453 | 0.665926428 | 0.824289729 |
| Zscan2 | 266.9103225 | -0.044995635 | 0.104201536 | -0.431813549 | 0.665876931 | 0.824289729 |
| Adat1 | 270.2829925 | 0.04759798 | 0.11036029 | 0.431296258 | 0.666252971 | 0.824514659 |
| BC005764 | 2301.52035 | 0.046697572 | 0.108266516 | 0.431320542 | 0.666235316 | 0.824514659 |
| Sobp | 3303.739988 | 0.029496516 | 0.068409305 | 0.431176966 | 0.666339701 | 0.824556132 |
| 2810468N07Rik | 896.4073398 | -0.042299047 | 0.098208688 | -0.430705756 | 0.666682333 | 0.824848364 |
| Ptk2 | 5572.36413 | -0.029974773 | 0.069590882 | -0.430728451 | 0.666665829 | 0.824848364 |
| Fam169a | 2410.100082 | 0.03360058 | 0.078069025 | 0.43039579 | 0.666907757 | 0.825012842 |
| Zfp429 | 61.47528034 | 0.061000359 | 0.141737189 | 0.430376528 | 0.666921766 | 0.825012842 |
| Ap1b1 | 9134.871198 | 0.032117106 | 0.074672945 | 0.430103641 | 0.667120251 | 0.825060833 |
| Olfr464 | 15.24344999 | 0.063291619 | 0.147173769 | 0.430046872 | 0.667161546 | 0.825060833 |
| Ptgr1 | 35.32123312 | -0.069072108 | 0.160621472 | -0.430030352 | 0.667173563 | 0.825060833 |
| Zfp507 | 1225.348798 | -0.034412417 | 0.079983671 | -0.430243025 | 0.667018867 | 0.825060833 |
| Meis1 | 245.0294757 | -0.067732913 | 0.157571093 | -0.429856214 | 0.667300238 | 0.825151627 |
| Zfp938 | 551.7235508 | -0.035297306 | 0.082184742 | -0.429487342 | 0.667568604 | 0.8254176 |
| Sephs1 | 1023.780118 | -0.036346273 | 0.084646927 | -0.429386801 | 0.667641757 | 0.825442179 |
| Tmod1 | 2687.42237 | -0.025700202 | 0.059866038 | -0.429295186 | 0.66770842 | 0.825458729 |
| 6430548M08Rik | 15780.44524 | -0.028368179 | 0.066103617 | -0.429147155 | 0.667816138 | 0.825526028 |
| H1f0 | 3398.65047 | -0.025889557 | 0.060416693 | -0.428516618 | 0.668275037 | 0.825961505 |
| Igf2bp2 | 40.46602135 | -0.068767341 | 0.160470482 | -0.428535766 | 0.6682611 | 0.825961505 |
| D430020J02Rik | 112.7601341 | -0.05669348 | 0.132384123 | -0.428249842 | 0.668469232 | 0.826003851 |
| Dnajc1 | 951.9734186 | -0.034814096 | 0.081282953 | -0.428307465 | 0.668427285 | 0.826003851 |
| Polr1c | 670.9800609 | -0.038910356 | 0.090839729 | -0.428340735 | 0.668403066 | 0.826003851 |
| Otud6b | 2493.777066 | -0.028985555 | 0.067718497 | -0.428030105 | 0.668629203 | 0.826069761 |
| Tcf15 | 10.56466275 | -0.06840767 | 0.159793378 | -0.428100782 | 0.668577748 | 0.826069761 |
| Larp4 | 1542.208037 | 0.031993358 | 0.074784684 | 0.427806294 | 0.668792154 | 0.826205202 |
| Snta1 | 1110.505937 | 0.029239656 | 0.068378216 | 0.427616547 | 0.668930317 | 0.82631 |
| Pih1d1 | 653.0168881 | -0.037998836 | 0.088914499 | -0.427363774 | 0.669114389 | 0.826471488 |
| Agtr1a | 26.77549627 | -0.071408016 | 0.167262577 | -0.426921654 | 0.669436393 | 0.826737404 |
| Egr1 | 11151.10528 | -0.071335836 | 0.167074078 | -0.426971301 | 0.669400231 | 0.826737404 |
| Rab1b | 5317.849902 | -0.027667973 | 0.064827471 | -0.426793969 | 0.6695294 | 0.826786365 |
| Eaf2 | 37.33129048 | 0.06731804 | 0.157770894 | 0.42668225 | 0.669610781 | 0.826820962 |
| Nsun3 | 181.6446103 | 0.05257777 | 0.12328435 | 0.426475619 | 0.66976131 | 0.82694093 |
| Zkscan8 | 827.7428554 | 0.043505039 | 0.1020363 | 0.426368254 | 0.66983953 | 0.826971608 |
| Ermp1 | 2702.450553 | 0.027285228 | 0.064016163 | 0.42622405 | 0.669944594 | 0.827035419 |
| 5730559C18Rik | 90.1446376 | 0.058978895 | 0.138449147 | 0.425996813 | 0.670110168 | 0.827173912 |
| Eif4a3 | 1548.534764 | -0.026211035 | 0.061553188 | -0.425827425 | 0.670233601 | 0.827194473 |
| Mpzl3 | 53.23578763 | 0.060940781 | 0.143103906 | 0.425849876 | 0.670217241 | 0.827194473 |
| Slc30a6 | 357.7358486 | -0.039407253 | 0.092561654 | -0.425740584 | 0.670296886 | 0.827206687 |
| Nog | 173.8712881 | -0.05419453 | 0.12740214 | -0.425381628 | 0.670558497 | 0.827463631 |
| Bves | 33.25624571 | -0.066537377 | 0.156512509 | -0.425124978 | 0.67074557 | 0.827527874 |
| Grk6 | 2871.794578 | -0.026419301 | 0.062149843 | -0.425090385 | 0.670770787 | 0.827527874 |
| Smad2 | 1640.372517 | -0.026269432 | 0.061779318 | -0.425214022 | 0.670680664 | 0.827527874 |
| Adamts18 | 91.11222444 | -0.064276276 | 0.151334397 | -0.424730118 | 0.671033425 | 0.827720077 |
| Rnf19b | 1702.067611 | -0.035175385 | 0.082807487 | -0.424785084 | 0.670993352 | 0.827720077 |
| Snai3 | 23.54896297 | 0.071232435 | 0.168073927 | 0.423816089 | 0.671699943 | 0.828476272 |
| Cd4 | 602.259641 | -0.052523266 | 0.12398252 | -0.42363444 | 0.671832434 | 0.828507779 |
| Tnni2 | 8.924590337 | 0.068047852 | 0.16061271 | 0.42367663 | 0.671801661 | 0.828507779 |
| Epor | 306.2094584 | 0.033669758 | 0.079574083 | 0.423124677 | 0.672204299 | 0.828834427 |
| Trappc10 | 5278.031982 | 0.033617909 | 0.079445059 | 0.423159217 | 0.6721791 | 0.828834427 |
| Adrb1 | 867.5492348 | 0.048751679 | 0.115258964 | 0.422975168 | 0.672313379 | 0.82890296 |
| Itm2b | 19310.33097 | 0.021495044 | 0.050829089 | 0.422888638 | 0.672376513 | 0.828914839 |
| Myzap | 30.33260973 | 0.068693778 | 0.162502756 | 0.422723773 | 0.672496809 | 0.82899718 |
| Maf1 | 2356.112032 | 0.027621725 | 0.065392652 | 0.422397999 | 0.672734538 | 0.829205242 |
| Pcdh1 | 4811.754715 | -0.039356691 | 0.093185939 | -0.422345806 | 0.672772628 | 0.829205242 |
| Slc33a1 | 1139.42915 | -0.024068338 | 0.057123965 | -0.421335207 | 0.673510325 | 0.82998242 |
| Slc41a2 | 951.0694831 | -0.036709122 | 0.087114985 | -0.421386998 | 0.673472512 | 0.82998242 |
| Hpse | 54.31243035 | 0.064145222 | 0.152470416 | 0.420706023 | 0.673969763 | 0.830416501 |
| Mphosph10 | 636.6394307 | 0.031681754 | 0.075302663 | 0.420725549 | 0.673955503 | 0.830416501 |
| Actrt3 | 29.59252788 | 0.070068244 | 0.166580949 | 0.420625791 | 0.674028358 | 0.830417276 |
| Pex1 | 893.831509 | 0.02559368 | 0.060867046 | 0.420485008 | 0.674131179 | 0.830417276 |
| Zfp119b | 82.76413208 | -0.064022862 | 0.152251017 | -0.420508599 | 0.674113949 | 0.830417276 |
| Nlrx1 | 359.4824604 | 0.039993616 | 0.095134884 | 0.420388549 | 0.674201632 | 0.830438038 |
| Arhgap30 | 167.2045975 | 0.050245828 | 0.119626063 | 0.420024087 | 0.674467858 | 0.830567874 |
| Eif3b | 3586.265112 | 0.028509209 | 0.067872563 | 0.420040264 | 0.67445604 | 0.830567874 |
| Trip12 | 8072.241072 | -0.03467379 | 0.082549776 | -0.420034944 | 0.674459927 | 0.830567874 |
| Galnt7 | 487.3457676 | -0.030760224 | 0.073274394 | -0.419795 | 0.674635218 | 0.830575929 |
| Gm14379 | 10.2125842 | 0.06381314 | 0.151991841 | 0.419845828 | 0.674598084 | 0.830575929 |
| Nup205 | 1307.013797 | 0.03330795 | 0.079326423 | 0.419884679 | 0.674569701 | 0.830575929 |
| Grb10 | 2062.083747 | -0.042115717 | 0.100461873 | -0.419220905 | 0.675054696 | 0.830976799 |
| Phldb1 | 4435.029842 | 0.051977994 | 0.123992543 | 0.419202577 | 0.675068089 | 0.830976799 |
| Ccdc129 | 63.36399785 | -0.069119003 | 0.164930974 | -0.419078366 | 0.675158861 | 0.831022513 |
| Znhit2 | 551.6008514 | -0.040613708 | 0.096932236 | -0.418990722 | 0.675222913 | 0.831035334 |
| 9130019O22Rik | 143.2438487 | -0.052760179 | 0.126005998 | -0.418711649 | 0.675426881 | 0.831055994 |
| Fam149a | 1223.616837 | -0.03123285 | 0.074609551 | -0.418617315 | 0.675495833 | 0.831055994 |
| Pcdha2 | 102.6958712 | -0.057005871 | 0.136201416 | -0.418540958 | 0.675551647 | 0.831055994 |
| Rpap2 | 674.5631901 | 0.031260253 | 0.074703838 | 0.41845578 | 0.675613911 | 0.831055994 |
| S100a13 | 427.7237441 | -0.039196569 | 0.093652105 | -0.418533773 | 0.675556899 | 0.831055994 |
| Sfxn5 | 6937.514704 | -0.050376441 | 0.120358517 | -0.418553193 | 0.675542703 | 0.831055994 |
| Zfp943 | 432.0137575 | 0.048637919 | 0.116232395 | 0.41845407 | 0.675615161 | 0.831055994 |
| Ppef1 | 24.01062762 | -0.069503469 | 0.166197912 | -0.418197004 | 0.675803086 | 0.831221165 |
| Pdcd6ip | 3498.002413 | 0.022819034 | 0.054648357 | 0.417561205 | 0.676267967 | 0.831660915 |
| Plxnb1 | 4677.240226 | -0.024892522 | 0.059611801 | -0.417577077 | 0.676256359 | 0.831660915 |
| Fam149b | 1069.503648 | -0.027126659 | 0.06498795 | -0.417410597 | 0.676378105 | 0.831730346 |
| Fbl | 988.5161814 | -0.030124828 | 0.072211959 | -0.417172283 | 0.676552397 | 0.831746635 |
| Gabrg2 | 6528.350616 | 0.038369908 | 0.091971985 | 0.417191262 | 0.676538516 | 0.831746635 |
| Irgq | 6052.776393 | -0.031840709 | 0.076318482 | -0.417208359 | 0.676526012 | 0.831746635 |
| Psmc5 | 5800.996883 | -0.033427916 | 0.080151177 | -0.417060819 | 0.676633923 | 0.831780863 |
| 4930539E08Rik | 431.6877385 | -0.048340777 | 0.11593116 | -0.416978291 | 0.676694287 | 0.831789074 |
| Gm1078 | 56.91073823 | 0.065940065 | 0.158189416 | 0.41684246 | 0.676793643 | 0.831845209 |
| Neurl3 | 45.36199933 | 0.068437687 | 0.164385587 | 0.416324132 | 0.677172836 | 0.832192166 |
| Slc6a7 | 3208.578681 | -0.038042387 | 0.091380003 | -0.416309761 | 0.677183351 | 0.832192166 |
| Ctdspl2 | 830.0826495 | -0.035385283 | 0.08503937 | -0.416104716 | 0.677333379 | 0.832310523 |
| Fam20a | 446.0180101 | -0.048977651 | 0.117757509 | -0.415919556 | 0.677468869 | 0.832362764 |
| Uqcrc1 | 8117.058384 | -0.028359583 | 0.068188502 | -0.415899786 | 0.677483336 | 0.832362764 |
| Calr4 | 15.88165734 | -0.069628797 | 0.167486492 | -0.415727836 | 0.67760917 | 0.832451354 |
| M6pr | 3902.498128 | -0.020148604 | 0.048492246 | -0.415501562 | 0.677774772 | 0.832588783 |
| Gm1141 | 13.95942987 | 0.069234781 | 0.16685795 | 0.414932469 | 0.67819134 | 0.832842534 |
| Naa10 | 758.6089307 | -0.02849518 | 0.06866557 | -0.41498949 | 0.678152906 | 0.832842534 |
| Nkx6-2 | 278.3828384 | -0.053523058 | 0.128940283 | -0.415099584 | 0.678069004 | 0.832842534 |
| Pip4k2a | 3801.823976 | -0.029766304 | 0.07173889 | -0.414925625 | 0.67819635 | 0.832842534 |
| Gzmm | 52.18773383 | -0.062038578 | 0.149550732 | -0.414832993 | 0.678264165 | 0.832859802 |
| Ptprk | 2786.966436 | 0.032125973 | 0.077489515 | 0.414584768 | 0.6784459 | 0.833016941 |
| Ing1 | 1196.328541 | -0.024370669 | 0.058809273 | -0.414401815 | 0.67857986 | 0.8331154 |
| Erich2 | 14.03996946 | 0.068119284 | 0.164536998 | 0.414005877 | 0.678869803 | 0.833141285 |
| Got1 | 13979.28938 | 0.025833826 | 0.062396673 | 0.414025704 | 0.678855283 | 0.833141285 |
| Pcm1 | 4244.235441 | -0.036043109 | 0.087054898 | -0.414027356 | 0.678854073 | 0.833141285 |
| Sgpp2 | 1175.770352 | 0.038099865 | 0.092011673 | 0.41407643 | 0.678818134 | 0.833141285 |
| Thsd4 | 429.2092741 | -0.060368752 | 0.145808002 | -0.414029076 | 0.678852813 | 0.833141285 |
| Polr3f | 971.9951725 | 0.03221143 | 0.077865489 | 0.413680448 | 0.679108148 | 0.833367784 |
| Ypel5 | 3604.148929 | -0.032937368 | 0.079650648 | -0.413522913 | 0.679223539 | 0.833443376 |
| Adamts15 | 250.6182692 | -0.038684573 | 0.093599321 | -0.413299719 | 0.679387035 | 0.833506033 |
| Dusp7 | 2576.178196 | -0.03368325 | 0.081642093 | -0.412572099 | 0.679920146 | 0.833506033 |
| Ecm2 | 150.5324112 | -0.056678243 | 0.137287779 | -0.412842593 | 0.679721943 | 0.833506033 |
| Esm1 | 11.91931579 | -0.068266928 | 0.165428 | -0.412668522 | 0.679849491 | 0.833506033 |
| Gabarap | 4629.714203 | -0.019388335 | 0.046992855 | -0.412580485 | 0.679914001 | 0.833506033 |
| Got2 | 11586.04955 | 0.031151311 | 0.075494702 | 0.412629104 | 0.679878375 | 0.833506033 |
| Metap1 | 1573.517111 | 0.027838949 | 0.067454133 | 0.412709309 | 0.679819603 | 0.833506033 |
| Rnf2 | 895.9670994 | 0.040274253 | 0.097537496 | 0.412910469 | 0.67967221 | 0.833506033 |
| Snora28 | 35.56134574 | 0.06649318 | 0.160855103 | 0.413373146 | 0.679333246 | 0.833506033 |
| Syf2 | 1426.741235 | 0.033282253 | 0.080560676 | 0.413132736 | 0.679509367 | 0.833506033 |
| Tmem248 | 2060.144527 | 0.03147079 | 0.076180287 | 0.413109371 | 0.679526484 | 0.833506033 |
| Unc5b | 1323.874436 | -0.068188771 | 0.165225327 | -0.412701684 | 0.679825191 | 0.833506033 |
| Ube4a | 3101.553318 | -0.02872725 | 0.069661652 | -0.412382557 | 0.680059046 | 0.833610353 |
| Gpatch11 | 1061.399188 | -0.025453797 | 0.061787654 | -0.411956038 | 0.680371646 | 0.833798326 |
| Pcdh8 | 1208.425982 | 0.048354583 | 0.117368133 | 0.41199073 | 0.680346218 | 0.833798326 |
| Phf19 | 40.98642829 | 0.063172588 | 0.153349001 | 0.411953049 | 0.680373837 | 0.833798326 |
| Gpr3 | 257.4173278 | -0.065311378 | 0.158602963 | -0.411791663 | 0.680492133 | 0.833877342 |
| Sfmbt2 | 109.2970527 | -0.061898236 | 0.150359849 | -0.411667323 | 0.680583279 | 0.833923079 |
| Ankmy1 | 15.39265554 | 0.068861152 | 0.16738576 | 0.411391939 | 0.680785164 | 0.834104485 |
| Gprc5a | 13.00459215 | 0.060073292 | 0.146059266 | 0.411293951 | 0.680857005 | 0.834126546 |
| Cnot11 | 1171.390626 | 0.02685076 | 0.06531239 | 0.411112806 | 0.68098982 | 0.834127781 |
| Fam76a | 2070.424221 | 0.020548532 | 0.04997343 | 0.411189139 | 0.680933851 | 0.834127781 |
| Rpl21 | 9906.652497 | 0.028882592 | 0.070261587 | 0.411072302 | 0.681019519 | 0.834127781 |
| 6030458C11Rik | 1663.573487 | 0.023457165 | 0.057079583 | 0.410955434 | 0.681105214 | 0.834166799 |
| Pacs1 | 4444.263963 | -0.026494292 | 0.064501462 | -0.410754906 | 0.681252262 | 0.834215012 |
| Spats2 | 1031.01526 | -0.034294694 | 0.08348009 | -0.410812859 | 0.681209763 | 0.834215012 |
| Dennd2d | 15.34345509 | -0.066004395 | 0.160824054 | -0.410413699 | 0.681502499 | 0.834389547 |
| Txnl4a | 4107.043261 | 0.029601396 | 0.07212004 | 0.410446192 | 0.681478668 | 0.834389547 |
| Aldh1l2 | 158.0304863 | -0.045691695 | 0.111426619 | -0.410060854 | 0.681761308 | 0.83457452 |
| Bphl | 788.699158 | -0.033124039 | 0.080775707 | -0.410074273 | 0.681751465 | 0.83457452 |
| 2010107G23Rik | 795.435503 | -0.03071787 | 0.07504445 | -0.409329008 | 0.68229823 | 0.834770118 |
| A930012L18Rik | 79.56746871 | 0.066324821 | 0.161962891 | 0.40950628 | 0.682168159 | 0.834770118 |
| Amn | 99.27541316 | -0.057702752 | 0.140947664 | -0.409391334 | 0.682252498 | 0.834770118 |
| Ano5 | 54.65094741 | -0.065587281 | 0.160172649 | -0.409478658 | 0.682188426 | 0.834770118 |
| Casp8ap2 | 573.727199 | 0.036309526 | 0.088690013 | 0.409398136 | 0.682247508 | 0.834770118 |
| Ppp1r1a | 2731.781984 | -0.03014624 | 0.073638491 | -0.409381559 | 0.682259671 | 0.834770118 |
| Yipf4 | 1095.404243 | -0.038188722 | 0.09326966 | -0.409444203 | 0.682213706 | 0.834770118 |
| Pcdha10 | 100.6843495 | -0.059437697 | 0.145255475 | -0.409194197 | 0.682397153 | 0.834825226 |
| 2310009B15Rik | 128.9386165 | 0.046550827 | 0.113800209 | 0.409057485 | 0.682497476 | 0.83487957 |
| Pld3 | 25469.9894 | -0.03587209 | 0.087709651 | -0.408986808 | 0.682549342 | 0.83487957 |
| C1rl | 20.98685579 | 0.064222249 | 0.15706913 | 0.408878873 | 0.682628554 | 0.834910548 |
| Amotl2 | 789.8739848 | 0.043092997 | 0.105482066 | 0.408533873 | 0.682881767 | 0.835099429 |
| Cad | 912.7019598 | -0.042352184 | 0.103705673 | -0.408388306 | 0.682988618 | 0.835099429 |
| Cmklr1 | 167.0508792 | 0.052072633 | 0.127482752 | 0.408468064 | 0.682930072 | 0.835099429 |
| Map1lc3b | 7861.185546 | 0.02381103 | 0.0583173 | 0.408301312 | 0.683052476 | 0.835099429 |
| Tex30 | 98.51765765 | 0.058672218 | 0.143676377 | 0.408363707 | 0.683006674 | 0.835099429 |
| Manea | 1372.564735 | -0.037703714 | 0.092416128 | -0.407977647 | 0.683290086 | 0.835324017 |
| A630033H20Rik | 10.0229666 | -0.06385784 | 0.156590368 | -0.40780184 | 0.683419162 | 0.835349992 |
| Tsacc | 26.71589533 | 0.067829382 | 0.166321004 | 0.407822107 | 0.683404282 | 0.835349992 |
| Gm11627 | 73.93308866 | -0.059754313 | 0.14670314 | -0.407314477 | 0.683777031 | 0.83565557 |
| Kdm3a | 2216.805962 | 0.041904546 | 0.102873567 | 0.407340267 | 0.683758092 | 0.83565557 |
| Rassf8 | 329.7976563 | 0.0619367 | 0.152106005 | 0.407194311 | 0.683865279 | 0.835697503 |
| Yy2 | 38.90033311 | -0.061939187 | 0.152196998 | -0.406967206 | 0.684032075 | 0.835835407 |
| 1700030C10Rik | 41.75334185 | 0.067525145 | 0.166045509 | 0.406666493 | 0.684252953 | 0.835958036 |
| Cxcr2 | 20.95648698 | -0.06740143 | 0.165764232 | -0.406610216 | 0.684294292 | 0.835958036 |
| Sytl3 | 8.126615697 | 0.063615322 | 0.156409218 | 0.40672361 | 0.684210998 | 0.835958036 |
| Bckdk | 1456.264235 | 0.029875973 | 0.07351349 | 0.406401241 | 0.684447808 | 0.836079655 |
| Nkap | 594.8974103 | -0.031904205 | 0.078518468 | -0.406327401 | 0.684502055 | 0.836080003 |
| Klhl25 | 410.6960091 | -0.038728145 | 0.095345092 | -0.406189179 | 0.684603604 | 0.836138124 |
| Aga | 181.2810359 | 0.045689568 | 0.112523673 | 0.406044053 | 0.684710232 | 0.836202439 |

| | | | | | | | |
|---|---|---|---|---|---|---|---|
| Ero1lb | 973.709698 | 0.0141192953522281 | 0.0623876621060495 | 0.226315506553642 | 0.820956038756699 | 0.957611693785944 | NM_026184 |
| Fut10 | 469.0794231 | 0.0155309168055484 | 0.0686788232363239 | 0.226138365127581 | 0.821093806295548 | 0.957611693785944 | NM_001012517,NM_134161 |
| Nosip | 1825.258263 | 0.015308625021468 | 0.0676332471033939 | 0.226347627492511 | 0.82093105805045 | 0.957611693785944 | NM_001163684,NM_025533 |
| Rbpms | 139.2830989 | -0.0212648690375412 | 0.0940380312769326 | -0.226130521330442 | 0.821099906750192 | 0.957611693785944 | NM_019733,NM_001042674,NM_001042675 |
| Rtkn2 | 133.2368028 | 0.0208201190252101 | 0.0921315302014015 | 0.225982559713237 | 0.821214984819738 | 0.957611693785944 | NM_001081346 |
| Coq2 | 2387.020376 | -0.0102139097735958 | 0.0452694899902491 | -0.225624582379786 | 0.821493419832825 | 0.957699646263451 | NM_027978 |
| Loxl1 | 73.28581121 | -0.0204341525304044 | 0.0905483137012073 | -0.22567126537368 | 0.821457108492563 | 0.957699646263451 | NM_010729 |
| Mis18a | 175.3863125 | 0.0198519195430333 | 0.087926910099007 | 0.225777518175945 | 0.82137446351308 | 0.957699646263451 | NM_025642 |
| Rabggtb | 1464.600151 | 0.0108236570572406 | 0.0479885765998191 | 0.225546532615459 | 0.821554129979958 | 0.957699646263451 | NR_028097,NM_001163478,NR_028095,NR_028094,NR_028098,NM_001163479,NM_011231,NR_028096 |
| Tjp2 | 1500.042857 | -0.0152377408350183 | 0.0675902643651547 | -0.225442825268358 | 0.821634791469255 | 0.957716817596084 | NM_011597,NM_001198985 |
| 2410018L13Rik | 108.1296907 | 0.0209078137833862 | 0.0929177424432709 | 0.225014224771453 | 0.821968208239126 | 0.957785689782947 | NR_015504 |
| 4930430F08Rik | 538.053771 | -0.014744161119921 | 0.0656076015342902 | -0.224732512317416 | 0.822187370311468 | 0.957785689782947 | NM_175128 |
| Barhl2 | 29.80961645 | -0.0189363957545013 | 0.0843623518373269 | -0.224465005326259 | 0.822395493892835 | 0.957785689782947 | NM_001005477 |
| C77370 | 1527.262193 | 0.0160711214950198 | 0.0715214665928937 | 0.22470346681365 | 0.822209967449343 | 0.957785689782947 | NM_001077354 |
| Cmtm5 | 1503.613946 | 0.01752401297708 | 0.0778075085381993 | 0.225222646335021 | 0.821806072761536 | 0.957785689782947 | NM_029390,NM_026066 |
| Dkkl1 | 728.1476348 | -0.0174620287219339 | 0.0777837043460009 | -0.224494691642076 | 0.822372396973676 | 0.957785689782947 | NM_015789 |
| Lmod1 | 49.87048321 | -0.0195066717664978 | 0.0869407780941499 | -0.224367347453155 | 0.822471475979952 | 0.957785689782947 | NM_053106 |
| Pno1 | 833.6434115 | 0.0169972148200814 | 0.0757266616845244 | 0.22445482795599 | 0.822403412253318 | 0.957785689782947 | NM_025443 |
| Rufy1 | 1572.564519 | -0.013193142021231 | 0.0588061167290905 | -0.224349825410335 | 0.822485109070114 | 0.957785689782947 | NM_172557 |
| Senp1 | 691.977563 | -0.0162349858031144 | 0.072176141365258 | -0.224935629641863 | 0.822029351003006 | 0.957785689782947 | NM_144851 |
| Slc18b1 | 867.929869 | -0.0140763854562199 | 0.0627224670134558 | -0.224423338661091 | 0.822427912173015 | 0.957785689782947 | NM_183116 |
| Zkscan4 | 221.3423296 | 0.0198183722037894 | 0.0882442793174996 | 0.224585348274914 | 0.822301864117872 | 0.957785689782947 | NM_001039115 |
| Nt5dc1 | 81.69760609 | 0.0213070590644701 | 0.0950608488142326 | 0.224141266675498 | 0.822647383078564 | 0.957821087415776 | NM_176968 |
| Tgfb1i1 | 510.4480104 | -0.0167736578353595 | 0.0748164562482225 | -0.224197438324379 | 0.822603676664807 | 0.957821087415776 | NM_009365 |
| Shroom3 | 190.1372097 | -0.0210496524761585 | 0.0939557071780986 | -0.224038039927235 | 0.822727703874756 | 0.957837831729158 | NM_001077596,NM_001077595,NM_015756 |
| Arhgap33 | 11187.42028 | 0.0165439250171443 | 0.0738891656615897 | 0.223901905902079 | 0.822833632683388 | 0.95788438438523 | NM_178252 |
| 1110059M19Rik | 40.01463741 | 0.006752450311462 | 0.0304258468818961 | 0.221931384118015 | 0.824367298688239 | 0.957903915785494 | NM_026841 |
| 1500004A13Rik | 1091.179059 | 0.0169345090545926 | 0.0767290112286129 | 0.220705425280882 | 0.825321806865957 | 0.957903915785494 | NR_015498 |
| 4632428C04Rik | 38.03352851 | -0.0205253966292321 | 0.0918084625985505 | -0.223567588959453 | 0.823093785573576 | 0.957903915785494 | NR_033631 |
| A2m | 207.2851278 | 0.0122194742426738 | 0.0548023487846475 | 0.222973549741302 | 0.82355609254041 | 0.957903915785494 | NM_175628 |
| Ap2a1 | 13082.83682 | 0.0122891168762638 | 0.0556922550520425 | 0.220661146954456 | 0.825356285968005 | 0.957903915785494 | NM_001077264,NM_007458 |
| Brd1 | 2334.702483 | -0.0127580221958368 | 0.0575161954479578 | -0.221816170149512 | 0.824456991053888 | 0.957903915785494 | NM_001033274 |
| Celf4 | 16709.70869 | 0.0117503644358853 | 0.0530303718479682 | 0.221578013248186 | 0.824642399934292 | 0.957903915785494 | NM_133195,NM_001174074,NM_001146295,NM_001146294,NM_001146293,NM_001146292 |
| Cnnm1 | 6011.652752 | 0.0093132547646519 | 0.0421102539224628 | 0.221163585992863 | 0.824965061369229 | 0.957903915785494 | NM_031396 |
| Crb2 | 298.8308377 | -0.0202683343222595 | 0.0907480323384049 | -0.223347369634172 | 0.823265162607443 | 0.957903915785494 | NM_001163566 |
| Diap1 | 2403.909993 | 0.0162177544971614 | 0.0726684400748385 | 0.223174661248532 | 0.823399572018146 | 0.957903915785494 | NM_007858 |
| E230025N22Rik | 23.23881003 | 0.0173041680720916 | 0.0779629790086237 | 0.221953654056467 | 0.82434996213591 | 0.957903915785494 | NM_172831 |
| Entpd4 | 7503.919256 | 0.0145504700695668 | 0.0655869671686489 | 0.221850021394526 | 0.824430638123865 | 0.957903915785494 | NM_026174 |
| Epb4.1l5 | 396.2296822 | -0.0202122685973416 | 0.0904388263865103 | -0.223491053620709 | 0.823153345236414 | 0.957903915785494 | NM_145506,NM_001113416 |
| Fundc1 | 1317.063268 | -0.0129097357182532 | 0.0581020393322681 | -0.222190750387027 | 0.82416539428892 | 0.957903915785494 | NM_028058 |
| Galnt6 | 835.4253515 | 0.0204297378136495 | 0.0922515424133198 | 0.22145686976286 | 0.824736715791311 | 0.957903915785494 | NM_001161767,NM_001161768,NM_172451 |
| Hspa12b | 329.6672846 | 0.0204160436403755 | 0.0920740423552846 | 0.221735063630599 | 0.824520132650776 | 0.957903915785494 | NM_028306 |
| Klhl20 | 931.7685458 | -0.0116781105171312 | 0.0528084648705355 | -0.22114088235212 | 0.824982738641973 | 0.957903915785494 | NM_001039482 |
| Mgll | 10334.47094 | -0.0112918327618739 | 0.0509592410336391 | -0.221585575703924 | 0.824636512293505 | 0.957903915785494 | NM_011844,NM_001166249,NM_001166251,NM_001166250 |
| Ndufa4 | 6566.228299 | 0.0139077634882817 | 0.0629827557572581 | 0.220818592661834 | 0.825233686058534 | 0.957903915785494 | NM_010886 |
| Nqo1 | 182.6227743 | 0.0204735845522869 | 0.0926536695745863 | 0.220968955134645 | 0.825116605705016 | 0.957903915785494 | NM_008706 |
| Pigg | 831.9668946 | 0.0171524798690943 | 0.0775737642536281 | 0.221111867319137 | 0.825005330154524 | 0.957903915785494 | NM_001081234 |
| Rab23 | 1300.573011 | -0.0151663216957973 | 0.0684627521818056 | -0.221526614289804 | 0.824682416113027 | 0.957903915785494 | NM_008999,NM_001159729 |
| Rab43 | 1010.413249 | -0.0151382081799687 | 0.0681544772604088 | -0.222116121911223 | 0.824223487843446 | 0.957903915785494 | NM_133717,NM_001039394 |
| Rft1 | 475.0497361 | 0.0167087919567373 | 0.0755975504512704 | 0.221022928084259 | 0.825074580396012 | 0.957903915785494 | NM_177815 |
| Rprd2 | 2354.08606 | 0.0135412694816308 | 0.0606849083205923 | 0.223140643306136 | 0.823426046917274 | 0.957903915785494 | NM_001081293 |
| Sec24b | 2723.814316 | -0.0117068761122368 | 0.0523122735427951 | -0.223788325748444 | 0.822922014311036 | 0.957903915785494 | NM_207209 |
| Sertad3 | 82.23851848 | -0.0209912139956222 | 0.0948267001604217 | -0.221363961417096 | 0.824809050988977 | 0.957903915785494 | NM_133210 |
| Slc6a7 | 3672.330954 | 0.0154510997744982 | 0.0691206700998734 | 0.223538049503465 | 0.823116773004308 | 0.957903915785494 | NM_201353 |
| Smc6 | 1751.248001 | -0.0165810848952426 | 0.0745674745579423 | -0.222363503572302 | 0.824030920542418 | 0.957903915785494 | NM_025695 |
| Sqrdl | 233.8218677 | 0.0200199841752168 | 0.0901359396046468 | 0.222108786606411 | 0.824229197966892 | 0.957903915785494 | NM_021507,NR_027888,NM_001162503 |
| Synpr | 4144.656626 | 0.0169383355250045 | 0.0762904938674509 | 0.222024195497195 | 0.824295048073882 | 0.957903915785494 | NM_001163032,NM_028052 |
| Tab1 | 998.4255223 | -0.015877768384607 | 0.0714198493974755 | -0.222315904031691 | 0.824067972244302 | 0.957903915785494 | NM_025609 |
| Tbc1d19 | 1099.698519 | -0.013366840744677 | 0.0605614223471722 | -0.220715436108002 | 0.825314011578963 | 0.957903915785494 | NM_144517 |
| Tti2 | 946.2853163 | -0.0121414683980545 | 0.0547542960300054 | -0.221744580396047 | 0.824512723769891 | 0.957903915785494 | NM_001112729,NM_144927,NM_001199988 |
| Ube3c | 4742.603854 | 0.0136094040448435 | 0.0613287303497268 | 0.221909111231163 | 0.824384637621521 | 0.957903915785494 | NM_133907 |
| Vps54 | 1302.855638 | -0.0140371587129275 | 0.0629420534125073 | -0.223017171380338 | 0.82352214221234 | 0.957903915785494 | NM_139061 |
| Zfp277 | 840.024259 | -0.015914405880467 | 0.0720472437761882 | -0.22088847603809 | 0.825179270598543 | 0.957903915785494 | NM_172575,NM_178845 |
| Zfp712 | 256.0083457 | 0.0204146815748685 | 0.0921232453371202 | 0.221601849784623 | 0.824623842376274 | 0.957903915785494 | NM_001166218 |
| Mrpl36 | 723.6044324 | -0.0139428885494809 | 0.0632475440582286 | -0.220449485542781 | 0.825521109303281 | 0.958018665314329 | NM_053163 |
| Btg1 | 978.7452275 | -0.015910227179571 | 0.0723812802689071 | -0.219811353439207 | 0.826018077111211 | 0.958041983654335 | NM_007569 |
| Chchd5 | 268.4690692 | -0.0174941852150814 | 0.0794806517491961 | -0.220106212393488 | 0.825788436659637 | 0.958041983654335 | NM_025395 |
| Fam102b | 2675.364313 | 0.0158736696344432 | 0.0721259128332332 | 0.22008275543279 | 0.825806704737156 | 0.958041983654335 | NM_001163567 |
| Med1 | 2082.332923 | -0.012996847439142 | 0.0590954360456477 | -0.219929800147386 | 0.82592582729866 | 0.958041983654335 | NM_134027,NM_013634,NM_001080118 |
| Ptpn7 | 77.37257228 | -0.0189308641522554 | 0.0859682524893048 | -0.220207618557915 | 0.825709463503759 | 0.958041983654335 | NM_177081 |
| Slc17a5 | 900.8355735 | -0.0140830109046167 | 0.0639640204270492 | -0.220170821199057 | 0.82573812036999 | 0.958041983654335 | NM_172773,NM_001276452 |
| Slc22a17 | 24948.50851 | 0.0122268247056629 | 0.055640668233663 | 0.219746187344774 | 0.82606883142438 | 0.958041983654335 | NM_021551 |
| Tmem209 | 1419.646656 | -0.0153421722206963 | 0.0697919446265872 | -0.219827263773528 | 0.826005685530853 | 0.958041983654335 | NM_178625 |
| Lrrc23 | 96.85106101 | -0.0205338674783597 | 0.0936624324125796 | -0.219232695003144 | 0.826468787985776 | 0.958429316164888 | NM_013588 |
| Frmd4a | 1991.289574 | -0.0119785162791028 | 0.0547620637460528 | -0.218737488321305 | 0.826854544592068 | 0.958455003409227 | NM_172475,NM_001177844,NM_001177843 |
| Pgp | 1058.514557 | -0.0166543510077939 | 0.0761529595621033 | -0.218696044166375 | 0.826886830698364 | 0.958455003409227 | NM_025954 |
| Prpf8 | 11075.16149 | 0.00995484917974843 | 0.0454686334242777 | 0.218938825076566 | 0.826697702034872 | 0.958455003409227 | NM_138659 |
| Snora28 | 39.00434522 | -0.0202287373872189 | 0.092482182051928 | -0.218731186250132 | 0.826859454055373 | 0.958455003409227 | NR_033168 |
| Vrk2 | 71.16682616 | 0.0207807428993781 | 0.0949778116594012 | 0.218795764361256 | 0.826809146491832 | 0.958455003409227 | NM_027260,NM_001252447 |
| Zfp882 | 392.5939632 | -0.0188768072828798 | 0.0862827047729293 | -0.218778575991075 | 0.826822536486176 | 0.958455003409227 | NM_001166645 |
| Poc1a | 353.8016106 | -0.0172422005278058 | 0.0788970550857035 | -0.218540483026598 | 0.827008019601359 | 0.95851898928663 | NM_027354 |
| Lrp4 | 1614.337485 | 0.0159998658779086 | 0.0732645608336294 | 0.218384791990242 | 0.827129313824664 | 0.958524924271472 | NM_172668,NM_001145857 |
| Lrrn4cl | 63.6927127 | 0.0205892825939512 | 0.0942885931625185 | 0.218364511584799 | 0.82714511398478 | 0.958524924271472 | NM_001013019 |
| Pkd2l1 | 55.08915914 | -0.0201291358030327 | 0.0922227248470674 | -0.218266548038055 | 0.827221436901941 | 0.958536901119783 | NM_181422 |
| 5330434G04Rik | 219.6216714 | 0.0197515950034767 | 0.0906186616261579 | 0.217963879062359 | 0.827457255118497 | 0.958657209112401 | NR_015552 |
| Samm50 | 3210.781337 | -0.0109938450959708 | 0.0504313548712263 | -0.217996227228932 | 0.827432050976646 | 0.958657209112401 | NM_178614 |
| Cox17 | 1040.03762 | -0.0142417191945533 | 0.065395730652634 | -0.217777507070023 | 0.827602470707917 | 0.958672527073621 | NM_001017429 |
| Lrrc46 | 35.97957151 | -0.0187895182698052 | 0.0862545442745028 | -0.217838009902505 | 0.827555328029266 | 0.958672527073621 | NM_027026 |
| Cdk5r2 | 10398.75056 | -0.0127029888320909 | 0.0583779857608474 | -0.217598957321519 | 0.827741596962659 | 0.958680787552191 | NM_009872 |
| Ftsj3 | 1534.944843 | -0.011740926211514 | 0.0539459734107228 | -0.217642308947201 | 0.827707816808292 | 0.958680787552191 | NM_025310 |
| Ankef1 | 33.97826602 | 0.0185157289701043 | 0.0851781071313623 | 0.217376619341273 | 0.827914850634899 | 0.958723519221682 | NM_175667 |
| Fstl3 | 75.66473761 | -0.0206807741798839 | 0.0951021084401926 | -0.217458629667391 | 0.827850944293048 | 0.958723519221682 | NM_031380 |
| Pkib | 328.963345 | -0.0182881713476002 | 0.084161898352474 | -0.217297514737709 | 0.827976493779155 | 0.958723519221682 | NM_001039050,NM_001039051,NM_001039052,NM_001190402,NM_001039053,NM_008863 |
| Siae | 1209.929236 | -0.0140843401035518 | 0.0648422246801309 | -0.21720939053264 | 0.828045166796146 | 0.958726613492812 | NM_011734 |
| Cenpj | 149.1597413 | -0.0196080029756078 | 0.0903736501649953 | -0.216965929115505 | 0.828234897097604 | 0.958819612652589 | NM_001014996 |
| U2af1 | 1460.892451 | -0.0127461484405309 | 0.0587550905205415 | -0.216936921168978 | 0.828257503756347 | 0.958819612652589 | NM_024187,NM_001163769 |
| Flii | 2811.669352 | -0.0122809260888364 | 0.056750334913679 | -0.216402706830126 | 0.828673856513108 | 0.959225152578644 | NM_022009 |
| Sco1 | 425.3247988 | -0.014817074279107 | 0.0684979230347132 | -0.216314212499524 | 0.828742831342351 | 0.959228555225418 | NM_001040026 |
| Megf9 | 3332.210653 | 0.0159900173142423 | 0.0739815984344971 | 0.216135061320685 | 0.828882470566088 | 0.959237314168498 | NM_172694 |
| Neil2 | 93.52987264 | 0.0205108975696477 | 0.0948950718761134 | 0.216142916214078 | 0.828876347963192 | 0.959237314168498 | NM_201610 |
| Igsf21 | 1807.409356 | 0.0149667662440499 | 0.0693494199289238 | 0.215816747213994 | 0.82913059354349 | 0.959448020537938 | NM_198610 |
| Gtf2a1 | 1339.992257 | -0.0137809148411342 | 0.06400630453303 | -0.215305584999406 | 0.829529075537986 | 0.959526943085267 | NM_031391,NM_175335 |
| Nt5c1a | 137.7849023 | 0.0194734471357963 | 0.0904397444444283 | 0.215319572776569 | 0.829518170631956 | 0.959526943085267 | NM_001085502 |
| Rps18 | 4312.937453 | 0.0120586383216441 | 0.0559687776338513 | 0.215452951296022 | 0.829414190054898 | 0.959526943085267 | NM_011296 |
| Sgol2 | 45.76606423 | -0.0195249098496472 | 0.0905733015817886 | -0.215570256451522 | 0.829322742583239 | 0.959526943085267 | NM_199007,NM_001177867 |
| Vps52 | 3489.843625 | 0.0111148459999969 | 0.0515728403396967 | 0.215517429848469 | 0.829363924283226 | 0.959526943085267 | NM_172620 |
| Abr | 26431.83324 | 0.0113376246239859 | 0.0527357251398644 | 0.214989451532458 | 0.829775542733674 | 0.959659198546922 | NM_198894,NM_198895,NM_198018 |
| Top2b | 4618.694173 | -0.0150647557496335 | 0.0700529007184919 | -0.215048279159364 | 0.829729677670291 | 0.959659198546922 | NM_009409 |
| Asb16 | 34.05496244 | 0.0190657748107037 | 0.0887331401082796 | 0.214866449980673 | 0.829871442985343 | 0.959693701202539 | NM_148953 |
| Abhd14a | 965.3864621 | -0.014325281683192 | 0.0668739716874479 | -0.214213113438884 | 0.830380870383304 | 0.960002080899847 | NM_145919,NM_001110272,NM_001110271 |
| Fam193a | 2099.228464 | -0.00996836717246871 | 0.0465305893882471 | -0.214232557625556 | 0.830365708098594 | 0.960002080899847 | NM_001243123 |
| H1fx | 526.7986928 | -0.0169411381552673 | 0.0790084721022661 | -0.214421791796445 | 0.830218149433734 | 0.960002080899847 | NM_198622 |
| Nphp1 | 609.2267704 | 0.0166058152151603 | 0.0775607793504369 | 0.214100675034885 | 0.830468549400212 | 0.960002080899847 | NM_016902 |
| Ptk7 | 526.0911573 | 0.016684900340175 | 0.0779150953588273 | 0.214142076876565 | 0.830436264164392 | 0.960002080899847 | NM_175168 |
| Gfra2 | 3117.68483 | 0.0137437856921054 | 0.0642556954147105 | 0.213892100978786 | 0.830631200201911 | 0.960113695721569 | NM_008115 |
| Mfsd8 | 776.047092 | 0.0163887903244524 | 0.0766625989393996 | 0.213778172814196 | 0.830720047045478 | 0.960118025690994 | NM_028140 |
| Ppig | 2515.492643 | -0.0160832771395003 | 0.0752547427852735 | -0.213717787666768 | 0.830767139261332 | 0.960118025690994 | NM_001081086 |
| Abca4 | 217.4940995 | -0.0133661935841957 | 0.0626341643117505 | -0.213401004564662 | 0.831014197021157 | 0.960162026441394 | NM_007378 |
| Csgalnact1 | 892.2368223 | -0.0129594786863918 | 0.0607968232184935 | -0.213160458068962 | 0.831201809341906 | 0.960162026441394 | NM_001252623,NM_172753 |
| Des | 77.99675585 | 0.0202621611767076 | 0.095021120489654 | 0.213238499738738 | 0.831140940309266 | 0.960162026441394 | NM_010043 |
| Kcnv1 | 3273.894911 | 0.018493541154992 | 0.0867247477700095 | 0.213244104255409 | 0.831136569074423 | 0.960162026441394 | NM_026200 |
| Mad2l1bp | 317.8831195 | 0.0173978870670657 | 0.0815289749636453 | 0.213395140498499 | 0.831018770538307 | 0.960162026441394 | NM_025649 |
| Mipol1 | 108.4291163 | -0.0201864054636727 | 0.0946146373483588 | -0.213353937925577 | 0.831050905513417 | 0.960162026441394 | NM_001164370 |
| Tyw5 | 366.2367048 | -0.0176724535324709 | 0.0829838075559531 | -0.212962673718665 | 0.831356076875701 | 0.960263865513394 | NR_004860,NM_001037742,NM_001114102 |
| Zfp688 | 265.4392461 | -0.0182130451663991 | 0.0855709553660047 | -0.212841437710939 | 0.831450641563479 | 0.960296733668688 | NM_026999 |
| Celsr1 | 352.2629801 | -0.0181975781043506 | 0.0855338745290602 | -0.212752879540935 | 0.831519718917951 | 0.960300161949607 | NM_009886 |
| Chst5 | 29.79830454 | 0.018952476772057 | 0.0891589976271186 | 0.212569424022914 | 0.831662822467765 | 0.960389073787006 | NM_019950 |
| A130010J15Rik | 230.549164 | -0.0175203559511968 | 0.0826148429423593 | -0.212072738108585 | 0.832050287800162 | 0.960519604033291 | NM_001160360,NM_181048,NM_001160359 |
| Arid2 | 2013.26159 | 0.0135753551933738 | 0.064254131355766 | 0.211275989682421 | 0.832671917502689 | 0.960519604033291 | NM_175251 |
| Becn1 | 4603.173043 | -0.00872901570298344 | 0.0412108338332818 | -0.211813615281279 | 0.832252446064124 | 0.960519604033291 | NM_019584 |
| C630043F03Rik | 234.3894136 | 0.0175594199482926 | 0.0828196949387311 | 0.212019857852445 | 0.832091542162892 | 0.960519604033291 | NR_027923 |
| Ccnk | 1268.21783 | -0.0116826211232654 | 0.0552518868206613 | -0.211442935174056 | 0.832541656572149 | 0.960519604033291 | NM_009832 |
| Decr2 | 1485.158105 | 0.012702843102442 | 0.0598765756353754 | 0.212150460637516 | 0.831989653657493 | 0.960519604033291 | NM_011933 |
| Fat1 | 1929.096596 | 0.0180018270843194 | 0.0852407866098892 | 0.211187951217603 | 0.832740612267559 | 0.960519604033291 | NM_001081286 |
| Ftsj1 | 858.0402353 | 0.0125309290629211 | 0.0591424554283342 | 0.211877051302097 | 0.83220295454736 | 0.960519604033291 | NM_133991 |
| Grin2d | 471.5495049 | 0.0182250541101793 | 0.0860998855548602 | 0.211673383929578 | 0.832361854124207 | 0.960519604033291 | NM_008172 |
| Nelfcd | 1829.640954 | 0.0111701741189323 | 0.0527202851756885 | 0.211876208212989 | 0.832203612304286 | 0.960519604033291 | NM_020580 |
| Rsl24d1 | 1374.753259 | 0.0135632375331868 | 0.0640853363929532 | 0.211643385157891 | 0.832385259493395 | 0.960519604033291 | NM_198609 |
| Rsph1 | 148.0019047 | -0.0199524499006646 | 0.094119397001164 | -0.211990838619778 | 0.832114181621026 | 0.960519604033291 | NM_025290 |
| Sorcs1 | 2350.759028 | 0.0131634463003385 | 0.0622746581823823 | 0.211377254962798 | 0.832592903681558 | 0.960519604033291 | NM_021377,NM_001252501 |
| Sphk2 | 1135.25874 | -0.0147591562848929 | 0.0698978537436754 | -0.211153211356342 | 0.832767719481981 | 0.960519604033291 | NM_001172561,NM_020011,NM_203280 |
| Tpm1 | 7665.884669 | 0.0119605758821038 | 0.0566125693163938 | 0.211270677634485 | 0.832676062357966 | 0.960519604033291 | [illegible]M_001164256,NM_001164255,NM_001164254,NM_001164253,NM_001164252,NM_001164251,NM_001164250,NM_024427,NM_001164248,NM_00116424[illegible] |
| A4galt | 98.42837766 | 0.0200526431719987 | 0.095047379753997 | 0.210975233866512 | 0.832906596902006 | 0.960603511440932 | NM_001170954,NM_001004150 |
| Oprd1 | 1066.366838 | 0.0169932080124463 | 0.0806715246872975 | 0.210646917587291 | 0.833162798697583 | 0.960810393705274 | NM_013622 |
| Smpd2 | 521.3697832 | -0.0176492114404961 | 0.0838140376049926 | -0.210575840811716 | 0.833218265834045 | 0.960810393705274 | NM_009213 |
| Dscr3 | 1426.670589 | -0.0130513065283666 | 0.0620438037881783 | -0.210356324588426 | 0.833389577892981 | 0.960817764718627 | NM_007834 |
| Fam194a | 92.12351764 | -0.0200008548133887 | 0.0951002484470389 | -0.21031338130023 | 0.833423092073158 | 0.960817764718627 | NM_001081262 |
| Tpp2 | 2504.815207 | 0.016091286249667 | 0.0764990881581659 | 0.210346118327547 | 0.833397543125797 | 0.960817764718627 | NM_009418 |
| Gucy1a3 | 5725.253391 | -0.0154856279407896 | 0.0737331320988026 | -0.210022651961113 | 0.833649993589496 | 0.960926821685528 | NM_021896 |
| Polr2l | 1011.955389 | 0.0145482804723941 | 0.0692580142841588 | 0.210059162434313 | 0.833621498003088 | 0.960926821685528 | NM_025593 |
| Lage3 | 355.6992567 | -0.0181572891542725 | 0.0865458142335691 | -0.209799738035507 | 0.8338239775308 | 0.961051017835981 | NM_025410 |
| Sytl5 | 154.6093884 | -0.0198718890123377 | 0.0947566085820875 | -0.209715072222353 | 0.833890061187161 | 0.961051017835981 | NM_177704 |
| Lias | 1124.591008 | 0.0122555522849971 | 0.0584637658256208 | 0.209626460285703 | 0.833959226142294 | 0.961054479884408 | NM_024471 |
| 4931403E22Rik | 24.28644521 | 0.0134426593888843 | 0.0642784611476471 | 0.209131630547388 | 0.834345483001174 | 0.961118357341399 | NR_045306 |
| 6430571L13Rik | 244.4606738 | -0.0191996958721779 | 0.0917782049709013 | -0.209196681044974 | 0.834294703249837 | 0.961118357341399 | NM_175486 |
| Ankrd35 | 704.4636499 | -0.015205177865776 | 0.0726923142372532 | -0.209171740167046 | 0.834314172528531 | 0.961118357341399 | NM_001081139 |
| Armcx6 | 306.0821621 | 0.0179425307249248 | 0.0857545173796476 | 0.209231318339659 | 0.834267664949791 | 0.961118357341399 | NM_001007578 |
| Nudt7 | 169.3910412 | 0.0181930955383453 | 0.086955105075328 | 0.209224007291864 | 0.834273372027686 | 0.961118357341399 | NM_024446,NM_024437 |
| Ankrd52 | 3743.396521 | -0.0134402608678331 | 0.0644045962052738 | -0.208684809155477 | 0.83469429959374 | 0.961348408066625 | NM_172790 |
| Dyx1c1 | 100.8995025 | -0.0198301007144366 | 0.0950892047657189 | -0.208542081756747 | 0.834805728343394 | 0.961348408066625 | NM_026314,NM_001163725 |
| Foxl2 | 26.68476654 | -0.0177406549293743 | 0.0849902009846574 | -0.20873765120966 | 0.834653046099713 | 0.961348408066625 | NM_012020 |
| Smim11 | 281.1019079 | 0.0186634412255363 | 0.089497145869462 | 0.20853671973806 | 0.834809914591244 | 0.961348408066625 | NM_138743 |
| Lrp6 | 2312.212666 | 0.0129038270945204 | 0.0619522130597457 | 0.208286781976233 | 0.835005051781329 | 0.96149361818145 | NM_008514 |
| Ufm1 | 2417.973957 | 0.0123665845324182 | 0.0593960071051099 | 0.208205654473267 | 0.835068393706263 | 0.96149361818145 | NM_026435 |
| Abhd11 | 900.5424293 | 0.0140199490843101 | 0.0676874226704694 | 0.207127831599751 | 0.83591002690691 | 0.961504005808573 | NR_033794,NM_001190437,NM_145215 |
| Acss3 | 52.00767089 | 0.0187764067247555 | 0.0903568782548573 | 0.207802738290664 | 0.835382994440573 | 0.961504005808573 | NM_198636,NM_001142804 |
| Clasp1 | 5500.032111 | 0.0109818878508865 | 0.0529590474688731 | 0.20736566036882 | 0.835724298766228 | 0.961504005808573 | NM_177548,NM_029709,NM_001081276 |
| Efemp2 | 377.4468579 | 0.0190393045104621 | 0.0919363994655874 | 0.207092126960972 | 0.835937910598683 | 0.961504005808573 | NM_001164352,NM_021474 |
| Ly6a | 935.7630607 | 0.016383482480416 | 0.0787303670486062 | 0.208096101854845 | 0.835153930807996 | 0.961504005808573 | NM_001271416,NM_001271417,NM_001271418,NM_001271419,NM_001271446,NM_010738 |
| Npm3 | 153.9901569 | 0.018344398681898 | 0.0883245055150746 | 0.20769319426042 | 0.835468532003269 | 0.961504005808573 | NM_008723 |
| Pogk | 3681.671189 | 0.0128940282514323 | 0.0621361530288976 | 0.20751249671726 | 0.835609634108177 | 0.961504005808573 | NM_175170,NM_001142948 |
| Polr2i | 462.4219065 | 0.015726382281446 | 0.0757192310688658 | 0.207693370091714 | 0.835468394703673 | 0.961504005808573 | NM_027259 |
| Rragd | 7845.945252 | 0.0146706118775694 | 0.0708190172890489 | 0.207156388766184 | 0.835887725212938 | 0.961504005808573 | NM_027491 |
| Slc35a3 | 523.7483129 | -0.0157723501127921 | 0.0760351135238597 | -0.207435083368985 | 0.835670085837516 | 0.961504005808573 | NM_144902 |
| Thap4 | 797.9404761 | 0.0135216935504588 | 0.0652669006607474 | 0.207175358620805 | 0.835872910794608 | 0.961504005808573 | NM_025920 |
| Tyw1 | 653.1093517 | 0.0125459927942156 | 0.0605528055967677 | 0.20719094137044 | 0.835860741563468 | 0.961504005808573 | NM_001015876,NM_178897 |
| Usp3 | 646.0074536 | 0.0154955361670409 | 0.0746114650929568 | 0.207683043721704 | 0.835476458159847 | 0.961504005808573 | NM_144937 |
| Zc3h12c | 601.2991478 | 0.0165323214427064 | 0.0798937185887364 | 0.206928926763426 | 0.83606536512188 | 0.961574465064167 | NM_001162921 |
| 4930539E08Rik | 500.2937884 | 0.0167406853437075 | 0.0810585252292031 | 0.206525905774514 | 0.836380131016492 | 0.961664000289241 | NM_172450 |
| Mast1 | 4013.927499 | 0.0126593415548063 | 0.0612728639508249 | 0.206606003678336 | 0.836317571173627 | 0.961664000289241 | NM_019945 |
| Ptrh1 | 97.3902771 | 0.0194502499239826 | 0.0940885056099936 | 0.206722912622354 | 0.836226262213405 | 0.961664000289241 | NM_178595 |
| Ror1 | 54.59995757 | -0.0193961405593144 | 0.0939325030596782 | -0.206490191653803 | 0.836408025585451 | 0.961664000289241 | NM_013845 |
| Kcnj9 | 7123.77755 | -0.0118787779308832 | 0.0576420467162244 | -0.206078350919133 | 0.836729709280285 | 0.961729368334026 | NM_008429 |
| Nckap5 | 156.1565278 | -0.0193307322041112 | 0.0937595232659181 | -0.206173533426422 | 0.836655360975445 | 0.961729368334026 | NM_176957,NM_172484,NM_001081756 |
| Ptgis | 47.70772622 | 0.0188286423962714 | 0.0913371183291525 | 0.206144475988595 | 0.836678057966557 | 0.961729368334026 | NM_008968 |
| Sema3f | 559.2375161 | -0.0161259662679048 | 0.0782488390000183 | -0.206085693717462 | 0.836723973672283 | 0.961729368334026 | NM_011349 |
| Ntsr1 | 503.2064413 | -0.0175581785108198 | 0.0852507302059399 | -0.205959274113014 | 0.836822723788726 | 0.96176017768455 | NM_018766 |
| Aacs | 1801.540069 | 0.0135526075725082 | 0.0658449450727049 | 0.205826089725546 | 0.836926760859819 | 0.961803649386846 | NM_030210 |
| Brsk2 | 9400.409769 | 0.00929022509145954 | 0.0452290150233134 | 0.205404099263954 | 0.837256417809595 | 0.961831963033541 | NM_001009930,NM_029426,NM_001009929 |
| Camkk2 | 13453.56029 | 0.0124815064043926 | 0.0608223349867026 | 0.205212549092721 | 0.837406065328821 | 0.961831963033541 | NM_001199676,NM_145358 |
| Col18a1 | 295.9233549 | -0.0190598613258614 | 0.092711817358096 | -0.205581789560261 | 0.837117603505598 | 0.961831963033541 | NM_009929,NM_001109991 |
| Fbxo8 | 717.0555845 | -0.0135402927806101 | 0.0660026530693825 | -0.205147704769634 | 0.837456725995606 | 0.961831963033541 | NM_015791 |
| Kcnk5 | 27.095239 | -0.0165124494338657 | 0.0804863145290928 | -0.20515847359239 | 0.837448312581424 | 0.961831963033541 | NM_021542 |
| Mnf1 | 1631.707752 | -0.0110061161387269 | 0.0536578810681445 | -0.205116488382189 | 0.837481114446387 | 0.961831963033541 | NM_026063 |
| Pcdh1 | 5096.426384 | 0.0162160434629154 | 0.0788773514335849 | 0.205585547285641 | 0.837114667969342 | 0.961831963033541 | NM_029357 |
| Prkx | 505.9890009 | 0.0168530715664879 | 0.082103473680287 | 0.205266242840274 | 0.837364116789858 | 0.961831963033541 | NM_016979 |
| 4930426L09Rik | 27.07507493 | 0.0167451755593566 | 0.0817845034928246 | 0.204747535831477 | 0.837769378849128 | 0.961858836323303 | NR_024323 |
| Cntnap5a | 1096.894068 | 0.0173542360633733 | 0.084725898642712 | 0.204827996414129 | 0.837706512750837 | 0.961858836323303 | NM_001077425 |
| Pxk | 1636.585095 | 0.00956474268533282 | 0.0466794339593087 | 0.20490271355198 | 0.837648135093984 | 0.961858836323303 | NM_145458,NM_178279 |
| Rad9a | 449.2376603 | -0.0155274066003599 | 0.0758314486474757 | -0.20476209906715 | 0.837758000109969 | 0.961858836323303 | NM_011237 |
| Ppp1r37 | 6378.701395 | 0.0126872041748117 | 0.0620232124731601 | 0.204555740809169 | 0.837919237827688 | 0.961954860403462 | NM_199149 |
| Ap1s3 | 119.2128339 | 0.0191018319750786 | 0.0936475411908092 | 0.203975798319768 | 0.838372411449897 | 0.962077671018172 | NM_183027 |
| Dnajc4 | 409.9737754 | 0.0144108406245423 | 0.0705600927069672 | 0.204235001283089 | 0.838169860687317 | 0.962077671018172 | NM_020566 |
| Gm7102 | 31.0553998 | -0.018065104897263 | 0.0885844033699069 | -0.203930987962153 | 0.838407429003366 | 0.962077671018172 | NM_001177513 |
| Krt80 | 165.6963024 | 0.0193832220271984 | 0.0949728999432792 | 0.204092136164892 | 0.838281499438321 | 0.962077671018172 | NM_028770 |
| Mrpl16 | 914.4551455 | -0.0139773927925798 | 0.0684378030694977 | -0.204234971984504 | 0.838169883581708 | 0.962077671018172 | NM_025606 |
| Scnm1 | 841.7269886 | -0.013813731567764 | 0.0677441603538461 | -0.203910292719123 | 0.838423601639752 | 0.962077671018172 | NM_027013,NM_001163573 |
| 2310009A05Rik | 130.8533915 | 0.0188177352890156 | 0.0923776406585085 | 0.203704437078869 | 0.838584474613772 | 0.962110266032671 | NR_040377 |
| 2310030G06Rik | 89.07557876 | -0.0193336211323641 | 0.0948739618646761 | -0.203782162696449 | 0.838523732462142 | 0.962110266032671 | NM_025865 |
| Map2k6 | 431.1362792 | 0.0161250990588034 | 0.0792325652818307 | 0.203516054307296 | 0.838731698724976 | 0.962130192139818 | NM_011943 |
| Wdr45 | 1514.940947 | 0.0109975471432246 | 0.054038625638311 | 0.203512709905558 | 0.838734312478763 | 0.962130192139818 | NM_172372 |
| Slc1a2 | 126585.6799 | -0.0129991809450105 | 0.0639296574193006 | -0.203335689095778 | 0.838872662275726 | 0.9622129099982 | NM_011393,NM_001077514,NM_001077515 |
| Brsk1 | 16089.54361 | 0.01198938325996 | 0.0591684193788789 | 0.202631460935051 | 0.839423097816561 | 0.962312360420122 | NM_001168572,NM_001003920 |
| Dclre1a | 400.8582844 | -0.0145395602638242 | 0.0717468583252423 | -0.202650828248306 | 0.839407958975577 | 0.962312360420122 | NM_018831 |
| Dhodh | 721.5631406 | 0.0144354917464563 | 0.0711426495659237 | 0.20290911056215 | 0.839206073211496 | 0.962312360420122 | NM_020046 |
| Fibp | 1282.158371 | -0.0149219640291836 | 0.0735494560091752 | -0.202883404430919 | 0.839226165874714 | 0.962312360420122 | NM_021438,NM_001253832 |
| Mll1 | 6452.190213 | 0.0178900348327709 | 0.0880754518339705 | 0.203121692370027 | 0.839039917077681 | 0.962312360420122 | NM_001081049 |
| Xpo5 | 2303.253858 | 0.00974718266870166 | 0.0480740764085783 | 0.202753404680331 | 0.839327779082641 | 0.962312360420122 | NM_028198 |
| Zfp820 | 25.81433504 | -0.0166762647267409 | 0.082289690706667 | -0.202653146263312 | 0.839406147057578 | 0.962312360420122 | NM_029281 |
| Gpld1 | 2151.630525 | -0.0140439435739468 | 0.0693559321861776 | -0.202490877582721 | 0.839532989336938 | 0.962362389765496 | NM_008156 |
| Etv3 | 1394.00133 | 0.0131085755293313 | 0.0648117923684395 | 0.202256025490118 | 0.839716576041628 | 0.962420939212616 | NM_001083318,NM_012051 |
| Kif2a | 3281.134747 | 0.014701647505362 | 0.0726728427973537 | 0.202299056146147 | 0.839682937807581 | 0.962420939212616 | NM_001145779,NM_008442 |
| Zmym2 | 3927.327832 | -0.0151149722667586 | 0.074860258320639 | -0.201909165234492 | 0.839987736822463 | 0.962655768448371 | NM_029498 |
| Il1r1 | 192.7285406 | 0.0183764165016901 | 0.0911186854708959 | 0.201675610295757 | 0.840170330965886 | 0.962746468032997 | NM_008362,NM_001123382 |
| Kank1 | 751.3449037 | 0.0142193518453173 | 0.0705190718961866 | 0.201638386084407 | 0.840199433791429 | 0.962746468032997 | NM_181404 |
| Capn7 | 2008.785801 | -0.0113724128797694 | 0.0564650903494305 | -0.201406086652691 | 0.840381056268833 | 0.962802683441996 | NM_009796 |
| Zfp322a | 1389.836772 | -0.0146757248751176 | 0.072845472416981 | -0.201463788869555 | 0.840335941203958 | 0.962802683441996 | NM_001111107,NM_172586 |
| Ndufaf4 | 1006.791975 | 0.0159669594229814 | 0.0793142775110025 | 0.20131254956923 | 0.840454190302243 | 0.962810533779999 | NM_026742 |
| Apoc1 | 35.58683875 | 0.0162100998481256 | 0.0808067485460913 | 0.200603292915314 | 0.84100878304051 | 0.96309494742331 | NM_001110009,NM_007469 |
| BC048403 | 526.6467308 | -0.0141234411745657 | 0.0705295785384196 | -0.200248483930358 | 0.841286250301568 | 0.96309494742331 | NM_173022 |
| Cdc42bpb | 8489.705522 | -0.00929197768246164 | 0.0463691096842496 | -0.20039154829013 | 0.841174368923073 | 0.96309494742331 | NM_183016 |
| D630023F18Rik | 158.2330122 | 0.0186150482035179 | 0.0928308950073132 | 0.200526432520676 | 0.841068887631912 | 0.96309494742331 | NM_175293 |
| Dip2a | 4555.945589 | 0.0105478689398924 | 0.0526782485523855 | 0.200231959675029 | 0.841299173058955 | 0.96309494742331 | NM_001081419 |
| Jmjd7 | 117.1452895 | -0.0187537192700361 | 0.0936319841206215 | -0.200291806759928 | 0.841252369981408 | 0.96309494742331 | NM_001114637 |
| Rnf135 | 163.6868104 | 0.0176575962955284 | 0.0879971818156123 | 0.200660929488944 | 0.840963712025812 | 0.96309494742331 | NM_028019 |

| | | | | | | |
|---|---|---|---|---|---|---|
| Kbtbd8 | 267.8250472 | -0.048316193 | 0.119084397 | -0.405730675 | 0.684940501 | 0.836417727 |
| Lrrc10b | 1828.373756 | 0.044464635 | 0.109643123 | 0.405539664 | 0.68508087 | 0.836523208 |
| Pllp | 1571.60785 | 0.067917795 | 0.167506536 | 0.405463554 | 0.685136803 | 0.836525582 |
| Rangap1 | 10335.5674 | -0.025702348 | 0.063450772 | -0.405075424 | 0.685422071 | 0.836796779 |
| Xrn2 | 1863.653306 | -0.025624681 | 0.063280051 | -0.404940912 | 0.685520945 | 0.836796779 |
| Zfp354a | 300.3412002 | 0.040246773 | 0.099374148 | 0.405002446 | 0.685475713 | 0.836796779 |
| Mfsd9 | 227.6935512 | -0.046850541 | 0.115767722 | -0.404694331 | 0.68570221 | 0.836952107 |
| Ccdc53 | 414.3496882 | 0.035158573 | 0.086908667 | 0.404546224 | 0.685811094 | 0.837009587 |
| Dtna | 5221.727049 | -0.017857933 | 0.044158009 | -0.404409837 | 0.685911367 | 0.837009587 |
| Fam179a | 146.8151778 | 0.053414648 | 0.132071751 | 0.404436583 | 0.685891703 | 0.837009587 |
| Arc | 9540.124169 | 0.060139412 | 0.148748453 | 0.404302777 | 0.685990084 | 0.83703972 |
| Psme2 | 772.3363048 | -0.028931852 | 0.071612342 | -0.4040065 | 0.686207939 | 0.83723961 |
| Smarca5 | 2757.44182 | 0.032067671 | 0.079436033 | 0.403691747 | 0.686439409 | 0.837456079 |
| Rnf220 | 5440.385001 | 0.029801739 | 0.073854293 | 0.403520746 | 0.686565176 | 0.837543567 |
| Mgst3 | 2227.095869 | -0.039387021 | 0.097649408 | -0.403351354 | 0.686689768 | 0.837599793 |
| Plagl2 | 418.5569075 | 0.037298967 | 0.092481878 | 0.403311088 | 0.686719386 | 0.837599793 |
| 1700003M07Rik | 660.3946791 | -0.038614474 | 0.095838055 | -0.402913794 | 0.687011643 | 0.83763747 |
| Bcan | 9766.838403 | 0.031938297 | 0.079270706 | 0.402901638 | 0.687020586 | 0.83763747 |
| Fam192a | 1543.191143 | 0.020741125 | 0.051478878 | 0.402905544 | 0.687017712 | 0.83763747 |
| Neurl2 | 60.00124942 | 0.057442313 | 0.142481912 | 0.403155125 | 0.68683411 | 0.83763747 |
| Wdr5 | 1106.548806 | 0.025343042 | 0.06288924 | 0.402978986 | 0.686963683 | 0.83763747 |
| Ptpla | 193.4111578 | -0.043673695 | 0.108511975 | -0.402478115 | 0.687332191 | 0.837951449 |
| Vit | 82.39122005 | -0.055692689 | 0.138524453 | -0.402042298 | 0.687652898 | 0.838276475 |
| Abcf2 | 4194.718615 | 0.031491521 | 0.078369054 | 0.401836171 | 0.6878046 | 0.83832949 |
| Dnmt3b | 25.11731642 | 0.067067145 | 0.166871628 | 0.401908617 | 0.687751281 | 0.83832949 |
| Cdc42ep1 | 821.9142013 | 0.053383909 | 0.132941511 | 0.401559368 | 0.688008338 | 0.838511853 |
| Pofut1 | 934.8873864 | -0.029263613 | 0.072893089 | -0.401459367 | 0.688081948 | 0.838535606 |
| Gpx3 | 386.4456488 | -0.067225304 | 0.167503861 | -0.401335845 | 0.688172876 | 0.838580459 |
| 1600002K03Rik | 71.03084095 | 0.058013124 | 0.144772723 | 0.40071861 | 0.688627308 | 0.838994182 |
| Hexb | 4530.336024 | 0.054426439 | 0.135842691 | 0.400657839 | 0.688672055 | 0.838994182 |
| Sfpi1 | 107.2398593 | 0.055718714 | 0.139069387 | 0.400654054 | 0.688674843 | 0.838994182 |
| Agpat6 | 3763.62684 | -0.023944907 | 0.059827772 | -0.400230637 | 0.688986651 | 0.839242074 |
| Asf1a | 355.9358031 | -0.035580053 | 0.088884471 | -0.400295486 | 0.688938892 | 0.839242074 |
| Ormdl1 | 278.3572034 | -0.039656793 | 0.099104302 | -0.400152078 | 0.689044509 | 0.83924657 |
| Ccdc79 | 37.12780319 | -0.065955654 | 0.164864214 | -0.400060466 | 0.689111982 | 0.839262777 |
| Fcrl1 | 17.80957779 | 0.066442633 | 0.166127649 | 0.399949278 | 0.689193876 | 0.839296542 |
| Rtn4rl2 | 832.2749924 | -0.04348449 | 0.108745749 | -0.399873011 | 0.689250052 | 0.839298986 |
| Chaf1a | 208.4733668 | 0.042383003 | 0.106101926 | 0.399455553 | 0.689557567 | 0.839342364 |
| Dmc1 | 25.06697163 | -0.067073538 | 0.167966988 | -0.399325716 | 0.689653221 | 0.839342364 |
| Efha1 | 825.3441301 | 0.029182917 | 0.073072996 | 0.399366638 | 0.689623072 | 0.839342364 |
| Nfatc1 | 273.447019 | -0.039985787 | 0.100063617 | -0.399603657 | 0.689448463 | 0.839342364 |
| Parp1 | 3060.247051 | 0.028112605 | 0.070389922 | 0.399383951 | 0.689610318 | 0.839342364 |
| Phkg1 | 546.0504954 | 0.060236702 | 0.150858144 | 0.399293674 | 0.689676828 | 0.839342364 |
| Scaf8 | 2646.538638 | 0.0296285 | 0.074219712 | 0.399199874 | 0.689745936 | 0.839342364 |
| Tgfb3 | 609.7779621 | 0.05113336 | 0.12810551 | 0.399150356 | 0.68978242 | 0.839342364 |
| Tnfrsf22 | 37.57004468 | -0.066271884 | 0.166057775 | -0.399089314 | 0.689827395 | 0.839342364 |
| Zfand5 | 5774.781522 | -0.029129811 | 0.072978563 | -0.399155727 | 0.689778463 | 0.839342364 |
| Mfi2 | 33.32889935 | 0.066366362 | 0.166363551 | 0.398923688 | 0.689949434 | 0.839424934 |
| Gpr179 | 36.0180086 | 0.064177778 | 0.161075798 | 0.398432159 | 0.690311657 | 0.839697927 |
| Tmem74b | 126.2159995 | -0.056264341 | 0.141226258 | -0.398398585 | 0.690336401 | 0.839697927 |
| Zfp784 | 196.996887 | -0.03802579 | 0.095421127 | -0.398504938 | 0.690258019 | 0.839697927 |
| Olfr287 | 149.0922863 | -0.052759501 | 0.13248547 | -0.398228584 | 0.690461698 | 0.839784406 |
| Gm16062 | 34.96471668 | 0.06516609 | 0.163853342 | 0.397709863 | 0.690844066 | 0.840183513 |
| Snora65 | 38.85525724 | 0.066231292 | 0.166665587 | 0.397390324 | 0.69107965 | 0.840404057 |
| Fam222a | 391.3833895 | -0.049039226 | 0.123476484 | -0.39715438 | 0.691253621 | 0.840515881 |
| Mllt1 | 3411.091759 | -0.035317246 | 0.088933779 | -0.397118471 | 0.6912801 | 0.840515881 |
| Top2b | 4331.857518 | -0.024492177 | 0.061692551 | -0.397003792 | 0.691364664 | 0.840552739 |
| Mettl25 | 177.0620277 | -0.053379306 | 0.134525808 | -0.396796024 | 0.691517883 | 0.840673055 |
| Pianp | 6769.683666 | 0.023412932 | 0.05906214 | 0.396411856 | 0.691801221 | 0.840951524 |
| Lsm14a | 2436.089014 | 0.029168993 | 0.073624027 | 0.396188504 | 0.691965971 | 0.841034242 |
| Mapk8ip2 | 11704.49379 | 0.027230757 | 0.068734609 | 0.396172427 | 0.691977831 | 0.841034242 |
| Aldh3a2 | 2160.572348 | 0.017036943 | 0.043053913 | 0.395711849 | 0.692317613 | 0.841315224 |
| Hfe | 154.6631992 | -0.05146739 | 0.130041332 | -0.395777165 | 0.692269424 | 0.841315224 |
| 2010107G12Rik | 15.92267058 | 0.065523624 | 0.165653057 | 0.395547328 | 0.692439 | 0.841396743 |
| 1700003M02Rik | 56.26096886 | 0.062651637 | 0.158493806 | 0.395293914 | 0.692625991 | 0.841530804 |
| Mtag2 | 32.39563154 | 0.063263813 | 0.160179311 | 0.394956207 | 0.692875207 | 0.841530804 |
| Pfkp | 11400.6714 | -0.020219374 | 0.051161958 | -0.395203287 | 0.692692867 | 0.841530804 |
| Rab3ip | 1317.752991 | -0.034754612 | 0.087990749 | -0.394980297 | 0.692857429 | 0.841530804 |
| Snora7a | 24.12184307 | -0.066360765 | 0.167950999 | -0.395119798 | 0.692754478 | 0.841530804 |
| Zfp91 | 3705.715971 | -0.033603239 | 0.085062009 | -0.395044027 | 0.692810396 | 0.841530804 |
| Wdr12 | 1248.92305 | 0.026173169 | 0.066348901 | 0.39447781 | 0.693228307 | 0.841893666 |
| Aif1 | 206.2758759 | 0.064627624 | 0.1638966 | 0.394319493 | 0.693345173 | 0.841969599 |
| 2310067B10Rik | 3429.968763 | 0.02238125 | 0.056827085 | 0.393848289 | 0.69369305 | 0.842326028 |
| Anp32b | 1811.987244 | -0.030006641 | 0.076231737 | -0.393623995 | 0.693858663 | 0.842461102 |
| Zfp418 | 141.8325229 | -0.04409291 | 0.112084624 | -0.393389467 | 0.694031848 | 0.842605348 |
| 4933412E12Rik | 168.805319 | -0.048073042 | 0.122264864 | -0.393187707 | 0.694180848 | 0.842644671 |
| Chd2 | 2865.842598 | -0.043940338 | 0.11176937 | -0.393133985 | 0.694220524 | 0.842644671 |
| Mat2b | 4187.440749 | 0.028264945 | 0.07189817 | 0.393124685 | 0.694227393 | 0.842644671 |
| Ube2r2 | 3125.27766 | 0.020939221 | 0.053289438 | 0.39293379 | 0.694368384 | 0.842749784 |
| Alg6 | 291.4031744 | 0.044408702 | 0.113040349 | 0.392857084 | 0.69442504 | 0.842752531 |
| Abhd3 | 1836.469352 | 0.029619715 | 0.075442838 | 0.39261136 | 0.694606548 | 0.842774773 |
| Bmi1 | 1313.96627 | 0.027527885 | 0.070110763 | 0.392634218 | 0.694589663 | 0.842774773 |
| Tug1 | 3967.912928 | -0.027551446 | 0.070159533 | -0.392697115 | 0.694543202 | 0.842774773 |
| Nup210l | 20.13740858 | -0.065542483 | 0.167019857 | -0.3924233 | 0.694745473 | 0.842877328 |
| AA986860 | 146.5955933 | 0.0460316 | 0.117407751 | 0.392066109 | 0.695009368 | 0.843085508 |
| C8g | 31.25616881 | 0.065244137 | 0.166467579 | 0.391932997 | 0.695107722 | 0.843085508 |
| Grb7 | 90.33636937 | 0.047807846 | 0.121973216 | 0.391953637 | 0.695092471 | 0.843085508 |
| Zfp773 | 85.10809361 | 0.049244226 | 0.125656218 | 0.391896457 | 0.695134721 | 0.843085508 |
| 2610100L16Rik | 185.5937378 | -0.045279267 | 0.115594281 | -0.391708539 | 0.69527358 | 0.843187919 |
| Mc5r | 12.8930283 | 0.063131737 | 0.161278913 | 0.391444462 | 0.695468733 | 0.843303626 |
| Zic5 | 49.68080126 | -0.061336979 | 0.156698886 | -0.391432132 | 0.695477845 | 0.843303626 |
| Sst | 5341.811177 | 0.030222106 | 0.077237779 | 0.391286574 | 0.695585422 | 0.843368067 |
| Arl5b | 366.6798005 | -0.041079447 | 0.10500875 | -0.391200222 | 0.695649244 | 0.843379451 |
| N6amt1 | 718.4268342 | -0.029724888 | 0.07601353 | -0.391047338 | 0.695762246 | 0.843384465 |
| Spry2 | 1909.683635 | -0.047083299 | 0.120395533 | -0.391071809 | 0.695744157 | 0.843384465 |
| Tstd3 | 704.4772325 | 0.025840903 | 0.0661138 | 0.390854896 | 0.695904495 | 0.843490905 |
| Smagp | 42.09226627 | -0.059086622 | 0.151201547 | -0.39078054 | 0.69595946 | 0.843491543 |
| Srgap3 | 16963.41267 | -0.028421465 | 0.072766935 | -0.390582139 | 0.69610613 | 0.843603315 |
| 9030025P20Rik | 58.63900224 | 0.064237722 | 0.16452512 | 0.390443248 | 0.696208813 | 0.843661766 |
| Dtx4 | 2073.992369 | 0.031096735 | 0.079693813 | 0.390202624 | 0.696386721 | 0.84377249 |
| Gpr133 | 269.2582667 | 0.056524612 | 0.144870879 | 0.390172358 | 0.6964091 | 0.84377249 |
| Fam73a | 4280.187357 | 0.025325969 | 0.064972193 | 0.389797042 | 0.696686631 | 0.843827087 |
| Fuz | 439.7195122 | -0.056198743 | 0.14422155 | -0.389669528 | 0.696780932 | 0.843827087 |
| Hsd17b4 | 2302.102311 | 0.023866663 | 0.061232895 | 0.389768651 | 0.696707627 | 0.843827087 |
| Plxnb3 | 1186.867762 | 0.056628476 | 0.145244253 | 0.389884454 | 0.69662199 | 0.843827087 |
| Rab26 | 1197.178635 | -0.032308076 | 0.082909254 | -0.389679976 | 0.696773205 | 0.843827087 |
| Zfp81 | 341.8961846 | -0.04418349 | 0.113335699 | -0.389846191 | 0.696650286 | 0.843827087 |
| Arl6ip1 | 9707.698232 | -0.028794815 | 0.073926905 | -0.389503861 | 0.696903455 | 0.843843555 |
| Commd5 | 425.3007849 | -0.033547374 | 0.086119502 | -0.389544444 | 0.69687344 | 0.843843555 |
| Clcn6 | 2520.924009 | -0.020796594 | 0.053445316 | -0.389119112 | 0.697188036 | 0.844056214 |
| Kcnmb1 | 45.78864986 | -0.065392905 | 0.168047908 | -0.389132516 | 0.697178121 | 0.844056214 |
| Usf1 | 1790.081057 | 0.026916393 | 0.06922647 | 0.388816495 | 0.697411898 | 0.844261266 |
| Ltn1 | 2659.250855 | -0.023786881 | 0.061191174 | -0.388730582 | 0.697475457 | 0.844272245 |
| 9130017N09Rik | 9.187396398 | 0.060423061 | 0.15556757 | 0.388403966 | 0.69771711 | 0.844441138 |
| Ankrd46 | 7994.453846 | 0.019788812 | 0.050950634 | 0.388391877 | 0.697726055 | 0.844441138 |
| Fbxo33 | 1961.716611 | 0.035248455 | 0.090788653 | 0.388247359 | 0.697832989 | 0.844441138 |
| Rasgrf2 | 8122.328737 | 0.029229992 | 0.075274904 | 0.388309927 | 0.697786692 | 0.844441138 |
| Mon1a | 738.4380455 | -0.026846981 | 0.06919922 | -0.387966523 | 0.698040808 | 0.844626652 |
| Jak2 | 2195.093135 | -0.036377935 | 0.09390351 | -0.387396973 | 0.698462345 | 0.845070714 |
| Fabp3 | 2156.524791 | 0.029039182 | 0.075006478 | 0.387155648 | 0.698640982 | 0.845220846 |
| Gpr139 | 41.83853061 | 0.062438676 | 0.161548223 | 0.386501779 | 0.699125085 | 0.845410448 |
| Ispd | 249.1273609 | -0.050746475 | 0.131277369 | -0.3865592 | 0.699082568 | 0.845410448 |
| Mark3 | 2723.965011 | -0.029445165 | 0.076142225 | -0.386712699 | 0.698968914 | 0.845410448 |
| Rasip1 | 334.2003882 | 0.039342935 | 0.101731422 | 0.386733366 | 0.698953612 | 0.845410448 |
| Rsu1 | 759.3770822 | -0.031148325 | 0.080576545 | -0.386568146 | 0.699075944 | 0.845410448 |
| Taf9b | 1138.169589 | -0.028338511 | 0.073254365 | -0.386850811 | 0.698866658 | 0.845410448 |
| Bid | 395.5447106 | 0.037935109 | 0.098230812 | 0.386183397 | 0.699360849 | 0.84548184 |
| Them6 | 1651.046142 | -0.035819261 | 0.092765421 | -0.386127293 | 0.699402398 | 0.84548184 |
| Tor1aip2 | 2140.205571 | 0.022300524 | 0.057744035 | 0.386196149 | 0.699351405 | 0.84548184 |
| Zfp868 | 448.0792393 | -0.034222656 | 0.08861437 | -0.386197584 | 0.699350343 | 0.84548184 |
| Pcmt1 | 4767.629079 | -0.025948313 | 0.067250235 | -0.385847172 | 0.699609857 | 0.845666651 |
| Brs3 | 9.319675715 | 0.048610902 | 0.126043657 | 0.385667179 | 0.699743173 | 0.845695844 |
| Krtcap2 | 711.2375452 | 0.025658781 | 0.06652565 | 0.385697562 | 0.699720669 | 0.845695844 |
| Nme7 | 607.0840906 | -0.026452246 | 0.068604265 | -0.385577285 | 0.699809758 | 0.84571035 |
| B230217O12Rik | 371.3139376 | -0.036975898 | 0.095999573 | -0.385167321 | 0.700113453 | 0.846011375 |
| Bhlhb9 | 2573.006799 | -0.029201478 | 0.075872705 | -0.384874617 | 0.700330313 | 0.846207429 |
| Gdap2 | 1286.869663 | 0.037675469 | 0.097912069 | 0.384788811 | 0.70039389 | 0.846218257 |
| Nol11 | 1013.98569 | -0.029049945 | 0.075546685 | -0.384529713 | 0.700585878 | 0.846318228 |
| Ppp3cb | 9947.016993 | -0.033040943 | 0.085916135 | -0.384572033 | 0.700554518 | 0.846318228 |
| Rbm27 | 1572.999398 | -0.026981311 | 0.070246418 | -0.384095187 | 0.700907899 | 0.846575765 |
| Zfp593 | 279.453477 | -0.03957743 | 0.103040844 | -0.384094583 | 0.700908346 | 0.846575765 |
| Gm10790 | 58.58123094 | 0.062580504 | 0.163030761 | 0.383857031 | 0.701084415 | 0.84672242 |
| Csnk1g1 | 1527.726187 | -0.021563721 | 0.05620739 | -0.383645649 | 0.701241102 | 0.846734721 |
| Itih3 | 1230.272969 | 0.064399527 | 0.167825911 | 0.383728155 | 0.701179943 | 0.846734721 |
| Kif24 | 32.77392906 | 0.060248353 | 0.157051448 | 0.383621758 | 0.701258811 | 0.846734721 |
| Tpr | 4758.115382 | -0.030136161 | 0.078571991 | -0.383548392 | 0.701313197 | 0.846734721 |
| Ndufaf7 | 1213.227337 | 0.020498002 | 0.053494447 | 0.383179999 | 0.701586307 | 0.84699846 |
| Stoml2 | 1499.572987 | 0.026807907 | 0.069978922 | 0.383085457 | 0.701656402 | 0.847017086 |
| Ppil3 | 491.9346333 | -0.03359692 | 0.087727436 | -0.382969358 | 0.701742484 | 0.847055005 |
| Adap1 | 7550.623652 | -0.033917203 | 0.088592482 | -0.382845159 | 0.701834575 | 0.847100172 |
| Col4a5 | 190.3006083 | -0.053208792 | 0.139131099 | -0.38243637 | 0.702137718 | 0.847268053 |
| Rsl24d1 | 1280.848223 | -0.025595047 | 0.066919368 | -0.382506784 | 0.702085498 | 0.847268053 |
| Srek1ip1 | 488.5347867 | -0.033013907 | 0.086305935 | -0.382521864 | 0.702074314 | 0.847268053 |
| Epx | 11.18929245 | 0.061630194 | 0.161225498 | 0.38226084 | 0.702267898 | 0.847271714 |
| Fam53a | 611.20584 | 0.030691363 | 0.080314977 | 0.382137483 | 0.70235939 | 0.847271714 |
| Ocln | 342.3514371 | -0.03117342 | 0.081576481 | -0.382137352 | 0.702359487 | 0.847271714 |
| Wdr37 | 2972.896414 | 0.027708478 | 0.072505721 | 0.382155749 | 0.702345843 | 0.847271714 |
| Ppp2r2d | 2126.69928 | -0.032222859 | 0.084359731 | -0.381969674 | 0.702483859 | 0.847289811 |
| Smndc1 | 865.4516355 | -0.029340821 | 0.076806056 | -0.382011812 | 0.702452603 | 0.847289811 |
| Otof | 1550.687555 | 0.050984659 | 0.133526591 | 0.381831503 | 0.70258635 | 0.847347467 |
| 2010320M18Rik | 135.5676894 | -0.045816035 | 0.120049253 | -0.381643651 | 0.702725702 | 0.847383611 |
| Fbln5 | 280.9131924 | -0.054149807 | 0.141866258 | -0.381696165 | 0.702686745 | 0.847383611 |
| Arl8a | 7663.810906 | 0.020542989 | 0.053858527 | 0.381425007 | 0.702887908 | 0.847513249 |
| Klf6 | 2067.493996 | 0.026483878 | 0.069494044 | 0.381095657 | 0.70313227 | 0.847741918 |
| Uba2 | 3359.460742 | -0.028045681 | 0.073650905 | -0.380792083 | 0.703357534 | 0.847947529 |
| Aff2 | 504.5553694 | 0.038694218 | 0.101760258 | 0.380248826 | 0.703760719 | 0.848206358 |
| Gpr101 | 348.2286257 | 0.055498738 | 0.145907217 | 0.380370071 | 0.703670728 | 0.848206358 |
| Trap1 | 2511.036339 | 0.029011226 | 0.076303616 | 0.380207751 | 0.703791206 | 0.848206358 |
| Uhrf1bp1l | 4180.839776 | 0.038827242 | 0.102102742 | 0.380276194 | 0.703740405 | 0.848206358 |
| Ankrd34a | 2536.322731 | 0.026177021 | 0.068863187 | 0.380130837 | 0.703848296 | 0.848209186 |
| Adprhl2 | 640.9688262 | 0.025645905 | 0.067504638 | 0.379913231 | 0.704009826 | 0.84827189 |
| Gng8 | 37.21611361 | 0.063615578 | 0.167416468 | 0.379983996 | 0.703957295 | 0.84827189 |
| Grhl3 | 47.40409557 | 0.059924792 | 0.157863917 | 0.37959778 | 0.704244009 | 0.848488078 |
| Bcs1l | 666.1490812 | -0.045269534 | 0.11942293 | -0.379069031 | 0.704636603 | 0.848723455 |
| Dbil5 | 12.64777304 | 0.062051921 | 0.163640243 | 0.379197193 | 0.704541436 | 0.848723455 |
| Stk39 | 3028.607662 | 0.033032404 | 0.087139605 | 0.379074516 | 0.70463253 | 0.848723455 |
| Ube3c | 4442.656102 | -0.022501689 | 0.059365012 | -0.379039567 | 0.704658482 | 0.848723455 |
| Eda2r | 17.08784489 | 0.063115673 | 0.166611154 | 0.378820214 | 0.704821376 | 0.848818156 |
| Snapc1 | 999.5470086 | 0.026354631 | 0.069576545 | 0.378786147 | 0.704846675 | 0.848818156 |
| Gorasp1 | 1363.655214 | -0.022381743 | 0.059146764 | -0.378410272 | 0.705125839 | 0.849088346 |
| Cuzd1 | 8.582796885 | 0.058118933 | 0.15361726 | 0.378335958 | 0.705181037 | 0.849088824 |
| Csf1r | 4813.85558 | 0.038719839 | 0.102387454 | 0.378169759 | 0.705304489 | 0.849105497 |
| Rin1 | 3215.854483 | 0.032166781 | 0.085057851 | 0.37817533 | 0.705300351 | 0.849105497 |
| Pdap1 | 3467.31506 | 0.029755502 | 0.078712587 | 0.378027241 | 0.705410357 | 0.849166971 |
| Ift20 | 1240.024748 | 0.022663745 | 0.059992209 | 0.377778141 | 0.705595413 | 0.849191808 |
| Tox4 | 3198.239775 | 0.021593571 | 0.057159243 | 0.377779159 | 0.705594657 | 0.849191808 |
| Zfp618 | 79.27789691 | -0.059639818 | 0.157835378 | -0.377860897 | 0.705533932 | 0.849191808 |
| Egfl6 | 96.39370506 | -0.057913811 | 0.153334885 | -0.377694947 | 0.705657222 | 0.849200233 |
| Pcbp4 | 4421.508976 | 0.030369633 | 0.080471144 | 0.3773978 | 0.705878002 | 0.84939995 |
| 4930579K19Rik | 14.1102877 | 0.062157446 | 0.164778785 | 0.377217526 | 0.706011957 | 0.849429202 |
| Tnfsf13 | 13.30771191 | 0.060548467 | 0.160487252 | 0.377280101 | 0.705965458 | 0.849429202 |
| Lrrn1 | 2747.226897 | -0.030951237 | 0.082088794 | -0.377045823 | 0.706139551 | 0.849450793 |
| Rab11fip1 | 161.312349 | 0.050893968 | 0.134977548 | 0.377055066 | 0.706132683 | 0.849450793 |
| Dhx8 | 1862.569677 | 0.023418723 | 0.062224038 | 0.37636135 | 0.706648276 | 0.84980222 |
| Nhej1 | 58.00484063 | 0.057226202 | 0.152052743 | 0.37635758 | 0.706651078 | 0.84980222 |
| Pcgf3 | 1380.565886 | 0.037299055 | 0.09906375 | 0.376515675 | 0.706533564 | 0.84980222 |
| Zswim3 | 332.9282698 | -0.038112889 | 0.101257182 | -0.376396892 | 0.706621856 | 0.84980222 |
| Lrrn2 | 4845.566684 | 0.032390796 | 0.086101701 | 0.37619229 | 0.706773947 | 0.849884015 |
| Fmod | 1225.307613 | 0.063018581 | 0.167606159 | 0.375992035 | 0.706922818 | 0.849997062 |
| Otub2 | 1393.675464 | 0.03696995 | 0.098409536 | 0.375674463 | 0.707158925 | 0.850192579 |
| Rab2b | 1609.713451 | 0.026732494 | 0.071167901 | 0.375625716 | 0.70719517 | 0.850192579 |
| Trim26 | 788.4999411 | -0.026460061 | 0.07048361 | -0.375407287 | 0.707357586 | 0.850321859 |
| Raly | 2723.950879 | -0.02349248 | 0.062591658 | -0.375329255 | 0.707415611 | 0.850325639 |
| Tmem129 | 995.40102 | 0.02389205 | 0.063679026 | 0.37519496 | 0.707515478 | 0.850379708 |
| AW551984 | 527.8791313 | 0.061020023 | 0.162798798 | 0.374818635 | 0.707795354 | 0.850403021 |
| Anapc2 | 3495.689469 | -0.021861754 | 0.058329138 | -0.374799879 | 0.707809304 | 0.850403021 |
| Aqr | 1995.205657 | 0.02482254 | 0.066180105 | 0.375075555 | 0.707604277 | 0.850403021 |
| Atp2a2 | 39242.75966 | -0.022093619 | 0.058942614 | -0.374832697 | 0.707784895 | 0.850403021 |
| Herc4 | 1218.566154 | -0.035370802 | 0.094353272 | -0.374876267 | 0.70775249 | 0.850403021 |
| Cpn1 | 12.60344936 | 0.06219576 | 0.166043579 | 0.374574917 | 0.707976631 | 0.850538103 |
| Pum1 | 4424.690301 | -0.027524402 | 0.07351877 | -0.374386056 | 0.708117116 | 0.85064092 |
| Mapk3 | 7065.011126 | -0.02758795 | 0.073740793 | -0.374120597 | 0.708314595 | 0.850812182 |
| Cyyr1 | 129.675924 | -0.049720038 | 0.13298096 | -0.373888401 | 0.708487347 | 0.850953717 |
| 2310033P09Rik | 408.34126 | -0.032738151 | 0.087627654 | -0.373605249 | 0.708698029 | 0.850973901 |
| 2810403D21Rik | 8.541697037 | -0.056505798 | 0.151286654 | -0.373501541 | 0.708775198 | 0.850973901 |
| Btc | 15.5488052 | 0.062617156 | 0.167534388 | 0.373757032 | 0.70858509 | 0.850973901 |
| Dnm1l | 8928.123971 | -0.022806773 | 0.061062843 | -0.373496747 | 0.708778766 | 0.850973901 |
| Prrg1 | 117.0713054 | 0.05829068 | 0.15602729 | 0.37359285 | 0.708707254 | 0.850973901 |
| Hdac1 | 934.2778922 | -0.024674485 | 0.066080342 | -0.373401287 | 0.708849802 | 0.850993246 |
| Gal3st3 | 1754.830002 | 0.033857842 | 0.090719147 | 0.373216055 | 0.708987648 | 0.851092788 |
| 1700113A16Rik | 35.5575411 | 0.062712493 | 0.168119141 | 0.373024112 | 0.709130498 | 0.851132384 |
| Mterfd1 | 494.2881501 | 0.034486468 | 0.092435419 | 0.373087161 | 0.709083574 | 0.851132384 |
| Cxx1c | 3927.90585 | 0.023049154 | 0.061887956 | 0.372433598 | 0.709570043 | 0.851593977 |
| Ccdc59 | 558.5149376 | 0.030063618 | 0.080753151 | 0.372290345 | 0.709676686 | 0.85160178 |
| Tmem173 | 169.5640962 | 0.051440882 | 0.138178974 | 0.372277202 | 0.709686471 | 0.85160178 |
| Crybb3 | 21.8635798 | -0.062506166 | 0.167964804 | -0.372138473 | 0.709789753 | 0.85163609 |
| Phf3 | 3743.114935 | 0.0327928 | 0.088131097 | 0.372091137 | 0.709824995 | 0.85163609 |
| Rnf113a2 | 289.042663 | 0.036400475 | 0.097851425 | 0.371997391 | 0.709894792 | 0.851653883 |
| Cxxc5 | 2718.412233 | -0.033295937 | 0.089564763 | -0.37175264 | 0.710077028 | 0.851806555 |
| Ubxn4 | 2507.749842 | 0.02162758 | 0.058190217 | 0.37167038 | 0.710138281 | 0.851814084 |
| Mir9-2 | 52.88681209 | -0.06008887 | 0.161983999 | -0.370955591 | 0.710670609 | 0.852386624 |
| Kctd7 | 611.7363298 | -0.031145514 | 0.084009329 | -0.370738754 | 0.710832122 | 0.852448367 |
| Ubtd1 | 312.8229857 | 0.034847745 | 0.093991076 | 0.370755897 | 0.710819353 | 0.852448367 |
| Dfna5 | 389.4453581 | -0.032274116 | 0.087145709 | -0.370346593 | 0.711124262 | 0.852732707 |
| Mc3r | 19.25439407 | -0.056691494 | 0.153114044 | -0.370256658 | 0.711191265 | 0.852747056 |

| | | | | | | | |
|---|---|---|---|---|---|---|---|
| Smagp | 43.30734737 | -0.0187735603219792 | 0.0936656875928529 | -0.200431564689775 | 0.841143075260213 | 0.96309494742331 | NM_001033872,NM_174992 |
| Tcte1 | 124.3127379 | -0.018712809177248 | 0.0932458469000786 | -0.20068249470993 | 0.840946848451042 | 0.96309494742331 | NM_013688 |
| Asxl1 | 2805.421772 | -0.0114614785006628 | 0.057266298813233 | -0.200143517883756 | 0.84136833949326 | 0.963098226909306 | NM_001039939 |
| Fbln7 | 94.99374781 | 0.0190197933727556 | 0.0950897505461161 | 0.2000193844607 | 0.841465420847716 | 0.963133457035216 | NM_024237 |
| Aldh1a7 | 27.90760048 | 0.0166533845325836 | 0.0837141950530445 | 0.198931429992624 | 0.842316383259643 | 0.963252575469256 | NM_011921 |
| Crispld1 | 203.4479672 | 0.0187457419383362 | 0.0943019687668849 | 0.198784205499207 | 0.842431551615133 | 0.963252575469256 | NM_031402 |
| Fam219b | 830.3210016 | 0.013028965624445 | 0.0655151193102059 | 0.198869600813127 | 0.842364749562674 | 0.963252575469256 | NM_175273,NM_001166364 |
| Got1l1 | 29.99478459 | -0.0174117795959838 | 0.087417492747168 | -0.199179581211998 | 0.842122271239931 | 0.963252575469256 | NM_029674 |
| Ilkap | 1318.278851 | 0.0120406261051477 | 0.0604387546636804 | 0.199220287912108 | 0.842090430040174 | 0.963252575469256 | NM_023343 |
| Med11 | 271.3910865 | 0.016726361709064 | 0.0839237901836876 | 0.199304174328332 | 0.842024814036496 | 0.963252575469256 | NM_025397 |
| Pdgfc | 303.49703 | 0.0168592811290902 | 0.0845148118623583 | 0.199483152805775 | 0.841884820634849 | 0.963252575469256 | NM_019971 |
| Rps14 | 6558.156253 | -0.00948602749298372 | 0.0475212320350331 | -0.199616615284522 | 0.841780432194072 | 0.963252575469256 | NM_020600 |
| Serhl | 194.4272864 | 0.0181947414694185 | 0.091443956556309 | 0.198971502925014 | 0.84228503624901 | 0.963252575469256 | NM_023475 |
| Slc25a24 | 112.104759 | 0.0181947609714904 | 0.0911532813151582 | 0.199606209551391 | 0.841788570997794 | 0.963252575469256 | NM_172685 |
| Wnt6 | 40.34068802 | 0.0171585090205515 | 0.0861237215963337 | 0.199230928512057 | 0.842082106895955 | 0.963252575469256 | NM_009526 |
| Zeb1 | 3401.171368 | 0.0105090102331744 | 0.0527578986874212 | 0.199193116000279 | 0.842111684160594 | 0.963252575469256 | NM_011546 |
| Zfp11 | 310.2885203 | 0.0167339438358164 | 0.0841767728549822 | 0.19879526463488 | 0.84242290033924 | 0.963252575469256 | NM_172462 |
| Hist1h2ac | 33.41247888 | -0.0147095940060689 | 0.0740459522451311 | -0.198654937374192 | 0.842532676137122 | 0.963292377297744 | NM_178189 |
| Nbea | 11600.90382 | 0.0134516972825335 | 0.0678193985341916 | 0.198345865243139 | 0.842774469169556 | 0.963458383294478 | NM_030595 |
| Pik3ip1 | 1459.328507 | -0.013681866163652 | 0.0689958718587334 | -0.198299779321074 | 0.842810524337253 | 0.963458383294478 | NM_178149 |
| Kcnj4 | 4840.263268 | -0.0141298512324803 | 0.0714242560165922 | -0.197829869298153 | 0.843178175674892 | 0.963802815537731 | NM_008427 |
| Itm2a | 1518.211123 | -0.0119437827100708 | 0.0604630764066832 | -0.19753845520091 | 0.843406191355271 | 0.963987594273874 | NM_008409 |
| Bin1 | 10429.83477 | -0.011230913451845 | 0.0569668025784668 | -0.197148390703083 | 0.843711416148606 | 0.964060034459129 | NM_001083334,NM_009668 |
| Cd83 | 707.0422949 | -0.0180753434994959 | 0.0917769683971214 | -0.196948578877474 | 0.843867777650271 | 0.964060034459129 | NM_009856 |
| Dusp23 | 155.4429452 | 0.0186244043089651 | 0.0943776355600871 | 0.19733917043416 | 0.843562128404291 | 0.964060034459129 | NM_026725 |
| Gba2 | 4240.566934 | 0.0125866513310961 | 0.0638459557803929 | 0.19714093363078 | 0.843717251523559 | 0.964060034459129 | NM_172692 |
| Gimap1 | 58.51092558 | -0.0180533464901794 | 0.0916495678365717 | -0.196982341721151 | 0.843841356314699 | 0.964060034459129 | NM_008376,NM_175860 |
| Gm11127 | 55.21100843 | 0.0186500407921082 | 0.0946770813360145 | 0.196985801937833 | 0.843838648509284 | 0.964060034459129 | NM_001199967 |
| Arap2 | 4513.038782 | -0.0141705439115113 | 0.0720659187450527 | -0.19663308479619 | 0.844114678107846 | 0.964266264092054 | NM_178407 |
| Chchd10 | 4162.673959 | 0.0113608660010236 | 0.0579944190496709 | 0.195895849759152 | 0.844691685747586 | 0.96432828511403 | NM_175329 |
| Commd1 | 602.4585136 | 0.0132604277133077 | 0.0676456080583882 | 0.196027918055848 | 0.844588314453375 | 0.96432828511403 | NM_144514 |
| Fam195a | 203.9150226 | 0.0171989884235543 | 0.087675282800989 | 0.196166899884357 | 0.844479534752507 | 0.96432828511403 | NM_026633 |
| Hnf1b | 27.00949256 | 0.015832501361885 | 0.0808254300648644 | 0.195885148389301 | 0.844700061943475 | 0.96432828511403 | NM_009330 |
| Mrpl4 | 1455.572116 | -0.0129528963135669 | 0.0660812007701391 | -0.196014844806211 | 0.844598546908935 | 0.96432828511403 | NM_023167 |
| Nkpd1 | 26.70778424 | -0.016424195219626 | 0.0837517832509231 | -0.196105618078824 | 0.844527499049419 | 0.96432828511403 | NM_027116 |
| Tbc1d23 | 1492.797589 | -0.0134731948425506 | 0.0686436284006889 | -0.196277428167177 | 0.844393027477081 | 0.96432828511403 | NM_026254 |
| Wnt2 | 175.1085015 | -0.0175239687713165 | 0.08926716763823 | -0.196309228072916 | 0.844368138962442 | 0.96432828511403 | NM_023653 |
| Abcc12 | 37.10064146 | -0.0170993379205845 | 0.0873523221822158 | -0.195751383516921 | 0.844804764111529 | 0.964372023849436 | NM_172912 |
| Phtf1 | 2268.797143 | 0.0100232519049907 | 0.051238837571383 | 0.195618253263979 | 0.844908972264211 | 0.964415191820676 | NM_001163469,NM_001163468,NM_013629,NM_001163467 |
| Alas1 | 2452.107328 | -0.0105054263531456 | 0.0537431682098666 | -0.195474638043705 | 0.845021390603164 | 0.964467723729202 | NM_020559 |
| D16Ertd472e | 473.1518093 | -0.0154594004910695 | 0.079210272371664 | -0.19516913688331 | 0.845260539629964 | 0.964664880473354 | NM_001252440,NM_001252438,NM_025967,NM_001252439 |
| Extl1 | 3390.34685 | 0.0126914021686866 | 0.0651290152972305 | 0.194865562004379 | 0.845498194871496 | 0.964815880977639 | NM_019578 |
| Hccs | 848.2960044 | 0.0123034557182957 | 0.0631495549363269 | 0.194830442284213 | 0.845525689442747 | 0.964815880977639 | NM_008222 |
| A830080D01Rik | 261.3394527 | 0.0158248171563111 | 0.0814599786809871 | 0.194264931228181 | 0.845968443340258 | 0.964996027829392 | NM_001033472 |
| Cxcl14 | 3390.507432 | -0.0104485057589548 | 0.0537603817479744 | -0.194353265717807 | 0.845899280670583 | 0.964996027829392 | NM_019568 |
| Fam179a | 162.2638716 | 0.0175726115615025 | 0.0903892112168903 | 0.19441049794468 | 0.845854470567205 | 0.964996027829392 | NM_177087 |
| Hmg20a | 4135.277945 | 0.00873891421158554 | 0.0450175654923587 | 0.194122319054967 | 0.846080105952588 | 0.964996027829392 | NM_025812 |
| Mepce | 1794.514287 | 0.0101809097871102 | 0.0524465564316604 | 0.1941196997438 | 0.846082156852205 | 0.964996027829392 | NM_144913 |
| Zfp428 | 733.8074719 | 0.0134666817355092 | 0.0693133738326708 | 0.194286917385079 | 0.845951228873802 | 0.964996027829392 | NM_146183 |
| Dcdc2c | 140.0306626 | -0.0179757349904708 | 0.0926453440037904 | -0.194027397531552 | 0.84615442940915 | 0.965002688356545 | NM_001177964 |
| Cdc27 | 2660.407192 | 0.0121193331268728 | 0.0625501186131593 | 0.193753959154333 | 0.846368539030131 | 0.965166054138925 | NM_145436 |
| Zxda | 48.27086049 | -0.0182998703334127 | 0.0944876410185173 | -0.193674750857907 | 0.846430563381232 | 0.965166054138925 | NR_003292 |
| Usp45 | 1891.477701 | 0.015768531164378 | 0.0814796554900202 | 0.193527219396617 | 0.846546090976983 | 0.965222018644556 | NM_152825 |
| 2310010J17Rik | 43.58973866 | 0.0179217207100565 | 0.0928872150874333 | 0.192940661351372 | 0.847005440123076 | 0.965275632665579 | NR_046006,NR_046007 |
| Alx4 | 65.34837582 | -0.0164828667034271 | 0.0854900926865546 | -0.19280440791965 | 0.847112151241921 | 0.965275632665579 | NM_007442 |
| Amigo1 | 5808.236522 | 0.00807724778549218 | 0.0418254708136576 | 0.193117916627364 | 0.84686662133167 | 0.965275632665579 | NM_001004293,NM_146137 |
| Bhlhe41 | 2044.277858 | 0.0140181867328822 | 0.0726364738184821 | 0.1929910139624 | 0.846966005617252 | 0.965275632665579 | NM_024469,NM_001271768 |
| Comtd1 | 274.5551962 | 0.015210586360805 | 0.078986051537548 | 0.192573069101629 | 0.847293337862101 | 0.965275632665579 | NM_026965 |
| Lenep | 250.0339852 | 0.0154846712626758 | 0.0804065445842769 | 0.192579737666078 | 0.847288114870071 | 0.965275632665579 | NM_020517 |
| Lphn1 | 28496.32144 | 0.0108277984084393 | 0.0562009309333862 | 0.192662260724351 | 0.847223481233131 | 0.965275632665579 | NM_181039 |
| Mcrs1 | 2815.126758 | 0.0133601699214122 | 0.0692971623245077 | 0.19279533927881 | 0.847119253729971 | 0.965275632665579 | NM_001164156,NM_016766 |
| Neurl1a | 8033.44638 | -0.0142991554663529 | 0.0741957464773931 | -0.192722037923154 | 0.847176663229949 | 0.965275632665579 | NM_021360,NM_001163480 |
| Snapin | 1946.885079 | -0.0119060927976131 | 0.0618389689488754 | -0.192533818076047 | 0.847324080415724 | 0.965275632665579 | NM_133854 |
| Syt14 | 261.208803 | 0.0168303039030772 | 0.0873541756716192 | 0.192667422864197 | 0.847219438180659 | 0.965275632665579 | NM_181546 |
| 1500011K16Rik | 551.2832864 | -0.0125255609479061 | 0.065170890522937 | -0.192195638994648 | 0.847588961802887 | 0.965398236624199 | NR_015476 |
| Gm10548 | 57.17279584 | -0.0182581325357547 | 0.0949634947810148 | -0.192264749504404 | 0.847534829062823 | 0.965398236624199 | NR_040534 |
| Hid1 | 5099.097141 | -0.0101526427966928 | 0.0528393061895234 | -0.192141864245481 | 0.847631082879322 | 0.965398236624199 | NM_175454 |
| Gls2 | 1126.437479 | -0.0137266124132384 | 0.0715691705260722 | -0.191795046838469 | 0.847902750988779 | 0.965542581936173 | NM_001033264 |
| Map3k5 | 2173.506186 | 0.0111184942786142 | 0.0579917341270845 | 0.191725500986897 | 0.847957229640628 | 0.965542581936173 | NM_008580 |
| S100b | 5442.853195 | 0.0112086602704113 | 0.0584533312649002 | 0.191754003199845 | 0.847934902381539 | 0.965542581936173 | NM_009115 |
| 1110008L16Rik | 288.1971638 | -0.0160697038131715 | 0.0840651233774184 | -0.191157797283245 | 0.848401966738384 | 0.9657461773248 | NM_025373 |
| Rps19bp1 | 512.1755314 | 0.013086324526951 | 0.0684138131101965 | 0.191281905369469 | 0.848304736751262 | 0.9657461773248 | NM_175109 |
| Syap1 | 942.2355852 | 0.0127257900868888 | 0.0665150308003481 | 0.191322020508216 | 0.848273309870044 | 0.9657461773248 | NM_025932 |
| Trnp1 | 4671.302256 | -0.0094371551339338 | 0.0493653779243725 | -0.191169510509804 | 0.848392790149498 | 0.9657461773248 | NM_001081156 |
| Acer3 | 369.1613193 | -0.0133806696384104 | 0.0700848056788532 | -0.190921120616701 | 0.848587392675436 | 0.965881559787132 | NM_025408 |
| 4833412C05Rik | 28.44906964 | -0.0157478350854838 | 0.082671307427507 | -0.190487311444697 | 0.848927285155454 | 0.965892061058607 | NR_045954 |
| Hnrnpr | 2770.350116 | 0.0107799863444651 | 0.0565936773345365 | 0.190478810570004 | 0.848933945930242 | 0.965892061058607 | NM_028871 |
| Mettl9 | 2079.787327 | 0.0112837867516342 | 0.0591856586300948 | 0.190650691616982 | 0.848799272362417 | 0.965892061058607 | NM_021554 |
| Tagln2 | 436.6942107 | -0.0151340257668641 | 0.0794465600970377 | -0.190493153490587 | 0.848922707685462 | 0.965892061058607 | NM_178598 |
| Tmem259 | 3615.411663 | -0.0119641184625315 | 0.0627168118206062 | -0.19076413668401 | 0.848710387423092 | 0.965892061058607 | NM_001003949 |
| Zpbp | 52.15200596 | 0.0179178924553278 | 0.0941065055545246 | 0.190400146618411 | 0.848995582789225 | 0.965892061058607 | NM_001185153,NM_015785 |
| Acot4 | 48.10058109 | 0.0176161252748601 | 0.0932276365222685 | 0.18895818806532 | 0.850125587460255 | 0.96626084768017 | NM_134247 |
| Acvr1c | 1078.606956 | -0.0168977678278675 | 0.0893600044398645 | -0.18909766101499 | 0.850016274655387 | 0.96626084768017 | NM_001033369,NM_001111030 |
| Adnp2 | 1231.28136 | -0.0113589805846277 | 0.0599672033825884 | -0.18941988193376 | 0.849763743003384 | 0.96626084768017 | NM_175028 |
| Arhgap25 | 461.894994 | 0.0158896246616095 | 0.083725008449893 | 0.189783494272431 | 0.849478790492346 | 0.96626084768017 | NM_001037727,NM_175476 |
| BB031773 | 32.74659619 | -0.0162584625421828 | 0.086179571908548 | -0.188657963623166 | 0.850360900034717 | 0.96626084768017 | NR_028391 |
| Cml1 | 315.2452292 | -0.0158931633936948 | 0.0841654605238726 | -0.18883237012868 | 0.850224200534158 | 0.96626084768017 | NM_023160 |
| Gm13476 | 162.933881 | -0.0166531598422958 | 0.0882855235991726 | -0.188628431518436 | 0.850384047689886 | 0.96626084768017 | NM_001177394,NM_001177395 |
| Ldb1 | 3951.470828 | -0.00982014200744084 | 0.0519864871753008 | -0.188897972165861 | 0.850172782942154 | 0.96626084768017 | NM_001113408,NM_010697 |
| Med17 | 1100.66529 | 0.0108701390876121 | 0.0576221996842051 | 0.188644986605601 | 0.850371071577227 | 0.96626084768017 | NM_144933 |
| Pced1b | 186.7548915 | -0.0166385173868395 | 0.0877005013234292 | -0.189719752290567 | 0.849528741826243 | 0.96626084768017 | NM_172293 |
| Rassf1 | 494.3528427 | 0.0139557069655011 | 0.0737249847900302 | 0.189294131497581 | 0.849862294565682 | 0.96626084768017 | NM_019713,NM_001243748 |
| Thoc7 | 1221.891325 | 0.0119902783479485 | 0.0633056224686799 | 0.189403055848327 | 0.849776929597044 | 0.96626084768017 | NM_025435,NM_001013578 |
| Ube3b | 6738.534981 | 0.00905360720478974 | 0.0479513411931458 | 0.188808216402586 | 0.850243131912092 | 0.96626084768017 | NM_054093 |
| Unc119 | 880.7050804 | 0.0131517655872358 | 0.0695304043839702 | 0.189151288616234 | 0.849974244451589 | 0.96626084768017 | NM_011676 |
| Vav2 | 354.0151408 | 0.015220832178114 | 0.0803100675198662 | 0.189525829676943 | 0.849680712782457 | 0.96626084768017 | NM_009500 |
| Wdfy1 | 948.7605502 | -0.0142718312707489 | 0.0753355318096649 | -0.189443559073913 | 0.849745187313188 | 0.96626084768017 | NM_001111279,NM_027057 |
| 2010003O02Rik | 55.07511054 | 0.0175461795116197 | 0.0932063644598629 | 0.18825087335292 | 0.850679994140533 | 0.966301569207939 | NR_045265 |
| Car4 | 1282.607159 | -0.0112069795248337 | 0.0594647928044477 | -0.188464114584377 | 0.85051284383518 | 0.966301569207939 | NM_007607 |
| Cluh | 4664.25772 | -0.00899430569926974 | 0.04778023109697 | -0.188243243968741 | 0.850685974599416 | 0.966301569207939 | NM_001081158 |
| Rasd2 | 8973.813095 | -0.0160954902751311 | 0.0854919358109509 | -0.18826910541274 | 0.850665702578313 | 0.966301569207939 | NM_029182 |
| Sumf1 | 958.1317009 | 0.0132356050506219 | 0.0704035303504571 | 0.18799632610378 | 0.850879531178665 | 0.966445857369715 | NM_145937 |
| Cacna2d4 | 108.3932638 | -0.0177797779229694 | 0.0948128501414392 | -0.1875249810173 | 0.851249039071253 | 0.966638800934101 | NM_001033382 |
| Dhx29 | 1188.454684 | 0.012421913319379 | 0.0662359778992759 | 0.187540272120218 | 0.851237051198259 | 0.966638800934101 | NM_172594 |
| Kptn | 755.9254643 | -0.0137300901386901 | 0.0731661948866751 | -0.187656200516594 | 0.851146167120327 | 0.966638800934101 | NM_133727 |
| 1110017D15Rik | 69.71163517 | 0.0177387696525374 | 0.0948007177776146 | 0.187116406588285 | 0.851569364747959 | 0.966650029598661 | NM_001048005,NM_001253788,NM_001253780,NM_028624,NR_045551,NM_001253777,NM_001253778,NR_045549 |
| Actr2 | 12059.73247 | 0.0125184608449576 | 0.0669028501391271 | 0.187114014110385 | 0.851571240542273 | 0.966650029598661 | NM_146243 |
| Cdc73 | 696.6687525 | 0.0148216810604817 | 0.0792499823774104 | 0.187024408281843 | 0.851641495548641 | 0.966650029598661 | NM_145991 |
| Dmtf1 | 2678.774545 | 0.0100500581269652 | 0.0537427362099837 | 0.187003097268766 | 0.851658204516293 | 0.966650029598661 | NM_001110327,NM_011806 |
| Ltc4s | 96.25697335 | 0.016868682756058 | 0.0901052588632868 | 0.187210857266968 | 0.851495312469163 | 0.966650029598661 | NM_008521 |
| Rab28 | 2084.5362 | 0.0109280417171167 | 0.0583393497665704 | 0.187318538188073 | 0.851410888864944 | 0.966650029598661 | NM_027295 |
| A930011G23Rik | 23.05016019 | 0.0147713930699992 | 0.0790575784905277 | 0.186843479803382 | 0.851783355214859 | 0.966691729721501 | NR_030692 |
| Tradd | 180.0105838 | -0.016369857168399 | 0.0876394080740941 | -0.186786487130986 | 0.851828042161311 | 0.966691729721501 | NM_001033161 |
| Bche | 190.9669227 | 0.0176991636652466 | 0.094819965898615 | 0.186660725908415 | 0.851926650994551 | 0.966728109706026 | NM_009738 |
| Htra3 | 183.7194423 | 0.0169554039961103 | 0.0908922833332643 | 0.186543932821468 | 0.852018230024399 | 0.966756507524951 | NM_030127,NM_001042615 |
| 5730559C18Rik | 96.96011841 | -0.017722703330178 | 0.0950929996699017 | -0.186372323848224 | 0.852152794547097 | 0.966758161011491 | NM_028872 |
| Cd1d1 | 33.71856723 | -0.0151271630922363 | 0.081139340877585 | -0.186434384709368 | 0.852104129985519 | 0.966758161011491 | NM_007639 |
| 4430402I18Rik | 144.9287642 | 0.0172202213956576 | 0.0932485881141802 | 0.184670049637341 | 0.853487839018966 | 0.966905493159456 | NM_198651 |
| 5830418P13Rik | 27.21353515 | 0.0161306998394742 | 0.0870258302131314 | 0.185355311175649 | 0.85295035751099 | 0.966905493159456 | NR_040466 |
| Abcc6 | 62.82198544 | 0.0173350971493149 | 0.0939582264244307 | 0.184497917947155 | 0.853622860359922 | 0.966905493159456 | NM_018795 |
| Ano5 | 60.85278973 | 0.0172447238843423 | 0.0931486717524926 | 0.185131183943917 | 0.853126143099339 | 0.966905493159456 | NM_177694,NR_073508,NM_001271879 |
| Bloc1s6 | 1255.94068 | -0.0119720950509417 | 0.0644729161866397 | -0.1856918495246 | 0.852686420296713 | 0.966905493159456 | NM_019788 |
| Bok | 988.0539052 | 0.0158700349660217 | 0.0853383472805247 | 0.185966045415124 | 0.852471388646983 | 0.966905493159456 | NM_016778 |
| Coa5 | 4942.837494 | -0.0104874005760928 | 0.0566273568282981 | -0.185200248846014 | 0.853071973917363 | 0.966905493159456 | NM_198006 |
| Creld2 | 559.9558172 | -0.0141421216165012 | 0.0764173269993439 | -0.185064332551472 | 0.853178576834757 | 0.966905493159456 | NM_029720 |
| Gpr162 | 8079.021073 | 0.0125280929698995 | 0.0680246592882747 | 0.184169874586332 | 0.8538801918112 | 0.966905493159456 | NM_013533 |
| Gtf3c1 | 6768.949666 | -0.00968263949443668 | 0.0525408047390933 | -0.184287993732084 | 0.853787532233587 | 0.966905493159456 | NM_207239 |
| Hpse | 59.38766555 | 0.0174143078921866 | 0.094315726995618 | 0.184638431435679 | 0.85351264024304 | 0.966905493159456 | NM_152803 |
| Nmb | 99.52427721 | -0.0175161091667052 | 0.0947843604577226 | -0.184799571175226 | 0.853386244218774 | 0.966905493159456 | NM_026523 |
| Nos1 | 1267.095665 | 0.0170397556152772 | 0.0919628706980423 | 0.185289513973817 | 0.853001962262974 | 0.966905493159456 | NM_008712 |
| Pop1 | 350.915863 | -0.0153563533193426 | 0.0831253750956385 | -0.184737251431042 | 0.853435126611318 | 0.966905493159456 | NM_026340,NM_152894 |
| Ptch2 | 85.12928962 | 0.0175106756376935 | 0.0950511625490161 | 0.184223687202811 | 0.853837977792317 | 0.966905493159456 | NM_008958 |
| Pttg1 | 1119.834693 | -0.011845281297412 | 0.0636544622936207 | -0.186087210080778 | 0.852376371612368 | 0.966905493159456 | NM_013917,NM_001131054 |
| Rap2b | 2231.760971 | -0.0123783354882566 | 0.0669974757433814 | -0.18475823182789 | 0.853418666035773 | 0.966905493159456 | NM_028712 |
| Rngtt | 931.8290965 | 0.0136087058783091 | 0.073327301043006 | 0.185588528211718 | 0.852767450436546 | 0.966905493159456 | NM_011884 |
| Rpp21 | 363.7612991 | -0.0146754564216689 | 0.0792911405229307 | -0.185083179846868 | 0.853163794218355 | 0.966905493159456 | NM_026308 |
| Rps16 | 4338.508106 | 0.0112957643058549 | 0.060942021801972 | 0.185352634715007 | 0.852952456647061 | 0.966905493159456 | NM_013647 |
| S1pr1 | 6522.570883 | 0.0107981508233658 | 0.0585463149232232 | 0.184437753896661 | 0.853670054487165 | 0.966905493159456 | NM_007901 |
| Slc52a2 | 558.3052121 | 0.0157129093019438 | 0.0847032773987053 | 0.185505328536248 | 0.852832701233696 | 0.966905493159456 | NM_029643 |
| Upf3a | 1194.792755 | -0.0103677292522688 | 0.0560441227071936 | -0.184992265940814 | 0.853235101763116 | 0.966905493159456 | NM_025924 |
| Zbtb9 | 746.450629 | 0.0109573869984684 | 0.0594598979803269 | 0.184281967690119 | 0.853792259365109 | 0.966905493159456 | NM_001005916 |
| Acap2 | 4210.641873 | -0.012907094592925 | 0.0707413813981925 | -0.182454658614495 | 0.855225934222107 | 0.967189922578937 | NM_030138 |
| Armc4 | 35.21768091 | 0.0159542113319843 | 0.0870614752888144 | 0.183252251114036 | 0.854600097979515 | 0.967189922578937 | NM_001081393 |
| B4galnt1 | 5780.755611 | -0.0103456922350413 | 0.0568023614128262 | -0.182134896819715 | 0.855476863033142 | 0.967189922578937 | NM_001244618,NM_001244617,NM_008080,NR_045166,NM_027739 |
| Dars2 | 891.88776 | 0.0120688597674973 | 0.0660219915154928 | 0.182800601594473 | 0.854954476561607 | 0.967189922578937 | NM_172644 |
| Egr4 | 797.8118027 | 0.0146565745769144 | 0.0805558878072187 | 0.181942933978825 | 0.85562751033852 | 0.967189922578937 | NM_020596 |
| Gm10754 | 80.68761794 | 0.0168253091576075 | 0.0917432324492184 | 0.183395643563361 | 0.854487593832442 | 0.967189922578937 | NR_033537 |
| Gpc6 | 517.8641111 | 0.0130158368880068 | 0.070851873349282 | 0.183704908179943 | 0.854244958271983 | 0.967189922578937 | NM_011821,NM_001079844 |
| Gpr27 | 236.6178855 | -0.0173015602347984 | 0.0946994802196604 | -0.18269963250766 | 0.855033704123591 | 0.967189922578937 | NM_008158 |
| Ncoa6 | 3654.581071 | -0.0100999978706404 | 0.0553437330669778 | -0.182495782465871 | 0.855193663889182 | 0.967189922578937 | NM_001242558,NM_019825 |
| Pcmtd1 | 4621.669766 | -0.0124385185236384 | 0.068072441149752 | -0.182724731382485 | 0.855014009616007 | 0.967189922578937 | NM_183028 |
| Pik3r2 | 5688.676322 | -0.0111202248429225 | 0.0609217854173778 | -0.182532812634057 | 0.85516460612115 | 0.967189922578937 | NM_008841 |
| Prkcb | 55081.02425 | 0.0111294931942675 | 0.0611853592426128 | 0.18189797905961 | 0.855662790517824 | 0.967189922578937 | NM_008855 |
| Prmt10 | 838.6071938 | 0.0120602004597644 | 0.065792849747451 | 0.183305640446615 | 0.854558208947671 | 0.967189922578937 | NM_001081240 |
| Qsox1 | 1167.708755 | 0.0120075364656314 | 0.0658876168714221 | 0.18224268892687 | 0.855392272978095 | 0.967189922578937 | NM_023268,NM_001024945 |
| Rps17 | 4301.080316 | -0.0127587329229558 | 0.0697055464085788 | -0.183037556985359 | 0.854768550170319 | 0.967189922578937 | NM_009092 |
| Rps19 | 4343.316174 | 0.00988851347276707 | 0.053943546179647 | 0.183312262042165 | 0.854553013701954 | 0.967189922578937 | NM_023133 |
| Serpinh1 | 554.2349355 | 0.0118294140368789 | 0.064814317050742 | 0.182512361082473 | 0.855180654537127 | 0.967189922578937 | NM_001111043,NM_009825,NM_001111044 |
| Spag6 | 254.2905091 | 0.015959854277176 | 0.0877402625929268 | 0.181898866102352 | 0.855662094372597 | 0.967189922578937 | NM_015773 |
| Syk | 205.6052481 | 0.0166737568915484 | 0.0916063414886497 | 0.182015312702062 | 0.855570708833356 | 0.967189922578937 | NM_001198977,NM_011518 |
| Trmt5 | 443.8409479 | -0.0143362767870336 | 0.0785702227968553 | -0.182464504703013 | 0.855218207867436 | 0.967189922578937 | NM_029580 |
| Ttll9 | 58.01424054 | 0.017323908703665 | 0.0950465830162939 | 0.18226755927349 | 0.855372756163573 | 0.967189922578937 | NM_029064,NM_001083618 |
| Wdr34 | 1137.36903 | 0.0128958642113274 | 0.0706719448303138 | 0.182475015259462 | 0.85520996011139 | 0.967189922578937 | NM_001008498 |
| Zfp937 | 500.9130632 | -0.0143500453010628 | 0.0785220955558148 | -0.182751685363039 | 0.854992859550371 | 0.967189922578937 | NM_001142411 |
| Mast2 | 6274.169317 | 0.00898824479970496 | 0.0494375200703118 | 0.181810187625139 | 0.855731689205554 | 0.967192539480227 | NM_008641,NM_001042743 |
| Agfg1 | 1836.920975 | -0.0124185274669489 | 0.0686597332517511 | -0.180870604629565 | 0.856469142382456 | 0.96759800120291 | NM_010472 |
| Dnd1 | 39.77503095 | -0.0146030363506576 | 0.0807873030855259 | -0.180759052387205 | 0.856556705043937 | 0.96759800120291 | NM_173383 |
| Eif4b | 9544.972267 | 0.0093887644572478 | 0.0518644636293168 | 0.181024998626241 | 0.85634795410502 | 0.96759800120291 | NM_145625 |
| Lifr | 1824.003833 | 0.0103709944737045 | 0.0573737970179793 | 0.180761863651006 | 0.856554498327557 | 0.96759800120291 | NM_001113386,NM_013584 |
| Oxsr1 | 2324.579861 | -0.0115273319789704 | 0.0637534833247374 | -0.180811014203793 | 0.856515917522577 | 0.96759800120291 | NM_133985,NR_045693 |
| Rbbp6 | 3436.528365 | -0.00907624281677766 | 0.0501870321871577 | -0.180848367022989 | 0.856486597596728 | 0.96759800120291 | NM_011247,NM_175023 |
| Tor1aip1 | 1899.511469 | 0.0120730874367203 | 0.0666778160167934 | 0.181066030022333 | 0.856315747954631 | 0.96759800120291 | NM_001160019,NM_001160018,NM_144791 |
| Angpt1 | 189.2961842 | 0.0162714249163988 | 0.0900626657849492 | 0.180667813622701 | 0.856628324003031 | 0.967603657423642 | NM_009640 |
| Limd2 | 2630.607001 | -0.0136274828974871 | 0.0754974881677491 | -0.180502467409352 | 0.856758117498117 | 0.967675018643779 | NM_172397 |
| Gtpbp1 | 2552.638021 | -0.0087579904379951 | 0.0485842791506373 | -0.180263875292677 | 0.856945414400304 | 0.967736071645714 | NM_013818 |
| Map1a | 59203.09631 | 0.0116533365477993 | 0.0646176020604971 | 0.180343067155124 | 0.856883247200713 | 0.967736071645714 | NM_032393,NM_001173506 |
| Lars | 2024.007247 | -0.00947305899074801 | 0.052614413000322 | -0.180046843145261 | 0.857115793145261 | 0.967853234703674 | NM_134137 |
| Kcns3 | 290.3425046 | 0.0154323673408734 | 0.0857909415737011 | 0.179883412604998 | 0.85724409760454 | 0.967922873051189 | NM_001168564,NM_173417 |
| 5730522E02Rik | 63.70918848 | 0.0170098696644966 | 0.0950992389364104 | 0.178864414213352 | 0.858044165473231 | 0.967971163250461 | NR_027973 |
| Capza2 | 7552.446365 | 0.0124136872268287 | 0.069118645083405 | 0.17959968995123 | 0.857466848106819 | 0.967971163250461 | NM_007604 |
| Dock10 | 2166.314931 | 0.0145309333019982 | 0.0814189968469986 | 0.178471043180555 | 0.858353060293899 | 0.967971163250461 | NM_175291 |
| Dut | 772.5311393 | 0.0105965366056983 | 0.0593731257678494 | 0.178473618639029 | 0.858351037843132 | 0.967971163250461 | NM_023595,NM_001159646 |
| Igfbp6 | 1288.365096 | -0.0140516244314798 | 0.0785893422966649 | -0.178798091711172 | 0.858096243732734 | 0.967971163250461 | NM_008344 |
| Il20ra | 29.0373385 | -0.0143658494386315 | 0.0801422571943862 | -0.179254365194468 | 0.857737977700393 | 0.967971163250461 | NM_172786 |
| Lmo1 | 206.3346478 | 0.0164982013203304 | 0.0921543732690715 | 0.179027871766422 | 0.85791581672845 | 0.967971163250461 | NM_057173 |
| Nckipsd | 3707.596194 | 0.00993503648880446 | 0.0554776493850912 | 0.179081785167963 | 0.857873484130957 | 0.967971163250461 | NM_030729 |
| Ncor1 | 9706.989055 | 0.0101412568145987 | 0.0567275948061965 | 0.178771140381417 | 0.858117406837315 | 0.967971163250461 | NM_011308,NM_177229,NM_001252313 |
| Pot1a | 761.8136483 | 0.0131951984482972 | 0.0739241820907257 | 0.178496373921364 | 0.858333168661913 | 0.967971163250461 | NM_133931 |
| Ppp3cc | 819.6609617 | -0.011465879597886 | 0.0641376532928259 | -0.178769864646241 | 0.858118408590673 | 0.967971163250461 | NM_008915 |
| Ptgr2 | 1948.812509 | 0.00951413355150228 | 0.0532906128501974 | 0.178533010649493 | 0.858304398864451 | 0.967971163250461 | NM_029880,NM_001252626,NM_001252625 |
| Scaf1 | 7342.311376 | -0.0103139063076359 | 0.0575031062811981 | -0.179362594034477 | 0.857653000689836 | 0.967971163250461 | NM_001008422 |
| Srrm4 | 2162.002461 | 0.0125548418319441 | 0.0702600641500409 | 0.178691010089787 | 0.858180328481154 | 0.967971163250461 | NM_026886 |
| Tgfbi | 183.6781946 | 0.01691863652084 | 0.0945663045955527 | 0.178907662652133 | 0.858010205938016 | 0.967971163250461 | NM_009369 |
| Zfp781 | 102.2134697 | 0.0168824902193308 | 0.094173990098567 | 0.179269140042391 | 0.857726376976163 | 0.967971163250461 | NM_199062 |
| Anks1 | 1608.176164 | -0.0122027154371482 | 0.0685709087569735 | -0.177957615822136 | 0.858756262028494 | 0.968350680191423 | NM_181413 |
| Fam174b | 1409.823147 | 0.0125407823147622 | 0.0705128611856086 | 0.177850992058762 | 0.858839999794632 | 0.968369932237586 | NM_001162532 |
| Btbd6 | 2015.053841 | -0.0101374345941221 | 0.0570470895633528 | -0.177702923527134 | 0.858956289151542 | 0.968425881420001 | NM_001145900,NM_201646 |
| E230008N13Rik | 35.43349274 | 0.0137853317968091 | 0.0778073406365155 | 0.17717263800608 | 0.859372787348866 | 0.968520191286962 | NM_198660 |
| Jph2 | 50.48283032 | -0.0165031068759248 | 0.0929929924158797 | -0.177466134245043 | 0.859142263946432 | 0.968520191286962 | NM_021566,NM_001205076 |
| Mrps30 | 745.8398605 | -0.0135306555293109 | 0.0764067898963976 | -0.177087082805829 | 0.859439988000064 | 0.968520191286962 | NM_021556 |
| Nid1 | 369.5252661 | -0.0155025242425668 | 0.0874331243538962 | -0.177307220314105 | 0.85926707966156 | 0.968520191286962 | NM_010917 |
| Sphk1 | 137.3583932 | 0.0167914660707068 | 0.0947864058972464 | 0.177150572508358 | 0.859390118936817 | 0.968520191286962 | NM_025367,NM_001172475,NM_011451,NM_001172472,NM_001172473 |
| Tmem243 | 213.485771 | -0.0163764343398224 | 0.0923384338318508 | -0.177352307811978 | 0.859231666245831 | 0.968520191286962 | NM_001081029 |
| Kdsr | 1443.447199 | -0.0100976378475394 | 0.0570565733681499 | -0.176975889918692 | 0.859527327689169 | 0.968543477000455 | NM_027534 |
| 3110057O12Rik | 341.3545372 | -0.0138919407528307 | 0.0786878582478941 | -0.176544908733775 | 0.859865870634946 | 0.968627980076635 | NM_026622 |
| Abcd2 | 2390.834851 | 0.0116070712770175 | 0.0660784067360382 | 0.175656040306541 | 0.860564172967789 | 0.968627980076635 | NM_011994 |
| Cdh7 | 704.2156165 | 0.0138212877967075 | 0.0784120459514541 | 0.176264853556614 | 0.860085872483358 | 0.968627980076635 | NM_172853 |
| Cenpa | 82.25759956 | -0.0167663450706567 | 0.0949910212339957 | -0.176504524878782 | 0.859897594150951 | 0.968627980076635 | NM_007681 |
| Crls1 | 888.2216621 | 0.0111303497683998 | 0.0632823230081445 | 0.17588402636495 | 0.860385054830553 | 0.968627980076635 | NM_025646,NM_001024385 |
| Fbxo48 | 31.33718863 | -0.0153137911506029 | 0.0869502762564923 | -0.17612124779718 | 0.860198688510578 | 0.968627980076635 | NM_176982 |
| Lgr5 | 126.4892576 | 0.0166555280557154 | 0.0945731773243783 | 0.176112599015134 | 0.860205483045282 | 0.968627980076635 | NM_010195 |
| Nfyb | 1627.295796 | -0.0109956227035626 | 0.0621960434459396 | -0.176789745687278 | 0.859673543939317 | 0.968627980076635 | NM_010914 |
| Pck2 | 950.8260782 | 0.0119609251423227 | 0.0679048338489662 | 0.176142469752981 | 0.860182016459517 | 0.968627980076635 | NM_028994 |
| Rad1 | 361.353771 | 0.012773151292457 5 | 0.0725646807069191 | 0.176024357414965 | 0.860274806763482 | 0.968627980076635 | NM_011232 |
| Spopl | 361.3453259 | -0.0140421634914845 | 0.0799369920807795 | -0.175665397533276 | 0.860556821284831 | 0.968627980076635 | NM_001165997,NM_029773,NM_001165998 |
| Stau1 | 2871.69108 | 0.00821100669012217 | 0.0467578051709983 | 0.175607188149521 | 0.860602554789278 | 0.968627980076635 | NM_001109906,NM_011490,NM_001109905 |
| Trmt2b | 666.3208489 | -0.0124284701768352 | 0.0707709840915152 | -0.175615336375198 | 0.860596152925363 | 0.968627980076635 | NM_001167995,NM_172540,NM_001167994 |
| Wdr73 | 1211.433357 | -0.0110641690024753 | 0.0629836438170978 | -0.175667337294826 | 0.860555297275828 | 0.968627980076635 | NM_028026,NR_027976 |
| Zfp407 | 740.275358 | 0.0118812151337877 | 0.0674707681265371 | 0.176094262207082 | 0.860219888585168 | 0.968627980076635 | NM_001033341 |
| Tshz3 | 1223.705709 | -0.0114319233749817 | 0.0652217738032276 | -0.175277713382521 | 0.860861422817308 | 0.96884427270816 | NM_172298 |
| Mfsd7c | 125.0329891 | -0.0160375889767754 | 0.0915480824910952 | -0.175182139706043 | 0.860936517767463 | 0.968853723046961 | NM_145447 |
| Pnp2 | 30.99786526 | -0.0151970883787835 | 0.0868761559536924 | -0.174928186128355 | 0.861136062408327 | 0.968928152017301 | NM_001123371 |
| Ptgdr | 26.59633905 | -0.0146825345037802 | 0.0839092679583912 | -0.174981082078573 | 0.861094498555556 | 0.968928152017301 | NM_008962 |
| Espnl | 28.12643965 | 0.0142831028650056 | 0.0817062957445608 | 0.174810309717859 | 0.8612286870974 | 0.968957316147226 | NM_001033292 |
| Npy1r | 1069.653817 | -0.0114929074436249 | 0.0660046728955485 | -0.174122633132539 | 0.861769086230206 | 0.969358561020486 | NM_010934 |
| Tcte2 | 76.09221761 | -0.0165730671882517 | 0.0951021269976501 | -0.174265999210104 | 0.861656419050694 | 0.969358561020486 | NM_022311 |
| Tmprss7 | 92.37218828 | 0.0165498671044039 | 0.0950586063778757 | 0.174101722453358 | 0.861785519552427 | 0.969358561020486 | NM_172455 |
| Rgs7 | 3681.694182 | -0.0113954661452224 | 0.0655686494043706 | -0.173794431465944 | 0.862027020843035 | 0.969505810682436 | NM_011880,NM_001199003 |
| Tbc1d2 | 199.5177719 | -0.0161311480644804 | 0.0928329638989303 | -0.173765302614304 | 0.862049914000712 | 0.969505810682436 | NM_198664 |
| Polr2k | 292.9442166 | 0.014302783683985 | 0.0824370047016109 | 0.173499557823036 | 0.862258775416777 | 0.969665352244647 | NM_023127,NM_001039368 |
| 2900092D14Rik | 622.805232 | -0.0135454231442263 | 0.0783726938154591 | -0.172833451101236 | 0.862782342523634 | 0.96975908749173 | NR_027891 |
| Acvr2b | 138.3148488 | 0.015275977053632 | 0.0884207391268056 | 0.172764638754314 | 0.862836433213623 | 0.96975908749173 | NM_007397 |
| Cdhr2 | 33.62460414 | -0.0123364886323796 | 0.071497432510279 | -0.172544498442039 | 0.863009481206498 | 0.96975908749173 | NM_001033364 |
| Dcp1a | 1255.323369 | 0.0110266289752534 | 0.0637060240106815 | 0.173086127198969 | 0.862583729262425 | 0.96975908749173 | NM_133761 |
| Lum | 77.3735883 | 0.0159169774074545 | 0.0920519750200048 | 0.172912937544201 | 0.862719862145673 | 0.96975908749173 | NM_008524 |

| | | | | | | | | | | | | | | | |
|---|---|---|---|---|---|---|---|---|---|---|---|---|---|---|---|
| Pvrl1 | 1996.919697 | -0.03682486 | 0.099535343 | -0.369967681 | 0.711406572 | 0.852939211 | | Nup153 | 2393.438854 | -0.0127896790584893 | 0.0740637949877829 | -0.172684630332526 | 0.862899325507913 | 0.96975908749173 | NM_175749 |
| 2410018L13Rik | 86.02727569 | 0.059582139 | 0.161161469 | 0.369704618 | 0.711602591 | 0.853108213 | | Pgpep1 | 513.6042383 | 0.0123270731939042 | 0.0712019100605154 | 0.173128406013649 | 0.862550497314171 | 0.96975908749173 | NM_023217 |
| 2700054A10Rik | 194.575206 | -0.038662399 | 0.104624809 | -0.369533756 | 0.711729917 | 0.853128836 | | Psd2 | 2795.734583 | -0.0122462084448272 | 0.0706681244082842 | -0.173291827784687 | 0.86242204699797 | 0.96975908749173 | NM_028707 |
| Ascl1 | 136.3006672 | 0.04774349 | 0.129198338 | 0.369536408 | 0.711727941 | 0.853128836 | | Slc27a2 | 274.154173 | 0.0160046094381434 | 0.0927470978933521 | 0.172561835374588 | 0.86299585274443 | 0.96975908749173 | NM_011978 |
| 1110008F13Rik | 1252.470317 | -0.025444399 | 0.068906263 | -0.369261053 | 0.711933152 | 0.853306433 | | Ube2g1 | 1710.930153 | 0.0146106922970727 | 0.0845499478746431 | 0.172805455997857 | 0.862804348300003 | 0.96975908749173 | NM_025985 |
| Atg4b | 2507.266747 | -0.018497429 | 0.05019876 | -0.368483793 | 0.712512527 | 0.853612884 | | Dcaf13 | 823.2454982 | 0.0108404050109189 | 0.0628860811528156 | 0.172381627415713 | 0.863137515165101 | 0.969773370907033 | NM_198606 |
| D130017N08Rik | 449.1845871 | 0.026694792 | 0.072412376 | 0.368649583 | 0.712388933 | 0.853612884 | | Rhbdl1 | 1853.68513 | -0.0135130772085915 | 0.0784010018183947 | -0.172358476233412 | 0.863155714749898 | 0.969773370907033 | NM_144816 |
| Mettl11b | 22.23890637 | -0.061892815 | 0.167970376 | -0.368474589 | 0.71251939 | 0.853612884 | | Rell1 | 587.6470193 | -0.0144479747353479 | 0.0838746256678477 | -0.172256801390249 | 0.863235644137906 | 0.96978816448161 | NM_145923 |
| Pus7l | 105.9122602 | -0.046440173 | 0.126011959 | -0.368537823 | 0.712472248 | 0.853612884 | | Msh2 | 1437.400382 | 0.00885722949529845 | 0.0514696156006797 | 0.172086567811505 | 0.863369472565477 | 0.969788505914485 | NM_008628 |
| Slc4a7 | 741.987174 | -0.037062304 | 0.100535721 | -0.368648111 | 0.71239003 | 0.853612884 | | Tns1 | 2737.700327 | 0.0108469607758987 | 0.0630173598975107 | 0.17212655042261 | 0.863338040047838 | 0.969788505914485 | NM_027884 |
| Zfp523 | 3718.260338 | 0.026338448 | 0.071423184 | 0.368766088 | 0.712302084 | 0.853612884 | | AF529169 | 155.478479 | 0.0161734801551353 | 0.0943825284819318 | 0.171360954355354 | 0.863939954523809 | 0.969836459474903 | NM_153509 |
| Pigx | 295.3677739 | -0.031127037 | 0.084498136 | -0.36837543 | 0.712593315 | 0.853635443 | | Acy1 | 454.7470637 | -0.0142808779370885 | 0.0833055855226072 | -0.171427616137612 | 0.86388754163893 | 0.969836459474903 | NM_025371,NM_001276442 |
| Ntan1 | 1315.933607 | -0.028264974 | 0.076780372 | -0.368127599 | 0.712778092 | 0.853724781 | | Erlec1 | 2081.462225 | 0.0110872349191864 | 0.0645861366366585 | 0.171665863551489 | 0.863700224297622 | 0.969836459474903 | NM_025745 |
| Pqlc3 | 34.36435913 | 0.059888839 | 0.162667131 | 0.368168042 | 0.712747938 | 0.853724781 | | Klhl22 | 3640.274215 | 0.00928489844836116 | 0.0542076942607961 | 0.171283773917611 | 0.864000638458684 | 0.969836459474903 | NM_145479 |
| Cacng6 | 16.23484161 | -0.060784381 | 0.165403313 | -0.36749192 | 0.713252117 | 0.853949516 | | Lama1 | 167.8008718 | -0.0161790344948849 | 0.0941476257653736 | -0.171847503995532 | 0.863557418218816 | 0.969836459474903 | NM_008480 |
| Cttnbp2nl | 1506.722623 | 0.033779742 | 0.091864595 | 0.367712302 | 0.713087767 | 0.853949516 | | Mgrn1 | 10738.52821 | 0.0063671069492221 | 0.0371762825767222 | 0.171267983453753 | 0.864013053976644 | 0.969836459474903 | NM_029657,NM_001252437 |
| Ddx28 | 471.9630016 | -0.030581987 | 0.083194637 | -0.367595653 | 0.713174756 | 0.853949516 | | Pde8a | 839.9144272 | 0.0152381171248544 | 0.0888699881013766 | 0.171465276978228 | 0.863857931034663 | 0.969836459474903 | NM_008803 |
| Pold2 | 434.3870496 | 0.027602677 | 0.075067935 | 0.367702621 | 0.713094986 | 0.853949516 | | Rnf170 | 1665.722527 | 0.00905112818048674 | 0.052683585152407 | 0.171801675119546 | 0.863593448552103 | 0.969836459474903 | NM_029965 |
| Ppp1cc | 8068.296299 | -0.040447791 | 0.110082227 | -0.367432525 | 0.713296414 | 0.853949516 | | Slc26a7 | 23.44703058 | 0.0119639863912783 | 0.0698426482168964 | 0.171299151689154 | 0.863988547459277 | 0.969836459474903 | NM_145947 |
| Tmem245 | 1256.132194 | 0.033636339 | 0.091509882 | 0.36757056 | 0.71319347 | 0.853949516 | | Ap2a2 | 10859.00859 | 0.00923676282912748 | 0.0539807378992997 | 0.171112200177006 | 0.864135542998972 | 0.969899003060043 | NM_007459 |
| Esyt3 | 96.54783528 | 0.057862833 | 0.157615956 | 0.36711279 | 0.713534888 | 0.854047893 | | 1700019D03Rik | 126.9518453 | 0.0156077508795139 | 0.0913500823125928 | 0.170856451186387 | 0.864336639979719 | 0.969963136487673 | NM_144953 |
| Gpr176 | 612.3331152 | 0.027193203 | 0.074075607 | 0.367100639 | 0.713543951 | 0.854047893 | | Dak | 479.2199325 | 0.0132961175707344 | 0.0778530641590577 | 0.170784768902221 | 0.864393005771233 | 0.969963136487673 | NM_145496 |
| Gstp2 | 10.1928301 | -0.058140685 | 0.158350232 | -0.367165139 | 0.713495841 | 0.854047893 | | Sfrp1 | 442.8883934 | 0.0157057841093836 | 0.0918741830908403 | 0.170948830030462 | 0.864264000920499 | 0.969963136487673 | NM_013834 |
| Nmnat3 | 112.4271343 | -0.041862457 | 0.114080914 | -0.366954079 | 0.713653272 | 0.85411276 | | 1700019G17Rik | 581.9797721 | -0.0130213227384822 | 0.0764364695489444 | -0.170354842594402 | 0.864731083361924 | 0.96997240313572 | NM_029331,NM_001145895 |
| Chchd7 | 403.0671475 | 0.03437352 | 0.093812768 | 0.366405557 | 0.714062473 | 0.854404507 | | Acsf3 | 748.3739755 | -0.0132122816216889 | 0.0774413485795894 | -0.170610169683579 | 0.864530300923063 | 0.96997240313572 | NM_144932 |
| Eif4e3 | 586.1100906 | -0.030614361 | 0.083538921 | -0.366468231 | 0.714015714 | 0.854404507 | | Efcab2 | 468.7590422 | -0.0135189003876684 | 0.079299804535987 | -0.170478356999397 | 0.864633953824312 | 0.96997240313572 | NM_026626 |
| Lynx1 | 18057.98361 | -0.033634925 | 0.091776444 | -0.36648756 | 0.714001294 | 0.854404507 | | Gm6313 | 30.06735782 | 0.0151468571986721 | 0.0888967764148497 | 0.170387024249194 | 0.864705776082388 | 0.96997240313572 | NR_045867 |
| Hyal2 | 318.0637515 | -0.029185294 | 0.079712516 | -0.366131887 | 0.714266665 | 0.854582834 | | Perp | 166.6477878 | 0.0161489645307913 | 0.0947989098447885 | 0.17034968605896 | 0.8647351384144 | 0.96997240313572 | NM_022032 |
| Pnpo | 2070.508823 | 0.026462011 | 0.072295345 | 0.366026479 | 0.714345318 | 0.854610945 | | Gtf2e1 | 715.0286773 | -0.0104003446025091 | 0.0611102126524899 | -0.170189959273287 | 0.864860747866423 | 0.970038392672972 | NM_028812 |
| Rmnd5b | 1848.063911 | 0.025085883 | 0.068581012 | 0.365784677 | 0.714525755 | 0.854760813 | | Adamts5 | 94.76126503 | 0.0156678692222789 | 0.0928917675713899 | 0.168668006131305 | 0.866057785583024 | 0.970182416393846 | NM_011782 |
| Trpc4 | 601.3768542 | -0.043220159 | 0.118307158 | -0.365321592 | 0.714871362 | 0.855108229 | | Cdk2ap2 | 576.5200048 | 0.0107675726301413 | 0.0636123100132788 | 0.169268693872203 | 0.865585299300546 | 0.970182416393846 | NM_026373 |
| Dab2 | 449.0587846 | -0.046981303 | 0.128689547 | -0.365074741 | 0.715055616 | 0.855109657 | | Cks2 | 34.14137276 | 0.015318560749787 | 0.090746108364262 | 0.168806806439535 | 0.865948604397014 | 0.970182416393846 | NM_025415 |
| Dock8 | 268.6816601 | -0.034356433 | 0.094140032 | -0.364950302 | 0.715148505 | 0.855109657 | | Cox5a | 4376.584947 | 0.0122116993027463 | 0.072286335708405 | 0.168935099325091 | 0.865847690689985 | 0.970182416393846 | NM_007747 |
| Kcnn1 | 682.6949707 | -0.032875071 | 0.0900674 | -0.365005214 | 0.715107514 | 0.855109657 | | Cyb5r1 | 851.7955579 | -0.0129222333039402 | 0.0761281351830803 | -0.169743200366902 | 0.865212098053447 | 0.970182416393846 | NM_028057 |
| Map3k7 | 2590.583368 | 0.021760282 | 0.059597422 | 0.365121192 | 0.715020942 | 0.855109657 | | Dgcr14 | 1179.190837 | -0.0088024531849225 | 0.0519027034326832 | -0.169595273516709 | 0.865328439889205 | 0.970182416393846 | NM_001081633,NM_022408 |
| Tnni3k | 22.62762996 | -0.061195781 | 0.167616339 | -0.365094366 | 0.715040966 | 0.855109657 | | Dram2 | 611.6657998 | 0.0120246207723991 | 0.0712452805700381 | 0.168777786769725 | 0.865971431238393 | 0.970182416393846 | NM_001025582,NM_026013 |
| Specc1 | 4106.055133 | 0.030698386 | 0.084155724 | 0.364780719 | 0.715275099 | 0.85515718 | | Gpr37 | 3001.565326 | -0.0140253890109861 | 0.0828217697114533 | -0.169344232317781 | 0.865525886004082 | 0.970182416393846 | NM_010338 |
| Usp4 | 3907.474046 | 0.015776828 | 0.043253909 | 0.364749191 | 0.715298635 | 0.85515718 | | Hus1 | 712.7593272 | 0.0125088500032833 | 0.0738559667065917 | 0.169368171064328 | 0.865507057606752 | 0.970182416393846 | NM_008316 |
| Cdca8 | 42.98309132 | -0.059380725 | 0.162924094 | -0.364468653 | 0.715508078 | 0.85529714 | | Krtcap2 | 784.6708661 | -0.0107767654713961 | 0.0636088346225139 | -0.169422463645981 | 0.865464355474259 | 0.970182416393846 | NM_025327 |
| Mlf2 | 14957.98408 | -0.029087926 | 0.079814419 | -0.364444503 | 0.715526109 | 0.85529714 | | Ldlrad3 | 582.5760565 | 0.0112462430579724 | 0.0665913957847806 | 0.168884326952983 | 0.865887627392043 | 0.970182416393846 | NM_178886 |
| Cyb5rl | 102.4248412 | 0.050103877 | 0.137530884 | 0.364310007 | 0.715626528 | 0.855351186 | | Map1lc3a | 5161.88873 | -0.0097722532499493 | 0.0576398301322957 | -0.169539938398845 | 0.865371960725142 | 0.970182416393846 | NR_046364,NM_025735 |
| Ctcf | 2628.789448 | 0.024850325 | 0.06826695 | 0.364016925 | 0.715845371 | 0.855546759 | | Mdm2 | 1799.722896 | -0.0112242649046742 | 0.0664685688551216 | -0.16886575261067 | 0.865902237745828 | 0.970182416393846 | NM_010786 |
| Ctdsp1 | 1122.964765 | 0.022321903 | 0.061347454 | 0.363860303 | 0.715962329 | 0.855620542 | | Myh10 | 14808.47377 | 0.00903504455813963 | 0.0531682097601599 | 0.169933210068506 | 0.865062663090183 | 0.970182416393846 | NM_175260 |
| 2610301B20Rik | 1402.338923 | 0.026178152 | 0.072011924 | 0.363525248 | 0.716212556 | 0.855787564 | | Pcolce | 261.0200388 | -0.0124860610281753 | 0.0738966513210369 | -0.168966533732777 | 0.865822965080266 | 0.970182416393846 | NM_008788 |
| Hat1 | 553.594975 | -0.03104853 | 0.085400261 | -0.36356482 | 0.716183001 | 0.855787564 | | Znrf1 | 6462.519745 | 0.0119641049708562 | 0.0709232015356578 | 0.168690988446722 | 0.866039707374098 | 0.970182416393846 | NM_001168622,NM_001168623,NM_133206,NM_001168621 |
| 2210016F16Rik | 306.7016654 | -0.043908192 | 0.120852222 | -0.363321348 | 0.716364848 | 0.855903528 | | 6330416G13Rik | 658.1153801 | -0.0119325736247228 | 0.0711901148027633 | -0.167615597443307 | 0.866885699823158 | 0.970212009833669 | NM_144905 |
| 3930402G23Rik | 13.26162826 | -0.05947501 | 0.163790112 | -0.363117219 | 0.716517322 | 0.855923913 | | Cwc15 | 1903.716914 | 0.0126111847287256 | 0.0750459413956284 | 0.168046192694709 | 0.866546939247221 | 0.970212009833669 | NM_023153 |
| Ccdc103 | 98.58619842 | 0.045043597 | 0.124077339 | 0.363028391 | 0.716583676 | 0.855923913 | | Hapln4 | 6144.371188 | 0.00952709054683034 | 0.0565538522473913 | 0.168460505663782 | 0.866221011508047 | 0.970212009833669 | NM_177900 |
| Slc35e1 | 3929.044079 | 0.02288207 | 0.063014093 | 0.363126235 | 0.716510588 | 0.855923913 | | Idh2 | 1334.678393 | 0.0122129886592703 | 0.0728574700535916 | 0.167628503299481 | 0.866875546091928 | 0.970212009833669 | NM_173011 |
| Slc48a1 | 5558.591535 | 0.025719875 | 0.070853126 | 0.363002685 | 0.716602878 | 0.855923913 | | Kif5b | 12321.13885 | 0.0108000023724166 | 0.0640895076572118 | 0.168514360106827 | 0.866178647482512 | 0.970212009833669 | NM_008448 |
| Paqr9 | 856.0129954 | -0.029183669 | 0.080413003 | -0.362922269 | 0.716662951 | 0.855929682 | | Lrrc1 | 428.9328634 | -0.0145950830058683 | 0.086798014461814 | -0.168149963986668 | 0.866465303313709 | 0.970212009833669 | NM_172528,NM_001146048 |
| Cdh13 | 2552.047051 | -0.03431623 | 0.094670345 | -0.362481303 | 0.716992393 | 0.85599323 | | Mapk14 | 4351.200155 | -0.00950884685373037 | 0.0566928163581925 | -0.167725780170317 | 0.866799013852217 | 0.970212009833669 | NM_001168513,NM_011951,NM_001168508,NM_001168514 |
| Fibp | 1137.313197 | 0.027596319 | 0.076091975 | 0.362670555 | 0.716850998 | 0.85599323 | | Otub2 | 1513.909759 | 0.0146757463304262 | 0.0872202364719662 | 0.168260795017945 | 0.866378115120498 | 0.970212009833669 | NM_001177841,NM_026580,NR_033587 |
| Gm11346 | 9.504323874 | 0.054617504 | 0.150651317 | 0.362542491 | 0.716946677 | 0.85599323 | | Rapgef4 | 25921.22962 | 0.00732067446271916 | 0.0436675119232466 | 0.167645788374354 | 0.866861947028211 | 0.970212009833669 | NM_001204166,NM_001204167,NM_001204165,NM_019688 |
| Gm1604b | 15.63049129 | -0.059469929 | 0.164043063 | -0.362526324 | 0.716958756 | 0.85599323 | | Tuft1 | 110.8998611 | 0.0158724451019758 | 0.0946692485230775 | 0.167662100941961 | 0.866849113122476 | 0.970212009833669 | NM_011656 |
| Pcdhgb6 | 563.6521109 | -0.029672174 | 0.081841495 | -0.362556603 | 0.716936134 | 0.85599323 | | Zfp623 | 532.4771775 | -0.0118030177569233 | 0.0703999932131743 | -0.167656518391746 | 0.86685350518757 | 0.970212009833669 | NM_030199 |
| Sumo3 | 5204.251865 | -0.030780368 | 0.084986419 | -0.362179837 | 0.717217646 | 0.856196179 | | Zfp868 | 494.6570626 | 0.0132025394485765 | 0.0784823171137738 | 0.168223109792199 | 0.866407761028256 | 0.970212009833669 | NM_172754,NM_001045553 |
| Kcnh2 | 1589.47047 | -0.031252237 | 0.086323324 | -0.362037 | 0.717324381 | 0.856257624 | | Atg9b | 232.9466722 | 0.0156629165078986 | 0.0941895605748398 | 0.166291427758105 | 0.867927611783762 | 0.970480832019932 | NM_001002897 |
| Mthfs | 49.43999889 | -0.055574692 | 0.153605961 | -0.361800361 | 0.717501222 | 0.856364623 | | Dleu2 | 128.8139356 | -0.0153965579927262 | 0.0924575907788019 | -0.166525623943212 | 0.867743319700436 | 0.970480832019932 | NR_028264 |
| Rapgef4 | 24482.07475 | -0.031759053 | 0.087788178 | -0.361769133 | 0.71752456 | 0.856364623 | | Fam161a | 114.3384175 | -0.0157620867764381 | 0.094452781168611 | -0.166877952998553 | 0.867466080837845 | 0.970480832019932 | NM_028672 |
| Dnajc10 | 3333.191 | 0.02695325 | 0.074743854 | 0.360608238 | 0.71839233 | 0.857031081 | | Fam175a | 166.4856219 | -0.0156180319705996 | 0.0936894121263156 | -0.166700074385596 | 0.86760604700448 | 0.970480832019932 | NM_172405 |
| Rab3gap1 | 4366.750005 | 0.02710996 | 0.075184707 | 0.360578118 | 0.718414849 | 0.857031081 | | Fgfbp3 | 142.76027 | -0.0153248460478094 | 0.0920023686547806 | -0.166570125007462 | 0.867708302035219 | 0.970480832019932 | NM_028263 |
| Snip1 | 505.3550265 | 0.024290437 | 0.067362621 | 0.360592224 | 0.718404303 | 0.857031081 | | Glcci1 | 2487.342573 | 0.0112580208307966 | 0.0673442392108348 | 0.167171252696925 | 0.867235303121006 | 0.970480832019932 | NM_133236 |
| Tada2b | 1455.415048 | -0.034171762 | 0.094763949 | -0.360598756 | 0.71839942 | 0.857031081 | | Kcnh6 | 117.5273428 | -0.0156035518378954 | 0.0934708559119336 | -0.166934941224854 | 0.867421239768024 | 0.970480832019932 | NM_001037712 |
| Tmem44 | 3010.115454 | -0.035398579 | 0.098101151 | -0.360837555 | 0.718220887 | 0.857031081 | | R3hcc1l | 626.4440671 | 0.0133869622269361 | 0.0801351510561023 | 0.167054807416086 | 0.867326924691958 | 0.970480832019932 | NM_177464 |
| Xpo7 | 4815.894444 | 0.025228833 | 0.069957581 | 0.360630436 | 0.718375734 | 0.857031081 | | Rab9 | 952.7854024 | -0.00932282415264199 | 0.0560623013393281 | -0.166293996677264 | 0.867925590228263 | 0.970480832019932 | NM_019773 |
| Opalin | 572.3601181 | -0.056439292 | 0.156618724 | -0.360361078 | 0.718577129 | 0.857158676 | | Rps10 | 4086.260649 | -0.0109050488269693 | 0.0655653116483702 | -0.166323449897616 | 0.867902412713777 | 0.970480832019932 | NM_025963 |
| Fam72a | 41.64150374 | -0.060425266 | 0.167869981 | -0.359952781 | 0.718882445 | 0.85745686 | | Slc9a9 | 838.132309 | 0.0109531752209116 | 0.0656251450197571 | 0.166905158344626 | 0.86744467431348 | 0.970480832019932 | NM_177909 |
| Fbxo11 | 5194.978154 | -0.025223322 | 0.070158906 | -0.359517041 | 0.719208332 | 0.857779532 | | Trpv2 | 894.4255517 | 0.0110955937854752 | 0.066618821886388 | 0.166553437471436 | 0.867721433343565 | 0.970480832019932 | NM_011706 |
| Pbld1 | 19.82299317 | -0.059896621 | 0.166650898 | -0.359413729 | 0.719285606 | 0.857805663 | | A930015D03Rik | 42.77334827 | -0.0153293894083253 | 0.0923453075702666 | -0.166000740174708 | 0.868156367679423 | 0.970658528480126 | NR_015618 |
| D930020B18Rik | 18.57144959 | -0.059397065 | 0.165334523 | -0.359253859 | 0.719405189 | 0.857816222 | | Atf7ip | 2559.128372 | -0.0104380118243337 | 0.0630366991378604 | -0.16558626906377 | 0.868482553810704 | 0.970658528480126 | NM_019426 |
| Nexn | 395.0169962 | -0.05487608 | 0.15273344 | -0.359293157 | 0.719375793 | 0.857816222 | | Bard1 | 23.9959028 | -0.0136836440788305 | 0.0826830637838059 | -0.165495126240237 | 0.868554285638488 | 0.970658528480126 | NM_007525 |
| Lrrc1 | 411.7669936 | -0.040987195 | 0.114212236 | -0.358868686 | 0.719693326 | 0.858093759 | | Dsg2 | 106.0236825 | -0.0156945808343273 | 0.0947431280186409 | -0.16565402855645 | 0.868429225988176 | 0.970658528480126 | NM_007883 |
| Tubb6 | 51.81741495 | -0.059736246 | 0.166538237 | -0.358693878 | 0.719824109 | 0.858134425 | | Gdpd5 | 1500.242776 | 0.01138548792691 8 | 0.0686366705105068 | 0.165880539400219 | 0.868250962617338 | 0.970658528480126 | NM_201352 |
| Vil1 | 23.85968561 | 0.060204957 | 0.167888419 | 0.358601014 | 0.719893588 | 0.858134425 | | Qser1 | 771.3344142 | 0.0116161520751598 | 0.0701876478899807 | 0.165501372739661 | 0.868549369442655 | 0.970658528480126 | NM_001123327 |
| Zfp719 | 661.2410681 | -0.032598889 | 0.090888023 | -0.3586709 | 0.7198413 | 0.858134425 | | Senp3 | 2855.01886 | -0.00879707588401201 | 0.0530986488461607 | -0.165674194639099 | 0.868413355068796 | 0.970658528480126 | NM_030702,NM_001163571 |
| Spg11 | 1122.138438 | 0.025845463 | 0.072306798 | 0.357441675 | 0.720761182 | 0.859102528 | | AU021092 | 94.19210239 | -0.0154388013654048 | 0.0933496855316182 | -0.165386752804599 | 0.868639579824675 | 0.970679170438744 | NM_001033220 |
| Slc25a26 | 562.0719877 | -0.030457347 | 0.085229937 | -0.357355037 | 0.720826033 | 0.859113735 | | 2010012O05Rik | 966.3414227 | -0.0115531329809311 | 0.0701525860144573 | -0.164685774784555 | 0.869191314171483 | 0.970788710570998 | NM_025563 |
| 1700040L02Rik | 70.02454988 | -0.051432668 | 0.144041018 | -0.357069596 | 0.721039705 | 0.859218836 | | Abcb9 | 2392.384255 | 0.0112832050924036 | 0.068467606174513 | 0.164796255088056 | 0.869104351748808 | 0.970788710570998 | NM_019875 |
| Ngly1 | 871.0853123 | -0.028469248 | 0.079742443 | -0.357014993 | 0.721080581 | 0.859218836 | | E330020D12Rik | 25.03249191 | -0.0140876357908403 | 0.0855518287519771 | -0.164667851013234 | 0.869205422668759 | 0.970788710570998 | NR_033736 |
| Tead4 | 29.75796076 | -0.059477577 | 0.166542792 | -0.357130901 | 0.720993812 | 0.859218836 | | Hnrpll | 1890.33587 | 0.0106639472520441 | 0.0646558852488099 | 0.164933899072094 | 0.868996010178366 | 0.970788710570998 | NM_144802 |
| Mzf1 | 252.6883157 | -0.036252273 | 0.101633065 | -0.356697629 | 0.721318181 | 0.859435858 | | Pigh | 428.4001969 | 0.011440955519291 1 | 0.0694438146950749 | 0.164751253506564 | 0.869139773677597 | 0.970788710570998 | NM_029988 |
| Dmtf1 | 2396.924026 | -0.028111131 | 0.078840759 | -0.356555813 | 0.721424362 | 0.859467364 | | Sypl2 | 127.1166887 | -0.0154707799492697 | 0.0937256337191409 | -0.165064554224617 | 0.868893171893655 | 0.970788710570998 | NM_008596 |
| Wiz | 1925.727432 | 0.024490863 | 0.068695349 | 0.356514139 | 0.721455566 | 0.859467364 | | Vegfb | 2093.009567 | 0.0109362321188247 | 0.0662307572154328 | 0.165123162992864 | 0.868847041825717 | 0.970788710570998 | NM_001185164,NM_011697 |
| Trim24 | 1084.178235 | 0.0271531 | 0.07619455 | 0.356365381 | 0.721566952 | 0.85953397 | | B4galt4 | 823.5341468 | 0.0095308717749213 | 0.0579272624304174 | 0.164531713998566 | 0.869312582764894 | 0.970833749307515 | NM_019804 |
| Bok | 970.617301 | -0.040649146 | 0.114149982 | -0.356102957 | 0.721763463 | 0.859604432 | | Cdkn1c | 239.0325539 | -0.0154544031311466 | 0.0940322615146128 | -0.164352137045486 | 0.869453940230138 | 0.970842334982165 | NM_001161624,NM_009876 |
| Htr1f | 101.581009 | -0.043492956 | 0.122140484 | -0.356089601 | 0.721773464 | 0.859604432 | | Slc7a10 | 1166.429157 | 0.0134183642483882 | 0.081607084713486 | 0.164426462426623 | 0.869395433065274 | 0.970842334982165 | NM_017394 |
| Taf9 | 2439.287394 | 0.028840178 | 0.080997147 | 0.356064125 | 0.721792543 | 0.859604432 | | Zfp804a | 930.0865896 | -0.0142391759314013 | 0.0867221144134992 | -0.164193136061093 | 0.869579104406962 | 0.970907461231017 | NM_175513 |
| Slc35e3 | 1352.309264 | -0.017858622 | 0.050192247 | -0.355804386 | 0.721987064 | 0.859770008 | | Clgn | 334.1718788 | -0.0138875792406368 | 0.0847241977136444 | -0.163915146031537 | 0.869797943562804 | 0.971077159956444 | NM_009904 |
| Hhipl1 | 183.5973184 | 0.042515692 | 0.119680718 | 0.355242619 | 0.722407839 | 0.860204968 | | Tnks | 3590.50213 | 0.0102421836069489 | 0.0625527191918658 | 0.163736824542086 | 0.8699383269572 | 0.971097166614577 | NM_175091 |
| Dcaf6 | 3831.216018 | -0.026486771 | 0.074617059 | -0.354969383 | 0.722612528 | 0.86038258 | | Wbscr16 | 503.4276578 | 0.0126914860658373 | 0.0775182552153625 | 0.163722545490448 | 0.869949568302044 | 0.971097166614577 | NM_033572 |
| Trim12c | 128.4595416 | 0.057152878 | 0.161044871 | 0.354887913 | 0.722673564 | 0.860389135 | | Ggnbp2 | 3949.430413 | 0.0098417715027456 | 0.0603648765835641 | 0.163038045627766 | 0.870488479342624 | 0.971477515718733 | NM_153144 |
| Itga2b | 90.21095094 | 0.056716037 | 0.159876871 | 0.354748232 | 0.722778214 | 0.86044761 | | Lect1 | 28.27493804 | -0.0139306920041081 | 0.0854460602574371 | -0.163034924748278 | 0.870490936568569 | 0.971477515718733 | NM_010701 |
| Ccdc111 | 151.9455901 | 0.052856415 | 0.149151515 | 0.35438068 | 0.723053611 | 0.860456111 | | Tmc7 | 838.5860981 | 0.0114723044592854 | 0.0703030301474285 | 0.163183641376873 | 0.870373845850669 | 0.971477515718733 | NM_172476 |
| Col6a1 | 4536.20896 | -0.05641591 | 0.159214182 | -0.354339728 | 0.723084298 | 0.860456111 | | 1110046J04Rik | 99.0606268 | 0.0154107397787069 | 0.0949031425749369 | 0.162383872236247 | 0.871003570372911 | 0.971532689510015 | NR_040707 |
| Cyp2e1 | 20.29962298 | -0.052300098 | 0.147486292 | -0.354609889 | 0.722881867 | 0.860456111 | | 1810043G02Rik | 1156.271975 | -0.00979010271884373 | 0.0609309917271176 | -0.160675256406283 | 0.872349178722579 | 0.971532689510015 | NM_026431 |
| Dlg3 | 8949.812473 | -0.02439447 | 0.068892181 | -0.354096346 | 0.723266681 | 0.860456111 | | 2410004N09Rik | 104.3897386 | -0.0153758802949183 | 0.0946112057367094 | -0.16251648179717 | 0.870899150209254 | 0.971532689510015 | NR_038152,NR_038151 |
| Eva1b | 48.05962262 | -0.057066099 | 0.161167594 | -0.354079241 | 0.723279499 | 0.860456111 | | 2610301G19Rik | 4163.4591 | -0.00797582272704586 | 0.0490245888392794 | -0.162690252297547 | 0.870762322274754 | 0.971532689510015 | NM_146055 |
| Gsn | 2395.39108 | 0.033125283 | 0.093524271 | 0.354189158 | 0.723197128 | 0.860456111 | | 4930520O04Rik | 26.40960403 | 0.0135974349463401 | 0.0847721726739453 | 0.160399745782607 | 0.872566189862645 | 0.971532689510015 | NR_040383,NR_040384 |
| Lrrc8b | 2659.863119 | 0.027352104 | 0.077170523 | 0.354437198 | 0.723011262 | 0.860456111 | | Akr1c18 | 68.34866461 | -0.0153429600473222 | 0.0947087695873777 | -0.162001471607831 | 0.871304695003611 | 0.971532689510015 | NM_134066 |
| Slc35g2 | 540.3725439 | -0.028387753 | 0.080175159 | -0.354071677 | 0.723285167 | 0.860456111 | | Armc8 | 3360.073184 | 0.0088247694490098 | 0.0550253230906288 | 0.160376513091528 | 0.872584489968721 | 0.971532689510015 | NM_028768,NM_001166138 |
| Tgfbi | 173.0605788 | 0.049507631 | 0.139675868 | 0.354446565 | 0.723004243 | 0.860456111 | | Atp1a2 | 55752.70747 | 0.00917647182299424 | 0.0571648975590675 | 0.160526340723559 | 0.87246647377738 | 0.971532689510015 | NM_178405 |
| Fsbp | 11.73931349 | -0.058832099 | 0.166317329 | -0.353734032 | 0.723538215 | 0.860691064 | | Atpaf2 | 814.5768439 | 0.00926762267814893 | 0.0572167892260341 | 0.161973833266619 | 0.871326459774114 | 0.971532689510015 | NM_145427 |
| Tmem163 | 589.0797543 | 0.032103602 | 0.090790117 | 0.353602383 | 0.723636888 | 0.860742357 | | Bckdk | 1630.444517 | 0.0114739762951557 | 0.07052123079080 4 | 0.162702439626904 | 0.870752726043616 | 0.971532689510015 | NM_009739 |
| Sgsh | 311.9617191 | 0.045524263 | 0.128948074 | 0.353043369 | 0.724055928 | 0.861174679 | | Chmp6 | 1049.256316 | 0.0107129060381988 | 0.0668493358896411 | 0.16025448713334 | 0.872680609440085 | 0.971532689510015 | NM_001085498 |
| 1700008O03Rik | 20.55105543 | 0.059141992 | 0.167704402 | 0.352656171 | 0.724346221 | 0.861321592 | | Crabp1 | 67.97053399 | 0.0153238231593321 | 0.0944728194680484 | 0.162203512561778 | 0.871145593807088 | 0.971532689510015 | NM_013496 |
| Abcg3 | 21.8414568 | 0.05898381 | 0.167225098 | 0.352721041 | 0.724297584 | 0.861321592 | | Deb1 | 790.5072205 | 0.0121382582922344 | 0.0753731587731962 | 0.16104218649983 | 0.872060174231843 | 0.971532689510015 | NM_026794 |
| Ncor2 | 6540.951016 | -0.031606109 | 0.089622101 | -0.352659772 | 0.724343522 | 0.861321592 | | Herc3 | 8361.88624 | 0.0112411407858902 | 0.0695741203928381 | 0.16157072087177 | 0.871643915678358 | 0.971532689510015 | NM_028705 |
| Mllt4 | 4058.749112 | -0.024267406 | 0.068828883 | -0.352575911 | 0.7244064 | 0.861327047 | | Hs3st4 | 2847.699485 | -0.0123208865431581 | 0.0765455583614694 | -0.160961482375966 | 0.872123737612386 | 0.971532689510015 | NM_001252072 |
| Adamts4 | 766.4677925 | -0.042240392 | 0.120035031 | -0.351900541 | 0.724912853 | 0.861686848 | | Igfbp5 | 8495.664644 | 0.0122941131298604 | 0.0763006902205932 | 0.161127154869987 | 0.871993253180735 | 0.971532689510015 | NM_010518 |
| Ap4b1 | 621.9431798 | 0.024394737 | 0.069362802 | 0.351697684 | 0.725064998 | 0.861686848 | | Kif1a | 67083.24583 | 0.00861985477847741 | 0.0535626158816767 | 0.160930429490584 | 0.872148195397576 | 0.971532689510015 | NM_001110315,NM_008440 |
| Btf3l4 | 2774.364396 | 0.025263375 | 0.071855291 | 0.351586842 | 0.725148135 | 0.861686848 | | Ncdn | 65284.40875 | -0.0104597967108294 | 0.0651091413867426 | -0.160650201923246 | 0.872368912961948 | 0.971532689510015 | NM_011986 |
| Lgals1 | 349.2201738 | -0.033445984 | 0.095130743 | -0.35157913 | 0.725153919 | 0.861686848 | | Nup214 | 2103.680493 | 0.00774564747364849 | 0.0483171260900978 | 0.160308530337774 | 0.872638039542459 | 0.971532689510015 | NM_172268 |
| Pgap2 | 322.7726639 | -0.026886116 | 0.076394656 | -0.351937126 | 0.724885416 | 0.861686848 | | Palmd | 636.2300338 | 0.0138020352805427 | 0.0852682867571974 | 0.161865985648852 | 0.871411389053465 | 0.971532689510015 | NM_023245 |
| Pou2f3 | 13.88675794 | 0.05353565 | 0.152246961 | 0.351636901 | 0.725110587 | 0.861686848 | | Plekhh1 | 2184.176064 | -0.0139792219848998 | 0.0865207705992279 | -0.161570705948203 | 0.871643927431231 | 0.971532689510015 | NM_181073 |
| Tspan18 | 146.0749015 | 0.055250208 | 0.157079863 | 0.351733232 | 0.725038336 | 0.861686848 | | Rnf43 | 148.7551884 | -0.0147365248269137 | 0.0915657359921843 | -0.160939293145326 | 0.872141214218593 | 0.971532689510015 | NM_172448 |
| Tspyl1 | 7061.744047 | -0.020636963 | 0.058651287 | -0.35185865 | 0.724944271 | 0.861686848 | | Samd5 | 271.5793756 | 0.0141099889318028 | 0.0866254430095948 | 0.16288504210293 | 0.870608948211899 | 0.971532689510015 | NM_177271 |
| Pccb | 1897.269827 | 0.021339117 | 0.060708664 | 0.351500353 | 0.725213008 | 0.861690977 | | Sgsm3 | 2898.781809 | -0.00953667393063292 | 0.0588422535467122 | -0.162071867676893 | 0.871249259617384 | 0.971532689510015 | NM_134091 |
| Dnd1 | 34.02841686 | 0.058986871 | 0.1679657 | 0.35118403 | 0.725450291 | 0.861896471 | | Shank2 | 6273.009678 | -0.0113757253350674 | 0.0703001262514626 | -0.161816570490592 | 0.87145030365621 | 0.971532689510015 | NM_001113373,NM_001081370 |
| Lhcgr | 17.49059571 | -0.05626589 | 0.160246217 | -0.351121485 | 0.72549721 | 0.861896471 | | Slc10a4 | 135.9234815 | 0.0152375975001288 | 0.0950392046580882 | 0.160329598242614 | 0.872621444428189 | 0.971532689510015 | NM_173403 |
| Ddx4 | 97.52253123 | -0.054436128 | 0.155191773 | -0.350766842 | 0.725763275 | 0.862146452 | | Srl | 390.3556637 | 0.0119663629605022 | 0.0739714280968019 | 0.161770068097678 | 0.871486924730683 | 0.971532689510015 | NM_175347 |
| Arpc2 | 6530.824311 | 0.033108411 | 0.094490643 | 0.350388248 | 0.726047345 | 0.862162734 | | Tcn2 | 1030.356807 | 0.0150633831037796 | 0.0929922428949554 | 0.161985372487416 | 0.871317372802578 | 0.971532689510015 | NM_001130458,NM_001130459,NM_015749 |
| Dnalc4 | 1614.750679 | 0.030024189 | 0.085690905 | 0.350377772 | 0.726055206 | 0.862162734 | | Tlcd1 | 589.0941234 | 0.0132174679376686 | 0.0818665237581623 | 0.161451437424089 | 0.871737856808085 | 0.971532689510015 | NM_026708 |
| Fmnl3 | 660.8887709 | 0.031787232 | 0.090689632 | 0.350505694 | 0.725959218 | 0.862162734 | | Tmem120a | 498.7335034 | -0.0115526742802766 | 0.0717116511928249 | -0.161098985842799 | 0.872015438992708 | 0.971532689510015 | NM_172541 |
| Rbm11 | 220.7235669 | -0.041650217 | 0.118856439 | -0.350424573 | 0.726020087 | 0.862162734 | | Wdr20a | 625.2335653 | -0.0104395665253337 | 0.0642526319655099 | -0.162476869911532 | 0.870930341386619 | 0.971532689510015 | NM_027149 |
| Zfp948 | 337.6644491 | 0.036261806 | 0.103428118 | 0.350599109 | 0.725889125 | 0.862162734 | | Zfyve19 | 625.0546276 | 0.0110707374963973 | 0.0687966126465683 | 0.160919805067605 | 0.872156563403424 | 0.971532689510015 | NM_028054,NM_001164827 |
| Sipa1 | 550.6994291 | 0.027270032 | 0.077933161 | 0.349915642 | 0.726402008 | 0.862508445 | | Mpl | 44.12021185 | -0.0143321957695884 | 0.0894939910929074 | -0.160147017632833 | 0.872765264391313 | 0.971552473794789 | NM_001122949,NM_010823 |
| Gm10069 | 38.61119159 | -0.057489845 | 0.164386269 | -0.349724131 | 0.726545742 | 0.862613006 | | Arsb | 5989.766881 | 0.00716547654915747 | 0.0448150898688448 | 0.159889817695956 | 0.872967869629157 | 0.97169109258184 | NM_009712 |
| Asb7 | 763.4842776 | 0.023269157 | 0.06658079 | 0.349487543 | 0.726723321 | 0.862625537 | | Gpr137b | 432.2786248 | -0.0114893556129297 | 0.071889751258495 | -0.159819103721993 | 0.87302357491984 | 0.97169109258184 | NM_031999 |
| Fam45a | 1038.915575 | 0.026436935 | 0.075628437 | 0.349563416 | 0.726666371 | 0.862625537 | | Pde5a | 572.2291674 | 0.0119752984791732 | 0.0750583068724281 | 0.159546610870492 | 0.873238238426676 | 0.971855550979612 | NM_153422 |
| Gstm4 | 343.9333416 | 0.030947479 | 0.088533548 | 0.349556524 | 0.726671544 | 0.862625537 | | 3110082I17Rik | 299.9144535 | 0.0127404184075717 | 0.07993996809922 | 0.159374824765485 | 0.873373572307075 | 0.97191269446355 | NM_028469 |
| Chd4 | 8806.354994 | 0.031293596 | 0.089582499 | 0.349327119 | 0.726843741 | 0.862694072 | | Arrb1 | 17918.51732 | 0.00797909125144708 | 0.0503060620355702 | 0.158610929350925 | 0.873975417431635 | 0.97191269446355 | NM_177231,NM_178220 |
| Scfd2 | 664.3973305 | 0.026703269 | 0.076456209 | 0.349262275 | 0.726892417 | 0.862694072 | | Ccdc87 | 160.9675473 | -0.0147709612853463 | 0.092732601371004 | -0.159285527063462 | 0.873443922909648 | 0.97191269446355 | NM_207268 |
| Adcy4 | 170.0421125 | 0.046029161 | 0.131996035 | 0.348716242 | 0.727302349 | 0.863081236 | | Derl2 | 902.739533 | -0.00984252637223207 | 0.0619637597686724 | -0.15884327240595 | 0.873792355144127 | 0.97191269446355 | NM_033562 |
| Cmtm7 | 71.68286398 | -0.052886774 | 0.151772255 | -0.348461411 | 0.727493689 | 0.863081236 | | Fbxw4 | 842.9562814 | 0.00999910153041279 | 0.0631131575903814 | 0.158431330520796 | 0.87411692735067 | 0.97191269446355 | NM_013907 |
| Gm20754 | 33.03341295 | -0.058059063 | 0.166565507 | -0.348565944 | 0.727415199 | 0.863081236 | | Gpr75 | 456.9022827 | 0.0121088307874347 | 0.0765711024146921 | 0.158138389099531 | 0.874347751060222 | 0.97191269446355 | NM_175490 |
| Mbtps2 | 1104.842195 | -0.02951621 | 0.084705508 | -0.348456794 | 0.727497156 | 0.863081236 | | Mettl13 | 501.4273588 | 0.0120482199764793 | 0.0762369750722382 | 0.158036437897268 | 0.874428086199195 | 0.97191269446355 | NM_144877 |
| Rac1 | 11034.29477 | 0.01604279 | 0.046015458 | 0.348639143 | 0.727360237 | 0.863081236 | | Pip5k1c | 22788.76989 | 0.00936079793673505 | 0.0589803676344097 | 0.158710403345704 | 0.873897041389595 | 0.97191269446355 | NM_008844,NM_001146687 |
| Ctcfl | 19.05918266 | -0.058482329 | 0.168098706 | -0.347904696 | 0.727911758 | 0.863374782 | | Psmg3 | 331.7637159 | 0.0126350720037873 | 0.0794401050736719 | 0.159051552009777 | 0.873628258108771 | 0.97191269446355 | NM_025604 |
| Ifnlr1 | 10.28598187 | -0.049561526 | 0.142421309 | -0.34799235 | 0.727845929 | 0.863374782 | | Rbms3 | 842.0536132 | 0.0117865062132447 | 0.0742143146000804 | 0.15881715376284 | 0.87381293359561 | 0.97191269446355 | NM_178660,NM_001172122,NM_001172123,NM_001172121,NM_001172126,NM_001172124 |
| Laptm4b | 3333.20333 | 0.019474058 | 0.055964278 | 0.347973012 | 0.727860451 | 0.863374782 | | Scn4b | 16662.21722 | 0.0149931961039417 | 0.0946520738446927 | 0.158403249870075 | 0.874139053085016 | 0.97191269446355 | NM_001013390 |
| Itch | 2810.052805 | 0.030512989 | 0.087741768 | 0.347758989 | 0.728021191 | 0.863398894 | | Setmar | 269.0860962 | -0.0135064218759059 | 0.0853765479972089 | -0.158198266331258 | 0.874300569819777 | 0.97191269446355 | NM_178391,NM_001276356 |
| Mlf1 | 30.96103818 | 0.058440718 | 0.16806386 | 0.347729238 | 0.728043536 | 0.863398894 | | Slc30a3 | 4867.657389 | -0.0129705328032674 | 0.0821164740312788 | -0.15795287067887 | 0.874493936158515 | 0.97191269446355 | NM_011773 |
| Gm13152 | 60.84356989 | -0.055428356 | 0.159443518 | -0.347636311 | 0.728113333 | 0.863415581 | | Smurf1 | 2607.484706 | -0.00924688853506733 | 0.0580741383899359 | -0.159225582874421 | 0.873491148760395 | 0.97191269446355 | NM_029438,NM_001038627 |
| Mylk | 828.7129317 | -0.031073492 | 0.089412036 | -0.347531425 | 0.728192114 | 0.863442919 | | Srrd | 566.4964554 | -0.0109172502323872 | 0.06894087017589 | -0.158356722282933 | 0.874175714027315 | 0.97191269446355 | NM_027323 |
| 4632415L05Rik | 928.7117235 | -0.023128487 | 0.066569749 | -0.347432396 | 0.728266499 | 0.863465039 | | Thra | 20814.97132 | -0.00966344887454163 | 0.0609932892509 | -0.158434624418932 | 0.874114331978692 | 0.97191269446355 | NM_178060 |
| Lpo | 30.84883934 | -0.058317567 | 0.16801885 | -0.34708943 | 0.728524134 | 0.863638327 | | Trpc1 | 1432.356263 | -0.0120213271016266 | 0.0760686979514121 | -0.15803250779059 | 0.87443118305633 | 0.97191269446355 | NM_011643 |
| Prrt3 | 1881.010398 | 0.02515609 | 0.072476868 | 0.347091297 | 0.728522732 | 0.863638327 | | Zfp462 | 1967.967147 | 0.0102913041492589 | 0.0648622654104307 | 0.158663964080477 | 0.873933630957536 | 0.97191269446355 | NM_172867 |
| Ccdc96 | 72.82090446 | 0.046784519 | 0.134824622 | 0.347002784 | 0.728589227 | 0.863649414 | | Adck1 | 1489.944413 | 0.00978632863925848 | 0.0621409953106927 | 0.157485868874948 | 0.874861943931116 | 0.971956739056508 | NM_028105 |
| Gnaz | 2385.102323 | -0.026481756 | 0.076351431 | -0.346840341 | 0.728711268 | 0.863727998 | | Frzb | 804.6025923 | 0.0136693807747986 | 0.0868078795335469 | 0.157467050782136 | 0.874876773573186 | 0.971956739056508 | NM_011356 |
| Tmem230 | 834.4622663 | 0.032450726 | 0.093626774 | 0.346596645 | 0.728894367 | 0.863878935 | | Pcp4 | 8286.438994 | 0.0133736875952481 | 0.0849699629268995 | 0.157393120281265 | 0.874935035102774 | 0.971956739056508 | NM_008791 |
| 4933417D19Rik | 14.66866737 | 0.057840978 | 0.167137561 | 0.346068099 | 0.729291537 | 0.864151352 | | Polr3gl | 703.5166762 | 0.0111641368290108 | 0.0708636255181602 | 0.157543969100901 | 0.874816158189453 | 0.971956739056508 | NM_027241 |
| Scamp4 | 1725.596758 | 0.02987764 | 0.086321262 | 0.346121443 | 0.729251449 | 0.864151352 | | Rpgrip1l | 1188.773251 | -0.0121719223407987 | 0.07716009438649 | -0.157748930163697 | 0.874654642470707 | 0.971956739056508 | NM_173431 |
| Smim4 | 111.5682331 | -0.047319331 | 0.136717545 | -0.34611016 | 0.729259928 | 0.864151352 | | Snapc3 | 611.681576 | 0.0122316209880653 | 0.0776064922598483 | 0.157610795590534 | 0.874763496255177 | 0.971956739056508 | NM_029949 |
| Rfx8 | 8.163543776 | -0.051202944 | 0.148051484 | -0.345845532 | 0.729458804 | 0.864253503 | | Kcng2 | 175.7111684 | -0.0148734234011182 | 0.0946560027522907 | -0.157131327846593 | 0.875141348198419 | 0.972003598230991 | NM_001190373 |
| Ubr5 | 7064.848839 | -0.027188539 | 0.078623916 | -0.345804948 | 0.729489306 | 0.864253503 | | Ldb3 | 405.0693128 | -0.0124317516195662 | 0.0791056370382048 | -0.157153801992166 | 0.875123636465136 | 0.972003598230991 | NM_001039072,NM_001039073,NM_001039071,NM_001039076,NM_001039074,NM_001039075,NM_011918 |
| Suz12 | 1498.348722 | -0.031044729 | 0.089796479 | -0.345723236 | 0.72955072 | 0.864260177 | | Pak7 | 2535.989718 | -0.0133696466995135 | 0.0851570043622032 | -0.156999964942962 | 0.875244875730509 | 0.972003598230991 | NM_172858 |
| Btbd10 | 3329.755175 | -0.037180141 | 0.107575007 | -0.34562062 | 0.729627847 | 0.864285464 | | Zfp961 | 458.4466749 | 0.0114715099908912 | 0.0730283427744052 | 0.157082983881043 | 0.875179447986674 | 0.972003598230991 | NM_001164581 |
| Lmtk2 | 9799.599148 | 0.025978649 | 0.075182621 | 0.345540612 | 0.729687985 | 0.864290622 | | Ccdc71l | 599.8711559 | 0.0117462950295742 | 0.0748606019430 12 | 0.156908904346189 | 0.875316642136486 | 0.972008985832475 | NM_001162903 |
| Apold1 | 269.2489309 | 0.050202694 | 0.145495579 | 0.345046182 | 0.730059652 | 0.864400447 | | 2010107E04Rik | 2019.914044 | -0.00953681751407566 | 0.0609017309789251 | -0.156593537831886 | 0.875565195774794 | 0.972151437342662 | NM_027360 |
| Cbfa2t2 | 2673.953218 | -0.02421978 | 0.070169906 | -0.345159069 | 0.729974788 | 0.864400447 | | AU041133 | 170.7801018 | -0.0137296772968493 | 0.0876932255863021 | -0.156564856692805 | 0.875587801198324 | 0.972151437342662 | NM_001163064 |
| Mettl8 | 430.5957936 | 0.027807078 | 0.080587964 | 0.3450525 | 0.730054902 | 0.864400447 | | Dnajc8 | 3113.662759 | 0.00921995879118441 | 0.0589475293459724 | 0.156409588213121 | 0.8757101798899 | 0.972151437342662 | NM_172400 |
| Negr1 | 9060.383222 | 0.025623179 | 0.074245733 | 0.345113155 | 0.730009304 | 0.864400447 | | Dusp2 | 44.0150948 | -0.0142478916947383 | 0.0912206439837889 | -0.156191527186218 | 0.875882055097588 | 0.972151437342662 | NM_010090 |
| Plekhg1 | 771.2916429 | -0.031657595 | 0.091739152 | -0.345082712 | 0.73003219 | 0.864400447 | | Gm14057 | 35.85260156 | -0.0131911597220588 | 0.0844765338448918 | -0.156151763355718 | 0.87591339749007 | 0.972151437342662 | NR_024097 |
| Nostrin | 151.2481392 | 0.053659323 | 0.155638012 | 0.344770033 | 0.730267263 | 0.864580192 | | Phlpp1 | 4909.866873 | -0.00995373467295851 | 0.0637098316821164 | -0.156235457073929 | 0.875847429188424 | 0.972151437342662 | NM_133821 |
| Slfn3 | 20.03007843 | -0.057864353 | 0.167887876 | -0.344660698 | 0.730349668 | 0.86461145 | | Rab30 | 442.0431365 | -0.0135744392606721 | 0.0868982277431346 | -0.15621077222423 | 0.875866885966557 | 0.972151437342662 | NM_029494 |
| Brd3 | 2160.017833 | 0.023675528 | 0.068736247 | 0.344440223 | 0.730515243 | 0.864620115 | | Pcm1 | 4428.357916 | 0.011681623003489 | 0.0749128904653543 | 0.15593608697947 | 0.87608339993489 | 0.972265831419834 | NM_023662 |
| Med7 | 455.5872962 | -0.029143654 | 0.084592937 | -0.344516394 | 0.730457969 | 0.864620115 | | Slitrk4 | 1127.917675 | 0.01266854860354 4 | 0.0813427470657981 | 0.155742817405668 | 0.876235745538555 | 0.97236061418587 | NM_178740 |
| Pcna | 1460.851695 | 0.027652107 | 0.08028407 | 0.344428315 | 0.730524198 | 0.864620115 | | Agtpbp1 | 8496.356879 | 0.00940832701265044 | 0.0607325571443164 | 0.154914060185113 | 0.876889069037969 | 0.972549750591973 | NM_023328,NM_001048008 |
| C2cd5 | 2221.632983 | 0.031169825 | 0.090636302 | 0.343900009 | 0.730921485 | 0.864877663 | | Aldh7a1 | 3588.296109 | -0.00937684170784259 | 0.0607115992924195 | -0.154448932611357 | 0.877255773842115 | 0.972549750591973 | NM_138600,NM_001127338 |
| Cstf1 | 841.2285045 | 0.021111964 | 0.061383625 | 0.343934788 | 0.730895329 | 0.864877663 | | Cdh11 | 1915.482287 | 0.0117512215870549 | 0.0760978280329238 | 0.15442256225724 | 0.877276564918533 | 0.972549750591973 | NM_009866 |
| Orai1 | 183.4613945 | 0.032373954 | 0.094129095 | 0.343931429 | 0.730897856 | 0.864877663 | | Cenpo | 147.6816724 | -0.0140007890053809 | 0.0904321927683949 | -0.154820850592866 | 0.876962553010389 | 0.972549750591973 | NM_134046 |
| Slc10a3 | 117.5178445 | 0.039476765 | 0.114810757 | 0.343842039 | 0.730965083 | 0.864877663 | | Gas2 | 275.6668239 | -0.0120868185548234 | 0.0781416129697536 | -0.154678385759735 | 0.877074870558279 | 0.972549750591973 | NM_008087 |
| Btrc | 5301.500974 | 0.019835283 | 0.057756053 | 0.343432107 | 0.731273409 | 0.864988796 | | Gm5918 | 153.8066863 | 0.0141479320139627 | 0.0915377226007118 | 0.154558488151121 | 0.877169398297981 | 0.972549750591973 | NM_001198789 |
| Coq9 | 2049.315332 | 0.02490255 | 0.072488804 | 0.343536502 | 0.731194885 | 0.864988796 | | Otud7a | 1901.645031 | 0.0095122814512817 | 0.0613854638266756 | 0.154959836715416 | 0.876852980423055 | 0.972549750591973 | NM_130880 |
| Dgkd | 4958.494059 | 0.022361845 | 0.065143259 | 0.343271828 | 0.731393973 | 0.864988796 | | Rbck1 | 3532.506644 | -0.00828050887536881 | 0.0535396396904714 | -0.154661273838242 | 0.877088361556346 | 0.972549750591973 | NM_019705,NM_001083921 |
| Pja2 | 24020.60627 | -0.025777128 | 0.075076657 | -0.343344117 | 0.731339595 | 0.864988796 | | Sftpc | 30.26068669 | -0.0127788271887461 | 0.08238891753655 1 | -0.15510371504853 | 0.876739553446186 | 0.972549750591973 | NM_011359 |
| Trp53rk | 111.367297 | 0.042657287 | 0.124213717 | 0.343418486 | 0.731283655 | 0.864988796 | | Tshz1 | 1627.556463 | -0.00776915303810539 | 0.0500447131767641 | -0.155244231506808 | 0.876628779258036 | 0.972549750591973 | NM_001081300 |
| Zfp354b | 99.15564212 | 0.047452333 | 0.138110591 | 0.343582147 | 0.731160553 | 0.864988796 | | Tspan7 | 48434.60493 | 0.00706702861065212 | 0.0457114689843475 | 0.154600776734434 | 0.877136057614832 | 0.972549750591973 | NM_019634 |
| Ppp1r10 | 2030.734903 | -0.016909313 | 0.049270695 | -0.343192094 | 0.731453952 | 0.864993706 | | Uck2 | 448.3182204 | 0.0110733427816777 | 0.0715257782294514 | 0.15481611044 | 0.876966290049633 | 0.972549750591973 | NM_030724 |
| Fam167a | 91.90246557 | -0.050243604 | 0.146504704 | -0.342948743 | 0.731637022 | 0.865144166 | | Zcrb1 | 1160.669351 | 0.0119126713162744 | 0.0768168147168303 | 0.155078954525621 | 0.876759073309 | 0.972549750591973 | NM_026025 |
| Aldh3b1 | 181.9754703 | -0.040863138 | 0.119343212 | -0.342400183 | 0.732049751 | 0.865272295 | | Ccng2 | 1605.189679 | -0.0126068318173455 | 0.0817318792436947 | -0.154246200307674 | 0.877415615464957 | 0.972570049191249 | NM_007635 |
| Ccdc102a | 54.89448219 | 0.051252741 | 0.149750864 | 0.342253389 | 0.73216021 | 0.865272295 | | Foxp1 | 4726.978309 | -0.0124668084005219 | 0.0809665760037697 | -0.1539747512596 | 0.87762964372841 | 0.972570049191249 | NM_053202,NM_001197321,NM_001197322 |
| Fam186b | 49.81298711 | 0.0565383 | 0.165075447 | 0.342499756 | 0.731974827 | 0.865272295 | | Rasal2 | 4602.60228 | -0.0100376369590653 | 0.0651199723943683 | -0.15414068203649 | 0.877498811938593 | 0.972570049191249 | NM_177644 |
| Fam199x | 365.0926378 | -0.037334102 | 0.109035833 | -0.342402131 | 0.732048285 | 0.865272295 | | Slmap | 5241.713447 | 0.0117352948735406 | 0.0762050922657762 | 0.153996203201383 | 0.877612729282388 | 0.972570049191249 | NM_032008 |
| Lmbrd2 | 1538.315373 | -0.032077711 | 0.093703573 | -0.342331783 | 0.73210122 | 0.865272295 | | Vezf1 | 1685.423193 | 0.0114616666234606 | 0.074396996011155 | 0.154060879309455 | 0.877561733745735 | 0.972570049191249 | NM_016686 |
| Mfap1a | 1101.280602 | -0.027407263 | 0.079998277 | -0.342598164 | 0.731900783 | 0.865272295 | | Sema4f | 2698.173373 | -0.0103569214171621 | 0.0674124128034991 | -0.153635228089989 | 0.877897358692097 | 0.972792511431947 | NM_011350,NM_001113481 |
| Msln | 50.56342227 | 0.056348289 | 0.16453837 | 0.342462912 | 0.73200255 | 0.865272295 | | Plvap | 377.1127685 | 0.0129730212468608 | 0.0845580724634737 | 0.15342144006966 | 0.878065938307925 | 0.972905096862007 | NM_032398 |
| Rsph3a | 163.4196193 | 0.034250786 | 0.100086762 | 0.342210953 | 0.732192143 | 0.865272295 | | Ednrb | 2032.682548 | 0.0117799315482014 | 0.0770287022254042 | 0.152929118729412 | 0.878454172574892 | 0.973035729400601 | NM_001136061,NM_007904,NM_001276296 |
| Gmds | 284.2111233 | -0.032797477 | 0.095863478 | -0.342126926 | 0.732255375 | 0.865281023 | | Ehd4 | 717.9992949 | 0.0124519384019777 | 0.0813524483635913 | 0.153061630626356 | 0.878349673597675 | 0.973035729400601 | NM_133838 |
| Med13l | 5431.808213 | 0.037757868 | 0.11040076 | 0.342007318 | 0.732345385 | 0.86532139 | | Rap1gap2 | 13380.93028 | 0.00829430590590763 | 0.0541821730577214 | 0.153081824478903 | 0.878333748897099 | 0.973035729400601 | NM_001015046 |
| Arhgap26 | 4977.694183 | 0.033116562 | 0.097091582 | 0.341085819 | 0.733038977 | 0.865332757 | | Wasf1 | 8929.277257 | -0.00830145902732422 | 0.0543121392011493 | -0.152847211496846 | 0.878518765736534 | 0.973035729400601 | NM_031877 |
| Btaf1 | 1672.089714 | -0.030148177 | 0.088266098 | -0.341560094 | 0.732681974 | 0.865332757 | | Zfp54 | 58.31056292 | 0.01452167270620 3 | 0.09492165926878 7 | 0.152985870854643 | 0.878409417548757 | 0.973035729400601 | NM_011760 |
| Btbd19 | 250.1686548 | 0.039139387 | 0.114783288 | 0.340985063 | 0.733114827 | 0.865332757 | | Gin1 | 229.621728 | -0.0134943032839714 | 0.0883358560652381 | -0.152761334808433 | 0.878586490133633 | 0.973036547398685 | NM_026250 |
| Ddhd1 | 2762.002079 | -0.020193397 | 0.059111933 | -0.341612875 | 0.732642247 | 0.865332757 | | Dlec1 | 146.9967857 | -0.0145196631062596 | 0.0951019030838081 | -0.152674790255924 | 0.878654742123076 | 0.973037949537227 | NM_177117 |
| Glod4 | 2752.528443 | -0.018482786 | 0.054107505 | -0.341593764 | 0.732656631 | 0.865332757 | | Chchd2 | 4089.995581 | -0.00902957085187181 | 0.0592043142281225 | -0.152515420026311 | 0.87878042928741 | 0.973102951351496 | NM_024166 |
| Gm14634 | 29.08526529 | -0.056070133 | 0.164173317 | -0.341530124 | 0.732704531 | 0.865332757 | | Itfg3 | 808.9742712 | 0.0112860590747861 | 0.0740433478210265 | 0.152425024082732 | 0.878851721311509 | 0.973107714290036 | NM_001206335,NM_207217 |
| Mad2l1 | 244.957894 | 0.030570847 | 0.089508863 | 0.341539889 | 0.732697181 | 0.865332757 | | Neurod1 | 282.2763105 | 0.0127812166250396 | 0.0840198703355019 | 0.152121356222077 | 0.879091220402613 | 0.973224530719333 | NM_010894 |
| Map2k2 | 2449.132867 | 0.024575821 | 0.072061438 | 0.341039833 | 0.733073595 | 0.865332757 | | Zfp558 | 137.6182573 | -0.0140509918572727 | 0.0923372448477374 | -0.152170360729764 | 0.879052570403441 | 0.973224530719333 | NM_028935 |
| Pik3r1 | 6156.615693 | 0.030501356 | 0.089398911 | 0.341182639 | 0.732966092 | 0.865332757 | | Arnt2 | 18677.36844 | -0.0063621320917926 8 | 0.0420129527531764 | -0.151432633863414 | 0.879634448188405 | 0.97341376813304 | NM_007488 |
| Qdpr | 5190.034086 | -0.052893157 | 0.155132031 | -0.340955744 | 0.733136899 | 0.865332757 | | Gh | 290.9693413 | -0.00520836124388798 | 0.0343885934309096 | -0.151456070872806 | 0.879615961381779 | 0.97341376813304 | NM_008117 |
| Snx6 | 1226.815068 | -0.02077683 | 0.060782896 | -0.341820338 | 0.732486103 | 0.865332757 | | March11 | 122.416473 | -0.0143094223670763 | 0.0943211629211417 | -0.15170956256169 | 0.879416014646674 | 0.97341376813304 | NM_177597 |
| Spdl1 | 21.23886745 | 0.057278649 | 0.167881303 | 0.341185397 | 0.732964016 | 0.865332757 | | Mark3 | 2814.894508 | 0.0084782028522818 1 | 0.05594269032307 | 0.151551575430499 | 0.879540629313806 | 0.97341376813304 | NM_022801,NM_021516 |
| Thbs4 | 231.5327989 | 0.055478779 | 0.162600348 | 0.341197172 | 0.732955152 | 0.865332757 | | Nup85 | 1270.562155 | 0.00828312000660924 | 0.0547120206587041 | 0.151394883736422 | 0.879664225133032 | 0.97341376813304 | NM_001002929 |
| Vmn2r57 | 22.07518669 | -0.055672752 | 0.16327707 | -0.340971036 | 0.733125386 | 0.865332757 | | Tspyl2 | 4704.129776 | -0.0113959709997568 | 0.0752426028817665 | -0.151456363327356 | 0.879615730697868 | 0.97341376813304 | NM_029836 |
| A330076C08Rik | 14.55609804 | -0.055819023 | 0.163990714 | -0.340379171 | 0.733571003 | 0.865510426 | | Vmn2r57 | 27.56865001 | 0.0125397869329216 | 0.082909937451144 | 0.1512458858181 | 0.879781754948714 | 0.973469665781918 | NM_177764 |
| Amica1 | 20.82008421 | 0.056868006 | 0.167008537 | 0.340509573 | 0.733472815 | 0.865510426 | | Rps5 | 5123.989623 | 0.0071292688559649 8 | 0.0471770214028996 | 0.151117400886331 | 0.879883106211959 | 0.973507654873556 | NM_009095 |
| Eml1 | 2424.6355 | 0.025614263 | 0.075276101 | 0.340270849 | 0.733652568 | 0.865510426 | | Trpm2 | 1830.208709 | -0.00978659751866496 | 0.0648827827444464 | -0.15083504598148 | 0.880105839834919 | 0.973679926080886 | NM_138301 |
| Inpp1 | 754.489519 | 0.023638965 | 0.069396811 | 0.340634742 | 0.733378572 | 0.865510426 | | Acaa1a | 1447.369919 | -0.0122777617978537 | 0.0816681303658371 | -0.150337245910427 | 0.880498548912156 | 0.973779118328922 | NM_130864 |
| Isca1 | 6903.527287 | -0.02069802 | 0.060790348 | -0.340482011 | 0.733493568 | 0.865510426 | | BC030336 | 2164.28034 | 0.0105434161978452 | 0.0700558590140357 | 0.150500134410355 | 0.880370044701402 | 0.973779118328922 | NM_001164580 |
| Psma8 | 8.264657935 | 0.051211141 | 0.150549059 | 0.340162476 | 0.733734175 | 0.865510426 | | Gprin3 | 354.6743416 | 0.0139418483894199 | 0.0927622879815409 | 0.150296512653873 | 0.880530684238633 | 0.973779118328922 | NM_183183 |
| Spag9 | 17926.11447 | -0.022076264 | 0.064848335 | -0.340429154 | 0.733533367 | 0.865510426 | | Tmem216 | 215.990843 | -0.0122009119652655 | 0.0811482849363346 | -0.150353294278959 | 0.880485888069278 | 0.973779118328922 | NM_026798 |
| Ttc30b | 404.298401 | -0.03701469 | 0.108814794 | -0.340162292 | 0.733734313 | 0.865510426 | | Zfp456 | 89.841218 | 0.0142724644609022 | 0.0949555626620894 | 0.150306775725109 | 0.880522587466414 | 0.973779118328922 | NM_001001186 |
| Atf2 | 4710.102518 | -0.021271527 | 0.06257795 | -0.33992049 | 0.733916406 | 0.865637095 | | Ccdc33 | 47.52935367 | -0.0128272947884026 | 0.0855017877319121 | -0.150023702763059 | 0.880745914792536 | 0.973868855771971 | NM_001166282,NM_029212 |
| Selm | 1826.322815 | 0.02759373 | 0.081188758 | 0.339871321 | 0.733953435 | 0.865637095 | | Mmp23 | 42.24978894 | -0.0139671847205238 | 0.0930700835846965 | -0.150071689876728 | 0.880708055214463 | 0.973868855771971 | NM_011985 |
| Edaradd | 12.02261461 | -0.053900075 | 0.158696585 | -0.339642313 | 0.734125909 | 0.865774611 | | 5730405O15Rik | 40.72667198 | -0.0135926515621854 | 0.0909584701829092 | -0.149437996646731 | 0.881208031339541 | 0.974259967609829 | NR_038158 |
| Faim | 520.295638 | -0.02384171 | 0.07028601 | -0.33920989 | 0.734451619 | 0.865899032 | | Kcna3 | 318.6664613 | -0.0137501793685282 | 0.0920850842603585 | -0.149320375595807 | 0.881300838110502 | 0.974259967609829 | NM_008418 |
| Rep15 | 23.39507524 | 0.05672322 | 0.167223783 | 0.339205461 | 0.734454956 | 0.865899032 | | Nmnat3 | 125.745363 | 0.0139753283590458 | 0.0935812480462009 | 0.149338982443857 | 0.881286156603738 | 0.974259967609829 | NM_144533 |
| Sgca | 12.2212277 | 0.056546062 | 0.166689708 | 0.339229473 | 0.734436868 | 0.865899032 | | Abhd2 | 2695.287489 | -0.0109683085378199 | 0.073591549329013 | -0.149043044178657 | 0.881519667918608 | 0.974279441237594 | NM_018811 |
| Ube2f | 1073.730474 | 0.019287664 | 0.056824703 | 0.339423932 | 0.734290393 | 0.865899032 | | Sh3rf1 | 1514.720633 | 0.00819511603880251 | 0.0549821008655352 | 0.149050616651491 | 0.881513692699039 | 0.974279441237594 | NM_021506 |
| Klk14 | 28.65982534 | -0.055709342 | 0.164333895 | -0.339000923 | 0.734609035 | 0.86601479 | | Sowaha | 3352.180346 | 0.011969214218526 4 | 0.0802307861627231 | 0.149184805371975 | 0.881407809374952 | 0.974279441237594 | NM_183173 |
| C230004F18Rik | 165.7937705 | 0.036333128 | 0.10722639 | 0.338845017 | 0.734726486 | 0.866021467 | | Prr3 | 1084.5584 | -0.0103551417109113 | 0.0695171882554976 | -0.148958005505817 | 0.88158676994187 | 0.974279475059008 | NM_145487,NM_001165892,NR_028516,NR_028507 |
| Men1 | 2415.984577 | 0.023460083 | 0.069227089 | 0.338885881 | 0.734695701 | 0.866021467 | | Nelfe | 769.0763749 | -0.00985019471340072 | 0.0661856821819274 | -0.148826670492344 | 0.881690404992172 | 0.974319880024061 | NM_001045863,NM_138580,NM_001045864 |
| Ppp4r2 | 2692.462702 | -0.026562473 | 0.078472675 | -0.33849328 | 0.73499149 | 0.866267926 | | Hhipl1 | 192.2454467 | 0.0126297090371026 | 0.0849738267021165 | 0.148630578700159 | 0.881845142691492 | 0.974354773869459 | NM_001044380 |
| Atp1b3 | 3771.220246 | 0.022618485 | 0.066894043 | 0.338124055 | 0.735269704 | 0.866398113 | | Mtus1 | 2389.715737 | -0.00940592340137976 | 0.0632898362648973 | -0.148616649314933 | 0.881856134659355 | 0.974354773869459 | NM_001005864,NM_001005865,NM_001005863 |
| Eif4ebp1 | 94.05779067 | -0.049550081 | 0.146542324 | -0.338128121 | 0.73526664 | 0.866398113 | | 3110039M20Rik | 119.8263547 | -0.0138888528568146 | 0.0936118618642204 | -0.148366377724222 | 0.882053633056986 | 0.974397329601898 | NR_026733 |
| Npas3 | 411.289914 | -0.040602758 | 0.120073518 | -0.338149152 | 0.735250792 | 0.866398113 | | Dapk2 | 71.35436094 | 0.0140934125805632 | 0.0950249038455313 | 0.148312831796945 | 0.882095889044813 | 0.974397329601898 | NM_010019 |
| Gm14057 | 40.20866927 | 0.056448889 | 0.167048917 | 0.337918319 | 0.735424743 | 0.866514902 | | Msc | 46.55535208 | 0.0135684835978864 | 0.0914464270201718 | 0.148376312120904 | 0.88204579332407 | 0.974397329601898 | NM_010827 |
| Cotl1 | 2145.843974 | 0.021474212 | 0.063619375 | 0.337542017 | 0.735708344 | 0.866717235 | | Rpl24 | 5179.205126 | -0.00883627185686455 | 0.0596742679750306 | -0.148075077528591 | 0.882283517850897 | 0.974530482875989 | NM_024218 |
| Eftud1 | 596.7551554 | 0.034461089 | 0.102071868 | 0.337615938 | 0.73565263 | 0.866717235 | | Gng8 | 46.3617978 | -0.0122802036057859 | 0.0830270189027081 | -0.147906112589396 | 0.882416864119878 | 0.974603662424373 | NM_010320 |
| A230072E10Rik | 24.84955612 | 0.056477327 | 0.167708804 | 0.33675827 | 0.736299133 | 0.867149496 | | Cox4i2 | 46.08836447 | 0.0137134736353966 | 0.09282011983620 3 | 0.147742468546651 | 0.882546014341261 | 0.974672196785612 | NM_053091 |
| Celf5 | 11000.68403 | -0.025039439 | 0.074341547 | -0.336816216 | 0.736255448 | 0.867149496 | | Vwa9 | 863.0864855 | 0.00838397383140928 | 0.0568679282615239 | 0.14742885995166 | 0.882793527457324 | 0.974871429211273 | NM_001077631,NM_175153 |
| Gli2 | 211.6036066 | -0.045321458 | 0.134562833 | -0.336805173 | 0.736263773 | 0.867149496 | | Lgi3 | 5428.266753 | 0.00891656295837052 | 0.060584984460753 | 0.147174469676503 | 0.882994311388108 | 0.974944919977475 | NM_145219 |
| Gosr2 | 4135.373615 | -0.018378971 | 0.054547849 | -0.336933008 | 0.736167402 | 0.867149496 | | P2ry12 | 1197.301845 | -0.01029239599685 | 0.0699142849743279 | -0.147214492726763 | 0.882962721689176 | 0.974944919977475 | NM_027571 |
| Ckap2 | 24.35405877 | -0.056368467 | 0.16753916 | -0.336449502 | 0.736531926 | 0.867298501 | | Irf6 | 61.74919424 | -0.0139001779088575 | 0.0945354356598974 | -0.147036693826272 | 0.88310305758643 | 0.974990880481909 | NM_016851 |

| | | | | | | | | | | | | | | |
|---|---|---|---|---|---|---|---|---|---|---|---|---|---|---|
| Cpne3 | 632.7993878 | 0.029129623 | 0.086581421 | 0.336441967 | 0.736537607 | 0.867298501 | | 1700030K09Rik | 360.389532 | 0.0116272648407206 | 0.079341803889557 | 0.146546514834798 | 0.883489972715068 | 0.975177866872649 | NM_028170 |
| Daam1 | 2839.69438 | -0.017914043 | 0.053271535 | -0.336277948 | 0.736661278 | 0.867312298 | | 2410004P03Rik | 105.2778305 | 0.0137654992949686 | 0.0938865565559032 | 0.146618427599613 | 0.883433207755784 | 0.975177866872649 | NM_001201332,NM_001201333 |
| Marveld2 | 38.81004968 | -0.052316338 | 0.155555897 | -0.336318577 | 0.736630642 | 0.867312298 | | Fam109b | 23.77420839 | -0.0119288563778833 | 0.0813082956545437 | -0.146711430633913 | 0.883359795898511 | 0.975177866872649 | NM_177391 |
| Rbks | 54.81880893 | -0.05290575 | 0.157362467 | -0.336203101 | 0.736717715 | 0.867312839 | | Gm15800 | 12714.29872 | 0.0108508323072801 | 0.0740762521122905 | 0.146481929064548 | 0.883540954558098 | 0.975177866872649 | NM_181421 |
| Arid3c | 8.671034986 | 0.053323558 | 0.158839878 | 0.335706367 | 0.737092304 | 0.867687902 | | E130308A19Rik | 368.6290194 | -0.0118663813620961 | 0.0811439032050553 | -0.146238730125035 | 0.883732931978192 | 0.975315648831121 | NM_001015681,NM_153158 |
| 1810043G02Rik | 1065.549107 | 0.022216711 | 0.06623775 | 0.335408608 | 0.737316875 | 0.867845457 | | Myrip | 3358.162033 | -0.00842373502352401 | 0.0576820221886948 | -0.146037442930962 | 0.883891830089912 | 0.975416905256102 | NM_144557 |
| Efemp2 | 333.4872041 | -0.036061168 | 0.107523191 | -0.335380376 | 0.737338169 | 0.867845457 | | Adprhl2 | 708.6101081 | 0.0109318182622124 | 0.07498815336559 | 0.145780603623034 | 0.884094588374141 | 0.97556654441832 | NM_133883 |
| Thoc3 | 1056.491454 | -0.024121314 | 0.071945362 | -0.33527268 | 0.7374194 | 0.867875138 | | Chrna7 | 535.4292273 | 0.0100972803440557 | 0.0693235001740084 | 0.145654508481403 | 0.884194135241097 | 0.975602279416041 | NM_007390 |
| D2Wsu81e | 1134.606289 | -0.023277702 | 0.069492757 | -0.334965872 | 0.737650831 | 0.867949724 | | A630019I02Rik | 26.66690767 | -0.0118884236972734 | 0.0819094827543916 | -0.145140993417347 | 0.884599552882288 | 0.975676134861822 | NR_046182 |
| Gm16432 | 23.58158421 | 0.055214976 | 0.164820902 | 0.334999841 | 0.737625206 | 0.867949724 | | Dusp18 | 1734.69552 | 0.0109853752714883 | 0.0757746926873056 | 0.144974197610025 | 0.884731243866166 | 0.975676134861822 | NM_173745 |
| Hist2h3c2 | 36.91056024 | -0.054266252 | 0.161935012 | -0.335111297 | 0.737541131 | 0.867949724 | | Haghl | 3028.070549 | 0.00994064581734188 | 0.0684728873771386 | 0.145176378536373 | 0.884571615532643 | 0.975676134861822 | NM_001271434,NM_001271435,NM_001271436,NM_026897,NM_001271433,NM_001271437 |
| Paqr4 | 2419.366756 | -0.030636098 | 0.091578893 | -0.334532307 | 0.737977917 | 0.868136804 | | Jakmip2 | 2492.858387 | 0.00776439883573117 | 0.0533696098940053 | 0.145483522385711 | 0.884329124568426 | 0.975676134861822 | NM_001163637 |
| Rhot1 | 3174.185746 | -0.020014936 | 0.059805509 | -0.334667095 | 0.737876226 | 0.868136804 | | Slc16a2 | 1878.895627 | -0.0110176191306851 | 0.0759651134951666 | -0.14503524873146 | 0.88468304155199 | 0.975676134861822 | NM_009197 |
| Tctex1d2 | 258.5524557 | 0.032193875 | 0.096229449 | 0.334553248 | 0.737962117 | 0.868136804 | | Wbp4 | 2302.876885 | -0.00842629815314724 | 0.0580774846790992 | -0.145087174482604 | 0.884642044420308 | 0.975676134861822 | NM_018765 |
| Arhgef16 | 27.12201044 | -0.055649893 | 0.1665622 | -0.334108776 | 0.738297478 | 0.86844679 | | Wdr85 | 890.5520037 | 0.00830350144754189 | 0.0571515955280905 | 0.145289057476281 | 0.884482653924221 | 0.975676134861822 | NM_026044 |
| Hp1bp3 | 10147.57499 | 0.021844245 | 0.065413012 | 0.333943427 | 0.738422249 | 0.868527619 | | Ddi2 | 529.5747245 | -0.0112702045671111 | 0.0779582224847141 | -0.144567233678538 | 0.885052569139176 | 0.975769680706692 | NM_001017966 |
| Ap3m2 | 6943.140625 | 0.019128639 | 0.057308271 | 0.33378497 | 0.738541826 | 0.868561308 | | Flt3 | 358.9529399 | -0.0123232297173214 | 0.0853164752600574 | -0.14444138344626 | 0.885151940153661 | 0.975769680706692 | NM_010229 |
| Cbx4 | 1613.814629 | 0.022332648 | 0.066912916 | 0.333756902 | 0.738563007 | 0.868561308 | | Kif6 | 100.3460042 | 0.0136374312940641 | 0.0942641278924689 | 0.144672545102427 | 0.884969416903468 | 0.975769680706692 | NM_177052 |
| BC049352 | 30.91641004 | 0.054707204 | 0.16395963 | 0.33366265 | 0.738634137 | 0.868579032 | | Mzf1 | 277.7978555 | -0.0134075998315998 | 0.0926344613220895 | -0.144736630841752 | 0.884918816436642 | 0.975769680706692 | NM_145819 |
| Pdgfb | 1146.216983 | -0.03379265 | 0.101341146 | -0.333454394 | 0.73879131 | 0.868697924 | | Rnf113a2 | 308.1940034 | 0.011337702275998 | 0.0784762851467856 | 0.144472973647917 | 0.885126996443132 | 0.975769680706692 | NM_025525 |
| Esrra | 1752.360015 | -0.021656955 | 0.064969626 | -0.333339697 | 0.738877877 | 0.868733786 | | Sppl2a | 1621.557235 | -0.00925964860699941 | 0.0641526604619924 | -0.144337717879765 | 0.885233795570903 | 0.975785863899145 | NM_023220 |
| Nacad | 1944.206602 | 0.025837314 | 0.077565102 | 0.333104882 | 0.739055115 | 0.868810315 | | Grm7 | 2323.287845 | -0.00815504403775292 | 0.0566373859721689 | -0.143986942507556 | 0.885510780558134 | 0.976017112020142 | NM_177328 |
| Snrpd1 | 817.1731181 | -0.020198387 | 0.060629992 | -0.333141842 | 0.739027216 | 0.868810315 | | Gatm | 4266.368844 | 0.0112946738383701 | 0.0786317633655439 | 0.143640093455152 | 0.885784678936657 | 0.976107811869013 | NM_025961 |
| 2410066E13Rik | 1786.320262 | -0.024874012 | 0.074900586 | -0.332093694 | 0.739818512 | 0.869641758 | | Leprot | 1059.170495 | 0.00774924719846047 | 0.0539537999531646 | 0.14362745914441 | 0.885794656206618 | 0.976107811869013 | NM_175036 |
| Ndufs1 | 5778.796976 | 0.019044253 | 0.057383635 | 0.331876039 | 0.739982864 | 0.869768965 | | Tjp3 | 24.45205681 | -0.00881364830376981 | 0.0613482786832025 | -0.143665779919968 | 0.88576439448313 | 0.976107811869013 | NM_013769 |
| 9330133O14Rik | 287.649951 | -0.029965612 | 0.090349306 | -0.331663996 | 0.74014299 | 0.869891185 | | Blm | 198.4191157 | -0.0130393145624956 | 0.0909723874570601 | -0.143332663096814 | 0.886027460726996 | 0.976290290110016 | NM_001042527,NM_007550 |
| Arhgap11a | 157.2080622 | 0.041561997 | 0.1255517 | 0.331034919 | 0.740618109 | 0.870317557 | | Ctbp2 | 733.2917133 | 0.011049754892143 | 0.0773145997185548 | 0.142919383044948 | 0.886353850263726 | 0.976378470899936 | NM_001170744,NM_009980 |
| Srp54a | 117.988052 | 0.038475709 | 0.116211698 | 0.331082929 | 0.740581846 | 0.870317557 | | Gale | 465.6589584 | 0.0122763429725921 | 0.0859653059421007 | 0.142805784706454 | 0.886443568375902 | 0.976378470899936 | NM_178389 |
| Oma1 | 240.2759288 | 0.031388434 | 0.094889988 | 0.330787628 | 0.740804907 | 0.870471047 | | Herc2 | 10721.02451 | 0.00983100063271346 | 0.0688155673270446 | 0.14286012619778 | 0.886400650172029 | 0.976378470899936 | NM_010418 |
| Fndc1 | 230.1010035 | 0.050388316 | 0.152442333 | 0.330540181 | 0.740991837 | 0.87062467 | | Psmg1 | 363.501582 | 0.0106380041428416 | 0.0744604916494011 | 0.142867766612808 | 0.886394615898213 | 0.976378470899936 | NM_019537 |
| AI847159 | 11.61781671 | -0.052631922 | 0.159306009 | -0.330382526 | 0.741110943 | 0.870698586 | | Rspo2 | 564.3439972 | -0.011854682792436 | 0.0829774998035944 | -0.142866232659404 | 0.886395827388792 | 0.976378470899936 | NM_172815 |
| Kbtbd7 | 1264.054909 | -0.029155222 | 0.088309601 | -0.330147813 | 0.741288277 | 0.870751603 | | Cbs | 1064.927887 | 0.0115687132073911 | 0.0811512811478507 | 0.142557369936204 | 0.886639767373576 | 0.976496358037897 | NM_001271353,NM_178224,NM_144855 |
| Rpl7 | 8331.632546 | -0.022929385 | 0.06946201 | -0.33009965 | 0.741324667 | 0.870751603 | | Cit | 4152.018881 | 0.00657891165650595 | 0.0461677869668994 | 0.142500043617486 | 0.886685044920822 | 0.976496358037897 | NM_007708 |
| Scamp3 | 1538.93972 | 0.021852688 | 0.066188159 | 0.330160083 | 0.741279006 | 0.870751603 | | Gemin5 | 1303.06476 | 0.00913903895206418 | 0.0642149378371406 | 0.142319517231991 | 0.886827630919955 | 0.976505318885936 | NM_001166669,NM_001166671,NM_001166670,NM_172558 |
| Aplp1 | 47280.54444 | -0.020483442 | 0.062105495 | -0.329816902 | 0.741538315 | 0.870758226 | | Nenf | 1168.917259 | -0.00854610866558904 | 0.060024763533314 | -0.142376382055148 | 0.886782716715351 | 0.976505318885936 | NM_025424 |
| C030016D13Rik | 15.16833982 | -0.055151078 | 0.167265933 | -0.329720926 | 0.741610841 | 0.870758226 | | Tceb1 | 1954.558891 | -0.0103838968860171 | 0.07303845428546 | -0.142170271641198 | 0.886945513029199 | 0.976561094691287 | NM_026456 |
| Exog | 1092.216366 | 0.026919149 | 0.081628163 | 0.32977772 | 0.741567924 | 0.870758226 | | Pnrc1 | 1089.528457 | -0.0111759657617909 | 0.0786749853476052 | -0.142052339919897 | 0.887038663547308 | 0.976589633673503 | NM_001033225 |
| Hsd17b11 | 1473.618405 | 0.019593955 | 0.059393064 | 0.329903079 | 0.741473197 | 0.870758226 | | Dazl | 130.4178549 | 0.0134646077092578 | 0.0949519122053866 | 0.141804492363808 | 0.887234435538373 | 0.97661226973329 | NM_010021 |
| Wbscr27 | 323.5035875 | 0.036585998 | 0.110960705 | 0.329720312 | 0.741611304 | 0.870758226 | | Kdelc2 | 405.2574782 | -0.0114789689225984 | 0.080968405823286 | -0.141770963697264 | 0.887260919982805 | 0.97661226973329 | NM_212445 |
| Slc35a1 | 593.5531464 | -0.031506146 | 0.095726714 | -0.329125954 | 0.742060488 | 0.871219611 | | Rmnd5b | 2019.534382 | 0.00937665563804328 | 0.0661271096089332 | 0.141797451809032 | 0.887239996893798 | 0.97661226973329 | NM_025346 |
| Hyi | 151.8503011 | -0.034566651 | 0.105106786 | -0.328871731 | 0.742252642 | 0.871313163 | | Mfsd3 | 197.7727406 | -0.0120057204478031 | 0.0848090557754451 | -0.141561774718863 | 0.887426162139077 | 0.9767201417815 | NM_027122 |
| Klhl12 | 1816.372597 | -0.017765545 | 0.0540135 | -0.328909341 | 0.742224213 | 0.871313163 | | Pnlip | 32.24304588 | -0.0118766520934139 | 0.0840384369260531 | -0.141324047993234 | 0.887613952717766 | 0.976852812878117 | NM_026925 |
| Loxl3 | 164.9779882 | 0.03919079 | 0.119288928 | 0.328536694 | 0.742505905 | 0.871467912 | | Tmem51 | 172.5953054 | -0.0129140779826951 | 0.0914453455320552 | -0.141221818426704 | 0.887694710195612 | 0.976867678810716 | NM_145402 |
| Rad51c | 122.6735446 | -0.036638465 | 0.111531654 | -0.328502836 | 0.7425315 | 0.871467912 | | Atxn1l | 2520.162697 | 0.00836661180331601 | 0.0593793914988044 | 0.140900935360452 | 0.88794820320957 | 0.97692460803016 | NM_001080930 |
| Rangrf | 283.6757771 | -0.042722945 | 0.130064871 | -0.328474124 | 0.742553206 | 0.871467912 | | Cpxm2 | 109.6520991 | 0.0120090009355406 | 0.0851570527147636 | 0.141021800927812 | 0.887852719800296 | 0.97692460803016 | NM_018867 |
| Ak2 | 679.7522014 | 0.032243224 | 0.098449797 | 0.327509298 | 0.743282712 | 0.871861787 | | Eid2b | 908.5997358 | 0.0103055848900045 | 0.0731322053837204 | 0.14091719012071 | 0.887935361906604 | 0.97692460803016 | NM_001177427 |
| H2-T10 | 50.77292073 | 0.052955554 | 0.161605304 | 0.327684504 | 0.743150221 | 0.871861787 | | C330013E15Rik | 39.68580651 | -0.0128846136505999 | 0.0915833436593094 | -0.140687303343398 | 0.88811697579188 | 0.977036291302094 | NR_045701 |
| Prps2 | 1343.299138 | 0.036272727 | 0.110750177 | 0.327518454 | 0.743275788 | 0.871861787 | | Cecr6 | 3681.137511 | 0.0106504074810753 | 0.0758187066699915 | 0.140472027931474 | 0.888287051812689 | 0.97703864986882 | NM_033567 |
| Ranbp1 | 1285.713858 | 0.022502492 | 0.068673698 | 0.327672637 | 0.743159194 | 0.871861787 | | Mrps10 | 688.1369642 | -0.00939150765980283 | 0.0668771806998588 | -0.140429180200514 | 0.888320903815056 | 0.97703864986882 | NM_183086,NM_001146212,NM_001146211 |
| Rasa3 | 3611.415396 | 0.022564779 | 0.068885732 | 0.327568256 | 0.743238127 | 0.871861787 | | Tfrc | 4169.980125 | 0.0106767114326344 | 0.0759465045456037 | 0.140581999086256 | 0.888200169643089 | 0.97703864986882 | NM_011638 |
| Rps6kl1 | 1396.054718 | -0.025842233 | 0.078884464 | -0.327595976 | 0.743217165 | 0.871861787 | | A230065H16Rik | 39.22237975 | 0.0114404714902447 | 0.0816226662124267 | 0.140162923133022 | 0.888531265722173 | 0.977089223453417 | NM_001101503 |
| Snapc2 | 1012.315525 | 0.021451455 | 0.065449948 | 0.327753584 | 0.743097985 | 0.871861787 | | Gm13399 | 43.88141948 | -0.0129425718812371 | 0.092370690727263 | -0.140115568903255 | 0.888568679734245 | 0.977089223453417 | NR_040524 |
| Gna13 | 2860.771725 | 0.021852536 | 0.066753159 | 0.327363317 | 0.743393108 | 0.871894162 | | Mrpl9 | 1432.886224 | 0.00785957158223095 | 0.0560409084684217 | 0.140247040903338 | 0.888464805887649 | 0.977089223453417 | NM_030116 |
| Igsf8 | 6794.012464 | 0.025219452 | 0.077063522 | 0.327255383 | 0.743474735 | 0.871894162 | | Ccdc42 | 64.57823312 | 0.012525893543543 | 0.0895434616979139 | 0.139886188293688 | 0.887499140129809 | 0.977140573164518 | NM_177779 |
| Ube2q2 | 2518.42581 | -0.018384086 | 0.056177571 | -0.32724957 | 0.743479132 | 0.871894162 | | Tesc | 1388.594384 | -0.011063970311279 | 0.0790656935320961 | -0.139933893159208 | 0.888712221874921 | 0.977140573164518 | NM_021344 |
| Cox6b2 | 56.03317897 | 0.049863796 | 0.152421585 | 0.327143928 | 0.743559028 | 0.871921863 | | Mterfd3 | 526.4987803 | -0.0100607088299461 | 0.0719749719978854 | -0.139780656395972 | 0.88883329719297 | 0.977158289186792 | NM_028832 |
| Btbd6 | 1870.13243 | 0.021724814 | 0.066480453 | 0.326784986 | 0.743830516 | 0.87217421 | | 5330413P13Rik | 94.94763417 | -0.0130426371480045 | 0.0945055073781017 | -0.138009281256202 | 0.890233082360416 | 0.977182599239554 | NR_029381 |
| Kcna2 | 13619.57716 | -0.024837525 | 0.076043288 | -0.326623497 | 0.74395267 | 0.872186452 | | Acacb | 276.8663701 | -0.0130824279685192 | 0.0946027335311038 | -0.138288054480137 | 0.890012765948111 | 0.977182599239554 | NM_133904 |
| Vtcn1 | 9.615922797 | 0.050647125 | 0.155063256 | 0.326622347 | 0.74395354 | 0.872186452 | | Akr7a5 | 677.3000956 | 0.00938071075215234 | 0.0680457813740493 | 0.137858814502935 | 0.890352000802429 | 0.977182599239554 | NM_025337 |
| Adsl | 920.1517154 | -0.023062421 | 0.070659821 | -0.326386632 | 0.744131851 | 0.872202723 | | Babam1 | 2575.35959 | 0.00704936738617968 | 0.0513670161462147 | 0.137235290562992 | 0.890844816958578 | 0.977182599239554 | NM_026636 |
| Exosc1 | 457.1513466 | 0.026632061 | 0.081598137 | 0.326380753 | 0.744136297 | 0.872202723 | | Cd320 | 414.2262987 | 0.0101324268700492 | 0.0726445467264485 | 0.139479524983535 | 0.889071234475028 | 0.977182599239554 | NM_019421 |
| Ncoa5 | 2082.659723 | -0.027723156 | 0.069591899 | -0.326520135 | 0.744030858 | 0.872202723 | | Chst7 | 55.0833245 | -0.0126105066725459 | 0.0914202905795748 | -0.137939910194984 | 0.890287908108977 | 0.977182599239554 | NM_021715 |
| Cdh12 | 445.41938 | 0.036423907 | 0.111630011 | 0.32629135 | 0.744203932 | 0.872216016 | | Cp | 764.3784287 | 0.0129790089826387 | 0.0929616727051543 | 0.139616775440392 | 0.888962785562209 | 0.977182599239554 | NM_007752,NM_001042611,NM_001276248,NM_001276250 |
| Atp2b2 | 30955.47886 | -0.023258484 | 0.07139818 | -0.325757388 | 0.744607921 | 0.872491502 | | Fuz | 526.351661 | -0.0111701537502139 | 0.0812614871809527 | -0.13745938128526 | 0.890667696956632 | 0.977182599239554 | NM_027376 |
| Gpr180 | 839.2537099 | -0.021200882 | 0.065074002 | -0.325796493 | 0.744578332 | 0.872491502 | | Gnpda1 | 889.1186941 | 0.00773191098086691 | 0.0560085689009222 | 0.138048715269702 | 0.890201916870949 | 0.977182599239554 | NM_011937 |
| Nck1 | 702.3564888 | 0.023186494 | 0.071145718 | 0.325901474 | 0.744498901 | 0.872491502 | | Gpbp1l1 | 1577.878183 | -0.0072330963719483 | 0.0522812449763714 | -0.13834973469391 | 0.889964020794276 | 0.977182599239554 | NM_029868 |
| Nrl | 11.43725739 | -0.054039002 | 0.16604577 | -0.325446422 | 0.744843226 | 0.872701221 | | Hsd17b10 | 840.7753128 | -0.0094791982068531 | 0.0682023030798497 | -0.138986482549644 | 0.889460830732597 | 0.977182599239554 | NM_016763 |
| Nup160 | 558.8627217 | 0.027497558 | 0.084578224 | 0.325113916 | 0.745094857 | 0.872930035 | | Med6 | 538.6247171 | -0.0102347413816599 | 0.0745132162530218 | -0.137354712309105 | 0.890750426027383 | 0.977182599239554 | NM_027213 |
| Klhl10 | 8.328962422 | -0.048219853 | 0.148382538 | -0.324969861 | 0.745203882 | 0.872982115 | | Nrf1 | 662.7632218 | -0.0101120998359821 | 0.072658172952142 | -0.13917360463554 | 0.889312965940951 | 0.977182599239554 | NM_010938,NM_001164230,NM_001164226,NM_001164227,NM_001164228,NM_001164229 |
| Prr18 | 1194.081594 | -0.054096886 | 0.166499965 | -0.324906289 | 0.745251997 | 0.872982115 | | Pdlim4 | 287.0967909 | -0.0117546353863522 | 0.0855493392155256 | -0.137401825591412 | 0.890713188121553 | 0.977182599239554 | NM_019417 |
| Nacc1 | 5961.913637 | 0.024233346 | 0.074701595 | 0.324401993 | 0.745633711 | 0.873297205 | | Ppil2 | 2620.639036 | 0.00608675307488063 | 0.043733191730546 | 0.139179255710012 | 0.889308500494015 | 0.977182599239554 | NM_001252444,NM_144954,NM_001252445 |
| Tpd52l2 | 1772.891706 | -0.020957439 | 0.064596899 | -0.324434136 | 0.745609379 | 0.873297205 | | Ppp1r3d | 333.7994168 | -0.0102719568722749 | 0.0748182307071003 | -0.137292164960272 | 0.890799863250015 | 0.977182599239554 | NM_001085501 |
| Lrrc42 | 686.7737961 | 0.026227612 | 0.080871001 | 0.324314172 | 0.745700191 | 0.873309052 | | Rnf144a | 1475.012214 | -0.00731326038093439 | 0.0533371453767162 | -0.137113831819858 | 0.890940819158701 | 0.977182599239554 | NM_080563,NM_001081977 |
| Ap4s1 | 1102.478813 | -0.024746341 | 0.076364259 | -0.32405658 | 0.745895199 | 0.873471409 | | Rps3a1 | 10782.13131 | 0.00869044820805715 | 0.0625829593581812 | 0.138862851759999 | 0.889558526500059 | 0.977182599239554 | NM_016959 |
| Tlr9 | 58.65205663 | 0.051752438 | 0.159878434 | 0.323698678 | 0.746166172 | 0.873722693 | | Ryk | 1223.159318 | 0.00760635351494679 | 0.0551791401160314 | 0.137848351731326 | 0.890360269940152 | 0.977182599239554 | NM_013649,NM_001042607 |
| Lef1 | 216.4999994 | 0.041776453 | 0.129152525 | 0.32346602 | 0.746342337 | 0.873862932 | | Sdsl | 91.01492069 | 0.0131251923920655 | 0.0950482213331449 | 0.138089826489878 | 0.890169426032142 | 0.977182599239554 | NM_133902 |
| A630072M18Rik | 248.651983 | 0.02833527 | 0.087703893 | 0.323078812 | 0.746635556 | 0.873899395 | | Sh3kbp1 | 1785.014709 | -0.00784710950088444 | 0.0562505732931174 | -0.139502747109683 | 0.889052885287516 | 0.977182599239554 | NM_021389,NM_001135728,NM_001135727 |
| A830080D01Rik | 234.7024779 | -0.037205585 | 0.115269352 | -0.322770836 | 0.746868802 | 0.873899395 | | Slc25a23 | 20296.77918 | 0.00673530039069019 | 0.04911963507959 | 0.137124992765606 | 0.890931997696226 | 0.977182599239554 | NM_025877 |
| Acsf2 | 692.3929229 | 0.026820628 | 0.083156693 | 0.322531195 | 0.747050309 | 0.873899395 | | Slitrk1 | 3948.262583 | -0.00953006506796882 | 0.0693810862034009 | -0.13735825697554 | 0.890747624347189 | 0.977182599239554 | NM_199065 |
| Akap13 | 1856.759863 | -0.033105212 | 0.102628709 | -0.322572627 | 0.747018927 | 0.873899395 | | Tcof1 | 1333.523405 | 0.00833411400635654 | 0.0606497186436367 | 0.137413893959275 | 0.890703649433162 | 0.977182599239554 | NM_001198984,NM_011552 |
| Fut10 | 425.7383717 | 0.026927149 | 0.083475278 | 0.322576336 | 0.747016118 | 0.873899395 | | Tmem245 | 1337.147633 | -0.0100075003996825 | 0.07283326597547 | -0.137402952844322 | 0.890712297154153 | 0.977182599239554 | NM_175518 |
| Gm13826 | 65.29880927 | -0.054234087 | 0.168048039 | -0.322729661 | 0.746899987 | 0.873899395 | | Tpm2 | 384.3063759 | -0.010630671346486 | 0.077462010739505 | -0.137237224350341 | 0.890843288481006 | 0.977182599239554 | NM_009416 |
| Gm6313 | 27.61326038 | -0.053263189 | 0.164769917 | -0.323257967 | 0.746499884 | 0.873899395 | | Urm1 | 1422.852831 | 0.00697420121837275 | 0.0508241938900416 | 0.137222072492905 | 0.890855264614413 | 0.977182599239554 | NM_026615 |
| Grap2 | 12.91169784 | -0.05373861 | 0.16627204 | -0.323196912 | 0.746546119 | 0.873899395 | | Usp38 | 1338.183907 | 0.00967235491972991 | 0.0698819860168581 | 0.138409845956533 | 0.889916515964467 | 0.977182599239554 | NM_027554 |
| Inppl1 | 1648.337644 | 0.020402154 | 0.063175901 | 0.322942034 | 0.746739142 | 0.873899395 | | Wdr31 | 180.9244038 | 0.0122896860250218 | 0.0891048596843683 | 0.137923858121262 | 0.890300594553843 | 0.977182599239554 | NM_023597 |
| Pmpca | 3970.37913 | 0.013781858 | 0.042723722 | 0.322580941 | 0.747012629 | 0.873899395 | | Wdr53 | 331.9328779 | -0.0108569003783422 | 0.0781121284210767 | -0.138991224510184 | 0.889457083564566 | 0.977182599239554 | NM_026898,NM_001185162 |
| Rims1 | 6072.698806 | 0.029864716 | 0.092372596 | 0.323307101 | 0.746462677 | 0.873899395 | | Zmiz2 | 14004.48304 | -0.00718195052309779 | 0.0522135211323707 | -0.137549630198091 | 0.890596366282052 | 0.977182599239554 | NM_001005867,NM_028601 |
| Rnf130 | 1861.552785 | -0.018543139 | 0.057456197 | -0.322735235 | 0.746895765 | 0.873899395 | | Rad51 | 52.33237969 | -0.0128677776085968 | 0.0939778280153212 | -0.136923547610602 | 0.891091225830579 | 0.977273774738769 | NM_011234 |
| Pak2 | 2141.887296 | 0.020434673 | 0.06339492 | 0.322339289 | 0.747195672 | 0.874003453 | | Chrm3 | 2096.367786 | 0.00783242436541839 | 0.0572924068732573 | 0.136709640821084 | 0.891260308812618 | 0.977385417923305 | NM_033269 |
| Ass1 | 1682.123934 | 0.02410641 | 0.07481603 | 0.322209159 | 0.747294246 | 0.874052771 | | Fam120a | 6251.138544 | -0.00757629912885317 | 0.0555254443716461 | -0.136447338955868 | 0.891467652512125 | 0.977391434854791 | NM_001033268 |
| Aqp4 | 4582.598944 | -0.028565737 | 0.08870562 | -0.322028487 | 0.747431114 | 0.874111615 | | Pgf | 83.36128123 | 0.0127744680283368 | 0.093570580864589 | 0.13652226918227 | 0.891408421102917 | 0.977391434854791 | NM_008827,NM_001271705 |
| P4ha2 | 417.7503252 | -0.033362311 | 0.103611654 | -0.321993803 | 0.747457389 | 0.874111615 | | Rab11fip1 | 164.6209819 | 0.0124725123025343 | 0.0912983427413355 | 0.136612691184013 | 0.891336944445364 | 0.977391434854791 | NM_001080813,NM_029423 |
| Rpia | 332.3970037 | 0.035781448 | 0.111169154 | 0.321864894 | 0.74755505 | 0.874159844 | | B3gnt3 | 41.5585744 | -0.0117171550752277 | 0.0860027707429848 | -0.136241599822916 | 0.891630289843312 | 0.977495969076525 | NM_028189 |
| Kdm5a | 2096.379063 | -0.024892692 | 0.077418943 | -0.32153231 | 0.747807031 | 0.87438851 | | Kcnc2 | 5076.347687 | 0.00726483373073646 | 0.0533910465545548 | 0.1360683897311 | 0.891767216415963 | 0.977572302894746 | NM_001025581 |
| Dnm3 | 7535.458429 | -0.023963957 | 0.07454962 | -0.321449755 | 0.747869583 | 0.874395662 | | Paox | 260.7971857 | 0.0124273057929144 | 0.0914586125814967 | 0.135879010649115 | 0.891916928641258 | 0.977662640012294 | NM_153783 |
| Arhgap40 | 9.341655097 | -0.050629512 | 0.157560375 | -0.321334043 | 0.747957261 | 0.874432189 | | Bcs1l | 778.5521498 | -0.00972516388449746 | 0.0716527282977302 | -0.135726358444966 | 0.892037609510864 | 0.977690693008285 | NM_025784 |
| B3galt5 | 2414.65666 | -0.022544355 | 0.070237643 | -0.32097255 | 0.748231193 | 0.874579799 | | Rassf2 | 3369.776528 | 0.00836483562838302 | 0.0616528582164542 | 0.135676363924853 | 0.892077133769228 | 0.977690693008285 | NM_175445 |
| Gm684 | 51.86542157 | -0.053146855 | 0.165595437 | -0.320943959 | 0.74825286 | 0.874579799 | | Asb6 | 1161.012773 | -0.00998629841345138 | 0.0736663886545908 | -0.135561123544083 | 0.892168240586959 | 0.977716775765083 | NM_133346 |
| Srpk2 | 9243.14002 | 0.015771759 | 0.049134068 | 0.320994358 | 0.748214667 | 0.874579799 | | 2300009A05Rik | 156.1417298 | -0.0121825683291466 | 0.0914211161645728 | -0.133257707193336 | 0.893989573996718 | 0.977844986642332 | NM_027090 |
| Mrps22 | 542.6896349 | -0.030971993 | 0.096542651 | -0.320811499 | 0.748353245 | 0.874631152 | | AU040320 | 3672.260992 | 0.00693698667742483 | 0.0527877019878771 | 0.131412931728264 | 0.895448659337654 | 0.977844986642332 | NM_001035525,NM_001035526,NM_133886 |
| Pus1 | 469.9164399 | 0.026375179 | 0.082439448 | 0.319933963 | 0.749018391 | 0.875342506 | | Ankrd39 | 315.1428888 | -0.00993865871998065 | 0.075196731504213 | -0.132168759481572 | 0.894850810632124 | 0.977844986642332 | NM_026241 |
| Epdr1 | 4923.01199 | -0.019700179 | 0.061590814 | -0.319855801 | 0.749077645 | 0.87534573 | | Appl1 | 4446.848321 | -0.00823779883252711 | 0.062905617865542 | -0.130954898974763 | 0.895810985470132 | 0.977844986642332 | NM_145221 |
| Pon2 | 2221.097596 | 0.029466126 | 0.092186793 | 0.31963501 | 0.749245031 | 0.875475303 | | Arhgef28 | 1464.236884 | 0.010095459593651 | 0.0763434973438273 | 0.1322373213816 | 0.894796582120629 | 0.977844986642332 | NM_012026 |
| C530005A16Rik | 249.0886477 | 0.045339465 | 0.141882302 | 0.319556875 | 0.74930427 | 0.875478498 | | Armc3 | 39.34785784 | -0.01111399393016 | 0.0840364902106296 | -0.132252000319193 | 0.894784971990875 | 0.977844986642332 | NM_001081083,NM_001271564,NM_001271565,NM_001271563 |
| Gemin6 | 164.8311605 | 0.03698991 | 0.115797044 | 0.319437428 | 0.749394833 | 0.875518288 | | B4galnt3 | 39.66004695 | -0.0114235954091241 | 0.0869376395347934 | -0.131399880077861 | 0.895458983524903 | 0.977844986642332 | NM_198884 |
| Ttc39b | 3043.251009 | 0.023872737 | 0.074771037 | 0.319277859 | 0.749515821 | 0.875593616 | | Calr3 | 47.605826 | -0.0124107742754156 | 0.0935989329363457 | -0.132595253878117 | 0.894513486133604 | 0.977844986642332 | NM_028500,NM_029782 |
| Fibin | 100.7398516 | 0.053105493 | 0.16643118 | 0.319083797 | 0.74966297 | 0.875699492 | | Cbx6 | 19377.43988 | -0.00798177582966997 | 0.0593143587491396 | -0.134567345883778 | 0.892953960931255 | 0.977844986642332 | NM_028763 |
| B930025P03Rik | 12.59728235 | -0.051352892 | 0.161340584 | -0.318288747 | 0.750265919 | 0.876337743 | | Ccdc120 | 518.3985206 | -0.00939128267289528 | 0.0705934282247254 | -0.133033384396623 | 0.894166978215078 | 0.977844986642332 | NM_207202 |
| Loxl2 | 98.41123997 | 0.046401348 | 0.145990742 | 0.317837608 | 0.750608122 | 0.876671358 | | Ctnnd2 | 9236.343425 | 0.00512537347756494 | 0.0379096528257203 | 0.135199694418926 | 0.892453988694297 | 0.977844986642332 | NM_008729 |
| Pou2f2 | 225.566791 | 0.038086189 | 0.119859099 | 0.317758014 | 0.750668501 | 0.876675794 | | Ctxn1 | 13355.9745 | -0.00558088512903422 | 0.0419396146971025 | -0.133069537460958 | 0.894138386442227 | 0.977844986642332 | NM_183315 |
| 3110082J24Rik | 30.79067243 | -0.053122968 | 0.167587477 | -0.316986504 | 0.751253843 | 0.876924335 | | Fam154b | 68.75033599 | -0.0127725689706511 | 0.0950956324700711 | -0.134312887341813 | 0.89315516295469 | 0.977844986642332 | NM_177894 |
| Fsd1l | 1063.504579 | 0.023534441 | 0.074240894 | 0.317001045 | 0.751242809 | 0.876924335 | | Fam173b | 213.1752957 | -0.0110986039625898 | 0.0840299292181046 | -0.132079177810358 | 0.894921665311642 | 0.977844986642332 | NM_026546 |
| Galnt9 | 2924.626317 | -0.044087415 | 0.139084771 | -0.316982335 | 0.751257006 | 0.876924335 | | Fzd10 | 54.28605642 | -0.0119376552224191 | 0.0901658187370748 | -0.132396681909244 | 0.894670539027593 | 0.977844986642332 | NM_175284 |
| Mpst | 698.6365587 | -0.035718466 | 0.112692432 | -0.316955323 | 0.751277503 | 0.876924335 | | Gltp | 1690.657045 | -0.00817721157915797 | 0.0606293672451172 | -0.134872124693278 | 0.892712979413973 | 0.977844986642332 | NM_019821 |
| Picalm | 5658.954887 | 0.020250563 | 0.063805089 | 0.317381636 | 0.75095404 | 0.876924335 | | Gtpbp8 | 394.891646 | -0.0103845629939123 | 0.0780425975598932 | -0.133062754426424 | 0.894143750817481 | 0.977844986642332 | NM_001159329,NM_025332 |
| Snrnp40 | 692.9643211 | -0.026335313 | 0.083073489 | -0.317012245 | 0.751234311 | 0.876924335 | | Gzf1 | 1909.848379 | 0.00736540326069981 | 0.0547941286780924 | 0.13441957082611 | 0.893070806792181 | 0.977844986642332 | NM_028986 |
| Tmem241 | 236.2524372 | 0.035792754 | 0.112853504 | 0.317161209 | 0.751121282 | 0.876924335 | | Hsbp1l1 | 40.9693029 | 0.0124561821362218 | 0.0927670317383931 | 0.134273802910378 | 0.893186067878279 | 0.977844986642332 | NM_001136181 |
| Trmt112 | 693.3137594 | -0.021366669 | 0.067443889 | -0.316806602 | 0.751390354 | 0.876989993 | | Igsf9b | 759.8492799 | -0.0123101316834975 | 0.0934568039467769 | -0.131720015703812 | 0.895205753220103 | 0.977844986642332 | NM_001129787,NM_001033323 |
| Atg9b | 194.579822 | 0.04119628 | 0.13024734 | 0.316292677 | 0.751780369 | 0.877114837 | | Itgb1 | 2658.92756 | -0.00801625662320502 | 0.0606818904291882 | -0.132102948120238 | 0.894902864088268 | 0.977844986642332 | NM_010578 |
| Col5a2 | 201.5184001 | -0.046561305 | 0.147089705 | -0.316550399 | 0.751584778 | 0.877114837 | | Leng1 | 299.2239346 | 0.0111795975269456 | 0.0836004331209915 | 0.133726550324995 | 0.893618809603352 | 0.977844986642332 | NM_027203 |
| Kctd12b | 176.0821419 | -0.036642688 | 0.115845014 | -0.316307859 | 0.751768847 | 0.877114837 | | Lrrc41 | 2012.319612 | 0.00704458827361943 | 0.0532232669766074 | 0.132359185630518 | 0.894700195736879 | 0.977844986642332 | NM_153521 |
| Prmt3 | 829.8614498 | 0.021365489 | 0.067537175 | 0.316351528 | 0.751735704 | 0.877114837 | | Lsm5 | 107.6937223 | -0.0124489064837204 | 0.0944519289999721 | -0.13180150596738 | 0.89514129535567 | 0.977844986642332 | NM_025520 |
| Tmem115 | 1095.15483 | -0.018385199 | 0.058118434 | -0.316340235 | 0.751744275 | 0.877114837 | | Mocs3 | 224.0218564 | -0.0115689682369543 | 0.0869447359323904 | -0.133061169407087 | 0.894145004333617 | 0.977844986642332 | NM_001160330 |
| Serbp1 | 5035.74888 | 0.024687245 | 0.078117149 | 0.3160285 | 0.751980876 | 0.877216661 | | Mpg | 458.6849364 | -0.011107417784395 | 0.0842900062863084 | -0.131776212552011 | 0.895161302083237 | 0.977844986642332 | NM_010822 |
| Zswim7 | 249.4677752 | 0.032268396 | 0.102083633 | 0.316097644 | 0.751928395 | 0.877216661 | | Mrpl40 | 730.5285936 | 0.00902614260416958 | 0.0681087003956407 | 0.132525544486051 | 0.894568619636863 | 0.977844986642332 | NM_010922 |
| 2900076A07Rik | 123.717829 | 0.040815082 | 0.129196393 | 0.31591503 | 0.752067003 | 0.877251084 | | Myo9a | 5087.245522 | 0.00982549805135236 | 0.0736727041501405 | 0.133366871281507 | 0.893903244208424 | 0.977844986642332 | NM_173018 |
| Acss1 | 1073.232393 | -0.040663143 | 0.128918297 | -0.315417935 | 0.752444351 | 0.87730433 | | Nol8 | 1145.609368 | 0.00998947355350253 | 0.0749968528450549 | 0.133198570000544 | 0.894036341744944 | 0.977844986642332 | NM_001271397,NR_073168,NR_073167 |
| Cep120 | 2177.01557 | 0.025628132 | 0.081302573 | 0.315219204 | 0.752595226 | 0.87730433 | | Npy | 2767.435272 | 0.00837864653554081 | 0.0633735111757742 | 0.13221054633223 | 0.894817759583057 | 0.977844986642332 | NM_023456 |
| Gpr37 | 2419.471675 | 0.049652545 | 0.157463483 | 0.315327362 | 0.752513112 | 0.87730433 | | Nup188 | 1042.157766 | -0.00678945073203921 | 0.0516826871733371 | -0.131367990005401 | 0.895484209457365 | 0.977844986642332 | NM_198304 |
| Pitpnm3 | 5699.723898 | 0.027544156 | 0.087377929 | 0.315230128 | 0.752586932 | 0.87730433 | | Pcyt2 | 2747.083534 | -0.00992623566107026 | 0.0749709001602651 | -0.13240118018926 | 0.894666981239556 | 0.977844986642332 | NM_024229 |
| Ptov1 | 4545.89114 | -0.017704385 | 0.056117269 | -0.315489068 | 0.75239035 | 0.87730433 | | Pick1 | 1828.081261 | 0.00726899102954839 | 0.0550519758515085 | 0.132038694653849 | 0.894953685764963 | 0.977844986642332 | NM_008837,NM_001045558 |
| Tmx1 | 1109.275501 | 0.028772074 | 0.091191597 | 0.315512338 | 0.752372685 | 0.87730433 | | Plekhm2 | 5995.415628 | -0.00735702180967073 | 0.0549391221953382 | -0.133912256251794 | 0.89347195849378 | 0.977844986642332 | NM_001033150 |
| Vkorc1 | 346.606721 | -0.037874492 | 0.120019323 | -0.315569947 | 0.752328951 | 0.87730433 | | Pramef8 | 756.7395568 | 0.00873352618248578 | 0.0665160880102058 | 0.131299456172855 | 0.895538421978709 | 0.977844986642332 | NM_172877 |
| Zfp605 | 555.0832218 | -0.023881988 | 0.075771669 | -0.315183608 | 0.752622251 | 0.87730433 | | Prelp | 1208.443865 | 0.012665708782251 | 0.094565739096896 | 0.133935491893879 | 0.893453584647753 | 0.977844986642332 | NM_054077 |
| Zfp647 | 95.11270836 | -0.04328718 | 0.137196941 | -0.31551126 | 0.752373503 | 0.87730433 | | Prg4 | 161.8328851 | 0.00785844308100834 | 0.0596505784871466 | 0.131741271925832 | 0.89518893972541 | 0.977844986642332 | NM_001110146,NM_021400 |
| Cdk12 | 1108.151734 | -0.026355387 | 0.083648909 | -0.315071496 | 0.75270737 | 0.877337546 | | Prkaa2 | 2244.601114 | 0.00978563225286151 | 0.0747472369645033 | 0.13091630741493 | 0.895841514260248 | 0.977844986642332 | NM_178143 |
| Ip6k1 | 11115.03151 | 0.022774788 | 0.072327915 | 0.314882408 | 0.752850939 | 0.877372881 | | Psg16 | 108.7579398 | 0.0124691943027794 | 0.0945818881700606 | 0.131834905646623 | 0.895114876792219 | 0.977844986642332 | NM_007676 |
| Phkb | 1570.296888 | 0.039519987 | 0.125486239 | 0.314934825 | 0.75281114 | 0.877372881 | | Psmg4 | 129.072841 | -0.0122235083989737 | 0.0933792566280334 | -0.130901753120151 | 0.895853027828456 | 0.977844986642332 | NM_001101430,NM_001110515 |
| 4930529M08Rik | 31.25026332 | 0.052703486 | 0.167437725 | 0.314764707 | 0.752940311 | 0.877408124 | | Rbm22 | 1321.373989 | 0.0068119940691952 | 0.0511496404489654 | 0.133178210086393 | 0.894052443158214 | 0.977844986642332 | NM_025776 |
| Rpl23 | 5647.02169 | -0.020945931 | 0.066559799 | -0.314693423 | 0.752994438 | 0.877408124 | | Scap | 4464.600711 | -0.00686395071432622 | 0.0523549991085945 | -0.131104017404127 | 0.895693023167386 | 0.977844986642332 | NM_001103162,NM_001001144 |
| BC049715 | 16.82902151 | -0.052830637 | 0.168117865 | -0.314247006 | 0.753332988 | 0.877736601 | | Shroom1 | 318.701116 | -0.0116729148426193 | 0.0891080268789288 | -0.130997343914699 | 0.895777408548319 | 0.977844986642332 | NM_027917 |
| Bcl9l | 1782.408814 | 0.02101478 | 0.067030286 | 0.313511717 | 0.75389192 | 0.878261353 | | Sirt3 | 2370.750974 | -0.00730459408278235 | 0.0543838426714466 | -0.134315519536054 | 0.89315308162637 | 0.977844986642332 | NM_001177804,NM_022433,NM_001127351 |
| Crtap | 197.589956 | 0.035452143 | 0.113083045 | 0.313505381 | 0.753896734 | 0.878261353 | | Smarcd2 | 663.4774802 | 0.00945786312958465 | 0.0701892251233319 | 0.134746733423153 | 0.892812122177299 | 0.977844986642332 | NM_001130187,NM_031878 |
| Exo5 | 304.325051 | -0.035581763 | 0.113611967 | -0.31318675 | 0.754138784 | 0.878477282 | | Spcs3 | 3894.241774 | -0.00949409190658177 | 0.0709300289541063 | -0.133851516016224 | 0.893519989790115 | 0.977844986642332 | NM_029701 |
| Dctn2 | 8988.493529 | 0.023062266 | 0.073728099 | 0.312801579 | 0.75443141 | 0.878686038 | | St8sia4 | 461.2523997 | 0.0102796693124793 | 0.0762473353248102 | 0.134820046742465 | 0.892754155537571 | 0.977844986642332 | NM_001159745,NM_009183 |
| Ttll1 | 1567.328126 | 0.020274857 | 0.064807801 | 0.312845938 | 0.754397712 | 0.878686038 | | Stoml1 | 2452.489191 | -0.0100195633862506 | 0.0740544708598029 | -0.135299911942107 | 0.89237475469841 | 0.977844986642332 | NM_026942,NR_028146 |
| 0610040B10Rik | 39.07994039 | -0.048487587 | 0.155195302 | -0.312429479 | 0.75471415 | 0.878883205 | | Taf8 | 722.8615797 | 0.00954271125208806 | 0.0721600042182608 | 0.132243773478965 | 0.894791478908774 | 0.977844986642332 | NM_022015 |
| Abi1 | 4114.116412 | -0.024630263 | 0.078826962 | -0.312459874 | 0.754691053 | 0.878883205 | | Thada | 625.2692457 | 0.00974770096412737 | 0.0721888373400051 | 0.135030585382838 | 0.892587692144122 | 0.977844986642332 | NM_183021 |
| Mir707 | 10.6946565 | 0.049148616 | 0.157352536 | 0.312347146 | 0.754776714 | 0.878890005 | | Ulk2 | 5447.263166 | 0.0080189745507627 | 0.0599321198236157 | 0.133800949713828 | 0.893559976186698 | 0.977844986642332 | NM_013881 |
| Man2a1 | 965.5910422 | -0.025500006 | 0.081795215 | -0.311754256 | 0.755227291 | 0.879348587 | | Zfp429 | 69.69011215 | 0.0127348627654112 | 0.0950184343040188 | 0.134025180047327 | 0.893382663265341 | 0.977844986642332 | NM_001080941 |
| Ttc7b | 8006.847535 | -0.021038767 | 0.067572737 | -0.311349938 | 0.755534607 | 0.879640307 | | Zfp568 | 195.2849814 | 0.0112382709001647 | 0.0858136038076227 | 0.130961416409228 | 0.89580583156965 | 0.977844986642332 | NM_001033355,NM_001167872,NM_001167873 |
| Abcd4 | 453.1293278 | 0.037973193 | 0.122207819 | 0.310727338 | 0.756008641 | 0.879680258 | | Dis3l | 1414.884636 | -0.00728484685475065 | 0.0557509210225628 | -0.130668128367035 | 0.896037846003427 | 0.97789975590126 | NM_001001295,NM_001177784,NM_172519 |
| Ccdc66 | 562.8360292 | 0.027725163 | 0.089130975 | 0.311060914 | 0.755754314 | 0.879680258 | | Rbbp9 | 1198.542945 | 0.00828198618859352 | 0.0634585840403642 | 0.130742397097751 | 0.895979092150907 | 0.97789975590126 | NM_015754 |
| Desi1 | 5137.662373 | -0.025033663 | 0.080535877 | -0.310838651 | 0.755923285 | 0.879680258 | | Gapvd1 | 3888.919729 | 0.00760074916266067 | 0.0583501337292876 | 0.130261041010188 | 0.896359902216174 | 0.978030610277611 | NM_025709 |
| Dlx1as | 574.1928372 | -0.027146056 | 0.087295312 | -0.31096809 | 0.75582488 | 0.879680258 | | Gp5 | 31.52605263 | 0.0114496562887317 | 0.0878993677206953 | 0.13025188846002 | 0.896361763367897 | 0.978030610277611 | NM_008148 |
| Ercc6 | 1268.420034 | -0.022121818 | 0.0712539 | -0.310464665 | 0.756207626 | 0.879680258 | | Hnrnpa3 | 7071.993316 | -0.00819317775147726 | 0.0629001919753126 | -0.130256800403613 | 0.896363257092972 | 0.978030610277611 | NM_053263,NM_146130,NM_198090 |
| Fbxo3 | 2782.695023 | -0.018039865 | 0.058079832 | -0.310604636 | 0.756101202 | 0.879680258 | | Ust | 969.0802248 | 0.00922844736807785 | 0.0708919989775645 | 0.130176148242038 | 0.896427064934333 | 0.978030610277611 | NM_177387 |
| Lrfn1 | 1371.266007 | 0.027613717 | 0.088935208 | 0.310492523 | 0.756186445 | 0.879680258 | | 1300018J18Rik | 927.7246237 | 0.00804616756936598 | 0.0623361398781092 | 0.129077090514416 | 0.897296649466261 | 0.978039388558981 | NM_027905 |
| Me3 | 1888.54766 | 0.022448348 | 0.072242867 | 0.310734456 | 0.756002501 | 0.879680258 | | 1810022K09Rik | 482.5878422 | -0.00957064585439482 | 0.0739835975100369 | -0.129361275794646 | 0.897071431548122 | 0.978039388558981 | NM_001099674 |
| Nutf2 | 476.9424126 | -0.023966927 | 0.07721089 | -0.310408635 | 0.756250228 | 0.879680258 | | Bgn | 793.8867094 | -0.0122555255616807 | 0.0942708183616079 | -0.130003385720812 | 0.896563748007883 | 0.978039388558981 | NM_007542 |
| Srpx | 19.20947324 | -0.052129024 | 0.167850824 | -0.310567577 | 0.756129379 | 0.879680258 | | Cdc16 | 4792.631404 | 0.00880901742882657 | 0.0684811137140186 | 0.12863426055989 | 0.897647055051186 | 0.978039388558981 | NM_001168304,NM_198164 |
| Trip13 | 21.2721915 | 0.052210029 | 0.167907146 | 0.310945844 | 0.755841792 | 0.879680258 | | Chat | 249.6982636 | -0.0123347764163167 | 0.0950648063103207 | -0.129751239128939 | 0.896763242218964 | 0.978039388558981 | NM_009891 |
| Xpa | 189.8406325 | 0.041275818 | 0.132667322 | 0.31112272 | 0.755707329 | 0.879680258 | | Fam101b | 569.9483183 | 0.00908491321383712 | 0.0706148873311173 | 0.128654361101469 | 0.897631149754425 | 0.978039388558981 | NM_029658 |
| Dnajb5 | 4907.142787 | -0.036929444 | 0.119020171 | -0.310278868 | 0.7563489 | 0.879728988 | | Fam160a1 | 133.9560094 | -0.0123132016167582 | 0.0947897761864146 | -0.129900192435974 | 0.896645392037468 | 0.978039388558981 | NM_172682 |
| Strip1 | 3124.122388 | -0.0220394 | 0.071064899 | -0.310130603 | 0.756461641 | 0.879794075 | | Fdx1l | 25.54956931 | 0.00935954349882795 | 0.072183277365001 | 0.129663598557608 | 0.896832583517944 | 0.978039388558981 | NM_001039824 |
| Efcab14 | 2396.544081 | 0.019323912 | 0.062372759 | 0.309813331 | 0.756702914 | 0.879942581 | | Fgd1 | 669.4661294 | 0.00790711187135347 | 0.0613690967374455 | 0.128845172761502 | 0.897480160712666 | 0.978039388558981 | NM_008001 |
| Meox1 | 21.57115546 | -0.052022422 | 0.167899381 | -0.309842843 | 0.756680469 | 0.879942581 | | Fra10ac1 | 480.2927714 | -0.0100593249432092 | 0.0779396446598694 | -0.129065573587208 | 0.897305762419436 | 0.978039388558981 | NM_001081075 |
| Tnfrsf21 | 2892.729433 | -0.01938768 | 0.062630311 | -0.309557468 | 0.756897504 | 0.880102809 | | Isoc2b | 178.6466474 | -0.0118522999895189 | 0.0916300304872969 | -0.129348951547894 | 0.897081092849455 | 0.978039388558981 | NM_026158 |
| Ptgds | 25568.77765 | 0.049043388 | 0.15857036 | 0.309284713 | 0.75710459 | 0.88027797 | | Mansc1 | 521.5091614 | -0.0102981853109433 | 0.0799767141232876 | -0.128764796401465 | 0.897543761979481 | 0.978039388558981 | NM_026345 |
| Prrg2 | 139.3613697 | 0.037818227 | 0.122436607 | 0.308880064 | 0.757412763 | 0.880569778 | | Me1 | 2524.02341 | 0.00641535199009721 | 0.0497195508471964 | 0.129030771211381 | 0.897333300439215 | 0.978039388558981 | NM_001198933,NM_008615 |
| 1700086O06Rik | 117.7000298 | -0.037700145 | 0.122097471 | -0.308770896 | 0.757495809 | 0.880600245 | | Metap1 | 1699.784029 | 0.00761913698618779 | 0.0591571818785043 | 0.1287947928594 | 0.897520025909402 | 0.978039388558981 | NM_175224 |
| Dhdh | 492.9445713 | -0.032732911 | 0.106069733 | -0.30859803 | 0.75762732 | 0.880620982 | | Npy2r | 413.8489335 | 0.0123263878421235 | 0.0950683675290921 | 0.129658141424923 | 0.896836901232306 | 0.978039388558981 | NM_008731,NM_001205099 |
| Ints5 | 876.266885 | 0.027967105 | 0.090622015 | 0.308612705 | 0.757616155 | 0.880620982 | | Rbm47 | 39.65496104 | 0.00887848617276626 | 0.0690017215212396 | 0.128670501721256 | 0.89761837753742 | 0.978039388558981 | NM_001127382,NM_139065,NM_178446 |
| Pfdn2 | 1709.104182 | -0.026293557 | 0.085274572 | -0.308339943 | 0.757823675 | 0.880783139 | | Rtn4rl1 | 3324.918469 | -0.00958700872883966 | 0.0741921251230079 | -0.129218665852048 | 0.897184626819269 | 0.978039388558981 | NM_177708 |
| A430105I19Rik | 343.7695633 | 0.034636466 | 0.11240313 | 0.308131698 | 0.757982129 | 0.880865123 | | Zfp398 | 589.8777824 | 0.00751921964963277 | 0.0579279230388535 | 0.129560010901371 | 0.896914465555515 | 0.978039388558981 | NM_027477,NM_173034 |
| Col4a3bp | 3236.718876 | 0.025935535 | 0.084179553 | 0.308097796 | 0.758007919 | 0.880865123 | | Phactr1 | 24972.06156 | 0.007107006268283 | 0.0552869945196586 | 0.128547534336596 | 0.897715658326434 | 0.978040802154671 | NM_198419,NM_001005748,NM_001005740 |
| Crebbp | 2235.255626 | 0.0329432 | 0.107050753 | 0.307734405 | 0.758284438 | 0.881093622 | | Csnk1g1 | 1640.976758 | -0.00796479651510601 | 0.0621823332681706 | -0.128087771823496 | 0.898079513005702 | 0.978273928543883 | NM_173185 |
| Mybl2 | 39.80392499 | 0.048790022 | 0.158568795 | 0.307689932 | 0.75831828 | 0.881093622 | | Oxnad1 | 650.8212551 | 0.00803954480484112 | 0.0628225102381938 | 0.127972358544077 | 0.898170847836978 | 0.978273928543883 | NM_145460 |
| Crnkl1 | 721.7406018 | 0.023464738 | 0.07637332 | 0.307382217 | 0.758552462 | 0.881299629 | | Reep3 | 1608.853786 | 0.00812801265044871 | 0.063531505647066 | 0.127936723168532 | 0.898199048946012 | 0.978273928543883 | NM_001204915,NM_178606 |
| Eif3l | 2483.004582 | 0.024322238 | 0.079197021 | 0.307110513 | 0.758759256 | 0.881435884 | | Tmem55b | 2638.203295 | -0.00614162042264527 | 0.0479795368553227 | -0.128004996021218 | 0.898145019316399 | 0.978273928543883 | NM_001033271 |
| Nudcd2 | 373.4802144 | -0.029636904 | 0.096512425 | -0.307078638 | 0.758783517 | 0.881435884 | | Ccdc174 | 858.9483871 | 0.00875112374763077 | 0.068452323713643 | 0.127842810336716 | 0.898273370260536 | 0.978281524696697 | NM_172730 |
| C2cd2l | 7997.75983 | -0.022200314 | 0.072372411 | -0.30675106 | 0.759032862 | 0.881581457 | | Rnps1 | 5681.723714 | 0.00558537671194559 | 0.043764061170113 2 | 0.127624735516088 | 0.898445955094181 | 0.978396127423201 | NM_009070,NM_001080127,NM_001080128 |
| Cklf | 46.78292553 | 0.047346221 | 0.154461724 | 0.306523971 | 0.759205731 | 0.881581457 | | Armc7 | 419.3541851 | -0.00987624645067195 | 0.0776045675251368 | -0.127263726422713 | 0.898731668940737 | 0.978597653736118 | NM_177778 |
| Gm4944 | 308.1623449 | -0.044267741 | 0.144427559 | -0.306504806 | 0.759220321 | 0.881581457 | | Exd1 | 41.40792365 | -0.0114939692339019 | 0.0904072053455671 | -0.127135544008556 | 0.898833119657668 | 0.978597653736118 | NM_172857 |
| Itpka | 4856.380867 | -0.032574978 | 0.106320755 | -0.306383999 | 0.75931229 | 0.881581457 | | Mob3b | 240.2037888 | 0.0111424053239087 | 0.0876193218783872 | 0.127168358360203 | 0.898807148390764 | 0.978597653736118 | NM_178061 |
| Mfsd6 | 4503.544478 | -0.015295309 | 0.049914136 | -0.30643242 | 0.759275427 | 0.881581457 | | Vps13d | 5244.282046 | 0.0091143817697691 6 | 0.071757335847045 | 0.127016724662089 | 0.898927161396848 | 0.978626691632362 | NM_001128198,NM_001276502,NM_001276465 |
| Tmem87b | 1557.173958 | 0.019801825 | 0.064610573 | 0.30647963 | 0.759239487 | 0.881581457 | | Mia | 31.20169135 | 0.0089410155381081 1 | 0.0704443258263351 | 0.126923147226225 | 0.899001225964549 | 0.978633978444322 | NM_019394 |
| Wdr55 | 389.7170651 | -0.024465578 | 0.079782875 | -0.306651999 | 0.759108327 | 0.881587457 | | Tmem215 | 639.278744 | -0.0114160708216733 | 0.0900160917138415 | -0.126822555882171 | 0.89908084287353 | 0.97864730787442 | NM_001166009,NM_177175 |
| Gins2 | 25.34136488 | -0.050593779 | 0.165175155 | -0.306303806 | 0.759373342 | 0.881592279 | | Gm16973 | 188.0957953 | -0.0107361408458182 | 0.0847709998893469 | -0.126648746149418 | 0.899218413701379 | 0.978708076765569 | NR_046210,NR_046209 |
| Akt1s1 | 1179.126679 | 0.025195572 | 0.082303722 | 0.306129196 | 0.759506279 | 0.881680549 | | Rgs3 | 1318.231437 | 0.00886627697863654 | 0.0700438676791925 | 0.12658177328592 | 0.899271423694758 | 0.978708076765569 | NM_134257,NM_019492,NM_001081650 |
| Gigyf2 | 2902.176205 | 0.021076429 | 0.068885586 | 0.305962827 | 0.759632949 | 0.881761531 | | Cops7b | 883.3743325 | 0.00774225581775533 | 0.0613094109419658 | 0.126281725085958 | 0.8995089217128 | 0.978889960032469 | NM_172974 |
| Ankrd49 | 338.8693129 | -0.023908916 | 0.078228826 | -0.305627948 | 0.759887938 | 0.881991439 | | Cuta | 1314.525452 | 0.00809168789771834 | 0.0642858531568659 | 0.125870507042877 | 0.899834428931663 | 0.978889960032469 | NM_026948,NM_026307 |
| Cpsf2 | 2200.8284 | -0.021293947 | 0.069703115 | -0.305494905 | 0.75998925 | 0.882042954 | | Gpn2 | 265.8036472 | -0.00995286271273173 | 0.0790861784201333 | -0.125848377934746 | 0.899851946113168 | 0.978889960032469 | NM_133884 |
| Fdx1l | 23.64457283 | -0.050463184 | 0.165248538 | -0.305377493 | 0.76007866 | 0.882080652 | | Jmjd6 | 821.3639332 | 0.00976807163418581 | 0.0777995268822451 | 0.125553696328451 | 0.90008521783445 | 0.978889960032469 | NM_033398 |
| Ly96 | 22.78309308 | 0.051133572 | 0.167641191 | 0.305017949 | 0.760352482 | 0.882312336 | | Kcnq2 | 7577.579728 | -0.00727174712256564 | 0.0578333335306524 | -0.125736257017805 | 0.899940700663443 | 0.978889960032469 | NM_001006669,NM_001006668,NM_001003824,NM_001003825,NM_001006671,NM_001006676,NM_001006677,NM_001006674,NM_001006675,NM_0010066 |
| Col13a1 | 55.00126042 | -0.046888151 | 0.153765654 | -0.304932539 | 0.760417533 | 0.88234174 | | Med28 | 2010.38411 | 0.0067843625315584 | 0.0538143190111097 | 0.126069838961593 | 0.89967664197115 | 0.978889960032469 | NM_025895 |
| Fxyd7 | 1809.908539 | 0.038480482 | 0.126565237 | 0.304036742 | 0.761099899 | 0.882736869 | | Prkcg | 12670.70839 | 0.00819015840125801 | 0.0650567844633548 | 0.125892456395095 | 0.899817054091346 | 0.978889960032469 | NM_011102 |
| Lap3 | 1822.310906 | -0.028173721 | 0.092567956 | -0.304357167 | 0.760855796 | 0.882736869 | | Sema6b | 4549.714818 | -0.0084498643745789 5 | 0.0669851310471516 | -0.126145373495366 | 0.899616851456195 | 0.978889960032469 | NM_013662,NM_001130456 |
| Pecr | 145.3310414 | 0.036576014 | 0.120282298 | 0.304084761 | 0.761063317 | 0.882736869 | | Tdp1 | 609.7884405 | 0.00879870506428189 | 0.0700079577289482 | 0.12568149892828 | 0.899984047442426 | 0.978889960032469 | NM_028354 |
| Pole2 | 14.41585235 | 0.048699786 | 0.160088962 | 0.304204519 | 0.760972082 | 0.882736869 | | Zfp248 | 507.7925851 | -0.00923261995831368 | 0.0735553707787971 | -0.12551931776782 | 0.900112432614188 | 0.978889960032469 | NM_028335 |
| Sdcbp | 6322.274343 | 0.016592555 | 0.054527766 | 0.304295524 | 0.760902755 | 0.882736869 | | Zfp236 | 2148.315324 | 0.00741783014288245 | 0.0591900724848026 | 0.125322000556677 | 0.900268477234591 | 0.978986367742901 | NM_177832 |
| Zfml | 4458.901212 | -0.020098389 | 0.066096661 | -0.304075712 | 0.761070211 | 0.882736869 | | Crip2 | 3881.458183 | 0.00807679731498616 | 0.0645991012519393 | 0.12502956168827 | 0.900500147122403 | 0.978986367742901 | NM_024223 |
| Spata24 | 110.5729162 | -0.040149415 | 0.132096936 | -0.30393903 | 0.761174342 | 0.882757129 | | D230025D16Rik | 1735.228881 | 0.0072701341232877 4 | 0.05814516334087 | 0.12503420242656 | 0.900496473178826 | 0.979053977320898 | NM_145604 |
| Slco2b1 | 1052.869527 | -0.020693403 | 0.068125059 | -0.303756117 | 0.761313081 | 0.882862667 | | Nkx3-1 | 63.68155376 | -0.0116499700489629 | 0.0933993746669865 | -0.124732848845342 | 0.900735051143914 | 0.979053977320898 | NM_010921 |
| Itpa | 1704.06511 | 0.020350795 | 0.06703579 | 0.303581039 | 0.761447099 | 0.882875204 | | Rpl5 | 9360.257815 | -0.00676010494316339 | 0.054104708439569 | -0.124944863172407 | 0.900567200958774 | 0.979053977320898 | NM_016980 |
| Wdr31 | 148.761649 | 0.040488688 | 0.133342828 | 0.303643534 | 0.761399481 | 0.882875204 | | Twf1 | 2050.430273 | -0.00621924567345726 | 0.0498099683837213 | -0.124859454617552 | 0.90063481466995 | 0.979053977320898 | NM_008971 |
| Cep170b | 12215.13351 | -0.026116892 | 0.086229387 | -0.302876934 | 0.76198365 | 0.883190125 | | Zfp229 | 97.82090343 | -0.0115735702823707 | 0.0927604122830908 | -0.124768341487632 | 0.900706951332915 | 0.979053977320898 | NM_001164676 |
| Rnf219 | 580.8321946 | 0.022988912 | 0.075886073 | 0.302939799 | 0.761935741 | 0.883190125 | | 3110062M04Rik | 155.911508 | 0.0108832544494818 | 0.0894464165726195 | 0.121673443968343 | 0.903157667668399 | 0.979150580904713 | NM_001135611,NM_199145 |
| Slc25a38 | 654.6583019 | -0.022640849 | 0.074721003 | -0.303005149 | 0.761885938 | 0.883190125 | | 9330117O12Rik | 109.6485205 | 0.0114187089249935 | 0.0931393738219773 | 0.122598085599316 | 0.90242537047875 | 0.979150580904713 | NR_045400 |
| Tcte1 | 107.0245045 | 0.041370885 | 0.136604928 | 0.302850604 | 0.762003717 | 0.883190125 | | Ascc3 | 1079.825933 | -0.00956178137741126 | 0.0775283018671273 | -0.123332784382777 | 0.901843580819085 | 0.979150580904713 | NM_198007 |
| Tmem218 | 502.1212783 | -0.023738924 | 0.078374926 | -0.302889274 | 0.761974246 | 0.883190125 | | Ash2l | 1558.297531 | -0.00603968690284565 | 0.0496048536756017 | -0.121759665697 1 | 0.903092288042852 | 0.979150580904713 | NM_001080793,NM_011791 |
| Sh2d5 | 4824.921479 | -0.027381039 | 0.090445762 | -0.302734357 | 0.762092313 | 0.8832674 | | Bcar3 | 634.2313261 | 0.00962573190125004 | 0.0775464094395285 | 0.124128660125216 | 0.901213406141635 | 0.979150580904713 | NM_013867 |
| Cwc25 | 678.065601 | -0.024772424 | 0.081877284 | -0.302555519 | 0.762228617 | 0.883300031 | | Bckdha | 1501.392653 | 0.00875456153084099 | 0.0718569302879402 | 0.121833224655712 | 0.903031100474007 | 0.979150580904713 | NM_007533 |
| Tbc1d10a | 871.9608594 | -0.024532393 | 0.081098341 | -0.302501789 | 0.76226967 | 0.883300031 | | Btf3 | 3570.803107 | 0.00746697177725234 | 0.0603652166739661 | 0.124135132450417 | 0.901208281607044 | 0.979150580904713 | NM_145455,NM_001170540 |
| Elmod2 | 585.4417674 | 0.025536226 | 0.084441749 | 0.302412334 | 0.762337753 | 0.883312978 | | Cadm3 | 24273.37534 | 0.00785181105136677 | 0.0633965966851376 | 0.123852077635420 | 0.902182535558444 | 0.979150580904713 | NM_053199 |
| Kcne4 | 64.94067904 | 0.049029375 | 0.162254771 | 0.302175246 | 0.762518473 | 0.883312978 | | Cav2 | 1089.292713 | 0.00960242616440797 | 0.0787520942992071 | 0.121932327639757 | 0.902952612886314 | 0.979150580904713 | NM_016900 |
| Ncoa7 | 3112.174392 | 0.022458226 | 0.074323222 | 0.302169702 | 0.762522699 | 0.883395148 | | Cdc20 | 39.91358251 | 0.0113050382025364 | 0.0916484791120875 | 0.123352163746331 | 0.901828235496193 | 0.979150580904713 | NM_023223 |
| Gpr1 | 26.14264596 | 0.050334655 | 0.166821374 | 0.301727851 | 0.762859535 | 0.883467684 | | Cdc42se1 | 2699.247703 | 0.00582952878053796 | 0.0475271559992284 | 0.122656798160458 | 0.902378875533834 | 0.979150580904713 | NM_172395,NM_001038708 |
| Nop58 | 978.3830433 | 0.023809763 | 0.078925464 | 0.301674028 | 0.762900569 | 0.883467684 | | Cers2 | 2715.639997 | -0.00966804870645999 | 0.0783133670010345 | -0.123453365328198 | 0.901748100800612 | 0.979150580904713 | NM_029789 |
| Psmg2 | 507.0191698 | 0.026247864 | 0.087008205 | 0.301671132 | 0.76290277 | 0.883467684 | | D330023K18Rik | 88.31312862 | 0.0116182717198275 | 0.094083376189238 | 0.123489099672795 | 0.901719805426438 | 0.979150580904713 | NR_040334 |
| Taf5 | 236.4150467 | 0.033557008 | 0.111197317 | 0.301778941 | 0.762820585 | 0.883467684 | | Fam89a | 29.29906413 | -0.0105718390603915 | 0.0855727589562522 | -0.123542108368811 | 0.901677832006015 | 0.979150580904713 | NM_001081120 |
| Tlr12 | 39.68963515 | 0.050643855 | 0.167895694 | 0.301638795 | 0.762927431 | 0.883467684 | | Fmr1 | 1989.725751 | 0.00767041237234495 | 0.0621370685189944 | 0.123219685812739 | 0.901933137333453 | 0.979150580904713 | NM_008031 |
| Ube2j1 | 4046.004225 | 0.022708143 | 0.075264338 | 0.301711852 | 0.762871733 | 0.883467684 | | Foxn3 | 569.6087551 | 0.00954744401843814 | 0.0766950832073121 | 0.124485737796656 | 0.9009306921299 | 0.979150580904713 | NM_183186 |
| Rnf24 | 2476.544661 | -0.024588548 | 0.081556228 | -0.301491973 | 0.76303937 | 0.883531276 | | Foxo3 | 1690.966981 | 0.0063572084058399 | 0.0516169794430468 | 0.123161185997998 | 0.901979460597258 | 0.979150580904713 | NM_019740 |
| Abca17 | 32.03836886 | 0.050585711 | 0.168009383 | 0.301088603 | 0.763346932 | 0.883545658 | | Gng12 | 2639.408872 | 0.00624765644898935 | 0.0511665051328821 | 0.122104420318758 | 0.902816321169154 | 0.979150580904713 | NM_001177556,NM_001177557,NM_001177558,NM_001177559,NM_025278,NM_001177560 |
| Bricd5 | 14.13602637 | -0.050140654 | 0.166388478 | -0.301346912 | 0.763149972 | 0.883545658 | | Kcnk4 | 599.8267932 | 0.00966967689421443 | 0.0778320947426899 | 0.12423765448151 | 0.901127109211766 | 0.979150580904713 | NM_008431 |
| Cct2 | 5035.270111 | -0.020173456 | 0.067043486 | -0.300901056 | 0.763489945 | 0.883545658 | | Krt10 | 222.8525473 | -0.0106208030404261 | 0.0866334562443882 | -0.122594705334916 | 0.902428047347248 | 0.979150580904713 | NM_010660 |
| Colec12 | 656.2896989 | 0.037987746 | 0.126319354 | 0.300727836 | 0.763622041 | 0.883545658 | | Lnx1 | 1127.340581 | 0.00866707229799283 | 0.0703876920228903 | 0.123133349722197 | 0.902001025977719 | 0.979150580904713 | NM_001159577,NM_001159580,NM_010727,NM_001159578,NM_001159579 |
| Gm16023 | 12.20423533 | -0.050303071 | 0.167231448 | -0.300799113 | 0.763567684 | 0.883545658 | | Mina | 435.313742 | -0.00859166389379648 | 0.0705595095054322 | -0.121764790515374 | 0.903085299502072 | 0.979150580904713 | NM_025910 |
| Il1a | 38.19105039 | 0.050353816 | 0.167343633 | 0.300900696 | 0.763490219 | 0.883545658 | | Nbr1 | 8592.516243 | 0.00587657577693442 | 0.0481008293184167 | 0.122172026141835 | 0.902762780373785 | 0.979150580904713 | NM_008676,NM_001252222,NM_001252223,NM_001252220 |
| Insr | 2606.667835 | -0.020184344 | 0.067083439 | -0.30088415 | 0.763502837 | 0.883545658 | | Piezo1 | 378.8623929 | -0.0109765213419664 | 0.0894150164815757 | -0.122759260959574 | 0.90229773522906 | 0.979150580904713 | NM_001037298 |
| Nudt16l1 | 1384.285795 | -0.019189135 | 0.063788804 | -0.300822926 | 0.763549525 | 0.883545658 | | Ppp1r14c | 409.9627382 | 0.00947480041507313 | 0.0765409606853631 | 0.123787320010155 | 0.901483671756991 | 0.979150580904713 | NM_133485 |
| Slamf6 | 8.921558944 | -0.047126082 | 0.156455033 | -0.301211671 | 0.763253091 | 0.883545658 | | Ptger3 | 64.66865513 | -0.0117193857089646 | 0.0950846758770902 | -0.123252097152999 | 0.901907472454632 | 0.979150580904713 | NM_011196 |
| Tmprss7 | 84.63888987 | 0.041436548 | 0.137647208 | 0.301034424 | 0.763388245 | 0.883545658 | | Ptk2 | 5983.627454 | 0.00904656199083111 | 0.0731622545793807 | 0.123650672643168 | 0.9015918693473 | 0.979150580904713 | NM_007982,NM_001130409 |
| Exosc9 | 1175.019969 | 0.020184691 | 0.067187655 | 0.300422609 | 0.76385482 | 0.88359866 | | Rbm4 | 461.3814006 | -0.00961882359100587 | 0.0784836008855694 | -0.12255838877711 | 0.902456807694829 | 0.979150580904713 | NM_009032 |

| | | | | | | | | | | | | | | |
|---|---|---|---|---|---|---|---|---|---|---|---|---|---|---|
| Kdelc2 | 364.5407453 | -0.026587043 | 0.088482565 | -0.30047776 | 0.763812758 | 0.88359866 | Reps1 | 1713.649796 | -0.00766165110958897 | 0.0627680730099478 | -0.122062869579805 | 0.902849227719804 | 0.979150580904713 | NM_009048,NM_001111065 |
| Klf13 | 8368.027943 | -0.03257404 | 0.108405842 | -0.300482335 | 0.763809269 | 0.88359866 | Tmem234 | 1293.503996 | -0.00823256909545335 | 0.0667679729821552 | -0.123301168625467 | 0.901868615466307 | 0.979150580904713 | NM_029748 |
| Setmar | 248.7700971 | 0.027867366 | 0.092777207 | 0.300368665 | 0.763895963 | 0.88359866 | Tmem74b | 146.2096044 | -0.0116847669591974 | 0.0950496878635057 | -0.122933249144143 | 0.902159956300035 | 0.979150580904713 | NM_001160363 |
| 4930511M06Rik | 8.185131572 | -0.044233494 | 0.147334258 | -0.300225452 | 0.764005192 | 0.883659036 | Ttf2 | 110.9686642 | 0.0115558542914126 | 0.0946402928282081 | 0.122102900847833 | 0.90281752452744 | 0.979150580904713 | NM_001013026 |
| Ncaph2 | 4285.153713 | -0.016732277 | 0.055769654 | -0.300024762 | 0.764158268 | 0.883770113 | Ubxn8 | 671.7578533 | -0.00862878285535703 | 0.0706287511770525 | -0.122170967368889 | 0.902763618871324 | 0.979150580904713 | NM_178648 |
| Prps1l3 | 550.3667741 | -0.025938107 | 0.086530518 | -0.299756751 | 0.764362706 | 0.883940571 | Wdr5 | 1195.502164 | -0.00719166023070411 | 0.0581697537283645 | -0.123632296335429 | 0.901606419874855 | 0.979150580904713 | NM_080848 |
| Krt80 | 162.7778352 | 0.049138404 | 0.163980649 | 0.299659774 | 0.764436685 | 0.883960146 | Zfp574 | 1373.21872 | 0.00822891275527662 | 0.0676490457509579 | 0.121641224409438 | 0.903183163537261 | 0.979150580904713 | NM_001168506,NM_175477 |
| Ltbp4 | 2584.522188 | 0.038631867 | 0.129009047 | 0.299450837 | 0.764596078 | 0.884012509 | Mastl | 48.39117928 | -0.011372096984252 | 0.0939142695056815 | -0.121090192620452 | 0.90361959716932 | 0.979550617050862 | NM_025979 |
| Olfr316 | 38.6680573 | -0.047331036 | 0.158046617 | -0.299475162 | 0.76457752 | 0.884012509 | Dgcr6 | 1511.894041 | 0.00725208592087503 | 0.0599513737958388 | 0.120966134080123 | 0.903717859258885 | 0.979584032803117 | NM_010047 |
| Isy1 | 638.6514619 | -0.024436665 | 0.081674545 | -0.299195603 | 0.764790804 | 0.884105702 | Gabbr2 | 11965.88807 | 0.00657271528419172 | 0.0545026043324658 | 0.120594517724294 | 0.904012211373277 | 0.979829979287287 | NM_001081141 |
| Klf8 | 204.687072 | 0.031901723 | 0.10660281 | 0.299257807 | 0.764743345 | 0.884105702 | 4930506C21Rik | 52.307608 | -0.0112131649079555 | 0.0940133859541514 | -0.119272003599827 | 0.905059863035047 | 0.980118791143422 | NR_073374 |
| Asrgl1 | 5807.253655 | 0.024514627 | 0.082033179 | 0.298837947 | 0.765063696 | 0.884223251 | 4932438A13Rik | 5373.247914 | 0.00855702205268501 | 0.0715996084904197 | 0.119512134676406 | 0.904869626857762 | 0.980118791143422 | NM_172679 |
| Idnk | 433.7869275 | 0.025092054 | 0.083935445 | 0.298944667 | 0.764982265 | 0.884223251 | 4933413G19Rik | 58.43830025 | -0.0110564308127642 | 0.0940758256526027 | -0.117526800706408 | 0.90644260731506 | 0.980118791143422 | NM_027697 |
| Pepd | 848.7736388 | -0.021987174 | 0.073574126 | -0.298843831 | 0.765059206 | 0.884223251 | Adamtsl2 | 271.6674647 | 0.0108310101160085 | 0.0906312572064723 | 0.119506343063672 | 0.904874215013801 | 0.980118791143422 | NM_029981 |
| Hnrpll | 1770.995578 | -0.019131344 | 0.064078454 | -0.298561267 | 0.765274822 | 0.884342751 | Adat2 | 211.0826864 | 0.0108687654441637 | 0.0907389770737202 | 0.119780559520011 | 0.904656982309785 | 0.980118791143422 | NM_025748 |
| Mfn2 | 9641.225098 | 0.015971496 | 0.053496377 | 0.298552851 | 0.765281245 | 0.884342751 | Aqp11 | 459.8841116 | -0.00848967915297683 | 0.0722926169868622 | -0.117434940203087 | 0.906515397340137 | 0.980118791143422 | NM_175105 |
| Psmc2 | 3680.664745 | 0.023923112 | 0.080198453 | 0.298298924 | 0.765475026 | 0.884500712 | Aste1 | 211.3921407 | -0.00997650203801161 | 0.0850058497886589 | -0.117362535199814 | 0.906572771430747 | 0.980118791143422 | NM_025651,NM_001164828 |
| Calm3 | 52351.66938 | -0.022042115 | 0.074055484 | -0.297643248 | 0.765975463 | 0.884503851 | Cck | 6296.987685 | 0.00757268265833232 | 0.0630250812171377 | 0.120153477188589 | 0.904361570524151 | 0.980118791143422 | NM_031161 |
| Casp2 | 475.7705935 | 0.025897173 | 0.086994429 | 0.297687718 | 0.765941519 | 0.884503851 | Ccsap | 2451.185702 | -0.00917651211055678 | 0.0773411628749035 | -0.118649781945992 | 0.905552823594806 | 0.980118791143422 | NM_028536 |
| Ddit3 | 844.3870488 | 0.024836372 | 0.083294828 | 0.298174244 | 0.765570179 | 0.884503851 | Celsr2 | 10760.62116 | 0.00723002993627715 | 0.0611320832454435 | 0.118268993177425 | 0.905854524798638 | 0.980118791143422 | NM_001004177,NM_017392 |
| Dnahc17 | 14.61769339 | -0.050077466 | 0.168004751 | -0.298071726 | 0.76564842 | 0.884503851 | Dck | 1024.592299 | -0.00899681546444122 | 0.0765633644608216 | -0.117508099700151 | 0.906457425880329 | 0.980118791143422 | NM_007832 |
| H2-Ke6 | 546.4619963 | -0.030992668 | 0.104167318 | -0.29752775 | 0.766063625 | 0.884503851 | Dpf1 | 1987.90373 | 0.00951072893982 | 0.0809202833269477 | 0.11753207661664 | 0.906438426721304 | 0.980118791143422 | NM_013874 |
| Isoc2a | 587.927983 | 0.029004883 | 0.097419879 | 0.297730641 | 0.765908755 | 0.884503851 | Endou | 192.6774268 | 0.0106459439140422 | 0.0888679066241976 | 0.119795146734597 | 0.904645426601783 | 0.980118791143422 | NM_008902,NM_001168693 |
| Jakmip1 | 2990.335259 | 0.019182597 | 0.064415792 | 0.297793388 | 0.765860861 | 0.884503851 | Fam13a | 944.0073635 | -0.00809862504032628 | 0.068880217353088 | -0.11757548613431 | 0.906404029426573 | 0.980118791143422 | NM_153574 |
| Mtpap | 897.0318741 | 0.019774764 | 0.066475908 | 0.297472634 | 0.766105698 | 0.884503851 | Gm3893 | 842.3859658 | -0.00530527131377601 | 0.0450809018485841 | -0.117683344747519 | 0.906318564033067 | 0.980118791143422 | NR_033506 |
| Ptpn18 | 32.03810674 | -0.049025207 | 0.164733536 | -0.297603073 | 0.766006129 | 0.884503851 | Gm608 | 3227.970725 | 0.00870627340185065 | 0.0740255256527045 | 0.117611774115549 | 0.906375275305246 | 0.980118791143422 | NM_001029889 |
| Rec8 | 182.2562181 | -0.04447448 | 0.149476264 | -0.297535402 | 0.766057784 | 0.884503851 | Hspa1a | 242.6632594 | 0.0105047202238345 | 0.0892882042236556 | 0.117649585577077 | 0.906345314128637 | 0.980118791143422 | NM_010479 |
| Rps15a | 5554.302223 | -0.019466631 | 0.065388361 | -0.297707896 | 0.765926116 | 0.884503851 | Kbtbd3 | 272.5478923 | 0.0105100350748983 | 0.087700138991024 | 0.119840574893205 | 0.904609439426464 | 0.980118791143422 | NM_001164574,NM_026962 |
| Adck4 | 847.6899211 | 0.02234379 | 0.075154905 | 0.297303157 | 0.766235071 | 0.884521397 | Klhl4 | 632.9310612 | -0.00796833961147916 | 0.067804801913747 | -0.117518809679815 | 0.906448939352651 | 0.980118791143422 | NM_172781 |
| Gm9776 | 120.3672513 | -0.03722929 | 0.125219056 | -0.29731329 | 0.766227336 | 0.884521397 | Msrb2 | 694.3106236 | -0.00857824218312129 | 0.0726123415384155 | -0.118137523200281 | 0.905958692400296 | 0.980118791143422 | NM_029619 |
| Dusp18 | 1576.996776 | 0.024465691 | 0.082354805 | 0.297076671 | 0.766407975 | 0.884589181 | Pabpc1 | 5746.709121 | -0.00572940675334552 | 0.0477260023240545 | -0.120047908359126 | 0.904445196961879 | 0.980118791143422 | NM_008774 |
| Mamstr | 87.92722086 | 0.0412269 | 0.138753276 | 0.297123792 | 0.766372001 | 0.884589181 | Pdzrn3 | 1433.470527 | -0.00997708664340807 | 0.0848373035183176 | -0.117602590247978 | 0.906382552470096 | 0.980118791143422 | NM_018884 |
| 2810001G20Rik | 246.0069339 | -0.038476869 | 0.129664648 | -0.296741399 | 0.766663948 | 0.884818712 | Rarres1 | 25.44891848 | 0.0100697001368077 | 0.0849683041795643 | 0.118511252331544 | 0.905662579856287 | 0.980118791143422 | NM_001164763 |
| Phf6 | 922.692861 | 0.027549908 | 0.092929726 | 0.296459588 | 0.766879125 | 0.884935217 | Rfx7 | 1702.931677 | -0.00825639628163356 | 0.0699558639921839 | -0.118022933467822 | 0.906049486598063 | 0.980118791143422 | NM_001033536 |
| Wrnip1 | 1939.183233 | 0.019826427 | 0.066864348 | 0.296517164 | 0.766835161 | 0.884935217 | Sipa1l1 | 22932.85322 | -0.00778942967464175 | 0.0655643128996401 | -0.118805937714395 | 0.905429104390242 | 0.980118791143422 | NM_172579,NM_001167983 |
| Slc16a13 | 180.7554453 | -0.030920351 | 0.104363351 | -0.29627595 | 0.767019351 | 0.884965212 | Slc16a7 | 420.5249418 | -0.00917312564448779 | 0.0770136807447547 | -0.119110339303093 | 0.905187939481222 | 0.980118791143422 | NM_011391 |
| Sox5 | 1524.364923 | -0.024077987 | 0.081267305 | -0.296281355 | 0.767015224 | 0.884965212 | Smim4 | 111.7624851 | -0.0113141994808594 | 0.0951000579466551 | -0.118971530881779 | 0.905297910623998 | 0.980118791143422 | NR_024069 |
| Tomm6 | 2195.278163 | -0.019270207 | 0.06507321 | -0.296131196 | 0.767129891 | 0.885026845 | Spats2 | 1059.499553 | 0.00710534138632657 | 0.0601315847009905 | 0.118163215249664 | 0.905938335692602 | 0.980118791143422 | NM_139140 |
| Ntmt1 | 595.9173927 | -0.024984962 | 0.084413003 | -0.295984753 | 0.767241726 | 0.885089963 | Stxbp4 | 1019.758238 | 0.00770332170810275 | 0.0644026921382688 | 0.119611796531179 | 0.904790674538325 | 0.980118791143422 | NM_011505 |
| Sgpp1 | 1187.96751 | -0.022171115 | 0.075022852 | -0.295524825 | 0.767592991 | 0.885429259 | Tada2a | 1123.870879 | -0.00665379828748846 | 0.0561656620135062 | -0.118467370435132 | 0.905697347620488 | 0.980118791143422 | NM_172562 |
| Clvs2 | 1077.803794 | -0.033500185 | 0.113612723 | -0.294862974 | 0.768098557 | 0.885915696 | Tfdp1 | 1738.840059 | 0.00662393027331305 | 0.0562069287243804 | 0.11784899875591 | 0.906187304630564 | 0.980118791143422 | NM_009361 |
| Ppp1r21 | 3008.228931 | 0.025294597 | 0.085795856 | 0.294823062 | 0.768129048 | 0.885915696 | Tmem167 | 2140.395816 | 0.00773188294069593 | 0.0657176795391242 | 0.117653012019282 | 0.906342599079101 | 0.980118791143422 | NM_025335 |
| Ppp2r1a | 22225.32532 | 0.021131643 | 0.071711206 | 0.294677005 | 0.768240631 | 0.88597844 | Tmem175 | 1518.97997 | 0.00769129730878214 | 0.0644021252909933 | 0.119426141203104 | 0.904937751814036 | 0.980118791143422 | NM_001163532,NM_001163531,NM_028223 |
| Ccdc34 | 239.4850173 | -0.034403027 | 0.116796471 | -0.294555363 | 0.768333565 | 0.886015592 | Zkscan5 | 641.5067351 | -0.00840436474189303 | 0.0715380413407652 | -0.117481057411952 | 0.906478854085003 | 0.980118791143422 | NM_016683,NM_001167944 |
| Pigyl | 452.9574557 | -0.02356638 | 0.0800257 | -0.294485143 | 0.768387215 | 0.886015592 | Cmtm7 | 83.70242574 | -0.0110828217165265 | 0.094644738705992 | -0.117099184466604 | 0.906781455992373 | 0.980201396200635 | NM_133978,NM_001252479 |
| Cited2 | 915.6787917 | 0.032755534 | 0.111349712 | 0.294168102 | 0.768629454 | 0.886163015 | Dip2b | 4319.887514 | -0.00661697221191063 | 0.0565501713277843 | -0.117010648359602 | 0.906851615278834 | 0.980201396200635 | NM_001159361,NM_172819 |
| Stx1b | 9293.042055 | -0.01554056 | 0.052825551 | -0.294186429 | 0.76861545 | 0.886163015 | Plekhg3 | 861.44023 | 0.0101700079004238 | 0.0868673801266783 | 0.11707510788967 | 0.906800535090633 | 0.980201396200635 | NM_153804 |
| Dolpp1 | 637.1222478 | 0.02545981 | 0.086717444 | 0.293595029 | 0.769067374 | 0.886601927 | Ankrd34c | 355.928771 | 0.00996902956233651 | 0.0853992737457251 | 0.116734359966797 | 0.907070561092881 | 0.980365101966904 | NM_207260 |
| Mir3061 | 40.65219261 | 0.048060105 | 0.16374666 | 0.293502811 | 0.769137852 | 0.886617206 | 0610010K14Rik | 636.0191122 | -0.00879313447609135 | 0.0755639074887965 | -0.116366857780549 | 0.907361800573796 | 0.980388092467641 | NM_145758,NM_026757,NM_001177601,NM_001177603,NM_001177606,NM_001177607 |
| Gm1976 | 851.3978434 | -0.02505219 | 0.085420464 | -0.29328089 | 0.769307459 | 0.886746747 | Bbx | 1566.452449 | -0.00785007259528052 | 0.0674496157865328 | -0.116384244798736 | 0.90734802136483 | 0.980388092467641 | NM_027444 |
| Bcat1 | 3867.723807 | 0.024863687 | 0.084845536 | 0.293046496 | 0.769486613 | 0.88688727 | Lpin1 | 1465.621247 | 0.00821626365644754 | 0.0704651275574859 | 0.116600422666441 | 0.907176702771156 | 0.980388092467641 | NM_172950,NM_001130412,NM_015763 |
| Lrif1 | 376.2870324 | -0.029724326 | 0.101482565 | -0.292900815 | 0.769597966 | 0.886949634 | Ppp1r3f | 1352.533368 | 0.007316786775082 | 0.0628701860817523 | 0.116379276586962 | 0.907351958668974 | 0.980388092467641 | NM_138605 |
| Cxx1a | 1045.486083 | -0.01721869 | 0.058824322 | -0.292713789 | 0.769740929 | 0.886958984 | Zscan20 | 311.6451596 | -0.00958500055810112 | 0.0825108271843711 | -0.116166579407614 | 0.907520523159481 | 0.980486658193724 | NM_177758 |
| Itga11 | 97.28021762 | 0.038419736 | 0.131228261 | 0.292770287 | 0.769697741 | 0.886958984 | Tmem223 | 784.9414655 | 0.00929992022463468 | 0.0801198973702727 | 0.116075039158566 | 0.907593070939657 | 0.980492112782199 | NM_025791 |
| Prss36 | 218.7301914 | 0.039431357 | 0.134760427 | 0.292603385 | 0.769825326 | 0.886958984 | Zfp677 | 75.08966222 | 0.0110191129671082 | 0.0950151409506759 | 0.115972179348009 | 0.907674590655775 | 0.980507258411972 | NM_172486 |
| Ttc39c | 873.6800332 | 0.040523694 | 0.138499621 | 0.292590652 | 0.769835059 | 0.886958984 | Cadm2 | 6497.704025 | 0.00921284287248226 | 0.0796844806719131 | 0.115615625155552 | 0.907956464480677 | 0.980520080517904 | NM_001145977,NM_178721 |
| Vdac2 | 6375.710804 | -0.016742134 | 0.057283502 | -0.292267988 | 0.770081733 | 0.887177215 | Chpt1 | 947.4007214 | -0.00735985633473662 | 0.063653413463441 | -0.115623906626213 | 0.907950615231757 | 0.980520080517904 | NM_144807,NM_001146690,NR_027477 |
| Gapvd1 | 3619.632255 | -0.014566427 | 0.049892494 | -0.291956276 | 0.770320055 | 0.887384277 | Dcun1d5 | 1066.683768 | -0.0081459202737173 | 0.070367599279831 | -0.115762372982534 | 0.907840871978455 | 0.980520080517904 | NM_029775 |
| Pced1b | 167.2704856 | 0.030531959 | 0.104603381 | 0.291883093 | 0.77037601 | 0.887384277 | Pdzd4 | 5808.908936 | -0.00587161743408645 | 0.050720260867358 | -0.115764732548236 | 0.907839001890201 | 0.980520080517904 | NM_001029868 |
| Gm14403 | 279.9965452 | -0.031924569 | 0.109432249 | -0.291729079 | 0.770493774 | 0.88745395 | Adck3 | 1739.548598 | -0.0067425064501687 | 0.0585486524160017 | -0.115160745327862 | 0.908317712303622 | 0.980523964515567 | NM_001163290,NM_023341 |
| E330011O21Rik | 14.93124897 | 0.047961373 | 0.164511075 | 0.291538872 | 0.770639219 | 0.887532087 | C2cd4c | 912.7639364 | -0.00907507214871701 | 0.0795101745382645 | -0.114137243458692 | 0.909128999112214 | 0.980523964515567 | NM_198614,NM_001168624 |
| Lysmd2 | 1116.330546 | 0.025968556 | 0.089111748 | 0.291415633 | 0.77073346 | 0.887532087 | Cox7a1 | 44.96097332 | -0.0103054878982353 | 0.0897932265384004 | -0.114769101139585 | 0.908628140902272 | 0.980523964515567 | NM_009944 |
| Pvrl2 | 205.3598162 | -0.031569922 | 0.108331698 | -0.291419064 | 0.770730836 | 0.887532087 | Cyp7b1 | 420.106057 | 0.00810051980527904 | 0.0710609597126032 | 0.113993954458827 | 0.909242585847769 | 0.980523964515567 | NM_007825 |
| Coprs | 609.4114605 | 0.024079612 | 0.082750322 | 0.290991162 | 0.771058079 | 0.887773937 | Dom3z | 585.7571396 | -0.00861811516327987 | 0.0752539185942493 | -0.114520483773697 | 0.908825209516165 | 0.980523964515567 | NM_001163770,NM_033613 |
| Pbx4 | 229.5874826 | -0.0316704 | 0.108826528 | -0.291017275 | 0.771038108 | 0.887773937 | Fbxo18 | 3934.543049 | -0.0058057537219552 | 0.0508531756277543 | -0.114166984662932 | 0.909105423171584 | 0.980523964515567 | NM_015792 |
| 3110043O21Rik | 940.9841575 | -0.020751245 | 0.071376733 | -0.290728415 | 0.771259038 | 0.887901166 | Gm5069 | 214.5950139 | -0.0104570451331907 | 0.091235144010432 | -0.114616415051584 | 0.908749168128853 | 0.980523964515567 | NR_003623 |
| Gm9958 | 44.37979541 | -0.046568666 | 0.160196674 | -0.290696835 | 0.771283193 | 0.887901166 | Gm6981 | 35.64135011 | 0.0106172929119036 | 0.0925489782757945 | 0.114720800917588 | 0.9086664260335 | 0.980523964515567 | NR_023357 |
| Dhx38 | 1559.743586 | 0.02271191 | 0.078226687 | 0.290334556 | 0.771560305 | 0.887979926 | Krt8 | 23.34132622 | 0.00459595602539325 | 0.040143256098283 | 0.114488869914811 | 0.908850268905564 | 0.980523964515567 | NM_031170 |
| Igfbp6 | 1164.660539 | 0.029998042 | 0.103302935 | 0.290389055 | 0.771518616 | 0.887979926 | Mgmt | 31.6669326 | -0.0103302404181533 | 0.0904026937735247 | -0.114269237251021 | 0.909024367853209 | 0.980523964515567 | NM_008598 |
| Kif23 | 54.25964979 | -0.044564784 | 0.153532047 | -0.290263724 | 0.771614489 | 0.887979926 | Nelfa | 814.8573388 | -0.007431257082613 | 0.0648565414283479 | -0.114579916211272 | 0.908778099391569 | 0.980523964515567 | NM_011914 |
| Tor2a | 742.4029251 | 0.0212208 | 0.07310215 | 0.290289689 | 0.771594626 | 0.887979926 | Rnf39 | 185.4180772 | 0.0109771243295504 | 0.0950931959472277 | 0.115435433841578 | 0.908099994532136 | 0.980523964515567 | NM_001099632 |
| Usp37 | 1062.485511 | -0.017575404 | 0.060571871 | -0.290157856 | 0.771695476 | 0.887979926 | Scrt1 | 6787.10549 | -0.00654519341043191 | 0.0573416242818706 | -0.114143843890292 | 0.909123766923638 | 0.980523964515567 | NM_130893 |
| Zfp281 | 1564.812668 | 0.021059528 | 0.072578314 | 0.290162819 | 0.771691679 | 0.887979926 | Slc12a5 | 29614.64337 | -0.00525905316658281 | 0.0460915594599178 | -0.114100135213611 | 0.909158415040144 | 0.980523964515567 | NM_020333 |
| Hmgn5 | 336.9339212 | 0.029442042 | 0.101574125 | 0.289857697 | 0.771925106 | 0.888178197 | Snhg10 | 46.33150997 | -0.0106249933859425 | 0.0927855993922823 | -0.114511232944907 | 0.908832542370561 | 0.980523964515567 | NR_003145 |
| Setd2 | 3149.6809 | 0.021556882 | 0.074408833 | 0.289708647 | 0.772039141 | 0.888243443 | Sptlc1 | 1249.912726 | 0.00631353188526859 | 0.0552218215223473 | 0.114330380838916 | 0.908975899967803 | 0.980523964515567 | NM_009269 |
| Farsb | 1985.79909 | 0.018547933 | 0.064057813 | 0.2895499 | 0.7721606 | 0.888308362 | Stk25 | 9744.835576 | -0.005720273496454 | 0.0499701144153678 | -0.114473892313018 | 0.908862141247945 | 0.980523964515567 | NM_021537 |
| Ybey | 245.789461 | 0.026698444 | 0.092227371 | 0.289485033 | 0.772210232 | 0.888308362 | Tmem219 | 371.9407676 | 0.00943876448809003 | 0.0821018846078937 | 0.114964041728983 | 0.908473623562193 | 0.980523964515567 | NM_026827,NM_028389 |
| Tmem184b | 3505.343241 | 0.01833948 | 0.063441345 | 0.289077723 | 0.772521901 | 0.888600915 | Zfp768 | 469.1098115 | 0.00866180310469447 | 0.0759600870224387 | 0.114030979218544 | 0.909213235743573 | 0.980523964515567 | NM_146202 |
| Snrpd3 | 1190.349532 | -0.021158553 | 0.073216494 | -0.28898615 | 0.772591976 | 0.888615551 | Cdh1 | 66.62953834 | 0.00874652131499382 | 0.0768238972913852 | 0.113851569932975 | 0.909355457434843 | 0.980572888033445 | NM_009864 |
| Fam32a | 2910.001655 | -0.015470884 | 0.053564328 | -0.288828494 | 0.772712626 | 0.888622387 | Alpl | 405.439408 | 0.0104431293227158 | 0.0923986247891553 | 0.113022562257241 | 0.910012667728689 | 0.980590354840512 | NM_007431 |
| Tph1 | 14.6630428 | -0.045322825 | 0.156881923 | -0.288897692 | 0.77265967 | 0.888622387 | Arl6ip6 | 261.8617282 | -0.00886052456915923 | 0.0787225091651585 | -0.112553889136968 | 0.910384243795893 | 0.980590354840512 | NM_022989 |
| Akr1e1 | 895.3758632 | 0.024601442 | 0.08523442 | 0.288632718 | 0.772862456 | 0.888688505 | En2 | 30.5336832 | 0.00980476616921984 | 0.0863147663798116 | 0.113514768098044 | 0.909622455507996 | 0.980590354840512 | NM_010134 |
| Trim33 | 2594.32056 | 0.023221288 | 0.080460873 | 0.288603478 | 0.772884833 | 0.888688505 | Gm16596 | 41.20961124 | 0.0105183959790286 | 0.0929082025890016 | 0.113212780851642 | 0.909861863160258 | 0.980590354840512 | NR_045751,NR_045753,NR_045752 |
| Becn1 | 4206.227103 | -0.013231618 | 0.0458968 | -0.288290638 | 0.773124273 | 0.888897854 | Hmcn1 | 152.7869976 | 0.0106199634336923 | 0.0937762101538444 | 0.113247948667042 | 0.909833982607314 | 0.980590354840512 | NM_001024720 |
| Ccs | 499.6885177 | 0.02225488 | 0.077234318 | 0.288147552 | 0.773233794 | 0.888918104 | Hspb1 | 188.5256052 | -0.0105378003995247 | 0.0935142389100764 | -0.112686586795171 | 0.910279035697159 | 0.980590354840512 | NM_013560 |
| Eif2s2 | 2471.138503 | 0.021482373 | 0.074693961 | 0.287605216 | 0.773648952 | 0.888918104 | Hspg2 | 398.8755258 | -0.00973158879407215 | 0.0862132978527044 | -0.112878048241451 | 0.910127240050909 | 0.980590354840512 | NM_008305 |
| Fhl4 | 32.35717745 | -0.047948763 | 0.166534851 | -0.287920294 | 0.773407752 | 0.888918104 | Ints9 | 696.7570001 | -0.00720335825333946 | 0.063616243969359 | -0.113231429645689 | 0.909847078642934 | 0.980590354840512 | NM_001253731,NM_153414 |
| Galnt2 | 3082.777041 | 0.016542961 | 0.057503781 | 0.287684747 | 0.773588067 | 0.888918104 | Larp6 | 1490.169747 | -0.00622482181918356 | 0.0551901989981498 | -0.112788537316059 | 0.910198206247569 | 0.980590354840512 | NM_026235 |
| Insrr | 22.66198715 | 0.048277034 | 0.16767866 | 0.287914004 | 0.773412567 | 0.888918104 | Macrod1 | 329.864913 | -0.0102562883527792 | 0.0904154425635958 | -0.113435139639616 | 0.909685582087644 | 0.980590354840512 | NM_134147 |
| Kcmf1 | 2148.387075 | 0.022882912 | 0.079538872 | 0.2876947 | 0.773580447 | 0.888918104 | Mtpap | 1000.318225 | 0.00768623735607075 | 0.0675754695243415 | 0.113743010743004 | 0.909441516106236 | 0.980590354840512 | NM_026157 |
| Mgme1 | 244.3784871 | 0.031820849 | 0.110577357 | 0.287770024 | 0.773522784 | 0.888918104 | Mtss1l | 8681.038097 | 0.00672841207502812 | 0.0594649379565651 | 0.113149232240732 | 0.909912243893056 | 0.980590354840512 | NM_198625 |
| P2rx3 | 101.2352895 | -0.035403594 | 0.123103073 | -0.287593095 | 0.773658231 | 0.888918104 | Nadk | 2918.481874 | 0.00516980768852969 | 0.0458919780093072 | 0.112651664033335 | 0.910306723730198 | 0.980590354840512 | NM_001159637,NM_138671 |
| Pdzk1ip1 | 9.49998641 | -0.042734176 | 0.148541135 | -0.287692535 | 0.773582105 | 0.888918104 | Ppp1r1b | 16180.97583 | -0.00977240401899429 | 0.0868177423768799 | -0.112562291433148 | 0.910377582067238 | 0.980590354840512 | NM_144828 |
| Sec31b | 15.42932273 | -0.047676552 | 0.165960972 | -0.287275687 | 0.773901237 | 0.889131379 | Tbc1d1 | 1104.140064 | 0.00823577418703358 | 0.0726946914577811 | 0.113292649323874 | 0.909798544711079 | 0.980590354840512 | NM_019636 |
| 1200011I18Rik | 1886.887827 | -0.019454601 | 0.067907005 | -0.286488863 | 0.774503722 | 0.889236585 | Rab3d | 576.8000482 | 0.00881772742295315 | 0.0784534975322901 | 0.11239431893172 | 0.91051075962327 | 0.980653910750176 | NM_031874 |
| Abt1 | 506.2398727 | -0.02212829 | 0.077133089 | -0.286884526 | 0.774200739 | 0.889236585 | Mpv17 | 1912.263767 | -0.00544747034568338 | 0.0486021613338688 | -0.112082882657469 | 0.910757689558134 | 0.980847138087296 | NM_008622 |
| Ccno | 67.12171089 | -0.045877911 | 0.159870634 | -0.28696897 | 0.77413608 | 0.889236585 | Sap30l | 755.7232547 | -0.00779555929056542 | 0.0696439594625691 | -0.111934464248191 | 0.910875369790372 | 0.980901150683886 | NM_001081168 |
| Cks1b | 58.90302479 | -0.048065384 | 0.167822204 | -0.286406584 | 0.774566732 | 0.889236585 | 2310039H08Rik | 68.30290355 | -0.0105690502732784 | 0.0951016072382536 | -0.111134297097632 | 0.911509852036587 | 0.980985114484439 | NM_025966 |
| Dlc1 | 3070.282061 | 0.019930228 | 0.069556795 | 0.286531718 | 0.774470904 | 0.889236585 | 3110043O21Rik | 1028.37232 | -0.00756807489792899 | 0.0681381163369823 | -0.111069623065312 | 0.911561136948296 | 0.980985114484439 | NM_001081343 |
| Ifi47 | 44.3159028 | -0.047952778 | 0.167298164 | -0.286630628 | 0.77439516 | 0.889236585 | Gnpat | 2533.631011 | 0.00556386004651155 | 0.0500600741209481 | 0.111143663772229 | 0.911502424525321 | 0.980985114484439 | NM_010322 |
| Pdlim1 | 319.4947746 | -0.036674986 | 0.128029121 | -0.286458157 | 0.774527237 | 0.889236585 | H3f3a | 4044.162289 | 0.00603802659463482 | 0.0541795300396106 | 0.111444794560241 | 0.911263640347344 | 0.980985114484439 | NM_008210 |
| Snord96a | 51.7638111 | 0.044284565 | 0.154479015 | 0.286670424 | 0.774364686 | 0.889236585 | Llgl2 | 52.53909829 | 0.0103113559830958 | 0.0927444787893912 | 0.111180267738755 | 0.911473398676677 | 0.980985114484439 | NM_001252532,NM_145438 |
| Tbc1d12 | 561.7835412 | 0.025605359 | 0.089401488 | 0.286408643 | 0.774565156 | 0.889236585 | Rasl10b | 12932.17956 | 0.00622666669432064 | 0.0558679006327428 | 0.111453386001609 | 0.911256827809497 | 0.980985114484439 | NM_001013386 |
| Wwp1 | 2080.366097 | -0.021718094 | 0.075688605 | -0.286940075 | 0.774158205 | 0.889236585 | Sec16a | 3347.12439 | -0.00712743082611711 | 0.0639548756293841 | -0.111444682770088 | 0.911263728990796 | 0.980985114484439 | NM_153125 |
| 6330409D20Rik | 59.57369671 | 0.043076654 | 0.150716293 | 0.285812856 | 0.775021461 | 0.889641698 | Sstr4 | 320.2458093 | -0.00993793129651522 | 0.0892846734308781 | -0.111306128080414 | 0.911373596098657 | 0.980985114484439 | NM_009219 |
| Arsk | 239.8758108 | -0.033071854 | 0.115734897 | -0.285755248 | 0.775065587 | 0.889641698 | Zfp575 | 344.613131 | -0.00851240961230672 | 0.0765144943058279 | -0.111252249518669 | 0.91141631962667 | 0.980985114484439 | NM_001033205 |
| Hace1 | 1923.585569 | 0.021595145 | 0.07560108 | 0.285645986 | 0.775149279 | 0.889641698 | Kcnmb4 | 1843.557377 | 0.00741663637369313 | 0.0670537012364299 | 0.110607412222365 | 0.911927669453058 | 0.981306861728656 | NM_021452 |
| Plxna1 | 5055.633147 | 0.026423678 | 0.092484258 | 0.285710001 | 0.775100244 | 0.889641698 | Usp13 | 1245.577465 | 0.00714379103411519 | 0.064706345731161 | 0.110403252623721 | 0.912089573679509 | 0.981408381279151 | NM_001013024 |
| Ephb3 | 942.4561403 | -0.036428702 | 0.127567146 | -0.28556492 | 0.775211375 | 0.889647066 | Bcat2 | 429.339478 | -0.00948852691388098 | 0.0861239066938152 | -0.110172973778515 | 0.91227219557239 | 0.98149575637022 | NM_001243053,NM_009737,NM_001243052 |
| Mir3064 | 37.45961149 | 0.047032251 | 0.164816995 | 0.285360444 | 0.77536801 | 0.889695026 | Itgav | 3789.224334 | -0.00729214547081765 | 0.0662137037802118 | -0.110130457209025 | 0.912305913707194 | 0.98149575637022 | NM_008402 |
| Unc50 | 1089.507123 | 0.020075875 | 0.070336492 | 0.285426157 | 0.77531767 | 0.889695026 | Ppcs | 186.2137886 | 0.0096495677743476 | 0.0877380800573478 | 0.109981524191553 | 0.912424027575357 | 0.981550131419658 | NM_026494 |
| Grwd1 | 602.3074049 | 0.022191332 | 0.077802292 | 0.285227229 | 0.775470061 | 0.889746232 | Ercc6 | 1376.60222 | 0.007698657015817 | 0.0700863690453917 | 0.109845282623087 | 0.912532077968047 | 0.981593673323745 | NM_001081221 |
| Rad54l2 | 1465.062182 | -0.020851055 | 0.073150555 | -0.285043022 | 0.775611181 | 0.889842254 | Prss57 | 35.92737339 | 0.0100490888080721 | 0.0916292975135047 | 0.109671132277217 | 0.912670195388595 | 0.98166481460904 | NM_001042710 |
| Cdk5 | 4007.767527 | 0.015029167 | 0.052740157 | 0.284966284 | 0.775669973 | 0.889843815 | Rundc3a | 12670.92831 | -0.00649752908562138 | 0.0592886373604241 | -0.109591472749187 | 0.912733373682342 | 0.98166481460904 | NM_016759,NM_001252347 |
| Zfp280d | 1450.307783 | -0.023831404 | 0.083675932 | -0.28480596 | 0.775792806 | 0.889918837 | 9130019O22Rik | 146.647588 | -0.00999041991161964 | 0.09268068023459 | -0.107793985610941 | 0.914159113878648 | 0.981708669866274 | NM_030226 |
| Aqp9 | 63.190852 | -0.040193331 | 0.141258987 | -0.284536454 | 0.775999302 | 0.890089812 | A730020E08Rik | 58.51039697 | -0.0100454996741353 | 0.0930771052314768 | -0.107926644787166 | 0.914053881085124 | 0.981708669866274 | NR_040287 |
| 2310057M21Rik | 586.4641659 | -0.021851674 | 0.076819893 | -0.284453322 | 0.776063001 | 0.890092881 | Apbb2 | 2301.215755 | 0.00520959452203117 | 0.0486602390343426 | 0.107060602771688 | 0.914740901907564 | 0.981708669866274 | NM_001201415,NM_001201414,NM_001201416,NM_009686,NM_001201413 |
| Angptl4 | 94.05376116 | 0.047341926 | 0.166483379 | 0.284364279 | 0.776131231 | 0.890092881 | Aurka | 47.22963877 | -0.0100723383541527 | 0.0928986599473788 | -0.108422859488587 | 0.91366026877219 | 0.981708669866274 | NM_011497 |
| Bend5 | 283.9789911 | -0.036051316 | 0.126803714 | -0.284308047 | 0.77617432 | 0.890092881 | Ckap2l | 33.46441539 | 0.00932295357840554 | 0.0859522572261629 | 0.108466652060974 | 0.913625532212088 | 0.981708669866274 | NM_181589 |
| Smg8 | 698.7915834 | 0.022498997 | 0.079180679 | 0.284147562 | 0.776297299 | 0.890168025 | Clec4a2 | 27.31727405 | -0.00926095154136053 | 0.0851700756983446 | -0.10873480462975 | 0.91341283540935 | 0.981708669866274 | NM_001170333,NM_011999,NM_001170332 |
| Chrm1 | 8391.945492 | 0.023766565 | 0.08377077 | 0.283709524 | 0.776632996 | 0.890235035 | Engase | 479.9984845 | 0.00877357115084031 | 0.0818531971142266 | 0.107186664176315 | 0.914640894922237 | 0.981708669866274 | NM_172573 |
| Dazap2 | 5009.593901 | -0.013163688 | 0.046419507 | -0.283580957 | 0.776731532 | 0.890235035 | Golga7b | 4796.407064 | 0.00626781984896256 | 0.058540566027355 | 0.107067974813119 | 0.914735053485302 | 0.981708669866274 | NM_001141983,NM_027694 |
| Gan | 204.9582929 | 0.034468743 | 0.121431073 | 0.283854389 | 0.776521972 | 0.890235035 | Gpr62 | 389.3153734 | 0.00963865427859612 | 0.0886475733174172 | 0.108730040968898 | 0.913416613858837 | 0.981708669866274 | NM_001159652 |
| Gid8 | 1707.407579 | 0.017710763 | 0.062445575 | 0.283619177 | 0.776702239 | 0.890235035 | Hecw2 | 2576.911207 | -0.00805637237186238 | 0.0749482468346473 | -0.107492472634304 | 0.914398296691464 | 0.981708669866274 | NM_001001883,NM_172655 |
| Myo5b | 1925.539543 | 0.028626117 | 0.100984092 | 0.283471545 | 0.776815391 | 0.890235035 | Herpud2 | 1698.247301 | 0.00643830429303333 | 0.0591699106559191 | 0.10881044472879 | 0.913352839315456 | 0.981708669866274 | NM_020586 |
| Sgk2 | 69.86783744 | 0.046668842 | 0.164515625 | 0.283674221 | 0.776660052 | 0.890235035 | Larp4b | 2789.322717 | 0.00558885020305855 | 0.0519594812195138 | 0.107561701385109 | 0.914343378543366 | 0.981708669866274 | NM_172585 |
| Stx1a | 9710.462948 | 0.036632172 | 0.129204129 | 0.283521687 | 0.77677696 | 0.890235035 | Lingo3 | 1955.788584 | 0.00848413241903583 | 0.0794322166277218 | 0.106809714990062 | 0.914939940121211 | 0.981708669866274 | NM_001013758 |
| Tnpo2 | 8398.545695 | 0.014249308 | 0.050260857 | 0.283507064 | 0.776788168 | 0.890235035 | Mum1 | 1480.728394 | -0.00798508044481741 | 0.0743416020521862 | -0.107410658694334 | 0.914463199011567 | 0.981708669866274 | NM_023431 |
| 5430435G22Rik | 71.78652017 | -0.043416699 | 0.153416178 | -0.28299948 | 0.777177236 | 0.890235241 | N28178 | 12763.90353 | 0.00694566443330481 | 0.0636776056289277 | 0.109075464831069 | 0.913142634988846 | 0.981708669866274 | NM_172690 |
| Als2 | 1988.688954 | 0.021486671 | 0.075907617 | 0.283063443 | 0.777128204 | 0.890235241 | Ndufaf3 | 1060.040343 | -0.00651212998543183 | 0.0600803850415533 | -0.108390283799411 | 0.913686108132797 | 0.981708669866274 | NM_023247 |
| Calcr | 39.84509933 | -0.0304389 | 0.10744218 | -0.283304939 | 0.776943092 | 0.890235241 | Ngrn | 1167.507569 | 0.00563086079714855 | 0.0517893336219742 | 0.108726264721803 | 0.91341960911101 | 0.981708669866274 | NR_028053,NM_031375 |
| Heca | 949.2197709 | -0.032996982 | 0.116619115 | -0.282946604 | 0.777217769 | 0.890235241 | Nipal4 | 256.3882201 | 0.0102525751174058 | 0.095083228512174 | 0.107827376897421 | 0.914132625871823 | 0.981708669866274 | NM_172524 |
| Setd3 | 6793.818741 | 0.01839059 | 0.064988296 | 0.282983111 | 0.777189784 | 0.890235241 | Nubp1 | 550.3504674 | -0.00812777837902515 | 0.0757024171823305 | -0.107364846216855 | 0.914499541919162 | 0.981708669866274 | NM_011955 |
| Sipa1l3 | 2536.48869 | 0.02458359 | 0.086878206 | 0.282966132 | 0.777202799 | 0.890235241 | Pcdhb11 | 135.1990926 | 0.00994939066370552 | 0.091322304080307 | 0.108948090654351 | 0.913243662805458 | 0.981708669866274 | NM_053136 |
| Sqrdl | 205.9951386 | -0.029694681 | 0.104840633 | -0.283236374 | 0.776995647 | 0.890235241 | Pnma3 | 1182.28157 | 0.00940941789370968 | 0.0867245172471954 | 0.108497783468653 | 0.913600838672499 | 0.981708669866274 | NM_153169 |
| Pln | 55.51473402 | 0.047517595 | 0.167984987 | 0.282868106 | 0.777277944 | 0.890238355 | Rplp1 | 11216.18111 | 0.00688023539140708 | 0.0643444719315397 | 0.106928150699992 | 0.914845980188515 | 0.981708669866274 | NM_018853 |
| Arap3 | 512.7888566 | -0.024704428 | 0.087381812 | -0.282718194 | 0.777392868 | 0.890304169 | Rusc1 | 7797.96964 | -0.00531133640925991 | 0.0497424756175144 | -0.106776680157627 | 0.91496614822812 | 0.981708669866274 | NM_028188,NM_001083807,NM_001083808 |
| Nptn | 20576.01339 | 0.019935845 | 0.070575451 | 0.282475637 | 0.777578825 | 0.890451316 | Sepw1 | 16381.45137 | -0.00624548120060365 | 0.0576264650192378 | -0.108378697157958 | 0.91369529879402 | 0.981708669866274 | NM_009156 |
| 2010012O05Rik | 861.5766989 | 0.022210538 | 0.078796253 | 0.281873026 | 0.778040873 | 0.890481986 | Sh3yl1 | 290.9273296 | 0.00840105977777554 | 0.0782972840521823 | 0.107296950072707 | 0.914553404065239 | 0.981708669866274 | NM_013709 |
| Asph | 3212.173557 | -0.023465136 | 0.0832443 | -0.28188279 | 0.778033386 | 0.890481986 | Smarcc2 | 13988.81522 | 0.00513190233638586 | 0.0480837176608471 | 0.106728484943347 | 0.915004383954254 | 0.981708669866274 | NM_198160,NM_001114096,NM_001114097 |
| B230120H23Rik | 408.8499808 | 0.026644062 | 0.094488216 | 0.281982909 | 0.777956615 | 0.890481986 | Snta1 | 1217.256665 | 0.00716430016534777 | 0.0659468936478101 | 0.10863741670091 | 0.913490082139036 | 0.981708669866274 | NM_009228 |
| D130020L05Rik | 86.87235258 | 0.035714752 | 0.126753187 | 0.281766107 | 0.778122861 | 0.890481986 | Syndig1 | 1759.862923 | -0.0064487662047731 | 0.0602765559494949 | -0.106986308411125 | 0.914799841713519 | 0.981708669866274 | NM_001085521 |
| Gabre | 22.44163423 | 0.044420205 | 0.157649102 | 0.281766306 | 0.778122708 | 0.890481986 | Tal1 | 116.0591691 | -0.0101105808115879 | 0.0946928251455942 | -0.106772406420892 | 0.914969538793899 | 0.981708669866274 | NM_011527 |
| Ipo5 | 6515.633211 | 0.020929028 | 0.074212876 | 0.282013437 | 0.777933207 | 0.890481986 | Tmem141 | 488.0438216 | -0.00708494201779967 | 0.0654866113717421 | -0.108189168280234 | 0.91384563698199 | 0.981708669866274 | NM_001040130,NM_001109993 |
| Lrriq1 | 89.36996211 | -0.046826666 | 0.165863823 | -0.28231995 | 0.77769819 | 0.890481986 | Tsc1 | 4187.291018 | 0.00617860198102823 | 0.0574152921183267 | 0.107612480108867 | 0.914303096789877 | 0.981708669866274 | NM_022887 |
| Parl | 770.8607167 | -0.021256311 | 0.075399011 | -0.281917643 | 0.778006661 | 0.890481986 | Xkrx | 83.54524752 | 0.0102461824369021 | 0.0950828187483336 | 0.107760608822734 | 0.914185590479837 | 0.981708669866274 | NM_183319 |
| Tfg | 1696.969997 | -0.020800307 | 0.073715003 | -0.282171965 | 0.777811654 | 0.890481986 | Zmym1 | 481.593906 | -0.00931452589682678 | 0.0862046748005991 | -0.108051285134737 | 0.913955010654718 | 0.981708669866274 | NM_026670 |
| Ccdc85a | 1705.205901 | -0.028826201 | 0.102409924 | -0.281478587 | 0.778343349 | 0.890668527 | Tube1 | 107.4196023 | -0.00984851003232829 | 0.092366512168456 | -0.106624249428914 | 0.915087079986565 | 0.981724883595631 | NM_028006 |
| Rnf17 | 24.30282666 | 0.047290321 | 0.168087401 | 0.281343639 | 0.778446842 | 0.890675607 | Dek | 1925.911067 | 0.00773162672815567 | 0.0728272434372273 | 0.106163934857975 | 0.915452285005951 | 0.982038598655695 | NM_025900 |
| Shroom1 | 292.2662869 | -0.029841284 | 0.106075715 | -0.281320606 | 0.778464507 | 0.890675607 | Ppp1r12a | 2907.392225 | -0.0069916587616253 | 0.0659077535873986 | -0.106082492287555 | 0.915516901887005 | 0.982038598655695 | NM_027892 |
| Palld | 87.26360503 | 0.037311905 | 0.132854023 | 0.280848894 | 0.7788263 | 0.890957966 | Rgs20 | 1474.378709 | 0.00827616244391705 | 0.0780769718315722 | 0.106000043928066 | 0.915582317334027 | 0.982038598655695 | NM_021374,NM_001177795 |
| Ppm1f | 3855.923264 | 0.019781586 | 0.070416823 | 0.280921304 | 0.77877076 | 0.890957966 | 1810013L24Rik | 1841.550695 | 0.00653659478785547 | 0.0647999571564887 | 0.100873443049814 | 0.919650924929444 | 0.982742917326017 | NM_001081400 |
| AI115009 | 80.29800702 | 0.042833063 | 0.152560279 | 0.280761569 | 0.778893282 | 0.890968808 | 9230112J17Rik | 37.72316541 | 0.0091458757617164 | 0.0906514736861147 | 0.100890535915439 | 0.919637356018433 | 0.982742917326017 | NR_040463,NR_040464 |
| Galnt18 | 1276.170404 | -0.022741001 | 0.081033612 | -0.280636647 | 0.778989105 | 0.890983874 | A230070E04Rik | 632.7449953 | -0.00686643369232277 | 0.0654828389284653 | -0.104858521784979 | 0.916488072280401 | 0.982742917326017 | NR_045897 |
| Lama1 | 177.1675749 | -0.03881344 | 0.138363909 | -0.280517085 | 0.779080819 | 0.890983874 | Akr1e1 | 1041.958752 | 0.00595118450451222 | 0.0594773019169029 | 0.100058077833234 | 0.920298217292153 | 0.982742917326017 | NM_018859 |
| Mill2 | 35.4384986 | -0.045517765 | 0.162436019 | -0.280219654 | 0.779308989 | 0.890983874 | Ap4s1 | 1209.96338 | -0.00745669468640328 | 0.0713507070420417 | -0.10450764954592 | 0.916766498062974 | 0.982742917326017 | NM_021710 |
| Mrps30 | 706.2789369 | -0.023091752 | 0.082416072 | -0.280185058 | 0.77933553 | 0.890983874 | Arl6ip4 | 1140.053383 | -0.00588001913437171 | 0.0584711500509689 | -0.100562741270629 | 0.919897574877995 | 0.982742917326017 | NM_144509 |
| Nipsnap1 | 2558.592551 | -0.014970008 | 0.053436698 | -0.280144698 | 0.779366493 | 0.890983874 | Asb2 | 119.4041281 | -0.00960797446915924 | 0.0944975002598707 | -0.101674377023065 | 0.919015140898462 | 0.982742917326017 | NM_023049 |
| Pigk | 2924.147807 | 0.013138905 | 0.04685452 | 0.280419166 | 0.779155935 | 0.890983874 | Atp6v0e | 615.6580127 | 0.00768430861315302 | 0.0749648183412923 | 0.10250553237078 | 0.918355421785822 | 0.982742917326017 | NM_025272 |
| Rrs1 | 620.4528669 | 0.01839026 | 0.065644993 | 0.280147185 | 0.779364585 | 0.890983874 | Cbfa2t2 | 2806.846059 | 0.0060966454819343 | 0.0602064569127471 | 0.101262319600865 | 0.919342226999252 | 0.982742917326017 | NM_009823,NM_172860 |
| Rtn1 | 37747.73286 | -0.019375575 | 0.069090361 | -0.280438174 | 0.779141353 | 0.890983874 | Ccbl2 | 245.1571875 | -0.00878743501405001 | 0.0875386120454302 | -0.100383531435129 | 0.920039843731936 | 0.982742917326017 | NM_173763 |
| Hmgn3 | 1100.811943 | 0.023432164 | 0.083674445 | 0.280039675 | 0.779447066 | 0.891010243 | Cdhr3 | 102.0095948 | -0.00973989805757473 | 0.0931920830790265 | -0.104514221978656 | 0.916761282581427 | 0.982742917326017 | NM_001024478 |
| Adat2 | 190.8223108 | 0.026897194 | 0.096151824 | 0.279736703 | 0.779679517 | 0.89114959 | Cspg5 | 11569.35997 | 0.00456745014709521 | 0.0437340601633041 | 0.104436910957735 | 0.916822632116565 | 0.982742917326017 | NM_013884,NM_001166273 |
| Lig4 | 504.5315982 | -0.02138826 | 0.076460137 | -0.279730865 | 0.779683997 | 0.89114959 | Cux1 | 3630.50548 | 0.00578542194638207 | 0.0554726130998057 | 0.104293301200234 | 0.916936593792094 | 0.982742917326017 | NM_198602,NM_009986 |
| Cntfr | 2746.717561 | 0.013828274 | 0.049557888 | 0.279032754 | 0.780219689 | 0.891299632 | Dbx2 | 565.1526707 | 0.00736626762821797 | 0.071444040550828 | 0.103105417490733 | 0.917879305405977 | 0.982742917326017 | NM_207533 |
| Dcaf17 | 561.4762993 | 0.027839182 | 0.099824824 | 0.278880349 | 0.78033665 | 0.891299632 | Dedd2 | 506.6341878 | -0.00770293738112003 | 0.0759388661134246 | -0.1014360336855 | 0.919204333233646 | 0.982742917326017 | NM_207677 |
| Fam132b | 25.52662574 | 0.046745028 | 0.167669639 | 0.278792443 | 0.780404114 | 0.891299632 | Dtx2 | 289.2863771 | 0.00871999672226203 | 0.0861847181157425 | 0.101177992025819 | 0.919409166872665 | 0.982742917326017 | NM_001256098,NM_023742,NM_001256097,NM_001256096 |
| Gad1 | 19339.09233 | -0.03235108 | 0.115992703 | -0.278906164 | 0.780316838 | 0.891299632 | Eral1 | 714.6234029 | -0.00643733896664239 | 0.0632579889099202 | -0.101763256745471 | 0.918944591065526 | 0.982742917326017 | NM_022313 |
| Isoc2b | 147.2186409 | 0.04069316 | 0.145821409 | 0.279061629 | 0.78019753 | 0.891299632 | Fat3 | 2981.612239 | 0.00865395005965291 | 0.0830716456756725 | 0.104174534996448 | 0.917030842128798 | 0.982742917326017 | NM_001080814 |
| Lrfn4 | 1316.480953 | -0.022984403 | 0.082355458 | -0.27908779 | 0.780177453 | 0.891299632 | Fign | 162.4140934 | 0.00931964500491441 | 0.0908439430894969 | 0.102589613439973 | 0.918288686616672 | 0.982742917326017 | NM_021716,NM_001267846,NM_001267847 |
| Mrpl24 | 1097.081974 | 0.02296467 | 0.082287299 | 0.279079161 | 0.780184075 | 0.891299632 | Fmnl1 | 5825.951168 | 0.00920134227543265 | 0.0904007456714271 | 0.101783920111412 | 0.918928189249094 | 0.982742917326017 | NM_001077698,NM_019679 |
| Ptpn12 | 1249.98716 | 0.026177429 | 0.093772698 | 0.27915832 | 0.780123328 | 0.891299632 | Fry | 6724.041142 | -0.00694419556369643 | 0.0674757298908287 | -0.102913974771843 | 0.918031246330813 | 0.982742917326017 | NM_172887 |
| Setd1b | 1984.839387 | -0.021854059 | 0.078314519 | -0.279055017 | 0.780202604 | 0.891299632 | Grrp1 | 57.03749983 | -0.00980079124402744 | 0.0945489677485978 | -0.103658363252441 | 0.917440469898889 | 0.982742917326017 | NM_001099296 |
| Srcin1 | 13775.71067 | -0.020315218 | 0.072883577 | -0.278735194 | 0.780448051 | 0.891299632 | Haus2 | 866.1103992 | 0.00670829890207111 | 0.0654523599808924 | 0.102491321994035 | 0.918366700622823 | 0.982742917326017 | NM_025475 |
| Vwc2l | 310.1385039 | -0.035906124 | 0.128705446 | -0.278979057 | 0.780260897 | 0.891299632 | Ippk | 790.5313762 | -0.00770521007631743 | 0.0760453491425387 | -0.101323883224927 | 0.919293357685022 | 0.982742917326017 | NR_075085,NM_001276399,NM_199056 |
| B230209E15Rik | 987.9569785 | 0.029120052 | 0.104536168 | 0.278564376 | 0.780579155 | 0.891305125 | Mcm8 | 228.7785018 | 0.00850912958155702 | 0.0822662623300384 | 0.103434012200771 | 0.917618519070799 | 0.982742917326017 | NM_025676 |
| Mnat1 | 379.8134247 | -0.028124889 | 0.100985557 | -0.278504073 | 0.780625439 | 0.891305125 | Mcu | 1151.10804 | 0.00641411900093281 | 0.0637429087525554 | 0.100624824415087 | 0.919848289688553 | 0.982742917326017 | NM_001033259 |
| Rd3 | 15.99540595 | -0.044133356 | 0.158456211 | -0.278520833 | 0.780612575 | 0.891305125 | Med8 | 866.1745062 | -0.00607871538057995 | 0.058309615255396 | -0.104248936542544 | 0.916971799758508 | 0.982742917326017 | NM_020000,NM_173719 |
| 5033404E19Rik | 15.44656618 | -0.046719549 | 0.167870615 | -0.278306893 | 0.780776785 | 0.89141224 | Ndufa3 | 2167.225618 | -0.00676571288674595 | 0.0668006658127457 | -0.10128211753541 | 0.919325651130371 | 0.982742917326017 | NM_025348 |
| Klhdc2 | 4523.563465 | 0.017986548 | 0.064718622 | 0.277919209 | 0.781074378 | 0.891554914 | Ngef | 17321.9276 | 0.00605019225823106 | 0.0587281961103285 | 0.103020229786472 | 0.917946915330279 | 0.982742917326017 | NM_019867,NM_001111314 |
| Osbpl10 | 1225.832075 | 0.031478805 | 0.113214773 | 0.278045032 | 0.780977791 | 0.891554914 | Nit1 | 1231.498558 | -0.00653703390361146 | 0.0634163419830361 | -0.103081220064067 | 0.917898509832923 | 0.982742917326017 | NM_001242580,NM_012049 |
| Rcan1 | 2503.90635 | 0.02593748 | 0.093306385 | 0.277981836 | 0.781026302 | 0.891554914 | Nol6 | 6319.547752 | 0.00489885391829718 | 0.0488879120671645 | 0.100205832304045 | 0.920180915813963 | 0.982742917326017 | NM_139236 |
| Gm13251 | 26.33329925 | -0.046710045 | 0.168121527 | -0.277835003 | 0.78113902 | 0.891563018 | Nrep | 2687.971195 | 0.00780810939237307 | 0.0758059845074866 | 0.103001226659116 | 0.917961997400046 | 0.982742917326017 | NM_001267717,NM_053078,NM_001109989,NM_001109988 |
| Aldh7a1 | 3120.493645 | -0.026565878 | 0.095735453 | -0.277492585 | 0.781401899 | 0.891666005 | Nsl1 | 68.51409678 | -0.00985085730939608 | 0.0948161829467552 | -0.103894261541069 | 0.917253261090688 | 0.982742917326017 | NM_198654 |
| Nsun2 | 2938.212673 | 0.019430205 | 0.069992962 | 0.277602265 | 0.781317693 | 0.891666005 | Pacrg | 417.2048372 | 0.00747283474146586 | 0.0741220265087199 | 0.100818003681898 | 0.919694934788174 | 0.982742917326017 | NM_027032 |
| Tdrd1 | 28.17556201 | -0.045070928 | 0.162389622 | -0.277548082 | 0.781359291 | 0.891666005 | Pcdhb17 | 533.7120901 | -0.00693986182980143 | 0.0675969481646657 | -0.102665312831816 | 0.918228604479697 | 0.982742917326017 | NM_053142 |
| Ndel1 | 3895.00482 | -0.023830187 | 0.085977988 | -0.277166143 | 0.781652536 | 0.891886323 | Pex12 | 300.6649488 | 0.00814328192645893 | 0.0813218700982969 | 0.100136429310047 | 0.920236014260773 | 0.982742917326017 | NM_134025 |
| Cope | 2528.779574 | -0.017523485 | 0.063295624 | -0.276851441 | 0.78189418 | 0.892042002 | Pld1 | 470.1117957 | -0.00718494174503015 | 0.0718034074075203 | -0.100064077798593 | 0.920293453918188 | 0.982742917326017 | NM_008875,NM_001164056 |
| Slc4a10 | 8714.103571 | 0.019312344 | 0.069760325 | 0.276838497 | 0.781904119 | 0.892042002 | Psmc3ip | 71.54118031 | -0.00958187574407103 | 0.0944920174358475 | -0.101404076281643 | 0.919229700751225 | 0.982742917326017 | NM_008949 |
| Thap11 | 1067.494443 | 0.020877111 | 0.075506101 | 0.276495682 | 0.782167376 | 0.89227664 | Ptk2b | 25393.54815 | -0.00637922424245372 | 0.0618034765787481 | -0.103217886688389 | 0.917790044208997 | 0.982742917326017 | NM_001162365,NM_172498,NM_001162366 |
| Atp7a | 422.9781785 | -0.030629624 | 0.11082083 | -0.276388691 | 0.782249542 | 0.892304675 | R3hcc1 | 737.7819684 | 0.00779999367987928 | 0.0762938724365053 | 0.102236174816932 | 0.918569214825335 | 0.982742917326017 | NM_001146012 |
| 2510039O18Rik | 931.4140059 | 0.020370448 | 0.07379391 | 0.276045114 | 0.782513416 | 0.892461248 | Rap1b | 1768.888707 | 0.00697538538977326 | 0.0686872921017738 | 0.101552778925072 | 0.919111662546485 | 0.982742917326017 | NM_024457 |
| Tssk4 | 14.31179391 | 0.046035621 | 0.166804809 | 0.275984977 | 0.782559605 | 0.892461248 | Rasal1 | 3960.031883 | 0.00790994949969239 | 0.0780591715852944 | 0.10133273693597 | 0.919286329618524 | 0.982742917326017 | NM_013832 |
| Vasp | 509.0329317 | 0.02547592 | 0.092265938 | 0.276114035 | 0.782460482 | 0.892461248 | Rgs10 | 1030.717594 | -0.00744339188408396 | 0.0726685954002245 | -0.102429279705893 | 0.918415944045546 | 0.982742917326017 | NM_026418 |
| Pdpr | 803.6551546 | 0.022809692 | 0.082686189 | 0.275858541 | 0.782656718 | 0.892506307 | Rgs4 | 22990.81809 | -0.00828787347730105 | 0.0803887943458958 | -0.103097372522346 | 0.917885690336125 | 0.982742917326017 | NM_009062 |
| H2-M3 | 117.089485 | -0.034741341 | 0.125977348 | -0.275774507 | 0.782721265 | 0.892514224 | Sfmbt1 | 1727.169171 | 0.00695596932949808 | 0.0695107278485687 | 0.100070443006321 | 0.920288400581342 | 0.982742917326017 | NM_001166531,NM_001166532,NM_019460 |
| Fam109a | 411.6674098 | 0.024045564 | 0.087339582 | 0.275311183 | 0.783077173 | 0.892809779 | Sh3bgrl3 | 3806.247994 | 0.00500695306520445 | 0.0493088909420247 | 0.101542601537954 | 0.919119741165877 | 0.982742917326017 | NM_080559 |
| Zfp786 | 193.0017087 | -0.031823069 | 0.115599584 | -0.275287057 | 0.783095708 | 0.892809779 | Shisa2 | 131.9794624 | 0.00979915023524229 | 0.0933810055302004 | 0.104937296183571 | 0.916425564257523 | 0.982742917326017 | NM_145463 |
| Blvra | 537.7480437 | -0.023260277 | 0.084523891 | -0.275191743 | 0.78316893 | 0.892827563 | Slc12a4 | 991.6542787 | 0.00790246854799097 | 0.0773319805530837 | 0.102188880867553 | 0.918606753334385 | 0.982742917326017 | NM_001253804,NR_045594,NM_009195 |
| Mbd5 | 2679.117595 | -0.028307745 | 0.10290354 | -0.275090102 | 0.783247016 | 0.892850888 | Slc22a6 | 332.1742669 | -0.00829569271474341 | 0.0815509928766509 | -0.101723994057203 | 0.918975756431853 | 0.982742917326017 | NM_008766 |
| Gm16894 | 34.25461438 | -0.045382331 | 0.165037607 | -0.274981754 | 0.783330257 | 0.892880085 | Slc38a4 | 46.00036715 | 0.0090057509509588 | 0.0894830220739772 | 0.100642007190074 | 0.919834649062728 | 0.982742917326017 | NM_027052 |
| Ajuba | 82.96797721 | 0.037989508 | 0.138483462 | 0.274325232 | 0.783834695 | 0.893282702 | Tm4sf1 | 353.308567 | -0.00821883262255861 | 0.0821265622603721 | -0.100075205832941 | 0.920284619376568 | 0.982742917326017 | NM_008536 |
| Gm16515 | 597.0403147 | -0.0199552 | 0.072730158 | -0.274373116 | 0.783797901 | 0.893282702 | Tmem161b | 650.7549454 | 0.00686719241789836 | 0.0669073416920575 | 0.102637352556985 | 0.918250796321745 | 0.982742917326017 | NR_029418,NR_029419,NM_175187 |
| Myo1g | 22.87287971 | 0.046037681 | 0.167838842 | 0.27429694 | 0.783856436 | 0.893282702 | Tsc22d2 | 1312.884419 | -0.00792854773554299 | 0.0767746965253866 | -0.103270323353493 | 0.917748428195335 | 0.982742917326017 | NM_001081229 |
| Hs3st1 | 658.363842 | -0.026559199 | 0.096855955 | -0.274213387 | 0.783920641 | 0.893290168 | Tubgcp3 | 1801.822961 | 0.00550781350846975 | 0.0537713363500294 | 0.10243028874373 | 0.918415143162251 | 0.982742917326017 | NM_198031 |
| Dlgap1 | 19578.01017 | -0.017585033 | 0.06416161 | -0.274074062 | 0.784027706 | 0.893346469 | Ubxn4 | 2709.824582 | -0.00586987157190894 | 0.0558989457292105 | -0.105008627539134 | 0.916368962785859 | 0.982742917326017 | NM_026390 |
| Thap7 | 590.8718038 | 0.021665473 | 0.079205091 | 0.273536368 | 0.78444094 | 0.893751594 | Uqcrq | 2238.614266 | -0.00534263205724728 | 0.052648268006703 | -0.101477831266303 | 0.919171154790738 | 0.982742917326017 | NM_025352 |
| Eda | 276.4629197 | -0.030135514 | 0.110304657 | -0.273202558 | 0.784697514 | 0.893758522 | Wwp1 | 2254.396443 | 0.00720400254044166 | 0.0689468536889871 | 0.104486313080192 | 0.916783429404841 | 0.982742917326017 | NM_177327,NM_001276292,NR_074089,NR_074090 |
| Ippk | 776.8478436 | 0.028593207 | 0.104658755 | 0.273204154 | 0.784696287 | 0.893758522 | Zc2hc1c | 132.7258129 | -0.00934018228458201 | 0.092434624115 | -0.101046359781392 | 0.919513658695494 | 0.982742917326017 | NM_172414 |
| Josd2 | 899.8417534 | 0.040030974 | 0.146515366 | 0.273220309 | 0.78468387 | 0.893758522 | Zfp786 | 200.4972397 | 0.0088283097681243 | 0.0869072440747908 | 0.101583128795648 | 0.919087571436894 | 0.982742917326017 | NM_177882 |
| Lrrc40 | 916.1603725 | 0.019681806 | 0.072054302 | 0.273152412 | 0.784736059 | 0.893758522 | Zfp954 | 460.6428495 | 0.00798566019501368 | 0.0781189600728562 | 0.102224353570068 | 0.918578597657256 | 0.982742917326017 | NM_172738 |
| Rpl30 | 8.556208069 | -0.043314411 | 0.158615427 | -0.273078172 | 0.784793125 | 0.893758522 | 6330403M23Rik | 1092.568648 | 0.00790966114001463 | 0.0795000956449778 | 0.0994924732586067 | 0.920747263719472 | 0.982762672073409 | NM_001267796 |
| Trim14 | 15.2114696 | 0.045611721 | 0.166921285 | 0.273252875 | 0.784658838 | 0.893758522 | Bfsp1 | 31.90639337 | 0.0081292873600207 | 0.0820408668246427 | 0.0990882675264335 | 0.921068187399579 | 0.982762672073409 | NM_009751 |
| Dazap1 | 1235.727374 | 0.020877227 | 0.076491626 | 0.272934798 | 0.784903337 | 0.893818337 | Brca1 | 88.21459617 | -0.00918847031816691 | 0.093363820843882 | -0.0984157485749366 | 0.921602169832299 | 0.982762672073409 | NM_009764 |
| Dbt | 780.5452209 | -0.018927367 | 0.069504889 | -0.272317051 | 0.785378247 | 0.894293419 | Cdk13 | 2274.56798 | 0.00736618739728969 | 0.0741184915595544 | 0.0993839356723948 | 0.920833437089566 | 0.982762672073409 | NM_027118,NM_001081058 |
| Entpd3 | 566.0440671 | -0.026299315 | 0.096688069 | -0.272001654 | 0.785620748 | 0.894503812 | Cep350 | 2463.311594 | -0.00781736170442117 | 0.0792063914998546 | -0.0986960970748873 | 0.921379567819488 | 0.982762672073409 | NM_001039184 |
| Guca1b | 19.17304051 | 0.045494177 | 0.167627148 | 0.271401009 | 0.786082626 | 0.894963936 | Crot | 1407.253973 | -0.00651043583905052 | 0.0651662258192493 | -0.0999050621269434 | 0.920419697464735 | 0.982762672073409 | NM_023733 |
| 4632427E13Rik | 12.35297793 | -0.043359733 | 0.159995362 | -0.271006189 | 0.786386272 | 0.895243858 | Gkn3 | 108.2477305 | 0.00903560783237198 | 0.0918095923040757 | 0.0984168168664317 | 0.921601321576972 | 0.982762672073409 | NM_026860 |
| Cpne1 | 1554.097881 | 0.017747336 | 0.065506244 | 0.27092587 | 0.786448047 | 0.895248405 | Gpr37l1 | 8218.292541 | 0.00572122020269789 | 0.0577695217787907 | 0.0990352702694235 | 0.921110266121186 | 0.982762672073409 | NM_134438 |
| 1700123M08Rik | 67.99566086 | -0.0364712 | 0.134717271 | -0.270724011 | 0.786603308 | 0.895359364 | Grasp | 1737.91751 | -0.00796529412621862 | 0.0798941039230513 | -0.099698147110959 | 0.920583971605715 | 0.982762672073409 | NM_019518 |
| Pigv | 246.8083063 | 0.027342101 | 0.101042158 | 0.270600926 | 0.786697983 | 0.895391868 | Il6st | 3903.252464 | -0.00649836601053945 | 0.0652384019410407 | -0.0996095216497233 | 0.920654334242365 | 0.982762672073409 | NM_010560 |
| Zfp846 | 392.1786724 | 0.029344886 | 0.108469182 | 0.270536626 | 0.786747443 | 0.895391868 | Lsm3 | 492.8204973 | 0.00695720353832707 | 0.0705314217995743 | 0.0986397744553769 | 0.921424288551094 | 0.982762672073409 | NM_026309 |
| Ska2 | 272.2157243 | -0.025847542 | 0.095609453 | -0.270345049 | 0.786894811 | 0.895493808 | Malsu1 | 215.5526816 | -0.00870794284330221 | 0.087962600633147 | -0.098995968520976 | 0.921141471035999 | 0.982762672073409 | NM_029353 |
| Fstl4 | 809.1844048 | 0.023367837 | 0.086517976 | 0.27009228 | 0.787089261 | 0.89564931 | Mrpl33 | 485.7496905 | 0.00773513777524683 | 0.0782598541948708 | 0.0988391539292415 | 0.921265980342921 | 0.982762672073409 | NM_025796 |
| Prom2 | 18.32913512 | 0.045055893 | 0.166904196 | 0.269950628 | 0.787198237 | 0.895707533 | Rab7l1 | 128.0619605 | -0.00930339626915268 | 0.093907109698056 | -0.0990702013837539 | 0.921082531516914 | 0.982762672073409 | NM_144875 |
| Cd300a | 58.37782299 | 0.039027132 | 0.144692528 | 0.269724587 | 0.787372143 | 0.895804021 | Rchy1 | 1256.626475 | -0.0059467884706709 | 0.0601623202080729 | -0.0988457301863323 | 0.921260758818022 | 0.982762672073409 | NR_073442,NM_001271797,NM_026557 |
| Hecw1 | 2700.168424 | -0.024387752 | 0.090428795 | -0.269690113 | 0.787398668 | 0.895804021 | Trp53i13 | 140.8800767 | 0.00935736869294031 | 0.0950470863537049 | 0.0984498215770459 | 0.921575114895788 | 0.982762672073409 | NM_001024920 |
| Ahi1 | 8857.041659 | 0.030570579 | 0.113464861 | 0.269427723 | 0.787600555 | 0.895932558 | Wdr4 | 906.7833943 | 0.00644659466279238 | 0.0650331453698399 | 0.0991278312948105 | 0.921036774733056 | 0.982762672073409 | NM_021322 |
| Cntnap1 | 11866.67093 | -0.021149221 | 0.078558474 | -0.269216286 | 0.787763248 | 0.895932558 | Wscd2 | 2396.084525 | 0.00747689905347707 | 0.0756307412423108 | 0.0988605184337068 | 0.921248966992557 | 0.982762672073409 | NM_177292 |
| Pi4k2b | 72.43232759 | 0.040240184 | 0.149416085 | 0.269316277 | 0.787686308 | 0.895932558 | Zfp963 | 161.8802626 | 0.00865176324084267 | 0.0870986766076182 | 0.099332889750083 | 0.920873965299175 | 0.982762672073409 | NM_001200023 |
| Tnfrsf1b | 234.8457863 | 0.029775882 | 0.110534531 | 0.269380819 | 0.787636645 | 0.895932558 | Ezh1 | 3365.78814 | -0.00456825389267947 | 0.0466129219119214 | -0.0980040234618101 | 0.921929098490033 | 0.982822698126098 | NM_007970 |
| Zfp39 | 476.6660527 | -0.026775409 | 0.099474888 | -0.269167521 | 0.787800772 | 0.895932558 | Med31 | 276.1023753 | -0.00814354203215717 | 0.0830472036781889 | -0.0980591961134984 | 0.921885288102872 | 0.982822698126098 | NM_026068 |
| Znf41-ps | 42.23565308 | -0.044237989 | 0.164458385 | -0.268991997 | 0.787935841 | 0.896020398 | Tspan15 | 599.8475083 | -0.00784202606212215 | 0.0799270013262566 | -0.0981148539541916 | 0.921841092686764 | 0.982822698126098 | NM_197996 |
| Ube2h | 5326.312223 | 0.013445897 | 0.050087268 | 0.268449398 | 0.78835342 | 0.896363682 | Ttl | 4101.575723 | 0.00587025843355048 | 0.0598890165692911 | 0.0980189485455758 | 0.921917247056976 | 0.982822698126098 | NM_027192 |
| Znrd1 | 440.343332 | 0.022391346 | 0.083390047 | 0.268513413 | 0.788304152 | 0.896363682 | Tmem63a | 1754.561001 | -0.00851816621031134 | 0.0870041027793464 | -0.0979053393828392 | 0.92200746014503 | 0.982834106264527 | NM_144794 |

| | | | | | | | | | | | | | | |
|---|---|---|---|---|---|---|---|---|---|---|---|---|---|---|
| Magel2 | 143.6801864 | 0.03961851 | 0.147864511 | 0.267937926 | 0.7887471 | 0.896745492 | | Cib1 | 387.0930004 | -0.00658843000876049 | 0.0674878478402533 | -0.0976239459340235 | 0.922230909222129 | 0.982928034878615 | NM_011870 |
| Creb3l4 | 11.23549125 | 0.043955628 | 0.164305323 | 0.267524065 | 0.789065688 | 0.896976066 | | Dscaml1 | 2779.667792 | -0.00639927515860577 | 0.0655356872330986 | -0.097645655806503 | 0.92221369614938 | 0.982928034878615 | NM_001081270 |
| Rpl6 | 9126.420504 | 0.018233499 | 0.068142323 | 0.267579653 | 0.789022895 | 0.896976066 | | A930016O22Rik | 32.53476756 | -0.0087749672812174 | 0.0899702054635971 | -0.0975319244410067 | 0.922303983036775 | 0.98293379728847 | NR_073015,NR_073014 |
| Pds5a | 1835.559765 | 0.022757815 | 0.085103977 | 0.26741188 | 0.789152053 | 0.89700843 | | Afmid | 141.7332787 | -0.008916470027662 | 0.092344775409133 | -0.0965563020556586 | 0.923078760186608 | 0.982966209989785 | NM_027827 |
| Trabd2b | 268.567508 | -0.04053713 | 0.151671312 | -0.267269591 | 0.789261598 | 0.897067136 | | Aldh1l1 | 3119.825443 | -0.00690846750469173 | 0.0714504228412387 | -0.0966889659987345 | 0.92297340262995 | 0.982966209989785 | NM_027406 |
| Nhlh2 | 15.12859802 | 0.042954127 | 0.160782349 | 0.267156978 | 0.789348299 | 0.897099872 | | Atg13 | 4909.815825 | 0.00372011742829323 | 0.0384035515152611 | 0.0968690988596433 | 0.922830348906204 | 0.982966209989785 | NM_145528 |
| Dner | 6628.236437 | -0.024096925 | 0.090352611 | -0.266698715 | 0.789701143 | 0.897435053 | | Cpe | 81611.37246 | 0.00504351802152769 | 0.0521646847262391 | 0.0966845299266379 | 0.922976925597913 | 0.982966209989785 | NM_013494 |
| 5031414D18Rik | 72.77979042 | 0.037618641 | 0.141197717 | 0.266425278 | 0.7899117 | 0.897568135 | | Dagla | 6605.384912 | -0.00737780086734955 | 0.0763918910128404 | -0.0965783248657825 | 0.923061270265744 | 0.982966209989785 | NM_198114 |
| Gcnt2 | 1771.547953 | -0.026611942 | 0.099952559 | -0.266245726 | 0.790049969 | 0.897568135 | | Eml3 | 481.1156758 | -0.00777405842689886 | 0.0802950976134514 | -0.0968185936372458 | 0.922870457716846 | 0.982966209989785 | NM_144872 |
| Plekhm3 | 2587.265647 | -0.022516267 | 0.084567366 | -0.266252437 | 0.790044801 | 0.897568135 | | Msln | 48.42781794 | 0.00772270572492 5 | 0.0799804500844894 | 0.0965574176785318 | 0.923077874188165 | 0.982966209989785 | NM_018857 |
| Prmt7 | 1457.117018 | 0.01831571 | 0.06877224 | 0.266324176 | 0.789989555 | 0.897568135 | | Nub1 | 4901.333861 | 0.00361161037089771 | 0.0372900571548131 | 0.0968518325381959 | 0.922844060963487 | 0.982966209989785 | NM_016736 |
| E330033B04Rik | 43.14432678 | 0.043079724 | 0.161927494 | 0.266043294 | 0.790205867 | 0.897679426 | | Ptprn | 27360.54396 | -0.00613528109250772 | 0.0635122402068506 | -0.0965999793508457 | 0.923044072894965 | 0.982966209989785 | NM_008985 |
| Nt5dc2 | 142.2024771 | 0.041921697 | 0.15775013 | 0.265747463 | 0.790433707 | 0.897872424 | | Rtn4rl2 | 894.8026959 | 0.00737731721788546 | 0.0762584337467499 | 0.096741001003212 | 0.922932078401018 | 0.982966209989785 | NM_199223 |
| Rab21 | 1524.996577 | 0.018047305 | 0.068017761 | 0.265332237 | 0.790753533 | 0.898169873 | | Ung | 77.58844354 | 0.00915093210694169 | 0.094018250751165 | 0.0973314439891161 | 0.922463185846828 | 0.982966209989785 | NM_011677,NM_001040691 |
| Cenpp | 20.92333691 | -0.044265274 | 0.16692753 | -0.265176595 | 0.790873424 | 0.898170716 | | 1700030L20Rik | 36.93925134 | -0.00849803350675543 | 0.0884014832902357 | -0.0961299877611224 | 0.923417334800758 | 0.98311053982086 | NR_040592 |
| Chst1 | 10613.64237 | 0.026885978 | 0.101444672 | 0.265030953 | 0.790985617 | 0.898170716 | | Tmem87a | 1307.516738 | 0.0061643993907197 | 0.0641016077228481 | 0.0961660652471044 | 0.923388681878411 | 0.98311053982086 | NM_001110496,NM_173734,NM_001110497 |
| Efhb | 10.095097 | 0.042613942 | 0.160788966 | 0.265030261 | 0.79098615 | 0.898170716 | | Tmub2 | 2659.444652 | 0.00524538494040344 | 0.0545325152345654 | 0.0961882084081582 | 0.923371095716825 | 0.98311053982086 | NM_028076 |
| Lysmd4 | 999.2058148 | -0.02020682 | 0.076241405 | -0.265037358 | 0.790980683 | 0.898170716 | | Tnrc6b | 3555.091005 | 0.00747880466531186 | 0.0779634499306744 | 0.0959270616162068 | 0.923578501624618 | 0.983210063363811 | NM_144812,NM_177124 |
| A330076H08Rik | 80.31321737 | 0.041396467 | 0.156361451 | 0.264748544 | 0.791203178 | 0.898211972 | | Ccdc114 | 115.2826705 | -0.00905868661957807 | 0.0946241844436948 | -0.0957333125018199 | 0.923732382848409 | 0.983229764289294 | NM_001033243 |
| Cry1 | 757.6069619 | -0.033695351 | 0.127297639 | -0.264697373 | 0.791242601 | 0.898211972 | | Daxx | 1021.624849 | 0.0060208408742357 6 | 0.0628655783910787 | 0.0957732518864438 | 0.923700661585124 | 0.983229764289294 | NM_007829,NM_001199733 |
| Ict1 | 929.3242243 | 0.016940804 | 0.064034156 | 0.264558873 | 0.791349306 | 0.898211972 | | Anp32b | 2022.265699 | -0.00520935572196716 | 0.0563858790048559 | -0.0923875944457393 | 0.926390095471324 | 0.983236500670538 | NM_130889 |
| Rnd2 | 2085.071817 | -0.020120338 | 0.076051455 | -0.264562173 | 0.791346763 | 0.898211972 | | Ap1s1 | 8898.84601 | -0.00693247390000304 | 0.0736996550328997 | -0.0940638581945652 | 0.925058430029212 | 0.983236500670538 | NM_007457 |
| Tceal8 | 915.333067 | -0.017550313 | 0.066287804 | -0.264759299 | 0.791194893 | 0.898211972 | | Arfip1 | 696.2051525 | 0.00666346856910507 | 0.071261276008492 | 0.0935075673962253 | 0.925500338124274 | 0.983236500670538 | NM_001081093 |
| Zfp759 | 264.2109664 | 0.02858673 | 0.108065461 | 0.264531607 | 0.791370313 | 0.898211972 | | Arx | 598.9354266 | -0.00643717549229788 | 0.0683434804832294 | -0.0941885816581653 | 0.924959354966758 | 0.983236500670538 | NM_007492 |
| Hmox2 | 2548.1047 | -0.016156485 | 0.061253954 | -0.263762326 | 0.791963066 | 0.898802261 | | Atp11a | 3442.988939 | 0.0057210501366312 6 | 0.06073218545886 | 0.0942012887138188 | 0.924949261082647 | 0.983236500670538 | NM_015804 |
| Ptcd1 | 1070.810211 | 0.021854635 | 0.082874976 | 0.263706085 | 0.792006407 | 0.898802261 | | Atp6v0a2 | 2101.520501 | 0.00443486998053219 | 0.0471730908000983 | 0.094012707357368 | 0.925099062434485 | 0.983236500670538 | NM_011596 |
| Cyth2 | 3027.412243 | -0.018298409 | 0.069461189 | -0.263433565 | 0.792216423 | 0.898898281 | | Atraid | 1825.743944 | -0.00557806552407406 | 0.0588535593202059 | -0.0947787285608608 | 0.924490582943956 | 0.983236500670538 | NM_027855 |
| Maea | 3095.757541 | -0.015422299 | 0.058557437 | -0.263370448 | 0.792265066 | 0.898898281 | | Baiap2l1 | 33.82250138 | 0.00735638537286915 | 0.0784790452620941 | 0.0937369376538827 | 0.925318127439606 | 0.983236500670538 | NM_025833 |
| Ssbp3 | 4795.561219 | 0.018747014 | 0.071142327 | 0.263514201 | 0.792154279 | 0.898898281 | | C2cd3 | 1854.783901 | -0.00561261638898337 | 0.0591158508648079 | -0.0949426643933261 | 0.924360368271869 | 0.983236500670538 | NM_001017985 |
| Lnp | 2270.636896 | 0.02603539 | 0.098917338 | 0.263203509 | 0.792393726 | 0.898978428 | | Camkv | 22608.97461 | 0.00581332125115801 | 0.0621056945433637 | 0.0936036750559003 | 0.925423990162316 | 0.983236500670538 | NM_145621 |
| Vmp1 | 3654.976116 | -0.019472616 | 0.07410662 | -0.262764865 | 0.792731818 | 0.899296148 | | Ccdc121 | 25.92657945 | 0.0075251010112816 | 0.0791479339908909 | 0.0950764048010105 | 0.924254139403112 | 0.983236500670538 | NM_207280 |
| Polr3gl | 628.0053753 | 0.020719883 | 0.078887467 | 0.262651142 | 0.792819478 | 0.899329745 | | Cmtm3 | 350.6673158 | 0.0078178114100208 | 0.0844793400687135 | 0.0925411041760267 | 0.926268134904108 | 0.983236500670538 | NM_024217 |
| Xrn1 | 733.6349894 | -0.027293602 | 0.104026468 | -0.262371703 | 0.793034886 | 0.899508238 | | Col14a1 | 87.66907545 | -0.00876911995165102 | 0.0946473810733084 | -0.0926504236272419 | 0.926181283730975 | 0.983236500670538 | NM_181277 |
| Cuedc1 | 1013.776411 | 0.023637883 | 0.09013445 | 0.262251367 | 0.793127654 | 0.899547608 | | Ctdspl | 1763.560267 | -0.0054399543765165 | 0.0575508337368974 | -0.094524336543691 | 0.924692651225676 | 0.983236500670538 | NM_133710 |
| 1110034G24Rik | 241.3123629 | 0.027853759 | 0.106393815 | 0.261798668 | 0.793476666 | 0.899577456 | | Ddx21 | 1248.518501 | -0.00604769098948288 | 0.0648318556793491 | -0.093282706874751 | 0.925678970088752 | 0.983236500670538 | NM_019553 |
| Adrb2 | 77.65653248 | 0.03476209 | 0.132812528 | 0.261738032 | 0.793523417 | 0.899577456 | | Dpm3 | 263.7164595 | -0.00887257258005292 | 0.0941403189935664 | -0.094248380236095 | 0.924911853909618 | 0.983236500670538 | NM_026767 |
| Dpp6 | 11489.48664 | -0.016651203 | 0.063566122 | -0.261950909 | 0.79335929 | 0.899577456 | | Ephb1 | 2013.756961 | -0.00636269175991412 | 0.0687440814176135 | -0.0925562118033318 | 0.926256132274544 | 0.983236500670538 | NM_173447,NM_001168296 |
| Eif3d | 2533.167786 | -0.01652823 | 0.06313186 | -0.261804895 | 0.793471865 | 0.899577456 | | Fam129c | 74.38265268 | -0.00832168772901945 | 0.0897164810838007 | -0.092755395981776 | 0.926097887029535 | 0.983236500670538 | NM_001166213 |
| Poll | 388.8964172 | -0.026884591 | 0.102796009 | -0.261533412 | 0.793681187 | 0.899577456 | | Fam78b | 1812.157363 | -0.00490427179268856 | 0.0517432138779909 | -0.0947809659495196 | 0.924488805722149 | 0.983236500670538 | NM_175461,NM_001160262,NM_001160261 |
| Tm7sf2 | 569.2492683 | 0.034528564 | 0.131990145 | 0.261599565 | 0.79363018 | 0.899577456 | | Fgfr1op2 | 3752.481763 | 0.00606660613423141 | 0.0696291001007009 | 0.0948828309525273 | 0.924407893902023 | 0.983236500670538 | NM_026218 |
| Txndc9 | 1551.095204 | -0.01941339 | 0.074107996 | -0.2619608 | 0.793351664 | 0.899577456 | | Gm16062 | 37.78092223 | 0.00784701007704967 | 0.0844573921093471 | 0.0929108735312373 | 0.92597436728777 | 0.983236500670538 | NR_045686 |
| Vamp2 | 48121.43284 | -0.011969191 | 0.045775287 | -0.261477138 | 0.793724577 | 0.899577456 | | Gpx7 | 103.0267194 | -0.00891826795894657 | 0.0937250625727624 | -0.0951535023198621 | 0.924192902113028 | 0.983236500670538 | NM_024198 |
| Vwa7 | 41.05197614 | 0.043776892 | 0.167425043 | 0.261471588 | 0.793728857 | 0.899577456 | | Lhx2 | 1633.697123 | 0.00684461954987152 | 0.0723783584138137 | 0.0945672118002223 | 0.924658594277008 | 0.983236500670538 | NM_010710 |
| Zcchc6 | 2626.625449 | 0.019331296 | 0.073934778 | 0.261464181 | 0.793734568 | 0.899577456 | | Lrrc38 | 101.4123467 | 0.00880448276969513 | 0.0947037026614075 | 0.0929687279617107 | 0.925928405066598 | 0.983236500670538 | NM_001162983 |
| Bcdin3d | 96.70941969 | -0.031976003 | 0.122481757 | -0.261067473 | 0.794040475 | 0.899794369 | | Mblac1 | 95.79886949 | 0.00871212696631927 | 0.0930826704255906 | 0.0935955847257698 | 0.92543041709723 | 0.983236500670538 | NM_177878 |
| Extl3 | 8191.856638 | 0.016141388 | 0.061840303 | 0.261017281 | 0.794079181 | 0.899794369 | | Med10 | 681.8334371 | 0.00632761383059033 | 0.0663553978157787 | 0.0953594438263721 | 0.924029328368215 | 0.983236500670538 | NM_138596 |
| Tmem88b | 2417.102433 | 0.042381573 | 0.162387697 | 0.26099005 | 0.794100181 | 0.899794369 | | Nptx1 | 19297.5259 | -0.00774364077654172 | 0.0835464070461954 | -0.0926867001265538 | 0.92615246328555 | 0.983236500670538 | NM_008730 |
| Chodl | 44.57409464 | 0.042111588 | 0.16149468 | 0.260761454 | 0.794276472 | 0.899928311 | | Olfml1 | 574.700076 | 0.00826700273697081 | 0.0886268446350215 | 0.093278766394263 | 0.92568210048836 | 0.983236500670538 | NM_172907 |
| 2210018M11Rik | 1964.574117 | 0.017925558 | 0.068764238 | 0.260681399 | 0.794338212 | 0.899932455 | | Rassf7 | 43.50245613 | 0.00857516529018809 | 0.0900279717385314 | 0.0952500109087537 | 0.92411624756273 | 0.983236500670538 | NM_025886 |
| Il1rapl2 | 143.6440427 | 0.034757001 | 0.133704315 | 0.259954222 | 0.794899086 | 0.900502042 | | Rhbdd1 | 389.8765516 | 0.00752004401308871 | 0.0805821283289891 | 0.0933214866501832 | 0.925648162690453 | 0.983236500670538 | NM_001122685,NM_029777 |
| Ndufs3 | 3573.91356 | -0.016146162 | 0.06215408 | -0.259776379 | 0.795036272 | 0.900528223 | | Rrp12 | 1764.319122 | 0.00560445890655159 | 0.0589116515678715 | 0.0951332844589283 | 0.924208960783709 | 0.983236500670538 | NM_199447 |
| Siglece | 38.91030855 | 0.041086077 | 0.15816111 | 0.259773572 | 0.795038438 | 0.900528223 | | Rxfp2 | 62.65206272 | -0.00880900471350848 | 0.0940660731155364 | -0.0936469911176688 | 0.925389580090614 | 0.983236500670538 | NM_080468 |
| Dek | 1760.98955 | 0.021167776 | 0.081592213 | 0.259433771 | 0.795300576 | 0.900638073 | | Slc38a9 | 954.1071972 | -0.00663574724621149 | 0.0715383798459028 | -0.0927578631289277 | 0.926095926981002 | 0.983236500670538 | NM_178746 |
| Iah1 | 302.8662738 | 0.022009542 | 0.08483482 | 0.259439957 | 0.795295804 | 0.900638073 | | Stard10 | 1909.920579 | 0.00539335686291768 | 0.0579231680479978 | 0.0931122561951117 | 0.925814380702672 | 0.983236500670538 | NM_019990 |
| Zdhhc9 | 2506.767678 | -0.018587327 | 0.071649052 | -0.259421809 | 0.795309805 | 0.900638073 | | Tfdp2 | 1790.74725 | -0.00618750309291453 | 0.0661735380237684 | -0.093504190310817 | 0.925503020894709 | 0.983236500670538 | NM_178667,NM_001184710,NM_001184706,NM_001184711,NM_001184709,NM_001184708 |
| Rbm14 | 1179.990966 | 0.01727389 | 0.066626544 | 0.259264381 | 0.79543126 | 0.900681926 | | Tgfb2 | 476.9701734 | -0.00807477689109823 | 0.0874679911297746 | -0.0923169354503391 | 0.926446233283605 | 0.983236500670538 | NM_009367 |
| Zfp869 | 1151.939719 | -0.019567954 | 0.075487557 | -0.259220919 | 0.795464791 | 0.900681926 | | Traf4 | 322.4643552 | -0.00815926867645178 | 0.0857347170906081 | -0.0951687828844028 | 0.924180765065486 | 0.983236500670538 | NM_009423 |
| Cnot4 | 1335.05416 | 0.02079997 | 0.080330411 | 0.258930207 | 0.795689091 | 0.900804236 | | Ucn3 | 28.53939441 | 0.00539424373940826 | 0.0584079508876297 | 0.0923546136687139 | 0.926416298298548 | 0.983236500670538 | NM_031250 |
| Socs1 | 21.6289523 | -0.0433299 | 0.167332917 | -0.258944268 | 0.795678242 | 0.900804236 | | Zfp646 | 1425.101555 | -0.00560326813787286 | 0.0595031179884406 | -0.0941676390632401 | 0.924975990800181 | 0.983236500670538 | NM_172749 |
| Pex6 | 2017.418013 | -0.030577602 | 0.118219747 | -0.258650546 | 0.79590488 | 0.90091781 | | Zfp97 | 58.34418934 | 0.0087500089487772 | 0.0942911850464272 | 0.0927977503355045 | 0.9260642382724 | 0.983236500670538 | NM_011765 |
| Psme3 | 4400.86656 | -0.01507503 | 0.058283627 | -0.258649477 | 0.795905704 | 0.90091781 | | Cpa2 | 47.28031565 | -0.00850258054352605 | 0.0923549970319456 | -0.0920641093257251 | 0.926647103919125 | 0.983377836903069 | NM_001024698 |
| Ccdc86 | 314.61713 | 0.025821007 | 0.099876944 | 0.258528206 | 0.795999283 | 0.900934512 | | Prkaa1 | 1285.024192 | -0.00554788123479965 | 0.0603584403476302 | -0.0919155830211487 | 0.926765110413295 | 0.983431221700893 | NM_001013367 |
| Gemin5 | 1196.74765 | 0.01592868 | 0.061624505 | 0.258476948 | 0.796036754 | 0.900934512 | | Bace2 | 171.1222896 | 0.00857215689072044 | 0.0950970864599685 | 0.0901411095736242 | 0.928175081455824 | 0.983501978429095 | NM_019517 |
| Cenpm | 60.43224162 | -0.040662624 | 0.157559368 | -0.25807811 | 0.796346625 | 0.901219386 | | Ccdc173 | 94.10101891 | -0.0085600063589284 | 0.0949766582416276 | -0.0901274766541305 | 0.928185916445303 | 0.983501978429095 | NM_001077684 |
| Dhx15 | 3385.917432 | 0.02240925 | 0.086833567 | 0.257975406 | 0.796425888 | 0.90124326 | | Cdc14a | 149.4671059 | 0.00788350444313608 | 0.0870167475217855 | 0.0905975535475211 | 0.927812375925613 | 0.983501978429095 | NM_001080818,NM_001173553 |
| Nudcd3 | 6683.59845 | 0.014681675 | 0.056937977 | 0.257853812 | 0.796519732 | 0.901283629 | | Clasp2 | 11216.05616 | 0.00495659352677016 | 0.0548042065291282 | 0.0904418445349766 | 0.927936105792896 | 0.983501978429095 | NM_001114347,NM_029633,NM_001081960 |
| Arfgap2 | 2877.693048 | 0.011207503 | 0.043543953 | 0.257383682 | 0.796882599 | 0.901493186 | | Dpm1 | 1407.069953 | 0.00599241646838029 | 0.0654173957977561 | 0.0916027976244483 | 0.927013628615372 | 0.983501978429095 | NM_010072 |
| Arsj | 101.7584186 | 0.039324169 | 0.152862332 | 0.257252182 | 0.796984103 | 0.901493186 | | Exoc6 | 1394.072023 | 0.00642849968168659 | 0.0709941691965485 | 0.0905496853394985 | 0.927850412889677 | 0.983501978429095 | NM_175353 |
| BC017612 | 54.75053544 | -0.041468195 | 0.161206203 | -0.257236968 | 0.796995848 | 0.901493186 | | Gnai2 | 7232.486432 | 0.00509520472841125 | 0.0559309111723615 | 0.0910981892054187 | 0.927414571466452 | 0.983501978429095 | NM_008138 |
| E430018J23Rik | 194.071015 | 0.029494142 | 0.114598264 | 0.257369077 | 0.796893872 | 0.901493186 | | Hs3st3b1 | 53.5085122 | -0.00865668974370342 | 0.0945211891819024 | -0.0915846469836933 | 0.927028050110427 | 0.983501978429095 | NM_018805 |
| Polm | 276.0519778 | 0.027912317 | 0.108444273 | 0.257388575 | 0.796878822 | 0.901493186 | | Hspa2 | 1398.296944 | -0.00682554921072218 | 0.0746960475185896 | -0.0913776489850217 | 0.927192520963631 | 0.983501978429095 | NM_008301,NM_001002012 |
| Chmp7 | 3678.19903 | -0.011082504 | 0.043159143 | -0.256782309 | 0.797346828 | 0.901824348 | | Huwe1 | 15499.57196 | 0.00657297278897755 | 0.072594329069185 | 0.0905438878388596 | 0.927855019702182 | 0.983501978429095 | NM_021523 |
| Echdc2 | 338.4211601 | 0.036500378 | 0.142244957 | 0.256602264 | 0.797485827 | 0.901915723 | | Ict1 | 1031.989255 | 0.00558070879739448 | 0.0618125828673022 | 0.0902843488901769 | 0.928061257130713 | 0.983501978429095 | NM_026729 |
| E230029C05Rik | 29.30619119 | 0.04291006 | 0.167325137 | 0.25644718 | 0.797605561 | 0.901985298 | | Ighmbp2 | 555.7961507 | -0.00618909723367673 | 0.0679206061234112 | -0.091122526533865 | 0.92739523351773 | 0.983501978429095 | NM_009212 |
| 4930579G24Rik | 57.1945659 | -0.03936036 | 0.153729779 | -0.256036013 | 0.79792303 | 0.902278457 | | Mpc2 | 1677.973958 | 0.006126684181808 | 0.0676016282947351 | 0.0911615381705347 | 0.927364235750097 | 0.983501978429095 | NM_027430 |
| Aaed1 | 193.5681617 | -0.027446668 | 0.107355124 | -0.255662391 | 0.798211538 | 0.902312383 | | Nedd4l | 7436.105284 | 0.00667036103288521 | 0.0730751916592257 | 0.0912807873839231 | 0.92726948369533 | 0.983501978429095 | NM_001114386,NM_031881 |
| Fbxw7 | 8382.801364 | -0.016151384 | 0.063141324 | -0.255797354 | 0.798107317 | 0.902312383 | | Nr3c1 | 4636.656443 | -0.00634206473776658 | 0.0698690285612152 | -0.0907707587806238 | 0.927674745220543 | 0.983501978429095 | NM_008173 |
| Myh4 | 22.2132619 | 0.042968763 | 0.168096195 | 0.255620079 | 0.798244213 | 0.902312383 | | Pfkfb1 | 30.58378175 | -0.00718397870842386 | 0.0796093857688822 | -0.0902403484091688 | 0.928096221711352 | 0.983501978429095 | NM_008824 |
| Ubap2 | 2014.848389 | 0.014360149 | 0.056147024 | 0.255759754 | 0.798136352 | 0.902312383 | | Pggt1b | 299.3668635 | 0.00727720420205337 | 0.0803313474613924 | 0.0905898435918909 | 0.927818502387914 | 0.983501978429095 | NM_172627 |
| Vps72 | 888.9115345 | 0.01636018 | 0.063981486 | 0.255701784 | 0.798181117 | 0.902312383 | | Pole4 | 962.9182221 | -0.00673774486445757 | 0.0739819215663013 | -0.0910728556627084 | 0.927434701032413 | 0.983501978429095 | NM_025882 |
| Spag7 | 1544.127887 | 0.013545848 | 0.053016735 | 0.255501365 | 0.79833589 | 0.90235018 | | Trip12 | 8214.272414 | -0.00576137024958379 | 0.0634101594884203 | -0.0908587881826084 | 0.927604796958323 | 0.983501978429095 | NM_133975 |
| Angptl1 | 32.38231405 | 0.042304591 | 0.165661392 | 0.255367833 | 0.798439013 | 0.902400909 | | Yif1a | 520.0652572 | 0.00739250626900892 | 0.0815754015442563 | 0.0906217576507832 | 0.927793142959965 | 0.983501978429095 | NM_026553 |
| Scamp5 | 19054.15493 | -0.011285818 | 0.044207569 | -0.255291529 | 0.798497942 | 0.902401686 | | Arhgap12 | 1678.59176 | 0.00649885498709495 | 0.0726786454083851 | 0.089419043937563 | 0.928748889222031 | 0.983596264569788 | NM_001039692,NM_029277 |
| Ubc | 5276.473282 | -0.022274722 | 0.087279673 | -0.255210881 | 0.798560228 | 0.902406255 | | Cped1 | 184.7211006 | -0.00852860402523227 | 0.0950593356938191 | -0.0897187421202104 | 0.928510721947552 | 0.983596264569788 | NM_001081351 |
| AU022252 | 561.5916079 | -0.021962725 | 0.086196332 | -0.254798832 | 0.798878477 | 0.902436799 | | Ntn4 | 289.8833626 | -0.00741826311471328 | 0.0827950607011957 | -0.0895978945107066 | 0.928606757615731 | 0.983596264569788 | NM_021320 |
| Dnajb13 | 12.90382355 | 0.041199476 | 0.161672786 | 0.254832472 | 0.798852494 | 0.902436799 | | Pdzd11 | 1121.799801 | -0.00587934394585521 | 0.0656935381032929 | -0.0894965336825192 | 0.928687308251456 | 0.983596264569788 | NM_028303 |
| Mospd2 | 347.0758614 | 0.020315138 | 0.079713638 | 0.254851479 | 0.798837813 | 0.902436799 | | Thbs1 | 107.9440572 | 0.00764458857531366 | 0.0854043522349494 | 0.0895105269844237 | 0.928676187843374 | 0.983596264569788 | NM_011580 |
| Spast | 1963.878183 | 0.018216461 | 0.071433175 | 0.255014023 | 0.798712269 | 0.902436799 | | Wbscr22 | 885.0188712 | 0.00636486617457508 | 0.0709589676338878 | 0.0896978406931532 | 0.928527331903898 | 0.983596264569788 | NM_025375 |
| Yif1a | 441.3166213 | -0.029719748 | 0.116556781 | -0.254980856 | 0.798737885 | 0.902436799 | | Zdhhc13 | 849.6433241 | 0.00727958259098255 | 0.0812485170413237 | 0.0895964979555268 | 0.928607867441943 | 0.983596264569788 | NM_028031 |
| BC068281 | 315.2953418 | 0.023221547 | 0.091283724 | 0.254388689 | 0.799195288 | 0.902499395 | | Anp32a | 7296.919244 | 0.00435407669972467 | 0.0487495088077477 | 0.0893152937580508 | 0.928831339984022 | 0.983611871736379 | NM_009672 |
| Eva1a | 307.9052506 | -0.024733133 | 0.097153023 | -0.254579135 | 0.799048176 | 0.902499395 | | Col8a2 | 168.5281401 | 0.00413010861077414 | 0.046322759249395 | 0.0891593825086766 | 0.928955244833259 | 0.983640460632105 | NM_199473 |
| Hdac8 | 440.9794588 | 0.020700658 | 0.08138649 | 0.254350056 | 0.799225132 | 0.902499395 | | Ddx50 | 2625.077459 | -0.00620797649334968 | 0.0697992116926821 | -0.0889404957850051 | 0.929129200078006 | 0.983640460632105 | NM_053183 |
| Lrrc17 | 251.7539354 | -0.034157285 | 0.134212402 | -0.254501705 | 0.799107987 | 0.902499395 | | Kcnh3 | 7377.584719 | -0.00618525017642861 | 0.0695205220377285 | -0.088970134215504 | 0.929105645411118 | 0.983640460632105 | NM_010601 |
| Lyn | 415.821327 | -0.029451314 | 0.115754644 | -0.254428786 | 0.799164314 | 0.902499395 | | Ppp1r16a | 3154.976095 | 0.00621016213214493 | 0.0696966005136577 | 0.0891027923092601 | 0.929000218281965 | 0.983640460632105 | NM_033371 |
| Lrrc28 | 848.8341289 | 0.017974266 | 0.070693934 | 0.254254715 | 0.799298783 | 0.902516786 | | Dpy19l4 | 672.3623002 | -0.00700581829186839 | 0.0791370415338247 | -0.0885276749810513 | 0.929457289231679 | 0.98368924828812 | NM_001081201 |
| Cog8 | 803.2138014 | 0.019063593 | 0.075071773 | 0.253938223 | 0.799543285 | 0.902575325 | | Ece2 | 1433.2077 | 0.00768603063342243 | 0.0867268863636815 | 0.0886233895161345 | 0.92938121906143 | 0.98368924828812 | NM_177942,NM_139293,NM_177941,NM_025462,NM_177940 |
| Fam229b | 261.6089568 | -0.02994843 | 0.11801689 | -0.253763933 | 0.799677939 | 0.902575325 | | Ikbkg | 1993.181154 | -0.00451729464537321 | 0.0510679955180599 | -0.0884564706240662 | 0.929513880077291 | 0.98368924828812 | NM_010547,NM_001161424,NM_001161423,NM_001161422,NM_001161421,NM_178590,NM_001136067 |
| Klhl18 | 1525.119764 | -0.015791033 | 0.062159328 | -0.254041235 | 0.799463702 | 0.902575325 | | Prr15 | 34.01909748 | -0.00799661092626463 | 0.0902471971154194 | -0.0886078555893244 | 0.929393564788261 | 0.98368924828812 | NM_030024 |
| Nsun4 | 482.1980431 | -0.019628913 | 0.077382833 | -0.253659792 | 0.799758399 | 0.902575325 | | Unc79 | 3242.497643 | 0.00610156763927996 | 0.0687758774887172 | 0.0887166818086898 | 0.929307074646434 | 0.98368924828812 | NM_001081017 |
| Nt5c1a | 130.7807405 | -0.029630497 | 0.116789331 | -0.253708937 | 0.799720429 | 0.902575325 | | Dnajc17 | 302.304977 | -0.00666949146302341 | 0.0754975239132841 | -0.0883405324747331 | 0.929606024610069 | 0.983715095322056 | NM_139139 |
| Tcf12 | 2004.053621 | -0.019747112 | 0.077773205 | -0.253906372 | 0.799567892 | 0.902575325 | | Agap3 | 10525.94679 | -0.00517568930117576 | 0.0586488340304981 | -0.0882488013058254 | 0.929678930752609 | 0.983720581884644 | NM_139153,NM_001256431 |
| Twf2 | 1461.227018 | 0.021322782 | 0.084005724 | 0.253825346 | 0.799630491 | 0.902575325 | | Sorcs3 | 2301.781588 | -0.00579954369068434 | 0.0658652219571996 | -0.0880516837011949 | 0.929835597973175 | 0.98381469124906 | NM_025696 |
| AA465934 | 36.49250901 | 0.042069891 | 0.166010957 | 0.253416352 | 0.799946493 | 0.902665401 | | Zdhhc17 | 5677.587165 | -0.0056226847544924 | 0.0639865411840416 | -0.0878729284384996 | 0.929977673328079 | 0.983893349072372 | NM_172554 |
| BC051226 | 95.50094421 | 0.03192256 | 0.125974126 | 0.253405689 | 0.799954732 | 0.902665401 | | Exoc3l4 | 79.95728257 | 0.00832305976761995 | 0.0949092901034245 | 0.0876948901266688 | 0.930119181066039 | 0.983918325794051 | NM_028807 |
| Ccdc162 | 26.54197927 | -0.042429274 | 0.167569588 | -0.25320659 | 0.800108575 | 0.902703835 | | Wdr96 | 64.28653905 | 0.00756848440520519 | 0.0863264976682458 | 0.0876727842509146 | 0.930136751328921 | 0.983918325794051 | NM_027559 |
| Nrsn2 | 3560.574223 | -0.023647204 | 0.093500758 | -0.252909224 | 0.800338362 | 0.902703835 | | Sorcs2 | 2725.897011 | 0.00542684071392467 | 0.0620353575220078 | 0.0874798007249242 | 0.930290140557717 | 0.983937278414256 | NM_030889 |
| Phgdh | 2506.51496 | -0.025155296 | 0.099460788 | -0.252916717 | 0.800332571 | 0.902703835 | | Srgap1 | 1164.257432 | 0.00733639432546783 | 0.0837841014120648 | 0.0875630841868932 | 0.930223943988353 | 0.983937278414256 | NM_001242411,NM_001081037 |
| Rassf2 | 3021.817833 | -0.01748886 | 0.069140323 | -0.252947334 | 0.800308911 | 0.902703835 | | 2810006K23Rik | 477.253085 | -0.00626122870477665 | 0.0721461812122271 | -0.0867853100499711 | 0.93084216445325 | 0.984208871023001 | NM_028310,NM_001134717 |
| Rb1 | 1193.018664 | -0.021210654 | 0.08377548 | -0.253184511 | 0.800125635 | 0.902703835 | | Aim1l | 58.45100518 | 0.00822547647542896 | 0.094996772806528 | 0.0865869042960133 | 0.930999875660148 | 0.984208871023001 | NM_001162970 |
| Smyd1 | 91.45201895 | -0.042465103 | 0.167812016 | -0.253051622 | 0.800228322 | 0.902703835 | | Aspa | 674.0575175 | -0.00752458624815737 | 0.0873247733471847 | -0.0861678302701252 | 0.931333003291001 | 0.984208871023001 | NM_023113 |
| Pick1 | 1636.407717 | -0.01567441 | 0.062063189 | -0.252555659 | 0.800611598 | 0.902893264 | | Cdk5rap3 | 1017.936141 | 0.00561008540427505 | 0.0650053269251877 | 0.0863019335435617 | 0.931226401467288 | 0.984208871023001 | NM_030248 |
| Tor1a | 923.1722241 | 0.019891716 | 0.078766257 | 0.252541091 | 0.800622857 | 0.902893264 | | Chst9 | 22.3502679 | -0.00660912279849547 | 0.0768068116269444 | -0.086048654520336 | 0.931427739911414 | 0.984208871023001 | NM_199055 |
| Fam129c | 61.2611326 | 0.041966813 | 0.166312257 | 0.252337464 | 0.800780233 | 0.903005016 | | Commd7 | 970.3313391 | -0.00474680451446179 | 0.0547975960199051 | -0.086624320394229 | 0.9309701336828 | 0.984208871023001 | NM_133850,NM_001195390 |
| Dnajb12 | 1329.805461 | -0.020525529 | 0.081533691 | -0.25174292 | 0.80123978 | 0.903335626 | | Echs1 | 1836.231642 | 0.00507229822405629 | 0.0586649920634966 | 0.0864620964524158 | 0.931090085551131 | 0.984208871023001 | NM_053119 |
| Gpc5 | 1535.771489 | 0.021982806 | 0.087311852 | 0.251773447 | 0.801216182 | 0.903335626 | | Gpnmb | 230.4196012 | 0.00615536158133193 | 0.0714829563046856 | 0.0861094993762654 | 0.931379372265255 | 0.984208871023001 | NM_053110 |
| Impdh1 | 2298.052411 | -0.021363981 | 0.084868005 | -0.251731869 | 0.801248322 | 0.903335626 | | Kifc1 | 46.64749451 | 0.0080236056906759 | 0.092766242518899 | 0.0864927313299477 | 0.931074739780 95 | 0.984208871023001 | NM_001195298 |
| Golim4 | 943.5162319 | -0.019794654 | 0.078742338 | -0.251385149 | 0.801516349 | 0.903572054 | | L1cam | 7584.437566 | -0.00502160526959718 | 0.0583561249856889 | -0.0860510404148436 | 0.931425843277921 | 0.984208871023001 | NM_008478 |
| Gins1 | 26.87842405 | 0.041720872 | 0.166053174 | 0.25125071 | 0.801620775 | 0.903624029 | | Nck1 | 764.9875855 | -0.00664497062606204 | 0.0765458420491235 | -0.0868103406801831 | 0.930822267981359 | 0.984208871023001 | NM_010878 |
| A4gnt | 8.878031344 | -0.039058858 | 0.155601257 | -0.251018913 | 0.801799487 | 0.903659159 | | Tma7 | 2248.119689 | 0.00575370269524936 | 0.066632610466718 | 0.0863496515437175 | 0.931188469612132 | 0.984208871023001 | NM_183250 |
| Cdh18 | 896.5594038 | -0.023279017 | 0.092755445 | -0.250971968 | 0.801835782 | 0.903659159 | | Zfp52 | 178.7123271 | 0.00794043087206558 | 0.0917522662806558 | 0.0865420680485106 | 0.931035516025383 | 0.984208871023001 | NM_144515 |
| Snhg8 | 336.7867829 | -0.028925283 | 0.115199979 | -0.251087571 | 0.801746405 | 0.903659159 | | Cdkl2 | 1634.048465 | 0.00593687696712614 | 0.0691434985527672 | 0.08586312651283 | 0.931575223961028 | 0.984221521802159 | NM_001276315,NM_016912,NM_177270 |
| Gm2a | 2334.528424 | -0.016786299 | 0.066972518 | -0.250644585 | 0.802088908 | 0.903888686 | | Hmgxb3 | 2623.005012 | 0.0047291028278487 | 0.0550496305963917 | 0.0859061682451813 | 0.931541008090051 | 0.984221521802159 | NM_178277,NM_134134 |
| Nop9 | 415.5390289 | 0.022137514 | 0.088367587 | 0.250516215 | 0.802188166 | 0.903934796 | | Mbnl1 | 3066.142725 | -0.0061561388186659 | 0.0718358426725145 | -0.085697314917446 | 0.931707366663134 | 0.984289193786814 | NM_001253713,NM_001253708,NM_001253709,NM_001253711,NM_001253710,NM_020007 |
| Cryl1 | 464.2802902 | 0.02511184 | 0.100321941 | 0.250312545 | 0.802345656 | 0.903980773 | | Brdt | 509.7457967 | 0.00680245629925258 | 0.0799940975983973 | 0.0850369777705808 | 0.932231993155597 | 0.984387567574209 | NM_054054,NM_001079873 |
| Polr3d | 684.2946357 | 0.015451801 | 0.061714412 | 0.250375897 | 0.802296668 | 0.903980773 | | Chd9 | 2653.391874 | 0.00618789457268466 | 0.073346201046039 | 0.0843655770092382 | 0.93276577525067 | 0.984387567574209 | NM_177224 |
| 2210404O09Rik | 153.6185303 | 0.033999007 | 0.135953232 | 0.250078694 | 0.802526492 | 0.904053037 | | Creg1 | 1178.401181 | 0.00571796095022404 | 0.0675173135126574 | 0.0846888102138735 | 0.932508792162412 | 0.984387567574209 | NM_011804 |
| Arl6ip5 | 4772.529839 | -0.013447536 | 0.05375837 | -0.250147766 | 0.802473078 | 0.904053037 | | Cys1 | 291.7728167 | -0.00790878858047078 | 0.0928070907263855 | -0.0852175035179966 | 0.932088475403574 | 0.984387567574209 | NM_001004455,NM_001161807,NM_138686 |
| Ryk | 1144.263596 | -0.015655842 | 0.062634724 | -0.249954677 | 0.802622399 | 0.904095344 | | Pcsk5 | 458.7280607 | -0.00669192809349282 | 0.0793765446336122 | -0.0843061253007875 | 0.932813042456852 | 0.984387567574209 | NM_001163144,NM_001190483 |
| Crispld2 | 123.8667687 | 0.036515541 | 0.14614793 | 0.249853287 | 0.802700809 | 0.904117937 | | Pik3r3 | 2766.769062 | -0.005937891656863 | 0.0705790040700984 | -0.0841310988150081 | 0.93295219900827 | 0.984387567574209 | NM_181585 |
| Plod2 | 164.0105132 | 0.034370283 | 0.137656364 | 0.249681758 | 0.802833467 | 0.904201623 | | Ppif | 1024.156637 | -0.00476328485597676 | 0.0562346780316311 | -0.0847036921469658 | 0.932496960610486 | 0.984387567574209 | NM_134084 |
| Fra10ac1 | 415.5731097 | -0.029349042 | 0.117605542 | -0.249554926 | 0.80293156 | 0.904246372 | | Prkcsh | 3163.912858 | 0.00478078695399463 | 0.056604142704405 | 0.0844600187474014 | 0.932690689635074 | 0.984387567574209 | NM_008925 |
| Hddc3 | 354.1106827 | 0.02265633 | 0.090965157 | 0.249066024 | 0.80330971 | 0.90453926 | | Rarb | 871.5478667 | -0.00651826405783037 | 0.0763212983213848 | -0.0854055709375163 | 0.931938964379505 | 0.984387567574209 | NM_011243 |
| Pcca | 1025.357392 | -0.017338339 | 0.069622287 | -0.249034318 | 0.803334235 | 0.90453926 | | Rps28 | 2111.927665 | -0.00687708485874123 | 0.0811274142665486 | -0.0847689393396194 | 0.932445087448521 | 0.984387567574209 | NM_016844 |
| Ube2t | 36.89201971 | -0.041783996 | 0.167812432 | -0.248992256 | 0.803366772 | 0.90453926 | | Slc12a2 | 3430.075009 | -0.00522808526066231 | 0.0616035847822226 | -0.0848666389029606 | 0.932367414299944 | 0.984387567574209 | NM_009194 |
| Lrrc51 | 87.81519744 | -0.040922811 | 0.164552793 | -0.248691077 | 0.803599752 | 0.904735834 | | Sp7 | 66.82767682 | -0.00786838522345039 | 0.0923858177661289 | -0.0851687565675415 | 0.93212722895457 | 0.984387567574209 | NM_130458 |
| Ift43 | 224.2838269 | -0.022297555 | 0.089701428 | -0.248575248 | 0.803689357 | 0.904770972 | | Synj2bp | 2114.156788 | 0.00585828187986062 | 0.0692753704313701 | 0.0845651469401281 | 0.932607108490843 | 0.984387567574209 | NM_025292 |
| Rtn3 | 47567.32987 | -0.013424615 | 0.054029055 | -0.248470293 | 0.803770552 | 0.904796639 | | Usp25 | 1541.275454 | -0.00498520943283529 | 0.0592136607097273 | -0.0841901914707385 | 0.932905216574217 | 0.984387567574209 | NM_013918 |
| Lmbr1 | 1351.937033 | -0.015728532 | 0.063392639 | -0.24811291 | 0.804047047 | 0.904978122 | | Ybey | 271.0944102 | -0.00680571273812389 | 0.0797649063617137 | -0.0853221428890257 | 0.932005288239963 | 0.984387567574209 | NM_172550 |
| Snrpb | 2121.638785 | 0.024755684 | 0.099776681 | 0.248110918 | 0.804048588 | 0.904978122 | | Zfp869 | 1279.975548 | -0.00475907482554298 | 0.0559883661340858 | -0.0850011378104074 | 0.932260486143089 | 0.984387567574209 | NM_181274,NM_001039965,NM_001039967 |
| Cep89 | 496.3590553 | 0.023484967 | 0.094721008 | 0.247938311 | 0.804182137 | 0.905062688 | | Zfp960 | 44.76405434 | -0.00761632577528068 | 0.0903747636462953 | -0.0842749177755984 | 0.932837854160397 | 0.984387567574209 | NM_001005358,NM_001163919 |
| Dync2li1 | 615.5838307 | -0.020819446 | 0.084151278 | -0.247404989 | 0.804594814 | 0.90542198 | | Rdh13 | 904.8739876 | 0.00460682619049433 | 0.0548223275371474 | 0.0840319336564607 | 0.933031042165264 | 0.984399253231597 | NM_175372 |
| Tsc22d2 | 1260.71676 | -0.021040171 | 0.085053852 | -0.247374697 | 0.804618255 | 0.90542198 | | Dstyk | 3392.921403 | 0.00497503023188153 | 0.0594454943203739 | 0.083690619259876 | 0.933302415725388 | 0.984471054449712 | NM_172516 |
| B230319C09Rik | 13.20432393 | -0.04096216 | 0.165836757 | -0.247002903 | 0.804905978 | 0.905484189 | | Prrc2a | 21973.51427 | -0.00423418329273836 | 0.050502022543195 | -0.0838418558210575 | 0.933182169021705 | 0.984471054449712 | NM_020027,NM_001199044 |
| Ccdc126 | 333.5876204 | -0.025793365 | 0.10442104 | -0.247013105 | 0.804898082 | 0.905484189 | | Rps4x | 9600.625649 | -0.00420751166266281 | 0.0502431105027145 | -0.0837430569199232 | 0.933260722884649 | 0.984471054449712 | NM_009094 |
| Ppp1r3e | 258.2518698 | 0.023143751 | 0.093698943 | 0.247001192 | 0.804907301 | 0.905484189 | | AI987944 | 316.6186107 | 0.00640562347070876 | 0.0766358390202914 | 0.0835852200823782 | 0.933386218475837 | 0.984481307127396 | NM_001199330,NM_183167 |
| Zmat2 | 7168.934951 | -0.014607992 | 0.05914127 | -0.247001658 | 0.804906941 | 0.905484189 | | Adpgk | 625.4500734 | -0.00625018782044205 | 0.0749219271246588 | -0.0834226782506899 | 0.933515456723995 | 0.984481307127396 | NM_028121 |
| Bckdha | 1363.433234 | 0.017454409 | 0.070781905 | 0.246594219 | 0.805222279 | 0.905535571 | | Wnt7b | 1236.344943 | -0.00579018998561523 | 0.0693824467630105 | -0.0834532400593015 | 0.933491156663359 | 0.984481307127396 | NM_001163634,NM_001163633,NM_009528 |
| Blnk | 399.808407 | 0.034324968 | 0.139213767 | 0.246563033 | 0.805246417 | 0.905535571 | | Henmt1 | 55.47743679 | 0.00782193671914148 | 0.0939996543933783 | 0.083212398701037 | 0.93368265419479 | 0.984586151349076 | NM_001078646,NM_025723 |
| Epn1 | 14098.95882 | 0.023637474 | 0.095896629 | 0.246489099 | 0.805303642 | 0.905535571 | | Osgin2 | 665.5196933 | 0.00650838493151631 | 0.0786915129436819 | 0.0827075841860386 | 0.934080454228961 | 0.984937933487942 | NM_145950 |
| Hcn2 | 4748.070076 | 0.022103975 | 0.089559807 | 0.246806861 | 0.8050577 | 0.905535571 | | Bola1 | 215.6643379 | -0.00717385184919661 | 0.0875485154694836 | -0.0819414448174986 | 0.934693277035188 | 0.985294233832594 | NM_026975 |
| Ier5 | 2676.015341 | -0.031114767 | 0.126087062 | -0.246772081 | 0.805084618 | 0.905535571 | | Nit2 | 322.2546995 | -0.00638253437876181 | 0.0777636675149233 | -0.0820760463430684 | 0.934586241093609 | 0.985294233832594 | NM_023175 |
| Mylpf | 19.17409249 | 0.041350805 | 0.167647144 | 0.246653798 | 0.805176166 | 0.905535571 | | Pclo | 9072.322961 | -0.00639092536286248 | 0.077769349776967 | -0.082169930773116 | 0.934511584330859 | 0.985294233832594 | NM_011995,NM_001110796 |
| Hsf5 | 15.43381094 | 0.040253905 | 0.163366625 | 0.246402255 | 0.805370861 | 0.905543596 | | Stmn4 | 9551.004108 | -0.00439282589400866 | 0.0535541356665072 | -0.0820258947201333 | 0.934626121827279 | 0.985294233832594 | NM_019675 |
| Wee1 | 978.6111253 | 0.020445547 | 0.08300102 | 0.246328863 | 0.805427669 | 0.905543596 | | 6720489N17Rik | 105.2280976 | 0.00772281528038137 | 0.0944641700303094 | 0.0817539102699305 | 0.9348424076016 | 0.985379929817926 | NM_173381 |
| Proca1 | 92.43109062 | 0.038367841 | 0.15582008 | 0.246231684 | 0.80550289 | 0.905562456 | | 1110032A03Rik | 1138.980879 | -0.00468603152236414 | 0.0640054114919426 | -0.0732130520394209 | 0.941636580236901 | 0.98541611256672 | NM_023483 |
| Pigq | 4166.684369 | -0.027546348 | 0.111912193 | -0.246142508 | 0.805571918 | 0.905574353 | | 1700052N19Rik | 1001.534053 | 0.00483909219193053 | 0.0639035844066219 | 0.0757248965647047 | 0.939637962864651 | 0.98541611256672 | NM_024261 |
| 5-Mar | 2731.516544 | -0.01481606 | 0.060214912 | -0.246053009 | 0.805641198 | 0.905583862 | | 1810011H11Rik | 23.44506111 | 0.00570310983824403 | 0.076920445862425 | 0.0741429638674254 | 0.940896629094565 | 0.98541611256672 | NM_001163616 |
| Osbpl9 | 1747.035349 | 0.018888632 | 0.076789122 | 0.245980571 | 0.805697272 | 0.905583862 | | 3110001I22Rik | 57.75753143 | 0.00711119058758734 | 0.0906194916790962 | 0.0784730796412944 | 0.937451743506723 | 0.98541611256672 | NM_025653 |
| Fbxw8 | 1947.195259 | -0.015298292 | 0.062226343 | -0.245849118 | 0.805799033 | 0.905632542 | | 3230401D17Rik | 719.853454 | 0.00551791835476013 | 0.0688375036042347 | 0.080158606367891 | 0.93611111512051 | 0.98541611256672 | NM_025699 |
| Tlcd2 | 10.17774983 | 0.038793143 | 0.157872424 | 0.245724633 | 0.805895402 | 0.905675155 | | Abhd3 | 2111.800781 | 0.00361696026388377 | 0.0451939618857454 | 0.0800319359702915 | 0.936211856196418 | 0.98541611256672 | NM_134130 |
| Enpp4 | 1379.939745 | -0.019523561 | 0.079573938 | -0.245351201 | 0.80618451 | 0.9057368 | | Acot8 | 286.4455354 | -0.0065881265884968 | 0.0906415194209487 | -0.0726833202993966 | 0.94205821911343 | 0.98541611256672 | NM_133240 |
| Gnao1 | 28590.1635 | 0.015708462 | 0.063988955 | 0.245487082 | 0.806079309 | 0.9057368 | | Agbl2 | 64.25407257 | -0.00731488778491018 | 0.0940059917193393 | -0.0778129952264025 | 0.937976809097896 | 0.98541611256672 | NM_178755 |
| Nop16 | 606.0944139 | -0.015899021 | 0.064797958 | -0.245363 | 0.806175375 | 0.9057368 | | Akt1s1 | 1318.408652 | 0.00537019429281561 | 0.0711542176946788 | 0.0754726067800928 | 0.939838691982583 | 0.98541611256672 | NM_001253920,NM_026270 |
| Prkag2 | 3259.603873 | -0.013320769 | 0.054300156 | -0.245317322 | 0.80621074 | 0.9057368 | | Alpk3 | 24.93399999 | -0.00618543670156269 | 0.0786050655989859 | -0.0786900634766415 | 0.937279149168887 | 0.98541611256672 | NM_054085 |
| Trio | 7767.073632 | -0.0234605 | 0.095649293 | -0.245276247 | 0.806242542 | 0.9057368 | | Anxa1 | 59.9237341 | -0.00678334043665856 | 0.0911332027365119 | -0.0744332497154833 | 0.94066565272641 | 0.98541611256672 | NM_010730 |
| Csgalnact2 | 462.2501273 | -0.021322509 | 0.087118622 | -0.244752595 | 0.806648002 | 0.906126596 | | Ap3b1 | 2269.20709 | 0.00372491476103522 | 0.0479206534021208 | 0.077730884213522 | 0.938042126373388 | 0.98541611256672 | NM_009680 |
| Sh2b1 | 3208.352808 | 0.01631594 | 0.066732215 | 0.244498697 | 0.806844611 | 0.906281746 | | Arhgap28 | 26.82185867 | 0.00613323229005549 | 0.0854277001920362 | 0.071794421203759 | 0.942765512599517 | 0.98541611256672 | NM_172964 |
| Lysmd3 | 247.8475825 | 0.030085765 | 0.12337949 | 0.243847381 | 0.807349023 | 0.906773168 | | Arhgap39 | 3216.402141 | -0.00557732324640296 | 0.0764748622875442 | -0.0726686270516527 | 0.942069814507096 | 0.98541611256672 | NM_001168288,NM_198420 |
| Maml2 | 429.1354292 | -0.025290163 | 0.103740621 | -0.243782644 | 0.807399163 | 0.906773168 | | Arhgef37 | 121.8525847 | -0.00703896298642424 | 0.0904392052495256 | -0.077830880612047 | 0.93796258177227 | 0.98541611256672 | NM_177828 |
| Nbeal1 | 594.7405356 | 0.026996131 | 0.110901803 | 0.243423731 | 0.807677162 | 0.90698013 | | Atp1a1 | 52519.46437 | 0.00518167856215672 | 0.0662682835611635 | 0.0781924366182533 | 0.937674978302782 | 0.98541611256672 | NM_144900 |
| Vcp | 13642.10763 | 0.01182067 | 0.048566072 | 0.243393578 | 0.807700519 | 0.90698013 | | Bag1 | 3709.71144 | -0.0040897319982875 | 0.0556931047681126 | -0.0734333633457098 | 0.941461269143258 | 0.98541611256672 | NM_009736,NM_001171739 |
| Mef2a | 3269.672315 | -0.018947969 | 0.077897149 | -0.243243426 | 0.807816828 | 0.907044998 | | Brap | 3024.454239 | 0.00435124291275 | 0.056058413450172 | 0.0776198012921028 | 0.938130490747412 | 0.98541611256672 | NM_028227 |
| Asb16 | 29.41225114 | 0.040242204 | 0.165565354 | 0.24305933 | 0.807959437 | 0.907139385 | | Brd2 | 8777.746503 | 0.00348586406830758 | 0.0448991394456572 | 0.077637658791359 | 0.938116285391309 | 0.98541611256672 | NM_010238,NM_001204973,NR_037970 |
| Ddx25 | 1680.872346 | 0.01705652 | 0.07025461 | 0.242781507 | 0.808174662 | 0.907315283 | | C1qtnf7 | 28.41916252 | -0.00634092567568592 | 0.0821847285000227 | -0.077154549165228 | 0.938500598326594 | 0.98541611256672 | NM_001135172,NM_175425 |
| D7Ertd715e | 141.6731363 | 0.034304345 | 0.141421018 | 0.242568933 | 0.80833935 | 0.907434422 | | Car2 | 7066.430313 | 0.00604666156029366 | 0.0763411430256004 | 0.0792058033276495 | 0.936868928709142 | 0.98541611256672 | NM_009801 |
| 1810022K09Rik | 422.6895378 | 0.024593865 | 0.101480661 | 0.242350265 | 0.808508768 | 0.907558854 | | Car5b | 179.658932 | 0.00626375614086113 | 0.0859642926584034 | 0.0728646272441448 | 0.941913842461533 | 0.98541611256672 | NM_181315 |
| C2cd2 | 632.7923645 | 0.019562623 | 0.080838367 | 0.241996764 | 0.808782671 | 0.907669035 | | Cdk17 | 8819.62247 | 0.00581887221851676 | 0.0786401737998645 | 0.0739936337542376 | 0.941015450903863 | 0.98541611256672 | NM_146239 |
| Gm14326 | 732.6571993 | 0.025491944 | 0.105281356 | 0.242131606 | 0.808678189 | 0.907669035 | | Cdt1 | 72.46638864 | -0.00683186395889988 | 0.0950873848679039 | -0.0718482684494519 | 0.942722659341401 | 0.98541611256672 | NM_026014 |
| Rbmx2 | 206.6897772 | 0.022295421 | 0.092124119 | 0.242015022 | 0.808768524 | 0.907669035 | | Chrna1 | 25.81485492 | 0.00666282993273179 | 0.0846539862929191 | 0.0787066294749206 | 0.937265972282931 | 0.98541611256672 | NM_007389 |
| P2rx7 | 460.5599437 | 0.027883889 | 0.115300254 | 0.241837189 | 0.808906321 | 0.907742054 | | Cnep1r1 | 885.8621043 | -0.00531939108200064 | 0.0695916690565769 | -0.0764371821241428 | 0.939071289062997 | 0.98541611256672 | NM_029074 |
| Gm10416 | 10.56428286 | 0.039244343 | 0.162400777 | 0.241651203 | 0.809050443 | 0.907838034 | | Cnot1 | 5741.366166 | -0.00501334091056246 | 0.0627470310790156 | -0.0798976592892387 | 0.936318651496344 | 0.98541611256672 | NM_001205226,NM_178078,NM_153164 |
| Ccnj | 335.8546819 | -0.02132529 | 0.088418029 | -0.241186905 | 0.80941026 | 0.907978755 | | Cnpy4 | 387.4161507 | -0.00545032418023995 | 0.0724812717079674 | -0.0751963100509567 | 0.940058520196972 | 0.98541611256672 | NM_178612 |
| Rprm | 1794.654288 | 0.021907808 | 0.090823312 | 0.241213488 | 0.809389658 | 0.907978755 | | Col4a3 | 32.70271373 | 0.00488088887846115 | 0.0672408051454037 | 0.072588198013194 | 0.942133818565235 | 0.98541611256672 | NM_007734 |
| Skap2 | 435.8208284 | 0.021055146 | 0.087253543 | 0.241309926 | 0.809314918 | 0.907978755 | | Cul7 | 1506.536895 | 0.00423267315028191 | 0.058662908910973 | 0.0721524593454004 | 0.942480578456734 | 0.98541611256672 | NM_025611 |
| Socs2 | 447.5557506 | 0.024599947 | 0.101938138 | 0.241322309 | 0.809305321 | 0.907978755 | | Cwf19l1 | 655.4172107 | -0.00434438610963218 | 0.0605060756102946 | -0.07180082439346 | 0.942760416744563 | 0.98541611256672 | NM_001081077 |
| 8430408G22Rik | 45.14930867 | 0.037977505 | 0.157594194 | 0.240982896 | 0.809568373 | 0.90802464 | | Cyp2d22 | 1367.153644 | -0.00555948954243003 | 0.0757770040128887 | -0.0733664468297196 | 0.941514517166761 | 0.98541611256672 | NM_019823,NM_001163472 |
| Rhbdf2 | 67.14118425 | 0.037041503 | 0.153681788 | 0.241027281 | 0.809533972 | 0.90802464 | | Dlgap2 | 2840.331024 | -0.00493531551173317 | 0.0669298811764323 | -0.0737385966474869 | 0.941218386687942 | 0.98541611256672 | NM_172910,NM_001145965 |
| 9130011E15Rik | 820.5687599 | -0.022410709 | 0.093116655 | -0.240673478 | 0.809808196 | 0.908187085 | | Dnajb5 | 4486.084487 | -0.00414072687953725 | 0.0517943294570174 | -0.0799455639825113 | 0.93628055095862 | 0.98541611256672 | NM_001159883,NM_001159885,NM_178055,NM_001159884,NM_020266 |
| Slc41a3 | 202.0083861 | 0.030241958 | 0.125670531 | 0.240644787 | 0.809830435 | 0.908187085 | | Dse | 253.579465 | 0.00645411729234271 | 0.0880451815118811 | 0.0733046054481899 | 0.941563726944679 | 0.98541611256672 | NM_172508 |
| Dbx2 | 546.0121797 | 0.026670073 | 0.11091535 | 0.240454301 | 0.809978087 | 0.908286928 | | Ebna1bp2 | 1525.109608 | 0.0036317622531744 | 0.0480511710784704 | 0.0755811392659611 | 0.939752342222196 | 0.98541611256672 | NM_026932 |
| Abcc3 | 144.0955549 | -0.029896562 | 0.1244421 | -0.240244755 | 0.81010452 | 0.908361319 | | Efha1 | 941.162954 | 0.0044717203231022 | 0.0565505036649938 | 0.079074809830037 | 0.936973119603365 | 0.98541611256672 | NM_028643 |
| Mgmt | 26.83399539 | -0.040187817 | 0.167359771 | -0.240128296 | 0.810230799 | 0.908361319 | | Eid2 | 1166.125216 | 0.00447052608095576 | 0.0594154031434694 | 0.075241870700782 | 0.940022270751168 | 0.98541611256672 | NM_198425 |
| Myt1 | 440.1319134 | -0.025879841 | 0.107913178 | -0.239820955 | 0.810469062 | 0.908361319 | | Epg5 | 2953.966204 | -0.0047256269443721 | 0.0620721205526995 | -0.0761253667878211 | 0.939319358915229 | 0.98541611256672 | NM_001195633 |
| Selplg | 1250.746319 | -0.033790217 | 0.140719857 | -0.240124018 | 0.810234115 | 0.908361319 | | Fam13c | 2682.423928 | 0.00502560570051425 | 0.0619354796501553 | 0.0811425975693021 | 0.935328549394408 | 0.98541611256672 | NM_001143777,NM_001143776,NM_024244 |
| Tmem240 | 728.5438775 | -0.023813098 | 0.099226077 | -0.239988308 | 0.810339321 | 0.908361319 | | Fancc | 369.8610948 | -0.00582874794684529 | 0.0784674706963745 | -0.074282347769298 | 0.940785722661827 | 0.98541611256672 | NM_001042673,NM_007985 |
| Tubg1 | 1406.213298 | 0.020160084 | 0.084040653 | 0.239884908 | 0.810419481 | 0.908361319 | | Ftl1 | 10727.26742 | 0.00392223387539344 | 0.0509100102851425 | 0.0770424883716453 | 0.938589744558803 | 0.98541611256672 | NM_010240 |
| Wdr75 | 1053.119343 | 0.0163205 | 0.0680691 | 0.239763713 | 0.81051344 | 0.908361319 | | Fxr1 | 2563.692165 | 0.00436400083228769 | 0.057988243107043 | 0.0752566485629163 | 0.940010513058976 | 0.98541611256672 | NM_001113189,NM_001113188,NM_008053 |
| Zfp667 | 688.2143782 | 0.025968258 | 0.108290284 | 0.23980229 | 0.810483532 | 0.908361319 | | Gfer | 436.7557836 | 0.00496093013405689 | 0.0692826972008524 | 0.0716041715246019 | 0.942916920203109 | 0.98541611256672 | NM_023040 |
| Gm7694 | 140.4494037 | -0.033977762 | 0.141938874 | -0.239383059 | 0.810808565 | 0.908618954 | | Gm10421 | 34.4608198 | 0.00664353906596925 | 0.0916240505767391 | 0.0725086811175741 | 0.942197097122986 | 0.98541611256672 | NR_033538 |
| Rnf114 | 2365.282758 | -0.014063509 | 0.058765451 | -0.239315936 | 0.81086061 | 0.908618954 | | Gm17762 | 83.33225226 | -0.00709901927949491 | 0.0950536074451007 | -0.0746843751679286 | 0.940465839759952 | 0.98541611256672 | NR_028378 |
| Zfyve9 | 2509.064972 | 0.019933558 | 0.083350735 | 0.239152759 | 0.810987133 | 0.908695013 | | Gm9833 | 23.11272387 | -0.00610613633559793 | 0.0852879084250959 | -0.0715943965370031 | 0.942924699548976 | 0.98541611256672 | NR_045710 |
| 5730405O15Rik | 36.13177291 | -0.038183137 | 0.160083456 | -0.238520196 | 0.811477653 | 0.909178882 | | Gmnn | 119.1149744 | 0.00705264953864685 | 0.0943470952386437 | 0.0747521640259058 | 0.940411902847154 | 0.98541611256672 | NM_020567 |
| Ccz1 | 1146.622472 | -0.015432467 | 0.064839978 | -0.238008524 | 0.811874483 | 0.909426203 | | Gpt | 563.9166865 | 0.0064246984792708 | 0.0889463296207531 | 0.0722311815075928 | 0.942417930721144 | 0.98541611256672 | NM_182805 |
| Fndc3a | 2365.846673 | 0.021832856 | 0.091717968 | 0.238043391 | 0.81184744 | 0.909426203 | | Gtf2h1 | 1834.280726 | 0.004617189228273 | 0.0595934963695296 | 0.0774780707552811 | 0.938243235828937 | 0.98541611256672 | NM_008186 |
| Kat5 | 2056.108397 | 0.012601636 | 0.052943257 | 0.238021542 | 0.811864386 | 0.909426203 | | Gtf2i | 10596.72743 | 0.00282540659829513 | 0.0347489898775691 | 0.0813090282120393 | 0.935196194290405 | 0.98541611256672 | NM_001080748,NM_001080749,NM_001080746,NM_010365,NM_001080747 |
| Fzd7 | 308.3830433 | 0.020487271 | 0.086165885 | 0.237765454 | 0.812063014 | 0.909490489 | | H2-Ke6 | 610.5704531 | -0.00579670894715041 | 0.0738113423873548 | -0.0785341217170912 | 0.937403188824415 | 0.98541611256672 | NM_013543 |
| Gm5535 | 13.29186154 | 0.038229895 | 0.160874472 | 0.237707493 | 0.812107972 | 0.909490489 | | Hapln1 | 668.0623773 | 0.00639887215827337 | 0.0868578786886487 | 0.0736706013879387 | 0.941272491896269 | 0.98541611256672 | NM_013500 |
| Zfp84 | 813.8602586 | -0.01678109 | 0.07058587 | -0.237740073 | 0.812082701 | 0.909490489 | | Hmgcl | 801.7499349 | 0.00455519126694886 | 0.0615899400020811 | 0.0739599887058656 | 0.941042222413729 | 0.98541611256672 | NM_008254 |
| Ccdc19 | 62.52335278 | -0.035124605 | 0.147920055 | -0.237456679 | 0.812302523 | 0.90957688 | | Homez | 768.9738387 | 0.00510351917974771 | 0.0642259738976843 | 0.0794619196880988 | 0.936665219488487 | 0.98541611256672 | NM_001177705,NM_183174 |
| Pon3 | 86.29999124 | 0.03987509 | 0.167873928 | 0.237529977 | 0.812245667 | 0.90957688 | | Ildr1 | 23.81969779 | 0.00644169009531601 | 0.0821466455405558 | 0.078416585127581 | 0.937496384126466 | 0.98541611256672 | NM_134109 |
| Acadl | 1250.297944 | -0.016070873 | 0.067704327 | -0.23736847 | 0.812370948 | 0.909587364 | | Kcnc3 | 6827.119155 | 0.00464708058793008 | 0.0637276987964894 | 0.0729208911617891 | 0.941869069455885 | 0.98541611256672 | NM_008422 |
| Eme2 | 545.1727203 | 0.029926144 | 0.126152539 | 0.237221893 | 0.812484653 | 0.909587364 | | Kcnk12 | 222.3240153 | 0.00642428679781242 | 0.0869596731564027 | 0.0738766207901669 | 0.941108558910217 | 0.98541611256672 | NM_199251 |
| Fam212a | 48.04433735 | 0.037541272 | 0.158349011 | 0.237079297 | 0.812595273 | 0.909587364 | | Kctd4 | 758.0290171 | 0.00604743711042493 | 0.0774626792280314 | 0.0780690413847259 | 0.937773133407141 | 0.98541611256672 | NM_026214 |
| Frg1 | 733.8853626 | 0.017680713 | 0.074581323 | 0.237066225 | 0.812605414 | 0.909587364 | | Kif1c | 4801.455113 | 0.00474216712921039 | 0.0615620008562004 | 0.0770307505158479 | 0.93859908226361 | 0.98541611256672 | NM_153103 |
| Glis1 | 18.42866847 | 0.039805058 | 0.167906716 | 0.237066502 | 0.812605199 | 0.909587364 | | Klhl26 | 2234.087364 | 0.00461516322512008 | 0.0566644825366446 | 0.0814471961715258 | 0.935086316658561 | 0.98541611256672 | NM_178771,NM_001122830 |
| Abcb8 | 1765.224281 | -0.022043598 | 0.093089013 | -0.236801285 | 0.812810954 | 0.909620291 | | Ksr1 | 1612.142272 | 0.00503792172651223 | 0.0689741071109176 | 0.0730407675797923 | 0.941773676292984 | 0.98541611256672 | NM_013571 |
| Prpf31 | 1207.418397 | -0.01403338 | 0.059251443 | -0.236844528 | 0.812777406 | 0.909620291 | | Lgals1 | 389.8960049 | -0.00659758382500261 | 0.0903886078498349 | -0.0729913202774775 | 0.941813024502751 | 0.98541611256672 | NM_008495 |
| Zfand3 | 2970.793702 | -0.014162939 | 0.059782326 | -0.236908466 | 0.812727802 | 0.909620291 | | Lrguk | 46.90375575 | -0.00720549076277671 | 0.0939636542015191 | -0.0766838074147633 | 0.938875086417074 | 0.98541611256672 | NM_028886 |
| Gjd3 | 9.349774015 | 0.037346362 | 0.157828569 | 0.236626118 | 0.812946856 | 0.909640976 | | Ly6g6e | 45.40872896 | 0.0065801069037107 | 0.0872307835473757 | 0.0754333119126117 | 0.939869955629577 | 0.98541611256672 | NM_027366 |
| Gpr183 | 64.07246037 | -0.035687294 | 0.150782364 | -0.23668082 | 0.812904416 | 0.909640976 | | Lyrm5 | 624.8592829 | -0.00540594767599395 | 0.0703317652074637 | -0.0768635290191787 | 0.938732111307037 | 0.98541611256672 | NM_133688,NM_001163628 |
| Rptor | 3441.702703 | -0.012604632 | 0.053286835 | -0.236543082 | 0.813011281 | 0.90964737 | | Mbd3 | 3490.526175 | -0.00410825639513331 | 0.0547472317144307 | -0.0750404407032406 | 0.940182535553746 | 0.98541611256672 | NM_013595 |
| Upp1 | 55.91329225 | -0.035120577 | 0.148742565 | -0.23611652 | 0.813342255 | 0.909951975 | | Mef2c | 21580.62981 | -0.005828243780976 | 0.078174372017463 | -0.0745544048588463 | 0.940569252757177 | 0.98541611256672 | NM_001170537,NM_025282 |
| Reep6 | 390.8513979 | -0.028720251 | 0.12169528 | -0.236001352 | 0.813431621 | 0.909960249 | | Mfi2 | 40.68885353 | -0.00748310562595945 | 0.0930823591332079 | -0.0803923073678285 | 0.935925244934584 | 0.98541611256672 | NM_013900 |
| Rimkla | 671.3257058 | -0.023916551 | 0.101360378 | -0.235955617 | 0.81346711 | 0.909960249 | | Mgea5 | 10812.80011 | -0.00562100007232725 | 0.0693961125589028 | -0.0809987744989856 | 0.935442927111345 | 0.98541611256672 | NM_023799 |
| Acaa1a | 1180.3101 | 0.031708261 | 0.134458038 | 0.235835351 | 0.813560436 | 0.909998455 | | Mrps24 | 518.7173079 | -0.00524153830162382 | 0.0724137751936201 | -0.0723831658770584 | 0.942296981334509 | 0.98541611256672 | NM_026080 |
| Dak | 407.2197754 | -0.027952226 | 0.118613702 | -0.235657644 | 0.813698339 | 0.910021804 | | Myb | 32.44200348 | 0.00670309295615409 | 0.0865481660275132 | 0.0774492778278308 | 0.93826614043708 | 0.98541611256672 | NM_010848,NM_001198914 |
| Etl4 | 4749.685033 | 0.020836305 | 0.088411378 | 0.23567447 | 0.813685282 | 0.910021804 | | Naa25 | 1537.312407 | 0.00577162579979549 | 0.0733080171857524 | 0.0787311677680625 | 0.937246454125208 | 0.98541611256672 | NM_172722 |
| Mtif2 | 927.2501011 | -0.014519696 | 0.061656082 | -0.235494947 | 0.8138246 | 0.910097319 | | Nek8 | 128.4052748 | -0.00727547591308887 | 0.0944483253048495 | -0.0770311918408836 | 0.938773169233394 | 0.98541611256672 | NM_080849 |
| BC065397 | 101.5208191 | -0.03940141 | 0.16745964 | -0.235288992 | 0.813984438 | 0.910144684 | | Nudt2 | 492.7057614 | -0.00540576473905901 | 0.0666232157247955 | -0.0811393548067227 | 0.935331128241236 | 0.98541611256672 | NM_025539 |
| Dph2 | 413.0690843 | 0.021890244 | 0.093012767 | 0.235346661 | 0.813939682 | 0.910144684 | | Numbl | 4094.789864 | -0.00580039019090232 | 0.0734833260706017 | -0.0789347801826761 | 0.937084498952531 | 0.98541611256672 | NM_010950 |
| 2610020H08Rik | 26.82693275 | 0.039173815 | 0.166666383 | 0.235043289 | 0.814175134 | 0.910226534 | | Nup133 | 1812.331158 | -0.00366570444038491 | 0.0463985495042754 | -0.0790047206119478 | 0.937028868297114 | 0.98541611256672 | NM_172288 |
| Timm50 | 771.7919499 | 0.016365104 | 0.069615794 | 0.235077458 | 0.814148614 | 0.910226534 | | Nup43 | 129.1605596 | -0.00662319412916074 | 0.0909554619234542 | -0.072817992334915 | 0.941950095315289 | 0.98541611256672 | NM_145706 |
| Stra13 | 247.5100693 | -0.026247974 | 0.111774783 | -0.234829123 | 0.814341363 | 0.910346686 | | Optc | 36.41011206 | 0.00658489993700904 | 0.0820566516100505 | 0.0802482164188443 | 0.936039842626604 | 0.98541611256672 | NM_001160420,NM_054076,NM_001160422,NM_001160421 |
| Gfra4 | 1661.769917 | -0.021220805 | 0.090498283 | -0.234488487 | 0.81460577 | 0.910517271 | | Pank2 | 2160.976897 | 0.004356744457333404 | 0.0600429157294288 | 0.0725605097688265 | 0.942155852484724 | 0.98541611256672 | NM_153501 |
| Pitpnm2 | 12354.86163 | -0.019189518 | 0.081864653 | -0.234405416 | 0.814670255 | 0.910517271 | | Pdgfrl | 29.14374952 | 0.00625324496845707 | 0.0871386349940603 | 0.0717620257522202 | 0.942791293930352 | 0.98541611256672 | NM_026840 |

| | | | | | | |
|---|---|---|---|---|---|---|
| Slc24a6 | 269.630771 | 0.030227373 | 0.128953061 | 0.234406014 | 0.81466979 | 0.910517271 |
| Fbxo45 | 2777.918009 | -0.013736659 | 0.058653458 | -0.234200321 | 0.814829466 | 0.910563847 |
| Vav3 | 301.1180343 | 0.027862823 | 0.118952268 | 0.234235324 | 0.814802293 | 0.910563847 |
| 8-Sep | 8313.202878 | 0.016290897 | 0.069754941 | 0.233544696 | 0.815338466 | 0.911066941 |
| Panx2 | 3515.223801 | -0.014902183 | 0.063829811 | -0.233467445 | 0.815398446 | 0.911068258 |
| 2010001M06Rik | 70.87226574 | -0.035731332 | 0.153551066 | -0.232699992 | 0.815994375 | 0.911208422 |
| Apex2 | 132.657123 | -0.030574111 | 0.131220613 | -0.232997778 | 0.815763131 | 0.911208422 |
| Bnip1 | 579.2204776 | 0.018705499 | 0.080353195 | 0.232790984 | 0.815923714 | 0.911208422 |
| Cbr1 | 1818.429086 | -0.014332999 | 0.061484219 | -0.233116704 | 0.815670784 | 0.911208422 |
| Fbln2 | 293.6164151 | 0.030466785 | 0.13081024 | 0.232908255 | 0.815832648 | 0.911208422 |
| Fbxo2 | 2319.33533 | 0.023006039 | 0.098835488 | 0.232771041 | 0.815939202 | 0.911208422 |
| Rc3h2 | 2340.179596 | 0.022122028 | 0.09500817 | 0.232843426 | 0.81588299 | 0.911208422 |
| Sh3glb2 | 5779.828244 | -0.015530388 | 0.06668869 | -0.232878884 | 0.815855456 | 0.911208422 |
| Jph3 | 10255.47169 | 0.017815535 | 0.076627597 | 0.23249502 | 0.816153555 | 0.911320494 |
| Qpctl | 655.664884 | 0.014854835 | 0.063923596 | 0.232384223 | 0.816239602 | 0.911350897 |
| Gm10432 | 109.6548149 | 0.030360571 | 0.131253851 | 0.231311849 | 0.817072542 | 0.912215158 |
| Klf4 | 283.8481897 | 0.035243143 | 0.152560268 | 0.231011281 | 0.817306038 | 0.912344363 |
| Nphs1as | 9.720336112 | -0.031418047 | 0.135980819 | -0.231047638 | 0.817277793 | 0.912344363 |
| Rp9 | 682.0548254 | -0.018334968 | 0.079452769 | -0.23076562 | 0.817496892 | 0.912425936 |
| Snapin | 1731.801023 | -0.017664978 | 0.076539436 | -0.230795766 | 0.81747347 | 0.912425936 |
| Ubxn1 | 1877.978809 | 0.018960181 | 0.082221736 | 0.230598155 | 0.817627001 | 0.912505421 |
| Ets2 | 3283.047716 | 0.025878867 | 0.11235705 | 0.230327043 | 0.817837648 | 0.912551433 |
| Fam185a | 620.8499388 | -0.020338325 | 0.088329985 | -0.230253912 | 0.817894471 | 0.912551433 |
| Psip1 | 5497.157328 | -0.014363534 | 0.062353425 | -0.230356781 | 0.817814542 | 0.912551433 |
| Zfand2b | 524.7684696 | -0.021338639 | 0.092709747 | -0.230166085 | 0.817962715 | 0.912551433 |
| Zfp953 | 48.12744804 | -0.036018179 | 0.156485086 | -0.230170045 | 0.817959638 | 0.912551433 |
| Mettl3 | 816.4399769 | 0.023155746 | 0.100841828 | 0.229624416 | 0.818383633 | 0.912870298 |
| Nme6 | 144.4761539 | -0.025111169 | 0.109383104 | -0.229570822 | 0.818425282 | 0.912870298 |
| Zfp120 | 515.5925515 | -0.021384604 | 0.09314998 | -0.229571749 | 0.818424561 | 0.912870298 |
| Gpr150 | 130.0591477 | 0.031305161 | 0.136411199 | 0.229491135 | 0.81848721 | 0.912873656 |
| Ykt6 | 3756.089337 | -0.015977197 | 0.069657963 | -0.229366418 | 0.818584135 | 0.912916043 |
| Txnrd2 | 468.1251824 | 0.017698934 | 0.077217039 | 0.229210215 | 0.818705534 | 0.912985716 |
| Ace2 | 73.33532919 | -0.032306672 | 0.141008883 | -0.229110903 | 0.81878272 | 0.913006079 |
| Casp6 | 92.26922438 | -0.031585424 | 0.13797199 | -0.228926351 | 0.818926159 | 0.913095376 |
| Kif7 | 97.70569441 | -0.034206736 | 0.149468236 | -0.228856224 | 0.818980666 | 0.913095376 |
| Adam17 | 682.3202812 | -0.021555372 | 0.094280579 | -0.228630031 | 0.819156481 | 0.913111003 |
| Smpd5 | 16.56457916 | -0.038388687 | 0.16792163 | -0.228610733 | 0.819171482 | 0.913111003 |
| Zbtb37 | 144.2714277 | -0.030481585 | 0.133333349 | -0.228611859 | 0.819170606 | 0.913111003 |
| A230001M10Rik | 132.6031211 | -0.037426568 | 0.164137176 | -0.228020058 | 0.819630648 | 0.913228629 |
| Aar2 | 1775.926419 | 0.015982571 | 0.070348458 | 0.227191488 | 0.820274846 | 0.913228629 |
| Acbd6 | 1269.956756 | 0.015438353 | 0.067998802 | 0.227038607 | 0.820393722 | 0.913228629 |
| BC030476 | 12.03320977 | 0.036225289 | 0.158928763 | 0.22793413 | 0.819697449 | 0.913228629 |
| Cgn | 207.4320808 | -0.030766322 | 0.134776445 | -0.228276703 | 0.819431135 | 0.913228629 |
| D11Wsu47e | 311.5929534 | 0.024077341 | 0.105895611 | 0.227368641 | 0.820137103 | 0.913228629 |
| Emr1 | 311.1617796 | 0.034716481 | 0.152914274 | 0.227032309 | 0.820398619 | 0.913228629 |
| Fancb | 93.61499575 | -0.029538151 | 0.129696617 | -0.227748047 | 0.819842118 | 0.913228629 |
| Gin1 | 202.448412 | 0.023495471 | 0.102990159 | 0.228133169 | 0.819542715 | 0.913228629 |
| Ift80 | 1001.891828 | -0.015087886 | 0.066478541 | -0.226958743 | 0.820455823 | 0.913228629 |
| Kbtbd13 | 9.75476809 | 0.034117553 | 0.15006488 | 0.227352015 | 0.82015003 | 0.913228629 |
| Lin7c | 2396.304023 | -0.017735803 | 0.07784205 | -0.227843477 | 0.819767926 | 0.913228629 |
| Psmd1 | 5930.389645 | -0.015180018 | 0.066836594 | -0.227121357 | 0.820329378 | 0.913228629 |
| Serpine1 | 73.92736112 | 0.037184821 | 0.163477217 | 0.227461793 | 0.820064676 | 0.913228629 |
| Slc29a1 | 865.8956978 | -0.022505962 | 0.099013553 | -0.227301836 | 0.820189045 | 0.913228629 |
| Snrpn | 52.58330523 | 0.03812647 | 0.167283391 | 0.227915454 | 0.819711968 | 0.913228629 |
| Ssh1 | 499.5685849 | 0.018447213 | 0.080983604 | 0.227789484 | 0.819809902 | 0.913228629 |
| Tmem131 | 3696.317512 | -0.012648549 | 0.055657091 | -0.227258539 | 0.82022271 | 0.913228629 |
| Zfp13 | 578.302945 | 0.020042739 | 0.088062531 | 0.227596676 | 0.819959804 | 0.913228629 |
| Zfp830 | 342.4723617 | -0.017693574 | 0.077660194 | -0.227833244 | 0.819775881 | 0.913228629 |
| Apex1 | 1189.181777 | -0.012294239 | 0.054218614 | -0.226753107 | 0.820615729 | 0.913341003 |
| Acot3 | 80.16038207 | -0.03229615 | 0.142685268 | -0.226345369 | 0.820932814 | 0.913628288 |
| 0610037L13Rik | 961.0195021 | 0.028150491 | 0.124414214 | 0.226264265 | 0.82099589 | 0.91363286 |
| Clstn3 | 18337.68636 | -0.012933957 | 0.057199697 | -0.226119325 | 0.821108615 | 0.91369268 |
| Gtf3c6 | 534.8002293 | -0.018870975 | 0.083519179 | -0.225947796 | 0.821242023 | 0.913775505 |
| Cnbd2 | 176.9133733 | 0.024963457 | 0.11054244 | 0.225826906 | 0.82133605 | 0.913814502 |
| Fam175b | 1221.178758 | -0.012544703 | 0.055570421 | -0.225744241 | 0.821400347 | 0.913820419 |
| Slc35a3 | 464.3128657 | -0.020108784 | 0.089131417 | -0.225608257 | 0.821506118 | 0.913872472 |
| AW011738 | 159.0116633 | -0.025185838 | 0.111723665 | -0.225429752 | 0.821644969 | 0.913961314 |
| Pskh1 | 442.4544604 | -0.01945868 | 0.086497008 | -0.224963623 | 0.822007574 | 0.914233389 |
| Zfp119a | 92.17451907 | 0.033852444 | 0.150430878 | 0.225036536 | 0.821950851 | 0.914233389 |
| Pkd2 | 1478.201243 | 0.017296769 | 0.077079106 | 0.224402825 | 0.822443872 | 0.914652983 |
| Cox7a1 | 36.86101703 | -0.037672863 | 0.168041956 | -0.224187246 | 0.822611607 | 0.914708213 |
| Cyp4f16 | 135.6896764 | -0.028713296 | 0.128060833 | -0.224216066 | 0.822589183 | 0.914708213 |
| Ezr | 2520.595334 | 0.02595503 | 0.115944696 | 0.223856985 | 0.822868587 | 0.91486265 |
| Ppapdc3 | 914.4584521 | 0.016735207 | 0.074754475 | 0.223868966 | 0.822859264 | 0.91486265 |
| Pkm | 50060.38061 | 0.015551355 | 0.069557631 | 0.223575109 | 0.823087933 | 0.915040858 |
| Gm10125 | 48.28660933 | -0.035837892 | 0.160397293 | -0.223432025 | 0.823199282 | 0.915098987 |
| Lrmp | 67.01253871 | -0.03374186 | 0.15113401 | -0.223257889 | 0.8233348 | 0.915183972 |
| Cd209b | 9.024613783 | 0.03069995 | 0.13761002 | 0.223093855 | 0.823462461 | 0.915205309 |
| Mog | 1972.154735 | 0.037406527 | 0.167681043 | 0.223081428 | 0.823472132 | 0.915205309 |
| Slc35b2 | 815.8139418 | -0.016605151 | 0.074482858 | -0.222939231 | 0.823582803 | 0.915262656 |
| A230072C01Rik | 299.285 | 0.027294379 | 0.122714084 | 0.22242255 | 0.823984959 | 0.915643904 |
| Ap2a2 | 10279.06416 | -0.015042637 | 0.067819235 | -0.221804881 | 0.82446578 | 0.915701468 |
| Apoa1bp | 1451.300584 | 0.014152412 | 0.06388748 | 0.2215209 | 0.824686865 | 0.915701468 |
| Calcrl | 194.14616 | -0.028580848 | 0.128728412 | -0.222024399 | 0.82429489 | 0.915701468 |
| Cetn3 | 1171.451407 | 0.015071566 | 0.067970755 | 0.221736047 | 0.824519367 | 0.915701468 |
| Dmrta1 | 9.124115221 | -0.032765252 | 0.14777958 | -0.22171705 | 0.824534157 | 0.915701468 |
| Eif2b5 | 2208.508654 | -0.017147889 | 0.077396606 | -0.22155867 | 0.824657459 | 0.915701468 |
| Elmo1 | 6678.457418 | -0.018879506 | 0.085186114 | -0.221626568 | 0.824604599 | 0.915701468 |
| Pex14 | 1775.004927 | 0.016930041 | 0.076253135 | 0.22202419 | 0.824295052 | 0.915701468 |
| Pnpla1 | 23.490394 | -0.036743955 | 0.16572521 | -0.221716145 | 0.824534861 | 0.915701468 |
| Slc31a1 | 884.4683802 | 0.019581862 | 0.088287573 | 0.221796352 | 0.82447242 | 0.915701468 |
| Zfp524 | 169.7678777 | -0.023540157 | 0.106180739 | -0.221698939 | 0.824548256 | 0.915701468 |
| Sos2 | 1631.220216 | -0.018247847 | 0.082407788 | -0.22143352 | 0.824754895 | 0.915711382 |
| 8430410A17Rik | 586.9472905 | -0.022126569 | 0.100051156 | -0.221152561 | 0.824973646 | 0.915793302 |
| Ech1 | 1305.745837 | 0.015069385 | 0.068199855 | 0.220959191 | 0.825124209 | 0.915793302 |
| Noc3l | 610.2383296 | 0.019216023 | 0.086960199 | 0.220974916 | 0.825111964 | 0.915793302 |
| Tmem192 | 554.5327578 | 0.019956735 | 0.090234149 | 0.221166106 | 0.824963099 | 0.915793302 |
| Zfp68 | 1415.394268 | -0.01553688 | 0.070306083 | -0.220989127 | 0.825100899 | 0.915793302 |
| Cacna1d | 2213.955075 | -0.019713415 | 0.089312035 | -0.220725181 | 0.825306423 | 0.915825236 |
| Dffa | 1658.795263 | 0.013582649 | 0.061546547 | 0.220689048 | 0.825334559 | 0.915825236 |
| Mastl | 43.73873704 | 0.033599777 | 0.152350442 | 0.220542697 | 0.825448523 | 0.915825236 |
| N4bp2l2 | 1212.088554 | -0.020065252 | 0.090951151 | -0.220615706 | 0.825391671 | 0.915825236 |
| Usp11 | 6887.169032 | 0.014261618 | 0.064633294 | 0.220654355 | 0.825361575 | 0.915825236 |
| Hhat | 26.66258117 | 0.036590543 | 0.16637129 | 0.219933036 | 0.825923307 | 0.91628639 |
| Fermt2 | 3400.889783 | -0.011483916 | 0.052293241 | -0.219606135 | 0.826177913 | 0.916503228 |
| Mphosph6 | 358.5510027 | -0.018627255 | 0.084953882 | -0.219263143 | 0.826445071 | 0.916733959 |
| Uros | 757.3493189 | -0.024888297 | 0.113645854 | -0.218998723 | 0.826651043 | 0.91689679 |
| A430005L14Rik | 456.9157829 | 0.022544573 | 0.103241431 | 0.218367496 | 0.827142789 | 0.916976744 |
| Anp32e | 4071.882198 | -0.011881593 | 0.054312982 | -0.21876157 | 0.826835784 | 0.916976744 |
| Arhgef9 | 18781.64348 | 0.012619225 | 0.057767821 | 0.218447317 | 0.827080602 | 0.916976744 |
| Arid3a | 279.5531646 | -0.018068853 | 0.082824181 | -0.21815915 | 0.827305112 | 0.916976744 |
| Clec10a | 29.32471976 | -0.036271345 | 0.165873006 | -0.218669365 | 0.826907615 | 0.916976744 |
| Fgfbp3 | 127.2092524 | -0.028315557 | 0.129568366 | -0.218537576 | 0.827010284 | 0.916976744 |
| Fus | 7776.590099 | 0.021780879 | 0.099845184 | 0.218146518 | 0.827314954 | 0.916976744 |
| Grtp1 | 90.68636162 | 0.029594553 | 0.13546735 | 0.218462622 | 0.827068678 | 0.916976744 |
| Kpna3 | 3486.819845 | -0.017076161 | 0.078247853 | -0.21823169 | 0.827248595 | 0.916976744 |
| Slc25a17 | 1080.913507 | -0.014701002 | 0.067318804 | -0.218378832 | 0.827133957 | 0.916976744 |
| H2afj | 594.9409748 | -0.0251402 | 0.115376466 | -0.217897126 | 0.827509267 | 0.917126508 |
| Ece1 | 2557.885601 | -0.011187121 | 0.051427615 | -0.217531392 | 0.827794245 | 0.917158757 |
| Fgfbp1 | 81.39850415 | 0.036497573 | 0.167899248 | 0.217377822 | 0.827913914 | 0.917158757 |
| Kif13a | 1513.942364 | 0.015240475 | 0.069983873 | 0.217771237 | 0.827607356 | 0.917158757 |
| Tchh | 250.5199436 | 0.022654972 | 0.104188056 | 0.217443082 | 0.82786306 | 0.917158757 |
| Tmem168 | 555.349853 | -0.017334725 | 0.079762956 | -0.217328016 | 0.827952725 | 0.917158757 |
| Tmem48 | 439.3963653 | 0.019272362 | 0.088631276 | 0.217444258 | 0.827862144 | 0.917158757 |
| Zfp746 | 1047.750062 | 0.015811235 | 0.072665163 | 0.217590304 | 0.82774834 | 0.917158757 |
| Zbtb26 | 319.0922871 | 0.020099973 | 0.092662154 | 0.216916748 | 0.828273226 | 0.917448195 |
| Ubash3b | 1253.519859 | 0.016701404 | 0.07708067 | 0.216674354 | 0.828462135 | 0.917591845 |
| Gfm1 | 1653.999973 | -0.014109718 | 0.065151547 | -0.216567663 | 0.828545288 | 0.917618349 |
| Zfp772 | 445.3973207 | -0.021030623 | 0.09741382 | -0.215889518 | 0.829073867 | 0.918138125 |
| 5930412G12Rik | 70.66179274 | 0.036231139 | 0.167991448 | 0.21567252 | 0.829243023 | 0.91825982 |
| 9030624J02Rik | 2154.927746 | -0.012903503 | 0.059884267 | -0.215474012 | 0.829397772 | 0.918365546 |
| Igf2as | 10.84695316 | -0.033791035 | 0.157088337 | -0.215108492 | 0.829682733 | 0.918615425 |
| Keap1 | 2720.706465 | 0.013782257 | 0.064141981 | 0.214871083 | 0.829867831 | 0.91875471 |
| Creb3l1 | 117.2056915 | 0.035009217 | 0.163265396 | 0.214431337 | 0.830210707 | 0.918979996 |
| Fcna | 10.64088507 | 0.029282414 | 0.136497986 | 0.214526344 | 0.830136625 | 0.918979996 |
| Ubr3 | 8552.910562 | -0.017065179 | 0.07960177 | -0.214381898 | 0.830249257 | 0.918979996 |
| Taf12 | 466.4672604 | -0.019114225 | 0.089223697 | -0.214228123 | 0.830369166 | 0.919047065 |
| Ghr | 352.1717997 | 0.022913285 | 0.107035289 | 0.214072249 | 0.830490716 | 0.91911594 |
| Ctr9 | 2314.529318 | -0.016072072 | 0.075189798 | -0.213753357 | 0.8307394 | 0.919325496 |
| Shprh | 1739.053546 | 0.019407841 | 0.090898692 | 0.213510672 | 0.830928666 | 0.919469273 |
| Dnahc7b | 165.7432123 | 0.035021406 | 0.164214317 | 0.213266462 | 0.831119131 | 0.919483028 |
| Oprd1 | 1006.069683 | -0.017656512 | 0.082779324 | -0.213296157 | 0.831095971 | 0.919483028 |
| Sacs | 299.8026036 | -0.027650661 | 0.129597284 | -0.213358334 | 0.831047477 | 0.919483028 |
| Bag5 | 1803.76061 | -0.020271723 | 0.095099351 | -0.213163635 | 0.831199331 | 0.9195061 |
| Atp6v1a | 20267.80659 | 0.015979285 | 0.074989406 | 0.213087229 | 0.831258925 | 0.919506375 |
| Uri1 | 1439.924126 | -0.019665783 | 0.092333171 | -0.212987191 | 0.831336953 | 0.919527039 |
| Cd3eap | 824.7792293 | 0.015131191 | 0.071068581 | 0.212909707 | 0.831397391 | 0.919528244 |
| 2310030G06Rik | 84.12084755 | 0.027802062 | 0.130817285 | 0.212525909 | 0.831696767 | 0.919597921 |
| 9430023L20Rik | 812.3570065 | -0.019856362 | 0.093497838 | -0.212372419 | 0.831816501 | 0.919597921 |
| Cml5 | 57.47623243 | -0.034976549 | 0.164406803 | -0.212743927 | 0.831526702 | 0.919597921 |
| Tbc1d13 | 2051.516885 | -0.01388035 | 0.06533844 | -0.21243773 | 0.831765553 | 0.919597921 |
| Wbscr16 | 437.0924488 | 0.017751642 | 0.083536779 | 0.212500916 | 0.831716263 | 0.919597921 |
| Zdhhc3 | 3233.997734 | 0.010844375 | 0.050996186 | 0.212650708 | 0.831599416 | 0.919597921 |
| Wbp1l | 1739.545726 | -0.014667344 | 0.069161712 | -0.212073175 | 0.832049947 | 0.919790374 |
| B3gnt9 | 128.2517424 | 0.026427958 | 0.124808178 | 0.211748607 | 0.832303165 | 0.919939024 |
| Exosc3 | 197.2737878 | -0.020182908 | 0.095296818 | -0.211789947 | 0.832270912 | 0.919939024 |
| Kif14 | 15.81632744 | -0.035101413 | 0.166075669 | -0.211357951 | 0.832607966 | 0.920210273 |
| 1810011O10Rik | 35.76285289 | 0.035498476 | 0.168100468 | 0.211174167 | 0.832751368 | 0.920237479 |
| Stk38l | 1409.986719 | 0.012957765 | 0.061354675 | 0.211194417 | 0.832735567 | 0.920237479 |
| Patz1 | 1092.522772 | -0.01448719 | 0.068720439 | -0.210813406 | 0.833032877 | 0.920482912 |
| Stat5a | 76.86898896 | 0.030553571 | 0.145030427 | 0.210670079 | 0.833144724 | 0.920488074 |
| Zbtb45 | 909.1845817 | -0.014657752 | 0.069581739 | -0.21065516 | 0.833156367 | 0.920488074 |
| Pcyt2 | 2424.828928 | 0.017239281 | 0.081895789 | 0.210502656 | 0.833275379 | 0.920520216 |
| Wdr26 | 5847.944787 | 0.016757262 | 0.07961995 | 0.21046562 | 0.833304282 | 0.920520216 |
| Eif2c4 | 663.3262626 | 0.017871374 | 0.085031281 | 0.210174108 | 0.833531787 | 0.920705889 |
| Dnph1 | 79.79991871 | -0.026495651 | 0.126241235 | -0.209881116 | 0.833760461 | 0.920864051 |
| Fhl3 | 34.28614244 | 0.034266301 | 0.163357842 | 0.209762203 | 0.833853274 | 0.920864051 |
| Smim11 | 251.8271336 | 0.025670403 | 0.122344002 | 0.209821506 | 0.833806988 | 0.920864051 |
| Prdm16 | 533.4657366 | -0.018464537 | 0.08817614 | -0.20940514 | 0.834131981 | 0.921106187 |
| Ccdc15 | 79.06884742 | 0.029632725 | 0.141722755 | 0.209089393 | 0.834378455 | 0.921120761 |
| Egln2 | 2669.206408 | 0.017156881 | 0.082087415 | 0.209007442 | 0.834442429 | 0.921120761 |
| Praf2 | 2754.444032 | -0.012605055 | 0.06028999 | -0.209073761 | 0.834390657 | 0.921120761 |
| Rnmtl1 | 350.6032545 | 0.017689499 | 0.084601629 | 0.20909171 | 0.834376646 | 0.921120761 |
| Whsc1 | 3905.255635 | -0.014087498 | 0.067305862 | -0.209305667 | 0.834209628 | 0.921120761 |
| Gm9866 | 938.384012 | 0.014150273 | 0.067736359 | 0.208902177 | 0.834524605 | 0.921145845 |
| Blzf1 | 819.136785 | 0.013063865 | 0.062564455 | 0.208806506 | 0.834599293 | 0.921162661 |
| Taf3 | 774.0498192 | 0.020160973 | 0.096642962 | 0.208612942 | 0.834750406 | 0.921258727 |
| Vrk1 | 609.9927974 | 0.018709646 | 0.089716144 | 0.208542695 | 0.834805249 | 0.921258727 |
| D3Ertd254e | 1196.647647 | 0.013372334 | 0.064330296 | 0.207869934 | 0.835330525 | 0.921732596 |
| Klf11 | 345.8746993 | -0.025068645 | 0.12061491 | -0.207840347 | 0.835353628 | 0.921732596 |
| Zhx1 | 3147.920426 | 0.011156568 | 0.053700954 | 0.20775363 | 0.83542134 | 0.921741668 |
| Ephx4 | 1872.042929 | 0.017384738 | 0.08389211 | 0.20722733 | 0.835832324 | 0.922129452 |
| Rfc5 | 333.0806462 | -0.019786364 | 0.095521352 | -0.207140746 | 0.835899942 | 0.922138391 |
| Zfp62 | 1244.622908 | 0.017404757 | 0.084112472 | 0.206922423 | 0.836070445 | 0.922260819 |
| P2ry14 | 93.23992548 | 0.027735606 | 0.134142432 | 0.206762364 | 0.83619545 | 0.922262177 |
| Pipox | 71.6765521 | -0.034739045 | 0.168071385 | -0.206692204 | 0.836250246 | 0.922262177 |
| Rab2a | 8426.314297 | -0.01244967 | 0.060195116 | -0.20682193 | 0.836148929 | 0.922262177 |
| Dpys | 11.14665628 | 0.033847309 | 0.16387341 | 0.206545458 | 0.83636486 | 0.922322928 |
| Kctd11 | 83.42781557 | -0.027842401 | 0.134898816 | -0.206394704 | 0.836482607 | 0.922387126 |
| Pop4 | 774.3721191 | -0.013433448 | 0.0651506 | -0.206190709 | 0.836641945 | 0.922497175 |
| Fam217a | 31.35323975 | 0.034145353 | 0.165813146 | 0.205926694 | 0.836848174 | 0.922658906 |
| Mogs | 1004.493087 | -0.012914657 | 0.062741777 | -0.205838244 | 0.836917266 | 0.922669427 |
| Cyb5d2 | 541.4193384 | -0.019348723 | 0.094058563 | -0.205709318 | 0.83701798 | 0.922697724 |
| Fezf2 | 845.3938196 | -0.02729931 | 0.132744576 | -0.205652919 | 0.837062038 | 0.922697724 |
| BC052688 | 25.97601984 | 0.034506101 | 0.168095561 | 0.205276693 | 0.837355952 | 0.922767037 |
| Gpr182 | 77.84557444 | 0.034045223 | 0.166275571 | 0.204751803 | 0.837766045 | 0.922767037 |
| Kdm3b | 2668.277346 | -0.011036058 | 0.053892585 | -0.204778786 | 0.837744962 | 0.922767037 |
| Lor | 122.8689931 | 0.0306024 | 0.149518053 | 0.204673612 | 0.837827138 | 0.922767037 |
| Nln | 1236.738217 | -0.014556002 | 0.07103154 | -0.204923078 | 0.837632224 | 0.922767037 |
| Nova1 | 2308.875951 | -0.014838123 | 0.072398483 | -0.204950739 | 0.837610613 | 0.922767037 |
| Prkcg | 11518.57231 | -0.017296681 | 0.084465682 | -0.204777614 | 0.837745878 | 0.922767037 |
| Rad17 | 536.5242237 | -0.013012783 | 0.063583169 | -0.20465767 | 0.837839594 | 0.922767037 |
| Rgs3 | 1143.772897 | -0.01932083 | 0.094237874 | -0.205021921 | 0.837554998 | 0.922767037 |
| Sftpc | 26.13468982 | -0.034442023 | 0.168091537 | -0.204900398 | 0.837649945 | 0.922767037 |
| Spty2d1 | 850.3143967 | 0.018193092 | 0.088763003 | 0.204962559 | 0.837601377 | 0.922767037 |
| Tiparp | 676.3074774 | 0.029035233 | 0.141764924 | 0.204812531 | 0.837718596 | 0.922767037 |
| Gpr107 | 2040.234263 | -0.010773115 | 0.052662226 | -0.204570069 | 0.837908042 | 0.922776829 |
| Cct8 | 6209.580887 | -0.011888253 | 0.058287215 | -0.203959869 | 0.83838486 | 0.922846414 |
| Fam13b | 3080.89765 | 0.016367692 | 0.080261258 | 0.203930176 | 0.838408063 | 0.922846414 |
| Pik3r5 | 168.5248051 | 0.020483429 | 0.100377886 | 0.204063167 | 0.838304137 | 0.922846414 |
| Prr15 | 29.20239371 | -0.033680989 | 0.165162709 | -0.203926113 | 0.838411238 | 0.922846414 |
| Rgmb | 1773.137858 | -0.014780339 | 0.072399842 | -0.204148769 | 0.838237244 | 0.922846414 |
| Stpg1 | 55.08842667 | 0.033551728 | 0.164566518 | 0.203879431 | 0.838447719 | 0.922846414 |
| Trub2 | 899.9603082 | -0.012421951 | 0.06091361 | -0.203927345 | 0.838410276 | 0.922846414 |
| Zhx3 | 1364.20454 | -0.024919567 | 0.122048912 | -0.20417689 | 0.83821527 | 0.922846414 |
| Hpgds | 243.5163283 | -0.030655793 | 0.150473297 | -0.203729124 | 0.838565182 | 0.922910138 |
| Nudt7 | 145.790949 | -0.024818126 | 0.12196453 | -0.203486426 | 0.838754854 | 0.922948524 |
| Rrp12 | 1550.121614 | -0.016532463 | 0.081255207 | -0.203463423 | 0.838772832 | 0.922948524 |
| Timp1 | 14.35168281 | 0.030730609 | 0.151043143 | 0.203455833 | 0.838778764 | 0.922948524 |
| Atoh7 | 12.08333944 | -0.032941739 | 0.162168381 | -0.203132933 | 0.839031132 | 0.923095104 |
| Dnajb6 | 5090.944604 | -0.009920257 | 0.048832859 | -0.203147176 | 0.839019999 | 0.923095104 |
| Ankrd34c | 318.3573853 | -0.022384585 | 0.110400978 | -0.202757121 | 0.839324874 | 0.923352712 |
| Amigo1 | 5297.982592 | -0.007814362 | 0.038659396 | -0.202133587 | 0.839812291 | 0.923775413 |
| Gm2897 | 9.349600119 | 0.032099766 | 0.158820852 | 0.202113044 | 0.839828351 | 0.923775413 |
| Camsap1 | 5447.76945 | -0.017366465 | 0.086078348 | -0.20175184 | 0.840110733 | 0.923777635 |
| Fggy | 407.6595333 | -0.028003081 | 0.138815249 | -0.201729141 | 0.840128479 | 0.923777635 |
| Gjc2 | 457.5489394 | -0.033103777 | 0.163944241 | -0.201920949 | 0.839978525 | 0.923777635 |
| Inpp5d | 752.5291971 | -0.017911914 | 0.088764424 | -0.201791593 | 0.840079654 | 0.923777635 |
| Ttc38 | 355.9054899 | 0.017619569 | 0.08729119 | 0.201848199 | 0.8400354 | 0.923777635 |
| Rab3il1 | 388.1476444 | -0.022510052 | 0.111777417 | -0.201382828 | 0.840399241 | 0.924009782 |
| Gipc2 | 48.14432263 | 0.029903241 | 0.148723883 | 0.201065493 | 0.840647364 | 0.924217005 |
| Dhx30 | 4709.26039 | 0.011286047 | 0.056246284 | 0.200654085 | 0.840969064 | 0.924293811 |
| Foxo4 | 785.9827068 | -0.017644205 | 0.087992941 | -0.200518419 | 0.841075154 | 0.924293811 |
| Gpr153 | 1197.687537 | -0.024332031 | 0.121209407 | -0.200743752 | 0.840898947 | 0.924293811 |
| Gstt2 | 74.08820043 | -0.02868563 | 0.142884172 | -0.200761423 | 0.840885129 | 0.924293811 |

| | | | | | | |
|---|---|---|---|---|---|---|
| Pex11a | 172.9826394 | -0.00642879284251083 | 0.0867900049794559 | -0.0740729631716531 | 0.940952328408718 | 0.9854611256672 | NM_011068 |
| Pias1 | 2157.218428 | 0.00507397104551521 | 0.0636631271981601 | 0.0797003111349179 | 0.936475611992441 | 0.9854611256672 | NM_019663 |
| Polr1e | 254.4927396 | -0.00643973982935362 | 0.0862753621384661 | -0.0746417015209773 | 0.940499793632881 | 0.9854611256672 | NM_022811 |
| Prmt3 | 930.193407 | -0.00431864548146415 | 0.0594904952379303 | -0.072593873427879 | 0.94212930215469 | 0.9854611256672 | NM_133740 |
| Pter | 78.19600695 | 0.00722592110767787 | 0.0950872429502436 | 0.0759925399399684 | 0.939425033302543 | 0.9854611256672 | NM_008961 |
| Rasip1 | 360.4600868 | -0.00614743408105094 | 0.0796791863137026 | -0.077152320015519 | 0.938502371644895 | 0.9854611256672 | NM_028544 |
| Rc3h2 | 2416.822941 | -0.00580674934308611 | 0.0767265083308458 | -0.0756811364078673 | 0.939672783911786 | 0.9854611256672 | NM_001100591 |
| Rev3l | 1797.269011 | -0.00520673398110247 | 0.0710996352710973 | -0.0732315146378685 | 0.941621888651971 | 0.9854611256672 | NM_011264 |
| Rnf144b | 1493.494485 | -0.00393613947408034 | 0.0496989119926346 | -0.0791997111458634 | 0.936873774344549 | 0.9854611256672 | NM_146042,NM_001170643 |
| Rpa3 | 153.9752133 | -0.00726466517611145 | 0.0901212929485961 | -0.0806098640890018 | 0.935752221328503 | 0.9854611256672 | NM_026632 |
| Rpl14 | 7111.608304 | 0.00487593274380683 | 0.0666833720181958 | 0.0731206685600172 | 0.941710094549801 | 0.9854611256672 | NM_025974 |
| Rpl32 | 4250.97326 | -0.00500792403308558 | 0.0631705129273233 | -0.0792762920707501 | 0.936812863127848 | 0.9854611256672 | NM_172086 |
| Scarf2 | 109.8188694 | -0.0072557062074567 | 0.0946880434588511 | -0.0766274805393971 | 0.938919896912485 | 0.9854611256672 | NM_153790 |
| Scg3 | 12711.69994 | -0.00505257786064258 | 0.0641292915975785 | -0.0787873643193864 | 0.937201754615775 | 0.9854611256672 | NM_001164790,NM_009130 |
| Slc10a3 | 129.4227104 | -0.00728237058496451 | 0.0911001698762668 | -0.0799380571392512 | 0.936286521444673 | 0.9854611256672 | NM_145406,NM_001256104 |
| Slc1a3 | 25730.02288 | -0.00417796479572791 | 0.0572190038373617 | -0.0730170837577544 | 0.941792522925206 | 0.9854611256672 | NM_148938 |
| Slc25a40 | 416.674286 | 0.00537454905648498 | 0.0744911458811729 | 0.0721501729220058 | 0.942482398016347 | 0.9854611256672 | NM_178766 |
| Slc6a20a | 469.4821467 | -0.00636526645058337 | 0.0883837357538878 | -0.0720185268962607 | 0.942587163804256 | 0.9854611256672 | NM_139142 |
| Smc3 | 2429.404374 | 0.00436201239423463 | 0.0606234819225066 | 0.0719525216286731 | 0.942639692113087 | 0.9854611256672 | NM_007790 |
| Snip1 | 555.3148445 | 0.00468857090625061 | 0.0649310522144369 | 0.0722084541425025 | 0.942436017306181 | 0.9854611256672 | NM_175246 |
| Snrpe | 620.005571 | -0.00560259199131472 | 0.0726872814580527 | -0.0770780235404448 | 0.938561475655359 | 0.9854611256672 | NM_009227 |
| Stim2 | 3107.196888 | 0.00526170826528658 | 0.0720034102617162 | 0.0730758202446446 | 0.941745782853123 | 0.9854611256672 | NM_001081103 |
| Thsd7a | 1988.324923 | 0.00538100823565264 | 0.0718796838618922 | 0.0748613230685818 | 0.940325049889373 | 0.9854611256672 | NM_001164805 |
| Tk2 | 1802.166996 | 0.00422713507405445 | 0.0570784090279908 | 0.074058389959354 | 0.940963924300221 | 0.9854611256672 | NM_021028,NR_045642 |
| Tll1 | 91.43329482 | -0.00709389205849877 | 0.0939734276004804 | -0.0754882762035436 | 0.939826225155774 | 0.9854611256672 | NM_009390 |
| Tmem39b | 314.4369258 | 0.00618463143542279 | 0.0768782913506613 | 0.0804470459315118 | 0.93588171088202 | 0.9854611256672 | NM_199305 |
| Tprgl | 5720.650099 | 0.00385186120915747 | 0.0497471230166844 | 0.0774288235294655 | 0.938282411744789 | 0.9854611256672 | NM_026388 |
| Tpst2 | 795.155637 | 0.0056235748335687 | 0.0723911142962517 | 0.0776832196636028 | 0.938080042532675 | 0.9854611256672 | NM_009419 |
| Tvp23b | 881.7163569 | -0.00462083334901021 | 0.0621282445690531 | -0.0743757268704789 | 0.940711422449591 | 0.9854611256672 | NM_026210 |
| Ube2d1 | 1693.881823 | -0.00531558496887545 | 0.0716601389943402 | -0.0741777094417202 | 0.940868982267302 | 0.9854611256672 | NM_145420 |
| Uspl1 | 1735.579214 | 0.00458428785620254 | 0.0639651039672799 | 0.0716685750803679 | 0.94286566528341 | 0.9854611256672 | NM_001013378,NM_001115150,NM_001115151,NM_001115149,NM_001115153 |
| Wiz | 2055.749603 | 0.00397711434572251 | 0.0538504052188237 | 0.073854863850352 | 0.941125871143053 | 0.9854611256672 | NR_033525,NM_011717,NM_212438 |
| Zfp191 | 1394.196608 | -0.00518075301186536 | 0.0720615231884658 | -0.0718934707821246 | 0.942686686166341 | 0.9854611256672 | NM_021559 |
| Zfp259 | 1524.184613 | -0.00428070634373208 | 0.0529721915578282 | -0.08081044445856 | 0.935592701761267 | 0.9854611256672 | NM_011752 |
| Zfp263 | 943.4552145 | -0.00437807609507952 | 0.0547933472604213 | -0.0799015996279883 | 0.936315517580256 | 0.9854611256672 | NM_148924 |
| Zfp472 | 192.274666 | -0.00698150209168099 | 0.0920330692810678 | -0.0758586250161838 | 0.939531574421142 | 0.9854611256672 | NM_153063 |
| Zkscan14 | 381.2526345 | 0.00571653513780581 | 0.0748754303087283 | 0.0763472759252969 | 0.939142814822134 | 0.9854611256672 | NM_023322 |
| Nop9 | 471.2768741 | 0.00506379906992428 | 0.0708784984736883 | 0.0714433739281888 | 0.943044890378269 | 0.985515832923362 | NM_026403 |
| Zkscan17 | 617.4866582 | 0.00549701448537889 | 0.0770820557159418 | 0.0713138023411851 | 0.943148010518554 | 0.985552694107799 | NM_001130529,NM_172941 |
| Nt5m | 1235.092451 | 0.00366463521073714 | 0.0516189553999856 | 0.0709939823915568 | 0.943402544763567 | 0.985747760411133 | NM_134029 |
| Rbl2 | 2418.098025 | -0.00379831549771621 | 0.0538232013640056 | -0.0705702262492387 | 0.943739807324816 | 0.986029234064611 | NM_011250 |
| Gltscr1l | 2953.863421 | -0.00292862874658077 | 0.0416857460095041 | -0.0702549198930748 | 0.943990762516362 | 0.98622049887174 | NM_001100452 |
| Tax1bp3 | 284.4462723 | 0.00557327789472045 | 0.0794469581841218 | 0.0701509286460551 | 0.944073531314306 | 0.986236038538052 | NM_029564 |
| Zbtb17 | 1407.887688 | 0.00527035522491865 | 0.0752433987537488 | 0.0700440877500376 | 0.944158568836044 | 0.98625394584441 | NM_009541 |
| Fn3k | 1288.393926 | 0.00438514689443267 | 0.062856717353524 | 0.0697641728531472 | 0.944381363591311 | 0.986415739377823 | NM_001038699,NM_022014 |
| Ankrd12 | 3121.170217 | -0.00522257444520142 | 0.075048858161343 | -0.0695889927328904 | 0.944520798227853 | 0.98648395068099 | NM_001025572 |
| Rbm6 | 2301.653597 | -0.00350826747893686 | 0.0504703287565108 | -0.069511484576654 | 0.944582491396247 | 0.98648395068099 | NM_029169,NM_011251 |
| 4931440F15Rik | 24.11256228 | -0.00575233974859534 | 0.0829580434819686 | -0.0693403497377037 | 0.944718708565487 | 0.986555281137474 | NM_176829 |
| Nyx | 22.71452184 | -0.00587867898804783 | 0.0850559444329862 | -0.0691154395761195 | 0.944897731412597 | 0.986671299439248 | NM_173415 |
| Zbtb12 | 216.3888816 | 0.00590097821794864 | 0.0855277923431504 | 0.0689948618604936 | 0.944993709396213 | 0.986700591050128 | NM_198886 |
| Osgep | 748.9787549 | -0.00404469535691562 | 0.0587283113551182 | -0.0688713035261336 | 0.945092060734223 | 0.986732356373559 | NM_133676 |
| Gatad2a | 1424.566333 | -0.00396407912142765 | 0.0578728376718822 | -0.0684963668777143 | 0.945390512097084 | 0.986781137082095 | NM_001113346,NM_001113345,NM_145596 |
| Ikzf5 | 1002.176369 | 0.00383293916455785 | 0.0562317608455439 | 0.0681632427461427 | 0.945655686928499 | 0.986781137082095 | NM_175115 |
| Il17d | 219.5708673 | -0.00612559946007442 | 0.0896942366280418 | -0.0682942370698468 | 0.945551411572103 | 0.986781137082095 | NM_145837 |
| Jph3 | 11191.90093 | 0.00384038108703632 | 0.0562009331038865 | 0.0683330484911601 | 0.945520516711781 | 0.986781137082095 | NM_020605 |
| Obfc1 | 483.7242922 | -0.00482497532327823 | 0.0708683216891133 | -0.0680836685316823 | 0.945719030811665 | 0.986781137082095 | NM_175360 |
| Pdcd7 | 898.1852476 | 0.00381765529544887 | 0.0558602018877469 | 0.0683430271720211 | 0.945512573445929 | 0.986781137082095 | NM_016688 |
| Pgd | 1425.006829 | -0.00444778740054795 | 0.0652628359393529 | -0.0681519173436037 | 0.945664702327673 | 0.986781137082095 | NM_001081274 |
| Ppt1 | 5056.973455 | 0.00379460901973883 | 0.0557665393378631 | 0.0680445490215753 | 0.945750171448226 | 0.986781137082095 | NM_008917 |
| Setdb2 | 218.4210756 | 0.00611440889445031 | 0.0890451234654234 | 0.0686664093045428 | 0.945255156560859 | 0.986781137082095 | NM_001081024 |
| Cebpg | 1461.664759 | -0.00465981258550408 | 0.0687030880006625 | -0.0678253732271706 | 0.945924645353046 | 0.986798748684979 | NM_009884 |
| Dhrs11 | 198.6231972 | -0.00626816208244912 | 0.0922817837378071 | -0.0679241539181595 | 0.94584601110695 | 0.986798748684979 | NM_177564 |
| Prom2 | 23.62464428 | 0.00525777664921896 | 0.0775857131088683 | 0.0677673303310526 | 0.945970850574028 | 0.986798748684979 | NM_178047,NM_138750 |
| 1190003J15Rik | 28.26943196 | -0.00494589686646826 | 0.0734014045399128 | -0.0673815017228842 | 0.946277995203032 | 0.986835677961034 | NM_029821 |
| Pura | 585.0067479 | -0.00513800927206089 | 0.0762221236142936 | -0.0674083721159585 | 0.946256604365881 | 0.986835677961034 | NM_008989 |
| Sntb2 | 448.8835523 | 0.00574313244740185 | 0.0850229928610432 | 0.0675479920683115 | 0.946145457093592 | 0.986835677961034 | NM_009229 |
| Zfp101 | 336.5926581 | -0.00543997105005155 | 0.0804420980199737 | -0.0676259220476921 | 0.946083419821203 | 0.986835677961034 | NM_009542 |
| 9630033F20Rik | 1558.318974 | -0.00373014026267127 | 0.0556957149217866 | -0.0669735592389737 | 0.946602752597109 | 0.987003270241239 | NM_177003 |
| Omg | 5472.185215 | 0.00474801526861403 | 0.0707759406737726 | 0.0670851594964884 | 0.946513908284913 | 0.987003270241239 | NM_019409 |
| Seh1l | 1604.97342 | 0.00358880315524831 | 0.0536253918896636 | 0.0669235790879144 | 0.946642541718363 | 0.987003270241239 | NM_001039088,NM_028112 |
| Pon3 | 110.0013073 | 0.0046830866568432 | 0.0702526008789266 | 0.0666606872664264 | 0.946851831681188 | 0.987150628507927 | NM_173006 |
| Klf5 | 206.8216571 | 0.00542446398573719 | 0.0816472999983961 | 0.0664377632309182 | 0.947029305892697 | 0.987216964782958 | NM_009769 |
| Rhot2 | 3512.82576 | 0.00482460527440681 | 0.0726487426746263 | 0.0664100312928317 | 0.947051384018347 | 0.987216964782958 | NM_145999 |
| Antxr1 | 891.1906019 | -0.00480299093685921 | 0.0724888680667378 | -0.0662583244152367 | 0.947172162574804 | 0.987272017333639 | NM_054041 |
| Ruvbl2 | 933.1314018 | 0.00429117655003837 | 0.0648963748069775 | 0.0661235171117293 | 0.94727948787433 | 0.987313040170949 | NM_011304 |
| Mkl1 | 3139.587201 | 0.00469949308808329 | 0.0712176899084979 | 0.0659877215074134 | 0.947387600965526 | 0.987335056413536 | NM_001082536,NM_153049 |
| Upb1 | 29.32443846 | -0.00537386749882792 | 0.0815133356785428 | -0.0659262371499601 | 0.947436551793217 | 0.987335056413536 | NM_133995 |
| Chrna10 | 25.02863998 | -0.00546602602295703 | 0.0831174179581533 | -0.0657627024182703 | 0.947566751085372 | 0.987347864318807 | NM_001081424 |
| Edem2 | 888.4636587 | 0.00457682745846736 | 0.0696200763730245 | 0.0657400522508006 | 0.947584784281185 | 0.987347864318807 | NM_145537 |
| Clstn2 | 3200.494602 | -0.00362793494621963 | 0.0555728026594398 | -0.0652825621995758 | 0.947949026073425 | 0.987367301863797 | NM_022319 |
| Dnahc1 | 581.1062147 | 0.0061580064407349 | 0.0945654207469713 | 0.0651190085349684 | 0.948079245931174 | 0.987367301863797 | NM_001033668 |
| Gtf2ird2 | 728.9228891 | -0.00497916814943468 | 0.0764402872670638 | -0.0651380093855308 | 0.948064117565084 | 0.987367301863797 | NM_053266 |
| Gulp1 | 87.09385818 | 0.00619934552952368 | 0.0947095636579471 | 0.0654563836014832 | 0.947810632665926 | 0.987367301863797 | NM_028450 |
| Mars | 2355.168809 | -0.00355585707028953 | 0.0543746356947619 | -0.0653955106982368 | 0.947859098374451 | 0.987367301863797 | NM_001171582,NM_001003913 |
| Tspan5 | 8940.87061 | -0.00336813785271501 | 0.0513533743566911 | -0.0655874690788681 | 0.947706265857832 | 0.987367301863797 | NM_019571 |
| Ydjc | 463.3549999 | 0.00537825375517449 | 0.0822956117581144 | 0.0653528619603973 | 0.947893054505737 | 0.987367301863797 | NM_026940 |
| Tsta3 | 770.5214808 | -0.00433669484911189 | 0.0666951195803642 | -0.0650226714697827 | 0.948155949226978 | 0.987376393897131 | NM_031201 |
| Dcaf10 | 979.813922 | 0.00483012144020825 | 0.0747086210945671 | 0.0646527986923253 | 0.948450445332077 | 0.987378592162887 | NM_153167 |
| Gm16515 | 654.4632133 | -0.00421054775238591 | 0.0650904091566213 | -0.0646876829772945 | 0.94842266984234 | 0.987378592162887 | NM_025294 |
| Gtf3c3 | 1298.514192 | -0.00421470030079655 | 0.065032573882287 | -0.0648090649914714 | 0.948326023886524 | 0.987378592162887 | NM_001033194 |
| Phrf1 | 2447.189144 | -0.00363669657951625 | 0.0563171498891675 | -0.0645752952106648 | 0.948512155210338 | 0.987378592162887 | NM_001081118 |
| St6galnac6 | 4872.693625 | -0.00314188834538546 | 0.0487055598772725 | -0.0645077965082907 | 0.948565899327625 | 0.987378592162887 | NM_016973,NM_001025310,NM_001025311 |
| Tub | 10736.14774 | -0.0037860288129508 | 0.0586238703406722 | -0.0645816932752752 | 0.948507060927003 | 0.987378592162887 | NM_021885 |
| Cars | 1726.344533 | 0.00324460730828969 | 0.0504233051966757 | 0.0643473746045429 | 0.948693632108527 | 0.987440792491292 | NM_013742,NM_001252593 |
| Prc1 | 43.57620233 | 0.00573888082741423 | 0.0895329956189065 | 0.0640979427499728 | 0.948892239927191 | 0.987576748383061 | NM_145150 |
| Bcl7a | 4411.218361 | -0.00344205325279396 | 0.054218866636534 | -0.0634844191020884 | 0.949380765970865 | 0.987872851467964 | NM_029850 |
| Msx1 | 116.7575214 | -0.00459488226366666 | 0.0722033942896302 | -0.0636380368107788 | 0.949258444114654 | 0.987872851467964 | NM_010835 |
| Tacc1 | 10441.74042 | -0.00453194570864842 | 0.0713298787788981 | -0.0635350260820733 | 0.949340468793901 | 0.987872851467964 | NM_199323,NM_177089 |
| Lamb1 | 790.3441974 | -0.00488373065441804 | 0.0770667344826068 | -0.0633701516900298 | 0.949471754965202 | 0.98789676331384 | NM_008482 |
| Htt | 8226.465366 | 0.00418743648973951 | 0.0664552447060816 | 0.0630113771796299 | 0.949757444646714 | 0.987981713043409 | NM_010414 |
| Il18 | 624.1089526 | 0.00446259519935554 | 0.0706893293866468 | 0.0631296864473936 | 0.94966323507284 | 0.987981713043409 | NM_008360 |
| Sema3b | 182.502565 | -0.00362331557024209 | 0.0574843643458631 | -0.0630313235863909 | 0.949741561289944 | 0.987981713043409 | NM_009153,NM_001042779 |
| Elp2 | 4317.915401 | 0.00248928328667958 | 0.0405562740999119 | 0.0613785004151795 | 0.951057774254637 | 0.988202090759861 | NM_021448 |
| Fam60a | 92.26203855 | 0.00581028275566063 | 0.0933538746980943 | 0.0622393315162443 | 0.950372241167685 | 0.988202090759861 | NM_019643 |
| Fhod3 | 3041.551438 | 0.00441725563626854 | 0.0709736275418874 | 0.0622379859851682 | 0.950373312668836 | 0.988202090759861 | NM_175276 |
| Ggt1 | 83.3602167 | -0.00586499686011221 | 0.0949655698123367 | -0.061759192007189 | 0.950754601479653 | 0.988202090759861 | NM_008116 |
| Gm7120 | 451.8071838 | 0.00389573283591831 | 0.0629176278985084 | 0.0619179865172676 | 0.950628143810917 | 0.988202090759861 | NM_001039244,NM_001177666 |
| Hn1 | 3004.893843 | -0.00307719703286598 | 0.0500309385124336 | -0.0615058826470193 | 0.95095632960346 | 0.988202090759861 | NM_008258 |
| Jmjd4 | 1709.149775 | 0.00351480309687136 | 0.0568374111869362 | 0.0618396057010989 | 0.950690563038364 | 0.988202090759861 | NR_037992,NM_001205068,NM_178659 |
| Mageh1 | 853.2570649 | -0.00383843960443175 | 0.0620424019392983 | -0.0618680045332101 | 0.950667947352684 | 0.988202090759861 | NM_023788 |
| Mir9-2 | 50.11727891 | 0.00570956071151693 | 0.0914831383708663 | 0.0624110717362009 | 0.950235478179176 | 0.988202090759861 | NR_029545 |
| Mrps33 | 1682.549137 | 0.00328994643420208 | 0.053579920548529 | 0.0614026000882601 | 0.951038581698113 | 0.988202090759861 | NM_010270,NM_001010930 |
| Nelfb | 2317.280311 | 0.00312622055029684 | 0.0507067719386786 | 0.0616529199310397 | 0.950839233051778 | 0.988202090759861 | NM_021393 |
| Rpl23a | 7434.902101 | 0.00427528972360462 | 0.0684764528027605 | 0.0624344508019298 | 0.950216860691405 | 0.988202090759861 | NM_207523 |
| Sod3 | 627.7371786 | 0.0047788011225228 | 0.0777955154974151 | 0.061427719733814 | 0.951018576883649 | 0.988202090759861 | NM_011435 |
| Tex10 | 876.8678412 | 0.00476768126854945 | 0.0774162881957164 | 0.0615849891497801 | 0.95089333117333 | 0.988202090759861 | NM_172304 |
| Tgs1 | 1418.371686 | -0.0044257418075848 | 0.0721023542490132 | -0.0613813772612754 | 0.951055483183422 | 0.988202090759861 | NM_054089 |
| Tmbim1 | 1668.979178 | 0.00466885563429373 | 0.0755989613408789 | 0.0617581981482744 | 0.950755392953502 | 0.988202090759861 | NM_027154 |
| Dhx37 | 758.1138608 | 0.00418708589158234 | 0.0685536450906658 | 0.0610775092417727 | 0.951297480728531 | 0.988239090686022 | NM_203319 |
| Megf8 | 9826.483825 | -0.00386701414292492 | 0.0632396157760241 | -0.0611486027464293 | 0.951240862147283 | 0.988239090686022 | NM_001160400 |
| Rab11fip2 | 2866.811064 | 0.00449041608211923 | 0.0733206979648771 | 0.061243498858539 | 0.95116528764876 | 0.988239090686022 | NM_001164367,NM_001033172 |
| Mid1ip1 | 3375.886461 | -0.00410278372861241 | 0.0673548790041068 | -0.0609129403730687 | 0.951428543656605 | 0.988304564155881 | NM_026524,NM_001166635 |
| Atpaf1 | 2225.955664 | -0.00405574844992857 | 0.0668688363076516 | -0.0606522959554552 | 0.951636124002739 | 0.988430624036573 | NM_181040 |
| Mroh5 | 91.62659939 | 0.00576007080966142 | 0.095066860236157 | 0.0605896817813562 | 0.95168599117276 | 0.988430624036573 | NM_001033365 |
| 5830417I10Rik | 385.1354286 | 0.00483948601931126 | 0.0799900198587562 | 0.0605011228632855 | 0.951756521574384 | 0.988433204574927 | NR_028359 |
| 1110007C09Rik | 182.3023159 | -0.00546029636565158 | 0.0905614287497747 | -0.0602938407778285 | 0.951921607370453 | 0.988463311792351 | NM_026738 |
| Cndp2 | 2066.915182 | 0.00310447845512235 | 0.0514285695833229 | 0.0603648610154824 | 0.951865044447126 | 0.988463311792351 | NM_023149 |
| Cntnap1 | 13343.88059 | -0.00300116839634086 | 0.0504614364605894 | -0.0594744939273537 | 0.95257418040403 | 0.988510896443608 | NM_016782 |
| Gm514 | 65.46510476 | -0.00516520170065205 | 0.0869044988719476 | -0.0594353775431452 | 0.952605335649198 | 0.988510896443608 | NM_001111145 |
| Gpatch11 | 1151.569811 | -0.00354200073433217 | 0.0590086932268259 | -0.0600250660814823 | 0.952135670834717 | 0.988510896443608 | NM_181649 |
| Hibch | 566.0517258 | 0.00404577124291089 | 0.0675844699694616 | 0.059862439473728 | 0.952265195178009 | 0.988510896443608 | NM_146108 |
| Hn1l | 226.3204416 | 0.00559734633503333 | 0.094133097794897 | 0.0594620432786475 | 0.952584097033936 | 0.988510896443608 | NM_198937 |
| Lrrc10b | 1986.978125 | -0.00561585338276583 | 0.0948319249431654 | -0.0592190170781783 | 0.952777662785096 | 0.988510896443608 | NM_001111140 |
| Prkrip1 | 576.9201059 | -0.00403227522415253 | 0.0680984628773532 | -0.0592124264451423 | 0.952782912137937 | 0.988510896443608 | NM_025774 |
| Rpap2 | 727.8441722 | 0.0040783779697979 | 0.0684027628846307 | 0.0596230005603218 | 0.952455899156346 | 0.988510896443608 | NM_001163461,NM_001163462,NM_144911 |
| Syde1 | 593.7899244 | 0.00447424407102557 | 0.0755644037868861 | 0.0592110020962285 | 0.952784046613449 | 0.988510896443608 | NM_027875 |
| Tmem184b | 3836.107586 | 0.00358406214842265 | 0.0596814380463622 | 0.060053213624619 | 0.952113252786209 | 0.988510896443608 | NM_001253819,NM_172608,NM_001253817,NM_001253820 |
| Vimp | 1236.997722 | -0.0038155690110708 | 0.064233250270252 | -0.0594017739257683 | 0.952632100167997 | 0.988510896443608 | NM_024439 |
| Zbtb45 | 987.4302567 | -0.00363950366995741 | 0.0611715574041553 | -0.0594966651888805 | 0.952556521567618 | 0.988510896443608 | NM_001024699 |
| Acta2 | 488.1541435 | 0.00530303974195974 | 0.0901446492735639 | 0.0588281144215947 | 0.953089015163159 | 0.988615469455874 | NM_007392 |
| Cmpk2 | 1116.615678 | -0.00379786474590898 | 0.0645452548075948 | -0.0588403401184202 | 0.95307927733659 | 0.988615469455874 | NM_020557 |
| March8 | 2873.753103 | 0.00358800547766918 | 0.0608496727494407 | 0.058965074347128 | 0.952979926321555 | 0.988615469455874 | NM_027920 |
| Gns | 5561.183148 | -0.0031208257603715 | 0.053310102794433 | -0.0585409818548951 | 0.953317719650023 | 0.988782091798374 | NM_029364 |
| 0610010F05Rik | 1691.971804 | -0.00384308158726365 | 0.0670235375451905 | -0.0573392830044588 | 0.954274928499552 | 0.988860025065572 | NM_027860 |
| Dand5 | 303.4825889 | 0.00435552675491392 | 0.0757465862623009 | 0.0575012943795418 | 0.95414587505069 | 0.988860025065572 | NM_201227,NR_033145 |
| Dnttip1 | 799.0132628 | -0.00322149317268269 | 0.0562760997277288 | -0.0572444285987958 | 0.954350487258113 | 0.988860025065572 | NM_133763 |
| Efcab1 | 134.4967677 | 0.0051735291548116 | 0.0903643888531486 | 0.0572518579550083 | 0.954344569195226 | 0.988860025065572 | NM_025769 |
| Efhc2 | 76.50053619 | 0.00544325432948647 | 0.0948126701309987 | 0.0574106216180364 | 0.954218102131158 | 0.988860025065572 | NM_028916 |
| Eml1 | 2756.897811 | -0.00344400663350596 | 0.0596724639679247 | -0.0577151738757962 | 0.953975506838989 | 0.988860025065572 | NM_001043335,NM_001043336 |
| Gnpda2 | 723.0288657 | 0.0039266944554513 | 0.0679041978087497 | 0.0578269765664668 | 0.95388644993594 | 0.988860025065572 | NM_001038015 |
| H6pd | 459.8175166 | 0.00498104436632303 | 0.0873962528420505 | 0.0569937978385089 | 0.95455013572005 | 0.988860025065572 | NM_173371 |
| Hcn3 | 688.2607413 | 0.00457611234767168 | 0.0801672153378317 | 0.0570820918300272 | 0.954479801810142 | 0.988860025065572 | NM_008227 |
| Med16 | 2452.075047 | 0.00378355916246947 | 0.065808268662396 | 0.0574936742657608 | 0.954151944980056 | 0.988860025065572 | NM_001163276,NM_198107 |
| Naaa | 1451.96377 | -0.00432519726730789 | 0.0754682167981018 | -0.0573115074240985 | 0.954297053822277 | 0.988860025065572 | NM_001163687,NM_025972 |
| Pars2 | 362.3155421 | -0.00442670653969861 | 0.076559627388091 | -0.0578203772761203 | 0.953891706612417 | 0.988860025065572 | NM_001083887,NM_172272 |
| Parvb | 998.8332647 | -0.00400567509008751 | 0.0695008844658344 | -0.0576348793382139 | 0.954039466144917 | 0.988860025065572 | NM_133167 |
| Pde3a | 90.33810263 | 0.00541937712206401 | 0.0939655576223526 | 0.0576740803672392 | 0.95400824019039 | 0.988860025065572 | NM_018779 |
| Qrsl1 | 528.5759043 | 0.00376421723443042 | 0.0651878913595986 | 0.0577441171346641 | 0.953952451909346 | 0.988860025065572 | NM_001081054 |
| Rltpr | 2153.53289 | 0.00342969329246039 | 0.06010503041206 | 0.0570616680325683 | 0.954496071125018 | 0.988860025065572 | NM_001033320 |
| Unc5b | 1589.670146 | -0.0047297256061118 | 0.0824902003683066 | -0.0573368180098275 | 0.954276892050321 | 0.988860025065572 | NM_029770 |
| Kxd1 | 1020.935424 | 0.00340647373105479 | 0.0599403721737687 | 0.0568310407079111 | 0.954679786979718 | 0.988923809860043 | NM_029366 |
| Dlx5 | 195.0280668 | 0.00482906986354155 | 0.0854117599580962 | 0.0565386998922717 | 0.954912666759787 | 0.988953471934455 | NM_010056,NM_198854 |
| Haus3 | 210.2873934 | -0.00502340436009902 | 0.088739246055264 | -0.0566085986010136 | 0.954856984837771 | 0.988953471934455 | NM_146159 |
| Zfp109 | 336.5568628 | -0.00446052311566367 | 0.0786695054409111 | -0.0566995189643588 | 0.954784557211777 | 0.988953471934455 | NM_020262 |
| Trip11 | 1172.032843 | 0.00473379567437838 | 0.0842843519441578 | 0.056164585301845 | 0.955210693444863 | 0.989191597132678 | NM_028446 |
| Abhd16a | 3019.27487 | -0.00342412536692883 | 0.0613015055447254 | -0.0558571169908804 | 0.955455633146349 | 0.989344398053186 | NM_178592 |
| Anpep | 194.6729138 | -0.00528844113023921 | 0.0950480890511334 | -0.0556396365569661 | 0.955628887989452 | 0.989344398053186 | NM_008486 |
| Aph1a | 2358.991537 | 0.00273048890193635 | 0.0499937138925117 | 0.0546166445606941 | 0.956443878109138 | 0.989344398053186 | NM_146104,NM_146134 |
| Cd200r1 | 24.3849909 | -0.00445766089777659 | 0.0799484213402055 | -0.055756709426542 | 0.95553562213534 | 0.989344398053186 | NM_021325 |
| Cul5 | 2679.123418 | 0.00386554532433463 | 0.0707826329898627 | 0.0546114938234672 | 0.956447981678431 | 0.989344398053186 | NM_027807,NM_001161618 |
| Gng7 | 12528.69263 | -0.00417405989234032 | 0.0763251281743888 | -0.054687885787802 | 0.956387120660836 | 0.989344398053186 | NM_001038655,NM_010319 |
| Il1rapl2 | 172.5155861 | -0.00499968250815088 | 0.0909247346872575 | -0.0549870453331282 | 0.956148784504837 | 0.989344398053186 | NM_030688 |
| Klk14 | 33.55137992 | 0.00499087095981279 | 0.0905543330489704 | 0.055114656491521 | 0.956047119701098 | 0.989344398053186 | NM_174866 |
| Lztfl1 | 1471.578493 | -0.00383825790325586 | 0.0690117014738181 | -0.055617494153684 | 0.955646527756585 | 0.989344398053186 | NM_033322 |
| Mapk15 | 53.84247939 | 0.00463652053933074 | 0.0847205390139846 | 0.0547272313572675 | 0.956355774381947 | 0.989344398053186 | NM_177922 |
| Mettl15 | 150.4015073 | 0.00512544434055454 | 0.0929033944644153 | 0.055169613232138 | 0.956003337180649 | 0.989344398053186 | NM_029790 |
| Msto1 | 749.0445541 | -0.00368988339561035 | 0.0675391985886796 | -0.0546332126041664 | 0.956430678430893 | 0.989344398053186 | NM_144898 |
| Pkdcc | 332.0798693 | -0.00491344762021854 | 0.0892277173885782 | -0.0550663825548842 | 0.956085578325227 | 0.989344398053186 | NM_134117 |
| Prkab1 | 607.2578945 | -0.0045607888720411 | 0.0824570570032692 | -0.0553110799462595 | 0.955890635159801 | 0.989344398053186 | NM_031869 |
| Spata21 | 27.45849805 | -0.00439707415413879 | 0.0803169285609295 | -0.0547465426395522 | 0.956340389273103 | 0.989344398053186 | NM_177867 |
| Syt3 | 2434.435512 | 0.00346220965566086 | 0.0633247751956966 | 0.0546738562427009 | 0.956398297895817 | 0.989344398053186 | NM_001114116,NM_016663 |
| A330009N23Rik | 498.2306525 | -0.00409680533369072 | 0.078228679903219 | -0.0523696084193048 | 0.958234189842925 | 0.989630473997713 | NR_045326,NR_045327,NR_045328 |
| A430035B10Rik | 84.03437181 | 0.00498550982913877 | 0.0950451399931504 | 0.0524541268443401 | 0.958166846456329 | 0.989630473997713 | NR_040452 |
| Alyref | 547.0065176 | 0.0037051575359417 | 0.0710871256088604 | 0.0521213581813453 | 0.958431994727027 | 0.989630473997713 | NM_011568 |
| Ankle2 | 1755.497957 | 0.00297514137087875 | 0.0570308181787329 | 0.0521672573862565 | 0.958395422214749 | 0.989630473997713 | NM_027922,NM_001253814 |
| Ankrd49 | 372.218217 | -0.00395763544077779 | 0.0732180023640277 | -0.0540527645209587 | 0.956893125643568 | 0.989630473997713 | NM_019683 |
| Atp1b3 | 4049.763391 | 0.00298899986351119 | 0.0567116557302337 | 0.0527052124474955 | 0.957966785867799 | 0.989630473997713 | NM_007502 |
| Disc1 | 35.72378749 | -0.00477441985099317 | 0.0889373421323635 | -0.0536829607960119 | 0.957187758539293 | 0.989630473997713 | NM_174854,NM_174853 |
| Gm19522 | 43.85926249 | -0.00465976388183316 | 0.0885847862232702 | -0.0526023043063922 | 0.95804878094359 | 0.989630473997713 | NR_040402 |
| Gpr63 | 179.8116388 | 0.004925608902756 | 0.0914450224443525 | 0.053864155435616 | 0.957043395005482 | 0.989630473997713 | NM_030733 |
| Kcna4 | 1113.829681 | 0.00361961287309045 | 0.0683989438259712 | 0.0529191339898449 | 0.957796339036893 | 0.989630473997713 | NM_021275 |
| Nme5 | 311.1559055 | 0.00399060347343522 | 0.076807837131946 | 0.0519556808581899 | 0.958564007238595 | 0.989630473997713 | NM_080637 |
| Npdc1 | 6559.044262 | 0.00304277618007267 | 0.0569464931528696 | 0.0534321959370618 | 0.957387553195621 | 0.989630473997713 | NM_008721 |
| Pecr | 161.7224913 | -0.00481547833700395 | 0.0900613832593345 | -0.0534688471654675 | 0.957358351490178 | 0.989630473997713 | NM_023523 |
| Plekha1 | 3911.807691 | -0.00338223016456532 | 0.0648935315520659 | -0.0521196810170775 | 0.958433331094116 | 0.989630473997713 | NM_133942 |
| Ptbp3 | 946.8823875 | -0.00436291482997548 | 0.0813154437685101 | -0.0536541968878124 | 0.95721067578931 | 0.989630473997713 | NM_178164,NM_144904 |
| Ptchd1 | 722.3166706 | 0.00427829953342037 | 0.0814797473471382 | 0.0525075208590548 | 0.958124302824333 | 0.989630473997713 | NM_001093750 |
| Ralgapa1 | 6907.960971 | -0.00327720938608622 | 0.0623381914739087 | -0.0525714543300135 | 0.958073361652437 | 0.989630473997713 | NM_019994,NM_001112714,NM_001003719 |
| Sh3d19 | 740.1844444 | 0.0039705085797551 | 0.074385202263285 | 0.0533776673175071 | 0.957430998739854 | 0.989630473997713 | NM_001082414 |
| Slc22a2 | 25.43934457 | 0.00366120524603829 | 0.0696404988871182 | 0.0525729324824742 | 0.958072183886111 | 0.989630473997713 | NM_013667 |
| Snrnp25 | 360.9419796 | -0.00388675566736259 | 0.0723036258257234 | -0.0537560270729854 | 0.957129544142709 | 0.989630473997713 | NM_030093 |
| Suv39h2 | 173.4107454 | -0.00478556174528034 | 0.0883606387624229 | -0.0541594290433706 | 0.956808144149447 | 0.989630473997713 | NR_027509,NM_022724 |
| Tbx6 | 39.51283578 | -0.00450829493410271 | 0.084181395319454 | -0.0535545284916519 | 0.957290085492757 | 0.989630473997713 | NM_011538 |
| Tecpr1 | 4367.512512 | 0.00312857272794109 | 0.0601213054009073 | 0.0520376712893841 | 0.958498676714727 | 0.989630473997713 | NM_027410 |
| Whsc1l1 | 3737.940638 | -0.00363862013859045 | 0.0694211706907923 | -0.0524136960293736 | 0.958199061264589 | 0.989630473997713 | NM_001081269,NM_001001735 |
| Wnk1 | 10331.97098 | 0.00381766282649341 | 0.0728837148954134 | 0.0523801898952554 | 0.95822575861854 | 0.989630473997713 | NM_198703,NM_001199084,NM_001185021,NM_001199083,NM_001185020 |
| Zfp113 | 405.7193929 | 0.00372388025541701 | 0.0712160072691103 | 0.0522899330953118 | 0.958297674570819 | 0.989630473997713 | NM_019747 |
| Zmat1 | 595.6323751 | -0.00385245351210371 | 0.0726469698245891 | -0.0530297894241937 | 0.957708172571628 | 0.989630473997713 | NM_175446 |
| Osbpl6 | 3314.779031 | 0.00337446070353988 | 0.0652513682806159 | 0.051714788401492 | 0.958755953567366 | 0.98975829589365 | NM_145525 |
| Fam221a | 39.44116953 | 0.00453697457494735 | 0.0880447011070832 | 0.0515303535351811 | 0.958902915350463 | 0.98976932767575 | NM_001172216,NM_172727 |
| Rlf | 1638.363162 | 0.00303046899356096 | 0.0587990852388712 | 0.0515393901325112 | 0.958895714757146 | 0.98976932767575 | NM_001081013 |
| Isyna1 | 1270.942009 | 0.00330783830802415 | 0.064601390666312 | 0.0512038250865257 | 0.959163103866792 | 0.989967546310148 | NM_023627 |
| Rffl | 690.8468133 | -0.00382245934992778 | 0.0751150195097063 | -0.0508880830342305 | 0.959414701568112 | 0.990016193434567 | NM_001164569,NM_001007465,NM_026097,NM_001164570,NM_001164571 |
| Tefm | 158.1871947 | -0.00470196385548837 | 0.0921278299047813 | -0.0510373885974313 | 0.95929572756453 | 0.990016193434567 | NM_183275 |
| Xpo4 | 357.536599 | 0.00426582434878088 | 0.0836894171306038 | 0.0509720881688509 | 0.959347762040849 | 0.990016193434567 | NM_020506 |
| 1110020A21Rik | 24.36474314 | 0.00404174733846394 | 0.0800818822801184 | 0.05047018405894 | 0.959747708788708 | 0.990191032543401 | NR_027930,NR_027929 |
| AA987161 | 436.5951786 | -0.00434466780291273 | 0.086171543413849 | -0.050418823091597 | 0.959788636803737 | 0.990191032543401 | NM_001163246 |
| Fam45a | 1135.876909 | 0.00285546464479577 | 0.0564655723191375 | 0.0505700115567231 | 0.959668159549295 | 0.990191032543401 | NM_001167829,NM_026437 |
| Pop4 | 858.73021 | -0.00324363000334392 | 0.0646857237722669 | -0.050144449411488 | 0.96000727876154 | 0.990205618922824 | NM_025390 |
| Psph | 507.969375 | 0.00392611385125387 | 0.0782148384748549 | 0.0501965346705417 | 0.959965773007001 | 0.990205618922824 | NM_133900 |
| Tagap1 | 846.0640037 | -0.00304933490863407 | 0.0607720824249365 | -0.0501765742913368 | 0.959981679041591 | 0.990205618922824 | NM_147155 |
| Rbmx2 | 229.9177574 | -0.00414197911010462 | 0.0831592507386777 | -0.0498077973684552 | 0.960275552995901 | 0.990412005312302 | NM_173376 |
| Cdc45 | 75.62735604 | 0.00464488616660512 | 0.0951018303116933 | 0.0488411858255688 | 0.961045859780352 | 0.990548518419332 | NM_009862,NM_001161623 |
| Cul4a | 2972.301681 | 0.0026133460391162 | 0.0534934115488147 | 0.0488536057703372 | 0.961035961913714 | 0.990548518419332 | NM_146207 |
| Klhl29 | 2811.520509 | 0.00353761889088608 | 0.0723879088662614 | 0.0488703009424118 | 0.961022656985854 | 0.990548518419332 | NM_001164493 |
| Nsa2 | 1784.717678 | -0.00326639203261761 | 0.0666189048041223 | -0.0490310076729975 | 0.96089458509958 | 0.990548518419332 | NM_021552 |
| Pigyl | 489.616155 | 0.00346443503288001 | 0.0703715372132993 | 0.0492306288887672 | 0.960735502529404 | 0.990548518419332 | NM_001082532 |
| Pla2g6 | 949.9523014 | 0.00338525192787274 | 0.0687043020827036 | 0.0492727795094652 | 0.960701911965465 | 0.990548518419332 | NM_016915,NM_001199023,NM_001199024,NM_001199025 |
| Star | 68.46508068 | 0.00460516056202628 | 0.093223514398022 | 0.0493991305923561 | 0.960601221004587 | 0.990548518419332 | NM_011485 |
| Stx1a | 11587.47922 | -0.00346584462708473 | 0.0700499473355071 | -0.0494767627802049 | 0.96053935513023 | 0.990548518419332 | NM_016801 |
| Szrd1 | 1386.992041 | 0.00308907751711267 | 0.0634301646814649 | 0.0487004492677179 | 0.961158017838392 | 0.990548518419332 | NM_174996,NM_001025608 |
| Tcp11l2 | 1157.458083 | 0.003023454487371 | 0.0617961764341795 | 0.0489262388360119 | 0.960978078330754 | 0.990548518419332 | NM_146008 |
| Zfp493 | 153.2149501 | -0.00447488459336167 | 0.0918043969697888 | -0.0487436848458825 | 0.961123561758987 | 0.990548518419332 | NM_028402 |
| Arf3 | 61608.71545 | -0.00258413256571669 | 0.0537646871188749 | -0.048063751584811 | 0.961665434845361 | 0.99057216324187 | NM_007478 |
| Btbd16 | 35.73183486 | 0.00446568007843594 | 0.092761360272762 | 0.0481415975930575 | 0.961603394535336 | 0.99057216324187 | NM_001081038 |

| | | | | | | | | | | | | | |
|---|---|---|---|---|---|---|---|---|---|---|---|---|---|
| Kcnrg | 18.47035517 | -0.033143681 | 0.165250239 | -0.200566618 | 0.841037463 | 0.924293811 | | Ccdc166 | 234.1979389 | 0.00440608533262513 | 0.0909185571471212 | 0.0484618923889802 | 0.961348134206248 | 0.99057216324187 | NM_146059,NM_001163518 |
| Tram1 | 866.7421984 | 0.01216284 | 0.060559817 | 0.20084011 | 0.840823599 | 0.924293811 | | Dtx4 | 2221.806757 | -0.002584838396871 | 0.0539614692679016 | -0.0479015570172505 | 0.961794698496719 | 0.99057216324187 | NM_172442 |
| Fkbpl | 252.9610019 | 0.026551754 | 0.132477973 | 0.200423916 | 0.841149057 | 0.924309467 | | Gpr52 | 416.5177017 | 0.00452756806827701 | 0.0943758633400979 | 0.04797379232401 | 0.961737129246403 | 0.99057216324187 | NM_001146330 |
| A330035P11Rik | 23.35424737 | 0.032955813 | 0.164923502 | 0.199824841 | 0.841617573 | 0.924555649 | | Pprc1 | 1234.624616 | -0.00351311413 4245 | 0.0732059657556153 | -0.0479894513784967 | 0.961724649502521 | 0.99057216324187 | NM_001081214 |
| BC020402 | 133.750866 | 0.025546446 | 0.127708838 | 0.200036631 | 0.841451933 | 0.924555649 | | Rnf145 | 5013.386771 | -0.00255334473252758 | 0.0530998623067728 | -0.048085712873909 | 0.96164793250907 | 0.99057216324187 | NM_028862,NM_001166553 |
| Camk1g | 1627.500206 | -0.027427099 | 0.137303035 | -0.199755956 | 0.841671449 | 0.924555649 | | Tro | 9626.730052 | -0.00280199349761436 | 0.0584359369124212 | -0.0479498343940946 | 0.961756222935211 | 0.99057216324187 | NM_019548,NM_001002272 |
| Ddx19b | 1465.477765 | -0.013979938 | 0.069910806 | -0.199968198 | 0.841505453 | 0.924555649 | | Uba6 | 852.204835 | 0.00432989588260635 | 0.0898494504300342 | 0.0481905661290387 | 0.961564368592318 | 0.99057216324187 | NM_172712 |
| Zfp398 | 526.993776 | 0.015029909 | 0.07520725 | 0.199846541 | 0.841600601 | 0.924555649 | | Acot5 | 94.24578925 | 0.00442556577853483 | 0.0926598660402923 | 0.047761409201805 | 0.961906392432166 | 0.990595298076336 | NM_145444 |
| Nt5dc1 | 68.25857394 | -0.031338288 | 0.15710204 | -0.199477284 | 0.841889411 | 0.924729515 | | Zfp862 | 604.5282903 | 0.0031226937764719 | 0.0654622070263757 | 0.0477022379525597 | 0.961953550507008 | 0.990595298076336 | NR_027808,NR_015597 |
| Snrpa1 | 659.1078367 | -0.015461882 | 0.07754719 | -0.199386747 | 0.841960227 | 0.924741742 | | Elmo2 | 7909.753245 | 0.00274371476160825 | 0.0578632010334519 | 0.0474172654226657 | 0.962180668680087 | 0.990758941890334 | NM_207706,NM_207705,NM_080287 |
| Rasal1 | 3560.212332 | -0.020937242 | 0.105057815 | -0.199292568 | 0.842033892 | 0.924757097 | | Gtf2h5 | 1750.806132 | 0.00208973879735396 | 0.0442183349113797 | 0.0472595542442846 | 0.962306363079141 | 0.990818133689226 | NM_181392 |
| Aqp6 | 12.38268249 | -0.031418642 | 0.15795969 | -0.198902912 | 0.842338692 | 0.924950886 | | Camk2g | 14345.41197 | 0.0021771066315428 | 0.0463192776475905 | 0.0470021714955665 | 0.962511496840062 | 0.99088887335923 | NM_178597,NM_001039139,NM_001039138 |
| Fam194a | 74.8162093 | 0.032338688 | 0.162638335 | 0.19883804 | 0.842389438 | 0.924950886 | | Smpd3 | 4814.58865 | 0.00335458930122046 | 0.0712705307562805 | 0.0470683923021696 | 0.962458718694194 | 0.99088887335923 | NM_021491 |
| Tesk1 | 3904.763772 | -0.014274921 | 0.07177418 | -0.198886573 | 0.842351473 | 0.924950886 | | 2310014L17Rik | 46.37805263 | -0.00404236793572561 | 0.0931165249644287 | -0.0434119286267376 | 0.965373168997424 | 0.99099371420617 | NM_029809 |
| Dnaja3 | 2653.465148 | 0.009887132 | 0.049745159 | 0.198755673 | 0.842453872 | 0.924956086 | | 4930481A15Rik | 71.31914043 | -0.00427534313572456 | 0.094155093032659 | -0.0454074548494323 | 0.96378253900766 | 0.99099371420617 | NR_027939,NR_027938,NR_027940,NR_015545 |
| Adrbk1 | 10165.95731 | 0.017915739 | 0.090288307 | 0.198428122 | 0.842710116 | 0.925040775 | | 6330408A02Rik | 648.1914189 | -0.00318771039321961 | 0.0702440292165817 | -0.0453805174442801 | 0.963804009814416 | 0.99099371420617 | NM_177312 |
| Tatdn1 | 255.6895049 | 0.022954524 | 0.115678283 | 0.198434173 | 0.842705383 | 0.925040775 | | Ccdc122 | 57.62077178 | -0.0041997713789843 | 0.0942540559120075 | -0.0445579910418399 | 0.964459627671757 | 0.99099371420617 | NM_175369 |
| Zfp609 | 2542.565259 | 0.019934032 | 0.10038408 | 0.198577628 | 0.842593156 | 0.925040775 | | Cenpv | 545.1090951 | -0.00326532995381138 | 0.0722957607043386 | -0.0451662714659758 | 0.963974778272316 | 0.99099371420617 | NM_028448 |
| Trnau1ap | 460.4308242 | -0.017390173 | 0.087773491 | -0.198125573 | 0.842946817 | 0.925235051 | | Cidea | 73.71269446 | -0.00399264227431882 | 0.0911734916114835 | -0.0437917008962717 | 0.965070442465036 | 0.99099371420617 | NM_007702 |
| Hnrnpul1 | 3451.958726 | 0.011843657 | 0.059803334 | 0.198043416 | 0.843011095 | 0.925240059 | | Dnmt1 | 2520.133156 | -0.00211972537478192 | 0.0488963482753591 | -0.0433514045434378 | 0.965421414809111 | 0.99099371420617 | NM_010066,NM_001199433,NM_001199432,NM_001199431 |
| Fitm2 | 572.6114104 | 0.012373242 | 0.062550445 | 0.197812211 | 0.843191992 | 0.925307509 | | Ecel1 | 1206.789129 | -0.00353915035707213 | 0.0789533190452041 | -0.0448258591262745 | 0.964246113203506 | 0.99099371420617 | NM_021306 |
| Spata21 | 24.57871801 | 0.033239722 | 0.168011559 | 0.197841876 | 0.843168782 | 0.925307509 | | Fam188b | 295.4877911 | 0.00350151547792255 | 0.0749700047238956 | 0.0467055523181326 | 0.962747905067198 | 0.99099371420617 | NM_177883,NM_001142781 |
| Dtd2 | 284.7222374 | -0.017055597 | 0.086361161 | -0.197491524 | 0.843442914 | 0.925451775 | | Fbxo41 | 6782.836369 | -0.00257636049656411 | 0.0595283038157024 | -0.0432795885557303 | 0.965478661947248 | 0.99099371420617 | NM_001001160 |
| Rfwd3 | 1455.186398 | -0.01192125 | 0.060344403 | -0.197553526 | 0.843394399 | 0.925451775 | | Frg1 | 811.0419143 | 0.00342921904854059 | 0.0787865573917614 | 0.0435254332980816 | 0.965282690893395 | 0.99099371420617 | NM_013522 |
| Necab3 | 2625.21566 | -0.017624998 | 0.089444064 | -0.197050507 | 0.843788014 | 0.92558548 | | Fst | 144.7548266 | -0.00408709700090655 | 0.0904066228000104 | -0.0452079380284747 | 0.963941567089245 | 0.99099371420617 | NM_008046 |
| Ttn | 17.95583686 | -0.032644632 | 0.165549319 | -0.197189768 | 0.843679038 | 0.92558548 | | Grik5 | 9775.990764 | 0.00293766668244591 | 0.0678612022970972 | 0.0432893403447933 | 0.965470888430742 | 0.99099371420617 | NM_008168 |
| Vcl | 1709.549346 | 0.01746234 | 0.08862763 | 0.19703043 | 0.843803725 | 0.92558548 | | Grpel2 | 910.0781272 | -0.00256912319244884 | 0.057095757983867 | -0.0449967437716612 | 0.964109904420991 | 0.99099371420617 | NM_021296 |
| Xpnpep1 | 1422.339839 | 0.011844306 | 0.060079331 | 0.197144439 | 0.843714509 | 0.92558548 | | Gtf3a | 740.8547991 | -0.00274007678174108 | 0.0627441577444927 | -0.0436706281547239 | 0.965166952208729 | 0.99099371420617 | NM_025652 |
| 4930455F23Rik | 835.7737628 | 0.015665281 | 0.079627398 | 0.196732303 | 0.84403703 | 0.925644801 | | Heyl | 280.6482672 | 0.00424786099638322 | 0.0945922960695682 | 0.0449070502872572 | 0.964181397199111 | 0.99099371420617 | NM_013905 |
| Bend3 | 274.8563567 | -0.019224806 | 0.097644908 | -0.196884875 | 0.84391763 | 0.925644801 | | Il1rap | 1460.17887 | 0.0034880281932175 | 0.0775242358181115 | 0.0449927452545442 | 0.964113091548218 | 0.99099371420617 | NM_134103,NM_008364,NM_001159318,NM_001159317 |
| Dennd1b | 470.9442163 | -0.018249141 | 0.092758931 | -0.196737295 | 0.844033124 | 0.925644801 | | Maneal | 2896.320105 | -0.00214063339871632 | 0.0491447068634557 | -0.0435577610558119 | 0.965256921513905 | 0.99099371420617 | NM_001007573 |
| Snx25 | 2484.099202 | 0.018655494 | 0.095098724 | 0.196169757 | 0.844477299 | 0.92606209 | | Mmab | 1529.747413 | -0.00284582720556893 | 0.0617202605414005 | -0.0461084768676573 | 0.963223789630913 | 0.99099371420617 | NM_029956 |
| Sap30bp | 1530.187915 | 0.015403719 | 0.078593656 | 0.195991888 | 0.844616516 | 0.926149207 | | Mrpl52 | 446.5201876 | 0.0033451844530923 | 0.0770426909851334 | 0.0434198807222064 | 0.965366830120243 | 0.99099371420617 | NM_026851 |
| Adamts7 | 14.25209264 | -0.031825256 | 0.163225321 | -0.194977445 | 0.845410605 | 0.926167851 | | Mrps28 | 262.6467706 | -0.00355541828075829 | 0.0801563510470126 | -0.0443560395940802 | 0.964620602457663 | 0.99099371420617 | NM_025434 |
| Arsb | 5349.719046 | -0.011769654 | 0.060233767 | -0.195399607 | 0.845080124 | 0.926167851 | | Mtg1 | 669.296821 | -0.00297249126012301 | 0.0657178162049024 | -0.0452311326179045 | 0.963923079395988 | 0.99099371420617 | NM_199301 |
| Asb6 | 1079.897126 | 0.015217288 | 0.077798323 | 0.195599175 | 0.844923906 | 0.926167851 | | Nampt | 2200.187764 | 0.00280569802497495 | 0.0628371787883094 | 0.0446502863285285 | 0.96438605990652 | 0.99099371420617 | NM_021524 |
| Cox11 | 507.0562127 | 0.015523617 | 0.079528095 | 0.195196642 | 0.845239008 | 0.926167851 | | Ndufab1 | 1399.7453 | -0.00346643007606388 | 0.0764418759146818 | -0.0453472659401088 | 0.963830513391555 | 0.99099371420617 | NM_028177 |
| Dnajb14 | 754.8567296 | -0.016525466 | 0.084711714 | -0.19507887 | 0.845331204 | 0.926167851 | | Pddc1 | 1100.293617 | -0.00320158058715284 | 0.0703973086513522 | -0.0454787356006591 | 0.963725723890818 | 0.99099371420617 | NM_172116 |
| Dok2 | 22.33949463 | 0.032811367 | 0.168004801 | 0.195300175 | 0.84515796 | 0.926167851 | | Ppil4 | 1682.786445 | 0.00291887335196235 | 0.0632956010144486 | 0.0461149480403236 | 0.963218631870197 | 0.99099371420617 | NM_026141 |
| Grem1 | 40.53111566 | -0.030539214 | 0.156584509 | -0.19503343 | 0.845366777 | 0.926167851 | | Prkci | 2820.941124 | 0.00290437193286113 | 0.065583402606962 | 0.0442851669387586 | 0.96467709514312 | 0.99099371420617 | NM_008857 |
| Ints4 | 1309.395563 | 0.012400662 | 0.063382579 | 0.195647797 | 0.844885847 | 0.926167851 | | Rad54l2 | 1548.374487 | -0.00276643217664907 | 0.0599718537370989 | -0.0461284207975596 | 0.963207557815436 | 0.99099371420617 | NM_030730 |
| Mdh1b | 47.13552877 | -0.032626121 | 0.167037662 | -0.195321941 | 0.845140921 | 0.926167851 | | Ranbp1 | 1419.521914 | -0.0033153135994824 | 0.0711832085607353 | -0.0465743771110528 | 0.962852453965857 | 0.99099371420617 | NM_011239 |
| Pex3 | 979.7850956 | -0.013991136 | 0.071603708 | -0.195396804 | 0.845082318 | 0.926167851 | | Rfc3 | 219.8446472 | 0.00362635010774649 | 0.0824295432498423 | 0.0439933301189732 | 0.964909720515769 | 0.99099371420617 | NM_027009 |
| Samd14 | 1743.403481 | 0.024022417 | 0.122813925 | 0.195600108 | 0.844923175 | 0.926167851 | | Rsph3a | 176.9169997 | 0.00414874520955571 | 0.0889675052561454 | 0.0466321405507674 | 0.962806415431004 | 0.99099371420617 | NM_025789 |
| Soat1 | 814.0589817 | -0.013771564 | 0.070549304 | -0.195204819 | 0.845232607 | 0.926167851 | | Slc35f4 | 503.3612859 | 0.00343304487389256 | 0.0776460921039483 | 0.0442140071814854 | 0.964733816856301 | 0.99099371420617 | NM_029238 |
| Taco1 | 418.6189808 | -0.015836213 | 0.080852296 | -0.195865962 | 0.84471508 | 0.926167851 | | Smg7 | 5717.168177 | -0.00256940847532393 | 0.0587739415755886 | -0.043716797043796 | 0.965130149912102 | 0.99099371420617 | NM_001160257,NM_001160256,NM_001005507 |
| A830019L24Rik | 13.27229272 | -0.032544112 | 0.166993928 | -0.194882011 | 0.845485317 | 0.926174459 | | Spag8 | 48.39261381 | -0.00422262985634897 | 0.0927123560008995 | -0.0455454918685056 | 0.963672515230868 | 0.99099371420617 | NM_001007463 |
| Cnpy4 | 356.6338269 | -0.015393897 | 0.079017205 | -0.19481703 | 0.84553619 | 0.926174459 | | Ssh1 | 534.160192 | 0.00307722939742698 | 0.0706560725600693 | 0.0435522282222923 | 0.965261331891038 | 0.99099371420617 | NM_198109 |
| Erf | 2011.426768 | -0.01359097 | 0.069828412 | -0.194633815 | 0.845679628 | 0.926203011 | | Thap2 | 483.5536159 | -0.00388639140487966 | 0.0854557311022064 | -0.0454784173601121 | 0.963725977547582 | 0.99099371420617 | NM_025780 |
| Thap4 | 757.8091901 | -0.015652824 | 0.080423067 | -0.194631026 | 0.845681812 | 0.926203011 | | Ticam1 | 283.0181989 | -0.00342329573657583 | 0.076967113323353 | -0.0444773824658583 | 0.964523880309704 | 0.99099371420617 | NM_174989 |
| Rdh10 | 309.1945007 | 0.019176912 | 0.098623471 | 0.194445726 | 0.845826889 | 0.926296425 | | Tmem160 | 657.3014426 | -0.00339041784997791 | 0.0782213852904469 | -0.0433438737678809 | 0.965427417856092 | 0.99099371420617 | NM_026938 |
| E330009J07Rik | 674.6469564 | 0.02037472 | 0.105092441 | 0.193874269 | 0.846274332 | 0.926589958 | | Tmem65 | 4215.340058 | 0.00300205777503421 | 0.0651296570581935 | 0.0460935603015975 | 0.963235678689706 | 0.99099371420617 | NM_175212 |
| Kcnq3 | 1884.500857 | -0.016938677 | 0.087312018 | -0.194001664 | 0.846174579 | 0.926589958 | | Wdr16 | 50.72828986 | -0.00380732733970436 | 0.0818228350011661 | -0.0465313544764135 | 0.962886743885629 | 0.99099371420617 | NM_027963 |
| Srsf4 | 2040.587141 | 0.011379428 | 0.05868343 | 0.193912119 | 0.846244694 | 0.926589958 | | Wnt9b | 26.07869138 | -0.00330860284025635 | 0.0739869934128473 | -0.0447187091627625 | 0.964331520859853 | 0.99099371420617 | NM_011719 |
| A330032B11Rik | 8.622986397 | 0.030440849 | 0.157459087 | 0.19332545 | 0.846704096 | 0.926969536 | | Zfp668 | 519.8427311 | 0.00328160881771367 | 0.0709359128890903 | 0.0462615998751446 | 0.96310174539504 | 0.99099371420617 | NM_146259 |
| Adam23 | 6781.288896 | 0.011789072 | 0.060995177 | 0.193278755 | 0.846740664 | 0.926969536 | | 3110007F17Rik | 58.06912255 | -0.00406120983395746 | 0.0946042989387971 | -0.0429283857024807 | 0.965758621083731 | 0.991035612699844 | NM_028426 |
| Tmed1 | 367.7612492 | -0.02036904 | 0.105567217 | -0.192948535 | 0.846999274 | 0.927187138 | | Amdhd2 | 468.3782464 | 0.00397450335691888 | 0.092374579872579 | 0.0430259424443531 | 0.965680853918194 | 0.991035612699844 | NM_172935 |
| Dhx32 | 1385.512775 | 0.017535956 | 0.091071875 | 0.192550727 | 0.847310837 | 0.927331646 | | Bid | 439.2711674 | -0.00334513068713229 | 0.0780004665286909 | -0.0428860343534208 | 0.965792381480036 | 0.991035612699844 | NM_007544 |
| Ncoa1 | 6428.528386 | -0.017226121 | 0.089445292 | -0.192588343 | 0.847281375 | 0.927331646 | | Ccne2 | 136.8195307 | -0.00407961004699735 | 0.0949693121441791 | -0.0429571401002023 | 0.965735699538073 | 0.991035612699844 | NM_001037134,NM_009830 |
| Nfil3 | 337.4089507 | 0.025167509 | 0.130664363 | 0.192611884 | 0.847262937 | 0.927331646 | | Cd7 | 39.383808 | -0.00381483851678355 | 0.0906328154550925 | -0.0420911399213209 | 0.966426043231535 | 0.991265690155436 | NM_009854 |
| Pxmp4 | 388.5847757 | -0.022598696 | 0.117422071 | -0.192456972 | 0.847384269 | 0.927346508 | | D330041H03Rik | 25.39932817 | -0.0036273889139046 | 0.0859697848341548 | -0.0421937651804321 | 0.966344232801145 | 0.991265690155436 | NR_033554 |
| Kpna4 | 2773.296383 | -0.014550523 | 0.075911714 | -0.191676906 | 0.847995297 | 0.927884119 | | Nrcam | 15978.63759 | 0.00221429032247799 | 0.0522441767412004 | 0.0423834857891785 | 0.966192992950483 | 0.991265690155436 | NM_176930,NM_001146031 |
| Zfp212 | 595.0481712 | 0.015553825 | 0.08112725 | 0.191721339 | 0.84796049 | 0.927884119 | | Rpl17 | 8761.145761 | 0.00306741419830843 | 0.0721610991087453 | 0.0425078641566407 | 0.966093842727963 | 0.991265690155436 | NM_001002239 |
| B230209K01Rik | 63.1950679 | 0.028309981 | 0.147807756 | 0.191532447 | 0.848108462 | 0.927924127 | | Rpl41 | 7104.953221 | 0.00293947567693402 | 0.0697158173091653 | 0.0421636837007431 | 0.966368212929214 | 0.991265690155436 | NM_018860 |
| Gpr83 | 1037.447373 | -0.019242718 | 0.100565316 | -0.191345471 | 0.848254938 | 0.927924127 | | Tecpr2 | 4888.496435 | 0.00219104509683912 | 0.0519884030190237 | 0.0421448817352086 | 0.96638320148389 | 0.991265690155436 | NM_001081057 |
| Npas4 | 1200.606961 | -0.03012002 | 0.157429025 | -0.191324437 | 0.848271417 | 0.927924127 | | Dtnb | 2713.301159 | -0.00212786634973967 | 0.0511676751137239 | -0.0415861448660768 | 0.966828618482532 | 0.991586616527937 | NM_007886,NR_027869,NM_001162465,NR_027870 |
| Zfp386 | 573.832646 | -0.017657632 | 0.092260678 | -0.191388496 | 0.848221233 | 0.927924127 | | Rbm11 | 248.6423534 | 0.00379637114109235 | 0.0914184748989882 | 0.0415273952588589 | 0.966875453428453 | 0.991586616527937 | NM_198302 |
| Gtf2b | 1081.669442 | -0.018729953 | 0.097991018 | -0.191139485 | 0.848416313 | 0.92801711 | | Scaf11 | 3013.903696 | 0.0026487149716232 | 0.0642292205070462 | 0.0412384729366695 | 0.967105782778389 | 0.991752813317253 | NM_028148 |
| Gss | 443.3035945 | 0.013777695 | 0.072338027 | 0.190462688 | 0.848946579 | 0.928531575 | | Bfar | 1537.489389 | -0.00232370084234855 | 0.0566451669068691 | -0.0410220495275257 | 0.967278317674564 | 0.991789711480251 | NM_025976,NM_001177552 |
| Apoa2 | 20.9844396 | -0.031848781 | 0.167511403 | -0.190129032 | 0.849208021 | 0.928542254 | | Zbtb40 | 674.7613357 | 0.00360058707535377 | 0.0876976682469698 | 0.0410568165303321 | 0.967250600970354 | 0.991789711480251 | NM_198248 |
| Ing2 | 979.5376707 | -0.014571501 | 0.076627813 | -0.190159428 | 0.849184203 | 0.928542254 | | Os9 | 6309.895424 | -0.00205492272763186 | 0.0502061887893236 | -0.0409296697714853 | 0.967351964202617 | 0.991795216827161 | NM_001171026,NM_177614 |
| Rrp9 | 619.674667 | 0.017565374 | 0.092270698 | 0.190367846 | 0.849020892 | 0.928542254 | | Actn4 | 7174.926895 | 0.00236631031080639 | 0.058318437129582 | 0.0405756811614912 | 0.967634171807982 | 0.991805208479729 | NM_021895 |
| Serinc2 | 225.1979403 | 0.030234624 | 0.159048416 | 0.190096981 | 0.849233136 | 0.928542254 | | Cbfa2t3 | 1426.230555 | 0.00376359296278985 | 0.0926049694042663 | 0.0406413714836394 | 0.967581801712478 | 0.991805208479729 | NM_001109873,NM_009824,NM_177289 |
| Suv39h1 | 1187.937113 | 0.01579404 | 0.083096864 | 0.190067817 | 0.849255989 | 0.928542254 | | Pcbp4 | 5016.420561 | -0.00236645918068996 | 0.0582500800307313 | -0.0406258528647767 | 0.967594173561136 | 0.991805208479729 | NM_021567 |
| Ido2 | 76.15362988 | 0.028316441 | 0.149081247 | 0.189939659 | 0.849356415 | 0.928586529 | | Pdcl | 861.4130015 | -0.0030350584117036 | 0.0748014393608334 | -0.0405748664415778 | 0.967634821325535 | 0.991805208479729 | NM_026176 |
| Armc10 | 818.6670109 | 0.012936698 | 0.068212668 | 0.18965243 | 0.8495815 | 0.928767075 | | Bcl9l | 1924.052684 | 0.00230577911597885 | 0.0569740132058006 | 0.0404707161429888 | 0.967717853039702 | 0.991820329729395 | NM_030256 |
| Hs6st1 | 2689.588529 | 0.016665209 | 0.087931425 | 0.189525065 | 0.849681312 | 0.928810656 | | Sla2 | 35.06453242 | 0.00336871215421914 | 0.0835348975270472 | 0.0403270040898584 | 0.96783242513512 | 0.991867772506897 | NM_029983 |
| Mios | 907.7136053 | 0.013856363 | 0.073218165 | 0.189247617 | 0.849898749 | 0.928982802 | | Inpp1 | 812.3228262 | 0.00246005904971946 | 0.0612287411731273 | 0.0401781745400174 | 0.967951077771363 | 0.991919390173321 | NM_008384 |
| Cenph | 19.03299071 | 0.031760951 | 0.168098096 | 0.188942957 | 0.850137525 | 0.929047173 | | Bloc1s5 | 609.9466861 | -0.00284079185901835 | 0.0714550827220519 | -0.0397563301419515 | 0.968287392182683 | 0.992194036318119 | NM_139063 |
| Isg20l2 | 552.0488136 | -0.014355722 | 0.075967855 | -0.188971006 | 0.850115541 | 0.929047173 | | Abcc3 | 154.2733847 | -0.00368984969245592 | 0.0941335267118379 | -0.0391980394376524 | 0.968732496738961 | 0.99223016835637 8 | NM_029600 |
| Myh14 | 1477.210071 | 0.024081433 | 0.127399218 | 0.189023395 | 0.850074481 | 0.929047173 | | Cbx7 | 2783.488223 | -0.00191793018175061 | 0.0487237184474666 | -0.0393633787170514 | 0.968600676817386 | 0.992230168356378 | NM_144811 |
| 4833412C05Rik | 27.76268487 | -0.031603568 | 0.167742076 | -0.188405731 | 0.850558608 | 0.929441785 | | Ccdc50 | 2397.839496 | 0.00236102710233554 | 0.0602220707555441 | 0.0392053456932678 | 0.968726671668033 | 0.992230168356378 | NM_001025615,NM_026202 |
| Zfp53 | 168.2957928 | -0.019424675 | 0.103144467 | -0.188324938 | 0.850621938 | 0.929445437 | | Glce | 1883.029285 | -0.00246192113987557 | 0.0624810883666968 | -0.039402660936806 | 0.968569358437904 | 0.992230168356378 | NM_033320 |
| Gbe1 | 453.2431497 | -0.017221079 | 0.09166574 | -0.187868216 | 0.850979959 | 0.929771065 | | Hic1 | 118.6124997 | -0.00360689998960739 | 0.0918001094745603 | -0.0392908027044013 | 0.968658539334381 | 0.992230168356378 | NM_001098203,NM_010430 |
| Prkag1 | 1877.766315 | 0.016572308 | 0.088275992 | 0.187732895 | 0.851086042 | 0.929821403 | | Raph1 | 2585.685301 | -0.0028592415459064 | 0.0729354188255627 | -0.0392023737156395 | 0.968729041141526 | 0.992230168356378 | NM_001045513 |
| Ccdc85b | 1943.503213 | -0.018104509 | 0.096518048 | -0.187576409 | 0.851208721 | 0.929876317 | | Csnk2a1 | 6078.6453 | -0.00224402140629021 | 0.0581981879904601 | -0.0385582693168737 | 0.969242573792884 | 0.992548842723171 | NM_007788 |
| Uxt | 170.7777474 | -0.019350028 | 0.103191524 | -0.187515672 | 0.851256337 | 0.929876317 | | Dusp22 | 1057.054804 | -0.00216977571249767 | 0.0564090992131025 | -0.0384649948814229 | 0.969316940855769 | 0.992548842723171 | NM_134068,NM_001037955 |
| Nisch | 17129.72234 | -0.010048584 | 0.053639938 | -0.187333999 | 0.851398767 | 0.929966338 | | Pias4 | 650.2752329 | 0.00283768111690836 | 0.0733789737198956 | 0.0386715836029602 | 0.969152229455449 | 0.992548842723171 | NM_021501 |
| Arfip1 | 623.7055757 | 0.014701619 | 0.07856456 | 0.187127874 | 0.851560373 | 0.930000411 | | Srek1ip1 | 527.6717307 | -0.0027636391897867 | 0.0718409709614525 | -0.0384688451840327 | 0.969313871030819 | 0.992548842723171 | NM_026075 |
| Ccdc115 | 589.8750615 | 0.015187064 | 0.0811403 | 0.187170413 | 0.851527022 | 0.930000411 | | Dhrs13 | 213.0520859 | 0.00332054017955178 | 0.0867024095266892 | 0.038298130325082 | 0.969449981478745 | 0.992567416518978 | NM_183286 |
| Cog4 | 2012.736954 | 0.010996807 | 0.058786166 | 0.187064538 | 0.851610032 | 0.930000411 | | Dram1 | 54.93842303 | 0.00359336771101958 | 0.0938930995024878 | 0.0382708391783825 | 0.969471740711225 | 0.992567416518978 | NM_027878 |
| Larp4b | 2626.189986 | -0.011652162 | 0.062320862 | -0.186970484 | 0.851683775 | 0.930015393 | | Tmtc1 | 5225.992893 | 0.00236657977127465 | 0.0620279376895888 | 0.0381534492266679 | 0.969565335983878 | 0.992593281450547 | NM_198967 |
| Galm | 100.0177695 | 0.025618158 | 0.137184164 | 0.186742825 | 0.851862277 | 0.930144758 | | Gm10638 | 26.23123298 | -0.00320242391685633 | 0.0844385149490179 | -0.0379261042048155 | 0.969746599872291 | 0.992708887226561 | NR_027829 |
| Zscan12 | 271.5163544 | 0.015552226 | 0.083318167 | 0.18666068 | 0.851926687 | 0.930149537 | | Gm5607 | 522.0120726 | -0.0031866755813334 | 0.0842791520162929 | -0.0378109592360083 | 0.96983840641503 | 0.992732907588241 | NR_027975 |
| Il11ra1 | 572.0125643 | -0.019851818 | 0.106421323 | -0.18653985 | 0.852021432 | 0.930187434 | | Traf1 | 34.71493907 | 0.00295107188336374 | 0.0783126753195295 | 0.0376831958724798 | 0.969940274231535 | 0.992740432068765 | NM_009421 |
| Chst5 | 26.48265673 | 0.03130002 | 0.168074849 | 0.186226675 | 0.852267006 | 0.930279 | | Ttc12 | 119.1031302 | 0.0035737953345727 | 0.0949711903424702 | 0.0376303089398333 | 0.969982441990361 | 0.992740432068765 | NM_172770 |
| Nell2 | 21824.50246 | 0.015467306 | 0.083064499 | 0.186208383 | 0.85228135 | 0.930279 | | Ccdc101 | 431.8118579 | 0.00278178021929413 | 0.0744365899420608 | 0.0373711399388309 | 0.970189083582742 | 0.992821600974353 | NM_029339 |
| Psen2 | 521.0037318 | 0.013517266 | 0.072615627 | 0.18614817 | 0.852328567 | 0.930279 | | Hcn2 | 5489.81836 | 0.00226903594972938 | 0.0612979352107484 | 0.0370165151881252 | 0.970471837548867 | 0.992821600974353 | NM_008226 |
| Zfp422 | 776.668348 | -0.013068961 | 0.070215431 | -0.186126619 | 0.852345468 | 0.930279 | | Ppp6r3 | 3753.013353 | 0.002056288702316 | 0.0553647128588778 | 0.0371407814858065 | 0.970372755522279 | 0.992821600974353 | NM_001164159,NM_028999,NM_029456 |
| Rpl3l | 12.82023564 | -0.03095389 | 0.166417576 | -0.18600133 | 0.852443718 | 0.9303207 | | Rras | 336.7052014 | 0.00295645799743116 | 0.0798283408376215 | 0.0370351928451686 | 0.97045694514628 | 0.992821600974353 | NM_009101 |
| Mccc2 | 625.639699 | 0.016437901 | 0.088455551 | 0.185832332 | 0.852576249 | 0.930345794 | | Tbc1d20 | 1526.55144 | 0.00213788668094016 | 0.0572879899071153 | 0.0373182351904204 | 0.970231266040059 | 0.992821600974353 | NM_024196 |
| Trmt10a | 214.4803079 | -0.022996337 | 0.123756736 | -0.185818874 | 0.852586803 | 0.930345794 | | Tmem165 | 848.7683778 | -0.00249157005697472 | 0.0669976232858253 | -0.0371889320065158 | 0.970334363488421 | 0.992821600974353 | NM_011626 |
| AA543186 | 8.427306637 | -0.027843815 | 0.150265941 | -0.185296913 | 0.852996159 | 0.930648992 | | 3632454L22Rik | 43.67757844 | -0.00337919072291068 | 0.0925967958906527 | -0.0364936031577287 | 0.970888779254253 | 0.992984126231971 | NR_040281,NR_040282,NR_040283 |
| Abra | 8.25901321 | -0.029367637 | 0.159033834 | -0.184662825 | 0.853493506 | 0.930648992 | | C8g | 32.38664308 | -0.00316647062555521 | 0.0867262513064853 | -0.0365110975956414 | 0.970874830008577 | 0.992984126231971 | NM_027062,NM_001271777 |
| Cpd | 4869.484081 | -0.010740822 | 0.058085609 | -0.184913647 | 0.853296767 | 0.930648992 | | Nsmce2 | 964.9906692 | -0.00227359957646235 | 0.0621784814401339 | -0.0365656982025484 | 0.970831294233979 | 0.992984126231971 | NM_001164604,NM_026746 |
| Elovl7 | 726.9286716 | 0.025928214 | 0.140450701 | 0.184607224 | 0.853537119 | 0.930648992 | | R74862 | 175.2880224 | -0.00306667110643671 | 0.084077499003118 | -0.0364743378763322 | 0.970904140497913 | 0.992984126231971 | NR_015529 |
| Fam118a | 699.0848997 | -0.013104741 | 0.070783137 | -0.185139304 | 0.853119774 | 0.930648992 | | B230217C12Rik | 1188.819909 | -0.00211109995088999 | 0.0580730336286854 | -0.0363524999294545 | 0.971001288687039 | 0.993013567971552 | NM_001080935 |
| Foxr2 | 21.14520136 | -0.030960789 | 0.167544743 | -0.184791166 | 0.853392837 | 0.930648992 | | 4921507P07Rik | 38.04780287 | 0.00317045592182045 | 0.0924663076054403 | 0.0342876881744753 | 0.972647742511624 | 0.99301409004836 | NM_027564 |
| Lsm7 | 651.2409455 | -0.014629177 | 0.079142634 | -0.184845713 | 0.853350052 | 0.930648992 | | Abca5 | 2270.303528 | -0.00254698833269565 | 0.0714322169555946 | -0.0356560168681167 | 0.97155664168676 | 0.99301409004836 | NM_147219 |
| Rgma | 2021.873058 | 0.01012398 | 0.054728917 | 0.184984111 | 0.853241498 | 0.930648992 | | Acadvl | 1905.657939 | -0.00255402166352346 | 0.0722813462042061 | -0.035334450693654 | 0.971813052792982 | 0.99301409004836 | NM_017366 |
| Shcbp1 | 26.20193717 | -0.030309907 | 0.164240729 | -0.184545621 | 0.853585441 | 0.930648992 | | Alkbh5 | 3295.265184 | 0.00213917081065864 | 0.0596729614333522 | 0.0358482428100681 | 0.971403365566939 | 0.99301409004836 | NM_172943 |
| Smarcad1 | 1025.307903 | 0.015363266 | 0.083023644 | 0.185046876 | 0.853192269 | 0.930648992 | | Arpc4 | 3055.208674 | 0.00160981564654416 | 0.0466338574717153 | 0.0345203192234432 | 0.972462239606109 | 0.99301409004836 | NM_026552,NM_001170486,NM_001170485 |
| Strn3 | 2711.490628 | 0.013184776 | 0.071372099 | 0.184732916 | 0.853438527 | 0.930648992 | | BC018242 | 7510.594424 | -0.00232021446908663 | 0.0668750313954744 | -0.0346947795114403 | 0.972323123769268 | 0.99301409004836 | NM_144935 |
| Vwa9 | 807.2230178 | 0.013534621 | 0.073335291 | 0.184558084 | 0.853575665 | 0.930648992 | | Bola3 | 393.4482145 | -0.00306288265387335 | 0.0872157659250373 | -0.0351184517850148 | 0.971985288242507 | 0.99301409004836 | NM_175277 |
| Foxf1 | 33.81081508 | 0.029991954 | 0.162748812 | 0.184283705 | 0.853790896 | 0.930807497 | | Cdk5 | 4402.63096 | 0.001877194787029 | 0.0538522946654675 | 0.0348582135392782 | 0.972192801112141 | 0.99301409004836 | NM_007668 |
| Gm15545 | 14.60392867 | -0.030572763 | 0.165970727 | -0.184205753 | 0.853852047 | 0.930808669 | | Cers6 | 1428.572648 | -0.00294033927006148 | 0.0863862530735754 | -0.0340371200908226 | 0.972847550287637 | 0.99301409004836 | NM_172856 |
| Adh5 | 1683.810814 | 0.009814553 | 0.05340946 | 0.183760575 | 0.854201286 | 0.930861916 | | Cyp4f14 | 377.2152851 | -0.0028588990618157 | 0.0830741338563218 | -0.0344138293004681 | 0.972547155817162 | 0.99301409004836 | NM_001204333,NM_022434,NM_001204336,NM_001204335,NM_001204334 |
| Coil | 784.7968806 | 0.011216755 | 0.061039823 | 0.183761266 | 0.854200744 | 0.930861916 | | Cyp4x1 | 378.5281887 | 0.00278084933529322 | 0.0793602003367143 | 0.0350408557878441 | 0.97204716265573 | 0.99301409004836 | NM_001003947 |
| Fstl3 | 69.54222957 | 0.024693306 | 0.134366951 | 0.183775145 | 0.854189856 | 0.930861916 | | Hif1an | 2936.210625 | 0.00205185680841125 | 0.0600027668719998 | 0.0341960365392542 | 0.972720827077738 | 0.99301409004836 | NM_176958 |
| Katnal1 | 3971.691127 | 0.008739849 | 0.047555446 | 0.183782303 | 0.85418424 | 0.930861916 | | Kif4 | 67.40690373 | -0.00336927040071479 | 0.0950951824917952 | -0.0354305056515927 | 0.971736459984117 | 0.99301409004836 | NM_008446 |
| Xrcc2 | 261.2427055 | 0.017718479 | 0.096356492 | 0.18388464 | 0.854103955 | 0.930861916 | | Lin54 | 546.1683159 | 0.00284453792642311 | 0.0834834488194552 | 0.0340730763600199 | 0.972818877966761 | 0.99301409004836 | NM_001115010,NM_172714 |
| Fam60a | 88.72269921 | -0.025180752 | 0.137140303 | -0.183613072 | 0.854317007 | 0.930922548 | | Mfap1b | 184.6184543 | -0.00314812460373995 | 0.0891243956417898 | -0.0353228157236874 | 0.971822330364351 | 0.99301409004836 | NM_001081975 |
| Cant1 | 1932.726305 | 0.010213039 | 0.05572207 | 0.18328534 | 0.854574136 | 0.931137248 | | Mfsd9 | 260.3968407 | 0.00289636755575975 | 0.08353470523281 | 0.0346726255925525 | 0.972340789436944 | 0.99301409004836 | NM_172499 |
| Gimap8 | 60.41812464 | 0.026956229 | 0.14727823 | 0.183029283 | 0.854775043 | 0.931269015 | | Pcp4l1 | 5781.254648 | 0.00297770893333282 | 0.0823212314411512 | 0.0361718220342887 | 0.971145354039771 | 0.99301409004836 | NM_025557 |
| Tmem160 | 593.9720746 | -0.016131478 | 0.088160752 | -0.182978002 | 0.854815279 | 0.931269015 | | Pfn1 | 3182.569188 | -0.00187049657547388 | 0.0543724080016838 | -0.0344015769067274 | 0.972556926027821 | 0.99301409004836 | NM_011072 |
| Ehmt2 | 7905.799161 | 0.011900013 | 0.065116166 | 0.182750515 | 0.854993778 | 0.931332508 | | Prrg1 | 146.4025335 | 0.00323071257778548 | 0.0936319790828233 | 0.0345043713636311 | 0.972474956581368 | 0.99301409004836 | NM_001164275,NM_027322 |
| Fam151b | 177.2680439 | -0.021716939 | 0.118828978 | -0.182757939 | 0.854987753 | 0.931332508 | | Ptchd4 | 172.9067352 | 0.00319918919004166 | 0.092096676446956 | 0.0347372925219974 | 0.972289223728888 | 0.99301409004836 | NM_028474 |
| Midn | 2183.873371 | -0.024589066 | 0.134666581 | -0.182592189 | 0.855118014 | 0.931402355 | | Rimklb | 890.8495223 | -0.00218564778674566 | 0.0631759949105486 | -0.0345961751744525 | 0.972401751444427 | 0.99301409004836 | NM_027664 |
| Mmp28 | 111.1680894 | -0.021838346 | 0.119806537 | -0.182280091 | 0.855362922 | 0.931603621 | | Rpe65 | 224.7034138 | 0.0026365439808179 | 0.0761977799945156 | 0.0346013227814205 | 0.972397646705875 | 0.99301409004836 | NM_029987 |
| Naif1 | 96.22778709 | 0.024516392 | 0.13462835 | 0.182104227 | 0.855500931 | 0.931642409 | | Socs4 | 209.4577036 | 0.0031408275139842 | 0.0912745169896079 | 0.0344107820843555 | 0.972549585704681 | 0.99301409004836 | NM_080843 |
| Senp2 | 4274.369935 | -0.010447225 | 0.057376653 | -0.182081465 | 0.855518794 | 0.931642409 | | Tnpo2 | 9274.185961 | 0.00158140115358024 | 0.0437273856964639 | 0.0361650057142136 | 0.971150789120196 | 0.99301409004836 | NM_001122843,NM_145390 |
| 4930481A15Rik | 64.08803779 | -0.026740508 | 0.147335511 | -0.181493981 | 0.855979857 | 0.931766735 | | Wdpcp | 396.1461506 | -0.00261726594487804 | 0.0765207781018748 | -0.0342033367903497 | 0.972715005725475 | 0.99301409004836 | NM_145425 |
| Fam189a1 | 3209.389555 | 0.012739998 | 0.070103856 | 0.181730339 | 0.855794355 | 0.931766735 | | Zfml | 4750.223563 | 0.00272159126694307 | 0.0768628264262523 | 0.0354084203441876 | 0.971754070459942 | 0.99301409004836 | NM_008717,NM_001166371 |
| Fgfr2 | 2224.543239 | -0.015813242 | 0.087136711 | -0.181476235 | 0.855993785 | 0.931766735 | | Zfp395 | 437.3462991 | 0.00315471788471491 | 0.0885678948153574 | 0.0356192036774921 | 0.971585995716886 | 0.99301409004836 | NM_199029 |
| Gjc3 | 1671.650198 | 0.023061021 | 0.127021183 | 0.181552559 | 0.855933883 | 0.931766735 | | Pcid2 | 1242.308043 | 0.00172152662180396 | 0.0507181583205873 | 0.0339430034293095 | 0.972922601152177 | 0.993020917955067 | NM_178708 |
| Pfkfb3 | 1926.590514 | 0.022420726 | 0.123319732 | 0.181809723 | 0.855732054 | 0.931766735 | | Shroom2 | 4670.04058 | -0.00196054748187467 | 0.057952218348054 | -0.0338304130154235 | 0.973012383741185 | 0.993042779893519 | NM_172441 |
| Snx2 | 2017.458977 | -0.012094036 | 0.066592035 | -0.181613851 | 0.855885778 | 0.931766735 | | Nfix | 4158.861074 | 0.0019782802731232 | 0.0589359718426676 | 0.0335666013687585 | 0.973227755498524 | 0.993070301460729 | NM_001081981,NM_001081982,NM_010906 |
| Aimp1 | 1379.377835 | -0.012430511 | 0.06875195 | -0.180802305 | 0.856522753 | 0.932277031 | | Rpl36 | 1896.91782 | -0.00207174369397077 | 0.0617704422398552 | -0.0335394019995221 | 0.973244445242664 | 0.993070301460729 | NM_018730 |
| H2-DMa | 504.0634724 | -0.024259737 | 0.134252294 | -0.180702591 | 0.856601025 | 0.932280843 | | Zfp385c | 22.64857211 | -0.00263424627442394 | 0.0781763508421355 | -0.0336962041083674 | 0.973119405938013 | 0.993070301460729 | NM_177790 |
| Slc35f5 | 675.4444576 | 0.012568278 | 0.069589548 | 0.180605835 | 0.856676976 | 0.932280843 | | Hoga1 | 34.56806819 | 0.00293520067575804 | 0.0890129202816327 | 0.0329749958373594 | 0.973694527226041 | 0.993459766979383 | NM_026152 |
| Tlr3 | 584.2735962 | 0.015604809 | 0.086420736 | 0.180567884 | 0.856706766 | 0.932280843 | | Anapc4 | 2212.225623 | -0.00179334183246801 | 0.0548461228922745 | -0.032697695623634 | 0.973915661536094 | 0.993518394767634 | NM_024213 |
| Ly6c1 | 1011.477383 | 0.020797984 | 0.115327064 | 0.180339144 | 0.856886327 | 0.932407044 | | Larp7 | 699.2378742 | 0.00198440044514928 | 0.0611709862457944 | 0.0324402231668041 | 0.974120985907161 | 0.993518394767634 | NM_138593 |
| Naa60 | 4317.511343 | -0.006978166 | 0.038710205 | -0.18026683 | 0.856943095 | 0.932407044 | | Prss36 | 264.0763371 | -0.00286341214586056 | 0.088408744117985 | -0.0323883364075303 | 0.974162363808028 | 0.993518394767634 | NM_001081374 |
| Ifrd1 | 1410.914111 | 0.020552304 | 0.114104536 | 0.180118206 | 0.85705977 | 0.932468512 | | Rnf114 | 2506.185198 | -0.00167320188232397 | 0.051572494060061 | -0.0324436875279945 | 0.974118223201113 | 0.993518394767634 | NM_030743 |
| Jmjd8 | 1302.631359 | -0.011971617 | 0.066574244 | -0.179823544 | 0.857291099 | 0.93252375 | | Shcbp1 | 28.9861601 | -0.00282949041129219 | 0.0864445507388505 | -0.032731853970068 | 0.97388842169956 | 0.993518394767634 | NM_011369 |
| Larp1 | 6294.587717 | -0.012919946 | 0.071840312 | -0.179842572 | 0.857276161 | 0.93252375 | | Zcchc6 | 2754.316261 | 0.00173125984682275 | 0.0532369447869062 | 0.0325198948540819 | 0.974057450620245 | 0.993518394767634 | NM_153538 |
| Trp63 | 22.58944417 | -0.029807007 | 0.16573154 | -0.179851143 | 0.857269432 | 0.93252375 | | Gli2 | 199.9130977 | -0.00294669188328376 | 0.0915401301687605 | -0.0321901648801606 | 0.974320399404754 | 0.993609809165506 | NM_001081125 |
| Hnrnpr | 2528.107511 | -0.010524565 | 0.058660523 | -0.179414783 | 0.857612024 | 0.932741872 | | Ipp | 424.7521678 | 0.00258802092160491 | 0.0807756520796407 | 0.032039616579675 | 0.974440457641283 | 0.993662484218833 | NM_008389 |
| Pgr | 974.4240757 | 0.019250236 | 0.107290461 | 0.179421691 | 0.857606601 | 0.932741872 | | Stxbp5l | 5984.349757 | 0.00223159009597832 | 0.0699182584864525 | 0.0319171292919247 | 0.974538138399273 | 0.993692334577273 | NM_001114611,NM_001114612,NM_001114613,NM_172440 |
| Nek3 | 57.00170093 | 0.026706281 | 0.148989758 | 0.179249107 | 0.857742106 | 0.932778042 | | Ctsf | 2509.109376 | -0.00197888200355666 | 0.0626986497844877 | -0.0315617961528456 | 0.97482151045386 | 0.993841750480964 | NM_019861 |
| Zzz3 | 2356.201211 | 0.010039344 | 0.05601716 | 0.179219076 | 0.857765686 | 0.932778042 | | Rdh14 | 801.3052661 | 0.00191259119403279 | 0.0605124501219605 | 0.0316065733609867 | 0.974785801226316 | 0.993841750480964 | NM_023697 |
| Ski | 6830.00348 | -0.009068559 | 0.050689819 | -0.178902971 | 0.85801389 | 0.932982469 | | Hmgb3 | 1248.657637 | -0.00211423047405299 | 0.067614521756923 | -0.0312688815821805 | 0.975055107167103 | 0.99394038503223 4 | NM_008253 |
| B230118H07Rik | 732.6164316 | 0.01576599 | 0.08820319 | 0.178746259 | 0.858136944 | 0.932987515 | | Rap1a | 1120.474605 | -0.00204091704829047 | 0.0651947810439727 | -0.0313049145285711 | 0.975026371103377 | 0.993940385032234 | NM_145541 |
| Rabac1 | 2860.612652 | 0.013428402 | 0.075191098 | 0.178590322 | 0.858259394 | 0.932987515 | | Npas2 | 2252.450966 | 0.0021051095149171 | 0.0677611860969188 | 0.0310665977998992 | 0.975216427898841 | 0.99403507344457 | NM_008719 |
| Sertad3 | 82.50069842 | -0.025859823 | 0.144798329 | -0.178592 | 0.858258077 | 0.932987515 | | Cdk5rap1 | 347.86108 | -0.00286034866118697 | 0.0946017122559115 | -0.0302356965109607 | 0.97587907983829 | 0.994349486251819 | NM_025876 |
| Znrd1as | 104.2594589 | 0.025216151 | 0.14111337 | 0.17869427 | 0.858177769 | 0.932987515 | | Foxj1 | 319.8285421 | -0.00286385008994454 | 0.0947852037401646 | -0.0302141049123577 | 0.975896299574163 | 0.994349486251819 | NM_008240 |
| Dip2b | 4044.149763 | -0.008806491 | 0.049362686 | -0.178403809 | 0.858405858 | 0.933081266 | | Mypn | 178.4685264 | -0.00290156862278651 | 0.0950289193541437 | -0.0305335327656758 | 0.975641550561654 | 0.994349486251819 | NM_182992 |
| Get4 | 1755.022801 | 0.01446647 | 0.081176228 | 0.178210665 | 0.858557534 | 0.933180669 | | Tm7sf3 | 1948.777735 | -0.00149619496967198 | 0.0496006628228742 | -0.0301648180592858 | 0.975935606876786 | 0.994349486251819 | NM_026281 |
| Fzd1 | 420.3180346 | -0.021516984 | 0.120814396 | -0.178099504 | 0.858644831 | 0.933210088 | | Vrk3 | 997.5237429 | -0.00171909511813237 | 0.0569493485217399 | -0.0301863877771337 | 0.975918404565646 | 0.994349486251819 | NM_133945 |
| Syap1 | 868.6686939 | 0.011848808 | 0.066631075 | 0.17782706 | 0.858858795 | 0.93337716 | | Zfp599 | 203.6688154 | -0.0028353710742152 | 0.0931047636605955 | -0.0304535553578254 | 0.975705333639134 | 0.994349486251819 | NM_181419 |
| Eml6 | 1171.415609 | 0.015660125 | 0.088202961 | 0.177546474 | 0.859079164 | 0.933420236 | | Zfp639 | 969.6446165 | 0.00170688857235096 | 0.0568451679146473 | 0.0300269774013835 | 0.976045538012434 | 0.994391743945056 | NM_144519,NM_001161818 |
| Golt1b | 1347.891973 | -0.0118331 | 0.066647134 | -0.177548528 | 0.85907755 | 0.933420236 | | Phf3 | 3899.7618 | -0.00217620475227432 | 0.0728535516420973 | -0.0298709493665486 | 0.976169974553758 | 0.994448772730759 | NM_001081080 |
| Kdm4a | 2273.818255 | -0.014287363 | 0.080425704 | -0.17764672 | 0.859000431 | 0.933420236 | | Exosc5 | 346.9434651 | 0.00230221428167074 | 0.0778042199748686 | 0.0295898382171864 | 0.976394169694438 | 0.994548466985672 | NM_138586 |
| Rhot2 | 3389.89882 | -0.016421563 | 0.092633898 | -0.177273796 | 0.859293332 | 0.933587469 | | Scn8a | 15640.99245 | 0.00171205393359197 | 0.0578855244089625 | 0.029576547005021 | 0.976404769907935 | 0.994548466985672 | NM_011323,NM_001077499 |
| Ube2i | 7013.49087 | -0.007844319 | 0.044317392 | -0.177003172 | 0.859505898 | 0.933752937 | | Cplx2 | 61181.34839 | -0.00174939753015902 | 0.0596646822129021 | -0.029320486848763 | 0.976608987783074 | 0.994686730930931 | NM_009946 |
| Atad2b | 704.6344387 | -0.017217593 | 0.097426468 | -0.176723976 | 0.859725207 | 0.933829543 | | Asxl3 | 409.8959446 | 0.00253525133905255 | 0.0870576114970036 | 0.0291215356757152 | 0.976767660095113 | 0.994708849589288 | NM_001167777 |
| Lsm11 | 1303.944447 | -0.019415971 | 0.109811988 | -0.176811036 | 0.85965682 | 0.933829543 | | Flrt2 | 2998.242836 | 0.00241385790993479 | 0.0827988944715151 | 0.0291532625567267 | 0.976742356450118 | 0.994708849589288 | NM_201518 |
| Sgta | 4380.261946 | 0.011842485 | 0.067026652 | 0.176683217 | 0.859757225 | 0.933829543 | | Hltf | 1063.162478 | 0.0018682459217397 | 0.0645323302347166 | 0.0289505417663446 | 0.976904035994422 | 0.994777989965298 | NM_144959,NM_009210 |
| E130012A19Rik | 2263.404472 | -0.022373602 | 0.126776155 | -0.176481152 | 0.859915955 | 0.933871017 | | Nf1 | 4622.201162 | 0.00201193893262482 | 0.0697819756802998 | 0.0288317851853657 | 0.976998750499507 | 0.994804699968866 | NM_010897 |
| Mal | 8265.278186 | -0.017999683 | 0.101958451 | -0.176539392 | 0.859870204 | 0.933871017 | | Atg10 | 409.4699528 | 0.00216042189909968 | 0.0752646444674106 | 0.0287043393931813 | 0.977100395459902 | 0.994838462497409 | NM_025770 |
| Acy1 | 402.5595259 | -0.018339215 | 0.104278165 | -0.175868217 | 0.860397475 | 0.93408313 | | Ythdc1 | 2610.998642 | 0.00216189510385037 | 0.0760427433973785 | 0.0284299888041318 | 0.977319198280755 | 0.994991496651685 | NM_177680 |
| Atp6v0e2 | 14821.55807 | 0.01059722 | 0.06026324 | 0.175848834 | 0.860412704 | 0.93408313 | | Wdr86 | 42.30540776 | -0.00116700084074132 | 0.041231709037322 | -0.0283034797244271 | 0.977420105294344 | 0.995024490118833 | NM_001081441 |
| B4galt3 | 1553.322516 | 0.01094893 | 0.062259175 | 0.175860507 | 0.860403533 | 0.93408313 | | Gpr173 | 529.7028759 | 0.00207451781559369 | 0.0737752251881067 | 0.0281194372542304 | 0.977566891515437 | 0.995104181230079 | NM_027543 |
| Fa2h | 1266.646751 | 0.024079557 | 0.136802506 | 0.176016928 | 0.860280643 | 0.93408313 | | Ankrd42 | 577.2417736 | 0.00204417247526868 | 0.0730356842950242 | 0.0279886810810362 | 0.977671178800485 | 0.995119468206711 | NM_028665 |
| Shroom4 | 180.2670878 | -0.02326056 | 0.132198268 | -0.175952079 | 0.86033159 | 0.93408313 | | Ccdc51 | 217.4783224 | -0.00236055865285887 | 0.0852168246700661 | -0.0277006173604595 | 0.977900931311226 | 0.995119468206711 | NM_025689 |
| Ehmt1 | 1302.023168 | 0.010291587 | 0.058621763 | 0.175559148 | 0.860640299 | 0.934210484 | | Frmd5 | 1412.093086 | 0.00170594650923154 | 0.0616516282661862 | 0.0276707453997155 | 0.977924756554766 | 0.995119468206711 | NM_172673 |
| Gm4925 | 12.29861451 | -0.029195699 | 0.166313636 | -0.175546033 | 0.860650603 | 0.934210484 | | Mad2l2 | 300.6812423 | 0.00230696713049254 | 0.0830546289967727 | 0.0277765027471519 | 0.977840406821897 | 0.995119468206711 | NM_027985 |
| Golph3l | 823.0960671 | -0.010964919 | 0.062550361 | -0.17529746 | 0.860845908 | 0.934292638 | | S100a1 | 1457.540538 | 0.00199080724705845 | 0.07216926489 1268 | 0.0275852504533314 | 0.977992945622953 | 0.995119468206711 | NM_011309 |
| Pcdha11 | 122.1904716 | 0.026130243 | 0.149178591 | 0.175160814 | 0.860953274 | 0.934292638 | | Triqk | 392.9499376 | 0.00215552177259928 | 0.078110167439953 | 0.0275959179610815 | 0.977984437422226 | 0.995119468206711 | NM_173746,NM_001171801 |
| Rad23b | 7500.137476 | 0.010797137 | 0.061625362 | 0.175206063 | 0.86091772 | 0.934292638 | | Wbp1 | 883.5808662 | -0.00202955743850291 | 0.0738363922602707 | -0.0274872237981074 | 0.978071129930647 | 0.995129315218364 | NM_016757,NM_001083923,NM_001083922 |
| Rrm2b | 1442.642542 | -0.014218375 | 0.08118165 | -0.175142723 | 0.860967489 | 0.934292638 | | Tbr1 | 4839.668549 | 0.0019606190242314 | 0.0717205006066457 | 0.0273369400331504 | 0.978190993983615 | 0.995181564547275 | NM_009322 |
| Gtpbp6 | 825.2574961 | 0.016903531 | 0.096694238 | 0.174814258 | 0.861225584 | 0.934441821 | | Cacna1s | 49.87721707 | -0.00246183865606002 | 0.0913277650207023 | -0.0269560812695019 | 0.978494763359673 | 0.995273516640162 | NM_014193,NM_001081023 |
| Slc35a2 | 930.2609508 | 0.015007002 | 0.085820179 | 0.174865652 | 0.8611852 | 0.934441821 | | Kazald1 | 207.2002742 | -0.00243851106554561 | 0.0907184071805483 | -0.0268800030923435 | 0.978555442974996 | 0.995273516640162 | NM_178929 |
| Ahcy | 1374.559168 | -0.016828835 | 0.096468134 | -0.174449684 | 0.86151207 | 0.934625736 | | Pard6g | 277.9627062 | 0.0023922158553416 | 0.0888925805001163 | 0.0269113107290036 | 0.978530472128364 | 0.995273516640162 | NM_053117 |
| Col22a1 | 107.1362851 | 0.023478159 | 0.134587726 | 0.174445024 | 0.861515733 | 0.934625736 | | Zmym4 | 2602.425746 | -0.00184082597690769 | 0.0683475220184474 | -0.0269333243187785 | 0.978512914189204 | 0.995273516640162 | NM_001114399 |
| Sigirr | 56.1650546 | -0.025129764 | 0.144159729 | -0.174318892 | 0.861614853 | 0.934667824 | | Cep57l1 | 176.988499 | 0.0024346708174194 | 0.0911297266333112 | 0.0267165381414569 | 0.978685822311205 | 0.995336431764514 | NM_001243075,NM_026643,NM_029132,NM_001243074 |
| Slc6a17 | 26027.30972 | -0.013439028 | 0.077156152 | -0.174179608 | 0.861724311 | 0.934721119 | | 1810010H24Rik | 98.41960669 | 0.00247806541366147 | 0.0943097794219819 | 0.0262758054238846 | 0.979037352722036 | 0.995367690830036 | NM_001163473 |
| Arl2 | 1502.057455 | -0.013260248 | 0.076176907 | -0.174071762 | 0.861809065 | 0.934747613 | | 2210404O09Rik | 177.035663 | -0.00222030185524256 | 0.0862823232597953 | -0.0257329861508743 | 0.97947031341901 | 0.995367690830036 | NM_001256493 |
| Fam117b | 3117.335118 | 0.013505229 | 0.077675446 | 0.173867408 | 0.861969667 | 0.93479093 | | Adamts4 | 888.6728786 | -0.00227085143507534 | 0.0874900271715194 | -0.0259555461175416 | 0.979292795552427 | 0.995367690830036 | NM_172845 |
| Mir3082 | 8.646482355 | 0.023990345 | 0.137928954 | 0.173932622 | 0.861918414 | 0.93479093 | | Arfip2 | 3887.100798 | 0.00115146924934032 | 0.0439946892127104 | 0.0261729147300728 | 0.979119419393865 | 0.995367690830036 | NM_029802 |
| Gm4787 | 15.75465411 | 0.028962348 | 0.1667884 | 0.173647257 | 0.862142691 | 0.934840346 | | Cyp26b1 | 708.288211 | -0.0021507758272596 | 0.0828920711719699 | -0.0259467014981149 | 0.979299850161812 | 0.995367690830036 | NM_175475,NM_001177713 |
| U2af1l4 | 872.8100903 | -0.015304611 | 0.08810341 | -0.173711907 | 0.862091879 | 0.934840346 | | Fbxl15 | 461.8507642 | 0.00208377899179412 | 0.0808881900430327 | 0.0257612265855565 | 0.979447788279534 | 0.995367690830036 | NM_133694 |
| Vim | 1536.529387 | -0.016590545 | 0.09557915 | -0.173579123 | 0.86219624 | 0.934840346 | | Ppt2 | 1109.02625 | 0.00185679244651565 | 0.0720844006415205 | 0.0257585889594835 | 0.979449892102519 | 0.995367690830036 | NM_019441 |
| Epas1 | 6564.44705 | -0.010289574 | 0.059322564 | -0.173451261 | 0.862296735 | 0.934842752 | | Sympk | 6114.783238 | 0.00150329543260825 | 0.0580129601822672 | 0.0259130964509506 | 0.979326654097651 | 0.995367690830036 | NM_026605 |
| Irs3 | 9.760925453 | 0.025720749 | 0.148312411 | 0.173422767 | 0.862319131 | 0.934842752 | | Tipin | 397.2249815 | 0.00230328526242981 | 0.0872412345198807 | 0.0264013373389957 | 0.978937227480629 | 0.995367690830036 | NM_025372 |
| F930015N05Rik | 66.58481139 | -0.024917239 | 0.143817773 | -0.17325563 | 0.862450499 | 0.934919753 | | Tll2 | 30.89567076 | -0.00229508627133718 | 0.0884865773016753 | -0.025937112060653 | 0.97930749885176 | 0.995367690830036 | NM_011904 |
| Trim25 | 349.6181088 | 0.021487266 | 0.124100372 | 0.173144248 | 0.862538045 | 0.934949242 | | Tmem120b | 287.0773651 | -0.00209515387717132 | 0.0805093735550413 | -0.026023725991843 | 0.979238414252705 | 0.995367690830036 | NM_001039723 |
| Dgkz | 20930.5799 | -0.016876264 | 0.097625907 | -0.172866651 | 0.862756245 | 0.93512034 | | Dcaf12l2 | 31.4662004 | 0.00221562208967824 | 0.0866152412778963 | 0.0255800486956983 | 0.979592299697382 | 0.995422017866679 | NM_175539 |
| 2310044G17Rik | 4038.125675 | -0.008549817 | 0.049585618 | -0.172425332 | 0.863103158 | 0.935401714 | | Adrb2 | 85.66519558 | -0.00240289309572287 | 0.094995611134351 | -0.025294780116994 | 0.979819837463805 | 0.995583586947344 | NM_007420 |
| Arl6ip6 | 235.7356215 | 0.015949672 | 0.092524729 | 0.172382806 | 0.863136589 | 0.935401714 | | B230118H07Rik | 848.9419204 | -0.00188320760496073 | 0.07520133944922 | -0.0250422082738616 | 0.980021296819766 | 0.9957186373088 | NM_026592 |
| Igf2 | 3141.66255 | 0.027784624 | 0.161361082 | 0.172189126 | 0.863288847 | 0.935501286 | | A630066F11Rik | 29.46303411 | 0.00202982403165392 | 0.0824724421358433 | 0.0246121489686279 | 0.980364328760183 | 0.995858198571358 | NR_030698 |
| Psmd6 | 2014.471549 | -0.01100942 | 0.064002729 | -0.17201486 | 0.863425847 | 0.935584311 | | Actr3 | 7768.162087 | -0.00149960821158043 | 0.0606818275003131 | -0.024712640890268 | 0.980284172188127 | 0.995858198571358 | NM_023735,NM_001205385,NM_001205386 |
| Nphs1 | 81.4127743 | 0.026257464 | 0.152759697 | 0.17188738 | 0.863526068 | 0.935627475 | | Ctla2a | 177.506956 | 0.00232687563889868 | 0.0944248062776984 | 0.0246426307940252 | 0.980340015566634 | 0.995858198571358 | NM_007796,NM_001145799 |
| Asf1b | 28.25143863 | -0.028591719 | 0.1664508 | -0.171772794 | 0.863616155 | 0.935659653 | | Fsd1 | 1599.295283 | 0.00151952484349918 | 0.0625347485297772 | 0.0242988878859188 | 0.980614200208776 | 0.996042365724962 | NM_183178 |
| Hnrnpa3 | 6121.153902 | -0.017976562 | 0.104766768 | -0.171586492 | 0.863762627 | 0.935752912 | | Man1c1 | 2247.063226 | 0.00160976364596218 | 0.0665586706156536 | 0.0241856339838553 | 0.980704537199787 | 0.99606447401511 | NM_207237 |
| Atmin | 4509.069956 | 0.014056039 | 0.082135603 | 0.171132108 | 0.864119889 | 0.935781262 | | Zfp105 | 191.0701573 | -0.00202338868623154 | 0.0851492260200376 | -0.0237628547058712 | 0.981041769327371 | 0.996337323725749 | NM_009544 |
| Dcdc2c | 124.5243508 | -0.021553907 | 0.125908439 | -0.171187154 | 0.864076608 | 0.935781262 | | Brms1 | 945.9424076 | 0.0015397889990629 | 0.0672830964666117 | 0.0228852279387439 | 0.981741823708041 | 0.996444753305681 | NM_134155 |
| Gm4349 | 20.24079729 | 0.028633696 | 0.167205481 | 0.171248548 | 0.864028336 | 0.935781262 | | Cbx8 | 270.6729617 | -0.00186589324077207 | 0.0832011693984585 | -0.0224262862440807 | 0.982107912231684 | 0.996444753305681 | NM_013926 |
| Psd2 | 2463.7829 | -0.015767126 | 0.092163331 | -0.171078085 | 0.864162368 | 0.935781262 | | Dag1 | 5225.080223 | 0.000999949491205885 | 0.0434861816812919 | 0.0229946491631403 | 0.981654541171351 | 0.996444753305681 | NM_010017,NM_001276481,NM_001276493,NM_001276492,NM_001276482,NM_001276485,NM_001276486,NM_001276494 |
| Tcp1 | 7167.527191 | -0.010465955 | 0.061198863 | -0.171015509 | 0.864211571 | 0.935781262 | | Gsg1l | 4885.554324 | 0.00190774738664255 | 0.0849202038460106 | 0.0224651767216899 | 0.982076889935708 | 0.996444753305681 | NM_001101488 |
| Wdr65 | 45.16773207 | 0.028799916 | 0.168086999 | 0.171339343 | 0.863956946 | 0.935781262 | | Htr3a | 297.9637017 | -0.00200427871500626 | 0.0857530107751763 | -0.0233726920709641 | 0.981352987624452 | 0.996444753305681 | NM_001099644,NM_013561 |
| Zfp526 | 403.2339095 | 0.015865082 | 0.092731292 | 0.171086605 | 0.864155668 | 0.935781262 | | Inpp5j | 3132.145556 | -0.00147322186616265 | 0.0648498510984456 | -0.0227174286634247 | 0.981875673360197 | 0.996444753305681 | NM_172439 |
| Rce1 | 521.4163589 | 0.015598805 | 0.091309845 | 0.170833769 | 0.864354476 | 0.935870597 | | Nisch | 18828.5405 | 0.00117527717736246 | 0.0512283111368412 | 0.0229419465776074 | 0.981696580660382 | 0.996444753305681 | NM_022656 |
| Abracl | 198.5400436 | 0.022395558 | 0.13117762 | 0.170726974 | 0.864438452 | 0.93589612 | | Ppp2r1b | 661.0791764 | 0.00182514653866798 | 0.0782661450956745 | 0.0233197449093331 | 0.981395221835891 | 0.996444753305681 | NM_001034085,NM_028614 |
| Clip4 | 2506.099618 | -0.018294823 | 0.10729707 | -0.17050627 | 0.864612004 | 0.936018613 | | Rapgef3 | 1594.753649 | -0.00177490875725793 | 0.0779248696244399 | -0.0227771796836122 | 0.98182801127643 | 0.996444753305681 | NM_001177811,NM_001177810,NM_144850 |
| Hk1 | 16890.39822 | -0.012041658 | 0.070721709 | -0.170268199 | 0.864799219 | 0.936113441 | | Rnf146 | 2310.574701 | 0.00135818487119143 | 0.0604446689411045 | 0.0224698868400588 | 0.982073132753396 | 0.996444753305681 | NM_001110198,NM_001110197,NM_001110196,NM_026518 |
| Tab3 | 1069.814722 | -0.014374682 | 0.084437138 | -0.170241223 | 0.864820433 | 0.936113441 | | Sbno1 | 7475.514196 | -0.00147079986011141 | 0.0640635067502142 | -0.0229584662894916 | 0.981683403308112 | 0.996444753305681 | NM_001081203 |
| 1810062O18Rik | 34.12228254 | -0.027184373 | 0.160096206 | -0.16980023 | 0.865167246 | 0.936273888 | | Scube3 | 222.3713692 | -0.00216452427326545 | 0.0948344351085356 | -0.0228242438602413 | 0.981790469256333 | 0.996444753305681 | NM_001004366 |
| 2610524H06Rik | 368.6887947 | -0.013268687 | 0.078428042 | -0.169182945 | 0.865652745 | 0.936273888 | | Smarcal1 | 3782.814542 | -0.00110032273715267 | 0.0486391501531009 | -0.0226221620585309 | 0.981951665583916 | 0.996444753305681 | NM_018817 |
| Ampd2 | 2899.248292 | -0.017107751 | 0.100912845 | -0.169529966 | 0.865379804 | 0.936273888 | | Xrn1 | 742.4843136 | 0.00167782399294668 | 0.0726094842729068 | 0.023107504615244 | 0.981564519468352 | 0.996444753305681 | NM_011916 |
| Cdkal1 | 421.2800841 | -0.014151287 | 0.083496878 | -0.169482832 | 0.865416875 | 0.936273888 | | S1pr2 | 124.4533986 | -0.00207345480720893 | 0.0929897892772714 | -0.0222976611015476 | 0.982210514590206 | 0.996479252335335 | NM_010333 |
| 9-Mar | 615.0578912 | -0.011291036 | 0.06667253 | -0.169350644 | 0.865520843 | 0.936273888 | | Ankra2 | 1161.372107 | 0.00115179158715523 | 0.052253378648417 | 0.0220424327947285 | 0.982414107274975 | 0.996546600717618 | NM_001271388,NM_001271389,NM_001271391,NM_001271390,NM_001271392,NM_023472 |
| Mdp1 | 1424.19307 | -0.010334746 | 0.061128224 | -0.169066688 | 0.865744187 | 0.936273888 | | Polh | 232.9251725 | -0.00186340394041515 | 0.084399620782359 | -0.0220783449397279 | 0.982385460500415 | 0.996546600717618 | NM_030715 |
| Mtrf1l | 239.6337421 | -0.015979038 | 0.094392104 | -0.169283632 | 0.86557355 | 0.936273888 | | A630072M18Rik | 275.5621077 | -0.00172962117781625 | 0.0843794112418781 | -0.0204981422880304 | 0.983645994005502 | 0.996711782404217 | NR_030699 |
| Ppp3r1 | 28685.26238 | -0.013233251 | 0.078046085 | -0.169556882 | 0.865358635 | 0.936273888 | | Atp5k | 2026.847485 | 0.00149053746466818 | 0.0709069748953207 | 0.0210210274358573 | 0.983228881912802 | 0.996711782404217 | NM_007507 |
| Set | 5879.256784 | -0.009164455 | 0.053941831 | -0.169895136 | 0.865092606 | 0.936273888 | | Brpf1 | 1552.828095 | -0.00103379772457294 | 0.0509510958572505 | -0.0202900047556023 | 0.983812029196728 | 0.996711782404217 | NM_030178 |
| Tlr6 | 21.81002346 | 0.028158953 | 0.166096093 | 0.16953411 | 0.865376545 | 0.936273888 | | C730002L08Rik | 149.2555482 | 0.00198585368120882 | 0.0948733017680156 | 0.0209316387666642 | 0.983300188062526 | 0.996711782404217 | NR_045778 |
| Toe1 | 438.9140672 | 0.013977515 | 0.082628399 | 0.169161146 | 0.865669891 | 0.936273888 | | Chmp1b | 968.5667817 | -0.00143052118009956 | 0.0702113977188561 | -0.0203744865730735 | 0.983744636390459 | 0.996711782404217 | NM_024190 |
| Ttyh1 | 19043.01706 | -0.007815355 | 0.046045622 | -0.169730687 | 0.865221939 | 0.936273888 | | Cnr1 | 6837.044198 | 0.00169123482042153 | 0.0782260454317739 | 0.0216198429958545 | 0.982751204808866 | 0.996711782404217 | NM_007726 |
| Zmym2 | 3695.228136 | -0.011353688 | 0.067160159 | -0.169053923 | 0.865754227 | 0.936273888 | | D630045J12Rik | 3009.074131 | -0.00164917795886883 | 0.0808814555495184 | -0.020390062934255 | 0.983732210833646 | 0.996711782404217 | NM_194061 |
| 1110054M08Rik | 111.5220471 | 0.022146323 | 0.131121525 | 0.16889922 | 0.865875912 | 0.93634013 | | Dab1 | 3881.395759 | 0.00142226374544479 | 0.066282129446015 | 0.0214577255940937 | 0.982880525779905 | 0.996711782404217 | NM_177259,NM_010014 |
| Fam53b | 1897.639704 | 0.019248602 | 0.114020447 | 0.16881711 | 0.865940499 | 0.936344623 | | Dgke | 3291.801127 | 0.00137315849607229 | 0.0671303267544541 | 0.0204551141408108 | 0.983680318303134 | 0.996711782404217 | NM_019505 |
| 4930556M19Rik | 9.227431491 | 0.026757754 | 0.159243386 | 0.168030548 | 0.866559247 | 0.936411385 | | Kctd11 | 95.94548898 | -0.00193329721630213 | 0.0931171864609574 | -0.0207619805728638 | 0.983435526304328 | 0.996711782404217 | NM_153143 |
| Abhd6 | 2074.904578 | -0.014446968 | 0.085792103 | -0.168395073 | 0.866272484 | 0.936411385 | | Lrrc9 | 97.99483767 | 0.00199238884733203 | 0.0938573015448179 | 0.0212278513715914 | 0.983063897103647 | 0.996711782404217 | NM_030070,NM_001142729,NM_001142728 |
| Diap1 | 2418.892902 | 0.019786781 | 0.117799254 | 0.167970341 | 0.866606612 | 0.936411385 | | Mir425 | 24.98596934 | -0.00177403130147174 | 0.0844343944400792 | -0.0210107659708595 | 0.983237067569424 | 0.996711782404217 | NR_029947 |

| | | | | | | | | | | | | | | | |
|---|---|---|---|---|---|---|---|---|---|---|---|---|---|---|---|
| Gorasp2 | 3845.741731 | 0.011352127 | 0.067424972 | 0.168366801 | 0.866294724 | 0.936411385 | | Ptplad2 | 45.96307611 | 0.00189073741248511 | 0.0918552322958449 | 0.0205838836310977 | 0.983577596742727 | 0.996711782404217 | NM_025760 |
| Ipo13 | 4119.97244 | 0.010286325 | 0.061215714 | 0.168034058 | 0.866556485 | 0.936411385 | | Samd10 | 964.6014628 | -0.00119583149195843 | 0.0562018886940956 | -0.021277425363181 | 0.983024351712978 | 0.996711782404217 | NM_172676 |
| Olfr78 | 44.40332962 | -0.026908616 | 0.159705959 | -0.168488489 | 0.866198999 | 0.936411385 | | Tbc1d16 | 2821.498093 | 0.0012725661475733 | 0.0614955194961351 | 0.0206936400895561 | 0.983490042408361 | 0.996711782404217 | NM_172443 |
| Riiad1 | 48.29344377 | 0.026391183 | 0.156876712 | 0.168228811 | 0.866403276 | 0.936411385 | | Tgfb3 | 613.8277586 | -0.00190805451897812 | 0.0880237407898761 | -0.0216765899183817 | 0.982705937923814 | 0.996711782404217 | NM_009368 |
| Sertad2 | 1327.933488 | -0.013266656 | 0.078713369 | -0.168543871 | 0.866155434 | 0.936411385 | | Tnrc6c | 4388.364021 | 0.00124581795950618 | 0.0584685248424749 | 0.02130749771544 | 0.983000362885915 | 0.996711782404217 | NM_198022 |
| Slc25a15 | 496.9165571 | -0.012732928 | 0.075719383 | -0.168159422 | 0.866457863 | 0.936411385 | | Zfp874a | 573.0016137 | 0.0017568875125319 | 0.0818024170072324 | 0.0214772078479854 | 0.982864984771782 | 0.996711782404217 | NM_177712 |
| Tgfb1 | 225.5138237 | -0.01894941 | 0.112569822 | -0.168334728 | 0.866319954 | 0.936411385 | | Egfl8 | 60.67257021 | 0.00187708049586609 | 0.0934911357427027 | 0.0200776306861006 | 0.983981444675275 | 0.996744383915834 | NM_152922 |
| Zbtb25 | 353.6152405 | 0.015267709 | 0.091038583 | 0.167705914 | 0.866814643 | 0.93656606 | | Zfp78 | 146.4260065 | 0.00180674167039435 | 0.0899406018896483 | 0.0200881652160957 | 0.983973041031299 | 0.996744383915834 | NM_177888,NM_001025163,NM_001112805 |
| Zfp874b | 447.417298 | 0.013014108 | 0.077633725 | 0.167634725 | 0.866870651 | 0.93656606 | | Bola2 | 544.9824701 | 0.00159452507101966 | 0.0811212778898477 | 0.0196560645060945 | 0.984317739446254 | 0.997015513785391 | NM_175103 |
| Ccdc28a | 585.1135855 | 0.011614931 | 0.069369334 | 0.1674361 | 0.867026923 | 0.936619688 | | Rrm1 | 864.704614 | 0.00133182711749067 | 0.0682337505831116 | 0.0195185975577944 | 0.984427401163218 | 0.997057065213841 | NM_009103 |
| Wdr34 | 1019.914335 | -0.01243662 | 0.074284857 | -0.167417967 | 0.867041189 | 0.936619688 | | Arpp19 | 14002.75836 | 0.00150964959333853 | 0.0816720975334239 | 0.0184842809903834 | 0.985252517377077 | 0.99736187093676 | NM_021548,NM_001142655 |
| Morc4 | 155.3713355 | -0.022746238 | 0.135991738 | -0.167261911 | 0.867163973 | 0.936621721 | | Asphd2 | 3533.728155 | 0.00128153187600573 | 0.0691592683238437 | 0.0185301537605178 | 0.985215922469756 | 0.99736187093676 | NM_028386 |
| Ric8 | 2169.884845 | 0.010002849 | 0.059792811 | 0.167291832 | 0.867140431 | 0.936621721 | | Dach2 | 24.33081698 | 0.00148179341266799 | 0.0806781098067413 | 0.0183667343746342 | 0.985346290087667 | 0.99736187093676 | NM_033605,NM_001142570 |
| Dpy19l4 | 635.531608 | -0.015182925 | 0.091012606 | -0.166822222 | 0.867509933 | 0.936799477 | | Noc4l | 859.9697702 | 0.00118511829514564 | 0.0631391381236515 | 0.0187699472999569 | 0.985024628178619 | 0.99736187093676 | NM_153570 |
| Dusp28 | 674.7630908 | 0.016658425 | 0.099802143 | 0.166914498 | 0.867437325 | 0.936799477 | | Nufip1 | 552.157216 | -0.00131497259171138 | 0.0693158866850253 | -0.0189707245279379 | 0.984864459648232 | 0.99736187093676 | NM_013745 |
| Zdhhc5 | 2646.557679 | -0.013466238 | 0.080710513 | -0.16684614 | 0.867491113 | 0.936799477 | | Prmt6 | 508.7583402 | 0.00134408023720844 | 0.0723074524409786 | 0.0185884053694957 | 0.985169452414239 | 0.99736187093676 | NR_024139,NM_178891 |
| Hexa | 2136.531987 | 0.017103872 | 0.102663372 | 0.166601507 | 0.867683608 | 0.936921724 | | Timm10b | 557.6468016 | 0.00143488766130692 | 0.0764891506014292 | 0.018759361949041 | 0.985033072579865 | 0.99736187093676 | NM_019502 |
| 9130227L01Rik | 9.932814981 | 0.026764769 | 0.160754641 | 0.166494535 | 0.867767783 | 0.936947319 | | Vstm5 | 562.772975 | 0.00119868954216366 | 0.0652524961763961 | 0.018370018197055 | 0.985343670418429 | 0.99736187093676 | NM_026955 |
| Pigp | 651.4541943 | -0.014333709 | 0.086135268 | -0.16640929 | 0.867834864 | 0.936954455 | | Zc3h18 | 1839.110378 | 0.000849208471815637 | 0.0457985923875982 | 0.018542239565546 | 0.985206281049013 | 0.99736187093676 | NM_001029994,NM_001029993 |
| Zfp444 | 1031.943148 | -0.012546151 | 0.075519971 | -0.166130247 | 0.868054451 | 0.937126231 | | Mei4 | 28.78520641 | -0.00157049805258038 | 0.0870431257265404 | -0.0180427579946329 | 0.985604743007355 | 0.997496291574243 | NM_175213 |
| Gnpnat1 | 320.5264742 | -0.015922907 | 0.095989505 | -0.165881748 | 0.868250011 | 0.93715637 | | Ppp1r36 | 50.68604427 | -0.00168386282854836 | 0.0934020864799576 | -0.018028107208394 | 0.985616430742469 | 0.997496291574243 | NM_001163103 |
| Mad2l1bp | 302.0338083 | 0.016423135 | 0.099015554 | 0.165864196 | 0.868263825 | 0.93715637 | | Atp7a | 432.5885375 | 0.00149278374079623 | 0.0857810587961945 | 0.01740225361805 | 0.98611571130122 | 0.997552088479139 | NM_009726,NM_001109757 |
| Zfp219 | 1163.620912 | 0.014398027 | 0.086752279 | 0.165967127 | 0.86818282 | 0.93715637 | | Cox14 | 1395.505238 | -0.000896909721668168 | 0.0515852024770875 | -0.0173869574722818 | 0.986127914013527 | 0.997552088479139 | NM_183256 |
| Cd79a | 12.35307453 | 0.026221886 | 0.158731072 | 0.165196929 | 0.868788982 | 0.937641239 | | Igfbpl1 | 22.49598629 | -0.00137810094255818 | 0.0797320354300647 | -0.0172841560500102 | 0.986209925357137 | 0.997552088479139 | NM_018741 |
| Fam167b | 11.85442749 | -0.025027567 | 0.151578676 | -0.16511272 | 0.868855261 | 0.937641239 | | Pid1 | 1712.766339 | -0.00128278507027662 | 0.074276792838622 | -0.017270334666491 | 0.986220951579851 | 0.997552088479139 | NM_001003948 |
| Leng9 | 221.9550056 | -0.015918412 | 0.09648146 | -0.164989338 | 0.868952374 | 0.937641239 | | Prr5l | 617.0124351 | 0.00156688867990609 | 0.0892617734690606 | 0.0175538600568944 | 0.98599476533931 | 0.997552088479139 | NM_175181,NM_001083810,NM_001110849 |
| Polg2 | 67.75887767 | -0.026285431 | 0.159393578 | -0.164908971 | 0.869015631 | 0.937641239 | | Slc25a18 | 2317.090168 | -0.00126991831952041 | 0.0715073720852606 | -0.017759264289649 | 0.985830902018089 | 0.997552088479139 | NM_001081048 |
| Ywhab | 24096.32406 | -0.007528205 | 0.045637418 | -0.164956856 | 0.868977941 | 0.937641239 | | Slc30a1 | 1622.57602 | 0.00126857185955322 | 0.0722062092372934 | 0.0175687364418242 | 0.985982897531606 | 0.997552088479139 | NM_009579 |
| Col6a4 | 215.4654053 | 0.027025995 | 0.164060037 | 0.164732349 | 0.869154654 | 0.937725939 | | Ube2r2 | 3400.369608 | -0.000912614872912714 | 0.0514951647894938 | -0.0177223410516963 | 0.985860357863811 | 0.997552088479139 | NM_026275 |
| Ing3 | 562.1509928 | 0.014055574 | 0.085463727 | 0.164462453 | 0.869367102 | 0.937824541 | | Dnajc15 | 722.1665811 | 0.00111935637294622 | 0.065827296890931 | 0.017004440799094 | 0.986433073039566 | 0.997581727802046 | NM_025384 |
| Snapc5 | 580.3158523 | 0.01448811 | 0.088092816 | 0.164464143 | 0.869365772 | 0.937824541 | | Mrps18b | 833.4781751 | -0.00115274468448462 | 0.0679025759628935 | -0.0169764499820236 | 0.986455403257027 | 0.997581727802046 | NM_025878 |
| Etv4 | 310.5063925 | 0.022191709 | 0.135089388 | 0.164274258 | 0.869515246 | 0.937919049 | | Osr1 | 71.98074577 | -0.00158560697664343 | 0.093406439577425 | -0.01697534970626 | 0.986456281023584 | 0.997581727802046 | NM_011859 |
| Ndufs2 | 6986.995156 | -0.0102067 | 0.062214235 | -0.164057305 | 0.869686032 | 0.938037966 | | Agl | 2103.422878 | -0.000960751241855346 | 0.0597627613165985 | -0.0160760851856507 | 0.987173692307525 | 0.997691963533785 | NM_001081326 |
| Itgb7 | 18.59006237 | 0.027544647 | 0.167982683 | 0.163973132 | 0.869752295 | 0.938044136 | | Atp13a2 | 6698.055681 | 0.00112340718477538 | 0.06801214566132 | 0.0165177436155243 | 0.986821346660788 | 0.997691963533785 | NM_001164366,NM_029097 |
| 1700011J10Rik | 278.0213912 | 0.014985983 | 0.091496951 | 0.163786693 | 0.869899068 | 0.93813713 | | Dhx16 | 1128.052173 | -0.00111117441921988 | 0.0669030282267555 | -0.0166087313036678 | 0.98674875894679 | 0.997691963533785 | NM_026987 |
| Hspa12a | 9771.55351 | -0.009691682 | 0.059310609 | -0.163405532 | 0.870199147 | 0.938395434 | | Galk2 | 935.1282756 | -0.00104998229039227 | 0.0650625569365859 | -0.0161380422139825 | 0.987124264163366 | 0.997691963533785 | NM_175154 |
| Ssr3 | 4804.285719 | -0.012905814 | 0.079089733 | -0.163179383 | 0.870377199 | 0.938522118 | | Gpaa1 | 1479.46611 | 0.00100379715890652 | 0.0619759399835396 | 0.0161965620718801 | 0.987077578174085 | 0.997691963533785 | NM_010331 |
| Ly6g6f | 13.0132176 | 0.024712933 | 0.151615148 | 0.162997781 | 0.870520182 | 0.938610974 | | Mamdc2 | 56.10741145 | 0.00155718903051999 | 0.0949286549675459 | 0.0164037827255886 | 0.986912261977461 | 0.997691963533785 | NM_174857 |
| Pdcd5 | 932.4693285 | 0.015109821 | 0.092783149 | 0.162850918 | 0.870635816 | 0.938670332 | | Pwwp2a | 900.8080263 | -0.00117886730019372 | 0.0733862499346762 | -0.0160638716550181 | 0.987183436036837 | 0.997691963533785 | NM_001164231,NM_027557 |
| 4931403E22Rik | 27.13302725 | 0.024627988 | 0.151418141 | 0.162648859 | 0.870794915 | 0.938711223 | | Smg5 | 4552.065266 | -0.000821289443617785 | 0.050388548569069 | -0.0162991288088408 | 0.986995752559635 | 0.997691963533785 | NM_178246 |
| Sde2 | 858.7217018 | -0.013972244 | 0.085883631 | -0.162688096 | 0.87076402 | 0.938711223 | | Tpd52 | 2731.114212 | -0.000966892855577619 | 0.0579008262833523 | -0.0166991201653057 | 0.986676649070293 | 0.997691963533785 | NM_001025262,NM_001025263,NM_001025261,NM_009412,NM_001025264 |
| Dbn1 | 9118.412048 | 0.010527404 | 0.064790162 | 0.162484608 | 0.870924248 | 0.938785328 | | Foxo4 | 809.6526748 | -0.0011285994754584 | 0.0708791183756824 | -0.0159228768828141 | 0.987295919201796 | 0.997717927614865 | NM_018789 |
| Fam175a | 146.504264 | 0.02111675 | 0.130151134 | 0.162247914 | 0.87111063 | 0.938790297 | | Mgst1 | 221.1813861 | 0.00148352003955855 | 0.0950901480459051 | 0.0156011960234027 | 0.987552551510845 | 0.997717927614865 | NM_019946 |
| Fbrsl1 | 1661.515585 | -0.012921422 | 0.079570505 | -0.162389588 | 0.87099907 | 0.938790297 | | Olfm1 | 38352.88984 | 0.000841136156182106 | 0.0531751280071383 | 0.0158182253189723 | 0.987379408553735 | 0.997717927614865 | NM_001038613,NM_001038612,NM_001038614,NM_019498 |
| Zfp273 | 195.1523405 | -0.019073015 | 0.117525479 | -0.16228834 | 0.871078797 | 0.938790297 | | Rnf169 | 2731.328973 | 0.000915695360086374 | 0.0580774077573018 | 0.0157668083932559 | 0.987420428209379 | 0.997717927614865 | NM_175388 |
| Frmd4b | 892.7113225 | 0.012038995 | 0.074399377 | 0.161815803 | 0.871450908 | 0.939091694 | | Xrcc1 | 1058.952446 | -0.000961346191146017 | 0.0615367757532874 | -0.0156223035636485 | 0.987535712182589 | 0.997717927614865 | NM_009532 |
| Zfp72 | 138.4753732 | 0.020322321 | 0.126156442 | 0.161088258 | 0.872023889 | 0.939643795 | | Tubgcp5 | 1336.507417 | 0.00104614325345638 | 0.0676749611911603 | 0.0154583502530774 | 0.987666512202468 | 0.997763666197444 | NM_146190 |
| Nsmce4a | 1049.429617 | 0.014735959 | 0.091580862 | 0.160906534 | 0.872167016 | 0.939732666 | | 2310008H04Rik | 124.5047635 | -0.00145183365784041 | 0.0946397394504979 | -0.0153406345608105 | 0.98776042459978 | 0.997789146574158 | NM_146068 |
| Cenpn | 35.67927986 | -0.02696999 | 0.167864943 | -0.160664814 | 0.872357404 | 0.939807093 | | Plcxd1 | 790.6206687 | 0.00103955410431824 | 0.0682174999089506 | 0.015238818568633 | 0.987841652512833 | 0.997801811028539 | NR_027665,NM_207279 |
| Tpst2 | 750.4762317 | 0.01416992 | 0.088188898 | 0.16067691 | 0.872347876 | 0.939807093 | | Strn4 | 10329.78007 | 0.00103353134101609 | 0.0684297593203301 | 0.0151035361117956 | 0.987949579875164 | 0.997841440499696 | NM_001039878,NM_133789 |
| Coq7 | 646.4501694 | 0.014124141 | 0.088114113 | 0.160293749 | 0.872649683 | 0.940056608 | | Aifm2 | 345.3009638 | -0.00126887076656936 | 0.0845287598699397 | -0.0150111130048721 | 0.988023314486073 | 0.99784653175448 | NM_178058,NM_001039194,NM_153779 |
| Mir682 | 56.23566017 | 0.026865879 | 0.167955249 | 0.159958556 | 0.872913721 | 0.940275667 | | Bdkrb2 | 24.44729177 | 0.00121708613011134 | 0.0823619178139436 | 0.0147772922536936 | 0.988209855760957 | 0.997965542601756 | NM_009747 |
| D830015G02Rik | 14.16509282 | 0.025456755 | 0.159235324 | 0.159868765 | 0.872984454 | 0.940286488 | | 2310040G24Rik | 38.75058803 | -0.00134278226289382 | 0.0926921998707565 | -0.014486464500423 | 0.988441877895972 | 0.997974805283964 | NR_040293,NR_040292 |
| Chmp4b | 6094.359911 | -0.009062981 | 0.056846709 | -0.159428426 | 0.873331345 | 0.940594735 | | AI462493 | 534.6690026 | -0.000960552933514798 | 0.0674114573857658 | -0.014249104985492 | 0.988631243841129 | 0.997974805283964 | NM_001160356 |
| Dtymk | 535.7868936 | -0.017931661 | 0.112630447 | -0.159207939 | 0.873505049 | 0.94068029 | | Actn3 | 46.09859252 | 0.0013062289344472 | 0.0915327269741541 | 0.0142706218598299 | 0.98861407760471 | 0.997974805283964 | NM_013456 |
| Phc2 | 3118.060063 | 0.010196562 | 0.064059432 | 0.159173468 | 0.873532207 | 0.94068029 | | Casp3 | 274.112759 | 0.00114631400398712 | 0.0794247563941002 | 0.0144327040589107 | 0.988484768038271 | 0.997974805283964 | NM_009810 |
| Paip1 | 2364.819852 | 0.009681235 | 0.060879272 | 0.159023494 | 0.873650364 | 0.940742146 | | Cdh23 | 68.05795678 | -0.0013446268352235 | 0.0932632360651149 | -0.0144175442752673 | 0.988496862537182 | 0.997974805283964 | NM_001252635,NM_023370,NR_045556 |
| A430033K04Rik | 669.2723637 | -0.011347727 | 0.071457435 | -0.158803999 | 0.873823298 | 0.940752848 | | Smek1 | 1629.277246 | 0.000938022585958552 | 0.0651927557109864 | 0.0143884481600534 | 0.988520075470446 | 0.997974805283964 | NM_001160214,NM_211355 |
| Ndufa4l2 | 80.71192522 | 0.02253701 | 0.142019313 | 0.158689757 | 0.873913308 | 0.940752848 | | Cdk2ap1 | 3152.297831 | 0.000843293797769938 | 0.0616929055948294 | 0.0136692183588872 | 0.989093881344482 | 0.998005843636446 | NM_013812 |
| Nupr1l | 12.20577834 | -0.025195334 | 0.15883527 | -0.158625562 | 0.873963888 | 0.940752848 | | Iqcb1 | 995.2776678 | -0.000915243733808122 | 0.0649962781784734 | -0.0140814791163111 | 0.988764976516212 | 0.998005843636446 | NM_177128 |
| Rab10 | 5226.402416 | -0.010594458 | 0.066759868 | -0.158695005 | 0.873909174 | 0.940752848 | | Lrrc51 | 109.1215858 | 0.00130318175670789 | 0.0941947379057525 | 0.0138349740726685 | 0.988961639925092 | 0.998005843636446 | NM_001162974,NM_027053,NM_001162973 |
| Tdo2 | 12.27823485 | -0.025298892 | 0.159335913 | -0.158777085 | 0.873844503 | 0.940752848 | | Optn | 811.1451742 | 0.000956118060125281 | 0.0702627831704624 | 0.0136077453380358 | 0.98914292515723 | 0.998005843636446 | NM_181848 |
| Tmem125 | 273.8593757 | 0.026370475 | 0.166491276 | 0.158389529 | 0.874149864 | 0.940887669 | | Spats2l | 1155.315904 | 0.000916191636808203 | 0.0667829310782698 | 0.0137189491688291 | 0.989054205619364 | 0.998005843636446 | NM_144882,NM_001164566 |
| 1700048O20Rik | 309.413122 | 0.021783935 | 0.137701808 | 0.158196437 | 0.874302012 | 0.940905765 | | Txlng | 509.8051626 | 0.000985945059029865 | 0.0703692127181613 | 0.0140110287005585 | 0.988821182270337 | 0.998005843636446 | NM_178935 |
| Arhgap25 | 417.2620577 | -0.022531239 | 0.142544686 | -0.158064392 | 0.874406059 | 0.940905765 | | Zbtb24 | 577.5114593 | -0.000911934559004978 | 0.0667466604164817 | -0.0136626245165637 | 0.989099141978214 | 0.998005843636446 | NM_153398 |
| Rqcd1 | 1239.515105 | -0.010165519 | 0.064277695 | -0.158150019 | 0.874338587 | 0.940905765 | | Adam10 | 3561.168864 | 0.000795099213217911 | 0.0663423798304641 | 0.0119847858224224 | 0.99043775334067 | 0.998028377581853 | NM_007399 |
| Ttc5 | 1312.680461 | 0.013246464 | 0.083806597 | 0.158059917 | 0.874409585 | 0.940905765 | | Arfgap2 | 3135.34028 | 0.000604339423845444 | 0.0452647493172185 | 0.013351215525578 | 0.989347587739343 | 0.998028377581853 | NM_001166024,NM_023854 |
| Noa1 | 666.1895431 | -0.012939446 | 0.082039366 | -0.157722401 | 0.874675548 | 0.941126593 | | Asap1 | 6955.515529 | -0.000798183577121354 | 0.0633747860952481 | -0.0125946551665158 | 0.989951184760824 | 0.998028377581853 | NM_001276462,NM_001276463,NM_001276461,NM_001276467,NM_010026 |
| Zfp939 | 236.7884263 | 0.016306482 | 0.103530082 | 0.157504769 | 0.874847049 | 0.941245759 | | Car15 | 213.5527915 | 0.00115850427376714 | 0.0913586729625258 | 0.0126808351763422 | 0.98988242855239 | 0.998028377581853 | NM_030558 |
| Fgd5 | 384.7542704 | 0.01753349 | 0.111466471 | 0.157298332 | 0.875009735 | 0.941355425 | | Cenpm | 71.83074623 | 0.00110026683064584 | 0.0940723996467543 | 0.0116959579512949 | 0.990668188485546 | 0.998028377581853 | NM_025639,NM_001080158,NM_178269 |
| Aste1 | 202.5166763 | -0.016480473 | 0.105029099 | -0.156913397 | 0.875313102 | 0.941551041 | | Chst3 | 136.6735208 | 0.00110310737819133 | 0.0933854110774645 | 0.011812416580533 | 0.990575274362129 | 0.998028377581853 | NM_016803 |
| Kctd15 | 416.5116248 | -0.016300842 | 0.103845649 | -0.156971839 | 0.875267043 | 0.941551041 | | Cnih2 | 5558.486328 | -0.000796694999475962 | 0.0672426553243362 | -0.0118480597714829 | 0.990546837200398 | 0.998028377581853 | NM_009920 |
| Il6st | 3669.446005 | 0.010612232 | 0.067673785 | 0.156814514 | 0.875391034 | 0.941569502 | | Fdxacb1 | 183.037781 | -0.00102978266506852 | 0.0857946133208495 | -0.0120028825261722 | 0.990423315298655 | 0.998028377581853 | NM_198675 |
| 4930426L09Rik | 23.66841967 | -0.026295945 | 0.167789507 | -0.156719839 | 0.875465651 | 0.941584395 | | Gm19345 | 47.51595982 | 0.00115002364535353 | 0.0926360666461409 | 0.0124144265510159 | 0.990094975148125 | 0.998028377581853 | NM_001270489 |
| Gna15 | 76.39278047 | -0.023150763 | 0.148036728 | -0.156385268 | 0.875729349 | 0.941802633 | | Gpr89 | 962.9290353 | -0.000677925571154355 | 0.0580159125278013 | -0.0116851660452551 | 0.990676798592368 | 0.998028377581853 | NM_026229 |
| Fbln7 | 79.98409254 | -0.022624109 | 0.14507332 | -0.155949484 | 0.87607284 | 0.942041265 | | Klk6 | 260.1341053 | 0.00111234385286247 | 0.0951020230259895 | 0.0116963216708702 | 0.990667898299162 | 0.998028377581853 | NM_001164697,NM_001164696,NM_011177,NM_001164698 |
| Hnrnpf | 3236.496776 | -0.010771316 | 0.069043957 | -0.156006645 | 0.876027784 | 0.942041265 | | Npc2 | 2502.742232 | 0.00095148531184052 | 0.0736911189544196 | 0.0129118043712845 | 0.989698156885608 | 0.998028377581853 | NM_023409 |
| Bag4 | 2823.315215 | -0.011408582 | 0.073296242 | -0.155650298 | 0.876308676 | 0.94205372 | | Ntan1 | 1448.107902 | -0.000805296228813928 | 0.0663066109947198 | -0.0121450367728499 | 0.99030990088718 | 0.998028377581853 | NM_010946 |
| Kcnk5 | 36.15723668 | -0.026138712 | 0.167944226 | -0.155639241 | 0.876317392 | 0.94205372 | | Nup98 | 1198.677622 | 0.00081594573450217 | 0.0639968193801782 | 0.0127497857300529 | 0.989827418417311 | 0.998028377581853 | NM_022979 |
| Med4 | 811.3447634 | 0.014993058 | 0.096411585 | 0.155510954 | 0.876418519 | 0.94205372 | | Pcnxl2 | 5259.152887 | -0.000659384190035605 | 0.0530104269909079 | -0.0124387639840875 | 0.990075558185272 | 0.998028377581853 | NM_175561 |
| Por | 2798.563722 | 0.012610459 | 0.080944767 | 0.155790916 | 0.876197831 | 0.94205372 | | Plce1 | 621.2339759 | -0.00104038189310219 | 0.0816255539033368 | -0.0127457866238095 | 0.989830608983186 | 0.998028377581853 | NM_019588 |
| Tpm3 | 6104.833985 | -0.008905319 | 0.057279245 | -0.155471995 | 0.87644923 | 0.94205372 | | Sdc1 | 38.32032578 | 0.00108235996359437 | 0.0834074928945301 | 0.0129767713431097 | 0.989646325084345 | 0.998028377581853 | NM_011519 |
| Zfp322a | 1271.928996 | -0.012624075 | 0.081188878 | -0.155490199 | 0.87643488 | 0.94205372 | | Sema3d | 206.5692842 | -0.00107975118084717 | 0.0855979860151777 | -0.0126142124495278 | 0.989935581546161 | 0.998028377581853 | NM_028882 |
| 2310047M10Rik | 785.9265885 | 0.011704978 | 0.075416122 | 0.155205254 | 0.876659506 | 0.942136042 | | Stox1 | 168.6806984 | 0.00116051401686726 | 0.087561079935454 | 0.0132537654597538 | 0.989425334763067 | 0.998028377581853 | NM_001033260 |
| Prob1 | 239.5822716 | -0.014272987 | 0.091998654 | -0.155143437 | 0.876708239 | 0.942136042 | | Tmem91 | 525.8558413 | 0.00108933539157689 | 0.0919141856237391 | 0.0118516568926173 | 0.990543967314482 | 0.998028377581853 | NM_177102 |
| Trp53i13 | 115.2111705 | -0.023514704 | 0.151426869 | -0.155287529 | 0.876594647 | 0.942136042 | | Tsr1 | 1584.168318 | 0.000714539741248637 | 0.0551125359171049 | 0.0129651036621392 | 0.989655633763749 | 0.998028377581853 | NM_177325 |
| Thumpd2 | 143.2234464 | -0.018081733 | 0.116691127 | -0.154953797 | 0.876857742 | 0.942231351 | | Vangl2 | 254.6860014 | -0.001123441269085 | 0.0882451699228802 | -0.0127309094557316 | 0.989842478282876 | 0.998028377581853 | NM_033509 |
| 1600014C10Rik | 1285.164816 | 0.010307636 | 0.066681858 | 0.154579324 | 0.877152971 | 0.942287189 | | Aldh9a1 | 1533.266815 | 0.000787937277086793 | 0.0719960450050652 | 0.0109441744616855 | 0.99126798647939 | 0.998279586921439 | NM_019993 |
| E130308A19Rik | 366.9387067 | 0.019118436 | 0.123641097 | 0.154628492 | 0.877114207 | 0.942287189 | | Gm16432 | 26.88371015 | -0.000955965891368745 | 0.0866691762495226 | -0.0110300562753302 | 0.991199466842107 | 0.998279586921439 | NM_001034899 |
| Sash3 | 90.12942851 | 0.023456062 | 0.15172531 | 0.154595579 | 0.877140156 | 0.942287189 | | Lig4 | 537.6282132 | -0.000903855238631598 | 0.0832601651193194 | -0.0108557944526808 | 0.991338499335107 | 0.998279586921439 | NM_176953 |
| Taf7 | 258.6372587 | -0.015061469 | 0.097316214 | -0.154768343 | 0.877003949 | 0.942287189 | | Prdm4 | 1595.927161 | 0.000580489228985798 | 0.0527688936588499 | 0.0110005950236261 | 0.991222972093907 | 0.998279586921439 | NM_181650 |
| Tbc1d9b | 11676.7819 | 0.011772268 | 0.07619679 | 0.154498221 | 0.877216914 | 0.942290548 | | Prex1 | 7686.81451 | 0.000768219168223128 | 0.0703503184399608 | 0.0109199103182276 | 0.991287345308019 | 0.998279586921439 | NM_177782 |
| Camsap2 | 7914.872972 | -0.013393235 | 0.086905114 | -0.154113316 | 0.877520389 | 0.942518136 | | Pwp2 | 1428.84286 | 0.000730462412099342 | 0.0649541579069945 | 0.0112458145196073 | 0.99102732734806 | 0.998279586921439 | NM_029546 |
| Gm16197 | 18.97442506 | 0.02583713 | 0.167859801 | 0.153920893 | 0.87767211 | 0.942518136 | | Adat1 | 307.1992107 | 0.000865875823856051 | 0.0832222459076214 | 0.0104043794350034 | 0.991698656055964 | 0.998293270403219 | NM_013925 |
| Osr1 | 73.55707676 | 0.023458631 | 0.152362272 | 0.153966141 | 0.877636433 | 0.942518136 | | Arhgef33 | 58.43782204 | -0.000948654120752249 | 0.0928579683755983 | -0.0102161843226536 | 0.991848806049477 | 0.998293270403219 | NM_001145452 |
| Tcf4 | 9297.539176 | 0.008420636 | 0.054702622 | 0.153934774 | 0.877661166 | 0.942518136 | | Atat1 | 2125.760381 | -0.000562476048119692 | 0.0549127913211491 | -0.0102430787906981 | 0.991827348491401 | 0.998293270403219 | NM_001142745,NM_001142744,NM_028476 |
| Slc26a5 | 10.66626192 | -0.024122216 | 0.157041592 | -0.153603998 | 0.877921984 | 0.942721133 | | Cacna1b | 4033.031622 | 0.000569997410425819 | 0.0616956837720342 | 0.00923885392910081 | 0.992628565957146 | 0.998293270403219 | NM_001042528,NM_007579 |
| Rnf185 | 1447.286638 | 0.011677565 | 0.076160135 | 0.153329101 | 0.878138752 | 0.942888554 | | Ccdc79 | 41.30852495 | 0.000923778945885732 | 0.0887338924811129 | 0.0104106663198885 | 0.991693640119241 | 0.998293270403219 | NM_180958 |
| Cers6 | 1315.767097 | -0.013630231 | 0.089339762 | -0.152566233 | 0.878740356 | 0.943469134 | | D7Ertd443e | 218.4410538 | -0.000913444613882423 | 0.0944849055841183 | -0.00966762477281834 | 0.992286471609197 | 0.998293270403219 | NM_001199941,NM_001081331 |
| Cyth4 | 487.6159804 | 0.01941896 | 0.127650407 | 0.152126109 | 0.879087472 | 0.943698723 | | Ei24 | 4491.384074 | 0.000608116175265164 | 0.0599772645099536 | 0.010139111548914 | 0.991910298040795 | 0.998293270403219 | NM_007915,NM_001199494 |
| Pde2a | 22668.44252 | -0.010128869 | 0.066609506 | -0.15206342 | 0.879136915 | 0.943698723 | | Fchsd2 | 1973.863046 | 0.000524471041214516 | 0.0567862317262647 | 0.00923588386253033 | 0.992630935626304 | 0.998293270403219 | NM_001146010,NM_199012 |
| Rpl14 | 6660.641903 | 0.010923351 | 0.07179587 | 0.152144564 | 0.879072917 | 0.943698723 | | Fyn | 4990.959839 | -0.000401076379141633 | 0.0419124068294899 | -0.00956939506655659 | 0.992364843949783 | 0.998293270403219 | NM_008054,NM_001122893,NM_001122892 |
| Gm10334 | 13.37430389 | -0.023400189 | 0.1542345 | -0.151718255 | 0.879409159 | 0.943925563 | | Gmds | 314.5227905 | 0.000800109234861509 | 0.0801859058555964 | 0.00997817791448779 | 0.992038698007196 | 0.998293270403219 | NM_146041 |
| Fzd6 | 395.1158759 | -0.017723871 | 0.117036379 | -0.151438987 | 0.879629437 | 0.943965819 | | Gtf2h4 | 545.0405969 | -0.000685658735672371 | 0.0764614388132667 | -0.00896737945707351 | 0.992845162271863 | 0.998293270403219 | NM_010364 |
| Hltf | 898.5618034 | -0.017884193 | 0.11806619 | -0.151475989 | 0.87960025 | 0.943965819 | | Ide | 2117.602129 | -0.000563464509962796 | 0.0568268980434287 | -0.00991545428948419 | 0.992088741743473 | 0.998293270403219 | NM_031156 |
| Tmem219 | 330.7484836 | -0.013630865 | 0.089947909 | -0.151541768 | 0.879548365 | 0.943965819 | | Lgr6 | 87.35356704 | 0.000876320910762289 | 0.0950801000805247 | 0.00921665953254277 | 0.992646273769556 | 0.998293270403219 | NM_001033409 |
| Gm15612 | 14.98544943 | -0.022533292 | 0.149110892 | -0.151117682 | 0.879882884 | 0.944172409 | | Med13l | 5432.756733 | 0.000685494137816467 | 0.0714965116640367 | 0.00958779836752897 | 0.992350160913666 | 0.998293270403219 | NM_172424 |
| Arap1 | 715.4586533 | 0.010510447 | 0.069615465 | 0.150978618 | 0.879992583 | 0.944181141 | | Minos1 | 2281.166637 | 0.000548472841152937 | 0.0553159121499627 | 0.00991528151368117 | 0.992088879591842 | 0.998293270403219 | NR_033745,NM_001163006 |
| Cnn3 | 2522.322828 | 0.010105897 | 0.066947367 | 0.150952863 | 0.880012899 | 0.944181141 | | Ppm1f | 4349.489281 | 0.000660905487538158 | 0.0627232296187207 | 0.0105368535956398 | 0.991592962762502 | 0.998293270403219 | NM_176833 |
| Mis12 | 984.6210667 | 0.012404985 | 0.082530873 | 0.150307209 | 0.880522246 | 0.944662213 | | Prkcdbp | 480.2667471 | -0.000727500732618219 | 0.079960880028017 | -0.00909820817834064 | 0.992740780313549 | 0.998293270403219 | NM_028444 |
| Cbln3 | 36.0143865 | 0.024832732 | 0.16553768 | 0.150012569 | 0.880754699 | 0.94471823 | | Rnf40 | 2864.912118 | 0.000421682368528538 | 0.0466067256660945 | 0.00904767203664135 | 0.992781100661185 | 0.998293270403219 | NM_172281 |
| Cytl1 | 25.07304251 | -0.025136332 | 0.167513636 | -0.150055441 | 0.880720875 | 0.94471823 | | Rps15 | 8610.970614 | 0.000508963201922546 | 0.0569072431271956 | 0.00894373323945676 | 0.992864028467252 | 0.998293270403219 | NM_009091 |
| Suox | 529.9425783 | 0.015083354 | 0.100549546 | 0.150009172 | 0.880757379 | 0.94471823 | | Slc24a4 | 1122.568292 | 0.000773367179320123 | 0.0765813157662569 | 0.0100986405310743 | 0.991942587587955 | 0.998293270403219 | NM_172152 |
| Gm20597 | 12.22818238 | 0.024518703 | 0.163584025 | 0.14988446 | 0.880855773 | 0.944758365 | | Vps4b | 1837.229103 | 0.000566594035802642 | 0.0623209789429634 | 0.00909154582313594 | 0.992746095884057 | 0.998293270403219 | NM_009190 |
| Prr22 | 34.40837143 | 0.025001901 | 0.167027748 | 0.14968711 | 0.881011479 | 0.94485996 | | Zbtbd6 | 27.58780106 | 0.000793402790495861 | 0.0838311940546358 | 0.0094642907027994 | 0.992448701301281 | 0.998293270403219 | NM_001034882 |
| Fam188a | 1158.275097 | 0.008587126 | 0.057468507 | 0.149423169 | 0.881219731 | 0.944968814 | | Ppp2r3a | 2003.602511 | 0.000650378562389081 | 0.0740677083828647 | 0.00878080651919835 | 0.99299397326458 | 0.998354821126889 | NM_001161362,NM_172144 |
| Olfm2 | 3697.667851 | -0.01704423 | 0.114140632 | -0.149326577 | 0.881295945 | 0.944968814 | | Slc15a4 | 886.3789451 | 0.000603503365072819 | 0.0720980209770906 | 0.00837059543235708 | 0.993321309132378 | 0.998545698024701 | NM_133895 |
| Rpl36a | 2433.186553 | 0.011617202 | 0.077772784 | 0.149373612 | 0.881258833 | 0.944968814 | | Tmem88b | 3043.769463 | -0.000734643125798951 | 0.0877189654407198 | -0.00837496340851649 | 0.993317824113797 | 0.998545698024701 | NM_001033394 |
| Mrm1 | 519.0977907 | -0.014576433 | 0.097724018 | -0.149159164 | 0.881428042 | 0.944987803 | | Lin7c | 2562.375208 | 0.000582382925717183 | 0.0708252541955371 | 0.00822281447955439 | 0.993439217214703 | 0.998595119317748 | NM_011699 |
| Nsun6 | 269.5786637 | 0.014842172 | 0.099563619 | 0.149072246 | 0.881496625 | 0.944987803 | | Ap5s1 | 600.7333367 | -0.000649907076882512 | 0.0825423923066716 | -0.00787361571091734 | 0.993717828495937 | 0.998598766982496 | NM_027129 |
| Vezf1 | 1580.460218 | -0.01176952 | 0.078915033 | -0.149141668 | 0.881441847 | 0.944987803 | | Bbs10 | 200.7019849 | 0.000726716961886902 | 0.0910310004251086 | 0.00798318109757318 | 0.993630410713132 | 0.998598766982496 | NM_027914 |
| 1700019L03Rik | 8.502020403 | 0.021972329 | 0.147567974 | 0.148896323 | 0.881635443 | 0.945021542 | | Cxxc5 | 3062.519806 | -0.00055135486873304 | 0.0699044167458372 | -0.0078872679924888 | 0.99370693588948 | 0.998598766982496 | NM_133687 |
| Myh1 | 24.65241061 | 0.025017723 | 0.168042031 | 0.148877769 | 0.881650084 | 0.945021542 | | Phyhipl | 7447.076843 | -0.000466921977024093 | 0.0584771772249818 | -0.00798468734610229 | 0.993629208938989 | 0.998598766982496 | NM_001162846,NM_178621 |
| Snd1 | 3064.39746 | -0.007636581 | 0.051324973 | -0.148788796 | 0.881720292 | 0.94503142 | | Pcdh8 | 1300.519623 | -0.000652918302740046 | 0.083855260676882 | -0.00778625333067563 | 0.993787531453602 | 0.998599728963408 | NM_021543,NM_001042726 |
| Trib1 | 859.966982 | -0.024983689 | 0.168014526 | -0.148699577 | 0.881790695 | 0.945041504 | | C5ar1 | 35.3715493 | -0.000690293094929621 | 0.0902503537577781 | -0.00764864697131593 | 0.993897322173554 | 0.998640969903371 | NM_001173550,NM_007577 |
| Lsm12 | 533.4127831 | -0.011556167 | 0.077882013 | -0.148380433 | 0.882042541 | 0.945246032 | | Batf3 | 37.74946985 | 0.000599020455029765 | 0.0849040607401738 | 0.00705526272604213 | 0.994370761499371 | 0.998727174362212 | NM_030060 |
| 4833422C13Rik | 133.6161986 | -0.018892619 | 0.127548957 | -0.148120533 | 0.882247645 | 0.945269692 | | Glp2r | 110.9693943 | -0.000621402117924617 | 0.0950611265370662 | -0.00653686886071479 | 0.994784370404613 | 0.998727174362212 | NM_175681 |
| Fez2 | 1431.770481 | -0.010151042 | 0.068491529 | -0.148208726 | 0.882178046 | 0.945269692 | | Gpr22 | 1945.48966 | 0.000624277530917521 | 0.0902181364029493 | 0.00691964560350986 | 0.994478965665997 | 0.998727174362212 | NM_175191 |
| Mrps15 | 821.8131202 | 0.013524767 | 0.091270395 | 0.148183508 | 0.882197947 | 0.945269692 | | Gpr30 | 141.0321922 | -0.000581497211022258 | 0.0893643329042092 | -0.0065070391298682 | 0.994808170580122 | 0.998727174362212 | NM_029771 |
| Asic2 | 2476.611449 | -0.009294476 | 0.062869443 | -0.147837733 | 0.88247083 | 0.945443441 | | Gse1 | 3019.740882 | -0.000577691827433339 | 0.0803158963108642 | -0.00719274581954949 | 0.994261068645347 | 0.998727174362212 | NM_198671,NM_001145896,NM_001145897 |
| Abcb9 | 2065.501332 | -0.014722898 | 0.100129344 | -0.147038795 | 0.883101399 | 0.945450326 | | Kit | 1895.80043 | -0.000365714779261351 | 0.053453129710878 | -0.00684178421805161 | 0.994541088592723 | 0.998727174362212 | NM_021099,NM_001122733 |
| B630005N14Rik | 710.6606425 | -0.014294388 | 0.097134512 | -0.147160749 | 0.883005141 | 0.945450326 | | Lrp10 | 1327.509155 | -0.000554396581140638 | 0.0817309317676857 | -0.0067831917384286 | 0.994587837542764 | 0.998727174362212 | NM_022993 |
| Bet1l | 955.4434053 | 0.012144761 | 0.08278807 | 0.146696992 | 0.883371193 | 0.945450326 | | Ptpn4 | 1688.941045 | 0.00053797117038933 | 0.08129597539351 | 0.00661743915101949 | 0.99472008600439 | 0.998727174362212 | NM_019933 |
| C2cd3 | 1778.371188 | -0.010931797 | 0.074472784 | -0.146789156 | 0.883298444 | 0.945450326 | | Purg | 490.1602947 | -0.000509172106891477 | 0.075071438889529 | -0.00678250096747884 | 0.994588388665562 | 0.998727174362212 | NM_152821,NM_001098233 |
| C2cd4c | 773.9773914 | -0.018946458 | 0.128810568 | -0.147087761 | 0.88306275 | 0.945450326 | | Setd5 | 5150.818357 | -0.000365140283920918 | 0.0545905105011639 | -0.00668871348827437 | 0.994663218569713 | 0.998727174362212 | NM_028385 |
| Chmp2a | 2398.925999 | 0.008172516 | 0.055610913 | 0.146958858 | 0.883164494 | 0.945450326 | | Snai2 | 24.08880296 | 0.000578675466293109 | 0.0824450962615219 | 0.00701891916600483 | 0.994399758746818 | 0.998727174362212 | NM_011415 |
| Cth | 118.8401007 | 0.019616487 | 0.13358341 | 0.14684823 | 0.883251815 | 0.945450326 | | Suv420h1 | 2258.553996 | -0.000336591317102737 | 0.0505154201204285 | -0.00666314001349104 | 0.994683622795715 | 0.998727174362212 | NM_001167887,NM_001167886,NM_001167885,NM_001167884,NM_001167889,NM_001167888,NM_144871 |
| Eya1 | 428.3649427 | 0.013572443 | 0.092537335 | 0.146669919 | 0.883392563 | 0.945450326 | | Rnh1 | 1420.347374 | 0.000449431215330027 | 0.0715601328416586 | 0.00628046927085065 | 0.994988943477144 | 0.998839626326814 | NM_145135,NM_001172100,NM_001172101 |
| Ifnar2 | 617.4991832 | -0.0128693 | 0.087360973 | -0.147311774 | 0.882885939 | 0.945450326 | | 9430038I01Rik | 46.22349184 | -0.000520972920183679 | 0.0941646545016543 | -0.00553257401028883 | 0.995585667135698 | 0.998915149237698 | NM_029886 |
| Rab5a | 3672.750555 | -0.009563287 | 0.065106855 | -0.146886022 | 0.883221985 | 0.945450326 | | Anapc10 | 308.6105478 | 0.000460099036325844 | 0.0871825234160453 | 0.00527742279413268 | 0.995789245377497 | 0.998915149237698 | NM_026904 |
| Rad21l | 17.42333812 | -0.024784342 | 0.167786821 | -0.14771328 | 0.882569051 | 0.945450326 | | Arid4a | 1714.846966 | -0.000400266827640428 | 0.0721515241299692 | -0.00554758658901526 | 0.995573689014735 | 0.998915149237698 | NM_001081195 |
| Rad51ap2 | 31.03090573 | 0.024612464 | 0.167346568 | 0.147074804 | 0.883072977 | 0.945450326 | | BC049352 | 34.0869798 | -0.000455903765305557 | 0.0899976843916202 | -0.00506572772830149 | 0.99595815134308 | 0.998915149237698 | NM_001198971 |
| Rhbdd2 | 3201.831224 | -0.008843938 | 0.060149514 | -0.147032583 | 0.883106302 | 0.945450326 | | Dbnl | 2681.366883 | -0.000345297283222312 | 0.0680812457522527 | -0.00507184143602258 | 0.995953273372745 | 0.998915149237698 | NM_013810,NM_001146309,NM_001146308 |
| Ubqln2 | 6647.014602 | 0.008731367 | 0.059348605 | 0.147120011 | 0.883037295 | 0.945450326 | | Ehhadh | 157.6520547 | 0.00049901350661022 | 0.0908910379649073 | 0.00549023887368335 | 0.995619445174554 | 0.998915149237698 | NM_023737 |
| Zfp612 | 2380.739916 | 0.010984234 | 0.074793392 | 0.146861026 | 0.883241715 | 0.945450326 | | Golt1a | 40.03716177 | -0.000523853133020574 | 0.0909977166930151 | -0.00575677228020804 | 0.995406785647759 | 0.998915149237698 | NM_026680 |
| Actr5 | 243.2720842 | -0.012804967 | 0.08759251 | -0.146187924 | 0.883773039 | 0.945726878 | | Iah1 | 330.2683267 | -0.000471838388137438 | 0.0792461221628638 | -0.00595408803938359 | 0.995249353149058 | 0.998915149237698 | NM_026347 |
| Psmd13 | 2530.165012 | -0.011518748 | 0.07876229 | -0.146246998 | 0.883726405 | 0.945726878 | | Mat2a | 8545.739586 | 0.000419903508110183 | 0.0697338541051811 | 0.00602151585479318 | 0.995195554500636 | 0.998915149237698 | NM_145569 |
| Cul9 | 3362.681816 | 0.009571489 | 0.06552039 | 0.146084127 | 0.883854977 | 0.945749241 | | Mrpl44 | 756.9796563 | -0.000350729539065924 | 0.0661509354790608 | -0.00530195886915224 | 0.995769668695944 | 0.998915149237698 | NM_001081210 |
| Pxdc1 | 140.863038 | -0.018766531 | 0.128782811 | -0.145722325 | 0.884140597 | 0.945858897 | | Olfr316 | 42.49757096 | 0.000496070990171054 | 0.0913006091252246 | 0.00543338094810146 | 0.995664810558869 | 0.998915149237698 | NM_001011818 |
| Rbms2 | 608.1929993 | -0.012202837 | 0.083673485 | -0.145838753 | 0.884048683 | 0.945858897 | | Sike1 | 1089.708568 | -0.000350014910255757 | 0.0678739409774265 | -0.00515683788528155 | 0.995885456905104 | 0.998915149237698 | NM_025679 |
| Slc52a2 | 493.9896092 | 0.012311651 | 0.084456003 | 0.145775914 | 0.88409829 | 0.945858897 | | Tmem144 | 454.2430676 | 0.000419811237313997 | 0.0759028005904374 | 0.00553090576432415 | 0.995586998183032 | 0.998915149237698 | NM_027495 |
| Lsm10 | 207.2721434 | 0.014712808 | 0.10115299 | 0.145451047 | 0.884354763 | 0.945957384 | | Bbs4 | 2679.620795 | -0.000258814106291697 | 0.0543787906583168 | -0.00475946785793615 | 0.996202508415655 | 0.998948537358147 | NM_175325 |
| Tpk1 | 85.35820468 | -0.018330993 | 0.12597976 | -0.145507443 | 0.884310239 | 0.945957384 | | Cacna1c | 2246.321271 | -0.000304330360353293 | 0.0662314075537359 | -0.00459495534813116 | 0.996333768971384 | 0.998948537358147 | NM_001255997,NM_001159533,NM_001255999,NM_001255998,NM_001159534,NM_001159535,NM_009781,NM_001256002,NM_001256000,NM_00125600 |
| Dqx1 | 29.89861834 | -0.023846921 | 0.165143384 | -0.14440131 | 0.885183582 | 0.946647876 | | Dusp28 | 686.1498932 | -0.000332126849442839 | 0.0736936372994728 | -0.00450685922982954 | 0.996404058776139 | 0.998948537358147 | NM_175118 |
| Gal3st4 | 161.6075657 | 0.021447541 | 0.148501714 | 0.144426217 | 0.885163915 | 0.946647876 | | Fau | 4848.646461 | -0.000235515046936034 | 0.0498840645572712 | -0.00472124813858427 | 0.996233002997019 | 0.998948537358147 | NM_007990,NM_001190436,NM_001160239 |
| Urm1 | 1298.505535 | -0.009146832 | 0.063287642 | -0.14452794 | 0.885083595 | 0.946647876 | | Sh3pxd2b | 1100.229319 | 0.000290424180630153 | 0.0644386252544277 | 0.00450698908431784 | 0.9964039551683 | 0.998948537358147 | NM_177364 |
| Ccl27a | 1563.130936 | 0.019039047 | 0.132012267 | 0.144221802 | 0.885325326 | 0.946712918 | | Six3os1 | 202.6254942 | 0.000445869913430518 | 0.0943055465536802 | 0.00472792884113897 | 0.996227672627087 | 0.998948537358147 | NR_038085,NR_038084,NR_038086,NR_038083,NR_038082,NR_015387,NR_015386,NR_015385 |
| F12 | 23.74888089 | 0.023963177 | 0.16623149 | 0.14415546 | 0.885377711 | 0.946712918 | | AA415398 | 255.6004113 | -0.000352940124264202 | 0.0814762617066987 | -0.00433181538856961 | 0.996543722190508 | 0.999019607214584 | NM_001004178 |
| Mafg | 973.5102133 | 0.01337885 | 0.092849266 | 0.144092146 | 0.885427707 | 0.946712918 | | Cntn3 | 1226.946852 | 0.000281335352191416 | 0.0726253636577091 | 0.00387403942292405 | 0.996908971488295 | 0.999132450481302 | NM_008779 |
| Anxa8 | 17.78290713 | 0.024111675 | 0.168038983 | 0.143488583 | 0.885904328 | 0.947157166 | | Mapk8ip1 | 16644.26024 | -0.000209300660717834 | 0.0544294246160404 | -0.00384535868593682 | 0.996931855235076 | 0.999132450481302 | NM_011162,NM_001202446,NM_001202445 |
| Cryzl1 | 1627.212727 | -0.009463941 | 0.066029748 | -0.143328444 | 0.886030792 | 0.947171527 | | Sesn2 | 488.8658519 | 0.000287630797056157 | 0.0748383531052299 | 0.00384336085872601 | 0.996933449258783 | 0.999132450481302 | NM_144907 |
| Npepl1 | 452.0506384 | -0.017050521 | 0.118970881 | -0.143316756 | 0.886040023 | 0.947171527 | | Spsb3 | 2099.193824 | 0.000234595284572947 | 0.0638615665828708 | 0.00367349717721625 | 0.997068979910296 | 0.999132450481302 | NM_027141,NM_001163751,NM_001163750 |
| Acot13 | 1414.828958 | -0.012606161 | 0.088094181 | -0.143098683 | 0.886212245 | 0.947254819 | | Stx6 | 3433.393598 | 0.000157980461283473 | 0.0427564536485895 | 0.00369489159652708 | 0.99705190974929 | 0.999132450481302 | NM_021433 |
| Tex264 | 2099.229709 | 0.009037454 | 0.063171032 | 0.14306327 | 0.886240213 | 0.947254819 | | Xrcc5 | 931.1260125 | 0.000227722422006989 | 0.0607552751428597 | 0.00374819176559604 | 0.997009382661805 | 0.999132450481302 | NM_009533 |
| Vezt | 1829.219662 | -0.007576155 | 0.053084241 | -0.142719478 | 0.886511733 | 0.947479671 | | Cyp2a5 | 32.93540652 | 0.000243204511649902 | 0.0872639069935097 | 0.00278700003276257 | 0.99777629858162 | 0.99928974856568 | NM_007812 |
| Nell1 | 1934.637275 | 0.010922752 | 0.076578643 | 0.14263444 | 0.886578896 | 0.947486096 | | Grip1 | 630.1448681 | 0.000211360002092078 | 0.0734026583542635 | 0.00287945977476715 | 0.997702526677097 | 0.99928974856568 | NM_133442,NM_130891,NM_028736 |
| Flywch2 | 228.3754321 | 0.014599729 | 0.10241519 | 0.142554328 | 0.88664217 | 0.947488363 | | Lyrm2 | 694.540003 | -0.000252561409612883 | 0.0779749076388834 | -0.00323900876910997 | 0.997415649429636 | 0.99928974856568 | NM_175364 |
| Fzd8 | 118.166857 | -0.019708393 | 0.138558891 | -0.142238385 | 0.886891713 | 0.947689669 | | Mettl10 | 495.2643526 | -0.000225671989408865 | 0.0784397250312367 | -0.0028770114800759 | 0.99770448012554 | 0.99928974856568 | NM_028095 |
| Axl | 2324.856205 | -0.008711061 | 0.061300293 | -0.142104722 | 0.886997289 | 0.947715232 | | Nek4 | 755.7873638 | 0.00022547331598731 | 0.069988864932354 | 0.00322155983248528 | 0.997429571594134 | 0.99928974856568 | NM_011849 |
| Slc13a4 | 581.6961566 | 0.019147704 | 0.13486599 | 0.141975782 | 0.887099135 | 0.947715232 | | Nle1 | 359.2750216 | 0.000247530687070899 | 0.086198646697953 | 0.00287163078021708 | 0.997708773285149 | 0.99928974856568 | NM_145431 |
| Zfp960 | 42.15163643 | -0.022600511 | 0.159156158 | -0.142002114 | 0.887078336 | 0.947715232 | | Phlpp2 | 2996.945819 | -0.000163590971498202 | 0.0557095737892086 | -0.00293649655474282 | 0.997657018103381 | 0.99928974856568 | NM_001122594 |
| Camk2a | 133864.2327 | 0.012878527 | 0.090792969 | 0.141844982 | 0.887202453 | 0.947760259 | | Stxbp5 | 5162.033456 | 0.000191039700148713 | 0.0596974106569722 | 0.00320013377542366 | 0.997446667026139 | 0.99928974856568 | NM_001081344 |
| Mplkip | 213.8935559 | 0.015674491 | 0.110569771 | 0.141761088 | 0.887268721 | 0.947765705 | | Pabpc4 | 1286.28693 | -0.000170983984824492 | 0.0641540960573902 | -0.00266520760687728 | 0.997873474516708 | 0.999318172411561 | NM_148917,NM_130881 |
| Tomm20 | 15656.25893 | -0.011693676 | 0.082545959 | -0.141662616 | 0.887346505 | 0.947783452 | | Ambra1 | 3647.260573 | -0.000107225462116868 | 0.0426023040686295 | -0.00251689349815763 | 0.997991811656865 | 0.9993677827194 | NM_172669,NM_001080754 |
| Slc25a25 | 2660.084245 | -0.013145657 | 0.092941241 | -0.141440518 | 0.887521947 | 0.947905498 | | Gm13889 | 524.4390432 | -0.000188978735564862 | 0.0843260265137073 | -0.00224104874115162 | 0.998211903306155 | 0.999381494756717 | NM_001145034 |
| Dcaf10 | 946.929872 | 0.012733841 | 0.090127444 | 0.14128705 | 0.88764318 | 0.947969633 | | Gypc | 77.59445086 | -0.000223523417456331 | 0.0932726850235715 | -0.00239645098026119 | 0.99808791059231 | 0.999381494756717 | NM_001048207 |
| Cmip | 13605.92313 | 0.007802127 | 0.05534535 | 0.140971672 | 0.887892322 | 0.948067112 | | Mkks | 1322.156651 | -0.000142072347974433 | 0.0633809406921608 | -0.00224156262786433 | 0.998211493284911 | 0.999381494756717 | NM_001141946,NM_021527 |
| Spata17 | 14.13120632 | -0.023027916 | 0.163389077 | -0.140939137 | 0.887918024 | 0.948067112 | | Syt7 | 15018.64047 | -0.000122316840124925 | 0.0594944725064757 | -0.00205593620670578 | 0.9983596013983 | 0.999460480352289 | NM_173068,NM_018801,NM_173067 |
| Syncrip | 2513.80012 | -0.013573633 | 0.096267255 | -0.140999483 | 0.887870351 | 0.948067112 | | Stard3 | 1373.756455 | -0.000112603586419158 | 0.0611133581149245 | -0.00184253639290129 | 0.998529869491221 | 0.999562048392907 | NM_021547 |
| Rmnd5a | 6258.636161 | -0.010292076 | 0.073143968 | -0.140709836 | 0.888099175 | 0.948129857 | | Fem1c | 842.9121598 | -0.000125697180339345 | 0.0744799947991497 | -0.00168766365623834 | 0.998653439864073 | 0.999616859665486 | NM_173423 |
| Zfp157 | 963.8884807 | 0.01145286 | 0.081391872 | 0.140712577 | 0.888097009 | 0.948129857 | | Ubap2l | 5705.227779 | 7.03492693203112e-05 | 0.0446571835907413 | 0.00157531809361342 | 0.998743078534621 | 0.99963770128808 | NM_001165984,NM_001165985,NM_001165986,NM_001165987,NM_001165983,NM_001165988,NM_028475,NM_153489 |
| Plaa | 1717.741828 | 0.013763256 | 0.097941519 | 0.140525249 | 0.888245005 | 0.948220209 | | 1110038F14Rik | 505.4804255 | 0.000102869377941928 | 0.0749900727937992 | 0.00137177327757487 | 0.998905483624171 | 0.999662490880104 | NM_054099 |
| Armcx4 | 3263.543913 | -0.012037519 | 0.085748317 | -0.140381984 | 0.888358192 | 0.948275702 | | Atic | 1283.481602 | 8.41740376719737e-05 | 0.0611064166250207 | 0.001377499161643 | 0.998900915033992 | 0.999662490880104 | NM_026195 |
| Phospho2 | 904.4462402 | 0.015476922 | 0.110323327 | 0.140286937 | 0.888433285 | 0.948290529 | | Tfcp2l1 | 403.8052919 | 9.08263416294951e-05 | 0.078739633820405 | 0.00115350221003895 | 0.999079638599858 | 0.99976789957988 | NM_023755 |
| Ddb1 | 10855.09087 | -0.008820483 | 0.062921797 | -0.140181678 | 0.888516448 | 0.948313966 | | Foxo6 | 358.5640331 | -9.38502255209411e-05 | 0.0899387735209896 | -0.00104349016388397 | 0.999167415459984 | 0.999786862090771 | NM_194060 |
| Cd53 | 252.0220639 | 0.017764764 | 0.126978142 | 0.139904107 | 0.888735756 | 0.948414813 | | Aatk | 13313.88148 | -4.29351687051473e-05 | 0.0554995197043343 | -0.000773613338167216 | 0.999382745923014 | 0.999917241493403 | NM_001198785,NM_007377,NM_001198787 |
| Tmem82 | 23.57419305 | 0.023456886 | 0.167753286 | 0.139829669 | 0.888794571 | 0.948414813 | | Ank3 | 10207.45761 | 4.81528937309087e-05 | 0.0719121546511126 | 0.0006696071611889 | 0.99946573082421 | 0.999917241493403 | NM_170689,NM_170730,NM_170687,NM_170690,NM_170729,NM_146005,NM_170728,NM_009670,NM_170688 |
| Trim36 | 268.6231078 | 0.019991874 | 0.142864123 | 0.139936282 | 0.888710334 | 0.948414813 | | Fastkd1 | 518.4055211 | -2.70310506107756e-05 | 0.0718062228324184 | -0.000376444401954698 | 0.999699640830774 | 0.999917241493403 | NM_177244 |
| N4bp2 | 260.751339 | 0.021026107 | 0.150590677 | 0.139624229 | 0.888956896 | 0.948457385 | | Gfra1 | 1810.269635 | 1.49375690395461e-05 | 0.0540688072617111 | 0.000276269623763721 | 0.999779568735384 | 0.999917241493403 | NM_010279 |
| Tead1 | 808.9096693 | -0.018800591 | 0.134618837 | -0.139657951 | 0.888930251 | 0.948457385 | | Il11ra1 | 646.1591078 | 3.3568816942196e-05 | 0.0826556162327169 | 0.000406128687585886 | 0.999675956199384 | 0.999917241493403 | NM_001172054,NM_010549,NM_001163401 |
| Amz1 | 624.5352566 | -0.015178556 | 0.108921084 | -0.139353697 | 0.88917066 | 0.948501164 | | Ints4 | 1463.528862 | -1.71324650154268e-05 | 0.0603684436988055 | -0.000283798354996615 | 0.999773561677207 | 0.999917241493403 | NM_027256 |
| Pigm | 545.2732085 | -0.012875428 | 0.092403041 | -0.139339877 | 0.88918158 | 0.948501164 | | Parp11 | 852.6964715 | -2.49157443259773e-05 | 0.0622600115979176 | -0.000400188559020613 | 0.99968069573587 | 0.999917241493403 | NM_181402 |
| Usp47 | 4655.399538 | -0.009196692 | 0.065945957 | -0.139458009 | 0.889088236 | 0.948501164 | | Sept1 | 28.48284513 | 6.74457254559182e-06 | 0.0856294075105761 | 7.87646760811558e-05 | 0.999937154881083 | 0.999947931801811 | NM_017461 |
| Prdm4 | 1471.746885 | 0.006626593 | 0.047662126 | 0.139032671 | 0.889424332 | 0.948694796 | | Snx7 | 548.7133995 | -4.71894747321351e-06 | 0.072312379925275 | -6.5257808940736e-05 | 0.999947931801811 | 0.999947931801811 | NM_001190156,NM_029655 |
| Gtf2h2 | 574.5250723 | 0.012275773 | 0.088458488 | 0.138774393 | 0.889628429 | 0.948727446 | | 0610005C13Rik | 0.546936012 | 0.0022560506561488 | 0.0178643354055514 | 0.126287970133371 | 0.899503978460615 | NA | NR_038166,NR_038165 |
| Sdhaf1 | 373.4587599 | -0.011329334 | 0.081646103 | -0.138761475 | 0.889638638 | 0.948727446 | | 0610008F07Rik | 0 | NA | NA | NA | NA | NA | NR_027970 |
| Tpx2 | 138.4942745 | -0.016017427 | 0.115400112 | -0.138799058 | 0.889608938 | 0.948727446 | | 0610009B14Rik | 0.725539079 | 0.00747045825642903 | 0.0180749991191235 | 0.41330338149368 | 0.679384352696264 | NA | NR_037995 |
| Hydin | 79.45290013 | -0.022211309 | 0.160529204 | -0.138363042 | 0.889953504 | 0.948997908 | | 0610010B08Rik | 9.730001072 | 0.0259912915669745 | 0.0648797610191647 | 0.400607079290829 | 0.688709432740887 | NA | NM_001177543 |
| Oxgr1 | 13.71520652 | -0.022313043 | 0.161497284 | -0.138163581 | 0.890111137 | 0.949100678 | | 0610012H03Rik | 0.406364829 | 0.0153428535423256 | 0.0145741015204257 | 1.05274781576226 | 0.292456587556691 | NA | NM_001159638,NM_028747 |
| Stk25 | 8821.653956 | -0.008145688 | 0.059005848 | -0.138048832 | 0.890201825 | 0.949132058 | | 0610031O16Rik | 0 | NA | NA | NA | NA | NA | NR_045760 |
| 4930513N10Rik | 26.34595695 | 0.022874549 | 0.165944722 | 0.137844389 | 0.890363401 | 0.949197578 | | 0610038B21Rik | 11.0614681 | -0.0845604013439811 | 0.0611449712935778 | -1.38294939968126 | 0.166680381663875 | NA | NR_028125 |
| Wwp2 | 1724.72234 | 0.010201599 | 0.0740233 | 0.137816049 | 0.8903858 | 0.949197578 | | 0610038L08Rik | 0.194331409 | -0.00840968081320881 | 0.00964081822647896 | -0.872299489073576 | 0.383045016303155 | NA | NR_045447 |
| S100a11 | 152.3316644 | -0.015323827 | 0.111355116 | -0.137612242 | 0.89054688 | 0.949238674 | | 0610039K10Rik | 1.256297036 | -0.0258215064455508 | 0.0227204453977424 | -1.13648768734598 | 0.255752515717239 | NA | NR_028113 |
| Vwc2 | 211.4320399 | -0.014005566 | 0.101764941 | -0.137626631 | 0.890535507 | 0.949238674 | | 0610043K17Rik | 15.21360369 | -0.00417388150002846 | 0.0679386955505608 | -0.061435997058881 | 0.951011984984013 | NA | NR_040640 |
| Fam168a | 14558.70205 | 0.006752528 | 0.049134441 | 0.137429639 | 0.890691205 | 0.949261902 | | 1100001G20Rik | 0 | NA | NA | NA | NA | NA | NM_183249 |
| Rhof | 1704.272908 | 0.01099219 | 0.079964174 | 0.13746394 | 0.890664094 | 0.949261902 | | 1110006O24Rik | 17.05743458 | 0.00125806440619661 | 0.0731444530043015 | 0.0171997240326977 | 0.986277282347329 | NA | NR_027810 |
| Fancd2 | 192.1573724 | -0.015553346 | 0.11344008 | -0.137106272 | 0.890946795 | 0.949468989 | | 1110015O18Rik | 11.89367556 | 0.0717339233349421 | 0.0572308063557063 | 1.25341451401357 | 0.210054892104038 | NA | NR_045272 |
| Cmtm5 | 1197.720742 | -0.022679606 | 0.166193847 | -0.136464778 | 0.891453867 | 0.949944032 | | 1110025L11Rik | 0 | NA | NA | NA | NA | NA | NM_001276278 |
| Crtc3 | 736.0602214 | -0.010951644 | 0.080322746 | -0.136345495 | 0.89154816 | 0.949979176 | | 1110028F11Rik | 0.342801831 | 0.00329783501100829 | 0.0128022060068578 | 0.257598964525467 | 0.796716428937095 | NA | NR_045139 |
| Slc35b1 | 1413.961795 | 0.007856621 | 0.05766349 | 0.136249495 | 0.891624049 | 0.949994706 | | 1110028F18Rik | 0 | NA | NA | NA | NA | NA | NR_045470 |
| Nhlh1 | 19.30096348 | -0.022767078 | 0.167314743 | -0.136073355 | 0.891763291 | 0.950074875 | | 1110036E04Rik | 0.120209049 | -0.00674972139751468 | 0.00863521114606269 | -0.781650996523957 | 0.434419707908266 | NA | NR_040713 |
| Zfp319 | 906.8293593 | -0.008973745 | 0.065983816 | -0.135999179 | 0.89182193 | 0.950074875 | | 1190002F15Rik | 1.531935788 | 0.0330025220492728 | 0.0261788813289373 | 1.26065440438789 | 0.20743338819763 | NA | NR_037956,NR_037955 |
| Cops2 | 2772.235842 | 0.01093053 | 0.080464827 | 0.135842334 | 0.891945923 | 0.950141639 | | 1190003K10Rik | 0.314698307 | 0.00249824212998287 | 0.0109738852417382 | 0.227653385738081 | 0.819915713647591 | NA | NM_001195435 |
| Akr1c13 | 9.949949216 | 0.020615761 | 0.152154401 | 0.135492374 | 0.892222593 | 0.950199203 | | 1300002K09Rik | 5.912161705 | 0.00231348929209209 | 0.0407088657637033 | 0.0568301093309957 | 0.954680528911897 | NA | NM_028788 |
| BC033916 | 15.55111826 | 0.021943659 | 0.162932534 | 0.134679421 | 0.892865344 | 0.950199203 | | 1300015D01Rik | 0.09572496 | 0.00498255289994511 | 0.00847999972823692 | 0.587565219295241 | 0.556824162160132 | NA | NR_040321 |
| Cd80 | 16.39149383 | 0.022685861 | 0.168116002 | 0.13494171 | 0.892657961 | 0.950199203 | | 1300017J02Rik | 2.471300781 | 0.0403932163374658 | 0.036901097276181 | 1.0946345588357 | 0.273676768966791 | NA | NM_027918 |
| Cstf3 | 924.6161184 | -0.00973183 | 0.072041229 | -0.135086947 | 0.89254313 | 0.950199203 | | 1500002O10Rik | 4.6156299 | -0.0800440080917262 | 0.047158702190621 | -1.69733271641317 | 0.0896337752372145 | NA | NR_040496 |
| Ddx56 | 1432.753155 | 0.007971567 | 0.059456586 | 0.134073749 | 0.893344257 | 0.950199203 | | 1500012K07Rik | 9.296915535 | 0.000743661313476433 | 0.0636143212886025 | 0.0116901555878058 | 0.990672817785303 | NA | NR_045814,NR_045812 |
| Ebf3 | 41.58639907 | 0.022091859 | 0.164392424 | 0.134384899 | 0.893098222 | 0.950199203 | | 1500015L24Rik | 21.48047971 | -0.0364471777400956 | 0.0761877836304338 | -0.47838611393253 | 0.632375415131794 | NA | NR_045817 |
| Eif4e2 | 2134.786825 | -0.007124623 | 0.052601597 | -0.135444987 | 0.892260057 | 0.950199203 | | 1500015O10Rik | 108.9475325 | 0.0109256272335657 | 0.0329894505523258 | 0.331185486591725 | NA | NA | NM_024283 |
| Fgd4 | 545.1506482 | -0.014466169 | 0.107424257 | -0.134663893 | 0.892877622 | 0.950199203 | | 1500016L03Rik | 3.555802094 | 0.0538734725869158 | 0.0372287622332001 | 1.44709276793715 | 0.147870943280932 | NA | NR_038057 |
| Galnt14 | 799.7241445 | 0.012044727 | 0.089740688 | 0.134217013 | 0.893230973 | 0.950199203 | | 1500017E21Rik | 12.72435476 | 0.0767130554866152 | 0.0675329140904612 | 1.13593581026071 | 0.255983429736871 | NA | NR_033510 |
| Gnai2 | 6623.334358 | -0.006613817 | 0.049255298 | -0.134276248 | 0.893184135 | 0.950199203 | | 1600002D24Rik | 0.309462921 | 0.00239536941478217 | 0.0109127344998192 | 0.219502216866162 | 0.826258853162846 | NA | NR_040484,NR_040485 |
| Gp5 | 26.03207002 | 0.02215906 | 0.165056146 | 0.13425165 | 0.893203585 | 0.950199203 | | 1600010M07Rik | 2.486319034 | -0.0224074279013278 | 0.0336747513754834 | -0.665407374548318 | 0.505789969410723 | NA | NR_037959 |
| Grik5 | 8809.341348 | 0.009723477 | 0.071796825 | 0.135430456 | 0.892271546 | 0.950199203 | | 1600014C23Rik | 2.553059075 | -0.0373114422648886 | 0.0348464679046625 | -1.07073814100672 | 0.284287187981184 | NA | NM_028164 |
| Lhx6 | 1025.496852 | -0.01556513 | 0.115676482 | -0.134557431 | 0.8929618 | 0.950199203 | | 1600014K23Rik | 0.291497113 | -0.00941849595073383 | 0.0102038556585987 | -0.923033044160808 | 0.355989985190414 | NA | NM_028046 |
| Mnd1 | 12.08354297 | -0.022266144 | 0.165249734 | -0.134742387 | 0.892815559 | 0.950199203 | | 1600015I10Rik | 0 | NA | NA | NA | NA | NA | NM_001081273 |
| Mrps2 | 1162.576365 | -0.00810478 | 0.059975345 | -0.135135187 | 0.89250499 | 0.950199203 | | 1600016N20Rik | 6.993231167 | -0.0504358582311813 | 0.0512102200710951 | -0.984878763675713 | 0.324683625804973 | NA | NM_028050 |
| Nnat | 4762.766555 | -0.0168729 | 0.12507643 | -0.13490072 | 0.892690371 | 0.950199203 | | 1600019K03Rik | 0 | NA | NA | NA | NA | NA | NR_040481 |
| Oasl1 | 10.02168297 | 0.021017553 | 0.155478588 | 0.135179725 | 0.892469777 | 0.950199203 | | 1600023N17Rik | 14.69097073 | 0.0391188776467615 | 0.0742497385606504 | 0.526855426094295 | 0.598293989566887 | NA | NR_073433 |
| Plce1 | 683.2283187 | 0.022339489 | 0.165414542 | 0.135051545 | 0.892571121 | 0.950199203 | | 1600025M17Rik | 0 | NA | NA | NA | NA | NA | NR_038168 |
| Ptpre | 1341.6084 | 0.006575796 | 0.048942296 | 0.134358149 | 0.893119374 | 0.950199203 | | 1600029D21Rik | 7.669103177 | -0.0371499150702197 | 0.0546001312098103 | -0.680399739836976 | 0.496251385716647 | NA | NM_029639 |
| R3hdm2 | 11071.28847 | 0.008816192 | 0.065296758 | 0.135017304 | 0.892598193 | 0.950199203 | | 1600029I14Rik | 3.540352257 | 0.00324340229653438 | 0.0414573691467817 | 0.0782346386971872 | 0.937641408751233 | NA | NR_028123 |
| Shank1 | 15649.75751 | -0.013244302 | 0.098822735 | -0.1340208 | 0.893386127 | 0.950199203 | | 1700001C02Rik | 12.45136468 | -0.0351976353520997 | 0.068428136439472 | -0.514373723785884 | 0.606990726621755 | NA | NM_029285 |
| Srsf9 | 1112.756146 | -0.011188332 | 0.083290734 | -0.134328645 | 0.893142703 | 0.950199203 | | 1700001C19Rik | 3.612210519 | 0.0295707775243651 | 0.0405243203539091 | 0.729704465518882 | 0.465570850963038 | NA | NM_029296,NM_001172091 |

| | | | | | | | | | | | | | | | |
|---|---|---|---|---|---|---|---|---|---|---|---|---|---|---|---|
| Stx12 | 4461.650201 | 0.007150024 | 0.053362367 | 0.133990002 | 0.893410481 | 0.950199203 | | 1700001D01Rik | 0 | NA | NA | NA | NA | NA | NR_045475 |
| Zfp738 | 559.3449046 | 0.014268906 | 0.106591608 | 0.133865193 | 0.893509175 | 0.950238942 | | 1700001F09Rik | 0 | NA | NA | NA | NA | NA | NM_027940 |
| Casp12 | 38.08740145 | 0.02235555 | 0.167277637 | 0.133643388 | 0.893684573 | 0.950305245 | | 1700001G11Rik | 7.729889812 | -0.0188206194022827 | 0.059925567096509 | -0.314066604859532 | 0.753470452620033 | NA | NR_038077 |
| Fam26e | 221.5847582 | -0.014632546 | 0.109563035 | -0.133553673 | 0.89375552 | 0.950305245 | | 1700001G17Rik | 8.904351366 | 0.0296355530785435 | 0.0610408775507951 | 0.485503391622806 | 0.627319307421784 | NA | NR_033199 |
| Gpt2 | 2410.923909 | 0.009898594 | 0.074112196 | 0.133562275 | 0.893748717 | 0.950305245 | | 1700001J03Rik | 3.185234464 | 0.0200856366124947 | 0.042613474182236 | 0.471344733043796 | 0.637394576000082 | NA | NM_001008547 |
| Ppp5c | 4285.859346 | -0.007256878 | 0.054396744 | -0.133406477 | 0.893871923 | 0.950328085 | | 1700001J11Rik | 2.449557285 | -0.00432860813858558 | 0.0344698511956898 | -0.125576641280274 | 0.90006705411627 | NA | NR_033613 |
| Slc25a40 | 348.0477285 | -0.017495065 | 0.131328292 | -0.133216272 | 0.894022342 | 0.950328085 | | 1700001K19Rik | 0.30309771 | -0.00396074557973908 | 0.0109135518926259 | -0.362919938321391 | 0.716664692123243 | NA | NM_025488 |
| Usp51 | 49.87343534 | -0.022266212 | 0.167125046 | -0.13323085 | 0.894010813 | 0.950328085 | | 1700001K23Rik | 0.209568245 | 0.00686197416234131 | 0.00977127153168884 | 0.702260104029194 | 0.482516968822431 | NA | NR_036590 |
| Zfp513 | 828.8889855 | 0.010045251 | 0.075362625 | 0.13329221 | 0.893962288 | 0.950328085 | | 1700001P01Rik | 6.991761568 | -0.0760869286077592 | 0.0549657282357322 | -1.38426126697429 | 0.166278469458771 | NA | NM_028156 |
| Car15 | 186.9197194 | -0.01556325 | 0.117063314 | -0.132947287 | 0.894235069 | 0.950489 | | 1700003C15Rik | 0 | NA | NA | NA | NA | NA | NR_045478 |
| Mtf1 | 1388.641237 | 0.011145645 | 0.083939875 | 0.132781287 | 0.894366354 | 0.950563334 | | 1700003F12Rik | 3.910003615 | -0.115568535510907 | 0.0426389440114871 | -2.71039863181819 | 0.00672023867282199 | NA | NM_029305 |
| Il17rb | 53.02902458 | -0.019307071 | 0.145631729 | -0.132574621 | 0.894529805 | 0.950606639 | | 1700003F17Rik | 0 | NA | NA | NA | NA | NA | NR_045932 |
| Tmem41b | 1435.130404 | -0.008409024 | 0.063423796 | -0.132584685 | 0.894521845 | 0.950606639 | | 1700003G13Rik | 0 | NA | NA | NA | NA | NA | NR_040720 |
| A330041J22Rik | 39.1580887 | -0.021289209 | 0.160684069 | -0.132491099 | 0.894595863 | 0.950611638 | | 1700003G18Rik | 0.106148608 | 0.00515554063321497 | 0.00854200771766808 | 0.603551390213729 | 0.546141939141592 | NA | NR_029433 |
| Rb1cc1 | 3929.369159 | 0.008998143 | 0.067978003 | 0.132368448 | 0.89469287 | 0.950638867 | | 1700003H04Rik | 0 | NA | NA | NA | NA | NA | NR_015460 |
| Thsd7b | 263.5779133 | -0.01626495 | 0.122936609 | -0.132303553 | 0.894744197 | 0.950638867 | | 1700003L19Rik | 1.16541587 | 0.0151444903870475 | 0.0226113304501365 | 0.669774404493569 | 0.503001612390762 | NA | NR_040507 |
| Iws1 | 1395.55833 | 0.010003335 | 0.075756688 | 0.132045569 | 0.894948249 | 0.950790466 | | 1700003P14Rik | 0 | NA | NA | NA | NA | NA | NR_045982 |
| Babam1 | 2320.239589 | 0.007418353 | 0.056253759 | 0.131873011 | 0.895084736 | 0.950796033 | | 1700006A11Rik | 0 | NA | NA | NA | NA | NA | NM_027939 |
| Hspa1b | 200.294678 | -0.019842128 | 0.150702695 | -0.131664053 | 0.89525002 | 0.950796033 | | 1700006E09Rik | 0 | NA | NA | NA | NA | NA | NM_029287 |
| Lox | 24.04041747 | -0.02182607 | 0.166016766 | -0.131469069 | 0.895404254 | 0.950796033 | | 1700006F04Rik | 0.345062444 | -0.0054408567472439 | 0.0129188235112097 | -0.421157293659353 | 0.673640226968649 | NA | NR_045621 |
| Malat1 | 9205.227744 | -0.018504491 | 0.140409697 | -0.131789269 | 0.895150975 | 0.950796033 | | 1700006H21Rik | 0 | NA | NA | NA | NA | NA | NR_045900 |
| Nupl1 | 1339.042615 | -0.011075942 | 0.083939253 | -0.131951873 | 0.895022358 | 0.950796033 | | 1700007B14Rik | 5.201223647 | -0.0023708711296342 | 0.045709053317712 | -0.0518687427883242 | 0.95863328037803 | NA | NM_001164235,NM_027944 |
| Pkib | 295.324103 | 0.013351484 | 0.101595316 | 0.131418301 | 0.895444412 | 0.950796033 | | 1700007F19Rik | 9.122245547 | 0.0500924968271985 | 0.0594182418676673 | 0.843049125195617 | 0.399200969920877 | NA | NR_040538 |
| Wfs1 | 6854.469288 | 0.017498895 | 0.133090224 | 0.131481446 | 0.895394463 | 0.950796033 | | 1700007G11Rik | 21.93590021 | 0.0475809698989979 | 0.0740976289235391 | 0.642138899587414 | 0.520783000685426 | NA | NM_001024614 |
| Zbtb21 | 749.7603584 | -0.009431491 | 0.071725267 | -0.131494676 | 0.895383998 | 0.950796033 | | 1700007J10Rik | 5.936779414 | 0.00210424845715142 | 0.0512201518702914 | 0.0410824329939544 | 0.967230179219619 | NA | NR_045476 |
| Whsc1l1 | 3505.082883 | 0.007970818 | 0.060736476 | 0.131236094 | 0.895588544 | 0.95088391 | | 1700007K09Rik | 4.725129896 | -0.0163551257250255 | 0.0435748536812231 | -0.375334036567818 | 0.707412055819365 | NA | NM_027037 |
| Efna4 | 8.988549763 | -0.020414968 | 0.155885272 | -0.130961491 | 0.895805771 | 0.950984215 | | 1700007L15Rik | 13.46687688 | -0.0434986804201415 | 0.0740897293846037 | -0.587108102316551 | 0.557131106305494 | NA | NR_045709 |
| Rpp38 | 178.2689245 | -0.015094259 | 0.115251105 | -0.130968451 | 0.895800265 | 0.950984215 | | 1700007P06Rik | 19.30882388 | 0.0830590533534366 | 0.0797639947727176 | 1.0413100997525 | 0.297731650804626 | NA | NR_040554 |
| Efcab11 | 9.084649627 | 0.020969714 | 0.160543606 | 0.130616935 | 0.896078346 | 0.951208405 | | 1700008A04Rik | 0 | NA | NA | NA | NA | NA | NM_027050 |
| Peo1 | 621.5958862 | -0.009333946 | 0.071635705 | -0.130297392 | 0.896331143 | 0.951411573 | | 1700008F21Rik | 0.108766301 | 0.00519171381361946 | 0.00856003371502575 | 0.606506234257728 | 0.544178643801544 | NA | NM_001168369,NM_029292 |
| Inip | 745.0784196 | 0.0089172 | 0.068534638 | 0.130112305 | 0.896477575 | 0.951501818 | | 1700008I05Rik | 0.235049888 | -0.00142616332698746 | 0.00985869345204131 | -0.144660480004291 | 0.884978943250011 | NA | NM_027952 |
| E130215H24Rik | 14.35097443 | -0.021325141 | 0.164313867 | -0.129782965 | 0.896738141 | 0.95158282 | | 1700008K24Rik | 0 | NA | NA | NA | NA | NA | NR_038141,NR_038142,NR_038143,NR_038144,NR_038145 |
| Mdh2 | 9992.289569 | 0.008592264 | 0.066199651 | 0.129793195 | 0.896730047 | 0.95158282 | | 1700008P02Rik | 0 | NA | NA | NA | NA | NA | NM_027048 |
| Shisa4 | 3170.291312 | -0.017621961 | 0.135666764 | -0.129891512 | 0.89665226 | 0.95158282 | | 1700009C05Rik | 1.62017637 | -0.000191074095932097 | 0.0253251746003369 | -0.00754482837522291 | 0.993980155038269 | NA | NR_046040 |
| Pdik1l | 543.787077 | -0.009762427 | 0.075267292 | -0.129703449 | 0.896801054 | 0.951584408 | | 1700009J07Rik | 1.072889252 | 0.00441242058576141 | 0.0254938642538106 | 0.17307774693677 | 0.862590316335949 | NA | NR_015547 |
| Cstb | 599.1638882 | 0.01352667 | 0.104529639 | 0.129405118 | 0.897037098 | 0.951704519 | | 1700009N14Rik | 0 | NA | NA | NA | NA | NA | NM_001081095 |
| Mettl1 | 183.5823193 | 0.013770763 | 0.106390783 | 0.12943568 | 0.897012916 | 0.951704519 | | 1700010B08Rik | 0.108766301 | 0.00519171381361946 | 0.00856003371502575 | 0.606506234257728 | 0.544178643801544 | NA | NM_029308 |
| Nfkbie | 96.30878828 | 0.016742832 | 0.129463243 | 0.129324982 | 0.897100505 | 0.951706622 | | 1700010D01Rik | 0 | NA | NA | NA | NA | NA | NM_029590 |
| 2900079G21Rik | 208.6325909 | 0.013409575 | 0.103761734 | 0.1292343 | 0.897172256 | 0.951717577 | | 1700010I02Rik | 0 | NA | NA | NA | NA | NA | NR_040587 |
| Pirt | 8.396537137 | -0.019791813 | 0.153287778 | -0.129115399 | 0.897266337 | 0.951752217 | | 1700010J16Rik | 0.09572496 | 0.00498255289994511 | 0.00847999972823692 | 0.587565219295241 | 0.556824162160132 | NA | NR_040579 |
| Trh | 268.6055651 | 0.016998232 | 0.131817065 | 0.128953198 | 0.897394682 | 0.951823193 | | 1700010K23Rik | 1.446992607 | 0.0190097171773459 | 0.0222041475984796 | 0.856133616164919 | 0.391923876350532 | NA | NR_040512 |
| Klhl15 | 209.882254 | 0.015603453 | 0.121099064 | 0.128848663 | 0.897477399 | 0.951845767 | | 1700011A15Rik | 0 | NA | NA | NA | NA | NA | NM_025487 |
| Serpinb9 | 363.2894913 | -0.016158566 | 0.125580035 | -0.128671456 | 0.897617622 | 0.951916874 | | 1700011B04Rik | 0 | NA | NA | NA | NA | NA | NR_045616 |
| Tspyl4 | 21133.37498 | -0.006284679 | 0.048866692 | -0.128608652 | 0.89766732 | 0.951916874 | | 1700011E24Rik | 0.940453988 | 0.0159158482497036 | 0.0215262403132148 | 0.739369626006311 | 0.459682580960081 | NA | NM_029298 |
| Hif1a | 2660.89268 | 0.009629944 | 0.075049869 | 0.128313941 | 0.897900533 | 0.952099019 | | 1700011H14Rik | 0.106347839 | -0.00653347234140708 | 0.00854389509723638 | -0.764694822098226 | 0.444453290197145 | NA | NM_025956 |
| 4632428N05Rik | 1314.845776 | -0.010116115 | 0.079104561 | -0.127882834 | 0.898241696 | 0.952125419 | | 1700011I03Rik | 3.312975103 | -0.000279252089561064 | 0.0423310441838256 | -0.00659686277400557 | 0.994736503219581 | NA | NM_029290 |
| Asap3 | 238.0782798 | -0.017095858 | 0.133630779 | -0.127933538 | 0.898201569 | 0.952125419 | | 1700011L22Rik | 0.732155478 | 0.00595239010997236 | 0.0176116097102367 | 0.337981036822123 | 0.735377478191037 | NA | NM_026315 |
| Hook3 | 2926.548077 | 0.009693743 | 0.075663759 | 0.128116058 | 0.898057128 | 0.952125419 | | 1700011M02Rik | 0 | NA | NA | NA | NA | NA | NR_073044 |
| Ly6a | 840.7973301 | -0.011866088 | 0.092693419 | -0.128014353 | 0.898137614 | 0.952125419 | | 1700012A03Rik | 0 | NA | NA | NA | NA | NA | NM_029587 |
| Prkaca | 7682.850179 | 0.006129347 | 0.047874688 | 0.128028971 | 0.898126046 | 0.952125419 | | 1700012B07Rik | 0 | NA | NA | NA | NA | NA | NM_001162428,NM_027038 |
| Rgs17 | 3085.962909 | 0.012160141 | 0.095137437 | 0.127816568 | 0.898294138 | 0.952125419 | | 1700012B09Rik | 12.24050919 | -0.0372325540685039 | 0.0589127978648246 | -0.63199432751325 | 0.527390583864667 | NA | NM_029306 |
| Erlin1 | 1135.659156 | -0.009407409 | 0.073695378 | -0.127652632 | 0.898423878 | 0.952132665 | | 1700012D01Rik | 11.53481532 | -0.0750272625286781 | 0.0655045731518975 | -1.14537442072477 | 0.252054089663715 | NA | NR_045171 |
| Mettl15 | 128.0784093 | 0.016147776 | 0.12645803 | 0.127692769 | 0.898392113 | 0.952132665 | | 1700012D14Rik | 17.19661905 | -0.0878658903477993 | 0.0708822169075664 | -1.23960415152337 | 0.215121844366991 | NA | NR_015573 |
| Acpl2 | 575.0811664 | -0.010419503 | 0.081676942 | -0.127569695 | 0.898489515 | 0.9521371 | | 1700012I11Rik | 0 | NA | NA | NA | NA | NA | NR_045140 |
| Ckb | 46198.40988 | -0.007916275 | 0.062138119 | -0.127398038 | 0.898625369 | 0.952215939 | | 1700012L04Rik | 0.237667581 | -0.00136938695604711 | 0.00989612066541175 | -0.13837613771559 | 0.889943154921195 | NA | NM_029588 |
| Bin2 | 244.5924536 | 0.019614738 | 0.154251452 | 0.1271608 | 0.898813131 | 0.952349767 | | 1700012P22Rik | 0.570373077 | -0.0091673637046035 | 0.0181304019377976 | -0.50563488531888 | 0.613112987054115 | NA | NM_027056 |
| Ngrn | 1082.374322 | -0.008031186 | 0.063287437 | -0.126900164 | 0.899019417 | 0.952438075 | | 1700013D24Rik | 0 | NA | NA | NA | NA | NA | NM_001177502 |
| Rnls | 22.99296527 | -0.021316199 | 0.167880893 | -0.126972154 | 0.898962438 | 0.952438075 | | 1700013G23Rik | 17.76692386 | 0.0829065378326902 | 0.0778547496388632 | 1.06488734749338 | 0.286926911316185 | NA | NR_045273 |
| Derl2 | 798.0939639 | -0.011523458 | 0.091171327 | -0.126393445 | 0.899420491 | 0.952724376 | | 1700013G24Rik | 0 | NA | NA | NA | NA | NA | NM_027063 |
| Letm1 | 3202.125936 | 0.008313104 | 0.065749121 | 0.126436726 | 0.899386232 | 0.952724376 | | 1700013H16Rik | 0.859859974 | 0.0087125691981874 | 0.0185923316855392 | 0.468610895370584 | 0.639347787996225 | NA | NM_001200013 |
| Mfap4 | 62.80055167 | -0.019970421 | 0.158086786 | -0.126325679 | 0.89947413 | 0.952724376 | | 1700015E13Rik | 0 | NA | NA | NA | NA | NA | NM_001039593 |
| Tmem143 | 703.0138949 | 0.014471242 | 0.114707176 | 0.126158123 | 0.89960676 | 0.952799722 | | 1700015F17Rik | 3.303416737 | 0.0151714079746832 | 0.0414065260123288 | 0.366401372821423 | 0.714065595494583 | NA | NM_001200025 |
| B3gnt6 | 13.87083927 | -0.020991617 | 0.166780011 | -0.125864108 | 0.899839494 | 0.952981075 | | 1700015G11Rik | 0 | NA | NA | NA | NA | NA | NM_001195601 |
| Rbm20 | 242.306444 | -0.018727476 | 0.148917871 | -0.125757072 | 0.899924223 | 0.952991929 | | 1700016C15Rik | 0 | NA | NA | NA | NA | NA | NM_027077 |
| Zfp71-rs1 | 494.0088994 | -0.010904564 | 0.086753633 | -0.125695761 | 0.899972758 | 0.952991929 | | 1700016D06Rik | 0.212297216 | 0.00681050119636477 | 0.00973439053551308 | 0.699633035218656 | 0.484156506270654 | NA | NM_024271 |
| Wnt10a | 304.5620225 | 0.017328798 | 0.138100367 | 0.125479738 | 0.900143765 | 0.953107872 | | 1700016G22Rik | 0.378011608 | -0.0142484740670214 | 0.0125319145197416 | -1.13697504436179 | 0.255548718289805 | NA | NR_045891 |
| Nudcd1 | 538.4244488 | -0.010131925 | 0.080885541 | -0.125262493 | 0.900315745 | 0.953224829 | | 1700016H13Rik | 0.321461979 | -0.00442641171345637 | 0.0111335060485946 | -0.397575722700138 | 0.690942958938875 | NA | NM_028824,NM_001163550 |
| Musk | 78.52697804 | 0.019019062 | 0.152092259 | 0.125049506 | 0.900484358 | 0.953338206 | | 1700016L04Rik | 1.66120297 | -0.00472872780314909 | 0.0307676577022534 | -0.153691511031168 | 0.877852978469984 | NA | NR_045824 |
| Acbd3 | 1366.151762 | 0.008282871 | 0.066344157 | 0.124847027 | 0.900644656 | 0.953358248 | | 1700016L21Rik | 0 | NA | NA | NA | NA | NA | NR_040460 |
| Calhm2 | 158.3646228 | 0.017919945 | 0.144086167 | 0.124369642 | 0.901022609 | 0.953358248 | | 1700016P04Rik | 0 | NA | NA | NA | NA | NA | NR_038149 |
| Fam49b | 5641.237353 | -0.008534813 | 0.06849423 | -0.1246063 | 0.90083524 | 0.953358248 | | 1700017D01Rik | 0 | NA | NA | NA | NA | NA | NM_027058 |
| Ikbkg | 1791.045074 | -0.007376092 | 0.05938506 | -0.124207869 | 0.901150692 | 0.953358248 | | 1700017G19Rik | 0.192890664 | -0.000903864895041124 | 0.00963528282151475 | -0.093807821917056 | 0.92526181809679 | NA | NR_040445 |
| Nfe2l3 | 145.5723182 | -0.017942915 | 0.144210292 | -0.124421875 | 0.900981254 | 0.953358248 | | 1700017J07Rik | 0.387222073 | -0.00720181929823146 | 0.0119779253649773 | -0.60125765345718 | 0.547668389729015 | NA | NR_040326 |
| Pcdhga3 | 587.4198588 | -0.012075166 | 0.097297211 | -0.124105984 | 0.90123136 | 0.953358248 | | 1700017N19Rik | 0 | NA | NA | NA | NA | NA | NM_001081246 |
| Rrbp1 | 2195.598937 | 0.005487427 | 0.043982308 | 0.124764407 | 0.900710066 | 0.953358248 | | 1700018A04Rik | 2.040357079 | 0.014071800998042 | 0.0294420616337052 | 0.477948900899408 | 0.632686575093083 | NA | NR_029439 |
| Ruvbl1 | 1283.71855 | 0.008354773 | 0.067184135 | 0.124356337 | 0.901033143 | 0.953358248 | | 1700018B08Rik | 0 | NA | NA | NA | NA | NA | NM_029597 |
| Ruvbl2 | 836.4055891 | 0.008114127 | 0.065217884 | 0.12441568 | 0.900986159 | 0.953358248 | | 1700018B24Rik | 0.09572496 | 0.00498255289994511 | 0.00847999972823692 | 0.587565219295241 | 0.556824162160132 | NA | NR_003617 |
| Skida1 | 289.4352532 | -0.014756732 | 0.118916736 | -0.124092978 | 0.901241658 | 0.953358248 | | 1700018C11Rik | 0 | NA | NA | NA | NA | NA | NM_029324 |
| Spry4 | 640.8021378 | 0.020415338 | 0.164436582 | 0.124153264 | 0.901193926 | 0.953358248 | | 1700018F24Rik | 0.106148608 | 0.00515554063321497 | 0.00854200771766808 | 0.603551390213729 | 0.546141939141592 | NA | NM_027069 |
| Top1 | 3006.688218 | 0.008854064 | 0.071071455 | 0.124579744 | 0.900856265 | 0.953358248 | | 1700018G05Rik | 0 | NA | NA | NA | NA | NA | NR_045422 |
| Gm20257 | 31.09443483 | -0.020544662 | 0.1657685 | -0.123935865 | 0.901366056 | 0.953424745 | | 1700019A02Rik | 1.655031696 | 0.00584944924316989 | 0.0310277120130202 | 0.18852338324899 | 0.85046638696099 | NA | NM_027070 |
| Dtx1 | 2745.514973 | 0.006678579 | 0.053948539 | 0.123795365 | 0.901477302 | 0.95343761 | | 1700019B03Rik | 5.966871673 | 0.00162714706538321 | 0.0478473875155235 | 0.0340070200249762 | 0.972871552769982 | NA | NM_029598 |
| Mafa | 14.5686718 | -0.020696634 | 0.167225167 | -0.12376507 | 0.90150129 | 0.95343761 | | 1700019B21Rik | 0 | NA | NA | NA | NA | NA | NR_045443,NR_045442 |
| Snap47 | 13389.23683 | 0.007507171 | 0.060813325 | 0.123446148 | 0.901753816 | 0.953639589 | | 1700019E08Rik | 0 | NA | NA | NA | NA | NA | NR_040497 |
| Mpg | 426.9234781 | -0.009584608 | 0.077731163 | -0.123304577 | 0.901865917 | 0.953693046 | | 1700019G24Rik | 4.022786096 | -0.004740066398078 | 0.0412231672233141 | -0.114985497654756 | 0.908456616989207 | NA | NR_040255 |
| 1700066M21Rik | 779.9621018 | -0.011243137 | 0.091277492 | -0.123175354 | 0.901968241 | 0.953736158 | | 1700019L03Rik | 12.15822791 | 0.0192699178453994 | 0.0644255006074257 | 0.299103889977043 | 0.764860778304797 | NA | NM_025619 |
| Btbd7 | 651.358974 | -0.009751146 | 0.079267978 | -0.123014947 | 0.902095262 | 0.953805376 | | 1700019M22Rik | 0 | NA | NA | NA | NA | NA | NM_027076 |
| Pfn1 | 2882.110852 | -0.00848593 | 0.069228154 | -0.12257918 | 0.902440342 | 0.953858521 | | 1700019N19Rik | 4.53142407 | -0.0546030584754443 | 0.0422118285673352 | -1.29354875940385 | 0.195821331483358 | NA | NM_026208 |
| Slc22a17 | 22120.49638 | -0.007807489 | 0.063586446 | -0.122785433 | 0.90227701 | 0.953858521 | | 1700019O17Rik | 0.106148608 | 0.00515554063321497 | 0.00854200771766808 | 0.603551390213729 | 0.546141939141592 | NA | NM_027966 |
| Spata2l | 2011.458395 | -0.011538775 | 0.094028553 | -0.122715645 | 0.902332275 | 0.953858521 | | 1700020A23Rik | 0 | NA | NA | NA | NA | NA | NM_029375,NM_001163483 |
| Trdmt1 | 157.4064013 | 0.015349987 | 0.125011144 | 0.122788951 | 0.902274224 | 0.953858521 | | 1700020G17Rik | 0.12890128 | -0.0068640067066208 | 0.00870804083610955 | -0.788237771940376 | 0.430557639008166 | NA | NR_045979 |
| Usp45 | 1768.588964 | -0.009774961 | 0.079754735 | -0.122562765 | 0.902453341 | 0.953858521 | | 1700020L24Rik | 8.261759608 | 0.00382833243263323 | 0.0534850762668394 | 0.0715775820068669 | 0.942938081271325 | NA | NM_025492 |
| Rpn2 | 5011.598066 | 0.006718546 | 0.054891845 | 0.122396073 | 0.902585349 | 0.953932974 | | 1700020M21Rik | 0 | NA | NA | NA | NA | NA | NR_040742 |
| Amer2 | 845.8173871 | -0.01184249 | 0.096838642 | -0.122290953 | 0.902668596 | 0.953955885 | | 1700020N01Rik | 1.304871827 | 0.00691241858892325 | 0.0245064517048548 | 0.282065256617867 | 0.777893473192189 | NA | NR_027968 |
| 4931429I11Rik | 14.8141693 | -0.020111119 | 0.16480017 | -0.122033366 | 0.902872594 | 0.9539997 | | 1700020N15Rik | 0 | NA | NA | NA | NA | NA | NM_029316 |
| Mapk12 | 228.8538536 | -0.014439364 | 0.118350244 | -0.122005361 | 0.902894772 | 0.9539997 | | 1700020N18Rik | 1.662577843 | -0.0150436382335009 | 0.0272347974414536 | -0.552368280536695 | 0.580696058038436 | NA | NR_026924 |
| Tfpi | 165.9017495 | 0.014867838 | 0.121730097 | 0.122137729 | 0.902789942 | 0.9539997 | | 1700021F07Rik | 0.756487453 | 0.010985648029265 | 0.0200867684793789 | 0.546909675418566 | 0.584440788202663 | NA | NM_028158 |
| Eif2c1 | 4114.239159 | -0.008885916 | 0.072887048 | -0.121913507 | 0.902967518 | 0.954011506 | | 1700021N21Rik | 1.886347427 | 0.0253713942835138 | 0.0321142237425148 | 0.790036044057499 | 0.429506718624935 | NA | NR_045800 |
| Dgke | 2985.479348 | -0.008015733 | 0.065832748 | -0.121759056 | 0.903089841 | 0.954075685 | | 1700022A21Rik | 0 | NA | NA | NA | NA | NA | NR_003953 |
| Chic1 | 1330.480205 | 0.008255748 | 0.067873933 | 0.121633553 | 0.903189239 | 0.954113462 | | 1700022A22Rik | 0 | NA | NA | NA | NA | NA | NR_045509 |
| Kbtbd2 | 2139.009392 | -0.009301334 | 0.076517409 | -0.121558403 | 0.903248758 | 0.954113462 | | 1700022E09Rik | 0 | NA | NA | NA | NA | NA | NR_040568 |
| Ppargc1a | 2193.101935 | -0.014012445 | 0.11543082 | -0.121392577 | 0.903380096 | 0.954122826 | | 1700022H16Rik | 0 | NA | NA | NA | NA | NA | NR_045488 |
| Rgs12 | 2446.854962 | -0.010629988 | 0.087567657 | -0.12139171 | 0.903380783 | 0.954122826 | | 1700022I11Rik | 0.127687691 | 0.00542298404407807 | 0.00869724871942359 | 0.62352868349814 | 0.532937159177571 | NA | NM_026088 |
| Pik3r6 | 172.9471594 | -0.014130104 | 0.116508989 | -0.121279092 | 0.90346998 | 0.954123446 | | 1700022P22Rik | 1.533755251 | 0.0296685445821269 | 0.0249994532325866 | 1.18676773872215 | 0.235319247409901 | NA | NM_029336 |
| Trim59 | 580.067647 | 0.016578397 | 0.136745438 | 0.12123547 | 0.90350453 | 0.954123446 | | 1700023A16Rik | 3.649312805 | -0.0168655119259229 | 0.0391439634790674 | -0.430858564819115 | 0.666571213135771 | NA | NM_027059 |
| B430212C06Rik | 9.053560191 | 0.018923953 | 0.156286665 | 0.121084888 | 0.903623799 | 0.954184362 | | 1700023C21Rik | 0 | NA | NA | NA | NA | NA | NR_045909 |
| Cyp27b1 | 12.6113855 | -0.019311519 | 0.159953851 | -0.12073182 | 0.903903455 | 0.954249731 | | 1700023E05Rik | 8.537211841 | 0.020296087299286 | 0.062655865704192 | 0.323929564633372 | 0.745991360419879 | NA | NM_027970 |
| Gm3414 | 68.06144855 | 0.018933623 | 0.156676822 | 0.120845081 | 0.903813743 | 0.954249731 | | 1700023F02Rik | 1.170791692 | -0.0410079582481214 | 0.0229986992338828 | -1.78305554723314 | 0.0745772616151429 | NA | NR_038039 |
| Pkn2 | 1091.404654 | 0.008720749 | 0.072177055 | 0.120824399 | 0.903830124 | 0.954249731 | | 1700023L04Rik | 6.222151013 | -0.0362121562920526 | 0.0535479208790615 | -0.676256999292992 | 0.498877489006918 | NA | NR_040264,NR_040263 |
| Pltp | 2624.837207 | -0.019890289 | 0.164903216 | -0.120617956 | 0.903993646 | 0.954249731 | | 1700024B18Rik | 0 | NA | NA | NA | NA | NA | NR_045479 |
| Pycard | 81.33847453 | 0.019726985 | 0.163505325 | 0.120650414 | 0.903967936 | 0.954249731 | | 1700024F13Rik | 14.51472345 | -0.0490852432925802 | 0.0760034956220662 | -0.645828759464705 | 0.518390260542136 | NA | NR_045363 |
| Cyb5d1 | 411.6726485 | 0.009079058 | 0.075511015 | 0.120234889 | 0.904297081 | 0.954439985 | | 1700024G13Rik | 7.267599535 | -0.0118093289505173 | 0.0514900713526737 | -0.229351574784797 | 0.81859567071818 | NA | NM_001034037 |
| Sh3bgrl3 | 3408.3619 | -0.009485502 | 0.078872594 | -0.120263601 | 0.904274337 | 0.954439985 | | 1700024P03Rik | 0 | NA | NA | NA | NA | NA | NR_040508 |
| Tmem87a | 1179.998807 | 0.007502279 | 0.062471502 | 0.120091227 | 0.904410882 | 0.954495075 | | 1700024P04Rik | 0 | NA | NA | NA | NA | NA | NM_027064 |
| Cyp4f17 | 74.3098625 | -0.017314129 | 0.144480974 | -0.119836742 | 0.904612475 | 0.95457779 | | 1700025B11Rik | 0 | NA | NA | NA | NA | NA | NR_040310 |
| Kdelr3 | 20.8035302 | -0.020146817 | 0.168056616 | -0.119881127 | 0.904577315 | 0.95457779 | | 1700025C18Rik | 0.127687691 | 0.00542298404407807 | 0.00869724871942359 | 0.62352868349814 | 0.532937159177571 | NA | NR_033448 |
| Npy1r | 1008.81523 | 0.008117226 | 0.067971882 | 0.119420351 | 0.904942339 | 0.954860842 | | 1700025E21Rik | 1.154524012 | -0.00569842988285404 | 0.0235620268140258 | -0.241848034883906 | 0.808897917032229 | NA | NM_027073,NM_029373 |
| Pole3 | 383.5262326 | -0.013832844 | 0.116116483 | -0.11912903 | 0.905173132 | 0.955010028 | | 1700025F22Rik | 0 | NA | NA | NA | NA | NA | NM_027074 |
| Zfp839 | 495.6389862 | 0.010819577 | 0.090854938 | 0.11908628 | 0.905207001 | 0.955010028 | | 1700025F24Rik | 0.240418098 | -0.0088029805258837 | 0.00986414519853158 | -0.892422034419582 | 0.372166765031741 | NA | NR_040578 |
| Aox4 | 36.63004758 | -0.01961695 | 0.164928067 | -0.118942457 | 0.905320945 | 0.955065208 | | 1700025K24Rik | 0 | NA | NA | NA | NA | NA | NR_045825 |
| C5ar2 | 46.08286919 | -0.018673785 | 0.157290577 | -0.118721574 | 0.905495944 | 0.955067686 | | 1700025M24Rik | 0 | NA | NA | NA | NA | NA | NR_040687,NR_040688,NR_040689 |
| Polr3h | 1344.222246 | -0.010183221 | 0.085699565 | -0.118824653 | 0.905414277 | 0.955067686 | | 1700025N23Rik | 0.474254397 | -0.00202179240680356 | 0.0141073909053298 | -0.143314410181951 | 0.886041875630184 | NA | NR_040523 |
| Tmem151a | 8629.333015 | -0.010133825 | 0.085369047 | -0.118706083 | 0.905508217 | 0.955067686 | | 1700026D11Rik | 0.192890664 | -0.000903864895041124 | 0.00963528282151475 | -0.093807821917056 | 0.92526181809679 | NA | NR_028286 |
| Agpat3 | 8408.979351 | 0.006970984 | 0.059136288 | 0.117879964 | 0.906162769 | 0.955432864 | | 1700026F02Rik | 0 | NA | NA | NA | NA | NA | NR_045487 |
| Fndc4 | 2970.414117 | -0.00816436 | 0.069257962 | -0.11788334 | 0.906160094 | 0.955432864 | | 1700026L06Rik | 8.457015926 | -0.0173303704717183 | 0.0524698322590387 | -0.330292088340589 | 0.741179270412783 | NA | NM_027283 |
| Hexim2 | 321.535283 | 0.013202657 | 0.111717412 | 0.118179044 | 0.905925794 | 0.955432864 | | 1700027A15Rik | 0.192890664 | -0.000903864895041124 | 0.00963528282151475 | -0.093807821917056 | 0.92526181809679 | NA | NR_038001,NR_038002 |
| Nrep | 2158.152474 | -0.019463996 | 0.165113836 | -0.117882283 | 0.906160931 | 0.955432864 | | 1700027F09Rik | 0 | NA | NA | NA | NA | NA | NR_040681 |
| Tmem161a | 942.0641698 | -0.008868627 | 0.075117872 | -0.118062814 | 0.906017888 | 0.955432864 | | 1700027H10Rik | 0.223412651 | 0.00718831964189475 | 0.00999993766105603 | 0.718836445339964 | 0.472241699104885 | NA | NR_040594 |
| Pdcd2l | 392.6584843 | 0.008908807 | 0.07567425 | 0.11772574 | 0.906284971 | 0.955461008 | | 1700027I24Rik | 0.12890128 | -0.0068640067066208 | 0.00870804083610955 | -0.788237771940376 | 0.430557639008166 | NA | NR_040741 |
| Zbtb22 | 1396.52152 | 0.009818406 | 0.083425554 | 0.117690626 | 0.906312794 | 0.955461008 | | 1700027J07Rik | 0.349184399 | -0.00508887990706381 | 0.0114407387221738 | -0.444803437141765 | 0.656461810238787 | NA | NR_040581 |
| Gemin8 | 343.2293741 | 0.010165546 | 0.086915963 | 0.116958334 | 0.906893072 | 0.956007705 | | 1700028B04Rik | 2.068070408 | 0.0202952848711236 | 0.0302732308761897 | 0.670403663029112 | 0.502600500194145 | NA | NR_033605 |
| Puf60 | 4792.285115 | -0.006761095 | 0.057893414 | -0.116785217 | 0.907030259 | 0.956029949 | | 1700028D13Rik | 0 | NA | NA | NA | NA | NA | NR_045377,NR_045378 |
| Seh1l | 1486.889284 | 0.007214505 | 0.061780732 | 0.11677598 | 0.907037579 | 0.956029949 | | 1700028E10Rik | 7.200830189 | -0.0153521387523475 | 0.059383140257925 | -0.258526893082227 | 0.796000296498594 | NA | NR_045700,NR_045699 |
| Trpm2 | 1559.0131 | 0.012006891 | 0.102942207 | 0.1166372 | 0.907147558 | 0.956080828 | | 1700028I16Rik | 0 | NA | NA | NA | NA | NA | NR_038042 |
| Tmtc3 | 547.1921766 | -0.010911905 | 0.093634459 | -0.116537278 | 0.907226744 | 0.95609925 | | 1700028K03Rik | 18.19629399 | 0.0154988861333146 | 0.079567935045048 | 0.194788090510829 | 0.845558846103258 | NA | NM_182745,NM_175241 |
| Adh1 | 14.13129829 | -0.018381701 | 0.157848268 | -0.116451714 | 0.907294552 | 0.956105679 | | 1700028M03Rik | 0 | NA | NA | NA | NA | NA | NR_036591 |
| Lonrf1 | 1321.721782 | -0.008894707 | 0.07647629 | -0.116306728 | 0.907409454 | 0.95616173 | | 1700028P15Rik | 0 | NA | NA | NA | NA | NA | NR_040509 |
| Mrpl16 | 843.8670443 | -0.008997183 | 0.077523798 | -0.116057044 | 0.907607332 | 0.956240175 | | 1700029B22Rik | 0 | NA | NA | NA | NA | NA | NR_040531 |
| Paqr8 | 5627.920049 | -0.00913367 | 0.078660804 | -0.116114629 | 0.907561695 | 0.956240175 | | 1700029F12Rik | 0 | NA | NA | NA | NA | NA | NM_001080777,NM_025585 |
| Slc6a12 | 47.73186695 | 0.019414982 | 0.167685241 | 0.115782292 | 0.907825085 | 0.95640456 | | 1700029H14Rik | 0.761934134 | 0.009675330791516 | 0.0198161699821609 | 0.488254329682578 | 0.625369706179624 | NA | NM_025601,NM_001080781 |
| Ahcyl2 | 7230.650512 | 0.007863996 | 0.06829816 | 0.115142138 | 0.908332461 | 0.956463738 | | 1700029J03Rik | 0 | NA | NA | NA | NA | NA | NR_040494 |
| Arhgef10l | 2156.661495 | 0.014215295 | 0.123328713 | 0.115263469 | 0.908236293 | 0.956463738 | | 1700029M20Rik | 0.771558907 | 0.0179230073090733 | 0.0189976784432228 | 0.943431449407816 | 0.345460263245835 | NA | NR_027909,NR_015613 |
| CK137956 | 81.90248776 | 0.015827107 | 0.138573029 | 0.114214913 | 0.90906743 | 0.956463738 | | 1700029N11Rik | 0 | NA | NA | NA | NA | NA | NR_045489 |
| Cat | 1780.542594 | -0.01294155 | 0.113410297 | -0.114112656 | 0.90914849 | 0.956463738 | | 1700029P11Rik | 0 | NA | NA | NA | NA | NA | NM_025503 |
| Coq4 | 613.7277962 | -0.011694159 | 0.101769923 | -0.114907814 | 0.908518192 | 0.956463738 | | 1700030A11Rik | 1.480134068 | 0.00728758890425572 | 0.0298254253713778 | 0.244341490976665 | 0.806966351928431 | NA | NR_045457 |
| Dcxr | 150.4381454 | 0.014865078 | 0.129187926 | 0.115065536 | 0.908393177 | 0.956463738 | | 1700030F04Rik | 0.925531857 | -0.00422483633719062 | 0.0224349404029688 | -0.188315023855894 | 0.850629708711975 | NA | NR_045731 |
| Gnb1l | 158.6264512 | 0.012256131 | 0.106536814 | 0.115041272 | 0.908412409 | 0.956463738 | | 1700030F18Rik | 0.21100899 | -0.000459554483117493 | 0.00972904351832043 | -0.0472353199214416 | 0.96232567770125 | NA | NM_028180 |
| Homez | 704.0498184 | 0.007922229 | 0.06862874 | 0.115436026 | 0.908099525 | 0.956463738 | | 1700030M09Rik | 0.217532603 | 0.00684685248201087 | 0.00976000296730706 | 0.701521557416086 | 0.482977586094298 | NA | NR_045903 |
| Hsp90ab1 | 62236.55288 | -0.006212361 | 0.053911814 | -0.115231909 | 0.908261307 | 0.956463738 | | 1700030N03Rik | 0 | NA | NA | NA | NA | NA | NR_045304 |
| Kcp | 63.51843393 | -0.018948426 | 0.164377715 | -0.115273695 | 0.908228188 | 0.956463738 | | 1700030O20Rik | 0 | NA | NA | NA | NA | NA | NR_045345 |
| Pnck | 2488.505235 | 0.01004438 | 0.087365191 | 0.11497005 | 0.908468861 | 0.956463738 | | 1700031A10Rik | 0 | NA | NA | NA | NA | NA | NR_045439 |
| Pno1 | 762.4529719 | -0.008465338 | 0.073423137 | -0.11529524 | 0.908211111 | 0.956463738 | | 1700031F05Rik | 0 | NA | NA | NA | NA | NA | NM_028496 |
| Slc16a14 | 209.0704238 | 0.013808256 | 0.120244563 | 0.114834762 | 0.908576095 | 0.956463738 | | 1700031M16Rik | 0.46293569 | -0.00199393361883231 | 0.0140214968410126 | -0.142205475024613 | 0.886917707298231 | NA | NR_015496 |
| Slc39a7 | 3402.003784 | -0.005738184 | 0.050301449 | -0.114075917 | 0.909177613 | 0.956463738 | | 1700031P21Rik | 0 | NA | NA | NA | NA | NA | NR_045910 |
| Sort1 | 27082.2747 | 0.008538676 | 0.074808586 | 0.114140317 | 0.909126563 | 0.956463738 | | 1700034E13Rik | 0 | NA | NA | NA | NA | NA | NM_030097 |
| St3gal5 | 6035.551493 | -0.007349071 | 0.064309106 | -0.114277306 | 0.909017972 | 0.956463738 | | 1700034F02Rik | 3.638767852 | 0.0213275244118548 | 0.0449779542827013 | 0.474177288673563 | 0.635373487445621 | NA | NM_028504,NM_001163521 |
| Tmem204 | 307.6590033 | -0.010375588 | 0.090561681 | -0.1145693 | 0.908786514 | 0.956463738 | | 1700034G24Rik | 0.09572496 | 0.00498255289994511 | 0.00847999972823692 | 0.587565219295241 | 0.556824162160132 | NA | NR_045397,NR_045396 |
| Uck1 | 1485.092848 | -0.007293255 | 0.063923156 | -0.114094106 | 0.909163194 | 0.956463738 | | 1700034I23Rik | 0.854866745 | -0.00481489060925801 | 0.0222282865355402 | -0.216610965562263 | 0.828511539344553 | NA | NR_045380 |
| Zbtb1 | 460.9006936 | -0.011269535 | 0.098670311 | -0.114214045 | 0.909068119 | 0.956463738 | | 1700034J05Rik | 0.565037534 | 0.00303603369225299 | 0.0165574101691361 | 0.183364044330575 | 0.85451238599073 | NA | NM_028509,NM_001164236 |
| Zfhx2 | 2551.685006 | -0.008441085 | 0.073724874 | -0.114494394 | 0.90884589 | 0.956463738 | | 1700034K08Rik | 0 | NA | NA | NA | NA | NA | NR_040756 |
| Zfp959 | 155.2584251 | 0.015049763 | 0.131141944 | 0.114759343 | 0.908635875 | 0.956463738 | | 1700034O15Rik | 0 | NA | NA | NA | NA | NA | NM_029671 |
| Cdkl1 | 507.6461099 | 0.00977873 | 0.085785355 | 0.113990665 | 0.909245193 | 0.956469891 | | 1700034P13Rik | 2.301102195 | -0.0383802071094812 | 0.0321999675914683 | -1.1919330974622 | 0.233287479194816 | NA | NR_040462 |
| Aldh18a1 | 903.8181963 | -0.007498777 | 0.06594477 | -0.113712988 | 0.909465317 | 0.956511805 | | 1700036G14Rik | 0.23403553 | -0.000407313579519725 | 0.00983856423584417 | -0.0413996971261099 | 0.966977254250032 | NA | NR_040542 |
| Fam134a | 8263.703757 | 0.006266921 | 0.05509244 | 0.11375283 | 0.909433732 | 0.956511805 | | 1700039E15Rik | 1.849439134 | -0.0268873224607724 | 0.0282450253821189 | -0.951931255044788 | 0.341131847203866 | NA | NM_001033176 |
| Mcm3 | 133.9981209 | -0.015427611 | 0.135678904 | -0.113706776 | 0.909470241 | 0.956511805 | | 1700039E22Rik | 0.108766301 | 0.00519171381361946 | 0.00856003371502575 | 0.606506234257728 | 0.544178643801544 | NA | NR_045315 |
| Armc4 | 26.7379031 | 0.018851512 | 0.165925659 | 0.113614207 | 0.909543625 | 0.956524056 | | 1700040N02Rik | 14.21524246 | -0.101309728959173 | 0.0676433382065586 | -1.49770445464724 | 0.134210053964819 | NA | NR_040610 |
| Bean1 | 566.6786218 | -0.010087987 | 0.089104443 | -0.113215312 | 0.909859857 | 0.956579732 | | 1700041C23Rik | 0 | NA | NA | NA | NA | NA | NR_033784 |
| Dedd | 1070.650126 | 0.00932881 | 0.082376704 | 0.113245731 | 0.909835741 | 0.956579732 | | 1700041M19Rik | 0 | NA | NA | NA | NA | NA | NR_040573 |
| Prr16 | 289.7261963 | 0.013257001 | 0.117146865 | 0.113165647 | 0.90989923 | 0.956579732 | | 1700042B14Rik | 0.108766301 | 0.00519171381361946 | 0.00856003371502575 | 0.606506234257728 | 0.544178643801544 | NA | NM_001081671 |
| Wdr82 | 6249.904103 | 0.005107405 | 0.045101436 | 0.113242628 | 0.909838201 | 0.956579732 | | 1700042G07Rik | 0 | NA | NA | NA | NA | NA | NM_001099295 |
| Zfp433 | 41.25529608 | 0.018182206 | 0.160679754 | 0.113158042 | 0.909905259 | 0.956579732 | | 1700042G15Rik | 0.106148608 | 0.00515554063321497 | 0.00854200771766808 | 0.603551390213729 | 0.546141939141592 | NA | NR_038179,NR_038178 |
| Klhdc10 | 4548.620363 | -0.00617608 | 0.054624484 | -0.113064313 | 0.909979568 | 0.956591808 | | 1700042O10Rik | 0.435709867 | 0.015952827522772 | 0.0148578726417494 | 1.07369526630251 | 0.282959283681906 | NA | NR_045178 |
| Taf4b | 227.0967711 | 0.010774342 | 0.09535845 | 0.112987802 | 0.910040226 | 0.956591808 | | 1700044C05Rik | 0 | NA | NA | NA | NA | NA | NR_045624 |
| Dkkl1 | 639.0562889 | -0.014680317 | 0.130055341 | -0.112877467 | 0.910127701 | 0.956618859 | | 1700044K03Rik | 0 | NA | NA | NA | NA | NA | NR_033785 |
| C130060C02Rik | 10.85089783 | -0.018507212 | 0.164266371 | -0.112665862 | 0.910295467 | 0.956675494 | | 1700045H11Rik | 0.097165704 | -0.00637828455464458 | 0.00848781691535724 | -0.75146349388194 | 0.452373763047372 | NA | NR_040649 |
| Gipr | 56.31207524 | 0.018313971 | 0.162568682 | 0.112653745 | 0.910305074 | 0.956675494 | | 1700046C09Rik | 0 | NA | NA | NA | NA | NA | NR_045918 |
| 2610044O15Rik8 | 274.3519795 | -0.011255253 | 0.100356178 | -0.112153065 | 0.910702043 | 0.956833082 | | 1700047E10Rik | 0.106347839 | -0.00653347234140708 | 0.00854389509723638 | -0.764694822098226 | 0.444453290197145 | NA | NR_073363 |
| C330027C09Rik | 110.9948917 | -0.016454939 | 0.146508223 | -0.112314101 | 0.910574362 | 0.956833082 | | 1700047G03Rik | 0 | NA | NA | NA | NA | NA | NR_040447 |
| Lipo1 | 172.7298145 | 0.015415541 | 0.137427483 | 0.112172185 | 0.910686883 | 0.956833082 | | 1700047I17Rik2 | 0 | NA | NA | NA | NA | NA | NM_001100116 |
| Tra2a | 1697.730144 | 0.013100424 | 0.116661043 | 0.112294765 | 0.910589693 | 0.956833082 | | 1700047L14Rik | 0 | NA | NA | NA | NA | NA | NR_040691 |
| Tmem109 | 1501.349548 | -0.010448655 | 0.093274268 | -0.112020768 | 0.910806939 | 0.956878405 | | 1700048M11Rik | 0 | NA | NA | NA | NA | NA | NR_045300 |
| Hnrnpl | 5373.954906 | 0.008185926 | 0.073251516 | 0.111750938 | 0.91102089 | 0.957038285 | | 1700049E15Rik | 0 | NA | NA | NA | NA | NA | NR_033636 |
| 0610043K17Rik | 13.43557615 | 0.018251914 | 0.163727531 | 0.111477364 | 0.911237814 | 0.957071495 | | 1700049E22Rik | 0 | NA | NA | NA | NA | NA | NR_040525 |
| Cenpl | 56.04421654 | -0.016430989 | 0.147354997 | -0.111506157 | 0.911214984 | 0.957071495 | | 1700049L16Rik | 0 | NA | NA | NA | NA | NA | NR_003644 |
| Rab13 | 158.6167318 | 0.011962421 | 0.107235387 | 0.111552927 | 0.911177898 | 0.957071495 | | 1700051A21Rik | 2.664904091 | -0.0263709161209457 | 0.0382223733136936 | -0.689934031686567 | 0.490235673344972 | NA | NR_045922 |
| Alyref2 | 81.42775452 | -0.016331463 | 0.147017587 | -0.1110851 | 0.911548864 | 0.957333295 | | 1700052I22Rik | 0 | NA | NA | NA | NA | NA | NR_033786 |
| Zfp317 | 1045.148324 | 0.009010826 | 0.081262943 | 0.110884809 | 0.911707692 | 0.957435203 | | 1700054A03Rik | 0 | NA | NA | NA | NA | NA | NR_045320 |
| Cxcl5 | 9.024761255 | 0.017767527 | 0.160579034 | 0.110646617 | 0.91189658 | 0.957533205 | | 1700054K19Rik | 0 | NA | NA | NA | NA | NA | NR_027865 |
| Gtf3c5 | 776.9756172 | 0.008577469 | 0.077546072 | 0.110611264 | 0.911924615 | 0.957533205 | | 1700054M17Rik | 0.703269883 | 0.0115284326264998 | 0.0160827339190083 | 0.716820453820616 | 0.473484889366507 | NA | NR_045919 |
| Nsa2 | 1669.867427 | 0.007514802 | 0.068217907 | 0.110158787 | 0.912283446 | 0.957845071 | | 1700054O13Rik | 0 | NA | NA | NA | NA | NA | NM_026096 |
| Gltscr1 | 828.6145941 | 0.008766322 | 0.079713801 | 0.109972453 | 0.912431222 | 0.957935313 | | 1700055C04Rik | 0 | NA | NA | NA | NA | NA | NR_040726 |
| 1200014J11Rik | 851.4867616 | -0.010445507 | 0.095216536 | -0.109702656 | 0.912645194 | 0.958095036 | | 1700055N04Rik | 0.769254926 | 0.00138643920361091 | 0.021287703408129 | 0.0651286414992741 | 0.948071576218993 | NA | NM_028545 |
| Ern2 | 13.93636741 | -0.018139584 | 0.166426952 | -0.108994268 | 0.913207036 | 0.958554964 | | 1700056E22Rik | 1.746807055 | -0.024661830632346 | 0.0294364138066327 | -0.837800106845525 | 0.402142978742772 | NA | NM_028516 |
| Nr2c2ap | 490.9915276 | -0.008931279 | 0.081923963 | -0.109019127 | 0.913187319 | 0.958554964 | | 1700057G04Rik | 0 | NA | NA | NA | NA | NA | NM_001033184 |
| Lca5 | 371.168883 | 0.013096488 | 0.120559642 | 0.108630783 | 0.913495344 | 0.958741946 | | 1700057H15Rik | 0.233836299 | 0.00743372775021133 | 0.0101683766597817 | 0.731063374118848 | 0.464740444385902 | NA | NR_040774 |
| Slc18b1 | 793.2573755 | -0.008703264 | 0.080130475 | -0.108613655 | 0.91350893 | 0.958741946 | | 1700058G18Rik | 4.316964816 | 0.00510134631973494 | 0.0406806139029013 | 0.12539993452191 | 0.90020693995008 | NA | NR_028107 |
| B930003M22Rik | 25.38807303 | -0.01800604 | 0.16661423 | -0.10807024 | 0.913939975 | 0.958938792 | | 1700060C16Rik | 0.127687691 | 0.00542298404407807 | 0.00869724871942359 | 0.62352868349814 | 0.532937159177571 | NA | NR_045732 |
| Dennd4c | 952.7512021 | -0.009799404 | 0.090680559 | -0.1080651 | 0.913944052 | 0.958938792 | | 1700060C20Rik | 0.120209049 | -0.00674972139751468 | 0.00863521114606269 | -0.781650996523957 | 0.434419707908266 | NA | NR_036606 |
| Nr3c2 | 1286.47972 | 0.009286106 | 0.085758376 | 0.108282207 | 0.913771836 | 0.958938792 | | 1700061F12Rik | 0 | NA | NA | NA | NA | NA | NR_038180 |
| Tmem69 | 467.3109662 | 0.009899799 | 0.091573342 | 0.108107866 | 0.913910128 | 0.958938792 | | 1700061G19Rik | 6.024746964 | 0.0240853010520719 | 0.0517178775996925 | 0.465705519443341 | 0.641426301062156 | NA | NM_030141 |
| Ckap5 | 7591.947189 | -0.006649455 | 0.061834647 | -0.107536074 | 0.914363708 | 0.959068549 | | 1700061I17Rik | 0.346765936 | -0.0134445992440418 | 0.0121756794716388 | -1.104217573677 | 0.26949876817541 | NA | NR_038029 |
| Fastkd2 | 695.0841194 | -0.009306513 | 0.086745735 | -0.10728496 | 0.914562916 | 0.959068549 | | 1700063A18Rik | 0 | NA | NA | NA | NA | NA | NR_040467 |
| Fkbp14 | 397.6364237 | -0.009241841 | 0.085970513 | -0.10750013 | 0.914392222 | 0.959068549 | | 1700063O14Rik | 0.892156306 | 0.0166009348859502 | 0.0227109157872756 | 0.730967216005058 | 0.464799177918741 | NA | NR_045383 |
| Gstm2 | 92.35188109 | -0.016121525 | 0.149929805 | -0.107527155 | 0.914370784 | 0.959068549 | | 1700064J06Rik | 0 | NA | NA | NA | NA | NA | NR_045348 |
| Kdelr2 | 745.2885152 | -0.006662552 | 0.062007873 | -0.107446865 | 0.914434477 | 0.959068549 | | 1700064M15Rik | 1.425541861 | -0.0215267105728621 | 0.0255480806007078 | -0.842596001997335 | 0.39945442782674 | NA | NR_045288 |
| Rpl28 | 4181.10371 | -0.007003841 | 0.065238825 | -0.107356943 | 0.914505811 | 0.959068549 | | 1700065D16Rik | 0.998756267 | -0.000597607076860035 | 0.0229044501883569 | -0.0260913085424691 | 0.979184509482611 | NA | NR_073361,NM_001271569 |
| Stk24 | 1889.851866 | -0.008784906 | 0.08165155 | -0.107590191 | 0.914320778 | 0.959068549 | | 1700065I16Rik | 0 | NA | NA | NA | NA | NA | NR_040315 |
| Taf15 | 1555.251379 | 0.013656929 | 0.127287823 | 0.107291715 | 0.914557557 | 0.959068549 | | 1700065I17Rik | 0.21491491 | 0.00699378003108193 | 0.00986423294990756 | 0.709003940458184 | 0.478322031551065 | NA | NM_026099 |
| Ncoa4 | 3930.470419 | 0.005568666 | 0.051950364 | 0.107192059 | 0.914636615 | 0.959080923 | | 1700065J11Rik | 0.106347839 | -0.00653347234140708 | 0.00854389509723638 | -0.764694822098226 | 0.444453290197145 | NA | NR_040526 |
| Arhgap27 | 705.1589356 | 0.009341777 | 0.087282287 | 0.107029465 | 0.914765604 | 0.959140337 | | 1700065J18Rik | 0.933090672 | 0.0287826424912678 | 0.0228035693916266 | 1.26219900038266 | 0.206877188328527 | NA | NR_040468 |
| Cnp | 10698.14211 | 0.013912745 | 0.130275889 | 0.106794471 | 0.914952034 | 0.959140337 | | 1700065O20Rik | 0 | NA | NA | NA | NA | NA | NR_045386 |
| Cntnap5a | 942.3713868 | -0.012237886 | 0.114629235 | -0.106760602 | 0.914978904 | 0.959140337 | | 1700066B17Rik | 0.539767185 | 0.0117536499441691 | 0.0167940487954241 | 0.699869941271792 | 0.484008530842235 | NA | NR_040465 |
| Lrch2 | 612.1794191 | 0.010787176 | 0.100897411 | 0.106912316 | 0.914858542 | 0.959140337 | | 1700066B19Rik | 1.782248378 | -0.00733161100187794 | 0.0273492726522974 | -0.268073344950987 | 0.788642862794807 | NA | NM_001033168 |
| Slk | 4355.104365 | -0.009469322 | 0.088721815 | -0.106730486 | 0.915002796 | 0.959140337 | | 1700066J24Rik | 3.248001682 | -0.00854074629086581 | 0.0344696050162269 | -0.247776157772773 | 0.804307603141091 | NA | NR_045726 |
| Atp6v1h | 6305.724868 | 0.006648692 | 0.062664232 | 0.106100279 | 0.91550279 | 0.959270321 | | 1700066N21Rik | 0 | NA | NA | NA | NA | NA | NR_045924 |
| Cnih4 | 1379.421505 | 0.007194052 | 0.067867299 | 0.106001743 | 0.915580969 | 0.959270321 | | 1700066O22Rik | 0 | NA | NA | NA | NA | NA | NR_015541 |
| Gm14420 | 1019.792743 | -0.008980748 | 0.084423044 | -0.106377918 | 0.915282513 | 0.959270321 | | 1700067G17Rik | 0 | NA | NA | NA | NA | NA | NR_040471 |
| Ndufa10 | 4163.439181 | -0.005384895 | 0.050732325 | -0.106143272 | 0.915468679 | 0.959270321 | | 1700067K01Rik | 1.205583332 | -0.00152956731310843 | 0.0243380238530726 | -0.0628468162551876 | 0.9498884855448 | NA | NM_183097 |
| Nfatc4 | 74.22616369 | -0.017285484 | 0.163147702 | -0.105949906 | 0.915622098 | 0.959270321 | | 1700067P10Rik | 1.662250936 | -0.00290593153422718 | 0.0290328604530801 | -0.100091120505451 | 0.920271984742096 | NA | NM_026625 |
| Tctex1d1 | 8.886442323 | 0.016591308 | 0.156567577 | 0.105968992 | 0.915606955 | 0.959270321 | | 1700069L16Rik | 0.21491491 | 0.00699378003108193 | 0.00986423294990756 | 0.709003940458184 | 0.478322031551065 | NA | NR_033216 |
| Thap2 | 453.9801629 | 0.009895664 | 0.093111503 | 0.106277569 | 0.915362128 | 0.959270321 | | 1700069P05Rik | 0 | NA | NA | NA | NA | NA | NR_040527 |
| Yod1 | 357.2769674 | 0.009250261 | 0.08689027 | 0.106459114 | 0.915218093 | 0.959270321 | | 1700071K01Rik | 0.09572496 | 0.00498255289994511 | 0.00847999972823692 | 0.587565219295241 | 0.556824162160132 | NA | NM_001033765 |
| Axin1 | 1125.106509 | 0.008208521 | 0.078036638 | 0.105188043 | 0.916226598 | 0.959773843 | | 1700071M16Rik | 0.506412023 | -0.00476029410615358 | 0.0144818618101892 | -0.328707328418527 | 0.742376914602839 | NA | NR_045445,NR_045444 |
| Ppp1r16b | 8477.186128 | 0.009375605 | 0.089082916 | 0.105245822 | 0.916180752 | 0.959773843 | | 1700072B07Rik | 0 | NA | NA | NA | NA | NA | NR_040727 |
| Ap1m1 | 2427.713154 | 0.007119125 | 0.067756201 | 0.105069719 | 0.916320487 | 0.959782428 | | 1700072O05Rik | 0 | NA | NA | NA | NA | NA | NR_045733 |
| Fgf2 | 21.90388538 | -0.017633163 | 0.168080348 | -0.104909128 | 0.916447916 | 0.959782428 | | 1700073E17Rik | 0.448266046 | 0.00696861170553033 | 0.0141123204466955 | 0.493796305990352 | 0.621450042256624 | NA | NR_003625 |
| Rras | 295.6393632 | 0.010860285 | 0.103564 | 0.104865449 | 0.916482576 | 0.959782428 | | 1700074P13Rik | 0 | NA | NA | NA | NA | NA | NM_028550 |
| Tmem216 | 189.8635601 | 0.012106077 | 0.115398251 | 0.104906934 | 0.916449657 | 0.959782428 | | 1700080E11Rik | 0 | NA | NA | NA | NA | NA | NM_028562 |
| Pcdhga9 | 438.3674225 | 0.010783577 | 0.102963523 | 0.104732015 | 0.916588458 | 0.959828438 | | 1700080N15Rik | 0 | NA | NA | NA | NA | NA | NR_040500 |
| Cxadr | 941.1407024 | 0.009268007 | 0.08856428 | 0.104647239 | 0.916655729 | 0.959834011 | | 1700080O16Rik | 0.120209049 | -0.00674972139751468 | 0.00863521114606269 | -0.781650996523957 | 0.434419707908266 | NA | NM_028851 |
| Gdpgp1 | 341.3943283 | 0.009358514 | 0.089682532 | 0.10435158 | 0.916890346 | 0.960014801 | | 1700081H04Rik | 0 | NA | NA | NA | NA | NA | NR_040693 |
| Atic | 1135.888063 | 0.006858914 | 0.06590389 | 0.104074497 | 0.91711023 | 0.96018014 | | 1700081H22Rik | 0 | NA | NA | NA | NA | NA | NR_045928 |
| Zfp27 | 727.8884507 | 0.008922646 | 0.085836934 | 0.103948794 | 0.917209985 | 0.960219696 | | 1700084E18Rik | 8.641373437 | -0.020684456721418 | 0.0555043825558112 | -0.372663486538549 | 0.709398915434901 | NA | NR_028299 |
| Espl1 | 51.19082012 | -0.017126746 | 0.165379119 | -0.103560511 | 0.917518127 | 0.960447004 | | 1700084F23Rik | 0 | NA | NA | NA | NA | NA | NR_045965 |
| Rprd2 | 2219.766922 | -0.006812227 | 0.065806551 | -0.103518979 | 0.917551087 | 0.960447004 | | 1700084J12Rik | 0 | NA | NA | NA | NA | NA | NR_033608 |

| | | | | | | | | | | | | | | | |
|---|---|---|---|---|---|---|---|---|---|---|---|---|---|---|---|
| Tmem132b | 7073.334779 | -0.009188575 | 0.088966727 | -0.103281028 | 0.917739932 | 0.960579782 | | 1700085B03Rik | 0 | NA | NA | NA | NA | NA | NR_038127,NR_038128,NR_038129 |
| Gdi2 | 8218.755862 | 0.005240927 | 0.050884374 | 0.102996789 | 0.917965519 | 0.960750997 | | 1700085C21Rik | 0.212297216 | 0.00681050119636477 | 0.00973439053551308 | 0.699633035218656 | 0.484156506270654 | NA | NR_046045 |
| Osgin2 | 605.3389432 | -0.008976945 | 0.08777697 | -0.102269935 | 0.918542418 | 0.96128985 | | 1700091E21Rik | 2.624174888 | -0.0302277382762113 | 0.0371254123246842 | -0.814206129533362 | 0.415526874606368 | NA | NR_045366 |
| Ccdc104 | 2972.470668 | -0.005938486 | 0.058213307 | -0.102012514 | 0.918746742 | 0.961396449 | | 1700091H14Rik | 9.518185756 | 0.0114212875294971 | 0.0639420928206048 | 0.178619232272246 | 0.858236692179302 | NA | NR_073362 |
| Spopl | 335.5293538 | 0.008211547 | 0.080516996 | 0.101985258 | 0.918768376 | 0.961396449 | | 1700092C02Rik | 0.106347839 | -0.00653347234140708 | 0.00854389509723638 | -0.764694822098226 | 0.444453290197145 | NA | NR_045467 |
| Mpc1 | 2852.030273 | -0.004863468 | 0.047799363 | -0.101747552 | 0.918957057 | 0.961504568 | | 1700092C10Rik | 0 | NA | NA | NA | NA | NA | NR_045931 |
| Nrn1 | 5992.384935 | 0.009717182 | 0.09554871 | 0.101698725 | 0.918995815 | 0.961504568 | | 1700092E19Rik | 0 | NA | NA | NA | NA | NA | NR_045933 |
| Trappc1 | 1036.212537 | -0.007871819 | 0.077476097 | -0.101603197 | 0.919071642 | 0.961518975 | | 1700092K14Rik | 2.606155627 | -0.0228289615344235 | 0.0363641047352998 | -0.627788356144039 | 0.530142594876859 | NA | NR_045930 |
| Clint1 | 2080.405397 | 0.007505495 | 0.074186746 | 0.101170298 | 0.919415274 | 0.961813535 | | 1700092M07Rik | 2.524417425 | 0.00568986871118847 | 0.0347750640784674 | 0.163619215721636 | 0.870030916613847 | NA | NM_001177347 |
| Mknk2 | 1738.556951 | 0.009457847 | 0.093587666 | 0.101058688 | 0.919503872 | 0.961841277 | | 1700093K21Rik | 4.259849704 | -0.0018530070265066 | 0.0454355804491435 | -0.0407831705502405 | 0.967468756151315 | NA | NM_001110133,NM_026105 |
| Dync1li1 | 2535.881072 | 0.006003032 | 0.059500814 | 0.100889922 | 0.919637843 | 0.961851542 | | 1700094J05Rik | 1.091849934 | -0.00395625916141542 | 0.0201804737179647 | -0.196043919320563 | 0.844575790270937 | NA | NR_040580 |
| Prss50 | 20.46506222 | 0.016958587 | 0.168081258 | 0.100895167 | 0.919633679 | 0.961851542 | | 1700094M24Rik | 0 | NA | NA | NA | NA | NA | NR_046049 |
| Dscr3 | 1303.525707 | -0.005959997 | 0.059183547 | -0.100703618 | 0.919785739 | 0.961876366 | | 1700095A21Rik | 0 | NA | NA | NA | NA | NA | NR_045468 |
| Rnf10 | 9304.627882 | 0.006490959 | 0.064417743 | 0.100763529 | 0.919738179 | 0.961876366 | | 1700095B10Rik | 0.113843286 | 0.00525947738444006 | 0.00859368002043176 | 0.612016897526494 | 0.540526577873086 | NA | NR_040675,NR_040676 |
| Pibf1 | 383.5208723 | -0.010104772 | 0.10067556 | -0.100369668 | 0.92005085 | 0.962082111 | | 1700096J18Rik | 0.43832756 | 0.0192156728843132 | 0.016298340988036 | 1.17899563510291 | 0.238399913921073 | NA | NR_027883 |
| Zfp560 | 265.8422889 | -0.010844311 | 0.108119445 | -0.100299356 | 0.920106669 | 0.962082111 | | 1700097N02Rik | 0.12890128 | -0.0068640067066208 | 0.00870804083610955 | -0.788237771940376 | 0.430557639008166 | NA | NR_045287 |
| Tspo | 101.1323323 | -0.012317522 | 0.123032321 | -0.100116144 | 0.920252119 | 0.962169265 | | 1700100L14Rik | 0 | NA | NA | NA | NA | NA | NR_045934 |
| Gt(ROSA)26Sor | 100.8166139 | 0.014085894 | 0.141035047 | 0.099875133 | 0.920443459 | 0.962304383 | | 1700101E01Rik | 10.5973199 | 0.0169239805149632 | 0.0581885167054611 | 0.290847429581838 | 0.771168009010344 | NA | NM_001085514,NM_001166705 |
| Rhobtb3 | 921.3101975 | 0.011464106 | 0.115027427 | 0.099664107 | 0.920610997 | 0.962411754 | | 1700101O22Rik | 0 | NA | NA | NA | NA | NA | NR_045045 |
| Sphk1 | 148.0795913 | 0.01553883 | 0.156029109 | 0.099589299 | 0.92067039 | 0.962411754 | | 1700102H20Rik | 4.676830309 | -0.0822583066673321 | 0.048456094869175 | -1.69758431605804 | 0.0895862450486202 | NA | NR_045302,NR_045303 |
| 3110082I17Rik | 277.8307144 | -0.009020973 | 0.090904605 | -0.099235602 | 0.920951208 | 0.962640357 | | 1700104A03Rik | 0.12890128 | -0.0068640067066208 | 0.00870804083610955 | -0.788237771940376 | 0.430557639008166 | NA | NR_045805 |
| Gm5077 | 9.747069564 | 0.015290097 | 0.154367877 | 0.099049733 | 0.921098783 | 0.962729664 | | 1700105P06Rik | 4.320698374 | -0.0638822725872685 | 0.0496547465083777 | -1.28652902450102 | 0.198258496033346 | NA | NR_045703 |
| Atg2b | 2937.258318 | -0.009752592 | 0.098692747 | -0.098817713 | 0.921283004 | 0.962792315 | | 1700106J16Rik | 0 | NA | NA | NA | NA | NA | NM_028859 |
| Rad21 | 4971.8104 | 0.006053248 | 0.061247519 | 0.098832538 | 0.921271234 | 0.962792315 | | 1700108F19Rik | 0.727270497 | 0.0079199612099441 | 0.0180887901047817 | 0.437838084474786 | 0.661503661118593 | NA | NR_015485 |
| Dsc3 | 20.36601679 | -0.016509183 | 0.167599826 | -0.098503581 | 0.921532429 | 0.962988025 | | 1700108J01Rik | 0 | NA | NA | NA | NA | NA | NR_015532 |
| 4930519G04Rik | 9.315846225 | -0.015921966 | 0.162048462 | -0.098254345 | 0.92173033 | 0.963129871 | | 1700109F18Rik | 0 | NA | NA | NA | NA | NA | NR_027899 |
| Ankrd11 | 8780.980855 | -0.007471755 | 0.07650391 | -0.097665012 | 0.922198299 | 0.963164169 | | 1700109G14Rik | 0.76626644 | 0.00898974185599649 | 0.0179364997362135 | 0.501198226421309 | 0.616231622388488 | NA | NR_033788 |
| Arsi | 19.21450399 | -0.016419574 | 0.167564068 | -0.097989825 | 0.921940373 | 0.963164169 | | 1700109G15Rik | 0 | NA | NA | NA | NA | NA | NR_046197 |
| Dtnbp1 | 1879.735553 | 0.009495668 | 0.097038617 | 0.09785453 | 0.922047806 | 0.963164169 | | 1700109I08Rik | 0 | NA | NA | NA | NA | NA | NR_045936 |
| Rad54l | 22.45863722 | 0.016456679 | 0.168055593 | 0.097924019 | 0.921992628 | 0.963164169 | | 1700110C19Rik | 19.55653517 | -0.08998780537591 | 0.0805497437179032 | -1.1171705982214 | 0.263921389353705 | NA | NR_045461 |
| Scrn2 | 107.1911142 | -0.013495625 | 0.137897583 | -0.097867014 | 0.922037893 | 0.963164169 | | 1700110I01Rik | 0 | NA | NA | NA | NA | NA | NR_038059 |
| Slc25a47 | 72.36580198 | 0.012846491 | 0.131500273 | 0.097691743 | 0.922177072 | 0.963164169 | | 1700110K17Rik | 1.139995443 | -0.0124468788676632 | 0.0204601067563447 | -0.608348676568044 | 0.542956243317571 | NA | NR_040728 |
| Zfp143 | 458.6307551 | -0.008637859 | 0.088418566 | -0.09769282 | 0.922176217 | 0.963164169 | | 1700110M21Rik | 0.120209049 | -0.00674972139751468 | 0.00863521114606269 | -0.781650996523957 | 0.434419707908266 | NA | NM_029680 |
| Alkbh6 | 922.1739909 | 0.007941385 | 0.081506005 | 0.097433125 | 0.92238244 | 0.963291556 | | 1700111N16Rik | 0 | NA | NA | NA | NA | NA | NR_033213 |
| Alkbh4 | 209.1106547 | -0.010273647 | 0.105537529 | -0.097345913 | 0.922451696 | 0.963298954 | | 1700112E06Rik | 14.45170484 | -0.0694443453889413 | 0.0659279946165641 | -1.05333623133584 | 0.292186919076533 | NA | NM_028275 |
| Ankrd52 | 3504.184953 | -0.005982595 | 0.06152692 | -0.097235404 | 0.922539453 | 0.96332567 | | 1700112H15Rik | 0 | NA | NA | NA | NA | NA | NR_040472 |
| Angpt2 | 70.5707085 | 0.015714696 | 0.161792786 | 0.097128533 | 0.922624322 | 0.963349366 | | 1700112J05Rik | 0 | NA | NA | NA | NA | NA | NR_077218 |
| Chek2 | 50.25845472 | -0.014473256 | 0.149273114 | -0.096958223 | 0.922759571 | 0.96342566 | | 1700113H08Rik | 0.530784281 | 0.00237221600747039 | 0.0147292791801344 | 0.161054453409359 | 0.872050512762703 | NA | NM_029685 |
| Zfp109 | 308.1359012 | 0.008482579 | 0.087601469 | 0.09683147 | 0.922860232 | 0.963465833 | | 1700119H24Rik | 0.242744565 | -0.00126183740302166 | 0.00996759682058903 | -0.126593944933168 | 0.899261789625903 | NA | NR_040536 |
| Rpl18a | 6791.102507 | -0.007235561 | 0.074803647 | -0.096727377 | 0.922942898 | 0.963487216 | | 1700120C14Rik | 16.1201867 | -0.0269953961597932 | 0.0649245011654205 | -0.415796743528485 | 0.677558741979808 | NA | NR_045627 |
| Ino80e | 909.081151 | -0.006361709 | 0.065851218 | -0.096607311 | 0.92303825 | 0.963521839 | | 1700120E14Rik | 0 | NA | NA | NA | NA | NA | NR_045368 |
| Avp | 98.95776403 | 0.010308915 | 0.107210402 | 0.09615592 | 0.923396739 | 0.963701272 | | 1700120G07Rik | 0.396404207 | -0.00976909470369045 | 0.013847504194659 | -0.705476926842774 | 0.480513485844527 | NA | NR_046050 |
| BC016423 | 1398.750865 | -0.007401878 | 0.076917132 | -0.096231855 | 0.923336431 | 0.963701272 | | 1700120K04Rik | 1.640092612 | -0.000247990671374761 | 0.0248365851114786 | -0.00998489406903804 | 0.992033339558113 | NA | NR_027915 |
| Det1 | 134.4439488 | -0.01234593 | 0.128268653 | -0.096250559 | 0.923321577 | 0.963701272 | | 1700121L16Rik | 0 | NA | NA | NA | NA | NA | NR_045453 |
| Tmem202 | 67.22361334 | 0.013716598 | 0.142861517 | 0.096013249 | 0.92351005 | 0.963754611 | | 1700121N20Rik | 0.459875358 | -0.0114023571436737 | 0.0144430867924611 | -0.789468159232103 | 0.429838432600178 | NA | NR_036593 |
| Hpdl | 38.20390556 | -0.01552926 | 0.161988137 | -0.095866648 | 0.923626483 | 0.963811202 | | 1700122O11Rik | 0.127687691 | 0.00542298404407807 | 0.00869724871942359 | 0.62352868349814 | 0.532937159177571 | NA | NM_029689 |
| Cenpe | 63.03364068 | -0.014939392 | 0.156007101 | -0.095760974 | 0.923710413 | 0.963833871 | | 1700123I01Rik | 1.677439366 | -0.0234061812894186 | 0.0284867152208706 | -0.821652517952306 | 0.411274690185843 | NA | NM_001165919 |
| Atl1 | 4497.646216 | 0.007135811 | 0.074894402 | 0.095278296 | 0.924093782 | 0.964025796 | | 1700123K08Rik | 1.287534481 | 0.0144890689248764 | 0.0284654339439756 | 0.50900572790821 | 0.610748209195957 | NA | NM_029693 |
| Egf | 53.65474773 | -0.015608856 | 0.16393095 | -0.095216044 | 0.924143227 | 0.964025796 | | 1700123L14Rik | 2.114527486 | -0.00357395973354053 | 0.0307633278418907 | -0.116175979136881 | 0.907513073698696 | NA | NR_003643 |
| Fsd1 | 1439.695119 | 0.006182816 | 0.064798681 | 0.095415774 | 0.923984587 | 0.964025796 | | 1700123O12Rik | 0 | NA | NA | NA | NA | NA | NR_045185 |
| Gtf3c1 | 6529.694508 | 0.007323265 | 0.076850026 | 0.095292947 | 0.924082145 | 0.964025796 | | 1700123O21Rik | 0.21100899 | -0.000459554483117493 | 0.00972904351832043 | -0.0472353199214416 | 0.96232567770125 | NA | NR_045799 |
| Akap12 | 1310.823654 | 0.011906066 | 0.125395172 | 0.094948362 | 0.924355843 | 0.964182673 | | 1700125G02Rik | 0.429381296 | -0.0100524906077422 | 0.0141519698525479 | -0.71032447867548 | 0.477502943458686 | NA | NR_040651 |
| Sec16a | 3243.065085 | 0.007692294 | 0.081173332 | 0.094763807 | 0.924502435 | 0.964270665 | | 1700125G22Rik | 0.481442367 | -0.00116566404370014 | 0.0141552163455238 | -0.0823487268047827 | 0.934369407578898 | NA | NR_040548 |
| Stx8 | 680.4846079 | 0.005976103 | 0.063116102 | 0.094684279 | 0.924565605 | 0.96427164 | | 1700125H03Rik | 0.620756173 | -0.015134952006537 | 0.0158687890016453 | -0.953755954847455 | 0.34020718921476 | NA | NR_038181,NR_038189 |
| Chn1 | 43142.48266 | -0.007424474 | 0.078715853 | -0.094319935 | 0.924855015 | 0.964378734 | | 1700125H20Rik | 3.248894681 | -0.0278417944611147 | 0.0406458233457028 | -0.684985372895841 | 0.493353151142199 | NA | NM_028589 |
| Fuca2 | 1472.079957 | -0.006831982 | 0.072358263 | -0.094418826 | 0.924776462 | 0.964378734 | | 1700126H18Rik | 0 | NA | NA | NA | NA | NA | NR_040695 |
| Gtpbp8 | 343.9583588 | -0.009479067 | 0.100424836 | -0.094389668 | 0.924799623 | 0.964378734 | | 1700128A07Rik | 0 | NA | NA | NA | NA | NA | NR_045938 |
| Coro2a | 1557.258025 | -0.008966294 | 0.095310079 | -0.094074985 | 0.925049592 | 0.964386918 | | 1700128F08Rik | 0 | NA | NA | NA | NA | NA | NR_033618 |
| Ggta1 | 207.905302 | -0.011497537 | 0.122152207 | -0.094124678 | 0.925010117 | 0.964386918 | | 1700129C05Rik | 0 | NA | NA | NA | NA | NA | NM_026461 |
| Rin3 | 145.4148716 | 0.01389528 | 0.147647147 | 0.094111402 | 0.925020663 | 0.964386918 | | 1810006J02Rik | 0 | NA | NA | NA | NA | NA | NR_040439 |
| 1810010H24Rik | 93.82252502 | 0.012389605 | 0.13213979 | 0.093761346 | 0.925298738 | 0.964581756 | | 1810007C17Rik | 0.108766301 | 0.00519171381361946 | 0.00856003371502575 | 0.606506234257728 | 0.544178643801544 | NA | NR_045472 |
| Akt1 | 6646.963061 | 0.006313688 | 0.06760388 | 0.093392397 | 0.92559183 | 0.964692796 | | 1810007D17Rik | 1.018675763 | -0.0209788678325802 | 0.023870443505993 | -0.878863764190773 | 0.379475147942568 | NA | NR_038136 |
| Gdf9 | 27.08459714 | 0.015665872 | 0.16788384 | 0.093313755 | 0.925654305 | 0.964692796 | | 1810007I06Rik | 0 | NA | NA | NA | NA | NA | NR_040574 |
| Poldip3 | 3166.60469 | -0.004885659 | 0.052246727 | -0.093511291 | 0.92549738 | 0.964692796 | | 1810008I18Rik | 0 | NA | NA | NA | NA | NA | NR_045301 |
| Tceanc2 | 729.536223 | -0.005796336 | 0.062102608 | -0.09333482 | 0.92563757 | 0.964692796 | | 1810009J06Rik | 0 | NA | NA | NA | NA | NA | NM_023707 |
| 1110008P14Rik | 2727.88039 | -0.011954374 | 0.12850458 | -0.093026834 | 0.925882243 | 0.964696902 | | 1810010D01Rik | 5.158765809 | -0.05816270497934 | 0.0502397483193793 | -1.15770295284112 | 0.246985278534204 | NA | NR_033626 |
| 3110001I22Rik | 56.60339567 | 0.014207755 | 0.153037229 | 0.092838556 | 0.92603182 | 0.964696902 | | 1810012K16Rik | 6.28149351 | -0.11494048632599 | 0.0546107204315004 | -2.10472386040325 | 0.0353153522501865 | NA | NR_045473 |
| Gid4 | 3346.752378 | -0.004587189 | 0.049339919 | -0.092971157 | 0.925926475 | 0.964696902 | | 1810013A23Rik | 0 | NA | NA | NA | NA | NA | NR_045427 |
| Heatr1 | 1104.786695 | -0.007506866 | 0.080692305 | -0.093030752 | 0.92587913 | 0.964696902 | | 1810018F18Rik | 0 | NA | NA | NA | NA | NA | NR_038140 |
| Map9 | 5159.779301 | -0.006459223 | 0.069366609 | -0.093117179 | 0.92581047 | 0.964696902 | | 1810019D21Rik | 2.908809968 | -0.0173084871049433 | 0.0314338995863524 | -0.550631239926022 | 0.581886489047297 | NA | NR_040345,NR_040344,NR_040346 |
| Tmem8c | 12.46530337 | -0.014719441 | 0.158524403 | -0.092852842 | 0.92602047 | 0.964696902 | | 1810019J16Rik | 4.635627769 | 0.029321900660787 | 0.0426298780984986 | 0.687825111604523 | 0.491562920547659 | NA | NM_133707,NM_001083916 |
| Ggps1 | 1556.760242 | 0.00655224 | 0.070637933 | 0.092758089 | 0.926095747 | 0.964698636 | | 1810024B03Rik | 17.46279312 | -0.0730487952834514 | 0.0741271599534903 | -0.98545250255486 | 0.324401851232447 | NA | NM_198630 |
| 3110040N11Rik | 200.2479906 | -0.010311439 | 0.111887495 | -0.092158995 | 0.926571716 | 0.964884184 | | 1810033B17Rik | 11.04505102 | -0.0449165581150988 | 0.0648160959421956 | -0.692984627694275 | 0.488319198602862 | NA | NM_026985 |
| Atp6v1g2 | 18419.25597 | 0.00563271 | 0.061011689 | 0.092321818 | 0.926442354 | 0.964884184 | | 1810034E14Rik | 8.11036596 | 0.0418968629359934 | 0.0527623435280911 | 0.794067513579778 | 0.427156128672949 | NA | NR_045798 |
| Fbxw11 | 5840.817051 | 0.005025535 | 0.054393946 | 0.092391446 | 0.926387036 | 0.964884184 | | 1810035I16Rik | 0 | NA | NA | NA | NA | NA | NR_040641 |
| Gnptab | 3552.563981 | -0.005160593 | 0.05597666 | -0.092191876 | 0.926545593 | 0.964884184 | | 1810044D09Rik | 15.16966339 | -0.00714603157065669 | 0.0743017767045741 | -0.0961757832396056 | 0.923380963816472 | NA | NR_038153 |
| Snhg10 | 48.81038557 | -0.014813644 | 0.160769776 | -0.092141968 | 0.926585245 | 0.964884184 | | 1810046K07Rik | 0.441471797 | -0.00252154620662118 | 0.0153626428894375 | -0.164134922927543 | 0.869624930091173 | NA | NM_027217 |
| Bdp1 | 1652.553161 | -0.008134477 | 0.088435951 | -0.091981567 | 0.926712685 | 0.964952039 | | 1810053B23Rik | 0 | NA | NA | NA | NA | NA | NR_040488,NR_040489,NR_040486,NR_040487,NR_040491,NR_040490 |
| Iqsec2 | 8981.57022 | -0.008630984 | 0.094373746 | -0.091455347 | 0.927130785 | 0.965037944 | | 1810062G17Rik | 3.298856266 | -0.0329616644761447 | 0.0401556674732087 | -0.820847131930661 | 0.411733348121287 | NA | NM_028183 |
| Mrpl4 | 1368.996757 | 0.006533102 | 0.071311346 | 0.091613784 | 0.927004899 | 0.965037944 | | 1810064F22Rik | 0.546021118 | 0.0187286890633388 | 0.0160811508703624 | 1.16463611431292 | 0.244166318845605 | NA | NM_027981 |
| Nup50 | 1760.151316 | 0.005786979 | 0.063208041 | 0.091554471 | 0.927052026 | 0.965037944 | | 1810065E05Rik | 0 | NA | NA | NA | NA | NA | NM_027239 |
| Pcsk1n | 7834.327333 | 0.014376645 | 0.1572758 | 0.091410405 | 0.927166495 | 0.965037944 | | 2010001E11Rik | 12.49285768 | -0.0704674654193762 | 0.0738138750361225 | -0.954664219767507 | 0.339747529577256 | NA | NM_001163503 |
| Prcc | 1597.469384 | -0.007103005 | 0.077514137 | -0.091634958 | 0.926988075 | 0.965037944 | | 2010003K11Rik | 0 | NA | NA | NA | NA | NA | NM_027237 |
| Xlr3a | 19.19988855 | 0.013191312 | 0.144313413 | 0.091407386 | 0.927168893 | 0.965037944 | | 2010005H15Rik | 1.140093182 | 0.0318778696170583 | 0.0233971874447357 | 1.36246588152333 | 0.173050904499492 | NA | NM_029733 |
| Haus8 | 237.5863527 | -0.009455107 | 0.103644752 | -0.091226103 | 0.927312934 | 0.965058208 | | 2010009K17Rik | 0 | NA | NA | NA | NA | NA | NR_040609 |
| Wsb2 | 12312.67374 | -0.008687331 | 0.095203949 | -0.091249691 | 0.927294192 | 0.965058208 | | 2010010A06Rik | 0.44014219 | 0.0124040145219905 | 0.0131092631396785 | 0.946202268565852 | 0.34404542930782 | NA | NR_045394,NR_045393 |
| Mipol1 | 93.46203294 | 0.012756516 | 0.140047371 | 0.091087148 | 0.927423344 | 0.965108288 | | 2010016I18Rik | 0.558660649 | 0.00129212023544911 | 0.0165122284328457 | 0.0782523231618368 | 0.937627341715388 | NA | NR_033207 |
| D19Bwg1357e | 1163.727505 | 0.006697117 | 0.073743056 | 0.090816912 | 0.927638072 | 0.965162524 | | 2010106C02Rik | 0.288615623 | 0.00197374456361263 | 0.0106648328298147 | 0.185070370544844 | 0.853173841012254 | NA | NR_045435 |
| Fancc | 326.8671602 | -0.009188663 | 0.101211927 | -0.09078637 | 0.927662341 | 0.965162524 | | 2010106E10Rik | 0.324852276 | 0.00269252691297653 | 0.0110865316558406 | 0.242864675496421 | 0.808110231248329 | NA | NM_001168590,NM_026333 |
| Syt12 | 1819.121522 | -0.007615453 | 0.083766944 | -0.090912385 | 0.927562209 | 0.965162524 | | 2010107G12Rik | 18.20194058 | -0.0870254732398891 | 0.0796845910875328 | -1.09212423697189 | 0.27477849759038 | NA | NM_001025573 |
| 1190007I07Rik | 143.3317598 | 0.011740821 | 0.130194697 | 0.090178949 | 0.928145012 | 0.96528002 | | 2010109A12Rik | 1.611207363 | 0.00903491503798267 | 0.0260531244783857 | 0.346788157615347 | 0.728750474543914 | NA | NM_029363 |
| Aamp | 4706.653545 | -0.007936662 | 0.087951122 | -0.090239464 | 0.928096924 | 0.96528002 | | 2010109I03Rik | 12.0968058 | 0.0203946796371138 | 0.0625206015685304 | 0.32620734806524 | 0.744267482264083 | NA | NM_025929 |
| Cbr4 | 341.7830142 | -0.00829273 | 0.091743701 | -0.090390189 | 0.927977153 | 0.96528002 | | 2010308F09Rik | 10.26049401 | -0.00472438236757157 | 0.0645133631178061 | -0.073231066235758 | 0.94162224546704 | NA | NR_045430,NR_045429 |
| Chst14 | 71.98985693 | 0.01309629 | 0.145058638 | 0.090282728 | 0.928062545 | 0.96528002 | | 2010310C07Rik | 0 | NA | NA | NA | NA | NA | NR_045169 |
| Fam213b | 1812.453392 | 0.007492157 | 0.082922882 | 0.090350898 | 0.928008375 | 0.96528002 | | 2200002D01Rik | 1.306873329 | 0.00402091073934755 | 0.0266929043888503 | 0.15063593982778 | 0.880262908987317 | NA | NM_028179 |
| Vrk2 | 61.55109073 | -0.012921861 | 0.143299449 | -0.090173839 | 0.928149073 | 0.96528002 | | 2200002J24Rik | 0.106148608 | 0.00515554063321497 | 0.00854200771766808 | 0.603551390213729 | 0.546141939141592 | NA | NM_026961 |
| Ctsk | 133.4660094 | -0.011386127 | 0.126539442 | -0.08998085 | 0.928302433 | 0.965309926 | | 2210010C04Rik | 0.453828193 | 0.016316890065186 | 0.0150244471794912 | 1.08602265828849 | 0.277468967825936 | NA | NM_023333 |
| Unc119 | 794.7407132 | 0.006478043 | 0.071961311 | 0.090021195 | 0.928270372 | 0.965309926 | | 2210019I11Rik | 0 | NA | NA | NA | NA | NA | NR_038157 |
| 4930539J05Rik | 20.49971388 | 0.015093239 | 0.167997421 | 0.089842088 | 0.928412702 | 0.965352267 | | 2210039B01Rik | 16.34856465 | -0.00410798147368475 | 0.0720512855607894 | -0.0570146867153239 | 0.95453349586519 | NA | NR_044985 |
| Ptchd2 | 1779.008469 | 0.006707008 | 0.074710907 | 0.089772803 | 0.928467761 | 0.965352267 | | 2210404O07Rik | 20.4099799 | 0.130286000696853 | 0.0810745053107136 | 1.60699100410837 | 0.108056341510561 | NA | NM_001099917 |
| Fam155a | 3259.964307 | -0.004650443 | 0.051887305 | -0.089625842 | 0.928584548 | 0.96540891 | | 2210407C18Rik | 1.070974855 | 0.0309279962700555 | 0.0237929824899231 | 1.29987891527067 | 0.193642472676736 | NA | NM_144544 |
| Nos1ap | 3410.102435 | 0.0093835 | 0.104803737 | 0.089534021 | 0.928657517 | 0.965419992 | | 2210409D07Rik | 1.583313647 | 0.027043396415274 | 0.0276684320073251 | 0.977409793518996 | 0.328366316449997 | NA | NR_045360 |
| Aftph | 3434.620288 | -0.005519066 | 0.061823566 | -0.089271233 | 0.928866356 | 0.965507535 | | 2210409E12Rik | 0 | NA | NA | NA | NA | NA | NR_003492 |
| Sox10 | 1522.089714 | 0.0114482 | 0.12813357 | 0.089345827 | 0.928807075 | 0.965507535 | | 2210414B05Rik | 0 | NA | NA | NA | NA | NA | NR_040643 |
| 9130221H12Rik | 277.362702 | 0.008514152 | 0.095823147 | 0.088852765 | 0.929198923 | 0.965521426 | | 2210416O15Rik | 2.824126492 | 0.0230294748350693 | 0.0393981458979351 | 0.584531944592762 | 0.558862479677697 | NA | NR_045499 |
| F730043M19Rik | 274.6463471 | 0.01003784 | 0.113015163 | 0.088818523 | 0.929226137 | 0.965521426 | | 2210417A02Rik | 0.430785401 | -0.0007334963684434 | 0.0137568789005558 | -0.0533185160489976 | 0.957478127511317 | NA | NR_028285 |
| Mcrs1 | 2498.203394 | -0.005879474 | 0.066222259 | -0.088783949 | 0.929253614 | 0.965521426 | | 2210417K05Rik | 13.07325707 | -0.0445201110103168 | 0.0690891254621721 | -0.644386663060202 | 0.51932473078872 | NA | NR_038038 |
| Pgm2 | 2626.346154 | -0.007514407 | 0.084429805 | -0.089001825 | 0.92908046 | 0.965521426 | | 2210420H20Rik | 0 | NA | NA | NA | NA | NA | NR_045389 |
| Synpo | 10628.55799 | -0.009221126 | 0.103665857 | -0.08895046 | 0.929121282 | 0.965521426 | | 2300002M23Rik | 1.063647385 | 0.00908224865538348 | 0.0212300305739807 | 0.427801958350196 | 0.668795311038778 | NA | NM_175148 |
| Wipi2 | 5049.482316 | 0.005763065 | 0.064846973 | 0.088871765 | 0.929183823 | 0.965521426 | | 2300003K06Rik | 0 | NA | NA | NA | NA | NA | NM_001195383 |
| Zfp809 | 377.5169647 | 0.008850475 | 0.100041454 | 0.08846808 | 0.929504653 | 0.965717501 | | 2300005B03Rik | 0.382899838 | 0.00820727460669677 | 0.0106737841638539 | 0.768918921415912 | 0.441941444022319 | NA | NM_001081961 |
| Henmt1 | 50.54880876 | 0.013676025 | 0.154734011 | 0.08838409 | 0.929571406 | 0.965722098 | | 2310001H17Rik | 6.814855266 | 0.00512749891530342 | 0.0535458582933893 | 0.0957590200012996 | 0.923711965034696 | NA | NR_040266,NR_040265,NR_040268 |
| 9330179D12Rik | 79.86706808 | -0.013741492 | 0.155928615 | -0.088126813 | 0.929775885 | 0.965740267 | | 2310001K24Rik | 0.543831507 | 0.0105450399825839 | 0.0120876531324133 | 0.87238108730198 | 0.383000514602336 | NA | NR_028122 |
| Mypop | 1387.588612 | 0.005771083 | 0.065476182 | 0.088140183 | 0.929765259 | 0.965740267 | | 2310002D06Rik | 0.489410913 | 0.00565929899362522 | 0.0126443813664561 | 0.447574209414358 | 0.654460519707199 | NA | NR_045490 |
| Ythdf2 | 1410.257702 | -0.006109565 | 0.069315558 | -0.088141326 | 0.929764351 | 0.965740267 | | 2310002F09Rik | 9.331062062 | -0.105987334101773 | 0.0596798508107032 | -1.77593161950005 | 0.0757441885517027 | NA | NR_077063 |
| Gpr156 | 47.35591155 | -0.013033328 | 0.148409143 | -0.087820252 | 0.930019541 | 0.965928594 | | 2310002J15Rik | 1.420358485 | 0.0159486233345523 | 0.028441098745837 | 0.560759746909804 | 0.574961331004079 | NA | NM_026415 |
| Kcng3 | 65.71152108 | 0.013937824 | 0.159380392 | 0.087450055 | 0.930313784 | 0.965975189 | | 2310002L09Rik | 0.12890128 | -0.0068640067066208 | 0.00870804083610955 | -0.788237771940376 | 0.430557639008166 | NA | NM_027104,NM_027981 |
| Mir132 | 12.86860148 | 0.012978124 | 0.148302017 | 0.087511447 | 0.930264987 | 0.965975189 | | 2310005A03Rik | 0 | NA | NA | NA | NA | NA | NR_040634 |
| Onecut2 | 208.9002343 | 0.011188251 | 0.127801619 | 0.087543893 | 0.930239197 | 0.965975189 | | 2310005E17Rik | 0 | NA | NA | NA | NA | NA | NR_045498 |
| Vgll4 | 419.589731 | -0.006364854 | 0.072625522 | -0.087639362 | 0.930163316 | 0.965975189 | | 2310005G13Rik | 1.636182147 | -0.028999592397335 | 0.0273072141487659 | -1.06197549993014 | 0.288246807755297 | NA | NM_183281 |
| Bcl6 | 1234.413767 | -0.011561643 | 0.132451699 | -0.0872895 | 0.9304414 | 0.965986374 | | 2310007B03Rik | 2.633311402 | 0.00354501546530127 | 0.0331744097656202 | 0.106859940850405 | 0.91490009372978 | NA | NM_172411,NM_001159940 |
| Spryd3 | 6716.414232 | 0.006870055 | 0.078713152 | 0.087279627 | 0.930449248 | 0.965986374 | | 2310007L24Rik | 6.803695725 | 0.0275666473466885 | 0.0561759074275982 | 0.490720107765372 | 0.623624422698185 | NA | NM_029345 |
| Actr1a | 8736.460976 | 0.004481038 | 0.051428965 | 0.087130625 | 0.930567683 | 0.966044602 | | 2310008N11Rik | 0.212695677 | -0.00857796107890413 | 0.00973699866180651 | -0.880965621629515 | 0.378336427605205 | NA | NR_045904 |
| Adnp | 5772.486053 | -0.004839853 | 0.055780969 | -0.086765304 | 0.930858067 | 0.966194882 | | 2310010M20Rik | 12.40052891 | -6.50966087391909e-05 | 0.0732776987118612 | -0.000888354982259479 | 0.999291195368371 | NA | NM_183283 |
| Lrp5 | 510.8987494 | -0.008287535 | 0.095574138 | -0.086713152 | 0.930899522 | 0.966194882 | | 2310014F07Rik | 1.934979665 | -0.0208204084104561 | 0.0330306615516811 | -0.63033579808505 | 0.528474906218574 | NA | NR_040714 |
| Pik3cd | 1209.048638 | 0.006330809 | 0.072911159 | 0.086829084 | 0.930807369 | 0.966194882 | | 2310015B20Rik | 3.427190836 | 0.00353826657113116 | 0.0403326682041745 | 0.087727064156022 | 0.930093608461971 | NA | NR_038041 |
| Lrriq3 | 16.26361541 | 0.01407704 | 0.163359225 | 0.086172298 | 0.931329452 | 0.966576363 | | 2310015D24Rik | 7.381563182 | -0.103422586001764 | 0.0593097783476294 | -1.74376955846266 | 0.0811992806648093 | NA | NR_037997 |
| D130040H23Rik | 98.53539653 | -0.012356729 | 0.144169568 | -0.085709691 | 0.931697199 | 0.966893262 | | 2310016D03Rik | 0.127687691 | 0.00542298404407807 | 0.00869724871942359 | 0.62352868349814 | 0.532937159177571 | NA | NR_045491 |
| Eri3 | 4486.789116 | 0.005750613 | 0.067439399 | 0.085270818 | 0.932046091 | 0.967125065 | | 2310020H05Rik | 0 | NA | NA | NA | NA | NA | NR_045495 |
| Kcnu1 | 127.0979844 | -0.012084503 | 0.141793856 | -0.085225855 | 0.932081836 | 0.967125065 | | 2310030A07Rik | 0.127687691 | 0.00542298404407807 | 0.00869724871942359 | 0.62352868349814 | 0.532937159177571 | NA | NR_040603 |
| Pafah2 | 310.1529266 | -0.008299969 | 0.09742529 | -0.085193168 | 0.932107822 | 0.967125065 | | 2310034C09Rik | 0 | NA | NA | NA | NA | NA | NM_054100 |
| Vps26b | 5657.925355 | -0.004826864 | 0.056807239 | -0.084969177 | 0.932285896 | 0.967245056 | | 2310034G01Rik | 16.25403758 | -0.0293697068039733 | 0.0772917345931556 | -0.379985091013523 | 0.703956482239361 | NA | NR_040418 |
| Prima1 | 173.4761502 | 0.009171923 | 0.108782303 | 0.084314478 | 0.932806401 | 0.96772028 | | 2310034O05Rik | 4.645129841 | -0.0305917181339266 | 0.0456439818831052 | -0.670224570070866 | 0.502714643317622 | NA | NR_040679 |
| 1600012H06Rik | 527.922023 | -0.00771546 | 0.09180318 | -0.084043492 | 0.933021852 | 0.967755144 | | 2310039L15Rik | 20.82079808 | -0.00101601899907666 | 0.0840070274815652 | -0.0120944524468458 | 0.990350258375785 | NA | NR_045337 |
| Ctnna3 | 70.19221622 | -0.013040146 | 0.155172356 | -0.084036531 | 0.933027387 | 0.967755144 | | 2310042E22Rik | 12.3423468 | 0.00210832425813311 | 0.0705968006682664 | 0.0298643031720389 | 0.976175275084982 | NA | NM_025634 |
| Eif4g1 | 11861.20377 | 0.004938552 | 0.058677818 | 0.084163872 | 0.932926142 | 0.967755144 | | 2310043L19Rik | 0.875876633 | 0.00645570492003864 | 0.0223130538603817 | 0.289324131086385 | 0.772333347653152 | NA | NR_037994 |
| 4932441J04Rik | 23.10635807 | 0.013840736 | 0.167442492 | 0.082659638 | 0.934122179 | 0.967854022 | | 2310043O21Rik | 0 | NA | NA | NA | NA | NA | NR_045757 |
| BC004004 | 2056.246026 | -0.005113492 | 0.061846985 | -0.082679723 | 0.934106208 | 0.967854022 | | 2310050B05Rik | 9.135076595 | -0.0680164657912292 | 0.0540827074268094 | -1.25763795910692 | 0.208522718814503 | NA | NR_015477 |
| Cyb5r1 | 738.7952058 | 0.008211656 | 0.098840666 | 0.083079727 | 0.933788145 | 0.967854022 | | 2310050C09Rik | 0 | NA | NA | NA | NA | NA | NM_025621 |
| Glt28d2 | 13.25781499 | -0.013801469 | 0.164878393 | -0.083706962 | 0.933289422 | 0.967854022 | | 2310057J18Rik | 0.09572496 | 0.00498255289994511 | 0.00847999972823692 | 0.587565219295241 | 0.556824162160132 | NA | NM_026336 |
| Mrgprf | 44.801966 | -0.013951182 | 0.167947905 | -0.083068507 | 0.933797067 | 0.967854022 | | 2310057N15Rik | 0 | NA | NA | NA | NA | NA | NM_027170 |
| Pank2 | 2071.923451 | 0.006394251 | 0.077202377 | 0.08282453 | 0.933991064 | 0.967854022 | | 2310061N02Rik | 0 | NA | NA | NA | NA | NA | NM_027155 |
| Papss1 | 2600.48444 | -0.004154017 | 0.049698383 | -0.083584556 | 0.933386747 | 0.967854022 | | 2310065F04Rik | 3.374394011 | 0.0121607501481583 | 0.0421507578826637 | 0.288506085276343 | 0.772959373080764 | NA | NR_038055 |
| Phip | 1864.166766 | -0.007299971 | 0.08769089 | -0.083246627 | 0.933655438 | 0.967854022 | | 2310068J16Rik | 5.449938812 | -0.00947740778138532 | 0.0456799757723145 | -0.207474010683021 | 0.835639687554797 | NA | NR_028124 |
| Sgol2 | 41.79432965 | -0.013154784 | 0.158906707 | -0.082783062 | 0.934024038 | 0.967854022 | | 2310069B03Rik | 0.237667581 | -0.00136938695604711 | 0.00989612066541175 | -0.13837613771559 | 0.889943154921195 | NA | NR_040520 |
| Slc4a1ap | 1124.905011 | 0.00564399 | 0.067781506 | 0.083267411 | 0.933638912 | 0.967854022 | | 2310079G19Rik | 0 | NA | NA | NA | NA | NA | NM_027173 |
| Smpd3 | 4604.49502 | 0.007164874 | 0.085597611 | 0.083704133 | 0.933291671 | 0.967854022 | | 2310081J21Rik | 0.192890664 | -0.000903864895041124 | 0.00963528282151475 | -0.093807821917056 | 0.92526181809679 | NA | NR_045474 |
| Stk4 | 2041.696668 | 0.006361011 | 0.076279282 | 0.083391066 | 0.933540592 | 0.967854022 | | 2410003L11Rik | 2.235867929 | 0.018966769357678 | 0.0343419099488934 | 0.552292210477046 | 0.580748166588827 | NA | NR_045496 |
| Tinagl1 | 328.052975 | -0.01167691 | 0.140865486 | -0.082894045 | 0.93393579 | 0.967854022 | | 2410004I01Rik | 14.04308908 | 0.0287578775603905 | 0.0744245871447072 | 0.386402916881155 | 0.699198290205441 | NA | NR_037963 |
| Ttc23 | 137.6945784 | -0.010284783 | 0.123023238 | -0.083600327 | 0.933374207 | 0.967854022 | | 2410007B07Rik | 0.322281847 | -0.00648722354265905 | 0.0126857541512189 | -0.511378627185182 | 0.609085957750446 | NA | NR_040539 |
| Urgcp | 3087.921149 | -0.00587621 | 0.070483796 | -0.083369661 | 0.933557611 | 0.967854022 | | 2410012E07Rik | 0 | NA | NA | NA | NA | NA | NR_045939 |
| Vmn2r-ps129 | 13.95260298 | -0.013731065 | 0.165510829 | -0.08296173 | 0.933881969 | 0.967854022 | | 2410012M07Rik | 0.424793663 | 0.00460829551054132 | 0.0121338282724052 | 0.379789082809217 | 0.704101986892093 | NA | NM_028033 |
| Dcaf5 | 4175.404969 | -0.006379166 | 0.077260758 | -0.0825667 | 0.93419608 | 0.967865869 | | 2410017I17Rik | 0.217532603 | 0.00684685248201087 | 0.00976000296730706 | 0.701521557416086 | 0.482977586094298 | NA | NR_033517 |
| Chp1 | 6534.915863 | -0.005715103 | 0.069487122 | -0.082246936 | 0.93445035 | 0.968064571 | | 2410021H03Rik | 0.194331409 | -0.00840968081320881 | 0.00964081822647896 | -0.872299489073576 | 0.383045016303155 | NA | NR_045428 |
| 7530420F21Rik | 16.91309614 | -0.013321429 | 0.162129686 | -0.082165269 | 0.934515291 | 0.968067121 | | 2410057H14Rik | 4.173405219 | 0.0428663371360472 | 0.0389025647563873 | 1.10188974440327 | 0.270509607702049 | NA | NR_040588 |
| 1700045I19Rik | 18.92302234 | -0.01375968 | 0.168048353 | -0.081879291 | 0.934742703 | 0.968108518 | | 2410076I21Rik | 4.783323044 | -0.0262905455640634 | 0.0472570531223907 | -0.55633061790742 | 0.57798485995458 | NA | NM_028598 |
| Ilk | 2682.990721 | -0.005732029 | 0.069883527 | -0.082022611 | 0.934628733 | 0.968108518 | | 2410088K16Rik | 0.113843286 | 0.00525947738444006 | 0.00859368002043176 | 0.612016897526494 | 0.540526577873086 | NA | NR_040493 |
| Wdtc1 | 3136.325127 | -0.007285369 | 0.088903965 | -0.081946498 | 0.934689259 | 0.968108518 | | 2410114N07Rik | 0 | NA | NA | NA | NA | NA | NR_040652 |
| Sdsl | 71.29409587 | 0.0133257 | 0.163461476 | 0.081521962 | 0.93502686 | 0.96833809 | | 2410124H12Rik | 0.855649974 | 0.0109583843546098 | 0.0188026796024985 | 0.582809715757409 | 0.560021403729732 | NA | NM_029740 |
| Nmrk1 | 179.478278 | 0.011208014 | 0.137754703 | 0.081362119 | 0.935153974 | 0.968405004 | | 2410137F16Rik | 1.144499055 | -0.0271139083213468 | 0.0211025359222329 | -1.28486492909037 | 0.198839488228039 | NA | NR_015595 |
| Uspl1 | 1583.709729 | 0.004329826 | 0.053481699 | 0.080959023 | 0.93547454 | 0.968672227 | | 2410137M14Rik | 13.90654139 | -0.0587987934655903 | 0.0691724948758537 | -0.850031411634329 | 0.395307622559344 | NA | NM_029747 |
| 1700030K09Rik | 326.6655629 | 0.00744176 | 0.092068422 | 0.080828587 | 0.935578273 | 0.9687149 | | 2410141K09Rik | 0 | NA | NA | NA | NA | NA | NM_183119 |
| 8430427H17Rik | 2717.094938 | 0.004995738 | 0.062041233 | 0.080522874 | 0.935821404 | 0.968837153 | | 2510003D18Rik | 0.959722939 | -0.0144735800355514 | 0.0196978545991067 | -0.734779514323744 | 0.462473785230667 | NA | NR_040373 |
| Clvs1 | 2400.068038 | -0.006471659 | 0.080292351 | -0.080601186 | 0.935759123 | 0.968837153 | | 2510049J12Rik | 2.521736267 | 0.0379692072989477 | 0.0354435729110757 | 1.07125789474466 | 0.284053486310497 | NA | NM_001101431 |
| 4930447C04Rik | 75.62695959 | -0.013416989 | 0.167294313 | -0.080199909 | 0.936078262 | 0.969038324 | | 2600006L11Rik | 0 | NA | NA | NA | NA | NA | NR_040743 |
| 1110017D15Rik | 61.5582291 | -0.011622887 | 0.145329946 | -0.079975857 | 0.936256458 | 0.969093297 | | 2610016A17Rik | 0.241530977 | 0.00760948419201657 | 0.0102872826258192 | 0.73969817577658 | 0.459483155250455 | NA | NR_045347 |
| Pi15 | 9.523208202 | -0.012417621 | 0.155265038 | -0.07997693 | 0.936255605 | 0.969093297 | | 2610020C07Rik | 21.50471589 | 0.0210021970511256 | 0.0808139183611442 | 0.259883414602794 | 0.794953705056864 | NA | NR_038156 |
| Rhoh | 30.89465425 | 0.01319374 | 0.165262233 | 0.079835181 | 0.936368343 | 0.969144363 | | 2610027K06Rik | 2.058034101 | -0.0173116242052387 | 0.0314767508969949 | -0.549981294508114 | 0.582332203509684 | NA | NR_077059 |
| Adcy10 | 8.387214761 | -0.011917851 | 0.149673539 | -0.07962564 | 0.936535002 | 0.969252109 | | 2610028E06Rik | 13.97994078 | 0.057779130223304 | 0.0717540336960654 | 0.805238775398244 | 0.420681904585106 | NA | NR_015560 |
| Vps52 | 3074.097243 | -0.005881734 | 0.074192647 | -0.079276512 | 0.936812688 | 0.969474739 | | 2610028H24Rik | 2.37094498 | -0.0556562261016392 | 0.0314830436647575 | -1.76781592956152 | 0.0770916809736959 | NA | NM_029816 |
| Scarf2 | 100.4106626 | 0.010525248 | 0.133099913 | 0.079077796 | 0.936970744 | 0.969573546 | | 2610035F20Rik | 6.052528907 | -0.0154574506135234 | 0.0519117665642039 | -0.297763910507762 | 0.765883360993908 | NA | NR_045046 |
| Eif3e | 2656.774389 | -0.006027796 | 0.076577872 | -0.078714592 | 0.937259638 | 0.969742961 | | 2610037D02Rik | 3.291275177 | -0.0493302018759803 | 0.0428495455801073 | -1.15124212423111 | 0.249632639766356 | NA | NR_040423 |
| H6pd | 416.8723981 | -0.008288047 | 0.10528719 | -0.078718473 | 0.937256552 | 0.969742961 | | 2610203C22Rik | 18.46878777 | 0.0149136411393082 | 0.0704074814389971 | 0.211818983359456 | 0.83224825797096 | NA | NR_015470 |
| Zfp184 | 114.8890868 | 0.009987746 | 0.127216188 | 0.078510024 | 0.937422357 | 0.969846554 | | 2610204G22Rik | 20.50866306 | -0.126199144556079 | 0.0829650577852902 | -1.5211119949157 | 0.128231734878411 | NA | NR_027971,NR_027972 |
| Tmsb15b1 | 10.78356615 | -0.012612815 | 0.161053046 | -0.078314666 | 0.937577751 | 0.969942557 | | 2610206C17Rik | 1.520360396 | 0.000705170457483921 | 0.0286944904180991 | 0.0245751169374011 | 0.980393867111713 | NA | NR_038175 |
| Gtf3a | 661.6806021 | 0.005399073 | 0.069260842 | 0.077952742 | 0.937865645 | 0.970068747 | | 2610305D13Rik | 11.03381714 | -0.0865964071168201 | 0.0688332200134203 | -1.25806125443407 | 0.208369604324312 | NA | NM_145078 |
| Oxsm | 419.4854133 | -0.007842087 | 0.100549571 | -0.077992247 | 0.937834221 | 0.970068747 | | 2610307P16Rik | 4.58676699 | 0.00895984059874709 | 0.0464706415370785 | 0.192806475279626 | 0.847110532103615 | NA | NR_045053,NR_045054 |
| Tmem136 | 871.5745593 | -0.007008244 | 0.089935536 | -0.077925194 | 0.937887559 | 0.970068747 | | 2610528A11Rik | 11.86244139 | -0.0640810057153511 | 0.0695176955310628 | -0.921794159398129 | 0.356635954555646 | NA | NM_001206684 |
| 1700019G17Rik | 486.5187861 | -0.010241584 | 0.13207937 | -0.077541132 | 0.938193071 | 0.970115459 | | 2610528J11Rik | 1.604340883 | 0.0271925760168648 | 0.0309916074952215 | 0.877417411183252 | 0.380259959536602 | NA | NM_025572 |
| Arl5c | 73.47917961 | -0.012190556 | 0.157335275 | -0.077481393 | 0.938240593 | 0.970115459 | | 2700069I18Rik | 11.41234745 | -0.0195717575186395 | 0.0656677879119897 | -0.298041979804015 | 0.765671123497911 | NA | NM_045905 |
| Fmnl2 | 1955.363534 | 0.00469223 | 0.060745927 | 0.077243536 | 0.938429808 | 0.970115459 | | 2700070H01Rik | 14.14887958 | 0.0110695380944154 | 0.0715460688002665 | 0.154719026216772 | 0.877042829862059 | NA | NR_046019 |
| Klhl24 | 2822.895878 | 0.006151223 | 0.079320336 | 0.077549135 | 0.938186705 | 0.970115459 | | 2700081L22Rik | 7.690517445 | 0.0539057293027272 | 0.0588619621294801 | 0.915799055154659 | 0.359772299175468 | NA | NR_045175 |
| Prkab1 | 546.0617654 | -0.006280806 | 0.081316775 | -0.077238745 | 0.93843362 | 0.970115459 | | 2700086A05Rik | 0 | NA | NA | NA | NA | NA | NR_015611 |
| Stil | 31.08527343 | -0.012647669 | 0.163289113 | -0.077455678 | 0.938261049 | 0.970115459 | | 2700089I24Rik | 0 | NA | NA | NA | NA | NA | NR_045308 |
| Tnfrsf13c | 12.39775922 | 0.012678673 | 0.163185513 | 0.077694846 | 0.938070794 | 0.970115459 | | 2700099C18Rik | 10.42749203 | 0.0148300068473195 | 0.0665199079166589 | 0.222940880584194 | 0.823581518866881 | NA | NR_024720 |
| Zfp579 | 404.6656401 | 0.010306945 | 0.133172238 | 0.0773956 | 0.938308841 | 0.970115459 | | 2810007J24Rik | 0 | NA | NA | NA | NA | NA | NM_001199306,NM_175250 |
| Olfml1 | 504.5213273 | 0.008217022 | 0.106817137 | 0.076926067 | 0.93868236 | 0.970307858 | | 2810403D21Rik | 9.585207386 | 0.0876337123437444 | 0.0625036396750285 | 1.40205774894667 | 0.160898002574195 | NA | NR_015493 |
| Fam198a | 297.4867065 | 0.008182682 | 0.106835247 | 0.076591595 | 0.938948445 | 0.97051816 | | 2810404M03Rik | 6.422898558 | -0.0185236066159007 | 0.0508653387609518 | -0.364169532084604 | 0.715731417042842 | NA | NR_045497 |
| Tprkb | 2298.747064 | 0.006542309 | 0.085789411 | 0.076260094 | 0.939212174 | 0.970725997 | | 2810405F15Rik | 2.827417771 | 0.00633678629749526 | 0.0363803150055384 | 0.174181732525696 | 0.861722641416205 | NA | NR_033447 |
| Mtfp1 | 2056.671375 | -0.006346638 | 0.083352079 | -0.076142533 | 0.939305702 | 0.970757908 | | 2810408A11Rik | 12.73600034 | -0.0424169227950101 | 0.058926168859879 | -0.719831674376686 | 0.471628640349086 | NA | NM_027419 |
| Rap1gap | 10353.71211 | 0.004801876 | 0.063256824 | 0.075910792 | 0.939490071 | 0.970883689 | | 2810408I11Rik | 12.03941746 | -0.0641735046489864 | 0.0649290592609748 | -0.988363382735124 | 0.322974714007038 | NA | NR_038009 |
| Med9 | 1749.001415 | 0.004111811 | 0.054289312 | 0.075738858 | 0.939626861 | 0.97096029 | | 2810417H13Rik | 8.190380104 | -0.0416435823447893 | 0.0570145653226677 | -0.730402522743301 | 0.465144176786847 | NA | NM_026515 |
| Gtf2i | 9835.058737 | -0.003246493 | 0.042922409 | -0.075636316 | 0.939708443 | 0.970979835 | | 2810433D01Rik | 7.44881634 | -0.0179083311099217 | 0.0515858634149049 | -0.347155788900635 | 0.728474283072824 | NA | NR_033474 |
| Hmgcl | 758.9093466 | -0.005756765 | 0.076278603 | -0.075470244 | 0.939840572 | 0.971043809 | | 2810442I21Rik | 2.063262406 | -0.00511153526040872 | 0.031694033022443 | -0.161277526807307 | 0.871874822554535 | NA | NR_015469 |
| Ppih | 342.7335673 | -0.00652199 | 0.086516524 | -0.075384329 | 0.939908928 | 0.971043809 | | 2810442N19Rik | 0.212297216 | 0.00681050119636477 | 0.00973439053551308 | 0.699633035218656 | 0.484156506270654 | NA | NR_040562 |
| Tm2d3 | 959.3000823 | 0.004836901 | 0.064216151 | 0.075322181 | 0.939958374 | 0.971043809 | | 2810454H06Rik | 19.41361911 | -0.101271172617363 | 0.0792696689150778 | -1.27755261253653 | 0.201407218240412 | NA | NR_029441 |
| Gcc1 | 896.1615592 | -0.004837321 | 0.064612797 | -0.074866293 | 0.940321095 | 0.97135376 | | 2810471M01Rik | 0.530790165 | 0.0141864695832445 | 0.014009315407773 | 1.01264545556403 | 0.311229539689781 | NA | NR_045906 |
| Lrpprc | 2867.53332 | -0.005056311 | 0.067643248 | -0.074749675 | 0.940413883 | 0.971384847 | | 2900002K06Rik | 9.141635964 | -0.0415383200162395 | 0.0645954116689363 | -0.643053723833069 | 0.520189240456689 | NA | NR_033561 |
| Fam154b | 56.91552099 | 0.011806756 | 0.158451531 | 0.074513357 | 0.940601913 | 0.971514302 | | 2900057B20Rik | 0 | NA | NA | NA | NA | NA | NR_045365 |
| Necab2 | 3048.566826 | -0.008009581 | 0.107909799 | -0.074224784 | 0.940831526 | 0.971686686 | | 2900060B14Rik | 0.574512993 | 0.0153233651922408 | 0.0145548017823169 | 1.05280480087731 | 0.292430464201114 | NA | NR_027901 |
| Gimap4 | 12.28994306 | -0.012333638 | 0.166836841 | -0.073926348 | 0.94106899 | 0.971737617 | | 2900092C05Rik | 0.12890128 | -0.0068640067066208 | 0.00870804083610955 | -0.788237771940376 | 0.430557639008166 | NA | NM_028434 |
| Psmc3 | 4912.858166 | -0.005200973 | 0.070219034 | -0.074067852 | 0.940956395 | 0.971737617 | | 2900093L17Rik | 3.222003427 | 0.0227839185374589 | 0.0321736102602709 | 0.708155483737964 | 0.47884870731168 | NA | NR_040666 |
| Tmem88 | 95.43750643 | -0.011243905 | 0.151957365 | -0.073993616 | 0.941015306 | 0.971737617 | | 3100003L05Rik | 1.028835615 | -0.00170557342712946 | 0.0246150675469716 | -0.0692898129925831 | 0.944758934304104 | NA | NR_045907 |
| Arhgap1 | 3366.549017 | -0.005084057 | 0.069304024 | -0.073358755 | 0.941520638 | 0.972074416 | | 3110009F21Rik | 1.18385265 | -0.0084058301902892 | 0.0242638420052558 | -0.346434426521093 | 0.729016256418651 | NA | NR_045466 |
| Pde12 | 466.9658067 | 0.005070808 | 0.06908819 | 0.073396158 | 0.941490875 | 0.972074416 | | 3110015C05Rik | 0.973123212 | -0.00975088329291321 | 0.0214919860122312 | -0.45369856872994 | 0.650045797226756 | NA | NR_045908 |
| Tnfaip2 | 43.05045635 | -0.011571267 | 0.157972542 | -0.073248594 | 0.941608297 | 0.972100144 | | 3110021A11Rik | 6.469588882 | -0.0898112674430694 | 0.0549430879813408 | -1.63462358492792 | 0.102127980379135 | NA | NR_030776 |
| Mak16 | 1080.603763 | -0.005316367 | 0.07277219 | -0.073054929 | 0.941762408 | 0.972175724 | | 3110021N24Rik | 15.14203202 | 0.00368630278046392 | 0.0721448271893277 | 0.0510958709595361 | 0.959249126193909 | NA | NM_001254730 |
| Naca | 5629.003993 | -0.004692384 | 0.064280201 | -0.072998894 | 0.941806997 | 0.972175724 | | 3110079O15Rik | 9.48931331 | 0.0359123901274371 | 0.0615362875409961 | 0.583596956568301 | 0.559491508264247 | NA | NM_028473 |
| Nin | 2260.93216 | -0.005293064 | 0.07265865 | -0.072848372 | 0.941926778 | 0.972234595 | | 3110099E03Rik | 6.644073915 | 0.00124774166707025 | 0.0488679066172512 | 0.025532946947021 | 0.979629869184466 | NA | NR_030712 |
| 4921524J17Rik | 260.9266079 | 0.010021695 | 0.13782981 | 0.072710651 | 0.942036373 | 0.972282944 | | 3200001D21Rik | 0.257802559 | -0.00892927589713349 | 0.00993479775307735 | -0.898787888698349 | 0.368765652383134 | NA | NR_045888 |
| Nat2 | 25.73834198 | 0.012070128 | 0.166313633 | 0.072574498 | 0.942144721 | 0.972330001 | | 3300005D01Rik | 0 | NA | NA | NA | NA | NA | NR_045079,NR_045081,NR_045080 |
| Ndufv1 | 5770.351763 | -0.005912573 | 0.081716599 | -0.072354614 | 0.942319703 | 0.972445816 | | 3425401B19Rik | 18.39658629 | -0.0656273175613306 | 0.0704331098524459 | -0.931767995177507 | 0.351456438004052 | NA | NM_001195097 |
| Cyp4f14 | 320.5134533 | 0.00866893 | 0.120258617 | 0.072085733 | 0.94253368 | 0.972601855 | | 3830403N18Rik | 3.073693115 | 0.00957534259670099 | 0.0360500107067743 | 0.265612753199589 | 0.790537462685931 | NA | NM_027510 |
| Hagh | 2275.049144 | 0.006813964 | 0.094629649 | 0.072006648 | 0.942596617 | 0.972602024 | | 3830417A13Rik | 0 | NA | NA | NA | NA | NA | NM_027512 |
| Gm10638 | 24.4786567 | 0.012088469 | 0.168104581 | 0.071910406 | 0.942673208 | 0.972616281 | | 3930402G23Rik | 11.29137687 | -0.00407174331021613 | 0.0669077552612265 | -0.0608560740727126 | 0.951473832380932 | NA | NR_030715 |
| Celf1 | 9904.330412 | 0.004293896 | 0.059792174 | 0.071813682 | 0.942750184 | 0.972629446 | | 4632427E13Rik | 14.29396869 | -0.0180151626898531 | 0.0668321572575102 | -0.269558299913006 | 0.787500084918362 | NA | NR_015510 |
| Zbtb40 | 650.5554704 | -0.008061655 | 0.112378522 | -0.071736614 | 0.942811517 | 0.972629446 | | 4732416N19Rik | 7.744872783 | -0.0324049744766374 | 0.0603190968304322 | -0.537225790494405 | 0.591111658497864 | NA | NR_015615 |
| 4932416H05Rik | 257.4662096 | -0.00660409 | 0.092408737 | -0.071466079 | 0.943026821 | 0.972722027 | | 4732456N10Rik | 6.092106704 | 0.0607604313400338 | 0.036773574964546 | 1.6522851367759 | 0.0984764378065017 | NA | NM_177717 |
| Muc2 | 8.862724349 | -0.010592989 | 0.148168998 | -0.071492612 | 0.943005704 | 0.972722027 | | 4732490B19Rik | 0.980038849 | -0.0178321388274387 | 0.0219170299243396 | -0.813620225413645 | 0.41586254760585 | NA | NR_040276 |
| 2810474O19Rik | 493.7560424 | 0.006040051 | 0.085117943 | 0.070960963 | 0.943428824 | 0.972853049 | | 4833403I15Rik | 2.28468946 | -0.0224331823143251 | 0.0314185275144919 | -0.714011256701247 | 0.475220218822806 | NA | NM_029008 |
| Fam220a | 1357.815134 | -0.006842247 | 0.096272255 | -0.071071852 | 0.94334057 | 0.972853049 | | 4833417C18Rik | 10.22409467 | -0.084900679082069 | 0.0617544029361252 | -1.37481175504012 | 0.169189812811293 | NA | NR_045187 |
| Gm2721 | 10.79135489 | 0.011624016 | 0.163297111 | 0.071183229 | 0.943251929 | 0.972853049 | | 4833419F23Rik | 5.963233664 | -0.0209740247533397 | 0.0532398435327395 | -0.393953538583221 | 0.693615341864737 | NA | NR_040328 |
| Rras2 | 402.9845218 | 0.008138882 | 0.11474421 | 0.070930652 | 0.943452948 | 0.972853049 | | 4833423E24Rik | 3.375788921 | 0.00391760473372324 | 0.0356690713923307 | 0.109831979942309 | 0.912542628137926 | NA | NM_001081664 |
| Sap30l | 739.2251498 | 0.007201341 | 0.101553197 | 0.070912006 | 0.943467788 | 0.972853049 | | 4833427F10Rik | 0.346765936 | -0.0134445992440418 | 0.0121756794716388 | -1.104217537677 | 0.26949876817541 | NA | NR_045459 |
| Ino80b | 314.043753 | 0.005961579 | 0.084357903 | 0.070670066 | 0.943660345 | 0.97298685 | | 4833427G06Rik | 8.604086648 | -0.0168559249890742 | 0.0627017892164409 | -0.268826858048485 | 0.788062924159334 | NA | NM_177702 |
| Eif2b1 | 892.1375464 | 0.004909488 | 0.069784713 | 0.07035191 | 0.943913567 | 0.973183178 | | 4833428L15Rik | 1.010476088 | -0.00904307485072493 | 0.0219108813500439 | -0.412720725663864 | 0.679811238195707 | NA | NR_040732 |
| Acot12 | 12.38480903 | -0.011311877 | 0.16104794 | -0.070239192 | 0.94400328 | 0.973210914 | | 4921501E09Rik | 0.403899654 | -0.00132471380270723 | 0.0135291380196343 | -0.097915610054737 | 0.921999304520122 | NA | NM_001009544 |
| Trim13 | 263.072493 | 0.007534811 | 0.107508434 | 0.070085768 | 0.944125394 | 0.973272046 | | 4921504E06Rik | 1.058845616 | 0.0151474797114503 | 0.025703770404822 | 0.589309641071515 | 0.555653578592418 | NA | NM_027600 |
| Sox7 | 86.25078052 | -0.009433982 | 0.135100449 | -0.069829391 | 0.944329453 | 0.97341764 | | 4921506M07Rik | 0.09572496 | 0.00498255289994511 | 0.00847999972823692 | 0.587565219295241 | 0.556824162160132 | NA | NM_001037743 |
| Gpr160 | 37.12839143 | -0.010870989 | 0.155902932 | -0.069729218 | 0.944409186 | 0.973435067 | | 4921507L20Rik | 0.409445549 | -0.00940499723891646 | 0.0139579455876461 | -0.673809564585253 | 0.500432393249778 | NA | NR_045827 |
| Marveld1 | 434.4174166 | -0.006443494 | 0.092539691 | -0.069629519 | 0.944488541 | 0.973452102 | | 4921508A21Rik | 19.14415816 | 0.00525567810436847 | 0.0800421494283967 | 0.0656613814334165 | 0.947647419181346 | NA | NR_046034 |
| 2700038G22Rik | 30.97042964 | -0.01168773 | 0.168108256 | -0.069525021 | 0.944571717 | 0.973473073 | | 4921508D12Rik | 1.059196812 | 0.004823055134659 | 0.023760646637228 | 0.202985011658015 | 0.839146747309004 | NA | NR_045802 |
| Hrc | 13.38579228 | -0.011367328 | 0.16463901 | -0.069043954 | 0.944954633 | 0.973738167 | | 4921509C19Rik | 0 | NA | NA | NA | NA | NA | NM_198655 |
| Sik1 | 768.448703 | 0.011449437 | 0.165700345 | 0.069097243 | 0.944912215 | 0.973738167 | | 4921509O07Rik | 1.00093551 | -0.00120810722131787 | 0.0245517842324889 | -0.0492064939100925 | 0.960754736145751 | NA | NR_045501 |
| Fam122b | 212.0208943 | -0.007688263 | 0.112300965 | -0.068461238 | 0.945418475 | 0.97415135 | | 4921511C10Rik | 0.46301088 | -0.00202035599165177 | 0.0155329105772313 | -0.130069376348132 | 0.896511538394695 | NA | NR_045502 |

| | | | | | | |
|---|---|---|---|---|---|---|
| 9430083A17Rik | 102.4786737 | -0.009589845 | 0.141621933 | -0.067714404 | 0.946012983 | 0.974313896 |
| Abhd14a | 847.5860223 | -0.006352051 | 0.093736824 | -0.067764734 | 0.945972917 | 0.974313896 |
| BC055402 | 10.74711078 | -0.011053039 | 0.162740628 | -0.06791813 | 0.945850806 | 0.974313896 |
| Harbi1 | 213.1063559 | 0.006355709 | 0.09386649 | 0.0677101 | 0.946016409 | 0.974313896 |
| Sec22b | 2119.657405 | -0.004162503 | 0.06115745 | -0.06806207 | 0.945736224 | 0.974313896 |
| Sema3g | 301.0340115 | -0.008717249 | 0.128605984 | -0.067782607 | 0.94595869 | 0.974313896 |
| Zfp954 | 430.6654951 | -0.005826521 | 0.085911312 | -0.067820189 | 0.945928773 | 0.974313896 |
| Lin52 | 1022.319749 | -0.004289962 | 0.063595218 | -0.067457301 | 0.946217654 | 0.974411039 |
| Tekt2 | 34.85591462 | 0.011294409 | 0.16768575 | 0.067354614 | 0.9462994 | 0.974411039 |
| Zfp654 | 567.9641119 | -0.006802611 | 0.100914843 | -0.067409417 | 0.946255772 | 0.974411039 |
| Cactin | 989.4322206 | 0.005168289 | 0.07716528 | 0.066976871 | 0.946600116 | 0.974464907 |
| Cd82 | 680.0478734 | -0.008735052 | 0.130093636 | -0.067144343 | 0.946466793 | 0.974464907 |
| Cul4b | 1963.434461 | -0.005430359 | 0.081082938 | -0.06697289 | 0.946603286 | 0.974464907 |
| S1pr2 | 112.6307831 | -0.007923385 | 0.118097703 | -0.067091783 | 0.946508636 | 0.974464907 |
| Bloc1s2a | 844.5482714 | -0.004421678 | 0.06620942 | -0.066783223 | 0.94675428 | 0.974490853 |
| Epb4.1l4a | 130.8769088 | -0.010586295 | 0.158419553 | -0.066824422 | 0.946721481 | 0.974490853 |
| Cacnb3 | 11473.69893 | -0.004923024 | 0.074158438 | -0.066385213 | 0.947071143 | 0.974575798 |
| Fzr1 | 2485.594993 | -0.004620119 | 0.069618341 | -0.066363527 | 0.947088407 | 0.974575798 |
| Mtrf1 | 265.3600913 | 0.005414439 | 0.081328258 | 0.066575121 | 0.946919952 | 0.974575798 |
| Trappc6a | 186.033769 | 0.007531103 | 0.113317436 | 0.066460231 | 0.947011419 | 0.974575798 |
| 1500015L24Rik | 19.24326038 | 0.010994645 | 0.166729753 | 0.065942911 | 0.947423276 | 0.974855641 |
| Rbfox2 | 11969.74879 | 0.005512853 | 0.0837307 | 0.065840287 | 0.947504981 | 0.97487497 |
| Anapc1 | 3795.769805 | -0.003851175 | 0.058664727 | -0.065647204 | 0.947658707 | 0.974968394 |
| Bfar | 1396.121496 | 0.003918516 | 0.059796093 | 0.065531312 | 0.947750976 | 0.974998581 |
| Zfp148 | 4125.639607 | 0.004838061 | 0.073999896 | 0.065379299 | 0.947872006 | 0.975058349 |
| A130010J15Rik | 218.2191523 | 0.006219038 | 0.09538217 | 0.065201265 | 0.948013754 | 0.975127361 |
| Ccdc58 | 190.2807117 | 0.007841222 | 0.120447414 | 0.065100788 | 0.948093753 | 0.975127361 |
| Ppp2r5d | 4658.48742 | -0.005506074 | 0.084633436 | -0.065057899 | 0.948127901 | 0.975127361 |
| Fam98b | 1571.510892 | 0.003411073 | 0.052649147 | 0.064788757 | 0.948342193 | 0.975218288 |
| Plp1 | 49858.16529 | 0.010048777 | 0.155038674 | 0.06481465 | 0.948321577 | 0.975218288 |
| Acad9 | 1584.367166 | 0.004156651 | 0.064318371 | 0.064626193 | 0.94847163 | 0.975243916 |
| Snord99 | 14.02318418 | 0.010449128 | 0.16175283 | 0.06459935 | 0.948493002 | 0.975243916 |
| Xrcc3 | 296.7279934 | -0.008774742 | 0.136019014 | -0.064511141 | 0.948563236 | 0.975251412 |
| Angptl2 | 173.1158367 | 0.010616632 | 0.165221774 | 0.064256859 | 0.948765704 | 0.97538782 |
| Ier5l | 317.6297975 | -0.009154008 | 0.14261604 | -0.064186387 | 0.948821817 | 0.97538782 |
| Hiatl1 | 1436.60328 | -0.003559058 | 0.055823757 | -0.063755258 | 0.949165105 | 0.975508052 |
| Lsm2 | 196.8088213 | 0.007301621 | 0.114583372 | 0.06372322 | 0.949190616 | 0.975508052 |
| Nrd1 | 6064.886265 | -0.003712292 | 0.058182628 | -0.063804126 | 0.949126193 | 0.975508052 |
| Pdgfd | 83.22977343 | 0.008639413 | 0.135285865 | 0.063860425 | 0.949081364 | 0.975508052 |
| Vwce | 14.9139325 | 0.010522902 | 0.166318202 | 0.063269696 | 0.949551746 | 0.975814469 |
| Amigo2 | 436.8699894 | -0.009431617 | 0.150147576 | -0.062815648 | 0.949913306 | 0.975997623 |
| Bmf | 171.1240867 | -0.008605235 | 0.137007358 | -0.062808564 | 0.949918946 | 0.975997623 |
| E030011O05Rik | 18.15322911 | -0.010538072 | 0.167702491 | -0.062837899 | 0.949895587 | 0.975997623 |
| 8030462N17Rik | 1074.09818 | -0.005646903 | 0.090393257 | -0.062470404 | 0.95018823 | 0.976122789 |
| Meaf6 | 1341.154369 | -0.003970438 | 0.063610212 | -0.062418242 | 0.950229768 | 0.976122789 |
| Trim23 | 1992.977334 | -0.005689204 | 0.091071525 | -0.062469621 | 0.950188854 | 0.976122789 |
| Fhl1 | 3417.754905 | -0.006312011 | 0.101329014 | -0.06229224 | 0.950330108 | 0.976161144 |
| Ctu1 | 286.7803078 | 0.006027457 | 0.096956558 | 0.062166574 | 0.950430181 | 0.976199219 |
| Foxj3 | 3362.595721 | -0.003815194 | 0.062118628 | -0.061417871 | 0.95102642 | 0.976746875 |
| Cubn | 36.03025949 | -0.010266064 | 0.167606967 | -0.061250816 | 0.95115946 | 0.976761844 |
| Hacl1 | 200.0825677 | 0.007474242 | 0.122203836 | 0.061162093 | 0.951230119 | 0.976761844 |
| Rab42 | 30.2532644 | 0.009790224 | 0.159938354 | 0.061212484 | 0.951189987 | 0.976761844 |
| Dynlt1a | 41.20297215 | -0.009665877 | 0.158450782 | -0.061002395 | 0.951357302 | 0.976827702 |
| Nek11 | 66.01732707 | 0.008994637 | 0.147751071 | 0.060876969 | 0.951457191 | 0.976847879 |
| Yrdc | 678.0361482 | 0.004914833 | 0.080810287 | 0.060819393 | 0.951503045 | 0.976847879 |
| Rnf34 | 2139.328822 | 0.005572195 | 0.091772486 | 0.060717492 | 0.9515842 | 0.976866468 |
| Lipa | 888.3373778 | -0.003984588 | 0.065838185 | -0.060520922 | 0.951740753 | 0.976913781 |
| Sfr1 | 2837.817735 | 0.004144172 | 0.068497258 | 0.060501287 | 0.951756391 | 0.976913781 |
| Btbd16 | 30.44447211 | -0.009825854 | 0.163092197 | -0.060247236 | 0.951958725 | 0.976992017 |
| Npas2 | 2211.395207 | -0.006027003 | 0.100032487 | -0.06025046 | 0.951956158 | 0.976992017 |
| Naa30 | 941.3673321 | 0.006539833 | 0.108916858 | 0.06004427 | 0.952120376 | 0.977093198 |
| A630066F11Rik | 25.74618602 | -0.009898697 | 0.16639836 | -0.059487951 | 0.952563463 | 0.977224279 |
| Ankef1 | 27.39198519 | 0.009975288 | 0.167203563 | 0.059659544 | 0.952426794 | 0.977224279 |
| Nubp2 | 794.0849247 | 0.004194273 | 0.070367914 | 0.059604908 | 0.952470309 | 0.977224279 |
| Pgrmc2 | 3100.645419 | 0.003647188 | 0.061154992 | 0.059638435 | 0.952443606 | 0.977224279 |
| R74862 | 163.6936304 | -0.006186852 | 0.103888555 | -0.059552776 | 0.952511831 | 0.977224279 |
| 2700046G09Rik | 96.18597442 | 0.00823954 | 0.139166834 | 0.059206207 | 0.952787866 | 0.977246133 |
| 4933411K20Rik | 1254.382468 | -0.004104786 | 0.070567674 | -0.058168085 | 0.953614742 | 0.977246133 |
| Abcc1 | 1059.609969 | 0.004113347 | 0.070681532 | 0.058195494 | 0.95359291 | 0.977246133 |
| Apbb1 | 16349.25817 | 0.004680395 | 0.079391335 | 0.058953472 | 0.952989168 | 0.977246133 |
| Cdk13 | 2154.752081 | -0.004412785 | 0.074680995 | -0.059088466 | 0.952881645 | 0.977246133 |
| Cep57l1 | 155.8028308 | -0.006931669 | 0.118580506 | -0.058455382 | 0.953385902 | 0.977246133 |
| Cops7b | 805.7245426 | 0.00462374 | 0.078701722 | 0.058750179 | 0.953151091 | 0.977246133 |
| Gemin7 | 632.4105373 | -0.003999334 | 0.067970227 | -0.058839501 | 0.953079945 | 0.977246133 |
| Hsd3b7 | 257.0368928 | 0.006980037 | 0.119659715 | 0.058332388 | 0.95348387 | 0.977246133 |
| Hsp90aa1 | 43613.85765 | 0.003880922 | 0.06598576 | 0.058814534 | 0.953099832 | 0.977246133 |
| Kcng2 | 168.8928808 | -0.007982291 | 0.137246599 | -0.058160209 | 0.953621015 | 0.977246133 |
| Nme5 | 270.0574552 | -0.005900529 | 0.100452812 | -0.058739308 | 0.95315975 | 0.977246133 |
| Sarnp | 1077.007126 | 0.004310534 | 0.073686096 | 0.058498605 | 0.953351473 | 0.977246133 |
| Sco1 | 378.8551667 | -0.005293503 | 0.090919272 | -0.058222013 | 0.953571787 | 0.977246133 |
| Scyl1 | 2810.376304 | 0.003201255 | 0.054071517 | 0.059204089 | 0.952789553 | 0.977246133 |
| Slc26a6 | 268.9945167 | 0.005626126 | 0.095199947 | 0.059097992 | 0.952874058 | 0.977246133 |
| Tmbim4 | 968.1389399 | -0.003457344 | 0.059491382 | -0.058115035 | 0.953656998 | 0.977246133 |
| Rabep1 | 6210.979818 | -0.003077299 | 0.053162054 | -0.057885251 | 0.953840031 | 0.977369052 |
| Pex16 | 626.9319964 | -0.004925009 | 0.08523702 | -0.057780159 | 0.953923743 | 0.977390191 |
| Dach2 | 21.94758897 | 0.009619436 | 0.166821333 | 0.057663107 | 0.954016981 | 0.977421087 |
| Stag2 | 2476.029157 | 0.004233482 | 0.073795191 | 0.057367998 | 0.954252055 | 0.977597285 |
| Pcbp1 | 4028.49714 | 0.003697711 | 0.064646813 | 0.057198654 | 0.95438695 | 0.977639006 |
| Slc7a14 | 6549.082324 | 0.006396928 | 0.111915705 | 0.057158451 | 0.954418975 | 0.977639006 |
| Agbl1 | 12.74074008 | 0.009210792 | 0.162521243 | 0.056674387 | 0.954804578 | 0.977713021 |
| Gpr25 | 251.9183422 | 0.005959122 | 0.105216433 | 0.056636803 | 0.954834517 | 0.977713021 |
| Mettl2 | 541.840372 | 0.004291354 | 0.07569776 | 0.056690632 | 0.954791637 | 0.977713021 |
| Olig2 | 877.0502211 | 0.004965607 | 0.087174494 | 0.056961699 | 0.954575705 | 0.977713021 |
| Pacsin1 | 20078.34289 | -0.003778743 | 0.066477968 | -0.056842033 | 0.954671031 | 0.977713021 |
| Psen1 | 2561.575049 | -0.003260898 | 0.057628242 | -0.05658507 | 0.954875728 | 0.977713021 |
| Snrpa | 1522.537082 | 0.00379388 | 0.067132597 | 0.056513236 | 0.954932952 | 0.977713021 |
| Far1 | 2281.220372 | 0.003728609 | 0.066288439 | 0.056248261 | 0.955144035 | 0.977864522 |
| Nap1l5 | 9994.425805 | -0.005011961 | 0.089596017 | -0.055939553 | 0.955389962 | 0.977922446 |
| Rpp30 | 284.6175693 | -0.005691787 | 0.101722529 | -0.055954046 | 0.955378416 | 0.977922446 |
| Siah1a | 806.2354318 | 0.003639596 | 0.064997774 | 0.055995699 | 0.955345234 | 0.977922446 |
| Mgat4b | 2072.041188 | 0.004750535 | 0.086566237 | 0.054877452 | 0.956236096 | 0.978659229 |
| Sox17 | 169.1801436 | 0.008991289 | 0.163767533 | 0.054902757 | 0.956215935 | 0.978659229 |
| Diap3 | 42.3746058 | 0.009159237 | 0.167149154 | 0.054796789 | 0.956300359 | 0.978660354 |
| Atp6ap1 | 15633.25101 | -0.002813573 | 0.051464297 | -0.054670388 | 0.956401061 | 0.978698768 |
| Zdhhc15 | 430.8799153 | -0.005830956 | 0.107325475 | -0.054329656 | 0.956672522 | 0.978911905 |
| Rgag4 | 1276.380522 | 0.005445396 | 0.100820139 | 0.054010999 | 0.956926401 | 0.979107024 |
| Nr4a1 | 3817.794586 | 0.008555188 | 0.158634597 | 0.053930151 | 0.956990815 | 0.979108273 |
| Tmed9 | 3613.415968 | 0.003549217 | 0.065950874 | 0.053816074 | 0.957081703 | 0.979136606 |
| Mterfd2 | 439.9342985 | -0.004534303 | 0.085155318 | -0.05324744 | 0.957534757 | 0.979524959 |
| Tgs1 | 1329.798506 | -0.003488217 | 0.0655915 | -0.053180933 | 0.957587747 | 0.979524959 |
| Gm5468 | 120.8048269 | -0.00718032 | 0.136221637 | -0.052710569 | 0.957962518 | 0.979843626 |
| Lama2 | 679.8431429 | -0.00447044 | 0.085047467 | -0.052564062 | 0.958079252 | 0.979898339 |
| Cpeb4 | 2154.009699 | 0.005145618 | 0.098334173 | 0.052327874 | 0.958267444 | 0.97996401 |
| Polr3b | 741.6605126 | 0.003302725 | 0.063119784 | 0.052324719 | 0.958269957 | 0.97996401 |
| Arfrp1 | 1179.621703 | 0.00513677 | 0.098600303 | 0.052096894 | 0.958451487 | 0.980084962 |
| Slc6a9 | 2037.336911 | -0.006062075 | 0.116584707 | -0.051997176 | 0.958530944 | 0.980101527 |
| Ropn1l | 15.23705801 | -0.008632749 | 0.16648075 | -0.05185434 | 0.958644757 | 0.980153218 |
| Cdadc1 | 2148.249051 | 0.002758667 | 0.05370501 | 0.051367028 | 0.959033058 | 0.980168304 |
| Chd1l | 367.5512368 | 0.004185183 | 0.081331214 | 0.051458509 | 0.958960163 | 0.980168304 |
| Efcab5 | 149.4599485 | -0.005906504 | 0.114962165 | -0.05137781 | 0.959024466 | 0.980168304 |
| Efhd2 | 9789.435918 | 0.005075965 | 0.098985112 | 0.051280081 | 0.95910234 | 0.980168304 |
| Ggcx | 588.9321664 | -0.006473137 | 0.126128802 | -0.051321637 | 0.959069227 | 0.980168304 |
| Pctp | 77.35417305 | -0.008259466 | 0.159696003 | -0.051719926 | 0.958751859 | 0.980168304 |
| Pdgfa | 738.0017226 | -0.004123908 | 0.080343637 | -0.051328373 | 0.959063859 | 0.980168304 |
| Clec14a | 166.7864288 | -0.006292612 | 0.122919662 | -0.051192882 | 0.959171824 | 0.980174663 |
| Fam104a | 1285.607952 | 0.003851724 | 0.075381455 | 0.051096434 | 0.959248677 | 0.980188551 |
| Nkrf | 1292.220283 | -0.003720215 | 0.073125956 | -0.050874069 | 0.959425869 | 0.98030496 |
| 2310039L15Rik | 17.82082552 | -0.008428966 | 0.168096725 | -0.050143544 | 0.960008 | 0.98049586 |
| Btbd2 | 2909.317488 | 0.003253568 | 0.064522285 | 0.050425497 | 0.959783319 | 0.98049586 |
| Cideb | 149.1887687 | 0.008304306 | 0.165237388 | 0.050256824 | 0.95991773 | 0.98049586 |
| Dusp27 | 27.1016467 | 0.008228596 | 0.163678439 | 0.050272938 | 0.959904889 | 0.98049586 |
| Dut | 693.1935947 | 0.003481451 | 0.069733771 | 0.049924889 | 0.960182244 | 0.98049586 |
| Man2a2 | 11034.22239 | 0.002270259 | 0.045385774 | 0.050021379 | 0.960105351 | 0.98049586 |
| Ror2 | 36.42642384 | -0.00834492 | 0.165972723 | -0.050278866 | 0.959900165 | 0.98049586 |
| Slc25a46 | 3359.500275 | 0.003826131 | 0.076156218 | 0.050240564 | 0.959930687 | 0.98049586 |
| Tsen15 | 546.9296642 | -0.003690376 | 0.073805512 | -0.050001356 | 0.960121308 | 0.98049586 |
| Axin2 | 1081.939312 | 0.003516955 | 0.071109375 | 0.049458383 | 0.960554002 | 0.980746208 |
| Ptafr | 36.16946345 | 0.008115902 | 0.164026793 | 0.049479122 | 0.960537475 | 0.980746208 |
| Suclg2 | 1009.37556 | -0.00297963 | 0.060735037 | -0.049059488 | 0.960871888 | 0.98100613 |
| Fastkd1 | 479.97532 | -0.004935354 | 0.100919764 | -0.048903741 | 0.960996008 | 0.981068205 |
| Fut11 | 719.7919859 | 0.003251783 | 0.066649446 | 0.048789354 | 0.961087166 | 0.981096624 |
| Pde11a | 99.96411309 | 0.007482275 | 0.154423856 | 0.048452646 | 0.961355344 | 0.981305731 |
| Samsn1 | 63.37936516 | 0.007661068 | 0.159379578 | 0.048068067 | 0.961661996 | 0.981554082 |
| Comp | 10.00674596 | 0.007257308 | 0.15378689 | 0.047190682 | 0.962361254 | 0.98200904 |
| Rrm1 | 777.3881959 | 0.00413322 | 0.087563626 | 0.047202479 | 0.962351852 | 0.98200904 |
| Rs1 | 8.459530939 | -0.007195348 | 0.152344108 | -0.04723089 | 0.962329209 | 0.98200904 |
| Stom | 570.183771 | 0.004323496 | 0.091157302 | 0.047428964 | 0.962171345 | 0.98200904 |
| 1110008J03Rik | 487.1465197 | -0.004010393 | 0.087640028 | -0.045759837 | 0.963501671 | 0.982137951 |
| AF529169 | 158.6245214 | -0.007075104 | 0.155685073 | -0.045444971 | 0.963752637 | 0.982137951 |
| Arl4d | 866.6304843 | 0.006698774 | 0.143216735 | 0.04677368 | 0.962693606 | 0.982137951 |
| Cbx8 | 255.483013 | -0.00443187 | 0.096445481 | -0.045952078 | 0.963348446 | 0.982137951 |
| Chd8 | 4609.269977 | 0.002865937 | 0.06215293 | 0.04611105 | 0.963221739 | 0.982137951 |
| Disp2 | 15021.49942 | 0.002665886 | 0.057100767 | 0.046687391 | 0.96276238 | 0.982137951 |
| Fbxo44 | 3575.105884 | -0.0028674 | 0.063232312 | -0.045347073 | 0.963830667 | 0.982137951 |
| Frmd8 | 889.5579279 | 0.005251624 | 0.116312405 | 0.045151019 | 0.963986936 | 0.982137951 |
| Fsip1 | 12.82421761 | 0.007634898 | 0.166329954 | 0.045902124 | 0.963388261 | 0.982137951 |
| Gse1 | 2850.179647 | -0.003778298 | 0.082246097 | -0.045938933 | 0.963358923 | 0.982137951 |
| Inhbe | 8.131688114 | -0.006905114 | 0.150797791 | -0.04579055 | 0.963477191 | 0.982137951 |
| Kctd4 | 690.0822562 | -0.003944364 | 0.085006513 | -0.04640073 | 0.962990855 | 0.982137951 |
| Lysmd1 | 827.4086473 | -0.003479754 | 0.076838085 | -0.045286839 | 0.963878678 | 0.982137951 |
| Mamdc2 | 49.91430138 | 0.006741999 | 0.144846045 | 0.046545968 | 0.962875096 | 0.982137951 |
| Mto1 | 815.1214825 | 0.003548518 | 0.077095049 | 0.046027834 | 0.963288065 | 0.982137951 |
| Obfc1 | 426.6985832 | 0.003915796 | 0.084021844 | 0.046604495 | 0.96282845 | 0.982137951 |
| Pde1a | 8720.519525 | -0.004357806 | 0.095476325 | -0.045642787 | 0.963594965 | 0.982137951 |
| Psmb10 | 1049.272001 | 0.00406844 | 0.089974962 | 0.045217467 | 0.963933972 | 0.982137951 |
| Rnf152 | 809.4292148 | -0.004721924 | 0.104644556 | -0.045123457 | 0.964008904 | 0.982137951 |
| Six3os1 | 188.1669923 | -0.007422247 | 0.162999395 | -0.045535426 | 0.963680538 | 0.982137951 |
| Spen | 3396.743117 | 0.004501472 | 0.098446264 | 0.045725169 | 0.963529303 | 0.982137951 |
| Ttc1 | 1397.041825 | -0.003385809 | 0.073702243 | -0.045939029 | 0.963358846 | 0.982137951 |
| Ube3b | 6260.927891 | 0.003159855 | 0.068567306 | 0.046083991 | 0.963243306 | 0.982137951 |
| Zmym5 | 1384.54329 | 0.003219723 | 0.070199185 | 0.045865531 | 0.963417427 | 0.982137951 |
| Zfp493 | 133.1399687 | 0.005725169 | 0.128558279 | 0.044533645 | 0.964479034 | 0.982552314 |
| Snx21 | 919.2078478 | -0.003390138 | 0.076275026 | -0.044446244 | 0.964548701 | 0.982558683 |
| Tmem206 | 387.6941321 | 0.003618075 | 0.081786304 | 0.044238154 | 0.96471457 | 0.982663043 |
| Gm5124 | 697.063281 | -0.003199907 | 0.072756585 | -0.043980989 | 0.964919557 | 0.98274263 |
| Pigs | 2204.547156 | 0.003321502 | 0.075456974 | 0.04401849 | 0.964889665 | 0.98274263 |
| 2210015D19Rik | 475.4594587 | -0.003065715 | 0.070105717 | -0.043729884 | 0.965119718 | 0.982881879 |
| Capsl | 39.03536385 | 0.006912649 | 0.159480618 | 0.04334476 | 0.965426712 | 0.983126192 |
| Evpl | 216.1698738 | 0.004426387 | 0.102297543 | 0.043269731 | 0.96548652 | 0.983126192 |
| Sv2a | 19617.77277 | 0.001596253 | 0.036984334 | 0.043160239 | 0.9655738 | 0.983128201 |
| Wdr47 | 6127.093932 | -0.002318444 | 0.05378215 | -0.043108056 | 0.965615398 | 0.983128201 |
| Senp6 | 2874.706153 | 0.002979048 | 0.069511364 | 0.04285699 | 0.965815534 | 0.983267355 |
| Dcun1d2 | 1702.580519 | -0.002520898 | 0.059217931 | -0.042569835 | 0.966044442 | 0.983435781 |
| Ncoa6 | 3536.951371 | 0.003325574 | 0.078669356 | 0.042272803 | 0.966281226 | 0.983612201 |
| Tspan2 | 3678.412705 | 0.004323105 | 0.102722713 | 0.042085195 | 0.966430782 | 0.983635193 |
| Zbbx | 11.49861351 | -0.006503412 | 0.154510209 | -0.042090503 | 0.966426551 | 0.983635193 |
| Mars2 | 337.7014098 | 0.003673713 | 0.08783402 | 0.041825622 | 0.96663771 | 0.983711068 |
| Rpp40 | 119.6578433 | 0.00526221 | 0.125917736 | 0.041790857 | 0.966665423 | 0.983711068 |
| Znrf1 | 6104.394776 | 0.003409104 | 0.081649794 | 0.041752753 | 0.9666958 | 0.983711068 |
| Wac | 4679.775774 | -0.003112016 | 0.074718909 | -0.041649651 | 0.966777992 | 0.983730098 |
| Gstm7 | 593.5406854 | 0.00455294 | 0.110112368 | 0.041348125 | 0.967018368 | 0.983852768 |
| Zfp36l2 | 996.9950571 | 0.0057473 | 0.139028106 | 0.041339121 | 0.967025545 | 0.983852768 |
| Adamts6 | 146.8056434 | 0.005662724 | 0.138581031 | 0.040862185 | 0.967405765 | 0.984110362 |
| Slc24a4 | 1145.890085 | 0.004733738 | 0.115628632 | 0.040939151 | 0.967344406 | 0.984110362 |
| Gm13399 | 40.15132026 | -0.006669718 | 0.164267058 | -0.040602892 | 0.967612478 | 0.984256023 |
| Acyp1 | 341.6497221 | -0.003122839 | 0.077612893 | -0.040236084 | 0.96790491 | 0.984424227 |
| Bre | 1266.240981 | -0.002829513 | 0.07023688 | -0.040285291 | 0.96786568 | 0.984424227 |
| Ap1s1 | 8167.578864 | -0.003501231 | 0.087715098 | -0.039915946 | 0.968160138 | 0.984489938 |
| Cldn11 | 3438.816519 | 0.004714588 | 0.118098054 | 0.039920965 | 0.968156137 | 0.984489938 |
| Sox2ot | 1187.128559 | -0.006427447 | 0.160837404 | -0.039962387 | 0.968123114 | 0.984489938 |
| Trove2 | 1663.904543 | 0.003225374 | 0.08121899 | 0.039712065 | 0.968322683 | 0.984590605 |
| Edem3 | 2190.712256 | -0.003613864 | 0.091825824 | -0.039355644 | 0.968606843 | 0.984750291 |
| Sec11c | 1295.041111 | -0.004041508 | 0.102511083 | -0.039425079 | 0.968551485 | 0.984750291 |
| Zfyve19 | 553.0646934 | 0.003093714 | 0.078945153 | 0.039188145 | 0.968740385 | 0.984821437 |
| Cnot1 | 5610.769466 | 0.00343527 | 0.088684359 | 0.03873592 | 0.969100935 | 0.985123336 |
| Actl6a | 433.003353 | -0.003766829 | 0.097850611 | -0.038495718 | 0.969292446 | 0.985253372 |
| Coro1a | 3037.449818 | -0.003785358 | 0.099312325 | -0.038115697 | 0.969595436 | 0.98536742 |
| Mab21l2 | 8.138141471 | -0.004338139 | 0.113370529 | -0.038265139 | 0.969476285 | 0.98536742 |
| Sec14l1 | 10262.44675 | 0.001754777 | 0.045963057 | 0.038177987 | 0.969545772 | 0.98536742 |
| Mapk14 | 3956.92498 | 0.001854529 | 0.049049087 | 0.037809646 | 0.969839454 | 0.985550764 |
| Rcsd1 | 247.4616877 | 0.004167505 | 0.110750411 | 0.037629702 | 0.969982926 | 0.985631916 |
| Vprbp | 1521.806675 | 0.003208061 | 0.085907482 | 0.037343203 | 0.970211358 | 0.985799382 |
| Agpat4 | 1777.268959 | 0.002914364 | 0.079260148 | 0.036769596 | 0.970668716 | 0.986190316 |
| Armc7 | 385.3887491 | -0.003305575 | 0.090067699 | -0.036701003 | 0.970723409 | 0.986190316 |
| Hccs | 747.401672 | -0.002930734 | 0.080197339 | -0.036544033 | 0.970848569 | 0.986252801 |
| Slc35d2 | 25.41426128 | -0.006025861 | 0.168005418 | -0.035867065 | 0.971388357 | 0.986736459 |
| Clec5a | 102.0897927 | 0.005342752 | 0.149441873 | 0.035751369 | 0.97148061 | 0.986765476 |
| 10-Sep | 178.4881779 | -0.004935324 | 0.138971798 | -0.035513132 | 0.971670575 | 0.986893732 |
| Vat1 | 1880.065325 | -0.003754861 | 0.10597752 | -0.035430733 | 0.971736279 | 0.986895772 |
| 2010300C02Rik | 11787.69074 | 0.003723328 | 0.106166639 | 0.035070599 | 0.972023446 | 0.98706365 |
| Adnp2 | 1096.30649 | -0.002758645 | 0.078892178 | -0.034967286 | 0.972105827 | 0.98706365 |
| Cryz | 594.1064462 | -0.003477279 | 0.099624496 | -0.03490386 | 0.972156403 | 0.98706365 |
| Sema4c | 657.1191632 | 0.003085459 | 0.088259259 | 0.03495904 | 0.972112402 | 0.98706365 |
| Xpnpep2 | 10.9238657 | 0.005762924 | 0.165646731 | 0.03479045 | 0.972246836 | 0.987090785 |
| Atp6v0d1 | 8909.064425 | 0.002169196 | 0.062519447 | 0.034696334 | 0.972321884 | 0.987102298 |
| Isg20 | 17.28502391 | -0.005781609 | 0.167831847 | -0.034448821 | 0.972519253 | 0.987237981 |
| Dapk1 | 5580.837138 | 0.003849414 | 0.115989933 | 0.033187479 | 0.973525083 | 0.987751553 |
| Gm16998 | 19.32956427 | -0.005611498 | 0.167908013 | -0.033420074 | 0.973339602 | 0.987751553 |
| Hhex | 73.58037285 | 0.004801122 | 0.144418124 | 0.033244597 | 0.973479534 | 0.987751553 |
| Map4 | 17264.18494 | -0.001629119 | 0.048301905 | -0.033727837 | 0.973094181 | 0.987751553 |
| Padi2 | 1366.575876 | 0.004144529 | 0.124929946 | 0.033174827 | 0.973535172 | 0.987751553 |
| Rsl1d1 | 1767.970769 | -0.002245538 | 0.06678787 | -0.033621946 | 0.973178622 | 0.987751553 |
| Rwdd4a | 933.008118 | 0.002073982 | 0.06228831 | 0.033296493 | 0.973438151 | 0.987751553 |
| Swap70 | 549.3918771 | -0.003226478 | 0.096667396 | -0.033377104 | 0.973373868 | 0.987751553 |
| Ttl | 3719.099822 | -0.002314237 | 0.070327071 | -0.032906774 | 0.973748931 | 0.987903741 |

| | | | | | | | |
|---|---|---|---|---|---|---|---|
| 4921511C20Rik | 0 | NA | NA | NA | NA | NA | NR_003646 |
| 4921511H03Rik | 0 | NA | NA | NA | NA | NA | NM_027603 |
| 4921511I17Rik | 0.106148608 | 0.00515554063321497 | 0.00854200771766808 | 0.603551390213729 | 0.546141939141592 | NA | NR_045829 |
| 4921511M17Rik | 0 | NA | NA | NA | NA | NA | NM_001201358 |
| 4921513I03Rik | 0 | NA | NA | NA | NA | NA | NR_038000,NR_038003 |
| 4921515E04Rik | 0.749695855 | 0.00940038764022294 | 0.0197663147619112 | 0.475576138164969 | 0.634376378513764 | NA | NR_045711 |
| 4921517D22Rik | 0.108766301 | 0.00519171381361946 | 0.00856003371502575 | 0.606506234257728 | 0.544178643801544 | NA | NM_183290 |
| 4921524L21Rik | 0 | NA | NA | NA | NA | NA | NM_027598 |
| 4921525B02Rik | 0.729403027 | 0.000619083492944162 | 0.0178928282305471 | 0.0345995325594891 | 0.972399074241543 | NA | NR_045148 |
| 4921525O09Rik | 8.890816197 | -0.0101626480712944 | 0.0602159493084826 | -0.168770038303835 | 0.865977526192325 | NA | NR_045061 |
| 4921530D09Rik | 0.297598527 | 0.00928691468127708 | 0.0113567208765242 | 0.81774614188045 | 0.413502157150213 | NA | NM_029859 |
| 4921530L18Rik | 8.357808954 | -0.0420929043267674 | 0.0622611456842005 | -0.676070185734617 | 0.498996084816089 | NA | NR_038162 |
| 4921530L21Rik | 1.996111418 | -0.0113264263823434 | 0.0260693533743541 | -0.434472854761556 | 0.663945102743292 | NA | NM_025733 |
| 4921531P14Rik | 0.326298904 | 0.00765111147689903 | 0.0103094497373689 | 0.742145475443356 | 0.457999199689495 | NA | NR_045361 |
| 4921533I20Rik | 0 | NA | NA | NA | NA | NA | NR_033607 |
| 4921536K21Rik | 20.06948696 | 0.0394733860440334 | 0.0822576191446956 | 0.479875134418826 | 0.631316183244626 | NA | NM_026150 |
| 4921539E11Rik | 0 | NA | NA | NA | NA | NA | NM_001163494,NM_027612 |
| 4922502D21Rik | 0 | NA | NA | NA | NA | NA | NM_199034 |
| 4922502H24Rik | 0 | NA | NA | NA | NA | NA | NR_046187 |
| 4922502N22Rik | 0.665186487 | -0.00424909791802404 | 0.0155908467193633 | -0.272537982991444 | 0.785208389955022 | NA | NR_045149 |
| 4922505E12Rik | 0.443441737 | -0.0212067781731997 | 0.0152604065834807 | -1.38966010225153 | 0.164632115633729 | NA | NM_001033547 |
| 4930401C15Rik | 0.113843286 | 0.00525947738444006 | 0.00859368002043176 | 0.612016897526494 | 0.540526577873086 | NA | NR_045349 |
| 4930401O10Rik | 0 | NA | NA | NA | NA | NA | NR_045942 |
| 4930401O12Rik | 0 | NA | NA | NA | NA | NA | NR_045957 |
| 4930402F06Rik | 1.028572868 | 0.000144484746001159 | 0.0218015440620265 | 0.00662727124235295 | 0.994712241302534 | NA | NM_001272040,NM_001080709 |
| 4930402F11Rik | 0 | NA | NA | NA | NA | NA | NR_045940 |
| 4930402K13Rik | 0 | NA | NA | NA | NA | NA | NM_001270700 |
| 4930404A05Rik | 0 | NA | NA | NA | NA | NA | NR_040495 |
| 4930404A10Rik | 0 | NA | NA | NA | NA | NA | NM_029105 |
| 4930404H11Rik | 1.828556031 | -0.0593106892782078 | 0.0304180715410496 | -1.94985041041038 | 0.051193951139142 | NA | NR_045941 |
| 4930404I05Rik | 0.441671028 | -0.0102554657446552 | 0.0142643784006446 | -0.718956372062575 | 0.472167801269588 | NA | NR_028368 |
| 4930404N11Rik | 3.326830391 | 0.02247463181181 | 0.0369058565273161 | 0.608971960728653 | 0.542543024417571 | NA | NM_001014836 |
| 4930405A10Rik | 0 | NA | NA | NA | NA | NA | NR_046307 |
| 4930405A21Rik | 8.906832054 | -0.0220144656415558 | 0.0605924089337207 | -0.363320521975556 | 0.71636546493559 | NA | NR_040505 |
| 4930405D11Rik | 0 | NA | NA | NA | NA | NA | NR_045956 |
| 4930405J17Rik | 0 | NA | NA | NA | NA | NA | NR_045350 |
| 4930405P13Rik | 0.108766301 | 0.00519171381361946 | 0.00856003371502575 | 0.606506234257728 | 0.544178643801544 | NA | NR_045955 |
| 4930406D18Rik | 0 | NA | NA | NA | NA | NA | NR_040543 |
| 4930407I10Rik | 0.09572496 | 0.00498255289994511 | 0.00847999972823692 | 0.587565219295241 | 0.556824162160132 | NA | NM_001166475 |
| 4930408F14Rik | 0 | NA | NA | NA | NA | NA | NM_001243002 |
| 4930412B13Rik | 0 | NA | NA | NA | NA | NA | NR_040631 |
| 4930412O13Rik | 1.602984943 | 0.000866859271773564 | 0.0212075992592023 | 0.0408749364404093 | 0.967395598567057 | NA | NR_024257 |
| 4930413E15Rik | 0 | NA | NA | NA | NA | NA | NR_040694 |
| 4930413F20Rik | 3.823996262 | -0.0378760410873697 | 0.0410348001288685 | -0.92302243384691 | 0.355995514401149 | NA | NR_045883 |
| 4930413G21Rik | 3.653778029 | 0.0147936052490789 | 0.044879363868521 | 0.329630457606715 | 0.741679205871118 | NA | NR_045614 |
| 4930413M19Rik | 0 | NA | NA | NA | NA | NA | NR_045759 |
| 4930414N06Rik | 0.427212126 | 0.012175076767397 | 0.0129887840374575 | 0.93735308342075 | 0.348576965801222 | NA | NR_040294 |
| 4930415F15Rik | 5.804862899 | 0.0113074528455194 | 0.0499433531114142 | 0.226405560321403 | 0.820886003693955 | NA | NM_028669 |
| 4930415L06Rik | 0 | NA | NA | NA | NA | NA | NM_001033880 |
| 4930415O20Rik | 1.535240012 | 0.0167598219597328 | 0.0262257322177115 | 0.639060210811353 | 0.522783763077004 | NA | NM_001201322 |
| 4930417O13Rik | 2.734422186 | -0.00115967651812203 | 0.0395763872694073 | -0.0293022329256027 | 0.976623546051284 | NA | NR_015527 |
| 4930417O22Rik | 1.812044272 | -0.0268251444095595 | 0.0302592298476184 | -0.886511142043188 | 0.375342152784878 | NA | NR_045806 |
| 4930419G24Rik | 1.563857722 | 0.00522603894827344 | 0.0271132482464829 | 0.192748537569686 | 0.847155908657191 | NA | NR_040595 |
| 4930423M02Rik | 0 | NA | NA | NA | NA | NA | NR_038183 |
| 4930425K10Rik | 0.212695677 | -0.00857796107890413 | 0.00973699866180651 | -0.880965621629515 | 0.378336427605205 | NA | NR_038182 |
| 4930425O10Rik | 0.431048149 | -0.0027962092323494 | 0.0152772432954657 | -0.183031007510322 | 0.854773689089455 | NA | NR_040545 |
| 4930428D18Rik | 0 | NA | NA | NA | NA | NA | NM_001033799 |
| 4930428E07Rik | 0.106148608 | 0.00515554063321497 | 0.00854200771766808 | 0.603551390213729 | 0.546141939141592 | NA | NR_045943 |
| 4930428G15Rik | 0.318844286 | -0.00448436457897182 | 0.0111065302755339 | -0.403759272042883 | 0.686389748741264 | NA | NR_040730 |
| 4930428O21Rik | 0 | NA | NA | NA | NA | NA | NR_045871 |
| 4930429D17Rik | 0.191449919 | 0.00665508079272897 | 0.00962405648474729 | 0.691504752001018 | 0.489248395458281 | NA | NR_040699 |
| 4930429F11Rik | 0 | NA | NA | NA | NA | NA | NR_033463 |
| 4930429F24Rik | 1.254331433 | -0.0223894742936397 | 0.027101622602135 | -0.826130398992273 | 0.408730125054296 | NA | NR_040734 |
| 4930430A15Rik | 0 | NA | NA | NA | NA | NA | NM_026248 |
| 4930430D24Rik | 0 | NA | NA | NA | NA | NA | NM_001034856 |
| 4930430F21Rik | 0.354244579 | -0.00570556803240082 | 0.0130207392936895 | -0.438190789609467 | 0.661247984684689 | NA | NR_045186 |
| 4930430J02Rik | 0 | NA | NA | NA | NA | NA | NR_040729 |
| 4930431F12Rik | 2.225650977 | -0.0367214366092281 | 0.0296102046118835 | -1.24016152845126 | 0.214915654867232 | NA | NR_073013 |
| 4930431P03Rik | 1.853216132 | 0.0275797924959527 | 0.0293166473087203 | 0.940755339637663 | 0.346830252359333 | NA | NR_045059 |
| 4930432J09Rik | 1.322596434 | -0.00496117619111256 | 0.0222177918802599 | -0.223297446382171 | 0.823304014588365 | NA | NR_045953 |
| 4930432K09Rik | 0 | NA | NA | NA | NA | NA | NM_028648,NM_001163551 |
| 4930432M17Rik | 0.255375383 | 0.00708304372802858 | 0.00992471678592415 | 0.713677163873753 | 0.475426830432214 | NA | NM_001033814 |
| 4930433B08Rik | 0 | NA | NA | NA | NA | NA | NR_040544 |
| 4930433I11Rik | 0.417516903 | -0.00258181724625811 | 0.0136578683727324 | -0.189035153641739 | 0.850065264847436 | NA | NM_207248 |
| 4930433N12Rik | 0 | NA | NA | NA | NA | NA | NR_027988 |
| 4930434J06Rik | 0.097165704 | -0.00637828455464458 | 0.00848781691535724 | -0.75146349388194 | 0.452373763047372 | NA | NR_046274 |
| 4930435E12Rik | 0 | NA | NA | NA | NA | NA | NM_029042 |
| 4930438E09Rik | 0 | NA | NA | NA | NA | NA | NR_045961 |
| 4930440C22Rik | 0 | NA | NA | NA | NA | NA | NR_040473 |
| 4930441J16Rik | 0.09572496 | 0.00498255289994511 | 0.00847999972823692 | 0.587565219295241 | 0.556824162160132 | NA | NR_040501 |
| 4930442J19Rik | 0 | NA | NA | NA | NA | NA | NR_040747 |
| 4930442L01Rik | 0 | NA | NA | NA | NA | NA | NR_015596 |
| 4930443O20Rik | 0.21511414 | -0.000824240786584897 | 0.00975135300262786 | -0.08452578697159 | 0.93263840116295 | NA | NR_040504 |
| 4930444F02Rik | 0 | NA | NA | NA | NA | NA | NR_038034,NR_038035 |
| 4930444G20Rik | 0.441181124 | -0.0125130927877208 | 0.0157590304717885 | -0.794026815933979 | 0.427179820341429 | NA | NM_053264 |
| 4930444M15Rik | 2.121325723 | -0.0430902368008292 | 0.0351739716659315 | -1.22506031477148 | 0.220552496173708 | NA | NR_045660 |
| 4930444P10Rik | 12.66172503 | -0.0803190573846954 | 0.0731048820673612 | -1.09868253820158 | 0.271906561709382 | NA | NM_001243238 |
| 4930447A16Rik | 0 | NA | NA | NA | NA | NA | NM_029113 |
| 4930447J18Rik | 0 | NA | NA | NA | NA | NA | NR_045959 |
| 4930447K03Rik | 7.223210312 | -0.0878875981855496 | 0.0599084082371994 | -1.46703277171996 | 0.14236714060777 | NA | NR_046184 |
| 4930447N08Rik | 5.358918605 | -0.0155491270970372 | 0.0391546402699837 | -0.397120928447331 | 0.691278287529702 | NA | NR_045168 |
| 4930448C13Rik | 0.108766301 | 0.00519171381361946 | 0.00856003371502575 | 0.606506234257728 | 0.544178643801544 | NA | NR_045960 |
| 4930448F12Rik | 0.106347839 | -0.00653347234140708 | 0.00854389509723638 | -0.764694822098226 | 0.444453290197145 | NA | NR_046032 |
| 4930448H16Rik | 0.127687691 | 0.00542298404407807 | 0.00869724871942359 | 0.62352868349814 | 0.532937159177571 | NA | NR_040700 |
| 4930448I06Rik | 0 | NA | NA | NA | NA | NA | NR_040475 |
| 4930448I18Rik | 0.108766301 | 0.00519171381361946 | 0.00856003371502575 | 0.606506234257728 | 0.544178643801544 | NA | NR_040696 |
| 4930448K20Rik | 0 | NA | NA | NA | NA | NA | NR_004448 |
| 4930449E01Rik | 0.33403441 | 0.00244344761020736 | 0.0111793636306143 | 0.218567683362231 | 0.826986829057954 | NA | NR_045921 |
| 4930449E18Rik | 0 | NA | NA | NA | NA | NA | NR_045319 |
| 4930449I24Rik | 0 | NA | NA | NA | NA | NA | NM_026136 |
| 4930451I11Rik | 2.797978986 | -0.00432641609920133 | 0.0388185505861946 | -0.111452283350836 | 0.911257702150212 | NA | NM_183131 |
| 4930452A19Rik | 0.491854894 | -0.0112205126393542 | 0.0146859494221809 | -0.764030456376714 | 0.444849093073699 | NA | NR_045433 |
| 4930452G13Rik | 13.10987468 | 0.00299067050616527 | 0.0639119073981468 | 0.0467936356136977 | 0.96267770152334 | NA | NR_045060 |
| 4930452N14Rik | 0.928930868 | -0.0202568164750673 | 0.0199773156883906 | -1.01399090803972 | 0.310587090694266 | NA | NR_040629 |
| 4930453H23Rik | 0.192890664 | -0.000903864895041124 | 0.00963528282151475 | -0.093807821917056 | 0.92526181809679 | NA | NM_001252013 |
| 4930453L07Rik | 0 | NA | NA | NA | NA | NA | NR_073360 |
| 4930455B14Rik | 0.212496447 | -0.000886988073087717 | 0.00973851166817767 | -0.0910804549309223 | 0.927428662789121 | NA | NR_045968 |
| 4930455C13Rik | 0.897270483 | -0.0226848763574602 | 0.0215039136422151 | -1.05491850157576 | 0.291462599744108 | NA | NR_045352 |
| 4930455D15Rik | 0.342602601 | 0.0130117013735539 | 0.0134333274893572 | 0.968613426856654 | 0.332738101639367 | NA | NR_045381 |
| 4930455F16Rik | 0.226556888 | -0.00933175792380659 | 0.010153338242423 | -0.919082739193727 | 0.358052296835844 | NA | NR_040572,NR_040570,NR_040571 |
| 4930455H04Rik | 0.439740379 | -0.00302812286801214 | 0.0153501872321072 | -0.197269441878752 | 0.843616691299491 | NA | NR_040596 |
| 4930455J16Rik | 0 | NA | NA | NA | NA | NA | NR_045469 |
| 4930456L15Rik | 0.217532603 | 0.00684685248201087 | 0.00976000296730706 | 0.701521557416086 | 0.482977586094298 | NA | NR_045887 |
| 4930459L07Rik | 0 | NA | NA | NA | NA | NA | NR_046190 |
| 4930461G14Rik | 0.694002734 | 0.000549708884025569 | 0.017735641584602 | 0.0309945868833313 | 0.975273856641709 | NA | NR_040736 |
| 4930465A12Rik | 0 | NA | NA | NA | NA | NA | NM_029137 |
| 4930465K10Rik | 1.907950431 | 0.0288217047872574 | 0.0322671871604603 | 0.893220243956532 | 0.371739239081312 | NA | NR_027978 |
| 4930465M20Rik | 0.758715399 | -0.000653546848989629 | 0.0182613232701632 | -0.0357885811077802 | 0.971450938191414 | NA | NR_045973 |
| 4930467D21Rik | 2.683043962 | -0.0307963045770054 | 0.0353682328537217 | -0.870733482907523 | 0.383899691870993 | NA | NR_045981 |
| 4930467E23Rik | 0 | NA | NA | NA | NA | NA | NM_001039553 |
| 4930467K11Rik | 2.337336422 | 0.00060046843827526 | 0.0289242386403269 | 0.0207600430124398 | 0.983437071920744 | NA | NR_045353 |
| 4930468A15Rik | 0.313257562 | 0.00962480958133652 | 0.0115601382610462 | 0.832586026567595 | 0.405078242417788 | NA | NM_001201395 |
| 4930469G21Rik | 0.290056368 | -0.00425303587871565 | 0.0107652157825088 | -0.395072051005791 | 0.692789714682806 | NA | NM_001195189 |
| 4930470P17Rik | 0 | NA | NA | NA | NA | NA | NR_027825 |
| 4930471C04Rik | 5.476811247 | -0.0524867561443182 | 0.0542566891859742 | -0.967378528468826 | 0.333354839767151 | NA | NR_046192 |
| 4930471G03Rik | 0.288615623 | 0.00197374456361263 | 0.0106648328298147 | 0.185070370544844 | 0.853173841012254 | NA | NR_015571 |
| 4930471I20Rik | 0.720323536 | -0.0117594621465869 | 0.0178152669716301 | -0.660077795371423 | 0.509203907186778 | NA | NR_040613 |
| 4930471M09Rik | 0.614955356 | 0.00634865703588795 | 0.0187883268995901 | 0.337904331227409 | 0.735435283502589 | NA | NR_045980 |
| 4930473A02Rik | 7.106418008 | -0.0470293686528887 | 0.0458921251811299 | -1.02478079773535 | 0.305466628869711 | NA | NR_040348 |
| 4930473O22Rik | 0.527723949 | -0.00725383165734869 | 0.0164320372881941 | -0.441444449652043 | 0.658891268855762 | NA | NR_045356 |
| 4930474G06Rik | 6.373690834 | -0.0233243439006809 | 0.0492544821162821 | -0.473547642742762 | 0.635822519135091 | NA | NR_045398 |
| 4930474H20Rik | 0 | NA | NA | NA | NA | NA | NR_045712 |
| 4930474M22Rik | 0 | NA | NA | NA | NA | NA | NR_027986 |
| 4930474N05Rik | 0 | NA | NA | NA | NA | NA | NM_175008 |
| 4930474N09Rik | 2.837501275 | -0.00334063101254588 | 0.0378048164658296 | -0.0883652223405279 | 0.929586401687542 | NA | NR_038130 |
| 4930478L05Rik | 0 | NA | NA | NA | NA | NA | NR_040569 |
| 4930478P22Rik | 0 | NA | NA | NA | NA | NA | NR_046060 |
| 4930479M11Rik | 0.220191124 | -0.000706246424282353 | 0.00977545482624743 | -0.072246911968337 | 0.942405412335504 | NA | NR_027849,NR_027848 |
| 4930480E11Rik | 0 | NA | NA | NA | NA | NA | NM_001177966 |
| 4930480G23Rik | 0.108766301 | 0.00519171381361946 | 0.00856003371502575 | 0.606506234257728 | 0.544178643801544 | NA | NR_040768 |
| 4930480K15Rik | 0 | NA | NA | NA | NA | NA | NR_045463 |
| 4930480M12Rik | 0 | NA | NA | NA | NA | NA | NR_046278 |
| 4930483J18Rik | 0.86768302 | -0.0195906577659667 | 0.0193616483327505 | -1.01182799260065 | 0.311620303242382 | NA | NR_015603 |
| 4930483K19Rik | 0.291497113 | -0.00941849595073383 | 0.0102038556585987 | -0.923033044160808 | 0.355989985190414 | NA | NR_045354 |
| 4930483O08Rik | 2.29962735 | 0.0182998797342594 | 0.031827150340865 | 0.574977009825571 | 0.565306845580103 | NA | NR_046279 |
| 4930486F22Rik | 0 | NA | NA | NA | NA | NA | NR_038024 |
| 4930486I03Rik | 1.199798456 | -0.0182318616933587 | 0.0246578303196898 | -0.739394401574749 | 0.459667540802587 | NA | NR_045734 |
| 4930486L24Rik | 8.588095533 | 0.0921642695113599 | 0.0597211043307139 | 1.54324456227379 | 0.122771448863965 | NA | NM_178098 |
| 4930487D11Rik | 0 | NA | NA | NA | NA | NA | NR_046191 |
| 4930487H11Rik | 0.097165704 | -0.00637828455464458 | 0.00848781691535724 | -0.75146349388194 | 0.452373763047372 | NA | NR_040601 |
| 4930488B22Rik | 0 | NA | NA | NA | NA | NA | NR_040627 |
| 4930488L21Rik | 10.01787041 | 0.0202727150306412 | 0.0631370170434794 | 0.321090795542025 | 0.748141585398868 | NA | NR_026888 |
| 4930500F04Rik | 0.106148608 | 0.00515554063321497 | 0.00854200771766808 | 0.603551390213729 | 0.546141939141592 | NA | NR_045758 |
| 4930500J02Rik | 10.65566619 | -0.0183080167555671 | 0.0524727695800447 | -0.348905096912011 | 0.727160558578827 | NA | NR_040323,NR_040322 |
| 4930500L23Rik | 0.318645055 | 0.00214885549292704 | 0.0110130133643737 | 0.195119666328422 | 0.845299266962597 | NA | NR_040701 |
| 4930502A04Rik | 0.09572496 | 0.00498255289994511 | 0.00847999972823692 | 0.587565219295241 | 0.556824162160132 | NA | NR_040737 |
| 4930502E09Rik | 0.120209049 | -0.00674972139751468 | 0.00863521114606269 | -0.781650996523957 | 0.434419707908266 | NA | NR_046281 |
| 4930502E18Rik | 8.998824987 | 0.00798615839975207 | 0.0601188746059768 | 0.132839452702565 | 0.894320352095702 | NA | NM_029142 |
| 4930503B20Rik | 2.296046608 | -0.0374514817918826 | 0.0339186897388527 | -1.10415473239767 | 0.269526022365898 | NA | NM_029144 |
| 4930503E14Rik | 1.497647165 | -0.0161060174029531 | 0.0299681416152465 | -0.537437976960141 | 0.590965116609815 | NA | NM_029131 |
| 4930503E24Rik | 0 | NA | NA | NA | NA | NA | NR_028310 |
| 4930503H13Rik | 0.097165704 | -0.00637828455464458 | 0.00848781691535724 | -0.75146349388194 | 0.452373763047372 | NA | NR_033598 |
| 4930503O07Rik | 0.332178952 | 0.0127721041124076 | 0.0133101933856661 | 0.95957314385545 | 0.337270090816746 | NA | NR_040477 |
| 4930504O13Rik | 0 | NA | NA | NA | NA | NA | NM_207527 |
| 4930505A04Rik | 4.92959076 | -0.0414465614583488 | 0.0483361957687458 | -0.857464283218337 | 0.391188345821494 | NA | NM_001100394 |
| 4930505G20Rik | 0 | NA | NA | NA | NA | NA | NR_045761 |
| 4930509E16Rik | 1.231859255 | -0.0125660170725749 | 0.0263021422799298 | -0.477756410061076 | 0.63282358943769 | NA | NR_045735 |
| 4930509J09Rik | 0 | NA | NA | NA | NA | NA | NR_040547 |
| 4930509K18Rik | 0 | NA | NA | NA | NA | NA | NR_040663 |
| 4930511A02Rik | 2.419896121 | -0.00822503326660879 | 0.029081856371921 | -0.282823529606253 | 0.777312116398267 | NA | NR_045966 |
| 4930511E03Rik | 0.113843286 | 0.00525947738444006 | 0.00859368002043176 | 0.612016897526494 | 0.540526577873086 | NA | NR_040761 |
| 4930511M06Rik | 6.010115187 | -0.0483822175704907 | 0.0556833546890206 | -0.86888115561095 | 0.384912137757207 | NA | NR_027944,NR_027945,NR_015494 |
| 4930511M11Rik | 0.204491261 | 0.00675251727497272 | 0.00969338978195025 | 0.696610517772262 | 0.486046571838551 | NA | NM_029154,NM_029141 |
| 4930512B01Rik | 1.839717732 | -0.0441288956722024 | 0.0291786610884766 | -1.51236876628414 | 0.130440076041335 | NA | NR_033573 |
| 4930513D17Rik | 0.910332818 | -0.0103592200788528 | 0.019241961685964 | -0.538366110894468 | 0.590324317910754 | NA | NR_045416 |
| 4930513O06Rik | 0 | NA | NA | NA | NA | NA | NM_029174 |
| 4930515B02Rik | 0 | NA | NA | NA | NA | NA | NR_040654 |
| 4930515G01Rik | 9.873490607 | 0.0618520382735823 | 0.0654909349191634 | 0.944436636153191 | 0.344946567311108 | NA | NR_027872 |
| 4930515G16Rik | 0 | NA | NA | NA | NA | NA | NR_003100 |
| 4930515L03Rik | 0.700811285 | -0.0114874470022257 | 0.0177217311170982 | -0.64821246447772 | 0.516847541179713 | NA | NR_040632 |
| 4930515L19Rik | 0 | NA | NA | NA | NA | NA | NR_045440 |
| 4930517E11Rik | 0 | NA | NA | NA | NA | NA | NR_040611 |
| 4930518P08Rik | 0 | NA | NA | NA | NA | NA | NR_045364 |
| 4930519D14Rik | 0.227686571 | 0.00691460391109036 | 0.00980755285755433 | 0.705028462402254 | 0.480792523671872 | NA | NR_045975 |
| 4930519F16Rik | 2.409221783 | -0.0663017629365639 | 0.0366088812055842 | -1.81108410727533 | 0.0701278332516997 | NA | NM_029170 |
| 4930519F24Rik | 0 | NA | NA | NA | NA | NA | NR_040763 |
| 4930519G04Rik | 8.420917434 | -0.054460160784701 | 0.0605316239269188 | -0.899697666305005 | 0.368281165656153 | NA | NM_026263 |
| 4930519H02Rik | 0 | NA | NA | NA | NA | NA | NR_045974 |
| 4930520P13Rik | 0.191449919 | 0.00665508079272897 | 0.00962405648474729 | 0.691504752001018 | 0.489248395458281 | NA | NR_036596 |
| 4930521E06Rik | 0 | NA | NA | NA | NA | NA | NR_040602 |
| 4930522H14Rik | 0 | NA | NA | NA | NA | NA | NM_001199090,NM_026291 |
| 4930522O17Rik | 0 | NA | NA | NA | NA | NA | NR_040665 |
| 4930523O13Rik | 0 | NA | NA | NA | NA | NA | NR_040451 |
| 4930524B15Rik | 5.603620684 | -0.0427688257248663 | 0.0520688278836866 | -0.821390214897961 | 0.411424035400043 | NA | NM_026262 |
| 4930524C18Rik | 0 | NA | NA | NA | NA | NA | NR_045661 |
| 4930524L23Rik | 17.06832118 | 0.0747903089083487 | 0.0761877666899667 | 0.981657714324339 | 0.326268504086013 | NA | NR_029473 |
| 4930524N10Rik | 0 | NA | NA | NA | NA | NA | NM_001256259 |
| 4930524O05Rik | 0 | NA | NA | NA | NA | NA | NR_045316 |
| 4930524O08Rik | 0.09572496 | 0.00498255289994511 | 0.00847999972823692 | 0.587565219295241 | 0.556824162160132 | NA | NR_040735 |
| 4930525D18Rik | 0.945476319 | -0.0360146609759424 | 0.0222936980041049 | -1.61546374985931 | 0.106210304433469 | NA | NR_040738 |
| 4930525M21Rik | 0 | NA | NA | NA | NA | NA | NM_001243943 |
| 4930526D03Rik | 0 | NA | NA | NA | NA | NA | NM_199023 |
| 4930526L06Rik | 0.579264538 | -0.0148713971815782 | 0.0158007516498203 | -0.941182895039514 | 0.346611140782892 | NA | NR_045317 |
| 4930527F14Rik | 0.108766301 | 0.00519171381361946 | 0.00856003371502575 | 0.606506234257728 | 0.544178643801544 | NA | NR_045809 |
| 4930527G23Rik | 0 | NA | NA | NA | NA | NA | NR_045395 |
| 4930528D03Rik | 2.222777668 | -0.0023344511561025 | 0.0354763990563131 | -0.065802934294344 | 0.947534720072803 | NA | NR_045977 |
| 4930528P14Rik | 0.83948556 | -0.0134917974640886 | 0.0168643326286169 | -0.800019648639668 | 0.423699413166397 | NA | NR_040516 |
| 4930529C04Rik | 0 | NA | NA | NA | NA | NA | NR_033593 |
| 4930529F24Rik | 0 | NA | NA | NA | NA | NA | NR_045713 |
| 4930529K09Rik | 0.097165704 | -0.00637828455464458 | 0.00848781691535724 | -0.75146349388194 | 0.452373763047372 | NA | NR_040457 |
| 4930529L06Rik | 0 | NA | NA | NA | NA | NA | NR_040537 |
| 4930533B01Rik | 0 | NA | NA | NA | NA | NA | NR_040614 |
| 4930533P14Rik | 0 | NA | NA | NA | NA | NA | NR_040478 |
| 4930539C22Rik | 0 | NA | NA | NA | NA | NA | NR_040600 |
| 4930539J05Rik | 20.63338835 | -0.0473542141234406 | 0.0787752199230399 | -0.60113083999897 | 0.547752843860089 | NA | NR_030689 |
| 4930539M17Rik | 5.21443376 | -0.00378117888223361 | 0.0454286107330683 | -0.0832334253946537 | 0.933665935318867 | NA | NR_040597 |
| 4930539N22Rik | 0 | NA | NA | NA | NA | NA | NR_040598 |
| 4930540M03Rik | 0 | NA | NA | NA | NA | NA | NR_040746 |
| 4930542C21Rik | 0 | NA | NA | NA | NA | NA | NR_040565 |
| 4930542D17Rik | 0.097165704 | -0.00637828455464458 | 0.00848781691535724 | -0.75146349388194 | 0.452373763047372 | NA | NR_040566 |
| 4930542H20Rik | 0 | NA | NA | NA | NA | NA | NR_045898 |
| 4930543E12Rik | 0.452133968 | -0.0170158559534382 | 0.0136860195687122 | -1.24330203299858 | 0.213756555340871 | NA | NR_045978 |
| 4930544D05Rik | 0.257802559 | -0.00892927589713349 | 0.00993479775307735 | -0.898787888698349 | 0.368765652383134 | NA | NM_001145537 |
| 4930544G11Rik | 0 | NA | NA | NA | NA | NA | NM_001161773 |
| 4930544M13Rik | 0 | NA | NA | NA | NA | NA | NR_045976 |
| 4930545E07Rik | 0 | NA | NA | NA | NA | NA | NR_045374 |
| 4930545H06Rik | 0.749629687 | -0.00725985925980626 | 0.0199354037785364 | -0.364169160577657 | 0.715731694444519 | NA | NR_045357 |
| 4930545L23Rik | 0.397844952 | -0.0152960324097223 | 0.0129817556207533 | -1.17827148011239 | 0.23868839386615 | NA | NR_040517 |
| 4930546C10Rik | 2.127829097 | -0.00732256130033827 | 0.034588654977054 | -0.211704135509057 | 0.832337861560425 | NA | NR_038051 |
| 4930546K05Rik | 1.171869033 | 0.0182238719473896 | 0.0251510157796093 | 0.72457796961681 | 0.468710978204706 | NA | NR_040754 |
| 4930547C10Rik | 0 | NA | NA | NA | NA | NA | NM_026652 |
| 4930547E08Rik | 0 | NA | NA | NA | NA | NA | NR_040514 |
| 4930547E14Rik | 0 | NA | NA | NA | NA | NA | NR_040564 |
| 4930548G14Rik | 0 | NA | NA | NA | NA | NA | NR_045811 |
| 4930548H24Rik | 0.438563431 | -0.012587699918621 | 0.0157371680443719 | -0.799870718996528 | 0.423785704269421 | NA | NM_026296 |

| | | | | | | |
|---|---|---|---|---|---|---|
| Acot6 | 129.0098936 | -0.004092168 | 0.125852082 | -0.032515693 | 0.974060801 | 0.988155441 |
| Lrp2bp | 17.00275331 | -0.005298453 | 0.165548536 | -0.032005437 | 0.974467715 | 0.988503521 |
| Sil1 | 693.5147739 | -0.002260879 | 0.070864069 | -0.031904441 | 0.974548257 | 0.988520503 |
| Abcf1 | 3444.169422 | 0.001334993 | 0.042315921 | 0.031548246 | 0.974832317 | 0.98858135 |
| Aqp1 | 91.21261106 | 0.003280536 | 0.103718952 | 0.031629091 | 0.974767844 | 0.98858135 |
| Ccdc50 | 2262.571162 | -0.00262221 | 0.083220457 | -0.031509195 | 0.97486346 | 0.98858135 |
| Nsf | 38819.19215 | -0.001928306 | 0.060856526 | -0.031686106 | 0.974722375 | 0.98858135 |
| Cd1d1 | 32.54327098 | -0.005093142 | 0.163182207 | -0.031211382 | 0.975100963 | 0.988757481 |
| Mrpl37 | 1369.966361 | 0.001892326 | 0.061511563 | 0.030763753 | 0.975457948 | 0.988883873 |
| Prickle1 | 3462.182822 | 0.002618281 | 0.085087885 | 0.030771495 | 0.975451773 | 0.988883873 |
| Tango6 | 248.1212418 | 0.003314154 | 0.107951024 | 0.030700537 | 0.975508363 | 0.988883873 |
| Usp15 | 2450.505048 | 0.002000606 | 0.064661848 | 0.03093951 | 0.97531778 | 0.988883873 |
| Zfp128 | 357.9806866 | -0.00254551 | 0.083037526 | -0.03065494 | 0.975544727 | 0.988883873 |
| Celf3 | 5659.13181 | 0.00269921 | 0.088477276 | 0.030507382 | 0.975662406 | 0.988938462 |
| Mrps26 | 1189.58237 | -0.001858848 | 0.06167794 | -0.030137965 | 0.975957023 | 0.989172376 |
| Fadd | 111.7609819 | 0.004711326 | 0.158478851 | 0.029728423 | 0.976283644 | 0.989438695 |
| Kif21a | 8666.123326 | 0.001674923 | 0.057035947 | 0.029366102 | 0.976572608 | 0.989551211 |
| Mgst1 | 211.8973778 | -0.00373632 | 0.126859479 | -0.029452431 | 0.976503757 | 0.989551211 |
| Paip2b | 3515.682181 | 0.002360915 | 0.080442824 | 0.02934898 | 0.976586264 | 0.989551211 |
| Lyve1 | 47.28689215 | -0.004872152 | 0.166685596 | -0.029229591 | 0.976681481 | 0.989582976 |
| Coq5 | 1367.132236 | 0.001647332 | 0.056608635 | 0.029100364 | 0.976784545 | 0.98960427 |
| Kctd10 | 2514.235073 | 0.001762763 | 0.060694797 | 0.029043073 | 0.976830238 | 0.98960427 |
| Aass | 199.9081355 | 0.003573214 | 0.124632008 | 0.028670113 | 0.977127693 | 0.989687009 |
| Kcnd1 | 267.568035 | 0.003421789 | 0.119139533 | 0.02872085 | 0.977087227 | 0.989687009 |
| Rnaseh2a | 653.4693222 | 0.002201751 | 0.076929657 | 0.028620313 | 0.977167411 | 0.989687009 |
| Tmem26 | 13.05834903 | -0.004356366 | 0.15095382 | -0.028858932 | 0.976977099 | 0.989687009 |
| Bco2 | 154.7218001 | -0.003870107 | 0.13595729 | -0.028465606 | 0.9772908 | 0.989703857 |
| Metap2 | 2494.62294 | -0.001629565 | 0.057337225 | -0.028420721 | 0.977326598 | 0.989703857 |
| Pcdh7 | 4754.351195 | 0.002609328 | 0.092009986 | 0.028359187 | 0.977375675 | 0.989703857 |
| Commd10 | 749.7467277 | 0.002324648 | 0.083314259 | 0.027902162 | 0.977740184 | 0.989854996 |
| Rab15 | 12222.68083 | -0.002641388 | 0.094788243 | -0.027866202 | 0.977768865 | 0.989854996 |
| Rfc2 | 1038.923328 | 0.002585927 | 0.092567043 | 0.027935716 | 0.977713422 | 0.989854996 |
| Vps37b | 1198.68308 | -0.002144972 | 0.077014194 | -0.027851643 | 0.977780477 | 0.989854996 |
| 5330413P13Rik | 90.78374376 | -0.00372425 | 0.135079343 | -0.027570836 | 0.978004443 | 0.989914466 |
| Arhgef4 | 8839.262913 | 0.002123502 | 0.077112625 | 0.027537674 | 0.978030892 | 0.989914466 |
| Nt5c | 854.1199201 | 0.002257567 | 0.081977984 | 0.027538704 | 0.97803007 | 0.989914466 |
| Nup155 | 886.5601017 | -0.00211878 | 0.078034319 | -0.027151899 | 0.97833858 | 0.990096536 |
| Rag1 | 10.12924035 | -0.004139332 | 0.152280533 | -0.027182278 | 0.978314351 | 0.990096536 |
| 1600023N17Rik | 13.30493131 | -0.004499118 | 0.167417626 | -0.026873619 | 0.978560535 | 0.990191805 |
| Rasd2 | 8500.339938 | -0.002628553 | 0.097619176 | -0.02692661 | 0.97851827 | 0.990191805 |
| Rbmxl1 | 1487.789283 | -0.002066028 | 0.077445511 | -0.026677186 | 0.97871721 | 0.990285669 |
| Plscr4 | 132.2827043 | 0.003571602 | 0.134521333 | 0.026550447 | 0.978818297 | 0.990323279 |
| Aldh9a1 | 1303.543604 | -0.002770205 | 0.106124073 | -0.026103459 | 0.979174818 | 0.990326427 |
| Cab39 | 7065.999796 | -0.001858733 | 0.07042627 | -0.026392606 | 0.978944192 | 0.990326427 |
| Il27ra | 15.85672355 | -0.004121346 | 0.158771101 | -0.025957787 | 0.979291008 | 0.990326427 |
| Nadkd1 | 1323.673688 | -0.00229767 | 0.088947468 | -0.025831768 | 0.979391523 | 0.990326427 |
| Neurl4 | 5312.826621 | -0.001665767 | 0.063877725 | -0.026077429 | 0.97919558 | 0.990326427 |
| Rarres1 | 21.18027367 | 0.004405938 | 0.16772645 | 0.026268594 | 0.979043105 | 0.990326427 |
| Rnase6 | 12.00093778 | -0.004047316 | 0.156718844 | -0.025825329 | 0.979396659 | 0.990326427 |
| Tmem236 | 9.708465889 | -0.0039695 | 0.151812044 | -0.026147466 | 0.979139718 | 0.990326427 |
| Uhrf2 | 1747.800848 | 0.001914713 | 0.073902783 | 0.025908531 | 0.979330296 | 0.990326427 |
| Myh13 | 23.58856819 | 0.004298491 | 0.168086438 | 0.025573096 | 0.979597845 | 0.990465219 |
| Kcnk12 | 195.6128008 | 0.002987966 | 0.120127797 | 0.02487323 | 0.98015608 | 0.990964979 |
| Dennd2c | 38.62710557 | -0.004107979 | 0.165768984 | -0.024781349 | 0.980229368 | 0.990974411 |
| Fcrls | 1049.299538 | 0.00386032 | 0.156356063 | 0.024689287 | 0.9803028 | 0.990983988 |
| Atp2a3 | 434.5558618 | 0.002851754 | 0.117091652 | 0.024354886 | 0.980569533 | 0.991124297 |
| Smarcd1 | 7756.527655 | -0.001173761 | 0.048118719 | -0.024393033 | 0.980539106 | 0.991124297 |
| Tmem120a | 473.5902356 | -0.002058137 | 0.085616241 | -0.024039093 | 0.980821426 | 0.991314232 |
| Gm16386 | 25.04830294 | 0.003967832 | 0.166506675 | 0.02382987 | 0.980988314 | 0.991418234 |
| Kirrel2 | 352.2542595 | -0.003447814 | 0.145494654 | -0.023697184 | 0.981094153 | 0.991460527 |
| Piezo1 | 341.5240115 | 0.002136649 | 0.09128698 | 0.023405847 | 0.981326541 | 0.991566025 |
| Slc43a1 | 8.342027049 | -0.003471896 | 0.148112058 | -0.02344101 | 0.981298492 | 0.991566025 |
| Pemt | 62.88058871 | 0.00323902 | 0.138915275 | 0.023316512 | 0.981397801 | 0.991573363 |
| Wbp5 | 1523.56789 | -0.002090061 | 0.09028048 | -0.023150747 | 0.981530026 | 0.991642294 |
| Ccdc6 | 2516.882976 | -0.0025616 | 0.112144761 | -0.022841908 | 0.981776379 | 0.991657986 |
| Ddx51 | 802.8650365 | -0.002052875 | 0.094147373 | -0.021804905 | 0.982603581 | 0.991657986 |
| E130307A14Rik | 547.485058 | -0.001841821 | 0.083705085 | -0.02200369 | 0.982445012 | 0.991657986 |
| Egfem1 | 187.2674802 | -0.0026555 | 0.116763893 | -0.022742475 | 0.981855695 | 0.991657986 |
| Fam221b | 45.74367774 | -0.00339704 | 0.156416615 | -0.0217179 | 0.982672985 | 0.991657986 |
| Hdac4 | 1274.619603 | 0.00222812 | 0.098441051 | 0.022634052 | 0.981942181 | 0.991657986 |
| Mir212 | 13.63421025 | 0.00357123 | 0.160901731 | 0.022195097 | 0.982292329 | 0.991657986 |
| Mmp19 | 27.56480974 | 0.003743919 | 0.167245996 | 0.022385704 | 0.982140284 | 0.991657986 |
| Ntng2 | 642.6352997 | 0.002780037 | 0.127449164 | 0.021812908 | 0.982597197 | 0.991657986 |
| Plcb2 | 170.3644098 | 0.003024491 | 0.134237082 | 0.022530964 | 0.982024413 | 0.991657986 |
| Pogz | 2689.165033 | -0.001430227 | 0.064274557 | -0.022251843 | 0.982247063 | 0.991657986 |
| Slc9a2 | 238.8313981 | 0.002851884 | 0.125619273 | 0.022702602 | 0.981887501 | 0.991657986 |
| Tbx1 | 44.81016938 | -0.003622845 | 0.158988405 | -0.022786848 | 0.981820299 | 0.991657986 |
| Tmem178 | 2281.969434 | 0.002656471 | 0.120365645 | 0.022070013 | 0.982392107 | 0.991657986 |
| Tmem41a | 507.4769731 | 0.001802434 | 0.083111142 | 0.021687034 | 0.982697607 | 0.991657986 |
| Tnpo1 | 1923.527351 | 0.002175728 | 0.095662554 | 0.022743775 | 0.981854658 | 0.991657986 |
| Zdhhc18 | 1163.638478 | 0.001246604 | 0.056438259 | 0.022087926 | 0.982377818 | 0.991657986 |
| Zmynd19 | 661.0664706 | 0.001650568 | 0.074941837 | 0.022024649 | 0.982428293 | 0.991657986 |
| Rgn | 10.15687085 | 0.003405989 | 0.158195522 | 0.021530252 | 0.982822672 | 0.991719601 |
| C030034I22Rik | 82.23014544 | -0.003228001 | 0.150816747 | -0.021403462 | 0.982923812 | 0.991757068 |
| Epha4 | 7563.200335 | -0.001738958 | 0.08179665 | -0.021259522 | 0.983038634 | 0.991808334 |
| Magix | 19.19766925 | 0.003203462 | 0.152765067 | 0.020969857 | 0.983269701 | 0.991976869 |
| Kcnk1 | 3415.837048 | -0.001354949 | 0.065733732 | -0.020612697 | 0.983554612 | 0.992174822 |
| Pdzk1 | 10.90663115 | 0.003238757 | 0.157501546 | 0.020563335 | 0.983593989 | 0.992174822 |
| Usp3 | 620.5620409 | 0.00204291 | 0.099945057 | 0.020440331 | 0.983692111 | 0.992209203 |
| Tm9sf4 | 4023.717384 | -0.001223899 | 0.060619961 | -0.020189709 | 0.983892037 | 0.992281665 |
| Zfp672 | 1071.752725 | -0.001572108 | 0.07758748 | -0.020262397 | 0.983834053 | 0.992281665 |
| Nme4 | 61.49123493 | 0.003113063 | 0.155693431 | 0.019994825 | 0.984047501 | 0.992373859 |
| Ola1 | 3219.105232 | 0.001461761 | 0.07422509 | 0.019693629 | 0.984287773 | 0.992487471 |
| Tmem28 | 278.9204815 | -0.002536698 | 0.128812145 | -0.019693004 | 0.984288271 | 0.992487471 |
| Bai2 | 17780.92237 | 0.001519213 | 0.078021277 | 0.019471773 | 0.984464755 | 0.992536238 |
| Pacs2 | 5988.197758 | -0.001769458 | 0.090853946 | -0.019475854 | 0.984461499 | 0.992536238 |
| Fbxo41 | 6372.723859 | 0.001401646 | 0.072975559 | 0.019207056 | 0.984675929 | 0.99268455 |
| Pcdhgb5 | 183.0302292 | 0.002506155 | 0.131344967 | 0.019080711 | 0.984776719 | 0.992699952 |
| Sgcb | 2422.764933 | 0.000868053 | 0.045652275 | 0.019014452 | 0.984829576 | 0.992699952 |
| Tmem98 | 347.7060939 | 0.003177439 | 0.167701803 | 0.018946961 | 0.984883417 | 0.992699952 |
| Spata3 | 11.59395914 | -0.003014432 | 0.16026466 | -0.018809086 | 0.984993405 | 0.992746232 |
| Arfgef2 | 3861.951108 | -0.001701558 | 0.092112444 | -0.01847262 | 0.98526182 | 0.992823012 |
| Scp2 | 3809.812125 | 0.000836352 | 0.044963602 | 0.018600644 | 0.985159689 | 0.992823012 |
| Tceanc | 71.20026113 | 0.002751489 | 0.148445861 | 0.018535303 | 0.985211815 | 0.992823012 |
| 1110004F10Rik | 2800.142859 | -0.001281422 | 0.07079693 | -0.018099968 | 0.985559104 | 0.993057993 |
| Golt1a | 33.23396233 | -0.003003961 | 0.166963962 | -0.017991675 | 0.985645495 | 0.993080459 |
| Zfp668 | 480.4279658 | 0.001305532 | 0.073554825 | 0.017749104 | 0.985839007 | 0.993210844 |
| Adora2b | 116.3628664 | -0.002579305 | 0.147382411 | -0.017500768 | 0.98603712 | 0.993345848 |
| Atxn1l | 2460.056662 | -0.00152341 | 0.088108897 | -0.017290077 | 0.986205201 | 0.993385996 |
| Kcnq2 | 7046.441661 | 0.001090031 | 0.063000029 | 0.017302079 | 0.986195627 | 0.993385996 |
| AU040320 | 3338.256367 | 0.000963815 | 0.059182281 | 0.016285525 | 0.987006605 | 0.993681113 |
| Erbb2 | 136.7994706 | 0.002106182 | 0.127113835 | 0.016569259 | 0.986780249 | 0.993681113 |
| Frat2 | 193.9293929 | 0.001966083 | 0.119653138 | 0.016431524 | 0.986890131 | 0.993681113 |
| Lztr1 | 6350.258598 | 0.001455694 | 0.087417554 | 0.016652188 | 0.98671409 | 0.993681113 |
| Mir5133 | 10.68092198 | 0.002350102 | 0.14507416 | 0.016199312 | 0.987075385 | 0.993681113 |
| Olfml3 | 870.0806096 | 0.002692105 | 0.162431527 | 0.016573781 | 0.986776641 | 0.993681113 |
| Osbpl3 | 1459.937824 | -0.00141628 | 0.086745438 | -0.016326849 | 0.986973638 | 0.993681113 |
| Rcor2 | 472.2407876 | -0.001624162 | 0.099837223 | -0.016268097 | 0.987020509 | 0.993681113 |
| Tsg101 | 2392.474888 | -0.001490636 | 0.091301186 | -0.016326583 | 0.98697385 | 0.993681113 |
| Git1 | 16622.79819 | 0.001599394 | 0.099729211 | 0.016037368 | 0.98720458 | 0.993746606 |
| Bckdhb | 700.0788937 | -0.001075777 | 0.06806251 | -0.015805714 | 0.98738939 | 0.993868071 |
| Tgfbr3 | 566.5736921 | 0.001678441 | 0.107970966 | 0.015545301 | 0.987597144 | 0.994012612 |
| Ncbp2 | 1875.304417 | -0.000829732 | 0.053664783 | -0.015461393 | 0.987664085 | 0.994015416 |
| Fnta | 1717.65578 | 0.00090605 | 0.059794542 | 0.015152724 | 0.987910338 | 0.994198673 |
| Gen1 | 37.2093745 | 0.002336947 | 0.156403349 | 0.014941798 | 0.988078614 | 0.994243474 |
| Mus81 | 699.6740013 | -0.001272845 | 0.086036196 | -0.014794299 | 0.988196288 | 0.994243474 |
| Ptcd3 | 1808.491483 | -0.001434729 | 0.097103948 | -0.014775189 | 0.988211534 | 0.994243474 |
| Ube2d1 | 1539.912066 | 0.001080004 | 0.072437366 | 0.01490949 | 0.988104389 | 0.994243474 |
| Ankrd40 | 5652.590667 | 0.001049128 | 0.072294241 | 0.014511911 | 0.988421577 | 0.994390229 |
| Hcrtr1 | 93.29354779 | -0.002152407 | 0.149276866 | -0.01441889 | 0.988495789 | 0.994400322 |
| Gm14325 | 796.1449138 | 0.001454586 | 0.101456665 | 0.014337022 | 0.988561103 | 0.994401463 |
| Slc12a5 | 27128.00226 | 0.000767148 | 0.054094762 | 0.014181557 | 0.988685134 | 0.994461664 |
| Akap2 | 3477.887049 | -0.000885137 | 0.064873061 | -0.013644136 | 0.989113892 | 0.994505822 |
| Irx1 | 11.85203082 | 0.002281948 | 0.163506049 | 0.013956351 | 0.988864805 | 0.994505822 |
| Kif3c | 11925.10338 | 0.000961997 | 0.070952256 | 0.013558367 | 0.98918232 | 0.994505822 |
| Prpf8 | 10564.95747 | -0.001020504 | 0.075688785 | -0.0134829 | 0.989242528 | 0.994505822 |
| Ptger2 | 17.25324598 | 0.002280266 | 0.1673403 | 0.013626522 | 0.989127945 | 0.994505822 |
| Rbl1 | 89.18921211 | 0.001790145 | 0.132524714 | 0.013508011 | 0.989222494 | 0.994505822 |
| Rbpms2 | 95.1584948 | -0.00175344 | 0.127388021 | -0.013764562 | 0.989017816 | 0.994505822 |
| Rnf41 | 3283.745133 | 0.000947558 | 0.069936905 | 0.013548749 | 0.989189993 | 0.994505822 |
| Cdc27 | 2473.86381 | -0.000936024 | 0.069933918 | -0.01338441 | 0.989321105 | 0.994520288 |
| Pabpc5 | 54.1662827 | 0.002040712 | 0.156791984 | 0.013015409 | 0.989615499 | 0.99475169 |
| Myh15 | 60.90675207 | 0.002057085 | 0.164094185 | 0.012536002 | 0.989997979 | 0.995071599 |
| 4632434I11Rik | 25.98516669 | -0.002020526 | 0.167527571 | -0.012060855 | 0.990377063 | 0.995108658 |
| Auh | 1526.627737 | 0.0007585 | 0.06385699 | 0.011878097 | 0.990522873 | 0.995108658 |
| Ccdc87 | 131.6507694 | -0.001811512 | 0.152924596 | -0.011845783 | 0.990548654 | 0.995108658 |
| Lgi4 | 1434.531332 | -0.001551298 | 0.128349622 | -0.012086499 | 0.990356604 | 0.995108658 |
| Lrrc6 | 247.0874761 | -0.001407613 | 0.118687475 | -0.011859829 | 0.990537447 | 0.995108658 |
| Lyst | 3153.545997 | -0.001600931 | 0.132486392 | -0.012083741 | 0.990358804 | 0.995108658 |
| Pla2g2d | 16.24653744 | -0.001999872 | 0.163386675 | -0.012240114 | 0.990234046 | 0.995108658 |
| Terf2ip | 2328.154726 | 0.000799079 | 0.067053076 | 0.011917116 | 0.990491742 | 0.995108658 |
| Hes1 | 248.3937809 | 0.001428428 | 0.122410824 | 0.011669133 | 0.99068959 | 0.9951212 |
| Rap1gap2 | 12501.18537 | -0.001025869 | 0.087634889 | -0.011706173 | 0.990660039 | 0.9951212 |
| Acad12 | 79.61071545 | -0.001665999 | 0.145267747 | -0.011468472 | 0.990849684 | 0.99521749 |
| Akna | 261.9427128 | -0.001149356 | 0.102197599 | -0.011246406 | 0.991026855 | 0.995252438 |
| Dpp8 | 8896.008392 | -0.0005846 | 0.052240346 | -0.011190587 | 0.99107139 | 0.995252438 |
| Gpr34 | 366.3839922 | 0.001867766 | 0.16811907 | 0.01110978 | 0.99113586 | 0.995252438 |
| Phc3 | 2744.1408 | -0.000797074 | 0.072185538 | -0.011042011 | 0.991189929 | 0.995252438 |
| Snrpb2 | 652.4126242 | 0.000950131 | 0.086200805 | 0.011022304 | 0.991205652 | 0.995252438 |
| Mterf | 59.80421933 | 0.001637926 | 0.151958411 | 0.010778777 | 0.991399946 | 0.995318523 |
| Stbd1 | 46.14457439 | 0.001736364 | 0.16040784 | 0.010824684 | 0.991363321 | 0.995318523 |
| Avil | 10.92911177 | -0.001614692 | 0.15745005 | -0.010255268 | 0.991817623 | 0.995593735 |
| Mkrn2 | 1157.719147 | 0.000732318 | 0.072349977 | 0.010121885 | 0.991924042 | 0.995593735 |
| Papola | 4072.027534 | -0.000830241 | 0.082096145 | -0.010113037 | 0.991931101 | 0.995593735 |
| 2-Sep | 2153.618269 | -0.000864322 | 0.084675078 | -0.010207519 | 0.99185572 | 0.995593735 |
| Gna11 | 5692.958845 | 0.000823935 | 0.083820209 | 0.009829794 | 0.992157085 | 0.995756048 |
| B3gnt7 | 40.69134208 | 0.001586532 | 0.168070111 | 0.009439702 | 0.992468319 | 0.995905917 |
| Fam101a | 17.36029988 | -0.001582826 | 0.168095498 | -0.00941623 | 0.992487047 | 0.995905917 |
| Gm11837 | 25.83735658 | 0.001558223 | 0.165751743 | 0.009400944 | 0.992499242 | 0.995905917 |
| Mxra8 | 218.7329288 | -0.001058822 | 0.113689213 | -0.009313306 | 0.992569164 | 0.995911581 |
| Mycl1 | 526.5835398 | -0.001164943 | 0.128205654 | -0.009086522 | 0.992750105 | 0.995992825 |
| Omd | 35.65791024 | 0.001509393 | 0.166771188 | 0.00905068 | 0.9927787 | 0.995992825 |
| Ppp1r17 | 18.66143946 | -0.001434873 | 0.16153546 | -0.008882714 | 0.992912713 | 0.996062776 |
| Caml | 843.456301 | 0.000472014 | 0.058882645 | 0.008016185 | 0.993604079 | 0.996562765 |
| Pde5a | 514.8273888 | -0.000758537 | 0.092836744 | -0.008170651 | 0.993480836 | 0.996562765 |
| Zfp942 | 457.6363565 | 0.001074722 | 0.132902675 | 0.008086537 | 0.993547947 | 0.996562765 |
| Golph3 | 2958.782088 | -0.00066526 | 0.085508761 | -0.007780025 | 0.993792501 | 0.996622719 |
| Slc35a4 | 2041.410042 | -0.000466242 | 0.059812484 | -0.007795061 | 0.993780504 | 0.996622719 |
| Acp1 | 1994.917265 | 0.000388099 | 0.05408669 | 0.007175502 | 0.994274827 | 0.996807126 |
| Bmx | 61.26489296 | -0.00116046 | 0.157502282 | -0.007367892 | 0.994121326 | 0.996807126 |
| Fam43b | 849.2272051 | -0.000613415 | 0.089245032 | -0.006873381 | 0.994515879 | 0.996807126 |
| Gm5918 | 130.1198713 | -0.00087513 | 0.129396702 | -0.006763156 | 0.994603824 | 0.996807126 |
| Jag1 | 352.4823198 | 0.000649683 | 0.095392869 | 0.006810608 | 0.994565963 | 0.996807126 |
| Mfsd3 | 175.8922193 | -0.000745592 | 0.108627142 | -0.006863774 | 0.994523544 | 0.996807126 |
| Plrg1 | 1346.263033 | -0.000587879 | 0.0858402 | -0.006848532 | 0.994535705 | 0.996807126 |
| Ppapdc1b | 598.3588504 | 0.000737008 | 0.099627551 | 0.00739763 | 0.994097599 | 0.996807126 |
| Samd9l | 278.0205214 | 0.00095588 | 0.133756259 | 0.007146429 | 0.994298023 | 0.996807126 |
| Timm9 | 828.7492188 | 0.000530912 | 0.078732755 | 0.006743214 | 0.994619735 | 0.996807126 |
| E2f1 | 613.2955741 | -0.000684424 | 0.111908831 | -0.006115907 | 0.995120243 | 0.99724423 |
| Nptxr | 128.1616818 | -0.00063963 | 0.11451604 | -0.005585502 | 0.995543437 | 0.997539288 |
| Pcdhac1 | 135.5874904 | 0.000832668 | 0.148084182 | 0.005622935 | 0.995513571 | 0.997539288 |
| Tmem56 | 2489.703399 | -0.000477591 | 0.098601722 | -0.004843642 | 0.996135348 | 0.99806784 |
| Psmb3 | 1875.521334 | 0.000353337 | 0.076583755 | 0.004613735 | 0.996318785 | 0.998122543 |
| Rps19-ps3 | 46.27803518 | -0.000694092 | 0.148410179 | -0.004676849 | 0.996268428 | 0.998122543 |
| Gstk1 | 299.8220455 | 0.000451616 | 0.099756995 | 0.004527161 | 0.99638786 | 0.998127207 |
| Hkdc1 | 547.9840648 | -0.000668757 | 0.15261185 | -0.004382081 | 0.996503617 | 0.998178629 |
| Ninj2 | 69.13077651 | -0.000681194 | 0.165557552 | -0.004114547 | 0.996717076 | 0.998327906 |
| Acaa1b | 12.80983927 | 0.000559127 | 0.15975484 | 0.003499907 | 0.997207484 | 0.998496366 |
| Adamts13 | 33.87468898 | -0.000591131 | 0.162275631 | -0.003642758 | 0.997093506 | 0.998496366 |
| Eif3k | 2610.413977 | 0.000187263 | 0.051557469 | 0.00363213 | 0.997101986 | 0.998496366 |
| Nr4a3 | 376.2466614 | -0.000576655 | 0.163903347 | -0.003518261 | 0.99719284 | 0.998496366 |
| Sf3b3 | 3715.511069 | 0.000158634 | 0.044406627 | 0.003572294 | 0.997149728 | 0.998496366 |
| Tbl1x | 1552.544523 | -0.000271419 | 0.086524969 | -0.003136891 | 0.997497127 | 0.998678719 |
| Tysnd1 | 512.4210779 | 0.000347951 | 0.111878406 | 0.003110086 | 0.997518514 | 0.998678719 |
| Isca2 | 801.6239039 | 0.000213654 | 0.070766964 | 0.003019118 | 0.997591096 | 0.998686854 |
| Srp54c | 562.3280563 | -0.000228141 | 0.079550465 | -0.002867874 | 0.997711771 | 0.998743131 |
| Sim1 | 10.55397087 | 0.000308483 | 0.110836533 | 0.002783224 | 0.997779311 | 0.998746214 |
| AF251705 | 117.7600714 | 0.00035095 | 0.139450625 | 0.002516661 | 0.997991997 | 0.998765534 |
| Rsg1 | 36.59120384 | -0.000413558 | 0.157190115 | -0.002630941 | 0.997900815 | 0.998765534 |
| Uso1 | 3796.483903 | -0.000201437 | 0.07785354 | -0.002587378 | 0.997935573 | 0.998765534 |
| 2810408I11Rik | 12.93392093 | -0.000273083 | 0.166876541 | -0.001636437 | 0.998694313 | 0.999065354 |
| 6530402F18Rik | 617.9998662 | -0.000170737 | 0.103713633 | -0.00164623 | 0.998686499 | 0.999065354 |
| Arhgap36 | 156.6084681 | -0.000217707 | 0.140817256 | -0.001546024 | 0.998766452 | 0.999065354 |
| Clcf1 | 15.12012217 | 0.000262736 | 0.168121392 | 0.001562776 | 0.998753085 | 0.999065354 |
| Hist1h4h | 8.883851178 | -0.000285968 | 0.151345079 | -0.001889511 | 0.998492389 | 0.999065354 |
| Magi2 | 4732.345226 | 0.000100652 | 0.073438427 | 0.001370561 | 0.998906451 | 0.999065354 |
| Mgat5b | 4226.267606 | -0.000137794 | 0.09467706 | -0.001455412 | 0.99883875 | 0.999065354 |
| Plekhg6 | 10.90760365 | -0.000225978 | 0.157194825 | -0.001437567 | 0.998852988 | 0.999065354 |
| Smcr8 | 2495.934167 | -0.00016619 | 0.081525538 | -0.002038508 | 0.998373507 | 0.999065354 |
| Zfp511 | 451.450827 | 0.000104494 | 0.083447117 | 0.00125222 | 0.999000874 | 0.999065354 |
| Zfp677 | 63.29793703 | -0.000204645 | 0.156325169 | -0.0013091 | 0.99895549 | 0.999065354 |
| Sumf1 | 916.2325874 | 3.95E-05 | 0.088585964 | 0.000445386 | 0.999644634 | 0.999644634 |
| 0610005C13Rik | 0.787035502 | -0.008368886 | 0.067921826 | -0.123213505 | 0.901938032 | NA |
| 0610008F07Rik | 0 | NA | NA | NA | NA | NA |
| 0610009B14Rik | 0.400957162 | -0.004374601 | 0.047713229 | -0.091685289 | 0.926948086 | NA |
| 0610012H03Rik | 0 | NA | NA | NA | NA | NA |
| 0610031O16Rik | 0 | NA | NA | NA | NA | NA |
| 0610038L08Rik | 0.159967308 | 0.0240345 | 0.030870969 | 0.778547006 | 0.436246606 | NA |
| 0610039K10Rik | 1.397022734 | 0.013049902 | 0.067191501 | 0.194219532 | 0.84600399 | NA |
| 1100001G20Rik | 0.503646335 | -0.052098212 | 0.044076498 | -1.181995275 | 0.237207576 | NA |
| 1110025L11Rik | 0 | NA | NA | NA | NA | NA |
| 1110028F11Rik | 0.2777807 | 0.011208478 | 0.038523646 | 0.290950592 | 0.771089108 | NA |
| 1110028F18Rik | 0 | NA | NA | NA | NA | NA |
| 1110036E04Rik | 0.099009884 | 0.018205726 | 0.027896865 | 0.652608315 | 0.514008823 | NA |
| 1190002F15Rik | 2.065659253 | -0.174155995 | 0.097164863 | -1.792376267 | 0.073072714 | NA |

| | | | | | | | |
|---|---|---|---|---|---|---|---|
| 4930548J01Rik | 0 | NA | NA | NA | NA | NA | NR_045462 |
| 4930548K13Rik | 0 | NA | NA | NA | NA | NA | NR_040656 |
| 4930549C01Rik | 0 | NA | NA | NA | NA | NA | NM_026300 |
| 4930549G23Rik | 0.577577851 | -0.00758998248925242 | 0.0167529983589172 | -0.453052183653588 | 0.650511165667378 | NA | NR_045376 |
| 4930550C14Rik | 13.4583258 | 0.0184094337946296 | 0.0742333995408532 | 0.24799394758283 | 0.804139089690942 | NA | NM_029247 |
| 4930550L24Rik | 0.194331409 | -0.00840968081320881 | 0.00964081822647896 | -0.872299489073576 | 0.383045016303155 | NA | NM_023774 |
| 4930552N02Rik | 0 | NA | NA | NA | NA | NA | NR_040661 |
| 4930552P12Rik | 1.178231564 | -7.61532774363492e-05 | 0.0221958235298877 | -0.00343097327899573 | 0.997262484762975 | NA | NR_045318 |
| 4930553E22Rik | 0.212496447 | -0.000886988073087717 | 0.00973851166817767 | -0.0910804549309223 | 0.927428662789121 | NA | NR_040567 |
| 4930554C24Rik | 0 | NA | NA | NA | NA | NA | NR_040739 |
| 4930555B11Rik | 0 | NA | NA | NA | NA | NA | NR_040633 |
| 4930555G01Rik | 9.366760938 | 0.115141832486402 | 0.0651304098348949 | 1.76786592896138 | 0.077083319883085 | NA | NM_175393 |
| 4930556C24Rik | 0.108766301 | 0.00519171381361946 | 0.00856003371502575 | 0.606506234257728 | 0.544178643801544 | NA | NR_040506 |
| 4930556G01Rik | 0 | NA | NA | NA | NA | NA | NR_040655 |
| 4930556J02Rik | 0 | NA | NA | NA | NA | NA | NR_045714 |
| 4930556M19Rik | 11.39173576 | -0.00226733052374292 | 0.0640657208218051 | -0.0353906971568987 | 0.97176820265994 | NA | NR_045065,NR_045064,NR_045063 |
| 4930556N09Rik | 0 | NA | NA | NA | NA | NA | NR_045358 |
| 4930557A04Rik | 0 | NA | NA | NA | NA | NA | NM_029229 |
| 4930557J02Rik | 0 | NA | NA | NA | NA | NA | NR_040703 |
| 4930558C23Rik | 0 | NA | NA | NA | NA | NA | NR_015490 |
| 4930558G05Rik | 0 | NA | NA | NA | NA | NA | NR_045441 |
| 4930558J18Rik | 1.241283012 | -0.0173854866394753 | 0.0222224037884548 | -0.782340506678562 | 0.434014487398897 | NA | NR_037999 |
| 4930558K02Rik | 1.104512261 | -0.0208402088271592 | 0.0222282883097322 | -0.937553469559541 | 0.348473933192933 | NA | NM_001204904,NM_001204905 |
| 4930562C15Rik | 0.097165704 | -0.00637828455464458 | 0.00848781691535724 | -0.75146349388194 | 0.452373763047372 | NA | NM_030192 |
| 4930562F07Rik | 0.974347369 | 0.0105967587980166 | 0.0242788972352495 | 0.436459642105639 | 0.662503274356058 | NA | NR_038030,NR_038031 |
| 4930563D23Rik | 0.106347839 | -0.00653347234140708 | 0.00854389509723638 | -0.764694822098226 | 0.444453290197145 | NA | NM_029252 |
| 4930563E18Rik | 2.611025974 | 0.0037354570310491 | 0.0323694641277838 | 0.115400644765164 | 0.908127567929016 | NA | NR_045379 |
| 4930563F08Rik | 0 | NA | NA | NA | NA | NA | NR_040704 |
| 4930563M20Rik | 0 | NA | NA | NA | NA | NA | NR_046193 |
| 4930564B18Rik | 0.237667581 | -0.00136938695604711 | 0.00989612066541175 | -0.13837613771559 | 0.889943154921195 | NA | NM_029230 |
| 4930564C03Rik | 0 | NA | NA | NA | NA | NA | NM_029257 |
| 4930564D02Rik | 0 | NA | NA | NA | NA | NA | NM_029228 |
| 4930564G21Rik | 0 | NA | NA | NA | NA | NA | NR_045359 |
| 4930564K09Rik | 0 | NA | NA | NA | NA | NA | NR_040381,NR_040382 |
| 4930565D16Rik | 0 | NA | NA | NA | NA | NA | NR_040752 |
| 4930565N06Rik | 0 | NA | NA | NA | NA | NA | NR_040476 |
| 4930567H12Rik | 1.65398349 | 0.0251064559467012 | 0.029726716401113 | 0.844575485833383 | 0.398347900841139 | NA | NR_015535 |
| 4930567H17Rik | 0 | NA | NA | NA | NA | NA | NM_001033807 |
| 4930567J20Rik | 0 | NA | NA | NA | NA | NA | NR_040576 |
| 4930567K20Rik | 0 | NA | NA | NA | NA | NA | NR_046010 |
| 4930568D16Rik | 0 | NA | NA | NA | NA | NA | NM_029463 |
| 4930568E12Rik | 0 | NA | NA | NA | NA | NA | NR_040755 |
| 4930568G15Rik | 1.578783041 | -0.0100788090233686 | 0.0300102456909153 | -0.335845601771254 | 0.736987300141941 | NA | NR_040480 |
| 4930568K20Rik | 0 | NA | NA | NA | NA | NA | NR_045369 |
| 4930571K23Rik | 0 | NA | NA | NA | NA | NA | NM_001145759 |
| 4930572K03Rik | 0 | NA | NA | NA | NA | NA | NR_045371 |
| 4930572O03Rik | 0.928072022 | -0.0141091509023587 | 0.0205681575508878 | -0.685970576968362 | 0.492731662647397 | NA | NR_073011 |
| 4930572O13Rik | 0.332414823 | -0.0166094830116705 | 0.0135191688281235 | -1.22858758721308 | 0.219226471765921 | NA | NR_045718 |
| 4930573O16Rik | 0 | NA | NA | NA | NA | NA | NR_040620 |
| 4930577N17Rik | 17.48180138 | -0.0157570637525743 | 0.076215824377788 | -0.206742679505367 | 0.836210823975927 | NA | NR_073429 |
| 4930578E11Rik | 0.097165704 | -0.00637828455464458 | 0.00848781691535724 | -0.75146349388194 | 0.452373763047372 | NA | NR_045391 |
| 4930578I06Rik | 0 | NA | NA | NA | NA | NA | NM_026359 |
| 4930578M01Rik | 20.50078066 | -0.0210397309048985 | 0.079509996910827 | -0.264617428277544 | 0.791304192780582 | NA | NR_045991 |
| 4930578N18Rik | 0 | NA | NA | NA | NA | NA | NR_040575 |
| 4930579F01Rik | 0 | NA | NA | NA | NA | NA | NM_001163385,NM_001163386 |
| 4930579G18Rik | 3.399227486 | -0.0166404992135831 | 0.0401350977200034 | -0.414612151430975 | 0.678425851253301 | NA | NR_038053 |
| 4930579K19Rik | 15.76751413 | 0.0764241484333672 | 0.0744016270000562 | 1.02718383340339 | 0.304333909114524 | NA | NR_029444 |
| 4930583K01Rik | 1.05463012 | -0.026022832145195 | 0.0212027729016247 | -1.2273315507332 | 0.219698001762876 | NA | NR_027879 |
| 4930583P06Rik | 0 | NA | NA | NA | NA | NA | NR_040612 |
| 4930584F24Rik | 0 | NA | NA | NA | NA | NA | NR_029445 |
| 4930590L20Rik | 0.321063518 | 0.00979019190676043 | 0.0116583710553074 | 0.839756417111422 | 0.401044974403149 | NA | NR_040604 |
| 4930591A17Rik | 0 | NA | NA | NA | NA | NA | NM_026596 |
| 4930592A05Rik | 0.7778633 | 0.00773788564091058 | 0.0160624563313481 | 0.481737380714867 | 0.629992517915888 | NA | NR_045070 |
| 4930592I03Rik | 0 | NA | NA | NA | NA | NA | NR_033307 |
| 4930593A02Rik | 2.444879053 | 0.0225128843399535 | 0.0332275681962163 | 0.677536321858097 | 0.498065732816751 | NA | NR_045167 |
| 4930593C16Rik | 0.667302911 | 0.0168496834363377 | 0.0196003650817975 | 0.859661713749694 | 0.389975545354716 | NA | NR_040753 |
| 4930594C11Rik | 0.617836846 | -0.0101602134386162 | 0.0172680300439502 | -0.588382890969995 | 0.556275318974247 | NA | NR_024017 |
| 4930595M18Rik | 0 | NA | NA | NA | NA | NA | NM_173435 |
| 4930596D02Rik | 0 | NA | NA | NA | NA | NA | NM_001033766 |
| 4930596M17Rik | 0.127687691 | 0.00542298404407807 | 0.00869724871942359 | 0.62352868349814 | 0.532937159177571 | NA | NR_040740 |
| 4930597G03Rik | 0 | NA | NA | NA | NA | NA | NR_045736 |
| 4930598F16Rik | 0 | NA | NA | NA | NA | NA | NR_040479 |
| 4930599N23Rik | 4.57635825 | 0.00683499122147687 | 0.0411412576691647 | 0.166134717524683 | 0.86805093315307 | NA | NR_045813 |
| 4931402G19Rik | 0.587554335 | 0.0194196800280105 | 0.0163709404560003 | 1.18622873745123 | 0.235531980343605 | NA | NR_040608 |
| 4931403G20Rik | 0 | NA | NA | NA | NA | NA | NR_038173,NR_038172 |
| 4931406B18Rik | 0 | NA | NA | NA | NA | NA | NM_028737 |
| 4931406H21Rik | 1.07426648 | -0.00854735005060804 | 0.02019350213211 | -0.423272297924824 | 0.672096603342298 | NA | NR_033492 |
| 4931408C20Rik | 0 | NA | NA | NA | NA | NA | NM_001033764 |
| 4931408D14Rik | 16.54639845 | -0.0267702986975508 | 0.0760118342086915 | -0.352185932312232 | 0.724698826349951 | NA | NR_040299,NR_040298 |
| 4931409K22Rik | 5.827644362 | -0.0628820784532509 | 0.0516816564461152 | -1.21671948573888 | 0.223710962629254 | NA | NR_033188,NM_177676 |
| 4931417E11Rik | 0 | NA | NA | NA | NA | NA | NM_025737 |
| 4931417G12Rik | 3.2449091 | 0.0263217757347866 | 0.0369918448416548 | 0.711556178056491 | 0.476739651467616 | NA | NM_027633 |
| 4931419H13Rik | 0.236453993 | 0.00749401214086028 | 0.0102093235778702 | 0.734036107652062 | 0.462926731928034 | NA | NR_040593 |
| 4931420L22Rik | 0 | NA | NA | NA | NA | NA | NR_040561 |
| 4931423N10Rik | 6.442202124 | -0.0357640195065553 | 0.0472089154474145 | -0.757569183015706 | 0.448708934668264 | NA | NM_027635 |
| 4931428L18Rik | 0.561390173 | -0.0205242846516337 | 0.0186466397331107 | -1.10069615466367 | 0.271028920405334 | NA | NR_033445 |
| 4931429I11Rik | 18.86230472 | 0.0387909564708113 | 0.0734499234589808 | 0.528127935932767 | 0.597410542531007 | NA | NM_001081121 |
| 4931429L15Rik | 4.334018639 | -0.0429431275099268 | 0.0410092108334374 | -1.04715810514726 | 0.295026664117275 | NA | NM_183104 |
| 4931429P17Rik | 0.738051225 | 0.0173715648208205 | 0.018838057476965 | 0.922152660488606 | 0.356448951891403 | NA | NR_038004 |
| 4931430N09Rik | 0.120209049 | -0.00674972139751468 | 0.00863521114606269 | -0.781650996523957 | 0.434419707908266 | NA | NR_046053 |
| 4931431B13Rik | 0.341529857 | 0.00771401917148477 | 0.0103511161961187 | 0.745235492031021 | 0.456129370356692 | NA | NR_045183 |
| 4931431C16Rik | 1.922618846 | 0.0209061610938172 | 0.0262322904504038 | 0.796962855124813 | 0.425472604368336 | NA | NR_045807 |
| 4931431F19Rik | 0 | NA | NA | NA | NA | NA | NM_027634 |
| 4931440J10Rik | 4.31903238 | 0.0538142277128026 | 0.0466411754144984 | 1.15379227119723 | 0.248585343580315 | NA | NR_045503 |
| 4931440L10Rik | 0.548592792 | -0.0138980954682334 | 0.0155684624275219 | -0.892708289783608 | 0.372013409853722 | NA | NM_001145300,NR_026896 |
| 4931440P22Rik | 9.681013394 | -0.0100266907680046 | 0.0654774006624722 | -0.153132083231143 | 0.878294115485757 | NA | NR_027955 |
| 4932411N23Rik | 0 | NA | NA | NA | NA | NA | NM_177705 |
| 4932412D23Rik | 2.915764801 | 0.0369453875526855 | 0.0407578156686052 | 0.906461422100786 | 0.36469167508212 | NA | NR_040521,NR_040522 |
| 4932413F04Rik | 0 | NA | NA | NA | NA | NA | NR_040764 |
| 4932414J04Rik | 0 | NA | NA | NA | NA | NA | NR_028259 |
| 4932414N04Rik | 0 | NA | NA | NA | NA | NA | NM_183113 |
| 4932415M13Rik | 0.714984696 | 3.31804448861758e-05 | 0.0158947881165835 | 0.00208750469920123 | 0.998334413439585 | NA | NR_073205 |
| 4932416K20Rik | 0 | NA | NA | NA | NA | NA | NM_001002775 |
| 4932429P05Rik | 0 | NA | NA | NA | NA | NA | NM_001085511 |
| 4932435O22Rik | 4.514933801 | -0.048638820108907 | 0.0475554673894614 | -1.0227808237184 | 0.306411486772814 | NA | NR_027643 |
| 4932443I19Rik | 20.11316877 | -0.0930395222411793 | 0.0763990995605388 | -1.21780914665695 | 0.22329650766996 | NA | NM_001101519 |
| 4932702P03Rik | 0.789727694 | 0.00345787351447456 | 0.0182591541064175 | 0.189377530542843 | 0.849796933844778 | NA | NR_045182 |
| 4933400A11Rik | 0 | NA | NA | NA | NA | NA | NR_003635 |
| 4933400B14Rik | 0.516150229 | 0.00784758350140605 | 0.0148035917613958 | 0.530113477046204 | 0.59603325549776 | NA | NR_045432 |
| 4933400C23Rik | 1.526865223 | 0.012416310961404 | 0.0283835540624994 | 0.437447365966356 | 0.661786939597487 | NA | NR_040770 |
| 4933400F21Rik | 0.127687691 | 0.00542298404407807 | 0.00869724871942359 | 0.62352868349814 | 0.532937159177571 | NA | NR_015540 |
| 4933400L20Rik | 0 | NA | NA | NA | NA | NA | NR_045730 |
| 4933401B06Rik | 0 | NA | NA | NA | NA | NA | NR_033580 |
| 4933401D09Rik | 0.106148608 | 0.00515554063321497 | 0.00854200771766808 | 0.603551390213729 | 0.546141939141592 | NA | NR_045431 |
| 4933401H06Rik | 0.57007369 | 0.0101860070818361 | 0.0153947996431531 | 0.661652461736742 | 0.508193977563616 | NA | NR_040540 |
| 4933401P06Rik | 0 | NA | NA | NA | NA | NA | NR_045505 |
| 4933402C06Rik | 0.53945737 | -0.00772964175130549 | 0.0165162376234508 | -0.468002575860887 | 0.639782747655422 | NA | NR_045504 |
| 4933402D24Rik | 9.614585055 | -0.0102716236407137 | 0.0610857942933877 | -0.168150774816488 | 0.866464665446857 | NA | NM_001256158 |
| 4933402E13Rik | 0 | NA | NA | NA | NA | NA | NM_001199996,NM_001199997 |
| 4933402J07Rik | 1.643947532 | -0.00373228614782902 | 0.0277643594217178 | -0.134427237853345 | 0.893064744410264 | NA | NM_177901 |
| 4933402J10Rik | 0 | NA | NA | NA | NA | NA | NR_040682,NR_040683 |
| 4933402J15Rik | 1.03580242 | -0.000969439091587504 | 0.0229836277784065 | -0.0421795506320508 | 0.966355564281951 | NA | NR_046059 |
| 4933402N03Rik | 0.106148608 | 0.00515554063321497 | 0.00854200771766808 | 0.603551390213729 | 0.546141939141592 | NA | NM_173409 |
| 4933402N22Rik | 0 | NA | NA | NA | NA | NA | NM_001177510 |
| 4933402P03Rik | 0 | NA | NA | NA | NA | NA | NM_175368 |
| 4933403O08Rik | 4.029635093 | 0.0273857532807568 | 0.0348630193067034 | 0.785524427469514 | 0.432146153970874 | NA | NM_001177389 |
| 4933404G15Rik | 0 | NA | NA | NA | NA | NA | NR_045920 |
| 4933404K08Rik | 0.256588971 | -0.000984450977364047 | 0.0101556058348068 | -0.0969367060298841 | 0.922776658860967 | NA | NR_046038 |
| 4933405D12Rik | 1.900325692 | -0.0552214682178813 | 0.0317251467503837 | -1.74062136425621 | 0.0817499702447958 | NA | NR_046036 |
| 4933405E24Rik | 0 | NA | NA | NA | NA | NA | NR_045506 |
| 4933405L10Rik | 10.60579032 | 0.000591300481139299 | 0.0670655377761457 | 0.00881675597850223 | 0.992965337668636 | NA | NM_027655 |
| 4933405O20Rik | 0.201873568 | 0.00673381845182595 | 0.00968011632508873 | 0.69563404257585 | 0.486658041207307 | NA | NM_172901 |
| 4933406D12Rik | 0 | NA | NA | NA | NA | NA | NR_040502 |
| 4933406F09Rik | 0.113843286 | 0.00525947738444006 | 0.00859368002043176 | 0.612016897526494 | 0.540526577873086 | NA | NR_015568 |
| 4933406G16Rik | 0 | NA | NA | NA | NA | NA | NR_046037 |
| 4933406I18Rik | 15.92333845 | -0.19876705445801 | 0.0730297830371401 | -2.72172593415654 | 0.00649419758134419 | NA | NR_029438,NR_029437 |
| 4933406J08Rik | 0 | NA | NA | NA | NA | NA | NM_028914 |
| 4933406J10Rik | 0 | NA | NA | NA | NA | NA | NR_046004 |
| 4933406K04Rik | 0 | NA | NA | NA | NA | NA | NR_015512 |
| 4933406M09Rik | 0.44803889 | 0.00538868503566239 | 0.0140821177917709 | 0.382661551007004 | 0.701970726278947 | NA | NM_173771 |
| 4933407E24Rik | 1.496508618 | 0.0117865722001179 | 0.0283964386835157 | 0.415072197309026 | 0.678089051737528 | NA | NR_045819 |
| 4933407G14Rik | 0 | NA | NA | NA | NA | NA | NR_045841 |
| 4933407I05Rik | 3.219468406 | 0.00763646684575858 | 0.0336239201184392 | 0.227114114560687 | 0.820335008973768 | NA | NR_040731 |
| 4933408J17Rik | 1.265031602 | -2.95821923166855e-05 | 0.025616749599433 | -0.00115479882417792 | 0.999078604052145 | NA | NR_045810 |
| 4933408N05Rik | 0.575450061 | -0.00902957083271323 | 0.0181609101915546 | -0.497198143566189 | 0.61904933044207 | NA | NR_045831 |
| 4933409G03Rik | 0 | NA | NA | NA | NA | NA | NM_177651 |
| 4933409K07Rik | 1.306977674 | 0.00223035423609889 | 0.0264929533030609 | 0.08418669714113 | 0.93290799478285 | NA | NR_033123 |
| 4933411E08Rik | 1.001886351 | -0.0114633533507541 | 0.0246882276748435 | -0.464324677402212 | 0.64241514629516 | NA | NR_045342 |
| 4933411G06Rik | 0 | NA | NA | NA | NA | NA | NR_045158 |
| 4933411G11Rik | 0.24789674 | -0.000769869626663926 | 0.0100368571301073 | -0.0767042528038554 | 0.938858821262798 | NA | NM_177880 |
| 4933411K16Rik | 1.923758284 | 0.0140365297081197 | 0.0290147591955786 | 0.483772055921757 | 0.628547648917798 | NA | NM_025752 |
| 4933412E24Rik | 0 | NA | NA | NA | NA | NA | NM_027668 |
| 4933413J09Rik | 0 | NA | NA | NA | NA | NA | NR_038005 |
| 4933413L06Rik | 0 | NA | NA | NA | NA | NA | NR_045508 |
| 4933415F23Rik | 0.451584937 | 0.00589671599968194 | 0.0141196736199318 | 0.417624100840258 | 0.676221972889845 | NA | NM_025746 |
| 4933416C03Rik | 0 | NA | NA | NA | NA | NA | NM_001161855 |
| 4933416E03Rik | 0 | NA | NA | NA | NA | NA | NR_040498 |
| 4933416I08Rik | 1.102862949 | 0.0123811483402552 | 0.0240852222925813 | 0.514055805250707 | 0.607212974961356 | NA | NM_027700 |
| 4933416M06Rik | 0 | NA | NA | NA | NA | NA | NR_045846 |
| 4933416M07Rik | 0 | NA | NA | NA | NA | NA | NR_045840 |
| 4933417A18Rik | 0 | NA | NA | NA | NA | NA | NM_025750 |
| 4933417D19Rik | 17.9403931 | 0.0348303926170066 | 0.0770166871977781 | 0.452244752199774 | 0.651092671773518 | NA | NR_045849 |
| 4933417E11Rik | 0.564827182 | 0.000975917913205956 | 0.017975469681116 | 0.0542916502610888 | 0.956702801869902 | NA | NR_040454 |
| 4933417G07Rik | 0 | NA | NA | NA | NA | NA | NR_033592 |
| 4933417O13Rik | 12.21227934 | 0.0258126886866 | 0.0657646048625698 | 0.392501235893404 | 0.69468789835581 | NA | NR_045842 |
| 4933421I07Rik | 0 | NA | NA | NA | NA | NA | NM_027702 |
| 4933422A05Rik | 1.083795384 | -0.0153830750659655 | 0.0248872196644872 | -0.618111435240652 | 0.536501884875252 | NA | NR_040715 |
| 4933422H20Rik | 1.80367557 | -0.0530893414245944 | 0.0313052392455271 | -1.69586122655746 | 0.0899121640792621 | NA | NM_001033775 |
| 4933424G05Rik | 0.574923518 | 0.012679746025291 | 0.0184110453820736 | 0.688703208435787 | 0.491010056720594 | NA | NR_045372 |
| 4933424G06Rik | 9.494021602 | 0.0383488347047232 | 0.0635186827326027 | 0.603741026339635 | 0.546015833323178 | NA | NR_040290 |
| 4933425B07Rik | 0 | NA | NA | NA | NA | NA | NR_046460 |
| 4933425H06Rik | 0.205932006 | -0.000576954122277892 | 0.00970424421944593 | -0.0594537925088239 | 0.952590668569529 | NA | NR_040519 |
| 4933425L06Rik | 0.47798497 | 0.0167779152807644 | 0.0152327156690434 | 1.10143953614661 | 0.270705406272229 | NA | NM_025751 |
| 4933426D04Rik | 0.106148608 | 0.00515554063321497 | 0.00854200771766808 | 0.603551390213729 | 0.546141939141592 | NA | NR_040684 |
| 4933427D06Rik | 1.027095484 | 0.0284053175386635 | 0.0230948845973133 | 1.22993979116778 | 0.218719652138007 | NA | NM_175017 |
| 4933427E11Rik | 3.383151085 | -0.0169547290096333 | 0.0341945606403247 | -0.495831169991378 | 0.62001353135832 | NA | NR_033197 |
| 4933427E13Rik | 0.562437198 | -0.0135432430921432 | 0.0156537553229193 | -0.865175340534037 | 0.386942557907326 | NA | NR_045853 |
| 4933427G17Rik | 15.23388054 | -0.0150994701424456 | 0.0740130444019631 | -0.204010931646599 | 0.838344956343853 | NA | NM_028955 |
| 4933427I22Rik | 20.81032013 | -0.0282096288160358 | 0.0817862862483006 | -0.34491881353302 | 0.730155406418331 | NA | NR_045160,NR_045159 |
| 4933428C19Rik | 0.226357657 | -0.0012264097261801 | 0.00980336873216358 | -0.125100846421945 | 0.900443713214317 | NA | NR_040644 |
| 4933429K18Rik | 0 | NA | NA | NA | NA | NA | NR_045307 |
| 4933429O19Rik | 2.211507293 | -0.00231967051311306 | 0.0207175251830617 | -0.111966583489884 | 0.910849902434564 | NA | NR_033783 |
| 4933430H16Rik | 0.12890128 | -0.0068640067066208 | 0.00870804083610955 | -0.788237771940376 | 0.430557639008166 | NA | NR_045855 |
| 4933430I17Rik | 15.56437843 | 0.100868527293321 | 0.073781715394261 | 1.36712092900414 | 0.171587404221123 | NA | NM_177607 |
| 4933430M04Rik | 2.325797149 | -0.0105457621572618 | 0.0329229836396895 | -0.320316113286545 | 0.748728709506493 | NA | NR_045857 |
| 4933430N04Rik | 0 | NA | NA | NA | NA | NA | NR_045854 |
| 4933431G14Rik | 0 | NA | NA | NA | NA | NA | NR_045163,NR_045165 |
| 4933432G23Rik | 0 | NA | NA | NA | NA | NA | NR_040716 |
| 4933432I03Rik | 0.217532603 | 0.00684685248201087 | 0.00976000296730706 | 0.701521557416086 | 0.482977586094298 | NA | NR_045657 |
| 4933433C11Rik | 1.021822652 | -0.0280155843502878 | 0.0224661992141043 | -1.24701041254453 | 0.212393679638605 | NA | NM_028961 |
| 4933433F19Rik | 0 | NA | NA | NA | NA | NA | NR_045856 |
| 4933433G15Rik | 6.836029445 | -0.0880732636923023 | 0.052278805914217 | -1.68468392022609 | 0.092049570476585 | NA | NR_040719 |
| 4933433G19Rik | 2.263605847 | 0.0036517514958335 | 0.03414208521353 | 0.1069574829128 | 0.914822709918294 | NA | NR_045851 |
| 4933433H22Rik | 0.342602601 | 0.0130117013735539 | 0.0134333274893572 | 0.968613426856654 | 0.332738101639367 | NA | NR_045458 |
| 4933434I20Rik | 0.461522871 | 0.0164668620078686 | 0.0150925202423685 | 1.09106111791999 | 0.275245990292358 | NA | NM_026233 |
| 4933436C20Rik | 16.16201907 | -0.0586270725376285 | 0.0738059980242554 | -0.794340217692897 | 0.426997396850208 | NA | NR_033641 |
| 4933436E23Rik | 0.217532603 | 0.00684685248201087 | 0.00976000296730706 | 0.701521557416086 | 0.482977586094298 | NA | NR_040455 |
| 4933436H12Rik | 0.309462921 | 0.00239536941478217 | 0.0109127344998192 | 0.219502216866162 | 0.826258853162846 | NA | NR_046208 |
| 4933436I01Rik | 0 | NA | NA | NA | NA | NA | NM_025763 |
| 4933438B17Rik | 0 | NA | NA | NA | NA | NA | NR_040685 |
| 4933438K21Rik | 0.335323189 | -0.0061413109930725 | 0.0128252480301014 | -0.478845397660815 | 0.632048617772247 | NA | NR_045446 |
| 4933439K11Rik | 4.146532363 | -0.0690314160625473 | 0.0462900269363571 | -1.49128053343881 | 0.135887857590343 | NA | NR_040558 |
| 4933440J02Rik | 0 | NA | NA | NA | NA | NA | NR_045344 |
| 4933440M02Rik | 0.313099713 | -0.00621682986300086 | 0.0125746085216642 | -0.494395499652349 | 0.621026891670966 | NA | NR_045804,NR_045803 |
| 5031410I06Rik | 2.236617333 | 0.0085471421337152 | 0.036830532819621 | 0.232066752212768 | 0.816486167876386 | NA | NM_207657 |
| 5031425F14Rik | 3.620920833 | 0.0161004504159944 | 0.0416940209904877 | 0.386157296262397 | 0.699380178117619 | NA | NR_015558 |
| 5031426D15Rik | 5.23184231 | -0.0475896109158114 | 0.0436113915736516 | -1.09121972949296 | 0.275176208493705 | NA | NR_027890 |
| 5031434C07Rik | 0.685151557 | 0.0142512902668939 | 0.0179647362160698 | 0.793292486763365 | 0.427607433181685 | NA | NR_045986,NR_045987 |
| 5033403H07Rik | 0.106148608 | 0.00515554063321497 | 0.00854200771766808 | 0.603551390213729 | 0.546141939141592 | NA | NR_040413 |
| 5033404E19Rik | 16.41084863 | -0.0481552475136654 | 0.0786959256741261 | -0.611915383180987 | 0.540593743223261 | NA | NR_033600 |
| 5033406O09Rik | 7.494689832 | 0.0452451945802717 | 0.0552514616606016 | 0.818895884749687 | 0.412845818043384 | NA | NR_029464 |
| 5330411J11Rik | 3.104631568 | -0.00778533882116634 | 0.0410595115173542 | -0.189611092130889 | 0.849613894633193 | NA | NR_040510 |
| 5330439B14Rik | 0 | NA | NA | NA | NA | NA | NR_037679 |
| 5430401F13Rik | 1.77566614 | -0.0222642925279734 | 0.0298684172148718 | -0.74541253283712 | 0.456022369701903 | NA | NM_001244628 |
| 5430402E10Rik | 2.930929908 | -0.00260991526020439 | 0.0115806195881553 | -0.225369224879283 | NA | NA | NM_027768 |
| 5430402O13Rik | 21.63667863 | -0.0452677372649591 | 0.0821637995641123 | -0.550945033032909 | 0.581671355554468 | NA | NR_015581,NR_027892 |
| 5430403N17Rik | 1.438699779 | 0.00603348175525568 | 0.0260605514174431 | 0.231517808606969 | 0.816912551949171 | NA | NR_046181 |
| 5430419D17Rik | 1.011123841 | 0.00805933578685623 | 0.0219086777149019 | 0.367860438303605 | 0.712977300583394 | NA | NM_175166 |
| 5430421F17Rik | 4.118278412 | -0.00148978562065869 | 0.0427679116282419 | -0.0348341914285781 | 0.972211956350157 | NA | NR_040352 |
| 5430421N21Rik | 0.106148608 | 0.00515554063321497 | 0.00854200771766808 | 0.603551390213729 | 0.546141939141592 | NA | NM_001201323 |
| 5430425J12Rik | 0 | NA | NA | NA | NA | NA | NM_027780 |
| 5430427M07Rik | 0 | NA | NA | NA | NA | NA | NR_045858 |
| 5430427O19Rik | 15.23594099 | -0.13460745756431 | 0.0752599703682242 | -1.78856644383084 | 0.0736846660600701 | NA | NM_001163539 |
| 5430428K19Rik | 0 | NA | NA | NA | NA | NA | NR_045426 |
| 5430434I15Rik | 0 | NA | NA | NA | NA | NA | NR_040541 |
| 5430437J10Rik | 0.448676571 | -0.00228182964137481 | 0.0154195577302497 | -0.147982820343698 | 0.882356326337823 | NA | NR_045274 |
| 5430440P10Rik | 1.973607392 | -0.0139692268944793 | 0.0302939879651514 | -0.461122085033795 | 0.644711017645273 | NA | NR_045859 |
| 5530400C23Rik | 0 | NA | NA | NA | NA | NA | NM_027784 |
| 5530401A14Rik | 0.763355183 | 0.00743895855830316 | 0.0182289350455689 | 0.408085197500959 | 0.683211127083107 | NA | NR_038010 |
| 5730403I07Rik | 2.007448435 | 0.00495050661391135 | 0.0310084242141785 | 0.159650376933625 | 0.873156492843507 | NA | NR_040378 |
| 5730408K05Rik | 11.00904932 | 0.00246588195014253 | 0.0620569274866381 | 0.0397358046879372 | 0.968303756194915 | NA | NR_027866 |
| 5730412P04Rik | 1.38199105 | -0.0104097805951466 | 0.025924072802956 | -0.401548810415299 | 0.68801610926895 | NA | NR_045425,NR_045424 |
| 5730416F02Rik | 0 | NA | NA | NA | NA | NA | NR_033596 |
| 5730420D15Rik | 3.190337572 | 0.0125227601923091 | 0.0400829520758804 | 0.312421105326834 | 0.754720512800642 | NA | NR_045339,NR_045338 |
| 5730422E09Rik | 6.063063076 | 0.0302209853217352 | 0.0481713068459899 | 0.627364863036738 | 0.530420094529315 | NA | NR_015478 |

| | | | | | | | | | | | | | | | |
|---|---|---|---|---|---|---|---|---|---|---|---|---|---|---|---|
| 1190003K10Rik | 0.17833704 | -0.002390466 | 0.031282281 | -0.076415987 | 0.939088151 | NA | | 5730435O14Rik | 1.298720366 | 0.023769142997922 | 0.0273093964705849 | 0.87036500508255 | 0.384100964667858 | NA | NR_045341 |
| 1300002K09Rik | 4.318427049 | 0.218488128 | 0.129086896 | 1.692566282 | 0.090538054 | NA | | 5730457N03Rik | 0 | NA | NA | NA | NA | NA | NR_038163 |
| 1300015D01Rik | 0.640586365 | -0.082316043 | 0.054836985 | -1.501104465 | 0.133328544 | NA | | 5730460C07Rik | 0 | NA | NA | NA | NA | NA | NR_045801 |
| 1300017J02Rik | 3.063965952 | -0.141407668 | 0.118448093 | -1.193836595 | 0.232541894 | NA | | 5730488B01Rik | 0.429829819 | 0.0124796205617181 | 0.0131493818755305 | 0.949065186473999 | 0.342587459210353 | NA | NR_073462 |
| 1300018J18Rik | 838.4683718 | 0.221761906 | 0.092208535 | 2.405004119 | NA | NA | | 5830403M04Rik | 2.249928154 | -0.023563789766575 | 0.0347219455893523 | -0.67864255204066 | 0.497364374252873 | NA | NR_045993 |
| 1500002O10Rik | 5.66066331 | -0.019385632 | 0.142637895 | -0.135908007 | 0.891894005 | NA | | 5830411N06Rik | 1.820817091 | -0.01941395218137 | 0.0246649863413558 | -0.787105734123947 | 0.431219972966004 | NA | NM_001128146,NM_001128145,NM_175533 |
| 1500016L03Rik | 2.023832634 | -0.027536726 | 0.096273574 | -0.286025798 | 0.774858363 | NA | | 5830416I19Rik | 7.346571602 | -0.0149944971749643 | 0.0587036722511625 | -0.255426902610294 | 0.798393394468488 | NA | NR_045384 |
| 1600002D24Rik | 0.496539307 | -0.049785487 | 0.052778288 | -0.943294857 | 0.345530106 | NA | | 5830428M24Rik | 0 | NA | NA | NA | NA | NA | NR_038061,NR_038060 |
| 1600010M07Rik | 1.65989692 | 0.123419835 | 0.094419236 | 1.307147157 | 0.191162738 | NA | | 5830432E09Rik | 2.993198669 | -0.028421723830361 | 0.0377335319087876 | -0.75322193265831 | 0.451316561709919 | NA | NR_015548 |
| 1600014C23Rik | 2.580700065 | 0.038362432 | 0.11349426 | 0.338012092 | 0.735354076 | NA | | 5830473C10Rik | 3.20689873 | 0.0372650687018995 | 0.0413288948867519 | 0.901671065824818 | 0.367231627566041 | NA | NM_001252661 |
| 1600014K23Rik | 0.347667565 | 0.012696751 | 0.035421431 | 0.358448274 | 0.720007871 | NA | | 5930438M14Rik | 20.46830921 | -0.120416576151801 | 0.0573613879989588 | -2.09926189641692 | 0.0357938203657862 | NA | NR_046158 |
| 1600015I10Rik | 0.107716603 | -0.020477583 | 0.027954224 | -0.732539849 | 0.46383913 | NA | | 6030407O03Rik | 12.91768032 | -0.0366676945813902 | 0.0641905329622927 | -0.571232125505638 | 0.567842304259998 | NA | NR_045311 |
| 1600016N20Rik | 5.956205954 | 0.142192876 | 0.143802093 | 0.988809503 | 0.322756354 | NA | | 6030408B16Rik | 6.045332183 | -0.0574920941643278 | 0.0469907951572905 | -1.22347566096481 | 0.221150091546285 | NA | NR_033803 |
| 1600019K03Rik | 0 | NA | NA | NA | NA | NA | | 6030440G07Rik | 0 | NA | NA | NA | NA | NA | NR_036598 |
| 1600025M17Rik | 0 | NA | NA | NA | NA | NA | | 6030466F02Rik | 1.590136553 | -0.0231153296542657 | 0.0272667524907769 | -0.847747807960795 | 0.396578435181234 | NA | NR_040702 |
| 1600029D21Rik | 5.514682936 | 0.320197518 | 0.141155291 | 2.268406061 | 0.023304469 | NA | | 6030468B19Rik | 3.236310119 | -0.0709846299652147 | 0.0413246116223639 | -1.71773253706272 | 0.0858454117177311 | NA | NM_029964 |
| 1600029I14Rik | 2.401394009 | 0.039527292 | 0.10482021 | 0.377096101 | 0.706102189 | NA | | 6030498E09Rik | 0.833729473 | 0.0148173274781542 | 0.0210919133494098 | 0.702512248779403 | 0.482359765690377 | NA | NM_183126 |
| 1700001C19Rik | 3.199909664 | -0.063890173 | 0.124721809 | -0.512261439 | 0.608468049 | NA | | 6230400D17Rik | 17.07324714 | -0.0565241594471918 | 0.0749585453516607 | -0.754072256632173 | 0.450805835279672 | NA | NR_029446 |
| 1700001D01Rik | 0 | NA | NA | NA | NA | NA | | 6330407A03Rik | 0 | NA | NA | NA | NA | NA | NR_028126 |
| 1700001F09Rik | 0.192142086 | -0.026384592 | 0.031527717 | -0.836869715 | 0.402665805 | NA | | 6330410L21Rik | 7.002595156 | -0.0142922647316427 | 0.0509528543674362 | -0.280499785715182 | 0.7790940898424 | NA | NR_040589 |
| 1700001G11Rik | 6.790179138 | 0.042434055 | 0.150812379 | 0.281369843 | 0.778426746 | NA | | 6330415B21Rik | 0 | NA | NA | NA | NA | NA | NR_045141 |
| 1700001G17Rik | 7.440824058 | 0.098663269 | 0.152781807 | 0.64577891 | 0.518422548 | NA | | 6430531B16Rik | 22.34841792 | 0.0107689176030811 | 0.0845559857871754 | 0.127358430072428 | 0.89865671624951 | NA | NM_001033465 |
| 1700001J03Rik | 2.254932462 | -0.035214803 | 0.100425809 | -0.350654906 | 0.725847259 | NA | | 6430562O15Rik | 2.337926215 | -0.00170355308112022 | 0.0310630977645166 | -0.0548416997568796 | 0.956264578770396 | NA | NR_015515 |
| 1700001J11Rik | 2.485277953 | -0.006079885 | 0.106896884 | -0.056876165 | 0.954643841 | NA | | 6430706D22Rik | 0 | NA | NA | NA | NA | NA | NR_040291 |
| 1700001K19Rik | 0.352713512 | 0.01988513 | 0.041853977 | 0.475107309 | 0.634710489 | NA | | 6530411M01Rik | 12.1557455 | -0.0483259149003368 | 0.0675966480645993 | -0.714915846924151 | 0.474661044318895 | NA | NR_027881 |
| 1700001K23Rik | 0.107716603 | -0.020477583 | 0.027954224 | -0.732539849 | 0.46383913 | NA | | 6720468P15Rik | 16.7774214 | -0.0497573637221371 | 0.0777492154527182 | -0.63997255062202 | 0.522190445056381 | NA | NR_040307,NR_040306 |
| 1700001P01Rik | 5.989583001 | 0.256745114 | 0.145391429 | 1.765888917 | 0.077414487 | NA | | 6720483E21Rik | 20.83510442 | -0.115411326316413 | 0.0739941297332626 | -1.55973624843556 | 0.118822222356041 | NA | NR_040492 |
| 1700003C15Rik | 0 | NA | NA | NA | NA | NA | | 7420426K07Rik | 0 | NA | NA | NA | NA | NA | NM_001033983 |
| 1700003F12Rik | 5.668743844 | 0.190346875 | 0.146811413 | 1.296540045 | 0.194789493 | NA | | 7420461P10Rik | 0.222609587 | 0.00717651388923987 | 0.00999169132085706 | 0.718248158273197 | 0.472604288901417 | NA | NM_001177581 |
| 1700003F17Rik | 0 | NA | NA | NA | NA | NA | | 7420700N18Rik | 0 | NA | NA | NA | NA | NA | NR_046272 |
| 1700003G13Rik | 0 | NA | NA | NA | NA | NA | | 7420701I03Rik | 0 | NA | NA | NA | NA | NA | NR_045860 |
| 1700003G18Rik | 0.094392817 | 0.017895942 | 0.027818137 | 0.643319236 | 0.520016977 | NA | | 7530416G11Rik | 0.322281847 | -0.00648722354265905 | 0.0126857541512189 | -0.511378627185182 | 0.609085957750446 | NA | NM_001256072 |
| 1700003H04Rik | 0 | NA | NA | NA | NA | NA | | 7530420F21Rik | 14.24436944 | 0.0264710138088646 | 0.0748579558165395 | 0.353616573150078 | 0.723626252306256 | NA | NR_015466 |
| 1700003L19Rik | 1.181434631 | 0.01240029 | 0.079220911 | 0.156527991 | 0.875616858 | NA | | 7630403G23Rik | 9.47790446 | -0.00205352192669275 | 0.0455639288975351 | -0.0450690266704335 | 0.964052289462129 | NA | NR_040744 |
| 1700003P14Rik | 0 | NA | NA | NA | NA | NA | | 8030411F24Rik | 0 | NA | NA | NA | NA | NA | NM_030135 |
| 1700006A11Rik | 0 | NA | NA | NA | NA | NA | | 8030423F21Rik | 2.702957672 | 0.0367122571561286 | 0.0380731830340774 | 0.964255001302865 | 0.334918095216808 | NA | NR_045738 |
| 1700006E09Rik | 0 | NA | NA | NA | NA | NA | | 8030423J24Rik | 1.830766549 | -0.0114658005628313 | 0.030118160545652 | -0.380693918722291 | 0.703430381848856 | NA | NM_029873 |
| 1700006F04Rik | 0.298391179 | 0.008861245 | 0.039220874 | 0.225931861 | 0.821254417 | NA | | 8030442B05Rik | 2.221277528 | -0.0358870383373318 | 0.0330567131849654 | -1.08562028343622 | 0.277647022139788 | NA | NR_040615 |
| 1700006H21Rik | 0 | NA | NA | NA | NA | NA | | 8030443G20Rik | 0.242744565 | -0.00126183740302166 | 0.00996759682058903 | -0.126593944933168 | 0.899261789625903 | NA | NR_040664 |
| 1700007B14Rik | 5.000454671 | 0.271909199 | 0.120059613 | 2.264784899 | 0.023525883 | NA | | 8430403D17Rik | 0 | NA | NA | NA | NA | NA | NR_045875 |
| 1700007J10Rik | 4.664008559 | 0.177386577 | 0.132557753 | 1.338183347 | 0.180836676 | NA | | 8430422H06Rik | 0 | NA | NA | NA | NA | NA | NR_045373 |
| 1700007K09Rik | 5.328206121 | 0.020225251 | 0.133870142 | 0.151081122 | 0.879911724 | NA | | 8430423G03Rik | 0 | NA | NA | NA | NA | NA | NR_040686 |
| 1700008A04Rik | 0 | NA | NA | NA | NA | NA | | 8430431K14Rik | 0.217532603 | 0.00684685248201087 | 0.00976000296730706 | 0.701521557416086 | 0.482977586094298 | NA | NR_002849 |
| 1700008F21Rik | 0 | NA | NA | NA | NA | NA | | 8430436N08Rik | 0.33403441 | 0.00244344761020736 | 0.0111793636306143 | 0.218567683362231 | 0.826986829057954 | NA | NR_040645 |
| 1700008I05Rik | 0.201994201 | 0.000227241 | 0.031727614 | 0.00716224 | 0.994285408 | NA | | 8430437L04Rik | 0 | NA | NA | NA | NA | NA | NR_040503 |
| 1700008K24Rik | 0 | NA | NA | NA | NA | NA | | 9030204H09Rik | 0.544628687 | 0.00309863725245576 | 0.0164300944981167 | 0.188595217928354 | 0.850410081096137 | NA | NR_040618 |
| 1700008P02Rik | 0 | NA | NA | NA | NA | NA | | 9030404E10Rik | 0.226066984 | -0.0093084363525918 | 0.0101406635841621 | -0.917931679257152 | 0.358654635211298 | NA | NR_045878 |
| 1700009C05Rik | 1.413023452 | 0.042920249 | 0.080989815 | 0.529946255 | 0.596149195 | NA | | 9030619P08Rik | 0.12890128 | -0.0068640067066208 | 0.00870804083610955 | -0.788237771940376 | 0.430557639008166 | NA | NM_001039720 |
| 1700009J07Rik | 1.022906055 | -0.048349521 | 0.077944357 | -0.620308166 | 0.53505492 | NA | | 9130008F23Rik | 3.572183108 | 0.00394823092756085 | 0.0321407750978445 | 0.122841808125083 | 0.902232366859569 | NA | NM_027834 |
| 1700009N14Rik | 0 | NA | NA | NA | NA | NA | | 9130015A21Rik | 0.713363127 | -0.0143745352822909 | 0.0180311098994497 | -0.797207457691201 | 0.425330555521743 | NA | NR_045050,NR_045051 |
| 1700010B08Rik | 0 | NA | NA | NA | NA | NA | | 9130015L21Rik | 0 | NA | NA | NA | NA | NA | NR_040499 |
| 1700010D01Rik | 0 | NA | NA | NA | NA | NA | | 9130017N09Rik | 10.90361137 | -0.0310533748155099 | 0.0641616371105612 | -0.48398663459911 | 0.628395354516106 | NA | NM_030218 |
| 1700010I02Rik | 0 | NA | NA | NA | NA | NA | | 9130019P16Rik | 7.618092065 | 0.00757122108683138 | 0.0538775603036477 | 0.140526427777369 | 0.888244073314361 | NA | NR_033635 |
| 1700010J16Rik | 0 | NA | NA | NA | NA | NA | | 9130204L05Rik | 0 | NA | NA | NA | NA | NA | NM_001101461 |
| 1700010K23Rik | 0.843194396 | 0.001915388 | 0.057336671 | 0.033405992 | 0.973350831 | NA | | 9130206I24Rik | 4.213188275 | -0.0651201832988439 | 0.0435450796199484 | -1.49546593707482 | 0.134792883301021 | NA | NR_030721 |
| 1700011A15Rik | 0.28821313 | -0.029561117 | 0.033427731 | -0.884329147 | 0.376518559 | NA | | 9130209A04Rik | 0 | NA | NA | NA | NA | NA | NR_033453 |
| 1700011B04Rik | 0 | NA | NA | NA | NA | NA | | 9130221F21Rik | 0.108766301 | 0.00519171381361946 | 0.00856003371502575 | 0.606506234257728 | 0.544178643801544 | NA | NR_046180 |
| 1700011E24Rik | 1.014018918 | -0.01008284 | 0.071659643 | -0.140704577 | 0.888103329 | NA | | 9130227L01Rik | 10.78565312 | -0.0472622064003937 | 0.0623855914093243 | -0.75758208478456 | 0.448701208503042 | NA | NR_045837 |
| 1700011H14Rik | 0.181709657 | 0.024794574 | 0.031325423 | 0.791516008 | 0.428642938 | NA | | 9130230L23Rik | 0 | NA | NA | NA | NA | NA | NR_027961 |
| 1700011I03Rik | 3.089749683 | -0.036099319 | 0.115858701 | -0.311580556 | 0.755359313 | NA | | 9130409I23Rik | 0.127687691 | 0.00542298404407807 | 0.00869724871942359 | 0.62352868349814 | 0.532937159177571 | NA | NM_001033819 |
| 1700011L22Rik | 0.287632815 | 0.04259931 | 0.040911109 | 1.041265103 | 0.297752528 | NA | | 9230009I02Rik | 8.788628684 | -0.0498523762212364 | 0.0624144987193465 | -0.798730699503058 | 0.424446581742498 | NA | NR_045865 |
| 1700011M02Rik | 0 | NA | NA | NA | NA | NA | | 9230102K24Rik | 1.866512249 | 0.00474597141793534 | 0.0315958257543768 | 0.150208810962249 | 0.880599874623351 | NA | NR_028438 |
| 1700012A03Rik | 0 | NA | NA | NA | NA | NA | | 9230102O04Rik | 0.491929936 | 0.0015689969597419 | 0.0144579934045654 | 0.108521073141896 | 0.913582365278159 | NA | NR_040511 |
| 1700012B07Rik | 0 | NA | NA | NA | NA | NA | | 9230104L09Rik | 0.997185341 | -0.0143594568740809 | 0.0197803361418445 | -0.725946049203079 | 0.467871845056282 | NA | NM_029960 |
| 1700012I11Rik | 0 | NA | NA | NA | NA | NA | | 9230105E05Rik | 0 | NA | NA | NA | NA | NA | NR_040626 |
| 1700012L04Rik | 0.333037401 | -0.016093237 | 0.040545167 | -0.396921218 | 0.691425557 | NA | | 9230110F15Rik | 0 | NA | NA | NA | NA | NA | NM_029863 |
| 1700012P22Rik | 0.292249882 | 0.042980062 | 0.04108841 | 1.046038582 | 0.29554322 | NA | | 9230112D13Rik | 0.09572496 | 0.00498255289994511 | 0.00847999972823692 | 0.587565219295241 | 0.556824162160132 | NA | NM_030062 |
| 1700013D24Rik | 0.188785634 | 0.025025111 | 0.031461603 | 0.79541754 | 0.42637066 | NA | | 9230115E21Rik | 0.30673395 | 0.00299118959348609 | 0.0123970371859449 | 0.241282618469303 | 0.809336081443423 | NA | NR_033560 |
| 1700013G24Rik | 0 | NA | NA | NA | NA | NA | | 9230116N13Rik | 12.64693172 | -0.146153256640757 | 0.0688452483883173 | -2.12292438566549 | 0.0337601886346593 | NA | NR_024328 |
| 1700013H16Rik | 0.653760176 | -0.030006487 | 0.054578099 | -0.549789885 | 0.582463497 | NA | | 9330111N05Rik | 9.001082341 | 0.0741431457335516 | 0.0632706047643966 | 1.17184190051038 | 0.241260536717713 | NA | NR_015587 |
| 1700015E13Rik | 0.094392817 | 0.017895942 | 0.027818137 | 0.643319236 | 0.520016977 | NA | | 9330158H04Rik | 21.24006886 | -0.0491308433792597 | 0.072291234719107 | -0.679623796303401 | 0.496742698217376 | NA | NR_015589 |
| 1700015F17Rik | 2.761763786 | 0.067473314 | 0.109707585 | 0.615028702 | 0.538535766 | NA | | 9330162B11Rik | 0.558930329 | 0.00131328921510046 | 0.0165135184962504 | 0.0795281281453529 | 0.936612559437325 | NA | NR_038007 |
| 1700015G11Rik | 0 | NA | NA | NA | NA | NA | | 9330175E14Rik | 2.187960253 | -0.0395105617141962 | 0.0303828866464502 | -1.30042158844089 | 0.193456514729249 | NA | NR_015514 |
| 1700016C15Rik | 0 | NA | NA | NA | NA | NA | | 9330175M20Rik | 12.09548082 | 0.0570337226861325 | 0.0660973676877103 | 0.86287434252449 | 0.388206558891148 | NA | NR_045151 |
| 1700016D06Rik | 0 | NA | NA | NA | NA | NA | | 9330178D15Rik | 0 | NA | NA | NA | NA | NA | NR_040553 |
| 1700016G22Rik | 0.499641834 | 0.064869181 | 0.050070714 | 1.295551338 | 0.195130103 | NA | | 9330182O14Rik | 0 | NA | NA | NA | NA | NA | NM_001256056 |
| 1700016H13Rik | 0.410728284 | -0.005925811 | 0.04256373 | -0.139222079 | 0.889274662 | NA | | 9430007A20Rik | 0.09572496 | 0.00498255289994511 | 0.00847999972823692 | 0.587565219295241 | 0.556824162160132 | NA | NM_198662 |
| 1700016L04Rik | 1.09910115 | 0.022207625 | 0.066299534 | 0.334958993 | 0.73765602 | NA | | 9430014N10Rik | 0.597814748 | 0.0201603017007698 | 0.0166745069430061 | 1.20904934518773 | 0.226643888277166 | NA | NR_045737 |
| 1700016L21Rik | 0 | NA | NA | NA | NA | NA | | 9430018G01Rik | 1.620084179 | 0.00615053706277432 | 0.0253936487787475 | 0.242207692024229 | 0.808619235104196 | NA | NR_045988 |
| 1700016P04Rik | 0 | NA | NA | NA | NA | NA | | 9430019J16Rik | 0 | NA | NA | NA | NA | NA | NR_040635 |
| 1700017D01Rik | 0 | NA | NA | NA | NA | NA | | 9430060I03Rik | 7.176976948 | -0.0545747428199059 | 0.0554827649870687 | -0.983634157970059 | 0.325295423896 | NA | NR_015525 |
| 1700017G19Rik | 0.402750607 | -0.037318681 | 0.048991399 | -0.76173946 | 0.446215519 | NA | | 9430069I07Rik | 0 | NA | NA | NA | NA | NA | NM_001256161 |
| 1700017J07Rik | 0.239950962 | 0.027417169 | 0.032831728 | 0.835081494 | 0.403671825 | NA | | 9430076C15Rik | 1.012800456 | -0.032898706521558 | 0.024612160161566 | -1.33668504940627 | 0.18132546409342 | NA | NR_015553 |
| 1700017N19Rik | 0 | NA | NA | NA | NA | NA | | 9430076G02Rik | 0.758139492 | 0.00103915156195008 | 0.019857698129089 | 0.0523299103045511 | 0.958265820983542 | NA | NR_040338 |
| 1700018A04Rik | 1.349866079 | 0.002120246 | 0.087195859 | 0.024315901 | 0.98060063 | NA | | 9530002B09Rik | 0 | NA | NA | NA | NA | NA | NM_023865 |
| 1700018B08Rik | 0.094392817 | 0.017895942 | 0.027818137 | 0.643319236 | 0.520016977 | NA | | 9530003J23Rik | 0.257802559 | -0.00892927589713349 | 0.00993479775307735 | -0.898787888698349 | 0.368765652383134 | NA | NM_029906 |
| 1700018B24Rik | 0 | NA | NA | NA | NA | NA | | 9530026F06Rik | 0 | NA | NA | NA | NA | NA | NR_040483 |
| 1700018C11Rik | 0 | NA | NA | NA | NA | NA | | 9530026P05Rik | 12.89476998 | 0.0492693221084623 | 0.0636920810357418 | 0.773554911493849 | 0.439194044381702 | NA | NR_015530 |
| 1700018F24Rik | 0 | NA | NA | NA | NA | NA | | 9530027J09Rik | 2.367267478 | -0.0180004418900075 | 0.0335395991433367 | -0.536692219041731 | 0.591480231628257 | NA | NR_045916 |
| 1700018G05Rik | 0 | NA | NA | NA | NA | NA | | 9530048J24Rik | 0.658477857 | 0.0211121211175937 | 0.0170573850724673 | 1.23771146795948 | 0.215823065378972 | NA | NR_046194 |
| 1700019A02Rik | 1.732172825 | 0.01584073 | 0.093920608 | 0.168660855 | 0.866063411 | NA | | 9530052E02Rik | 13.79833074 | 0.0462477069463801 | 0.0661159377225566 | 0.699494078726526 | 0.484243312200862 | NA | NR_046017 |
| 1700019B03Rik | 5.325715112 | 0.11904574 | 0.130617208 | 0.91140931 | 0.36207975 | NA | | 9530053A07Rik | 7.95183268 | 0.0604333029110478 | 0.051743253714098 | 1.16794554986754 | 0.242828729086991 | NA | NM_001164655 |
| 1700019B21Rik | 0 | NA | NA | NA | NA | NA | | 9530059O14Rik | 2.865997635 | -0.0524531886352628 | 0.0413662211577873 | -1.26801982794574 | 0.204790869838282 | NA | NR_015610 |
| 1700019E08Rik | 0 | NA | NA | NA | NA | NA | | 9630028B13Rik | 14.62226931 | 0.0360456516511586 | 0.0648196878869577 | 0.556091101734714 | 0.578148577773077 | NA | NR_038006 |
| 1700019G24Rik | 2.199194789 | 0.093608401 | 0.091510596 | 1.022924175 | 0.306343698 | NA | | 9630028H03Rik | 0.682786739 | 0.0153351655445467 | 0.0182097640510302 | 0.842139716998645 | 0.399709752113727 | NA | NR_015544 |
| 1700019M22Rik | 0 | NA | NA | NA | NA | NA | | 9830107B12Rik | 0.191449919 | 0.00665508079272897 | 0.00962405648474729 | 0.691504752001018 | 0.489248395458281 | NA | NM_177824,NM_001177897,NM_001177896 |
| 1700019N19Rik | 6.154910495 | -0.063797621 | 0.146511642 | -0.43544404 | 0.66324015 | NA | | 9830132P13Rik | 0.343816189 | 0.00220638102164924 | 0.0128049404084123 | 0.172307012081036 | 0.863196171959189 | NA | NR_040552 |
| 1700019O17Rik | 0 | NA | NA | NA | NA | NA | | 9930111J21Rik2 | 0.437549073 | -0.00919688418782782 | 0.0142039622589292 | -0.647487230687783 | 0.517316656204546 | NA | NM_173434 |
| 1700020A23Rik | 0.094392817 | 0.017895942 | 0.027818137 | 0.643319236 | 0.520016977 | NA | | A1bg | 0.09572496 | 0.00498255289994511 | 0.00847999972823692 | 0.587565219295241 | 0.556824162160132 | NA | NM_001081067 |
| 1700020G17Rik | 0.204276544 | 2.70E-05 | 0.031767541 | 0.000850493 | 0.999321405 | NA | | A1cf | 0.644943959 | 0.0116957533623467 | 0.0160737826258774 | 0.72762918564779 | 0.466840609703208 | NA | NM_001081074 |
| 1700020L24Rik | 6.022953304 | 0.19663498 | 0.139041814 | 1.414214722 | 0.157298867 | NA | | A230020J21Rik | 0.097165704 | -0.00637828455464458 | 0.00848781691535724 | -0.75146349388194 | 0.452373763047372 | NA | NR_027298 |
| 1700020M21Rik | 0 | NA | NA | NA | NA | NA | | A230028O05Rik | 7.763369906 | 0.00132075709519351 | 0.0528952743012262 | 0.0249692834122025 | 0.980079464252447 | NA | NR_040374 |
| 1700020N01Rik | 0.620554997 | 0.021634778 | 0.053065255 | 0.40770138 | 0.683492924 | NA | | A230108P19Rik | 0.726181501 | -0.00864880776890885 | 0.019853072907064 | -0.43564075996676 | 0.663097392919228 | NA | NR_040333 |
| 1700020N15Rik | 0 | NA | NA | NA | NA | NA | | A330032B11Rik | 10.25026493 | 0.00936423534246792 | 0.0657066105932312 | 0.142515878660047 | 0.886672538030031 | NA | NR_045329 |
| 1700020N18Rik | 1.495483865 | 0.062587367 | 0.087569837 | 0.714713756 | 0.474785936 | NA | | A330040F15Rik | 10.12829481 | 0.0623049422658107 | 0.0620781453759737 | 1.00365340956086 | 0.315545701219314 | NA | NR_015503 |
| 1700021F07Rik | 1.755178943 | -0.177768483 | 0.087435428 | -2.033140181 | 0.042038362 | NA | | A330070K13Rik | 13.90318997 | -0.00352313145581737 | 0.0693419426367935 | -0.0508080870227591 | 0.959478446689893 | NA | NM_198665 |
| 1700021N21Rik | 2.167429004 | -0.162674127 | 0.106199892 | -1.531772994 | 0.125578462 | NA | | A330076C08Rik | 18.65219595 | 0.0685032728725322 | 0.0726059344829808 | 0.943494128411634 | 0.345428217188667 | NA | NR_045088 |
| 1700022A21Rik | 0 | NA | NA | NA | NA | NA | | A330093E20Rik | 5.245324324 | -0.0675812452406373 | 0.0486891671770043 | -1.38801399077033 | 0.165132784876428 | NA | NR_040342 |
| 1700022A22Rik | 0 | NA | NA | NA | NA | NA | | A3galt2 | 7.479995603 | 0.00138295813885313 | 0.0554942094805114 | 0.0249207647392257 | 0.980118164509949 | NA | NM_001009819 |
| 1700022E09Rik | 0 | NA | NA | NA | NA | NA | | A430071A18Rik | 0.520471911 | 0.00216599178832626 | 0.0146598204825785 | 0.147750225925365 | 0.882539892036391 | NA | NR_045313,NR_045312 |
| 1700022H16Rik | 0.119397641 | -0.020477583 | 0.027954224 | -0.732539845 | 0.463839132 | NA | | A430088P11Rik | 0.097165704 | -0.00637828455464458 | 0.00848781691535724 | -0.75146349388194 | 0.452373763047372 | NA | NR_045310,NR_045309 |
| 1700022I11Rik | 0 | NA | NA | NA | NA | NA | | A430089I19Rik | 0 | NA | NA | NA | NA | NA | NM_177913 |
| 1700022P22Rik | 0.294090812 | 0.009484222 | 0.033616198 | 0.282132504 | 0.77784191 | NA | | A430090L17Rik | 0.345220294 | 0.0102849425769615 | 0.0119473016652568 | 0.860859034544216 | 0.389315686243366 | NA | NR_045836 |
| 1700023A16Rik | 2.179656538 | 0.082062822 | 0.086572956 | 0.947903666 | 0.343178498 | NA | | A430092G05Rik | 0.426726963 | 0.00796577679624491 | 0.015424583208532 | 0.516433843855095 | 0.605551433285488 | NA | NR_040622 |
| 1700023C21Rik | 0 | NA | NA | NA | NA | NA | | A430093F15Rik | 4.098456256 | 0.036858254591699 | 0.0411507447440041 | 0.895688639925999 | 0.370419079324913 | NA | NR_027805 |
| 1700023E05Rik | 7.707835974 | -0.111299066 | 0.158723178 | -0.701214954 | 0.483168879 | NA | | A430107P09Rik | 17.62972331 | 0.00384520950442947 | 0.0751695332387362 | 0.0511538297333469 | 0.959202942179327 | NA | NM_001205242 |
| 1700023F02Rik | 0.978055737 | 0.13187226 | 0.070074519 | 1.881886059 | 0.059851488 | NA | | A4gnt | 9.347932523 | 0.0643867106153567 | 0.0637672568207489 | 1.0097142926557 | 0.312632192808243 | NA | NM_001077424 |
| 1700023L04Rik | 4.933102782 | 0.176106676 | 0.142383379 | 1.23684855 | 0.216143314 | NA | | A530006G24Rik | 0 | NA | NA | NA | NA | NA | NR_046014 |
| 1700024B18Rik | 0 | NA | NA | NA | NA | NA | | A530013C23Rik | 2.257348453 | -0.0111950817461078 | 0.0296464075313825 | -0.377620179924232 | 0.705712771408032 | NA | NR_015500 |
| 1700024G13Rik | 5.754891192 | -0.009231793 | 0.13331388 | -0.069248554 | 0.944791775 | NA | | A530016L24Rik | 0.544512808 | 0.00114718639850937 | 0.0146457089524885 | 0.0783291817576672 | 0.937566205082552 | NA | NM_177039 |
| 1700024P03Rik | 0 | NA | NA | NA | NA | NA | | A530032D15Rik | 0.194331409 | -0.00840968081320881 | 0.00964081822647896 | -0.872299489073576 | 0.383045016303155 | NA | NM_213615 |
| 1700024P04Rik | 0 | NA | NA | NA | NA | NA | | A530046M15Rik | 0 | NA | NA | NA | NA | NA | NR_046131 |
| 1700025B11Rik | 0 | NA | NA | NA | NA | NA | | A530050N04Rik | 6.075994793 | 0.0255812912976618 | 0.0498071792463668 | 0.513606505823712 | 0.607527130105561 | NA | NR_045419 |
| 1700025C18Rik | 0 | NA | NA | NA | NA | NA | | A530053G22Rik | 1.720483642 | -0.0157290183925281 | 0.0271061672587773 | -0.580274527282528 | 0.561729502350024 | NA | NR_015565 |
| 1700025E21Rik | 0.785077497 | 0.025108046 | 0.067164331 | 0.373830062 | 0.708530752 | NA | | A530064D06Rik | 2.518090489 | -0.0496322765824622 | 0.0338792375811745 | -1.46497619562847 | 0.142927412430819 | NA | NM_001113556,NM_178796 |
| 1700025F22Rik | 0 | NA | NA | NA | NA | NA | | A530065N20Rik | 1.025780582 | 0.0131097453202344 | 0.0194406196980617 | 0.674348118724916 | 0.500090018626307 | NA | NM_046142 |
| 1700025F24Rik | 0.425134012 | -0.003256152 | 0.042864108 | -0.075964537 | 0.939447312 | NA | | A530088E08Rik | 1.545027225 | -0.0196233544486569 | 0.0283626425727514 | -0.691873276558142 | 0.489016914145406 | NA | NR_029458 |
| 1700025K24Rik | 0 | NA | NA | NA | NA | NA | | A530099J19Rik | 0.220191124 | -0.000706246424282353 | 0.00977545482624743 | -0.072246911968337 | 0.942405412335504 | NA | NM_175688 |
| 1700025M24Rik | 0.116696964 | -0.020477583 | 0.027954224 | -0.732539847 | 0.463839131 | NA | | A630001G21Rik | 21.48783318 | -0.201825658041601 | 0.0828445276317206 | -2.43619782514546 | 0.0148425647455428 | NA | NM_177055 |
| 1700025N23Rik | 0.198019769 | 0.025308392 | 0.031628095 | 0.80018706 | 0.423602426 | NA | | A630010A05Rik | 0 | NA | NA | NA | NA | NA | NR_033556 |
| 1700026D11Rik | 0.199381295 | -0.004055078 | 0.031639813 | -0.128163771 | 0.89801937 | NA | | A630012P03Rik | 0 | NA | NA | NA | NA | NA | NR_045367 |
| 1700026F02Rik | 0 | NA | NA | NA | NA | NA | | A630020A06 | 0.883981836 | -0.00389200043149929 | 0.0208567869087587 | -0.186605945034845 | 0.85196960516556 | NA | NR_045740 |
| 1700026L06Rik | 4.384532908 | 0.260504994 | 0.122357671 | 2.12904505 | 0.033250533 | NA | | A630023A22Rik | 0 | NA | NA | NA | NA | NA | NM_001251843 |
| 1700027A15Rik | 0.079983654 | 0.016740032 | 0.027674178 | 0.604897175 | 0.54524732 | NA | | A630033H20Rik | 10.2089395 | -0.0249191854573942 | 0.0635855652451465 | -0.391900038339225 | 0.695132074655225 | NA | NM_001122596,NM_175442,NM_001122595 |
| 1700027F09Rik | 0 | NA | NA | NA | NA | NA | | A630073D07Rik | 0 | NA | NA | NA | NA | NA | NM_001142969 |
| 1700027H10Rik | 0.690975005 | 0.005249197 | 0.047660626 | 0.110136963 | 0.912300754 | NA | | A630075F10Rik | 0.097165704 | -0.00637828455464458 | 0.00848781691535724 | -0.75146349388194 | 0.452373763047372 | NA | NR_033634,NR_033632 |
| 1700027I24Rik | 0.222620122 | -0.001447027 | 0.032059667 | -0.04513545 | 0.963999345 | NA | | A630076J17Rik | 0.194331409 | -0.00840968081320881 | 0.00964081822647896 | -0.872299489073576 | 0.383045016303155 | NA | NM_001256174 |
| 1700027J07Rik | 0.510163613 | 0.009578335 | 0.050846372 | 0.188377952 | 0.850580382 | NA | | A630077J23Rik | 2.550528828 | 0.0317363811137837 | 0.037780484729149 | 0.840020485213572 | 0.400896900814044 | NA | NR_040667 |
| 1700028B04Rik | 1.206882925 | 0.053053473 | 0.077520621 | 0.684378841 | 0.493735974 | NA | | A630095E13Rik | 0 | NA | NA | NA | NA | NA | NM_001033325 |
| 1700028D13Rik | 0 | NA | NA | NA | NA | NA | | A630095N17Rik | 1.165960768 | -0.0162736633746058 | 0.0237247934967768 | -0.685934879762673 | 0.492754173904241 | NA | NM_001243091,NM_001243090 |
| 1700028E10Rik | 7.104082621 | 0.015533361 | 0.156379048 | 0.099331474 | 0.92087509 | NA | | A730008H23Rik | 0 | NA | NA | NA | NA | NA | NM_172505 |
| 1700028I16Rik | 0.544515322 | -0.071444103 | 0.051231676 | -1.394529886 | 0.163157651 | NA | | A730017C20Rik | 1673.97398 | 0.0941309431796579 | 0.0950643404498672 | 0.99018141538886 | NA | NA | NM_001167925,NR_033765,NR_033764,NM_173759 |
| 1700028M03Rik | 0.116696964 | -0.020477583 | 0.027954224 | -0.732539847 | 0.463839131 | NA | | A730018C14Rik | 6.521334774 | -0.0350144865700841 | 0.0518459202467742 | -0.675356641437235 | 0.499449205656958 | NA | NR_036459 |
| 1700028P15Rik | 0 | NA | NA | NA | NA | NA | | A730020M07Rik | 19.82716672 | 0.168763197468622 | 0.0779013071371508 | 2.16637183213758 | 0.0302827846290138 | NA | NR_036458,NR_036456,NR_036457 |
| 1700029B22Rik | 0 | NA | NA | NA | NA | NA | | A730043L09Rik | 8.732907499 | -0.0682567088576155 | 0.0603389895112036 | -1.13122061556801 | 0.257962247023806 | NA | NR_040769 |
| 1700029F12Rik | 0 | NA | NA | NA | NA | NA | | A730082K24Rik | 0 | NA | NA | NA | NA | NA | NR_040319,NR_040317 |
| 1700029H14Rik | 0.571399219 | 0.021750442 | 0.053110482 | 0.409532001 | 0.682149287 | NA | | A730085A09Rik | 5.000880027 | -0.0241197498542583 | 0.0396289423225714 | -0.608639757728797 | 0.542763245637167 | NA | NR_040437 |
| 1700029J03Rik | 0 | NA | NA | NA | NA | NA | | A730085K08Rik | 6.659131101 | 0.0325262629110486 | 0.0553702566025928 | 0.587432042161162 | 0.556913579212255 | NA | NR_045967 |
| 1700029M20Rik | 1.763654503 | -0.237893836 | 0.094151439 | -2.526714805 | 0.011513496 | NA | | A730090H04Rik | 17.38644728 | 0.0643037163999628 | 0.0796267216575065 | 0.807564534385182 | 0.419341310067009 | NA | NR_040279 |
| 1700029N11Rik | 0 | NA | NA | NA | NA | NA | | A730090N16Rik | 2.43617978 | -0.0191701228754629 | 0.035018617018169 | -0.547426612122253 | 0.584085676431672 | NA | NR_040390 |
| 1700029P11Rik | 0 | NA | NA | NA | NA | NA | | A730098P11Rik | 8.658001106 | 0.0631277616852846 | 0.0623875412963284 | 1.01186487515897 | 0.311602665680784 | NA | NR_038091 |
| 1700030A11Rik | 1.17684774 | 0.072940977 | 0.07688226 | 0.948736124 | 0.342754837 | NA | | A830019L24Rik | 13.43938145 | -0.0298970663438666 | 0.0720201345431245 | -0.415120945462338 | 0.678053366989142 | NA | NR_040551 |
| 1700030F04Rik | 0.880480036 | -0.025404946 | 0.074833042 | -0.339488353 | 0.734241869 | NA | | A930001A20Rik | 5.182807737 | 0.00969523839659097 | 0.0505799810332882 | 0.191681337167621 | 0.847991825687811 | NA | NR_040549,NR_040550 |
| 1700030F18Rik | 0.276690427 | -0.012033946 | 0.033317809 | -0.361186582 | 0.717959972 | NA | | A930001C03Rik | 1.794854418 | -0.0131820367633516 | 0.0242493134489022 | -0.543604535078017 | 0.586713638332199 | NA | NR_045990,NR_045989 |
| 1700030M09Rik | 0.098353386 | -0.020212831 | 0.02786466 | -0.725393061 | 0.468210929 | NA | | A930003A15Rik | 9.866450946 | 0.0130297317669189 | 0.0667775746917162 | 0.195121368619206 | 0.845297934341833 | NA | NR_015488,NR_027896 |
| 1700030N03Rik | 0 | NA | NA | NA | NA | NA | | A930003O13Rik | 11.43787874 | 0.0124911184536566 | 0.0627865317348765 | 0.198945826573154 | 0.842305121520392 | NA | NR_027362 |
| 1700030O20Rik | 0 | NA | NA | NA | NA | NA | | A930006I01Rik | 0.097165704 | -0.00637828455464458 | 0.00848781691535724 | -0.75146349388194 | 0.452373763047372 | NA | NR_040332 |
| 1700031A10Rik | 0 | NA | NA | NA | NA | NA | | A930006K02Rik | 19.83852871 | -0.0518496859879413 | 0.0784651817860914 | -0.66079864734516 | 0.50874145039826 | NA | NR_077219 |
| 1700031F05Rik | 0 | NA | NA | NA | NA | NA | | A930007I19Rik | 14.63490669 | -0.0536523639926317 | 0.0713988180206886 | -0.751446109053029 | 0.452384222063693 | NA | NR_015567 |
| 1700031M16Rik | 0.395349417 | 0.027211441 | 0.048374467 | 0.562516604 | 0.573764094 | NA | | A930009A15Rik | 14.81798236 | 0.0173189966537876 | 0.0724167363736094 | 0.239157376057879 | 0.810983552663362 | NA | NM_029982 |
| 1700031P21Rik | 0 | NA | NA | NA | NA | NA | | A930018P22Rik | 1.744636652 | -0.0349721474084709 | 0.0260149520325642 | -1.34430950957336 | 0.178848321733107 | NA | NM_026634 |
| 1700034E13Rik | 0.119397641 | -0.020477583 | 0.027954224 | -0.732539845 | 0.463839132 | NA | | A930019D19Rik | 0.703384099 | -0.00596706768149885 | 0.0159608783635798 | -0.373855845873418 | 0.708511568121965 | NA | NR_040619 |
| 1700034F02Rik | 3.160626376 | -0.105399317 | 0.124784832 | -0.844648464 | 0.398307142 | NA | | A930041C12Rik | 1.028348205 | -0.0221623296676117 | 0.0225951960086746 | -0.980842549854548 | 0.326670391760056 | NA | NR_046195 |
| 1700034G24Rik | 0.188785634 | 0.025025111 | 0.031461603 | 0.79541754 | 0.42637066 | NA | | AA467197 | 5.030761406 | 0.0387443750340386 | 0.047348363173868 | 0.818283303517068 | 0.413195437654716 | NA | NM_001004174 |
| 1700034I23Rik | 0.46990427 | 0.035927024 | 0.051562175 | 0.696770909 | 0.485946174 | NA | | AA536875 | 0.334833285 | -0.00610342348777058 | 0.012817048366707 | -0.47619571317407 | 0.633934953261501 | NA | NR_045143,NR_045144,NR_045145 |
| 1700034J05Rik | 0.567794913 | -0.037435068 | 0.058745955 | -0.637236527 | 0.523970788 | NA | | AA543186 | 9.438754294 | -0.0478256783692797 | 0.0675438221944185 | -0.7080688775891 | 0.478902485503748 | NA | NR_027448 |
| 1700034K08Rik | 0 | NA | NA | NA | NA | NA | | AA545190 | 0 | NA | NA | NA | NA | NA | NR_033776 |
| 1700034O15Rik | 0 | NA | NA | NA | NA | NA | | AA619741 | 0.194331409 | -0.00840968081320881 | 0.00964081822647896 | -0.872299489073576 | 0.383045016303155 | NA | NR_033627 |
| 1700034P13Rik | 2.790014232 | 0.094678183 | 0.115412086 | 0.820348946 | 0.412017211 | NA | | AA792892 | 0.106148608 | 0.00515554063321497 | 0.00854200771766808 | 0.603551390213729 | 0.546141939141592 | NA | NM_178894 |
| 1700036G14Rik | 0.08731684 | 0.017365056 | 0.027733412 | 0.626142072 | 0.531221759 | NA | | AB099516 | 0.427411356 | 0.0060983518828938 | 0.0139232555105122 | 0.43799755583667 | 0.661388055218398 | NA | NM_001024672 |
| 1700039E15Rik | 1.781638371 | 0.01592076 | 0.072687075 | 0.219031518 | 0.826625496 | NA | | AB124611 | 12.63636353 | -0.12735234247559 | 0.0720127930413812 | -1.76846831093482 | 0.0769826453419399 | NA | NM_206536,NM_001198794 |
| 1700039E22Rik | 0 | NA | NA | NA | NA | NA | | AF357355 | 5.026700563 | -0.0299523149423338 | 0.0405188255849561 | -0.739219720954956 | 0.4597735875975 | NA | NR_028433 |
| 1700041C23Rik | 0 | NA | NA | NA | NA | NA | | AF357399 | 7.506786207 | -0.0702296182699029 | 0.058391639542298 | -1.20273413831838 | 0.229079218760261 | NA | NR_028129 |
| 1700041M19Rik | 0.098353386 | -0.020212831 | 0.02786466 | -0.725393061 | 0.468210929 | NA | | AF357425 | 6.577827015 | -0.0527533930930175 | 0.0556096864870034 | -0.94863676502378 | 0.342805385518008 | NA | NR_046302 |
| 1700042B14Rik | 0.096071043 | -0.020098795 | 0.027825948 | -0.722304036 | 0.470107572 | NA | | AF357426 | 7.577226937 | -0.0216451519109026 | 0.0517480522748862 | -0.418279547912709 | 0.675742741128282 | NA | NR_046303 |
| 1700042G07Rik | 0 | NA | NA | NA | NA | NA | | AF366264 | 0 | NA | NA | NA | NA | NA | NM_153093 |
| 1700042G15Rik | 0 | NA | NA | NA | NA | NA | | AI182371 | 11.8628424 | -0.0272641203321508 | 0.0681405266190181 | -0.400116079005198 | 0.68907102190133 | NA | NM_178885,NM_001243103,NM_001243102 |
| 1700042O10Rik | 0 | NA | NA | NA | NA | NA | | AI314831 | 0.120209049 | -0.00674972139751468 | 0.00863521114606269 | -0.781650996523957 | 0.434419707908266 | NA | NR_015462 |
| 1700044C05Rik | 0 | NA | NA | NA | NA | NA | | AI427809 | 4.927230866 | -0.0458241210456548 | 0.0490383357498021 | -0.934455061433029 | 0.350069206505353 | NA | NR_033140,NR_033139 |
| 1700044K03Rik | 0 | NA | NA | NA | NA | NA | | AI463170 | 1.011354418 | 0.0066952313593629 | 0.0210943631674164 | 0.317394334506611 | 0.750944405567597 | NA | NR_046044 |
| 1700045H11Rik | 0.079983654 | 0.016740032 | 0.027674178 | 0.604897175 | 0.54524732 | NA | | AI506816 | 1.858313118 | -0.000606898089826422 | 0.0288861224675708 | -0.0210100227369651 | 0.983237660453399 | NA | NR_015554,NR_038090 |
| 1700046C09Rik | 0 | NA | NA | NA | NA | NA | | AI507597 | 2.008287894 | 0.0199085559854652 | 0.0330746160570179 | 0.601928559084238 | 0.547221692964838 | NA | NR_033566 |
| 1700047E10Rik | 0.30236719 | -0.01694275 | 0.039582453 | -0.428036895 | 0.668624259 | NA | | AI606473 | 21.79203477 | -0.0126509840337338 | 0.0775556056957918 | -0.163121465176311 | 0.870422799384952 | NA | NR_040387 |
| 1700047G03Rik | 0 | NA | NA | NA | NA | NA | | AI607873 | 16.16421211 | -0.0694303890566175 | 0.0659169177263689 | -1.05330151122711 | 0.29220282652325 | NA | NM_001204910 |
| 1700047I17Rik2 | 0 | NA | NA | NA | NA | NA | | AI646519 | 0 | NA | NA | NA | NA | NA | NR_040330 |
| 1700047L14Rik | 0 | NA | NA | NA | NA | NA | | AI662270 | 4.04762712 | -0.0524746937903383 | 0.0391961149369126 | -1.33877283181759 | 0.180644637821764 | NA | NR_015519 |
| 1700048M11Rik | 0 | NA | NA | NA | NA | NA | | AI747448 | 0.635802653 | 0.00568284268458012 | 0.0171635211184364 | 0.331100049073021 | 0.740568914569294 | NA | NM_001033199 |
| 1700049E15Rik | 0 | NA | NA | NA | NA | NA | | AI847159 | 12.35379395 | -0.0313331893218891 | 0.0697292909749424 | -0.449354767326529 | 0.653175755645191 | NA | NR_045264 |
| 1700049E22Rik | 0 | NA | NA | NA | NA | NA | | AU015228 | 0.360627147 | -0.0100772056525624 | 0.010553896215922 | -0.954832741045871 | 0.339662287230353 | NA | NM_001033197 |
| 1700049L16Rik | 0 | NA | NA | NA | NA | NA | | AU015836 | 0.097165704 | -0.00637828455464458 | 0.00848781691535724 | -0.75146349388194 | 0.452373763047372 | NA | NR_028320 |
| 1700051A21Rik | 1.76604708 | 0.096531081 | 0.095244804 | 1.013504957 | 0.31081903 | NA | | AU016765 | 0 | NA | NA | NA | NA | NA | NR_045899 |
| 1700052I22Rik | 0 | NA | NA | NA | NA | NA | | AU018091 | 0 | NA | NA | NA | NA | NA | NM_001004153 |
| 1700054A03Rik | 0.119397641 | -0.020477583 | 0.027954224 | -0.732539845 | 0.463839132 | NA | | AU018829 | 0 | NA | NA | NA | NA | NA | NM_001200055 |
| 1700054K19Rik | 0 | NA | NA | NA | NA | NA | | AU019990 | 0.384276662 | 0.0023869237934625 | 0.0116652479776256 | 0.204618349994849 | 0.837870317514098 | NA | NR_033468,NR_033469 |
| 1700054M17Rik | 0.651627236 | -0.059904957 | 0.056388213 | -1.062366654 | 0.288069266 | NA | | AU021063 | 1.630926438 | -0.0262761091164492 | 0.0268214619192103 | -0.979667297613988 | 0.327250373641236 | NA | NR_045996 |
| 1700054O13Rik | 0 | NA | NA | NA | NA | NA | | AU022751 | 0.665083679 | -0.00345676881088412 | 0.0189776069938744 | -0.182149878643808 | 0.85546510591964 | NA | NM_001033211,NM_001166433 |
| 1700055C04Rik | 0 | NA | NA | NA | NA | NA | | AU022793 | 7.69148397 | -0.118792464953515 | 0.0563529263271984 | -2.10800880621137 | 0.035030227711814 | NA | NR_045719 |
| 1700055N04Rik | 2.253040213 | -0.154269086 | 0.094692377 | -1.629160568 | 0.103279032 | NA | | AU023871 | 1.603182284 | 0.0103159310476967 | 0.0235491844348347 | 0.43805895173325 | 0.661343549518274 | NA | NM_001033221,NM_001191012 |
| 1700056E22Rik | 2.635905444 | 0.037144121 | 0.112075612 | 0.331420196 | 0.740327112 | NA | | AU040972 | 8.472841476 | -0.0519505572470685 | 0.0522136185607056 | -0.994961825652223 | 0.319754831222753 | NA | NR_045305 |
| 1700057G04Rik | 0 | NA | NA | NA | NA | NA | | AV039307 | 19.59031109 | -0.0120767633089205 | 0.0766601465262066 | -0.157536397413381 | 0.874822125015629 | NA | NR_038350,NR_038349 |
| 1700057H15Rik | 0.098353386 | -0.020212831 | 0.02786466 | -0.725393061 | 0.468210929 | NA | | AV051173 | 4.227758262 | -0.0212673512587954 | 0.042464467588347 | -0.500826984691351 | 0.616492892619942 | NA | NR_040442 |
| 1700058G18Rik | 5.410811809 | 0.122361825 | 0.125293175 | 0.976604074 | 0.328765202 | NA | | AV320801 | 0 | NA | NA | NA | NA | NA | NM_177918 |
| 1700060C16Rik | 0 | NA | NA | NA | NA | NA | | AW046200 | 16.02101879 | -0.00887601547777514 | 0.069443753287944 | -0.12781589498729 | 0.898294670859872 | NA | NR_040698 |
| 1700060C20Rik | 0.197363271 | -0.000561558 | 0.031650166 | -0.017742642 | 0.985844163 | NA | | AY074887 | 1.794163629 | -0.0169043489637137 | 0.0291532479669398 | -0.579844447619814 | 0.562019520814455 | NA | NM_145229 |
| 1700061F12Rik | 0 | NA | NA | NA | NA | NA | | AY358078 | 0.491638859 | -0.00244243873529541 | 0.0142369039247984 | -0.171556874176911 | 0.863785914098802 | NA | NM_194347 |
| 1700061G19Rik | 5.80828273 | -0.169261873 | 0.140020175 | -1.208839177 | 0.226724637 | NA | | AY512915 | 0.428629686 | -0.0106292681563651 | 0.0141635099071664 | -0.750468508585358 | 0.452972583388393 | NA | NR_033559 |
| 1700061I17Rik | 0.285336609 | 0.031742875 | 0.03528833 | 0.899528977 | 0.368370968 | NA | | AY512931 | 5.63855703 | 0.0239873679366072 | 0.0429585670459345 | 0.558383800627198 | 0.576582332221375 | NA | NR_033588 |
| 1700063A18Rik | 0 | NA | NA | NA | NA | NA | | AY761184 | 0 | NA | NA | NA | NA | NA | NM_001007582 |
| 1700063O14Rik | 1.386926346 | -0.16451203 | 0.088094712 | -1.867445009 | 0.061839469 | NA | | AY761185 | 0 | NA | NA | NA | NA | NA | NM_001012640 |
| 1700064J06Rik | 0 | NA | NA | NA | NA | NA | | Aadac | 0 | NA | NA | NA | NA | NA | NM_023383 |
| 1700064M15Rik | 1.360944942 | 0.053265395 | 0.088665645 | 0.600744463 | 0.5480102 | NA | | Aadacl2 | 0 | NA | NA | NA | NA | NA | NM_001128091 |
| 1700065D16Rik | 1.437427219 | 0.011840258 | 0.087890685 | 0.134715731 | 0.892836635 | NA | | Aadacl3 | 0.106347839 | -0.00653347234140708 | 0.00854389509723638 | -0.764694822098226 | 0.444453290197145 | NA | NM_001085503 |
| 1700065I16Rik | 0 | NA | NA | NA | NA | NA | | Aadat | 4.590899464 | -0.0737982530075065 | 0.0495001993357892 | -1.49086779442825 | 0.135996208273427 | NA | NM_011834 |
| 1700065I17Rik | 0 | NA | NA | NA | NA | NA | | Abca12 | 9.920506747 | -0.0114156521569779 | 0.0607595461356664 | -0.187882446183659 | 0.85096880359936 | NA | NM_175210 |
| 1700065J11Rik | 0.08731684 | 0.017365056 | 0.027733412 | 0.626142072 | 0.531221759 | NA | | Abca13 | 9.487931622 | -0.140900299279444 | 0.0636352887828528 | -2.21418495891875 | 0.0268160581190012 | NA | NM_178259 |
| 1700065J18Rik | 0.282093074 | 0.009461846 | 0.038656194 | 0.244769212 | 0.806635134 | NA | | Abca14 | 0.356663042 | 0.00289666753272811 | 0.0129429520293053 | 0.223802694019843 | 0.822910833613774 | NA | NM_026458 |
| 1700065O20Rik | 0 | NA | NA | NA | NA | NA | | Abca15 | 1.86664755 | -0.00427633257264839 | 0.032011625378521 | -0.133586861712986 | 0.893729273897017 | NA | NM_177213 |
| 1700066B17Rik | 0.199381295 | -0.004055078 | 0.031639813 | -0.128163771 | 0.89801937 | NA | | Abca16 | 0.308221406 | 0.00253133711108074 | 0.0124107148809387 | 0.203963843772494 | 0.838381753432823 | NA | NR_024051 |
| 1700066B19Rik | 1.988627642 | -0.036136824 | 0.088810508 | -0.406898072 | 0.684082852 | NA | | Abcb11 | 1.083999946 | 0.0147278207928579 | 0.0253869098833607 | 0.580134441746728 | 0.561823959210836 | NA | NM_021022 |
| 1700066J24Rik | 4.91723478 | 0.019788423 | 0.106371591 | 0.186031088 | 0.852420382 | NA | | Abcb5 | 0 | NA | NA | NA | NA | NA | NM_029961 |
| 1700066N21Rik | 0 | NA | NA | NA | NA | NA | | Abcc2 | 10.12131111 | -0.0644695528197231 | 0.0603135220606656 | -1.06890711430974 | 0.28511152591965 | NA | NM_013806 |
| 1700066O22Rik | 0 | NA | NA | NA | NA | NA | | Abcg5 | 1.070025063 | 0.0213610437441172 | 0.0225705863293443 | 0.946410670614502 | 0.343939164961787 | NA | NM_031884 |
| 1700067G17Rik | 0 | NA | NA | NA | NA | NA | | Abcg8 | 0.127687691 | 0.00542298404407807 | 0.00869724871942359 | 0.62352868349814 | 0.532937159177571 | NA | NM_026180 |
| 1700067K01Rik | 1.22238232 | -0.038266874 | 0.075601477 | -0.506165695 | 0.612740335 | NA | | Abhd12b | 14.96552665 | 0.0155990633351248 | 0.044521332382888 | 0.35037278761047 | 0.726058946001471 | NA | NM_001195033 |
| 1700067P10Rik | 1.352373962 | 0.01319769 | 0.082785853 | 0.159419633 | 0.873338272 | NA | | Abhd16b | 0 | NA | NA | NA | NA | NA | NM_183181 |
| 1700069L16Rik | 0 | NA | NA | NA | NA | NA | | Abo | 0.09572496 | 0.00498255289994511 | 0.00847999972823692 | 0.587565219295241 | 0.556824162160132 | NA | NM_030718 |
| 1700069P05Rik | 0 | NA | NA | NA | NA | NA | | Abp1 | 2.477613783 | -0.0121281870286346 | 0.0377060499076554 | -0.321650956765222 | 0.747717135635165 | NA | NM_001161621,NM_029638,NM_001161622 |
| 1700071K01Rik | 0 | NA | NA | NA | NA | NA | | Abra | 9.294690511 | -0.00209285154548801 | 0.0617099632863474 | -0.0339143216756869 | 0.972945472712433 | NA | NM_175456 |
| 1700071M16Rik | 1.025937788 | -0.019215963 | 0.066067457 | -0.290853686 | 0.771163224 | NA | | Acaa1b | 13.06777948 | -0.0434825315716432 | 0.0657618784234078 | -0.661211823842393 | 0.508476479622871 | NA | NM_146230 |
| 1700072B07Rik | 0 | NA | NA | NA | NA | NA | | Acap1 | 2.194760634 | -0.0448651763090205 | 0.0328261583976363 | -1.36675074084365 | 0.171703447507759 | NA | NM_153788 |
| 1700072O05Rik | 0 | NA | NA | NA | NA | NA | | Acbd7 | 5.681388768 | -0.0749467534029375 | 0.053553659785779 | -1.39947024540869 | 0.161672014926704 | NA | NM_030063 |
| 1700073E17Rik | 0.079983654 | 0.016740032 | 0.027674178 | 0.604897175 | 0.54524732 | NA | | Accsl | 14.8359031 | -0.013306477704119 | 0.0728243220703208 | -0.18272024134011 | 0.855017532841957 | NA | NM_001033452 |
| 1700074P13Rik | 0 | NA | NA | NA | NA | NA | | Ace3 | 0 | NA | NA | NA | NA | NA | NM_001101453 |
| 1700080E11Rik | 0 | NA | NA | NA | NA | NA | | Acer1 | 0.108766301 | 0.00519171381361946 | 0.00856003371502575 | 0.606506234257728 | 0.544178643801544 | NA | NM_175731 |
| 1700080N15Rik | 0 | NA | NA | NA | NA | NA | | Acmsd | 0.723721657 | 0.01180888712453 | 0.0213044017315752 | 0.554293299258813 | 0.579378135381561 | NA | NM_001033041 |

| | | | | | | | | | | | | | | | |
|---|---|---|---|---|---|---|---|---|---|---|---|---|---|---|---|
| 1700080O16Rik | 0.099009884 | 0.018205726 | 0.027896865 | 0.652608315 | 0.514008823 | NA | | Acnat1 | 6.584851945 | 0.0116789172626695 | 0.0408971826536954 | 0.285567770317162 | 0.775209191459065 | NA | NM_001164565 |
| 1700081H04Rik | 0 | NA | NA | NA | NA | NA | | Acnat2 | 0.255375383 | 0.00708304372802858 | 0.00992471678592415 | 0.713677163873753 | 0.475426830432214 | NA | NM_145368 |
| 1700081H22Rik | 0 | NA | NA | NA | NA | NA | | Acot10 | 0.202072798 | -0.00115107594702963 | 0.00968429294483289 | -0.11886009165427 | 0.905386199803646 | NA | NM_022816 |
| 1700084F23Rik | 0 | NA | NA | NA | NA | NA | | Acot12 | 10.04418092 | -0.0112099098467599 | 0.0636325109790273 | -0.176166391586442 | 0.860163223448779 | NA | NM_028790 |
| 1700084J12Rik | 0.098353386 | -0.020212831 | 0.02786466 | -0.725393061 | 0.468210929 | NA | | Acox2 | 8.518887032 | 0.00126042591186088 | 0.0414733004487242 | 0.0303912613229136 | 0.975755014092292 | NA | NM_053115,NM_001161667 |
| 1700085B03Rik | 0 | NA | NA | NA | NA | NA | | Acoxl | 1.027057486 | -0.0104360012283847 | 0.0231147221884016 | -0.451487201244459 | 0.651638447427263 | NA | NM_028765 |
| 1700085C21Rik | 0.394855623 | -0.002402222 | 0.042167018 | -0.056969207 | 0.954569725 | NA | | Acp5 | 11.66981932 | 0.0104257345303603 | 0.0705234565839572 | 0.147833572478805 | 0.882474113428185 | NA | NM_007388,NM_001102404,NM_001102405 |
| 1700091E21Rik | 1.870113224 | 0.15100164 | 0.099934872 | 1.511000488 | 0.130788327 | NA | | Acpp | 11.63108551 | -0.0339727761890324 | 0.0574964423521971 | -0.590867448474996 | 0.554609238016789 | NA | NM_207668,NM_019807 |
| 1700091H14Rik | 7.58037232 | 0.00378325 | 0.151503259 | 0.024971409 | 0.980077769 | NA | | Acpt | 0 | NA | NA | NA | NA | NA | NM_001195034 |
| 1700092C02Rik | 0.08731684 | 0.017365056 | 0.027733412 | 0.626142072 | 0.531221759 | NA | | Acrv1 | 0 | NA | NA | NA | NA | NA | NM_007391 |
| 1700092C10Rik | 0 | NA | NA | NA | NA | NA | | Acsbg2 | 1.700927331 | -0.0196610643466672 | 0.0312786893165741 | -0.628576988878145 | 0.529626029102695 | NA | NM_001039114 |
| 1700092E19Rik | 0 | NA | NA | NA | NA | NA | | Acsm1 | 0 | NA | NA | NA | NA | NA | NM_054094 |
| 1700092K14Rik | 2.149771734 | 0.043087493 | 0.091281036 | 0.472031155 | 0.636904551 | NA | | Acsm2 | 0.113843286 | 0.00525947738444006 | 0.00859368002043176 | 0.612016897526494 | 0.540526577873086 | NA | NM_001177977,NM_001177978,NM_146197 |
| 1700092M07Rik | 1.382630839 | 0.038598511 | 0.077139913 | 0.500370169 | 0.616814454 | NA | | Acsm3 | 5.070424228 | -0.0398928218040261 | 0.0488384099824547 | -0.816832935764242 | 0.414023907471209 | NA | NM_016870,NM_212442,NM_212441 |
| 1700093K21Rik | 5.207369283 | 0.198452885 | 0.126825587 | 1.564770083 | 0.117636829 | NA | | Acsm4 | 2.126352761 | 0.00540439346408994 | 0.0329118255798371 | 0.164208255509256 | 0.869567202369676 | NA | NM_178414 |
| 1700094J05Rik | 1.235265611 | 0.066574415 | 0.071141067 | 0.935808499 | 0.349371795 | NA | | Acsm5 | 8.175990469 | 0.0502230956734052 | 0.0577838519862777 | 0.86915451197909 | 0.384762623916191 | NA | NM_178758 |
| 1700094M24Rik | 0 | NA | NA | NA | NA | NA | | Actbl2 | 0.774573665 | -0.00944649768394838 | 0.0200622431252253 | -0.470859495869173 | 0.637741074271472 | NA | NM_175497 |
| 1700095A21Rik | 0 | NA | NA | NA | NA | NA | | Actc1 | 0.108766301 | 0.00519171381361946 | 0.00856003371502575 | 0.606506234257728 | 0.544178643801544 | NA | NM_009608 |
| 1700095B10Rik | 0.352791569 | -0.038810225 | 0.038158703 | -1.017074017 | 0.30911822 | NA | | Actg2 | 3.014386258 | 0.0418971603409291 | 0.0328500948936331 | 1.27540454530161 | 0.202166090713363 | NA | NM_009610 |
| 1700096J18Rik | 0.096071043 | -0.020098795 | 0.027825948 | -0.722304036 | 0.470107572 | NA | | Actl10 | 0.120209049 | -0.00674972139751468 | 0.00863521114606269 | -0.781650996523957 | 0.434419707908266 | NA | NM_001171640 |
| 1700097N02Rik | 0.311993147 | -0.014421208 | 0.039860783 | -0.361789373 | 0.717509434 | NA | | Actl11 | 0 | NA | NA | NA | NA | NA | NM_026338 |
| 1700100L14Rik | 0 | NA | NA | NA | NA | NA | | Actl7a | 0 | NA | NA | NA | NA | NA | NM_009611 |
| 1700101E01Rik | 6.980296523 | 0.342701974 | 0.153761258 | 2.228792724 | 0.025827699 | NA | | Actl7b | 0.664630415 | -0.0223659551732091 | 0.0186081101603519 | -1.20194662329891 | 0.229384209039899 | NA | NM_025271 |
| 1700101O22Rik | 0 | NA | NA | NA | NA | NA | | Actl9 | 4.933858128 | -0.046020337228154 | 0.0421887884076184 | -1.09081912434925 | 0.275352479605337 | NA | NM_183282 |
| 1700102H20Rik | 5.830630321 | -0.068945968 | 0.136910925 | -0.503582663 | 0.614554678 | NA | | Actrt1 | 0 | NA | NA | NA | NA | NA | NM_028514 |
| 1700104A03Rik | 0.105923158 | 0.018524693 | 0.027977518 | 0.662127842 | 0.507889294 | NA | | Actrt2 | 0 | NA | NA | NA | NA | NA | NM_028513 |
| 1700105P06Rik | 3.909002408 | 0.146085372 | 0.133234882 | 1.096449896 | 0.272881938 | NA | | Adad1 | 0.127687691 | 0.00542298404407807 | 0.00869724871942359 | 0.62352868349814 | 0.532937159177571 | NA | NM_009350 |
| 1700106J16Rik | 0 | NA | NA | NA | NA | NA | | Adad2 | 3.406875155 | 0.0266603955676753 | 0.0423705913808239 | 0.629219340557547 | 0.5292054686945 | NA | NM_029428 |
| 1700108F19Rik | 0.495097952 | -0.018652806 | 0.05081183 | -0.367095744 | 0.713547602 | NA | | Adam1b | 0.235249118 | -0.00959082599542929 | 0.0102925929774586 | -0.93181825186654 | 0.351430460444064 | NA | NM_172125 |
| 1700108J01Rik | 0.119397641 | -0.020477583 | 0.027954224 | -0.732539845 | 0.463839132 | NA | | Adam2 | 0.684756679 | -0.00192347900033274 | 0.0175468996051627 | -0.109619308459873 | 0.912711297004945 | NA | NM_009618 |
| 1700109F18Rik | 0.323149808 | -0.030167563 | 0.033779321 | -0.893077861 | 0.371815478 | NA | | Adam20 | 0 | NA | NA | NA | NA | NA | NM_001009548 |
| 1700109G14Rik | 1.172532083 | -0.100619872 | 0.080734232 | -1.246309893 | 0.212650647 | NA | | Adam21 | 18.57972341 | -0.0465026166086472 | 0.0806260542514932 | -0.576769098281726 | 0.564095447239493 | NA | NM_020330 |
| 1700109G15Rik | 0 | NA | NA | NA | NA | NA | | Adam24 | 0 | NA | NA | NA | NA | NA | NM_010086 |
| 1700109I08Rik | 0 | NA | NA | NA | NA | NA | | Adam25 | 0.520319392 | 0.0140235985383759 | 0.0139295254941114 | 1.00675349955777 | 0.314053245642838 | NA | NM_011781 |
| 1700110I01Rik | 0 | NA | NA | NA | NA | NA | | Adam26a | 0.127687691 | 0.00542298404407807 | 0.00869724871942359 | 0.62352868349814 | 0.532937159177571 | NA | NM_010085 |
| 1700110K17Rik | 1.860880301 | 0.07121156 | 0.08778855 | 0.811171393 | 0.417267251 | NA | | Adam26b | 0 | NA | NA | NA | NA | NA | NM_001009547 |
| 1700110M21Rik | 0.099009884 | 0.018205726 | 0.027896865 | 0.652608315 | 0.514008823 | NA | | Adam28 | 1.873391939 | -0.0170177261372877 | 0.0342006934089856 | -0.497584242921128 | 0.61877711212635 | NA | NM_010082,NM_001048175,NM_183366 |
| 1700111N16Rik | 0.188785634 | 0.025025111 | 0.031461603 | 0.79541754 | 0.42637066 | NA | | Adam29 | 0 | NA | NA | NA | NA | NA | NM_175939 |
| 1700112H15Rik | 0 | NA | NA | NA | NA | NA | | Adam3 | 5.039618237 | -0.0586536793559516 | 0.0473825513933317 | -1.23787507492064 | 0.215762385659604 | NA | NM_009619 |
| 1700112J05Rik | 0 | NA | NA | NA | NA | NA | | Adam30 | 0 | NA | NA | NA | NA | NA | NM_027665 |
| 1700113H08Rik | 0.66421776 | -0.022294278 | 0.060705401 | -0.367253607 | 0.713429856 | NA | | Adam32 | 13.84849078 | 0.0820761639499564 | 0.0717345789559831 | 1.14416457368934 | 0.252555391947022 | NA | NM_153397 |
| 1700119H24Rik | 0.496818212 | 0.030597582 | 0.045890909 | 0.666746028 | 0.504934373 | NA | | Adam34 | 0.09572496 | 0.00498255289994511 | 0.00847999972823692 | 0.587565219295241 | 0.556824162160132 | NA | NM_145745 |
| 1700120E14Rik | 0 | NA | NA | NA | NA | NA | | Adam39 | 0 | NA | NA | NA | NA | NA | NM_001025380 |
| 1700120G07Rik | 0.247284148 | 0.028796741 | 0.033643376 | 0.855940884 | 0.392030479 | NA | | Adam4 | 5.892466854 | 0.00143494627570475 | 0.0550557223607255 | 0.0260635264451345 | 0.979206668853323 | NA | NM_009620 |
| 1700120K04Rik | 1.267252396 | 0.099744843 | 0.074046513 | 1.347056583 | 0.17796201 | NA | | Adam5 | 10.4070559 | 0.0538883063668454 | 0.0694589168916306 | 0.775829926212665 | 0.437849411653269 | NA | NM_001272057,NM_007401,NM_001272059,NM_001272058 |
| 1700121L16Rik | 0 | NA | NA | NA | NA | NA | | Adam6a | 0 | NA | NA | NA | NA | NA | NM_174885 |
| 1700121N20Rik | 1.017859841 | -0.064033937 | 0.073737633 | -0.868402388 | 0.385174088 | NA | | Adam6b | 0 | NA | NA | NA | NA | NA | NM_001009545 |
| 1700122O11Rik | 0 | NA | NA | NA | NA | NA | | Adam7 | 0.219991894 | 0.00711449016834187 | 0.00994861810862167 | 0.715123456410123 | 0.474532761039468 | NA | NM_007402 |
| 1700123I01Rik | 2.289901513 | 0.025548849 | 0.106337215 | 0.240262538 | 0.810126735 | NA | | Adamdec1 | 0.227686571 | 0.00691460391109036 | 0.00980755285755433 | 0.705028462402254 | 0.480792523671872 | NA | NM_021475 |
| 1700123K08Rik | 0.569095902 | 0.022553468 | 0.05830359 | 0.386828113 | 0.698883463 | NA | | Adamts12 | 6.87992501 | -0.0247146947166477 | 0.056122648937812 | -0.440369355053668 | 0.659669616365811 | NA | NM_175501 |
| 1700123L14Rik | 1.304710972 | 0.030808095 | 0.071729445 | 0.429504154 | 0.667556372 | NA | | Adamts14 | 12.1249835 | -0.0937607226852841 | 0.0680622815220492 | -1.37757243202183 | 0.168335337333912 | NA | NM_001081127 |
| 1700123O12Rik | 0 | NA | NA | NA | NA | NA | | Adamts19 | 9.335933272 | 0.0381506373457873 | 0.0497707731763646 | 0.76652691752646 | 0.443362841847049 | NA | NM_175506 |
| 1700123O21Rik | 0.199381295 | -0.004055078 | 0.031639813 | -0.128163771 | 0.89801937 | NA | | Adamts7 | 12.51700345 | -0.0145483928155017 | 0.0665076284294556 | -0.218747731035592 | 0.82684656528851 | NA | NM_001003911 |
| 1700125G02Rik | 0.38258743 | 0.019046329 | 0.042646787 | 0.446606435 | 0.655159248 | NA | | Adcy10 | 8.954459603 | 0.0181840384130583 | 0.0605613992205218 | 0.300257897722027 | 0.763980445136419 | NA | NM_173029 |
| 1700125G22Rik | 0.391976801 | -0.003620343 | 0.047492484 | -0.076229803 | 0.939236272 | NA | | Adh1 | 12.84448624 | -0.107770893917615 | 0.0610921083336802 | -1.76407226493116 | 0.0777198133066705 | NA | NM_007409 |
| 1700125H03Rik | 1.302823544 | -0.056690712 | 0.074863899 | -0.757250325 | 0.448899905 | NA | | Adh4 | 0 | NA | NA | NA | NA | NA | NM_011996 |
| 1700125H20Rik | 2.701148872 | 0.116217923 | 0.115881161 | 1.002906098 | 0.31590617 | NA | | Adh6-ps1 | 0 | NA | NA | NA | NA | NA | NR_033581 |
| 1700126H18Rik | 0 | NA | NA | NA | NA | NA | | Adh6a | 0 | NA | NA | NA | NA | NA | NM_026945 |
| 1700128A07Rik | 0 | NA | NA | NA | NA | NA | | Adh7 | 4.741622799 | 0.0161259148435909 | 0.0337670679734565 | 0.477563371989156 | 0.632961005956073 | NA | NM_009626 |
| 1700128F08Rik | 0 | NA | NA | NA | NA | NA | | Adig | 0.525532643 | -0.0123294079548737 | 0.0151790883024921 | -0.81226274656097 | 0.41664088193826 | NA | NM_145635 |
| 1700129C05Rik | 0 | NA | NA | NA | NA | NA | | Adipoq | 1.233807261 | -0.0393619938063293 | 0.0250831789099168 | -1.56925858351898 | 0.116587698568262 | NA | NM_009605 |
| 1810006J02Rik | 0 | NA | NA | NA | NA | NA | | Adm2 | 0 | NA | NA | NA | NA | NA | NM_182928 |
| 1810007C17Rik | 0 | NA | NA | NA | NA | NA | | Adprhl1 | 4.816383266 | 0.0170720471544634 | 0.0493690274845707 | 0.345804809701769 | 0.729489410291821 | NA | NM_172750 |
| 1810007D17Rik | 1.365135949 | 0.016537606 | 0.084668701 | 0.195321361 | 0.845141375 | NA | | Adtrp | 6.474368897 | 0.0144910507070286 | 0.0522875180757164 | 0.277141681998454 | 0.781671317211262 | NA | NM_175417,NM_001145875 |
| 1810007I06Rik | 0 | NA | NA | NA | NA | NA | | Afm | 0 | NA | NA | NA | NA | NA | NM_145146 |
| 1810008I18Rik | 0 | NA | NA | NA | NA | NA | | Afp | 3.52164106 | -0.0247957527831361 | 0.0408876445735475 | -0.606436321821724 | 0.544225055357846 | NA | NM_007423 |
| 1810009J06Rik | 0 | NA | NA | NA | NA | NA | | Agbl1 | 11.5398979 | 0.0282990557193912 | 0.0671372074328806 | 0.421510765810191 | 0.673382151472777 | NA | NM_001199224 |
| 1810010D01Rik | 4.821765732 | 0.036112285 | 0.121594504 | 0.296989449 | 0.766474564 | NA | | Ager | 19.84716907 | 0.000938345284217964 | 0.0830170748848578 | 0.0113030395917878 | 0.99098167124831 | NA | NM_001271424,NM_001271423,NM_001271422,NR_073175,NR_073176,NR_073173,NM_007425 |
| 1810012K16Rik | 5.815315053 | 0.209559767 | 0.143749318 | 1.45781399 | 0.144891821 | NA | | Agr2 | 0.410265417 | -0.01013448436559 | 0.0139857084979753 | -0.724631459826089 | 0.468678153504599 | NA | NM_011783 |
| 1810013A23Rik | 0 | NA | NA | NA | NA | NA | | Agr3 | 0.106148608 | 0.00515554063321497 | 0.00854200771766808 | 0.603551390213729 | 0.546141939141592 | NA | NM_207531 |
| 1810018F18Rik | 0 | NA | NA | NA | NA | NA | | Agrp | 6.924577723 | 0.0349066470583858 | 0.0417358523654364 | 0.836370771890447 | NA | NA | NM_007427,NM_001271806 |
| 1810019D21Rik | 2.62439483 | 0.164416764 | 0.105783488 | 1.554276254 | 0.120118535 | NA | | Agtr1b | 3.092091181 | -0.0290157498208965 | 0.04158378027806 | -0.697766043079193 | 0.48532351835035 | NA | NM_175086 |
| 1810019J16Rik | 4.780621242 | -0.028542864 | 0.133833793 | -0.213270977 | 0.83111561 | NA | | Agtr2 | 21.22505221 | -0.0764619527588244 | 0.0802958407578632 | -0.95225296898503 | 0.340968703453143 | NA | NM_007429 |
| 1810034E14Rik | 6.08985207 | -0.097671016 | 0.140143226 | -0.696937117 | 0.485842148 | NA | | Agxt | 0 | NA | NA | NA | NA | NA | NM_016702 |
| 1810035I16Rik | 0 | NA | NA | NA | NA | NA | | Agxt2 | 0.364150398 | -0.0138946944620142 | 0.0123764228912162 | -1.12267450653093 | 0.261575759058949 | NA | NM_001031851 |
| 1810046K07Rik | 0.294043327 | 0.010508917 | 0.039092051 | 0.2688249 | 0.788064431 | NA | | Ahsg | 0.106347839 | -0.00653347234140708 | 0.00854389509723638 | -0.764694822098226 | 0.444453290197145 | NA | NM_013465,NR_075099,NM_001276449,NM_001276450 |
| 1810053B23Rik | 0 | NA | NA | NA | NA | NA | | Aicda | 0 | NA | NA | NA | NA | NA | NM_009645 |
| 1810062G17Rik | 3.223892963 | 0.062676836 | 0.125600841 | 0.499016055 | 0.617768073 | NA | | Aipl1 | 1.061152941 | 0.00996751159764588 | 0.0220705976535041 | 0.451619469220099 | 0.651543141841376 | NA | NM_053245 |
| 1810064F22Rik | 0.284856677 | 0.008904946 | 0.033416532 | 0.266483248 | 0.789867059 | NA | | Aire | 5.872841891 | 0.0307017884390307 | 0.0491619877713359 | 0.624502584839164 | 0.532297573317468 | NA | 271549,NR_073358,NM_001271559,NM_001271558,NM_009646,NM_001271552,NM_001271551,NM_001271553,NM_001271557,NM_001271556,NM_00 |
| 1810065E05Rik | 0.308183275 | 0.007047773 | 0.033991235 | 0.207340891 | 0.835743642 | NA | | Airn | 16.28575355 | -0.119249801237776 | 0.0780988477067774 | -1.52690858750567 | 0.126783743882578 | NA | NR_027773,NR_027784,NR_002853,NR_027772 |
| 2010003K11Rik | 0.308183275 | 0.007047773 | 0.033991235 | 0.207340891 | 0.835743642 | NA | | Akap14 | 9.170735326 | -0.0294135737230075 | 0.0595424358303178 | -0.493993457150954 | 0.621310800271971 | NA | NM_001033785 |
| 2010005H15Rik | 1.375106598 | -0.141443519 | 0.075358259 | -1.876947794 | 0.060525249 | NA | | Akap3 | 13.62895301 | -0.0171506799256466 | 0.0696710497151287 | -0.246166521040983 | 0.80555333057786 | NA | NM_009650 |
| 2010009K17Rik | 0 | NA | NA | NA | NA | NA | | Akap4 | 0 | NA | NA | NA | NA | NA | NM_009651,NM_001042542 |
| 2010010A06Rik | 0 | NA | NA | NA | NA | NA | | Aknad1 | 0.450693223 | -0.0111640731504787 | 0.0143617019199425 | -0.777350289868945 | 0.436952132210492 | NA | NM_177859 |
| 2010016I18Rik | 0.408673203 | -0.002822473 | 0.042495157 | -0.066418693 | 0.947044488 | NA | | Akp3 | 0.204491261 | 0.00675251727497272 | 0.00969338978195025 | 0.696610517772262 | 0.486046571838551 | NA | NM_007432 |
| 2010106C02Rik | 0.079983654 | 0.016740032 | 0.027674178 | 0.604897175 | 0.54524732 | NA | | Akr1b3 | 0.09572496 | 0.00498255289994511 | 0.00847999972823692 | 0.587565219295241 | 0.556824162160132 | NA | NM_009658 |
| 2010106E10Rik | 0.398790038 | -0.006260381 | 0.047666372 | -0.131337473 | 0.895508349 | NA | | Akr1b7 | 1.471065161 | -0.00104255905676359 | 0.0262071841655204 | -0.0397814221542824 | 0.968267387479056 | NA | NM_009731 |
| 2010109A12Rik | 1.249209019 | -0.013485133 | 0.089181369 | -0.151210208 | 0.879809898 | NA | | Akr1c12 | 1.349922685 | -0.0275135176105035 | 0.0230250734567912 | -1.19493723492936 | 0.232111553628913 | NA | NM_013777 |
| 2010308F09Rik | 6.56425722 | 0.305770883 | 0.151325555 | 2.020616298 | 0.0433195 | NA | | Akr1c13 | 10.89633378 | -0.00780200092895575 | 0.0614758443060934 | -0.126911651511591 | 0.899010324641015 | NA | NM_013778 |
| 2010310C07Rik | 0 | NA | NA | NA | NA | NA | | Akr1c19 | 0.576654935 | 0.0131665485119237 | 0.0184212881884473 | 0.714746350919199 | 0.474765791144485 | NA | NM_001013785 |
| 2200002D01Rik | 2.751391874 | -0.1236684 | 0.111516433 | -1.10897019 | 0.267443042 | NA | | Akr1c20 | 0 | NA | NA | NA | NA | NA | NM_054080 |
| 2200002J24Rik | 0 | NA | NA | NA | NA | NA | | Akr1c21 | 0.506611253 | -0.0128144177033946 | 0.0150562545197539 | -0.851102622274483 | 0.394712348897175 | NA | NM_029901 |
| 2210010C04Rik | 0.430100701 | -0.046155006 | 0.041541744 | -1.111051218 | 0.266546304 | NA | | Akr1c6 | 0 | NA | NA | NA | NA | NA | NM_030611 |
| 2210019I11Rik | 0 | NA | NA | NA | NA | NA | | Akr1cl | 0.108766301 | 0.00519171381361946 | 0.00856003371502575 | 0.606506234257728 | 0.544178643801544 | NA | NM_027582 |
| 2210407C18Rik | 1.119479332 | -0.053604557 | 0.078463285 | -0.683180128 | 0.494493028 | NA | | Akr1d1 | 0.106347839 | -0.00653347234140708 | 0.00854389509723638 | -0.764694822098226 | 0.444453290197145 | NA | NM_145364 |
| 2210409D07Rik | 1.094514179 | -0.05351922 | 0.067126426 | -0.79728987 | 0.425282702 | NA | | Alas2 | 11.20152991 | 0.0279642971023577 | 0.0428929114767583 | 0.651956142392207 | 0.514429465531393 | NA | NM_001102446,NM_009653 |
| 2210409E12Rik | 0.188785634 | 0.025025111 | 0.031461603 | 0.79541754 | 0.42637066 | NA | | Alb | 1.271591302 | 0.00787116196619805 | 0.0244997144401497 | 0.321275661617465 | 0.748001498776945 | NA | NM_009654 |
| 2210414B05Rik | 0 | NA | NA | NA | NA | NA | | Aldh8a1 | 2.991591206 | -0.00108846792888505 | 0.0392239533886861 | -0.027750082152581 | 0.977861479283496 | NA | NM_178713 |
| 2210416O15Rik | 1.648555063 | 0.089082826 | 0.096799845 | 0.920278602 | 0.357427188 | NA | | Aldoart1 | 0.291497113 | -0.00941849595073383 | 0.0102038556585987 | -0.923033044160808 | 0.355989985190414 | NA | NM_001199270 |
| 2210417A02Rik | 0.159967308 | 0.0240345 | 0.030870969 | 0.778547006 | 0.436246606 | NA | | Aldoart2 | 0.494547629 | 0.00162457911534448 | 0.0144772416068025 | 0.112216067084294 | 0.910652089942672 | NA | NR_003959 |
| 2210420H20Rik | 0.098353386 | -0.020212831 | 0.02786466 | -0.725393061 | 0.468210929 | NA | | Allc | 21.18265081 | -0.0428853308168852 | 0.0809450882457354 | -0.529807697370008 | 0.596245267812789 | NA | NM_053156 |
| 2300002M23Rik | 0.824582689 | -0.009713019 | 0.062308342 | -0.155886342 | 0.876122611 | NA | | Alms1-ps2 | 3.798375822 | -0.0458055431494962 | 0.0466320439103901 | -0.982276119775447 | 0.325963835846348 | NA | NR_040440 |
| 2300003K06Rik | 0 | NA | NA | NA | NA | NA | | Alox12e | 5.988396507 | 0.0557175789501324 | 0.0491132594441765 | 1.13447121165848 | 0.256596942863222 | NA | NM_145684 |
| 2300005B03Rik | 0.768098354 | -0.017953171 | 0.049913376 | -0.359686576 | 0.719081532 | NA | | Alox15 | 3.143714617 | 0.0143442726689035 | 0.036312199026034 | 0.395026273639317 | 0.692823497979988 | NA | NM_009660 |
| 2310001H17Rik | 5.601893641 | 0.034433326 | 0.127895942 | 0.26922923 | 0.787753288 | NA | | Alpi | 0 | NA | NA | NA | NA | NA | NM_001081082 |
| 2310001K24Rik | 0.107716603 | -0.020477583 | 0.027954224 | -0.732539849 | 0.46383913 | NA | | Alpk2 | 19.38808774 | 0.0603084653211317 | 0.081858255830328 | 0.736742613306302 | 0.461278887333129 | NA | NM_001037294 |
| 2310002D06Rik | 0.20671448 | -0.003314027 | 0.03178208 | -0.10427343 | 0.916952363 | NA | | Alppl2 | 0 | NA | NA | NA | NA | NA | NM_007433 |
| 2310002F09Rik | 6.146738874 | 0.343511533 | 0.134253687 | 2.558674865 | 0.010507196 | NA | | Als2cr11 | 0.548112961 | 0.0106380473538722 | 0.0168343100001557 | 0.631926544882077 | 0.527434876908273 | NA | NM_175200 |
| 2310002J15Rik | 0.867184421 | 0.09414792 | 0.068136439 | 1.381755799 | 0.167046695 | NA | | Als2cr12 | 1.38811635 | 0.00172890108292559 | 0.0271479307457159 | 0.06368445164825 | 0.949221485400666 | NA | NM_175370 |
| 2310002L09Rik | 0.105923158 | 0.018524693 | 0.027977518 | 0.662127842 | 0.507889294 | NA | | Alx1 | 4.480073271 | -0.0287824941387418 | 0.0479670558109835 | -0.6000471292664 | 0.548474826682082 | NA | NM_172553 |
| 2310005A03Rik | 0.094392817 | 0.017895942 | 0.027818137 | 0.643319236 | 0.520016977 | NA | | Ambn | 0.290056368 | -0.00425303587871565 | 0.0107652157825088 | -0.395072051005791 | 0.692789714682806 | NA | NM_009664 |
| 2310005E17Rik | 0 | NA | NA | NA | NA | NA | | Ambp | 0 | NA | NA | NA | NA | NA | NM_007443 |
| 2310005G13Rik | 1.542146306 | 0.042012054 | 0.090068075 | 0.466447787 | 0.640895013 | NA | | Amd1 | 0 | NA | NA | NA | NA | NA | NM_009665 |
| 2310007B03Rik | 1.318607928 | 0.094192747 | 0.080628132 | 1.168236742 | 0.242711284 | NA | | Amd2 | 0 | NA | NA | NA | NA | NA | NM_007444 |
| 2310007L24Rik | 3.804890523 | 0.232236128 | 0.131254596 | 1.769356159 | 0.076834457 | NA | | Amdhd1 | 7.286250525 | -0.0636738774541834 | 0.0558411042747749 | -1.14026895207634 | 0.254174268677397 | NA | NM_027908 |
| 2310008N11Rik | 0.17463368 | 0.024560283 | 0.031186242 | 0.787535833 | 0.430968261 | NA | | Amelx | 0.337583802 | -0.0131877509542174 | 0.0120595734966011 | -1.0935503612905 | 0.274152229987901 | NA | NM_009666,NM_001081978 |
| 2310014F07Rik | 1.484926722 | 0.099947604 | 0.090094341 | 1.109366068 | 0.267272294 | NA | | Amica1 | 21.70011885 | 0.0734386168619275 | 0.0785987491668414 | 0.9343484169963 | 0.350124196740237 | NA | NM_001005421 |
| 2310015B20Rik | 2.653928998 | 0.017448554 | 0.106482186 | 0.163863598 | 0.869838524 | NA | | Ampd1 | 0.113843286 | 0.00525947738444006 | 0.00859368002043176 | 0.612016897526494 | 0.540526577873086 | NA | NM_001033303 |
| 2310015D24Rik | 5.901268509 | 0.262936099 | 0.142809491 | 1.841166836 | 0.065597114 | NA | | Amtn | 0 | NA | NA | NA | NA | NA | NM_027793 |
| 2310016D03Rik | 0 | NA | NA | NA | NA | NA | | Amy2a2 | 0 | NA | NA | NA | NA | NA | NM_001160152 |
| 2310020H05Rik | 0 | NA | NA | NA | NA | NA | | Amy2a3 | 0 | NA | NA | NA | NA | NA | NM_001160151 |
| 2310030A07Rik | 0.096071043 | -0.020098795 | 0.027825948 | -0.722304036 | 0.470107572 | NA | | Amy2a4 | 0 | NA | NA | NA | NA | NA | NM_001160150 |
| 2310034C09Rik | 0 | NA | NA | NA | NA | NA | | Amy2a5 | 0 | NA | NA | NA | NA | NA | NM_001042711 |
| 2310034O05Rik | 4.304087238 | 0.013054971 | 0.128955331 | 0.101236377 | 0.91936282 | NA | | Amy2b | 0.097165704 | -0.00637828455464458 | 0.00848781691535724 | -0.75146349388194 | 0.452373763047372 | NA | NM_001190404,NM_001190403 |
| 2310043L19Rik | 1.683619574 | -0.027326437 | 0.085849419 | -0.318306603 | 0.750252376 | NA | | Ang2 | 0 | NA | NA | NA | NA | NA | NM_007449 |
| 2310043O21Rik | 0.094392817 | 0.017895942 | 0.027818137 | 0.643319236 | 0.520016977 | NA | | Ang3 | 0 | NA | NA | NA | NA | NA | NM_001123394 |
| 2310050C09Rik | 0 | NA | NA | NA | NA | NA | | Ang4 | 0 | NA | NA | NA | NA | NA | NM_177544 |
| 2310057J18Rik | 0 | NA | NA | NA | NA | NA | | Ang5 | 0 | NA | NA | NA | NA | NA | NM_007448 |
| 2310057N15Rik | 0 | NA | NA | NA | NA | NA | | Ang6 | 0 | NA | NA | NA | NA | NA | NM_001011876 |
| 2310061N02Rik | 0 | NA | NA | NA | NA | NA | | Angpt4 | 0.340400173 | -0.006017291565113 | 0.0128784657053825 | -0.467236680422116 | 0.640330553117361 | NA | NM_009641 |
| 2310065F04Rik | 3.373993267 | 0.025467206 | 0.12625687 | 0.201709471 | 0.840143858 | NA | | Angptl3 | 5.866712144 | -0.0426500943846253 | 0.0524059690804809 | -0.813840391332651 | 0.41573639257191 | NA | NM_013913 |
| 2310068J16Rik | 6.350782555 | -0.130231293 | 0.150385883 | -0.865980837 | 0.386500671 | NA | | Angptl7 | 8.152985034 | 0.045766872500469 | 0.059600793652441 | 0.767890319839635 | 0.442552348900333 | NA | NM_001039554 |
| 2310069B03Rik | 0.222620122 | -0.001447027 | 0.032059667 | -0.04513545 | 0.963999345 | NA | | Ankar | 0.907937044 | -0.00444098824676889 | 0.0186114986597066 | -0.238615295198312 | 0.811403903829943 | NA | NM_176980 |
| 2310079G19Rik | 0 | NA | NA | NA | NA | NA | | Ankdd1b | 11.30595212 | 0.00153326756075571 | 0.0612864847043769 | 0.0250180373071096 | 0.980040576420607 | NA | NM_001042714 |
| 2310081J21Rik | 0.079983654 | 0.016740032 | 0.027674178 | 0.604897175 | 0.54524732 | NA | | Ankk1 | 0.756170104 | 0.000906820833292873 | 0.0195727681323753 | 0.046330740095618 | 0.963046638555713 | NA | NM_172922 |
| 2410003L11Rik | 1.778094025 | 0.009087409 | 0.098055755 | 0.092675939 | 0.926161013 | NA | | Ankle1 | 8.188618525 | -0.0146103426687452 | 0.0537352162552665 | -0.271895112496421 | 0.78570266938426 | NA | NM_172756 |
| 2410007B07Rik | 0.494509999 | 0.03737204 | 0.052072961 | 0.717686096 | 0.472950858 | NA | | Ankmy1 | 15.58967802 | -0.0612472075300096 | 0.0710157717766755 | -0.862445144194374 | 0.388442607286498 | NA | NM_172850 |
| 2410012E07Rik | 0 | NA | NA | NA | NA | NA | | Ankrd1 | 1.191895526 | 0.00701349904399481 | 0.0256254794989486 | 0.273692402293685 | 0.784321016993066 | NA | NM_013468 |
| 2410012M07Rik | 0.08731684 | 0.017365056 | 0.027733412 | 0.626142072 | 0.531221759 | NA | | Ankrd2 | 12.40681504 | -0.0936264220747082 | 0.0677975198073356 | -1.38097119689293 | 0.167287816978472 | NA | NM_020033 |
| 2410017I17Rik | 0 | NA | NA | NA | NA | NA | | Ankrd22 | 1.25525968 | -0.00260977422407211 | 0.0243884696470306 | -0.107008527465759 | 0.914782214663392 | NA | NM_024204 |
| 2410021H03Rik | 0.159967308 | 0.0240345 | 0.030870969 | 0.778547006 | 0.436246606 | NA | | Ankrd33 | 1.131216723 | 0.020249090575647 | 0.0214336408395393 | 0.944734062086777 | 0.344794662643495 | NA | NM_144790 |
| 2410057H14Rik | 3.283049985 | -0.087052309 | 0.12333788 | -0.705803515 | 0.480310336 | NA | | Ankrd36 | 0 | NA | NA | NA | NA | NA | NM_023816 |
| 2410076I21Rik | 4.680900391 | -0.06182659 | 0.121138272 | -0.510380318 | 0.609785043 | NA | | Ankrd53 | 0 | NA | NA | NA | NA | NA | NM_029245 |
| 2410088K16Rik | 0 | NA | NA | NA | NA | NA | | Ankrd60 | 0.255375383 | 0.00708304372802858 | 0.00992471678592415 | 0.713677163873753 | 0.475426830432214 | NA | NM_027303,NM_001199955 |
| 2410114N07Rik | 0 | NA | NA | NA | NA | NA | | Ankrd61 | 3.345195297 | 0.0214018968295124 | 0.0370345416339448 | 0.577890150256269 | 0.56333828568003 | NA | NM_025732 |
| 2410124H12Rik | 1.206912129 | -0.098297715 | 0.071099962 | -1.382528372 | 0.166809525 | NA | | Ankrd66 | 11.15524748 | -0.0466357889216099 | 0.0678013900784262 | -0.687829392106357 | 0.491560224664814 | NA | NM_001254953 |
| 2410137F16Rik | 1.577581099 | 0.043788493 | 0.086383756 | 0.506906564 | 0.61222038 | NA | | Ankrd7 | 1.827834041 | -0.0214863697900761 | 0.0330018088426435 | -0.651066427677512 | 0.515003607109214 | NA | NM_001167757,NM_029202 |
| 2410141K09Rik | 0 | NA | NA | NA | NA | NA | | Anks4b | 8.388243669 | 0.0323427917014694 | 0.0516718803070833 | 0.625926355094064 | 0.531363246984331 | NA | NM_028085 |
| 2510003D18Rik | 0.639497534 | 0.043360211 | 0.050060413 | 0.866157682 | 0.386403697 | NA | | Ano7 | 16.69086913 | 0.051310865400205 | 0.0718508658049688 | 0.714130091897329 | 0.475146739958127 | NA | NM_207031,NM_001271884 |
| 2510049J12Rik | 1.751491653 | 0.009023362 | 0.079900997 | 0.112931778 | 0.910084642 | NA | | Ano9 | 0.397645721 | -0.00694727832916359 | 0.0120596122586279 | -0.576078084450291 | 0.564562403875035 | NA | NM_178381 |
| 2600006L11Rik | 0 | NA | NA | NA | NA | NA | | Antxrl | 0 | NA | NA | NA | NA | NA | NM_172808 |
| 2610016A17Rik | 0.660333954 | -0.101834242 | 0.060586068 | -1.680819444 | 0.092797991 | NA | | Anxa10 | 3.186994858 | 0.0798987270609642 | 0.0416224455287061 | 1.91960674213291 | 0.0549075917935621 | NA | NM_011922,NM_001136089 |
| 2610027K06Rik | 2.452697712 | 0.02012729 | 0.114137613 | 0.176342303 | 0.86002503 | NA | | Anxa13 | 0.830491695 | -0.00782861969203861 | 0.0169001158611265 | -0.463228758688331 | 0.643200404419513 | NA | NM_027211 |
| 2610028H24Rik | 2.231059021 | 0.134326748 | 0.102585819 | 1.309408549 | 0.190395997 | NA | | Aoah | 11.32795662 | -0.00944004869879284 | 0.0637315748456518 | -0.148122005797836 | 0.882246482891769 | NA | NM_012054 |
| 2610035F20Rik | 6.484895822 | -0.102606829 | 0.148692803 | -0.690059149 | 0.490156992 | NA | | Aox3l1 | 1.006701342 | -0.000516357338594926 | 0.0190279616265529 | -0.0271367658149134 | 0.978350650670216 | NA | NM_001008419 |
| 2610037D02Rik | 2.54646873 | 0.128129051 | 0.112391189 | 1.140027538 | 0.254274829 | NA | | Apcs | 0 | NA | NA | NA | NA | NA | NM_011318 |
| 2610206C17Rik | 1.068033753 | 0.068003709 | 0.080072483 | 0.84927689 | 0.395727237 | NA | | Aplnr | 6.431248792 | -0.0544465926988939 | 0.0534468321111192 | -1.01870570337445 | 0.308342704112123 | NA | NM_011784 |
| 2610307P16Rik | 7.718447562 | -0.385415747 | 0.147331387 | -2.615978546 | 0.008897215 | NA | | Apoa1 | 13.39905657 | -0.110550187080672 | 0.0681331435038203 | -1.62256108254382 | 0.104683265560041 | NA | NM_009692 |
| 2610528J11Rik | 1.22007994 | -0.050163986 | 0.086589842 | -0.57932876 | 0.562367363 | NA | | Apoa4 | 0 | NA | NA | NA | NA | NA | NM_007468 |
| 2700086A05Rik | 0 | NA | NA | NA | NA | NA | | Apoa5 | 0.241530977 | 0.00760948419201657 | 0.0102872826258192 | 0.73969817577658 | 0.459483155250455 | NA | NM_080434 |
| 2700089I24Rik | 0.107716603 | -0.020477583 | 0.027954224 | -0.732539849 | 0.46383913 | NA | | Apob | 2.070968059 | 0.00527556226992189 | 0.0320693019918049 | 0.164505054437107 | 0.869333568089459 | NA | NM_009693 |
| 2810007J24Rik | 0 | NA | NA | NA | NA | NA | | Apobec2 | 4.194527813 | 0.0267946204443387 | 0.0480124828131494 | 0.558076126756777 | 0.576792401769655 | NA | NM_009694 |
| 2810404M03Rik | 4.904431192 | 0.047550992 | 0.130091949 | 0.365518331 | 0.714724526 | NA | | Apobec4 | 0.405586341 | -0.0100214244197644 | 0.0139407229051103 | -0.718859738334718 | 0.472227345659372 | NA | NM_001081197 |
| 2810405F15Rik | 2.621914376 | -0.036617563 | 0.107992482 | -0.339075109 | 0.734553149 | NA | | Apoc2 | 0.840692375 | -0.00431066191165442 | 0.0218609206727126 | -0.197185744195811 | 0.843682186088316 | NA | NM_009695 |
| 2810442I21Rik | 1.913210347 | -0.036894559 | 0.089449939 | -0.412460413 | 0.680001991 | NA | | Apoc3 | 13.81388057 | 0.0455691234676547 | 0.0701392141418623 | 0.649695381181308 | 0.51588900800746 | NA | NM_023114 |
| 2810442N19Rik | 0.096071043 | -0.020098795 | 0.027825948 | -0.722304036 | 0.470107572 | NA | | Apoc4 | 0.227686571 | 0.00691460391109036 | 0.00980755285755433 | 0.705028462402254 | 0.480792523671872 | NA | NM_007385 |
| 2810471M01Rik | 0 | NA | NA | NA | NA | NA | | Apof | 2.483368486 | -0.0182475688667038 | 0.0363666621970373 | -0.501766391642902 | 0.615831856681224 | NA | NM_133997 |
| 2900057B20Rik | 0 | NA | NA | NA | NA | NA | | Apoh | 15.83451376 | 0.0874386601115528 | 0.0766042869596296 | 1.14143298739446 | 0.25368978382489 | NA | NM_013475 |
| 2900060B14Rik | 0.096071043 | -0.020098795 | 0.027825948 | -0.722304036 | 0.470107572 | NA | | Apol10a | 0 | NA | NA | NA | NA | NA | NM_177744 |
| 2900092C05Rik | 0.105923158 | 0.018524693 | 0.027977518 | 0.662127842 | 0.507889294 | NA | | Apol10b | 17.35188762 | -0.0827643055623687 | 0.073835271310207 | -1.12093182694011 | 0.262316882137912 | NA | NM_177820 |
| 2900093L17Rik | 1.048127784 | 0.10735569 | 0.071642996 | 1.498481295 | 0.134008249 | NA | | Apol11a | 0 | NA | NA | NA | NA | NA | NM_001177533 |
| 3100003L05Rik | 1.071449219 | -0.042018488 | 0.076873758 | -0.546590785 | 0.584659902 | NA | | Apol11b | 0.9347476 | -0.00437476682407002 | 0.0190490564376407 | -0.229657927592967 | 0.818357589984352 | NA | NM_001143686 |
| 3110009F21Rik | 0.902191072 | 0.013251232 | 0.063848263 | 0.207542558 | 0.835586159 | NA | | Apol6 | 6.750461165 | -0.0235666376850459 | 0.0476360629000338 | -0.494722616654979 | 0.620795934513895 | NA | NM_028010,NM_001163621 |
| 3110015C05Rik | 1.176732261 | 0.066978284 | 0.07561092 | 0.885828185 | 0.375710119 | NA | | Apol7a | 16.29096818 | -0.00248660314337062 | 0.0757571583604641 | -0.0328233423373536 | 0.973815463746316 | NA | NM_001164640,NM_029419 |
| 3110021A11Rik | 5.658365749 | 0.139867743 | 0.140350995 | 0.99655683 | 0.318979669 | NA | | Apol7b | 5.474585189 | -0.0860815531765837 | 0.0474810485855156 | -1.81296655699477 | 0.0698369795689749 | NA | NM_001024848 |
| 3110079O15Rik | 6.442475272 | -0.011307627 | 0.141434464 | -0.07994959 | 0.936277349 | NA | | Apol7c | 0.113843286 | 0.00525947738444006 | 0.00859368002043176 | 0.612016897526494 | 0.540526577873086 | NA | NM_175391 |
| 3110099E03Rik | 7.584837601 | -0.210768864 | 0.133039148 | -1.584261978 | 0.113134114 | NA | | Apol7d | 0.752792827 | 0.000899543766135771 | 0.019838776277957 | 0.0453427042843999 | 0.963834149326249 | NA | NR_040308 |
| 3200001D21Rik | 0.211846316 | 0.025707621 | 0.031860375 | 0.806883811 | 0.419733426 | NA | | Apol7e | 0 | NA | NA | NA | NA | NA | NM_001134802 |
| 3300005D01Rik | 0.313786592 | -0.036039695 | 0.036798757 | -0.979372609 | 0.327395906 | NA | | Apol9a | 3.542994518 | -0.0219648866030416 | 0.0442687039199363 | -0.496171892512755 | 0.619773139986776 | NA | NM_001162883,NM_173786 |
| 3830403N18Rik | 2.284420513 | -0.010745656 | 0.097777774 | -0.109898762 | 0.912489664 | NA | | Apol9b | 6.63364543 | 0.0553405777707215 | 0.0536371458324542 | 1.03175843740061 | 0.302185294160347 | NA | NM_001168660,NM_173743 |
| 3830417A13Rik | 0 | NA | NA | NA | NA | NA | | Apom | 1.152755946 | 0.0157828842599894 | 0.024146480436276 | 0.653630838732019 | 0.513349671005954 | NA | NM_018816 |
| 4732416N19Rik | 7.177832134 | 0.12671669 | 0.146110089 | 0.867268586 | 0.385794865 | NA | | Apon | 0.750337725 | 0.0108779074749183 | 0.0200592805016097 | 0.542288018458355 | 0.587620108272692 | NA | NM_133996 |
| 4732456N10Rik | 4.55523015 | 0.089755881 | 0.10153235 | 0.884012637 | 0.376689391 | NA | | Apoo-ps | 0 | NA | NA | NA | NA | NA | NR_004438 |
| 4732490B19Rik | 0.539953942 | 0.069964435 | 0.051895658 | 1.348175111 | 0.177602068 | NA | | Aqp1 | 129.0974644 | 0.00439966927135023 | 0.0267398936521472 | 0.164535780455392 | NA | NA | NM_007472 |
| 4833403I15Rik | 3.239762966 | 0.126166008 | 0.101478368 | 1.243279829 | 0.213764734 | NA | | Aqp12 | 0.212297216 | 0.00681050119636477 | 0.00973439053551308 | 0.699633035218656 | 0.484156506270654 | NA | NM_177587,NM_001159658 |
| 4833419F23Rik | 5.458823561 | 0.033687057 | 0.141094359 | 0.238755518 | 0.811295164 | NA | | Aqp2 | 0.892364251 | -0.0283633641540671 | 0.0232386828340015 | -1.22052374296221 | 0.222266394532471 | NA | NM_009699 |
| 4833423E24Rik | 5.246093872 | -0.129244546 | 0.134098976 | -0.963799647 | 0.335146384 | NA | | Aqp3 | 1.912165725 | -0.00363122350905033 | 0.0324457447810266 | -0.111916787041171 | 0.910889386091874 | NA | NM_016689 |
| 4833427F10Rik | 0.285336609 | 0.031742875 | 0.03528833 | 0.899528977 | 0.368370968 | NA | | Aqp5 | 7.539287504 | -0.0231443445685017 | 0.0615843679121579 | -0.375815249115069 | 0.707054250319758 | NA | NM_009701 |
| 4833427G06Rik | 5.338523982 | 0.267809996 | 0.144546026 | 1.852766231 | 0.063915872 | NA | | Aqp6 | 16.20689406 | 0.0144309171006465 | 0.0728042648085762 | 0.198215271297494 | 0.842876639764183 | NA | NM_175087 |
| 4833428L15Rik | 0.969874391 | -0.007538203 | 0.071551649 | -0.105353313 | 0.916095461 | NA | | Aqp7 | 1.536631908 | 0.0220610470901532 | 0.0289752335031093 | 0.761375989870309 | 0.446432524239767 | NA | NM_007473 |
| 4921501E09Rik | 0.279364949 | 0.005060019 | 0.033327057 | 0.151829152 | 0.879321689 | NA | | Aqp8 | 0.249110328 | -0.010000816291709 | 0.0105091939847586 | -0.951625434473194 | 0.341286977626714 | NA | NM_001109045,NM_007474 |
| 4921504E06Rik | 0.502404983 | 0.007339048 | 0.055675222 | 0.13181893 | 0.895127513 | NA | | Areg | 3.383278858 | 0.0631976097252283 | 0.041909278693829 | 1.50796223879019 | 0.131564191788256 | NA | NM_009704 |
| 4921506M07Rik | 0 | NA | NA | NA | NA | NA | | Arg1 | 3.22235964 | 0.0326171836650069 | 0.0403973416684751 | 0.807409159065049 | 0.419430791822337 | NA | NM_007482 |
| 4921507L20Rik | 0.345637534 | 0.019330126 | 0.041637051 | 0.464253003 | 0.642466491 | NA | | Arhgap40 | 12.03655567 | 0.039420714216144 | 0.0626772275314659 | 0.628947957156426 | 0.529383128072843 | NA | NM_001145015 |
| 4921508D12Rik | 0.740533532 | 0.01739619 | 0.062889035 | 0.276617211 | 0.782074048 | NA | | Arhgap8 | 6.446520908 | 0.00277901413239657 | 0.0536259845024158 | 0.0518221559600716 | 0.958670401365795 | NA | NM_001164628,NM_001205334,NM_001164627,NM_028455 |
| 4921509C19Rik | 0.238795281 | -0.027601606 | 0.032267774 | -0.855392297 | 0.392334007 | NA | | Arhgef38 | 0.387222073 | -0.00720181929823146 | 0.0119779253649773 | -0.60125765345718 | 0.547668389729015 | NA | NM_029953 |
| 4921509O07Rik | 1.096585181 | -0.028447705 | 0.083638729 | -0.340125984 | 0.733761655 | NA | | Arid3c | 8.899705224 | -0.0811179752481474 | 0.0622382869989164 | -1.30334524228727 | 0.19245692181383 | NA | NM_001252622,NM_001017362 |
| 4921511C10Rik | 0.496152747 | 0.00873411 | 0.050596247 | 0.172623666 | 0.862947248 | NA | | Arl13a | 0.403747135 | 0.0119836939206955 | 0.0128877677856313 | 0.929850236288107 | 0.35244863171373 | NA | NM_028947 |
| 4921511C20Rik | 0 | NA | NA | NA | NA | NA | | Arl14 | 0 | NA | NA | NA | NA | NA | NM_027843 |
| 4921511H03Rik | 0 | NA | NA | NA | NA | NA | | Arl14epl | 0.108766301 | 0.00519171381361946 | 0.00856003371502575 | 0.606506234257728 | 0.544178643801544 | NA | NM_001033446 |
| 4921511I17Rik | 0.283178451 | 0.028390488 | 0.033368195 | 0.850824814 | 0.394866675 | NA | | Arl5a | 4048.754827 | 0.0816637886021927 | 0.0571711798928235 | 1.42840831263732 | NA | NA | NM_182994 |
| 4921511M17Rik | 0 | NA | NA | NA | NA | NA | | Arl9 | 2.454849386 | -0.0490639305476849 | 0.0324602257421052 | -1.5115092217009 | 0.13065876088444 | NA | NM_206935 |
| 4921513I03Rik | 0 | NA | NA | NA | NA | NA | | Armc12 | 0.805688471 | -0.0124238225780724 | 0.0166971554392408 | -0.744068211096278 | 0.456835209781343 | NA | NM_026290 |
| 4921515E04Rik | 0.302603776 | 0.00963845 | 0.039370813 | 0.244812063 | 0.806601953 | NA | | Arrdc5 | 0 | NA | NA | NA | NA | NA | NM_029799 |
| 4921517D22Rik | 0.238795281 | -0.027601606 | 0.032267774 | -0.855392297 | 0.392334007 | NA | | Arsi | 19.93692438 | -0.0647712381964069 | 0.0802387383752397 | -0.807231513206273 | 0.419533113095013 | NA | NM_001038499 |
| 4921524L21Rik | 0 | NA | NA | NA | NA | NA | | Art1 | 1.724394973 | -0.00893713451464244 | 0.0314881343953219 | -0.283825469062092 | 0.77654413533209 | NA | NM_009710 |
| 4921525B02Rik | 0.470066974 | 0.035966715 | 0.05156661 | 0.697480692 | 0.485502018 | NA | | Art2a-ps | 0 | NA | NA | NA | NA | NA | NR_033804,NM_007490 |
| 4921530D09Rik | 0.107716603 | -0.020477583 | 0.027954224 | -0.732539849 | 0.46383913 | NA | | Art2b | 0 | NA | NA | NA | NA | NA | NM_019915 |
| 4921530L18Rik | 7.312735488 | 0.072810329 | 0.155593381 | 0.467952615 | 0.639818476 | NA | | Art4 | 3.398784008 | 0.0231189867914923 | 0.0402257312145729 | 0.574731299927663 | 0.56547303555469 | NA | NM_026639 |
| 4921530L21Rik | 1.817384064 | 0.014184023 | 0.080545876 | 0.176098685 | 0.860216414 | NA | | Art5 | 0.871835 | 0.0156832015936662 | 0.0226385090250168 | 0.692766541132873 | 0.488456072824408 | NA | NM_007491 |
| 4921531P14Rik | 0 | NA | NA | NA | NA | NA | | Artn | 16.38987981 | 0.0385723298245062 | 0.0678440831893041 | 0.568543755199382 | 0.569665803815772 | NA | NM_009711 |
| 4921533I20Rik | 0 | NA | NA | NA | NA | NA | | Asb10 | 11.10414919 | -0.0598008606988107 | 0.0601074934072257 | -0.994898594317725 | 0.319785586621975 | NA | NM_080444 |
| 4921539E11Rik | 0 | NA | NA | NA | NA | NA | | Asb12 | 0.989058214 | 0.0234309775542899 | 0.0209549498985171 | 1.11815955980635 | 0.263498852558756 | NA | NM_080858 |
| 4922502D21Rik | 0 | NA | NA | NA | NA | NA | | Asb14 | 3.707845727 | -0.0300582249452385 | 0.0416846315031632 | -0.721086497860909 | 0.47085629780616 | NA | NM_001170748,NM_080856 |
| 4922502H24Rik | 0 | NA | NA | NA | NA | NA | | Asb15 | 2.048915888 | -0.0162763231944938 | 0.0341175288125066 | -0.477066298791469 | 0.633314911908832 | NA | NM_080847 |
| 4922502N22Rik | 0.430253841 | 0.022125686 | 0.043852981 | 0.504542347 | 0.613880312 | NA | | Asb17 | 0.120209049 | -0.00674972139751468 | 0.00863521114606269 | -0.781650996523957 | 0.434419707908266 | NA | NM_025758 |
| 4922505E12Rik | 0.555364211 | 0.060011321 | 0.059945413 | 1.001099473 | 0.31677872 | NA | | Asb18 | 13.53016113 | 0.0413292931383664 | 0.0755407245628238 | 0.547112744516962 | 0.58430127708124 | NA | NM_139152 |
| 4930401C15Rik | 0 | NA | NA | NA | NA | NA | | Asb4 | 58.97699677 | 0.0577322167719021 | 0.0400423180509491 | 1.4417800862189 | NA | NA | NM_023048 |
| 4930401O10Rik | 0 | NA | NA | NA | NA | NA | | Asb5 | 19.10611198 | 0.0996656970244506 | 0.077990770455991 | 1.27791655912273 | 0.20127884872674 | NA | NM_029569 |
| 4930401O12Rik | 0 | NA | NA | NA | NA | NA | | Asb9 | 0 | NA | NA | NA | NA | NA | NM_027027 |
| 4930402F06Rik | 0.56395571 | 0.017428757 | 0.047045094 | 0.370469173 | 0.711032941 | NA | | Ascl2 | 5.693798609 | -0.0297885510845628 | 0.0500442796841597 | -0.595243877473406 | 0.55168046457455 | NA | NM_008554 |
| 4930402F11Rik | 0 | NA | NA | NA | NA | NA | | Ascl3 | 3.112209689 | 0.0173212238893964 | 0.0354996650985296 | 0.487926402722991 | 0.625601972220027 | NA | NM_020051 |
| 4930402K13Rik | 0 | NA | NA | NA | NA | NA | | Ascl4 | 3.919982356 | -0.0223255599693746 | 0.0356662404719816 | -0.625957759324618 | 0.531342647840101 | NA | NM_001163614 |
| 4930404A05Rik | 0 | NA | NA | NA | NA | NA | | Ascl5 | 11.76699077 | -0.0660387974214677 | 0.0699578528002961 | -0.943979764644636 | 0.345179988913784 | NA | NM_001270609 |
| 4930404A10Rik | 0.192746203 | -0.000975302 | 0.031567365 | -0.030895906 | 0.975352555 | NA | | Asgr2 | 2.938424935 | -0.0600018171216046 | 0.0303507617020558 | -1.97694600585726 | 0.0480477337608217 | NA | NM_007493 |
| 4930404H11Rik | 2.137369726 | 0.104910205 | 0.10463188 | 1.002660038 | 0.316024917 | NA | | Asic3 | 12.07581403 | -0.117363324817083 | 0.0641424540243706 | -1.82972925813677 | 0.0672904335840258 | NA | NM_183000 |
| 4930404I05Rik | 0.761449927 | -0.003542071 | 0.055175041 | -0.064196988 | 0.948813375 | NA | | Asic5 | 0.606475725 | 0.0152833423794426 | 0.014534742771663 | 1.05150415246695 | 0.293027103254472 | NA | NM_021370 |
| 4930404N11Rik | 4.061198555 | -0.06188911 | 0.102513511 | -0.60371662 | 0.546032062 | NA | | Astl | 0.315916084 | 0.00270415235986604 | 0.0125014265294581 | 0.216307503267249 | 0.828748060749853 | NA | NM_172539 |
| 4930405A10Rik | 0 | NA | NA | NA | NA | NA | | Asz1 | 0.12890128 | -0.0068640067066208 | 0.00870804083610955 | -0.788237771940376 | 0.430557639008166 | NA | NM_023729 |
| 4930405D11Rik | 0.107716603 | -0.020477583 | 0.027954224 | -0.732539849 | 0.46383913 | NA | | Atf7ip2 | 0.854022854 | 0.0036786923731306 | 0.0184721231882335 | 0.199148323971436 | 0.842146721150413 | NA | NM_029253,NM_153123 |
| 4930405J17Rik | 0 | NA | NA | NA | NA | NA | | Atoh1 | 0 | NA | NA | NA | NA | NA | NM_007500 |
| 4930405P13Rik | 0.20210942 | -0.001755803 | 0.031727492 | -0.055340098 | 0.955867518 | NA | | Atoh7 | 11.25117341 | -0.0579656561205465 | 0.0688463313203484 | -0.841957080484463 | 0.399811977884925 | NA | NM_016864 |
| 4930406D18Rik | 0 | NA | NA | NA | NA | NA | | Atp12a | 4.121024504 | 0.0386649857566918 | 0.0396626904398648 | 0.974845259559844 | 0.329637022804074 | NA | NM_138652 |
| 4930407I10Rik | 0.094392817 | 0.017895942 | 0.027818137 | 0.643319236 | 0.520016977 | NA | | Atp1a4 | 4.351165372 | -0.0262118242632099 | 0.0469148040567367 | -0.558711152912639 | 0.576358866524913 | NA | NM_013734 |
| 4930408F14Rik | 0 | NA | NA | NA | NA | NA | | Atp1b4 | 0.108766301 | 0.00519171381361946 | 0.00856003371502575 | 0.606506234257728 | 0.544178643801544 | NA | NM_133690 |

| | | | | | | | | | | | | | | | |
|---|---|---|---|---|---|---|---|---|---|---|---|---|---|---|---|
| 4930412B13Rik | 0 | NA | NA | NA | NA | NA | | Atp2a1 | 18.87376361 | -0.0372223545872005 | 0.0676135319973593 | -0.550516346175411 | 0.581965268380206 | NA | NM_007504 |
| 4930412O13Rik | 0.66557215 | 0.04559458 | 0.049811598 | 0.915340648 | 0.360012826 | NA | | Atp4b | 0 | NA | NA | NA | NA | NA | NM_009724 |
| 4930413E15Rik | 0 | NA | NA | NA | NA | NA | | Atp6v0a4 | 13.88961335 | 0.0663592206411948 | 0.0684824421053704 | 0.9689961193132 | 0.33254712540672 | NA | NM_080467 |
| 4930413F20Rik | 3.863764201 | 0.021671767 | 0.125973555 | 0.172034259 | 0.863410596 | NA | | Atp6v0c | 2.952785157 | 0.00765883027450603 | 0.040992193095117 | 0.186836314337579 | 0.851788973503716 | NA | NM_009729 |
| 4930413G21Rik | 3.735394276 | -0.053166696 | 0.126238523 | -0.421160631 | 0.67363779 | NA | | Atp6v0c-ps2 | 0 | NA | NA | NA | NA | NA | NR_037854 |
| 4930413M19Rik | 0 | NA | NA | NA | NA | NA | | Atp6v0d2 | 2.790396319 | 0.0329774690109271 | 0.0242586592743192 | 1.3594102063933 | 0.174016638696674 | NA | NM_175406 |
| 4930414N06Rik | 0.574278041 | 0.012588346 | 0.03989752 | 0.315517008 | 0.75236914 | NA | | Atp6v1b1 | 1.454548221 | 0.00821297157931775 | 0.0294045003192386 | 0.27931002024014 | 0.780006917139159 | NA | NM_134157 |
| 4930415F15Rik | 6.800379707 | 0.161059409 | 0.128369336 | 1.254656401 | 0.209603519 | NA | | Atp6v1e2 | 1.289684196 | -0.0253530470229735 | 0.026514746557605 | -0.956186662689471 | 0.338977936994691 | NA | NM_029121 |
| 4930415L06Rik | 0 | NA | NA | NA | NA | NA | | Atp6v1g3 | 0.12890128 | -0.0068640067066208 | 0.00870804083610955 | -0.788237771940376 | 0.430557639008166 | NA | NM_177397 |
| 4930415O20Rik | 1.339143676 | -0.02737891 | 0.081591324 | -0.335561533 | 0.737201536 | NA | | Atp8b3 | 5.832107403 | 0.0809207389034759 | 0.0540034727469681 | 1.49843583731417 | 0.134020051242341 | NA | NM_026094 |
| 4930417O13Rik | 4.964691473 | -0.356244184 | 0.141475704 | -2.518059094 | 0.011800352 | NA | | Atp8b4 | 3.323783005 | -0.00934204181103063 | 0.0372819299960169 | -0.250578277788427 | 0.802140177816594 | NA | NM_001080944 |
| 4930417O22Rik | 1.440912637 | 0.05008834 | 0.082749727 | 0.605299152 | 0.544980245 | NA | | Atp8b5 | 18.7828216 | -0.0772745693319553 | 0.0800686027715497 | -0.965104506100022 | 0.334492470198551 | NA | NM_177195 |
| 4930419G24Rik | 1.09495344 | 0.103030006 | 0.078097338 | 1.319251196 | 0.187085148 | NA | | Aurkb | 8.930552684 | -0.0337553710771544 | 0.0644531914760412 | -0.523719156555686 | 0.600473889623136 | NA | NM_011496 |
| 4930423M02Rik | 0 | NA | NA | NA | NA | NA | | Aurkc | 5.238413299 | 0.00160145681979409 | 0.0478858352118812 | 0.0334432262214514 | 0.973321139386329 | NA | NM_020572,NM_001080965,NM_001080966 |
| 4930425K10Rik | 0.17463368 | 0.024560283 | 0.031186242 | 0.787535833 | 0.430968261 | NA | | Avil | 12.63040505 | 0.0313715494683348 | 0.0670412878648694 | 0.467943717482997 | 0.639824839053468 | NA | NM_009635 |
| 4930425O10Rik | 0.593519161 | -0.035580971 | 0.059082103 | -0.60222925 | 0.547021548 | NA | | Avp | 265.6136494 | 0.0276705594061733 | 0.0267014227876163 | 1.03629531752917 | NA | NA | NM_009732 |
| 4930428D18Rik | 0 | NA | NA | NA | NA | NA | | Avpr1a | 13.19834505 | 0.0681485030474093 | 0.0534233232175856 | 1.27563204501244 | NA | NA | NM_016847 |
| 4930428E07Rik | 0.116696964 | -0.020477583 | 0.027954224 | -0.732539847 | 0.463839131 | NA | | Avpr1b | 2.241994274 | -0.0343709268746935 | 0.0288553433704982 | -1.19114600139655 | 0.233596273604877 | NA | NM_011924 |
| 4930428G15Rik | 0.712869317 | -0.061164802 | 0.067168122 | -0.91062248 | 0.362494321 | NA | | Avpr2 | 0 | NA | NA | NA | NA | NA | NM_001276299,NM_019404,NM_001276298 |
| 4930428O21Rik | 0 | NA | NA | NA | NA | NA | | Awat1 | 0.120209049 | -0.00674972139751468 | 0.00863521114606269 | -0.781650996523957 | 0.434419707908266 | NA | NM_001081136 |
| 4930429D17Rik | 0 | NA | NA | NA | NA | NA | | Awat2 | 7.800138389 | -0.0638268567714299 | 0.0612834460783412 | -1.04150240979982 | 0.297642435370483 | NA | NM_177746 |
| 4930429F11Rik | 0 | NA | NA | NA | NA | NA | | Aym1 | 0.424594432 | 0.00882658952495119 | 0.0110666225409165 | 0.79758657099912 | 0.425110446545159 | NA | NM_001012726 |
| 4930429F24Rik | 1.878795999 | -0.006604545 | 0.109082731 | -0.060546206 | 0.951720616 | NA | | Azgp1 | 0.949664049 | 0.0277265436537805 | 0.0223438841533915 | 1.24090079698932 | 0.214642398313254 | NA | NM_013478 |
| 4930430A15Rik | 0 | NA | NA | NA | NA | NA | | B020004C17Rik | 0 | NA | NA | NA | NA | NA | NM_001256060 |
| 4930430D24Rik | 0 | NA | NA | NA | NA | NA | | B020004J07Rik | 9.358360774 | 0.101338769973961 | 0.0650324113759681 | 1.55820983740551 | 0.119166654783711 | NA | NM_001033790 |
| 4930430F21Rik | 0.401377075 | -0.004225889 | 0.047720475 | -0.08855505 | 0.929435533 | NA | | B020014A21Rik | 0 | NA | NA | NA | NA | NA | NR_045946 |
| 4930430J02Rik | 0 | NA | NA | NA | NA | NA | | B020018J22Rik | 1.256833248 | -0.024017592040233 | 0.0271129149453777 | -0.885835849395736 | 0.375705988727776 | NA | NR_045950 |
| 4930431F12Rik | 2.357898175 | 0.060875398 | 0.098157387 | 0.620181528 | 0.535138281 | NA | | B130006D01Rik | 0.106347839 | -0.00653347234140708 | 0.00854389509723638 | -0.764694822098226 | 0.444453290197145 | NA | NR_028263 |
| 4930431P03Rik | 1.064313701 | -0.021011496 | 0.076941904 | -0.273082613 | 0.784789711 | NA | | B130024G19Rik | 16.98105827 | 0.0813867123882867 | 0.0689765465165124 | 1.17991863174541 | 0.238032578753896 | NA | NR_045850 |
| 4930432J09Rik | 0.649580461 | 0.072845422 | 0.063527461 | 1.146676111 | 0.251515507 | NA | | B230119M05Rik | 0.437122686 | -0.00309378938484024 | 0.015328819040773 | -0.201828293269761 | 0.840050961859409 | NA | NR_045454 |
| 4930432K09Rik | 0 | NA | NA | NA | NA | NA | | B230208H11Rik | 4.696593207 | -0.0482291789911342 | 0.0481763828379615 | -1.00109589284339 | 0.316780450494506 | NA | NR_038028,NR_038027 |
| 4930432M17Rik | 0 | NA | NA | NA | NA | NA | | B230214G05Rik | 0.22897535 | -0.00116364198856673 | 0.00981572003153595 | -0.118548816065269 | 0.905632818135836 | NA | NR_045281 |
| 4930433B08Rik | 0 | NA | NA | NA | NA | NA | | B230319C09Rik | 15.10490354 | 0.00867855882623351 | 0.0719920869875553 | 0.12054878792073 | 0.904048434223004 | NA | NR_028382 |
| 4930433I11Rik | 0.185906812 | 0.024895454 | 0.031385107 | 0.793225068 | 0.427646705 | NA | | B230323A14Rik | 3.064303286 | 0.0443905785543935 | 0.0364549689229998 | 1.21768252355818 | 0.223344640820302 | NA | NR_040765 |
| 4930433N12Rik | 0.094392817 | 0.017895942 | 0.027818137 | 0.643319236 | 0.520016977 | NA | | B230378P21Rik | 11.50972309 | -0.17924463447065 | 0.0686784262221064 | -2.60991179225587 | 0.00905655716466637 | NA | NR_040277 |
| 4930434J06Rik | 0.17833704 | -0.002390466 | 0.031282281 | -0.076415987 | 0.939088151 | NA | | B3gnt6 | 14.01018673 | 0.0266716236014017 | 0.0740680497939137 | 0.360096204444597 | 0.718775191085712 | NA | NM_001081167 |
| 4930435E12Rik | 0 | NA | NA | NA | NA | NA | | B3gnt8 | 13.88971931 | 0.00735730305859026 | 0.0724358972471128 | 0.101569847799235 | 0.919098113612417 | NA | NM_001036740,NM_146184 |
| 4930438E09Rik | 0 | NA | NA | NA | NA | NA | | B430010I23Rik | 0.523136316 | 0.0108872515365422 | 0.0166665021953125 | 0.653241538563756 | 0.513600574882611 | NA | NR_015457 |
| 4930440C22Rik | 0 | NA | NA | NA | NA | NA | | B430212C06Rik | 8.203070453 | -0.0219476917143918 | 0.0545743938621344 | -0.402160979924687 | 0.687565556963263 | NA | NR_033214 |
| 4930441J16Rik | 0 | NA | NA | NA | NA | NA | | B430306N03Rik | 8.761835432 | -0.122281211012283 | 0.0649684672299572 | -1.88216247398089 | 0.0598139590426028 | NA | NM_177083 |
| 4930442J19Rik | 0 | NA | NA | NA | NA | NA | | B4galnt2 | 6.160238756 | -0.0282621202321181 | 0.054084621169595 | -0.522553724532814 | 0.601284850547472 | NA | NM_008081 |
| 4930442L01Rik | 0 | NA | NA | NA | NA | NA | | B930018H19Rik | 7.539884275 | -0.0660814641604827 | 0.0510096593024442 | -1.29546962407013 | 0.195158273502812 | NA | NR_040706 |
| 4930443O20Rik | 0.185670226 | -0.001645086 | 0.031432796 | -0.052336624 | 0.958260472 | NA | | B930025P03Rik | 16.51731414 | 0.0245071832108689 | 0.0644429805209454 | 0.380292516155481 | 0.703728290210616 | NA | NR_040705 |
| 4930444F02Rik | 0 | NA | NA | NA | NA | NA | | B930092H01Rik | 0.106148608 | 0.00515554063321497 | 0.00854200771766808 | 0.603551390213729 | 0.546141939141592 | NA | NR_045334 |
| 4930444G20Rik | 0.380940254 | 0.025976956 | 0.047990576 | 0.54129287 | 0.588305734 | NA | | BB014433 | 0 | NA | NA | NA | NA | NA | NR_037972 |
| 4930444M15Rik | 2.513581065 | 0.181080302 | 0.110869587 | 1.633272988 | 0.102411595 | NA | | BB019430 | 0.224626239 | -0.00168521210467683 | 0.00978955093367521 | -0.172143964119932 | 0.863324350255464 | NA | NR_033565 |
| 4930447A16Rik | 0 | NA | NA | NA | NA | NA | | BB123696 | 0.735831993 | 0.00054648811743211 | 0.0179149977808566 | 0.0305045037748244 | 0.975664701561217 | NA | NR_027893 |
| 4930447J18Rik | 0 | NA | NA | NA | NA | NA | | BB283400 | 0 | NA | NA | NA | NA | NA | NR_038124 |
| 4930447K03Rik | 7.44066142 | 0.025083196 | 0.155849513 | 0.160944977 | 0.872136737 | NA | | BB287469 | 0 | NA | NA | NA | NA | NA | NM_001177573 |
| 4930448C13Rik | 0 | NA | NA | NA | NA | NA | | BB557941 | 9.030406223 | -0.0409117037573289 | 0.0635092009542251 | -0.644185458841096 | 0.519455179063966 | NA | NR_040356 |
| 4930448F12Rik | 0.08731684 | 0.017365056 | 0.027733412 | 0.626142072 | 0.531221759 | NA | | BC002163 | 19.3039701 | -0.0231067696548582 | 0.0772377849018924 | -0.299164064378704 | 0.764814866828257 | NA | NR_002445 |
| 4930448H16Rik | 0 | NA | NA | NA | NA | NA | | BC016579 | 7.121960708 | -0.0744846144089118 | 0.0584902652733899 | -1.27345318166643 | 0.202857276040802 | NA | NM_145389 |
| 4930448I06Rik | 0 | NA | NA | NA | NA | NA | | BC018473 | 9.663879306 | -0.0880112867734139 | 0.0611449961944675 | -1.43938657700624 | 0.150041025321771 | NA | NR_003364 |
| 4930448I18Rik | 0 | NA | NA | NA | NA | NA | | BC021614 | 0 | NA | NA | NA | NA | NA | NM_144869 |
| 4930448K20Rik | 0 | NA | NA | NA | NA | NA | | BC021767 | 3.833913218 | -0.0419368253678627 | 0.0441509411015903 | -0.949851222228016 | 0.342187854640059 | NA | NR_033629 |
| 4930449E01Rik | 0.08731684 | 0.017365056 | 0.027733412 | 0.626142072 | 0.531221759 | NA | | BC024386 | 0 | NA | NA | NA | NA | NA | NR_015583 |
| 4930449E18Rik | 0 | NA | NA | NA | NA | NA | | BC024582 | 2.964245369 | -0.0336982852306968 | 0.0382035201810876 | -0.882072779444519 | 0.377737451282654 | NA | NR_026843 |
| 4930449I24Rik | 0 | NA | NA | NA | NA | NA | | BC027072 | 16.12087067 | 0.0701971491402768 | 0.0754236242989681 | 0.930705064795423 | 0.352006149060345 | NA | NM_146082 |
| 4930451I11Rik | 2.411114941 | -0.043380949 | 0.11076614 | -0.391644498 | 0.695320905 | NA | | BC030476 | 15.05141573 | 0.0153855110880321 | 0.0667153296993796 | 0.230614330429895 | 0.817614433603962 | NA | NM_173421 |
| 4930452A19Rik | 0.430253841 | 0.022125686 | 0.043852981 | 0.504542347 | 0.613880312 | NA | | BC030870 | 0 | NA | NA | NA | NA | NA | NR_033217 |
| 4930452N14Rik | 1.232026643 | -0.00778314 | 0.066195024 | -0.117578928 | 0.906401302 | NA | | BC033916 | 17.06183678 | 0.0913502887189756 | 0.0719269326749979 | 1.27004287992847 | 0.204069356405772 | NA | NR_040470 |
| 4930453H23Rik | 0.079983654 | 0.016740032 | 0.027674178 | 0.604897175 | 0.54524732 | NA | | BC039771 | 6.458923014 | -0.0120099578723411 | 0.0536975119405842 | -0.223659485110409 | 0.823022273500792 | NA | NR_033220 |
| 4930453L07Rik | 0.212768007 | -0.02694223 | 0.031868764 | -0.845411806 | 0.397880955 | NA | | BC039966 | 16.51023395 | 0.0511146702659277 | 0.0789969223879257 | 0.647046349665646 | 0.51760194630012 | NA | NR_040670 |
| 4930455B14Rik | 0.832005321 | -0.09663103 | 0.066726197 | -1.4481723 | 0.147568869 | NA | | BC048502 | 0 | NA | NA | NA | NA | NA | NM_177631 |
| 4930455C13Rik | 0.543893889 | 0.070201746 | 0.051980176 | 1.350548462 | 0.17684012 | NA | | BC048507 | 0.571777182 | 0.00200539641477231 | 0.0165938352331473 | 0.120851893886857 | 0.903808345910362 | NA | NM_001001185 |
| 4930455D15Rik | 0.119397641 | -0.020477583 | 0.027954224 | -0.732539845 | 0.463839132 | NA | | BC048562 | 0 | NA | NA | NA | NA | NA | NM_001004192 |
| 4930455F16Rik | 0.410740291 | -0.005062362 | 0.047943713 | -0.105589704 | 0.915907894 | NA | | BC048602 | 0 | NA | NA | NA | NA | NA | NR_045280 |
| 4930455H04Rik | 0.861910856 | 0.04703845 | 0.058004879 | 0.810939544 | 0.41740039 | NA | | BC048609 | 1.850648943 | 0.0190006576015673 | 0.0264311684525877 | 0.718873160513156 | 0.472219074848059 | NA | NM_001111317 |
| 4930455J16Rik | 0 | NA | NA | NA | NA | NA | | BC048644 | 4.505176405 | 0.00182999066865317 | 0.0463594496575305 | 0.0394739515281522 | 0.968512520994854 | NA | NM_001033485 |
| 4930456L15Rik | 0 | NA | NA | NA | NA | NA | | BC048671 | 0 | NA | NA | NA | NA | NA | NM_177738 |
| 4930459L07Rik | 0 | NA | NA | NA | NA | NA | | BC048679 | 0 | NA | NA | NA | NA | NA | NM_183143,NM_001193274 |
| 4930461G14Rik | 0.462752832 | 0.005884822 | 0.049969805 | 0.117767555 | 0.906251838 | NA | | BC048943 | 9.172351895 | -0.00640940131728418 | 0.0524246470777658 | -0.122259312643089 | 0.902693654013519 | NA | NM_001127685 |
| 4930465A12Rik | 0 | NA | NA | NA | NA | NA | | BC049635 | 2.650237588 | 0.00387657193826076 | 0.0340965555026063 | 0.113693945946078 | 0.909480411837634 | NA | NM_177785 |
| 4930465K10Rik | 1.060764143 | -0.020894344 | 0.066152052 | -0.315853302 | 0.752113858 | NA | | BC049715 | 18.03857757 | -0.032805443822294 | 0.0774082732508178 | -0.423797643902972 | 0.671713396556698 | NA | NM_178776 |
| 4930465M20Rik | 0.386222787 | 0.053439715 | 0.045634083 | 1.171048305 | 0.241579361 | NA | | BC049730 | 0 | NA | NA | NA | NA | NA | NM_199150 |
| 4930467D21Rik | 2.016869231 | 0.118619644 | 0.094075112 | 1.260903567 | 0.207343593 | NA | | BC049762 | 3.522296248 | -0.0194567392913503 | 0.0427206583131369 | -0.455440998795827 | 0.648792005387122 | NA | NM_177567 |
| 4930467E23Rik | 0 | NA | NA | NA | NA | NA | | BC051019 | 9.741549949 | -0.0505830635034222 | 0.0565796481469856 | -0.894015165524091 | 0.371313777014837 | NA | NM_001040700 |
| 4930467K11Rik | 2.686659219 | -0.053715893 | 0.107438707 | -0.499967794 | 0.617097755 | NA | | BC051070 | 1.022498331 | 0.000419229489371584 | 0.0233039322427881 | 0.0179896459105662 | 0.985647113442288 | NA | NM_178412 |
| 4930468A15Rik | 0.38092772 | -0.000766687 | 0.035977 | -0.021310467 | 0.982997994 | NA | | BC051537 | 2.72092569 | -0.0152206707569613 | 0.0328305372095237 | -0.463613210463885 | 0.642924887925015 | NA | NR_046183 |
| 4930469G21Rik | 0.258320694 | 0.006870708 | 0.033097595 | 0.207589347 | 0.835549623 | NA | | BC051628 | 3.610209457 | 0.01685601347053 | 0.0349223703073968 | 0.482670944788641 | 0.629329396707844 | NA | NM_199312 |
| 4930470P17Rik | 0 | NA | NA | NA | NA | NA | | BC051665 | 1.560338262 | 0.012777045249529 | 0.0220552023966744 | 0.579321151523669 | 0.562372496078968 | NA | NM_199148 |
| 4930471C04Rik | 6.068235747 | 0.148690851 | 0.13758957 | 1.080684028 | 0.27983769 | NA | | BC053393 | 2.691327572 | 0.012226567319393 | 0.0374638639028917 | 0.326356281644757 | 0.744154810464221 | NA | NM_001025435 |
| 4930471G03Rik | 0.079983654 | 0.016740032 | 0.027674178 | 0.604897175 | 0.54524732 | NA | | BC055111 | 5.303674951 | 0.0365529884724447 | 0.0456765189651457 | 0.800257753887442 | 0.423561474316627 | NA | NM_183182 |
| 4930471I20Rik | 0.815331231 | 0.023730175 | 0.06737475 | 0.352211695 | 0.724679507 | NA | | BC055402 | 10.23975924 | -0.0871056983245603 | 0.0662246169501421 | -1.31530694077308 | 0.188406766411555 | NA | NR_037990 |
| 4930471M09Rik | 0.820973202 | -0.049337952 | 0.072961054 | -0.676223119 | 0.498898996 | NA | | BC061212 | 0 | NA | NA | NA | NA | NA | NM_198667 |
| 4930473O22Rik | 0.265890466 | 0.030259369 | 0.03447172 | 0.877802699 | 0.3800508 | NA | | BC061237 | 0 | NA | NA | NA | NA | NA | NM_198677 |
| 4930474G06Rik | 6.829476927 | -0.008029662 | 0.144632649 | -0.05551763 | 0.955726085 | NA | | BC080695 | 0 | NA | NA | NA | NA | NA | NM_001007579 |
| 4930474H20Rik | 0 | NA | NA | NA | NA | NA | | BC089597 | 0 | NA | NA | NA | NA | NA | NM_145424 |
| 4930474M22Rik | 0 | NA | NA | NA | NA | NA | | BC090627 | 0 | NA | NA | NA | NA | NA | NR_004843 |
| 4930474N05Rik | 0 | NA | NA | NA | NA | NA | | BC094916 | 0.227686571 | 0.00691460391109036 | 0.00980755285755433 | 0.705028462402254 | 0.480792523671872 | NA | NM_001024721 |
| 4930474N09Rik | 3.418441142 | -0.06720669 | 0.121752223 | -0.551995593 | 0.580951372 | NA | | BC096441 | 0 | NA | NA | NA | NA | NA | NM_001034097,NM_001159503 |
| 4930478L05Rik | 0 | NA | NA | NA | NA | NA | | BC100451 | 6.913574096 | 0.0479167148772217 | 0.0553788469872924 | 0.865253025008213 | 0.386899927564353 | NA | NM_021440 |
| 4930478P22Rik | 0 | NA | NA | NA | NA | NA | | BC100530 | 0 | NA | NA | NA | NA | NA | NM_001082546 |
| 4930479M11Rik | 0.08731684 | 0.017365056 | 0.027733412 | 0.626142072 | 0.531221759 | NA | | BC107364 | 1.925270583 | 0.00550854360611597 | 0.0342363163433095 | 0.160897672251835 | 0.872173995693186 | NA | NM_001256181,NM_001256180 |
| 4930480E11Rik | 0 | NA | NA | NA | NA | NA | | BC117090 | 0 | NA | NA | NA | NA | NA | NM_001001332 |
| 4930480G23Rik | 0.094392817 | 0.017895942 | 0.027818137 | 0.643319236 | 0.520016977 | NA | | BC125332 | 5.227761903 | 0.0391853876251055 | 0.0497124215234764 | 0.788241377592125 | 0.430555530353008 | NA | NM_172521 |
| 4930480K15Rik | 0 | NA | NA | NA | NA | NA | | Baat | 5.211806029 | -0.014096740080793 | 0.0453339308807396 | -0.310953403045447 | 0.755836045667379 | NA | NM_007519 |
| 4930480M12Rik | 0 | NA | NA | NA | NA | NA | | Baat1 | 16.37996782 | -0.0815984023076013 | 0.0732508151623659 | -1.1139589658727 | 0.265296790853973 | NA | NM_172724 |
| 4930483J18Rik | 1.04385259 | 0.004672836 | 0.066845675 | 0.069904832 | 0.944269407 | NA | | Banf2 | 0 | NA | NA | NA | NA | NA | NM_001044750,NM_207275 |
| 4930483K19Rik | 0.239950962 | 0.027417169 | 0.032831728 | 0.835081494 | 0.403671825 | NA | | Barhl1 | 4.660516633 | 0.0240798392900405 | 0.0355627975539427 | 0.677107565947689 | 0.498337708841732 | NA | NM_001164186,NM_019446 |
| 4930483O08Rik | 1.313476444 | 0.109374984 | 0.086680888 | 1.261811991 | 0.207016446 | NA | | Barx1 | 0 | NA | NA | NA | NA | NA | NM_007526 |
| 4930486F22Rik | 0 | NA | NA | NA | NA | NA | | Batf | 5.354196921 | -0.0377590490447373 | 0.0454715249310753 | -0.830388888474087 | 0.406318945891404 | NA | NM_016767 |
| 4930486I03Rik | 2.188966278 | -0.12387563 | 0.109669576 | -1.12953505 | 0.258672193 | NA | | Batf2 | 10.06878781 | -0.101019291765189 | 0.0559533417748123 | -1.80542016903562 | 0.0710089573837405 | NA | NM_028967 |
| 4930486L24Rik | 6.815623883 | 0.105972792 | 0.134994638 | 0.785014825 | 0.432444876 | NA | | Bc1 | 0.127687691 | 0.00542298404407807 | 0.00869724871942359 | 0.62352868349814 | 0.532937159177571 | NA | NR_038088 |
| 4930487D11Rik | 0.096071043 | -0.020098795 | 0.027825948 | -0.722304036 | 0.470107572 | NA | | Bcl2a1a | 6.520767286 | -0.161997023345862 | 0.0561694362963068 | -2.88407778371302 | 0.00392561776621373 | NA | NM_009742 |
| 4930487H11Rik | 0.187700257 | -0.003168524 | 0.031449734 | -0.100748845 | 0.919749836 | NA | | Bcl2a1c | 0 | NA | NA | NA | NA | NA | NM_007535 |
| 4930488B22Rik | 0 | NA | NA | NA | NA | NA | | Bcl2a1d | 1.238160007 | -0.0102640616226104 | 0.0249057921866823 | -0.4121154446996 | 0.680254808828002 | NA | NM_007536 |
| 4930488L21Rik | 8.031833972 | 0.012865558 | 0.153101826 | 0.084032689 | 0.933030442 | NA | | Bcl2l10 | 1.015389671 | -0.0214626428177984 | 0.0235150845239417 | -0.912718080853435 | 0.361390832034588 | NA | NM_013479 |
| 4930500F04Rik | 0 | NA | NA | NA | NA | NA | | Bcl2l14 | 0.203513543 | -0.00860782669689374 | 0.00975338121443284 | -0.882547960306941 | 0.377480556050336 | NA | NM_025778 |
| 4930500J02Rik | 7.694952451 | 0.155917232 | 0.145144368 | 1.074221712 | 0.282723323 | NA | | Bcl2l15 | 11.77644069 | -0.0420359217448798 | 0.0658190647656937 | -0.638658751754094 | 0.523044951866882 | NA | NM_001142960,NM_001142959 |
| 4930500L23Rik | 0.40780145 | -0.036172458 | 0.049100092 | -0.736708546 | 0.461299609 | NA | | Bcl3 | 16.53613191 | -0.252931909900107 | 0.0746477117931848 | -3.38834110013804 | 0.000703167585209072 | NA | NM_033601 |
| 4930502A04Rik | 0 | NA | NA | NA | NA | NA | | Bcmo1 | 4.339483503 | 0.0500209414880188 | 0.0376500446753638 | 1.3285758866775 | 0.183987931833423 | NA | NM_021486,NM_001163028 |
| 4930502E09Rik | 0.099009884 | 0.018205726 | 0.027896865 | 0.652608315 | 0.514008823 | NA | | Bdkrb1 | 3.190715645 | 0.031466221793565 | 0.0404174024780916 | 0.778531520194089 | 0.436255732061912 | NA | NM_007539 |
| 4930503B20Rik | 2.495604498 | 0.020978341 | 0.111466346 | 0.18820336 | 0.850717239 | NA | | Best2 | 1.645174031 | -0.0348109431398753 | 0.0273503815444639 | -1.27277724017424 | 0.203097099726711 | NA | NM_001130194,NR_052011 |
| 4930503E14Rik | 1.459175472 | 0.008307894 | 0.078779715 | 0.10545728 | 0.916012967 | NA | | Best3 | 9.729391705 | 0.00125011679871401 | 0.0544618273219835 | 0.0229540002637664 | 0.981686965742281 | NA | NM_001007583 |
| 4930503E24Rik | 0 | NA | NA | NA | NA | NA | | Bet3l | 3.839309565 | 0.0384329534912733 | 0.0475276374906271 | 0.808644307196052 | 0.41871977135892 | NA | NM_001162937 |
| 4930503H13Rik | 0.176054697 | -0.002190495 | 0.031239146 | -0.070120197 | 0.944097991 | NA | | Bex6 | 0 | NA | NA | NA | NA | NA | NM_001033539 |
| 4930503O07Rik | 0 | NA | NA | NA | NA | NA | | Bglap | 1.582125684 | -0.00250385071746596 | 0.0245392023861064 | -0.102034722973865 | 0.918729113955719 | NA | NM_007541,NM_001037939 |
| 4930504O13Rik | 0 | NA | NA | NA | NA | NA | | Bglap2 | 1.208833268 | -0.0263899732558993 | 0.0233947504815673 | -1.12802969523833 | 0.259307378349755 | NA | NM_001032298 |
| 4930505A04Rik | 6.08102324 | -0.060985046 | 0.15135395 | -0.402929993 | 0.686999726 | NA | | Bglap3 | 11.55314369 | -0.0361892151691387 | 0.0663871720744308 | -0.545123614070572 | 0.585668502685847 | NA | NM_031368 |
| 4930505G20Rik | 0 | NA | NA | NA | NA | NA | | Bhlha15 | 4.674872679 | -0.0207021688192609 | 0.0432666889234332 | -0.478478231969551 | 0.632309863982652 | NA | NM_010800 |
| 4930509E16Rik | 1.005958322 | 0.045463035 | 0.072765671 | 0.624786867 | 0.532110951 | NA | | Bhlhe23 | 2.888036141 | 0.00198653816933398 | 0.0373100533958345 | 0.0532440451976348 | 0.957537462371317 | NA | NM_080641 |
| 4930509J09Rik | 0 | NA | NA | NA | NA | NA | | Bhmt | 4.343513701 | 0.0330045793018902 | 0.036840182606381 | 0.895885334080117 | 0.370314007780955 | NA | NM_016668 |
| 4930509K18Rik | 0 | NA | NA | NA | NA | NA | | Bhmt2 | 14.77101316 | -0.0303297472907511 | 0.0731550998831566 | -0.41459511830609 | 0.678438322435157 | NA | NM_022884 |
| 4930511A02Rik | 2.134322429 | 0.023974154 | 0.087649941 | 0.273521619 | 0.784452276 | NA | | Bik | 2.99264569 | 0.0229827596857887 | 0.0401663461856461 | 0.572189453816984 | 0.567193631476543 | NA | NM_007546 |
| 4930511E03Rik | 0 | NA | NA | NA | NA | NA | | Birc7 | 8.494131837 | 0.0844611529416281 | 0.0631404800959032 | 1.33767042653685 | 0.181003895469745 | NA | NM_001163247 |
| 4930511M11Rik | 0 | NA | NA | NA | NA | NA | | Blk | 0.120209049 | -0.00674972139751468 | 0.00863521114606269 | -0.781650996523957 | 0.434419707908266 | NA | NM_007549 |
| 4930512B01Rik | 1.54174701 | 0.130241365 | 0.088845497 | 1.465930961 | 0.142667096 | NA | | Bmp10 | 0 | NA | NA | NA | NA | NA | NM_009756 |
| 4930513D17Rik | 0.929186171 | -0.000153963 | 0.058545282 | -0.002629812 | 0.997901716 | NA | | Bmp15 | 4.826487318 | 0.0191334228717443 | 0.0501583225993816 | 0.381460580820543 | 0.702861515882878 | NA | NM_009757 |
| 4930513O06Rik | 0 | NA | NA | NA | NA | NA | | Bmp8a | 5.143282247 | -0.0467630053614616 | 0.0514806276886329 | -0.908361211994061 | 0.363687411750655 | NA | NM_001256019,NM_007558 |
| 4930515B02Rik | 0 | NA | NA | NA | NA | NA | | Bmp8b | 6.66869795 | -0.0369318822539249 | 0.0517932642468861 | -0.713063422260459 | 0.475806512434563 | NA | NM_007559 |
| 4930515G01Rik | 7.402560496 | -0.000999662 | 0.149758177 | -0.006675172 | 0.994674023 | NA | | Bnc1 | 0.776462932 | 0.00764764447012096 | 0.0179430948670789 | 0.426216576726263 | 0.669950039542914 | NA | NM_007562 |
| 4930515G16Rik | 0 | NA | NA | NA | NA | NA | | Bnc2 | 15.57552781 | 0.00145008481676018 | 0.069451874608626 | 0.0208789874273613 | 0.983342188574467 | NA | NM_172870 |
| 4930515L03Rik | 1.09530653 | 0.005333132 | 0.072668172 | 0.073390207 | 0.94149561 | NA | | Bnipl | 6.133403478 | -0.0803809402086582 | 0.0512604255719759 | -1.56808959956436 | 0.116860223407589 | NA | NM_001168356,NM_134253 |
| 4930515L19Rik | 0 | NA | NA | NA | NA | NA | | Boll | 0 | NA | NA | NA | NA | NA | NM_029267,NM_001113367 |
| 4930517E11Rik | 0 | NA | NA | NA | NA | NA | | Bpi | 0.309462921 | 0.00239536941478217 | 0.0109127344998192 | 0.219502216866162 | 0.826258853162846 | NA | NM_177850 |
| 4930518P08Rik | 0 | NA | NA | NA | NA | NA | | Bpifa1 | 11.17320953 | 0.0357796518559142 | 0.0417040646118951 | 0.857941598472127 | 0.390924712643541 | NA | NM_011126 |
| 4930519D14Rik | 0.094392817 | 0.017895942 | 0.027818137 | 0.643319236 | 0.520016977 | NA | | Bpifa2 | 0.217532603 | 0.00684685248201087 | 0.00976000296730706 | 0.701521557416086 | 0.482977586094298 | NA | NM_008953 |
| 4930519F16Rik | 1.681417607 | 0.226274572 | 0.095808368 | 2.361741218 | 0.018189334 | NA | | Bpifa3 | 0 | NA | NA | NA | NA | NA | NM_028528 |
| 4930519F24Rik | 0.119397641 | -0.020477583 | 0.027954224 | -0.732539845 | 0.463839132 | NA | | Bpifa5 | 0.747274741 | -0.00748289856866147 | 0.0180925780158936 | -0.413589404566229 | 0.679174834170192 | NA | NM_025990 |
| 4930519H02Rik | 0.107716603 | -0.020477583 | 0.027954224 | -0.732539849 | 0.46383913 | NA | | Bpifa6 | 0.12890128 | -0.0068640067066208 | 0.00870804083610955 | -0.788237771940376 | 0.430557639008166 | NA | NM_001080811 |
| 4930520P13Rik | 0 | NA | NA | NA | NA | NA | | Bpifb1 | 6.436800035 | -0.00273423339920367 | 0.0502608882868531 | -0.0544008172636867 | 0.956615827738526 | NA | NM_153418,NM_001012392 |
| 4930521E06Rik | 0 | NA | NA | NA | NA | NA | | Bpifb2 | 0 | NA | NA | NA | NA | NA | NM_025631 |
| 4930522H14Rik | 0 | NA | NA | NA | NA | NA | | Bpifb3 | 1.950685126 | -0.000613196374556278 | 0.0257826727176767 | -0.0237832741884773 | 0.981025481540702 | NA | NM_194357 |
| 4930522O17Rik | 0.119397641 | -0.020477583 | 0.027954224 | -0.732539845 | 0.463839132 | NA | | Bpifb4 | 6.177014064 | -0.0769606802170624 | 0.0495459408550002 | -1.5533195835819 | 0.12034680393976 | NA | NM_001034875 |
| 4930523O13Rik | 0 | NA | NA | NA | NA | NA | | Bpifb5 | 1.734144193 | -0.0260843419352703 | 0.0294667450987205 | -0.885212867857704 | 0.376041833421946 | NA | NM_144890 |
| 4930524B15Rik | 6.554640627 | 0.05191124 | 0.144854754 | 0.358367525 | 0.720068291 | NA | | Bpifb6 | 1.327498615 | -0.00931378059104271 | 0.0253662463203806 | -0.367172204882341 | 0.713490570860533 | NA | NM_199303 |
| 4930524C18Rik | 0 | NA | NA | NA | NA | NA | | Bpifb9a | 4.120746202 | -0.0620463313576399 | 0.0323822683952369 | -1.91605883196145 | 0.0553576076314567 | NA | NM_175167 |
| 4930524N10Rik | 0 | NA | NA | NA | NA | NA | | Bpifb9b | 12.61121309 | -0.0130670288028303 | 0.0198515001218454 | -0.658238859664355 | 0.51038465821605 | NA | NM_001025574 |
| 4930524O05Rik | 0 | NA | NA | NA | NA | NA | | Bpifc | 7.364241138 | -0.0513314726492913 | 0.0573277914232051 | -0.895402934160714 | 0.370571732727168 | NA | NM_177772 |
| 4930524O08Rik | 0 | NA | NA | NA | NA | NA | | Brat1 | 0.631880482 | -0.0161216627837558 | 0.0164156601117484 | -0.982090435231279 | 0.326055297142213 | NA | NR_074085,NM_001276287,NM_181066 |
| 4930525D18Rik | 0.882749213 | 0.107936301 | 0.072744355 | 1.483775627 | 0.137868455 | NA | | Bricd5 | 15.29271671 | -0.0248894180418813 | 0.0704980033782063 | -0.353051389389782 | 0.724049914787013 | NA | NM_175682 |
| 4930525M21Rik | 0 | NA | NA | NA | NA | NA | | Brs3 | 13.73253843 | 0.0531724633124602 | 0.051206816685792 | 1.03838642497013 | 0.299090186987693 | NA | NM_009766 |
| 4930526D03Rik | 0 | NA | NA | NA | NA | NA | | Bsnd | 0.623897432 | 0.0213829806133127 | 0.0171670258447213 | 1.24558445980832 | 0.212916990412024 | NA | NM_080458 |
| 4930526L06Rik | 0.978497497 | 0.043411735 | 0.06751127 | 0.643029462 | 0.520204983 | NA | | Bsph1 | 1.069612833 | -0.035848219801572 | 0.0217295206379416 | -1.64974738278294 | 0.0989946145975716 | NA | NM_001033418 |
| 4930527F14Rik | 0.107716603 | -0.020477583 | 0.027954224 | -0.732539849 | 0.46383913 | NA | | Bsph2 | 0 | NA | NA | NA | NA | NA | NM_001080942 |
| 4930527G23Rik | 0.098353386 | -0.020212831 | 0.02786466 | -0.725393061 | 0.468210929 | NA | | Bspry | 13.74744501 | -0.0312658282277685 | 0.0684085070982249 | -0.457045907797333 | 0.647638048767425 | NA | NM_138653 |
| 4930528D03Rik | 1.522758917 | 0.143613366 | 0.093881166 | 1.529735638 | 0.126082178 | NA | | Bst1 | 5.469989599 | 0.0630120933465474 | 0.0467106649963554 | 1.3489872891226 | 0.177341048857485 | NA | NM_009763 |
| 4930528P14Rik | 0.618726074 | 0.039228334 | 0.048427554 | 0.810041617 | 0.417916257 | NA | | Bsx | 3.740478274 | 0.0345132507069862 | 0.0250588125126805 | 1.37728995296651 | NA | NA | NM_178245 |
| 4930529C04Rik | 0 | NA | NA | NA | NA | NA | | Btc | 18.96012651 | 0.0422332602009408 | 0.0791392311766377 | 0.533657701408253 | 0.593578378958857 | NA | NM_007568 |
| 4930529F24Rik | 0 | NA | NA | NA | NA | NA | | Btg3 | 0 | NA | NA | NA | NA | NA | NM_009770 |
| 4930529K09Rik | 0.079983654 | 0.016740032 | 0.027674178 | 0.604897175 | 0.54524732 | NA | | Btg4 | 0.21100899 | -0.000459554483117493 | 0.00972904351832043 | -0.0472353199214416 | 0.96232567770125 | NA | NM_019493 |
| 4930529L06Rik | 0 | NA | NA | NA | NA | NA | | Btla | 13.64167669 | 0.00806804851785076 | 0.0681257016468693 | 0.118428850240275 | 0.905727867440694 | NA | NM_177584,NM_001037719 |
| 4930533B01Rik | 0 | NA | NA | NA | NA | NA | | Btn1a1 | 3.336976282 | 0.048394921445514 | 0.037465379588958 | 1.29172377209217 | 0.196452821918978 | NA | NM_013483 |
| 4930533P14Rik | 0.323149808 | -0.030167563 | 0.033779321 | -0.893077861 | 0.371815478 | NA | | Btnl1 | 0 | NA | NA | NA | NA | NA | NM_001111094 |
| 4930539C22Rik | 0 | NA | NA | NA | NA | NA | | Btnl10 | 0.120209049 | -0.00674972139751468 | 0.00863521114606269 | -0.781650996523957 | 0.434419707908266 | NA | NM_138678 |
| 4930539M17Rik | 5.329181811 | 0.014030299 | 0.140677651 | 0.099733673 | 0.920555766 | NA | | Btnl2 | 0.347878816 | 0.00340886725081251 | 0.0128564130570379 | 0.265149169965912 | 0.790894550259911 | NA | NM_079835 |
| 4930539N22Rik | 0 | NA | NA | NA | NA | NA | | Btnl4 | 0 | NA | NA | NA | NA | NA | NM_030746 |
| 4930540M03Rik | 0 | NA | NA | NA | NA | NA | | Btnl5 | 0.638578749 | 0.00457849037287473 | 0.0152692283897995 | 0.299850801624879 | 0.764290963053352 | NA | NR_004051 |
| 4930542C21Rik | 0 | NA | NA | NA | NA | NA | | Btnl6 | 0.240418098 | -0.0088029805258837 | 0.00986414519853158 | -0.892422034419582 | 0.372166765031741 | NA | NR_030747 |
| 4930542D17Rik | 0.079983654 | 0.016740032 | 0.027674178 | 0.604897175 | 0.54524732 | NA | | Btnl9 | 0.981826011 | 0.0207600109109852 | 0.0234779098667829 | 0.884235906380956 | 0.376568879109931 | NA | NM_172793 |
| 4930542H20Rik | 0 | NA | NA | NA | NA | NA | | Bub1 | 7.457294122 | 0.01435164500734 | 0.0583554365897748 | 0.245935012160542 | 0.805732540234863 | NA | NM_009772,NM_001113179 |
| 4930543E12Rik | 0.488510588 | 0.029306206 | 0.046230877 | 0.633909798 | 0.526139691 | NA | | C030007H22Rik | 1.111433229 | 0.00402764479295851 | 0.0255580670971919 | 0.157588004509192 | 0.874781456472301 | NA | NR_040482 |
| 4930544D05Rik | 0.331243956 | 0.008596723 | 0.034710864 | 0.247666645 | 0.804392342 | NA | | C030016D13Rik | 17.49752904 | 0.0165150506821659 | 0.0795695464250853 | 0.207554917982532 | 0.83557650794815 | NA | NR_027987 |
| 4930544G11Rik | 0 | NA | NA | NA | NA | NA | | C030034L19Rik | 3.85451335 | -0.0418696312778326 | 0.043562181346121 | -0.961146342630542 | 0.336478591130379 | NA | NR_015499 |
| 4930544M13Rik | 0 | NA | NA | NA | NA | NA | | C130021I20Rik | 7.322352946 | 0.080744317224494 | 0.0525883178706316 | 1.5354040686969 | 0.124684604931428 | NA | NR_046275 |
| 4930545E07Rik | 0 | NA | NA | NA | NA | NA | | C130026I21Rik | 1.872426285 | -0.0252554625693569 | 0.0320370461148243 | -0.788320573589668 | 0.430509216507266 | NA | NM_175219,NM_001037909 |
| 4930545H06Rik | 0.813882095 | 0.031557003 | 0.065495559 | 0.481818976 | 0.629934548 | NA | | C130050O18Rik | 7.584762748 | 0.0430019601095982 | 0.0571712118486689 | 0.752161074063351 | 0.45195419939601 | NA | NM_177000 |
| 4930545L23Rik | 0.425621188 | 0.027403281 | 0.044342756 | 0.617987764 | 0.536583404 | NA | | C130060C02Rik | 10.75026041 | -0.0594768056682147 | 0.0650594053605713 | -0.914192273024679 | 0.360615823617191 | NA | NR_045355 |
| 4930546C10Rik | 2.10939122 | -0.057982471 | 0.099752708 | -0.581262119 | 0.561063807 | NA | | C130079G13Rik | 0 | NA | NA | NA | NA | NA | NM_177661 |
| 4930546K05Rik | 0.565738727 | -0.007881918 | 0.057869715 | -0.13620108 | 0.891662321 | NA | | C130080G10Rik | 6.221132432 | 0.0419101447170543 | 0.0510905871864903 | 0.820310492108347 | 0.412039126625556 | NA | NR_028422 |
| 4930547C10Rik | 0 | NA | NA | NA | NA | NA | | C130083M11Rik | 1.069576044 | -0.00713366330650953 | 0.0232914006477897 | -0.306278845758745 | 0.759392344419655 | NA | NR_040717 |
| 4930547E08Rik | 0 | NA | NA | NA | NA | NA | | C1ql4 | 0.655896803 | 0.00512107898138762 | 0.0174866649142207 | 0.292856242543025 | 0.769632036515633 | NA | NM_001024702 |
| 4930547E14Rik | 0 | NA | NA | NA | NA | NA | | C1qtnf3 | 4.104191035 | 0.0305045689609286 | 0.0356859878166434 | 0.854805228249889 | 0.392658983340752 | NA | NM_030888,NM_001204134 |
| 4930548G14Rik | 0 | NA | NA | NA | NA | NA | | C1qtnf5 | 0 | NA | NA | NA | NA | NA | NM_001040631,NM_001040632,NM_001190319,NM_145613,NM_001190313 |
| 4930548H24Rik | 0.380940254 | 0.025976956 | 0.047990576 | 0.54129287 | 0.588305734 | NA | | C1qtnf9 | 16.28389737 | 0.0602125202852838 | 0.0770835281389446 | 0.78113342420899 | 0.434724025237835 | NA | NM_183175 |
| 4930548J01Rik | 0 | NA | NA | NA | NA | NA | | C1rb | 0.794369964 | 0.00768744715138506 | 0.0166305698590855 | 0.462247969644006 | 0.643903506869677 | NA | NM_001113356 |
| 4930548K13Rik | 0 | NA | NA | NA | NA | NA | | C1rl | 20.83768123 | -0.124544366855504 | 0.0697796993603523 | -1.78482234800611 | 0.0742901372172414 | NA | NM_181344 |
| 4930549C01Rik | 0 | NA | NA | NA | NA | NA | | C230024C17Rik | 1.505577713 | -0.0409669450109451 | 0.0272962509514515 | -1.50082680159293 | 0.133400364605902 | NA | NR_046171 |
| 4930549G23Rik | 1.00591031 | 0.019068594 | 0.071772183 | 0.265682233 | 0.790483948 | NA | | C230035I16Rik | 21.02954182 | -0.121583285879725 | 0.0746978053121624 | -1.6276687831942 | 0.10359513570756 | NA | NR_015492 |
| 4930550L24Rik | 0.159967308 | 0.0240345 | 0.030870969 | 0.778547006 | 0.436246606 | NA | | C230079O03Rik | 0 | NA | NA | NA | NA | NA | NR_040459 |
| 4930552N02Rik | 0 | NA | NA | NA | NA | NA | | C2cd4a | 3.283019704 | -0.00341154703912722 | 0.0421462505585114 | -0.0809454457731889 | 0.935485338017692 | NA | NM_001163143 |
| 4930552P12Rik | 0.931403597 | -0.00929293 | 0.064794173 | -0.143422307 | 0.885956667 | NA | | C2cd4d | 3.727446145 | 0.0311159579667253 | 0.0403977440649436 | 0.770239989557415 | 0.441157546551316 | NA | NM_001136117 |
| 4930553E22Rik | 0.08731684 | 0.017365056 | 0.027733412 | 0.626142072 | 0.531221759 | NA | | C330008G21Rik | 0 | NA | NA | NA | NA | NA | NR_040617 |
| 4930554C24Rik | 0 | NA | NA | NA | NA | NA | | C330011F03Rik | 0 | NA | NA | NA | NA | NA | NR_046166 |
| 4930555B11Rik | 0 | NA | NA | NA | NA | NA | | C330013F16Rik | 0.106347839 | -0.00653347234140708 | 0.00854389509723638 | -0.764694822098226 | 0.444453290197145 | NA | NR_045456,NR_045455 |
| 4930555G01Rik | 5.980794058 | -0.133563684 | 0.15019306 | -0.889279997 | 0.373852619 | NA | | C330021F23Rik | 2.613490235 | -0.0118551179338984 | 0.0344439774142582 | -0.344185510033198 | 0.730706778684002 | NA | NM_001024728 |
| 4930556C24Rik | 0 | NA | NA | NA | NA | NA | | C330022C24Rik | 1.677133784 | 0.0263581853622308 | 0.0305836436893034 | 0.86183927690243 | 0.388775967964088 | NA | NR_045717 |
| 4930556G01Rik | 0 | NA | NA | NA | NA | NA | | C330024C12Rik | 0.108766301 | 0.00519171381361946 | 0.00856003371502575 | 0.606506234257728 | 0.544178643801544 | NA | NR_046016 |
| 4930556J02Rik | 0 | NA | NA | NA | NA | NA | | C330024D21Rik | 3.579928947 | -0.0204044366302017 | 0.0450192578925141 | -0.453237960495005 | 0.65037740065867 | NA | NR_015582 |
| 4930556N09Rik | 0.32048461 | -0.048989879 | 0.042627894 | -1.149244659 | 0.25045511 | NA | | C330046G13Rik | 0 | NA | NA | NA | NA | NA | NR_040658 |
| 4930557A04Rik | 0 | NA | NA | NA | NA | NA | | C430002E04Rik | 0 | NA | NA | NA | NA | NA | NR_040385 |
| 4930557J02Rik | 0 | NA | NA | NA | NA | NA | | C430049B03Rik | 6.089537161 | -0.0039221265222234 | 0.0526756752586918 | -0.0744580207650252 | 0.940645942981377 | NA | NR_038184,NR_038185 |
| 4930558C23Rik | 0 | NA | NA | NA | NA | NA | | C4bp | 0.660042781 | -0.00273650593056747 | 0.0155456188198701 | -0.176030685061551 | 0.860269835648373 | NA | NM_007576 |
| 4930558G05Rik | 0 | NA | NA | NA | NA | NA | | C4bp-ps1 | 5.12714124 | -0.0368857702782682 | 0.0452770431100109 | -0.814668267727771 | 0.415262221843606 | NA | NR_028304 |
| 4930558J18Rik | 1.445046221 | 0.012865077 | 0.080275315 | 0.160261929 | 0.872674748 | NA | | C530044C16Rik | 7.76552431 | 0.050984082005173 | 0.0518022626057925 | 0.984205697599627 | 0.325014384949695 | NA | NR_045984 |
| 4930558K02Rik | 0.6359905 | 0.05770125 | 0.047097028 | 1.225156918 | 0.220516103 | NA | | C6 | 0.113843286 | 0.00525947738444006 | 0.00859368002043176 | 0.612016897526494 | 0.540526577873086 | NA | NM_016704 |
| 4930562C15Rik | 0.17833704 | -0.002390466 | 0.031282281 | -0.076415987 | 0.939088151 | NA | | C630028M04Rik | 0 | NA | NA | NA | NA | NA | NR_040668 |
| 4930562F07Rik | 0.795601132 | -0.041375411 | 0.068722503 | -0.602064955 | 0.547130901 | NA | | C630031E19Rik | 11.87871877 | -0.0731761792738364 | 0.0592769140610445 | -1.23448024299103 | 0.21702400432709 | NA | NR_046080 |
| 4930563D23Rik | 0.08731684 | 0.017365056 | 0.027733412 | 0.626142072 | 0.531221759 | NA | | C730027H18Rik | 0 | NA | NA | NA | NA | NA | NR_038040 |
| 4930563E18Rik | 1.896076537 | 0.04518369 | 0.096495718 | 0.468245545 | 0.639609006 | NA | | C730036E19Rik | 1.614645163 | -0.00595333399914109 | 0.0303531540763004 | -0.196135597117053 | 0.844504034859806 | NA | NR_038011 |
| 4930563F08Rik | 0 | NA | NA | NA | NA | NA | | C86187 | 1.91680046 | 0.00140817780503804 | 0.0298441521806956 | 0.0471843795900795 | 0.962366276936766 | NA | NR_015609 |
| 4930563M20Rik | 0 | NA | NA | NA | NA | NA | | C86695 | 0.106347839 | -0.00653347234140708 | 0.00854389509723638 | -0.764694822098226 | 0.444453290197145 | NA | NM_001081662 |
| 4930564B18Rik | 0.105923158 | 0.018524693 | 0.027977518 | 0.662127842 | 0.507889294 | NA | | C87198 | 0.108766301 | 0.00519171381361946 | 0.00856003371502575 | 0.606506234257728 | 0.544178643801544 | NA | NR_046002,NR_046003 |
| 4930564C03Rik | 0 | NA | NA | NA | NA | NA | | C87414 | 0 | NA | NA | NA | NA | NA | NM_001164284,NM_001164285 |
| 4930564D02Rik | 0 | NA | NA | NA | NA | NA | | C87499 | 0.256588971 | -0.000984450977364047 | 0.0101556058348068 | -0.0969367060298841 | 0.922776658860967 | NA | NM_198663,NR_027385 |
| 4930564G21Rik | 0 | NA | NA | NA | NA | NA | | C87977 | 0 | NA | NA | NA | NA | NA | NM_001177542 |
| 4930564K09Rik | 0 | NA | NA | NA | NA | NA | | C8a | 1.74505306 | -0.0150247996610112 | 0.0296296545515532 | -0.507086562041056 | 0.612094083360284 | NA | NM_146148 |
| 4930565D16Rik | 0 | NA | NA | NA | NA | NA | | C8b | 3.119695849 | 0.02396860511282 | 0.0424191678385965 | 0.565041850986322 | 0.57204529848465 | NA | NM_133882 |
| 4930565N06Rik | 0 | NA | NA | NA | NA | NA | | C9 | 0.106148608 | 0.00515554063321497 | 0.00854200771766808 | 0.603551390213729 | 0.546141939141592 | NA | NM_013485 |
| 4930567H12Rik | 0.688757862 | 0.006342159 | 0.055170474 | 0.114955679 | 0.908480252 | NA | | C920009B18Rik | 1.38569564 | -0.0204572677756911 | 0.0265281344458665 | -0.771153652641364 | 0.44061586205351 | NA | NR_015465 |
| 4930567H17Rik | 0 | NA | NA | NA | NA | NA | | C920021L13Rik | 21.98077923 | -0.171321102493531 | 0.0839302424803014 | -2.04123206880691 | 0.0412277655809184 | NA | NR_040446 |
| 4930567J20Rik | 0 | NA | NA | NA | NA | NA | | Cabp2 | 0.342801831 | 0.00329783501100829 | 0.0128022060068578 | 0.257598964525467 | 0.796716428937095 | NA | NM_001160253,NM_001160252,NM_013878 |
| 4930567K20Rik | 0.096071043 | -0.020098795 | 0.027825948 | -0.722304036 | 0.470107572 | NA | | Cabp4 | 1.908898629 | 0.0056697483415585 | 0.031541692356528 | 0.179754094278491 | 0.857345623943981 | NA | NM_144532 |
| 4930568D16Rik | 0 | NA | NA | NA | NA | NA | | Cabp5 | 8.288640541 | 0.0546727005908052 | 0.0572943472823152 | 0.954242489602125 | 0.33996091152346 | NA | NM_013877 |
| 4930568E12Rik | 0 | NA | NA | NA | NA | NA | | Cabs1 | 0.346765936 | -0.0134445992440418 | 0.0121756794716388 | -1.104217573677 | 0.269498768617541 | NA | NM_027631 |
| 4930568G15Rik | 1.328994491 | 0.001593344 | 0.086013489 | 0.018524352 | 0.985220551 | NA | | Cabyr | 17.34649615 | -0.0986985590586621 | 0.0740658220596349 | -1.3325789995174 | 0.182670000594186 | NA | NM_001042420,NM_181731,NM_027687,NM_001042418,NM_001042419 |
| 4930568K20Rik | 0 | NA | NA | NA | NA | NA | | Cacna1f | 10.87853332 | 0.0147820669698553 | 0.0521344394689165 | 0.28353746813886 | 0.776764864119625 | NA | NM_019582 |
| 4930571K23Rik | 0 | NA | NA | NA | NA | NA | | Cacng1 | 11.14023722 | -0.000381004923620321 | 0.0615338703131236 | -0.00619179196240257 | 0.995059696356646 | NA | NM_007582 |
| 4930572K03Rik | 0.238795281 | -0.027601606 | 0.032267774 | -0.855392297 | 0.392334007 | NA | | Cacng6 | 15.49176918 | -0.125872882951838 | 0.0757609795848687 | -1.66144740526794 | 0.0966236236820599 | NA | NM_133183 |
| 4930572O03Rik | 0.705750341 | 0.031598865 | 0.054521254 | 0.579569665 | 0.562204855 | NA | | Calca | 20.57864512 | 0.161712199363833 | 0.0834755737413966 | 1.93723974710027 | 0.0527160436871647 | NA | NM_007587,NM_001033954 |
| 4930572O13Rik | 0.380940254 | 0.025976956 | 0.047990576 | 0.54129287 | 0.588305734 | NA | | Calcb | 2.173775682 | -0.0291690539409661 | 0.0344279380748261 | -0.847249518038802 | 0.39685605808535 | NA | NM_054084 |
| 4930573O16Rik | 0 | NA | NA | NA | NA | NA | | Calcoco2 | 2.121417314 | -0.0373605942396154 | 0.0343047887025827 | -1.08907810403749 | 0.276119444696403 | NA | NM_001271018 |
| 4930578E11Rik | 0.176054697 | -0.002190495 | 0.031239146 | -0.070120197 | 0.944097991 | NA | | Calhm1 | 0 | NA | NA | NA | NA | NA | NM_001081271 |
| 4930578I06Rik | 0 | NA | NA | NA | NA | NA | | Calm4 | 0 | NA | NA | NA | NA | NA | NM_020036 |

| | | | | | | | | | | | | | | | |
|---|---|---|---|---|---|---|---|---|---|---|---|---|---|---|---|
| 4930578N18Rik | 0 | NA | NA | NA | NA | NA | | Calm5 | 0.127687691 | 0.00542298404407807 | 0.00869724871942359 | 0.62352868349814 | 0.532937159177571 | NA | NM_001008706 |
| 4930579F01Rik | 0 | NA | NA | NA | NA | NA | | Calml3 | 0.106347839 | -0.00653347234140708 | 0.00854389509723638 | -0.764694822098226 | 0.444453290197145 | NA | NM_027416 |
| 4930579G18Rik | 3.406179514 | 0.084716588 | 0.115538925 | 0.733229847 | 0.463418254 | NA | | Calr4 | 18.68087361 | 0.0437033749159445 | 0.0743603061455729 | 0.587724515689698 | 0.556717217448788 | NA | NM_001033226 |
| 4930583K01Rik | 0.974763109 | 0.077267739 | 0.069555094 | 1.110885411 | 0.266617676 | NA | | Camp | 4.154289592 | -0.0271926050863757 | 0.0339699009710313 | -0.800491149784775 | 0.423426289188727 | NA | NM_009921 |
| 4930583P06Rik | 0 | NA | NA | NA | NA | NA | | Capn13 | 0.680543483 | 0.00539322587076443 | 0.0173797421979938 | 0.310316793502512 | 0.756320061711495 | NA | NM_001033444 |
| 4930584F24Rik | 0 | NA | NA | NA | NA | NA | | Capn8 | 2.165906263 | 0.00501773708086392 | 0.0319194568654493 | 0.15719995180417 | 0.875087266283638 | NA | NM_130890,NM_001145806 |
| 4930590L20Rik | 0 | NA | NA | NA | NA | NA | | Capn9 | 7.405676436 | -0.0973199432079045 | 0.059192526750092 | -1.64412550960672 | 0.10015027965695 | NA | NM_023709 |
| 4930591A17Rik | 0 | NA | NA | NA | NA | NA | | Capns2 | 4.162440638 | -0.00752311824500813 | 0.0394159104446838 | -0.190865012634075 | 0.848631352083363 | NA | NM_027112 |
| 4930592A05Rik | 0.68657355 | 0.032306041 | 0.060686154 | 0.532346161 | 0.594486266 | NA | | Caps2 | 8.470756141 | -0.0390672260844637 | 0.0536252670646096 | -0.728522732341717 | 0.466293658317821 | NA | NM_178278 |
| 4930592I03Rik | 0 | NA | NA | NA | NA | NA | | Capza3 | 0 | NA | NA | NA | NA | NA | NM_007605 |
| 4930593A02Rik | 1.28787035 | 0.023814336 | 0.075742054 | 0.314413659 | 0.753206884 | NA | | Car1 | 6.121786073 | -0.0382133624185541 | 0.0553339742832659 | -0.690594935815239 | 0.489820130843685 | NA | NM_009799,NM_001083957 |
| 4930593C16Rik | 0.49242343 | -0.024792866 | 0.039435663 | -0.6286915 | 0.529551044 | NA | | Car5a | 7.286279045 | -0.0155027261832992 | 0.0490553358931878 | -0.316025278413231 | 0.751983321239202 | NA | NM_007608 |
| 4930594C11Rik | 0.751088818 | -0.011157514 | 0.061845162 | -0.180410466 | 0.856830339 | NA | | Car6 | 1.200717788 | -0.00209521311649233 | 0.0261117435547681 | -0.080240260942274 | 0.936046169775015 | NA | NM_009802 |
| 4930595M18Rik | 0 | NA | NA | NA | NA | NA | | Card11 | 12.35078047 | 0.00337536542436738 | 0.0587683967256618 | 0.0574350435340955 | 0.954198648361096 | NA | NM_175362 |
| 4930596D02Rik | 0 | NA | NA | NA | NA | NA | | Card14 | 8.506194928 | -0.127714737869583 | 0.0618171399778273 | -2.06600851989257 | 0.0388276751190881 | NA | NM_130886 |
| 4930596M17Rik | 0.605945404 | -0.004798112 | 0.052429254 | -0.091515938 | 0.927082643 | NA | | Casc5 | 4.943653599 | -0.00640690660912214 | 0.0465857019773152 | -0.137529463702017 | 0.890612305342061 | NA | NM_029617 |
| 4930597G03Rik | 0 | NA | NA | NA | NA | NA | | Casp14 | 0.120209049 | -0.00674972139751468 | 0.00863521114606269 | -0.781650996523957 | 0.434419707908266 | NA | NM_009809 |
| 4930598F16Rik | 0 | NA | NA | NA | NA | NA | | Casp4 | 12.76275709 | -0.230922901836436 | 0.0698805119280152 | -3.30453935532574 | 0.00095132618310958 | NA | NM_007609 |
| 4930599N23Rik | 5.22008009 | 0.087699406 | 0.134255255 | 0.653228851 | 0.513608753 | NA | | Cass4 | 17.83253685 | -0.131366541075031 | 0.0794349611139015 | -1.65376226327681 | 0.0981758262099779 | NA | NM_001080820,NM_001276422,NM_001033538,NM_001276423 |
| 4931402G19Rik | 0.997312756 | -0.104805702 | 0.06156002 | -1.702496218 | 0.088662387 | NA | | Catsper1 | 0 | NA | NA | NA | NA | NA | NM_139301 |
| 4931403G20Rik | 0.19442443 | -0.026451352 | 0.031568725 | -0.837897386 | 0.402088337 | NA | | Catsper3 | 7.038414255 | -0.0193171498398089 | 0.0550645309224572 | -0.350809305304201 | 0.725731415721284 | NA | NM_001252487,NM_029772,NM_001252488 |
| 4931406B18Rik | 0 | NA | NA | NA | NA | NA | | Catsper4 | 0.991618214 | -0.00254423527945597 | 0.0210686364877148 | -0.120759370495548 | 0.903881632170047 | NA | NM_177866,NM_001130030 |
| 4931406H21Rik | 0.789628455 | 0.016746707 | 0.050267784 | 0.333149899 | 0.739021135 | NA | | Catsperb | 0 | NA | NA | NA | NA | NA | NM_173023 |
| 4931408C20Rik | 0.096071043 | -0.020098795 | 0.027825948 | -0.722304036 | 0.470107572 | NA | | Catsperg2 | 10.77627923 | 0.187644679633437 | 0.056784309653626 | 3.30451634928797 | 0.000951404265804529 | NA | NM_029714 |
| 4931409K22Rik | 5.124992282 | 0.145853595 | 0.141286002 | 1.032328699 | 0.301918161 | NA | | Cav3 | 4.642390565 | 0.0470614132652847 | 0.0457829351859082 | 1.02792477315369 | 0.303985215361358 | NA | NM_007617 |
| 4931417E11Rik | 0 | NA | NA | NA | NA | NA | | Cblc | 0.955625209 | -0.000482163945000589 | 0.0231467627334633 | -0.0208307291413812 | 0.983380684743392 | NA | NM_001161844,NM_023224 |
| 4931417G12Rik | 2.927860015 | -0.140760812 | 0.117456152 | -1.198411569 | 0.230756831 | NA | | Ccdc105 | 13.01339065 | 0.0454975986985328 | 0.070564234524046 | 0.644768543234587 | 0.519077189997772 | NA | NM_027630 |
| 4931419H13Rik | 0.430483556 | -0.061355922 | 0.047577744 | -1.289592921 | 0.197192035 | NA | | Ccdc11 | 16.47753607 | 0.0309836432342553 | 0.0718949856176802 | 0.43095694321463 | 0.666499677972489 | NA | NM_028948,NR_029424 |
| 4931420L22Rik | 0 | NA | NA | NA | NA | NA | | Ccdc125 | 11.15303906 | -0.0160181427115761 | 0.0644278267168133 | -0.24862149676385 | 0.803653578511766 | NA | NM_001168386,NM_183115 |
| 4931423N10Rik | 7.255305665 | 0.135750534 | 0.130148345 | 1.043044647 | 0.296927615 | NA | | Ccdc144b | 0.127687691 | 0.00542298404407807 | 0.00869724871942359 | 0.62352868349814 | 0.532937159177571 | NA | NM_178418 |
| 4931428L18Rik | 0.466626353 | 0.089501577 | 0.058821838 | 1.521570567 | 0.128116717 | NA | | Ccdc147 | 12.97084452 | 0.139521866959006 | 0.072608141469603 | 1.92157331306181 | 0.054659469259432 | NA | NM_001163267 |
| 4931429L15Rik | 3.548963464 | 0.10799846 | 0.122005073 | 0.885196474 | 0.376050673 | NA | | Ccdc150 | 4.821910895 | -0.0766308524912794 | 0.0481069726419762 | -1.59292610369779 | 0.1111767695315 | NA | NM_030025 |
| 4931429P17Rik | 0.185670226 | -0.001645086 | 0.031432796 | -0.052336624 | 0.958260472 | NA | | Ccdc152 | 0.435709867 | 0.015952827522772 | 0.0148578726417494 | 1.07369526630251 | 0.282959283681906 | NA | NM_001166063 |
| 4931430N09Rik | 0.287795519 | 0.031925978 | 0.035387605 | 0.902179657 | 0.366961439 | NA | | Ccdc154 | 0.9994637 | -0.0279596293898284 | 0.0223959066178655 | -1.24842587830424 | 0.21187513696654 | NA | NM_001079929 |
| 4931431B13Rik | 0.094392817 | 0.017895942 | 0.027818137 | 0.643319236 | 0.520016977 | NA | | Ccdc158 | 14.2416676 | 0.0168069395687077 | 0.0736974314728332 | 0.228053260918642 | 0.819604835035124 | NA | NM_177230 |
| 4931431C16Rik | 0.900243631 | 0.028123536 | 0.058318278 | 0.482242223 | 0.629633886 | NA | | Ccdc169 | 5.252289339 | 0.0190125352580516 | 0.0477045100154507 | 0.398547962276392 | 0.690226311322208 | NA | NM_177203 |
| 4931431F19Rik | 0 | NA | NA | NA | NA | NA | | Ccdc170 | 20.28776809 | 0.0980892967497833 | 0.0714687221423012 | 1.37247867052216 | 0.169914474392788 | NA | NM_001195672 |
| 4931440J10Rik | 5.291001609 | -0.206325516 | 0.135621765 | -1.521330421 | 0.12817694 | NA | | Ccdc172 | 1.646965382 | 0.00934786783058715 | 0.0283172365753899 | 0.330112290643192 | 0.74131511672896 | NA | NM_029372 |
| 4931440L10Rik | 0.456340463 | 0.030209904 | 0.045608803 | 0.662370032 | 0.507734105 | NA | | Ccdc175 | 1.565342685 | 0.0230122985379643 | 0.0259424274840443 | 0.887052630372308 | 0.375050565585517 | NA | NM_028687 |
| 4931440P22Rik | 7.476089336 | 0.288538908 | 0.157216927 | 1.835291613 | 0.066462493 | NA | | Ccdc178 | 0 | NA | NA | NA | NA | NA | NM_027616 |
| 4932411N23Rik | 0.096071043 | -0.020098795 | 0.027825948 | -0.722304036 | 0.470107572 | NA | | Ccdc27 | 2.030132062 | -0.0271875811414532 | 0.0293424627498764 | -0.926560983418737 | 0.354154520411794 | NA | NM_001033455 |
| 4932412D23Rik | 3.061015359 | -0.208674211 | 0.121981904 | -1.710698097 | 0.087136859 | NA | | Ccdc38 | 3.356248116 | 0.0113583157661424 | 0.0372941479578595 | 0.304560269857264 | 0.760701083273134 | NA | NM_175488 |
| 4932413F04Rik | 0 | NA | NA | NA | NA | NA | | Ccdc42b | 10.72285132 | -0.00821757730211165 | 0.0655406056280522 | -0.125381467311227 | 0.900221559270896 | NA | NM_001195094 |
| 4932414J04Rik | 0 | NA | NA | NA | NA | NA | | Ccdc54 | 0.106148608 | 0.00515554063321497 | 0.00854200771766808 | 0.603551390213729 | 0.546141939141592 | NA | NM_027046 |
| 4932414N04Rik | 0 | NA | NA | NA | NA | NA | | Ccdc63 | 10.85506877 | -0.0263681258057752 | 0.0636186649853837 | -0.414471536173123 | 0.678528808484365 | NA | NM_183307 |
| 4932415M13Rik | 0.450736154 | 0.045589387 | 0.042063495 | 1.083823084 | 0.278443248 | NA | | Ccdc64b | 0.22897535 | -0.00116364198856673 | 0.00981572003153595 | -0.118548816065269 | 0.905632818135836 | NA | NM_153784 |
| 4932416K20Rik | 0.107716603 | -0.020477583 | 0.027954224 | -0.732539849 | 0.46383913 | NA | | Ccdc67 | 16.39275236 | 0.0574287189133909 | 0.0656120007716753 | 0.875277666249479 | 0.381422843076537 | NA | NM_181816 |
| 4932429P05Rik | 0 | NA | NA | NA | NA | NA | | Ccdc68 | 3.152226755 | -0.000297175371846468 | 0.0371671164779703 | -0.0079956531473893 | 0.993620459774736 | NA | NM_201362 |
| 4932435O22Rik | 5.460629182 | -0.040521896 | 0.148776287 | -0.272367973 | 0.785339096 | NA | | Ccdc69 | 14.75189616 | -0.0609341284105937 | 0.0721835048674356 | -0.844155856971737 | 0.398582318082337 | NA | NM_177471 |
| 4932702P03Rik | 1.398967995 | -0.087155862 | 0.079085499 | -1.10204605 | 0.270441652 | NA | | Ccdc7 | 0 | NA | NA | NA | NA | NA | NM_001197041,NM_029061 |
| 4933400A11Rik | 0 | NA | NA | NA | NA | NA | | Ccdc70 | 0 | NA | NA | NA | NA | NA | NM_026459 |
| 4933400B14Rik | 0.197363271 | -0.000561558 | 0.031650166 | -0.017742642 | 0.985844163 | NA | | Ccdc73 | 1.47116782 | 0.00919180331401171 | 0.0255961290536515 | 0.359109117427287 | 0.719513461205874 | NA | NM_177600 |
| 4933400C23Rik | 0.62365482 | 0.069517633 | 0.062256133 | 1.116639103 | 0.264148665 | NA | | Ccdc78 | 18.43796605 | -0.0346341411719657 | 0.0734039718733744 | -0.471829252396742 | 0.637048669375117 | NA | NM_001165929 |
| 4933400F21Rik | 0.096071043 | -0.020098795 | 0.027825948 | -0.722304036 | 0.470107572 | NA | | Ccdc83 | 0.780074698 | -0.0261002232209602 | 0.0192895200812179 | -1.35307789468406 | 0.176030746774021 | NA | NM_183293,NM_029256 |
| 4933400L20Rik | 0.295060159 | -0.029687054 | 0.033501022 | -0.886153701 | 0.375534709 | NA | | Ccer1 | 0.240418098 | -0.0088029805258837 | 0.00986414519853158 | -0.892422034419582 | 0.372166765031741 | NA | NM_025724 |
| 4933401B06Rik | 0 | NA | NA | NA | NA | NA | | Ccin | 0.322082616 | 0.0021341954732379 | 0.0125665209615259 | 0.16983184763484 | 0.86514237995411 | NA | NM_001002787 |
| 4933401D09Rik | 0 | NA | NA | NA | NA | NA | | Ccl1 | 0 | NA | NA | NA | NA | NA | NM_011329 |
| 4933401H06Rik | 0.174376471 | 0.024538273 | 0.031173126 | 0.787161136 | 0.431187544 | NA | | Ccl11 | 3.20107698 | -0.0212280027285613 | 0.0395469605412169 | -0.536779627006654 | 0.591419845927653 | NA | NM_011330 |
| 4933401P06Rik | 0 | NA | NA | NA | NA | NA | | Ccl19 | 0.510493435 | -0.00472572262369758 | 0.0128382628052112 | -0.368096735157918 | 0.712801105220293 | NA | NM_011888 |
| 4933402C06Rik | 0.947736072 | -0.037611715 | 0.058791758 | -0.639744696 | 0.522338592 | NA | | Ccl2 | 5.620617041 | -0.0753317995767074 | 0.0484729249589608 | -1.55410055490743 | 0.120160432880679 | NA | NM_011333 |
| 4933402D24Rik | 7.163047344 | 0.223585585 | 0.155171886 | 1.440889781 | 0.149615824 | NA | | Ccl20 | 0 | NA | NA | NA | NA | NA | NM_001159738,NM_016960 |
| 4933402E13Rik | 0.096071043 | -0.020098795 | 0.027825948 | -0.722304036 | 0.470107572 | NA | | Ccl21a | 0 | NA | NA | NA | NA | NA | NM_011124 |
| 4933402J07Rik | 1.042779188 | 0.116034349 | 0.074231793 | 1.563135483 | 0.118020731 | NA | | Ccl21b | 0 | NA | NA | NA | NA | NA | NM_011335 |
| 4933402J10Rik | 0 | NA | NA | NA | NA | NA | | Ccl21c | 0 | NA | NA | NA | NA | NA | NM_023052 |
| 4933402J15Rik | 1.208531101 | -0.036599839 | 0.079628192 | -0.45963418 | 0.645778821 | NA | | Ccl22 | 9.872632942 | -0.0765794000013239 | 0.0652654355596035 | -1.17335308260355 | 0.240654243509412 | NA | NM_009137 |
| 4933402N03Rik | 0 | NA | NA | NA | NA | NA | | Ccl24 | 4.457229419 | 0.0547460251965233 | 0.0468371864565936 | 1.16885810908516 | 0.242460803679039 | NA | NM_019577 |
| 4933402N22Rik | 0 | NA | NA | NA | NA | NA | | Ccl26 | 0 | NA | NA | NA | NA | NA | NM_001013412 |
| 4933402P03Rik | 0 | NA | NA | NA | NA | NA | | Ccl27b | 0 | NA | NA | NA | NA | NA | NM_001199961,NM_001199960,NM_001199959 |
| 4933403O08Rik | 2.848565718 | -0.00935093 | 0.116946144 | -0.079959287 | 0.936269636 | NA | | Ccl3 | 22.66162113 | -0.23196032410088 | 0.0648678868574497 | -3.5758883993034 | NA | NA | NM_011337 |
| 4933404G15Rik | 0 | NA | NA | NA | NA | NA | | Ccl4 | 11.02313557 | -0.148997265500968 | 0.049825302399724 | -2.99039360174146 | 0.00278618175005588 | NA | NM_013652 |
| 4933404K08Rik | 0.222620122 | -0.001447027 | 0.032059667 | -0.04513545 | 0.963999345 | NA | | Ccl5 | 4.50238755 | -0.0297656523154657 | 0.0391316675458858 | -0.760653817795076 | 0.446863865790826 | NA | NM_013653 |
| 4933405D12Rik | 1.532017869 | 0.163082275 | 0.090868546 | 1.794705448 | 0.072700642 | NA | | Ccl7 | 2.706960573 | 0.00130278769932201 | 0.0351144548040066 | 0.0371011797447405 | 0.970404331377312 | NA | NM_013654 |
| 4933405E24Rik | 0 | NA | NA | NA | NA | NA | | Ccl8 | 4.761070116 | -0.0493548186840495 | 0.0445071263813571 | -1.10891946294522 | 0.267464926793617 | NA | NM_021443 |
| 4933405L10Rik | 7.40056411 | 0.127715269 | 0.153739476 | 0.830725274 | 0.406128845 | NA | | Ccnb1 | 11.94177566 | -0.03175357005182 | 0.0695657761300729 | -0.456453903316592 | 0.648063611598563 | NA | NM_172301 |
| 4933405O20Rik | 0.212768007 | -0.02694223 | 0.031868764 | -0.845411806 | 0.397880955 | NA | | Ccnb1ip1 | 0.12890128 | -0.0068640067066208 | 0.00870804083610955 | -0.788237771940376 | 0.430557639008166 | NA | NM_001111119 |
| 4933406D12Rik | 0 | NA | NA | NA | NA | NA | | Ccnb3 | 0.194331409 | -0.00840968081320881 | 0.00964081822647896 | -0.872299489073576 | 0.383045016303155 | NA | NM_183015 |
| 4933406F09Rik | 0 | NA | NA | NA | NA | NA | | Ccr1 | 5.885725369 | -0.0734338874073456 | 0.0477815801636845 | -1.53686602987562 | 0.12432611971925 | NA | NM_009912 |
| 4933406G16Rik | 0 | NA | NA | NA | NA | NA | | Ccr10 | 5.212091725 | -0.0434720228315962 | 0.0468132141971827 | -0.928627174551335 | 0.353082333811841 | NA | NM_007721 |
| 4933406J08Rik | 0 | NA | NA | NA | NA | NA | | Ccr1l1 | 1.039507966 | -0.00925915393369021 | 0.0231253889299715 | -0.40038911179954 | 0.688869942575022 | NA | NM_007718 |
| 4933406J10Rik | 0.119397641 | -0.020477583 | 0.027954224 | -0.732539845 | 0.463839132 | NA | | Ccr2 | 6.074415662 | 0.0124551188988551 | 0.0467196448738188 | 0.266592756269747 | 0.789782733510691 | NA | NM_009915 |
| 4933406K04Rik | 0 | NA | NA | NA | NA | NA | | Ccr3 | 1.122568499 | -0.0165085270052448 | 0.025492783772702 | -0.647576473108534 | 0.517258918155039 | NA | NM_009914 |
| 4933406M09Rik | 0.105923158 | 0.018524693 | 0.027977518 | 0.662127842 | 0.507889294 | NA | | Ccr4 | 0.194331409 | -0.00840968081320881 | 0.00964081822647896 | -0.872299489073576 | 0.383045016303155 | NA | NM_009916 |
| 4933407E24Rik | 0.896399808 | -0.000816113 | 0.069363515 | -0.011765745 | 0.99061251 | NA | | Ccr6 | 6.608129769 | -0.06653363259234 | 0.0568866048019362 | -1.16958346914871 | 0.242168632985179 | NA | NM_009835,NM_001190338,NM_001190337,NM_001190336,NM_001190335,NM_001190334,NM_001190333 |
| 4933407G14Rik | 0.096071043 | -0.020098795 | 0.027825948 | -0.722304036 | 0.470107572 | NA | | Ccr7 | 3.375673998 | 0.00598644956000355 | 0.0431673959472034 | 0.138679886257799 | 0.889703112938998 | NA | NM_007719 |
| 4933407I05Rik | 2.918431594 | -0.065131308 | 0.103673378 | -0.628235609 | 0.529849606 | NA | | Ccr8 | 0 | NA | NA | NA | NA | NA | NM_007720 |
| 4933408J17Rik | 1.306557241 | -0.07162614 | 0.091615212 | -0.781814924 | 0.434323349 | NA | | Ccr9 | 11.70624472 | -0.0537624348959557 | 0.0691534766441234 | -0.777436471815107 | 0.436901301580402 | NA | NM_001166625,NM_009913 |
| 4933408N05Rik | 0.496037528 | 0.012714628 | 0.055562508 | 0.228834661 | 0.818997426 | NA | | Ccrl1 | 9.099144152 | 0.0116778930868462 | 0.0430268540915584 | 0.271409410086 | 0.786076165376725 | NA | NM_145700 |
| 4933409G03Rik | 0 | NA | NA | NA | NA | NA | | Cct6b | 5.993991038 | 0.00733342063489557 | 0.0503622923189139 | 0.145613320943722 | 0.884226651484317 | NA | NM_009839 |
| 4933409K07Rik | 1.09986382 | 0.036652755 | 0.083827004 | 0.437242813 | 0.661935264 | NA | | Cct8l1 | 3.099425143 | -0.064101177772705 | 0.0395411765571325 | -1.62112469466067 | 0.104990898843846 | NA | NM_198621 |
| 4933411E08Rik | 1.335811433 | -0.057819597 | 0.082562911 | -0.700309579 | 0.483733991 | NA | | Cd101 | 10.97252196 | -0.0418069261146056 | 0.0678559599899648 | -0.61611280896753 | 0.53782006875398 | NA | NM_001167906,NM_001099332 |
| 4933411G06Rik | 0 | NA | NA | NA | NA | NA | | Cd160 | 0.097165704 | -0.00637828455464458 | 0.00848781691535724 | -0.75146349388194 | 0.452373763047372 | NA | NM_018767,NM_001163496,NM_001163497 |
| 4933411G11Rik | 0.099009884 | 0.018205726 | 0.027896865 | 0.652608315 | 0.514008823 | NA | | Cd163l1 | 2.240102337 | -0.0643204871034609 | 0.034650467146372 | -1.8562660881816 | 0.0634156402252877 | NA | NM_172909 |
| 4933411K16Rik | 1.808784137 | -0.094933916 | 0.092210675 | -1.029532823 | 0.303229364 | NA | | Cd177 | 1.205319222 | 0.0149994492529565 | 0.0228148765685992 | 0.657441613057013 | 0.510897001970147 | NA | NM_026862 |
| 4933412E24Rik | 0 | NA | NA | NA | NA | NA | | Cd19 | 1.011132555 | -0.0203747850965182 | 0.0223944708740059 | -0.909813195013596 | 0.362921034639273 | NA | NM_009844 |
| 4933413J09Rik | 0 | NA | NA | NA | NA | NA | | Cd1d2 | 0 | NA | NA | NA | NA | NA | NM_007640,NR_038188 |
| 4933413L06Rik | 0 | NA | NA | NA | NA | NA | | Cd2 | 1.20151611 | 0.00788722970152225 | 0.0265971484168724 | 0.296544185034469 | 0.766814528719377 | NA | NM_013486 |
| 4933415F23Rik | 0.489904938 | 0.03017109 | 0.045748741 | 0.659495523 | 0.50957762 | NA | | Cd200r2 | 1.823617806 | 0.00991932844754334 | 0.0280555428442226 | 0.353560382082786 | 0.723668369414965 | NA | NM_206535 |
| 4933416C03Rik | 0 | NA | NA | NA | NA | NA | | Cd200r3 | 0 | NA | NA | NA | NA | NA | NM_027578,NM_001128132,NM_001128133,NM_029018 |
| 4933416E03Rik | 0 | NA | NA | NA | NA | NA | | Cd200r4 | 9.428398792 | -0.0695607903447389 | 0.0639049573494262 | -1.08850382239342 | 0.276372750055577 | NA | NM_207244 |
| 4933416I08Rik | 0.723371454 | 0.006844428 | 0.061016294 | 0.112173772 | 0.910685625 | NA | | Cd207 | 0.106347839 | -0.00653347234140708 | 0.00854389509723638 | -0.764694822098226 | 0.444453290197145 | NA | NM_144943 |
| 4933416M06Rik | 0 | NA | NA | NA | NA | NA | | Cd209b | 14.13743795 | 0.0342313168833996 | 0.0691288400918107 | 0.495181415425698 | 0.62047206794792 | NA | NM_026972,NM_001037800 |
| 4933416M07Rik | 0.094392817 | 0.017895942 | 0.027818137 | 0.643319236 | 0.520016977 | NA | | Cd209d | 2.432853711 | -0.0084124858018285 | 0.0340893457866133 | -0.246777566647584 | 0.805080372857917 | NA | NM_130904 |
| 4933417A18Rik | 0 | NA | NA | NA | NA | NA | | Cd209e | 0.798890638 | 0.0111014457697147 | 0.0202552307697946 | 0.548077970371465 | 0.58363836630687 | NA | NM_130905 |
| 4933417E11Rik | 0.408720688 | -0.001528354 | 0.047884645 | -0.03191741 | 0.974537915 | NA | | Cd209f | 16.9247893 | -0.164645656715883 | 0.077759551968728 | -2.11736889613379 | 0.0342285518401889 | NA | NM_026956 |
| 4933417G07Rik | 0 | NA | NA | NA | NA | NA | | Cd209g | 5.318352073 | -0.0695212535350655 | 0.0527117084251453 | -1.31889585088654 | 0.187203933462868 | NA | NM_027343 |
| 4933421I07Rik | 0 | NA | NA | NA | NA | NA | | Cd22 | 29.40777017 | -0.439350672195134 | 0.0793769303496595 | -5.53499197134194 | NA | NA | NM_001043317,NM_009845 |
| 4933422A05Rik | 0.524867658 | 0.066447841 | 0.050604724 | 1.313075849 | 0.189157391 | NA | | Cd226 | 1.020220713 | -0.017675267286604 | 0.0220074050686853 | -0.803150904499614 | 0.421887518307939 | NA | NM_178687,NM_001039149 |
| 4933422H20Rik | 2.50170239 | 0.023268841 | 0.107765362 | 0.215921334 | 0.829049067 | NA | | Cd244 | 2.377071555 | 0.00426843582508349 | 0.0289572028269919 | 0.147404977289614 | 0.882812377129672 | NA | NM_018729 |
| 4933424G05Rik | 0.832944429 | -0.043859678 | 0.073013575 | -0.600705798 | 0.548035956 | NA | | Cd247 | 1.982945442 | -0.0135933260297587 | 0.0334979004845628 | -0.405796358372462 | 0.684892234621236 | NA | NM_001113391,NM_031162,NM_001113393,NM_001113394,NM_001113392 |
| 4933424G06Rik | 7.064976964 | 0.003371442 | 0.153226087 | 0.022003052 | 0.982445521 | NA | | Cd27 | 0.335323189 | -0.0061431309930725 | 0.0128252480301014 | -0.478845397660815 | 0.632048617772247 | NA | NM_001033126,NM_001042564 |
| 4933425B07Rik | 0 | NA | NA | NA | NA | NA | | Cd28 | 10.59661273 | -0.0320403418474722 | 0.059531330150754 | -0.53820974210277 | 0.590432255045003 | NA | NM_007642 |
| 4933425H06Rik | 0.079983654 | 0.016740032 | 0.027674178 | 0.604897175 | 0.54524732 | NA | | Cd300c | 0.127687691 | 0.00542298404407807 | 0.00869724871942359 | 0.62352868349814 | 0.532937159177571 | NA | NM_199225 |
| 4933425L06Rik | 0.531154764 | -0.070672053 | 0.050962944 | -1.38673411 | 0.165522855 | NA | | Cd300e | 7.623722105 | 0.0119481842712648 | 0.0515777446815329 | 0.23165387213107 | 0.816806861502045 | NA | NM_172050 |
| 4933426D04Rik | 0.116696964 | -0.020477583 | 0.027954224 | -0.732539847 | 0.463839131 | NA | | Cd300lb | 2.725536848 | 0.00742761236816128 | 0.0333395988272491 | 0.222786495021966 | 0.823701679283965 | NA | NM_199221 |
| 4933427D06Rik | 2.117011045 | -0.056698448 | 0.085191871 | -0.665538244 | 0.505706291 | NA | | Cd300ld | 3.823994927 | 0.00649205155007086 | 0.0434867945196236 | 0.149287884328685 | 0.88132647504678 | NA | NM_145437 |
| 4933427E11Rik | 2.800614345 | 0.195179288 | 0.104946691 | 1.859794581 | 0.062914594 | NA | | Cd300lf | 3.804090961 | -0.0244101349001593 | 0.0454744999339035 | -0.53678731895104 | 0.591414532091592 | NA | NM_145634,NM_001169153 |
| 4933427E13Rik | 0.55073328 | 0.049633073 | 0.054814118 | 0.905479733 | 0.365211292 | NA | | Cd300lg | 9.523451092 | 0.0885665921033479 | 0.0583542387439706 | 1.5177405105383 | 0.129079827279041 | NA | NM_027987,NM_001160712,NM_001160713,NM_001160711 |
| 4933428C19Rik | 0.195080928 | -0.000361156 | 0.031609476 | -0.011425558 | 0.990883922 | NA | | Cd300lh | 0.724429695 | 0.0183631535391828 | 0.0159140421839553 | 1.1538962467844 | 0.248542708319589 | NA | NM_199201 |
| 4933429K18Rik | 0 | NA | NA | NA | NA | NA | | Cd3d | 0.589409793 | 0.0100492600676131 | 0.015506378103943 | 0.648072683398435 | 0.516937940950164 | NA | NM_013487 |
| 4933429O19Rik | 1.152370116 | 0.065394523 | 0.061053146 | 1.071108167 | 0.284120797 | NA | | Cd3e | 2.054425479 | -0.0285476912843924 | 0.0315586385955298 | -0.904591977184848 | 0.365681587179484 | NA | NM_007648 |
| 4933430H16Rik | 0.105923158 | 0.018524693 | 0.027977518 | 0.662127842 | 0.507889294 | NA | | Cd3g | 0.403899654 | -0.00132471380270723 | 0.0135291380196343 | -0.097915610054737 | 0.921999304520122 | NA | NM_009850 |
| 4933430M04Rik | 1.602568339 | 0.04362618 | 0.087617457 | 0.497916531 | 0.618542875 | NA | | Cd40 | 15.69028893 | -0.0947225378285509 | 0.0731067892189981 | -1.29567361445461 | 0.195087955447646 | NA | NM_170704,NM_011611,NR_027852,NM_170702,NM_170703 |
| 4933430N04Rik | 0 | NA | NA | NA | NA | NA | | Cd40lg | 0.240418098 | -0.0088029805258837 | 0.00986414519853158 | -0.892422034419582 | 0.372166765031741 | NA | NM_011616 |
| 4933431G14Rik | 0 | NA | NA | NA | NA | NA | | Cd5 | 5.492453844 | -0.00380321168602303 | 0.049817996633239 | -0.0763421241930371 | 0.939146913348188 | NA | NM_007650 |
| 4933432G23Rik | 0 | NA | NA | NA | NA | NA | | Cd5l | 0 | NA | NA | NA | NA | NA | NM_009690 |
| 4933432I03Rik | 0.19046386 | -0.000774927 | 0.031526133 | -0.024580468 | 0.980389599 | NA | | Cd69 | 0.841087458 | -0.0277488738209466 | 0.0198072420278735 | -1.40094586524956 | 0.161230261951153 | NA | NM_001033122 |
| 4933433C11Rik | 1.372710149 | 0.073171619 | 0.085220248 | 0.85861777 | 0.390551431 | NA | | Cd70 | 1.700390045 | -0.0353899684182915 | 0.0290966214434341 | -1.21629133083688 | 0.223873962759114 | NA | NM_011617 |
| 4933433F19Rik | 0 | NA | NA | NA | NA | NA | | Cd74 | 332.7995493 | -0.13157187026483 | 0.0503120842679549 | -2.61511468227189 | NA | NA | NM_010545,NM_001042605 |
| 4933433G15Rik | 5.462890164 | 0.181224023 | 0.127936922 | 1.416510729 | 0.156626025 | NA | | Cd79a | 10.11619103 | -0.0346544195162693 | 0.0635134978793903 | -0.545622909669952 | 0.585325173076284 | NA | NM_007655 |
| 4933433G19Rik | 3.338064257 | 0.013275869 | 0.109267928 | 0.121498311 | 0.903296353 | NA | | Cd79b | 9.467615619 | 0.00216318339777681 | 0.0650068287286091 | 0.0332762486662391 | 0.973454294086787 | NA | NM_008339 |
| 4933433H22Rik | 0.421503194 | -0.044552504 | 0.040840123 | -1.090900344 | 0.275316736 | NA | | Cd80 | 17.99361608 | 0.0347912618709413 | 0.0784439227831336 | 0.443517619167585 | 0.657391377953568 | NA | NM_009855 |
| 4933434I20Rik | 0 | NA | NA | NA | NA | NA | | Cd8a | 1.121830405 | -0.0186408668218757 | 0.0204963955918302 | -0.909470484132628 | 0.363101831066608 | NA | NM_009857,NM_001081110 |
| 4933436E23Rik | 0 | NA | NA | NA | NA | NA | | Cd8b1 | 0.405586341 | -0.0100214244197644 | 0.0139407229051103 | -0.718859738334718 | 0.472227345659372 | NA | NM_009858 |
| 4933436H12Rik | 0.079983654 | 0.016740032 | 0.027674178 | 0.604897175 | 0.54524732 | NA | | Cd96 | 2.226441281 | 0.0104300223110242 | 0.0321088207559128 | 0.324833552446909 | 0.745307049210358 | NA | NM_032465 |
| 4933436I01Rik | 0 | NA | NA | NA | NA | NA | | Cdc25c | 18.29826254 | 0.0089210783537029 | 0.0686812227851898 | 0.129891082190031 | 0.89665259989617 | NA | NM_009860 |
| 4933438B17Rik | 0.566356902 | 0.035306671 | 0.036986662 | 0.954578446 | 0.339790921 | NA | | Cdc6 | 17.96821184 | -0.0566703550303294 | 0.0809355193772223 | -0.700191405039382 | 0.483807778501482 | NA | NM_001025779,NM_011799 |
| 4933438K21Rik | 0.294043327 | 0.010508917 | 0.039092051 | 0.2688249 | 0.788064431 | NA | | Cdca2 | 12.33035596 | -0.0364068235684558 | 0.0663071475547976 | -0.549063334965034 | 0.582961986030403 | NA | NM_001110162,NM_175384 |
| 4933439K11Rik | 4.788981835 | 0.192743155 | 0.141262722 | 1.364430423 | 0.172432138 | NA | | Cdca5 | 3.983261408 | -0.0530659817496789 | 0.046755424897309 | -1.13496951137178 | 0.256388093084623 | NA | NM_026410 |
| 4933440J02Rik | 0 | NA | NA | NA | NA | NA | | Cdcp1 | 16.5790013 | 0.0150831657069133 | 0.0741848677568351 | 0.203318630375581 | 0.83888599470291 | NA | NM_133974 |
| 4933440M02Rik | 0.617197563 | -0.009986935 | 0.063001754 | -0.158518368 | 0.874048348 | NA | | Cdcp2 | 0.217532603 | 0.00684685248201087 | 0.00976000296730706 | 0.701525557416086 | 0.482977586094298 | NA | NM_172873 |
| 5031410I06Rik | 1.406075504 | 0.071746676 | 0.086475795 | 0.82965038 | 0.406736484 | NA | | Cdh16 | 0.383063074 | 0.00808564447825504 | 0.0105955130109064 | 0.763119678106395 | 0.44539202539159 | NA | NM_007663,NM_001252628,NM_001252627 |
| 5031425F14Rik | 3.624529577 | -0.156543277 | 0.119421821 | -1.310843152 | 0.18991076 | NA | | Cdh17 | 1.969219946 | -0.0274453540102579 | 0.0311022987118829 | -0.882422044251414 | 0.377548619082304 | NA | NM_019753 |
| 5031426D15Rik | 7.783855974 | -0.041969229 | 0.14812144 | -0.283343376 | 0.77691363 | NA | | Cdh26 | 8.450385051 | -0.0420119184366638 | 0.0591679699992544 | -0.710044952314456 | 0.477676260769974 | NA | NM_198656 |
| 5031434C07Rik | 0.529876295 | -0.026705656 | 0.04018514 | -0.664565453 | 0.506328472 | NA | | Cdh3 | 15.66940397 | 0.0397058884123429 | 0.0550263714359999 | 0.721579260564619 | 0.470553193858538 | NA | NM_001037809,NM_007665 |
| 5033403H07Rik | 0 | NA | NA | NA | NA | NA | | Cdhr5 | 1.733664362 | 0.000325129658427082 | 0.031398081209005 | 0.0103550804987992 | 0.991737988797133 | NA | NM_028069,NM_001114322 |
| 5033406O09Rik | 6.163381864 | -0.067236878 | 0.148473129 | -0.452855535 | 0.650652771 | NA | | Cdk1 | 13.55456425 | -0.0225716590492435 | 0.0587102663916466 | -0.384458467598692 | 0.700638672982512 | NA | NM_007659 |
| 5330411J11Rik | 3.284653191 | -0.067891171 | 0.124746327 | -0.544233826 | 0.586280577 | NA | | Cdk6 | 22.03415341 | -0.111838789937058 | 0.0854724739574732 | -1.30847727647036 | 0.190711477018367 | NA | NM_009873 |
| 5330439B14Rik | 0 | NA | NA | NA | NA | NA | | Cdkn2a | 9.82797659 | 0.0355063703463339 | 0.0494169213287241 | 0.718506321147438 | 0.472445151753173 | NA | NM_001040654,NM_009877 |
| 5430401F13Rik | 1.104555619 | 0.076855026 | 0.069738205 | 1.102050531 | 0.270439704 | NA | | Cdkn3 | 11.08123189 | 0.0795763395040981 | 0.0692448403164304 | 1.1492024079495 | 0.250472514664695 | NA | NM_028222 |
| 5430402E10Rik | 1.709813011 | 0.036245416 | 0.04818324 | 0.75224115 | NA | NA | | Cdr1 | 0.617638168 | -0.0161328761400947 | 0.0160571498532969 | -1.00471604783474 | 0.315033598516922 | NA | NM_001166658 |
| 5430403N17Rik | 0.990847688 | -0.007537041 | 0.064922126 | -0.11609356 | 0.907578393 | NA | | Cdrt4 | 2.250340267 | 0.00987550254709289 | 0.0307908048214251 | 0.320728951528452 | 0.748415805284325 | NA | NM_025496 |
| 5430419D17Rik | 0.402033573 | 0.020557702 | 0.043166232 | 0.476245001 | 0.633899843 | NA | | Cdx1 | 0 | NA | NA | NA | NA | NA | NM_009880 |
| 5430421F17Rik | 4.448672424 | 0.010685712 | 0.125815844 | 0.084931372 | 0.932315951 | NA | | Cdx2 | 0 | NA | NA | NA | NA | NA | NM_007673 |
| 5430421N21Rik | 0.331747315 | -0.03734372 | 0.03744515 | -0.99729124 | 0.318623164 | NA | | Cdx4 | 0 | NA | NA | NA | NA | NA | NM_007674 |
| 5430425J12Rik | 0 | NA | NA | NA | NA | NA | | Ceacam-ps1 | 1.781110721 | -0.0236518955233236 | 0.0280420393011952 | -0.843444204227885 | 0.398980058457449 | NA | NR_003247 |
| 5430427M07Rik | 0.233393928 | -0.027475306 | 0.032191697 | -0.85349047 | 0.39338737 | NA | | Ceacam10 | 4.579360345 | -0.030140182136695 | 0.045871221396129 | -0.657060815460179 | 0.511141813110275 | NA | NM_007675 |
| 5430428K19Rik | 0 | NA | NA | NA | NA | NA | | Ceacam11 | 0 | NA | NA | NA | NA | NA | NM_023289 |
| 5430434I15Rik | 0 | NA | NA | NA | NA | NA | | Ceacam12 | 0 | NA | NA | NA | NA | NA | NM_001162523,NM_001162524,NM_026087 |
| 5430437J10Rik | 0.923168337 | -0.053697833 | 0.07081308 | -0.758303882 | 0.448269084 | NA | | Ceacam13 | 0 | NA | NA | NA | NA | NA | NM_028171,NM_027210 |
| 5430440P10Rik | 2.025330369 | -0.039732542 | 0.096551468 | -0.411516702 | 0.680693697 | NA | | Ceacam14 | 0 | NA | NA | NA | NA | NA | NM_025957 |
| 5530400C23Rik | 0 | NA | NA | NA | NA | NA | | Ceacam15 | 1.241349419 | -0.0245626244918859 | 0.0267015441843945 | -0.919895281046754 | 0.357627485120715 | NA | NM_175315 |
| 5530401A14Rik | 1.195750858 | -0.076546543 | 0.064444899 | -1.1877828 | 0.234918993 | NA | | Ceacam16 | 0.724629857 | -0.0196962196529729 | 0.01689181684908245 | -1.16558310231477 | 0.243783043026436 | NA | NM_001033419 |
| 5730403I07Rik | 2.921794355 | -0.096465799 | 0.092218981 | -1.046051453 | 0.295537277 | NA | | Ceacam18 | 0.437366647 | -0.00211524396583317 | 0.0153267415786962 | -0.13801002352472 | 0.890232495729196 | NA | NM_028236 |
| 5730412P04Rik | 1.225651752 | -0.000777335 | 0.074383427 | -0.010450375 | 0.991661959 | NA | | Ceacam19 | 0 | NA | NA | NA | NA | NA | NM_177036 |
| 5730416F02Rik | 0 | NA | NA | NA | NA | NA | | Ceacam2 | 17.50371015 | -0.0345886474571203 | 0.0742181747801646 | -0.466040125071418 | 0.641186779723595 | NA | NM_007543,NM_001113369,NM_001113368 |
| 5730420D15Rik | 2.378564255 | 0.06379947 | 0.106297844 | 0.60019533 | 0.548376066 | NA | | Ceacam20 | 0 | NA | NA | NA | NA | NA | NM_027839 |
| 5730422E09Rik | 5.36260156 | -0.101837742 | 0.131950107 | -0.771789769 | 0.440238952 | NA | | Ceacam3 | 1.105533145 | -0.00401860330965244 | 0.0238506579456762 | -0.168490249569025 | 0.866197613733427 | NA | NM_054059 |
| 5730435O14Rik | 1.274846761 | -0.063631618 | 0.08842494 | -0.71961166 | 0.471764131 | NA | | Ceacam5 | 0 | NA | NA | NA | NA | NA | NM_028480 |
| 5730457N03Rik | 0 | NA | NA | NA | NA | NA | | Ceacam9 | 11.85566558 | -0.00865236562121012 | 0.0681147051815881 | -0.127026397576612 | 0.898919505539116 | NA | NM_011927 |
| 5730460C07Rik | 0 | NA | NA | NA | NA | NA | | Cebpe | 0.475335162 | -0.00223769088993525 | 0.0124701600518868 | -0.179443638303317 | 0.857589369010932 | NA | NM_207131 |
| 5730488B01Rik | 0 | NA | NA | NA | NA | NA | | Cel | 0.301656965 | 0.00287749821367573 | 0.0123358006508999 | 0.233264000862871 | 0.81555641073489 | NA | NM_009885 |
| 5830403M04Rik | 1.966094221 | 0.013819393 | 0.098611424 | 0.140139876 | 0.888549475 | NA | | Cela2a | 0.423306759 | -0.00976599639876241 | 0.0140919448469068 | -0.693019771568724 | 0.488297143743493 | NA | NM_007919 |
| 5830411N06Rik | 1.463700863 | 0.137456779 | 0.093695528 | 1.467058059 | 0.142360262 | NA | | Cela3b | 0 | NA | NA | NA | NA | NA | NM_026419 |
| 5830416I19Rik | 4.743051969 | 0.209804681 | 0.137766789 | 1.522897369 | 0.127784384 | NA | | Cenph | 21.44171775 | -0.048653396826853 | 0.0776662881486054 | -0.626441638793918 | 0.531025305396289 | NA | NM_021886 |
| 5830428M24Rik | 0 | NA | NA | NA | NA | NA | | Cenpi | 17.90266986 | -0.0490320841950002 | 0.0783229309141017 | -0.626024634455708 | 0.531298783428223 | NA | NM_145924 |
| 5830432E09Rik | 3.127986928 | 0.075015536 | 0.116468975 | 0.644081707 | 0.519522452 | NA | | Cep55 | 8.296040583 | -0.0519588252861347 | 0.0582352228415859 | -0.892223344409198 | 0.372273231983985 | NA | NM_028760,NM_001164362,NM_028293 |
| 5830473C10Rik | 2.529883987 | -0.019438374 | 0.111860035 | -0.173774073 | 0.862043021 | NA | | Cer1 | 5.52947549 | 0.00726054708276441 | 0.0530674706298378 | 0.136817281784711 | 0.891175223213105 | NA | NM_009887 |
| 6030408B16Rik | 6.325265018 | 0.100117471 | 0.144425714 | 0.693210841 | 0.488177246 | NA | | Cers1 | 0 | NA | NA | NA | NA | NA | NM_138647 |
| 6030440G07Rik | 0 | NA | NA | NA | NA | NA | | Cers3 | 8.631064728 | 0.0181856165406787 | 0.057920048091683 | 0.313977925430798 | 0.753537804552706 | NA | NM_001164201 |
| 6030466F02Rik | 1.43368476 | 0.060713864 | 0.090463583 | 0.671141489 | 0.502130399 | NA | | Ces1a | 0.405340399 | -0.00677209336026796 | 0.0121196852483227 | -0.558768088569395 | 0.576320003803731 | NA | NM_001013764 |
| 6030468B19Rik | 2.58199959 | 0.13739291 | 0.106765721 | 1.286863507 | 0.198141867 | NA | | Ces1b | 0 | NA | NA | NA | NA | NA | NM_001081372 |
| 6030498E09Rik | 0.558783501 | -0.006176017 | 0.052494503 | -0.117650732 | 0.906344406 | NA | | Ces1c | 0 | NA | NA | NA | NA | NA | NM_007954 |
| 6330407A03Rik | 0.107716603 | -0.020477583 | 0.027954224 | -0.732539849 | 0.46383913 | NA | | Ces1d | 7.94638497 | 0.0474895454687101 | 0.0506073747886608 | 0.938391798962681 | 0.348043098954607 | NA | NM_053200 |
| 6330410L21Rik | 6.399064506 | 0.07381682 | 0.137644779 | 0.536284921 | 0.59176165 | NA | | Ces1e | 0.191449919 | 0.00665508079272897 | 0.00962405648474729 | 0.691504752001018 | 0.489248395458281 | NA | NM_133660 |
| 6330415B21Rik | 0 | NA | NA | NA | NA | NA | | Ces1f | 3.035197234 | -0.0305722798189084 | 0.0372047390930143 | -0.821730794630105 | 0.411230128708017 | NA | NM_144930 |
| 6430562O15Rik | 1.492400801 | 0.032328451 | 0.083519906 | 0.3870748 | 0.698700833 | NA | | Ces1g | 0.719463989 | 0.0238008131970467 | 0.0180951363112226 | 1.31531549625771 | 0.18840389225459 | NA | NM_021456 |
| 6430706D22Rik | 0 | NA | NA | NA | NA | NA | | Ces2a | 0 | NA | NA | NA | NA | NA | NM_133960,NM_001190330 |
| 7420426K07Rik | 0 | NA | NA | NA | NA | NA | | Ces2b | 1.605141492 | -0.000451970045122661 | 0.027333577689253 | -0.0165353416322213 | 0.986807307392345 | NA | NM_198171 |
| 7420461P10Rik | 0.096071043 | -0.020098795 | 0.027825948 | -0.722304036 | 0.470107572 | NA | | Ces2c | 0 | NA | NA | NA | NA | NA | NM_145603 |
| 7420700N18Rik | 0 | NA | NA | NA | NA | NA | | Ces2d-ps | 0 | NA | NA | NA | NA | NA | NR_033726 |
| 7420701I03Rik | 0 | NA | NA | NA | NA | NA | | Ces2e | 0.779254277 | -0.007543178618867 | 0.0184685079825491 | -0.408434651353242 | 0.682954597991981 | NA | NM_172759,NM_001163756 |
| 7530416G11Rik | 0.186326724 | 0.024939385 | 0.031411053 | 0.793968451 | 0.427213798 | NA | | Ces2f | 0 | NA | NA | NA | NA | NA | NM_001079865 |
| 7630403G23Rik | 6.137286725 | 0.0464188 | 0.104719735 | 0.443266975 | 0.65757264 | NA | | Ces2g | 10.11002416 | -0.0573881121184171 | 0.0624395673536414 | -0.919098490759646 | 0.358044058612955 | NA | NM_197999 |
| 8030411F24Rik | 0 | NA | NA | NA | NA | NA | | Ces2h | 0 | NA | NA | NA | NA | NA | NM_001272045 |
| 8030423F21Rik | 2.141433242 | 0.034766227 | 0.097894192 | 0.355140854 | 0.722484072 | NA | | Ces3a | 0 | NA | NA | NA | NA | NA | NM_198672,NM_001164681 |
| 8030423J24Rik | 1.331013364 | 0.035204865 | 0.084182414 | 0.418197386 | 0.675802807 | NA | | Ces3b | 0 | NA | NA | NA | NA | NA | NM_144511,NM_001159415 |
| 8030442B05Rik | 2.133971828 | 0.051578361 | 0.100061993 | 0.515464057 | 0.60622878 | NA | | Ces4a | 0 | NA | NA | NA | NA | NA | NM_146213 |
| 8030443G20Rik | 0.663524823 | -0.038052693 | 0.059997452 | -0.634238477 | 0.5259252 | NA | | Ces5a | 16.58951276 | -0.0176175647808098 | 0.0771934322044144 | -0.228226214040556 | 0.819470383289988 | NA | NM_001003951 |
| 8430403D17Rik | 0.098353386 | -0.020212831 | 0.02786466 | -0.725393061 | 0.468210929 | NA | | Cetn1 | 0 | NA | NA | NA | NA | NA | NM_007593 |
| 8430422H06Rik | 0 | NA | NA | NA | NA | NA | | Cfc1 | 0.209568245 | 0.00686197416234131 | 0.00977127153168884 | 0.702260104029194 | 0.482516968822431 | NA | NM_007685 |
| 8430423G03Rik | 0 | NA | NA | NA | NA | NA | | Cfd | 0.108766301 | 0.00519171381361946 | 0.00856003371502575 | 0.606506234257728 | 0.544178643801544 | NA | NM_013459 |
| 8430431K14Rik | 0 | NA | NA | NA | NA | NA | | Cfhr1 | 0 | NA | NA | NA | NA | NA | NM_015780 |
| 8430436N08Rik | 0.302785524 | -0.016957309 | 0.039595455 | -0.428264025 | 0.668458907 | NA | | Cfhr2 | 0.725209114 | 0.00832586729883459 | 0.0183396579494299 | 0.453981602153786 | 0.649842068725904 | NA | NM_001025575 |
| 8430437L04Rik | 0.098353386 | -0.020212831 | 0.02786466 | -0.725393061 | 0.468210929 | NA | | Cfi | 0.736588345 | -0.00763592360411818 | 0.0198865451885206 | -0.383974367177963 | 0.70099744598282 | NA | NM_007686 |
| 9030204H09Rik | 0.362076728 | 0.018991228 | 0.042095091 | 0.451150657 | 0.65188097 | NA | | Cga | 9.106314582 | -0.0509164528646532 | 0.0488753231565295 | -1.0417619685416 | 0.297522050618395 | NA | NM_009889 |
| 9030404E10Rik | 0.185906812 | 0.024895454 | 0.031385107 | 0.793225068 | 0.427646705 | NA | | Ch25h | 15.55986639 | 0.02410877230944 | 0.0735700743319486 | 0.32769808279194 | 0.743139953132727 | NA | NM_009890 |
| 9030619P08Rik | 0.309710804 | -0.014226733 | 0.039782345 | -0.357614233 | 0.720632025 | NA | | Chad | 5.366179878 | 0.0484336821034721 | 0.0497158153722158 | 0.974210756493792 | 0.329951904924108 | NA | NM_007689 |
| 9130008F23Rik | 4.619969587 | -0.064091833 | 0.121275204 | -0.528482583 | 0.597164432 | NA | | Chaf1b | 17.65788618 | 0.0685155331362223 | 0.0789748393622453 | 0.867561538453434 | 0.38563440948119 | NA | NM_028083 |
| 9130015A21Rik | 1.12994131 | 0.036923979 | 0.07408221 | 0.498418966 | 0.618188772 | NA | | Chi3l3 | 4.938303949 | -0.0333715800007313 | 0.0417060965169287 | -0.800160714805464 | 0.423617687709857 | NA | NM_009892 |
| 9130015L21Rik | 0.283178451 | 0.028390488 | 0.033368195 | 0.850824814 | 0.394866675 | NA | | Chi3l4 | 0 | NA | NA | NA | NA | NA | NM_145126 |
| 9130019P16Rik | 7.620208972 | 0.006877866 | 0.155372474 | 0.044266952 | 0.964691614 | NA | | Chit1 | 5.977157792 | -0.0226884159264708 | 0.0489381409553254 | -0.463614176664018 | 0.642924195561721 | NA | NM_027979 |
| 9130204L05Rik | 0 | NA | NA | NA | NA | NA | | Chkb | 22.29064703 | -0.191897995701263 | 0.0854370107911407 | -2.24607572203547 | 0.0246991581189929 | NA | NM_007692,NR_037154,NR_037153 |
| 9130206I24Rik | 5.39408971 | -0.015742662 | 0.133937251 | -0.117537595 | 0.906434054 | NA | | Chmp4c | 3.678402829 | -0.00939125512983161 | 0.0396927184284059 | -0.236598940603443 | 0.812967942147175 | NA | NM_025519 |
| 9130209A04Rik | 0 | NA | NA | NA | NA | NA | | Chrdl2 | 6.42185293 | -0.00785205330506828 | 0.054054598834926 | -0.145261522133337 | 0.88450439333047 | NA | NM_133709 |
| 9130221F21Rik | 0 | NA | NA | NA | NA | NA | | Chrna6 | 29.03269317 | 0.048802235969985 | 0.063078839922562 | 0.773670473805423 | NA | NA | NM_021369 |
| 9130230L23Rik | 0 | NA | NA | NA | NA | NA | | Chrna9 | 0.212297216 | 0.00681050119636477 | 0.00973439053551308 | 0.699633035218656 | 0.484156506270654 | NA | NM_001081104 |
| 9130409I23Rik | 0.192142086 | -0.026384592 | 0.031527717 | -0.836869715 | 0.402665805 | NA | | Chrnb4 | 18.62219421 | -0.00410422293891301 | 0.0628378956654649 | -0.0653144554802246 | 0.947923633111417 | NA | NM_148944 |
| 9230009I02Rik | 6.460502141 | 0.126379548 | 0.1389964 | 0.909228929 | 0.363229297 | NA | | Chrnd | 0 | NA | NA | NA | NA | NA | NM_021600 |
| 9230102K24Rik | 1.840445218 | -0.054323261 | 0.094912351 | -0.572351865 | 0.567083619 | NA | | Chrne | 3.430925882 | 0.0516124406186159 | 0.0349798279264088 | 1.47549155265141 | 0.140080437681896 | NA | NM_009603 |
| 9230102O04Rik | 0.159967308 | 0.0240345 | 0.030870969 | 0.778547006 | 0.436246606 | NA | | Chrng | 0 | NA | NA | NA | NA | NA | NM_009604 |
| 9230104L09Rik | 0.833822031 | 0.048595634 | 0.059165953 | 0.821344557 | 0.411450034 | NA | | Chst13 | 0.429381296 | -0.0100524906077422 | 0.0141519698525479 | -0.71032447867548 | 0.477502943458686 | NA | NM_027928 |
| 9230105E05Rik | 0 | NA | NA | NA | NA | NA | | Chst4 | 6.704431706 | 0.0423943801265064 | 0.0507811646273164 | 0.834844581404135 | 0.403805221012355 | NA | NM_011998 |
| 9230110F15Rik | 0.094392817 | 0.017895942 | 0.027818137 | 0.643319236 | 0.520016977 | NA | | Cib3 | 0.534690201 | 0.0116522952639082 | 0.0167582688787382 | 0.69531616590135 | 0.48685718543538 | NA | NM_001080812 |
| 9230112D13Rik | 0 | NA | NA | NA | NA | NA | | Cib4 | 0 | NA | NA | NA | NA | NA | NM_028483 |
| 9230115E21Rik | 0.389426822 | -0.00542245 | 0.0474348 | -0.114313744 | 0.908989088 | NA | | Cidec | 0.674591037 | 0.00389701139981464 | 0.0175660161813603 | 0.221849471136765 | 0.824431066493655 | NA | NM_178373 |
| 9330111N05Rik | 6.798952139 | -0.025466348 | 0.155174401 | -0.164114364 | 0.869641115 | NA | | Ciita | 9.678908858 | -0.0460845543528973 | 0.0266228541263504 | -1.73101479406313 | NA | NA | NM_007575,NM_001243761,NM_001243760 |
| 9330162B11Rik | 0.289311041 | 0.012154199 | 0.038946437 | 0.312074719 | 0.754983739 | NA | | Cilp | 5.825388647 | 0.0764277830402489 | 0.044317341142706 | 1.72455704854098 | 0.0846073295829954 | NA | NM_173385 |
| 9330175E14Rik | 1.46230452 | 0.156422717 | 0.081557834 | 1.917936127 | 0.055119111 | NA | | Cilp2 | 16.9315897 | -0.0285927661402095 | 0.074416334047789 | -0.384227018249081 | 0.700810194723957 | NA | NM_026818 |
| 9330178D15Rik | 0 | NA | NA | NA | NA | NA | | Ckap2 | 22.17471522 | -0.0227549724027093 | 0.0820744093536881 | -0.277248079905759 | 0.781589623579319 | NA | NM_001004140 |
| 9330182O14Rik | 0 | NA | NA | NA | NA | NA | | Ckm | 2.457932771 | 0.034789493695203 | 0.0348753108673023 | 0.99753931449025 | 0.318502800698898 | NA | NM_007710 |
| 9430007A20Rik | 0 | NA | NA | NA | NA | NA | | Ckmt2 | 1.074382203 | -0.0124409809366344 | 0.0216622125687854 | -0.574317184688765 | 0.565753182390687 | NA | NM_198415 |
| 9430014N10Rik | 0.423821016 | -0.060879619 | 0.047397786 | -1.284440147 | 0.198987993 | NA | | Clca1 | 9.099916493 | 0.0323926742051003 | 0.0597612862529736 | 0.542034421213424 | 0.587794793667132 | NA | NM_009899 |
| 9430018G01Rik | 1.1451679 | 0.008926323 | 0.072644483 | 0.122876814 | 0.902204646 | NA | | Clca2 | 2.910918961 | 0.0144554989049153 | 0.0390514644340299 | 0.370165347559121 | 0.711259294257472 | NA | NM_030601 |
| 9430019J16Rik | 0 | NA | NA | NA | NA | NA | | Clca3 | 1.376225769 | 0.00873259919326466 | 0.0197179340685925 | 0.442875970823653 | 0.657855448851006 | NA | NM_017474 |
| 9430060I03Rik | 7.122908069 | -0.012088913 | 0.143328609 | -0.084344032 | 0.932782905 | NA | | Clca4 | 0.106148608 | 0.00515554063321497 | 0.00854200771766808 | 0.603551390213729 | 0.546141939141592 | NA | NM_139148 |

| | | | | | | |
|---|---|---|---|---|---|---|
| 9430069I07Rik | 0 | NA | NA | NA | NA | NA |
| 9430076C15Rik | 1.2997562 | -0.001626253 | 0.081439436 | -0.01996886 | 0.984068214 | NA |
| 9430076G02Rik | 1.27944664 | -0.029819558 | 0.086373056 | -0.345241433 | 0.729912872 | NA |
| 9530002B09Rik | 0 | NA | NA | NA | NA | NA |
| 9530003J23Rik | 0.211846316 | 0.025707621 | 0.031860375 | 0.806883811 | 0.419733426 | NA |
| 9530026F06Rik | 0.203787646 | -0.02671144 | 0.031728022 | -0.841887971 | 0.399850664 | NA |
| 9530027J09Rik | 2.461712552 | 0.106928818 | 0.10883103 | 0.982521413 | 0.325843039 | NA |
| 9530048J24Rik | 0.521607058 | -0.053101502 | 0.044487442 | -1.193629028 | 0.232623114 | NA |
| 9530053A07Rik | 5.088403885 | 0.305295635 | 0.117933637 | 2.588707018 | 0.009633702 | NA |
| 9530059O14Rik | 3.90566549 | 0.047944508 | 0.131037513 | 0.365883834 | 0.71445176 | NA |
| 9630028H03Rik | 0.492226933 | -0.036718726 | 0.046334526 | -0.792470094 | 0.428086623 | NA |
| 9830107B12Rik | 0 | NA | NA | NA | NA | NA |
| 9830132P13Rik | 0.105923158 | 0.018524693 | 0.027977518 | 0.662127842 | 0.507889294 | NA |
| 9930111J21Rik2 | 0.254617334 | 0.029389883 | 0.033981526 | 0.864878265 | 0.387105608 | NA |
| A1bg | 0 | NA | NA | NA | NA | NA |
| A1cf | 0.08731684 | 0.017365056 | 0.027733412 | 0.626142072 | 0.531221759 | NA |
| A230020J21Rik | 0.187700257 | -0.003168524 | 0.031449734 | -0.100748845 | 0.919749836 | NA |
| A230028O05Rik | 7.252867673 | -0.008809902 | 0.133754343 | -0.065866289 | 0.94748428 | NA |
| A230108P19Rik | 0.346294032 | 0.050460286 | 0.044390535 | 1.136735257 | 0.255648976 | NA |
| A330093E20Rik | 6.749109249 | 0.016053548 | 0.142900097 | 0.11234106 | 0.910552987 | NA |
| A3galt2 | 7.237645065 | 0.021751122 | 0.145357671 | 0.149638626 | 0.881049732 | NA |
| A430071A18Rik | 0.483152594 | -0.021244376 | 0.05063691 | -0.419543292 | 0.674819123 | NA |
| A430088P11Rik | 0.174376471 | 0.024538273 | 0.031173126 | 0.787161136 | 0.431187544 | NA |
| A430089I19Rik | 0 | NA | NA | NA | NA | NA |
| A430090L17Rik | 0 | NA | NA | NA | NA | NA |
| A430092G05Rik | 0.379842343 | -0.027920334 | 0.043022777 | -0.648966337 | 0.516360135 | NA |
| A430093F15Rik | 2.465706733 | 0.010129674 | 0.111331717 | 0.090986418 | 0.927503383 | NA |
| A530006G24Rik | 0 | NA | NA | NA | NA | NA |
| A530013C23Rik | 2.442407716 | 0.087283208 | 0.097467933 | 0.895506915 | 0.370516171 | NA |
| A530016L24Rik | 0.520737658 | -0.0194622 | 0.051269202 | -0.379608015 | 0.70423641 | NA |
| A530032D15Rik | 0.506479192 | -0.018468102 | 0.045386828 | -0.406904433 | 0.68407818 | NA |
| A530046M15Rik | 0 | NA | NA | NA | NA | NA |
| A530050N04Rik | 5.660482064 | 0.045064192 | 0.12986375 | 0.347011328 | 0.728582809 | NA |
| A530053G22Rik | 1.112233245 | 0.118486707 | 0.073791026 | 1.605706178 | 0.108338483 | NA |
| A530064D06Rik | 1.664664629 | 0.131011716 | 0.079362674 | 1.650797653 | 0.098779899 | NA |
| A530065N20Rik | 0.198019769 | 0.025308392 | 0.031628095 | 0.80018706 | 0.423602426 | NA |
| A530088E08Rik | 1.489254681 | 0.099355415 | 0.093643733 | 1.060993744 | 0.288692744 | NA |
| A530099J19Rik | 0.204013804 | -0.003115899 | 0.031740777 | -0.098167077 | 0.921799625 | NA |
| A630010A05Rik | 0 | NA | NA | NA | NA | NA |
| A630012P03Rik | 0.096071043 | -0.020098795 | 0.027825948 | -0.722304036 | 0.470107572 | NA |
| A630020A06 | 1.158197331 | -0.063360987 | 0.07979942 | -0.794003111 | 0.42719362 | NA |
| A630023A22Rik | 0 | NA | NA | NA | NA | NA |
| A630073D07Rik | 0 | NA | NA | NA | NA | NA |
| A630075F10Rik | 0.079983654 | 0.016740032 | 0.027674178 | 0.604897175 | 0.54524732 | NA |
| A630076J17Rik | 0.501077839 | -0.018122177 | 0.045283609 | -0.400192852 | 0.689014479 | NA |
| A630077J23Rik | 3.174756188 | -0.231080716 | 0.12488884 | -1.850291157 | 0.064271597 | NA |
| A630095E13Rik | 0.192142086 | -0.026384592 | 0.031527717 | -0.836869715 | 0.402665805 | NA |
| A630095N17Rik | 1.318866669 | -0.014521743 | 0.072030855 | -0.201604475 | 0.840225947 | NA |
| A730008H23Rik | 0 | NA | NA | NA | NA | NA |
| A730018C14Rik | 8.063125415 | -0.195203337 | 0.14620121 | -1.335169092 | 0.18182101 | NA |
| A730082K24Rik | 0 | NA | NA | NA | NA | NA |
| A730085A09Rik | 3.397070152 | 0.259675453 | 0.108332692 | 2.397018369 | 0.016529095 | NA |
| A730090N16Rik | 1.353239558 | 0.100290753 | 0.081106574 | 1.236530504 | 0.216261434 | NA |
| A730098P11Rik | 7.212616839 | -0.145312751 | 0.154259555 | -0.942001623 | 0.346191809 | NA |
| A930001A20Rik | 4.837739006 | 0.012436999 | 0.141469758 | 0.087912777 | 0.929946002 | NA |
| A930001C03Rik | 2.223242678 | 0.121634856 | 0.098371412 | 1.236485818 | 0.216278034 | NA |
| A930006I01Rik | 0.079983654 | 0.016740032 | 0.027674178 | 0.604897175 | 0.54524732 | NA |
| A930018P22Rik | 1.843354373 | 0.069817574 | 0.092858753 | 0.751868533 | 0.452130123 | NA |
| A930019D19Rik | 0.536358603 | 0.018999954 | 0.051864556 | 0.366337928 | 0.714112931 | NA |
| A930041C12Rik | 1.176855315 | 0.004953334 | 0.066942803 | 0.073993531 | 0.941015533 | NA |
| AA467197 | 3.381262845 | 0.031271476 | 0.118097646 | 0.26479339 | 0.791168629 | NA |
| AA536875 | 0.482409049 | 0.03844785 | 0.05183958 | 0.741669784 | 0.458287431 | NA |
| AA545190 | 0 | NA | NA | NA | NA | NA |
| AA619741 | 0.159967308 | 0.0240345 | 0.030870969 | 0.778547006 | 0.436246606 | NA |
| AA792892 | 0 | NA | NA | NA | NA | NA |
| AB099516 | 0.195033442 | -0.002424765 | 0.031596358 | -0.076741901 | 0.93882887 | NA |
| AF357355 | 3.574150081 | 0.111350073 | 0.106999133 | 1.040663321 | 0.298031832 | NA |
| AF357399 | 6.262553216 | 0.142496222 | 0.145564736 | 0.97891994 | 0.327619539 | NA |
| AF357425 | 4.230150771 | 0.244130117 | 0.130024704 | 1.877567183 | 0.060440399 | NA |
| AF357426 | 5.609738731 | 0.096073778 | 0.137180076 | 0.7003479 | 0.483710064 | NA |
| AF366264 | 0 | NA | NA | NA | NA | NA |
| AI314831 | 0.099009884 | 0.018205726 | 0.027896865 | 0.652608315 | 0.514008823 | NA |
| AI427809 | 5.07471216 | -0.044295103 | 0.136950375 | -0.32343908 | 0.746362737 | NA |
| AI463170 | 0.48794879 | 0.064131883 | 0.049799661 | 1.287797577 | 0.197816436 | NA |
| AI506816 | 2.116057961 | -0.06180638 | 0.092132999 | -0.670838684 | 0.502323301 | NA |
| AI507597 | 2.297210766 | -0.060771891 | 0.104072412 | -0.583938528 | 0.559261671 | NA |
| AI646519 | 0 | NA | NA | NA | NA | NA |
| AI662270 | 4.790869034 | -0.039136686 | 0.117567014 | -0.33288832 | 0.739218587 | NA |
| AI747448 | 0.263371537 | 0.009921338 | 0.037961873 | 0.261350055 | 0.793822569 | NA |
| AU015228 | 0.297029653 | 0.02866449 | 0.033517577 | 0.855207712 | 0.392436167 | NA |
| AU015836 | 0.196680618 | -0.003857632 | 0.031597546 | -0.122086447 | 0.902830556 | NA |
| AU016765 | 0 | NA | NA | NA | NA | NA |
| AU018091 | 0 | NA | NA | NA | NA | NA |
| AU018829 | 0 | NA | NA | NA | NA | NA |
| AU019990 | 0.200315975 | 0.025370485 | 0.031664208 | 0.801235428 | 0.422995365 | NA |
| AU021063 | 1.577856553 | 0.091196744 | 0.083203315 | 1.096071045 | 0.273047685 | NA |
| AU022751 | 0.722087843 | -0.018583788 | 0.054979351 | -0.33801396 | 0.735352668 | NA |
| AU022793 | 5.942553399 | 0.331664192 | 0.139439668 | 2.378549788 | 0.01738089 | NA |
| AU023871 | 0.824694589 | 0.002268418 | 0.057226179 | 0.039639519 | 0.96838052 | NA |
| AV051173 | 4.482025111 | -0.090561204 | 0.134030697 | -0.675675093 | 0.499246953 | NA |
| AV320801 | 0 | NA | NA | NA | NA | NA |
| AY074887 | 1.746146875 | 0.066616651 | 0.093763116 | 0.710478217 | 0.477407634 | NA |
| AY358078 | 0.600695175 | -0.01924069 | 0.047747217 | -0.40296987 | 0.68697039 | NA |
| AY512915 | 0.381360167 | 0.020495541 | 0.042644778 | 0.480610819 | 0.630793125 | NA |
| AY512931 | 3.937876758 | 0.045757036 | 0.124981246 | 0.366111214 | 0.71428209 | NA |
| AY761184 | 0 | NA | NA | NA | NA | NA |
| AY761185 | 0 | NA | NA | NA | NA | NA |
| Aadac | 0 | NA | NA | NA | NA | NA |
| Aadacl2 | 0 | NA | NA | NA | NA | NA |
| Aadacl3 | 0.08731684 | 0.017365056 | 0.027733412 | 0.626142072 | 0.531221759 | NA |
| Aadat | 6.836992621 | -0.119528564 | 0.150494808 | -0.794237129 | 0.427057397 | NA |
| Abca12 | 6.783017849 | 0.064861459 | 0.135527712 | 0.478584473 | 0.632234266 | NA |
| Abca14 | 1.030459691 | -0.007957983 | 0.064045404 | -0.124255332 | 0.901113113 | NA |
| Abca15 | 1.021931072 | 0.119306829 | 0.075489238 | 1.580448199 | 0.114004261 | NA |
| Abca16 | 0.08731684 | 0.017365056 | 0.027733412 | 0.626142072 | 0.531221759 | NA |
| Abcb11 | 0.781051818 | -0.037914647 | 0.067597685 | -0.56088677 | 0.57487473 | NA |
| Abcb5 | 0.192142086 | -0.026384592 | 0.031527717 | -0.836869715 | 0.402665805 | NA |
| Abcg5 | 1.062502086 | -0.033374209 | 0.06048551 | -0.551771973 | 0.581104592 | NA |
| Abcg8 | 0.215468684 | -0.027008432 | 0.031909031 | -0.846419662 | 0.397318674 | NA |
| Abhd12b | 6.56843143 | 0.215727634 | 0.116561549 | 1.850761555 | 0.064203864 | NA |
| Abhd16b | 0.098353386 | -0.020212831 | 0.02786466 | -0.725393061 | 0.468210929 | NA |
| Abo | 0 | NA | NA | NA | NA | NA |
| Abp1 | 1.087822263 | 0.131401943 | 0.077071413 | 1.704937519 | 0.088206078 | NA |
| Acap1 | 3.34557331 | -0.091379074 | 0.111964918 | -0.816140234 | 0.414419935 | NA |
| Acbd7 | 7.53369653 | 0.006552191 | 0.146003894 | 0.044876826 | 0.964205488 | NA |
| Ace3 | 0 | NA | NA | NA | NA | NA |
| Acer1 | 0.609905974 | -0.029829418 | 0.054035901 | -0.552029618 | 0.580928061 | NA |
| Acmsd | 0.87601496 | 0.003546103 | 0.064153208 | 0.055275543 | 0.955918946 | NA |
| Acnat1 | 4.202123611 | 0.108514166 | 0.124992797 | 0.868163353 | 0.385304913 | NA |
| Acnat2 | 0 | NA | NA | NA | NA | NA |
| Acot10 | 0.08731684 | 0.017365056 | 0.027733412 | 0.626142072 | 0.531221759 | NA |
| Acox2 | 5.121583048 | 0.190673171 | 0.12464736 | 1.529700845 | 0.126090794 | NA |
| Acoxl | 0.876552974 | 0.032384074 | 0.064824785 | 0.499563152 | 0.617382709 | NA |
| Acpt | 0 | NA | NA | NA | NA | NA |
| Acrv1 | 0 | NA | NA | NA | NA | NA |
| Acsbg2 | 1.704303594 | 0.033296316 | 0.090317104 | 0.368660143 | 0.71238106 | NA |
| Acsm1 | 0 | NA | NA | NA | NA | NA |
| Acsm2 | 0.341110531 | -0.038008058 | 0.037770309 | -1.006294602 | 0.314273876 | NA |
| Acsm3 | 4.98918845 | 0.251118734 | 0.130025547 | 1.931303044 | 0.053445587 | NA |
| Acsm4 | 1.474061535 | 0.040944629 | 0.079840068 | 0.512833097 | 0.608068076 | NA |
| Actbl2 | 0.571288896 | 0.022514058 | 0.053122995 | 0.423810033 | 0.671704361 | NA |
| Actc1 | 0 | NA | NA | NA | NA | NA |
| Actg2 | 3.660722762 | -0.190788708 | 0.123651213 | -1.542958637 | 0.122840812 | NA |
| Actl10 | 0.197363271 | -0.000561558 | 0.031650166 | -0.017742642 | 0.985844163 | NA |
| Actl11 | 0 | NA | NA | NA | NA | NA |
| Actl7a | 0 | NA | NA | NA | NA | NA |
| Actl7b | 0.910275419 | 0.002696959 | 0.070454943 | 0.038279207 | 0.969465069 | NA |
| Actl9 | 5.064570303 | -0.018445694 | 0.121375821 | -0.151971732 | 0.879209231 | NA |
| Actr8 | 1805.598313 | 0.1420983 | 0.080864198 | 1.757246147 | NA | NA |
| Actrt1 | 0 | NA | NA | NA | NA | NA |
| Actrt2 | 0 | NA | NA | NA | NA | NA |
| Ada | 20.83374261 | 0.2270041 | 0.157922884 | 1.437436386 | NA | NA |
| Adad1 | 0.215050351 | -0.027006021 | 0.031907566 | -0.846382992 | 0.397339123 | NA |
| Adad2 | 3.510446752 | -0.083303528 | 0.112342153 | -0.741516216 | 0.458380504 | NA |
| Adam1b | 0.193239998 | 0.025145508 | 0.031532422 | 0.797449311 | 0.42519013 | NA |
| Adam2 | 0.746849799 | -0.00504642 | 0.061759293 | -0.081711098 | 0.934876453 | NA |
| Adam20 | 0 | NA | NA | NA | NA | NA |
| Adam24 | 0 | NA | NA | NA | NA | NA |
| Adam25 | 0 | NA | NA | NA | NA | NA |
| Adam26a | 0 | NA | NA | NA | NA | NA |
| Adam26b | 0 | NA | NA | NA | NA | NA |
| Adam28 | 1.418099254 | 0.061945414 | 0.090504709 | 0.684444096 | 0.493694779 | NA |
| Adam29 | 0 | NA | NA | NA | NA | NA |
| Adam3 | 3.516109624 | 0.16018811 | 0.115274822 | 1.389619243 | 0.164644529 | NA |
| Adam30 | 0 | NA | NA | NA | NA | NA |
| Adam34 | 0 | NA | NA | NA | NA | NA |
| Adam39 | 0 | NA | NA | NA | NA | NA |
| Adam4 | 7.268415126 | -0.239545171 | 0.157467127 | -1.521239231 | 0.128199814 | NA |
| Adam5 | 7.351368263 | 0.003698338 | 0.156765848 | 0.023591476 | 0.981178471 | NA |
| Adam6a | 0 | NA | NA | NA | NA | NA |
| Adam6b | 0 | NA | NA | NA | NA | NA |
| Adam7 | 0 | NA | NA | NA | NA | NA |
| Adamdec1 | 0.098353386 | -0.020212831 | 0.02786466 | -0.725393061 | 0.468210929 | NA |
| Adamts12 | 8.020384222 | -0.177224349 | 0.150919601 | -1.174296437 | 0.24027631 | NA |
| Adh4 | 0 | NA | NA | NA | NA | NA |
| Adh6-ps1 | 0 | NA | NA | NA | NA | NA |
| Adh6a | 0 | NA | NA | NA | NA | NA |
| Adh7 | 3.457528501 | -0.027422988 | 0.106834914 | -0.256685635 | 0.797421461 | NA |
| Adig | 0.639411646 | 0.032045962 | 0.060168178 | 0.532606496 | 0.594306005 | NA |
| Adipoq | 1.743847353 | 0.074828092 | 0.098163715 | 0.762278525 | 0.445893788 | NA |
| Adipor2 | 3159.946284 | 0.27510772 | 0.160253584 | 1.716702445 | NA | NA |
| Adm2 | 0 | NA | NA | NA | NA | NA |
| Adprhl1 | 4.519450566 | 0.05331919 | 0.137129957 | 0.388822334 | 0.697407578 | NA |
| Afm | 0 | NA | NA | NA | NA | NA |
| Afp | 4.10379429 | -0.083586349 | 0.117267397 | -0.712784212 | 0.475979297 | NA |
| Agr2 | 1.226054931 | -0.039562708 | 0.075085596 | -0.526901426 | 0.598262043 | NA |
| Agr3 | 0.977524795 | -0.093181919 | 0.064611519 | -1.442187401 | 0.149249514 | NA |
| Agrp | 4.094995815 | 0.004568953 | 0.130892431 | 0.03490617 | 0.972154561 | NA |
| Agt | 964.1615949 | 0.346699071 | 0.142928413 | 2.425683343 | NA | NA |
| Agtr1b | 4.917559474 | -0.162501733 | 0.141827691 | -1.145768723 | 0.25189086 | NA |
| Agxt | 0 | NA | NA | NA | NA | NA |
| Agxt2 | 0.299163155 | 0.032734233 | 0.035823547 | 0.913763036 | 0.360841373 | NA |
| Ahsg | 0.20671448 | -0.003314027 | 0.03178208 | -0.10427343 | 0.916952363 | NA |
| Aicda | 0.19442443 | -0.026451352 | 0.031568725 | -0.837897386 | 0.402088337 | NA |
| Aipl1 | 0.958438028 | -0.024468712 | 0.063760441 | -0.383760081 | 0.701156278 | NA |
| Aire | 6.525529669 | 0.251428125 | 0.119637062 | 2.101590597 | 0.035589154 | NA |
| Akap14 | 6.612560473 | 0.229692455 | 0.150132845 | 1.529928086 | 0.126034531 | NA |
| Akap4 | 0 | NA | NA | NA | NA | NA |
| Aknad1 | 0.915478062 | -0.025627691 | 0.074998556 | -0.341709124 | 0.732569805 | NA |
| Akp3 | 0.295060159 | -0.029687054 | 0.033501022 | -0.886153701 | 0.375534709 | NA |
| Akr1b3 | 0 | NA | NA | NA | NA | NA |
| Akr1b7 | 1.01527448 | 0.083273158 | 0.071468685 | 1.165169853 | 0.243950246 | NA |
| Akr1c12 | 1.073983606 | 0.05470851 | 0.060038048 | 0.911230649 | 0.362173858 | NA |
| Akr1c19 | 0.305079873 | -0.015054679 | 0.039642215 | -0.379763828 | 0.704120735 | NA |
| Akr1c20 | 0 | NA | NA | NA | NA | NA |
| Akr1c21 | 0.327267802 | 0.035667551 | 0.037337881 | 0.955264473 | 0.339443969 | NA |
| Akr1c6 | 0 | NA | NA | NA | NA | NA |
| Akr1cl | 0.206069989 | -0.026776453 | 0.031767726 | -0.842882251 | 0.399294301 | NA |
| Akr1d1 | 0.08731684 | 0.017365056 | 0.027733412 | 0.626142072 | 0.531221759 | NA |
| Alas2 | 24.17344701 | -0.175998604 | 0.091829495 | -1.916580334 | NA | NA |
| Alb | 0.702296617 | 0.005938661 | 0.054546958 | 0.10887246 | 0.913303651 | NA |
| Aldh8a1 | 1.984465515 | 0.031198082 | 0.097224269 | 0.320887796 | 0.748295423 | NA |
| Aldoart1 | 0.239950962 | 0.027417169 | 0.032831728 | 0.835081494 | 0.403671825 | NA |
| Aldoart2 | 0.977385348 | 0.0095598 | 0.058332584 | 0.163884395 | 0.869822152 | NA |
| Alms1-ps2 | 3.595120074 | 0.020417395 | 0.128248318 | 0.159202046 | 0.873509692 | NA |
| Alox12e | 4.988882264 | 0.033035474 | 0.128125116 | 0.257837613 | 0.796532235 | NA |
| Alox15 | 2.286331333 | 0.01030815 | 0.102305164 | 0.100758844 | 0.919741898 | NA |
| Alpi | 0.094392817 | 0.017895942 | 0.027818137 | 0.643319236 | 0.520016977 | NA |
| Alppl2 | 0 | NA | NA | NA | NA | NA |
| Als2cr11 | 0.488462861 | -0.011555605 | 0.044402921 | -0.26024426 | 0.794675365 | NA |
| Als2cr12 | 0.810586661 | 0.022431908 | 0.063397743 | 0.353828177 | 0.723467655 | NA |
| Alx1 | 4.136739917 | 0.074635799 | 0.129062468 | 0.578292049 | 0.563066962 | NA |
| Ambn | 0.256038351 | 0.007085343 | 0.03307033 | 0.214250763 | 0.830351512 | NA |
| Ambp | 0 | NA | NA | NA | NA | NA |
| Amd1 | 0 | NA | NA | NA | NA | NA |
| Amd2 | 0 | NA | NA | NA | NA | NA |
| Amelx | 0.376356809 | 0.021703132 | 0.042541541 | 0.510163288 | 0.60993707 | NA |
| Ampd1 | 0.116696964 | -0.020477583 | 0.027954224 | -0.732539847 | 0.463839131 | NA |
| Amtn | 0 | NA | NA | NA | NA | NA |
| Amy2a2 | 0 | NA | NA | NA | NA | NA |
| Amy2a3 | 0 | NA | NA | NA | NA | NA |
| Amy2a4 | 0 | NA | NA | NA | NA | NA |
| Amy2a5 | 0 | NA | NA | NA | NA | NA |
| Amy2b | 0.079983654 | 0.016740032 | 0.027674178 | 0.604897175 | 0.54524732 | NA |
| Ang2 | 0 | NA | NA | NA | NA | NA |
| Ang3 | 0 | NA | NA | NA | NA | NA |
| Ang4 | 0.098353386 | -0.020212831 | 0.02786466 | -0.725393061 | 0.468210929 | NA |
| Ang5 | 0 | NA | NA | NA | NA | NA |
| Ang6 | 0 | NA | NA | NA | NA | NA |
| Angpt4 | 1.123832365 | -0.079502269 | 0.075748052 | -1.049561891 | 0.293919586 | NA |
| Angptl3 | 6.915324161 | -0.035852231 | 0.154708243 | -0.231740923 | 0.816739244 | NA |
| Ankar | 0.599827183 | 0.017101474 | 0.046253359 | 0.369734744 | 0.711580142 | NA |
| Ankk1 | 0.472380367 | 0.010114997 | 0.055121332 | 0.183504213 | 0.854402413 | NA |
| Ankrd1 | 1.798583869 | 0.034259767 | 0.089266409 | 0.383792377 | 0.701132339 | NA |
| Ankrd22 | 1.287552525 | 0.078051246 | 0.076533001 | 1.019837784 | 0.3078054 | NA |

| | | | | | | | |
|---|---|---|---|---|---|---|---|
| Clca5 | 0.411864012 | -0.000821182903821995 | 0.0119636778998887 | -0.0686396700658109 | 0.945276441167731 | NA | NM_178697 |
| Clca6 | 0.12890128 | -0.0068640067066208 | 0.00870804083610955 | -0.788237771940376 | 0.430557639008166 | NA | NM_207208 |
| Clcf1 | 17.25641078 | 0.0148717552381357 | 0.0786654571395248 | 0.189050642796856 | 0.850053125158675 | NA | NM_019952 |
| Clcnka | 2.341993474 | 0.0274669247093893 | 0.0251901851894437 | 1.09038200802508 | 0.275544903976427 | NA | NM_024412,NM_001146307 |
| Clcnkb | 1.359549035 | -0.0240792979563096 | 0.0258974249653355 | -0.929795066055425 | 0.35247720142502 | NA | NM_019701 |
| Cldn13 | 0.09572496 | 0.00498255289994511 | 0.00847999972823692 | 0.587565219295241 | 0.556824162160132 | NA | NM_020504 |
| Cldn15 | 4.06319975 | 0.00348421754864906 | 0.04444817293942 | 0.0783883187594195 | 0.937519165218383 | NA | NM_021719 |
| Cldn16 | 0.12890128 | -0.0068640067066208 | 0.00870804083610955 | -0.788237771940376 | 0.430557639008166 | NA | NM_053241 |
| Cldn17 | 0 | NA | NA | NA | NA | NA | NM_181490 |
| Cldn18 | 0.212496447 | -0.000886988073087717 | 0.00973851166817767 | -0.0910804549309223 | 0.927428662789121 | NA | NM_001194922,NM_001194923,NM_001194921,NM_019815 |
| Cldn19 | 18.47388086 | -0.0535818689092221 | 0.0752271433974717 | -0.712267759871139 | 0.476298984369399 | NA | NM_001038590,NM_153105 |
| Cldn2 | 72.39906469 | 0.00837142738093306 | 0.0268996154084545 | 0.311209928239417 | NA | NA | NM_016675 |
| Cldn20 | 8.758740003 | 0.00107860250607579 | 0.060398488813996 | 0.0178581041886241 | 0.985752051693389 | NA | NM_001101560 |
| Cldn22 | 20.96914417 | -0.0985152309500032 | 0.0779296102267431 | -1.26415659802948 | 0.206173825427487 | NA | NM_029383 |
| Cldn24 | 0 | NA | NA | NA | NA | NA | NM_001111318 |
| Cldn3 | 6.534962883 | 0.0510827846942461 | 0.048784933751805 | 1.04710165138548 | 0.295052697771231 | NA | NM_009902 |
| Cldn4 | 0.410423267 | 0.00622751620557347 | 0.0137781518810749 | 0.451984871361979 | 0.65127988120664 | NA | NM_009903 |
| Cldn6 | 1.00649821 | -0.0126453213368404 | 0.0246989511738075 | -0.511978069346135 | 0.608666358218408 | NA | NM_018777 |
| Cldn7 | 0.191449919 | 0.00665508079272897 | 0.00962405648474729 | 0.691504752001018 | 0.489248395458281 | NA | NM_001193619,NM_016887 |
| Cldn8 | 0 | NA | NA | NA | NA | NA | NM_018778 |
| Cldn9 | 4.861988429 | 0.00399138372141647 | 0.0373680144487141 | 0.106812839277143 | 0.914937461480162 | NA | NM_020293 |
| Cldnd2 | 0.855261623 | 0.0106366777616144 | 0.01889571866718 | 0.562914697713472 | 0.57349297137794 | NA | NM_028849 |
| Clec12b | 0.966868033 | 0.0283616791679565 | 0.0225523527524119 | 1.25759292076182 | 0.208539014894539 | NA | NM_027709,NM_001204223 |
| Clec1b | 1.078493332 | -0.0334379273506072 | 0.023613554054819 | -1.41604805752581 | 0.156761434455351 | NA | NM_001204253,NM_001204239,NM_019985 |
| Clec2e | 0.430558245 | -0.00274495969186159 | 0.0152720675302497 | -0.179737267820784 | 0.857358834377078 | NA | NM_153506 |
| Clec2g | 0 | NA | NA | NA | NA | NA | NM_027562,NM_001168224,NM_001168223 |
| Clec2h | 0 | NA | NA | NA | NA | NA | NM_053165 |
| Clec2i | 0.753793022 | -0.0342940459171646 | 0.0193468272712849 | -1.77259275830021 | 0.0762962125703219 | NA | NM_020257 |
| Clec3a | 1.614509012 | 0.0114097174155453 | 0.0259072338089033 | 0.440406625412252 | 0.659642627181868 | NA | NM_001007223 |
| Clec4a1 | 8.570775218 | 0.00202063128140246 | 0.0586050900928911 | 0.0344787675985088 | 0.972495373282093 | NA | NM_199311 |
| Clec4a4 | 0.097165704 | -0.00637828455464458 | 0.00848781691535724 | -0.75146349388194 | 0.452373763047372 | NA | NM_001005860 |
| Clec4b1 | 0.108766301 | 0.00519171381361946 | 0.00856003371502575 | 0.606506234257728 | 0.544178643801544 | NA | NM_001190310,NM_027218 |
| Clec4b2 | 0 | NA | NA | NA | NA | NA | NM_001004159 |
| Clec4d | 1.960996837 | 0.019232504726383 | 0.0242304961746413 | 0.793731361824565 | 0.427351838521415 | NA | NM_010819,NM_001163161 |
| Clec4e | 0.21491491 | 0.00699378003108193 | 0.00986423294990756 | 0.709003940458184 | 0.478322031551065 | NA | NM_019948 |
| Clec4f | 0.201873568 | 0.00673381845182595 | 0.00968011632508873 | 0.69563404257585 | 0.486658041207307 | NA | NM_016751 |
| Clec4g | 4.159828353 | 0.0489539863622731 | 0.0430771750583703 | 1.1364251786692 | 0.255778663055836 | NA | NM_029465 |
| Clec4n | 4.67078998 | 0.0442537224724719 | 0.0479329560570215 | 0.9232420888006 | 0.355881059601299 | NA | NM_020001,NM_001190320,NM_001190321 |
| Clec7a | 82.32206052 | -0.140446556454903 | 0.0437941460533508 | -3.20697100210171 | NA | NA | NM_020008 |
| Clec9a | 0.636822305 | -0.0223358741738281 | 0.0184472944296263 | -1.21079404131788 | 0.225974349464334 | NA | NM_001205365,NM_001205364,NM_001205363,NM_172732 |
| Clhc1 | 6.822282124 | -0.000705403635738279 | 0.0503419246877822 | -0.01401225003043 | 0.988820207885743 | NA | NM_001081099 |
| Clic3 | 5.476162645 | -0.0212163627217718 | 0.0527122189863437 | -0.402494205893105 | 0.687320351682312 | NA | NM_027085 |
| Clnk | 2.851749329 | -0.0397291678888369 | 0.0352748899102037 | -1.12627333465737 | 0.260049840956887 | NA | NM_013748 |
| Clps | 0.108766301 | 0.00519171381361946 | 0.00856003371502575 | 0.606506234257728 | 0.544178643801544 | NA | NM_025469 |
| Clpsl2 | 0 | NA | NA | NA | NA | NA | NM_001034871 |
| Clrn1 | 13.58346615 | -0.00670159906457478 | 0.0733312001897715 | -0.0913881001160751 | 0.927184216912865 | NA | NM_153386,NM_153385,NM_153384 |
| Clrn2 | 8.262242196 | -0.0598543846420894 | 0.0559357176328787 | -1.07005661453975 | 0.284593826154432 | NA | NM_001163317 |
| Clrn3 | 0.106148608 | 0.00515554063321497 | 0.00854200771766808 | 0.603551390213729 | 0.546141939141592 | NA | NM_178669 |
| Cma1 | 0.892192393 | 0.013190493865958 | 0.020530657924501 | 0.642477894009263 | 0.520562938039777 | NA | NM_010780 |
| Cma2 | 0 | NA | NA | NA | NA | NA | NM_001024714 |
| Cmah | 8.722428804 | -0.0214562061274512 | 0.0617374575559463 | -0.347539516152048 | 0.728186036732311 | NA | NM_007717,NM_001111110 |
| Cml2 | 13.95577759 | -0.00344645556447712 | 0.070977999453567 | -0.0485566737722969 | 0.961272598529589 | NA | NM_053096 |
| Cmtm1 | 0.90694462 | 0.00518926559409092 | 0.0224060069612201 | 0.231601534493513 | 0.816847515562142 | NA | NM_181990 |
| Cmtm2a | 0.326298904 | 0.00765111147689903 | 0.0103094497373689 | 0.742145475443356 | 0.457999199689495 | NA | NM_027022 |
| Cmtm2b | 0 | NA | NA | NA | NA | NA | NM_028524 |
| Cndp1 | 16.30639992 | 0.00685694252175968 | 0.0569213650812734 | 0.12046342374202 | 0.90411605223154 | NA | NM_177450 |
| Cnfn | 0.397844952 | -0.0152960324097223 | 0.0129817556207533 | -1.17827148011239 | 0.23868839386615 | NA | NM_028219,NM_001081375 |
| Cnga1 | 0 | NA | NA | NA | NA | NA | NM_007723 |
| Cnga2 | 20.97599604 | -0.00361572177070046 | 0.0789193579730329 | -0.0458153976865342 | 0.963457386118863 | NA | NM_007724 |
| Cnga3 | 6.836651156 | 0.0700174427252926 | 0.0406709944914703 | 1.72155718346098 | 0.085149765019045 | NA | NM_009918 |
| Cngb1 | 23.85274932 | 0.0770387936985594 | 0.0535013133062905 | 1.43994210492589 | NA | NA | NM_001195413,NM_145601 |
| Cngb3 | 0.855971059 | 0.019608972768116 | 0.0164360009307566 | 1.19305011302487 | 0.23284974744188 | NA | NM_013927 |
| Cnksr1 | 10.54540088 | -0.0472124652175851 | 0.0632402328344259 | -0.746557422411705 | 0.455330757567101 | NA | NM_001081047 |
| Cnn1 | 13.66317086 | -0.00220548926834625 | 0.0620245684919762 | -0.0355583170019977 | 0.971634545499874 | NA | NM_009922 |
| Cnpy1 | 3.608850288 | 0.0532832370285067 | 0.0388913726086224 | 1.37005287945773 | 0.170670395422516 | NA | NM_175651 |
| Cnr2 | 17.00650197 | -0.159607055162752 | 0.0693911688141531 | -2.30010616466513 | 0.0214422061166775 | NA | NM_009924 |
| Cntd1 | 20.50529426 | -0.0242517462767264 | 0.0789667993924298 | -0.307113197740307 | 0.758757212484146 | NA | NM_026562 |
| Cntf | 3.457793133 | -0.0184957157129453 | 0.0371737829921843 | -0.49754730953355 | 0.618803149650936 | NA | NM_170786 |
| Col10a1 | 16.75788107 | 0.0410103178462107 | 0.0759613607979167 | 0.539883928031677 | 0.58927708234819 | NA | NM_009925 |
| Col6a6 | 20.99624879 | -0.0974121150785768 | 0.0813162900910533 | -1.19794096569704 | 0.230940000118855 | NA | NM_172927,NM_001102607 |
| Col9a3 | 607.5346762 | -0.0130060341722018 | 0.0693556322418507 | -0.187526719197777 | NA | NA | NM_009936 |
| Colec10 | 0 | NA | NA | NA | NA | NA | NM_173422 |
| Colec11 | 18.23717655 | -0.073169504550434 | 0.0781670256007056 | -0.936066122359574 | 0.349239144296859 | NA | NM_027866 |
| Comp | 12.94748285 | 0.0334558344592859 | 0.0685601772859871 | 0.487977653846059 | 0.625565669321488 | NA | NM_016685 |
| Corin | 10.86330466 | 0.0556587715734955 | 0.0584287946115045 | 0.952591473837052 | 0.340797098846981 | NA | NM_001122756,NM_016869 |
| Cox7b2 | 4.835529098 | -0.0400788734922588 | 0.048866016532208 | -0.820178855091299 | 0.412114154240779 | NA | NM_030052 |
| Cox8b | 21.55887925 | 0.104178120277575 | 0.0592189377255903 | 1.75920278678954 | 0.0785430711032991 | NA | NM_007751 |
| Cox8c | 0 | NA | NA | NA | NA | NA | NM_001039049 |
| Cpa1 | 0.108766301 | 0.00519171381361946 | 0.00856003371502575 | 0.606506234257728 | 0.544178643801544 | NA | NM_025350 |
| Cpa3 | 0.212297216 | 0.00681050119636477 | 0.00973439053551308 | 0.699633035218656 | 0.484156506270654 | NA | NM_007753 |
| Cpa4 | 15.08362295 | 0.122429777388703 | 0.0738138072073823 | 1.65862976075373 | 0.0971904209659764 | NA | NM_027926 |
| Cpa5 | 2.351345804 | 0.0218561120517943 | 0.0326542769528777 | 0.669318511732296 | 0.503292320592815 | NA | NM_144537 |
| Cpa6 | 9.879624675 | 0.06816136312888 | 0.061984076452519 | 1.09965925169657 | 0.271480615236968 | NA | NM_177834 |
| Cpb1 | 3.899067489 | -0.0472149668702747 | 0.0424089047111296 | -1.113326722109 | 0.265568133793728 | NA | NM_029706 |
| Cpb2 | 0.108766301 | 0.00519171381361946 | 0.00856003371502575 | 0.606506234257728 | 0.544178643801544 | NA | NM_019775 |
| Cphx1 | 0 | NA | NA | NA | NA | NA | NM_175342 |
| Cphx2 | 0 | NA | NA | NA | NA | NA | NM_001270506 |
| Cplx4 | 2.211179868 | 0.0536267047678334 | 0.0361651643525926 | 1.48282762508693 | 0.138120214421046 | NA | NM_145493 |
| Cpn1 | 14.71494729 | -0.0529764768210039 | 0.0742921193881013 | -0.713083396426684 | 0.475794153100729 | NA | NM_030703 |
| Cpn2 | 0 | NA | NA | NA | NA | NA | NM_027904 |
| Cps1 | 17.74640388 | 0.0286945491029881 | 0.0710109499789348 | 0.404086258689685 | 0.686149289361053 | NA | NM_001080809 |
| Cpsf4l | 1.224820421 | 0.0346728917640153 | 0.0245568460104342 | 1.41194401550113 | 0.157966446184744 | NA | NM_001164532,NM_029794,NM_026682 |
| Cpt1b | 2.278823929 | -0.0204707659245476 | 0.0341381593886135 | -0.599644687679777 | 0.548743058351709 | NA | NM_009948 |
| Cpvl | 1.058559131 | 0.0138828046554868 | 0.0240159208324937 | 0.578066723000822 | 0.563219072651097 | NA | NM_027749 |
| Cpxcr1 | 0 | NA | NA | NA | NA | NA | NM_001033471 |
| Cpz | 7.936509421 | -0.0272907171991859 | 0.0419513751733935 | -0.650532124069542 | 0.515348558331862 | NA | NM_153107 |
| Cr2 | 11.59745735 | -0.00200274585791913 | 0.0642760974736533 | -0.0311584856056336 | 0.975143147513279 | NA | NM_007758 |
| Crb3 | 18.80099779 | 0.024579631780534 | 0.0682889673220923 | 0.359935619828624 | 0.718895278979618 | NA | NM_177638 |
| Crct1 | 0.732979872 | -0.0299239625492877 | 0.0180910298233121 | -1.65407734338748 | 0.0981117988672218 | NA | NM_028798 |
| Creb3l3 | 21.70365303 | -0.0617244228693033 | 0.0792251792785366 | -0.77910108164344 | 0.435920173026188 | NA | NM_145365 |
| Creb3l4 | 12.6386198 | -0.000373165379278152 | 0.0712732445237699 | -0.00523570074256547 | 0.995822534298371 | NA | NM_030080 |
| Crisp1 | 0.09572496 | 0.00498255289994511 | 0.00847999972823692 | 0.587565219295241 | 0.556824162160132 | NA | NM_009638 |
| Crisp2 | 0.09572496 | 0.00498255289994511 | 0.00847999972823692 | 0.587565219295241 | 0.556824162160132 | NA | NM_001204071,NM_009420 |
| Crisp3 | 0 | NA | NA | NA | NA | NA | NM_009639 |
| Crisp4 | 0 | NA | NA | NA | NA | NA | NM_030033 |
| Crnn | 0 | NA | NA | NA | NA | NA | NM_001081200 |
| Crp | 0 | NA | NA | NA | NA | NA | NM_007768 |
| Crtam | 3.510511234 | -0.000116080497733719 | 0.0441774921004274 | -0.00262759365040091 | 0.997903486006757 | NA | NM_019465 |
| Crx | 1.113887639 | 0.019695619373771 | 0.0222744145753216 | 0.884226128914396 | 0.376574156104941 | NA | NM_007770,NM_001113330 |
| Crxos1 | 5.224075751 | -0.0248203675938443 | 0.0463410435625051 | -0.535602258511215 | 0.592233467517969 | NA | NM_001205274,NM_001033638,NR_038092,NM_001145190 |
| Cryaa | 3.18609695 | -0.0245920547867287 | 0.0374850283105617 | -0.656050052383171 | 0.511791920101087 | NA | NM_013501 |
| Cryba1 | 0.860872084 | -0.00354288055139853 | 0.0207833891619826 | -0.170466930286772 | 0.864642939502643 | NA | NM_009965 |
| Cryba2 | 10.37114946 | -0.0177746836571972 | 0.0620080889528398 | -0.286651047586965 | 0.774379523526174 | NA | NM_021541 |
| Cryba4 | 20.49017498 | 0.0430573585244259 | 0.0768289001191533 | 0.560431796598007 | 0.575184948490693 | NA | NM_021351 |
| Crybb2 | 0.127687691 | 0.00542298404407807 | 0.00869724871942359 | 0.62352868349814 | 0.532937159177571 | NA | NM_007773 |
| Cryga | 13.75682525 | -0.0688274108766363 | 0.0742905004977249 | -0.926463146909935 | 0.354205340623916 | NA | NM_007774 |
| Crygb | 0.108766301 | 0.00519171381361946 | 0.00856003371502575 | 0.606506234257728 | 0.544178643801544 | NA | NM_144761 |
| Crygc | 0 | NA | NA | NA | NA | NA | NM_001082573,NM_007775 |
| Crygd | 0 | NA | NA | NA | NA | NA | NM_007776 |
| Cryge | 0.097165704 | -0.00637828455464458 | 0.00848781691535724 | -0.75146349388194 | 0.452373763047372 | NA | NM_007777 |
| Crygf | 0 | NA | NA | NA | NA | NA | NM_027010 |
| Crygn | 11.07150858 | -0.0570748448542705 | 0.0611443781060299 | -0.933443868793588 | 0.350590839144942 | NA | NM_153076 |
| Crygs | 0.530631763 | 0.0226984517873837 | 0.0176918220882368 | 1.28299118508973 | 0.199495164357502 | NA | NM_009967 |
| Csf2 | 0.323681211 | 0.00985023388331581 | 0.0116938364156777 | 0.842344080520041 | 0.399595383885569 | NA | NM_009969 |
| Csf3 | 0.097165704 | -0.00637828455464458 | 0.00848781691535724 | -0.75146349388194 | 0.452373763047372 | NA | NM_009971 |
| Csl | 0 | NA | NA | NA | NA | NA | NM_027945 |
| Csn1s1 | 0.323880442 | 0.00225021938481196 | 0.0110697885432795 | 0.203275733408482 | 0.838919521495622 | NA | NM_007784 |
| Csn1s2a | 0 | NA | NA | NA | NA | NA | NM_007785 |
| Csn1s2b | 0.108766301 | 0.00519171381361946 | 0.00856003371502575 | 0.606506234257728 | 0.544178643801544 | NA | NM_009973 |
| Csn2 | 0.53223091 | 0.00711496316961995 | 0.0130511675608932 | 0.545159131274919 | 0.585644076975072 | NA | NM_009972 |
| Csn3 | 0 | NA | NA | NA | NA | NA | NM_007786 |
| Csnka2ip | 1.260580072 | 0.0162669137153747 | 0.0248376018168337 | 0.654930932355547 | 0.51251222362967 | NA | NM_173861 |
| Csrp3 | 0.224626239 | -0.00168521210467683 | 0.00978955093367521 | -0.172143964119932 | 0.863324350255464 | NA | NM_001198841,NM_013808 |
| Cst10 | 0 | NA | NA | NA | NA | NA | NM_021405 |
| Cst11 | 0.91583343 | -0.0147855486074796 | 0.0223722397795399 | -0.660888170034788 | 0.50868403320743 | NA | NM_030059 |
| Cst12 | 0.346765936 | -0.0134445992440418 | 0.0121756794716388 | -1.104217573677 | 0.26949876817541 | NA | NM_027054 |
| Cst13 | 0 | NA | NA | NA | NA | NA | NM_027024 |
| Cst8 | 0 | NA | NA | NA | NA | NA | NM_009978 |
| Cst9 | 0 | NA | NA | NA | NA | NA | NM_009979 |
| Csta | 0 | NA | NA | NA | NA | NA | NM_001033239 |
| Cstl1 | 8.062861158 | 0.00818906295178319 | 0.0486818651270913 | 0.168215883479493 | 0.866413445786805 | NA | NM_177655 |
| Ctag2 | 0 | NA | NA | NA | NA | NA | NM_027302 |
| Ctcfl | 18.0840661 | -0.0640379958779518 | 0.076517055840602 | -0.836911394125694 | 0.402642375230371 | NA | NM_001081387 |
| Ctf2 | 0 | NA | NA | NA | NA | NA | NM_198858 |
| Ctla4 | 3.128608248 | 0.018038706604781 | 0.0407541979088865 | 0.442622049515239 | 0.658039133369348 | NA | NM_009843 |
| Ctrb1 | 0.567655227 | 0.00309433163276112 | 0.0165734891223922 | 0.186703693465513 | 0.851892960057586 | NA | NM_025583 |
| Ctrc | 0 | NA | NA | NA | NA | NA | NM_001033875 |
| Cts3 | 0 | NA | NA | NA | NA | NA | NM_054092,NM_026906 |
| Cts6 | 0 | NA | NA | NA | NA | NA | NM_021445 |
| Cts7 | 0 | NA | NA | NA | NA | NA | NM_019539 |
| Cts8 | 0 | NA | NA | NA | NA | NA | NM_019541 |
| Cts8-ps | 0 | NA | NA | NA | NA | NA | NR_027871 |
| Ctse | 2.249497915 | -0.021800667019664 | 0.025160026807829 | -0.86648027786999 | 0.38622683718935 | NA | NM_007799 |
| Ctsg | 0.203513543 | -0.00860782669689374 | 0.00975338121443284 | -0.882547960306941 | 0.377480556050336 | NA | NM_007800 |
| Ctsj | 0 | NA | NA | NA | NA | NA | NM_012007 |
| Ctsll3 | 0.127687691 | 0.00542298404407807 | 0.00869724871942359 | 0.62352868349814 | 0.532937159177571 | NA | NM_027344 |
| Ctsm | 0 | NA | NA | NA | NA | NA | NM_022326 |
| Ctsq | 0 | NA | NA | NA | NA | NA | NM_029636 |
| Ctsr | 0 | NA | NA | NA | NA | NA | NM_020284 |
| Ctsw | 4.950156864 | 0.00134912259941364 | 0.0452648549020124 | 0.0298050795111166 | 0.976222507703896 | NA | NM_009985 |
| Cuzd1 | 10.4711266 | -0.044970971517572 | 0.0644729200698387 | -0.697517212945502 | 0.485479171042842 | NA | NM_008411 |
| Cxcl1 | 3.190413697 | -0.0745578868854331 | 0.040010848764895 | -1.86344177109408 | 0.0624001377496979 | NA | NM_008176 |
| Cxcl11 | 0.938461913 | 0.000340360790670743 | 0.0215905221289374 | 0.0157643612617669 | 0.987422380495174 | NA | NM_019494,NR_038116 |
| Cxcl13 | 0.737471969 | -0.0143732124242958 | 0.0181357300241403 | -0.792535641254235 | 0.428048418433195 | NA | NM_018866 |
| Cxcl17 | 0.09572496 | 0.00498255289994511 | 0.00847999972823692 | 0.587565219295241 | 0.556824162160132 | NA | NM_153576 |
| Cxcl2 | 0 | NA | NA | NA | NA | NA | NM_009140 |
| Cxcl3 | 0 | NA | NA | NA | NA | NA | NM_203320 |
| Cxcl5 | 7.958610589 | -0.124850849537058 | 0.0606005988643981 | -2.06022468220865 | 0.0393770663608805 | NA | NM_009141 |
| Cxcl9 | 13.8710099 | -0.0216308666048536 | 0.0159947797461164 | -1.3523703951038 | NA | NA | NM_008599 |
| Cxcr1 | 0 | NA | NA | NA | NA | NA | NM_178241 |
| Cxcr2 | 19.7530006 | -0.0448419287783039 | 0.0740727158732955 | -0.605377138527065 | 0.544928437652292 | NA | NM_009909 |
| Cxcr3 | 2.753435642 | -0.0360883312413011 | 0.0348229454371525 | -1.03633770171545 | 0.300044623103032 | NA | NM_009910 |
| Cxcr5 | 0 | NA | NA | NA | NA | NA | NM_007551 |
| Cxcr6 | 1.574834332 | -0.0257964342698038 | 0.0259738937934911 | -0.993167773569178 | 0.320628200362407 | NA | NM_030712 |
| Cyb5r2 | 16.1171627 | -0.0599562896763565 | 0.0759211611281847 | -0.789717764920992 | 0.429692614211455 | NA | NM_001205227,NM_177216 |
| Cyct | 0.227686571 | 0.00691460391109036 | 0.00980755285755433 | 0.705028462402254 | 0.480792523671872 | NA | NM_009989 |
| Cylc1 | 0 | NA | NA | NA | NA | NA | NM_026134 |
| Cylc2 | 0 | NA | NA | NA | NA | NA | NM_001162865 |
| Cym | 1.889006502 | -0.00474098199211666 | 0.0327500769159704 | -0.144762468933462 | 0.88489841547055 | NA | NM_001111143 |
| Cyp11b1 | 0.565118408 | -0.00667244984124853 | 0.014900288684848 | -0.447806749411082 | 0.654292671926396 | NA | NM_001033229 |
| Cyp11b2 | 0.48121521 | -0.00269058588389411 | 0.0141642517458129 | -0.189956090316559 | 0.849343538826259 | NA | NM_009991 |
| Cyp17a1 | 0.375584431 | 0.00259421861314003 | 0.0115885238266724 | 0.223861006970458 | 0.822865457651262 | NA | NM_007809 |
| Cyp19a1 | 13.20762524 | 0.0529173456332002 | 0.064472185674928 | 0.82077790723603 | 0.411772784933843 | NA | NM_007810 |
| Cyp1a1 | 4.659616567 | 0.0234060996550871 | 0.0481055594676434 | 0.486557061472913 | 0.626572258729862 | NA | NM_009992,NM_001136059 |
| Cyp1a2 | 0.205932006 | -0.000576954122277892 | 0.00970424421944593 | -0.0594537925088239 | 0.952590668569529 | NA | NM_009993 |
| Cyp21a1 | 1.873735045 | -0.0437625893430573 | 0.0287666401649714 | -1.5212965119349 | 0.128185445192809 | NA | NM_009995 |
| Cyp24a1 | 2.725777577 | 0.00190085858948859 | 0.0337313727879135 | 0.0563528380964589 | 0.955060726959779 | NA | NM_009996 |
| Cyp26a1 | 5.824360881 | 0.0166719038681681 | 0.0437147322606712 | 0.381379525985735 | 0.702921650996075 | NA | NM_007811 |
| Cyp26c1 | 0.394963462 | -0.00200109807023417 | 0.0134533619193562 | -0.148743346252736 | 0.881756156306212 | NA | NM_001105201 |
| Cyp27b1 | 13.11496418 | -0.0635950575893194 | 0.0684544754915312 | -0.929012414932418 | 0.35288265242173 | NA | NM_010009 |
| Cyp2a12 | 0 | NA | NA | NA | NA | NA | NM_133657 |
| Cyp2a22 | 0 | NA | NA | NA | NA | NA | NM_001101467 |
| Cyp2a4 | 0.240418098 | -0.0088029805258837 | 0.00986414519853158 | -0.892422034419582 | 0.372166765031741 | NA | NM_009997 |
| Cyp2ab1 | 0.680584311 | -0.00304766953677561 | 0.0190500482702324 | -0.159982247474821 | 0.872895058553006 | NA | NM_183158 |
| Cyp2b10 | 2.206759211 | 0.0380750768847593 | 0.0309883836464235 | 1.22868870216642 | 0.219188543720473 | NA | NM_009999 |
| Cyp2b13 | 0 | NA | NA | NA | NA | NA | NM_007813 |
| Cyp2b19 | 0 | NA | NA | NA | NA | NA | NM_007814 |
| Cyp2b23 | 2.310920538 | 0.0709973299411223 | 0.0325493412920558 | 2.18122171211036 | 0.0291670216511208 | NA | NM_001081148 |
| Cyp2b9 | 0 | NA | NA | NA | NA | NA | NM_010000 |
| Cyp2c29 | 0 | NA | NA | NA | NA | NA | NM_007815 |
| Cyp2c37 | 0 | NA | NA | NA | NA | NA | NM_010001 |
| Cyp2c38 | 0 | NA | NA | NA | NA | NA | NM_010002 |
| Cyp2c39 | 0 | NA | NA | NA | NA | NA | NM_010003 |
| Cyp2c40 | 0 | NA | NA | NA | NA | NA | NM_010004 |
| Cyp2c44 | 8.971566109 | -0.00214359792344029 | 0.0622114333290058 | -0.0344566554527663 | 0.972513005744788 | NA | NM_001167905,NM_001001446 |
| Cyp2c50 | 0 | NA | NA | NA | NA | NA | NM_001167877,NM_134144,NM_001167875 |
| Cyp2c53-ps | 0 | NA | NA | NA | NA | NA | NR_033614 |
| Cyp2c54 | 0 | NA | NA | NA | NA | NA | NM_206537 |
| Cyp2c55 | 0.884519814 | -0.0131100391088587 | 0.0208438272750339 | -0.628965061736119 | 0.529371929775454 | NA | NM_028089 |
| Cyp2c65 | 0.201873568 | 0.00673381845182595 | 0.00968011632508873 | 0.69563404257585 | 0.486658041207307 | NA | NM_028191 |
| Cyp2c66 | 0 | NA | NA | NA | NA | NA | NM_001011707 |
| Cyp2c67 | 0 | NA | NA | NA | NA | NA | NM_001024719 |
| Cyp2c68 | 0 | NA | NA | NA | NA | NA | NM_001039555 |
| Cyp2c69 | 0 | NA | NA | NA | NA | NA | NM_001104525 |
| Cyp2c70 | 0.108766301 | 0.00519171381361946 | 0.00856003371502575 | 0.606506234257728 | 0.544178643801544 | NA | NM_145499 |
| Cyp2d10 | 0 | NA | NA | NA | NA | NA | NM_010005 |
| Cyp2d11 | 0 | NA | NA | NA | NA | NA | NM_001104531 |
| Cyp2d12 | 0 | NA | NA | NA | NA | NA | NM_201360 |
| Cyp2d13 | 0 | NA | NA | NA | NA | NA | NR_003552 |
| Cyp2d26 | 2.185797528 | -0.0228254030174323 | 0.0283712761317233 | -0.804525073579969 | 0.421093794632035 | NA | NM_029562 |
| Cyp2d34 | 0 | NA | NA | NA | NA | NA | NM_145474 |
| Cyp2d37-ps | 0 | NA | NA | NA | NA | NA | NR_033515 |
| Cyp2d40 | 0 | NA | NA | NA | NA | NA | NM_023623 |
| Cyp2d9 | 0 | NA | NA | NA | NA | NA | NM_010006 |
| Cyp2e1 | 21.62497288 | -0.0870980068138807 | 0.0723345556676982 | -1.2040995622342 | 0.228551099122422 | NA | NM_021282 |
| Cyp2f2 | 8.192871199 | -0.00738230388376702 | 0.0546423058128284 | -0.13510234925031 | 0.892530952776582 | NA | NM_007817 |
| Cyp2g1 | 4.907342637 | -9.30636236406112e-05 | 0.0138761663341821 | -0.00670672442224631 | 0.994648848245897 | NA | NM_013809 |
| Cyp2j11 | 0 | NA | NA | NA | NA | NA | NM_001004141 |
| Cyp2j13 | 2.577506032 | 0.0382991483358695 | 0.0306421754911226 | 1.24988346036218 | 0.21134212215109 | NA | NM_145548 |
| Cyp2j5 | 0 | NA | NA | NA | NA | NA | NM_010007 |
| Cyp2j8 | 3.484359187 | 0.0395538712261838 | 0.0426562206815825 | 0.927270878530075 | 0.353785910365012 | NA | NM_001104927 |
| Cyp2r1 | 8.969278955 | -0.0327048428134396 | 0.0602002508931317 | -0.543267550022302 | 0.586945603184656 | NA | NM_177382 |
| Cyp2t4 | 0 | NA | NA | NA | NA | NA | NM_001100184 |
| Cyp2w1 | 0.127687691 | 0.00542298404407807 | 0.00869724871942359 | 0.62352868349814 | 0.532937159177571 | NA | NM_001160265 |
| Cyp3a11 | 0 | NA | NA | NA | NA | NA | NM_007818 |

| | | | | | | | | | | | | | | | |
|---|---|---|---|---|---|---|---|---|---|---|---|---|---|---|---|
| Ankrd33 | 0.563199176 | -0.006483013 | 0.052564167 | -0.123335213 | 0.901841657 | NA | | Cyp3a13 | 20.35255979 | -0.0966025421805359 | 0.0697705343446434 | -1.3845750657914 | 0.166182440107021 | NA | NM_007819 |
| Ankrd36 | 0.116696964 | -0.020477583 | 0.027954224 | -0.732539847 | 0.463839131 | NA | | Cyp3a16 | 0 | NA | NA | NA | NA | NA | NM_007820 |
| Ankrd53 | 0 | NA | NA | NA | NA | NA | | Cyp3a25 | 0 | NA | NA | NA | NA | NA | NR_030782,NM_019792 |
| Ankrd60 | 0.096071043 | -0.020098795 | 0.027825948 | -0.722304036 | 0.470107572 | NA | | Cyp3a41a | 0 | NA | NA | NA | NA | NA | NM_017396 |
| Ankrd61 | 2.998291199 | -0.07531032 | 0.118007854 | -0.638180576 | 0.52335614 | NA | | Cyp3a41b | 0 | NA | NA | NA | NA | NA | NM_001105159 |
| Ankrd7 | 2.336743905 | -0.076060354 | 0.113097251 | -0.672521688 | 0.501251638 | NA | | Cyp3a44 | 0 | NA | NA | NA | NA | NA | NM_177380 |
| Anks4b | 7.515308668 | 0.121738445 | 0.144283321 | 0.843745797 | 0.39881147 | NA | | Cyp3a57 | 0.406628625 | -0.000921993720139942 | 0.0119199762705297 | -0.0773486204347094 | 0.938346213215504 | NA | NM_001100180 |
| Ano9 | 0.54939413 | 0.01643428 | 0.052034848 | 0.315832191 | 0.752129883 | NA | | Cyp3a59 | 0 | NA | NA | NA | NA | NA | NM_001105160 |
| Antxrl | 0 | NA | NA | NA | NA | NA | | Cyp4a10 | 0 | NA | NA | NA | NA | NA | NM_010011 |
| Anxa10 | 2.311589427 | -0.178605716 | 0.108068424 | -1.65270955 | 0.09838999 | NA | | Cyp4a12a | 0 | NA | NA | NA | NA | NA | NM_177406 |
| Anxa13 | 1.378542077 | -0.040044204 | 0.077213795 | -0.518614634 | 0.604029502 | NA | | Cyp4a12b | 3.071682626 | 0.0145344933265169 | 0.0319292354143028 | 0.455209563834597 | 0.648958480691893 | NA | NM_172306 |
| Aox3l1 | 0.48472991 | 0.029813289 | 0.045640658 | 0.653217767 | 0.513615898 | NA | | Cyp4a14 | 0 | NA | NA | NA | NA | NA | NM_007822 |
| Apcs | 0 | NA | NA | NA | NA | NA | | Cyp4a29-ps | 0 | NA | NA | NA | NA | NA | NM_001100183 |
| Apoa4 | 0.215050351 | -0.027006021 | 0.031907566 | -0.846382992 | 0.397339123 | NA | | Cyp4a30b-ps | 0 | NA | NA | NA | NA | NA | NM_001100185 |
| Apoa5 | 0.107716603 | -0.020477583 | 0.027954224 | -0.732539849 | 0.46383913 | NA | | Cyp4a31 | 0 | NA | NA | NA | NA | NA | NM_001252539,NM_201640 |
| Apob | 1.711512134 | -0.016972912 | 0.088064362 | -0.192733039 | 0.847168047 | NA | | Cyp4a32 | 0 | NA | NA | NA | NA | NA | NM_001100181 |
| Apobec2 | 2.468537556 | 0.048974147 | 0.110988758 | 0.441253219 | 0.659029689 | NA | | Cyp4f18 | 1.396185696 | -0.0149232372878576 | 0.0231267602717999 | -0.645280061386487 | 0.518745711303114 | NA | NM_024444 |
| Apobec4 | 0.493412615 | 0.00310454 | 0.044827045 | 0.069255952 | 0.944785887 | NA | | Cyp4f39 | 0.745705075 | 0.022826690232968 | 0.0177236619796887 | 1.28792177706431 | 0.197773194015859 | NA | NM_177307 |
| Apoc2 | 0.912251645 | 0.081212828 | 0.06270407 | 1.295176341 | 0.195259404 | NA | | Cyp4f40 | 0.113843286 | 0.00525947738444006 | 0.00859368002043176 | 0.612016897526494 | 0.540526577873086 | NA | NM_001101588 |
| Apoc4 | 0.405897065 | -0.027138935 | 0.04364914 | -0.621751886 | 0.534105028 | NA | | Cyp4f41-ps | 4.719036762 | 0.0306598921851193 | 0.0501055678277781 | 0.611905892185534 | 0.540600023002553 | NA | NR_033585 |
| Apod | 10116.30964 | 0.302350263 | 0.144000193 | 2.099651796 | NA | NA | | Cyp7a1 | 0 | NA | NA | NA | NA | NA | NM_007824 |
| Apof | 3.089470404 | -0.107454105 | 0.121800419 | -0.882214582 | 0.377660778 | NA | | Cyp8b1 | 0 | NA | NA | NA | NA | NA | NM_010012 |
| Apol10a | 0.507828167 | -0.069254262 | 0.050465732 | -1.372302744 | 0.169969212 | NA | | Cypt1 | 0 | NA | NA | NA | NA | NA | NM_025738 |
| Apol11a | 0 | NA | NA | NA | NA | NA | | Cypt10 | 0 | NA | NA | NA | NA | NA | NM_001039944 |
| Apol11b | 1.366416063 | -0.020011337 | 0.07350953 | -0.272227785 | 0.785446879 | NA | | Cypt12 | 0.314540458 | -0.0125494430549186 | 0.0117659602535877 | -1.06658893829698 | 0.286157497745293 | NA | NM_029289 |
| Apol6 | 5.585729203 | 0.268061819 | 0.136635685 | 1.961872689 | 0.049777308 | NA | | Cypt14 | 0 | NA | NA | NA | NA | NA | NM_001191032 |
| Apol7b | 4.031407462 | 0.29226632 | 0.12256045 | 2.384670752 | 0.017094413 | NA | | Cypt15 | 0 | NA | NA | NA | NA | NA | NM_001177380 |
| Apol7c | 0 | NA | NA | NA | NA | NA | | Cypt2 | 0 | NA | NA | NA | NA | NA | NM_173436 |
| Apol7d | 0.781403623 | -0.009451213 | 0.067897486 | -0.139198272 | 0.889293474 | NA | | Cypt3 | 0 | NA | NA | NA | NA | NA | NM_173367 |
| Apol7e | 0 | NA | NA | NA | NA | NA | | Cypt4 | 0 | NA | NA | NA | NA | NA | NM_173412 |
| Apol9a | 3.435832313 | 0.074800035 | 0.131540197 | 0.568647736 | 0.569595223 | NA | | Cypt7 | 0 | NA | NA | NA | NA | NA | NM_001039943 |
| Apol9b | 5.046188569 | -0.118540898 | 0.135451484 | -0.875153922 | 0.381490161 | NA | | Cypt8 | 0 | NA | NA | NA | NA | NA | NM_001039941 |
| Apom | 1.537810199 | -0.145311625 | 0.097659617 | -1.487939734 | 0.136766788 | NA | | Cypt9 | 0 | NA | NA | NA | NA | NA | NM_001039942 |
| Apon | 0.668229972 | 0.034229313 | 0.060509279 | 0.565687011 | 0.571606567 | NA | | Cysltr1 | 21.7105504 | -0.0301867502485388 | 0.0845454841166595 | -0.357047458701471 | 0.721056277221643 | NA | NM_021476 |
| Apoo-ps | 0 | NA | NA | NA | NA | NA | | Cysltr2 | 7.910038257 | -0.0563369723260185 | 0.0602358653460656 | -0.935272897672387 | 0.349647678830253 | NA | NM_133720,NM_001162412 |
| Aqp12 | 0 | NA | NA | NA | NA | NA | | Cytip | 6.595034199 | -0.081414396764822 | 0.0489152247911395 | -1.66439788659766 | 0.096032942141842 | NA | NM_139200 |
| Aqp2 | 1.788037816 | -0.082425166 | 0.10200462 | -0.808053268 | 0.419059918 | NA | | D030018L15Rik | 0.354244579 | -0.00570556803240082 | 0.0130207392936895 | -0.438190789609467 | 0.661247984684689 | NA | NR_003627 |
| Aqp3 | 1.584931511 | -0.02089577 | 0.09401094 | -0.22226956 | 0.824104047 | NA | | D030024E09Rik | 0 | NA | NA | NA | NA | NA | NR_040350 |
| Aqp5 | 7.883463462 | -0.118305929 | 0.157776108 | -0.74983425 | 0.453354538 | NA | | D030025E07Rik | 0.333619697 | 0.00358748059577537 | 0.0127060559346679 | 0.282344152601052 | 0.77767963325269 | NA | NR_045704 |
| Aqp7 | 0.967893051 | 0.018368493 | 0.072823062 | 0.252234557 | 0.800859769 | NA | | D030025P21Rik | 0.22897535 | -0.00116364198856673 | 0.00981572003153595 | -0.118548816065269 | 0.905632818135836 | NA | NR_028577 |
| Aqp8 | 0.312649645 | 0.012040436 | 0.039723828 | 0.303103628 | 0.761810889 | NA | | D030040B21Rik | 0.300679247 | -0.0121548274246918 | 0.0115806462900197 | -1.04958109593304 | 0.293910752236529 | NA | NR_037998 |
| Areg | 5.148644673 | -0.121385393 | 0.115016147 | -1.055376966 | 0.291252953 | NA | | D030045P18Rik | 1.073374414 | 0.0108962625077468 | 0.0221101494955679 | 0.492817224502759 | 0.622141739056699 | NA | NR_040624 |
| Arg1 | 1.344317085 | 0.139473702 | 0.085254256 | 1.635973488 | 0.101845136 | NA | | D030047H15Rik | 4.749439604 | -0.0487491691048526 | 0.0470518672709482 | -1.03607299629854 | 0.300168089059046 | NA | NR_033548 |
| Arhgap8 | 5.170951952 | 0.051176054 | 0.139942827 | 0.365692586 | 0.714594479 | NA | | D130009I18Rik | 0 | NA | NA | NA | NA | NA | NR_015593 |
| Arhgef11 | 4184.845402 | 0.070485036 | 0.09767587 | 0.721621794 | NA | NA | | D130058E03 | 0 | NA | NA | NA | NA | NA | NR_073373 |
| Arhgef38 | 0.336022005 | 0.013628574 | 0.035131903 | 0.387925872 | 0.698070891 | NA | | D16Ertd519e | 0.12890128 | -0.0068640067066208 | 0.00870804083610955 | -0.788237771940376 | 0.430557639008166 | NA | NR_040474 |
| Arid5b | 941.6196678 | 0.286301178 | 0.133880563 | 2.138482028 | NA | NA | | D17Ertd648e | 0.451410283 | -0.0118078003715272 | 0.0158420142559 | -0.745347162348983 | 0.456061876943773 | NA | NR_045808 |
| Arl13a | 0.107716603 | -0.020477583 | 0.027954224 | -0.732539849 | 0.46383913 | NA | | D1Pas1 | 0 | NA | NA | NA | NA | NA | NM_033077 |
| Arl14 | 0 | NA | NA | NA | NA | NA | | D230030E09Rik | 0.106347839 | -0.00653347234140708 | 0.00854389509723638 | -0.764694822098226 | 0.444453290197145 | NA | NR_045947 |
| Arl14epl | 0.211089781 | -0.002448206 | 0.031868916 | -0.076821119 | 0.93876585 | NA | | D330022K07Rik | 1.990296802 | -0.0448012978301017 | 0.0312144280064401 | -1.43527530989382 | 0.151208660581962 | NA | NR_045739 |
| Arl9 | 3.583664441 | -0.020384064 | 0.129789218 | -0.157055144 | 0.875201389 | NA | | D330045A20Rik | 10.59280126 | 0.0315536878502971 | 0.0618926929641518 | 0.509812812129098 | 0.610182608836098 | NA | NM_175326 |
| Armc12 | 0.664808826 | 0.07862935 | 0.054816974 | 1.434397841 | 0.151458764 | NA | | D4Ertd617e | 17.53492791 | -0.0469213352648357 | 0.0713302298828021 | -0.657804346655394 | 0.510663860991653 | NA | NR_029469 |
| Arrdc5 | 0.098353386 | -0.020212831 | 0.02786466 | -0.725393061 | 0.468210929 | NA | | D530049I02Rik | 0 | NA | NA | NA | NA | NA | NR_040605 |
| Art1 | 1.104411432 | 0.039149796 | 0.074125616 | 0.528154746 | 0.597391936 | NA | | D5Ertd577e | 0 | NA | NA | NA | NA | NA | NM_177187 |
| Art2a-ps | 0 | NA | NA | NA | NA | NA | | D5Ertd605e | 1.902378653 | 0.000644154829876656 | 0.0291804546638594 | 0.0220748729688045 | 0.982388230057419 | NA | NR_033625 |
| Art2b | 0 | NA | NA | NA | NA | NA | | D630010B17Rik | 1.834930761 | -0.00509731421444458 | 0.0290841434100557 | -0.175260936606515 | 0.860874604715359 | NA | NR_045629 |
| Art4 | 3.142597282 | -0.072302224 | 0.091704501 | -0.788426123 | 0.430447496 | NA | | D630013N20Rik | 4.141915698 | -0.0330008722833067 | 0.0398080008962221 | -0.829000993276169 | 0.407103845174123 | NA | NR_045291,NR_045292 |
| Art5 | 0.685174529 | -0.047276997 | 0.061327002 | -0.77090018 | 0.440766101 | NA | | D630029K05Rik | 2.002195999 | 0.0365516117024279 | 0.0311797178137021 | 1.17228808550554 | 0.241081413465289 | NA | NR_027847,NR_027846 |
| Asb12 | 0.634307402 | -0.006355127 | 0.053469976 | -0.118854125 | 0.905390927 | NA | | D630032N06Rik | 6.672860791 | -0.0746900123620799 | 0.0555428195404005 | -1.34472849920325 | 0.178712927719529 | NA | NR_028329 |
| Asb14 | 3.916359886 | 0.143258216 | 0.136141293 | 1.052276007 | 0.292672936 | NA | | D630033O11Rik | 3.62288829 | -0.0542734428743486 | 0.0403838924396574 | -1.3439378820515 | 0.178968474797252 | NA | NM_001243261 |
| Asb15 | 2.114224913 | 0.009215635 | 0.096415217 | 0.095582784 | 0.923851939 | NA | | D630039A03Rik | 12.6205393 | 0.0724318535481491 | 0.0643702836962879 | 1.12523744481067 | 0.260488429692097 | NA | NM_178727 |
| Asb17 | 0.099009884 | 0.018205726 | 0.027896865 | 0.652608315 | 0.514008823 | NA | | D630045M09Rik | 0.226556888 | -0.00933175792380659 | 0.010153338242423 | -0.919082739193727 | 0.358052296835844 | NA | NR_045293 |
| Asb9 | 0.498228802 | 0.006953757 | 0.044429446 | 0.156512357 | 0.875629179 | NA | | D6Ertd474e | 20.77966601 | 0.0428980749094522 | 0.0763065181990336 | 0.562181002644614 | 0.573992703448005 | NA | NR_027803 |
| Ascl2 | 7.976169801 | 0.042936466 | 0.148797997 | 0.288555405 | 0.772921626 | NA | | D6Ertd527e | 2.380519081 | -0.0126046812670561 | 0.0321176822047053 | -0.392453016588148 | 0.694723519840731 | NA | NM_001167938,NM_001167937 |
| Ascl3 | 3.163888827 | 0.043878647 | 0.11099558 | 0.395318869 | 0.692607576 | NA | | D730001G18Rik | 0.106347839 | -0.00653347234140708 | 0.00854389509723638 | -0.764694822098226 | 0.444453290197145 | NA | NR_027836,NR_027837 |
| Ascl4 | 5.229341579 | 0.086470426 | 0.133779624 | 0.6463647 | 0.518043196 | NA | | D730005E14Rik | 5.125161974 | -0.0359037725001626 | 0.0417565641932443 | -0.859835410164597 | 0.38987977684852 | NA | NR_030675 |
| Asgr2 | 3.05986852 | 0.106795161 | 0.108520943 | 0.984097239 | 0.325067704 | NA | | D730039F16Rik | 21.61974239 | 0.13497307795917 | 0.0834285622277553 | 1.61782816765679 | 0.105699634202447 | NA | NM_030021 |
| Asic5 | 0.20210942 | -0.001755803 | 0.031727492 | -0.055340098 | 0.955867518 | NA | | D730045A05Rik | 0.781166032 | 0.00116510353360099 | 0.0213324113912195 | 0.0546165884500312 | 0.956443922812246 | NA | NR_045390 |
| Astl | 0.183387883 | -0.001444816 | 0.031390672 | -0.046026918 | 0.963288796 | NA | | D730048I06Rik | 0.108766301 | 0.00519171381361946 | 0.00856003371502575 | 0.606506234257728 | 0.544178643801544 | NA | NR_026593 |
| Asz1 | 0.105923158 | 0.018524693 | 0.027977518 | 0.662127842 | 0.507889294 | NA | | D730050B12Rik | 0 | NA | NA | NA | NA | NA | NR_046196 |
| Atf7ip2 | 0.776419309 | -0.033338817 | 0.062710421 | -0.531631219 | 0.594981438 | NA | | D7Ertd143e | 0 | NA | NA | NA | NA | NA | NR_028425 |
| Atoh1 | 0 | NA | NA | NA | NA | NA | | D830005E20Rik | 0 | NA | NA | NA | NA | NA | NR_040657 |
| Atp12a | 4.689740524 | -0.232568477 | 0.128532634 | -1.809411894 | 0.070387037 | NA | | D830013O20Rik | 0.41620922 | -0.0157695646389328 | 0.0131796358585653 | -1.19650988905618 | 0.231497641828105 | NA | NR_046013 |
| Atp1a4 | 3.813822435 | 0.092473371 | 0.132323449 | 0.698843416 | 0.484649892 | NA | | D830015G02Rik | 12.83432344 | -0.0620044191608555 | 0.068817871501052 | -0.900992980579292 | 0.367592051865354 | NA | NR_033497 |
| Atp1b4 | 0 | NA | NA | NA | NA | NA | | D830046C22Rik | 1.849948845 | -0.0108265967391137 | 0.0221914741771179 | -0.487871903087773 | 0.625640577145218 | NA | NR_033147 |
| Atp4b | 0 | NA | NA | NA | NA | NA | | D930007P13Rik | 0.192890664 | -0.000903864895041124 | 0.00963528282151475 | -0.093807821917056 | 0.92526181809679 | NA | NR_045743 |
| Atp6v0c | 2.314865729 | -0.012024494 | 0.114414417 | -0.10509597 | 0.916299657 | NA | | D930015M05Rik | 4.88896531 | -0.0403506949106179 | 0.0470673377800811 | -0.857297157938988 | 0.391280678811942 | NA | NR_040621 |
| Atp6v0c-ps2 | 0 | NA | NA | NA | NA | NA | | D930020B18Rik | 16.84389861 | -0.0108807559860489 | 0.0737400342906198 | -0.147555613320796 | 0.882693486978748 | NA | NM_177335 |
| Atp6v0d2 | 0.451040743 | 0.02612425 | 0.044851076 | 0.582466505 | 0.560252497 | NA | | D930028M14Rik | 4.488474599 | -0.0221910055070826 | 0.0446497720884481 | -0.497001540413774 | 0.619187965051371 | NA | NR_045848,NR_045847 |
| Atp6v1b1 | 0.652296579 | 0.083224305 | 0.067807775 | 1.227356387 | 0.219688671 | NA | | D930032P07Rik | 0.804933061 | -0.00813400335156511 | 0.018321366296201 | -0.443962705622654 | 0.657069547611335 | NA | NR_045330 |
| Atp6v1e2 | 1.495931226 | 0.071170774 | 0.091244895 | 0.779997327 | 0.435392449 | NA | | DQ267100 | 10.80658683 | -0.0371226579264213 | 0.0631592205192463 | -0.587763079107492 | 0.55669132914273 | NA | NR_046304 |
| Atp6v1g3 | 0.317012939 | 0.010888466 | 0.039852335 | 0.273220278 | 0.784683893 | NA | | DQ267101 | 1.199012197 | -0.0173001737450995 | 0.0236385780423927 | -0.731861862167592 | 0.464252886094916 | NA | NR_046305 |
| Atp8b3 | 6.755804481 | -0.138494199 | 0.141631099 | -0.977851615 | 0.328147719 | NA | | DQ267102 | 11.08398629 | 0.0202230753702788 | 0.0673953961601165 | 0.300066125024819 | 0.76412671756489 | NA | NR_046306 |
| Atp8b4 | 3.429688571 | -0.093214292 | 0.117465375 | -0.79354697 | 0.427459215 | NA | | DXBay18 | 0 | NA | NA | NA | NA | NA | NM_001025384 |
| Aurkc | 4.002984267 | 0.171843579 | 0.1313428 | 1.308359338 | 0.190751458 | NA | | Daf2 | 0 | NA | NA | NA | NA | NA | NM_007827 |
| Avpr1b | 1.493018595 | 0.124975862 | 0.073750772 | 1.694570211 | 0.090156982 | NA | | Dao | 6.392253965 | 0.048947031055581 | 0.0518821956652116 | 0.943426360970325 | 0.345462864907662 | NA | NM_010018 |
| Avpr2 | 0 | NA | NA | NA | NA | NA | | Dapl1 | 8.95994435 | -0.0442562196832856 | 0.057810263755832 | -0.765542601054488 | 0.443948508905021 | NA | NM_029723 |
| Awat1 | 0.099009884 | 0.018205726 | 0.027896865 | 0.652608315 | 0.514008823 | NA | | Dbh | 17.48410389 | -0.00538533796895495 | 0.0740442714680919 | -0.0727313249516631 | 0.942019920846209 | NA | NM_138942 |
| Aym1 | 0.299858689 | -0.034980473 | 0.036264327 | -0.964597324 | 0.33474654 | NA | | Dbil5 | 14.9798487 | -0.0517646288806886 | 0.0729685293270226 | -0.709410335635181 | 0.47806987571583 | NA | NM_021294 |
| Azgp1 | 0.695983529 | -0.04154153 | 0.056086079 | -0.740674517 | 0.458890814 | NA | | Dbx1 | 0 | NA | NA | NA | NA | NA | NM_001005232 |
| B020004C17Rik | 0 | NA | NA | NA | NA | NA | | Dcpp1 | 0 | NA | NA | NA | NA | NA | NM_019910 |
| B020004J07Rik | 7.203025663 | -0.048641878 | 0.151001513 | -0.322128417 | 0.747355411 | NA | | Dcpp2 | 0 | NA | NA | NA | NA | NA | NM_001039238 |
| B020014A21Rik | 0 | NA | NA | NA | NA | NA | | Dcpp3 | 0 | NA | NA | NA | NA | NA | NM_001077633 |
| B020018J22Rik | 0.999349742 | 0.048801405 | 0.07392027 | 0.660189745 | 0.509132072 | NA | | Dcst1 | 1.494023757 | -0.00214355843669974 | 0.0276822366086693 | -0.0774344380839676 | 0.938277945387898 | NA | NM_029974 |
| B130006D01Rik | 0.419064155 | -0.029347008 | 0.044028886 | -0.666539874 | 0.505066086 | NA | | Dcstamp | 4.176112139 | 0.0451177024197393 | 0.039746609230758 | 1.13513336842894 | 0.256319442324325 | NA | NM_029422 |
| B230119M05Rik | 0.818183462 | -0.07350399 | 0.07059028 | -1.041276359 | 0.297747306 | NA | | Dct | 17.66044213 | -0.0257558223724127 | 0.0732793748117055 | -0.351474373772884 | 0.725232494439565 | NA | NM_010024 |
| B230208H11Rik | 4.25069826 | 0.178552036 | 0.136488432 | 1.308184389 | 0.190810776 | NA | | Ddi1 | 0 | NA | NA | NA | NA | NA | NM_027942 |
| B230214G05Rik | 0.099009884 | 0.018205726 | 0.027896865 | 0.652608315 | 0.514008823 | NA | | Ddx3y | 0.223412651 | 0.00718831964189475 | 0.00999993766105603 | 0.718836445339964 | 0.472241699104885 | NA | NM_012008 |
| B230323A14Rik | 1.848492479 | -0.086778593 | 0.095571815 | -0.907993563 | 0.363881622 | NA | | Ddx43 | 6.361504066 | -0.0108055003628656 | 0.0532116756392359 | -0.203066342735091 | 0.839083178232751 | NA | NM_001191044 |
| B430010I23Rik | 1.388852301 | -0.075926941 | 0.076505134 | -0.992442438 | 0.320981746 | NA | | Dear1 | 0.526837674 | -0.00550876317531243 | 0.01480074035604 | -0.372195109352376 | 0.709747587128406 | NA | NM_001040461 |
| B4galnt2 | 4.739173048 | 0.147341103 | 0.138913892 | 1.060664996 | 0.288842173 | NA | | Defa-ps1 | 0 | NA | NA | NA | NA | NA | NR_003146 |
| B930018H19Rik | 7.905492961 | 0.171055224 | 0.150025173 | 1.140176817 | 0.254212644 | NA | | Defa-ps12 | 0 | NA | NA | NA | NA | NA | NR_002878 |
| B930092H01Rik | 0.094392817 | 0.017895942 | 0.027818137 | 0.643319236 | 0.520016977 | NA | | Defa-ps13 | 0 | NA | NA | NA | NA | NA | NR_002881 |
| BB014433 | 0 | NA | NA | NA | NA | NA | | Defa-rs1 | 0 | NA | NA | NA | NA | NA | NM_007844 |
| BB019430 | 0.21363976 | -0.000753674 | 0.031922632 | -0.02360938 | 0.98116419 | NA | | Defa-rs7 | 0 | NA | NA | NA | NA | NA | NM_007848 |
| BB123696 | 0.639521788 | 0.057858035 | 0.047157191 | 1.226918608 | 0.219853184 | NA | | Defa17 | 0 | NA | NA | NA | NA | NA | NM_001167790 |
| BB283400 | 0 | NA | NA | NA | NA | NA | | Defa20 | 0 | NA | NA | NA | NA | NA | NM_183268 |
| BB287469 | 0 | NA | NA | NA | NA | NA | | Defa21 | 0 | NA | NA | NA | NA | NA | NM_183253 |
| BB557941 | 7.23918054 | 0.27121741 | 0.15688831 | 1.728729245 | 0.083857565 | NA | | Defa22 | 0 | NA | NA | NA | NA | NA | NM_207658 |
| BC016579 | 6.725059 | 0.237332762 | 0.145391178 | 1.632373889 | 0.102600746 | NA | | Defa23 | 0 | NA | NA | NA | NA | NA | NM_001012307 |
| BC021614 | 0.096071043 | -0.020098795 | 0.027825948 | -0.722304036 | 0.470107572 | NA | | Defa24 | 0 | NA | NA | NA | NA | NA | NM_001024225 |
| BC021767 | 3.096620928 | 0.227057757 | 0.119923008 | 1.893362757 | 0.058309645 | NA | | Defa25 | 0 | NA | NA | NA | NA | NA | NM_007849 |
| BC024386 | 0 | NA | NA | NA | NA | NA | | Defa26 | 0 | NA | NA | NA | NA | NA | NM_001079933 |
| BC024582 | 2.943682564 | 0.085967715 | 0.117368464 | 0.732460082 | 0.463887799 | NA | | Defa3 | 0 | NA | NA | NA | NA | NA | NM_007850 |
| BC030870 | 0 | NA | NA | NA | NA | NA | | Defa4 | 0 | NA | NA | NA | NA | NA | NM_010039 |
| BC039771 | 6.94654834 | -0.064269477 | 0.142735937 | -0.450268362 | 0.652516949 | NA | | Defa5 | 0 | NA | NA | NA | NA | NA | NM_007851 |
| BC048502 | 0 | NA | NA | NA | NA | NA | | Defa6 | 0 | NA | NA | NA | NA | NA | NM_007852 |
| BC048507 | 0.186326724 | 0.024939385 | 0.031411053 | 0.793968451 | 0.427213798 | NA | | Defb10 | 0.106148608 | 0.00515554063321497 | 0.00854200771766808 | 0.603551390213729 | 0.546141939141592 | NA | NM_139225 |
| BC048562 | 0.119397641 | -0.020477583 | 0.027954224 | -0.732539845 | 0.463839132 | NA | | Defb11 | 1.616575638 | -0.00886170705986563 | 0.0183428664972429 | -0.483114624488905 | 0.629014350655317 | NA | NM_139221 |
| BC048602 | 0 | NA | NA | NA | NA | NA | | Defb12 | 0.59690524 | 0.0156587704108518 | 0.0147112737758238 | 1.0644061588049 | 0.287144744982081 | NA | NM_152802 |
| BC048609 | 0.609009071 | 0.027901577 | 0.054375796 | 0.513124931 | 0.607863933 | NA | | Defb13 | 0 | NA | NA | NA | NA | NA | NM_139223 |
| BC048644 | 3.888042553 | -0.052750898 | 0.125030752 | -0.42190339 | 0.673095536 | NA | | Defb14 | 0.35787663 | -0.00674720089483589 | 0.0130723048459276 | -0.516144702434617 | 0.605753348373098 | NA | NM_183026 |
| BC048671 | 0.119397641 | -0.020477583 | 0.027954224 | -0.732539845 | 0.463839132 | NA | | Defb15 | 0 | NA | NA | NA | NA | NA | NM_139222 |
| BC048679 | 0.096071043 | -0.020098795 | 0.027825948 | -0.722304036 | 0.470107572 | NA | | Defb18 | 0 | NA | NA | NA | NA | NA | NM_001039123 |
| BC049635 | 2.342311452 | 0.138256711 | 0.104499118 | 1.3230419 | 0.185821444 | NA | | Defb19 | 0 | NA | NA | NA | NA | NA | NM_145157 |
| BC049730 | 0 | NA | NA | NA | NA | NA | | Defb2 | 0 | NA | NA | NA | NA | NA | NM_010030 |
| BC049762 | 3.134227431 | 0.205648659 | 0.113506461 | 1.811779322 | 0.070020301 | NA | | Defb20 | 0.235249118 | -0.00959082599542929 | 0.0102925929774586 | -0.93181825186654 | 0.351430460444064 | NA | NM_176950 |
| BC051019 | 6.67735047 | 0.156031214 | 0.127020367 | 1.228395233 | 0.219298637 | NA | | Defb21 | 0 | NA | NA | NA | NA | NA | NM_207276 |
| BC051070 | 0.579950701 | 0.023389586 | 0.058461528 | 0.400085095 | 0.689093842 | NA | | Defb22 | 0 | NA | NA | NA | NA | NA | NM_001002791 |
| BC051537 | 2.482475864 | 0.067703871 | 0.086602696 | 0.781775561 | 0.434346486 | NA | | Defb23 | 0 | NA | NA | NA | NA | NA | NM_001037933 |
| BC051628 | 3.40388302 | -0.099261099 | 0.127053494 | -0.78125438 | 0.434652896 | NA | | Defb25 | 0.385490251 | -0.00508296712693953 | 0.0117912533198291 | -0.431079461111366 | 0.666410594179892 | NA | NM_001039122 |
| BC051665 | 0.461343821 | 0.036894045 | 0.051391044 | 0.717908062 | 0.472813976 | NA | | Defb26 | 0 | NA | NA | NA | NA | NA | NM_001039120 |
| BC053393 | 1.735623088 | 0.023509006 | 0.087789743 | 0.26778762 | 0.7888628 | NA | | Defb28 | 0.33403441 | 0.00244344761020736 | 0.0111793636306143 | 0.218567683362231 | 0.826986829057954 | NA | NM_001037502 |
| BC055111 | 3.288159444 | 0.158698338 | 0.119825494 | 1.324412139 | 0.185366205 | NA | | Defb29 | 0 | NA | NA | NA | NA | NA | NM_001001444 |
| BC061212 | 0 | NA | NA | NA | NA | NA | | Defb3 | 0 | NA | NA | NA | NA | NA | NM_013756 |
| BC061237 | 0.098353386 | -0.020212831 | 0.02786466 | -0.725393061 | 0.468210929 | NA | | Defb30 | 0.770412418 | 0.00126807230863351 | 0.018053434755678 | 0.0702399474556876 | 0.944002679353667 | NA | NM_001039566 |
| BC080695 | 0 | NA | NA | NA | NA | NA | | Defb34 | 0 | NA | NA | NA | NA | NA | NM_183035 |
| BC089597 | 0.288817247 | -0.014517254 | 0.039073354 | -0.37153846 | 0.710236516 | NA | | Defb35 | 0 | NA | NA | NA | NA | NA | NM_139224 |
| BC090627 | 0 | NA | NA | NA | NA | NA | | Defb36 | 0 | NA | NA | NA | NA | NA | NM_001037247 |
| BC094916 | 0 | NA | NA | NA | NA | NA | | Defb37 | 0 | NA | NA | NA | NA | NA | NM_181683 |
| BC096441 | 0 | NA | NA | NA | NA | NA | | Defb38 | 0 | NA | NA | NA | NA | NA | NM_183036 |
| BC100451 | 5.237384986 | -0.033081727 | 0.145372682 | -0.227564947 | 0.819984473 | NA | | Defb39 | 0 | NA | NA | NA | NA | NA | NM_183038 |
| BC100530 | 0 | NA | NA | NA | NA | NA | | Defb4 | 0 | NA | NA | NA | NA | NA | NM_019728 |
| BC107364 | 1.651366412 | -0.041334664 | 0.086431294 | -0.478237244 | 0.632481357 | NA | | Defb40 | 0 | NA | NA | NA | NA | NA | NM_183039 |
| BC117090 | 0 | NA | NA | NA | NA | NA | | Defb41 | 0.23403553 | -0.000407313579519725 | 0.00983856423584417 | -0.0413996971261099 | 0.966977254250032 | NA | NM_183124 |
| BC125332 | 2.627048511 | 0.091651976 | 0.112830304 | 0.812299298 | 0.416619914 | NA | | Defb42 | 0.332904726 | -0.0130643184082516 | 0.0120034188534089 | -1.08838311549392 | 0.276426011853847 | NA | NM_001034910 |
| Baat | 5.476431584 | 0.084660008 | 0.138559451 | 0.611001325 | 0.541198703 | NA | | Defb43 | 0 | NA | NA | NA | NA | NA | NM_001039121 |
| Bag3 | 372.7242188 | 0.393999397 | 0.12888778 | 3.056918177 | NA | NA | | Defb44-ps | 0 | NA | NA | NA | NA | NA | NR_002879 |
| Banf2 | 0 | NA | NA | NA | NA | NA | | Defb45 | 0 | NA | NA | NA | NA | NA | NM_001037752 |
| Barhl1 | 2.010134074 | 0.116289232 | 0.087097976 | 1.335154244 | 0.181825869 | NA | | Defb46 | 0 | NA | NA | NA | NA | NA | NM_001025351 |
| Barx1 | 0 | NA | NA | NA | NA | NA | | Defb47 | 0.917323538 | -0.00406203540923328 | 0.019292472583573 | -0.210550275069044 | 0.833238217118502 | NA | NM_001039125 |
| Batf | 5.419938029 | -0.01380056 | 0.125152616 | -0.110269848 | 0.912195369 | NA | | Defb48 | 0 | NA | NA | NA | NA | NA | NM_001037751 |
| Bc1 | 0 | NA | NA | NA | NA | NA | | Defb5 | 0 | NA | NA | NA | NA | NA | NM_030734 |
| Bcl2a1a | 6.646647669 | 0.301929753 | 0.147171219 | 2.051554339 | 0.040212994 | NA | | Defb50 | 0 | NA | NA | NA | NA | NA | NM_199067 |
| Bcl2a1c | 0 | NA | NA | NA | NA | NA | | Defb6 | 0.354045348 | 0.00283843574875446 | 0.0129156729755236 | 0.219766771281183 | 0.826052799641565 | NA | NM_054074 |
| Bcl2a1d | 1.446188375 | 0.02523781 | 0.079787286 | 0.316313679 | 0.751764429 | NA | | Defb7 | 0 | NA | NA | NA | NA | NA | NM_139220 |
| Bcl2l10 | 1.091020553 | 0.03623669 | 0.078890562 | 0.459328585 | 0.645998224 | NA | | Defb8 | 0 | NA | NA | NA | NA | NA | NM_153108 |
| Bcl2l14 | 0.167300494 | 0.024299763 | 0.031030539 | 0.783091891 | 0.433573153 | NA | | Defb9 | 0 | NA | NA | NA | NA | NA | NM_139219 |
| Bcmo1 | 2.427182447 | 0.127250925 | 0.090455439 | 1.40678025 | 0.159492561 | NA | | Dennd2d | 20.14053706 | 0.0810892849887403 | 0.0820874469565663 | 0.987840260541248 | 0.32323088652587 | NA | NM_028110,NM_001093754,NR_051982 |
| Bdkrb1 | 2.029057989 | -0.014098727 | 0.10124509 | -0.139253442 | 0.889249879 | NA | | Depdc1a | 1.717669478 | -0.0169079963743609 | 0.0299402131912907 | -0.564725316627981 | 0.572260611683238 | NA | NM_001172092,NM_001172093,NM_029523 |
| Best2 | 2.085187786 | 0.009796423 | 0.099723293 | 0.098236055 | 0.921744853 | NA | | Depdc1b | 14.49786466 | -0.173950211376683 | 0.0760319683852003 | -2.28785621457806 | 0.0221458958855567 | NA | NM_178683 |
| Best3 | 7.278845561 | 0.256466239 | 0.1341041 | 1.912441446 | 0.055819596 | NA | | Dfnb59 | 2.522778796 | 0.0154064844820212 | 0.0350780585891776 | 0.439205734343988 | 0.660512470392779 | NA | NM_001080711 |
| Bet3l | 5.266578649 | -0.028364686 | 0.134085205 | -0.211542249 | 0.832464168 | NA | | Dhh | 6.653340462 | -0.0114645822349368 | 0.0579112406333903 | -0.197968168347728 | 0.84306996878928 | NA | NM_007857 |
| Beta-s | 296.0418766 | -0.188675608 | 0.093899957 | -2.009325826 | NA | NA | | Dhrs2 | 2.342921304 | 0.0359560126311314 | 0.0362596361825006 | 0.991626403810534 | 0.321379804226573 | NA | NM_027790 |
| Bex6 | 0 | NA | NA | NA | NA | NA | | Dhrs7c | 0.466157216 | 0.00712081961299664 | 0.0157450460764072 | 0.452257781809015 | 0.651083286259444 | NA | NM_001013013 |
| Bglap | 1.768971194 | -0.020394106 | 0.084638305 | -0.240955985 | 0.809589231 | NA | | Dhrs9 | 2.156142019 | -0.0729512546935573 | 0.0349874870514315 | -2.0850670008486 | 0.0370632191537505 | NA | NM_175512 |
| Bglap2 | 1.256570555 | 0.021432703 | 0.069188086 | 0.309774477 | 0.756732462 | NA | | Dio1 | 0 | NA | NA | NA | NA | NA | NM_007860 |
| Bglap3 | 7.422566327 | 0.535619198 | 0.147779456 | 3.624449656 | 0.000289577 | NA | | Dio3os | 8.385694236 | 0.0607159705465111 | 0.0560886678109791 | 1.08249977965471 | 0.27903051325693 | NA | NR_002866 |
| Bhlha15 | 4.158081394 | 0.118235459 | 0.129962801 | 0.90976386 | 0.362947058 | NA | | Dkk1 | 0.73024527 | 0.0172385779990236 | 0.018800214012851 | 0.916935200165282 | 0.359176597186361 | NA | NM_010051 |
| Bhlhe23 | 2.453154771 | 0.019244711 | 0.097268501 | 0.197851415 | 0.843161318 | NA | | Dkk2 | 20.60244448 | 0.0443476684461397 | 0.0811981686519172 | 0.546165870270432 | 0.584951924338559 | NA | NM_020265 |
| Bhmt | 2.953675465 | 0.089629847 | 0.093638558 | 0.95718952 | 0.338471606 | NA | | Dkk4 | 0 | NA | NA | NA | NA | NA | NM_145592 |
| Bik | 2.824761723 | -0.008820612 | 0.11210762 | -0.078679862 | 0.937287264 | NA | | Dlgap5 | 7.278894875 | 0.0226085697222002 | 0.0567263904322325 | 0.398554703550357 | 0.690221343229589 | NA | NM_144553 |
| Birc7 | 3.402648078 | 0.145769176 | 0.115999903 | 1.256631883 | 0.208886964 | NA | | Dlx3 | 2.450194858 | 0.0020259615906316 | 0.0316455004638296 | 0.0640205261707663 | 0.948953882812207 | NA | NM_010055 |
| Blk | 0.421776838 | -0.034999817 | 0.049427494 | -0.708104206 | 0.478880548 | NA | | Dlx4 | 14.29464018 | 0.0455844251729236 | 0.0651223057283718 | 0.699981744550912 | 0.483938705190555 | NA | NM_007867 |
| Bmp10 | 0 | NA | NA | NA | NA | NA | | Dlx6as2 | 1.100519677 | 0.00456730392173208 | 0.0276267815631954 | 0.165321606908301 | 0.868690852961063 | NA | NR_002839 |
| Bmp15 | 7.566656856 | -0.346971736 | 0.152251678 | -2.278935381 | 0.022670907 | NA | | Dmbt1 | 0.592959185 | -0.00509395626934084 | 0.0152022007437956 | -0.335080187085401 | 0.737564598206156 | NA | NM_007769 |
| Bmp8a | 5.134289876 | 0.071577255 | 0.141882678 | 0.504481985 | 0.613922718 | NA | | Dmbx1 | 8.299835162 | -0.00029672533069034 | 0.0516198442772915 | -0.00574828023688701 | 0.995413561205907 | NA | NM_130865,NM_001025567 |
| Bmp8b | 7.181411658 | -0.076495296 | 0.15240289 | -0.501928117 | 0.615718086 | NA | | Dmgdh | 14.27196448 | -0.0203474695691898 | 0.0661952944427608 | -0.307385437899733 | 0.758550010893191 | NA | NM_028772 |
| Bnc1 | 0.193239998 | 0.025145508 | 0.031532422 | 0.797449311 | 0.42519013 | NA | | Dmrt1 | 0 | NA | NA | NA | NA | NA | NM_015826 |
| Boll | 0 | NA | NA | NA | NA | NA | | Dmrt3 | 6.176869938 | -0.0143731832858999 | 0.0467529192469251 | -0.307428573817777 | 0.75851718179937 | NA | NM_177360 |
| Bpi | 0.482559746 | 0.027836625 | 0.0455364 | 0.611304907 | 0.540997743 | NA | | Dmrta1 | 9.207022633 | -0.00607201723248048 | 0.0586024715740975 | -0.103613671392733 | 0.917475937860914 | NA | NM_175647 |
| Bpifa2 | 0.119397641 | -0.020477583 | 0.027954224 | -0.732539845 | 0.463839132 | NA | | Dmrtb1 | 18.30721047 | 0.00783638601919865 | 0.0813104168622929 | 0.0963761633699447 | 0.923221822859955 | NA | NM_019872 |
| Bpifa3 | 0 | NA | NA | NA | NA | NA | | Dmrtc1b | 0 | NA | NA | NA | NA | NA | NM_001039116 |
| Bpifa5 | 0.360960404 | 0.037959713 | 0.038487497 | 0.986286873 | 0.32399236 | NA | | Dmrtc1c1 | 0 | NA | NA | NA | NA | NA | NM_001142691 |
| Bpifa6 | 0.105923158 | 0.018524693 | 0.027977518 | 0.662127842 | 0.507889294 | NA | | Dmrtc1c2 | 0 | NA | NA | NA | NA | NA | NM_001142690 |
| Bpifb1 | 7.364245161 | -0.20212543 | 0.150245184 | -1.345303886 | 0.178527119 | NA | | Dmrtc2 | 0.552498712 | -0.00736465969491979 | 0.0165944977873112 | -0.443801300244898 | 0.657186248260847 | NA | NM_027732 |
| Bpifb2 | 0.288817247 | -0.014517254 | 0.039073354 | -0.37153846 | 0.710236516 | NA | | Dnaaf1 | 19.37044385 | -0.0348189018246627 | 0.0801683522945837 | -0.434322283395802 | 0.664054424473226 | NA | NM_026648 |
| Bpifb3 | 2.614841149 | -0.051131894 | 0.089706022 | -0.569994002 | 0.568681766 | NA | | Dnahc17 | 15.48493669 | 0.0112755120123625 | 0.0760556668946034 | 0.148253410597107 | 0.882142781920872 | NA | NM_001167746 |
| Bpifb4 | 4.235457987 | 0.272341675 | 0.126416023 | 2.154328774 | 0.031214398 | NA | | Dnahc8 | 21.46121323 | 0.0921692818445744 | 0.0743764862513695 | 1.23922608461323 | 0.215261783130583 | NA | NM_013811 |
| Bpifb5 | 1.33467163 | 0.081170829 | 0.078213963 | 1.037804842 | 0.299360922 | NA | | Dnajb13 | 15.90774647 | 0.00475471143688795 | 0.0712090390242104 | 0.0667711782386418 | 0.946763868620872 | NA | NM_153527 |
| Bpifb6 | 1.204479727 | 0.022259603 | 0.079788357 | 0.278983096 | 0.780257797 | NA | | Dnajb7 | 0.536670213 | 0.0190472850833179 | 0.0162169355655615 | 1.17453047811123 | 0.240182611472615 | NA | NM_021317 |
| Bpifb9a | 2.943751015 | 0.132924612 | 0.075548193 | 1.759467792 | 0.078498086 | NA | | Dnajb8 | 0 | NA | NA | NA | NA | NA | NM_019964 |
| Bpifb9b | 7.716286422 | 0.193258552 | 0.094622319 | 2.042420371 | 0.041109851 | NA | | Dnajc22 | 1.00246825 | -0.0114853152145707 | 0.0243488214011263 | -0.471699020883178 | 0.637141636105064 | NA | NM_176835 |
| Brat1 | 0.953203133 | -0.012833519 | 0.065301315 | -0.196527732 | 0.844197129 | NA | | Dnajc5b | 0.689569867 | 0.0142940141213119 | 0.0182330003215352 | 0.783963904417262 | 0.433061290475319 | NA | NM_025489,NM_001163537,NM_001163536 |
| Bsnd | 0.292777816 | -0.03443077 | 0.035983808 | -0.956840623 | 0.338647705 | NA | | Dnajc5g | 0.235249118 | -0.00959082599542929 | 0.0102925929774586 | -0.93181825186654 | 0.351430460444064 | NA | NM_177677 |
| Bsph1 | 1.117230264 | 0.051571758 | 0.06768252 | 0.761965688 | 0.446080483 | NA | | Dnase1 | 0.385781328 | -0.00133843355935467 | 0.01173576040662 | -0.11404745095168 | 0.909200178383767 | NA | NM_010061 |
| Bsph2 | 0 | NA | NA | NA | NA | NA | | Dnase1l3 | 10.63385292 | -0.0360759045240652 | 0.0512891813970606 | -0.703382341877906 | 0.481817507964678 | NA | NM_007870 |
| Bst1 | 6.492475 | 0.055254644 | 0.115978935 | 0.47641965 | 0.633775438 | NA | | Dnase2b | 1.879966503 | -0.00209406991007555 | 0.0264073214513927 | -0.0792988381623655 | 0.936794930405315 | NA | NM_019957 |
| Bsx | 0.675759004 | 0.004073513 | 0.060574347 | 0.06724815 | 0.946384154 | NA | | Dnm3os | 1.421425898 | -0.0256015188697277 | 0.0294293333825333 | -0.869932000733751 | 0.384337566148203 | NA | NR_002870 |
| Btg3 | 0 | NA | NA | NA | NA | NA | | Dnmt3l | 3.785301734 | 0.00299142190116933 | 0.0458013418941329 | 0.0653129750670587 | 0.947924811793484 | NA | NM_019448,NM_001081695 |
| Btg4 | 0.297734681 | -0.017963657 | 0.03944387 | -0.455423286 | 0.648804746 | NA | | Dntt | 1.069428219 | 0.0167237254292122 | 0.0218507062325352 | 0.765363153540133 | 0.44405532753605 | NA | NM_001043228,NM_009345 |
| Btn1a1 | 1.634812729 | -0.045320037 | 0.090787953 | -0.499185585 | 0.617648649 | NA | | Doxl2 | 0 | NA | NA | NA | NA | NA | NM_001029987 |
| Btnl1 | 0.119397641 | -0.020477583 | 0.027954224 | -0.732539845 | 0.463839132 | NA | | Dpcr1 | 0.778605831 | 0.00107214574328684 | 0.0160726636343245 | 0.0667061644342001 | 0.946815626736596 | NA | NM_001033366 |
| Btnl10 | 0.314060235 | -0.015766464 | 0.039947878 | -0.394675873 | 0.693082111 | NA | | Dpep1 | 3.082210665 | 0.0191685576180554 | 0.023987792625074 | 0.799096353618584 | NA | NA | NM_007876 |
| Btnl2 | 0.08731684 | 0.017365056 | 0.027733412 | 0.626142072 | 0.531221759 | NA | | Dpep2 | 2.742759324 | 0.047008329414003 | 0.0378850814089176 | 1.24081373632572 | 0.214674565613874 | NA | NM_176913 |
| Btnl4 | 0 | NA | NA | NA | NA | NA | | Dpep3 | 0 | NA | NA | NA | NA | NA | NM_027960 |
| Btnl5 | 0.462360653 | -0.013923057 | 0.043980692 | -0.316572026 | 0.751568365 | NA | | Dpp8 | 9723.264526 | 0.027330192479708 | 0.0579074685741233 | 0.471963170773465 | NA | NA | NM_028906 |
| Btnl6 | 0.198019769 | 0.025308392 | 0.031628095 | 0.80018706 | 0.423602426 | NA | | Dppa1 | 1.29059397 | -0.00764928525627544 | 0.026733246869332 | -0.286133790394559 | 0.774775651985112 | NA | NM_178247,NM_001163358 |
| Btnl9 | 0.762823459 | -0.031698507 | 0.061350021 | -0.516682909 | 0.605377529 | NA | | Dppa2 | 0.677494072 | -0.0222487452219733 | 0.0186801957681695 | -1.19103383594537 | 0.233640302057453 | NA | NM_028615 |
| C030007H22Rik | 2.060017668 | -0.091560523 | 0.094327218 | -0.970669174 | 0.331713047 | NA | | Dppa3 | 0 | NA | NA | NA | NA | NA | NM_139218 |
| C030034L19Rik | 3.921997651 | 0.236487143 | 0.118383829 | 1.997630463 | 0.045756738 | NA | | Dppa4 | 0 | NA | NA | NA | NA | NA | NM_028610,NM_001018002 |
| C130021I20Rik | 5.811273544 | -0.191888635 | 0.147708656 | -1.299102166 | 0.19390887 | NA | | Dppa5a | 2.26022915 | 0.00216735032237925 | 0.0310047489705318 | 0.069903817780921 | 0.94427021441566 | NA | NM_025274 |
| C130026I21Rik | 1.924063848 | 0.102343837 | 0.102984531 | 0.99377873 | 0.320330604 | NA | | Dpt | 4.74169612 | 0.0351370423603621 | 0.0489347677075607 | 0.718038401047386 | 0.47273360949667 | NA | NM_019759 |
| C130050O18Rik | 6.790989308 | -0.127379167 | 0.14490064 | -0.879079397 | 0.379358228 | NA | | Dpy19l2 | 10.85011542 | 0.0126168792455956 | 0.063192241536544 | 0.199658675476787 | 0.841747535150457 | NA | NM_001166207 |
| C130079G13Rik | 0 | NA | NA | NA | NA | NA | | Dpys | 14.05175839 | 0.0273598475404371 | 0.0733705548123905 | 0.37289955909965 | 0.709223200176741 | NA | NM_022722,NM_001164466 |
| C130080G10Rik | 4.632132215 | 0.016732535 | 0.137320041 | 0.121850641 | 0.903017307 | NA | | Drd3 | 5.093873114 | -0.0522662475474396 | 0.0510171265143691 | -1.02448434708918 | 0.305606560429321 | NA | NM_007877 |
| C130083M11Rik | 1.351018946 | -0.009805109 | 0.088731714 | -0.110502868 | 0.912010575 | NA | | Drd4 | 8.488410245 | -0.0808091106620268 | 0.0548005551259856 | -1.47460386991058 | 0.14031907388177 | NA | NM_007878 |
| C1ql4 | 0.497028206 | -0.021472216 | 0.055798694 | -0.384815747 | 0.700373932 | NA | | Dsc1 | 3.090523456 | 0.0069350000676749 | 0.0373291192707011 | 0.185779900602103 | 0.852617367125049 | NA | NM_013504 |
| C1qtnf3 | 2.839363405 | 0.018647936 | 0.10561413 | 0.176566674 | 0.859848773 | NA | | Dscc1 | 5.028209847 | -0.0460891197544582 | 0.0454081669697481 | -1.01499626234998 | 0.310107609182016 | NA | NM_183089 |
| C1qtnf5 | 0 | NA | NA | NA | NA | NA | | Dsg1a | 1.346430562 | -0.0193338188960549 | 0.0241979993423665 | -0.798984189664173 | 0.424299579616666 | NA | NM_010079 |
| C1rb | 0.269026497 | 0.030520093 | 0.03461589 | 0.881678683 | 0.377950592 | NA | | Dsg1b | 1.315972644 | -0.0371891067040679 | 0.0239354287550581 | -1.55372636457198 | 0.120249701473485 | NA | NM_181682 |
| C230024C17Rik | 1.292691633 | 0.079787348 | 0.077758798 | 1.026087726 | 0.304850235 | NA | | Dsg1c | 10.17379232 | 0.0712231503935011 | 0.0624809298440042 | 1.13991822098877 | 0.254320373123972 | NA | NM_181680 |
| C230079O03Rik | 0 | NA | NA | NA | NA | NA | | Dsg3 | 0.299039272 | 0.00281759532524203 | 0.012303739144426 | 0.229003174739646 | 0.818866448605973 | NA | NM_030596 |
| C2cd4a | 2.754249344 | -0.038932739 | 0.112024442 | -0.347537893 | 0.728187256 | NA | | Dsg4 | 0 | NA | NA | NA | NA | NA | NM_181564 |
| C2cd4d | 3.12534758 | -0.095787147 | 0.118604197 | -0.807620217 | 0.419309245 | NA | | Dspp | 9.237751076 | 0.0243862230548031 | 0.0607369569406688 | 0.401505513004627 | 0.688047980044088 | NA | NM_010080 |
| C330008G21Rik | 0 | NA | NA | NA | NA | NA | | Duox1 | 3.647264892 | 0.0308295596495433 | 0.0453598171226185 | 0.679666753642406 | 0.496715491679582 | NA | NM_001099297 |
| C330011F03Rik | 0 | NA | NA | NA | NA | NA | | Duox2 | 1.901967724 | 0.0122088289587582 | 0.0278576424120666 | 0.438257795766302 | 0.661199416295548 | NA | NM_177610 |
| C330013F16Rik | 0.300084847 | -0.016747215 | 0.039501486 | -0.423964169 | 0.671591945 | NA | | Duoxa1 | 1.99090577 | 0.0259364573802457 | 0.0293119675094652 | 0.884841912159955 | 0.376241900869603 | NA | NM_145395 |
| C330021F23Rik | 2.771310574 | 0.021981985 | 0.112445875 | 0.195489471 | 0.845009779 | NA | | Duoxa2 | 0.465271493 | -0.00989654861615473 | 0.0144508438382702 | -0.68484226436284 | 0.493443462071998 | NA | NM_025777 |
| C330022C24Rik | 1.325156643 | 0.001301153 | 0.081950265 | 0.015877345 | 0.987332244 | NA | | Dupd1 | 2.199804773 | 0.00856333949653228 | 0.035047063520291 | 0.244338287901764 | 0.806968832450158 | NA | NM_001013826 |
| C330024C12Rik | 0.094392817 | 0.017895942 | 0.027818137 | 0.643319236 | 0.520016977 | NA | | Dusp13 | 9.34228095 | -0.00947054619386417 | 0.0619285073353385 | -0.152927086431808 | 0.878455775262737 | NA | NM_001007268,NM_013849 |
| C330024D21Rik | 2.92332916 | 0.087083422 | 0.12020982 | 0.724428521 | 0.468802695 | NA | | Dusp21 | 0 | NA | NA | NA | NA | NA | NM_028568 |
| C330046G13Rik | 0.119397641 | -0.020477583 | 0.027954224 | -0.732539845 | 0.463839132 | NA | | Dux | 1.160395969 | -0.0395209645145706 | 0.0256867933365873 | -1.53857135831269 | 0.12390897532773 | NA | NM_001081954 |
| C430002E04Rik | 0 | NA | NA | NA | NA | NA | | Duxbl1 | 0 | NA | NA | NA | NA | NA | NM_183389 |
| C430049B03Rik | 6.112321871 | -0.081808695 | 0.149951652 | -0.545567146 | 0.585363513 | NA | | Duxbl2 | 0 | NA | NA | NA | NA | NA | NM_001177538 |

| | | | | | | |
|---|---|---|---|---|---|---|
| C4bp | 0.765844708 | 0.066775506 | 0.059510282 | 1.122083506 | 0.261826936 | NA |
| C4bp-ps1 | 4.112967842 | 0.187839373 | 0.131127314 | 1.432496158 | 0.152001877 | NA |
| C530044C16Rik | 6.273837178 | -0.074356482 | 0.133433124 | -0.557256545 | 0.577352161 | NA |
| C6 | 0.652231925 | -0.101419571 | 0.060468106 | -1.677240726 | 0.093495418 | NA |
| C630028M04Rik | 0 | NA | NA | NA | NA | NA |
| C730027H18Rik | 0.096071043 | -0.020098795 | 0.027825948 | -0.722304036 | 0.470107572 | NA |
| C730036E19Rik | 2.532110895 | -0.136872893 | 0.11569686 | -1.183030317 | 0.236797132 | NA |
| C86187 | 1.036144045 | 0.047346346 | 0.07129148 | 0.664123476 | 0.506611286 | NA |
| C86695 | 0.410466648 | -0.020907232 | 0.037449064 | -0.55828449 | 0.576650134 | NA |
| C87198 | 0.119397641 | -0.020477583 | 0.027954224 | -0.732539845 | 0.463839132 | NA |
| C87414 | 0 | NA | NA | NA | NA | NA |
| C87499 | 0.105923158 | 0.018524693 | 0.027977518 | 0.662127842 | 0.507889294 | NA |
| C87977 | 0 | NA | NA | NA | NA | NA |
| C8a | 1.560028588 | 0.060622267 | 0.078764365 | 0.769666169 | 0.441497943 | NA |
| C8b | 3.447200476 | 0.113633108 | 0.103216963 | 1.10091505 | 0.270933631 | NA |
| C9 | 0 | NA | NA | NA | NA | NA |
| C920009B18Rik | 1.47656618 | -0.00521879 | 0.078807297 | -0.06622217 | 0.947200946 | NA |
| Cabp2 | 0.08731684 | 0.017365056 | 0.027733412 | 0.626142072 | 0.531221759 | NA |
| Cabp4 | 1.249089198 | 0.115590482 | 0.08163889 | 1.415875226 | 0.15681204 | NA |
| Cabp5 | 6.461264216 | -0.095840601 | 0.144840321 | -0.661698343 | 0.508164567 | NA |
| Cabs1 | 0.797272051 | 0.015791619 | 0.056720173 | 0.27841275 | 0.780695533 | NA |
| Cacna1f | 7.407490114 | 0.167424236 | 0.137324519 | 1.219186769 | 0.222773313 | NA |
| Calcb | 3.460229114 | -0.11511207 | 0.101596372 | -1.133033276 | 0.257200279 | NA |
| Calcoco2 | 1.659453276 | 0.080678432 | 0.092747754 | 0.869869383 | 0.384371789 | NA |
| Calhm1 | 0.098353386 | -0.020212831 | 0.02786466 | -0.725393061 | 0.468210929 | NA |
| Calm4 | 0 | NA | NA | NA | NA | NA |
| Calm5 | 0 | NA | NA | NA | NA | NA |
| Calml3 | 0.181709657 | 0.024794574 | 0.031325423 | 0.791516008 | 0.428642938 | NA |
| Calml4 | 24.27553477 | 0.414091445 | 0.148607148 | 2.78648404 | NA | NA |
| Camp | 4.92462669 | -0.038932099 | 0.130681482 | -0.297915958 | 0.765767308 | NA |
| Capn13 | 0.577221778 | 0.048337644 | 0.054475647 | 0.887325731 | 0.374903556 | NA |
| Capn8 | 1.130778681 | 0.149419225 | 0.081821028 | 1.826171453 | 0.06782444 | NA |
| Capn9 | 7.137506979 | 0.094543834 | 0.150215766 | 0.629386891 | 0.529095798 | NA |
| Capns2 | 5.140823299 | -0.151525387 | 0.133278981 | -1.136903852 | 0.255578482 | NA |
| Capza3 | 0 | NA | NA | NA | NA | NA |
| Car1 | 4.365296727 | 0.287287806 | 0.138705328 | 2.07120959 | 0.038339214 | NA |
| Car6 | 0.686664619 | 0.040716927 | 0.066216252 | 0.614908356 | 0.538615244 | NA |
| Casc5 | 6.936931086 | -0.305600687 | 0.147197613 | -2.076125285 | 0.037882365 | NA |
| Casp14 | 0.099009884 | 0.018205726 | 0.027896865 | 0.652608315 | 0.514008823 | NA |
| Catsper1 | 0.283178451 | 0.028390488 | 0.033368195 | 0.850824814 | 0.394866675 | NA |
| Catsper4 | 0.711136086 | 0.059233718 | 0.058127284 | 1.019034662 | 0.308186511 | NA |
| Catsperb | 0 | NA | NA | NA | NA | NA |
| Catsperg2 | 2.428626099 | 0.037121406 | 0.102317898 | 0.362804624 | 0.716750837 | NA |
| Cav3 | 3.428642594 | 0.017420055 | 0.115867918 | 0.150344072 | 0.880493164 | NA |
| Cblc | 0.448020035 | 0.075829036 | 0.05417878 | 1.399607666 | 0.161630837 | NA |
| Ccdc144b | 0.098353386 | -0.020212831 | 0.02786466 | -0.725393061 | 0.468210929 | NA |
| Ccdc147 | 6.580716721 | 0.0815182 | 0.152400081 | 0.534896042 | 0.592721745 | NA |
| Ccdc150 | 4.960270646 | 0.201929436 | 0.139347613 | 1.449105807 | 0.147308036 | NA |
| Ccdc152 | 0.332165648 | -0.049934083 | 0.043026356 | -1.160546414 | 0.245826408 | NA |
| Ccdc154 | 0.937815091 | 0.042109344 | 0.066319649 | 0.634945222 | 0.525464142 | NA |
| Ccdc169 | 6.182358922 | -0.138730546 | 0.136232021 | -1.018340219 | 0.308516301 | NA |
| Ccdc172 | 1.914837824 | 0.003450926 | 0.094072603 | 0.036683645 | 0.970737249 | NA |
| Ccdc175 | 1.012519264 | -0.002321965 | 0.06532011 | -0.035547483 | 0.971643185 | NA |
| Ccdc178 | 0.432801378 | -0.061508142 | 0.047634613 | -1.291248906 | 0.196617382 | NA |
| Ccdc27 | 2.025826332 | 0.009484835 | 0.082766829 | 0.114597055 | 0.908764514 | NA |
| Ccdc38 | 3.560846977 | -0.036810944 | 0.127471875 | -0.288776988 | 0.772752043 | NA |
| Ccdc42b | 6.692989405 | 0.266806993 | 0.144128746 | 1.851171264 | 0.064144918 | NA |
| Ccdc54 | 0.19442443 | -0.026451352 | 0.031568725 | -0.837897386 | 0.402088337 | NA |
| Ccdc64b | 0.197363271 | -0.000561558 | 0.031650166 | -0.017742642 | 0.985844163 | NA |
| Ccdc68 | 3.776624343 | -0.052750183 | 0.117658022 | -0.448334772 | 0.653911609 | NA |
| Ccdc7 | 0 | NA | NA | NA | NA | NA |
| Ccdc70 | 0 | NA | NA | NA | NA | NA |
| Ccdc73 | 1.245860835 | -0.055379176 | 0.0814382 | -0.680014731 | 0.496495133 | NA |
| Ccdc83 | 0.874461425 | 0.034875448 | 0.068401208 | 0.509865971 | 0.610145364 | NA |
| Ccer1 | 0.198019769 | 0.025308392 | 0.031628095 | 0.80018706 | 0.423602426 | NA |
| Ccin | 1.246725701 | 0.063284555 | 0.059794406 | 1.058369155 | NA | NA |
| Ccl1 | 0 | NA | NA | NA | NA | NA |
| Ccl11 | 2.177316376 | 0.111290404 | 0.095260493 | 1.168274494 | 0.24269606 | NA |
| Ccl19 | 0.545128971 | 0.052193958 | 0.044892258 | 1.162649418 | 0.244971772 | NA |
| Ccl2 | 7.657881003 | 0.100684772 | 0.142001661 | 0.709039393 | 0.478300032 | NA |
| Ccl20 | 0.331747315 | -0.03734372 | 0.03744515 | -0.99729124 | 0.318623164 | NA |
| Ccl21a | 0 | NA | NA | NA | NA | NA |
| Ccl21b | 0 | NA | NA | NA | NA | NA |
| Ccl21c | 0 | NA | NA | NA | NA | NA |
| Ccl22 | 7.745376831 | 0.233001805 | 0.14669812 | 1.588308044 | 0.112216699 | NA |
| Ccl24 | 5.011820588 | -0.017888037 | 0.129120674 | -0.138537356 | 0.889815748 | NA |
| Ccl26 | 0 | NA | NA | NA | NA | NA |
| Ccl27b | 0 | NA | NA | NA | NA | NA |
| Ccl5 | 3.853689555 | 0.197422328 | 0.11401529 | 1.731542575 | 0.083355038 | NA |
| Ccl7 | 3.246695258 | -0.094285733 | 0.117179106 | -0.804629228 | 0.421033671 | NA |
| Ccl8 | 3.889292092 | 0.087408529 | 0.112510266 | 0.776893814 | 0.437221421 | NA |
| Ccnb1ip1 | 0.323674185 | -0.015335151 | 0.04024759 | -0.381020351 | 0.703188147 | NA |
| Ccnb3 | 0.159967308 | 0.0240345 | 0.030870969 | 0.778547006 | 0.436246606 | NA |
| Ccr10 | 4.373914362 | 0.133965387 | 0.127212233 | 1.053085725 | 0.292301705 | NA |
| Ccr1l1 | 1.053006083 | 0.006999787 | 0.0772405 | 0.090623268 | 0.927791942 | NA |
| Ccr3 | 1.577672267 | -0.049590578 | 0.095854562 | -0.517352299 | 0.604910252 | NA |
| Ccr4 | 0.159967308 | 0.0240345 | 0.030870969 | 0.778547006 | 0.436246606 | NA |
| Ccr6 | 7.210458705 | 0.070182679 | 0.153807148 | 0.456303104 | 0.648172032 | NA |
| Ccr7 | 3.227672923 | -0.017707268 | 0.125379273 | -0.141229627 | 0.887688542 | NA |
| Ccr8 | 0 | NA | NA | NA | NA | NA |
| Ccrl1 | 4.164982232 | 0.168830964 | 0.106907222 | 1.579228797 | 0.114283587 | NA |
| Cct6b | 5.668538221 | 0.041738134 | 0.133284779 | 0.313150038 | 0.754166675 | NA |
| Cct8l1 | 2.531438489 | 0.135460055 | 0.100688233 | 1.345341474 | 0.178514986 | NA |
| Cd160 | 0.079983654 | 0.016740032 | 0.027674178 | 0.604897175 | 0.54524732 | NA |
| Cd163l1 | 3.295479378 | 0.039348761 | 0.123032794 | 0.319823357 | 0.74910224 | NA |
| Cd177 | 3.22070232 | -0.182423077 | 0.106474589 | -1.71330154 | 0.086657079 | NA |
| Cd19 | 1.431574452 | -0.044563126 | 0.090335411 | -0.493307389 | 0.621795408 | NA |
| Cd1d2 | 0 | NA | NA | NA | NA | NA |
| Cd2 | 2.21153379 | -0.207226095 | 0.10275758 | -2.016650212 | 0.043732029 | NA |
| Cd200r2 | 1.931964708 | -0.008143172 | 0.08424728 | -0.096657978 | 0.922998012 | NA |
| Cd200r3 | 0 | NA | NA | NA | NA | NA |
| Cd200r4 | 6.705642094 | 0.412995008 | 0.140600688 | 2.937361216 | 0.003310183 | NA |
| Cd207 | 0.185670226 | -0.001645086 | 0.031432796 | -0.052336624 | 0.958260472 | NA |
| Cd209d | 2.214390628 | -0.033180086 | 0.100762333 | -0.329290567 | 0.741936073 | NA |
| Cd209e | 0.275064582 | 0.010932281 | 0.038417727 | 0.284563456 | 0.775978612 | NA |
| Cd209g | 5.493189024 | -0.037289074 | 0.130204764 | -0.286387935 | 0.774581015 | NA |
| Cd226 | 0.976978344 | 0.024011131 | 0.080907063 | 0.296774229 | 0.766638882 | NA |
| Cd244 | 2.627008601 | 0.11305792 | 0.09932635 | 1.138246996 | 0.25501736 | NA |
| Cd247 | 3.053224201 | -0.165473618 | 0.118748421 | -1.393480581 | 0.163474513 | NA |
| Cd27 | 0.488467756 | -0.020554715 | 0.055630496 | -0.369486469 | 0.711765157 | NA |
| Cd300c | 0.098353386 | -0.020212831 | 0.02786466 | -0.725393061 | 0.468210929 | NA |
| Cd300e | 4.939095036 | 0.126572946 | 0.134377331 | 0.941921865 | 0.346232645 | NA |
| Cd300lb | 2.653118307 | 0.042060645 | 0.113232116 | 0.37145508 | 0.710298607 | NA |
| Cd300ld | 3.963134501 | -0.112237914 | 0.124950874 | -0.898256334 | 0.369048907 | NA |
| Cd300lf | 6.429006114 | 0.235143999 | 0.104052941 | 2.259849607 | NA | NA |
| Cd300lg | 5.582805756 | 0.094471637 | 0.150070218 | 0.62951622 | 0.529011153 | NA |
| Cd300lh | 0.116696964 | -0.020477583 | 0.027954224 | -0.732539847 | 0.463839131 | NA |
| Cd3d | 0.879765535 | 0.027449169 | 0.061992718 | 0.442780535 | 0.657924484 | NA |
| Cd3e | 1.473692792 | 0.064687221 | 0.076260439 | 0.848240873 | 0.396303839 | NA |
| Cd3g | 0.588168521 | -0.033702544 | 0.053860702 | -0.625735339 | 0.53148855 | NA |
| Cd40lg | 0.198019769 | 0.025308392 | 0.031628095 | 0.80018706 | 0.423602426 | NA |
| Cd5 | 5.295899881 | 0.160072967 | 0.128744657 | 1.243336774 | 0.213743758 | NA |
| Cd5l | 0.48156345 | -0.012077111 | 0.044778632 | -0.269707011 | 0.787385666 | NA |
| Cd69 | 0.810921301 | 0.051807665 | 0.063085565 | 0.821228515 | 0.411516117 | NA |
| Cd70 | 1.912147608 | 0.005744978 | 0.094550521 | 0.060760932 | 0.951549604 | NA |
| Cd8a | 1.366151416 | -0.005040101 | 0.076024171 | -0.066296029 | 0.947142145 | NA |
| Cd8b1 | 0.884508339 | -0.04816716 | 0.06562523 | -0.733973201 | 0.462965071 | NA |
| Cd96 | 1.779697318 | -0.024466514 | 0.09673155 | -0.252932101 | 0.800320683 | NA |
| Cdcp2 | 0.098353386 | -0.020212831 | 0.02786466 | -0.725393061 | 0.468210929 | NA |
| Cdh16 | 0 | NA | NA | NA | NA | NA |
| Cdh17 | 2.472933667 | 0.014869444 | 0.112928582 | 0.131671217 | 0.895244353 | NA |
| Cdhr5 | 0.846679412 | 0.067107966 | 0.071303602 | 0.941158145 | 0.346623822 | NA |
| Cdkn1a | 1019.537859 | 0.11224457 | 0.152161752 | 0.737666122 | NA | NA |
| Cdkn2a | 4.153287324 | 0.340939665 | 0.125446185 | 2.717816134 | 0.006571434 | NA |
| Cdr1 | 1.191371392 | 0.085765618 | 0.070050215 | 1.224344827 | 0.220822173 | NA |
| Cdrt4 | 1.650567937 | -0.033895378 | 0.092427697 | -0.366723165 | 0.713825525 | NA |
| Cdx1 | 0 | NA | NA | NA | NA | NA |
| Cdx2 | 0 | NA | NA | NA | NA | NA |
| Cdx4 | 0 | NA | NA | NA | NA | NA |
| Ceacam-ps1 | 1.014480811 | 0.116186393 | 0.073507767 | 1.580600225 | 0.113969474 | NA |
| Ceacam10 | 4.137747377 | 0.061850514 | 0.123778286 | 0.499687921 | 0.617294839 | NA |
| Ceacam11 | 0 | NA | NA | NA | NA | NA |
| Ceacam12 | 0 | NA | NA | NA | NA | NA |
| Ceacam13 | 0 | NA | NA | NA | NA | NA |
| Ceacam14 | 0 | NA | NA | NA | NA | NA |
| Ceacam15 | 1.645313989 | -0.050038391 | 0.097491763 | -0.513257626 | 0.607771121 | NA |
| Ceacam16 | 0.730358311 | 0.059975352 | 0.057700588 | 1.039423589 | 0.298607778 | NA |
| Ceacam18 | 0.412387467 | -0.005395598 | 0.042599376 | -0.126659095 | 0.899210223 | NA |
| Ceacam19 | 0 | NA | NA | NA | NA | NA |
| Ceacam20 | 0 | NA | NA | NA | NA | NA |
| Ceacam3 | 1.08876445 | 0.018664533 | 0.066257674 | 0.281696177 | 0.778176486 | NA |
| Ceacam5 | 0 | NA | NA | NA | NA | NA |
| Cebpe | 0.307917359 | 0.010394288 | 0.03404619 | 0.305299606 | 0.760137976 | NA |
| Cel | 0.393805724 | -0.036573061 | 0.048755423 | -0.750133191 | 0.453174491 | NA |
| Cela2a | 0.258977192 | 0.029727007 | 0.03417262 | 0.869907163 | 0.38435114 | NA |
| Cela3b | 0 | NA | NA | NA | NA | NA |
| Cep55 | 8.057431714 | -0.029397394 | 0.156868106 | -0.187401985 | 0.851345466 | NA |
| Cer1 | 4.428946437 | -0.038589098 | 0.132362777 | -0.291540408 | 0.770638044 | NA |
| Cers1 | 0 | NA | NA | NA | NA | NA |
| Ces1a | 2.303102152 | -0.246464794 | 0.102371746 | -2.407547029 | 0.016060094 | NA |
| Ces1b | 0 | NA | NA | NA | NA | NA |
| Ces1c | 0 | NA | NA | NA | NA | NA |
| Ces1d | 4.568624031 | 0.036060239 | 0.140350017 | 0.256930775 | 0.797232213 | NA |
| Ces1e | 0 | NA | NA | NA | NA | NA |
| Ces1f | 3.2535107 | -0.019126911 | 0.109824549 | -0.174158798 | 0.861740665 | NA |
| Ces1g | 0.213790458 | -0.002646789 | 0.031909369 | -0.082947074 | 0.933893623 | NA |
| Ces2a | 0 | NA | NA | NA | NA | NA |
| Ces2b | 0.838780774 | 0.045559078 | 0.062686379 | 0.726777954 | 0.46736199 | NA |
| Ces2c | 0 | NA | NA | NA | NA | NA |
| Ces2d-ps | 0 | NA | NA | NA | NA | NA |
| Ces2e | 0.763784651 | -0.00317115 | 0.056229574 | -0.056396483 | 0.955025958 | NA |
| Ces2f | 0 | NA | NA | NA | NA | NA |
| Ces2h | 0 | NA | NA | NA | NA | NA |
| Ces3a | 0 | NA | NA | NA | NA | NA |
| Ces3b | 0 | NA | NA | NA | NA | NA |
| Ces4a | 0 | NA | NA | NA | NA | NA |
| Cetn1 | 0 | NA | NA | NA | NA | NA |
| Cfc1 | 0.203787646 | -0.02671144 | 0.031728022 | -0.841887971 | 0.399850664 | NA |
| Cfd | 0 | NA | NA | NA | NA | NA |
| Cfhr1 | 0 | NA | NA | NA | NA | NA |
| Cfhr2 | 0.572845755 | -0.029822974 | 0.053532279 | -0.557102627 | 0.577457312 | NA |
| Cfi | 0.871541253 | 0.00020978 | 0.064142277 | 0.003270542 | 0.99739049 | NA |
| Cga | 4.56743289 | 0.289719723 | 0.105259322 | 2.752437679 | 0.00591534 | NA |
| Chad | 3.328561322 | -0.006873845 | 0.126534448 | -0.054323906 | 0.956677103 | NA |
| Chi3l3 | 7.451930034 | -0.057053337 | 0.137564507 | -0.41473879 | 0.678333133 | NA |
| Chi3l4 | 0.098353386 | -0.020212831 | 0.02786466 | -0.725393061 | 0.468210929 | NA |
| Chit1 | 5.854022429 | 0.04452478 | 0.147620979 | 0.30161553 | 0.762945168 | NA |
| Chmp4c | 3.675516495 | 0.040397073 | 0.114601444 | 0.352500558 | 0.7244629 | NA |
| Chrdl2 | 5.33060177 | 0.084383174 | 0.143046078 | 0.589902044 | 0.555256323 | NA |
| Chrna9 | 0.217751027 | -0.027071875 | 0.031947577 | -0.847384294 | 0.396780956 | NA |
| Chrnd | 0 | NA | NA | NA | NA | NA |
| Chrne | 1.690479618 | -0.035285547 | 0.098151323 | -0.359501494 | 0.719219961 | NA |
| Chrng | 0.096071043 | -0.020098795 | 0.027825948 | -0.722304036 | 0.470107572 | NA |
| Chst13 | 1.103949313 | -0.080511457 | 0.075243076 | -1.070018145 | 0.284611141 | NA |
| Chst4 | 4.187077356 | -0.01092005 | 0.114141695 | -0.095670996 | 0.923781877 | NA |
| Cib3 | 0.470878708 | 0.028927796 | 0.04534639 | 0.637929415 | 0.523519629 | NA |
| Cib4 | 0 | NA | NA | NA | NA | NA |
| Cidec | 0.331243956 | 0.008596723 | 0.034710864 | 0.247666645 | 0.804392342 | NA |
| Ciita | 9.936993295 | 0.141291351 | 0.109518408 | 1.290115088 | NA | NA |
| Cilp | 5.520717217 | -0.435128263 | 0.144845084 | -3.004094107 | 0.002663729 | NA |
| Ckm | 0.969645158 | 0.020592352 | 0.076153894 | 0.270404457 | 0.786849112 | NA |
| Ckmt2 | 1.03630432 | 0.003079144 | 0.064486102 | 0.047748952 | 0.961916321 | NA |
| Clca2 | 2.012024816 | 0.049548493 | 0.09627614 | 0.514649766 | 0.606797782 | NA |
| Clca3 | 0.7238653 | 0.106446607 | 0.063481683 | 1.676808206 | 0.093579993 | NA |
| Clca4 | 0.192142086 | -0.026384592 | 0.031527717 | -0.836869715 | 0.402665805 | NA |
| Clca5 | 0.352713512 | 0.01988513 | 0.041853977 | 0.475107309 | 0.634710489 | NA |
| Clca6 | 0.105923158 | 0.018524693 | 0.027977518 | 0.662127842 | 0.507889294 | NA |
| Clcnka | 6.973684801 | 0.073460396 | 0.063858896 | 1.150354928 | NA | NA |
| Clcnkb | 1.171397964 | 0.084597537 | 0.085157829 | 0.993420544 | 0.320505054 | NA |
| Cldn13 | 0.236094605 | -0.028257939 | 0.032632891 | -0.865934257 | 0.386526215 | NA |
| Cldn15 | 5.074131718 | -0.051908619 | 0.133040948 | -0.390170243 | 0.696410664 | NA |
| Cldn16 | 0.105923158 | 0.018524693 | 0.027977518 | 0.662127842 | 0.507889294 | NA |
| Cldn17 | 0 | NA | NA | NA | NA | NA |
| Cldn18 | 0.195033442 | -0.002424765 | 0.031596358 | -0.076741901 | 0.93882887 | NA |
| Cldn2 | 30.84619758 | 0.280923164 | 0.117259105 | 2.395747128 | NA | NA |
| Cldn20 | 7.192744513 | 0.061794147 | 0.156725465 | 0.39428275 | 0.693372298 | NA |
| Cldn24 | 0.119397641 | -0.020477583 | 0.027954224 | -0.732539845 | 0.463839132 | NA |
| Cldn3 | 3.452382754 | 0.224953466 | 0.121639461 | 1.849346131 | 0.064407849 | NA |
| Cldn4 | 0.079983654 | 0.016740032 | 0.027674178 | 0.604897175 | 0.54524732 | NA |
| Cldn6 | 0.79738299 | 0.056420574 | 0.069491893 | 0.811901524 | 0.41684814 | NA |
| Cldn7 | 0.358192922 | -0.030696978 | 0.034083535 | -0.900639514 | 0.367780018 | NA |
| Cldn8 | 0 | NA | NA | NA | NA | NA |
| Cldn9 | 3.68848754 | 0.142388042 | 0.12567826 | 1.132956819 | 0.257232387 | NA |
| Cldnd2 | 0.68889895 | 0.008020532 | 0.060682319 | 0.132172472 | 0.894847874 | NA |
| Clec12b | 1.117520908 | -0.112944557 | 0.069043422 | -1.635848203 | 0.101871361 | NA |
| Clec1b | 2.16508377 | -0.053897296 | 0.107256072 | -0.502510441 | 0.61530851 | NA |
| Clec2e | 0.724560782 | -0.05162733 | 0.062704856 | -0.823338625 | 0.410315455 | NA |
| Clec2g | 0.500981137 | -0.05018208 | 0.04329235 | -1.159144275 | 0.246397381 | NA |
| Clec2h | 0.236094605 | -0.028257939 | 0.032632891 | -0.865934257 | 0.386526215 | NA |
| Clec2i | 1.473017754 | -0.004436983 | 0.091300537 | -0.048597552 | 0.96124002 | NA |
| Clec3a | 1.276319948 | 0.026881504 | 0.082439033 | 0.326077382 | 0.744365809 | NA |
| Clec4a4 | 0.079983654 | 0.016740032 | 0.027674178 | 0.604897175 | 0.54524732 | NA |
| Clec4b1 | 0.298180463 | -0.015304498 | 0.039412854 | -0.388312359 | 0.697784893 | NA |
| Clec4b2 | 0 | NA | NA | NA | NA | NA |
| Clec4d | 1.084798614 | 0.060484599 | 0.072840762 | 0.830367456 | 0.40633106 | NA |

| | | | | | | | |
|---|---|---|---|---|---|---|---|
| Duxbl3 | 0 | NA | NA | NA | NA | NA | NM_001177539 |
| Dydc1 | 1.328186792 | 0.00296812948729158 | 0.0246783380222547 | 0.120272665226278 | 0.904267156893977 | NA | NM_027094 |
| Dydc2 | 15.28233037 | 0.0101533599707385 | 0.068766804239148 | 0.147649146751513 | 0.882619666636351 | NA | NM_027717 |
| Dynap | 0 | NA | NA | NA | NA | NA | NM_029346 |
| Dynlt1c | 0 | NA | NA | NA | NA | NA | NM_001166630 |
| Dynlt1f | 0 | NA | NA | NA | NA | NA | NM_001166627,NM_001199948 |
| Dyrk4 | 8.664852141 | -0.09605529065802 | 0.0588154825545159 | -1.63316335233647 | 0.102434645356725 | NA | NM_207210 |
| Dytn | 3.982354247 | -0.0102020625049651 | 0.0461012102647552 | -0.221297064575432 | 0.824861135458024 | NA | NM_001081658 |
| E030002O03Rik | 2.31887147 | -0.0459475312048293 | 0.0325282359134077 | -1.41254297734266 | 0.157790145546769 | NA | NM_172905 |
| E030011O05Rik | 16.68083249 | -0.0754921559462913 | 0.0698878162409548 | -1.08019051111877 | 0.280057352970746 | NA | NR_015511 |
| E030018B13Rik | 0 | NA | NA | NA | NA | NA | NM_001256311 |
| E030019B13Rik | 19.86825102 | 0.024744717307597 | 0.0755428756892435 | 0.327558582881964 | 0.743245441685269 | NA | NR_045082 |
| E030024N20Rik | 0.905180643 | -0.0108831505862658 | 0.0192382192696352 | -0.565704675351282 | 0.571594556438066 | NA | NR_033228 |
| E030025P04Rik | 0.217374753 | -0.00904456656736547 | 0.00999660569474087 | -0.90476376117583 | 0.365590553873005 | NA | NR_037978 |
| E030030I06Rik | 20.21601571 | -0.0941887715489643 | 0.0794155370240079 | -1.18602448687705 | 0.23561262947407 | NA | NM_001254745,NM_001254744 |
| E030044B06Rik | 0.657073752 | 0.0133411052068515 | 0.0178061472719538 | 0.749241540187911 | 0.4537116359395 | NA | NR_045343 |
| E130006D01Rik | 16.47743923 | -0.0165112843068237 | 0.0762175727182412 | -0.216633562549442 | 0.828493927659038 | NA | NR_045834,NR_045832,NR_045833 |
| E130018N17Rik | 1.953163287 | 0.00377544668580255 | 0.0228669943992227 | 0.165104631587739 | 0.868861627563907 | NA | NR_040327 |
| E130112N10Rik | 21.87284141 | 0.0323362631345587 | 0.0818336566204327 | 0.395146257297817 | 0.692734952384498 | NA | NR_015604 |
| E130201H02Rik | 3.035542264 | -0.0019845396360008 | 0.0358334085725827 | -0.0553823851833965 | 0.955833828884879 | NA | NR_024324 |
| E130215H24Rik | 14.56544889 | -0.018453165890621 | 0.0707400363209767 | -0.260858869323892 | 0.794201344413823 | NA | NR_040331 |
| E130218I03Rik | 0.526995524 | 0.00942377320457171 | 0.0150941210165823 | 0.624334016814811 | 0.532408248420893 | NA | NR_040436,NR_040435 |
| E130304I02Rik | 3.724907869 | -0.000246464158636182 | 0.044546988075622 | -0.00553267839831932 | 0.995585583847375 | NA | NR_033567 |
| E130310I04Rik | 2.526221834 | -0.00706362802949866 | 0.0363769488526077 | -0.194178683267778 | 0.846035973479127 | NA | NR_045722,NR_045723 |
| E230016K23Rik | 3.566196814 | 0.0235383581108071 | 0.043084847197304 | 0.546325672295293 | 0.584842092316135 | NA | NR_036452,NR_036453 |
| E230019M04Rik | 1.855420341 | -0.00589850747867529 | 0.0276466567197569 | -0.213353373554933 | 0.831051345682953 | NA | NM_177921 |
| E2f7 | 16.08061732 | -0.0526080820765697 | 0.0764495307998018 | -0.688141333585609 | 0.491363783627998 | NA | NM_178609 |
| E2f8 | 17.80085121 | -0.0784626997023253 | 0.0745062931760485 | -1.05310164225897 | 0.292294410273729 | NA | NM_001013368 |
| E330011O21Rik | 15.51602597 | -0.0915560778445175 | 0.0714565464318162 | -1.28128327517032 | 0.200094185216017 | NA | NR_045698 |
| E330012B07Rik | 0.882475236 | -0.0155605875899721 | 0.0222774358371339 | -0.69849096205383 | 0.484870208076359 | NA | NR_033640 |
| E330013P04Rik | 6.045918369 | 0.0206740587554231 | 0.0491981555246424 | 0.420220200024935 | 0.674324599436898 | NA | NR_026942 |
| E330014E10Rik | 0 | NA | NA | NA | NA | NA | NM_001122668 |
| E330017A01Rik | 0.106148608 | 0.00515554063321497 | 0.00854200771766808 | 0.603551390213729 | 0.546141939141592 | NA | NM_175011 |
| E330017L17Rik | 2.423488993 | -0.0370331052268538 | 0.037982640153531 | -0.975000818193811 | 0.329559854106012 | NA | NR_045190 |
| E330021D16Rik | 0.212297216 | 0.00681050119636477 | 0.00973439053551308 | 0.699633035218656 | 0.484156506270654 | NA | NM_001201390 |
| E330023G01Rik | 0.760890561 | -0.0208141952908415 | 0.0175374649033995 | -1.18684173599155 | 0.235290052800064 | NA | NR_045332 |
| E330034G19Rik | 6.454303816 | 0.0571075339367344 | 0.0583644295760472 | 0.97846469761732 | 0.327844543173712 | NA | NM_001033214 |
| E430016F16Rik | 5.605175975 | -0.0764918358416614 | 0.0486590300123542 | -1.57199672542261 | 0.115951310642808 | NA | NR_015542 |
| EU599041 | 0 | NA | NA | NA | NA | NA | NM_001177525 |
| Ear1 | 0.106347839 | -0.00653347234140708 | 0.00854389509723638 | -0.764694822098226 | 0.444453290197145 | NA | NM_007894 |
| Ear10 | 0 | NA | NA | NA | NA | NA | NM_053112 |
| Ear11 | 0.576646845 | -0.0149496061681509 | 0.0157865123101243 | -0.946986001370507 | 0.343645912207582 | NA | NM_053113 |
| Ear12 | 0 | NA | NA | NA | NA | NA | NM_001012766 |
| Ear14 | 0 | NA | NA | NA | NA | NA | NM_017389 |
| Ear2 | 0.364141684 | 0.0106460894690162 | 0.0121537141469398 | 0.875953584254473 | 0.381055267460423 | NA | NM_007895 |
| Ear3 | 0 | NA | NA | NA | NA | NA | NM_017388 |
| Ear4 | 0 | NA | NA | NA | NA | NA | NM_001017422 |
| Ear5 | 0 | NA | NA | NA | NA | NA | NM_019398 |
| Ear6 | 0 | NA | NA | NA | NA | NA | NM_053111 |
| Ear7 | 0 | NA | NA | NA | NA | NA | NM_017385 |
| Eci3 | 3.102882784 | 0.0100947022073126 | 0.0323331667713061 | 0.312208893076043 | 0.754881773978464 | NA | NM_026947 |
| Ect2 | 20.38607892 | -0.155305497362072 | 0.0808257594761444 | -1.92148515978883 | 0.0546705715160091 | NA | NM_001177626,NM_007900,NM_001177625 |
| Ect2l | 14.96070415 | 0.112604493511607 | 0.0683848073121702 | 1.64663026683073 | 0.0996340653574421 | NA | NM_001195036 |
| Eda2r | 20.20779828 | -0.0683243870828128 | 0.080001275518825 | -0.854039971734394 | 0.393082842294262 | NA | NM_175540,NM_001161432,NM_001161433 |
| Edar | 9.896441771 | -0.00875373143166108 | 0.0630655538958066 | -0.138803687447564 | 0.889605280033684 | NA | NM_010100 |
| Edaradd | 14.00363609 | 0.136016050593327 | 0.0696810647062305 | 1.95198008478716 | 0.0509405690968704 | NA | NM_133643 |
| Eddm3b | 0 | NA | NA | NA | NA | NA | NM_203508 |
| Edn2 | 1.743607036 | 0.00168044210280648 | 0.0316486482761846 | 0.0530968048980154 | 0.957654777185736 | NA | NM_007902 |
| Efcab11 | 8.751759098 | 0.0118225717774251 | 0.0586183022071943 | 0.201687379747653 | 0.840161129357559 | NA | NM_030172 |
| Efcab3 | 12.93209589 | -0.0483804685017254 | 0.0591952733243403 | -0.817302899112251 | 0.413755350422808 | NA | NM_001081046 |
| Efcab8 | 0.423505437 | -0.00171692463125296 | 0.015208307437769 | -0.112893866610631 | 0.910114698980405 | NA | NR_036629 |
| Efcab9 | 7.328702053 | 0.0159752590022789 | 0.053916424858555 | 0.296296704467861 | 0.767003502478569 | NA | NM_027031 |
| Efhb | 12.24538815 | -0.000622962041176636 | 0.0621994856816005 | -0.0100155497163687 | 0.992008881113415 | NA | NM_172497 |
| Efna4 | 7.90850905 | -0.0699528231030411 | 0.0597156956023157 | -1.17143110194849 | 0.241425536735511 | NA | NM_007910 |
| Egfbp2 | 0 | NA | NA | NA | NA | NA | NM_010115 |
| Ehf | 8.295292148 | -0.0164055547127515 | 0.04879492564921 | -0.336214360294186 | 0.73670922466165 | NA | NM_007914 |
| Eid3 | 4.163133451 | -0.0227755170292985 | 0.0420027732318381 | -0.542238411344578 | 0.587654277252063 | NA | NM_025499 |
| Eif2s3y | 0.300216221 | 0.00934385809732901 | 0.0113912670188907 | 0.820265040037567 | 0.412065031496518 | NA | NM_012011 |
| Eif3j1 | 0 | NA | NA | NA | NA | NA | NM_144545 |
| Eif3j2 | 0 | NA | NA | NA | NA | NA | NM_001256055 |
| Eif4e1b | 0.191449919 | 0.00665508079272897 | 0.00962405648474729 | 0.691504752001018 | 0.489248395458281 | NA | NM_001033269,NM_001039683 |
| Elane | 0.980505671 | -0.00920314633300144 | 0.0214995462809833 | -0.428062351303748 | 0.66860572602786 | NA | NM_015779 |
| Elf3 | 3.9942423 | -0.0351946201802625 | 0.0408789436665342 | -0.860947397940588 | 0.389267014955284 | NA | NM_007921,NM_001163131 |
| Elf5 | 13.05146958 | -0.0170425936162181 | 0.0731778071542921 | -0.232892925860494 | 0.815844551768932 | NA | NM_010125,NM_001145813 |
| Ell3 | 2.349348143 | -0.0141618026899924 | 0.0365417216760836 | -0.38755159966262 | 0.698347892559491 | NA | NM_145973 |
| Elovl3 | 0.845218932 | 0.0128293511123917 | 0.0207928797093263 | 0.617006941402025 | 0.537230147247559 | NA | NM_007703 |
| Eme1 | 2.806630389 | 0.00552200705017815 | 0.0402208763234677 | 0.137292062106469 | 0.890799944545679 | NA | NM_177752 |
| Emilin3 | 8.02964709 | -0.0111055474886674 | 0.0503636038072782 | -0.220507403147002 | 0.825476007390453 | NA | NM_182840 |
| Emr4 | 1.538712315 | 0.0015484591416534 | 0.0285327649462673 | 0.0542695089161338 | 0.956720442100622 | NA | NM_139138 |
| En1 | 3.673177089 | 0.0128441296976608 | 0.0303206676381277 | 0.423609725582344 | 0.671850461334341 | NA | NM_010133 |
| Enam | 0 | NA | NA | NA | NA | NA | NM_017468 |
| Eno1 | 2.300857202 | -0.0398136064389617 | 0.0326745911896115 | -1.21848828063134 | 0.223038475586472 | NA | NM_023119 |
| Enpp2 | 17413.19297 | 0.0154524055306892 | 0.0596177094627807 | 0.259191533353627 | NA | NA | NM_015744,NM_001136077 |
| Enpp3 | 20.61971984 | -0.0436717299485718 | 0.0721981750624341 | -0.604886895143904 | 0.545254151174968 | NA | NM_134005 |
| Enpp7 | 0 | NA | NA | NA | NA | NA | NM_001030291 |
| Enthd1 | 1.245331172 | -0.00829646054887064 | 0.0245020775622383 | -0.338602329855361 | 0.734909327213527 | NA | NM_001163189 |
| Entpd8 | 0.097165704 | -0.00637828455464458 | 0.00848781691535724 | -0.75146349388194 | 0.452373763047372 | NA | NM_028093 |
| Eomes | 8.104965014 | 0.014502886546257 | 0.0519410953246528 | 0.279217957488345 | 0.780077563379226 | NA | NM_010136,NM_001164789 |
| Epb4.2 | 0.688850708 | -0.0152035927138371 | 0.019432969118874 | -0.782360771575084 | 0.434002581159999 | NA | NM_013513 |
| Epcam | 5.809712952 | 0.000676377130091887 | 0.0525599412332858 | 0.0128686812469939 | 0.989732561302281 | NA | NM_008532 |
| Epgn | 2.744144728 | -0.00118104907033327 | 0.0346173845029002 | -0.0341172242586475 | 0.972783673508232 | NA | NM_053087 |
| Epha1 | 12.27091342 | 0.00184044524476699 | 0.0709277674103017 | 0.0259481626444043 | 0.979298684728138 | NA | NM_023580 |
| Ephx3 | 12.14611267 | -0.0812550618053063 | 0.0710120498026722 | -1.14424329435775 | 0.252522752778033 | NA | NM_001033163 |
| Epo | 0.561306268 | 0.0121683135725874 | 0.016940050114277 | 0.718316267691089 | 0.472562301925324 | NA | NM_007942 |
| Eppin | 0.09572496 | 0.00498255289994511 | 0.00847999972823692 | 0.587565219295241 | 0.556824162160132 | NA | NM_029325 |
| Eppk1 | 6.822632887 | -0.0721498171550582 | 0.0531357734558523 | -1.35783884307252 | 0.174514826595876 | NA | NM_144848 |
| Eps8l3 | 5.085628937 | -0.027453278332267 | 0.0446462161887184 | -0.614907167411964 | 0.538616029167188 | NA | NM_133867 |
| Epx | 12.83652883 | -0.0518542124549119 | 0.0744537337480562 | -0.696462216796021 | 0.486139411229709 | NA | NM_007946 |
| Epyc | 3.0075103 | -0.0258149095752076 | 0.0395545625762302 | -0.652640501976394 | 0.513988068144861 | NA | NM_007884 |
| Eras | 0 | NA | NA | NA | NA | NA | NM_181548 |
| Ercc6l | 5.57820661 | -0.0709070940574881 | 0.0458296411325388 | -1.54718850737726 | 0.121817796436168 | NA | NM_146235 |
| Ereg | 0.240418098 | -0.0088029805258837 | 0.00986414519853158 | -0.892422034419582 | 0.372166765031741 | NA | NM_007950 |
| Erich2 | 16.79315303 | 0.0165252766817685 | 0.07547972282106 | 0.218936637074636 | 0.826699406464539 | NA | NM_025744 |
| Ern2 | 15.78449249 | 0.0194662678679282 | 0.0574846791552159 | 0.338634017863557 | 0.734885452596811 | NA | NM_012016 |
| Erp27 | 3.529799523 | -0.0306716297785055 | 0.0403590583641486 | -0.759968914580808 | 0.447273166242724 | NA | NM_026983 |
| Erv3 | 17.79198645 | -0.0989510495953285 | 0.0798286168546527 | -1.23954358091276 | 0.215144259734309 | NA | NM_001166206 |
| Esco2 | 6.97862036 | -0.0175973710040662 | 0.0535071415250656 | -0.328878921626239 | 0.74224720698928 | NA | NM_028039 |
| Esm1 | 10.29384499 | -0.0523370941339855 | 0.0584211149743522 | -0.895859213864067 | 0.370327959805934 | NA | NM_023612 |
| Esrp1 | 4.255649014 | 0.0198315685420842 | 0.0445578430996609 | 0.445074697572943 | 0.656265773395379 | NA | NM_194055 |
| Esrp2 | 3.655329606 | 0.0687490422381653 | 0.0420300637756672 | 1.63571110919814 | 0.10190006334359 | NA | NM_176838 |
| Esx1 | 0 | NA | NA | NA | NA | NA | NM_007957 |
| Etd | 0 | NA | NA | NA | NA | NA | NR_034074 |
| Etv2 | 0 | NA | NA | NA | NA | NA | NM_007959 |
| Evx1 | 0.12890128 | -0.0068640067066208 | 0.00870804083610955 | -0.788237771940376 | 0.430557639008166 | NA | NM_007966 |
| Evx2 | 0 | NA | NA | NA | NA | NA | NM_007967 |
| F10 | 0.698589963 | -0.0144374447495517 | 0.0194736370359372 | -0.741384093937275 | 0.458460586007868 | NA | NM_007972,NM_001242368 |
| F11 | 0.89196579 | -0.00625981745031726 | 0.0223463230867447 | -0.280127402884926 | 0.779379761535478 | NA | NM_028066 |
| F13b | 0 | NA | NA | NA | NA | NA | NM_031164 |
| F2 | 2.526762571 | -0.0243696469889379 | 0.0300423780859011 | -0.81117569718539 | 0.417264779937834 | NA | NM_010168 |
| F2rl1 | 15.75543931 | -0.163090073471495 | 0.07283695460609 | -2.23911164811192 | 0.0251486525601896 | NA | NM_007974 |
| F2rl3 | 6.835461511 | 0.0110381138054626 | 0.0537616914094247 | 0.20531559770695 | 0.837325558433517 | NA | NM_007975 |
| F420014N23Rik | 11.88504573 | -0.0109613610125546 | 0.0712303926246605 | -0.153886011415297 | 0.877699613992795 | NA | NR_045715,NR_045716 |
| F5 | 171.4930781 | -0.000919091501415032 | 0.0311921351533225 | -0.0294654885565643 | NA | NA | NM_007976 |
| F630028O10Rik | 1.975428467 | -0.0187546529927891 | 0.026539363973798 | -0.706673039011236 | 0.479769687918487 | NA | NR_030718 |
| F630111L10Rik | 0 | NA | NA | NA | NA | NA | NR_045641 |
| F630206G17Rik | 0.328758195 | 0.0127026590537822 | 0.0132741836670923 | 0.95694465078655 | 0.338595192596037 | NA | NR_045876 |
| F7 | 2.656525822 | -0.0122785607777526 | 0.0353796939474885 | -0.347051073872396 | 0.728552949061255 | NA | NM_010172 |
| F730035M05Rik | 0.882616281 | 0.00455182326906565 | 0.0205688653981807 | 0.2212967599792 | 0.824861372611985 | NA | NR_045174 |
| F830002L21Rik | 0 | NA | NA | NA | NA | NA | NR_033558 |
| F830016B08Rik | 11.40428454 | -0.108075122035293 | 0.0432007651219775 | -2.50169462809612 | 0.0123600484442559 | NA | NM_001101475 |
| F830045P16Rik | 1.441765289 | 0.00839437432097663 | 0.0258918173576338 | 0.32420954485459 | 0.745779396288565 | NA | NM_177653 |
| F9 | 9.353768716 | -0.122744505867291 | 0.0671055318211071 | -1.82912648981769 | 0.0673806614593992 | NA | NM_007979 |
| Fabp1 | 0 | NA | NA | NA | NA | NA | NM_017399 |
| Fabp12 | 0.255375383 | 0.00708304372802858 | 0.00992471678592415 | 0.713677163873753 | 0.475426830432214 | NA | NM_029310 |
| Fabp2 | 0 | NA | NA | NA | NA | NA | NM_007980 |
| Fabp4 | 4.619292328 | 0.0416379703126382 | 0.046247626914181 | 0.900326634919092 | 0.367946450775821 | NA | NM_024406 |
| Fabp6 | 0 | NA | NA | NA | NA | NA | NM_008375 |
| Fabp9 | 0 | NA | NA | NA | NA | NA | NM_011598 |
| Faim3 | 2.3892318 | -0.00470538286190562 | 0.0336519597537594 | -0.139824928364832 | 0.888798316791153 | NA | NM_026976 |
| Fam101a | 16.22956798 | -0.0452756480787423 | 0.0774037987807388 | -0.584927985343385 | 0.558596140343389 | NA | NM_028443 |
| Fam110c | 1.549691887 | 0.030214095715753 | 0.0274006473698998 | 1.10267817062395 | 0.270166949818879 | NA | NM_027828 |
| Fam115e | 0 | NA | NA | NA | NA | NA | NM_203396 |
| Fam122c | 0 | NA | NA | NA | NA | NA | NM_028671 |
| Fam124b | 0.67239973 | -0.00319004940147154 | 0.0190122711351518 | -0.167788970544054 | 0.866749299714491 | NA | NM_173425 |
| Fam150a | 1.619597959 | 0.0155055171159973 | 0.0267790018208326 | 0.579017740083758 | 0.562577203100415 | NA | NM_001195732 |
| Fam151a | 1.957808024 | -0.0199240901152225 | 0.0283959186768662 | -0.70165330243231 | 0.482895401773012 | NA | NM_146149 |
| Fam154a | 1.466440785 | -0.0230401269379799 | 0.0275858908491462 | -0.835214170315367 | 0.403597132619808 | NA | NM_001081096 |
| Fam159a | 2.640132242 | 0.0315697440620897 | 0.0289909445657021 | 1.08895189636003 | 0.27617509906612 | NA | NM_001099303 |
| Fam159b | 18.45272737 | 0.0212744932114464 | 0.0821361010030967 | 0.259015134047381 | 0.795623563584992 | NA | NM_029984 |
| Fam162b | 0 | NA | NA | NA | NA | NA | NM_029894 |
| Fam166a | 3.554082237 | -0.0145159005238286 | 0.0380377408616693 | -0.381618366259399 | 0.702744459164507 | NA | NM_026624 |
| Fam167b | 10.99419472 | -0.121181218310297 | 0.0629020035058064 | -1.92650808489926 | 0.0540409633164469 | NA | NM_182783 |
| Fam170a | 0.106347839 | -0.00653347234140708 | 0.00854389509723638 | -0.764694822098226 | 0.444453290197145 | NA | NM_001004061 |
| Fam170b | 0.554225534 | 0.0015352068880447 | 0.0130113578911368 | 0.11798975179143 | 0.906075778045834 | NA | NM_001164485 |
| Fam177a | 0 | NA | NA | NA | NA | NA | NM_028527 |
| Fam178b | 7.044463249 | 0.046700274248172 | 0.0471942658325465 | 0.989532804978316 | 0.322402527742016 | NA | NM_001126046,NM_201365,NM_027957 |
| Fam187a | 3.830482669 | 0.0336605811320045 | 0.0438390766040757 | 0.767821399068317 | 0.442593299437994 | NA | NM_025766 |
| Fam187b | 3.085768913 | 0.0632342404356452 | 0.0375829463435336 | 1.68252483074747 | 0.092467114102165 | NA | NM_001242647,NM_175240,NR_038860 |
| Fam19a3 | 8.242350781 | 0.00782372551790491 | 0.0577570420945003 | 0.135459248503481 | 0.892248782210369 | NA | NM_183224 |
| Fam19a4 | 21.40448009 | 0.10662718451892 | 0.0778554662715416 | 1.36955296301264 | 0.170826491173858 | NA | NM_177233 |
| Fam209 | 1.629426551 | 0.017575569331208 | 0.0224807597958978 | 0.781804951913373 | 0.434329210409221 | NA | NM_029608 |
| Fam227b | 2.743729671 | 0.0572525103681588 | 0.0349457782106958 | 1.63832409234016 | 0.101354104482701 | NA | NM_029455 |
| Fam229a | 4.049355045 | -0.0140372892462629 | 0.0447623574474405 | -0.31359584362252 | 0.753828016724078 | NA | NM_001085491 |
| Fam24a | 0 | NA | NA | NA | NA | NA | NM_183272 |
| Fam25c | 0.348893173 | 0.00231788807319956 | 0.0128591005169735 | 0.180252737750983 | 0.856954157665002 | NA | NM_183278 |
| Fam26d | 0.80879701 | -0.0187453513438165 | 0.020679607618227 | -0.906465523421942 | 0.364689505184929 | NA | NM_001081165 |
| Fam26f | 8.113123751 | -0.0380617745695285 | 0.0611192767647619 | -0.622745827245666 | 0.53345156253807 | NA | NM_175449 |
| Fam3b | 0.09572496 | 0.00498255289994511 | 0.00847999972823692 | 0.587565219295241 | 0.556824162160132 | NA | NM_020622 |
| Fam46b | 3.493317917 | 0.0430283738956046 | 0.0409671181176325 | 1.05031488356231 | 0.293573364469661 | NA | NM_175307 |
| Fam46d | 0.235249118 | -0.00959082599542929 | 0.0102925929774586 | -0.93181825186654 | 0.351430460444064 | NA | NM_001271008,NM_001163104 |
| Fam47c | 0 | NA | NA | NA | NA | NA | NM_001164739 |
| Fam47e | 9.509144535 | -0.00700423818266531 | 0.0547912411847939 | -0.127834997550835 | 0.898279553231299 | NA | NM_001033478,NM_001170571 |
| Fam50b | 0 | NA | NA | NA | NA | NA | NM_138746 |
| Fam64a | 7.351570161 | 0.0335289272582822 | 0.0563073190037496 | 0.59546303840269 | 0.551533998757137 | NA | NM_144526 |
| Fam65c | 15.81082549 | -0.066306412094855 | 0.0654391673148687 | -1.01325268666414 | 0.310939480448881 | NA | NM_001080708 |
| Fam71a | 0 | NA | NA | NA | NA | NA | NM_001109759 |
| Fam71b | 0.106347839 | -0.00653347234140708 | 0.00854389509723638 | -0.764694822098226 | 0.444453290197145 | NA | NM_001013783,NR_075102 |
| Fam71d | 4.179820251 | -0.0498792102216805 | 0.0460370090713475 | -1.08345896546795 | 0.278604755173078 | NA | NM_029069,NM_027597 |
| Fam71e2 | 0.120209049 | -0.00674972139751468 | 0.00863521114606269 | -0.781650996523957 | 0.434419707908266 | NA | NM_172895 |
| Fam71f1 | 0 | NA | NA | NA | NA | NA | NM_207258 |
| Fam71f2 | 20.72549595 | -0.0982246122626713 | 0.0827501190308479 | -1.1870026703654 | 0.235226567197071 | NA | NM_001101486 |
| Fam83a | 0.36293681 | -0.00593061268544428 | 0.0131125568226957 | -0.452284994119481 | 0.651063684812157 | NA | NM_173862 |
| Fam83b | 3.387348643 | -0.0130917461865086 | 0.0437173845762674 | -0.29946316124354 | 0.764586676099671 | NA | NM_001045518 |
| Fam83c | 0 | NA | NA | NA | NA | NA | NM_027788 |
| Fam83e | 2.281906748 | -0.0200565090086367 | 0.0348571779563521 | -0.575391072500227 | 0.565026840526275 | NA | NM_001033170 |
| Fam83g | 5.166910194 | 0.00899613328027243 | 0.0521178711610709 | 0.172611295892531 | 0.862956972342395 | NA | NM_178618 |
| Fam92b | 10.41981396 | -0.0251523018566383 | 0.0658771641540417 | -0.381806080750899 | 0.702605208077967 | NA | NM_001033980 |
| Fasl | 3.826065304 | -0.0261270565731219 | 0.0423440534418394 | -0.617018316609866 | 0.537222644325207 | NA | NM_010177,NM_001205243 |
| Fate1 | 3.345602043 | 0.00755617914347053 | 0.0435837139853406 | 0.173371621014493 | 0.862359330448163 | NA | NR_003243 |
| Fbp1 | 10.72618356 | 0.0438719041711772 | 0.0647491951982321 | 0.677566787306957 | 0.498046410440608 | NA | NM_019395 |
| Fbp2 | 1.191606103 | 0.00840161885292042 | 0.02618780905057 | 0.320821754759875 | 0.748345472242931 | NA | NM_007994 |
| Fbxl13 | 13.50234993 | -0.0296555500548065 | 0.0700681718470799 | -0.423238530035124 | 0.67212123780577 | NA | NM_001199632,NM_177076 |
| Fbxl22 | 9.732944358 | -0.0116884947146882 | 0.0631005281542806 | -0.185236083699805 | 0.853043868095121 | NA | NM_175206 |
| Fbxo15 | 6.795287828 | 0.0370682197402322 | 0.0525430710083405 | 0.70548255038896 | 0.48050998739361 | NA | NM_015798 |
| Fbxo24 | 8.470166489 | 0.0720532084194725 | 0.0589148684679309 | 1.22300550426745 | 0.221327617342334 | NA | NM_027708 |
| Fbxo39 | 0.108766301 | 0.00519171381361946 | 0.00856003371502575 | 0.606506234257728 | 0.544178643801544 | NA | NM_001099688 |
| Fbxo40 | 8.094448538 | 0.0464337424720491 | 0.0528852625958631 | 0.878009112422963 | 0.37993877359785 | NA | NM_001037321 |
| Fbxo43 | 5.554942104 | -0.0253989295629613 | 0.0491699136095434 | -0.516554284895706 | 0.605467335020931 | NA | NM_001081253 |
| Fbxw13 | 0.319043516 | -0.00957618023178583 | 0.0102890835921761 | -0.930712647632453 | 0.352002225550712 | NA | NM_177598 |
| Fbxw14 | 0 | NA | NA | NA | NA | NA | NM_015793 |
| Fbxw15 | 0.581325921 | -0.0182437040905998 | 0.0174236536372216 | -1.04706535554784 | 0.295069436410656 | NA | NM_199036 |
| Fbxw16 | 0 | NA | NA | NA | NA | NA | NM_177070 |
| Fbxw18 | 0.12890128 | -0.0068640067066208 | 0.00870804083610955 | -0.788237771940376 | 0.430557639008166 | NA | NM_001033794 |
| Fbxw19 | 0.106148608 | 0.00515554063321497 | 0.00854200771766808 | 0.603551390213729 | 0.546141939141592 | NA | NM_177703 |
| Fbxw20 | 0 | NA | NA | NA | NA | NA | NM_001008428 |
| Fbxw21 | 0 | NA | NA | NA | NA | NA | NM_177069 |
| Fbxw22 | 0 | NA | NA | NA | NA | NA | NM_001014395 |
| Fbxw24 | 0.212496447 | -0.000886988073087717 | 0.00973851166817767 | -0.0910804549309223 | 0.927428662789121 | NA | NM_001013776 |
| Fbxw26 | 0 | NA | NA | NA | NA | NA | NM_198674 |
| Fbxw28 | 0 | NA | NA | NA | NA | NA | NM_001177420,NM_001177419 |
| Fcamr | 0.812623213 | 0.0306215266680823 | 0.0204900577835628 | 1.49445779956009 | 0.135056004055277 | NA | NM_001170632,NM_144960 |
| Fcer1a | 0.212496447 | -0.000886988073087717 | 0.00973851166817767 | -0.0910804549309223 | 0.927428662789121 | NA | NM_010184 |
| Fcer2a | 0.489410913 | 0.00565929899362522 | 0.0126443813664561 | 0.447574209414358 | 0.654460519707199 | NA | NM_013517,NM_001253746,NM_001253747,NM_001253745,NM_001253743,NM_001253739,NM_001253737 |
| Fcgbp | 20.98965864 | 0.0837798786956976 | 0.0734543033786155 | 1.14057141436438 | 0.254048318352478 | NA | NM_001122603 |
| Fcgr4 | 21.22068383 | -0.154311857790745 | 0.0549865325554109 | -2.80635731367936 | 0.00501050975926208 | NA | NM_144559 |
| Fcna | 8.27766832 | 0.0849242887423712 | 0.0614888872475213 | 1.38113230770483 | 0.167238283484929 | NA | NM_007995 |
| Fcnb | 0.55890461 | 0.00181148496239832 | 0.0147357984933791 | 0.122930899415612 | 0.902161816999182 | NA | NM_010190 |
| Fcrl1 | 16.69972909 | -0.0615351501536632 | 0.072603922013894 | -0.847545813597885 | 0.396690962483565 | NA | NM_178165,NM_001136236,NM_153090 |
| Fcrl5 | 0 | NA | NA | NA | NA | NA | NM_183222,NM_001113238 |
| Fcrl6 | 1.096054655 | -0.00290625451315608 | 0.0197436239525898 | -0.147199648865621 | 0.882974437743611 | NA | NM_001164725 |
| Fcrla | 0.21511414 | -0.000824240786584897 | 0.00975135300262786 | -0.08452578697159 | 0.93263840116295 | NA | NM_001160215,NR_027666,NM_145141 |
| Fcrlb | 14.66400033 | 0.0389541067281356 | 0.0749816885080178 | 0.519514930955046 | 0.603401703583629 | NA | NM_001029984 |
| Fer1l4 | 0.713561805 | -0.00748950297278271 | 0.0195095236227617 | -0.383889587342088 | 0.701060284266191 | NA | NM_001136556 |
| Ferd3l | 1.323332537 | 0.0102798722395146 | 0.0248546472398139 | 0.41359960333887 | 0.679167363790519 | NA | NM_033522 |
| Fermt1 | 16.43977992 | -0.0168183127293506 | 0.0767562585203759 | -0.219113243057385 | 0.826561834955325 | NA | NM_198029 |
| Fetub | 4.099065369 | 0.00830269790536544 | 0.0377002307383757 | 0.220229365782475 | 0.825692527416297 | NA | NM_021564,NM_001083905,NM_001083904 |
| Fev | 1.697717137 | 0.061628796185564 | 0.0309106497472732 | 1.99377226585153 | 0.0461769477665121 | NA | NM_153111 |
| Fezf1 | 31.45713106 | 0.0786240083671335 | 0.0424947227728112 | 1.85020640768689 | NA | NA | NM_028462 |
| Ffar1 | 0 | NA | NA | NA | NA | NA | NM_194057 |
| Ffar2 | 2.783664535 | 0.0610264788112055 | 0.0379325088234475 | 1.60881736283899 | 0.107656283151142 | NA | NM_001168510,NM_001168511,NM_001168509,NM_146187,NM_001168512 |
| Ffar3 | 2.000452318 | -0.0214290815015587 | 0.0301853984192335 | -0.709915476480991 | 0.47775655252545 | NA | NM_001033316 |
| Ffar4 | 2.557899846 | 0.0263169108299804 | 0.032399495309106 | 0.812262986781277 | 0.416640744127751 | NA | NM_181748 |
| Fga | 0.257802559 | -0.00892927589713349 | 0.00993479775307735 | -0.898787888698349 | 0.368765652383134 | NA | NM_010196,NM_001111048 |
| Fgb | 0.12890128 | -0.0068640067066208 | 0.00870804083610955 | -0.788237771940376 | 0.430557639008166 | NA | NM_181849 |
| Fgf15 | 1.309825175 | 0.0143391093461124 | 0.0273097385106611 | 0.525054801989945 | 0.599545094976334 | NA | NM_008003 |

| | | | | | | | | | | | | | | |
|---|---|---|---|---|---|---|---|---|---|---|---|---|---|---|
| Clec4e | 0 | NA | NA | NA | NA | NA | | Fgf16 | 19.04434867 | 0.0425107767716839 | 0.0721829680047055 | 0.588930850957981 | 0.555907661233541 | NA | NM_030614 |
| Clec4f | 0.094392817 | 0.017895942 | 0.027818137 | 0.643319236 | 0.520016977 | NA | | Fgf20 | 0.315716853 | 0.0123948421219712 | 0.0131137196252776 | 0.945181266349427 | 0.344566341855147 | NA | NM_030610 |
| Clec4g | 3.369466128 | -0.135668073 | 0.107077407 | -1.267009323 | 0.205151956 | NA | | Fgf21 | 0.192890664 | -0.000903864895041124 | 0.00963528282151475 | -0.093807821917056 | 0.92526181809679 | NA | NM_020013 |
| Clec4n | 6.992929713 | -0.475495151 | 0.148866014 | -3.194114892 | 0.001402603 | NA | | Fgf23 | 17.94398926 | 0.0321309681005133 | 0.0657186593423724 | 0.488916974601104 | 0.624900477639294 | NA | NM_022657 |
| Clec9a | 1.090837823 | -0.027094434 | 0.075375517 | -0.35945934 | 0.71925149 | NA | | Fgf3 | 19.48548822 | 0.00767776597280639 | 0.0812072590648008 | 0.0945453160373234 | 0.924675986642532 | NA | NM_008007 |
| Clhc1 | 5.619247619 | 0.084229664 | 0.141310262 | 0.596061905 | 0.551133872 | NA | | Fgf4 | 0 | NA | NA | NA | NA | NA | NM_010202 |
| Clic3 | 6.26346902 | 0.164607011 | 0.14159867 | 1.162489806 | 0.245036563 | NA | | Fgf6 | 0 | NA | NA | NA | NA | NA | NM_010204 |
| Clnk | 2.97680312 | 0.080992202 | 0.116628595 | 0.694445493 | 0.487402872 | NA | | Fgf7 | 21.65813229 | -0.0421396139126694 | 0.0798207675693344 | -0.52792794652176 | 0.597549347009448 | NA | NM_008008 |
| Clps | 0.192746203 | -0.000975302 | 0.031567365 | -0.030895906 | 0.975352555 | NA | | Fgf8 | 0.475177312 | -0.022041534045028 | 0.01555480462569 | -1.41702416555105 | 0.156475862316816 | NA | NM_001166361,NM_001166363,NM_001166362,NM_010205 |
| Clpsl2 | 0 | NA | NA | NA | NA | NA | | Fgfr4 | 7.884167257 | -0.0335565863218129 | 0.0603226501721599 | -0.556285014435589 | 0.578016029867635 | NA | NM_008011 |
| Clrn2 | 7.996239079 | 0.25158323 | 0.156610172 | 1.606429689 | 0.108179532 | NA | | Fgg | 0 | NA | NA | NA | NA | NA | NM_133862 |
| Clrn3 | 0.116696964 | -0.020477583 | 0.027954224 | -0.732539847 | 0.463839131 | NA | | Fgl1 | 9.77194487 | 0.0641705867650605 | 0.0639590136946143 | 1.00330794767187 | 0.31571230263738 | NA | NM_145594 |
| Cma1 | 0.609727828 | -0.037051079 | 0.059330555 | -0.624485625 | 0.532308708 | NA | | Fgr | 14.5826347 | -0.246762036003808 | 0.0560920259919274 | -4.39923557119011 | 1.08632855016735e-05 | NA | NM_010208 |
| Cma2 | 0 | NA | NA | NA | NA | NA | | Fhl5 | 3.349380727 | 0.0314340511175992 | 0.0413162400572636 | 0.760815869837918 | 0.446767054131816 | NA | NM_021318 |
| Cmah | 6.407250863 | 0.081367871 | 0.142703912 | 0.570186691 | 0.568551082 | NA | | Figla | 0.855438572 | -0.0236316782330062 | 0.0182692787594406 | -1.29352004226191 | 0.195831256785303 | NA | NM_012013 |
| Cmtm1 | 0.392633299 | 0.026946 | 0.048301387 | 0.557872169 | 0.576931677 | NA | | Fitm1 | 0 | NA | NA | NA | NA | NA | NM_026808 |
| Cmtm2a | 0.096071043 | -0.020098795 | 0.027825948 | -0.722304036 | 0.470107572 | NA | | Fkbp11 | 19.21931376 | 0.0610021238004903 | 0.0701720735152386 | 0.869321950237698 | 0.38467106013663 | NA | NM_024169 |
| Cmtm2b | 0 | NA | NA | NA | NA | NA | | Fkbp6 | 2.363767144 | 0.00669287120015257 | 0.0289164380224643 | 0.231455589203383 | 0.816960883400124 | NA | NM_033571 |
| Cnfn | 0.612124479 | 0.05749822 | 0.056841187 | 1.011559104 | 0.311748908 | NA | | Flg2 | 0 | NA | NA | NA | NA | NA | NM_001013804 |
| Cnga1 | 0 | NA | NA | NA | NA | NA | | Fmo3 | 0 | NA | NA | NA | NA | NA | NM_008030 |
| Cnga3 | 7.394195149 | -0.432416454 | 0.135990071 | -3.179764904 | 0.001473946 | NA | | Fmo4 | 2.521405816 | 0.0202059470908335 | 0.033438938631343 | 0.604264008304798 | 0.545668131215877 | NA | NM_144878 |
| Cngb3 | 0.233393928 | -0.027475306 | 0.032191697 | -0.85349047 | 0.39338737 | NA | | Fmo6 | 0.95380211 | -0.00233859089827368 | 0.0187688548474476 | -0.124599551612586 | 0.900840583392775 | NA | NM_001178038 |
| Cnpy1 | 1.353565994 | 0.029979155 | 0.07482703 | 0.400646063 | 0.688680727 | NA | | Fmo9 | 0.113843286 | 0.00525947738444006 | 0.00859368002043176 | 0.612016897526494 | 0.540526577873086 | NA | NM_172844 |
| Cntf | 3.262999941 | 0.107163825 | 0.098631722 | 1.086504656 | 0.277255782 | NA | | Fmr1nb | 0.880555709 | 0.00633574565904043 | 0.0223290441724765 | 0.28374459784759 | 0.776606114580096 | NA | NM_001166620,NM_001166619,NM_174993 |
| Col8a2 | 118.1302481 | 0.154562689 | 0.161210953 | 0.95876047 | NA | NA | | Fnd3c2 | 0 | NA | NA | NA | NA | NA | NM_001033424 |
| Colec10 | 0.094392817 | 0.017895942 | 0.027818137 | 0.643319236 | 0.520016977 | NA | | Fndc3c1 | 0.819350582 | -0.00662082177968036 | 0.0164143511050961 | -0.40335568170129 | 0.68668658470651 | NA | NM_001007580 |
| Cox7b2 | 5.399662925 | -0.04210873 | 0.145802866 | -0.288805913 | 0.772729907 | NA | | Fndc7 | 8.500584234 | 0.0485540747733775 | 0.0591879309005138 | 0.820337424110834 | 0.412023777504499 | NA | NM_177091,NM_001165965 |
| Cox8c | 0 | NA | NA | NA | NA | NA | | Fndc8 | 0.335123958 | 0.00243718958797335 | 0.0127138988834395 | 0.191694901014818 | 0.847981200318863 | NA | NM_030224 |
| Cp | 859.6799956 | -0.035654034 | 0.143638879 | -0.248219943 | NA | NA | | Folr1 | 129.7458546 | 0.0141046857916944 | 0.0273438019206446 | 0.515827529493816 | NA | NA | NM_008034,NM_001252553,NM_001252554,NM_001252552 |
| Cpa1 | 0 | NA | NA | NA | NA | NA | | Folr2 | 14.23968427 | -0.0644922220073963 | 0.0642389263054274 | -1.00394302514903 | 0.315406076576052 | NA | NM_008035 |
| Cpa3 | 1.006239153 | 0.013014933 | 0.062058454 | 0.209720547 | 0.833885788 | NA | | Folr4 | 0.22897535 | -0.00113636419885667 | 0.00981572003153595 | -0.118548816065269 | 0.905632818135836 | NA | NM_176807,NM_022888 |
| Cpa5 | 1.71620555 | 0.041323327 | 0.096663668 | 0.427495957 | 0.669018129 | NA | | Fosl1 | 3.543819132 | 0.0341286291886489 | 0.0456726452290041 | 0.747244417693061 | 0.454916036747418 | NA | NM_010235 |
| Cpb1 | 5.316919271 | -0.100469728 | 0.146862974 | -0.684105226 | 0.493908722 | NA | | Foxa1 | 1.201777077 | 0.0356654615038406 | 0.0245161353903467 | 1.45477502616028 | 0.145731554179399 | NA | NM_008259 |
| Cpb2 | 0 | NA | NA | NA | NA | NA | | Foxa2 | 0.106148608 | 0.00515554063321497 | 0.00854200771766808 | 0.603551390213729 | 0.546141939141592 | NA | NM_010446 |
| Cphx1 | 0 | NA | NA | NA | NA | NA | | Foxa3 | 0 | NA | NA | NA | NA | NA | NM_008260 |
| Cphx2 | 0 | NA | NA | NA | NA | NA | | Foxb1 | 4.825874593 | 0.0572329182774995 | 0.0439672247739233 | 1.30171778118331 | 0.193012878856142 | NA | NM_022378 |
| Cplx4 | 2.388063154 | 0.001138628 | 0.084311275 | 0.013505048 | 0.989224859 | NA | | Foxb2 | 1.134372229 | 0.00379908300905239 | 0.0218232192133099 | 0.174084445191997 | 0.861799097482844 | NA | NM_008023 |
| Cpn2 | 0 | NA | NA | NA | NA | NA | | Foxd2 | 9.614899422 | 0.0334817308896199 | 0.0555433337610267 | 0.60280376820149 | 0.546639239683555 | NA | NM_008593 |
| Cpsf4l | 0.295716657 | -0.010253415 | 0.03372058 | -0.304069933 | 0.761074613 | NA | | Foxd3 | 0.097165704 | -0.00637828455464458 | 0.00848781691535724 | -0.75146349388194 | 0.452373763047372 | NA | NM_010425 |
| Cpt1b | 2.04416705 | 0.107359693 | 0.101147295 | 1.061419315 | 0.288499383 | NA | | Foxd4 | 0.097165704 | -0.00637828455464458 | 0.00848781691535724 | -0.75146349388194 | 0.452373763047372 | NA | NM_008022 |
| Cpvl | 0.350688377 | 0.02015342 | 0.04180392 | 0.482094028 | 0.629739152 | NA | | Foxe1 | 0 | NA | NA | NA | NA | NA | NM_183298 |
| Cpxcr1 | 0 | NA | NA | NA | NA | NA | | Foxe3 | 0 | NA | NA | NA | NA | NA | NM_015758 |
| Cpz | 7.517777699 | 0.086076124 | 0.150796912 | 0.57080827 | 0.568129616 | NA | | Foxh1 | 1.867489414 | 0.043831541764575 | 0.0318041838512913 | 1.37816904749139 | 0.168151101328433 | NA | NM_007989 |
| Crct1 | 0.698757213 | 0.06007153 | 0.05689194 | 1.055888217 | 0.291019288 | NA | | Foxi1 | 3.10567469 | -0.0276347185965674 | 0.0409319613539803 | -0.675137904034989 | 0.499588153848432 | NA | NM_023907 |
| Crisp1 | 0 | NA | NA | NA | NA | NA | | Foxi2 | 0.21511414 | -0.000824240786584897 | 0.00975135300262786 | -0.08452578697159 | 0.93263840116295 | NA | NM_183193 |
| Crisp2 | 0 | NA | NA | NA | NA | NA | | Foxi3 | 0 | NA | NA | NA | NA | NA | NM_001101464 |
| Crisp3 | 0 | NA | NA | NA | NA | NA | | Foxl1 | 0.765234524 | 0.00136147124924975 | 0.0160239832153894 | 0.0849645953162377 | 0.932289537738774 | NA | NM_008024 |
| Crisp4 | 0 | NA | NA | NA | NA | NA | | Foxl2os | 19.71469933 | -0.161191697372493 | 0.08009179068251 | -2.01258700796776 | 0.0441580948804861 | NA | NR_003248 |
| Crnn | 0 | NA | NA | NA | NA | NA | | Foxn1 | 0.5396566 | -0.0178265475807991 | 0.0171376156815873 | -1.04019998534288 | 0.298246998331762 | NA | NM_008238 |
| Crp | 0 | NA | NA | NA | NA | NA | | Foxn4 | 0.551567013 | 0.00939524269285595 | 0.0152551110071042 | 0.615875078751027 | 0.537976970668696 | NA | NM_148935 |
| Crtam | 2.957490911 | 0.055928676 | 0.120427015 | 0.464419677 | 0.642347095 | NA | | Foxp3 | 3.475135377 | -0.0162000882931096 | 0.0409465460893515 | -0.395639921808265 | 0.692370681802368 | NA | NM_001199348,NM_001199347,NM_054039 |
| Crx | 0.508673765 | -0.022465145 | 0.056012253 | -0.401075549 | 0.688364502 | NA | | Foxr1 | 0 | NA | NA | NA | NA | NA | NM_001033469 |
| Crxos1 | 3.825199017 | 0.261274637 | 0.129334325 | 2.020149226 | 0.043367911 | NA | | Foxs1 | 13.39738723 | 0.0211910982386642 | 0.0732008469616827 | 0.289492527999802 | 0.772204497022903 | NA | NM_010226 |
| Cryaa | 3.527884846 | 0.054035705 | 0.114948909 | 0.470084542 | 0.638294618 | NA | | Fpr-rs3 | 0 | NA | NA | NA | NA | NA | NM_008040 |
| Cryba1 | 1.263743685 | -9.22E-05 | 0.086134906 | -0.001070538 | 0.999145834 | NA | | Fpr-rs4 | 0 | NA | NA | NA | NA | NA | NM_008041 |
| Cryba2 | 6.796622269 | 0.184559996 | 0.13483829 | 1.368750641 | 0.171077235 | NA | | Fpr-rs6 | 0 | NA | NA | NA | NA | NA | NM_177316 |
| Crybb2 | 0.098353386 | -0.020212831 | 0.02786466 | -0.725393061 | 0.468210929 | NA | | Fpr1 | 0.86200232 | -0.0022923037749072 | 0.018566172482088 | -0.123466685291151 | 0.901737553696376 | NA | NM_013521 |
| Crygb | 0 | NA | NA | NA | NA | NA | | Fpr2 | 6.806976057 | -0.0792447540320594 | 0.0528527552686338 | -1.499349534935 | 0.133782978094408 | NA | NM_008039 |
| Crygc | 0 | NA | NA | NA | NA | NA | | Fpr3 | 0 | NA | NA | NA | NA | NA | NM_008042 |
| Crygd | 0 | NA | NA | NA | NA | NA | | Frk | 15.55729792 | -0.117683808986212 | 0.0747727108098259 | -1.57388715363716 | 0.115513540342089 | NA | NM_010237,NM_001159544 |
| Cryge | 0.079983654 | 0.016740032 | 0.027674178 | 0.604897175 | 0.54524732 | NA | | Fsbp | 9.941548812 | -0.108627340953143 | 0.0675640008336574 | -1.60776951650012 | 0.107885667059782 | NA | NM_001256142 |
| Crygf | 0 | NA | NA | NA | NA | NA | | Fscb | 0.106148608 | 0.00515554063321497 | 0.00854200771766808 | 0.603551390213729 | 0.546141939141592 | NA | NM_001163271 |
| Crygs | 0.495946458 | 0.007147691 | 0.044404124 | 0.160969084 | 0.87211775 | NA | | Fscn2 | 7.360289024 | -0.00107329440603088 | 0.0586188334061466 | -0.0183097196526319 | 0.985391773605603 | NA | NM_172802 |
| Csf2 | 0 | NA | NA | NA | NA | NA | | Fscn3 | 0.861412953 | -0.0128519535542059 | 0.017014756248769 | -0.75534162031476 | 0.450044032004319 | NA | NM_019569 |
| Csf3 | 0.295416859 | -0.013215976 | 0.033806277 | -0.390932592 | 0.695847062 | NA | | Fsd2 | 8.460259195 | -0.0389943446608733 | 0.0608788735191575 | -0.640523426383742 | 0.521832364061887 | NA | NM_172904 |
| Csl | 0 | NA | NA | NA | NA | NA | | Fshb | 0.09572496 | 0.00498255289994511 | 0.00847999972823692 | 0.587565219295241 | 0.556824162160132 | NA | NM_008045 |
| Csn1s1 | 0.276102474 | 0.031058433 | 0.034913626 | 0.889579115 | 0.373691925 | NA | | Fshr | 0.204491261 | 0.00675251727497272 | 0.00969338978195025 | 0.696610517772262 | 0.486046571838551 | NA | NM_013523 |
| Csn1s2a | 0 | NA | NA | NA | NA | NA | | Fsip1 | 19.92612328 | 0.154773220222623 | 0.0800881514880334 | 1.93253580394784 | 0.0532934072904119 | NA | NM_027759 |
| Csn1s2b | 0 | NA | NA | NA | NA | NA | | Ftcd | 2.8407767 | 0.000610461642076193 | 0.0369796891455998 | 0.0165080252479308 | 0.986829099739138 | NA | NM_080845 |
| Csn2 | 0.079983654 | 0.016740032 | 0.027674178 | 0.604897175 | 0.54524732 | NA | | Fthl17 | 0 | NA | NA | NA | NA | NA | NM_031261 |
| Csn3 | 0 | NA | NA | NA | NA | NA | | Ftmt | 0 | NA | NA | NA | NA | NA | NM_026286 |
| Csnka2ip | 0.48794879 | 0.064131883 | 0.049799661 | 1.287797577 | 0.197816436 | NA | | Fut1 | 1.713121285 | -0.0238270278662139 | 0.0302549469056127 | -0.787541552809456 | 0.430964913764111 | NA | NM_008051,NM_001271981 |
| Cspg5 | 10656.73371 | -0.232533607 | 0.113845531 | -2.042536101 | NA | NA | | Fut2 | 8.902867462 | 0.0564200386091912 | 0.0612287962783392 | 0.921462482337756 | 0.356809020214175 | NA | NM_001271993,NM_018876 |
| Csrp2bp | 1388.527159 | 0.203391239 | 0.102418302 | 1.985887628 | NA | NA | | Fut4 | 13.954792 | 0.0457901230031045 | 0.0615136269615181 | 0.744389906186967 | 0.456640623752456 | NA | NM_010242 |
| Csrp3 | 0.321356363 | -0.01100297 | 0.034551013 | -0.318455781 | 0.750139232 | NA | | Fut4-ps1 | 0.217374753 | -0.00904456656736547 | 0.00999660569474087 | -0.90476376117583 | 0.365590553873005 | NA | NR_033644 |
| Cst10 | 0 | NA | NA | NA | NA | NA | | Fut7 | 9.175731575 | -0.0331138907441378 | 0.0625654582345011 | -0.529267932794863 | 0.596619597363986 | NA | NM_013524,NM_001177367,NM_001177366 |
| Cst11 | 0.691297128 | 0.034620722 | 0.06074969 | 0.569891329 | 0.568751406 | NA | | Fxyd3 | 2.825558303 | -0.0523910465636507 | 0.038372883680014 | -1.36531429330494 | 0.172154288477575 | NA | NM_008557 |
| Cst12 | 0.800245648 | -0.038051497 | 0.068702165 | -0.553861688 | 0.579673506 | NA | | Fxyd4 | 0.09572496 | 0.00498255289994511 | 0.00847999972823692 | 0.587565219295241 | 0.556824162160132 | NA | NM_033648,NM_001173372 |
| Cst13 | 0 | NA | NA | NA | NA | NA | | G630055G22Rik | 0 | NA | NA | NA | NA | NA | NR_045404 |
| Cst8 | 0 | NA | NA | NA | NA | NA | | G630071F17Rik | 0 | NA | NA | NA | NA | NA | NR_045401 |
| Cst9 | 0 | NA | NA | NA | NA | NA | | G630090E17Rik | 1.088190654 | 0.016635693857436 | 0.0265161677823743 | 0.627379265132497 | 0.530410656147677 | NA | NM_001173500 |
| Csta | 0 | NA | NA | NA | NA | NA | | G630093K05Rik | 9.26186484 | 0.0583519856371121 | 0.0594901008250961 | 0.980868830743283 | 0.32665742989336 | NA | NR_045156 |
| Cstl1 | 5.864419947 | 0.061379307 | 0.14608311 | 0.420167034 | 0.674363436 | NA | | G6b | 0 | NA | NA | NA | NA | NA | NR_001462 |
| Ctag2 | 0 | NA | NA | NA | NA | NA | | G6pc | 0.988647175 | 0.00891844052596055 | 0.0210778504637456 | 0.423119071904437 | 0.672208388125198 | NA | NM_008061 |
| Ctf2 | 0 | NA | NA | NA | NA | NA | | G6pc2 | 0.23403553 | -0.000407313579519725 | 0.00983856423584417 | -0.0413996971261099 | 0.966977254250032 | NA | NM_021331 |
| Ctla2a | 199.18628 | 0.282252577 | 0.162631217 | 1.735537511 | NA | NA | | G6pd2 | 0 | NA | NA | NA | NA | NA | NM_019468 |
| Ctla4 | 2.082501219 | 0.057465537 | 0.099780127 | 0.575921667 | 0.564668129 | NA | | G730013B05Rik | 7.247731364 | 0.00536019892183303 | 0.0558711447588759 | 0.0959385912883321 | 0.923569344511283 | NA | NR_040380,NR_040379 |
| Ctrb1 | 0.167300494 | 0.024299763 | 0.031030539 | 0.783091891 | 0.433573153 | NA | | Gabra6 | 12.95759981 | 0.0609326614961248 | 0.0429378468295152 | 1.41908982390426 | 0.155872830308786 | NA | NM_008068,NM_001099641 |
| Ctrc | 0 | NA | NA | NA | NA | NA | | Gabrp | 0.432488341 | 0.00620036638839739 | 0.0139681256348908 | 0.443893944718651 | 0.657119262665896 | NA | NM_146017 |
| Cts3 | 0 | NA | NA | NA | NA | NA | | Gabrr1 | 5.897516715 | -0.126424480505751 | 0.0518678418780411 | -2.4374347558747 | 0.0147918832676072 | NA | NM_008075 |
| Cts6 | 0 | NA | NA | NA | NA | NA | | Gabrr3 | 2.528714734 | -0.00725473604090748 | 0.0339285237459568 | -0.213824099605041 | 0.830684230791586 | NA | NM_001081190 |
| Cts7 | 0 | NA | NA | NA | NA | NA | | Gal | 47.98962653 | 0.0573874099358012 | 0.0340866017938761 | 1.68357674029305 | NA | NA | NM_010253 |
| Cts8 | 0 | NA | NA | NA | NA | NA | | Gal3st2 | 3.365335519 | -0.00798245342187244 | 0.0391017309417338 | -0.204145781519678 | 0.83823957909398 | NA | NM_199366 |
| Cts8-ps | 0 | NA | NA | NA | NA | NA | | Galnt5 | 1.457884469 | -0.0126109693148204 | 0.0265929951089116 | -0.474221473104934 | 0.635341982371487 | NA | NM_172855 |
| Ctse | 3.374962363 | 0.129652481 | 0.101517341 | 1.277146145 | 0.201550656 | NA | | Galntl5 | 0.09572496 | 0.00498255289994511 | 0.00847999972823692 | 0.587565219295241 | 0.556824162160132 | NA | NM_026449 |
| Ctsg | 0.286698134 | 0.007861574 | 0.038801625 | 0.202609399 | 0.839440343 | NA | | Galp | 0.106148608 | 0.00515554063321497 | 0.00854200771766808 | 0.603551390213729 | 0.546141939141592 | NA | NM_178028 |
| Ctsj | 0 | NA | NA | NA | NA | NA | | Galr2 | 0.108766301 | 0.00519171381361946 | 0.00856003371502575 | 0.606506234257728 | 0.544178643801544 | NA | NM_010254 |
| Ctsll3 | 0 | NA | NA | NA | NA | NA | | Galr3 | 3.259397857 | -0.0313783745635622 | 0.0401808133305081 | -0.780929303383153 | 0.43484407612986 | NA | NM_015738 |
| Ctsm | 0 | NA | NA | NA | NA | NA | | Gapdhs | 0.649109394 | 0.0160670453296311 | 0.0195022451001213 | 0.823856189230807 | 0.410021276282451 | NA | NM_008085 |
| Ctsq | 0 | NA | NA | NA | NA | NA | | Gapt | 0.21511414 | -0.000824240786584897 | 0.00975135300262786 | -0.08452578697159 | 0.93263840116295 | NA | NM_177713 |
| Ctsr | 0 | NA | NA | NA | NA | NA | | Gas2l2 | 16.23990206 | -0.0144301261807721 | 0.0727788126038696 | -0.19827372369091 | 0.84283090901858 | NA | NM_001013759 |
| Ctsw | 4.625655994 | -0.057633096 | 0.132025276 | -0.436530771 | 0.662451679 | NA | | Gast | 2.16200003 | -0.000890185007463129 | 0.0321395142548531 | -0.0276975252458494 | 0.977903397515469 | NA | NM_010257 |
| Cxcl1 | 3.061675514 | 0.281468208 | 0.11865801 | 2.372096143 | 0.017687488 | NA | | Gata1 | 0 | NA | NA | NA | NA | NA | NM_008089 |
| Cxcl11 | 0.946603521 | -0.013609344 | 0.077083244 | -0.176553859 | 0.85985884 | NA | | Gata3 | 8.562112109 | 0.00836351340931149 | 0.0347293041226096 | 0.240820068832495 | 0.80969457462403 | NA | NM_008091 |
| Cxcl13 | 2.081694735 | -0.132821048 | 0.089437611 | -1.48506928 | 0.137525473 | NA | | Gata4 | 0 | NA | NA | NA | NA | NA | NM_008092 |
| Cxcl17 | 0 | NA | NA | NA | NA | NA | | Gata5 | 0 | NA | NA | NA | NA | NA | NM_008093 |
| Cxcl2 | 0.379249494 | 0.016397727 | 0.036387388 | 0.450643137 | 0.652246771 | NA | | Gata6 | 3.150551789 | -0.0139535889643603 | 0.0381680006600049 | -0.365583439611022 | 0.71467593421712 | NA | NM_010258 |
| Cxcl3 | 0.096071043 | -0.020098795 | 0.027825948 | -0.722304036 | 0.470107572 | NA | | Gbp1 | 0 | NA | NA | NA | NA | NA | NM_010259 |
| Cxcl9 | 11.7388471 | 0.0620493 | 0.049980769 | 1.241463479 | NA | NA | | Gbp10 | 16.74372577 | -0.0418426765126077 | 0.0238424074327736 | -1.75496860501977 | NA | NA | NM_001039646 |
| Cxcr1 | 0 | NA | NA | NA | NA | NA | | Gbp6 | 147.0475794 | -0.1194222953971 | 0.0500267774531421 | -2.38716746264454 | NA | NA | NM_194336 |
| Cxcr3 | 2.801578802 | 0.029349885 | 0.103400683 | 0.283846142 | 0.776528292 | NA | | Gbp8 | 10.71689967 | -0.10110349208363 | 0.04063928169757 | -2.48782674940032 | NA | NA | NM_029509 |
| Cxcr5 | 0 | NA | NA | NA | NA | NA | | Gbx1 | 13.72742537 | -0.0159892903509532 | 0.0676996973670174 | -0.236179642935052 | 0.813293275313777 | NA | NM_015739 |
| Cxcr6 | 1.83817932 | -0.007441211 | 0.084213542 | -0.088361215 | 0.929589587 | NA | | Gbx2 | 8.666318005 | 0.106056394520401 | 0.0620390236884198 | 1.70951101766286 | 0.0873563326247969 | NA | NM_010262 |
| Cyct | 0.107716603 | -0.020477583 | 0.027954224 | -0.732539849 | 0.46383913 | NA | | Gc | 0.22897535 | -0.00116364198856673 | 0.00981572003153595 | -0.118548816065269 | 0.905632818135836 | NA | NM_008096 |
| Cylc1 | 0 | NA | NA | NA | NA | NA | | Gcg | 1.149091228 | 0.0102640672188071 | 0.0202924555683681 | 0.505807056431688 | 0.612992104358162 | NA | NM_008100 |
| Cylc2 | 0 | NA | NA | NA | NA | NA | | Gcgr | 0.868302965 | 0.0132450885416623 | 0.0208745228369232 | 0.634509763175723 | 0.525748197265887 | NA | NM_008101 |
| Cym | 1.731090764 | -0.014411747 | 0.092176278 | -0.156349844 | 0.87575727 | NA | | Gchfr | 6.732028513 | -0.0169207745351889 | 0.0570820744153957 | -0.296428865076866 | 0.766902584195136 | NA | NM_177157 |
| Cyp11b1 | 0.491454083 | 0.009574229 | 0.044346165 | 0.215897555 | 0.829067603 | NA | | Gckr | 4.520779335 | 0.0143523196152407 | 0.0441222986689136 | 0.325284947707238 | 0.744965421954984 | NA | NM_144909 |
| Cyp11b2 | 0.211846316 | 0.025707621 | 0.031860375 | 0.806883811 | 0.419733426 | NA | | Gcm1 | 0.531587345 | 0.00279591970985478 | 0.0163446605787419 | 0.171060126723658 | 0.864176487957744 | NA | NM_008103 |
| Cyp17a1 | 0.289473745 | 0.012204471 | 0.038954522 | 0.313300481 | 0.754052385 | NA | | Gcm2 | 0.23403553 | -0.000407313579519725 | 0.00983856423584417 | -0.0413996971261099 | 0.966977254250032 | NA | NM_008104 |
| Cyp1a1 | 3.824364194 | 0.007771535 | 0.133483278 | 0.058221041 | 0.95357256 | NA | | Gcnt3 | 0 | NA | NA | NA | NA | NA | NM_028087 |
| Cyp1a2 | 0.196680618 | -0.003857632 | 0.031597546 | -0.122086447 | 0.902830556 | NA | | Gcnt7 | 0 | NA | NA | NA | NA | NA | NM_001039560 |
| Cyp21a1 | 1.666562999 | 0.129478863 | 0.086451318 | 1.497708378 | 0.134209034 | NA | | Gcsam | 0.106347839 | -0.00653347234140708 | 0.00854389509723638 | -0.764694822098226 | 0.444453290197145 | NA | NM_001159297,NM_008099 |
| Cyp24a1 | 1.70862315 | 0.086528132 | 0.092714498 | 0.933275096 | 0.35067795 | NA | | Gdf1 | 0 | NA | NA | NA | NA | NA | NM_001163282,NM_008107 |
| Cyp26a1 | 4.792384107 | 0.049591066 | 0.128326215 | 0.386445325 | 0.699166888 | NA | | Gdf15 | 2.014736869 | 0.00994511842064368 | 0.0347125429806273 | 0.286499275670871 | 0.774495748161129 | NA | NM_011819 |
| Cyp26c1 | 0.167300494 | 0.024299763 | 0.031030539 | 0.783091891 | 0.433573153 | NA | | Gdf2 | 0.191449919 | 0.00665508079272897 | 0.00962405648474729 | 0.691504752001018 | 0.489248395458281 | NA | NM_019506 |
| Cyp2a12 | 0 | NA | NA | NA | NA | NA | | Gdf3 | 0.360627147 | -0.0100772056525624 | 0.010553896215922 | -0.954832741045871 | 0.339662287230353 | NA | NM_008108 |
| Cyp2a22 | 0 | NA | NA | NA | NA | NA | | Gdf5 | 17.1008736 | 0.134404831211077 | 0.0728731010197266 | 1.84436821447592 | 0.0651294963346806 | NA | NM_008109 |
| Cyp2a4 | 0.198019769 | 0.025308392 | 0.031628095 | 0.80018706 | 0.423602426 | NA | | Gdf6 | 3.393105803 | 0.0169680483063502 | 0.0374512122446905 | 0.45307073628186 | 0.65049780670139 | NA | NM_013526 |
| Cyp2ab1 | 0.556822015 | 0.070925085 | 0.052229705 | 1.357945369 | 0.17448102 | NA | | Gdf7 | 7.559636392 | 0.0346718728364811 | 0.0581056426536343 | 0.596704059245311 | 0.550704981984354 | NA | NM_013527 |
| Cyp2b10 | 1.93622122 | -0.10604424 | 0.079189005 | -1.339128328 | 0.1805289 | NA | | Gdpd3 | 17.07200613 | -0.11389996381642 | 0.0742262315235023 | -1.534497461054 | 0.124907317394739 | NA | NM_024228 |
| Cyp2b13 | 0 | NA | NA | NA | NA | NA | | Gdpd4 | 3.261757732 | -0.0502103243775241 | 0.040650531107647 | -1.23517019358398 | 0.21676716980656 | NA | NM_177696 |
| Cyp2b19 | 0 | NA | NA | NA | NA | NA | | Gfi1 | 0.945824685 | 0.00362436441659644 | 0.0190896911260764 | 0.189859772620709 | 0.84941901584295 | NA | NM_010278,NM_001267621 |
| Cyp2b23 | 0.389457872 | -0.034711543 | 0.048622122 | -0.713904319 | 0.475286347 | NA | | Gfi1b | 0 | NA | NA | NA | NA | NA | NM_001160406,NM_008114 |
| Cyp2b9 | 0 | NA | NA | NA | NA | NA | | Gfra3 | 0.35787663 | -0.00674720089483589 | 0.0130723048459276 | -0.516144702434617 | 0.605753348373098 | NA | NM_010280 |
| Cyp2c29 | 0 | NA | NA | NA | NA | NA | | Gfral | 0.215934008 | -0.00149029401866511 | 0.00975123948835942 | -0.152831239602324 | 0.878531361484804 | NA | NM_205844 |
| Cyp2c37 | 0 | NA | NA | NA | NA | NA | | Ggt6 | 3.887703818 | 0.0144961561872703 | 0.0459583740826429 | 0.315419256590826 | 0.752443347857866 | NA | NM_027819 |
| Cyp2c38 | 0 | NA | NA | NA | NA | NA | | Ghrh | 7.211534777 | 0.0838290068545169 | 0.0439867836335348 | 1.90577714326462 | 0.0566791256181563 | NA | NM_010285 |
| Cyp2c39 | 0 | NA | NA | NA | NA | NA | | Ghrhr | 1.167604397 | -0.0274704629583764 | 0.0190379823527601 | -1.44292932146739 | NA | NA | NM_001003685 |
| Cyp2c40 | 0 | NA | NA | NA | NA | NA | | Ghrl | 3.032253809 | -0.0208113070457979 | 0.0348653861965165 | -0.596904532435014 | 0.550571121046601 | NA | NM_021488 |
| Cyp2c50 | 0 | NA | NA | NA | NA | NA | | Ghsr | 15.88990149 | 0.0885721757747463 | 0.044265631221258 | 2.00092426858268 | NA | NA | NM_177330 |
| Cyp2c53-ps | 0 | NA | NA | NA | NA | NA | | Gif | 0 | NA | NA | NA | NA | NA | NM_008118 |
| Cyp2c54 | 0 | NA | NA | NA | NA | NA | | Gimap3 | 2.701455704 | -0.0150511027709085 | 0.0355229012242585 | -0.423701394091936 | 0.671783598348119 | NA | NM_031247 |
| Cyp2c55 | 0.755177424 | 0.04798083 | 0.063384139 | 0.756984806 | 0.449058965 | NA | | Gimap4 | 12.90805057 | -0.0302584096405576 | 0.0654144774759768 | -0.462564417053854 | 0.64367661899157 | NA | NM_001243199,NM_175048,NM_174990,NM_001243200,NM_001243201 |
| Cyp2c65 | 0.404910094 | -0.043499547 | 0.040365123 | -1.077651771 | 0.281189189 | NA | | Gimap5 | 20.05676061 | 0.110020859961949 | 0.0721519077099198 | 1.52485032556974 | 0.127296434208186 | NA | NM_175035 |
| Cyp2c66 | 0 | NA | NA | NA | NA | NA | | Gimap7 | 0.215934008 | -0.00149029401866511 | 0.00975123948835942 | -0.152831239602324 | 0.878531361484804 | NA | NM_146167 |
| Cyp2c67 | 0 | NA | NA | NA | NA | NA | | Gimap9 | 16.85501736 | -0.0653100399565625 | 0.0754579889411697 | -0.86551524726535 | 0.386756050926774 | NA | NM_174960 |
| Cyp2c68 | 0 | NA | NA | NA | NA | NA | | Gip | 0 | NA | NA | NA | NA | NA | NM_008119 |
| Cyp2c69 | 0 | NA | NA | NA | NA | NA | | Gja10 | 0.735229523 | -0.0202023741119362 | 0.0173018086236863 | -1.16764521856282 | 0.242949902396816 | NA | NM_010289 |
| Cyp2c70 | 0.116696964 | -0.020477583 | 0.027954224 | -0.732539847 | 0.463839131 | NA | | Gja3 | 4.366468414 | 0.00919455433555953 | 0.0400690794024988 | 0.229467571320995 | 0.81850552255259 | NA | NM_016975,NM_001271623 |
| Cyp2d10 | 0 | NA | NA | NA | NA | NA | | Gja5 | 19.88664392 | 0.00957299470881115 | 0.0743447039285795 | 0.128764985304234 | 0.897543612501233 | NA | NM_008121,NM_001271628 |
| Cyp2d11 | 0 | NA | NA | NA | NA | NA | | Gja6 | 7.776498753 | -0.0653268255309061 | 0.0571178290417561 | -1.14372038690667 | 0.252739615799264 | NA | NM_001001496 |
| Cyp2d12 | 0 | NA | NA | NA | NA | NA | | Gja8 | 0.106347839 | -0.00653347234140708 | 0.00854389509723638 | -0.764694822098226 | 0.444453290197145 | NA | NM_008123 |
| Cyp2d13 | 0 | NA | NA | NA | NA | NA | | Gjb3 | 3.431121622 | 0.00967394560243806 | 0.0368952063696417 | 0.262200609627109 | 0.793166783415796 | NA | NM_001160012,NM_008126 |
| Cyp2d26 | 1.875993474 | 0.071241832 | 0.093101132 | 0.765209084 | 0.444147052 | NA | | Gjb4 | 0.21100899 | -0.000459554483117493 | 0.00972904351832043 | -0.0472353199214416 | 0.96232567770125 | NA | NM_008127 |
| Cyp2d34 | 0 | NA | NA | NA | NA | NA | | Gjb5 | 0.527194754 | 0.002237243785892 | 0.0163124918614922 | 0.137149112771257 | 0.890912932833979 | NA | NM_010291 |
| Cyp2d37-ps | 0 | NA | NA | NA | NA | NA | | Gjd3 | 10.78553117 | -0.0097893286881423 | 0.0582722094541518 | -0.167993092759671 | 0.866588713034821 | NA | NM_178596 |
| Cyp2d40 | 0 | NA | NA | NA | NA | NA | | Gjd4 | 0.905989442 | 0.0167808334636018 | 0.0213047413340349 | 0.787657226177813 | 0.430897231600891 | NA | NM_153086 |
| Cyp2d9 | 0 | NA | NA | NA | NA | NA | | Gje1 | 0.87416202 | 0.0058361348514583 | 0.0235370486296303 | 0.247955253154014 | 0.804169028589126 | NA | NM_029722 |
| Cyp2g1 | 3.774254895 | -0.005822804 | 0.074314066 | -0.078353992 | 0.93754647 | NA | | Gk2 | 0.21511414 | -0.000824240786584897 | 0.00975135300262786 | -0.08452578697159 | 0.93263840116295 | NA | NM_010294 |
| Cyp2j11 | 0 | NA | NA | NA | NA | NA | | Gkn1 | 2.912255501 | -0.00363976401288982 | 0.0384330096512854 | -0.0947041110210867 | 0.924549852534798 | NA | NM_025466 |
| Cyp2j13 | 2.94176132 | -0.210737144 | 0.115223812 | -1.828937432 | 0.067408982 | NA | | Gkn2 | 0 | NA | NA | NA | NA | NA | NM_025467 |
| Cyp2j5 | 0 | NA | NA | NA | NA | NA | | Glb1l3 | 0.67401178 | -0.0299631922056688 | 0.0181025651064097 | -1.65519041249351 | 0.0978858794123746 | NA | NM_001113323 |
| Cyp2j8 | 3.210243929 | -0.128843303 | 0.115225143 | -1.118187402 | 0.263486964 | NA | | Glipr1l1 | 0.215934008 | -0.00149029401866511 | 0.00975123948835942 | -0.152831239602324 | 0.878531361484804 | NA | NM_027018 |
| Cyp2t4 | 0.094392817 | 0.017895942 | 0.027818137 | 0.643319236 | 0.520016977 | NA | | Glipr1l2 | 0 | NA | NA | NA | NA | NA | NM_026223 |
| Cyp2w1 | 0 | NA | NA | NA | NA | NA | | Glis1 | 21.09973864 | -0.0274166741349588 | 0.0818093212197441 | -0.335128977067494 | 0.737527794957734 | NA | NM_147221 |
| Cyp3a11 | 0 | NA | NA | NA | NA | NA | | Glod5 | 6.494854489 | 0.0311132334325403 | 0.0543230071490352 | 0.572745049757299 | 0.566817330021039 | NA | NM_027227 |
| Cyp3a16 | 0 | NA | NA | NA | NA | NA | | Glp1r | 24.55941584 | 0.0616838879984628 | 0.0546942191538667 | 1.12779538592429 | NA | NA | NM_021332 |
| Cyp3a25 | 0 | NA | NA | NA | NA | NA | | Glra4 | 6.424763043 | 0.110699916803219 | 0.0580655786106598 | 1.90646368213227 | 0.0565900733013834 | NA | NM_010297 |
| Cyp3a41a | 0 | NA | NA | NA | NA | NA | | Glrp1 | 6.339215949 | -0.0158127227806244 | 0.0465361067343254 | -0.339794707599827 | 0.734011133757908 | NA | NM_008132 |
| Cyp3a41b | 0 | NA | NA | NA | NA | NA | | Glt28d2 | 14.31486856 | -0.000984270653825848 | 0.0729940208156046 | -0.0134842640921547 | 0.989241439897946 | NA | NM_177130 |
| Cyp3a44 | 0 | NA | NA | NA | NA | NA | | Glt6d1 | 0 | NA | NA | NA | NA | NA | NM_001039095 |
| Cyp3a57 | 1.037445604 | 0.04466278 | 0.065037715 | 0.686721228 | 0.492258417 | NA | | Gltpd2 | 0.191449919 | 0.00665508079272897 | 0.00962405648474729 | 0.691504752001018 | 0.489248395458281 | NA | NM_146020 |
| Cyp3a59 | 0 | NA | NA | NA | NA | NA | | Glyat | 0.793813653 | 0.0110035824521579 | 0.0202348768917179 | 0.543792903265038 | 0.586583992937947 | NA | NM_145935 |
| Cyp4a10 | 0 | NA | NA | NA | NA | NA | | Glyatl3 | 0.217374753 | -0.00904456656736547 | 0.00999660569474087 | -0.90476376117583 | 0.365590553873005 | NA | NM_001145060 |
| Cyp4a12a | 0 | NA | NA | NA | NA | NA | | Glycam1 | 2.191804102 | -0.0431279964576192 | 0.0327415391406091 | -1.31722568912858 | 0.187762985410331 | NA | NM_008134 |
| Cyp4a12b | 3.14207922 | -0.096682315 | 0.116253188 | -0.831653017 | 0.405604827 | NA | | Gm10007 | 2.190538874 | 0.0260462274337775 | 0.0327234685742955 | 0.795949468945874 | 0.426061406434977 | NA | NR_040449 |
| Cyp4a14 | 0 | NA | NA | NA | NA | NA | | Gm10012 | 0.745256552 | 0.000594825147775844 | 0.0179590771362528 | 0.0331211422092013 | 0.973577982953178 | NA | NR_028042 |
| Cyp4a29-ps | 0 | NA | NA | NA | NA | NA | | Gm10024 | 0 | NA | NA | NA | NA | NA | NM_001081452 |
| Cyp4a30b-ps | 0 | NA | NA | NA | NA | NA | | Gm10046 | 2.985689659 | 0.0170599740867179 | 0.0406409570123075 | 0.419772941900741 | 0.674651333300805 | NA | NR_033484 |
| Cyp4a31 | 0 | NA | NA | NA | NA | NA | | Gm10052 | 0 | NA | NA | NA | NA | NA | NR_002885 |
| Cyp4a32 | 0 | NA | NA | NA | NA | NA | | Gm10057 | 0 | NA | NA | NA | NA | NA | NM_001243016 |
| Cyp4f18 | 3.549067979 | 0.149919138 | 0.092241097 | 1.625296564 | 0.104099383 | NA | | Gm10058 | 0 | NA | NA | NA | NA | NA | NM_001109969 |
| Cyp4f39 | 0.765385359 | -0.115948354 | 0.064415651 | -1.800002827 | 0.071860192 | NA | | Gm10081 | 0 | NA | NA | NA | NA | NA | NM_001162940 |
| Cyp4f40 | 0 | NA | NA | NA | NA | NA | | Gm10094 | 0 | NA | NA | NA | NA | NA | NM_001142441 |
| Cyp4f41-ps | 3.218777658 | 0.102065351 | 0.119120128 | 0.856827075 | 0.39154046 | NA | | Gm10096 | 0 | NA | NA | NA | NA | NA | NM_001102678 |
| Cyp7a1 | 0.206069989 | -0.026776453 | 0.031767726 | -0.842882251 | 0.399294301 | NA | | Gm101 | 14.92818003 | 0.0316402064761256 | 0.0617499710557183 | 0.51239224788585 | 0.608376515345107 | NA | NM_001115074 |
| Cyp8b1 | 0 | NA | NA | NA | NA | NA | | Gm10100 | 0 | NA | NA | NA | NA | NA | NM_001205033 |
| Cypt1 | 0 | NA | NA | NA | NA | NA | | Gm10104 | 0 | NA | NA | NA | NA | NA | NM_001177481 |
| Cypt10 | 0 | NA | NA | NA | NA | NA | | Gm10142 | 0 | NA | NA | NA | NA | NA | NM_001205035 |
| Cypt12 | 0.258977192 | 0.029727007 | 0.03417262 | 0.869907163 | 0.38435114 | NA | | Gm10147 | 0 | NA | NA | NA | NA | NA | NM_001099919 |
| Cypt14 | 0 | NA | NA | NA | NA | NA | | Gm10190 | 6.175972794 | 0.0182765617871371 | 0.0512787829743471 | 0.356415669932731 | 0.721529295977156 | NA | NR_028385 |
| Cypt15 | 0 | NA | NA | NA | NA | NA | | Gm10220 | 2.603009291 | -0.0337011475892121 | 0.0374765060758977 | -0.899260660024184 | 0.368513836514775 | NA | NM_001134299 |
| Cypt2 | 0 | NA | NA | NA | NA | NA | | Gm10228 | 0 | NA | NA | NA | NA | NA | NM_001270487 |
| Cypt3 | 0 | NA | NA | NA | NA | NA | | Gm10229 | 0 | NA | NA | NA | NA | NA | NM_001199334 |
| Cypt4 | 0 | NA | NA | NA | NA | NA | | Gm10230 | 0 | NA | NA | NA | NA | NA | NM_001099347 |
| Cypt7 | 0 | NA | NA | NA | NA | NA | | Gm10248 | 0.72587044 | 0.0141246289585777 | 0.0167923334326615 | 0.841135570301679 | 0.40027199001691 | NA | NR_033550 |
| Cypt8 | 0 | NA | NA | NA | NA | NA | | Gm10256 | 0 | NA | NA | NA | NA | NA | NM_001270510 |
| Cypt9 | 0 | NA | NA | NA | NA | NA | | Gm10267 | 0 | NA | NA | NA | NA | NA | NM_001165950 |
| Cytip | 7.628773967 | 0.111026357 | 0.147352558 | 0.753474241 | 0.451164985 | NA | | Gm10272 | 0 | NA | NA | NA | NA | NA | NR_026831 |
| D030018L15Rik | 0.401795408 | -0.00423014 | 0.047730791 | -0.088624972 | 0.929379961 | NA | | Gm10280 | 0.215934008 | -0.00149029401866511 | 0.00975123948835942 | -0.152831239602324 | 0.878531361484804 | NA | NR_033584 |
| D030024E09Rik | 0 | NA | NA | NA | NA | NA | | Gm10318 | 0 | NA | NA | NA | NA | NA | NM_001162944 |
| D030025E07Rik | 0.46655017 | -0.036459544 | 0.045831149 | -0.795518874 | 0.426311736 | NA | | Gm10319 | 0 | NA | NA | NA | NA | NA | NR_003624 |
| D030025P21Rik | 0.412413621 | -0.026778838 | 0.043795168 | -0.611456443 | 0.540897446 | NA | | Gm10324 | 0.21511414 | -0.000824240786584897 | 0.00975135300262786 | -0.08452578697159 | 0.93263840116295 | NA | NM_001177832 |
| D030040B21Rik | 0.462334498 | 0.005887023 | 0.049959785 | 0.117835246 | 0.906198202 | NA | | Gm10334 | 16.67583771 | 0.0136873703316756 | 0.0747748212979249 | 0.183047850788451 | 0.854760473348918 | NA | NM_001103153 |
| D030045P18Rik | 1.457485358 | -0.095836437 | 0.083616539 | -1.146142118 | 0.251736353 | NA | | Gm10352 | 0 | NA | NA | NA | NA | NA | NM_001270511 |
| D030047H15Rik | 5.035610353 | 0.160455316 | 0.132056806 | 1.21504768 | 0.224347907 | NA | | Gm10364 | 1.424602839 | 0.0215148164167072 | 0.0253057297731043 | 0.850195454136785 | 0.395216428680378 | NA | NR_073535 |
| D130009I18Rik | 0 | NA | NA | NA | NA | NA | | Gm10373 | 1.147038825 | 0.000442870942116817 | 0.02010128234679 | 0.0220319746012393 | 0.982422449680152 | NA | NR_046064 |
| D130058E03 | 0.098353386 | -0.020212831 | 0.02786466 | -0.725393061 | 0.468210929 | NA | | Gm10375 | 0 | NA | NA | NA | NA | NA | NM_001098269 |
| D16Ertd519e | 0.317012939 | 0.010888466 | 0.039852335 | 0.273220278 | 0.784683893 | NA | | Gm10377 | 0 | NA | NA | NA | NA | NA | NM_001244671 |
| D17Ertd648e | 0.266310378 | 0.0407635 | 0.040043874 | 1.017970943 | 0.308691765 | NA | | Gm10389 | 1.539676372 | -0.026954810104748 | 0.0269343371864399 | -1.00076010477505 | 0.316942801457473 | NA | NR_033541 |
| D1Pas1 | 0 | NA | NA | NA | NA | NA | | Gm10390 | 0.746784837 | -0.009022343265935891 | 0.0199441129530133 | -0.452381245564283 | 0.650994355346521 | NA | NR_045793 |
| D230030E09Rik | 0.08731684 | 0.017365056 | 0.027733412 | 0.626142072 | 0.531221759 | NA | | Gm10400 | 0 | NA | NA | NA | NA | NA | NR_033555 |
| D330022K07Rik | 2.055724125 | 0.162640569 | 0.098066497 | 1.6584723 | 0.097222174 | NA | | Gm10408 | 0.97901975 | -0.00940412552189008 | 0.0229864393837812 | -0.40911623435361 | 0.682454362993824 | NA | NM_001256501 |
| D530049I02Rik | 0 | NA | NA | NA | NA | NA | | Gm10409 | 0 | NA | NA | NA | NA | NA | NR_033121 |
| D5Ertd577e | 0 | NA | NA | NA | NA | NA | | Gm10413 | 0 | NA | NA | NA | NA | NA | NM_029288 |
| D5Ertd605e | 2.405648674 | -0.064213337 | 0.106402337 | -0.60349555 | 0.546179075 | NA | | Gm10415 | 0.217374753 | -0.00904456656736547 | 0.00999660569474087 | -0.90476376117583 | 0.365590553873005 | NA | NR_045480 |
| D630010B17Rik | 2.410794075 | -0.077905693 | 0.106244561 | -0.733267585 | 0.463395241 | NA | | Gm10416 | 11.58903031 | -0.0273067865445747 | 0.0680189489860219 | -0.401458519304472 | 0.688082572246612 | NA | NR_027964 |
| D630013N20Rik | 3.010989214 | 0.126214459 | 0.110023053 | 1.147163757 | 0.251313947 | NA | | Gm10433 | 12.71088226 | -0.0362790618718231 | 0.0679355829500082 | -0.53402149943307 | 0.593326659522199 | NA | NR_045282 |
| D630029K05Rik | 1.675520834 | -0.144985409 | 0.094226611 | -1.538688569 | 0.123880344 | NA | | Gm10436 | 0 | NA | NA | NA | NA | NA | NM_001105254 |
| D630032N06Rik | 6.094611392 | 0.209464017 | 0.150812663 | 1.388902047 | 0.164862538 | NA | | Gm10439 | 0 | NA | NA | NA | NA | NA | NM_001037716 |
| D630033O11Rik | 4.324882926 | 0.083293511 | 0.13742743 | 0.606090869 | 0.544454414 | NA | | Gm10440 | 2.329855727 | -0.0275417613264784 | 0.0328757547960842 | -0.837752973195883 | 0.402169455335716 | NA | NR_038045 |
| D630045M09Rik | 0.383033497 | -0.001876918 | 0.041875907 | -0.044820955 | 0.964250022 | NA | | Gm10445 | 0 | NA | NA | NA | NA | NA | NR_046063 |
| D6Ertd527e | 2.420643215 | -0.058708771 | 0.110519336 | -0.531208139 | 0.595274554 | NA | | Gm1045 | 1.053020722 | 0.00425323751001939 | 0.0199868983066863 | 0.212801278355258 | 0.831481966598336 | NA | NM_001177577 |
| D730001G18Rik | 0.08731684 | 0.017365056 | 0.027733412 | 0.626142072 | 0.531221759 | NA | | Gm10466 | 0.09572496 | 0.00498255289994511 | 0.00847999972823692 | 0.587565219295241 | 0.556824162160132 | NA | NR_033491 |
| D730005E14Rik | 4.188776461 | 0.22374399 | 0.126599061 | 1.767343206 | 0.077170768 | NA | | Gm10471 | 3.388938125 | 0.0148502068314364 | 0.0387897729036706 | 0.382838199860437 | 0.701839735910677 | NA | NM_001177579 |
| D730045A05Rik | 0.566001468 | -0.004543743 | 0.057856841 | -0.078534241 | 0.937403094 | NA | | Gm10474 | 3.271741785 | -0.0312526725104627 | 0.0370750945602536 | -0.842955975733834 | 0.399253065876382 | NA | NR_033481 |
| D730048I06Rik | 0 | NA | NA | NA | NA | NA | | Gm10485 | 0.249110328 | -0.010000816291709 | 0.0105091939847586 | -0.951625434473194 | 0.341286977626714 | NA | NR_045884 |
| D730050B12Rik | 0 | NA | NA | NA | NA | NA | | Gm10486 | 0 | NA | NA | NA | NA | NA | NM_001109970 |
| D7Ertd143e | 0 | NA | NA | NA | NA | NA | | Gm10487 | 0 | NA | NA | NA | NA | NA | NM_001100609 |
| D830005E20Rik | 0 | NA | NA | NA | NA | NA | | Gm10488 | 0 | NA | NA | NA | NA | NA | NM_001099325 |
| D830013O20Rik | 0.341934174 | 0.0366924 | 0.037856746 | 0.969243346 | 0.332423789 | NA | | Gm10494 | 1.127933422 | -0.0186090767810054 | 0.0216093937813799 | 0.86115667422731 | 0.389151758618021 | NA | NR_033462 |
| D830031N03Rik | 392.919938 | -0.083108005 | 0.157079395 | -0.52908279 | NA | NA | | Gm10510 | 0.194331409 | -0.00840968081320881 | 0.00964081821647896 | -0.872294489037576 | 0.383045016303155 | NA | NR_033511 |
| D830046C22Rik | 2.078483941 | 0.00507376 | 0.091802309 | 0.055268325 | 0.955924697 | NA | | Gm1051 | 0 | NA | NA | NA | NA | NA | NR_033125 |
| D930007P13Rik | 0.079983654 | 0.016740032 | 0.027674178 | 0.604897175 | 0.54524732 | NA | | Gm10532 | 7.833005902 | -0.0923703795670599 | 0.0519682443654314 | -1.7774389089908 | 0.0754960538216541 | NA | NR_045879 |

| | | | | | | | | | | | | | | | |
|---|---|---|---|---|---|---|---|---|---|---|---|---|---|---|---|
| D930015M05Rik | 4.259944431 | 0.137513506 | 0.130806399 | 1.051275065 | 0.293132276 | NA | | Gm10536 | 1.774559936 | -0.0399491545598507 | 0.0315541407410229 | -1.26605109889472 | 0.205494788473218 | NA | NR_033455 |
| D930028M14Rik | 5.369676831 | -0.050589606 | 0.134747415 | -0.375440274 | 0.707333057 | NA | | Gm10538 | 0 | NA | NA | NA | NA | NA | NR_045892 |
| D930032P07Rik | 0.397753127 | 0.04024206 | 0.039595247 | 1.016335617 | 0.309469593 | NA | | Gm10549 | 1.600590109 | -0.0167542763413988 | 0.0289888227057493 | -0.577956425187145 | 0.563293538756556 | NA | NR_045415 |
| DQ267100 | 7.292888916 | 0.171924681 | 0.142912322 | 1.203008098 | 0.228973187 | NA | | Gm10556 | 0.591637739 | 0.000660690444770769 | 0.0150958543075647 | 0.0437663501057831 | 0.965090650095086 | NA | NR_045881 |
| DQ267101 | 0.619937596 | 0.079772058 | 0.055246832 | 1.443920948 | 0.148761213 | NA | | Gm10578 | 0 | NA | NA | NA | NA | NA | NR_045885 |
| DQ267102 | 7.417180309 | 0.118441321 | 0.155041872 | 0.763931187 | 0.444908251 | NA | | Gm10584 | 6.330093869 | -0.0127568360161649 | 0.0564860198299662 | -0.225840589486839 | 0.821325406665125 | NA | NR_028578 |
| DXBay18 | 0 | NA | NA | NA | NA | NA | | Gm10591 | 0 | NA | NA | NA | NA | NA | NM_001193668 |
| Dab1 | 4052.864631 | -0.051749828 | 0.111027227 | -0.466100337 | NA | NA | | Gm10619 | 0 | NA | NA | NA | NA | NA | NR_077222 |
| Daf2 | 0.188785634 | 0.025025111 | 0.031461603 | 0.79541754 | 0.42637066 | NA | | Gm10636 | 3.494092062 | 0.0175618859383246 | 0.0412278284677501 | 0.42597164563402 | 0.670128507212912 | NA | NR_033542 |
| Dao | 3.940688519 | -0.018999632 | 0.131076522 | -0.144950685 | 0.884749808 | NA | | Gm10637 | 0.568632945 | 0.0196170899458736 | 0.0164542588165949 | 1.19221960493832 | 0.233175148150358 | NA | NR_040697 |
| Dbx1 | 0 | NA | NA | NA | NA | NA | | Gm10639 | 0.332904726 | -0.0130643184082516 | 0.0120034188534089 | -1.08838311549392 | 0.276420011853847 | NA | NM_001122660 |
| Dcaf8 | 6303.052294 | 0.145013089 | 0.073143629 | 1.982579889 | NA | NA | | Gm10640 | 5.773040383 | -0.0846917206149967 | 0.0541007823514595 | -1.56544354691225 | 0.117478944483927 | NA | NR_046062 |
| Dcbld2 | 1685.706633 | 0.162802744 | 0.113311397 | 1.436772894 | NA | NA | | Gm10649 | 5.231603004 | -0.0120003162679166 | 0.0485368549598014 | -0.247241323688058 | 0.804721466109382 | NA | NR_028579 |
| Dclre1b | 341.6925532 | 0.17318999 | 0.128943557 | 1.343145746 | NA | NA | | Gm10658 | 13.27176259 | -0.064262471321436 | 0.0674976668108316 | -0.95206952118119 | 0.341061725200485 | NA | NR_045886 |
| Dcpp1 | 0 | NA | NA | NA | NA | NA | | Gm10662 | 0 | NA | NA | NA | NA | NA | NM_001201364 |
| Dcpp2 | 0 | NA | NA | NA | NA | NA | | Gm10664 | 0.12890128 | -0.0068640067066208 | 0.00870804083610955 | -0.788237771940376 | 0.430557639008166 | NA | NR_040692 |
| Dcpp3 | 0 | NA | NA | NA | NA | NA | | Gm10665 | 0 | NA | NA | NA | NA | NA | NM_001167160 |
| Dcst1 | 1.016765751 | 0.015991548 | 0.071806909 | 0.222702077 | 0.823767385 | NA | | Gm10666 | 0 | NA | NA | NA | NA | NA | NM_001167573 |
| Dcstamp | 2.851231242 | 0.075215276 | 0.092361064 | 0.81436129 | 0.415438008 | NA | | Gm10670 | 0 | NA | NA | NA | NA | NA | NM_001167161 |
| Dct | 24.09630804 | -0.344028829 | 0.157984581 | -2.177610165 | NA | NA | | Gm10677 | 3.695166467 | -0.000214748357885154 | 0.0394410936125703 | -0.00544478710439995 | 0.995655709897476 | NA | NR_046048 |
| Ddi1 | 0 | NA | NA | NA | NA | NA | | Gm10681 | 0 | NA | NA | NA | NA | NA | NM_001270429 |
| Ddx43 | 5.904419629 | -0.033536154 | 0.145966234 | -0.229752819 | 0.818283849 | NA | | Gm10684 | 7.558916914 | 0.0283133282253674 | 0.0538319364437286 | 0.525957825332249 | 0.598917510313521 | NA | NR_033547 |
| Dear1 | 0.954295622 | 0.01559776 | 0.066390676 | 0.234939012 | 0.81425607 | NA | | Gm10696 | 0 | NA | NA | NA | NA | NA | NM_001146107 |
| Defa-ps1 | 0 | NA | NA | NA | NA | NA | | Gm10697 | 0 | NA | NA | NA | NA | NA | NM_001163731 |
| Defa-ps12 | 0 | NA | NA | NA | NA | NA | | Gm10714 | 9.183686047 | -0.0183988917360033 | 0.056197378830703 | -0.327397685066964 | 0.743367117123373 | NA | NR_040530 |
| Defa-ps13 | 0 | NA | NA | NA | NA | NA | | Gm10731 | 0.839618383 | -0.0278054792658696 | 0.0199681836460941 | -1.39248915968922 | 0.163774322026961 | NA | NR_045392 |
| Defa-rs1 | 0.116696964 | -0.020477583 | 0.027954224 | -0.732539847 | 0.463839131 | NA | | Gm10745 | 0 | NA | NA | NA | NA | NA | NR_040751 |
| Defa-rs7 | 0 | NA | NA | NA | NA | NA | | Gm10768 | 1.528836905 | 0.00637944559564524 | 0.0254558627953565 | 0.250608107331917 | 0.802117112974568 | NA | NR_033472 |
| Defa17 | 0 | NA | NA | NA | NA | NA | | Gm10778 | 14.8010387 | 0.069627164447342 | 0.0700753750831583 | 0.993603878177114 | 0.320415755371452 | NA | NM_001142963 |
| Defa20 | 0 | NA | NA | NA | NA | NA | | Gm10782 | 0 | NA | NA | NA | NA | NA | NR_046061 |
| Defa21 | 0 | NA | NA | NA | NA | NA | | Gm10785 | 12.54066838 | -0.0361557562243082 | 0.0676357703233576 | -0.534565601181924 | 0.59295027544374 | NA | NR_040389 |
| Defa22 | 0 | NA | NA | NA | NA | NA | | Gm10787 | 0 | NA | NA | NA | NA | NA | NR_045882 |
| Defa23 | 0 | NA | NA | NA | NA | NA | | Gm10789 | 9.734770095 | 0.0474398973004676 | 0.0616372164866 | 0.769663200329318 | 0.441499704621206 | NA | NR_033476 |
| Defa24 | 0 | NA | NA | NA | NA | NA | | Gm10791 | 9.762593256 | 0.0138181618107218 | 0.0583252389256856 | 0.236915648615311 | 0.812722229712768 | NA | NR_045889 |
| Defa25 | 0 | NA | NA | NA | NA | NA | | Gm10804 | 0 | NA | NA | NA | NA | NA | NR_040532,NR_040533 |
| Defa26 | 0 | NA | NA | NA | NA | NA | | Gm10818 | 2.115918774 | -0.0340041314719318 | 0.0316229186482241 | -1.07530022292365 | 0.282240336445582 | NA | NR_033493 |
| Defa3 | 0 | NA | NA | NA | NA | NA | | Gm10823 | 0 | NA | NA | NA | NA | NA | NR_033475 |
| Defa4 | 0 | NA | NA | NA | NA | NA | | Gm10825 | 6.489731019 | -0.0271124271297508 | 0.0557600989747554 | -0.486233482871426 | 0.626801634257743 | NA | NR_028580 |
| Defa5 | 0 | NA | NA | NA | NA | NA | | Gm10857 | 1.053336509 | -0.0102906017955657 | 0.0216696909861686 | -0.474884565826711 | 0.634869252948079 | NA | NR_033471,NR_033470 |
| Defa6 | 0 | NA | NA | NA | NA | NA | | Gm10863 | 13.75417552 | -0.14784590284371 | 0.0710552763548663 | -2.08071673812559 | 0.0374598410911349 | NA | NR_029470 |
| Defb10 | 0 | NA | NA | NA | NA | NA | | Gm10865 | 13.77090466 | -0.0913843821661115 | 0.0699215903233303 | -1.30695514423418 | 0.191227946030201 | NA | NR_045746 |
| Defb11 | 0.804457876 | 0.06189952 | 0.048640448 | 1.272593545 | 0.20316231 | NA | | Gm10872 | 0.120209049 | -0.00674972139751468 | 0.00863521114606269 | -0.781650996523957 | 0.434419707908266 | NA | NR_045747 |
| Defb12 | 0 | NA | NA | NA | NA | NA | | Gm10921 | 0 | NA | NA | NA | NA | NA | NM_001085553 |
| Defb13 | 0 | NA | NA | NA | NA | NA | | Gm10922 | 0 | NA | NA | NA | NA | NA | NM_001085554 |
| Defb14 | 0.204933042 | 0.025509582 | 0.031745334 | 0.803569494 | 0.421645647 | NA | | Gm10941 | 12.52539075 | -0.149826866296215 | 0.068769957174304 | -2.17866743636999 | 0.0293563825077247 | NA | NR_026944 |
| Defb15 | 0 | NA | NA | NA | NA | NA | | Gm1110 | 2.910855909 | 0.0186773391985077 | 0.0333493536320824 | 0.560051010420183 | 0.575444644417062 | NA | NM_001177509 |
| Defb18 | 0.098353386 | -0.020212831 | 0.02786466 | -0.725393061 | 0.468210929 | NA | | Gm11110 | 21.02024724 | 0.00453520487123301 | 0.0814703373526335 | 0.0556669460150015 | 0.955607131913051 | NA | NR_033508 |
| Defb19 | 0 | NA | NA | NA | NA | NA | | Gm11128 | 7.7388057 | 0.00380759232293408 | 0.0560900138535176 | 0.0678836046087959 | 0.945878290270398 | NA | NM_001201389 |
| Defb2 | 0 | NA | NA | NA | NA | NA | | Gm11149 | 4.221256969 | -0.00236170944544232 | 0.0327958622192577 | -0.072012421251591 | 0.942592022787622 | NA | NR_029465 |
| Defb20 | 0.407030455 | 0.026116724 | 0.048660926 | 0.536708323 | 0.591469106 | NA | | Gm11166 | 1.445909434 | -0.0141236509123426 | 0.0290852338818142 | -0.485595232609545 | 0.627254177227443 | NA | NR_024558 |
| Defb21 | 0 | NA | NA | NA | NA | NA | | Gm11186 | 0.113843286 | 0.00525947738444006 | 0.00859368002043176 | 0.612016897526494 | 0.540526577873086 | NA | NR_046047 |
| Defb22 | 0 | NA | NA | NA | NA | NA | | Gm11190 | 7.402993694 | -0.0445896680540204 | 0.0596928849165477 | -0.746984638392968 | 0.455072833596387 | NA | NR_033549 |
| Defb23 | 0.094392817 | 0.017895942 | 0.027818137 | 0.643319236 | 0.520016977 | NA | | Gm11194 | 0 | NA | NA | NA | NA | NA | NR_003282 |
| Defb25 | 0.644647693 | -0.028208895 | 0.054433229 | -0.518229311 | 0.604298287 | NA | | Gm11201 | 2.149941643 | 0.0100267045932246 | 0.0319384085791816 | 0.313938766496973 | 0.753567546318599 | NA | NR_045873 |
| Defb26 | 0 | NA | NA | NA | NA | NA | | Gm11202 | 0.354968264 | -0.0136522058293636 | 0.0122686640796521 | -1.11277036690622 | 0.265807065332778 | NA | NR_003283 |
| Defb28 | 0.419482488 | -0.03706176 | 0.049390016 | -0.75038972 | 0.453020021 | NA | | Gm11213 | 0 | NA | NA | NA | NA | NA | NR_028583,NR_028584 |
| Defb29 | 0 | NA | NA | NA | NA | NA | | Gm1123 | 2.291310793 | -0.00770669732685002 | 0.034422472818187 | -0.223885639115915 | 0.822846290435048 | NA | NM_001080776 |
| Defb3 | 0 | NA | NA | NA | NA | NA | | Gm11240 | 0.209568245 | 0.00686197416234131 | 0.00977127153168884 | 0.702260104029194 | 0.482516968822431 | NA | NR_046041,NR_046042 |
| Defb30 | 0.498045124 | 0.003767638 | 0.044410484 | 0.084836674 | 0.932391237 | NA | | Gm11346 | 7.025365611 | -0.0239724944349239 | 0.0577579505857861 | -0.415050987644 | 0.678104577916417 | NA | NR_024599 |
| Defb34 | 0 | NA | NA | NA | NA | NA | | Gm11351 | 0.240418098 | -0.0088029805258837 | 0.00986414519853158 | -0.892422034419582 | 0.372166765031741 | NA | NR_045062 |
| Defb35 | 0 | NA | NA | NA | NA | NA | | Gm1140 | 0 | NA | NA | NA | NA | NA | NM_001126317 |
| Defb36 | 0 | NA | NA | NA | NA | NA | | Gm1141 | 12.15427643 | -0.11962055653174 | 0.0708962827089454 | -1.68726133389567 | 0.0915531124063562 | NA | NR_027801 |
| Defb37 | 0 | NA | NA | NA | NA | NA | | Gm11413 | 7.667104193 | -0.0321791578065375 | 0.054998895806035 | -0.585087342844555 | 0.558488989051739 | NA | NR_045450 |
| Defb38 | 0 | NA | NA | NA | NA | NA | | Gm11426 | 0.220191124 | -0.000706246424282353 | 0.00977545482624743 | -0.072246911968337 | 0.942405412335504 | NA | NR_033582 |
| Defb39 | 0 | NA | NA | NA | NA | NA | | Gm11435 | 0 | NA | NA | NA | NA | NA | NM_001045543 |
| Defb4 | 0 | NA | NA | NA | NA | NA | | Gm11437 | 0.428231225 | 0.00541815605275971 | 0.0139174168418424 | 0.389307593092286 | 0.697048620028689 | NA | NM_001037932 |
| Defb40 | 0 | NA | NA | NA | NA | NA | | Gm11468 | 1.224102218 | 0.00929240672655605 | 0.0266311889566623 | 0.348929472945345 | 0.727142257913413 | NA | NR_033467 |
| Defb41 | 0.08731684 | 0.017365056 | 0.027733412 | 0.626142072 | 0.531221759 | NA | | Gm11487 | 0 | NA | NA | NA | NA | NA | NM_001013393 |
| Defb42 | 0.556822015 | 0.070925085 | 0.052229705 | 1.357945369 | 0.17448102 | NA | | Gm11517 | 17.14904064 | -0.0371875584917328 | 0.077409672906907 | -0.480399375107224 | 0.630943438574518 | NA | NR_033523 |
| Defb43 | 0 | NA | NA | NA | NA | NA | | Gm11529 | 2.860514319 | 0.0370271560665695 | 0.0397794610014227 | 0.930810904281615 | 0.351951388123985 | NA | NR_033524 |
| Defb44-ps | 0 | NA | NA | NA | NA | NA | | Gm11541 | 0 | NA | NA | NA | NA | NA | NM_001007584 |
| Defb45 | 0 | NA | NA | NA | NA | NA | | Gm11544 | 0 | NA | NA | NA | NA | NA | NM_001205037 |
| Defb46 | 0 | NA | NA | NA | NA | NA | | Gm11545 | 6.98125267 | -0.18960547878619 | 0.0556321317939305 | -3.40820084854048 | 0.000653927430351452 | NA | NM_001105561 |
| Defb47 | 0.688621028 | 0.009994152 | 0.060688229 | 0.164680234 | 0.869195675 | NA | | Gm11548 | 0 | NA | NA | NA | NA | NA | NR_040590 |
| Defb48 | 0 | NA | NA | NA | NA | NA | | Gm11554 | 0 | NA | NA | NA | NA | NA | NM_001099313 |
| Defb5 | 0 | NA | NA | NA | NA | NA | | Gm11559 | 0 | NA | NA | NA | NA | NA | NM_001177484 |
| Defb50 | 0 | NA | NA | NA | NA | NA | | Gm11562 | 0 | NA | NA | NA | NA | NA | NM_001177537 |
| Defb6 | 0.509507115 | -0.017094169 | 0.051034812 | -0.334951159 | 0.73766193 | NA | | Gm11563 | 0 | NA | NA | NA | NA | NA | NM_001126320 |
| Defb7 | 0 | NA | NA | NA | NA | NA | | Gm11564 | 0 | NA | NA | NA | NA | NA | NM_001100614 |
| Defb8 | 0 | NA | NA | NA | NA | NA | | Gm11565 | 0 | NA | NA | NA | NA | NA | NM_001126323 |
| Defb9 | 0 | NA | NA | NA | NA | NA | | Gm11567 | 0 | NA | NA | NA | NA | NA | NM_001101613 |
| Depdc1a | 1.971358144 | -0.047190598 | 0.092630525 | -0.509449751 | 0.610437011 | NA | | Gm11568 | 0 | NA | NA | NA | NA | NA | NM_001205030 |
| Dfnb59 | 2.167006197 | -0.005730924 | 0.109633413 | -0.052273516 | 0.958310755 | NA | | Gm11569 | 0 | NA | NA | NA | NA | NA | NM_001099312 |
| Dhh | 5.817106699 | 0.118128211 | 0.138605369 | 0.85226288 | 0.394068203 | NA | | Gm11570 | 0.204491261 | 0.00675251727497272 | 0.00969338978195025 | 0.696610517772262 | 0.486046571838551 | NA | NM_001256057 |
| Dhrs2 | 1.268245534 | 0.002510205 | 0.091751101 | 0.027358849 | 0.97817352 | NA | | Gm11595 | 0 | NA | NA | NA | NA | NA | NM_001126322 |
| Dhrs7c | 0.21363976 | -0.000753674 | 0.031922632 | -0.02360938 | 0.98116419 | NA | | Gm11596 | 0 | NA | NA | NA | NA | NA | NM_001099311 |
| Dhrs9 | 2.54759367 | 0.067456978 | 0.112489165 | 0.599675332 | 0.548722631 | NA | | Gm11648 | 4.37029336 | -0.0503430861994002 | 0.0437348225808144 | -1.15109844349717 | 0.249691738284424 | NA | NR_045346 |
| Dio1 | 0.196706773 | -0.026517754 | 0.031609464 | -0.838918169 | 0.401515231 | NA | | Gm11710 | 0 | NA | NA | NA | NA | NA | NM_001101656 |
| Dkk1 | 0.392799438 | 0.019887619 | 0.042925294 | 0.463307699 | 0.643143828 | NA | | Gm11711 | 0 | NA | NA | NA | NA | NA | NM_001101657 |
| Dkk4 | 0 | NA | NA | NA | NA | NA | | Gm11747 | 0.212496447 | -0.000886988073087717 | 0.00973851166817767 | -0.0910804549309223 | 0.927428662789121 | NA | NR_045902 |
| Dlx3 | 3.760782338 | -0.211117866 | 0.118229402 | -1.785662982 | 0.074153843 | NA | | Gm11756 | 0 | NA | NA | NA | NA | NA | NM_001126326 |
| Dlx6as2 | 0.997141522 | -0.047699308 | 0.081438456 | -0.585709869 | 0.558070501 | NA | | Gm11757 | 0 | NA | NA | NA | NA | NA | NM_001085538 |
| Dmbt1 | 0.278003423 | 0.03119216 | 0.03498768 | 0.891518392 | 0.372651129 | NA | | Gm11758 | 0 | NA | NA | NA | NA | NA | NM_001097978 |
| Dmrt1 | 0 | NA | NA | NA | NA | NA | | Gm11937 | 0 | NA | NA | NA | NA | NA | NM_001099346 |
| Dmrt3 | 3.573628683 | 0.294315984 | 0.113503582 | 2.593010534 | 0.009513986 | NA | | Gm11938 | 0 | NA | NA | NA | NA | NA | NM_001127354 |
| Dmrtc1b | 0 | NA | NA | NA | NA | NA | | Gm11944 | 21.89968336 | -0.00806713919699806 | 0.0819335683452971 | -0.0984595125040848 | 0.921567420039604 | NA | NR_045708 |
| Dmrtc1c1 | 0 | NA | NA | NA | NA | NA | | Gm11961 | 1.327998927 | -0.0079360143768446 | 0.0257554569203751 | -0.308129434526414 | 0.757983844617993 | NA | NR_027797,NR_027798 |
| Dmrtc1c2 | 0 | NA | NA | NA | NA | NA | | Gm11978 | 3.812917842 | -0.000152161903686009 | 0.0422434956196048 | -0.00360201970632829 | 0.9971260103034 | NA | NR_028586 |
| Dmrtc2 | 0.484073412 | 0.009324742 | 0.050383912 | 0.185073807 | 0.853171145 | NA | | Gm11981 | 0 | NA | NA | NA | NA | NA | NR_046025 |
| Dnajb7 | 0.19442443 | -0.026451352 | 0.031568725 | -0.837897386 | 0.402088337 | NA | | Gm11985 | 0 | NA | NA | NA | NA | NA | NR_045101 |
| Dnajb8 | 0 | NA | NA | NA | NA | NA | | Gm12 | 0 | NA | NA | NA | NA | NA | NM_001195544 |
| Dnajc22 | 0.977141333 | -0.006109669 | 0.063937801 | -0.095556438 | 0.923872864 | NA | | Gm12060 | 3.897726663 | -0.0338670593733281 | 0.038535073458816 | -0.878863236358513 | 0.379475434169987 | NA | NR_004857 |
| Dnajc5b | 1.34728483 | -0.129015236 | 0.080200045 | -1.608667881 | 0.107688983 | NA | | Gm12130 | 0 | NA | NA | NA | NA | NA | NR_040295 |
| Dnajc5g | 0.193239998 | 0.025145508 | 0.031532422 | 0.797449311 | 0.42519013 | NA | | Gm12159 | 0 | NA | NA | NA | NA | NA | NR_045100 |
| Dnase1 | 0.566503969 | 0.017827893 | 0.052996707 | 0.336396242 | 0.736572083 | NA | | Gm12169 | 0 | NA | NA | NA | NA | NA | NM_001163357 |
| Dnase1l3 | 6.793814045 | 0.252224576 | 0.117630774 | 2.144205689 | 0.032016409 | NA | | Gm12171 | 0.321262748 | 0.00283574568300987 | 0.0125641879413187 | 0.2257006737128 | 0.821434234059046 | NA | NM_001163353 |
| Dnase2b | 1.242544597 | 0.02098239 | 0.072567323 | 0.289143781 | 0.772471351 | NA | | Gm12185 | 7.932670599 | -0.0548381596839503 | 0.0484822934867176 | -1.13109664869658 | 0.258014414700675 | NA | NM_001045540 |
| Dnm3os | 1.628968287 | 0.006649644 | 0.090978817 | 0.073090022 | 0.941734482 | NA | | Gm12191 | 0 | NA | NA | NA | NA | NA | NR_028101 |
| Dnmt3l | 3.539878472 | -0.008370061 | 0.128629357 | -0.065071158 | 0.948117344 | NA | | Gm12228 | 0 | NA | NA | NA | NA | NA | NR_040284 |
| Dntt | 0.312649645 | 0.012040436 | 0.039723828 | 0.303103628 | 0.761810889 | NA | | Gm12238 | 0.624360458 | -0.00217922033612074 | 0.0172426565232261 | -0.126385417072207 | 0.899426845076867 | NA | NR_028480 |
| Doxl2 | 0 | NA | NA | NA | NA | NA | | Gm12250 | 22.8306853 | -0.140796375415865 | 0.0533303686073167 | -2.64007879736554 | NA | NA | NM_001135115 |
| Dpcr1 | 0.364243852 | 0.020744286 | 0.042194559 | 0.491634149 | 0.622977998 | NA | | Gm12253 | 0 | NA | NA | NA | NA | NA | NM_001045542 |
| Dpep1 | 0.611217879 | 0.037217074 | 0.037942619 | 0.980877835 | NA | NA | | Gm12295 | 2.113712748 | 0.00271004379580579 | 0.0319771800698917 | 0.084749305282158 | 0.932460696988693 | NA | NR_040280 |
| Dpep2 | 2.906442131 | 0.012397415 | 0.100063396 | 0.123895602 | 0.901397935 | NA | | Gm12298 | 0 | NA | NA | NA | NA | NA | NR_033539 |
| Dpep3 | 0 | NA | NA | NA | NA | NA | | Gm12359 | 13.00483454 | 0.00644187100452926 | 0.0657910863313454 | 0.0979140391767647 | 0.922000551905511 | NA | NR_033551 |
| Dppa1 | 1.738467432 | -0.069286381 | 0.097356143 | -0.711679603 | 0.476663201 | NA | | Gm12360 | 3.066196279 | -0.0299276093793545 | 0.036524542923546 | -0.819383542786551 | 0.412567621424785 | NA | NR_040625 |
| Dppa2 | 0.871485526 | 0.006170912 | 0.065128794 | 0.09474937 | 0.924513903 | NA | | Gm12409 | 0.428541593 | 0.00660357460510141 | 0.0139432791970574 | 0.47360269501704 | 0.635783253288317 | NA | NR_046068 |
| Dppa3 | 0 | NA | NA | NA | NA | NA | | Gm12504 | 3.437001253 | -0.0125084433498902 | 0.0398051050615194 | -0.314242189049826 | 0.753337102146952 | NA | NR_040414 |
| Dppa4 | 0 | NA | NA | NA | NA | NA | | Gm12505 | 0 | NA | NA | NA | NA | NA | NR_040674 |
| Dppa5a | 3.237291537 | 0.04849488 | 0.110106154 | 0.440437504 | 0.659620267 | NA | | Gm12530 | 5.800533179 | -0.0292577972414213 | 0.0543412062004539 | -0.538409050647407 | 0.590294679354421 | NA | NR_040669 |
| Dpt | 4.263485727 | -0.116786472 | 0.130039205 | -0.89808664 | 0.369139361 | NA | | Gm12597 | 0 | NA | NA | NA | NA | NA | NM_206870 |
| Dpy19l2 | 7.935042318 | 0.072922031 | 0.153326317 | 0.475600223 | 0.634359216 | NA | | Gm12603 | 0.12890128 | -0.0068640067066208 | 0.00870804083610955 | -0.788237771940376 | 0.430557639008166 | NA | NR_033533 |
| Drap1 | 3856.029312 | -0.055052089 | 0.116475557 | -0.472649291 | NA | NA | | Gm12633 | 0.326298904 | 0.00765111147689903 | 0.0103094497373689 | 0.742145475443356 | 0.457999199689495 | NA | NR_033610 |
| Dsc1 | 2.589115619 | -0.063544332 | 0.107979268 | -0.588486412 | 0.556205852 | NA | | Gm12657 | 4.44738203 | 0.0128736313119974 | 0.0403633609994599 | 0.318943492147981 | 0.749769363258368 | NA | NM_001081019 |
| Dscc1 | 3.842157774 | 0.129346656 | 0.123408192 | 1.048120503 | 0.294583091 | NA | | Gm12669 | 0 | NA | NA | NA | NA | NA | NR_033611 |
| Dsg1a | 1.78476557 | -0.039501569 | 0.082623023 | -0.478093976 | 0.632583319 | NA | | Gm12695 | 3.397062918 | 0.0210487334723748 | 0.0413659084087237 | 0.508842529563205 | 0.610862605991059 | NA | NM_001081284 |
| Dsg1b | 1.318872376 | 0.046211558 | 0.068456056 | 0.675054338 | 0.499641243 | NA | | Gm12709 | 1.004606608 | 0.0159173757340332 | 0.0200662102802637 | 0.793242745476899 | 0.427636407422682 | NA | NR_040444 |
| Dsg3 | 1.289828268 | -0.073754435 | 0.078075202 | -0.944658915 | 0.344833038 | NA | | Gm12718 | 0.127687691 | 0.00542298404407807 | 0.00869724871942359 | 0.62352868349814 | 0.532937159177571 | NA | NR_040673 |
| Dsg4 | 0 | NA | NA | NA | NA | NA | | Gm12789 | 0 | NA | NA | NA | NA | NA | NM_001085520 |
| Duox1 | 2.139866676 | 0.06955864 | 0.111811441 | 0.622106643 | 0.533871747 | NA | | Gm12794 | 0.240418098 | -0.0088029805258837 | 0.00986414519853158 | -0.892422034419582 | 0.372166765031741 | NA | NM_001085516 |
| Duox2 | 2.520612473 | -0.124688883 | 0.100292205 | -1.243255967 | 0.213773525 | NA | | Gm128 | 4.982652806 | -0.0221172389914617 | 0.0426008567327156 | -0.519173572734198 | 0.603639706242362 | NA | NM_001024841 |
| Duoxa1 | 1.486072896 | -0.093221287 | 0.08844968 | -1.053947138 | 0.29190712 | NA | | Gm12830 | 0.414768476 | -0.00783927838205996 | 0.0122486160043821 | -0.640013400636883 | 0.522163887340352 | NA | NR_033617 |
| Duoxa2 | 0.466789057 | 0.062805705 | 0.049307408 | 1.273757982 | 0.202749201 | NA | | Gm12839 | 0.108766301 | 0.00519171381361946 | 0.00856003371502575 | 0.606506234257728 | 0.544178643801544 | NA | NR_033575 |
| Dupd1 | 2.002378788 | -0.028313667 | 0.110639196 | -0.255909914 | 0.798020399 | NA | | Gm12886 | 0 | NA | NA | NA | NA | NA | NM_001144948 |
| Dusp13 | 7.723204011 | 0.200993409 | 0.150781543 | 1.333010691 | 0.182528295 | NA | | Gm12887 | 0 | NA | NA | NA | NA | NA | NM_001099309 |
| Dusp21 | 0 | NA | NA | NA | NA | NA | | Gm12888 | 0 | NA | NA | NA | NA | NA | NM_001033791 |
| Dux | 1.104854567 | 0.068644114 | 0.08014215 | 0.856529481 | 0.391704973 | NA | | Gm13003 | 16.52187612 | -0.00616135683686721 | 0.065544911711891 | -0.0940020617305895 | 0.925107518966226 | NA | NR_040443 |
| Duxbl1 | 0 | NA | NA | NA | NA | NA | | Gm13011 | 0 | NA | NA | NA | NA | NA | NM_001126318 |
| Duxbl2 | 0 | NA | NA | NA | NA | NA | | Gm13023 | 0 | NA | NA | NA | NA | NA | NM_001007077 |
| Duxbl3 | 0 | NA | NA | NA | NA | NA | | Gm13032 | 0.120209049 | -0.00674972139751468 | 0.00863521114606269 | -0.781650996523957 | 0.434419707908266 | NA | NR_045944 |
| Dydc1 | 1.163652955 | 0.016304391 | 0.079213808 | 0.205827638 | 0.836925551 | NA | | Gm13034 | 0.319043516 | -0.00957618023178583 | 0.0102890835921761 | -0.930712647632453 | 0.352002225550712 | NA | NR_030771 |
| Dynap | 0 | NA | NA | NA | NA | NA | | Gm13037 | 0 | NA | NA | NA | NA | NA | NR_040672 |
| Dynlt1c | 0 | NA | NA | NA | NA | NA | | Gm13040 | 0 | NA | NA | NA | NA | NA | NM_001113736 |
| Dynlt1f | 0 | NA | NA | NA | NA | NA | | Gm13043 | 0 | NA | NA | NA | NA | NA | NM_001039595 |
| Dyrk4 | 8.052629629 | 0.081786825 | 0.14085509 | 0.580645156 | 0.561479632 | NA | | Gm13051 | 3.538946092 | -0.0162945844654342 | 0.0372810829750049 | -0.437073796283196 | 0.662057830127222 | NA | NM_001037926 |
| Dytn | 2.675503349 | 0.146201412 | 0.116647499 | 1.253360879 | 0.210074402 | NA | | Gm13057 | 0 | NA | NA | NA | NA | NA | NM_001113735 |
| E030002O03Rik | 5.963835012 | -0.376582793 | 0.14013703 | -2.687246855 | 0.007204369 | NA | | Gm13078 | 0 | NA | NA | NA | NA | NA | NM_001085412 |
| E030018B13Rik | 0 | NA | NA | NA | NA | NA | | Gm13083 | 0 | NA | NA | NA | NA | NA | NM_001126324 |
| E030024N20Rik | 1.254731069 | 0.00474194 | 0.081515214 | 0.058172459 | 0.953611258 | NA | | Gm13084 | 0 | NA | NA | NA | NA | NA | NM_001005371 |
| E030025P04Rik | 0.91606928 | -0.07312839 | 0.066468253 | -1.100200264 | 0.271244876 | NA | | Gm13088 | 0 | NA | NA | NA | NA | NA | NM_001126325 |
| E030044B06Rik | 0.527043497 | -0.046865061 | 0.053304187 | -0.879200371 | 0.379292643 | NA | | Gm13102 | 0 | NA | NA | NA | NA | NA | NM_001085419 |
| E130018N17Rik | 1.949237493 | 0.041858869 | 0.102735546 | 0.407442908 | 0.683682718 | NA | | Gm13103 | 0 | NA | NA | NA | NA | NA | NM_177571 |
| E130201H02Rik | 4.146769221 | -0.170755899 | 0.130373563 | -1.309743281 | 0.190282697 | NA | | Gm13109 | 0 | NA | NA | NA | NA | NA | NM_001126315 |
| E130218I03Rik | 0.865351476 | 0.022878361 | 0.056461578 | 0.405202304 | 0.685328812 | NA | | Gm13119 | 0 | NA | NA | NA | NA | NA | NM_001034101 |
| E130304I02Rik | 2.753772459 | 0.080587187 | 0.115470125 | 0.697905081 | 0.485236557 | NA | | Gm13124 | 0 | NA | NA | NA | NA | NA | NM_001085542 |
| E130310I04Rik | 2.721459421 | 0.06797549 | 0.106170778 | 0.640246701 | 0.522012225 | NA | | Gm13125 | 0 | NA | NA | NA | NA | NA | NM_001115077 |
| E230016K23Rik | 3.351906977 | 0.021074134 | 0.123095519 | 0.171201472 | 0.86406535 | NA | | Gm13128 | 0 | NA | NA | NA | NA | NA | NM_001085541 |
| E230019M04Rik | 0.976884471 | 0.068046414 | 0.068004534 | 1.00061584 | 0.317012569 | NA | | Gm13139 | 6.598793456 | -0.0331407089084731 | 0.0531166547324299 | -0.623923119319472 | 0.532678076277797 | NA | NM_001083918 |
| E330012B07Rik | 0.583727788 | 0.046764408 | 0.054560797 | 0.857106391 | 0.391386089 | NA | | Gm13154 | 0.564763665 | -0.0062993601387677 | 0.0166593544926572 | -0.378127504372646 | 0.705335877003366 | NA | NM_001014397 |
| E330013P04Rik | 4.000472248 | 0.030423815 | 0.115649079 | 0.263070104 | 0.792496546 | NA | | Gm13177 | 0 | NA | NA | NA | NA | NA | NM_001081248 |
| E330014E10Rik | 0 | NA | NA | NA | NA | NA | | Gm13178 | 0.227686571 | 0.00691460391109036 | 0.00980755285755433 | 0.705028462402254 | 0.480792523671872 | NA | NM_001085536 |
| E330017A01Rik | 0 | NA | NA | NA | NA | NA | | Gm13212 | 6.238532706 | 0.00804055778092145 | 0.0401906364598144 | 0.200060474010184 | 0.841433285568123 | NA | NM_001205101 |
| E330017L17Rik | 2.113500612 | 0.087607488 | 0.101099921 | 0.866543582 | 0.386192137 | NA | | Gm1322 | 0 | NA | NA | NA | NA | NA | NM_001033477 |
| E330021D16Rik | 0 | NA | NA | NA | NA | NA | | Gm13238 | 14.33041661 | -0.0575303415627335 | 0.0707846031036659 | -0.812752195254649 | 0.416360148923696 | NA | NR_033612 |
| E330023G01Rik | 0.825193745 | 0.019755774 | 0.050840231 | 0.388585451 | 0.697582831 | NA | | Gm13242 | 0.097165704 | -0.00637828455464458 | 0.00848781691535724 | -0.75146349388194 | 0.452373763047372 | NA | NM_001103158 |
| E330034G19Rik | 7.070014016 | -0.077931341 | 0.150930311 | -0.516339898 | 0.605617035 | NA | | Gm13247 | 2.196852926 | 0.0200671662506139 | 0.0309026072117019 | 0.649368065067825 | 0.516100500463062 | NA | NM_001243138,NM_001243139 |
| E430016F16Rik | 7.844773586 | -0.092996683 | 0.149346694 | -0.622689931 | 0.533488301 | NA | | Gm13271 | 0 | NA | NA | NA | NA | NA | NM_001085528 |
| EU599041 | 0 | NA | NA | NA | NA | NA | | Gm13272 | 0 | NA | NA | NA | NA | NA | NM_001161608 |
| Ear1 | 0.08731684 | 0.017365056 | 0.027733412 | 0.626142072 | 0.531221759 | NA | | Gm13275 | 0 | NA | NA | NA | NA | NA | NM_001085533 |
| Ear10 | 0 | NA | NA | NA | NA | NA | | Gm13276 | 0 | NA | NA | NA | NA | NA | NM_001085532 |
| Ear11 | 0.601913201 | 0.016398223 | 0.047539851 | 0.344936356 | 0.730142218 | NA | | Gm13277 | 0 | NA | NA | NA | NA | NA | NM_001098840 |
| Ear12 | 0 | NA | NA | NA | NA | NA | | Gm13278 | 0 | NA | NA | NA | NA | NA | NM_001098841 |
| Ear14 | 0 | NA | NA | NA | NA | NA | | Gm13279 | 0 | NA | NA | NA | NA | NA | NM_001243166 |
| Ear2 | 0.19442443 | -0.026451352 | 0.031568725 | -0.837897386 | 0.402088337 | NA | | Gm13280 | 0 | NA | NA | NA | NA | NA | NM_206867 |
| Ear3 | 0 | NA | NA | NA | NA | NA | | Gm13283 | 0 | NA | NA | NA | NA | NA | NM_001085531 |
| Ear4 | 0 | NA | NA | NA | NA | NA | | Gm13285 | 0 | NA | NA | NA | NA | NA | NM_001161609 |
| Ear5 | 0 | NA | NA | NA | NA | NA | | Gm13286 | 0 | NA | NA | NA | NA | NA | NM_001243150 |
| Ear6 | 0 | NA | NA | NA | NA | NA | | Gm13287 | 0 | NA | NA | NA | NA | NA | NM_001161610 |
| Ear7 | 0 | NA | NA | NA | NA | NA | | Gm13288 | 0 | NA | NA | NA | NA | NA | NM_001243167 |
| Eci3 | 2.696487227 | 0.082874488 | 0.113218419 | 0.731987681 | 0.464176087 | NA | | Gm13289 | 0 | NA | NA | NA | NA | NA | NM_001243165 |
| Ect2l | 7.825503896 | 0.182843732 | 0.152792908 | 1.196676826 | 0.231432543 | NA | | Gm13290 | 0 | NA | NA | NA | NA | NA | NM_001243155 |
| Eddm3b | 0 | NA | NA | NA | NA | NA | | Gm13298 | 0.839767923 | 0.0123379582262517 | 0.0191585146787882 | 0.643993463643193 | 0.519579672549587 | NA | NM_001085530 |
| Edn2 | 1.77512903 | 0.006815807 | 0.101457163 | 0.067179155 | 0.946439079 | NA | | Gm13304 | 0 | NA | NA | NA | NA | NA | NM_001193666 |
| Efcab8 | 0.378390275 | 0.024206488 | 0.047912756 | 0.50522012 | 0.61340424 | NA | | Gm13305 | 0 | NA | NA | NA | NA | NA | NM_001099348 |
| Efcab9 | 7.840917801 | -0.055421878 | 0.158129876 | -0.350483285 | 0.725976032 | NA | | Gm13306 | 0 | NA | NA | NA | NA | NA | NM_001164046,NM_001199963,NM_001199962 |
| Egfbp2 | 0 | NA | NA | NA | NA | NA | | Gm13308 | 0 | NA | NA | NA | NA | NA | NM_001177580 |
| Ehf | 5.217180314 | 0.307076261 | 0.116888929 | 2.627077364 | 0.008612175 | NA | | Gm13315 | 0.332414823 | -0.0166094830116705 | 0.0135191688281235 | -1.22858758721308 | 0.219226471765921 | NA | NR_028497 |
| Eid3 | 3.601856976 | 0.083050739 | 0.130042195 | 0.638644548 | 0.523054194 | NA | | Gm1332 | 1.557541471 | 0.0215880344773298 | 0.030197968803121 | 0.71488366048973 | 0.474680934235318 | NA | NM_001177439 |
| Eif3j1 | 0 | NA | NA | NA | NA | NA | | Gm13363 | 0 | NA | NA | NA | NA | NA | NR_002688 |
| Eif3j2 | 0.119397641 | -0.020477583 | 0.027954224 | -0.732539845 | 0.463839132 | NA | | Gm13483 | 16.75208756 | -0.0708496425147506 | 0.0772738846001168 | -0.916863994626246 | 0.359213913359254 | NA | NR_040362,NR_040361 |
| Eif4e1b | 0.390895054 | 0.015442627 | 0.036628692 | 0.421599181 | 0.673317604 | NA | | Gm13490 | 0.106347839 | -0.00653347234140708 | 0.00854389509723638 | -0.764694822098226 | 0.444453290197145 | NA | NR_040639 |
| Elane | 1.329531748 | -0.068571592 | 0.08965843 | -0.764809201 | 0.444385168 | NA | | Gm13497 | 1.027545051 | -0.0158744260324277 | 0.0194726277894952 | -0.81521745313652 | 0.414947849272224 | NA | NR_040636 |
| Elf3 | 5.286250829 | -0.126652234 | 0.133251651 | -0.950474034 | 0.341871442 | NA | | Gm13498 | 0.308174694 | -0.0038530932116974 | 0.0109687422478024 | -0.351279401470974 | 0.725378746751542 | NA | NR_033595 |
| Ell3 | 3.148015308 | -0.034543619 | 0.123897159 | -0.278808803 | 0.780391558 | NA | | Gm13539 | 0 | NA | NA | NA | NA | NA | NR_045340 |
| Elovl3 | 0.766232536 | -0.054016814 | 0.063110901 | -0.855903078 | 0.392051392 | NA | | Gm13544 | 2.086943085 | 0.0125144941409119 | 0.0280902518535976 | 0.445510214936328 | 0.655951079432322 | NA | NR_040365 |
| Elovl4 | 1842.92971 | 0.113001034 | 0.121353031 | 0.931176039 | NA | NA | | Gm13546 | 1.402126726 | 0.00690562499907936 | 0.0256251350825778 | 0.269486384240542 | 0.787555418595751 | NA | NR_045896,NR_045895 |
| Eme1 | 4.324249447 | -0.274760079 | 0.13995426 | -1.963213398 | 0.049621383 | NA | | Gm13547 | 4.466372608 | -0.0429588096012272 | 0.0471446320542127 | -0.911213169546596 | 0.36218306636819 | NA | NM_001177392 |
| Emilin3 | 5.724997593 | 0.106658926 | 0.135393543 | 0.787769665 | 0.430831448 | NA | | Gm13580 | 0.106347839 | -0.00653347234140708 | 0.00854389509723638 | -0.764694822098226 | 0.444453290197145 | NA | NR_046065 |
| Emr4 | 1.817799692 | -0.086568146 | 0.101525858 | -0.852670915 | 0.393841825 | NA | | Gm13582 | 0.203314312 | -0.000639443352387801 | 0.00969102748600992 | -0.0659830294889689 | 0.947391336513325 | NA | NR_045335 |
| En1 | 1.78813753 | 0.088671736 | 0.086840617 | 1.021085979 | 0.307213706 | NA | | Gm136 | 4.76045503 | -0.00523599983361608 | 0.0433677598439394 | -0.120734846633952 | 0.903901057255619 | NA | NM_001033255 |
| Enam | 0 | NA | NA | NA | NA | NA | | Gm13704 | 15.73702858 | -0.105689095406252 | 0.0588348452137265 | -1.79636905684583 | 0.072435841668037 | NA | NR_040623 |
| Eno1 | 2.611817104 | 0.013854512 | 0.098205679 | 0.141076485 | 0.88780952 | NA | | Gm13710 | 7.330188777 | -0.0459165285045239 | 0.0555269780691838 | -0.826922877872342 | 0.408280775387452 | NA | NR_046046 |
| Enpp2 | 12168.30993 | 0.298143001 | 0.166153531 | 1.794382581 | NA | NA | | Gm13718 | 0 | NA | NA | NA | NA | NA | NM_001145100 |
| Enpp7 | 0 | NA | NA | NA | NA | NA | | Gm13749 | 16.83532029 | 0.0622132745913425 | 0.0744124961126227 | 0.836059504000285 | 0.403121428711476 | NA | NR_027824 |
| Enthd1 | 0.955421901 | 0.066721921 | 0.069484429 | 0.960242778 | 0.336933042 | NA | | Gm13752 | 0 | NA | NA | NA | NA | NA | NR_040370 |
| Entpd8 | 0.276690427 | -0.012033946 | 0.033317809 | -0.361186582 | 0.717959972 | NA | | Gm13769 | 0 | NA | NA | NA | NA | NA | NM_001270423 |
| Ep300 | 3273.437727 | -0.059880036 | 0.09823326 | -0.609569872 | NA | NA | | Gm13807 | 0 | NA | NA | NA | NA | NA | NR_040529 |
| Epb4.2 | 0.494244083 | 0.041448004 | 0.052104748 | 0.795474608 | 0.426337476 | NA | | Gm13871 | 0 | NA | NA | NA | NA | NA | NM_001177578 |
| Epcam | 6.107276653 | -0.068676491 | 0.146376 | -0.469178627 | 0.638941961 | NA | | Gm13939 | 0 | NA | NA | NA | NA | NA | NR_033473 |
| Epgn | 2.711438282 | 0.025508325 | 0.10108538 | 0.252344351 | 0.80077491 | NA | | Gm13944 | 0.318844286 | -0.00448436457897182 | 0.0111065302755339 | -0.403759272042883 | 0.686389748741264 | NA | NR_040368 |
| Epo | 0.59261035 | -0.06016431 | 0.055570839 | -1.082659739 | 0.278959481 | NA | | Gm14005 | 6.409563778 | 0.0167648445105365 | 0.0576676593610617 | 0.290714842535406 | 0.77126941895867 | NA | NR_028589,NR_028590,NR_028591 |
| Eppin | 0 | NA | NA | NA | NA | NA | | Gm14015 | 16.0461305 | 0.0827686254033913 | 0.0707728131152527 | 1.16949746322226 | 0.242203262709665 | NA | NR_040637 |
| Eppk1 | 6.623047455 | 0.15723547 | 0.151422488 | 1.038389157 | 0.299088916 | NA | | Gm14023 | 9.967836739 | 0.0580674410606521 | 0.0664961969984719 | 0.873244541518465 | 0.382529801675622 | NA | NR_040371 |
| Eps8l3 | 5.852289433 | 0.163762554 | 0.117493252 | 1.393803913 | 0.163376826 | NA | | Gm14047 | 2.629237912 | 0.0109287059251646 | 0.0363790932335645 | 0.300411718758328 | 0.763863125989505 | NA | NR_015474 |
| Epyc | 1.96563487 | 0.131724268 | 0.100098432 | 1.315947373 | 0.188191707 | NA | | Gm14085 | 0.538437718 | 0.0143037664246173 | 0.0140664739385652 | 1.01686936520755 | 0.309215578500882 | NA | NM_001085518 |
| Eras | 0 | NA | NA | NA | NA | NA | | Gm14092 | 0.933509117 | -0.0234738955680579 | 0.0216376897901449 | -1.08486145220315 | 0.277983023154311 | NA | NM_001037929 |
| Ereg | 0.198019769 | 0.025308392 | 0.031628095 | 0.80018706 | 0.423602426 | NA | | Gm14124 | 5.134910273 | -0.0102199342221013 | 0.0395285047518066 | -0.258545935047904 | 0.795985602584924 | NA | NM_001142410 |
| Erp27 | 3.119599062 | 0.148250721 | 0.119277426 | 1.242906769 | 0.213902191 | NA | | Gm14137 | 2.972374042 | -0.0550101981056792 | 0.0376735033940012 | -1.46018270534506 | 0.144239867156234 | NA | NM_001039223 |
| Esco2 | 6.581219312 | 0.002982056 | 0.151701691 | 0.019657365 | 0.984316702 | NA | | Gm14139 | 0.288615623 | 0.00197374456361263 | 0.0106648328298147 | 0.185070370544844 | 0.853173841012254 | NA | NM_001145863 |
| Esrp1 | 3.052494327 | -0.035666884 | 0.108028346 | -0.330162273 | 0.741277352 | NA | | Gm14151 | 0.335323189 | -0.0061413109930725 | 0.0128252480301014 | -0.478845397660815 | 0.632048617772247 | NA | NM_001097977 |
| Esrp2 | 2.340204657 | 0.01422642 | 0.0984252 | 0.144540421 | 0.885073741 | NA | | Gm14164 | 1.766862212 | 0.0026789595756827 | 0.0239318595830626 | 0.111941137143337 | 0.91087007884239 | NA | NR_033505 |
| Esx1 | 0 | NA | NA | NA | NA | NA | | Gm14207 | 2.709108448 | 0.0272450984806449 | 0.0353424206502216 | 0.7708894291731 | 0.440772473324439 | NA | NR_030683 |
| Etd | 0 | NA | NA | NA | NA | NA | | Gm14288 | 0 | NA | NA | NA | NA | NA | NM_001033123 |
| Etv2 | 0.096071043 | -0.020098795 | 0.027825948 | -0.722304036 | 0.470107572 | NA | | Gm14305 | 0 | NA | NA | NA | NA | NA | NM_001099327 |
| Evx1 | 0.105923158 | 0.018524693 | 0.027977518 | 0.662127842 | 0.507889294 | NA | | Gm14306 | 0 | NA | NA | NA | NA | NA | NM_001242944 |
| Evx2 | 0 | NA | NA | NA | NA | NA | | Gm14308 | 2.098716026 | -0.0518177459807515 | 0.0317761784892288 | -1.63071043921522 | 0.102951432810131 | NA | NM_001099349 |
| F10 | 0.608590881 | 0.025239389 | 0.058857007 | 0.428825565 | 0.668050172 | NA | | Gm14318 | 0.108766301 | 0.00519171381361946 | 0.00856003371502575 | 0.606506234257728 | 0.544178643801544 | NA | NR_045877 |
| F11 | 1.014523425 | -0.008946388 | 0.077241892 | -0.115823006 | 0.907792817 | NA | | Gm14327 | 4.458747783 | -0.0699782668012742 | 0.0438310637485462 | -1.5965450257546 | 0.110367161503196 | NA | NR_038101 |
| F13b | 0 | NA | NA | NA | NA | NA | | Gm14345 | 0 | NA | NA | NA | NA | NA | NM_001085545 |
| F2 | 2.575664693 | 0.015768548 | 0.099068314 | 0.159168429 | 0.873536176 | NA | | Gm14346 | 0 | NA | NA | NA | NA | NA | NM_001085551 |
| F2rl3 | 4.338555771 | 0.074491482 | 0.116644866 | 0.638617748 | 0.523071632 | NA | | Gm14347 | 0 | NA | NA | NA | NA | NA | NM_001085543 |
| F5 | 90.66509134 | 0.306242572 | 0.130779031 | 2.34167948 | NA | NA | | Gm14351 | 0 | NA | NA | NA | NA | NA | NM_001085552 |
| F630028O10Rik | 3.168600947 | -0.113463967 | 0.101133298 | -1.121924925 | 0.261894362 | NA | | Gm14354 | 0 | NA | NA | NA | NA | NA | NM_029106 |
| F630111L10Rik | 0 | NA | NA | NA | NA | NA | | Gm14374 | 0 | NA | NA | NA | NA | NA | NM_001085523 |
| F630206G17Rik | 0 | NA | NA | NA | NA | NA | | Gm14379 | 14.30840983 | 0.0208337381670456 | 0.0623360492495854 | 0.334216531490951 | 0.738216170222258 | NA | NR_026741 |

| | | | | | | |
|---|---|---|---|---|---|---|
| F7 | 3.810119778 | 0.055875058 | 0.107332469 | 0.520579268 | 0.602659895 | NA |
| F730035M05Rik | 0.862304613 | 0.072123711 | 0.060987028 | 1.18260741 | 0.236964775 | NA |
| F830002L21Rik | 0 | NA | NA | NA | NA | NA |
| F830045P16Rik | 3.177521128 | -0.054521823 | 0.100498841 | -0.542511952 | 0.587465876 | NA |
| Fabp1 | 0 | NA | NA | NA | NA | NA |
| Fabp12 | 0 | NA | NA | NA | NA | NA |
| Fabp2 | 0 | NA | NA | NA | NA | NA |
| Fabp4 | 2.614357751 | 0.013335283 | 0.118686773 | 0.112356947 | 0.910540391 | NA |
| Fabp6 | 0.238795281 | -0.027601606 | 0.032267774 | -0.855392297 | 0.392334007 | NA |
| Fabp9 | 0 | NA | NA | NA | NA | NA |
| Faim3 | 2.164581991 | 0.008983142 | 0.104935757 | 0.085606108 | 0.931779543 | NA |
| Fam110c | 3.070669514 | -0.266438682 | 0.110310311 | -2.415356089 | 0.015719833 | NA |
| Fam115e | 0.116696964 | -0.020477583 | 0.027954224 | -0.732539847 | 0.463839131 | NA |
| Fam122c | 0 | NA | NA | NA | NA | NA |
| Fam124b | 1.211753037 | -0.015175223 | 0.08484861 | -0.178850585 | 0.858055024 | NA |
| Fam150a | 1.362713392 | -0.078483362 | 0.086579937 | -0.906484396 | 0.364679521 | NA |
| Fam151a | 3.3190959 | 0.04965642 | 0.112951216 | 0.439627135 | 0.660207184 | NA |
| Fam154a | 1.591349109 | -0.010914459 | 0.086180799 | -0.126646068 | 0.899220533 | NA |
| Fam159a | 1.405242769 | -0.067305686 | 0.08552105 | -0.787007247 | 0.431277624 | NA |
| Fam162b | 0 | NA | NA | NA | NA | NA |
| Fam166a | 2.776405486 | 0.138982001 | 0.109737877 | 1.266490704 | 0.205337455 | NA |
| Fam170a | 0.08731684 | 0.017365056 | 0.027733412 | 0.626142072 | 0.531221759 | NA |
| Fam170b | 0.38968403 | -0.004310943 | 0.042055996 | -0.102504825 | 0.918355984 | NA |
| Fam177a | 0 | NA | NA | NA | NA | NA |
| Fam178b | 3.829658447 | -0.006411603 | 0.120401891 | -0.053251678 | 0.957531381 | NA |
| Fam187a | 2.566500152 | -0.003690655 | 0.106798931 | -0.034557039 | 0.972432959 | NA |
| Fam187b | 2.035303952 | -0.085013408 | 0.105238561 | -0.807816139 | 0.419196433 | NA |
| Fam209 | 1.449700838 | -0.062223199 | 0.083969511 | -0.741021334 | 0.458680505 | NA |
| Fam213a | 3923.705842 | 0.107790065 | 0.084980665 | 1.268406934 | NA | NA |
| Fam227b | 1.101352946 | -0.029980011 | 0.083655862 | -0.3583731 | 0.72006412 | NA |
| Fam229a | 2.800884478 | 0.091358648 | 0.11708871 | 0.780251553 | 0.435242823 | NA |
| Fam24a | 0 | NA | NA | NA | NA | NA |
| Fam25c | 0.333037401 | -0.016093237 | 0.040545167 | -0.396921218 | 0.691425557 | NA |
| Fam26d | 0.665247938 | 0.039121182 | 0.061388665 | 0.63727045 | 0.523948696 | NA |
| Fam3b | 0 | NA | NA | NA | NA | NA |
| Fam46b | 3.030819446 | -0.069823169 | 0.110165979 | -0.633799742 | 0.526211521 | NA |
| Fam46d | 0.670842878 | 0.03854673 | 0.060574172 | 0.636355866 | 0.5245445 | NA |
| Fam47c | 0 | NA | NA | NA | NA | NA |
| Fam47e | 7.995450745 | 0.077393978 | 0.155229753 | 0.498576958 | 0.618077441 | NA |
| Fam50b | 0 | NA | NA | NA | NA | NA |
| Fam71a | 0 | NA | NA | NA | NA | NA |
| Fam71b | 0.08731684 | 0.017365056 | 0.027733412 | 0.626142072 | 0.531221759 | NA |
| Fam71d | 4.862649671 | 0.064585519 | 0.140295324 | 0.460354042 | 0.645262117 | NA |
| Fam71e2 | 0.099009884 | 0.018205726 | 0.027896865 | 0.652608315 | 0.514008823 | NA |
| Fam71f1 | 0 | NA | NA | NA | NA | NA |
| Fam83a | 0.426633926 | -0.004170814 | 0.042905384 | -0.097209576 | 0.922559963 | NA |
| Fam83b | 3.412064704 | -0.092500129 | 0.119400519 | -0.774704576 | 0.438514247 | NA |
| Fam83c | 0.238795281 | -0.027601606 | 0.032267774 | -0.855392297 | 0.392334007 | NA |
| Fam83e | 3.100974519 | -0.146460362 | 0.126535299 | -1.15746644 | 0.247081843 | NA |
| Fam83g | 7.944149917 | -0.110072069 | 0.148975254 | -0.738861431 | 0.459991145 | NA |
| Fasl | 3.499930511 | 0.167464601 | 0.12045149 | 1.390307425 | 0.164435544 | NA |
| Fate1 | 3.933553786 | 0.091768208 | 0.121420309 | 0.755789615 | 0.449775344 | NA |
| Fbp1 | 6.741081777 | 0.104716587 | 0.141920064 | 0.737856119 | 0.460601887 | NA |
| Fbp2 | 0.678304087 | 0.007077915 | 0.060597108 | 0.116802858 | 0.907016279 | NA |
| Fbxo15 | 4.952400966 | -0.052408258 | 0.130119018 | -0.402771698 | 0.687116184 | NA |
| Fbxo24 | 6.798809193 | -0.16018688 | 0.15314045 | -1.046012856 | 0.295555097 | NA |
| Fbxo39 | 0 | NA | NA | NA | NA | NA |
| Fbxo40 | 6.1891596 | 0.056783087 | 0.136295278 | 0.416618152 | 0.67695773 | NA |
| Fbxo43 | 4.772613506 | 0.165549102 | 0.135142861 | 1.224993319 | 0.220577738 | NA |
| Fbxw13 | 1.218222965 | -0.060677382 | 0.06307571 | -0.961976984 | 0.336061165 | NA |
| Fbxw14 | 0 | NA | NA | NA | NA | NA |
| Fbxw15 | 0.869516844 | 0.009508591 | 0.062816444 | 0.151371049 | 0.879683026 | NA |
| Fbxw16 | 0.094392817 | 0.017895942 | 0.027818137 | 0.643319236 | 0.520016977 | NA |
| Fbxw18 | 0.204276544 | 2.70E-05 | 0.031767541 | 0.000850493 | 0.999321405 | NA |
| Fbxw19 | 0 | NA | NA | NA | NA | NA |
| Fbxw20 | 0 | NA | NA | NA | NA | NA |
| Fbxw21 | 0 | NA | NA | NA | NA | NA |
| Fbxw22 | 0 | NA | NA | NA | NA | NA |
| Fbxw24 | 0.592206887 | -0.00501336 | 0.052942776 | -0.094693948 | 0.924557925 | NA |
| Fbxw26 | 0 | NA | NA | NA | NA | NA |
| Fbxw28 | 0 | NA | NA | NA | NA | NA |
| Fcamr | 0.772426891 | 0.028456539 | 0.048674378 | 0.584630763 | 0.558796018 | NA |
| Fcer1a | 0.314431083 | -0.017921647 | 0.039992366 | -0.448126696 | 0.654061762 | NA |
| Fcer2a | 0.722610492 | -0.050589987 | 0.061769794 | -0.819008519 | 0.412781553 | NA |
| Fcnb | 0.373417968 | -0.003176841 | 0.047020903 | -0.067562317 | 0.946134054 | NA |
| Fcrl5 | 0.203787646 | -0.02671144 | 0.031728022 | -0.841887971 | 0.399850664 | NA |
| Fcrl6 | 1.1676572 | 0.012223131 | 0.075074963 | 0.162812343 | 0.870666189 | NA |
| Fcrla | 0.440108409 | -0.030798726 | 0.044545461 | -0.691399874 | 0.489314283 | NA |
| Fer1l4 | 0.421660619 | 0.058159795 | 0.047506763 | 1.224242436 | 0.220860785 | NA |
| Ferd3l | 0.709143477 | 0.005126568 | 0.054605503 | 0.09388372 | 0.925201526 | NA |
| Fetub | 2.839570631 | 0.077230581 | 0.10495259 | 0.735861596 | 0.461814931 | NA |
| Fev | 0.975805528 | 0.028297693 | 0.057303592 | 0.493820588 | 0.621432892 | NA |
| Ffar1 | 0 | NA | NA | NA | NA | NA |
| Ffar2 | 1.906626708 | -0.141934789 | 0.096254684 | -1.474575388 | 0.140326736 | NA |
| Ffar3 | 3.190905903 | -0.133988432 | 0.107408912 | -1.247461028 | 0.212228501 | NA |
| Ffar4 | 1.74008905 | 0.008127778 | 0.099620747 | 0.081587199 | 0.934974981 | NA |
| Fga | 0.61964287 | -0.002240474 | 0.053268353 | -0.042060124 | 0.966450769 | NA |
| Fgb | 0.200315975 | 0.025370485 | 0.031664208 | 0.801235428 | 0.422995365 | NA |
| Fgf15 | 1.659710916 | -0.101587859 | 0.087624442 | -1.159355271 | 0.246311401 | NA |
| Fgf20 | 0.203787646 | -0.02671144 | 0.031728022 | -0.841887971 | 0.399850664 | NA |
| Fgf21 | 0.079983654 | 0.016740032 | 0.027674178 | 0.604897175 | 0.54524732 | NA |
| Fgf4 | 0 | NA | NA | NA | NA | NA |
| Fgf6 | 0 | NA | NA | NA | NA | NA |
| Fgf8 | 0.390839854 | 0.053765426 | 0.045767491 | 1.174751441 | 0.240094173 | NA |
| Fgg | 0.098353386 | -0.020212831 | 0.02786466 | -0.725393061 | 0.468210929 | NA |
| Fgl1 | 7.191908138 | -0.035068286 | 0.157767221 | -0.222278658 | 0.824096965 | NA |
| Fhl5 | 5.232462433 | -0.306394061 | 0.143279146 | -2.138441427 | 0.032480933 | NA |
| Figla | 0.841377836 | 0.042306321 | 0.058185242 | 0.727097102 | 0.467166474 | NA |
| Fitm1 | 0.533252617 | -0.072906618 | 0.051718649 | -1.409677544 | 0.158634918 | NA |
| Fkbp5 | 2046.344257 | 0.210444595 | 0.157993516 | 1.331982481 | NA | NA |
| Fkbp6 | 1.41373376 | 0.013835613 | 0.076512541 | 0.180828044 | 0.85650255 | NA |
| Flg2 | 0 | NA | NA | NA | NA | NA |
| Fmo3 | 0.116696964 | -0.020477583 | 0.027954224 | -0.732539847 | 0.463839131 | NA |
| Fmo4 | 3.176147293 | 0.010691655 | 0.107144884 | 0.099786893 | 0.920513514 | NA |
| Fmo6 | 0.601929635 | 0.059495052 | 0.0606896 | 0.980317088 | 0.326929622 | NA |
| Fmo9 | 0 | NA | NA | NA | NA | NA |
| Fmr1nb | 0.976635247 | -0.01263563 | 0.077170442 | -0.163736661 | 0.869938456 | NA |
| Fnd3c2 | 0 | NA | NA | NA | NA | NA |
| Fndc3c1 | 0.501157356 | 0.032769269 | 0.046039771 | 0.711760042 | 0.47661338 | NA |
| Fndc7 | 7.809630633 | -0.118504735 | 0.149768535 | -0.79125255 | 0.428796631 | NA |
| Fndc8 | 0.287795519 | 0.031925978 | 0.035387605 | 0.902179657 | 0.366961439 | NA |
| Folr1 | 46.57336198 | 0.422003245 | 0.12823039 | 3.29097686 | NA | NA |
| Folr4 | 0.30279753 | -0.014859216 | 0.039561964 | -0.375593495 | 0.707219128 | NA |
| Fosl1 | 2.366274081 | 0.256716804 | 0.098691656 | 2.601200693 | 0.009289809 | NA |
| Foxa1 | 0.268769288 | 0.03049543 | 0.034602455 | 0.881308274 | 0.378150989 | NA |
| Foxa2 | 0 | NA | NA | NA | NA | NA |
| Foxa3 | 0 | NA | NA | NA | NA | NA |
| Foxb1 | 2.808150522 | 0.01249967 | 0.113219953 | 0.11040165 | 0.912090845 | NA |
| Foxb2 | 0.770230356 | -0.005970955 | 0.061934977 | -0.096406831 | 0.923197467 | NA |
| Foxd3 | 3.520537522 | -0.178102128 | 0.080494249 | -2.212606862 | 0.026924762 | NA |
| Foxd4 | 0.079983654 | 0.016740032 | 0.027674178 | 0.604897175 | 0.54524732 | NA |
| Foxe1 | 0 | NA | NA | NA | NA | NA |
| Foxe3 | 0 | NA | NA | NA | NA | NA |
| Foxh1 | 0.626277961 | -0.006339789 | 0.058658962 | -0.108078782 | 0.913933199 | NA |
| Foxi1 | 2.438620476 | 0.109743235 | 0.113148543 | 0.96990409 | 0.332094301 | NA |
| Foxi2 | 0.185670226 | -0.001645086 | 0.031432796 | -0.052336624 | 0.958260472 | NA |
| Foxi3 | 0 | NA | NA | NA | NA | NA |
| Foxl1 | 1.495943527 | -0.128616887 | 0.092341476 | -1.392839847 | 0.163668226 | NA |
| Foxn1 | 0.700186587 | 0.005332081 | 0.060807798 | 0.087687447 | 0.930125097 | NA |
| Foxn4 | 0.08731684 | 0.017365056 | 0.027733412 | 0.626142072 | 0.531221759 | NA |
| Foxp3 | 3.104468821 | 0.137387151 | 0.116806182 | 1.176197596 | 0.239515925 | NA |
| Foxr1 | 0.528836942 | -0.072640212 | 0.051627513 | -1.407005851 | 0.159425654 | NA |
| Fpr-rs3 | 0 | NA | NA | NA | NA | NA |
| Fpr-rs4 | 0 | NA | NA | NA | NA | NA |
| Fpr-rs6 | 0 | NA | NA | NA | NA | NA |
| Fpr1 | 1.538364536 | 0.048392463 | 0.076911902 | 0.629193423 | 0.529222434 | NA |
| Fpr2 | 6.236296654 | 0.227658796 | 0.135529912 | 1.679767908 | 0.093002481 | NA |
| Fpr3 | 0 | NA | NA | NA | NA | NA |
| Frmpd1 | 622.6516966 | 0.204622233 | 0.136374933 | 1.50043874 | NA | NA |
| Fscb | 0 | NA | NA | NA | NA | NA |
| Fscn2 | 6.00751901 | 0.036516432 | 0.135683288 | 0.269129916 | 0.78782971 | NA |
| Fscn3 | 0.498928155 | 0.04751468 | 0.042883458 | 1.107995533 | 0.267863745 | NA |
| Fshb | 0.192142086 | -0.026384592 | 0.031527717 | -0.836869715 | 0.402665805 | NA |
| Fshr | 0.431541485 | -0.028919407 | 0.044298965 | -0.652823532 | 0.513870051 | NA |
| Ftcd | 3.16605085 | -0.077390674 | 0.122079584 | -0.633936248 | 0.526122428 | NA |
| Fthl17 | 0 | NA | NA | NA | NA | NA |
| Ftmt | 0 | NA | NA | NA | NA | NA |
| Fut1 | 1.049546946 | 0.098197456 | 0.077943846 | 1.259848741 | 0.207723933 | NA |
| Fut2 | 4.395458818 | 0.292207881 | 0.128666096 | 2.271055773 | 0.023143602 | NA |
| Fut4-ps1 | 0.820184021 | -0.034797393 | 0.063024181 | -0.552127644 | 0.580860904 | NA |
| Fxyd3 | 3.789960698 | 0.00572286 | 0.129084969 | 0.044334053 | 0.964638128 | NA |
| Fxyd4 | 0.211089781 | -0.002448206 | 0.031868916 | -0.076821119 | 0.93876585 | NA |
| G630055G22Rik | 0.19046386 | -0.000774927 | 0.031526133 | -0.024580468 | 0.980389599 | NA |
| G630071F17Rik | 0 | NA | NA | NA | NA | NA |
| G630090E17Rik | 1.842049084 | -0.193652633 | 0.103772397 | -1.866128554 | 0.062023382 | NA |
| G630093K05Rik | 7.183520563 | -0.128233229 | 0.148141392 | -0.865613768 | 0.386702003 | NA |
| G6b | 0 | NA | NA | NA | NA | NA |
| G6pc | 0.843897386 | 0.005955343 | 0.070126262 | 0.084923151 | 0.932322486 | NA |
| G6pc2 | 0.08731684 | 0.017365056 | 0.027733412 | 0.626142072 | 0.531221759 | NA |
| G6pd2 | 0 | NA | NA | NA | NA | NA |
| G730013B05Rik | 6.028322006 | 0.036204347 | 0.150148774 | 0.241123161 | 0.809459662 | NA |
| Gabra6 | 6.479264846 | 0.055393262 | 0.120871636 | 0.458281726 | 0.646750051 | NA |
| Gabrp | 0.185670226 | -0.001645086 | 0.031432796 | -0.052336624 | 0.958260472 | NA |
| Gabrr1 | 6.273610202 | 0.381262814 | 0.1369402 | 2.784155518 | 0.00536673 | NA |
| Gabrr2 | 39.57276487 | 0.578255091 | 0.155960107 | 3.70771156 | NA | NA |
| Gabrr3 | 2.518757579 | -0.016978466 | 0.114334698 | -0.148497932 | 0.881949818 | NA |
| Gal3st2 | 1.582316677 | 0.152511966 | 0.085650141 | 1.780638826 | 0.074971473 | NA |
| Galnt16 | 6936.344997 | 0.097446571 | 0.165043944 | 0.590428033 | NA | NA |
| Galnt5 | 1.004129514 | 0.045353969 | 0.071174536 | 0.63722184 | 0.523980354 | NA |
| Galntl5 | 0 | NA | NA | NA | NA | NA |
| Galp | 0.192142086 | -0.026384592 | 0.031527717 | -0.836869715 | 0.402665805 | NA |
| Galr2 | 0.573673924 | -0.001525252 | 0.046178452 | -0.033029522 | 0.973651045 | NA |
| Galr3 | 3.99998227 | -0.081628314 | 0.127764679 | -0.638895783 | 0.522890732 | NA |
| Gapdhs | 0.105923158 | 0.018524693 | 0.027977518 | 0.662127842 | 0.507889294 | NA |
| Gapt | 0.183387883 | -0.001444816 | 0.031390672 | -0.046026918 | 0.963288796 | NA |
| Gast | 3.825093049 | -0.166486861 | 0.119860133 | -1.389009481 | 0.164829867 | NA |
| Gata1 | 0.30883905 | -0.035660532 | 0.036608049 | -0.97411725 | 0.329998325 | NA |
| Gata3 | 3.030947112 | 0.230762804 | 0.092189626 | 2.503132016 | 0.012309962 | NA |
| Gata4 | 0.094392817 | 0.017895942 | 0.027818137 | 0.643319236 | 0.520016977 | NA |
| Gata5 | 0 | NA | NA | NA | NA | NA |
| Gata6 | 2.593282027 | 0.063245836 | 0.115025439 | 0.549842158 | 0.58242764 | NA |
| Gatsl2 | 4780.810923 | 0.006269933 | 0.122850168 | 0.05103724 | NA | NA |
| Gbp1 | 0 | NA | NA | NA | NA | NA |
| Gbp10 | 13.05975424 | 0.090124964 | 0.061635827 | 1.462217151 | NA | NA |
| Gbp6 | 126.1925848 | 0.32540605 | 0.138247608 | 2.353791538 | NA | NA |
| Gbp8 | 9.717919696 | 0.15023243 | 0.102796668 | 1.461452329 | NA | NA |
| Gc | 1.827451186 | -0.108507875 | 0.088999008 | -1.219203195 | 0.22276708 | NA |
| Gcg | 0.707079824 | 0.006315794 | 0.06085822 | 0.103778824 | 0.917344871 | NA |
| Gcgr | 0.593519161 | -0.035580971 | 0.059082103 | -0.60222925 | 0.547021548 | NA |
| Gckr | 3.499604695 | 0.086851506 | 0.123832684 | 0.701361734 | 0.483077296 | NA |
| Gcm1 | 0.159967308 | 0.0240345 | 0.030870969 | 0.778547006 | 0.436246606 | NA |
| Gcm2 | 0.08731684 | 0.017365056 | 0.027733412 | 0.626142072 | 0.531221759 | NA |
| Gcnt3 | 0 | NA | NA | NA | NA | NA |
| Gcnt7 | 0.327786745 | -0.013119639 | 0.034805201 | -0.376944789 | 0.706214636 | NA |
| Gcsam | 0.195033442 | -0.002424765 | 0.031596358 | -0.076741901 | 0.93882887 | NA |
| Gdf1 | 0.098353386 | -0.020212831 | 0.02786466 | -0.725393061 | 0.468210929 | NA |
| Gdf15 | 1.598617737 | -0.019268878 | 0.101151324 | -0.190495563 | 0.84892082 | NA |
| Gdf2 | 0.098353386 | -0.020212831 | 0.02786466 | -0.725393061 | 0.468210929 | NA |
| Gdf3 | 0.297029653 | 0.02866449 | 0.033517577 | 0.855207712 | 0.392436167 | NA |
| Gdf6 | 3.613165193 | -0.133304603 | 0.102818013 | -1.2965102 | 0.194799768 | NA |
| Gdf7 | 5.801625888 | -0.027578161 | 0.144280865 | -0.191142191 | 0.848414193 | NA |
| Gem | 89.20727141 | 0.181509619 | 0.152133927 | 1.193091 | NA | NA |
| Gfi1 | 0.615267569 | -0.003068902 | 0.053224915 | -0.057659123 | 0.954020155 | NA |
| Gfi1b | 0.107716603 | -0.020477583 | 0.027954224 | -0.732539849 | 0.46383913 | NA |
| Gfra3 | 0.513772093 | -0.015418338 | 0.051085511 | -0.301814299 | 0.76279363 | NA |
| Gfral | 0.099009884 | 0.018205726 | 0.027896865 | 0.652608315 | 0.514008823 | NA |
| Ggt6 | 3.368657239 | -0.096339354 | 0.116431997 | -0.82743023 | 0.407993252 | NA |
| Ghrh | 3.987779931 | -0.145342207 | 0.12010446 | -1.210131639 | 0.226228384 | NA |
| Ghrhr | 1.075850375 | 0.055512784 | 0.058551877 | 0.948095722 | 0.343080726 | NA |
| Ghrl | 2.238825209 | 0.169029556 | 0.097527166 | 1.733153566 | 0.083068378 | NA |
| Ghsr | 6.587346289 | 0.075959234 | 0.132407323 | 0.573678494 | 0.566185384 | NA |
| Gif | 0 | NA | NA | NA | NA | NA |
| Gimap3 | 3.510398947 | -0.113363313 | 0.105957268 | -1.069896525 | 0.284665888 | NA |
| Gimap6 | 136.4938202 | 0.192641666 | 0.161529907 | 1.1926068 | NA | NA |
| Gimap7 | 0.295716657 | -0.010253415 | 0.03372058 | -0.304069933 | 0.761074613 | NA |
| Gip | 0 | NA | NA | NA | NA | NA |
| Gja10 | 0.517954385 | 0.0503077 | 0.044100896 | 1.140741003 | 0.253977718 | NA |
| Gja3 | 4.961887536 | -0.029290092 | 0.129110669 | -0.22686035 | 0.820532334 | NA |
| Gja6 | 5.048565387 | 0.244324266 | 0.127284965 | 1.919506094 | 0.054920316 | NA |
| Gja8 | 0.08731684 | 0.017365056 | 0.027733412 | 0.626142072 | 0.531221759 | NA |
| Gjb3 | 3.015169002 | -0.044134795 | 0.111096007 | -0.397267158 | 0.691170463 | NA |
| Gjb4 | 0.079983654 | 0.016740032 | 0.027674178 | 0.604897175 | 0.54524732 | NA |
| Gjb5 | 0.400694422 | -0.006405997 | 0.042335824 | -0.151313856 | 0.879728139 | NA |
| Gjd4 | 0.721947876 | -0.045751712 | 0.056502661 | -0.809726681 | 0.41809728 | NA |
| Gje1 | 0.982925089 | 0.044350041 | 0.071094565 | 0.623817603 | 0.532747378 | NA |
| Gk2 | 0.08731684 | 0.017365056 | 0.027733412 | 0.626142072 | 0.531221759 | NA |
| Gkn1 | 2.904403723 | -0.02022624 | 0.119027508 | -0.169929123 | 0.865065877 | NA |
| Gkn2 | 0.119397641 | -0.020477583 | 0.027954224 | -0.732539845 | 0.463839132 | NA |
| Glb1l3 | 1.079110674 | 0.030086877 | 0.074340139 | 0.404719133 | 0.685683976 | NA |
| Glipr1l1 | 0.218407525 | -0.002234018 | 0.031987735 | -0.069839845 | 0.944321133 | NA |
| Glipr1l2 | 0 | NA | NA | NA | NA | NA |
| Glod5 | 3.668959318 | 0.230269405 | 0.12459466 | 1.84814827 | 0.064580896 | NA |
| Glra4 | 6.164400178 | -0.030199606 | 0.126484026 | -0.238762215 | 0.811289971 | NA |
| Glrp1 | 4.989901877 | 0.157099965 | 0.132768472 | 1.183262583 | 0.236705097 | NA |
| Glt6d1 | 0 | NA | NA | NA | NA | NA |

| | | | | | | | |
|---|---|---|---|---|---|---|---|
| Gm14391 | 0 | NA | NA | NA | NA | NA | NM_001177480,NM_001099308 |
| Gm14405 | 11.98265158 | -0.0459006385141887 | 0.0730435797385055 | -0.62840072568338 | 0.529741461820167 | NA | NR_040256 |
| Gm14430 | 0 | NA | NA | NA | NA | NA | NM_001100415 |
| Gm14431 | 0 | NA | NA | NA | NA | NA | NM_001177406,NM_001177404 |
| Gm14434 | 0 | NA | NA | NA | NA | NA | NM_001101804 |
| Gm14436 | 0 | NA | NA | NA | NA | NA | NM_001242943 |
| Gm14440 | 0 | NA | NA | NA | NA | NA | NM_001199308 |
| Gm14458 | 0 | NA | NA | NA | NA | NA | NM_001099326 |
| Gm14459 | 0 | NA | NA | NA | NA | NA | NM_001126491 |
| Gm14461 | 1.417282238 | -0.0294186993799203 | 0.0230642093805722 | -1.27551302082354 | 0.202127718421343 | NA | NM_177843 |
| Gm14474 | 0 | NA | NA | NA | NA | NA | NM_001242947 |
| Gm14475 | 0 | NA | NA | NA | NA | NA | NM_001242954 |
| Gm14476 | 0 | NA | NA | NA | NA | NA | NM_001242950 |
| Gm14477 | 0 | NA | NA | NA | NA | NA | NM_001242949 |
| Gm14478 | 0 | NA | NA | NA | NA | NA | NM_001242953 |
| Gm14479 | 0 | NA | NA | NA | NA | NA | NM_001242951 |
| Gm14482 | 0 | NA | NA | NA | NA | NA | NM_001242952 |
| Gm14483 | 0 | NA | NA | NA | NA | NA | NM_001111037 |
| Gm14484 | 0 | NA | NA | NA | NA | NA | NM_001025260 |
| Gm14492 | 3.577989769 | 0.0548369553550813 | 0.0402251781781508 | 1.36324953272344 | 0.172803881331961 | NA | NR_033261 |
| Gm14496 | 0 | NA | NA | NA | NA | NA | NM_001205282 |
| Gm14501 | 0.106148608 | 0.00515554063321497 | 0.00854200771766808 | 0.603551390213729 | 0.546141939141592 | NA | NM_001085537 |
| Gm14511 | 0 | NA | NA | NA | NA | NA | NM_001085525 |
| Gm14525 | 0 | NA | NA | NA | NA | NA | NM_001162364 |
| Gm14548 | 0 | NA | NA | NA | NA | NA | NM_001166672 |
| Gm14625 | 0 | NA | NA | NA | NA | NA | NM_001220498 |
| Gm14632 | 0 | NA | NA | NA | NA | NA | NM_001100610 |
| Gm14635 | 0 | NA | NA | NA | NA | NA | NR_045321 |
| Gm14685 | 0 | NA | NA | NA | NA | NA | NM_001025383 |
| Gm14692 | 0 | NA | NA | NA | NA | NA | NM_001163195 |
| Gm14718 | 0 | NA | NA | NA | NA | NA | NR_038463 |
| Gm14725 | 0 | NA | NA | NA | NA | NA | NM_001081476 |
| Gm14743 | 4.735802286 | -0.00862736390553638 | 0.0132258906114535 | -0.652308729823089 | NA | NA | NM_001126321 |
| Gm14744 | 2.03804561 | -0.0067387430277643 | 0.0119694624551868 | -0.562994625113191 | 0.573438543940529 | NA | NM_001085544 |
| Gm14781 | 0 | NA | NA | NA | NA | NA | NM_001205268 |
| Gm14812 | 0.097165704 | -0.00637828455464458 | 0.00848781691535724 | -0.75146349388194 | 0.452373763047372 | NA | NR_033544 |
| Gm14819 | 0 | NA | NA | NA | NA | NA | NM_001110250 |
| Gm14850 | 0 | NA | NA | NA | NA | NA | NM_001177522 |
| Gm14851 | 0 | NA | NA | NA | NA | NA | NM_001177482 |
| Gm14858 | 0.936484148 | 0.00473704306912293 | 0.0210326058617932 | 0.225223783503119 | 0.821805188155645 | NA | NR_040285 |
| Gm14873 | 0.876617123 | -0.00539609469719791 | 0.0222968866369469 | -0.242011128506898 | 0.808771540210163 | NA | NR_045913,NR_045912 |
| Gm14920 | 0 | NA | NA | NA | NA | NA | NM_001102665 |
| Gm15008 | 0.685014685 | 0.00315810812178894 | 0.0157090153829008 | 0.20103794189587 | 0.840668906327281 | NA | NR_045917 |
| Gm15023 | 0 | NA | NA | NA | NA | NA | NM_001099321 |
| Gm15056 | 0 | NA | NA | NA | NA | NA | NM_001177471 |
| Gm15070 | 0.980770624 | -5.95080759934087e-05 | 0.0213565299059627 | -0.00278641128757506 | 0.997776768330492 | NA | NM_001195579 |
| Gm15080 | 0 | NA | NA | NA | NA | NA | NM_001198988 |
| Gm15085 | 0 | NA | NA | NA | NA | NA | NM_001122734 |
| Gm15091 | 0 | NA | NA | NA | NA | NA | NM_001122735 |
| Gm15093 | 0 | NA | NA | NA | NA | NA | NM_001099920 |
| Gm15097 | 0 | NA | NA | NA | NA | NA | NM_001198987 |
| Gm15104 | 4.151745343 | -0.0502740450590804 | 0.0449632960224189 | -1.11811298339903 | 0.263518741988436 | NA | NM_001101501 |
| Gm15107 | 0 | NA | NA | NA | NA | NA | NM_001081648 |
| Gm15114 | 0 | NA | NA | NA | NA | NA | NM_001082966 |
| Gm15127 | 0 | NA | NA | NA | NA | NA | NM_001114400 |
| Gm15133 | 1.124665711 | -0.0313231331325413 | 0.0228983371958475 | -1.36792173443151 | 0.171336575735923 | NA | NR_040749 |
| Gm15140 | 0 | NA | NA | NA | NA | NA | NM_001243017 |
| Gm15179 | 0.323523361 | -0.00593920960320982 | 0.0126896719225698 | -0.468034921584249 | 0.639759616748484 | NA | NR_037976 |
| Gm15217 | 0 | NA | NA | NA | NA | NA | NR_037981 |
| Gm1527 | 0 | NA | NA | NA | NA | NA | NM_001033479 |
| Gm15284 | 0 | NA | NA | NA | NA | NA | NM_001177485 |
| Gm15292 | 0 | NA | NA | NA | NA | NA | NM_001177487 |
| Gm15293 | 0 | NA | NA | NA | NA | NA | NM_001177486 |
| Gm15299 | 0 | NA | NA | NA | NA | NA | NM_001170955 |
| Gm15308 | 0 | NA | NA | NA | NA | NA | NM_001177521 |
| Gm15315 | 0 | NA | NA | NA | NA | NA | NM_001177528 |
| Gm15319 | 0 | NA | NA | NA | NA | NA | NM_001177408 |
| Gm15348 | 0.666283812 | 0.0220619021746092 | 0.0174328938476098 | 1.26553298422305 | 0.205680332185405 | NA | NR_033546 |
| Gm15350 | 0.534316316 | 0.00242828786212454 | 0.0147532078991331 | 0.164593888917355 | 0.869263641550438 | NA | NR_045775 |
| Gm15386 | 0 | NA | NA | NA | NA | NA | NM_001040027 |
| Gm15401 | 1.323608543 | 0.00273748833772807 | 0.0246799719289916 | 0.110919426715892 | 0.911680240367948 | NA | NR_040421,NR_040422 |
| Gm15408 | 9.047503379 | 0.00763867089031013 | 0.0550906896040157 | 0.138656294651889 | 0.889721756208469 | NA | NR_040429 |
| Gm15412 | 0 | NA | NA | NA | NA | NA | NR_046043 |
| Gm15413 | 2.971967653 | 0.025199159015596 | 0.0377416750568988 | 0.667674632289268 | 0.504341306310484 | NA | NR_045874 |
| Gm15421 | 20.34535197 | -0.0668144741593864 | 0.0719441051105798 | -0.928699774035564 | 0.353044697905877 | NA | NR_004442 |
| Gm15441 | 2.796897423 | 0.0269520284803878 | 0.0376209990320844 | 0.716409164397847 | 0.47373873781004 | NA | NR_040409 |
| Gm15455 | 2.783679713 | -0.0304733017808147 | 0.0351574623168688 | -0.866766250253319 | 0.386070097570487 | NA | NM_001161816 |
| Gm15471 | 1.946779206 | 0.0146832632816997 | 0.031192624778076 | 0.470728686225212 | 0.637834496403391 | NA | NR_040412 |
| Gm1553 | 0.288615623 | 0.00197374456361263 | 0.0106648328298147 | 0.185070370544844 | 0.853173841012254 | NA | NM_001255990 |
| Gm15545 | 13.93130976 | 0.0021357632385991 | 0.0712328756478343 | 0.0299828305283929 | 0.976080746268335 | NA | NR_045266 |
| Gm156 | 0 | NA | NA | NA | NA | NA | NM_001014997 |
| Gm15612 | 14.00544087 | 0.0176777026874791 | 0.0548132381232343 | 0.322507906716533 | 0.747067948800069 | NA | NR_045880 |
| Gm1564 | 0.257802559 | -0.00892927589713349 | 0.00993479775307735 | -0.898787888698349 | 0.368765652383134 | NA | NM_001127576 |
| Gm15645 | 14.50455922 | -0.0946966317182601 | 0.0707610378833749 | -1.33825950764508 | 0.180811856674276 | NA | NR_033578 |
| Gm15698 | 0 | NA | NA | NA | NA | NA | NR_003564 |
| Gm15708 | 10.22986784 | -0.0835487508520329 | 0.067648500953892 | -1.235042161673 | 0.216814813227502 | NA | NR_040432 |
| Gm15713 | 0 | NA | NA | NA | NA | NA | NR_046026 |
| Gm15760 | 12.39661844 | -0.116669836092161 | 0.0705908428742103 | -1.65276162377124 | 0.0983793872219077 | NA | NR_030670 |
| Gm15772 | 0 | NA | NA | NA | NA | NA | NR_003373 |
| Gm15787 | 6.628577017 | -0.00690880258720503 | 0.0577086712516016 | -0.119718621783607 | 0.904706048406115 | NA | NR_040431,NR_040430 |
| Gm15850 | 1.523124282 | 0.015897864082014 | 0.027378102879621 | 0.580678075172535 | 0.561457441040953 | NA | NR_046170,NR_046167,NR_046169,NR_046168 |
| Gm1587 | 0.217532603 | 0.00684685248201087 | 0.00976000296730706 | 0.701521557416086 | 0.482977586094298 | NA | NM_001033440 |
| Gm15880 | 2.079917153 | -0.0144947073729877 | 0.0332438378470912 | -0.436011853976 | 0.662828129061534 | NA | NR_040343 |
| Gm15881 | 0 | NA | NA | NA | NA | NA | NM_001177534 |
| Gm15910 | 9.910299235 | -0.0394557119056075 | 0.0627213312550888 | -0.62906368720301 | 0.529307362441366 | NA | NR_038023 |
| Gm15915 | 1.165605285 | 0.011836952642705 | 0.0221301867782153 | 0.534878117448025 | 0.592734140828908 | NA | NR_038017 |
| Gm15941 | 0.523892668 | 0.00262461874340838 | 0.0162928159600756 | 0.161090553642772 | 0.872022080178953 | NA | NR_045283 |
| Gm15987 | 0.515605119 | -0.0132903332792225 | 0.012110641568181 | -1.09740951413681 | 0.272462416240765 | NA | NR_045009 |
| Gm15997 | 17.74214006 | -0.0427271355872812 | 0.0812981236009096 | -0.525561153133467 | 0.599193153538449 | NA | NR_045423 |
| Gm16023 | 11.88668495 | -0.0412774567951358 | 0.0671902901281708 | -0.614336635790615 | 0.538992897996162 | NA | NR_040441 |
| Gm1604b | 12.90969182 | -0.0114475992383267 | 0.0728462015600625 | -0.157147510689188 | 0.875128594595281 | NA | NM_001033442 |
| Gm16063 | 0 | NA | NA | NA | NA | NA | NR_046178 |
| Gm16065 | 1.077925408 | 0.0129236131404148 | 0.0235172952405581 | 0.549536543561635 | 0.58263729315782 | NA | NR_040416 |
| Gm16119 | 13.71066085 | -0.0843021019357958 | 0.0666219448589431 | -1.26538038050805 | 0.205735004792722 | NA | NR_027289,NR_027290 |
| Gm16130 | 0.192890664 | -0.000903864895041124 | 0.00963528282151475 | -0.093807821917056 | 0.92526181809679 | NA | NM_001243265 |
| Gm16157 | 0 | NA | NA | NA | NA | NA | NR_040341,NR_040340 |
| Gm16197 | 21.06422548 | -0.0520850061799952 | 0.076459644956778 | -0.681209103304605 | 0.49573918867458 | NA | NR_036469 |
| Gm16291 | 11.91751746 | 0.00363134034905995 | 0.0676759863087345 | 0.0536577381006899 | 0.957207854374527 | NA | NR_045788 |
| Gm16294 | 6.976440534 | 0.01707755226438 | 0.0563035553144643 | 0.303312147323542 | 0.76165198974178 | NA | NR_046185 |
| Gm1631 | 0 | NA | NA | NA | NA | NA | NR_037979 |
| Gm16325 | 1.500677145 | 0.0284008392830901 | 0.0276491624497439 | 1.02718624243004 | 0.304332774972001 | NA | NR_045949 |
| Gm16336 | 0 | NA | NA | NA | NA | NA | NR_045267 |
| Gm16367 | 0 | NA | NA | NA | NA | NA | NM_001031622 |
| Gm16381 | 0 | NA | NA | NA | NA | NA | NM_001166062 |
| Gm16390 | 0 | NA | NA | NA | NA | NA | NM_001097980 |
| Gm16404 | 0 | NA | NA | NA | NA | NA | NM_001220497 |
| Gm16405 | 0 | NA | NA | NA | NA | NA | NM_001166646 |
| Gm16430 | 0 | NA | NA | NA | NA | NA | NM_001166601 |
| Gm16442 | 0 | NA | NA | NA | NA | NA | NM_001167148 |
| Gm16445 | 0 | NA | NA | NA | NA | NA | NM_001243032 |
| Gm16451 | 0 | NA | NA | NA | NA | NA | NM_001167149 |
| Gm1647 | 0.760714661 | 0.00206428918130612 | 0.0160854521112801 | 0.128332680177421 | 0.897885703879568 | NA | NM_001243000 |
| Gm16497 | 0.106148608 | 0.00515554063321497 | 0.00854200771766808 | 0.603551390213729 | 0.546141939141592 | NA | NR_045003 |
| Gm16501 | 0 | NA | NA | NA | NA | NA | NM_001113395 |
| Gm16523 | 0 | NA | NA | NA | NA | NA | NR_033526 |
| Gm1653 | 21.2467651 | 0.0274813623960505 | 0.0798304038011032 | 0.344246816845874 | 0.730660676642325 | NA | NR_040591 |
| Gm16548 | 2.31402558 | 0.016383149544412 | 0.0282777330788558 | 0.57936573270303 | 0.56234242087578 | NA | NR_037987 |
| Gm16551 | 2.257490815 | 0.0260383234024326 | 0.0281346155855911 | 0.925490640638712 | 0.354710749906639 | NA | NR_045284 |
| Gm1661 | 19.82913182 | 0.0720433020977637 | 0.0779960170582628 | 0.923679244338177 | 0.355653341629421 | NA | NM_001145637,NM_001033774 |
| Gm16617 | 0.226066984 | -0.0093084363525918 | 0.0101406635841621 | -0.917931679257152 | 0.358654635211298 | NA | NR_045728 |
| Gm16675 | 12.20743184 | -0.0496191695643733 | 0.0678674603660134 | -0.731118702494155 | 0.464706651595199 | NA | NR_045750 |
| Gm16796 | 8.423860254 | 0.0386746920488047 | 0.0623422584963231 | 0.620360779054639 | 0.535020288149993 | NA | NR_040367 |
| Gm16833 | 0.515605119 | -0.0132903332792225 | 0.012110641568181 | -1.09740951413681 | 0.272462416240765 | NA | NR_045754 |
| Gm16853 | 8.67318582 | 0.0154871145609518 | 0.056905293540009 | 0.272155955931642 | 0.785502106053427 | NA | NR_045742 |
| Gm16863 | 0.309861382 | -0.0124208717281794 | 0.0117059258046019 | -1.0610755557067 | 0.28865556572345 | NA | NR_046162 |
| Gm16880 | 0.411706162 | -0.0156488736868625 | 0.013129479323261 | -1.191888368272 | 0.233305019639917 | NA | NR_037986 |
| Gm16897 | 14.79302018 | 0.0630183301075663 | 0.0743355119506328 | 0.847755379009398 | 0.396574217865499 | NA | NR_033564 |
| Gm16907 | 9.282788243 | 0.0421276439599499 | 0.05862678384932 | 0.718573341294391 | 0.472403843918739 | NA | NR_045794 |
| Gm16982 | 1.676041641 | -0.00901591253659639 | 0.0231777971715995 | -0.388989189518142 | 0.697284143385399 | NA | NR_040337 |
| Gm16998 | 16.51142448 | -0.164956754790496 | 0.0769140552325556 | -2.14468934568248 | 0.0319776933207104 | NA | NR_038016 |
| Gm17019 | 1.772317822 | 0.00472334685394943 | 0.0302630933189709 | 0.156076142123533 | 0.875973003711604 | NA | NM_182957 |
| Gm1715 | 4.341144704 | -0.0459331677821162 | 0.0471729203249412 | -0.973718978297608 | 0.330196091517354 | NA | NR_045333 |
| Gm1720 | 0.611119886 | 0.00580156497226671 | 0.0172573406772955 | 0.336179547054981 | 0.736735475349771 | NA | NR_046015 |
| Gm17252 | 0 | NA | NA | NA | NA | NA | NM_001199953 |
| Gm17359 | 0.113843286 | 0.00525947738444006 | 0.00859368002043176 | 0.612016897526494 | 0.540526577873086 | NA | NM_001142953 |
| Gm17365 | 0 | NA | NA | NA | NA | NA | NM_001167592 |
| Gm17384 | 8.315182849 | 0.00988006138626329 | 0.0608050615394681 | 0.162487482721323 | 0.870921984631895 | NA | NM_001170705 |
| Gm17455 | 13.17172147 | 0.103929398887438 | 0.0639041962442971 | 1.6263313678202 | 0.103879182319788 | NA | NM_001164374 |
| Gm17495 | 0 | NA | NA | NA | NA | NA | NM_001177584 |
| Gm17660 | 7.425864102 | -0.0782589495590593 | 0.0592563431027117 | -1.3206847648936 | 0.186606496048422 | NA | NM_001163772 |
| Gm17677 | 0.226556888 | -0.00933175792380659 | 0.010153338242423 | -0.919082739193727 | 0.358052296835844 | NA | NM_001167588 |
| Gm17689 | 0 | NA | NA | NA | NA | NA | NM_001167584 |
| Gm17727 | 0.217532603 | 0.00684685248201087 | 0.00976000296730706 | 0.701521557416086 | 0.482977586094298 | NA | NM_001167673 |
| Gm17745 | 1.317769838 | 0.0206078208253472 | 0.0257238282375178 | 0.801117960945295 | 0.423063359184659 | NA | NR_038014 |
| Gm17746 | 0.855988487 | -0.0214035359126361 | 0.0178722428531702 | -1.19758533321628 | 0.231078488746131 | NA | NR_045844 |
| Gm17751 | 7.00940474 | 0.0733189447555433 | 0.0534018147757346 | 1.37296728703795 | 0.169762516656958 | NA | NR_038012,NR_038013 |
| Gm17757 | 0 | NA | NA | NA | NA | NA | NR_040453 |
| Gm17801 | 4.682773056 | 0.0573149241669679 | 0.0494255700885111 | 1.15962090198107 | 0.246203187061323 | NA | NR_027452 |
| Gm17830 | 0.330974078 | -0.00671789744880547 | 0.01278556667705 | -0.525428212803741 | 0.599285545196596 | NA | NR_033486 |
| Gm18409 | 0.801396501 | 0.0304684228345504 | 0.0204400036985651 | 1.49062707051708 | 0.136059432994227 | NA | NR_038018 |
| Gm18853 | 0 | NA | NA | NA | NA | NA | NR_040456 |
| Gm19276 | 0.548733837 | 0.00269688126639513 | 0.0164543653001976 | 0.163900655977459 | 0.869809350688666 | NA | NR_040357 |
| Gm19277 | 0 | NA | NA | NA | NA | NA | NR_040360 |
| Gm19299 | 0.224853396 | -0.000161157395207779 | 0.00979162641959728 | -0.0164586952465051 | 0.986868454038988 | NA | NR_045748 |
| Gm19303 | 0.436125133 | -0.00267289099106318 | 0.0153191773454902 | -0.174480060566049 | 0.861488199773005 | NA | NR_045092 |
| Gm19395 | 0 | NA | NA | NA | NA | NA | NR_038015 |
| Gm19402 | 0 | NA | NA | NA | NA | NA | NM_001205032 |
| Gm19424 | 0.319043516 | -0.00957618023178583 | 0.0102890835921761 | -0.930712647632453 | 0.352002225550712 | NA | NR_040320 |
| Gm19434 | 1.26643968 | -0.0103275938126033 | 0.0249420603814085 | -0.414063379475313 | 0.67882769114058 | NA | NR_040296 |
| Gm19466 | 0.648160155 | -0.0100749839718643 | 0.0174310369773315 | -0.577991084808467 | 0.563270138261572 | NA | NR_040349 |
| Gm19510 | 0.21100899 | -0.000459554483117493 | 0.00972904351832043 | -0.0472353199214416 | 0.96232567770125 | NA | NR_045076 |
| Gm19557 | 3.52462533 | -0.0422473044871625 | 0.0377213165719822 | -1.11998488723328 | 0.262720202268585 | NA | NR_045322 |
| Gm19583 | 0.12890128 | -0.0068640067066208 | 0.00870804083610955 | -0.788237771940376 | 0.430557639008166 | NA | NR_045792 |
| Gm19589 | 0.321063518 | 0.00979019190676043 | 0.0116583710553074 | 0.839756417111422 | 0.401044974403149 | NA | NR_037971 |
| Gm19619 | 11.12975644 | 0.00394487857698426 | 0.0625650192150077 | 0.0630524632850747 | 0.949724727734636 | NA | NR_040428 |
| Gm1965 | 0 | NA | NA | NA | NA | NA | NM_001033491 |
| Gm19668 | 0 | NA | NA | NA | NA | NA | NM_001205034 |
| Gm1968 | 0.384276662 | 0.0023869237934625 | 0.0116652479776256 | 0.204618349994849 | 0.837870317514098 | NA | NR_037689 |
| Gm19689 | 2.214672762 | -0.0444391112782302 | 0.034353052529822 | -1.29360007352047 | 0.195803597067436 | NA | NR_045094 |
| Gm19705 | 17.32564883 | -0.0150307290197231 | 0.076289646187355 | -0.197021873490016 | 0.84381042069207 | NA | NR_045324,NR_045325 |
| Gm19710 | 1.661338131 | -0.00279545349543736 | 0.0310053036954271 | -0.0901604939238075 | 0.928159677703876 | NA | NR_045749 |
| Gm19782 | 0.120209049 | -0.00674972139751468 | 0.00863521114606269 | -0.781650996523957 | 0.434419707908266 | NA | NR_045071 |
| Gm19784 | 0 | NA | NA | NA | NA | NA | NR_040461 |
| Gm1987 | 0 | NA | NA | NA | NA | NA | NM_001193667 |
| Gm19897 | 2.301024175 | 0.00266543640953584 | 0.030858258629843 | 0.0863767603191354 | 0.931166920452119 | NA | NR_040339 |
| Gm1993 | 0 | NA | NA | NA | NA | NA | NM_001102677 |
| Gm1995 | 0 | NA | NA | NA | NA | NA | NR_033562 |
| Gm19990 | 1.254337168 | 0.0121837722926657 | 0.0254284082351629 | 0.479140187619679 | 0.631838901675509 | NA | NR_045048 |
| Gm2002 | 0 | NA | NA | NA | NA | NA | NM_001100596 |
| Gm20098 | 0.113843286 | 0.00525947738444006 | 0.00859368002043176 | 0.612016897526494 | 0.540526577873086 | NA | NR_045095 |
| Gm20110 | 0.561481615 | 0.000827798194506355 | 0.0149136418996772 | 0.0555061064275837 | 0.955735265226365 | NA | NR_038019 |
| Gm2012 | 0 | NA | NA | NA | NA | NA | NM_001104946 |
| Gm20125 | 2.036085988 | -0.0328451863527454 | 0.0333267099737537 | -0.985551420425613 | 0.324353286795802 | NA | NR_038020 |
| Gm20139 | 0.604369265 | -0.0155706206868308 | 0.0159775092716347 | -0.97453366617497 | 0.329791631588806 | NA | NR_038022,NR_038021 |
| Gm2016 | 0 | NA | NA | NA | NA | NA | NM_001122662 |
| Gm20172 | 0 | NA | NA | NA | NA | NA | NM_001204913 |
| Gm20187 | 12.6754678 | 0.0802356291443277 | 0.0735970183167 | 1.0902021709502 | 0.275624097262933 | NA | NR_045052 |
| Gm20199 | 0.097165704 | -0.00637828455464458 | 0.00848781691535724 | -0.75146349388194 | 0.452373763047372 | NA | NR_045640 |
| Gm2022 | 0 | NA | NA | NA | NA | NA | NM_001177574 |
| Gm20268 | 0 | NA | NA | NA | NA | NA | NR_037989 |
| Gm2027 | 6.231627958 | -0.0311317354340493 | 0.0526292445381251 | -0.591529209800782 | 0.554165889642402 | NA | NR_045113 |
| Gm2030 | 0 | NA | NA | NA | NA | NA | NM_001100445 |
| Gm20324 | 8.18426936 | -0.00307427115086829 | 0.0572646405062185 | -0.0536853304882696 | 0.95718587052101 | NA | NR_045068 |
| Gm20337 | 11.50094285 | -0.0529532195040789 | 0.0680254847639238 | -0.778432078622419 | 0.436314333658602 | NA | NR_045057 |
| Gm20356 | 0 | NA | NA | NA | NA | NA | NR_040450 |
| Gm20362 | 1.793146381 | -0.0111989977449068 | 0.032159961121742 | -0.348227962792393 | 0.727668989155138 | NA | NR_040301,NR_040302 |
| Gm20408 | 0 | NA | NA | NA | NA | NA | NM_001177587 |
| Gm20410 | 0 | NA | NA | NA | NA | NA | NM_001177585 |
| Gm2042 | 0 | NA | NA | NA | NA | NA | NM_001270792,NM_001270793 |
| Gm20516 | 0 | NA | NA | NA | NA | NA | NM_001177586 |
| Gm20554 | 1.409755441 | 0.00588158137615079 | 0.0289704865942293 | 0.203019764856913 | 0.839119583778458 | NA | NR_030701 |
| Gm20556 | 0 | NA | NA | NA | NA | NA | NR_040347 |
| Gm20579 | 0 | NA | NA | NA | NA | NA | NM_001244763 |
| Gm20580 | 0 | NA | NA | NA | NA | NA | NM_001177583 |
| Gm20581 | 0 | NA | NA | NA | NA | NA | NM_001177582 |
| Gm20594 | 1.939887592 | 0.0141434378157084 | 0.0307846427258361 | 0.459431604961863 | 0.645924256740647 | NA | NM_001190732 |
| Gm20597 | 13.66357099 | -0.021193041762081 | 0.0648224128685332 | -0.32694003237835 | 0.743713241950883 | NA | NR_037973 |
| Gm20604 | 0 | NA | NA | NA | NA | NA | NM_001142939 |
| Gm20611 | 0.212496447 | -0.000886988073087717 | 0.00973851166817767 | -0.0910804549309223 | 0.927428662789121 | NA | NR_040638 |
| Gm20735 | 1.107523121 | -0.0176106458168386 | 0.0258345803554138 | -0.681669513286606 | 0.495447949092388 | NA | NR_015497 |
| Gm20740 | 0 | NA | NA | NA | NA | NA | NR_045279 |
| Gm20741 | 0 | NA | NA | NA | NA | NA | NR_045150 |
| Gm20744 | 0.684132458 | 0.0265637158685231 | 0.0191107587783643 | 1.38998750267292 | 0.164532672346648 | NA | NR_045744 |
| Gm20745 | 0.194331409 | -0.00840968081320881 | 0.00964081822647896 | -0.872299489073576 | 0.383045016303155 | NA | NR_045745 |
| Gm20750 | 0.240418098 | -0.0088029805258837 | 0.00986414519853158 | -0.892422034419582 | 0.372166765031741 | NA | NR_040555 |
| Gm20752 | 6.029504315 | -0.0349970695554339 | 0.0463755088915225 | -0.754645509924127 | 0.450461709357438 | NA | NR_040750 |
| Gm20753 | 12.33641579 | -0.00324304941242737 | 0.0687137996377355 | -0.0471964791573888 | 0.962356633622322 | NA | NR_040630 |
| Gm20755 | 1.09377793 | -0.0219573406403266 | 0.0230913840878788 | -0.950888892444194 | 0.341660780974032 | NA | NR_040559 |
| Gm20756 | 0.127687691 | 0.00542298404407807 | 0.00869724871942359 | 0.62352868349814 | 0.532937159177571 | NA | NR_040771 |
| Gm20757 | 0.366568861 | -0.00548477312881077 | 0.011623202474977 | -0.471881406231944 | 0.637011440575618 | NA | NR_046027 |
| Gm20758 | 0.191449919 | 0.00665508079272897 | 0.00962405648474729 | 0.691504752001018 | 0.489248395458281 | NA | NR_046029 |
| Gm20759 | 0 | NA | NA | NA | NA | NA | NR_046030 |
| Gm20765 | 0 | NA | NA | NA | NA | NA | NM_001270644 |
| Gm2083 | 0 | NA | NA | NA | NA | NA | NM_001134644 |
| Gm20878 | 0 | NA | NA | NA | NA | NA | NM_001270431 |

| | | | | | | | | | | | | | | | |
|---|---|---|---|---|---|---|---|---|---|---|---|---|---|---|---|
| Gltpd2 | 0.119397641 | -0.020477583 | 0.027954224 | -0.732539845 | 0.463839132 | NA | | Gm21002 | 0 | NA | NA | NA | NA | NA | NM_001270555 |
| Glyat | 0.477174001 | 0.011008699 | 0.05521377 | 0.199383206 | 0.841962996 | NA | | Gm21057 | 0.972280655 | -0.026018801039846 | 0.0222773117662254 | -1.16795066266896 | 0.242826666616079 | NA | NR_045695 |
| Glyatl3 | 0.178993538 | 0.024685672 | 0.031260828 | 0.789667872 | 0.429721759 | NA | | Gm2109 | 3.233702143 | -0.0329453272863177 | 0.0374639859083022 | -0.879386602561604 | 0.379191693766218 | NA | NR_046066 |
| Glycam1 | 2.218913329 | 0.217720313 | 0.097647593 | 2.229653655 | 0.025770445 | NA | | Gm21119 | 0 | NA | NA | NA | NA | NA | NM_001270553 |
| Gm10007 | 1.665083702 | -0.004579291 | 0.09434744 | -0.048536461 | 0.961288707 | NA | | Gm21276 | 0 | NA | NA | NA | NA | NA | NR_073038 |
| Gm10012 | 0.814547248 | -0.010378377 | 0.06227327 | -0.166658623 | 0.867638664 | NA | | Gm21293 | 0 | NA | NA | NA | NA | NA | NM_001270898 |
| Gm10024 | 0 | NA | NA | NA | NA | NA | | Gm21304 | 0 | NA | NA | NA | NA | NA | NM_001270901 |
| Gm10046 | 2.368722812 | 0.040270498 | 0.110488348 | 0.364477329 | 0.7155016 | NA | | Gm21312 | 0 | NA | NA | NA | NA | NA | NM_001270642 |
| Gm10052 | 0 | NA | NA | NA | NA | NA | | Gm21319 | 0 | NA | NA | NA | NA | NA | NM_001270722 |
| Gm10057 | 0 | NA | NA | NA | NA | NA | | Gm21498 | 0 | NA | NA | NA | NA | NA | NM_001270613 |
| Gm10058 | 0 | NA | NA | NA | NA | NA | | Gm21541 | 0 | NA | NA | NA | NA | NA | NM_001270360 |
| Gm10081 | 0 | NA | NA | NA | NA | NA | | Gm21586 | 0 | NA | NA | NA | NA | NA | NM_001270432 |
| Gm10094 | 0 | NA | NA | NA | NA | NA | | Gm21637 | 0 | NA | NA | NA | NA | NA | NM_001270685 |
| Gm10096 | 0 | NA | NA | NA | NA | NA | | Gm21677 | 0 | NA | NA | NA | NA | NA | NM_001270513 |
| Gm10100 | 0 | NA | NA | NA | NA | NA | | Gm21693 | 0 | NA | NA | NA | NA | NA | NM_001270514 |
| Gm10104 | 0 | NA | NA | NA | NA | NA | | Gm21704 | 0 | NA | NA | NA | NA | NA | NM_001270515 |
| Gm10142 | 0 | NA | NA | NA | NA | NA | | Gm21708 | 0 | NA | NA | NA | NA | NA | NM_001270516 |
| Gm10147 | 0 | NA | NA | NA | NA | NA | | Gm2176 | 0 | NA | NA | NA | NA | NA | NR_028424 |
| Gm10190 | 5.185794867 | -0.105169283 | 0.13768921 | -0.763816446 | 0.444976635 | NA | | Gm21945 | 0 | NA | NA | NA | NA | NA | NM_001251916 |
| Gm10220 | 2.282862464 | 0.093317574 | 0.110217915 | 0.846664303 | 0.397182261 | NA | | Gm21946 | 0 | NA | NA | NA | NA | NA | NM_001256051 |
| Gm10228 | 0 | NA | NA | NA | NA | NA | | Gm21947 | 0 | NA | NA | NA | NA | NA | NM_001256050 |
| Gm10229 | 0 | NA | NA | NA | NA | NA | | Gm21949 | 0 | NA | NA | NA | NA | NA | NM_001113419 |
| Gm10230 | 0 | NA | NA | NA | NA | NA | | Gm21950 | 0 | NA | NA | NA | NA | NA | NM_001270667 |
| Gm10248 | 0.414849978 | -0.030430887 | 0.043968299 | -0.692109732 | 0.488868421 | NA | | Gm21951 | 0 | NA | NA | NA | NA | NA | NM_001270669 |
| Gm10256 | 0 | NA | NA | NA | NA | NA | | Gm2381 | 0.958745221 | -0.0324820250824193 | 0.0210869469027087 | -1.54038539729272 | 0.12346643875281 | NA | NR_046214 |
| Gm10267 | 0 | NA | NA | NA | NA | NA | | Gm2447 | 0.587881648 | 0.00934991935365208 | 0.0154798745688504 | 0.60400485236919 | 0.545840416096457 | NA | NR_038079 |
| Gm10272 | 0 | NA | NA | NA | NA | NA | | Gm2506 | 0 | NA | NA | NA | NA | NA | NM_001199965,NM_001199966 |
| Gm10280 | 0.099009884 | 0.018205726 | 0.027896865 | 0.652608315 | 0.514008823 | NA | | Gm2516 | 0.127687691 | 0.00542298404407807 | 0.00869724871942359 | 0.62352868349814 | 0.532937159177571 | NA | NR_046067 |
| Gm10318 | 0 | NA | NA | NA | NA | NA | | Gm2518 | 12.24174935 | 0.046312221990813 | 0.0692707817040708 | 0.668567913505896 | 0.503771146292441 | NA | NR_015538 |
| Gm10319 | 0 | NA | NA | NA | NA | NA | | Gm2663 | 0 | NA | NA | NA | NA | NA | NM_001102660 |
| Gm10324 | 0.08731684 | 0.017365056 | 0.027733412 | 0.626142072 | 0.531221759 | NA | | Gm2696 | 0 | NA | NA | NA | NA | NA | NM_001205009 |
| Gm10352 | 0 | NA | NA | NA | NA | NA | | Gm2721 | 12.50821608 | 0.0225360243079977 | 0.0711723756983228 | 0.316640045900965 | 0.751516745863545 | NA | NR_045085 |
| Gm10364 | 1.021338241 | -0.011381811 | 0.07725403 | -0.147329678 | 0.882871809 | NA | | Gm2762 | 20.46052548 | 0.0488173120248981 | 0.0782443622919057 | 0.623908363426565 | 0.532687767453434 | NA | NR_037991 |
| Gm10373 | 0.806298578 | 0.016681995 | 0.049256587 | 0.338675417 | 0.734854262 | NA | | Gm2799 | 0 | NA | NA | NA | NA | NA | NM_001168334 |
| Gm10375 | 0 | NA | NA | NA | NA | NA | | Gm2825 | 0 | NA | NA | NA | NA | NA | NM_001168337 |
| Gm10377 | 0 | NA | NA | NA | NA | NA | | Gm2837 | 0 | NA | NA | NA | NA | NA | NR_040388 |
| Gm10389 | 2.031687473 | 0.01096503 | 0.109148051 | 0.100460152 | 0.919979017 | NA | | Gm2848 | 0.120209049 | -0.00674972139751468 | 0.00863521114606269 | -0.781650996523957 | 0.434419707908266 | NA | NR_046069 |
| Gm10390 | 0.844386285 | 0.031879823 | 0.064648562 | 0.493125017 | 0.621924255 | NA | | Gm2863 | 0 | NA | NA | NA | NA | NA | NM_001099333 |
| Gm10400 | 0.096071043 | -0.020098795 | 0.027825948 | -0.722304036 | 0.470107572 | NA | | Gm2897 | 9.877661889 | -0.0229286541454176 | 0.0597258209855554 | -0.383898517710838 | 0.70105366503643 | NA | NM_001177715 |
| Gm10408 | 0.440944058 | 0.059506558 | 0.048029448 | 1.238959851 | 0.215360367 | NA | | Gm2913 | 0 | NA | NA | NA | NA | NA | NM_001243015 |
| Gm10409 | 0 | NA | NA | NA | NA | NA | | Gm2933 | 0 | NA | NA | NA | NA | NA | NM_001145038 |
| Gm10413 | 0 | NA | NA | NA | NA | NA | | Gm3020 | 0 | NA | NA | NA | NA | NA | NR_033117 |
| Gm10415 | 0.273386355 | 0.041384988 | 0.040339907 | 1.025906866 | 0.304935486 | NA | | Gm3139 | 0 | NA | NA | NA | NA | NA | NM_001243937,NM_001243938 |
| Gm10436 | 0 | NA | NA | NA | NA | NA | | Gm3143 | 0.097165704 | -0.00637828455464458 | 0.00848781691535724 | -0.75146349388194 | 0.452373763047372 | NA | NR_038348,NR_038347 |
| Gm10439 | 0 | NA | NA | NA | NA | NA | | Gm3238 | 0 | NA | NA | NA | NA | NA | NM_001101630 |
| Gm10440 | 2.269802738 | 0.113465519 | 0.106587171 | 1.064532609 | 0.28708749 | NA | | Gm3258 | 0.212297216 | 0.00681050119636477 | 0.00973439053551308 | 0.699633035218656 | 0.484156506270654 | NA | NM_011509 |
| Gm10445 | 0 | NA | NA | NA | NA | NA | | Gm3259 | 0 | NA | NA | NA | NA | NA | NM_001270456 |
| Gm1045 | 0.678127538 | -0.001548773 | 0.04101571 | -0.037760472 | 0.969878661 | NA | | Gm3264 | 1.285375627 | -0.0270957763128255 | 0.0272353519936807 | -0.994875201874111 | 0.319796965077772 | NA | NM_001242945 |
| Gm10466 | 0 | NA | NA | NA | NA | NA | | Gm3279 | 10.6191946 | -0.0835216780417994 | 0.0630498240076163 | -1.32469327799726 | 0.185272903585325 | NA | NR_046071 |
| Gm10471 | 3.549274368 | 0.013462761 | 0.114199705 | 0.117887877 | 0.906156499 | NA | | Gm3285 | 0 | NA | NA | NA | NA | NA | NM_001101631 |
| Gm10474 | 3.710584375 | 0.001827684 | 0.119330919 | 0.015316094 | 0.987780003 | NA | | Gm3286 | 0 | NA | NA | NA | NA | NA | NM_001167793,NM_001167792,NM_001122678 |
| Gm10485 | 0.301004085 | 0.013136417 | 0.039354285 | 0.333798913 | 0.738531304 | NA | | Gm3317 | 0 | NA | NA | NA | NA | NA | NM_001242941 |
| Gm10486 | 0 | NA | NA | NA | NA | NA | | Gm3336 | 1.205649186 | 0.0213446253948177 | 0.0276944249655935 | 0.770719212308449 | 0.440873381635667 | NA | NM_001195253 |
| Gm10487 | 0 | NA | NA | NA | NA | NA | | Gm3376 | 0 | NA | NA | NA | NA | NA | NM_001270512 |
| Gm10488 | 0 | NA | NA | NA | NA | NA | | Gm3383 | 0 | NA | NA | NA | NA | NA | NM_001205306 |
| Gm10494 | 0.613100444 | -0.006928698 | 0.062938259 | -0.110087217 | 0.912340206 | NA | | Gm3402 | 0 | NA | NA | NA | NA | NA | NM_001243111 |
| Gm10510 | 0.256038351 | 0.007085343 | 0.03307033 | 0.214250763 | 0.830351512 | NA | | Gm3404 | 0 | NA | NA | NA | NA | NA | NM_001243109 |
| Gm10512 | 0.107716603 | -0.020477583 | 0.027954224 | -0.732539849 | 0.46383913 | NA | | Gm3409 | 0 | NA | NA | NA | NA | NA | NM_001243113 |
| Gm10536 | 1.848793241 | 0.060591749 | 0.106909603 | 0.566756847 | 0.570879392 | NA | | Gm3415 | 0 | NA | NA | NA | NA | NA | NM_001243114 |
| Gm10538 | 0 | NA | NA | NA | NA | NA | | Gm3417 | 0 | NA | NA | NA | NA | NA | NM_001123368 |
| Gm10549 | 0.945542035 | 0.081901904 | 0.07456633 | 1.098376493 | 0.272040123 | NA | | Gm3428 | 0 | NA | NA | NA | NA | NA | NR_030730 |
| Gm10556 | 0.615078183 | -0.001857088 | 0.053211811 | -0.034899925 | 0.97215954 | NA | | Gm3434 | 0 | NA | NA | NA | NA | NA | NR_030729 |
| Gm10578 | 0 | NA | NA | NA | NA | NA | | Gm3435 | 19.50219111 | -0.023923014164352 | 0.0769161079713986 | -0.311027362087117 | 0.755779820606132 | NA | NM_001123372 |
| Gm10584 | 5.185747891 | -0.037390519 | 0.135246782 | -0.276461433 | 0.782193678 | NA | | Gm3448 | 0 | NA | NA | NA | NA | NA | NM_001123367 |
| Gm10591 | 0 | NA | NA | NA | NA | NA | | Gm3494 | 0 | NA | NA | NA | NA | NA | NM_001256885 |
| Gm10619 | 0 | NA | NA | NA | NA | NA | | Gm3500 | 0 | NA | NA | NA | NA | NA | NM_001256886 |
| Gm10636 | 2.154172627 | 0.072376572 | 0.110402682 | 0.655568967 | 0.512101498 | NA | | Gm362 | 0 | NA | NA | NA | NA | NA | NM_001195271 |
| Gm10637 | 0.482637559 | -0.049175986 | 0.042867026 | -1.14717514 | 0.251309244 | NA | | Gm364 | 0 | NA | NA | NA | NA | NA | NM_001128625 |
| Gm10639 | 0.479713553 | 0.008997466 | 0.050300534 | 0.178874167 | 0.858036507 | NA | | Gm3646 | 0 | NA | NA | NA | NA | NA | NM_001177348 |
| Gm10640 | 6.972240599 | -0.029134361 | 0.151803251 | -0.191921853 | 0.84780342 | NA | | Gm3696 | 2.099391675 | 0.0166104004975075 | 0.0309933328895953 | 0.535934633318632 | 0.5920037275718 | NA | NM_001024712 |
| Gm10649 | 6.142534515 | -0.083052187 | 0.142983268 | -0.580852493 | 0.561339873 | NA | | Gm3701 | 0 | NA | NA | NA | NA | NA | NM_001243011 |
| Gm10662 | 0 | NA | NA | NA | NA | NA | | Gm3706 | 0 | NA | NA | NA | NA | NA | NM_001243001 |
| Gm10664 | 0.394136287 | -0.017186747 | 0.03688538 | -0.465950104 | 0.641251216 | NA | | Gm3716 | 4.784573863 | -0.0124383581162384 | 0.0440109358273555 | -0.28261971445077 | 0.77746836625481 | NA | NR_045078 |
| Gm10665 | 0 | NA | NA | NA | NA | NA | | Gm3750 | 0 | NA | NA | NA | NA | NA | NM_001099643 |
| Gm10666 | 0 | NA | NA | NA | NA | NA | | Gm3763 | 0 | NA | NA | NA | NA | NA | NM_001243024 |
| Gm10670 | 0 | NA | NA | NA | NA | NA | | Gm3776 | 0.386703839 | -0.0106102602379525 | 0.0108279273240121 | -0.979897622181403 | 0.327136656825014 | NA | NM_001243092 |
| Gm10677 | 6.693545595 | -0.04670418 | 0.125735101 | -0.371449016 | 0.710303123 | NA | | Gm382 | 0 | NA | NA | NA | NA | NA | NM_001033241 |
| Gm10681 | 0 | NA | NA | NA | NA | NA | | Gm3833 | 0 | NA | NA | NA | NA | NA | NR_030680 |
| Gm10684 | 3.96246455 | 0.214059037 | 0.12896362 | 1.659840473 | 0.096946549 | NA | | Gm3925 | 0.113843286 | 0.00525947738444006 | 0.00859368002043176 | 0.612016897526494 | 0.540526577873086 | NA | NR_040424,NR_040425,NR_040426 |
| Gm10696 | 0.107716603 | -0.020477583 | 0.027954224 | -0.732539849 | 0.46383913 | NA | | Gm3985 | 0.521227213 | 0.00794086785200512 | 0.0148411354916805 | 0.535057971572089 | 0.59260977103426 | NA | NM_001177589 |
| Gm10697 | 0 | NA | NA | NA | NA | NA | | Gm4013 | 7.653573207 | -0.01418182407006 | 0.0523301521890864 | -0.271006742323553 | 0.78638584662649 | NA | NR_033452 |
| Gm10714 | 5.966882616 | 0.222710908 | 0.141662646 | 1.572121616 | 0.115922349 | NA | | Gm4027 | 0 | NA | NA | NA | NA | NA | NM_001177564 |
| Gm10731 | 1.168665658 | 0.004111664 | 0.072666983 | 0.056582281 | 0.954877949 | NA | | Gm4064 | 0 | NA | NA | NA | NA | NA | NM_001270518 |
| Gm10745 | 0 | NA | NA | NA | NA | NA | | Gm4070 | 0 | NA | NA | NA | NA | NA | NM_001243040,NM_001243039 |
| Gm10768 | 1.119424133 | -0.021548962 | 0.077527442 | -0.277952703 | 0.781048666 | NA | | Gm41 | 0 | NA | NA | NA | NA | NA | NR_036689 |
| Gm10782 | 0 | NA | NA | NA | NA | NA | | Gm4133 | 0 | NA | NA | NA | NA | NA | NM_001167158 |
| Gm10787 | 0.116696964 | -0.020477583 | 0.027954224 | -0.732539847 | 0.463839131 | NA | | Gm4141 | 0 | NA | NA | NA | NA | NA | NM_001167163 |
| Gm10789 | 6.078012854 | 0.015137487 | 0.129637506 | 0.116767803 | 0.907044058 | NA | | Gm4172 | 0 | NA | NA | NA | NA | NA | NM_001166839 |
| Gm10791 | 7.070179103 | 0.046711269 | 0.141089308 | 0.331075893 | 0.740587161 | NA | | Gm4175 | 0 | NA | NA | NA | NA | NA | NM_001167165 |
| Gm10804 | 0 | NA | NA | NA | NA | NA | | Gm4177 | 0 | NA | NA | NA | NA | NA | NM_001167167 |
| Gm10818 | 2.069276574 | 0.069858037 | 0.094961905 | 0.735642754 | 0.461948137 | NA | | Gm4179 | 0 | NA | NA | NA | NA | NA | NM_001167169 |
| Gm1082 | 2.166970131 | 0.011149646 | 0.097126817 | 0.114794721 | 0.908607833 | NA | | Gm4187 | 0 | NA | NA | NA | NA | NA | NM_001167159 |
| Gm10823 | 0 | NA | NA | NA | NA | NA | | Gm4201 | 0 | NA | NA | NA | NA | NA | NM_001167162 |
| Gm10825 | 5.723088238 | -0.040102144 | 0.128624986 | -0.311775695 | 0.755210997 | NA | | Gm4214 | 0 | NA | NA | NA | NA | NA | NM_001167164 |
| Gm10857 | 0.607229355 | 0.041512175 | 0.049138713 | 0.844795719 | 0.398224905 | NA | | Gm4216 | 0 | NA | NA | NA | NA | NA | NM_001167166 |
| Gm10872 | 0.506202321 | -0.046639326 | 0.052906751 | -0.881538272 | 0.378026549 | NA | | Gm4220 | 0 | NA | NA | NA | NA | NA | NM_001167168 |
| Gm10921 | 0 | NA | NA | NA | NA | NA | | Gm4224 | 0 | NA | NA | NA | NA | NA | NR_046074 |
| Gm10922 | 0 | NA | NA | NA | NA | NA | | Gm4251 | 0.236453993 | 0.00749401214086028 | 0.0102093235778702 | 0.734036107652062 | 0.462926731928034 | NA | NR_046075,NR_046076 |
| Gm1110 | 1.351077591 | 0.096155621 | 0.083031192 | 1.158066254 | 0.246837 | NA | | Gm4262 | 0.555591603 | 0.0193863533460825 | 0.0163586421030564 | 1.18508328649482 | 0.235984518583227 | NA | NR_040518 |
| Gm11128 | 6.173766686 | -0.039183861 | 0.133097611 | -0.294399433 | 0.7684527 | NA | | Gm4265 | 0.205932006 | -0.000576954122277892 | 0.00970424421944593 | -0.0594537925088239 | 0.952590668569529 | NA | NR_046077 |
| Gm11149 | 4.404330934 | 0.067551017 | 0.101183254 | 0.667610641 | 0.504382164 | NA | | Gm4278 | 0 | NA | NA | NA | NA | NA | NR_046078 |
| Gm11166 | 1.589607904 | 0.014514141 | 0.097236105 | 0.14926699 | 0.881342961 | NA | | Gm428 | 0 | NA | NA | NA | NA | NA | NM_001081644 |
| Gm11186 | 0 | NA | NA | NA | NA | NA | | Gm4297 | 0 | NA | NA | NA | NA | NA | NM_001100446 |
| Gm11194 | 0 | NA | NA | NA | NA | NA | | Gm4301 | 0 | NA | NA | NA | NA | NA | NM_001166637 |
| Gm11201 | 2.19009043 | -0.047749419 | 0.091201009 | -0.523562398 | 0.600582941 | NA | | Gm4302 | 0 | NA | NA | NA | NA | NA | NM_001166634 |
| Gm11202 | 0.29182997 | 0.032201186 | 0.035537591 | 0.906116166 | 0.36487437 | NA | | Gm4303 | 0 | NA | NA | NA | NA | NA | NM_001166638 |
| Gm11213 | 0 | NA | NA | NA | NA | NA | | Gm4305 | 0 | NA | NA | NA | NA | NA | NM_001166639 |
| Gm1123 | 2.051606762 | 0.064583559 | 0.094182112 | 0.685730627 | 0.49288299 | NA | | Gm4307 | 0 | NA | NA | NA | NA | NA | NM_001166641 |
| Gm11240 | 0.215468684 | -0.027008432 | 0.031909031 | -0.846419662 | 0.397318674 | NA | | Gm4312 | 0 | NA | NA | NA | NA | NA | NM_001166636 |
| Gm11351 | 0.198019769 | 0.025308392 | 0.031628095 | 0.80018706 | 0.423602426 | NA | | Gm4340 | 0 | NA | NA | NA | NA | NA | NM_001177535 |
| Gm1140 | 0 | NA | NA | NA | NA | NA | | Gm4345 | 0 | NA | NA | NA | NA | NA | NM_001255977 |
| Gm11413 | 7.376766201 | 0.069026574 | 0.148567712 | 0.46461356 | 0.64220822 | NA | | Gm436 | 0 | NA | NA | NA | NA | NA | NM_001085504 |
| Gm11426 | 0.08731684 | 0.017365056 | 0.027733412 | 0.626142072 | 0.531221759 | NA | | Gm4371 | 13.50497348 | -0.0234930985225305 | 0.063305112080495 | -0.371109026592641 | 0.710556328097123 | NA | NR_028311 |
| Gm11435 | 0.098353386 | -0.020212831 | 0.02786466 | -0.725393061 | 0.468210929 | NA | | Gm438 | 0 | NA | NA | NA | NA | NA | NM_001126316 |
| Gm11437 | 0.293434314 | -0.014075032 | 0.039227749 | -0.358802961 | 0.719742498 | NA | | Gm44 | 0 | NA | NA | NA | NA | NA | NM_001101450 |
| Gm11468 | 2.177079743 | -0.213911707 | 0.103844481 | -2.059923699 | 0.039405836 | NA | | Gm4461 | 1.501906774 | -0.0368971904902418 | 0.0286225126634429 | -1.28909683521138 | 0.197364423676222 | NA | NM_001199062 |
| Gm11487 | 0 | NA | NA | NA | NA | NA | | Gm4477 | 2.814227903 | 0.045446152085541 | 0.0395169278199077 | 1.15004264229889 | 0.250126308513092 | NA | NM_001253910 |
| Gm11529 | 2.633445258 | -0.176207923 | 0.113559765 | -1.551675658 | 0.12073985 | NA | | Gm4489 | 0.309662151 | -0.00555052144305575 | 0.0125234542884447 | -0.443210101239973 | 0.657613773219227 | NA | NR_027637 |
| Gm11541 | 0 | NA | NA | NA | NA | NA | | Gm4498 | 0 | NA | NA | NA | NA | NA | NM_001167151 |
| Gm11544 | 0 | NA | NA | NA | NA | NA | | Gm4541 | 0 | NA | NA | NA | NA | NA | NR_033693 |
| Gm11545 | 6.538721738 | 0.519609241 | 0.148984531 | 3.48767243 | 0.000487245 | NA | | Gm4559 | 0 | NA | NA | NA | NA | NA | NM_001199309 |
| Gm11548 | 0 | NA | NA | NA | NA | NA | | Gm4566 | 0.765868818 | 0.000972757989970912 | 0.0163003512738302 | 0.0596771182184673 | 0.952412796263542 | NA | NR_028023 |
| Gm11554 | 0 | NA | NA | NA | NA | NA | | Gm4567 | 0 | NA | NA | NA | NA | NA | NM_001037248 |
| Gm11559 | 0 | NA | NA | NA | NA | NA | | Gm4598 | 2.894554109 | -0.0240303760383631 | 0.0390474830881828 | -0.615414212078514 | 0.538281208052563 | NA | NR_030681 |
| Gm11562 | 0 | NA | NA | NA | NA | NA | | Gm4710 | 0 | NA | NA | NA | NA | NA | NR_033456 |
| Gm11563 | 0 | NA | NA | NA | NA | NA | | Gm4719 | 0 | NA | NA | NA | NA | NA | NR_045952 |
| Gm11564 | 0 | NA | NA | NA | NA | NA | | Gm4724 | 0 | NA | NA | NA | NA | NA | NM_001256480 |
| Gm11565 | 0 | NA | NA | NA | NA | NA | | Gm4736 | 0 | NA | NA | NA | NA | NA | NM_053251 |
| Gm11567 | 0 | NA | NA | NA | NA | NA | | Gm4745 | 0.127687691 | 0.00542298404407807 | 0.00869724871942359 | 0.62352868349814 | 0.532937159177571 | NA | NM_001038676 |
| Gm11568 | 0 | NA | NA | NA | NA | NA | | Gm4759 | 2.177775734 | -0.0332982231912218 | 0.0315499284033175 | -1.05541358970946 | 0.291236210293006 | NA | NR_003967 |
| Gm11569 | 0 | NA | NA | NA | NA | NA | | Gm4763 | 0 | NA | NA | NA | NA | NA | NM_177593 |
| Gm11570 | 0 | NA | NA | NA | NA | NA | | Gm4776 | 0 | NA | NA | NA | NA | NA | NR_037691,NR_037690 |
| Gm11595 | 0 | NA | NA | NA | NA | NA | | Gm4787 | 17.8510807 | -0.026534139049661 | 0.0772529068927453 | -0.343471075936338 | 0.731244096866397 | NA | NM_001038995 |
| Gm11596 | 0.098353386 | -0.020212831 | 0.02786466 | -0.725393061 | 0.468210929 | NA | | Gm4788 | 3.384828324 | 0.0237515318734475 | 0.0409421836987822 | 0.580123719051997 | 0.561831189624653 | NA | NM_001029977,NM_001160303,NM_001160304 |
| Gm11648 | 4.544781541 | 0.068948765 | 0.138959469 | 0.496178959 | 0.619768155 | NA | | Gm4791 | 0.455532238 | -0.010604440074692 | 0.0143879871610707 | -0.73703430201719 | 0.461101489736641 | NA | NM_001243258 |
| Gm11710 | 0 | NA | NA | NA | NA | NA | | Gm4792 | 0.707957873 | 0.0146284583399266 | 0.018328843114901 | 0.798111383693057 | 0.42480585594714 | NA | NR_033209 |
| Gm11711 | 0 | NA | NA | NA | NA | NA | | Gm4794 | 0 | NA | NA | NA | NA | NA | NM_001101452 |
| Gm11747 | 0.2777807 | 0.011208478 | 0.038523646 | 0.290950592 | 0.771089108 | NA | | Gm4814 | 3.082110864 | -0.0453458894531308 | 0.0395698051754669 | -1.14597201709866 | 0.251806730680277 | NA | NR_036451 |
| Gm11756 | 0 | NA | NA | NA | NA | NA | | Gm4827 | 1.402403799 | 0.00937308615751376 | 0.0293881420212173 | 0.318941093681481 | 0.749771182056887 | NA | NR_045935 |
| Gm11757 | 0 | NA | NA | NA | NA | NA | | Gm4832 | 0.349184399 | -0.00508887990706381 | 0.0114407387221738 | -0.444803437141765 | 0.656461810238787 | NA | NM_001190356 |
| Gm11758 | 0 | NA | NA | NA | NA | NA | | Gm4836 | 0 | NA | NA | NA | NA | NA | NM_009529 |
| Gm11937 | 0 | NA | NA | NA | NA | NA | | Gm4841 | 10.56741595 | -0.103605972877816 | 0.03838121388906 | -2.69939281173562 | NA | NA | NM_001034859 |
| Gm11938 | 0 | NA | NA | NA | NA | NA | | Gm4846 | 0 | NA | NA | NA | NA | NA | NM_001164306 |
| Gm11961 | 1.284420953 | -0.023135237 | 0.082292133 | -0.281135459 | 0.778606505 | NA | | Gm4847 | 4.393942755 | 0.0569086912006442 | 0.0295016370867535 | 1.92900112740512 | NA | NA | NM_001164312 |
| Gm11978 | 2.445186259 | 0.100952169 | 0.097897036 | 1.031207611 | 0.302443472 | NA | | Gm4850 | 0 | NA | NA | NA | NA | NA | NR_015347 |
| Gm11981 | 0 | NA | NA | NA | NA | NA | | Gm4858 | 0 | NA | NA | NA | NA | NA | NM_001034860,NM_001177694 |
| Gm11985 | 0 | NA | NA | NA | NA | NA | | Gm4861 | 1.818954947 | -0.0313090104248024 | 0.0244292776511835 | -1.28161834630774 | 0.199976561177942 | NA | NM_177665 |
| Gm12 | 0 | NA | NA | NA | NA | NA | | Gm4871 | 0 | NA | NA | NA | NA | NA | NM_001101463 |
| Gm12060 | 3.907643635 | 0.041831464 | 0.122959699 | 0.340204669 | 0.733702402 | NA | | Gm4872 | 0 | NA | NA | NA | NA | NA | NR_073372,NR_073371 |
| Gm12130 | 0 | NA | NA | NA | NA | NA | | Gm4884 | 0 | NA | NA | NA | NA | NA | NM_183166 |
| Gm12159 | 0 | NA | NA | NA | NA | NA | | Gm4890 | 20.3708262 | -0.0397414622539801 | 0.0837467120895903 | -0.474543552366159 | 0.635112348327665 | NA | NR_045823,NR_045822 |
| Gm12169 | 0.107716603 | -0.020477583 | 0.027954224 | -0.732539849 | 0.46383913 | NA | | Gm4894 | 0.643456503 | 0.0147615494184696 | 0.0177555332453006 | 0.831377419902418 | 0.405760450893113 | NA | NM_177701 |
| Gm12171 | 0.20671448 | -0.003314027 | 0.03178208 | -0.10427343 | 0.916952363 | NA | | Gm4906 | 0 | NA | NA | NA | NA | NA | NM_001114529 |
| Gm12185 | 8.060386651 | 0.00437008 | 0.133305673 | 0.032782402 | 0.973848112 | NA | | Gm4907 | 0 | NA | NA | NA | NA | NA | NM_001034864 |
| Gm12191 | 0 | NA | NA | NA | NA | NA | | Gm4922 | 0 | NA | NA | NA | NA | NA | NM_177706 |
| Gm12228 | 0 | NA | NA | NA | NA | NA | | Gm4925 | 13.03643043 | -0.00545647630800483 | 0.0728099340797107 | -0.0749413713523103 | 0.940261359508759 | NA | NM_001037166 |
| Gm12238 | 0.462717353 | 0.004293417 | 0.044257148 | 0.097010698 | 0.922717899 | NA | | Gm4926 | 6.073344593 | 0.00981471269913036 | 0.0536207255547223 | 0.183039535507851 | 0.854766997762057 | NA | NR_028066 |
| Gm12250 | 22.08773812 | 0.203168811 | 0.139805838 | 1.453221225 | NA | NA | | Gm4937 | 0 | NA | NA | NA | NA | NA | NM_001013760 |
| Gm12253 | 0 | NA | NA | NA | NA | NA | | Gm4952 | 6.999717798 | -0.0570155187284096 | 0.0594566763701175 | -0.958942245165005 | 0.337587841251874 | NA | NM_001013762,NM_001167907 |
| Gm12295 | 1.534685539 | 0.015731773 | 0.084450203 | 0.186284614 | 0.852221573 | NA | | Gm4956 | 0.222609587 | 0.00717651388923987 | 0.00999169132085706 | 0.718248158273197 | 0.472604288901417 | NA | NR_002858 |
| Gm12298 | 0 | NA | NA | NA | NA | NA | | Gm4961 | 5.437016421 | -0.0196302822578228 | 0.0512824819270122 | -0.382787289541905 | 0.701877486514586 | NA | NR_045694 |
| Gm12360 | 4.13031607 | 0.095576326 | 0.09404658 | 1.016265824 | 0.309502819 | NA | | Gm4971 | 0 | NA | NA | NA | NA | NA | NR_033604 |
| Gm12409 | 0.368800901 | -0.003607529 | 0.046901082 | -0.07691782 | 0.938688921 | NA | | Gm4975 | 1.006182018 | 0.00958660557334846 | 0.0215786138705495 | 0.444264197453027 | 0.656851582897926 | NA | NM_001195687 |
| Gm12504 | 3.0697094 | -0.033102735 | 0.107900842 | -0.30678848 | 0.759004377 | NA | | Gm4981 | 1.121076854 | -0.0253891651823065 | 0.0247033140870962 | -1.02776352568697 | 0.30406107745199 | NA | NM_001034869 |
| Gm12505 | 0 | NA | NA | NA | NA | NA | | Gm4984 | 0 | NA | NA | NA | NA | NA | NM_001101484 |
| Gm12530 | 5.056708818 | 0.044885282 | 0.144270955 | 0.311117939 | 0.755710964 | NA | | Gm5 | 7.162955226 | 0.0521528936202508 | 0.0572496050294446 | 0.910973859006147 | 0.362309147422969 | NA | NR_024513 |
| Gm12597 | 0 | NA | NA | NA | NA | NA | | Gm5039 | 0 | NA | NA | NA | NA | NA | NR_003647 |
| Gm12603 | 0.105923158 | 0.018524693 | 0.027977518 | 0.662127842 | 0.507889294 | NA | | Gm5065 | 0.12890128 | -0.0068640067066208 | 0.00870804083610955 | -0.788237771940376 | 0.430557639008166 | NA | NR_003622 |
| Gm12633 | 0 | NA | NA | NA | NA | NA | | Gm5071 | 0 | NA | NA | NA | NA | NA | NM_001256004 |
| Gm12657 | 2.130322593 | 0.099612284 | 0.090831579 | 1.096670184 | 0.272785595 | NA | | Gm5072 | 0 | NA | NA | NA | NA | NA | NM_001114678 |
| Gm12669 | 0 | NA | NA | NA | NA | NA | | Gm5077 | 7.921354853 | -0.116788682949593 | 0.0610162751317219 | -1.91405789188982 | 0.0556127590615698 | NA | NM_173864 |
| Gm12695 | 2.027333788 | 0.006431673 | 0.090511717 | 0.07105901 | 0.943350791 | NA | | Gm5082 | 0 | NA | NA | NA | NA | NA | NM_177808,NM_001145878 |
| Gm12709 | 0.649215748 | 0.028699483 | 0.054876546 | 0.522982684 | 0.600986303 | NA | | Gm5084 | 2.683582072 | -0.0460549291918173 | 0.0378104865112814 | -1.21804645856847 | 0.223206318447003 | NA | NR_036449 |
| Gm12718 | 0.107716603 | -0.020477583 | 0.027954224 | -0.732539849 | 0.46383913 | NA | | Gm5087 | 2.76313607 | 0.0268403656215734 | 0.0356152630442459 | 0.753619749718786 | 0.451077581817775 | NA | NM_177813 |
| Gm12789 | 0.119397641 | -0.020477583 | 0.027954224 | -0.732539845 | 0.463839132 | NA | | Gm5088 | 0.226066984 | -0.0093084363525918 | 0.0101406635841621 | -0.917931679257152 | 0.358654635211298 | NA | NR_002862 |
| Gm12794 | 0.198019769 | 0.025308392 | 0.031628095 | 0.80018706 | 0.423602426 | NA | | Gm5091 | 0 | NA | NA | NA | NA | NA | NR_046164 |
| Gm128 | 4.202001019 | 0.079275767 | 0.133426887 | 0.594151365 | 0.552410878 | NA | | Gm5095 | 0 | NA | NA | NA | NA | NA | NR_033454 |
| Gm12830 | 1.402624803 | -0.010824009 | 0.069768248 | -0.155142337 | 0.876709106 | NA | | Gm5105 | 8.235451455 | -0.0465533205366993 | 0.0523695893815133 | -0.888938047567218 | 0.374036375980395 | NA | NR_037975 |
| Gm12839 | 0 | NA | NA | NA | NA | NA | | Gm5108 | 0.321262748 | 0.00283574568300987 | 0.0125641879413187 | 0.2257006737128 | 0.821434234059046 | NA | NM_001256184 |
| Gm12886 | 0 | NA | NA | NA | NA | NA | | Gm5111 | 16.59091814 | -0.0642325970829381 | 0.0693521701222866 | -0.926180059970419 | 0.354352413298218 | NA | NM_183309 |
| Gm12887 | 0 | NA | NA | NA | NA | NA | | Gm5114 | 0 | NA | NA | NA | NA | NA | NM_177890 |
| Gm12888 | 0 | NA | NA | NA | NA | NA | | Gm5122 | 0.710417717 | -0.00299628299833557 | 0.0191905029122542 | -0.156133636103006 | 0.875927685701998 | NA | NR_040767 |
| Gm13011 | 0 | NA | NA | NA | NA | NA | | Gm5127 | 4.841476578 | 0.0417075601394018 | 0.0471691663780536 | 0.884212364601105 | 0.376581584917719 | NA | NM_001033541 |
| Gm13023 | 0 | NA | NA | NA | NA | NA | | Gm5128 | 0 | NA | NA | NA | NA | NA | NM_183320 |
| Gm13032 | 0.316760911 | -0.015973748 | 0.040037625 | -0.398968415 | 0.689916477 | NA | | Gm5129 | 0.451410283 | -0.0118078003715272 | 0.0158420142559 | -0.745347162348983 | 0.456061876943773 | NA | NR_028426 |
| Gm13034 | 0.261950519 | 0.027936351 | 0.033119325 | 0.843506059 | 0.398945478 | NA | | Gm5132 | 0.226357657 | -0.0012264097261801 | 0.00980336873216358 | -0.125100846421945 | 0.900443713214317 | NA | NM_001085517 |
| Gm13037 | 0.192746203 | -0.000975302 | 0.031567365 | -0.030895906 | 0.975352555 | NA | | Gm5134 | 3.136669825 | -0.0387389660456726 | 0.0391165323775498 | -0.990347653308504 | 0.322004222035941 | NA | NM_198635 |
| Gm13040 | 0 | NA | NA | NA | NA | NA | | Gm5136 | 0 | NA | NA | NA | NA | NA | NM_203660 |
| Gm13043 | 0 | NA | NA | NA | NA | NA | | Gm5142 | 0 | NA | NA | NA | NA | NA | NM_001004158 |
| Gm13051 | 3.005711473 | 0.048107853 | 0.097982051 | 0.490986386 | 0.623436076 | NA | | Gm5150 | 0.097165704 | -0.00637828455464458 | 0.00848781691535724 | -0.75146349388194 | 0.452373763047372 | NA | NM_001081687 |
| Gm13057 | 0 | NA | NA | NA | NA | NA | | Gm5166 | 0 | NA | NA | NA | NA | NA | NR_027707 |
| Gm13078 | 0 | NA | NA | NA | NA | NA | | Gm5168 | 0 | NA | NA | NA | NA | NA | NM_001025607 |
| Gm13083 | 0 | NA | NA | NA | NA | NA | | Gm5169 | 0 | NA | NA | NA | NA | NA | NM_001040669 |
| Gm13084 | 0 | NA | NA | NA | NA | NA | | Gm5176 | 14.50623841 | -0.0393552650381123 | 0.075838563458816 | -0.518934737727254 | 0.603806252484865 | NA | NR_033603 |
| Gm13088 | 0 | NA | NA | NA | NA | NA | | Gm5177 | 2.247010132 | 0.00183176551922502 | 0.0332970213281556 | 0.0550128944319743 | 0.956128191079047 | NA | NR_033630 |
| Gm13102 | 0 | NA | NA | NA | NA | NA | | Gm525 | 2.11527068 | 0.0294150038135387 | 0.0307275130032904 | 0.957285537895268 | 0.338423152879882 | NA | NM_001033266 |
| Gm13103 | 0 | NA | NA | NA | NA | NA | | Gm5294 | 0.445061325 | -0.00152805503338926 | 0.0138785295966984 | -0.110102084139575 | 0.912328415304257 | NA | NM_001195128 |
| Gm13109 | 0 | NA | NA | NA | NA | NA | | Gm53 | 0 | NA | NA | NA | NA | NA | NR_037977 |
| Gm13119 | 0 | NA | NA | NA | NA | NA | | Gm5334 | 0 | NA | NA | NA | NA | NA | NR_003648 |
| Gm13124 | 0 | NA | NA | NA | NA | NA | | Gm5346 | 0 | NA | NA | NA | NA | NA | NM_001025240 |
| Gm13125 | 0 | NA | NA | NA | NA | NA | | Gm5347 | 0 | NA | NA | NA | NA | NA | NM_001079931 |
| Gm13128 | 0 | NA | NA | NA | NA | NA | | Gm5382 | 0 | NA | NA | NA | NA | NA | NM_001034100 |
| Gm13139 | 4.772703669 | 0.22596694 | 0.138142542 | 1.635752005 | 0.101891501 | NA | | Gm5409 | 0 | NA | NA | NA | NA | NA | NM_001003664 |
| Gm13154 | 0.853962284 | -0.033059725 | 0.056950476 | -0.58049954 | 0.561577797 | NA | | Gm5414 | 0 | NA | NA | NA | NA | NA | NM_001003670 |
| Gm13177 | 0 | NA | NA | NA | NA | NA | | Gm5415 | 0.217374753 | -0.00904456656736547 | 0.00999660569474087 | -0.90476376117583 | 0.365590553873005 | NA | NM_001164286 |
| Gm13178 | 0.098353386 | -0.020212831 | 0.02786466 | -0.725393061 | 0.468210929 | NA | | Gm5416 | 0 | NA | NA | NA | NA | NA | NM_001082542 |
| Gm13212 | 3.161633673 | 0.139904147 | 0.096489302 | 1.449944645 | 0.147073956 | NA | | Gm5420 | 0.106148608 | 0.00515554063321497 | 0.00854200771766808 | 0.603551390213729 | 0.546141939141592 | NA | NR_045843 |
| Gm1322 | 0 | NA | NA | NA | NA | NA | | Gm5424 | 0.354261383 | -0.00497680888617611 | 0.0114881291174227 | -0.433213174687285 | 0.664859908514983 | NA | NR_002687 |
| Gm13242 | 0.079983654 | 0.016740032 | 0.027674178 | 0.604897175 | 0.54524732 | NA | | Gm5431 | 8.342219603 | -0.0603022653309729 | 0.0537628612858089 | -1.12163422646722 | 0.262017992856438 | NA | NM_001024230 |
| Gm13247 | 2.081103138 | -0.06677113 | 0.092140514 | -0.72466635 | 0.468656743 | NA | | Gm5434 | 0.09572496 | 0.00498255289994511 | 0.00847999972823692 | 0.587565219295241 | 0.556824162160132 | NA | NR_003649 |
| Gm13271 | 0 | NA | NA | NA | NA | NA | | Gm5441 | 0 | NA | NA | NA | NA | NA | NR_044987 |
| Gm13272 | 0 | NA | NA | NA | NA | NA | | Gm5458 | 3.15857681 | -0.0376991529796076 | 0.0376425287880696 | -1.00150426109406 | 0.31658308110082 | NA | NM_001024706 |
| Gm13275 | 0 | NA | NA | NA | NA | NA | | Gm5460 | 0 | NA | NA | NA | NA | NA | NM_001034880 |
| Gm13276 | 0 | NA | NA | NA | NA | NA | | Gm5464 | 4.561012357 | -0.0670840833480953 | 0.044770371342303 | -1.49840354986532 | 0.134028434666421 | NA | NM_001034881 |
| Gm13277 | 0 | NA | NA | NA | NA | NA | | Gm5475 | 1.341305976 | 0.0139027556836409 | 0.0269049642820273 | 0.516735704902163 | 0.605340667913755 | NA | NR_040351 |
| Gm13278 | 0 | NA | NA | NA | NA | NA | | Gm5476 | 0 | NA | NA | NA | NA | NA | NR_002868 |
| Gm13279 | 0 | NA | NA | NA | NA | NA | | Gm5477 | 0 | NA | NA | NA | NA | NA | NR_002869 |
| Gm13280 | 0 | NA | NA | NA | NA | NA | | Gm5478 | 0 | NA | NA | NA | NA | NA | NR_003960 |
| Gm13283 | 0 | NA | NA | NA | NA | NA | | Gm5480 | 3.566236658 | 0.00183250343285165 | 0.027742889163071 | 0.0660530856062002 | 0.947335561496008 | NA | NR_045596,NR_045595,NM_001253796,NM_001253797,NM_001253803,NM_001253802 |
| Gm13285 | 0 | NA | NA | NA | NA | NA | | Gm5483 | 0.097165704 | -0.00637828455464458 | 0.00848781691535724 | -0.75146349388194 | 0.452373763047372 | NA | NM_001082547 |
| Gm13286 | 0 | NA | NA | NA | NA | NA | | Gm5485 | 0 | NA | NA | NA | NA | NA | NR_015373 |
| Gm13287 | 0 | NA | NA | NA | NA | NA | | Gm5494 | 0.191449919 | 0.00665508079272897 | 0.00962405648474729 | 0.691504752001018 | 0.489248395458281 | NA | NM_001177529 |
| Gm13288 | 0 | NA | NA | NA | NA | NA | | Gm5512 | 1.105671852 | 0.00438280228130223 | 0.0255547225321469 | 0.171506549358493 | 0.863825481099159 | NA | NR_002891 |
| Gm13289 | 0 | NA | NA | NA | NA | NA | | Gm5523 | 2.312306773 | -0.0156618855251871 | 0.0305042935395424 | -0.513432166684497 | 0.607649049493319 | NA | NR_004447 |
| Gm13290 | 0 | NA | NA | NA | NA | NA | | Gm5531 | 1.375280419 | -0.0141126785978278 | 0.0239629929051448 | -0.588936392615543 | 0.555903943623883 | NA | NM_001008426 |
| Gm13298 | 0.482734261 | -0.021234089 | 0.050628366 | -0.4194109 | 0.67491586 | NA | | Gm5535 | 19.99808624 | 0.0523969398262957 | 0.0745227269315642 | 0.703100141174556 | 0.481993344195576 | NA | NM_001033778 |
| Gm13304 | 0 | NA | NA | NA | NA | NA | | Gm5538 | 0.12890128 | -0.0068640067066208 | 0.00870804083610955 | -0.788237771940376 | 0.430557639008166 | NA | NM_001101531 |
| Gm13305 | 0 | NA | NA | NA | NA | NA | | Gm5544 | 1.788290623 | 0.0202444991689265 | 0.0266017408273389 | 0.761021592546342 | 0.446644170334043 | NA | NM_001033779 |
| Gm13306 | 0 | NA | NA | NA | NA | NA | | Gm5547 | 0.522557059 | -0.0187239880719638 | 0.0143512744871933 | -1.30469165569041 | 0.191997863962332 | NA | NR_045845 |
| Gm13308 | 0 | NA | NA | NA | NA | NA | | Gm5549 | 0 | NA | NA | NA | NA | NA | NM_001270430 |
| Gm13315 | 0.585367496 | 0.027251914 | 0.062924522 | 0.433088935 | 0.664950161 | NA | | Gm5591 | 0 | NA | NA | NA | NA | NA | NM_001013810 |
| Gm1332 | 0.496526426 | 0.041177769 | 0.052144109 | 0.78969169 | 0.429707846 | NA | | Gm5592 | 0 | NA | NA | NA | NA | NA | NM_001033782 |
| Gm13363 | 0 | NA | NA | NA | NA | NA | | Gm5615 | 0 | NA | NA | NA | NA | NA | NM_001033783 |
| Gm13490 | 0.37445586 | 0.021594193 | 0.042489861 | 0.508219911 | 0.611299129 | NA | | Gm5622 | 2.835227714 | -0.0342704652648238 | 0.0383677723402157 | -0.893209669848432 | 0.371744900647404 | NA | NM_001013816 |
| Gm13497 | 0.899153043 | 0.057189445 | 0.063719345 | 0.897520924 | 0.369441013 | NA | | Gm5627 | 0 | NA | NA | NA | NA | NA | NR_033301 |

| | | | | | | |
|---|---|---|---|---|---|---|
| Gm13498 | 0.276664272 | 0.005276976 | 0.033299614 | 0.158469592 | 0.87408678 | NA |
| Gm13539 | 0 | NA | NA | NA | NA | NA |
| Gm13544 | 3.018081471 | -0.154871573 | 0.111627587 | -1.387395155 | 0.165321302 | NA |
| Gm13546 | 0.744732266 | 0.0208816 | 0.067843496 | 0.307790742 | 0.758241567 | NA |
| Gm13547 | 3.314472048 | 0.200455762 | 0.113211871 | 1.770624943 | 0.076623091 | NA |
| Gm13580 | 0.08731684 | 0.017365056 | 0.027733412 | 0.626142072 | 0.531221759 | NA |
| Gm13582 | 0.079983654 | 0.016740032 | 0.027674178 | 0.604897175 | 0.54524732 | NA |
| Gm136 | 7.486667715 | -0.332955286 | 0.140212455 | -2.374648435 | 0.017565672 | NA |
| Gm13710 | 4.935292295 | 0.197781296 | 0.13092614 | 1.510632606 | 0.130882082 | NA |
| Gm13718 | 0 | NA | NA | NA | NA | NA |
| Gm13752 | 0 | NA | NA | NA | NA | NA |
| Gm13769 | 0 | NA | NA | NA | NA | NA |
| Gm13807 | 0 | NA | NA | NA | NA | NA |
| Gm13871 | 0 | NA | NA | NA | NA | NA |
| Gm13939 | 0 | NA | NA | NA | NA | NA |
| Gm13944 | 0.17463368 | 0.024560283 | 0.031186242 | 0.787535833 | 0.430968261 | NA |
| Gm14005 | 6.259972334 | -0.189276401 | 0.148707537 | -1.272809737 | 0.203085565 | NA |
| Gm14023 | 6.950434306 | -0.065195302 | 0.14628259 | -0.445680525 | 0.655828034 | NA |
| Gm14047 | 2.932336519 | -0.067815572 | 0.119467639 | -0.567648046 | 0.570273976 | NA |
| Gm14085 | 0 | NA | NA | NA | NA | NA |
| Gm14092 | 1.280276179 | -0.017309126 | 0.073661637 | -0.23498155 | 0.814223053 | NA |
| Gm14124 | 4.114478889 | 0.012894769 | 0.104654401 | 0.123212866 | 0.901938538 | NA |
| Gm14137 | 5.393404973 | -0.045281017 | 0.114638313 | -0.394990257 | 0.692850079 | NA |
| Gm14139 | 0.491740777 | -0.038091983 | 0.046390976 | -0.821107608 | 0.411584977 | NA |
| Gm14151 | 0.186326724 | 0.024939385 | 0.031411053 | 0.793968451 | 0.427213798 | NA |
| Gm14164 | 1.100526109 | 0.060038741 | 0.072132852 | 0.832335606 | 0.405219538 | NA |
| Gm14207 | 1.481295904 | 0.039395746 | 0.088746968 | 0.443910889 | 0.657107012 | NA |
| Gm14288 | 0 | NA | NA | NA | NA | NA |
| Gm14305 | 0 | NA | NA | NA | NA | NA |
| Gm14306 | 0 | NA | NA | NA | NA | NA |
| Gm14308 | 1.402071703 | 0.146343382 | 0.078928163 | 1.854133891 | 0.063720007 | NA |
| Gm14318 | 0 | NA | NA | NA | NA | NA |
| Gm14327 | 4.815060157 | 0.072283916 | 0.136988002 | 0.527666036 | 0.59773115 | NA |
| Gm14345 | 0 | NA | NA | NA | NA | NA |
| Gm14346 | 0 | NA | NA | NA | NA | NA |
| Gm14347 | 0 | NA | NA | NA | NA | NA |
| Gm14351 | 0 | NA | NA | NA | NA | NA |
| Gm14354 | 0 | NA | NA | NA | NA | NA |
| Gm14374 | 0 | NA | NA | NA | NA | NA |
| Gm14391 | 0 | NA | NA | NA | NA | NA |
| Gm14430 | 0 | NA | NA | NA | NA | NA |
| Gm14431 | 0 | NA | NA | NA | NA | NA |
| Gm14434 | 0 | NA | NA | NA | NA | NA |
| Gm14436 | 0 | NA | NA | NA | NA | NA |
| Gm14440 | 0 | NA | NA | NA | NA | NA |
| Gm14458 | 0 | NA | NA | NA | NA | NA |
| Gm14459 | 0 | NA | NA | NA | NA | NA |
| Gm14461 | 2.338466806 | -0.020661496 | 0.099826237 | -0.206974609 | 0.836029688 | NA |
| Gm14474 | 0 | NA | NA | NA | NA | NA |
| Gm14475 | 0 | NA | NA | NA | NA | NA |
| Gm14476 | 0 | NA | NA | NA | NA | NA |
| Gm14477 | 0 | NA | NA | NA | NA | NA |
| Gm14478 | 0 | NA | NA | NA | NA | NA |
| Gm14479 | 0 | NA | NA | NA | NA | NA |
| Gm14482 | 0 | NA | NA | NA | NA | NA |
| Gm14483 | 0.098353386 | -0.020212831 | 0.02786466 | -0.725393061 | 0.468210929 | NA |
| Gm14484 | 0 | NA | NA | NA | NA | NA |
| Gm14492 | 3.401484765 | -0.077609153 | 0.117692894 | -0.659420886 | 0.509625534 | NA |
| Gm14496 | 0 | NA | NA | NA | NA | NA |
| Gm14501 | 0 | NA | NA | NA | NA | NA |
| Gm14511 | 0 | NA | NA | NA | NA | NA |
| Gm14525 | 0 | NA | NA | NA | NA | NA |
| Gm14548 | 0 | NA | NA | NA | NA | NA |
| Gm14625 | 0 | NA | NA | NA | NA | NA |
| Gm14632 | 0 | NA | NA | NA | NA | NA |
| Gm14635 | 0 | NA | NA | NA | NA | NA |
| Gm14685 | 0 | NA | NA | NA | NA | NA |
| Gm14692 | 0 | NA | NA | NA | NA | NA |
| Gm14718 | 0 | NA | NA | NA | NA | NA |
| Gm14725 | 0 | NA | NA | NA | NA | NA |
| Gm14743 | 2.913104147 | 0.132434134 | 0.069315834 | 1.910589925 | NA | NA |
| Gm14744 | 1.229087554 | 0.083549556 | 0.056066624 | 1.490183475 | 0.136176 | NA |
| Gm14781 | 0 | NA | NA | NA | NA | NA |
| Gm14812 | 0.079983654 | 0.016740032 | 0.027674178 | 0.604897175 | 0.54524732 | NA |
| Gm14819 | 0 | NA | NA | NA | NA | NA |
| Gm14850 | 0 | NA | NA | NA | NA | NA |
| Gm14851 | 0 | NA | NA | NA | NA | NA |
| Gm14858 | 0.386222787 | 0.053439715 | 0.045634083 | 1.171048305 | 0.241579361 | NA |
| Gm14873 | 1.789798875 | -0.027703176 | 0.081284736 | -0.340816457 | 0.73324176 | NA |
| Gm14920 | 0 | NA | NA | NA | NA | NA |
| Gm15008 | 0.310199702 | 0.010205801 | 0.034113052 | 0.299175856 | 0.76480587 | NA |
| Gm15023 | 0 | NA | NA | NA | NA | NA |
| Gm15056 | 0 | NA | NA | NA | NA | NA |
| Gm15070 | 0.575738104 | 0.018512958 | 0.053137856 | 0.348394908 | 0.727543626 | NA |
| Gm15080 | 0 | NA | NA | NA | NA | NA |
| Gm15085 | 0 | NA | NA | NA | NA | NA |
| Gm15091 | 0 | NA | NA | NA | NA | NA |
| Gm15093 | 0 | NA | NA | NA | NA | NA |
| Gm15097 | 0 | NA | NA | NA | NA | NA |
| Gm15104 | 4.111783717 | 0.20070567 | 0.12065897 | 1.663412759 | 0.096229841 | NA |
| Gm15107 | 0 | NA | NA | NA | NA | NA |
| Gm15114 | 0 | NA | NA | NA | NA | NA |
| Gm15127 | 0 | NA | NA | NA | NA | NA |
| Gm15133 | 1.055623674 | 0.111406771 | 0.072854512 | 1.529167761 | 0.126222861 | NA |
| Gm15140 | 0 | NA | NA | NA | NA | NA |
| Gm15179 | 0.582829523 | 0.023716638 | 0.058499643 | 0.405415095 | 0.685172417 | NA |
| Gm15217 | 0 | NA | NA | NA | NA | NA |
| Gm1527 | 0 | NA | NA | NA | NA | NA |
| Gm15284 | 0 | NA | NA | NA | NA | NA |
| Gm15292 | 0 | NA | NA | NA | NA | NA |
| Gm15293 | 0 | NA | NA | NA | NA | NA |
| Gm15299 | 0 | NA | NA | NA | NA | NA |
| Gm15308 | 0 | NA | NA | NA | NA | NA |
| Gm15315 | 0 | NA | NA | NA | NA | NA |
| Gm15319 | 0 | NA | NA | NA | NA | NA |
| Gm15348 | 0 | NA | NA | NA | NA | NA |
| Gm15350 | 0.537538576 | 0.051328891 | 0.044528031 | 1.152732117 | 0.249020355 | NA |
| Gm15386 | 0 | NA | NA | NA | NA | NA |
| Gm15401 | 0.927537728 | 0.084647147 | 0.074477453 | 1.136547297 | 0.255727583 | NA |
| Gm15408 | 7.199789303 | -0.054936747 | 0.14384481 | -0.381916781 | 0.702523092 | NA |
| Gm15412 | 0 | NA | NA | NA | NA | NA |
| Gm15413 | 2.044128532 | 0.119959044 | 0.106424412 | 1.127176011 | 0.259668071 | NA |
| Gm15417 | 73.02544883 | 0.176601473 | 0.16747169 | 1.054515379 | NA | NA |
| Gm15441 | 2.092731098 | 0.10037557 | 0.098003676 | 1.024202087 | 0.305739833 | NA |
| Gm15455 | 2.552545577 | 0.143903929 | 0.108635775 | 1.324645856 | 0.185288639 | NA |
| Gm15471 | 2.881483846 | 0.074193832 | 0.092582574 | 0.801380094 | 0.422911636 | NA |
| Gm1553 | 0.079983654 | 0.016740032 | 0.027674178 | 0.604897175 | 0.54524732 | NA |
| Gm156 | 0 | NA | NA | NA | NA | NA |
| Gm1564 | 0.211846316 | 0.025707621 | 0.031860375 | 0.806883811 | 0.419733426 | NA |
| Gm15698 | 0.098353386 | -0.020212831 | 0.02786466 | -0.725393061 | 0.468210929 | NA |
| Gm15713 | 0 | NA | NA | NA | NA | NA |
| Gm15772 | 0 | NA | NA | NA | NA | NA |
| Gm15787 | 6.597693041 | 0.006260648 | 0.151751078 | 0.04125604 | 0.967091778 | NA |
| Gm15850 | 1.979993066 | -0.122100993 | 0.091664009 | -1.332049459 | 0.182843937 | NA |
| Gm1587 | 0.309826022 | -0.01193339 | 0.034225532 | -0.348669245 | 0.727337636 | NA |
| Gm15880 | 3.002832699 | -0.056348994 | 0.112379008 | -0.501419215 | 0.616076119 | NA |
| Gm15881 | 0 | NA | NA | NA | NA | NA |
| Gm15915 | 0.587717662 | 0.024545414 | 0.058572002 | 0.419063942 | 0.675169402 | NA |
| Gm15941 | 0.901370294 | -0.023544105 | 0.064578436 | -0.364581526 | 0.715423807 | NA |
| Gm15987 | 0.659787236 | 0.016087308 | 0.046903158 | 0.342989874 | 0.731606078 | NA |
| Gm16063 | 0 | NA | NA | NA | NA | NA |
| Gm16065 | 0.651198668 | 0.010138487 | 0.065005828 | 0.155962742 | 0.876062389 | NA |
| Gm16130 | 0.079983654 | 0.016740032 | 0.027674178 | 0.604897175 | 0.54524732 | NA |
| Gm16157 | 0.652614779 | -0.082917553 | 0.055030012 | -1.506769682 | 0.131869703 | NA |
| Gm16294 | 5.002996032 | 0.039285375 | 0.137260593 | 0.286210149 | 0.774717171 | NA |
| Gm1631 | 0.29109959 | -0.010630118 | 0.033578644 | -0.316573778 | 0.751567036 | NA |
| Gm16325 | 1.21199048 | -0.080292872 | 0.085386543 | -0.940345746 | 0.347040242 | NA |
| Gm16336 | 0 | NA | NA | NA | NA | NA |
| Gm16367 | 0 | NA | NA | NA | NA | NA |
| Gm16381 | 0 | NA | NA | NA | NA | NA |
| Gm16390 | 0 | NA | NA | NA | NA | NA |
| Gm16404 | 0 | NA | NA | NA | NA | NA |
| Gm16405 | 0 | NA | NA | NA | NA | NA |
| Gm16430 | 0 | NA | NA | NA | NA | NA |
| Gm16442 | 0 | NA | NA | NA | NA | NA |
| Gm16445 | 0 | NA | NA | NA | NA | NA |
| Gm16451 | 0 | NA | NA | NA | NA | NA |
| Gm1647 | 0.840912053 | 0.002766328 | 0.063990717 | 0.043230145 | 0.965518076 | NA |
| Gm16497 | 0.227114243 | -0.027513412 | 0.032207244 | -0.854261616 | 0.39296005 | NA |
| Gm16501 | 0 | NA | NA | NA | NA | NA |
| Gm16523 | 0.404218839 | -0.002314653 | 0.036541701 | -0.063342793 | 0.94949354 | NA |
| Gm16548 | 1.429623161 | 0.050192426 | 0.084987252 | 0.590587707 | 0.554796704 | NA |
| Gm16551 | 0.946752622 | -0.006409306 | 0.071468774 | -0.089679802 | 0.928541667 | NA |
| Gm16617 | 0.401340017 | -0.003486686 | 0.042331768 | -0.082365706 | 0.934355906 | NA |
| Gm16796 | 7.151077362 | 0.113291412 | 0.154698788 | 0.732335482 | 0.463963827 | NA |
| Gm16833 | 0.423692631 | 0.031061933 | 0.034784099 | 0.892992313 | 0.371861289 | NA |
| Gm16853 | 6.300706512 | 0.243538079 | 0.148707516 | 1.637698518 | 0.1014846 | NA |
| Gm16863 | 0.539474011 | 0.04933016 | 0.054620124 | 0.903149904 | 0.366446341 | NA |
| Gm16880 | 1.013259476 | -0.016392334 | 0.071729171 | -0.228530927 | 0.819233516 | NA |
| Gm16982 | 1.937057278 | 0.072024504 | 0.085701883 | 0.840407481 | 0.400679956 | NA |
| Gm17019 | 0.813807001 | 0.049577646 | 0.06258422 | 0.792174864 | 0.428258723 | NA |
| Gm1715 | 3.988540077 | 0.003802252 | 0.116258024 | 0.032705289 | 0.973909606 | NA |
| Gm1720 | 0.159967308 | 0.0240345 | 0.030870969 | 0.778547006 | 0.436246606 | NA |
| Gm17252 | 0 | NA | NA | NA | NA | NA |
| Gm17359 | 0.19442443 | -0.026451352 | 0.031568725 | -0.837897386 | 0.402088337 | NA |
| Gm17365 | 0 | NA | NA | NA | NA | NA |
| Gm17384 | 5.411198423 | 0.176646725 | 0.142620572 | 1.238578157 | 0.215501762 | NA |
| Gm17495 | 0.116696964 | -0.020477583 | 0.027954224 | -0.732539847 | 0.463839131 | NA |
| Gm17660 | 6.825038994 | 0.286873783 | 0.151756259 | 1.890358815 | 0.058709988 | NA |
| Gm17677 | 0.186326724 | 0.024939385 | 0.031411053 | 0.793968451 | 0.427213798 | NA |
| Gm17689 | 0 | NA | NA | NA | NA | NA |
| Gm17727 | 0 | NA | NA | NA | NA | NA |
| Gm17745 | 1.856371848 | -0.137630911 | 0.089636171 | -1.535439431 | 0.124675924 | NA |
| Gm17746 | 0.914430724 | 0.026377469 | 0.058012971 | 0.454682264 | 0.649337842 | NA |
| Gm17751 | 7.248427673 | -0.336861661 | 0.152608484 | -2.207358666 | 0.027289012 | NA |
| Gm17757 | 0 | NA | NA | NA | NA | NA |
| Gm17801 | 3.177281092 | -0.082888088 | 0.124350382 | -0.666568823 | 0.505047589 | NA |
| Gm17830 | 0.426633926 | -0.004170814 | 0.042905384 | -0.097209576 | 0.922559963 | NA |
| Gm18409 | 1.082787731 | -0.073424906 | 0.057334288 | -1.280645646 | 0.20031816 | NA |
| Gm18853 | 0 | NA | NA | NA | NA | NA |
| Gm19276 | 0.275017096 | 0.008844542 | 0.038389071 | 0.2303922 | 0.817787022 | NA |
| Gm19277 | 0 | NA | NA | NA | NA | NA |
| Gm19299 | 0.176054697 | -0.002190495 | 0.031239146 | -0.070120197 | 0.944097991 | NA |
| Gm19303 | 0.282397767 | 0.011595411 | 0.038695572 | 0.299657307 | 0.764438567 | NA |
| Gm19395 | 0 | NA | NA | NA | NA | NA |
| Gm19402 | 0.098353386 | -0.020212831 | 0.02786466 | -0.725393061 | 0.468210929 | NA |
| Gm19424 | 0.360303906 | 0.01488513 | 0.035847896 | 0.415230234 | 0.677973368 | NA |
| Gm19434 | 1.594634258 | -0.015377979 | 0.093891344 | -0.163784838 | 0.869900528 | NA |
| Gm19466 | 0.338960846 | 0.036479516 | 0.037750094 | 0.966342396 | 0.333872878 | NA |
| Gm19510 | 0.187700257 | -0.003168524 | 0.031449734 | -0.100748845 | 0.919749836 | NA |
| Gm19557 | 2.938296458 | 0.070952002 | 0.1066624 | 0.665201628 | 0.505921539 | NA |
| Gm19583 | 0.21363976 | -0.000753674 | 0.031922632 | -0.02360938 | 0.98116419 | NA |
| Gm19589 | 0.107716603 | -0.020477583 | 0.027954224 | -0.732539849 | 0.46383913 | NA |
| Gm19619 | 8.065104231 | 0.164940638 | 0.156455409 | 1.054234169 | 0.29177572 | NA |
| Gm1965 | 0.188785634 | 0.025025111 | 0.031461603 | 0.79541754 | 0.42637066 | NA |
| Gm19668 | 0 | NA | NA | NA | NA | NA |
| Gm1968 | 0.105923158 | 0.018524693 | 0.027977518 | 0.662127842 | 0.507889294 | NA |
| Gm19689 | 2.262677967 | 0.052066007 | 0.10442818 | 0.498581957 | 0.618073919 | NA |
| Gm19710 | 2.050198435 | -0.037167857 | 0.104183566 | -0.356753552 | 0.721276311 | NA |
| Gm19782 | 0.305079873 | -0.015054679 | 0.039642215 | -0.379763828 | 0.704120735 | NA |
| Gm19784 | 0 | NA | NA | NA | NA | NA |
| Gm1987 | 0 | NA | NA | NA | NA | NA |
| Gm19897 | 2.557211408 | 0.060659439 | 0.097579969 | 0.621638224 | 0.534179781 | NA |
| Gm1993 | 0 | NA | NA | NA | NA | NA |
| Gm1995 | 0 | NA | NA | NA | NA | NA |
| Gm19990 | 1.264991571 | -0.0324123 | 0.085073853 | -0.380990153 | 0.703210555 | NA |
| Gm2002 | 0 | NA | NA | NA | NA | NA |
| Gm20098 | 0.215433205 | -0.027029776 | 0.031922004 | -0.84674434 | 0.397137638 | NA |
| Gm20110 | 0.294090812 | 0.009484222 | 0.033616198 | 0.282132504 | 0.77784191 | NA |
| Gm2012 | 0 | NA | NA | NA | NA | NA |
| Gm20125 | 1.627670774 | 0.215002718 | 0.093678218 | 2.295119629 | 0.021726269 | NA |
| Gm20139 | 1.140243808 | -0.052958208 | 0.079629872 | -0.66505455 | 0.506015603 | NA |
| Gm2016 | 0 | NA | NA | NA | NA | NA |
| Gm20172 | 0 | NA | NA | NA | NA | NA |
| Gm20199 | 0.176054697 | -0.002190495 | 0.031239146 | -0.070120197 | 0.944097991 | NA |
| Gm2022 | 0 | NA | NA | NA | NA | NA |
| Gm20268 | 0 | NA | NA | NA | NA | NA |
| Gm2027 | 5.893859806 | -0.051956358 | 0.138081687 | -0.376272616 | 0.706714235 | NA |
| Gm2030 | 0 | NA | NA | NA | NA | NA |
| Gm20324 | 7.74618319 | 0.049083581 | 0.157553918 | 0.311535132 | 0.755393839 | NA |
| Gm20356 | 0 | NA | NA | NA | NA | NA |
| Gm20362 | 1.746981493 | -0.005619916 | 0.100535985 | -0.055899543 | 0.955421835 | NA |
| Gm20408 | 0 | NA | NA | NA | NA | NA |
| Gm20410 | 0 | NA | NA | NA | NA | NA |
| Gm2042 | 0 | NA | NA | NA | NA | NA |
| Gm20516 | 0 | NA | NA | NA | NA | NA |
| Gm20554 | 1.055791187 | 0.040993609 | 0.080037269 | 0.51218151 | 0.608523982 | NA |
| Gm20556 | 0.096071043 | -0.020098795 | 0.027825948 | -0.722304036 | 0.470107572 | NA |
| Gm20579 | 0 | NA | NA | NA | NA | NA |
| Gm20580 | 0 | NA | NA | NA | NA | NA |
| Gm20581 | 0 | NA | NA | NA | NA | NA |
| Gm20594 | 1.362017742 | -0.003296715 | 0.073343854 | -0.044948753 | 0.964148157 | NA |
| Gm20604 | 0 | NA | NA | NA | NA | NA |
| Gm20611 | 0.281741269 | -0.015233555 | 0.038833149 | -0.392282259 | 0.69484967 | NA |
| Gm20735 | 1.169054187 | -0.010860347 | 0.079141224 | -0.137227436 | 0.890851025 | NA |

| | | | | | | | |
|---|---|---|---|---|---|---|---|
| Gm5634 | 0 | NA | NA | NA | NA | NA | NM_001085524 |
| Gm5635 | 0 | NA | NA | NA | NA | NA | NM_001038697 |
| Gm5640 | 2.472465543 | 0.0118717038363346 | 0.0326191027967974 | 0.363949428967746 | 0.715895773097993 | NA | NM_001099302 |
| Gm5643 | 6.414334197 | 0.0546842630891696 | 0.0501955313443948 | 1.08942492737008 | 0.275966543836272 | NA | NR_002883 |
| Gm5662 | 0 | NA | NA | NA | NA | NA | NM_001013824 |
| Gm5712 | 0 | NA | NA | NA | NA | NA | NR_033594 |
| Gm572 | 3.214112978 | 0.0215619134925446 | 0.040544600832567 | 0.531807270259897 | 0.594859486870527 | NA | NM_001085505 |
| Gm5725 | 0 | NA | NA | NA | NA | NA | NM_001166711 |
| Gm5726 | 0 | NA | NA | NA | NA | NA | NM_001167146 |
| Gm5728 | 0 | NA | NA | NA | NA | NA | NM_001166713 |
| Gm5741 | 3.458912933 | -0.0362193132690103 | 0.0443228360269467 | -0.817170481757762 | 0.413831008897134 | NA | NM_001195531 |
| Gm5766 | 0 | NA | NA | NA | NA | NA | NR_003628 |
| Gm5771 | 0.194331409 | -0.00840968081320881 | 0.00964081822647896 | -0.872299489073576 | 0.383045016303155 | NA | NM_001038997 |
| Gm5779 | 0.106148608 | 0.00515554063321497 | 0.00854200771766808 | 0.603551390213729 | 0.546141939141592 | NA | NR_033602 |
| Gm5795 | 0 | NA | NA | NA | NA | NA | NM_001270806 |
| Gm5796 | 5.296437332 | 0.0161135757156587 | 0.0518006531547816 | 0.31106896794353 | 0.755748191616188 | NA | NM_001029930 |
| Gm5797 | 0.879810723 | -0.00524696272514982 | 0.0186548511857417 | -0.28126532197481 | 0.778506906152088 | NA | NM_001025085 |
| Gm5800 | 0 | NA | NA | NA | NA | NA | NM_001034102 |
| Gm5801 | 0.09572496 | 0.00498255289994511 | 0.00847999972823692 | 0.587565219295241 | 0.556824162160132 | NA | NR_002889 |
| Gm5803 | 2.699114828 | -0.0202661658770619 | 0.0323429360947425 | -0.626602539042714 | 0.530919803567946 | NA | NM_001165971 |
| Gm581 | 16.4940275 | -0.0842426515343294 | 0.0616889230337355 | -1.36560418615608 | 0.172063231944283 | NA | NM_001195255 |
| Gm5833 | 0 | NA | NA | NA | NA | NA | NR_040304 |
| Gm5860 | 10.1522238 | -0.0778870373768622 | 0.0585074882879617 | -1.3312319440807 | 0.183112703517995 | NA | NR_040659 |
| Gm5868 | 2.147800347 | 0.0259628257271159 | 0.0257534681591855 | 1.0081292960869 | 0.313392395104517 | NA | NM_001024147 |
| Gm5878 | 1.080185468 | 0.0276065141118769 | 0.0262813849788947 | 1.05042082576874 | 0.293524674840422 | NA | NM_001034902 |
| Gm5885 | 0 | NA | NA | NA | NA | NA | NM_001185040 |
| Gm5886 | 0 | NA | NA | NA | NA | NA | NM_001177652 |
| Gm5891 | 0 | NA | NA | NA | NA | NA | NM_001034904 |
| Gm590 | 0 | NA | NA | NA | NA | NA | NM_001195437 |
| Gm5901 | 0.43199899 | -0.00998173168171151 | 0.0141728176617135 | -0.70428703169421 | 0.481254040757148 | NA | NM_001195727 |
| Gm5916 | 0.106148608 | 0.00515554063321497 | 0.00854200771766808 | 0.603551390213729 | 0.546141939141592 | NA | NM_001167587 |
| Gm5925 | 0 | NA | NA | NA | NA | NA | NR_040410 |
| Gm5934 | 0 | NA | NA | NA | NA | NA | NM_001100444 |
| Gm5935 | 0 | NA | NA | NA | NA | NA | NM_001081657 |
| Gm5936 | 1.386410611 | -0.00160448425721149 | 0.0276655642811772 | -0.0579957177415365 | 0.953752039534521 | NA | NM_001081670 |
| Gm5938 | 0.191449919 | 0.00665508079272897 | 0.00962405648474729 | 0.691504752001018 | 0.489248395458281 | NA | NM_001085534 |
| Gm5941 | 0 | NA | NA | NA | NA | NA | NM_001034103 |
| Gm595 | 0 | NA | NA | NA | NA | NA | NM_001085499 |
| Gm597 | 0.30673395 | 0.00299118959348609 | 0.0123970371859449 | 0.241282618469303 | 0.809336081443423 | NA | NM_001013750 |
| Gm6026 | 0 | NA | NA | NA | NA | NA | NM_001177569 |
| Gm6034 | 1.685180203 | 0.00159327820396201 | 0.0314393319941742 | 0.0506778644106447 | 0.959582215620766 | NA | NM_001034909 |
| Gm6040 | 0 | NA | NA | NA | NA | NA | NM_001025353 |
| Gm6042 | 0 | NA | NA | NA | NA | NA | NR_002872 |
| Gm6084 | 0 | NA | NA | NA | NA | NA | NM_001038500 |
| Gm6086 | 3.976320993 | -0.0376436337096724 | 0.0442771443524264 | -0.850182058039826 | 0.395223875310368 | NA | NM_001039219 |
| Gm609 | 14.43189301 | -0.0740387005403338 | 0.0713680718649856 | -1.03742049638668 | 0.299539930188414 | NA | NM_001005854 |
| Gm6116 | 0 | NA | NA | NA | NA | NA | NR_045866 |
| Gm6121 | 0 | NA | NA | NA | NA | NA | NM_001114754 |
| Gm614 | 0.674103222 | -0.0122544577310835 | 0.0178407673803168 | -0.686879519801567 | 0.492158653799437 | NA | NM_001033362 |
| Gm6150 | 1.389506481 | -0.0358110838372315 | 0.0269466019567603 | -1.32896473903075 | 0.183859603370072 | NA | NR_038036 |
| Gm6164 | 0 | NA | NA | NA | NA | NA | NM_001167153 |
| Gm6194 | 0.485489131 | -0.00204318730788208 | 0.012539019635632 | -0.162946336097599 | 0.870560687406046 | NA | NR_033512 |
| Gm6213 | 0.226066984 | -0.0093084363525918 | 0.0101406635841621 | -0.917931679257152 | 0.358654635211298 | NA | NR_044988 |
| Gm6225 | 18.44304911 | -0.0183278634837225 | 0.0737300292649389 | -0.248580716248786 | 0.803685126561992 | NA | NR_033457 |
| Gm6249 | 2.438780768 | -0.0578705852063816 | 0.0371508105364136 | -1.55772066263963 | 0.119299480172224 | NA | NR_046021 |
| Gm6251 | 0.202072798 | -0.00115107594702963 | 0.00968429294483289 | -0.11886009165427 | 0.905386199803646 | NA | NM_001101561 |
| Gm6260 | 0.927811108 | -0.00146770345071052 | 0.0207904108622603 | -0.070595211438306 | 0.943719921624443 | NA | NR_040405 |
| Gm6268 | 0 | NA | NA | NA | NA | NA | NR_044989 |
| Gm6289 | 0.12890128 | -0.0068640067066208 | 0.00870804083610955 | -0.788237771940376 | 0.430557639008166 | NA | NM_001039222 |
| Gm6300 | 0.926794364 | 0.00538358145984031 | 0.0210069974878708 | 0.256275627345066 | 0.79773801553336 | NA | NR_033591 |
| Gm6307 | 0.23403553 | -0.000407313579519725 | 0.00983856423584417 | -0.0413996971261099 | 0.966977254250032 | NA | NR_045331 |
| Gm6329 | 0 | NA | NA | NA | NA | NA | NR_040690 |
| Gm6367 | 0 | NA | NA | NA | NA | NA | NR_044992 |
| Gm6370 | 0 | NA | NA | NA | NA | NA | NM_001243110 |
| Gm6377 | 0.959495378 | -0.0123856036127272 | 0.0190500658464001 | -0.650160672020343 | 0.515588441788402 | NA | NM_001037917 |
| Gm6402 | 10.61500856 | 0.015641491564072 | 0.06697812667937 | 0.233531338356911 | 0.815348837692081 | NA | NR_030688 |
| Gm6406 | 0.106347839 | -0.00653347234140708 | 0.00854389509723638 | -0.764694822098226 | 0.444453290197145 | NA | NM_001134661 |
| Gm6408 | 1.171846807 | -0.0147654289472348 | 0.0237028112094904 | -0.622939988709982 | 0.533323958488662 | NA | NM_001243104 |
| Gm6416 | 0.227686571 | 0.00691460391109036 | 0.00980755285755433 | 0.705028462402254 | 0.480792523671872 | NA | NR_046023 |
| Gm6432 | 0 | NA | NA | NA | NA | NA | NM_001244762 |
| Gm6455 | 0 | NA | NA | NA | NA | NA | NR_003596 |
| Gm6460 | 0 | NA | NA | NA | NA | NA | NM_001037919 |
| Gm648 | 1.803776798 | 0.0276706228223134 | 0.0295755914581227 | 0.935589838042407 | 0.349484408650378 | NA | NM_001033372 |
| Gm6484 | 0 | NA | NA | NA | NA | NA | NM_001080940 |
| Gm6498 | 0.865688855 | -0.0227720756554356 | 0.0213994738643991 | -1.06414184758626 | 0.28726444590683 | NA | NR_003630 |
| Gm6524 | 0 | NA | NA | NA | NA | NA | NR_028307 |
| Gm6525 | 0.097165704 | -0.0063782845546458 | 0.00848781691535724 | -0.75146349388194 | 0.452373763047372 | NA | NR_036654 |
| Gm6531 | 0.120209049 | -0.00674972139751468 | 0.00863521114606269 | -0.781650996523957 | 0.434419707908266 | NA | NM_001037922 |
| Gm6567 | 0.21511414 | -0.000824240786584897 | 0.00975135300262786 | -0.08452578697159 | 0.93263840116295 | NA | NR_046024 |
| Gm6568 | 0.336143057 | -0.00539628300464043 | 0.011316386701961 | -0.47685565602891 | 0.633464910573091 | NA | NR_040420 |
| Gm6578 | 0 | NA | NA | NA | NA | NA | NR_003631 |
| Gm6583 | 2.559359672 | -0.00530846012460831 | 0.0347246263977379 | -0.152873066618627 | 0.878498375947209 | NA | NM_001039228 |
| Gm6588 | 0 | NA | NA | NA | NA | NA | NM_001177504 |
| Gm6592 | 0 | NA | NA | NA | NA | NA | NM_001081564 |
| Gm6602 | 0.562699945 | -0.0148439742075509 | 0.0155126138524171 | -0.956897035455956 | 0.338619227682003 | NA | NR_045362 |
| Gm6607 | 0.360627147 | -0.0100772056525624 | 0.010553896215922 | -0.954832741045871 | 0.339662287230353 | NA | NR_033622 |
| Gm6614 | 0 | NA | NA | NA | NA | NA | NM_001081318 |
| Gm6623 | 1.469736246 | -0.0115614049249475 | 0.0278258524679818 | -0.4154914908088 | 0.677782143044108 | NA | NR_033619 |
| Gm6634 | 0 | NA | NA | NA | NA | NA | NR_040556 |
| Gm6639 | 0 | NA | NA | NA | NA | NA | NR_040748 |
| Gm6642 | 0.212496447 | -0.000886988073087717 | 0.00973851166817767 | -0.0910804549309223 | 0.927428662789121 | NA | NR_033643 |
| Gm6644 | 0 | NA | NA | NA | NA | NA | NR_037965 |
| Gm6654 | 1.087101106 | 0.0144290513509944 | 0.0240711690969368 | 0.599432927120713 | 0.548884225050553 | NA | NR_038089 |
| Gm6696 | 0 | NA | NA | NA | NA | NA | NM_001177523 |
| Gm6710 | 0 | NA | NA | NA | NA | NA | NM_001164689 |
| Gm6756 | 0.222609587 | 0.00717651388923987 | 0.00999169132085706 | 0.718248158273197 | 0.472604288901417 | NA | NR_076393 |
| Gm6760 | 0.120209049 | -0.00674972139751468 | 0.00863521114606269 | -0.781650996523957 | 0.434419707908266 | NA | NM_001177377 |
| Gm6763 | 0 | NA | NA | NA | NA | NA | NM_001270899 |
| Gm6772 | 0 | NA | NA | NA | NA | NA | NM_001195767 |
| Gm6787 | 5.971818024 | -0.00808522160219894 | 0.0521098083995498 | -0.155157384963035 | 0.876697243233284 | NA | NR_003632 |
| Gm6792 | 0.223412651 | 0.00718831964189475 | 0.00999993766105603 | 0.718836445339964 | 0.472241699104885 | NA | NM_001177417,NM_001177416 |
| Gm6793 | 0 | NA | NA | NA | NA | NA | NR_033513 |
| Gm6812 | 0 | NA | NA | NA | NA | NA | NM_001098842 |
| Gm6878 | 0.212695677 | -0.00857796107890413 | 0.00973699866180651 | -0.880965621629515 | 0.378336427605205 | NA | NM_001037931 |
| Gm6880 | 0.217532603 | 0.00684685248201087 | 0.00976000296730706 | 0.701521557416086 | 0.482977586094298 | NA | NM_001099305 |
| Gm6890 | 0.097165704 | -0.0063782845546458 | 0.00848781691535724 | -0.75146349388194 | 0.452373763047372 | NA | NM_001099306 |
| Gm6902 | 0 | NA | NA | NA | NA | NA | NM_001270494 |
| Gm6904 | 0.203314312 | -0.000639443352387801 | 0.00969102748600992 | -0.0659830294889689 | 0.947391336513325 | NA | NM_001164329 |
| Gm6927 | 0.452312811 | 0.00549325758014353 | 0.0141190566598134 | 0.389066898199991 | 0.697226659524734 | NA | NM_001101585 |
| Gm6936 | 3.216180835 | -0.0277064163420695 | 0.0415992496229688 | -0.666031637425777 | 0.505390878608447 | NA | NR_045001 |
| Gm6938 | 1.234382615 | -0.0229172019679422 | 0.0253583405229904 | -0.903734293936349 | 0.366136309983333 | NA | NR_033482 |
| Gm6994 | 0 | NA | NA | NA | NA | NA | NR_033141 |
| Gm7008 | 17.73926759 | -0.075244522297611 | 0.0785788204972517 | -0.957567469471531 | 0.338280909391479 | NA | NR_045157 |
| Gm7030 | 0 | NA | NA | NA | NA | NA | NM_001177467 |
| Gm7056 | 1.476431974 | -0.0246723427501514 | 0.0292310892645509 | -0.844044589883688 | 0.398644489138927 | NA | NR_037571 |
| Gm7073 | 0 | NA | NA | NA | NA | NA | NM_001039240 |
| Gm7092 | 1.453518001 | 0.00673687964419086 | 0.0298164561441135 | 0.225945015451505 | 0.82124418571535 | NA | NM_001039243 |
| Gm7104 | 0 | NA | NA | NA | NA | NA | NR_033570 |
| Gm711 | 13.40238324 | -0.0179426832622443 | 0.0643185411259104 | -0.278965955199754 | 0.78027095178931 | NA | NM_198628 |
| Gm7134 | 0 | NA | NA | NA | NA | NA | NR_033597 |
| Gm7157 | 0 | NA | NA | NA | NA | NA | NM_001199311 |
| Gm7168 | 0.931597681 | 0.021623398874696 | 0.01990988384695 | 1.08606353713151 | 0.277450882941307 | NA | NM_001122977 |
| Gm7173 | 8.556949004 | 0.0373975596512877 | 0.0604140911612775 | 0.619020479037806 | 0.535902867874779 | NA | NM_001099307 |
| Gm7244 | 7.385878191 | 0.00266012286307812 | 0.0497331731666198 | 0.0534878973872424 | 0.957343173332153 | NA | NM_001101597 |
| Gm7257 | 0 | NA | NA | NA | NA | NA | NM_001167586 |
| Gm7271 | 3.470884289 | 0.0770602583627903 | 0.0427599670544916 | 1.80215897417759 | 0.0715203973266339 | NA | NR_033501 |
| Gm732 | 0 | NA | NA | NA | NA | NA | NM_001033252 |
| Gm7325 | 0 | NA | NA | NA | NA | NA | NM_001177468,NM_001177469,NM_001177470 |
| Gm7334 | 0 | NA | NA | NA | NA | NA | NR_002700 |
| Gm7337 | 0 | NA | NA | NA | NA | NA | NR_003652 |
| Gm7367 | 0 | NA | NA | NA | NA | NA | NR_003376 |
| Gm7444 | 2.927048557 | -0.0311476692956477 | 0.0317062105705803 | -0.982383852725345 | 0.325910778251156 | NA | NR_033529 |
| Gm7457 | 2.248518919 | -0.0388281424118015 | 0.0336297065532319 | -1.15457868626778 | 0.248262998925762 | NA | NR_045707 |
| Gm7534 | 0 | NA | NA | NA | NA | NA | NM_001080712 |
| Gm7538 | 0.55289662 | -0.00712573376272161 | 0.0165938993615511 | -0.429418885065213 | 0.667618412796209 | NA | NR_046033 |
| Gm7550 | 0 | NA | NA | NA | NA | NA | NR_033689 |
| Gm757 | 0 | NA | NA | NA | NA | NA | NM_001033410 |
| Gm7609 | 5.832558768 | 0.00105505176432003 | 0.0498034280509527 | 0.0211843201484169 | 0.983098622185749 | NA | NM_001081746 |
| Gm7616 | 0.226066984 | -0.0093084363525918 | 0.0101406635841621 | -0.917931679257152 | 0.358654635211298 | NA | NM_001101600 |
| Gm766 | 3.536026832 | 0.0196838541781103 | 0.0424513705599292 | 0.463680063057619 | 0.642876983175829 | NA | NM_001145390 |
| Gm7714 | 0.108766301 | 0.00519171381361946 | 0.00856003371502575 | 0.606506234257728 | 0.544178643801544 | NA | NM_001110779 |
| Gm773 | 0 | NA | NA | NA | NA | NA | NM_001033423 |
| Gm7849 | 0 | NA | NA | NA | NA | NA | NM_001177518 |
| Gm7854 | 11.39097677 | -0.0554298706711619 | 0.066594521610214 | -0.832348807843381 | 0.405212088466438 | NA | NR_028417 |
| Gm7861 | 0 | NA | NA | NA | NA | NA | NM_001177526 |
| Gm7903 | 0 | NA | NA | NA | NA | NA | NM_001242937 |
| Gm7904 | 0.33403441 | 0.00244344761020736 | 0.0111793636306143 | 0.218567683362231 | 0.826986829057954 | NA | NR_003372 |
| Gm7977 | 0 | NA | NA | NA | NA | NA | NR_040408 |
| Gm7978 | 0 | NA | NA | NA | NA | NA | NM_001270457 |
| Gm805 | 0.337583802 | -0.0131877509542174 | 0.0120595734966011 | -1.0935503612905 | 0.274152229987901 | NA | NR_046081 |
| Gm806 | 0 | NA | NA | NA | NA | NA | NM_001033400 |
| Gm8096 | 0 | NA | NA | NA | NA | NA | NR_033590 |
| Gm813 | 0.523864742 | -0.00787665053554553 | 0.0164156490859341 | -0.479825713519588 | 0.631351327241889 | NA | NM_001033404 |
| Gm815 | 0 | NA | NA | NA | NA | NA | NM_001033407 |
| Gm8179 | 0.257802559 | -0.00892927589713349 | 0.00993479775307735 | -0.898787888698349 | 0.368765652383134 | NA | NR_046039 |
| Gm8221 | 0 | NA | NA | NA | NA | NA | NR_033577 |
| Gm8234 | 19.98504479 | -0.133703062151211 | 0.0702367160954032 | -1.90360639824904 | 0.0569614649085856 | NA | NR_004433 |
| Gm826 | 3.661728426 | 0.0763956112153656 | 0.0420954648151498 | 1.81481809384539 | 0.06955186884966 | NA | NM_001033411 |
| Gm8267 | 1.730092743 | -0.0133523087717547 | 0.0291487560262129 | -0.458074737726278 | 0.646898747629183 | NA | NM_001162954 |
| Gm829 | 0 | NA | NA | NA | NA | NA | NM_001033412 |
| Gm8298 | 0 | NA | NA | NA | NA | NA | NM_001243003 |
| Gm8300 | 0 | NA | NA | NA | NA | NA | NM_001177565 |
| Gm833 | 1.855722136 | 0.0115280720536848 | 0.0285496133939325 | 0.403790828780007 | 0.686366541169238 | NA | NR_033138 |
| Gm8363 | 0.700284741 | 0.0052126192686363 | 0.0176992507084066 | 0.294510731245842 | 0.768367664500615 | NA | NR_073378 |
| Gm8369 | 1.146326815 | 0.00512656809237268 | 0.0264100937719887 | 0.194113967812188 | 0.84608664491228 | NA | NM_001164202 |
| Gm839 | 0 | NA | NA | NA | NA | NA | NR_033190 |
| Gm8439 | 0 | NA | NA | NA | NA | NA | NM_001101603 |
| Gm8453 | 0 | NA | NA | NA | NA | NA | NM_001167150 |
| Gm853 | 0.751133396 | 0.00677378319489913 | 0.0161600893112329 | 0.419167435553136 | 0.67509376940704 | NA | NM_001034872 |
| Gm8579 | 0 | NA | NA | NA | NA | NA | NR_036696 |
| Gm8580 | 2.328062488 | -0.00677896783588147 | 0.0345450771485097 | -0.19623542326273 | 0.84442590327426 | NA | NR_027478 |
| Gm8615 | 2.940016543 | -0.026596031090951 | 0.0392511255961742 | -0.677586456107726 | 0.498033935931744 | NA | NR_028061 |
| Gm8633 | 0 | NA | NA | NA | NA | NA | NR_045179 |
| Gm8660 | 0 | NA | NA | NA | NA | NA | NM_001166751 |
| Gm8677 | 0 | NA | NA | NA | NA | NA | NM_001167147 |
| Gm8693 | 0 | NA | NA | NA | NA | NA | NM_001167154 |
| Gm8709 | 0 | NA | NA | NA | NA | NA | NR_033633 |
| Gm8720 | 0 | NA | NA | NA | NA | NA | NM_001166756 |
| Gm8764 | 0 | NA | NA | NA | NA | NA | NM_001270900 |
| Gm8765 | 0 | NA | NA | NA | NA | NA | NM_001244649 |
| Gm8773 | 3.473818726 | 0.0369816607562456 | 0.0423769161156379 | 0.872684096580558 | 0.38283528839115 | NA | NR_033499 |
| Gm8787 | 9.866530148 | -0.0701917980768314 | 0.0649903895438296 | -1.08003350294575 | 0.280127261351267 | NA | NM_001099310 |
| Gm8801 | 0.557041062 | -0.0182495569332047 | 0.0172627764149361 | -1.05716232977534 | 0.290437508715972 | NA | NR_028278 |
| Gm8817 | 0.226066984 | -0.0093084363525918 | 0.0101406635841621 | -0.917931679257152 | 0.358654635211298 | NA | NM_001101606 |
| Gm884 | 2.463864808 | -0.0334398426630587 | 0.0358667652789818 | -0.9323350573422 | 0.351163395531092 | NA | NM_001033434 |
| Gm8882 | 0 | NA | NA | NA | NA | NA | NM_001177588 |
| Gm8883 | 0 | NA | NA | NA | NA | NA | NR_027658,NR_027659 |
| Gm8884 | 0 | NA | NA | NA | NA | NA | NR_026561 |
| Gm889 | 20.21979035 | 0.0165389158712936 | 0.0759284148357659 | 0.217822483283333 | 0.827567426022084 | NA | NM_001033437 |
| Gm8898 | 0 | NA | NA | NA | NA | NA | NM_001177407,NM_001177405 |
| Gm8909 | 0.297797758 | 0.00172794575940152 | 0.010770200141429 | 0.160437664733338 | 0.872536321716291 | NA | NM_001081032 |
| Gm8979 | 1.305524027 | 0.0344605020536603 | 0.0274215025975024 | 1.25669634372257 | 0.208863612596778 | NA | NR_030719 |
| Gm8989 | 0 | NA | NA | NA | NA | NA | NR_030720 |
| Gm8994 | 0 | NA | NA | NA | NA | NA | NM_001142734 |
| Gm9 | 0 | NA | NA | NA | NA | NA | NM_001033234 |
| Gm904 | 0.120209049 | -0.00674972139751468 | 0.00863521114606269 | -0.781650996523957 | 0.434419707908266 | NA | NM_001033770 |
| Gm9041 | 0 | NA | NA | NA | NA | NA | NM_001244651 |
| Gm9047 | 0.120209049 | -0.00674972139751468 | 0.00863521114606269 | -0.781650996523957 | 0.434419707908266 | NA | NM_001145360 |
| Gm9054 | 3.103595811 | 0.00445054209106099 | 0.0405841063049987 | 0.109662192820366 | 0.91267728528048 | NA | NR_045872 |
| Gm906 | 0 | NA | NA | NA | NA | NA | NM_001033438 |
| Gm9112 | 0 | NA | NA | NA | NA | NA | NM_001177365 |
| Gm9125 | 0 | NA | NA | NA | NA | NA | NM_001163730 |
| Gm9159 | 10.53945768 | -0.0363146134576422 | 0.0608726474198428 | -0.596567013213304 | 0.550796500246142 | NA | NR_033398 |
| Gm9199 | 0 | NA | NA | NA | NA | NA | NR_027860 |
| Gm9268 | 0 | NA | NA | NA | NA | NA | NM_001105061 |
| Gm933 | 0 | NA | NA | NA | NA | NA | NM_001256309 |
| Gm9376 | 0 | NA | NA | NA | NA | NA | NM_001101609 |
| Gm94 | 0 | NA | NA | NA | NA | NA | NM_001033280 |
| Gm9513 | 0 | NA | NA | NA | NA | NA | NM_001128510 |
| Gm9573 | 2.489867221 | -0.0440560168185671 | 0.0328936979324536 | -1.33934521162792 | 0.180458316652438 | NA | NM_001244654 |
| Gm9696 | 0 | NA | NA | NA | NA | NA | NR_037189 |
| Gm97 | 0 | NA | NA | NA | NA | NA | NM_001033283 |
| Gm9705 | 0.479198558 | 0.0074134133596442 | 0.0158498358413752 | 0.467728084621031 | 0.639979054507854 | NA | NM_001100187 |
| Gm9731 | 0 | NA | NA | NA | NA | NA | NR_003107 |
| Gm9733 | 0 | NA | NA | NA | NA | NA | NM_001076679 |
| Gm9758 | 0.09572496 | 0.00498255289994511 | 0.00847999972823692 | 0.587565219295241 | 0.556824162160132 | NA | NM_198666 |
| Gm9767 | 13.84558561 | 0.0323238379291751 | 0.0724243166501924 | 0.446311949138552 | 0.65537192601403 | NA | NR_028030 |
| Gm9839 | 0 | NA | NA | NA | NA | NA | NM_001199956 |
| Gm9846 | 0 | NA | NA | NA | NA | NA | NM_001190258 |
| Gm9855 | 0 | NA | NA | NA | NA | NA | NR_037190 |
| Gm9871 | 2.197402966 | 0.012564886074405 | 0.0342411295692466 | 0.366953024986947 | 0.713654057945592 | NA | NR_027989 |
| Gm9895 | 13.59746244 | 0.0954817821048458 | 0.0705540519044044 | 1.35331394197199 | 0.175955357032616 | NA | NR_045687 |
| Gm9920 | 3.907953278 | -0.00735030756314674 | 0.0479485752124244 | -0.15329564081066 | 0.878165138175533 | NA | NR_045093 |
| Gm9926 | 0.685637272 | 0.00400573241083254 | 0.0159185285465727 | 0.251639616005526 | 0.801319634574229 | NA | NR_040528 |
| Gm9961 | 0 | NA | NA | NA | NA | NA | NR_033509 |
| Gm9992 | 0.353527519 | -0.00579453045469275 | 0.0115029412048641 | -0.503743377584378 | 0.614441721879931 | NA | NM_001142539 |
| Gm9994 | 0.108766301 | 0.00519171381361946 | 0.00856003371502575 | 0.606506234257728 | 0.544178643801544 | NA | NM_001205249 |
| Gm9999 | 0.406628625 | -0.000921993720139942 | 0.0119199762705297 | -0.0773486204347094 | 0.938346213215504 | NA | NR_033461 |
| Gmcl1l | 0 | NA | NA | NA | NA | NA | NM_027955 |
| Gml | 0.09572496 | 0.00498255289994511 | 0.00847999972823692 | 0.587565219295241 | 0.556824162160132 | NA | NM_001177524 |
| Gmnc | 21.8116485 | -0.0575780666273209 | 0.0813649807779168 | -0.707651695813442 | 0.479161581229434 | NA | NM_001013761 |
| Gna14 | 12.45963215 | 0.00617550992592354 | 0.06706847289487 | 0.0920776880607326 | 0.926636315480222 | NA | NM_008137 |
| Gnat1 | 6.869559469 | 0.00472668946596844 | 0.0563100588335005 | 0.0839404107167509 | 0.933103809812951 | NA | NM_008140 |
| Gnat2 | 2.682671864 | -0.0343625683689969 | 0.0359771026856727 | -0.955123281305283 | 0.339515356787131 | NA | NM_008141 |
| Gnat3 | 0 | NA | NA | NA | NA | NA | NM_001081143 |

| | | | | | | | | | | | | | | | |
|---|---|---|---|---|---|---|---|---|---|---|---|---|---|---|---|
| Gm20740 | 0 | NA | NA | NA | NA | NA | | Gnb3 | 10.72234035 | 0.00852464514311103 | 0.0577438828627072 | 0.147628540383739 | 0.882635929922381 | NA | NM_013530 |
| Gm20741 | 0 | NA | NA | NA | NA | NA | | Gngt1 | 0 | NA | NA | NA | NA | NA | NM_010314 |
| Gm20744 | 0.224413567 | -0.027287719 | 0.032077135 | -0.850690659 | 0.394941214 | NA | | Gnmt | 19.04769443 | -0.0599431621832146 | 0.0750590896510691 | -0.798612965623156 | 0.424514867220388 | NA | NM_010321 |
| Gm20745 | 0.159967308 | 0.0240345 | 0.030870969 | 0.778547006 | 0.436246606 | NA | | Gnrh1 | 10.58063559 | 0.0108228611026773 | 0.0618207262421918 | 0.175068488524012 | 0.861025818363274 | NA | NM_008145 |
| Gm20750 | 0.296373155 | 0.009272162 | 0.033639931 | 0.275629621 | 0.782832556 | NA | | Gnrhr | 0 | NA | NA | NA | NA | NA | NM_010323 |
| Gm20752 | 4.973363912 | 0.084306828 | 0.122115078 | 0.690388352 | 0.48995 | NA | | Gp1ba | 0.332178952 | 0.0127721041124076 | 0.0133101933856661 | 0.95957314385545 | 0.337270090816746 | NA | NM_010326 |
| Gm20755 | 0.619937596 | 0.079772058 | 0.055246832 | 1.443920948 | 0.148761213 | NA | | Gp2 | 2.485826243 | -0.0468059380616066 | 0.0291501101227355 | -1.60568649190456 | 0.108342810741047 | NA | NM_025989 |
| Gm20756 | 0.403835985 | -0.003083065 | 0.042384291 | -0.072740739 | 0.94201243 | NA | | Gp49a | 5.613307118 | 0.0258586969277399 | 0.0343848551717247 | 0.752037395492768 | 0.452028570542081 | NA | NM_008147 |
| Gm20757 | 0.32854328 | 0.00879128 | 0.034635881 | 0.253820007 | 0.799634616 | NA | | Gp6 | 0.108766301 | 0.00519171381361946 | 0.00856003371502575 | 0.606506234257728 | 0.544178643801544 | NA | NM_001163014 |
| Gm20758 | 0 | NA | NA | NA | NA | NA | | Gp9 | 9.686344736 | 0.0324433328032413 | 0.0625378870591543 | 0.518778844775381 | 0.603914972077994 | NA | NM_018762 |
| Gm20759 | 0 | NA | NA | NA | NA | NA | | Gpa33 | 0 | NA | NA | NA | NA | NA | NM_021610 |
| Gm20765 | 0 | NA | NA | NA | NA | NA | | Gpat2 | 18.46041544 | 0.0229289171050916 | 0.0739137575992464 | 0.31021176368018 | 0.756399925378435 | NA | NM_001081089 |
| Gm2083 | 0 | NA | NA | NA | NA | NA | | Gpbar1 | 0.340383369 | -0.00403721153616492 | 0.0113238420937375 | -0.356523121988573 | 0.7214488393547 | NA | NM_174985 |
| Gm20878 | 0 | NA | NA | NA | NA | NA | | Gpha2 | 0.212297216 | 0.00681050119636477 | 0.00973439053551308 | 0.699633035218656 | 0.484156506270654 | NA | NM_130453 |
| Gm21002 | 0 | NA | NA | NA | NA | NA | | Gphb5 | 0.191449919 | 0.00665508079272897 | 0.00962405648474729 | 0.691504752001018 | 0.489248395458281 | NA | NM_175644 |
| Gm21057 | 1.653400267 | -0.008468466 | 0.100130603 | -0.084574206 | 0.932599906 | NA | | Gpihbp1 | 0.203513543 | -0.00860782669689374 | 0.00975338121443284 | -0.882547960306941 | 0.377480556050336 | NA | NM_026730 |
| Gm2109 | 2.980264325 | 0.06341751 | 0.11607839 | 0.546333468 | 0.584836734 | NA | | Gpr110 | 0.337741652 | 0.00190180888516019 | 0.0112128429016166 | 0.169609875198198 | 0.865316955796633 | NA | NM_133776 |
| Gm21119 | 0 | NA | NA | NA | NA | NA | | Gpr113 | 0 | NA | NA | NA | NA | NA | NM_001014394 |
| Gm21276 | 0 | NA | NA | NA | NA | NA | | Gpr114 | 19.48681293 | -0.162459497220154 | 0.0685376343094088 | -2.37036919726089 | 0.0177703305256657 | NA | NM_001145972,NM_001033468 |
| Gm21293 | 0 | NA | NA | NA | NA | NA | | Gpr119 | 0.233836299 | 0.00743372775021133 | 0.0101683766597817 | 0.731063374118848 | 0.464740444385902 | NA | NM_181751 |
| Gm21304 | 0 | NA | NA | NA | NA | NA | | Gpr128 | 0.097165704 | -0.00637828455464458 | 0.00848781691535724 | -0.75146349388194 | 0.452373763047372 | NA | NM_172825 |
| Gm21312 | 0 | NA | NA | NA | NA | NA | | Gpr132 | 1.569175784 | 0.0114770529608671 | 0.0253367332952042 | 0.452980770138964 | 0.650562588578504 | NA | NM_019925 |
| Gm21319 | 0 | NA | NA | NA | NA | NA | | Gpr141 | 0.57694017 | 0.000332632151082112 | 0.015006458564943 | 0.0221659327310697 | 0.982315592652382 | NA | NM_181754 |
| Gm21498 | 0 | NA | NA | NA | NA | NA | | Gpr142 | 0.45890573 | -0.00161511075132098 | 0.0154987944447753 | -0.104208798760179 | 0.917003651590928 | NA | NM_181749 |
| Gm21541 | 0 | NA | NA | NA | NA | NA | | Gpr143 | 3.680114979 | -0.0076753041556262 | 0.042228749999103 | -0.181755419134813 | 0.855774672055083 | NA | NM_010951 |
| Gm21586 | 0 | NA | NA | NA | NA | NA | | Gpr15 | 0.800262629 | 0.000744585656389101 | 0.019741164776912 | 0.0377174125642231 | 0.969912992656191 | NA | NM_001162955 |
| Gm21637 | 0 | NA | NA | NA | NA | NA | | Gpr152 | 2.709964921 | -0.00488233811576248 | 0.0347973838907147 | -0.140307619995114 | 0.888416943996503 | NA | NM_206973 |
| Gm21677 | 0 | NA | NA | NA | NA | NA | | Gpr171 | 0.621742765 | -0.00224171440648176 | 0.0172290991614795 | -0.130112107746976 | 0.896477730956858 | NA | NM_173398 |
| Gm21693 | 0 | NA | NA | NA | NA | NA | | Gpr174 | 5.433178473 | -0.0199704392841116 | 0.0510133906895438 | -0.391474454337828 | 0.695446567567095 | NA | NM_001177781,NM_001177782,NM_001033251 |
| Gm21704 | 0 | NA | NA | NA | NA | NA | | Gpr18 | 0.458888926 | -0.000592276211756187 | 0.0139816623856329 | -0.0423609292958461 | 0.966210974279136 | NA | NM_182806 |
| Gm21708 | 0 | NA | NA | NA | NA | NA | | Gpr20 | 7.753135342 | 0.0532757139982597 | 0.0570464041829481 | 0.933901352088805 | 0.350354781331944 | NA | NM_173365 |
| Gm2176 | 0 | NA | NA | NA | NA | NA | | Gpr31b | 0.127687691 | 0.00542298404407807 | 0.00869724871942359 | 0.62352868349814 | 0.532937159177571 | NA | NM_001013832 |
| Gm21945 | 0 | NA | NA | NA | NA | NA | | Gpr33 | 0.097165704 | -0.00637828455464458 | 0.00848781691535724 | -0.75146349388194 | 0.452373763047372 | NA | NM_008159 |
| Gm21946 | 0 | NA | NA | NA | NA | NA | | Gpr35 | 13.90372398 | -0.0552977351594324 | 0.073377567970845 | -0.753605450393283 | 0.451086170583518 | NA | NM_001104529,NM_001271766,NM_022320 |
| Gm21947 | 0 | NA | NA | NA | NA | NA | | Gpr39 | 10.44618563 | -0.00748446336558742 | 0.0569194657639147 | -0.131492157650087 | 0.895385989999562 | NA | NM_027677 |
| Gm21949 | 0 | NA | NA | NA | NA | NA | | Gpr44 | 4.393186002 | 0.0814769983974767 | 0.0495359594652315 | 1.64480509264516 | 0.100010011633139 | NA | NM_009962 |
| Gm21950 | 0 | NA | NA | NA | NA | NA | | Gpr50 | 27.82497676 | 0.0484585498882722 | 0.0404620011215321 | 1.19763107471431 | NA | NA | NM_010340 |
| Gm21951 | 0 | NA | NA | NA | NA | NA | | Gpr55 | 5.296343738 | 0.03672903566751 | 0.0516745698797116 | 0.71077583718661 | 0.477223156114853 | NA | NM_001033290 |
| Gm2381 | 1.105618087 | 0.098496357 | 0.077202422 | 1.275819518 | 0.202019326 | NA | | Gpr65 | 8.840092391 | -0.154945880280633 | 0.0633505909433035 | -2.44584743367752 | 0.0144512137775544 | NA | NM_008152 |
| Gm2447 | 0.505546546 | 0.009202456 | 0.05076589 | 0.181272426 | 0.856153749 | NA | | Gpr82 | 0.817903953 | -0.0160291740708363 | 0.0188524077718122 | -0.850245457495508 | 0.395188633534042 | NA | NM_175669 |
| Gm2506 | 0 | NA | NA | NA | NA | NA | | Gpr87 | 0 | NA | NA | NA | NA | NA | NM_032399 |
| Gm2516 | 0 | NA | NA | NA | NA | NA | | Gpr97 | 4.195243604 | -0.0162569815750132 | 0.0408291545345184 | -0.398170909007409 | 0.69050420817088 | NA | NM_173036 |
| Gm2663 | 0 | NA | NA | NA | NA | NA | | Gprc5a | 9.704609962 | 0.0248512252173959 | 0.0625539428291237 | 0.397276719794963 | 0.691163412735365 | NA | NM_181444 |
| Gm2696 | 0 | NA | NA | NA | NA | NA | | Gprc5d | 4.113543059 | -0.00802817224175721 | 0.0414163338448173 | -0.193840726507516 | 0.846300596624007 | NA | NM_001205396,NM_053118 |
| Gm2799 | 0 | NA | NA | NA | NA | NA | | Gprc6a | 0.315716853 | 0.0123948421219712 | 0.0131137196252776 | 0.945181266349427 | 0.344566341855147 | NA | NM_153071 |
| Gm2825 | 0 | NA | NA | NA | NA | NA | | Gprin2 | 19.43680601 | 0.0293258781088077 | 0.07829207107751 | 0.374570217714317 | 0.707980125867868 | NA | NM_183209 |
| Gm2837 | 0 | NA | NA | NA | NA | NA | | Gpx2 | 2.931517556 | 0.0234323309529987 | 0.0392214684138915 | 0.597436350565076 | 0.550216090461215 | NA | NM_030677 |
| Gm2848 | 0.099009884 | 0.018205726 | 0.027896865 | 0.652608315 | 0.514008823 | NA | | Gpx2-ps1 | 9.033993797 | -0.106805607476586 | 0.0580679347112833 | -1.83932161540839 | 0.0658678955970058 | NA | NR_033563 |
| Gm2863 | 0 | NA | NA | NA | NA | NA | | Gpx3 | 477.309003 | 0.0925237147782256 | 0.0659399890374937 | 1.40315029057127 | NA | NA | NM_008161 |
| Gm2913 | 0 | NA | NA | NA | NA | NA | | Gpx5 | 0 | NA | NA | NA | NA | NA | NM_010343 |
| Gm2933 | 0 | NA | NA | NA | NA | NA | | Grap2 | 14.15510344 | -0.014150929576857 | 0.074568738323858 | -0.189770269618864 | 0.849489153927879 | NA | NM_010815 |
| Gm3020 | 0 | NA | NA | NA | NA | NA | | Grhl2 | 3.606149882 | -0.0082350388646645 | 0.0424472391295989 | -0.194006466227909 | 0.8461708187829 | NA | NM_026496 |
| Gm3139 | 0 | NA | NA | NA | NA | NA | | Grifin | 2.926006549 | 0.0149521931757254 | 0.0383409024127758 | 0.389980210031339 | 0.69655118078307 | NA | NM_030022 |
| Gm3143 | 0.079983654 | 0.016740032 | 0.027674178 | 0.604897175 | 0.54524732 | NA | | Grin3b | 8.094519872 | 0.0385630962496114 | 0.058457335542296 | 0.659679335225767 | 0.509459630554525 | NA | NM_130455 |
| Gm3238 | 0 | NA | NA | NA | NA | NA | | Grm6 | 0.434749506 | -0.0165846961888169 | 0.0135129615593836 | -1.22731764727772 | 0.21970322534406 | NA | NM_173372 |
| Gm3258 | 0.295060159 | -0.029687054 | 0.033501022 | -0.886153701 | 0.375534709 | NA | | Grxcr1 | 0.427940552 | -0.00281048961418964 | 0.0152501962985345 | -0.184292028717015 | 0.853784366990649 | NA | NM_001018019 |
| Gm3259 | 0 | NA | NA | NA | NA | NA | | Grxcr2 | 11.53202877 | 0.0437985386639976 | 0.0470589537638625 | 0.930716370869073 | 0.35200029908469 | NA | NM_001033426 |
| Gm3264 | 1.613866166 | 0.026648896 | 0.096497914 | 0.276160333 | 0.782424922 | NA | | Gsc | 0.536376888 | 0.00160239556588434 | 0.014760441545214 | 0.108560137647367 | 0.913551379374449 | NA | NM_010351 |
| Gm3285 | 0 | NA | NA | NA | NA | NA | | Gsc2 | 0.754611645 | 0.00878900532872483 | 0.0178892490254901 | 0.491300966083121 | 0.623213596672045 | NA | NM_029469 |
| Gm3286 | 0 | NA | NA | NA | NA | NA | | Gsdma | 1.055462159 | 0.0161779229921083 | 0.0202231025998394 | 0.799972354006492 | 0.423726814965496 | NA | NM_021347 |
| Gm3317 | 0 | NA | NA | NA | NA | NA | | Gsdma2 | 0.106148608 | 0.00515554063321497 | 0.00854200771766808 | 0.603551390213729 | 0.546141939141592 | NA | NM_029727 |
| Gm3336 | 0.589328065 | -0.006539609 | 0.058189931 | -0.112383855 | 0.910519056 | NA | | Gsdma3 | 0.106148608 | 0.00515554063321497 | 0.00854200771766808 | 0.603551390213729 | 0.546141939141592 | NA | NM_001007461 |
| Gm3376 | 0 | NA | NA | NA | NA | NA | | Gsdmc | 0.423704667 | -0.00952366370425414 | 0.0140901230955794 | -0.675910610549747 | 0.49909740061161 | NA | NM_031378 |
| Gm3383 | 0 | NA | NA | NA | NA | NA | | Gsdmc2 | 0 | NA | NA | NA | NA | NA | NM_001168274,NM_177912 |
| Gm3402 | 0 | NA | NA | NA | NA | NA | | Gsdmc3 | 0 | NA | NA | NA | NA | NA | NM_183194 |
| Gm3404 | 0 | NA | NA | NA | NA | NA | | Gsdmc4 | 2.289432896 | 0.0109779895459366 | 0.035003421593795 | 0.313626184129456 | 0.753804970166888 | NA | NM_028992 |
| Gm3409 | 0 | NA | NA | NA | NA | NA | | Gsdmcl-ps | 0 | NA | NA | NA | NA | NA | NR_029414 |
| Gm3415 | 0 | NA | NA | NA | NA | NA | | Gsdmcl1 | 0 | NA | NA | NA | NA | NA | NM_001167581 |
| Gm3417 | 0 | NA | NA | NA | NA | NA | | Gsg1 | 3.354542912 | 0.0171374079616408 | 0.035942765820637 | 0.476797140408186 | 0.633506582208017 | NA | NM_010352,NM_001080553,NM_001080552 |
| Gm3428 | 0 | NA | NA | NA | NA | NA | | Gsg2 | 6.536854203 | 0.0208109113939815 | 0.0524068800590725 | 0.397102658477736 | 0.691291759597167 | NA | NM_010353 |
| Gm3434 | 0.116696964 | -0.020477583 | 0.027954224 | -0.732539847 | 0.463839131 | NA | | Gsta1 | 0.113843286 | 0.00525947738444006 | 0.00859368002043176 | 0.612016897526494 | 0.540526577873086 | NA | NM_008181 |
| Gm3448 | 0 | NA | NA | NA | NA | NA | | Gsta2 | 2.645002416 | 0.0362145141939124 | 0.0369865074175283 | 0.979127706898594 | 0.327516883125412 | NA | NM_008182 |
| Gm3494 | 0 | NA | NA | NA | NA | NA | | Gstm3 | 1.40269462 | 0.00208077771565643 | 0.0227273474112172 | 0.0915539186341408 | 0.927052465211743 | NA | NM_010359 |
| Gm3500 | 0 | NA | NA | NA | NA | NA | | Gstp2 | 11.1338592 | 0.0100556673374551 | 0.0613853887716612 | 0.163812065683247 | 0.869879092952307 | NA | NM_181796 |
| Gm362 | 0 | NA | NA | NA | NA | NA | | Gstt4 | 1.223288234 | 0.0269309030890852 | 0.0253074206133644 | 1.06415045217463 | 0.28726054854167 | NA | NM_029472 |
| Gm364 | 0.094392817 | 0.017895942 | 0.027818137 | 0.643319236 | 0.520016977 | NA | | Gsx1 | 12.17077174 | -0.0282144295557687 | 0.0577928882769097 | -0.488198987747101 | 0.625408901484268 | NA | NM_008178 |
| Gm3646 | 0 | NA | NA | NA | NA | NA | | Gsx2 | 7.42049852 | -0.00597333552671963 | 0.0534196040865391 | -0.111819165058635 | 0.910966791304421 | NA | NM_133256 |
| Gm3696 | 1.101142027 | 0.008536422 | 0.072641135 | 0.117514982 | 0.906451972 | NA | | Gtf2a1l | 6.597662945 | -0.0119373325551696 | 0.0561366408152063 | -0.212647789069987 | 0.831601693525728 | NA | NM_023630 |
| Gm3701 | 0 | NA | NA | NA | NA | NA | | Gtsf1 | 2.826755307 | 0.0290558965462175 | 0.0353596021722318 | 0.821725776344717 | 0.411232985439917 | NA | NM_028797 |
| Gm3706 | 0 | NA | NA | NA | NA | NA | | Gtsf1l | 1.933802046 | -0.0195903944031033 | 0.022522449309321 | -0.869816338980314 | 0.384400780927561 | NA | NM_026630 |
| Gm3716 | 4.910453018 | -0.097938929 | 0.120135335 | -0.815238321 | 0.414935907 | NA | | Guca1a | 3.943673694 | -0.0452402049951359 | 0.0437909507799158 | -1.03309483327968 | 0.301559521025853 | NA | NM_008189 |
| Gm3750 | 0 | NA | NA | NA | NA | NA | | Guca1b | 19.60695319 | -0.0622788087288454 | 0.0787989800928879 | -0.790350441787843 | 0.429323136005788 | NA | NM_146079 |
| Gm3763 | 0 | NA | NA | NA | NA | NA | | Guca2a | 0.315716853 | 0.0123948421219712 | 0.0131137196252776 | 0.945181266349427 | 0.344566341855147 | NA | NM_008190 |
| Gm3776 | 0.413840517 | 0.018406376 | 0.037259486 | 0.494005092 | 0.621302583 | NA | | Guca2b | 0.334833285 | -0.00610342348777058 | 0.012817048366707 | -0.47619571317407 | 0.633934953261501 | NA | NM_008191 |
| Gm382 | 0 | NA | NA | NA | NA | NA | | Gucy1b2 | 8.7577185 | -0.0340567326009944 | 0.0556044900161347 | -0.612481700508577 | 0.540219101680731 | NA | NM_001204340,NM_172810 |
| Gm3833 | 0 | NA | NA | NA | NA | NA | | Gucy2c | 13.83646313 | 0.0291305225503541 | 0.0712456644506062 | 0.408874319230329 | 0.682631896083366 | NA | NM_001127318,NM_145067 |
| Gm3925 | 0 | NA | NA | NA | NA | NA | | Gucy2d | 0.680293638 | -0.0129639980077018 | 0.0193652009949206 | -0.669448151408406 | 0.503209644497719 | NA | NM_001130693 |
| Gm3985 | 0.197363271 | -0.000561558 | 0.031650166 | -0.017742642 | 0.985844163 | NA | | Gulo | 3.444514516 | 0.0275583122132898 | 0.043003824395328 | 0.640833986297361 | 0.521630549173349 | NA | NM_178747 |
| Gm4013 | 7.753198208 | -0.02975479 | 0.146804805 | -0.202682671 | 0.839383069 | NA | | Gvin1 | 0 | NA | NA | NA | NA | NA | NM_001039160,NM_029000 |
| Gm4027 | 0 | NA | NA | NA | NA | NA | | Gxylt2 | 22.14229852 | -0.0929546165382185 | 0.0843851076051764 | -1.10155238496747 | 0.270656318447935 | NA | NM_198612 |
| Gm4064 | 0 | NA | NA | NA | NA | NA | | Gykl1 | 0.097165704 | -0.00637828455464458 | 0.00848781691535724 | -0.75146349388194 | 0.452373763047372 | NA | NM_010293 |
| Gm4070 | 0 | NA | NA | NA | NA | NA | | Gyltl1b | 12.73194804 | 0.013610424993269 | 0.044315487418253 | 0.307125697723073 | 0.758747698385796 | NA | NM_172670,NM_001166633 |
| Gm41 | 0 | NA | NA | NA | NA | NA | | Gypa | 1.368677131 | -0.00734037064467297 | 0.022631033781944 | -0.324349771884015 | 0.745673242089403 | NA | NM_010369 |
| Gm4133 | 0 | NA | NA | NA | NA | NA | | Gys2 | 2.767438859 | 0.0401561963569912 | 0.0387474259135665 | 1.03635778145798 | 0.30003525873549 | NA | NM_145572 |
| Gm4141 | 0 | NA | NA | NA | NA | NA | | Gzma | 0.763547837 | 0.00600973649828535 | 0.0141975776579579 | 0.423293088657051 | 0.672081436193138 | NA | NM_010370 |
| Gm4172 | 0 | NA | NA | NA | NA | NA | | Gzmb | 1.142123786 | -0.0278550429745613 | 0.0214429084263202 | -1.29903287468086 | 0.193932647387751 | NA | NM_013542 |
| Gm4175 | 0 | NA | NA | NA | NA | NA | | Gzmc | 0.233836299 | 0.00743372775021133 | 0.0101683766597817 | 0.731063374118848 | 0.464740444385902 | NA | NM_010371 |
| Gm4177 | 0 | NA | NA | NA | NA | NA | | Gzmd | 0 | NA | NA | NA | NA | NA | NM_010372 |
| Gm4179 | 0 | NA | NA | NA | NA | NA | | Gzme | 0 | NA | NA | NA | NA | NA | NM_010373 |
| Gm4187 | 0 | NA | NA | NA | NA | NA | | Gzmf | 0 | NA | NA | NA | NA | NA | NM_010374 |
| Gm4201 | 0 | NA | NA | NA | NA | NA | | Gzmg | 0.127687691 | 0.00542298404407807 | 0.00869724871942359 | 0.62352868349814 | 0.532937159177571 | NA | NM_010375 |
| Gm4214 | 0 | NA | NA | NA | NA | NA | | Gzmk | 3.16118579 | -0.0141715542117707 | 0.0424335257836371 | -0.333970697698549 | 0.738401670251633 | NA | NM_008196 |
| Gm4216 | 0 | NA | NA | NA | NA | NA | | Gzmn | 0 | NA | NA | NA | NA | NA | NM_153052 |
| Gm4220 | 0 | NA | NA | NA | NA | NA | | H19 | 8.285261359 | 0.0199973681010476 | 0.0517207400158486 | 0.38664118291656 | 0.699021865031459 | NA | NR_001592 |
| Gm4224 | 0 | NA | NA | NA | NA | NA | | H1fnt | 0 | NA | NA | NA | NA | NA | NM_027304 |
| Gm4251 | 0 | NA | NA | NA | NA | NA | | H1foo | 0 | NA | NA | NA | NA | NA | NM_138311 |
| Gm4262 | 0.487806363 | -0.025218575 | 0.039384805 | -0.640312313 | 0.521969578 | NA | | H2-Aa | 122.9707825 | -0.100363112214454 | 0.0455014617020422 | -2.20571182683455 | NA | NA | NM_010378 |
| Gm4265 | 0.371083244 | -0.002928723 | 0.041579259 | -0.070437114 | 0.943845752 | NA | | H2-Ab1 | 96.19247695 | -0.0842756534160819 | 0.04060080592789 | -2.07571380641462 | NA | NA | NM_207105 |
| Gm4278 | 0 | NA | NA | NA | NA | NA | | H2-DMb2 | 15.73232806 | -0.0133998298968647 | 0.0703994321741827 | -0.190340028080208 | 0.849042689078015 | NA | NM_010388 |
| Gm428 | 0 | NA | NA | NA | NA | NA | | H2-Ea-ps | 0.097165704 | -0.00637828455464458 | 0.00848781691535724 | -0.75146349388194 | 0.452373763047372 | NA | NM_010381 |
| Gm4297 | 0 | NA | NA | NA | NA | NA | | H2-Eb1 | 106.1703908 | -0.0978102059917765 | 0.0505944587322878 | -1.93321973280361 | NA | NA | NM_010382 |
| Gm4301 | 0 | NA | NA | NA | NA | NA | | H2-Eb2 | 0.237667581 | -0.00136938695604711 | 0.00989612066541175 | -0.13837613771559 | 0.889943154921195 | NA | NM_001033978 |
| Gm4302 | 0 | NA | NA | NA | NA | NA | | H2-L | 0 | NA | NA | NA | NA | NA | NM_001267808 |
| Gm4303 | 0 | NA | NA | NA | NA | NA | | H2-M1 | 0 | NA | NA | NA | NA | NA | NM_177636 |
| Gm4305 | 0 | NA | NA | NA | NA | NA | | H2-M10.1 | 0 | NA | NA | NA | NA | NA | NM_013544 |
| Gm4307 | 0 | NA | NA | NA | NA | NA | | H2-M10.2 | 0 | NA | NA | NA | NA | NA | NM_177923 |
| Gm4312 | 0 | NA | NA | NA | NA | NA | | H2-M10.3 | 0.120209049 | -0.00674972139751468 | 0.00863521114606269 | -0.781650996523957 | 0.434419707908266 | NA | NM_201608 |
| Gm4340 | 0 | NA | NA | NA | NA | NA | | H2-M10.4 | 0 | NA | NA | NA | NA | NA | NM_177634 |
| Gm4345 | 0 | NA | NA | NA | NA | NA | | H2-M10.5 | 0 | NA | NA | NA | NA | NA | NM_177637 |
| Gm436 | 0 | NA | NA | NA | NA | NA | | H2-M10.6 | 0 | NA | NA | NA | NA | NA | NM_201611 |
| Gm438 | 0 | NA | NA | NA | NA | NA | | H2-M11 | 0.108766301 | 0.00519171381361946 | 0.00856003371502575 | 0.606506234257728 | 0.544178643801544 | NA | NM_177635 |
| Gm44 | 0 | NA | NA | NA | NA | NA | | H2-M2 | 1.035380625 | 0.0136129200795333 | 0.0247588310645384 | 0.5498207909755 | 0.582442296901335 | NA | NM_008204 |
| Gm4461 | 2.006666071 | 0.020792586 | 0.103606388 | 0.200688264 | 0.840942337 | NA | | H2-M5 | 20.93562416 | -0.0983529827213608 | 0.0775333023419186 | -1.26852564963154 | 0.204610296862967 | NA | NM_001115075 |
| Gm4477 | 1.064432019 | 0.028665342 | 0.074319474 | 0.385704324 | 0.69971566 | NA | | H2-M9 | 0.09572496 | 0.00498255289994511 | 0.00847999972823692 | 0.587565219295241 | 0.556824162160132 | NA | NM_008205 |
| Gm4489 | 0.479444338 | 0.008662674 | 0.055255961 | 0.156773568 | 0.875423305 | NA | | H2-Q10 | 3.534138123 | 0.0396821421964533 | 0.042124201679249 | 0.942027162879181 | 0.346178733288136 | NA | NM_010391 |
| Gm4498 | 0 | NA | NA | NA | NA | NA | | H2-Q5 | 20.2984589 | -0.131607808058521 | 0.0776393256371219 | -1.69511786686095 | 0.0900530635031841 | NA | NR_051981 |
| Gm4541 | 0 | NA | NA | NA | NA | NA | | H2-Q6 | 0 | NA | NA | NA | NA | NA | NM_207648 |
| Gm4559 | 0.094392817 | 0.017895942 | 0.027818137 | 0.643319236 | 0.520016977 | NA | | H2-Q7 | 0 | NA | NA | NA | NA | NA | NM_001198560,NM_010394,NM_001198561 |
| Gm4566 | 0.38134816 | 0.013258311 | 0.036322589 | 0.365015595 | 0.715099765 | NA | | H2-Q8 | 0 | NA | NA | NA | NA | NA | NM_023124 |
| Gm4567 | 0 | NA | NA | NA | NA | NA | | H2-Q9 | 0 | NA | NA | NA | NA | NA | NM_001201460 |
| Gm4598 | 3.777051371 | -0.147144992 | 0.120949555 | -1.216581515 | 0.223763479 | NA | | H2-T22 | 1.762293184 | 0.0326042815493356 | 0.0317875781735851 | 1.02569253219892 | 0.305036535497908 | NA | NM_010397 |
| Gm4710 | 0.283178451 | 0.028390488 | 0.033368195 | 0.850824814 | 0.394866675 | NA | | H2-T3 | 0.23403553 | -0.000407313579519725 | 0.00983856423584417 | -0.0413996971261099 | 0.966977254250032 | NA | NM_008208 |
| Gm4719 | 0 | NA | NA | NA | NA | NA | | H2-T9 | 0 | NA | NA | NA | NA | NA | NM_010399 |
| Gm4724 | 0 | NA | NA | NA | NA | NA | | H2afb1 | 0.120209049 | -0.00674972139751468 | 0.00863521114606269 | -0.781650996523957 | 0.434419707908266 | NA | NM_026627 |
| Gm4736 | 0 | NA | NA | NA | NA | NA | | H2afy3 | 0.106437839 | -0.00653347234140708 | 0.00854389509723638 | -0.764694822098226 | 0.444453290197145 | NA | NR_003523 |
| Gm4745 | 0 | NA | NA | NA | NA | NA | | H2bfm | 0 | NA | NA | NA | NA | NA | NM_027067 |
| Gm4759 | 2.439005598 | 0.006430841 | 0.091774583 | 0.07007213 | 0.944136249 | NA | | H60c | 0.631545537 | 0.00489766623125536 | 0.0171330903077031 | 0.285860060461677 | 0.774985304925285 | NA | NM_001204916 |
| Gm4763 | 0.227114243 | -0.027513412 | 0.032207244 | -0.854261616 | 0.39296005 | NA | | Haao | 8.508164124 | -0.00516146304522586 | 0.0572006584184159 | -0.0902343292531771 | 0.92810100478952 | NA | NM_025325 |
| Gm4776 | 0.098353386 | -0.020212831 | 0.02786466 | -0.725393061 | 0.468210929 | NA | | Habp2 | 0.097165704 | -0.00637828455464458 | 0.00848781691535724 | -0.75146349388194 | 0.452373763047372 | NA | NM_146101 |
| Gm4788 | 3.340403103 | -0.141652066 | 0.126845024 | -1.116733335 | 0.26410836 | NA | | Hal | 1.272710417 | -0.00269688510211617 | 0.0261913301646256 | -0.102968619202038 | 0.917987876785079 | NA | NM_010401 |
| Gm4791 | 0.285336609 | 0.031742875 | 0.03528833 | 0.899528977 | 0.368370968 | NA | | Hamp | 0.108766301 | 0.00519171381361946 | 0.00856003371502575 | 0.606506234257728 | 0.544178643801544 | NA | NM_032541 |
| Gm4792 | 0.309710804 | -0.014226733 | 0.039782345 | -0.357614233 | 0.720632025 | NA | | Hamp2 | 0.12890128 | -0.0068640067066208 | 0.00870804083610955 | -0.788237771940376 | 0.430557639008166 | NA | NM_183257 |
| Gm4794 | 0 | NA | NA | NA | NA | NA | | Hand1 | 0.106148608 | 0.00515554063321497 | 0.00854200771766808 | 0.603551390213729 | 0.546141939141592 | NA | NM_008213 |
| Gm4814 | 3.46518233 | 0.066414662 | 0.130114548 | 0.510432251 | 0.609748667 | NA | | Hand2 | 0 | NA | NA | NA | NA | NA | NM_010402 |
| Gm4827 | 2.326590804 | -0.057497865 | 0.096373257 | -0.596616389 | 0.550763527 | NA | | Hao1 | 3.93938632 | 0.050835103798083 | 0.0439067891763032 | 1.1577959753322 | 0.246947306192677 | NA | NM_010403 |
| Gm4832 | 0.296373155 | 0.009272162 | 0.033639931 | 0.275629621 | 0.782832556 | NA | | Hao2 | 0 | NA | NA | NA | NA | NA | NM_019545 |
| Gm4836 | 0 | NA | NA | NA | NA | NA | | Has2 | 12.96911832 | -0.0126885033874063 | 0.0673809379107271 | -0.188309984705426 | 0.8506336587111 | NA | NM_008216 |
| Gm4841 | 9.063191654 | 0.269685464 | 0.111962688 | 2.408708371 | NA | NA | | Has2as | 2.026683379 | -0.0428695506352461 | 0.0318615215366282 | -1.34549602679718 | 0.178465103525682 | NA | NR_002874 |
| Gm4846 | 0 | NA | NA | NA | NA | NA | | Havcr1 | 0.217374753 | -0.00904456656736547 | 0.00999660569474087 | -0.90476376117583 | 0.365590553873005 | NA | NM_001166632,NM_001166631,NM_134248 |
| Gm4847 | 0.927575411 | 0.00625276 | 0.069430007 | 0.090058465 | 0.928240755 | NA | | Hba-a1 | 0 | NA | NA | NA | NA | NA | NM_008218 |
| Gm4850 | 0 | NA | NA | NA | NA | NA | | Hba-a2 | 0 | NA | NA | NA | NA | NA | NM_001083955 |
| Gm4858 | 0 | NA | NA | NA | NA | NA | | Hba-x | 0 | NA | NA | NA | NA | NA | NM_010405 |
| Gm4861 | 1.64668743 | 0.047849013 | 0.068404387 | 0.699502111 | 0.484238294 | NA | | Hbb-b1 | 0 | NA | NA | NA | NA | NA | NM_008220 |
| Gm4871 | 0 | NA | NA | NA | NA | NA | | Hbb-b2 | 0 | NA | NA | NA | NA | NA | NM_016956 |
| Gm4872 | 0 | NA | NA | NA | NA | NA | | Hbb-bh1 | 0 | NA | NA | NA | NA | NA | NM_008219 |
| Gm4884 | 0 | NA | NA | NA | NA | NA | | Hbb-bh2 | 0 | NA | NA | NA | NA | NA | NM_001127686 |
| Gm4894 | 0.409448625 | -0.030061717 | 0.043829456 | -0.685879317 | 0.492789214 | NA | | Hbb-y | 1.266060517 | -0.0164513592252827 | 0.0267476182869241 | -0.615058845569258 | 0.538515859292922 | NA | NM_008221 |
| Gm4906 | 0 | NA | NA | NA | NA | NA | | Hbq1a | 14.55619515 | -0.0115520114654523 | 0.0745982937206309 | -0.154856242539733 | 0.87693465081247 | NA | NM_175000 |
| Gm4907 | 0 | NA | NA | NA | NA | NA | | Hbq1b | 4.426962402 | 0.0122771413376169 | 0.0473410332814492 | 0.259334038288254 | 0.795377518748198 | NA | NM_001033981 |
| Gm4922 | 0 | NA | NA | NA | NA | NA | | Hc | 10.69454291 | 0.00746938816378054 | 0.0620852688547391 | 0.12030854180976 | 0.904238737876668 | NA | NM_010406 |
| Gm4937 | 0 | NA | NA | NA | NA | NA | | Hcrt | 21.47296838 | 0.0266110661111678 | 0.021465924640888 | 1.23968878845682 | NA | NA | NM_010410 |
| Gm4952 | 5.288185368 | 0.333357223 | 0.146925878 | 2.268880256 | 0.023275609 | NA | | Hcst | 12.81806759 | -0.0541670963285187 | 0.0700606509645094 | -0.773145775593179 | 0.439436112857494 | NA | NM_011827 |
| Gm4956 | 0.188785634 | 0.025025111 | 0.031461603 | 0.79541754 | 0.42637066 | NA | | Hdgfl1 | 0.223412651 | 0.00718831964189475 | 0.00999993766105603 | 0.718836445339964 | 0.472241699104885 | NA | NM_008232 |
| Gm4961 | 5.368778638 | 0.018221312 | 0.147291842 | 0.123708901 | 0.901545764 | NA | | Hdhd1a | 0.106148608 | 0.00515554063321497 | 0.00854200771766808 | 0.603551390213729 | 0.546141939141592 | NA | NM_026108 |
| Gm4971 | 0.215050351 | -0.027006021 | 0.031907566 | -0.846382992 | 0.397339123 | NA | | Helt | 4.770978134 | -0.0281954823194092 | 0.0454951118694373 | -0.619747510464973 | 0.535424030958458 | NA | NM_173789 |
| Gm4975 | 0.561778158 | 0.022427986 | 0.058194018 | 0.385400203 | 0.699940932 | NA | | Hemgn | 0.342818636 | 0.00260767036841674 | 0.0127977035531239 | 0.203760804240556 | 0.83854042388457 | NA | NM_053149 |
| Gm4981 | 0.844193991 | 0.068484887 | 0.07127605 | 0.960840099 | 0.336632573 | NA | | Hemt1 | 0 | NA | NA | NA | NA | NA | NM_010416 |
| Gm4984 | 0.283178451 | 0.028390488 | 0.033368195 | 0.850824814 | 0.394866675 | NA | | Hepacam2 | 17.986423 | 0.00820384367120136 | 0.0749652174903253 | 0.109435334757218 | 0.9128572089547 | NA | NM_178899 |
| Gm5 | 5.809474054 | 0.011902801 | 0.145173819 | 0.081989999 | 0.934654667 | NA | | Hephl1 | 0.346566706 | -0.0051482901046312 | 0.0114161257772475 | -0.450966484172047 | 0.652013705001052 | NA | NM_001164797 |
| Gm5039 | 0 | NA | NA | NA | NA | NA | | Hes2 | 0.455333007 | -0.00242452963031724 | 0.0139658608669154 | -0.173604022940029 | 0.862176670184748 | NA | NM_008236 |
| Gm5065 | 0.105923158 | 0.018524693 | 0.027977518 | 0.662127842 | 0.507889294 | NA | | Hes3 | 19.36480714 | 0.00486165803408813 | 0.0820114684863071 | 0.059280221703381 | 0.95272891420112 | NA | NM_008237 |
| Gm5071 | 0 | NA | NA | NA | NA | NA | | Hes7 | 15.3203102 | -0.126554046509009 | 0.0634620022546254 | -1.99417040138826 | 0.0461334356117353 | NA | NM_033041 |
| Gm5072 | 0 | NA | NA | NA | NA | NA | | Hesx1 | 0.12890128 | -0.0068640067066208 | 0.00870804083610955 | -0.788237771940376 | 0.430557639008166 | NA | NM_010420 |
| Gm5082 | 0.096071043 | -0.020098795 | 0.027825948 | -0.722304036 | 0.470107572 | NA | | Hgd | 1.960486095 | 0.0186027130035286 | 0.0341233659911837 | 0.545160550935535 | 0.585643100663138 | NA | NM_013547,NR_027778 |
| Gm5084 | 2.61041684 | 0.186756539 | 0.111605932 | 1.673356747 | 0.094257087 | NA | | Hgfac | 6.076266127 | -0.0502753832610736 | 0.0476532116998748 | -1.05502612452889 | 0.291413376827451 | NA | NM_019447 |
| Gm5087 | 1.871052239 | 0.05152163 | 0.100714692 | 0.511560222 | 0.608958831 | NA | | Hhla1 | 0.903806998 | -0.0386023359078386 | 0.0229724239618271 | -1.68037713268672 | 0.0928839623221594 | NA | NM_001145096 |
| Gm5088 | 0.185906812 | 0.024895454 | 0.031385107 | 0.793225068 | 0.427646705 | NA | | Higd1c | 0 | NA | NA | NA | NA | NA | NM_001002900 |
| Gm5091 | 0 | NA | NA | NA | NA | NA | | Hils1 | 0 | NA | NA | NA | NA | NA | NM_018792 |
| Gm5095 | 0.116696964 | -0.020477583 | 0.027954224 | -0.732539847 | 0.463839131 | NA | | Hipk4 | 5.714803124 | -0.0415715046337251 | 0.0461024418683728 | -0.901720233223567 | 0.367205502064654 | NA | NM_001033315 |
| Gm5105 | 7.748551166 | -0.000784878 | 0.133317147 | -0.005887296 | 0.995302644 | NA | | Hist1h1a | 0.660023499 | -0.00368992069631123 | 0.0174374856096879 | -0.211608529974138 | 0.832412454074931 | NA | NM_030609 |
| Gm5108 | 0.183387883 | -0.001444816 | 0.031390672 | -0.046026918 | 0.963288796 | NA | | Hist1h1b | 1.847495584 | -0.00301814880137018 | 0.0289065369235306 | -0.104410597829633 | 0.91684351279948 | NA | NM_020034 |
| Gm5114 | 0 | NA | NA | NA | NA | NA | | Hist1h1d | 3.194053724 | 0.00167111898398685 | 0.0298030113170427 | 0.0560721521127375 | 0.955284328419739 | NA | NM_145713 |
| Gm5122 | 0.913584581 | 0.006419397 | 0.078297819 | 0.081986924 | 0.934657112 | NA | | Hist1h1t | 0 | NA | NA | NA | NA | NA | NM_010377 |
| Gm5127 | 5.072092974 | -0.017973835 | 0.13035956 | -0.137878914 | 0.890336116 | NA | | Hist1h2aa | 0 | NA | NA | NA | NA | NA | NM_175658 |
| Gm5128 | 0 | NA | NA | NA | NA | NA | | Hist1h2ab | 0.097165704 | -0.00637828455464458 | 0.00848781691535724 | -0.75146349388194 | 0.452373763047372 | NA | NM_175660 |
| Gm5129 | 0.266310378 | 0.0407635 | 0.040043874 | 1.017970943 | 0.308691765 | NA | | Hist1h2ad | 1.924128283 | -0.00226403559348797 | 0.0245173052726569 | -0.0923443897406193 | 0.926424421101589 | NA | NM_178188 |
| Gm5132 | 0.215706849 | -0.002035226 | 0.0319478 | -0.063704747 | 0.949205325 | NA | | Hist1h2ae | 0.120209049 | -0.00674972139751468 | 0.00863521114606269 | -0.781650996523957 | 0.434419707908266 | NA | NM_178187 |
| Gm5134 | 2.979985258 | 0.011748176 | 0.101498697 | 0.11574706 | 0.907853008 | NA | | Hist1h2af | 0.113843286 | 0.00525947738444006 | 0.00859368002043176 | 0.612016897526494 | 0.540526577873086 | NA | NM_175661 |
| Gm5136 | 0.096071043 | -0.020098795 | 0.027825948 | -0.722304036 | 0.470107572 | NA | | Hist1h2ag | 0 | NA | NA | NA | NA | NA | NM_178186 |
| Gm5142 | 0 | NA | NA | NA | NA | NA | | Hist1h2ah | 0 | NA | NA | NA | NA | NA | NM_175659 |
| Gm5150 | 0.079983654 | 0.016740032 | 0.027674178 | 0.604897175 | 0.54524732 | NA | | Hist1h2ai | 0.113843286 | 0.00525947738444006 | 0.00859368002043176 | 0.612016897526494 | 0.540526577873086 | NA | NM_178182 |
| Gm5166 | 0 | NA | NA | NA | NA | NA | | Hist1h2ak | 0.113843286 | 0.00525947738444006 | 0.00859368002043176 | 0.612016897526494 | 0.540526577873086 | NA | NM_178183 |
| Gm5168 | 0 | NA | NA | NA | NA | NA | | Hist1h2an | 0 | NA | NA | NA | NA | NA | NM_178184 |
| Gm5169 | 0 | NA | NA | NA | NA | NA | | Hist1h2ao | 0 | NA | NA | NA | NA | NA | NM_001177544 |
| Gm5177 | 1.83737654 | 0.049376459 | 0.100508986 | 0.491264122 | 0.623239652 | NA | | Hist1h2ap | 0 | NA | NA | NA | NA | NA | NM_178185 |
| Gm525 | 1.273412036 | -0.047122374 | 0.076601304 | -0.615164125 | 0.538446337 | NA | | Hist1h2ba | 0 | NA | NA | NA | NA | NA | NM_175663 |
| Gm5294 | 0.178993538 | 0.024685672 | 0.031260828 | 0.789667872 | 0.429721759 | NA | | Hist1h2bb | 0.319043516 | -0.00957618023178583 | 0.0102890835921761 | -0.930712647632453 | 0.352002225550712 | NA | NM_175664 |
| Gm53 | 0 | NA | NA | NA | NA | NA | | Hist1h2bf | 0 | NA | NA | NA | NA | NA | NM_178195 |
| Gm5334 | 0 | NA | NA | NA | NA | NA | | Hist1h2bg | 10.16125153 | -0.0664974717249454 | 0.0664911967869335 | -1.00009437246305 | 0.317264839271426 | NA | NM_178196 |
| Gm5346 | 0.116696964 | -0.020477583 | 0.027954224 | -0.732539847 | 0.463839131 | NA | | Hist1h2bh | 1.658423627 | -0.00898891016483143 | 0.023555224064638 | -0.381610047103136 | 0.702750630743902 | NA | NM_178197 |
| Gm5347 | 0 | NA | NA | NA | NA | NA | | Hist1h2bj | 0.677809771 | 0.0107739313623017 | 0.0159463824365314 | 0.675634828474939 | 0.499272523024431 | NA | NM_178198 |
| Gm5382 | 0 | NA | NA | NA | NA | NA | | Hist1h2bk | 0 | NA | NA | NA | NA | NA | NM_175665 |
| Gm5409 | 0 | NA | NA | NA | NA | NA | | Hist1h2bl | 0 | NA | NA | NA | NA | NA | NM_178199 |
| Gm5414 | 0 | NA | NA | NA | NA | NA | | Hist1h2bm | 0 | NA | NA | NA | NA | NA | NM_178200 |
| Gm5415 | 0.178993538 | 0.024685672 | 0.031260828 | 0.789667872 | 0.429721759 | NA | | Hist1h2bn | 2.378402104 | -0.0205387543785259 | 0.0378568863823225 | -0.542536810109049 | 0.587448756458835 | NA | NM_178201 |
| Gm5416 | 0 | NA | NA | NA | NA | NA | | Hist1h2bp | 0.223412651 | 0.00718831964189475 | 0.00999993766105603 | 0.718836445339964 | 0.472241699104885 | NA | NM_178202 |
| Gm5420 | 0.098353386 | -0.020212831 | 0.02786466 | -0.725393061 | 0.468210929 | NA | | Hist1h2bq | 0 | NA | NA | NA | NA | NA | NM_001097979 |
| Gm5424 | 0.525806514 | 0.006325424 | 0.04539272 | 0.139348856 | 0.889174485 | NA | | Hist1h2br | 0 | NA | NA | NA | NA | NA | NM_001110555 |
| Gm5434 | 0 | NA | NA | NA | NA | NA | | Hist1h3a | 0 | NA | NA | NA | NA | NA | NM_013550 |
| Gm5441 | 0.094392817 | 0.017895942 | 0.027818137 | 0.643319236 | 0.520016977 | NA | | Hist1h3b | 0.438854104 | -0.00212522163852104 | 0.0138327359250247 | -0.153637114887469 | 0.87789587090842 | NA | NM_178203 |
| Gm5458 | 2.847557002 | 0.108818506 | 0.112722406 | 0.965367134 | 0.334360957 | NA | | Hist1h3c | 0.240418098 | -0.0088029805258837 | 0.00986414519853158 | -0.892422034419582 | 0.372166765031741 | NA | NM_175653 |
| Gm5460 | 0 | NA | NA | NA | NA | NA | | Hist1h3d | 2.104481424 | -0.00524843840777111 | 0.0306179995223253 | -0.171416764310293 | 0.863896073856886 | NA | NM_178204 |
| Gm5464 | 5.830661967 | -0.028250892 | 0.142275042 | -0.198565339 | 0.842602769 | NA | | Hist1h3e | 0.192890664 | -0.000903864895041124 | 0.00963528282151475 | -0.093807821917056 | 0.92526181809679 | NA | NM_178205 |
| Gm5475 | 0.650129455 | 0.071216437 | 0.063544202 | 1.120738548 | 0.262399169 | NA | | Hist1h3f | 0.364141684 | 0.0106460894690162 | 0.0121537141469398 | 0.875953584254473 | 0.381055267460423 | NA | NM_013548 |
| Gm5476 | 0 | NA | NA | NA | NA | NA | | Hist1h3g | 0.603271003 | 0.00961714697636968 | 0.0155766124128925 | 0.617409403369998 | 0.536964721194581 | NA | NM_145073 |
| Gm5477 | 0 | NA | NA | NA | NA | NA | | Hist1h3h | 0 | NA | NA | NA | NA | NA | NM_178206 |
| Gm5478 | 0 | NA | NA | NA | NA | NA | | Hist1h3i | 0.916997476 | 0.0114933310665482 | 0.0189930988650839 | 0.605131955990452 | 0.5450913227602 | NA | NM_178207 |
| Gm5480 | 2.052333237 | 0.069070467 | 0.078997461 | 0.874337808 | 0.381934315 | NA | | Hist1h4a | 0.205932006 | -0.000576954122277892 | 0.00970424421944593 | -0.0594537925088239 | 0.952590668569529 | NA | NM_178192 |
| Gm5483 | 0.176054697 | -0.002190495 | 0.031239146 | -0.070120197 | 0.944097991 | NA | | Hist1h4b | 0.589493697 | -0.0173320451977439 | 0.0174706335897436 | -0.992067351691177 | 0.321164671490061 | NA | NM_178193 |
| Gm5485 | 0 | NA | NA | NA | NA | NA | | Hist1h4c | 0.864229324 | -0.00829133471530604 | 0.0186793554387384 | -0.44387691762163 | 0.657131573719512 | NA | NM_178208 |
| Gm5494 | 0 | NA | NA | NA | NA | NA | | Hist1h4d | 1.970355761 | -0.0168987914710144 | 0.0303742302190961 | -0.556352913279436 | 0.577969621384924 | NA | NM_175654 |
| Gm5512 | 0.779778063 | -0.006753362 | 0.062007213 | -0.108912518 | 0.913271878 | NA | | Hist1h4f | 0.332414823 | -0.0166094830116705 | 0.0135191688281235 | -1.22858758721308 | 0.219226471765921 | NA | NM_175655 |
| Gm5523 | 2.538534456 | -0.010241435 | 0.113260938 | -0.090423363 | 0.927950792 | NA | | Hist1h4h | 11.056691 | -0.0322745875535834 | 0.0692545644287729 | -0.466028309033367 | 0.641195237382983 | NA | NM_153173 |
| Gm5531 | 1.678235804 | 0.061666331 | 0.095355726 | 0.646697731 | 0.517827592 | NA | | Hist1h4i | 21.00121644 | -0.0178610320215568 | 0.0743843436499226 | -0.240118163919235 | 0.810238653306725 | NA | NM_175656 |
| Gm5538 | 0.105923158 | 0.018524693 | 0.027977518 | 0.662127842 | 0.507889294 | NA | | Hist1h4j | 0.723986006 | -0.0241577472732078 | 0.0189778213271058 | -1.27294629119011 | 0.203037101161474 | NA | NM_178210 |
| Gm5544 | 1.970400815 | -0.026780741 | 0.093228206 | -0.287260072 | 0.773913193 | NA | | Hist1h4k | 10.32971507 | 0.0234736885527303 | 0.064974980573535 | 0.361272729064753 | 0.717895578040459 | NA | NM_178211 |
| Gm5547 | 0.644719697 | 0.02588267 | 0.053786651 | 0.481209923 | 0.630367309 | NA | | Hist1h4m | 0.257802559 | -0.00892927589713349 | 0.00993479775307735 | -0.898787888698349 | 0.368765652383134 | NA | NM_001195421 |
| Gm5549 | 0 | NA | NA | NA | NA | NA | | Hist1h4n | 0 | NA | NA | NA | NA | NA | NM_175657 |
| Gm5591 | 0 | NA | NA | NA | NA | NA | | Hist2h2aa1 | 0 | NA | NA | NA | NA | NA | NM_013549 |
| Gm5592 | 0 | NA | NA | NA | NA | NA | | Hist2h2aa2 | 0 | NA | NA | NA | NA | NA | NM_178212 |
| Gm5615 | 0 | NA | NA | NA | NA | NA | | Hist2h2ab | 2.007974036 | -0.0116439359706569 | 0.0339680363845049 | -0.342790964978137 | 0.731755723506476 | NA | NM_178213 |
| Gm5622 | 2.363747454 | 0.040331076 | 0.100621713 | 0.400818821 | 0.688553521 | NA | | Hist2h2ac | 0.120209049 | -0.00674972139751468 | 0.00863521114606269 | -0.781650996523957 | 0.434419707908266 | NA | NM_175662 |
| Gm5627 | 0 | NA | NA | NA | NA | NA | | Hist2h2bb | 0.201873568 | 0.00673381845182595 | 0.00968011632508873 | 0.69563404257585 | 0.486658041207307 | NA | NM_175666 |
| Gm5634 | 0 | NA | NA | NA | NA | NA | | Hist2h3b | 0 | NA | NA | NA | NA | NA | NM_178215 |
| Gm5635 | 0 | NA | NA | NA | NA | NA | | Hist2h4 | 4.663266761 | -0.0697028171438835 | 0.048870722615803 | -1.42626941884719 | 0.153790588385471 | NA | NM_033596 |

| | | | | | | |
|---|---|---|---|---|---|---|
| Gm5640 | 2.475794889 | -0.075222178 | 0.106993061 | -0.703056604 | 0.482020475 | NA |
| Gm5643 | 6.117196228 | -0.192416509 | 0.125489112 | -1.533332305 | 0.125193999 | NA |
| Gm5662 | 0 | NA | NA | NA | NA | NA |
| Gm5712 | 0 | NA | NA | NA | NA | NA |
| Gm572 | 2.4021855 | -0.043518809 | 0.112342109 | -0.387377537 | 0.698476731 | NA |
| Gm5725 | 0 | NA | NA | NA | NA | NA |
| Gm5726 | 0 | NA | NA | NA | NA | NA |
| Gm5728 | 0 | NA | NA | NA | NA | NA |
| Gm5741 | 1.964853267 | 0.273286686 | 0.102867309 | 2.656691307 | 0.007891166 | NA |
| Gm5766 | 0.096071043 | -0.020098795 | 0.027825948 | -0.722304036 | 0.470107572 | NA |
| Gm5771 | 0.159967308 | 0.0240345 | 0.030870969 | 0.778547006 | 0.436246606 | NA |
| Gm5779 | 0.215433205 | -0.027029776 | 0.031922004 | -0.84674434 | 0.397137638 | NA |
| Gm5795 | 0 | NA | NA | NA | NA | NA |
| Gm5796 | 4.550372248 | -0.03313484 | 0.138826635 | -0.238677831 | 0.811355408 | NA |
| Gm5797 | 1.07331165 | -0.036005461 | 0.071304466 | -0.5049538 | 0.613591286 | NA |
| Gm5800 | 0 | NA | NA | NA | NA | NA |
| Gm5801 | 0.107716603 | -0.020477583 | 0.027954224 | -0.732539849 | 0.46383913 | NA |
| Gm5803 | 1.760068487 | 0.136834324 | 0.097766945 | 1.399597007 | 0.161634031 | NA |
| Gm5833 | 0 | NA | NA | NA | NA | NA |
| Gm5868 | 1.537306715 | -0.052823434 | 0.069287568 | -0.762379681 | 0.44583343 | NA |
| Gm5878 | 0.680997126 | -0.019373436 | 0.059986914 | -0.322961042 | 0.746724746 | NA |
| Gm5885 | 0 | NA | NA | NA | NA | NA |
| Gm5886 | 0 | NA | NA | NA | NA | NA |
| Gm5891 | 0 | NA | NA | NA | NA | NA |
| Gm590 | 0.318806384 | -0.016982193 | 0.040116674 | -0.423320068 | 0.672061754 | NA |
| Gm5901 | 0.4546761 | 0.062003256 | 0.049007249 | 1.265185398 | 0.205804876 | NA |
| Gm5916 | 0 | NA | NA | NA | NA | NA |
| Gm5925 | 0 | NA | NA | NA | NA | NA |
| Gm5934 | 0 | NA | NA | NA | NA | NA |
| Gm5935 | 0 | NA | NA | NA | NA | NA |
| Gm5936 | 1.368957027 | -0.018483132 | 0.083396809 | -0.221628774 | 0.824602881 | NA |
| Gm5938 | 0 | NA | NA | NA | NA | NA |
| Gm5941 | 0 | NA | NA | NA | NA | NA |
| Gm595 | 0 | NA | NA | NA | NA | NA |
| Gm597 | 0.199381295 | -0.004055078 | 0.031639813 | -0.128163771 | 0.89801937 | NA |
| Gm6026 | 0 | NA | NA | NA | NA | NA |
| Gm6034 | 1.095546289 | 0.039378448 | 0.075289599 | 0.523026405 | 0.600955879 | NA |
| Gm6040 | 0 | NA | NA | NA | NA | NA |
| Gm6042 | 0 | NA | NA | NA | NA | NA |
| Gm6084 | 0 | NA | NA | NA | NA | NA |
| Gm6086 | 3.953030292 | -0.013715926 | 0.120487487 | -0.113836931 | 0.909367063 | NA |
| Gm6116 | 0 | NA | NA | NA | NA | NA |
| Gm6121 | 0 | NA | NA | NA | NA | NA |
| Gm614 | 0.61296065 | 0.018665908 | 0.053631683 | 0.348038835 | 0.727811019 | NA |
| Gm6150 | 1.559676214 | 0.015996788 | 0.094049439 | 0.170089135 | 0.864940038 | NA |
| Gm6164 | 0 | NA | NA | NA | NA | NA |
| Gm6194 | 0.521672338 | 0.008656229 | 0.045342019 | 0.190909656 | 0.848596375 | NA |
| Gm6213 | 0.39969727 | 0.025482209 | 0.048473027 | 0.525698742 | 0.599097538 | NA |
| Gm6249 | 3.350400091 | -0.001580262 | 0.125806947 | -0.012561006 | 0.989978031 | NA |
| Gm6251 | 0.506154836 | -0.058553951 | 0.057703036 | -1.014746452 | 0.310226705 | NA |
| Gm6260 | 1.226944435 | -0.07725345 | 0.082825485 | -0.9327256 | 0.350961664 | NA |
| Gm6268 | 0 | NA | NA | NA | NA | NA |
| Gm6289 | 0.105923158 | 0.018524693 | 0.027977518 | 0.662127842 | 0.507889294 | NA |
| Gm6300 | 0.999400785 | -0.066967927 | 0.073744079 | -0.908112592 | 0.363818738 | NA |
| Gm6307 | 0.08731684 | 0.017365056 | 0.027733412 | 0.626142072 | 0.531221759 | NA |
| Gm6329 | 0 | NA | NA | NA | NA | NA |
| Gm6367 | 0 | NA | NA | NA | NA | NA |
| Gm6370 | 0 | NA | NA | NA | NA | NA |
| Gm6377 | 0.621055902 | 0.040831999 | 0.04851525 | 0.841632255 | 0.399993829 | NA |
| Gm6406 | 0.08731684 | 0.017365056 | 0.027733412 | 0.626142072 | 0.531221759 | NA |
| Gm6408 | 1.368388281 | 0.045329629 | 0.082036148 | 0.552556767 | 0.580566953 | NA |
| Gm6416 | 0.396533849 | -0.033875382 | 0.048794365 | -0.694247834 | 0.4875268 | NA |
| Gm6432 | 0 | NA | NA | NA | NA | NA |
| Gm6455 | 0 | NA | NA | NA | NA | NA |
| Gm6460 | 0 | NA | NA | NA | NA | NA |
| Gm648 | 1.514310546 | -0.053729152 | 0.086623841 | -0.620258251 | 0.535087777 | NA |
| Gm6484 | 0.107716603 | -0.020477583 | 0.027954224 | -0.732539849 | 0.46383913 | NA |
| Gm6498 | 0.720587189 | 0.082389711 | 0.065783028 | 1.252446319 | 0.210407277 | NA |
| Gm6524 | 0 | NA | NA | NA | NA | NA |
| Gm6525 | 0.17833704 | -0.002390466 | 0.031282281 | -0.076415987 | 0.939088151 | NA |
| Gm6531 | 0.099009884 | 0.018205726 | 0.027896865 | 0.652608315 | 0.514008823 | NA |
| Gm6567 | 0.183387883 | -0.001444816 | 0.031390672 | -0.046026918 | 0.963288796 | NA |
| Gm6568 | 0.198019769 | 0.025308392 | 0.031628095 | 0.80018706 | 0.423602426 | NA |
| Gm6578 | 0 | NA | NA | NA | NA | NA |
| Gm6583 | 1.648508042 | 0.087776321 | 0.088978852 | 0.986485205 | 0.323895072 | NA |
| Gm6588 | 0 | NA | NA | NA | NA | NA |
| Gm6592 | 0 | NA | NA | NA | NA | NA |
| Gm6602 | 0.384346493 | 0.039441134 | 0.039210102 | 1.005892153 | 0.314467451 | NA |
| Gm6607 | 0.601453029 | -0.00149993 | 0.046549396 | -0.032222328 | 0.97429475 | NA |
| Gm6614 | 0 | NA | NA | NA | NA | NA |
| Gm6623 | 1.254307926 | 0.091666933 | 0.083938858 | 1.092067902 | 0.274803256 | NA |
| Gm6634 | 0 | NA | NA | NA | NA | NA |
| Gm6639 | 0 | NA | NA | NA | NA | NA |
| Gm6642 | 0.08731684 | 0.017365056 | 0.027733412 | 0.626142072 | 0.531221759 | NA |
| Gm6644 | 0 | NA | NA | NA | NA | NA |
| Gm6654 | 0.645748536 | -0.016484187 | 0.054188787 | -0.304199228 | 0.760976113 | NA |
| Gm6696 | 0 | NA | NA | NA | NA | NA |
| Gm6710 | 0 | NA | NA | NA | NA | NA |
| Gm6756 | 0.094392817 | 0.017895942 | 0.027818137 | 0.643319236 | 0.520016977 | NA |
| Gm6760 | 0.099009884 | 0.018205726 | 0.027896865 | 0.652608315 | 0.514008823 | NA |
| Gm6763 | 0 | NA | NA | NA | NA | NA |
| Gm6772 | 0.224413567 | -0.027287719 | 0.032077135 | -0.850690659 | 0.394941214 | NA |
| Gm6787 | 6.609826932 | -0.066410947 | 0.14908206 | -0.445465716 | 0.65598323 | NA |
| Gm6792 | 0.116696964 | -0.020477583 | 0.027954224 | -0.732539847 | 0.463839131 | NA |
| Gm6793 | 0 | NA | NA | NA | NA | NA |
| Gm6812 | 0 | NA | NA | NA | NA | NA |
| Gm6878 | 0.69449026 | -0.032567002 | 0.049908573 | -0.65253322 | 0.51405725 | NA |
| Gm6880 | 0 | NA | NA | NA | NA | NA |
| Gm6890 | 0.176054697 | -0.002190495 | 0.031239146 | -0.070120197 | 0.944097991 | NA |
| Gm6902 | 0 | NA | NA | NA | NA | NA |
| Gm6904 | 0.176054697 | -0.002190495 | 0.031239146 | -0.070120197 | 0.944097991 | NA |
| Gm6927 | 0.330336725 | -0.015889236 | 0.040460155 | -0.392713161 | 0.694531349 | NA |
| Gm6936 | 2.865676108 | -0.001987056 | 0.106925254 | -0.018583596 | 0.985173289 | NA |
| Gm6938 | 1.53743108 | 0.000985347 | 0.094385048 | 0.01043965 | 0.991670516 | NA |
| Gm6994 | 0.211089781 | -0.002448206 | 0.031868916 | -0.076821119 | 0.93876585 | NA |
| Gm7030 | 0 | NA | NA | NA | NA | NA |
| Gm7056 | 1.62794826 | 0.077392097 | 0.087061263 | 0.888938362 | 0.374036207 | NA |
| Gm7073 | 0 | NA | NA | NA | NA | NA |
| Gm7092 | 2.727667465 | -0.206376365 | 0.114401215 | -1.803970045 | 0.071236003 | NA |
| Gm7104 | 0 | NA | NA | NA | NA | NA |
| Gm7134 | 0 | NA | NA | NA | NA | NA |
| Gm7157 | 0 | NA | NA | NA | NA | NA |
| Gm7168 | 0.199381295 | -0.004055078 | 0.031639813 | -0.128163771 | 0.89801937 | NA |
| Gm7173 | 7.355165769 | -0.168024644 | 0.156136567 | -1.076138967 | 0.281865111 | NA |
| Gm7257 | 0 | NA | NA | NA | NA | NA |
| Gm7271 | 1.876384503 | -0.043772021 | 0.089319141 | -0.49006317 | 0.624089199 | NA |
| Gm732 | 0 | NA | NA | NA | NA | NA |
| Gm7325 | 0 | NA | NA | NA | NA | NA |
| Gm7334 | 0 | NA | NA | NA | NA | NA |
| Gm7337 | 0 | NA | NA | NA | NA | NA |
| Gm7367 | 0 | NA | NA | NA | NA | NA |
| Gm7444 | 4.761500957 | -0.161747097 | 0.120763461 | -1.339371161 | 0.180449873 | NA |
| Gm7457 | 3.116500362 | 0.000662869 | 0.119346788 | 0.005554141 | 0.99556846 | NA |
| Gm7534 | 0 | NA | NA | NA | NA | NA |
| Gm7538 | 0.491394591 | 0.007996605 | 0.050513606 | 0.158305954 | 0.874215717 | NA |
| Gm7550 | 0 | NA | NA | NA | NA | NA |
| Gm757 | 0 | NA | NA | NA | NA | NA |
| Gm7609 | 5.468562067 | 0.066313116 | 0.143798188 | 0.461154045 | 0.64468809 | NA |
| Gm7616 | 0.41930074 | -0.004836851 | 0.042752125 | -0.113137094 | 0.909921867 | NA |
| Gm766 | 3.5986732 | 0.034657506 | 0.114361721 | 0.303051632 | 0.761850514 | NA |
| Gm7714 | 0 | NA | NA | NA | NA | NA |
| Gm773 | 0 | NA | NA | NA | NA | NA |
| Gm7849 | 0 | NA | NA | NA | NA | NA |
| Gm7861 | 0 | NA | NA | NA | NA | NA |
| Gm7903 | 0 | NA | NA | NA | NA | NA |
| Gm7904 | 0.20671448 | -0.003314027 | 0.03178208 | -0.10427343 | 0.916952363 | NA |
| Gm7977 | 0 | NA | NA | NA | NA | NA |
| Gm7978 | 0 | NA | NA | NA | NA | NA |
| Gm805 | 0.590147267 | 0.023835975 | 0.058603081 | 0.406735863 | 0.684201998 | NA |
| Gm806 | 0.094392817 | 0.017895942 | 0.027818137 | 0.643319236 | 0.520016977 | NA |
| Gm8096 | 0 | NA | NA | NA | NA | NA |
| Gm813 | 0.389920616 | 0.025130846 | 0.048220387 | 0.521166418 | 0.602250846 | NA |
| Gm815 | 0 | NA | NA | NA | NA | NA |
| Gm8179 | 0.211846316 | 0.025707621 | 0.031860375 | 0.806883811 | 0.419733426 | NA |
| Gm8221 | 0 | NA | NA | NA | NA | NA |
| Gm826 | 2.326774602 | -0.071440128 | 0.101492335 | -0.703896777 | 0.481497059 | NA |
| Gm8267 | 1.622612839 | 0.01422999 | 0.092979317 | 0.15304468 | 0.878363041 | NA |
| Gm829 | 0.107716603 | -0.020477583 | 0.027954224 | -0.732539849 | 0.46383913 | NA |
| Gm8298 | 0 | NA | NA | NA | NA | NA |
| Gm8300 | 0 | NA | NA | NA | NA | NA |
| Gm833 | 0.743993901 | 0.030682617 | 0.050374558 | 0.60908956 | 0.542465077 | NA |
| Gm8363 | 0.575591037 | 0.053180578 | 0.045290032 | 1.174222573 | 0.240305887 | NA |
| Gm8369 | 1.154714898 | 0.047998337 | 0.078758791 | 0.609434669 | 0.542236364 | NA |
| Gm839 | 0 | NA | NA | NA | NA | NA |
| Gm8439 | 0.116696964 | -0.020477583 | 0.027954224 | -0.732539847 | 0.463839131 | NA |
| Gm8453 | 0 | NA | NA | NA | NA | NA |
| Gm853 | 0.17463368 | 0.024560283 | 0.031186242 | 0.787535833 | 0.430968261 | NA |
| Gm8579 | 0.81708555 | -0.12266306 | 0.06615006 | -1.854315171 | 0.063694083 | NA |
| Gm8580 | 3.072772518 | 0.010377089 | 0.120374847 | 0.086206454 | 0.9313023 | NA |
| Gm8615 | 3.091924734 | 0.043490516 | 0.121885067 | 0.356815789 | 0.721229716 | NA |
| Gm8633 | 0.107716603 | -0.020477583 | 0.027954224 | -0.732539849 | 0.46383913 | NA |
| Gm8660 | 0 | NA | NA | NA | NA | NA |
| Gm8677 | 0 | NA | NA | NA | NA | NA |
| Gm8693 | 0 | NA | NA | NA | NA | NA |
| Gm8709 | 0 | NA | NA | NA | NA | NA |
| Gm8720 | 0 | NA | NA | NA | NA | NA |
| Gm8764 | 0 | NA | NA | NA | NA | NA |
| Gm8765 | 0 | NA | NA | NA | NA | NA |
| Gm8773 | 2.689334559 | -0.061088421 | 0.108456042 | -0.563255123 | 0.573261172 | NA |
| Gm8787 | 7.939486688 | 0.178328192 | 0.154002115 | 1.15795937 | 0.246880617 | NA |
| Gm8801 | 0.567932444 | 0.090023784 | 0.05875239 | 1.532257406 | 0.125458928 | NA |
| Gm8817 | 0.281977855 | 0.011495295 | 0.038675761 | 0.297222198 | 0.766296876 | NA |
| Gm884 | 1.742690937 | 0.105767348 | 0.087959738 | 1.202451825 | 0.22918852 | NA |
| Gm8882 | 0 | NA | NA | NA | NA | NA |
| Gm8883 | 0 | NA | NA | NA | NA | NA |
| Gm8884 | 0 | NA | NA | NA | NA | NA |
| Gm8898 | 0 | NA | NA | NA | NA | NA |
| Gm8909 | 0.08731684 | 0.017365056 | 0.027733412 | 0.626142072 | 0.531221759 | NA |
| Gm8979 | 0.286710141 | 0.009849273 | 0.038825804 | 0.25367852 | 0.79974393 | NA |
| Gm8989 | 0 | NA | NA | NA | NA | NA |
| Gm8994 | 0 | NA | NA | NA | NA | NA |
| Gm9 | 0 | NA | NA | NA | NA | NA |
| Gm904 | 0.099009884 | 0.018205726 | 0.027896865 | 0.652608315 | 0.514008823 | NA |
| Gm9041 | 0 | NA | NA | NA | NA | NA |
| Gm9047 | 0.626586934 | -0.036676868 | 0.059529235 | -0.616115222 | 0.537818476 | NA |
| Gm9054 | 2.778805595 | 0.021395882 | 0.111773788 | 0.191421283 | 0.848195547 | NA |
| Gm906 | 0 | NA | NA | NA | NA | NA |
| Gm9112 | 0 | NA | NA | NA | NA | NA |
| Gm9125 | 0 | NA | NA | NA | NA | NA |
| Gm9199 | 0.119397641 | -0.020477583 | 0.027954224 | -0.732539845 | 0.463839132 | NA |
| Gm9268 | 0 | NA | NA | NA | NA | NA |
| Gm933 | 0 | NA | NA | NA | NA | NA |
| Gm9376 | 0 | NA | NA | NA | NA | NA |
| Gm94 | 0 | NA | NA | NA | NA | NA |
| Gm9513 | 0 | NA | NA | NA | NA | NA |
| Gm9573 | 2.274881693 | 0.061073732 | 0.095381592 | 0.64030942 | 0.521971458 | NA |
| Gm9696 | 0 | NA | NA | NA | NA | NA |
| Gm97 | 0 | NA | NA | NA | NA | NA |
| Gm9705 | 0.320973508 | -0.015129019 | 0.040159764 | -0.376720802 | 0.706381102 | NA |
| Gm9731 | 0.096071043 | -0.020098795 | 0.027825948 | -0.722304036 | 0.470107572 | NA |
| Gm9733 | 0 | NA | NA | NA | NA | NA |
| Gm9758 | 0.116696964 | -0.020477583 | 0.027954224 | -0.732539847 | 0.463839131 | NA |
| Gm9839 | 0 | NA | NA | NA | NA | NA |
| Gm9846 | 0 | NA | NA | NA | NA | NA |
| Gm9855 | 0 | NA | NA | NA | NA | NA |
| Gm9871 | 1.567935623 | 0.049808998 | 0.08930336 | 0.557750549 | 0.577014735 | NA |
| Gm9895 | 7.430599815 | 0.18773158 | 0.155673391 | 1.205932359 | 0.227843572 | NA |
| Gm9920 | 3.84398225 | 0.065978439 | 0.13475695 | 0.48961066 | 0.624409432 | NA |
| Gm9926 | 2.168183447 | -0.258871212 | 0.102067534 | -2.536273797 | 0.01120391 | NA |
| Gm9961 | 0.096071043 | -0.020098795 | 0.027825948 | -0.722304036 | 0.470107572 | NA |
| Gm9992 | 0.331243956 | 0.008596723 | 0.034710864 | 0.247666645 | 0.804392342 | NA |
| Gm9994 | 0 | NA | NA | NA | NA | NA |
| Gm9999 | 0.159967308 | 0.0240345 | 0.030870969 | 0.778547006 | 0.436246606 | NA |
| Gmcl1l | 0 | NA | NA | NA | NA | NA |
| Gml | 0 | NA | NA | NA | NA | NA |
| Gnat1 | 5.119184197 | 0.067078729 | 0.141623744 | 0.473640413 | 0.635756351 | NA |
| Gnat2 | 2.442187145 | 0.050275045 | 0.104274793 | 0.482139964 | 0.629706522 | NA |
| Gnat3 | 0 | NA | NA | NA | NA | NA |
| Gnb3 | 5.962343211 | 0.272265562 | 0.138601293 | 1.964379668 | 0.049486079 | NA |
| Gngt1 | 0 | NA | NA | NA | NA | NA |
| Gnrhr | 0 | NA | NA | NA | NA | NA |
| Gp1ba | 0.439846772 | -0.045662679 | 0.041334256 | -1.104717565 | 0.26928199 | NA |
| Gp2 | 5.904305298 | -0.208975532 | 0.132330794 | -1.579190494 | 0.11429237 | NA |
| Gp49a | 1.928387987 | 0.153676866 | 0.088963825 | 1.727408476 | 0.084094329 | NA |
| Gp6 | 0.098353386 | -0.020212831 | 0.02786466 | -0.725393061 | 0.468210929 | NA |
| Gpa33 | 0 | NA | NA | NA | NA | NA |
| Gpbar1 | 0.17463368 | 0.024560283 | 0.031186242 | 0.787535833 | 0.430968261 | NA |
| Gpha2 | 0.547215998 | -0.090429391 | 0.05723373 | -1.58000172 | 0.114106473 | NA |
| Gphb5 | 0 | NA | NA | NA | NA | NA |
| Gpihbp1 | 0.356086128 | 0.051237183 | 0.044718313 | 1.145776311 | 0.251887719 | NA |
| Gpr110 | 0.407193159 | 0.02051662 | 0.04329096 | 0.473923893 | 0.63555418 | NA |
| Gpr113 | 0.094392817 | 0.017895942 | 0.027818137 | 0.643319236 | 0.520016977 | NA |
| Gpr119 | 0.38488829 | -0.02558324 | 0.043075085 | -0.593921989 | 0.552564291 | NA |
| Gpr128 | 0.929519008 | 0.066206034 | 0.050206807 | 1.318666504 | NA | NA |
| Gpr132 | 0.655039105 | 0.023132371 | 0.053842275 | 0.429632135 | 0.667463257 | NA |

| | | | | | | | |
|---|---|---|---|---|---|---|---|
| Hist3h2bb-ps | 5.18475471 | -0.0103193941420658 | 0.0442002755035323 | -0.233469000464513 | 0.815397238521398 | NA | NM_206882 |
| Hmga1 | 15.5043637 | 0.0293369610646621 | 0.0783189084757412 | 0.374583375019189 | 0.707970339094017 | NA | 01166537,NM_001166535,NM_001166539,NM_001166543,NM_001166544,NM_001166545,NM_001039356,NM_001166542,NM_001166541,NM_0011665 |
| Hmga1-rs1 | 0 | NA | NA | NA | NA | NA | NM_001166477,NM_001166476 |
| Hmga2 | 3.565496224 | 0.00804095613253912 | 0.0419606862915651 | 0.191630710629142 | 0.848031484713103 | NA | NM_010441 |
| Hmgb1-rs17 | 1.496993781 | 0.0205241149305303 | 0.0281556681570805 | 0.728951442957285 | 0.466031365394327 | NA | NR_033589 |
| Hmgb4 | 0.12890128 | -0.0068640067066208 | 0.00870804083610955 | -0.788237771940376 | 0.430557639008166 | NA | NM_027036 |
| Hmx1 | 0 | NA | NA | NA | NA | NA | NM_010445 |
| Hmx2 | 1.815946383 | 0.0094944773776442 | 0.0114632172052221 | 0.828255908238307 | 0.407525587464867 | NA | NM_145998 |
| Hmx3 | 1.451804699 | 0.0257082222792187 | 0.0187699085159417 | 1.36965090998627 | 0.170795899430051 | NA | NM_008257 |
| Hnf1a | 0.323722592 | -0.0163509104806049 | 0.013414304634911 | -1.21891599494851 | 0.222876078142229 | NA | NM_009327 |
| Hnf4a | 0.12890128 | -0.0068640067066208 | 0.00870804083610955 | -0.788237771940376 | 0.430557639008166 | NA | NM_008261 |
| Hnf4g | 1.074416698 | 0.0157795681053513 | 0.0199625544032011 | 0.790458364527787 | 0.429260128447807 | NA | NM_013920 |
| Hormad1 | 0.127687691 | 0.00542298404407807 | 0.00869724871942359 | 0.62352868349814 | 0.532937159177571 | NA | NM_026489 |
| Hormad2 | 0.985901806 | -0.00624884779514409 | 0.0194733618202687 | -0.32089209109441 | 0.748292167645687 | NA | NM_029458 |
| Hotair | 0 | NA | NA | NA | NA | NA | NR_047528 |
| Hoxa1 | 0.807392905 | -0.027153622605454 | 0.0198194017258183 | -1.3700525869094 | 0.170670486737627 | NA | NM_010449 |
| Hoxa10 | 0.09572496 | 0.00498255289994511 | 0.00847999972823692 | 0.587565219295241 | 0.556824162160132 | NA | NM_008263,NM_001122950 |
| Hoxa11 | 0 | NA | NA | NA | NA | NA | NM_010450 |
| Hoxa11as | 0 | NA | NA | NA | NA | NA | NR_015348 |
| Hoxa13 | 0 | NA | NA | NA | NA | NA | NM_008264 |
| Hoxa2 | 0.533360734 | 0.0142262365174673 | 0.0140287124632757 | 1.01407998451096 | 0.310544587895273 | NA | NM_010451 |
| Hoxa3 | 0 | NA | NA | NA | NA | NA | NM_010452 |
| Hoxa4 | 0.415611529 | 0.0048600098249294 | 0.0120675527377329 | 0.402733671901271 | 0.68714416024218 | NA | NM_008265 |
| Hoxa5 | 0.219991894 | 0.00711449016834187 | 0.00994861810862167 | 0.715123456410123 | 0.474532761039468 | NA | NM_010453 |
| Hoxa6 | 0 | NA | NA | NA | NA | NA | NM_010454 |
| Hoxa7 | 0.52184809 | 0.00243956128556291 | 0.017699154548832 | 0.137834904985555 | 0.890370897446906 | NA | NM_010455 |
| Hoxa9 | 0 | NA | NA | NA | NA | NA | NM_010456 |
| Hoxb1 | 0 | NA | NA | NA | NA | NA | NM_008266 |
| Hoxb13 | 0.226556888 | -0.00933175792380659 | 0.010153338242423 | -0.919082739193727 | 0.358052296835844 | NA | NM_008267 |
| Hoxb2 | 0.106347839 | -0.00653347234140708 | 0.00854389509723638 | -0.764694822098226 | 0.444453290197145 | NA | NM_134032 |
| Hoxb3 | 0.710417164 | 0.0146703204675107 | 0.0183413920405657 | 0.799847712489017 | 0.423799035352418 | NA | NM_001079869,NM_010458 |
| Hoxb4 | 1.65841631 | 0.0252376323198093 | 0.0280500181558199 | 0.899736755235325 | 0.368260358386409 | NA | NM_010459 |
| Hoxb5 | 0.521648859 | 0.0133137968174299 | 0.0180533220562906 | 0.737470742277642 | 0.460836129175685 | NA | NM_008268 |
| Hoxb6 | 0.318445824 | 0.0076171880905019 | 0.0102869072557308 | 0.740474070693932 | 0.459012388854512 | NA | NM_008269 |
| Hoxb7 | 0.530743041 | 0.0103814776656352 | 0.0119942749221342 | 0.865536077256095 | 0.386744623285174 | NA | NM_010460 |
| Hoxb8 | 1.835394799 | -0.00418526520678957 | 0.0262017534306856 | -0.159732256769048 | 0.873091989907674 | NA | NM_010461 |
| Hoxb9 | 0.212297216 | 0.00681050119636477 | 0.00973439053551308 | 0.699633035218656 | 0.484156506270654 | NA | NM_008270 |
| Hoxc10 | 0 | NA | NA | NA | NA | NA | NM_010462 |
| Hoxc11 | 0 | NA | NA | NA | NA | NA | NM_001024842 |
| Hoxc12 | 0 | NA | NA | NA | NA | NA | NM_010463 |
| Hoxc13 | 0.343816189 | 0.00220638102164924 | 0.0128049404084123 | 0.172307012081036 | 0.863196171959189 | NA | NM_010464 |
| Hoxc4 | 1.088186466 | 0.00300942833822286 | 0.021894932618312 | 0.137448622961548 | 0.89067620014844 | NA | NM_013553 |
| Hoxc5 | 0.106347839 | -0.00653347234140708 | 0.00854389509723638 | -0.764694822098226 | 0.444453290197145 | NA | NM_175730 |
| Hoxc6 | 0.106148608 | 0.00515554063321497 | 0.00854200771766808 | 0.603551390213729 | 0.546141939141592 | NA | NM_010465 |
| Hoxc8 | 0 | NA | NA | NA | NA | NA | NM_010466 |
| Hoxc9 | 0 | NA | NA | NA | NA | NA | NM_008272 |
| Hoxd1 | 2.579380224 | -0.0284724862523747 | 0.0364292270303472 | -0.781583595739096 | 0.434459330621135 | NA | NM_010467 |
| Hoxd10 | 0 | NA | NA | NA | NA | NA | NM_013554 |
| Hoxd11 | 0 | NA | NA | NA | NA | NA | NR_073086,NM_008273 |
| Hoxd12 | 0 | NA | NA | NA | NA | NA | NM_008274 |
| Hoxd13 | 0 | NA | NA | NA | NA | NA | NM_008275 |
| Hoxd3 | 0.249110328 | -0.010000816291709 | 0.0105091939847586 | -0.951625434473194 | 0.341286977626714 | NA | NM_010468 |
| Hoxd4 | 0.215934008 | -0.00149029401866511 | 0.00975123948835942 | -0.152831239602324 | 0.878531361484804 | NA | NM_010469 |
| Hoxd8 | 0.203314312 | -0.000639443352387801 | 0.00969102748600992 | -0.0659830294889689 | 0.947391336513325 | NA | NM_008276 |
| Hoxd9 | 0 | NA | NA | NA | NA | NA | NM_013555 |
| Hp | 3.412429754 | -0.0480016768493988 | 0.0373354768809162 | -1.28568538182874 | 0.198552884976407 | NA | NM_017370 |
| Hpd | 2.333305631 | -0.0115296382521685 | 0.0313092510905212 | -0.368250208822756 | 0.712686675392033 | NA | NM_008277 |
| Hpse2 | 1.951739169 | -0.0344585813550881 | 0.0304673705341233 | -1.13099951689282 | 0.258055294769214 | NA | NM_001081257 |
| Hpx | 19.63998885 | -0.0684819455842567 | 0.0820331920376307 | -0.834807763579921 | 0.403825953944475 | NA | NM_017371 |
| Hrasls5 | 0.730710545 | 0.00403870488601952 | 0.0161183001464204 | 0.250566427559449 | 0.802149340715342 | NA | NM_025731 |
| Hrc | 14.59806848 | -0.038118595242862 | 0.0730032484029588 | -0.522149302623594 | 0.601566381318717 | NA | NM_010473 |
| Hrg | 0.783267985 | -0.0254222372681514 | 0.0192886511667334 | -1.31798937356472 | 0.187507205199555 | NA | NM_053176 |
| Hrh4 | 0 | NA | NA | NA | NA | NA | NM_153087 |
| Hrnr | 0.425192124 | -0.00757776848570139 | 0.0123226000471034 | -0.614948830338989 | 0.538588513594647 | NA | NM_133698 |
| Hs3st6 | 8.415898699 | 0.0157882933980114 | 0.0586335450087539 | 0.269270660603145 | 0.787721408070857 | NA | NM_001012402 |
| Hsd11b2 | 1.075834258 | -0.00620348645789856 | 0.0241810310531343 | -0.2565435048765 | 0.797531192065825 | NA | NM_008289 |
| Hsd17b1 | 9.165575409 | -0.0114244529661705 | 0.0457024210760436 | -0.249974786831567 | 0.802606846952315 | NA | NM_010475 |
| Hsd17b13 | 9.089481088 | 0.0459560119743974 | 0.0518527341471618 | 0.886279435988291 | 0.375466967443539 | NA | NM_001163486,NM_198030 |
| Hsd17b14 | 8.133562054 | -0.0876899521085256 | 0.05622386272339 | -1.55965719644597 | 0.118840912334086 | NA | NM_025330 |
| Hsd17b2 | 5.157178834 | -0.0171394902940181 | 0.0482076403118397 | -0.355534728170643 | 0.722189032777002 | NA | NM_008290 |
| Hsd17b3 | 0 | NA | NA | NA | NA | NA | NM_008291 |
| Hsd17b6 | 0 | NA | NA | NA | NA | NA | NM_013786 |
| Hsd3b1 | 0 | NA | NA | NA | NA | NA | NM_008293 |
| Hsd3b2 | 0 | NA | NA | NA | NA | NA | NM_153193 |
| Hsd3b3 | 0 | NA | NA | NA | NA | NA | NM_001161742,NM_001012306,NM_001161743,NM_001161744,NM_001161745 |
| Hsd3b4 | 0 | NA | NA | NA | NA | NA | NM_001111336,NM_008294 |
| Hsd3b5 | 0 | NA | NA | NA | NA | NA | NM_008295 |
| Hsd3b6 | 0 | NA | NA | NA | NA | NA | NM_013821 |
| Hsf2bp | 14.17981283 | -0.0287711569844946 | 0.0735927060299794 | -0.390951203408312 | 0.6958333050162 | NA | NM_028902 |
| Hsf3 | 2.457946924 | 0.0300776806177825 | 0.038160812836294 | 0.788182388745561 | 0.430590028928933 | NA | NM_172931 |
| Hsf5 | 18.94364384 | 0.132464253314117 | 0.0787475789380691 | 1.6821374714046 | 0.0925421857081575 | NA | NM_001045527 |
| Hsfy2 | 0 | NA | NA | NA | NA | NA | NM_027661 |
| Hsh2d | 1.857906822 | -0.0377229902530265 | 0.0293410454144422 | -1.28567301267523 | 0.198557203566154 | NA | NM_197944 |
| Hspa1l | 19.2298203 | -0.0769101515378165 | 0.0782252056971403 | -0.983188869270408 | 0.32551449193887 | NA | NM_013558 |
| Hspb7 | 13.96690146 | -0.052873545453428 | 0.0727783915365786 | -0.726500604603959 | 0.467531936987795 | NA | NM_013868 |
| Hspb9 | 0 | NA | NA | NA | NA | NA | NM_029307 |
| Htr2b | 3.676363584 | 0.037941788868021 | 0.0454066709420955 | 0.835599441245229 | 0.403380283203057 | NA | NM_008311 |
| Htr3b | 1.303480001 | 0.0116606365944307 | 0.025973996342298 | 0.448935021040319 | 0.653478531725009 | NA | NM_020274 |
| Hus1b | 0.330974078 | -0.00671789744880547 | 0.01278556667705 | -0.525428212803741 | 0.599285545196596 | NA | NM_153072 |
| Hyal4 | 0 | NA | NA | NA | NA | NA | NM_029848 |
| Hyal5 | 0 | NA | NA | NA | NA | NA | NM_028957 |
| Hyal6 | 0.12890128 | -0.0068640067066208 | 0.00870804083610955 | -0.788237771940376 | 0.430557639008166 | NA | NM_028920 |
| I730028E13Rik | 1.635074717 | 0.0128515817553484 | 0.0287942556205837 | 0.446324500438253 | 0.65536286090483 | NA | NR_045705 |
| I730030J21Rik | 10.31155572 | -0.0592987897784426 | 0.0639145280090709 | -0.927782643971183 | 0.35352032877602 | NA | NR_045782,NR_045781 |
| Iapp | 1.033043189 | 0.0289774377128288 | 0.0199244466886594 | 1.45436599397875 | 0.145844862749635 | NA | NM_010491 |
| Ibsp | 5.979585314 | 0.00493510236536707 | 0.0462493786232864 | 0.106706349626117 | 0.915021945098382 | NA | NM_008318 |
| Icos | 3.462577356 | -0.0178915175479478 | 0.0427653247384759 | -0.418365057610585 | 0.675680230182897 | NA | NM_017480 |
| Idi2 | 3.793146691 | -0.00677412894139484 | 0.0415502578711371 | -0.163034582418332 | 0.87049120610229 | NA | NM_177197 |
| Ifi202b | 0.517564097 | -0.0186052918188176 | 0.0143061080534879 | -1.30051386088067 | 0.193424908760123 | NA | NM_011940,NM_008327 |
| Ifi205 | 0.418957648 | -0.0103528989990422 | 0.0140690420066446 | -0.735863820305088 | 0.461813577149106 | NA | NM_172648 |
| Ifi27l2b | 2.267211278 | 0.00685212853695722 | 0.0298707129231478 | 0.229392869014763 | 0.818563578113521 | NA | NM_145449 |
| Ifi44l | 0.22897535 | -0.00116364198856673 | 0.00981572003153595 | -0.118548816065269 | 0.905632818135836 | NA | NM_031367 |
| Ifi47 | 41.93980667 | -0.165874551478518 | 0.079724019663446 | -2.08060948480465 | NA | NA | NM_001271676,NM_001271677,NM_008330 |
| Ifitm5 | 1.33181971 | -0.0142632151781434 | 0.0227827014061656 | -0.626054607127643 | 0.531279124494749 | NA | NM_053088 |
| Ifitm6 | 5.926722942 | -0.040301752208806 | 0.0474881253671981 | -0.848670102202937 | 0.396064887064239 | NA | NM_001033632 |
| Ifitm7 | 20.43010166 | -0.135744140265602 | 0.0800820245658502 | -1.69506379242425 | 0.0900633199220655 | NA | NM_001270718 |
| Ifltd1 | 20.98962547 | 0.025096019497353 | 0.0772848574038853 | 0.324721042910165 | 0.745392207051531 | NA | NM_028742 |
| Ifna1 | 0 | NA | NA | NA | NA | NA | NM_010502 |
| Ifna11 | 0 | NA | NA | NA | NA | NA | NM_008333 |
| Ifna12 | 0 | NA | NA | NA | NA | NA | NM_177361 |
| Ifna13 | 0 | NA | NA | NA | NA | NA | NM_177347 |
| Ifna14 | 0 | NA | NA | NA | NA | NA | NM_206975 |
| Ifna2 | 0 | NA | NA | NA | NA | NA | NM_010503 |
| Ifna4 | 0 | NA | NA | NA | NA | NA | NM_010504 |
| Ifna5 | 0 | NA | NA | NA | NA | NA | NM_010505 |
| Ifna6 | 0 | NA | NA | NA | NA | NA | NM_206871 |
| Ifna7 | 0 | NA | NA | NA | NA | NA | NM_008334 |
| Ifna9 | 0 | NA | NA | NA | NA | NA | NM_010507 |
| Ifnab | 0 | NA | NA | NA | NA | NA | NM_008336 |
| Ifnb1 | 0 | NA | NA | NA | NA | NA | NM_010510 |
| Ifne | 0.09572496 | 0.00498255289994511 | 0.00847999972823692 | 0.587565219295241 | 0.556824162160132 | NA | NM_177348 |
| Ifng | 0.751621763 | 0.00975522122263679 | 0.0200538040384948 | 0.486452405933104 | 0.626646442070338 | NA | NM_008337 |
| Ifnk | 0.439983788 | 0.0126652380404026 | 0.0132459535836759 | 0.956159023236422 | 0.338991898718962 | NA | NM_199157 |
| Ifnl2 | 0 | NA | NA | NA | NA | NA | NM_001024673 |
| Ifnl3 | 0 | NA | NA | NA | NA | NA | NM_177396 |
| Ifnlr1 | 12.89042552 | 0.0317399943910071 | 0.0615736198298822 | 0.515480403437372 | 0.606217360034459 | NA | NM_174851 |
| Ifnz | 0.09572496 | 0.00498255289994511 | 0.00847999972823692 | 0.587565219295241 | 0.556824162160132 | NA | NM_197889 |
| Igbp1b | 10.62473705 | 0.00406495350519887 | 0.055722642841196 | 0.0729497614961226 | 0.941846095449017 | NA | NM_015777 |
| Igf2as | 10.66763695 | 0.0019010120252624 | 0.058978247138404 | 0.0322324266572606 | 0.974286696874159 | NA | NR_002855 |
| Igf2bp1 | 1.152334539 | -0.00111152334921972 | 0.023739555988848 | -0.0468215728104551 | 0.962655435270884 | NA | NM_009951 |
| Igf2bp3 | 0.613315933 | 0.00450825894462919 | 0.0170318739200569 | 0.264695415536175 | 0.791244109440564 | NA | NM_023670 |
| Igfals | 1.813150625 | -0.0318580951652922 | 0.0321026218580922 | -0.992382968161261 | 0.32101074409394 | NA | NM_008340 |
| Igfbp1 | 0.437122686 | -0.00309378938484024 | 0.015328819040773 | -0.201828293269761 | 0.840050961859409 | NA | NM_008341 |
| Igfl3 | 0 | NA | NA | NA | NA | NA | NM_001003393 |
| Igj | 15.31356732 | -0.149842656565642 | 0.0770312069319407 | -1.94522015886409 | 0.0517484866106758 | NA | NM_152839 |
| Igll1 | 0.440787404 | -0.00207676726209905 | 0.0153548750617872 | -0.135251329219043 | 0.892413165045382 | NA | NM_001190325 |
| Igsf23 | 0.205932006 | -0.000576954122277892 | 0.00970424421944593 | -0.0594537925088239 | 0.952590668569529 | NA | NM_027308 |
| Igsf5 | 12.94797278 | -0.0134318459111593 | 0.0735459303817657 | -0.182632075513039 | 0.855086714986146 | NA | NM_001177886,NM_001177887,NM_028078 |
| Igtp | 411.4049186 | -0.122454345536283 | 0.0599463661674123 | -2.04273175115076 | NA | NA | NM_018738 |
| Ihh | 15.05506724 | -0.039638108025211 | 0.0669218152702741 | -0.59230473448944 | 0.553646546098741 | NA | NM_010544 |
| Iigp1 | 128.6605486 | -0.103450512182519 | 0.0438098705460764 | -2.36135169752936 | NA | NA | NM_021792,NM_001146275 |
| Ikzf3 | 8.304615533 | -0.0523058937137965 | 0.0597609864033469 | -0.875251512094638 | 0.381437070518889 | NA | NM_011771 |
| Il10 | 0.388662817 | -0.0108564151646332 | 0.0109525618994156 | -0.991221530116391 | 0.321577419693911 | NA | NM_010548 |
| Il11 | 15.82387403 | 0.040153796026308 | 0.0751959364699654 | 0.533988908328126 | 0.593349207993341 | NA | NM_008350 |
| Il11ra2 | 0 | NA | NA | NA | NA | NA | NM_010550 |
| Il12b | 0.120209049 | -0.00674972139751468 | 0.00863521114606269 | -0.781650996523957 | 0.434419707908266 | NA | NM_008352 |
| Il12rb2 | 10.15642188 | 0.00674988306666967 | 0.0512880107144403 | 0.131607425841713 | 0.895294811648766 | NA | NM_008354 |
| Il13 | 0.09572496 | 0.00498255289994511 | 0.00847999972823692 | 0.587565219295241 | 0.556824162160132 | NA | NM_008355 |
| Il13ra2 | 4.470941776 | 0.0219753306187088 | 0.0387882607552991 | 0.566545913397434 | 0.571022730652623 | NA | NM_008356 |
| Il15 | 15.95075998 | -0.0775071234797623 | 0.0745244200545833 | -1.04002316855327 | 0.298329136820587 | NA | NM_008357,NM_001254747 |
| Il17a | 0 | NA | NA | NA | NA | NA | NM_010552 |
| Il17b | 8.761059008 | 0.00403538505661083 | 0.0634754566316241 | 0.0635739429182832 | 0.949309480298315 | NA | NM_019508 |
| Il17c | 1.497785723 | 0.00826037769674001 | 0.0274853444267853 | 0.300537536240223 | 0.763767169155336 | NA | NM_145834 |
| Il17f | 0.681515317 | 0.00705864756381606 | 0.0191356397277724 | 0.368874396896778 | 0.712221347979209 | NA | NM_145856 |
| Il17re | 11.41325271 | 0.0388955407489599 | 0.0598562226787614 | 0.649816159594732 | 0.515810979445461 | NA | NM_001034029,NM_145826,NM_001034031 |
| Il18r1 | 1.299042347 | -0.0261460460847111 | 0.0268392436399955 | -0.974172239554048 | 0.329971025829274 | NA | NM_008365,NM_001161843,NM_001161842 |
| Il18rap | 1.756437712 | 0.0017683433503801 | 0.029573094401418 | 0.0597956820607624 | 0.952318364641429 | NA | NM_010553 |
| Il19 | 0.863694892 | 0.00336837038561511 | 0.0184922592123728 | 0.182150290396178 | 0.855464782793929 | NA | NM_001009940 |
| Il1b | 14.5472279 | -0.109734552413049 | 0.075912973534917 | -1.44553094554484 | 0.148308807291266 | NA | NM_008361 |
| Il1f10 | 0 | NA | NA | NA | NA | NA | NM_153077 |
| Il1f5 | 4.376965801 | -0.0120522796726826 | 0.0444550774156213 | -0.271111431434545 | 0.786305329714989 | NA | NM_001146087,NM_019451,NM_001146088 |
| Il1f6 | 0 | NA | NA | NA | NA | NA | NM_019450 |
| Il1f8 | 0.233836299 | 0.00743372775021133 | 0.0101683766597817 | 0.731063374118848 | 0.464740444385902 | NA | NM_027163 |
| Il1r2 | 12.3452988 | -0.135565718654944 | 0.0723340026151039 | -1.87416310108404 | 0.0609079482172305 | NA | NM_010555 |
| Il1rl1 | 12.5048087 | 0.00267934212492771 | 0.0677677593633848 | 0.0395371213405555 | 0.968462158092548 | NA | NM_010743,NM_001025602 |
| Il1rn | 3.319060698 | -0.0535121017104085 | 0.0330064507164987 | -1.62126192149645 | 0.104961477803695 | NA | NM_001159562,NM_001039701,NM_031167 |
| Il2 | 0 | NA | NA | NA | NA | NA | NM_008366 |
| Il20 | 0 | NA | NA | NA | NA | NA | NM_021380 |
| Il20rb | 12.44698962 | -0.108455034452365 | 0.0730346851766485 | -1.484979831022 | 0.137549166943818 | NA | NM_001033543 |
| Il21 | 2.000708935 | -0.00734587002093875 | 0.0300918808576051 | -0.244114685143792 | 0.807141999447707 | NA | NM_021782 |
| Il22 | 1.243066132 | 0.00796886032985169 | 0.0266486593404766 | 0.299034192603746 | 0.764913956589602 | NA | NM_016971 |
| Il22ra1 | 4.127623594 | 0.0466346226290686 | 0.0465367143253194 | 1.00210389377868 | 0.316293417509969 | NA | NM_178257 |
| Il22ra2 | 0 | NA | NA | NA | NA | NA | NM_178258 |
| Il23a | 2.386181236 | -0.00318542707283732 | 0.0350162757724225 | -0.0909698990703643 | 0.927516508920242 | NA | NM_031252 |
| Il23r | 0.528166588 | 0.00231952184738176 | 0.0147118373261157 | 0.157663641594532 | 0.874721851892895 | NA | NM_144548 |
| Il24 | 0 | NA | NA | NA | NA | NA | NM_053095 |
| Il25 | 3.995688887 | 0.0682920947106752 | 0.0380798680831154 | 1.79339105276354 | 0.0729104172280796 | NA | NM_080729 |
| Il27 | 1.691845501 | -0.0454737907416162 | 0.0300343736573795 | -1.5140582340875 | 0.130011070386082 | NA | NM_145636 |
| Il27ra | 17.35649141 | -0.0599764816809169 | 0.0710606681380341 | -0.844017981429807 | 0.39865935761426 | NA | NM_016671 |
| Il2ra | 14.92368204 | -0.0182793943510069 | 0.0663773008792059 | -0.275386225545265 | 0.783019525548055 | NA | NM_008367 |
| Il2rb | 18.81958276 | -0.00488577256647869 | 0.0692443914051492 | -0.0705583870019452 | 0.943749230188482 | NA | NM_008368 |
| Il3 | 0.212297216 | 0.00681050119636477 | 0.00973439053551308 | 0.699633035218656 | 0.484156506270654 | NA | NM_010556 |
| Il31 | 0 | NA | NA | NA | NA | NA | NM_029594 |
| Il31ra | 7.699513683 | 0.0250797363289927 | 0.0487034945704228 | 0.514947367744395 | 0.606589799439224 | NA | NM_139299 |
| Il4 | 0.255375383 | 0.00708304372802858 | 0.00992471678592415 | 0.713677163873753 | 0.475426830432214 | NA | NM_021283,NR_027491 |
| Il4i1 | 1.064684283 | 0.0471358590993255 | 0.0253015010148219 | 1.86296690744603 | 0.0624669222599845 | NA | NM_010215 |
| Il5 | 4.173938155 | -0.00738281953369447 | 0.0442998357447682 | -0.166655686405482 | 0.867640974927785 | NA | NM_010558 |
| Il5ra | 0.330974078 | -0.00671789744880547 | 0.01278556667705 | -0.525428212803741 | 0.599285545196596 | NA | NM_008370 |
| Il6 | 2.857597098 | -0.064739139995109 | 0.0342024613659462 | -1.89282108391084 | 0.0583816667258552 | NA | NM_031168 |
| Il7 | 4.27378977 | -0.0165959268059401 | 0.0473132707287483 | -0.350766847235867 | 0.725763270884253 | NA | NM_008371 |
| Il7r | 11.8053372 | 0.0166666514552977 | 0.0707738177296681 | 0.235491767859106 | 0.813827067519064 | NA | NM_008372 |
| Il8 | 0 | NA | NA | NA | NA | NA | NM_011339 |
| Il9 | 0 | NA | NA | NA | NA | NA | NM_008373 |
| Il9r | 0.449986343 | -0.00255968786282767 | 0.013925376636135 | -0.1838146234541 | 0.854158883753557 | NA | NM_008374,NM_001134458 |
| Iltifb | 2.99550181 | 0.0212790517663531 | 0.0395498920729119 | 0.53803058999818 | 0.590555930047978 | NA | NM_054079 |
| Inhbc | 0 | NA | NA | NA | NA | NA | NM_010565 |
| Inhbe | 9.166563898 | -0.0179346693705876 | 0.0580464936400657 | -0.308970761986018 | 0.75734376793099 | NA | NM_008382 |
| Ins1 | 0 | NA | NA | NA | NA | NA | NM_008386 |
| Ins2 | 1.539320337 | 0.0192654564324905 | 0.0272730357406897 | 0.706392079549386 | 0.479944345068951 | NA | NM_001185084,NM_001185083,NM_008387 |
| Insl3 | 3.262886568 | 0.0134649109987139 | 0.0381051692415127 | 0.353361794914819 | 0.723817223928908 | NA | NM_013564 |
| Insl5 | 15.60169665 | -0.0597286501139268 | 0.0747721978976723 | -0.798808270898591 | 0.424401593955765 | NA | NM_011831 |
| Insl6 | 11.67445781 | 0.0457379435913184 | 0.0625126416200604 | 0.73165910775783 | 0.464376661340504 | NA | NM_013754 |
| Insm2 | 5.212682721 | 0.0349804203395756 | 0.0432773573332926 | 0.80828457408294 | 0.418926780649366 | NA | NM_020287 |
| Iqcf1 | 0 | NA | NA | NA | NA | NA | NM_001146701,NM_028843 |
| Iqcf3 | 0.574236473 | 0.00239212947531775 | 0.018035069611814 | 0.132637662443552 | 0.89447994523857 | NA | NM_026645 |
| Iqcf4 | 0.513177942 | -0.00154094418427993 | 0.0127186663577136 | -0.121156113458812 | 0.903567384360659 | NA | NM_026090 |
| Iqcf5 | 2.986798258 | 0.0337319080632353 | 0.0347130700364196 | 0.971735085022589 | 0.331182357613949 | NA | NM_029300 |
| Iqcf6 | 0 | NA | NA | NA | NA | NA | NM_001101628 |
| Iqch | 4.297471241 | -0.0315111438078007 | 0.0492566858485618 | -0.639733332946531 | 0.522345980970812 | NA | NM_030068 |
| Iqcj | 1.351053393 | 0.00286782640592304 | 0.0271352610248914 | 0.105686339383003 | 0.915831219237367 | NA | NM_177585 |
| Irg1 | 1.573195958 | -0.0424736494360563 | 0.0270123697454748 | -1.57237776012494 | 0.115862968898949 | NA | NM_008392 |
| Irgc1 | 4.411005129 | 0.0229563772431568 | 0.0367398443148959 | 0.62483599675596 | 0.53207870254053 | NA | NM_199013 |
| Irgm2 | 303.3758548 | -0.151037486178652 | 0.0681455010260691 | -2.21639703141771 | NA | NA | NM_019440 |
| Irs3 | 10.91409598 | 0.0125196117424736 | 0.0573464319823239 | 0.218315443693735 | 0.827183342333192 | NA | NM_010571 |
| Irs4 | 205.7260439 | 0.0747606018111001 | 0.0452942747995529 | 1.65055301452443 | NA | NA | NM_010572 |
| Irx1 | 19.36873148 | 0.126711049071712 | 0.0714949351431877 | 1.7723080497652 | 0.0763434357284229 | NA | NM_010573 |
| Irx2 | 9.65513574 | 0.0107582710412495 | 0.0378391106300708 | 0.284316170811368 | 0.776168094764269 | NA | NM_010574 |
| Irx3 | 11.42235705 | 0.0180506377498018 | 0.0551433741577524 | 0.32734010251464 | 0.743410664221228 | NA | NM_008393,NM_001253822 |
| Irx4 | 0.300216221 | 0.00934385809732901 | 0.0113912670188907 | 0.820265040037567 | 0.412065031496518 | NA | NM_018885 |
| Irx5 | 6.328190972 | -0.0174275590704197 | 0.0495757181756678 | -0.351534172609793 | 0.72518764023949 | NA | NM_018826 |
| Irx6 | 8.569213613 | 0.0216017554537378 | 0.0589083844365541 | 0.366700863728548 | 0.71384216221595 | NA | NM_022428 |
| Isg20 | 18.33046176 | -0.058405161969769 | 0.0806696914068009 | -0.72400378569993 | 0.469063410932455 | NA | NM_001113527,NM_020583 |
| Isl2 | 0.815821339 | -0.00760613633732843 | 0.0163602588453808 | -0.464915403186053 | 0.641992039478835 | NA | NM_027397 |
| Isx | 2.760722691 | 0.0223499747519286 | 0.039740812999438 | 0.562393495881545 | 0.573847949508446 | NA | NM_027837 |
| Itga2 | 2.495017543 | -0.0330230127602088 | 0.0353289525918232 | -0.934729459481679 | 0.349927740833657 | NA | NM_008396 |
| Itgad | 13.78773318 | 0.208233106756763 | 0.0744595943505208 | 2.79659201172249 | 0.00516447032148052 | NA | NM_001029872 |
| Itgae | 9.535240201 | -0.0554800444368398 | 0.0642949055234419 | -0.862899540564871 | 0.388192703312539 | NA | NM_172944,NM_008399 |
| Itgb1bp2 | 11.78536701 | -0.0839072480307203 | 0.0623191033910804 | -1.34641295309024 | 0.178169375848314 | NA | NM_013712 |
| Itgb2l | 0.212695677 | -0.00857796107890413 | 0.00973699866180651 | -0.880965621629515 | 0.378336427605205 | NA | NM_008405 |
| Itgb6 | 1.451383634 | 0.0158011587703787 | 0.026279577218267 | 0.601271422258473 | 0.547659220489216 | NA | NM_001159564,NM_021359 |
| Itgb7 | 21.00765731 | 0.0485909947602335 | 0.0749735071589313 | 0.648108866739137 | 0.51691453953185 | NA | NM_013566 |
| Itih1 | 0.108766301 | 0.00519171381361946 | 0.00856003371502575 | 0.606506234257728 | 0.544178643801544 | NA | NM_008406 |
| Itih3 | 1389.227926 | 0.0821099373736827 | 0.0757337852082634 | 1.08419164772876 | NA | NA | NM_008407 |
| Itih4 | 4.422393775 | -0.016545823197931 | 0.0467981740129496 | -0.353557025394893 | 0.723670885392753 | NA | NM_001159299,NM_018746 |
| Itln1 | 0 | NA | NA | NA | NA | NA | NM_010584 |

| | | | | | | | | | | | | | | | |
|---|---|---|---|---|---|---|---|---|---|---|---|---|---|---|---|
| Gpr141 | 0.445240244 | -0.001897651 | 0.037393919 | -0.050747581 | 0.959526662 | NA | | Ivl | 0 | NA | NA | NA | NA | NA | NM_008412 |
| Gpr142 | 0.577809731 | -0.005504678 | 0.058021926 | -0.094872373 | 0.924416201 | NA | | Iyd | 8.970726697 | -0.0717196347509128 | 0.0445675432892792 | -1.60923464606058 | NA | NA | NM_027391 |
| Gpr143 | 2.696545174 | 0.098010003 | 0.112966934 | 0.86759904 | 0.385613872 | NA | | Izumo1 | 2.447151125 | -0.0258923012847694 | 0.0380379726266405 | -0.680696143795932 | 0.496063776878904 | NA | NM_001018013 |
| Gpr15 | 0.999636632 | 0.044679399 | 0.071154745 | 0.627918755 | 0.530057164 | NA | | Izumo2 | 0 | NA | NA | NA | NA | NA | NM_029317 |
| Gpr152 | 2.458535501 | 0.05356629 | 0.111667445 | 0.479694778 | 0.631444441 | NA | | Izumo3 | 0 | NA | NA | NA | NA | NA | NM_027034 |
| Gpr171 | 0.692149418 | -0.025692469 | 0.061038031 | -0.420925582 | 0.673809424 | NA | | Kap | 0 | NA | NA | NA | NA | NA | NM_010594 |
| Gpr174 | 4.064567381 | 0.10918384 | 0.128744314 | 0.848067281 | 0.396400502 | NA | | Kbtbd12 | 11.911247 | -0.0286345344901684 | 0.0659509929733352 | -0.434178974405218 | 0.664158480025683 | NA | NM_029013,NM_001142724 |
| Gpr18 | 0.369409914 | 0.025046824 | 0.047675316 | 0.52536252 | 0.599331203 | NA | | Kbtbd13 | 11.48134195 | -0.0673018119560868 | 0.0660984824162094 | -1.01820510087206 | 0.308580495869688 | NA | NM_028974 |
| Gpr20 | 7.823481412 | -0.207879345 | 0.154900744 | -1.342016442 | 0.179590665 | NA | | Kcna10 | 4.924735604 | 0.00784044068468617 | 0.0461169804495283 | 0.170012013107991 | 0.865000689094597 | NA | NM_001081140 |
| Gpr31b | 0.094392817 | 0.017895942 | 0.027818137 | 0.643319236 | 0.520016977 | NA | | Kcna7 | 0.850958727 | -0.0381728471828127 | 0.0203927603410246 | -1.87188230256497 | 0.0612228885050994 | NA | NM_010596 |
| Gpr33 | 0.079983654 | 0.016740032 | 0.027674178 | 0.604897175 | 0.54524732 | NA | | Kcne1 | 0.910229902 | 0.00482227140965231 | 0.02096028599058 | 0.230060008403875 | 0.818045140077278 | NA | NM_008424 |
| Gpr44 | 3.09601992 | -0.16617475 | 0.125894881 | -1.319948434 | 0.186852235 | NA | | Kcne2 | 59.05984924 | -0.000364193367031491 | 0.0250946169295162 | -0.014512808386532 | NA | NA | NM_134110 |
| Gpr55 | 6.356275839 | -0.303130177 | 0.142468495 | -2.127699726 | 0.033361988 | NA | | Kcne3 | 6.355423897 | -0.0636292703101446 | 0.0538192525009646 | -1.18227711001754 | 0.237095764958083 | NA | NM_001190871,NM_001190870,NM_020574,NM_001190869,NM_001190950 |
| Gpr65 | 12.8400628 | 0.507655199 | 0.144868566 | 3.504246718 | NA | NA | | Kcnj1 | 0.120209049 | -0.00674972139751468 | 0.00863521114606269 | -0.781650996523957 | 0.434419707908266 | NA | NM_001168354,NM_019659 |
| Gpr82 | 0.485069967 | 0.062268854 | 0.049079739 | 1.268728307 | 0.204537983 | NA | | Kcnj13 | 155.011273 | 0.00121225328839355 | 0.0306552302250255 | 0.0395447458556004 | NA | NA | NM_001110227 |
| Gpr87 | 0 | NA | NA | NA | NA | NA | | Kcnj15 | 3.032868895 | -0.010374757244618 | 0.0394226969267555 | -0.263167110659465 | 0.792421779426804 | NA | NM_001039056,NM_001039057,NM_019664,NM_001271687,NM_001271695,NR_073406,NM_001271690,NM_001271691,NM_001271692,NM_001271693 |
| Gpr97 | 3.673893711 | 0.1026279 | 0.117888708 | 0.870549025 | 0.38400044 | NA | | Kcnk15 | 1.461269618 | 0.0172858339952477 | 0.0229742474668986 | 0.752400444025566 | 0.45181027965498 | NA | NM_001030292 |
| Gprc5d | 4.99614278 | 0.030720847 | 0.13167471 | 0.233308637 | 0.815521753 | NA | | Kcnk16 | 0.212695677 | -0.00857796107890413 | 0.00973699866180651 | -0.880965621629515 | 0.378336427605205 | NA | NM_029006 |
| Gprc6a | 0.579680688 | 0.02709434 | 0.040889424 | 0.662624645 | 0.507570982 | NA | | Kcnk18 | 0.217532603 | 0.00684685248201087 | 0.00976000296730706 | 0.701521557416086 | 0.482977586094298 | NA | NM_207261 |
| Gpx2 | 1.839163825 | 0.064538389 | 0.095235325 | 0.67767279 | 0.497979182 | NA | | Kcnk7 | 3.976956415 | 0.0239630714701151 | 0.0409993927384708 | 0.584473814599453 | 0.558901577571778 | NA | NM_010609 |
| Gpx2-ps1 | 7.184999987 | 0.196824021 | 0.134395797 | 1.464510242 | 0.143054587 | NA | | Kcnmb3 | 0.503272123 | 0.00529663843074171 | 0.0127240585271732 | 0.416269574635353 | 0.677212753630587 | NA | NM_001195074 |
| Gpx5 | 0 | NA | NA | NA | NA | NA | | Kcnn4 | 8.696782589 | -0.0216371488083591 | 0.064138373900495 | -0.337351066023708 | 0.735852268341611 | NA | NM_008433,NM_001163510 |
| Grhl2 | 3.923822387 | -0.099916815 | 0.132772092 | -0.752543804 | 0.451724098 | NA | | Kcnq1 | 22.32249441 | 0.011126706204095 | 0.0809327162446823 | 0.137480943682353 | 0.890650654451135 | NA | NM_008434 |
| Grifin | 3.414847239 | -0.120549396 | 0.126160407 | -0.955524786 | 0.339312377 | NA | | Kcnrg | 20.48787797 | 0.108423082197982 | 0.080441899324463 | 1.34784338893661 | 0.17770875924829 | NA | NM_206974,NM_001039105 |
| Grin3b | 5.485007072 | 0.123063841 | 0.137212765 | 0.896883331 | 0.369781176 | NA | | Kcnv2 | 0.636891649 | 0.011752306170045 | 0.012754461277781 | 0.921427092378906 | 0.356827489452057 | NA | NM_183179 |
| Grm6 | 0.477384717 | 0.028178173 | 0.045982715 | 0.612799235 | 0.540009097 | NA | | Kctd19 | 0.430848918 | 0.00680112998966961 | 0.0154512079789059 | 0.440168173191025 | 0.659815309070607 | NA | NM_177791 |
| Grxcr1 | 0.469085263 | 0.061308216 | 0.048717378 | 1.258446538 | 0.20823031 | NA | | Kdelr3 | 16.03923588 | -0.139623215743587 | 0.0716379376486687 | -1.94901221791638 | 0.0512939656501513 | NA | NM_134090 |
| Grxcr2 | 6.163260754 | 0.192086484 | 0.118781075 | 1.61714721 | 0.105846508 | NA | | Kdm5d | 0 | NA | NA | NA | NA | NA | NM_011419 |
| Gsc | 0.17463368 | 0.024560283 | 0.031186242 | 0.787535833 | 0.430968261 | NA | | Keg1 | 0 | NA | NA | NA | NA | NA | NM_029550 |
| Gsc2 | 0.474461318 | 0.009131132 | 0.05516185 | 0.165533453 | 0.868524121 | NA | | Kel | 0.829413177 | 0.0015915661270765 | 0.0182600459029313 | 0.0871611241032535 | 0.930543440182331 | NA | NM_032540 |
| Gsdma | 2.521827192 | -0.235250413 | 0.104617938 | -2.248662297 | 0.02453399 | NA | | Kera | 0.86531832 | 0.00450441083425911 | 0.016856405785879 | 0.267222496389625 | 0.789297856138724 | NA | NM_008438 |
| Gsdma2 | 0 | NA | NA | NA | NA | NA | | Khdc1a | 0 | NA | NA | NA | NA | NA | NM_183322 |
| Gsdma3 | 0 | NA | NA | NA | NA | NA | | Khdc1b | 0.106148608 | 0.00515554063321497 | 0.00854200771766808 | 0.603551390213729 | 0.546141939141592 | NA | NM_001113187 |
| Gsdmc | 1.176063034 | -0.050141483 | 0.086599856 | -0.579001921 | 0.562587877 | NA | | Khdc1c | 0 | NA | NA | NA | NA | NA | NM_001033904 |
| Gsdmc2 | 0 | NA | NA | NA | NA | NA | | Khdc3 | 8.979998024 | -0.0550851964169312 | 0.0551627891173531 | -0.998593386925072 | 0.317991704986153 | NA | NM_025890 |
| Gsdmc3 | 0 | NA | NA | NA | NA | NA | | Kif12 | 0.46281165 | 0.00749841064087392 | 0.015721197953068 | 0.476961785180665 | 0.633389334144131 | NA | NM_010616 |
| Gsdmc4 | 2.780089505 | 0.074454858 | 0.109044961 | 0.682790451 | 0.494739265 | NA | | Kif14 | 15.47479433 | -0.0524818936188718 | 0.0736506179343637 | -0.712579134986251 | 0.476106226776446 | NA | NM_001081258 |
| Gsdmcl-ps | 0 | NA | NA | NA | NA | NA | | Kif15 | 14.90649256 | -0.0156330013583822 | 0.0780155047008 | -0.200383262510918 | 0.841180848607364 | NA | NM_010620 |
| Gsdmcl1 | 0 | NA | NA | NA | NA | NA | | Kif18b | 5.320192453 | -0.0549564577292601 | 0.0443435459778621 | -1.23933385383064 | 0.215221886443438 | NA | NM_197959 |
| Gsg1 | 3.269141262 | 0.006816792 | 0.115749724 | 0.05889251 | 0.953037724 | NA | | Kif2b | 0 | NA | NA | NA | NA | NA | NM_028547 |
| Gsta1 | 0 | NA | NA | NA | NA | NA | | Kif2c | 7.480792864 | 0.0794768134131515 | 0.055501624063382 | 1.43197725312863 | 0.152151619220279 | NA | NM_134471 |
| Gsta2 | 1.303579416 | 0.095315121 | 0.082125978 | 1.160596481 | 0.245806037 | NA | | Kif4-ps | 0 | NA | NA | NA | NA | NA | NR_033653 |
| Gstm3 | 0.787779753 | 0.017407436 | 0.055623753 | 0.312949687 | 0.754318887 | NA | | Kir3dl1 | 0 | NA | NA | NA | NA | NA | NM_177749 |
| Gstt4 | 0.489742234 | 0.039007481 | 0.051990224 | 0.750284927 | 0.453083119 | NA | | Kir3dl2 | 9.851000786 | 0.0264110483045525 | 0.0617459329675042 | 0.427737456302622 | 0.668842276485471 | NA | NM_177748 |
| Gtf2a1l | 4.33086103 | 0.15690267 | 0.132253029 | 1.186382428 | 0.235471308 | NA | | Kis2 | 0.341198496 | 0.00159525481779074 | 0.0112445641955764 | 0.141868976871359 | 0.887183499289969 | NA | NR_003188 |
| Gtsf1 | 2.568829213 | -0.032771883 | 0.11391905 | -0.287676935 | 0.773594047 | NA | | Kiss1 | 15.76579921 | -0.0341850953820401 | 0.0705819865076548 | -0.484331726457325 | 0.628150463268469 | NA | NM_178260 |
| Gtsf1l | 2.368506006 | -0.015161177 | 0.082039574 | -0.184803214 | 0.853383387 | NA | | Kl | 662.9426602 | 0.00978480391490694 | 0.0364448491072125 | 0.26848249216569 | NA | NA | NM_013823 |
| Guca1a | 3.181988315 | 0.157189115 | 0.118609329 | 1.325267722 | 0.185082371 | NA | | Klb | 7.817126689 | -0.0372181683706313 | 0.0585326686004072 | -0.635852922147008 | 0.524872289764552 | NA | NM_031180 |
| Guca2a | 0 | NA | NA | NA | NA | NA | | Klf1 | 7.439810736 | 0.0245707117453698 | 0.056059247850358 | 0.438298990577929 | 0.66116955757361 | NA | NM_010635 |
| Guca2b | 0.378048898 | -0.001605613 | 0.041752189 | -0.03845579 | 0.96932428 | NA | | Klf17 | 0.825057078 | -0.0024642921022481 | 0.0184513341553949 | -0.133556309884919 | 0.893753434237974 | NA | NM_029416 |
| Gucy2d | 0.759356684 | 0.023263342 | 0.067947642 | 0.342371591 | 0.732071265 | NA | | Klhdc7b | 16.61368745 | -0.00340148048965268 | 0.0756701005874992 | -0.0449514466512367 | 0.964146009760973 | NA | NM_001160178 |
| Gulo | 2.04977914 | -0.006919657 | 0.105252772 | -0.065743226 | 0.947582258 | NA | | Klhl10 | 9.834012777 | -0.000162095954088188 | 0.063507667322688 | -0.00255238400214835 | 0.997963494422637 | NA | NM_025727 |
| Gvin1 | 0 | NA | NA | NA | NA | NA | | Klhl30 | 7.677939076 | 0.0474062606681715 | 0.0538098125888538 | 0.880996576412371 | 0.378319673014025 | NA | NM_027551 |
| Gykl1 | 0.079983654 | 0.016740032 | 0.027674178 | 0.604897175 | 0.54524732 | NA | | Klhl31 | 2.798314717 | -0.0136757944025861 | 0.0336559847880236 | -0.406340640118562 | 0.684492328488097 | NA | NM_172925 |
| Gyltl1b | 7.647445691 | 0.212429596 | 0.14326102 | 1.482815046 | 0.138123557 | NA | | Klhl34 | 4.821208803 | -0.0471407236545412 | 0.0467806478614112 | -1.00769711001431 | 0.313599892825028 | NA | NM_001081667 |
| Gypa | 1.66986205 | -0.064037491 | 0.08945648 | -0.715850776 | 0.474083496 | NA | | Klhl41 | 17.48058856 | -0.103141200571693 | 0.0777285033389638 | -1.32694180565793 | 0.184527933817204 | NA | NM_001081087 |
| Gys2 | 3.691120737 | -0.087120239 | 0.122240439 | -0.712695727 | 0.476034061 | NA | | Klk1 | 0.09572496 | 0.00498255289994511 | 0.00847999972823692 | 0.587565219295241 | 0.556824162160132 | NA | NM_010639 |
| Gzma | 0.546533824 | -0.023452341 | 0.046571786 | -0.50357401 | 0.61456076 | NA | | Klk10 | 4.156257411 | -0.0294387201507149 | 0.03908897037663 | -0.75312088978213 | 0.451377272412585 | NA | NM_133712 |
| Gzmb | 0.877155726 | 0.074701388 | 0.061603867 | 1.212608758 | 0.225279439 | NA | | Klk11 | 0.443931641 | -0.0212290374277822 | 0.0152683528378383 | -1.39039473696023 | 0.164409043779599 | NA | NM_019974,NM_001177373 |
| Gzmc | 0.107716603 | -0.020477583 | 0.027954224 | -0.732539849 | 0.46383913 | NA | | Klk12 | 0 | NA | NA | NA | NA | NA | NM_027097 |
| Gzmd | 0 | NA | NA | NA | NA | NA | | Klk13 | 0.598202733 | -0.0182433857947567 | 0.0175333329581267 | -1.04049731094058 | 0.298108912726659 | NA | NM_001039042 |
| Gzme | 0 | NA | NA | NA | NA | NA | | Klk15 | 0 | NA | NA | NA | NA | NA | NM_174865 |
| Gzmf | 0 | NA | NA | NA | NA | NA | | Klk1b1 | 0 | NA | NA | NA | NA | NA | NM_010645 |
| Gzmg | 0 | NA | NA | NA | NA | NA | | Klk1b11 | 0 | NA | NA | NA | NA | NA | NM_010640 |
| Gzmk | 2.414602173 | 0.018978304 | 0.108979724 | 0.174145273 | 0.861751294 | NA | | Klk1b16 | 0 | NA | NA | NA | NA | NA | NM_008454 |
| Gzmn | 0 | NA | NA | NA | NA | NA | | Klk1b21 | 0.255375383 | 0.00708304372802858 | 0.00992471678592415 | 0.713677163873753 | 0.475426830432214 | NA | NM_010642 |
| H19 | 7.225761096 | -0.030305485 | 0.145582921 | -0.208166486 | 0.835098976 | NA | | Klk1b22 | 0 | NA | NA | NA | NA | NA | NM_010114 |
| H1fnt | 0 | NA | NA | NA | NA | NA | | Klk1b24 | 0 | NA | NA | NA | NA | NA | NM_010643 |
| H1foo | 0.308183275 | 0.007047773 | 0.033991235 | 0.207340891 | 0.835743642 | NA | | Klk1b26 | 0 | NA | NA | NA | NA | NA | NM_010644 |
| H2-Ab1 | 94.76645758 | 0.108464792 | 0.114756139 | 0.945176378 | NA | NA | | Klk1b27 | 0 | NA | NA | NA | NA | NA | NM_020268 |
| H2-Ea-ps | 0.079983654 | 0.016740032 | 0.027674178 | 0.604897175 | 0.54524732 | NA | | Klk1b3 | 0.424992894 | -0.00135729073984099 | 0.0120779861810505 | -0.112377239011126 | 0.910524301635971 | NA | NM_008693 |
| H2-Eb2 | 0.536442192 | -0.057169132 | 0.05827754 | -0.980980534 | 0.326602341 | NA | | Klk1b4 | 0.106148608 | 0.00515554063321497 | 0.00854200771766808 | 0.603551390213729 | 0.546141939141592 | NA | NM_010915 |
| H2-L | 0 | NA | NA | NA | NA | NA | | Klk1b5 | 0 | NA | NA | NA | NA | NA | NM_008456 |
| H2-M1 | 0 | NA | NA | NA | NA | NA | | Klk1b7-ps | 0 | NA | NA | NA | NA | NA | NR_033120 |
| H2-M10.1 | 0 | NA | NA | NA | NA | NA | | Klk1b8 | 0 | NA | NA | NA | NA | NA | NM_008457 |
| H2-M10.2 | 0 | NA | NA | NA | NA | NA | | Klk1b9 | 0 | NA | NA | NA | NA | NA | NM_010116 |
| H2-M10.3 | 0.099009884 | 0.018205726 | 0.027896865 | 0.652608315 | 0.514008823 | NA | | Klk4 | 0.215934008 | -0.00149029401866511 | 0.00975123948835942 | -0.152831239602324 | 0.878531361484804 | NA | NM_019928 |
| H2-M10.4 | 0 | NA | NA | NA | NA | NA | | Klk5 | 0 | NA | NA | NA | NA | NA | NM_026806 |
| H2-M10.5 | 0 | NA | NA | NA | NA | NA | | Klk7 | 1.088407242 | -0.0154742467966074 | 0.0240981292245203 | -0.642134775377587 | 0.520785678264974 | NA | NM_011872 |
| H2-M10.6 | 0 | NA | NA | NA | NA | NA | | Klk9 | 2.071874923 | 0.0216059060926366 | 0.0348687679355368 | 0.619634916053824 | 0.535498173696449 | NA | NM_028660 |
| H2-M11 | 0.233393928 | -0.027475306 | 0.032191697 | -0.85349047 | 0.39338737 | NA | | Klkb1 | 3.754403219 | -0.0325091507686063 | 0.0458590974832781 | -0.708892074914041 | 0.478391453470503 | NA | NM_008455 |
| H2-M2 | 0.480100836 | 0.043799064 | 0.056601691 | 0.773811928 | 0.439042018 | NA | | Klra1 | 0 | NA | NA | NA | NA | NA | NM_016659 |
| H2-M9 | 0.119397641 | -0.020477583 | 0.027954224 | -0.732539845 | 0.463839132 | NA | | Klra10 | 0 | NA | NA | NA | NA | NA | NM_008459 |
| H2-Q10 | 1.376818378 | 0.222349937 | 0.089057002 | 2.496714807 | 0.012534972 | NA | | Klra12 | 0 | NA | NA | NA | NA | NA | NM_010646 |
| H2-Q6 | 0 | NA | NA | NA | NA | NA | | Klra13-ps | 0 | NA | NA | NA | NA | NA | NR_033451 |
| H2-Q7 | 0 | NA | NA | NA | NA | NA | | Klra14 | 0 | NA | NA | NA | NA | NA | NM_008461,NM_001252149 |
| H2-Q8 | 0 | NA | NA | NA | NA | NA | | Klra15 | 0 | NA | NA | NA | NA | NA | NM_013793 |
| H2-Q9 | 0 | NA | NA | NA | NA | NA | | Klra17 | 0 | NA | NA | NA | NA | NA | NM_133203 |
| H2-T22 | 1.100885548 | -0.034925214 | 0.086748596 | -0.40260264 | 0.687240568 | NA | | Klra18 | 0 | NA | NA | NA | NA | NA | NM_053153 |
| H2-T3 | 0.370495291 | 0.0385767 | 0.038790203 | 0.994495936 | 0.319981483 | NA | | Klra19 | 0 | NA | NA | NA | NA | NA | NM_053154 |
| H2-T9 | 0 | NA | NA | NA | NA | NA | | Klra2 | 4.23552351 | -0.0253198863105485 | 0.0449592000815367 | -0.563174750988208 | 0.573315894234286 | NA | NM_001170851,NM_008462 |
| H2afb1 | 0.099009884 | 0.018205726 | 0.027896865 | 0.652608315 | 0.514008823 | NA | | Klra21 | 0 | NA | NA | NA | NA | NA | NM_053151 |
| H2afy3 | 0.08731684 | 0.017365056 | 0.027733412 | 0.626142072 | 0.531221759 | NA | | Klra22 | 0 | NA | NA | NA | NA | NA | NM_053152 |
| H2bfm | 0 | NA | NA | NA | NA | NA | | Klra23 | 0 | NA | NA | NA | NA | NA | NM_024470 |
| H60c | 0.178993538 | 0.024685672 | 0.031260828 | 0.789667872 | 0.429721759 | NA | | Klra3 | 0 | NA | NA | NA | NA | NA | NM_010648 |
| Haao | 6.298059786 | 0.074869019 | 0.145676105 | 0.513941658 | 0.607292781 | NA | | Klra33 | 0 | NA | NA | NA | NA | NA | NM_001039118 |
| Habp2 | 0.079983654 | 0.016740032 | 0.027674178 | 0.604897175 | 0.54524732 | NA | | Klra4 | 0 | NA | NA | NA | NA | NA | NM_010649,NM_001252577 |
| Hal | 0.721176579 | 0.036635741 | 0.061061229 | 0.599983687 | 0.548517108 | NA | | Klra5 | 0 | NA | NA | NA | NA | NA | NM_008463 |
| Hamp | 0.283178451 | 0.028390488 | 0.033368195 | 0.850824814 | 0.394866675 | NA | | Klra6 | 0 | NA | NA | NA | NA | NA | NM_008464 |
| Hamp2 | 0.505798573 | 0.029734949 | 0.046020057 | 0.646130196 | 0.518195041 | NA | | Klra7 | 0 | NA | NA | NA | NA | NA | NM_014194,NM_001110323 |
| Hand1 | 0 | NA | NA | NA | NA | NA | | Klra8 | 0.097165704 | -0.00637828455464458 | 0.00848781691535724 | -0.75146349388194 | 0.452373763047372 | NA | NM_010650,NM_001101620 |
| Hand2 | 0 | NA | NA | NA | NA | NA | | Klra9 | 0.308221406 | 0.00253133711108074 | 0.0124107148809387 | 0.203963843772494 | 0.838381753432823 | NA | NM_010651 |
| Hao1 | 3.010711308 | 0.088411103 | 0.097130101 | 0.910233815 | 0.362699215 | NA | | Klrb1 | 0.233836299 | 0.00743372775021133 | 0.0101683766597817 | 0.731063374118848 | 0.464740444385902 | NA | NM_001099918 |
| Hao2 | 0 | NA | NA | NA | NA | NA | | Klrb1-ps1 | 0 | NA | NA | NA | NA | NA | NR_073569 |
| Has2as | 2.140560232 | 0.089250519 | 0.104967094 | 0.850271409 | 0.395174208 | NA | | Klrb1a | 1.525437933 | -0.0213265388760635 | 0.0307071470583896 | -0.694513848372533 | 0.487360019211565 | NA | NM_001159902,NM_010737 |
| Havcr1 | 0.275064582 | 0.010932281 | 0.038417727 | 0.284563456 | 0.775978612 | NA | | Klrb1b | 2.350758835 | -0.0476484247248414 | 0.02934471017639 | -1.62374834981939 | 0.104429527451778 | NA | NM_030599 |
| Hba-a1 | 0 | NA | NA | NA | NA | NA | | Klrb1c | 2.538109108 | -0.0591076911157267 | 0.0323701303825515 | -1.82599484207168 | 0.0678510387841022 | NA | NM_008527,NM_001159904 |
| Hba-a2 | 0 | NA | NA | NA | NA | NA | | Klrb1f | 0.538795351 | 0.00894919074331237 | 0.0149864819206075 | 0.597150871747065 | 0.550406656091267 | NA | NM_153094,NR_024262,NR_024263 |
| Hba-x | 0 | NA | NA | NA | NA | NA | | Klrc1 | 0.113843286 | 0.00525947738444006 | 0.00859368002043176 | 0.612016897526494 | 0.540526577873086 | NA | NM_001136068,NM_010652 |
| Hbb-b1 | 0 | NA | NA | NA | NA | NA | | Klrc2 | 0 | NA | NA | NA | NA | NA | NM_010653,NM_001098669 |
| Hbb-b2 | 0 | NA | NA | NA | NA | NA | | Klrc3 | 0 | NA | NA | NA | NA | NA | NM_021378 |
| Hbb-bh1 | 0 | NA | NA | NA | NA | NA | | Klrd1 | 0.51821185 | -0.00504928813924851 | 0.0147324435090702 | -0.34273256409501 | 0.731799662266437 | NA | NM_010654 |
| Hbb-bh2 | 0 | NA | NA | NA | NA | NA | | Klre1 | 0.48193227 | -0.00120568868924353 | 0.0141586618979216 | -0.0851555533945277 | 0.932137725428612 | NA | NM_153590 |
| Hbb-y | 1.966967144 | 0.046089237 | 0.090247736 | 0.510696876 | 0.609563329 | NA | | Klrg1 | 7.076320156 | 0.0250310113153765 | 0.0584790062944813 | 0.428034142531911 | 0.66862626304525 | NA | NM_016970 |
| Hbq1b | 3.15856341 | 0.03774795 | 0.120402931 | 0.313513551 | 0.753890527 | NA | | Klrg2 | 9.928711501 | 0.0486590791970757 | 0.0579408337723974 | 0.839806333961603 | 0.401016981503537 | NA | NM_001033171 |
| Hcrt | 2.286357048 | 0.205823382 | 0.106548396 | 1.931736086 | 0.053392088 | NA | | Klri1 | 0.63302697 | -0.000739290844147518 | 0.0134895523394399 | -0.0548046981504365 | 0.956294057447289 | NA | NM_001012520 |
| Hdgfl1 | 0.094392817 | 0.017895942 | 0.027818137 | 0.643319236 | 0.520016977 | NA | | Klri2 | 4.297419383 | -0.0196799646802307 | 0.0430253936650727 | -0.457403477430738 | 0.647381065350997 | NA | NM_177155 |
| Hdhd1a | 0 | NA | NA | NA | NA | NA | | Klrk1 | 1.238867292 | -0.0232703998484142 | 0.0250709310833713 | -0.928182514284388 | 0.35331290309943 | NA | NM_033078,NM_001083322 |
| Helt | 5.602517392 | -0.02684434 | 0.147888845 | -0.181517001 | 0.85596179 | NA | | Kmo | 2.916864259 | -0.014711536181673 | 0.0399967766332233 | -0.367818044853467 | 0.713008912990261 | NA | NM_133809 |
| Hemgn | 1.339731961 | -0.144898562 | 0.077259302 | -1.875483705 | 0.060726209 | NA | | Kncn | 4.642396435 | -0.0180380058997546 | 0.0491574241939664 | -0.366943675254013 | 0.713661032238039 | NA | NM_001039124 |
| Hemt1 | 0 | NA | NA | NA | NA | NA | | Kng1 | 0 | NA | NA | NA | NA | NA | NM_023125,NM_001102411,NM_001102412 |
| Hephl1 | 5.853980551 | -0.080719501 | 0.058643611 | -1.376441522 | 0.16868498 | NA | | Kng2 | 0.397381925 | 0.00594993753234186 | 0.0136539817993225 | 0.435765743633633 | 0.663006700565674 | NA | NM_001102409,NM_001102410,NM_201375 |
| Herpud1 | 2018.258476 | 0.136115423 | 0.117697413 | 1.156486104 | NA | NA | | Kntc1 | 17.95387604 | 0.0432655993680273 | 0.0783431806069386 | 0.552257376236719 | 0.580772029030485 | NA | NM_001042421 |
| Hes2 | 0.494522006 | 0.031199547 | 0.04639328 | 0.672501424 | 0.501264534 | NA | | Kpna7 | 0.108766301 | 0.00519171381361946 | 0.00856003371502575 | 0.606506234257728 | 0.544178643801544 | NA | NM_001013774 |
| Hesx1 | 0.486850878 | 0.009782776 | 0.044755885 | 0.218580767 | 0.826976636 | NA | | Kprp | 0.240418098 | -0.0088029805258837 | 0.00986414519853158 | -0.892422034419582 | 0.372166765031741 | NA | NM_028629 |
| Hexim1 | 1888.243271 | 0.159838936 | 0.138901855 | 1.150732915 | NA | NA | | Krt13 | 0.106347839 | -0.00653347234140708 | 0.00854389509723638 | -0.764694822098226 | 0.444453290197145 | NA | NM_010662 |
| Hgd | 1.956849132 | -0.01272358 | 0.100747789 | -0.126291406 | 0.899501259 | NA | | Krt14 | 1.247129162 | 0.00328403185712699 | 0.0225013676807154 | 0.145948099854461 | 0.883962359899648 | NA | NM_016958 |
| Hgfac | 4.516198336 | 0.12697407 | 0.117480587 | 1.080808945 | 0.279782109 | NA | | Krt15 | 3.28384213 | -0.00271420857583465 | 0.0273898257232093 | -0.0990955036831327 | 0.921062442058557 | NA | NM_008469 |
| Hhla1 | 1.092684631 | 0.036603875 | 0.080081372 | 0.457083514 | 0.64761102 | NA | | Krt16 | 0.794999316 | -0.0114342328407381 | 0.0215197252237376 | -0.531337306673668 | 0.595185057461477 | NA | NM_008470 |
| Hibch | 504.4762041 | -0.154015475 | 0.11809773 | -1.304135786 | NA | NA | | Krt17 | 3.364229612 | -0.0152541894995539 | 0.0428564930124183 | -0.355936485403362 | 0.72188813134641 | NA | NM_010663 |
| Higd1c | 0 | NA | NA | NA | NA | NA | | Krt19 | 0.640926902 | -0.00345313849404909 | 0.0173394395742869 | -0.199149371538503 | 0.842145901724335 | NA | NM_008471 |
| Hils1 | 0 | NA | NA | NA | NA | NA | | Krt20 | 3.144813577 | -0.00736023455090807 | 0.0420662254775097 | -0.174967791080831 | 0.861104942136909 | NA | NM_023256 |
| Hipk1 | 6342.8212 | 0.009305169 | 0.097045344 | 0.095884753 | NA | NA | | Krt23 | 1.465321169 | 0.0380953128121441 | 0.0248845152142111 | 1.53088426614751 | 0.125797998462844 | NA | NM_033373 |
| Hipk4 | 7.761176972 | -0.056312021 | 0.141812906 | -0.397086714 | 0.691303517 | NA | | Krt24 | 0.784472859 | -0.00655563145152798 | 0.0182208948037237 | -0.359786471638498 | 0.719006820738146 | NA | NM_029393 |
| Hist1h1a | 0.735221671 | 0.004778139 | 0.061109148 | 0.078190247 | 0.93767672 | NA | | Krt25 | 2.46567565 | 0.0304200168253751 | 0.0317191573060355 | 0.959042402415487 | 0.337537384480137 | NA | NM_133730 |
| Hist1h1b | 0.892063337 | 0.057228649 | 0.062091197 | 0.921687 | 0.356691864 | NA | | Krt26 | 1.845863774 | 0.0510049446186743 | 0.0281370043436923 | 1.81273542825139 | 0.0698726374139353 | NA | NM_001033397 |
| Hist1h1d | 3.359303027 | -0.074191839 | 0.098423094 | -0.753805192 | 0.450966206 | NA | | Krt27 | 0.09572496 | 0.00498255289994511 | 0.00847999972823692 | 0.587565219295241 | 0.556824162160132 | NA | NM_010666 |
| Hist1h1t | 0 | NA | NA | NA | NA | NA | | Krt28 | 4.287144234 | 0.030535663470545 | 0.0457235647559659 | 0.667832082505353 | 0.504240784905114 | NA | NM_027574 |
| Hist1h2aa | 0 | NA | NA | NA | NA | NA | | Krt31 | 0.865465048 | 0.0156859484054229 | 0.0226204267827649 | 0.693441753158009 | 0.48803236729368 | NA | NM_010659 |
| Hist1h2ab | 0.411730969 | -0.0302294 | 0.043889081 | -0.688768129 | 0.490969195 | NA | | Krt32 | 0.995237888 | -0.00829226226837843 | 0.0210679269150701 | -0.393596498687628 | 0.693878966640475 | NA | NM_001159374 |
| Hist1h2ad | 0.850310816 | 0.055425758 | 0.053806532 | 1.030093493 | 0.302966119 | NA | | Krt33a | 0.715278666 | 0.00610176527515277 | 0.017545607969377 | 0.347765964325796 | 0.728015952544506 | NA | NM_027983 |
| Hist1h2ae | 0.401150917 | -0.033366739 | 0.048906957 | -0.682249333 | 0.495081305 | NA | | Krt33b | 0.113843286 | 0.00525947738444006 | 0.00859368002043176 | 0.612016897526494 | 0.540526577873086 | NA | NM_013570 |
| Hist1h2af | 0.096071043 | -0.020098795 | 0.027825948 | -0.722304036 | 0.470107572 | NA | | Krt34 | 0.127687691 | 0.00542298404407807 | 0.00869724871942359 | 0.62352868349814 | 0.532937159177571 | NA | NM_027563 |
| Hist1h2ag | 0 | NA | NA | NA | NA | NA | | Krt35 | 0 | NA | NA | NA | NA | NA | NM_016880 |
| Hist1h2ah | 0 | NA | NA | NA | NA | NA | | Krt36 | 0 | NA | NA | NA | NA | NA | NM_001174099 |
| Hist1h2ai | 0 | NA | NA | NA | NA | NA | | Krt39 | 0 | NA | NA | NA | NA | NA | NM_213730 |
| Hist1h2ak | 0.116696964 | -0.020477583 | 0.027954224 | -0.732539847 | 0.463839131 | NA | | Krt4 | 0 | NA | NA | NA | NA | NA | NM_008475 |
| Hist1h2an | 0 | NA | NA | NA | NA | NA | | Krt40 | 0 | NA | NA | NA | NA | NA | NM_001039666 |
| Hist1h2ao | 0 | NA | NA | NA | NA | NA | | Krt42 | 0.226357657 | -0.0012264097261801 | 0.00980336873216358 | -0.125100846421945 | 0.900443713214317 | NA | NM_212483 |
| Hist1h2ap | 0 | NA | NA | NA | NA | NA | | Krt5 | 0 | NA | NA | NA | NA | NA | NM_027011 |
| Hist1h2ba | 0 | NA | NA | NA | NA | NA | | Krt6a | 0.54329922 | 0.00949149198305067 | 0.0150322747613217 | 0.631407563642495 | 0.527774071765445 | NA | NM_008476 |
| Hist1h2bb | 1.107346981 | -0.060832732 | 0.079704043 | -0.763232701 | 0.44532463 | NA | | Krt6b | 0 | NA | NA | NA | NA | NA | NM_010669 |
| Hist1h2bf | 0.215050351 | -0.027006021 | 0.031907566 | -0.846382992 | 0.397339123 | NA | | Krt7 | 4.008040747 | -0.0199181702269446 | 0.0455583212102509 | -0.437201584646251 | 0.661965160622737 | NA | NM_033073 |
| Hist1h2bh | 1.279162208 | 0.0237592 | 0.074111338 | 0.320587925 | 0.74852269 | NA | | Krt71 | 0.106347839 | -0.00653347234140708 | 0.00854389509723638 | -0.764694822098226 | 0.444453290197145 | NA | NM_019956 |
| Hist1h2bj | 0.75689519 | -0.059168597 | 0.067594424 | -0.875347296 | 0.381384967 | NA | | Krt72 | 0 | NA | NA | NA | NA | NA | NM_213728 |
| Hist1h2bk | 0 | NA | NA | NA | NA | NA | | Krt74 | 0.108766301 | 0.00519171381361946 | 0.00856003371502575 | 0.606506234257728 | 0.544178643801544 | NA | NR_033444 |
| Hist1h2bl | 0 | NA | NA | NA | NA | NA | | Krt75 | 1.20445185 | 0.0476832619066961 | 0.0254314828652817 | 1.87496978289818 | 0.0607968806920877 | NA | NM_133357 |
| Hist1h2bm | 0 | NA | NA | NA | NA | NA | | Krt76 | 0.314698307 | 0.00249824212998287 | 0.0109738852417382 | 0.227653385738081 | 0.819915713647591 | NA | NM_001033177 |
| Hist1h2bn | 3.186758882 | -0.141833258 | 0.11571051 | -1.225759514 | 0.220289187 | NA | | Krt78 | 1.146559101 | 0.0123583451898674 | 0.024150255895207 | 0.511727297776588 | 0.608841878482975 | NA | NM_212487 |
| Hist1h2bp | 0.646335094 | -0.079163576 | 0.053841353 | -1.470311779 | 0.141477332 | NA | | Krt79 | 0.443603775 | 0.00781161574515981 | 0.0155660529051266 | 0.501836643673948 | 0.615782434690321 | NA | NM_146063 |
| Hist1h2bq | 0 | NA | NA | NA | NA | NA | | Krt81 | 0 | NA | NA | NA | NA | NA | NM_001166157 |
| Hist1h2br | 0 | NA | NA | NA | NA | NA | | Krt82 | 0.097165704 | -0.00637828455464458 | 0.00848781691535724 | -0.75146349388194 | 0.452373763047372 | NA | NM_053249 |
| Hist1h3a | 0 | NA | NA | NA | NA | NA | | Krt83 | 1.898568809 | 0.0153874881280809 | 0.029578209770744 | 0.520230542935049 | 0.602902899753147 | NA | NM_001003668 |
| Hist1h3b | 0.303023688 | 0.009739077 | 0.039388711 | 0.247255542 | 0.804710463 | NA | | Krt84 | 0.108766301 | 0.00519171381361946 | 0.00856003371502575 | 0.606506234257728 | 0.544178643801544 | NA | NM_008474 |
| Hist1h3c | 0.390161855 | -0.000140598 | 0.036193922 | -0.003884577 | 0.996900564 | NA | | Krt85 | 0.551209933 | 0.00205829740722053 | 0.0164676731494798 | 0.124990178547815 | 0.900531325746378 | NA | NM_016879 |
| Hist1h3d | 2.527979752 | -0.088721967 | 0.106429093 | -0.833625136 | 0.404492257 | NA | | Krt86 | 0.337255936 | 0.012887312700231 | 0.0133695761693565 | 0.963928290394809 | 0.335081879137593 | NA | NM_010667 |
| Hist1h3e | 0.079983654 | 0.016740032 | 0.027674178 | 0.604897175 | 0.54524732 | NA | | Krtap1-3 | 0 | NA | NA | NA | NA | NA | NM_001085526 |
| Hist1h3f | 0.094392817 | 0.017895942 | 0.027818137 | 0.643319236 | 0.520016977 | NA | | Krtap1-4 | 0 | NA | NA | NA | NA | NA | NM_001039502 |
| Hist1h3g | 0.517847881 | -0.057108045 | 0.05793189 | -0.985779063 | 0.324241542 | NA | | Krtap1-5 | 0 | NA | NA | NA | NA | NA | NM_027157 |
| Hist1h3h | 0.116696964 | -0.020477583 | 0.027954224 | -0.732539847 | 0.463839131 | NA | | Krtap10-10 | 0.384340583 | 0.00429661131557121 | 0.0117774169097335 | 0.364817799055773 | 0.715247417762362 | NA | NM_001024709 |
| Hist1h3i | 0.497448118 | -0.02127213 | 0.055798949 | -0.381228152 | 0.703033961 | NA | | Krtap10-4 | 0 | NA | NA | NA | NA | NA | NM_001135991 |
| Hist1h4a | 0.286053643 | -0.017027416 | 0.03902234 | -0.436350468 | 0.662582471 | NA | | Krtap11-1 | 0.217374753 | -0.00904456656736547 | 0.00999660569474087 | -0.90476376117583 | 0.365590553873005 | NA | NM_001113406 |
| Hist1h4b | 0.496263686 | 0.037917571 | 0.052109423 | 0.727652863 | 0.466826112 | NA | | Krtap12-1 | 0 | NA | NA | NA | NA | NA | NM_010670 |
| Hist1h4c | 1.888179012 | -0.122173901 | 0.088073245 | -1.387185192 | 0.1653853 | NA | | Krtap13 | 0 | NA | NA | NA | NA | NA | NM_010671 |
| Hist1h4d | 1.393488188 | 0.116951635 | 0.087710608 | 1.333380738 | 0.18240689 | NA | | Krtap13-1 | 0.191449919 | 0.00665508079272897 | 0.00962405648474729 | 0.691504752001018 | 0.489248395458281 | NA | NM_183189 |
| Hist1h4f | 0.392621292 | 0.024891049 | 0.048289092 | 0.515459033 | 0.60623229 | NA | | Krtap14 | 0 | NA | NA | NA | NA | NA | NM_013707 |
| Hist1h4j | 1.149130049 | -0.015105454 | 0.073409579 | -0.205769517 | 0.836970954 | NA | | Krtap15 | 0 | NA | NA | NA | NA | NA | NM_013713 |
| Hist1h4k | 7.871869586 | -0.063463666 | 0.135785275 | -0.467382534 | 0.640226217 | NA | | Krtap16-1 | 0 | NA | NA | NA | NA | NA | NM_130870 |
| Hist1h4m | 0.211846316 | 0.025707621 | 0.031860375 | 0.806883811 | 0.419733426 | NA | | Krtap16-3 | 0 | NA | NA | NA | NA | NA | NM_183296 |
| Hist1h4n | 0 | NA | NA | NA | NA | NA | | Krtap17-1 | 0.895073387 | -0.00624204304225022 | 0.0223531770478874 | -0.279246347348202 | 0.780055777644527 | NA | NM_001099774 |
| Hist2h2aa1 | 0 | NA | NA | NA | NA | NA | | Krtap19-1 | 0 | NA | NA | NA | NA | NA | NM_130876 |
| Hist2h2aa2 | 0 | NA | NA | NA | NA | NA | | Krtap19-3 | 0 | NA | NA | NA | NA | NA | NM_130857 |
| Hist2h2ab | 1.662834137 | 0.028157252 | 0.095539667 | 0.294717924 | 0.768209369 | NA | | Krtap19-4 | 0 | NA | NA | NA | NA | NA | NM_130873 |
| Hist2h2ac | 0.099009884 | 0.018205726 | 0.027896865 | 0.652608315 | 0.514008823 | NA | | Krtap19-5 | 0 | NA | NA | NA | NA | NA | NM_010676 |
| Hist2h2bb | 0.094392817 | 0.017895942 | 0.027818137 | 0.643319236 | 0.520016977 | NA | | Krtap19-9b | 0 | NA | NA | NA | NA | NA | NM_133359 |
| Hist2h3b | 0 | NA | NA | NA | NA | NA | | Krtap2-4 | 0 | NA | NA | NA | NA | NA | NM_027800 |
| Hist2h4 | 4.721943855 | 0.038976049 | 0.137838291 | 0.282766486 | 0.777355847 | NA | | Krtap20-2 | 0 | NA | NA | NA | NA | NA | NM_001163615 |
| Hist3h2bb-ps | 4.869970795 | 0.030168454 | 0.132748238 | 0.227260677 | 0.820221048 | NA | | Krtap21-1 | 0 | NA | NA | NA | NA | NA | NM_028621 |
| Hmga1-rs1 | 0 | NA | NA | NA | NA | NA | | Krtap22-2 | 0 | NA | NA | NA | NA | NA | NM_001191018 |
| Hmga2 | 2.380176363 | 0.038319994 | 0.11059335 | 0.346494555 | 0.728971076 | NA | | Krtap24-1 | 0 | NA | NA | NA | NA | NA | NM_001163141 |
| Hmgb1-rs17 | 0.683694727 | 0.038873006 | 0.066191532 | 0.587280648 | 0.557015236 | NA | | Krtap26-1 | 0 | NA | NA | NA | NA | NA | NM_027105 |
| Hmgb4 | 0.105923158 | 0.018524693 | 0.027977518 | 0.662127842 | 0.507889294 | NA | | Krtap27-1 | 0 | NA | NA | NA | NA | NA | NM_001163105 |
| Hmx1 | 0 | NA | NA | NA | NA | NA | | Krtap3-1 | 0 | NA | NA | NA | NA | NA | NM_023511 |
| Hmx2 | 0.489484589 | -0.049559653 | 0.043029867 | -1.15175008 | 0.249423787 | NA | | Krtap3-2 | 0 | NA | NA | NA | NA | NA | NM_025720 |
| Hmx3 | 1.269676182 | -0.121350778 | 0.065944701 | -1.840189982 | 0.06574035 | NA | | Krtap3-3 | 0 | NA | NA | NA | NA | NA | NM_025524 |
| Hnf1a | 0.362381421 | 0.026644729 | 0.047496261 | 0.560985827 | 0.574807199 | NA | | Krtap31-1 | 0 | NA | NA | NA | NA | NA | NM_027568 |
| Hnf4a | 0.311993147 | -0.014421208 | 0.039860783 | -0.361789373 | 0.717509434 | NA | | Krtap31-2 | 0 | NA | NA | NA | NA | NA | NM_001025244 |
| Hnf4g | 0.824562923 | -0.076709381 | 0.070664541 | -1.085542766 | 0.277681333 | NA | | Krtap4-1 | 0 | NA | NA | NA | NA | NA | NM_001048196 |
| Hormad1 | 0 | NA | NA | NA | NA | NA | | Krtap4-13 | 0 | NA | NA | NA | NA | NA | NM_027087 |
| Hormad2 | 1.272288978 | -0.024683576 | 0.075761823 | -0.325804936 | 0.744571944 | NA | | Krtap4-16 | 0.366568861 | -0.00548477312881077 | 0.011623202474977 | -0.471881406231944 | 0.637011440575618 | NA | NM_001013823 |
| Hotair | 0 | NA | NA | NA | NA | NA | | Krtap4-2 | 0 | NA | NA | NA | NA | NA | NM_026807 |
| Hoxa1 | 1.25874131 | 0.084597018 | 0.078748822 | 1.07426391 | 0.282704415 | NA | | Krtap4-6 | 0 | NA | NA | NA | NA | NA | NM_026834 |
| Hoxa10 | 0 | NA | NA | NA | NA | NA | | Krtap4-7 | 0 | NA | NA | NA | NA | NA | NM_029613 |
| Hoxa11 | 0 | NA | NA | NA | NA | NA | | Krtap4-8 | 0 | NA | NA | NA | NA | NA | NM_001085547 |
| Hoxa11as | 0 | NA | NA | NA | NA | NA | | Krtap4-9 | 0 | NA | NA | NA | NA | NA | NM_001085548 |
| Hoxa13 | 0 | NA | NA | NA | NA | NA | | Krtap5-1 | 0.12890128 | -0.0068640067066208 | 0.00870804083610955 | -0.788237771940376 | 0.430557639008166 | NA | NM_015808 |
| Hoxa2 | 0.098353386 | -0.020212831 | 0.02786466 | -0.725393061 | 0.468210929 | NA | | Krtap5-2 | 0.257802559 | -0.00892927589713349 | 0.00993479775307735 | -0.898787888698349 | 0.368765652383134 | NA | NM_027844 |
| Hoxa3 | 0 | NA | NA | NA | NA | NA | | Krtap5-3 | 0 | NA | NA | NA | NA | NA | NM_023860 |
| Hoxa4 | 0.176054697 | -0.002190495 | 0.031239146 | -0.070120197 | 0.944097991 | NA | | Krtap5-4 | 0 | NA | NA | NA | NA | NA | NM_015809 |
| Hoxa5 | 0.192142086 | -0.026384592 | 0.031527717 | -0.836869715 | 0.402665805 | NA | | Krtap5-5 | 0 | NA | NA | NA | NA | NA | NM_001037822 |
| Hoxa6 | 0 | NA | NA | NA | NA | NA | | Krtap6-1 | 0 | NA | NA | NA | NA | NA | NM_010672 |
| Hoxa7 | 0.815988888 | -0.065845293 | 0.064875033 | -1.014955824 | 0.310126886 | NA | | Krtap6-2 | 0 | NA | NA | NA | NA | NA | NM_010673 |
| Hoxa9 | 0 | NA | NA | NA | NA | NA | | Krtap6-5 | 0 | NA | NA | NA | NA | NA | NM_130856 |
| Hoxb1 | 0 | NA | NA | NA | NA | NA | | Krtap7-1 | 0.608701423 | -0.00256536330197423 | 0.0171609609923798 | -0.149488324291009 | 0.88116832171686 | NA | NM_027771 |
| Hoxb13 | 0.186326724 | 0.024939385 | 0.031411053 | 0.793968451 | 0.427213798 | NA | | Krtap8-1 | 0 | NA | NA | NA | NA | NA | NM_010675 |
| Hoxb2 | 0.183387883 | -0.001444816 | 0.031390672 | -0.046026918 | 0.963288796 | NA | | Krtap9-1 | 0 | NA | NA | NA | NA | NA | NM_015741 |
| Hoxb3 | 0.200315975 | 0.025370485 | 0.031664208 | 0.801235428 | 0.422995365 | NA | | Krtap9-3 | 0 | NA | NA | NA | NA | NA | NM_029351 |
| Hoxb4 | 1.229976046 | -0.070115861 | 0.075895558 | -0.923846696 | 0.355566139 | NA | | Krtap9-5 | 0 | NA | NA | NA | NA | NA | NM_001085527 |
| Hoxb5 | 0.482559746 | 0.027836625 | 0.0455364 | 0.611304907 | 0.540997743 | NA | | Krtdap | 0.212695677 | -0.00857796107890413 | 0.00973699866180651 | -0.880965621629515 | 0.378336427605205 | NA | NM_001033131 |
| Hoxb6 | 0.096071043 | -0.020098795 | 0.027825948 | -0.722304036 | 0.470107572 | NA | | Kynu | 4.395592164 | -0.014050820847474 | 0.037781318615131 | -0.371898635688348 | 0.709968320960979 | NA | NM_027552 |
| Hoxb7 | 0 | NA | NA | NA | NA | NA | | L1td1 | 3.977013092 | -0.0309272664086137 | 0.0451660813294103 | -0.684745399607538 | 0.493504595149195 | NA | NM_001081202 |
| Hoxb8 | 0.964034454 | 0.053194504 | 0.061906427 | 0.859272723 | 0.390190069 | NA | | LOC100038947 | 0.255375383 | 0.00708304372802858 | 0.00992471678592415 | 0.713677163873753 | 0.475426830432214 | NA | NM_001173459 |
| Hoxb9 | 0.352791569 | -0.038810225 | 0.038158703 | -1.017074017 | 0.30911822 | NA | | LOC100043315 | 0.548750089 | 0.0192720871638031 | 0.0163109611450132 | 1.18154209261269 | 0.237387442149662 | NA | NR_015482 |
| Hoxc10 | 0.238795281 | -0.027601606 | 0.032267774 | -0.855392297 | 0.392334007 | NA | | LOC100048884 | 0.227686571 | 0.00691460391109036 | 0.00980755285755433 | 0.705028462402254 | 0.480792523671872 | NA | NM_001199333 |
| Hoxc11 | 0 | NA | NA | NA | NA | NA | | LOC100502924 | 0 | NA | NA | NA | NA | NA | NM_001256481 |
| Hoxc12 | 0 | NA | NA | NA | NA | NA | | LOC100503280 | 0.340400173 | -0.006017291565113 | 0.0128784657053825 | -0.467236680422116 | 0.640330553117361 | NA | NM_001251890 |
| Hoxc13 | 0.321356363 | -0.01100297 | 0.034551013 | -0.318455781 | 0.750139232 | NA | | LOC100504608 | 2.507285137 | 0.0296512175370918 | 0.0306773457214701 | 0.966550946301064 | 0.333768566893965 | NA | NM_001205036 |
| Hoxc4 | 0.990867558 | 0.0197887 | 0.077313249 | 0.255954836 | 0.797985711 | NA | | LOC100861615 | 6.141076021 | 0.0636428046318823 | 0.0564273641187605 | 1.12787130190834 | 0.259374275331748 | NA | NM_001270812 |
| Hoxc5 | 0.40113891 | -0.035620199 | 0.048927249 | -0.728023742 | 0.466599053 | NA | | LOC100861978 | 0 | NA | NA | NA | NA | NA | NM_001267703,NM_001267702,NM_001267704 |
| Hoxc6 | 0 | NA | NA | NA | NA | NA | | LOC171588 | 6.615725779 | -0.00218628843326067 | 0.0529848691191081 | -0.041262505119074 | 0.967086624178187 | NA | NR_038065 |
| Hoxc8 | 0.116696964 | -0.020477583 | 0.027954224 | -0.732539847 | 0.463839131 | NA | | LOC547349 | 0.12890128 | -0.0068640067066208 | 0.00870804083610955 | -0.788237771940376 | 0.430557639008166 | NA | NM_001025208 |
| Hoxc9 | 0 | NA | NA | NA | NA | NA | | LOC666331 | 0 | NA | NA | NA | NA | NA | NM_001256318 |
| Hoxd1 | 2.362826339 | 0.059516032 | 0.105533821 | 0.563952218 | 0.572786651 | NA | | Lactbl1 | 1.097861156 | 0.0142384788110028 | 0.0254115965922406 | 0.560314215571582 | 0.575265132761939 | NA | NM_001243262 |
| Hoxd10 | 0 | NA | NA | NA | NA | NA | | Lalba | 0.556711097 | -0.014001777383149 | 0.0154439683508995 | -0.9066178500899 | 0.36460891901238 | NA | NM_010679 |
| Hoxd11 | 0 | NA | NA | NA | NA | NA | | Lamp3 | 0.255375383 | 0.00708304372802858 | 0.00992471678592415 | 0.713677163873753 | 0.475426830432214 | NA | NM_177356 |

| | | | | | | |
|---|---|---|---|---|---|---|
| Hoxd12 | 0 | NA | NA | NA | NA | NA |
| Hoxd13 | 0 | NA | NA | NA | NA | NA |
| Hoxd3 | 0.301004085 | 0.013136417 | 0.039354285 | 0.333798913 | 0.738531304 | NA |
| Hoxd4 | 0.099009884 | 0.018205726 | 0.027896865 | 0.652608315 | 0.514008823 | NA |
| Hoxd8 | 0.079983654 | 0.016740032 | 0.027674178 | 0.604897175 | 0.54524732 | NA |
| Hoxd9 | 0 | NA | NA | NA | NA | NA |
| Hp | 6.869078324 | -0.214982673 | 0.130941795 | -1.641818585 | 0.100627606 | NA |
| Hpd | 2.115674275 | 0.049641122 | 0.095095014 | 0.522016028 | 0.601659171 | NA |
| Hpse2 | 2.599510969 | -0.011149662 | 0.107279551 | -0.103930919 | 0.91722417 | NA |
| Hrasls5 | 0.936130138 | -0.042000539 | 0.065679436 | -0.63947777 | 0.522512171 | NA |
| Hrg | 1.017409249 | 0.01373711 | 0.070249651 | 0.195547015 | 0.844964735 | NA |
| Hrh4 | 0 | NA | NA | NA | NA | NA |
| Hrnr | 0.570133794 | 0.036094382 | 0.048404861 | 0.745676794 | 0.45586268 | NA |
| Hs3st6 | 7.689334033 | -0.138802215 | 0.156066491 | -0.889378713 | 0.373799581 | NA |
| Hsd11b2 | 0.67038813 | 0.035419061 | 0.061414033 | 0.576725864 | 0.564124657 | NA |
| Hsd17b1 | 7.248374535 | 0.160981282 | 0.134076167 | 1.200670373 | 0.229879091 | NA |
| Hsd17b13 | 5.879659122 | 0.06002237 | 0.135413755 | 0.443251651 | 0.657583723 | NA |
| Hsd17b14 | 6.812879294 | 0.279941837 | 0.146770905 | 1.907338764 | 0.056476734 | NA |
| Hsd17b2 | 5.158410381 | 0.056811922 | 0.131521664 | 0.431958663 | 0.665771457 | NA |
| Hsd17b3 | 0 | NA | NA | NA | NA | NA |
| Hsd17b6 | 0 | NA | NA | NA | NA | NA |
| Hsd3b1 | 0 | NA | NA | NA | NA | NA |
| Hsd3b2 | 0.098353386 | -0.020212831 | 0.02786466 | -0.725393061 | 0.468210929 | NA |
| Hsd3b3 | 0 | NA | NA | NA | NA | NA |
| Hsd3b4 | 0 | NA | NA | NA | NA | NA |
| Hsd3b5 | 0 | NA | NA | NA | NA | NA |
| Hsd3b6 | 0 | NA | NA | NA | NA | NA |
| Hsf3 | 1.66649381 | 0.031307164 | 0.098873152 | 0.316639691 | 0.751517015 | NA |
| Hsfy2 | 0 | NA | NA | NA | NA | NA |
| Hsh2d | 1.200692057 | 0.095444877 | 0.066470454 | 1.435899284 | 0.151031002 | NA |
| Hspb9 | 0 | NA | NA | NA | NA | NA |
| Htr2b | 3.310832292 | 0.007602766 | 0.119997645 | 0.063357624 | 0.949481731 | NA |
| Htr3b | 0.794161071 | 0.023670767 | 0.072294609 | 0.327420914 | 0.743349551 | NA |
| Hus1b | 0.429334602 | -0.005411584 | 0.048367196 | -0.111885422 | 0.910914255 | NA |
| Hyal4 | 0 | NA | NA | NA | NA | NA |
| Hyal5 | 0 | NA | NA | NA | NA | NA |
| Hyal6 | 0.105923158 | 0.018524693 | 0.027977518 | 0.662127842 | 0.507889294 | NA |
| I730028E13Rik | 2.004273009 | -0.160314373 | 0.102328518 | -1.566663683 | 0.117193324 | NA |
| I730030J21Rik | 7.334782236 | 0.115656824 | 0.132390552 | 0.87360331 | 0.382334323 | NA |
| Iapp | 0 | NA | NA | NA | NA | NA |
| Ibsp | 7.028872326 | 0.085146592 | 0.127034714 | 0.670262395 | 0.502690535 | NA |
| Icos | 3.419283575 | -0.07026302 | 0.11921927 | -0.589359586 | 0.555620081 | NA |
| Idi2 | 3.00797442 | 0.093299555 | 0.120094986 | 0.776881349 | 0.437228776 | NA |
| Ifi202b | 0.803429042 | 0.090552676 | 0.058568157 | 1.546107653 | 0.122078571 | NA |
| Ifi205 | 0.801425426 | -0.042094233 | 0.068780797 | -0.612005601 | 0.540534052 | NA |
| Ifi27l2b | 2.796705782 | -0.154592643 | 0.113204604 | -1.365603845 | 0.172063339 | NA |
| Ifi44l | 0.493865508 | 0.010205312 | 0.05056179 | 0.201838429 | 0.840043037 | NA |
| Ifitm2 | 259.4752542 | 0.308139079 | 0.162140184 | 1.900448565 | NA | NA |
| Ifitm5 | 0.977458471 | 0.100649779 | 0.069659942 | 1.444873139 | 0.148493522 | NA |
| Ifitm6 | 5.42448444 | 0.205485975 | 0.137368359 | 1.49587559 | 0.134686078 | NA |
| Ifna1 | 0 | NA | NA | NA | NA | NA |
| Ifna11 | 0 | NA | NA | NA | NA | NA |
| Ifna12 | 0 | NA | NA | NA | NA | NA |
| Ifna13 | 0 | NA | NA | NA | NA | NA |
| Ifna14 | 0 | NA | NA | NA | NA | NA |
| Ifna2 | 0.107716603 | -0.020477583 | 0.027954224 | -0.732539849 | 0.46383913 | NA |
| Ifna4 | 0 | NA | NA | NA | NA | NA |
| Ifna5 | 0 | NA | NA | NA | NA | NA |
| Ifna6 | 0 | NA | NA | NA | NA | NA |
| Ifna7 | 0 | NA | NA | NA | NA | NA |
| Ifna9 | 0 | NA | NA | NA | NA | NA |
| Ifnab | 0 | NA | NA | NA | NA | NA |
| Ifnb1 | 0 | NA | NA | NA | NA | NA |
| Ifne | 0 | NA | NA | NA | NA | NA |
| Ifng | 0.293623414 | 0.010408489 | 0.039073364 | 0.266383235 | 0.789944076 | NA |
| Ifnk | 0 | NA | NA | NA | NA | NA |
| Ifnl2 | 0 | NA | NA | NA | NA | NA |
| Ifnl3 | 0 | NA | NA | NA | NA | NA |
| Ifnz | 0 | NA | NA | NA | NA | NA |
| Igf2bp1 | 0.948524069 | 0.061319841 | 0.066722407 | 0.919029207 | 0.358080295 | NA |
| Igf2bp3 | 1.220477769 | -0.013382445 | 0.077957831 | -0.171662608 | 0.863702784 | NA |
| Igfals | 2.102523036 | 0.133804318 | 0.096411606 | 1.38784451 | 0.165184398 | NA |
| Igfbp1 | 0.505383842 | 0.011024618 | 0.05572892 | 0.197825804 | 0.843181356 | NA |
| Igfl3 | 0 | NA | NA | NA | NA | NA |
| Igll1 | 0.669316088 | -0.01851675 | 0.059854163 | -0.309364442 | 0.757044316 | NA |
| Igsf23 | 0.079983654 | 0.016740032 | 0.027674178 | 0.604897175 | 0.54524732 | NA |
| Igtp | 342.6334407 | 0.368037063 | 0.154679316 | 2.379355383 | NA | NA |
| Iigp1 | 113.2435397 | 0.197562664 | 0.118672532 | 1.664771619 | NA | NA |
| Il10 | 0.319934616 | 0.029547171 | 0.033991092 | 0.869262165 | 0.384703752 | NA |
| Il11ra2 | 0 | NA | NA | NA | NA | NA |
| Il12b | 0.71227231 | -0.086203635 | 0.065414181 | -1.317812664 | 0.187566367 | NA |
| Il12rb2 | 7.465627489 | -0.000413354 | 0.120699625 | -0.003424651 | 0.99726753 | NA |
| Il13 | 0 | NA | NA | NA | NA | NA |
| Il13ra2 | 7.424376002 | -0.369589113 | 0.140048974 | -2.638999081 | 0.008315121 | NA |
| Il17a | 0 | NA | NA | NA | NA | NA |
| Il17b | 6.968546564 | 0.036197373 | 0.151297694 | 0.239246034 | 0.810914809 | NA |
| Il17c | 1.051226501 | 0.034639185 | 0.07267005 | 0.476663841 | 0.633601515 | NA |
| Il17f | 0.298391179 | 0.008861245 | 0.039220874 | 0.225931861 | 0.821254417 | NA |
| Il17re | 7.915955225 | 0.214883859 | 0.156594606 | 1.372230273 | 0.169991764 | NA |
| Il18r1 | 2.089358884 | -0.051005697 | 0.089413523 | -0.570447237 | 0.568374399 | NA |
| Il18rap | 2.655103376 | -0.088742223 | 0.109395688 | -0.811204029 | 0.417248512 | NA |
| Il19 | 0.57410637 | 0.02462778 | 0.058376336 | 0.421879507 | 0.673112969 | NA |
| Il1f10 | 0.096071043 | -0.020098795 | 0.027825948 | -0.722304036 | 0.470107572 | NA |
| Il1f5 | 3.248080251 | 0.065925601 | 0.109009998 | 0.604766556 | 0.545334118 | NA |
| Il1f6 | 0 | NA | NA | NA | NA | NA |
| Il1f8 | 0 | NA | NA | NA | NA | NA |
| Il1rn | 3.015185515 | 0.148202274 | 0.099959021 | 1.482630301 | 0.138172662 | NA |
| Il2 | 0 | NA | NA | NA | NA | NA |
| Il20 | 0 | NA | NA | NA | NA | NA |
| Il21 | 2.68038864 | 0.036686079 | 0.089529743 | 0.409764148 | 0.681978968 | NA |
| Il22 | 1.313344777 | 0.030458062 | 0.081741128 | 0.372616127 | 0.709434168 | NA |
| Il22ra1 | 3.056421495 | 0.079193051 | 0.116960709 | 0.677091061 | 0.49834818 | NA |
| Il22ra2 | 0.119397641 | -0.020477583 | 0.027954224 | -0.732539845 | 0.463839132 | NA |
| Il23a | 3.721289891 | -0.281273911 | 0.130513745 | -2.155128646 | 0.031151769 | NA |
| Il23r | 0.451066898 | 0.006365339 | 0.044041609 | 0.144530111 | 0.885081881 | NA |
| Il24 | 0 | NA | NA | NA | NA | NA |
| Il25 | 2.540226355 | -0.015535638 | 0.103527021 | -0.150063604 | 0.880714435 | NA |
| Il27 | 1.241576115 | 0.158271463 | 0.083179902 | 1.902760871 | 0.057071755 | NA |
| Il3 | 0 | NA | NA | NA | NA | NA |
| Il31 | 0.096071043 | -0.020098795 | 0.027825948 | -0.722304036 | 0.470107572 | NA |
| Il4 | 0 | NA | NA | NA | NA | NA |
| Il4i1 | 0.330487422 | -0.017886856 | 0.040493356 | -0.441723219 | 0.658689506 | NA |
| Il5 | 2.601214813 | 0.125010189 | 0.108609286 | 1.151008288 | 0.249728826 | NA |
| Il5ra | 0.385382084 | -0.000943527 | 0.041930191 | -0.022502333 | 0.982047251 | NA |
| Il6 | 2.260663149 | 0.142455921 | 0.090968713 | 1.565988088 | 0.117351406 | NA |
| Il7 | 3.796125002 | 0.072403111 | 0.130125431 | 0.556410156 | 0.577930498 | NA |
| Il8 | 0 | NA | NA | NA | NA | NA |
| Il9 | 0 | NA | NA | NA | NA | NA |
| Il9r | 0.305736371 | 0.008542183 | 0.033941149 | 0.251676297 | 0.80129128 | NA |
| Iltifb | 1.699873486 | 0.097147322 | 0.096393182 | 1.007823587 | 0.313539161 | NA |
| Inhbc | 0.107716603 | -0.020477583 | 0.027954224 | -0.732539849 | 0.46383913 | NA |
| Ins1 | 0 | NA | NA | NA | NA | NA |
| Ins2 | 0.468677007 | 0.028797021 | 0.045304689 | 0.635630028 | 0.525017593 | NA |
| Insl3 | 1.825258177 | 0.037173765 | 0.093714226 | 0.396671532 | 0.691609696 | NA |
| Insm2 | 3.910246587 | -0.113580811 | 0.130822248 | -0.868207148 | 0.385280942 | NA |
| Iqcf1 | 0 | NA | NA | NA | NA | NA |
| Iqcf3 | 0.186326724 | 0.024939385 | 0.031411053 | 0.793968451 | 0.427213798 | NA |
| Iqcf4 | 0.211846316 | 0.025707621 | 0.031860375 | 0.806883811 | 0.419733426 | NA |
| Iqcf5 | 2.293894031 | 0.022767951 | 0.095841592 | 0.237558149 | 0.812223814 | NA |
| Iqcf6 | 0 | NA | NA | NA | NA | NA |
| Iqch | 2.970735406 | 0.202007729 | 0.124224906 | 1.626145151 | 0.103918781 | NA |
| Iqcj | 1.623363065 | -0.102028201 | 0.091583758 | -1.114042529 | 0.265260942 | NA |
| Irg1 | 1.562964034 | 0.074159782 | 0.078666255 | 0.942714026 | 0.345827197 | NA |
| Irgc1 | 4.580136633 | -0.23723407 | 0.128670826 | -1.84372851 | 0.065222716 | NA |
| Irx2 | 5.247987084 | 0.188455073 | 0.112781148 | 1.670980273 | 0.094725574 | NA |
| Irx3 | 6.141851399 | 0.334152361 | 0.142946491 | 2.337604503 | 0.019407774 | NA |
| Irx4 | 0.094392817 | 0.017895942 | 0.027818137 | 0.643319236 | 0.520016977 | NA |
| Irx5 | 3.390072367 | 0.181013927 | 0.111149641 | 1.628560606 | 0.103406069 | NA |
| Isl2 | 0.852951773 | 0.017080043 | 0.056027866 | 0.304849064 | 0.760481112 | NA |
| Isx | 2.802752224 | -0.096197048 | 0.12116548 | -0.793931144 | 0.427235518 | NA |
| Itga2 | 5.231023707 | -0.105884164 | 0.138144283 | -0.766475176 | 0.443393617 | NA |
| Itgad | 11.56699328 | 0.276335662 | 0.13545839 | 2.040004035 | NA | NA |
| Itgae | 7.832432227 | 0.064394042 | 0.150255071 | 0.428564854 | 0.668239927 | NA |
| Itgam | 1131.312302 | -0.111903595 | 0.166502778 | -0.672082447 | NA | NA |
| Itgb2l | 0.36509754 | 0.021286825 | 0.04223302 | 0.504032755 | 0.61423836 | NA |
| Itgb6 | 0.910840021 | -0.026687877 | 0.070723857 | -0.377353252 | 0.705911103 | NA |
| Itih1 | 0 | NA | NA | NA | NA | NA |
| Itih4 | 4.566125969 | 0.206095701 | 0.126725605 | 1.626314596 | 0.103882748 | NA |
| Itln1 | 0 | NA | NA | NA | NA | NA |
| Ivl | 0.096071043 | -0.020098795 | 0.027825948 | -0.722304036 | 0.470107572 | NA |
| Izumo1 | 1.858657362 | 0.082709532 | 0.099163 | 0.834076536 | 0.404237857 | NA |
| Izumo2 | 0 | NA | NA | NA | NA | NA |
| Izumo3 | 0 | NA | NA | NA | NA | NA |
| Kap | 0 | NA | NA | NA | NA | NA |
| Kcna10 | 4.083566732 | 0.206485294 | 0.124999434 | 1.651889826 | 0.098557013 | NA |
| Kcna7 | 1.007082075 | 0.091882827 | 0.076574122 | 1.199920073 | 0.230170383 | NA |
| Kcne1 | 0.38827344 | 0.020990736 | 0.04283191 | 0.490072386 | 0.624082678 | NA |
| Kcne2 | 26.11502812 | 0.212993065 | 0.090336557 | 2.357772676 | NA | NA |
| Kcne3 | 5.558619948 | 0.146664536 | 0.140124964 | 1.046669567 | 0.295252005 | NA |
| Kcnj1 | 0.099009884 | 0.018205726 | 0.027896865 | 0.652608315 | 0.514008823 | NA |
| Kcnj15 | 2.982287336 | -0.067712734 | 0.10854698 | -0.623810392 | 0.532752114 | NA |
| Kcnk15 | 3.251105874 | -0.183021399 | 0.090816472 | -2.015288578 | 0.043874421 | NA |
| Kcnk16 | 0.17463368 | 0.024560283 | 0.031186242 | 0.787535833 | 0.430968261 | NA |
| Kcnk18 | 0 | NA | NA | NA | NA | NA |
| Kcnk7 | 3.266791574 | 0.049750173 | 0.12016794 | 0.41400538 | 0.678870167 | NA |
| Kcnmb3 | 0.099009884 | 0.018205726 | 0.027896865 | 0.652608315 | 0.514008823 | NA |
| Kcnv2 | 0.531537619 | -0.072805061 | 0.051684011 | -1.408657322 | 0.15893652 | NA |
| Kctd19 | 0.602691698 | -0.043058929 | 0.048770142 | -0.882895305 | 0.37729284 | NA |
| Keg1 | 0 | NA | NA | NA | NA | NA |
| Kel | 0.612869747 | -0.006355786 | 0.053223994 | -0.119415802 | 0.904945943 | NA |
| Kera | 0.239950962 | 0.027417169 | 0.032831728 | 0.835081494 | 0.403671825 | NA |
| Khdc1a | 0 | NA | NA | NA | NA | NA |
| Khdc1b | 0 | NA | NA | NA | NA | NA |
| Khdc1c | 0 | NA | NA | NA | NA | NA |
| Kif12 | 0.197363271 | -0.000561558 | 0.031650166 | -0.017742642 | 0.985844163 | NA |
| Kif2b | 0 | NA | NA | NA | NA | NA |
| Kif4-ps | 0 | NA | NA | NA | NA | NA |
| Kir3dl1 | 0 | NA | NA | NA | NA | NA |
| Kir3dl2 | 5.239499727 | 0.173088669 | 0.128892893 | 1.342887615 | 0.179308367 | NA |
| Kis2 | 0.105923158 | 0.018524693 | 0.027977518 | 0.662127842 | 0.507889294 | NA |
| Kl | 348.806466 | 0.383400764 | 0.150019669 | 2.555669971 | NA | NA |
| Klb | 6.73275033 | 0.035676134 | 0.148306622 | 0.24055658 | 0.809898806 | NA |
| Klf1 | 4.867912659 | 0.223518709 | 0.142719895 | 1.5661356 | 0.117316875 | NA |
| Klf17 | 0.639411646 | 0.032045962 | 0.060168178 | 0.532606496 | 0.594306005 | NA |
| Klhl31 | 2.227178724 | 0.106658768 | 0.105957151 | 1.006621709 | 0.314116598 | NA |
| Klhl34 | 6.350984302 | 0.181332575 | 0.136217171 | 1.331202032 | 0.183122543 | NA |
| Klk1 | 0.233393928 | -0.027475306 | 0.032191697 | -0.85349047 | 0.39338737 | NA |
| Klk10 | 3.584123508 | 0.024521382 | 0.112562779 | 0.21784627 | 0.827548892 | NA |
| Klk11 | 0.365320263 | 0.05192513 | 0.045006765 | 1.15371833 | 0.248615666 | NA |
| Klk12 | 0 | NA | NA | NA | NA | NA |
| Klk13 | 1.002455104 | -0.033529498 | 0.073335503 | -0.457206902 | 0.647522338 | NA |
| Klk15 | 0 | NA | NA | NA | NA | NA |
| Klk1b1 | 0 | NA | NA | NA | NA | NA |
| Klk1b11 | 0 | NA | NA | NA | NA | NA |
| Klk1b16 | 0 | NA | NA | NA | NA | NA |
| Klk1b21 | 0 | NA | NA | NA | NA | NA |
| Klk1b22 | 0 | NA | NA | NA | NA | NA |
| Klk1b24 | 0 | NA | NA | NA | NA | NA |
| Klk1b26 | 0 | NA | NA | NA | NA | NA |
| Klk1b27 | 0.638908139 | -0.060041808 | 0.056221338 | -1.067954093 | 0.28554122 | NA |
| Klk1b3 | 0.65992155 | 0.044480374 | 0.050358557 | 0.883273402 | 0.377088571 | NA |
| Klk1b4 | 0 | NA | NA | NA | NA | NA |
| Klk1b5 | 0 | NA | NA | NA | NA | NA |
| Klk1b7-ps | 0 | NA | NA | NA | NA | NA |
| Klk1b8 | 0 | NA | NA | NA | NA | NA |
| Klk1b9 | 0 | NA | NA | NA | NA | NA |
| Klk4 | 0.218407525 | -0.002234018 | 0.031987735 | -0.069839845 | 0.944321133 | NA |
| Klk5 | 0.116696964 | -0.020477583 | 0.027954224 | -0.732539847 | 0.463839131 | NA |
| Klk7 | 0.830848704 | 0.060916436 | 0.066285696 | 0.918998219 | 0.358096504 | NA |
| Klk9 | 1.340873711 | 0.000938095 | 0.090559374 | 0.010358894 | 0.991734946 | NA |
| Klkb1 | 3.41490234 | 0.061658413 | 0.124287637 | 0.496094495 | 0.619827743 | NA |
| Klra1 | 0 | NA | NA | NA | NA | NA |
| Klra10 | 0 | NA | NA | NA | NA | NA |
| Klra12 | 0 | NA | NA | NA | NA | NA |
| Klra13-ps | 0 | NA | NA | NA | NA | NA |
| Klra14 | 0.196706773 | -0.026517754 | 0.031609464 | -0.838918169 | 0.401515231 | NA |
| Klra15 | 0 | NA | NA | NA | NA | NA |
| Klra17 | 0 | NA | NA | NA | NA | NA |
| Klra18 | 0 | NA | NA | NA | NA | NA |
| Klra19 | 0 | NA | NA | NA | NA | NA |
| Klra2 | 3.456128443 | 0.225087158 | 0.125761906 | 1.789788064 | 0.073487989 | NA |
| Klra21 | 0 | NA | NA | NA | NA | NA |
| Klra22 | 0 | NA | NA | NA | NA | NA |
| Klra23 | 0 | NA | NA | NA | NA | NA |
| Klra3 | 0 | NA | NA | NA | NA | NA |
| Klra33 | 0 | NA | NA | NA | NA | NA |
| Klra4 | 0 | NA | NA | NA | NA | NA |
| Klra5 | 0 | NA | NA | NA | NA | NA |
| Klra6 | 0 | NA | NA | NA | NA | NA |
| Klra7 | 0 | NA | NA | NA | NA | NA |
| Klra8 | 0.079983654 | 0.016740032 | 0.027674178 | 0.604897175 | 0.54524732 | NA |
| Klra9 | 0.08731684 | 0.017365056 | 0.027733412 | 0.626142072 | 0.531221759 | NA |
| Klrb1 | 0.098353386 | -0.020212831 | 0.02786466 | -0.725393061 | 0.468210929 | NA |
| Klrb1-ps1 | 0 | NA | NA | NA | NA | NA |
| Klrb1a | 1.517537733 | 0.114815172 | 0.095204409 | 1.205985873 | 0.227822937 | NA |

| | | | | | | | | |
|---|---|---|---|---|---|---|---|---|
| | Lao1 | 8.050104979 | 0.0612854503874838 | 0.0480971877852881 | 1.27420028507841 | 0.202592444216184 | NA | NM_133892 |
| | Lat | 17.16880597 | -0.0532265551245093 | 0.0756474746920428 | -0.703613112548594 | 0.48167374317344 | NA | NM_010689 |
| | Lax1 | 14.63061844 | -0.0560077820005496 | 0.0734596250169467 | -0.762429456829231 | 0.445803730919775 | NA | NM_001159649,NM_172842 |
| | Lbx1 | 0.205932006 | -0.000576954122277892 | 0.00970424421944593 | -0.0594537925088239 | 0.952590668569529 | NA | NM_010691 |
| | Lbx2 | 1.972697308 | 0.0227945993180183 | 0.0301798531231816 | 0.755291923555103 | 0.450073843577124 | NA | NM_010692 |
| | Lce1a1 | 0 | NA | NA | NA | NA | NA | NM_025984 |
| | Lce1a2 | 0 | NA | NA | NA | NA | NA | NM_028625 |
| | Lce1b | 0 | NA | NA | NA | NA | NA | NM_026822 |
| | Lce1c | 0 | NA | NA | NA | NA | NA | NM_028622 |
| | Lce1d | 0 | NA | NA | NA | NA | NA | NM_027137 |
| | Lce1e | 0 | NA | NA | NA | NA | NA | NM_026811 |
| | Lce1f | 0 | NA | NA | NA | NA | NA | NM_026394 |
| | Lce1g | 0.212695677 | -0.00857796107890413 | 0.00973699866180651 | -0.880965621629515 | 0.378336427605205 | NA | NM_025413 |
| | Lce1h | 0 | NA | NA | NA | NA | NA | NM_026335 |
| | Lce1i | 0 | NA | NA | NA | NA | NA | NM_029667 |
| | Lce1k | 0 | NA | NA | NA | NA | NA | NM_001254760 |
| | Lce1l | 0 | NA | NA | NA | NA | NA | NM_028628 |
| | Lce1m | 0 | NA | NA | NA | NA | NA | NM_025420 |
| | Lce3a | 0.108766301 | 0.00519171381361946 | 0.00856003371502575 | 0.606506234257728 | 0.544178643801544 | NA | NM_001039594 |
| | Lce3b | 0 | NA | NA | NA | NA | NA | NM_025501 |
| | Lce3c | 0.953969319 | -0.0163245560836365 | 0.0174714448690956 | -0.934356385859777 | 0.350120087478754 | NA | NM_033175 |
| | Lce3d | 0.106148608 | 0.00515554063321497 | 0.00854200771766808 | 0.603551390213729 | 0.546141939141592 | NA | NM_001270426 |
| | Lce3e | 0 | NA | NA | NA | NA | NA | NM_001254725 |
| | Lce3f | 0.097165704 | -0.00637828455464458 | 0.00848781691535724 | -0.75146349388194 | 0.452373763047372 | NA | NM_001018079 |
| | Lce6a | 0 | NA | NA | NA | NA | NA | NM_001166172 |
| | Lcn10 | 0 | NA | NA | NA | NA | NA | NM_178036 |
| | Lcn11 | 1.23620877 | -0.0139864000568279 | 0.0246090368603567 | -0.568344065482669 | 0.569801363111582 | NA | NM_001100455 |
| | Lcn12 | 0.432447513 | 0.0122685117249055 | 0.0130381092331998 | 0.940973227442017 | 0.346718579229735 | NA | NM_029958 |
| | Lcn2 | 11.95296539 | 0.0835298235688088 | 0.054922835056476 | 1.52085782685684 | 0.128295519160553 | NA | NM_008491 |
| | Lcn3 | 0.226066984 | -0.0093084363525918 | 0.0101406635841621 | -0.917931679257152 | 0.358654635211298 | NA | NM_010694 |
| | Lcn4 | 0 | NA | NA | NA | NA | NA | NM_010695 |
| | Lcn5 | 0 | NA | NA | NA | NA | NA | NM_007947,NM_001276257,NM_001042630 |
| | Lcn6 | 0 | NA | NA | NA | NA | NA | NM_001276448,NM_177840 |
| | Lcn8 | 0.106148608 | 0.00515554063321497 | 0.00854200771766808 | 0.603551390213729 | 0.546141939141592 | NA | NM_033145 |
| | Lcn9 | 0 | NA | NA | NA | NA | NA | NM_029959 |
| | Lctl | 11.45639405 | -0.0053855569525568 | 0.0605283298149832 | -0.088975806354129 | 0.929101137577074 | NA | NM_145835 |
| | Ldhal6b | 6.757425588 | -0.0105885287239021 | 0.058082609413491 | -0.182301188442176 | 0.85534636607104 | NA | NM_175349 |
| | Ldhc | 1.080858687 | -0.0314745771756955 | 0.0239010948656669 | -1.31686758922946 | 0.187883012170781 | NA | NM_013580 |
| | Ldlrad1 | 0.291497113 | -0.00941849595073383 | 0.0102038556585987 | -0.923033044160808 | 0.355989985190414 | NA | NM_001081272 |
| | Ldlrad2 | 3.900073275 | 0.0192640915749363 | 0.0436369503607731 | 0.441462829452297 | 0.658877965447819 | NA | NM_001033979 |
| | Leap2 | 0.120209049 | -0.00674972139751468 | 0.00863521114606269 | -0.781650996523957 | 0.434419707908266 | NA | NM_153069 |
| | Lect2 | 0.217532603 | 0.00684685248201087 | 0.00976000296730706 | 0.701521557416086 | 0.482977586094298 | NA | NM_010702 |
| | Lefty2 | 17.65535745 | 0.0669366619560358 | 0.0736242290519159 | 0.909166219028734 | 0.363262392733809 | NA | NM_177099 |
| | Lelp1 | 0 | NA | NA | NA | NA | NA | NM_027042 |
| | Lep | 0 | NA | NA | NA | NA | NA | NM_008493 |
| | Lgals2 | 3.661322689 | 0.0279705010534392 | 0.0416517109435029 | 0.671533063584804 | 0.501881003713362 | NA | NM_025622 |
| | Lgals4 | 8.303348144 | -0.144458784636031 | 0.0609421590924142 | -2.37042446128254 | 0.0177676742097265 | NA | NM_010706 |
| | Lgals6 | 0.224626239 | -0.00168521210467683 | 0.00978955093367521 | -0.172143964119932 | 0.863324350255464 | NA | NM_010707 |
| | Lgals7 | 3.58848747 | -0.0411154409074805 | 0.0430885151647118 | -0.954208812958882 | 0.339977954528684 | NA | NM_008496 |
| | Lgsn | 0.226066984 | -0.0093084363525918 | 0.0101406635841621 | -0.917931679257152 | 0.358654635211298 | NA | NM_153601 |
| | Lhb | 5.764121871 | -0.000497881671822517 | 0.0447296217826633 | -0.0111309162022803 | 0.991118997203693 | NA | NM_008497 |
| | Lhcgr | 19.33113077 | -0.0621251399421435 | 0.0796688943894946 | -0.77979166672527 | 0.435513512195815 | NA | NM_013582 |
| | Lhfpl1 | 4.541119856 | -0.00148868652265725 | 0.047776120012369 | -0.0311596362842323 | 0.975142229850173 | NA | NM_178358 |
| | Lhx1 | 8.55942623 | 0.107078106438613 | 0.0477411499819788 | 2.24288913189215 | 0.024903964771731 | NA | NM_008498 |
| | Lhx3 | 7.304399762 | -0.00844125574372068 | 0.0477410859052215 | -0.176813232955756 | 0.859655094430063 | NA | NM_001039653 |
| | Lhx4 | 1.064075119 | -0.00442591548194255 | 0.0234149302393488 | -0.189021083415606 | 0.850076292475604 | NA | NM_010712 |
| | Lhx5 | 11.93645414 | 0.11508654224135 | 0.0568916351883087 | 2.02290796986972 | 0.043082636708001 | NA | NM_008499 |
| | Lilra5 | 18.63702593 | -0.119953530689062 | 0.0757956115370837 | -1.58259202949201 | 0.113514481357957 | NA | NM_001081239 |
| | Lilra6 | 0.291497113 | -0.00941849595073383 | 0.0102038556585987 | -0.923033044160808 | 0.355989985190414 | NA | NM_011090,NR_028115 |
| | Lilrb4 | 20.31163142 | -0.117255690868773 | 0.0565608894764777 | -2.07308781658281 | 0.0381641076489764 | NA | NM_013532 |
| | Lim2 | 0 | NA | NA | NA | NA | NA | NM_177693 |
| | Lin28a | 0.560996453 | -0.00719580254083206 | 0.016648491668553 | -0.432219487752399 | 0.665581896938222 | NA | NM_145833 |
| | Lipc | 5.837605561 | -0.0279534246784002 | 0.0557497589121828 | -0.501408889004031 | 0.616083385182047 | NA | NM_008280 |
| | Lipf | 0.411706162 | -0.0156488736868625 | 0.013129479323261 | -1.191888368272 | 0.233305019639917 | NA | NM_026334 |
| | Lipi | 0.12890128 | -0.0068640067066208 | 0.00870804083610955 | -0.788237771940376 | 0.430557639008166 | NA | NM_177142,NM_001252513 |
| | Lipk | 0.222609587 | 0.00717651388923987 | 0.00999169132085706 | 0.718248158273197 | 0.472604288901417 | NA | NM_172837,NM_001205349 |
| | Lipm | 4.350341414 | -0.0154732219205039 | 0.0450298149545798 | -0.343621707886458 | 0.731130797221949 | NA | NM_023903 |
| | Lipn | 0.108766301 | 0.00519171381361946 | 0.00856003371502575 | 0.606506234257728 | 0.544178643801544 | NA | NM_027340 |
| | Lman1l | 2.659868646 | -0.0192730433675281 | 0.034997014947624 | -0.550705350052623 | 0.581835676513833 | NA | NM_199222 |
| | Lmod2 | 0.233836299 | 0.00743372775021133 | 0.0101683766597817 | 0.731063374118848 | 0.464740444385902 | NA | NM_053098 |
| | Lmod3 | 1.750447079 | -0.0258349073704444 | 0.0292128609458633 | -0.884367587903189 | 0.376497813740182 | NA | NM_001081157 |
| | Lmx1a | 11.2885041 | 0.00554688288553377 | 0.0417026372731981 | 0.133010362131191 | NA | NA | NM_033652 |
| | Lmx1b | 1.240293532 | 0.0217826869131236 | 0.0258700090309931 | 0.842005384962458 | 0.399784939251317 | NA | NM_010725 |
| | Lox | 15.78146708 | -0.140451421793049 | 0.0694678483244297 | -2.02181908869713 | 0.0431950449968129 | NA | NM_010728 |
| | Loxhd1 | 10.15196697 | 0.106617134531021 | 0.0683043608869167 | 1.56091255589865 | 0.118544384372006 | NA | NM_172834 |
| | Loxl4 | 5.853579617 | 0.00371535808598743 | 0.045422118303752 | 0.0817962311035708 | 0.93480875317647 | NA | NM_053083,NM_001164311 |
| | Lpar3 | 13.00001555 | -0.0189725848228113 | 0.0704270142783646 | -0.269393570311267 | 0.787626833488164 | NA | NM_022983 |
| | Lpar5 | 16.66724293 | -0.0967364585114065 | 0.0710974236867524 | -1.360618338825 | 0.173634333096376 | NA | NM_001163269,NM_001163268 |
| | Lpcat2b | 1.246483544 | 0.0316358659773357 | 0.0272527194026731 | 1.16083336528364 | 0.245709671434249 | NA | NM_027599 |
| | Lrat | 6.48853038 | 0.0132745347098385 | 0.0466467015891809 | 0.284576063421327 | 0.77596895192449 | NA | NM_023624 |
| | Lrch4 | 0 | NA | NA | NA | NA | NA | NM_146164,NM_001168652 |
| | Lrcol1 | 0 | NA | NA | NA | NA | NA | NM_001033459 |
| | Lrit1 | 0.194331409 | -0.00840968081320881 | 0.00964081822647896 | -0.872299489073576 | 0.383045016303155 | NA | NM_146245 |
| | Lrit2 | 0.321791944 | -0.0064493982079258 | 0.012677176873412 | -0.508740887054451 | 0.610933858924308 | NA | NM_173418 |
| | Lrit3 | 2.276302754 | -0.045270601709444 | 0.034288011832043 | -1.32030407394859 | 0.186733515949295 | NA | NM_001205102 |
| | Lrp2 | 22.17130481 | -0.0226020349859218 | 0.0699472140935933 | -0.323129881279889 | 0.746596881104631 | NA | NM_001081088 |
| | Lrp2bp | 19.68937987 | -0.0307492700916103 | 0.0794346049888516 | -0.387101693222064 | 0.698680924141362 | NA | NM_026278 |
| | Lrr1 | 1.050075312 | 0.0174016079199067 | 0.0216854702241303 | 0.802454719222238 | 0.422289971296264 | NA | NM_001081406 |
| | Lrrc10 | 1.044785728 | 0.00376596121953573 | 0.0237330836492202 | 0.158679810647339 | 0.873921145393016 | NA | NM_146242 |
| | Lrrc14b | 9.877003295 | 0.0489066653813225 | 0.058985798351874 | 0.82912610743309 | 0.407033052038764 | NA | NM_001033042 |
| | Lrrc15 | 1.192965478 | 0.00736651504714934 | 0.0242411806503884 | 0.303884334405606 | 0.761216013088892 | NA | NM_028973 |
| | Lrrc19 | 0 | NA | NA | NA | NA | NA | NM_175305 |
| | Lrrc26 | 0 | NA | NA | NA | NA | NA | NM_146117 |
| | Lrrc30 | 0.361523991 | 0.0105955510260098 | 0.0121250525610889 | 0.873856090324303 | 0.382196629856021 | NA | NM_001033340 |
| | Lrrc34 | 7.556988423 | 0.0813201629562312 | 0.0523160069583768 | 1.5544030915993 | 0.120088296258356 | NA | NM_027941 |
| | Lrrc39 | 10.78513989 | 0.0102508032955727 | 0.0601155462645797 | 0.170518342301291 | 0.864602510530169 | NA | NM_027321,NM_175413 |
| | Lrrc52 | 0 | NA | NA | NA | NA | NA | NM_001013382 |
| | Lrrc63 | 0.74940253 | -0.0102063864539079 | 0.0213296713146236 | -0.47850650407868 | 0.632289746143334 | NA | NM_027581 |
| | Lrrc66 | 0.794511502 | -0.0269120401284838 | 0.0197582908288356 | -1.36206316435063 | 0.173177951976176 | NA | NM_153568 |
| | Lrrc69 | 2.651490013 | 0.00388792900191091 | 0.035607240330044 | 0.109189281895301 | 0.91305236128862 | NA | NM_028499 |
| | Lrrc72 | 0.257802559 | -0.00892927589713349 | 0.00993479775307735 | -0.898787888698349 | 0.368765652383134 | NA | NM_027699,NM_001177877 |
| | Lrrc8e | 0.865135084 | 0.0118261255425992 | 0.0192252540657293 | 0.615134941892927 | 0.538465607988394 | NA | NM_028175 |
| | Lrrd1 | 13.68845613 | -0.0794920886358245 | 0.0709958616469986 | -1.11967214414652 | 0.262853499671042 | NA | NM_172879 |
| | Lrriq3 | 17.7069029 | -0.0700927903671263 | 0.080875270911079 | -0.866677657799652 | 0.386118650369993 | NA | NM_028938 |
| | Lrriq4 | 0.203513543 | -0.00860782669689374 | 0.00975338121443284 | -0.882547960306941 | 0.377480556050336 | NA | NM_026668 |
| | Lrtm1 | 0.24789674 | -0.000769869626663926 | 0.0100368571301073 | -0.0767042528038554 | 0.938858821262798 | NA | NM_176920 |
| | Lta | 5.631046331 | 0.0314685276971805 | 0.0514767398411228 | 0.611315475577991 | 0.540990747704982 | NA | NM_010735 |
| | Ltb | 13.45619781 | 0.031256432602233 | 0.070548630710125 | 0.443048040587231 | 0.657730986785606 | NA | NM_008518 |
| | Ltb4r1 | 4.873109598 | 0.00373284511744738 | 0.0501603396057682 | 0.0744182584644645 | 0.940677580932038 | NA | NM_008519 |
| | Ltb4r2 | 2.557732746 | -0.0270692565159877 | 0.0336448655283428 | -0.804558320888048 | 0.421074601763495 | NA | NM_020490 |
| | Ltf | 8.409557156 | -0.0232805912314546 | 0.0435646120358804 | -0.534392254251694 | 0.593070176844876 | NA | NM_008522 |
| | Luzp4 | 0 | NA | NA | NA | NA | NA | NM_001114383 |
| | Ly6c2 | 1.642337728 | 0.0210497166034852 | 0.0250925975307032 | 0.838881529810888 | 0.401535793463324 | NA | NM_001099217 |
| | Ly6d | 16.91919721 | 0.0500325649767039 | 0.0671382586127908 | 0.745216900325918 | 0.456140607707513 | NA | NM_010742 |
| | Ly6f | 0 | NA | NA | NA | NA | NA | NM_008530 |
| | Ly6g5b | 8.822887049 | -0.0297848203357946 | 0.0566588666113882 | -0.525686836273706 | 0.599105811460907 | NA | NM_148939 |
| | Ly6g5c | 0 | NA | NA | NA | NA | NA | NM_148947 |
| | Ly6g6c | 0.108766301 | 0.00519171381361946 | 0.00856003371502575 | 0.606506234257728 | 0.544178643801544 | NA | NM_023463 |
| | Ly6g6d | 10.93927542 | 0.0699862878808363 | 0.0627644427681844 | 1.11506268189658 | 0.26482356184201 | NA | NM_033478 |
| | Ly6g6f | 17.55738223 | -0.00557730262895243 | 0.0744408704248425 | -0.074922587513044 | 0.940276304827651 | NA | NM_001163192 |
| | Ly6i | 0.443441737 | -0.0212067781731997 | 0.0152604065834807 | -1.38966010225153 | 0.164632115633729 | NA | NM_020498 |
| | Ly6k | 1.231991385 | 0.00877732775309191 | 0.0262403010860342 | 0.334497981723367 | 0.738003813631272 | NA | NM_029627 |
| | Ly9 | 47.20549678 | -0.461462823151847 | 0.0811384037869452 | -5.6873539731391 | NA | NA | NM_008534 |
| | Lyg1 | 0 | NA | NA | NA | NA | NA | NM_027111 |
| | Lyg2 | 4.663625196 | 0.0614511450448745 | 0.0472189177356281 | 1.30140943485682 | 0.19311834587907 | NA | NM_001033427 |
| | Lypd2 | 4.052908143 | -0.0390561374915625 | 0.0418323734216875 | -0.933634271664688 | 0.350492580498547 | NA | NM_026671 |
| | Lypd3 | 1.255514117 | 0.0170414539838759 | 0.0230676440834445 | 0.738760053789213 | 0.460052712201488 | NA | NM_133743 |
| | Lypd4 | 0 | NA | NA | NA | NA | NA | NM_182785 |
| | Lypd5 | 0 | NA | NA | NA | NA | NA | NM_029806 |
| | Lypd8 | 0.120209049 | -0.00674972139751468 | 0.00863521114606269 | -0.781650996523957 | 0.434419707908266 | NA | NM_001083884,NM_027339 |
| | Lyzl1 | 0 | NA | NA | NA | NA | NA | NM_026092 |
| | Lyzl4 | 21.02682536 | 0.0598772291689541 | 0.0776618337508778 | 0.770999425033243 | 0.44070727239511 | NA | NM_026915 |
| | Lyzl6 | 0.861411351 | 0.0150225755760322 | 0.0222813556491598 | 0.674221793887962 | 0.500170315862861 | NA | NM_027083 |
| | M1ap | 1.833683077 | -0.0154512115385654 | 0.0322189538363577 | -0.479569002055223 | 0.631533892304172 | NA | NM_033079 |
| | Mab21l2 | 12.94346964 | 0.0327763683940737 | 0.0526870701701803 | 0.62209510394496 | 0.533879334191458 | NA | NM_011839 |
| | Mab21l3 | 4.987563243 | 0.0144078431283286 | 0.0509840431258314 | 0.282595146343518 | 0.777487201395602 | NA | NM_172295 |
| | Macc1 | 4.16186299 | 0.0217745255459758 | 0.0438451561613137 | 0.496623286409647 | 0.619454728763784 | NA | NM_001163136 |
| | Madcam1 | 1.746014756 | -0.029949741599017 | 0.0312620822652162 | -0.958021328999591 | 0.338052003416038 | NA | NM_013591 |
| | Mael | 3.052938119 | 0.00451638688280381 | 0.0372267169999696 | 0.121321116842173 | 0.903436694737355 | NA | NM_175296 |
| | Mafa | 12.6026177 | -0.113653215774767 | 0.0698111160820597 | -1.62801029625674 | 0.103522702576896 | NA | NM_194350 |
| | Magea1 | 0 | NA | NA | NA | NA | NA | NM_020015 |
| | Magea10 | 0.224853396 | -0.000161157395207779 | 0.00979162641959728 | -0.0164586952465051 | 0.986868454038988 | NA | NM_001085506 |
| | Magea2 | 0 | NA | NA | NA | NA | NA | NM_020016 |
| | Magea3 | 0 | NA | NA | NA | NA | NA | NM_020017 |
| | Magea4 | 0 | NA | NA | NA | NA | NA | NM_020280 |
| | Magea5 | 0 | NA | NA | NA | NA | NA | NM_020018 |
| | Magea6 | 0 | NA | NA | NA | NA | NA | NM_020019 |
| | Magea8 | 0 | NA | NA | NA | NA | NA | NM_020020 |
| | Mageb1 | 0 | NA | NA | NA | NA | NA | NM_010759 |
| | Mageb16 | 0 | NA | NA | NA | NA | NA | NM_028025,NM_001113734 |
| | Mageb16-ps1 | 0.428167708 | -0.000792272155220925 | 0.0137354739435619 | -0.0576807293637128 | 0.95400294387567 | NA | NR_033647 |
| | Mageb18 | 0.346566706 | -0.0051482901046312 | 0.0114161257772475 | -0.450966484172047 | 0.652013705001052 | NA | NM_173783 |
| | Mageb2 | 0 | NA | NA | NA | NA | NA | NM_031171 |
| | Mageb3 | 4.439427723 | -0.0352461537721979 | 0.0355926600960513 | -0.990264669094182 | 0.322044770824372 | NA | NM_008545 |
| | Mageb4 | 0 | NA | NA | NA | NA | NA | NM_001033492 |
| | Mageb5 | 0 | NA | NA | NA | NA | NA | NM_028847 |
| | Magix | 20.85908421 | -0.1663362021609 | 0.0824240327903243 | -2.01805464413561 | 0.0435855706250569 | NA | NR_037581,NM_018832 |
| | Mansc4 | 7.526702359 | -0.02135321343574 | 0.0469477465806789 | -0.454829357976635 | 0.649232007633881 | NA | NM_001034903 |
| | Marc1 | 5.298693257 | 0.0153540156920383 | 0.0460895029488756 | 0.333134764093023 | 0.739032558724351 | NA | NM_001081361 |
| | Marcksl1-ps4 | 0.915593775 | 0.000329591085993255 | 0.019199722177854 | 0.0171664507923673 | 0.986303826633073 | NA | NR_028405 |
| | Marco | 0 | NA | NA | NA | NA | NA | NM_010766 |
| | Marveld3 | 18.21266796 | 0.0115430075610985 | 0.0801272005364285 | 0.144058540468423 | 0.885454243067668 | NA | NM_028584,NM_212447 |
| | Mat1a | 1.880538289 | 0.0353174754016489 | 0.0326900820557221 | 1.08037279751847 | 0.279976204252721 | NA | NM_133653 |
| | Matn1 | 13.4832559 | -0.0472605668394741 | 0.0712023802957594 | -0.663749816272487 | 0.506850449616684 | NA | NM_010769 |
| | Matn3 | 2.811696213 | 0.0461277912508327 | 0.0404757313474023 | 1.1396407109959 | 0.254436017178205 | NA | NM_010770 |
| | Mb | 3.941334226 | -0.0151878321191911 | 0.0357672850815296 | -0.424629157191306 | 0.671107034416425 | NA | NM_001164047,NM_013593,NM_001164048 |
| | Mbd3l1 | 0 | NA | NA | NA | NA | NA | NM_028557 |
| | Mbd3l2 | 0.120209049 | -0.00674972139751468 | 0.00863521114606269 | -0.781650996523957 | 0.434419707908266 | NA | NM_144934 |
| | Mbl1 | 0 | NA | NA | NA | NA | NA | NM_010775 |
| | Mbl2 | 0.795913517 | 0.000216250178044455 | 0.0200046477095122 | 0.0108099968159713 | 0.991375038418332 | NA | NM_010776 |
| | Mbnl3 | 4.206337952 | 0.0276220150752195 | 0.0411627146007735 | 0.671044544635072 | 0.502192152924293 | NA | NM_134163 |
| | Mboat4 | 7.333695251 | -0.0399123212742626 | 0.0502436174582213 | -0.794375948496356 | 0.426976601707745 | NA | NM_001126314 |
| | Mc1r | 2.042746324 | 0.00366533070634582 | 0.0330726021157346 | 0.110826801396495 | 0.911753691826133 | NA | NM_008559 |
| | Mc2r | 1.086318744 | -0.0229185914621276 | 0.0227446460892226 | -1.00764775025395 | 0.313623596789435 | NA | NM_008560,NM_001271717,NM_001271716 |
| | Mc5r | 11.14756877 | -0.00267608536481605 | 0.0680091022924828 | -0.0393489294022316 | 0.968612196777467 | NA | NM_013596 |
| | Mcin | 10.43921798 | 0.123115488643494 | 0.0656070984607508 | 1.87655743863063 | 0.0605787753481238 | NA | NM_001037914 |
| | Mcoln2 | 1.739256276 | 0.02851510671744 | 0.0299254886024216 | 0.952870213625603 | 0.340655833600221 | NA | NM_026656,NM_001005846 |
| | Mcoln3 | 0.84596859 | -0.0100828465457381 | 0.0186645548563993 | -0.540213609342048 | 0.589049728235551 | NA | NM_134160 |
| | Mcpt-ps1 | 0.12890128 | -0.0068640067066208 | 0.00870804083610955 | -0.788237771940376 | 0.430557639008166 | NA | NR_028284 |
| | Mcpt1 | 0 | NA | NA | NA | NA | NA | NM_008570 |
| | Mcpt2 | 0 | NA | NA | NA | NA | NA | NM_008571 |
| | Mcpt4 | 0.554788539 | 0.0231539205280358 | 0.0178655474153742 | 1.29600957584489 | 0.194972185828537 | NA | NM_010779 |
| | Mcpt8 | 0.108766301 | 0.00519171381361946 | 0.00856003371502575 | 0.606506234257728 | 0.544178643801544 | NA | NM_008572 |
| | Mcpt9 | 0 | NA | NA | NA | NA | NA | NM_010782 |
| | Mdfi | 10.30369588 | -0.0422785763429792 | 0.0652508723176485 | -0.647938867961219 | 0.517024490269797 | NA | NM_001109973,NM_001276390,NM_001276391,NM_010783 |
| | Mef2b | 5.651830588 | 0.000462771529786009 | 0.042875931702016 | 0.0107932705230111 | 0.991388383290713 | NA | NM_001045484,NM_008578 |
| | Mefv | 0.980017856 | -0.0283740305814941 | 0.0226929478380431 | -1.25034573665776 | 0.211173277654577 | NA | NM_001161791,NM_019453,NM_001161790 |
| | Meiob | 0.318334547 | 0.012456712416834 | 0.0131461389967144 | 0.947556725206336 | 0.343355165469973 | NA | NM_029197 |
| | Meox1 | 22.00125762 | 0.00480745541035737 | 0.0822623214399395 | 0.0584405512293661 | 0.953397714740035 | NA | NM_010791 |
| | Meox2 | 0.898895954 | 0.0105508889655561 | 0.0189145469999084 | 0.557818750066135 | 0.576968157776341 | NA | NM_008584 |
| | Mep1a | 2.830565892 | 0.0406974167663268 | 0.0354895955534622 | 1.14674219673818 | 0.251488185240611 | NA | NM_008585 |
| | Mep1b | 0.120209049 | -0.00674972139751468 | 0.00863521114606269 | -0.781650996523957 | 0.434419707908266 | NA | NM_008586 |
| | Mepe | 0.106148608 | 0.00515554063321497 | 0.00854200771766808 | 0.603551390213729 | 0.546141939141592 | NA | NM_053172 |
| | Mesp1 | 0.922719674 | -0.00318865764883762 | 0.0206553152078563 | -0.154374678708597 | 0.877314317776235 | NA | NM_008588 |
| | Mettl21c | 1.547359131 | 0.0105813718573631 | 0.0288199081138862 | 0.367154947737837 | 0.713503442550626 | NA | NM_001013799 |
| | Mettl21e | 0.960856407 | 0.0112008708815201 | 0.0194888404740873 | 0.574732544833179 | 0.565472193482985 | NA | NM_207281 |
| | Mettl7a2 | 0.108766301 | 0.00519171381361946 | 0.00856003371502575 | 0.606506234257728 | 0.544178643801544 | NA | NM_199477 |
| | Mettl7a3 | 4.393208858 | -0.0114750144466977 | 0.046581038643813 | -0.246345182090992 | 0.805415037095701 | NA | NM_001081471 |
| | Mettl7b | 0.431200114 | -0.0011657596226791 | 0.0137630292631745 | -0.0847022556144893 | 0.932498102693247 | NA | NM_027853 |
| | Mfap5 | 4.943196057 | -0.00606202698902843 | 0.0468234370383506 | -0.129465655929174 | 0.896989198802725 | NA | NM_015776 |
| | Mfrp | 0 | NA | NA | NA | NA | NA | NM_147126,NM_001190314 |
| | Mfsd6l | 2.19348567 | -0.018584552743233 | 0.0335137990510703 | -0.554534349117293 | 0.579213204729468 | NA | NM_146004 |
| | Mgam | 0.965269439 | 0.0197418899809618 | 0.0234264921977314 | 0.842716434638627 | 0.399387053468819 | NA | NM_001171003 |
| | Mgarp | 3.184530062 | 0.0244765657480765 | 0.0371827375522502 | 0.658277667524651 | 0.510359725533686 | NA | NR_028121,NM_026358 |
| | Mgl2 | 14.7431898 | -0.042356848884247 | 0.0688328704727986 | -0.615357874708792 | 0.538318404629218 | NA | NM_145137 |
| | Mgst2 | 3.205764872 | 0.00991345082207775 | 0.0378962835575689 | 0.261594274990529 | 0.793634258259574 | NA | NM_174995 |
| | Mia2 | 0 | NA | NA | NA | NA | NA | NM_177321 |
| | Mill1 | 0 | NA | NA | NA | NA | NA | NM_153749 |
| | Milr1 | 14.08035286 | -0.224805488481666 | 0.06836710500317 | -3.28821131845853 | 0.00100826128311654 | NA | NM_001271373,NM_001271372,NR_073165,NM_001271374,NM_001033435,NM_001271375 |
| | Miox | 0 | NA | NA | NA | NA | NA | NM_019977 |
| | Mip | 0 | NA | NA | NA | NA | NA | NM_008600 |
| | Mir100 | 0.120209049 | -0.00674972139751468 | 0.00863521114606269 | -0.781650996523957 | 0.434419707908266 | NA | NR_029790 |
| | Mir101a | 0 | NA | NA | NA | NA | NA | NR_029537 |
| | Mir101b | 0 | NA | NA | NA | NA | NA | NR_029778 |
| | Mir101c | 0 | NA | NA | NA | NA | NA | NR_039546 |
| | Mir103-1 | 0 | NA | NA | NA | NA | NA | NR_029754 |
| | Mir103-2 | 3.148041587 | -0.0487586502328889 | 0.0375417257859957 | -1.29878553028794 | 0.194017541756219 | NA | NR_029755 |
| | Mir105 | 0 | NA | NA | NA | NA | NA | NR_030546 |
| | Mir106a | 0 | NA | NA | NA | NA | NA | NR_029657 |
| | Mir106b | 0.705747355 | -0.0240071728049076 | 0.0162288114837101 | -1.47929334375504 | 0.139061931083352 | NA | NR_029658 |
| | Mir107 | 0 | NA | NA | NA | NA | NA | NR_029783 |
| | Mir10a | 0 | NA | NA | NA | NA | NA | NR_029784 |
| | Mir10b | 0 | NA | NA | NA | NA | NA | NR_029566 |
| | Mir1186 | 0 | NA | NA | NA | NA | NA | NR_035414 |
| | Mir1186b | 0 | NA | NA | NA | NA | NA | NR_037310 |
| | Mir1187 | 0 | NA | NA | NA | NA | NA | NR_035415 |
| | Mir1188 | 18.60809112 | -0.178101307138374 | 0.0736812205878554 | -2.41718725229329 | 0.0156409685526962 | NA | NR_035419 |
| | Mir1190 | 0 | NA | NA | NA | NA | NA | NR_035421 |
| | Mir1192 | 0.670958985 | 0.00585915373888986 | 0.0190788768857453 | 0.307101606345995 | 0.758766035062188 | NA | NR_035423 |
| | Mir1193 | 0.120209049 | -0.00674972139751468 | 0.00863521114606269 | -0.781650996523957 | 0.434419707908266 | NA | NR_035424 |
| | Mir1196 | 0 | NA | NA | NA | NA | NA | NR_035428 |
| | Mir1197 | 0 | NA | NA | NA | NA | NA | NR_035429 |
| | Mir1199 | 0 | NA | NA | NA | NA | NA | NR_035431 |
| | Mir1224 | 0 | NA | NA | NA | NA | NA | NR_035407 |
| | Mir122a | 0 | NA | NA | NA | NA | NA | NR_029600 |

| | | | | | | | | | | | | | | | |
|---|---|---|---|---|---|---|---|---|---|---|---|---|---|---|---|
| Klrb1b | 2.323347446 | 0.06007905 | 0.086588676 | 0.693844191 | 0.487779926 | NA | | Mir1247 | 0 | NA | NA | NA | NA | NA | NR_037204 |
| Klrb1c | 2.511132967 | 0.176184219 | 0.101887222 | 1.729208193 | 0.08377184 | NA | | Mir124a-1 | 0 | NA | NA | NA | NA | NA | NR_029813 |
| Klrb1f | 0.478448042 | -0.035850381 | 0.046036495 | -0.778738272 | 0.436133906 | NA | | Mir124a-2 | 6.577125017 | -0.0250786412403344 | 0.0480941982748828 | -0.521448368823973 | 0.602054464119644 | NA | NR_029814 |
| Klrc1 | 0.096071043 | -0.020098795 | 0.027825948 | -0.722304036 | 0.470107572 | NA | | Mir124a-3 | 0.427411356 | 0.0060983518828938 | 0.0139232555105122 | 0.43799755583667 | 0.661388055218398 | NA | NR_029538 |
| Klrc2 | 0 | NA | NA | NA | NA | NA | | Mir1251 | 0.108766301 | 0.00519171381361946 | 0.00856003371502575 | 0.606506234257728 | 0.544178643801544 | NA | NR_037219 |
| Klrc3 | 0 | NA | NA | NA | NA | NA | | Mir125a | 5.686836025 | -0.0616765833185484 | 0.0460759927493615 | -1.33858392707997 | 0.180706161468518 | NA | NR_029539 |
| Klrd1 | 1.248728633 | -0.11829835 | 0.084904083 | -1.393317575 | 0.163523778 | NA | | Mir125b-1 | 0 | NA | NA | NA | NA | NA | NR_029822 |
| Klre1 | 1.200119047 | -0.119395654 | 0.078267184 | -1.525488039 | 0.127137415 | NA | | Mir125b-2 | 0 | NA | NA | NA | NA | NA | NR_029540 |
| Klrg1 | 4.883981353 | 0.018926089 | 0.135593325 | 0.139579796 | 0.888992005 | NA | | Mir126 | 3.261818363 | 0.0291155784668866 | 0.0416877864448262 | 0.698419871859136 | 0.484914652436865 | NA | NR_029541 |
| Klri1 | 0.661489635 | -0.024113131 | 0.059861226 | -0.402817203 | 0.687082704 | NA | | Mir1264 | 0 | NA | NA | NA | NA | NA | NR_037205 |
| Klri2 | 3.935442842 | 0.05164906 | 0.13296822 | 0.388431613 | 0.697696654 | NA | | Mir127 | 0 | NA | NA | NA | NA | NA | NR_029542 |
| Klrk1 | 1.188686129 | 0.022501498 | 0.075096263 | 0.29963539 | 0.764455287 | NA | | Mir128-1 | 6.764855657 | -0.061405509436413 | 0.0546441107299699 | -1.1237351768767 | 0.261125388598151 | NA | NR_029543 |
| Kmo | 3.029795415 | -0.081131485 | 0.111186097 | -0.729690915 | 0.465579136 | NA | | Mir128-2 | 0 | NA | NA | NA | NA | NA | NR_029823 |
| Kncn | 2.897474947 | 0.218710898 | 0.121601032 | 1.798594092 | 0.072082913 | NA | | Mir129-1 | 6.227782697 | 0.000743050312746701 | 0.0510878863277268 | 0.0145445499150241 | 0.988395537322696 | NA | NR_029567 |
| Kng1 | 0 | NA | NA | NA | NA | NA | | Mir129-2 | 0 | NA | NA | NA | NA | NA | NR_029752 |
| Kng2 | 0.176054697 | -0.002190495 | 0.031239146 | -0.070120197 | 0.944097991 | NA | | Mir1298 | 0 | NA | NA | NA | NA | NA | NR_037210 |
| Kpna7 | 0 | NA | NA | NA | NA | NA | | Mir1306 | 0 | NA | NA | NA | NA | NA | NR_035467 |
| Kprp | 0.292412586 | 0.032260595 | 0.035568585 | 0.906996868 | 0.364408454 | NA | | Mir130a | 0 | NA | NA | NA | NA | NA | NR_029544 |
| Krt13 | 0.195033442 | -0.002424765 | 0.031596358 | -0.076741901 | 0.93882887 | NA | | Mir130b | 0 | NA | NA | NA | NA | NA | NR_029659 |
| Krt14 | 0.501644273 | 0.06509729 | 0.050114127 | 1.298980819 | 0.193950512 | NA | | Mir132 | 10.9902922 | 0.0144546376239703 | 0.055090001529876 | 0.262382233119586 | 0.793026768904007 | NA | NR_029546 |
| Krt15 | 1.894032703 | 0.063778806 | 0.073840837 | 0.86373353 | 0.38773429 | NA | | Mir133a-1 | 0.106148608 | 0.00515554063321497 | 0.00854200771766808 | 0.603551390213729 | 0.546141939141592 | NA | NR_029547 |
| Krt16 | 1.016873591 | -0.007411404 | 0.076118005 | -0.097367296 | 0.922434715 | NA | | Mir133a-2 | 0 | NA | NA | NA | NA | NA | NR_029901 |
| Krt17 | 3.942048268 | -0.116151483 | 0.131116868 | -0.88586224 | 0.375691766 | NA | | Mir133b | 0 | NA | NA | NA | NA | NA | NR_029902 |
| Krt18 | 18.54729659 | 0.509573785 | 0.144505286 | 3.52633319 | NA | NA | | Mir134 | 5.481220332 | 0.00289260076841043 | 0.0517755966280183 | 0.0558680335292379 | 0.955446936588885 | NA | NR_029548 |
| Krt19 | 2.742119319 | -0.119798582 | 0.10298342 | -1.163280286 | 0.244715802 | NA | | Mir135a-1 | 0.12890128 | -0.0068640067066208 | 0.00870804083610955 | -0.788237771940376 | 0.430557639008166 | NA | NR_029549 |
| Krt20 | 4.468513605 | -0.0499318 | 0.135565812 | -0.368321479 | 0.712633539 | NA | | Mir135a-2 | 0.634600579 | -0.0147616782762498 | 0.0159374417382655 | -0.926226336615071 | 0.354328368471606 | NA | NR_029812 |
| Krt23 | 0.748066559 | 0.059406784 | 0.047704345 | 1.24531181 | 0.213017156 | NA | | Mir135b | 0.351510867 | 0.00237641026287115 | 0.0128867963543482 | 0.184406597072461 | 0.853694494854753 | NA | NR_029777 |
| Krt24 | 1.105906919 | 0.000490973 | 0.072693744 | 0.006753991 | 0.994611136 | NA | | Mir136 | 0.556841831 | -0.0081599366863108 | 0.0166339239489947 | -0.490559937109968 | 0.623737727854109 | NA | NR_029550 |
| Krt25 | 2.731886909 | -0.173978101 | 0.104059339 | -1.671912429 | 0.094541591 | NA | | Mir137 | 1.258010048 | 0.00245344295263711 | 0.0270740764490826 | 0.0906196359920613 | 0.927794828862064 | NA | NR_029551 |
| Krt26 | 2.268360061 | -0.257914606 | 0.105798543 | -2.43778978 | 0.014777365 | NA | | Mir138-1 | 3.040431364 | -0.0466591403177232 | 0.0360152178780606 | -1.29553958206502 | 0.19513415600691 | NA | NR_029819 |
| Krt27 | 0.116696964 | -0.020477583 | 0.027954224 | -0.732539847 | 0.463839131 | NA | | Mir138-2 | 7.804399111 | 0.00920870591406584 | 0.0589048577579561 | 0.156331858942857 | 0.875771445252787 | NA | NR_029552 |
| Krt28 | 3.243874308 | -0.020074229 | 0.11958889 | -0.167860318 | 0.866693169 | NA | | Mir139 | 0 | NA | NA | NA | NA | NA | NR_029791 |
| Krt31 | 0.280299629 | 0.041967777 | 0.040615819 | 1.033286501 | 0.301469842 | NA | | Mir140 | 1.58693622 | 0.0041573334776022 | 0.0279916769508623 | 0.148520343561415 | 0.881932132156747 | NA | NR_029553 |
| Krt32 | 1.128431217 | -0.017479721 | 0.059519 | -0.293683037 | 0.769000118 | NA | | Mir141 | 0 | NA | NA | NA | NA | NA | NR_029554 |
| Krt33a | 0.185906812 | 0.024895454 | 0.031385107 | 0.793225068 | 0.427646705 | NA | | Mir142 | 0.550080249 | -0.0174816736561517 | 0.0172048016522707 | -1.01609271699127 | 0.309585236886294 | NA | NR_029555 |
| Krt33b | 0.224413567 | -0.027287719 | 0.032077135 | -0.850690659 | 0.394941214 | NA | | Mir143 | 0 | NA | NA | NA | NA | NA | NR_029601 |
| Krt34 | 0 | NA | NA | NA | NA | NA | | Mir144 | 0 | NA | NA | NA | NA | NA | NR_029556 |
| Krt35 | 0.116696964 | -0.020477583 | 0.027954224 | -0.732539847 | 0.463839131 | NA | | Mir145 | 1.595195537 | -0.00943968369435271 | 0.0280630975536558 | -0.336373548084074 | 0.736589194097885 | NA | NR_029557 |
| Krt36 | 0 | NA | NA | NA | NA | NA | | Mir146 | 0 | NA | NA | NA | NA | NA | NR_029558 |
| Krt39 | 0 | NA | NA | NA | NA | NA | | Mir146b | 0 | NA | NA | NA | NA | NA | NR_030468 |
| Krt4 | 0.119397641 | -0.020477583 | 0.027954224 | -0.732539845 | 0.463839132 | NA | | Mir147 | 0 | NA | NA | NA | NA | NA | NR_030547 |
| Krt40 | 0 | NA | NA | NA | NA | NA | | Mir148a | 0 | NA | NA | NA | NA | NA | NR_029719 |
| Krt42 | 0.622661843 | -0.005421728 | 0.053328618 | -0.101666391 | 0.91902148 | NA | | Mir148b | 0 | NA | NA | NA | NA | NA | NR_029766 |
| Krt5 | 0.213790458 | -0.002646789 | 0.031909369 | -0.082947074 | 0.933893623 | NA | | Mir149 | 4.708733545 | 0.00262269917262966 | 0.0413736530301366 | 0.0633905633307092 | 0.94945550151068 | NA | NR_029559 |
| Krt6a | 0.363162105 | 0.038102764 | 0.038558083 | 0.988191333 | 0.323058952 | NA | | Mir150 | 0 | NA | NA | NA | NA | NA | NR_029560 |
| Krt6b | 0 | NA | NA | NA | NA | NA | | Mir152 | 0 | NA | NA | NA | NA | NA | NR_029562 |
| Krt7 | 4.331912649 | -0.085230925 | 0.133233088 | -0.639712899 | 0.522359268 | NA | | Mir153 | 0 | NA | NA | NA | NA | NA | NR_029563 |
| Krt71 | 0.183387883 | -0.001444816 | 0.031390672 | -0.046026918 | 0.963288796 | NA | | Mir154 | 1.947616703 | -0.0189168158420936 | 0.0341364207900639 | -0.554153464372565 | 0.579473823028684 | NA | NR_029564 |
| Krt72 | 0 | NA | NA | NA | NA | NA | | Mir155 | 0 | NA | NA | NA | NA | NA | NR_029565 |
| Krt74 | 0 | NA | NA | NA | NA | NA | | Mir15a | 0.12890128 | -0.0068640067066208 | 0.00870804083610955 | -0.788237771940376 | 0.430557639008166 | NA | NR_029733 |
| Krt75 | 0.533252617 | -0.072906618 | 0.051718649 | -1.409677544 | 0.158634918 | NA | | Mir15b | 0 | NA | NA | NA | NA | NA | NR_029529 |
| Krt76 | 0.187700257 | -0.003168524 | 0.031449734 | -0.100748845 | 0.919749836 | NA | | Mir16-1 | 0.674396547 | 0.00517986742673555 | 0.0190921663358927 | 0.271308521809678 | 0.786153752737112 | NA | NR_029734 |
| Krt78 | 1.113675977 | 0.006779323 | 0.078429329 | 0.086438619 | 0.931117748 | NA | | Mir16-2 | 0 | NA | NA | NA | NA | NA | NR_029735 |
| Krt79 | 0.185670226 | -0.001645086 | 0.031432796 | -0.052336624 | 0.958260472 | NA | | Mir17 | 0 | NA | NA | NA | NA | NA | NR_029785 |
| Krt8 | 10.70122885 | 0.38933577 | 0.129842208 | 2.998530106 | NA | NA | | Mir17hg | 13.80863866 | -0.185596340011579 | 0.073168709608764 | -2.53655341202503 | 0.0111949663798089 | NA | NR_029382 |
| Krt81 | 0 | NA | NA | NA | NA | NA | | Mir18 | 0 | NA | NA | NA | NA | NA | NR_029736 |
| Krt82 | 0.079983654 | 0.016740032 | 0.027674178 | 0.604897175 | 0.54524732 | NA | | Mir181a-1 | 0.127687691 | 0.00542298404407807 | 0.00869724871942359 | 0.62352868349814 | 0.532937159177571 | NA | NR_029795 |
| Krt83 | 2.889392607 | -0.217218845 | 0.112683449 | -1.927690773 | 0.0538936 | NA | | Mir181a-2 | 0 | NA | NA | NA | NA | NA | NR_029568 |
| Krt84 | 0.094392817 | 0.017895942 | 0.027818137 | 0.643319236 | 0.520016977 | NA | | Mir181b-1 | 0.913601056 | 0.00313390167757523 | 0.0190019154554998 | 0.16492556684164 | 0.869002568527539 | NA | NR_029820 |
| Krt85 | 0.467810785 | 0.011930807 | 0.055036386 | 0.216780353 | 0.828379524 | NA | | Mir181b-2 | 0 | NA | NA | NA | NA | NA | NR_029904 |
| Krt86 | 0.32546763 | -0.049399607 | 0.042801411 | -1.154158374 | 0.248435245 | NA | | Mir181c | 1.084659312 | -0.0136700021667274 | 0.0221821550055361 | -0.616261231756594 | 0.537722121382489 | NA | NR_029821 |
| Krtap1-3 | 0 | NA | NA | NA | NA | NA | | Mir181d | 0.968508701 | -0.0182553499626338 | 0.0222374921967457 | -0.820926649512457 | 0.411688050290064 | NA | NR_030534 |
| Krtap1-4 | 0 | NA | NA | NA | NA | NA | | Mir182 | 0 | NA | NA | NA | NA | NA | NR_029569 |
| Krtap1-5 | 0 | NA | NA | NA | NA | NA | | Mir183 | 0 | NA | NA | NA | NA | NA | NR_030713 |
| Krtap10-10 | 0.079983654 | 0.016740032 | 0.027674178 | 0.604897175 | 0.54524732 | NA | | Mir1839 | 14.16468634 | -0.126981755944828 | 0.0690499526466971 | -1.83898396852706 | 0.0659175440773251 | NA | NR_035501 |
| Krtap10-4 | 0.116696964 | -0.020477583 | 0.027954224 | -0.732539847 | 0.463839131 | NA | | Mir184 | 0 | NA | NA | NA | NA | NA | NR_029570 |
| Krtap11-1 | 0.178993538 | 0.024685672 | 0.031260828 | 0.789667872 | 0.429721759 | NA | | Mir1843 | 0 | NA | NA | NA | NA | NA | NR_037207 |
| Krtap12-1 | 0 | NA | NA | NA | NA | NA | | Mir1843b | 0 | NA | NA | NA | NA | NA | NR_039543 |
| Krtap13 | 0 | NA | NA | NA | NA | NA | | Mir185 | 0 | NA | NA | NA | NA | NA | NR_029571 |
| Krtap13-1 | 0 | NA | NA | NA | NA | NA | | Mir187 | 0 | NA | NA | NA | NA | NA | NR_029573 |
| Krtap14 | 0 | NA | NA | NA | NA | NA | | Mir188 | 0 | NA | NA | NA | NA | NA | NR_029574 |
| Krtap15 | 0 | NA | NA | NA | NA | NA | | Mir1892 | 0 | NA | NA | NA | NA | NA | NR_035439 |
| Krtap16-1 | 0 | NA | NA | NA | NA | NA | | Mir1893 | 0 | NA | NA | NA | NA | NA | NR_035446 |
| Krtap16-3 | 0 | NA | NA | NA | NA | NA | | Mir1894 | 3.493832003 | 0.049049937645392 | 0.0442419736674083 | 1.10867426517017 | 0.267570727641674 | NA | NR_035445 |
| Krtap17-1 | 0.912211048 | 0.041259068 | 0.075318195 | 0.547796821 | 0.583831422 | NA | | Mir1895 | 0.587590975 | 0.00239791249935951 | 0.0166905740213935 | 0.143668665696334 | 0.885762115606808 | NA | NR_035435 |
| Krtap19-1 | 0 | NA | NA | NA | NA | NA | | Mir1896 | 0.856955244 | 0.00605930071278908 | 0.0222508058372452 | 0.272318259262616 | 0.785377317700854 | NA | NR_035441 |
| Krtap19-3 | 0 | NA | NA | NA | NA | NA | | Mir1897 | 0 | NA | NA | NA | NA | NA | NR_035433 |
| Krtap19-4 | 0 | NA | NA | NA | NA | NA | | Mir1898 | 5.217000903 | -0.0833560267042121 | 0.0509051369845395 | -1.63747770150443 | 0.10153069441132 | NA | NR_035443 |
| Krtap19-5 | 0 | NA | NA | NA | NA | NA | | Mir1899 | 0 | NA | NA | NA | NA | NA | NR_035437 |
| Krtap19-9b | 0 | NA | NA | NA | NA | NA | | Mir18b | 0 | NA | NA | NA | NA | NA | NR_030548 |
| Krtap2-4 | 0 | NA | NA | NA | NA | NA | | Mir190 | 0.127687691 | 0.00542298404407807 | 0.00869724871942359 | 0.62352868349814 | 0.532937159177571 | NA | NR_029576 |
| Krtap20-2 | 0 | NA | NA | NA | NA | NA | | Mir1900 | 0 | NA | NA | NA | NA | NA | NR_035438 |
| Krtap21-1 | 0 | NA | NA | NA | NA | NA | | Mir1901 | 6.376823963 | -0.0291646363198481 | 0.0555835009148504 | -0.524699521257684 | 0.599792090175091 | NA | NR_035447 |
| Krtap22-2 | 0 | NA | NA | NA | NA | NA | | Mir1902 | 0 | NA | NA | NA | NA | NA | NR_035432 |
| Krtap24-1 | 0 | NA | NA | NA | NA | NA | | Mir1903 | 0 | NA | NA | NA | NA | NA | NR_035436 |
| Krtap26-1 | 0 | NA | NA | NA | NA | NA | | Mir1904 | 0 | NA | NA | NA | NA | NA | NR_035442 |
| Krtap27-1 | 0 | NA | NA | NA | NA | NA | | Mir1905 | 1.607791299 | 0.0198128910063879 | 0.0281385939138883 | 0.704118019081577 | 0.481359279487467 | NA | NR_035434 |
| Krtap3-1 | 0 | NA | NA | NA | NA | NA | | Mir1906-1 | 0 | NA | NA | NA | NA | NA | NR_035440 |
| Krtap3-2 | 0 | NA | NA | NA | NA | NA | | Mir1906-2 | 0 | NA | NA | NA | NA | NA | NR_037313 |
| Krtap3-3 | 0 | NA | NA | NA | NA | NA | | Mir1907 | 0.531833288 | 0.00147671265859378 | 0.0147317495456928 | 0.100240141472235 | 0.920153678198582 | NA | NR_035444 |
| Krtap31-1 | 0 | NA | NA | NA | NA | NA | | Mir190b | 0 | NA | NA | NA | NA | NA | NR_030543 |
| Krtap31-2 | 0 | NA | NA | NA | NA | NA | | Mir191 | 5.066423869 | -0.0190160646359713 | 0.0380926020665296 | -0.4992062396462 | 0.617634099314954 | NA | NR_029577 |
| Krtap4-1 | 0 | NA | NA | NA | NA | NA | | Mir1912 | 0 | NA | NA | NA | NA | NA | NR_037300 |
| Krtap4-13 | 0 | NA | NA | NA | NA | NA | | Mir192 | 0.120209049 | -0.00674972139751468 | 0.00863521114606269 | -0.781650996523957 | 0.434419707908266 | NA | NR_029720 |
| Krtap4-16 | 0.211846316 | 0.025707621 | 0.031860375 | 0.806883811 | 0.419733426 | NA | | Mir1929 | 0.347878816 | 0.00340886725081251 | 0.0128564130570379 | 0.265149169965912 | 0.790894550259911 | NA | NR_035450 |
| Krtap4-2 | 0 | NA | NA | NA | NA | NA | | Mir193 | 0 | NA | NA | NA | NA | NA | NR_029579 |
| Krtap4-6 | 0 | NA | NA | NA | NA | NA | | Mir1930 | 1.093290274 | -0.00828655717979403 | 0.0213836661788225 | -0.387518076203448 | 0.698372705429972 | NA | NR_035451 |
| Krtap4-7 | 0 | NA | NA | NA | NA | NA | | Mir1931 | 0.203513543 | -0.00860782669689374 | 0.00975338121443284 | -0.882547960306941 | 0.377480556050336 | NA | NR_035452 |
| Krtap4-8 | 0 | NA | NA | NA | NA | NA | | Mir1932 | 0 | NA | NA | NA | NA | NA | NR_035453 |
| Krtap4-9 | 0 | NA | NA | NA | NA | NA | | Mir1933 | 0 | NA | NA | NA | NA | NA | NR_035454 |
| Krtap5-1 | 0.105923158 | 0.018524693 | 0.027977518 | 0.662127842 | 0.507889294 | NA | | Mir1934 | 0.657115133 | -0.0163160432289019 | 0.0205334600145153 | -0.794607592552247 | 0.426841800384946 | NA | NR_035455 |
| Krtap5-2 | 0.211846316 | 0.025707621 | 0.031860375 | 0.806883811 | 0.419733426 | NA | | Mir1936 | 0 | NA | NA | NA | NA | NA | NR_035457 |
| Krtap5-3 | 0 | NA | NA | NA | NA | NA | | Mir1938 | 0.648422349 | 0.00582902956687644 | 0.0189632857398013 | 0.307384998932021 | 0.758550344976929 | NA | NR_035459 |
| Krtap5-4 | 0 | NA | NA | NA | NA | NA | | Mir193b | 0 | NA | NA | NA | NA | NA | NR_030549 |
| Krtap5-5 | 0.107716603 | -0.020477583 | 0.027954224 | -0.732539849 | 0.46383913 | NA | | Mir194-1 | 0 | NA | NA | NA | NA | NA | NR_029580 |
| Krtap6-1 | 0 | NA | NA | NA | NA | NA | | Mir194-2 | 0 | NA | NA | NA | NA | NA | NR_029830 |
| Krtap6-2 | 0 | NA | NA | NA | NA | NA | | Mir1940 | 0.113843286 | 0.00525947738444006 | 0.00859368002043176 | 0.612016897526494 | 0.540526577873086 | NA | NR_035461 |
| Krtap6-5 | 0 | NA | NA | NA | NA | NA | | Mir1941 | 0 | NA | NA | NA | NA | NA | NR_035462 |
| Krtap7-1 | 0.341676965 | 0.050095582 | 0.044235127 | 1.132484193 | 0.257430926 | NA | | Mir1942 | 0 | NA | NA | NA | NA | NA | NR_035463 |
| Krtap8-1 | 0 | NA | NA | NA | NA | NA | | Mir1943 | 7.513756175 | -0.124400412107558 | 0.0551124488934055 | -2.25721075011861 | 0.0239949090086439 | NA | NR_035464 |
| Krtap9-1 | 0 | NA | NA | NA | NA | NA | | Mir1945 | 0 | NA | NA | NA | NA | NA | NR_035466 |
| Krtap9-3 | 0 | NA | NA | NA | NA | NA | | Mir1946a | 0 | NA | NA | NA | NA | NA | NR_035468 |
| Krtap9-5 | 0 | NA | NA | NA | NA | NA | | Mir1947 | 0.427212126 | 0.012175076767397 | 0.0129887840374575 | 0.93735308342075 | 0.348576965801222 | NA | NR_035469 |
| Krtdap | 0.17463368 | 0.024560283 | 0.031186242 | 0.787535833 | 0.430968261 | NA | | Mir1948 | 0 | NA | NA | NA | NA | NA | NR_035471 |
| Kynu | 4.014358091 | -0.026246882 | 0.125619344 | -0.208939807 | 0.834495228 | NA | | Mir1949 | 3.139771678 | -0.0676863296819005 | 0.0397180680768449 | -1.70416973834034 | 0.0883493809815636 | NA | NR_035472 |
| L1td1 | 4.493681656 | -0.106457821 | 0.123886038 | -0.859320573 | 0.390163677 | NA | | Mir195 | 0.097165704 | -0.0063782845546458 | 0.00848781691535724 | -0.75146349388194 | 0.452373763047372 | NA | NR_029581 |
| LOC100038947 | 0.203787646 | -0.02671144 | 0.031728022 | -0.841887971 | 0.399850664 | NA | | Mir1951 | 0 | NA | NA | NA | NA | NA | NR_035476 |
| LOC100043315 | 0.41925633 | -0.060548742 | 0.047272363 | -1.280848629 | 0.20024684 | NA | | Mir1952 | 0 | NA | NA | NA | NA | NA | NR_035477 |
| LOC100048884 | 0 | NA | NA | NA | NA | NA | | Mir1953 | 0 | NA | NA | NA | NA | NA | NR_035478 |
| LOC100502924 | 0 | NA | NA | NA | NA | NA | | Mir1955 | 0 | NA | NA | NA | NA | NA | NR_035480 |
| LOC100503280 | 0.305724365 | 0.009518016 | 0.039476503 | 0.241105846 | 0.809473082 | NA | | Mir1956 | 0.300216221 | 0.00934385809732901 | 0.0113912670188907 | 0.820265040037567 | 0.412065031496518 | NA | NR_035481 |
| LOC100504608 | 1.228183666 | -0.022943163 | 0.073840375 | -0.31071298 | 0.756018829 | NA | | Mir1960 | 0 | NA | NA | NA | NA | NA | NR_035486 |
| LOC100861615 | 6.091513533 | -0.048265545 | 0.135024574 | -0.357457486 | 0.720749348 | NA | | Mir1961 | 0 | NA | NA | NA | NA | NA | NR_035487 |
| LOC100861978 | 0 | NA | NA | NA | NA | NA | | Mir1962 | 0 | NA | NA | NA | NA | NA | NR_035488 |
| LOC171588 | 5.818998857 | -0.020013229 | 0.148150586 | -0.135087074 | 0.89254303 | NA | | Mir1963 | 0 | NA | NA | NA | NA | NA | NR_035489 |
| LOC547349 | 0.105923158 | 0.018524693 | 0.027977518 | 0.662127842 | 0.507889294 | NA | | Mir1964 | 0.57283292 | -0.0311343653719056 | 0.0184476705313912 | -1.68771256614358 | 0.0914664184018825 | NA | NR_035490 |
| LOC666331 | 0 | NA | NA | NA | NA | NA | | Mir1965 | 0 | NA | NA | NA | NA | NA | NR_035491 |
| Lactbl1 | 0.593755747 | -0.004769304 | 0.058238563 | -0.081892538 | 0.934732168 | NA | | Mir1966 | 0 | NA | NA | NA | NA | NA | NR_035492 |
| Lalba | 0.798357233 | -0.007830999 | 0.062135221 | -0.126031561 | 0.899706942 | NA | | Mir1967 | 0.106347839 | -0.00653347234140708 | 0.00854389509723638 | -0.764694822098226 | 0.444453290197145 | NA | NR_035493 |
| Lamp3 | 0 | NA | NA | NA | NA | NA | | Mir1968 | 3.172473764 | 0.00116464094543287 | 0.0406226062752974 | 0.0286697741041074 | 0.97712796322465 | NA | NR_035494 |
| Lao1 | 6.420233442 | 0.088157996 | 0.116576444 | 0.756224782 | 0.449514437 | NA | | Mir1969 | 2.054516533 | -0.0337304637541791 | 0.0338971274739933 | -0.995083249459941 | 0.319695776726002 | NA | NR_035495 |
| Lbx1 | 0.079983654 | 0.016740032 | 0.027674178 | 0.604897175 | 0.54524732 | NA | | Mir196a-1 | 0 | NA | NA | NA | NA | NA | NR_029721 |
| Lbx2 | 1.310361776 | -0.029823595 | 0.087516462 | -0.340776974 | 0.733271486 | NA | | Mir196a-2 | 0 | NA | NA | NA | NA | NA | NR_029722 |
| Lce1a1 | 0 | NA | NA | NA | NA | NA | | Mir196b | 0 | NA | NA | NA | NA | NA | NR_029912 |
| Lce1a2 | 0 | NA | NA | NA | NA | NA | | Mir1971 | 0 | NA | NA | NA | NA | NA | NR_035499 |
| Lce1b | 0 | NA | NA | NA | NA | NA | | Mir1981 | 0 | NA | NA | NA | NA | NA | NR_035502 |
| Lce1c | 0 | NA | NA | NA | NA | NA | | Mir1982 | 0 | NA | NA | NA | NA | NA | NR_035503 |
| Lce1d | 0 | NA | NA | NA | NA | NA | | Mir1983 | 0.226556888 | -0.00933175792380659 | 0.010153338242423 | -0.919082739193727 | 0.358052296835844 | NA | NR_035500 |
| Lce1e | 0 | NA | NA | NA | NA | NA | | Mir199a-1 | 0.240418098 | -0.0088029805258837 | 0.00986414519853158 | -0.892422034419582 | 0.372166765031741 | NA | NR_029585 |
| Lce1f | 0 | NA | NA | NA | NA | NA | | Mir199a-2 | 0 | NA | NA | NA | NA | NA | NR_029810 |
| Lce1g | 0.17463368 | 0.024560283 | 0.031186242 | 0.787535833 | 0.430968261 | NA | | Mir199b | 0 | NA | NA | NA | NA | NA | NR_029811 |
| Lce1h | 0 | NA | NA | NA | NA | NA | | Mir19a | 0 | NA | NA | NA | NA | NA | NR_029786 |
| Lce1i | 0 | NA | NA | NA | NA | NA | | Mir19b-1 | 0 | NA | NA | NA | NA | NA | NR_029815 |
| Lce1k | 0 | NA | NA | NA | NA | NA | | Mir19b-2 | 0 | NA | NA | NA | NA | NA | NR_029715 |
| Lce1l | 0 | NA | NA | NA | NA | NA | | Mir1a-1 | 0 | NA | NA | NA | NA | NA | NR_029528 |
| Lce1m | 0 | NA | NA | NA | NA | NA | | Mir1a-2 | 0 | NA | NA | NA | NA | NA | NR_029781 |
| Lce3a | 0.302141032 | -0.047437531 | 0.04196367 | -1.130442867 | 0.25828966 | NA | | Mir1b | 0.41913649 | 0.0122773187407356 | 0.0130433482186402 | 0.94127048783304 | 0.346566262520942 | NA | NR_035413 |
| Lce3b | 0 | NA | NA | NA | NA | NA | | Mir200a | 0 | NA | NA | NA | NA | NA | NR_029723 |
| Lce3c | 0.717316046 | 0.045366621 | 0.050111211 | 0.905318791 | 0.365296524 | NA | | Mir200b | 0 | NA | NA | NA | NA | NA | NR_029587 |
| Lce3d | 0 | NA | NA | NA | NA | NA | | Mir200c | 0 | NA | NA | NA | NA | NA | NR_029792 |
| Lce3e | 0 | NA | NA | NA | NA | NA | | Mir201 | 0 | NA | NA | NA | NA | NA | NR_029588 |
| Lce3f | 1.181425567 | -0.106516642 | 0.066956276 | -1.590838815 | 0.111645855 | NA | | Mir202 | 0 | NA | NA | NA | NA | NA | NR_029589 |
| Lce6a | 0 | NA | NA | NA | NA | NA | | Mir203 | 0 | NA | NA | NA | NA | NA | NR_029590 |
| Lcn10 | 0 | NA | NA | NA | NA | NA | | Mir204 | 0 | NA | NA | NA | NA | NA | NR_029591 |
| Lcn11 | 0.593158268 | 0.071669397 | 0.052434572 | 1.366834783 | 0.171677098 | NA | | Mir205 | 0 | NA | NA | NA | NA | NA | NR_029592 |
| Lcn12 | 0 | NA | NA | NA | NA | NA | | Mir206 | 0 | NA | NA | NA | NA | NA | NR_029593 |
| Lcn3 | 0.293623414 | 0.010408489 | 0.039073364 | 0.266383235 | 0.789944076 | NA | | Mir207 | 0 | NA | NA | NA | NA | NA | NR_029594 |
| Lcn4 | 0 | NA | NA | NA | NA | NA | | Mir208a | 0 | NA | NA | NA | NA | NA | NR_029724 |
| Lcn5 | 0 | NA | NA | NA | NA | NA | | Mir208b | 0 | NA | NA | NA | NA | NA | NR_030607 |
| Lcn6 | 0 | NA | NA | NA | NA | NA | | Mir20a | 0 | NA | NA | NA | NA | NA | NR_029737 |
| Lcn8 | 0.238795281 | -0.027601606 | 0.032267774 | -0.855392297 | 0.392334007 | NA | | Mir20b | 0 | NA | NA | NA | NA | NA | NR_030273 |
| Lcn9 | 0 | NA | NA | NA | NA | NA | | Mir21 | 0 | NA | NA | NA | NA | NA | NR_029738 |
| Ldhal6b | 7.216227394 | -0.168309871 | 0.154698833 | -1.087984105 | 0.276602124 | NA | | Mir210 | 0.09572496 | 0.00498255289994511 | 0.00847999972823692 | 0.587565219295241 | 0.556824162160132 | NA | NR_029793 |
| Ldhc | 0.994426385 | 0.049832203 | 0.071188917 | 0.699999454 | 0.483927646 | NA | | Mir211 | 0 | NA | NA | NA | NA | NA | NR_029805 |
| Ldlrad1 | 0.239950962 | 0.027417169 | 0.032831728 | 0.835081494 | 0.403671825 | NA | | Mir212 | 11.84910035 | -0.00251781027824436 | 0.0664632315627762 | -0.0378827544048355 | 0.96978116317015 | NA | NR_029794 |
| Ldlrad2 | 2.087130886 | 0.110319212 | 0.101415702 | 1.087792229 | 0.276686841 | NA | | Mir2136 | 0 | NA | NA | NA | NA | NA | NR_035516 |
| Leap2 | 0.305079873 | -0.015054679 | 0.039642215 | -0.379763828 | 0.704120735 | NA | | Mir2137 | 0 | NA | NA | NA | NA | NA | NR_035517 |
| Lect2 | 0 | NA | NA | NA | NA | NA | | Mir214 | 0 | NA | NA | NA | NA | NA | NR_029796 |
| Lelp1 | 0 | NA | NA | NA | NA | NA | | Mir215 | 0 | NA | NA | NA | NA | NA | NR_029908 |
| Lep | 0 | NA | NA | NA | NA | NA | | Mir216a | 0 | NA | NA | NA | NA | NA | NR_029797 |
| Lgals2 | 2.826545916 | -0.059761391 | 0.122620378 | -0.487369159 | 0.625996745 | NA | | Mir216b | 0 | NA | NA | NA | NA | NA | NR_030419 |
| Lgals4 | 7.737277884 | 0.425836631 | 0.156769947 | 2.716315469 | 0.006601298 | NA | | Mir217 | 0 | NA | NA | NA | NA | NA | NR_029828 |
| Lgals6 | 0.60415196 | 0.020563279 | 0.052843793 | 0.389133293 | 0.697177546 | NA | | Mir218-1 | 2.133570485 | -0.0225264804452168 | 0.0277817229760147 | -0.8108381350093 | 0.417458631409438 | NA | NR_029798 |
| Lgals7 | 3.880974019 | 0.009929444 | 0.131055042 | 0.075765448 | 0.939605706 | NA | | Mir218-2 | 0.12890128 | -0.0068640067066208 | 0.00870804083610955 | -0.788237771940376 | 0.430557639008166 | NA | NR_029799 |
| Lgsn | 0.473045832 | 0.03947874 | 0.051663738 | 0.764147964 | 0.444779072 | NA | | Mir219-1 | 0 | NA | NA | NA | NA | NA | NR_029800 |
| Lhb | 2.559019543 | 0.252937408 | 0.107532361 | 2.352198041 | 0.018662836 | NA | | Mir219-2 | 2.773660428 | -0.000467624905406704 | 0.0380404528228738 | -0.0122928322537086 | 0.990191985957771 | NA | NR_029838 |
| Lhfpl1 | 4.535166914 | -0.100523214 | 0.120567251 | -0.833752223 | 0.404420623 | NA | | Mir22 | 0 | NA | NA | NA | NA | NA | NR_029739 |
| Lhx1 | 5.917210998 | -0.18460631 | 0.124027708 | -1.488427972 | 0.136638064 | NA | | Mir221 | 3.040170242 | 0.0198843849412809 | 0.0401329253703764 | 0.495463133020408 | 0.620273239635804 | NA | NR_029806 |
| Lhx3 | 5.555580945 | 0.322154567 | 0.119185858 | 2.702959659 | 0.006872509 | NA | | Mir222 | 0 | NA | NA | NA | NA | NA | NR_029807 |
| Lhx4 | 0.426277686 | 0.058476664 | 0.047630606 | 1.227711936 | 0.219555124 | NA | | Mir223 | 0 | NA | NA | NA | NA | NA | NR_029801 |
| Lilra6 | 0.990536273 | -0.061771697 | 0.067614392 | -0.913587997 | 0.360933376 | NA | | Mir224 | 0.127687691 | 0.00542298404407807 | 0.00869724871942359 | 0.62352868349814 | 0.532937159177571 | NA | NR_029808 |
| Lim2 | 0 | NA | NA | NA | NA | NA | | Mir23a | 0.57007369 | 0.0101860070818361 | 0.0153947996431531 | 0.661652461736742 | 0.508193977563616 | NA | NR_029740 |
| Lin28a | 0.514098028 | 0.006684456 | 0.05090618 | 0.131309328 | 0.895530613 | NA | | Mir23b | 19.35736512 | -0.166922683436461 | 0.0712396228381102 | -2.3431157660083 | 0.019123448378073 | NA | NR_029530 |
| Lipf | 0.338960846 | 0.036479516 | 0.037750094 | 0.966342396 | 0.333872878 | NA | | Mir24-2 | 1.008052884 | -0.0108393263515856 | 0.0234210500632704 | -0.462802748907665 | 0.643505760677094 | NA | NR_029741 |
| Lipi | 0.300347587 | -0.013446771 | 0.039455705 | -0.340806761 | 0.73324906 | NA | | Mir25 | 3.192235021 | -0.0537627104825282 | 0.0392673450148684 | -1.36914554478209 | 0.17095378400829 | NA | NR_029787 |
| Lipk | 0.238795281 | -0.027601606 | 0.032267774 | -0.855392297 | 0.392334007 | NA | | Mir26a-1 | 2.532465486 | -0.0545237547169071 | 0.0360495567601137 | -1.51246671574153 | 0.130415173831674 | NA | NR_029742 |
| Lipm | 3.142511058 | 0.230372281 | 0.109191951 | 2.109791775 | 0.034876295 | NA | | Mir26a-2 | 0.728599411 | 0.0182583361895325 | 0.0190855331832144 | 0.956658428887415 | 0.338739687094635 | NA | NR_029803 |
| Lipn | 0 | NA | NA | NA | NA | NA | | Mir26b | 0.108766301 | 0.00519171381361946 | 0.00856003371502575 | 0.606506234257728 | 0.544178643801544 | NA | NR_029743 |
| Lman1l | 1.393655212 | 0.138459737 | 0.077590734 | 1.784488049 | 0.074344395 | NA | | Mir27a | 1.083360502 | -0.0238814563782348 | 0.0202985482878307 | -1.17651055827239 | 0.239390915794086 | NA | NR_029746 |
| Lmod2 | 0.094392817 | 0.017895942 | 0.027818137 | 0.643319236 | 0.520016977 | NA | | Mir2861 | 0.856803614 | -0.010795564536427 | 0.0148291637517496 | -0.72799550380265 | 0.466616338629313 | NA | NR_037217 |
| Lmod3 | 1.843987541 | -0.004747097 | 0.087084654 | -0.054511294 | 0.956527811 | NA | | Mir28b | 0 | NA | NA | NA | NA | NA | NR_039551 |
| Lmx1a | 5.016530087 | 0.264831897 | 0.10694634 | 2.476306323 | NA | NA | | Mir290 | 0 | NA | NA | NA | NA | NA | NR_029640 |
| Lmx1b | 0.819267527 | -0.011522036 | 0.055657027 | -0.207018536 | 0.835995383 | NA | | Mir291a | 0 | NA | NA | NA | NA | NA | NR_029641 |
| Loxl4 | 6.44770556 | 0.002642171 | 0.124206831 | 0.021272352 | 0.983028399 | NA | | Mir291b | 0 | NA | NA | NA | NA | NA | NR_030276 |
| Lpcat2b | 1.298488313 | -0.022962054 | 0.080392725 | -0.285623537 | 0.775166474 | NA | | Mir292 | 0 | NA | NA | NA | NA | NA | NR_029642 |
| Lrat | 5.053319707 | -0.021652086 | 0.106831773 | -0.202674591 | 0.839389384 | NA | | Mir293 | 0 | NA | NA | NA | NA | NA | NR_029643 |
| Lrch4 | 0 | NA | NA | NA | NA | NA | | Mir294 | 0 | NA | NA | NA | NA | NA | NR_029644 |
| Lrcol1 | 0.107716603 | -0.020477583 | 0.027954224 | -0.732539849 | 0.46383913 | NA | | Mir295 | 0 | NA | NA | NA | NA | NA | NR_029645 |
| Lrg1 | 49.01134731 | 0.271078349 | 0.16780799 | 1.615407876 | NA | NA | | Mir296 | 0 | NA | NA | NA | NA | NA | NR_029646 |
| Lrit1 | 0.515459553 | -0.019052845 | 0.045554446 | -0.418243369 | 0.67576919 | NA | | Mir297-1 | 0 | NA | NA | NA | NA | NA | NR_029647 |
| Lrit2 | 0.293623414 | 0.010408489 | 0.039073364 | 0.266383235 | 0.789944076 | NA | | Mir297a-3 | 0 | NA | NA | NA | NA | NA | NR_030551 |
| Lrit3 | 2.173935032 | 0.14230611 | 0.107900473 | 1.318864558 | 0.187214397 | NA | | Mir297b | 0 | NA | NA | NA | NA | NA | NR_030474 |
| Lrr1 | 1.045045392 | -0.104975171 | 0.076446864 | -1.37317825 | 0.169696939 | NA | | Mir297c | 0 | NA | NA | NA | NA | NA | NR_030555 |
| Lrrc10 | 1.328786961 | 0.04053153 | 0.081204073 | 0.499131732 | 0.617686584 | NA | | Mir298 | 0 | NA | NA | NA | NA | NA | NR_029649 |
| Lrrc15 | 1.154903561 | -0.039247414 | 0.080272433 | -0.488927671 | 0.624892905 | NA | | Mir299 | 0 | NA | NA | NA | NA | NA | NR_029650 |
| Lrrc19 | 0 | NA | NA | NA | NA | NA | | Mir29a | 0 | NA | NA | NA | NA | NA | NR_029744 |
| Lrrc26 | 0.287139021 | 0.008693067 | 0.033441491 | 0.259948559 | 0.794903454 | NA | | Mir29b-1 | 0 | NA | NA | NA | NA | NA | NR_029532 |
| Lrrc30 | 0 | NA | NA | NA | NA | NA | | Mir29b-2 | 0 | NA | NA | NA | NA | NA | NR_029809 |
| Lrrc34 | 4.067439929 | 0.197196167 | 0.127441899 | 1.547341726 | 0.121780865 | NA | | Mir29c | 0 | NA | NA | NA | NA | NA | NR_029745 |
| Lrrc52 | 0.603663061 | -0.004610385 | 0.052400067 | -0.087984343 | 0.92988912 | NA | | Mir300 | 0.12890128 | -0.0068640067066208 | 0.00870804083610955 | -0.788237771940376 | 0.430557639008166 | NA | NR_029651 |
| Lrrc58 | 4206.490262 | -0.008980542 | 0.135290053 | -0.066379912 | NA | NA | | Mir301 | 0 | NA | NA | NA | NA | NA | NR_029652 |
| Lrrc63 | 0.903760442 | 0.00255845 | 0.070410694 | 0.036336097 | 0.971014367 | NA | | Mir301b | 0 | NA | NA | NA | NA | NA | NR_030415 |
| Lrrc66 | 1.166819058 | 0.045440474 | 0.077118192 | 0.58923158 | 0.555705935 | NA | | Mir302a | 0 | NA | NA | NA | NA | NA | NR_029653 |
| Lrrc69 | 2.091909967 | 0.054846023 | 0.107070421 | 0.512242529 | 0.608481282 | NA | | Mir302b | 0 | NA | NA | NA | NA | NA | NR_030403 |
| Lrrc72 | 0.211846316 | 0.025707621 | 0.031860375 | 0.806883811 | 0.419733426 | NA | | Mir302c | 0 | NA | NA | NA | NA | NA | NR_030404 |
| Lrrc8e | 0.789559445 | 0.014020811 | 0.056659565 | 0.247457092 | 0.804554494 | NA | | Mir302d | 0 | NA | NA | NA | NA | NA | NR_030405 |
| Lrriq4 | 0.275017096 | 0.008844542 | 0.038389071 | 0.2303922 | 0.817787022 | NA | | Mir3057 | 1.389777318 | -0.00747472914918147 | 0.0271920816325053 | -0.274886242627567 | 0.783403636941381 | NA | NR_037218 |
| Lrtm1 | 0.295716657 | -0.010253415 | 0.03372058 | -0.304069933 | 0.761074613 | NA | | Mir3058 | 1.091337543 | 0.0045488242174623 | 0.0266588412497298 | 0.170630980350971 | 0.86451393635248 | NA | NR_037212 |
| Lta | 5.73735428 | -0.10244084 | 0.13750598 | -0.744991892 | 0.456276622 | NA | | Mir3059 | 0 | NA | NA | NA | NA | NA | NR_037211 |
| Ltb4r1 | 4.365710893 | 0.055336194 | 0.135407208 | 0.408665052 | 0.682785484 | NA | | Mir3060 | 16.29616498 | -0.116685761787835 | 0.0793172738654421 | -1.47112673067643 | 0.141256847256985 | NA | NR_037220 |
| Ltb4r2 | 2.065653484 | 0.051845857 | 0.096794205 | 0.535629757 | 0.592214459 | NA | | Mir3062 | 0 | NA | NA | NA | NA | NA | NR_037222 |
| Luzp4 | 0 | NA | NA | NA | NA | NA | | Mir3063 | 0 | NA | NA | NA | NA | NA | NR_037223 |
| Ly6c2 | 1.112398198 | -0.030243582 | 0.06703571 | -0.451156285 | 0.651876914 | NA | | Mir3065 | 0 | NA | NA | NA | NA | NA | NR_037225 |
| Ly6f | 0 | NA | NA | NA | NA | NA | | Mir3066 | 0 | NA | NA | NA | NA | NA | NR_037226 |
| Ly6g5c | 0 | NA | NA | NA | NA | NA | | Mir3067 | 0 | NA | NA | NA | NA | NA | NR_037227 |
| Ly6g6c | 0 | NA | NA | NA | NA | NA | | Mir3068 | 3.229482682 | -0.10066055434372 | 0.0412363427977544 | -2.44106405937633 | 0.0146440568747901 | NA | NR_037228 |
| Ly6i | 0.484297991 | 0.036542974 | 0.051863038 | 0.704605351 | 0.481055868 | NA | | Mir3069 | 0 | NA | NA | NA | NA | NA | NR_037229 |
| Ly6k | 0.982137931 | 0.021872688 | 0.07729474 | 0.282977698 | 0.777193933 | NA | | Mir3070a | 5.906870762 | -0.0742647855950334 | 0.0456202044456527 | -1.62789243269405 | 0.103547696270391 | NA | NR_037230 |
| Ly9 | 43.63071773 | 0.8464765 | 0.167386621 | 5.057014072 | NA | NA | | Mir3070b | 5.700176491 | -0.0958236775313728 | 0.0536014960646896 | -1.78770528001171 | 0.0738235694336588 | NA | NR_037231 |
| Lyg1 | 0 | NA | NA | NA | NA | NA | | Mir3071 | 0 | NA | NA | NA | NA | NA | NR_037232 |
| Lyg2 | 2.377896396 | 0.067007288 | 0.102529546 | 0.653541248 | 0.513407407 | NA | | Mir3073 | 0 | NA | NA | NA | NA | NA | NR_037234 |
| Lypd2 | 4.849282279 | -0.099581706 | 0.134744639 | -0.73904021 | 0.459882581 | NA | | Mir3074-1 | 0.097165704 | -0.0063782845546458 | 0.00848781691535724 | -0.75146349388194 | 0.452373763047372 | NA | NR_037235 |
| Lypd3 | 0.362333936 | 0.019030129 | 0.042103636 | 0.451983023 | 0.651281213 | NA | | Mir3074-2 | 0 | NA | NA | NA | NA | NA | NR_037294 |
| Lypd4 | 0 | NA | NA | NA | NA | NA | | Mir3075 | 0 | NA | NA | NA | NA | NA | NR_037236 |
| Lypd5 | 0 | NA | NA | NA | NA | NA | | Mir3076 | 0 | NA | NA | NA | NA | NA | NR_037237 |

| | | | | | | | | | | | | | | | |
|---|---|---|---|---|---|---|---|---|---|---|---|---|---|---|---|
| Lypd8 | 0.287795519 | 0.031925978 | 0.035387605 | 0.902179657 | 0.366961439 | NA | | Mir3077 | 0 | NA | NA | NA | NA | NA | NR_037238 |
| Lyzl1 | 0 | NA | NA | NA | NA | NA | | Mir3078 | 0 | NA | NA | NA | NA | NA | NR_037239 |
| Lyzl6 | 0.907724086 | -0.087049886 | 0.072820354 | -1.19540596 | 0.231928458 | NA | | Mir3081 | 0 | NA | NA | NA | NA | NA | NR_037242 |
| M1ap | 1.325353327 | 0.073222765 | 0.086063402 | 0.850800263 | 0.394880316 | NA | | Mir3082 | 10.21800983 | -0.0434678058285585 | 0.0558277094180833 | -0.778606292137766 | 0.436211671433385 | NA | NR_037243 |
| Mab21l3 | 3.198626589 | 0.014856769 | 0.105264105 | 0.141138032 | 0.887760899 | NA | | Mir3083 | 0 | NA | NA | NA | NA | NA | NR_037244 |
| Macc1 | 4.742395832 | -0.142333648 | 0.13380872 | -1.063709807 | 0.287460181 | NA | | Mir3085 | 0 | NA | NA | NA | NA | NA | NR_037246 |
| Madcam1 | 2.160123571 | 0.099281291 | 0.100597065 | 0.986920356 | 0.323681684 | NA | | Mir3086 | 0 | NA | NA | NA | NA | NA | NR_037247 |
| Mael | 2.721507992 | 0.073588115 | 0.09825567 | 0.748945223 | 0.453890222 | NA | | Mir3087 | 0 | NA | NA | NA | NA | NA | NR_037270 |
| Maff | 69.17769177 | 0.257166854 | 0.117863747 | 2.181899534 | NA | NA | | Mir3088 | 0 | NA | NA | NA | NA | NA | NR_037271 |
| Magea1 | 0 | NA | NA | NA | NA | NA | | Mir3089 | 0 | NA | NA | NA | NA | NA | NR_037272 |
| Magea10 | 0.17833704 | -0.002390466 | 0.031282281 | -0.076415987 | 0.939088151 | NA | | Mir3091 | 2.273203209 | 0.0242821795917378 | 0.0305608696531592 | 0.794551328784835 | 0.426874539847387 | NA | NR_037274 |
| Magea2 | 0 | NA | NA | NA | NA | NA | | Mir3092 | 0 | NA | NA | NA | NA | NA | NR_037275 |
| Magea3 | 0 | NA | NA | NA | NA | NA | | Mir3093 | 1.86405858 | -0.0411749626623215 | 0.02989543047833 | -1.3772995405491 | 0.168419657232523 | NA | NR_037276 |
| Magea4 | 0 | NA | NA | NA | NA | NA | | Mir3094 | 0 | NA | NA | NA | NA | NA | NR_037277 |
| Magea5 | 0 | NA | NA | NA | NA | NA | | Mir3095 | 0 | NA | NA | NA | NA | NA | NR_037278 |
| Magea6 | 0 | NA | NA | NA | NA | NA | | Mir3096 | 0 | NA | NA | NA | NA | NA | NR_037279 |
| Magea8 | 0 | NA | NA | NA | NA | NA | | Mir3096b | 0 | NA | NA | NA | NA | NA | NR_039541 |
| Mageb1 | 0 | NA | NA | NA | NA | NA | | Mir3097 | 1.174215799 | -0.00747269171651829 | 0.0239690915339142 | -0.311763660543624 | 0.755220143180104 | NA | NR_037280 |
| Mageb16 | 0 | NA | NA | NA | NA | NA | | Mir3099 | 0 | NA | NA | NA | NA | NA | NR_037213 |
| Mageb16-ps1 | 0.258320694 | 0.006870708 | 0.033097595 | 0.207589347 | 0.835549623 | NA | | Mir30a | 0 | NA | NA | NA | NA | NA | NR_029533 |
| Mageb18 | 0.413070119 | -0.002318447 | 0.042591244 | -0.054434824 | 0.956588734 | NA | | Mir30b | 0.631190298 | 0.00397774684543735 | 0.0171215330860167 | 0.232324221519977 | 0.816286200988879 | NA | NR_029534 |
| Mageb2 | 0 | NA | NA | NA | NA | NA | | Mir30c-1 | 0.113843286 | 0.00525947738444006 | 0.00859368002043176 | 0.612016897526494 | 0.540526577873086 | NA | NR_029716 |
| Mageb3 | 3.848484583 | 0.123354358 | 0.116432672 | 1.059447962 | 0.289395816 | NA | | Mir30c-2 | 0.097165704 | -0.00637828455464458 | 0.00848781691535724 | -0.75146349388194 | 0.452373763047372 | NA | NR_029717 |
| Mageb4 | 0.188785634 | 0.025025111 | 0.031461603 | 0.79541754 | 0.42637066 | NA | | Mir30d | 0.718908469 | -0.00610196463357164 | 0.0179451349621561 | -0.340034479898862 | 0.733830562153634 | NA | NR_029718 |
| Mageb5 | 0 | NA | NA | NA | NA | NA | | Mir30e | 0.108766301 | 0.00519171381361946 | 0.00856003371502575 | 0.606506234257728 | 0.544178643801544 | NA | NR_029602 |
| Mansc4 | 6.531528774 | 0.118896125 | 0.134343551 | 0.885015499 | 0.376148272 | NA | | Mir31 | 0 | NA | NA | NA | NA | NA | NR_029747 |
| 1-Mar | 3.674522383 | 0.051839156 | 0.132232611 | 0.392030044 | 0.695036015 | NA | | Mir3100 | 1.134515865 | -0.00779123174154452 | 0.0256267372816398 | -0.304027456008867 | 0.761106973753223 | NA | NR_037282 |
| Marcksl1-ps4 | 0.793177326 | -0.012109046 | 0.055799279 | -0.217010791 | 0.828199935 | NA | | Mir3101 | 16.4571339 | -0.051582716717099 | 0.0726376374781511 | -0.710137588555447 | 0.477618818854162 | NA | NR_037283 |
| Marco | 0.19442443 | -0.026451352 | 0.031568725 | -0.837897386 | 0.402088337 | NA | | Mir3102 | 0 | NA | NA | NA | NA | NA | NR_037289 |
| Mat1a | 1.361257865 | -0.000613543 | 0.081854933 | -0.007495494 | 0.994019517 | NA | | Mir3103 | 0 | NA | NA | NA | NA | NA | NR_037290 |
| Matn3 | 2.865118358 | -0.089187609 | 0.118298064 | -0.753922813 | 0.450895572 | NA | | Mir3104 | 1.491974438 | 0.00617083239213001 | 0.0275164948555058 | 0.224259391486241 | 0.822555472370921 | NA | NR_037291 |
| Mb | 5.31222965 | 0.13556448 | 0.121250778 | 1.118050393 | 0.263545471 | NA | | Mir3106 | 0 | NA | NA | NA | NA | NA | NR_037214 |
| Mbd3l1 | 0 | NA | NA | NA | NA | NA | | Mir3107 | 0 | NA | NA | NA | NA | NA | NR_037293 |
| Mbd3l2 | 0.099009884 | 0.018205726 | 0.027896865 | 0.652608315 | 0.514008823 | NA | | Mir3108 | 0 | NA | NA | NA | NA | NA | NR_037295 |
| Mbl1 | 0 | NA | NA | NA | NA | NA | | Mir3109 | 0 | NA | NA | NA | NA | NA | NR_037296 |
| Mbl2 | 0.681282946 | 0.008839375 | 0.054361526 | 0.162603511 | 0.870830622 | NA | | Mir3110 | 0 | NA | NA | NA | NA | NA | NR_037297 |
| Mbnl3 | 3.407290376 | -0.09714654 | 0.106540491 | -0.911827407 | 0.361859581 | NA | | Mir3112 | 6.611335474 | -0.0385175950407623 | 0.0582436034462554 | -0.661318887597754 | 0.508407831264887 | NA | NR_037299 |
| Mboat4 | 5.240030806 | 0.305571521 | 0.136591577 | 2.237118329 | 0.025278607 | NA | | Mir32 | 0.323681211 | 0.00985023388331581 | 0.0116938364156777 | 0.842344080520041 | 0.399595383885569 | NA | NR_029789 |
| Mc1r | 1.717398057 | 0.107937639 | 0.088991717 | 1.21289534 | 0.225169837 | NA | | Mir320 | 1.307155672 | -6.7087055730712e-05 | 0.0230475138104751 | -0.00291081529584423 | 0.997677508695783 | NA | NR_029802 |
| Mc2r | 0.831689407 | 0.122187568 | 0.067723391 | 1.804215155 | 0.071197584 | NA | | Mir322 | 0 | NA | NA | NA | NA | NA | NR_029756 |
| Mcm7 | 492.3551671 | 0.026997463 | 0.11970892 | 0.225525907 | NA | NA | | Mir323 | 0.240418098 | -0.0088029805258837 | 0.00986414519853158 | -0.892422034419582 | 0.372166765031741 | NA | NR_029757 |
| Mcoln2 | 0.962157358 | -0.03636776 | 0.072161417 | -0.503977905 | 0.614276904 | NA | | Mir324 | 0.576364262 | 0.00251948358971321 | 0.0180487455074274 | 0.139593280246342 | 0.888981350221907 | NA | NR_029758 |
| Mcoln3 | 0.760998328 | 0.011477899 | 0.05537348 | 0.207281514 | 0.83579001 | NA | | Mir326 | 0 | NA | NA | NA | NA | NA | NR_029760 |
| Mcpt-ps1 | 0.21363976 | -0.000753674 | 0.031922632 | -0.02360938 | 0.98116419 | NA | | Mir328 | 0.106347839 | -0.00653347234140708 | 0.00854389509723638 | -0.764694822098226 | 0.444453290197145 | NA | NR_029761 |
| Mcpt1 | 0 | NA | NA | NA | NA | NA | | Mir329 | 0 | NA | NA | NA | NA | NA | NR_029762 |
| Mcpt2 | 0 | NA | NA | NA | NA | NA | | Mir33 | 0 | NA | NA | NA | NA | NA | NR_029804 |
| Mcpt4 | 0.670575502 | -0.080313968 | 0.054217216 | -1.481336999 | 0.138516794 | NA | | Mir330 | 0.364141684 | 0.0106460894690162 | 0.0121537141469398 | 0.875935584254473 | 0.381055267460423 | NA | NR_029763 |
| Mcpt8 | 0 | NA | NA | NA | NA | NA | | Mir331 | 21.87748658 | -0.0830408995344233 | 0.0821707396116466 | -1.01058965669397 | 0.312212868349161 | NA | NR_029764 |
| Mcpt9 | 0 | NA | NA | NA | NA | NA | | Mir335 | 0.926546215 | -0.0053628285814382 | 0.0224419654200202 | -0.238964301079178 | 0.811133264477604 | NA | NR_029900 |
| Mef2b | 6.58410113 | 0.046991247 | 0.134334315 | 0.349808215 | 0.726482634 | NA | | Mir337 | 1.809274879 | -0.0109069346917537 | 0.0277862918639426 | -0.39252933587397 | 0.694667140113092 | NA | NR_029765 |
| Mefv | 0.958981039 | 0.044863195 | 0.072618989 | 0.617788754 | 0.536714597 | NA | | Mir338 | 4.992204301 | -0.048650461942182 | 0.0450685816472489 | -1.07947621522613 | 0.280375491176205 | NA | NR_029767 |
| Meiob | 0 | NA | NA | NA | NA | NA | | Mir339 | 1.360556778 | 0.03130609307843 | 0.0261428622169713 | 1.19750059571162 | 0.231111495505483 | NA | NR_029768 |
| Meox2 | 1.368920632 | -0.072981926 | 0.07790776 | -0.936773509 | 0.348875075 | NA | | Mir340 | 0.361523991 | 0.0105955510260098 | 0.0121250525610889 | 0.873856090324303 | 0.382196629856021 | NA | NR_029769 |
| Mep1a | 1.300725613 | 0.045436634 | 0.070135083 | 0.64784458 | 0.517085479 | NA | | Mir341 | 11.39807248 | -0.0770561998710391 | 0.0668833443738338 | -1.15209848718607 | 0.249280604728542 | NA | NR_029770 |
| Mep1b | 0.215706849 | -0.002035226 | 0.0319478 | -0.063704747 | 0.949205325 | NA | | Mir343 | 9.5040228 | -0.064424213377319 | 0.0610762762434438 | -1.05481567213644 | 0.291509635498709 | NA | NR_030759 |
| Mepe | 0.192746203 | -0.000975302 | 0.031567365 | -0.030895906 | 0.975352555 | NA | | Mir344 | 0 | NA | NA | NA | NA | NA | NR_029772 |
| Mesp1 | 0.686827323 | 0.040767534 | 0.066213896 | 0.615694537 | 0.538096144 | NA | | Mir344-2 | 6.431452148 | -0.166122191707496 | 0.054362663577348 | -3.05581406016162 | 0.00224450401142833 | NA | NR_030557 |
| Mettl21c | 1.186620711 | 0.016888272 | 0.076282014 | 0.221392583 | 0.824786767 | NA | | Mir344b | 0.633678068 | -0.00271054942267143 | 0.0172996419783739 | -0.156682399905146 | 0.875495158587407 | NA | NR_037285 |
| Mettl21e | 1.304414302 | -0.084831958 | 0.067412641 | -1.258398364 | 0.208247723 | NA | | Mir344c | 0 | NA | NA | NA | NA | NA | NR_037286 |
| Mettl7a2 | 0 | NA | NA | NA | NA | NA | | Mir344d-1 | 0.4298712 | -0.0123579986757156 | 0.015660003200329 | -0.789144070893677 | 0.430027806374149 | NA | NR_037215 |
| Mettl7a3 | 5.440438738 | 0.081190936 | 0.12005675 | 0.676271311 | 0.498868404 | NA | | Mir344d-2 | 0.12890128 | -0.0068640067066208 | 0.00870804083610955 | -0.788237771940376 | 0.430557639008166 | NA | NR_037216 |
| Mettl7b | 0.609883421 | -0.040524199 | 0.059374544 | -0.682518065 | 0.494911424 | NA | | Mir344d-3 | 0.212695677 | -0.00857796107890413 | 0.00973699866180651 | -0.880965621629515 | 0.378336427605205 | NA | NR_037209 |
| Mfap5 | 5.888324268 | -0.088737789 | 0.137057818 | -0.647447843 | 0.51734214 | NA | | Mir344e | 0.106347839 | -0.00653347234140708 | 0.00854389509723638 | -0.764694822098226 | 0.444453290197145 | NA | NR_037284 |
| Mfrp | 0 | NA | NA | NA | NA | NA | | Mir344f | 0.23403553 | -0.000407313579519725 | 0.00983856423584417 | -0.0413996971261099 | 0.966977254250032 | NA | NR_037288 |
| Mfsd6l | 1.847446006 | 0.041419523 | 0.093546088 | 0.442771303 | 0.657931162 | NA | | Mir344g | 1.309514955 | -0.00209985079277275 | 0.0228235685382164 | -0.0920036141226192 | 0.926695168115733 | NA | NR_037287 |
| Mgam | 4.339769707 | -0.286992334 | 0.103171518 | -2.781701188 | 0.00540748 | NA | | Mir345 | 2.597795488 | 0.0174760340291461 | 0.0326526460421809 | 0.535210347319798 | 0.592504412003594 | NA | NR_029773 |
| Mgarp | 2.090185046 | 0.056731167 | 0.100188161 | 0.566246213 | 0.571226419 | NA | | Mir346 | 0 | NA | NA | NA | NA | NA | NR_029774 |
| Mgst2 | 1.910121513 | 0.051605332 | 0.095108525 | 0.542594181 | 0.587409246 | NA | | Mir3471-1 | 0 | NA | NA | NA | NA | NA | NR_037304 |
| Mia2 | 0 | NA | NA | NA | NA | NA | | Mir3473 | 0 | NA | NA | NA | NA | NA | NR_037311 |
| Mill1 | 0 | NA | NA | NA | NA | NA | | Mir3473c | 0.469085418 | -0.00231634314321046 | 0.0155809327962042 | -0.148665241324625 | 0.881817789791702 | NA | NR_039566 |
| Miox | 0 | NA | NA | NA | NA | NA | | Mir3473d | 0.816830525 | -0.0124208601950181 | 0.0165083147879886 | -0.75240025129976 | 0.45181039551978 | NA | NR_039583 |
| Mip | 0 | NA | NA | NA | NA | NA | | Mir3474 | 0 | NA | NA | NA | NA | NA | NR_037312 |
| Mir100 | 0.099009884 | 0.018205726 | 0.027896865 | 0.652608315 | 0.514008823 | NA | | Mir3475 | 0 | NA | NA | NA | NA | NA | NR_037208 |
| Mir101a | 0 | NA | NA | NA | NA | NA | | Mir34a | 0.406564059 | 0.00565656510264079 | 0.013731356404231 | 0.41194510841608 | 0.680379657052235 | NA | NR_029751 |
| Mir101b | 0.188785634 | 0.025025111 | 0.031461603 | 0.79541754 | 0.42637066 | NA | | Mir34b | 0 | NA | NA | NA | NA | NA | NR_029655 |
| Mir101c | 0 | NA | NA | NA | NA | NA | | Mir34c | 0 | NA | NA | NA | NA | NA | NR_029654 |
| Mir103-1 | 0 | NA | NA | NA | NA | NA | | Mir350 | 0 | NA | NA | NA | NA | NA | NR_029775 |
| Mir103-2 | 1.920915016 | 0.167017145 | 0.088579629 | 1.885502865 | 0.059361978 | NA | | Mir351 | 0 | NA | NA | NA | NA | NA | NR_029776 |
| Mir105 | 0 | NA | NA | NA | NA | NA | | Mir3572 | 0 | NA | NA | NA | NA | NA | NR_039587 |
| Mir106a | 0 | NA | NA | NA | NA | NA | | Mir362 | 0 | NA | NA | NA | NA | NA | NR_029851 |
| Mir106b | 0.909184964 | 0.024388887 | 0.058249921 | 0.418693912 | 0.675439846 | NA | | Mir363 | 0.106347839 | -0.00653347234140708 | 0.00854389509723638 | -0.764694822098226 | 0.444453290197145 | NA | NR_029853 |
| Mir107 | 0 | NA | NA | NA | NA | NA | | Mir365-1 | 1.79566016 | 0.0239824369710766 | 0.0271522180394064 | 0.88325885333826 | 0.377096429752129 | NA | NR_029855 |
| Mir10a | 0 | NA | NA | NA | NA | NA | | Mir365-2 | 0 | NA | NA | NA | NA | NA | NR_029959 |
| Mir10b | 0 | NA | NA | NA | NA | NA | | Mir367 | 0 | NA | NA | NA | NA | NA | NR_030268 |
| Mir1186 | 0 | NA | NA | NA | NA | NA | | Mir369 | 5.775608065 | -0.00223312837891032 | 0.0522287524177346 | -0.0427566862223583 | 0.965895491778284 | NA | NR_030272 |
| Mir1186b | 0 | NA | NA | NA | NA | NA | | Mir370 | 2.217855882 | 0.00705012553782974 | 0.0332056271003215 | 0.212317193002552 | 0.831859583110827 | NA | NR_029918 |
| Mir1187 | 0 | NA | NA | NA | NA | NA | | Mir374 | 22.22510651 | -0.133373361401732 | 0.0751246610049719 | -1.77536057557591 | 0.0758383692516108 | NA | NR_030418 |
| Mir1190 | 0 | NA | NA | NA | NA | NA | | Mir374c | 0 | NA | NA | NA | NA | NA | NR_037298 |
| Mir1192 | 0.383703858 | 0.028442486 | 0.048086527 | 0.591485549 | 0.554195135 | NA | | Mir375 | 0 | NA | NA | NA | NA | NA | NR_029876 |
| Mir1193 | 0.099009884 | 0.018205726 | 0.027896865 | 0.652608315 | 0.514008823 | NA | | Mir376a | 0 | NA | NA | NA | NA | NA | NR_029877 |
| Mir1196 | 0 | NA | NA | NA | NA | NA | | Mir376b | 0 | NA | NA | NA | NA | NA | NR_029915 |
| Mir1197 | 0 | NA | NA | NA | NA | NA | | Mir376c | 0.191449919 | 0.00665508079272897 | 0.00962405648474729 | 0.691504752001018 | 0.489248395458281 | NA | NR_030270 |
| Mir1199 | 0.116696964 | -0.020477583 | 0.027954224 | -0.732539847 | 0.463839131 | NA | | Mir377 | 15.9916664 | -0.0346546061388394 | 0.0663073744746262 | -0.522635776388653 | 0.601227738936174 | NA | NR_029878 |
| Mir1224 | 0 | NA | NA | NA | NA | NA | | Mir378b | 0 | NA | NA | NA | NA | NA | NR_039545 |
| Mir122a | 0 | NA | NA | NA | NA | NA | | Mir379 | 0 | NA | NA | NA | NA | NA | NR_029880 |
| Mir1247 | 0 | NA | NA | NA | NA | NA | | Mir380 | 0 | NA | NA | NA | NA | NA | NR_029881 |
| Mir124a-1 | 0 | NA | NA | NA | NA | NA | | Mir381 | 0.113843286 | 0.00525947738444006 | 0.00859368002043176 | 0.612016897526494 | 0.540526577873086 | NA | NR_029882 |
| Mir124a-2 | 6.188467794 | 0.159161527 | 0.132752934 | 1.198930394 | 0.230555012 | NA | | Mir382 | 9.292188856 | -0.0516677603669342 | 0.0598190254062155 | -0.863734573007031 | 0.38773371669834 | NA | NR_029883 |
| Mir124a-3 | 0.398821088 | -0.027919467 | 0.043501191 | -0.641809245 | 0.520997046 | NA | | Mir383 | 0 | NA | NA | NA | NA | NA | NR_029884 |
| Mir1251 | 0.119397641 | -0.020477583 | 0.027954224 | -0.732539845 | 0.463839132 | NA | | Mir384 | 0.74312821 | 0.0204532679252645 | 0.0204060644267776 | 1.00231320932344 | 0.316192344760037 | NA | NR_029910 |
| Mir125a | 4.436323189 | 0.310515415 | 0.135676455 | 2.288646281 | 0.022099913 | NA | | Mir3960 | 1.394163851 | -0.0302408130950407 | 0.0197258785462942 | -1.53305278769051 | NA | NA | NR_039536 |
| Mir125b-1 | 0 | NA | NA | NA | NA | NA | | Mir3966 | 0 | NA | NA | NA | NA | NA | NR_039547 |
| Mir125b-2 | 0 | NA | NA | NA | NA | NA | | Mir3967 | 0 | NA | NA | NA | NA | NA | NR_039548 |
| Mir126 | 1.690534935 | 0.064467984 | 0.088732415 | 0.726543777 | 0.467505481 | NA | | Mir3968 | 0 | NA | NA | NA | NA | NA | NR_039549 |
| Mir1264 | 0 | NA | NA | NA | NA | NA | | Mir3969 | 0 | NA | NA | NA | NA | NA | NR_039550 |
| Mir127 | 0 | NA | NA | NA | NA | NA | | Mir3971 | 0.09572496 | 0.00498255289994511 | 0.00847999972823692 | 0.587565219295241 | 0.556824162160132 | NA | NR_039553 |
| Mir128-1 | 6.461650651 | 0.179996805 | 0.137606822 | 1.308051461 | 0.190855855 | NA | | Mir409 | 10.07523633 | -0.096074052064963 | 0.0588946533704587 | -1.63128648471091 | 0.102829883828566 | NA | NR_029913 |
| Mir128-2 | 0 | NA | NA | NA | NA | NA | | Mir410 | 0 | NA | NA | NA | NA | NA | NR_029914 |
| Mir129-1 | 5.202608281 | 0.20374429 | 0.134992744 | 1.509298087 | 0.131222623 | NA | | Mir411 | 0 | NA | NA | NA | NA | NA | NR_029916 |
| Mir129-2 | 0 | NA | NA | NA | NA | NA | | Mir412 | 0.127687691 | 0.00542298404407807 | 0.00869724871942359 | 0.62352868349814 | 0.532937159177571 | NA | NR_029917 |
| Mir1298 | 0 | NA | NA | NA | NA | NA | | Mir421 | 18.63963674 | -0.0398489636140579 | 0.0681077676489167 | -0.585086914307222 | 0.558489277184985 | NA | NR_030558 |
| Mir1306 | 0 | NA | NA | NA | NA | NA | | Mir423 | 0 | NA | NA | NA | NA | NA | NR_030756 |
| Mir130a | 0 | NA | NA | NA | NA | NA | | Mir429 | 0 | NA | NA | NA | NA | NA | NR_029958 |
| Mir130b | 0 | NA | NA | NA | NA | NA | | Mir431 | 5.9531147 | -0.00789798937881747 | 0.0529838837517247 | -0.14906399492771 | 0.881503136304214 | NA | NR_029951 |
| Mir133a-1 | 0 | NA | NA | NA | NA | NA | | Mir432 | 2.11306047 | -0.000380596747701731 | 0.0312786347306419 | -0.012167946298784 | 0.990291623080742 | NA | NR_035526 |
| Mir133a-2 | 0 | NA | NA | NA | NA | NA | | Mir433 | 0 | NA | NA | NA | NA | NA | NR_029952 |
| Mir133b | 0 | NA | NA | NA | NA | NA | | Mir434 | 5.092605892 | -0.0243461545374858 | 0.0471053271268038 | -0.516845036909474 | 0.605264338245878 | NA | NR_029953 |
| Mir134 | 5.490815863 | -0.122742067 | 0.141703851 | -0.866187235 | 0.386387492 | NA | | Mir448 | 0 | NA | NA | NA | NA | NA | NR_029956 |
| Mir135a-1 | 0.105923158 | 0.018524693 | 0.027977518 | 0.662127842 | 0.507889294 | NA | | Mir449a | 0 | NA | NA | NA | NA | NA | NR_029961 |
| Mir135a-2 | 0.416779358 | 0.041366775 | 0.04012708 | 1.03089422 | 0.302590428 | NA | | Mir449b | 0 | NA | NA | NA | NA | NA | NR_030602 |
| Mir135b | 0.222620122 | -0.001447027 | 0.032059667 | -0.04513545 | 0.963999345 | NA | | Mir449c | 0 | NA | NA | NA | NA | NA | NR_030452 |
| Mir136 | 0.525223898 | 0.006204767 | 0.045384814 | 0.136714611 | 0.89125638 | NA | | Mir450-1 | 0 | NA | NA | NA | NA | NA | NR_029963 |
| Mir137 | 1.72245743 | 0.012298194 | 0.092857863 | 0.132441065 | 0.894635436 | NA | | Mir450-2 | 0 | NA | NA | NA | NA | NA | NR_030274 |
| Mir138-1 | 4.336868572 | 0.16737562 | 0.128531445 | 1.302215339 | 0.192842783 | NA | | Mir450b | 0 | NA | NA | NA | NA | NA | NR_030498 |
| Mir138-2 | 7.888519922 | -0.090004968 | 0.158504068 | -0.567840113 | 0.570143539 | NA | | Mir451 | 0 | NA | NA | NA | NA | NA | NR_029971 |
| Mir139 | 0 | NA | NA | NA | NA | NA | | Mir452 | 0 | NA | NA | NA | NA | NA | NR_029974 |
| Mir140 | 0.813705528 | 0.030122804 | 0.06445444 | 0.467350329 | 0.640249254 | NA | | Mir453 | 8.759935517 | -0.0491573009887648 | 0.0575080553426903 | -0.85478983241281 | 0.392667508028476 | NA | NR_030559 |
| Mir141 | 0 | NA | NA | NA | NA | NA | | Mir455 | 0 | NA | NA | NA | NA | NA | NR_030477 |
| Mir142 | 0.598714191 | 0.017606709 | 0.053437106 | 0.3294847 | 0.741789357 | NA | | Mir463 | 0 | NA | NA | NA | NA | NA | NR_030147 |
| Mir143 | 0.212768007 | -0.02694223 | 0.031868764 | -0.845411806 | 0.397880955 | NA | | Mir465 | 0 | NA | NA | NA | NA | NA | NR_030149 |
| Mir144 | 0 | NA | NA | NA | NA | NA | | Mir465b-1 | 0 | NA | NA | NA | NA | NA | NR_030560 |
| Mir145 | 1.422393584 | -0.006857031 | 0.075784459 | -0.090480703 | 0.927905228 | NA | | Mir465b-2 | 0 | NA | NA | NA | NA | NA | NR_030561 |
| Mir146 | 0 | NA | NA | NA | NA | NA | | Mir465c-1 | 0 | NA | NA | NA | NA | NA | NR_030562 |
| Mir146b | 0.119397641 | -0.020477583 | 0.027954224 | -0.732539845 | 0.463839132 | NA | | Mir465c-2 | 0 | NA | NA | NA | NA | NA | NR_030563 |
| Mir147 | 0 | NA | NA | NA | NA | NA | | Mir466 | 0 | NA | NA | NA | NA | NA | NR_030150 |
| Mir148a | 0 | NA | NA | NA | NA | NA | | Mir4660 | 0 | NA | NA | NA | NA | NA | NR_037248 |
| Mir148b | 0 | NA | NA | NA | NA | NA | | Mir466g | 0 | NA | NA | NA | NA | NA | NR_030569 |
| Mir149 | 2.972681264 | 0.073213464 | 0.110445894 | 0.662889864 | 0.507401094 | NA | | Mir466i | 0 | NA | NA | NA | NA | NA | NR_035412 |
| Mir150 | 0 | NA | NA | NA | NA | NA | | Mir467a-1 | 0 | NA | NA | NA | NA | NA | NR_035406 |
| Mir152 | 0 | NA | NA | NA | NA | NA | | Mir467a-10 | 0 | NA | NA | NA | NA | NA | NR_037267 |
| Mir153 | 0.298180463 | -0.015304498 | 0.039412854 | -0.388312359 | 0.697784893 | NA | | Mir467a-2 | 0 | NA | NA | NA | NA | NA | NR_037249 |
| Mir154 | 1.201797363 | 0.081344257 | 0.081416452 | 0.999113264 | 0.317739826 | NA | | Mir467a-3 | 0 | NA | NA | NA | NA | NA | NR_037252 |
| Mir155 | 0.119397641 | -0.020477583 | 0.027954224 | -0.732539845 | 0.463839132 | NA | | Mir467a-4 | 0 | NA | NA | NA | NA | NA | NR_037254 |
| Mir15a | 0.321391842 | -0.01514286 | 0.04017217 | -0.376949019 | 0.706211492 | NA | | Mir467a-5 | 0 | NA | NA | NA | NA | NA | NR_037256 |
| Mir15b | 0 | NA | NA | NA | NA | NA | | Mir467a-6 | 0 | NA | NA | NA | NA | NA | NR_037258 |
| Mir16-1 | 0.303286429 | 0.012912943 | 0.039427608 | 0.32751017 | 0.743282052 | NA | | Mir467a-7 | 0 | NA | NA | NA | NA | NA | NR_037261 |
| Mir16-2 | 0 | NA | NA | NA | NA | NA | | Mir467a-8 | 0 | NA | NA | NA | NA | NA | NR_037263 |
| Mir17 | 0 | NA | NA | NA | NA | NA | | Mir467a-9 | 0 | NA | NA | NA | NA | NA | NR_037265 |
| Mir18 | 0 | NA | NA | NA | NA | NA | | Mir467b | 0 | NA | NA | NA | NA | NA | NR_030472 |
| Mir181a-1 | 0 | NA | NA | NA | NA | NA | | Mir467d | 0 | NA | NA | NA | NA | NA | NR_030572 |
| Mir181a-2 | 0.215468684 | -0.027008432 | 0.031909031 | -0.846419662 | 0.397318674 | NA | | Mir467f | 0 | NA | NA | NA | NA | NA | NR_035420 |
| Mir181b-1 | 0.599170685 | -0.002006017 | 0.052323611 | -0.038338665 | 0.969417664 | NA | | Mir468 | 0.956550331 | -0.00561414710530632 | 0.0190195582230234 | -0.295177576654243 | 0.767858230482831 | NA | NR_030151 |
| Mir181b-2 | 0 | NA | NA | NA | NA | NA | | Mir470 | 0 | NA | NA | NA | NA | NA | NR_030153 |
| Mir181c | 0.859830262 | 0.09472059 | 0.069693768 | 1.359096978 | 0.17411586 | NA | | Mir471 | 0 | NA | NA | NA | NA | NA | NR_030154 |
| Mir181d | 1.161288641 | 0.052504258 | 0.086764318 | 0.605136528 | 0.545088285 | NA | | Mir483 | 0 | NA | NA | NA | NA | NA | NR_030251 |
| Mir182 | 0 | NA | NA | NA | NA | NA | | Mir484 | 0.108766301 | 0.00519171381361946 | 0.00856003371502575 | 0.606506234257728 | 0.544178643801544 | NA | NR_030252 |
| Mir183 | 0 | NA | NA | NA | NA | NA | | Mir485 | 0.428430455 | -0.00286174811964168 | 0.0152554242063721 | -0.18758889172327 | 0.851198934776155 | NA | NR_030253 |
| Mir184 | 0 | NA | NA | NA | NA | NA | | Mir486 | 1.485314722 | -0.000893977406700569 | 0.0255761285287312 | -0.0349535859462197 | 0.972116751285645 | NA | NR_030254 |
| Mir1843 | 0 | NA | NA | NA | NA | NA | | Mir487b | 0.12890128 | -0.0068640067066208 | 0.00870804083610955 | -0.788237771940376 | 0.430557639008166 | NA | NR_030271 |
| Mir1843b | 0 | NA | NA | NA | NA | NA | | Mir488 | 0 | NA | NA | NA | NA | NA | NR_030441 |
| Mir185 | 0.116696964 | -0.020477583 | 0.027954224 | -0.732539847 | 0.463839131 | NA | | Mir489 | 0 | NA | NA | NA | NA | NA | NR_030250 |
| Mir187 | 0 | NA | NA | NA | NA | NA | | Mir490 | 0 | NA | NA | NA | NA | NA | NR_030524 |
| Mir188 | 0 | NA | NA | NA | NA | NA | | Mir491 | 2.72934465 | 0.0204866234698223 | 0.0400590480655639 | 0.511410641518296 | 0.609063544997526 | NA | NR_030478 |
| Mir1892 | 0 | NA | NA | NA | NA | NA | | Mir493 | 0.834263005 | 0.000290881808641226 | 0.0201386721824231 | 0.0144439417855515 | 0.988475802564229 | NA | NR_030573 |
| Mir1893 | 0 | NA | NA | NA | NA | NA | | Mir494 | 0.42201798 | -0.000629897072052632 | 0.0120467086091002 | -0.0522878980883462 | 0.95829929605327 | NA | NR_030269 |
| Mir1894 | 3.582350746 | -0.174863184 | 0.118121118 | -1.480371907 | 0.138774023 | NA | | Mir495 | 0 | NA | NA | NA | NA | NA | NR_030446 |
| Mir1895 | 1.373629463 | 0.032823391 | 0.073130882 | 0.448830776 | 0.653553736 | NA | | Mir496 | 7.118165946 | -0.0304391592851897 | 0.0520384845819711 | -0.584935543948093 | 0.558591057744752 | NA | NR_030437 |
| Mir1896 | 1.097917236 | -0.026940225 | 0.074160089 | -0.363271205 | 0.716402301 | NA | | Mir497 | 0 | NA | NA | NA | NA | NA | NR_030444 |
| Mir1897 | 0 | NA | NA | NA | NA | NA | | Mir499 | 0 | NA | NA | NA | NA | NA | NR_030757 |
| Mir1898 | 6.583395593 | 0.084719094 | 0.148226174 | 0.571552861 | 0.567624939 | NA | | Mir500 | 0 | NA | NA | NA | NA | NA | NR_030495 |
| Mir1899 | 0 | NA | NA | NA | NA | NA | | Mir501 | 0 | NA | NA | NA | NA | NA | NR_030496 |
| Mir18b | 0 | NA | NA | NA | NA | NA | | Mir503 | 0 | NA | NA | NA | NA | NA | NR_030275 |
| Mir190 | 0 | NA | NA | NA | NA | NA | | Mir504 | 0 | NA | NA | NA | NA | NA | NR_030574 |
| Mir1900 | 0 | NA | NA | NA | NA | NA | | Mir5046 | 0 | NA | NA | NA | NA | NA | NR_039555 |
| Mir1901 | 6.378478339 | 0.020424364 | 0.153466177 | 0.133087073 | 0.894124518 | NA | | Mir505 | 0 | NA | NA | NA | NA | NA | NR_030499 |
| Mir1902 | 0 | NA | NA | NA | NA | NA | | Mir509 | 0 | NA | NA | NA | NA | NA | NR_030575 |
| Mir1903 | 0 | NA | NA | NA | NA | NA | | Mir5098 | 0 | NA | NA | NA | NA | NA | NR_039557 |
| Mir1904 | 0 | NA | NA | NA | NA | NA | | Mir5100 | 0.354244579 | -0.00570556803240082 | 0.0130207392936895 | -0.438190789609467 | 0.661247984684689 | NA | NR_039559 |
| Mir1905 | 1.115578782 | 0.005731202 | 0.083430484 | 0.068694338 | 0.945232925 | NA | | Mir5101 | 0 | NA | NA | NA | NA | NA | NR_039560 |
| Mir1906-1 | 0 | NA | NA | NA | NA | NA | | Mir5103 | 2.300549262 | -0.0398385317446347 | 0.0365385641700577 | -1.09031464835942 | 0.275574564746966 | NA | NR_039562 |
| Mir1906-2 | 0 | NA | NA | NA | NA | NA | | Mir5104 | 0 | NA | NA | NA | NA | NA | NR_039563 |
| Mir1907 | 0.518444887 | -0.021172618 | 0.051250408 | -0.413120974 | 0.679517983 | NA | | Mir5106 | 0 | NA | NA | NA | NA | NA | NR_039565 |
| Mir190b | 0 | NA | NA | NA | NA | NA | | Mir5107 | 0.097165704 | -0.00637828455464458 | 0.00848781691535724 | -0.75146349388194 | 0.452373763047372 | NA | NR_039567 |
| Mir191 | 4.417109132 | 0.032452265 | 0.123406214 | 0.262971074 | 0.792572875 | NA | | Mir5108 | 0 | NA | NA | NA | NA | NA | NR_039568 |
| Mir1912 | 0 | NA | NA | NA | NA | NA | | Mir5109 | 0 | NA | NA | NA | NA | NA | NR_039569 |
| Mir192 | 0.099009884 | 0.018205726 | 0.027896865 | 0.652608315 | 0.514008823 | NA | | Mir511 | 0 | NA | NA | NA | NA | NA | NR_030609 |
| Mir1929 | 0.527163612 | -0.038923887 | 0.04705911 | -0.827127563 | 0.408164763 | NA | | Mir5112 | 1.845075032 | 0.00431641016216521 | 0.0279042154327466 | 0.154686669925138 | 0.877068339354874 | NA | NR_039572 |
| Mir193 | 0 | NA | NA | NA | NA | NA | | Mir5113 | 0 | NA | NA | NA | NA | NA | NR_039573 |
| Mir1930 | 0.707852793 | 0.070978149 | 0.063325329 | 1.120849269 | 0.262352028 | NA | | Mir5114 | 0 | NA | NA | NA | NA | NA | NR_039574 |
| Mir1931 | 0.167300494 | 0.024299763 | 0.031030539 | 0.783091891 | 0.433573153 | NA | | Mir5116 | 0 | NA | NA | NA | NA | NA | NR_039576 |
| Mir1932 | 0.096071043 | -0.020098795 | 0.027825948 | -0.722304036 | 0.470107572 | NA | | Mir5117 | 0 | NA | NA | NA | NA | NA | NR_039577 |
| Mir1933 | 0 | NA | NA | NA | NA | NA | | Mir5119 | 4.803802496 | -0.0290459653884623 | 0.0444898684108608 | -0.652866965580226 | 0.513842047331301 | NA | NR_039579 |
| Mir1934 | 0.659372557 | 0.083679911 | 0.06789324 | 1.23252199 | 0.217754157 | NA | | Mir5121 | 0 | NA | NA | NA | NA | NA | NR_039581 |
| Mir1936 | 0 | NA | NA | NA | NA | NA | | Mir5122 | 0 | NA | NA | NA | NA | NA | NR_039582 |
| Mir1938 | 0.392396713 | -0.003471664 | 0.047500251 | -0.073087276 | 0.941736667 | NA | | Mir5123 | 0 | NA | NA | NA | NA | NA | NR_039584 |
| Mir193b | 0.296502237 | 0.007885408 | 0.033590428 | 0.234751624 | 0.814401517 | NA | | Mir5126 | 0 | NA | NA | NA | NA | NA | NR_039588 |
| Mir194-1 | 0 | NA | NA | NA | NA | NA | | Mir5127 | 0 | NA | NA | NA | NA | NA | NR_039589 |
| Mir194-2 | 0 | NA | NA | NA | NA | NA | | Mir5128 | 0.120209049 | -0.00674972139751468 | 0.00863521114606269 | -0.781650996523957 | 0.434419707908266 | NA | NR_039590 |
| Mir1940 | 0 | NA | NA | NA | NA | NA | | Mir5129 | 0 | NA | NA | NA | NA | NA | NR_039591 |
| Mir1941 | 0 | NA | NA | NA | NA | NA | | Mir5130 | 0 | NA | NA | NA | NA | NA | NR_039592 |
| Mir1942 | 0 | NA | NA | NA | NA | NA | | Mir5131 | 0 | NA | NA | NA | NA | NA | NR_039593 |
| Mir1943 | 5.355910632 | 0.236561267 | 0.11430196 | 2.069616889 | 0.038488235 | NA | | Mir5132 | 0 | NA | NA | NA | NA | NA | NR_039594 |
| Mir1945 | 0 | NA | NA | NA | NA | NA | | Mir5133 | 9.959030639 | -0.0761158970104896 | 0.0532055421617723 | -1.43060090956423 | 0.152544626649066 | NA | NR_039595 |
| Mir1946a | 0 | NA | NA | NA | NA | NA | | Mir5134 | 0 | NA | NA | NA | NA | NA | NR_039596 |
| Mir1947 | 0.19046386 | -0.000774927 | 0.031526133 | -0.024580468 | 0.980389599 | NA | | Mir5135 | 0 | NA | NA | NA | NA | NA | NR_039597 |
| Mir1948 | 0.094392817 | 0.017895942 | 0.027818137 | 0.643319236 | 0.520016977 | NA | | Mir5136 | 0 | NA | NA | NA | NA | NA | NR_039598 |
| Mir1949 | 3.481742175 | 0.174092853 | 0.131864775 | 1.32023774 | 0.186755655 | NA | | Mir532 | 0 | NA | NA | NA | NA | NA | NR_030242 |
| Mir195 | 0.079983654 | 0.016740032 | 0.027674178 | 0.604897175 | 0.54524732 | NA | | Mir539 | 0 | NA | NA | NA | NA | NA | NR_030262 |
| Mir1951 | 0 | NA | NA | NA | NA | NA | | Mir540 | 0.891101612 | -0.0243941801002644 | 0.0182958067490843 | -1.33332082235102 | 0.18242654334025 | NA | NR_030260 |
| Mir1952 | 0 | NA | NA | NA | NA | NA | | Mir541 | 12.86242223 | -0.0131762570338687 | 0.0670781328488584 | -0.196431481829669 | 0.844272457282634 | NA | NR_030263 |
| Mir1953 | 0 | NA | NA | NA | NA | NA | | Mir542 | 0 | NA | NA | NA | NA | NA | NR_030264 |
| Mir1955 | 0.098353386 | -0.020212831 | 0.02786466 | -0.725393061 | 0.468210929 | NA | | Mir543 | 0 | NA | NA | NA | NA | NA | NR_030261 |
| Mir1956 | 0 | NA | NA | NA | NA | NA | | Mir546 | 0 | NA | NA | NA | NA | NA | NR_030259 |
| Mir1960 | 0 | NA | NA | NA | NA | NA | | Mir547 | 0 | NA | NA | NA | NA | NA | NR_030265 |
| Mir1961 | 0 | NA | NA | NA | NA | NA | | Mir551b | 0 | NA | NA | NA | NA | NA | NR_030422 |
| Mir1962 | 0 | NA | NA | NA | NA | NA | | Mir568 | 18.79264089 | 0.0146397669000657 | 0.0758576547514423 | 0.192989975079441 | 0.846966819232282 | NA | NR_030576 |
| Mir1963 | 0 | NA | NA | NA | NA | NA | | Mir574 | 0 | NA | NA | NA | NA | NA | NR_030577 |
| Mir1964 | 1.098050894 | -0.021040418 | 0.072437663 | -0.290462406 | 0.771462507 | NA | | Mir582 | 0 | NA | NA | NA | NA | NA | NR_030644 |
| Mir1965 | 0 | NA | NA | NA | NA | NA | | Mir592 | 0.209568245 | 0.00686197416234131 | 0.00977127153168884 | 0.702260104029194 | 0.482516968822431 | NA | NR_030420 |
| Mir1966 | 0 | NA | NA | NA | NA | NA | | Mir598 | 0 | NA | NA | NA | NA | NA | NR_030611 |
| Mir1967 | 0.185670226 | -0.001645086 | 0.031432796 | -0.052336624 | 0.958260472 | NA | | Mir599 | 0 | NA | NA | NA | NA | NA | NR_035527 |
| Mir1968 | 2.217277567 | 0.039560172 | 0.105632057 | 0.374509154 | 0.708025547 | NA | | Mir615 | 0 | NA | NA | NA | NA | NA | NR_030526 |
| Mir1969 | 1.549136228 | 0.168408375 | 0.091425718 | 1.842024088 | 0.065471627 | NA | | Mir653 | 0 | NA | NA | NA | NA | NA | NR_030612 |
| Mir196a-1 | 0 | NA | NA | NA | NA | NA | | Mir654 | 0.097165704 | -0.00637828455464458 | 0.00848781691535724 | -0.75146349388194 | 0.452373763047372 | NA | NR_030578 |
| Mir196a-2 | 0 | NA | NA | NA | NA | NA | | Mir664 | 0 | NA | NA | NA | NA | NA | NR_035529 |
| Mir196b | 0 | NA | NA | NA | NA | NA | | Mir665 | 0.598294728 | -0.0149228688602199 | 0.0157322564925091 | -0.948552349583507 | 0.342848335797541 | NA | NR_030425 |
| Mir1971 | 0 | NA | NA | NA | NA | NA | | Mir666 | 0 | NA | NA | NA | NA | NA | NR_030435 |
| Mir1981 | 0 | NA | NA | NA | NA | NA | | Mir667 | 0.113843286 | 0.00525947738444006 | 0.00859368002043176 | 0.612016897526494 | 0.540526577873086 | NA | NR_030426 |
| Mir1982 | 0 | NA | NA | NA | NA | NA | | Mir668 | 5.995332248 | -0.0516767821390411 | 0.0515001834213965 | -1.00342908910051 | 0.315653874758717 | NA | NR_030424 |
| Mir1983 | 0.186326724 | 0.024939385 | 0.031411053 | 0.793968451 | 0.427213798 | NA | | Mir669a-1 | 0 | NA | NA | NA | NA | NA | NR_035408 |
| Mir199a-1 | 0.198019769 | 0.025308392 | 0.031628095 | 0.80018706 | 0.423602426 | NA | | Mir669a-10 | 0 | NA | NA | NA | NA | NA | NR_037264 |
| Mir199a-2 | 0 | NA | NA | NA | NA | NA | | Mir669a-11 | 0 | NA | NA | NA | NA | NA | NR_037266 |
| Mir199b | 0 | NA | NA | NA | NA | NA | | Mir669a-12 | 0 | NA | NA | NA | NA | NA | NR_037268 |
| Mir19a | 0 | NA | NA | NA | NA | NA | | Mir669a-2 | 0.217532603 | 0.00684685248201087 | 0.00976000296730706 | 0.701521557416086 | 0.482977586094298 | NA | NR_030470 |
| Mir19b-1 | 0 | NA | NA | NA | NA | NA | | Mir669a-3 | 0 | NA | NA | NA | NA | NA | NR_030471 |

| | | | | | | |
|---|---|---|---|---|---|---|
| Mir19b-2 | 0 | NA | NA | NA | NA | NA |
| Mir1a-1 | 0 | NA | NA | NA | NA | NA |
| Mir1a-2 | 0 | NA | NA | NA | NA | NA |
| Mir1b | 0.096071043 | -0.020098795 | 0.027825948 | -0.722304036 | 0.470107572 | NA |
| Mir200a | 0 | NA | NA | NA | NA | NA |
| Mir200b | 0 | NA | NA | NA | NA | NA |
| Mir200c | 0 | NA | NA | NA | NA | NA |
| Mir201 | 0 | NA | NA | NA | NA | NA |
| Mir202 | 0 | NA | NA | NA | NA | NA |
| Mir203 | 0 | NA | NA | NA | NA | NA |
| Mir204 | 0 | NA | NA | NA | NA | NA |
| Mir205 | 0 | NA | NA | NA | NA | NA |
| Mir206 | 0 | NA | NA | NA | NA | NA |
| Mir207 | 0 | NA | NA | NA | NA | NA |
| Mir208a | 0 | NA | NA | NA | NA | NA |
| Mir208b | 0 | NA | NA | NA | NA | NA |
| Mir20a | 0 | NA | NA | NA | NA | NA |
| Mir20b | 0 | NA | NA | NA | NA | NA |
| Mir21 | 0 | NA | NA | NA | NA | NA |
| Mir210 | 0 | NA | NA | NA | NA | NA |
| Mir211 | 0 | NA | NA | NA | NA | NA |
| Mir2136 | 0 | NA | NA | NA | NA | NA |
| Mir2137 | 0 | NA | NA | NA | NA | NA |
| Mir214 | 0 | NA | NA | NA | NA | NA |
| Mir215 | 0 | NA | NA | NA | NA | NA |
| Mir216a | 0 | NA | NA | NA | NA | NA |
| Mir216b | 0 | NA | NA | NA | NA | NA |
| Mir217 | 0 | NA | NA | NA | NA | NA |
| Mir218-1 | 1.62510428 | 0.082863464 | 0.092234695 | 0.898397979 | 0.368973413 | NA |
| Mir218-2 | 0.200315975 | 0.025370485 | 0.031664208 | 0.801235428 | 0.422995365 | NA |
| Mir219-1 | 0 | NA | NA | NA | NA | NA |
| Mir219-2 | 1.863088764 | 0.058447026 | 0.102840709 | 0.568325775 | 0.56981378 | NA |
| Mir22 | 0 | NA | NA | NA | NA | NA |
| Mir221 | 1.732378575 | 0.016218783 | 0.08655574 | 0.187379638 | 0.851362987 | NA |
| Mir222 | 0.331747315 | -0.03734372 | 0.03744515 | -0.99729124 | 0.318623164 | NA |
| Mir223 | 0 | NA | NA | NA | NA | NA |
| Mir224 | 0 | NA | NA | NA | NA | NA |
| Mir23a | 0.389426822 | -0.00542245 | 0.0474348 | -0.114313744 | 0.908989088 | NA |
| Mir24-2 | 0.898641866 | 0.005814992 | 0.070393532 | 0.082606915 | 0.934164103 | NA |
| Mir25 | 3.069236884 | 0.051379836 | 0.102669605 | 0.500438625 | 0.616766262 | NA |
| Mir26a-1 | 1.967147038 | 0.165729845 | 0.101439177 | 1.633785391 | 0.102303921 | NA |
| Mir26a-2 | 0.176054697 | -0.002190495 | 0.031239146 | -0.070120197 | 0.944097991 | NA |
| Mir26b | 0.116696964 | -0.020477583 | 0.027954224 | -0.732539847 | 0.463839131 | NA |
| Mir27a | 1.218550218 | 0.080075385 | 0.079375039 | 1.008823258 | 0.313059405 | NA |
| Mir2861 | 0.927827864 | 0.009771901 | 0.056235838 | 0.173766435 | 0.862049024 | NA |
| Mir28b | 0 | NA | NA | NA | NA | NA |
| Mir290 | 0 | NA | NA | NA | NA | NA |
| Mir291a | 0 | NA | NA | NA | NA | NA |
| Mir291b | 0 | NA | NA | NA | NA | NA |
| Mir292 | 0 | NA | NA | NA | NA | NA |
| Mir293 | 0 | NA | NA | NA | NA | NA |
| Mir294 | 0 | NA | NA | NA | NA | NA |
| Mir295 | 0 | NA | NA | NA | NA | NA |
| Mir296 | 0.238795281 | -0.027601606 | 0.032267774 | -0.855392297 | 0.392334007 | NA |
| Mir297-1 | 0 | NA | NA | NA | NA | NA |
| Mir297a-3 | 0 | NA | NA | NA | NA | NA |
| Mir297b | 0 | NA | NA | NA | NA | NA |
| Mir297c | 0 | NA | NA | NA | NA | NA |
| Mir298 | 0 | NA | NA | NA | NA | NA |
| Mir299 | 0 | NA | NA | NA | NA | NA |
| Mir29a | 0 | NA | NA | NA | NA | NA |
| Mir29b-1 | 0 | NA | NA | NA | NA | NA |
| Mir29b-2 | 0 | NA | NA | NA | NA | NA |
| Mir29c | 0 | NA | NA | NA | NA | NA |
| Mir300 | 0.201994201 | 0.000227241 | 0.031727614 | 0.00716224 | 0.994285408 | NA |
| Mir301 | 0.096071043 | -0.020098795 | 0.027825948 | -0.722304036 | 0.470107572 | NA |
| Mir301b | 0 | NA | NA | NA | NA | NA |
| Mir302a | 0 | NA | NA | NA | NA | NA |
| Mir302b | 0 | NA | NA | NA | NA | NA |
| Mir302c | 0 | NA | NA | NA | NA | NA |
| Mir302d | 0 | NA | NA | NA | NA | NA |
| Mir3057 | 0.791376948 | 0.082635319 | 0.066510581 | 1.24243869 | 0.214074748 | NA |
| Mir3058 | 0.676440077 | 0.007297889 | 0.060582559 | 0.120461886 | 0.90411727 | NA |
| Mir3059 | 0 | NA | NA | NA | NA | NA |
| Mir3062 | 0 | NA | NA | NA | NA | NA |
| Mir3063 | 0 | NA | NA | NA | NA | NA |
| Mir3065 | 0 | NA | NA | NA | NA | NA |
| Mir3066 | 0 | NA | NA | NA | NA | NA |
| Mir3067 | 0 | NA | NA | NA | NA | NA |
| Mir3068 | 3.272901237 | 0.09265432 | 0.107422202 | 0.862524867 | 0.388398755 | NA |
| Mir3069 | 0 | NA | NA | NA | NA | NA |
| Mir3070a | 4.880662248 | 0.266296126 | 0.130175445 | 2.045670945 | 0.040788756 | NA |
| Mir3070b | 6.499130046 | 0.18487701 | 0.149589309 | 1.235897218 | 0.216496771 | NA |
| Mir3071 | 0 | NA | NA | NA | NA | NA |
| Mir3073 | 0 | NA | NA | NA | NA | NA |
| Mir3074-1 | 0.17833704 | -0.002390466 | 0.031282281 | -0.076415987 | 0.939088151 | NA |
| Mir3074-2 | 0.217751027 | -0.027071875 | 0.031947577 | -0.847384294 | 0.396780956 | NA |
| Mir3075 | 0 | NA | NA | NA | NA | NA |
| Mir3076 | 0.096071043 | -0.020098795 | 0.027825948 | -0.722304036 | 0.470107572 | NA |
| Mir3077 | 0 | NA | NA | NA | NA | NA |
| Mir3078 | 0 | NA | NA | NA | NA | NA |
| Mir3081 | 0 | NA | NA | NA | NA | NA |
| Mir3083 | 0 | NA | NA | NA | NA | NA |
| Mir3085 | 0 | NA | NA | NA | NA | NA |
| Mir3086 | 0 | NA | NA | NA | NA | NA |
| Mir3087 | 0 | NA | NA | NA | NA | NA |
| Mir3088 | 0 | NA | NA | NA | NA | NA |
| Mir3089 | 0 | NA | NA | NA | NA | NA |
| Mir3091 | 2.100454759 | -0.029999523 | 0.092428139 | -0.324571322 | 0.745505535 | NA |
| Mir3092 | 0 | NA | NA | NA | NA | NA |
| Mir3093 | 2.001486636 | 0.026368064 | 0.089586297 | 0.294331446 | 0.768504646 | NA |
| Mir3094 | 0.316104413 | -0.036204367 | 0.036880698 | -0.981661674 | 0.326266553 | NA |
| Mir3095 | 0 | NA | NA | NA | NA | NA |
| Mir3096 | 0 | NA | NA | NA | NA | NA |
| Mir3096b | 0 | NA | NA | NA | NA | NA |
| Mir3097 | 0.706253102 | 0.062939234 | 0.058126208 | 1.082803018 | 0.278895866 | NA |
| Mir3099 | 0 | NA | NA | NA | NA | NA |
| Mir30a | 0 | NA | NA | NA | NA | NA |
| Mir30b | 0.383703858 | 0.028442486 | 0.048086527 | 0.591485549 | 0.554195135 | NA |
| Mir30c-1 | 0 | NA | NA | NA | NA | NA |
| Mir30c-2 | 0.079983654 | 0.016740032 | 0.027674178 | 0.604897175 | 0.54524732 | NA |
| Mir30d | 0.327267802 | 0.035667551 | 0.037337881 | 0.955264473 | 0.339443969 | NA |
| Mir30e | 0 | NA | NA | NA | NA | NA |
| Mir31 | 0.309826022 | -0.01193339 | 0.034225532 | -0.348669245 | 0.727337636 | NA |
| Mir3100 | 0.70093952 | 0.072751279 | 0.064177266 | 1.133598915 | 0.256962828 | NA |
| Mir3102 | 0 | NA | NA | NA | NA | NA |
| Mir3103 | 0 | NA | NA | NA | NA | NA |
| Mir3104 | 0.588203125 | 0.0520691 | 0.055442676 | 0.939151999 | 0.34765271 | NA |
| Mir3106 | 0 | NA | NA | NA | NA | NA |
| Mir3107 | 0 | NA | NA | NA | NA | NA |
| Mir3108 | 0 | NA | NA | NA | NA | NA |
| Mir3109 | 0 | NA | NA | NA | NA | NA |
| Mir3110 | 0 | NA | NA | NA | NA | NA |
| Mir3112 | 4.820484465 | 0.195429931 | 0.137986327 | 1.416299248 | 0.156687908 | NA |
| Mir32 | 0.107716603 | -0.020477583 | 0.027954224 | -0.732539849 | 0.46383913 | NA |
| Mir320 | 1.168957922 | 0.046872974 | 0.075889022 | 0.617651578 | 0.536805037 | NA |
| Mir322 | 0 | NA | NA | NA | NA | NA |
| Mir323 | 0.198019769 | 0.025308392 | 0.031628095 | 0.80018706 | 0.423602426 | NA |
| Mir324 | 0.671194683 | 0.056551227 | 0.056497169 | 1.000956827 | 0.316847681 | NA |
| Mir326 | 0 | NA | NA | NA | NA | NA |
| Mir328 | 0.389457872 | -0.034711543 | 0.048622122 | -0.713904319 | 0.475286347 | NA |
| Mir329 | 0 | NA | NA | NA | NA | NA |
| Mir33 | 0 | NA | NA | NA | NA | NA |
| Mir330 | 0.203787646 | -0.02671144 | 0.031728022 | -0.841887971 | 0.399850664 | NA |
| Mir335 | 0.681939444 | 0.039928515 | 0.061568448 | 0.648522352 | 0.516647159 | NA |
| Mir337 | 1.634991426 | 0.155901605 | 0.086358302 | 1.805287994 | 0.071029627 | NA |
| Mir338 | 3.325487554 | 0.251093364 | 0.116348254 | 2.158118885 | 0.03091859 | NA |
| Mir339 | 0.422001417 | -0.006176072 | 0.048208622 | -0.128111359 | 0.898060847 | NA |
| Mir340 | 0 | NA | NA | NA | NA | NA |
| Mir344 | 0.192142086 | -0.026384592 | 0.031527717 | -0.836869715 | 0.402665805 | NA |
| Mir344-2 | 6.856383389 | 0.192744819 | 0.143224784 | 1.34575046 | 0.178383007 | NA |
| Mir344b | 0.811177505 | -0.011323829 | 0.062251274 | -0.181905174 | 0.855657144 | NA |
| Mir344c | 0 | NA | NA | NA | NA | NA |
| Mir344d-1 | 0.266310378 | 0.0407635 | 0.040043874 | 1.017970943 | 0.308691765 | NA |
| Mir344d-2 | 0.105923158 | 0.018524693 | 0.027977518 | 0.662127842 | 0.507889294 | NA |
| Mir344d-3 | 0.272987066 | 0.007834485 | 0.033279985 | 0.2354113 | 0.813889516 | NA |
| Mir344e | 0.08731684 | 0.017365056 | 0.027733412 | 0.626142072 | 0.531221759 | NA |
| Mir344f | 0.20671448 | -0.003314027 | 0.03178208 | -0.10427343 | 0.916952363 | NA |
| Mir344g | 1.23487479 | -0.007358988 | 0.080815105 | -0.091059565 | 0.927445261 | NA |
| Mir345 | 2.224772481 | 0.004362455 | 0.102982554 | 0.042361109 | 0.966210831 | NA |
| Mir346 | 0 | NA | NA | NA | NA | NA |
| Mir3471-1 | 0 | NA | NA | NA | NA | NA |
| Mir3473 | 0 | NA | NA | NA | NA | NA |
| Mir3473c | 0.193239998 | 0.025145508 | 0.031532422 | 0.797449311 | 0.42519013 | NA |
| Mir3473d | 0.988148424 | -0.001676897 | 0.057931531 | -0.028946194 | 0.976907504 | NA |
| Mir3474 | 0 | NA | NA | NA | NA | NA |
| Mir3475 | 0 | NA | NA | NA | NA | NA |
| Mir34a | 0.20671448 | -0.003314027 | 0.03178208 | -0.10427343 | 0.916952363 | NA |
| Mir34b | 0 | NA | NA | NA | NA | NA |
| Mir34c | 0 | NA | NA | NA | NA | NA |
| Mir350 | 0 | NA | NA | NA | NA | NA |
| Mir351 | 0 | NA | NA | NA | NA | NA |
| Mir3572 | 0 | NA | NA | NA | NA | NA |
| Mir362 | 0 | NA | NA | NA | NA | NA |
| Mir363 | 0.08731684 | 0.017365056 | 0.027733412 | 0.626142072 | 0.531221759 | NA |
| Mir365-1 | 0.984004379 | -0.031039372 | 0.065916481 | -0.4708894 | 0.637719718 | NA |
| Mir365-2 | 0 | NA | NA | NA | NA | NA |
| Mir367 | 0 | NA | NA | NA | NA | NA |
| Mir369 | 3.201080901 | 0.165684723 | 0.118335717 | 1.400124379 | 0.161476076 | NA |
| Mir370 | 2.605066898 | -0.025067966 | 0.111088766 | -0.225657072 | 0.821468148 | NA |
| Mir374c | 0 | NA | NA | NA | NA | NA |
| Mir375 | 0 | NA | NA | NA | NA | NA |
| Mir376a | 0 | NA | NA | NA | NA | NA |
| Mir376b | 0 | NA | NA | NA | NA | NA |
| Mir376c | 0.119397641 | -0.020477583 | 0.027954224 | -0.732539845 | 0.463839132 | NA |
| Mir378b | 0 | NA | NA | NA | NA | NA |
| Mir379 | 0 | NA | NA | NA | NA | NA |
| Mir380 | 0 | NA | NA | NA | NA | NA |
| Mir381 | 0.096071043 | -0.020098795 | 0.027825948 | -0.722304036 | 0.470107572 | NA |
| Mir382 | 7.259339403 | 0.137739379 | 0.146864577 | 0.937866586 | 0.348312977 | NA |
| Mir383 | 0 | NA | NA | NA | NA | NA |
| Mir384 | 0.794367966 | -0.094767332 | 0.066368305 | -1.427900438 | 0.153320518 | NA |
| Mir3960 | 1.285712233 | 0.048271091 | 0.053775144 | 0.89764688 | 0.369373837 | NA |
| Mir3966 | 0 | NA | NA | NA | NA | NA |
| Mir3967 | 0 | NA | NA | NA | NA | NA |
| Mir3968 | 0 | NA | NA | NA | NA | NA |
| Mir3969 | 0 | NA | NA | NA | NA | NA |
| Mir3971 | 0.096071043 | -0.020098795 | 0.027825948 | -0.722304036 | 0.470107572 | NA |
| Mir409 | 7.371817728 | 0.159969339 | 0.123064958 | 1.299877251 | 0.193643043 | NA |
| Mir410 | 0 | NA | NA | NA | NA | NA |
| Mir411 | 0 | NA | NA | NA | NA | NA |
| Mir412 | 0.094392817 | 0.017895942 | 0.027818137 | 0.643319236 | 0.520016977 | NA |
| Mir423 | 0.681390527 | -0.014222642 | 0.054510777 | -0.26091431 | 0.794158589 | NA |
| Mir429 | 0 | NA | NA | NA | NA | NA |
| Mir431 | 3.907237645 | 0.135417483 | 0.122655561 | 1.10404683 | 0.269572824 | NA |
| Mir432 | 1.196030218 | 0.044886332 | 0.075085309 | 0.597804446 | 0.549970424 | NA |
| Mir433 | 0 | NA | NA | NA | NA | NA |
| Mir434 | 3.581388194 | 0.238684672 | 0.128963735 | 1.850789073 | 0.064199904 | NA |
| Mir448 | 0 | NA | NA | NA | NA | NA |
| Mir449a | 0 | NA | NA | NA | NA | NA |
| Mir449b | 0 | NA | NA | NA | NA | NA |
| Mir449c | 0 | NA | NA | NA | NA | NA |
| Mir450-1 | 0 | NA | NA | NA | NA | NA |
| Mir450-2 | 0 | NA | NA | NA | NA | NA |
| Mir450b | 0 | NA | NA | NA | NA | NA |
| Mir451 | 0 | NA | NA | NA | NA | NA |
| Mir452 | 0 | NA | NA | NA | NA | NA |
| Mir455 | 0 | NA | NA | NA | NA | NA |
| Mir463 | 0 | NA | NA | NA | NA | NA |
| Mir465 | 0 | NA | NA | NA | NA | NA |
| Mir465b-1 | 0 | NA | NA | NA | NA | NA |
| Mir465b-2 | 0 | NA | NA | NA | NA | NA |
| Mir465c-1 | 0 | NA | NA | NA | NA | NA |
| Mir465c-2 | 0 | NA | NA | NA | NA | NA |
| Mir466 | 0 | NA | NA | NA | NA | NA |
| Mir4660 | 0 | NA | NA | NA | NA | NA |
| Mir466g | 0 | NA | NA | NA | NA | NA |
| Mir466i | 0 | NA | NA | NA | NA | NA |
| Mir467a-1 | 0 | NA | NA | NA | NA | NA |
| Mir467a-10 | 0 | NA | NA | NA | NA | NA |
| Mir467a-2 | 0 | NA | NA | NA | NA | NA |
| Mir467a-3 | 0 | NA | NA | NA | NA | NA |
| Mir467a-4 | 0 | NA | NA | NA | NA | NA |
| Mir467a-5 | 0 | NA | NA | NA | NA | NA |
| Mir467a-6 | 0 | NA | NA | NA | NA | NA |
| Mir467a-7 | 0 | NA | NA | NA | NA | NA |
| Mir467a-8 | 0 | NA | NA | NA | NA | NA |
| Mir467a-9 | 0 | NA | NA | NA | NA | NA |
| Mir467b | 0 | NA | NA | NA | NA | NA |
| Mir467d | 0 | NA | NA | NA | NA | NA |
| Mir467f | 0 | NA | NA | NA | NA | NA |
| Mir468 | 0.62501449 | 0.025957127 | 0.053549922 | 0.48472764 | 0.627869557 | NA |
| Mir470 | 0 | NA | NA | NA | NA | NA |
| Mir471 | 0 | NA | NA | NA | NA | NA |
| Mir483 | 0 | NA | NA | NA | NA | NA |

| | | | | | | | |
|---|---|---|---|---|---|---|---|
| Mir669a-4 | 0 | NA | NA | NA | NA | NA | NR_037250 |
| Mir669a-5 | 0 | NA | NA | NA | NA | NA | NR_037251 |
| Mir669a-6 | 0 | NA | NA | NA | NA | NA | NR_037253 |
| Mir669a-7 | 0 | NA | NA | NA | NA | NA | NR_037255 |
| Mir669a-8 | 0 | NA | NA | NA | NA | NA | NR_037259 |
| Mir669a-9 | 0 | NA | NA | NA | NA | NA | NR_037260 |
| Mir669b | 0 | NA | NA | NA | NA | NA | NR_030469 |
| Mir669c | 0 | NA | NA | NA | NA | NA | NR_030473 |
| Mir669e | 0 | NA | NA | NA | NA | NA | NR_035426 |
| Mir669g | 0 | NA | NA | NA | NA | NA | NR_035411 |
| Mir669h | 0 | NA | NA | NA | NA | NA | NR_035418 |
| Mir669i | 0 | NA | NA | NA | NA | NA | NR_035417 |
| Mir669j | 0 | NA | NA | NA | NA | NA | NR_035416 |
| Mir669k | 0 | NA | NA | NA | NA | NA | NR_035410 |
| Mir669m-1 | 0 | NA | NA | NA | NA | NA | NR_035474 |
| Mir669m-2 | 0 | NA | NA | NA | NA | NA | NR_035475 |
| Mir669p-1 | 0 | NA | NA | NA | NA | NA | NR_037257 |
| Mir669p-2 | 0 | NA | NA | NA | NA | NA | NR_037262 |
| Mir670 | 1.166737156 | -0.0151457629878309 | 0.022349249079344 | -0.677685542546 | 0.49797109504299 | NA | NR_030431 |
| Mir671 | 0 | NA | NA | NA | NA | NA | NR_030423 |
| Mir672 | 0 | NA | NA | NA | NA | NA | NR_030430 |
| Mir673 | 0 | NA | NA | NA | NA | NA | NR_030438 |
| Mir674 | 3.033230905 | -0.0236230985658337 | 0.0420397037215797 | -0.561923526442636 | 0.574168123705738 | NA | NR_030440 |
| Mir675 | 0 | NA | NA | NA | NA | NA | NR_030416 |
| Mir676 | 0 | NA | NA | NA | NA | NA | NR_030525 |
| Mir677 | 10.36118702 | -0.0447353691272842 | 0.065022960944045 | -0.687993417675656 | 0.491456926467778 | NA | NR_030442 |
| Mir678 | 0 | NA | NA | NA | NA | NA | NR_030443 |
| Mir679 | 0 | NA | NA | NA | NA | NA | NR_030445 |
| Mir680-1 | 0 | NA | NA | NA | NA | NA | NR_030447 |
| Mir680-2 | 0 | NA | NA | NA | NA | NA | NR_030448 |
| Mir680-3 | 0 | NA | NA | NA | NA | NA | NR_030449 |
| Mir681 | 0 | NA | NA | NA | NA | NA | NR_030450 |
| Mir683-1 | 0 | NA | NA | NA | NA | NA | NR_030453 |
| Mir683-2 | 0 | NA | NA | NA | NA | NA | NR_030647 |
| Mir684-1 | 0.432488341 | 0.00620036638839739 | 0.0139681256348908 | 0.443893944718651 | 0.657119262665896 | NA | NR_030454 |
| Mir684-2 | 0 | NA | NA | NA | NA | NA | NR_030455 |
| Mir686 | 0 | NA | NA | NA | NA | NA | NR_030457 |
| Mir687 | 0 | NA | NA | NA | NA | NA | NR_030459 |
| Mir688 | 1.225983948 | -0.0237877750383443 | 0.0270772768171749 | -0.878514305517455 | 0.379664677833096 | NA | NR_030460 |
| Mir690 | 0 | NA | NA | NA | NA | NA | NR_030463 |
| Mir691 | 0 | NA | NA | NA | NA | NA | NR_030464 |
| Mir692-1 | 0 | NA | NA | NA | NA | NA | NR_030465 |
| Mir693 | 0.360627147 | -0.0100772056525624 | 0.010553896215922 | -0.954832741045871 | 0.339662287230353 | NA | NR_030466 |
| Mir694 | 0 | NA | NA | NA | NA | NA | NR_030467 |
| Mir695 | 0 | NA | NA | NA | NA | NA | NR_030475 |
| Mir697 | 7.55552617 | -0.0368908540220187 | 0.0520399879492035 | -0.708894361352046 | 0.478390034489515 | NA | NR_030479 |
| Mir698 | 0 | NA | NA | NA | NA | NA | NR_030480 |
| Mir7-1 | 6.711561705 | -0.111510616401086 | 0.0547920246733388 | -2.03516145033688 | 0.0418346305098005 | NA | NR_029825 |
| Mir7-2 | 0 | NA | NA | NA | NA | NA | NR_029826 |
| Mir700 | 0.570173846 | 0.000854631162947742 | 0.0165809204867965 | 0.0515430469392994 | 0.958892800920339 | NA | NR_030481 |
| Mir701 | 0 | NA | NA | NA | NA | NA | NR_030482 |
| Mir702 | 0 | NA | NA | NA | NA | NA | NR_030483 |
| Mir703 | 11.36090892 | 0.0645823862618396 | 0.0580324517028793 | 1.11286675587127 | 0.265765659664068 | NA | NR_030484 |
| Mir704 | 0.255375383 | 0.00708304372802858 | 0.00992471678592415 | 0.713677163873753 | 0.475426830432214 | NA | NR_030485 |
| Mir705 | 0 | NA | NA | NA | NA | NA | NR_030486 |
| Mir706 | 0 | NA | NA | NA | NA | NA | NR_030487 |
| Mir707 | 10.40869857 | -0.0264734129000659 | 0.0581982190418058 | -0.454883557193547 | 0.649193012849281 | NA | NR_030488 |
| Mir708 | 0.108766301 | 0.00519171381361946 | 0.00856003371502575 | 0.606506234257728 | 0.544178643801544 | NA | NR_030489 |
| Mir709 | 3.225708945 | -0.0439699229280778 | 0.0426102545812057 | -1.03190941617776 | 0.302114554435122 | NA | NR_030490 |
| Mir710 | 0 | NA | NA | NA | NA | NA | NR_030491 |
| Mir711 | 0.097165704 | -0.00637828455464458 | 0.00848781691535724 | -0.75146349388194 | 0.452373763047372 | NA | NR_030492 |
| Mir713 | 0 | NA | NA | NA | NA | NA | NR_030493 |
| Mir717 | 0 | NA | NA | NA | NA | NA | NR_030497 |
| Mir718 | 0 | NA | NA | NA | NA | NA | NR_030758 |
| Mir719 | 0 | NA | NA | NA | NA | NA | NR_030458 |
| Mir721 | 0 | NA | NA | NA | NA | NA | NR_030500 |
| Mir741 | 0 | NA | NA | NA | NA | NA | NR_030530 |
| Mir742 | 0 | NA | NA | NA | NA | NA | NR_030531 |
| Mir743 | 0 | NA | NA | NA | NA | NA | NR_030532 |
| Mir743b | 0 | NA | NA | NA | NA | NA | NR_030535 |
| Mir744 | 0.290056368 | -0.00425303587871565 | 0.0107652157825088 | -0.395072051005791 | 0.692789714682806 | NA | NR_030417 |
| Mir758 | 0 | NA | NA | NA | NA | NA | NR_030421 |
| Mir759 | 0 | NA | NA | NA | NA | NA | NR_030436 |
| Mir760 | 3.651487852 | -0.00328704582403397 | 0.036317831537871 | -0.0905077667042526 | 0.927883722348503 | NA | NR_030439 |
| Mir761 | 7.789700109 | -0.0370940614055055 | 0.057390794087032 | -0.646341664993397 | 0.518058110346253 | NA | NR_030432 |
| Mir762 | 0.310639869 | 0.0122746672672506 | 0.0130505022290595 | 0.940551332953199 | 0.346934831808109 | NA | NR_030428 |
| Mir763 | 0 | NA | NA | NA | NA | NA | NR_030434 |
| Mir764 | 0 | NA | NA | NA | NA | NA | NR_030433 |
| Mir767 | 0 | NA | NA | NA | NA | NA | NR_035528 |
| Mir770 | 0 | NA | NA | NA | NA | NA | NR_030427 |
| Mir7b | 0.120209049 | -0.00674972139751468 | 0.00863521114606269 | -0.781650996523957 | 0.434419707908266 | NA | NR_029827 |
| Mir802 | 0 | NA | NA | NA | NA | NA | NR_030429 |
| Mir804 | 0.461806006 | -0.021704566626653 | 0.0154366531524305 | -1.40604096058449 | 0.159711962093329 | NA | NR_030529 |
| Mir871 | 0 | NA | NA | NA | NA | NA | NR_030536 |
| Mir872 | 0.097165704 | -0.00637828455464458 | 0.00848781691535724 | -0.75146349388194 | 0.452373763047372 | NA | NR_030604 |
| Mir873 | 0.3436217 | 0.002597311771164 | 0.0128044321936811 | 0.202844744060245 | 0.839256384147266 | NA | NR_030605 |
| Mir874 | 0.762249383 | -0.00942113202710957 | 0.0213796959178191 | -0.440657905674768 | 0.659460675128982 | NA | NR_030544 |
| Mir875 | 0 | NA | NA | NA | NA | NA | NR_030606 |
| Mir876 | 0 | NA | NA | NA | NA | NA | NR_030545 |
| Mir877 | 0 | NA | NA | NA | NA | NA | NR_030608 |
| Mir878 | 0 | NA | NA | NA | NA | NA | NR_030603 |
| Mir879 | 0 | NA | NA | NA | NA | NA | NR_030537 |
| Mir880 | 0 | NA | NA | NA | NA | NA | NR_030538 |
| Mir881 | 0 | NA | NA | NA | NA | NA | NR_030539 |
| Mir882 | 0 | NA | NA | NA | NA | NA | NR_030540 |
| Mir883a | 0 | NA | NA | NA | NA | NA | NR_030541 |
| Mir883b | 0 | NA | NA | NA | NA | NA | NR_030542 |
| Mir9-1 | 3.818828355 | -0.0741986602835388 | 0.0398144142473951 | -1.86361300765321 | 0.0623760696458489 | NA | NR_029817 |
| Mir9-3 | 0.665410991 | -0.00985345899576219 | 0.0160794716351147 | -0.612797436343865 | 0.54001028621475 | NA | NR_029818 |
| Mir92-1 | 0 | NA | NA | NA | NA | NA | NR_029816 |
| Mir92-2 | 0 | NA | NA | NA | NA | NA | NR_029748 |
| Mir92b | 0 | NA | NA | NA | NA | NA | NR_030579 |
| Mir93 | 0.668067976 | -0.0230565794173973 | 0.0186392402847743 | -1.23699137224125 | 0.216090285712039 | NA | NR_029749 |
| Mir96 | 0 | NA | NA | NA | NA | NA | NR_029750 |
| Mir98 | 0 | NA | NA | NA | NA | NA | NR_029753 |
| Mir99a | 1.917235761 | -0.00588033782209659 | 0.0306298994955745 | -0.19198031723696 | 0.847757623211622 | NA | NR_029535 |
| Mir99b | 0.582654835 | -0.00609107891811043 | 0.015161808814571 | -0.401738274938325 | 0.687876652915776 | NA | NR_029536 |
| Mira | 1.511422223 | 0.00207193334260678 | 0.0295180518684099 | 0.0701920760842676 | 0.944040781139563 | NA | NR_045199 |
| Mirlet7a-1 | 1.214889506 | -0.0130091408329704 | 0.0266731129244 | -0.487724881225616 | 0.625744725493531 | NA | NR_029725 |
| Mirlet7a-2 | 0 | NA | NA | NA | NA | NA | NR_029726 |
| Mirlet7b | 15.86373004 | -0.0477790082016767 | 0.0597154272835235 | -0.80011163572231 | 0.423646120203595 | NA | NR_029727 |
| Mirlet7c-1 | 0.224853396 | -0.000161157395207779 | 0.00979162641959728 | -0.0164586952465051 | 0.986868454038988 | NA | NR_029728 |
| Mirlet7c-2 | 15.04408271 | -0.0776752093972307 | 0.0691898719944428 | -1.1226384318715 | 0.261591086128819 | NA | NR_029729 |
| Mirlet7d | 16.2469985 | -0.075852849184658 | 0.0641097365167977 | -1.18317206255844 | 0.236740962695633 | NA | NR_029656 |
| Mirlet7e | 1.760403072 | -0.0186783346110686 | 0.0292174274842641 | -0.63928744654636 | 0.522635953468987 | NA | NR_029730 |
| Mirlet7f-1 | 3.533763314 | -0.0526809018164898 | 0.0426015477075483 | -1.23659596074148 | 0.21623712024964 | NA | NR_029731 |
| Mirlet7f-2 | 0 | NA | NA | NA | NA | NA | NR_029732 |
| Mirlet7g | 0.191449919 | 0.00665508079272897 | 0.00962405648474729 | 0.691504752001018 | 0.489248395458281 | NA | NR_029526 |
| Mirlet7i | 1.373352305 | -0.0228632718224534 | 0.023317232945198 | -0.980531089438803 | 0.32682403105001 | NA | NR_029527 |
| Mis18bp1 | 8.049298573 | 0.0213473608172307 | 0.0599676539300083 | 0.355981256864683 | 0.721854601827891 | NA | NM_172578 |
| Mixl1 | 0 | NA | NA | NA | NA | NA | NM_013729 |
| Mkln1 | 2524.590958 | 0.0601465587463357 | 0.0777651162624542 | 0.773438806975398 | NA | NA | NM_013791 |
| Mlana | 0.407027086 | -0.0158737350575048 | 0.0132224722193719 | -1.20051188568569 | 0.229940599374159 | NA | NM_029993 |
| Mlkl | 10.4517768 | 0.0270829815494539 | 0.0638692555919777 | 0.424037845727697 | 0.671538212876518 | NA | NM_029005 |
| Mmp10 | 0.097165704 | -0.00637828455464458 | 0.00848781691535724 | -0.75146349388194 | 0.452373763047372 | NA | NM_019471 |
| Mmp12 | 13.41825555 | 0.0751917453317415 | 0.0400622116214386 | 1.87687454807173 | 0.0605352899958333 | NA | NM_008605 |
| Mmp13 | 2.536290848 | 0.0313258199941081 | 0.0323686783900555 | 0.967781866674303 | 0.333153321837747 | NA | NM_008607 |
| Mmp1a | 0 | NA | NA | NA | NA | NA | NM_032006 |
| Mmp1b | 0 | NA | NA | NA | NA | NA | NM_032007 |
| Mmp20 | 0 | NA | NA | NA | NA | NA | NM_013903 |
| Mmp21 | 3.995002897 | -0.0606506010829709 | 0.0440098323906685 | -1.37811479363486 | 0.168167848764045 | NA | NM_152944 |
| Mmp25 | 12.11157119 | -0.115029031240157 | 0.0679219113765171 | -1.6935482071832 | 0.0903511669992106 | NA | NM_001033339 |
| Mmp27 | 0 | NA | NA | NA | NA | NA | NM_001030289 |
| Mmp3 | 0 | NA | NA | NA | NA | NA | NM_010809 |
| Mmp7 | 0 | NA | NA | NA | NA | NA | NM_010810 |
| Mmp8 | 0.872738459 | -0.00868259111810392 | 0.0167723602428986 | -0.517672587063596 | 0.604686727609731 | NA | NM_008611 |
| Mmrn1 | 1.014005331 | -0.00776182986303447 | 0.0218936720394162 | -0.354523893893199 | 0.722946300603283 | NA | NM_001163507,NM_027613 |
| Mnd1 | 13.68509732 | -0.0155430916902024 | 0.0725412032816448 | -0.214265699865159 | 0.830339864423533 | NA | NM_029797 |
| Mnda | 5.302428134 | -0.0561172003105667 | 0.0414743029525361 | -1.35305951675157 | 0.176036617403178 | NA | NM_001033450 |
| Mnx1 | 0 | NA | NA | NA | NA | NA | NM_019944 |
| Moap1 | 3.697028797 | 0.0123386474573751 | 0.0389589772859836 | 0.316708710467464 | 0.751464638795065 | NA | NM_001142937,NM_022323 |
| Mogat1 | 5.053378446 | 0.0606387583736537 | 0.0449793762606361 | 1.3481458262622 | 0.177611484640461 | NA | NM_026713 |
| Mogat2 | 0.556118146 | 0.00194738650019528 | 0.017922158543124 | 0.108658033322801 | 0.913473729257516 | NA | NM_177448 |
| Morc1 | 1.430629264 | -0.0273439313124201 | 0.0263195498670678 | -1.03892093331862 | 0.298841509888305 | NA | NM_010816 |
| Morn3 | 5.489750998 | -0.00371155258456075 | 0.0489564746433325 | -0.0758133140018945 | 0.939567623569665 | NA | NM_029112 |
| Morn5 | 20.7103893 | -0.0889443284197085 | 0.0732523955987788 | -1.21421733299861 | 0.224664743339193 | NA | NM_029309 |
| Mos | 4.28031773 | 0.0109781060536233 | 0.0476335345131795 | 0.230470112407589 | 0.817726485260826 | NA | NM_020021 |
| Mospd4 | 1.004578682 | 0.0110561930748441 | 0.0246951197611183 | 0.447707611130995 | 0.65436422797541 | NA | NR_045438 |
| Mov10l1 | 0.342801831 | 0.00329783501100829 | 0.0128022060068578 | 0.257598964525467 | 0.796716428937095 | NA | NM_031260 |
| Moxd2 | 0 | NA | NA | NA | NA | NA | NM_139296 |
| Mpo | 4.007323397 | 0.0253085254333219 | 0.0361638462161881 | 0.699829472839453 | 0.484033806355271 | NA | NM_010824 |
| Mptx1 | 0 | NA | NA | NA | NA | NA | NM_025470 |
| Mptx2 | 0.127687691 | 0.00542298404407807 | 0.00869724871942359 | 0.62352868349814 | 0.532937159177571 | NA | NM_001205011 |
| Mpz | 5.789201278 | -0.0298248511558671 | 0.0495344070409343 | -0.602103728247326 | 0.547105092987313 | NA | NM_008623 |
| Mrap | 11.84095858 | 0.0998147188176803 | 0.0688546611723447 | 1.44964359882393 | 0.147157931337884 | NA | NM_029844 |
| Mrgpra1 | 0.127687691 | 0.00542298404407807 | 0.00869724871942359 | 0.62352868349814 | 0.532937159177571 | NA | NM_153095 |
| Mrgpra2a | 0 | NA | NA | NA | NA | NA | NM_001172588 |
| Mrgpra2b | 0.192890664 | -0.000903864895041124 | 0.00963528282151475 | -0.093807821917056 | 0.92526181809679 | NA | NM_153101 |
| Mrgpra3 | 0 | NA | NA | NA | NA | NA | NM_153067 |
| Mrgpra4 | 0 | NA | NA | NA | NA | NA | NM_153524 |
| Mrgpra6 | 0 | NA | NA | NA | NA | NA | NM_205821 |
| Mrgpra9 | 0 | NA | NA | NA | NA | NA | NM_001177566 |
| Mrgprb1 | 0.106148608 | 0.00515554063321497 | 0.00854200771766808 | 0.603551390213729 | 0.546141939141592 | NA | NM_205810 |
| Mrgprb2 | 0 | NA | NA | NA | NA | NA | NM_175531 |
| Mrgprb3 | 0 | NA | NA | NA | NA | NA | NM_207537 |
| Mrgprb4 | 0 | NA | NA | NA | NA | NA | NM_205795 |
| Mrgprb5 | 0 | NA | NA | NA | NA | NA | NM_207538 |
| Mrgprb8 | 0 | NA | NA | NA | NA | NA | NM_207539 |
| Mrgprd | 0 | NA | NA | NA | NA | NA | NM_203490 |
| Mrgprg | 0 | NA | NA | NA | NA | NA | NM_203492 |
| Mrgprh | 4.081931266 | -0.0418957311228213 | 0.0475566937651533 | -0.880963914979303 | 0.378337351360287 | NA | NM_030726 |
| Mrgprx1 | 0 | NA | NA | NA | NA | NA | NM_207540 |
| Mrgprx2 | 0 | NA | NA | NA | NA | NA | NM_001034868 |
| Mroh2a | 0.941360207 | -0.0160046082660133 | 0.0213076793226173 | -0.751119257225964 | 0.452580887255788 | NA | NM_001177364 |
| Mroh2b | 0.212695677 | -0.00857796107890413 | 0.00973699866180651 | -0.880965621629515 | 0.378336427605205 | NA | NM_001166066 |
| Mroh4 | 4.86843983 | -0.0431306528685344 | 0.0486508515326254 | -0.886534387576153 | 0.375329632372231 | NA | NM_001177437 |
| Mroh9 | 0 | NA | NA | NA | NA | NA | NM_030071 |
| Ms4a1 | 0.695642305 | -0.0137884799616017 | 0.0194502279957286 | -0.708910968273981 | 0.47837972817522 | NA | NM_007641 |
| Ms4a10 | 0 | NA | NA | NA | NA | NA | NM_023529 |
| Ms4a13 | 0 | NA | NA | NA | NA | NA | NM_198224 |
| Ms4a15 | 1.341935732 | -0.0123159658460062 | 0.0265676158666228 | -0.463570608210986 | 0.642955416314544 | NA | NM_001034898 |
| Ms4a18 | 0 | NA | NA | NA | NA | NA | NM_001251849 |
| Ms4a2 | 0.471719916 | -0.00295777152446865 | 0.0156034558093928 | -0.189558746511024 | 0.849654916517588 | NA | NM_001276330,NM_001276329,NM_001276328,NM_013516 |
| Ms4a3 | 0.203513543 | -0.00860782669689374 | 0.00975338121443284 | -0.882547960306941 | 0.377480556050336 | NA | NM_133246 |
| Ms4a4b | 0.737538137 | -0.000507005013514291 | 0.0179235492786713 | -0.0282870878770434 | 0.977433178861746 | NA | NM_021718 |
| Ms4a4c | 2.845534898 | -0.0103126307923706 | 0.0302380223311637 | -0.341048454803945 | 0.733067104393303 | NA | NM_029499 |
| Ms4a4d | 1.866797768 | 0.0244185894258404 | 0.027747486063259 | 0.880028892353372 | 0.378843657957223 | NA | NM_025658 |
| Ms4a5 | 0 | NA | NA | NA | NA | NA | NM_183190 |
| Ms4a7 | 13.34998502 | -0.0148965951284294 | 0.0634486364100189 | -0.234781958624995 | 0.814377971320506 | NA | NM_027836,NM_001276398,NM_001025610 |
| Ms4a8a | 1.04183065 | -0.0112142622169364 | 0.0213687657402594 | -0.524796909341768 | 0.599724380291146 | NA | NM_022430 |
| Msgn1 | 0 | NA | NA | NA | NA | NA | NM_019544 |
| Msh4 | 0.97245336 | -0.00094240878197984 | 0.0192282724464867 | -0.0490116199779575 | 0.960910035666419 | NA | NM_031870 |
| Msh5 | 11.27344889 | -0.0549172448221389 | 0.0688360969867542 | -0.797797191097374 | 0.42498819160882 | NA | NM_013600,NM_001146215 |
| Msmb | 0 | NA | NA | NA | NA | NA | NM_020597 |
| Msmp | 1.001665186 | -0.0102503923726675 | 0.0230474953472166 | -0.44475081644416 | 0.656499841361145 | NA | NM_001099314 |
| Msr1 | 10.21934489 | -0.00287045295145848 | 0.047970084642635 | -0.0598383966349575 | 0.952284344260428 | NA | NM_031195,NM_001113326 |
| Mss51 | 15.61185022 | -0.0986170785565545 | 0.0737296351752094 | -1.33755006819447 | 0.181043150576351 | NA | NM_029104 |
| Mst1 | 0.657513041 | -0.0117054395187441 | 0.0175155144510865 | -0.668289792539781 | 0.503948627765929 | NA | NM_008243 |
| Mstn | 10.98835454 | -0.112347657328816 | 0.065579403328506 | -1.71315461298168 | 0.0866840990175734 | NA | NM_010834 |
| Msx1as | 0.226066984 | -0.0093084363525918 | 0.0101406635841621 | -0.917931679257152 | 0.358654635211298 | NA | NR_027920 |
| Msx2 | 13.81086082 | 0.00861597592806846 | 0.0658252446241461 | 0.130891665914264 | 0.895861007598145 | NA | NM_013601 |
| Msx3 | 1.765766832 | 0.0250949079940181 | 0.0285202172231309 | 0.879898908121403 | 0.378914076284849 | NA | NM_010836 |
| Mt4 | 0 | NA | NA | NA | NA | NA | NM_008631 |
| Mtap7d3 | 2.698113209 | -0.0300272850041412 | 0.0356275518355298 | -0.84281078707733 | 0.399334274092508 | NA | NM_177293 |
| Mtfr2 | 14.09236121 | -0.0333843265371818 | 0.0704573059309963 | -0.473823489218809 | 0.635625782864882 | NA | NM_027930 |
| Mtl5 | 5.816483074 | -0.0623729196407998 | 0.0474122677255677 | -1.31554390103902 | 0.188327173176466 | NA | NM_001039657,NM_001039658 |
| Mtnr1a | 0.751690259 | -0.00399961159961934 | 0.0141116826538377 | -0.283425562899236 | 0.77685063505499 | NA | NM_008639 |
| Mtnr1b | 0 | NA | NA | NA | NA | NA | NM_145712 |
| Muc13 | 1.978809442 | -0.018956190006976 | 0.0330107970989086 | -0.574242116910372 | 0.565803972493337 | NA | NM_010739 |
| Muc15 | 6.353283851 | 0.000373539491467449 | 0.0544563818932652 | 0.00685942544254203 | 0.994527013262355 | NA | NM_172979 |
| Muc19 | 1.319024518 | 0.00495306291341426 | 0.0206483558792164 | 0.239876866825981 | 0.810425714942896 | NA | NM_207243 |
| Muc2 | 6.903675111 | -0.0347507883845086 | 0.0563171721321298 | -0.617054924259642 | 0.537198498807017 | NA | NM_023566 |
| Muc20 | 0.552207486 | 0.00165345284777613 | 0.0164733503744343 | 0.100371376204211 | 0.920049493466764 | NA | NM_146071,NM_001145874 |
| Muc4 | 1.156918588 | 0.0158647693869235 | 0.0241520484418768 | 0.656870551792033 | 0.511264154758842 | NA | NM_080457 |
| Muc5ac | 0.874635119 | 0.00432932536049868 | 0.0189073158429283 | 0.228976201406078 | 0.818887413288574 | NA | NM_010844 |
| Muc5b | 0.205932006 | -0.000576954122277892 | 0.00970424421944593 | -0.0594537925088239 | 0.952590668569529 | NA | NM_028801 |
| Muc6 | 0.212496447 | -0.000886988073087717 | 0.00973851166817767 | -0.0910804549309223 | 0.927428662789121 | NA | NM_181729 |
| Mug-ps1 | 0 | NA | NA | NA | NA | NA | NR_027619 |
| Mug1 | 0 | NA | NA | NA | NA | NA | NM_008645 |
| Mug2 | 0.106347839 | -0.00653347234140708 | 0.00854389509723638 | -0.764694822098226 | 0.444453290197145 | NA | NM_008646 |
| Mup1 | 0 | NA | NA | NA | NA | NA | NM_001163010,NM_001163011,NM_031188 |
| Mup10 | 0 | NA | NA | NA | NA | NA | NM_001122647 |
| Mup11 | 0.106347839 | -0.00653347234140708 | 0.00854389509723638 | -0.764694822098226 | 0.444453290197145 | NA | NM_001164526 |
| Mup12 | 0 | NA | NA | NA | NA | NA | NM_001199995 |
| Mup13 | 0 | NA | NA | NA | NA | NA | NM_001134674 |
| Mup14 | 0 | NA | NA | NA | NA | NA | NM_001199999 |
| Mup15 | 0 | NA | NA | NA | NA | NA | NM_001200004 |
| Mup16 | 0 | NA | NA | NA | NA | NA | NM_001199936 |
| Mup17 | 0 | NA | NA | NA | NA | NA | NM_001200006 |
| Mup19 | 0 | NA | NA | NA | NA | NA | NM_001135127 |
| Mup2 | 0 | NA | NA | NA | NA | NA | NM_001045550,NM_008647 |
| Mup20 | 0 | NA | NA | NA | NA | NA | NM_001012323 |
| Mup21 | 0 | NA | NA | NA | NA | NA | NM_001009550 |
| Mup3 | 0.455241012 | -0.00262957968178519 | 0.0154738539002529 | -0.169936959385548 | 0.865059714452143 | NA | NM_001039544 |
| Mup4 | 0.73332599 | -0.00499566941928792 | 0.0162591179667226 | -0.307253408795761 | 0.758650495888469 | NA | NM_008648 |

| | | | | | | | | | | | | | | | |
|---|---|---|---|---|---|---|---|---|---|---|---|---|---|---|---|
| Mir484 | 0 | NA | NA | NA | NA | NA | | Mup5 | 10.4859403 | 0.0442757379291814 | 0.0621192299573051 | 0.712754133617117 | 0.475997912179095 | NA | NM_008649 |
| Mir485 | 0.590545564 | 0.020004006 | 0.053347245 | 0.374977293 | 0.707677354 | NA | | Mup6 | 0.108766301 | 0.00519171381361946 | 0.00856003371502575 | 0.606506234257728 | 0.544178643801544 | NA | NM_001081285 |
| Mir486 | 0.94480822 | 0.019528546 | 0.071623939 | 0.272653891 | 0.785119282 | NA | | Mup7 | 0 | NA | NA | NA | NA | NA | NM_001134675 |
| Mir487b | 0.222620122 | -0.001447027 | 0.032059667 | -0.04513545 | 0.963999345 | NA | | Mup8 | 0 | NA | NA | NA | NA | NA | NM_001134676 |
| Mir488 | 0 | NA | NA | NA | NA | NA | | Mup9 | 0 | NA | NA | NA | NA | NA | NM_001126319 |
| Mir489 | 0 | NA | NA | NA | NA | NA | | Mx1 | 18.93959695 | -0.215532059471882 | 0.066524841747769 | -3.23987331362739 | 0.00119582816670126 | NA | NM_010846,NR_003520 |
| Mir490 | 0 | NA | NA | NA | NA | NA | | Mybpc2 | 9.587488879 | -0.120206882000515 | 0.0643516492169352 | -1.86796894039633 | 0.0617664003492435 | NA | NM_146189 |
| Mir491 | 2.293765607 | -0.007964255 | 0.104865042 | -0.075947666 | 0.939460735 | NA | | Mybph | 9.203483394 | -0.0129951409160481 | 0.0572280196480761 | -0.227076543902126 | 0.82036422291081 | NA | NM_016749 |
| Mir493 | 0.635690241 | -0.003851216 | 0.053467645 | -0.072028909 | 0.942578901 | NA | | Mybphl | 1.603244846 | -0.0398578647216726 | 0.0290347418669443 | -1.37276456268586 | 0.169825550729227 | NA | NM_026831 |
| Mir494 | 0.256038351 | 0.007085343 | 0.03307033 | 0.214250763 | 0.830351512 | NA | | Mycs | 0.407027086 | -0.0158737350575048 | 0.0132224722193719 | -1.20051188568569 | 0.229940599374159 | NA | NM_010850 |
| Mir495 | 0 | NA | NA | NA | NA | NA | | Myf5 | 0 | NA | NA | NA | NA | NA | NM_008656 |
| Mir496 | 4.52837712 | 0.163733886 | 0.122973017 | 1.33146189 | 0.183037077 | NA | | Myf6 | 0 | NA | NA | NA | NA | NA | NM_008657 |
| Mir497 | 0 | NA | NA | NA | NA | NA | | Myh4 | 22.32417301 | -0.00592901271010406 | 0.0771808490192828 | -0.0768197394229591 | 0.938766947350722 | NA | NM_010855 |
| Mir499 | 0 | NA | NA | NA | NA | NA | | Myh6 | 1.149044516 | 0.00709666476957806 | 0.0243612440336453 | 0.29130962112513 | 0.770814529795199 | NA | NM_001164171,NM_010856 |
| Mir500 | 0 | NA | NA | NA | NA | NA | | Myh8 | 19.65741406 | 0.0131694848978525 | 0.0735022042681526 | 0.179171291922175 | 0.857803204671142 | NA | NM_177369 |
| Mir501 | 0 | NA | NA | NA | NA | NA | | Myl1 | 10.27370433 | 0.09073441198769 | 0.0626708111999865 | 1.44779380145808 | 0.147674726839457 | NA | NM_001113387,NM_021285 |
| Mir503 | 0 | NA | NA | NA | NA | NA | | Myl10 | 0.106347839 | -0.00653347234140708 | 0.00854389509723638 | -0.764694822098226 | 0.444453290197145 | NA | NM_001085387,NM_021611 |
| Mir504 | 0 | NA | NA | NA | NA | NA | | Myl2 | 11.47400791 | 0.041087333671082 | 0.0708027465145412 | 0.580307060018408 | 0.561707567278728 | NA | NM_010861 |
| Mir5046 | 0 | NA | NA | NA | NA | NA | | Myl3 | 2.185320775 | -0.0177105657630958 | 0.0297493856691387 | -0.595325428231223 | 0.551625961759104 | NA | NM_010859 |
| Mir505 | 0 | NA | NA | NA | NA | NA | | Myl7 | 0.822840093 | -0.00817134397086926 | 0.0202305442043767 | -0.403911228898206 | 0.686277998786826 | NA | NM_022879 |
| Mir509 | 0 | NA | NA | NA | NA | NA | | Mylk2 | 0.846191468 | -0.0155828612770763 | 0.018586259390956 | -0.838407607969726 | 0.401801818041628 | NA | NM_001081044 |
| Mir5098 | 0 | NA | NA | NA | NA | NA | | Mylk4 | 15.44037163 | -0.00872088509625615 | 0.0749455663585113 | -0.116362922051169 | 0.907364919642532 | NA | NM_001166030 |
| Mir5100 | 0.502431138 | -0.017833366 | 0.050924809 | -0.350190145 | 0.726196002 | NA | | Mylpf | 20.55906142 | 0.00423731557085853 | 0.0787135665449846 | 0.0538320871083501 | 0.957068944759541 | NA | NM_016754 |
| Mir5101 | 0 | NA | NA | NA | NA | NA | | Myo15 | 20.83593343 | -0.0808231249893992 | 0.0799920870470782 | -1.01038900187505 | 0.312308955001998 | NA | NM_001103171,NM_010862,NM_182698 |
| Mir5103 | 3.393330191 | -0.011278602 | 0.127968635 | -0.088135673 | 0.929768843 | NA | | Myo3a | 1.785826984 | 0.00898513536889501 | 0.0291528069701425 | 0.308208241425854 | 0.757923881769411 | NA | NM_148413 |
| Mir5104 | 0 | NA | NA | NA | NA | NA | | Myo7b | 6.303919108 | -0.0142459859141397 | 0.0549415833865029 | -0.259293326403101 | 0.795408928139137 | NA | NM_032394 |
| Mir5106 | 0 | NA | NA | NA | NA | NA | | Myocd | 10.17793859 | 0.0496094507419904 | 0.0677533145202843 | 0.732206993757304 | 0.464042236259533 | NA | NM_145136,NM_146386 |
| Mir5107 | 0.079983654 | 0.016740032 | 0.027674178 | 0.604897175 | 0.54524732 | NA | | Myod1 | 7.074066916 | 0.0617119460315655 | 0.0560257879393966 | 1.10149180049587 | 0.270682671176041 | NA | NM_010866 |
| Mir5108 | 0 | NA | NA | NA | NA | NA | | Myog | 0.570488403 | 0.0118280525685891 | 0.0169931929338767 | 0.69604650606946 | 0.486399705610745 | NA | NM_031189 |
| Mir5109 | 0 | NA | NA | NA | NA | NA | | Myom2 | 7.308155604 | 0.0580561459053118 | 0.0581789735508442 | 0.997888796621256 | 0.318333285191439 | NA | NM_008664 |
| Mir511 | 0 | NA | NA | NA | NA | NA | | Myot | 1.669919046 | 0.0331343250311391 | 0.0254869358570364 | 1.30005133677109 | 0.193583374737979 | NA | NM_001033621 |
| Mir5112 | 0.813522201 | 0.05673995 | 0.064692596 | 0.877070229 | 0.380448494 | NA | | Myoz1 | 0.354244579 | -0.00570556803240082 | 0.0130207392936895 | -0.438190789609467 | 0.661247984684689 | NA | NM_021508 |
| Mir5113 | 0 | NA | NA | NA | NA | NA | | Myoz2 | 4.160797024 | 0.0470516444794629 | 0.0479114508240561 | 0.982054261981116 | 0.326073116680258 | NA | NM_021503 |
| Mir5114 | 0 | NA | NA | NA | NA | NA | | Myoz3 | 1.221922479 | 0.00458976692823096 | 0.0228187599146842 | 0.2011400683206 | 0.840589052184378 | NA | NM_133363 |
| Mir5116 | 0 | NA | NA | NA | NA | NA | | Myrfl | 4.6581879 | 0.0203724567712768 | 0.0480799357361168 | 0.423720549109914 | 0.671769627011167 | NA | NM_001033333 |
| Mir5117 | 0 | NA | NA | NA | NA | NA | | Mzb1 | 3.107203627 | -0.0500257848341933 | 0.0362910119353381 | -1.37846210856086 | 0.168060658967703 | NA | NM_027222 |
| Mir5119 | 3.771806447 | 0.167827703 | 0.123225056 | 1.361960864 | 0.173210236 | NA | | Naa11 | 2.131511852 | 0.0468366634810982 | 0.0344198323123431 | 1.36074641666114 | 0.173593840524157 | NA | NM_001033191 |
| Mir5121 | 0 | NA | NA | NA | NA | NA | | Naaladl1 | 5.169334223 | -0.0238055181469687 | 0.0498439053281763 | -0.477601383563974 | 0.632933945948642 | NA | NM_001009546 |
| Mir5122 | 0 | NA | NA | NA | NA | NA | | Nags | 3.325266733 | -0.0197209791531854 | 0.0379489750382989 | -0.519670930065505 | 0.603292951530956 | NA | NM_145829 |
| Mir5123 | 0.094392817 | 0.017895942 | 0.027818137 | 0.643319236 | 0.520016977 | NA | | Naip1 | 14.56683364 | -0.0663436333328982 | 0.0746875920528477 | -0.888281861945081 | 0.374389153826343 | NA | NM_008670 |
| Mir5126 | 0 | NA | NA | NA | NA | NA | | Naip6 | 10.87268139 | -0.119973797368877 | 0.0647345431513146 | -1.85331959613034 | 0.0638365640561625 | NA | NM_010871 |
| Mir5127 | 0 | NA | NA | NA | NA | NA | | Naip7 | 0.219991894 | 0.00711449016834187 | 0.00994861810862167 | 0.715123456410123 | 0.474532761039468 | NA | NM_021545 |
| Mir5128 | 0.197363271 | -0.000561558 | 0.031650166 | -0.017742642 | 0.985844163 | NA | | Nanog | 1.837764489 | -0.0164917135159476 | 0.0318034694813743 | -0.51855076772696 | 0.604074048632095 | NA | NM_028016 |
| Mir5129 | 0 | NA | NA | NA | NA | NA | | Nanos2 | 18.81761408 | -0.121144678286241 | 0.0734999858732545 | -1.64822723224936 | 0.0993060506060568 | NA | NM_194064 |
| Mir5130 | 0 | NA | NA | NA | NA | NA | | Napsa | 2.854951619 | 0.049456753849453 | 0.0403973958482948 | 1.22425599004399 | 0.220855673522326 | NA | NM_008437 |
| Mir5131 | 0 | NA | NA | NA | NA | NA | | Nat3 | 0 | NA | NA | NA | NA | NA | NM_008674 |
| Mir5132 | 0 | NA | NA | NA | NA | NA | | Ncapg | 10.90842384 | -0.0181574775193581 | 0.0632268180793289 | -0.287179998471161 | 0.773974500484294 | NA | NM_019438 |
| Mir5134 | 0 | NA | NA | NA | NA | NA | | Nccrp1 | 13.81151309 | 0.276137571935732 | 0.0642863343398193 | 4.29543191055289 | 1.74353546652607e-05 | NA | NM_001081115 |
| Mir5135 | 0 | NA | NA | NA | NA | NA | | Ncr1 | 0.429580527 | -0.0208285968975787 | 0.0151249693245128 | -1.37710010848234 | 0.168481299181666 | NA | NM_010746 |
| Mir5136 | 0 | NA | NA | NA | NA | NA | | Nctc1 | 0 | NA | NA | NA | NA | NA | NR_002452 |
| Mir532 | 0 | NA | NA | NA | NA | NA | | Ndc80 | 10.09392329 | -0.0559151852786983 | 0.0664132736307068 | -0.841927859024334 | 0.399828335253308 | NA | NM_023294 |
| Mir539 | 0 | NA | NA | NA | NA | NA | | Ndufs5 | 0 | NA | NA | NA | NA | NA | NM_001030274 |
| Mir540 | 1.078837746 | 0.02419156 | 0.065681076 | 0.368318573 | 0.712635705 | NA | | Neil3 | 4.045797931 | 0.0213811053382692 | 0.0429498267363886 | 0.497815869421293 | 0.618613829576632 | NA | NM_146208 |
| Mir542 | 0 | NA | NA | NA | NA | NA | | Nepn | 0.754372279 | 0.000145829848151853 | 0.0198413924614785 | 0.00734977892478956 | 0.994135777667459 | NA | NM_025684 |
| Mir543 | 0.238795281 | -0.027601606 | 0.032267774 | -0.855392297 | 0.392334007 | NA | | Nespas | 1.69239165 | -0.0489840017895266 | 0.0245347856204404 | -1.99651232121291 | 0.0458781859671877 | NA | NR_002846 |
| Mir546 | 0 | NA | NA | NA | NA | NA | | Neurod4 | 20.20722562 | 0.0888934197236461 | 0.0794462834198408 | 1.11891224985165 | 0.263177576456611 | NA | NM_007501 |
| Mir547 | 0 | NA | NA | NA | NA | NA | | Neurog1 | 5.445158285 | 0.0466104536191636 | 0.0446093882430972 | 1.0448574942378 | 0.296088837542505 | NA | NM_010896 |
| Mir551b | 0 | NA | NA | NA | NA | NA | | Neurog2 | 10.58013349 | 0.0332838492273146 | 0.0646031096088164 | 0.515205064103793 | 0.60640973066058 | NA | NM_009718 |
| Mir574 | 0 | NA | NA | NA | NA | NA | | Neurog3 | 2.524382914 | -0.0115634758380826 | 0.0331223468790645 | -0.349114025050907 | 0.727003707757856 | NA | NM_009719 |
| Mir582 | 0 | NA | NA | NA | NA | NA | | Nfe2 | 1.736274808 | 0.000771200041338343 | 0.028053534866153 | 0.027490298282118 | 0.978068677773962 | NA | NM_008685 |
| Mir592 | 0 | NA | NA | NA | NA | NA | | Ngp | 15.15045116 | -0.00382520895479672 | 0.0524005269895111 | -0.0729994367339551 | 0.941806565737582 | NA | NM_008694 |
| Mir598 | 0.096071043 | -0.020098795 | 0.027825948 | -0.722304036 | 0.470107572 | NA | | Nhlh1 | 20.90783013 | -0.0657096980068828 | 0.0759949580531411 | -0.864658652234981 | 0.387226170083695 | NA | NM_010916 |
| Mir599 | 0 | NA | NA | NA | NA | NA | | Nhlh2 | 17.57108475 | 0.0282797652196848 | 0.0673603653863262 | 0.419827966453185 | 0.674611133102431 | NA | NM_178777 |
| Mir615 | 0 | NA | NA | NA | NA | NA | | Nhlrc4 | 18.70080914 | -0.0186465751481813 | 0.070706295375951 | -0.263718740305032 | 0.791996653919251 | NA | NM_001039038 |
| Mir653 | 0 | NA | NA | NA | NA | NA | | Niacr1 | 9.219611668 | -0.121749002047878 | 0.0450806660154041 | -2.70069217713591 | 0.00691953481367829 | NA | NM_030701 |
| Mir654 | 0.176054697 | -0.002190495 | 0.031239146 | -0.070120197 | 0.944097991 | NA | | Nipal1 | 0.731486784 | 0.0177782311105617 | 0.0188179351324125 | 0.944749303548189 | 0.344786879504999 | NA | NM_001081205 |
| Mir664 | 0 | NA | NA | NA | NA | NA | | Nipsnap3a | 0 | NA | NA | NA | NA | NA | NM_028529 |
| Mir665 | 1.005556649 | -0.004527101 | 0.071578148 | -0.063246976 | 0.949569839 | NA | | Nkapl | 9.529810526 | -0.0146476995037261 | 0.0525945032694995 | -0.278502478266024 | 0.780626662886153 | NA | NM_025719 |
| Mir666 | 0 | NA | NA | NA | NA | NA | | Nkg7 | 1.387827618 | -0.0129232761607913 | 0.0288430562433643 | -0.448055020652135 | 0.654113488596902 | NA | NM_024253 |
| Mir667 | 0 | NA | NA | NA | NA | NA | | Nkx1-2 | 2.45143039 | 0.0158710763741104 | 0.0317720744783481 | 0.499529119035843 | 0.617406678195334 | NA | NM_009123 |
| Mir668 | 5.45249312 | 0.092449238 | 0.139097008 | 0.664638584 | 0.506281684 | NA | | Nkx2-3 | 0 | NA | NA | NA | NA | NA | NM_008699 |
| Mir669a-1 | 0 | NA | NA | NA | NA | NA | | Nkx2-4 | 0.543831507 | 0.0105450399825839 | 0.0120876531324133 | 0.87238108730198 | 0.383000514602336 | NA | NM_023504 |
| Mir669a-10 | 0 | NA | NA | NA | NA | NA | | Nkx2-5 | 0 | NA | NA | NA | NA | NA | NM_008700 |
| Mir669a-11 | 0 | NA | NA | NA | NA | NA | | Nkx2-6 | 0 | NA | NA | NA | NA | NA | NM_010920 |
| Mir669a-12 | 0 | NA | NA | NA | NA | NA | | Nkx2-9 | 16.94863923 | -0.00482692760986338 | 0.0769618242244553 | -0.0627184667009176 | 0.949990692042209 | NA | NM_008701 |
| Mir669a-2 | 0 | NA | NA | NA | NA | NA | | Nkx3-2 | 0 | NA | NA | NA | NA | NA | NM_007524 |
| Mir669a-3 | 0 | NA | NA | NA | NA | NA | | Nkx6-1 | 3.397899967 | -0.0111809859512937 | 0.0368877883415899 | -0.303108059712201 | 0.761807512150902 | NA | NM_144955 |
| Mir669a-4 | 0 | NA | NA | NA | NA | NA | | Nkx6-3 | 2.878226646 | 0.00525541927509563 | 0.0404958208365473 | 0.129776830461296 | 0.896742994482468 | NA | NM_029002 |
| Mir669a-5 | 0 | NA | NA | NA | NA | NA | | Nlrc3 | 17.73668817 | -0.0410869813051267 | 0.0711295318238039 | -0.577636043027865 | 0.563509867439507 | NA | NM_001081280 |
| Mir669a-6 | 0 | NA | NA | NA | NA | NA | | Nlrc4 | 13.46760797 | -0.051635807090807 | 0.0650315886210518 | -0.794011159587321 | 0.427188934708275 | NA | NM_001033367 |
| Mir669a-7 | 0 | NA | NA | NA | NA | NA | | Nlrc5 | 83.70274765 | -0.131428114603678 | 0.0532114630012876 | -2.46992108825306 | NA | NA | NM_001033207 |
| Mir669a-8 | 0 | NA | NA | NA | NA | NA | | Nlrp10 | 10.63405449 | 0.0674052042459938 | 0.0621137380100453 | 1.08518995000901 | 0.277837534454254 | NA | NM_175532 |
| Mir669a-9 | 0 | NA | NA | NA | NA | NA | | Nlrp12 | 1.279829732 | -0.0164868780311294 | 0.026761560470015 | -0.616065645708593 | 0.537851194672231 | NA | NM_001033431 |
| Mir669b | 0 | NA | NA | NA | NA | NA | | Nlrp14 | 0 | NA | NA | NA | NA | NA | NM_001002894 |
| Mir669c | 0 | NA | NA | NA | NA | NA | | Nlrp1c-ps | 12.07922935 | -0.075483529416031 | 0.0644639289779273 | -1.17094211620078 | 0.241622044699104 | NA | NR_027858 |
| Mir669e | 0 | NA | NA | NA | NA | NA | | Nlrp2 | 3.004186556 | 0.0126330363632748 | 0.0390391718543771 | 0.323598984384151 | 0.746241656974147 | NA | NM_177690 |
| Mir669g | 0 | NA | NA | NA | NA | NA | | Nlrp3 | 53.12146834 | -0.231327021763424 | 0.0872246231192296 | -2.65208393560173 | NA | NA | NM_145827 |
| Mir669h | 0 | NA | NA | NA | NA | NA | | Nlrp4a | 0 | NA | NA | NA | NA | NA | NM_172896 |
| Mir669i | 0 | NA | NA | NA | NA | NA | | Nlrp4b | 0 | NA | NA | NA | NA | NA | NM_172481 |
| Mir669j | 0 | NA | NA | NA | NA | NA | | Nlrp4c | 0.617098604 | 0.00791652526967432 | 0.0135563267159957 | 0.583972740958896 | 0.559238651844108 | NA | NM_031389 |
| Mir669k | 0 | NA | NA | NA | NA | NA | | Nlrp4e | 0 | NA | NA | NA | NA | NA | NM_001004194 |
| Mir669m-1 | 0 | NA | NA | NA | NA | NA | | Nlrp4f | 6.434465653 | -0.00348928971685708 | 0.0552788133444465 | -0.0631216465359893 | 0.949669637225495 | NA | NM_175290 |
| Mir669m-2 | 0 | NA | NA | NA | NA | NA | | Nlrp5 | 0.203513543 | -0.00860782669689374 | 0.00975338121443284 | -0.882547960306941 | 0.377480556050336 | NA | NM_011860,NM_001039143 |
| Mir669p-1 | 0 | NA | NA | NA | NA | NA | | Nlrp9a | 0 | NA | NA | NA | NA | NA | NM_001048219,NM_001048220 |
| Mir669p-2 | 0 | NA | NA | NA | NA | NA | | Nlrp9b | 0.120209049 | -0.00674972139751468 | 0.00863521114606269 | -0.781650996523957 | 0.434419707908266 | NA | NM_194058 |
| Mir670 | 1.415539497 | -0.013598678 | 0.061933765 | -0.219568086 | 0.826207549 | NA | | Nlrp9c | 0 | NA | NA | NA | NA | NA | NM_001042612 |
| Mir671 | 0 | NA | NA | NA | NA | NA | | Nme8 | 0.893333255 | 0.0208110055293381 | 0.0201827998196838 | 1.03112579598801 | 0.3024818323469 | NA | NM_181591,NM_001167909 |
| Mir672 | 0 | NA | NA | NA | NA | NA | | Nme9 | 10.8260449 | 0.0269863449520396 | 0.049904818347152 | 0.540756300610394 | 0.588675566842315 | NA | NM_001165957 |
| Mir673 | 0 | NA | NA | NA | NA | NA | | Nmrk2 | 5.98511608 | -0.0157476278731883 | 0.0403104858561731 | -0.390658349526608 | 0.696049789363295 | NA | NM_027120 |
| Mir674 | 2.601193737 | 0.090443513 | 0.115514443 | 0.782962812 | 0.43364895 | NA | | Nms | 1.115474863 | -0.00430260108975698 | 0.0238609980888214 | -0.180319409680213 | 0.856901818699801 | NA | NM_001011684 |
| Mir675 | 0 | NA | NA | NA | NA | NA | | Nmu | 0.763449516 | 0.0183000292137685 | 0.0189724951459777 | 0.964555746250816 | 0.334767373931112 | NA | NM_019515 |
| Mir676 | 0 | NA | NA | NA | NA | NA | | Nmur1 | 3.599724091 | 0.0219844639615454 | 0.0428017811806878 | 0.513634324439395 | 0.607507676914773 | NA | NM_010341 |
| Mir677 | 7.452538808 | 0.21603724 | 0.150460123 | 1.435843828 | 0.151046785 | NA | | Nmur2 | 3.957705342 | 0.0540649884165738 | 0.0311121283606631 | 1.73774637947725 | 0.0822555125544276 | NA | NM_153079 |
| Mir678 | 0 | NA | NA | NA | NA | NA | | Nnmt | 17.5621366 | -0.0708950946211514 | 0.0779039143163471 | -0.910032509191583 | 0.362805365467034 | NA | NM_010924 |
| Mir679 | 0 | NA | NA | NA | NA | NA | | Nobox | 1.318592192 | 0.0140906469941846 | 0.0231153106220788 | 0.609580689810234 | 0.542139606464758 | NA | NM_130869 |
| Mir680-1 | 0 | NA | NA | NA | NA | NA | | Nodal | 21.64464367 | -0.0424283675812828 | 0.0781001027803121 | -0.54325623233339 | 0.586953394493007 | NA | NM_013611 |
| Mir680-2 | 0 | NA | NA | NA | NA | NA | | Noto | 0 | NA | NA | NA | NA | NA | NM_001007472 |
| Mir680-3 | 0 | NA | NA | NA | NA | NA | | Nox1 | 0.77388662 | -0.0216990625757548 | 0.0218835207855849 | -0.991570908007108 | 0.32140688657332 | NA | NM_172203 |
| Mir681 | 0.422179562 | -0.00343537 | 0.036938882 | -0.093001469 | 0.925902394 | NA | | Nox3 | 1.159403599 | 0.0154752401942027 | 0.0245260496370002 | 0.630971576068924 | 0.528059109692272 | NA | NM_198958 |
| Mir683-1 | 0 | NA | NA | NA | NA | NA | | Nox4 | 5.141287849 | 0.00552215956122006 | 0.0437751301102324 | 0.126148330052119 | 0.899614511160081 | NA | NM_015760 |
| Mir683-2 | 0 | NA | NA | NA | NA | NA | | Noxa1 | 0.192890664 | -0.000903864895041124 | 0.00963528282151475 | -0.093807821917056 | 0.92526181809679 | NA | NM_001163626,NM_172204 |
| Mir684-1 | 0.08731684 | 0.017365056 | 0.027733412 | 0.626142072 | 0.531221759 | NA | | Npb | 6.081205805 | -0.0209037388111477 | 0.0525814854250549 | -0.397549415771871 | 0.690962353893974 | NA | NM_153288 |
| Mir684-2 | 0 | NA | NA | NA | NA | NA | | Npcd | 0 | NA | NA | NA | NA | NA | NM_001013362,NM_001013360 |
| Mir686 | 0 | NA | NA | NA | NA | NA | | Npffr1 | 21.71576102 | 0.0735392301924757 | 0.0741968854557828 | 0.991136349467134 | 0.321619005756976 | NA | NM_001177511 |
| Mir687 | 0 | NA | NA | NA | NA | NA | | Npffr2 | 3.968499126 | -0.0166166836549255 | 0.0448295189884875 | -0.370663884642455 | 0.710887892967945 | NA | NM_133192 |
| Mir688 | 1.329130058 | 0.089015312 | 0.091802701 | 0.969637186 | 0.33222737 | NA | | Nphs1as | 10.63824656 | -0.0511574117133979 | 0.0592656715646915 | -0.863187919123753 | 0.388034154824583 | NA | NR_004443 |
| Mir690 | 0 | NA | NA | NA | NA | NA | | Nphs2 | 12.9140783 | -0.0213922459024353 | 0.0513057437590832 | -0.416956159974739 | 0.676710474491637 | NA | NM_130456 |
| Mir691 | 0 | NA | NA | NA | NA | NA | | Npm2 | 11.20811721 | 0.00609074517596897 | 0.0681249940536082 | 0.089405441579578 | 0.928759699037182 | NA | NM_181345 |
| Mir692-1 | 0 | NA | NA | NA | NA | NA | | Npm3-ps1 | 0.35787663 | -0.00674720089483589 | 0.0130723048459276 | -0.516144702434617 | 0.605753348373098 | NA | NR_002702 |
| Mir693 | 0.297029653 | 0.02866449 | 0.033517577 | 0.855207712 | 0.392436167 | NA | | Nppa | 5.52399073 | -0.0320914246562865 | 0.0492208252379572 | -0.651988756814641 | 0.514408425451053 | NA | NM_008725 |
| Mir694 | 0 | NA | NA | NA | NA | NA | | Nppb | 0.191449919 | 0.00665508079272897 | 0.00962405648474729 | 0.691504752001018 | 0.489248395458281 | NA | NM_008726 |
| Mir695 | 0 | NA | NA | NA | NA | NA | | Nps | 0.428139782 | -0.0129079788277846 | 0.0156498226724372 | -0.824800325087285 | 0.409484962090028 | NA | NM_001163611 |
| Mir697 | 4.83199621 | 0.242706135 | 0.128600216 | 1.887291823 | 0.059121086 | NA | | Npvf | 5.076898678 | 0.00665288331197042 | 0.0116614327667567 | 0.570503080113435 | NA | NA | NM_021892 |
| Mir698 | 0 | NA | NA | NA | NA | NA | | Npw | 2.562940129 | 0.00973990111415233 | 0.033424344988308 | 0.291401405698732 | 0.77074433958332 | NA | NM_001099664 |
| Mir7-1 | 6.288786782 | 0.176047001 | 0.141922735 | 1.240442562 | 0.214811747 | NA | | Npy6r | 0.343816189 | 0.00220638102164924 | 0.0128049404084123 | 0.172307012081036 | 0.863196171959189 | NA | NM_010935 |
| Mir7-2 | 0 | NA | NA | NA | NA | NA | | Nr0b1 | 1.195404933 | 0.0274127739153162 | 0.0254569667628448 | 1.07682797289604 | 0.281557125941616 | NA | NM_007430 |
| Mir700 | 0.299325859 | 0.043567584 | 0.041359933 | 1.053376566 | 0.29216844 | NA | | Nr0b2 | 1.012840553 | 0.0358229473902898 | 0.0221185214894235 | 1.61959050506244 | 0.105320270577563 | NA | NM_011850 |
| Mir701 | 0 | NA | NA | NA | NA | NA | | Nr1h4 | 3.319054382 | -0.0105681209718854 | 0.0432456436790847 | -0.244374232241954 | 0.806940996585286 | NA | NM_001163504,NM_001163700,NM_009108 |
| Mir702 | 0 | NA | NA | NA | NA | NA | | Nr1h5 | 0.506912888 | -0.0178698354858196 | 0.0140226761995497 | -1.27435271495418 | 0.202538442112223 | NA | NM_198658,NM_201622,NM_201619 |
| Mir703 | 7.726868215 | 0.02662373 | 0.152955264 | 0.174062199 | 0.86181658 | NA | | Nr1i2 | 0.428430455 | -0.00286174811964168 | 0.0152554242063721 | -0.18758889172327 | 0.851198934776155 | NA | NM_010936,NM_001098404 |
| Mir704 | 0 | NA | NA | NA | NA | NA | | Nr1i3 | 4.819949339 | 0.0416655505971613 | 0.0477736544880193 | 0.872144931001881 | 0.383129317220729 | NA | NM_001243062,NM_001243063,NM_009803 |
| Mir705 | 0 | NA | NA | NA | NA | NA | | Nr2e3 | 0.516150229 | 0.00784758350140605 | 0.0148035917613958 | 0.530113477046204 | 0.59603325549776 | NA | NM_013708 |
| Mir706 | 0 | NA | NA | NA | NA | NA | | Nr5a1 | 9.910560834 | 0.0355945370646387 | 0.0261096601626266 | 1.36327079107636 | NA | NA | NM_139051 |
| Mir708 | 0 | NA | NA | NA | NA | NA | | Nr5a2 | 2.530492686 | 0.0158596275898664 | 0.0254600777492036 | 0.622921412341817 | 0.533336166318827 | NA | NM_030676,NM_001159769 |
| Mir709 | 3.257913912 | -0.000278494 | 0.107607283 | -0.002588055 | 0.997935034 | NA | | Nrg4 | 14.91259462 | -0.00549419522249883 | 0.071411683437336 | -0.0769369234562297 | 0.938673723915512 | NA | NM_032002 |
| Mir710 | 0 | NA | NA | NA | NA | NA | | Nrk | 0.32976049 | 0.00299720392154094 | 0.0126582099054095 | 0.236779445430122 | 0.812827898340701 | NA | NM_013724 |
| Mir711 | 0.079983654 | 0.016740032 | 0.027674178 | 0.604897175 | 0.54524732 | NA | | Nrl | 12.95202956 | 0.0290513667778818 | 0.0737716002332555 | 0.3938014993036 | 0.693727597396101 | NA | NM_001271917,NM_008736,NM_001136074,NM_001271916 |
| Mir713 | 0 | NA | NA | NA | NA | NA | | Nron | 0.340383369 | -0.00403721153616492 | 0.0113238420937375 | -0.356523121988573 | 0.7214488393547 | NA | NR_045729 |
| Mir717 | 0 | NA | NA | NA | NA | NA | | Nrp | 5.007925122 | -0.00438271013267595 | 0.0478308376644574 | -0.0916293827722924 | 0.926992505570861 | NA | NM_001013372 |
| Mir718 | 0 | NA | NA | NA | NA | NA | | Nrtn | 15.67546915 | 0.00616158535023936 | 0.0690095984581154 | 0.0892859180158695 | 0.928854685165126 | NA | NM_008738 |
| Mir719 | 0 | NA | NA | NA | NA | NA | | Nt5c1b | 0.72527368 | 0.00103413122405449 | 0.0158688355062153 | 0.0651674298123168 | 0.948040693230286 | NA | NM_027588 |
| Mir721 | 0 | NA | NA | NA | NA | NA | | Ntf5 | 4.827497343 | -0.0423280402062349 | 0.0479525105646063 | -0.882707489302493 | 0.377394334642912 | NA | NM_198190 |
| Mir741 | 0 | NA | NA | NA | NA | NA | | Nuggc | 2.543681082 | -0.00217949098921892 | 0.0329816719547522 | -0.0660818830594454 | 0.947312634544385 | NA | NM_001195674 |
| Mir742 | 0 | NA | NA | NA | NA | NA | | Nup210l | 19.23016232 | -0.0338701684745234 | 0.0774305936670072 | -0.437426175759199 | 0.661802304292278 | NA | NM_029937 |
| Mir743 | 0 | NA | NA | NA | NA | NA | | Nup62-il4i1 | 3.85146023 | -0.0394092607025459 | 0.0405885249669286 | -0.970945870406879 | 0.33157523429647 | NA | NM_001171024 |
| Mir743b | 0 | NA | NA | NA | NA | NA | | Nup62cl | 9.359482255 | 0.110626574353964 | 0.0629024886107523 | 1.75869948546128 | 0.0786285648446502 | NA | NM_001081668 |
| Mir744 | 1.087655921 | -0.092254674 | 0.079640209 | -1.158393175 | 0.246703623 | NA | | Nupr1l | 13.38088614 | -0.0822074991605568 | 0.0735484044244018 | -1.1177332778858 | 0.263680925514822 | NA | NM_026916 |
| Mir758 | 0 | NA | NA | NA | NA | NA | | Nusap1 | 13.37826019 | 0.0025178344554902 | 0.0671564081438431 | 0.0374920953201848 | 0.970092642738031 | NA | NM_133851,NM_001042652 |
| Mir759 | 0 | NA | NA | NA | NA | NA | | Nutf2-ps1 | 0 | NA | NA | NA | NA | NA | NR_033574 |
| Mir760 | 2.69608766 | 0.077396299 | 0.110639458 | 0.699536143 | 0.484217034 | NA | | Nxf2 | 2.781649977 | 0.0409687160546798 | 0.0379904744603859 | 1.07839443009324 | 0.280857771628712 | NA | NM_031259 |
| Mir761 | 6.735877293 | 0.133968576 | 0.146888601 | 0.912042016 | 0.3617466 | NA | | Nxf3 | 5.389266607 | -0.0549291738528326 | 0.0458614933116783 | -1.19771882436382 | 0.231026498373389 | NA | NM_001024141 |
| Mir762 | 0.196706773 | -0.026517754 | 0.031609464 | -0.838918169 | 0.401515231 | NA | | Nxnl1 | 4.600733284 | -0.00825456699645173 | 0.0464505648558376 | -0.177706493388624 | 0.858953485434751 | NA | NM_145598 |
| Mir763 | 0 | NA | NA | NA | NA | NA | | Nxnl2 | 5.24656118 | -0.0374919341745788 | 0.0469507182091646 | -0.798538033168159 | 0.424558331278265 | NA | NM_029173 |
| Mir764 | 0 | NA | NA | NA | NA | NA | | Nxpe2 | 9.965410442 | 0.0558914978247966 | 0.0617227792403023 | 0.905524646049993 | 0.365187509084929 | NA | NM_030069 |
| Mir767 | 0 | NA | NA | NA | NA | NA | | Nxpe5 | 0.127687691 | 0.00542298404407807 | 0.00869724871942359 | 0.62352868349814 | 0.532937159177571 | NA | NM_001013773 |
| Mir770 | 0 | NA | NA | NA | NA | NA | | ORF63 | 18.62215235 | -0.115071945273814 | 0.0753001634440908 | -1.52817656709674 | 0.12646870514319 | NA | NM_144854 |
| Mir7b | 0.412413621 | -0.026778838 | 0.043795168 | -0.611456443 | 0.540897446 | NA | | OTTMUSG00000016609 | 0 | NA | NA | NA | NA | NA | NM_001100416 |
| Mir802 | 0 | NA | NA | NA | NA | NA | | Oas1d | 0.658320007 | 0.00654853700682598 | 0.0190198514551877 | 0.344300113082105 | 0.730620599255655 | NA | NM_133893 |
| Mir804 | 0.581676142 | 0.059713211 | 0.060372883 | 0.989073377 | 0.322627242 | NA | | Oas1e | 0 | NA | NA | NA | NA | NA | NM_145210 |
| Mir871 | 0 | NA | NA | NA | NA | NA | | Oas1f | 0.217532603 | 0.00684685248201087 | 0.00976000296730706 | 0.701521557416086 | 0.482977586094298 | NA | NM_145153 |
| Mir872 | 0.079983654 | 0.016740032 | 0.027674178 | 0.604897175 | 0.54524732 | NA | | Oas1g | 2.947954269 | -0.0806371486351498 | 0.0375240031269108 | -2.14894845740272 | 0.0316384859687255 | NA | NM_011852 |
| Mir873 | 0.389505357 | -0.025109253 | 0.043180094 | -0.581500646 | 0.560903083 | NA | | Oas1h | 0.640107034 | -0.00217022306698216 | 0.0154415665662979 | -0.140544228959178 | 0.88823000959471 | NA | NM_001159934,NM_145228 |
| Mir874 | 0.87355605 | 0.004156921 | 0.070283927 | 0.059144693 | 0.952836861 | NA | | Oas3 | 5.989800439 | 0.00978628929132801 | 0.040681957798307 | 0.24055600617469 | 0.80989925092143 | NA | NM_145226 |
| Mir875 | 0 | NA | NA | NA | NA | NA | | Oasl1 | 9.526263273 | -0.0724405943837207 | 0.0567891126230365 | -1.2756070844878 | 0.202094448498997 | NA | NM_145209 |
| Mir876 | 0 | NA | NA | NA | NA | NA | | Oaz1 | 0 | NA | NA | NA | NA | NA | NM_008753 |
| Mir877 | 0 | NA | NA | NA | NA | NA | | Oaz1-ps | 0 | NA | NA | NA | NA | NA | NR_027656 |
| Mir878 | 0 | NA | NA | NA | NA | NA | | Oaz3 | 2.700948297 | 0.00172103789654641 | 0.0339870056818182 | 0.0506381148330198 | 0.959613890526387 | NA | NM_016901 |
| Mir879 | 0 | NA | NA | NA | NA | NA | | Obox1 | 0 | NA | NA | NA | NA | NA | NM_027802 |
| Mir880 | 0 | NA | NA | NA | NA | NA | | Obox2 | 0 | NA | NA | NA | NA | NA | NM_145708 |
| Mir881 | 0 | NA | NA | NA | NA | NA | | Obox3 | 0 | NA | NA | NA | NA | NA | NM_145707 |
| Mir882 | 0 | NA | NA | NA | NA | NA | | Obox5 | 0.452914545 | -0.0106764464178018 | 0.0143684267899376 | -0.743049087689871 | 0.457451961798949 | NA | NM_145709 |
| Mir883a | 0 | NA | NA | NA | NA | NA | | Obox6 | 0 | NA | NA | NA | NA | NA | NM_145710 |
| Mir883b | 0 | NA | NA | NA | NA | NA | | Obp1a | 2.584336722 | -0.0160903923663518 | 0.025384687690089 | -0.633862136213479 | 0.526170797890223 | NA | NM_008754 |
| Mir9-1 | 4.354946689 | 0.154274195 | 0.13631842 | 1.131719364 | 0.257752438 | NA | | Obp2a | 1.181446803 | -0.0201409239344619 | 0.0209055903891301 | -0.963422872043559 | 0.335335352778839 | NA | NM_153558 |
| Mir9-3 | 0.574762868 | 0.049560631 | 0.055201391 | 0.897814902 | 0.369284239 | NA | | Obp2b | 2.096591503 | -0.0344758552922573 | 0.0317167956843972 | -1.08699049031669 | 0.277041012447552 | NA | NM_001099301 |
| Mir92-1 | 0 | NA | NA | NA | NA | NA | | Oc90 | 1.495963119 | -0.0469196337858909 | 0.0294655949341481 | -1.59235317972539 | 0.111305370093083 | NA | NM_010953 |
| Mir92-2 | 0 | NA | NA | NA | NA | NA | | Oca2 | 27.14640978 | 0.0139871195005607 | 0.035920952395599 | 0.389386098300512 | NA | NA | NM_021879 |
| Mir92b | 0 | NA | NA | NA | NA | NA | | Ocstamp | 0.542256936 | 0.00204518341975332 | 0.0164107438536312 | 0.124624662842494 | 0.900820702488011 | NA | NM_029021 |
| Mir93 | 0.685873859 | 0.036908102 | 0.061591586 | 0.599239353 | 0.549013284 | NA | | Odam | 0 | NA | NA | NA | NA | NA | NM_027128 |
| Mir96 | 0 | NA | NA | NA | NA | NA | | Odf1 | 1.368277527 | -0.0169678796954989 | 0.0260628733797717 | -0.651036416754733 | 0.51502297925167 | NA | NM_008757 |
| Mir98 | 0 | NA | NA | NA | NA | NA | | Odf3 | 4.861169343 | 0.000438262300846886 | 0.0488067996206821 | 0.00897953367672095 | 0.992835464998092 | NA | NM_027019 |
| Mir99a | 1.058404621 | 0.052246972 | 0.067612117 | 0.772745691 | 0.4396729 | NA | | Odf3l1 | 2.937249825 | 0.0189678142414106 | 0.0364195042884143 | 0.520814728591587 | 0.602495841799664 | NA | NM_198673 |
| Mir99b | 0.29182997 | 0.032201186 | 0.035537591 | 0.906116166 | 0.36487437 | NA | | Odf3l2 | 10.26046263 | 0.0460685854281369 | 0.0668253265950171 | 0.689388107406152 | 0.490579065372184 | NA | NM_001033473 |
| Mira | 0.957081083 | 0.088239072 | 0.078769759 | 1.120215085 | 0.262622117 | NA | | Odf4 | 12.46616216 | -0.0922539997638723 | 0.071744800047997 | -1.28586322217296 | 0.198490801052843 | NA | NM_145746 |
| Mirlet7a-1 | 1.968685763 | -0.017095067 | 0.101609264 | -0.168243196 | 0.866391959 | NA | | Ofcc1 | 0.310639869 | 0.0122746672672506 | 0.0130505022290595 | 0.940551332953199 | 0.346934831808109 | NA | NM_172143 |
| Mirlet7a-2 | 0 | NA | NA | NA | NA | NA | | Oit1 | 0 | NA | NA | NA | NA | NA | NM_146050 |
| Mirlet7c-1 | 0.079983654 | 0.016740032 | 0.027674178 | 0.604897175 | 0.54524732 | NA | | Oit3 | 7.775550235 | -0.0259037296590817 | 0.0410469090987376 | -0.631076254652227 | 0.527990666259841 | NA | NM_010959 |
| Mirlet7e | 1.250852019 | 0.088818589 | 0.081499644 | 1.089803398 | 0.275799757 | NA | | Olah | 2.174097225 | 0.0252744776513626 | 0.0328546881524461 | 0.769280704601081 | 0.44172668931506 | NA | NM_145921 |
| Mirlet7f-1 | 2.284652334 | 0.146834136 | 0.097290874 | 1.50922826 | 0.131240461 | NA | | Olfr1 | 0 | NA | NA | NA | NA | NA | NM_146921 |
| Mirlet7f-2 | 0 | NA | NA | NA | NA | NA | | Olfr10 | 0.127687691 | 0.00542298404407807 | 0.00869724871942359 | 0.62352868349814 | 0.532937159177571 | NA | NM_206822 |
| Mirlet7g | 0 | NA | NA | NA | NA | NA | | Olfr100 | 0 | NA | NA | NA | NA | NA | NM_207673 |
| Mirlet7i | 1.93374864 | -0.039596899 | 0.087544269 | -0.45230715 | 0.651047726 | NA | | Olfr1000 | 0 | NA | NA | NA | NA | NA | NM_001011695 |
| Mixl1 | 0 | NA | NA | NA | NA | NA | | Olfr1002 | 0 | NA | NA | NA | NA | NA | NM_146573 |
| Mlana | 1.605273873 | -0.035104211 | 0.061353749 | -0.572160804 | 0.567213039 | NA | | Olfr1006 | 0 | NA | NA | NA | NA | NA | NM_146570 |
| Mmp10 | 0.079983654 | 0.016740032 | 0.027674178 | 0.604897175 | 0.54524732 | NA | | Olfr1008 | 0 | NA | NA | NA | NA | NA | NM_146866 |
| Mmp12 | 2.696035589 | 0.289662924 | 0.117370765 | 2.467930788 | 0.013589659 | NA | | Olfr1009 | 0 | NA | NA | NA | NA | NA | NM_146572 |
| Mmp13 | 1.151436786 | 0.019515823 | 0.079789089 | 0.244592625 | 0.806771875 | NA | | Olfr101 | 0 | NA | NA | NA | NA | NA | NM_146834 |
| Mmp1a | 0 | NA | NA | NA | NA | NA | | Olfr1010 | 0 | NA | NA | NA | NA | NA | NM_207149 |
| Mmp1b | 0 | NA | NA | NA | NA | NA | | Olfr1012 | 0 | NA | NA | NA | NA | NA | NM_146568 |
| Mmp20 | 0.098353386 | -0.020212831 | 0.02786466 | -0.725393061 | 0.468210929 | NA | | Olfr1013 | 0 | NA | NA | NA | NA | NA | NM_146762 |
| Mmp21 | 3.056002172 | 0.345337112 | 0.123704552 | 2.791628179 | 0.005244358 | NA | | Olfr1014 | 0 | NA | NA | NA | NA | NA | NM_146569 |
| Mmp27 | 0 | NA | NA | NA | NA | NA | | Olfr1015 | 0 | NA | NA | NA | NA | NA | NM_146571 |
| Mmp3 | 0.456546308 | -0.046631236 | 0.041759841 | -1.116652618 | 0.264142884 | NA | | Olfr1016 | 0 | NA | NA | NA | NA | NA | NM_001011758 |
| Mmp7 | 0 | NA | NA | NA | NA | NA | | Olfr1018 | 0 | NA | NA | NA | NA | NA | NM_146586 |
| Mmp8 | 2.287495582 | -0.025068287 | 0.091182534 | -0.274924221 | 0.783374459 | NA | | Olfr1019 | 0 | NA | NA | NA | NA | NA | NM_147015 |
| Mmrn1 | 0.947272025 | 0.019086986 | 0.072770315 | 0.262290825 | 0.793097235 | NA | | Olfr102 | 0 | NA | NA | NA | NA | NA | NM_001011721 |
| Mnda | 5.093501151 | 0.270803598 | 0.126623554 | 2.138651066 | 0.032463938 | NA | | Olfr1020 | 0 | NA | NA | NA | NA | NA | NM_146580 |
| Mnx1 | 0.098353386 | -0.020212831 | 0.02786466 | -0.725393061 | 0.468210929 | NA | | Olfr1022 | 0 | NA | NA | NA | NA | NA | NM_146589 |
| Moap1 | 3.153424689 | -0.001762253 | 0.11992441 | -0.014694701 | 0.988275747 | NA | | Olfr1023 | 0 | NA | NA | NA | NA | NA | NM_146587 |
| Mogat1 | 3.653524889 | -0.057343705 | 0.122701267 | -0.467344033 | 0.640253757 | NA | | Olfr1024 | 0 | NA | NA | NA | NA | NA | NM_001005230 |
| Mogat2 | 0.745606615 | -0.061062738 | 0.067492909 | -0.904728193 | 0.365609401 | NA | | Olfr1026 | 0 | NA | NA | NA | NA | NA | NM_146584 |
| Morc1 | 1.391559479 | 0.015745358 | 0.061596964 | 0.255619067 | 0.798244994 | NA | | Olfr1028 | 0 | NA | NA | NA | NA | NA | NM_001011774 |
| Morn3 | 4.154172981 | 0.041431942 | 0.128366215 | 0.322763604 | 0.746874279 | NA | | Olfr1029 | 0.201873568 | 0.00673381845182595 | 0.00968011632508873 | 0.69563404257585 | 0.486658041207307 | NA | NM_001011852 |
| Mos | 3.193792803 | 0.102481655 | 0.121620599 | 0.842634027 | 0.399433154 | NA | | Olfr103 | 0 | NA | NA | NA | NA | NA | NM_146833 |
| Mospd4 | 1.28926284 | -0.115640977 | 0.084031933 | -1.37615514 | 0.168773607 | NA | | Olfr1030 | 0.127687691 | 0.00542298404407807 | 0.00869724871942359 | 0.62352868349814 | 0.532937159177571 | NA | NM_146588 |
| Mov10l1 | 0.195033442 | -0.002424765 | 0.031596358 | -0.076741901 | 0.93882887 | NA | | Olfr1031 | 0 | NA | NA | NA | NA | NA | NM_001011759 |
| Moxd2 | 0.098353386 | -0.020212831 | 0.02786466 | -0.725393061 | 0.468210929 | NA | | Olfr1032 | 0 | NA | NA | NA | NA | NA | NM_146579 |
| Mpo | 3.785362582 | -0.217414719 | 0.12679216 | -1.714733145 | 0.086394164 | NA | | Olfr1033 | 2.444877895 | -0.00269495668074595 | 0.0379671951009035 | -0.0709811897766928 | 0.943412726108008 | NA | NM_146578 |
| Mptx1 | 0 | NA | NA | NA | NA | NA | | Olfr1034 | 0.113843286 | 0.00525947738444006 | 0.00859368002043176 | 0.612016897526494 | 0.540526577873086 | NA | NM_001011872 |
| Mptx2 | 0 | NA | NA | NA | NA | NA | | Olfr1036 | 0 | NA | NA | NA | NA | NA | NM_207142 |
| Mpz | 7.078823613 | -0.169653088 | 0.131092866 | -1.294144324 | 0.195615574 | NA | | Olfr1037 | 0 | NA | NA | NA | NA | NA | NM_001011532 |
| Mrgpra1 | 0 | NA | NA | NA | NA | NA | | Olfr1038-ps | 0 | NA | NA | NA | NA | NA | NM_147013 |
| Mrgpra2a | 0 | NA | NA | NA | NA | NA | | Olfr1039 | 0 | NA | NA | NA | NA | NA | NM_001011784 |
| Mrgpra2b | 0.079983654 | 0.016740032 | 0.027674178 | 0.604897175 | 0.54524732 | NA | | Olfr1040 | 0 | NA | NA | NA | NA | NA | NM_207561 |
| Mrgpra3 | 0 | NA | NA | NA | NA | NA | | Olfr1042 | 0 | NA | NA | NA | NA | NA | NM_001011777 |
| Mrgpra4 | 0 | NA | NA | NA | NA | NA | | Olfr1043 | 0 | NA | NA | NA | NA | NA | NM_146577 |
| Mrgpra6 | 0 | NA | NA | NA | NA | NA | | Olfr1044 | 0 | NA | NA | NA | NA | NA | NM_147011 |
| Mrgpra9 | 0 | NA | NA | NA | NA | NA | | Olfr1045 | 0 | NA | NA | NA | NA | NA | NM_147017 |
| Mrgprb1 | 0.119397641 | -0.020477583 | 0.027954224 | -0.732539845 | 0.463839132 | NA | | Olfr1046 | 0 | NA | NA | NA | NA | NA | NM_146582 |
| Mrgprb2 | 0 | NA | NA | NA | NA | NA | | Olfr1047 | 0 | NA | NA | NA | NA | NA | NM_147012 |
| Mrgprb3 | 0 | NA | NA | NA | NA | NA | | Olfr1048 | 0 | NA | NA | NA | NA | NA | NM_147014 |
| Mrgprb4 | 0 | NA | NA | NA | NA | NA | | Olfr1049 | 0 | NA | NA | NA | NA | NA | NM_147016 |
| Mrgprb5 | 0 | NA | NA | NA | NA | NA | | Olfr1051 | 0 | NA | NA | NA | NA | NA | NM_207562 |
| Mrgprb8 | 0 | NA | NA | NA | NA | NA | | Olfr1052 | 0 | NA | NA | NA | NA | NA | NM_147010 |
| Mrgprd | 0.196706773 | -0.026517754 | 0.031609464 | -0.838918169 | 0.401515231 | NA | | Olfr1053 | 0 | NA | NA | NA | NA | NA | NM_001177857 |

| | | | | | | | | | | | | | | | |
|---|---|---|---|---|---|---|---|---|---|---|---|---|---|---|---|
| Mrgprg | 0 | NA | NA | NA | NA | NA | | Olfr1054 | 0 | NA | NA | NA | NA | NA | NM_147019 |
| Mrgprh | 5.56264731 | -0.119494366 | 0.146952059 | -0.813152035 | 0.416130896 | NA | | Olfr1055 | 0 | NA | NA | NA | NA | NA | NM_147021 |
| Mrgprx1 | 0 | NA | NA | NA | NA | NA | | Olfr1056 | 0 | NA | NA | NA | NA | NA | NM_147018 |
| Mrgprx2 | 0 | NA | NA | NA | NA | NA | | Olfr1057 | 0 | NA | NA | NA | NA | NA | NM_207563 |
| Mroh2a | 1.308692255 | 0.012060884 | 0.082918141 | 0.145455316 | 0.884351393 | NA | | Olfr1058 | 0 | NA | NA | NA | NA | NA | NM_146391 |
| Mroh2b | 0.408027608 | -0.004440697 | 0.036642628 | -0.121189371 | 0.903541043 | NA | | Olfr1061 | 0 | NA | NA | NA | NA | NA | NM_207134 |
| Mroh4 | 3.747974313 | 0.275704126 | 0.122883014 | 2.243630889 | 0.02485616 | NA | | Olfr1062 | 0 | NA | NA | NA | NA | NA | NM_147078 |
| Mroh9 | 0.098353386 | -0.020212831 | 0.02786466 | -0.725393061 | 0.468210929 | NA | | Olfr1065 | 0 | NA | NA | NA | NA | NA | NM_146408 |
| Mrps6 | 615.8638723 | 0.063948532 | 0.126733364 | 0.504591134 | NA | NA | | Olfr1066 | 0 | NA | NA | NA | NA | NA | NM_001011735 |
| Ms4a1 | 0.828011001 | -0.008119904 | 0.062313387 | -0.130307538 | 0.896323116 | NA | | Olfr107 | 0 | NA | NA | NA | NA | NA | NM_146511 |
| Ms4a10 | 0 | NA | NA | NA | NA | NA | | Olfr1076 | 0 | NA | NA | NA | NA | NA | NM_146406 |
| Ms4a13 | 0 | NA | NA | NA | NA | NA | | Olfr1077-ps1 | 0 | NA | NA | NA | NA | NA | NR_033507 |
| Ms4a18 | 0 | NA | NA | NA | NA | NA | | Olfr1079 | 0.108766301 | 0.00519171381361946 | 0.00856003371502575 | 0.606506234257728 | 0.544178643801544 | NA | NM_146407 |
| Ms4a2 | 0.58646488 | 0.050588692 | 0.054651257 | 0.925663823 | 0.354620714 | NA | | Olfr108 | 0 | NA | NA | NA | NA | NA | NM_146465 |
| Ms4a3 | 0.849765129 | -0.040631658 | 0.062482766 | -0.65028583 | 0.515507608 | NA | | Olfr1080 | 0 | NA | NA | NA | NA | NA | NM_146409 |
| Ms4a4b | 0.698099462 | 0.00568318 | 0.054502766 | 0.104273234 | 0.916952518 | NA | | Olfr1082 | 0 | NA | NA | NA | NA | NA | NM_207674 |
| Ms4a4c | 2.191486038 | 0.112669057 | 0.097010559 | 1.16141025 | 0.245475102 | NA | | Olfr1084 | 0 | NA | NA | NA | NA | NA | NM_207135 |
| Ms4a4d | 0.68531206 | 0.075058946 | 0.063967246 | 1.17339654 | 0.240636824 | NA | | Olfr1085 | 0 | NA | NA | NA | NA | NA | NM_146590 |
| Ms4a5 | 0 | NA | NA | NA | NA | NA | | Olfr1086 | 0 | NA | NA | NA | NA | NA | NM_146592 |
| Ms4a8a | 0.838007076 | 0.022662155 | 0.061762684 | 0.366923086 | 0.713676391 | NA | | Olfr1087 | 0 | NA | NA | NA | NA | NA | NM_146846 |
| Msgn1 | 0 | NA | NA | NA | NA | NA | | Olfr1089 | 0 | NA | NA | NA | NA | NA | NM_001011771 |
| Msh4 | 0.757306031 | 0.042257997 | 0.058686886 | 0.720058609 | 0.471488911 | NA | | Olfr109 | 0 | NA | NA | NA | NA | NA | NM_146835 |
| Msmb | 0 | NA | NA | NA | NA | NA | | Olfr1090 | 0 | NA | NA | NA | NA | NA | NM_146847 |
| Msmp | 0.897514379 | 0.004682035 | 0.069324716 | 0.067537749 | 0.946153612 | NA | | Olfr1093 | 0 | NA | NA | NA | NA | NA | NM_146366 |
| Mst1 | 0.594987921 | 0.017868075 | 0.05269327 | 0.339095951 | 0.734537448 | NA | | Olfr1094 | 0 | NA | NA | NA | NA | NA | NM_146365 |
| Msx1as | 0.185906812 | 0.024895454 | 0.031385107 | 0.793225068 | 0.427646705 | NA | | Olfr1095 | 0 | NA | NA | NA | NA | NA | NM_146730 |
| Msx3 | 3.894539214 | -0.350889337 | 0.126935976 | -2.764301712 | 0.005704477 | NA | | Olfr1097 | 0 | NA | NA | NA | NA | NA | NM_146843 |
| Mt4 | 0 | NA | NA | NA | NA | NA | | Olfr1098 | 0 | NA | NA | NA | NA | NA | NM_146845 |
| Mtap7d3 | 2.627510154 | 0.073243988 | 0.112918892 | 0.648642458 | 0.516569505 | NA | | Olfr1099 | 0 | NA | NA | NA | NA | NA | NM_146768 |
| Mtl5 | 4.998152067 | 0.213281802 | 0.1428941 | 1.492586483 | 0.135545464 | NA | | Olfr11 | 0 | NA | NA | NA | NA | NA | NM_146542 |
| Mtnr1a | 0.638084606 | 0.0288786 | 0.053770188 | 0.537074568 | 0.591216107 | NA | | Olfr110 | 0.960368879 | -0.0248313253657796 | 0.0182612482128621 | -1.35978247906898 | 0.173898768483643 | NA | NM_146328 |
| Mtnr1b | 0 | NA | NA | NA | NA | NA | | Olfr1100 | 0 | NA | NA | NA | NA | NA | NM_146594 |
| Muc13 | 1.682814255 | 0.013843078 | 0.087901789 | 0.157483463 | 0.87486384 | NA | | Olfr1101 | 0 | NA | NA | NA | NA | NA | NM_146591 |
| Muc15 | 5.511925791 | 0.004302542 | 0.147961621 | 0.029078767 | 0.97680177 | NA | | Olfr1102 | 0 | NA | NA | NA | NA | NA | NM_207154 |
| Muc19 | 0.498271657 | 0.024449561 | 0.039505462 | 0.618890637 | 0.535988407 | NA | | Olfr1104 | 0 | NA | NA | NA | NA | NA | NM_146767 |
| Muc20 | 0.633091198 | -0.008942341 | 0.058752393 | -0.152203852 | 0.879026156 | NA | | Olfr1105 | 0 | NA | NA | NA | NA | NA | NM_001011825 |
| Muc4 | 0.699494611 | -0.026379251 | 0.065754389 | -0.401178553 | 0.68828867 | NA | | Olfr1106 | 0 | NA | NA | NA | NA | NA | NM_146752 |
| Muc5ac | 0.894578973 | -0.028540078 | 0.073878633 | -0.386310311 | 0.699266865 | NA | | Olfr1107 | 0 | NA | NA | NA | NA | NA | NM_146844 |
| Muc5b | 0.079983654 | 0.016740032 | 0.027674178 | 0.604897175 | 0.54524732 | NA | | Olfr1109 | 0 | NA | NA | NA | NA | NA | NM_146766 |
| Muc6 | 0.181709657 | 0.024794574 | 0.031325423 | 0.791516008 | 0.428642938 | NA | | Olfr111 | 4.028216858 | -0.0794718487316535 | 0.0381923607268569 | -2.08083101487279 | 0.0374493762434598 | NA | NM_001005485 |
| Mug-ps1 | 0 | NA | NA | NA | NA | NA | | Olfr1110 | 0 | NA | NA | NA | NA | NA | NM_146769 |
| Mug1 | 0 | NA | NA | NA | NA | NA | | Olfr1111 | 0 | NA | NA | NA | NA | NA | NM_146593 |
| Mug2 | 0.08731684 | 0.017365056 | 0.027733412 | 0.626142072 | 0.531221759 | NA | | Olfr1112 | 0 | NA | NA | NA | NA | NA | NM_146661 |
| Mup1 | 0.098353386 | -0.020212831 | 0.02786466 | -0.725393061 | 0.468210929 | NA | | Olfr1113 | 0 | NA | NA | NA | NA | NA | NM_207565 |
| Mup10 | 0 | NA | NA | NA | NA | NA | | Olfr1115 | 0 | NA | NA | NA | NA | NA | NM_146297 |
| Mup11 | 0.08731684 | 0.017365056 | 0.027733412 | 0.626142072 | 0.531221759 | NA | | Olfr1116-ps | 0 | NA | NA | NA | NA | NA | NM_001011734 |
| Mup12 | 0 | NA | NA | NA | NA | NA | | Olfr1118 | 0 | NA | NA | NA | NA | NA | NM_207632 |
| Mup13 | 0 | NA | NA | NA | NA | NA | | Olfr112 | 0 | NA | NA | NA | NA | NA | NM_001013575 |
| Mup14 | 0 | NA | NA | NA | NA | NA | | Olfr1120 | 0 | NA | NA | NA | NA | NA | NM_147029 |
| Mup15 | 0 | NA | NA | NA | NA | NA | | Olfr1121 | 0 | NA | NA | NA | NA | NA | NM_146348 |
| Mup16 | 0 | NA | NA | NA | NA | NA | | Olfr1122 | 0 | NA | NA | NA | NA | NA | NM_147031 |
| Mup17 | 0 | NA | NA | NA | NA | NA | | Olfr1123 | 0 | NA | NA | NA | NA | NA | NM_146350 |
| Mup19 | 0 | NA | NA | NA | NA | NA | | Olfr1124 | 0 | NA | NA | NA | NA | NA | NM_147028 |
| Mup2 | 0 | NA | NA | NA | NA | NA | | Olfr1126 | 0.120209049 | -0.00674972139751468 | 0.00863521114606269 | -0.781650996523957 | 0.434419707908266 | NA | NM_146837 |
| Mup20 | 0.119397641 | -0.020477583 | 0.027954224 | -0.732539845 | 0.463839132 | NA | | Olfr1128 | 0 | NA | NA | NA | NA | NA | NM_146349 |
| Mup21 | 0 | NA | NA | NA | NA | NA | | Olfr1129 | 0 | NA | NA | NA | NA | NA | NM_001011836 |
| Mup3 | 0.480379018 | 0.040041947 | 0.051818222 | 0.772738725 | 0.439677024 | NA | | Olfr113 | 0 | NA | NA | NA | NA | NA | NM_146289 |
| Mup4 | 1.151479737 | -0.029335583 | 0.052330001 | -0.56058823 | 0.575078277 | NA | | Olfr1130 | 0 | NA | NA | NA | NA | NA | NM_146838 |
| Mup5 | 6.084986267 | 0.067297747 | 0.135854788 | 0.495365297 | 0.620342286 | NA | | Olfr1131 | 0.106148608 | 0.00515554063321497 | 0.00854200771766808 | 0.603551390213729 | 0.546141939141592 | NA | NM_146658 |
| Mup6 | 0.116696964 | -0.020477583 | 0.027954224 | -0.732539847 | 0.463839131 | NA | | Olfr1132 | 0 | NA | NA | NA | NA | NA | NM_146836 |
| Mup7 | 0 | NA | NA | NA | NA | NA | | Olfr1133 | 0 | NA | NA | NA | NA | NA | NM_146351 |
| Mup8 | 0 | NA | NA | NA | NA | NA | | Olfr1134 | 0 | NA | NA | NA | NA | NA | NM_147030 |
| Mup9 | 0 | NA | NA | NA | NA | NA | | Olfr1135 | 0 | NA | NA | NA | NA | NA | NM_146660 |
| Mybphl | 1.896426157 | 0.002717667 | 0.090605352 | 0.029994549 | 0.9760714 | NA | | Olfr1136 | 0 | NA | NA | NA | NA | NA | NM_146659 |
| Mycs | 0.453998628 | 0.026589972 | 0.045454031 | 0.584986 | 0.55855713 | NA | | Olfr1137 | 0 | NA | NA | NA | NA | NA | NM_001011833 |
| Myf5 | 0 | NA | NA | NA | NA | NA | | Olfr1138 | 0 | NA | NA | NA | NA | NA | NM_146639 |
| Myf6 | 0 | NA | NA | NA | NA | NA | | Olfr114 | 0 | NA | NA | NA | NA | NA | NM_146287 |
| Myh6 | 1.403832196 | -0.044467476 | 0.085158715 | -0.522171763 | 0.601550744 | NA | | Olfr1140 | 0.309462921 | 0.00239536941478217 | 0.0109127344998192 | 0.219502216866162 | 0.826258853162846 | NA | NM_146642 |
| Myl1 | 6.913453372 | 0.158164195 | 0.151182927 | 1.046177625 | 0.295479031 | NA | | Olfr1141 | 0 | NA | NA | NA | NA | NA | NM_146637 |
| Myl10 | 0.08731684 | 0.017365056 | 0.027733412 | 0.626142072 | 0.531221759 | NA | | Olfr1143 | 0 | NA | NA | NA | NA | NA | NM_146293 |
| Myl3 | 2.212555087 | 0.002935246 | 0.090917319 | 0.032284788 | 0.97424494 | NA | | Olfr1145 | 0 | NA | NA | NA | NA | NA | NM_146320 |
| Myl7 | 0.59419716 | 0.024548688 | 0.058662177 | 0.418475568 | 0.675599446 | NA | | Olfr1148 | 0 | NA | NA | NA | NA | NA | NM_001011519 |
| Mylk2 | 0.69074833 | 0.079811121 | 0.055198329 | 1.445897412 | 0.148205978 | NA | | Olfr115 | 0 | NA | NA | NA | NA | NA | NM_001011753 |
| Myo3a | 1.316026059 | 0.023699925 | 0.077516712 | 0.305739554 | 0.759802955 | NA | | Olfr1151 | 0 | NA | NA | NA | NA | NA | NM_146638 |
| Myo7b | 4.585168708 | 0.099602232 | 0.132212434 | 0.753349961 | 0.451239644 | NA | | Olfr1152 | 0 | NA | NA | NA | NA | NA | NM_001011834 |
| Myod1 | 3.454667478 | 0.054640306 | 0.11792442 | 0.463350219 | 0.643113355 | NA | | Olfr1153 | 0 | NA | NA | NA | NA | NA | NM_146640 |
| Myog | 0.535761119 | -0.039416426 | 0.047223402 | -0.834679942 | 0.403897938 | NA | | Olfr1154 | 0 | NA | NA | NA | NA | NA | NM_146647 |
| Myom2 | 6.623072989 | -0.070833386 | 0.152056726 | -0.465835267 | 0.641333419 | NA | | Olfr1155 | 0 | NA | NA | NA | NA | NA | NM_146643 |
| Myot | 2.399949825 | -0.322176302 | 0.110278779 | -2.921471435 | 0.003483822 | NA | | Olfr1156 | 0 | NA | NA | NA | NA | NA | NM_146817 |
| Myoz1 | 1.091107073 | -0.088395639 | 0.080114811 | -1.103362008 | 0.269869988 | NA | | Olfr1157 | 0 | NA | NA | NA | NA | NA | NM_146849 |
| Myoz2 | 2.173228614 | 0.010710975 | 0.10458328 | 0.102415745 | 0.918426686 | NA | | Olfr1158 | 0 | NA | NA | NA | NA | NA | NM_146645 |
| Myoz3 | 0.863460983 | -0.006944181 | 0.062492514 | -0.111120205 | 0.911521027 | NA | | Olfr116 | 0 | NA | NA | NA | NA | NA | NM_146632 |
| Myrfl | 5.680828004 | -0.212031363 | 0.143188575 | -1.480784078 | 0.138664121 | NA | | Olfr1160 | 0 | NA | NA | NA | NA | NA | NM_146649 |
| Mzb1 | 3.839294082 | 0.012605687 | 0.119707059 | 0.105304458 | 0.916134226 | NA | | Olfr1161 | 0 | NA | NA | NA | NA | NA | NM_146848 |
| Naa11 | 0.789799633 | 0.020365278 | 0.068182092 | 0.298689542 | 0.765176937 | NA | | Olfr1162 | 0 | NA | NA | NA | NA | NA | NM_001011835 |
| Naaladl1 | 6.947671097 | -0.206554886 | 0.150085579 | -1.376247385 | 0.168745056 | NA | | Olfr1163 | 0 | NA | NA | NA | NA | NA | NM_146644 |
| Nags | 3.153450993 | -0.008749018 | 0.125862701 | -0.069512399 | 0.944581763 | NA | | Olfr1164 | 0 | NA | NA | NA | NA | NA | NM_146641 |
| Naip7 | 0.334447991 | -0.050120769 | 0.043104878 | -1.162763289 | 0.244925556 | NA | | Olfr1166 | 0 | NA | NA | NA | NA | NA | NM_146650 |
| Nanog | 1.060817981 | 0.072334201 | 0.068997377 | 1.048361611 | 0.294472033 | NA | | Olfr1167 | 0 | NA | NA | NA | NA | NA | NM_146294 |
| Napsa | 4.658449301 | -0.41834785 | 0.140281831 | -2.982195533 | 0.002861892 | NA | | Olfr1168 | 0 | NA | NA | NA | NA | NA | NM_146531 |
| Nat3 | 0 | NA | NA | NA | NA | NA | | Olfr117 | 0 | NA | NA | NA | NA | NA | NM_207155 |
| Nccrp1 | 4.830353489 | -0.190264658 | 0.120531099 | -1.578552428 | 0.114438754 | NA | | Olfr1170 | 0 | NA | NA | NA | NA | NA | NM_146532 |
| Ncr1 | 0.472604946 | 0.035675895 | 0.051617383 | 0.691160469 | 0.489464703 | NA | | Olfr1173 | 0 | NA | NA | NA | NA | NA | NM_207566 |
| Nctc1 | 0 | NA | NA | NA | NA | NA | | Olfr1176 | 0 | NA | NA | NA | NA | NA | NM_146771 |
| Ndufs5 | 0 | NA | NA | NA | NA | NA | | Olfr1178 | 0 | NA | NA | NA | NA | NA | NM_001011868 |
| Neil3 | 5.018182292 | -0.212206768 | 0.13556121 | -1.565394466 | 0.117490445 | NA | | Olfr1179 | 0 | NA | NA | NA | NA | NA | NM_146917 |
| Nepn | 1.19780767 | 0.01177341 | 0.067964026 | 0.173230031 | 0.862470619 | NA | | Olfr118 | 0 | NA | NA | NA | NA | NA | NM_213721 |
| Nespas | 2.015159464 | 0.063619504 | 0.078921413 | 0.806112077 | 0.420178229 | NA | | Olfr1180 | 0 | NA | NA | NA | NA | NA | NM_146918 |
| Neurog1 | 3.385506913 | -0.031957493 | 0.117260483 | -0.272534212 | 0.785211289 | NA | | Olfr1181 | 0 | NA | NA | NA | NA | NA | NM_001011816 |
| Neurog2 | 7.943221544 | 0.06639644 | 0.154118602 | 0.430813927 | 0.666603672 | NA | | Olfr1182 | 0 | NA | NA | NA | NA | NA | NM_001011535 |
| Neurog3 | 2.230123776 | -0.011449681 | 0.09433143 | -0.12137716 | 0.903392307 | NA | | Olfr1183 | 0 | NA | NA | NA | NA | NA | NM_146529 |
| Nfe2 | 2.013152324 | 0.001533329 | 0.094740235 | 0.016184557 | 0.987087155 | NA | | Olfr1184 | 0 | NA | NA | NA | NA | NA | NM_146823 |
| Nipal1 | 0.378164117 | -0.004452483 | 0.047147495 | -0.094437328 | 0.924761765 | NA | | Olfr1186 | 0 | NA | NA | NA | NA | NA | NM_146530 |
| Nipsnap3a | 0 | NA | NA | NA | NA | NA | | Olfr1188 | 0 | NA | NA | NA | NA | NA | NM_146919 |
| Nkapl | 7.805105387 | 0.041705858 | 0.154498785 | 0.269942955 | 0.78720414 | NA | | Olfr1189 | 0 | NA | NA | NA | NA | NA | NM_146772 |
| Nkg7 | 1.097492992 | 0.10270995 | 0.078095624 | 1.315181875 | 0.188448785 | NA | | Olfr119 | 0 | NA | NA | NA | NA | NA | NM_001011830 |
| Nkx1-2 | 1.490843325 | 0.055405805 | 0.081854543 | 0.676881241 | 0.498481307 | NA | | Olfr1193 | 0 | NA | NA | NA | NA | NA | NM_001011517 |
| Nkx2-3 | 0.28821313 | -0.029561117 | 0.033427731 | -0.884329147 | 0.376518559 | NA | | Olfr1195 | 0 | NA | NA | NA | NA | NA | NM_146753 |
| Nkx2-4 | 0 | NA | NA | NA | NA | NA | | Olfr1196 | 0 | NA | NA | NA | NA | NA | NM_146464 |
| Nkx2-5 | 0 | NA | NA | NA | NA | NA | | Olfr1197 | 0 | NA | NA | NA | NA | NA | NM_001005225 |
| Nkx2-6 | 0 | NA | NA | NA | NA | NA | | Olfr1198 | 0 | NA | NA | NA | NA | NA | NM_207567 |
| Nkx3-2 | 0 | NA | NA | NA | NA | NA | | Olfr1199 | 0 | NA | NA | NA | NA | NA | NM_146458 |
| Nkx6-1 | 3.139826244 | 0.08464138 | 0.108929772 | 0.777027055 | 0.437142809 | NA | | Olfr12 | 0.09572496 | 0.00498255289994511 | 0.00847999972823692 | 0.587565219295241 | 0.556824162160132 | NA | NM_206896 |
| Nkx6-3 | 1.619407367 | 0.059613199 | 0.095760045 | 0.622526851 | 0.533595494 | NA | | Olfr120 | 0 | NA | NA | NA | NA | NA | NM_146631 |
| Nlrc5 | 74.26540573 | 0.334813377 | 0.144531556 | 2.316541709 | NA | NA | | Olfr1200 | 0 | NA | NA | NA | NA | NA | NM_001005227 |
| Nlrp12 | 1.391324294 | 0.029897338 | 0.087895529 | 0.340146295 | 0.73374636 | NA | | Olfr1201 | 0 | NA | NA | NA | NA | NA | NM_146895 |
| Nlrp14 | 0 | NA | NA | NA | NA | NA | | Olfr1202 | 0 | NA | NA | NA | NA | NA | NM_146462 |
| Nlrp2 | 2.613462047 | -0.065946138 | 0.114997382 | -0.573457735 | 0.566334808 | NA | | Olfr1204 | 0 | NA | NA | NA | NA | NA | NM_146463 |
| Nlrp3 | 53.20753095 | 0.210094283 | 0.164305906 | 1.278677605 | NA | NA | | Olfr1205 | 0 | NA | NA | NA | NA | NA | NM_146896 |
| Nlrp4a | 0 | NA | NA | NA | NA | NA | | Olfr1206 | 0 | NA | NA | NA | NA | NA | NM_001001810 |
| Nlrp4b | 0 | NA | NA | NA | NA | NA | | Olfr1208 | 0.106347839 | -0.00653347234140708 | 0.00854389509723638 | -0.764694822098226 | 0.444453290197145 | NA | NM_146778 |
| Nlrp4c | 0.08731684 | 0.017365056 | 0.027733412 | 0.626142072 | 0.531221759 | NA | | Olfr1209 | 0 | NA | NA | NA | NA | NA | NM_146461 |
| Nlrp4e | 0 | NA | NA | NA | NA | NA | | Olfr121 | 0.106347839 | -0.00653347234140708 | 0.00854389509723638 | -0.764694822098226 | 0.444453290197145 | NA | NM_146629 |
| Nlrp4f | 4.652014542 | 0.111635616 | 0.134249304 | 0.831554525 | 0.40566044 | NA | | Olfr1211 | 0.120209049 | -0.00674972139751468 | 0.00863521114606269 | -0.781650996523957 | 0.434419707908266 | NA | NM_001011804 |
| Nlrp5 | 0.167300494 | 0.024299763 | 0.031030539 | 0.783091891 | 0.433573153 | NA | | Olfr1212 | 0 | NA | NA | NA | NA | NA | NM_207140 |
| Nlrp9a | 0 | NA | NA | NA | NA | NA | | Olfr1213 | 0 | NA | NA | NA | NA | NA | NM_146898 |
| Nlrp9b | 0.099009884 | 0.018205726 | 0.027896865 | 0.652608315 | 0.514008823 | NA | | Olfr1214 | 0 | NA | NA | NA | NA | NA | NM_146897 |
| Nlrp9c | 0 | NA | NA | NA | NA | NA | | Olfr1215 | 0 | NA | NA | NA | NA | NA | NM_146459 |
| Nme8 | 0.195080928 | -0.000361156 | 0.031609476 | -0.011425558 | 0.990883922 | NA | | Olfr1216 | 0 | NA | NA | NA | NA | NA | NM_146893 |
| Nme9 | 5.450462961 | 0.225083681 | 0.119367065 | 1.885643088 | 0.059343067 | NA | | Olfr1217 | 0 | NA | NA | NA | NA | NA | NM_146901 |
| Nmrk2 | 3.515925956 | 0.172381982 | 0.107719335 | 1.600288213 | 0.10953466 | NA | | Olfr1218 | 0 | NA | NA | NA | NA | NA | NM_146818 |
| Nms | 1.56081153 | -0.040459704 | 0.086896442 | -0.465608296 | 0.641495904 | NA | | Olfr1219 | 0 | NA | NA | NA | NA | NA | NM_146899 |
| Nmu | 0.698265872 | -0.052029371 | 0.062441752 | -0.833246491 | 0.404705728 | NA | | Olfr122 | 0 | NA | NA | NA | NA | NA | NM_146288 |
| Nmur1 | 4.628174789 | -0.19431463 | 0.124295954 | -1.563322252 | 0.117976817 | NA | | Olfr1220 | 0 | NA | NA | NA | NA | NA | NM_146900 |
| Nmur2 | 1.441198444 | -0.071728574 | 0.083144879 | -0.862693826 | 0.388305828 | NA | | Olfr1221 | 0 | NA | NA | NA | NA | NA | NM_146902 |
| Nobox | 0.609036663 | 0.027795583 | 0.058866341 | 0.472181252 | 0.63679742 | NA | | Olfr1222 | 0 | NA | NA | NA | NA | NA | NM_001011860 |
| Noto | 0 | NA | NA | NA | NA | NA | | Olfr1223 | 0 | NA | NA | NA | NA | NA | NM_146892 |
| Nox1 | 1.646020596 | -0.126685703 | 0.099899233 | -1.26813489 | 0.204749784 | NA | | Olfr1225 | 0 | NA | NA | NA | NA | NA | NM_146891 |
| Nox3 | 0.592096564 | -0.004291432 | 0.052941551 | -0.081059815 | 0.935394384 | NA | | Olfr1226 | 0 | NA | NA | NA | NA | NA | NM_146967 |
| Nox4 | 4.143342874 | -0.018443496 | 0.12100948 | -0.152413645 | 0.878860696 | NA | | Olfr1228 | 0 | NA | NA | NA | NA | NA | NM_146971 |
| Noxa1 | 0.079983654 | 0.016740032 | 0.027674178 | 0.604897175 | 0.54524732 | NA | | Olfr1229 | 0 | NA | NA | NA | NA | NA | NM_001011761 |
| Npb | 4.868376212 | 0.13055971 | 0.141309944 | 0.923924435 | 0.35552566 | NA | | Olfr123 | 0.12890128 | -0.0068640067066208 | 0.00870804083610955 | -0.788237771940376 | 0.430557639008166 | NA | NM_146630 |
| Npcd | 0 | NA | NA | NA | NA | NA | | Olfr1230 | 0 | NA | NA | NA | NA | NA | NM_146789 |
| Npm2 | 7.918720856 | 0.251925001 | 0.155487655 | 1.620225092 | 0.105183933 | NA | | Olfr1231 | 0 | NA | NA | NA | NA | NA | NM_146454 |
| Npm3-ps1 | 0.832578476 | -0.037757824 | 0.068913294 | -0.547903343 | 0.583758273 | NA | | Olfr1232 | 0 | NA | NA | NA | NA | NA | NM_146323 |
| Nppb | 0.215468684 | -0.027008432 | 0.031909031 | -0.846419662 | 0.397318674 | NA | | Olfr1233 | 0 | NA | NA | NA | NA | NA | NM_146972 |
| Nps | 0.556402103 | 0.086806466 | 0.057717414 | 1.503990893 | 0.132583712 | NA | | Olfr1234 | 0 | NA | NA | NA | NA | NA | NM_146973 |
| Npvf | 1.236846178 | 0.019351132 | 0.073871169 | 0.261957839 | 0.793353947 | NA | | Olfr1238 | 0 | NA | NA | NA | NA | NA | NM_146790 |
| Npw | 1.693065885 | -0.004491479 | 0.082043825 | -0.054744878 | 0.956341715 | NA | | Olfr1239 | 0 | NA | NA | NA | NA | NA | NM_146970 |
| Npy6r | 0.513115595 | -0.045873581 | 0.053031465 | -0.86502571 | 0.387024678 | NA | | Olfr124 | 0 | NA | NA | NA | NA | NA | NM_147062 |
| Nr0b1 | 0.294043327 | 0.010508917 | 0.039092051 | 0.2688249 | 0.788064431 | NA | | Olfr1240 | 0 | NA | NA | NA | NA | NA | NM_146808 |
| Nr0b2 | 0.094392817 | 0.017895942 | 0.027818137 | 0.643319236 | 0.520016977 | NA | | Olfr1241 | 0 | NA | NA | NA | NA | NA | NM_146455 |
| Nr1h4 | 4.708145276 | -0.080325303 | 0.139081023 | -0.577543231 | 0.563572543 | NA | | Olfr1242 | 0 | NA | NA | NA | NA | NA | NM_146968 |
| Nr1h5 | 0.895456348 | 0.009786724 | 0.056538074 | 0.17309971 | 0.862573053 | NA | | Olfr1243 | 0 | NA | NA | NA | NA | NA | NM_146969 |
| Nr1i2 | 0.47910454 | -0.016023567 | 0.050473258 | -0.317466479 | 0.750889671 | NA | | Olfr1245 | 0 | NA | NA | NA | NA | NA | NM_146788 |
| Nr1i3 | 3.697592198 | -0.084714827 | 0.126150329 | -0.671538691 | 0.50187742 | NA | | Olfr1246 | 0 | NA | NA | NA | NA | NA | NM_146792 |
| Nr2e3 | 0.197363271 | -0.000561558 | 0.031650166 | -0.017742642 | 0.985844163 | NA | | Olfr1247 | 0 | NA | NA | NA | NA | NA | NM_146966 |
| Nr5a1 | 2.325661084 | 0.022814063 | 0.099116043 | 0.230175278 | 0.817955571 | NA | | Olfr1248 | 0 | NA | NA | NA | NA | NA | NM_146791 |
| Nr5a2 | 1.262916509 | -0.002348899 | 0.080299129 | -0.029251859 | 0.976663722 | NA | | Olfr1249 | 0 | NA | NA | NA | NA | NA | NM_001011796 |
| Nrk | 1.510314622 | -0.114129947 | 0.068192908 | -1.673633672 | 0.094202617 | NA | | Olfr125 | 0 | NA | NA | NA | NA | NA | NM_146290 |
| Nron | 0.623077958 | -0.025656387 | 0.047703389 | -0.537831528 | 0.590693363 | NA | | Olfr1250 | 0 | NA | NA | NA | NA | NA | NM_146965 |
| Nrp | 4.814916409 | -0.049398025 | 0.144472626 | -0.34191962 | 0.732411384 | NA | | Olfr1251 | 0 | NA | NA | NA | NA | NA | NM_001011529 |
| Nt5c1b | 0.345637534 | 0.019330126 | 0.041637051 | 0.464253003 | 0.642466491 | NA | | Olfr1252 | 0 | NA | NA | NA | NA | NA | NM_207568 |
| Ntf5 | 4.669376945 | 0.011842635 | 0.12996755 | 0.091119935 | 0.927397292 | NA | | Olfr1253 | 0 | NA | NA | NA | NA | NA | NM_146373 |
| Ntrk2 | 24371.76408 | -0.034132687 | 0.095398685 | -0.357789908 | NA | NA | | Olfr1254 | 0 | NA | NA | NA | NA | NA | NM_146476 |
| Nuggc | 2.196805493 | 0.065764734 | 0.105089153 | 0.625799448 | 0.531446494 | NA | | Olfr1255 | 0 | NA | NA | NA | NA | NA | NM_146977 |
| Nup62-il4i1 | 3.424502979 | 0.13945278 | 0.114487382 | 1.218062442 | 0.223200245 | NA | | Olfr1256 | 0 | NA | NA | NA | NA | NA | NM_146983 |
| Nup62cl | 5.384119915 | 0.217158713 | 0.140382346 | 1.546908989 | 0.121885193 | NA | | Olfr1257 | 0 | NA | NA | NA | NA | NA | NM_146982 |
| Nutf2-ps1 | 0 | NA | NA | NA | NA | NA | | Olfr1258 | 0 | NA | NA | NA | NA | NA | NM_146978 |
| Nxf2 | 2.731299807 | -0.210425303 | 0.114537431 | -1.837174986 | 0.066184067 | NA | | Olfr1259 | 0 | NA | NA | NA | NA | NA | NM_146341 |
| Nxf3 | 4.464267893 | 0.280771482 | 0.126178935 | 2.225185069 | 0.026068817 | NA | | Olfr126 | 0 | NA | NA | NA | NA | NA | NM_146890 |
| Nxnl1 | 4.194395197 | 0.014482128 | 0.128491486 | 0.112708856 | 0.91026138 | NA | | Olfr1260 | 0 | NA | NA | NA | NA | NA | NM_146981 |
| Nxnl2 | 4.299733494 | 0.067201784 | 0.125121139 | 0.537093769 | 0.591202845 | NA | | Olfr1261 | 0 | NA | NA | NA | NA | NA | NM_146474 |
| Nxpe5 | 0.482637559 | -0.049175986 | 0.042867026 | -1.14717514 | 0.251309244 | NA | | Olfr1262 | 0 | NA | NA | NA | NA | NA | NM_146974 |
| OTTMUSG00000016609 | 0 | NA | NA | NA | NA | NA | | Olfr1263 | 0 | NA | NA | NA | NA | NA | NM_146794 |
| Oas1d | 0.748706893 | 0.039722983 | 0.056595821 | 0.701871312 | 0.482759421 | NA | | Olfr1264 | 0 | NA | NA | NA | NA | NA | NM_021368 |
| Oas1e | 0.384284173 | -0.031469349 | 0.034522933 | -0.911549113 | 0.362006121 | NA | | Olfr1265 | 0 | NA | NA | NA | NA | NA | NM_146343 |
| Oas1f | 0.19046386 | -0.000774927 | 0.031526133 | -0.024580468 | 0.980389599 | NA | | Olfr1269 | 0 | NA | NA | NA | NA | NA | NM_146342 |
| Oas1g | 3.14379104 | 0.04901949 | 0.09227846 | 0.531212698 | 0.595271395 | NA | | Olfr127 | 0 | NA | NA | NA | NA | NA | NM_146377 |
| Oas1h | 0.575354256 | -0.004736431 | 0.046231168 | -0.102451031 | 0.91839868 | NA | | Olfr1270 | 0 | NA | NA | NA | NA | NA | NM_146985 |
| Oas3 | 3.864437366 | 0.060347289 | 0.120403974 | 0.501206783 | 0.616225601 | NA | | Olfr1271 | 0 | NA | NA | NA | NA | NA | NM_146793 |
| Oaz1 | 0 | NA | NA | NA | NA | NA | | Olfr1272 | 0 | NA | NA | NA | NA | NA | NM_146980 |
| Oaz1-ps | 0 | NA | NA | NA | NA | NA | | Olfr1273-ps | 0 | NA | NA | NA | NA | NA | NM_146975 |
| Oaz3 | 3.082645033 | 0.125802511 | 0.103314688 | 1.217663369 | 0.223351923 | NA | | Olfr1274-ps | 0 | NA | NA | NA | NA | NA | NM_146263 |
| Obox1 | 0 | NA | NA | NA | NA | NA | | Olfr1275 | 0 | NA | NA | NA | NA | NA | NM_001011795 |
| Obox2 | 0 | NA | NA | NA | NA | NA | | Olfr1276 | 0 | NA | NA | NA | NA | NA | NM_146395 |
| Obox3 | 0 | NA | NA | NA | NA | NA | | Olfr1277 | 0 | NA | NA | NA | NA | NA | NM_146396 |
| Obox5 | 0.393053211 | 0.021363521 | 0.042959842 | 0.497290494 | 0.618984214 | NA | | Olfr1278 | 0.120209049 | -0.00674972139751468 | 0.00863521114606269 | -0.781650996523957 | 0.434419707908266 | NA | NM_146394 |
| Obox6 | 0 | NA | NA | NA | NA | NA | | Olfr1279 | 0 | NA | NA | NA | NA | NA | NM_146393 |
| Obp1a | 1.264166687 | 0.110742242 | 0.064169661 | 1.725772584 | 0.084388334 | NA | | Olfr128 | 0 | NA | NA | NA | NA | NA | NM_206816 |
| Obp2a | 0.914950978 | 0.096606065 | 0.069296257 | 1.39410221 | 0.163286742 | NA | | Olfr1280 | 0 | NA | NA | NA | NA | NA | NM_146908 |
| Obp2b | 1.992791227 | 0.094284877 | 0.089429586 | 1.05429178 | 0.291749351 | NA | | Olfr1281 | 0.120209049 | -0.00674972139751468 | 0.00863521114606269 | -0.781650996523957 | 0.434419707908266 | NA | NM_001005568 |
| Oc90 | 2.799342917 | 0.002494517 | 0.12108851 | 0.020600777 | 0.983564121 | NA | | Olfr1282 | 0 | NA | NA | NA | NA | NA | NM_146907 |
| Oca2 | 16.29520498 | 0.102392568 | 0.148485906 | 0.68957769 | NA | NA | | Olfr1283 | 0 | NA | NA | NA | NA | NA | NM_207236 |
| Ociad2 | 3605.454997 | -0.003095451 | 0.103416732 | -0.029931821 | NA | NA | | Olfr1284 | 0 | NA | NA | NA | NA | NA | NM_146381 |
| Ocstamp | 0.582208971 | -0.023855297 | 0.047658271 | -0.500548931 | 0.616688612 | NA | | Olfr1286 | 0 | NA | NA | NA | NA | NA | NM_207254 |
| Odam | 0 | NA | NA | NA | NA | NA | | Olfr1287 | 0 | NA | NA | NA | NA | NA | NM_001011773 |
| Odf1 | 1.331330041 | 0.041256347 | 0.071099425 | 0.580262741 | 0.561737449 | NA | | Olfr1288 | 0 | NA | NA | NA | NA | NA | NM_146400 |
| Odf3 | 2.868109075 | 0.091449229 | 0.112579873 | 0.81230531 | 0.416616464 | NA | | Olfr1289 | 0 | NA | NA | NA | NA | NA | NM_146404 |
| Odf3l1 | 1.819315222 | 0.04220695 | 0.094135522 | 0.448363688 | 0.653890744 | NA | | Olfr129 | 0 | NA | NA | NA | NA | NA | NM_146327 |
| Odf3l2 | 7.508541522 | -0.01851026 | 0.156537286 | -0.118248251 | 0.905870959 | NA | | Olfr1290 | 0 | NA | NA | NA | NA | NA | NM_146262 |
| Ofcc1 | 0.116696964 | -0.020477583 | 0.027954224 | -0.732539847 | 0.463839131 | NA | | Olfr1294 | 0 | NA | NA | NA | NA | NA | NM_146885 |
| Oit1 | 0.098353386 | -0.020212831 | 0.02786466 | -0.725393061 | 0.468210929 | NA | | Olfr1295 | 0 | NA | NA | NA | NA | NA | NM_146403 |
| Oit3 | 5.117316794 | 0.212135524 | 0.124677522 | 1.701473695 | 0.088854073 | NA | | Olfr1297 | 0 | NA | NA | NA | NA | NA | NM_146888 |
| Olah | 1.314458349 | 0.119766747 | 0.080139039 | 1.49448693 | 0.135048395 | NA | | Olfr1298 | 0 | NA | NA | NA | NA | NA | NM_146886 |
| Olfr1 | 0 | NA | NA | NA | NA | NA | | Olfr1299 | 0 | NA | NA | NA | NA | NA | NM_146884 |
| Olfr10 | 0.188785634 | 0.025025111 | 0.031461603 | 0.79541754 | 0.42637066 | NA | | Olfr13 | 0 | NA | NA | NA | NA | NA | NM_146652 |
| Olfr100 | 0 | NA | NA | NA | NA | NA | | Olfr130 | 0 | NA | NA | NA | NA | NA | NM_146487 |
| Olfr1000 | 0 | NA | NA | NA | NA | NA | | Olfr1300-ps1 | 0 | NA | NA | NA | NA | NA | NR_033534 |
| Olfr1002 | 0 | NA | NA | NA | NA | NA | | Olfr1301 | 0 | NA | NA | NA | NA | NA | NM_146887 |
| Olfr1006 | 0 | NA | NA | NA | NA | NA | | Olfr1302 | 0 | NA | NA | NA | NA | NA | NM_146889 |
| Olfr1008 | 0 | NA | NA | NA | NA | NA | | Olfr1303 | 0 | NA | NA | NA | NA | NA | NM_146402 |
| Olfr1009 | 0 | NA | NA | NA | NA | NA | | Olfr1305 | 0 | NA | NA | NA | NA | NA | NM_146401 |
| Olfr101 | 0 | NA | NA | NA | NA | NA | | Olfr1306 | 0 | NA | NA | NA | NA | NA | NM_001011803 |
| Olfr1010 | 0 | NA | NA | NA | NA | NA | | Olfr1307 | 0 | NA | NA | NA | NA | NA | NM_001011787 |
| Olfr1012 | 0 | NA | NA | NA | NA | NA | | Olfr1308 | 0 | NA | NA | NA | NA | NA | NM_207151 |
| Olfr1013 | 0 | NA | NA | NA | NA | NA | | Olfr1309 | 0 | NA | NA | NA | NA | NA | NM_146447 |
| Olfr1014 | 0 | NA | NA | NA | NA | NA | | Olfr131 | 0 | NA | NA | NA | NA | NA | NM_146867 |
| Olfr1015 | 0 | NA | NA | NA | NA | NA | | Olfr1310 | 0 | NA | NA | NA | NA | NA | NM_146449 |
| Olfr1016 | 0 | NA | NA | NA | NA | NA | | Olfr1311 | 0 | NA | NA | NA | NA | NA | NM_146274 |
| Olfr1018 | 0 | NA | NA | NA | NA | NA | | Olfr1312 | 0 | NA | NA | NA | NA | NA | NM_146362 |
| Olfr1019 | 0 | NA | NA | NA | NA | NA | | Olfr1313 | 0 | NA | NA | NA | NA | NA | NM_207150 |
| Olfr102 | 0 | NA | NA | NA | NA | NA | | Olfr1314 | 0 | NA | NA | NA | NA | NA | NM_146450 |
| Olfr1020 | 0 | NA | NA | NA | NA | NA | | Olfr1316 | 0 | NA | NA | NA | NA | NA | NM_146742 |
| Olfr1022 | 0 | NA | NA | NA | NA | NA | | Olfr1317 | 0 | NA | NA | NA | NA | NA | NM_146448 |
| Olfr1023 | 0 | NA | NA | NA | NA | NA | | Olfr1318 | 0 | NA | NA | NA | NA | NA | NM_001011802 |
| Olfr1024 | 0 | NA | NA | NA | NA | NA | | Olfr132 | 0 | NA | NA | NA | NA | NA | NM_001005481 |
| Olfr1026 | 0 | NA | NA | NA | NA | NA | | Olfr1320 | 0 | NA | NA | NA | NA | NA | NM_207240 |
| Olfr1028 | 0 | NA | NA | NA | NA | NA | | Olfr1321 | 0 | NA | NA | NA | NA | NA | NM_207631 |
| Olfr1029 | 0.098353386 | -0.020212831 | 0.02786466 | -0.725393061 | 0.468210929 | NA | | Olfr1322 | 0 | NA | NA | NA | NA | NA | NM_001011794 |
| Olfr103 | 0 | NA | NA | NA | NA | NA | | Olfr1323 | 0 | NA | NA | NA | NA | NA | NM_146390 |
| Olfr1030 | 0 | NA | NA | NA | NA | NA | | Olfr1324 | 0 | NA | NA | NA | NA | NA | NM_146292 |
| Olfr1031 | 0 | NA | NA | NA | NA | NA | | Olfr1325 | 0 | NA | NA | NA | NA | NA | NM_146398 |
| Olfr1032 | 0 | NA | NA | NA | NA | NA | | Olfr1328 | 0 | NA | NA | NA | NA | NA | NM_146399 |
| Olfr1033 | 5.192019563 | -0.140129054 | 0.131726217 | -1.063790171 | 0.287423765 | NA | | Olfr1329 | 0 | NA | NA | NA | NA | NA | NM_001011870 |

| | | | | | | | | | | | | | | | |
|---|---|---|---|---|---|---|---|---|---|---|---|---|---|---|---|
| Olfr1034 | 0.188785634 | 0.025025111 | 0.031461603 | 0.79541754 | 0.42637066 | NA | | Olfr133 | 0 | NA | NA | NA | NA | NA | NM_146831 |
| Olfr1036 | 0 | NA | NA | NA | NA | NA | | Olfr1330 | 0 | NA | NA | NA | NA | NA | NM_146334 |
| Olfr1037 | 0 | NA | NA | NA | NA | NA | | Olfr1331 | 0 | NA | NA | NA | NA | NA | NM_001011856 |
| Olfr1038-ps | 0 | NA | NA | NA | NA | NA | | Olfr1333 | 0 | NA | NA | NA | NA | NA | NM_207157 |
| Olfr1039 | 0 | NA | NA | NA | NA | NA | | Olfr1335 | 0.108766301 | 0.00519171381361946 | 0.00856003371502575 | 0.606506234257728 | 0.544178643801544 | NA | NM_207703 |
| Olfr1040 | 0 | NA | NA | NA | NA | NA | | Olfr1336 | 0 | NA | NA | NA | NA | NA | NM_146915 |
| Olfr1042 | 0 | NA | NA | NA | NA | NA | | Olfr1337 | 0 | NA | NA | NA | NA | NA | NM_146309 |
| Olfr1043 | 0 | NA | NA | NA | NA | NA | | Olfr1338 | 0 | NA | NA | NA | NA | NA | NM_207152 |
| Olfr1044 | 0 | NA | NA | NA | NA | NA | | Olfr1339 | 0 | NA | NA | NA | NA | NA | NM_146852 |
| Olfr1045 | 0 | NA | NA | NA | NA | NA | | Olfr134 | 0 | NA | NA | NA | NA | NA | NM_146832 |
| Olfr1046 | 0 | NA | NA | NA | NA | NA | | Olfr1340 | 0.09572496 | 0.00498255289994511 | 0.00847999972823692 | 0.587565219295241 | 0.556824162160132 | NA | NM_146304 |
| Olfr1047 | 0 | NA | NA | NA | NA | NA | | Olfr1341 | 0 | NA | NA | NA | NA | NA | NM_146853 |
| Olfr1048 | 0 | NA | NA | NA | NA | NA | | Olfr1342 | 0.106347839 | -0.00653347234140708 | 0.00854389509723638 | -0.764694822098226 | 0.444453290197145 | NA | NM_146713 |
| Olfr1049 | 0 | NA | NA | NA | NA | NA | | Olfr1346 | 0.326538963 | -0.00431616909593923 | 0.0111843034107492 | -0.385913090643711 | 0.699561035292352 | NA | NM_146916 |
| Olfr1051 | 0 | NA | NA | NA | NA | NA | | Olfr1347 | 0.427100848 | 0.0157809700572465 | 0.0147784426861069 | 1.06783714579629 | 0.285593978988929 | NA | NM_146385 |
| Olfr1052 | 0 | NA | NA | NA | NA | NA | | Olfr1348 | 0 | NA | NA | NA | NA | NA | NM_146913 |
| Olfr1053 | 0 | NA | NA | NA | NA | NA | | Olfr1349 | 0 | NA | NA | NA | NA | NA | NM_207136 |
| Olfr1054 | 0 | NA | NA | NA | NA | NA | | Olfr135 | 0 | NA | NA | NA | NA | NA | NM_146332 |
| Olfr1055 | 0 | NA | NA | NA | NA | NA | | Olfr1350 | 0.24789674 | -0.000769869626663926 | 0.0100368571301073 | -0.0767042528038554 | 0.938858821262798 | NA | NM_146389 |
| Olfr1056 | 0 | NA | NA | NA | NA | NA | | Olfr1351 | 0 | NA | NA | NA | NA | NA | NM_147040 |
| Olfr1057 | 0 | NA | NA | NA | NA | NA | | Olfr1352 | 0 | NA | NA | NA | NA | NA | NM_147071 |
| Olfr1058 | 0 | NA | NA | NA | NA | NA | | Olfr1353 | 0 | NA | NA | NA | NA | NA | NM_147042 |
| Olfr1061 | 0 | NA | NA | NA | NA | NA | | Olfr1354 | 0 | NA | NA | NA | NA | NA | NM_001199840 |
| Olfr1062 | 0 | NA | NA | NA | NA | NA | | Olfr1355 | 0 | NA | NA | NA | NA | NA | NM_207571 |
| Olfr1065 | 0.098353386 | -0.020212831 | 0.02786466 | -0.725393061 | 0.468210929 | NA | | Olfr1356 | 0 | NA | NA | NA | NA | NA | NM_146308 |
| Olfr1066 | 0 | NA | NA | NA | NA | NA | | Olfr1357 | 0 | NA | NA | NA | NA | NA | NM_001011737 |
| Olfr107 | 0 | NA | NA | NA | NA | NA | | Olfr1359 | 0 | NA | NA | NA | NA | NA | NM_001011820 |
| Olfr1076 | 0 | NA | NA | NA | NA | NA | | Olfr136 | 0 | NA | NA | NA | NA | NA | NM_146807 |
| Olfr1077-ps1 | 0 | NA | NA | NA | NA | NA | | Olfr1360 | 0 | NA | NA | NA | NA | NA | NM_146543 |
| Olfr1079 | 0 | NA | NA | NA | NA | NA | | Olfr1361 | 0 | NA | NA | NA | NA | NA | NM_146541 |
| Olfr108 | 0 | NA | NA | NA | NA | NA | | Olfr1362 | 0 | NA | NA | NA | NA | NA | NM_146744 |
| Olfr1080 | 0 | NA | NA | NA | NA | NA | | Olfr1364 | 0 | NA | NA | NA | NA | NA | NM_146540 |
| Olfr1082 | 0 | NA | NA | NA | NA | NA | | Olfr1366 | 0 | NA | NA | NA | NA | NA | NM_146283 |
| Olfr1084 | 0 | NA | NA | NA | NA | NA | | Olfr1367 | 0 | NA | NA | NA | NA | NA | NM_146533 |
| Olfr1085 | 0 | NA | NA | NA | NA | NA | | Olfr1368 | 0 | NA | NA | NA | NA | NA | NM_146534 |
| Olfr1086 | 0 | NA | NA | NA | NA | NA | | Olfr137 | 0 | NA | NA | NA | NA | NA | NM_146488 |
| Olfr1087 | 0 | NA | NA | NA | NA | NA | | Olfr1370 | 0 | NA | NA | NA | NA | NA | NM_146535 |
| Olfr1089 | 0 | NA | NA | NA | NA | NA | | Olfr1371 | 0.097165704 | -0.00637828455464458 | 0.00848781691535724 | -0.75146349388194 | 0.452373763047372 | NA | NM_207253 |
| Olfr109 | 0 | NA | NA | NA | NA | NA | | Olfr1372-ps1 | 9.365635562 | 0.0761493515233942 | 0.0621619238516489 | 1.22501600344814 | 0.220569190846135 | NA | NR_034155 |
| Olfr1090 | 0 | NA | NA | NA | NA | NA | | Olfr1373 | 0 | NA | NA | NA | NA | NA | NM_207227 |
| Olfr1093 | 0 | NA | NA | NA | NA | NA | | Olfr1377 | 0.908957928 | 0.015486899935371 | 0.0224371785098518 | 0.690233842395601 | 0.490047144309203 | NA | NM_146911 |
| Olfr1094 | 0 | NA | NA | NA | NA | NA | | Olfr1378 | 0.753219214 | -0.00528043076947627 | 0.018076444703034 | -0.292116666536202 | 0.770197424120193 | NA | NM_146910 |
| Olfr1095 | 0 | NA | NA | NA | NA | NA | | Olfr138 | 0 | NA | NA | NA | NA | NA | NM_130868 |
| Olfr1097 | 0 | NA | NA | NA | NA | NA | | Olfr1380 | 0 | NA | NA | NA | NA | NA | NM_207573 |
| Olfr1098 | 0 | NA | NA | NA | NA | NA | | Olfr1381 | 0 | NA | NA | NA | NA | NA | NM_146469 |
| Olfr1099 | 0 | NA | NA | NA | NA | NA | | Olfr1382 | 0 | NA | NA | NA | NA | NA | NM_001011790 |
| Olfr11 | 0 | NA | NA | NA | NA | NA | | Olfr1383 | 0 | NA | NA | NA | NA | NA | NM_207574 |
| Olfr110 | 1.747658581 | -0.023432014 | 0.068517071 | -0.341987971 | 0.732359945 | NA | | Olfr1384 | 0 | NA | NA | NA | NA | NA | NM_146472 |
| Olfr1100 | 0 | NA | NA | NA | NA | NA | | Olfr1385 | 0 | NA | NA | NA | NA | NA | NM_001011805 |
| Olfr1101 | 0 | NA | NA | NA | NA | NA | | Olfr1386 | 0 | NA | NA | NA | NA | NA | NM_001011741 |
| Olfr1102 | 0 | NA | NA | NA | NA | NA | | Olfr1387 | 0 | NA | NA | NA | NA | NA | NM_146473 |
| Olfr1104 | 0 | NA | NA | NA | NA | NA | | Olfr1388 | 0 | NA | NA | NA | NA | NA | NM_146467 |
| Olfr1105 | 0 | NA | NA | NA | NA | NA | | Olfr1389 | 0.240418098 | -0.0088029805258837 | 0.00986414519853158 | -0.892422034419582 | 0.372166765031741 | NA | NM_147066 |
| Olfr1106 | 0 | NA | NA | NA | NA | NA | | Olfr139 | 0 | NA | NA | NA | NA | NA | NM_147003 |
| Olfr1107 | 0 | NA | NA | NA | NA | NA | | Olfr1390 | 0 | NA | NA | NA | NA | NA | NM_147065 |
| Olfr1109 | 0 | NA | NA | NA | NA | NA | | Olfr1391 | 0 | NA | NA | NA | NA | NA | NM_146468 |
| Olfr111 | 3.236110612 | 0.119609163 | 0.086359032 | 1.385022039 | 0.166045728 | NA | | Olfr1392 | 2.685136898 | 0.0174313168694868 | 0.0386092190988816 | 0.451480689750385 | 0.651643139426204 | NA | NM_146470 |
| Olfr1110 | 0 | NA | NA | NA | NA | NA | | Olfr1393 | 19.97134508 | -0.0236066158679452 | 0.0755566891507087 | -0.312435816514649 | 0.754709334089754 | NA | NM_146471 |
| Olfr1111 | 0 | NA | NA | NA | NA | NA | | Olfr1394 | 0 | NA | NA | NA | NA | NA | NM_146276 |
| Olfr1112 | 0 | NA | NA | NA | NA | NA | | Olfr1395 | 0.097165704 | -0.00637828455464458 | 0.00848781691535724 | -0.75146349388194 | 0.452373763047372 | NA | NM_146877 |
| Olfr1113 | 0 | NA | NA | NA | NA | NA | | Olfr1396 | 0.545002712 | 0.00110646631130581 | 0.0146471586700009 | 0.0755413617230752 | 0.939783989634957 | NA | NM_146337 |
| Olfr1115 | 0 | NA | NA | NA | NA | NA | | Olfr140 | 0 | NA | NA | NA | NA | NA | NM_020515 |
| Olfr1116-ps | 0 | NA | NA | NA | NA | NA | | Olfr1402 | 0 | NA | NA | NA | NA | NA | NM_146275 |
| Olfr1118 | 0 | NA | NA | NA | NA | NA | | Olfr1404 | 0 | NA | NA | NA | NA | NA | NM_146881 |
| Olfr112 | 0 | NA | NA | NA | NA | NA | | Olfr1406 | 0 | NA | NA | NA | NA | NA | NM_146763 |
| Olfr1120 | 0 | NA | NA | NA | NA | NA | | Olfr1408 | 0 | NA | NA | NA | NA | NA | NM_146764 |
| Olfr1121 | 0 | NA | NA | NA | NA | NA | | Olfr141 | 0 | NA | NA | NA | NA | NA | NM_181818 |
| Olfr1122 | 0 | NA | NA | NA | NA | NA | | Olfr1410 | 0 | NA | NA | NA | NA | NA | NM_146491 |
| Olfr1123 | 0 | NA | NA | NA | NA | NA | | Olfr1411 | 0 | NA | NA | NA | NA | NA | NM_146490 |
| Olfr1124 | 0 | NA | NA | NA | NA | NA | | Olfr1412 | 0 | NA | NA | NA | NA | NA | NM_146277 |
| Olfr1126 | 0.099009884 | 0.018205726 | 0.027896865 | 0.652608315 | 0.514008823 | NA | | Olfr1413 | 0 | NA | NA | NA | NA | NA | NM_147037 |
| Olfr1128 | 0 | NA | NA | NA | NA | NA | | Olfr1414 | 0 | NA | NA | NA | NA | NA | NM_147039 |
| Olfr1129 | 0 | NA | NA | NA | NA | NA | | Olfr1415 | 0 | NA | NA | NA | NA | NA | NM_001011525 |
| Olfr113 | 0 | NA | NA | NA | NA | NA | | Olfr1416 | 0 | NA | NA | NA | NA | NA | NM_147038 |
| Olfr1130 | 0 | NA | NA | NA | NA | NA | | Olfr1417 | 0 | NA | NA | NA | NA | NA | NM_146936 |
| Olfr1131 | 0 | NA | NA | NA | NA | NA | | Olfr1418 | 0 | NA | NA | NA | NA | NA | NM_001011524 |
| Olfr1132 | 0 | NA | NA | NA | NA | NA | | Olfr1419 | 0.599454723 | 0.00491859512176551 | 0.016959373639411 | 0.290022215816711 | 0.771799241909812 | NA | NM_001011775 |
| Olfr1133 | 0 | NA | NA | NA | NA | NA | | Olfr142 | 0 | NA | NA | NA | NA | NA | NM_146984 |
| Olfr1134 | 0 | NA | NA | NA | NA | NA | | Olfr1420 | 0 | NA | NA | NA | NA | NA | NM_146410 |
| Olfr1135 | 0 | NA | NA | NA | NA | NA | | Olfr1423 | 0 | NA | NA | NA | NA | NA | NM_146680 |
| Olfr1136 | 0 | NA | NA | NA | NA | NA | | Olfr1424 | 0 | NA | NA | NA | NA | NA | NM_146681 |
| Olfr1137 | 0 | NA | NA | NA | NA | NA | | Olfr1425 | 0 | NA | NA | NA | NA | NA | NM_001011853 |
| Olfr1138 | 0 | NA | NA | NA | NA | NA | | Olfr1426 | 0 | NA | NA | NA | NA | NA | NM_146809 |
| Olfr114 | 0 | NA | NA | NA | NA | NA | | Olfr1427 | 0 | NA | NA | NA | NA | NA | NM_146679 |
| Olfr1140 | 0.079983654 | 0.016740032 | 0.027674178 | 0.604897175 | 0.54524732 | NA | | Olfr1428 | 0.09572496 | 0.00498255289994511 | 0.00847999972823692 | 0.587565219295241 | 0.556824162160132 | NA | NM_146678 |
| Olfr1141 | 0 | NA | NA | NA | NA | NA | | Olfr143 | 0 | NA | NA | NA | NA | NA | NM_146806 |
| Olfr1143 | 0 | NA | NA | NA | NA | NA | | Olfr1431 | 0 | NA | NA | NA | NA | NA | NM_146414 |
| Olfr1145 | 0 | NA | NA | NA | NA | NA | | Olfr1433 | 0 | NA | NA | NA | NA | NA | NM_146685 |
| Olfr1148 | 0 | NA | NA | NA | NA | NA | | Olfr1434 | 0 | NA | NA | NA | NA | NA | NM_207626 |
| Olfr115 | 0 | NA | NA | NA | NA | NA | | Olfr1436 | 0 | NA | NA | NA | NA | NA | NM_146687 |
| Olfr1151 | 0 | NA | NA | NA | NA | NA | | Olfr1437 | 0.106347839 | -0.00653347234140708 | 0.00854389509723638 | -0.764694822098226 | 0.444453290197145 | NA | NM_001011839 |
| Olfr1152 | 0 | NA | NA | NA | NA | NA | | Olfr1440 | 0 | NA | NA | NA | NA | NA | NM_146684 |
| Olfr1153 | 0 | NA | NA | NA | NA | NA | | Olfr1441 | 0 | NA | NA | NA | NA | NA | NM_146683 |
| Olfr1154 | 0 | NA | NA | NA | NA | NA | | Olfr1442 | 0 | NA | NA | NA | NA | NA | NM_146697 |
| Olfr1155 | 0 | NA | NA | NA | NA | NA | | Olfr1443 | 0 | NA | NA | NA | NA | NA | NM_146698 |
| Olfr1156 | 0 | NA | NA | NA | NA | NA | | Olfr1444 | 0.120209049 | -0.00674972139751468 | 0.00863521114606269 | -0.781650996523957 | 0.434419707908266 | NA | NM_146702 |
| Olfr1157 | 0 | NA | NA | NA | NA | NA | | Olfr1445 | 0 | NA | NA | NA | NA | NA | NM_146699 |
| Olfr1158 | 0 | NA | NA | NA | NA | NA | | Olfr1446 | 0 | NA | NA | NA | NA | NA | NM_146704 |
| Olfr116 | 0 | NA | NA | NA | NA | NA | | Olfr1447 | 0 | NA | NA | NA | NA | NA | NM_146703 |
| Olfr1160 | 0 | NA | NA | NA | NA | NA | | Olfr1448 | 0 | NA | NA | NA | NA | NA | NM_146701 |
| Olfr1161 | 0 | NA | NA | NA | NA | NA | | Olfr1449 | 0 | NA | NA | NA | NA | NA | NM_146303 |
| Olfr1162 | 0 | NA | NA | NA | NA | NA | | Olfr145 | 0 | NA | NA | NA | NA | NA | NM_146313 |
| Olfr1163 | 0 | NA | NA | NA | NA | NA | | Olfr1450 | 0 | NA | NA | NA | NA | NA | NM_146371 |
| Olfr1164 | 0 | NA | NA | NA | NA | NA | | Olfr1451 | 0 | NA | NA | NA | NA | NA | NM_146705 |
| Olfr1166 | 0 | NA | NA | NA | NA | NA | | Olfr1453 | 0 | NA | NA | NA | NA | NA | NM_146700 |
| Olfr1167 | 0 | NA | NA | NA | NA | NA | | Olfr1454 | 0 | NA | NA | NA | NA | NA | NM_146692 |
| Olfr1168 | 0 | NA | NA | NA | NA | NA | | Olfr1457 | 0 | NA | NA | NA | NA | NA | NM_146575 |
| Olfr117 | 0 | NA | NA | NA | NA | NA | | Olfr1459 | 0 | NA | NA | NA | NA | NA | NM_146689 |
| Olfr1170 | 0 | NA | NA | NA | NA | NA | | Olfr146 | 0 | NA | NA | NA | NA | NA | NM_146747 |
| Olfr1173 | 0 | NA | NA | NA | NA | NA | | Olfr1461 | 0 | NA | NA | NA | NA | NA | NM_146302 |
| Olfr1176 | 0 | NA | NA | NA | NA | NA | | Olfr1462 | 0 | NA | NA | NA | NA | NA | NM_146693 |
| Olfr1178 | 0 | NA | NA | NA | NA | NA | | Olfr1463 | 0 | NA | NA | NA | NA | NA | NM_001011840 |
| Olfr1179 | 0 | NA | NA | NA | NA | NA | | Olfr1465 | 0 | NA | NA | NA | NA | NA | NM_001011841 |
| Olfr118 | 0 | NA | NA | NA | NA | NA | | Olfr1466 | 0 | NA | NA | NA | NA | NA | NM_146694 |
| Olfr1180 | 0 | NA | NA | NA | NA | NA | | Olfr1467 | 0 | NA | NA | NA | NA | NA | NM_146691 |
| Olfr1181 | 0 | NA | NA | NA | NA | NA | | Olfr1469 | 0 | NA | NA | NA | NA | NA | NM_146695 |
| Olfr1182 | 0 | NA | NA | NA | NA | NA | | Olfr147 | 0 | NA | NA | NA | NA | NA | NM_146869 |
| Olfr1183 | 0 | NA | NA | NA | NA | NA | | Olfr1471 | 0 | NA | NA | NA | NA | NA | NM_207132 |
| Olfr1184 | 0 | NA | NA | NA | NA | NA | | Olfr1472 | 0 | NA | NA | NA | NA | NA | NM_146690 |
| Olfr1186 | 0 | NA | NA | NA | NA | NA | | Olfr1474 | 0 | NA | NA | NA | NA | NA | NM_001011842 |
| Olfr1188 | 0 | NA | NA | NA | NA | NA | | Olfr1475 | 0 | NA | NA | NA | NA | NA | NM_146301 |
| Olfr1189 | 0 | NA | NA | NA | NA | NA | | Olfr1477 | 0 | NA | NA | NA | NA | NA | NM_146696 |
| Olfr119 | 0.098353386 | -0.020212831 | 0.02786466 | -0.725393061 | 0.468210929 | NA | | Olfr148 | 0 | NA | NA | NA | NA | NA | NM_146505 |
| Olfr1193 | 0.233393928 | -0.027475306 | 0.032191697 | -0.85349047 | 0.39338737 | NA | | Olfr1480 | 0 | NA | NA | NA | NA | NA | NM_207575 |
| Olfr1195 | 0.098353386 | -0.020212831 | 0.02786466 | -0.725393061 | 0.468210929 | NA | | Olfr1484 | 0 | NA | NA | NA | NA | NA | NM_146291 |
| Olfr1196 | 0 | NA | NA | NA | NA | NA | | Olfr1487 | 0 | NA | NA | NA | NA | NA | NM_146636 |
| Olfr1197 | 0 | NA | NA | NA | NA | NA | | Olfr1489 | 0 | NA | NA | NA | NA | NA | NM_146635 |
| Olfr1198 | 0 | NA | NA | NA | NA | NA | | Olfr149 | 0 | NA | NA | NA | NA | NA | NM_207138 |
| Olfr1199 | 0 | NA | NA | NA | NA | NA | | Olfr1490 | 0 | NA | NA | NA | NA | NA | NM_001011832 |
| Olfr12 | 0.215050351 | -0.027006021 | 0.031907566 | -0.846382992 | 0.397339123 | NA | | Olfr1491 | 0 | NA | NA | NA | NA | NA | NM_146345 |
| Olfr120 | 0 | NA | NA | NA | NA | NA | | Olfr1494 | 0 | NA | NA | NA | NA | NA | NM_146990 |
| Olfr1200 | 0 | NA | NA | NA | NA | NA | | Olfr1495 | 0 | NA | NA | NA | NA | NA | NM_146344 |
| Olfr1201 | 0 | NA | NA | NA | NA | NA | | Olfr1496 | 0 | NA | NA | NA | NA | NA | NM_146989 |
| Olfr1202 | 0 | NA | NA | NA | NA | NA | | Olfr1497 | 0 | NA | NA | NA | NA | NA | NM_146741 |
| Olfr1204 | 0 | NA | NA | NA | NA | NA | | Olfr1499 | 0 | NA | NA | NA | NA | NA | NM_146796 |
| Olfr1205 | 0.098353386 | -0.020212831 | 0.02786466 | -0.725393061 | 0.468210929 | NA | | Olfr15 | 0 | NA | NA | NA | NA | NA | NM_008762 |
| Olfr1206 | 0 | NA | NA | NA | NA | NA | | Olfr150 | 0.097165704 | -0.00637828455464458 | 0.00848781691535724 | -0.75146349388194 | 0.452373763047372 | NA | NM_146609 |
| Olfr1208 | 0.08731684 | 0.017365056 | 0.027733412 | 0.626142072 | 0.531221759 | NA | | Olfr1500 | 0 | NA | NA | NA | NA | NA | NM_001011831 |
| Olfr1209 | 0 | NA | NA | NA | NA | NA | | Olfr1501 | 0 | NA | NA | NA | NA | NA | NM_146633 |
| Olfr121 | 0.08731684 | 0.017365056 | 0.027733412 | 0.626142072 | 0.531221759 | NA | | Olfr1502 | 0 | NA | NA | NA | NA | NA | NM_146797 |
| Olfr1211 | 0.099009884 | 0.018205726 | 0.027896865 | 0.652608315 | 0.514008823 | NA | | Olfr1504 | 0 | NA | NA | NA | NA | NA | NM_146634 |
| Olfr1212 | 0 | NA | NA | NA | NA | NA | | Olfr1505 | 0 | NA | NA | NA | NA | NA | NM_001011850 |
| Olfr1213 | 0 | NA | NA | NA | NA | NA | | Olfr1506 | 0 | NA | NA | NA | NA | NA | NM_146265 |
| Olfr1214 | 0 | NA | NA | NA | NA | NA | | Olfr1507 | 2.123502352 | 0.00867673309324299 | 0.0249793599237675 | 0.347356101986713 | 0.728323808085531 | NA | NM_001170918,NM_020512 |
| Olfr1215 | 0 | NA | NA | NA | NA | NA | | Olfr1508 | 0.108766301 | 0.00519171381361946 | 0.00856003371502575 | 0.606506234257728 | 0.544178643801544 | NA | NM_020513 |
| Olfr1216 | 0 | NA | NA | NA | NA | NA | | Olfr1509 | 0 | NA | NA | NA | NA | NA | NM_020514 |
| Olfr1217 | 0 | NA | NA | NA | NA | NA | | Olfr151 | 0 | NA | NA | NA | NA | NA | NM_207664 |
| Olfr1218 | 0 | NA | NA | NA | NA | NA | | Olfr1510 | 0 | NA | NA | NA | NA | NA | NM_146431 |
| Olfr1219 | 0 | NA | NA | NA | NA | NA | | Olfr1511 | 0 | NA | NA | NA | NA | NA | NM_146271 |
| Olfr122 | 0 | NA | NA | NA | NA | NA | | Olfr1512 | 0 | NA | NA | NA | NA | NA | NM_146432 |
| Olfr1220 | 0 | NA | NA | NA | NA | NA | | Olfr1513 | 0 | NA | NA | NA | NA | NA | NM_001012269 |
| Olfr1221 | 0 | NA | NA | NA | NA | NA | | Olfr152 | 0 | NA | NA | NA | NA | NA | NM_146646 |
| Olfr1222 | 0 | NA | NA | NA | NA | NA | | Olfr153 | 0.443599587 | -0.00196617251786174 | 0.0138699370766132 | -0.141757854199425 | 0.887271275268317 | NA | NM_206823 |
| Olfr1223 | 0.098353386 | -0.020212831 | 0.02786466 | -0.725393061 | 0.468210929 | NA | | Olfr1532-ps1 | 0 | NA | NA | NA | NA | NA | NM_001011542 |
| Olfr1225 | 0 | NA | NA | NA | NA | NA | | Olfr1535 | 0 | NA | NA | NA | NA | NA | NM_207572 |
| Olfr1226 | 0.098353386 | -0.020212831 | 0.02786466 | -0.725393061 | 0.468210929 | NA | | Olfr1537 | 0 | NA | NA | NA | NA | NA | NM_207665 |
| Olfr1228 | 0.295060159 | -0.029687054 | 0.033501022 | -0.886153701 | 0.375534709 | NA | | Olfr154 | 0 | NA | NA | NA | NA | NA | NM_013728 |
| Olfr1229 | 0 | NA | NA | NA | NA | NA | | Olfr155 | 0 | NA | NA | NA | NA | NA | NM_019473 |
| Olfr123 | 0.105923158 | 0.018524693 | 0.027977518 | 0.662127842 | 0.507889294 | NA | | Olfr156 | 0 | NA | NA | NA | NA | NA | NM_019474 |
| Olfr1230 | 0 | NA | NA | NA | NA | NA | | Olfr157 | 0 | NA | NA | NA | NA | NA | NM_019475 |
| Olfr1231 | 0 | NA | NA | NA | NA | NA | | Olfr159 | 0 | NA | NA | NA | NA | NA | NM_019476 |
| Olfr1232 | 0 | NA | NA | NA | NA | NA | | Olfr16 | 0 | NA | NA | NA | NA | NA | NM_008763 |
| Olfr1233 | 0 | NA | NA | NA | NA | NA | | Olfr160 | 0 | NA | NA | NA | NA | NA | NM_030553 |
| Olfr1234 | 0 | NA | NA | NA | NA | NA | | Olfr161 | 0 | NA | NA | NA | NA | NA | NM_146860 |
| Olfr1238 | 0 | NA | NA | NA | NA | NA | | Olfr164 | 0 | NA | NA | NA | NA | NA | NM_146451 |
| Olfr1239 | 0 | NA | NA | NA | NA | NA | | Olfr165 | 0 | NA | NA | NA | NA | NA | NM_146466 |
| Olfr124 | 0 | NA | NA | NA | NA | NA | | Olfr166 | 0.255375383 | 0.00708304372802858 | 0.00992471678592415 | 0.713677163873753 | 0.475426830432214 | NA | NM_147068 |
| Olfr1240 | 0 | NA | NA | NA | NA | NA | | Olfr167 | 0 | NA | NA | NA | NA | NA | NM_146935 |
| Olfr1241 | 0 | NA | NA | NA | NA | NA | | Olfr168 | 0 | NA | NA | NA | NA | NA | NM_146357 |
| Olfr1242 | 0 | NA | NA | NA | NA | NA | | Olfr169 | 0 | NA | NA | NA | NA | NA | NM_001011855 |
| Olfr1243 | 0 | NA | NA | NA | NA | NA | | Olfr17 | 0 | NA | NA | NA | NA | NA | NM_020598 |
| Olfr1245 | 0 | NA | NA | NA | NA | NA | | Olfr170 | 0 | NA | NA | NA | NA | NA | NM_146957 |
| Olfr1246 | 0 | NA | NA | NA | NA | NA | | Olfr171 | 0 | NA | NA | NA | NA | NA | NM_146958 |
| Olfr1247 | 0 | NA | NA | NA | NA | NA | | Olfr172 | 0 | NA | NA | NA | NA | NA | NM_147001 |
| Olfr1248 | 0 | NA | NA | NA | NA | NA | | Olfr173 | 0 | NA | NA | NA | NA | NA | NM_147000 |
| Olfr1249 | 0 | NA | NA | NA | NA | NA | | Olfr175-ps1 | 0 | NA | NA | NA | NA | NA | NM_147002 |
| Olfr125 | 0 | NA | NA | NA | NA | NA | | Olfr176 | 0 | NA | NA | NA | NA | NA | NM_146993 |
| Olfr1250 | 0 | NA | NA | NA | NA | NA | | Olfr177 | 0 | NA | NA | NA | NA | NA | NM_146996 |
| Olfr1251 | 0 | NA | NA | NA | NA | NA | | Olfr178 | 0 | NA | NA | NA | NA | NA | NM_146997 |
| Olfr1252 | 0.192142086 | -0.026384592 | 0.031527717 | -0.836869715 | 0.402665805 | NA | | Olfr179 | 0 | NA | NA | NA | NA | NA | NM_207693 |
| Olfr1253 | 0 | NA | NA | NA | NA | NA | | Olfr18 | 0.24789674 | -0.000769869626663926 | 0.0100368571301073 | -0.0767042528038554 | 0.938858821262798 | NA | NM_146563 |
| Olfr1254 | 0 | NA | NA | NA | NA | NA | | Olfr180 | 0 | NA | NA | NA | NA | NA | NM_001011662 |
| Olfr1255 | 0 | NA | NA | NA | NA | NA | | Olfr181 | 0 | NA | NA | NA | NA | NA | NM_146999 |
| Olfr1256 | 0 | NA | NA | NA | NA | NA | | Olfr183 | 0 | NA | NA | NA | NA | NA | NM_146485 |
| Olfr1257 | 0 | NA | NA | NA | NA | NA | | Olfr186 | 0 | NA | NA | NA | NA | NA | NM_146321 |
| Olfr1258 | 0 | NA | NA | NA | NA | NA | | Olfr187 | 0 | NA | NA | NA | NA | NA | NM_146322 |
| Olfr1259 | 0.196706773 | -0.026517754 | 0.031609464 | -0.838918169 | 0.401515231 | NA | | Olfr19 | 0.255375383 | 0.00708304372802858 | 0.00992471678592415 | 0.713677163873753 | 0.475426830432214 | NA | NM_146335 |
| Olfr126 | 0 | NA | NA | NA | NA | NA | | Olfr190 | 0 | NA | NA | NA | NA | NA | NM_146397 |
| Olfr1260 | 0 | NA | NA | NA | NA | NA | | Olfr191 | 0 | NA | NA | NA | NA | NA | NM_001011807 |
| Olfr1261 | 0.196706773 | -0.026517754 | 0.031609464 | -0.838918169 | 0.401515231 | NA | | Olfr192 | 0 | NA | NA | NA | NA | NA | NM_207549 |
| Olfr1262 | 0 | NA | NA | NA | NA | NA | | Olfr193 | 0 | NA | NA | NA | NA | NA | NM_001011791 |
| Olfr1263 | 0 | NA | NA | NA | NA | NA | | Olfr194 | 0 | NA | NA | NA | NA | NA | NM_001005524 |
| Olfr1264 | 0 | NA | NA | NA | NA | NA | | Olfr195 | 0 | NA | NA | NA | NA | NA | NM_146998 |
| Olfr1265 | 0 | NA | NA | NA | NA | NA | | Olfr196 | 0 | NA | NA | NA | NA | NA | NM_146779 |
| Olfr1269 | 0 | NA | NA | NA | NA | NA | | Olfr197 | 0 | NA | NA | NA | NA | NA | NM_146484 |
| Olfr127 | 0 | NA | NA | NA | NA | NA | | Olfr198 | 0 | NA | NA | NA | NA | NA | NM_001011808 |
| Olfr1270 | 0.107716603 | -0.020477583 | 0.027954224 | -0.732539849 | 0.46383913 | NA | | Olfr199 | 0 | NA | NA | NA | NA | NA | NM_207550 |
| Olfr1271 | 0 | NA | NA | NA | NA | NA | | Olfr2 | 0 | NA | NA | NA | NA | NA | NM_010983 |
| Olfr1272 | 0 | NA | NA | NA | NA | NA | | Olfr20 | 0.113843286 | 0.00525947738444006 | 0.00859368002043176 | 0.612016897526494 | 0.540526577873086 | NA | NM_146923 |
| Olfr1273-ps | 0 | NA | NA | NA | NA | NA | | Olfr201 | 0 | NA | NA | NA | NA | NA | NM_146994 |
| Olfr1274-ps | 0 | NA | NA | NA | NA | NA | | Olfr202 | 0 | NA | NA | NA | NA | NA | NM_146995 |
| Olfr1275 | 0 | NA | NA | NA | NA | NA | | Olfr203 | 0 | NA | NA | NA | NA | NA | NM_146486 |
| Olfr1276 | 0 | NA | NA | NA | NA | NA | | Olfr204 | 0 | NA | NA | NA | NA | NA | NM_146992 |
| Olfr1277 | 0 | NA | NA | NA | NA | NA | | Olfr205 | 0 | NA | NA | NA | NA | NA | NM_001011736 |
| Olfr1278 | 0.099009884 | 0.018205726 | 0.027896865 | 0.652608315 | 0.514008823 | NA | | Olfr206 | 0 | NA | NA | NA | NA | NA | NM_146991 |
| Olfr1279 | 0 | NA | NA | NA | NA | NA | | Olfr207 | 0 | NA | NA | NA | NA | NA | NM_001011792 |
| Olfr128 | 0 | NA | NA | NA | NA | NA | | Olfr208 | 0 | NA | NA | NA | NA | NA | NM_146384 |
| Olfr1280 | 0 | NA | NA | NA | NA | NA | | Olfr209 | 0 | NA | NA | NA | NA | NA | NM_207551 |
| Olfr1281 | 0.099009884 | 0.018205726 | 0.027896865 | 0.652608315 | 0.514008823 | NA | | Olfr211 | 0 | NA | NA | NA | NA | NA | NM_146912 |
| Olfr1282 | 0 | NA | NA | NA | NA | NA | | Olfr212 | 2.192704541 | -0.00863898796430754 | 0.0330511914598851 | -0.261382043512496 | 0.793797903051081 | NA | NM_001011800 |
| Olfr1283 | 0 | NA | NA | NA | NA | NA | | Olfr213 | 0 | NA | NA | NA | NA | NA | NM_001011801 |
| Olfr1284 | 0 | NA | NA | NA | NA | NA | | Olfr214 | 0.097165704 | -0.00637828455464458 | 0.00848781691535724 | -0.75146349388194 | 0.452373763047372 | NA | NM_146759 |
| Olfr1286 | 0 | NA | NA | NA | NA | NA | | Olfr215 | 0 | NA | NA | NA | NA | NA | NM_146446 |
| Olfr1287 | 0 | NA | NA | NA | NA | NA | | Olfr218 | 0 | NA | NA | NA | NA | NA | NM_001001809 |
| Olfr1288 | 0 | NA | NA | NA | NA | NA | | Olfr220 | 0 | NA | NA | NA | NA | NA | NM_207694 |
| Olfr1289 | 0 | NA | NA | NA | NA | NA | | Olfr221 | 1.208201025 | -0.00151600269294522 | 0.0247326050716605 | -0.0612957142425047 | 0.951123703925274 | NA | NM_001001808 |
| Olfr129 | 0 | NA | NA | NA | NA | NA | | Olfr222 | 0 | NA | NA | NA | NA | NA | NM_001011789 |
| Olfr1290 | 0.096071043 | -0.020098795 | 0.027825948 | -0.722304036 | 0.470107572 | NA | | Olfr223 | 0 | NA | NA | NA | NA | NA | NM_146429 |
| Olfr1294 | 0 | NA | NA | NA | NA | NA | | Olfr224 | 0 | NA | NA | NA | NA | NA | NM_207695 |
| Olfr1295 | 0 | NA | NA | NA | NA | NA | | Olfr225 | 0 | NA | NA | NA | NA | NA | NM_001011740 |
| Olfr1297 | 0 | NA | NA | NA | NA | NA | | Olfr228 | 0 | NA | NA | NA | NA | NA | NM_146405 |
| Olfr1298 | 0 | NA | NA | NA | NA | NA | | Olfr229 | 0 | NA | NA | NA | NA | NA | NM_146613 |
| Olfr1299 | 0 | NA | NA | NA | NA | NA | | Olfr23 | 0 | NA | NA | NA | NA | NA | NM_010970 |
| Olfr13 | 0 | NA | NA | NA | NA | NA | | Olfr231 | 0 | NA | NA | NA | NA | NA | NM_001005520 |
| Olfr130 | 0 | NA | NA | NA | NA | NA | | Olfr235 | 0.217532603 | 0.00684685248201087 | 0.00976000296730706 | 0.701521557416086 | 0.482977586094298 | NA | NM_146686 |
| Olfr1300-ps1 | 0 | NA | NA | NA | NA | NA | | Olfr237-ps1 | 0 | NA | NA | NA | NA | NA | NM_146654 |
| Olfr1301 | 0 | NA | NA | NA | NA | NA | | Olfr239 | 0 | NA | NA | NA | NA | NA | NM_207175 |
| Olfr1302 | 0 | NA | NA | NA | NA | NA | | Olfr24 | 0 | NA | NA | NA | NA | NA | NM_146606 |
| Olfr1303 | 0 | NA | NA | NA | NA | NA | | Olfr242 | 0 | NA | NA | NA | NA | NA | NM_010974 |
| Olfr1305 | 0 | NA | NA | NA | NA | NA | | Olfr243 | 0 | NA | NA | NA | NA | NA | NM_001025386 |
| Olfr1306 | 0 | NA | NA | NA | NA | NA | | Olfr247 | 0 | NA | NA | NA | NA | NA | NM_146269 |
| Olfr1307 | 0 | NA | NA | NA | NA | NA | | Olfr248 | 0 | NA | NA | NA | NA | NA | NM_146714 |
| Olfr1308 | 0 | NA | NA | NA | NA | NA | | Olfr25 | 0 | NA | NA | NA | NA | NA | NM_146870 |
| Olfr1309 | 0 | NA | NA | NA | NA | NA | | Olfr250 | 0 | NA | NA | NA | NA | NA | NM_207552 |
| Olfr131 | 0 | NA | NA | NA | NA | NA | | Olfr251 | 0 | NA | NA | NA | NA | NA | NM_207553 |

| | | | | | | |
|---|---|---|---|---|---|---|
| Olfr1310 | 0 | NA | NA | NA | NA | NA |
| Olfr1311 | 0 | NA | NA | NA | NA | NA |
| Olfr1312 | 0 | NA | NA | NA | NA | NA |
| Olfr1313 | 0 | NA | NA | NA | NA | NA |
| Olfr1314 | 0 | NA | NA | NA | NA | NA |
| Olfr1316 | 0 | NA | NA | NA | NA | NA |
| Olfr1317 | 0 | NA | NA | NA | NA | NA |
| Olfr1318 | 0 | NA | NA | NA | NA | NA |
| Olfr132 | 0.098353386 | -0.020212831 | 0.02786466 | -0.725393061 | 0.468210929 | NA |
| Olfr1320 | 0 | NA | NA | NA | NA | NA |
| Olfr1321 | 0 | NA | NA | NA | NA | NA |
| Olfr1322 | 0 | NA | NA | NA | NA | NA |
| Olfr1323 | 0 | NA | NA | NA | NA | NA |
| Olfr1324 | 0 | NA | NA | NA | NA | NA |
| Olfr1325 | 0 | NA | NA | NA | NA | NA |
| Olfr1328 | 0 | NA | NA | NA | NA | NA |
| Olfr1329 | 0 | NA | NA | NA | NA | NA |
| Olfr133 | 0 | NA | NA | NA | NA | NA |
| Olfr1330 | 0 | NA | NA | NA | NA | NA |
| Olfr1331 | 0 | NA | NA | NA | NA | NA |
| Olfr1333 | 0 | NA | NA | NA | NA | NA |
| Olfr1335 | 0 | NA | NA | NA | NA | NA |
| Olfr1336 | 0 | NA | NA | NA | NA | NA |
| Olfr1337 | 0 | NA | NA | NA | NA | NA |
| Olfr1338 | 0 | NA | NA | NA | NA | NA |
| Olfr1339 | 0.233393928 | -0.027475306 | 0.032191697 | -0.85349047 | 0.39338737 | NA |
| Olfr134 | 0 | NA | NA | NA | NA | NA |
| Olfr1340 | 0 | NA | NA | NA | NA | NA |
| Olfr1341 | 0 | NA | NA | NA | NA | NA |
| Olfr1342 | 0.08731684 | 0.017365056 | 0.027733412 | 0.626142072 | 0.531221759 | NA |
| Olfr1346 | 0.29403132 | 0.006032586 | 0.033500653 | 0.180073692 | 0.857094716 | NA |
| Olfr1347 | 0.096071043 | -0.020098795 | 0.027825948 | -0.722304036 | 0.470107572 | NA |
| Olfr1348 | 0.094392817 | 0.017895942 | 0.027818137 | 0.643319236 | 0.520016977 | NA |
| Olfr1349 | 0.212768007 | -0.02694223 | 0.031868764 | -0.845411806 | 0.397880955 | NA |
| Olfr135 | 0 | NA | NA | NA | NA | NA |
| Olfr1350 | 0.099009884 | 0.018205726 | 0.027896865 | 0.652608315 | 0.514008823 | NA |
| Olfr1351 | 0 | NA | NA | NA | NA | NA |
| Olfr1352 | 0 | NA | NA | NA | NA | NA |
| Olfr1353 | 0 | NA | NA | NA | NA | NA |
| Olfr1354 | 0 | NA | NA | NA | NA | NA |
| Olfr1355 | 0 | NA | NA | NA | NA | NA |
| Olfr1356 | 0 | NA | NA | NA | NA | NA |
| Olfr1357 | 0 | NA | NA | NA | NA | NA |
| Olfr1359 | 0 | NA | NA | NA | NA | NA |
| Olfr136 | 0.098353386 | -0.020212831 | 0.02786466 | -0.725393061 | 0.468210929 | NA |
| Olfr1360 | 0 | NA | NA | NA | NA | NA |
| Olfr1361 | 0 | NA | NA | NA | NA | NA |
| Olfr1362 | 0 | NA | NA | NA | NA | NA |
| Olfr1364 | 0 | NA | NA | NA | NA | NA |
| Olfr1366 | 0 | NA | NA | NA | NA | NA |
| Olfr1367 | 0 | NA | NA | NA | NA | NA |
| Olfr1368 | 0 | NA | NA | NA | NA | NA |
| Olfr137 | 0 | NA | NA | NA | NA | NA |
| Olfr1370 | 0 | NA | NA | NA | NA | NA |
| Olfr1371 | 0.196680618 | -0.003857632 | 0.031597546 | -0.122086447 | 0.902830556 | NA |
| Olfr1372-ps1 | 7.366762932 | 0.011213609 | 0.149911666 | 0.074801442 | 0.940372694 | NA |
| Olfr1373 | 0 | NA | NA | NA | NA | NA |
| Olfr1377 | 0.406008005 | -0.00339866 | 0.047828467 | -0.071059364 | 0.943350509 | NA |
| Olfr1378 | 0.612124479 | 0.05749822 | 0.056841187 | 1.011559104 | 0.311748908 | NA |
| Olfr138 | 0 | NA | NA | NA | NA | NA |
| Olfr1380 | 0 | NA | NA | NA | NA | NA |
| Olfr1381 | 0 | NA | NA | NA | NA | NA |
| Olfr1382 | 0 | NA | NA | NA | NA | NA |
| Olfr1383 | 0 | NA | NA | NA | NA | NA |
| Olfr1384 | 0 | NA | NA | NA | NA | NA |
| Olfr1385 | 0 | NA | NA | NA | NA | NA |
| Olfr1386 | 0 | NA | NA | NA | NA | NA |
| Olfr1387 | 0 | NA | NA | NA | NA | NA |
| Olfr1388 | 0 | NA | NA | NA | NA | NA |
| Olfr1389 | 0.198019769 | 0.025308392 | 0.031628095 | 0.80018706 | 0.423602426 | NA |
| Olfr139 | 0 | NA | NA | NA | NA | NA |
| Olfr1390 | 0.192142086 | -0.026384592 | 0.031527717 | -0.836869715 | 0.402665805 | NA |
| Olfr1391 | 0.096071043 | -0.020098795 | 0.027825948 | -0.722304036 | 0.470107572 | NA |
| Olfr1392 | 2.880441076 | -0.095750908 | 0.116126803 | -0.824537533 | 0.409634198 | NA |
| Olfr1394 | 0 | NA | NA | NA | NA | NA |
| Olfr1395 | 0.079983654 | 0.016740032 | 0.027674178 | 0.604897175 | 0.54524732 | NA |
| Olfr1396 | 0.284680111 | 0.011373957 | 0.038774397 | 0.293336791 | 0.769264735 | NA |
| Olfr140 | 0 | NA | NA | NA | NA | NA |
| Olfr1402 | 0 | NA | NA | NA | NA | NA |
| Olfr1404 | 0 | NA | NA | NA | NA | NA |
| Olfr1406 | 0 | NA | NA | NA | NA | NA |
| Olfr1408 | 0.116696964 | -0.020477583 | 0.027954224 | -0.732539847 | 0.463839131 | NA |
| Olfr141 | 0 | NA | NA | NA | NA | NA |
| Olfr1410 | 0 | NA | NA | NA | NA | NA |
| Olfr1411 | 0 | NA | NA | NA | NA | NA |
| Olfr1412 | 0 | NA | NA | NA | NA | NA |
| Olfr1413 | 0 | NA | NA | NA | NA | NA |
| Olfr1414 | 0.096071043 | -0.020098795 | 0.027825948 | -0.722304036 | 0.470107572 | NA |
| Olfr1415 | 0 | NA | NA | NA | NA | NA |
| Olfr1416 | 0 | NA | NA | NA | NA | NA |
| Olfr1417 | 0 | NA | NA | NA | NA | NA |
| Olfr1418 | 0 | NA | NA | NA | NA | NA |
| Olfr1419 | 0.263371537 | 0.009921338 | 0.037961873 | 0.261350055 | 0.793822569 | NA |
| Olfr142 | 0 | NA | NA | NA | NA | NA |
| Olfr1420 | 0 | NA | NA | NA | NA | NA |
| Olfr1423 | 0 | NA | NA | NA | NA | NA |
| Olfr1424 | 0 | NA | NA | NA | NA | NA |
| Olfr1425 | 0 | NA | NA | NA | NA | NA |
| Olfr1426 | 0 | NA | NA | NA | NA | NA |
| Olfr1427 | 0 | NA | NA | NA | NA | NA |
| Olfr1428 | 0.096071043 | -0.020098795 | 0.027825948 | -0.722304036 | 0.470107572 | NA |
| Olfr143 | 0 | NA | NA | NA | NA | NA |
| Olfr1431 | 0.107716603 | -0.020477583 | 0.027954224 | -0.732539849 | 0.46383913 | NA |
| Olfr1433 | 0 | NA | NA | NA | NA | NA |
| Olfr1434 | 0 | NA | NA | NA | NA | NA |
| Olfr1436 | 0 | NA | NA | NA | NA | NA |
| Olfr1437 | 0.08731684 | 0.017365056 | 0.027733412 | 0.626142072 | 0.531221759 | NA |
| Olfr1440 | 0 | NA | NA | NA | NA | NA |
| Olfr1441 | 0 | NA | NA | NA | NA | NA |
| Olfr1442 | 0 | NA | NA | NA | NA | NA |
| Olfr1443 | 0 | NA | NA | NA | NA | NA |
| Olfr1444 | 0.099009884 | 0.018205726 | 0.027896865 | 0.652608315 | 0.514008823 | NA |
| Olfr1445 | 0.094392817 | 0.017895942 | 0.027818137 | 0.643319236 | 0.520016977 | NA |
| Olfr1446 | 0 | NA | NA | NA | NA | NA |
| Olfr1447 | 0 | NA | NA | NA | NA | NA |
| Olfr1448 | 0 | NA | NA | NA | NA | NA |
| Olfr1449 | 0 | NA | NA | NA | NA | NA |
| Olfr145 | 0 | NA | NA | NA | NA | NA |
| Olfr1450 | 0 | NA | NA | NA | NA | NA |
| Olfr1451 | 0 | NA | NA | NA | NA | NA |
| Olfr1453 | 0 | NA | NA | NA | NA | NA |
| Olfr1454 | 0 | NA | NA | NA | NA | NA |
| Olfr1457 | 0 | NA | NA | NA | NA | NA |
| Olfr1459 | 0 | NA | NA | NA | NA | NA |
| Olfr146 | 0 | NA | NA | NA | NA | NA |
| Olfr1461 | 0 | NA | NA | NA | NA | NA |
| Olfr1462 | 0 | NA | NA | NA | NA | NA |
| Olfr1463 | 0 | NA | NA | NA | NA | NA |
| Olfr1465 | 0 | NA | NA | NA | NA | NA |
| Olfr1466 | 0 | NA | NA | NA | NA | NA |
| Olfr1467 | 0 | NA | NA | NA | NA | NA |
| Olfr1469 | 0 | NA | NA | NA | NA | NA |
| Olfr147 | 0 | NA | NA | NA | NA | NA |
| Olfr1471 | 0 | NA | NA | NA | NA | NA |
| Olfr1472 | 0 | NA | NA | NA | NA | NA |
| Olfr1474 | 0 | NA | NA | NA | NA | NA |
| Olfr1475 | 0 | NA | NA | NA | NA | NA |
| Olfr1477 | 0 | NA | NA | NA | NA | NA |
| Olfr148 | 0 | NA | NA | NA | NA | NA |
| Olfr1480 | 0 | NA | NA | NA | NA | NA |
| Olfr1484 | 0 | NA | NA | NA | NA | NA |
| Olfr1487 | 0.098353386 | -0.020212831 | 0.02786466 | -0.725393061 | 0.468210929 | NA |
| Olfr1489 | 0 | NA | NA | NA | NA | NA |
| Olfr149 | 0 | NA | NA | NA | NA | NA |
| Olfr1490 | 0 | NA | NA | NA | NA | NA |
| Olfr1491 | 0 | NA | NA | NA | NA | NA |
| Olfr1494 | 0 | NA | NA | NA | NA | NA |
| Olfr1495 | 0 | NA | NA | NA | NA | NA |
| Olfr1496 | 0 | NA | NA | NA | NA | NA |
| Olfr1497 | 0 | NA | NA | NA | NA | NA |
| Olfr1499 | 0 | NA | NA | NA | NA | NA |
| Olfr15 | 0.098353386 | -0.020212831 | 0.02786466 | -0.725393061 | 0.468210929 | NA |
| Olfr150 | 0.079983654 | 0.016740032 | 0.027674178 | 0.604897175 | 0.54524732 | NA |
| Olfr1500 | 0 | NA | NA | NA | NA | NA |
| Olfr1501 | 0 | NA | NA | NA | NA | NA |
| Olfr1502 | 0 | NA | NA | NA | NA | NA |
| Olfr1504 | 0 | NA | NA | NA | NA | NA |
| Olfr1505 | 0 | NA | NA | NA | NA | NA |
| Olfr1506 | 0 | NA | NA | NA | NA | NA |
| Olfr1507 | 2.68215537 | -0.062068599 | 0.070310921 | -0.882773226 | 0.377358809 | NA |
| Olfr1508 | 0.096071043 | -0.020098795 | 0.027825948 | -0.722304036 | 0.470107572 | NA |
| Olfr1509 | 0 | NA | NA | NA | NA | NA |
| Olfr151 | 0 | NA | NA | NA | NA | NA |
| Olfr1510 | 0 | NA | NA | NA | NA | NA |
| Olfr1511 | 0 | NA | NA | NA | NA | NA |
| Olfr1512 | 0 | NA | NA | NA | NA | NA |
| Olfr1513 | 0 | NA | NA | NA | NA | NA |
| Olfr152 | 0 | NA | NA | NA | NA | NA |
| Olfr153 | 0.49741106 | -0.017678863 | 0.050840067 | -0.347734855 | 0.728039318 | NA |
| Olfr1532-ps1 | 0 | NA | NA | NA | NA | NA |
| Olfr1535 | 0 | NA | NA | NA | NA | NA |
| Olfr1537 | 0 | NA | NA | NA | NA | NA |
| Olfr154 | 0 | NA | NA | NA | NA | NA |
| Olfr155 | 0 | NA | NA | NA | NA | NA |
| Olfr156 | 0.107716603 | -0.020477583 | 0.027954224 | -0.732539849 | 0.46383913 | NA |
| Olfr157 | 0 | NA | NA | NA | NA | NA |
| Olfr159 | 0 | NA | NA | NA | NA | NA |
| Olfr16 | 0.094392817 | 0.017895942 | 0.027818137 | 0.643319236 | 0.520016977 | NA |
| Olfr160 | 0 | NA | NA | NA | NA | NA |
| Olfr161 | 0 | NA | NA | NA | NA | NA |
| Olfr164 | 0 | NA | NA | NA | NA | NA |
| Olfr165 | 0 | NA | NA | NA | NA | NA |
| Olfr166 | 0.116696964 | -0.020477583 | 0.027954224 | -0.732539847 | 0.463839131 | NA |
| Olfr167 | 0 | NA | NA | NA | NA | NA |
| Olfr168 | 0.116696964 | -0.020477583 | 0.027954224 | -0.732539847 | 0.463839131 | NA |
| Olfr169 | 0 | NA | NA | NA | NA | NA |
| Olfr17 | 0 | NA | NA | NA | NA | NA |
| Olfr170 | 0 | NA | NA | NA | NA | NA |
| Olfr171 | 0 | NA | NA | NA | NA | NA |
| Olfr172 | 0 | NA | NA | NA | NA | NA |
| Olfr173 | 0 | NA | NA | NA | NA | NA |
| Olfr175-ps1 | 0 | NA | NA | NA | NA | NA |
| Olfr176 | 0 | NA | NA | NA | NA | NA |
| Olfr177 | 0 | NA | NA | NA | NA | NA |
| Olfr178 | 0 | NA | NA | NA | NA | NA |
| Olfr179 | 0 | NA | NA | NA | NA | NA |
| Olfr18 | 0.478259379 | 0.031371149 | 0.045569182 | 0.688429064 | 0.491182626 | NA |
| Olfr180 | 0 | NA | NA | NA | NA | NA |
| Olfr181 | 0 | NA | NA | NA | NA | NA |
| Olfr183 | 0.188785634 | 0.025025111 | 0.031461603 | 0.79541754 | 0.42637066 | NA |
| Olfr186 | 0 | NA | NA | NA | NA | NA |
| Olfr187 | 0.233393928 | -0.027475306 | 0.032191697 | -0.85349047 | 0.39338737 | NA |
| Olfr19 | 0 | NA | NA | NA | NA | NA |
| Olfr190 | 0 | NA | NA | NA | NA | NA |
| Olfr191 | 0.098353386 | -0.020212831 | 0.02786466 | -0.725393061 | 0.468210929 | NA |
| Olfr192 | 0 | NA | NA | NA | NA | NA |
| Olfr193 | 0 | NA | NA | NA | NA | NA |
| Olfr194 | 0 | NA | NA | NA | NA | NA |
| Olfr195 | 0 | NA | NA | NA | NA | NA |
| Olfr196 | 0 | NA | NA | NA | NA | NA |
| Olfr197 | 0 | NA | NA | NA | NA | NA |
| Olfr198 | 0 | NA | NA | NA | NA | NA |
| Olfr199 | 0 | NA | NA | NA | NA | NA |
| Olfr2 | 0 | NA | NA | NA | NA | NA |
| Olfr20 | 0.116696964 | -0.020477583 | 0.027954224 | -0.732539847 | 0.463839131 | NA |
| Olfr201 | 0 | NA | NA | NA | NA | NA |
| Olfr202 | 0 | NA | NA | NA | NA | NA |
| Olfr203 | 0 | NA | NA | NA | NA | NA |
| Olfr204 | 0 | NA | NA | NA | NA | NA |
| Olfr205 | 0 | NA | NA | NA | NA | NA |
| Olfr206 | 0 | NA | NA | NA | NA | NA |
| Olfr207 | 0 | NA | NA | NA | NA | NA |
| Olfr208 | 0 | NA | NA | NA | NA | NA |
| Olfr209 | 0 | NA | NA | NA | NA | NA |
| Olfr211 | 0 | NA | NA | NA | NA | NA |
| Olfr212 | 3.176389577 | 0.095130843 | 0.092764089 | 1.025513684 | 0.305120872 | NA |
| Olfr213 | 0 | NA | NA | NA | NA | NA |
| Olfr214 | 0.079983654 | 0.016740032 | 0.027674178 | 0.604897175 | 0.54524732 | NA |
| Olfr215 | 0 | NA | NA | NA | NA | NA |
| Olfr218 | 0 | NA | NA | NA | NA | NA |
| Olfr220 | 0 | NA | NA | NA | NA | NA |
| Olfr221 | 0.979208136 | 0.017547049 | 0.071719569 | 0.244661946 | 0.806718195 | NA |
| Olfr222 | 0.290495473 | -0.034250308 | 0.035891155 | -0.954282697 | 0.339940564 | NA |
| Olfr223 | 0 | NA | NA | NA | NA | NA |
| Olfr224 | 0.192142086 | -0.026384592 | 0.031527717 | -0.836869715 | 0.402665805 | NA |

| | | | | | | | |
|---|---|---|---|---|---|---|---|
| Olfr257 | 0 | NA | NA | NA | NA | NA | NM_207554 |
| Olfr259 | 0 | NA | NA | NA | NA | NA | NM_146770 |
| Olfr26 | 0 | NA | NA | NA | NA | NA | NM_146783 |
| Olfr262 | 0 | NA | NA | NA | NA | NA | NM_146688 |
| Olfr263 | 0 | NA | NA | NA | NA | NA | NM_010984 |
| Olfr266 | 0 | NA | NA | NA | NA | NA | NM_146489 |
| Olfr267 | 0.628998992 | -0.00260283260632676 | 0.0172734575095016 | -0.150683938342687 | 0.88022504389996 | NA | NM_146920 |
| Olfr27 | 0 | NA | NA | NA | NA | NA | NM_146829 |
| Olfr270 | 0 | NA | NA | NA | NA | NA | NM_146607 |
| Olfr272 | 0 | NA | NA | NA | NA | NA | NM_146839 |
| Olfr273 | 0 | NA | NA | NA | NA | NA | NM_146824 |
| Olfr275 | 0 | NA | NA | NA | NA | NA | NM_146858 |
| Olfr279 | 0.127687691 | 0.00542298404407807 | 0.00869724871942359 | 0.62352868349814 | 0.532937159177571 | NA | NM_001001807 |
| Olfr281 | 0.326298904 | 0.00765111147689903 | 0.0103094497373689 | 0.742145475443356 | 0.457999199689495 | NA | NM_146280 |
| Olfr282 | 0 | NA | NA | NA | NA | NA | NM_146457 |
| Olfr283 | 0.985661481 | 0.0183970091163279 | 0.0223622236997987 | 0.822682456060639 | 0.410688594574762 | NA | NM_147036 |
| Olfr284 | 0 | NA | NA | NA | NA | NA | NM_146281 |
| Olfr285 | 0 | NA | NA | NA | NA | NA | NM_001011778 |
| Olfr286 | 0.544628687 | 0.00309863725245576 | 0.0164300944981167 | 0.188595217928354 | 0.850410081096137 | NA | NM_001011779 |
| Olfr288 | 10.8552333 | 0.0016263051404404 | 0.052641865793484 | 0.0308937594807232 | 0.975354266762277 | NA | NM_001011733 |
| Olfr29-ps1 | 0 | NA | NA | NA | NA | NA | NR_033638 |
| Olfr290 | 0 | NA | NA | NA | NA | NA | NM_146416 |
| Olfr291 | 0 | NA | NA | NA | NA | NA | NM_146415 |
| Olfr292 | 0 | NA | NA | NA | NA | NA | NM_146620 |
| Olfr293 | 0 | NA | NA | NA | NA | NA | NM_001011752 |
| Olfr294 | 0 | NA | NA | NA | NA | NA | NM_001011750 |
| Olfr295 | 0 | NA | NA | NA | NA | NA | NM_146851 |
| Olfr297 | 0 | NA | NA | NA | NA | NA | NM_146618 |
| Olfr298 | 0 | NA | NA | NA | NA | NA | NM_001011751 |
| Olfr299 | 0 | NA | NA | NA | NA | NA | NM_001011767 |
| Olfr3 | 0.120209049 | -0.00674972139751468 | 0.00863521114606269 | -0.781650996523957 | 0.434419707908266 | NA | NM_206903 |
| Olfr30 | 0 | NA | NA | NA | NA | NA | NM_146878 |
| Olfr301 | 0 | NA | NA | NA | NA | NA | NM_212436 |
| Olfr303 | 0 | NA | NA | NA | NA | NA | NM_146619 |
| Olfr304 | 0 | NA | NA | NA | NA | NA | NM_001011828 |
| Olfr305 | 0 | NA | NA | NA | NA | NA | NM_146616 |
| Olfr307 | 0 | NA | NA | NA | NA | NA | NM_146617 |
| Olfr308 | 0 | NA | NA | NA | NA | NA | NM_146621 |
| Olfr309 | 0 | NA | NA | NA | NA | NA | NM_001011866 |
| Olfr31 | 0 | NA | NA | NA | NA | NA | NM_147027 |
| Olfr310 | 0 | NA | NA | NA | NA | NA | NM_001011520 |
| Olfr311 | 0 | NA | NA | NA | NA | NA | NM_146537 |
| Olfr312 | 0 | NA | NA | NA | NA | NA | NM_001011819 |
| Olfr313 | 0 | NA | NA | NA | NA | NA | NM_146536 |
| Olfr314 | 0 | NA | NA | NA | NA | NA | NM_001011760 |
| Olfr315 | 0 | NA | NA | NA | NA | NA | NM_146538 |
| Olfr317 | 1.810214737 | -0.0416211946927376 | 0.0272382108534562 | -1.52804436813648 | 0.126501522446303 | NA | NM_001011769 |
| Olfr318 | 0 | NA | NA | NA | NA | NA | NM_146501 |
| Olfr319 | 0 | NA | NA | NA | NA | NA | NM_146500 |
| Olfr32 | 0 | NA | NA | NA | NA | NA | NM_010980 |
| Olfr320 | 0.097165704 | -0.00637828455464458 | 0.00848781691535724 | -0.75146349388194 | 0.452373763047372 | NA | NM_207230 |
| Olfr323 | 0 | NA | NA | NA | NA | NA | NM_146376 |
| Olfr324 | 2.253404505 | 0.00379110307958257 | 0.0299300149115196 | 0.126665592743271 | 0.899205079448631 | NA | NM_001011743 |
| Olfr325 | 0 | NA | NA | NA | NA | NA | NM_207153 |
| Olfr328 | 0.12890128 | -0.0068640067066208 | 0.00870804083610955 | -0.788237771940376 | 0.430557639008166 | NA | NM_146502 |
| Olfr329-ps | 0.203513543 | -0.00860782669689374 | 0.00975338121443284 | -0.882547960306941 | 0.377480556050336 | NA | NM_001011531 |
| Olfr33 | 0.333392541 | 0.00196325193564656 | 0.0126902925975457 | 0.154705017284334 | 0.877053874398876 | NA | NM_147073 |
| Olfr330 | 0 | NA | NA | NA | NA | NA | NM_146879 |
| Olfr331 | 0 | NA | NA | NA | NA | NA | NM_001011861 |
| Olfr332 | 0 | NA | NA | NA | NA | NA | NM_001011770 |
| Olfr338 | 0 | NA | NA | NA | NA | NA | NM_146947 |
| Olfr339 | 0 | NA | NA | NA | NA | NA | NM_146949 |
| Olfr340 | 0 | NA | NA | NA | NA | NA | NM_146951 |
| Olfr341 | 0 | NA | NA | NA | NA | NA | NM_146950 |
| Olfr342 | 0 | NA | NA | NA | NA | NA | NM_146948 |
| Olfr344 | 0 | NA | NA | NA | NA | NA | NM_146628 |
| Olfr345 | 0 | NA | NA | NA | NA | NA | NM_146945 |
| Olfr346 | 0 | NA | NA | NA | NA | NA | NM_146938 |
| Olfr347 | 0 | NA | NA | NA | NA | NA | NM_146943 |
| Olfr348 | 0 | NA | NA | NA | NA | NA | NM_146944 |
| Olfr350 | 0 | NA | NA | NA | NA | NA | NM_146627 |
| Olfr351 | 0 | NA | NA | NA | NA | NA | NM_146942 |
| Olfr352 | 0 | NA | NA | NA | NA | NA | NM_146940 |
| Olfr353 | 0 | NA | NA | NA | NA | NA | NM_146941 |
| Olfr354 | 0 | NA | NA | NA | NA | NA | NM_146939 |
| Olfr355 | 0 | NA | NA | NA | NA | NA | NM_146625 |
| Olfr356 | 0 | NA | NA | NA | NA | NA | NM_146624 |
| Olfr357 | 0 | NA | NA | NA | NA | NA | NM_146623 |
| Olfr358 | 0 | NA | NA | NA | NA | NA | NM_207235 |
| Olfr360 | 0 | NA | NA | NA | NA | NA | NM_146622 |
| Olfr361 | 0 | NA | NA | NA | NA | NA | NM_146368 |
| Olfr362 | 0 | NA | NA | NA | NA | NA | NM_147051 |
| Olfr365 | 0 | NA | NA | NA | NA | NA | NM_146662 |
| Olfr366 | 0 | NA | NA | NA | NA | NA | NM_001005569 |
| Olfr367-ps | 0 | NA | NA | NA | NA | NA | NM_001081010 |
| Olfr368 | 0 | NA | NA | NA | NA | NA | NM_146374 |
| Olfr370 | 0 | NA | NA | NA | NA | NA | NM_146270 |
| Olfr371 | 0 | NA | NA | NA | NA | NA | NM_146859 |
| Olfr372 | 0 | NA | NA | NA | NA | NA | NM_207555 |
| Olfr373 | 0 | NA | NA | NA | NA | NA | NM_146539 |
| Olfr374 | 0 | NA | NA | NA | NA | NA | NM_146338 |
| Olfr376 | 0 | NA | NA | NA | NA | NA | NM_146922,NM_001172686 |
| Olfr378 | 0 | NA | NA | NA | NA | NA | NM_147024 |
| Olfr38 | 0 | NA | NA | NA | NA | NA | NM_146986 |
| Olfr380 | 0 | NA | NA | NA | NA | NA | NM_147025 |
| Olfr381 | 0 | NA | NA | NA | NA | NA | NM_147022 |
| Olfr382 | 0 | NA | NA | NA | NA | NA | NM_146443 |
| Olfr384 | 0 | NA | NA | NA | NA | NA | NM_207224 |
| Olfr385 | 0 | NA | NA | NA | NA | NA | NM_147023 |
| Olfr389 | 0 | NA | NA | NA | NA | NA | NM_147009 |
| Olfr39 | 0 | NA | NA | NA | NA | NA | NM_146825 |
| Olfr390 | 0 | NA | NA | NA | NA | NA | NM_146347 |
| Olfr391-ps | 0 | NA | NA | NA | NA | NA | NM_001159775 |
| Olfr392 | 0 | NA | NA | NA | NA | NA | NM_147006 |
| Olfr393 | 0 | NA | NA | NA | NA | NA | NM_147008 |
| Olfr394 | 0 | NA | NA | NA | NA | NA | NM_147007 |
| Olfr395 | 0 | NA | NA | NA | NA | NA | NM_147005 |
| Olfr397 | 0 | NA | NA | NA | NA | NA | NM_146346 |
| Olfr398 | 0 | NA | NA | NA | NA | NA | NM_146710 |
| Olfr399 | 0 | NA | NA | NA | NA | NA | NM_147004 |
| Olfr401 | 0 | NA | NA | NA | NA | NA | NM_146706 |
| Olfr402 | 0 | NA | NA | NA | NA | NA | NM_146708 |
| Olfr403 | 0 | NA | NA | NA | NA | NA | NM_207622 |
| Olfr406 | 0 | NA | NA | NA | NA | NA | NM_001011863 |
| Olfr410 | 0 | NA | NA | NA | NA | NA | NM_146707 |
| Olfr411 | 0.108766301 | 0.00519171381361946 | 0.00856003371502575 | 0.606506234257728 | 0.544178643801544 | NA | NM_146709 |
| Olfr412 | 0 | NA | NA | NA | NA | NA | NM_001011851 |
| Olfr414 | 0 | NA | NA | NA | NA | NA | NM_146761 |
| Olfr417 | 0 | NA | NA | NA | NA | NA | NM_207137 |
| Olfr418-ps1 | 0 | NA | NA | NA | NA | NA | NM_146651 |
| Olfr419 | 0 | NA | NA | NA | NA | NA | NM_146715 |
| Olfr420 | 0 | NA | NA | NA | NA | NA | NM_146305 |
| Olfr421-ps1 | 0 | NA | NA | NA | NA | NA | NR_047667,NM_146720 |
| Olfr424 | 0 | NA | NA | NA | NA | NA | NM_146721 |
| Olfr426 | 0 | NA | NA | NA | NA | NA | NM_001206926 |
| Olfr427 | 0 | NA | NA | NA | NA | NA | NM_207158 |
| Olfr429 | 0 | NA | NA | NA | NA | NA | NM_146722 |
| Olfr43 | 0 | NA | NA | NA | NA | NA | NM_146711 |
| Olfr430 | 0 | NA | NA | NA | NA | NA | NM_146718 |
| Olfr432 | 0 | NA | NA | NA | NA | NA | NM_146716 |
| Olfr433 | 0 | NA | NA | NA | NA | NA | NM_146717 |
| Olfr434 | 0 | NA | NA | NA | NA | NA | NM_146369 |
| Olfr435 | 0 | NA | NA | NA | NA | NA | NM_146653 |
| Olfr437 | 0.212695677 | -0.00857796107890413 | 0.00973699866180651 | -0.880965621629515 | 0.378336427605205 | NA | NM_146296 |
| Olfr44 | 0 | NA | NA | NA | NA | NA | NM_146830 |
| Olfr441 | 0 | NA | NA | NA | NA | NA | NM_146655 |
| Olfr444 | 0 | NA | NA | NA | NA | NA | NM_146656 |
| Olfr446 | 0.543673105 | 0.0191808252780552 | 0.0162727917602594 | 1.17870526217251 | 0.23851555954191 | NA | NM_146295 |
| Olfr447 | 0 | NA | NA | NA | NA | NA | NM_146988 |
| Olfr448 | 0 | NA | NA | NA | NA | NA | NM_146273 |
| Olfr449 | 0 | NA | NA | NA | NA | NA | NM_147064 |
| Olfr45 | 0 | NA | NA | NA | NA | NA | NM_146963 |
| Olfr450 | 0 | NA | NA | NA | NA | NA | NM_146445 |
| Olfr452 | 0 | NA | NA | NA | NA | NA | NM_001011869 |
| Olfr453 | 0 | NA | NA | NA | NA | NA | NM_001011799 |
| Olfr455 | 0 | NA | NA | NA | NA | NA | NM_001081301 |
| Olfr456 | 0 | NA | NA | NA | NA | NA | NM_001011528 |
| Olfr457 | 0 | NA | NA | NA | NA | NA | NM_146987 |
| Olfr458 | 0 | NA | NA | NA | NA | NA | NM_146444 |
| Olfr459 | 0.113843286 | 0.00525947738444006 | 0.00859368002043176 | 0.612016897526494 | 0.540526577873086 | NA | NM_146576 |
| Olfr46 | 0.22897535 | -0.00116364198856673 | 0.00981572003153595 | -0.118548816065269 | 0.905632818135836 | NA | NM_146934 |
| Olfr460 | 0.106347839 | -0.00653347234140708 | 0.00854389509723638 | -0.764694822098226 | 0.444453290197145 | NA | NM_146383 |
| Olfr461 | 0 | NA | NA | NA | NA | NA | NM_146382 |
| Olfr462 | 0 | NA | NA | NA | NA | NA | NM_146411 |
| Olfr463 | 0 | NA | NA | NA | NA | NA | NM_146413 |
| Olfr464 | 18.3386413 | -0.11741316149378 | 0.0777025109541861 | -1.51106006809752 | 0.13077314737123 | NA | NM_146412 |
| Olfr466 | 0 | NA | NA | NA | NA | NA | NM_146819 |
| Olfr467 | 0 | NA | NA | NA | NA | NA | NM_001005488 |
| Olfr469 | 0 | NA | NA | NA | NA | NA | NM_146426 |
| Olfr47 | 0 | NA | NA | NA | NA | NA | NM_146370 |
| Olfr470 | 0 | NA | NA | NA | NA | NA | NM_146425 |
| Olfr472 | 0 | NA | NA | NA | NA | NA | NM_146774 |
| Olfr473 | 0 | NA | NA | NA | NA | NA | NM_146775 |
| Olfr474 | 0 | NA | NA | NA | NA | NA | NM_146495 |
| Olfr476 | 0 | NA | NA | NA | NA | NA | NM_146924 |
| Olfr477 | 0 | NA | NA | NA | NA | NA | NM_146926 |
| Olfr478 | 0 | NA | NA | NA | NA | NA | NM_146734 |
| Olfr479 | 0 | NA | NA | NA | NA | NA | NM_001011742 |
| Olfr48 | 0 | NA | NA | NA | NA | NA | NM_010990 |
| Olfr480 | 0 | NA | NA | NA | NA | NA | NM_020291 |
| Olfr481 | 0 | NA | NA | NA | NA | NA | NM_146925 |
| Olfr482 | 0 | NA | NA | NA | NA | NA | NM_146733 |
| Olfr483 | 0 | NA | NA | NA | NA | NA | NM_146735 |
| Olfr484 | 0 | NA | NA | NA | NA | NA | NM_146499 |
| Olfr485 | 0 | NA | NA | NA | NA | NA | NM_001011810 |
| Olfr486 | 0 | NA | NA | NA | NA | NA | NM_146496 |
| Olfr487 | 0 | NA | NA | NA | NA | NA | NM_001011811 |
| Olfr488 | 0 | NA | NA | NA | NA | NA | NM_146732 |
| Olfr49 | 0 | NA | NA | NA | NA | NA | NM_010991 |
| Olfr490 | 0 | NA | NA | NA | NA | NA | NM_146498 |
| Olfr491 | 0 | NA | NA | NA | NA | NA | NM_146736 |
| Olfr492 | 0 | NA | NA | NA | NA | NA | NM_146497 |
| Olfr493 | 0 | NA | NA | NA | NA | NA | NM_146310 |
| Olfr494 | 0 | NA | NA | NA | NA | NA | NM_146737 |
| Olfr495 | 0 | NA | NA | NA | NA | NA | NM_146364 |
| Olfr497 | 0 | NA | NA | NA | NA | NA | NM_146738 |
| Olfr498 | 0 | NA | NA | NA | NA | NA | NM_146307 |
| Olfr5 | 0.204491261 | 0.00675251727497272 | 0.00969338978195025 | 0.696610517772262 | 0.486046571838551 | NA | NM_146914 |
| Olfr50 | 0 | NA | NA | NA | NA | NA | NM_146946 |
| Olfr502 | 0 | NA | NA | NA | NA | NA | NM_146739 |
| Olfr503 | 0 | NA | NA | NA | NA | NA | NM_001011527 |
| Olfr504 | 0 | NA | NA | NA | NA | NA | NM_001011858 |
| Olfr506 | 0 | NA | NA | NA | NA | NA | NM_001011871 |
| Olfr507 | 0 | NA | NA | NA | NA | NA | NM_146743 |
| Olfr508 | 0 | NA | NA | NA | NA | NA | NM_146773 |
| Olfr509 | 0 | NA | NA | NA | NA | NA | NM_146372 |
| Olfr51 | 0 | NA | NA | NA | NA | NA | NM_146909 |
| Olfr510 | 0 | NA | NA | NA | NA | NA | NM_146311 |
| Olfr512 | 0 | NA | NA | NA | NA | NA | NM_146724 |
| Olfr513 | 0 | NA | NA | NA | NA | NA | NM_146723 |
| Olfr514 | 0 | NA | NA | NA | NA | NA | NM_146726 |
| Olfr516 | 0 | NA | NA | NA | NA | NA | NM_146725 |
| Olfr517 | 0 | NA | NA | NA | NA | NA | NM_001011846 |
| Olfr518 | 0 | NA | NA | NA | NA | NA | NM_146306 |
| Olfr519 | 0 | NA | NA | NA | NA | NA | NM_207160 |
| Olfr52 | 0 | NA | NA | NA | NA | NA | NM_146583 |
| Olfr520 | 0.393522717 | 0.00404856176060657 | 0.0118516400212778 | 0.34160350410053 | 0.732649299794451 | NA | NM_147063 |
| Olfr521 | 0 | NA | NA | NA | NA | NA | NM_146356 |
| Olfr522 | 0 | NA | NA | NA | NA | NA | NM_146952 |
| Olfr523 | 0.09572496 | 0.00498255289994511 | 0.00847999972823692 | 0.587565219295241 | 0.556824162160132 | NA | NM_146518 |
| Olfr524 | 0 | NA | NA | NA | NA | NA | NM_001011814 |
| Olfr525 | 0 | NA | NA | NA | NA | NA | NM_146956 |
| Olfr527 | 0 | NA | NA | NA | NA | NA | NM_001011776 |
| Olfr53 | 0.113843286 | 0.00525947738444006 | 0.00859368002043176 | 0.612016897526494 | 0.540526577873086 | NA | NM_146960 |
| Olfr530 | 0 | NA | NA | NA | NA | NA | NM_146519 |
| Olfr531 | 0 | NA | NA | NA | NA | NA | NM_146953 |
| Olfr532 | 0.097165704 | -0.00637828455464458 | 0.00848781691535724 | -0.75146349388194 | 0.452373763047372 | NA | NM_147026 |
| Olfr533 | 0 | NA | NA | NA | NA | NA | NM_001011815 |
| Olfr535 | 0 | NA | NA | NA | NA | NA | NM_146954 |
| Olfr536 | 0.900494549 | 0.0149940335432652 | 0.0171526935043628 | 0.874150379906892 | 0.382036364314131 | NA | NM_146520 |
| Olfr538 | 0 | NA | NA | NA | NA | NA | NM_001011867 |
| Olfr539 | 0 | NA | NA | NA | NA | NA | NM_146961 |
| Olfr54 | 0.127687691 | 0.00542298404407807 | 0.00869724871942359 | 0.62352868349814 | 0.532937159177571 | NA | NM_010997 |
| Olfr541 | 0 | NA | NA | NA | NA | NA | NM_146962 |
| Olfr543 | 0 | NA | NA | NA | NA | NA | NM_001011782 |
| Olfr544 | 0 | NA | NA | NA | NA | NA | NM_020289 |
| Olfr545 | 0.09572496 | 0.00498255289994511 | 0.00847999972823692 | 0.587565219295241 | 0.556824162160132 | NA | NM_146840 |
| Olfr547 | 0 | NA | NA | NA | NA | NA | NM_147079 |
| Olfr549 | 0 | NA | NA | NA | NA | NA | NM_147101 |
| Olfr55 | 0 | NA | NA | NA | NA | NA | NM_010998 |
| Olfr550 | 3.7527994 | 0.0136328195689966 | 0.0370264239941596 | 0.368191634470209 | 0.712730347614329 | NA | NM_147104 |
| Olfr551 | 0 | NA | NA | NA | NA | NA | NM_146755 |
| Olfr552 | 0 | NA | NA | NA | NA | NA | NM_147102 |
| Olfr553 | 0 | NA | NA | NA | NA | NA | NM_207621 |
| Olfr554 | 0 | NA | NA | NA | NA | NA | NM_146325 |

| | | | | | | |
|---|---|---|---|---|---|---|
| Olfr225 | 0 | NA | NA | NA | NA | NA |
| Olfr228 | 0 | NA | NA | NA | NA | NA |
| Olfr229 | 0 | NA | NA | NA | NA | NA |
| Olfr23 | 0 | NA | NA | NA | NA | NA |
| Olfr231 | 0 | NA | NA | NA | NA | NA |
| Olfr235 | 0 | NA | NA | NA | NA | NA |
| Olfr237-ps1 | 0 | NA | NA | NA | NA | NA |
| Olfr239 | 0 | NA | NA | NA | NA | NA |
| Olfr24 | 0 | NA | NA | NA | NA | NA |
| Olfr242 | 0 | NA | NA | NA | NA | NA |
| Olfr243 | 0 | NA | NA | NA | NA | NA |
| Olfr247 | 0 | NA | NA | NA | NA | NA |
| Olfr248 | 0 | NA | NA | NA | NA | NA |
| Olfr25 | 0 | NA | NA | NA | NA | NA |
| Olfr250 | 0 | NA | NA | NA | NA | NA |
| Olfr251 | 0 | NA | NA | NA | NA | NA |
| Olfr257 | 0 | NA | NA | NA | NA | NA |
| Olfr259 | 0 | NA | NA | NA | NA | NA |
| Olfr26 | 0 | NA | NA | NA | NA | NA |
| Olfr262 | 0 | NA | NA | NA | NA | NA |
| Olfr263 | 0 | NA | NA | NA | NA | NA |
| Olfr266 | 0.098353386 | -0.020212831 | 0.02786466 | -0.725393061 | 0.468210929 | NA |
| Olfr267 | 0.587805432 | 0.015924212 | 0.053274974 | 0.29890606 | 0.765011723 | NA |
| Olfr27 | 0 | NA | NA | NA | NA | NA |
| Olfr270 | 0 | NA | NA | NA | NA | NA |
| Olfr272 | 0.238795281 | -0.027601606 | 0.032267774 | -0.855392297 | 0.392334007 | NA |
| Olfr273 | 0 | NA | NA | NA | NA | NA |
| Olfr275 | 0 | NA | NA | NA | NA | NA |
| Olfr279 | 0 | NA | NA | NA | NA | NA |
| Olfr281 | 0.098353386 | -0.020212831 | 0.02786466 | -0.725393061 | 0.468210929 | NA |
| Olfr282 | 0 | NA | NA | NA | NA | NA |
| Olfr283 | 0.577209771 | 0.036509886 | 0.048527698 | 0.752351489 | 0.451839712 | NA |
| Olfr284 | 0 | NA | NA | NA | NA | NA |
| Olfr285 | 0 | NA | NA | NA | NA | NA |
| Olfr286 | 0.159967308 | 0.0240345 | 0.030870969 | 0.778547006 | 0.436246606 | NA |
| Olfr288 | 7.316914825 | 0.078441901 | 0.128301048 | 0.6113894 | 0.540941818 | NA |
| Olfr29-ps1 | 0 | NA | NA | NA | NA | NA |
| Olfr290 | 0 | NA | NA | NA | NA | NA |
| Olfr291 | 0 | NA | NA | NA | NA | NA |
| Olfr292 | 0 | NA | NA | NA | NA | NA |
| Olfr293 | 0 | NA | NA | NA | NA | NA |
| Olfr294 | 0 | NA | NA | NA | NA | NA |
| Olfr295 | 0 | NA | NA | NA | NA | NA |
| Olfr297 | 0 | NA | NA | NA | NA | NA |
| Olfr298 | 0 | NA | NA | NA | NA | NA |
| Olfr299 | 0 | NA | NA | NA | NA | NA |
| Olfr3 | 0.099009884 | 0.018205726 | 0.027896865 | 0.652608315 | 0.514008823 | NA |
| Olfr30 | 0 | NA | NA | NA | NA | NA |
| Olfr301 | 0 | NA | NA | NA | NA | NA |
| Olfr303 | 0 | NA | NA | NA | NA | NA |
| Olfr304 | 0 | NA | NA | NA | NA | NA |
| Olfr305 | 0 | NA | NA | NA | NA | NA |
| Olfr307 | 0 | NA | NA | NA | NA | NA |
| Olfr308 | 0 | NA | NA | NA | NA | NA |
| Olfr309 | 0.290495473 | -0.034250308 | 0.035891155 | -0.954282697 | 0.339940564 | NA |
| Olfr31 | 0.094392817 | 0.017895942 | 0.027818137 | 0.643319236 | 0.520016977 | NA |
| Olfr310 | 0 | NA | NA | NA | NA | NA |
| Olfr311 | 0 | NA | NA | NA | NA | NA |
| Olfr312 | 0 | NA | NA | NA | NA | NA |
| Olfr313 | 0 | NA | NA | NA | NA | NA |
| Olfr314 | 0 | NA | NA | NA | NA | NA |
| Olfr315 | 0 | NA | NA | NA | NA | NA |
| Olfr317 | 1.721536622 | 0.125837949 | 0.096092464 | 1.309550652 | 0.190347892 | NA |
| Olfr318 | 0 | NA | NA | NA | NA | NA |
| Olfr319 | 0 | NA | NA | NA | NA | NA |
| Olfr32 | 0.775415375 | -0.069517603 | 0.050698856 | -1.371186808 | 0.17031673 | NA |
| Olfr320 | 0.079983654 | 0.016740032 | 0.027674178 | 0.604897175 | 0.54524732 | NA |
| Olfr323 | 0 | NA | NA | NA | NA | NA |
| Olfr324 | 1.583113224 | 0.018576751 | 0.093897186 | 0.197841397 | 0.843169156 | NA |
| Olfr325 | 0 | NA | NA | NA | NA | NA |
| Olfr328 | 0.105923158 | 0.018524693 | 0.027977518 | 0.662127842 | 0.507889294 | NA |
| Olfr329-ps | 0.167300494 | 0.024299763 | 0.031030539 | 0.783091891 | 0.433573153 | NA |
| Olfr33 | 0.298065244 | -0.009392943 | 0.033782767 | -0.278039479 | 0.780982053 | NA |
| Olfr330 | 0 | NA | NA | NA | NA | NA |
| Olfr331 | 0 | NA | NA | NA | NA | NA |
| Olfr332 | 0 | NA | NA | NA | NA | NA |
| Olfr338 | 0 | NA | NA | NA | NA | NA |
| Olfr339 | 0 | NA | NA | NA | NA | NA |
| Olfr340 | 0 | NA | NA | NA | NA | NA |
| Olfr341 | 0 | NA | NA | NA | NA | NA |
| Olfr342 | 0.098353386 | -0.020212831 | 0.02786466 | -0.725393061 | 0.468210929 | NA |
| Olfr344 | 0 | NA | NA | NA | NA | NA |
| Olfr345 | 0 | NA | NA | NA | NA | NA |
| Olfr346 | 0 | NA | NA | NA | NA | NA |
| Olfr347 | 0 | NA | NA | NA | NA | NA |
| Olfr348 | 0 | NA | NA | NA | NA | NA |
| Olfr350 | 0 | NA | NA | NA | NA | NA |
| Olfr351 | 0 | NA | NA | NA | NA | NA |
| Olfr352 | 0 | NA | NA | NA | NA | NA |
| Olfr353 | 0 | NA | NA | NA | NA | NA |
| Olfr354 | 0 | NA | NA | NA | NA | NA |
| Olfr355 | 0 | NA | NA | NA | NA | NA |
| Olfr356 | 0 | NA | NA | NA | NA | NA |
| Olfr357 | 0 | NA | NA | NA | NA | NA |
| Olfr358 | 0 | NA | NA | NA | NA | NA |
| Olfr360 | 0 | NA | NA | NA | NA | NA |
| Olfr361 | 0 | NA | NA | NA | NA | NA |
| Olfr362 | 0 | NA | NA | NA | NA | NA |
| Olfr365 | 0 | NA | NA | NA | NA | NA |
| Olfr366 | 0 | NA | NA | NA | NA | NA |
| Olfr367-ps | 0 | NA | NA | NA | NA | NA |
| Olfr368 | 0 | NA | NA | NA | NA | NA |
| Olfr370 | 0 | NA | NA | NA | NA | NA |
| Olfr371 | 0 | NA | NA | NA | NA | NA |
| Olfr372 | 0 | NA | NA | NA | NA | NA |
| Olfr373 | 0 | NA | NA | NA | NA | NA |
| Olfr374 | 0 | NA | NA | NA | NA | NA |
| Olfr376 | 0 | NA | NA | NA | NA | NA |
| Olfr378 | 0 | NA | NA | NA | NA | NA |
| Olfr38 | 0 | NA | NA | NA | NA | NA |
| Olfr380 | 0 | NA | NA | NA | NA | NA |
| Olfr381 | 0 | NA | NA | NA | NA | NA |
| Olfr382 | 0 | NA | NA | NA | NA | NA |
| Olfr384 | 0 | NA | NA | NA | NA | NA |
| Olfr385 | 0 | NA | NA | NA | NA | NA |
| Olfr389 | 0 | NA | NA | NA | NA | NA |
| Olfr39 | 0 | NA | NA | NA | NA | NA |
| Olfr390 | 0 | NA | NA | NA | NA | NA |
| Olfr391-ps | 0 | NA | NA | NA | NA | NA |
| Olfr392 | 0 | NA | NA | NA | NA | NA |
| Olfr393 | 0 | NA | NA | NA | NA | NA |
| Olfr394 | 0 | NA | NA | NA | NA | NA |
| Olfr395 | 0 | NA | NA | NA | NA | NA |
| Olfr397 | 0 | NA | NA | NA | NA | NA |
| Olfr398 | 0 | NA | NA | NA | NA | NA |
| Olfr399 | 0 | NA | NA | NA | NA | NA |
| Olfr401 | 0 | NA | NA | NA | NA | NA |
| Olfr402 | 0 | NA | NA | NA | NA | NA |
| Olfr403 | 0 | NA | NA | NA | NA | NA |
| Olfr406 | 0 | NA | NA | NA | NA | NA |
| Olfr410 | 0 | NA | NA | NA | NA | NA |
| Olfr411 | 0 | NA | NA | NA | NA | NA |
| Olfr412 | 0 | NA | NA | NA | NA | NA |
| Olfr414 | 0.098353386 | -0.020212831 | 0.02786466 | -0.725393061 | 0.468210929 | NA |
| Olfr417 | 0 | NA | NA | NA | NA | NA |
| Olfr418-ps1 | 0 | NA | NA | NA | NA | NA |
| Olfr419 | 0 | NA | NA | NA | NA | NA |
| Olfr420 | 0 | NA | NA | NA | NA | NA |
| Olfr421-ps1 | 0 | NA | NA | NA | NA | NA |
| Olfr424 | 0 | NA | NA | NA | NA | NA |
| Olfr426 | 0 | NA | NA | NA | NA | NA |
| Olfr427 | 0 | NA | NA | NA | NA | NA |
| Olfr429 | 0 | NA | NA | NA | NA | NA |
| Olfr43 | 0.096071043 | -0.020098795 | 0.027825948 | -0.722304036 | 0.470107572 | NA |
| Olfr430 | 0 | NA | NA | NA | NA | NA |
| Olfr432 | 0 | NA | NA | NA | NA | NA |
| Olfr433 | 0 | NA | NA | NA | NA | NA |
| Olfr434 | 0 | NA | NA | NA | NA | NA |
| Olfr435 | 0 | NA | NA | NA | NA | NA |
| Olfr437 | 0.17463368 | 0.024560283 | 0.031186242 | 0.787535833 | 0.430968261 | NA |
| Olfr44 | 0 | NA | NA | NA | NA | NA |
| Olfr441 | 0 | NA | NA | NA | NA | NA |
| Olfr444 | 0 | NA | NA | NA | NA | NA |
| Olfr446 | 0.19442443 | -0.026451352 | 0.031568725 | -0.837897386 | 0.402088337 | NA |
| Olfr447 | 0 | NA | NA | NA | NA | NA |
| Olfr448 | 0 | NA | NA | NA | NA | NA |
| Olfr449 | 0.119397641 | -0.020477583 | 0.027954224 | -0.732539845 | 0.463839132 | NA |
| Olfr45 | 0 | NA | NA | NA | NA | NA |
| Olfr450 | 0 | NA | NA | NA | NA | NA |
| Olfr452 | 0 | NA | NA | NA | NA | NA |
| Olfr453 | 0 | NA | NA | NA | NA | NA |
| Olfr455 | 0 | NA | NA | NA | NA | NA |
| Olfr456 | 0 | NA | NA | NA | NA | NA |
| Olfr457 | 0 | NA | NA | NA | NA | NA |
| Olfr458 | 0 | NA | NA | NA | NA | NA |
| Olfr459 | 0 | NA | NA | NA | NA | NA |
| Olfr46 | 0.099009884 | 0.018205726 | 0.027896865 | 0.652608315 | 0.514008823 | NA |
| Olfr460 | 0.08731684 | 0.017365056 | 0.027733412 | 0.626142072 | 0.531221759 | NA |
| Olfr461 | 0 | NA | NA | NA | NA | NA |
| Olfr462 | 0 | NA | NA | NA | NA | NA |
| Olfr463 | 0.346511884 | -0.038380278 | 0.037950969 | -1.011312187 | 0.311867035 | NA |
| Olfr466 | 0 | NA | NA | NA | NA | NA |
| Olfr467 | 0 | NA | NA | NA | NA | NA |
| Olfr469 | 0 | NA | NA | NA | NA | NA |
| Olfr47 | 0 | NA | NA | NA | NA | NA |
| Olfr470 | 0 | NA | NA | NA | NA | NA |
| Olfr472 | 0 | NA | NA | NA | NA | NA |
| Olfr473 | 0 | NA | NA | NA | NA | NA |
| Olfr474 | 0 | NA | NA | NA | NA | NA |
| Olfr476 | 0 | NA | NA | NA | NA | NA |
| Olfr477 | 0 | NA | NA | NA | NA | NA |
| Olfr478 | 0 | NA | NA | NA | NA | NA |
| Olfr479 | 0 | NA | NA | NA | NA | NA |
| Olfr48 | 0.098353386 | -0.020212831 | 0.02786466 | -0.725393061 | 0.468210929 | NA |
| Olfr480 | 0 | NA | NA | NA | NA | NA |
| Olfr481 | 0 | NA | NA | NA | NA | NA |
| Olfr482 | 0 | NA | NA | NA | NA | NA |
| Olfr483 | 0 | NA | NA | NA | NA | NA |
| Olfr484 | 0 | NA | NA | NA | NA | NA |
| Olfr485 | 0 | NA | NA | NA | NA | NA |
| Olfr486 | 0 | NA | NA | NA | NA | NA |
| Olfr487 | 0 | NA | NA | NA | NA | NA |
| Olfr488 | 0 | NA | NA | NA | NA | NA |
| Olfr49 | 0 | NA | NA | NA | NA | NA |
| Olfr490 | 0 | NA | NA | NA | NA | NA |
| Olfr491 | 0 | NA | NA | NA | NA | NA |
| Olfr492 | 0 | NA | NA | NA | NA | NA |
| Olfr493 | 0 | NA | NA | NA | NA | NA |
| Olfr494 | 0 | NA | NA | NA | NA | NA |
| Olfr495 | 0 | NA | NA | NA | NA | NA |
| Olfr497 | 0 | NA | NA | NA | NA | NA |
| Olfr498 | 0 | NA | NA | NA | NA | NA |
| Olfr5 | 0 | NA | NA | NA | NA | NA |
| Olfr50 | 0 | NA | NA | NA | NA | NA |
| Olfr502 | 0 | NA | NA | NA | NA | NA |
| Olfr503 | 0 | NA | NA | NA | NA | NA |
| Olfr504 | 0 | NA | NA | NA | NA | NA |
| Olfr506 | 0 | NA | NA | NA | NA | NA |
| Olfr507 | 0 | NA | NA | NA | NA | NA |
| Olfr508 | 0 | NA | NA | NA | NA | NA |
| Olfr509 | 0 | NA | NA | NA | NA | NA |
| Olfr51 | 0 | NA | NA | NA | NA | NA |
| Olfr510 | 0 | NA | NA | NA | NA | NA |
| Olfr512 | 0 | NA | NA | NA | NA | NA |
| Olfr513 | 0 | NA | NA | NA | NA | NA |
| Olfr514 | 0 | NA | NA | NA | NA | NA |
| Olfr516 | 0 | NA | NA | NA | NA | NA |
| Olfr517 | 0 | NA | NA | NA | NA | NA |
| Olfr518 | 0 | NA | NA | NA | NA | NA |
| Olfr519 | 0 | NA | NA | NA | NA | NA |
| Olfr52 | 0 | NA | NA | NA | NA | NA |
| Olfr520 | 0.185670226 | -0.001645086 | 0.031432796 | -0.052336624 | 0.958260472 | NA |
| Olfr521 | 0 | NA | NA | NA | NA | NA |
| Olfr522 | 0.096071043 | -0.020098795 | 0.027825948 | -0.722304036 | 0.470107572 | NA |
| Olfr523 | 0.215050351 | -0.027006021 | 0.031907566 | -0.846382992 | 0.397339123 | NA |
| Olfr524 | 0 | NA | NA | NA | NA | NA |
| Olfr525 | 0 | NA | NA | NA | NA | NA |
| Olfr527 | 0 | NA | NA | NA | NA | NA |
| Olfr53 | 0 | NA | NA | NA | NA | NA |
| Olfr530 | 0 | NA | NA | NA | NA | NA |
| Olfr531 | 0 | NA | NA | NA | NA | NA |
| Olfr532 | 0.079983654 | 0.016740032 | 0.027674178 | 0.604897175 | 0.54524732 | NA |
| Olfr533 | 0.116696964 | -0.020477583 | 0.027954224 | -0.732539847 | 0.463839131 | NA |
| Olfr535 | 0 | NA | NA | NA | NA | NA |
| Olfr536 | 0.215706849 | -0.002035226 | 0.0319478 | -0.063704747 | 0.949205325 | NA |

| | | | | | | | |
|---|---|---|---|---|---|---|---|
| Olfr555 | 0 | NA | NA | NA | NA | NA | NM_147103 |
| Olfr556 | 0 | NA | NA | NA | NA | NA | NM_146754 |
| Olfr557 | 0 | NA | NA | NA | NA | NA | NM_146361 |
| Olfr558 | 12.55436411 | -0.0536144615040809 | 0.0615068262284891 | -0.871683108878205 | 0.383381277491496 | NA | NM_147093 |
| Olfr559 | 0 | NA | NA | NA | NA | NA | NM_147112 |
| Olfr56 | 0.603776958 | -0.0158364197761598 | 0.0159657748266675 | -0.991897978525192 | 0.321247295325773 | NA | NM_010999 |
| Olfr560 | 0.313257562 | 0.00962480958133652 | 0.0115601382610462 | 0.832586026567595 | 0.405078242417788 | NA | NM_147113 |
| Olfr561 | 0.240418098 | -0.0088029805258837 | 0.00986414519853158 | -0.892422034419582 | 0.372166765031741 | NA | NM_147092 |
| Olfr564 | 0.488314838 | -0.00797063425337698 | 0.0128168213807136 | -0.621888533561916 | 0.534015165389952 | NA | NM_146359 |
| Olfr566 | 0 | NA | NA | NA | NA | NA | NM_001011536 |
| Olfr568 | 0 | NA | NA | NA | NA | NA | NM_147091 |
| Olfr569 | 0 | NA | NA | NA | NA | NA | NM_147088 |
| Olfr57 | 0 | NA | NA | NA | NA | NA | NM_147041 |
| Olfr570 | 0.628307206 | -0.00200236215886405 | 0.0153904808700936 | -0.130103937347077 | 0.896484195048209 | NA | NM_147110 |
| Olfr571 | 0 | NA | NA | NA | NA | NA | NM_147085 |
| Olfr572 | 0 | NA | NA | NA | NA | NA | NM_147089 |
| Olfr574 | 0.427411356 | 0.0060983518828938 | 0.0139232555105122 | 0.43799755583667 | 0.661388055218398 | NA | NM_146360 |
| Olfr575 | 0 | NA | NA | NA | NA | NA | NM_147114 |
| Olfr576 | 0 | NA | NA | NA | NA | NA | NM_001001805 |
| Olfr577 | 0.217532603 | 0.00684685248201087 | 0.00976000296730706 | 0.701521557416086 | 0.482977586094298 | NA | NM_147109 |
| Olfr578 | 0 | NA | NA | NA | NA | NA | NM_147115 |
| Olfr58 | 0.393522717 | 0.00404856176060657 | 0.0118516400212778 | 0.34160350410053 | 0.732649299794451 | NA | NM_011001 |
| Olfr582 | 0 | NA | NA | NA | NA | NA | NM_147053 |
| Olfr583 | 0 | NA | NA | NA | NA | NA | NM_146757 |
| Olfr584 | 0 | NA | NA | NA | NA | NA | NM_147054 |
| Olfr585 | 0 | NA | NA | NA | NA | NA | NM_147087 |
| Olfr586 | 0 | NA | NA | NA | NA | NA | NM_147111 |
| Olfr589 | 0 | NA | NA | NA | NA | NA | NM_147052 |
| Olfr59 | 0 | NA | NA | NA | NA | NA | NM_011002 |
| Olfr591 | 0 | NA | NA | NA | NA | NA | NM_001011847 |
| Olfr592 | 0 | NA | NA | NA | NA | NA | NM_207556 |
| Olfr593 | 0 | NA | NA | NA | NA | NA | NM_146380 |
| Olfr594 | 0.106148608 | 0.00515554063321497 | 0.00854200771766808 | 0.603551390213729 | 0.546141939141592 | NA | NM_207143 |
| Olfr596 | 0 | NA | NA | NA | NA | NA | NM_001190381 |
| Olfr597 | 0 | NA | NA | NA | NA | NA | NM_001011845 |
| Olfr598 | 0 | NA | NA | NA | NA | NA | NM_001011793 |
| Olfr599 | 0 | NA | NA | NA | NA | NA | NM_146731 |
| Olfr6 | 0 | NA | NA | NA | NA | NA | NM_206897 |
| Olfr60 | 0 | NA | NA | NA | NA | NA | NM_146955 |
| Olfr600 | 0 | NA | NA | NA | NA | NA | NM_147046 |
| Olfr601 | 0 | NA | NA | NA | NA | NA | NM_146314 |
| Olfr603 | 0 | NA | NA | NA | NA | NA | NM_147070 |
| Olfr605 | 0 | NA | NA | NA | NA | NA | NM_001011854 |
| Olfr606 | 0 | NA | NA | NA | NA | NA | NM_147094 |
| Olfr608 | 0 | NA | NA | NA | NA | NA | NM_146756 |
| Olfr609 | 0.240418098 | -0.0088029805258837 | 0.00986414519853158 | -0.892422034419582 | 0.372166765031741 | NA | NM_147082 |
| Olfr61 | 0 | NA | NA | NA | NA | NA | NM_146964 |
| Olfr610 | 0 | NA | NA | NA | NA | NA | NM_147081 |
| Olfr611 | 0 | NA | NA | NA | NA | NA | NM_146727 |
| Olfr612 | 0 | NA | NA | NA | NA | NA | NM_001200027 |
| Olfr615 | 0 | NA | NA | NA | NA | NA | NM_147080 |
| Olfr616 | 0 | NA | NA | NA | NA | NA | NM_147099 |
| Olfr617 | 0 | NA | NA | NA | NA | NA | NM_146841 |
| Olfr618 | 0 | NA | NA | NA | NA | NA | NM_147047 |
| Olfr619 | 0 | NA | NA | NA | NA | NA | NM_147076 |
| Olfr62 | 0.108766301 | 0.00519171381361946 | 0.00856003371502575 | 0.606506234257728 | 0.544178643801544 | NA | NM_146315 |
| Olfr620 | 0 | NA | NA | NA | NA | NA | NM_146812 |
| Olfr622 | 0.300480017 | -0.00401601363468843 | 0.0108825526077113 | -0.369032319847708 | 0.712103634607305 | NA | NM_147083 |
| Olfr623 | 0 | NA | NA | NA | NA | NA | NM_147122 |
| Olfr624 | 0 | NA | NA | NA | NA | NA | NM_001011865 |
| Olfr628 | 0 | NA | NA | NA | NA | NA | NM_147097 |
| Olfr629 | 0 | NA | NA | NA | NA | NA | NM_146821 |
| Olfr63 | 0 | NA | NA | NA | NA | NA | NM_146937 |
| Olfr630 | 0 | NA | NA | NA | NA | NA | NM_147098 |
| Olfr631 | 0 | NA | NA | NA | NA | NA | NM_001271020,NM_146959 |
| Olfr632 | 0 | NA | NA | NA | NA | NA | NM_147119 |
| Olfr633 | 0 | NA | NA | NA | NA | NA | NM_146354 |
| Olfr635 | 0 | NA | NA | NA | NA | NA | NM_147118 |
| Olfr638 | 0 | NA | NA | NA | NA | NA | NM_147120 |
| Olfr639 | 0 | NA | NA | NA | NA | NA | NM_147084 |
| Olfr64 | 0 | NA | NA | NA | NA | NA | NM_013617 |
| Olfr640 | 0 | NA | NA | NA | NA | NA | NM_146822 |
| Olfr641 | 0 | NA | NA | NA | NA | NA | NM_147072 |
| Olfr642 | 0 | NA | NA | NA | NA | NA | NM_146329 |
| Olfr643 | 0 | NA | NA | NA | NA | NA | NM_147077 |
| Olfr644 | 0 | NA | NA | NA | NA | NA | NM_147121 |
| Olfr645 | 0 | NA | NA | NA | NA | NA | NM_207144 |
| Olfr646 | 0 | NA | NA | NA | NA | NA | NM_147056 |
| Olfr648 | 0 | NA | NA | NA | NA | NA | NM_146751 |
| Olfr649 | 0 | NA | NA | NA | NA | NA | NM_147055 |
| Olfr65 | 0 | NA | NA | NA | NA | NA | NM_013616 |
| Olfr651 | 0 | NA | NA | NA | NA | NA | NM_146813 |
| Olfr652 | 0 | NA | NA | NA | NA | NA | NM_147048 |
| Olfr653 | 0 | NA | NA | NA | NA | NA | NM_147074 |
| Olfr654 | 0 | NA | NA | NA | NA | NA | NM_146379 |
| Olfr655 | 0 | NA | NA | NA | NA | NA | NM_146820 |
| Olfr656 | 0 | NA | NA | NA | NA | NA | NM_147075 |
| Olfr657 | 0 | NA | NA | NA | NA | NA | NM_146312 |
| Olfr658 | 4.357951792 | 0.0481403792238574 | 0.0470676548792279 | 1.02279111520177 | 0.306406619762875 | NA | NM_147049 |
| Olfr659 | 0 | NA | NA | NA | NA | NA | NM_147050 |
| Olfr66 | 0 | NA | NA | NA | NA | NA | NM_013618 |
| Olfr661 | 0.108766301 | 0.00519171381361946 | 0.00856003371502575 | 0.606506234257728 | 0.544178643801544 | NA | NM_146748 |
| Olfr663 | 0 | NA | NA | NA | NA | NA | NM_001011757 |
| Olfr665 | 0 | NA | NA | NA | NA | NA | NM_146814 |
| Olfr666 | 0 | NA | NA | NA | NA | NA | NM_147096 |
| Olfr667 | 0 | NA | NA | NA | NA | NA | NM_147060 |
| Olfr668 | 0 | NA | NA | NA | NA | NA | NM_147059 |
| Olfr669 | 0 | NA | NA | NA | NA | NA | NM_147043 |
| Olfr67 | 0 | NA | NA | NA | NA | NA | NM_013619 |
| Olfr670 | 0 | NA | NA | NA | NA | NA | NM_207146 |
| Olfr671 | 0 | NA | NA | NA | NA | NA | NM_001011755 |
| Olfr672 | 0 | NA | NA | NA | NA | NA | NM_146760 |
| Olfr675 | 0 | NA | NA | NA | NA | NA | NM_001011848 |
| Olfr676 | 0 | NA | NA | NA | NA | NA | NM_147095 |
| Olfr677 | 0 | NA | NA | NA | NA | NA | NM_146358 |
| Olfr678 | 0 | NA | NA | NA | NA | NA | NM_146758 |
| Olfr679 | 0 | NA | NA | NA | NA | NA | NM_147044 |
| Olfr68 | 0 | NA | NA | NA | NA | NA | NM_013620 |
| Olfr681 | 0.120209049 | -0.00674972139751468 | 0.00863521114606269 | -0.781650996523957 | 0.434419707908266 | NA | NM_207557 |
| Olfr683 | 0.315716853 | 0.0123948421219712 | 0.0131137196252776 | 0.945181266349427 | 0.344566341855147 | NA | NM_147045 |
| Olfr684 | 0 | NA | NA | NA | NA | NA | NM_207249 |
| Olfr685 | 0 | NA | NA | NA | NA | NA | NM_001011857 |
| Olfr686 | 0 | NA | NA | NA | NA | NA | NM_147069 |
| Olfr688 | 0 | NA | NA | NA | NA | NA | NM_001011533 |
| Olfr689 | 0 | NA | NA | NA | NA | NA | NM_146750 |
| Olfr69 | 0 | NA | NA | NA | NA | NA | NM_013621 |
| Olfr690 | 1.72318098 | 0.00760413980817141 | 0.0318533165751858 | 0.238723644058313 | 0.811319881441241 | NA | NM_020290 |
| Olfr691 | 0.755522351 | -0.0180977557915085 | 0.0201615859975957 | -0.89763552300234 | 0.369379894229839 | NA | NM_147061 |
| Olfr692 | 0 | NA | NA | NA | NA | NA | NM_146355 |
| Olfr693 | 0 | NA | NA | NA | NA | NA | NM_146453 |
| Olfr694 | 0 | NA | NA | NA | NA | NA | NM_146452 |
| Olfr695 | 0 | NA | NA | NA | NA | NA | NM_146598 |
| Olfr697 | 0.354968264 | -0.0136522058293636 | 0.0122686640796521 | -1.11277036690622 | 0.265807065332778 | NA | NM_146599 |
| Olfr698 | 0 | NA | NA | NA | NA | NA | NM_146602 |
| Olfr699 | 0 | NA | NA | NA | NA | NA | NM_001011862 |
| Olfr70 | 0 | NA | NA | NA | NA | NA | NM_019485 |
| Olfr700 | 0.223412651 | 0.00718831964189475 | 0.00999993766105603 | 0.718836445339964 | 0.472241699104885 | NA | NM_146600 |
| Olfr701 | 0.209568245 | 0.00686197416234131 | 0.00977127153168884 | 0.702260104029194 | 0.482516968822431 | NA | NM_028910 |
| Olfr702 | 0 | NA | NA | NA | NA | NA | NM_146597 |
| Olfr703 | 0 | NA | NA | NA | NA | NA | NM_146596 |
| Olfr704 | 0 | NA | NA | NA | NA | NA | NM_001011749 |
| Olfr705 | 0 | NA | NA | NA | NA | NA | NM_147032 |
| Olfr706 | 0 | NA | NA | NA | NA | NA | NM_146353 |
| Olfr707 | 0.106347839 | -0.00653347234140708 | 0.00854389509723638 | -0.764694822098226 | 0.444453290197145 | NA | NM_001005570 |
| Olfr71 | 0 | NA | NA | NA | NA | NA | NM_019486 |
| Olfr710 | 0 | NA | NA | NA | NA | NA | NM_146601 |
| Olfr711 | 0 | NA | NA | NA | NA | NA | NM_147035 |
| Olfr713 | 0 | NA | NA | NA | NA | NA | NM_147034 |
| Olfr714 | 0 | NA | NA | NA | NA | NA | NM_147033 |
| Olfr715 | 0 | NA | NA | NA | NA | NA | NM_146780 |
| Olfr716 | 0 | NA | NA | NA | NA | NA | NM_146604 |
| Olfr720 | 0 | NA | NA | NA | NA | NA | NM_146392 |
| Olfr722 | 0 | NA | NA | NA | NA | NA | NM_146494 |
| Olfr723 | 0 | NA | NA | NA | NA | NA | NM_001011530 |
| Olfr724 | 0 | NA | NA | NA | NA | NA | NM_146492 |
| Olfr725 | 0 | NA | NA | NA | NA | NA | NM_146317 |
| Olfr726 | 0 | NA | NA | NA | NA | NA | NM_146316 |
| Olfr727 | 0.205932006 | -0.000576954122277892 | 0.00970424421944593 | -0.0594537925088239 | 0.952590668569529 | NA | NM_146319 |
| Olfr728 | 0 | NA | NA | NA | NA | NA | NM_001011809 |
| Olfr729 | 0 | NA | NA | NA | NA | NA | NM_146278 |
| Olfr73 | 0 | NA | NA | NA | NA | NA | NM_054090 |
| Olfr730 | 0 | NA | NA | NA | NA | NA | NM_146493 |
| Olfr731 | 0 | NA | NA | NA | NA | NA | NM_146363 |
| Olfr732 | 0 | NA | NA | NA | NA | NA | NM_146665 |
| Olfr733 | 0 | NA | NA | NA | NA | NA | NM_146663 |
| Olfr734 | 0 | NA | NA | NA | NA | NA | NM_146664 |
| Olfr735 | 0 | NA | NA | NA | NA | NA | NM_001011754 |
| Olfr736 | 0 | NA | NA | NA | NA | NA | NM_146666 |
| Olfr738 | 0 | NA | NA | NA | NA | NA | NM_146420 |
| Olfr739 | 0 | NA | NA | NA | NA | NA | NM_146668 |
| Olfr74 | 0 | NA | NA | NA | NA | NA | NM_054091 |
| Olfr740 | 0 | NA | NA | NA | NA | NA | NM_146667 |
| Olfr741 | 0.12890128 | -0.0068640067066208 | 0.00870804083610955 | -0.788237771940376 | 0.430557639008166 | NA | NM_207133 |
| Olfr742 | 0.194331409 | -0.00840968081320881 | 0.00964081822647896 | -0.872299489073576 | 0.383045016303155 | NA | NM_146430 |
| Olfr743 | 0 | NA | NA | NA | NA | NA | NM_001177508 |
| Olfr744 | 0 | NA | NA | NA | NA | NA | NM_001011738 |
| Olfr745 | 0 | NA | NA | NA | NA | NA | NM_146299 |
| Olfr746 | 0 | NA | NA | NA | NA | NA | NM_146298 |
| Olfr747 | 0 | NA | NA | NA | NA | NA | NM_207156 |
| Olfr748 | 0 | NA | NA | NA | NA | NA | NM_001011837 |
| Olfr749 | 0 | NA | NA | NA | NA | NA | NM_020288 |
| Olfr75-ps1 | 0 | NA | NA | NA | NA | NA | NR_002859,NR_033528 |
| Olfr750 | 0 | NA | NA | NA | NA | NA | NM_207558 |
| Olfr76 | 0 | NA | NA | NA | NA | NA | NM_146682 |
| Olfr761 | 0 | NA | NA | NA | NA | NA | NM_001011829 |
| Olfr763 | 0 | NA | NA | NA | NA | NA | NM_146862 |
| Olfr765 | 0 | NA | NA | NA | NA | NA | NM_001085477 |
| Olfr767 | 0 | NA | NA | NA | NA | NA | NM_146318 |
| Olfr768 | 0 | NA | NA | NA | NA | NA | NM_146864 |
| Olfr769 | 0 | NA | NA | NA | NA | NA | NM_146267 |
| Olfr77 | 0 | NA | NA | NA | NA | NA | NM_146339 |
| Olfr770 | 0 | NA | NA | NA | NA | NA | NM_146863 |
| Olfr771 | 0 | NA | NA | NA | NA | NA | NM_146547 |
| Olfr772 | 0 | NA | NA | NA | NA | NA | NM_146266 |
| Olfr773 | 0 | NA | NA | NA | NA | NA | NM_207008 |
| Olfr774 | 0 | NA | NA | NA | NA | NA | NM_207620 |
| Olfr775 | 0 | NA | NA | NA | NA | NA | NM_146545 |
| Olfr776 | 0 | NA | NA | NA | NA | NA | NM_207559 |
| Olfr777 | 0 | NA | NA | NA | NA | NA | NM_146544 |
| Olfr780 | 0 | NA | NA | NA | NA | NA | NM_146284 |
| Olfr781 | 0 | NA | NA | NA | NA | NA | NM_146728 |
| Olfr782 | 0 | NA | NA | NA | NA | NA | NM_001011797 |
| Olfr784 | 0 | NA | NA | NA | NA | NA | NM_146729 |
| Olfr786 | 0 | NA | NA | NA | NA | NA | NM_146549 |
| Olfr787 | 0 | NA | NA | NA | NA | NA | NM_001011822 |
| Olfr788 | 0 | NA | NA | NA | NA | NA | NM_146551 |
| Olfr790 | 0 | NA | NA | NA | NA | NA | NM_146933 |
| Olfr791 | 0 | NA | NA | NA | NA | NA | NM_146930 |
| Olfr792 | 0 | NA | NA | NA | NA | NA | NM_001011849 |
| Olfr794 | 0 | NA | NA | NA | NA | NA | NM_146378 |
| Olfr796 | 0 | NA | NA | NA | NA | NA | NM_146931 |
| Olfr798 | 0 | NA | NA | NA | NA | NA | NM_146556 |
| Olfr799 | 0 | NA | NA | NA | NA | NA | NM_146927 |
| Olfr8 | 0 | NA | NA | NA | NA | NA | NM_207201 |
| Olfr800 | 0 | NA | NA | NA | NA | NA | NM_146548 |
| Olfr801 | 0 | NA | NA | NA | NA | NA | NM_146285 |
| Olfr802 | 0 | NA | NA | NA | NA | NA | NM_146932 |
| Olfr803 | 0 | NA | NA | NA | NA | NA | NM_146554 |
| Olfr804 | 0 | NA | NA | NA | NA | NA | NM_001011821 |
| Olfr805 | 0 | NA | NA | NA | NA | NA | NM_146555 |
| Olfr806 | 0 | NA | NA | NA | NA | NA | NM_146553 |
| Olfr807 | 0 | NA | NA | NA | NA | NA | NM_146929 |
| Olfr808 | 0 | NA | NA | NA | NA | NA | NM_146928 |
| Olfr809 | 0 | NA | NA | NA | NA | NA | NM_146324 |
| Olfr810 | 0 | NA | NA | NA | NA | NA | NM_146550 |
| Olfr811 | 0 | NA | NA | NA | NA | NA | NM_146552 |
| Olfr812 | 0 | NA | NA | NA | NA | NA | NM_146795 |
| Olfr813 | 0 | NA | NA | NA | NA | NA | NM_207147 |
| Olfr814 | 0 | NA | NA | NA | NA | NA | NM_207159 |
| Olfr815 | 0 | NA | NA | NA | NA | NA | NM_146670 |
| Olfr816 | 0 | NA | NA | NA | NA | NA | NM_146672 |
| Olfr818 | 0 | NA | NA | NA | NA | NA | NM_146777 |
| Olfr819 | 0 | NA | NA | NA | NA | NA | NM_001165944 |
| Olfr820 | 0 | NA | NA | NA | NA | NA | NM_146675 |
| Olfr821 | 0 | NA | NA | NA | NA | NA | NM_146776 |
| Olfr822 | 0 | NA | NA | NA | NA | NA | NM_146671 |
| Olfr823 | 0 | NA | NA | NA | NA | NA | NM_146673 |
| Olfr824 | 0 | NA | NA | NA | NA | NA | NM_146674 |
| Olfr825 | 0 | NA | NA | NA | NA | NA | NM_146677 |
| Olfr826 | 0 | NA | NA | NA | NA | NA | NM_146676 |
| Olfr827 | 0 | NA | NA | NA | NA | NA | NM_146300 |
| Olfr828 | 0 | NA | NA | NA | NA | NA | NM_146605 |
| Olfr829 | 0 | NA | NA | NA | NA | NA | NM_147067 |
| Olfr830 | 0 | NA | NA | NA | NA | NA | NM_146566 |
| Olfr832 | 0 | NA | NA | NA | NA | NA | NM_001011824 |

| | | | | | | | | | | | | | | | |
|---|---|---|---|---|---|---|---|---|---|---|---|---|---|---|---|
| Olfr538 | 0 | NA | NA | NA | NA | NA | | Olfr834 | 0 | NA | NA | NA | NA | NA | NM_001011823 |
| Olfr539 | 0 | NA | NA | NA | NA | NA | | Olfr835 | 0 | NA | NA | NA | NA | NA | NM_001012266 |
| Olfr54 | 0 | NA | NA | NA | NA | NA | | Olfr836 | 0.120209049 | -0.00674972139751468 | 0.00863521114606269 | -0.781650996523957 | 0.434419707908266 | NA | NM_146564 |
| Olfr541 | 0 | NA | NA | NA | NA | NA | | Olfr837 | 0 | NA | NA | NA | NA | NA | NM_146565 |
| Olfr543 | 0 | NA | NA | NA | NA | NA | | Olfr843 | 0 | NA | NA | NA | NA | NA | NM_146567 |
| Olfr544 | 0 | NA | NA | NA | NA | NA | | Olfr845 | 0 | NA | NA | NA | NA | NA | NM_207145 |
| Olfr545 | 0 | NA | NA | NA | NA | NA | | Olfr846 | 0 | NA | NA | NA | NA | NA | NM_146282 |
| Olfr547 | 0 | NA | NA | NA | NA | NA | | Olfr847 | 0 | NA | NA | NA | NA | NA | NM_146525 |
| Olfr549 | 0 | NA | NA | NA | NA | NA | | Olfr849 | 0 | NA | NA | NA | NA | NA | NM_146527 |
| Olfr55 | 0 | NA | NA | NA | NA | NA | | Olfr850 | 0 | NA | NA | NA | NA | NA | NM_146523 |
| Olfr550 | 2.860648009 | 0.150094874 | 0.097489485 | 1.53960065 | 0.123657727 | NA | | Olfr851 | 0 | NA | NA | NA | NA | NA | NM_146905 |
| Olfr551 | 0 | NA | NA | NA | NA | NA | | Olfr853 | 0 | NA | NA | NA | NA | NA | NM_146906 |
| Olfr552 | 0 | NA | NA | NA | NA | NA | | Olfr854 | 0 | NA | NA | NA | NA | NA | NM_146522 |
| Olfr553 | 0 | NA | NA | NA | NA | NA | | Olfr855 | 0 | NA | NA | NA | NA | NA | NM_146524 |
| Olfr554 | 0 | NA | NA | NA | NA | NA | | Olfr857 | 0.458678574 | -0.00329924320639459 | 0.0155038389300741 | -0.212801695197876 | 0.831481641452064 | NA | NM_001012265 |
| Olfr555 | 0 | NA | NA | NA | NA | NA | | Olfr859 | 0.690783455 | 0.00419901188792458 | 0.017647408750934 | 0.237939288831984 | 0.811928182360044 | NA | NM_146526 |
| Olfr556 | 0 | NA | NA | NA | NA | NA | | Olfr860 | 0 | NA | NA | NA | NA | NA | NM_146528 |
| Olfr557 | 0 | NA | NA | NA | NA | NA | | Olfr862 | 0 | NA | NA | NA | NA | NA | NM_146562 |
| Olfr559 | 0 | NA | NA | NA | NA | NA | | Olfr866 | 0 | NA | NA | NA | NA | NA | NM_146558 |
| Olfr56 | 1.12784763 | -0.033726047 | 0.068604092 | -0.491604007 | 0.62299931 | NA | | Olfr867 | 0 | NA | NA | NA | NA | NA | NM_001011748 |
| Olfr560 | 0.669314316 | -0.102290746 | 0.060715769 | -1.684747598 | 0.092037279 | NA | | Olfr868 | 0 | NA | NA | NA | NA | NA | NM_146559 |
| Olfr561 | 0.292412586 | 0.032260595 | 0.035568585 | 0.906996868 | 0.364408454 | NA | | Olfr869 | 0 | NA | NA | NA | NA | NA | NM_146557 |
| Olfr564 | 0.510820111 | 0.029651871 | 0.046114588 | 0.643004131 | 0.520221419 | NA | | Olfr870 | 0 | NA | NA | NA | NA | NA | NM_146904 |
| Olfr566 | 0 | NA | NA | NA | NA | NA | | Olfr871 | 0 | NA | NA | NA | NA | NA | NM_146903 |
| Olfr568 | 0 | NA | NA | NA | NA | NA | | Olfr872 | 0 | NA | NA | NA | NA | NA | NM_146560 |
| Olfr569 | 0 | NA | NA | NA | NA | NA | | Olfr873 | 0 | NA | NA | NA | NA | NA | NM_146561 |
| Olfr57 | 0 | NA | NA | NA | NA | NA | | Olfr874 | 0.215934008 | -0.00149029401866511 | 0.00975123948835942 | -0.152831239602324 | 0.878531361484804 | NA | NM_146882 |
| Olfr570 | 0.254617334 | 0.029389883 | 0.033981526 | 0.864878265 | 0.387105608 | NA | | Olfr875 | 0.09572496 | 0.00498255289994511 | 0.00847999972823692 | 0.587565219295241 | 0.556824162160132 | NA | NM_146749 |
| Olfr571 | 0 | NA | NA | NA | NA | NA | | Olfr876 | 0.127687691 | 0.00542298404407807 | 0.00869724871942359 | 0.62352868349814 | 0.532937159177571 | NA | NM_146883 |
| Olfr572 | 0 | NA | NA | NA | NA | NA | | Olfr877 | 0.444845289 | 0.00837525398380596 | 0.0155822522737987 | 0.537486740468757 | 0.590931441532687 | NA | NM_146417 |
| Olfr574 | 0.195033442 | -0.002424765 | 0.031596358 | -0.076741901 | 0.93882887 | NA | | Olfr878 | 0.650179346 | 0.00867115731878333 | 0.0138177707927729 | 0.627536630099451 | 0.530307532921215 | NA | NM_146798 |
| Olfr575 | 0 | NA | NA | NA | NA | NA | | Olfr881 | 0.097165704 | -0.00637828455464458 | 0.00848781691535724 | -0.75146349388194 | 0.452373763047372 | NA | NM_146418 |
| Olfr576 | 0 | NA | NA | NA | NA | NA | | Olfr883 | 0.12890128 | -0.0068640067066208 | 0.00870804083610955 | -0.788237771940376 | 0.430557639008166 | NA | NM_146419 |
| Olfr577 | 0 | NA | NA | NA | NA | NA | | Olfr884 | 0 | NA | NA | NA | NA | NA | NM_001011798 |
| Olfr578 | 0 | NA | NA | NA | NA | NA | | Olfr885 | 0 | NA | NA | NA | NA | NA | NM_001011739 |
| Olfr58 | 0.08731684 | 0.017365056 | 0.027733412 | 0.626142072 | 0.531221759 | NA | | Olfr887 | 0 | NA | NA | NA | NA | NA | NM_146423 |
| Olfr582 | 0 | NA | NA | NA | NA | NA | | Olfr888 | 0 | NA | NA | NA | NA | NA | NM_146424 |
| Olfr583 | 0 | NA | NA | NA | NA | NA | | Olfr889 | 0 | NA | NA | NA | NA | NA | NM_146482 |
| Olfr584 | 0 | NA | NA | NA | NA | NA | | Olfr890 | 0 | NA | NA | NA | NA | NA | NM_146481 |
| Olfr585 | 0 | NA | NA | NA | NA | NA | | Olfr891 | 0 | NA | NA | NA | NA | NA | NM_146478 |
| Olfr586 | 0 | NA | NA | NA | NA | NA | | Olfr893 | 0 | NA | NA | NA | NA | NA | NM_146336 |
| Olfr589 | 0 | NA | NA | NA | NA | NA | | Olfr894 | 0 | NA | NA | NA | NA | NA | NM_146868 |
| Olfr59 | 0 | NA | NA | NA | NA | NA | | Olfr895 | 0 | NA | NA | NA | NA | NA | NM_146875 |
| Olfr591 | 0 | NA | NA | NA | NA | NA | | Olfr898 | 0.678454432 | 0.0166244498836941 | 0.0196617984455118 | 0.845520308315893 | 0.39782039880612 | NA | NM_146871 |
| Olfr592 | 0 | NA | NA | NA | NA | NA | | Olfr899 | 0 | NA | NA | NA | NA | NA | NM_146479 |
| Olfr593 | 0 | NA | NA | NA | NA | NA | | Olfr9 | 0 | NA | NA | NA | NA | NA | NM_146861 |
| Olfr594 | 0 | NA | NA | NA | NA | NA | | Olfr90 | 0 | NA | NA | NA | NA | NA | NM_146477 |
| Olfr596 | 0 | NA | NA | NA | NA | NA | | Olfr900 | 0 | NA | NA | NA | NA | NA | NM_146874 |
| Olfr597 | 0 | NA | NA | NA | NA | NA | | Olfr901 | 0 | NA | NA | NA | NA | NA | NM_001011806 |
| Olfr598 | 0 | NA | NA | NA | NA | NA | | Olfr902 | 0 | NA | NA | NA | NA | NA | NM_146802 |
| Olfr599 | 0 | NA | NA | NA | NA | NA | | Olfr904 | 0 | NA | NA | NA | NA | NA | NM_146801 |
| Olfr6 | 0.098353386 | -0.020212831 | 0.02786466 | -0.725393061 | 0.468210929 | NA | | Olfr905 | 0 | NA | NA | NA | NA | NA | NM_146804 |
| Olfr60 | 0 | NA | NA | NA | NA | NA | | Olfr906 | 0 | NA | NA | NA | NA | NA | NM_146803 |
| Olfr600 | 0 | NA | NA | NA | NA | NA | | Olfr907 | 0 | NA | NA | NA | NA | NA | NM_146805 |
| Olfr601 | 0 | NA | NA | NA | NA | NA | | Olfr908 | 0 | NA | NA | NA | NA | NA | NM_146872 |
| Olfr603 | 0 | NA | NA | NA | NA | NA | | Olfr91 | 0 | NA | NA | NA | NA | NA | NM_182714 |
| Olfr605 | 0 | NA | NA | NA | NA | NA | | Olfr910 | 0 | NA | NA | NA | NA | NA | NM_146811 |
| Olfr606 | 0 | NA | NA | NA | NA | NA | | Olfr911-ps1 | 0 | NA | NA | NA | NA | NA | NM_146873 |
| Olfr608 | 0 | NA | NA | NA | NA | NA | | Olfr912 | 0 | NA | NA | NA | NA | NA | NM_146810 |
| Olfr609 | 0.198019769 | 0.025308392 | 0.031628095 | 0.80018706 | 0.423602426 | NA | | Olfr913 | 0 | NA | NA | NA | NA | NA | NM_001011523 |
| Olfr61 | 0.098353386 | -0.020212831 | 0.02786466 | -0.725393061 | 0.468210929 | NA | | Olfr914 | 0 | NA | NA | NA | NA | NA | NM_146786 |
| Olfr610 | 0.094392817 | 0.017895942 | 0.027818137 | 0.643319236 | 0.520016977 | NA | | Olfr915 | 0 | NA | NA | NA | NA | NA | NM_146785 |
| Olfr611 | 0 | NA | NA | NA | NA | NA | | Olfr916 | 0 | NA | NA | NA | NA | NA | NM_146784 |
| Olfr612 | 0 | NA | NA | NA | NA | NA | | Olfr917 | 0 | NA | NA | NA | NA | NA | NM_001011864 |
| Olfr615 | 0 | NA | NA | NA | NA | NA | | Olfr918 | 0 | NA | NA | NA | NA | NA | NM_146375 |
| Olfr616 | 0 | NA | NA | NA | NA | NA | | Olfr919 | 0 | NA | NA | NA | NA | NA | NM_146440 |
| Olfr617 | 0 | NA | NA | NA | NA | NA | | Olfr92 | 0 | NA | NA | NA | NA | NA | NM_146456 |
| Olfr618 | 0 | NA | NA | NA | NA | NA | | Olfr920 | 3.984140183 | -0.0292923687588997 | 0.0424046468257506 | -0.690782047525783 | 0.489702519067108 | NA | NM_146787 |
| Olfr619 | 0 | NA | NA | NA | NA | NA | | Olfr921 | 0.106148608 | 0.00515554063321497 | 0.00854200771766808 | 0.603551390213729 | 0.546141939141592 | NA | NM_146782 |
| Olfr62 | 0 | NA | NA | NA | NA | NA | | Olfr922 | 0 | NA | NA | NA | NA | NA | NM_146781 |
| Olfr620 | 0 | NA | NA | NA | NA | NA | | Olfr923 | 0 | NA | NA | NA | NA | NA | NM_146816 |
| Olfr622 | 0.159967308 | 0.0240345 | 0.030870969 | 0.778547006 | 0.436246606 | NA | | Olfr924 | 0 | NA | NA | NA | NA | NA | NM_207560 |
| Olfr623 | 0 | NA | NA | NA | NA | NA | | Olfr926 | 0 | NA | NA | NA | NA | NA | NM_146815 |
| Olfr624 | 0 | NA | NA | NA | NA | NA | | Olfr93 | 0 | NA | NA | NA | NA | NA | NM_001011813 |
| Olfr628 | 0 | NA | NA | NA | NA | NA | | Olfr930 | 0 | NA | NA | NA | NA | NA | NM_146272 |
| Olfr629 | 0 | NA | NA | NA | NA | NA | | Olfr933 | 0 | NA | NA | NA | NA | NA | NM_146441 |
| Olfr63 | 0 | NA | NA | NA | NA | NA | | Olfr934 | 0 | NA | NA | NA | NA | NA | NM_146442 |
| Olfr630 | 0 | NA | NA | NA | NA | NA | | Olfr935 | 0 | NA | NA | NA | NA | NA | NM_146746 |
| Olfr631 | 0 | NA | NA | NA | NA | NA | | Olfr936 | 0 | NA | NA | NA | NA | NA | NM_207139 |
| Olfr632 | 0 | NA | NA | NA | NA | NA | | Olfr937 | 0 | NA | NA | NA | NA | NA | NM_146439 |
| Olfr633 | 0 | NA | NA | NA | NA | NA | | Olfr938 | 0 | NA | NA | NA | NA | NA | NM_146438 |
| Olfr635 | 0 | NA | NA | NA | NA | NA | | Olfr94 | 0 | NA | NA | NA | NA | NA | NM_001011518 |
| Olfr638 | 0 | NA | NA | NA | NA | NA | | Olfr943 | 0 | NA | NA | NA | NA | NA | NM_146326 |
| Olfr639 | 0 | NA | NA | NA | NA | NA | | Olfr944 | 0 | NA | NA | NA | NA | NA | NM_146507 |
| Olfr64 | 0 | NA | NA | NA | NA | NA | | Olfr945 | 0 | NA | NA | NA | NA | NA | NM_146506 |
| Olfr640 | 0.098353386 | -0.020212831 | 0.02786466 | -0.725393061 | 0.468210929 | NA | | Olfr947-ps1 | 0 | NA | NA | NA | NA | NA | NR_033620 |
| Olfr641 | 0 | NA | NA | NA | NA | NA | | Olfr948 | 0 | NA | NA | NA | NA | NA | NM_001011756 |
| Olfr642 | 0 | NA | NA | NA | NA | NA | | Olfr95 | 0.097165704 | -0.00637828455464458 | 0.00848781691535724 | -0.75146349388194 | 0.452373763047372 | NA | NM_146513 |
| Olfr643 | 0 | NA | NA | NA | NA | NA | | Olfr951 | 0 | NA | NA | NA | NA | NA | NM_001011812 |
| Olfr644 | 0 | NA | NA | NA | NA | NA | | Olfr952 | 0 | NA | NA | NA | NA | NA | NM_146503 |
| Olfr645 | 0 | NA | NA | NA | NA | NA | | Olfr954 | 0 | NA | NA | NA | NA | NA | NM_146331 |
| Olfr646 | 0 | NA | NA | NA | NA | NA | | Olfr955 | 0 | NA | NA | NA | NA | NA | NM_207141 |
| Olfr648 | 0 | NA | NA | NA | NA | NA | | Olfr957 | 0 | NA | NA | NA | NA | NA | NM_146745 |
| Olfr649 | 0 | NA | NA | NA | NA | NA | | Olfr958 | 0 | NA | NA | NA | NA | NA | NM_146330 |
| Olfr65 | 0 | NA | NA | NA | NA | NA | | Olfr959 | 0 | NA | NA | NA | NA | NA | NM_146508 |
| Olfr651 | 0 | NA | NA | NA | NA | NA | | Olfr96 | 0 | NA | NA | NA | NA | NA | NM_146514 |
| Olfr652 | 0 | NA | NA | NA | NA | NA | | Olfr960 | 0 | NA | NA | NA | NA | NA | NM_146279 |
| Olfr653 | 0 | NA | NA | NA | NA | NA | | Olfr961 | 0 | NA | NA | NA | NA | NA | NM_146504 |
| Olfr654 | 0 | NA | NA | NA | NA | NA | | Olfr963 | 0 | NA | NA | NA | NA | NA | NM_001011827 |
| Olfr655 | 0 | NA | NA | NA | NA | NA | | Olfr965 | 0 | NA | NA | NA | NA | NA | NM_001011859 |
| Olfr656 | 0 | NA | NA | NA | NA | NA | | Olfr967 | 0 | NA | NA | NA | NA | NA | NM_001011826 |
| Olfr657 | 0 | NA | NA | NA | NA | NA | | Olfr968 | 0 | NA | NA | NA | NA | NA | NM_146612 |
| Olfr658 | 2.070826888 | 0.058282488 | 0.102596741 | 0.568073485 | 0.569985071 | NA | | Olfr969 | 0 | NA | NA | NA | NA | NA | NM_146826 |
| Olfr659 | 0 | NA | NA | NA | NA | NA | | Olfr97 | 0 | NA | NA | NA | NA | NA | NM_146512 |
| Olfr66 | 0 | NA | NA | NA | NA | NA | | Olfr970 | 0.240418098 | -0.0088029805258837 | 0.00986414519853158 | -0.892422034419582 | 0.372166765031741 | NA | NM_146611 |
| Olfr661 | 0 | NA | NA | NA | NA | NA | | Olfr971 | 0 | NA | NA | NA | NA | NA | NM_146614 |
| Olfr663 | 0 | NA | NA | NA | NA | NA | | Olfr972 | 0 | NA | NA | NA | NA | NA | NM_146610 |
| Olfr665 | 0 | NA | NA | NA | NA | NA | | Olfr974 | 0 | NA | NA | NA | NA | NA | NM_147107 |
| Olfr666 | 0 | NA | NA | NA | NA | NA | | Olfr975 | 0 | NA | NA | NA | NA | NA | NM_146828 |
| Olfr667 | 0 | NA | NA | NA | NA | NA | | Olfr976 | 0 | NA | NA | NA | NA | NA | NM_146367 |
| Olfr668 | 0 | NA | NA | NA | NA | NA | | Olfr978 | 0 | NA | NA | NA | NA | NA | NM_147105 |
| Olfr669 | 0 | NA | NA | NA | NA | NA | | Olfr979 | 0 | NA | NA | NA | NA | NA | NM_147108 |
| Olfr67 | 0 | NA | NA | NA | NA | NA | | Olfr98 | 0 | NA | NA | NA | NA | NA | NM_146510 |
| Olfr670 | 0 | NA | NA | NA | NA | NA | | Olfr980 | 0 | NA | NA | NA | NA | NA | NM_147106 |
| Olfr671 | 0 | NA | NA | NA | NA | NA | | Olfr981 | 0 | NA | NA | NA | NA | NA | NM_146286 |
| Olfr672 | 0 | NA | NA | NA | NA | NA | | Olfr982 | 0 | NA | NA | NA | NA | NA | NM_146854 |
| Olfr675 | 0 | NA | NA | NA | NA | NA | | Olfr983 | 0 | NA | NA | NA | NA | NA | NM_146827 |
| Olfr676 | 0 | NA | NA | NA | NA | NA | | Olfr984 | 0 | NA | NA | NA | NA | NA | NM_146608 |
| Olfr677 | 0 | NA | NA | NA | NA | NA | | Olfr985 | 0 | NA | NA | NA | NA | NA | NM_146855 |
| Olfr678 | 0 | NA | NA | NA | NA | NA | | Olfr986 | 0 | NA | NA | NA | NA | NA | NM_146615 |
| Olfr679 | 0.094392817 | 0.017895942 | 0.027818137 | 0.643319236 | 0.520016977 | NA | | Olfr987 | 0 | NA | NA | NA | NA | NA | NM_001011785 |
| Olfr68 | 0 | NA | NA | NA | NA | NA | | Olfr988 | 0 | NA | NA | NA | NA | NA | NM_001011534 |
| Olfr681 | 0.099009884 | 0.018205726 | 0.027896865 | 0.652608315 | 0.514008823 | NA | | Olfr99 | 0 | NA | NA | NA | NA | NA | NM_146515 |
| Olfr683 | 0 | NA | NA | NA | NA | NA | | Olfr992 | 0 | NA | NA | NA | NA | NA | NM_146865 |
| Olfr684 | 0 | NA | NA | NA | NA | NA | | Olfr993 | 0 | NA | NA | NA | NA | NA | NM_146435 |
| Olfr685 | 0 | NA | NA | NA | NA | NA | | Olfr994 | 0 | NA | NA | NA | NA | NA | NM_146433 |
| Olfr686 | 0 | NA | NA | NA | NA | NA | | Olfr995 | 0 | NA | NA | NA | NA | NA | NM_146434 |
| Olfr688 | 0 | NA | NA | NA | NA | NA | | Olfr996 | 0 | NA | NA | NA | NA | NA | NM_146437 |
| Olfr689 | 0 | NA | NA | NA | NA | NA | | Olfr998 | 0 | NA | NA | NA | NA | NA | NM_146436 |
| Olfr69 | 0 | NA | NA | NA | NA | NA | | Olig3 | 3.202049419 | 0.00311104832025895 | 0.0336502928120781 | 0.0924523402406265 | 0.926338655955027 | NA | NM_053008 |
| Olfr690 | 1.599022041 | 0.012544975 | 0.091741038 | 0.136743326 | 0.891233682 | NA | | Olr1 | 2.740774214 | -0.0860153455474191 | 0.039975589087875 | -2.1516967606991 | 0.0314212454451118 | NA | NM_138648 |
| Olfr691 | 1.723081843 | -0.107161713 | 0.100602915 | -1.065194912 | 0.286787736 | NA | | Omp | 19.31650379 | -0.0303188356306833 | 0.053830925989413 | -0.563223371573547 | 0.573282790070658 | NA | NM_011010 |
| Olfr692 | 0 | NA | NA | NA | NA | NA | | Omt2a | 0 | NA | NA | NA | NA | NA | NM_001111286,NM_001164523 |
| Olfr693 | 0 | NA | NA | NA | NA | NA | | Omt2b | 0 | NA | NA | NA | NA | NA | NM_205822 |
| Olfr694 | 0 | NA | NA | NA | NA | NA | | Onecut1 | 4.667185566 | -0.0133338869374813 | 0.0386864893457308 | -0.344665209042876 | 0.730346076346337 | NA | NM_008262 |
| Olfr695 | 0 | NA | NA | NA | NA | NA | | Onecut3 | 6.492253781 | 0.0259196323963202 | 0.0191481942414224 | 1.35363324966954 | NA | NA | NM_139226 |
| Olfr697 | 0.695061837 | 0.009378663 | 0.060757132 | 0.154363163 | 0.877323397 | NA | | Ooep | 5.386247955 | -0.0265298839434957 | 0.0445400318287243 | -0.595641333295732 | 0.551414857916813 | NA | NM_026480 |
| Olfr698 | 0 | NA | NA | NA | NA | NA | | Oog1 | 0 | NA | NA | NA | NA | NA | NM_178657 |
| Olfr699 | 0 | NA | NA | NA | NA | NA | | Oog2 | 0 | NA | NA | NA | NA | NA | NM_198661 |
| Olfr70 | 0 | NA | NA | NA | NA | NA | | Oog3 | 0 | NA | NA | NA | NA | NA | NM_201258 |
| Olfr700 | 0.119397641 | -0.020477583 | 0.027954224 | -0.732539845 | 0.463839132 | NA | | Oog4 | 0 | NA | NA | NA | NA | NA | NM_173773 |
| Olfr701 | 0 | NA | NA | NA | NA | NA | | Oosp1 | 0.09572496 | 0.00498255289994511 | 0.00847999972823692 | 0.587565219295241 | 0.556824162160132 | NA | NM_133353 |
| Olfr702 | 0 | NA | NA | NA | NA | NA | | Opn1mw | 16.8867866 | 9.16558074680783e-05 | 0.0684358114584057 | 0.00133929598429304 | 0.998931396731249 | NA | NM_008106 |
| Olfr703 | 0 | NA | NA | NA | NA | NA | | Opn1sw | 1.754973435 | -0.0323889422969295 | 0.0285769573760712 | -1.13339365946811 | 0.257048975568632 | NA | NM_007538 |
| Olfr704 | 0 | NA | NA | NA | NA | NA | | Opn4 | 1.651225453 | -0.0185342179590576 | 0.0233252638385062 | -0.794598427155222 | 0.426847133560641 | NA | NM_013887,NM_001128599 |
| Olfr705 | 0 | NA | NA | NA | NA | NA | | Opn5 | 1.046073401 | 0.0253517053006967 | 0.0212326688494456 | 1.19399522878908 | 0.23247983442001 | NA | NM_181753 |
| Olfr706 | 0 | NA | NA | NA | NA | NA | | Orc1 | 6.230791539 | 0.0262351597923845 | 0.0490558005712926 | 0.534802398225202 | 0.592786504531828 | NA | NM_011015 |
| Olfr707 | 0.08731684 | 0.017365056 | 0.027733412 | 0.626142072 | 0.531221759 | NA | | Orm1 | 0.106347839 | -0.00653347234140708 | 0.00854389509723638 | -0.764694822098226 | 0.444453290197145 | NA | NM_008768 |
| Olfr71 | 0 | NA | NA | NA | NA | NA | | Orm2 | 0 | NA | NA | NA | NA | NA | NM_011016 |
| Olfr710 | 0 | NA | NA | NA | NA | NA | | Orm3 | 0.097165704 | -0.00637828455464458 | 0.00848781691535724 | -0.75146349388194 | 0.452373763047372 | NA | NM_013623 |
| Olfr711 | 0 | NA | NA | NA | NA | NA | | Osm | 4.925129957 | -0.094616335111715 | 0.0491062653553796 | -1.92676707192007 | 0.0540086648282711 | NA | NM_001013365 |
| Olfr713 | 0 | NA | NA | NA | NA | NA | | Osr2 | 0.212695677 | -0.00857796107890413 | 0.00973699866180651 | -0.880965621629515 | 0.378336427605205 | NA | NM_054049 |
| Olfr714 | 0 | NA | NA | NA | NA | NA | | Ostn | 1.400948497 | 0.0210407191400323 | 0.0220287285319693 | 0.955149050454585 | 0.339502326921059 | NA | NM_198112 |
| Olfr715 | 0 | NA | NA | NA | NA | NA | | Otc | 0.440787404 | -0.00207676726209905 | 0.0153548750617872 | -0.135251329219043 | 0.892413165045382 | NA | NM_008769 |
| Olfr716 | 0 | NA | NA | NA | NA | NA | | Otoa | 4.725076963 | -0.0191123776894087 | 0.0465847160326933 | -0.410271421983029 | 0.681606853412878 | NA | NM_139310 |
| Olfr720 | 0 | NA | NA | NA | NA | NA | | Otog | 7.592172514 | -0.0354352918983094 | 0.0556357844943864 | -0.636915471226703 | 0.524179904674843 | NA | NM_013624 |
| Olfr722 | 0 | NA | NA | NA | NA | NA | | Otogl | 15.62130939 | -0.153559297566809 | 0.0710653584470091 | -2.16081788543025 | 0.030709409922246 | NA | NM_001177567 |
| Olfr723 | 0 | NA | NA | NA | NA | NA | | Otol1 | 0.217532603 | 0.00684685248201087 | 0.00976000296730706 | 0.701521557416086 | 0.482977586094298 | NA | NM_001018031 |
| Olfr724 | 0 | NA | NA | NA | NA | NA | | Otop1 | 1.489672776 | -0.0221788961813212 | 0.0202569199639198 | -1.09487998278241 | NA | NA | NM_172709 |
| Olfr725 | 0 | NA | NA | NA | NA | NA | | Otop3 | 4.88749934 | -0.033499242873159 | 0.0452845873381879 | -0.739749324046717 | 0.459452113232439 | NA | NM_027132 |
| Olfr726 | 0 | NA | NA | NA | NA | NA | | Otor | 0.360627147 | -0.0100772056525624 | 0.010553896215922 | -0.954832741045871 | 0.339662287230353 | NA | NM_020595 |
| Olfr727 | 0.274408084 | -0.016030385 | 0.038579263 | -0.415518171 | 0.677762616 | NA | | Otos | 2.434381313 | -0.0224732008166085 | 0.0340438188850213 | -0.660125730679888 | 0.509173147842098 | NA | NM_153114 |
| Olfr728 | 0 | NA | NA | NA | NA | NA | | Otp | 15.53315368 | 0.0269764963586002 | 0.0319029437019936 | 0.845580163717444 | NA | NA | NM_011021 |
| Olfr729 | 0 | NA | NA | NA | NA | NA | | Ott | 0 | NA | NA | NA | NA | NA | NM_011022 |
| Olfr73 | 0 | NA | NA | NA | NA | NA | | Otud6a | 0 | NA | NA | NA | NA | NA | NM_001163191 |
| Olfr730 | 0 | NA | NA | NA | NA | NA | | Otx2os1 | 1.819187122 | 0.0277375572994228 | 0.0252839755259623 | 1.09704098040045 | 0.272623478224786 | NA | NR_029384 |
| Olfr731 | 0 | NA | NA | NA | NA | NA | | Ovch2 | 0 | NA | NA | NA | NA | NA | NM_172908 |
| Olfr732 | 0 | NA | NA | NA | NA | NA | | Ovol1 | 0.909119122 | 0.0103566161653807 | 0.0193319060455882 | 0.535726593174927 | 0.592147521681499 | NA | NM_019935 |
| Olfr733 | 0 | NA | NA | NA | NA | NA | | Oxct2a | 1.406187698 | -0.0211515696058832 | 0.0267981323904871 | -0.789292675238504 | 0.429940966734354 | NA | NM_022033 |
| Olfr734 | 0 | NA | NA | NA | NA | NA | | Oxct2b | 3.174094448 | -0.0129488168257541 | 0.0396754593267213 | -0.326368416282785 | 0.744145630562802 | NA | NM_181859 |
| Olfr735 | 0 | NA | NA | NA | NA | NA | | Oxgr1 | 13.54600362 | 0.0657596599168352 | 0.0664065565444197 | 0.990258542811931 | 0.322047764457387 | NA | NM_001001490 |
| Olfr736 | 0 | NA | NA | NA | NA | NA | | P2rx1 | 13.13994985 | -0.0229645597394484 | 0.0631982762110814 | -0.363373198071844 | 0.716326120287856 | NA | NM_008771 |
| Olfr738 | 0 | NA | NA | NA | NA | NA | | P2rx2 | 4.341453551 | 0.0461196301247678 | 0.0478396144955689 | 0.964046859722073 | 0.335022433011737 | NA | NM_153400,NM_001164834,NM_001164833 |
| Olfr739 | 0 | NA | NA | NA | NA | NA | | P2ry10 | 0.12890128 | -0.0068640067066208 | 0.00870804083610955 | -0.788237771940376 | 0.430557639008166 | NA | NM_172435 |
| Olfr74 | 0 | NA | NA | NA | NA | NA | | P2ry4 | 1.112289045 | 0.0139176846494259 | 0.0249082456501932 | 0.558758125517279 | 0.576326804221211 | NA | NM_020621 |
| Olfr740 | 0.096071043 | -0.020098795 | 0.027825948 | -0.722304036 | 0.470107572 | NA | | Pabpc1l | 14.03903673 | -0.0606633032646081 | 0.0718468057549808 | -0.844342384148409 | 0.398478108164451 | NA | NM_001114079 |
| Olfr741 | 0.222620122 | -0.001447027 | 0.032059667 | -0.04513545 | 0.963999345 | NA | | Pabpc2 | 0 | NA | NA | NA | NA | NA | NM_011033 |
| Olfr742 | 0.159967308 | 0.0240345 | 0.030870969 | 0.778547006 | 0.436246606 | NA | | Pabpc4l | 10.41113065 | -0.119262048262762 | 0.0594861372488189 | -2.00487128226047 | 0.0449768095311758 | NA | NM_001101479 |
| Olfr743 | 0 | NA | NA | NA | NA | NA | | Pabpc6 | 0 | NA | NA | NA | NA | NA | NM_001163836 |
| Olfr744 | 0 | NA | NA | NA | NA | NA | | Pabpn1l | 0.762737197 | 0.00995187591412606 | 0.0201031108857983 | 0.495041586879794 | 0.620570765283153 | NA | NM_001007462 |
| Olfr745 | 0 | NA | NA | NA | NA | NA | | Padi1 | 3.123640962 | 0.0479506612111127 | 0.0376725670969288 | 1.27282701727597 | 0.203079431797421 | NA | NM_011059 |
| Olfr746 | 0 | NA | NA | NA | NA | NA | | Padi3 | 0.715248492 | -0.0151610035213367 | 0.0184023520037972 | -0.823862271420976 | 0.410017819972652 | NA | NM_011060 |
| Olfr747 | 0 | NA | NA | NA | NA | NA | | Padi4 | 2.337343682 | -0.00814001061408853 | 0.0311538109300424 | -0.261284586735388 | 0.793873051844783 | NA | NM_011061 |
| Olfr748 | 0 | NA | NA | NA | NA | NA | | Padi6 | 17.00937294 | 0.0329349186794422 | 0.073024051197932 | 0.451014674468991 | 0.651978972701093 | NA | NM_153106 |
| Olfr749 | 0 | NA | NA | NA | NA | NA | | Pah | 5.588214659 | 0.0345585876285542 | 0.0506742898135087 | 0.681974779631575 | 0.495254898443624 | NA | NM_008777 |
| Olfr75-ps1 | 0 | NA | NA | NA | NA | NA | | Panx3 | 0 | NA | NA | NA | NA | NA | NM_172454 |
| Olfr750 | 0.116696964 | -0.020477583 | 0.027954224 | -0.732539847 | 0.463839131 | NA | | Papolb | 4.859969328 | -0.0187011884352679 | 0.0486876383178343 | -0.384105474847352 | 0.700900273827198 | NA | NM_019943 |
| Olfr76 | 0 | NA | NA | NA | NA | NA | | Parpbp | 20.05051672 | -0.0650330905042144 | 0.0762872939526581 | -0.852476043318199 | 0.393949930021342 | NA | NM_029249 |
| Olfr761 | 0 | NA | NA | NA | NA | NA | | Pate2 | 8.388622204 | 0.0231498617977608 | 0.0642053409789259 | 0.360559751646818 | 0.718428581604378 | NA | NM_001033421 |
| Olfr763 | 0 | NA | NA | NA | NA | NA | | Pate4 | 0 | NA | NA | NA | NA | NA | NM_020264 |
| Olfr765 | 0 | NA | NA | NA | NA | NA | | Patl2 | 2.566150773 | -0.00912690875536417 | 0.0392508150647795 | -0.23252787847338 | 0.816128036637853 | NA | NM_026251 |
| Olfr767 | 0 | NA | NA | NA | NA | NA | | Pax1 | 3.185137048 | -0.0355696471141701 | 0.037628955378519 | -0.945273307652743 | 0.344519362006355 | NA | NM_008780 |
| Olfr768 | 0 | NA | NA | NA | NA | NA | | Pax2 | 9.250346322 | 0.0145116655866135 | 0.0575535119709157 | 0.252142138501415 | 0.800931200082415 | NA | NM_011037 |
| Olfr769 | 0 | NA | NA | NA | NA | NA | | Pax3 | 0.795913517 | 0.000216250178044455 | 0.0200046477095122 | 0.0108099968159713 | 0.991375038418332 | NA | NM_001159520,NM_008781 |
| Olfr77 | 0 | NA | NA | NA | NA | NA | | Pax4 | 0.766701302 | -0.00997533422416961 | 0.0200383513424227 | -0.497812123048819 | 0.618616470390866 | NA | NM_011038,NM_001159925,NM_001159926 |
| Olfr770 | 0 | NA | NA | NA | NA | NA | | Pax5 | 1.391655813 | 0.0109546377886272 | 0.020934906967077 | 0.523271386199846 | 0.600785410231382 | NA | NM_008782 |
| Olfr771 | 0 | NA | NA | NA | NA | NA | | Pax6os1 | 5.713588866 | -0.0252398734872618 | 0.0444679737505648 | -0.567596662461852 | 0.570308873927693 | NA | NR_002867 |
| Olfr772 | 0 | NA | NA | NA | NA | NA | | Pax7 | 8.389591352 | -0.011877764459883 | 0.0201080875754643 | -0.590695878725742 | 0.554724209835615 | NA | NM_011039 |
| Olfr773 | 0 | NA | NA | NA | NA | NA | | Pax8 | 1.20695477 | 0.00635402392173369 | 0.0244210691389875 | 0.26018614850853 | 0.794720188304121 | NA | NM_011040 |
| Olfr774 | 0 | NA | NA | NA | NA | NA | | Pax9 | 0.127687691 | 0.00542298404407807 | 0.00869724871942359 | 0.62352868349814 | 0.532937159177571 | NA | NM_011041 |
| Olfr775 | 0 | NA | NA | NA | NA | NA | | Pbk | 12.14453337 | -0.116272208646551 | 0.0701498188066822 | -1.65748409082812 | 0.0974216427123755 | NA | NM_023209 |
| Olfr776 | 0 | NA | NA | NA | NA | NA | | Pbld1 | 19.6262789 | -0.0898283202623059 | 0.0697738166079635 | -1.28742162360162 | 0.197947371434488 | NA | NM_026701 |
| Olfr777 | 0 | NA | NA | NA | NA | NA | | Pbp2 | 4.41781871 | 0.00993937588661403 | 0.0481366058957034 | 0.206482690286671 | 0.836413884565941 | NA | NM_029595 |
| Olfr780 | 0 | NA | NA | NA | NA | NA | | Pbsn | 0 | NA | NA | NA | NA | NA | NM_017471 |
| Olfr781 | 0 | NA | NA | NA | NA | NA | | Pcdh12 | 12.56836415 | 0.0140336288432518 | 0.0681969538697942 | 0.205780875052655 | 0.836962080979571 | NA | NM_017378 |
| Olfr782 | 0 | NA | NA | NA | NA | NA | | Pcdha4 | 0.220191124 | -0.000706246424282353 | 0.00977545482624743 | -0.072246911968337 | 0.942405412335504 | NA | NM_007766 |
| Olfr784 | 0 | NA | NA | NA | NA | NA | | Pcdha4-g | 0 | NA | NA | NA | NA | NA | NM_001174154 |
| Olfr786 | 0 | NA | NA | NA | NA | NA | | Pcdhb1 | 7.912004206 | 0.121863947999345 | 0.0569464033981856 | 2.13997620090662 | 0.0323566901196086 | NA | NM_053126 |
| Olfr787 | 0 | NA | NA | NA | NA | NA | | Pck1 | 0.800526829 | -0.0100887621095731 | 0.014650583637018 | -0.688625269786632 | 0.491059114628679 | NA | NM_011044 |
| Olfr788 | 0.096071043 | -0.020098795 | 0.027825948 | -0.722304036 | 0.470107572 | NA | | Pcp2 | 11.80046778 | 0.0810164990324996 | 0.0486418961698662 | 1.66557032952777 | 0.0957990248651924 | NA | NM_001129804,NM_008790,NM_001129803 |
| Olfr790 | 0 | NA | NA | NA | NA | NA | | Pdc | 0.235049888 | -0.00142616332698746 | 0.00985869345204131 | -0.144660480004291 | 0.884978943250011 | NA | NM_001159730,NM_024458 |
| Olfr791 | 0 | NA | NA | NA | NA | NA | | Pdcd1 | 10.8440444 | -0.28325805071313 | 0.0620314252136553 | -4.5663637380167 | 4.9625736400795e-06 | NA | NM_008798 |
| Olfr792 | 0 | NA | NA | NA | NA | NA | | Pdcd1lg2 | 5.153699133 | 0.00395507424794044 | 0.0491788339412587 | 0.0804222859912569 | 0.93590140265266 | NA | NM_021396 |
| Olfr794 | 0 | NA | NA | NA | NA | NA | | Pdcl2 | 0.106148608 | 0.00515554063321497 | 0.00854200771766808 | 0.603551390213729 | 0.546141939141592 | NA | NM_023508 |
| Olfr796 | 0.098353386 | -0.020212831 | 0.02786466 | -0.725393061 | 0.468210929 | NA | | Pde6a | 3.357753918 | 0.000983371307837189 | 0.0401036939537232 | 0.024520716445022 | 0.980437259348621 | NA | NM_146086 |
| Olfr798 | 0 | NA | NA | NA | NA | NA | | Pde6b | 10.76250585 | 0.0269825657618467 | 0.0620965103134152 | 0.434526282163999 | 0.663906313713146 | NA | NM_008806 |
| Olfr799 | 0 | NA | NA | NA | NA | NA | | Pde6c | 8.805477227 | -0.0213052435511784 | 0.0620351876601007 | -0.343438044677365 | 0.731268942511369 | NA | NM_033614,NM_001170959 |
| Olfr8 | 0 | NA | NA | NA | NA | NA | | Pde6g | 8.757120053 | -0.0132494356812478 | 0.0572669513550443 | -0.231362685942609 | 0.817033051079931 | NA | NM_012065 |
| Olfr800 | 0 | NA | NA | NA | NA | NA | | Pde6h | 5.379326757 | -0.00176193719558362 | 0.0524625489162335 | -0.0335846662425207 | 0.973208349936807 | NA | NM_023898 |
| Olfr801 | 0 | NA | NA | NA | NA | NA | | Pdha2 | 0 | NA | NA | NA | NA | NA | NM_008811 |
| Olfr802 | 0 | NA | NA | NA | NA | NA | | Pdia2 | 19.75583382 | -0.0438751738698401 | 0.0790008347428207 | -0.555376079413229 | 0.578637450645878 | NA | NM_001081070 |
| Olfr803 | 0 | NA | NA | NA | NA | NA | | Pdilt | 1.764357048 | -0.0218854908614086 | 0.024800470672936 | -0.882462722180978 | 0.377526630088606 | NA | NM_027943 |
| Olfr804 | 0 | NA | NA | NA | NA | NA | | Pdx1 | 0 | NA | NA | NA | NA | NA | NM_008814 |
| Olfr805 | 0 | NA | NA | NA | NA | NA | | Pdxk-ps | 2.898917696 | 0.0137004240807732 | 0.041107974224351 | 0.333278988791852 | 0.738923698171965 | NA | NR_027316 |
| Olfr806 | 0 | NA | NA | NA | NA | NA | | Pdzd9 | 2.025972155 | -0.0118553651311532 | 0.0335325079107172 | -0.353548418231019 | 0.723677336835251 | NA | NM_001040137,NM_001040136 |
| Olfr807 | 0 | NA | NA | NA | NA | NA | | Pdzk1 | 10.24380282 | -0.034914955365071 | 0.0621501004749159 | -0.561784375218555 | 0.574262938931015 | NA | NM_001146001,NM_021517 |
| Olfr808 | 0 | NA | NA | NA | NA | NA | | Pdzk1ip1 | 7.921058025 | 0.0152609199197592 | 0.0544875292677775 | 0.280080967605631 | 0.779415386226659 | NA | NM_001164558,NM_026018,NM_001164557 |
| Olfr809 | 0 | NA | NA | NA | NA | NA | | Pea15b | 0 | NA | NA | NA | NA | NA | NR_027806 |
| Olfr810 | 0 | NA | NA | NA | NA | NA | | Pebp4 | 3.24573145 | -0.0294075412810046 | 0.0375565767095049 | -0.783019749336261 | 0.433615514701353 | NA | NM_028526,NM_028560 |
| Olfr811 | 0 | NA | NA | NA | NA | NA | | Peg12 | 2.648344487 | -0.00474274800475251 | 0.0397870424855026 | -0.119203331247369 | 0.90511426751573 | NA | NM_013788 |
| Olfr812 | 0 | NA | NA | NA | NA | NA | | Peg3as | 3.422907728 | 0.00721221079544745 | 0.0400017237805728 | 0.180297500052988 | 0.856919018180947 | NA | NR_023846 |
| Olfr813 | 0 | NA | NA | NA | NA | NA | | Per1 | 5332.737938 | 0.110837857598238 | 0.0913693237719121 | 1.2130751659597 | NA | NA | NM_001159367,NM_011065 |
| Olfr814 | 0 | NA | NA | NA | NA | NA | | Pet117 | 0 | NA | NA | NA | NA | NA | NM_001164813 |
| Olfr815 | 0 | NA | NA | NA | NA | NA | | Pet2 | 0 | NA | NA | NA | NA | NA | NM_008821 |
| Olfr816 | 0 | NA | NA | NA | NA | NA | | Pex11c | 17.59566597 | 0.0834248406570277 | 0.0728372593855206 | 1.14535941303706 | 0.252060303873339 | NA | NM_026951 |

| | | | | | | | | | | | | | | | |
|---|---|---|---|---|---|---|---|---|---|---|---|---|---|---|---|
| Olfr818 | 0 | NA | NA | NA | NA | NA | | Pf4 | 18.28679974 | 0.00314085008680948 | 0.0735077719873824 | 0.0427281361125815 | 0.965918250671319 | NA | NM_019932 |
| Olfr819 | 0 | NA | NA | NA | NA | NA | | Pfn3 | 1.452028206 | -0.00480184967229497 | 0.024893287328074 | -0.19289737064496 | 0.847039344284183 | NA | NM_029303 |
| Olfr820 | 0 | NA | NA | NA | NA | NA | | Pfpl | 0.423464609 | 0.00499580379007427 | 0.0121362222011964 | 0.411644060833179 | 0.680600331734096 | NA | NM_019540 |
| Olfr821 | 0 | NA | NA | NA | NA | NA | | Pga5 | 0 | NA | NA | NA | NA | NA | NM_021453 |
| Olfr822 | 0 | NA | NA | NA | NA | NA | | Pgc | 1.981941967 | 0.0168604031958518 | 0.0319897699808494 | 0.527056093430659 | 0.598154635776538 | NA | NM_025973 |
| Olfr823 | 0 | NA | NA | NA | NA | NA | | Pgk2 | 0 | NA | NA | NA | NA | NA | NM_031190 |
| Olfr824 | 0 | NA | NA | NA | NA | NA | | Pglyrp2 | 0.353527519 | -0.00579453045469275 | 0.0115029412048641 | -0.503743377584378 | 0.614441721879931 | NA | NR_073184,NM_001271478,NM_001271479,NM_021319,NM_001271476,NM_001271477 |
| Olfr825 | 0 | NA | NA | NA | NA | NA | | Pglyrp3 | 0.435065206 | 0.00898360253900676 | 0.0111639713759994 | 0.804695948819784 | 0.42099515794143 | NA | NM_207247 |
| Olfr826 | 0 | NA | NA | NA | NA | NA | | Pglyrp4 | 0 | NA | NA | NA | NA | NA | NM_001165968,NM_207263 |
| Olfr827 | 0 | NA | NA | NA | NA | NA | | Pgpep1l | 22.29078339 | -0.0348965493890602 | 0.0759527036514013 | -0.459451049290151 | 0.645910296394347 | NA | NM_030101 |
| Olfr828 | 0 | NA | NA | NA | NA | NA | | Pgr15l | 26.80586984 | 0.0504571928486785 | 0.0384830375953315 | 1.31115410844802 | NA | NA | NM_001033361 |
| Olfr829 | 0 | NA | NA | NA | NA | NA | | Phf11a | 3.506920076 | -0.0353456444875148 | 0.0382140011213242 | -0.924939641240321 | 0.354997305005362 | NA | NM_172603 |
| Olfr830 | 0 | NA | NA | NA | NA | NA | | Phf11b | 16.35940056 | -0.137402534933852 | 0.0771229798693903 | -1.78160303409628 | 0.0748139893118476 | NA | NM_001164327 |
| Olfr832 | 0 | NA | NA | NA | NA | NA | | Phf11c | 13.39382483 | -0.0169335562255089 | 0.0671533706101658 | -0.252162416743166 | 0.800915526655298 | NA | NM_001164289 |
| Olfr834 | 0 | NA | NA | NA | NA | NA | | Phgr1 | 0 | NA | NA | NA | NA | NA | NM_001145644 |
| Olfr835 | 0 | NA | NA | NA | NA | NA | | Phlda2 | 0.459676127 | -0.00315679208243135 | 0.0140053422032237 | -0.225399139601511 | 0.821668780922467 | NA | NM_009434 |
| Olfr836 | 0.099009884 | 0.018205726 | 0.027896865 | 0.652608315 | 0.514008823 | NA | | Phldb3 | 3.56803334 | -0.0348748423000513 | 0.0445724126236876 | -0.782431110348229 | 0.433961256470578 | NA | NM_001102613 |
| Olfr837 | 0 | NA | NA | NA | NA | NA | | Phox2a | 1.091611411 | 0.01574243856984 | 0.0257716039059316 | 0.610844347418235 | 0.541302631127072 | NA | NM_008887 |
| Olfr843 | 0 | NA | NA | NA | NA | NA | | Phox2b | 0 | NA | NA | NA | NA | NA | NM_008888 |
| Olfr845 | 0 | NA | NA | NA | NA | NA | | Phxr4 | 0 | NA | NA | NA | NA | NA | NR_028271 |
| Olfr846 | 0 | NA | NA | NA | NA | NA | | Pi15 | 9.752901974 | 0.00891314484642831 | 0.0641723605445145 | 0.138893828601575 | 0.889534047761422 | NA | NM_053191 |
| Olfr847 | 0 | NA | NA | NA | NA | NA | | Pif1 | 9.454403623 | -0.115770381615784 | 0.0592560781405651 | -1.95373006868862 | 0.0507331483344848 | NA | NM_172453 |
| Olfr849 | 0.098353386 | -0.020212831 | 0.02786466 | -0.725393061 | 0.468210929 | NA | | Pigr | 4.08118176 | 0.0474775489330387 | 0.0447657495968121 | 1.06057754780498 | 0.28888193057977 | NA | NM_011082 |
| Olfr850 | 0 | NA | NA | NA | NA | NA | | Pih1d3 | 0 | NA | NA | NA | NA | NA | NM_029062 |
| Olfr851 | 0 | NA | NA | NA | NA | NA | | Pik3c2g | 8.458390493 | -0.0271985863079351 | 0.056202010805759 | -0.483943295230854 | 0.628426112789859 | NA | NM_207683,NM_011084 |
| Olfr853 | 0 | NA | NA | NA | NA | NA | | Pilra | 11.21593858 | -0.0993936094580348 | 0.0660720066054079 | -1.50432254996626 | 0.132498335562708 | NA | NM_153510 |
| Olfr854 | 0 | NA | NA | NA | NA | NA | | Pilrb1 | 0.674595778 | -0.00442354080162154 | 0.0190302132440242 | -0.232448304435611 | 0.816189834648375 | NA | NM_133209 |
| Olfr855 | 0 | NA | NA | NA | NA | NA | | Pilrb2 | 2.780893249 | -0.0289600467167043 | 0.0340329438781379 | -0.850941570626443 | 0.394801810941677 | NA | NM_001024932 |
| Olfr857 | 0.299325859 | 0.043567584 | 0.041359933 | 1.053376566 | 0.29216844 | NA | | Pin1-ps1 | 0.209568245 | 0.00686197416234131 | 0.00977127153168884 | 0.702260104029194 | 0.482516968822431 | NA | NM_001033768 |
| Olfr859 | 0.500059445 | -0.006790017 | 0.038562195 | -0.176079638 | 0.860231377 | NA | | Pinc | 0.120209049 | -0.00674972139751468 | 0.00863521114606269 | -0.781650996523957 | 0.434419707908266 | NA | NR_003202 |
| Olfr860 | 0 | NA | NA | NA | NA | NA | | Pinlyp | 1.005770336 | -0.00767200141426159 | 0.0195402451146192 | -0.392625648719305 | 0.694595992853922 | NA | NM_001037143 |
| Olfr862 | 0 | NA | NA | NA | NA | NA | | Pip | 0.209568245 | 0.00686197416234131 | 0.00977127153168884 | 0.702260104029194 | 0.482516968822431 | NA | NM_008843 |
| Olfr866 | 0 | NA | NA | NA | NA | NA | | Pira1 | 0 | NA | NA | NA | NA | NA | NM_011087 |
| Olfr867 | 0 | NA | NA | NA | NA | NA | | Pira11 | 0 | NA | NA | NA | NA | NA | NM_011088 |
| Olfr868 | 0 | NA | NA | NA | NA | NA | | Pira2 | 0.21511414 | -0.000824240786584897 | 0.00975135300262786 | -0.08452578697159 | 0.93263840116295 | NA | NM_011089 |
| Olfr869 | 0 | NA | NA | NA | NA | NA | | Pira4 | 0 | NA | NA | NA | NA | NA | NM_011091 |
| Olfr870 | 0 | NA | NA | NA | NA | NA | | Pira6 | 1.327112512 | 0.00246479470910847 | 0.0250838376557129 | 0.0982622652458091 | 0.92172404110213 | NA | NM_011093,NM_008848 |
| Olfr871 | 0 | NA | NA | NA | NA | NA | | Pira7 | 0 | NA | NA | NA | NA | NA | NM_011094 |
| Olfr872 | 0 | NA | NA | NA | NA | NA | | Pirt | 7.287741437 | -0.0149946460778901 | 0.0518771661669674 | -0.289041348743485 | 0.772549735592254 | NA | NM_178656 |
| Olfr873 | 0 | NA | NA | NA | NA | NA | | Pisd-ps1 | 0 | NA | NA | NA | NA | NA | NR_003517 |
| Olfr874 | 0.099009884 | 0.018205726 | 0.027896865 | 0.652608315 | 0.514008823 | NA | | Pisd-ps3 | 0 | NA | NA | NA | NA | NA | NR_003518 |
| Olfr875 | 0 | NA | NA | NA | NA | NA | | Pitx1 | 0.290056368 | -0.00425303587871565 | 0.0107652157825088 | -0.395072051005791 | 0.692789714682806 | NA | NM_011097 |
| Olfr876 | 0 | NA | NA | NA | NA | NA | | Pitx2 | 3.032052365 | 0.0151724180975051 | 0.030316381067024 | 0.500469302848572 | 0.616744665709944 | NA | NM_011098,NM_001042504,NM_001042502 |
| Olfr877 | 0.520888355 | -0.021443664 | 0.051299307 | -0.418010783 | 0.675939233 | NA | | Pitx3 | 1.506925887 | -0.00982612374885568 | 0.0298003982739993 | -0.329731289444843 | 0.741603009177721 | NA | NM_008852 |
| Olfr878 | 0.831003462 | -0.009820528 | 0.06234679 | -0.157514569 | 0.874839327 | NA | | Piwil1 | 7.404292387 | 0.00170946299978076 | 0.0451337789115745 | 0.0378754680198598 | 0.969786972694899 | NA | NM_021311 |
| Olfr881 | 0.375043813 | -0.019924845 | 0.036566726 | -0.544890048 | 0.585829141 | NA | | Piwil4 | 1.063707118 | 0.00473163077282997 | 0.0254788871289687 | 0.185707905878285 | 0.85267382817869 | NA | NM_177905 |
| Olfr883 | 0.105923158 | 0.018524693 | 0.027977518 | 0.662127842 | 0.507889294 | NA | | Pkd1l2 | 19.65486971 | -0.0792029272352239 | 0.0752524415620046 | -1.05249644518131 | 0.292571840617704 | NA | NM_029686 |
| Olfr884 | 0 | NA | NA | NA | NA | NA | | Pkd2l2 | 20.05317315 | 0.0175353596210881 | 0.0707241433981007 | 0.247940219259821 | 0.804180660786307 | NA | NM_016927,NM_001163004 |
| Olfr885 | 0 | NA | NA | NA | NA | NA | | Pkhd1 | 2.570534169 | -0.0247404045457233 | 0.0373825289969385 | -0.661817303686153 | 0.508088315148205 | NA | NM_153179 |
| Olfr887 | 0 | NA | NA | NA | NA | NA | | Pklr | 4.148702938 | 0.0226253587821321 | 0.0449777372414434 | 0.50303461600742 | 0.614939934821887 | NA | NM_013631,NM_001099779 |
| Olfr888 | 0 | NA | NA | NA | NA | NA | | Pkp3 | 2.691068741 | 0.0172697534211523 | 0.0317024933181406 | 0.54474432808319 | 0.585929372798654 | NA | NM_019762,NM_001162924 |
| Olfr889 | 0 | NA | NA | NA | NA | NA | | Pla2g10 | 0 | NA | NA | NA | NA | NA | NM_011987 |
| Olfr890 | 0 | NA | NA | NA | NA | NA | | Pla2g12b | 0 | NA | NA | NA | NA | NA | NM_023530 |
| Olfr891 | 0 | NA | NA | NA | NA | NA | | Pla2g1b | 0 | NA | NA | NA | NA | NA | NM_011107 |
| Olfr893 | 0 | NA | NA | NA | NA | NA | | Pla2g2a | 0.12890128 | -0.0068640067066208 | 0.00870804083610955 | -0.788237771940376 | 0.430557639008166 | NA | NM_001082531,NR_002926 |
| Olfr894 | 0 | NA | NA | NA | NA | NA | | Pla2g2c | 8.447316024 | 0.0102588917332926 | 0.062111099619642 | 0.165170022687028 | 0.868810159596014 | NA | NM_008868 |
| Olfr895 | 0 | NA | NA | NA | NA | NA | | Pla2g2d | 17.46767024 | -0.0621104560814888 | 0.0718023046506589 | -0.865020369244079 | 0.387027608932413 | NA | NM_011109 |
| Olfr898 | 0.201994201 | 0.000227241 | 0.031727614 | 0.00716224 | 0.994285408 | NA | | Pla2g2e | 0 | NA | NA | NA | NA | NA | NM_012044 |
| Olfr899 | 0 | NA | NA | NA | NA | NA | | Pla2g2f | 5.010397569 | -0.0752967482028152 | 0.0449977210450861 | -1.6733458151663 | 0.0942592381053928 | NA | NM_012045 |
| Olfr9 | 0 | NA | NA | NA | NA | NA | | Pla2g4c | 7.286950046 | -0.0500376980207207 | 0.0576808568191915 | -0.867492280455727 | 0.385672339612089 | NA | NM_001004762,NM_001168504 |
| Olfr90 | 0 | NA | NA | NA | NA | NA | | Pla2g4d | 3.894030992 | 0.0324392364174731 | 0.0453822696572051 | 0.714799781115022 | 0.474732770451327 | NA | NM_001024137 |
| Olfr900 | 0 | NA | NA | NA | NA | NA | | Pla2g4f | 1.516743762 | -0.0135706852200216 | 0.0272196568023405 | -0.498561951701563 | 0.618088015110777 | NA | NM_001024145 |
| Olfr901 | 0.098353386 | -0.020212831 | 0.02786466 | -0.725393061 | 0.468210929 | NA | | Plac1 | 0.120209049 | -0.00674972139751468 | 0.00863521114606269 | -0.781650996523957 | 0.434419707908266 | NA | NM_019538 |
| Olfr902 | 0 | NA | NA | NA | NA | NA | | Plac1l | 0.204491261 | 0.00675251727497272 | 0.00969338978195025 | 0.69661051777262 | 0.486046571838551 | NA | NM_001037634 |
| Olfr904 | 0 | NA | NA | NA | NA | NA | | Plac8 | 2.612746898 | -0.00264190261785469 | 0.0315537558947617 | -0.0837270411378595 | 0.933273456908864 | NA | NM_139198 |
| Olfr905 | 0 | NA | NA | NA | NA | NA | | Plac8l1 | 1.941924685 | 0.00795004837313799 | 0.0324670530724736 | 0.244865105415996 | 0.806560881128653 | NA | NM_027072 |
| Olfr906 | 0 | NA | NA | NA | NA | NA | | Plac9a | 0 | NA | NA | NA | NA | NA | NM_207229 |
| Olfr907 | 0 | NA | NA | NA | NA | NA | | Plac9b | 0 | NA | NA | NA | NA | NA | NM_001270503 |
| Olfr908 | 0 | NA | NA | NA | NA | NA | | Plbd1 | 6.918319111 | 0.0342744266749496 | 0.0455386786697648 | 0.752644294392009 | 0.451663692748173 | NA | NM_025806 |
| Olfr91 | 0 | NA | NA | NA | NA | NA | | Pldi | 0.21491491 | 0.00699378003108193 | 0.00986423294990756 | 0.709003940458184 | 0.478322031551065 | NA | NR_033616 |
| Olfr910 | 0 | NA | NA | NA | NA | NA | | Plek2 | 15.16473187 | -0.0368980617184178 | 0.0745151969219489 | -0.495174987688306 | 0.620476604786924 | NA | NM_013738 |
| Olfr911-ps1 | 0 | NA | NA | NA | NA | NA | | Plekhg6 | 11.07522905 | 0.0157443123175333 | 0.0614990212431283 | 0.256009152654483 | 0.797943769980176 | NA | NM_198604 |
| Olfr912 | 0 | NA | NA | NA | NA | NA | | Plekhs1 | 21.03565159 | -0.0940716504852567 | 0.0805047388031407 | -1.16852314390202 | 0.242595809291905 | NA | NM_001164263,NM_172641 |
| Olfr913 | 0 | NA | NA | NA | NA | NA | | Plg | 3.127638071 | -0.0168874229269748 | 0.0420573626866723 | -0.401533093094455 | 0.688027678561823 | NA | NM_008877 |
| Olfr914 | 0 | NA | NA | NA | NA | NA | | Plin1 | 0.883295104 | -0.0117639965825542 | 0.0188272125224723 | -0.624840058957887 | 0.532076036158417 | NA | NM_001113471,NM_175640 |
| Olfr915 | 0.096071043 | -0.020098795 | 0.027825948 | -0.722304036 | 0.470107572 | NA | | Plscr5 | 0.574236473 | 0.00239212947531775 | 0.018035069611814 | 0.132637662443552 | 0.89447994523857 | NA | NM_001195693 |
| Olfr916 | 0 | NA | NA | NA | NA | NA | | Pmch | 252.5295557 | 0.0259701990312754 | 0.0218275766688132 | 1.18978846920653 | NA | NA | NM_029971 |
| Olfr917 | 0 | NA | NA | NA | NA | NA | | Pmp2 | 1.538947633 | -0.00693531470754725 | 0.0261252046683756 | -0.26546451197538 | 0.790651644735881 | NA | NM_001030305 |
| Olfr918 | 0 | NA | NA | NA | NA | NA | | Pnliprp1 | 0.39882267 | -0.00143678238067353 | 0.0134841982730002 | -0.106553044651564 | 0.915143571362664 | NA | NM_018874 |
| Olfr919 | 0 | NA | NA | NA | NA | NA | | Pnliprp2 | 2.650133983 | -0.0162780513323571 | 0.0370944130437873 | -0.438827575277766 | 0.660786477966729 | NA | NM_011128 |
| Olfr92 | 0 | NA | NA | NA | NA | NA | | Pnma5 | 6.092390332 | -0.0103028838283403 | 0.0541585118230159 | -0.190235726232822 | 0.849124416777461 | NA | NM_001100461 |
| Olfr920 | 3.481590909 | 0.274191145 | 0.115275251 | 2.378577742 | 0.017379572 | NA | | Pnmt | 8.644115552 | -0.0237036804334023 | 0.0548397976950026 | -0.432235008692644 | 0.665570617419868 | NA | NM_008890 |
| Olfr921 | 0.295060159 | -0.029687054 | 0.033501022 | -0.886153701 | 0.375534709 | NA | | Pnpla5 | 0.337583802 | -0.0131877509542174 | 0.0120595734966011 | -1.0935503612905 | 0.274152229987901 | NA | NM_029427 |
| Olfr922 | 0 | NA | NA | NA | NA | NA | | Podnl1 | 1.355888624 | -0.00996834611364508 | 0.0256989755584502 | -0.387888851482539 | 0.698098288712727 | NA | NM_001013384 |
| Olfr923 | 0 | NA | NA | NA | NA | NA | | Pof1b | 0.290056368 | -0.00425303587871565 | 0.0107652157825088 | -0.395072051005791 | 0.692789714682806 | NA | NM_181579 |
| Olfr924 | 0 | NA | NA | NA | NA | NA | | Pole2 | 16.64288983 | -0.074004742756428 | 0.0772202358362356 | -0.958359450149481 | 0.337881535262158 | NA | NM_011133 |
| Olfr926 | 0 | NA | NA | NA | NA | NA | | Pom121l12 | 0 | NA | NA | NA | NA | NA | NM_001164166 |
| Olfr93 | 0.098353386 | -0.020212831 | 0.02786466 | -0.725393061 | 0.468210929 | NA | | Pom121l2 | 1.769221581 | -0.0627666584098966 | 0.0298356654615064 | -2.10374588396151 | 0.0354006196472828 | NA | NM_001162929,NM_001162928 |
| Olfr930 | 0 | NA | NA | NA | NA | NA | | Pon1 | 5.488229233 | 0.0348644269865574 | 0.0285476156648654 | 1.22127281647085 | 0.221982742152234 | NA | NM_011134 |
| Olfr933 | 0 | NA | NA | NA | NA | NA | | Popdc2 | 16.58293519 | -0.0873795726082457 | 0.0732615292011065 | -1.19270746271736 | 0.232983961928085 | NA | NM_022318,NM_001081984 |
| Olfr934 | 0 | NA | NA | NA | NA | NA | | Poteg | 0 | NA | NA | NA | NA | NA | NM_026256,NM_027618 |
| Olfr935 | 0 | NA | NA | NA | NA | NA | | Pou1f1 | 0.249110328 | -0.010000816291709 | 0.0105091939847586 | -0.951625434473194 | 0.341286977626714 | NA | NM_008849 |
| Olfr936 | 0 | NA | NA | NA | NA | NA | | Pou2af1 | 1.621032158 | -0.0214247911983268 | 0.0282606213622588 | -0.758114654440649 | 0.448382347504923 | NA | NM_011136 |
| Olfr937 | 0 | NA | NA | NA | NA | NA | | Pou2f3 | 11.64641492 | 0.0299296062455528 | 0.06023458386795 | 0.496884087572787 | 0.619270793324536 | NA | NM_011139 |
| Olfr938 | 0 | NA | NA | NA | NA | NA | | Pou4f1 | 5.002682619 | -0.0275226802645664 | 0.0476300766065604 | -0.577842452194912 | 0.563370491097116 | NA | NM_011143 |
| Olfr94 | 0 | NA | NA | NA | NA | NA | | Pou4f2 | 1.061619024 | -0.0039423909973558 | 0.0151879531264497 | -0.259573555734127 | 0.795192736745619 | NA | NM_138944 |
| Olfr943 | 0 | NA | NA | NA | NA | NA | | Pou4f3 | 0.337583802 | -0.0131877509542174 | 0.0120595734966011 | -1.0935503612905 | 0.274152229987901 | NA | NM_138945 |
| Olfr944 | 0 | NA | NA | NA | NA | NA | | Pou5f1 | 0.201873568 | 0.00673381845182595 | 0.00968011632508873 | 0.69563404257585 | 0.486658041207307 | NA | NM_013633,NM_001252452 |
| Olfr945 | 0 | NA | NA | NA | NA | NA | | Pou5f2 | 0 | NA | NA | NA | NA | NA | NM_029315 |
| Olfr947-ps1 | 0.098353386 | -0.020212831 | 0.02786466 | -0.725393061 | 0.468210929 | NA | | Pp2d1 | 6.503277534 | 0.0995462014948701 | 0.0564560347174292 | 1.76325174081236 | 0.077858040710355 | NA | NM_173449 |
| Olfr948 | 0 | NA | NA | NA | NA | NA | | Ppbp | 0.999363745 | 0.00458646785158276 | 0.0168420274070329 | 0.272322787556297 | 0.785373836161249 | NA | NM_023785 |
| Olfr95 | 0.187700257 | -0.003168524 | 0.031449734 | -0.100748845 | 0.919749836 | NA | | Ppifos | 0.24789674 | -0.000769869626663926 | 0.0100368571301073 | -0.0767042528038554 | 0.938858821262798 | NA | NR_028021 |
| Olfr951 | 0 | NA | NA | NA | NA | NA | | Ppm1n | 1.343699343 | -0.0375533040371918 | 0.0269291015108697 | -1.3945249536838 | 0.163159139483383 | NA | NM_177691 |
| Olfr952 | 0 | NA | NA | NA | NA | NA | | Ppp1r14d | 0.361523991 | 0.0105955510260098 | 0.0121250525610889 | 0.873856090324303 | 0.382196629856021 | NA | NM_028104 |
| Olfr954 | 0 | NA | NA | NA | NA | NA | | Ppp1r1c | 1.449683927 | 0.0147880647150002 | 0.0254879352832726 | 0.580198613604665 | 0.561780688464085 | NA | NM_172420 |
| Olfr955 | 0 | NA | NA | NA | NA | NA | | Ppp1r2-ps3 | 1.280040084 | -0.00992540937301732 | 0.0229917730644941 | -0.431693951796392 | 0.665963863833991 | NA | NR_003650 |
| Olfr957 | 0 | NA | NA | NA | NA | NA | | Ppp1r2-ps7 | 0 | NA | NA | NA | NA | NA | NR_033731 |
| Olfr958 | 0 | NA | NA | NA | NA | NA | | Ppp1r2-ps9 | 0 | NA | NA | NA | NA | NA | NR_033171 |
| Olfr959 | 0 | NA | NA | NA | NA | NA | | Ppp1r27 | 0 | NA | NA | NA | NA | NA | NM_026814 |
| Olfr96 | 0 | NA | NA | NA | NA | NA | | Ppp1r3a | 2.564106344 | -0.0154985678470531 | 0.0299499293836142 | -0.517482617355766 | 0.604819300152218 | NA | NM_080464 |
| Olfr960 | 0 | NA | NA | NA | NA | NA | | Ppp2ca | 14304.24727 | 0.10567994955415 | 0.055870011254003 | 1.89153263409407 | NA | NA | NM_019411 |
| Olfr961 | 0 | NA | NA | NA | NA | NA | | Ppp3r2 | 5.322600447 | 0.0584542246892499 | 0.0520938840381933 | 1.1220938075263 | 0.261822556546363 | NA | NM_001004025 |
| Olfr963 | 0 | NA | NA | NA | NA | NA | | Ppy | 0 | NA | NA | NA | NA | NA | NM_008918 |
| Olfr965 | 0 | NA | NA | NA | NA | NA | | Ppyr1 | 1.131070143 | 0.00433270898118638 | 0.0223213357622698 | 0.194106169421547 | 0.846092751002515 | NA | NM_008919 |
| Olfr967 | 0 | NA | NA | NA | NA | NA | | Pram1 | 17.99966471 | -0.11538212377866 | 0.0757372388923527 | -1.52345299968824 | 0.127645410927822 | NA | NR_028279,NM_001002842 |
| Olfr968 | 0 | NA | NA | NA | NA | NA | | Prame | 0.300679247 | -0.0121548274246918 | 0.0115806462900197 | -1.04958109593304 | 0.293910752236529 | NA | NM_029459 |
| Olfr969 | 0 | NA | NA | NA | NA | NA | | Pramef12 | 0 | NA | NA | NA | NA | NA | NM_029948 |
| Olfr97 | 0 | NA | NA | NA | NA | NA | | Pramef17 | 0 | NA | NA | NA | NA | NA | NM_001085540 |
| Olfr970 | 0.431413697 | -0.00280547 | 0.037124934 | -0.075568363 | 0.939762507 | NA | | Pramef6 | 0 | NA | NA | NA | NA | NA | NM_001085414 |
| Olfr971 | 0 | NA | NA | NA | NA | NA | | Pramel1 | 1.630984956 | -0.00912663681694123 | 0.0280438964962013 | -0.325441110445457 | 0.744847245752354 | NA | NM_031377 |
| Olfr972 | 0 | NA | NA | NA | NA | NA | | Pramel3 | 0.097165704 | -0.00637828455464458 | 0.00848781691535724 | -0.75146349388194 | 0.452373763047372 | NA | NM_031390 |
| Olfr974 | 0 | NA | NA | NA | NA | NA | | Pramel4 | 0 | NA | NA | NA | NA | NA | NM_001001319 |
| Olfr975 | 0 | NA | NA | NA | NA | NA | | Pramel5 | 0 | NA | NA | NA | NA | NA | NM_001085418 |
| Olfr976 | 0 | NA | NA | NA | NA | NA | | Pramel6 | 0 | NA | NA | NA | NA | NA | NM_178249 |
| Olfr978 | 0 | NA | NA | NA | NA | NA | | Pramel7 | 0 | NA | NA | NA | NA | NA | NM_178250 |
| Olfr979 | 0 | NA | NA | NA | NA | NA | | Prap1 | 0.345220294 | 0.0102849425769615 | 0.0119473016652568 | 0.860859034544216 | 0.389315686243366 | NA | NM_009475 |
| Olfr98 | 0 | NA | NA | NA | NA | NA | | Prb1 | 0 | NA | NA | NA | NA | NA | NM_198669 |
| Olfr980 | 0 | NA | NA | NA | NA | NA | | Prdm12 | 8.685178865 | -0.00348326530078251 | 0.021702956122747 | -0.16049727424605 | 0.872489368631209 | NA | NM_001123362 |
| Olfr981 | 0 | NA | NA | NA | NA | NA | | Prdm13 | 3.048187347 | 0.038599400413962 | 0.0276390725293525 | 1.39655194192822 | 0.1625483505977 | NA | NM_001080771 |
| Olfr982 | 0 | NA | NA | NA | NA | NA | | Prdm14 | 0.257802559 | -0.00892927589713349 | 0.00993479775307735 | -0.898787888698349 | 0.368765652383134 | NA | NM_001081209 |
| Olfr983 | 0.096071043 | -0.020098795 | 0.027825948 | -0.722304036 | 0.470107572 | NA | | Prdm6 | 14.69580203 | -0.102949839907877 | 0.0690737188959651 | -1.49043430053237 | 0.136110079258695 | NA | NM_001033281 |
| Olfr984 | 0 | NA | NA | NA | NA | NA | | Prdx6b | 0.351100342 | 0.010390520699914 | 0.0120080616872527 | 0.8652954132426 | 0.38687666769005 | NA | NM_177256 |
| Olfr985 | 0.107716603 | -0.020477583 | 0.027954224 | -0.732539849 | 0.46383913 | NA | | Prelid2 | 7.729893033 | -0.0344824227886069 | 0.0572312621699756 | -0.602510262419075 | 0.546834534099772 | NA | NM_029942 |
| Olfr986 | 0 | NA | NA | NA | NA | NA | | Prf1 | 0.106148608 | 0.00515554063321497 | 0.00854200771766808 | 0.603551390213729 | 0.546141939141592 | NA | NM_011073 |
| Olfr987 | 0 | NA | NA | NA | NA | NA | | Prg2 | 0.840452317 | 0.0134375081512177 | 0.0203697227798464 | 0.659680462834408 | 0.509458906784074 | NA | NM_008920 |
| Olfr988 | 0 | NA | NA | NA | NA | NA | | Prg3 | 0 | NA | NA | NA | NA | NA | NM_016914 |
| Olfr99 | 0 | NA | NA | NA | NA | NA | | Prh1 | 0 | NA | NA | NA | NA | NA | NM_011174 |
| Olfr992 | 0 | NA | NA | NA | NA | NA | | Prhoxnb | 0 | NA | NA | NA | NA | NA | NM_001039678 |
| Olfr993 | 0 | NA | NA | NA | NA | NA | | Prkag3 | 4.307395596 | 0.0184829145846756 | 0.0421022184674316 | 0.439000966159852 | 0.66066083626015 | NA | NM_153744 |
| Olfr994 | 0.116696964 | -0.020477583 | 0.027954224 | -0.732539847 | 0.463839131 | NA | | Prl2a1 | 0 | NA | NA | NA | NA | NA | NM_019991 |
| Olfr995 | 0 | NA | NA | NA | NA | NA | | Prl2b1 | 0 | NA | NA | NA | NA | NA | NM_025532 |
| Olfr996 | 0 | NA | NA | NA | NA | NA | | Prl2c1 | 0 | NA | NA | NA | NA | NA | NM_001045532 |
| Olfr998 | 0 | NA | NA | NA | NA | NA | | Prl2c2 | 0 | NA | NA | NA | NA | NA | NM_031191 |
| Olig3 | 1.495177282 | 0.133257063 | 0.069937737 | 1.905367104 | 0.056732368 | NA | | Prl2c3 | 0 | NA | NA | NA | NA | NA | NM_011118 |
| Olr1 | 2.329184167 | 0.193705392 | 0.109915037 | 1.762319309 | 0.078015363 | NA | | Prl2c4 | 0 | NA | NA | NA | NA | NA | NM_011954 |
| Omt2a | 0 | NA | NA | NA | NA | NA | | Prl2c5 | 0 | NA | NA | NA | NA | NA | NM_181852 |
| Omt2b | 0 | NA | NA | NA | NA | NA | | Prl3a1 | 0 | NA | NA | NA | NA | NA | NM_025896 |
| Onecut1 | 3.472786976 | 0.220068871 | 0.112243424 | 1.960639328 | 0.049921111 | NA | | Prl3b1 | 0 | NA | NA | NA | NA | NA | NM_008865 |
| Onecut3 | 2.239489832 | -0.17551623 | 0.081101217 | -2.164162712 | 0.030451864 | NA | | Prl3c1 | 0 | NA | NA | NA | NA | NA | NM_013766,NM_001163218 |
| Ooep | 5.206067247 | 0.001695608 | 0.127415209 | 0.013307738 | 0.989382275 | NA | | Prl3d1 | 0 | NA | NA | NA | NA | NA | NM_001205322,NM_008864 |
| Oog1 | 0 | NA | NA | NA | NA | NA | | Prl3d2 | 0 | NA | NA | NA | NA | NA | NM_172155 |
| Oog2 | 0 | NA | NA | NA | NA | NA | | Prl3d3 | 0 | NA | NA | NA | NA | NA | NM_172156 |
| Oog3 | 0 | NA | NA | NA | NA | NA | | Prl4a1 | 0 | NA | NA | NA | NA | NA | NM_011165 |
| Oog4 | 0 | NA | NA | NA | NA | NA | | Prl5a1 | 0 | NA | NA | NA | NA | NA | NM_023746 |
| Oosp1 | 0.206069989 | -0.026776453 | 0.031767726 | -0.842882251 | 0.399294301 | NA | | Prl6a1 | 0 | NA | NA | NA | NA | NA | NM_011166 |
| Opn1sw | 2.057131987 | 0.017078223 | 0.104042247 | 0.164147005 | 0.869615419 | NA | | Prl7a1 | 0 | NA | NA | NA | NA | NA | NM_001164058,NM_008930 |
| Opn4 | 1.291219282 | 0.041594807 | 0.065863348 | 0.631531924 | 0.527692782 | NA | | Prl7a2 | 0 | NA | NA | NA | NA | NA | NM_011168 |
| Opn5 | 1.030044237 | -0.128301444 | 0.074244241 | -1.728099602 | 0.083970369 | NA | | Prl7b1 | 0 | NA | NA | NA | NA | NA | NM_029355 |
| Orc1 | 4.243598391 | 0.064110754 | 0.139122119 | 0.460823586 | 0.644925179 | NA | | Prl7c1 | 0 | NA | NA | NA | NA | NA | NM_026206 |
| Orm1 | 0.08731684 | 0.017365056 | 0.027733412 | 0.626142072 | 0.531221759 | NA | | Prl7d1 | 0 | NA | NA | NA | NA | NA | NM_011120 |
| Orm2 | 0 | NA | NA | NA | NA | NA | | Prl8a1 | 0 | NA | NA | NA | NA | NA | NM_028477 |
| Orm3 | 0.079983654 | 0.016740032 | 0.027674178 | 0.604897175 | 0.54524732 | NA | | Prl8a2 | 0 | NA | NA | NA | NA | NA | NM_010088 |
| Osm | 5.813465361 | 0.176672858 | 0.148990891 | 1.185796377 | 0.235702723 | NA | | Prl8a6 | 0 | NA | NA | NA | NA | NA | NM_011167,NM_001271379,NM_001271378,NR_073166 |
| Osr2 | 0.698905935 | -0.042900743 | 0.056927169 | -0.753607537 | 0.451084917 | NA | | Prl8a8 | 0 | NA | NA | NA | NA | NA | NM_023741 |
| Ostn | 0.654429243 | -0.028855515 | 0.054562931 | -0.528848329 | 0.596910669 | NA | | Prl8a9 | 0 | NA | NA | NA | NA | NA | NM_023332 |
| Otc | 0.820565582 | -0.063884022 | 0.064857528 | -0.98499009 | 0.324628939 | NA | | Prlh | 0 | NA | NA | NA | NA | NA | NM_001101647 |
| Otoa | 5.009308354 | 0.127446385 | 0.137663904 | 0.925779245 | 0.354560715 | NA | | Prlhr | 7.949361486 | 0.0163000452554565 | 0.0461172328742849 | 0.353448033187295 | 0.723752581227441 | NA | NM_201615 |
| Otog | 5.739167022 | 0.202669602 | 0.145976704 | 1.388369492 | 0.165024561 | NA | | Prm1 | 0 | NA | NA | NA | NA | NA | NM_013637 |
| Otol1 | 0 | NA | NA | NA | NA | NA | | Prm2 | 0.106148608 | 0.00515554063321497 | 0.00854200771766808 | 0.603551390213729 | 0.546141939141592 | NA | NM_008933 |
| Otop1 | 1.24152155 | 0.028934276 | 0.045989539 | 0.629149084 | 0.529251459 | NA | | Prm3 | 0 | NA | NA | NA | NA | NA | NM_013638 |
| Otop3 | 5.621215564 | -0.09303327 | 0.115766404 | -0.803629259 | 0.42161112 | NA | | Prnd | 7.235977917 | 0.0237255258149851 | 0.0589213251458518 | 0.402664498061708 | 0.687195054399056 | NA | NM_001126338,NM_023043 |
| Otor | 0.39538304 | 0.017028994 | 0.036795472 | 0.462801355 | 0.64350676 | NA | | Proc | 0.106148608 | 0.00515554063321497 | 0.00854200771766808 | 0.603551390213729 | 0.546141939141592 | NA | NM_001042768,NR_045173,NM_008934,NM_001042767 |
| Otos | 2.505579875 | -0.008737251 | 0.110786601 | -0.07886559 | 0.937139533 | NA | | Prodh2 | 0 | NA | NA | NA | NA | NA | NM_019546 |
| Otp | 6.230244395 | 0.303533233 | 0.119398721 | 2.542181609 | 0.011016292 | NA | | Prok1 | 0.220191124 | -0.000706246424282353 | 0.00977545482624743 | -0.072246911968337 | 0.942405412335504 | NA | NM_001044382 |
| Ott | 0 | NA | NA | NA | NA | NA | | Prok2 | 7.155140366 | 0.0462302484629691 | 0.0564380502704374 | 0.819132628456245 | 0.412710747846611 | NA | NM_001037539,NM_001170419,NM_015768 |
| Otud6a | 0 | NA | NA | NA | NA | NA | | Prokr1 | 8.034121145 | -0.043282973775097 | 0.0595939504409617 | -0.726298113396199 | 0.467656035673223 | NA | NM_021381 |
| Otx2os1 | 0.273223652 | 0.041357205 | 0.040327385 | 1.025536483 | 0.30511012 | NA | | Prol1 | 0 | NA | NA | NA | NA | NA | NM_008644 |
| Ovch2 | 0 | NA | NA | NA | NA | NA | | Prop1 | 0.733507625 | 0.0172814210666845 | 0.0188167074751345 | 0.918408339478159 | 0.35840512633898 | NA | NM_008936 |
| Ovol1 | 0.406270745 | 0.000396321 | 0.042421082 | 0.009342549 | 0.992545833 | NA | | Proz | 1.138838504 | -0.0311260278531602 | 0.0224988306449869 | -1.38345091548549 | 0.166526647878555 | NA | NM_025834 |
| Oxct2a | 0.838197859 | 0.08496648 | 0.066223674 | 1.283022739 | 0.19948411 | NA | | Prp2 | 0.106148608 | 0.00515554063321497 | 0.00854200771766808 | 0.603551390213729 | 0.546141939141592 | NA | NM_031499 |
| Oxct2b | 3.565518149 | -0.121942891 | 0.121289381 | -1.00538803 | 0.314710041 | NA | | Prph2 | 1.192608397 | -0.0020225764203795 | 0.0276614826790204 | -0.0731188723268804 | 0.941711523910339 | NA | NM_008938 |
| P2rx2 | 3.147652957 | -0.067471213 | 0.126862922 | -0.53184344 | 0.594834434 | NA | | Prpmp5 | 0 | NA | NA | NA | NA | NA | NM_001024705 |
| P2ry10 | 0.105923158 | 0.018524693 | 0.027977518 | 0.662127842 | 0.507889294 | NA | | Prps1l1 | 0 | NA | NA | NA | NA | NA | NM_029294 |
| P2ry4 | 0.284916696 | 0.042355674 | 0.040798016 | 1.038179738 | 0.299186384 | NA | | Prr15l | 0.415611529 | 0.0048600098249294 | 0.0120675527377329 | 0.402733671901271 | 0.68714416024218 | NA | NM_146026 |
| P4hb | 4610.350828 | -0.056078986 | 0.065364279 | -0.857945462 | NA | NA | | Prr19 | 5.274557021 | -0.0558322668668877 | 0.0496627104574277 | -1.12422915206669 | 0.260915825267022 | NA | NM_001081294 |
| Pabpc2 | 0 | NA | NA | NA | NA | NA | | Prr23a | 0.097165704 | -0.00637828455464458 | 0.00848781691535724 | -0.75146349388194 | 0.452373763047372 | NA | NM_001134660 |
| Pabpc6 | 0.098353386 | -0.020212831 | 0.02786466 | -0.725393061 | 0.468210929 | NA | | Prr9 | 0.097165704 | -0.00637828455464458 | 0.00848781691535724 | -0.75146349388194 | 0.452373763047372 | NA | NM_175424 |
| Pabpn1l | 0.474724059 | 0.012514508 | 0.055169993 | 0.226835404 | 0.820551733 | NA | | Prrx2 | 19.50130942 | -0.0823585879517754 | 0.0790487769690725 | -1.04187048945739 | 0.297471727677905 | NA | NM_009116 |
| Padi1 | 4.905138669 | -0.245018244 | 0.115233128 | -2.126283034 | 0.0334797 | NA | | Prrxl1 | 0.962850976 | 0.00928228210521239 | 0.0229616465808904 | 0.404251588513581 | 0.686027721236885 | NA | NM_001001796 |
| Padi3 | 1.366927568 | -0.06609889 | 0.080328433 | -0.822857947 | 0.41058878 | NA | | Prss1 | 0 | NA | NA | NA | NA | NA | NM_053243 |
| Padi4 | 2.314169231 | -0.052098663 | 0.100979433 | -0.515933412 | 0.605900917 | NA | | Prss2 | 0 | NA | NA | NA | NA | NA | NM_009430 |
| Pah | 3.588377053 | 0.128634487 | 0.125737589 | 1.023039238 | 0.306289294 | NA | | Prss21 | 0 | NA | NA | NA | NA | NA | NM_020487 |
| Panx3 | 0.094392817 | 0.017895942 | 0.027818137 | 0.643319236 | 0.520016977 | NA | | Prss27 | 0.314540458 | -0.0125494430549186 | 0.0117659602535877 | -1.06658893829698 | 0.286157497745293 | NA | NM_175440 |
| Papolb | 6.672834408 | -0.203629957 | 0.152389182 | -1.336249424 | 0.181467761 | NA | | Prss28 | 0 | NA | NA | NA | NA | NA | NM_053259 |
| Papss2 | 731.0706084 | -0.159608637 | 0.121798581 | -1.310431009 | NA | NA | | Prss29 | 0.212496447 | -0.000886988073087717 | 0.00973851166817767 | -0.0910804549309223 | 0.927428662789121 | NA | NM_053260 |
| Pate2 | 6.935396093 | -0.033564293 | 0.152523321 | -0.220060074 | 0.825824369 | NA | | Prss3 | 0.255375383 | 0.00708304372802858 | 0.00992471678592415 | 0.713677163873753 | 0.475426830432214 | NA | NM_011645 |
| Pate4 | 0.096071043 | -0.020098795 | 0.027825948 | -0.722304036 | 0.470107572 | NA | | Prss30 | 20.4621217 | -0.0743762259294248 | 0.0811956784202385 | -0.916012124986274 | 0.359660535446301 | NA | NM_013921 |
| Patl2 | 3.437888617 | -0.10195791 | 0.129845326 | -0.785225873 | 0.432321148 | NA | | Prss32 | 1.21667189 | -0.0234230796590811 | 0.0230590313293217 | -1.01578766794494 | 0.309730509777423 | NA | NM_027220 |
| Pax1 | 2.52653858 | 0.128005298 | 0.100384222 | 1.275153561 | 0.202254895 | NA | | Prss33 | 1.34024055 | -0.0271201809207276 | 0.0259533995151715 | -1.04495678513615 | 0.296042942959589 | NA | NM_001081399 |
| Pax2 | 5.556533266 | 0.171593911 | 0.137104289 | 1.251557565 | 0.210731126 | NA | | Prss34 | 0.217532603 | 0.00684685248201087 | 0.00976000296730706 | 0.701521557416086 | 0.482977586094298 | NA | NM_178372 |
| Pax3 | 0.516243536 | 0.008771227 | 0.050948872 | 0.172157442 | 0.863313755 | NA | | Prss37 | 0 | NA | NA | NA | NA | NA | NM_026317 |
| Pax4 | 0.721075248 | 0.006675597 | 0.060994402 | 0.109446069 | 0.912848696 | NA | | Prss38 | 1.921803007 | 0.0132654109487022 | 0.0321204549779503 | 0.412989509576015 | 0.679614299650459 | NA | NM_001045521 |
| Pax5 | 0.637093824 | 0.026258985 | 0.054685418 | 0.480182574 | 0.631097576 | NA | | Prss39 | 0 | NA | NA | NA | NA | NA | NM_009355 |
| Pax6os1 | 7.078914153 | -0.034704897 | 0.147094445 | -0.235936147 | 0.813482219 | NA | | Prss40 | 0 | NA | NA | NA | NA | NA | NM_009356 |
| Pax7 | 4.52595865 | 0.175388583 | 0.082573036 | 2.124041836 | 0.033666645 | NA | | Prss41 | 4.238815633 | -0.0114957095969619 | 0.0470504120161116 | -0.244327501170986 | 0.806977185912568 | NA | NM_027644 |
| Pax8 | 0.580633354 | 0.022423036 | 0.053248904 | 0.421098549 | 0.673683121 | NA | | Prss42 | 0.24789674 | -0.000769869626663926 | 0.0100368571301073 | -0.0767042528038554 | 0.938858821262798 | NA | NM_153099 |
| Pax9 | 0 | NA | NA | NA | NA | NA | | Prss43 | 8.025508264 | 0.0155933368931184 | 0.0572009933817231 | 0.272606050546331 | 0.78515606038722 | NA | NM_199471 |
| Pbp2 | 5.181955143 | -0.216329473 | 0.13401829 | -1.614178726 | 0.106488665 | NA | | Prss44 | 0.127687691 | 0.00542298404407807 | 0.00869724871942359 | 0.62352868349814 | 0.532937159177571 | NA | NM_148940 |
| Pbsn | 0 | NA | NA | NA | NA | NA | | Prss45 | 12.94196397 | -0.0613996596031257 | 0.0727820476919241 | -0.843609949846721 | 0.398887402441878 | NA | NM_153172 |
| Pcdha4 | 0.08731684 | 0.017365056 | 0.027733412 | 0.626142072 | 0.531221759 | NA | | Prss46 | 0.308221406 | 0.00253133711108074 | 0.0124107148809387 | 0.203963843772494 | 0.838381753432823 | NA | NM_183103 |
| Pcdha4-g | 0 | NA | NA | NA | NA | NA | | Prss48 | 0 | NA | NA | NA | NA | NA | NM_001001650 |
| Pcdhb1 | 6.652721741 | -0.066901165 | 0.136739181 | -0.48926112 | 0.624656843 | NA | | Prss51 | 0 | NA | NA | NA | NA | NA | NM_001193631 |
| Pck1 | 1.585148945 | -0.004545358 | 0.0723927 | -0.062787513 | 0.949935709 | NA | | Prss52 | 0 | NA | NA | NA | NA | NA | NM_028525 |
| Pcnt | 1554.201552 | 0.130183332 | 0.118868667 | 1.095186277 | NA | NA | | Prss54 | 9.870856792 | 0.0412945102991698 | 0.0651077618432136 | 0.634248653772055 | 0.525918559957795 | NA | NM_027640 |
| Pcp2 | 6.44973494 | -0.096827672 | 0.137231645 | -0.705578304 | 0.480450421 | NA | | Prss55 | 1.590962535 | -0.0172234897073177 | 0.0261951774736781 | -0.657506127783425 | 0.51085553213843 | NA | NM_001081063 |
| Pdc | 0.298669361 | 0.012533675 | 0.03926893 | 0.319175364 | 0.749593537 | NA | | Prss56 | 8.186573763 | 0.0511982894802336 | 0.0622397742001353 | 0.822597609618615 | 0.410736858451813 | NA | NM_027084 |
| Pdcd1lg2 | 4.391468329 | 0.02915534 | 0.134584049 | 0.216632954 | 0.828494402 | NA | | Prss58 | 0 | NA | NA | NA | NA | NA | NM_175020 |
| Pdcl2 | 0.206069989 | -0.026776453 | 0.031767726 | -0.842882251 | 0.399294301 | NA | | Prss8 | 4.439235943 | 0.0480742657095115 | 0.0481297316704194 | 0.99884757385959 | 0.317868535997076 | NA | NM_133351 |
| Pde6a | 6.060335561 | -0.171546757 | 0.135696943 | -1.264190284 | 0.206161737 | NA | | Psapl1 | 0.655475157 | 0.00478692383777398 | 0.0189945617116142 | 0.25201549319493 | 0.801029088388892 | NA | NM_175249 |
| Pde6c | 7.108978609 | 0.117915475 | 0.146447249 | 0.805173707 | 0.420719447 | NA | | Psca | 1.500426011 | -0.0165592730361935 | 0.0279753012419277 | -0.591924744366128 | 0.553900983279666 | NA | NM_028216 |
| Pde6g | 5.982113432 | 0.068252227 | 0.129232659 | 0.528134512 | 0.597405979 | NA | | Psg-ps1 | 0 | NA | NA | NA | NA | NA | NR_002857 |
| Pde6h | 4.163190705 | 0.054426054 | 0.135918579 | 0.400431303 | 0.688838872 | NA | | Psg17 | 1.326564174 | 0.00173200533140248 | 0.0256878760372062 | 0.0674250112735616 | 0.946243358374737 | NA | NM_007677 |
| Pdha2 | 0 | NA | NA | NA | NA | NA | | Psg18 | 0 | NA | NA | NA | NA | NA | NM_001163685,NM_011963 |
| Pdilt | 1.620847932 | 0.034163249 | 0.084139753 | 0.406029824 | 0.684720687 | NA | | Psg19 | 0.255375383 | 0.00708304372802858 | 0.00992471678592415 | 0.713677163873753 | 0.475426830432214 | NA | NM_011964 |
| Pdx1 | 0 | NA | NA | NA | NA | NA | | Psg20 | 0 | NA | NA | NA | NA | NA | NM_054058 |
| Pdxk-ps | 2.905076689 | 0.015946435 | 0.110471739 | 0.144348549 | 0.885225244 | NA | | Psg21 | 0 | NA | NA | NA | NA | NA | NM_027403 |
| Pdzd9 | 2.126268668 | -0.040047852 | 0.109038784 | -0.367280803 | 0.713409572 | NA | | Psg22 | 0.191449919 | 0.00665508079272897 | 0.00962405648474729 | 0.691504752001018 | 0.489248395458281 | NA | NM_001004152 |
| Pea15b | 0 | NA | NA | NA | NA | NA | | Psg25 | 0 | NA | NA | NA | NA | NA | NM_054060 |
| Pebp4 | 3.5279941 | -0.032343569 | 0.11958746 | -0.270459539 | 0.78680674 | NA | | Psg26 | 0 | NA | NA | NA | NA | NA | NM_001029893 |
| Peg12 | 3.499186594 | -0.204394151 | 0.13043656 | -1.567000475 | 0.11711458 | NA | | Psg27 | 0 | NA | NA | NA | NA | NA | NM_001037168 |
| Peg3as | 6.231941421 | 0.154966317 | 0.098504051 | 1.573197401 | NA | NA | | Psg28 | 0 | NA | NA | NA | NA | NA | NM_054063 |
| Pet117 | 0 | NA | NA | NA | NA | NA | | Psg29 | 3.458559647 | -0.00616049761915238 | 0.0448091556333277 | -0.137483010605324 | 0.890649020797477 | NA | NM_054064 |
| Pet2 | 0 | NA | NA | NA | NA | NA | | Psma8 | 12.68257091 | 0.0519264322669377 | 0.0690890881063267 | 0.751586591894569 | 0.452299709044859 | NA | NM_001163609 |
| Pfn3 | 1.414062082 | -0.006063038 | 0.080297207 | -0.07550746 | 0.939810962 | NA | | Psmb11 | 8.45064709 | 0.00303764590675885 | 0.063984613919372 | 0.0474746305508171 | 0.96213494941871 | NA | NM_175204 |
| Pfpl | 0.57091032 | -0.005729398 | 0.057927547 | -0.098906271 | 0.92121269 | NA | | Psmb9 | 70.11715032 | -0.161087122965678 | 0.0759356660714005 | -2.12136313934762 | NA | NA | NM_013585 |

| | | | | | | | | | | | | | | | |
|---|---|---|---|---|---|---|---|---|---|---|---|---|---|---|---|
| Pga5 | 0.096071043 | -0.020098795 | 0.027825948 | -0.722304036 | 0.470107572 | NA | | Psors1c2 | 5.424324711 | -0.0425425031090018 | 0.0424118185181828 | -1.00308132486144 | 0.315821624447142 | NA | NM_020576 |
| Pgc | 0.784043763 | 0.094117759 | 0.070872186 | 1.327992891 | 0.184180455 | NA | | Pspn | 0.329777294 | 0.00230606622174634 | 0.0126536401242093 | 0.182245282709938 | 0.855390237522515 | NA | NM_008954 |
| Pgk2 | 0 | NA | NA | NA | NA | NA | | Ptchd3 | 0 | NA | NA | NA | NA | NA | NM_029049 |
| Pglyrp2 | 0.310199702 | 0.010205801 | 0.034113052 | 0.299175856 | 0.76480587 | NA | | Ptcra | 9.517173675 | 0.0631051158470472 | 0.0628233891045483 | 1.00448442445584 | 0.315145175023801 | NA | NM_011195 |
| Pglyrp3 | 0 | NA | NA | NA | NA | NA | | Ptf1a | 0.297598527 | 0.00928691468127708 | 0.0113567208765242 | 0.817746141888045 | 0.413502157150213 | NA | NM_018809 |
| Pglyrp4 | 0 | NA | NA | NA | NA | NA | | Ptger2 | 21.50792901 | 0.0543153992890872 | 0.0802084016140359 | 0.677178427647191 | 0.498292753148438 | NA | NM_008964 |
| Phf11a | 3.333410359 | 0.10281034 | 0.119701353 | 0.858890377 | 0.390400999 | NA | | Ptgir | 4.940020943 | -0.025605643909362 | 0.0445220660597072 | -0.575122544291251 | 0.565208421995152 | NA | NM_008967 |
| Phf2 | 2545.963116 | 0.026272047 | 0.097650381 | 0.269041926 | NA | NA | | Pth | 0.202072798 | -0.00115107594702963 | 0.00968429294483289 | -0.11886009165427 | 0.905386199803646 | NA | NM_020623 |
| Phgr1 | 0 | NA | NA | NA | NA | NA | | Pth2 | 0.344835288 | -0.007099476111916 | 0.0129416197693129 | -0.548577089921174 | 0.583295711774425 | NA | NM_053256 |
| Phlda2 | 0.740683258 | -0.038907871 | 0.057196921 | -0.680244161 | 0.496349875 | NA | | Ptk6 | 3.018710889 | -0.0184404814797424 | 0.0375918912362609 | -0.490544127291867 | 0.623748912245103 | NA | NM_009184 |
| Phldb3 | 4.528429979 | 0.1785205 | 0.130187157 | 1.37126045 | 0.17029378 | NA | | Ptpn20 | 16.78955382 | -0.0018111356386089 | 0.0733947100035287 | -0.0246766509265017 | 0.980312879270068 | NA | NM_008978 |
| Phox2a | 0.266310378 | 0.0407635 | 0.040043874 | 1.017970943 | 0.308691765 | NA | | Ptprcap | 5.423080402 | -0.00104219412184352 | 0.0528166270091786 | -0.0197323112220401 | 0.984256915165632 | NA | NM_016933 |
| Phox2b | 0.098353386 | -0.020212831 | 0.02786466 | -0.725393061 | 0.468210929 | NA | | Ptprq | 4.939503204 | -0.0406742486214803 | 0.0474212727876647 | -0.85772157157411 | 0.391046225604942 | NA | NM_001081432 |
| Phxr4 | 0 | NA | NA | NA | NA | NA | | Ptx3 | 4.226919148 | -0.0458382894062882 | 0.0390642137311434 | -1.17340872957963 | 0.240631938193981 | NA | NM_008987 |
| Pigr | 3.885978194 | -0.070447289 | 0.116903789 | -0.602609111 | 0.546768758 | NA | | Ptx4 | 6.286111793 | 0.0756198175692217 | 0.0546288553223366 | 1.38424678904634 | 0.166282901030528 | NA | NM_001163416,NM_026747 |
| Pih1d3 | 0.196706773 | -0.026517754 | 0.031609464 | -0.838918169 | 0.401515231 | NA | | Pxt1 | 1.617534884 | -0.00346589244617452 | 0.0267149863774263 | -0.129735886712004 | 0.896775389008121 | NA | NM_153390 |
| Pik3c2g | 7.169320049 | 0.004016622 | 0.138447562 | 0.029011863 | 0.97685513 | NA | | Pydc3 | 2.743227701 | -0.0336868158258564 | 0.034710764069473 | -0.970500555920718 | 0.331797047945733 | NA | NM_001162938 |
| Pilrb1 | 0.932836247 | -0.05196266 | 0.070817098 | -0.733758678 | 0.463095829 | NA | | Pydc4 | 0.410265417 | -0.01013448436559 | 0.0139857084979753 | -0.724631459826089 | 0.468678153504599 | NA | NM_001177349,NM_001177350 |
| Pilrb2 | 2.07725609 | 0.103125854 | 0.097169719 | 1.061296197 | 0.288555313 | NA | | Pyhin1 | 13.546221 | -0.0783534676071344 | 0.0645425399341449 | -1.2139817814279 | 0.224754681259184 | NA | NM_175026 |
| Pin1-ps1 | 0 | NA | NA | NA | NA | NA | | Pyy | 0.217532603 | 0.00684685248201087 | 0.00976000296730706 | 0.701521557416086 | 0.482977586094298 | NA | NM_145435 |
| Pinc | 0.40343326 | -0.026145848 | 0.043560066 | -0.600225165 | 0.548356185 | NA | | Pzp | 0.297598527 | 0.00928691468127708 | 0.0113567208765242 | 0.817746141888045 | 0.413502157150213 | NA | NM_007376 |
| Pinlyp | 0.876573426 | 0.005823502 | 0.0575016 | 0.101275475 | 0.919331784 | NA | | Qprt | 15.72880647 | 0.0844576950483143 | 0.0708432794511945 | 1.19217652969466 | 0.233192034219494 | NA | NM_133686 |
| Pip | 0 | NA | NA | NA | NA | NA | | Qrfp | 0.877224048 | 0.0143423644501641 | 0.0170477521165224 | 0.841305314163017 | 0.40017691432478 | NA | NM_183424 |
| Pira1 | 0 | NA | NA | NA | NA | NA | | Qrich2 | 1.982759282 | 0.000745286888249584 | 0.0305434028644165 | 0.024400912090835 | 0.9805328208001 | NA | NM_001033267 |
| Pira11 | 0 | NA | NA | NA | NA | NA | | R3hdml | 0.394764232 | 0.00589493804908813 | 0.0136286083697903 | 0.432541451712348 | 0.665347931870215 | NA | NM_001099331 |
| Pira2 | 0.08731684 | 0.017365056 | 0.027733412 | 0.626142072 | 0.531221759 | NA | | Rab17 | 0.617637615 | -0.00138456702327747 | 0.0153163529566188 | -0.0903979574771515 | 0.927970979746458 | NA | NM_008998,NM_001159725 |
| Pira4 | 0 | NA | NA | NA | NA | NA | | Rab19 | 2.520500328 | -0.0184148172711509 | 0.0368801846071568 | -0.499314671748616 | 0.617557720731271 | NA | NM_011226 |
| Pira6 | 1.055108535 | 0.037618857 | 0.080056788 | 0.469902153 | 0.638424926 | NA | | Rab20 | 16.76572115 | 0.0439962949102857 | 0.0662625638425505 | 0.663969100483756 | 0.506710087923641 | NA | NM_011227 |
| Pira7 | 0 | NA | NA | NA | NA | NA | | Rab25 | 2.848920711 | 0.0312837215663126 | 0.0388235195660879 | 0.805793032572935 | 0.420362196130757 | NA | NM_016899 |
| Pisd-ps1 | 0.094392817 | 0.017895942 | 0.027818137 | 0.643319236 | 0.520016977 | NA | | Rab38 | 12.27904393 | 0.0135013418534371 | 0.0702230240029535 | 0.192263748893373 | 0.847535612811185 | NA | NM_028238 |
| Pisd-ps3 | 0 | NA | NA | NA | NA | NA | | Rab44 | 2.78164657 | -0.0251638977220598 | 0.0367717719345971 | -0.684326492800421 | 0.493769021400784 | NA | NM_001002786 |
| Pitx1 | 0.159967308 | 0.0240345 | 0.030870969 | 0.778547006 | 0.436246606 | NA | | Rad51b | 9.388619497 | 0.0165532643632186 | 0.0625264857661888 | 0.264740040326556 | 0.791209729940709 | NA | NM_001252562,NM_009014 |
| Pitx2 | 1.539596081 | 0.097937816 | 0.069972898 | 1.399653567 | 0.161617085 | NA | | Raet1a | 0.12890128 | -0.0068640067066208 | 0.00870804083610955 | -0.788237771940376 | 0.430557639008166 | NA | NM_009016 |
| Pitx3 | 1.584338663 | 0.042659999 | 0.094421335 | 0.451804664 | 0.651409709 | NA | | Raet1b | 0 | NA | NA | NA | NA | NA | NM_009017 |
| Piwil1 | 7.162601587 | -0.106265545 | 0.145177836 | -0.731968106 | 0.464188035 | NA | | Raet1c | 0 | NA | NA | NA | NA | NA | NM_009018 |
| Piwil4 | 0.956200006 | 0.0198166 | 0.077208804 | 0.256662444 | 0.797439366 | NA | | Raet1d | 0.584133578 | 0.0236872078946436 | 0.0180665615514324 | 1.31110769623817 | 0.189821380953494 | NA | NM_020030 |
| Pkhd1 | 3.419336713 | 0.122665716 | 0.117696724 | 1.042218617 | 0.297310334 | NA | | Raet1e | 0 | NA | NA | NA | NA | NA | NM_198193 |
| Pklr | 2.107931662 | 0.125979004 | 0.105215142 | 1.1973467 | 0.231171449 | NA | | Rag1 | 8.967626615 | -0.0513217038941634 | 0.0543326304571395 | -0.944583456798557 | 0.344871576110936 | NA | NM_009019 |
| Pkp3 | 2.513033096 | -0.038644634 | 0.113222479 | -0.341315911 | 0.732865771 | NA | | Rag2 | 0 | NA | NA | NA | NA | NA | NM_009020 |
| Pla2g10 | 0.107716603 | -0.020477583 | 0.027954224 | -0.732539849 | 0.46383913 | NA | | Rasef | 4.299766666 | 0.0236918418000903 | 0.0425590546318443 | 0.55668158057165 | 0.577745003899559 | NA | NM_001017427 |
| Pla2g12b | 0 | NA | NA | NA | NA | NA | | Rasl2-9 | 4.7721782 | 0.0235749487018577 | 0.0499485389043731 | 0.471984751085354 | 0.636937672967648 | NA | NM_009028 |
| Pla2g1b | 0 | NA | NA | NA | NA | NA | | Raver1-fdx1l | 0 | NA | NA | NA | NA | NA | NR_038081 |
| Pla2g2a | 0.105923158 | 0.018524693 | 0.027977518 | 0.662127842 | 0.507889294 | NA | | Rax | 5.135484584 | -0.00629379327225052 | 0.0423803007004707 | -0.148507518073854 | 0.881942253180907 | NA | NM_013833 |
| Pla2g2c | 6.13330658 | 0.093089917 | 0.146138441 | 0.636998152 | 0.524126048 | NA | | Rbbp8nl | 0 | NA | NA | NA | NA | NA | NM_173031 |
| Pla2g2e | 0.29109959 | -0.010630118 | 0.033578644 | -0.316573778 | 0.751567036 | NA | | Rbm44 | 1.034513641 | 0.0104190478856975 | 0.0247588875062922 | 0.420820518815702 | 0.67388614724443 | NA | NM_001033408 |
| Pla2g2f | 5.340863401 | 0.15590213 | 0.141126347 | 1.104698963 | 0.269290053 | NA | | Rbm46 | 11.52924913 | -0.0303283807040695 | 0.0673233562016201 | -0.450488246801631 | 0.652358426989235 | NA | NM_001146328 |
| Pla2g3 | 119.7263356 | 0.381232508 | 0.15684787 | 2.430587733 | NA | NA | | Rbmxl2 | 0.120209049 | -0.00674972139751468 | 0.00863521114606269 | -0.781650996523957 | 0.434419707908266 | NA | NM_029660 |
| Pla2g4c | 6.523544008 | 0.100244289 | 0.146846293 | 0.682647735 | 0.494829463 | NA | | Rbmy | 0 | NA | NA | NA | NA | NA | NM_001166384,NM_011253 |
| Pla2g4d | 2.085772536 | 0.117154982 | 0.095012777 | 1.233044501 | 0.217559162 | NA | | Rbp2 | 1.38813166 | 0.0206486794887591 | 0.0250774789196552 | 0.823395348269042 | 0.410283207499563 | NA | NM_009034 |
| Pla2g4f | 2.029914565 | -0.058556496 | 0.094340198 | -0.620695075 | 0.534800271 | NA | | Rbp3 | 4.887548185 | -0.0365800426479061 | 0.0448960875676983 | -0.814771278070668 | 0.415203244468788 | NA | NM_015745 |
| Plac1 | 0.206726487 | -0.001342295 | 0.031808201 | -0.04219965 | 0.966339542 | NA | | Rbp7 | 2.274294225 | 0.000358956010586974 | 0.035314828010554 | 0.010164455861988 | 0.991890077246682 | NA | NM_022020 |
| Plac1l | 0.407192437 | -0.059678651 | 0.046941173 | -1.2713498 | 0.203604233 | NA | | Rbpjl | 17.68455356 | -0.0516394793436321 | 0.0769365425426264 | -0.671195736603597 | 0.502095844300186 | NA | NM_009036 |
| Plac8 | 1.839029156 | 0.020541468 | 0.096883849 | 0.212021595 | 0.832090187 | NA | | Rcvrn | 8.638259164 | -0.0807202104426048 | 0.061997807217515 | -1.30198492600559 | 0.192921538647421 | NA | NM_009038 |
| Plac8l1 | 1.457471117 | 0.027500759 | 0.085429105 | 0.321913228 | 0.747518431 | NA | | Rd3 | 19.33013986 | 0.0143611531231708 | 0.0673479561773735 | 0.213238143193953 | 0.831141218396043 | NA | NM_023727,NM_001177900 |
| Plac9a | 0 | NA | NA | NA | NA | NA | | Rdh1 | 4.068035654 | -0.0444923363256051 | 0.0421311596334801 | -1.05604347738505 | 0.290948351864451 | NA | NM_080436 |
| Plac9b | 0 | NA | NA | NA | NA | NA | | Rdh16 | 6.818557127 | 0.0300783547051449 | 0.0564806048699025 | 0.53254306986314 | 0.594349919929976 | NA | NM_009040 |
| Plaur | 55.7093842 | 0.254864934 | 0.161760859 | 1.575566151 | NA | NA | | Rdh18-ps | 4.712843211 | -0.0577050049874016 | 0.0469114437752241 | -1.230083756618 | 0.218665742056165 | NA | NR_037604 |
| Plbd1 | 5.892752642 | -0.185260901 | 0.144428705 | -1.282715241 | 0.199591858 | NA | | Rdh19 | 0.202072798 | -0.00115107594702963 | 0.00968429294483289 | -0.11886009165427 | 0.905386199803646 | NA | NM_147222 |
| Pldi | 0 | NA | NA | NA | NA | NA | | Rdh5 | 151.8046017 | 0.0157525334658692 | 0.0488981971938399 | 0.322149575441886 | NA | NA | NM_134006 |
| Plg | 3.298327254 | 0.142602649 | 0.126589991 | 1.126492292 | 0.259957201 | NA | | Rdh7 | 0 | NA | NA | NA | NA | NA | NM_017473,NM_001150749 |
| Plin1 | 1.10321352 | -0.016789173 | 0.066089342 | -0.254037532 | 0.799466563 | NA | | Rdh8 | 0 | NA | NA | NA | NA | NA | NM_001030290 |
| Plscr5 | 1.185070533 | -0.13680812 | 0.084077097 | -1.627174639 | 0.103700013 | NA | | Reg1 | 0 | NA | NA | NA | NA | NA | NM_009042 |
| Pmp2 | 1.392519306 | -0.014294857 | 0.082638399 | -0.172980812 | 0.86266651 | NA | | Reg2 | 0 | NA | NA | NA | NA | NA | NM_009043 |
| Pnliprp1 | 0.796903708 | -0.040192877 | 0.050691198 | -0.792896579 | 0.427838081 | NA | | Reg3a | 0.423306759 | -0.00976599639876241 | 0.0140919448469068 | -0.693019771568724 | 0.488297143743493 | NA | NM_011259 |
| Pnliprp2 | 3.244481446 | -0.137778797 | 0.121076856 | -1.13794495 | 0.25514347 | NA | | Reg3b | 1.322442219 | 0.00347238023829701 | 0.0287166446426094 | 0.120918731332028 | 0.903755405576791 | NA | NM_011036 |
| Pnma5 | 5.469156186 | -0.062300479 | 0.132487665 | -0.470236068 | 0.638186368 | NA | | Reg3d | 0 | NA | NA | NA | NA | NA | NM_013893,NM_001161741 |
| Pnpla2 | 849.433174 | 0.395853393 | 0.167492267 | 2.363412946 | NA | NA | | Reg3g | 0.528213853 | -0.00859352304702679 | 0.0178531968357867 | -0.481343656605024 | 0.630272274096989 | NA | NM_011260 |
| Pnpla5 | 0.394700387 | 0.01999451 | 0.042974513 | 0.465264371 | 0.641742146 | NA | | Reg4 | 0.12890128 | -0.0068640067066208 | 0.00870804083610955 | -0.788237771940376 | 0.430557639008166 | NA | NM_026328 |
| Podnl1 | 1.495842343 | -0.0117407 | 0.074261612 | -0.158099182 | 0.874378645 | NA | | Rem1 | 16.10280817 | 0.0572845411365371 | 0.0738513092025144 | 0.77567400977819 | 0.437941489398001 | NA | NM_009047 |
| Pof1b | 0.384380875 | -0.006409867 | 0.041941234 | -0.152829724 | 0.878532557 | NA | | Ren1 | 0 | NA | NA | NA | NA | NA | NM_031192 |
| Pom121l12 | 0 | NA | NA | NA | NA | NA | | Ren2 | 0 | NA | NA | NA | NA | NA | NM_031193 |
| Pom121l2 | 1.548674272 | 0.114262852 | 0.082902948 | 1.378272486 | 0.168119175 | NA | | Rergl | 0.353527519 | -0.00579453045469275 | 0.0115029412048641 | -0.503743377584378 | 0.614441721879931 | NA | NM_001128090 |
| Pon1 | 0.949583441 | 0.083272253 | 0.063202749 | 1.31754164 | 0.187657133 | NA | | Retn | 20.75461809 | -0.107435056902834 | 0.070593285905753 | -1.52188774788394 | 0.128037209646712 | NA | NM_001204959,NM_022984 |
| Poteg | 0 | NA | NA | NA | NA | NA | | Retnla | 0.543403425 | 0.0186829485926561 | 0.0160617774961266 | 1.16319308975371 | 0.24475116957772 | NA | NM_020509 |
| Pou1f1 | 0.516054436 | -0.019319368 | 0.056115458 | -0.344278905 | 0.730636547 | NA | | Retnlb | 0 | NA | NA | NA | NA | NA | NM_023881 |
| Pou2af1 | 3.088322072 | -0.118224156 | 0.116071805 | -1.018543273 | 0.308419847 | NA | | Retnlg | 2.251450064 | -0.00819499183687602 | 0.0255101044863398 | -0.321244934189285 | 0.748024782632821 | NA | NM_181596 |
| Pou4f1 | 3.232891575 | 0.254519548 | 0.117724772 | 2.16198803 | 0.030619099 | NA | | Rex2 | 0 | NA | NA | NA | NA | NA | NM_001177767 |
| Pou4f2 | 0.587618527 | 0.02809576 | 0.041193526 | 0.68204309 | 0.495211705 | NA | | Rfpl3s | 0 | NA | NA | NA | NA | NA | NM_183111 |
| Pou4f3 | 0.376356809 | 0.021703132 | 0.042541541 | 0.510163288 | 0.60993707 | NA | | Rfpl4 | 0 | NA | NA | NA | NA | NA | NM_001145013,NM_138954 |
| Pou5f1 | 0 | NA | NA | NA | NA | NA | | Rfpl4b | 0 | NA | NA | NA | NA | NA | NM_001177783 |
| Pou5f2 | 0 | NA | NA | NA | NA | NA | | Rfx6 | 0.108766301 | 0.00519171381361946 | 0.00856003371502575 | 0.606506234257728 | 0.544178643801544 | NA | NM_001159389,NM_177306 |
| Pp2d1 | 5.414729412 | -0.026397436 | 0.137055751 | -0.192603638 | 0.847269395 | NA | | Rfx8 | 9.193314498 | -0.0204563021385362 | 0.0599732809310977 | -0.341090262546051 | 0.733035631519246 | NA | NM_001145660 |
| Ppbp | 1.959774784 | -0.166789816 | 0.090069715 | -1.851785756 | 0.064056594 | NA | | Rgn | 8.798153655 | -0.0434044884472704 | 0.0607610985196912 | -0.714346670891805 | 0.475012839621713 | NA | NM_009060 |
| Ppifos | 0.669327356 | 0.044675377 | 0.049579597 | 0.901083916 | 0.367543704 | NA | | Rgr | 0.525396929 | 0.00146880074099759 | 0.0162984279665609 | 0.0901191663399132 | 0.928192518654153 | NA | NM_021340 |
| Ppm1n | 1.551335448 | 0.050106203 | 0.096567402 | 0.518872845 | 0.603849415 | NA | | Rgs1 | 7.654425543 | 0.0137065674402862 | 0.0618641143271936 | 0.221559260798489 | 0.824656999425966 | NA | NM_015811 |
| Ppp1r14d | 0.608846751 | -0.058874571 | 0.04678185 | -1.258491744 | 0.208213971 | NA | | Rgs13 | 11.54510417 | -0.0823206328111377 | 0.058441722978108 | -1.40859352900965 | 0.158955393382329 | NA | NM_153171 |
| Ppp1r1c | 3.493365301 | -0.244963968 | 0.112490805 | -2.177635481 | 0.029433186 | NA | | Rgs18 | 2.147296486 | -0.0177666845055225 | 0.0339365404382828 | -0.523526684690596 | 0.600607786262249 | NA | NM_022881 |
| Ppp1r2-ps3 | 1.174269328 | -0.001670633 | 0.060912306 | -0.027426856 | 0.978119278 | NA | | Rgs9bp | 13.93581596 | -0.0146325076225666 | 0.0628922611394173 | -0.232659906918114 | 0.816025504521785 | NA | NM_145840 |
| Ppp1r2-ps7 | 0 | NA | NA | NA | NA | NA | | Rhag | 0.237667581 | -0.00136938695604711 | 0.00989612066541175 | -0.13837613771559 | 0.889943154921195 | NA | NM_011269 |
| Ppp1r2-ps9 | 0 | NA | NA | NA | NA | NA | | Rhbdl2 | 9.318544184 | 0.0322487547602896 | 0.0580422935410844 | 0.555607864418086 | 0.578478954033641 | NA | NM_183163 |
| Ppp1r27 | 0 | NA | NA | NA | NA | NA | | Rhbg | 0.645700311 | 0.00650051685141044 | 0.017450379515449 | 0.372514353951755 | 0.709509926862719 | NA | NM_021375 |
| Ppp1r3a | 1.744453034 | 0.119676409 | 0.090430701 | 1.323404638 | 0.18570085 | NA | | Rhd | 4.142573185 | -0.00618522668197482 | 0.045455621866487 | -0.136071764679453 | 0.891764548410606 | NA | NM_011270 |
| Ppp3r2 | 5.521500261 | -0.120183725 | 0.14070536 | -0.854151718 | 0.393020931 | NA | | Rho | 15.60699633 | -0.0327024569545678 | 0.0731380177642025 | -0.447133487538598 | 0.654778681462298 | NA | NM_145383 |
| Ppy | 0.107716603 | -0.020477583 | 0.027954224 | -0.732539849 | 0.46383913 | NA | | Rhod | 18.90698801 | -0.0215982227604898 | 0.0770658686038355 | -0.280256657736741 | 0.779280600920335 | NA | NM_007485 |
| Ppyr1 | 0.573420282 | 0.014545625 | 0.047116924 | 0.308713384 | 0.757539562 | NA | | Rhox1 | 0 | NA | NA | NA | NA | NA | NM_001025084 |
| Pqbp1 | 2003.471158 | 0.092169247 | 0.075802169 | 1.215918333 | NA | NA | | Rhox10 | 0 | NA | NA | NA | NA | NA | NM_001024850 |
| Prame | 0.247284148 | 0.028796741 | 0.033643376 | 0.855940884 | 0.392030479 | NA | | Rhox11 | 0 | NA | NA | NA | NA | NA | NM_198598 |
| Pramef12 | 0 | NA | NA | NA | NA | NA | | Rhox12 | 0 | NA | NA | NA | NA | NA | NM_001025083 |
| Pramef17 | 0 | NA | NA | NA | NA | NA | | Rhox13 | 0 | NA | NA | NA | NA | NA | NM_001185002 |
| Pramef6 | 0 | NA | NA | NA | NA | NA | | Rhox2a | 0 | NA | NA | NA | NA | NA | NM_029203 |
| Pramel1 | 1.116174192 | 0.037311252 | 0.077241891 | 0.483044253 | 0.629064315 | NA | | Rhox2b | 0 | NA | NA | NA | NA | NA | NM_001099316 |
| Pramel3 | 0.079983654 | 0.016740032 | 0.027674178 | 0.604897175 | 0.54524732 | NA | | Rhox2c | 0 | NA | NA | NA | NA | NA | NM_001099318 |
| Pramel4 | 0 | NA | NA | NA | NA | NA | | Rhox2d | 0 | NA | NA | NA | NA | NA | NM_001081669 |
| Pramel5 | 0 | NA | NA | NA | NA | NA | | Rhox2e | 0 | NA | NA | NA | NA | NA | NM_001085348 |
| Pramel6 | 0 | NA | NA | NA | NA | NA | | Rhox2f | 0 | NA | NA | NA | NA | NA | NM_001085356 |
| Pramel7 | 0 | NA | NA | NA | NA | NA | | Rhox2g | 0 | NA | NA | NA | NA | NA | NM_001114153 |
| Prap1 | 0.119397641 | -0.020477583 | 0.027954224 | -0.732539845 | 0.463839132 | NA | | Rhox2h | 0 | NA | NA | NA | NA | NA | NM_001100465 |
| Prb1 | 0 | NA | NA | NA | NA | NA | | Rhox3a | 0 | NA | NA | NA | NA | NA | NM_194063 |
| Prdm13 | 0.734759766 | 0.030213253 | 0.050271506 | 0.601001553 | 0.547838952 | NA | | Rhox3c | 0 | NA | NA | NA | NA | NA | NM_001102457 |
| Prdm14 | 0.445240244 | -0.001897651 | 0.037393919 | -0.050747581 | 0.959526662 | NA | | Rhox3e | 0 | NA | NA | NA | NA | NA | NM_001184969 |
| Prdx6b | 0 | NA | NA | NA | NA | NA | | Rhox3f | 0 | NA | NA | NA | NA | NA | NM_001040089 |
| Prelid2 | 6.692044179 | -0.015802065 | 0.133551379 | -0.118321988 | 0.905812535 | NA | | Rhox3g-ps | 0 | NA | NA | NA | NA | NA | NM_001145406 |
| Prf1 | 0.323149808 | -0.030167563 | 0.033779321 | -0.893077861 | 0.371815478 | NA | | Rhox3h | 0 | NA | NA | NA | NA | NA | NM_001114157 |
| Prg2 | 1.141916517 | -0.093944404 | 0.069903023 | -1.34392476 | 0.178972718 | NA | | Rhox4a | 0.097165704 | -0.00637828455464458 | 0.00848781691535724 | -0.75146349388194 | 0.452373763047372 | NA | NM_001039688 |
| Prg3 | 0.096071043 | -0.020098795 | 0.027825948 | -0.722304036 | 0.470107572 | NA | | Rhox4b | 0 | NA | NA | NA | NA | NA | NM_021300 |
| Prh1 | 0 | NA | NA | NA | NA | NA | | Rhox4c | 0 | NA | NA | NA | NA | NA | NM_001039689 |
| Prhoxnb | 0 | NA | NA | NA | NA | NA | | Rhox4d | 0 | NA | NA | NA | NA | NA | NM_001039695 |
| Prkag3 | 3.219630112 | -0.001288579 | 0.108444624 | -0.011882372 | 0.990519462 | NA | | Rhox4e | 0 | NA | NA | NA | NA | NA | NM_201236 |
| Prl2a1 | 0 | NA | NA | NA | NA | NA | | Rhox4f | 0 | NA | NA | NA | NA | NA | NM_001039696 |
| Prl2b1 | 0 | NA | NA | NA | NA | NA | | Rhox4g | 0.108766301 | 0.00519171381361946 | 0.00856003371502575 | 0.606506234257728 | 0.544178643801544 | NA | NM_001039698 |
| Prl2c1 | 0 | NA | NA | NA | NA | NA | | Rhox6 | 0 | NA | NA | NA | NA | NA | NM_008955 |
| Prl2c2 | 0 | NA | NA | NA | NA | NA | | Rhox7 | 0 | NA | NA | NA | NA | NA | NM_001025086 |
| Prl2c3 | 0 | NA | NA | NA | NA | NA | | Rhox8 | 0.553189945 | 0.0107391262364186 | 0.0168692373309267 | 0.636610062787506 | 0.524378868944667 | NA | NM_001004193 |
| Prl2c4 | 0 | NA | NA | NA | NA | NA | | Rhox9 | 0 | NA | NA | NA | NA | NA | NM_023894 |
| Prl2c5 | 0 | NA | NA | NA | NA | NA | | Ribc1 | 13.05053646 | 0.0241733522571395 | 0.0691027480806179 | 0.349817524318107 | 0.726475646497532 | NA | NM_025660 |
| Prl3a1 | 0 | NA | NA | NA | NA | NA | | Ribc2 | 7.833225282 | -0.0234618345711521 | 0.0473662671644134 | -0.4953279195448 | 0.62036866602088 | NA | NM_026357 |
| Prl3b1 | 0 | NA | NA | NA | NA | NA | | Ripk3 | 10.38469083 | 0.00412818802761226 | 0.066432975370733 | 0.0621406463367727 | 0.950450828429858 | NA | NM_001164108,NM_001164107,NM_019955 |
| Prl3c1 | 0 | NA | NA | NA | NA | NA | | Ripply1 | 0.308420637 | -0.00472633378880091 | 0.0109973596733179 | -0.42976986560402 | 0.667363055492239 | NA | NM_001037915 |
| Prl3d1 | 0 | NA | NA | NA | NA | NA | | Ripply3 | 11.10919622 | 0.0854185103515661 | 0.0677325807201574 | 1.2611140671648 | 0.207267753085126 | NA | NM_133229 |
| Prl3d2 | 0 | NA | NA | NA | NA | NA | | Rln1 | 12.29038666 | 0.0167035570614852 | 0.0633569213289397 | 0.263642183223563 | 0.792055650623358 | NA | NM_011272 |
| Prl3d3 | 0 | NA | NA | NA | NA | NA | | Rln3 | 0.108766301 | 0.00519171381361946 | 0.00856003371502575 | 0.606506234257728 | 0.544178643801544 | NA | NM_173184 |
| Prl4a1 | 0 | NA | NA | NA | NA | NA | | Rmi2 | 9.294668307 | 0.0261303460959429 | 0.057289402258091 | 0.45611134112073 | 0.648309915208771 | NA | NM_001162932 |
| Prl5a1 | 0 | NA | NA | NA | NA | NA | | Rmnd1 | 0.23403553 | -0.000407313579519725 | 0.00983856423584417 | -0.0413996971261099 | 0.966977254250032 | NA | NM_025343 |
| Prl6a1 | 0 | NA | NA | NA | NA | NA | | Rmrp | 4.131041172 | 0.0699532420685211 | 0.0436003846593147 | 1.60441800261908 | 0.108621945403697 | NA | NR_001460 |
| Prl7a1 | 0 | NA | NA | NA | NA | NA | | Rn4.5s | 0.535907978 | 0.011145936732061 | 0.0167583139908443 | 0.665098931679552 | 0.505987217919985 | NA | NR_002841 |
| Prl7a2 | 0 | NA | NA | NA | NA | NA | | Rnase1 | 17.0044005 | 0.0377673708138555 | 0.0683510005595689 | 0.552550372411019 | 0.580571332658944 | NA | NM_011271 |
| Prl7b1 | 0 | NA | NA | NA | NA | NA | | Rnase10 | 1.582225195 | -0.00437818779702935 | 0.0295550925575149 | -0.14813649419332 | 0.882235048958603 | NA | NM_001162863,NM_029145 |
| Prl7c1 | 0 | NA | NA | NA | NA | NA | | Rnase11 | 0 | NA | NA | NA | NA | NA | NM_001011877 |
| Prl7d1 | 0 | NA | NA | NA | NA | NA | | Rnase12 | 0.432646743 | 0.00474316701276897 | 0.0122006285717131 | 0.388764151362323 | 0.697450621870491 | NA | NM_001011875 |
| Prl8a1 | 0 | NA | NA | NA | NA | NA | | Rnase13 | 1.801764817 | 0.000492263004610089 | 0.0316286005072149 | 0.0155638566587161 | 0.987582340396621 | NA | NM_001011687 |
| Prl8a2 | 0 | NA | NA | NA | NA | NA | | Rnase6 | 13.18818965 | 0.0373198649732714 | 0.0630588969774051 | 0.59182552759595 | 0.553967427155109 | NA | NM_030098 |
| Prl8a6 | 0 | NA | NA | NA | NA | NA | | Rnase9 | 0 | NA | NA | NA | NA | NA | NM_183032 |
| Prl8a8 | 0 | NA | NA | NA | NA | NA | | Rnaset2a | 0 | NA | NA | NA | NA | NA | NM_001083938 |
| Prl8a9 | 0 | NA | NA | NA | NA | NA | | Rnaset2b | 14.40300816 | -0.0399076919779763 | 0.0679162520985605 | -0.587601505455011 | 0.556799800320989 | NA | NM_026611 |
| Prlh | 0 | NA | NA | NA | NA | NA | | Rnf133 | 0.192890664 | -0.000903864895041124 | 0.00963528282151475 | -0.093807821917056 | 0.92526181809679 | NA | NM_198251 |
| Prm1 | 0 | NA | NA | NA | NA | NA | | Rnf148 | 0.191449919 | 0.00665508079272897 | 0.00962405648474729 | 0.691504752001018 | 0.489248395458281 | NA | NM_027754 |
| Prm2 | 0.094392817 | 0.017895942 | 0.027818137 | 0.643319236 | 0.520016977 | NA | | Rnf151 | 2.090118535 | -0.0390269421649181 | 0.0325496065009389 | -1.19899889308316 | 0.230528376016983 | NA | NM_026205 |
| Prm3 | 0 | NA | NA | NA | NA | NA | | Rnf183 | 0.976674389 | 3.06767982168194e-05 | 0.0244932651899801 | 0.00125245850150551 | 0.999000682959866 | NA | NM_153504 |
| Prnd | 7.866505926 | -0.00404365 | 0.148905287 | -0.027155852 | 0.978335428 | NA | | Rnf186 | 0.106347839 | -0.00653347234140708 | 0.00854389509723638 | -0.764694822098226 | 0.444453290197145 | NA | NM_025786 |
| Proc | 0 | NA | NA | NA | NA | NA | | Rnf222 | 1.495121871 | -0.0169959119576182 | 0.0268828467007605 | -0.632221436472254 | 0.527242191725796 | NA | NM_177060 |
| Prodh2 | 0.415899877 | -0.004022046 | 0.042665509 | -0.094269271 | 0.924895259 | NA | | Rnu11 | 0.236453993 | 0.00749401214086028 | 0.0102093235778702 | 0.734036107652062 | 0.462926731928034 | NA | NR_002865 |
| Prok1 | 0.614929368 | -0.038004199 | 0.059400239 | -0.639798746 | 0.522303448 | NA | | Rnu12 | 0.257802559 | -0.00892927589713349 | 0.00993479775307735 | -0.898787888698349 | 0.368765652383134 | NA | NR_004432 |
| Prokr1 | 7.357789906 | -0.019095274 | 0.148106986 | -0.12892892 | 0.897413893 | NA | | Rnu73b | 5.881157751 | -0.0680503949218533 | 0.0498065209311317 | -1.36629488769046 | 0.171846424979715 | NA | NR_004418 |
| Prol1 | 0 | NA | NA | NA | NA | NA | | Ropn1 | 3.393929158 | 0.0697096499314512 | 0.0432163456574086 | 1.61303897567056 | 0.106736040330974 | NA | NM_030744 |
| Prop1 | 0.403421253 | -0.028229169 | 0.04362283 | -0.647119161 | 0.517554825 | NA | | Ropn1l | 15.60194401 | -0.0577382480050938 | 0.072679531078329 | -0.794422406810351 | 0.426949564092387 | NA | NM_145852 |
| Proz | 1.66627929 | 0.075725797 | 0.085845564 | 0.882116601 | 0.377713756 | NA | | Ros1 | 0.097165704 | -0.00637828455464458 | 0.00848781691535724 | -0.75146349388194 | 0.452373763047372 | NA | NM_011282 |
| Prp2 | 0 | NA | NA | NA | NA | NA | | Rp1 | 2.692349903 | 0.0102608065585265 | 0.0370521723816005 | 0.276928609012459 | 0.781834924532491 | NA | NM_011283,NM_001195662 |
| Prph2 | 1.475740698 | 0.033545409 | 0.097006331 | 0.345806383 | 0.729488228 | NA | | Rpl1l1 | 0.300480017 | -0.00401601363468843 | 0.0108825526077113 | -0.369032319847708 | 0.712103634607305 | NA | NM_146246 |
| Prpmp5 | 0 | NA | NA | NA | NA | NA | | Rpl10l | 0 | NA | NA | NA | NA | NA | NM_001162933 |
| Prps1l1 | 0 | NA | NA | NA | NA | NA | | Rpl26 | 0 | NA | NA | NA | NA | NA | NM_009080 |
| Prr15l | 0.282093074 | 0.009461846 | 0.038656194 | 0.244769212 | 0.806635134 | NA | | Rpl30 | 8.953789965 | -0.016141320735998 | 0.0651245218048021 | -0.247853193983956 | 0.804247995833524 | NA | NM_001163485,NM_009083 |
| Prr19 | 4.462012358 | 0.208689708 | 0.138380097 | 1.508090485 | 0.13153137 | NA | | Rpl31-ps12 | 1.855502249 | -0.0082843713720054 | 0.0257368693985776 | -0.321887298867175 | 0.747538075571742 | NA | NM_001258458 |
| Prr23a | 0.174376471 | 0.024538273 | 0.031173126 | 0.787161136 | 0.431187544 | NA | | Rpl34 | 7.955198875 | 0.00687736794323134 | 0.0566574861392978 | 0.121384982142036 | 0.903386111434354 | NA | NM_026724,NM_001005859 |
| Prr9 | 0.17833704 | -0.002390466 | 0.031282281 | -0.076415987 | 0.939088151 | NA | | Rpl34-ps1 | 0 | NA | NA | NA | NA | NA | NM_001199350 |
| Prrxl1 | 0.651198668 | 0.010138487 | 0.065005828 | 0.155962742 | 0.876062389 | NA | | Rpl39l | 0.981815442 | -0.0116301130511015 | 0.0233639625909847 | -0.497779989409378 | 0.61863912156366 | NA | NM_026594 |
| Prss1 | 0 | NA | NA | NA | NA | NA | | Rpl3l | 11.96112832 | -0.0511124243579378 | 0.0678946855856443 | -0.752819221667402 | 0.451558554470352 | NA | NM_001163945,NM_025425 |
| Prss2 | 0 | NA | NA | NA | NA | NA | | Rplp2-ps1 | 0 | NA | NA | NA | NA | NA | NR_038063 |
| Prss21 | 0 | NA | NA | NA | NA | NA | | Rpph1 | 6.638601367 | -0.00864826959197949 | 0.0518477307973716 | -0.166801313364671 | 0.867526385250056 | NA | NR_002142 |
| Prss27 | 0.258977192 | 0.029727007 | 0.03417262 | 0.869907163 | 0.38435114 | NA | | Rprl1 | 0 | NA | NA | NA | NA | NA | NR_004434 |
| Prss28 | 0 | NA | NA | NA | NA | NA | | Rprl2 | 0 | NA | NA | NA | NA | NA | NR_004439 |
| Prss29 | 0.08731684 | 0.017365056 | 0.027733412 | 0.626142072 | 0.531221759 | NA | | Rprl3 | 0 | NA | NA | NA | NA | NA | NR_024198 |
| Prss3 | 0 | NA | NA | NA | NA | NA | | Rps15a-ps4 | 14.50369155 | 0.0817936287313761 | 0.0742158560399507 | 1.10210449755301 | 0.270416244335406 | NA | NR_036572 |
| Prss32 | 1.242096834 | 0.027552611 | 0.083794036 | 0.328813513 | 0.742296649 | NA | | Rps15a-ps6 | 20.72409232 | -0.0344923982094224 | 0.0717774407254124 | -0.480546504038428 | 0.630838844091229 | NA | NR_029471 |
| Prss33 | 0.985455347 | 0.079724183 | 0.073074396 | 1.091000244 | 0.275272775 | NA | | Rptn | 0.120209049 | -0.00674972139751468 | 0.00863521114606269 | -0.781650996523957 | 0.434419707908266 | NA | NM_009100 |
| Prss34 | 0.119397641 | -0.020477583 | 0.027954224 | -0.732539845 | 0.463839132 | NA | | Rrh | 9.490046542 | -0.00761827489239815 | 0.0423733043763559 | -0.179789492571392 | NA | NA | NM_009102 |
| Prss37 | 0 | NA | NA | NA | NA | NA | | Rs1 | 10.24727284 | 0.00202986885984947 | 0.0649346063256793 | 0.0312602012194957 | 0.97506202971035 | NA | NM_011302 |
| Prss38 | 1.430750791 | 0.046143255 | 0.087228565 | 0.528992478 | 0.596810668 | NA | | Rsc1a1 | 0.106347839 | -0.00653347234140708 | 0.00854389509723638 | -0.764694822098226 | 0.444453290197145 | NA | NM_023544 |
| Prss39 | 0 | NA | NA | NA | NA | NA | | Rsph6a | 6.740492765 | -0.0493413045113803 | 0.0524529937380761 | -0.9406766133839 | 0.346870607228706 | NA | NM_001159671,NM_031255 |
| Prss40 | 0 | NA | NA | NA | NA | NA | | Rspo4 | 7.316038993 | 0.0104570586110562 | 0.0566871583633203 | 0.184469620862535 | 0.853645057207973 | NA | NM_001040689 |
| Prss41 | 4.056226237 | 0.027637124 | 0.136255618 | 0.202832913 | 0.839265632 | NA | | Rtdr1 | 8.886900332 | 0.0171292586959352 | 0.0583687458078566 | 0.29346627992184 | 0.769165770135182 | NA | NM_001163533,NM_027730,NM_001163535,NM_001163534 |
| Prss42 | 0.099009884 | 0.018205726 | 0.027896865 | 0.652608315 | 0.514008823 | NA | | Rtp2 | 4.62875023 | 0.020450334506513 | 0.0416379158926805 | 0.491146928660471 | 0.623322531929829 | NA | NM_001008230 |
| Prss43 | 5.199971063 | 0.076795151 | 0.134078207 | 0.572763858 | 0.566804593 | NA | | Rtp3 | 7.045666317 | -0.0429713465969922 | 0.05142248538889 | -0.835652852483017 | 0.40335022624186 | NA | NM_153100 |
| Prss44 | 0.196706773 | -0.026517754 | 0.031609464 | -0.838918169 | 0.401515231 | NA | | Rubie | 1.420532821 | -0.0175021012012443 | 0.0233839021433833 | -0.748467945765701 | 0.454177952471756 | NA | NR_046459 |
| Prss46 | 0.185670226 | -0.001645086 | 0.031432796 | -0.052336624 | 0.958260472 | NA | | Rufy4 | 3.038122281 | 0.0836638871429468 | 0.0385368574049081 | 2.17100959385161 | 0.0299304471718492 | NA | NM_001170641,NM_001034060 |
| Prss48 | 0 | NA | NA | NA | NA | NA | | Runx3 | 0.952960804 | -0.00911083934139455 | 0.0229215505897717 | -0.397479189102507 | 0.6910141299592 | NA | NM_019732 |
| Prss51 | 0 | NA | NA | NA | NA | NA | | Rxfp4 | 0 | NA | NA | NA | NA | NA | NM_181817 |
| Prss52 | 0.119397641 | -0.020477583 | 0.027954224 | -0.732539845 | 0.463839132 | NA | | S100a14 | 2.465526828 | -0.0137756772313317 | 0.0359127555933641 | -0.383587307732999 | 0.701284348662261 | NA | NM_001163525,NM_001163526,NM_025393 |
| Prss54 | 4.806953644 | 0.385914742 | 0.142130464 | 2.715214822 | 0.006623279 | NA | | S100a2 | 0.215934008 | -0.00149029401866511 | 0.00975123948835942 | -0.152831239602324 | 0.878531361484804 | NA | NM_001195760 |
| Prss55 | 1.471134817 | 0.035870757 | 0.08525442 | 0.420749531 | 0.673937989 | NA | | S100a5 | 3.864879878 | -0.0267098240160817 | 0.0267467375070509 | -0.998619888073568 | 0.31797886211208 | NA | NM_011312 |
| Prss58 | 0 | NA | NA | NA | NA | NA | | S100a7a | 16.0240126 | -0.0914026809244281 | 0.0770963716205866 | -1.18556397665829 | 0.235794535409817 | NA | NM_199422 |
| Prss8 | 2.008251573 | 0.049147042 | 0.101145204 | 0.485905811 | 0.627033948 | NA | | S100a8 | 9.604427418 | -0.0472813055361664 | 0.052907520546136 | -0.893659446674249 | 0.371504129350405 | NA | NM_013650 |
| Psapl1 | 0.803193456 | -0.004987786 | 0.068001862 | -0.07334778 | 0.941529371 | NA | | S100g | 0 | NA | NA | NA | NA | NA | NM_009789 |
| Psca | 2.727767399 | 0.033442762 | 0.092677388 | 0.360851366 | 0.718210562 | NA | | S100z | 0.106347839 | -0.00653347234140708 | 0.00854389509723638 | -0.764694822098226 | 0.444453290197145 | NA | NM_001081159 |
| Psg-ps1 | 0 | NA | NA | NA | NA | NA | | S1pr4 | 21.09466036 | 0.150494290049176 | 0.0834928855122103 | 1.8024804044791 | 0.0714698549940377 | NA | NM_010102 |
| Psg17 | 1.455416154 | 0.025154363 | 0.077252925 | 0.325610491 | 0.744719073 | NA | | Saa1 | 1.718321026 | 0.0363250350193237 | 0.0294290256256177 | 1.23432679971922 | 0.217081153422116 | NA | NM_009117 |
| Psg18 | 0 | NA | NA | NA | NA | NA | | Saa2 | 2.191075973 | -0.0116009511182782 | 0.0310864710151574 | -0.37318327682231 | 0.70901204178908 | NA | NM_011314 |
| Psg19 | 0 | NA | NA | NA | NA | NA | | Saa3 | 0.223412651 | 0.00718831964189475 | 0.00999993766105603 | 0.718836445339964 | 0.472241699104885 | NA | NM_011315 |
| Psg20 | 0 | NA | NA | NA | NA | NA | | Saa4 | 0 | NA | NA | NA | NA | NA | NM_011316 |
| Psg21 | 0 | NA | NA | NA | NA | NA | | Sall4 | 7.013722063 | -0.00440322588147659 | 0.0592710820154431 | -0.0742896153022705 | 0.940779939987083 | NA | NM_175303,NM_201396,NM_201395 |
| Psg22 | 0.094392817 | 0.017895942 | 0.027818137 | 0.643319236 | 0.520016977 | NA | | Samd7 | 2.24010455 | 0.0375204394433921 | 0.0302296273413351 | 1.2411810115862 | 0.214538887807428 | NA | NR_028060,NM_029489 |
| Psg25 | 0 | NA | NA | NA | NA | NA | | Samt1 | 0 | NA | NA | NA | NA | NA | NM_030036 |
| Psg26 | 0 | NA | NA | NA | NA | NA | | Samt2 | 0 | NA | NA | NA | NA | NA | NM_001037167 |
| Psg27 | 0 | NA | NA | NA | NA | NA | | Samt3 | 0 | NA | NA | NA | NA | NA | NM_028554 |
| Psg28 | 0 | NA | NA | NA | NA | NA | | Samt4 | 0 | NA | NA | NA | NA | NA | NM_029199 |
| Psg29 | 3.212231462 | 0.057291191 | 0.125186582 | 0.457646421 | 0.647206487 | NA | | Sap25 | 0 | NA | NA | NA | NA | NA | NM_001081962 |
| Psmb11 | 6.886808652 | 0.039236154 | 0.149717425 | 0.26206805 | 0.793268979 | NA | | Sapcd1 | 6.115335559 | -0.0124993117361689 | 0.0558390429096717 | -0.223845379233817 | 0.822877618246753 | NA | NR_028258,NM_023893 |
| Psmb2 | 1840.493634 | -0.021000219 | 0.080518345 | -0.26081285 | NA | NA | | Satl1 | 0 | NA | NA | NA | NA | NA | NM_028655 |
| Psors1c2 | 4.538298468 | 0.233022611 | 0.141324948 | 1.648842717 | 0.099179861 | NA | | Sbk2 | 5.76904807 | 0.00228620449449653 | 0.0484582944242123 | 0.0471788064698006 | 0.962370718696732 | NA | NM_001146329 |
| Pspn | 0.481615831 | -0.011066428 | 0.044268784 | -0.249982642 | 0.802600772 | NA | | Sbp | 0 | NA | NA | NA | NA | NA | NM_011321 |
| Ptchd3 | 0 | NA | NA | NA | NA | NA | | Sbpl | 0 | NA | NA | NA | NA | NA | NM_001077421 |
| Ptf1a | 0.381531838 | 0.016204264 | 0.036434783 | 0.444747101 | 0.656502526 | NA | | Sbspon | 15.19441133 | 0.0717789075818683 | 0.0763725893683008 | 0.939851695164089 | 0.347293637846146 | NA | NM_001033288 |
| Ptgir | 5.488531867 | -0.101224414 | 0.138770167 | -0.729439306 | 0.465732982 | NA | | Scarna10 | 0.355458167 | -0.0172950524609032 | 0.0137931659887683 | -1.25388561806524 | 0.209883583198022 | NA | NR_028517 |
| Pth | 0.20671448 | -0.003314027 | 0.03178208 | -0.10427343 | 0.916952363 | NA | | Scarna13 | 4.927500618 | -0.0160546888662327 | 0.0486769007244647 | -0.329821509325545 | 0.741534833885168 | NA | NR_028576 |
| Pth2 | 0.731905974 | -0.048379781 | 0.062701219 | -0.771592351 | 0.440355906 | NA | | Scarna17 | 1.791971035 | 0.0043238144546753 | 0.0263621347895644 | 0.164016096920455 | 0.869718471906415 | NA | NR_028560 |
| Ptk6 | 2.504663572 | 0.060484835 | 0.109573857 | 0.552000604 | 0.580947939 | NA | | Scarna2 | 0 | NA | NA | NA | NA | NA | NR_028538 |
| Ptpn11 | 8646.390569 | 0.016434726 | 0.100931217 | 0.162830953 | NA | NA | | Scarna3a | 0 | NA | NA | NA | NA | NA | NR_028518 |
| Ptprcap | 4.998701277 | -0.029774673 | 0.145182462 | -0.205084505 | 0.837506103 | NA | | Scarna3b | 0 | NA | NA | NA | NA | NA | NR_028544 |
| Ptprq | 6.32643376 | -0.159113074 | 0.149571784 | -1.063790715 | 0.287423519 | NA | | Scarna6 | 3.620121666 | -0.0238190715583407 | 0.0443471171383865 | -0.537105297826071 | 0.591194881530442 | NA | NR_028519 |
| Ptx3 | 5.661119263 | -0.018929939 | 0.127926135 | -0.147975538 | 0.882362074 | NA | | Scarna8 | 0.120209049 | -0.00674972139751468 | 0.00863521114606269 | -0.781650996523957 | 0.434419707908266 | NA | NR_028545 |
| Ptx4 | 4.32173572 | 0.270517178 | 0.124909831 | 2.165699663 | 0.030334145 | NA | | Scarna9 | 0.191449919 | 0.00665508079272897 | 0.00962405648474729 | 0.691504752001018 | 0.489248395458281 | NA | NR_028568 |
| Pxt1 | 1.172185394 | 0.038541959 | 0.075817592 | 0.508351143 | 0.611207109 | NA | | Scd4 | 17.22780539 | -0.0231868868184192 | 0.0777330551084241 | -0.298288633916132 | 0.765482878485967 | NA | NM_183216 |
| Pydc3 | 1.79767076 | 0.152674599 | 0.085799484 | 1.77943494 | 0.075168482 | NA | | Scgb1a1 | 0 | NA | NA | NA | NA | NA | NM_011681 |
| Pydc4 | 0.92872216 | 0.017464004 | 0.065307653 | 0.267411287 | 0.78915251 | NA | | Scgb1b2 | 0.191449919 | 0.00665508079272897 | 0.00962405648474729 | 0.691504752001018 | 0.489248395458281 | NA | NM_020563 |
| Pyy | 0 | NA | NA | NA | NA | NA | | Scgb1b20 | 0 | NA | NA | NA | NA | NA | NM_001270543 |
| Pzp | 0 | NA | NA | NA | NA | NA | | Scgb1b24 | 0 | NA | NA | NA | NA | NA | NM_001099329 |
| Qrfp | 0.105923158 | 0.018524693 | 0.027977518 | 0.662127842 | 0.507889294 | NA | | Scgb1b27 | 0 | NA | NA | NA | NA | NA | NM_009596 |
| Qrich2 | 0.922304787 | 0.065687929 | 0.067719607 | 0.969998679 | 0.332047151 | NA | | Scgb1b29 | 0 | NA | NA | NA | NA | NA | NM_001256066 |
| R3hdml | 0.682018854 | -0.0685167 | 0.057665156 | -1.188181988 | 0.234761718 | NA | | Scgb1b3 | 0 | NA | NA | NA | NA | NA | NM_001256073 |
| Rab17 | 1.031810426 | -0.040174341 | 0.05338871 | -0.752487574 | 0.4517579 | NA | | Scgb1b30 | 0 | NA | NA | NA | NA | NA | NM_001099330 |
| Rab19 | 2.959517648 | -0.064979996 | 0.123729994 | -0.525175778 | 0.599461002 | NA | | Scgb1b7 | 0 | NA | NA | NA | NA | NA | NM_001270542 |
| Rab25 | 1.493587948 | -3.10E-05 | 0.090496353 | -0.000342893 | 0.999726411 | NA | | Scgb1c1 | 3.743693204 | -0.0202794619017168 | 0.0410061082733186 | -0.494547343204281 | 0.620919679649755 | NA | NM_001099742 |
| Rab44 | 2.617581641 | 0.01035885 | 0.103891834 | 0.099708028 | 0.920576127 | NA | | Scgb2b19 | 0 | NA | NA | NA | NA | NA | NM_001199336 |
| Rad51b | 7.232306618 | 0.070867641 | 0.154712864 | 0.458059138 | 0.646909955 | NA | | Scgb2b2 | 0.09572496 | 0.00498255289994511 | 0.00847999972823692 | 0.587565219295241 | 0.556824162160132 | NA | NM_207262 |
| Raet1a | 0.105923158 | 0.018524693 | 0.027977518 | 0.662127842 | 0.507889294 | NA | | Scgb2b20 | 0 | NA | NA | NA | NA | NA | NM_001009952 |
| Raet1b | 0 | NA | NA | NA | NA | NA | | Scgb2b23-ps | 0 | NA | NA | NA | NA | NA | NR_045685 |

| | | | | | | |
|---|---|---|---|---|---|---|
| Raet1c | 0 | NA | NA | NA | NA | NA |
| Raet1d | 0.307160824 | -0.016020939 | 0.039725656 | -0.403289483 | 0.686735278 | NA |
| Raet1e | 0 | NA | NA | NA | NA | NA |
| Rag2 | 0.116696964 | -0.020477583 | 0.027954224 | -0.732539847 | 0.463839131 | NA |
| Rasef | 3.455393393 | -0.069929101 | 0.125673393 | -0.556435212 | 0.577913373 | NA |
| Rasl2-9 | 3.856998568 | -0.089202206 | 0.122030712 | -0.730981611 | 0.464790385 | NA |
| Raver1-fdx1l | 0 | NA | NA | NA | NA | NA |
| Rax | 2.36282512 | 0.180793304 | 0.098567262 | 1.834212496 | 0.066622457 | NA |
| Rbbp8nl | 0 | NA | NA | NA | NA | NA |
| Rbm44 | 1.102006092 | -0.063204017 | 0.070650799 | -0.894597346 | 0.371002371 | NA |
| Rbmxl2 | 0.099009884 | 0.018205726 | 0.027896865 | 0.652608315 | 0.514008823 | NA |
| Rbmy | 0 | NA | NA | NA | NA | NA |
| Rbp2 | 0.991301619 | -0.011024931 | 0.077182441 | -0.142842476 | 0.88641459 | NA |
| Rbp3 | 3.562431764 | 0.214066409 | 0.117492438 | 1.82195904 | 0.068461201 | NA |
| Rbp7 | 2.043720657 | -0.027469301 | 0.098803711 | -0.278018918 | 0.780997837 | NA |
| Rcvrn | 6.934324495 | 0.106488628 | 0.137276122 | 0.77572579 | 0.437910909 | NA |
| Rdh1 | 4.02785589 | 0.134807819 | 0.120160741 | 1.121895709 | 0.261906786 | NA |
| Rdh16 | 5.69937055 | 0.241813316 | 0.140816375 | 1.717224401 | 0.085938179 | NA |
| Rdh18-ps | 6.089229591 | 0.105723181 | 0.132679412 | 0.796831845 | 0.425548698 | NA |
| Rdh19 | 0.08731684 | 0.017365056 | 0.027733412 | 0.626142072 | 0.531221759 | NA |
| Rdh5 | 91.15875645 | 0.306568967 | 0.166926322 | 1.836552579 | NA | NA |
| Rdh7 | 0 | NA | NA | NA | NA | NA |
| Rdh8 | 0.283178451 | 0.028390488 | 0.033368195 | 0.850824814 | 0.394866675 | NA |
| Reg1 | 0 | NA | NA | NA | NA | NA |
| Reg2 | 0 | NA | NA | NA | NA | NA |
| Reg3a | 0.357330579 | 0.020235566 | 0.041992186 | 0.481888834 | 0.629884919 | NA |
| Reg3b | 0.79644831 | 0.019239514 | 0.068219575 | 0.282023363 | 0.777925596 | NA |
| Reg3d | 0 | NA | NA | NA | NA | NA |
| Reg3g | 0.383007342 | 0.024584967 | 0.04803763 | 0.511785598 | 0.608801071 | NA |
| Reg4 | 0.201994201 | 0.000227241 | 0.031727614 | 0.00716224 | 0.994285408 | NA |
| Ren1 | 0 | NA | NA | NA | NA | NA |
| Ren2 | 0 | NA | NA | NA | NA | NA |
| Rergl | 1.981022479 | -0.21192552 | 0.094872729 | -2.233787539 | 0.025497056 | NA |
| Retnla | 2.229168003 | -0.225033942 | 0.087173379 | -2.581452553 | 0.009838551 | NA |
| Retnlb | 0 | NA | NA | NA | NA | NA |
| Retnlg | 3.754170542 | -0.033909857 | 0.118334712 | -0.28655883 | 0.774450142 | NA |
| Rex2 | 0 | NA | NA | NA | NA | NA |
| Rfpl3s | 0 | NA | NA | NA | NA | NA |
| Rfpl4 | 0 | NA | NA | NA | NA | NA |
| Rfpl4b | 0.119397641 | -0.020477583 | 0.027954224 | -0.732539845 | 0.463839132 | NA |
| Rfx6 | 0.19046386 | -0.000774927 | 0.031526133 | -0.024580468 | 0.980389599 | NA |
| Rgr | 0.5855302 | 0.02347199 | 0.058537891 | 0.400970889 | 0.688441557 | NA |
| Rgs1 | 6.11725672 | 0.096417133 | 0.138892139 | 0.694187115 | 0.487564872 | NA |
| Rgs18 | 4.329224149 | -0.117003015 | 0.130286346 | -0.898045103 | 0.369161504 | NA |
| Rhag | 0.296387018 | 0.012756677 | 0.039194327 | 0.325472542 | 0.74482346 | NA |
| Rhbg | 0.686557386 | -0.05415787 | 0.062368571 | -0.868351935 | 0.385201699 | NA |
| Rhd | 3.913436963 | 0.107411919 | 0.122906442 | 0.87393237 | 0.382155085 | NA |
| Rhou | 1799.611557 | 0.154249562 | 0.12017974 | 1.28349056 | NA | NA |
| Rhox1 | 0 | NA | NA | NA | NA | NA |
| Rhox10 | 0 | NA | NA | NA | NA | NA |
| Rhox11 | 0 | NA | NA | NA | NA | NA |
| Rhox12 | 0 | NA | NA | NA | NA | NA |
| Rhox13 | 0 | NA | NA | NA | NA | NA |
| Rhox2a | 0 | NA | NA | NA | NA | NA |
| Rhox2b | 0 | NA | NA | NA | NA | NA |
| Rhox2c | 0 | NA | NA | NA | NA | NA |
| Rhox2d | 0 | NA | NA | NA | NA | NA |
| Rhox2e | 0 | NA | NA | NA | NA | NA |
| Rhox2f | 0 | NA | NA | NA | NA | NA |
| Rhox2g | 0 | NA | NA | NA | NA | NA |
| Rhox2h | 0 | NA | NA | NA | NA | NA |
| Rhox3a | 0 | NA | NA | NA | NA | NA |
| Rhox3c | 0 | NA | NA | NA | NA | NA |
| Rhox3e | 0 | NA | NA | NA | NA | NA |
| Rhox3f | 0 | NA | NA | NA | NA | NA |
| Rhox3g-ps | 0 | NA | NA | NA | NA | NA |
| Rhox3h | 0 | NA | NA | NA | NA | NA |
| Rhox4a | 0.174376471 | 0.024538273 | 0.031173126 | 0.787161136 | 0.431187544 | NA |
| Rhox4b | 0 | NA | NA | NA | NA | NA |
| Rhox4c | 0 | NA | NA | NA | NA | NA |
| Rhox4d | 0 | NA | NA | NA | NA | NA |
| Rhox4e | 0 | NA | NA | NA | NA | NA |
| Rhox4f | 0 | NA | NA | NA | NA | NA |
| Rhox4g | 0 | NA | NA | NA | NA | NA |
| Rhox6 | 0 | NA | NA | NA | NA | NA |
| Rhox7 | 0 | NA | NA | NA | NA | NA |
| Rhox8 | 0.099009884 | 0.018205726 | 0.027896865 | 0.652608315 | 0.514008823 | NA |
| Rhox9 | 0 | NA | NA | NA | NA | NA |
| Ribc2 | 6.357543594 | 0.015626106 | 0.135185025 | 0.115590513 | 0.907977082 | NA |
| Ripk3 | 7.602369188 | 0.043477767 | 0.151801714 | 0.286411571 | 0.774562913 | NA |
| Ripply1 | 0.462846809 | -0.009373068 | 0.037958243 | -0.246931029 | 0.804961602 | NA |
| Rln3 | 0.331747315 | -0.03734372 | 0.03744515 | -0.99729124 | 0.318623164 | NA |
| Rmi2 | 7.68667395 | -0.104345867 | 0.146692338 | -0.711324593 | 0.476883115 | NA |
| Rmnd1 | 0.440108409 | -0.030798726 | 0.044545461 | -0.691399874 | 0.489314283 | NA |
| Rmrp | 3.751697272 | -0.24950485 | 0.123210895 | -2.025022636 | 0.04286504 | NA |
| Rn4.5s | 0.284023613 | -0.011266173 | 0.033405616 | -0.337253873 | 0.735925529 | NA |
| Rnase10 | 1.441550775 | -0.032974422 | 0.091699494 | -0.359592186 | 0.719152128 | NA |
| Rnase11 | 0.119397641 | -0.020477583 | 0.027954224 | -0.732539845 | 0.463839132 | NA |
| Rnase12 | 0.412784469 | -0.036563067 | 0.049225712 | -0.742763586 | 0.457624825 | NA |
| Rnase13 | 1.366069526 | 0.075899547 | 0.086602413 | 0.876413765 | 0.380805138 | NA |
| Rnase9 | 0 | NA | NA | NA | NA | NA |
| Rnaset2a | 0 | NA | NA | NA | NA | NA |
| Rnf125 | 56.35163046 | 0.178217722 | 0.168039044 | 1.060573292 | NA | NA |
| Rnf133 | 0.079983654 | 0.016740032 | 0.027674178 | 0.604897175 | 0.54524732 | NA |
| Rnf148 | 0 | NA | NA | NA | NA | NA |
| Rnf151 | 1.976521841 | 0.040251082 | 0.100397971 | 0.400915297 | 0.688482487 | NA |
| Rnf183 | 0.472604946 | 0.035675895 | 0.051617383 | 0.691160469 | 0.489464703 | NA |
| Rnf186 | 0.204013804 | -0.003115899 | 0.031740777 | -0.098167077 | 0.921799625 | NA |
| Rnf222 | 1.535178008 | 0.014924316 | 0.078164365 | 0.190935034 | 0.848576492 | NA |
| Rnu11 | 0 | NA | NA | NA | NA | NA |
| Rnu12 | 0.211846316 | 0.025707621 | 0.031860375 | 0.806883811 | 0.419733426 | NA |
| Rnu73b | 4.901059966 | 0.180714247 | 0.135132054 | 1.337315922 | 0.181119536 | NA |
| Ropn1 | 2.88413121 | -0.113528915 | 0.114095572 | -0.995033484 | 0.319719979 | NA |
| Ros1 | 0.079983654 | 0.016740032 | 0.027674178 | 0.604897175 | 0.54524732 | NA |
| Rp1 | 1.924705671 | 0.029416796 | 0.095110125 | 0.309291945 | 0.757099458 | NA |
| Rp1l1 | 0.159967308 | 0.0240345 | 0.030870969 | 0.778547006 | 0.436246606 | NA |
| Rpl10l | 0 | NA | NA | NA | NA | NA |
| Rpl26 | 0 | NA | NA | NA | NA | NA |
| Rpl31-ps12 | 2.541509973 | -0.092173001 | 0.102889947 | -0.895840684 | 0.370337857 | NA |
| Rpl34 | 7.220905668 | -0.044252212 | 0.1528405 | -0.289531975 | 0.772174314 | NA |
| Rpl34-ps1 | 0 | NA | NA | NA | NA | NA |
| Rpl39l | 1.060129594 | -0.013374988 | 0.071765407 | -0.186370961 | 0.852153863 | NA |
| Rplp2-ps1 | 0 | NA | NA | NA | NA | NA |
| Rpph1 | 6.871012262 | -0.10230378 | 0.151463495 | -0.675435222 | 0.499399294 | NA |
| Rprl1 | 0 | NA | NA | NA | NA | NA |
| Rprl2 | 0 | NA | NA | NA | NA | NA |
| Rprl3 | 0 | NA | NA | NA | NA | NA |
| Rptn | 0.745603014 | -0.017866445 | 0.055148533 | -0.323969547 | 0.74596109 | NA |
| Rrh | 5.27185439 | 0.443849627 | 0.138547587 | 3.203589726 | 0.001357257 | NA |
| Rsc1a1 | 0.08731684 | 0.017365056 | 0.027733412 | 0.626142072 | 0.531221759 | NA |
| Rsph6a | 6.251109469 | 0.08820279 | 0.136193845 | 0.647626847 | 0.517226329 | NA |
| Rspo4 | 8.088924366 | -0.116627937 | 0.154057832 | -0.757039972 | 0.449025915 | NA |
| Rtdr1 | 5.549808354 | 0.098825807 | 0.13748284 | 0.718822851 | 0.472250076 | NA |
| Rtp1 | 51.04770619 | 0.597057864 | 0.155319259 | 3.844068461 | NA | NA |
| Rtp2 | 4.043698216 | -0.114263572 | 0.123126775 | -0.928015634 | 0.35339946 | NA |
| Rubie | 1.345916069 | 0.052699266 | 0.07091647 | 0.743117448 | 0.457410577 | NA |
| Rufy4 | 2.09458199 | -0.114836471 | 0.093463254 | -1.228680432 | 0.219191646 | NA |
| Runx3 | 0.842833339 | 0.002721243 | 0.069092262 | 0.039385634 | 0.968582934 | NA |
| Rxfp4 | 0 | NA | NA | NA | NA | NA |
| S100a14 | 1.85227154 | 0.054955778 | 0.096856315 | 0.567394886 | 0.570445924 | NA |
| S100a2 | 0.099009884 | 0.018205726 | 0.027896865 | 0.652608315 | 0.514008823 | NA |
| S100g | 0.094392817 | 0.017895942 | 0.027818137 | 0.643319236 | 0.520016977 | NA |
| S100z | 0.400720577 | -0.028041581 | 0.043551246 | -0.643875506 | 0.519656166 | NA |
| Saa1 | 0.842506355 | -0.012771225 | 0.067335073 | -0.189666756 | 0.849570272 | NA |
| Saa2 | 1.548906565 | 0.122461294 | 0.087530823 | 1.399064815 | 0.161793548 | NA |
| Saa3 | 0.233393928 | -0.027475306 | 0.032191697 | -0.85349047 | 0.39338737 | NA |
| Saa4 | 0 | NA | NA | NA | NA | NA |
| Sall4 | 5.625156988 | 0.100267113 | 0.150266578 | 0.667261572 | 0.504605068 | NA |
| Samd7 | 1.823176112 | -0.07817623 | 0.097817348 | -0.799206193 | 0.424170861 | NA |
| Samt1 | 0 | NA | NA | NA | NA | NA |
| Samt2 | 0 | NA | NA | NA | NA | NA |
| Samt3 | 0 | NA | NA | NA | NA | NA |
| Samt4 | 0 | NA | NA | NA | NA | NA |
| Sap25 | 0 | NA | NA | NA | NA | NA |
| Sapcd1 | 7.798835615 | 0.12963868 | 0.140046685 | 0.925681896 | 0.354611319 | NA |
| Sat1 | 1335.670063 | 0.349017909 | 0.168019537 | 2.07724598 | NA | NA |
| Satl1 | 0 | NA | NA | NA | NA | NA |
| Sbk2 | 4.261273317 | 0.00978297 | 0.126437501 | 0.077373956 | 0.938326059 | NA |
| Sbp | 0 | NA | NA | NA | NA | NA |
| Sbpl | 0.096071043 | -0.020098795 | 0.027825948 | -0.722304036 | 0.470107572 | NA |
| Scarna10 | 0.743813171 | -0.024893294 | 0.066152898 | -0.376299372 | 0.706694346 | NA |
| Scarna13 | 3.647477527 | 0.087140605 | 0.124696778 | 0.698820024 | 0.484664513 | NA |
| Scarna17 | 2.04761523 | -0.070980183 | 0.088615341 | -0.800992045 | 0.423136251 | NA |
| Scarna2 | 0.098353386 | -0.020212831 | 0.02786466 | -0.725393061 | 0.468210929 | NA |
| Scarna3a | 0 | NA | NA | NA | NA | NA |
| Scarna3b | 0 | NA | NA | NA | NA | NA |
| Scarna6 | 2.707959894 | 0.110666663 | 0.11640052 | 0.950740281 | 0.341736235 | NA |
| Scarna8 | 0.099009884 | 0.018205726 | 0.027896865 | 0.652608315 | 0.514008823 | NA |
| Scarna9 | 0 | NA | NA | NA | NA | NA |
| Sccpdh | 2502.747246 | -0.404563707 | 0.071706625 | -5.641929248 | NA | NA |
| Scgb1a1 | 0 | NA | NA | NA | NA | NA |
| Scgb1b2 | 0 | NA | NA | NA | NA | NA |
| Scgb1b20 | 0 | NA | NA | NA | NA | NA |
| Scgb1b24 | 0 | NA | NA | NA | NA | NA |
| Scgb1b27 | 0 | NA | NA | NA | NA | NA |
| Scgb1b29 | 0 | NA | NA | NA | NA | NA |
| Scgb1b3 | 0 | NA | NA | NA | NA | NA |
| Scgb1b30 | 0 | NA | NA | NA | NA | NA |
| Scgb1b7 | 0 | NA | NA | NA | NA | NA |
| Scgb1c1 | 2.319617599 | 0.129513312 | 0.100420196 | 1.289713797 | 0.197150048 | NA |
| Scgb2b19 | 0.107716603 | -0.020477583 | 0.027954224 | -0.732539849 | 0.46383913 | NA |
| Scgb2b2 | 0 | NA | NA | NA | NA | NA |
| Scgb2b20 | 0 | NA | NA | NA | NA | NA |
| Scgb2b23-ps | 0 | NA | NA | NA | NA | NA |
| Scgb2b24 | 0 | NA | NA | NA | NA | NA |
| Scgb2b26 | 0 | NA | NA | NA | NA | NA |
| Scgb2b27 | 0 | NA | NA | NA | NA | NA |
| Scgb2b3 | 0 | NA | NA | NA | NA | NA |
| Scgb2b7 | 0 | NA | NA | NA | NA | NA |
| Scgb3a1 | 1.612864918 | 0.084380987 | 0.098480193 | 0.856832057 | 0.391537706 | NA |
| Scgb3a2 | 0.08731684 | 0.017365056 | 0.027733412 | 0.626142072 | 0.531221759 | NA |
| Scimp | 1.164911991 | 0.079176849 | 0.08200121 | 0.965557078 | 0.334265862 | NA |
| Scin | 6.758425107 | 0.05192602 | 0.137656258 | 0.377215107 | 0.706013754 | NA |
| Scn10a | 40.82910494 | 0.315661711 | 0.125490269 | 2.515427796 | NA | NA |
| Scn11a | 16.77910324 | 0.403017476 | 0.15671383 | 2.571677794 | NA | NA |
| Scn4a | 0.671056634 | 0.056541089 | 0.056494479 | 1.000825029 | 0.316911407 | NA |
| Scnn1b | 0.919709443 | -0.056949659 | 0.070846114 | -0.803850137 | 0.42148353 | NA |
| Scnn1g | 6.319470388 | 0.070038414 | 0.13893941 | 0.504093217 | 0.614195873 | NA |
| Scp2d1 | 0 | NA | NA | NA | NA | NA |
| Sct | 1.116057616 | -0.017872692 | 0.060353447 | -0.296133734 | 0.767127953 | NA |
| Sdhc | 2822.274744 | -0.132054187 | 0.065111067 | -2.02813736 | NA | NA |
| Sdr16c5 | 0 | NA | NA | NA | NA | NA |
| Sdr16c6 | 0 | NA | NA | NA | NA | NA |
| Sdr9c7 | 6.742021621 | 0.010525215 | 0.141459066 | 0.074404668 | 0.940688395 | NA |
| Sds | 4.559517294 | 0.065796938 | 0.123047332 | 0.534728688 | 0.592837481 | NA |
| Sectm1a | 0.096071043 | -0.020098795 | 0.027825948 | -0.722304036 | 0.470107572 | NA |
| Sectm1b | 0 | NA | NA | NA | NA | NA |
| Sel1l2 | 0.098353386 | -0.020212831 | 0.02786466 | -0.725393061 | 0.468210929 | NA |
| Sele | 1.286676614 | -0.024345638 | 0.085698685 | -0.284084155 | 0.776345889 | NA |
| Selenbp2 | 0.789636402 | -0.004616457 | 0.062054785 | -0.074393247 | 0.940697482 | NA |
| Sell | 2.355369266 | -0.081621642 | 0.101007089 | -0.808078355 | 0.419045477 | NA |
| Selp | 0.626781319 | -0.049557557 | 0.050238808 | -0.986439744 | 0.32391737 | NA |
| 12-Sep | 0 | NA | NA | NA | NA | NA |
| 14-Sep | 0.97104811 | -0.031165945 | 0.065495028 | -0.475852076 | 0.634179766 | NA |
| Serpina10 | 0.650108395 | -0.016118184 | 0.054231577 | -0.297210319 | 0.766305944 | NA |
| Serpina11 | 0.096071043 | -0.020098795 | 0.027825948 | -0.722304036 | 0.470107572 | NA |
| Serpina12 | 0.606904295 | -0.042153114 | 0.04878701 | -0.864023309 | 0.387575086 | NA |
| Serpina1a | 0.119397641 | -0.020477583 | 0.027954224 | -0.732539845 | 0.463839132 | NA |
| Serpina1b | 0.676114142 | -0.009201815 | 0.047624202 | -0.193217197 | 0.846788871 | NA |
| Serpina1c | 0.098353386 | -0.020212831 | 0.02786466 | -0.725393061 | 0.468210929 | NA |
| Serpina1d | 0.38092772 | -0.000766687 | 0.035977 | -0.021310467 | 0.982997994 | NA |
| Serpina1e | 0.08731684 | 0.017365056 | 0.027733412 | 0.626142072 | 0.531221759 | NA |
| Serpina1f | 0 | NA | NA | NA | NA | NA |
| Serpina3a | 0 | NA | NA | NA | NA | NA |
| Serpina3b | 0.413840517 | 0.018406376 | 0.037259486 | 0.494005092 | 0.621302583 | NA |
| Serpina3c | 0.986744043 | -0.004546526 | 0.066284846 | -0.068590737 | 0.945315393 | NA |
| Serpina3f | 4.307982241 | 0.119560657 | 0.090160649 | 1.326084693 | NA | NA |
| Serpina3h | 1.700969836 | -0.033251273 | 0.091200833 | -0.364593967 | 0.715414519 | NA |
| Serpina3i | 1.437030694 | 0.085119104 | 0.08647326 | 0.984340174 | 0.324948283 | NA |
| Serpina3j | 0 | NA | NA | NA | NA | NA |
| Serpina3k | 0 | NA | NA | NA | NA | NA |
| Serpina3m | 0.188785634 | 0.025025111 | 0.031461603 | 0.79541754 | 0.42637066 | NA |
| Serpina4-ps1 | 0.096071043 | -0.020098795 | 0.027825948 | -0.722304036 | 0.470107572 | NA |
| Serpina5 | 0.079983654 | 0.016740032 | 0.027674178 | 0.604897175 | 0.54524732 | NA |
| Serpina6 | 0 | NA | NA | NA | NA | NA |
| Serpina7 | 0.098353386 | -0.020212831 | 0.02786466 | -0.725393061 | 0.468210929 | NA |
| Serpinb10 | 0.725213963 | -0.019866562 | 0.06124627 | -0.324371786 | 0.745656578 | NA |
| Serpinb11 | 2.096562876 | -0.010921561 | 0.099007949 | -0.110309938 | 0.912163576 | NA |
| Serpinb12 | 0.079983654 | 0.016740032 | 0.027674178 | 0.604897175 | 0.54524732 | NA |
| Serpinb13 | 0 | NA | NA | NA | NA | NA |
| Serpinb1c | 0.341110531 | -0.038008058 | 0.037770309 | -1.006294602 | 0.314273876 | NA |
| Serpinb2 | 2.01169938 | -0.025748391 | 0.084858552 | -0.303427176 | 0.761564337 | NA |
| Serpinb3a | 0.192142086 | -0.026384592 | 0.031527717 | -0.836869715 | 0.402665805 | NA |
| Serpinb3b | 0.519763674 | 0.02504685 | 0.039752317 | 0.630072711 | 0.528647013 | NA |
| Serpinb3c | 0.211846316 | 0.025707621 | 0.031860375 | 0.806883811 | 0.419733426 | NA |

| | | | | | | | |
|---|---|---|---|---|---|---|---|
| Scgb2b24 | 0 | NA | NA | NA | NA | NA | NM_177446 |
| Scgb2b26 | 0 | NA | NA | NA | NA | NA | NM_178308 |
| Scgb2b27 | 0 | NA | NA | NA | NA | NA | NM_001276475,NM_001100464 |
| Scgb2b3 | 0 | NA | NA | NA | NA | NA | NM_001270541 |
| Scgb2b7 | 0 | NA | NA | NA | NA | NA | NM_001198871 |
| Scgb3a1 | 2.59611355 | 0.0230336288421057 | 0.0394309569723312 | 0.584150895913292 | 0.559118794978467 | NA | NM_054037,NM_170727 |
| Scgb3a2 | 0.393522717 | 0.00404856176060657 | 0.0118516400212778 | 0.34160350410053 | 0.732649299794451 | NA | NM_054038 |
| Scgn | 10.22539503 | -0.00375342116571476 | 0.0582603962142233 | -0.0644249165747765 | 0.948631890677295 | NA | NM_145399 |
| Scimp | 1.422092934 | -0.00305128241275488 | 0.0283060043237635 | -0.107796295720667 | 0.914157281355522 | NA | NM_001045526 |
| Scin | 7.860382517 | -0.0142994995581932 | 0.0589033393309638 | -0.24276212046057 | 0.808189681341322 | NA | NM_001146196,NM_009132 |
| Scn11a | 20.10733417 | 0.0935618028797475 | 0.079409973284816 | 1.17821224475387 | 0.238712002213252 | NA | NM_011887 |
| Scn4a | 1.554855131 | 0.000990897284622514 | 0.0297176426715846 | 0.0333437377780301 | 0.973400475432156 | NA | NM_133199 |
| Scnn1b | 1.824694551 | 0.0445326292016584 | 0.0285668590528901 | 1.55889134045883 | 0.119022100250681 | NA | NR_073548,NM_011325,NM_001272023 |
| Scnn1g | 6.224211241 | 0.0357638806273726 | 0.0518897616888425 | 0.689228076278923 | 0.490679751110633 | NA | NM_011326 |
| Scp2d1 | 0.108766301 | 0.00519171381361946 | 0.00856003371502575 | 0.606506234257728 | 0.544178643801544 | NA | NM_025490 |
| Sct | 1.70644656 | 0.0157238168442326 | 0.0276232223975801 | 0.56922456829693 | 0.569203749775241 | NA | NM_011328 |
| Sctr | 13.84736701 | -0.0831568847041834 | 0.0646294258091593 | -1.28667218783179 | 0.198208570915502 | NA | NM_001012322 |
| Sdr16c5 | 0 | NA | NA | NA | NA | NA | NM_181989 |
| Sdr16c6 | 0 | NA | NA | NA | NA | NA | NM_001080710 |
| Sdr9c7 | 5.880920262 | -0.0177653422620132 | 0.0445038261595086 | -0.39918685189762 | 0.689755530456932 | NA | NM_027301 |
| Sds | 6.973786394 | 0.0241591273558949 | 0.0499514403092365 | 0.483652267208552 | 0.628632674262658 | NA | NM_145565 |
| Sec14l4 | 13.83826438 | 0.00615951404023258 | 0.0673015076648674 | 0.0915211895534988 | 0.9270784700624 | NA | NM_146013 |
| Sec31b | 15.52319493 | -0.0170248033524949 | 0.0692456353641749 | -0.245861031716417 | 0.805789810211487 | NA | NM_001033343 |
| Sectm1a | 0.108766301 | 0.00519171381361946 | 0.00856003371502575 | 0.606506234257728 | 0.544178643801544 | NA | NM_145373 |
| Sectm1b | 0 | NA | NA | NA | NA | NA | NM_026907 |
| Sel1l2 | 0 | NA | NA | NA | NA | NA | NM_001033296 |
| Sele | 1.450016684 | 0.015971174456354 | 0.0280143299249838 | 0.570107316474152 | 0.56860491306777 | NA | NM_011345 |
| Selenbp2 | 0.93937571 | 0.0140455164805025 | 0.0211038507182479 | 0.665542827610969 | 0.505703360277796 | NA | NM_019414 |
| Sell | 1.346184353 | -0.0306963458747803 | 0.0240737030950072 | -1.2750986316329 | 0.202274333667572 | NA | NM_001164059,NM_011346 |
| Selp | 0.428541593 | 0.00660357460510141 | 0.0139432791970574 | 0.47360269501704 | 0.635783253288317 | NA | NM_011347 |
| Sept12 | 0 | NA | NA | NA | NA | NA | NM_029374,NM_027669 |
| Sept14 | 0.314540458 | -0.0125494430549186 | 0.0117659602535877 | -1.06658893829698 | 0.286157497745293 | NA | NM_028826 |
| Serinc4 | 15.76722809 | -0.0571767139582241 | 0.0734219073891865 | -0.778741876796367 | 0.436131782429959 | NA | NM_001025371 |
| Serpina10 | 0.505990377 | -0.00603384872883753 | 0.0146708271423194 | -0.411282109066114 | 0.680865686765499 | NA | NM_144834 |
| Serpina11 | 0.106148608 | 0.00515554063321497 | 0.00854200771766808 | 0.603551390213729 | 0.546141939141592 | NA | NM_001166350,NM_199314 |
| Serpina12 | 0.763755743 | 0.0196194243006595 | 0.0205021192720206 | 0.956946159582354 | 0.338594431009715 | NA | NM_026535 |
| Serpina1a | 0 | NA | NA | NA | NA | NA | NM_001252569,NM_009243 |
| Serpina1b | 0.468811549 | -0.01064851353293 | 0.0144942261566833 | -0.734672787482343 | 0.46253879711408 | NA | NM_009244 |
| Serpina1c | 0 | NA | NA | NA | NA | NA | NM_009245 |
| Serpina1d | 0.342602601 | 0.0130117013735539 | 0.0134333274893572 | 0.968613426856654 | 0.332738101639367 | NA | NM_009246 |
| Serpina1e | 0.515330361 | 0.00871678476311381 | 0.0149920422012791 | 0.581427443045093 | 0.560952406163085 | NA | NM_009247 |
| Serpina1f | 0 | NA | NA | NA | NA | NA | NM_026687,NM_001164742 |
| Serpina3a | 0 | NA | NA | NA | NA | NA | NM_001167705 |
| Serpina3b | 0.609313426 | -0.00678649988491643 | 0.0151510491169669 | -0.447922769738537 | 0.654208934783781 | NA | NM_173024 |
| Serpina3c | 1.816701476 | 0.0415003216533617 | 0.0280925806195345 | 1.47726982491968 | 0.139603323245933 | NA | NM_008458 |
| Serpina3f | 3.959729982 | -0.0553384064560567 | 0.0264254955623478 | -2.09412937310835 | NA | NA | NM_001168295,NM_001168294,NM_001033335 |
| Serpina3g | 16.21186766 | 0.00322836990132908 | 0.0459584757065592 | 0.070245365010405 | 0.943998367421137 | NA | NM_009251 |
| Serpina3h | 2.85656553 | 0.0296263939077669 | 0.0355629990212686 | 0.833067927990235 | 0.404806420311009 | NA | NR_033450 |
| Serpina3i | 1.433623348 | -0.00696212676933366 | 0.020665618683181 | -0.336894185268205 | 0.736196668845365 | NA | NM_001199940 |
| Serpina3j | 0 | NA | NA | NA | NA | NA | NM_001101472 |
| Serpina3k | 0 | NA | NA | NA | NA | NA | NM_011458 |
| Serpina3m | 0.419406171 | 0.0156214382132428 | 0.0147043726426825 | 1.06236686139865 | 0.288069171767418 | NA | NM_009253 |
| Serpina4-ps1 | 0 | NA | NA | NA | NA | NA | NR_002861 |
| Serpina5 | 0.097165704 | -0.00637828455464458 | 0.00848781691535724 | -0.75146349388194 | 0.452373763047372 | NA | NM_172953 |
| Serpina6 | 0 | NA | NA | NA | NA | NA | NM_007618 |
| Serpina7 | 0.217532603 | 0.00684685248201087 | 0.00976000296730706 | 0.701521557416086 | 0.482977586094298 | NA | NM_177920 |
| Serpinb10 | 0.67286561 | 0.00367924125592136 | 0.0156599513053686 | 0.234945893775546 | 0.814250728270062 | NA | NM_001160307,NM_198028 |
| Serpinb11 | 1.772136732 | -0.0390430081444346 | 0.0310311682186057 | -1.25818686133206 | 0.208324185450199 | NA | NM_025867 |
| Serpinb12 | 0.192890664 | -0.000903864895041124 | 0.00963528282151475 | -0.093807821917056 | 0.92526181809679 | NA | NM_027971,NM_001199213 |
| Serpinb13 | 0 | NA | NA | NA | NA | NA | NM_172852 |
| Serpinb1c | 0.223412651 | 0.00718831964189475 | 0.00999993766105603 | 0.718836445339964 | 0.472241699104885 | NA | NM_173051 |
| Serpinb2 | 1.563712549 | 0.011284524077885 | 0.0232754646887219 | 0.484824867249714 | 0.627800581500042 | NA | NM_001174170,NM_011111 |
| Serpinb3a | 0 | NA | NA | NA | NA | NA | NM_009126 |
| Serpinb3b | 0.62437142 | -0.0118744809041844 | 0.0139186896889702 | -0.853132095731271 | 0.393586054463293 | NA | NM_198680 |
| Serpinb3c | 0.363951167 | -0.00554477873176065 | 0.0115995521256745 | -0.478016622683891 | 0.632638373947178 | NA | NM_201363 |
| Serpinb3d | 0.09572496 | 0.00498255289994511 | 0.00847999972823692 | 0.587565219295241 | 0.556824162160132 | NA | NM_201376 |
| Serpinb5 | 3.883766842 | 0.0181077321706523 | 0.0409850652630929 | 0.441812939772439 | 0.658624574146887 | NA | NM_009257 |
| Serpinb6c | 5.890520457 | -0.0238546822113531 | 0.0529902722176527 | -0.450170969369099 | 0.652587167183965 | NA | NM_148942 |
| Serpinb6d | 0 | NA | NA | NA | NA | NA | NM_001076790 |
| Serpinb6e | 0.727020108 | 0.00387817822096635 | 0.0158986451074279 | 0.243931366148582 | 0.807283976125905 | NA | NM_001045535 |
| Serpinb7 | 0.24789674 | -0.000769869626663926 | 0.0100368571301073 | -0.0767042528038554 | 0.938858821262798 | NA | NM_027548 |
| Serpinb9b | 1.032804552 | -0.0246888085875893 | 0.0205864076457675 | -1.19927716444813 | 0.230420191375217 | NA | NM_011452 |
| Serpinb9c | 0.201873568 | 0.00673381845182595 | 0.00968011632508873 | 0.69563404257585 | 0.486658041207307 | NA | NM_011453,NM_001164524 |
| Serpinb9d | 0 | NA | NA | NA | NA | NA | NM_011460 |
| Serpinb9e | 0.108766301 | 0.00519171381361946 | 0.00856003371502575 | 0.606506234257728 | 0.544178643801544 | NA | NM_011456 |
| Serpinb9f | 0 | NA | NA | NA | NA | NA | NM_183197 |
| Serpinb9g | 0 | NA | NA | NA | NA | NA | NM_011455 |
| Serpinc1 | 0.428167708 | -0.000792272155220925 | 0.0137354739435619 | -0.0576807293637128 | 0.95400294387567 | NA | NM_080844 |
| Serpine3 | 0.108766301 | 0.00519171381361946 | 0.00856003371502575 | 0.606506234257728 | 0.544178643801544 | NA | NM_001199945 |
| Serpinf2 | 6.813703967 | -0.0184463112486517 | 0.0461609856022049 | -0.399608262432131 | 0.689445069876961 | NA | NM_008878 |
| Serpini2 | 0 | NA | NA | NA | NA | NA | NM_026460 |
| Sfta2 | 0.427940552 | -0.00281048961418964 | 0.0152501962985345 | -0.184292028717015 | 0.853784366990649 | NA | NM_001163194 |
| Sftpa1 | 0.360627147 | -0.0100772056525624 | 0.010553896215922 | -0.954832741045871 | 0.339662287230353 | NA | NM_023134 |
| Sftpb | 0.427411356 | 0.0060983518828938 | 0.0139232555105122 | 0.43799755583667 | 0.661388055218398 | NA | NM_147779 |
| Sftpd | 0 | NA | NA | NA | NA | NA | NM_009160 |
| Sgca | 11.34513306 | -0.0622901331546761 | 0.0691106732817912 | -0.901309887413409 | 0.367423578187164 | NA | NM_009161 |
| Sgcg | 5.675724316 | 0.0964146816497962 | 0.0532407942192255 | 1.81091741894001 | 0.0701536357825269 | NA | NM_011892 |
| Sgol1 | 6.616605979 | -0.0561247011399216 | 0.0572196899191402 | -0.980863426894381 | 0.326660095071568 | NA | NM_028232 |
| Sh2d1a | 0.636717407 | -0.00100888991360931 | 0.0154151701670083 | -0.0654478609498939 | 0.947817418207905 | NA | NM_011364 |
| Sh2d1b1 | 1.511807921 | -0.00572035864808812 | 0.0258079226884796 | -0.22165126256526 | 0.824585373150537 | NA | NM_012009 |
| Sh2d1b2 | 0 | NA | NA | NA | NA | NA | NM_001033499 |
| Sh2d2a | 0.46293569 | -0.00199393361883231 | 0.0140214968410126 | -0.142205475024613 | 0.886917707298231 | NA | NM_021309,NM_001025571 |
| Sh2d4a | 10.66255757 | 0.0651948425090181 | 0.0644459455502057 | 1.01162054420055 | 0.311719519142441 | NA | NM_028182 |
| Sh2d4b | 13.63584079 | -2.47324229961451e-05 | 0.0715026809656306 | -0.000345895044243632 | 0.999724015690043 | NA | NM_177816 |
| Sh2d7 | 3.390627198 | -0.0569388861456441 | 0.040839175002376 | -1.39422224230366 | 0.163250503306918 | NA | NM_173778 |
| Shbg | 1.358219832 | -0.00924255251157604 | 0.0256931719124812 | -0.359727967533904 | 0.719050575171194 | NA | NM_011367 |
| Shox2 | 21.8812246 | 0.0222985101859449 | 0.03098215848542 | 0.719721003184412 | 0.47169679185341 | NA | NM_013665 |
| Siglec15 | 0.927348726 | 0.0106629671647229 | 0.0193879712508652 | 0.549978490619385 | 0.582334126683772 | NA | NM_001101038 |
| Siglecg | 3.809185785 | 0.0178320340027034 | 0.0418910377077686 | 0.425676588083099 | 0.670343523973285 | NA | NM_172900 |
| Sim1 | 21.02888898 | 0.035297961007772 | 0.0320877112135317 | 1.1000460822175 | 0.271312044178843 | NA | NM_011376 |
| Sim2 | 5.442816465 | 0.0195782937852742 | 0.0508694075608266 | 0.384873634745276 | 0.700331040492041 | NA | NM_011377 |
| Sirpb1a | 0.476780095 | -0.0021277247032961 | 0.0156388378995265 | -0.136053888208696 | 0.891778680350359 | NA | NM_001002898 |
| Sirpb1b | 0 | NA | NA | NA | NA | NA | NM_001173460 |
| Sis | 4.516395738 | 0.00731414164480021 | 0.0491960888972601 | 0.148673234168572 | 0.881811482514925 | NA | NM_001081137 |
| Sit1 | 0.572749016 | 0.00251543981871357 | 0.0166019927512964 | 0.151514330622578 | 0.879570007137095 | NA | NM_019436 |
| Six1 | 8.538975441 | -0.0499446418146776 | 0.0591052268892835 | -0.845012267836047 | 0.398103989569387 | NA | NM_009189 |
| Six2 | 13.25143872 | 0.00349631914563039 | 0.0533934613433916 | 0.0654821593817335 | 0.947790110597092 | NA | NM_011380 |
| Six6 | 5.085214279 | 0.0464341068396488 | 0.0305188791246548 | 1.52148795013041 | NA | NA | NM_011384 |
| Ska1 | 1.165949806 | -0.000899481777423793 | 0.0272485688573612 | -0.0330102392581546 | 0.973666422345145 | NA | NM_025581,NM_001164355 |
| Skint1 | 0 | NA | NA | NA | NA | NA | NM_001102662 |
| Skint10 | 0 | NA | NA | NA | NA | NA | NM_177668 |
| Skint11 | 0 | NA | NA | NA | NA | NA | NM_177669,NM_001166027 |
| Skint2 | 0 | NA | NA | NA | NA | NA | NM_177865 |
| Skint3 | 0.768228894 | 0.00779222605114639 | 0.0182570496502471 | 0.426806422747552 | 0.669520328010908 | NA | NM_177578,NM_001102474 |
| Skint4 | 0.191449919 | 0.00665508079272897 | 0.00962405648474729 | 0.691504752001018 | 0.489248395458281 | NA | NM_178786 |
| Skint5 | 0.313257562 | 0.00962480958133652 | 0.0115601382610462 | 0.832586026567595 | 0.405078242417788 | NA | NM_001167876,NM_001167878 |
| Skint6 | 0 | NA | NA | NA | NA | NA | NM_001103199 |
| Skint7 | 0.408982522 | 0.0120846716222426 | 0.0129414896798793 | 0.933792934288797 | 0.350410714955054 | NA | NM_177818,NM_001142775 |
| Skint8 | 0 | NA | NA | NA | NA | NA | NM_001100466 |
| Skint9 | 0 | NA | NA | NA | NA | NA | NM_177864 |
| Skor1 | 12.57378579 | -0.0224099426940542 | 0.0627385803080943 | -0.357195565854443 | 0.720945405177861 | NA | NM_172446,NM_001163757,NM_001163755,NM_001163758 |
| Skor2 | 0.800420478 | 0.012971778913312 | 0.0166366452329611 | 0.779711217716653 | 0.435560874471577 | NA | NM_001109743 |
| Slamf1 | 11.13686847 | 0.00166465076861333 | 0.0647230129392136 | 0.0257196118199369 | 0.979480981060442 | NA | NM_013730 |
| Slamf6 | 9.123214006 | -0.0192818346630765 | 0.0607251756225393 | -0.317526206641709 | 0.750844357483177 | NA | NM_030710 |
| Slamf7 | 3.210977104 | -0.0410124336699479 | 0.0303433706090247 | -1.35161100585675 | 0.176499787667945 | NA | NM_144539 |
| Slamf8 | 15.18104501 | -0.306207076000792 | 0.0618122326096049 | -4.95382650121606 | 7.2768094880382e-07 | NA | NM_029084 |
| Slc10a1 | 0 | NA | NA | NA | NA | NA | NM_001177561,NM_011387 |
| Slc10a2 | 0 | NA | NA | NA | NA | NA | NM_011388 |
| Slc10a5 | 0.41774406 | -0.00103676765343045 | 0.0136489312371981 | -0.0759596217031919 | 0.939451222559279 | NA | NM_001010834 |
| Slc10a6 | 1.22971801 | 0.0228730516205554 | 0.023106703035799 | 0.989888154321213 | 0.32222879005765 | NA | NM_029415 |
| Slc12a1 | 5.520072124 | -0.0314258532459228 | 0.0465380482031536 | -0.675272265582321 | 0.499502801157696 | NA | NM_183354,NM_001079690 |
| Slc12a3 | 6.185878677 | -0.0174835501028333 | 0.0539530647234868 | -0.32405110242463 | 0.745899345656814 | NA | NM_019415,NM_001205311 |
| Slc13a1 | 6.266668738 | 0.0264646751977698 | 0.054372160052132 | 0.486732091798367 | 0.626448199833734 | NA | NM_019481 |
| Slc13a2 | 0.385490251 | -0.00508296712693953 | 0.0117912533198291 | -0.431079461111366 | 0.666410594179892 | NA | NM_022411 |
| Slc15a1 | 2.574388517 | -0.0130642312518734 | 0.0297151054531814 | -0.439649499896852 | 0.660190983519164 | NA | NM_053079 |
| Slc15a5 | 2.056374142 | 0.0373617705346304 | 0.0348428046592829 | 1.07229515247064 | 0.283587483388982 | NA | NM_177787 |
| Slc16a5 | 0.612759861 | -0.0102980728652475 | 0.0172427561673979 | -0.59724053192371 | 0.550346801747742 | NA | NM_001080934 |
| Slc16a8 | 20.07484796 | 0.00947501891385382 | 0.0233804980283875 | 0.405253083246973 | NA | NA | NM_020516 |
| Slc17a1 | 0.120209049 | -0.00674972139751468 | 0.00863521114606269 | -0.781650996523957 | 0.434419707908266 | NA | NM_001170638,NM_009198 |
| Slc17a2 | 0.593945617 | -0.0159087475407923 | 0.0159262665175918 | -0.998899994748919 | 0.317843138785385 | NA | NM_144836 |
| Slc17a3 | 0.742907396 | -0.0235934473102465 | 0.0190659822692268 | -1.23746298392017 | 0.215915248421502 | NA | NM_134069,NM_001164743 |
| Slc17a4 | 0 | NA | NA | NA | NA | NA | NM_177016 |
| Slc17a9 | 21.07770029 | 0.00175740692477341 | 0.0764936492057066 | 0.0229745468156107 | 0.981670576287911 | NA | NM_183161 |
| Slc18a1 | 5.995618548 | 0.0376994215041173 | 0.0551182641164409 | 0.683973309182503 | 0.493992020155418 | NA | NM_153054 |
| Slc1a7 | 4.297115518 | 0.0231014469990179 | 0.0489660205964018 | 0.471785264917269 | 0.637080069524132 | NA | NM_146255 |
| Slc22a1 | 6.326155981 | -0.0408062637221477 | 0.055310762103033 | -0.7377635413183 | 0.460658151944499 | NA | NM_009202 |
| Slc22a12 | 7.750617388 | -0.0558981101332528 | 0.0547059637298229 | -1.02179189108737 | 0.306879408092737 | NA | NM_009203 |
| Slc22a13 | 1.715931041 | 0.0427015970388442 | 0.0287043119257898 | 1.48763701945694 | 0.136846645097257 | NA | NM_133980 |
| Slc22a13b-ps | 20.0097511 | -0.072860630076235 | 0.0722917426054805 | -1.0078693285049 | 0.31351719782809 | NA | NR_033303 |
| Slc22a14 | 1.375413303 | 0.0119515953584962 | 0.0249139362595676 | 0.4797152579174 | 0.631429877011962 | NA | NM_001037749 |
| Slc22a16 | 0 | NA | NA | NA | NA | NA | NM_027572 |
| Slc22a19 | 0 | NA | NA | NA | NA | NA | NM_144785 |
| Slc22a20 | 0.346276033 | -0.0170215305728547 | 0.0136845010181804 | -1.24385467546394 | 0.213553053235879 | NA | NM_198650 |
| Slc22a22 | 0 | NA | NA | NA | NA | NA | NM_172378 |
| Slc22a26 | 0.106148608 | 0.00515554063321497 | 0.00854200771766808 | 0.603551390213729 | 0.546141939141592 | NA | NM_146232 |
| Slc22a27 | 0 | NA | NA | NA | NA | NA | NM_134256 |
| Slc22a28 | 0 | NA | NA | NA | NA | NA | NM_001013820 |
| Slc22a29 | 1.63290074 | -0.0179772105938536 | 0.0269230667737939 | -0.667725216629187 | 0.504309010449039 | NA | NM_172776 |
| Slc22a30 | 0.120209049 | -0.00674972139751468 | 0.00863521114606269 | -0.781650996523957 | 0.434419707908266 | NA | NM_177002 |
| Slc22a7 | 7.143797858 | -0.055069994169811 | 0.0471509748496684 | -1.16795027770668 | 0.242826821906937 | NA | NM_144856 |
| Slc23a1 | 9.653164508 | 0.0454698276362909 | 0.0675651201003149 | 0.67297782596673 | 0.500961397955973 | NA | NM_011397 |
| Slc24a1 | 9.098563725 | 0.0326322400525305 | 0.0571237232853326 | 0.571255481536677 | 0.567826474303194 | NA | NM_144813 |
| Slc24a5 | 16.70125483 | 0.00275882602669955 | 0.0455095109315996 | 0.0606208673797011 | 0.951661154320103 | NA | NM_175034 |
| Slc25a2 | 3.952268908 | -0.00960502130226802 | 0.04425609236027 | -0.217032747131799 | 0.828182824594393 | NA | NM_001159275 |
| Slc25a21 | 20.22036376 | -0.0679647421179226 | 0.0780020301655309 | -0.871320169150626 | 0.383579360835884 | NA | NM_001167976,NM_172577 |
| Slc25a31 | 2.145967408 | 0.00619034436811983 | 0.0255272077963821 | 0.242499862009866 | 0.808392863765404 | NA | NM_178386 |
| Slc25a41 | 3.406738091 | -0.0200611988600472 | 0.0421572184354719 | -0.475866283510948 | 0.634169644026467 | NA | NM_175333 |
| Slc25a43 | 7.277357917 | -0.0466542818263313 | 0.0569629183916776 | -0.819028995416561 | 0.412769870736301 | NA | NR_075097,NR_075096,NM_001085497 |
| Slc25a48 | 6.135086484 | 0.0597565556633013 | 0.0493593765594095 | 1.21064243166397 | 0.226032474616105 | NA | NM_177809 |
| Slc25a54 | 0 | NA | NA | NA | NA | NA | NM_029054 |
| Slc26a3 | 6.939871194 | 0.0553380003781379 | 0.0584271376387095 | 0.947128382710213 | 0.343573363444227 | NA | NM_021353 |
| Slc26a5 | 10.2863615 | -0.0318778717185697 | 0.0601165640861571 | -0.530267692492927 | 0.5959263432645 | NA | NM_030727 |
| Slc26a9 | 1.289261018 | 0.0219084246174841 | 0.021491627161564 | 1.01939348066979 | 0.308016199948962 | NA | NM_177243 |
| Slc27a5 | 16.30839846 | 0.0395396316231584 | 0.0793147666383178 | 0.498515387474599 | 0.618120826340597 | NA | NM_009512 |
| Slc27a6 | 4.94496703 | -0.0247918471045877 | 0.0475772669205459 | -0.521085987263416 | 0.60230687206246 | NA | NM_001081072 |
| Slc28a1 | 22.07906159 | -0.0323305200021327 | 0.0778718291634826 | -0.415176070081244 | 0.678013015394024 | NA | NM_001004184 |
| Slc28a2 | 4.19160987 | 0.00345292492835952 | 0.0459624363130589 | 0.0751249325610373 | 0.940115310559644 | NA | NM_172980 |
| Slc28a3 | 7.164824504 | 0.0708124693973903 | 0.0458639278720432 | 1.54396870662608 | 0.122595913818762 | NA | NM_022317 |
| Slc2a12 | 180.4482565 | 0.0187733661457439 | 0.0584944514120283 | 0.320942682469255 | NA | NA | NM_178934 |
| Slc2a2 | 1.690456458 | -0.00437343343385443 | 0.024656890427336 | -0.177371653848362 | 0.859216471228428 | NA | NM_031197 |
| Slc2a7 | 0 | NA | NA | NA | NA | NA | NM_001085529 |
| Slc30a2 | 3.227866408 | 0.0515218630477874 | 0.0428873635134727 | 1.20132968844313 | 0.22962333847187 | NA | NM_001039677 |
| Slc30a8 | 0.533274033 | -0.00646714190403665 | 0.016459981318383 | -0.392900926127658 | 0.694392657523221 | NA | NM_172816 |
| Slc34a1 | 2.340456278 | 0.00118096292134792 | 0.0372254773162828 | 0.0317245877417226 | 0.974691686570664 | NA | NM_011392 |
| Slc34a2 | 2.042086693 | 0.0286279162627946 | 0.0335618422007717 | 0.852990014419898 | 0.393664841744222 | NA | NM_011402 |
| Slc34a3 | 10.13532032 | -0.0544125088232586 | 0.0632804159070436 | -0.859863324905898 | 0.389864387231059 | NA | NM_080854 |
| Slc35g3 | 4.371261137 | -5.48736779878465e-06 | 0.039279625812609 | -0.000139700103686405 | 0.999888535444489 | NA | NM_019871 |
| Slc36a2 | 14.61858557 | -0.0050664863164046 | 0.0748272515158911 | -0.0677091061580502 | 0.946017200283614 | NA | NM_153170 |
| Slc36a3 | 0.212695677 | -0.00857796107890413 | 0.00973699866180651 | -0.880965621629515 | 0.378336427605205 | NA | NM_172258 |
| Slc38a8 | 16.58338342 | -0.0857554873499693 | 0.0714919492898382 | -1.19951250737764 | 0.230328724362811 | NA | NM_001009950 |
| Slc39a5 | 9.35167071 | -0.0957472036685538 | 0.0623693056824063 | -1.53516545712586 | 0.124743190945655 | NA | NM_028092,NM_001136237,NM_028051 |
| Slc3a1 | 10.17573481 | -0.0377638665470863 | 0.0590159895426477 | -0.639892118047032 | 0.522242738510689 | NA | NM_009205 |
| Slc43a1 | 8.121991202 | -0.0127276938655894 | 0.0620636558592763 | -0.205074833078611 | 0.837513658843743 | NA | NM_024497,NM_001081349,NM_001083809 |
| Slc44a3 | 4.415748996 | -0.0322100390664738 | 0.0378695177428207 | -0.850553188588733 | 0.395017602424069 | NA | NM_145394 |
| Slc44a4 | 21.47195614 | -0.0258817378922208 | 0.0813062134526891 | -0.318324231238256 | 0.750239005656443 | NA | NM_023557 |
| Slc45a2 | 2.705179238 | 0.00252872162025064 | 0.0374274502652842 | 0.0675632885042173 | 0.946133280121691 | NA | NM_053077 |
| Slc46a2 | 0 | NA | NA | NA | NA | NA | NM_021053 |
| Slc47a2 | 3.465496222 | -0.00755757157861871 | 0.0416536506478808 | -0.18143839642068 | 0.856023482999715 | NA | NM_001033542 |
| Slc4a1 | 4.388138427 | -0.00275743494626639 | 0.0326852857006358 | -0.0843631893421922 | 0.932767673565822 | NA | NM_011403 |
| Slc4a5 | 123.835318 | 0.00320727505881369 | 0.0245593885667392 | 0.130592626526432 | NA | NA | NM_001166067 |
| Slc4a9 | 7.77462547 | -0.0524906947635958 | 0.0595566723850866 | -0.881357078249722 | 0.37812458137361 | NA | NM_172830,NM_001271544,NM_001271545,NM_001271546,NM_001271547 |
| Slc51a | 0.097165704 | -0.00637828455464458 | 0.00848781691535724 | -0.75146349388194 | 0.452373763047372 | NA | NM_145932 |
| Slc51b | 0 | NA | NA | NA | NA | NA | NM_178933 |
| Slc5a1 | 2.53745628 | 0.0102237411118493 | 0.0375250825325174 | 0.272450862779312 | 0.785275368406998 | NA | NM_019810 |
| Slc5a10 | 4.514881584 | -0.0392559910769978 | 0.0461649162226899 | -0.850342517413766 | 0.395134684368668 | NA | NM_001033227 |
| Slc5a12 | 1.498691249 | 0.00273536459537942 | 0.0273685600994365 | 0.0999455062831655 | 0.920387588403236 | NA | NM_001003915,NM_001177624 |
| Slc5a2 | 0.768725731 | 0.0127518289364427 | 0.0214989796716453 | 0.593136471181508 | 0.553089824280234 | NA | NM_133254 |
| Slc5a4a | 1.34506611 | 0.0102631842184408 | 0.0248708282739284 | 0.412659526470193 | 0.679856082029379 | NA | NM_133184 |
| Slc5a4b | 1.402686381 | -0.0143223091766242 | 0.0294871938095222 | -0.485712857898305 | 0.627170766012346 | NA | NM_023219 |
| Slc5a8 | 3.665361673 | 0.0371576947474878 | 0.0428190516058126 | 0.867784160414326 | 0.385512502823585 | NA | NM_145423 |
| Slc5a9 | 0.598202733 | -0.0182433857947567 | 0.0175333329581267 | -1.04049731094058 | 0.298108912726659 | NA | NM_145551 |
| Slc6a14 | 0.434194484 | 0.00512053343735348 | 0.0139630860849759 | 0.366719320227003 | 0.71382839366562 | NA | NM_020049 |
| Slc6a18 | 0.315716853 | 0.0123948421219712 | 0.0131137196252776 | 0.945181266349427 | 0.344566341855147 | NA | NM_001040692,NM_001136087,NM_001168644,NM_001168645,NM_001168646,NM_001168643 |
| Slc6a19 | 12.95992685 | -0.00687973289463438 | 0.0622567691166677 | -0.110505781013819 | 0.912008265364775 | NA | NM_028878 |
| Slc6a2 | 2.885109541 | -0.0269272879076346 | 0.0409036723572972 | -0.658309788725628 | 0.510339089276015 | NA | NM_009209 |
| Slc6a20b | 2.037485204 | 0.00231831175480515 | 0.0283629075732053 | 0.0817374505354057 | 0.93485549676329 | NA | NM_011731 |
| Slc6a3 | 67.01662748 | 0.0553046081538969 | 0.0296028960340285 | 1.86821613974269 | NA | NA | NM_010020 |
| Slc6a4 | 6.531249875 | 0.127359554139645 | 0.0496145944020802 | 2.56697763378884 | 0.0102589229334639 | NA | NM_010484 |
| Slc6a5 | 13.15472404 | -0.0271397270386925 | 0.020923007268937 | -1.29712362519632 | NA | NA | NM_148931,NM_001146013 |
| Slc7a12 | 0 | NA | NA | NA | NA | NA | NM_080852 |
| Slc7a13 | 0.318445824 | 0.0076171880905019 | 0.0102869072557308 | 0.740474070693932 | 0.459012388854512 | NA | NM_028746 |
| Slc7a15 | 0.120209049 | -0.00674972139751468 | 0.00863521114606269 | -0.781650996523957 | 0.434419707908266 | NA | NM_177802,NM_001038660 |
| Slc7a9 | 0.332904726 | -0.0130643184082516 | 0.0120034188534089 | -1.08838311549392 | 0.276426011853847 | NA | NM_001199015,NM_021291,NM_001199016 |
| Slc9a3 | 16.719281 | -0.0143691997331825 | 0.068912283890182 | -0.208514344932771 | 0.83482738315241 | NA | NM_001081060 |
| Slc9b1 | 2.204987207 | -0.0312962043303701 | 0.0336705921351949 | -0.929481851839995 | 0.352639426085144 | NA | NM_028946 |
| Slc9c1 | 0 | NA | NA | NA | NA | NA | NM_198106 |
| Slco1a1 | 0 | NA | NA | NA | NA | NA | NM_013797 |
| Slco1a5 | 8.12752002 | -0.0109477278526532 | 0.0351778643845711 | -0.311210701507361 | 0.755640448091412 | NA | NM_130861,NM_001267707 |
| Slco1a6 | 0 | NA | NA | NA | NA | NA | NM_023718 |
| Slco1b2 | 4.555797743 | -0.0180893021172922 | 0.0423560078503485 | -0.427077598559453 | 0.669322808717531 | NA | NM_020495 |
| Slco6b1 | 0 | NA | NA | NA | NA | NA | NM_001039475 |
| Slco6c1 | 0.846373745 | -0.0131836247062292 | 0.0210300570346576 | -0.626894386662982 | 0.530728467070438 | NA | NM_028942 |
| Slco6d1 | 0.097165704 | -0.00637828455464458 | 0.00848781691535724 | -0.75146349388194 | 0.452373763047372 | NA | NM_001164233,NM_027584 |
| Slfn1 | 1.376489281 | -0.0341366597702136 | 0.0254326108284947 | -1.34223969377013 | 0.179518290008799 | NA | NM_011407 |
| Slfn10-ps | 5.822556287 | -0.00497850879599769 | 0.0453670453089388 | -0.109738440361175 | 0.912616813505273 | NA | NR_073523,NR_073524 |

| | | | | | | |
|---|---|---|---|---|---|---|
| Serpinb3d | 0 | NA | NA | NA | NA | NA |
| Serpinb5 | 4.934441871 | -0.232121113 | 0.139900385 | -1.659188523 | 0.09707781 | NA |
| Serpinb6c | 5.590896087 | 0.015753888 | 0.142647351 | 0.110439399 | 0.912060908 | NA |
| Serpinb6d | 0 | NA | NA | NA | NA | NA |
| Serpinb6e | 0.898256634 | 0.026555098 | 0.057925231 | 0.458437501 | 0.646638154 | NA |
| Serpinb7 | 0.649309414 | -0.061653223 | 0.057177405 | -1.078279488 | 0.280909048 | NA |
| Serpinb9b | 1.33413296 | 0.004628374 | 0.074655478 | 0.061996436 | 0.950565671 | NA |
| Serpinb9c | 0.094392817 | 0.017895942 | 0.027818137 | 0.643319236 | 0.520016977 | NA |
| Serpinb9d | 0 | NA | NA | NA | NA | NA |
| Serpinb9e | 0.098353386 | -0.020212831 | 0.02786466 | -0.725393061 | 0.468210929 | NA |
| Serpinb9f | 0 | NA | NA | NA | NA | NA |
| Serpinb9g | 0 | NA | NA | NA | NA | NA |
| Serpinc1 | 0.569860421 | -0.032391892 | 0.053568423 | -0.604682578 | 0.545389926 | NA |
| Serpine3 | 0.098353386 | -0.020212831 | 0.02786466 | -0.725393061 | 0.468210929 | NA |
| Serpinf2 | 5.348155506 | 0.082448973 | 0.115921372 | 0.711249119 | 0.476929876 | NA |
| Serpini2 | 0 | NA | NA | NA | NA | NA |
| Sfta2 | 0.376370672 | 0.027798864 | 0.047885955 | 0.580522285 | 0.561562463 | NA |
| Sftpa1 | 0.297029653 | 0.02866449 | 0.033517577 | 0.855207712 | 0.392436167 | NA |
| Sftpb | 0.185670226 | -0.001645086 | 0.031432796 | -0.052336624 | 0.958260472 | NA |
| Sftpd | 0 | NA | NA | NA | NA | NA |
| Sgcg | 3.714718138 | -0.005543271 | 0.109092518 | -0.050812566 | 0.959474878 | NA |
| Sh2d1a | 0.551455211 | -0.006423304 | 0.04591486 | -0.139895971 | 0.888742185 | NA |
| Sh2d1b1 | 1.607386356 | -0.052718474 | 0.092387153 | -0.570625599 | 0.568253462 | NA |
| Sh2d1b2 | 0 | NA | NA | NA | NA | NA |
| Sh2d2a | 0.610399768 | -0.006711683 | 0.062901991 | -0.106700643 | 0.915026472 | NA |
| Sh2d7 | 3.465215393 | 0.089890027 | 0.121590735 | 0.739283524 | 0.459734852 | NA |
| Shbg | 1.281199613 | 0.033476079 | 0.077403559 | 0.432487592 | 0.665387069 | NA |
| Shox2 | 6.221846719 | 0.259673605 | 0.104389761 | 2.487539038 | 0.012863034 | NA |
| Siglec15 | 0.932205047 | -0.017555772 | 0.06557263 | -0.267730174 | 0.788907022 | NA |
| Siglecg | 2.662740426 | 0.018562599 | 0.11140283 | 0.166625918 | 0.867664399 | NA |
| Sim2 | 4.214574456 | -0.077544818 | 0.129059375 | -0.600846068 | 0.547942517 | NA |
| Sirpb1a | 0.582088857 | -0.027172096 | 0.053599983 | -0.506942252 | 0.612195339 | NA |
| Sirpb1b | 0.299858689 | -0.034980473 | 0.036264327 | -0.964597324 | 0.33474654 | NA |
| Sis | 4.599129682 | 0.152244774 | 0.135595123 | 1.122789455 | 0.261526925 | NA |
| Sit1 | 0.286710141 | 0.009849273 | 0.038825804 | 0.25367852 | 0.79974393 | NA |
| Six6 | 1.810013564 | -0.049986132 | 0.089873706 | -0.556181931 | 0.57808649 | NA |
| Ska1 | 2.185053297 | -0.081721159 | 0.091527124 | -0.892862748 | 0.371930678 | NA |
| Skint1 | 0 | NA | NA | NA | NA | NA |
| Skint10 | 0.318806384 | -0.016982193 | 0.040116674 | -0.423320068 | 0.672061754 | NA |
| Skint11 | 0 | NA | NA | NA | NA | NA |
| Skint2 | 0 | NA | NA | NA | NA | NA |
| Skint3 | 1.581200634 | -0.05378995 | 0.084321719 | -0.637913348 | 0.523530089 | NA |
| Skint4 | 0.107716603 | -0.020477583 | 0.027954224 | -0.732539849 | 0.46383913 | NA |
| Skint5 | 0.107716603 | -0.020477583 | 0.027954224 | -0.732539849 | 0.46383913 | NA |
| Skint6 | 0 | NA | NA | NA | NA | NA |
| Skint7 | 0 | NA | NA | NA | NA | NA |
| Skint8 | 0 | NA | NA | NA | NA | NA |
| Skint9 | 0 | NA | NA | NA | NA | NA |
| Skor2 | 0.499336704 | -0.024198502 | 0.039525929 | -0.612218432 | 0.540393248 | NA |
| Slamf7 | 4.005037745 | -0.019113612 | 0.111189675 | -0.171900959 | 0.863515392 | NA |
| Slc10a1 | 0.096071043 | -0.020098795 | 0.027825948 | -0.722304036 | 0.470107572 | NA |
| Slc10a2 | 0 | NA | NA | NA | NA | NA |
| Slc10a5 | 0.159967308 | 0.0240345 | 0.030870969 | 0.778547006 | 0.436246606 | NA |
| Slc10a6 | 1.560736871 | 0.063371788 | 0.0726491 | 0.872299694 | 0.383044904 | NA |
| Slc12a1 | 3.49259856 | 0.19811565 | 0.106583684 | 1.858780267 | 0.063058291 | NA |
| Slc12a3 | 7.353508885 | 0.111521512 | 0.133253346 | 0.836913415 | 0.402641239 | NA |
| Slc13a1 | 4.86241085 | 0.079856996 | 0.131174675 | 0.608783641 | 0.542667858 | NA |
| Slc13a2 | 0.427314999 | -0.00123242 | 0.042901703 | -0.02872661 | 0.977082634 | NA |
| Slc15a1 | 2.06578724 | 0.083811144 | 0.090251847 | 0.928636337 | 0.353077584 | NA |
| Slc15a5 | 2.432967051 | -0.124535988 | 0.102797106 | -1.211473678 | 0.225713917 | NA |
| Slc16a5 | 1.269995199 | -0.081054214 | 0.078926637 | -1.026956382 | 0.304441003 | NA |
| Slc16a8 | 6.573843282 | 0.326076484 | 0.107915393 | 3.021593808 | NA | NA |
| Slc17a1 | 0.099009884 | 0.018205726 | 0.027896865 | 0.652608315 | 0.514008823 | NA |
| Slc17a2 | 0.600329945 | 0.053285905 | 0.055639064 | 0.957706721 | 0.338210667 | NA |
| Slc17a3 | 0.816087363 | 0.053149322 | 0.064635028 | 0.82229905 | 0.410906717 | NA |
| Slc17a4 | 0 | NA | NA | NA | NA | NA |
| Slc18a1 | 5.787397128 | -0.130868944 | 0.148436108 | -0.881651674 | 0.377965202 | NA |
| Slc1a7 | 3.047235252 | 0.010976726 | 0.122253463 | 0.08978663 | 0.928456773 | NA |
| Slc22a1 | 7.124424939 | -0.00445354 | 0.148188891 | -0.03005313 | 0.976024681 | NA |
| Slc22a12 | 7.10112088 | 0.112817447 | 0.147475559 | 0.764990805 | 0.44427702 | NA |
| Slc22a13 | 0.831476197 | -0.075869228 | 0.070717418 | -1.072850644 | 0.283338134 | NA |
| Slc22a14 | 0.56700977 | 0.058647955 | 0.060141457 | 0.97516684 | 0.329477508 | NA |
| Slc22a16 | 0 | NA | NA | NA | NA | NA |
| Slc22a19 | 0 | NA | NA | NA | NA | NA |
| Slc22a20 | 0.284916696 | 0.042355674 | 0.040798016 | 1.038179738 | 0.299186384 | NA |
| Slc22a22 | 0 | NA | NA | NA | NA | NA |
| Slc22a26 | 0 | NA | NA | NA | NA | NA |
| Slc22a27 | 0 | NA | NA | NA | NA | NA |
| Slc22a28 | 0 | NA | NA | NA | NA | NA |
| Slc22a29 | 1.288823074 | 0.094167423 | 0.072361247 | 1.301351585 | 0.193138137 | NA |
| Slc22a30 | 0.218407525 | -0.002234018 | 0.031987735 | -0.069839845 | 0.944321133 | NA |
| Slc23a1 | 6.475352322 | 0.04140952 | 0.152293342 | 0.271906303 | 0.785694064 | NA |
| Slc24a1 | 7.99271006 | -0.135920025 | 0.156082007 | -0.870824429 | 0.383850024 | NA |
| Slc25a2 | 5.289379588 | -0.123422725 | 0.137415725 | -0.898170313 | 0.369094758 | NA |
| Slc25a31 | 1.120484986 | 0.025598664 | 0.065749371 | 0.389337017 | 0.697026856 | NA |
| Slc25a41 | 3.529875707 | 0.169315375 | 0.112299314 | 1.507715135 | 0.13162745 | NA |
| Slc25a43 | 6.904239365 | 0.056192651 | 0.147297934 | 0.381489742 | 0.702839881 | NA |
| Slc25a48 | 6.14951664 | -0.118222169 | 0.138160763 | -0.855685553 | 0.392171733 | NA |
| Slc25a54 | 0 | NA | NA | NA | NA | NA |
| Slc26a3 | 5.006784055 | 0.026397753 | 0.138252049 | 0.190939327 | 0.848573129 | NA |
| Slc26a9 | 0.869904894 | -0.055282602 | 0.070581208 | -0.78324817 | 0.433481393 | NA |
| Slc27a6 | 7.372042461 | -0.334043524 | 0.149942228 | -2.227814868 | 0.025892863 | NA |
| Slc28a2 | 3.445389332 | 0.018368091 | 0.126347099 | 0.145378018 | 0.884412419 | NA |
| Slc28a3 | 2.271840048 | 0.068000825 | 0.091578763 | 0.742539238 | 0.457760688 | NA |
| Slc2a12 | 130.0413045 | 0.240041497 | 0.165165798 | 1.453336588 | NA | NA |
| Slc2a2 | 1.415985396 | -0.006921476 | 0.076339664 | -0.090666837 | 0.927757322 | NA |
| Slc2a5 | 204.357231 | -0.218856896 | 0.166955716 | -1.310867944 | NA | NA |
| Slc2a7 | 0 | NA | NA | NA | NA | NA |
| Slc30a2 | 3.854484238 | -0.122897272 | 0.120445105 | -1.020359213 | 0.30755813 | NA |
| Slc30a8 | 1.516755232 | -0.087377064 | 0.087474128 | -0.99889037 | 0.317847802 | NA |
| Slc34a1 | 1.880407214 | 0.061162988 | 0.102707221 | 0.595508155 | 0.55150385 | NA |
| Slc34a2 | 2.449511793 | -0.144336872 | 0.105649065 | -1.366191658 | 0.171878815 | NA |
| Slc34a3 | 6.109588583 | 0.228385488 | 0.131853278 | 1.732118389 | 0.083252486 | NA |
| Slc35g3 | 2.437441128 | 0.13187174 | 0.099519648 | 1.325082454 | 0.185143805 | NA |
| Slc36a3 | 0.17463368 | 0.024560283 | 0.031186242 | 0.787535833 | 0.430968261 | NA |
| Slc38a2 | 5998.012261 | 0.24738027 | 0.166917212 | 1.482053686 | NA | NA |
| Slc39a14 | 1535.394057 | 0.06927232 | 0.085215767 | 0.812904963 | NA | NA |
| Slc44a3 | 4.487310293 | 0.161364895 | 0.130852994 | 1.233176944 | 0.217509755 | NA |
| Slc45a2 | 2.017695165 | 0.01360999 | 0.094633027 | 0.143818603 | 0.885643713 | NA |
| Slc46a2 | 0 | NA | NA | NA | NA | NA |
| Slc47a2 | 3.802909098 | 0.076984124 | 0.127448864 | 0.604039314 | 0.545817504 | NA |
| Slc4a1 | 6.459954439 | -0.196756786 | 0.129307359 | -1.521620947 | 0.128104086 | NA |
| Slc4a5 | 49.33279172 | 0.303545429 | 0.103950937 | 2.920083635 | NA | NA |
| Slc4a9 | 5.511057126 | 0.123489862 | 0.12628237 | 0.977886794 | 0.328130317 | NA |
| Slc51a | 0.079983654 | 0.016740032 | 0.027674178 | 0.604897175 | 0.54524732 | NA |
| Slc51b | 0 | NA | NA | NA | NA | NA |
| Slc5a1 | 1.776605274 | 0.044468868 | 0.099695053 | 0.44604889 | 0.655561931 | NA |
| Slc5a10 | 4.65089161 | 0.130441909 | 0.128306044 | 1.016646643 | 0.309321557 | NA |
| Slc5a12 | 1.361654606 | -0.017884823 | 0.082788648 | -0.216029898 | 0.828964443 | NA |
| Slc5a2 | 0.357764354 | 0.026260735 | 0.047363929 | 0.554445872 | 0.57927374 | NA |
| Slc5a4a | 1.07156002 | 0.007911469 | 0.078408129 | 0.10090113 | 0.919628946 | NA |
| Slc5a4b | 1.63457504 | -0.055961005 | 0.095701821 | -0.584743374 | 0.558720284 | NA |
| Slc5a8 | 3.000929632 | -0.014338427 | 0.115992831 | -0.123614772 | 0.901620296 | NA |
| Slc5a9 | 0.983728672 | -0.024520894 | 0.065357323 | -0.375182046 | 0.707525081 | NA |
| Slc6a14 | 1.598951379 | -0.127632625 | 0.077648368 | -1.643725785 | 0.100232857 | NA |
| Slc6a18 | 0.334830846 | -0.037566726 | 0.037554766 | -1.000318476 | 0.317156409 | NA |
| Slc6a2 | 2.51436945 | 0.119831659 | 0.103380424 | 1.159132978 | 0.246401986 | NA |
| Slc6a20b | 1.580178914 | 0.056601605 | 0.074818355 | 0.756520312 | 0.449337299 | NA |
| Slc6a4 | 8.2200236 | -0.236383931 | 0.116077468 | -2.036432522 | NA | NA |
| Slc6a5 | 9.398780647 | 0.052891473 | 0.047345844 | 1.117130217 | NA | NA |
| Slc7a12 | 0 | NA | NA | NA | NA | NA |
| Slc7a13 | 0.096071043 | -0.020098795 | 0.027825948 | -0.722304036 | 0.470107572 | NA |
| Slc7a15 | 0.412796476 | -0.02682719 | 0.043806178 | -0.612406537 | 0.540268818 | NA |
| Slc7a9 | 0.369714607 | 0.021630852 | 0.042365317 | 0.510579254 | 0.609645706 | NA |
| Slc9b1 | 2.688258334 | 0.11790608 | 0.114244178 | 1.032053288 | 0.302047155 | NA |
| Slc9c1 | 0 | NA | NA | NA | NA | NA |
| Slco1a1 | 0 | NA | NA | NA | NA | NA |
| Slco1a5 | 5.505525164 | 0.190158591 | 0.118893587 | 1.599401576 | 0.109731402 | NA |
| Slco1a6 | 0.096071043 | -0.020098795 | 0.027825948 | -0.722304036 | 0.470107572 | NA |
| Slco1b2 | 4.756397274 | 0.232516176 | 0.126333168 | 1.840499847 | 0.065694887 | NA |
| Slco6b1 | 0 | NA | NA | NA | NA | NA |
| Slco6c1 | 0.951824705 | 0.018973528 | 0.077202833 | 0.245762072 | 0.805866419 | NA |
| Slco6d1 | 0.079983654 | 0.016740032 | 0.027674178 | 0.604897175 | 0.54524732 | NA |
| Slfn1 | 1.843410828 | -0.010768272 | 0.084241161 | -0.127826724 | 0.898286101 | NA |
| Slfn10-ps | 5.851185142 | 0.055919566 | 0.132030294 | 0.423535875 | 0.67190433 | NA |
| Slfn14-ps | 0.828450308 | -0.004796219 | 0.063212146 | -0.075874957 | 0.939518581 | NA |
| Slfn2 | 90.02144944 | 0.474883087 | 0.163292962 | 2.908166286 | NA | NA |
| Slfn4 | 1.779416325 | -0.11518335 | 0.077243412 | -1.491173767 | 0.135915879 | NA |
| Slfnl1 | 1.853738839 | -0.020551781 | 0.082403985 | -0.249402763 | 0.803049248 | NA |
| Sln | 7.139820866 | 0.053572088 | 0.148022269 | 0.361919112 | 0.717412477 | NA |
| Slpi | 1.524041911 | -0.026465255 | 0.085757308 | -0.308606411 | 0.757620943 | NA |
| Slurp1 | 0 | NA | NA | NA | NA | NA |
| Slx | 0 | NA | NA | NA | NA | NA |
| Slxl1 | 0 | NA | NA | NA | NA | NA |
| Smarca5-ps | 6.142828382 | -0.140467067 | 0.14601476 | -0.962005944 | 0.336046618 | NA |
| Smc1b | 3.729585714 | 0.190189383 | 0.122260053 | 1.555613457 | 0.119800037 | NA |
| Smcp | 0 | NA | NA | NA | NA | NA |
| Smgc | 0 | NA | NA | NA | NA | NA |
| Smim6 | 0.73395846 | 0.057529561 | 0.068965195 | 0.834182529 | 0.404178135 | NA |
| Smim9 | 2.064742545 | 0.007331969 | 0.090417284 | 0.081090348 | 0.935370101 | NA |
| Smlr1 | 1.861110957 | -0.034383872 | 0.092834115 | -0.3703797 | 0.711099597 | NA |
| Smok2a | 0.203787646 | -0.02671144 | 0.031728022 | -0.841887971 | 0.399850664 | NA |
| Smok2b | 0 | NA | NA | NA | NA | NA |
| Smok3a | 0 | NA | NA | NA | NA | NA |
| Smok3b | 0 | NA | NA | NA | NA | NA |
| Smok4a | 0 | NA | NA | NA | NA | NA |
| Smpx | 2.002337781 | -0.078651516 | 0.090071661 | -0.873210455 | 0.382548377 | NA |
| Smr2 | 0 | NA | NA | NA | NA | NA |
| Smr3a | 0.498722596 | 0.038106329 | 0.05215842 | 0.730588245 | 0.465030694 | NA |
| Smtnl1 | 0.297029653 | 0.02866449 | 0.033517577 | 0.855207712 | 0.392436167 | NA |
| Snhg9 | 6.228410644 | 0.120158539 | 0.144461314 | 0.831769668 | 0.405538968 | NA |
| Snora15 | 0.176054697 | -0.002190495 | 0.031239146 | -0.070120197 | 0.944097991 | NA |
| Snora17 | 7.419739404 | 0.163646196 | 0.148316452 | 1.103358354 | 0.269871574 | NA |
| Snora19 | 0 | NA | NA | NA | NA | NA |
| Snora20 | 0.776311287 | 0.056253754 | 0.069333946 | 0.811345061 | 0.41716754 | NA |
| Snora21 | 0.756517709 | -0.003487157 | 0.060621506 | -0.057523435 | 0.954128239 | NA |
| Snora23 | 4.454482807 | 0.052035229 | 0.127935034 | 0.406731663 | 0.684205083 | NA |
| Snora24 | 0 | NA | NA | NA | NA | NA |
| Snora26 | 0 | NA | NA | NA | NA | NA |
| Snora2b | 4.287580726 | 0.091161089 | 0.124264218 | 0.733606909 | 0.463188349 | NA |
| Snora30 | 5.453059429 | 0.04289294 | 0.143507137 | 0.298890639 | 0.76502349 | NA |
| Snora33 | 4.880171593 | 0.107710074 | 0.138614036 | 0.777050271 | 0.437129112 | NA |
| Snora35 | 0 | NA | NA | NA | NA | NA |
| Snora36b | 0.098353386 | -0.020212831 | 0.02786466 | -0.725393061 | 0.468210929 | NA |
| Snora44 | 7.757554327 | 0.236738768 | 0.152725308 | 1.550095202 | 0.121118668 | NA |
| Snora47 | 1.661032965 | 0.030399447 | 0.102875619 | 0.295497095 | 0.767614171 | NA |
| Snora5c | 4.709046317 | 0.12076528 | 0.124500499 | 0.969998363 | 0.332047308 | NA |
| Snora62 | 1.647731421 | -0.020662579 | 0.097009475 | -0.212995475 | 0.831330492 | NA |
| Snora68 | 5.229687796 | 0.143930448 | 0.137887133 | 1.043827977 | 0.296564985 | NA |
| Snora69 | 1.614440914 | 0.111076425 | 0.082010488 | 1.354417306 | 0.175603279 | NA |
| Snora74a | 2.927036111 | 0.079155726 | 0.112849439 | 0.701427737 | 0.483036117 | NA |
| Snora75 | 5.714637261 | 0.140444187 | 0.136321732 | 1.03024063 | 0.302897061 | NA |
| Snora78 | 4.210737126 | 0.08472242 | 0.12842511 | 0.659702924 | 0.50944449 | NA |
| Snord11 | 0 | NA | NA | NA | NA | NA |
| Snord110 | 0.836563683 | 0.120566208 | 0.067280645 | 1.791989478 | 0.073134651 | NA |
| Snord111 | 0.17463368 | 0.024560283 | 0.031186242 | 0.787535833 | 0.430968261 | NA |
| Snord116 | 0 | NA | NA | NA | NA | NA |
| Snord116l1 | 0 | NA | NA | NA | NA | NA |
| Snord116l2 | 0 | NA | NA | NA | NA | NA |
| Snord123 | 0.079983654 | 0.016740032 | 0.027674178 | 0.604897175 | 0.54524732 | NA |
| Snord15a | 0.185670226 | -0.001645086 | 0.031432796 | -0.052336624 | 0.958260472 | NA |
| Snord15b | 4.508321953 | 0.021899669 | 0.129521653 | 0.169081139 | 0.86573282 | NA |
| Snord16a | 4.669576605 | 0.108281837 | 0.123963081 | 0.873500687 | 0.382390232 | NA |
| Snord17 | 0.856084225 | 0.046257005 | 0.078427052 | 0.589809308 | 0.555318502 | NA |
| Snord1a | 0 | NA | NA | NA | NA | NA |
| Snord1b | 0.944323739 | 0.070449576 | 0.067910535 | 1.037388016 | 0.299555061 | NA |
| Snord1c | 1.966396074 | -0.024856255 | 0.087490008 | -0.28410393 | 0.776330735 | NA |
| Snord23 | 1.729293513 | 0.137780731 | 0.089556602 | 1.538476528 | 0.123932143 | NA |
| Snord33 | 0.079983654 | 0.016740032 | 0.027674178 | 0.604897175 | 0.54524732 | NA |
| Snord34 | 2.761218143 | 0.148125241 | 0.114082284 | 1.298407041 | 0.194147501 | NA |
| Snord35a | 3.313797802 | 0.112011181 | 0.107819687 | 1.03887503 | 0.298862861 | NA |
| Snord35b | 2.7360309 | -0.033000222 | 0.111615197 | -0.29566065 | 0.767489251 | NA |
| Snord38a | 0.97346973 | 0.040727718 | 0.065993421 | 0.61714815 | 0.537137012 | NA |
| Snord42a | 1.601975893 | 0.101605255 | 0.089815358 | 1.131268163 | 0.25794224 | NA |
| Snord43 | 1.06343616 | 0.106829216 | 0.082129948 | 1.300734002 | 0.193349519 | NA |
| Snord45b | 7.299069619 | 0.155278879 | 0.139179939 | 1.115669977 | 0.264563426 | NA |
| Snord45c | 0 | NA | NA | NA | NA | NA |
| Snord47 | 1.659266406 | 0.051411949 | 0.083406692 | 0.616400772 | 0.537630044 | NA |
| Snord49b | 2.026158894 | 0.010474814 | 0.104496421 | 0.100240892 | 0.920153082 | NA |
| Snord4a | 5.077063052 | -0.0371771 | 0.140092135 | -0.265376069 | 0.79071977 | NA |
| Snord55 | 1.899069637 | -0.072711789 | 0.092540098 | -0.785732788 | 0.43202405 | NA |
| Snord57 | 4.032632855 | 0.095197874 | 0.124156007 | 0.766760116 | 0.443224154 | NA |
| Snord58b | 4.051392012 | 0.111357297 | 0.130557126 | 0.852939244 | 0.393692997 | NA |
| Snord61 | 0 | NA | NA | NA | NA | NA |
| Snord64 | 2.774180084 | 0.094359898 | 0.117405762 | 0.803707559 | 0.421565888 | NA |
| Snord65 | 5.091137723 | 0.093665193 | 0.135875982 | 0.689343261 | 0.49060728 | NA |
| Snord66 | 2.34916592 | 0.050203948 | 0.100213218 | 0.500971322 | 0.616391306 | NA |
| Snord67 | 0 | NA | NA | NA | NA | NA |
| Snord68 | 5.346287061 | 0.215082612 | 0.133920108 | 1.606051659 | 0.10826256 | NA |
| Snord7 | 0 | NA | NA | NA | NA | NA |
| Snord70 | 0 | NA | NA | NA | NA | NA |
| Snord71 | 1.914277662 | -0.041711868 | 0.106185353 | -0.392821299 | 0.694451472 | NA |
| Snord72 | 2.028357065 | -0.093775826 | 0.088327722 | -1.061680569 | 0.288380723 | NA |
| Snord73a | 1.37279849 | 0.010632801 | 0.082723032 | 0.128534959 | 0.897725635 | NA |
| Snord8 | 0.921924053 | -0.061698577 | 0.076084357 | -0.810923289 | 0.417409725 | NA |

| | | | | | | | |
|---|---|---|---|---|---|---|---|
| Slfn14-ps | 0.57694017 | 0.000332632151082112 | 0.015006458564943 | 0.0221659327310697 | 0.982315592652382 | NA | NM_001166028 |
| Slfn3 | 20.78917277 | -0.0350531937585675 | 0.0794870981298746 | -0.440992243814132 | 0.659218612099536 | NA | NM_011409 |
| Slfn4 | 0.97778493 | 0.00911140280930583 | 0.0233447931009467 | 0.390297004128785 | 0.696316938352903 | NA | NM_011410 |
| Slfnl1 | 2.250463993 | 0.00102903454849566 | 0.029188087886172 | 0.0352552915596491 | 0.971876173316088 | NA | NM_177570 |
| Sln | 8.029300378 | 0.0120135156087125 | 0.0531975542486586 | 0.22582834452423 | 0.821334930738958 | NA | NM_025540 |
| Slpi | 1.457270812 | -0.0119161870102559 | 0.0259558950571442 | -0.459093665775014 | 0.646166905149441 | NA | NM_011414 |
| Slurp1 | 0 | NA | NA | NA | NA | NA | NM_020519 |
| Slx | 0 | NA | NA | NA | NA | NA | NM_001136476 |
| Slxl1 | 0 | NA | NA | NA | NA | NA | NM_029181 |
| Smarca5-ps | 6.232461676 | 0.0088159307896464 | 0.0527316814729708 | 0.167184708383806 | 0.867224716021397 | NA | NR_002888 |
| Smc1b | 5.73928136 | 0.00192795164054942 | 0.0456234553050946 | 0.0422579050108493 | 0.966293102224506 | NA | NM_080470 |
| Smcp | 0 | NA | NA | NA | NA | NA | NM_008574 |
| Smgc | 0 | NA | NA | NA | NA | NA | NM_198927 |
| Smim5 | 13.16704612 | -0.00134053204607646 | 0.0629223816461349 | -0.0213045344280736 | 0.983002726710568 | NA | NM_183259 |
| Smim6 | 1.367906726 | -0.0032965362304074 | 0.0255754141273253 | -0.128894735154466 | 0.897440942656606 | NA | NM_001162998 |
| Smim9 | 1.829118544 | 0.03301907085085 | 0.0285869834432979 | 1.15503865304092 | 0.248074598170091 | NA | NM_001033786 |
| Smlr1 | 1.895823843 | 0.0141193243992007 | 0.0271395902321022 | 0.520248252772056 | 0.60289055779881 | NA | NM_001195596 |
| Smok2a | 0 | NA | NA | NA | NA | NA | NM_013741 |
| Smok2b | 0 | NA | NA | NA | NA | NA | NM_001167913 |
| Smok3a | 0 | NA | NA | NA | NA | NA | NM_001126045,NM_001168602 |
| Smok3b | 0 | NA | NA | NA | NA | NA | NM_001039889 |
| Smok4a | 0 | NA | NA | NA | NA | NA | NR_030763 |
| Smpd5 | 18.08520383 | -0.0572783562882163 | 0.0778940187806283 | -0.735337028245113 | 0.462134262538612 | NA | NM_001195537 |
| Smpx | 2.231234673 | 0.0139385373809166 | 0.0326409334404815 | 0.427026310578176 | 0.669360164152029 | NA | NM_001252591,NM_025357,NR_045569 |
| Smr2 | 0 | NA | NA | NA | NA | NA | NM_001252679,NM_021289,NM_001252680,NM_001252681 |
| Smr3a | 0.330974078 | -0.00671789744880547 | 0.01278556667705 | -0.525428212803741 | 0.599285545196596 | NA | NM_011422 |
| Smtnl1 | 0.456352106 | -0.00873813301279707 | 0.012619114628994 | -0.692452146580877 | 0.488653427703681 | NA | NM_024230 |
| Snai1 | 17.18609486 | -0.00728666566148664 | 0.0727594693281881 | -0.100147317301333 | 0.92022737035174 | NA | NM_011427 |
| Snhg9 | 5.808687506 | -0.0340826958938153 | 0.0481564509437307 | -0.7077493300667 | 0.479100937452343 | NA | NR_027900 |
| Snora15 | 0.309462921 | 0.00239536941478217 | 0.0109127344998192 | 0.21950221686162 | 0.826258853162846 | NA | NR_003681 |
| Snora16a | 19.09954969 | -0.15306488573547 | 0.0748605476858863 | -2.04466692359409 | 0.0408877066887843 | NA | NR_029412 |
| Snora17 | 9.102667517 | -0.0358610765608566 | 0.0577555075005184 | -0.620911807597478 | 0.534657651798733 | NA | NR_028571 |
| Snora19 | 0 | NA | NA | NA | NA | NA | NR_034047 |
| Snora20 | 0.997626583 | -0.0184396377241903 | 0.021972078878983 | -0.83923045360211 | 0.401340001104809 | NA | NR_028479 |
| Snora21 | 0.776305083 | -0.010185648641066 | 0.0214378462820974 | -0.475124623389613 | 0.634698148752287 | NA | NR_028078 |
| Snora23 | 5.043238112 | -0.0309045863923364 | 0.048832168558105 | -0.632873519748837 | 0.526816241388956 | NA | NR_033336 |
| Snora24 | 0 | NA | NA | NA | NA | NA | NR_028573 |
| Snora26 | 0 | NA | NA | NA | NA | NA | NR_031758 |
| Snora2b | 4.81953814 | -0.0501207905549401 | 0.0434176016854388 | -1.15438874118534 | 0.24834082895338 | NA | NR_034052 |
| Snora3 | 11.79861909 | -0.0548230506773751 | 0.0685578767020595 | -0.799660860496401 | 0.423907315519476 | NA | NR_028079 |
| Snora30 | 5.702490234 | -0.0511315773007479 | 0.0507185754743191 | -1.00814300919469 | 0.313385812756358 | NA | NR_034045 |
| Snora31 | 21.05596023 | -0.0687398443240219 | 0.0797400227206714 | -0.862049469998485 | 0.388660295662306 | NA | NR_028481 |
| Snora33 | 7.139059484 | 0.0209963486572029 | 0.0574487120091317 | 0.365479884977498 | 0.714753219324922 | NA | NR_037680 |
| Snora35 | 0 | NA | NA | NA | NA | NA | NR_028446 |
| Snora36b | 0 | NA | NA | NA | NA | NA | NR_034044 |
| Snora41 | 11.92568077 | -0.00563825335728269 | 0.0582125940930985 | -0.096856246403751 | 0.922840555687903 | NA | NR_028558 |
| Snora43 | 11.65684916 | -0.118742816240896 | 0.0682881333539658 | -1.73884993495726 | 0.0820611624860577 | NA | NR_028572 |
| Snora44 | 9.864701051 | -0.0597652641017037 | 0.0624257271402814 | -0.957381945546278 | 0.338374507819652 | NA | NR_034050 |
| Snora47 | 0.890172945 | -0.035470780608832 | 0.0225829882923497 | -1.57068586980794 | 0.116255632949667 | NA | NR_034043 |
| Snora52 | 17.10556745 | -0.0897868223804326 | 0.0685710441184397 | -1.30939850099625 | 0.190399398984295 | NA | NR_034049 |
| Snora5c | 6.630578333 | 0.00403107562468774 | 0.049025874544646 | 0.0822234312417373 | 0.934469041093296 | NA | NR_034042 |
| Snora62 | 1.117309089 | -0.0076325295824109 | 0.026723222405166 | -0.285614117440246 | 0.775173689539248 | NA | NR_002902 |
| Snora64 | 19.91761423 | -0.0370816383895246 | 0.0613080062956189 | -0.604841694096557 | 0.545284187230101 | NA | NR_002897 |
| Snora68 | 5.310437037 | -0.0404379298529499 | 0.0480648613609953 | -0.841320014412136 | 0.400168681167363 | NA | NR_002901 |
| Snora69 | 2.138970587 | -0.0270848766101407 | 0.0298000341843973 | -0.908887434240655 | 0.363409547354629 | NA | NR_002900 |
| Snora70 | 18.562294 | -0.058383519946702 | 0.0786616717468339 | -0.742210515619405 | 0.457959798353242 | NA | NR_002899 |
| Snora74a | 3.933764526 | -0.0116745488314467 | 0.0391888843882436 | -0.297904597532993 | 0.765775978865164 | NA | NR_002905 |
| Snora75 | 7.830322142 | -0.0556811684960785 | 0.0573032936062469 | -0.971692288381979 | 0.331203654364653 | NA | NR_028478 |
| Snora78 | 4.772865748 | -0.0378199813398652 | 0.045249887388728 | -0.835802772611681 | 0.403265866454461 | NA | NR_028515 |
| Snora81 | 21.4831359 | -0.0673856950524219 | 0.0790027368238264 | -0.852953932503501 | 0.39368485149838 | NA | NR_034048 |
| Snord100 | 13.99743965 | -0.118758371566307 | 0.0704799672956072 | -1.68499470307939 | 0.0919895934649661 | NA | NR_037681 |
| Snord11 | 0 | NA | NA | NA | NA | NA | NR_028521 |
| Snord110 | 1.250600861 | -0.0430526831848107 | 0.0264067795874896 | -1.63036477212871 | 0.103024425430704 | NA | NR_028547 |
| Snord111 | 0.212695677 | -0.00857796107890413 | 0.00973699866180651 | -0.880965621629515 | 0.378336427605205 | NA | NR_028559 |
| Snord116 | 0 | NA | NA | NA | NA | NA | NR_002895 |
| Snord116l1 | 0 | NA | NA | NA | NA | NA | NR_033778 |
| Snord116l2 | 0 | NA | NA | NA | NA | NA | NR_033779 |
| Snord12 | 12.69521876 | -0.0051959471164429 | 0.0712998472167046 | -0.0728745897680627 | 0.941905914593888 | NA | NR_028540 |
| Snord123 | 0.205932006 | -0.000576954122277892 | 0.00970424421944593 | -0.0594537925088239 | 0.952590668569529 | NA | NR_028575 |
| Snord15a | 0.435106034 | 0.00765078179755927 | 0.0154951963355762 | 0.493751846176577 | 0.621481444770458 | NA | NR_002172 |
| Snord15b | 2.918914568 | -0.0371062679551138 | 0.0359059501413663 | -1.0334294959198 | 0.301402948977441 | NA | NR_002173 |
| Snord16a | 6.229619863 | -0.00754592527213209 | 0.0412954229748062 | -0.182730305892151 | 0.855009635439236 | NA | NR_028548 |
| Snord17 | 1.198369775 | -0.00235545658040324 | 0.0276633957969551 | -0.0851470512764201 | 0.932144484588442 | NA | NR_030762 |
| Snord19 | 14.63033988 | -0.149635685592514 | 0.066095228021913 | -2.26394083310983 | 0.0235777540719619 | NA | NR_028523 |
| Snord1a | 0 | NA | NA | NA | NA | NA | NR_028570 |
| Snord1b | 1.248136422 | -0.00855447387676706 | 0.0223955279087346 | -0.381972414833351 | 0.702481825828163 | NA | NR_028567 |
| Snord1c | 0.804933061 | -0.00813400335156511 | 0.018321366296201 | -0.443962705622654 | 0.657069547611335 | NA | NR_028569 |
| Snord22 | 15.68998674 | -0.110809559795714 | 0.0714209250764459 | -1.55149992354633 | 0.120781925651641 | NA | NR_004445 |
| Snord23 | 2.158748485 | -0.0436725584128486 | 0.0300312890494582 | -1.45423522583146 | 0.145881101877064 | NA | NR_028539 |
| Snord32a | 10.48661788 | -0.112497914081325 | 0.0620612697331249 | -1.81269114481684 | 0.0698794710342045 | NA | NR_000002 |
| Snord33 | 0.288615623 | 0.00197374456361263 | 0.0106648328298147 | 0.185070370544844 | 0.853173841012254 | NA | NR_001277 |
| Snord34 | 3.651696567 | -0.00822003348708651 | 0.0382663311843284 | -0.214811120707933 | 0.829914582220357 | NA | NR_002455 |
| Snord35a | 4.997169502 | 0.0134021466324863 | 0.0455520818702298 | 0.294215897105796 | 0.768592933834266 | NA | NR_000003 |
| Snord35b | 2.180108994 | -0.00663986173857889 | 0.0302444286793875 | -0.219539995579555 | 0.826229427720222 | NA | NR_000004 |
| Snord37 | 8.287161125 | -0.0761541948574068 | 0.0602070114222058 | -1.2648725299346 | 0.205917026042543 | NA | NR_028549 |
| Snord38a | 1.130209386 | -0.0222887562821523 | 0.0226499309582066 | -0.984054049580958 | 0.325088938302355 | NA | NR_028524 |
| Snord42a | 1.670153154 | -0.0383227857575299 | 0.0291089746627 | -1.3165281911024 | 0.187996822788432 | NA | NR_037682 |
| Snord42b | 21.8074519 | 0.00638717893058693 | 0.0793655394156048 | 0.0804779880237427 | 0.935857102453573 | NA | NR_037683 |
| Snord43 | 0.917895365 | -0.0360121107417158 | 0.0226902637034238 | -1.58711732981234 | 0.112486073637572 | NA | NR_028281 |
| Snord45b | 7.342379586 | -0.131414920314582 | 0.0548608174290043 | -2.39542402890091 | 0.016601154139376 | NA | NR_028561 |
| Snord45c | 0 | NA | NA | NA | NA | NA | NR_028525 |
| Snord47 | 2.540570966 | -0.00350485101171367 | 0.0308682332146394 | -0.113542326421568 | 0.909600608392563 | NA | NR_028543 |
| Snord49a | 8.36987764 | -0.0758257348819998 | 0.0571527657552959 | -1.32672030618175 | 0.18460122120623 | NA | NR_028550 |
| Snord49b | 2.042164714 | -0.0232989436071368 | 0.0338978588104434 | -0.687327885145323 | 0.4918761304236 | NA | NR_028526 |
| Snord4a | 4.97117751 | -0.0415783507351734 | 0.0505062436885027 | -0.823231895676264 | 0.41037613401115 | NA | NR_030702 |
| Snord52 | 14.614972 | -0.140878376453144 | 0.0755797467235941 | -1.86396994645082 | 0.0623259249041553 | NA | NR_028527 |
| Snord53 | 8.567590711 | -0.127130182319258 | 0.0629374973269213 | -2.0199433997017 | 0.0433892588587486 | NA | NR_028551 |
| Snord55 | 0.68199713 | -0.021730672623881 | 0.0184435835091904 | -1.17822399389212 | 0.23870731944437 | NA | NR_030704 |
| Snord57 | 5.688914333 | 0.0233232261219096 | 0.0530198594786002 | 0.439896038036905 | 0.660012405549794 | NA | NR_028528 |
| Snord58b | 5.082388009 | -0.048073682327742 | 0.0490071841574427 | -0.9809517350211 | 0.326616543275462 | NA | NR_028552 |
| Snord61 | 0 | NA | NA | NA | NA | NA | NR_002903 |
| Snord64 | 3.637874294 | -0.0189822894285476 | 0.0432177171708066 | -0.439224713177819 | 0.660498719854234 | NA | NR_028529 |
| Snord65 | 6.448611832 | -0.0096077150039081 | 0.0492275685961113 | -0.195169399543878 | 0.845260334010806 | NA | NR_028541 |
| Snord66 | 1.834900853 | -0.0167304196822951 | 0.0329865085664578 | -0.507189769677756 | 0.612021672453319 | NA | NR_028530 |
| Snord67 | 0 | NA | NA | NA | NA | NA | NR_028553 |
| Snord68 | 7.883669935 | -0.0409523957342026 | 0.059243997793929 | -0.691249700546022 | 0.489408635462957 | NA | NR_028128 |
| Snord69 | 15.83057544 | -0.133074880523273 | 0.0763231153783384 | -1.7435724401921 | 0.0812336724688184 | NA | NR_028531 |
| Snord7 | 0 | NA | NA | NA | NA | NA | NR_028362 |
| Snord70 | 0 | NA | NA | NA | NA | NA | NR_028554 |
| Snord71 | 2.224623015 | 0.00869344853035657 | 0.035810070492217 | 0.242765468229 | 0.808187087770813 | NA | NR_028532 |
| Snord72 | 2.134308492 | 0.0165429768539008 | 0.0292359097876532 | 0.565844434945109 | 0.571499536952406 | NA | NR_028091 |
| Snord73a | 2.36658174 | 0.027186087951129 | 0.0309534402622986 | 0.878289706111982 | 0.379786520670981 | NA | NR_004417 |
| Snord8 | 0.848365821 | 0.00377701463346565 | 0.0219052781786686 | 0.172424865032927 | 0.863103525446227 | NA | NR_028542 |
| Snord82 | 0 | NA | NA | NA | NA | NA | NR_002851 |
| Snord83b | 5.184917167 | -0.036001299401044 | 0.0528721676209216 | -0.680912113518081 | 0.495927102713527 | NA | NR_028282 |
| Snord85 | 0.517988882 | 0.00121152938701394 | 0.0146377493610679 | 0.0827674635717048 | 0.934036440640215 | NA | NR_028565 |
| Snord87 | 1.970019805 | -0.0173915611166994 | 0.0302394859172236 | -0.575127539016583 | 0.565205044255774 | NA | NR_004410 |
| Snord88a | 2.941289526 | 0.00577831687920662 | 0.0346683087529708 | 0.166674322661081 | 0.867626310437499 | NA | NR_028533 |
| Snord88c | 12.56359718 | -0.0777911468743662 | 0.0689016323919098 | -1.12901747279213 | 0.258890462899919 | NA | NR_028534 |
| Snord89 | 7.849671122 | -0.0542969474233245 | 0.0550857932675004 | -0.985679686224264 | 0.324290320833423 | NA | NR_028555 |
| Snord90 | 0 | NA | NA | NA | NA | NA | NR_028535 |
| Snord91a | 1.473685646 | -0.00279891843262591 | 0.0293598021275286 | -0.0953316517757305 | 0.924051402651799 | NA | NR_028562 |
| Snord92 | 0 | NA | NA | NA | NA | NA | NR_028556 |
| Snord93 | 0 | NA | NA | NA | NA | NA | NR_028536 |
| Snord95 | 7.156057332 | -0.0807247190207964 | 0.0564311694759098 | -1.43049877878674 | 0.152573916068631 | NA | NR_028564 |
| Snord99 | 14.46468841 | -0.0624878978115906 | 0.0685022065889147 | -0.912202700076273 | 0.361662022843218 | NA | NR_028537 |
| Sntn | 13.3729447 | 0.0205928276728257 | 0.0642489248529786 | 0.320516299999548 | 0.748576975979694 | NA | NM_177624 |
| Snurf | 0 | NA | NA | NA | NA | NA | NM_033174 |
| Snx31 | 8.724798734 | 0.0656333184581377 | 0.0625962957866771 | 1.04851761008048 | 0.294400193132694 | NA | NM_025712 |
| Soat2 | 7.281236281 | -0.0442067210291892 | 0.0584381241926467 | -0.75647056848467 | 0.449367111888965 | NA | NM_146064 |
| Sohlh1 | 2.479969315 | -0.0260492697626117 | 0.0358410488720266 | -0.726799872839178 | 0.467348560996944 | NA | NM_001001714 |
| Sohlh2 | 0.108766301 | 0.00519171381361946 | 0.00856003371502575 | 0.606506234257728 | 0.544178643801544 | NA | NM_028937 |
| Sost | 3.510119443 | -0.0306883909047394 | 0.0381666054066946 | -0.804063934366994 | 0.421360052108567 | NA | NM_024449 |
| Sostdc1 | 94.29098429 | 0.0126543399106236 | 0.0385353340020544 | 0.32838277488263 | NA | NA | NM_025312 |
| Sowahd | 0.618148956 | 0.00280000028365849 | 0.015137723011585 | 0.184968391977819 | 0.853253827300672 | NA | NM_173779 |
| Sox14 | 4.945473281 | 0.00553866247583784 | 0.0280795308506874 | 0.197249110225154 | 0.843632601047418 | NA | NM_011440 |
| Sox15 | 2.792802749 | 0.053512230689959 | 0.038829415226122 | 1.37813640453589 | 0.168161177619583 | NA | NM_009235 |
| Sox30 | 5.508660513 | -0.0218735055999959 | 0.051288325892179 | -0.426481177139209 | 0.669757260866825 | NA | NM_173384 |
| Sp5 | 5.734526696 | -0.0128267842520984 | 0.0457874227393418 | -0.280137720900313 | 0.779371845718117 | NA | NM_022435 |
| Spaca1 | 0.794049928 | -0.0172110328882211 | 0.0186039579351498 | -0.925127488904018 | 0.354899595735662 | NA | NM_026293 |
| Spaca3 | 0.09572496 | 0.00498255289994511 | 0.00847999972823692 | 0.587565219295241 | 0.556824162160132 | NA | NM_029367 |
| Spaca4 | 0 | NA | NA | NA | NA | NA | NM_027055 |
| Spaca5 | 0.322281847 | -0.00648722354265905 | 0.0126857541512189 | -0.511378627185182 | 0.609085957750446 | NA | NM_001085393 |
| Spaca7 | 0 | NA | NA | NA | NA | NA | NM_024279 |
| Spag11a | 0 | NA | NA | NA | NA | NA | NM_153115 |
| Spag11b | 0.224626239 | -0.00168521210467683 | 0.00978955093367521 | -0.172143964119932 | 0.863324350255464 | NA | NM_001034905,NM_001039563 |
| Spam1 | 0 | NA | NA | NA | NA | NA | NM_001079875,NM_009241 |
| Spata16 | 2.526488474 | 0.0142739472804577 | 0.0354597754100879 | 0.402539133860305 | 0.68728729382458 | NA | NM_029150,NM_027583 |
| Spata17 | 16.63550244 | 0.0173797601771008 | 0.0722919425422957 | 0.240410750713089 | 0.810011844706034 | NA | NM_029067,NM_028848 |
| Spata19 | 6.891230852 | -0.0266866346137352 | 0.0563932462892893 | -0.473223947364841 | 0.636053414460352 | NA | NM_029299 |
| Spata20 | 5.922937722 | -0.0156918183354248 | 0.0532114297458279 | -0.294895634460098 | 0.768073606929036 | NA | NM_144827 |
| Spata22 | 2.45023949 | 0.0489099443193318 | 0.0360917247478194 | 1.35515674745599 | 0.17536762184228 | NA | NM_001045531 |
| Spata25 | 2.732054541 | -0.010575297886664 | 0.034336272931314 | -0.307992014969673 | 0.758088408337915 | NA | NM_029370 |
| Spata3 | 13.25343049 | -0.0488608648782356 | 0.0705401264902628 | -0.692667667458468 | 0.488518134159585 | NA | NM_001122732,NM_027029,NM_028647,NM_027300 |
| Spata31 | 0 | NA | NA | NA | NA | NA | NM_030047 |
| Spata31d1a | 2.514100685 | 0.00410686698922663 | 0.0323531175009216 | 0.126938833301293 | 0.898988810705437 | NA | NM_028157 |
| Spata31d1b | 2.465217617 | 0.00629608301921066 | 0.025642611277497 | 0.245532054090601 | 0.806044492028428 | NA | NM_001167593 |
| Spata31d1c | 0.36293681 | -0.00593061268544428 | 0.0131125568226957 | -0.452284994119481 | 0.651063684812157 | NA | NM_001083890 |
| Spata31d1d | 1.486257525 | 0.00114328523887236 | 0.0269943517565053 | 0.0423527576874242 | 0.966217488433182 | NA | NM_177711 |
| Spata4 | 1.415146138 | -0.0119381841468809 | 0.0253468757946333 | -0.470992332294011 | 0.637646210533829 | NA | NM_133711 |
| Spata9 | 2.786785386 | -0.00467503985649199 | 0.0373177746771591 | -0.125276490812659 | 0.900304663157615 | NA | NM_029343 |
| Spatc1 | 0.880929594 | 0.0159450642115349 | 0.0223519863098274 | 0.713362293199172 | 0.475621599666033 | NA | NM_028852 |
| Spatc1l | 1.945154612 | -0.0118007000053265 | 0.0329861470023023 | -0.357747147749715 | 0.720532546201107 | NA | NM_029661 |
| Spc25 | 12.54616424 | -0.000281864602367953 | 0.0704086531723217 | -0.0040032664973452 | 0.996805864000604 | NA | NM_001199124,NM_025565,NM_001199123 |
| Spdef | 8.416982083 | 0.00326341590731912 | 0.057490557075562 | 0.0567643813753602 | 0.95473288771192 | NA | NM_013891 |
| Spdl1 | 20.0072745 | -0.12517456043572 | 0.0810996064770284 | -1.54346692756365 | 0.122717525888289 | NA | NM_027411 |
| Spdyb | 0.226556888 | -0.00933175792380659 | 0.010153338242423 | -0.919082739193727 | 0.358052296835844 | NA | NM_029048 |
| Speer1-ps1 | 0 | NA | NA | NA | NA | NA | NR_001586 |
| Speer2 | 0.67765611 | 0.0064431620234376 | 0.0191126188554661 | 0.337115602637307 | 0.736029755971199 | NA | NM_173069 |
| Speer3 | 0 | NA | NA | NA | NA | NA | NM_027650 |
| Speer4a | 2.761868621 | -0.00103109806696027 | 0.0369313359159739 | -0.0279193276221098 | 0.977726493238287 | NA | NM_029376 |
| Speer4b | 3.764123392 | 0.0263804633739005 | 0.0434193124863686 | 0.607574414776434 | 0.543469774143998 | NA | NM_028561 |
| Speer4d | 1.221072351 | 0.0493750222111172 | 0.025867672702111 | 1.90875394086333 | 0.0562938411007422 | NA | NM_025759 |
| Speer4e | 1.330708063 | 0.0112582275839268 | 0.0272987551548205 | 0.412408094071599 | 0.68004033151391 | NA | NM_001122661 |
| Speer4f | 0 | NA | NA | NA | NA | NA | NM_027609 |
| Speer5-ps1 | 0.106148608 | 0.00515554063321497 | 0.00854200771766808 | 0.603551390213729 | 0.546141939141592 | NA | NR_001582,NR_027506 |
| Speer6-ps1 | 0 | NA | NA | NA | NA | NA | NR_001581 |
| Speer7-ps1 | 5.297765712 | 0.0696304769266094 | 0.0510151299206146 | 1.3648985513702 | 0.172284937642466 | NA | NR_001585 |
| Speer8-ps1 | 0.346433883 | 0.00169660250222162 | 0.0112993204793637 | 0.150150843612249 | 0.880645607231878 | NA | NR_001584 |
| Speer9-ps1 | 0.108766301 | 0.00519171381361946 | 0.00856003371502575 | 0.606506234257728 | 0.544178643801544 | NA | NR_001583 |
| Spem1 | 5.799838121 | -0.0431444589848804 | 0.0548093904587827 | -0.787172756780164 | 0.431180742828467 | NA | NM_028855 |
| Spert | 0 | NA | NA | NA | NA | NA | NM_026457,NM_001164140,NM_001164141,NM_001164139 |
| Spesp1 | 1.5692159 | -0.0310576058441488 | 0.0292919747184634 | -1.0602769578581 | 0.289018618818341 | NA | NM_025721 |
| Spib | 0.826632487 | -0.0105657121388604 | 0.0189791274978208 | -0.556701678729621 | 0.577731269768818 | NA | NM_019866 |
| Spic | 0.583236734 | -0.00622433695838341 | 0.0149930010060907 | -0.415149505816404 | 0.678032460500875 | NA | NM_011461 |
| Spink11 | 0.097165704 | -0.00637828455464458 | 0.00848781691535724 | -0.75146349388194 | 0.452373763047372 | NA | NM_001048217 |
| Spink12 | 0.106148608 | 0.00515554063321497 | 0.00854200771766808 | 0.603551390213729 | 0.546141939141592 | NA | NM_030061 |
| Spink13 | 1.549054206 | 0.0332027343454432 | 0.0255613655068105 | 1.29894212171868 | 0.193963792754714 | NA | NM_001168423 |
| Spink14 | 0 | NA | NA | NA | NA | NA | NM_001039218 |
| Spink2 | 0.410423267 | 0.00622751620557347 | 0.0137781518810749 | 0.451984871361979 | 0.65127988120664 | NA | NM_183284 |
| Spink3 | 0 | NA | NA | NA | NA | NA | NM_009258 |
| Spink4 | 0.120209049 | -0.00674972139751468 | 0.00863521114606269 | -0.781650996523957 | 0.434419707908266 | NA | NM_011463 |
| Spink5 | 0.106148608 | 0.00515554063321497 | 0.00854200771766808 | 0.603551390213729 | 0.546141939141592 | NA | NM_001081180 |
| Spink6 | 0 | NA | NA | NA | NA | NA | NM_001013797 |
| Spink7 | 0 | NA | NA | NA | NA | NA | NM_001001803 |
| Spinkl | 0 | NA | NA | NA | NA | NA | NM_183123 |
| Spint3 | 0 | NA | NA | NA | NA | NA | NM_001177401 |
| Spint4 | 0.41620922 | -0.0157695646389328 | 0.0131796358585653 | -1.19650988905618 | 0.231497641828105 | NA | NM_030058 |
| Spint5 | 0 | NA | NA | NA | NA | NA | NM_001040055 |
| Spn | 7.032922148 | 0.0364178161148597 | 0.0577892836859524 | 0.63018286076649 | 0.52857495154929 | NA | NM_009259,NM_001037810 |
| Spn-ps | 0.656915902 | -0.00484699711628104 | 0.020240907535725 | -0.239465404786132 | 0.810744719146985 | NA | NR_033583 |
| Spns3 | 8.623028829 | 0.0106009722208489 | 0.0570868099230921 | 0.185699152486023 | 0.852680692979746 | NA | NM_029932 |
| Spo11 | 11.69815952 | -0.0984243544495669 | 0.0642928329175109 | -1.5308759932207 | 0.125800043467395 | NA | NM_001083959,NM_012046,NM_001083960 |
| Spon2 | 2.607883662 | 0.0398892851090181 | 0.0348560996887407 | 1.14439898511947 | 0.252458208920435 | NA | NM_133903 |
| Spp2 | 3.047594687 | 0.0142951794730619 | 0.0412023073432211 | 0.346950945100745 | 0.728628172349777 | NA | NM_029269 |
| Sppl2c | 0 | NA | NA | NA | NA | NA | NM_001082535,NM_199019 |
| Sprr1a | 2.480314828 | -0.0257816879953169 | 0.0387530604116877 | -0.665281340916788 | 0.505870562739618 | NA | NM_009264 |
| Sprr1b | 0 | NA | NA | NA | NA | NA | NM_009265 |
| Sprr2a1 | 0 | NA | NA | NA | NA | NA | NM_011468 |
| Sprr2a2 | 0 | NA | NA | NA | NA | NA | NM_001164787 |
| Sprr2b | 0 | NA | NA | NA | NA | NA | NM_011469 |
| Sprr2d | 0 | NA | NA | NA | NA | NA | NM_011470 |
| Sprr2e | 0 | NA | NA | NA | NA | NA | NM_011471 |
| Sprr2f | 0 | NA | NA | NA | NA | NA | NM_011472 |
| Sprr2g | 0.212297216 | 0.00681050119636477 | 0.00973439053551308 | 0.699633035218656 | 0.484156506270654 | NA | NR_003548 |
| Sprr2h | 0 | NA | NA | NA | NA | NA | NM_011474 |
| Sprr2i | 0 | NA | NA | NA | NA | NA | NM_011475 |
| Sprr2j-ps | 0 | NA | NA | NA | NA | NA | NR_003185 |
| Sprr2k | 0 | NA | NA | NA | NA | NA | NM_011477 |
| Sprr3 | 0 | NA | NA | NA | NA | NA | NM_011478,NM_001204427 |
| Sprr4 | 0 | NA | NA | NA | NA | NA | NM_173070 |
| Spta1 | 0.959331758 | 0.0215868888696522 | 0.019696336904451 | 1.09598495265249 | 0.273085359581418 | NA | NM_011465 |
| Sptlc3 | 6.629747119 | 0.0560163395550842 | 0.0411026984271822 | 1.36283849232729 | 0.172933416920777 | NA | NM_175467 |
| Spz1 | 0 | NA | NA | NA | NA | NA | NM_030237 |
| Srd5a2 | 0.742151045 | -0.0183357104060177 | 0.0203891045339888 | -0.899289636553285 | 0.368498406001784 | NA | NM_053188 |
| Srms | 14.63049548 | -0.0120120388839412 | 0.0724483843570497 | -0.165801335537613 | 0.868313295064208 | NA | NM_011481 |
| Srp54b | 8.459657433 | 0.00600448635039461 | 0.0617873511257708 | 0.0971798635318129 | 0.922583558920901 | NA | NM_001100109 |
| Sry | 0 | NA | NA | NA | NA | NA | NM_011564 |
| Ssu2 | 4.906319756 | -0.0065433639420745 | 0.0483877929148655 | -0.135227575962951 | 0.892431944875208 | NA | NM_175525 |
| Ssx9 | 0 | NA | NA | NA | NA | NA | NM_199063 |
| Ssxb1 | 0 | NA | NA | NA | NA | NA | NM_026492 |
| Ssxb10 | 0 | NA | NA | NA | NA | NA | NM_199064 |
| Ssxb2 | 0.127687691 | 0.00542298404407807 | 0.00869724871942359 | 0.62352868349814 | 0.532937159177571 | NA | NM_001134226,NM_001001450 |
| Ssxb3 | 0 | NA | NA | NA | NA | NA | NM_198898 |
| Ssxb5 | 0 | NA | NA | NA | NA | NA | NM_199319 |

| | | | | | | | | | | | | | | | |
|---|---|---|---|---|---|---|---|---|---|---|---|---|---|---|---|
| Snord82 | 0 | NA | NA | NA | NA | NA | | Ssxb6 | 0.106347839 | -0.00653347234140708 | 0.00854389509723638 | -0.764694822098226 | 0.444453290197145 | NA | NM_001205108 |
| Snord83b | 4.547536993 | 0.227758917 | 0.140969012 | 1.615666541 | 0.106166429 | NA | | Ssxb8 | 0 | NA | NA | NA | NA | NA | NM_001081565 |
| Snord85 | 0.29403132 | 0.006032586 | 0.033500653 | 0.180073692 | 0.857094716 | NA | | Ssxb9 | 0.12890128 | -0.0068640067066208 | 0.00870804083610955 | -0.788237771940376 | 0.430557639008166 | NA | NM_199066 |
| Snord87 | 1.214169242 | 0.070065494 | 0.07329384 | 0.955953374 | 0.339095791 | NA | | St6galnac1 | 0.458888926 | -0.000592276211756187 | 0.0139816623856329 | -0.0423609292958461 | 0.966210974279136 | NA | NM_011371 |
| Snord88a | 1.991792652 | 0.069812155 | 0.093519149 | 0.746501171 | 0.455364725 | NA | | Stap1 | 1.355862946 | -0.00852181165795662 | 0.0267764046479711 | -0.318258249006643 | 0.750289051501087 | NA | NM_019992 |
| Snord89 | 5.937936274 | 0.329545783 | 0.147550232 | 2.233448083 | 0.02551941 | NA | | Steap1 | 21.20695591 | 0.00396143372905228 | 0.0257936840961807 | 0.153581540127448 | NA | NA | NM_027399 |
| Snord90 | 0 | NA | NA | NA | NA | NA | | Steap4 | 20.18423418 | 0.0203806775391837 | 0.0657944234102539 | 0.309762993317871 | 0.756741195349054 | NA | NM_054098 |
| Snord91a | 0.855062497 | 0.11313603 | 0.075230074 | 1.503867056 | 0.132615601 | NA | | Stfa1 | 0 | NA | NA | NA | NA | NA | NM_001082543 |
| Snord92 | 0 | NA | NA | NA | NA | NA | | Stfa2 | 0 | NA | NA | NA | NA | NA | NM_001082545 |
| Snord93 | 0 | NA | NA | NA | NA | NA | | Stfa2l1 | 0 | NA | NA | NA | NA | NA | NM_173869 |
| Snord95 | 5.746609284 | 0.198417805 | 0.141353351 | 1.403700749 | 0.160407979 | NA | | Stfa3 | 0 | NA | NA | NA | NA | NA | NM_025288 |
| Sntn | 7.934930054 | 0.172702571 | 0.142813564 | 1.209286894 | 0.226552644 | NA | | Stk31 | 2.276189835 | -0.0229101860090932 | 0.0336118801928168 | -0.681609772427706 | 0.495485733953998 | NA | NM_029916 |
| Snurf | 0 | NA | NA | NA | NA | NA | | Stmn1-rs1 | 0.097165704 | -0.00637828455464458 | 0.00848781691535724 | -0.75146349388194 | 0.452373763047372 | NA | NR_029430 |
| Soat2 | 7.165783843 | 0.234555454 | 0.151916526 | 1.54397589 | 0.122594173 | NA | | Stmnd1 | 10.55987753 | 0.00691385859495877 | 0.0677550174958297 | 0.102042016229784 | 0.918723324994839 | NA | NM_001005422 |
| Sohlh1 | 2.407972676 | 0.13330053 | 0.112137835 | 1.188720381 | 0.234549718 | NA | | Stoml3 | 18.31726264 | -0.0183731071791311 | 0.0697614346956712 | -0.263370546481482 | 0.79226498995024 | NA | NM_153156 |
| Sohlh2 | 0.570960929 | -0.072956975 | 0.051753376 | -1.409704638 | 0.158626915 | NA | | Stpg2 | 3.085868376 | 0.0215752202942747 | 0.0345053406168901 | 0.625271911783819 | 0.531792611594387 | NA | NM_198659 |
| Sorbs1 | 6155.148757 | 0.252065664 | 0.102485928 | 2.459514869 | NA | NA | | Stra8 | 0 | NA | NA | NA | NA | NA | NM_009292 |
| Sost | 4.074268789 | -0.034610803 | 0.110089933 | -0.314386632 | 0.753227408 | NA | | Strc | 17.72364808 | -0.0984942815181961 | 0.0751190257596323 | -1.3111762369411 | 0.189798228753888 | NA | NM_080459 |
| Sostdc1 | 49.54682155 | 0.385153684 | 0.157084879 | 2.451882618 | NA | NA | | Stx11 | 8.714227986 | -0.0407818922065554 | 0.0640672344011124 | -0.636548347806429 | 0.524419079068941 | NA | NM_001163591,NM_001163590,NM_029075 |
| Sowahd | 0.528070844 | -0.018641267 | 0.051372038 | -0.362867963 | 0.71670352 | NA | | Stx19 | 0 | NA | NA | NA | NA | NA | NM_026588 |
| Sox14 | 1.838380592 | 0.081284516 | 0.062738168 | 1.295615074 | NA | NA | | Styk1 | 13.01413797 | 0.0492117912232519 | 0.0611905322661503 | 0.804238652626904 | 0.421259159796719 | NA | NM_172891 |
| Sox15 | 2.038582862 | -0.089736471 | 0.105899275 | -0.847375685 | 0.396785753 | NA | | Styxl1 | 6.073328644 | 0.0193628436309074 | 0.0516031137941433 | 0.375226264603919 | 0.707492198503741 | NA | NM_029659 |
| Sox30 | 4.138357827 | 0.087630711 | 0.127423461 | 0.687712532 | 0.491633826 | NA | | Sucnr1 | 0.12890128 | -0.0068640067066208 | 0.00870804083610955 | -0.788237771940376 | 0.430557639008166 | NA | NM_032400 |
| Spaca1 | 3.615942245 | -0.296482065 | 0.114712299 | -2.584570858 | 0.009750027 | NA | | Sult1b1 | 0.09572496 | 0.00498255289994511 | 0.00847999972823692 | 0.587565219295241 | 0.556824162160132 | NA | NM_019878 |
| Spaca3 | 0.107716603 | -0.020477583 | 0.027954224 | -0.732539849 | 0.46383913 | NA | | Sult1c1 | 8.933841044 | 0.037704026121874 | 0.035922000740159 | 1.04960818843598 | 0.293898290769618 | NA | NM_018751 |
| Spaca4 | 0 | NA | NA | NA | NA | NA | | Sult1c2 | 8.112205283 | 0.0489651008619896 | 0.0392912041598657 | 1.24621023735397 | 0.212687221630782 | NA | NM_026935 |
| Spaca5 | 0.303023688 | 0.009739077 | 0.039388711 | 0.247255542 | 0.804710463 | NA | | Sult1d1 | 0.205932006 | -0.000576954122277892 | 0.00970424421944593 | -0.0594537925088239 | 0.952590668569529 | NA | NM_016771 |
| Spaca7 | 0 | NA | NA | NA | NA | NA | | Sult1e1 | 0 | NA | NA | NA | NA | NA | NM_023135 |
| Spag11a | 0 | NA | NA | NA | NA | NA | | Sult2a1 | 0 | NA | NA | NA | NA | NA | NM_001111296 |
| Spag11b | 0.105923158 | 0.018524693 | 0.027977518 | 0.662127842 | 0.507889294 | NA | | Sult2a2 | 0 | NA | NA | NA | NA | NA | NM_009286 |
| Spam1 | 0 | NA | NA | NA | NA | NA | | Sult2a3 | 0 | NA | NA | NA | NA | NA | NM_001101586 |
| Spata16 | 3.264970887 | -0.101572799 | 0.120909337 | -0.84007407 | 0.400866858 | NA | | Sult2a4 | 0 | NA | NA | NA | NA | NA | NM_001101534 |
| Spata19 | 5.673169361 | 0.135287959 | 0.147839571 | 0.915099783 | 0.360139249 | NA | | Sult2a5 | 1.16921559 | -0.00558792844507372 | 0.0215639150232303 | -0.259133299266574 | 0.79553239296192 | NA | NM_001184980 |
| Spata20 | 4.280226671 | 0.07261472 | 0.129458911 | 0.560909398 | 0.574859303 | NA | | Sult2a6 | 0 | NA | NA | NA | NA | NA | NM_001081325 |
| Spata22 | 0.745115121 | 0.01765305 | 0.062917093 | 0.280576371 | 0.779035342 | NA | | Sult2a7 | 0 | NA | NA | NA | NA | NA | NM_001184981 |
| Spata25 | 2.817128849 | -0.047127984 | 0.110879361 | -0.425038376 | 0.6708087 | NA | | Sult3a1 | 0.12890128 | -0.0068640067066208 | 0.00870804083610955 | -0.788237771940376 | 0.430557639008166 | NA | NM_020565 |
| Spata31 | 0 | NA | NA | NA | NA | NA | | Sult5a1 | 3.848142728 | -0.0600700082263538 | 0.0462431898456923 | -1.29900226231797 | 0.193943152786264 | NA | NM_020564 |
| Spata31d1a | 1.725001558 | 0.047203527 | 0.086973592 | 0.54273402 | 0.587312948 | NA | | Sult6b1 | 1.667044668 | 0.00658010969880745 | 0.0305800198530571 | 0.21517676346929 | 0.829629506668506 | NA | NM_001163625 |
| Spata31d1b | 2.590111147 | 0.079996456 | 0.072614066 | 1.101666118 | 0.270606852 | NA | | Sun3 | 2.085080895 | -0.00872484584628018 | 0.0338189737981157 | -0.257986711789768 | 0.796417162636479 | NA | NM_177576 |
| Spata31d1c | 0.193239998 | 0.025145508 | 0.031532422 | 0.797449311 | 0.42519013 | NA | | Sun5 | 0 | NA | NA | NA | NA | NA | NM_029599 |
| Spata31d1d | 2.074998987 | 0.041834143 | 0.097289076 | 0.42999836 | 0.667196834 | NA | | Sva | 0 | NA | NA | NA | NA | NA | NM_009299 |
| Spata4 | 1.825730401 | -0.045155613 | 0.094649565 | -0.477082103 | 0.633303659 | NA | | Sval1 | 0 | NA | NA | NA | NA | NA | NM_027832 |
| Spata9 | 3.203432809 | 0.007843334 | 0.124095971 | 0.063203774 | 0.949604239 | NA | | Sval2 | 0.108766301 | 0.00519171381361946 | 0.00856003371502575 | 0.606506234257728 | 0.544178643801544 | NA | NM_032542 |
| Spatc1 | 0.913368603 | -0.058414623 | 0.059789942 | -0.976997495 | 0.328570393 | NA | | Sval3 | 0 | NA | NA | NA | NA | NA | NM_001003952 |
| Spatc1l | 1.20437504 | 0.045352351 | 0.06979633 | 0.64978132 | 0.515833487 | NA | | Svopl | 2.202772154 | 0.0108951733254635 | 0.0270471443406576 | 0.402821576586396 | 0.687079487045578 | NA | NM_177200 |
| Spdef | 7.113733933 | 0.199837831 | 0.136978785 | 1.458896217 | 0.144593675 | NA | | Svs1 | 0 | NA | NA | NA | NA | NA | NM_172888 |
| Spdyb | 0.186326724 | 0.024939385 | 0.031411053 | 0.793968451 | 0.427213798 | NA | | Svs2 | 0 | NA | NA | NA | NA | NA | NM_017390 |
| Speer1-ps1 | 0 | NA | NA | NA | NA | NA | | Svs3a | 0 | NA | NA | NA | NA | NA | NM_021363 |
| Speer2 | 0.714780812 | -0.040502242 | 0.056225019 | -0.720359779 | 0.471303507 | NA | | Svs3b | 0 | NA | NA | NA | NA | NA | NM_173377 |
| Speer3 | 0 | NA | NA | NA | NA | NA | | Svs4 | 0 | NA | NA | NA | NA | NA | NM_009300 |
| Speer4a | 1.675574735 | 0.048176622 | 0.088345027 | 0.545323528 | 0.585531025 | NA | | Svs5 | 0 | NA | NA | NA | NA | NA | NM_009301 |
| Speer4b | 2.754380874 | -0.022620124 | 0.111163425 | -0.203485315 | 0.838755723 | NA | | Svs6 | 0 | NA | NA | NA | NA | NA | NM_013679 |
| Speer4d | 0.38488829 | -0.02558324 | 0.043075085 | -0.593921989 | 0.552564291 | NA | | Syce1 | 7.089492757 | 0.0062365939912011 | 0.0529452296494247 | 0.117793312683626 | 0.906231428363599 | NA | NM_001143765 |
| Speer4e | 0.688668513 | 0.012091754 | 0.060689169 | 0.199240727 | 0.842074443 | NA | | Syce1l | 6.117300296 | 0.0128303608848872 | 0.0520275788197292 | 0.246606918406548 | 0.80521244977827 | NA | NM_001048145,NM_029139 |
| Speer4f | 0 | NA | NA | NA | NA | NA | | Syce3 | 0.328957426 | 0.00300711674092362 | 0.0126513499961959 | 0.237691372211489 | 0.812120475900523 | NA | NM_001162880,NM_001162882 |
| Speer5-ps1 | 0 | NA | NA | NA | NA | NA | | Sycn | 0 | NA | NA | NA | NA | NA | NM_026716 |
| Speer6-ps1 | 0.507592018 | 0.0062035 | 0.044582043 | 0.139147943 | 0.889333244 | NA | | Sycp1 | 1.193554861 | 0.0101740382579327 | 0.0199650742904684 | 0.509591805665852 | 0.610337465876378 | NA | NM_011516 |
| Speer7-ps1 | 4.087694419 | -0.202015384 | 0.130756363 | -1.544975555 | 0.122352176 | NA | | Sycp1-ps1 | 0 | NA | NA | NA | NA | NA | NR_024208 |
| Speer8-ps1 | 0.302629931 | -0.009711995 | 0.033929987 | -0.286236321 | 0.774697127 | NA | | Sycp3 | 10.03624008 | 0.0958534555583181 | 0.0589496772243078 | 1.62602172007811 | 0.103945034919942 | NA | NM_011517 |
| Speer9-ps1 | 0 | NA | NA | NA | NA | NA | | Syna | 18.41737594 | -0.0909047184518164 | 0.0673766845349155 | -1.34920142003586 | 0.177272278800474 | NA | NM_001013751 |
| Spem1 | 4.868364845 | 0.031948161 | 0.129093631 | 0.247480533 | 0.804536356 | NA | | Synb | 4.064533027 | 0.00665430231137565 | 0.0414569545856858 | 0.160511122388938 | 0.872478460821883 | NA | NM_173420 |
| Spert | 0 | NA | NA | NA | NA | NA | | Syngr4 | 1.849443373 | -0.0378663311329901 | 0.0304528491886505 | -1.24344132460034 | 0.213705250163417 | NA | NM_021482 |
| Spesp1 | 1.511865256 | 0.02825672 | 0.085173065 | 0.331756519 | 0.740073119 | NA | | Synpo2l | 9.88536195 | 0.0750875517038907 | 0.0635770256540234 | 1.18104851448235 | 0.237583451138469 | NA | NM_175132 |
| Spib | 1.633440668 | -0.056844572 | 0.090236294 | -0.629952426 | 0.528725711 | NA | | Syt8 | 1.193591353 | 0.00863312276594158 | 0.0223363148262202 | 0.386506137342196 | 0.699121858148968 | NA | NM_018802 |
| Spic | 1.367443798 | -0.0486333 | 0.078163966 | -0.622195902 | 0.533813061 | NA | | Sytl3 | 7.65893679 | -0.0391434561267998 | 0.0516156079730562 | -0.75836472074945 | 0.448232671790161 | NA | NM_183368,NM_031395,NM_183370 |
| Spink11 | 0.079983654 | 0.016740032 | 0.027674178 | 0.604897175 | 0.54524732 | NA | | Sytl4 | 63.80699916 | 0.0725919998043172 | 0.0442327757595782 | 1.64113598022611 | NA | NA | NM_013757 |
| Spink12 | 0.192746203 | -0.000975302 | 0.031567365 | -0.030895906 | 0.975352555 | NA | | T | 4.115627506 | -0.00380363170979711 | 0.0414510864291218 | -0.0917619304454427 | 0.926887191506175 | NA | NM_009309 |
| Spink13 | 1.634970793 | -0.19230465 | 0.091671591 | -2.09775622 | 0.035926686 | NA | | Taar1 | 0 | NA | NA | NA | NA | NA | NM_053205 |
| Spink14 | 0 | NA | NA | NA | NA | NA | | Taar2 | 0 | NA | NA | NA | NA | NA | NM_001007266 |
| Spink2 | 0.17833704 | -0.002390466 | 0.031282281 | -0.076415987 | 0.939088151 | NA | | Taar3 | 0.55367566 | 0.000661977171951082 | 0.014865518203506 | 0.044531052526306 | 0.964481100183787 | NA | NM_001008429 |
| Spink3 | 0 | NA | NA | NA | NA | NA | | Taar4 | 0 | NA | NA | NA | NA | NA | NM_001008499 |
| Spink4 | 0.289473745 | 0.012204471 | 0.038954522 | 0.313300481 | 0.754052385 | NA | | Taar5 | 0 | NA | NA | NA | NA | NA | NM_001009574 |
| Spink5 | 0.472189209 | -0.048684432 | 0.042637861 | -1.141812248 | 0.25353207 | NA | | Taar6 | 0 | NA | NA | NA | NA | NA | NM_001010828 |
| Spink6 | 0 | NA | NA | NA | NA | NA | | Taar7a | 0 | NA | NA | NA | NA | NA | NM_001010829 |
| Spink7 | 0 | NA | NA | NA | NA | NA | | Taar7b | 0 | NA | NA | NA | NA | NA | NM_001010827 |
| Spinkl | 0 | NA | NA | NA | NA | NA | | Taar7d | 0 | NA | NA | NA | NA | NA | NM_001010838 |
| Spint3 | 0 | NA | NA | NA | NA | NA | | Taar7e | 0 | NA | NA | NA | NA | NA | NM_001010835 |
| Spint4 | 0.458631138 | 0.026681677 | 0.045037875 | 0.592427534 | 0.553564333 | NA | | Taar7f | 0 | NA | NA | NA | NA | NA | NM_001010839 |
| Spint5 | 0 | NA | NA | NA | NA | NA | | Taar8a | 0 | NA | NA | NA | NA | NA | NM_001010830 |
| Spire1 | 8371.158319 | 0.165101596 | 0.112973923 | 1.461413315 | NA | NA | | Taar8b | 0 | NA | NA | NA | NA | NA | NM_001010837 |
| Spn | 6.57032193 | -0.126585441 | 0.153581259 | -0.824224529 | 0.409811992 | NA | | Taar8c | 0 | NA | NA | NA | NA | NA | NM_001010840 |
| Spn-ps | 0.518310624 | 0.007452973 | 0.050979418 | 0.146195733 | 0.883766874 | NA | | Taar9 | 0 | NA | NA | NA | NA | NA | NM_001010831 |
| Spns3 | 7.270725561 | 0.056149556 | 0.155121155 | 0.361972269 | 0.717372753 | NA | | Tac4 | 0.192890664 | -0.000903864895041124 | 0.00963528282151475 | -0.093807821917056 | 0.92526181809679 | NA | NM_053093 |
| Spon2 | 3.084358915 | -0.086554282 | 0.112880987 | -0.766774676 | 0.443215496 | NA | | Tacr2 | 21.16480622 | 0.0173855544007552 | 0.0805914924647977 | 0.215724437766792 | 0.829202551115105 | NA | NM_009314 |
| Spp2 | 3.712950175 | -0.240209296 | 0.127036008 | -1.890875661 | 0.058640944 | NA | | Tacstd2 | 7.87253979 | 0.0566090167130939 | 0.0585290194202469 | 0.967195713747275 | 0.333446204509173 | NA | NM_020047 |
| Sppl2c | 0 | NA | NA | NA | NA | NA | | Taf7l | 2.813960406 | 0.0428450107590602 | 0.0333590364178607 | 1.2843599623915 | 0.199016035129311 | NA | NM_028958 |
| Sprr1a | 4.524972019 | -0.15121316 | 0.133439168 | -1.133199209 | 0.257130606 | NA | | Tal2 | 4.767440794 | -0.0556299505958576 | 0.0470129277543293 | -1.18329049589418 | 0.236694038156917 | NA | NM_009317 |
| Sprr1b | 0 | NA | NA | NA | NA | NA | | Tap1 | 208.1511572 | -0.146568135452932 | 0.0687891364609928 | -2.13068724210609 | NA | NA | NM_013683,NM_001161730 |
| Sprr2a1 | 0 | NA | NA | NA | NA | NA | | Tarm1 | 0.774374434 | -0.000259785533563383 | 0.0199221389247857 | -0.0130400422637439 | 0.989595846464774 | NA | NM_177363 |
| Sprr2a2 | 0 | NA | NA | NA | NA | NA | | Tarsl2 | 1988.905996 | 0.0779881292475198 | 0.0698017853476592 | 1.11727986410501 | NA | NA | NM_172310 |
| Sprr2b | 0.096071043 | -0.020098795 | 0.027825948 | -0.722304036 | 0.470107572 | NA | | Tas1r1 | 6.482190712 | 0.0449191664558125 | 0.0517131124802091 | 0.868622372575298 | 0.385053713356256 | NA | NM_031867 |
| Sprr2d | 0 | NA | NA | NA | NA | NA | | Tas1r2 | 0.489247677 | 0.00603938983428819 | 0.0127232473563935 | 0.474673616343187 | 0.635019626078808 | NA | NM_031873 |
| Sprr2e | 0 | NA | NA | NA | NA | NA | | Tas1r3 | 14.58962066 | 0.0289794145357411 | 0.0710658514798477 | 0.407782555647825 | 0.683433321716303 | NA | NM_031872 |
| Sprr2f | 0 | NA | NA | NA | NA | NA | | Tas2r102 | 0 | NA | NA | NA | NA | NA | NM_199153 |
| Sprr2g | 0 | NA | NA | NA | NA | NA | | Tas2r103 | 0.097165704 | -0.00637828455464458 | 0.00848781691535724 | -0.75146349388194 | 0.452373763047372 | NA | NM_053211 |
| Sprr2h | 0 | NA | NA | NA | NA | NA | | Tas2r104 | 0 | NA | NA | NA | NA | NA | NM_207011 |
| Sprr2i | 0 | NA | NA | NA | NA | NA | | Tas2r105 | 0 | NA | NA | NA | NA | NA | NM_020501 |
| Sprr2j-ps | 0 | NA | NA | NA | NA | NA | | Tas2r106 | 0 | NA | NA | NA | NA | NA | NM_207016 |
| Sprr2k | 0 | NA | NA | NA | NA | NA | | Tas2r107 | 0 | NA | NA | NA | NA | NA | NM_199154 |
| Sprr3 | 0 | NA | NA | NA | NA | NA | | Tas2r108 | 0.706969034 | -0.0146327631434963 | 0.0195125242231779 | -0.749916462684765 | 0.453305018755021 | NA | NM_020502 |
| Sprr4 | 0.323149808 | -0.030167563 | 0.033779321 | -0.893077861 | 0.371815478 | NA | | Tas2r109 | 0 | NA | NA | NA | NA | NA | NM_207017 |
| Spta1 | 0.583851251 | -0.037236951 | 0.058974167 | -0.631411228 | 0.527771676 | NA | | Tas2r110 | 0 | NA | NA | NA | NA | NA | NM_199155 |
| Sptlc3 | 2.853528592 | 0.009473679 | 0.113370541 | 0.083563854 | 0.933403207 | NA | | Tas2r113 | 0 | NA | NA | NA | NA | NA | NM_207018 |
| Spz1 | 0 | NA | NA | NA | NA | NA | | Tas2r114 | 0 | NA | NA | NA | NA | NA | NM_207019 |
| Srd5a2 | 1.432144443 | -0.05491518 | 0.086347828 | -0.63597639 | 0.524791811 | NA | | Tas2r115 | 0 | NA | NA | NA | NA | NA | NM_207020 |
| Sry | 0 | NA | NA | NA | NA | NA | | Tas2r116 | 0 | NA | NA | NA | NA | NA | NM_053212 |
| Ssu2 | 4.484801144 | 0.02334115 | 0.135817984 | 0.171856106 | 0.863550656 | NA | | Tas2r117 | 0 | NA | NA | NA | NA | NA | NM_207021 |
| Ssx9 | 0 | NA | NA | NA | NA | NA | | Tas2r118 | 0 | NA | NA | NA | NA | NA | NM_207022 |
| Ssxb1 | 0 | NA | NA | NA | NA | NA | | Tas2r119 | 0 | NA | NA | NA | NA | NA | NM_020503 |
| Ssxb10 | 0 | NA | NA | NA | NA | NA | | Tas2r120 | 0 | NA | NA | NA | NA | NA | NM_207023 |
| Ssxb2 | 0 | NA | NA | NA | NA | NA | | Tas2r121 | 0 | NA | NA | NA | NA | NA | NM_207024 |
| Ssxb3 | 0 | NA | NA | NA | NA | NA | | Tas2r122 | 0 | NA | NA | NA | NA | NA | NM_001039128 |
| Ssxb5 | 0 | NA | NA | NA | NA | NA | | Tas2r123 | 0 | NA | NA | NA | NA | NA | NM_207025 |
| Ssxb6 | 0.08731684 | 0.017365056 | 0.027733412 | 0.626142072 | 0.531221759 | NA | | Tas2r124 | 0 | NA | NA | NA | NA | NA | NM_207026 |
| Ssxb8 | 0.192142086 | -0.026384592 | 0.031527717 | -0.836869715 | 0.402665805 | NA | | Tas2r125 | 0 | NA | NA | NA | NA | NA | NM_207027 |
| Ssxb9 | 0.225320798 | -0.001646023 | 0.032098875 | -0.051279768 | 0.95910259 | NA | | Tas2r126 | 0 | NA | NA | NA | NA | NA | NM_207028 |
| St6galnac1 | 0.167300494 | 0.024299763 | 0.031030539 | 0.783091891 | 0.433573153 | NA | | Tas2r129 | 0 | NA | NA | NA | NA | NA | NM_207029 |
| Stap1 | 1.094190362 | 0.00690199 | 0.071121416 | 0.097045169 | 0.922690524 | NA | | Tas2r130 | 0 | NA | NA | NA | NA | NA | NM_199156 |
| Steap1 | 10.8170662 | 0.206857191 | 0.121166028 | 1.707221036 | NA | NA | | Tas2r131 | 0 | NA | NA | NA | NA | NA | NM_207030 |
| Stfa1 | 0 | NA | NA | NA | NA | NA | | Tas2r134 | 0 | NA | NA | NA | NA | NA | NM_199158 |
| Stfa2 | 0 | NA | NA | NA | NA | NA | | Tas2r135 | 0.194331409 | -0.00840968081320881 | 0.00964081822647896 | -0.872299489073576 | 0.383045016303155 | NA | NM_199159 |
| Stfa2l1 | 0.188785634 | 0.025025111 | 0.031461603 | 0.79541754 | 0.42637066 | NA | | Tas2r136 | 0 | NA | NA | NA | NA | NA | NM_181276 |
| Stfa3 | 0 | NA | NA | NA | NA | NA | | Tas2r137 | 0.44803889 | 0.00538868503566239 | 0.0140821177917709 | 0.382661551007004 | 0.701970726278947 | NA | NM_001025385 |
| Stk31 | 1.559446096 | 0.112133069 | 0.092909366 | 1.206908134 | 0.227467525 | NA | | Tas2r138 | 0 | NA | NA | NA | NA | NA | NM_001001451 |
| Stmn1-rs1 | 0.079983654 | 0.016740032 | 0.027674178 | 0.604897175 | 0.54524732 | NA | | Tas2r139 | 0 | NA | NA | NA | NA | NA | NM_181275 |
| Stmnd1 | 6.707037581 | 0.368062238 | 0.150432828 | 2.446688291 | 0.014417547 | NA | | Tas2r140 | 0 | NA | NA | NA | NA | NA | NM_021562 |
| Ston1 | 177.4926521 | 0.163835912 | 0.157622854 | 1.039417238 | NA | NA | | Tas2r143 | 0.224853396 | -0.000161157395207779 | 0.00979162641959728 | -0.0164586952465051 | 0.986868454038988 | NA | NM_001001452 |
| Stpg2 | 2.114023909 | 0.045308749 | 0.097303285 | 0.4656446 | 0.641469913 | NA | | Tas2r144 | 0 | NA | NA | NA | NA | NA | NM_001001453 |
| Stra8 | 0.116696964 | -0.020477583 | 0.027954224 | -0.732539847 | 0.463839131 | NA | | Tat | 2.810170396 | 0.00369368911530158 | 0.0346264138939888 | 0.106672586038221 | 0.915048731658314 | NA | NM_146214 |
| Stx19 | 0.192142086 | -0.026384592 | 0.031527717 | -0.836869715 | 0.402665805 | NA | | Tbc1d21 | 0 | NA | NA | NA | NA | NA | NM_028854 |
| Styxl1 | 5.612173477 | 0.051955929 | 0.138398518 | 0.375408132 | 0.707356958 | NA | | Tbpl2 | 0.212297216 | 0.00681050119636477 | 0.00973439053551308 | 0.699633035218656 | 0.484156506270654 | NA | NM_199059 |
| Sucnr1 | 0.311993147 | -0.014421208 | 0.039860783 | -0.361789373 | 0.717509434 | NA | | Tbrg3 | 1.057718031 | 0.0223223215988425 | 0.0240295662415205 | 0.9289523320763 | 0.352913790425741 | NA | NR_027799 |
| Sult1b1 | 0 | NA | NA | NA | NA | NA | | Tbx10 | 0.57283292 | -0.0311343653719056 | 0.0184476705313912 | -1.68771256614358 | 0.0914664184018825 | NA | NM_001001320,NM_011533 |
| Sult1c1 | 4.124745454 | 0.239600029 | 0.107349233 | 2.23196778 | 0.025617093 | NA | | Tbx19 | 1.58698693 | 0.0102041178991794 | 0.0247890434516056 | 0.411638227150652 | 0.680604608225396 | NA | NM_032005 |
| Sult1c2 | 3.003228312 | 0.177472826 | 0.086205761 | 2.058711889 | 0.039521847 | NA | | Tbx20 | 1.078307322 | 0.0112098010525195 | 0.0204261874028589 | 0.548795564802785 | 0.583145754359592 | NA | NM_020496,NM_194263,NM_001205085 |
| Sult1d1 | 0.898933214 | -0.093259539 | 0.064431186 | -1.447428561 | 0.147776931 | NA | | Tbx21 | 6.367014925 | 0.00837138300021578 | 0.051116596396318 | 0.163770352300271 | 0.86991193195381 | NA | NM_019507 |
| Sult1e1 | 0 | NA | NA | NA | NA | NA | | Tbx22 | 3.023359733 | -0.0684240966246243 | 0.0381297172069591 | -1.794508368715 | 0.0727320640739599 | NA | NM_145224,NM_181319 |
| Sult2a1 | 0 | NA | NA | NA | NA | NA | | Tbx4 | 10.09789491 | -0.0679026737693276 | 0.067318953471835 | -1.00867096512035 | 0.313132640933283 | NA | NM_172798,NM_011536 |
| Sult2a2 | 0 | NA | NA | NA | NA | NA | | Tbx5 | 0.338696681 | 0.00369895467940622 | 0.0127616983847258 | 0.289848150919584 | 0.77193240913126 | NA | NM_011537 |
| Sult2a3 | 0 | NA | NA | NA | NA | NA | | Tc2n | 8.119880446 | 0.0306454476198553 | 0.0364942741590543 | 0.839733035552158 | 0.401058086965 | NA | NM_001082976,NM_028924 |
| Sult2a4 | 0 | NA | NA | NA | NA | NA | | Tcam1 | 8.84232474 | -0.0183005913100507 | 0.0591997907040292 | -0.309132702876348 | 0.75722058339128 | NA | NM_029467 |
| Sult2a5 | 0.707463931 | 0.031633586 | 0.054543609 | 0.579968699 | 0.561935726 | NA | | Tcea3 | 16.16589703 | 0.0632379088118966 | 0.072027409122673 | 0.877970061427497 | 0.379959966030299 | NA | NM_011542 |
| Sult2a6 | 0 | NA | NA | NA | NA | NA | | Tcf15 | 9.759064435 | -0.0190082949442079 | 0.0636690758836286 | -0.298548309055882 | 0.76528471086561 | NA | NM_009328 |
| Sult2a7 | 0 | NA | NA | NA | NA | NA | | Tcf21 | 0.743239487 | 0.0100732369116114 | 0.0143507286637083 | 0.701932086353615 | 0.482721517642827 | NA | NM_011545 |
| Sult3a1 | 0.105923158 | 0.018524693 | 0.027977518 | 0.662127842 | 0.507889294 | NA | | Tcfl5 | 21.50036819 | 0.143983820961933 | 0.0790837604572048 | 1.82064965208436 | 0.0686601300904631 | NA | NM_178254 |
| Sult5a1 | 5.699542381 | 0.026347987 | 0.145060757 | 0.18163415 | 0.855869847 | NA | | Tchhl1 | 0.127687691 | 0.00542298404407807 | 0.00869724871942359 | 0.62352868349814 | 0.532937159177571 | NA | NM_027762 |
| Sult6b1 | 1.912376999 | -0.065649506 | 0.100216088 | -0.655079509 | 0.512416564 | NA | | Tcl1 | 0 | NA | NA | NA | NA | NA | NM_009337 |
| Sun3 | 2.050813791 | 0.090764811 | 0.10590399 | 0.857048079 | 0.391418314 | NA | | Tcl1b1 | 0.194331409 | -0.00840968081320881 | 0.00964081822647896 | -0.872299489073576 | 0.383045016303155 | NA | NM_013773 |
| Sun5 | 0.096071043 | -0.020098795 | 0.027825948 | -0.722304036 | 0.470107572 | NA | | Tcl1b2 | 0.769338278 | -0.00959303185972974 | 0.0200434402845632 | -0.478612040824049 | 0.632214650815219 | NA | NM_013775 |
| Sva | 0 | NA | NA | NA | NA | NA | | Tcl1b3 | 0 | NA | NA | NA | NA | NA | NM_013772 |
| Sval1 | 0 | NA | NA | NA | NA | NA | | Tcl1b4 | 0 | NA | NA | NA | NA | NA | NM_013774 |
| Sval2 | 0 | NA | NA | NA | NA | NA | | Tcl1b5 | 0 | NA | NA | NA | NA | NA | NM_013776 |
| Sval3 | 0 | NA | NA | NA | NA | NA | | Tcp10a | 0.913990035 | 0.0024137841660322 | 0.0189950482159907 | 0.127074390050782 | 0.898881520895737 | NA | NR_046291,NM_011553 |
| Svopl | 0.957201742 | 0.038953393 | 0.065902215 | 0.591078666 | 0.554467713 | NA | | Tcp10b | 5.998320282 | -0.00225642942838785 | 0.0488541385999629 | -0.0461870681389839 | 0.963161149605757 | NA | NM_009341 |
| Svs1 | 0 | NA | NA | NA | NA | NA | | Tcp10c | 0.692298838 | 0.014348034420441 | 0.0182474700244136 | 0.786302671068619 | 0.431690188005767 | NA | NM_001167578 |
| Svs2 | 0 | NA | NA | NA | NA | NA | | Tcp11 | 18.89066489 | 0.0171356438235792 | 0.0725461877850312 | 0.236203229235911 | 0.813274973841269 | NA | NM_013687,NM_001085555 |
| Svs3a | 0.094392817 | 0.017895942 | 0.027818137 | 0.643319236 | 0.520016977 | NA | | Tcte3 | 0 | NA | NA | NA | NA | NA | NM_198104,NM_011560 |
| Svs3b | 0 | NA | NA | NA | NA | NA | | Tctex1d1 | 10.45028329 | 0.0118166103422851 | 0.0667272154554842 | 0.177088317886849 | 0.85943901787939 | NA | NM_001163767,NM_026100,NM_001163768 |
| Svs4 | 0 | NA | NA | NA | NA | NA | | Tctex1d4 | 14.12539754 | 0.0113527935078267 | 0.0701036972113251 | 0.161942864063276 | 0.871350847663297 | NA | NM_175030 |
| Svs5 | 0 | NA | NA | NA | NA | NA | | Tdg | 1.347988832 | -0.013306905632876 | 0.0224332300682961 | -0.593178315934183 | 0.553061822804725 | NA | NM_172552,NM_011561 |
| Svs6 | 0 | NA | NA | NA | NA | NA | | Tdgf1 | 21.73456736 | 0.0789638549880002 | 0.0770434509650553 | 1.02492624614928 | 0.305397989398113 | NA | NM_011562 |
| Syce1 | 5.775800117 | 0.047463816 | 0.142759344 | 0.332474325 | 0.739531124 | NA | | Tdh | 1.758036472 | 0.021237433232715 | 0.0285750586946568 | 0.743215734380491 | 0.457351078771363 | NA | NM_021480 |
| Syce1l | 4.444955211 | 0.146339423 | 0.128739573 | 1.136708939 | 0.255659981 | NA | | Tdo2 | 15.73434334 | 0.0119280304193538 | 0.0726320466898382 | 0.164225448172902 | 0.869553668332011 | NA | NM_019911 |
| Syce3 | 0.314431083 | -0.017921647 | 0.039992366 | -0.448126696 | 0.654061762 | NA | | Tdpoz1 | 0 | NA | NA | NA | NA | NA | NM_148949 |
| Sycn | 0 | NA | NA | NA | NA | NA | | Tdpoz2 | 0 | NA | NA | NA | NA | NA | NM_001007222 |
| Sycp1 | 0.37239624 | 0.052452486 | 0.04522608 | 1.159784039 | 0.246136744 | NA | | Tdpoz3 | 0 | NA | NA | NA | NA | NA | NM_207271 |
| Sycp1-ps1 | 0 | NA | NA | NA | NA | NA | | Tdpoz4 | 0 | NA | NA | NA | NA | NA | NM_207272 |
| Sycp3 | 6.175903059 | 0.070594094 | 0.138200365 | 0.510809751 | 0.60948428 | NA | | Tdpoz5 | 0 | NA | NA | NA | NA | NA | NM_207273 |
| Synb | 2.827455305 | 0.171654527 | 0.099670056 | 1.722227658 | 0.085028286 | NA | | Tdrd12 | 10.59576165 | 0.0856043452134638 | 0.0687290496146057 | 1.24553366725548 | 0.212935647939884 | NA | NM_028034 |
| Syne1 | 19737.15193 | -0.169729761 | 0.097243582 | -1.745408354 | NA | NA | | Tecrl | 10.5325273 | -0.0429761360527148 | 0.0660460900632559 | -0.65069917101155 | 0.515240698433342 | NA | NM_153801 |
| Syngr4 | 2.701203929 | 0.102966466 | 0.114471088 | 0.899497578 | 0.368387685 | NA | | Tecta | 12.76454034 | -0.0955738199085814 | 0.0713315088139382 | -1.33985417521277 | 0.180292758673716 | NA | NM_009347 |
| Syt8 | 0.607073637 | 0.043174474 | 0.049620922 | 0.870086075 | 0.384253367 | NA | | Tectb | 3.508855062 | -0.0426797244239562 | 0.0366419906455508 | -1.16477635827186 | 0.244109531028839 | NA | NM_009348 |
| T | 2.95375557 | 0.132193287 | 0.110911914 | 1.191876349 | 0.233309733 | NA | | Teddm1 | 0 | NA | NA | NA | NA | NA | NM_178244 |
| Taar1 | 0 | NA | NA | NA | NA | NA | | Tekt3 | 4.135558479 | 0.0114713249551037 | 0.0455280759010058 | 0.251961558402918 | 0.801070777305313 | NA | NM_027660 |
| Taar2 | 0.116696964 | -0.020477583 | 0.027954224 | -0.732539847 | 0.463839131 | NA | | Tepp | 4.168158204 | -0.0504915641655651 | 0.0418879211382281 | -1.20539675385048 | 0.228050174052767 | NA | NM_199455,NM_028532 |
| Taar3 | 0.305736371 | 0.008542183 | 0.033941149 | 0.251676297 | 0.80129128 | NA | | Terc | 0.364150398 | -0.0138946944620142 | 0.0123764228912162 | -1.12267450653093 | 0.261575759058949 | NA | NR_001579 |
| Taar4 | 0 | NA | NA | NA | NA | NA | | Tescl | 0.224626239 | -0.00168521210467683 | 0.00978955093367521 | -0.172143964119932 | 0.863324350255464 | NA | NM_001163810 |
| Taar5 | 0.094392817 | 0.017895942 | 0.027818137 | 0.643319236 | 0.520016977 | NA | | Tespa1 | 0.226066984 | -0.0093084363525918 | 0.0101406635841621 | -0.917931679257152 | 0.358654635211298 | NA | NM_183264 |
| Taar6 | 0 | NA | NA | NA | NA | NA | | Tex101 | 0 | NA | NA | NA | NA | NA | NM_019981 |
| Taar7a | 0 | NA | NA | NA | NA | NA | | Tex11 | 13.47697597 | -0.0902361474250281 | 0.0707338613399385 | -1.27571357926247 | 0.202056786563106 | NA | NM_031384,NM_001167997 |
| Taar7b | 0 | NA | NA | NA | NA | NA | | Tex12 | 0 | NA | NA | NA | NA | NA | NM_025687 |
| Taar7d | 0 | NA | NA | NA | NA | NA | | Tex13 | 0 | NA | NA | NA | NA | NA | NM_031381 |
| Taar7e | 0 | NA | NA | NA | NA | NA | | Tex13a | 0 | NA | NA | NA | NA | NA | NM_026469 |
| Taar7f | 0 | NA | NA | NA | NA | NA | | Tex14 | 18.90110364 | -0.0397914980216568 | 0.0772751372714461 | -0.514932738092465 | 0.60660002282114 | NA | NM_031386,NM_001199293 |
| Taar8a | 0 | NA | NA | NA | NA | NA | | Tex16 | 0.212695677 | -0.00857796107890413 | 0.00973699866180651 | -0.880965621629515 | 0.378336427605205 | NA | NM_031382 |
| Taar8b | 0 | NA | NA | NA | NA | NA | | Tex19.1 | 0 | NA | NA | NA | NA | NA | NM_028602 |
| Taar8c | 0 | NA | NA | NA | NA | NA | | Tex19.2 | 0.120209049 | -0.00674972139751468 | 0.00863521114606269 | -0.781650996523957 | 0.434419707908266 | NA | NM_027622 |
| Taar9 | 0 | NA | NA | NA | NA | NA | | Tex21 | 0.710575567 | 0.0116546194246954 | 0.0163895928770841 | 0.711098775430284 | 0.477023028782178 | NA | NM_001159532,NM_019784 |
| Tac4 | 0.783678328 | -0.040057892 | 0.068613333 | -0.583820817 | 0.559340871 | NA | | Tex22 | 1.628583663 | 0.0214413510896183 | 0.0284096628218785 | 0.75472036482975 | 0.450416784685627 | NA | NM_029381 |
| Tacstd2 | 5.654824625 | -0.017623005 | 0.145771708 | -0.120894548 | 0.903774561 | NA | | Tex24 | 0.12890128 | -0.0068640067066208 | 0.00870804083610955 | -0.788237771940376 | 0.430557639008166 | NA | NM_001013609 |
| Taf7l | 3.465168167 | -0.270614826 | 0.115410054 | -2.344811532 | 0.0190367 | NA | | Tex26 | 10.85960582 | -0.110364360609887 | 0.0594451692706705 | -1.85657408270413 | 0.0633717740780522 | NA | NM_029464 |
| Tal2 | 3.541156078 | 0.236510616 | 0.121988916 | 1.938787752 | 0.052527188 | NA | | Tex28 | 0 | NA | NA | NA | NA | NA | NM_001126488 |
| Tap1 | 175.3532398 | 0.407157652 | 0.164556707 | 2.474269568 | NA | NA | | Tex29 | 0.665008488 | -0.0033027686652756 | 0.017459755046754 | -0.189164662186348 | 0.849963763089328 | NA | NM_029326 |
| Tarm1 | 0.858186872 | 0.089817948 | 0.05829794 | 1.54067106 | 0.123396864 | NA | | Tex33 | 0.767812082 | -0.0159660870503751 | 0.018915894716357 | -0.844056667146111 | 0.398637740624054 | NA | NM_001163612,NM_001163613,NM_028522 |
| Tas1r1 | 4.508631947 | 0.036943638 | 0.128455738 | 0.28759819 | 0.773654331 | NA | | Tex35 | 1.012063008 | 0.0076842395979362 | 0.0234178170690105 | 0.328136460170106 | 0.742808487575273 | NA | NM_028540 |
| Tas1r2 | 0.181709657 | 0.024794574 | 0.031325423 | 0.791516008 | 0.428642938 | NA | | Tex36 | 0.800350573 | -0.00359413574856419 | 0.0186324335826513 | -0.192896742801794 | 0.847039835996788 | NA | NM_028654 |
| Tas2r102 | 0 | NA | NA | NA | NA | NA | | Tex37 | 0 | NA | NA | NA | NA | NA | NM_028825 |
| Tas2r103 | 0.079983654 | 0.016740032 | 0.027674178 | 0.604897175 | 0.54524732 | NA | | Tex38 | 5.739947728 | 0.0774072809515102 | 0.0537014850421112 | 1.44143650572809 | 0.149461403167561 | NA | NM_029196 |
| Tas2r104 | 0 | NA | NA | NA | NA | NA | | Tfap2a | 7.57410212 | 0.0215030174178642 | 0.0582162327383111 | 0.369364632619271 | 0.711855956117155 | NA | NM_001122948,NM_011547 |
| Tas2r105 | 0 | NA | NA | NA | NA | NA | | Tfap2c | 5.078949204 | 0.00786123334764579 | 0.0510746120230024 | 0.153916653230872 | 0.877675453193995 | NA | NM_001159696,NM_009335 |
| Tas2r106 | 0 | NA | NA | NA | NA | NA | | Tfap2e | 0.516571875 | 0.0112766580422993 | 0.0166265449728752 | 0.678232191997567 | 0.497624484335967 | NA | NM_198960 |
| Tas2r107 | 0 | NA | NA | NA | NA | NA | | Tfec | 1.101029416 | -0.0171326671157214 | 0.0254555158564971 | -0.673043406871229 | 0.500919676248094 | NA | NM_031198 |
| Tas2r108 | 0.631566968 | 0.019997227 | 0.053874878 | 0.371179071 | 0.71050416 | NA | | Tff1 | 0 | NA | NA | NA | NA | NA | NM_009362 |
| Tas2r109 | 0 | NA | NA | NA | NA | NA | | Tff2 | 0 | NA | NA | NA | NA | NA | NM_009363 |
| Tas2r110 | 0 | NA | NA | NA | NA | NA | | Tff3 | 0 | NA | NA | NA | NA | NA | NM_011575 |
| Tas2r113 | 0 | NA | NA | NA | NA | NA | | Tfpi2 | 0.674790267 | -0.00481617710132038 | 0.0190312105893828 | -0.253067301142012 | 0.80021620648252 | NA | NM_009364 |
| Tas2r114 | 0 | NA | NA | NA | NA | NA | | Tg | 7.946859883 | 0.017841691550639 | 0.0557230458973787 | 0.320185145361522 | 0.748827983224163 | NA | NM_009375 |
| Tas2r115 | 0 | NA | NA | NA | NA | NA | | Tgif2lx1 | 0 | NA | NA | NA | NA | NA | NM_153109 |
| Tas2r116 | 0 | NA | NA | NA | NA | NA | | Tgif2lx2 | 0 | NA | NA | NA | NA | NA | NM_001142750 |
| Tas2r117 | 0 | NA | NA | NA | NA | NA | | Tgm1 | 4.393657582 | -0.0755393319449775 | 0.0426799561687849 | -1.7699018163525 | 0.0767434982046553 | NA | NM_019984,NM_001161715,NM_001161714 |
| Tas2r118 | 0 | NA | NA | NA | NA | NA | | Tgm4 | 15.52487044 | 0.0928539312289177 | 0.0771609718015684 | 1.20337949433434 | 0.228829499349468 | NA | NM_177911 |
| Tas2r119 | 0 | NA | NA | NA | NA | NA | | Tgm5 | 1.242644486 | 0.00446563195577435 | 0.0226854940390307 | 0.196849667372955 | 0.84394518251042 | NA | NM_028799 |
| Tas2r120 | 0 | NA | NA | NA | NA | NA | | Tgm6 | 7.202611723 | -0.0227941421974742 | 0.0563828543996346 | -0.404274356809113 | 0.686010980193479 | NA | NM_177726 |
| Tas2r121 | 0 | NA | NA | NA | NA | NA | | Tgm7 | 13.6494803 | -0.0740567177716291 | 0.0721546045815775 | -1.02636163278951 | 0.304721155810512 | NA | NM_001160424 |
| Tas2r122 | 0 | NA | NA | NA | NA | NA | | Tgoln2 | 0 | NA | NA | NA | NA | NA | NM_009444 |
| Tas2r123 | 0 | NA | NA | NA | NA | NA | | Tgtp1 | 0 | NA | NA | NA | NA | NA | NM_011579 |
| Tas2r124 | 0 | NA | NA | NA | NA | NA | | Tgtp2 | 31.30998525 | -0.168143785433552 | 0.0676307491468121 | -2.48620320719126 | NA | NA | NM_001145164 |
| Tas2r125 | 0 | NA | NA | NA | NA | NA | | Theg | 15.05706019 | 0.0815077760794726 | 0.0722464706589859 | 1.1281904200442 | 0.259239508892167 | NA | NM_199226,NM_011583 |
| Tas2r126 | 0 | NA | NA | NA | NA | NA | | Them5 | 0.323722592 | -0.0163509104806049 | 0.013414304634911 | -1.21891599494851 | 0.222876078142229 | NA | NM_025416 |
| Tas2r129 | 0 | NA | NA | NA | NA | NA | | Themis | 0.886846282 | -0.00527703506239931 | 0.0235620656091726 | -0.223963176655658 | 0.822785956194466 | NA | NM_178666 |
| Tas2r130 | 0 | NA | NA | NA | NA | NA | | Themis3 | 0.50399356 | -0.0128887585200878 | 0.0150396090110343 | -0.856987605903288 | 0.39145173435571 | NA | NM_028998 |
| Tas2r131 | 0 | NA | NA | NA | NA | NA | | Ticrr | 5.776853749 | -0.0232811385417349 | 0.04346194805969 | -0.535667165902479 | 0.592188599830555 | NA | NM_029835 |
| Tas2r134 | 0 | NA | NA | NA | NA | NA | | Tigd4 | 2.814086805 | -0.0195505016373029 | 0.0387467230730464 | -0.504571744052928 | 0.613859659948012 | NA | NM_207278 |
| Tas2r135 | 0.254360125 | 0.029364299 | 0.033967282 | 0.864487756 | 0.387320003 | NA | | Tigit | 0.442291665 | -0.00273282415188234 | 0.0138652338129813 | -0.197099031198719 | 0.843750041549212 | NA | NM_001146325 |
| Tas2r136 | 0 | NA | NA | NA | NA | NA | | Timd2 | 0.454027424 | 0.00804978036414047 | 0.0156544059541427 | 0.514218194399783 | 0.607099448513707 | NA | NM_134249,NM_001161355,NM_001161356 |
| Tas2r137 | 0.725227826 | -0.022946998 | 0.065969169 | -0.347844275 | 0.727957137 | NA | | Timd4 | 3.421866688 | -0.0267592232464983 | 0.0349312338545436 | -0.766054338587803 | 0.443643970605386 | NA | NM_178759 |
| Tas2r138 | 0 | NA | NA | NA | NA | NA | | Timm8a2 | 0.4259239 | 0.00797407930705397 | 0.0154197355711079 | 0.517134633747876 | 0.605062178310543 | NA | NM_001037744 |
| Tas2r139 | 0 | NA | NA | NA | NA | NA | | Timp1 | 10.32580293 | 0.0376347659353952 | 0.0666508270260732 | 0.564655648168819 | 0.572308006774778 | NA | NM_011593,NM_001044384 |
| Tas2r140 | 0 | NA | NA | NA | NA | NA | | Tinag | 1.311727898 | -0.00797650444152448 | 0.028756156247003 | -0.277384236370458 | 0.781485084473367 | NA | NM_012033 |
| Tas2r143 | 0.293774112 | 0.008475859 | 0.039057974 | 0.217007122 | 0.828202795 | NA | | Tk1 | 14.33553245 | 0.0179925579019854 | 0.0707556010949244 | 0.254291640853237 | 0.799270257234743 | NA | NM_009387,NM_001271729 |
| Tas2r144 | 0 | NA | NA | NA | NA | NA | | Tktl1 | 3.448760418 | -0.0542677980510563 | 0.0426513509801923 | -1.27235824432053 | 0.203245862862264 | NA | NM_031379 |
| Tat | 1.611244111 | 0.079196092 | 0.091588384 | 0.864695809 | 0.38720577 | NA | | Tktl2 | 2.358548777 | 0.0191380953543542 | 0.0351470147386201 | 0.544515529887238 | 0.586086764583331 | NA | NM_001271574,NM_028927 |
| Tbc1d21 | 0 | NA | NA | NA | NA | NA | | Tlcd2 | 9.851044355 | -0.0269200785990535 | 0.0564891635859121 | -0.476552968572669 | 0.63368048062408 | NA | NM_027249 |
| Tbpl2 | 0 | NA | NA | NA | NA | NA | | Tln2 | 5780.046072 | 0.054491449781895 | 0.0619946135451001 | 0.878970714806591 | NA | NA | NM_001081242 |
| Tbrg3 | 0.494745862 | -0.017414199 | 0.05079001 | -0.342866614 | 0.73169881 | NA | | Tlr11 | 0.12890128 | -0.0068640067066208 | 0.00870804083610955 | -0.788237771940376 | 0.430557639008166 | NA | NM_205819 |

| | | | | | | |
|---|---|---|---|---|---|---|
| Tbx10 | 0.47124342 | 0.077409868 | 0.054708132 | 1.414960895 | 0.157079962 | NA |
| Tbx19 | 1.19880358 | -0.039255453 | 0.076895544 | -0.510503613 | 0.609698684 | NA |
| Tbx20 | 0.656506615 | -0.019769438 | 0.054376061 | -0.363568787 | 0.716180038 | NA |
| Tbx22 | 6.96165136 | -0.312816501 | 0.152173296 | -2.055659632 | 0.039815336 | NA |
| Tbx5 | 0.079983654 | 0.016740032 | 0.027674178 | 0.604897175 | 0.54524732 | NA |
| Tc2n | 4.428694762 | 0.144517302 | 0.086317465 | 1.674253308 | 0.094080827 | NA |
| Tcam1 | 7.072622623 | 0.138219208 | 0.146330259 | 0.944570241 | 0.344878326 | NA |
| Tcf21 | 0.08731684 | 0.017365056 | 0.027733412 | 0.626142072 | 0.531221759 | NA |
| Tchhl1 | 0.487806363 | -0.025218575 | 0.039384805 | -0.640312313 | 0.521969578 | NA |
| Tcl1 | 0 | NA | NA | NA | NA | NA |
| Tcl1b1 | 0.159967308 | 0.0240345 | 0.030870969 | 0.778547006 | 0.436246606 | NA |
| Tcl1b2 | 0.904023182 | 0.00650544 | 0.074708519 | 0.087077624 | 0.930609811 | NA |
| Tcl1b3 | 0 | NA | NA | NA | NA | NA |
| Tcl1b4 | 0 | NA | NA | NA | NA | NA |
| Tcl1b5 | 0 | NA | NA | NA | NA | NA |
| Tcp10a | 0.406927243 | 0.024331845 | 0.043361066 | 0.561144991 | 0.574698699 | NA |
| Tcp10b | 4.719536598 | 0.079476574 | 0.120586515 | 0.659083432 | 0.509842194 | NA |
| Tcp10c | 0.487454995 | 0.031555684 | 0.045749554 | 0.689748452 | 0.49035239 | NA |
| Tcte3 | 0 | NA | NA | NA | NA | NA |
| Tdg | 2.028800312 | 0.008983561 | 0.08875971 | 0.101212147 | 0.919382055 | NA |
| Tdh | 1.156995207 | -0.018693609 | 0.079249747 | -0.23588226 | 0.813524034 | NA |
| Tdpoz1 | 0 | NA | NA | NA | NA | NA |
| Tdpoz2 | 0 | NA | NA | NA | NA | NA |
| Tdpoz3 | 0.096071043 | -0.020098795 | 0.027825948 | -0.722304036 | 0.470107572 | NA |
| Tdpoz4 | 0 | NA | NA | NA | NA | NA |
| Tdpoz5 | 0 | NA | NA | NA | NA | NA |
| Tecrl | 6.731038187 | 0.354808408 | 0.14026961 | 2.529474539 | 0.011423346 | NA |
| Tectb | 3.84552122 | 0.021939578 | 0.126013754 | 0.174104628 | 0.861783236 | NA |
| Teddm1 | 0 | NA | NA | NA | NA | NA |
| Tekt3 | 2.135990805 | 0.130476131 | 0.106453419 | 1.225664071 | 0.220325116 | NA |
| Tepp | 3.447898974 | 0.095804654 | 0.109689984 | 0.873412965 | 0.382438027 | NA |
| Terc | 0.729646711 | -0.019163599 | 0.061280247 | -0.312720663 | 0.754492896 | NA |
| Tescl | 0.105923158 | 0.018524693 | 0.027977518 | 0.662127842 | 0.507889294 | NA |
| Tespa1 | 0.284260198 | 0.011273774 | 0.038754816 | 0.290899955 | 0.771127836 | NA |
| Tex101 | 0 | NA | NA | NA | NA | NA |
| Tex12 | 0 | NA | NA | NA | NA | NA |
| Tex13 | 0.098353386 | -0.020212831 | 0.02786466 | -0.725393061 | 0.468210929 | NA |
| Tex13a | 0 | NA | NA | NA | NA | NA |
| Tex16 | 0.272987066 | 0.007834485 | 0.033279985 | 0.2354113 | 0.813889516 | NA |
| Tex19.1 | 0 | NA | NA | NA | NA | NA |
| Tex19.2 | 0.099009884 | 0.018205726 | 0.027896865 | 0.652608315 | 0.514008823 | NA |
| Tex21 | 0.320973508 | -0.015129019 | 0.040159764 | -0.376720802 | 0.706381102 | NA |
| Tex22 | 0.913014257 | -0.020655493 | 0.069611272 | -0.296726267 | 0.766675501 | NA |
| Tex24 | 0.105923158 | 0.018524693 | 0.027977518 | 0.662127842 | 0.507889294 | NA |
| Tex28 | 0 | NA | NA | NA | NA | NA |
| Tex29 | 0.573334654 | -0.00282487 | 0.052673623 | -0.053629686 | 0.957230204 | NA |
| Tex33 | 1.096528261 | 0.001265473 | 0.079649808 | 0.015887959 | 0.987323776 | NA |
| Tex35 | 1.371065569 | -0.028309733 | 0.086162026 | -0.328563914 | 0.742485327 | NA |
| Tex36 | 0.453998628 | 0.026589972 | 0.045454031 | 0.584986 | 0.55855713 | NA |
| Tex37 | 0 | NA | NA | NA | NA | NA |
| Tex38 | 4.355556834 | -0.059106171 | 0.136627621 | -0.432607776 | 0.665299739 | NA |
| Tfap2a | 6.742375519 | -0.037887232 | 0.151565133 | -0.24997327 | 0.80260802 | NA |
| Tfap2c | 7.507192167 | -0.203723485 | 0.140626394 | -1.448685976 | 0.147425299 | NA |
| Tfap2e | 0.272729858 | 0.010322181 | 0.038323464 | 0.26934363 | 0.78766526 | NA |
| Tfec | 1.883608677 | -0.038493707 | 0.105120215 | -0.36618748 | 0.714225184 | NA |
| Tff1 | 0 | NA | NA | NA | NA | NA |
| Tff2 | 0 | NA | NA | NA | NA | NA |
| Tff3 | 0 | NA | NA | NA | NA | NA |
| Tfpi2 | 1.029540879 | -0.073577205 | 0.078163571 | -0.941323481 | 0.346539113 | NA |
| Tg | 7.738540515 | 0.082199922 | 0.150480033 | 0.546251359 | 0.584893167 | NA |
| Tgif2lx1 | 0 | NA | NA | NA | NA | NA |
| Tgif2lx2 | 0 | NA | NA | NA | NA | NA |
| Tgm1 | 4.589164824 | 0.094308232 | 0.124129342 | 0.759757765 | 0.447399393 | NA |
| Tgm5 | 0.505937128 | 0.03385261 | 0.046683903 | 0.725145244 | 0.46836293 | NA |
| Tgm6 | 7.479841179 | -0.135278741 | 0.1533584 | -0.882108458 | 0.377718159 | NA |
| Tgoln2 | 0 | NA | NA | NA | NA | NA |
| Tgtp1 | 0 | NA | NA | NA | NA | NA |
| Them5 | 0.463017151 | 0.009204469 | 0.04998486 | 0.184145136 | 0.853899599 | NA |
| Themis | 0.674726373 | 0.035983769 | 0.061458617 | 0.585495914 | 0.558214313 | NA |
| Themis3 | 0.63671097 | 0.032300813 | 0.060136437 | 0.537125483 | 0.591180939 | NA |
| Tigd4 | 2.461664092 | 0.017037577 | 0.108471884 | 0.157069061 | 0.875190421 | NA |
| Tigit | 0.400129188 | 0.02188733 | 0.04314476 | 0.507299852 | 0.611944442 | NA |
| Timd2 | 0.400720577 | -0.028041581 | 0.043551246 | -0.643875506 | 0.519656166 | NA |
| Timd4 | 2.710522476 | 0.14887368 | 0.112577536 | 1.322410185 | 0.186031599 | NA |
| Timm8a2 | 0.176054697 | -0.002190495 | 0.031239146 | -0.070120197 | 0.944097991 | NA |
| Tinag | 1.018815137 | 0.073477377 | 0.081170573 | 0.905221854 | 0.365347867 | NA |
| Tktl1 | 3.655482423 | 0.05044423 | 0.125631359 | 0.401525783 | 0.68803306 | NA |
| Tktl2 | 6.943658428 | -0.559927057 | 0.146636401 | -3.818472457 | 0.000134281 | NA |
| Tlr11 | 0.201994201 | 0.000227241 | 0.031727614 | 0.00716224 | 0.994285408 | NA |
| Tlr8 | 6.668945966 | -0.181774026 | 0.140074093 | -1.297699115 | 0.194390748 | NA |
| Tlx1 | 0 | NA | NA | NA | NA | NA |
| Tlx2 | 0 | NA | NA | NA | NA | NA |
| Tlx3 | 0 | NA | NA | NA | NA | NA |
| Tm4sf19 | 1.642775276 | 0.067198221 | 0.100893972 | 0.666028104 | 0.505393137 | NA |
| Tm4sf20 | 0 | NA | NA | NA | NA | NA |
| Tm4sf4 | 0.55073328 | 0.049633073 | 0.054814118 | 0.905479733 | 0.365211292 | NA |
| Tmc1 | 7.74719123 | 0.208245389 | 0.146201061 | 1.424376731 | 0.154337446 | NA |
| Tmc2 | 0.291593384 | 0.011931959 | 0.039023223 | 0.305765587 | 0.759783132 | NA |
| Tmc5 | 5.999963146 | 0.112682217 | 0.126481366 | 0.890899749 | 0.372982955 | NA |
| Tmco2 | 0.837310763 | -0.033554311 | 0.064018926 | -0.524131119 | 0.600187346 | NA |
| Tmco5b | 2.087200257 | 0.036995967 | 0.104147049 | 0.355228187 | 0.72241865 | NA |
| Tmed11 | 0 | NA | NA | NA | NA | NA |
| Tmed6 | 0.105923158 | 0.018524693 | 0.027977518 | 0.662127842 | 0.507889294 | NA |
| Tmem102 | 3.998689575 | 0.03048839 | 0.107871677 | 0.282635732 | 0.777456087 | NA |
| Tmem139 | 0 | NA | NA | NA | NA | NA |
| Tmem171 | 4.538185253 | 0.227651133 | 0.129760802 | 1.754390608 | 0.0793636 | NA |
| Tmem181c-ps | 0.206069989 | -0.026776453 | 0.031767726 | -0.842882251 | 0.399294301 | NA |
| Tmem182 | 3.632067248 | -0.316780394 | 0.122653805 | -2.582719657 | 0.009802494 | NA |
| Tmem184a | 4.528773212 | 0.212630843 | 0.110302079 | 1.927713838 | 0.05389073 | NA |
| Tmem190 | 0 | NA | NA | NA | NA | NA |
| Tmem207 | 0 | NA | NA | NA | NA | NA |
| Tmem210 | 0.283178451 | 0.028390488 | 0.033368195 | 0.850824814 | 0.394866675 | NA |
| Tmem211 | 1.154859085 | 0.086358106 | 0.072904567 | 1.184536298 | 0.236200836 | NA |
| Tmem213 | 0.116696964 | -0.020477583 | 0.027954224 | -0.732539847 | 0.463839131 | NA |
| Tmem217 | 1.887823986 | 0.002348191 | 0.084237717 | 0.027875769 | 0.977761235 | NA |
| Tmem221 | 7.859498067 | 0.085292255 | 0.156803724 | 0.543942789 | 0.586480842 | NA |
| Tmem225 | 0 | NA | NA | NA | NA | NA |
| Tmem233 | 0 | NA | NA | NA | NA | NA |
| Tmem235 | 0.99394839 | 0.049346079 | 0.073902494 | 0.667718726 | 0.504313154 | NA |
| Tmem239 | 0.094392817 | 0.017895942 | 0.027818137 | 0.643319236 | 0.520016977 | NA |
| Tmem247 | 0 | NA | NA | NA | NA | NA |
| Tmem252 | 47.3771998 | 0.204360485 | 0.13008139 | 1.571020154 | NA | NA |
| Tmem253 | 4.460908769 | 0.063004086 | 0.110064042 | 0.572431143 | 0.567029923 | NA |
| Tmem254a | 0 | NA | NA | NA | NA | NA |
| Tmem254b | 0 | NA | NA | NA | NA | NA |
| Tmem254c | 0 | NA | NA | NA | NA | NA |
| Tmem27 | 4.323201814 | 0.210631168 | 0.120882324 | 1.742448037 | 0.081430077 | NA |
| Tmem30b | 3.576165431 | 0.007358545 | 0.115199793 | 0.06387637 | 0.949068668 | NA |
| Tmem30c | 7.202564556 | 0.17144171 | 0.149584489 | 1.14611957 | 0.251745681 | NA |
| Tmem45b | 1.274413075 | -0.036644497 | 0.07236703 | -0.506370065 | 0.612596885 | NA |
| Tmem51as1 | 3.567674565 | 0.150962731 | 0.130016648 | 1.161103083 | 0.245599981 | NA |
| Tmem52b | 0.099009884 | 0.018205726 | 0.027896865 | 0.652608315 | 0.514008823 | NA |
| Tmem54 | 2.143751428 | -0.021525632 | 0.102567875 | -0.209867196 | 0.833771326 | NA |
| Tmem72 | 24.87395896 | 0.215595491 | 0.110048562 | 1.959094128 | NA | NA |
| Tmem89 | 1.997515811 | -0.080396475 | 0.102131164 | -0.78718847 | 0.431171546 | NA |
| Tmem92 | 0.079983654 | 0.016740032 | 0.027674178 | 0.604897175 | 0.54524732 | NA |
| Tmem95 | 1.153741678 | -0.001538765 | 0.058017103 | -0.026522614 | 0.978840496 | NA |
| Tmigd1 | 0.119397641 | -0.020477583 | 0.027954224 | -0.732539845 | 0.463839132 | NA |
| Tmod4 | 7.563143088 | 0.054579957 | 0.142124937 | 0.384028009 | 0.700957688 | NA |
| Tmprss11a | 5.172311587 | 0.185875555 | 0.102622617 | 1.811253316 | 0.070101649 | NA |
| Tmprss11bnl | 0 | NA | NA | NA | NA | NA |
| Tmprss11c | 0 | NA | NA | NA | NA | NA |
| Tmprss11d | 0.19442443 | -0.026451352 | 0.031568725 | -0.837897386 | 0.402088337 | NA |
| Tmprss11e | 0.116696964 | -0.020477583 | 0.027954224 | -0.732539847 | 0.463839131 | NA |
| Tmprss11f | 0.176054697 | -0.002190495 | 0.031239146 | -0.070120197 | 0.944097991 | NA |
| Tmprss11g | 0 | NA | NA | NA | NA | NA |
| Tmprss12 | 0 | NA | NA | NA | NA | NA |
| Tmprss13 | 0.512850401 | 0.033489004 | 0.046267694 | 0.723809661 | 0.469182597 | NA |
| Tmprss15 | 0.871501254 | -0.017336007 | 0.064315405 | -0.269546729 | 0.787508988 | NA |
| Tmprss2 | 1.442318935 | 0.123906339 | 0.090922286 | 1.362771921 | 0.172954403 | NA |
| Tmprss3 | 1.795150573 | 0.108706306 | 0.096860485 | 1.122297767 | 0.261735855 | NA |
| Tmsb15a | 0 | NA | NA | NA | NA | NA |
| Tmsb15b2 | 1.342234934 | 0.003684634 | 0.084180385 | 0.043770692 | 0.965087189 | NA |
| Tmsb15l | 0 | NA | NA | NA | NA | NA |
| Tnf | 2.607046118 | 0.104075028 | 0.112160005 | 0.927915685 | 0.353451308 | NA |
| Tnfrsf13b | 4.072251826 | 0.03083063 | 0.131797657 | 0.233923957 | 0.815044014 | NA |
| Tnfrsf17 | 7.474252658 | -0.267831735 | 0.154031663 | -1.738809612 | 0.082068257 | NA |
| Tnfrsf18 | 2.22352369 | -0.044768391 | 0.100717773 | -0.444493456 | 0.656685859 | NA |
| Tnfrsf26 | 1.509362283 | -0.131811912 | 0.085485001 | -1.541930285 | 0.123090536 | NA |
| Tnfsf11 | 0.953514938 | -0.009882079 | 0.072650881 | -0.136021457 | 0.891804318 | NA |
| Tnfsf14 | 0.280299629 | 0.041967777 | 0.040615819 | 1.033286501 | 0.301469842 | NA |
| Tnfsf15 | 1.036246275 | -0.043177279 | 0.073581656 | -0.586794062 | 0.557342025 | NA |
| Tnfsf18 | 0.895316464 | 0.045505672 | 0.078674053 | 0.57840762 | 0.562988951 | NA |
| Tnfsf4 | 0 | NA | NA | NA | NA | NA |
| Tnfsf8 | 3.460287471 | 0.198018841 | 0.119834684 | 1.652433447 | 0.098446222 | NA |
| Tnip3 | 2.977704263 | 0.030418475 | 0.119087463 | 0.255429701 | 0.798391234 | NA |
| Tnk1 | 6.239772721 | 0.102355081 | 0.144356066 | 0.709045932 | 0.478295974 | NA |
| Tnnc2 | 4.409021046 | 0.169022739 | 0.137940729 | 1.225328738 | 0.220451385 | NA |
| Tnnt3 | 2.97399399 | -0.03655763 | 0.10425375 | -0.350660099 | 0.725843363 | NA |
| Tnp1 | 0 | NA | NA | NA | NA | NA |
| Tnp2 | 0 | NA | NA | NA | NA | NA |
| Tomm20l | 0.116696964 | -0.020477583 | 0.027954224 | -0.732539847 | 0.463839131 | NA |
| Tomt | 2.604535377 | 0.178115516 | 0.106927723 | 1.665756194 | 0.095761985 | NA |
| Topaz1 | 0.754584575 | 0.085968027 | 0.057162772 | 1.503916326 | 0.132602913 | NA |
| Tpbpa | 0 | NA | NA | NA | NA | NA |
| Tpbpb | 0 | NA | NA | NA | NA | NA |
| Tppp2 | 0 | NA | NA | NA | NA | NA |
| Tprg | 0.427430218 | -0.003973439 | 0.048318641 | -0.082234071 | 0.93446058 | NA |
| Tpsab1 | 0 | NA | NA | NA | NA | NA |
| Tpsb2 | 1.029588275 | -0.068296389 | 0.07386073 | -0.924664408 | 0.355140499 | NA |
| Tpsg1 | 0.215433205 | -0.027029776 | 0.031922004 | -0.84674434 | 0.397137638 | NA |
| Tpte | 0.523233625 | -0.018953331 | 0.051293849 | -0.369504955 | 0.71175138 | NA |
| Trap1a | 0.079983654 | 0.016740032 | 0.027674178 | 0.604897175 | 0.54524732 | NA |
| Trat1 | 4.656008607 | -0.029134172 | 0.140489697 | -0.207375865 | 0.83571633 | NA |
| Trcg1 | 0 | NA | NA | NA | NA | NA |
| Treh | 1.119991334 | -0.008487325 | 0.066260281 | -0.128090685 | 0.898077207 | NA |
| Trem1 | 0.079983654 | 0.016740032 | 0.027674178 | 0.604897175 | 0.54524732 | NA |
| Trem3 | 1.974725285 | -0.1263264 | 0.101647741 | -1.242786103 | 0.213946665 | NA |
| Treml1 | 3.581138038 | 0.081416692 | 0.120772832 | 0.674130851 | 0.500228127 | NA |
| Treml4 | 2.56864326 | 0.116636441 | 0.109414127 | 1.066008965 | 0.286419589 | NA |
| Trex2 | 0 | NA | NA | NA | NA | NA |
| Trim10 | 1.496180113 | -0.092272274 | 0.090491902 | -1.019674371 | 0.30788292 | NA |
| Trim15 | 2.241585677 | 0.049665234 | 0.106436454 | 0.466618646 | 0.640772744 | NA |
| Trim30b | 3.204279456 | 0.174378247 | 0.114578223 | 1.521914401 | 0.12803053 | NA |
| Trim30e-ps1 | 3.224145527 | 0.026776696 | 0.113177776 | 0.236589702 | 0.81297511 | NA |
| Trim31 | 0.238795281 | -0.027601606 | 0.032267774 | -0.855392297 | 0.392334007 | NA |
| Trim34b | 1.334494249 | 0.003706998 | 0.081910033 | 0.045256947 | 0.963902504 | NA |
| Trim38 | 1.494402856 | 0.00830225 | 0.084687017 | 0.098034505 | 0.921904895 | NA |
| Trim40 | 5.231497624 | -0.138444575 | 0.141706871 | -0.976978558 | 0.328579768 | NA |
| Trim42 | 0.225320798 | -0.001646023 | 0.032098875 | -0.051279768 | 0.95910259 | NA |
| Trim43b | 2.192713984 | 0.070831641 | 0.087001845 | 0.814139527 | 0.415565024 | NA |
| Trim43c | 0.484476137 | 0.03706576 | 0.05187087 | 0.714577567 | 0.474870111 | NA |
| Trim50 | 0.451324107 | 0.006406988 | 0.044046195 | 0.145460652 | 0.884347181 | NA |
| Trim52 | 0 | NA | NA | NA | NA | NA |
| Trim55 | 0.536358603 | 0.018999954 | 0.051864556 | 0.366337928 | 0.714112931 | NA |
| Trim58 | 4.134495169 | -0.159624254 | 0.128512812 | -1.242088251 | 0.214204003 | NA |
| Trim60 | 0.079983654 | 0.016740032 | 0.027674178 | 0.604897175 | 0.54524732 | NA |
| Trim61 | 0.755112314 | -0.0351974 | 0.062581738 | -0.562422859 | 0.573827948 | NA |
| Trim63 | 5.998641539 | -0.103461712 | 0.145400001 | -0.711566103 | 0.476733503 | NA |
| Trim69 | 1.187744795 | 0.035950389 | 0.074039548 | 0.48555658 | 0.627281588 | NA |
| Trim75 | 0.388436144 | 0.026670728 | 0.048195357 | 0.553387917 | 0.579997811 | NA |
| Triml1 | 0 | NA | NA | NA | NA | NA |
| Triml2 | 1.481389728 | -0.116726324 | 0.091785309 | -1.271732097 | 0.203468322 | NA |
| Troap | 7.196862209 | -0.097420481 | 0.141967602 | -0.686216293 | 0.492576725 | NA |
| Trp53tg5 | 0.373757766 | 0.017971539 | 0.042393413 | 0.423922914 | 0.671622033 | NA |
| Trpd52l3 | 0 | NA | NA | NA | NA | NA |
| Trpm5 | 0.961060696 | 0.01515042 | 0.07168587 | 0.211344577 | 0.832618401 | NA |
| Trpm8 | 5.072425431 | 0.007314258 | 0.130694487 | 0.055964546 | 0.955370051 | NA |
| Trpv1 | 4.831119748 | 0.266247382 | 0.138219288 | 1.926267933 | 0.054070927 | NA |
| Trpv5 | 0 | NA | NA | NA | NA | NA |
| Try10 | 0.08731684 | 0.017365056 | 0.027733412 | 0.626142072 | 0.531221759 | NA |
| Try4 | 0 | NA | NA | NA | NA | NA |
| Try5 | 0 | NA | NA | NA | NA | NA |
| Tsc1 | 3905.656923 | 0.187964682 | 0.099547126 | 1.888197976 | NA | NA |
| Tsga13 | 3.634188703 | 0.122436544 | 0.127197137 | 0.962573116 | 0.335761794 | NA |
| Tsga8 | 0 | NA | NA | NA | NA | NA |
| Tshb | 1.259273622 | 0.134503086 | 0.085858944 | 1.566558828 | 0.117217848 | NA |
| Tsix | 5.271486292 | 0.212349726 | 0.13748138 | 1.544570806 | 0.122450112 | NA |
| Tslp | 5.126907474 | -0.201091064 | 0.141880112 | -1.417330883 | 0.15638621 | NA |
| Tspan1 | 4.514927444 | -0.191255442 | 0.136863564 | -1.39741679 | 0.162288273 | NA |
| Tspan10 | 2.59523789 | -0.088338573 | 0.099418998 | -0.888548208 | 0.374245936 | NA |
| Tspan32 | 6.197854676 | 0.050745529 | 0.151579969 | 0.334777273 | 0.737793106 | NA |
| Tspo2 | 1.044963501 | 0.077922586 | 0.069093725 | 1.127780945 | 0.259412443 | NA |
| Tspy-ps | 0.215050351 | -0.027006021 | 0.031907566 | -0.846382992 | 0.397339123 | NA |
| Tssk1 | 2.886031018 | 0.103274677 | 0.115846973 | 0.89147497 | 0.372674414 | NA |
| Tssk2 | 0 | NA | NA | NA | NA | NA |
| Tssk3 | 1.824711376 | 0.081781891 | 0.097352299 | 0.840061223 | 0.40087406 | NA |
| Tssk5 | 0.233393928 | -0.027475306 | 0.032191697 | -0.85349047 | 0.39338737 | NA |
| Tssk6 | 2.3286089 | -0.077148753 | 0.116204333 | -0.663905995 | 0.506750479 | NA |
| Tsx | 2.84739085 | -0.031909003 | 0.106203304 | -0.300452074 | 0.763832348 | NA |
| Ttc23l | 0.099009884 | 0.018205726 | 0.027896865 | 0.652608315 | 0.514008823 | NA |
| Ttc24 | 0.588839167 | -0.044984229 | 0.06340616 | -0.709461493 | 0.47803814 | NA |
| Ttc29 | 7.994592152 | 0.177986267 | 0.142321928 | 1.250589206 | 0.211084391 | NA |
| Ttc30a2 | 2.517119421 | -0.12207371 | 0.093403509 | -1.306949929 | 0.191229717 | NA |
| Ttc36 | 1.787386729 | -0.123140353 | 0.095388043 | -1.290941185 | 0.196724073 | NA |

| | | | | | | | |
|---|---|---|---|---|---|---|---|
| Tlr5 | 18.58821657 | 0.0608009149790552 | 0.0806724535556252 | 0.753676283530062 | 0.451043626181076 | NA | NM_016928 |
| Tlr6 | 20.98609501 | -0.0670042350082575 | 0.0802421263371358 | -0.835025666278339 | 0.403703257384908 | NA | NM_011604 |
| Tlr8 | 5.224068675 | 0.0119584364829806 | 0.0438216200120559 | 0.27288896393357 | 0.784938570176451 | NA | NM_133212 |
| Tlx1 | 0 | NA | NA | NA | NA | NA | NM_021901 |
| Tlx2 | 0.106148608 | 0.00515554063321497 | 0.00854200771766808 | 0.603551390213729 | 0.546141939141592 | NA | NM_009392 |
| Tlx3 | 0.127687691 | 0.00542298404407807 | 0.00869724871942359 | 0.62352868349814 | 0.532937159177571 | NA | NM_019916 |
| Tm4sf19 | 1.316398883 | -0.0381504551368867 | 0.0269156396208848 | -1.4174084537558 | 0.156363542487162 | NA | NM_001160402 |
| Tm4sf20 | 0 | NA | NA | NA | NA | NA | NM_025453 |
| Tm4sf4 | 1.419526267 | 0.0114108749217619 | 0.0225586397551406 | 0.505831692230542 | 0.612974808230347 | NA | NM_145539 |
| Tm4sf5 | 10.04134989 | 0.0393057034589181 | 0.0584442656488479 | 0.672533105216508 | 0.501244372018671 | NA | NM_029360 |
| Tmc1 | 14.99237296 | 0.0660501665351074 | 0.0720764439103703 | 0.916390473109955 | 0.359462130332422 | NA | NM_028953 |
| Tmc2 | 0.235249118 | -0.00959082599542929 | 0.0102925929774586 | -0.93181825186654 | 0.351430460444064 | NA | NM_138655 |
| Tmc3 | 14.66022472 | -0.0262325220771926 | 0.0617051372977636 | -0.425127035219212 | 0.670744070631327 | NA | NM_177695 |
| Tmc5 | 8.186954316 | -0.0647113043416162 | 0.0599055317095549 | -1.08022251860415 | 0.280043102992448 | NA | NM_028930,NM_001105252 |
| Tmco2 | 1.041925876 | 0.0146426742961957 | 0.0204069918732304 | 0.717532225580182 | 0.473045759591868 | NA | NM_001081312 |
| Tmco5b | 2.30639267 | -0.0180417145290287 | 0.036543107447856 | -0.493710463861692 | 0.621510674228373 | NA | NM_029232 |
| Tmed11 | 0 | NA | NA | NA | NA | NA | NM_026109 |
| Tmed6 | 0.351510867 | 0.00237641026287115 | 0.0128867963543482 | 0.184406597072461 | 0.853694494854753 | NA | NM_025458 |
| Tmem102 | 4.029851536 | 0.0155698322918328 | 0.0369025334662621 | 0.421917706708874 | 0.673085085229292 | NA | NM_001033433 |
| Tmem139 | 0.108766301 | 0.00519171381361946 | 0.00856003371502575 | 0.606506234257728 | 0.544178643801544 | NA | NM_175408 |
| Tmem150b | 8.431272126 | -0.027454441957602 | 0.0445668618897108 | -0.616028160689062 | 0.537875933973605 | NA | NM_001142792,NM_177887 |
| Tmem154 | 20.74783051 | -0.0850894264587221 | 0.0814977729685251 | -1.04407057223004 | 0.296452739773051 | NA | NM_177260 |
| Tmem171 | 6.384764122 | 0.015431312100142 | 0.0475515289207235 | 0.324517685348639 | 0.745546135149407 | NA | NM_001025606 |
| Tmem174 | 14.07253166 | -0.0222931339240418 | 0.0638889821036291 | -0.348935500144327 | 0.727137732929163 | NA | NM_026685 |
| Tmem181c-ps | 0.127687691 | 0.00542298404407807 | 0.00869724871942359 | 0.62352868349814 | 0.532937159177571 | NA | NR_028305 |
| Tmem182 | 2.06983075 | 0.0163755787189436 | 0.0358350575287635 | 0.456970906375677 | 0.64769195722478 | NA | NM_001081198 |
| Tmem184a | 11.51737104 | 0.0244851515098844 | 0.0438464551563163 | 0.558429442530594 | 0.576551172511864 | NA | NM_001161548,NM_144914 |
| Tmem190 | 0 | NA | NA | NA | NA | NA | NM_030028 |
| Tmem207 | 0.106148608 | 0.00515554063321497 | 0.00854200771766808 | 0.603551390213729 | 0.546141939141592 | NA | NM_001101640 |
| Tmem210 | 0.217532603 | 0.00684685248201087 | 0.00976000296730706 | 0.701521557416086 | 0.482977586094298 | NA | NM_030055 |
| Tmem211 | 1.510697141 | 0.000914641263813424 | 0.0257681475649741 | 0.0354950336071759 | 0.971685006488965 | NA | NM_001033428 |
| Tmem213 | 0.127687691 | 0.00542298404407807 | 0.00869724871942359 | 0.62352868349814 | 0.532937159177571 | NA | NM_029921 |
| Tmem217 | 2.99097361 | 0.0104970432602377 | 0.0333129298706698 | 0.315104174294791 | 0.752682559222265 | NA | NM_001162901,NM_001162902 |
| Tmem221 | 8.635703972 | -0.0215831158940957 | 0.0633823612945215 | -0.340522433265062 | 0.733463131808075 | NA | NM_001100462 |
| Tmem225 | 0 | NA | NA | NA | NA | NA | NM_029379 |
| Tmem233 | 0.106148608 | 0.00515554063321497 | 0.00854200771766808 | 0.603551390213729 | 0.546141939141592 | NA | NM_001101546 |
| Tmem235 | 1.030691626 | -0.0339622790235373 | 0.0246591474491256 | -1.37726898683764 | 0.168429099903118 | NA | NM_001085535 |
| Tmem236 | 9.695423514 | -0.0953195194131807 | 0.0617415736653178 | -1.54384661346468 | 0.122625495863726 | NA | NM_001081310 |
| Tmem239 | 0.127687691 | 0.00542298404407807 | 0.00869724871942359 | 0.62352868349814 | 0.532937159177571 | NA | NM_025753 |
| Tmem247 | 0 | NA | NA | NA | NA | NA | NM_030104 |
| Tmem253 | 6.609024931 | -0.0128474739546885 | 0.0521153352041595 | -0.246520029169133 | 0.805279701730543 | NA | NM_001033805 |
| Tmem254a | 0 | NA | NA | NA | NA | NA | NM_026679,NM_025311 |
| Tmem254b | 0 | NA | NA | NA | NA | NA | NM_001270495,NM_001270496 |
| Tmem254c | 0 | NA | NA | NA | NA | NA | NM_001270499,NM_001270498 |
| Tmem26 | 21.56230327 | 0.0973230375955919 | 0.060125832994861 | 1.61865595448649 | 0.105521308676302 | NA | NM_177794 |
| Tmem27 | 7.579073774 | -0.00140079130270354 | 0.0365008218501637 | -0.0383769797966141 | 0.969387114919411 | NA | NM_020626 |
| Tmem30b | 2.858747816 | -0.045925701342944 | 0.0356962279109863 | -1.28657015126266 | 0.198244153022555 | NA | NM_178715 |
| Tmem30c | 8.193614122 | -0.038668885171115 | 0.0613709213934184 | -0.630084807155298 | 0.528639099279304 | NA | NM_027651 |
| Tmem45a | 21.2810456 | -0.0545187329425297 | 0.0776775754341068 | -0.701859354361253 | 0.48276687902789 | NA | NM_019631 |
| Tmem45b | 1.661960464 | 0.0129474810073386 | 0.0284133762383854 | 0.455682594659307 | 0.648618239882679 | NA | NM_144936 |
| Tmem51as1 | 4.350639713 | -0.0368766362849725 | 0.0475813798709685 | -0.775022422320975 | 0.438326410291092 | NA | NR_027137 |
| Tmem52 | 20.46076827 | 0.0439611734908513 | 0.0788333224091413 | 0.557647098300574 | 0.577085388436082 | NA | NM_001253856,NM_001253858,NM_027161 |
| Tmem52b | 0.120209049 | -0.00674972139751468 | 0.00863521114606269 | -0.781650996523957 | 0.434419707908266 | NA | NM_001081186 |
| Tmem54 | 1.897081997 | 0.00605105868147167 | 0.0298894371645545 | 0.202448063781126 | 0.839566456750027 | NA | NM_133881,NM_025452 |
| Tmem71 | 21.72496814 | -0.0169215741595175 | 0.0813560656597138 | -0.207994007850564 | 0.835233646087008 | NA | NM_172514 |
| Tmem72 | 56.84427618 | 0.00676799128820042 | 0.0267875268415774 | 0.252654609297328 | NA | NA | NM_178768 |
| Tmem79 | 11.98766079 | -0.0494164432752239 | 0.0712532698750684 | -0.693532287877707 | 0.487975570496097 | NA | NM_024246 |
| Tmem82 | 20.56700936 | -0.0469752551191863 | 0.0759492421166458 | -0.618508543469595 | 0.536240167845143 | NA | NM_145987 |
| Tmem89 | 1.303856917 | 0.00200247786989859 | 0.0218466223540653 | 0.0916607536599801 | 0.926967580116191 | NA | NM_027066 |
| Tmem8c | 18.75197626 | 0.0862808903345064 | 0.0726680004399048 | 1.18732990879334 | 0.235097515149536 | NA | NM_001159602,NM_025376 |
| Tmem92 | 0.557032348 | 0.0120663252741177 | 0.0169105036814138 | 0.71354026476335 | 0.475511506599363 | NA | NM_001034896,NM_001163172 |
| Tmem95 | 1.699291406 | 0.00369743384902356 | 0.0253370347398922 | 0.145930014580676 | 0.88397663700953 | NA | NM_001195710 |
| Tmigd1 | 0.219991894 | 0.00711449016834187 | 0.00994861810862167 | 0.715123456410123 | 0.474532761039468 | NA | NM_025655 |
| Tmod4 | 7.516796618 | 0.0123864769047604 | 0.0541026801868908 | 0.228943868621164 | 0.818912543707626 | NA | NM_016712 |
| Tmprss11a | 12.47182721 | 0.00970158373891771 | 0.0249983131067517 | 0.388089536181442 | NA | NA | NM_001033233 |
| Tmprss11bnl | 0.191449919 | 0.00665508079272897 | 0.00962405648474729 | 0.691504752001018 | 0.489248395458281 | NA | NM_177024 |
| Tmprss11c | 0 | NA | NA | NA | NA | NA | NM_001030297 |
| Tmprss11d | 0.09572496 | 0.00498255289994511 | 0.00847999972823692 | 0.587565219295241 | 0.556824162160132 | NA | NM_145561 |
| Tmprss11e | 0 | NA | NA | NA | NA | NA | NM_172880 |
| Tmprss11f | 0.097165704 | -0.00637828455464458 | 0.00848781691535724 | -0.75146349388194 | 0.452373763047372 | NA | NM_178730 |
| Tmprss11g | 0 | NA | NA | NA | NA | NA | NM_177162 |
| Tmprss12 | 0 | NA | NA | NA | NA | NA | NM_183109 |
| Tmprss13 | 1.046042392 | -0.0023596610533475 | 0.0230130847801396 | -0.102535625966315 | 0.91833153642415 | NA | NM_001013373 |
| Tmprss15 | 0.761135951 | -0.008023821618461 | 0.0184076240300595 | -0.435896648332134 | 0.662911717008221 | NA | NM_178855,NM_008941 |
| Tmprss2 | 3.008733408 | 0.0113824035142301 | 0.04076500256257 | 0.279219987702915 | 0.780076005432867 | NA | NM_015775 |
| Tmprss3 | 1.608423898 | -0.0128109753444729 | 0.0294698637002407 | -0.434714441667683 | 0.663769714492241 | NA | NM_080727,NM_001163776 |
| Tmprss4 | 11.77360598 | -0.0706197791196979 | 0.0682317631107806 | -1.03499859742801 | 0.30066956621328 | NA | NM_145403 |
| Tmsb15a | 0 | NA | NA | NA | NA | NA | NM_030106 |
| Tmsb15b1 | 10.88147611 | -0.0778616508639077 | 0.0682706169637556 | -1.14048553135596 | 0.254084077177249 | NA | NM_001081983 |
| Tmsb15b2 | 1.609564543 | -0.00927496692439041 | 0.0280651178458115 | -0.330480241534943 | 0.741037119731379 | NA | NM_001080967 |
| Tmsb15l | 0 | NA | NA | NA | NA | NA | NM_207267 |
| Tnf | 2.197373991 | -0.0400957102110267 | 0.0330263499141403 | -1.21405212247992 | 0.224727821058814 | NA | NM_013693 |
| Tnfrsf13b | 4.191239149 | -0.0150104595539176 | 0.0438347616286135 | -0.342432786131986 | 0.732025218449508 | NA | NM_021349 |
| Tnfrsf13c | 12.79127943 | -0.0515835695670093 | 0.0679442626050006 | -0.759204200461994 | 0.447730413747827 | NA | NM_028075 |
| Tnfrsf17 | 6.656281683 | 0.00270276272654405 | 0.053299637660359 | 0.050708838656031 | 0.959557533483249 | NA | NM_011608 |
| Tnfrsf18 | 2.229469499 | 0.0311527327862977 | 0.0366291961224698 | 0.850489120267299 | 0.395053206693788 | NA | NM_009400,NM_021985 |
| Tnfrsf26 | 1.803338207 | 0.0366130302135887 | 0.0293337291384692 | 1.24815464275809 | 0.211974430652063 | NA | NM_175649 |
| Tnfsf11 | 1.635285069 | 0.0162463035519696 | 0.0297768600183503 | 0.545601636369907 | 0.585339799285022 | NA | NM_011613 |
| Tnfsf13 | 18.44835723 | 0.0194239915257509 | 0.0774236255000248 | 0.250879384687878 | 0.801907363428195 | NA | NM_023517,NM_001159505 |
| Tnfsf14 | 0.555628243 | 0.002012195892949 | 0.0179195962833076 | 0.112290247008712 | 0.910593274655051 | NA | NM_019418 |
| Tnfsf15 | 1.180454581 | 0.00607713374144627 | 0.0220485787454874 | 0.275624738065715 | 0.782836307160581 | NA | NM_177371 |
| Tnfsf18 | 0.902588669 | -0.0162981466257828 | 0.02266293661602 | -0.719154225329383 | 0.472045899681462 | NA | NM_183391 |
| Tnfsf4 | 0.113843286 | 0.00525947738444006 | 0.00859368002043176 | 0.612016897526494 | 0.540526577873086 | NA | NM_009452 |
| Tnfsf8 | 3.625124784 | -0.12078917231047 | 0.0433046837660188 | -2.78928655761836 | 0.00528243029171323 | NA | NM_009403 |
| Tnfsf9 | 11.76975247 | -0.0160844048041395 | 0.0686022733799413 | -0.234458772452903 | 0.81462883603974 | NA | NM_009404 |
| Tnip3 | 2.50517225 | -0.00316215042489371 | 0.0358508238721197 | -0.0882030057711682 | 0.929715328370576 | NA | NM_001001495 |
| Tnk1 | 5.597180968 | -0.0597892149787799 | 0.0454326752325559 | -1.31599591423435 | 0.188175414014839 | NA | NM_031880 |
| Tnn | 18.93388326 | 0.0428131048027512 | 0.0797921477067648 | 0.536557869830611 | 0.591573052162629 | NA | NM_177839 |
| Tnnc2 | 6.708040353 | -0.00656808270083586 | 0.0534699269387918 | -0.122836949232313 | 0.902236214555377 | NA | NM_009394 |
| Tnni2 | 9.29915725 | -0.046089514623191 | 0.0574352682155996 | -0.802460161762994 | 0.422286824182781 | NA | NM_009405 |
| Tnni3 | 14.4201095 | -0.0329032541862876 | 0.0695632070143435 | -0.472997948175437 | 0.636214643034575 | NA | NM_009406 |
| Tnnt3 | 2.988541184 | 0.0203321546068686 | 0.0356140014455827 | 0.570903402638865 | 0.568065124309835 | NA | NM_001163669,NM_001163668,NM_001163670,NM_011620,NM_001163667,NM_001163666,NM_001163665,NM_001163664 |
| Tnp1 | 0 | NA | NA | NA | NA | NA | NM_009407 |
| Tnp2 | 0 | NA | NA | NA | NA | NA | NM_013694 |
| Tns4 | 20.98582626 | -0.129304848959411 | 0.0754632599657208 | -1.71348082521412 | 0.0866241180057848 | NA | NM_172564 |
| Tomm20l | 0 | NA | NA | NA | NA | NA | NM_029227 |
| Tomt | 3.476529954 | -0.0461625578163512 | 0.0411113981691283 | -1.12286518756776 | 0.26149475456363 | NA | NM_001081679 |
| Topaz1 | 0.900394332 | -0.00434642921001173 | 0.0205972264470803 | -0.211020120654538 | 0.832871570849724 | NA | NM_001199736 |
| Tpbpa | 0 | NA | NA | NA | NA | NA | NM_009411 |
| Tpbpb | 0 | NA | NA | NA | NA | NA | NM_026429 |
| Tph1 | 17.61168378 | -0.013073211892193 | 0.0736178328351876 | -0.177582134500763 | 0.859051156127076 | NA | NM_001136084,NM_009414,NM_001276372 |
| Tpo | 14.35344157 | -0.0252148266135632 | 0.0704093317856023 | -0.358117680911144 | 0.720255247912487 | NA | NM_009417 |
| Tppp2 | 0 | NA | NA | NA | NA | NA | NM_001128634 |
| Tprg | 0.12890128 | -0.0068640067066208 | 0.00870804083610955 | -0.788237771940376 | 0.430557639008166 | NA | NM_175165 |
| Tpsab1 | 0 | NA | NA | NA | NA | NA | NM_031187,NR_033521 |
| Tpsb2 | 0.767280798 | 0.00795016384280367 | 0.0179184750925473 | 0.443685291395714 | 0.657270131100974 | NA | NM_010781 |
| Tpsg1 | 0 | NA | NA | NA | NA | NA | NM_012034 |
| Tpte | 0.212297216 | 0.00681050119636477 | 0.00973439053551308 | 0.699633035218656 | 0.484156506270654 | NA | NM_199257 |
| Tram2 | 17.75232845 | -0.0580981526187101 | 0.0787682624011203 | -0.737583270821063 | 0.460767724408822 | NA | NM_177409 |
| Trap1a | 0.415500251 | 0.00773121617951768 | 0.0153252904708914 | 0.504474365050519 | 0.613928071896657 | NA | NM_011635 |
| Trat1 | 5.257152409 | -0.0139254516980722 | 0.0522143339353235 | -0.266697871035209 | 0.789701793235211 | NA | NM_198297 |
| Trcg1 | 0 | NA | NA | NA | NA | NA | NM_001014398 |
| Trdn | 11.93532316 | 0.0129863490308343 | 0.0655494327602022 | 0.198115353314222 | 0.842954812706667 | NA | NM_029726 |
| Treh | 1.629630927 | 0.00616532106675359 | 0.0273264362067706 | 0.225617457765167 | 0.821498961591281 | NA | NM_021481 |
| Trem1 | 0.309462921 | 0.00239536941478217 | 0.0109127344998192 | 0.219502216866162 | 0.826258853162846 | NA | NM_021406 |
| Trem3 | 1.486938282 | -0.00365340857309925 | 0.0297723449556297 | -0.122711482032873 | 0.902335571240771 | NA | NM_021407 |
| Treml1 | 4.460721264 | -0.0161031700864379 | 0.0436917676435252 | -0.36856302582723 | 0.71245345933798 | NA | NM_027763 |
| Treml4 | 2.573964946 | -0.034251947831922 | 0.0332785423507927 | -1.029250243922 | 0.303362097619675 | NA | NM_001163796,NM_001163795,NM_001033922,NM_172623 |
| Trex2 | 0 | NA | NA | NA | NA | NA | NM_011907 |
| Trim10 | 2.150013696 | 0.0476522817290277 | 0.0341412808799513 | 1.39573796005441 | 0.162793418765846 | NA | NM_011280 |
| Trim14 | 20.6128322 | 0.0434845366813352 | 0.0787048182536928 | 0.552501583081858 | 0.580604750094735 | NA | NM_029077 |
| Trim15 | 2.092455193 | 0.00312023219971213 | 0.0306223314413786 | 0.101894011750389 | 0.918840803155313 | NA | NM_001024134,NM_001177872 |
| Trim29 | 6.863682406 | 0.0305294514451679 | 0.0548601950320578 | 0.556495496002664 | 0.577872172591521 | NA | NM_023655 |
| Trim30b | 3.061838828 | -0.0631692119092405 | 0.0324416167796108 | -1.94716596088212 | 0.0515148411377185 | NA | NM_175648 |
| Trim30e-ps1 | 3.819995155 | 0.0263929124622511 | 0.0389713090010478 | 0.677239567743066 | 0.498253966715513 | NA | NR_033673 |
| Trim31 | 0.217532603 | 0.00684685248201087 | 0.00976000296730706 | 0.701521557416086 | 0.482977586094298 | NA | NM_146077 |
| Trim34b | 1.138745056 | -0.0150539147414276 | 0.0226603702761308 | -0.664327835687867 | 0.506480509066667 | NA | NM_001243916 |
| Trim38 | 1.904837943 | 0.000688206377388729 | 0.0305737611443293 | 0.022509706088823 | 0.982041369623354 | NA | NM_001029935 |
| Trim40 | 4.971032975 | 0.023786151736576 | 0.0496277025182691 | 0.479291817464646 | 0.631731042442549 | NA | NM_001033235,NM_001172168 |
| Trim42 | 0.338469525 | 0.00207548790044268 | 0.0127461301155165 | 0.162832787805616 | 0.870650091848121 | NA | NM_030219 |
| Trim43b | 1.552173921 | -0.00770735974481781 | 0.0300061507202258 | -0.256859329164891 | 0.797287368019325 | NA | NM_001170884 |
| Trim43c | 0.217374753 | -0.00904456656736547 | 0.00999660569474087 | -0.90476376117583 | 0.365590553873005 | NA | NM_001177858 |
| Trim44 | 16237.15466 | 0.0480817428698805 | 0.0548777918772072 | 0.876160305018587 | NA | NA | NM_020267 |
| Trim50 | 0.309861382 | -0.0124208717281794 | 0.0117059258046019 | -1.0610755557067 | 0.28865556572345 | NA | NM_178240 |
| Trim52 | 0 | NA | NA | NA | NA | NA | NR_073203,NM_198601 |
| Trim54 | 20.64769469 | 0.0917690084131082 | 0.0795913311523637 | 1.15300255799759 | 0.24890933456782 | NA | NM_021447 |
| Trim55 | 0.41620922 | -0.0157695646389328 | 0.0131796358585653 | -1.19650988905618 | 0.231497641828105 | NA | NM_001081281 |
| Trim58 | 3.416377328 | 0.0160158501877575 | 0.0373344518669027 | 0.428983134528238 | 0.667935498407379 | NA | NM_001039047 |
| Trim6 | 9.202660641 | -0.104673659123742 | 0.0633999718641821 | -1.6510048198125 | 0.0987375907306784 | NA | NM_001013616 |
| Trim60 | 0.433618577 | 0.00670518071861956 | 0.0139879821382268 | 0.479352965449921 | 0.631687548112578 | NA | NM_153097 |
| Trim61 | 0.642209104 | -0.00129876803483414 | 0.0154459766184788 | -0.0840845526905927 | 0.932989206313358 | NA | NM_153110,NM_001177551 |
| Trim63 | 4.982828573 | 0.0127022360579182 | 0.0518522380336848 | 0.244969870917943 | 0.806479760207378 | NA | NM_001039048 |
| Trim69 | 1.167652745 | 0.00765696984418116 | 0.0247775255604443 | 0.309028834437164 | 0.757299592905365 | NA | NM_080510 |
| Trim71 | 13.87673465 | 0.0128981982045148 | 0.0690519526419966 | 0.186789767863425 | 0.851825469784123 | NA | NM_001042503 |
| Trim75 | 0.226556888 | -0.00933175792380659 | 0.010153338242423 | -0.919082739193727 | 0.358052296835844 | NA | NM_001033429 |
| Triml1 | 0 | NA | NA | NA | NA | NA | NM_177742 |
| Triml2 | 0.561190942 | -0.0088950510062541 | 0.018070301943092 | -0.492246949401669 | 0.622544778159548 | NA | NM_001160412 |
| Trip13 | 22.18870218 | -0.103057784802671 | 0.0808763730780028 | -1.27426318565591 | 0.202570158838935 | NA | NM_027182 |
| Troap | 7.899980241 | -0.016137573800203 | 0.0532246151158075 | -0.303197566860568 | 0.761739303164264 | NA | NM_001162506,NM_030159 |
| Trp53tg5 | 0.30309771 | -0.00396074557973908 | 0.0109135518926259 | -0.362919938321391 | 0.716664692123243 | NA | NM_001271575 |
| Trpa1 | 11.44852968 | -0.0218820468429422 | 0.0640137240725846 | -0.341833679573623 | 0.732476062068109 | NA | NM_177781 |
| Trpd52l3 | 0 | NA | NA | NA | NA | NA | NM_025741 |
| Trpm1 | 9.884618947 | -0.0419372645386526 | 0.0600467582328401 | -0.698410135248847 | 0.484920739769082 | NA | NM_018752,NM_001039104 |
| Trpm5 | 0.95029838 | -0.017977732682905 | 0.0218415765000685 | -0.823096843895341 | 0.410452923450678 | NA | NM_020277 |
| Trpm8 | 4.905015686 | -0.00271600317138418 | 0.0461918902481921 | -0.0587982686309418 | 0.953112787508981 | NA | NM_134252 |
| Trpv1 | 5.72608507 | -0.00184190765051625 | 0.0494275059293839 | -0.037264830905038 | 0.970273846876846 | NA | NM_001001445 |
| Trpv3 | 18.14646162 | -0.186273546223275 | 0.0767068014178319 | -2.42838369975328 | 0.0151662884217426 | NA | NM_145099 |
| Trpv4 | 120.8252734 | -0.00926024789679106 | 0.0584808351397692 | -0.158346710929471 | NA | NA | NM_022017 |
| Trpv5 | 0.127687691 | 0.00542298404407807 | 0.00869724871942359 | 0.62352868349814 | 0.532937159177571 | NA | NM_001007572 |
| Try10 | 0.106347839 | -0.00653347234140708 | 0.00854389509723638 | -0.764694822098226 | 0.444453290197145 | NA | NM_001038996 |
| Try4 | 0 | NA | NA | NA | NA | NA | NM_011646 |
| Try5 | 0 | NA | NA | NA | NA | NA | NM_001003405 |
| Tsga13 | 5.079378828 | -0.00397208623179497 | 0.050049102411019 | -0.0793637855715162 | 0.936743272679162 | NA | NM_054073 |
| Tsga8 | 0 | NA | NA | NA | NA | NA | NM_021898 |
| Tshb | 1.921619034 | -0.0228011919393758 | 0.0294856447070141 | -0.773298063038513 | 0.439346001843284 | NA | NM_001165940,NM_009432,NM_001165939 |
| Tshr | 20.99114888 | 0.00846706000908423 | 0.0824638761160766 | 0.10267598866158 | 0.918220131177505 | NA | NM_001113404,NM_011648 |
| Tsix | 6.039149835 | -0.0289673691651911 | 0.0534119506992753 | -0.542338723561807 | 0.587585183956822 | NA | NR_002844 |
| Tslp | 4.162648845 | 0.030112507216972 | 0.0462659253894079 | 0.650857125703703 | 0.515138720058348 | NA | NR_033206,NM_021367 |
| Tsnaxip1 | 15.30696427 | -0.045605788486688 | 0.0603500098407417 | -0.755688169845168 | 0.449836178605013 | NA | NM_024445 |
| Tspan1 | 4.926805836 | 0.0411320586830536 | 0.047784146656416 | 0.860788808865978 | 0.389354369767105 | NA | NM_133681 |
| Tspan10 | 1.515838434 | -0.0253748913102298 | 0.0275545960938984 | -0.920895055901354 | 0.357105220123721 | NA | NM_145363 |
| Tspan32 | 7.120293943 | -0.0196940380068017 | 0.0559164904198096 | -0.352204472400591 | 0.724684923091369 | NA | NM_001128082,NM_001128080,NM_001128081,NM_020286 |
| Tspan8 | 11.48362848 | 0.0555376904519845 | 0.0621458525122649 | 0.893666885348846 | 0.371500148145921 | NA | NM_001168680,NM_001168679,NM_146010 |
| Tspo2 | 1.715561293 | -0.024327256304394 | 0.0304154855641694 | -0.799831265329278 | 0.423808565782495 | NA | NM_027292 |
| Tspy-ps | 0 | NA | NA | NA | NA | NA | NR_027507 |
| Tssk1 | 4.772582658 | -0.00671054344858671 | 0.0479433119506352 | -0.139968291208088 | 0.888685043670681 | NA | NM_009435 |
| Tssk2 | 0 | NA | NA | NA | NA | NA | NM_009436 |
| Tssk3 | 3.501972258 | 0.0316848475266036 | 0.0440425244547927 | 0.719414881840535 | 0.471885330160239 | NA | NM_080442 |
| Tssk4 | 13.40366221 | -0.0972574959920158 | 0.0652855875769084 | -1.48972383648142 | 0.136296864442576 | NA | NM_027673,NM_001253889,NM_001253888 |
| Tssk5 | 0.09572496 | 0.00498255289994511 | 0.00847999972823692 | 0.587565219295241 | 0.556824162160132 | NA | NM_183099 |
| Tssk6 | 1.812640886 | -0.0122140653744158 | 0.0336413276199039 | -0.363067281779609 | 0.716554624737259 | NA | NM_032004 |
| Tstd1 | 19.84431139 | -0.0498631429820654 | 0.0804116047787235 | -0.620098841694289 | 0.535192714731135 | NA | NM_001164525 |
| Tsx | 2.677119502 | 0.0185747861624972 | 0.0398332577619958 | 0.466313508010863 | 0.640991111252381 | NA | NM_009440 |
| Ttc23l | 0.120209049 | -0.00674972139751468 | 0.00863521114606269 | -0.781650996523957 | 0.434419707908266 | NA | NM_029430 |
| Ttc24 | 0.203513543 | -0.00860782669689374 | 0.00975338121443284 | -0.882547960306941 | 0.377480556050336 | NA | NM_172526 |
| Ttc29 | 12.7599653 | -0.0186521197746578 | 0.0652529651447188 | -0.285843252230622 | 0.774998179079763 | NA | NM_183096 |
| Ttc30a2 | 1.986946704 | 0.0447705996141338 | 0.0305794268213727 | 1.46407582704731 | 0.143173231231322 | NA | NM_001081228 |
| Ttc36 | 2.108329158 | 0.0321905020489225 | 0.0331484395773318 | 0.971101580025374 | 0.331497697085507 | NA | NM_138951 |
| Ttc39d | 3.452478373 | 0.0769524450058001 | 0.0396313051823742 | 1.94170857234407 | 0.0521723903579713 | NA | NM_026351 |
| Ttk | 12.28877061 | -0.0463320127169239 | 0.0647258506581511 | -0.715819293926718 | 0.474102937318638 | NA | NM_001110265,NM_009445 |
| Ttll10 | 4.728755573 | -0.0653081338504363 | 0.0500318213903249 | -1.30533192747337 | 0.191779846858345 | NA | NM_029264 |
| Ttll2 | 0.194331409 | -0.00840968081320881 | 0.00964081822647896 | -0.872299489073576 | 0.383045016303155 | NA | NM_001098267 |
| Ttll6 | 17.64200936 | -0.0320400313346652 | 0.0655834943073989 | -0.488538033433979 | 0.625168792814758 | NA | NM_172799 |
| Ttn | 20.25288244 | -0.0416182694713894 | 0.0805072996777367 | -0.516950259640846 | 0.605190881524188 | NA | NM_028004,NM_011652 |
| Ttr | 10626.60361 | 0.0118980928188254 | 0.0240149631814495 | 0.495444974407296 | NA | NA | NM_013697 |
| Tuba3a | 0.34789562 | 0.0020947884034485 | 0.0113191368529813 | 0.185066090343873 | 0.853177198132332 | NA | NM_009446 |
| Tuba3b | 0 | NA | NA | NA | NA | NA | NM_009449 |
| Tubal3 | 0.227686571 | 0.00691460391109036 | 0.00980755285755433 | 0.705028462402254 | 0.480792523671872 | NA | NM_001033879 |
| Tubb1 | 1.798813337 | 0.0312288771432793 | 0.0296850143493846 | 1.05200815386928 | 0.292795808660617 | NA | NM_001080971 |
| Tubb2a-ps2 | 0.524155415 | 0.000932839619984173 | 0.0162880999266465 | 0.0572712363127204 | 0.954329132830581 | NA | NR_003964 |
| Tulp1 | 2.148672343 | -0.0107192022145517 | 0.0327762650352939 | -0.327041601689794 | 0.743636420140702 | NA | NM_021478 |
| Tulp2 | 2.634622341 | 0.0185504112579618 | 0.0374347318671369 | 0.495540112957154 | 0.620218914200224 | NA | NM_001045555,NM_008807 |
| Twist1 | 13.81463304 | 0.0359661121908392 | 0.0727894862659508 | 0.494111361899573 | 0.62122753414407 | NA | NM_011658 |
| Txk | 6.024010165 | -0.0163096511763336 | 0.0559450935402289 | -0.291529607768118 | 0.770646303043731 | NA | NM_001122754,NM_013698 |
| Txndc2 | 3.130159782 | 0.0254794708855725 | 0.0387562393765752 | 0.657428875851475 | 0.510905189607507 | NA | NM_001146002,NM_153519 |
| Txndc8 | 0 | NA | NA | NA | NA | NA | NM_026132 |
| Tymp | 11.62648812 | -0.0161070010507815 | 0.0613604063829663 | -0.262498278617216 | 0.792937312308786 | NA | NM_138302 |
| Tyms-ps | 0.35231393 | 0.00236581429106576 | 0.0128934416322417 | 0.183489742967443 | 0.854413766077074 | NA | NR_000040 |
| Tyr | 3.876374708 | 0.00777427050869211 | 0.042367657380721 | 0.183495406385856 | 0.854409322758363 | NA | NM_011661 |
| Tyrp1 | 4.059861114 | 0.024808276108767 | 0.0486093792245073 | 0.510359862737342 | 0.609799370945708 | NA | NM_031202 |
| U05342 | 0 | NA | NA | NA | NA | NA | NR_024093 |
| U90926 | 0.308420637 | -0.00472633378880091 | 0.0109973596733179 | -0.42976986560402 | 0.667363055492239 | NA | NR_033483 |
| Uba1y | 0.212695677 | -0.00857796107890413 | 0.00973699866180651 | -0.880965621629515 | 0.378336427605205 | NA | NM_011667 |
| Ubald2 | 0 | NA | NA | NA | NA | NA | NM_176902 |
| Ubash3a | 2.263418031 | -0.0443874078030514 | 0.0275236671716147 | -1.612699627789 | 0.106809781462927 | NA | NM_177823 |
| Ubd | 0.291497113 | -0.00941849595073383 | 0.0102038556585987 | -0.923033044160808 | 0.355989985190414 | NA | NM_023137 |
| Ube2c | 9.170607398 | -0.0119602701583054 | 0.0570398335553379 | -0.20968276751197 | 0.833915276081893 | NA | NM_026785 |
| Ube2d2b | 0 | NA | NA | NA | NA | NA | NM_001276397,NR_003639 |
| Ube2dnl1 | 0 | NA | NA | NA | NA | NA | NM_001276396,NR_003645 |
| Ube2dnl2 | 0 | NA | NA | NA | NA | NA | NM_001081661 |
| Ube2u | 1.825540927 | 0.00105131594763288 | 0.0286405616752827 | 0.0367072391789082 | 0.970718436504657 | NA | NM_001033773 |
| Ubl4b | 0.697055124 | -0.0304562192195744 | 0.0182487681311098 | -1.66894658317533 | 0.0951279665118085 | NA | NM_026261 |
| Ubqln3 | 0.106347839 | -0.00653347234140708 | 0.00854389509723638 | -0.764694822098226 | 0.444453290197145 | NA | NM_198623 |
| Ubqlnl | 0 | NA | NA | NA | NA | NA | NM_198624 |
| Ubtfl1 | 0 | NA | NA | NA | NA | NA | NM_001033793 |

| | | | | | | | | | | | | | | | |
|---|---|---|---|---|---|---|---|---|---|---|---|---|---|---|---|
| Ttc39d | 2.53430735 | -0.142404698 | 0.107101953 | -1.329618124 | 0.183644124 | NA | | Uchl4 | 0 | NA | NA | NA | NA | NA | NM_033607 |
| Ttll10 | 3.342846972 | 0.231011973 | 0.122705588 | 1.882652425 | 0.059747487 | NA | | Ucma | 19.97908135 | -0.0218238724965006 | 0.0758199382306872 | -0.287838172989538 | 0.773470615510581 | NA | NM_026754,NM_001113558,NM_001165932 |
| Ttll2 | 0.834746954 | -0.043086269 | 0.051353324 | -0.839016171 | 0.401460235 | NA | | Ucn | 7.337988093 | -0.0150363382205176 | 0.0503470726348302 | -0.298653674059202 | 0.765204307348931 | NA | NM_021290 |
| Ttr | 3242.384781 | 0.264883478 | 0.09876165 | 2.682047932 | NA | NA | | Ucn2 | 1.463835896 | -0.0248363023229376 | 0.0237755932637078 | -1.04461335822265 | 0.296201703184377 | NA | NM_145077 |
| Tuba3a | 0.099009884 | 0.018205726 | 0.027896865 | 0.652608315 | 0.514008823 | NA | | Ucp1 | 2.143707167 | 0.00888028299620792 | 0.0302710853879935 | 0.293358592279949 | 0.769248072352563 | NA | NM_009463 |
| Tuba3b | 0 | NA | NA | NA | NA | NA | | Ugt1a1 | 0.408982522 | 0.0120846716222426 | 0.0129414896798793 | 0.933792934288797 | 0.350410714955054 | NA | NM_201645 |
| Tubal3 | 0 | NA | NA | NA | NA | NA | | Ugt1a10 | 0 | NA | NA | NA | NA | NA | NM_201641 |
| Tubb1 | 1.56694682 | -0.108455904 | 0.092576313 | -1.171529743 | 0.241385909 | NA | | Ugt1a2 | 0 | NA | NA | NA | NA | NA | NM_013701 |
| Tubb2a-ps2 | 0.280719541 | 0.042027364 | 0.040642707 | 1.034069012 | 0.301103902 | NA | | Ugt1a5 | 0 | NA | NA | NA | NA | NA | NM_201643 |
| Tulp1 | 1.443701084 | 0.038351081 | 0.076630828 | 0.500465438 | 0.616747387 | NA | | Ugt1a6b | 0 | NA | NA | NA | NA | NA | NM_201410 |
| Tulp2 | 2.623988091 | -0.046266979 | 0.11508258 | -0.402032863 | 0.687659841 | NA | | Ugt1a9 | 0 | NA | NA | NA | NA | NA | NM_201644 |
| Txk | 5.499631949 | 0.067900859 | 0.13909452 | 0.488163436 | 0.625434081 | NA | | Ugt2a1 | 1.387563536 | 0.00931204555241746 | 0.0208030568080035 | 0.447628713335766 | 0.654421177112573 | NA | NM_053184 |
| Txndc2 | 2.185929593 | -0.022596257 | 0.0896969 | -0.251917929 | 0.801104501 | NA | | Ugt2a2 | 0.108766301 | 0.00519171381361946 | 0.00856003371502575 | 0.606506234257728 | 0.544178643801544 | NA | NM_001024148 |
| Txndc8 | 0 | NA | NA | NA | NA | NA | | Ugt2a3 | 0 | NA | NA | NA | NA | NA | NM_028094 |
| Tyms-ps | 0.105923158 | 0.018524693 | 0.027977518 | 0.662127842 | 0.507889294 | NA | | Ugt2b1 | 0 | NA | NA | NA | NA | NA | NM_152811 |
| Tyr | 4.464869866 | -0.147637378 | 0.137195215 | -1.076111708 | 0.2818773 | NA | | Ugt2b34 | 0 | NA | NA | NA | NA | NA | NM_153598 |
| U05342 | 0 | NA | NA | NA | NA | NA | | Ugt2b35 | 0 | NA | NA | NA | NA | NA | NM_172881 |
| U90926 | 0.291330644 | 0.006248311 | 0.033473087 | 0.186666694 | 0.851921971 | NA | | Ugt2b36 | 0 | NA | NA | NA | NA | NA | NM_001029867 |
| Ubald2 | 0.305482598 | 0.007240588 | 0.03390939 | 0.21352751 | 0.830915534 | NA | | Ugt2b37 | 0 | NA | NA | NA | NA | NA | NM_053215 |
| Ubap1 | 1953.538303 | 0.116286704 | 0.102450721 | 1.13505013 | NA | NA | | Ugt2b38 | 0 | NA | NA | NA | NA | NA | NM_133894 |
| Ubash3a | 1.943408164 | 0.070357288 | 0.070260689 | 1.001374872 | 0.316645608 | NA | | Ugt2b5 | 0.778565143 | 0.000269352352265916 | 0.0196602487899377 | 0.01370035319206 | 0.989069041667805 | NA | NM_009467 |
| Ubb | 29757.92344 | -0.021607675 | 0.076180919 | -0.283636317 | NA | NA | | Ugt3a1 | 0.319043516 | -0.00957618023178583 | 0.0102890835921761 | -0.930712647632453 | 0.352002225550712 | NA | NM_207216 |
| Ubd | 0.239950962 | 0.027417169 | 0.032831728 | 0.835081494 | 0.403671825 | NA | | Ugt3a2 | 0 | NA | NA | NA | NA | NA | NM_144845 |
| Ube2d2b | 0.59705218 | -0.055901067 | 0.045635394 | -1.224949792 | 0.220594138 | NA | | Umod | 0 | NA | NA | NA | NA | NA | NM_009470 |
| Ube2dnl1 | 0 | NA | NA | NA | NA | NA | | Umodl1 | 16.17612813 | 0.102219662043129 | 0.0708233554203326 | 1.44330442177529 | 0.148934734728718 | NA | NM_177465 |
| Ube2dnl2 | 0 | NA | NA | NA | NA | NA | | Unc5cl | 8.797076338 | 0.095903334316356 | 0.0600708473001496 | 1.59650377224024 | 0.110376364230396 | NA | NM_152823 |
| Ube2u | 0.886445463 | 0.057284504 | 0.065158193 | 0.879160416 | 0.379314303 | NA | | Unc93a | 0.599001055 | -0.00559145445160231 | 0.0133839057705952 | -0.417774493293793 | 0.676112001468125 | NA | NM_199252 |
| Ubl4b | 0.963937922 | 0.019238703 | 0.06542135 | 0.294073771 | 0.768701533 | NA | | Uncx | 6.865153869 | 0.0267207450307415 | 0.0496430037490411 | 0.538258022536714 | 0.59039892740191 | NA | NM_013702 |
| Ubqln3 | 0.08731684 | 0.017365056 | 0.027733412 | 0.626142072 | 0.531221759 | NA | | Upk1a | 20.20476212 | 0.0250933117608768 | 0.0729085229095364 | 0.344175286502679 | 0.730714466759956 | NA | NM_026815 |
| Ubqlnl | 0.096071043 | -0.020098795 | 0.027825948 | -0.722304036 | 0.470107572 | NA | | Upk2 | 1.281566481 | 0.0219545951758633 | 0.026367978686843 | 0.832623366265772 | 0.405057176627485 | NA | NM_009476 |
| Ubtfl1 | 0 | NA | NA | NA | NA | NA | | Upk3a | 1.885251996 | -0.0156664939469468 | 0.0297309145347193 | -0.526942887298396 | 0.598233250165769 | NA | NM_023478 |
| Uchl4 | 0 | NA | NA | NA | NA | NA | | Upk3b | 0.313099713 | -0.00621682986300086 | 0.0125746085216642 | -0.494395499652349 | 0.621026891670966 | NA | NM_175309 |
| Ucn | 7.068471288 | -0.019200273 | 0.150350244 | -0.127703638 | 0.898383511 | NA | | Upk3bl | 4.960631809 | 0.0302636835262629 | 0.0528707192840197 | 0.572409150775638 | 0.567044817873516 | NA | NM_027158 |
| Ucn2 | 1.776156778 | 0.013873087 | 0.084654647 | 0.163878627 | 0.869826693 | NA | | Uroc1 | 0.326298904 | 0.00765111147689903 | 0.0103094497373689 | 0.742145475443356 | 0.457999199689495 | NA | NM_144940 |
| Ucp1 | 2.024471489 | -0.032467156 | 0.089660064 | -0.3621139 | 0.717266917 | NA | | Ush1c | 6.059260386 | -0.0259367703423533 | 0.0530917102582997 | -0.488527685700211 | 0.625176120385193 | NA | NM_153677,NM_023649,NM_001163733 |
| Ugt1a1 | 0.284856677 | 0.008904946 | 0.033416532 | 0.266483248 | 0.789867059 | NA | | Ush2a | 5.553128838 | 0.0675702258597732 | 0.0470493901443541 | 1.43615519037459 | 0.150958186456707 | NA | NM_021408 |
| Ugt1a10 | 0 | NA | NA | NA | NA | NA | | Usp17la | 4.114706727 | -0.0479931284556416 | 0.0430516939123065 | -1.11477909680861 | 0.264945096146632 | NA | NM_007887 |
| Ugt1a2 | 0 | NA | NA | NA | NA | NA | | Usp17lb | 0 | NA | NA | NA | NA | NA | NM_201409 |
| Ugt1a5 | 0 | NA | NA | NA | NA | NA | | Usp17lc | 0 | NA | NA | NA | NA | NA | NM_010089 |
| Ugt1a6a | 56.13477206 | 0.811262311 | 0.166727719 | 4.865791446 | NA | NA | | Usp17ld | 0.120209049 | -0.00674972139751468 | 0.00863521114606269 | -0.781650996523957 | 0.434419707908266 | NA | NM_001001559 |
| Ugt1a6b | 0 | NA | NA | NA | NA | NA | | Usp17le | 0 | NA | NA | NA | NA | NA | NM_001256973 |
| Ugt1a9 | 0 | NA | NA | NA | NA | NA | | Usp26 | 0.127687691 | 0.00542298404407807 | 0.00869724871942359 | 0.62352868349814 | 0.532937159177571 | NA | NM_031388 |
| Ugt2a1 | 1.36967311 | -0.06198634 | 0.069773716 | -0.888390979 | 0.374330476 | NA | | Usp44 | 0.481442367 | -0.00116566404370014 | 0.0141552163455238 | -0.0823487268047827 | 0.934369407578898 | NA | NM_001206851,NM_183199 |
| Ugt2a2 | 0 | NA | NA | NA | NA | NA | | Usp50 | 4.627395278 | 0.0358530903761424 | 0.0378202381634114 | 0.947986901119726 | 0.343136122664917 | NA | NM_029163 |
| Ugt2a3 | 0 | NA | NA | NA | NA | NA | | Usp9y | 0 | NA | NA | NA | NA | NA | NM_148943 |
| Ugt2b1 | 0 | NA | NA | NA | NA | NA | | Utf1 | 0.529654598 | -0.0168018233714002 | 0.0170496396091865 | -0.985465016066804 | 0.324395707372624 | NA | NM_009482 |
| Ugt2b34 | 0 | NA | NA | NA | NA | NA | | Uts2 | 0.563966486 | 0.00627263724169609 | 0.0131957962212362 | 0.47535117521756 | 0.634536688845777 | NA | NM_011910 |
| Ugt2b35 | 0 | NA | NA | NA | NA | NA | | Uts2d | 0.24789674 | -0.000769869626663926 | 0.0100368571301073 | -0.0767042528038554 | 0.938858821262798 | NA | NM_198166 |
| Ugt2b36 | 0 | NA | NA | NA | NA | NA | | Uts2r | 6.509905286 | -0.00465011940583881 | 0.0514670422801456 | -0.09035140159264 | 0.928007974579832 | NA | NM_145440 |
| Ugt2b37 | 0 | NA | NA | NA | NA | NA | | Uty | 0.108766301 | 0.00519171381361946 | 0.00856003371502575 | 0.606506234257728 | 0.544178643801544 | NA | NM_009484 |
| Ugt2b38 | 0 | NA | NA | NA | NA | NA | | V1ra8 | 0 | NA | NA | NA | NA | NA | NM_053223 |
| Ugt2b5 | 0.284916696 | 0.042355674 | 0.040798016 | 1.038179738 | 0.299186384 | NA | | V1rd18 | 0 | NA | NA | NA | NA | NA | NM_207618 |
| Ugt3a1 | 0.458657292 | 0.005116849 | 0.037753393 | 0.135533495 | 0.892190083 | NA | | V1rd19 | 0 | NA | NA | NA | NA | NA | NM_207619 |
| Ugt3a2 | 0 | NA | NA | NA | NA | NA | | Vaultrc5 | 1.009678759 | 0.0168372573583483 | 0.0203379837795214 | 0.827872494190008 | 0.407742713709324 | NA | NR_027885 |
| Umod | 0 | NA | NA | NA | NA | NA | | Vax1 | 16.70707024 | 0.0432022407011467 | 0.0640953221034428 | 0.674031103727399 | 0.500291539287419 | NA | NM_009501 |
| Unc5cl | 5.534235381 | -0.059142708 | 0.146730175 | -0.403071203 | 0.686895844 | NA | | Vax2 | 0 | NA | NA | NA | NA | NA | NM_011912 |
| Unc93a | 0.508233334 | 0.033209451 | 0.046178565 | 0.719152935 | 0.472046695 | NA | | Vax2os | 1.610696021 | 0.00426991393245931 | 0.0267395304892976 | 0.159685449008476 | 0.873128863815716 | NA | NR_002871,NR_075094,NR_002873,NR_075095 |
| Uncx | 4.103913735 | 0.121382362 | 0.126181977 | 0.961962753 | 0.336068314 | NA | | Veph1 | 5.046815941 | 0.0110459816230239 | 0.0463278963885687 | 0.238430459487676 | 0.811547245265336 | NA | NM_145820 |
| Upk2 | 0.481360729 | 0.009303061 | 0.05529186 | 0.168253724 | 0.866383678 | NA | | Vgll1 | 0 | NA | NA | NA | NA | NA | NM_133251 |
| Upk3a | 2.505005272 | -0.068714024 | 0.099994133 | -0.687180562 | 0.491968952 | NA | | Vgll2 | 4.274707362 | 0.0382689005359402 | 0.0478530423048653 | 0.799717190228664 | 0.423874670905537 | NA | NM_153786 |
| Upk3b | 0.178993538 | 0.024685672 | 0.031260828 | 0.789667872 | 0.429721759 | NA | | Vmn1r-ps103 | 0 | NA | NA | NA | NA | NA | NM_134211 |
| Upk3bl | 2.866502003 | 0.097853592 | 0.1187077 | 0.824323884 | 0.409755551 | NA | | Vmn1r-ps79 | 0 | NA | NA | NA | NA | NA | NR_030707 |
| Uroc1 | 0 | NA | NA | NA | NA | NA | | Vmn1r1 | 0 | NA | NA | NA | NA | NA | NM_001166728 |
| Ush1c | 6.951718787 | 0.176395112 | 0.126009543 | 1.399855189 | 0.161556687 | NA | | Vmn1r10 | 0 | NA | NA | NA | NA | NA | NM_053231 |
| Ush2a | 3.868210486 | 0.043616446 | 0.109289363 | 0.399091406 | 0.689825855 | NA | | Vmn1r100 | 0 | NA | NA | NA | NA | NA | NM_001166844 |
| Usp17la | 3.144878331 | 0.185500021 | 0.123682925 | 1.499802997 | 0.133665441 | NA | | Vmn1r101 | 0 | NA | NA | NA | NA | NA | NM_001166836 |
| Usp17lb | 0 | NA | NA | NA | NA | NA | | Vmn1r103 | 0 | NA | NA | NA | NA | NA | NM_001166737 |
| Usp17lc | 0 | NA | NA | NA | NA | NA | | Vmn1r104 | 0 | NA | NA | NA | NA | NA | NM_001166738 |
| Usp17ld | 0.099009884 | 0.018205726 | 0.027896865 | 0.652608315 | 0.514008823 | NA | | Vmn1r107 | 0 | NA | NA | NA | NA | NA | NM_001166759 |
| Usp17le | 0 | NA | NA | NA | NA | NA | | Vmn1r11 | 0 | NA | NA | NA | NA | NA | NM_053233 |
| Usp26 | 0 | NA | NA | NA | NA | NA | | Vmn1r111 | 0 | NA | NA | NA | NA | NA | NM_001166746 |
| Usp44 | 0.480966971 | -0.012302917 | 0.044293682 | -0.277757829 | 0.781198266 | NA | | Vmn1r112 | 0 | NA | NA | NA | NA | NA | NM_001166847 |
| Usp50 | 2.553787272 | 0.13238994 | 0.087912352 | 1.505931046 | 0.132084876 | NA | | Vmn1r113 | 0 | NA | NA | NA | NA | NA | NM_001166716 |
| Usp53 | 924.941957 | 0.237369371 | 0.166002668 | 1.429912988 | NA | NA | | Vmn1r114 | 0 | NA | NA | NA | NA | NA | NM_001166837 |
| Usp9y | 0 | NA | NA | NA | NA | NA | | Vmn1r115 | 0 | NA | NA | NA | NA | NA | NM_001166745 |
| Utf1 | 0.465691673 | 0.035179317 | 0.051468901 | 0.683506283 | 0.494286982 | NA | | Vmn1r116 | 0 | NA | NA | NA | NA | NA | NM_001166744 |
| Uts2 | 0.200315975 | 0.025370485 | 0.031664208 | 0.801235428 | 0.422995365 | NA | | Vmn1r117 | 0 | NA | NA | NA | NA | NA | NM_001166743 |
| Uts2d | 0.099009884 | 0.018205726 | 0.027896865 | 0.652608315 | 0.514008823 | NA | | Vmn1r118 | 0 | NA | NA | NA | NA | NA | NM_001166742 |
| Uts2r | 6.559290417 | 0.091557576 | 0.14883831 | 0.615147914 | 0.538457042 | NA | | Vmn1r119 | 0 | NA | NA | NA | NA | NA | NM_001166708 |
| V1ra8 | 0 | NA | NA | NA | NA | NA | | Vmn1r12 | 0 | NA | NA | NA | NA | NA | NM_001101579 |
| V1rd18 | 0 | NA | NA | NA | NA | NA | | Vmn1r120 | 0 | NA | NA | NA | NA | NA | NM_001166715 |
| V1rd19 | 0 | NA | NA | NA | NA | NA | | Vmn1r121 | 0 | NA | NA | NA | NA | NA | NM_001166741 |
| Vaultrc5 | 1.329536028 | -0.091579739 | 0.08414314 | -1.088380347 | 0.276427234 | NA | | Vmn1r122 | 0 | NA | NA | NA | NA | NA | NM_001166714 |
| Vax2 | 0 | NA | NA | NA | NA | NA | | Vmn1r123 | 0 | NA | NA | NA | NA | NA | NM_001166707 |
| Vax2os | 1.1654809 | 0.04742536 | 0.081762314 | 0.580039354 | 0.561888079 | NA | | Vmn1r124 | 0 | NA | NA | NA | NA | NA | NM_001166757 |
| Veph1 | 4.757637665 | -0.114894081 | 0.13399422 | -0.857455502 | 0.391193197 | NA | | Vmn1r125 | 0 | NA | NA | NA | NA | NA | NM_001166740 |
| Vgll1 | 0 | NA | NA | NA | NA | NA | | Vmn1r126 | 0 | NA | NA | NA | NA | NA | NM_001166838 |
| Vgll2 | 3.827524474 | 0.066210044 | 0.125171202 | 0.528955889 | 0.59683605 | NA | | Vmn1r127 | 0 | NA | NA | NA | NA | NA | NM_001166726 |
| Vmn1r-ps103 | 0 | NA | NA | NA | NA | NA | | Vmn1r128 | 0 | NA | NA | NA | NA | NA | NM_001166739 |
| Vmn1r-ps79 | 0 | NA | NA | NA | NA | NA | | Vmn1r129 | 0 | NA | NA | NA | NA | NA | NM_001166725 |
| Vmn1r1 | 0 | NA | NA | NA | NA | NA | | Vmn1r13 | 0 | NA | NA | NA | NA | NA | NM_053235 |
| Vmn1r10 | 0 | NA | NA | NA | NA | NA | | Vmn1r130 | 0 | NA | NA | NA | NA | NA | NM_001166848 |
| Vmn1r100 | 0 | NA | NA | NA | NA | NA | | Vmn1r132 | 0 | NA | NA | NA | NA | NA | NM_001122682 |
| Vmn1r101 | 0 | NA | NA | NA | NA | NA | | Vmn1r135 | 0 | NA | NA | NA | NA | NA | NM_001166747 |
| Vmn1r103 | 0 | NA | NA | NA | NA | NA | | Vmn1r137 | 0 | NA | NA | NA | NA | NA | NM_001166849 |
| Vmn1r104 | 0 | NA | NA | NA | NA | NA | | Vmn1r139 | 0 | NA | NA | NA | NA | NA | NM_001166748 |
| Vmn1r107 | 0 | NA | NA | NA | NA | NA | | Vmn1r14 | 0 | NA | NA | NA | NA | NA | NM_053237 |
| Vmn1r11 | 0 | NA | NA | NA | NA | NA | | Vmn1r142 | 0 | NA | NA | NA | NA | NA | NM_001166749 |
| Vmn1r111 | 0 | NA | NA | NA | NA | NA | | Vmn1r143 | 0 | NA | NA | NA | NA | NA | NM_001166750 |
| Vmn1r112 | 0 | NA | NA | NA | NA | NA | | Vmn1r148 | 0 | NA | NA | NA | NA | NA | NM_030736 |
| Vmn1r113 | 0 | NA | NA | NA | NA | NA | | Vmn1r149 | 0 | NA | NA | NA | NA | NA | NM_001166840 |
| Vmn1r114 | 0 | NA | NA | NA | NA | NA | | Vmn1r15 | 0.191449919 | 0.00665508079272897 | 0.00962405648474729 | 0.691504752001018 | 0.489248395458281 | NA | NM_053236 |
| Vmn1r115 | 0 | NA | NA | NA | NA | NA | | Vmn1r151 | 0 | NA | NA | NA | NA | NA | NM_001166712 |
| Vmn1r116 | 0 | NA | NA | NA | NA | NA | | Vmn1r152 | 0 | NA | NA | NA | NA | NA | NM_001166752 |
| Vmn1r117 | 0 | NA | NA | NA | NA | NA | | Vmn1r155 | 0 | NA | NA | NA | NA | NA | NM_001166753 |
| Vmn1r118 | 0 | NA | NA | NA | NA | NA | | Vmn1r157 | 0 | NA | NA | NA | NA | NA | NM_001166754 |
| Vmn1r119 | 0 | NA | NA | NA | NA | NA | | Vmn1r158 | 0 | NA | NA | NA | NA | NA | NM_001166841 |
| Vmn1r12 | 0 | NA | NA | NA | NA | NA | | Vmn1r159 | 0 | NA | NA | NA | NA | NA | NM_001166758 |
| Vmn1r120 | 0 | NA | NA | NA | NA | NA | | Vmn1r16 | 0 | NA | NA | NA | NA | NA | NM_134184 |
| Vmn1r121 | 0 | NA | NA | NA | NA | NA | | Vmn1r160 | 0 | NA | NA | NA | NA | NA | NM_001166724 |
| Vmn1r122 | 0 | NA | NA | NA | NA | NA | | Vmn1r163 | 0 | NA | NA | NA | NA | NA | NM_001166755 |
| Vmn1r123 | 0 | NA | NA | NA | NA | NA | | Vmn1r165 | 0 | NA | NA | NA | NA | NA | NM_001166850 |
| Vmn1r124 | 0 | NA | NA | NA | NA | NA | | Vmn1r167 | 0 | NA | NA | NA | NA | NA | NM_001101562 |
| Vmn1r125 | 0 | NA | NA | NA | NA | NA | | Vmn1r168 | 0 | NA | NA | NA | NA | NA | NM_001166842 |
| Vmn1r126 | 0 | NA | NA | NA | NA | NA | | Vmn1r169 | 0 | NA | NA | NA | NA | NA | NM_001166843 |
| Vmn1r127 | 0 | NA | NA | NA | NA | NA | | Vmn1r17 | 0 | NA | NA | NA | NA | NA | NM_134171 |
| Vmn1r128 | 0 | NA | NA | NA | NA | NA | | Vmn1r170 | 0 | NA | NA | NA | NA | NA | NM_001166722 |
| Vmn1r129 | 0 | NA | NA | NA | NA | NA | | Vmn1r171 | 0 | NA | NA | NA | NA | NA | NM_030737 |
| Vmn1r13 | 0 | NA | NA | NA | NA | NA | | Vmn1r172 | 0 | NA | NA | NA | NA | NA | NM_030735 |
| Vmn1r130 | 0 | NA | NA | NA | NA | NA | | Vmn1r173 | 0 | NA | NA | NA | NA | NA | NM_001166718 |
| Vmn1r132 | 0 | NA | NA | NA | NA | NA | | Vmn1r174 | 0 | NA | NA | NA | NA | NA | NM_207548 |
| Vmn1r135 | 0 | NA | NA | NA | NA | NA | | Vmn1r175 | 0 | NA | NA | NA | NA | NA | NM_001166727 |
| Vmn1r137 | 0 | NA | NA | NA | NA | NA | | Vmn1r176 | 0 | NA | NA | NA | NA | NA | NM_001166721 |
| Vmn1r139 | 0 | NA | NA | NA | NA | NA | | Vmn1r177 | 0 | NA | NA | NA | NA | NA | NM_206872 |
| Vmn1r14 | 0 | NA | NA | NA | NA | NA | | Vmn1r178 | 0 | NA | NA | NA | NA | NA | NM_206868 |
| Vmn1r142 | 0 | NA | NA | NA | NA | NA | | Vmn1r179 | 0 | NA | NA | NA | NA | NA | NM_207545 |
| Vmn1r143 | 0 | NA | NA | NA | NA | NA | | Vmn1r18 | 0 | NA | NA | NA | NA | NA | NM_134181 |
| Vmn1r148 | 0 | NA | NA | NA | NA | NA | | Vmn1r180 | 0 | NA | NA | NA | NA | NA | NM_206869 |
| Vmn1r149 | 0 | NA | NA | NA | NA | NA | | Vmn1r181 | 0 | NA | NA | NA | NA | NA | NM_207546 |
| Vmn1r15 | 0 | NA | NA | NA | NA | NA | | Vmn1r183 | 0 | NA | NA | NA | NA | NA | NM_203489 |
| Vmn1r151 | 0 | NA | NA | NA | NA | NA | | Vmn1r184 | 0 | NA | NA | NA | NA | NA | NM_001167540 |
| Vmn1r152 | 0 | NA | NA | NA | NA | NA | | Vmn1r185 | 0 | NA | NA | NA | NA | NA | NM_134231 |
| Vmn1r155 | 0 | NA | NA | NA | NA | NA | | Vmn1r186 | 0 | NA | NA | NA | NA | NA | NM_001167567 |
| Vmn1r157 | 0 | NA | NA | NA | NA | NA | | Vmn1r187 | 0 | NA | NA | NA | NA | NA | NM_001167568 |
| Vmn1r158 | 0 | NA | NA | NA | NA | NA | | Vmn1r188 | 0 | NA | NA | NA | NA | NA | NM_145850 |
| Vmn1r159 | 0 | NA | NA | NA | NA | NA | | Vmn1r189 | 0 | NA | NA | NA | NA | NA | NM_145844 |
| Vmn1r16 | 0 | NA | NA | NA | NA | NA | | Vmn1r19 | 0 | NA | NA | NA | NA | NA | NM_134182 |
| Vmn1r160 | 0 | NA | NA | NA | NA | NA | | Vmn1r191 | 0 | NA | NA | NA | NA | NA | NM_145849 |
| Vmn1r163 | 0 | NA | NA | NA | NA | NA | | Vmn1r192 | 0 | NA | NA | NA | NA | NA | NM_145845 |
| Vmn1r165 | 0 | NA | NA | NA | NA | NA | | Vmn1r193 | 0 | NA | NA | NA | NA | NA | NM_134225 |
| Vmn1r167 | 0 | NA | NA | NA | NA | NA | | Vmn1r194 | 0 | NA | NA | NA | NA | NA | NM_001080972 |
| Vmn1r168 | 0 | NA | NA | NA | NA | NA | | Vmn1r195 | 0 | NA | NA | NA | NA | NA | NM_134223 |
| Vmn1r169 | 0 | NA | NA | NA | NA | NA | | Vmn1r196 | 0 | NA | NA | NA | NA | NA | NM_001167541 |
| Vmn1r17 | 0 | NA | NA | NA | NA | NA | | Vmn1r197 | 0 | NA | NA | NA | NA | NA | NM_134244 |
| Vmn1r170 | 0 | NA | NA | NA | NA | NA | | Vmn1r198 | 0 | NA | NA | NA | NA | NA | NM_134220 |
| Vmn1r171 | 0 | NA | NA | NA | NA | NA | | Vmn1r199 | 0 | NA | NA | NA | NA | NA | NM_134213 |
| Vmn1r172 | 0 | NA | NA | NA | NA | NA | | Vmn1r2 | 0 | NA | NA | NA | NA | NA | NM_001167534 |
| Vmn1r173 | 0 | NA | NA | NA | NA | NA | | Vmn1r20 | 0 | NA | NA | NA | NA | NA | NM_001101533 |
| Vmn1r174 | 0 | NA | NA | NA | NA | NA | | Vmn1r200 | 0.770647357 | 0.0167659012500455 | 0.0189781296399724 | 0.883432749596808 | 0.377002502774113 | NA | NM_134212 |
| Vmn1r175 | 0 | NA | NA | NA | NA | NA | | Vmn1r201 | 0 | NA | NA | NA | NA | NA | NM_134221 |
| Vmn1r176 | 0 | NA | NA | NA | NA | NA | | Vmn1r202 | 0 | NA | NA | NA | NA | NA | NM_134224 |
| Vmn1r177 | 0 | NA | NA | NA | NA | NA | | Vmn1r203 | 0 | NA | NA | NA | NA | NA | NM_134236 |
| Vmn1r178 | 0 | NA | NA | NA | NA | NA | | Vmn1r204 | 0 | NA | NA | NA | NA | NA | NM_001045544 |
| Vmn1r179 | 0 | NA | NA | NA | NA | NA | | Vmn1r205 | 0 | NA | NA | NA | NA | NA | NM_134217 |
| Vmn1r18 | 0 | NA | NA | NA | NA | NA | | Vmn1r206 | 3.215882515 | 0.023567429936152 | 0.0367869539002837 | 0.640646409594958 | 0.521752439598556 | NA | NM_134216 |
| Vmn1r180 | 0 | NA | NA | NA | NA | NA | | Vmn1r207-ps | 0.212297216 | 0.00681050119636477 | 0.00973439053551308 | 0.699633035218656 | 0.484156506270654 | NA | NM_001166709 |
| Vmn1r181 | 0 | NA | NA | NA | NA | NA | | Vmn1r208 | 0.106148608 | 0.00515554063321497 | 0.00854200771766808 | 0.603551390213729 | 0.546141939141592 | NA | NM_134218 |
| Vmn1r183 | 0 | NA | NA | NA | NA | NA | | Vmn1r209 | 0 | NA | NA | NA | NA | NA | NM_001013787 |
| Vmn1r184 | 0 | NA | NA | NA | NA | NA | | Vmn1r21 | 0 | NA | NA | NA | NA | NA | NM_134183 |
| Vmn1r185 | 0 | NA | NA | NA | NA | NA | | Vmn1r210 | 0 | NA | NA | NA | NA | NA | NM_134235 |
| Vmn1r186 | 0 | NA | NA | NA | NA | NA | | Vmn1r211 | 0 | NA | NA | NA | NA | NA | NM_134243 |
| Vmn1r187 | 0 | NA | NA | NA | NA | NA | | Vmn1r212 | 0 | NA | NA | NA | NA | NA | NM_134241 |
| Vmn1r188 | 0 | NA | NA | NA | NA | NA | | Vmn1r213 | 0 | NA | NA | NA | NA | NA | NM_134215 |
| Vmn1r189 | 0 | NA | NA | NA | NA | NA | | Vmn1r214 | 0 | NA | NA | NA | NA | NA | NM_134214 |
| Vmn1r19 | 0 | NA | NA | NA | NA | NA | | Vmn1r215 | 0 | NA | NA | NA | NA | NA | NM_134219 |
| Vmn1r191 | 0 | NA | NA | NA | NA | NA | | Vmn1r216 | 0 | NA | NA | NA | NA | NA | NM_134245 |
| Vmn1r192 | 0 | NA | NA | NA | NA | NA | | Vmn1r217 | 0 | NA | NA | NA | NA | NA | NM_134239 |
| Vmn1r193 | 0 | NA | NA | NA | NA | NA | | Vmn1r218 | 0 | NA | NA | NA | NA | NA | NM_134222 |
| Vmn1r194 | 0 | NA | NA | NA | NA | NA | | Vmn1r219 | 0 | NA | NA | NA | NA | NA | NM_134238 |
| Vmn1r195 | 0 | NA | NA | NA | NA | NA | | Vmn1r22 | 0 | NA | NA | NA | NA | NA | NM_134178 |
| Vmn1r196 | 0 | NA | NA | NA | NA | NA | | Vmn1r220 | 0 | NA | NA | NA | NA | NA | NM_134237 |
| Vmn1r197 | 0 | NA | NA | NA | NA | NA | | Vmn1r221 | 0 | NA | NA | NA | NA | NA | NM_001167542 |
| Vmn1r198 | 0 | NA | NA | NA | NA | NA | | Vmn1r222 | 0.224853396 | -0.000161157395207779 | 0.00979162641959728 | -0.0164586952465051 | 0.986868454038988 | NA | NM_134240 |
| Vmn1r199 | 0 | NA | NA | NA | NA | NA | | Vmn1r223 | 0.097165704 | -0.00637828455464458 | 0.00848781691535724 | -0.75146349388194 | 0.452373763047372 | NA | NM_001083311 |
| Vmn1r2 | 0 | NA | NA | NA | NA | NA | | Vmn1r224 | 0 | NA | NA | NA | NA | NA | NM_001166735 |
| Vmn1r20 | 0 | NA | NA | NA | NA | NA | | Vmn1r225 | 0 | NA | NA | NA | NA | NA | NM_134194 |
| Vmn1r200 | 0.486850878 | 0.009782776 | 0.044755885 | 0.218580767 | 0.826976636 | NA | | Vmn1r226 | 0 | NA | NA | NA | NA | NA | NM_134191 |
| Vmn1r201 | 0 | NA | NA | NA | NA | NA | | Vmn1r227 | 0 | NA | NA | NA | NA | NA | NM_134195 |
| Vmn1r202 | 0 | NA | NA | NA | NA | NA | | Vmn1r228 | 0 | NA | NA | NA | NA | NA | NM_134192 |
| Vmn1r203 | 0.116696964 | -0.020477583 | 0.027954224 | -0.732539847 | 0.463839131 | NA | | Vmn1r229 | 0 | NA | NA | NA | NA | NA | NM_134190 |
| Vmn1r204 | 0 | NA | NA | NA | NA | NA | | Vmn1r23 | 0 | NA | NA | NA | NA | NA | NM_134179 |
| Vmn1r205 | 0 | NA | NA | NA | NA | NA | | Vmn1r230 | 0 | NA | NA | NA | NA | NA | NM_134197 |
| Vmn1r206 | 3.545874401 | -0.118193148 | 0.108800615 | -1.086327933 | 0.277333933 | NA | | Vmn1r231 | 0 | NA | NA | NA | NA | NA | NM_134196 |
| Vmn1r207-ps | 0.39257328 | -0.00221316 | 0.042111238 | -0.052555094 | 0.958086398 | NA | | Vmn1r232 | 0.120209049 | -0.00674972139751468 | 0.00863521114606269 | -0.781650996523957 | 0.434419707908266 | NA | NM_134193 |
| Vmn1r208 | 0 | NA | NA | NA | NA | NA | | Vmn1r233 | 0 | NA | NA | NA | NA | NA | NM_134202 |
| Vmn1r209 | 0 | NA | NA | NA | NA | NA | | Vmn1r234 | 0 | NA | NA | NA | NA | NA | NM_134198 |
| Vmn1r21 | 0 | NA | NA | NA | NA | NA | | Vmn1r235 | 0 | NA | NA | NA | NA | NA | NM_134199 |
| Vmn1r210 | 0 | NA | NA | NA | NA | NA | | Vmn1r236 | 0 | NA | NA | NA | NA | NA | NM_134201 |
| Vmn1r211 | 0.094392817 | 0.017895942 | 0.027818137 | 0.643319236 | 0.520016977 | NA | | Vmn1r237 | 0 | NA | NA | NA | NA | NA | NM_134200 |
| Vmn1r212 | 0 | NA | NA | NA | NA | NA | | Vmn1r238 | 0 | NA | NA | NA | NA | NA | NM_001167539 |
| Vmn1r213 | 0 | NA | NA | NA | NA | NA | | Vmn1r24 | 0 | NA | NA | NA | NA | NA | NM_134173 |
| Vmn1r214 | 0 | NA | NA | NA | NA | NA | | Vmn1r25 | 0 | NA | NA | NA | NA | NA | NM_053238 |
| Vmn1r215 | 0 | NA | NA | NA | NA | NA | | Vmn1r26 | 0 | NA | NA | NA | NA | NA | NM_134172 |
| Vmn1r216 | 0 | NA | NA | NA | NA | NA | | Vmn1r27 | 0 | NA | NA | NA | NA | NA | NM_134436 |
| Vmn1r217 | 0 | NA | NA | NA | NA | NA | | Vmn1r28 | 0 | NA | NA | NA | NA | NA | NM_134180 |
| Vmn1r218 | 0 | NA | NA | NA | NA | NA | | Vmn1r29 | 0 | NA | NA | NA | NA | NA | NM_053232 |
| Vmn1r219 | 0 | NA | NA | NA | NA | NA | | Vmn1r3 | 0 | NA | NA | NA | NA | NA | NM_001167535 |
| Vmn1r22 | 0 | NA | NA | NA | NA | NA | | Vmn1r30 | 0 | NA | NA | NA | NA | NA | NM_134177 |
| Vmn1r220 | 0 | NA | NA | NA | NA | NA | | Vmn1r31 | 0.191449919 | 0.00665508079272897 | 0.00962405648474729 | 0.691504752001018 | 0.489248395458281 | NA | NM_001166729 |
| Vmn1r221 | 0 | NA | NA | NA | NA | NA | | Vmn1r32 | 0 | NA | NA | NA | NA | NA | NM_134170 |
| Vmn1r222 | 0.295034005 | -0.017733924 | 0.039348288 | -0.45069111 | 0.652212191 | NA | | Vmn1r33 | 0 | NA | NA | NA | NA | NA | NM_134169 |
| Vmn1r223 | 0.079983654 | 0.016740032 | 0.027674178 | 0.604897175 | 0.54524732 | NA | | Vmn1r34 | 0 | NA | NA | NA | NA | NA | NM_001166719 |
| Vmn1r224 | 0 | NA | NA | NA | NA | NA | | Vmn1r35 | 0 | NA | NA | NA | NA | NA | NM_134167 |
| Vmn1r225 | 0 | NA | NA | NA | NA | NA | | Vmn1r36 | 0 | NA | NA | NA | NA | NA | NM_134166 |
| Vmn1r226 | 0 | NA | NA | NA | NA | NA | | Vmn1r37 | 0 | NA | NA | NA | NA | NA | NM_134165 |
| Vmn1r227 | 0 | NA | NA | NA | NA | NA | | Vmn1r38 | 0 | NA | NA | NA | NA | NA | NM_134168 |
| Vmn1r228 | 0 | NA | NA | NA | NA | NA | | Vmn1r39 | 0 | NA | NA | NA | NA | NA | NM_001166720 |
| Vmn1r229 | 0 | NA | NA | NA | NA | NA | | Vmn1r4 | 0.113843286 | 0.00525947738444006 | 0.00859368002043176 | 0.612016897526494 | 0.540526577873086 | NA | NM_134176 |
| Vmn1r23 | 0 | NA | NA | NA | NA | NA | | Vmn1r40 | 0 | NA | NA | NA | NA | NA | NM_053228 |
| Vmn1r230 | 0 | NA | NA | NA | NA | NA | | Vmn1r41 | 0 | NA | NA | NA | NA | NA | NM_053230 |
| Vmn1r231 | 0 | NA | NA | NA | NA | NA | | Vmn1r42 | 0 | NA | NA | NA | NA | NA | NM_053221 |
| Vmn1r232 | 0.099009884 | 0.018205726 | 0.027896865 | 0.652608315 | 0.514008823 | NA | | Vmn1r43 | 0 | NA | NA | NA | NA | NA | NM_053220 |
| Vmn1r233 | 0 | NA | NA | NA | NA | NA | | Vmn1r44 | 0 | NA | NA | NA | NA | NA | NM_053227 |
| Vmn1r234 | 0 | NA | NA | NA | NA | NA | | Vmn1r45 | 0 | NA | NA | NA | NA | NA | NM_011684 |
| Vmn1r235 | 0 | NA | NA | NA | NA | NA | | Vmn1r46 | 0 | NA | NA | NA | NA | NA | NM_053229 |
| Vmn1r236 | 0 | NA | NA | NA | NA | NA | | Vmn1r47 | 0 | NA | NA | NA | NA | NA | NM_053219 |
| Vmn1r237 | 0 | NA | NA | NA | NA | NA | | Vmn1r48 | 0 | NA | NA | NA | NA | NA | NM_053218 |
| Vmn1r238 | 0 | NA | NA | NA | NA | NA | | Vmn1r49 | 0 | NA | NA | NA | NA | NA | NM_011911 |
| Vmn1r24 | 0 | NA | NA | NA | NA | NA | | Vmn1r5 | 0.108766301 | 0.00519171381361946 | 0.00856003371502575 | 0.606506234257728 | 0.544178643801544 | NA | NM_134174 |
| Vmn1r25 | 0 | NA | NA | NA | NA | NA | | Vmn1r50 | 0 | NA | NA | NA | NA | NA | NM_053225 |
| Vmn1r26 | 0 | NA | NA | NA | NA | NA | | Vmn1r51 | 0 | NA | NA | NA | NA | NA | NM_011683 |
| Vmn1r27 | 0 | NA | NA | NA | NA | NA | | Vmn1r52 | 0 | NA | NA | NA | NA | NA | NM_053222 |
| Vmn1r28 | 0 | NA | NA | NA | NA | NA | | Vmn1r53 | 0.09572496 | 0.00498255289994511 | 0.00847999972823692 | 0.587565219295241 | 0.556824162160132 | NA | NM_053226 |
| Vmn1r29 | 0 | NA | NA | NA | NA | NA | | Vmn1r54 | 0 | NA | NA | NA | NA | NA | NM_053224 |
| Vmn1r3 | 0 | NA | NA | NA | NA | NA | | Vmn1r55 | 0 | NA | NA | NA | NA | NA | NM_001166706 |
| Vmn1r30 | 0 | NA | NA | NA | NA | NA | | Vmn1r56 | 0 | NA | NA | NA | NA | NA | NM_030740 |
| Vmn1r31 | 0 | NA | NA | NA | NA | NA | | Vmn1r57 | 0 | NA | NA | NA | NA | NA | NM_001166734 |
| Vmn1r32 | 0.28821313 | -0.029561117 | 0.033427731 | -0.884329147 | 0.376518559 | NA | | Vmn1r59 | 0 | NA | NA | NA | NA | NA | NM_207543 |
| Vmn1r33 | 0 | NA | NA | NA | NA | NA | | Vmn1r6 | 0 | NA | NA | NA | NA | NA | NM_134175 |
| Vmn1r34 | 0 | NA | NA | NA | NA | NA | | Vmn1r60 | 0 | NA | NA | NA | NA | NA | NM_001166732 |
| Vmn1r35 | 0 | NA | NA | NA | NA | NA | | Vmn1r61 | 0 | NA | NA | NA | NA | NA | NM_001166733 |
| Vmn1r36 | 0 | NA | NA | NA | NA | NA | | Vmn1r62 | 0 | NA | NA | NA | NA | NA | NM_030741 |
| Vmn1r37 | 0 | NA | NA | NA | NA | NA | | Vmn1r63 | 0 | NA | NA | NA | NA | NA | NM_030742 |
| Vmn1r38 | 0 | NA | NA | NA | NA | NA | | Vmn1r64 | 0 | NA | NA | NA | NA | NA | NM_207544 |
| Vmn1r39 | 0 | NA | NA | NA | NA | NA | | Vmn1r65 | 1.277796275 | -0.0039982139465171 | 0.0250734191361799 | -0.159460260477513 | 0.873306265198408 | NA | NM_030738 |

| | | | | | | | | | | | | | | | |
|---|---|---|---|---|---|---|---|---|---|---|---|---|---|---|---|
| Vmn1r4 | 0.107716603 | -0.020477583 | 0.027954224 | -0.732539849 | 0.46383913 | NA | | Vmn1r66 | 0 | NA | NA | NA | NA | NA | NM_134230 |
| Vmn1r40 | 0 | NA | NA | NA | NA | NA | | Vmn1r67 | 0 | NA | NA | NA | NA | NA | NM_134229 |
| Vmn1r41 | 0 | NA | NA | NA | NA | NA | | Vmn1r68 | 0 | NA | NA | NA | NA | NA | NM_001172072 |
| Vmn1r42 | 0 | NA | NA | NA | NA | NA | | Vmn1r69 | 0 | NA | NA | NA | NA | NA | NM_145842 |
| Vmn1r43 | 0 | NA | NA | NA | NA | NA | | Vmn1r7 | 0 | NA | NA | NA | NA | NA | NM_001166710 |
| Vmn1r44 | 0 | NA | NA | NA | NA | NA | | Vmn1r70 | 0 | NA | NA | NA | NA | NA | NM_134228 |
| Vmn1r45 | 0 | NA | NA | NA | NA | NA | | Vmn1r71 | 0 | NA | NA | NA | NA | NA | NM_145848 |
| Vmn1r46 | 0 | NA | NA | NA | NA | NA | | Vmn1r72 | 0 | NA | NA | NA | NA | NA | NM_145843 |
| Vmn1r47 | 0 | NA | NA | NA | NA | NA | | Vmn1r73 | 0 | NA | NA | NA | NA | NA | NM_134203 |
| Vmn1r48 | 0 | NA | NA | NA | NA | NA | | Vmn1r74 | 0 | NA | NA | NA | NA | NA | NM_134206 |
| Vmn1r49 | 0 | NA | NA | NA | NA | NA | | Vmn1r75 | 0 | NA | NA | NA | NA | NA | NM_134207 |
| Vmn1r5 | 0 | NA | NA | NA | NA | NA | | Vmn1r76 | 0 | NA | NA | NA | NA | NA | NM_134205 |
| Vmn1r50 | 0 | NA | NA | NA | NA | NA | | Vmn1r77 | 0 | NA | NA | NA | NA | NA | NM_001166731 |
| Vmn1r51 | 0.098353386 | -0.020212831 | 0.02786466 | -0.725393061 | 0.468210929 | NA | | Vmn1r78 | 0 | NA | NA | NA | NA | NA | NM_134208 |
| Vmn1r52 | 0 | NA | NA | NA | NA | NA | | Vmn1r79 | 0 | NA | NA | NA | NA | NA | NM_001166835 |
| Vmn1r53 | 0 | NA | NA | NA | NA | NA | | Vmn1r8 | 0 | NA | NA | NA | NA | NA | NM_134187 |
| Vmn1r54 | 0 | NA | NA | NA | NA | NA | | Vmn1r80 | 0 | NA | NA | NA | NA | NA | NM_134204 |
| Vmn1r55 | 0 | NA | NA | NA | NA | NA | | Vmn1r81 | 0 | NA | NA | NA | NA | NA | NM_134210 |
| Vmn1r56 | 0 | NA | NA | NA | NA | NA | | Vmn1r82 | 0 | NA | NA | NA | NA | NA | NM_134234 |
| Vmn1r57 | 0 | NA | NA | NA | NA | NA | | Vmn1r83 | 0 | NA | NA | NA | NA | NA | NM_134209 |
| Vmn1r59 | 0 | NA | NA | NA | NA | NA | | Vmn1r84 | 0 | NA | NA | NA | NA | NA | NM_134233 |
| Vmn1r6 | 0 | NA | NA | NA | NA | NA | | Vmn1r85 | 0 | NA | NA | NA | NA | NA | NM_145847 |
| Vmn1r60 | 0 | NA | NA | NA | NA | NA | | Vmn1r86 | 0 | NA | NA | NA | NA | NA | NM_001167536 |
| Vmn1r61 | 0 | NA | NA | NA | NA | NA | | Vmn1r87 | 0 | NA | NA | NA | NA | NA | NM_134227 |
| Vmn1r62 | 0 | NA | NA | NA | NA | NA | | Vmn1r88 | 0 | NA | NA | NA | NA | NA | NM_001167537 |
| Vmn1r63 | 0 | NA | NA | NA | NA | NA | | Vmn1r89 | 0 | NA | NA | NA | NA | NA | NM_134226 |
| Vmn1r64 | 0 | NA | NA | NA | NA | NA | | Vmn1r9 | 0 | NA | NA | NA | NA | NA | NM_134185 |
| Vmn1r65 | 0.927369673 | 0.009575659 | 0.064866054 | 0.147622033 | 0.882641066 | NA | | Vmn1r90 | 0 | NA | NA | NA | NA | NA | NM_001244031 |
| Vmn1r66 | 0 | NA | NA | NA | NA | NA | | Vmn1r91 | 0 | NA | NA | NA | NA | NA | NM_001166736 |
| Vmn1r67 | 0 | NA | NA | NA | NA | NA | | Vmn1r93 | 0 | NA | NA | NA | NA | NA | NM_207547 |
| Vmn1r68 | 0 | NA | NA | NA | NA | NA | | Vmn1r94 | 0 | NA | NA | NA | NA | NA | NM_001166723 |
| Vmn1r69 | 0 | NA | NA | NA | NA | NA | | Vmn1r95 | 0 | NA | NA | NA | NA | NA | NM_001167538 |
| Vmn1r7 | 0 | NA | NA | NA | NA | NA | | Vmn2r-ps11 | 0.738960096 | -0.00495053255431637 | 0.0176884420139608 | -0.279873860592646 | 0.779574282337881 | NA | NR_003962 |
| Vmn1r70 | 0 | NA | NA | NA | NA | NA | | Vmn2r-ps129 | 14.36537776 | -0.0596866605524025 | 0.0718504888974699 | -0.830705628310996 | 0.406139946034753 | NA | NR_033648 |
| Vmn1r71 | 0 | NA | NA | NA | NA | NA | | Vmn2r-ps54 | 1.634148256 | 0.0279501968368006 | 0.0297657431523035 | 0.939005510253406 | 0.347727915246104 | NA | NR_004441 |
| Vmn1r72 | 0 | NA | NA | NA | NA | NA | | Vmn2r-ps60 | 0 | NA | NA | NA | NA | NA | NR_028441 |
| Vmn1r73 | 0 | NA | NA | NA | NA | NA | | Vmn2r1 | 17.31394169 | -0.0932750180859636 | 0.0755894339640895 | -1.23396899797234 | 0.217214456749086 | NA | NM_019918 |
| Vmn1r74 | 0 | NA | NA | NA | NA | NA | | Vmn2r10 | 0 | NA | NA | NA | NA | NA | NM_009491 |
| Vmn1r75 | 0 | NA | NA | NA | NA | NA | | Vmn2r100 | 0 | NA | NA | NA | NA | NA | NM_001104562 |
| Vmn1r76 | 0 | NA | NA | NA | NA | NA | | Vmn2r101 | 0 | NA | NA | NA | NA | NA | NM_001104563 |
| Vmn1r77 | 0 | NA | NA | NA | NA | NA | | Vmn2r102 | 0 | NA | NA | NA | NA | NA | NM_001104564 |
| Vmn1r78 | 0.116696964 | -0.020477583 | 0.027954224 | -0.732539847 | 0.463839131 | NA | | Vmn2r103 | 0 | NA | NA | NA | NA | NA | NM_001104565 |
| Vmn1r79 | 0 | NA | NA | NA | NA | NA | | Vmn2r104 | 0 | NA | NA | NA | NA | NA | NM_001104566 |
| Vmn1r8 | 0 | NA | NA | NA | NA | NA | | Vmn2r105 | 0 | NA | NA | NA | NA | NA | NM_001104567 |
| Vmn1r80 | 0 | NA | NA | NA | NA | NA | | Vmn2r106 | 0 | NA | NA | NA | NA | NA | NM_001104568 |
| Vmn1r81 | 0 | NA | NA | NA | NA | NA | | Vmn2r107 | 0 | NA | NA | NA | NA | NA | NM_001104569 |
| Vmn1r82 | 0 | NA | NA | NA | NA | NA | | Vmn2r108 | 0 | NA | NA | NA | NA | NA | NM_001104570 |
| Vmn1r83 | 0 | NA | NA | NA | NA | NA | | Vmn2r109 | 0.120209049 | -0.00674972139751468 | 0.00863521114606269 | -0.781650996523957 | 0.434419707908266 | NA | NM_001104571 |
| Vmn1r84 | 0 | NA | NA | NA | NA | NA | | Vmn2r11 | 0 | NA | NA | NA | NA | NA | NM_001104622 |
| Vmn1r85 | 0 | NA | NA | NA | NA | NA | | Vmn2r110 | 0 | NA | NA | NA | NA | NA | NM_001104572 |
| Vmn1r86 | 0 | NA | NA | NA | NA | NA | | Vmn2r111 | 0.113843286 | 0.00525947738444006 | 0.00859368002043176 | 0.612016897526494 | 0.540526577873086 | NA | NM_001104573 |
| Vmn1r87 | 0 | NA | NA | NA | NA | NA | | Vmn2r112 | 0 | NA | NA | NA | NA | NA | NM_001104575 |
| Vmn1r88 | 0 | NA | NA | NA | NA | NA | | Vmn2r113 | 0 | NA | NA | NA | NA | NA | NM_001104578 |
| Vmn1r89 | 0 | NA | NA | NA | NA | NA | | Vmn2r114 | 0 | NA | NA | NA | NA | NA | NM_001102584 |
| Vmn1r9 | 0 | NA | NA | NA | NA | NA | | Vmn2r115 | 0 | NA | NA | NA | NA | NA | NM_001104579 |
| Vmn1r90 | 0 | NA | NA | NA | NA | NA | | Vmn2r116 | 0 | NA | NA | NA | NA | NA | NM_001104580 |
| Vmn1r91 | 0 | NA | NA | NA | NA | NA | | Vmn2r117 | 0 | NA | NA | NA | NA | NA | NM_001104581 |
| Vmn1r93 | 0 | NA | NA | NA | NA | NA | | Vmn2r118 | 0 | NA | NA | NA | NA | NA | NM_001104582 |
| Vmn1r94 | 0 | NA | NA | NA | NA | NA | | Vmn2r12 | 0 | NA | NA | NA | NA | NA | NM_001104623 |
| Vmn1r95 | 0 | NA | NA | NA | NA | NA | | Vmn2r120 | 0 | NA | NA | NA | NA | NA | NM_001104591 |
| Vmn2r-ps11 | 0.864449938 | -0.020465133 | 0.056685193 | -0.361031362 | 0.718076002 | NA | | Vmn2r121 | 0 | NA | NA | NA | NA | NA | NM_001100616 |
| Vmn2r-ps54 | 1.387477171 | -0.021122606 | 0.087481186 | -0.241453125 | 0.809203943 | NA | | Vmn2r124 | 0 | NA | NA | NA | NA | NA | NM_001271883 |
| Vmn2r-ps60 | 0 | NA | NA | NA | NA | NA | | Vmn2r13 | 0.406628625 | -0.000921993720139942 | 0.0119199762705297 | -0.0773486204347094 | 0.938346213215504 | NA | NM_001104624 |
| Vmn2r10 | 0 | NA | NA | NA | NA | NA | | Vmn2r14 | 0 | NA | NA | NA | NA | NA | NM_001104625 |
| Vmn2r100 | 0 | NA | NA | NA | NA | NA | | Vmn2r15 | 0.191449919 | 0.00665508079272897 | 0.00962405648474729 | 0.691504752001018 | 0.489248395458281 | NA | NM_001104626 |
| Vmn2r101 | 0 | NA | NA | NA | NA | NA | | Vmn2r16 | 0 | NA | NA | NA | NA | NA | NM_001104627 |
| Vmn2r102 | 0 | NA | NA | NA | NA | NA | | Vmn2r17 | 0 | NA | NA | NA | NA | NA | NM_001104628 |
| Vmn2r103 | 0 | NA | NA | NA | NA | NA | | Vmn2r18 | 0.337741652 | 0.00190180888516019 | 0.0112128429016166 | 0.169609875198198 | 0.865316955796633 | NA | NM_001102582 |
| Vmn2r104 | 0 | NA | NA | NA | NA | NA | | Vmn2r19 | 0 | NA | NA | NA | NA | NA | NM_001104632 |
| Vmn2r105 | 0 | NA | NA | NA | NA | NA | | Vmn2r2 | 0.106347839 | -0.00653347234140708 | 0.00854389509723638 | -0.764694822098226 | 0.444453290197145 | NA | NM_001104592 |
| Vmn2r106 | 0 | NA | NA | NA | NA | NA | | Vmn2r20 | 0 | NA | NA | NA | NA | NA | NM_001104634 |
| Vmn2r107 | 0 | NA | NA | NA | NA | NA | | Vmn2r21 | 0 | NA | NA | NA | NA | NA | NM_001104635 |
| Vmn2r108 | 0 | NA | NA | NA | NA | NA | | Vmn2r22 | 0 | NA | NA | NA | NA | NA | NM_001104637 |
| Vmn2r109 | 0.099009884 | 0.018205726 | 0.027896865 | 0.652608315 | 0.514008823 | NA | | Vmn2r23 | 0 | NA | NA | NA | NA | NA | NM_001104638 |
| Vmn2r11 | 0 | NA | NA | NA | NA | NA | | Vmn2r24 | 0 | NA | NA | NA | NA | NA | NM_001104639 |
| Vmn2r110 | 0 | NA | NA | NA | NA | NA | | Vmn2r25 | 0 | NA | NA | NA | NA | NA | NM_001104641 |
| Vmn2r111 | 0 | NA | NA | NA | NA | NA | | Vmn2r26 | 0 | NA | NA | NA | NA | NA | NM_019917 |
| Vmn2r112 | 0 | NA | NA | NA | NA | NA | | Vmn2r27 | 1.293232342 | 0.00435840269573583 | 0.0266810047860555 | 0.163352269927019 | 0.870241081095853 | NA | NM_001104642 |
| Vmn2r113 | 0 | NA | NA | NA | NA | NA | | Vmn2r28 | 1.074723478 | 0.00201427539600421 | 0.0237855026075909 | 0.084685004527143 | 0.932511817791907 | NA | NM_001081405 |
| Vmn2r114 | 0 | NA | NA | NA | NA | NA | | Vmn2r3 | 0 | NA | NA | NA | NA | NA | NM_001104614 |
| Vmn2r115 | 0 | NA | NA | NA | NA | NA | | Vmn2r30 | 0.291497113 | -0.00941849595073383 | 0.0102038556585987 | -0.923033044160808 | 0.355989985190414 | NA | NM_009490 |
| Vmn2r116 | 0 | NA | NA | NA | NA | NA | | Vmn2r31 | 0 | NA | NA | NA | NA | NA | NM_001105062 |
| Vmn2r117 | 0 | NA | NA | NA | NA | NA | | Vmn2r32 | 0 | NA | NA | NA | NA | NA | NM_001105063 |
| Vmn2r118 | 0.116696964 | -0.020477583 | 0.027954224 | -0.732539847 | 0.463839131 | NA | | Vmn2r33 | 0 | NA | NA | NA | NA | NA | NM_001105065 |
| Vmn2r12 | 0 | NA | NA | NA | NA | NA | | Vmn2r34 | 0 | NA | NA | NA | NA | NA | NM_001105066 |
| Vmn2r120 | 0 | NA | NA | NA | NA | NA | | Vmn2r35 | 0 | NA | NA | NA | NA | NA | NM_001105067 |
| Vmn2r121 | 0 | NA | NA | NA | NA | NA | | Vmn2r36 | 0 | NA | NA | NA | NA | NA | NM_001105068 |
| Vmn2r124 | 0 | NA | NA | NA | NA | NA | | Vmn2r37 | 0.120209049 | -0.00674972139751468 | 0.00863521114606269 | -0.781650996523957 | 0.434419707908266 | NA | NM_009489 |
| Vmn2r13 | 0.159967308 | 0.0240345 | 0.030870969 | 0.778547006 | 0.436246606 | NA | | Vmn2r38 | 0 | NA | NA | NA | NA | NA | NM_001105070 |
| Vmn2r14 | 0 | NA | NA | NA | NA | NA | | Vmn2r39 | 0 | NA | NA | NA | NA | NA | NM_001105071 |
| Vmn2r15 | 0.107716603 | -0.020477583 | 0.027954224 | -0.732539849 | 0.46383913 | NA | | Vmn2r4 | 0 | NA | NA | NA | NA | NA | NM_001104615 |
| Vmn2r16 | 0 | NA | NA | NA | NA | NA | | Vmn2r40 | 0 | NA | NA | NA | NA | NA | NM_001105072 |
| Vmn2r17 | 0 | NA | NA | NA | NA | NA | | Vmn2r41 | 0 | NA | NA | NA | NA | NA | NM_001105073 |
| Vmn2r18 | 0.287795519 | 0.031925978 | 0.035387605 | 0.902179657 | 0.366961439 | NA | | Vmn2r42 | 0 | NA | NA | NA | NA | NA | NM_009493 |
| Vmn2r19 | 0 | NA | NA | NA | NA | NA | | Vmn2r43 | 0 | NA | NA | NA | NA | NA | NM_198961 |
| Vmn2r2 | 0.279458926 | -0.011015785 | 0.033355472 | -0.330254219 | 0.741207882 | NA | | Vmn2r44 | 0 | NA | NA | NA | NA | NA | NM_001105074 |
| Vmn2r20 | 0 | NA | NA | NA | NA | NA | | Vmn2r45 | 0 | NA | NA | NA | NA | NA | NM_001105075 |
| Vmn2r21 | 0 | NA | NA | NA | NA | NA | | Vmn2r46 | 0 | NA | NA | NA | NA | NA | NM_001105076 |
| Vmn2r22 | 0 | NA | NA | NA | NA | NA | | Vmn2r47 | 0 | NA | NA | NA | NA | NA | NM_001105151 |
| Vmn2r23 | 0 | NA | NA | NA | NA | NA | | Vmn2r48 | 0 | NA | NA | NA | NA | NA | NM_001105152 |
| Vmn2r24 | 0 | NA | NA | NA | NA | NA | | Vmn2r49 | 0 | NA | NA | NA | NA | NA | NM_001105156 |
| Vmn2r25 | 0 | NA | NA | NA | NA | NA | | Vmn2r5 | 0.641969186 | 0.00403054676387996 | 0.0152746337548859 | 0.263871908718642 | 0.791878622279925 | NA | NM_001104618 |
| Vmn2r26 | 0 | NA | NA | NA | NA | NA | | Vmn2r50 | 0 | NA | NA | NA | NA | NA | NM_001105178 |
| Vmn2r27 | 1.71778417 | -0.096222303 | 0.083292515 | -1.155233491 | 0.247994823 | NA | | Vmn2r51 | 0.106347839 | -0.00653347234140708 | 0.00854389509723638 | -0.764694822098226 | 0.444453290197145 | NA | NM_001105179 |
| Vmn2r28 | 1.080958715 | 0.007275374 | 0.079606288 | 0.091391953 | 0.927181156 | NA | | Vmn2r52 | 0 | NA | NA | NA | NA | NA | NM_001105191 |
| Vmn2r3 | 0 | NA | NA | NA | NA | NA | | Vmn2r53 | 0 | NA | NA | NA | NA | NA | NM_001104644 |
| Vmn2r30 | 0.239950962 | 0.027417169 | 0.032831728 | 0.835081494 | 0.403671825 | NA | | Vmn2r54 | 0.826762817 | -0.000274930933430262 | 0.0201115482302361 | -0.0136703017730343 | 0.989093016985822 | NA | NM_001081449 |
| Vmn2r31 | 0 | NA | NA | NA | NA | NA | | Vmn2r55 | 0.538596121 | 0.0147687205475919 | 0.0142918540542707 | 1.0333663142312 | 0.301432504409536 | NA | NM_001104645 |
| Vmn2r32 | 0 | NA | NA | NA | NA | NA | | Vmn2r56 | 0 | NA | NA | NA | NA | NA | NM_001104648 |
| Vmn2r33 | 0 | NA | NA | NA | NA | NA | | Vmn2r58 | 0 | NA | NA | NA | NA | NA | NM_001105055 |
| Vmn2r34 | 0 | NA | NA | NA | NA | NA | | Vmn2r59 | 0 | NA | NA | NA | NA | NA | NM_001105056 |
| Vmn2r35 | 0 | NA | NA | NA | NA | NA | | Vmn2r6 | 0 | NA | NA | NA | NA | NA | NM_001104619 |
| Vmn2r36 | 0 | NA | NA | NA | NA | NA | | Vmn2r60 | 0 | NA | NA | NA | NA | NA | NM_001105057 |
| Vmn2r37 | 0.099009884 | 0.018205726 | 0.027896865 | 0.652608315 | 0.514008823 | NA | | Vmn2r61 | 0 | NA | NA | NA | NA | NA | NM_001105058 |
| Vmn2r38 | 0 | NA | NA | NA | NA | NA | | Vmn2r62 | 0 | NA | NA | NA | NA | NA | NM_001105059 |
| Vmn2r39 | 0 | NA | NA | NA | NA | NA | | Vmn2r63 | 0 | NA | NA | NA | NA | NA | NM_001105060 |
| Vmn2r4 | 0 | NA | NA | NA | NA | NA | | Vmn2r65 | 0 | NA | NA | NA | NA | NA | NM_001105180 |
| Vmn2r40 | 0 | NA | NA | NA | NA | NA | | Vmn2r66 | 0 | NA | NA | NA | NA | NA | NM_001033878 |
| Vmn2r41 | 0 | NA | NA | NA | NA | NA | | Vmn2r67 | 0 | NA | NA | NA | NA | NA | NM_001102579 |
| Vmn2r42 | 0 | NA | NA | NA | NA | NA | | Vmn2r68 | 0 | NA | NA | NA | NA | NA | NM_001105181 |
| Vmn2r43 | 0 | NA | NA | NA | NA | NA | | Vmn2r69 | 0 | NA | NA | NA | NA | NA | NM_001105182 |
| Vmn2r44 | 0 | NA | NA | NA | NA | NA | | Vmn2r7 | 0 | NA | NA | NA | NA | NA | NM_175674 |
| Vmn2r45 | 0 | NA | NA | NA | NA | NA | | Vmn2r70 | 0 | NA | NA | NA | NA | NA | NM_001105183 |
| Vmn2r46 | 0 | NA | NA | NA | NA | NA | | Vmn2r71 | 0 | NA | NA | NA | NA | NA | NM_001105184 |
| Vmn2r47 | 0 | NA | NA | NA | NA | NA | | Vmn2r72 | 0 | NA | NA | NA | NA | NA | NM_001105185 |
| Vmn2r48 | 0 | NA | NA | NA | NA | NA | | Vmn2r73 | 0.106148608 | 0.00515554063321497 | 0.00854200771766808 | 0.603551390213729 | 0.546141939141592 | NA | NM_001105186 |
| Vmn2r49 | 0 | NA | NA | NA | NA | NA | | Vmn2r74 | 0 | NA | NA | NA | NA | NA | NM_001105187 |
| Vmn2r5 | 0.492397275 | -0.018455921 | 0.05075977 | -0.363593463 | 0.716161609 | NA | | Vmn2r75 | 0 | NA | NA | NA | NA | NA | NM_001102578 |
| Vmn2r50 | 0 | NA | NA | NA | NA | NA | | Vmn2r76 | 0 | NA | NA | NA | NA | NA | NM_001102580 |
| Vmn2r51 | 0.08731684 | 0.017365056 | 0.027733412 | 0.626142072 | 0.531221759 | NA | | Vmn2r77 | 0.09572496 | 0.00498255289994511 | 0.00847999972823692 | 0.587565219295241 | 0.556824162160132 | NA | NM_001105188 |
| Vmn2r52 | 0 | NA | NA | NA | NA | NA | | Vmn2r78 | 0.385490251 | -0.00508296712693953 | 0.0117912533198291 | -0.431079461111366 | 0.666410594179892 | NA | NM_001105189 |
| Vmn2r53 | 0 | NA | NA | NA | NA | NA | | Vmn2r79 | 0 | NA | NA | NA | NA | NA | NM_001105190 |
| Vmn2r54 | 0.715075039 | 0.032315171 | 0.056394524 | 0.573019664 | 0.56663138 | NA | | Vmn2r8 | 0 | NA | NA | NA | NA | NA | NM_001104620 |
| Vmn2r55 | 0.215050351 | -0.027006021 | 0.031907566 | -0.846382992 | 0.397339123 | NA | | Vmn2r80 | 0 | NA | NA | NA | NA | NA | NM_001103368 |
| Vmn2r56 | 0.313786592 | -0.036039695 | 0.036798757 | -0.979372609 | 0.327395906 | NA | | Vmn2r81 | 0 | NA | NA | NA | NA | NA | NM_175936 |
| Vmn2r58 | 0 | NA | NA | NA | NA | NA | | Vmn2r82 | 0 | NA | NA | NA | NA | NA | NM_001101572 |
| Vmn2r59 | 0 | NA | NA | NA | NA | NA | | Vmn2r83 | 0 | NA | NA | NA | NA | NA | NM_001104537 |
| Vmn2r6 | 0 | NA | NA | NA | NA | NA | | Vmn2r85 | 9.407297401 | 0.0591845461811216 | 0.0574861216967215 | 1.02954494814175 | 0.303223669506745 | NA | NM_001102602 |
| Vmn2r60 | 0 | NA | NA | NA | NA | NA | | Vmn2r86 | 2.729100944 | 0.0170277772116209 | 0.0302793691814751 | 0.562355744915535 | 0.573873664833134 | NA | NM_001103365 |
| Vmn2r61 | 0 | NA | NA | NA | NA | NA | | Vmn2r88 | 0 | NA | NA | NA | NA | NA | NM_011686 |
| Vmn2r62 | 0 | NA | NA | NA | NA | NA | | Vmn2r89 | 0 | NA | NA | NA | NA | NA | NM_009486,NM_009487 |
| Vmn2r63 | 0 | NA | NA | NA | NA | NA | | Vmn2r9 | 0 | NA | NA | NA | NA | NA | NM_001104621 |
| Vmn2r65 | 0 | NA | NA | NA | NA | NA | | Vmn2r90 | 0 | NA | NA | NA | NA | NA | NM_001104539 |
| Vmn2r66 | 0 | NA | NA | NA | NA | NA | | Vmn2r91 | 0 | NA | NA | NA | NA | NA | NM_001104540 |
| Vmn2r67 | 0 | NA | NA | NA | NA | NA | | Vmn2r92 | 0 | NA | NA | NA | NA | NA | NM_001104541 |
| Vmn2r68 | 0 | NA | NA | NA | NA | NA | | Vmn2r93 | 0 | NA | NA | NA | NA | NA | NM_001104542 |
| Vmn2r69 | 0 | NA | NA | NA | NA | NA | | Vmn2r94 | 0 | NA | NA | NA | NA | NA | NM_001104543 |
| Vmn2r7 | 0 | NA | NA | NA | NA | NA | | Vmn2r95 | 0 | NA | NA | NA | NA | NA | NM_001102581 |
| Vmn2r70 | 0 | NA | NA | NA | NA | NA | | Vmn2r96 | 0 | NA | NA | NA | NA | NA | NM_001104547 |
| Vmn2r71 | 0 | NA | NA | NA | NA | NA | | Vmn2r97 | 0 | NA | NA | NA | NA | NA | NM_001104549 |
| Vmn2r72 | 0 | NA | NA | NA | NA | NA | | Vmn2r98 | 0 | NA | NA | NA | NA | NA | NM_001104550 |
| Vmn2r73 | 0 | NA | NA | NA | NA | NA | | Vmn2r99 | 0 | NA | NA | NA | NA | NA | NM_001104551 |
| Vmn2r74 | 0 | NA | NA | NA | NA | NA | | Vmo1 | 4.232400386 | 0.0229887750424863 | 0.03459026434125 | 0.664602467783731 | 0.506304790432655 | NA | NM_001013607 |
| Vmn2r75 | 0 | NA | NA | NA | NA | NA | | Vnn1 | 5.79495185 | -0.0307953194607168 | 0.0504221078640692 | -0.610750338794574 | 0.541364874898259 | NA | NM_011704 |
| Vmn2r76 | 0 | NA | NA | NA | NA | NA | | Vnn3 | 2.054711154 | -0.0177192791643115 | 0.0242786760546873 | -0.729828888708723 | 0.465494783735512 | NA | NM_011979 |
| Vmn2r77 | 0.098353386 | -0.020212831 | 0.02786466 | -0.725393061 | 0.468210929 | NA | | Vpreb1 | 3.016205768 | 0.0719212764838457 | 0.0405501879511744 | 1.77363608204343 | 0.0761233653345621 | NA | NM_016982 |
| Vmn2r78 | 0.32854328 | 0.00879128 | 0.034635881 | 0.253820007 | 0.799634616 | NA | | Vpreb2 | 0 | NA | NA | NA | NA | NA | NM_016983 |
| Vmn2r79 | 0 | NA | NA | NA | NA | NA | | Vpreb3 | 1.561088071 | 0.000656472005209961 | 0.0292909208827333 | 0.0224121326822792 | 0.982119202302456 | NA | NM_009514 |
| Vmn2r8 | 0 | NA | NA | NA | NA | NA | | Vrtn | 4.278817104 | 0.0452361449550812 | 0.0488173593553634 | 0.926640554762232 | 0.354113191257074 | NA | NM_001033776,NM_001168589,NM_001168588 |
| Vmn2r80 | 0 | NA | NA | NA | NA | NA | | Vsig1 | 0.414170784 | 0.011937776581977 | 0.0128627015871451 | 0.928092477392742 | 0.353359601435565 | NA | NM_030181,NM_026103 |
| Vmn2r81 | 0 | NA | NA | NA | NA | NA | | Vsig4 | 0 | NA | NA | NA | NA | NA | NM_177789 |
| Vmn2r82 | 0 | NA | NA | NA | NA | NA | | Vsig8 | 6.936310254 | -0.0435954945403891 | 0.0582856615814539 | -0.747962592471642 | 0.454482721246978 | NA | NM_177723,NR_027644 |
| Vmn2r83 | 0 | NA | NA | NA | NA | NA | | Vsx1 | 3.067340106 | -0.00529433563713992 | 0.0311270284701825 | -0.17008805200315 | 0.864940889724457 | NA | NM_054068 |
| Vmn2r85 | 6.843460148 | -0.070462639 | 0.139289425 | -0.505872138 | 0.612946413 | NA | | Vsx2 | 13.5261872 | -0.0498656841787177 | 0.0731244660225723 | -0.681928865823444 | 0.495283931800487 | NA | NM_007701 |
| Vmn2r86 | 2.533008175 | -0.032033396 | 0.105125763 | -0.30471499 | 0.760583232 | NA | | Vtcn1 | 9.503312626 | 0.0418577527795574 | 0.0622067990913265 | 0.672880672064569 | 0.501023209373771 | NA | NM_178594 |
| Vmn2r88 | 0 | NA | NA | NA | NA | NA | | Vwa2 | 1.923373596 | 0.0576920028876384 | 0.0318614335568437 | 1.81071585447374 | 0.0701848473715047 | NA | NM_172840 |
| Vmn2r89 | 0 | NA | NA | NA | NA | NA | | Vwce | 14.49241638 | -0.0298455599751306 | 0.0659944698025623 | -0.452243348032351 | 0.651093683229493 | NA | NM_027913 |
| Vmn2r9 | 0 | NA | NA | NA | NA | NA | | Vwde | 0.106347839 | -0.00653347234140708 | 0.00854389509723638 | -0.764694822098226 | 0.444453290197145 | NA | NM_001013757 |
| Vmn2r90 | 0 | NA | NA | NA | NA | NA | | Wap | 0 | NA | NA | NA | NA | NA | NM_011709 |
| Vmn2r91 | 0 | NA | NA | NA | NA | NA | | Wbp2nl | 1.975287132 | -0.0136812973652254 | 0.0288530090052693 | -0.474172290409186 | 0.6353770514278 | NA | NM_029066 |
| Vmn2r92 | 0 | NA | NA | NA | NA | NA | | Wbscr25 | 11.80264077 | -0.0316096720574289 | 0.0692004401755907 | -0.456784262892285 | 0.647826118198283 | NA | NR_026907 |
| Vmn2r93 | 0 | NA | NA | NA | NA | NA | | Wbscr28 | 0.318445824 | 0.0076171880905019 | 0.0102869072557308 | 0.740474070693932 | 0.459012388854512 | NA | NM_029681,NM_194065 |
| Vmn2r94 | 0 | NA | NA | NA | NA | NA | | Wdr20b | 0.108766301 | 0.00519171381361946 | 0.00856003371502575 | 0.606506234257728 | 0.544178643801544 | NA | NM_027614 |
| Vmn2r95 | 0 | NA | NA | NA | NA | NA | | Wdr38 | 5.834878009 | -0.0153443656978009 | 0.0491359003343878 | -0.312284207542282 | 0.754824540905833 | NA | NM_029687 |
| Vmn2r96 | 0 | NA | NA | NA | NA | NA | | Wdr64 | 0.627996698 | 0.00754850179329448 | 0.0201536382472231 | 0.374547845937175 | 0.707996766736385 | NA | NM_029453 |
| Vmn2r97 | 0 | NA | NA | NA | NA | NA | | Wdr69 | 17.55190969 | -0.0565634726915898 | 0.071846771168645 | -0.78727925794771 | 0.43111840912901 | NA | NM_027725,NR_027636 |
| Vmn2r98 | 0 | NA | NA | NA | NA | NA | | Wdr72 | 12.7707595 | 0.00646356512584668 | 0.0255192045036964 | 0.253282390715214 | 0.800050002730889 | NA | NM_001033500 |
| Vmn2r99 | 0 | NA | NA | NA | NA | NA | | Wdr93 | 16.28104093 | -0.0440997703789037 | 0.0683045942996459 | -0.645634028443857 | 0.518516394355714 | NA | NM_001037927 |
| Vmo1 | 3.032667467 | -0.050000583 | 0.121282702 | -0.412264755 | 0.680145379 | NA | | Wdr95 | 17.5805258 | -0.0533174472925255 | 0.0786487833614409 | -0.67791827176141 | 0.497823514136265 | NA | NM_029440 |
| Vnn3 | 3.56905524 | -0.106108741 | 0.118010439 | -0.899147072 | 0.368574328 | NA | | Wee2 | 7.266115142 | -0.0516358440274018 | 0.0560029302170964 | -0.922020398347632 | 0.356517935763055 | NA | NM_201370 |
| Vpreb1 | 1.717552634 | -0.10940848 | 0.103708828 | -1.054958217 | 0.291444435 | NA | | Wfdc10 | 0.619517067 | 0.0154621408254966 | 0.0146184656184032 | 1.05771297953673 | 0.290186316004987 | NA | NM_001039501 |
| Vpreb2 | 0.212768007 | -0.02694223 | 0.031868764 | -0.845411806 | 0.397880955 | NA | | Wfdc11 | 0 | NA | NA | NA | NA | NA | NM_001161806 |
| Vpreb3 | 1.674477876 | -0.028956127 | 0.096876275 | -0.298898024 | 0.765017855 | NA | | Wfdc12 | 0.355458167 | -0.0172950524609032 | 0.0137931659887683 | -1.25388561806524 | 0.209883583198022 | NA | NM_138684 |
| Vrtn | 4.096327904 | -0.199447721 | 0.128421976 | -1.55306535 | 0.120407523 | NA | | Wfdc13 | 0 | NA | NA | NA | NA | NA | NM_001012704 |
| Vsig1 | 0.233393928 | -0.027475306 | 0.032191697 | -0.85349047 | 0.39338737 | NA | | Wfdc15a | 0.767122395 | 0.00978525436011461 | 0.0198376941896509 | 0.493265712565499 | 0.621824851892585 | NA | NM_183271 |
| Vsig4 | 0 | NA | NA | NA | NA | NA | | Wfdc15b | 0 | NA | NA | NA | NA | NA | NM_001045554,NM_138685 |
| Vsig8 | 7.590406203 | 0.020972284 | 0.15625742 | 0.134216246 | 0.89323158 | NA | | Wfdc16 | 0 | NA | NA | NA | NA | NA | NM_001012723 |
| Vsx1 | 6.03869358 | -0.068781925 | 0.118307491 | -0.581382669 | 0.560982575 | NA | | Wfdc17 | 5.58392691 | 0.0135466028146099 | 0.0540311752550742 | 0.250718270529156 | 0.802031933924929 | NA | NM_001081957 |
| Vwa2 | 0.589190971 | -0.008579232 | 0.052937852 | -0.162062328 | 0.871256772 | NA | | Wfdc2 | 18.47947389 | 0.0267429478884293 | 0.0352288522896132 | 0.759120611383476 | NA | NA | NM_026323 |
| Vwde | 0.658277769 | -0.062529329 | 0.056532976 | -1.106068236 | 0.268696986 | NA | | Wfdc3 | 17.37265845 | -0.113235999181945 | 0.0756462857545635 | -1.49691419813185 | 0.134415585333105 | NA | NM_027961 |
| Wap | 0 | NA | NA | NA | NA | NA | | Wfdc5 | 0.227686571 | 0.00691460391109036 | 0.00980755285755433 | 0.705028462402254 | 0.480792523671872 | NA | NM_145369 |
| Wbp2nl | 1.525674379 | 0.031567476 | 0.086285278 | 0.36585008 | 0.714476948 | NA | | Wfdc6a | 1.279167853 | -0.00386048791146289 | 0.0201600487685401 | -0.19149199269236 | 0.848140153370325 | NA | NM_001177848,NM_001033240 |
| Wbscr28 | 0 | NA | NA | NA | NA | NA | | Wfdc6b | 0 | NA | NA | NA | NA | NA | NM_001012725 |
| Wdr20b | 0.233393928 | -0.027475306 | 0.032191697 | -0.85349047 | 0.39338737 | NA | | Wfdc8 | 0.127687691 | 0.00542298404407807 | 0.00869724871942359 | 0.62352868349814 | 0.532937159177571 | NA | NM_001276431,NM_001276432,NM_001080550 |
| Wdr38 | 5.02981735 | -0.002470608 | 0.120626277 | -0.020481509 | 0.983659263 | NA | | Wfdc9 | 0.09572496 | 0.00498255289994511 | 0.00847999972823692 | 0.587565219295241 | 0.556824162160132 | NA | NM_001160414 |
| Wdr61 | 1237.717804 | -0.009051068 | 0.078275304 | -0.115631212 | NA | NA | | Wfikkn1 | 6.71714547 | 0.022654795594217 | 0.0557244310459913 | 0.406550505208734 | 0.684338155612662 | NA | NM_001100454 |
| Wdr64 | 0.492768124 | -0.024633558 | 0.055739517 | -0.44194064 | 0.658532161 | NA | | Wisp2 | 6.363977806 | -0.00818600718396916 | 0.0510787432624496 | -0.160262501798612 | 0.87267429625525 | NA | NM_016873 |
| Wdr72 | 4.569572229 | 0.167982415 | 0.083677933 | 2.007487623 | NA | NA | | Wisp3 | 5.556606437 | 0.0119219312487944 | 0.0455971289366499 | 0.261462322887875 | 0.793736001167321 | NA | NM_001127376 |
| Wee2 | 6.406454477 | 0.172377554 | 0.144930786 | 1.189378448 | 0.234290778 | NA | | Wnt1 | 3.788024226 | -0.00490990883576857 | 0.0389861125954019 | -0.125939943967321 | 0.899779463670922 | NA | NM_021279 |
| Wfdc10 | 0 | NA | NA | NA | NA | NA | | Wnt11 | 10.78524007 | -6.63956122867783e-05 | 0.0662813786449192 | -0.00100172346508468 | 0.999200740446684 | NA | NM_009519 |
| Wfdc11 | 0 | NA | NA | NA | NA | NA | | Wnt16 | 14.92581312 | 0.0218843801890118 | 0.0728885413905057 | 0.300244452303752 | 0.763990700200729 | NA | NM_053116 |
| Wfdc12 | 0.603371276 | -0.005189871 | 0.062807235 | -0.082631732 | 0.934144369 | NA | | Wnt3a | 0.575251383 | -0.0179565793531345 | 0.0173808499060826 | -1.03312435526242 | 0.301545706977181 | NA | NM_009522 |
| Wfdc13 | 0 | NA | NA | NA | NA | NA | | Wnt8a | 0.212695677 | -0.00857796107890413 | 0.00973699866180651 | -0.880965621629515 | 0.378336427605205 | NA | NM_009290 |
| Wfdc15a | 0.280299629 | 0.041967777 | 0.040615819 | 1.033286501 | 0.301469842 | NA | | Wnt8b | 13.53837413 | -0.0612617809331323 | 0.0714968219661579 | -0.856846210061333 | 0.3915298834682 | NA | NM_011720 |
| Wfdc15b | 0.098353386 | -0.020212831 | 0.02786466 | -0.725393061 | 0.468210929 | NA | | Wt1 | 5.622697468 | 0.0407793818133721 | 0.0430187885940023 | 0.947943518313241 | 0.343158208861549 | NA | NM_144783 |
| Wfdc16 | 0 | NA | NA | NA | NA | NA | | Xcl1 | 0.612959092 | -0.0166396230903786 | 0.0163182047198189 | -1.01969691985598 | 0.307872222935911 | NA | NM_008510 |
| Wfdc17 | 7.863379813 | -0.357414337 | 0.142113425 | -2.514993479 | 0.011903459 | NA | | Xcr1 | 2.937892694 | 0.0107111312733522 | 0.0380404108447588 | 0.28157243929525 | 0.778271375708697 | NA | NM_011798 |
| Wfdc2 | 7.400308631 | 0.323755539 | 0.13551065 | 2.389151985 | NA | NA | | Xirp1 | 5.012100265 | 0.00557196688561463 | 0.0511049637401689 | 0.109029856941959 | 0.913178809078756 | NA | NM_011724 |
| Wfdc5 | 0 | NA | NA | NA | NA | NA | | Xirp2 | 12.85751988 | -0.0151781501471778 | 0.0707956146614805 | -0.214393931315581 | 0.830239873756768 | NA | NM_001024618,NM_001083919 |
| Wfdc6a | 0.818336458 | 0.030833894 | 0.064495082 | 0.478081325 | 0.632592323 | NA | | Xkr5 | 9.957613239 | -0.000477946785855315 | 0.0583842580442097 | -0.00818622693626431 | 0.993468408868088 | NA | NM_176951,NM_001113350 |
| Wfdc6b | 0 | NA | NA | NA | NA | NA | | Xkr6 | 20.50614761 | -0.050811308431766 | 0.0809913480611076 | -0.627367115724869 | 0.530418618229499 | NA | NM_173393 |
| Wfdc8 | 0 | NA | NA | NA | NA | NA | | Xkr9 | 0.297598527 | 0.00928691468127708 | 0.0113567208765242 | 0.817746141888045 | 0.413502157150213 | NA | NM_001011873 |
| Wfdc9 | 0 | NA | NA | NA | NA | NA | | Xlr3a | 22.13421947 | -0.0606219208294041 | 0.0649728633884315 | -0.933034464973232 | 0.350802173812672 | NA | NM_001110784 |
| Wfikkn1 | 7.145698986 | 0.082598747 | 0.141746252 | 0.582722615 | 0.560080047 | NA | | Xlr3c | 0.226556888 | -0.00933175792380659 | 0.010153338242423 | -0.919082739193727 | 0.358052296835844 | NA | NM_011727 |
| Wisp2 | 7.165727003 | 0.110055098 | 0.138609263 | 0.793995262 | 0.427198189 | NA | | Xlr4a | 5.050836268 | -0.0130036711370575 | 0.0339188837977925 | -0.383375561960673 | 0.701441320436655 | NA | NM_001081642 |
| Wisp3 | 4.077437145 | 0.056489555 | 0.122511343 | 0.461096523 | 0.644729356 | NA | | Xlr4b | 6.538480856 | -0.0542697265736147 | 0.0492565696819966 | -1.10177641122765 | 0.270558887909022 | NA | NM_021365 |
| Wnt1 | 3.810729552 | -0.106960266 | 0.12112162 | -0.88308153 | 0.377192223 | NA | | Xlr4c | 1.614146089 | -0.00560590416174734 | 0.0201695698037095 | -0.277938707483802 | 0.781059409790409 | NA | NM_183094 |
| Wnt3 | 101.8637554 | 0.341800524 | 0.167404788 | 2.041760736 | NA | NA | | Xlr5a | 0.439983788 | 0.0126652380404026 | 0.0132459535836759 | 0.956159023236422 | 0.338991898718962 | NA | NM_001045539 |
| Wnt3a | 0.791119018 | -0.005831258 | 0.067937947 | -0.085832117 | 0.931599875 | NA | | Xlr5b | 0 | NA | NA | NA | NA | NA | NM_001111293 |
| Wnt8a | 0.17463368 | 0.024560283 | 0.031186242 | 0.787535833 | 0.430968261 | NA | | Xlr5c | 0 | NA | NA | NA | NA | NA | NM_031493 |
| Wt1 | 4.825393899 | -0.087355315 | 0.131370515 | -0.664953737 | 0.506080083 | NA | | Xpnpep2 | 13.02434224 | 0.0280553919907877 | 0.0719478645997154 | 0.389940579152347 | 0.69658048650192 | NA | NM_133213 |
| Xcl1 | 0.414584642 | 0.043381126 | 0.041065838 | 1.056379903 | 0.290794684 | NA | | Yipf7 | 0.773030112 | 0.00111062887160337 | 0.0161222948787903 | 0.0688877656656966 | 0.945078956968815 | NA | NM_023784 |
| Xcr1 | 2.082470675 | -0.017091836 | 0.099124538 | -0.172427904 | 0.863101136 | NA | | Zar1 | 4.72705282 | -0.0174970376215544 | 0.0450536219397117 | -0.388360288657103 | 0.697749427509066 | NA | NM_174877 |
| Xirp1 | 6.475558845 | -0.315428202 | 0.149592908 | -2.108577251 | 0.034981088 | NA | | Zar1l | 16.45912498 | 0.0181982692179993 | 0.0766239591201764 | 0.23750103005585 | 0.812268120191322 | NA | NM_001159693 |
| Xkr5 | 7.521844008 | 0.031874979 | 0.141376145 | 0.225462215 | 0.821619717 | NA | | Zbbx | 16.07888367 | 0.0338847963111059 | 0.0727405490752392 | 0.465830911945099 | 0.641336536751642 | NA | NM_172515 |
| Xkr9 | 0 | NA | NA | NA | NA | NA | | Zbp1 | 17.60772859 | -0.129992317407601 | 0.0478756588120125 | -2.7152068636387 | 0.00662343837317395 | NA | NM_001139519,NM_021394 |
| Xlr3c | 0.303023688 | 0.009739077 | 0.039388711 | 0.247255542 | 0.804710463 | NA | | Zbtb32 | 0.300480017 | -0.00401601363468843 | 0.0108825526077113 | -0.369032319847708 | 0.712103634607305 | NA | NM_021397 |
| Xlr4a | 2.900418569 | 0.142513721 | 0.096346356 | 1.479181231 | 0.139091884 | NA | | Zc2hc1b | 0 | NA | NA | NA | NA | NA | NM_029172 |
| Xlr4b | 4.791741083 | 0.143096307 | 0.115648122 | 1.237342246 | 0.21596005 | NA | | Zc3h12d | 2.491325265 | -0.00273619062612749 | 0.0310067169733424 | -0.0882450931028876 | 0.929681877972338 | NA | NM_172785 |
| Xlr4c | 0.696916283 | 0.076469279 | 0.054048147 | 1.414836276 | 0.157116505 | NA | | Zcchc13 | 0.097165704 | -0.00637828455464458 | 0.00848781691535724 | -0.75146349388194 | 0.452373763047372 | NA | NM_029158 |
| Xlr5a | 0.119397641 | -0.020477583 | 0.027954224 | -0.732539845 | 0.463839132 | NA | | Zcchc5 | 5.024948625 | -0.0592738566807834 | 0.0406002667149418 | -1.45993761806913 | 0.144307219330654 | NA | NM_199468 |
| Xlr5b | 0 | NA | NA | NA | NA | NA | | Zdhhc11 | 0.106148608 | 0.00515554063321497 | 0.00854200771766808 | 0.603551390213729 | 0.546141939141592 | NA | NM_027704 |
| Xlr5c | 0 | NA | NA | NA | NA | NA | | Zdhhc19 | 0.108766301 | 0.00519171381361946 | 0.00856003371502575 | 0.606506234257728 | 0.544178643801544 | NA | NM_199309 |
| Yipf7 | 0.550767766 | 0.020121298 | 0.052098809 | 0.386214157 | 0.69933807 | NA | | Zdhhc25 | 0 | NA | NA | NA | NA | NA | NM_027306 |
| Zbtb32 | 1.152464342 | 0.040710781 | 0.060016863 | 0.678322367 | 0.49756732 | NA | | Zfa-ps | 0.712046018 | -0.0123354953002287 | 0.0180230784534131 | -0.68442776477471 | 0.493705088613528 | NA | NR_037920 |

| | | | | | | |
|---|---|---|---|---|---|---|
| Zc2hc1b | 0 | NA | NA | NA | NA | NA |
| Zc3h12d | 2.241074628 | 0.00248366 | 0.097761553 | 0.025405284 | 0.979731696 | NA |
| Zcchc13 | 0.174376471 | 0.024538273 | 0.031173126 | 0.787161136 | 0.431187544 | NA |
| Zcchc5 | 4.805242689 | 0.139991527 | 0.104361919 | 1.34140431 | 0.179789219 | NA |
| Zdhhc11 | 0.188785634 | 0.025025111 | 0.031461603 | 0.79541754 | 0.42637066 | NA |
| Zdhhc19 | 0.4144578 | -0.044107517 | 0.040639925 | -1.085324756 | 0.277777845 | NA |
| Zdhhc25 | 0 | NA | NA | NA | NA | NA |
| Zfa-ps | 0.706356315 | 0.072659511 | 0.06424429 | 1.130987858 | 0.258060202 | NA |
| Zfp345 | 0.08731684 | 0.017365056 | 0.027733412 | 0.626142072 | 0.531221759 | NA |
| Zfp36l3 | 1.492030096 | 0.035174536 | 0.094780073 | 0.371117415 | 0.710550081 | NA |
| Zfp42 | 0.079983654 | 0.016740032 | 0.027674178 | 0.604897175 | 0.54524732 | NA |
| Zfp534 | 2.000606159 | 0.054859949 | 0.102743157 | 0.533952343 | 0.593374507 | NA |
| Zfp572 | 0.17463368 | 0.024560283 | 0.031186242 | 0.787535833 | 0.430968261 | NA |
| Zfp600 | 0.183387883 | -0.001444816 | 0.031390672 | -0.046026918 | 0.963288796 | NA |
| Zfp616 | 0 | NA | NA | NA | NA | NA |
| Zfp663 | 2.048067755 | -0.192161354 | 0.095886602 | -2.004048002 | 0.045064919 | NA |
| Zfp735 | 0 | NA | NA | NA | NA | NA |
| Zfp750 | 6.930758443 | 0.164315773 | 0.145321112 | 1.130708199 | 0.258177929 | NA |
| Zfp819 | 4.045034739 | 0.172203602 | 0.129765696 | 1.327034857 | 0.184497152 | NA |
| Zfp872 | 2.2216008 | 0.06052693 | 0.079721162 | 0.759232918 | 0.447713238 | NA |
| Zfp936 | 0.293434314 | -0.014075032 | 0.039227749 | -0.358802961 | 0.719742498 | NA |
| Zfp957 | 0.17463368 | 0.024560283 | 0.031186242 | 0.787535833 | 0.430968261 | NA |
| Zfy1 | 0 | NA | NA | NA | NA | NA |
| Zfy2 | 0 | NA | NA | NA | NA | NA |
| Zg16 | 0 | NA | NA | NA | NA | NA |
| Zim3 | 4.2945162 | 0.135609547 | 0.100124632 | 1.354407449 | 0.175606422 | NA |
| Znrf4 | 0 | NA | NA | NA | NA | NA |
| Zp1 | 4.558993664 | -0.002736762 | 0.136103234 | -0.020107986 | 0.983957229 | NA |
| Zp3 | 0.930606196 | -0.065800364 | 0.076152546 | -0.864059935 | 0.387554967 | NA |
| Zp3r | 3.360176357 | 0.036943991 | 0.120062992 | 0.307705068 | 0.758306763 | NA |
| Zp4-ps | 0 | NA | NA | NA | NA | NA |
| Zpbp2 | 1.182448374 | 0.052326943 | 0.085574332 | 0.611479415 | 0.540882242 | NA |
| Zpld1 | 6.036963971 | 0.025764405 | 0.135165346 | 0.19061398 | 0.848828037 | NA |
| Zscan10 | 1.096775958 | 0.146845934 | 0.073590533 | 1.995445999 | 0.045994258 | NA |
| Zscan4a | 0 | NA | NA | NA | NA | NA |
| Zscan4b | 0 | NA | NA | NA | NA | NA |
| Zscan4c | 0 | NA | NA | NA | NA | NA |
| Zscan4d | 0 | NA | NA | NA | NA | NA |
| Zscan4e | 0 | NA | NA | NA | NA | NA |
| Zscan4f | 0 | NA | NA | NA | NA | NA |
| Zscan5b | 0.256038351 | 0.007085343 | 0.03307033 | 0.214250763 | 0.830351512 | NA |
| Zswim2 | 0.686664619 | 0.040716927 | 0.066216252 | 0.614908356 | 0.538615244 | NA |
| Zyg11a | 0.270704723 | 0.00804778 | 0.033253949 | 0.242009739 | 0.808772617 | NA |
| a | 0.521693954 | 0.010491765 | 0.051043378 | 0.205546051 | 0.837145523 | NA |

| | | | | | | | |
|---|---|---|---|---|---|---|---|
| Zfp345 | 0.43002905 | 0.00615178557924862 | 0.0139465098832267 | 0.441098570951238 | 0.659141638025682 | NA | NM_001034900 |
| Zfp352 | 21.84462636 | -0.0136136352601502 | 0.0777077989106166 | -0.17519007681338 7 | 0.860930281309073 | NA | NM_153102 |
| Zfp36l3 | 1.972252113 | -0.011724936905281 | 0.0304417821841174 | -0.385159345611451 | 0.700119361669558 | NA | NM_001009549 |
| Zfp389 | 19.70221992 | -0.144989231241677 | 0.0811008913348126 | -1.78776372066136 | 0.0738141363501815 | NA | NR_026798 |
| Zfp42 | 0.203314312 | -0.000639443352387801 | 0.00969102748600992 | -0.0659830294889689 | 0.947391336513325 | NA | NM_009556 |
| Zfp457 | 9.808236318 | -0.0709559934922646 | 0.0590291070858609 | -1.20205093715979 | 0.229343793580863 | NA | NM_001003666 |
| Zfp474 | 18.97248162 | 0.0504806261204911 | 0.069627891386137 | 0.725005814703472 | 0.468448463215201 | NA | NM_025749 |
| Zfp534 | 2.683628588 | -0.00359058979553474 | 0.03809391919346 | -0.0942562453944403 | 0.924905606233539 | NA | NM_001127188 |
| Zfp572 | 0.531141502 | 0.0011504769157983 | 0.0128694827144426 | 0.089395738844048 | 0.928767409824307 | NA | NR_045613 |
| Zfp600 | 0.21511414 | -0.000824240786584897 | 0.00975135300262786 | -0.08452578697159 | 0.93263840116295 | NA | NM_001177546,NM_001177545 |
| Zfp616 | 0 | NA | NA | NA | NA | NA | NM_001177570 |
| Zfp648 | 15.93789832 | 0.03138086250125 | 0.0685307805815568 | 0.457909018910187 | 0.647017807066218 | NA | NM_001204908 |
| Zfp663 | 0.805095652 | -7.04241472584986e-05 | 0.0200366852474587 | -0.00351476037022793 | 0.997195632739652 | NA | NM_001005425 |
| Zfp735 | 0 | NA | NA | NA | NA | NA | NM_001126489 |
| Zfp750 | 8.85906099 | -0.0199552807019229 | 0.0607066500691789 | -0.328716552127035 | 0.742369942198933 | NA | NM_178763 |
| Zfp819 | 5.326630336 | -0.0259755042748643 | 0.0489459068874922 | -0.530698191670491 | 0.595627939352151 | NA | NM_028913 |
| Zfp872 | 3.395008597 | -0.0211361353755769 | 0.0359401803237025 | -0.588092079260872 | 0.556470489366933 | NA | NM_001033813 |
| Zfp936 | 0.332506265 | 0.00179938841223566 | 0.0111545909305638 | 0.1613137069245 | 0.871846328078977 | NA | NM_001034893 |
| Zfp957 | 0.212695677 | -0.00857796107890413 | 0.00973699866180651 | -0.880965621629515 | 0.378336427605205 | NA | NM_001033215 |
| Zfy1 | 0 | NA | NA | NA | NA | NA | NM_009570 |
| Zfy2 | 0 | NA | NA | NA | NA | NA | NM_009571 |
| Zg16 | 0.127687691 | 0.00542298404407807 | 0.00869724871942359 | 0.62352868349814 | 0.532937159177571 | NA | NM_026918 |
| Zim3 | 3.605831694 | 0.00845116722043532 | 0.0336910049414631 | 0.250843429429277 | 0.801935162934825 | NA | NR_036631 |
| Zmynd12 | 16.02131086 | 0.016705454799322 | 0.0712550944249293 | 0.234445760463093 | 0.814638936650975 | NA | NM_001014900 |
| Zmynd15 | 18.15214063 | -0.183571876790859 | 0.0751212420736146 | -2.44367467474737 | 0.0145385298328676 | NA | NM_001029929 |
| Znrf4 | 0 | NA | NA | NA | NA | NA | NM_011483 |
| Zp1 | 3.752133087 | -0.0612996690266118 | 0.0419921262298899 | -1.45978959700733 | 0.144347908516177 | NA | NM_009580 |
| Zp2 | 7.580646502 | -0.0817644497203866 | 0.0550208289009532 | -1.48606357544297 | 0.137262305673213 | NA | NM_011775 |
| Zp3 | 0.892355537 | 0.00607350812159236 | 0.02236225972905 | 0.271596350063964 | 0.785932406226564 | NA | NM_011776 |
| Zp3r | 3.036695988 | -0.00560822134858653 | 0.0385594371842742 | -0.145443547886476 | 0.88436068393922 | NA | NM_009581 |
| Zp4-ps | 0 | NA | NA | NA | NA | NA | NR_027813 |
| Zpbp2 | 1.880278259 | 0.0112193244977294 | 0.0280042642003512 | 0.400629147670613 | 0.688693182511645 | NA | NM_199419,NM_027061,NM_001166495,NM_001166494 |
| Zpld1 | 5.571485153 | -0.00158721811033654 | 0.0514344810930566 | -0.0308590283523013 | 0.9753819649872 | NA | NM_178720 |
| Zscan10 | 3.183903651 | 0.0181015072658075 | 0.0289382201054957 | 0.62552248202611 | 0.531628197681323 | NA | NM_001033425 |
| Zscan4a | 0 | NA | NA | NA | NA | NA | NR_033707 |
| Zscan4b | 0 | NA | NA | NA | NA | NA | NM_001185173 |
| Zscan4c | 0 | NA | NA | NA | NA | NA | NM_001013765 |
| Zscan4d | 0 | NA | NA | NA | NA | NA | NM_001100186 |
| Zscan4e | 0 | NA | NA | NA | NA | NA | NM_001161802 |
| Zscan4f | 0 | NA | NA | NA | NA | NA | NM_001110316 |
| Zscan5b | 0.194331409 | -0.00840968081320881 | 0.00964081822647896 | -0.872299489073576 | 0.383045016303155 | NA | NM_133204 |
| Zswim2 | 0.55897171 | -0.0307918840007222 | 0.0183473164462484 | -1.6782772614694 | 0.0932929849016459 | NA | NM_027964 |
| Zyg11a | 0.308420637 | -0.00472633378880091 | 0.0109973596733179 | -0.42976986560402 | 0.667363055492239 | NA | NM_001167936 |
| a | 0.591928964 | -0.00798522787408491 | 0.01684326960 5146 | -0.474090129843036 | 0.635435636733042 | NA | NM_015770 |
