## Supplementary table 3 for "DAP12 deficiency alters microglia-oligodendrocyte communication and enhances resilience against tau toxicity"

| | | | | | | | | | |
|---|---|---|---|---|---|---|---|---|---|
| Figure3B_snRNA_microglia_cell_ratio | | | | | | | | | |
| Microglia cell ratio of different clusters | | | | | | | | | |
| Raw counts | | | | | | | | | |
| | WT_1 | WT_2 | WT_3 | DAP12WT_1 | DAP12WT_2 | DAP12WT_3 | DAP12KO_1 | DAP12KO_2 | DAP12KO_3 |
| 1 | 215 | 242 | 185 | 219 | 244 | 139 | 193 | 252 | 191 |
| 2 | 80 | 46 | 14 | 160 | 382 | 199 | 106 | 131 | 19 |
| 3 | 40 | 46 | 37 | 40 | 60 | 47 | 35 | 29 | 30 |
| 4 | 32 | 33 | 27 | 55 | 65 | 62 | 32 | 18 | 24 |
| sum | 367 | 367 | 263 | 474 | 751 | 447 | 366 | 430 | 264 |
| | | 997 | | | 1672 | | | 1060 | |
| Cell Ratio | | | | | | | | | |
| | WT_1 | WT_2 | WT_3 | DAP12WT_1 | DAP12WT_2 | DAP12WT_3 | DAP12KO_1 | DAP12KO_2 | DAP12KO_3 |
| 1 | 0.585831063 | 0.659400545 | 0.703422053 | 0.462025316 | 0.324900133 | 0.310961969 | 0.527322404 | 0.586046512 | 0.723484848 |
| 2 | 0.217983651 | 0.125340599 | 0.053231939 | 0.337552743 | 0.508655126 | 0.445190157 | 0.289617486 | 0.304651163 | 0.071969697 |
| 3 | 0.108991826 | 0.125340599 | 0.140684411 | 0.084388186 | 0.079893475 | 0.105145414 | 0.095628415 | 0.06744186 | 0.113636364 |
| 4 | 0.08719346 | 0.089918256 | 0.102661597 | 0.116033755 | 0.086551265 | 0.138702461 | 0.087431694 | 0.041860465 | 0.090909091 |
| Figure3D_MG2 vs MG1 upregulation genes involved pathways | | | | | | | | | |
| Ingenuity Canonical Pathways | -log(p-value) | Ratio | z-score | Molecules | | | | | |
| STAT3 Pathway | 1.44 | 0.037 | 2.236 | FGFR2,INSR,RAP1A,RAP1B,TNFRSF11A | | | | | |
| IL-8 Signaling | 4.02 | 0.0547 | 2.828 | AKT3,FNBP1,GNAQ,GNAS,NCF1,NFKB1,PIK3CB,PRKCD,RAP1A,RAP1B,RHOH | | | | | |
| NF-κB Activation by Viruses | 3.18 | 0.0769 | 2.449 | AKT3,NFKB1,PIK3CB,PRKCD,RAP1A,RAP1B | | | | | |
| Chemokine Signaling | 2.43 | 0.0658 | 1.342 | CAMK1,GNAQ,PPP1R12A,RAP1A,RAP1B | | | | | |
| Neuroinflammation Signaling Pathway | 1.79 | 0.0315 | 2.121 | AKT3,APP,BIRC6,CREB1,GLS,GSK3B,NCF1,NFKB1,PIK3CB | | | | | |
| Figure3F_MG4 vs MG1 upregulation genes involved pathways | | | | | | | | | |
| Ingenuity Canonical Pathways | -log(p-value) | Ratio | z-score | Molecules | | | | | |
| Natural Killer Cell Signaling | 4.08 | 0.0765 | 3.051 | 1G,HLA-A,LCP2,MAP3K5,NFATC2,NFATC3,NFKB1,RAP1A,RAP1B,ROCK1,SYK,VAV3,WIPF1 | | | | | |
| NF-κB Activation by Viruses | 3.58 | 0.0641 | 2.236 | NFKB1,PRKCB,PRKCH,RAP1A,RAP1B | | | | | |
| STAT3 Pathway | 2.479 | 0.0296 | 2 | INSR,RAP1A,RAP1B,TNFRSF11A | | | | | |
| Actin Cytoskeleton Signaling | 2.28 | 0.0508 | 1.732 | ,ARHGEF12,ARPC2,DOCK1,IQGAP1,IQGAP2,ITGA4,PPP1R12A,RAP1A,RAP1B,ROCK1,VAV3 | | | | | |
| JAK/STAT Signaling | 2.71 | 0.0854 | 1.89 | BCL2L1,GNAQ,JAK1,NFKB1,PIAS1,RAP1A,RAP1B | | | | | |
| Figure3LN_p2ry12_staining | | | | | | | | | |
| P2ry12 positive area | | | | Branch length | | | | | |
| WTB6 | Dap12+/+ tau+ | Dap12-/- tau+ | | WTB6 | Dap12+/+ tau+ | Dap12-/- tau+ | | | |
| 0.94604194 | 0.80168195 | 0.71617034 | | 33.2732109 | 31.1440405 | 48.2460696 | | | |
| 0.7769186 | 0.42531653 | 0.85325061 | | 50.1761757 | 29.6887081 | 29.4262373 | | | |
| 1.02624906 | 0.66390186 | 0.61512389 | | 29.5808664 | 28.2865919 | 18.5927243 | | | |
| 1.2507904 | 0.58005752 | 0.68111718 | | 39.6176607 | 29.8686508 | 41.1610963 | | | |
| 1.03624906 | 0.81099703 | 0.75495388 | | 36.0233478 | 5.35781864 | 35.0988598 | | | |
| | 0.66697935 | 0.80873368 | | | 17.1927392 | 45.5057695 | | | |
| | 0.36011095 | 0.63197113 | | | 19.6585207 | 49.0356837 | | | |
| | 0.52395392 | 0.67495691 | | | 31.2958897 | 67.7860757 | | | |
| | 0.4310466 | 0.74223948 | | | 27.9993222 | 25.9640894 | | | |
| | 0.6 | 0.75828183 | | | 17.0051871 | 37.6253403 | | | |
| | 0.53 | 0.82218095 | | | 12.1885481 | 34.8679948 | | | |
| | | 0.57160218 | | | | 39.3498193 | | | |
| sFigure4F_IHC_p2ry12_microglia_branchpoints | | | | | | | | | |
| Branch points | | | | | | | | | |
| DAP12+/+ Tau- | DAP12+/+ Tau+ | DAP12-/- Tau+ | | | | | | | |
| 2.84662577 | 2.71794872 | 4.54117647 | | | | | | | |
| 3.46564885 | 2.26984127 | 2.41935484 | | | | | | | |
| 4.34939759 | 2.11811024 | 1.20253165 | | | | | | | |
| 2.55223881 | 2.45535714 | 3.37179487 | | | | | | | |
| 3.02325581 | 0.16949153 | 2.91428571 | | | | | | | |
| | 1.12878788 | 4.29896907 | | | | | | | |
| | 1.5 | 4.58955224 | | | | | | | |
| | 2.46153846 | 6.94174757 | | | | | | | |
| | 2.13157895 | 2.01550388 | | | | | | | |
| | 0.97087379 | 3.01265823 | | | | | | | |
| | 0.57894737 | 3.13924051 | | | | | | | |
| | | 3.72727273 | | | | | | | |
| Figure3B_pseudobulk_MG2_vs_MG1 | | | | | | | | | |
| MG2_versus_MG1_DEGs | | | | | | | | | |
| | p_val | avg_log2FC | pct.1 | pct.2 | p_val_adj | | | | |
| Kcnip4 | 5.76E-24 | 1.108671357 | 0.316 | 0.226 | 1.55E-19 | | | | |
| Lsamp | 1.67E-19 | 1.083246045 | 0.349 | 0.273 | 4.50E-15 | | | | |
| Tenm2 | 7.49E-16 | 0.986858787 | 0.253 | 0.163 | 2.01E-11 | | | | |
| Nrg3 | 2.39E-18 | 0.950062819 | 0.261 | 0.186 | 6.44E-14 | | | | |
| Snhg11 | 1.67E-43 | 0.873990665 | 0.584 | 0.509 | 4.50E-39 | | | | |
| Apoe | 5.86E-36 | 0.844332454 | 0.303 | 0.291 | 1.58E-19 | | | | |
| Nkain2 | 5.80E-27 | 0.843340873 | 0.255 | 0.215 | 1.56E-22 | | | | |
| Pcdh9 | 2.57E-26 | 0.841434882 | 0.257 | 0.223 | 6.92E-22 | | | | |
| Dlg2 | 6.48E-21 | 0.794404103 | 0.285 | 0.205 | 1.74E-16 | | | | |
| Ptprd | 5.02E-25 | 0.783946541 | 0.283 | 0.231 | 1.35E-20 | | | | |
| Meg3 | 5.87E-23 | 0.716479428 | 0.638 | 0.532 | 1.58E-18 | | | | |
| Gm26848 | 1.48E-12 | 0.585388632 | 0.311 | 0.246 | 3.99E-08 | | | | |
| Gria2 | 7.24E-30 | 0.558963715 | 0.287 | 0.27 | 1.95E-25 | | | | |
| Rbfox1 | 4.73E-24 | 0.515005223 | 0.297 | 0.315 | 1.27E-19 | | | | |
| Cd9 | 2.59E-35 | 0.449134399 | 0.276 | 0.299 | 6.98E-31 | | | | |
| Gm20388 | 1.34E-17 | 0.441719097 | 0.749 | 0.668 | 3.61E-13 | | | | |
| Rap1a | 1.15E-40 | 0.411283053 | 0.3 | 0.407 | 3.10E-19 | | | | |
| Cacna1a | 2.79E-22 | 0.399782346 | 0.534 | 0.522 | 7.51E-18 | | | | |
| Gm26733 | 8.37E-12 | 0.396899247 | 0.254 | 0.23 | 2.25E-07 | | | | |
| Gm26804 | 8.11E-17 | 0.381118248 | 0.361 | 0.346 | 2.18E-12 | | | | |
| Fgfr2 | 2.06E-35 | 0.378972476 | 0.422 | 0.464 | 5.55E-31 | | | | |
| Apbb2 | 3.45E-30 | 0.36393249 | 0.279 | 0.305 | 9.28E-26 | | | | |
| Ctsz | 5.01E-31 | 0.331066194 | 0.232 | 0.268 | 1.35E-26 | | | | |
| Cadm1 | 1.15E-32 | 0.300227206 | 0.409 | 0.472 | 3.10E-28 | | | | |
| Gm20275 | 7.99E-22 | 0.29133445 | 0.267 | 0.299 | 2.15E-17 | | | | |
| Gm44686 | 1.70E-22 | 0.289571191 | 0.235 | 0.255 | 4.58E-18 | | | | |
| Fgf13 | 1.56E-11 | 0.274844085 | 0.394 | 0.433 | 4.21E-07 | | | | |
| Arhgap45 | 3.20E-42 | 0.271461214 | 0.67 | 0.77 | 8.61E-38 | | | | |
| Fam105a | 4.06E-12 | 0.266635923 | 0.468 | 0.462 | 1.09E-07 | | | | |
| Kcnma1 | 5.27E-36 | 0.260033893 | 0.303 | 0.397 | 1.42E-31 | | | | |
| Malat1 | 4.48E-46 | 0.25042968 | 1 | 1 | 1.21E-41 | | | | |
| Srsf11 | 1.44E-29 | 0.243411619 | 0.302 | 0.351 | 3.87E-25 | | | | |
| Nbea | 9.27E-21 | 0.231197388 | 0.277 | 0.324 | 2.49E-16 | | | | |
| Glg1 | 1.04E-24 | 0.227657607 | 0.245 | 0.292 | 2.80E-20 | | | | |
| Ctsb | 7.81E-30 | 0.226441652 | 0.232 | 0.284 | 2.10E-25 | | | | |
| Arglu1 | 6.41E-41 | 0.220640457 | 0.401 | 0.498 | 1.72E-36 | | | | |
| Mir142hg | 8.58E-29 | 0.193420461 | 0.235 | 0.287 | 2.31E-24 | | | | |
| Rab10 | 4.61E-23 | 0.191044307 | 0.214 | 0.255 | 1.24E-18 | | | | |
| Myo9a | 2.27E-35 | 0.185544414 | 0.299 | 0.371 | 6.10E-31 | | | | |
| Plek | 1.08E-29 | 0.177913615 | 0.228 | 0.29 | 2.91E-25 | | | | |
| Eml6 | 4.94E-17 | 0.17378053 | 0.255 | 0.297 | 1.33E-12 | | | | |
| Cmss1 | 3.84E-33 | 0.172506713 | 0.509 | 0.611 | 1.03E-28 | | | | |
| Rsrc2 | 1.37E-27 | 0.171417643 | 0.231 | 0.284 | 3.68E-23 | | | | |
| Gm37240 | 4.47E-23 | 0.169896708 | 0.363 | 0.439 | 1.20E-18 | | | | |
| Gm17231 | 5.92E-23 | 0.169792331 | 0.223 | 0.279 | 1.59E-18 | | | | |
| Mknk1 | 1.02E-29 | 0.16908808 | 0.282 | 0.344 | 2.75E-25 | | | | |
| Tpr | 1.53E-26 | 0.16723785 | 0.205 | 0.26 | 4.12E-22 | | | | |
| Gm48678 | 1.14E-32 | 0.166391832 | 0.268 | 0.351 | 3.07E-28 | | | | |
| Lrrc4 | 6.77E-20 | 0.157581612 | 0.237 | 0.288 | 1.82E-15 | | | | |
| Cd180 | 3.95E-22 | 0.157141001 | 0.245 | 0.295 | 1.06E-17 | | | | |
| Asph | 1.66E-31 | 0.155812573 | 0.309 | 0.389 | 4.46E-27 | | | | |
| Arhgap17 | 4.11E-31 | 0.154073893 | 0.345 | 0.42 | 1.11E-26 | | | | |
| Zfp950 | 3.76E-27 | 0.153035596 | 0.214 | 0.277 | 1.01E-22 | | | | |
| Man2b1 | 2.39E-38 | 0.152571195 | 0.306 | 0.395 | 6.44E-34 | | | | |
| Vps13b | 1.43E-28 | 0.151770714 | 0.308 | 0.38 | 3.85E-24 | | | | |
| Abcc5 | 2.12E-32 | 0.149974236 | 0.195 | 0.263 | 5.70E-28 | | | | |
| Fnip2 | 5.69E-27 | 0.14943671 | 0.205 | 0.264 | 1.53E-22 | | | | |
| Hnrnpu | 2.54E-36 | 0.144773017 | 0.261 | 0.341 | 6.83E-32 | | | | |
| Gls | 6.94E-21 | 0.143081268 | 0.246 | 0.298 | 1.87E-16 | | | | |
| Kpna4 | 3.28E-29 | 0.141185974 | 0.26 | 0.335 | 8.83E-25 | | | | |
| Rsrp1 | 1.08E-33 | 0.137033515 | 0.631 | 0.746 | 2.89E-29 | | | | |
| Srsf7 | 1.10E-30 | 0.135898458 | 0.355 | 0.44 | 2.97E-26 | | | | |
| Celf1 | 3.63E-33 | 0.135484315 | 0.323 | 0.409 | 9.76E-29 | | | | |

| | | | | | |
|---|---|---|---|---|---|
| Med13l | 3.53E-44 | 0.135424072 | 0.267 | 0.361 | 9.51E-40 |
| Ttc3 | 8.71E-23 | 0.134045728 | 0.23 | 0.294 | 2.34E-18 |
| Ptprc | 8.85E-44 | 0.132057969 | 0.443 | 0.565 | 2.38E-39 |
| Aff4 | 1.83E-26 | 0.131469 | 0.191 | 0.252 | 4.92E-22 |
| Nfkb1 | 1.29E-36 | 0.129014222 | 0.297 | 0.406 | 3.46E-32 |
| Cd81 | 7.64E-33 | 0.128663627 | 0.335 | 0.431 | 2.06E-28 |
| Luc7l2 | 3.58E-40 | 0.12778485 | 0.51 | 0.632 | 9.64E-36 |
| Dmxl1 | 1.29E-28 | 0.127619761 | 0.239 | 0.316 | 3.47E-24 |
| Strbp | 1.13E-34 | 0.127552048 | 0.298 | 0.407 | 3.04E-30 |
| Gm16599 | 7.24E-23 | 0.125543756 | 0.256 | 0.319 | 1.95E-18 |
| Asb3 | 1.99E-20 | 0.124898335 | 0.197 | 0.251 | 5.35E-16 |
| Thoc2 | 5.25E-32 | 0.122374283 | 0.207 | 0.279 | 1.41E-27 |
| Arid4b | 1.15E-26 | 0.122033564 | 0.216 | 0.278 | 3.10E-22 |
| Akt3 | 4.54E-12 | 0.119284055 | 0.266 | 0.302 | 1.22E-07 |
| Basp1 | 4.90E-33 | 0.117933069 | 0.388 | 0.489 | 1.32E-28 |
| Frrs1 | 1.49E-29 | 0.117641821 | 0.21 | 0.28 | 4.02E-25 |
| St6gal1 | 1.54E-31 | 0.11682613 | 0.196 | 0.262 | 4.15E-27 |
| Usp50 | 1.29E-24 | 0.11487151 | 0.27 | 0.334 | 3.47E-20 |
| Tnrc6c | 3.10E-32 | 0.112984941 | 0.29 | 0.37 | 8.35E-28 |
| Marcks | 4.02E-28 | 0.112887361 | 0.277 | 0.351 | 1.08E-23 |
| Ttc14 | 1.67E-30 | 0.111679275 | 0.286 | 0.372 | 4.49E-26 |
| Ppp1r12a | 8.71E-37 | 0.110182303 | 0.381 | 0.481 | 2.34E-32 |
| Sltm | 2.00E-22 | 0.109904244 | 0.274 | 0.337 | 5.38E-18 |
| Plgrkt | 1.27E-30 | 0.109859005 | 0.246 | 0.321 | 3.42E-26 |
| Tmsb4x | 1.67E-29 | 0.108678368 | 0.346 | 0.444 | 4.50E-25 |
| Pik3cb | 2.92E-30 | 0.108284974 | 0.2 | 0.267 | 7.86E-26 |
| Paxbp1 | 2.46E-23 | 0.107897631 | 0.2 | 0.257 | 6.62E-19 |
| Pou2f2 | 1.44E-28 | 0.107672774 | 0.326 | 0.417 | 3.88E-24 |
| Hk2 | 8.70E-30 | 0.107060269 | 0.223 | 0.291 | 2.34E-25 |
| Rb1 | 1.79E-30 | 0.10534408 | 0.258 | 0.339 | 4.83E-26 |
| Celf2 | 5.62E-33 | 0.105208899 | 0.663 | 0.802 | 1.51E-28 |
| Son | 1.06E-33 | 0.104584045 | 0.512 | 0.618 | 2.85E-29 |
| Capza1 | 4.30E-26 | 0.104287626 | 0.218 | 0.286 | 1.16E-21 |
| Srek1 | 1.18E-29 | 0.102805724 | 0.22 | 0.29 | 3.18E-25 |
| Iffo1 | 3.46E-25 | 0.102381884 | 0.202 | 0.264 | 9.31E-21 |
| Nktr | 8.90E-25 | 0.101748406 | 0.249 | 0.327 | 2.39E-20 |
| Gnas | 7.41E-19 | 0.100276531 | 0.219 | 0.278 | 1.99E-14 |
| Ccnt2 | 1.65E-37 | 0.100257013 | 0.188 | 0.265 | 4.45E-33 |
| Tns3 | 9.94E-35 | 0.099853322 | 0.412 | 0.519 | 2.68E-30 |
| 9930021J03Rik | 3.57E-15 | 0.097997227 | 0.204 | 0.253 | 9.60E-11 |
| Cst3 | 5.22E-16 | 0.097773135 | 0.858 | 0.939 | 1.40E-11 |
| Plekho1 | 1.01E-35 | 0.0975004 | 0.374 | 0.476 | 2.73E-31 |
| Anxa3 | 5.44E-37 | 0.097385319 | 0.364 | 0.477 | 1.46E-32 |
| Spop | 5.27E-24 | 0.096992068 | 0.215 | 0.279 | 1.42E-19 |
| Snx30 | 8.12E-26 | 0.096374728 | 0.191 | 0.256 | 2.19E-21 |
| Fam13b | 3.45E-29 | 0.095118114 | 0.227 | 0.297 | 9.28E-25 |
| App | 3.23E-22 | 0.094938815 | 0.204 | 0.272 | 8.68E-18 |
| Smg1 | 8.32E-35 | 0.094785719 | 0.331 | 0.441 | 2.24E-30 |
| Senp2 | 1.97E-33 | 0.094283232 | 0.25 | 0.336 | 5.31E-29 |
| Atg16l2 | 3.69E-26 | 0.092196502 | 0.196 | 0.265 | 9.92E-22 |
| Dync1i2 | 1.09E-19 | 0.091455419 | 0.201 | 0.257 | 2.93E-15 |
| Prrc2c | 2.46E-23 | 0.091180429 | 0.265 | 0.329 | 6.61E-19 |
| Kdm7a | 3.26E-29 | 0.090918544 | 0.192 | 0.264 | 8.77E-25 |
| Smad7 | 6.72E-28 | 0.089915043 | 0.225 | 0.301 | 1.81E-23 |
| Ppcdc | 6.85E-40 | 0.089368752 | 0.31 | 0.415 | 1.84E-35 |
| Cacna1d | 2.51E-20 | 0.088133203 | 0.335 | 0.403 | 6.76E-16 |
| Fmnl2 | 3.36E-20 | 0.086703215 | 0.198 | 0.264 | 9.03E-16 |
| Myo1f | 5.06E-43 | 0.085816379 | 0.442 | 0.579 | 1.36E-38 |
| Wapl | 2.89E-28 | 0.084032469 | 0.275 | 0.351 | 7.79E-24 |
| Mafb | 6.61E-30 | 0.082748911 | 0.259 | 0.345 | 1.78E-25 |
| Gsap | 5.13E-29 | 0.082412708 | 0.231 | 0.314 | 1.38E-24 |
| Zfp644 | 1.36E-30 | 0.081352486 | 0.25 | 0.333 | 3.66E-26 |
| Pnisr | 1.29E-34 | 0.081189498 | 0.379 | 0.498 | 3.48E-30 |
| Slc38a9 | 3.80E-23 | 0.080791578 | 0.215 | 0.284 | 1.02E-18 |
| Birc6 | 5.60E-37 | 0.080338731 | 0.26 | 0.364 | 1.51E-32 |
| Lpp | 7.21E-25 | 0.080288614 | 0.179 | 0.25 | 1.94E-20 |
| Rab2a | 4.07E-32 | 0.07686758 | 0.23 | 0.312 | 1.09E-27 |
| Canx | 9.66E-32 | 0.07613853 | 0.199 | 0.277 | 2.60E-27 |
| Bclaf1 | 1.08E-21 | 0.074499753 | 0.201 | 0.263 | 2.92E-17 |
| Rbm25 | 2.23E-29 | 0.07429774 | 0.336 | 0.44 | 6.01E-25 |
| Fam91a1 | 1.62E-23 | 0.074241807 | 0.185 | 0.252 | 4.37E-19 |
| Osbpl11 | 8.12E-36 | 0.073114707 | 0.281 | 0.385 | 2.19E-31 |
| Rtn4 | 2.11E-31 | 0.070934195 | 0.326 | 0.434 | 5.69E-27 |
| Cdkal1 | 9.00E-36 | 0.070752156 | 0.247 | 0.349 | 2.42E-31 |
| Spag9 | 2.90E-34 | 0.070610428 | 0.297 | 0.401 | 7.80E-30 |
| Park2 | 6.15E-05 | 0.070516894 | 0.477 | 0.502 | 1 |
| Sh3kbp1 | 1.61E-35 | 0.06850073 | 0.419 | 0.534 | 4.32E-31 |
| Zfr | 2.22E-23 | 0.06810363 | 0.299 | 0.379 | 5.96E-19 |
| Ankrd12 | 7.33E-19 | 0.067259379 | 0.231 | 0.294 | 1.97E-14 |
| Dnmt3a | 6.75E-38 | 0.066698286 | 0.318 | 0.436 | 1.82E-33 |
| 4930517O19Rik | 4.61E-25 | 0.066496667 | 0.214 | 0.286 | 1.24E-20 |
| Smad3 | 5.28E-33 | 0.066022405 | 0.193 | 0.279 | 1.42E-28 |
| Arid2 | 2.62E-28 | 0.06601041 | 0.177 | 0.251 | 7.06E-24 |
| Ptpn1 | 1.40E-30 | 0.065362964 | 0.282 | 0.372 | 3.77E-26 |
| Strn3 | 1.31E-33 | 0.065304789 | 0.32 | 0.424 | 3.53E-29 |
| Mitf | 3.96E-32 | 0.064957919 | 0.381 | 0.496 | 1.06E-27 |
| Osbpl8 | 1.58E-17 | 0.064344679 | 0.215 | 0.271 | 4.25E-13 |
| Dock11 | 5.51E-27 | 0.064075973 | 0.286 | 0.373 | 1.48E-22 |
| C1qb | 3.74E-34 | 0.063797959 | 0.487 | 0.628 | 1.01E-29 |
| Gdi2 | 4.08E-30 | 0.063692837 | 0.319 | 0.411 | 1.10E-25 |
| Ube2d3 | 2.08E-32 | 0.063338181 | 0.182 | 0.261 | 5.59E-28 |
| Rabep1 | 9.02E-24 | 0.063205776 | 0.271 | 0.353 | 2.43E-19 |
| Ndfip1 | 1.01E-20 | 0.061843962 | 0.196 | 0.26 | 2.73E-16 |
| Phf3 | 2.04E-24 | 0.06056857 | 0.22 | 0.295 | 5.49E-20 |
| R3hdm1 | 2.75E-27 | 0.059298446 | 0.265 | 0.356 | 7.41E-23 |
| Dip2c | 4.43E-20 | 0.0587988 | 0.27 | 0.345 | 1.19E-15 |
| Pnn | 1.35E-30 | 0.057149041 | 0.215 | 0.299 | 3.64E-26 |
| Srsf5 | 2.60E-31 | 0.057007821 | 0.261 | 0.355 | 7.01E-27 |
| Prpf4b | 6.79E-30 | 0.056832772 | 0.36 | 0.465 | 1.83E-25 |
| Ubac2 | 5.61E-29 | 0.055397318 | 0.238 | 0.322 | 1.51E-24 |
| Luc7l3 | 2.05E-20 | 0.054912195 | 0.182 | 0.253 | 5.52E-16 |
| Gm42418 | 1.20E-12 | 0.054424523 | 0.827 | 0.907 | 3.22E-08 |
| Gm26542 | 1.77E-31 | 0.054280966 | 0.332 | 0.447 | 4.77E-27 |
| Synj1 | 1.97E-21 | 0.053765477 | 0.188 | 0.251 | 5.31E-17 |
| Wwox | 4.81E-15 | 0.052956037 | 0.399 | 0.472 | 1.29E-10 |
| Pnpla7 | 1.96E-38 | 0.052738488 | 0.354 | 0.477 | 5.28E-34 |
| Gm2245 | 5.36E-25 | 0.052620459 | 0.335 | 0.43 | 1.44E-20 |
| Dock1 | 2.35E-23 | 0.052564298 | 0.225 | 0.301 | 6.32E-19 |
| Phip | 3.04E-22 | 0.052446921 | 0.239 | 0.311 | 8.17E-18 |
| Gm35188 | 1.92E-14 | 0.052446636 | 0.744 | 0.837 | 5.17E-10 |
| Tsc22d4 | 3.15E-28 | 0.052441077 | 0.257 | 0.347 | 8.46E-24 |
| Sema4d | 8.84E-27 | 0.051470852 | 0.186 | 0.261 | 2.38E-22 |
| Disc1 | 2.85E-32 | 0.051387032 | 0.202 | 0.289 | 7.68E-28 |
| Lyst | 2.84E-27 | 0.049715808 | 0.193 | 0.274 | 7.63E-23 |
| Gphn | 2.36E-25 | 0.049423896 | 0.351 | 0.446 | 6.34E-21 |
| Eml4 | 8.41E-28 | 0.048287047 | 0.324 | 0.422 | 2.26E-23 |
| Nsd3 | 1.30E-35 | 0.047361666 | 0.29 | 0.404 | 3.50E-31 |
| Btbd1 | 3.21E-19 | 0.046138695 | 0.201 | 0.262 | 8.65E-15 |
| Klf12 | 8.45E-20 | 0.045380454 | 0.42 | 0.515 | 2.27E-15 |
| Camk1 | 1.19E-36 | 0.045002439 | 0.257 | 0.356 | 3.21E-32 |
| Serpine2 | 2.36E-23 | 0.0448089 | 0.185 | 0.256 | 6.35E-19 |
| Rasa1 | 8.80E-26 | 0.043995443 | 0.225 | 0.305 | 2.37E-21 |
| Spi1 | 4.88E-36 | 0.043041007 | 0.255 | 0.354 | 1.31E-31 |
| Map2k5 | 1.45E-30 | 0.042142789 | 0.177 | 0.253 | 3.89E-26 |
| Top1 | 2.92E-26 | 0.041988538 | 0.208 | 0.282 | 7.85E-22 |
| Prkcd | 7.46E-32 | 0.041751796 | 0.224 | 0.311 | 2.01E-27 |
| Snrnp70 | 1.01E-41 | 0.041544417 | 0.409 | 0.547 | 2.71E-37 |
| Mtdh | 3.29E-28 | 0.040845254 | 0.471 | 0.582 | 8.84E-24 |
| Znrf2 | 2.50E-30 | 0.040658141 | 0.19 | 0.272 | 6.72E-26 |
| Skap2 | 1.06E-39 | 0.040135527 | 0.421 | 0.556 | 2.86E-35 |
| Nufip2 | 2.00E-26 | 0.038809454 | 0.207 | 0.284 | 5.38E-22 |

| | | | | | |
|---|---|---|---|---|---|
| Tnfrsf11a | 9.63E-21 | 0.037777756 | 0.257 | 0.331 | 2.59E-16 |
| Itch | 1.67E-35 | 0.037035189 | 0.337 | 0.451 | 4.48E-31 |
| Fmnl3 | 1.52E-47 | 0.036962194 | 0.281 | 0.405 | 4.09E-43 |
| Tbc1d22a | 1.36E-34 | 0.036724819 | 0.284 | 0.386 | 3.66E-30 |
| Syngr1 | 1.19E-22 | 0.036167446 | 0.214 | 0.287 | 3.21E-18 |
| Trpm7 | 2.22E-28 | 0.036089605 | 0.221 | 0.309 | 5.98E-24 |
| Rnf169 | 7.42E-27 | 0.035335856 | 0.364 | 0.465 | 2.00E-22 |
| Acin1 | 3.26E-26 | 0.035026235 | 0.232 | 0.316 | 8.76E-22 |
| Dnajc1 | 2.93E-22 | 0.034361901 | 0.266 | 0.353 | 7.89E-18 |
| Larp1 | 7.36E-32 | 0.034286417 | 0.185 | 0.272 | 1.98E-27 |
| Tbc1d16 | 5.25E-40 | 0.034107632 | 0.236 | 0.345 | 1.41E-35 |
| Wdr26 | 2.01E-33 | 0.033952323 | 0.246 | 0.344 | 5.40E-29 |
| Atp6v0a1 | 9.87E-31 | 0.033941941 | 0.275 | 0.379 | 2.66E-26 |
| Insr | 2.00E-27 | 0.032849318 | 0.2 | 0.286 | 5.37E-23 |
| Fubp1 | 1.29E-27 | 0.032712175 | 0.224 | 0.316 | 3.46E-23 |
| Ubl3 | 1.08E-28 | 0.031828413 | 0.195 | 0.279 | 2.91E-24 |
| Atp6v0b | 4.70E-37 | 0.031773999 | 0.427 | 0.564 | 1.26E-32 |
| Mbtd1 | 2.00E-22 | 0.03110173 | 0.236 | 0.313 | 5.39E-18 |
| Rabgap1 | 2.87E-29 | 0.029523551 | 0.196 | 0.284 | 7.71E-25 |
| Zeb2 | 1.50E-17 | 0.028946399 | 0.755 | 0.853 | 4.02E-13 |
| Mlxipl | 3.17E-21 | 0.028612612 | 0.192 | 0.261 | 8.54E-17 |
| Ehmt1 | 3.24E-22 | 0.02860876 | 0.193 | 0.266 | 8.73E-18 |
| Nrf1 | 2.34E-27 | 0.028296212 | 0.192 | 0.274 | 6.30E-23 |
| Pcmtd1 | 3.94E-20 | 0.027985915 | 0.199 | 0.267 | 1.06E-15 |
| Sat1 | 1.51E-22 | 0.02756796 | 0.281 | 0.368 | 4.07E-18 |
| Rbm39 | 1.42E-26 | 0.027282509 | 0.641 | 0.765 | 3.83E-22 |
| Oxct1 | 1.30E-21 | 0.025905708 | 0.255 | 0.332 | 3.48E-17 |
| Fryl | 1.72E-20 | 0.025752224 | 0.188 | 0.254 | 4.64E-16 |
| Fnbp1 | 7.30E-37 | 0.025244341 | 0.327 | 0.445 | 1.96E-32 |
| Skil | 1.09E-25 | 0.025116599 | 0.199 | 0.276 | 2.92E-21 |
| Ewsr1 | 2.98E-30 | 0.024912132 | 0.216 | 0.311 | 8.03E-26 |
| Wipf1 | 3.65E-23 | 0.024872617 | 0.289 | 0.382 | 9.83E-19 |
| Ago3 | 3.38E-18 | 0.024687972 | 0.222 | 0.289 | 9.10E-14 |
| Stx16 | 4.63E-25 | 0.024389305 | 0.214 | 0.292 | 1.24E-20 |
| Cdk8 | 1.43E-25 | 0.023946577 | 0.178 | 0.253 | 3.86E-21 |
| Tmem243 | 7.58E-34 | 0.023596532 | 0.273 | 0.384 | 2.04E-29 |
| Capza2 | 1.24E-33 | 0.023590262 | 0.23 | 0.325 | 3.34E-29 |
| Tcf4 | 1.18E-39 | 0.023075726 | 0.474 | 0.623 | 3.16E-35 |
| Scaf8 | 7.49E-27 | 0.02269525 | 0.175 | 0.255 | 2.02E-22 |
| Dpy19l4 | 1.59E-22 | 0.022219602 | 0.207 | 0.282 | 4.27E-18 |
| Susd6 | 9.09E-22 | 0.021852889 | 0.206 | 0.282 | 2.45E-17 |
| Psap | 4.31E-31 | 0.021386741 | 0.371 | 0.493 | 1.16E-26 |
| Rasal2 | 1.25E-18 | 0.020461056 | 0.218 | 0.295 | 3.37E-14 |
| Rtn3 | 3.74E-27 | 0.020423003 | 0.376 | 0.489 | 1.01E-22 |
| Usp34 | 6.07E-33 | 0.020312947 | 0.347 | 0.463 | 1.63E-28 |
| Hps3 | 1.17E-35 | 0.020221933 | 0.222 | 0.322 | 3.14E-31 |
| Lrp1 | 9.71E-26 | 0.020148012 | 0.317 | 0.416 | 2.61E-21 |
| Sfpq | 7.86E-44 | 0.020045629 | 0.26 | 0.388 | 2.11E-39 |
| Creb1 | 1.40E-29 | 0.019844223 | 0.22 | 0.318 | 3.78E-25 |
| Gpatch8 | 1.44E-33 | 0.019514121 | 0.264 | 0.365 | 3.87E-29 |
| Comt | 4.68E-30 | 0.019396344 | 0.29 | 0.394 | 1.26E-25 |
| Supt20 | 2.07E-28 | 0.017767391 | 0.172 | 0.253 | 5.56E-24 |
| Elk3 | 5.73E-24 | 0.016798191 | 0.277 | 0.363 | 1.54E-19 |
| Mbd5 | 3.00E-42 | 0.016570028 | 0.273 | 0.401 | 8.08E-38 |
| Zfc3h1 | 7.60E-22 | 0.016321341 | 0.185 | 0.255 | 2.05E-17 |
| Dlg1 | 2.11E-18 | 0.016075639 | 0.186 | 0.253 | 5.67E-14 |
| Vgll4 | 3.69E-31 | 0.014354479 | 0.207 | 0.3 | 9.94E-27 |
| Ube3a | 2.49E-31 | 0.014226019 | 0.268 | 0.389 | 6.71E-27 |
| Nptn | 6.30E-28 | 0.01279571 | 0.27 | 0.376 | 1.69E-23 |
| Itm2b | 1.85E-26 | 0.012725519 | 0.453 | 0.58 | 4.99E-22 |
| Csnk1a1 | 6.83E-29 | 0.01189793 | 0.195 | 0.289 | 1.84E-24 |
| Rhoh | 6.11E-37 | 0.011748248 | 0.269 | 0.38 | 1.64E-32 |
| Adap2os | 8.28E-45 | 0.011671753 | 0.361 | 0.508 | 2.23E-40 |
| Asap1 | 1.25E-33 | 0.011333482 | 0.515 | 0.658 | 3.36E-29 |
| Trappc8 | 1.02E-25 | 0.011313921 | 0.193 | 0.277 | 2.75E-21 |
| Rap1b | 2.25E-42 | 0.010714123 | 0.227 | 0.344 | 6.05E-38 |
| Tcf12 | 1.43E-31 | 0.009952051 | 0.339 | 0.453 | 3.86E-27 |
| Zranb2 | 2.52E-29 | 0.009773809 | 0.172 | 0.267 | 6.78E-25 |
| Clk1 | 1.34E-34 | 0.009257077 | 0.275 | 0.385 | 3.60E-30 |
| Lrmda | 5.26E-12 | 0.008597204 | 0.71 | 0.803 | 1.41E-07 |
| Oxr1 | 1.07E-25 | 0.008542037 | 0.369 | 0.488 | 2.89E-21 |
| Fam193a | 1.33E-33 | 0.00720611 | 0.254 | 0.361 | 3.58E-29 |
| Gnaq | 1.95E-35 | 0.006723337 | 0.498 | 0.643 | 5.25E-31 |
| Slc16a6 | 8.63E-33 | 0.006271275 | 0.235 | 0.332 | 2.32E-28 |
| Rab3gap1 | 4.65E-26 | 0.005901229 | 0.172 | 0.253 | 1.25E-21 |
| Phf20l1 | 1.03E-24 | 0.005716564 | 0.231 | 0.322 | 2.76E-20 |
| Trim2 | 1.08E-22 | 0.005711448 | 0.18 | 0.262 | 2.91E-18 |
| Gsk3b | 1.86E-37 | 0.005524953 | 0.245 | 0.353 | 5.01E-33 |
| Ncf1 | 8.06E-31 | 0.004979433 | 0.215 | 0.316 | 2.17E-26 |
| Rbm26 | 2.46E-34 | 0.004608445 | 0.395 | 0.523 | 6.61E-30 |
| Atp9b | 6.05E-28 | 0.004471043 | 0.237 | 0.332 | 1.63E-23 |
| Clint1 | 1.14E-27 | 0.004089051 | 0.266 | 0.363 | 3.07E-23 |
| Cog5 | 1.66E-32 | 0.00408622 | 0.241 | 0.346 | 4.46E-28 |
| Nedd4l | 4.59E-30 | 0.003395619 | 0.17 | 0.262 | 1.23E-25 |
| Sft2d1 | 8.13E-31 | 0.003225523 | 0.305 | 0.41 | 2.19E-26 |
| Srpk2 | 4.80E-21 | 0.003014369 | 0.19 | 0.263 | 1.29E-16 |
| Lrrk1 | 1.81E-41 | 0.002936427 | 0.269 | 0.392 | 4.87E-37 |
| P2ry6 | 7.35E-32 | 0.002910257 | 0.285 | 0.389 | 1.98E-27 |
| Asxl2 | 9.66E-30 | 0.002827387 | 0.218 | 0.318 | 2.60E-25 |
| Erbin | 3.61E-38 | 0.002627348 | 0.252 | 0.372 | 9.71E-34 |
| Nf1 | 3.15E-26 | 0.002350044 | 0.399 | 0.508 | 8.46E-22 |
| Trip12 | 2.82E-26 | 0.002178692 | 0.261 | 0.36 | 7.59E-22 |
| 4930444A19Rik | 2.48E-30 | 0.001711198 | 0.312 | 0.427 | 6.66E-26 |
| Ubr3 | 1.44E-28 | 0.000639743 | 0.223 | 0.319 | 3.89E-24 |
| Rapgef6 | 2.63E-30 | 8.20E-05 | 0.247 | 0.348 | 7.07E-26 |
| Sf3b1 | 2.61E-32 | -8.62E-05 | 0.324 | 0.443 | 7.02E-28 |
| Huwe1 | 6.77E-20 | -0.000217063 | 0.237 | 0.314 | 1.82E-15 |
| Nr6a1 | 3.96E-27 | -0.001066996 | 0.429 | 0.557 | 1.06E-22 |
| R3hdm2 | 2.89E-27 | -0.001075651 | 0.259 | 0.361 | 7.77E-23 |
| Abi1 | 2.26E-32 | -0.00130968 | 0.314 | 0.431 | 6.08E-28 |
| Smap2 | 1.07E-40 | -0.001404318 | 0.479 | 0.637 | 2.88E-36 |
| Krcc1 | 2.05E-29 | -0.001637723 | 0.173 | 0.26 | 5.52E-25 |
| Gnb1 | 1.55E-25 | -0.002094435 | 0.208 | 0.296 | 4.16E-21 |
| Fndc3a | 8.35E-23 | -0.002804566 | 0.191 | 0.269 | 2.25E-18 |
| Ralgapa1 | 3.90E-26 | -0.002885941 | 0.188 | 0.276 | 1.05E-21 |
| Nsmce2 | 1.46E-17 | -0.002953101 | 0.202 | 0.269 | 3.93E-13 |
| Soga1 | 5.47E-54 | -0.002965324 | 0.366 | 0.527 | 1.47E-49 |
| Usp15 | 2.44E-27 | -0.003147878 | 0.19 | 0.277 | 6.57E-23 |
| Vps54 | 2.39E-24 | -0.003508574 | 0.208 | 0.289 | 6.42E-20 |
| Gfm2 | 1.47E-31 | -0.00360649 | 0.2 | 0.295 | 3.96E-27 |
| Snx5 | 2.38E-45 | -0.003902624 | 0.232 | 0.354 | 6.40E-41 |
| Pcm1 | 2.57E-32 | -0.004486854 | 0.222 | 0.324 | 6.91E-28 |
| Tbc1d23 | 5.80E-31 | -0.005056844 | 0.194 | 0.289 | 1.56E-26 |
| C1qc | 1.30E-37 | -0.005172677 | 0.511 | 0.674 | 3.50E-33 |
| Clip1 | 3.90E-27 | -0.005574395 | 0.175 | 0.259 | 1.05E-22 |
| Map4k3 | 2.76E-24 | -0.006089425 | 0.211 | 0.294 | 7.44E-20 |
| Gns | 4.09E-42 | -0.006541631 | 0.334 | 0.47 | 1.10E-37 |
| Gtf2i | 1.81E-35 | -0.006824276 | 0.215 | 0.319 | 4.87E-31 |
| Ssbp2 | 2.85E-21 | -0.007028365 | 0.261 | 0.349 | 7.66E-17 |
| Apba1 | 7.48E-26 | -0.007695622 | 0.352 | 0.469 | 2.01E-21 |
| Zcchc7 | 1.12E-23 | -0.008074398 | 0.458 | 0.571 | 3.01E-19 |
| Atp2b1 | 2.34E-33 | -0.008606835 | 0.318 | 0.447 | 6.29E-29 |
| Igf1r | 6.66E-21 | -0.00964664 | 0.229 | 0.312 | 1.79E-16 |
| Pde4d | 3.75E-26 | -0.010213449 | 0.294 | 0.411 | 1.01E-21 |
| Man1a2 | 4.84E-40 | -0.010396025 | 0.377 | 0.522 | 1.30E-35 |
| Kmt2e | 5.05E-37 | -0.010696544 | 0.266 | 0.388 | 1.36E-32 |
| Fkbp15 | 3.93E-22 | -0.01107424 | 0.183 | 0.257 | 1.06E-17 |
| Trem2 | 1.83E-36 | -0.011135386 | 0.275 | 0.396 | 4.92E-32 |
| Cdk12 | 8.86E-23 | -0.011769968 | 0.258 | 0.348 | 2.38E-18 |
| Tra2a | 9.14E-24 | -0.01186012 | 0.303 | 0.399 | 2.46E-19 |
| Phkb | 1.03E-28 | -0.011901617 | 0.217 | 0.31 | 2.78E-24 |

| | | | | | |
|---|---|---|---|---|---|
| Snx24 | 5.31E-33 | -0.01310827 | 0.309 | 0.435 | 1.43E-28 |
| Ccnl1 | 1.77E-23 | -0.013997382 | 0.184 | 0.263 | 4.77E-19 |
| Kdm2a | 1.16E-27 | -0.014187593 | 0.272 | 0.373 | 3.11E-23 |
| Rhoq | 8.13E-23 | -0.014800394 | 0.193 | 0.272 | 2.19E-18 |
| Lamp2 | 2.48E-25 | -0.014910785 | 0.231 | 0.321 | 6.68E-21 |
| Dcakd | 8.69E-30 | -0.0150492 | 0.199 | 0.289 | 2.34E-25 |
| AC149090.1 | 1.87E-26 | -0.016012603 | 0.534 | 0.662 | 5.02E-22 |
| Cpsf6 | 6.25E-30 | -0.016780332 | 0.208 | 0.306 | 1.68E-25 |
| Dnm2 | 6.38E-35 | -0.017103188 | 0.296 | 0.412 | 1.72E-30 |
| Rrbp1 | 3.60E-42 | -0.018090034 | 0.449 | 0.618 | 9.69E-38 |
| Lhfpl2 | 4.20E-26 | -0.018490338 | 0.337 | 0.454 | 1.13E-21 |
| Senp6 | 2.46E-23 | -0.018512374 | 0.212 | 0.297 | 6.61E-19 |
| Tmcc1 | 7.42E-25 | -0.01854732 | 0.32 | 0.432 | 2.00E-20 |
| Btbd7 | 1.03E-28 | -0.018551757 | 0.184 | 0.277 | 2.76E-24 |
| Mycbp2 | 3.02E-27 | -0.018821877 | 0.646 | 0.782 | 8.12E-23 |
| Kmt2c | 2.63E-32 | -0.018968077 | 0.337 | 0.462 | 7.07E-28 |
| Rere | 6.80E-38 | -0.019596408 | 0.471 | 0.63 | 1.83E-33 |
| Ncor1 | 1.33E-34 | -0.019666807 | 0.325 | 0.447 | 3.57E-30 |
| Rtn4rl1 | 6.80E-27 | -0.020544147 | 0.215 | 0.31 | 1.83E-22 |
| Wsb1 | 2.32E-37 | -0.020630842 | 0.247 | 0.37 | 6.26E-33 |
| Ambra1 | 3.75E-27 | -0.021188188 | 0.298 | 0.401 | 1.01E-22 |
| Bin1 | 1.27E-29 | -0.021234283 | 0.504 | 0.641 | 3.42E-25 |
| Khdrbs1 | 2.63E-37 | -0.021420545 | 0.22 | 0.331 | 7.08E-33 |
| Sesn1 | 6.26E-30 | -0.021786355 | 0.19 | 0.285 | 1.69E-25 |
| Arih1 | 4.13E-33 | -0.021982949 | 0.297 | 0.419 | 1.11E-28 |
| Cdc42 | 2.27E-35 | -0.022436787 | 0.303 | 0.429 | 6.11E-31 |
| Cmip | 1.63E-28 | -0.022463194 | 0.284 | 0.395 | 4.38E-24 |
| Arrb2 | 1.06E-28 | -0.022690221 | 0.168 | 0.254 | 2.84E-24 |
| Peli1 | 3.93E-26 | -0.022871817 | 0.244 | 0.344 | 1.06E-21 |
| Ctsh | 2.50E-32 | -0.022989153 | 0.219 | 0.323 | 6.72E-28 |
| Zmiz1 | 3.99E-32 | -0.023052715 | 0.31 | 0.431 | 1.07E-27 |
| Wdr33 | 2.33E-25 | -0.023547082 | 0.2 | 0.29 | 6.27E-21 |
| Daglb | 1.60E-33 | -0.024113618 | 0.227 | 0.334 | 4.30E-29 |
| Papola | 6.14E-25 | -0.024496106 | 0.245 | 0.343 | 1.65E-20 |
| Fus | 5.16E-38 | -0.024735915 | 0.395 | 0.54 | 1.39E-33 |
| Taok1 | 7.08E-30 | -0.024755838 | 0.233 | 0.341 | 1.90E-25 |
| Rin2 | 4.09E-27 | -0.024953931 | 0.22 | 0.319 | 1.10E-22 |
| Fermt3 | 1.22E-33 | -0.025531937 | 0.26 | 0.379 | 3.29E-29 |
| Tec | 7.02E-30 | -0.025538575 | 0.175 | 0.266 | 1.89E-25 |
| Wac | 2.05E-31 | -0.026096693 | 0.232 | 0.336 | 5.52E-27 |
| Pvt1 | 1.20E-26 | -0.026353747 | 0.237 | 0.339 | 3.24E-22 |
| Ap3b1 | 3.41E-36 | -0.026515443 | 0.256 | 0.371 | 9.18E-32 |
| Ifngr2 | 2.46E-21 | -0.026801238 | 0.186 | 0.266 | 6.62E-17 |
| Ptp4a2 | 3.32E-29 | -0.026918657 | 0.214 | 0.319 | 8.94E-25 |
| Pak2 | 4.52E-33 | -0.027400422 | 0.26 | 0.377 | 1.22E-28 |
| Stxbp5 | 1.68E-26 | -0.027687964 | 0.175 | 0.262 | 4.51E-22 |
| Rock1 | 2.29E-33 | -0.028138246 | 0.266 | 0.382 | 6.16E-29 |
| Rptor | 2.07E-36 | -0.028661366 | 0.407 | 0.55 | 5.58E-32 |
| Smyd3 | 3.72E-31 | -0.028880733 | 0.382 | 0.517 | 1.00E-26 |
| C1qa | 1.44E-35 | -0.028899423 | 0.525 | 0.693 | 3.88E-31 |
| Mbnl2 | 2.27E-35 | -0.029474999 | 0.405 | 0.546 | 6.11E-31 |
| Hnrnpdl | 7.34E-20 | -0.029571401 | 0.174 | 0.25 | 1.98E-15 |
| Fcer1g | 1.74E-26 | -0.029735743 | 0.184 | 0.279 | 4.69E-22 |
| Rsrc1 | 2.01E-24 | -0.029896769 | 0.233 | 0.327 | 5.41E-20 |
| Clta | 1.26E-28 | -0.029928351 | 0.219 | 0.315 | 3.38E-24 |
| Gm16337 | 2.70E-28 | -0.030035687 | 0.317 | 0.436 | 7.27E-24 |
| Pkig | 4.22E-26 | -0.030897687 | 0.243 | 0.343 | 1.14E-21 |
| Xist | 3.19E-16 | -0.031634587 | 0.763 | 0.868 | 8.59E-12 |
| Sfmbt1 | 1.27E-28 | -0.031885914 | 0.221 | 0.326 | 3.41E-24 |
| Srrm1 | 4.82E-34 | -0.032108502 | 0.176 | 0.276 | 1.30E-29 |
| Arid1b | 5.50E-33 | -0.032556003 | 0.318 | 0.443 | 1.48E-28 |
| Sipa1l2 | 4.27E-32 | -0.032628494 | 0.31 | 0.434 | 1.15E-27 |
| Smg6 | 5.44E-27 | -0.032701807 | 0.326 | 0.445 | 1.46E-22 |
| Tra2b | 1.72E-23 | -0.032916669 | 0.193 | 0.279 | 4.62E-19 |
| Zmynd8 | 6.85E-20 | -0.033304082 | 0.198 | 0.279 | 1.84E-15 |
| Lrch3 | 4.33E-25 | -0.033536067 | 0.213 | 0.304 | 1.16E-20 |
| Acer3 | 2.93E-27 | -0.034077592 | 0.318 | 0.43 | 7.87E-23 |
| Mark2 | 3.10E-29 | -0.034285833 | 0.206 | 0.31 | 8.33E-25 |
| B430010I23Rik | 2.11E-27 | -0.034489063 | 0.215 | 0.316 | 5.68E-23 |
| Babam2 | 1.62E-26 | -0.034638421 | 0.246 | 0.351 | 4.37E-22 |
| Ddx17 | 2.00E-27 | -0.034837694 | 0.281 | 0.391 | 5.39E-23 |
| Ppp2r5c | 3.79E-29 | -0.035401826 | 0.188 | 0.285 | 1.02E-24 |
| Gm26827 | 2.25E-19 | -0.035555786 | 0.225 | 0.312 | 6.05E-15 |
| Pias2 | 1.05E-20 | -0.035613869 | 0.181 | 0.259 | 2.82E-16 |
| Calm2 | 1.43E-20 | -0.035855411 | 0.214 | 0.303 | 3.85E-16 |
| Ly86 | 6.27E-27 | -0.036012419 | 0.622 | 0.772 | 1.69E-22 |
| Nsd1 | 3.23E-34 | -0.036434732 | 0.469 | 0.619 | 8.70E-30 |
| Ube2h | 6.81E-21 | -0.03652087 | 0.277 | 0.37 | 1.83E-16 |
| Trappc9 | 1.48E-17 | -0.036549298 | 0.188 | 0.264 | 3.98E-13 |
| Erp29 | 1.50E-24 | -0.036772819 | 0.168 | 0.253 | 4.03E-20 |
| Dis3l2 | 4.11E-25 | -0.037220229 | 0.17 | 0.255 | 1.11E-20 |
| Ubn2 | 2.56E-32 | -0.037320469 | 0.353 | 0.481 | 6.90E-28 |
| Ptprj | 1.28E-24 | -0.037513237 | 0.609 | 0.746 | 3.43E-20 |
| Tpm3 | 2.01E-35 | -0.037516494 | 0.227 | 0.339 | 5.42E-31 |
| Fndc3b | 2.13E-30 | -0.037840915 | 0.191 | 0.289 | 5.73E-26 |
| P2rx7 | 3.55E-38 | -0.038199639 | 0.238 | 0.361 | 9.56E-34 |
| Fbxl17 | 1.15E-35 | -0.038235335 | 0.428 | 0.581 | 3.09E-31 |
| Mylip | 2.04E-27 | -0.038258485 | 0.34 | 0.458 | 5.48E-23 |
| Plekhm3 | 2.47E-21 | -0.038406297 | 0.205 | 0.288 | 6.64E-17 |
| Abhd12 | 2.59E-24 | -0.039013113 | 0.494 | 0.626 | 6.96E-20 |
| Ankrd11 | 1.59E-33 | -0.039086215 | 0.397 | 0.536 | 4.28E-29 |
| Stard9 | 2.26E-34 | -0.039790472 | 0.306 | 0.432 | 6.07E-30 |
| Rapgef2 | 1.71E-18 | -0.040014377 | 0.189 | 0.266 | 4.60E-14 |
| Cerk | 1.09E-41 | -0.040280754 | 0.32 | 0.472 | 2.95E-37 |
| Map4k4 | 7.77E-37 | -0.040347128 | 0.4 | 0.548 | 2.09E-32 |
| Bmpr2 | 3.89E-27 | -0.040519433 | 0.195 | 0.296 | 1.05E-22 |
| Nisch | 1.52E-21 | -0.040633908 | 0.204 | 0.286 | 4.10E-17 |
| Dock7 | 1.72E-26 | -0.040976524 | 0.184 | 0.273 | 4.63E-22 |
| Arpc2 | 8.81E-32 | -0.041521979 | 0.205 | 0.311 | 2.37E-27 |
| Evl | 9.32E-32 | -0.041676609 | 0.268 | 0.385 | 2.51E-27 |
| Pibf1 | 3.76E-19 | -0.041770673 | 0.186 | 0.264 | 1.01E-14 |
| Ubr5 | 4.29E-26 | -0.041915933 | 0.167 | 0.252 | 1.15E-21 |
| Srrm2 | 4.07E-36 | -0.042646412 | 0.533 | 0.695 | 1.09E-31 |
| Kdm3b | 2.03E-36 | -0.042909533 | 0.253 | 0.379 | 5.47E-32 |
| Arhgap25 | 1.80E-33 | -0.043267956 | 0.209 | 0.319 | 4.84E-29 |
| Dhx9 | 2.98E-25 | -0.043712638 | 0.21 | 0.302 | 8.01E-21 |
| Map3k5 | 1.15E-34 | -0.044335998 | 0.243 | 0.369 | 3.09E-30 |
| Zdhhc20 | 7.72E-25 | -0.045149486 | 0.214 | 0.305 | 2.08E-20 |
| Irak2 | 1.94E-38 | -0.046440772 | 0.315 | 0.455 | 5.22E-34 |
| Ascc3 | 2.42E-25 | -0.04678229 | 0.215 | 0.314 | 6.52E-21 |
| Pitpnc1 | 7.69E-26 | -0.046788556 | 0.431 | 0.564 | 2.07E-21 |
| Fto | 2.10E-30 | -0.047015783 | 0.219 | 0.324 | 5.65E-26 |
| Rbm33 | 2.94E-26 | -0.047352358 | 0.17 | 0.261 | 7.91E-22 |
| Fbxo11 | 1.93E-26 | -0.047656376 | 0.244 | 0.35 | 5.18E-22 |
| Slc29a3 | 5.69E-38 | -0.047715634 | 0.154 | 0.252 | 1.53E-33 |
| Helz | 7.50E-28 | -0.048000882 | 0.193 | 0.287 | 2.02E-23 |
| Clasp2 | 1.44E-31 | -0.048149128 | 0.409 | 0.55 | 3.88E-27 |
| Hps4 | 2.35E-32 | -0.048154042 | 0.207 | 0.313 | 6.31E-28 |
| Chd6 | 1.14E-29 | -0.048395772 | 0.215 | 0.316 | 3.06E-25 |
| Chd7 | 2.33E-29 | -0.048767654 | 0.328 | 0.455 | 6.27E-25 |
| Zcchc11 | 2.41E-14 | -0.04888213 | 0.228 | 0.295 | 6.50E-10 |
| Zfp407 | 1.31E-20 | -0.049241194 | 0.193 | 0.28 | 3.52E-16 |
| Ext1 | 8.23E-14 | -0.049941784 | 0.252 | 0.329 | 2.22E-09 |
| Nrd1 | 2.16E-21 | -0.050281824 | 0.199 | 0.285 | 5.82E-17 |
| Mia2 | 1.37E-26 | -0.051844916 | 0.179 | 0.269 | 3.67E-22 |
| Sipa1 | 9.38E-37 | -0.051999806 | 0.218 | 0.336 | 2.52E-32 |
| Garnl3 | 1.33E-20 | -0.052185485 | 0.169 | 0.251 | 3.58E-16 |
| Crebbp | 5.92E-35 | -0.052347718 | 0.332 | 0.466 | 1.59E-30 |
| Wdr44 | 5.05E-28 | -0.053223124 | 0.233 | 0.334 | 1.36E-23 |
| Mdm4 | 9.93E-31 | -0.053233432 | 0.194 | 0.3 | 2.67E-26 |
| Cd84 | 4.33E-34 | -0.053419663 | 0.323 | 0.457 | 1.16E-29 |

| | | | | | |
|---|---|---|---|---|---|
| Brd4 | 5.18E-31 | -0.053475278 | 0.235 | 0.344 | 1.39E-26 |
| Rsbn1l | 2.20E-27 | -0.053862766 | 0.213 | 0.315 | 5.93E-23 |
| Trio | 5.82E-35 | -0.054035177 | 0.269 | 0.399 | 1.57E-30 |
| Arid4a | 3.01E-26 | -0.054317277 | 0.222 | 0.323 | 8.09E-22 |
| Cnot4 | 1.90E-31 | -0.054880275 | 0.299 | 0.415 | 5.12E-27 |
| Atxn7 | 7.32E-24 | -0.054897358 | 0.208 | 0.299 | 1.97E-19 |
| Zfp292 | 2.77E-35 | -0.055565889 | 0.255 | 0.384 | 7.44E-31 |
| Tbl1xr1 | 2.63E-34 | -0.056008045 | 0.275 | 0.403 | 7.08E-30 |
| Tubgcp5 | 3.22E-39 | -0.056481278 | 0.244 | 0.374 | 8.66E-35 |
| Jak1 | 1.16E-32 | -0.056812976 | 0.228 | 0.342 | 3.11E-28 |
| Ankhd1 | 2.80E-27 | -0.057110118 | 0.273 | 0.386 | 7.54E-23 |
| Rbm5 | 8.15E-30 | -0.057252736 | 0.281 | 0.399 | 2.19E-25 |
| Lcorl | 2.53E-19 | -0.057653908 | 0.217 | 0.303 | 6.81E-15 |
| Btk | 1.93E-22 | -0.057773821 | 0.179 | 0.261 | 5.18E-18 |
| Rsf1 | 2.54E-26 | -0.058272469 | 0.202 | 0.298 | 6.83E-22 |
| Kif21b | 3.54E-30 | -0.058595748 | 0.185 | 0.283 | 9.52E-26 |
| Ankfy1 | 3.39E-24 | -0.058845475 | 0.181 | 0.266 | 9.12E-20 |
| Elf1 | 2.16E-24 | -0.058875059 | 0.198 | 0.291 | 5.82E-20 |
| Ep300 | 1.31E-33 | -0.059190562 | 0.213 | 0.328 | 3.53E-29 |
| Thada | 4.78E-29 | -0.059682223 | 0.181 | 0.285 | 1.29E-24 |
| Magi1 | 2.48E-21 | -0.059705451 | 0.459 | 0.581 | 6.68E-17 |
| Pds5a | 1.16E-23 | -0.059921232 | 0.205 | 0.299 | 3.13E-19 |
| Parp8 | 6.24E-31 | -0.060599069 | 0.309 | 0.44 | 1.68E-26 |
| Osbpl9 | 1.53E-21 | -0.062010552 | 0.206 | 0.3 | 4.12E-17 |
| Lncpint | 2.69E-35 | -0.062446342 | 0.308 | 0.449 | 7.23E-31 |
| Cyfip1 | 6.34E-34 | -0.063188004 | 0.426 | 0.578 | 1.71E-29 |
| Sgms1 | 4.14E-22 | -0.063446553 | 0.174 | 0.26 | 1.12E-17 |
| Itgav | 6.57E-26 | -0.063549212 | 0.259 | 0.369 | 1.77E-21 |
| Slmap | 4.21E-21 | -0.064176075 | 0.222 | 0.31 | 1.13E-16 |
| Pias1 | 1.73E-36 | -0.064251984 | 0.291 | 0.425 | 4.66E-32 |
| Dmtf1 | 1.50E-31 | -0.064276025 | 0.166 | 0.268 | 4.05E-27 |
| Nr3c1 | 2.95E-27 | -0.06464977 | 0.245 | 0.349 | 7.94E-23 |
| Ddx5 | 2.62E-21 | -0.065363209 | 0.618 | 0.747 | 7.05E-17 |
| Slc24a3 | 2.38E-29 | -0.065532535 | 0.388 | 0.537 | 6.39E-25 |
| Ash1l | 2.77E-37 | -0.065670223 | 0.339 | 0.482 | 7.46E-33 |
| Peak1 | 3.83E-24 | -0.066127545 | 0.22 | 0.321 | 1.03E-19 |
| Rnpep | 1.67E-31 | -0.066407652 | 0.153 | 0.251 | 4.50E-27 |
| Nuak1 | 6.82E-28 | -0.066479887 | 0.225 | 0.334 | 1.83E-23 |
| Epc1 | 2.67E-24 | -0.066799215 | 0.192 | 0.282 | 7.19E-20 |
| Ccny | 8.61E-26 | -0.067323077 | 0.197 | 0.292 | 2.32E-21 |
| Ktn1 | 3.69E-29 | -0.067357322 | 0.188 | 0.3 | 9.93E-25 |
| Arhgap31 | 7.26E-20 | -0.067377225 | 0.225 | 0.319 | 1.95E-15 |
| Ppp3ca | 6.93E-33 | -0.068585426 | 0.538 | 0.702 | 1.86E-28 |
| Rnf111 | 6.27E-34 | -0.068873121 | 0.235 | 0.356 | 1.69E-29 |
| Ptbp3 | 3.81E-35 | -0.069017827 | 0.277 | 0.413 | 1.03E-30 |
| Bod1l | 2.64E-31 | -0.069583759 | 0.167 | 0.264 | 7.09E-27 |
| Atp2c1 | 1.55E-27 | -0.069596804 | 0.315 | 0.435 | 4.17E-23 |
| Snd1 | 1.92E-31 | -0.069751037 | 0.21 | 0.32 | 5.16E-27 |
| Ank2 | 3.53E-20 | -0.069755008 | 0.487 | 0.609 | 9.49E-16 |
| Med12l | 2.27E-36 | -0.069852143 | 0.413 | 0.566 | 6.11E-32 |
| Sh3glb1 | 1.16E-37 | -0.070451338 | 0.217 | 0.341 | 3.13E-33 |
| Scmh1 | 1.36E-26 | -0.071083505 | 0.21 | 0.314 | 3.65E-22 |
| Retreg1 | 1.21E-31 | -0.071508303 | 0.285 | 0.409 | 3.24E-27 |
| Aim2 | 5.40E-23 | -0.071987241 | 0.253 | 0.353 | 1.45E-18 |
| Fnip1 | 1.99E-29 | -0.072166448 | 0.242 | 0.357 | 5.36E-25 |
| Atxn7l1 | 1.32E-28 | -0.072222662 | 0.255 | 0.37 | 3.55E-24 |
| Asah1 | 1.01E-20 | -0.07239704 | 0.176 | 0.257 | 2.70E-16 |
| Rtn1 | 2.53E-28 | -0.072518901 | 0.318 | 0.444 | 6.80E-24 |
| Dip2b | 1.76E-30 | -0.072961039 | 0.652 | 0.813 | 4.73E-26 |
| Znrf1 | 1.55E-23 | -0.073734133 | 0.172 | 0.263 | 4.16E-19 |
| Mysm1 | 8.44E-27 | -0.073831884 | 0.164 | 0.255 | 2.27E-22 |
| Ube2k | 7.10E-21 | -0.074112936 | 0.193 | 0.279 | 1.91E-16 |
| Dgkd | 2.99E-33 | -0.07480316 | 0.285 | 0.414 | 8.05E-29 |
| Atad2b | 2.12E-28 | -0.075580747 | 0.229 | 0.338 | 5.71E-24 |
| Chd2 | 4.74E-32 | -0.075715698 | 0.288 | 0.422 | 1.28E-27 |
| Uvrag | 9.93E-36 | -0.075986978 | 0.324 | 0.466 | 2.67E-31 |
| Zbtb20 | 1.62E-29 | -0.075992177 | 0.595 | 0.756 | 4.35E-25 |
| Fam172a | 3.95E-31 | -0.077069882 | 0.405 | 0.546 | 1.06E-26 |
| Capzb | 1.67E-27 | -0.077380568 | 0.166 | 0.264 | 4.50E-23 |
| Tnrc18 | 7.34E-25 | -0.077430754 | 0.214 | 0.312 | 1.97E-20 |
| Hectd1 | 6.20E-27 | -0.077604887 | 0.164 | 0.259 | 1.67E-22 |
| Cul1 | 1.06E-28 | -0.077814278 | 0.259 | 0.373 | 2.84E-24 |
| Itsn2 | 6.59E-32 | -0.078450932 | 0.245 | 0.363 | 1.77E-27 |
| Crebrf | 2.02E-25 | -0.078641272 | 0.177 | 0.269 | 5.44E-21 |
| Hmg20a | 1.07E-21 | -0.078792391 | 0.164 | 0.254 | 2.88E-17 |
| Eif4g3 | 9.31E-31 | -0.079778653 | 0.325 | 0.459 | 2.51E-26 |
| Arhgef6 | 2.09E-24 | -0.079862558 | 0.18 | 0.272 | 5.63E-20 |
| Spred1 | 6.03E-27 | -0.080254626 | 0.158 | 0.252 | 1.62E-22 |
| Gapvd1 | 1.00E-22 | -0.080340948 | 0.208 | 0.31 | 2.70E-18 |
| Akap13 | 4.32E-34 | -0.080951913 | 0.418 | 0.568 | 1.16E-29 |
| Sparc | 5.65E-39 | -0.081054355 | 0.318 | 0.472 | 1.52E-34 |
| Ulk2 | 5.81E-31 | -0.081940481 | 0.422 | 0.572 | 1.56E-26 |
| Stag1 | 2.83E-30 | -0.081949207 | 0.405 | 0.547 | 7.61E-26 |
| Ptpra | 1.93E-27 | -0.082236416 | 0.31 | 0.434 | 5.18E-23 |
| Phf21a | 1.84E-29 | -0.08229096 | 0.341 | 0.482 | 4.94E-25 |
| Cds1 | 1.41E-24 | -0.08233203 | 0.167 | 0.256 | 3.80E-20 |
| Ogt | 5.31E-26 | -0.082590059 | 0.231 | 0.334 | 1.43E-21 |
| Phc2 | 1.82E-21 | -0.083754174 | 0.238 | 0.336 | 4.90E-17 |
| Fam168a | 8.71E-30 | -0.083887752 | 0.253 | 0.373 | 2.34E-25 |
| Rbm6 | 2.95E-27 | -0.083956958 | 0.28 | 0.4 | 7.94E-23 |
| Ankrd17 | 3.51E-28 | -0.084204666 | 0.267 | 0.384 | 9.44E-24 |
| Mapk14 | 1.18E-34 | -0.08468132 | 0.266 | 0.401 | 3.17E-30 |
| Prkcb | 1.31E-24 | -0.084846511 | 0.378 | 0.508 | 3.53E-20 |
| Tmcc3 | 1.06E-10 | -0.085011162 | 0.795 | 0.88 | 2.87E-06 |
| Pip4k2a | 3.14E-26 | -0.085122865 | 0.615 | 0.759 | 8.45E-22 |
| Ccdc50 | 4.97E-28 | -0.085187463 | 0.211 | 0.319 | 1.34E-23 |
| Cltc | 3.20E-32 | -0.08569935 | 0.248 | 0.368 | 8.62E-28 |
| Anks1 | 3.39E-31 | -0.085725172 | 0.184 | 0.29 | 9.13E-27 |
| Myh9 | 5.40E-33 | -0.086000291 | 0.193 | 0.309 | 1.45E-28 |
| Morc3 | 1.18E-37 | -0.086595681 | 0.194 | 0.318 | 3.18E-33 |
| Vti1a | 4.59E-32 | -0.086825995 | 0.294 | 0.429 | 1.23E-27 |
| Rgs10 | 1.55E-36 | -0.087120723 | 0.363 | 0.523 | 4.18E-32 |
| Atrx | 1.14E-33 | -0.087407268 | 0.289 | 0.422 | 3.07E-29 |
| Dst | 1.88E-18 | -0.087413477 | 0.685 | 0.813 | 5.05E-14 |
| Map3k3 | 3.99E-38 | -0.087461783 | 0.304 | 0.46 | 1.07E-33 |
| Dennd5a | 2.13E-36 | -0.087678837 | 0.181 | 0.304 | 5.72E-32 |
| Gatad2b | 1.67E-18 | -0.087692483 | 0.193 | 0.279 | 4.48E-14 |
| Setd5 | 5.25E-26 | -0.087863991 | 0.278 | 0.395 | 1.41E-21 |
| Kcnk13 | 1.59E-18 | -0.088488842 | 0.23 | 0.323 | 4.29E-14 |
| Gm32036 | 1.07E-30 | -0.088511524 | 0.356 | 0.492 | 2.87E-26 |
| Baz2b | 8.86E-40 | -0.088697905 | 0.351 | 0.515 | 2.39E-35 |
| Epb41 | 1.35E-18 | -0.09004502 | 0.177 | 0.259 | 3.64E-14 |
| Cryl1 | 2.80E-27 | -0.090135483 | 0.186 | 0.289 | 7.54E-23 |
| Tgfbr2 | 1.34E-36 | -0.09019983 | 0.371 | 0.532 | 3.60E-32 |
| Map4 | 2.59E-33 | -0.090468388 | 0.341 | 0.488 | 6.96E-29 |
| Akap8l | 5.98E-41 | -0.090693181 | 0.289 | 0.441 | 1.61E-36 |
| Usp9x | 2.31E-26 | -0.090897215 | 0.216 | 0.326 | 6.21E-22 |
| Ltc4s | 3.65E-30 | -0.091348513 | 0.154 | 0.257 | 9.82E-26 |
| Pwwp2a | 2.34E-33 | -0.092032616 | 0.325 | 0.466 | 6.31E-29 |
| Sik2 | 2.93E-19 | -0.092241089 | 0.237 | 0.334 | 7.89E-15 |
| Ppt1 | 4.17E-26 | -0.092517866 | 0.161 | 0.256 | 1.12E-21 |
| Grb2 | 2.41E-31 | -0.092651403 | 0.259 | 0.388 | 6.48E-27 |
| Cep170 | 8.13E-37 | -0.09305533 | 0.216 | 0.345 | 2.19E-32 |
| Fbxl20 | 3.13E-26 | -0.093291248 | 0.224 | 0.332 | 8.42E-22 |
| Kif13b | 1.05E-28 | -0.094062983 | 0.179 | 0.28 | 2.82E-24 |
| Asxl1 | 4.81E-26 | -0.094114954 | 0.221 | 0.332 | 1.29E-21 |
| Taf15 | 4.64E-30 | -0.094127342 | 0.232 | 0.347 | 1.25E-25 |
| Xrn1 | 3.10E-24 | -0.094233124 | 0.17 | 0.258 | 8.33E-20 |
| Apc | 7.03E-35 | -0.09424216 | 0.275 | 0.415 | 1.89E-30 |
| Ppp6r3 | 1.18E-29 | -0.094299162 | 0.176 | 0.281 | 3.18E-25 |
| Nr2c2 | 6.65E-20 | -0.094773094 | 0.173 | 0.26 | 1.79E-15 |

| | | | | | |
|---|---|---|---|---|---|
| Hdac9 | 6.90E-16 | -0.095141175 | 0.38 | 0.493 | 1.86E-11 |
| Dennd1a | 9.81E-27 | -0.095618716 | 0.318 | 0.441 | 2.64E-22 |
| Mon2 | 9.28E-24 | -0.095638061 | 0.209 | 0.307 | 2.50E-19 |
| Psme4 | 2.78E-29 | -0.096593348 | 0.177 | 0.283 | 7.48E-25 |
| Ski | 4.07E-28 | -0.097024589 | 0.303 | 0.424 | 1.09E-23 |
| Rnf216 | 1.04E-32 | -0.097039664 | 0.465 | 0.629 | 2.81E-28 |
| Git2 | 2.30E-37 | -0.097452734 | 0.334 | 0.488 | 6.19E-33 |
| Herc1 | 3.30E-27 | -0.097580257 | 0.246 | 0.36 | 8.89E-23 |
| Lysmd4 | 2.16E-30 | -0.097745239 | 0.695 | 0.856 | 5.81E-26 |
| Snx13 | 2.88E-32 | -0.098240741 | 0.237 | 0.361 | 7.74E-28 |
| Prkce | 4.62E-32 | -0.098343581 | 0.358 | 0.512 | 1.24E-27 |
| Ppp1r21 | 7.82E-34 | -0.098871591 | 0.22 | 0.344 | 2.10E-29 |
| Dnajc7 | 4.16E-34 | -0.099903101 | 0.21 | 0.333 | 1.12E-29 |
| Zmym2 | 3.34E-24 | -0.100287967 | 0.159 | 0.252 | 8.98E-20 |
| Btbd9 | 2.68E-22 | -0.100528248 | 0.407 | 0.536 | 7.21E-18 |
| Specc1 | 3.18E-28 | -0.100978246 | 0.467 | 0.619 | 8.56E-24 |
| Lcor | 6.67E-22 | -0.101031999 | 0.247 | 0.35 | 1.79E-17 |
| Scamp2 | 9.34E-41 | -0.101256275 | 0.208 | 0.339 | 2.51E-36 |
| Atf6 | 2.50E-25 | -0.101296331 | 0.164 | 0.258 | 6.73E-21 |
| Cask | 1.98E-42 | -0.101509898 | 0.339 | 0.508 | 5.34E-38 |
| Zmym4 | 1.40E-29 | -0.101679125 | 0.237 | 0.357 | 3.76E-25 |
| Crlf3 | 1.83E-25 | -0.102063927 | 0.317 | 0.44 | 4.92E-21 |
| Ptk2b | 1.85E-29 | -0.102260947 | 0.212 | 0.326 | 4.98E-25 |
| Pros1 | 1.47E-29 | -0.102322009 | 0.274 | 0.401 | 3.96E-25 |
| Rb1cc1 | 1.24E-26 | -0.102861558 | 0.179 | 0.279 | 3.35E-22 |
| Acss1 | 5.73E-25 | -0.103798531 | 0.163 | 0.259 | 1.54E-20 |
| Stk3 | 1.30E-23 | -0.103860104 | 0.173 | 0.268 | 3.50E-19 |
| Tacc1 | 3.26E-33 | -0.104227649 | 0.426 | 0.591 | 8.77E-29 |
| Pacsin2 | 5.53E-27 | -0.105366464 | 0.195 | 0.301 | 1.49E-22 |
| Nrros | 2.27E-36 | -0.105661771 | 0.271 | 0.413 | 6.10E-32 |
| Cdk13 | 9.74E-34 | -0.106363283 | 0.234 | 0.362 | 2.62E-29 |
| Bach2 | 6.47E-21 | -0.106800629 | 0.347 | 0.469 | 1.74E-16 |
| Nipbl | 9.12E-34 | -0.106984152 | 0.399 | 0.556 | 2.45E-29 |
| Scaper | 6.36E-21 | -0.107755404 | 0.202 | 0.304 | 1.71E-16 |
| Frmd4a | 4.21E-21 | -0.107838396 | 0.77 | 0.897 | 1.13E-16 |
| Mark3 | 5.38E-30 | -0.108060514 | 0.254 | 0.379 | 1.45E-25 |
| Lgmn | 7.03E-32 | -0.108665561 | 0.596 | 0.769 | 1.89E-27 |
| Zdhhc14 | 3.01E-22 | -0.109074929 | 0.253 | 0.361 | 8.09E-18 |
| Larp4b | 1.93E-30 | -0.109590952 | 0.2 | 0.316 | 5.20E-26 |
| Ppm1h | 1.72E-31 | -0.110084137 | 0.439 | 0.597 | 4.63E-27 |
| Pum1 | 9.37E-26 | -0.110191412 | 0.185 | 0.288 | 2.52E-21 |
| Fry | 1.86E-27 | -0.110218 | 0.195 | 0.302 | 5.01E-23 |
| Mgat4a | 2.74E-39 | -0.11079997 | 0.412 | 0.585 | 7.37E-35 |
| Hivep3 | 2.35E-25 | -0.111326978 | 0.381 | 0.52 | 6.33E-21 |
| Lims1 | 3.97E-23 | -0.111385938 | 0.383 | 0.513 | 1.07E-18 |
| Abca1 | 6.01E-32 | -0.111460757 | 0.288 | 0.426 | 1.62E-27 |
| Arhgap39 | 4.04E-33 | -0.11191431 | 0.369 | 0.522 | 1.09E-28 |
| Cux1 | 2.60E-39 | -0.111986921 | 0.368 | 0.536 | 7.01E-35 |
| Nrip1 | 2.63E-29 | -0.112098405 | 0.465 | 0.621 | 7.09E-25 |
| Prex1 | 3.12E-43 | -0.112513126 | 0.312 | 0.471 | 8.40E-39 |
| Irf2 | 5.16E-37 | -0.112621666 | 0.282 | 0.427 | 1.39E-32 |
| Jmjd1c | 1.19E-35 | -0.11268135 | 0.524 | 0.702 | 3.20E-31 |
| Ncoa2 | 7.50E-27 | -0.112904853 | 0.259 | 0.381 | 2.02E-22 |
| Gna15 | 5.01E-33 | -0.113169279 | 0.227 | 0.351 | 1.35E-28 |
| Tg | 4.54E-26 | -0.113282826 | 0.193 | 0.293 | 1.22E-21 |
| Diaph2 | 1.02E-24 | -0.113348307 | 0.591 | 0.743 | 2.76E-20 |
| Med13 | 4.12E-23 | -0.113459785 | 0.176 | 0.271 | 1.11E-18 |
| Atp8a1 | 2.21E-31 | -0.113747204 | 0.397 | 0.555 | 5.94E-27 |
| Plcg2 | 1.78E-34 | -0.113773362 | 0.307 | 0.452 | 4.78E-30 |
| Stag2 | 8.98E-33 | -0.113831926 | 0.228 | 0.356 | 2.42E-28 |
| Nlk | 2.37E-25 | -0.114352368 | 0.21 | 0.318 | 6.39E-21 |
| Snx29 | 2.88E-26 | -0.115471862 | 0.622 | 0.78 | 7.75E-22 |
| Lrba | 3.48E-32 | -0.116704557 | 0.307 | 0.453 | 9.37E-28 |
| St3gal3 | 3.61E-22 | -0.117065328 | 0.186 | 0.277 | 9.72E-18 |
| Ccdc93 | 3.64E-27 | -0.117236999 | 0.16 | 0.262 | 9.80E-23 |
| Foxn3 | 9.28E-30 | -0.117238335 | 0.717 | 0.874 | 2.50E-25 |
| Mctp1 | 3.86E-29 | -0.117510198 | 0.436 | 0.589 | 1.04E-24 |
| Bptf | 2.28E-32 | -0.117590775 | 0.238 | 0.366 | 6.13E-28 |
| Man1c1 | 8.17E-29 | -0.118155827 | 0.221 | 0.339 | 2.20E-24 |
| Zfp710 | 2.05E-31 | -0.118258193 | 0.536 | 0.71 | 5.50E-27 |
| Gtdc1 | 3.17E-25 | -0.118633349 | 0.29 | 0.421 | 8.52E-21 |
| Spire1 | 2.18E-35 | -0.119369966 | 0.285 | 0.427 | 5.86E-31 |
| Rps6ka1 | 5.31E-33 | -0.119546736 | 0.197 | 0.316 | 1.43E-28 |
| Il6st | 1.71E-24 | -0.119608861 | 0.184 | 0.278 | 4.61E-20 |
| Usp8 | 1.50E-26 | -0.119703525 | 0.179 | 0.279 | 4.04E-22 |
| Pid1 | 1.54E-23 | -0.119885683 | 0.425 | 0.56 | 4.13E-19 |
| Rnf13 | 5.18E-37 | -0.119947166 | 0.251 | 0.388 | 1.39E-32 |
| Hipk2 | 1.31E-30 | -0.119971923 | 0.187 | 0.306 | 3.54E-26 |
| Trim44 | 8.34E-26 | -0.120110699 | 0.232 | 0.345 | 2.24E-21 |
| Hpgd | 2.00E-33 | -0.120434139 | 0.228 | 0.364 | 5.37E-29 |
| Hpgds | 2.53E-28 | -0.120481103 | 0.55 | 0.714 | 6.80E-24 |
| Cmtm7 | 1.94E-35 | -0.12049885 | 0.184 | 0.302 | 5.22E-31 |
| Ints6l | 3.22E-39 | -0.121302956 | 0.347 | 0.516 | 8.66E-35 |
| Ppp1r9a | 1.80E-25 | -0.121490145 | 0.454 | 0.605 | 4.84E-21 |
| Lrch1 | 1.36E-38 | -0.122002936 | 0.534 | 0.723 | 3.67E-34 |
| Cep68 | 1.20E-26 | -0.122038722 | 0.192 | 0.299 | 3.22E-22 |
| Fars2 | 9.50E-21 | -0.122043419 | 0.195 | 0.293 | 2.56E-16 |
| Ube3c | 7.53E-33 | -0.122173993 | 0.169 | 0.279 | 2.02E-28 |
| Nek7 | 1.63E-39 | -0.122531966 | 0.333 | 0.493 | 4.39E-35 |
| Zfand6 | 5.22E-28 | -0.122564061 | 0.192 | 0.301 | 1.41E-23 |
| Pik3r5 | 2.98E-24 | -0.122626974 | 0.182 | 0.287 | 8.03E-20 |
| Il6ra | 1.53E-42 | -0.124115369 | 0.319 | 0.484 | 4.13E-38 |
| P4ha1 | 2.37E-21 | -0.124267911 | 0.2 | 0.298 | 6.37E-17 |
| Dnajc13 | 1.18E-19 | -0.124335635 | 0.164 | 0.256 | 3.18E-15 |
| Hsp90b1 | 1.48E-21 | -0.125384352 | 0.161 | 0.255 | 3.98E-17 |
| Cd86 | 2.66E-27 | -0.125439706 | 0.245 | 0.365 | 7.16E-23 |
| Chd9 | 1.51E-24 | -0.125828022 | 0.748 | 0.889 | 4.06E-20 |
| Lair1 | 4.63E-45 | -0.125946756 | 0.465 | 0.663 | 1.25E-40 |
| Inpp5d | 1.19E-29 | -0.125961255 | 0.859 | 0.972 | 3.20E-25 |
| Zmynd11 | 1.26E-35 | -0.127392674 | 0.206 | 0.335 | 3.40E-31 |
| Sipa1l1 | 3.59E-30 | -0.12819349 | 0.254 | 0.386 | 9.66E-26 |
| Tulp4 | 8.29E-22 | -0.128521638 | 0.182 | 0.282 | 2.23E-17 |
| Zfp638 | 1.15E-33 | -0.12860278 | 0.286 | 0.429 | 3.08E-29 |
| Tmem164 | 1.01E-24 | -0.129592595 | 0.241 | 0.36 | 2.71E-20 |
| Tsix | 6.06E-23 | -0.129849615 | 0.215 | 0.321 | 1.63E-18 |
| Ntpcr | 4.52E-34 | -0.130787875 | 0.196 | 0.32 | 1.22E-29 |
| Rfx3 | 8.37E-20 | -0.13087449 | 0.168 | 0.26 | 2.25E-15 |
| Kansl1l | 6.44E-28 | -0.131173223 | 0.251 | 0.377 | 1.73E-23 |
| Fgd2 | 2.30E-37 | -0.131475343 | 0.428 | 0.605 | 6.19E-33 |
| Ddx6 | 1.61E-27 | -0.131503291 | 0.187 | 0.298 | 4.34E-23 |
| Nav2 | 8.09E-09 | -0.13216345 | 0.912 | 0.962 | 0.00021769 |
| Stab1 | 7.58E-29 | -0.132247422 | 0.185 | 0.297 | 2.04E-24 |
| Rps6ka3 | 5.97E-27 | -0.13240158 | 0.22 | 0.34 | 1.61E-22 |
| Sh3pxd2a | 2.11E-32 | -0.132709293 | 0.217 | 0.344 | 5.69E-28 |
| Heatr5a | 2.77E-29 | -0.132746613 | 0.244 | 0.37 | 7.46E-25 |
| Dennd1b | 1.51E-25 | -0.133084407 | 0.156 | 0.256 | 4.07E-21 |
| Tbc1d5 | 1.53E-30 | -0.133157486 | 0.542 | 0.711 | 4.11E-26 |
| Tm6sf1 | 1.22E-32 | -0.133905879 | 0.229 | 0.367 | 3.29E-28 |
| Fam117b | 1.89E-33 | -0.134386813 | 0.189 | 0.313 | 5.09E-29 |
| Pan3 | 1.76E-47 | -0.134695702 | 0.428 | 0.632 | 4.72E-43 |
| Rnf180 | 8.63E-32 | -0.134981923 | 0.359 | 0.515 | 2.32E-27 |
| Fcho2 | 1.67E-41 | -0.135879372 | 0.331 | 0.496 | 4.50E-37 |
| Acap2 | 1.16E-38 | -0.136191426 | 0.353 | 0.524 | 3.11E-34 |
| Rock2 | 8.35E-38 | -0.13735825 | 0.368 | 0.536 | 2.25E-33 |
| Msi2 | 5.31E-37 | -0.138039855 | 0.199 | 0.332 | 1.43E-32 |
| Atxn2 | 3.90E-23 | -0.138113529 | 0.155 | 0.25 | 1.05E-18 |
| Zfp148 | 1.60E-31 | -0.139103143 | 0.244 | 0.38 | 4.30E-27 |
| Sirpa | 3.03E-35 | -0.139512806 | 0.478 | 0.655 | 8.14E-31 |
| Klf3 | 6.82E-30 | -0.140308855 | 0.179 | 0.295 | 1.84E-25 |
| Tnpo1 | 3.23E-24 | -0.141060893 | 0.185 | 0.289 | 8.70E-20 |
| Sppl3 | 2.47E-32 | -0.141462474 | 0.281 | 0.422 | 6.65E-28 |

| | | | | | |
|---|---|---|---|---|---|
| Pmepa1 | 2.03E-35 | -0.141994391 | 0.303 | 0.462 | 5.47E-31 |
| Camk1d | 2.34E-27 | -0.142308782 | 0.622 | 0.756 | 6.31E-33 |
| Atf7ip | 3.10E-36 | -0.142727042 | 0.21 | 0.347 | 8.33E-32 |
| Tmem135 | 8.23E-34 | -0.142730958 | 0.364 | 0.524 | 2.21E-29 |
| Zc3h7a | 1.23E-26 | -0.142837833 | 0.191 | 0.299 | 3.32E-22 |
| Taok3 | 8.68E-21 | -0.143409545 | 0.202 | 0.304 | 2.34E-16 |
| Akap10 | 7.75E-26 | -0.144492921 | 0.206 | 0.323 | 2.08E-21 |
| Sdccag8 | 2.94E-31 | -0.144569451 | 0.208 | 0.332 | 7.92E-27 |
| Rab14 | 2.50E-34 | -0.145060596 | 0.169 | 0.288 | 6.74E-30 |
| Xiap | 1.63E-31 | -0.146475473 | 0.226 | 0.353 | 4.38E-27 |
| Zfp532 | 2.65E-23 | -0.146667195 | 0.159 | 0.255 | 7.13E-19 |
| Cbl | 1.15E-33 | -0.147419438 | 0.235 | 0.374 | 3.10E-29 |
| Kansl1 | 1.81E-37 | -0.147607088 | 0.475 | 0.661 | 4.87E-33 |
| Tnks2 | 1.12E-26 | -0.147681503 | 0.152 | 0.255 | 3.01E-22 |
| Arsb | 8.10E-22 | -0.147779244 | 0.25 | 0.363 | 2.18E-17 |
| Hnrnph1 | 4.07E-39 | -0.148288371 | 0.237 | 0.384 | 1.10E-34 |
| Cacnb2 | 2.61E-30 | -0.149026202 | 0.405 | 0.576 | 7.03E-26 |
| Dyrk1a | 4.90E-26 | -0.149197469 | 0.188 | 0.302 | 1.32E-21 |
| E2f3 | 2.56E-29 | -0.149544103 | 0.186 | 0.299 | 6.88E-25 |
| Ctss | 1.67E-36 | -0.149841024 | 0.559 | 0.751 | 4.50E-32 |
| Pitpna | 5.31E-36 | -0.150017279 | 0.169 | 0.295 | 1.43E-31 |
| Tab2 | 5.58E-34 | -0.150414522 | 0.203 | 0.332 | 1.50E-29 |
| Mef2c | 6.05E-19 | -0.151466531 | 0.718 | 0.848 | 1.63E-24 |
| Tspan14 | 1.53E-26 | -0.151599502 | 0.197 | 0.312 | 4.11E-22 |
| Thrap3 | 3.41E-28 | -0.151757415 | 0.15 | 0.254 | 9.17E-24 |
| Fyb | 6.03E-43 | -0.152041088 | 0.582 | 0.785 | 1.62E-38 |
| Tpp2 | 3.27E-22 | -0.152612694 | 0.162 | 0.262 | 8.80E-18 |
| Ttc28 | 5.27E-37 | -0.152717131 | 0.354 | 0.527 | 1.42E-32 |
| Arl15 | 2.64E-28 | -0.153003608 | 0.355 | 0.506 | 7.09E-24 |
| Gab1 | 4.56E-32 | -0.153778449 | 0.211 | 0.342 | 1.23E-27 |
| Adap2 | 1.07E-47 | -0.154461009 | 0.529 | 0.741 | 2.88E-43 |
| Sdk1 | 1.46E-17 | -0.155023324 | 0.222 | 0.322 | 3.94E-13 |
| Abi3 | 1.99E-28 | -0.155179664 | 0.263 | 0.395 | 5.34E-24 |
| Rcbtb2 | 5.52E-28 | -0.155812772 | 0.259 | 0.39 | 1.49E-23 |
| Rab31 | 1.17E-30 | -0.156367999 | 0.252 | 0.388 | 3.14E-26 |
| Large1 | 2.36E-32 | -0.156819875 | 0.529 | 0.706 | 6.34E-28 |
| Smurf2 | 1.02E-30 | -0.157740353 | 0.212 | 0.342 | 2.75E-26 |
| Mapkap1 | 4.38E-25 | -0.157859476 | 0.166 | 0.27 | 1.18E-20 |
| Nckap1l | 9.20E-42 | -0.158170047 | 0.309 | 0.477 | 2.47E-37 |
| Nr6a1os | 1.16E-26 | -0.158211776 | 0.16 | 0.265 | 3.12E-22 |
| Herc2 | 2.65E-32 | -0.159609312 | 0.386 | 0.548 | 7.12E-28 |
| Exoc6b | 2.50E-23 | -0.159679963 | 0.283 | 0.409 | 6.73E-19 |
| Phactr2 | 8.78E-23 | -0.16069023 | 0.203 | 0.312 | 2.36E-18 |
| Ythdf3 | 1.33E-25 | -0.160918644 | 0.169 | 0.277 | 3.58E-21 |
| Syk | 4.88E-34 | -0.1609612 | 0.178 | 0.306 | 1.31E-29 |
| Tbc1d14 | 7.28E-32 | -0.16104938 | 0.179 | 0.299 | 1.96E-27 |
| Cradd | 1.64E-22 | -0.161166465 | 0.193 | 0.301 | 4.41E-18 |
| Adam10 | 3.28E-38 | -0.16124752 | 0.245 | 0.394 | 8.83E-34 |
| Rapgef1 | 7.30E-32 | -0.161679255 | 0.201 | 0.329 | 1.96E-27 |
| Apbb1ip | 3.08E-27 | -0.162573355 | 0.675 | 0.837 | 8.28E-23 |
| Setd2 | 5.57E-26 | -0.163066403 | 0.24 | 0.364 | 1.50E-21 |
| Rhobtb1 | 1.83E-35 | -0.16340907 | 0.247 | 0.388 | 4.92E-31 |
| Lcp1 | 8.68E-22 | -0.163624473 | 0.192 | 0.296 | 2.33E-17 |
| Gna12 | 9.68E-38 | -0.163790754 | 0.254 | 0.41 | 2.61E-33 |
| Pum2 | 1.49E-37 | -0.164362747 | 0.202 | 0.345 | 4.01E-33 |
| A630001G21Rik | 1.88E-33 | -0.16537627 | 0.157 | 0.274 | 5.05E-29 |
| Slc7a8 | 4.41E-28 | -0.165444189 | 0.291 | 0.434 | 1.19E-23 |
| Herc4 | 2.26E-21 | -0.165641016 | 0.184 | 0.284 | 6.09E-17 |
| Cflar | 3.58E-38 | -0.165690013 | 0.281 | 0.435 | 9.64E-34 |
| Slc7a7 | 2.94E-34 | -0.166266829 | 0.199 | 0.329 | 7.90E-30 |
| Etv6 | 1.48E-36 | -0.167943565 | 0.416 | 0.597 | 3.98E-32 |
| Itga6 | 3.56E-31 | -0.169355525 | 0.283 | 0.43 | 9.59E-27 |
| Zfand3 | 3.89E-39 | -0.169560404 | 0.328 | 0.498 | 1.05E-34 |
| Mtmr3 | 5.94E-30 | -0.170095038 | 0.247 | 0.382 | 1.60E-25 |
| Aftph | 4.19E-33 | -0.170884473 | 0.316 | 0.471 | 1.13E-28 |
| Bin2 | 5.87E-41 | -0.171323165 | 0.274 | 0.44 | 1.58E-26 |
| Cep350 | 1.96E-24 | -0.171434849 | 0.16 | 0.266 | 5.29E-20 |
| F11r | 2.33E-35 | -0.171645308 | 0.215 | 0.357 | 6.27E-31 |
| Tbl1x | 2.26E-31 | -0.172175582 | 0.186 | 0.312 | 6.09E-27 |
| Bcas3 | 4.67E-23 | -0.172829228 | 0.236 | 0.357 | 1.26E-18 |
| Pacs1 | 1.45E-31 | -0.173189414 | 0.207 | 0.339 | 3.89E-27 |
| Mllt10 | 7.39E-38 | -0.173701971 | 0.292 | 0.455 | 1.99E-33 |
| Washc4 | 2.41E-28 | -0.173983912 | 0.165 | 0.276 | 6.47E-24 |
| Cyth4 | 9.11E-40 | -0.174277619 | 0.433 | 0.624 | 2.45E-35 |
| Pik3cd | 3.04E-31 | -0.174321033 | 0.211 | 0.34 | 8.17E-27 |
| Notch2 | 2.93E-26 | -0.174458619 | 0.145 | 0.251 | 7.89E-22 |
| Immp2l | 5.53E-11 | -0.175386262 | 0.235 | 0.32 | 1.49E-06 |
| Hexb | 1.35E-19 | -0.177494897 | 0.843 | 0.946 | 3.63E-17 |
| Sbf2 | 2.92E-31 | -0.17810698 | 0.337 | 0.493 | 7.86E-27 |
| Frmd4b | 2.66E-27 | -0.179450415 | 0.561 | 0.731 | 7.14E-23 |
| Cd2ap | 2.32E-30 | -0.179523497 | 0.229 | 0.357 | 6.24E-26 |
| Hnrnpa2b1 | 1.90E-36 | -0.180030529 | 0.471 | 0.658 | 5.10E-32 |
| Lyn | 2.45E-33 | -0.180432291 | 0.674 | 0.853 | 6.58E-29 |
| Bmp2k | 1.27E-42 | -0.180494226 | 0.47 | 0.671 | 3.41E-38 |
| Nfat5 | 5.59E-36 | -0.180993007 | 0.239 | 0.391 | 1.50E-31 |
| Gab2 | 1.29E-26 | -0.181094477 | 0.645 | 0.809 | 3.48E-22 |
| Mlph | 1.74E-31 | -0.181138629 | 0.138 | 0.254 | 4.69E-27 |
| Csf3r | 1.88E-37 | -0.181359454 | 0.415 | 0.603 | 5.05E-33 |
| Tet3 | 3.17E-36 | -0.181447599 | 0.179 | 0.315 | 8.54E-32 |
| P2ry12 | 1.17E-19 | -0.181709361 | 0.66 | 0.804 | 3.15E-10 |
| Ophn1 | 3.13E-29 | -0.181910166 | 0.624 | 0.798 | 8.43E-25 |
| Wasf2 | 3.07E-27 | -0.182299088 | 0.449 | 0.613 | 8.27E-23 |
| Sort1 | 1.61E-32 | -0.182802423 | 0.177 | 0.307 | 4.32E-18 |
| Pik3r1 | 1.57E-36 | -0.182863723 | 0.38 | 0.557 | 4.23E-32 |
| Rnf130 | 2.54E-25 | -0.183022703 | 0.157 | 0.269 | 6.82E-21 |
| Hdac8 | 5.41E-19 | -0.183346871 | 0.193 | 0.296 | 1.46E-14 |
| Golm1 | 6.32E-33 | -0.183622949 | 0.274 | 0.42 | 1.70E-28 |
| Tcf20 | 1.44E-30 | -0.184132121 | 0.218 | 0.358 | 3.89E-26 |
| Wnk1 | 1.10E-33 | -0.185481699 | 0.563 | 0.752 | 2.95E-29 |
| Nlrp1b | 1.25E-22 | -0.185617556 | 0.185 | 0.297 | 3.36E-18 |
| Capn3 | 1.97E-35 | -0.185910639 | 0.247 | 0.394 | 5.31E-31 |
| Pafah1b1 | 2.49E-30 | -0.186127313 | 0.286 | 0.436 | 6.71E-26 |
| 1-Mar | 1.99E-40 | -0.186236041 | 0.424 | 0.618 | 5.35E-36 |
| Phf14 | 2.20E-39 | -0.186393579 | 0.418 | 0.609 | 5.93E-35 |
| Ifngr1 | 1.58E-25 | -0.187268404 | 0.206 | 0.324 | 4.25E-21 |
| Dock4 | 2.25E-17 | -0.187543904 | 0.871 | 0.956 | 6.05E-13 |
| Zfp608 | 4.14E-32 | -0.188369122 | 0.175 | 0.3 | 1.11E-27 |
| Slc9a9 | 1.47E-28 | -0.188413562 | 0.617 | 0.791 | 3.95E-24 |
| Snx2 | 2.50E-27 | -0.189013626 | 0.144 | 0.251 | 6.72E-23 |
| Wdfy2 | 4.38E-37 | -0.189583161 | 0.258 | 0.412 | 1.18E-32 |
| Ptgs1 | 1.21E-31 | -0.18967557 | 0.247 | 0.386 | 3.26E-27 |
| Exoc4 | 6.02E-32 | -0.190510562 | 0.41 | 0.582 | 1.62E-27 |
| Scamp5 | 3.14E-30 | -0.191175441 | 0.142 | 0.252 | 8.44E-26 |
| Eef2k | 2.44E-35 | -0.191473106 | 0.198 | 0.336 | 6.56E-31 |
| Abl1 | 9.26E-38 | -0.191702847 | 0.187 | 0.328 | 2.49E-33 |
| Il10ra | 2.85E-41 | -0.192962919 | 0.23 | 0.389 | 7.68E-37 |
| Zfp652 | 1.78E-30 | -0.193194662 | 0.399 | 0.571 | 4.78E-26 |
| Lnpep | 2.16E-35 | -0.193336202 | 0.198 | 0.334 | 5.82E-31 |
| Extl3 | 1.09E-33 | -0.193420651 | 0.299 | 0.456 | 2.93E-29 |
| Cdk14 | 1.11E-25 | -0.194796508 | 0.274 | 0.419 | 2.99E-21 |
| Nav3 | 2.20E-13 | -0.19589031 | 0.84 | 0.928 | 5.91E-09 |
| Kat6a | 1.71E-30 | -0.197500914 | 0.19 | 0.32 | 4.60E-26 |
| Plxdc2 | 4.43E-17 | -0.199308881 | 0.936 | 0.982 | 1.19E-12 |
| Qk | 8.38E-31 | -0.199505477 | 0.807 | 0.945 | 2.25E-26 |
| Ivns1abp | 9.39E-41 | -0.199549005 | 0.549 | 0.758 | 2.53E-36 |
| Ncoa3 | 2.83E-33 | -0.199987234 | 0.221 | 0.364 | 7.61E-29 |
| Arhgap12 | 6.04E-43 | -0.200698122 | 0.287 | 0.464 | 1.63E-38 |
| Selplg | 4.89E-43 | -0.201227005 | 0.559 | 0.771 | 1.31E-38 |
| Kdm6a | 1.43E-26 | -0.201295254 | 0.15 | 0.264 | 3.84E-22 |
| St3gal6 | 6.50E-34 | -0.201500312 | 0.467 | 0.654 | 1.75E-29 |
| Ldlrad4 | 3.27E-21 | -0.201604193 | 0.808 | 0.927 | 8.81E-17 |

| | | | | | |
|---|---|---|---|---|---|
| Cdh23 | 9.59E-39 | -0.20259172 | 0.287 | 0.458 | 2.58E-34 |
| Whrn | 2.73E-37 | -0.202661944 | 0.171 | 0.301 | 7.34E-33 |
| Pkn1 | 1.98E-32 | -0.203100584 | 0.202 | 0.338 | 5.33E-28 |
| Blnk | 9.66E-34 | -0.203477241 | 0.377 | 0.551 | 2.60E-29 |
| Dapp1 | 1.19E-40 | -0.204014537 | 0.281 | 0.452 | 3.20E-36 |
| Cd37 | 9.28E-47 | -0.20438798 | 0.319 | 0.51 | 2.50E-42 |
| Mkln1 | 1.78E-36 | -0.204409221 | 0.281 | 0.444 | 4.78E-32 |
| Pten | 8.35E-37 | -0.20454492 | 0.335 | 0.509 | 2.25E-32 |
| Evi5 | 4.49E-30 | -0.204594517 | 0.147 | 0.266 | 1.21E-25 |
| Cdk19 | 3.76E-33 | -0.204786175 | 0.236 | 0.381 | 1.01E-28 |
| Mertk | 2.10E-28 | -0.205125953 | 0.784 | 0.927 | 5.66E-24 |
| Havcr2 | 5.95E-39 | -0.205139116 | 0.227 | 0.381 | 1.60E-34 |
| Csad | 1.13E-30 | -0.205242439 | 0.165 | 0.293 | 3.05E-26 |
| Itpr2 | 4.57E-32 | -0.205301447 | 0.361 | 0.53 | 1.23E-27 |
| Supt3 | 3.02E-20 | -0.205438256 | 0.164 | 0.269 | 8.13E-16 |
| Atg10 | 1.66E-18 | -0.205863621 | 0.161 | 0.256 | 4.46E-14 |
| Plxna4 | 9.79E-30 | -0.206165095 | 0.431 | 0.603 | 2.63E-25 |
| Vrk2 | 5.05E-36 | -0.206281643 | 0.249 | 0.4 | 1.36E-31 |
| Hck | 1.83E-34 | -0.207767913 | 0.223 | 0.372 | 4.92E-30 |
| Cx3cr1 | 3.09E-30 | -0.209260415 | 0.652 | 0.829 | 8.32E-22 |
| Mtus1 | 9.16E-31 | -0.210081908 | 0.26 | 0.411 | 2.47E-26 |
| Gng2 | 2.75E-30 | -0.210404444 | 0.258 | 0.403 | 7.39E-26 |
| Cnot2 | 2.37E-33 | -0.210911397 | 0.269 | 0.429 | 6.39E-29 |
| Ankrd44 | 2.81E-27 | -0.210937955 | 0.588 | 0.762 | 7.55E-23 |
| Ptprm | 1.76E-30 | -0.211258957 | 0.391 | 0.57 | 4.73E-26 |
| Macf1 | 2.05E-30 | -0.211476984 | 0.709 | 0.878 | 5.50E-26 |
| Camk2d | 1.24E-45 | -0.21182767 | 0.383 | 0.593 | 3.34E-41 |
| Arid1a | 1.03E-32 | -0.212391145 | 0.23 | 0.377 | 2.78E-28 |
| Gbf1 | 4.25E-22 | -0.212576219 | 0.162 | 0.269 | 1.14E-17 |
| Mgat5 | 4.41E-37 | -0.212613544 | 0.361 | 0.541 | 1.19E-32 |
| Dennd4a | 2.73E-31 | -0.214309417 | 0.599 | 0.786 | 7.36E-27 |
| Tanc2 | 2.78E-17 | -0.216599841 | 0.952 | 0.984 | 7.47E-13 |
| Hook3 | 3.60E-29 | -0.217384477 | 0.186 | 0.312 | 9.69E-25 |
| Laptm5 | 1.73E-43 | -0.217876605 | 0.233 | 0.406 | 4.65E-39 |
| Vav1 | 1.32E-25 | -0.219155802 | 0.174 | 0.292 | 3.55E-21 |
| Fchsd2 | 1.24E-36 | -0.221192103 | 0.544 | 0.746 | 3.33E-32 |
| Stambpl1 | 3.28E-29 | -0.222683326 | 0.162 | 0.279 | 8.84E-25 |
| Fhit | 5.89E-14 | -0.222859207 | 0.399 | 0.52 | 1.59E-09 |
| Pld1 | 1.23E-39 | -0.22397637 | 0.302 | 0.481 | 3.32E-35 |
| Tcf7l2 | 5.71E-29 | -0.225140702 | 0.203 | 0.338 | 1.54E-24 |
| Unc93b1 | 1.53E-41 | -0.227990708 | 0.484 | 0.695 | 4.11E-37 |
| Slc1a3 | 2.64E-27 | -0.228386873 | 0.214 | 0.352 | 7.10E-23 |
| Mbnl1 | 9.76E-31 | -0.229438186 | 0.719 | 0.887 | 2.63E-26 |
| Runx1 | 1.42E-38 | -0.22964744 | 0.518 | 0.729 | 3.82E-34 |
| Atp8a2 | 9.22E-33 | -0.23006873 | 0.497 | 0.686 | 2.48E-28 |
| Galnt1 | 5.35E-25 | -0.230338829 | 0.165 | 0.285 | 1.44E-20 |
| Tgfbr1 | 6.13E-21 | -0.231016375 | 0.876 | 0.964 | 1.65E-16 |
| Rffl | 5.88E-41 | -0.231665065 | 0.193 | 0.346 | 1.58E-36 |
| Cttnbp2nl | 2.92E-30 | -0.231755635 | 0.406 | 0.582 | 7.85E-26 |
| Gpr34 | 7.96E-43 | -0.232080982 | 0.432 | 0.641 | 2.14E-38 |
| Tbc1d9 | 8.73E-32 | -0.232143593 | 0.184 | 0.325 | 2.35E-27 |
| Pla2g4a | 1.03E-33 | -0.232598171 | 0.188 | 0.325 | 2.76E-29 |
| Tnrc6b | 5.80E-32 | -0.232831447 | 0.364 | 0.539 | 1.56E-27 |
| AW554918 | 2.74E-25 | -0.233647004 | 0.182 | 0.306 | 7.37E-21 |
| Rab8b | 7.45E-28 | -0.233677771 | 0.269 | 0.42 | 2.01E-23 |
| Cd300c2 | 6.28E-31 | -0.233939451 | 0.144 | 0.267 | 1.69E-26 |
| Snta1 | 5.84E-27 | -0.234656557 | 0.142 | 0.253 | 1.57E-22 |
| Fer | 2.05E-30 | -0.235475905 | 0.17 | 0.303 | 5.50E-26 |
| Sall1 | 5.49E-34 | -0.236559997 | 0.235 | 0.387 | 1.48E-29 |
| Gm5086 | 1.98E-36 | -0.237985575 | 0.245 | 0.404 | 5.34E-32 |
| Maml2 | 6.26E-27 | -0.23979712 | 0.483 | 0.66 | 1.68E-22 |
| Usp24 | 2.24E-46 | -0.243191419 | 0.258 | 0.441 | 6.03E-42 |
| Pard3b | 1.41E-28 | -0.243556049 | 0.319 | 0.482 | 3.80E-24 |
| Fkbp5 | 1.73E-24 | -0.244307163 | 0.361 | 0.523 | 4.66E-20 |
| Smarca2 | 1.65E-29 | -0.24502102 | 0.221 | 0.364 | 4.44E-25 |
| Itgb5 | 5.11E-37 | -0.245125708 | 0.563 | 0.769 | 1.37E-32 |
| Plcl1 | 2.60E-15 | -0.2459127 | 0.709 | 0.836 | 6.99E-11 |
| Ikzf1 | 1.00E-44 | -0.246181262 | 0.573 | 0.795 | 2.70E-40 |
| Adgre1 | 1.37E-21 | -0.246850497 | 0.145 | 0.254 | 3.68E-17 |
| Arhgap5 | 5.43E-30 | -0.247674424 | 0.617 | 0.799 | 1.46E-25 |
| Ssh2 | 3.02E-27 | -0.247953572 | 0.776 | 0.92 | 8.13E-23 |
| Ccr5 | 4.44E-31 | -0.249693433 | 0.209 | 0.359 | 1.19E-26 |
| 0610040J01Rik | 5.65E-42 | -0.250314267 | 0.378 | 0.581 | 1.52E-37 |
| Atxn1 | 5.82E-21 | -0.250562933 | 0.3 | 0.443 | 1.57E-16 |
| Serinc3 | 1.86E-50 | -0.252541736 | 0.487 | 0.718 | 5.02E-46 |
| Mvb12b | 1.21E-45 | -0.253050335 | 0.221 | 0.391 | 3.25E-41 |
| St3gal5 | 4.68E-33 | -0.253294966 | 0.214 | 0.368 | 1.26E-28 |
| Prkca | 1.70E-32 | -0.255401947 | 0.408 | 0.601 | 4.58E-28 |
| Fam49b | 3.24E-30 | -0.256787053 | 0.615 | 0.801 | 8.73E-26 |
| Fat3 | 2.60E-18 | -0.257030245 | 0.304 | 0.445 | 6.99E-14 |
| Srgap2 | 1.17E-26 | -0.258665337 | 0.909 | 0.982 | 3.14E-22 |
| Fli1 | 1.26E-38 | -0.258781541 | 0.507 | 0.721 | 3.40E-34 |
| Peli2 | 7.04E-25 | -0.261025056 | 0.188 | 0.314 | 1.89E-20 |
| Casp8 | 1.76E-29 | -0.26155052 | 0.164 | 0.298 | 4.72E-25 |
| Cdk6 | 4.13E-24 | -0.263283582 | 0.177 | 0.301 | 1.11E-19 |
| Zeb1 | 2.40E-35 | -0.266467091 | 0.374 | 0.565 | 6.45E-31 |
| Tbxas1 | 2.61E-30 | -0.268150525 | 0.4 | 0.584 | 7.02E-26 |
| Fam102b | 1.01E-34 | -0.27073263 | 0.37 | 0.562 | 2.71E-30 |
| Numb | 1.04E-25 | -0.27093393 | 0.701 | 0.863 | 2.79E-21 |
| B4galt1 | 2.27E-26 | -0.271048439 | 0.231 | 0.373 | 6.12E-22 |
| Cd33 | 2.93E-28 | -0.271579897 | 0.216 | 0.338 | 7.88E-24 |
| Epb41l2 | 1.07E-22 | -0.274004684 | 0.755 | 0.894 | 2.89E-18 |
| Plcl2 | 1.59E-30 | -0.274022573 | 0.558 | 0.753 | 4.27E-26 |
| Ccnd3 | 1.96E-39 | -0.274314265 | 0.594 | 0.809 | 5.27E-35 |
| Mlxip | 6.37E-25 | -0.276437552 | 0.194 | 0.324 | 1.71E-20 |
| Rreb1 | 3.24E-37 | -0.277024166 | 0.619 | 0.822 | 8.71E-33 |
| Hnrnpa3 | 3.15E-32 | -0.278918064 | 0.142 | 0.275 | 8.48E-28 |
| Dido1 | 4.34E-25 | -0.279923062 | 0.153 | 0.274 | 1.17E-20 |
| Csf1r | 9.37E-35 | -0.280913822 | 0.626 | 0.826 | 2.52E-30 |
| Wdfy3 | 1.27E-39 | -0.282297943 | 0.355 | 0.556 | 3.42E-35 |
| Wdfy4 | 1.82E-33 | -0.28505999 | 0.179 | 0.324 | 4.91E-29 |
| Vsir | 2.43E-50 | -0.285871661 | 0.324 | 0.543 | 6.55E-46 |
| Pik3ap1 | 1.29E-37 | -0.285976741 | 0.26 | 0.437 | 3.47E-33 |
| Actr2 | 1.14E-37 | -0.286510927 | 0.251 | 0.425 | 3.08E-33 |
| Nos1ap | 3.84E-32 | -0.28729591 | 0.219 | 0.378 | 1.03E-27 |
| Ubash3b | 2.40E-32 | -0.287611292 | 0.456 | 0.655 | 6.44E-28 |
| Agps | 6.20E-33 | -0.288413311 | 0.201 | 0.358 | 1.67E-28 |
| Pde3b | 8.07E-28 | -0.288815004 | 0.696 | 0.865 | 2.17E-23 |
| Dock2 | 4.99E-40 | -0.290860128 | 0.538 | 0.76 | 1.34E-35 |
| Foxp1 | 5.32E-34 | -0.292525158 | 0.237 | 0.408 | 1.43E-29 |
| Picalm | 6.23E-32 | -0.292678167 | 0.536 | 0.739 | 1.68E-37 |
| Maml3 | 1.03E-24 | -0.294006608 | 0.728 | 0.881 | 2.77E-20 |
| Chn2 | 3.10E-42 | -0.294119144 | 0.499 | 0.723 | 8.35E-38 |
| Dock8 | 4.80E-39 | -0.295786983 | 0.642 | 0.849 | 1.29E-34 |
| Tmem119 | 2.02E-31 | -0.298012529 | 0.158 | 0.291 | 5.44E-27 |
| Slc8a1 | 3.38E-26 | -0.300058493 | 0.809 | 0.935 | 9.10E-22 |
| Rbm47 | 1.93E-38 | -0.303124772 | 0.251 | 0.43 | 5.20E-34 |
| Slco2b1 | 3.74E-40 | -0.306312669 | 0.543 | 0.763 | 1.01E-35 |
| Rassf2 | 6.06E-38 | -0.306398494 | 0.16 | 0.309 | 1.63E-33 |
| Bank1 | 3.37E-23 | -0.30657017 | 0.183 | 0.311 | 9.07E-19 |
| Rcsd1 | 2.04E-37 | -0.308762868 | 0.225 | 0.401 | 5.49E-33 |
| Col27a1 | 3.75E-46 | -0.319292184 | 0.206 | 0.387 | 1.01E-41 |
| Lpcat2 | 2.69E-52 | -0.321425277 | 0.462 | 0.713 | 7.24E-48 |
| Ralgps1 | 3.75E-31 | -0.323089133 | 0.128 | 0.263 | 1.01E-26 |
| Rap1gds1 | 2.44E-36 | -0.325718424 | 0.514 | 0.729 | 6.56E-32 |
| Tjp1 | 3.67E-36 | -0.32694029 | 0.257 | 0.432 | 9.89E-32 |
| Mef2a | 3.39E-28 | -0.328346748 | 0.836 | 0.947 | 9.12E-24 |
| Maf | 1.23E-41 | -0.332053858 | 0.288 | 0.496 | 3.30E-37 |
| Man1a | 1.80E-35 | -0.333422268 | 0.296 | 0.491 | 4.83E-31 |
| Nfia | 5.18E-26 | -0.333448192 | 0.492 | 0.68 | 1.39E-21 |
| Cables1 | 1.58E-23 | -0.335897618 | 0.199 | 0.334 | 4.24E-19 |

| | | | | | |
|---|---|---|---|---|---|
| AY036118 | 1.60E-27 | -0.336804794 | 0.41 | 0.597 | 4.30E-23 |
| P3h2 | 2.42E-34 | -0.338564096 | 0.349 | 0.551 | 6.50E-30 |
| Inpp4b | 6.32E-33 | -0.3393674 | 0.605 | 0.807 | 1.70E-28 |
| Zfhx3 | 1.94E-43 | -0.339894057 | 0.803 | 0.958 | 5.23E-39 |
| Pag1 | 2.05E-43 | -0.343116392 | 0.722 | 0.916 | 5.51E-39 |
| Fcrls | 1.37E-27 | -0.344124419 | 0.37 | 0.555 | 3.67E-23 |
| Itga9 | 7.53E-39 | -0.348283682 | 0.232 | 0.419 | 2.03E-34 |
| 4933406I18Rik | 1.37E-29 | -0.353430588 | 0.585 | 0.782 | 3.67E-25 |
| Cfh | 1.37E-41 | -0.357991729 | 0.454 | 0.687 | 3.67E-37 |
| Entpd1 | 7.86E-51 | -0.364889105 | 0.491 | 0.745 | 2.11E-46 |
| Rapgef5 | 3.47E-31 | -0.365541919 | 0.614 | 0.812 | 9.35E-27 |
| Lcp2 | 1.40E-39 | -0.376965194 | 0.177 | 0.35 | 3.77E-35 |
| Dock10 | 7.76E-44 | -0.377771355 | 0.601 | 0.831 | 2.09E-39 |
| Rasgrp3 | 1.09E-40 | -0.385683237 | 0.345 | 0.566 | 2.92E-36 |
| Arhgap22 | 2.58E-44 | -0.386383068 | 0.363 | 0.595 | 6.94E-40 |
| Csmd3 | 4.38E-30 | -0.39347453 | 0.93 | 0.98 | 1.18E-25 |
| Tlr7 | 1.03E-34 | -0.394758248 | 0.176 | 0.339 | 2.78E-30 |
| Abca9 | 4.14E-49 | -0.397015778 | 0.335 | 0.576 | 1.11E-44 |
| Gtf2h2 | 8.77E-45 | -0.397099757 | 0.274 | 0.49 | 2.36E-40 |
| A830008E24Rik | 4.27E-23 | -0.403786821 | 0.193 | 0.336 | 1.15E-18 |
| Bbs9 | 3.29E-32 | -0.405858909 | 0.363 | 0.57 | 8.85E-28 |
| Elmo1 | 9.23E-52 | -0.40601433 | 0.887 | 0.984 | 2.48E-47 |
| Siglech | 2.12E-47 | -0.451521043 | 0.505 | 0.761 | 5.70E-43 |
| Agmo | 3.35E-42 | -0.493840754 | 0.456 | 0.704 | 9.02E-38 |
| 8030442B05Rik | 1.00E-61 | -0.568336826 | 0.294 | 0.576 | 2.70E-57 |
| Ifitm10 | 4.46E-38 | -0.596982038 | 0.112 | 0.281 | 1.20E-33 |
| Figure3E_pseudobulk_MG4_vs_MG1 | | | | | |
| MG4_versus_MG1_DEGs | | | | | |
| | p_val | avg_log2FC | pct.1 | pct.2 | p_val_adj |
| Apoe | 2.29E-223 | 5.532149363 | 0.914 | 0.422 | 3.30E-190 |
| F13a1 | 3.60E-290 | 4.932857683 | 0.853 | 0.65 | 5.20E-287 |
| Mrc1 | 3.60E-290 | 4.760821622 | 0.868 | 0.648 | 5.20E-287 |
| Dab2 | 3.60E-290 | 4.070073574 | 0.885 | 0.664 | 5.20E-287 |
| Arhgap15 | 3.60E-290 | 4.036103101 | 0.986 | 0.633 | 5.20E-287 |
| Cd74 | 1.96E-254 | 3.500117691 | 0.589 | 0.419 | 2.83E-251 |
| Aoah | 3.60E-290 | 3.216272711 | 0.813 | 0.803 | 5.20E-287 |
| P2rx7 | 5.82E-87 | 3.122552021 | 0.856 | 0.643 | 8.40E-112 |
| Cd163 | 3.60E-290 | 2.872449725 | 0.796 | 0.752 | 5.20E-287 |
| Rbpj | 2.45E-245 | 2.829169526 | 0.802 | 0.665 | 3.53E-242 |
| Colec12 | 3.60E-290 | 2.602485696 | 0.632 | 0.796 | 5.20E-287 |
| Ebf1 | 1.11E-119 | 2.570911238 | 0.816 | 0.461 | 1.61E-116 |
| Trps1 | 2.87E-230 | 2.509303113 | 0.767 | 0.573 | 4.14E-227 |
| Mndal | 3.60E-290 | 2.425578598 | 0.799 | 0.744 | 5.20E-287 |
| Pde7b | 5.17E-171 | 2.408871126 | 0.578 | 0.451 | 7.46E-168 |
| Itga4 | 3.27E-265 | 2.402188733 | 0.756 | 0.442 | 4.72E-262 |
| F630028O10Rik | 3.60E-290 | 2.351936148 | 0.56 | 0.705 | 5.20E-287 |
| Cd38 | 3.60E-290 | 2.321963309 | 0.853 | 0.718 | 5.20E-287 |
| Gm30489 | 3.60E-290 | 2.295403189 | 0.713 | 0.518 | 5.20E-287 |
| H2-Aa | 3.60E-290 | 2.171692332 | 0.483 | 0.778 | 5.20E-287 |
| Samd4 | 1.02E-121 | 2.13139521 | 0.819 | 0.468 | 1.47E-118 |
| H2-Ab1 | 3.60E-290 | 2.117785746 | 0.454 | 0.82 | 5.20E-287 |
| Tnfaip8 | 8.69E-46 | 2.049510548 | 0.489 | 0.445 | 1.25E-52 |
| Mdfic | 8.98E-284 | 2.04627745 | 0.652 | 0.655 | 1.30E-280 |
| H2-Eb1 | 3.60E-290 | 2.03216091 | 0.319 | 0.642 | 5.20E-287 |
| Vav3 | 3.60E-290 | 1.995259433 | 0.569 | 0.644 | 5.20E-287 |
| Stab1 | 4.40E-73 | 1.990283143 | 0.681 | 0.41 | 6.35E-70 |
| Iqgap2 | 8.65E-274 | 1.965799495 | 0.667 | 0.834 | 1.25E-270 |
| 5033421B08Rik | 6.08E-133 | 1.864612864 | 0.753 | 0.665 | 8.78E-130 |
| Klra2 | 3.60E-290 | 1.862385916 | 0.425 | 0.772 | 5.20E-287 |
| Pstpip2 | 6.38E-204 | 1.839959453 | 0.664 | 0.725 | 9.20E-201 |
| Neat1 | 8.00E-76 | 1.837982603 | 0.612 | 0.32 | 1.15E-72 |
| Mcc | 1.96E-151 | 1.819028468 | 0.799 | 0.723 | 2.82E-148 |
| Gm26740 | 2.63E-77 | 1.805454277 | 0.799 | 0.663 | 3.79E-74 |
| Ifi207 | 4.00E-168 | 1.80167307 | 0.48 | 0.221 | 5.77E-165 |
| Ms4a4a | 3.60E-290 | 1.745041712 | 0.603 | 0.719 | 5.20E-287 |
| Adam33 | 3.60E-290 | 1.742376554 | 0.56 | 0.625 | 5.20E-287 |
| Hdac9 | 1.03E-55 | 1.695429198 | 0.922 | 0.85 | 1.49E-52 |
| Igf1 | 1.47E-215 | 1.659695897 | 0.747 | 0.731 | 2.12E-212 |
| Ms4a7 | 3.60E-290 | 1.618874055 | 0.428 | 0.583 | 5.20E-287 |
| Vcam1 | 3.73E-196 | 1.618696407 | 0.667 | 0.626 | 5.39E-193 |
| Lyz2 | 1.09E-210 | 1.605657789 | 0.466 | 0.6 | 1.57E-207 |
| Man2a1 | 6.44E-82 | 1.56297401 | 0.739 | 0.701 | 9.30E-79 |
| Gm4951 | 2.51E-157 | 1.552409547 | 0.511 | 0.614 | 3.63E-154 |
| Skap1 | 3.21E-68 | 1.546135135 | 0.615 | 0.26 | 4.64E-65 |
| Ccr2 | 1.56E-95 | 1.539702921 | 0.497 | 0.599 | 2.25E-92 |
| Kcnq5 | 6.89E-30 | 1.535782642 | 0.862 | 0.888 | 9.95E-27 |
| Ms4a6c | 4.28E-223 | 1.520997375 | 0.477 | 0.531 | 6.17E-220 |
| Ctsc | 8.70E-44 | 1.514515397 | 0.693 | 0.475 | 1.26E-50 |
| Rad51b | 4.26E-27 | 1.498707852 | 0.615 | 0.46 | 6.14E-24 |
| Bank1 | 8.59E-41 | 1.466646081 | 0.793 | 0.685 | 1.24E-37 |
| Slc16a10 | 1.41E-71 | 1.385023439 | 0.782 | 0.307 | 2.04E-68 |
| Mctp1 | 1.10E-44 | 1.380735476 | 0.868 | 0.789 | 1.58E-41 |
| Bcl2 | 1.66E-47 | 1.372581918 | 0.667 | 0.657 | 2.39E-44 |
| Ms4a6b | 7.64E-79 | 1.366442021 | 0.664 | 0.568 | 1.10E-75 |
| Apobec1 | 2.47E-34 | 1.348122736 | 0.578 | 0.463 | 3.57E-31 |
| Eps8 | 1.85E-89 | 1.316187892 | 0.618 | 0.579 | 2.66E-86 |
| Cbr2 | 9.34E-260 | 1.306547057 | 0.695 | 0.753 | 1.35E-256 |
| Ikzf3 | 5.55E-177 | 1.253411742 | 0.693 | 0.066 | 8.01E-174 |
| Gm20663 | 6.15E-20 | 1.235131563 | 0.566 | 0.541 | 8.88E-17 |
| Tmem163 | 8.71E-79 | 1.223715166 | 0.828 | 0.549 | 1.26E-75 |
| Fam129a | 1.52E-27 | 1.22245198 | 0.701 | 0.592 | 2.19E-24 |
| Pla2g7 | 1.20E-99 | 1.208854078 | 0.626 | 0.601 | 1.73E-96 |
| Tox | 2.29E-50 | 1.207801903 | 0.784 | 0.849 | 3.30E-47 |
| Cpq | 4.58E-247 | 1.205397986 | 0.328 | 0.757 | 6.61E-244 |
| Ms4a4b | 2.91E-76 | 1.195073437 | 0.33 | 0.025 | 4.19E-73 |
| Arhgap18 | 2.64E-42 | 1.17996691 | 0.601 | 0.51 | 3.81E-39 |
| Rbms1 | 6.25E-114 | 1.152873191 | 0.583 | 0.806 | 9.01E-111 |
| Cp | 9.87E-45 | 1.151792752 | 0.595 | 0.616 | 1.42E-41 |
| Ctsb | 7.81E-30 | 1.126441652 | 0.232 | 0.284 | 2.10E-25 |
| Mob3b | 1.77E-75 | 1.098967957 | 0.509 | 0.428 | 2.56E-72 |
| Atp8b4 | 3.60E-290 | 1.047621224 | 0.184 | 0.694 | 5.20E-287 |
| 3-Mar | 2.79E-38 | 1.032989146 | 0.687 | 0.366 | 4.02E-35 |
| Clec2d | 2.19E-249 | 1.013044493 | 0.284 | 0.647 | 3.16E-246 |
| Cenpf | 3.79E-167 | 1.00321175 | 0.434 | 0.66 | 5.47E-164 |
| Xylt1 | 1.88E-18 | 1.002490477 | 0.796 | 0.786 | 2.71E-15 |
| Arhgap6 | 1.64E-76 | 0.982928844 | 0.825 | 0.723 | 2.37E-73 |
| Creb5 | 1.80E-113 | 0.974220458 | 0.434 | 0.775 | 2.59E-110 |
| Alcam | 3.96E-279 | 0.921677116 | 0.164 | 0.778 | 5.72E-276 |
| Ptprc | 2.01E-16 | 0.919230917 | 0.833 | 0.66 | 2.90E-13 |
| Dpyd | 1.12E-39 | 0.909755448 | 0.695 | 0.853 | 1.61E-36 |
| Egfr | 3.60E-290 | 0.898461185 | 0.382 | 0.793 | 5.20E-287 |
| Ifi213 | 3.64E-228 | 0.887006719 | 0.307 | 0.668 | 5.25E-225 |
| Oasl2 | 9.50E-94 | 0.886128521 | 0.537 | 0.771 | 1.37E-90 |
| Myof | 1.38E-239 | 0.878116386 | 0.293 | 0.598 | 2.00E-236 |
| Tnfrsf11a | 9.63E-21 | 0.877777556 | 0.257 | 0.331 | 2.59E-08 |
| Pde4d | 4.38E-07 | 0.872056215 | 0.779 | 0.718 | 0.000631841 |
| Rbm47 | 1.33E-14 | 0.863148666 | 0.721 | 0.598 | 1.92E-11 |
| Ahnak | 7.26E-124 | 0.854244477 | 0.73 | 0.345 | 1.05E-120 |
| Bcar3 | 1.96E-81 | 0.851606882 | 0.601 | 0.152 | 2.83E-78 |
| Rasgef1b | 1.66E-26 | 0.841172672 | 0.638 | 0.585 | 2.40E-23 |
| Trpv4 | 1.30E-49 | 0.831883758 | 0.549 | 0.623 | 1.88E-46 |
| Runx2 | 2.40E-21 | 0.825594642 | 0.744 | 0.537 | 3.46E-18 |
| Themis | 5.26E-273 | 0.821504436 | 0.066 | 0.419 | 7.59E-270 |
| Xdh | 7.09E-243 | 0.816924597 | 0.368 | 0.678 | 1.02E-239 |
| Sntb1 | 2.01E-68 | 0.78386825 | 0.761 | 0.656 | 2.90E-65 |
| Cpne2 | 9.25E-75 | 0.774491751 | 0.675 | 0.58 | 1.33E-71 |
| Myo1e | 2.33E-29 | 0.771607419 | 0.767 | 0.46 | 3.36E-26 |
| Cobll1 | 3.84E-107 | 0.767911858 | 0.79 | 0.709 | 5.55E-104 |
| Galnt10 | 1.14E-19 | 0.763204448 | 0.638 | 0.646 | 1.65E-16 |
| Sema6d | 1.90E-30 | 0.757622212 | 0.583 | 0.672 | 2.75E-27 |

| | | | | | |
|---|---|---|---|---|---|
| Epsti1 | 3.98E-38 | 0.755480935 | 0.451 | 0.577 | 5.74E-35 |
| Ednrb | 3.57E-67 | 0.732274294 | 0.405 | 0.424 | 5.14E-64 |
| Usp18 | 5.70E-55 | 0.711603905 | 0.514 | 0.607 | 8.23E-52 |
| Gm16083 | 1.49E-78 | 0.703698364 | 0.767 | 0.496 | 2.15E-75 |
| Cd36 | 7.53E-136 | 0.702864068 | 0.336 | 0.562 | 1.09E-132 |
| Utrn | 3.85E-12 | 0.687570753 | 0.595 | 0.618 | 5.55E-09 |
| Tlr7 | 1.19E-07 | 0.671116057 | 0.75 | 0.627 | 0.000172372 |
| Notch2 | 1.38E-15 | 0.649425195 | 0.603 | 0.601 | 1.99E-12 |
| Prkch | 1.09E-29 | 0.636775165 | 0.437 | 0.548 | 1.57E-26 |
| Etv1 | 4.39E-27 | 0.634407631 | 0.661 | 0.821 | 6.34E-24 |
| Clic4 | 1.19E-46 | 0.631426596 | 0.621 | 0.257 | 1.72E-43 |
| Thrb | 3.40E-17 | 0.623873395 | 0.805 | 0.747 | 4.90E-14 |
| Esr1 | 1.62E-08 | 0.615402508 | 0.578 | 0.535 | 2.34E-05 |
| Ptgds | 2.70E-22 | 0.611059997 | 0.428 | 0.643 | 3.90E-19 |
| Syne2 | 1.51E-30 | 0.605094154 | 0.422 | 0.393 | 2.18E-27 |
| Osbpl3 | 4.10E-14 | 0.599235748 | 0.586 | 0.4 | 5.91E-11 |
| Gm30382 | 1.73E-51 | 0.582384888 | 0.468 | 0.736 | 2.50E-48 |
| Etv5 | 1.13E-115 | 0.577069929 | 0.724 | 0.184 | 1.63E-112 |
| Rasgrp1 | 1.51E-28 | 0.572711368 | 0.739 | 0.87 | 2.18E-25 |
| Adgrg6 | 1.30E-246 | 0.571977732 | 0.279 | 0.835 | 1.88E-243 |
| Mctp2 | 3.91E-126 | 0.568196176 | 0.221 | 0.455 | 5.64E-123 |
| Pltp | 3.17E-30 | 0.56750128 | 0.586 | 0.573 | 4.57E-27 |
| Maf | 1.95E-11 | 0.551263073 | 0.73 | 0.714 | 2.82E-08 |
| Dach1 | 9.99E-20 | 0.548753996 | 0.681 | 0.747 | 1.44E-16 |
| Calcrl | 1.94E-11 | 0.546033799 | 0.739 | 0.584 | 2.80E-08 |
| Il16 | 1.37E-16 | 0.544658777 | 0.54 | 0.586 | 1.97E-13 |
| Lcp1 | 6.17E-10 | 0.534992539 | 0.592 | 0.613 | 8.90E-07 |
| St3gal4 | 1.47E-43 | 0.527444848 | 0.44 | 0.697 | 2.12E-40 |
| Tgfbr2 | 4.44E-05 | 0.527317034 | 0.716 | 0.62 | 0.064101868 |
| Gm15261 | 6.35E-54 | 0.523994101 | 0.649 | 0.264 | 9.16E-51 |
| Hmga2 | 2.62E-41 | 0.521593947 | 0.365 | 0.136 | 3.77E-38 |
| Slc25a21 | 3.16E-134 | 0.518158148 | 0.773 | 0.15 | 4.56E-131 |
| Pparg | 2.31E-61 | 0.517797738 | 0.397 | 0.066 | 3.33E-58 |
| Acsl3 | 4.35E-14 | 0.511920329 | 0.767 | 0.656 | 6.27E-11 |
| Lrrk2 | 8.95E-48 | 0.502444359 | 0.816 | 0.745 | 1.29E-44 |
| Gm5086 | 3.56E-05 | 0.486607412 | 0.606 | 0.635 | 0.05136456 |
| Col14a1 | 6.90E-95 | 0.482207361 | 0.319 | 0.737 | 9.96E-92 |
| Camk4 | 4.43E-24 | 0.477171167 | 0.799 | 0.791 | 6.40E-21 |
| Frmpd4 | 7.60E-05 | 0.475975494 | 0.848 | 0.904 | 0.109621508 |
| Eda | 6.63E-62 | 0.463212334 | 0.468 | 0.748 | 9.57E-59 |
| Slit2 | 1.71E-30 | 0.45748438 | 0.79 | 0.526 | 2.46E-27 |
| Satb1 | 5.83E-37 | 0.456503642 | 0.782 | 0.438 | 8.41E-34 |
| Dock2 | 0.000215179 | 0.453821712 | 0.876 | 0.809 | 0.310503892 |
| Nhsl2 | 2.12E-35 | 0.448291097 | 0.319 | 0.525 | 3.06E-32 |
| Cpne8 | 1.37E-52 | 0.447954144 | 0.509 | 0.69 | 1.97E-49 |
| Sh3d19 | 1.34E-11 | 0.438789648 | 0.739 | 0.712 | 1.93E-08 |
| Abtb2 | 1.07E-76 | 0.435506441 | 0.451 | 0.804 | 1.54E-73 |
| Mill2 | 7.74E-48 | 0.430047143 | 0.491 | 0.395 | 1.12E-44 |
| Dpp4 | 3.60E-290 | 0.429569399 | 0.078 | 0.461 | 5.20E-287 |
| Prkcb | 4.95E-08 | 0.415067626 | 0.779 | 0.825 | 7.15E-05 |
| Ifitm3 | 1.15E-92 | 0.412613224 | 0.747 | 0.813 | 1.67E-89 |
| Atp10a | 9.88E-44 | 0.411558411 | 0.566 | 0.211 | 1.43E-40 |
| Man1a | 1.81E-10 | 0.399763274 | 0.796 | 0.626 | 2.61E-07 |
| Cnksr3 | 1.66E-07 | 0.399738662 | 0.601 | 0.499 | 0.000239241 |
| Stk39 | 6.45E-42 | 0.399313979 | 0.345 | 0.573 | 9.30E-39 |
| Tox2 | 1.93E-89 | 0.396086655 | 0.833 | 0.625 | 2.78E-86 |
| Grk5 | 0.001705932 | 0.388094498 | 0.603 | 0.634 | 1 |
| St18 | 5.91E-08 | 0.387672086 | 0.491 | 0.527 | 8.52E-05 |
| Airn | 3.90E-28 | 0.383306138 | 0.618 | 0.759 | 5.63E-25 |
| Asic2 | 2.08E-06 | 0.382494809 | 0.813 | 0.896 | 0.003004487 |
| Pdzd2 | 7.41E-52 | 0.377859193 | 0.779 | 0.398 | 1.07E-48 |
| Tbc1d1 | 0.048307824 | 0.376436551 | 0.356 | 0.332 | 1 |
| Pde8a | 1.81E-12 | 0.3715239 | 0.71 | 0.781 | 2.61E-09 |
| Plekhg1 | 4.91E-20 | 0.368582943 | 0.474 | 0.528 | 7.08E-17 |
| Btbd11 | 1.84E-48 | 0.366756533 | 0.586 | 0.804 | 2.65E-45 |
| Sgcz | 1.86E-12 | 0.365890543 | 0.701 | 0.857 | 2.68E-09 |
| Kitl | 9.02E-06 | 0.363002769 | 0.411 | 0.468 | 0.013010781 |
| Epha6 | 7.81E-17 | 0.362002687 | 0.871 | 0.921 | 1.13E-13 |
| Ntng1 | 6.23E-16 | 0.361294852 | 0.787 | 0.793 | 8.99E-13 |
| Gm6994 | 1.44E-42 | 0.360939028 | 0.451 | 0.795 | 2.07E-39 |
| Sugct | 8.69E-07 | 0.360908521 | 0.764 | 0.669 | 0.001253715 |
| Atp2b4 | 4.96E-09 | 0.358254288 | 0.359 | 0.302 | 7.16E-06 |
| Tshz3 | 1.35E-26 | 0.357104772 | 0.595 | 0.78 | 1.95E-23 |
| 9530059O14Rik | 1.63E-28 | 0.34316701 | 0.825 | 0.703 | 2.35E-25 |
| Klf2 | 2.82E-35 | 0.342476749 | 0.54 | 0.249 | 4.07E-32 |
| Ezr | 2.84E-88 | 0.336427129 | 0.647 | 0.169 | 4.10E-85 |
| Kit | 5.56E-06 | 0.33557424 | 0.468 | 0.422 | 0.008021666 |
| Spats2l | 1.70E-59 | 0.332560769 | 0.376 | 0.625 | 2.45E-56 |
| Etl4 | 1.08E-07 | 0.331860094 | 0.848 | 0.85 | 0.000155994 |
| Prkcq | 2.03E-92 | 0.330596191 | 0.086 | 0.5 | 2.93E-89 |
| Lin7a | 6.67E-42 | 0.32805663 | 0.828 | 0.779 | 9.63E-39 |
| Hs3st3a1 | 2.56E-113 | 0.325144134 | 0.356 | 0.77 | 3.70E-110 |
| Grik4 | 0.000269456 | 0.309723663 | 0.75 | 0.721 | 0.388824924 |
| Stxbp6 | 1.09E-124 | 0.308828458 | 0.161 | 0.664 | 1.57E-121 |
| Caln1 | 1.19E-25 | 0.308044103 | 0.805 | 0.784 | 1.72E-22 |
| Dlc1 | 0.003116045 | 0.307054671 | 0.796 | 0.729 | 1 |
| Rsad2 | 1.48E-55 | 0.305845198 | 0.644 | 0.804 | 2.14E-52 |
| Agbl1 | 1.82E-35 | 0.305748396 | 0.825 | 0.652 | 2.63E-32 |
| Slc7a2 | 8.73E-37 | 0.304696785 | 0.664 | 0.571 | 1.26E-33 |
| Runx1 | 0.066731531 | 0.30444341 | 0.874 | 0.866 | 1 |
| Plp1 | 2.23E-05 | 0.299451037 | 0.693 | 0.571 | 0.032152664 |
| Svil | 0.001352091 | 0.297823546 | 0.491 | 0.481 | 1 |
| Sphkap | 3.91E-05 | 0.296971544 | 0.813 | 0.748 | 0.056361318 |
| Edaradd | 8.36E-49 | 0.292730461 | 0.555 | 0.762 | 1.21E-45 |
| Cdh18 | 3.92E-05 | 0.289622124 | 0.83 | 0.871 | 0.056504444 |
| Sox5 | 4.16E-17 | 0.286584177 | 0.828 | 0.874 | 6.01E-14 |
| Heg1 | 1.21E-191 | 0.275916039 | 0.101 | 0.665 | 1.74E-188 |
| Lrrc4c | 1.61E-11 | 0.275149644 | 0.848 | 0.877 | 2.32E-08 |
| Crim1 | 7.79E-12 | 0.27502956 | 0.615 | 0.767 | 1.12E-08 |
| Rhoj | 3.03E-34 | 0.271920539 | 0.388 | 0.522 | 4.37E-31 |
| Samd3 | 1.31E-160 | 0.268519797 | 0.793 | 0.122 | 1.89E-157 |
| Tenm2 | 0.060099233 | 0.26767479 | 0.914 | 0.946 | 1 |
| Maml2 | 0.198855034 | 0.263154085 | 0.851 | 0.851 | 1 |
| Anln | 6.81E-15 | 0.26185134 | 0.664 | 0.534 | 9.83E-12 |
| Grip1 | 7.40E-18 | 0.259059322 | 0.822 | 0.846 | 1.07E-14 |
| Pip5k1b | 3.71E-26 | 0.258917422 | 0.517 | 0.753 | 5.36E-23 |
| Lingo2 | 0.001139841 | 0.25064724 | 0.885 | 0.92 | 1 |
| Cadps2 | 8.84E-30 | 0.248743432 | 0.563 | 0.827 | 1.28E-26 |
| Plcb4 | 0.008852011 | 0.246169841 | 0.305 | 0.245 | 1 |
| Fap | 3.76E-15 | 0.245843427 | 0.422 | 0.304 | 5.43E-12 |
| Dner | 0.006624273 | 0.244277874 | 0.443 | 0.399 | 1 |
| Zfp385b | 1.16E-20 | 0.243811983 | 0.802 | 0.879 | 1.68E-17 |
| Gm15155 | 2.73E-06 | 0.242903655 | 0.595 | 0.471 | 0.003945319 |
| Myo3b | 5.86E-43 | 0.24243353 | 0.509 | 0.168 | 8.46E-40 |
| Lama3 | 7.39E-80 | 0.240030269 | 0.448 | 0.06 | 1.07E-76 |
| Lrp4 | 4.08E-18 | 0.239525608 | 0.75 | 0.535 | 5.89E-15 |
| Adgre1 | 1.94E-24 | 0.239304045 | 0.497 | 0.654 | 2.80E-21 |
| Postn | 0.001810046 | 0.238325119 | 0.672 | 0.575 | 1 |
| Mbp | 1.53E-71 | 0.236441188 | 0.307 | 0.716 | 2.21E-68 |
| Lhfpl3 | 9.18E-06 | 0.235291181 | 0.626 | 0.638 | 0.013253807 |
| Gm28153 | 1.86E-34 | 0.228791958 | 0.813 | 0.481 | 2.69E-31 |
| Gja1 | 8.58E-56 | 0.228418184 | 0.81 | 0.385 | 1.24E-52 |
| Corin | 3.57E-120 | 0.225154354 | 0.776 | 0.216 | 5.14E-117 |
| Grm1 | 1.56E-13 | 0.225001306 | 0.753 | 0.856 | 2.25E-10 |
| Cobl | 6.57E-16 | 0.223263986 | 0.526 | 0.316 | 9.49E-13 |
| Sytl2 | 1.50E-101 | 0.219302962 | 0.17 | 0.621 | 2.17E-98 |
| Shisa6 | 1.54E-05 | 0.218782162 | 0.822 | 0.783 | 0.022259886 |
| Sh3rf1 | 0.000667272 | 0.217608228 | 0.494 | 0.444 | 0.9628737 |
| Hcn1 | 0.037752088 | 0.216695022 | 0.79 | 0.833 | 1 |
| Shisa9 | 3.54E-09 | 0.214741339 | 0.773 | 0.735 | 5.10E-06 |
| Ak9 | 1.13E-90 | 0.214508583 | 0.81 | 0.789 | 1.63E-87 |

| | | | | | |
|---|---|---|---|---|---|
| Itga8 | 1.60E-29 | 0.213735885 | 0.552 | 0.782 | 2.31E-26 |
| Mbnl3 | 3.11E-49 | 0.212461212 | 0.543 | 0.194 | 4.49E-46 |
| Apod | 3.34E-06 | 0.209689158 | 0.431 | 0.524 | 0.004814401 |
| Mid1 | 0.016497516 | 0.205881101 | 0.402 | 0.333 | 1 |
| Fmn1 | 1.36E-20 | 0.205413617 | 0.374 | 0.599 | 1.96E-17 |
| Zmat4 | 4.87E-15 | 0.203855713 | 0.517 | 0.726 | 7.03E-12 |
| Pde1c | 4.63E-33 | 0.2027036 | 0.707 | 0.368 | 6.68E-30 |
| Tes | 1.28E-09 | 0.198353108 | 0.374 | 0.523 | 1.85E-06 |
| Cit | 1.64E-26 | 0.198190122 | 0.675 | 0.383 | 2.36E-23 |
| Ctnna3 | 5.49E-15 | 0.19528823 | 0.744 | 0.534 | 7.92E-12 |
| Vsnl1 | 4.37E-22 | 0.194289279 | 0.618 | 0.343 | 6.30E-19 |
| Gfra1 | 3.75E-07 | 0.190557318 | 0.601 | 0.521 | 0.000540987 |
| A330008L17Rik | 0.242006607 | 0.190012633 | 0.621 | 0.665 | 1 |
| Agbl4 | 4.39E-11 | 0.189836491 | 0.839 | 0.87 | 6.34E-08 |
| Gm20754 | 0.000479197 | 0.18913187 | 0.842 | 0.9 | 0.691481175 |
| Klhl14 | 5.59E-86 | 0.187047167 | 0.819 | 0.321 | 8.07E-83 |
| Cntn4 | 1.00E-07 | 0.18636265 | 0.704 | 0.79 | 0.000144444 |
| Thsd4 | 1.48E-08 | 0.183824018 | 0.764 | 0.848 | 2.14E-05 |
| Cdh9 | 2.38E-13 | 0.179560184 | 0.767 | 0.642 | 3.44E-10 |
| Dgkb | 0.03608727 | 0.177173354 | 0.83 | 0.878 | 1 |
| Cachd1 | 8.31E-22 | 0.17716008 | 0.779 | 0.879 | 1.20E-18 |
| Tmtc2 | 4.49E-101 | 0.172511665 | 0.175 | 0.645 | 6.48E-98 |
| 6530403H02Rik | 6.91E-05 | 0.172308593 | 0.621 | 0.512 | 0.09976573 |
| Zfp462 | 0.000122022 | 0.172222809 | 0.621 | 0.667 | 0.176078226 |
| Hpgd | 0.078530757 | 0.169789054 | 0.687 | 0.627 | 1 |
| Mamdc2 | 4.06E-272 | 0.168644815 | 0.721 | 0.008 | 5.86E-269 |
| Tle4 | 5.81E-14 | 0.167072328 | 0.397 | 0.525 | 8.39E-11 |
| Eya2 | 1.61E-147 | 0.164375058 | 0.138 | 0.753 | 2.32E-144 |
| Ptprg | 1.64E-11 | 0.16384309 | 0.793 | 0.618 | 2.37E-08 |
| Igsf9b | 2.74E-20 | 0.162687348 | 0.81 | 0.755 | 3.96E-17 |
| Chsy3 | 0.003213175 | 0.161213614 | 0.83 | 0.876 | 1 |
| Pbx3 | 0.000869177 | 0.160374169 | 0.532 | 0.567 | 1 |
| Car2 | 1.30E-07 | 0.159448421 | 0.322 | 0.39 | 0.000188305 |
| Mal | 2.75E-20 | 0.15884335 | 0.586 | 0.325 | 3.97E-17 |
| Mpp7 | 9.79E-29 | 0.158606117 | 0.753 | 0.439 | 1.41E-25 |
| Ar | 2.71E-21 | 0.15848937 | 0.744 | 0.799 | 3.91E-18 |
| Sorcs2 | 4.65E-30 | 0.15815384 | 0.448 | 0.744 | 6.71E-27 |
| Sema3c | 1.48E-65 | 0.157353181 | 0.747 | 0.286 | 2.13E-62 |
| Kcnt2 | 0.001284144 | 0.1545316 | 0.833 | 0.841 | 1 |
| Camk2d | 0.699917345 | 0.15377515 | 0.773 | 0.762 | 1 |
| Tenm3 | 5.21E-21 | 0.153430271 | 0.69 | 0.891 | 7.52E-18 |
| Nrg1 | 0.415651649 | 0.152986325 | 0.816 | 0.789 | 1 |
| Npsr1 | 9.21E-78 | 0.151414125 | 0.799 | 0.357 | 1.33E-74 |
| Il1r1 | 1.29E-06 | 0.151088769 | 0.658 | 0.56 | 0.001858874 |
| Khdrbs2 | 0.001623215 | 0.150755826 | 0.819 | 0.791 | 1 |
| Enpp2 | 3.59E-08 | 0.146587831 | 0.825 | 0.686 | 5.18E-05 |
| Robo1 | 3.93E-09 | 0.146305918 | 0.618 | 0.764 | 5.67E-06 |
| Cntnap5a | 6.68E-08 | 0.145065978 | 0.842 | 0.719 | 9.65E-05 |
| Pam | 8.29E-118 | 0.145002465 | 0.264 | 0.814 | 1.20E-114 |
| Slc1a2 | 4.47E-09 | 0.144862642 | 0.641 | 0.468 | 6.45E-06 |
| Gm20319 | 9.04E-48 | 0.142742391 | 0.704 | 0.301 | 1.30E-44 |
| Fbn2 | 3.59E-07 | 0.142590968 | 0.828 | 0.767 | 0.000517558 |
| Ntm | 2.62E-05 | 0.142070628 | 0.845 | 0.892 | 0.037759545 |
| 4930473D10Rik | 1.34E-16 | 0.14178342 | 0.468 | 0.359 | 1.93E-13 |
| Cntn5 | 7.07E-05 | 0.140937969 | 0.721 | 0.818 | 0.10202645 |
| Lrig1 | 8.92E-30 | 0.140126193 | 0.81 | 0.672 | 1.29E-26 |
| Gpc6 | 6.65E-11 | 0.139852923 | 0.704 | 0.832 | 9.60E-08 |
| Ppfibp1 | 0.247985368 | 0.139767544 | 0.534 | 0.513 | 1 |
| Prr5l | 7.96E-53 | 0.138686881 | 0.256 | 0.639 | 1.15E-49 |
| Pde1a | 2.36E-07 | 0.138600933 | 0.825 | 0.876 | 0.000340882 |
| Sema6a | 1.67E-11 | 0.136733641 | 0.566 | 0.375 | 2.41E-08 |
| Mapk4 | 7.28E-08 | 0.136536996 | 0.684 | 0.783 | 0.000105054 |
| Scml4 | 0.001170829 | 0.136301901 | 0.261 | 0.257 | 1 |
| Cdh20 | 7.34E-05 | 0.134729859 | 0.589 | 0.559 | 0.105901343 |
| Il1rapl2 | 1.27E-16 | 0.132547395 | 0.678 | 0.861 | 1.83E-13 |
| Prlr | 1.62E-07 | 0.132156537 | 0.299 | 0.418 | 0.000233411 |
| Npy | 8.74E-66 | 0.131135444 | 0.753 | 0.299 | 1.26E-62 |
| Met | 5.40E-178 | 0.130592352 | 0.583 | 0.019 | 7.79E-175 |
| Meis2 | 1.72E-19 | 0.130433612 | 0.626 | 0.832 | 2.48E-16 |
| Nyap2 | 9.39E-09 | 0.130331819 | 0.805 | 0.69 | 1.36E-05 |
| Uaca | 1.21E-07 | 0.129273531 | 0.376 | 0.385 | 0.000175134 |
| Ldb2 | 4.32E-55 | 0.127954493 | 0.319 | 0.731 | 6.23E-52 |
| Mgp | 0.109687648 | 0.125738951 | 0.813 | 0.778 | 1 |
| Cyth4 | 9.54E-08 | 0.125714169 | 0.759 | 0.671 | 0.000137734 |
| Hs6st3 | 8.30E-06 | 0.125062555 | 0.833 | 0.915 | 0.011975779 |
| Ptchd4 | 1.01E-24 | 0.122775067 | 0.79 | 0.51 | 1.45E-21 |
| Ptchd1 | 3.44E-64 | 0.121669296 | 0.261 | 0.699 | 4.96E-61 |
| Sh3rf3 | 6.52E-08 | 0.121516877 | 0.764 | 0.785 | 9.40E-05 |
| Kcnh7 | 0.090307923 | 0.121271382 | 0.853 | 0.827 | 1 |
| Gm16168 | 8.54E-74 | 0.121134672 | 0.767 | 0.271 | 1.23E-70 |
| Dkk3 | 0.000275981 | 0.120045528 | 0.73 | 0.66 | 0.39823993 |
| Ptn | 0.125729262 | 0.119454768 | 0.52 | 0.5 | 1 |
| Rmst | 1.92E-26 | 0.118684361 | 0.69 | 0.39 | 2.77E-23 |
| Pdgfd | 7.11E-06 | 0.118631953 | 0.632 | 0.729 | 0.010263201 |
| Ly6a | 2.20E-81 | 0.118297105 | 0.437 | 0.049 | 3.17E-78 |
| Errfi1 | 1.21E-09 | 0.118229944 | 0.348 | 0.506 | 1.75E-06 |
| Lin28b | 0.000499593 | 0.118072115 | 0.534 | 0.526 | 0.720913359 |
| Gm31698 | 3.26E-118 | 0.117832155 | 0.825 | 0.218 | 4.70E-115 |
| Ghr | 3.70E-16 | 0.116597567 | 0.747 | 0.531 | 5.34E-13 |
| Rreb1 | 0.09413133 | 0.116527319 | 0.79 | 0.829 | 1 |
| Rspo2 | 1.40E-36 | 0.116168364 | 0.756 | 0.421 | 2.02E-33 |
| Aox3 | 2.13E-68 | 0.115480579 | 0.776 | 0.329 | 3.08E-65 |
| Clstn2 | 9.21E-05 | 0.114196665 | 0.842 | 0.888 | 0.132921753 |
| Ano3 | 0.001863262 | 0.11343583 | 0.787 | 0.819 | 1 |
| Cacng2 | 0.000128049 | 0.11308878 | 0.549 | 0.476 | 0.184775307 |
| 4930445B16Rik | 2.48E-53 | 0.112010834 | 0.621 | 0.223 | 3.58E-50 |
| Plpp3 | 0.0294067 | 0.111851595 | 0.575 | 0.5 | 1 |
| Rgcc | 1.16E-37 | 0.111068222 | 0.664 | 0.309 | 1.67E-34 |
| A830018L16Rik | 7.42E-05 | 0.11063396 | 0.773 | 0.847 | 0.107024324 |
| Slc4a4 | 5.35E-05 | 0.109178729 | 0.741 | 0.628 | 0.077260667 |
| 4930447C04Rik | 3.11E-15 | 0.107140028 | 0.767 | 0.625 | 4.49E-12 |
| Palld | 3.97E-90 | 0.105450354 | 0.224 | 0.735 | 5.73E-87 |
| Ptprr | 5.95E-05 | 0.105160407 | 0.782 | 0.818 | 0.085813001 |
| Pkp2 | 0.092444903 | 0.104876801 | 0.629 | 0.673 | 1 |
| Ros1 | 4.69E-22 | 0.104736484 | 0.543 | 0.47 | 6.76E-19 |
| Sox6 | 2.11E-12 | 0.103190157 | 0.813 | 0.72 | 3.04E-09 |
| Dcn | 1.18E-16 | 0.102717564 | 0.411 | 0.2 | 1.71E-13 |
| Cdh1 | 3.14E-33 | 0.101826844 | 0.802 | 0.524 | 4.53E-30 |
| Gnal | 0.004809894 | 0.101635939 | 0.721 | 0.672 | 1 |
| Kirrel3 | 3.72E-06 | 0.101601682 | 0.848 | 0.779 | 0.005361511 |
| Mkx | 9.10E-11 | 0.101590419 | 0.787 | 0.618 | 1.31E-07 |
| Emp1 | 1.19E-85 | 0.100683009 | 0.402 | 0.034 | 1.71E-82 |
| Igsf21 | 4.21E-54 | 0.100508859 | 0.362 | 0.764 | 6.07E-51 |
| Mgat4c | 3.27E-08 | 0.098556603 | 0.75 | 0.862 | 4.71E-05 |
| Lmo3 | 1.34E-77 | 0.098175645 | 0.776 | 0.266 | 1.93E-74 |
| Stard5 | 4.63E-55 | 0.096904118 | 0.365 | 0.743 | 6.67E-52 |
| Adam12 | 1.56E-147 | 0.096708136 | 0.747 | 0.105 | 2.25E-144 |
| Ryr3 | 0.002068019 | 0.09632022 | 0.851 | 0.873 | 1 |
| Gm48321 | 1.39E-06 | 0.096219256 | 0.307 | 0.353 | 0.002004 |
| Syndig1 | 0.00029082 | 0.095606236 | 0.81 | 0.746 | 0.419652983 |
| Mrvi1 | 1.43E-115 | 0.09453212 | 0.805 | 0.197 | 2.06E-112 |
| Vat1l | 2.33E-66 | 0.094008259 | 0.807 | 0.819 | 3.37E-63 |
| Greb1l | 0.088815027 | 0.093320924 | 0.322 | 0.345 | 1 |
| Grip1os2 | 1.59E-22 | 0.090419098 | 0.796 | 0.558 | 2.30E-19 |
| Ccdc114 | 4.59E-25 | 0.090417312 | 0.494 | 0.746 | 6.62E-22 |
| Rgs20 | 8.36E-07 | 0.090402977 | 0.782 | 0.744 | 0.00120608 |
| Parm1 | 1.37E-47 | 0.089029001 | 0.635 | 0.242 | 1.98E-44 |
| 4933413L06Rik | 1.03E-07 | 0.088375892 | 0.658 | 0.783 | 0.000148876 |
| Flrt2 | 0.00192508 | 0.087068047 | 0.79 | 0.744 | 1 |
| Phka1 | 0.000357802 | 0.086514038 | 0.466 | 0.561 | 0.516308894 |
| Nell1 | 1.06E-07 | 0.085971831 | 0.825 | 0.686 | 0.000153542 |

| | | | | | |
|---|---|---|---|---|---|
| Coro6 | 1.14E-93 | 0.085791346 | 0.103 | 0.584 | 1.65E-90 |
| Ttr | 0.005724641 | 0.085732374 | 0.796 | 0.718 | 1 |
| Cpne4 | 0.595106664 | 0.085541279 | 0.563 | 0.581 | 1 |
| Mylk | 0.000614799 | 0.085435521 | 0.483 | 0.418 | 0.887155548 |
| Kcnn2 | 1.95E-79 | 0.085090659 | 0.345 | 0.813 | 2.81E-76 |
| Galnt9 | 1.08E-47 | 0.084442966 | 0.167 | 0.53 | 1.56E-44 |
| Ank1 | 1.73E-27 | 0.082734666 | 0.149 | 0.4 | 2.49E-24 |
| Ephb1 | 0.093349846 | 0.082524271 | 0.624 | 0.6 | 1 |
| Gabra3 | 5.66E-13 | 0.082156266 | 0.359 | 0.18 | 8.17E-10 |
| Sgms2 | 2.79E-06 | 0.081565601 | 0.333 | 0.212 | 0.004025951 |
| Cntn6 | 3.06E-08 | 0.081255098 | 0.727 | 0.678 | 4.42E-05 |
| Tenm4 | 9.98E-88 | 0.080905513 | 0.276 | 0.781 | 1.44E-84 |
| Synpr | 0.012636277 | 0.080527188 | 0.575 | 0.636 | 1 |
| Ebf2 | 4.19E-44 | 0.080354785 | 0.805 | 0.659 | 6.05E-41 |
| Adgrl4 | 1.87E-214 | 0.080053385 | 0.644 | 0.015 | 2.70E-211 |
| Cd86 | 7.71E-05 | 0.079653936 | 0.46 | 0.548 | 0.111248572 |
| Ednra | 5.79E-165 | 0.079648401 | 0.431 | 0.002 | 8.35E-162 |
| Zfp521 | 3.48E-69 | 0.079198503 | 0.667 | 0.201 | 5.03E-66 |
| Il31ra | 3.22E-49 | 0.078521493 | 0.138 | 0.461 | 4.65E-46 |
| Cacng3 | 0.160093713 | 0.078307753 | 0.437 | 0.419 | 1 |
| Pawr | 1.13E-51 | 0.07730576 | 0.767 | 0.345 | 1.63E-48 |
| AU020206 | 7.27E-20 | 0.077224419 | 0.503 | 0.72 | 1.05E-16 |
| Sema3a | 4.38E-11 | 0.077004038 | 0.408 | 0.232 | 6.32E-08 |
| Luzp2 | 0.000786374 | 0.076899234 | 0.698 | 0.789 | 1 |
| Rspo3 | 6.15E-20 | 0.076866338 | 0.376 | 0.633 | 8.88E-17 |
| Fli1 | 0.005175968 | 0.076698703 | 0.793 | 0.817 | 1 |
| Prrg1 | 4.09E-118 | 0.076446967 | 0.098 | 0.455 | 5.90E-115 |
| Rai14 | 0.053407711 | 0.076240226 | 0.253 | 0.285 | 1 |
| A630012P03Rik | 2.32E-39 | 0.075859216 | 0.422 | 0.121 | 3.35E-36 |
| Homer1 | 7.12E-23 | 0.075303815 | 0.451 | 0.707 | 1.03E-19 |
| Dpf3 | 7.22E-11 | 0.075115102 | 0.767 | 0.587 | 1.04E-07 |
| 4930587E11Rik | 6.35E-10 | 0.075079225 | 0.672 | 0.819 | 9.17E-07 |
| Arhgap28 | 4.60E-179 | 0.074444631 | 0.543 | 0.011 | 6.63E-176 |
| Fn1 | 2.29E-124 | 0.074334559 | 0.805 | 0.178 | 3.30E-121 |
| Rab27b | 1.17E-08 | 0.074276751 | 0.724 | 0.562 | 1.69E-05 |
| Elavl2 | 7.16E-06 | 0.073944957 | 0.724 | 0.776 | 0.01033451 |
| Olfr112 | 0.007329486 | 0.07367687 | 0.546 | 0.519 | 1 |
| Atp1a2 | 4.60E-05 | 0.073307028 | 0.506 | 0.382 | 0.066333946 |
| Car13 | 9.32E-60 | 0.072195784 | 0.425 | 0.823 | 1.34E-56 |
| Phactr2 | 6.45E-24 | 0.071444775 | 0.578 | 0.804 | 9.30E-21 |
| Syt17 | 1.73E-07 | 0.071369789 | 0.695 | 0.547 | 0.000250237 |
| Id3 | 7.85E-37 | 0.071232967 | 0.529 | 0.198 | 1.13E-33 |
| Ttn | 0.000129141 | 0.070928989 | 0.448 | 0.334 | 0.186349829 |
| Gm48893 | 0.009605818 | 0.070642906 | 0.445 | 0.422 | 1 |
| Gm43154 | 4.58E-33 | 0.070535009 | 0.566 | 0.242 | 6.61E-30 |
| Tgfb2 | 2.78E-60 | 0.070194983 | 0.213 | 0.626 | 4.02E-57 |
| 4930467D21Rik | 4.77E-37 | 0.070054492 | 0.431 | 0.767 | 6.89E-34 |
| Tmtc1 | 3.53E-07 | 0.06899265 | 0.497 | 0.641 | 0.000509668 |
| Cftr | 0.00055024 | 0.068134079 | 0.736 | 0.646 | 0.793996208 |
| Kcnh8 | 7.57E-14 | 0.067330256 | 0.601 | 0.789 | 1.09E-10 |
| Kcnj13 | 2.51E-39 | 0.065882137 | 0.701 | 0.337 | 3.63E-36 |
| Wwtr1 | 0.661788191 | 0.065361476 | 0.644 | 0.621 | 1 |
| Fyb2 | 2.64E-45 | 0.064765565 | 0.675 | 0.287 | 3.81E-42 |
| Col19a1 | 4.71E-20 | 0.064594974 | 0.718 | 0.884 | 6.80E-17 |
| Mirt1 | 3.60E-114 | 0.064374899 | 0.236 | 0.814 | 5.19E-111 |
| Rorb | 6.78E-10 | 0.06408159 | 0.819 | 0.858 | 9.79E-07 |
| Trpc6 | 1.88E-18 | 0.063784202 | 0.402 | 0.182 | 2.72E-15 |
| Ralyl | 2.95E-07 | 0.06366686 | 0.787 | 0.892 | 0.000426366 |
| Gm26713 | 4.69E-40 | 0.063638492 | 0.543 | 0.197 | 6.77E-37 |
| Adgrv1 | 6.21E-10 | 0.06334954 | 0.253 | 0.118 | 8.96E-07 |
| Cntnap4 | 0.000378764 | 0.063241532 | 0.621 | 0.722 | 0.546556988 |
| Nnat | 2.04E-28 | 0.062891624 | 0.759 | 0.449 | 2.94E-25 |
| Slc2a13 | 0.977291569 | 0.062432809 | 0.655 | 0.653 | 1 |
| Nkain3 | 0.000619412 | 0.06183404 | 0.753 | 0.836 | 0.893811754 |
| Igfbp7 | 6.45E-91 | 0.061676082 | 0.698 | 0.169 | 9.30E-88 |
| Gpnmb | 1.99E-13 | 0.061615744 | 0.807 | 0.627 | 2.88E-10 |
| Ntng2 | 3.84E-67 | 0.059327698 | 0.092 | 0.463 | 5.54E-64 |
| Nwd1 | 1.24E-19 | 0.058645742 | 0.489 | 0.735 | 1.78E-16 |
| Slc22a2 | 1.09E-207 | 0.05792009 | 0.73 | 0.041 | 1.58E-204 |
| Fgf1 | 2.02E-32 | 0.057917529 | 0.782 | 0.45 | 2.92E-29 |
| Coch | 2.20E-36 | 0.057904786 | 0.675 | 0.324 | 3.18E-33 |
| Dach2 | 6.54E-12 | 0.057341994 | 0.256 | 0.109 | 9.44E-09 |
| Rcsd1 | 0.000463423 | 0.057190708 | 0.514 | 0.571 | 0.66871874 |
| Grm4 | 4.88E-06 | 0.057069734 | 0.471 | 0.464 | 0.007041789 |
| Ly6c1 | 6.34E-90 | 0.057013521 | 0.33 | 0.012 | 9.15E-87 |
| Rdh5 | 1.01E-05 | 0.056750546 | 0.457 | 0.573 | 0.014583673 |
| Dchs2 | 2.41E-167 | 0.056250584 | 0.695 | 0.06 | 3.47E-164 |
| Abca4 | 3.71E-118 | 0.05597891 | 0.736 | 0.142 | 5.35E-115 |
| Ptprt | 0.349101588 | 0.055919825 | 0.741 | 0.706 | 1 |
| Strip2 | 1.79E-51 | 0.055655486 | 0.402 | 0.789 | 2.59E-48 |
| Ndst4 | 3.74E-05 | 0.055395333 | 0.592 | 0.703 | 0.054007977 |
| Gm973 | 4.78E-43 | 0.054317787 | 0.394 | 0.092 | 6.89E-40 |
| Car12 | 1.78E-85 | 0.053979248 | 0.48 | 0.062 | 2.57E-82 |
| Myo16 | 0.093640475 | 0.053795181 | 0.471 | 0.413 | 1 |
| Cnih3 | 1.98E-07 | 0.053658008 | 0.609 | 0.687 | 0.000285132 |
| Kcnip1 | 0.000180076 | 0.053637843 | 0.764 | 0.685 | 0.259850177 |
| D630024D03Rik | 3.39E-105 | 0.053070069 | 0.724 | 0.158 | 4.90E-102 |
| Efr3b | 0.000245722 | 0.052331727 | 0.793 | 0.801 | 0.354577335 |
| Bmper | 4.17E-41 | 0.052143464 | 0.721 | 0.345 | 6.02E-38 |
| Ahcyl2 | 1.01E-21 | 0.051377443 | 0.448 | 0.675 | 1.45E-18 |
| Gm28175 | 1.39E-24 | 0.051154251 | 0.819 | 0.559 | 2.01E-21 |
| Grm8 | 3.62E-27 | 0.050524477 | 0.56 | 0.83 | 5.23E-24 |
| Matn2 | 0.000152444 | 0.050035412 | 0.572 | 0.666 | 0.219976758 |
| Neu4 | 2.03E-06 | 0.049897176 | 0.678 | 0.747 | 0.002930673 |
| Zfpm2 | 0.014013983 | 0.049761519 | 0.782 | 0.812 | 1 |
| Flt1 | 6.61E-19 | 0.049518423 | 0.661 | 0.408 | 9.54E-16 |
| Rgs5 | 8.09E-272 | 0.048692749 | 0.652 | 0.002 | 1.17E-268 |
| Eya1 | 1.31E-39 | 0.048339114 | 0.448 | 0.777 | 1.89E-36 |
| Egfem1 | 0.052065366 | 0.047831214 | 0.862 | 0.904 | 1 |
| Rtl4 | 4.51E-24 | 0.047663322 | 0.543 | 0.803 | 6.51E-21 |
| Trpm3 | 1.16E-16 | 0.047631413 | 0.73 | 0.902 | 1.67E-13 |
| Neto1 | 1.53E-16 | 0.046675697 | 0.624 | 0.826 | 2.20E-13 |
| Slc6a13 | 1.10E-11 | 0.046201748 | 0.718 | 0.569 | 1.59E-08 |
| Dcc | 0.013606949 | 0.045916742 | 0.856 | 0.908 | 1 |
| Bicc1 | 0.69745761 | 0.045812187 | 0.727 | 0.727 | 1 |
| Raver2 | 2.28E-05 | 0.045602825 | 0.629 | 0.747 | 0.032905305 |
| Gm13052 | 4.95E-56 | 0.045479503 | 0.279 | 0.701 | 7.14E-53 |
| Tnr | 0.010131467 | 0.045193103 | 0.67 | 0.687 | 1 |
| Cald1 | 2.38E-56 | 0.045098125 | 0.434 | 0.833 | 3.43E-53 |
| Fgfr3 | 3.71E-23 | 0.045030838 | 0.486 | 0.72 | 5.36E-20 |
| Nos1 | 3.23E-06 | 0.044662063 | 0.825 | 0.733 | 0.004654456 |
| Tspan2 | 7.47E-103 | 0.044607185 | 0.46 | 0.039 | 1.08E-99 |
| Dnah6 | 4.53E-55 | 0.043784239 | 0.796 | 0.365 | 6.54E-52 |
| Sorcs3 | 0.004300902 | 0.042919354 | 0.56 | 0.474 | 1 |
| Tead1 | 8.37E-32 | 0.042417525 | 0.687 | 0.36 | 1.21E-28 |
| Pamr1 | 8.16E-152 | 0.042060838 | 0.718 | 0.085 | 1.18E-148 |
| Angpt1 | 1.56E-10 | 0.041507158 | 0.241 | 0.418 | 2.25E-07 |
| Ror2 | 0.058387219 | 0.041338814 | 0.647 | 0.675 | 1 |
| Gatm | 4.69E-06 | 0.040477396 | 0.506 | 0.632 | 0.006765677 |
| Plcxd3 | 1.13E-52 | 0.040225489 | 0.523 | 0.145 | 1.64E-49 |
| Adra1a | 9.79E-40 | 0.039894899 | 0.56 | 0.211 | 1.41E-36 |
| Gm16070 | 4.70E-75 | 0.039092016 | 0.759 | 0.258 | 6.79E-72 |
| Grid2 | 0.00080711 | 0.03869745 | 0.805 | 0.857 | 1 |
| Col5a2 | 1.61E-10 | 0.038570561 | 0.382 | 0.214 | 2.33E-07 |
| Htr7 | 1.77E-155 | 0.037873567 | 0.583 | 0.032 | 2.56E-152 |
| Tbx18 | 1.37E-41 | 0.037381066 | 0.063 | 0.357 | 1.97E-38 |
| Sox5os4 | 2.16E-18 | 0.03699566 | 0.601 | 0.35 | 3.11E-15 |
| Alkal2 | 1.05E-10 | 0.036675621 | 0.58 | 0.391 | 1.52E-07 |
| Nxn | 1.16E-18 | 0.036530772 | 0.101 | 0.286 | 1.67E-15 |
| Fras1 | 0.279659448 | 0.036197266 | 0.802 | 0.82 | 1 |
| Gm48619 | 0.008269595 | 0.036161907 | 0.207 | 0.274 | 1 |

| | | | | | |
|---|---|---|---|---|---|
| Plppr1 | 4.81E-11 | 0.035945676 | 0.661 | 0.82 | 6.95E-08 |
| Sntg2 | 8.30E-11 | 0.034938806 | 0.664 | 0.824 | 1.20E-07 |
| Kl | 4.87E-24 | 0.034133034 | 0.19 | 0.437 | 7.03E-21 |
| Cacna2d2 | 1.89E-84 | 0.034069755 | 0.756 | 0.228 | 2.72E-81 |
| Enpp1 | 4.00E-47 | 0.033931798 | 0.072 | 0.35 | 5.77E-44 |
| Cntnap3 | 0.031223324 | 0.033766 | 0.394 | 0.324 | 1 |
| Fgd5 | 2.34E-104 | 0.033003932 | 0.695 | 0.141 | 3.38E-101 |
| Scel | 7.74E-47 | 0.03282723 | 0.376 | 0.075 | 1.12E-43 |
| Elovl7 | 4.10E-49 | 0.03262622 | 0.385 | 0.779 | 5.91E-46 |
| Ndrg2 | 7.15E-10 | 0.032365249 | 0.511 | 0.333 | 1.03E-06 |
| Paqr8 | 4.38E-24 | 0.032051599 | 0.534 | 0.26 | 6.32E-21 |
| Cep112 | 0.095562919 | 0.031906706 | 0.563 | 0.611 | 1 |
| 1700007F19Rik | 1.15E-81 | 0.031608364 | 0.784 | 0.26 | 1.66E-78 |
| Adamts12 | 6.21E-112 | 0.031333648 | 0.247 | 0.831 | 8.96E-109 |
| Lrriq1 | 6.84E-29 | 0.031260355 | 0.342 | 0.65 | 9.88E-26 |
| Gm26633 | 7.28E-46 | 0.031207937 | 0.25 | 0.025 | 1.05E-42 |
| Inpp4b | 1.40E-17 | 0.031078444 | 0.693 | 0.882 | 2.01E-14 |
| Col12a1 | 5.75E-212 | 0.030713992 | 0.632 | 0.014 | 8.29E-209 |
| 4930488N15Rik | 5.92E-16 | 0.03010029 | 0.454 | 0.238 | 8.55E-13 |
| Shc4 | 1.06E-06 | 0.029864176 | 0.523 | 0.38 | 0.001533561 |
| Fam20a | 1.27E-87 | 0.029333785 | 0.739 | 0.208 | 1.84E-84 |
| C130073E24Rik | 1.14E-07 | 0.029331549 | 0.247 | 0.37 | 0.000164617 |
| Rasgrf2 | 9.16E-09 | 0.029149299 | 0.537 | 0.701 | 1.32E-05 |
| Socs2 | 5.43E-26 | 0.029052653 | 0.276 | 0.551 | 7.83E-23 |
| Ptprb | 0.001498001 | 0.029020986 | 0.77 | 0.797 | 1 |
| Myl9 | 2.78E-69 | 0.028957935 | 0.81 | 0.335 | 4.02E-66 |
| Dcdc2a | 2.33E-236 | 0.028945214 | 0.672 | 0.014 | 3.36E-233 |
| Clmp | 1.18E-32 | 0.028744432 | 0.414 | 0.723 | 1.71E-29 |
| Gm13963 | 1.76E-24 | 0.02845225 | 0.816 | 0.547 | 2.55E-21 |
| Gpc4 | 0.01254673 | 0.028432082 | 0.701 | 0.623 | 1 |
| Gm46102 | 4.80E-90 | 0.028362552 | 0.066 | 0.584 | 6.93E-87 |
| Col1a2 | 1.39E-43 | 0.028319019 | 0.052 | 0.31 | 2.00E-40 |
| Thbs1 | 0.632067291 | 0.028271826 | 0.342 | 0.345 | 1 |
| Mecom | 0.001456581 | 0.028212076 | 0.589 | 0.51 | 1 |
| Baiap2l1 | 1.16E-24 | 0.027817821 | 0.641 | 0.356 | 1.67E-21 |
| Npr3 | 0.323687941 | 0.027661023 | 0.776 | 0.761 | 1 |
| Lama2 | 0.006383721 | 0.02765437 | 0.741 | 0.783 | 1 |
| Frk | 5.55E-125 | 0.02740522 | 0.33 | 0.001 | 8.01E-122 |
| Gm29260 | 9.05E-150 | 0.026964809 | 0.411 | 0.002 | 1.31E-146 |
| Slc26a7 | 1.51E-252 | 0.026860617 | 0.713 | 0.014 | 2.18E-249 |
| Col24a1 | 6.84E-11 | 0.02656617 | 0.385 | 0.214 | 9.86E-08 |
| Hapln1 | 1.09E-40 | 0.026198447 | 0.307 | 0.055 | 1.57E-37 |
| Gm32828 | 0.002110305 | 0.025773744 | 0.316 | 0.39 | 1 |
| Mei4 | 2.48E-12 | 0.025660547 | 0.681 | 0.477 | 3.58E-09 |
| Vtn | 1.00E-60 | 0.02542913 | 0.67 | 0.225 | 1.44E-57 |
| Pdgfra | 2.11E-66 | 0.02541658 | 0.336 | 0.033 | 3.05E-63 |
| Baiap3 | 2.87E-27 | 0.02530603 | 0.44 | 0.734 | 4.14E-24 |
| 5530401A14Rik | 0.139985712 | 0.024986041 | 0.305 | 0.351 | 1 |
| Fgf12 | 0.000158018 | 0.024917279 | 0.851 | 0.895 | 0.228019621 |
| Gm42303 | 0.008927977 | 0.024828185 | 0.618 | 0.625 | 1 |
| Gm14964 | 0.008275302 | 0.024662122 | 0.69 | 0.609 | 1 |
| Sgpp2 | 7.93E-185 | 0.024438377 | 0.103 | 0.833 | 1.14E-181 |
| St8sia2 | 1.18E-113 | 0.024174096 | 0.397 | 0.013 | 1.70E-110 |
| Efemp1 | 6.47E-17 | 0.024148736 | 0.572 | 0.335 | 9.34E-14 |
| Cldn14 | 1.69E-29 | 0.02397481 | 0.126 | 0.402 | 2.43E-26 |
| Cfap65 | 1.36E-42 | 0.023821689 | 0.103 | 0.455 | 1.96E-39 |
| Ankrd33b | 1.85E-46 | 0.023660357 | 0.264 | 0.656 | 2.67E-43 |
| Gm45341 | 8.77E-18 | 0.023554389 | 0.67 | 0.422 | 1.26E-14 |
| Plpp4 | 2.79E-15 | 0.023106679 | 0.253 | 0.474 | 4.03E-12 |
| Adarb2 | 4.56E-07 | 0.021697104 | 0.782 | 0.888 | 0.000657859 |
| Cped1 | 5.79E-59 | 0.021603775 | 0.325 | 0.752 | 8.35E-56 |
| Crispld1 | 0.458142946 | 0.021584386 | 0.402 | 0.437 | 1 |
| Eln | 1.39E-58 | 0.021522111 | 0.739 | 0.292 | 2.01E-55 |
| Tm4sf1 | 1.43E-145 | 0.021127118 | 0.753 | 0.112 | 2.06E-142 |
| Lhfp | 0.59246009 | 0.021096564 | 0.667 | 0.646 | 1 |
| Gm2164 | 8.89E-34 | 0.020893884 | 0.333 | 0.665 | 1.28E-30 |
| Sel1l3 | 1.47E-44 | 0.020688237 | 0.428 | 0.796 | 2.12E-41 |
| Nek10 | 6.19E-11 | 0.02065776 | 0.471 | 0.625 | 8.93E-08 |
| Slc35f1 | 3.71E-09 | 0.020515618 | 0.606 | 0.761 | 5.35E-06 |
| Ano2 | 7.22E-09 | 0.019941543 | 0.468 | 0.304 | 1.04E-05 |
| Gm10475 | 1.16E-102 | 0.019888314 | 0.284 | 0.001 | 1.67E-99 |
| Kctd8 | 0.000258869 | 0.019819506 | 0.477 | 0.368 | 0.373548485 |
| Foxp2 | 4.98E-10 | 0.019761105 | 0.618 | 0.435 | 7.19E-07 |
| Top2a | 2.34E-31 | 0.019727326 | 0.132 | 0.419 | 3.37E-28 |
| Rab38 | 0.018639249 | 0.019537921 | 0.537 | 0.46 | 1 |
| Sema5a | 0.244176235 | 0.019326866 | 0.796 | 0.801 | 1 |
| Aebp1 | 0.113795292 | 0.019234452 | 0.69 | 0.64 | 1 |
| Serpini1 | 5.02E-52 | 0.018803944 | 0.264 | 0.669 | 7.24E-49 |
| 2600014E21Rik | 4.98E-05 | 0.018758858 | 0.181 | 0.278 | 0.071922854 |
| Elfn1 | 9.75E-108 | 0.018224617 | 0.089 | 0.651 | 1.41E-104 |
| Gm16226 | 1.23E-35 | 0.017661234 | 0.42 | 0.131 | 1.78E-32 |
| Lockd | 2.52E-51 | 0.017624747 | 0.316 | 0.718 | 3.64E-48 |
| Gm36855 | 1.66E-32 | 0.017594085 | 0.086 | 0.368 | 2.39E-29 |
| Gad1 | 1.81E-77 | 0.017389652 | 0.23 | 0.731 | 2.62E-74 |
| Neurod6 | 3.59E-14 | 0.017297504 | 0.747 | 0.537 | 5.18E-11 |
| Lrtm1 | 4.75E-32 | 0.016658618 | 0.402 | 0.724 | 6.85E-29 |
| Steap2 | 5.00E-68 | 0.016651501 | 0.733 | 0.256 | 7.21E-65 |
| Ccdc60 | 6.41E-14 | 0.016404848 | 0.422 | 0.224 | 9.25E-11 |
| Kcnc1 | 6.79E-33 | 0.016359717 | 0.77 | 0.435 | 9.80E-30 |
| Teddm2 | 2.60E-71 | 0.01633997 | 0.247 | 0.734 | 3.75E-68 |
| Vwc2l | 0.166267356 | 0.015890375 | 0.739 | 0.715 | 1 |
| D5Ertd615e | 2.25E-19 | 0.015462217 | 0.29 | 0.101 | 3.24E-16 |
| Unc5c | 0.014911617 | 0.015455177 | 0.787 | 0.847 | 1 |
| Igsf5 | 4.42E-101 | 0.015362654 | 0.466 | 0.039 | 6.38E-98 |
| Cldn5 | 3.83E-22 | 0.015112524 | 0.81 | 0.554 | 5.52E-19 |
| Tmem132c | 3.11E-08 | 0.014967418 | 0.138 | 0.27 | 4.49E-05 |
| Ankfn1 | 1.86E-12 | 0.014837148 | 0.701 | 0.497 | 2.68E-09 |
| Slc6a1 | 4.54E-50 | 0.014630032 | 0.101 | 0.481 | 6.55E-47 |
| Pex5l | 8.02E-08 | 0.014295398 | 0.836 | 0.698 | 0.000115704 |
| Nr1h4 | 1.53E-93 | 0.014280391 | 0.276 | 0.002 | 2.21E-90 |
| Galnt14 | 2.01E-19 | 0.014162853 | 0.595 | 0.821 | 2.90E-16 |
| Smoc2 | 1.25E-21 | 0.013637357 | 0.784 | 0.519 | 1.81E-18 |
| Trpc3 | 8.61E-82 | 0.013578503 | 0.615 | 0.136 | 1.24E-78 |
| Slc6a20a | 3.70E-16 | 0.01354864 | 0.451 | 0.233 | 5.34E-13 |
| Wdr95 | 1.83E-232 | 0.012933646 | 0.595 | 0.004 | 2.65E-229 |
| 6330576A10Rik | 2.04E-40 | 0.012869082 | 0.497 | 0.787 | 2.95E-37 |
| Osbpl10 | 6.45E-43 | 0.012732513 | 0.17 | 0.536 | 9.31E-40 |
| Fbxw15 | 1.62E-14 | 0.012646532 | 0.19 | 0.391 | 2.34E-11 |
| Aspa | 1.21E-13 | 0.012364036 | 0.371 | 0.581 | 1.74E-10 |
| Lbp | 7.38E-76 | 0.012268328 | 0.672 | 0.186 | 1.07E-72 |
| A230006K03Rik | 0.062923927 | 0.012111016 | 0.793 | 0.742 | 1 |
| Fbln5 | 3.18E-66 | 0.012109673 | 0.802 | 0.335 | 4.59E-63 |
| Foxd1 | 6.36E-116 | 0.011999537 | 0.787 | 0.182 | 9.17E-113 |
| Gm13832 | 0.102505313 | 0.011798975 | 0.632 | 0.576 | 1 |
| Fa2h | 0.822984375 | 0.011768817 | 0.724 | 0.722 | 1 |
| Igfbp5 | 5.60E-16 | 0.011742753 | 0.733 | 0.509 | 8.08E-13 |
| Lrrc3b | 0.018256115 | 0.011731711 | 0.397 | 0.323 | 1 |
| Wnt5b | 6.82E-05 | 0.011032851 | 0.578 | 0.455 | 0.098363504 |
| F5 | 3.49E-111 | 0.010616671 | 0.526 | 0.05 | 5.04E-108 |
| Gna14 | 0.00187954 | 0.01021193 | 0.727 | 0.643 | 1 |
| Tox3 | 3.28E-08 | 0.010157863 | 0.555 | 0.687 | 4.74E-05 |
| Fmo2 | 1.31E-66 | 0.009216076 | 0.81 | 0.335 | 1.89E-63 |
| Gm38505 | 6.70E-110 | 0.008679206 | 0.739 | 0.159 | 9.67E-107 |
| Tmtc4 | 7.07E-11 | 0.008578471 | 0.739 | 0.554 | 1.02E-07 |
| Car8 | 9.12E-20 | 0.008538318 | 0.103 | 0.316 | 1.32E-16 |
| Htr2a | 0.002185744 | 0.008497127 | 0.483 | 0.569 | 1 |
| Gm1715 | 3.40E-21 | 0.008180301 | 0.468 | 0.724 | 4.90E-18 |
| Hs3st5 | 0.027822607 | 0.008165787 | 0.776 | 0.833 | 1 |
| Tmem232 | 1.19E-42 | 0.007731552 | 0.466 | 0.135 | 1.71E-39 |
| Gm48727 | 6.16E-109 | 0.00772674 | 0.784 | 0.194 | 8.89E-106 |
| Adamtsl1 | 2.67E-39 | 0.007690581 | 0.782 | 0.415 | 3.85E-36 |

| | | | | | |
|---|---|---|---|---|---|
| Agtr1b | 7.63E-184 | 0.007127493 | 0.756 | 0.069 | 1.10E-180 |
| Gm32647 | 1.08E-16 | 0.006658577 | 0.641 | 0.838 | 1.56E-13 |
| Pappa2 | 1.79E-134 | 0.006491535 | 0.56 | 0.041 | 2.58E-131 |
| Hcrtr2 | 3.16E-15 | 0.006276602 | 0.632 | 0.827 | 4.56E-12 |
| Spock1 | 1.05E-119 | 0.006073782 | 0.299 | 0.874 | 1.51E-116 |
| Mpped1 | 4.20E-34 | 0.00605935 | 0.282 | 0.619 | 6.07E-31 |
| Emid1 | 2.34E-18 | 0.005989403 | 0.132 | 0.319 | 3.38E-15 |
| L3mbtl4 | 3.23E-12 | 0.00570518 | 0.319 | 0.509 | 4.66E-09 |
| Reln | 4.90E-34 | 0.005624852 | 0.721 | 0.38 | 7.07E-31 |
| Ndnf | 0.10579392 | 0.005581516 | 0.526 | 0.5 | 1 |
| Satb2 | 4.45E-20 | 0.005455132 | 0.563 | 0.799 | 6.42E-17 |
| Epha10 | 0.663818994 | 0.005425638 | 0.615 | 0.592 | 1 |
| Col9a3 | 0.211983073 | 0.005395198 | 0.29 | 0.248 | 1 |
| Ltbp1 | 7.90E-09 | 0.005378602 | 0.79 | 0.636 | 1.14E-05 |
| Pdlim3 | 6.15E-178 | 0.005132883 | 0.667 | 0.041 | 8.87E-175 |
| Ror1 | 0.016492322 | 0.004904912 | 0.747 | 0.809 | 1 |
| A330093E20Rik | 1.42E-14 | 0.004896737 | 0.69 | 0.467 | 2.05E-11 |
| Fhod3 | 0.009855333 | 0.004609838 | 0.698 | 0.766 | 1 |
| Slc22a8 | 1.11E-100 | 0.004432292 | 0.563 | 0.077 | 1.60E-97 |
| Gpr176 | 1.10E-27 | 0.004274112 | 0.52 | 0.23 | 1.58E-24 |
| Gm49156 | 3.21E-183 | 0.004143838 | 0.644 | 0.034 | 4.63E-180 |
| Gm11418 | 4.36E-14 | 0.003947673 | 0.368 | 0.179 | 6.29E-11 |
| Nid1 | 3.73E-74 | 0.003883532 | 0.753 | 0.254 | 5.38E-71 |
| Spata17 | 6.36E-46 | 0.003768886 | 0.638 | 0.251 | 9.17E-43 |
| Rnf152 | 1.75E-89 | 0.003628199 | 0.733 | 0.2 | 2.53E-86 |
| Ptpn13 | 6.39E-68 | 0.003423137 | 0.678 | 0.211 | 9.22E-65 |
| Gm30624 | 4.06E-231 | 0.003322424 | 0.782 | 0.042 | 5.86E-228 |
| Nwd2 | 0.782719228 | 0.003238829 | 0.822 | 0.825 | 1 |
| Lgr5 | 4.66E-17 | 0.003219977 | 0.454 | 0.231 | 6.72E-14 |
| Unc5d | 1.88E-06 | 0.00276163 | 0.764 | 0.871 | 0.0027126 |
| Slc9a2 | 0.469909111 | 0.002740338 | 0.483 | 0.517 | 1 |
| Gad2 | 2.69E-11 | 0.002442722 | 0.71 | 0.517 | 3.88E-08 |
| Chgb | 0.000887576 | 0.002406676 | 0.477 | 0.582 | 1 |
| Akap13 | 0.016393879 | 0.0022057 | 0.805 | 0.78 | 1 |
| Vsx1 | 3.35E-235 | 0.002195141 | 0.782 | 0.041 | 4.83E-232 |
| Rarb | 0.000714154 | 0.001994576 | 0.77 | 0.674 | 1 |
| Slc13a1 | 1.12E-07 | 0.001923065 | 0.644 | 0.486 | 0.000161153 |
| Sema3d | 0.175440586 | 0.001804625 | 0.629 | 0.679 | 1 |
| Folr1 | 0.932513397 | 0.001688509 | 0.629 | 0.639 | 1 |
| Rab3b | 5.75E-09 | 0.00162506 | 0.417 | 0.258 | 8.30E-06 |
| Prkcg | 8.09E-39 | 0.001281411 | 0.422 | 0.769 | 1.17E-35 |
| Adamts19 | 5.71E-11 | 0.001258259 | 0.319 | 0.161 | 8.24E-08 |
| Ntf3 | 7.82E-05 | 0.001071722 | 0.256 | 0.161 | 0.112875226 |
| Osr1 | 3.18E-34 | 0.001026297 | 0.313 | 0.075 | 4.59E-31 |
| Tnfrsf11b | 2.60E-13 | 0.000908001 | 0.632 | 0.419 | 3.75E-10 |
| Hunk | 8.62E-44 | 0.000868159 | 0.224 | 0.604 | 1.24E-40 |
| Gm43507 | 6.90E-54 | 0.000782229 | 0.747 | 0.316 | 9.95E-51 |
| Gm816 | 2.19E-55 | 0.000668219 | 0.299 | 0.033 | 3.15E-52 |
| Lepr | 5.34E-147 | 0.000606774 | 0.707 | 0.085 | 7.70E-144 |
| Cyp26b1 | 0.349567488 | 0.000569615 | 0.388 | 0.352 | 1 |
| Gm35552 | 8.46E-28 | 0.000552994 | 0.52 | 0.79 | 1.22E-24 |
| Serpina1b | 2.76E-183 | 0.000494894 | 0.678 | 0.041 | 3.98E-180 |
| Otc | 4.06E-60 | 0.000330699 | 0.638 | 0.203 | 5.86E-57 |
| Gm12128 | 5.95E-276 | 0.000210467 | 0.661 | 0 | 8.59E-273 |
| Atp13a4 | 5.04E-17 | 0.000158032 | 0.29 | 0.111 | 7.27E-14 |
| Ano1 | 6.02E-65 | 0.000156605 | 0.807 | 0.336 | 8.69E-62 |
| Gm33819 | 6.51E-156 | 3.70E-05 | 0.422 | 0.002 | 9.39E-153 |
| Grpr | 1.55E-159 | 2.78E-05 | 0.425 | 0.002 | 2.24E-156 |
| Gm45829 | 2.17E-161 | 2.53E-05 | 0.44 | 0.002 | 3.13E-158 |
| 4930438E09Rik | 3.72E-141 | 1.24E-05 | 0.385 | 0.002 | 5.37E-138 |
| Krt73 | 4.02E-154 | 6.53E-06 | 0.391 | 0 | 5.80E-151 |
| 4930455M05Rik | 7.81E-174 | 2.03E-06 | 0.457 | 0.001 | 1.13E-170 |
| Gm26725 | 1.25E-99 | 5.52E-07 | 0.276 | 0.001 | 1.81E-96 |
| Skint1 | 1.09E-115 | -1.57E-06 | 0.299 | 0 | 1.57E-112 |
| Rpe65 | 9.47E-111 | -3.76E-05 | 0.287 | 0.001 | 1.37E-107 |
| Adamts9 | 3.14E-05 | -7.86E-05 | 0.724 | 0.602 | 0.045272523 |
| Gm12153 | 6.29E-129 | -9.62E-05 | 0.356 | 0.002 | 9.08E-126 |
| Svep1 | 1.02E-29 | -9.74E-05 | 0.477 | 0.187 | 1.48E-26 |
| Gpc3 | 8.02E-10 | -0.000120409 | 0.509 | 0.685 | 1.16E-06 |
| Ttc21a | 1.42E-106 | -0.00015274 | 0.293 | 0.001 | 2.05E-103 |
| Slc16a2 | 1.06E-05 | -0.000283014 | 0.75 | 0.854 | 0.015308442 |
| Paqr5 | 1.19E-119 | -0.00029325 | 0.5 | 0.034 | 1.72E-116 |
| Lmntd1 | 1.11E-193 | -0.000310287 | 0.727 | 0.05 | 1.60E-190 |
| Cgnl1 | 2.53E-60 | -0.000316943 | 0.296 | 0.025 | 3.65E-57 |
| Eps8l2 | 1.89E-05 | -0.000409295 | 0.256 | 0.154 | 0.027249194 |
| Chrna7 | 8.89E-06 | -0.00051394 | 0.776 | 0.871 | 0.012831309 |
| Fam180a | 1.57E-212 | -0.000625258 | 0.744 | 0.042 | 2.27E-209 |
| Defb9 | 2.52E-27 | -0.000631633 | 0.385 | 0.132 | 3.64E-24 |
| B020031H02Rik | 9.36E-136 | -0.000740211 | 0.368 | 0.001 | 1.35E-132 |
| Hectd2 | 5.66E-08 | -0.000934433 | 0.615 | 0.461 | 8.17E-05 |
| Maob | 1.75E-78 | -0.001269395 | 0.543 | 0.102 | 2.53E-75 |
| Sphk1 | 5.26E-90 | -0.001405609 | 0.33 | 0.012 | 7.60E-87 |
| 4933436I20Rik | 3.62E-16 | -0.001544179 | 0.385 | 0.184 | 5.23E-13 |
| Gap43 | 0.003055715 | -0.001600466 | 0.466 | 0.549 | 1 |
| A830019P07Rik | 1.66E-80 | -0.001649574 | 0.543 | 0.098 | 2.39E-77 |
| Trp63 | 1.16E-101 | -0.001746717 | 0.402 | 0.021 | 1.67E-98 |
| A2m | 8.03E-10 | -0.001767308 | 0.181 | 0.344 | 1.16E-06 |
| 1700042O10Rik | 2.45E-21 | -0.001784794 | 0.273 | 0.083 | 3.53E-18 |
| Slc6a12 | 2.37E-203 | -0.001806415 | 0.724 | 0.041 | 3.43E-200 |
| Mdga1 | 5.43E-31 | -0.002083782 | 0.293 | 0.617 | 7.83E-28 |
| Igfbp2 | 5.01E-36 | -0.002182813 | 0.638 | 0.29 | 7.23E-33 |
| Gm37459 | 0.094234556 | -0.002190886 | 0.773 | 0.821 | 1 |
| Gm16223 | 4.06E-160 | -0.002195902 | 0.624 | 0.041 | 5.85E-157 |
| 4930470O06Rik | 2.79E-90 | -0.002238371 | 0.256 | 0.001 | 4.02E-87 |
| Susd5 | 6.53E-90 | -0.002467346 | 0.747 | 0.209 | 9.42E-87 |
| Carmn | 7.47E-54 | -0.002558721 | 0.764 | 0.333 | 1.08E-50 |
| Stra6 | 4.02E-126 | -0.002569295 | 0.733 | 0.127 | 5.80E-123 |
| Rerg | 4.64E-08 | -0.002581078 | 0.178 | 0.318 | 6.70E-05 |
| Htr3a | 0.140033037 | -0.002646351 | 0.437 | 0.49 | 1 |
| Prr16 | 0.607580664 | -0.002664961 | 0.802 | 0.798 | 1 |
| Ptgfr | 0.046059119 | -0.002711658 | 0.305 | 0.37 | 1 |
| Ccbe1 | 4.55E-32 | -0.002827881 | 0.46 | 0.774 | 6.56E-29 |
| Adgrf5 | 7.69E-28 | -0.003029323 | 0.618 | 0.311 | 1.11E-24 |
| Rcan2 | 6.16E-32 | -0.003128359 | 0.756 | 0.425 | 8.89E-29 |
| Spo11 | 1.04E-162 | -0.003147431 | 0.359 | 0.001 | 1.51E-159 |
| Acvr1c | 1.79E-163 | -0.003180775 | 0.149 | 0.845 | 2.59E-160 |
| Grin3a | 8.60E-09 | -0.00320595 | 0.704 | 0.841 | 1.24E-05 |
| Penk | 1.02E-17 | -0.003487288 | 0.098 | 0.252 | 1.47E-14 |
| Cacng4 | 0.140267619 | -0.003587692 | 0.299 | 0.351 | 1 |
| Mme | 4.14E-89 | -0.003592354 | 0.526 | 0.078 | 5.98E-86 |
| Npas1 | 0.530320403 | -0.003619973 | 0.523 | 0.553 | 1 |
| Chrm2 | 5.20E-52 | -0.003968343 | 0.333 | 0.049 | 7.50E-49 |
| Gfap | 3.67E-17 | -0.004000547 | 0.379 | 0.177 | 5.30E-14 |
| Frem1 | 0.625354181 | -0.004134658 | 0.816 | 0.834 | 1 |
| Popdc3 | 7.06E-09 | -0.00420414 | 0.25 | 0.125 | 1.02E-05 |
| Htr4 | 7.61E-06 | -0.004210459 | 0.667 | 0.783 | 0.010980462 |
| Itih5 | 5.79E-25 | -0.004258096 | 0.135 | 0.401 | 8.36E-22 |
| Gm36431 | 8.67E-39 | -0.004283114 | 0.181 | 0.541 | 1.25E-35 |
| Slco1b2 | 1.33E-163 | -0.004405183 | 0.612 | 0.035 | 1.91E-160 |
| Mlip | 9.48E-13 | -0.004460356 | 0.632 | 0.807 | 1.37E-09 |
| Gli2 | 1.82E-129 | -0.004480764 | 0.661 | 0.086 | 2.63E-126 |
| 4930544I03Rik | 7.88E-47 | -0.004557966 | 0.532 | 0.168 | 1.14E-43 |
| Ntn4 | 1.40E-33 | -0.004948363 | 0.799 | 0.463 | 2.02E-30 |
| Tfcp2l1 | 4.69E-11 | -0.005000289 | 0.701 | 0.516 | 6.77E-08 |
| Oca2 | 6.16E-42 | -0.005082208 | 0.443 | 0.125 | 8.89E-39 |
| Aldh1a2 | 9.99E-213 | -0.00512165 | 0.598 | 0.009 | 1.44E-209 |
| Slc13a4 | 8.40E-64 | -0.005176961 | 0.724 | 0.261 | 1.21E-60 |
| Synpo2 | 1.10E-45 | -0.005191253 | 0.615 | 0.233 | 1.59E-42 |
| Mog | 3.43E-05 | -0.005384722 | 0.316 | 0.403 | 0.049559911 |
| Acss3 | 1.67E-89 | -0.005386148 | 0.253 | 0.785 | 2.41E-86 |
| 7630403G23Rik | 3.54E-47 | -0.005594579 | 0.17 | 0.563 | 5.11E-44 |

| | | | | | |
|---|---|---|---|---|---|
| Gm26691 | 2.76E-43 | -0.005601274 | 0.17 | 0.529 | 3.98E-40 |
| Bmp4 | 2.51E-250 | -0.005629076 | 0.638 | 0.004 | 3.62E-247 |
| Plekhh1 | 1.63E-30 | -0.005650574 | 0.672 | 0.349 | 2.35E-27 |
| Egln3 | 9.94E-59 | -0.005694651 | 0.037 | 0.418 | 1.43E-55 |
| Cdh6 | 0.026615868 | -0.005855732 | 0.753 | 0.791 | 1 |
| Cspg4 | 6.60E-62 | -0.005912906 | 0.681 | 0.234 | 9.52E-59 |
| Zbbx | 2.21E-40 | -0.005995546 | 0.523 | 0.182 | 3.18E-37 |
| Pdzph1 | 1.35E-44 | -0.006042268 | 0.336 | 0.062 | 1.95E-41 |
| Slc26a4 | 5.22E-53 | -0.006118724 | 0.368 | 0.779 | 7.54E-50 |
| Rlbp1 | 0.856607359 | -0.006300204 | 0.494 | 0.504 | 1 |
| Platr22 | 1.32E-160 | -0.006463771 | 0.721 | 0.077 | 1.91E-157 |
| Fam107a | 0.027221823 | -0.006509495 | 0.687 | 0.753 | 1 |
| F3 | 2.31E-07 | -0.006991607 | 0.747 | 0.607 | 0.000333606 |
| Hes5 | 5.75E-41 | -0.00703597 | 0.652 | 0.282 | 8.30E-38 |
| Gabrg1 | 5.29E-14 | -0.007072859 | 0.376 | 0.191 | 7.64E-11 |
| C230072F16Rik | 3.50E-26 | -0.007096014 | 0.043 | 0.271 | 5.05E-23 |
| Ceacam16 | 2.24E-17 | -0.007164787 | 0.083 | 0.278 | 3.23E-14 |
| Gm5149 | 1.81E-39 | -0.007173452 | 0.391 | 0.099 | 2.61E-36 |
| A330049N07Rik | 7.61E-62 | -0.007336228 | 0.71 | 0.257 | 1.10E-58 |
| 9630002D21Rik | 8.82E-127 | -0.007396864 | 0.629 | 0.075 | 1.27E-123 |
| Slc4a5 | 0.007633952 | -0.007475106 | 0.325 | 0.259 | 1 |
| 4930567K20Rik | 3.55E-05 | -0.007628678 | 0.385 | 0.512 | 0.051207724 |
| Fstl4 | 1.36E-29 | -0.007781141 | 0.802 | 0.49 | 1.96E-26 |
| Thsd7a | 0.75774023 | -0.007833732 | 0.704 | 0.718 | 1 |
| Pcp4l1 | 2.04E-159 | -0.007841724 | 0.767 | 0.101 | 2.94E-156 |
| Rfx4 | 0.639345775 | -0.007882292 | 0.744 | 0.764 | 1 |
| Sorcs1 | 0.103733668 | -0.007889401 | 0.73 | 0.676 | 1 |
| Slc47a1 | 2.10E-46 | -0.008202952 | 0.302 | 0.044 | 3.03E-43 |
| Gm17171 | 1.08E-37 | -0.008232015 | 0.115 | 0.451 | 1.56E-34 |
| Wdr72 | 1.23E-25 | -0.008238736 | 0.466 | 0.75 | 1.77E-22 |
| Cabcoco1 | 1.56E-52 | -0.008272721 | 0.71 | 0.288 | 2.25E-49 |
| Trpc5 | 8.93E-25 | -0.008528163 | 0.172 | 0.438 | 1.29E-21 |
| Tbx3os1 | 8.62E-86 | -0.008571997 | 0.422 | 0.04 | 1.24E-82 |
| Myo5b | 0.000336226 | -0.0086129 | 0.348 | 0.247 | 0.485174455 |
| Gm26911 | 3.87E-06 | -0.008738003 | 0.67 | 0.532 | 0.005583462 |
| Ntn1 | 0.039424527 | -0.00875727 | 0.782 | 0.724 | 1 |
| Rxfp2 | 7.51E-06 | -0.008928023 | 0.664 | 0.78 | 0.010840151 |
| Kcnk5 | 1.29E-102 | -0.008960248 | 0.29 | 0.001 | 1.86E-99 |
| 9330158H04Rik | 6.39E-25 | -0.009049641 | 0.46 | 0.198 | 9.22E-22 |
| Cort | 3.87E-13 | -0.009119745 | 0.42 | 0.227 | 5.59E-10 |
| Clu | 0.0596057 | -0.009185032 | 0.468 | 0.525 | 1 |
| Klf5 | 6.13E-09 | -0.009382265 | 0.675 | 0.819 | 8.85E-06 |
| Oprd1 | 2.57E-210 | -0.010072334 | 0.736 | 0.041 | 3.71E-207 |
| Ankrd55 | 3.01E-17 | -0.010291524 | 0.606 | 0.816 | 4.34E-14 |
| Kif26b | 2.91E-26 | -0.010430068 | 0.316 | 0.095 | 4.20E-23 |
| Gm28905 | 2.67E-17 | -0.01047523 | 0.675 | 0.863 | 3.85E-14 |
| Kank1 | 0.45586774 | -0.010478887 | 0.448 | 0.419 | 1 |
| Gm11351 | 2.83E-32 | -0.010539622 | 0.526 | 0.82 | 4.09E-29 |
| Cdh7 | 0.183790967 | -0.010622197 | 0.463 | 0.424 | 1 |
| Adamts18 | 9.54E-13 | -0.010638986 | 0.526 | 0.338 | 1.38E-09 |
| Scara5 | 2.03E-31 | -0.010688477 | 0.098 | 0.392 | 2.93E-28 |
| Vgll3 | 8.12E-125 | -0.010732712 | 0.33 | 0.002 | 1.17E-121 |
| Etnppl | 1.92E-16 | -0.010821846 | 0.296 | 0.117 | 2.76E-13 |
| Col4a3 | 0.002202991 | -0.010836564 | 0.414 | 0.51 | 1 |
| Pde5a | 1.47E-27 | -0.011115928 | 0.075 | 0.329 | 2.12E-24 |
| Prdm16 | 7.23E-30 | -0.011197571 | 0.767 | 0.449 | 1.04E-26 |
| Cpne5 | 2.24E-21 | -0.011259906 | 0.293 | 0.095 | 3.24E-18 |
| Trabd2b | 0.077415701 | -0.011326838 | 0.282 | 0.241 | 1 |
| Rassf9 | 6.58E-19 | -0.011331346 | 0.575 | 0.803 | 9.50E-16 |
| Gm20647 | 6.72E-22 | -0.011362136 | 0.201 | 0.466 | 9.70E-19 |
| Pik3ap1 | 3.60E-20 | -0.011428609 | 0.56 | 0.767 | 5.19E-17 |
| 9130410C08Rik | 3.29E-37 | -0.011442432 | 0.695 | 0.338 | 4.74E-34 |
| Myoc | 7.69E-91 | -0.011671211 | 0.641 | 0.135 | 1.11E-87 |
| Erg | 1.12E-15 | -0.011810187 | 0.761 | 0.539 | 1.61E-12 |
| Wdr86 | 1.63E-39 | -0.011852015 | 0.135 | 0.487 | 2.36E-36 |
| Gm44593 | 0.259120499 | -0.012037268 | 0.42 | 0.462 | 1 |
| Vwc2 | 8.71E-35 | -0.012062329 | 0.784 | 0.445 | 1.26E-31 |
| Crtac1 | 2.04E-32 | -0.012071149 | 0.376 | 0.704 | 2.94E-29 |
| Atp10b | 0.833170747 | -0.012074776 | 0.586 | 0.57 | 1 |
| Unc13c | 6.30E-53 | -0.012184084 | 0.218 | 0.641 | 9.09E-50 |
| Pappa | 1.22E-07 | -0.012498472 | 0.445 | 0.586 | 0.000176085 |
| Gm45680 | 2.90E-109 | -0.012505826 | 0.305 | 0.003 | 4.19E-106 |
| Siah3 | 2.42E-146 | -0.012565572 | 0.46 | 0.01 | 3.50E-143 |
| Gm42397 | 1.50E-10 | -0.012624751 | 0.612 | 0.426 | 2.17E-07 |
| Gm39185 | 1.76E-12 | -0.012756399 | 0.48 | 0.678 | 2.54E-09 |
| Adamtsl3 | 3.59E-12 | -0.01288786 | 0.253 | 0.109 | 5.18E-09 |
| Fgf10 | 3.67E-31 | -0.012897921 | 0.227 | 0.552 | 5.30E-28 |
| Otx2 | 0.0354825 | -0.012931413 | 0.718 | 0.671 | 1 |
| Lpar1 | 1.21E-114 | -0.012961406 | 0.06 | 0.654 | 1.74E-111 |
| Gm48508 | 2.45E-108 | -0.012985039 | 0.718 | 0.149 | 3.54E-105 |
| Stk33 | 0.043889314 | -0.013010982 | 0.713 | 0.727 | 1 |
| Cwh43 | 3.37E-07 | -0.01304105 | 0.138 | 0.264 | 0.000486097 |
| Pth2r | 7.68E-72 | -0.013044819 | 0.066 | 0.534 | 1.11E-68 |
| Frmpd1 | 1.78E-32 | -0.013345225 | 0.661 | 0.329 | 2.57E-29 |
| Gm40293 | 3.49E-08 | -0.013493077 | 0.727 | 0.844 | 5.04E-05 |
| Slco1a4 | 0.159436103 | -0.013737173 | 0.405 | 0.353 | 1 |
| Igfbpl1 | 2.26E-71 | -0.013940693 | 0.138 | 0.627 | 3.26E-68 |
| Lypd6b | 2.59E-205 | -0.014255716 | 0.724 | 0.041 | 3.73E-202 |
| Cyyr1 | 3.73E-73 | -0.014313346 | 0.282 | 0.012 | 5.38E-70 |
| Gm29571 | 3.79E-42 | -0.014568841 | 0.095 | 0.442 | 5.46E-39 |
| Sulf1 | 2.87E-51 | -0.014609464 | 0.698 | 0.285 | 4.14E-48 |
| Gm30613 | 8.81E-36 | -0.014686378 | 0.172 | 0.514 | 1.27E-32 |
| A230001M10Rik | 2.44E-21 | -0.014734932 | 0.698 | 0.431 | 3.52E-18 |
| Dkk2 | 9.92E-09 | -0.014921857 | 0.77 | 0.617 | 1.43E-05 |
| Otof | 1.22E-08 | -0.015002118 | 0.351 | 0.207 | 1.77E-05 |
| Piezo2 | 5.53E-136 | -0.015049704 | 0.181 | 0.833 | 7.98E-133 |
| Cdh13 | 0.006524291 | -0.015060518 | 0.555 | 0.622 | 1 |
| Stk32b | 7.89E-32 | -0.015086209 | 0.477 | 0.18 | 1.14E-28 |
| Slc17a8 | 5.62E-67 | -0.015171911 | 0.023 | 0.437 | 8.12E-64 |
| Gm10561 | 1.64E-06 | -0.015417343 | 0.287 | 0.426 | 0.002369371 |
| 6030407O03Rik | 1.32E-05 | -0.01565259 | 0.319 | 0.212 | 0.019021123 |
| Bmp5 | 0.153029824 | -0.015659965 | 0.33 | 0.375 | 1 |
| Hhip | 0.01091431 | -0.015661036 | 0.598 | 0.523 | 1 |
| Slc2a1 | 1.59E-55 | -0.015809727 | 0.244 | 0.672 | 2.29E-52 |
| Mettl24 | 8.55E-24 | -0.015955942 | 0.164 | 0.434 | 1.23E-20 |
| Pecam1 | 9.78E-15 | -0.016098426 | 0.635 | 0.827 | 1.41E-11 |
| Dpp10 | 7.76E-53 | -0.016164128 | 0.511 | 0.876 | 1.12E-49 |
| Slc44a5 | 0.00033222 | -0.016280392 | 0.563 | 0.672 | 0.479393837 |
| Cdh3 | 4.15E-14 | -0.016304178 | 0.555 | 0.755 | 5.99E-11 |
| Col8a2 | 1.12E-08 | -0.016965542 | 0.675 | 0.815 | 1.62E-05 |
| Ptpru | 2.85E-27 | -0.0171221 | 0.098 | 0.361 | 4.11E-24 |
| Ddah1 | 0.011967804 | -0.017176617 | 0.802 | 0.731 | 1 |
| Rgs6 | 2.60E-13 | -0.017250695 | 0.434 | 0.645 | 3.75E-10 |
| Gm14051 | 6.53E-39 | -0.017293066 | 0.351 | 0.71 | 9.42E-36 |
| Spag16 | 0.022809399 | -0.017726948 | 0.534 | 0.47 | 1 |
| Prdm6 | 3.30E-20 | -0.017761533 | 0.71 | 0.447 | 4.76E-17 |
| Dnah11 | 9.57E-45 | -0.017859792 | 0.583 | 0.213 | 1.38E-41 |
| Ntsr2 | 8.75E-05 | -0.018192875 | 0.181 | 0.285 | 0.126203444 |
| Gm12239 | 9.16E-19 | -0.018197547 | 0.466 | 0.712 | 1.32E-15 |
| Lama4 | 2.85E-88 | -0.018245245 | 0.71 | 0.185 | 4.11E-85 |
| Gm2516 | 1.28E-170 | -0.01876606 | 0.767 | 0.089 | 1.85E-167 |
| 4930555F03Rik | 7.81E-07 | -0.019116492 | 0.152 | 0.275 | 0.001127284 |
| Gpr149 | 2.00E-22 | -0.019508508 | 0.572 | 0.818 | 2.88E-19 |
| Plch2 | 0.016749549 | -0.019511486 | 0.46 | 0.399 | 1 |
| Adgrd1 | 3.11E-17 | -0.019552085 | 0.652 | 0.835 | 4.49E-14 |
| Tcerg1l | 0.38623197 | -0.019595369 | 0.54 | 0.51 | 1 |
| Fign | 5.36E-12 | -0.019869666 | 0.454 | 0.264 | 7.73E-09 |
| Lama1 | 1.82E-44 | -0.019921108 | 0.647 | 0.263 | 2.63E-41 |
| Ccdc141 | 5.77E-35 | -0.019923892 | 0.644 | 0.302 | 8.33E-32 |
| Phldb2 | 1.93E-53 | -0.020009415 | 0.776 | 0.348 | 2.78E-50 |
| Gulp1 | 0.003363269 | -0.020061399 | 0.195 | 0.276 | 1 |

| | | | | | |
|---|---|---|---|---|---|
| Grin2c | 5.51E-23 | -0.020103678 | 0.049 | 0.261 | 7.95E-20 |
| Usp29 | 9.55E-14 | -0.020210674 | 0.635 | 0.82 | 1.38E-10 |
| Gm14507 | 1.73E-12 | -0.020389959 | 0.647 | 0.445 | 2.49E-09 |
| Pde11a | 1.33E-24 | -0.020713945 | 0.474 | 0.752 | 1.92E-21 |
| Drc7 | 2.10E-43 | -0.020772396 | 0.466 | 0.821 | 3.03E-40 |
| Tcf7l1 | 9.21E-07 | -0.021081917 | 0.784 | 0.651 | 0.001328322 |
| Nell1os | 2.71E-19 | -0.02134843 | 0.629 | 0.376 | 3.91E-16 |
| Hmcn1 | 3.15E-85 | -0.0213754 | 0.293 | 0.812 | 4.55E-82 |
| Slc38a11 | 4.65E-07 | -0.021600708 | 0.721 | 0.576 | 0.000671589 |
| Pde3a | 0.246781719 | -0.021612327 | 0.787 | 0.763 | 1 |
| Epb41l4a | 8.93E-90 | -0.021690551 | 0.733 | 0.2 | 1.29E-86 |
| Stac | 4.75E-17 | -0.0220167 | 0.282 | 0.106 | 6.86E-14 |
| Dnaaf3 | 1.92E-11 | -0.022494546 | 0.103 | 0.259 | 2.78E-08 |
| Gm45323 | 1.53E-09 | -0.02292531 | 0.348 | 0.526 | 2.21E-06 |
| Lrrtm4 | 6.42E-05 | -0.023018894 | 0.885 | 0.915 | 0.092578994 |
| Qrfpr | 2.98E-20 | -0.023205719 | 0.489 | 0.244 | 4.31E-17 |
| Moxd1 | 7.40E-18 | -0.023307026 | 0.716 | 0.49 | 1.07E-14 |
| Ddr2 | 1.74E-22 | -0.023416936 | 0.126 | 0.375 | 2.50E-19 |
| Gm21847 | 1.23E-102 | -0.023726504 | 0.132 | 0.717 | 1.77E-99 |
| Gm33680 | 0.846147212 | -0.023762561 | 0.489 | 0.487 | 1 |
| Exoc3l2 | 0.003602087 | -0.024328878 | 0.29 | 0.221 | 1 |
| Nupr1 | 7.39E-15 | -0.024457166 | 0.178 | 0.369 | 1.07E-11 |
| Cpa6 | 4.74E-48 | -0.024544657 | 0.04 | 0.385 | 6.84E-45 |
| Lrrtm3 | 3.67E-06 | -0.024694592 | 0.667 | 0.787 | 0.005299229 |
| Gm13481 | 3.82E-15 | -0.024890029 | 0.216 | 0.432 | 5.51E-12 |
| Glra2 | 8.73E-32 | -0.024972518 | 0.193 | 0.517 | 1.26E-28 |
| Gm19410 | 7.34E-50 | -0.025397277 | 0.095 | 0.483 | 1.06E-46 |
| Gm13629 | 0.30035048 | -0.025504087 | 0.684 | 0.694 | 1 |
| Rab3c | 0.007339693 | -0.025562495 | 0.606 | 0.688 | 1 |
| Gm30551 | 1.31E-11 | -0.026091958 | 0.626 | 0.792 | 1.89E-08 |
| Egflam | 0.011410457 | -0.026191027 | 0.678 | 0.674 | 1 |
| Bmp6 | 2.95E-13 | -0.026512216 | 0.707 | 0.505 | 4.26E-10 |
| Htra1 | 4.23E-06 | -0.026803972 | 0.523 | 0.392 | 0.006109361 |
| Shroom3 | 1.04E-85 | -0.02684806 | 0.175 | 0.713 | 1.51E-82 |
| Grik3 | 1.20E-27 | -0.027274692 | 0.382 | 0.689 | 1.74E-24 |
| Abcc9 | 5.81E-140 | -0.027425467 | 0.374 | 0.005 | 8.38E-137 |
| Acsbg1 | 8.09E-40 | -0.027459445 | 0.049 | 0.35 | 1.17E-36 |
| Kcns3 | 0.038397495 | -0.027465634 | 0.379 | 0.432 | 1 |
| Sgcd | 1.14E-08 | -0.027867582 | 0.805 | 0.655 | 1.64E-05 |
| Zbtb7c | 0.000321657 | -0.027926198 | 0.52 | 0.417 | 0.464150812 |
| Zic4 | 1.26E-89 | -0.028110799 | 0.187 | 0.736 | 1.81E-86 |
| Sec14l5 | 1.15E-73 | -0.028240574 | 0.356 | 0.826 | 1.66E-70 |
| Gm15584 | 5.05E-20 | -0.02824667 | 0.27 | 0.53 | 7.29E-17 |
| Gm11728 | 2.06E-15 | -0.02828845 | 0.468 | 0.69 | 2.98E-12 |
| Prss12 | 6.86E-77 | -0.028364785 | 0.121 | 0.617 | 9.90E-74 |
| Smoc1 | 6.13E-07 | -0.028488767 | 0.509 | 0.655 | 0.000885091 |
| 4933432K03Rik | 1.57E-45 | -0.028951614 | 0.034 | 0.361 | 2.26E-42 |
| Trp73 | 1.87E-68 | -0.028986852 | 0.33 | 0.796 | 2.70E-65 |
| Cdh4 | 1.92E-37 | -0.030071976 | 0.526 | 0.197 | 2.77E-34 |
| Pou6f2 | 4.86E-27 | -0.030284265 | 0.509 | 0.791 | 7.02E-24 |
| Scrg1 | 3.95E-29 | -0.030320772 | 0.598 | 0.295 | 5.71E-26 |
| Zfhx4 | 1.98E-24 | -0.030928807 | 0.463 | 0.742 | 2.86E-21 |
| Prkg2 | 2.58E-07 | -0.031431108 | 0.489 | 0.637 | 0.000371777 |
| Tmem72 | 4.83E-120 | -0.031476315 | 0.184 | 0.804 | 6.97E-117 |
| Tll1 | 1.48E-26 | -0.031598819 | 0.701 | 0.401 | 2.14E-23 |
| Dsp | 2.80E-21 | -0.031933841 | 0.626 | 0.847 | 4.04E-18 |
| Slc47a2 | 0.020892812 | -0.031994929 | 0.624 | 0.571 | 1 |
| Gm43948 | 1.08E-19 | -0.032144215 | 0.46 | 0.705 | 1.56E-16 |
| Fam189a1 | 0.013884638 | -0.032461208 | 0.828 | 0.868 | 1 |
| Trpc7 | 1.89E-16 | -0.032588871 | 0.626 | 0.819 | 2.73E-13 |
| Gm29683 | 0.10762953 | -0.032592267 | 0.739 | 0.715 | 1 |
| C030029H02Rik | 7.52E-153 | -0.032969923 | 0.626 | 0.05 | 1.08E-149 |
| Gm29514 | 0.010480176 | -0.033232915 | 0.23 | 0.306 | 1 |
| Gm1604a | 0.156294148 | -0.033524284 | 0.756 | 0.739 | 1 |
| Gjc3 | 4.64E-16 | -0.033765903 | 0.626 | 0.394 | 6.70E-13 |
| Atp13a5 | 0.000285186 | -0.033836273 | 0.629 | 0.517 | 0.411523526 |
| Slc16a9 | 6.65E-06 | -0.034155993 | 0.408 | 0.545 | 0.009602908 |
| Tac1 | 2.34E-14 | -0.034271721 | 0.497 | 0.706 | 3.37E-11 |
| Hs3st4 | 0.10380748 | -0.034308693 | 0.83 | 0.864 | 1 |
| Tmem108 | 0.027980878 | -0.034672801 | 0.615 | 0.675 | 1 |
| Gm47448 | 1.38E-20 | -0.034773842 | 0.276 | 0.091 | 1.99E-17 |
| Zdbf2 | 0.214431941 | -0.035285322 | 0.52 | 0.486 | 1 |
| Tenm1 | 1.37E-10 | -0.03550281 | 0.727 | 0.871 | 1.98E-07 |
| Adamts2 | 1.49E-13 | -0.035563213 | 0.376 | 0.193 | 2.15E-10 |
| Enpp6 | 1.47E-13 | -0.03569014 | 0.371 | 0.189 | 2.12E-10 |
| B230110G15Rik | 1.01E-06 | -0.036250361 | 0.486 | 0.352 | 0.001461932 |
| Sv2c | 2.88E-72 | -0.036473511 | 0.287 | 0.774 | 4.15E-69 |
| Plch1 | 1.45E-12 | -0.038315097 | 0.678 | 0.834 | 2.09E-09 |
| 4930419G24Rik | 1.15E-44 | -0.038318604 | 0.46 | 0.821 | 1.66E-41 |
| Syt2 | 0.005019318 | -0.038536093 | 0.675 | 0.741 | 1 |
| Slc7a11 | 4.81E-05 | -0.038912791 | 0.615 | 0.518 | 0.069341479 |
| Col11a1 | 6.97E-26 | -0.039306938 | 0.526 | 0.799 | 1.01E-22 |
| Olfm2 | 0.00977778 | -0.039473509 | 0.727 | 0.752 | 1 |
| Adra1b | 1.53E-33 | -0.039601309 | 0.474 | 0.787 | 2.21E-30 |
| Nr4a3 | 0.072958688 | -0.039631816 | 0.658 | 0.704 | 1 |
| Adgra1 | 3.03E-10 | -0.039899643 | 0.259 | 0.437 | 4.37E-07 |
| Fbln1 | 5.18E-32 | -0.040584035 | 0.279 | 0.609 | 7.48E-29 |
| Prrx1 | 5.99E-12 | -0.040684793 | 0.716 | 0.828 | 8.65E-09 |
| C1qtnf7 | 8.31E-55 | -0.04081222 | 0.069 | 0.467 | 1.20E-51 |
| Arhgef26 | 7.40E-13 | -0.041427078 | 0.138 | 0.317 | 1.07E-09 |
| Mageb18 | 0.049802529 | -0.041448726 | 0.779 | 0.783 | 1 |
| Itgad | 0.186164752 | -0.041520904 | 0.221 | 0.26 | 1 |
| Col8a1 | 1.67E-05 | -0.041607451 | 0.638 | 0.747 | 0.024050769 |
| Il34 | 4.21E-16 | -0.042182346 | 0.73 | 0.51 | 6.07E-13 |
| Cemip | 5.69E-05 | -0.042329335 | 0.718 | 0.808 | 0.082042854 |
| 1700047M11Rik | 1.38E-15 | -0.042401016 | 0.158 | 0.366 | 1.99E-12 |
| Arsj | 0.040298454 | -0.042415509 | 0.624 | 0.576 | 1 |
| Glra3 | 0.000994183 | -0.042832881 | 0.652 | 0.738 | 1 |
| Nr4a2 | 2.30E-14 | -0.043032571 | 0.572 | 0.357 | 3.33E-11 |
| 9330111N05Rik | 0.000365311 | -0.043084546 | 0.658 | 0.563 | 0.527143585 |
| 9630013A20Rik | 1.13E-81 | -0.043262928 | 0.098 | 0.614 | 1.64E-78 |
| Kcnk2 | 0.232987706 | -0.043265512 | 0.716 | 0.757 | 1 |
| Rfx3 | 0.143871103 | -0.043521679 | 0.759 | 0.795 | 1 |
| Cdh8 | 3.13E-09 | -0.043998934 | 0.756 | 0.881 | 4.51E-06 |
| Trhr | 0.00187422 | -0.044921834 | 0.69 | 0.623 | 1 |
| Pard3b | 0.000107496 | -0.045219911 | 0.842 | 0.786 | 0.155117159 |
| Alk | 1.78E-06 | -0.045261136 | 0.305 | 0.446 | 0.002569658 |
| Tbx15 | 0.003470791 | -0.045618146 | 0.33 | 0.259 | 1 |
| 2010300C02Rik | 1.18E-24 | -0.0459757 | 0.635 | 0.869 | 1.70E-21 |
| Vwa3a | 0.002304624 | -0.046174823 | 0.71 | 0.652 | 1 |
| Slc32a1 | 3.01E-17 | -0.046410976 | 0.216 | 0.449 | 4.34E-14 |
| Nectin3 | 3.73E-196 | -0.047084915 | 0.066 | 0.813 | 5.38E-193 |
| Fam163a | 4.06E-17 | -0.048099309 | 0.73 | 0.509 | 5.86E-14 |
| Stmn2 | 0.091347236 | -0.048195675 | 0.695 | 0.737 | 1 |
| Pgm5 | 5.43E-101 | -0.048433608 | 0.221 | 0.794 | 7.84E-98 |
| Gm6999 | 6.06E-88 | -0.048828585 | 0.069 | 0.597 | 8.75E-85 |
| Ninj2 | 1.11E-63 | -0.048928799 | 0.187 | 0.652 | 1.61E-60 |
| Slco1c1 | 0.078794484 | -0.04911797 | 0.572 | 0.633 | 1 |
| Rph3a | 5.37E-77 | -0.049473461 | 0.158 | 0.664 | 7.74E-74 |
| Cntnap5c | 2.94E-22 | -0.050981997 | 0.517 | 0.775 | 4.25E-19 |
| St8sia6 | 5.45E-05 | -0.051400779 | 0.764 | 0.847 | 0.078610546 |
| 4921539H07Rik | 1.37E-124 | -0.051630204 | 0.057 | 0.666 | 1.97E-121 |
| Grik1 | 0.535978709 | -0.052317636 | 0.621 | 0.65 | 1 |
| Snrpn | 6.18E-54 | -0.052810438 | 0.382 | 0.793 | 8.92E-51 |
| Otx2os1 | 1.10E-71 | -0.053724357 | 0.316 | 0.792 | 1.59E-68 |
| Cntnap2 | 0.000298016 | -0.053728784 | 0.862 | 0.925 | 0.43003775 |
| Gm15398 | 2.01E-26 | -0.05440143 | 0.106 | 0.374 | 2.90E-23 |
| Gm29266 | 1.15E-19 | -0.054715373 | 0.307 | 0.567 | 1.66E-16 |
| Samd5 | 5.70E-90 | -0.054888001 | 0.305 | 0.83 | 8.23E-87 |
| Npas2 | 3.03E-118 | -0.055254595 | 0.224 | 0.83 | 4.38E-115 |
| Cpne9 | 1.25E-07 | -0.055466798 | 0.42 | 0.576 | 0.000179917 |

| | | | | | |
|---|---|---|---|---|---|
| Zic1 | 2.44E-102 | -0.056204344 | 0.348 | 0.879 | 3.52E-99 |
| Brinp3 | 0.003964777 | -0.056358191 | 0.578 | 0.668 | 1 |
| Frmpd3 | 4.22E-22 | -0.056632622 | 0.517 | 0.773 | 6.09E-19 |
| Nipal2 | 1.33E-96 | -0.056863257 | 0.239 | 0.795 | 1.92E-93 |
| Id4 | 8.84E-27 | -0.057362096 | 0.526 | 0.79 | 1.28E-23 |
| Tmem196 | 5.92E-138 | -0.057724895 | 0.115 | 0.782 | 8.55E-135 |
| Plce1 | 1.52E-35 | -0.057813176 | 0.405 | 0.742 | 2.19E-32 |
| Nr2f2 | 1.33E-12 | -0.058298344 | 0.175 | 0.361 | 1.92E-09 |
| D030068K23Rik | 7.09E-115 | -0.0585226 | 0.27 | 0.854 | 1.02E-111 |
| Abca8a | 3.60E-34 | -0.058621292 | 0.132 | 0.455 | 5.19E-31 |
| Il33 | 0.000280727 | -0.060018638 | 0.474 | 0.584 | 0.405088619 |
| Col4a4 | 2.47E-24 | -0.060194911 | 0.06 | 0.289 | 3.56E-21 |
| Pld5 | 1.10E-07 | -0.060245676 | 0.678 | 0.787 | 0.000159155 |
| Galnt17 | 2.64E-45 | -0.060419985 | 0.563 | 0.889 | 3.81E-42 |
| C530008M17Rik | 7.47E-13 | -0.06044638 | 0.603 | 0.789 | 1.08E-09 |
| Gm48749 | 5.93E-14 | -0.06065749 | 0.112 | 0.292 | 8.56E-11 |
| Thsd7b | 4.94E-23 | -0.060689747 | 0.52 | 0.781 | 7.13E-20 |
| Galnt13 | 7.27E-14 | -0.060692308 | 0.615 | 0.805 | 1.05E-10 |
| Sdk2 | 5.63E-61 | -0.060892036 | 0.144 | 0.595 | 8.12E-58 |
| Dlx6os1 | 7.71E-34 | -0.060953726 | 0.388 | 0.723 | 1.11E-30 |
| Gm49171 | 4.91E-39 | -0.061820917 | 0.055 | 0.364 | 7.09E-36 |
| Phkg1 | 0.083257115 | -0.063366974 | 0.368 | 0.426 | 1 |
| Gli3 | 2.20E-67 | -0.063442507 | 0.313 | 0.779 | 3.18E-64 |
| Gria4 | 0.541657724 | -0.063682133 | 0.767 | 0.791 | 1 |
| Sema5b | 2.76E-17 | -0.063922728 | 0.629 | 0.824 | 3.98E-14 |
| Tprkb | 1.60E-08 | -0.064557571 | 0.716 | 0.559 | 2.31E-05 |
| Sema3e | 1.25E-146 | -0.065071121 | 0.147 | 0.806 | 1.80E-143 |
| Klhl1 | 3.33E-39 | -0.065814234 | 0.353 | 0.717 | 4.81E-36 |
| Lurap1l | 0.000338256 | -0.066208525 | 0.761 | 0.82 | 0.488102928 |
| Bmpr1b | 0.005892914 | -0.068399327 | 0.58 | 0.612 | 1 |
| Aqp4 | 0.18011801 | -0.068511153 | 0.552 | 0.576 | 1 |
| Car10 | 0.653839869 | -0.068534292 | 0.802 | 0.805 | 1 |
| Junb | 3.39E-16 | -0.069277369 | 0.385 | 0.221 | 4.90E-13 |
| Dnah9 | 5.78E-74 | -0.069915748 | 0.267 | 0.762 | 8.34E-71 |
| Nrxn3 | 5.42E-07 | -0.070162607 | 0.882 | 0.938 | 0.000781433 |
| Neto2 | 8.59E-05 | -0.070267141 | 0.391 | 0.512 | 0.123961694 |
| Arhgap29 | 1.90E-19 | -0.071653299 | 0.721 | 0.472 | 2.74E-16 |
| Gabra5 | 2.69E-06 | -0.07294042 | 0.652 | 0.773 | 0.003876621 |
| Spag6l | 4.29E-20 | -0.073245174 | 0.129 | 0.36 | 6.19E-17 |
| Erbb4 | 1.53E-12 | -0.073637911 | 0.767 | 0.899 | 2.20E-09 |
| Hs3st2 | 1.79E-16 | -0.074013603 | 0.532 | 0.74 | 2.58E-13 |
| 9330185C12Rik | 2.47E-16 | -0.074811392 | 0.672 | 0.839 | 3.57E-13 |
| Mag | 2.89E-05 | -0.075319882 | 0.422 | 0.307 | 0.041694049 |
| Cnr1 | 1.63E-20 | -0.07588645 | 0.149 | 0.392 | 2.36E-17 |
| Fnbp1l | 0.448804343 | -0.076550874 | 0.497 | 0.514 | 1 |
| Bcas1 | 1.23E-10 | -0.077339526 | 0.181 | 0.35 | 1.77E-07 |
| Emcn | 5.19E-175 | -0.077659709 | 0.043 | 0.778 | 7.49E-172 |
| Gm10754 | 3.16E-13 | -0.078011506 | 0.681 | 0.846 | 4.57E-10 |
| Tmem132d | 9.83E-23 | -0.078419969 | 0.552 | 0.799 | 1.42E-19 |
| Ccnd3 | 0.252528192 | -0.079392709 | 0.776 | 0.809 | 1 |
| Calb1 | 3.07E-25 | -0.080269823 | 0.451 | 0.734 | 4.43E-22 |
| Cyp7b1 | 1.62E-106 | -0.080363061 | 0.282 | 0.844 | 2.34E-103 |
| Pdzrn4 | 0.001505177 | -0.080491922 | 0.687 | 0.599 | 1 |
| Abi3bp | 2.06E-31 | -0.080498187 | 0.066 | 0.343 | 2.97E-28 |
| Pcdh7 | 0.002051336 | -0.080900045 | 0.819 | 0.886 | 1 |
| Cfap44 | 9.36E-40 | -0.081215633 | 0.434 | 0.783 | 1.35E-36 |
| Gm2115 | 3.00E-76 | -0.08161366 | 0.325 | 0.813 | 4.33E-73 |
| Syt10 | 1.53E-28 | -0.082705835 | 0.724 | 0.887 | 2.21E-25 |
| Cldn11 | 5.25E-55 | -0.083309267 | 0.06 | 0.439 | 7.57E-52 |
| Cacna2d3 | 0.08421301 | -0.083627365 | 0.816 | 0.846 | 1 |
| Vcan | 1.63E-27 | -0.08424409 | 0.328 | 0.631 | 2.35E-24 |
| Kcnh5 | 1.78E-124 | -0.084355396 | 0.043 | 0.666 | 2.57E-121 |
| Spon1 | 0.001015349 | -0.085590146 | 0.532 | 0.632 | 1 |
| Rbpms | 9.36E-20 | -0.08591842 | 0.598 | 0.37 | 1.35E-16 |
| 9530026P05Rik | 0.014117102 | -0.086192004 | 0.693 | 0.748 | 1 |
| Gpc5 | 0.33844836 | -0.0868091 | 0.822 | 0.841 | 1 |
| 2900026A02Rik | 2.26E-104 | -0.086983483 | 0.147 | 0.726 | 3.26E-101 |
| Rgs12 | 3.32E-05 | -0.087752409 | 0.664 | 0.552 | 0.047955356 |
| Mobp | 4.86E-95 | -0.088694452 | 0.141 | 0.693 | 7.01E-92 |
| Hs6st2 | 2.11E-07 | -0.088756422 | 0.704 | 0.797 | 0.000303907 |
| Spp1 | 4.46E-83 | -0.089536247 | 0.256 | 0.777 | 6.44E-80 |
| Gsg1l | 0.00088939 | -0.090147737 | 0.29 | 0.377 | 1 |
| Pdzrn3 | 1.51E-08 | -0.090654947 | 0.718 | 0.566 | 2.17E-05 |
| Epha3 | 1.67E-60 | -0.092010965 | 0.391 | 0.819 | 2.41E-57 |
| Sst | 2.36E-59 | -0.094198396 | 0.204 | 0.656 | 3.40E-56 |
| 9430041J12Rik | 0.072052467 | -0.094266524 | 0.575 | 0.583 | 1 |
| Bcan | 2.86E-32 | -0.094679858 | 0.394 | 0.717 | 4.13E-29 |
| Rnf17 | 3.61E-29 | -0.094965982 | 0.302 | 0.619 | 5.21E-26 |
| Tmem200a | 5.46E-94 | -0.095522688 | 0.273 | 0.817 | 7.88E-91 |
| Chrm3 | 4.32E-07 | -0.097170941 | 0.695 | 0.818 | 0.000623569 |
| Serpine1 | 1.94E-15 | -0.097360971 | 0.124 | 0.32 | 2.80E-12 |
| Necab1 | 2.92E-50 | -0.098336166 | 0.284 | 0.699 | 4.21E-47 |
| Kcnc2 | 0.009705559 | -0.099074237 | 0.557 | 0.634 | 1 |
| Neb | 0.000431935 | -0.09912161 | 0.698 | 0.755 | 0.623281618 |
| Astn2 | 0.19198722 | -0.099467518 | 0.569 | 0.526 | 1 |
| Mef2c | 8.57E-07 | -0.099788611 | 0.802 | 0.9 | 1.23634E-16 |
| Nckap5 | 3.31E-34 | -0.099904911 | 0.566 | 0.857 | 4.78E-31 |
| Pde10a | 2.29E-05 | -0.100160229 | 0.652 | 0.768 | 0.033077078 |
| Ptprz1 | 2.29E-05 | -0.102161782 | 0.454 | 0.582 | 0.033069526 |
| Pcdh19 | 4.14E-24 | -0.102918727 | 0.586 | 0.829 | 5.98E-21 |
| Prox1 | 0.124269583 | -0.103009124 | 0.451 | 0.486 | 1 |
| Tmem178 | 1.43E-26 | -0.103839984 | 0.353 | 0.653 | 2.06E-23 |
| Ranbp3l | 1.50E-38 | -0.104284901 | 0.756 | 0.401 | 2.16E-35 |
| E130114P18Rik | 1.38E-22 | -0.105124278 | 0.382 | 0.659 | 1.99E-19 |
| 4930578G10Rik | 7.12E-20 | -0.10558439 | 0.569 | 0.799 | 1.03E-16 |
| 4930509J09Rik | 5.69E-108 | -0.105732007 | 0.316 | 0.872 | 8.21E-105 |
| Bdnf | 1.49E-39 | -0.106519288 | 0.563 | 0.864 | 2.16E-36 |
| Col23a1 | 1.60E-31 | -0.107619343 | 0.282 | 0.611 | 2.32E-28 |
| Brinp2 | 1.86E-62 | -0.10821966 | 0.279 | 0.737 | 2.69E-59 |
| Itgbl1 | 1.17E-06 | -0.108384922 | 0.787 | 0.852 | 0.001694935 |
| Scube1 | 1.35E-18 | -0.111805919 | 0.086 | 0.292 | 1.95E-15 |
| Tcf7l2 | 0.080092318 | -0.112029687 | 0.497 | 0.559 | 1 |
| Wdr17 | 5.51E-06 | -0.11211161 | 0.655 | 0.777 | 0.007951357 |
| Cdh19 | 7.86E-138 | -0.114103258 | 0.115 | 0.781 | 1.13E-134 |
| Cntnap5b | 1.17E-23 | -0.115105131 | 0.457 | 0.733 | 1.68E-20 |
| C1qb | 0.006774818 | -0.115930704 | 0.621 | 0.702 | 1 |
| Fhad1 | 0.000257712 | -0.117230374 | 0.445 | 0.344 | 0.371878844 |
| Oprm1 | 1.90E-06 | -0.11744464 | 0.328 | 0.469 | 0.002748462 |
| Trpc4 | 2.01E-36 | -0.11811889 | 0.371 | 0.719 | 2.90E-33 |
| Efcab6 | 2.21E-17 | -0.118830926 | 0.385 | 0.628 | 3.19E-14 |
| Tll2 | 2.63E-06 | -0.119375059 | 0.379 | 0.493 | 0.003788659 |
| Rnf182 | 2.06E-05 | -0.120517955 | 0.773 | 0.841 | 0.029674807 |
| Htr2c | 1.40E-17 | -0.121946253 | 0.448 | 0.681 | 2.02E-14 |
| Slc13a3 | 6.36E-19 | -0.122723136 | 0.342 | 0.597 | 9.18E-16 |
| Adamts17 | 6.66E-69 | -0.123527018 | 0.368 | 0.823 | 9.61E-66 |
| Ptger3 | 3.08E-07 | -0.124229233 | 0.305 | 0.2 | 0.000444678 |
| Vstm2a | 1.08E-12 | -0.124756541 | 0.471 | 0.668 | 1.56E-09 |
| Dock8 | 0.002036826 | -0.125984301 | 0.954 | 0.962 | 1 |
| Cntn3 | 4.60E-19 | -0.126444337 | 0.603 | 0.826 | 6.63E-16 |
| Blnk | 0.111263865 | -0.129056195 | 0.782 | 0.788 | 1 |
| Syt6 | 1.97E-46 | -0.130340185 | 0.089 | 0.458 | 2.84E-43 |
| Prex2 | 7.83E-52 | -0.130472532 | 0.218 | 0.642 | 1.13E-48 |
| Gabrg3 | 2.72E-15 | -0.133084321 | 0.744 | 0.889 | 3.93E-12 |
| Pcsk5 | 1.47E-69 | -0.134237153 | 0.457 | 0.882 | 2.12E-66 |
| Spock3 | 0.000268125 | -0.135860656 | 0.816 | 0.835 | 0.386904143 |
| Fstl5 | 2.88E-33 | -0.138434461 | 0.368 | 0.7 | 4.16E-30 |
| Htr1f | 5.65E-48 | -0.138583991 | 0.31 | 0.714 | 8.15E-45 |
| Rxfp1 | 6.05E-09 | -0.140480567 | 0.632 | 0.48 | 8.73E-06 |
| Tshz2 | 2.54E-31 | -0.140564877 | 0.42 | 0.739 | 3.67E-28 |
| P2ry6 | 4.58E-09 | -0.141606364 | 0.46 | 0.594 | 6.61E-06 |
| Zfp536 | 0.003920442 | -0.143874393 | 0.701 | 0.715 | 1 |

| | | | | | |
|---|---|---|---|---|---|
| Ipcef1 | 4.00E-49 | -0.145523929 | 0.161 | 0.514 | 5.77E-46 |
| L3mbtl3 | 0.041979802 | -0.146483832 | 0.601 | 0.553 | 1 |
| Galntl6 | 0.015395445 | -0.150067955 | 0.862 | 0.906 | 1 |
| Csf2rb2 | 6.40E-28 | -0.151932159 | 0.141 | 0.429 | 9.24E-25 |
| Gm48530 | 0.000125482 | -0.152114941 | 0.629 | 0.718 | 0.181071243 |
| Nxph1 | 4.17E-23 | -0.156049955 | 0.649 | 0.864 | 6.02E-20 |
| Mir670hg | 1.53E-110 | -0.159489963 | 0.287 | 0.859 | 2.21E-107 |
| Dgkh | 2.96E-09 | -0.160534858 | 0.466 | 0.639 | 4.27E-06 |
| Efr3a | 3.41E-11 | -0.160883441 | 0.486 | 0.671 | 4.92E-08 |
| Rbms3 | 4.58E-69 | -0.161778111 | 0.233 | 0.718 | 6.61E-66 |
| Kcnmb2 | 1.11E-69 | -0.162115228 | 0.399 | 0.844 | 1.61E-66 |
| Arhgap12 | 7.21E-10 | -0.162181796 | 0.658 | 0.809 | 1.04E-06 |
| Tbxas1 | 0.003098662 | -0.163037312 | 0.704 | 0.786 | 1 |
| Sv2b | 2.67E-92 | -0.171914846 | 0.247 | 0.791 | 3.85E-89 |
| Cd84 | 4.19E-05 | -0.172061654 | 0.58 | 0.696 | 0.060446273 |
| Afap1 | 1.64E-07 | -0.174450213 | 0.701 | 0.595 | 0.000237249 |
| Efna5 | 9.37E-07 | -0.180011211 | 0.629 | 0.518 | 0.001352487 |
| Ndst3 | 2.35E-24 | -0.183896115 | 0.641 | 0.857 | 3.39E-21 |
| Adgrl2 | 2.64E-05 | -0.185451142 | 0.471 | 0.401 | 0.03803394 |
| Zfp804a | 6.33E-36 | -0.185655225 | 0.546 | 0.84 | 9.13E-33 |
| Megf11 | 1.74E-52 | -0.188235753 | 0.129 | 0.542 | 2.51E-49 |
| Olfr111 | 5.62E-09 | -0.189868864 | 0.448 | 0.617 | 8.11E-06 |
| Pcdh11x | 2.82E-15 | -0.193297304 | 0.411 | 0.633 | 4.07E-12 |
| Dnah7b | 2.15E-36 | -0.196175781 | 0.368 | 0.713 | 3.10E-33 |
| Rnf220 | 1.81E-09 | -0.197482122 | 0.313 | 0.489 | 2.61E-15 |
| C1ql3 | 2.41E-15 | -0.201579239 | 0.506 | 0.718 | 3.48E-12 |
| Npas3 | 0.006482897 | -0.203209858 | 0.681 | 0.709 | 1 |
| Slc35f4 | 3.42E-24 | -0.204993541 | 0.494 | 0.749 | 4.94E-21 |
| Chil1 | 3.52E-05 | -0.206599271 | 0.44 | 0.351 | 0.050823585 |
| Hck | 1.01E-05 | -0.207872933 | 0.589 | 0.538 | 0.014568765 |
| Esrrg | 1.18E-29 | -0.209792434 | 0.287 | 0.608 | 1.70E-26 |
| Elavl4 | 9.11E-26 | -0.212599594 | 0.167 | 0.45 | 1.31E-22 |
| Pard3bos1 | 7.91E-103 | -0.213446876 | 0.747 | 0.261 | 1.14E-99 |
| Adamts6 | 1.82E-10 | -0.213663482 | 0.328 | 0.513 | 2.63E-07 |
| Trhde | 6.52E-46 | -0.214614324 | 0.5 | 0.851 | 9.41E-43 |
| Slc38a2 | 0.006235385 | -0.229316514 | 0.402 | 0.491 | 1 |
| Col25a1 | 0.000570182 | -0.231488162 | 0.517 | 0.603 | 0.822773086 |
| Runx1t1 | 1.25E-30 | -0.231932282 | 0.543 | 0.817 | 1.80E-27 |
| Bnc2 | 0.000379375 | -0.240557037 | 0.661 | 0.675 | 0.547438738 |
| Tbc1d9 | 0.000489345 | -0.242448511 | 0.276 | 0.38 | 0.706125103 |
| Zfp804b | 1.35E-18 | -0.244953738 | 0.672 | 0.86 | 1.95E-15 |
| Cdc14a | 3.70E-16 | -0.247491479 | 0.385 | 0.619 | 5.34E-13 |
| Prkca | 7.63E-05 | -0.249639002 | 0.876 | 0.905 | 0.110122658 |
| Gm4876 | 1.15E-18 | -0.253396898 | 0.095 | 0.299 | 1.66E-15 |
| Tmem44 | 0.001984197 | -0.254060296 | 0.365 | 0.425 | 1 |
| Sfmbt2 | 7.68E-79 | -0.263032542 | 0.089 | 0.592 | 1.11E-75 |
| Pcdh15 | 1.47E-69 | -0.284236845 | 0.296 | 0.772 | 2.12E-66 |
| Cdh12 | 4.62E-37 | -0.300412865 | 0.638 | 0.894 | 6.67E-34 |
| Epha7 | 6.94E-12 | -0.309461583 | 0.727 | 0.863 | 1.00E-08 |
| Col6a3 | 1.72E-13 | -0.314144093 | 0.756 | 0.822 | 2.47E-10 |
| Nrp2 | 2.03E-09 | -0.327262261 | 0.264 | 0.431 | 2.93E-06 |
| Fbxl7 | 3.95E-36 | -0.330163552 | 0.514 | 0.819 | 5.71E-33 |
| Glul | 8.89E-14 | -0.334016094 | 0.359 | 0.285 | 1.28E-10 |
| Fyb | 1.00E-05 | -0.336423374 | 0.741 | 0.846 | 0.014428787 |
| Lrrk1 | 4.72E-13 | -0.348017198 | 0.569 | 0.761 | 6.81E-10 |
| Lypd6 | 2.16E-17 | -0.373126231 | 0.701 | 0.519 | 3.11E-14 |
| Lcp2 | 1.10E-44 | -0.378667957 | 0.491 | 0.833 | 1.59E-41 |
| Cfh | 4.79E-13 | -0.383678846 | 0.652 | 0.826 | 6.92E-10 |
| Unc93b1 | 1.39E-06 | -0.397978222 | 0.621 | 0.723 | 0.002000946 |
| Arhgap25 | 7.09E-37 | -0.407658844 | 0.42 | 0.752 | 1.02E-33 |
| Adipor2 | 4.03E-46 | -0.409924685 | 0.233 | 0.62 | 5.82E-43 |
| Fcer1g | 5.08E-28 | -0.418574431 | 0.451 | 0.733 | 7.33E-25 |
| Pla2g4a | 1.11E-07 | -0.429193969 | 0.603 | 0.741 | 0.000160601 |
| Ccdc162 | 7.29E-40 | -0.431893663 | 0.118 | 0.466 | 1.05E-36 |
| Slc39a12 | 2.95E-25 | -0.442620391 | 0.365 | 0.64 | 4.26E-22 |
| Gm21798 | 3.57E-06 | -0.450113189 | 0.305 | 0.35 | 0.005156561 |
| Arhgap31 | 4.19E-65 | -0.45127516 | 0.371 | 0.801 | 6.05E-62 |
| Arhgap45 | 6.23E-13 | -0.46185485 | 0.635 | 0.766 | 8.99E-10 |
| Glis3 | 0.000173241 | -0.471059782 | 0.644 | 0.65 | 0.249987204 |
| Ptpro | 4.13E-81 | -0.477234129 | 0.302 | 0.781 | 5.95E-78 |
| Eya4 | 1.89E-06 | -0.497255152 | 0.388 | 0.429 | 0.002726332 |
| D7Ertd443e | 1.12E-18 | -0.499928259 | 0.437 | 0.625 | 1.61E-15 |
| Gm13269 | 1.14E-16 | -0.50077152 | 0.368 | 0.279 | 1.65E-13 |
| Rnf144b | 1.99E-27 | -0.514831322 | 0.342 | 0.644 | 1.98E-38 |
| Khdrbs3 | 6.22E-39 | -0.53949651 | 0.693 | 0.441 | 8.98E-36 |
| Cux2 | 1.64E-26 | -0.546982232 | 0.408 | 0.686 | 2.36E-23 |
| Myo1f | 3.38E-16 | -0.561413056 | 0.503 | 0.724 | 4.87E-13 |
| Chst9 | 8.54E-51 | -0.606866494 | 0.391 | 0.767 | 1.23E-47 |
| Gm10848 | 2.24E-13 | -0.607771889 | 0.368 | 0.318 | 3.24E-10 |
| Gm2245 | 3.39E-07 | -0.627548404 | 0.397 | 0.549 | 0.000489689 |
| Ubash3b | 1.30E-12 | -0.630361984 | 0.721 | 0.813 | 1.88E-09 |
| Arl15 | 5.50E-14 | -0.640196928 | 0.578 | 0.754 | 7.93E-11 |
| Arhgap24 | 3.32E-13 | -0.671081058 | 0.506 | 0.679 | 4.79E-10 |
| Nckap1l | 1.41E-17 | -0.693177707 | 0.601 | 0.811 | 2.03E-14 |
| Abca9 | 1.52E-27 | -0.697597819 | 0.606 | 0.863 | 2.20E-24 |
| Adap2os | 2.12E-11 | -0.726859291 | 0.569 | 0.651 | 3.06E-08 |
| Adamts16 | 3.41E-21 | -0.729511129 | 0.759 | 0.641 | 4.92E-18 |
| Entpd1 | 2.40E-23 | -0.746989569 | 0.624 | 0.851 | 3.47E-20 |
| Elmo1 | 3.06E-49 | -0.753243509 | 0.905 | 0.972 | 4.42E-46 |
| Cd37 | 4.25E-19 | -0.758831754 | 0.647 | 0.68 | 6.14E-16 |
| Spi1 | 4.23E-24 | -0.774597667 | 0.405 | 0.69 | 6.11E-21 |
| Npnt | 4.70E-17 | -0.80632727 | 0.805 | 0.791 | 6.78E-14 |
| Itpr2 | 1.88E-22 | -0.806797063 | 0.437 | 0.701 | 2.71E-19 |
| Tspan18 | 2.42E-20 | -0.81330055 | 0.428 | 0.653 | 3.50E-17 |
| Fcrls | 4.04E-19 | -0.815427775 | 0.529 | 0.745 | 5.83E-16 |
| Mrc2 | 9.23E-13 | -0.834340948 | 0.526 | 0.61 | 1.33E-09 |
| Laptm5 | 5.45E-45 | -0.837490535 | 0.328 | 0.713 | 7.86E-42 |
| Adcy8 | 1.51E-31 | -0.848632149 | 0.638 | 0.851 | 2.18E-28 |
| Oxr1 | 3.12E-15 | -0.853927371 | 0.675 | 0.83 | 4.50E-12 |
| Adap2 | 7.87E-28 | -0.863786268 | 0.655 | 0.858 | 1.13E-24 |
| C1qa | 2.08E-18 | -0.865482516 | 0.534 | 0.729 | 3.01E-15 |
| Itga9 | 7.52E-45 | -0.87476821 | 0.313 | 0.703 | 1.08E-41 |
| Myo1b | 2.86E-43 | -0.878480536 | 0.164 | 0.539 | 4.13E-40 |
| Lair1 | 1.53E-25 | -0.887945899 | 0.69 | 0.782 | 2.21E-22 |
| Ikzf1 | 2.48E-40 | -0.893577847 | 0.724 | 0.947 | 3.58E-37 |
| Zfhx3 | 1.10E-53 | -0.899933078 | 0.839 | 0.966 | 1.59E-50 |
| Gab1 | 2.25E-17 | -0.908148272 | 0.741 | 0.735 | 3.25E-14 |
| Chst8 | 2.65E-15 | -0.910168873 | 0.322 | 0.406 | 3.82E-12 |
| Irak2 | 4.07E-20 | -0.913733439 | 0.319 | 0.579 | 5.88E-17 |
| C1qc | 2.96E-32 | -0.977470008 | 0.474 | 0.787 | 4.28E-29 |
| Rgs10 | 1.02E-19 | -0.994996259 | 0.779 | 0.834 | 1.47E-16 |
| Slco2b1 | 7.68E-34 | -0.99849101 | 0.672 | 0.857 | 1.11E-30 |
| Ebf3 | 8.22E-20 | -1.000076126 | 0.77 | 0.714 | 1.19E-16 |
| Inpp5d | 8.50E-65 | -1.023095922 | 0.836 | 0.976 | 1.23E-61 |
| Csf1r | 7.06E-40 | -1.04398946 | 0.764 | 0.945 | 1.02E-36 |
| Havcr2 | 3.13E-22 | -1.072534308 | 0.486 | 0.585 | 4.52E-19 |
| Csf3r | 1.19E-36 | -1.082362833 | 0.497 | 0.637 | 1.72E-33 |
| Nhs | 4.30E-13 | -1.086779305 | 0.434 | 0.459 | 6.21E-10 |
| Sncaip | 1.30E-73 | -1.088135591 | 0.178 | 0.678 | 1.88E-70 |
| Il10ra | 2.68E-55 | -1.091525317 | 0.466 | 0.85 | 3.86E-52 |
| Cdh23 | 4.62E-37 | -1.09847885 | 0.402 | 0.489 | 6.67E-34 |
| Arhgap22 | 1.45E-31 | -1.116360532 | 0.408 | 0.724 | 2.10E-28 |
| Ctss | 1.07E-32 | -1.130766635 | 0.641 | 0.869 | 1.55E-29 |
| Ccr5 | 4.41E-55 | -1.137039782 | 0.227 | 0.661 | 6.36E-52 |
| Lrmda | 3.93E-35 | -1.151706522 | 0.741 | 0.935 | 5.66E-32 |
| Trem2 | 4.92E-28 | -1.185919308 | 0.664 | 0.665 | 7.10E-25 |
| Lgmn | 3.28E-41 | -1.19698019 | 0.511 | 0.811 | 4.73E-38 |
| Rasgrp3 | 8.63E-32 | -1.198752859 | 0.44 | 0.72 | 1.24E-28 |
| Maml3 | 1.16E-65 | -1.198824635 | 0.701 | 0.956 | 1.67E-62 |
| Fat3 | 6.93E-38 | -1.202671748 | 0.787 | 0.668 | 1.00E-34 |
| P3h2 | 8.67E-41 | -1.202715615 | 0.491 | 0.82 | 1.25E-37 |

| | | | | | | | | | |
|---|---|---|---|---|---|---|---|---|---|
| Agmo | 6.59E-34 | -1.206870087 | 0.601 | 0.829 | 9.51E-31 | | | | |
| Vsir | 7.45E-33 | -1.24874556 | 0.287 | 0.604 | 1.08E-29 | | | | |
| Anxa3 | 1.99E-35 | -1.300448467 | 0.405 | 0.666 | 2.87E-32 | | | | |
| Ly86 | 1.14E-16 | -1.313148219 | 0.733 | 0.831 | 1.64E-13 | | | | |
| Cacnb2 | 2.10E-44 | -1.326821824 | 0.851 | 0.795 | 3.02E-41 | | | | |
| Fgd2 | 1.38E-37 | -1.3829367 | 0.474 | 0.712 | 1.99E-34 | | | | |
| Garnl3 | 1.74E-42 | -1.399417946 | 0.414 | 0.731 | 2.50E-39 | | | | |
| Serpine2 | 1.74E-35 | -1.431826051 | 0.175 | 0.447 | 2.51E-32 | | | | |
| Fgf13 | 4.24E-34 | -1.485646636 | 0.56 | 0.792 | 6.13E-31 | | | | |
| Slc1a3 | 1.43E-42 | -1.532405038 | 0.351 | 0.442 | 2.06E-39 | | | | |
| A830008E24Rik | 2.94E-94 | -1.542606928 | 0.756 | 0.455 | 4.25E-91 | | | | |
| Apbb1ip | 1.28E-68 | -1.596260181 | 0.5 | 0.862 | 1.85E-65 | | | | |
| Itgb5 | 7.97E-83 | -1.617116392 | 0.46 | 0.873 | 1.15E-79 | | | | |
| Ptprm | 9.11E-66 | -1.64189155 | 0.431 | 0.817 | 1.31E-62 | | | | |
| 0610040J01Rik | 3.23E-57 | -1.655754318 | 0.658 | 0.912 | 4.67E-54 | | | | |
| Gpr34 | 2.27E-48 | -1.661945521 | 0.299 | 0.644 | 3.27E-45 | | | | |
| Sdk1 | 2.10E-34 | -1.664287982 | 0.681 | 0.694 | 3.03E-31 | | | | |
| Abi3 | 1.02E-43 | -1.664943561 | 0.457 | 0.732 | 1.47E-40 | | | | |
| Whrn | 4.88E-64 | -1.730929663 | 0.351 | 0.744 | 7.04E-61 | | | | |
| F11r | 3.06E-114 | -1.781939701 | 0.121 | 0.734 | 4.42E-111 | | | | |
| Tgfbr1 | 1.73E-163 | -1.80826365 | 0.575 | 0.972 | 2.49E-160 | | | | |
| Bin2 | 5.33E-126 | -1.845853711 | 0.193 | 0.822 | 7.69E-123 | | | | |
| Chn2 | 8.75E-85 | -1.933929038 | 0.376 | 0.789 | 1.26E-81 | | | | |
| Cst3 | 6.73E-125 | -2.079802324 | 0.822 | 0.953 | 9.71E-122 | | | | |
| Lpcat2 | 1.68E-97 | -2.095526551 | 0.471 | 0.881 | 2.43E-94 | | | | |
| Pag1 | 3.79E-114 | -2.150063659 | 0.471 | 0.919 | 5.47E-111 | | | | |
| Cx3cr1 | 6.67E-94 | -2.275930115 | 0.379 | 0.836 | 9.63E-141 | | | | |
| Hexb | 1.96E-241 | -2.311890662 | 0.474 | 0.93 | 2.83E-180 | | | | |
| Epb41l2 | 1.26E-157 | -2.32950844 | 0.779 | 0.975 | 1.81E-154 | | | | |
| 8030442B05Rik | 1.98E-86 | -2.361080215 | 0.279 | 0.716 | 2.86E-83 | | | | |
| Mertk | 5.23E-80 | -2.467976935 | 0.799 | 0.968 | 7.55E-87 | | | | |
| Siglech | 7.29E-80 | -2.532924419 | 0.491 | 0.847 | 1.05E-112 | | | | |
| Selplg | 7.42E-144 | -2.722272828 | 0.477 | 0.889 | 1.07E-140 | | | | |
| Plxdc2 | 3.60E-290 | -3.96124252 | 0.279 | 0.988 | 5.20E-287 | | | | ≈≈ |

Figure3H_Venndiagram_pseudobulk_DEGs

| Dap12+/+ tau+ vs tau- | | p_val | avg_log2FC | pct.1 | pct.2 | p_val_adj | Upregulation | Downregulation | Dap12-/- tau+ vs Dap12+/+ tau+ | | p_val | avg_log2FC | pct.1 | pct.2 | p_val_adj | Upregulation | Downregulation | Dap12+/+ tau+ vs tau- | | Names | total | elements | | Names | total | elements | | | Names | total | elements |
|---|---|---|---|---|---|---|---|---|---|---|---|---|---|---|---|---|---|---|---|---|---|---|---|---|---|---|---|---|---|---|---|
| | Scoc | 2.68E-137 | -1.9474858 | 0.015 | 0.357 | 7.22E-133 | 419 | 514 | | Kcnip4 | 9.73E-63 | -1.4925335 | 0.106 | 0.386 | 2.62E-58 | 629 | 347 | | | tau+ up Dap12ko down | 229 | Hps3 | | tau+ up only | 190 | Man1c1 | | | Dap12ko down only | 118 | Dhx9 |
| | Cacna1a | 1.34E-128 | 1.92370081 | 0.61 | 0.215 | 3.60E-124 | Cacna1a | Scoc | | Lsamp | 5.51E-60 | -0.9005729 | 0.139 | 0.428 | 1.48E-55 | Ifitm10 | Kcnip4 | | | | | Dmxl1 | | | | Dip2c | | | | | Fat3 |
| | Ptprd | 1.99E-27 | 0.98606268 | 0.353 | 0.169 | 5.37E-23 | Ptprd | Ifitm10 | | Csmd1 | 6.15E-58 | -1.3635235 | 0.068 | 0.313 | 1.66E-53 | Ppp2r5c | Lsamp | | | | | Rbm39 | | | | Snd1 | | | | | Wdr33 |
| | Csmd1 | 2.65E-26 | 0.91103201 | 0.313 | 0.139 | 7.12E-22 | Csmd1 | Cdh23 | | Cmss1 | 1.45E-57 | -0.9262552 | 0.478 | 0.62 | 3.90E-53 | Gm32036 | Csmd1 | | | | | Luc7l2 | | | | Erbin | | | | | Strbp |
| | Gm20388 | 1.92E-25 | 0.34625319 | 0.797 | 0.605 | 5.18E-21 | Gm20388 | Kcnrg | | Nrxn1 | 1.51E-39 | -1.0430957 | 0.093 | 0.298 | 4.06E-35 | Morc3 | Cmss1 | | | | | Vps54 | | | | Strn3 | | | | | C1qb |
| | Ifitm10 | 1.64E-23 | -0.8143352 | 0.11 | 0.266 | 4.42E-19 | Kcnip4 | Tbc1d9 | | Gm42418 | 5.09E-35 | -0.7250106 | 0.81 | 0.904 | 1.37E-30 | Lrp1 | Nrxn1 | | | | | Nlgn1 | | | | Map4k4 | | | | | Daglb |
| | Cdh23 | 7.87E-21 | -0.7010148 | 0.279 | 0.462 | 2.12E-16 | Meg3 | Fkbp5 | | Ptprd | 9.79E-32 | -1.0048374 | 0.185 | 0.353 | 2.63E-27 | Pla2g4a | Gm42418 | | | | | Gphn | | | | Ppcdc | | | | | Jak1 |
| | Kcnip4 | 1.46E-22 | 1.04838109 | 0.386 | 0.215 | 3.92E-18 | Cadm1 | Entpd1 | | Dlg2 | 2.35E-31 | -1.1187522 | 0.148 | 0.319 | 6.32E-27 | Dnajc7 | Ptprd | | | | | Osbpl8 | | | | Nav3 | | | | | Havcr2 |
| | Kcnrg | 4.15E-16 | -0.6107131 | 0.144 | 0.277 | 1.12E-11 | Apbb2 | AY036118 | | Ifitm10 | 8.52E-31 | 0.90897154 | 0.29 | 0.11 | 2.29E-26 | Elmo1 | Dlg2 | | | | | Sh3kbp1 | | | | Etv6 | | | | | Gm2245 |
| | Meg3 | 9.27E-21 | 0.69212651 | 0.661 | 0.528 | 2.50E-16 | Cadm2 | Gm32036 | | Nrg3 | 2.13E-29 | -0.9779618 | 0.124 | 0.301 | 5.72E-25 | Csmd3 | Nrg3 | | | | | AW554918 | | | | Rap1a | | | | | Rhoh |
| | Cadm1 | 1.02E-20 | 0.58316789 | 0.469 | 0.403 | 2.74E-16 | Ly86 | Lysmd4 | | Cst3 | 3.69E-29 | -0.7907851 | 0.861 | 0.86 | 9.93E-25 | Son | Cst3 | | | | | Runx1 | | | | Celf1 | | | | | Stx16 |
| | Apbb2 | 1.44E-18 | 0.70317654 | 0.305 | 0.209 | 3.87E-14 | Nrxn1 | Tm6sf1 | | Gm20388 | 8.33E-27 | -0.0828425 | 0.628 | 0.797 | 2.24E-22 | Wdfy2 | Gm20388 | | | | | Ptpra | | | | Scmh1 | | | | | Zfp148 |
| | Cadm2 | 1.82E-18 | 0.8653272 | 0.281 | 0.183 | 4.89E-14 | Nkain2 | Gm26827 | | Fkbp5 | 3.15E-26 | -0.5672726 | 0.415 | 0.435 | 8.49E-22 | Srrm1 | Fkbp5 | | | | | Acer3 | | | | Pacs1 | | | | | Rasal2 |
| | Ly86 | 2.43E-18 | 0.47319716 | 0.705 | 0.653 | 6.55E-14 | Dlg2 | Fat3 | | Tenm2 | 5.61E-26 | -0.717812 | 0.114 | 0.281 | 1.51E-21 | Usp8 | Tenm2 | | | | | Ly86 | | | | Capza1 | | | | | Cnot4 |
| | Nrxn1 | 3.92E-18 | 0.92668892 | 0.298 | 0.198 | 1.06E-13 | Man2b1 | P4ha1 | | Pcdh9 | 9.18E-26 | -0.8496976 | 0.151 | 0.306 | 2.47E-21 | Cbl | Pcdh9 | | | | | Tcf4 | | | | Ddx6 | | | | | Trappc8 |
| | Tbc1d9 | 2.71E-17 | -0.5584933 | 0.205 | 0.357 | 7.28E-13 | Arhgap17 | Picalm | | Nlgn1 | 5.87E-25 | -0.8932817 | 0.144 | 0.264 | 1.58E-20 | Tspan14 | Nlgn1 | | | | | Vps13b | | | | Hdac9 | | | | | C1qa |
| | Fkbp5 | 2.11E-17 | -0.5448861 | 0.435 | 0.608 | 5.68E-13 | Pcdh9 | Hsp90b1 | | Meg3 | 1.43E-23 | -0.7400831 | 0.487 | 0.661 | 3.84E-19 | Abl1 | Meg3 | | | | | Nrg3 | | | | Pwwp2a | | | | | Rock1 |
| | Entpd1 | 5.54E-18 | -0.5131055 | 0.559 | 0.719 | 1.49E-13 | AU020206 | Sort1 | | Nkain2 | 1.65E-23 | -0.8376583 | 0.179 | 0.3 | 4.44E-19 | Rapgef1 | Nkain2 | | | | | Insr | | | | Mllt10 | | | | | Mbd5 |
| | AY036118 | 2.65E-17 | -0.51083 | 0.452 | 0.62 | 7.13E-13 | Park2 | Ddx5 | | Ppp2r5c | 1.07E-21 | 0.14158511 | 0.31 | 0.206 | 2.87E-17 | Cdh23 | Snhg11 | | | | | Nrp1 | | | | Pip4k2a | | | | | Zfp950 |
| | Nkain2 | 3.98E-17 | 0.87232419 | 0.3 | 0.163 | 1.07E-12 | Arid1b | Peli1 | | Snhg11 | 1.75E-21 | -0.7078613 | 0.459 | 0.621 | 4.70E-17 | Zfp710 | Tmsb4x | | | | | Med13l | | | | Zmiz1 | | | | | Trim44 |
| | Gm32036 | 2.99E-15 | -0.490176 | 0.324 | 0.481 | 8.03E-11 | Tenm2 | E2f3 | | Gm32036 | 5.41E-21 | 0.56590006 | 0.508 | 0.324 | 1.45E-16 | Map2k5 | Cadm2 | | | | | Immp2l | | | | Plxdc2 | | | | | Creb1 |
| | Dlg2 | 5.21E-17 | 0.88096162 | 0.319 | 0.203 | 1.40E-12 | Cd9 | Bank1 | | Morc3 | 1.14E-20 | 0.21308747 | 0.337 | 0.218 | 3.06E-16 | Nrros | Cadm1 | | | | | Stk3 | | | | Pik3ap1 | | | | | Clip1 |
| | Lysmd4 | 2.83E-24 | -0.4648452 | 0.697 | 0.851 | 7.61E-20 | Rap1a | Gm16337 | | Tmsb4x | 4.14E-20 | -0.2832805 | 0.423 | 0.389 | 1.11E-15 | Gm16337 | C1qb | | | | | Bmpr2 | | | | Tbl1xr1 | | | | | Man1a |
| | Tm6sf1 | 1.03E-16 | -0.4617918 | 0.257 | 0.41 | 2.77E-12 | Znrf2 | Csad | | Lrp1 | 8.50E-20 | 0.32137997 | 0.444 | 0.292 | 2.29E-15 | Ptgs1 | Osbpl11 | | | | | Lnpep | | | | Fli1 | | | | | Lpp |
| | Man2b1 | 1.96E-16 | 0.44354069 | 0.359 | 0.313 | 5.26E-12 | Tns3 | Tcf7l2 | | Cadm2 | 7.41E-19 | -0.655589 | 0.159 | 0.281 | 1.99E-14 | Dido1 | C1qc | | | | | Camk1d | | | | Apc | | | | | Nr3c1 |
| | Gm26827 | 2.70E-10 | -0.4420799 | 0.211 | 0.323 | 7.28E-06 | Tgfbr2 | Stab1 | | Cadm1 | 7.59E-19 | -0.5853467 | 0.38 | 0.469 | 2.04E-14 | Ankhd1 | Cd81 | | | | | Cog5 | | | | Atxn7 | | | | | Capza2 |
| | Fat3 | 1.32E-08 | -0.4415668 | 0.327 | 0.442 | 0.00035394 | Lair1 | Sppl3 | | Pla2g4a | 3.37E-18 | 0.25887149 | 0.341 | 0.221 | 9.07E-14 | Tbxas1 | Pou2f2 | | | | | Tnpo1 | | | | Nfkb1 | | | | | Senp6 |
| | P4ha1 | 4.04E-12 | -0.4268208 | 0.217 | 0.336 | 1.09E-07 | Itsn2 | Tpp2 | | C1qb | 4.08E-18 | -0.4851518 | 0.538 | 0.563 | 1.10E-13 | Il10ra | Malat1 | | | | | Dock11 | | | | Chd6 | | | | | Malat1 |
| | Arhgap17 | 1.77E-15 | 0.38652519 | 0.389 | 0.357 | 4.76E-11 | Nrg3 | Tlr7 | | Dnajc7 | 4.31E-18 | 0.03622201 | 0.323 | 0.243 | 1.16E-13 | Tpr | Plek | | | | | Spag9 | | | | Asap1 | | | | | Myo9a |
| | Picalm | 7.25E-16 | -0.4185003 | 0.583 | 0.735 | 1.95E-11 | Nfkb1 | Atg16l2 | | Elmo1 | 9.56E-18 | 0.27187302 | 0.957 | 0.91 | 2.57E-13 | Atp6v0a1 | Cd9 | | | | | Gls | | | | Large1 | | | | | Mark3 |
| | Hsp90b1 | 7.82E-11 | -0.4080533 | 0.182 | 0.289 | 2.10E-06 | Tnpo1 | Tet3 | | Csmd3 | 1.13E-17 | 0.38064488 | 0.904 | 0.87 | 3.04E-13 | Ogt | Rhoh | | | | | Rapgef6 | | | | Ubl3 | | | | | Ppp1r21 |
| | Pcdh9 | 3.15E-15 | 0.74812709 | 0.306 | 0.216 | 8.49E-11 | Lsamp | Mlph | | Son | 1.25E-17 | 0.1880059 | 0.654 | 0.508 | 3.38E-13 | Brd4 | Gns | | | | | Nbea | | | | Hdac8 | | | | | Hnrnpm |
| | Sort1 | 1.54E-10 | -0.4037467 | 0.197 | 0.308 | 4.14E-06 | Wsb1 | Clk1 | | Osbpl11 | 2.16E-17 | -0.004027 | 0.387 | 0.313 | 5.80E-13 | Hps4 | Ptpra | | | | | Bach2 | | | | Tra2a | | | | | Btbd7 |
| | Ddx5 | 4.53E-15 | -0.3997017 | 0.628 | 0.771 | 1.22E-10 | Snhg11 | Tpm3 | | Wdfy2 | 2.65E-17 | 0.2636916 | 0.428 | 0.294 | 7.14E-13 | Gfm2 | Apoe | | | | | Heatr5a | | | | Nek7 | | | | | Phip |
| | AU020206 | 6.06E-15 | 0.66031263 | 0.263 | 0.171 | 1.63E-10 | Pou2f2 | Sfpq | | C1qc | 7.83E-17 | -0.2872827 | 0.602 | 0.578 | 2.11E-12 | Slc29a3 | Atp9b | | | | | Clta | | | | Rtn4 | | | | | Ep300 |
| | Peli1 | 6.31E-15 | -0.023561 | 0.28 | 0.355 | 1.70E-10 | Myo1f | Rapgef1 | | Srrm1 | 8.29E-17 | 0.12154902 | 0.278 | 0.196 | 2.23E-12 | Smad7 | Gsap | | | | | Atp2b1 | | | | Fus | | | | | B4galt1 |
| | Park2 | 7.15E-15 | 0.5083433 | 0.564 | 0.405 | 1.92E-10 | Ash1l | Capza2 | | Cd81 | 9.71E-17 | -0.1857875 | 0.396 | 0.349 | 2.61E-12 | Siglech | Tnpo1 | | | | | Zbtb20 | | | | Scaper | | | | | Fkbp5 |
| | E2f3 | 7.65E-14 | -0.3994275 | 0.19 | 0.308 | 2.06E-09 | Ncor1 | Dnajc7 | | Usp8 | 1.38E-16 | 0.08944185 | 0.29 | 0.214 | 3.72E-12 | Cnot2 | Fam13b | | | | | Ssbp2 | | | | Ube2k | | | | | Papola |
| | Arid1b | 1.56E-14 | 0.2760076 | 0.398 | 0.401 | 4.20E-10 | Nlgn1 | Pten | | Pou2f2 | 1.49E-16 | -0.0938409 | 0.425 | 0.366 | 4.01E-12 | Hnrnpa3 | Dhx9 | | | | | Ctsh | | | | Ubr3 | | | | | Gm44686 |
| | Tenm2 | 1.80E-14 | 0.78449454 | 0.281 | 0.154 | 4.85E-10 | Rabgap1 | Cables1 | | Cbl | 1.57E-16 | 0.08386235 | 0.377 | 0.285 | 4.24E-12 | Taok1 | Kmt2e | | | | | Cacna1d | | | | Git2 | | | | | Elk3 |
| | Bank1 | 1.35E-08 | -0.3886626 | 0.227 | 0.332 | 0.00036288 | Atxn7 | Bin2 | | Tspan14 | 2.87E-16 | 0.15481911 | 0.308 | 0.218 | 7.73E-12 | Cerk | Wdr33 | | | | | Srek1 | | | | Itga6 | | | | | Dst |
| | Cd9 | 3.38E-14 | 0.57392528 | 0.287 | 0.232 | 9.10E-10 | Fam13b | Nr6a1 | | Malat1 | 2.91E-16 | -0.1437644 | 1 | 1 | 7.83E-12 | Selplg | Clint1 | | | | | Mbtd1 | | | | Frrs1 | | | | | Itch |
| | Gm16337 | 4.61E-14 | -0.3818373 | 0.294 | 0.435 | 1.24E-09 | Cacna1d | Nckap1l | | Abl1 | 3.47E-16 | 0.22771044 | 0.325 | 0.219 | 9.34E-12 | Tsc22d4 | Ap3b1 | | | | | Clint1 | | | | Lyst | | | | | Pcmtd1 |
| | Rap1a | 4.69E-14 | 0.10653923 | 0.331 | 0.379 | 1.26E-09 | Anxa3 | Arid2 | | Rapgef1 | 3.96E-16 | 0.30306303 | 0.341 | 0.222 | 1.07E-11 | Zfhx3 | C1qa | | | | | Tmcc1 | | | | Mtus1 | | | | | Sfpq |
| | Csad | 3.25E-10 | -0.3810079 | 0.185 | 0.288 | 8.74E-06 | Slmap | Pag1 | | Cdh23 | 4.76E-16 | 0.4478552 | 0.428 | 0.279 | 1.28E-11 | Zfand3 | Fth1 | | | | | Ppp1r12a | | | | Ncoa3 | | | | | Larp4b |
| | Znrf2 | 1.43E-13 | 0.02347377 | 0.21 | 0.262 | 3.83E-09 | Lrch3 | Pacsin2 | | Plek | 8.33E-16 | -0.2811262 | 0.267 | 0.26 | 2.24E-11 | Cd33 | Hnrnpm | | | | | Map3k5 | | | | Nsd1 | | | | | Fyb |
| | Tcf7l2 | 1.04E-10 | -0.3761858 | 0.239 | 0.353 | 2.79E-06 | Ankrd17 | Irf2 | | Cd9 | 9.08E-16 | -0.4375205 | 0.273 | 0.287 | 2.44E-11 | Bin2 | Myo1f | | | | | Ankrd12 | | | | Dync1i2 | | | | | Top1 |
| | Tns3 | 1.56E-13 | 0.35429439 | 0.467 | 0.433 | 4.21E-09 | Gnb1 | Cyfip1 | | Rhoh | 9.31E-16 | -0.107024 | 0.341 | 0.295 | 2.51E-11 | Pkn1 | Trem2 | | | | | Marcks | | | | Wac | | | | | Tbc1d5 |
| | Tgfbr2 | 1.57E-13 | 0.40021561 | 0.477 | 0.427 | 4.23E-09 | Fars2 | Rassf2 | | Zfp710 | 9.65E-16 | 0.41074449 | 0.723 | 0.569 | 2.60E-11 | Tubgcp5 | Pitpna | | | | | Pcdh9 | | | | Mapk14 | | | | | Lncpint |
| | Stab1 | 3.02E-08 | -0.3758313 | 0.241 | 0.342 | 0.00081187 | Khdrbs1 | B4galt1 | | Gns | 1.03E-15 | -0.0923386 | 0.436 | 0.383 | 2.77E-11 | Arsb | Wdfy3 | | | | | Gnaq | | | | Znrf2 | | | | | Fubp1 |
| | Sppl3 | 2.37E-13 | -0.0334078 | 0.317 | 0.391 | 6.39E-09 | Mtmr3 | Sipa1 | | Map2k5 | 1.19E-15 | 0.01234948 | 0.26 | 0.198 | 3.21E-11 | Ubash3b | Tns3 | | | | | Park2 | | | | Rsrc2 | | | | | Rnf169 |
| | Lair1 | 2.61E-13 | 0.20221034 | 0.539 | 0.564 | 7.01E-09 | Gsap | Lpcat2 | | Nrros | 1.27E-15 | 0.17318112 | 0.451 | 0.334 | 3.41E-11 | Tet3 | Peli1 | | | | | Magi1 | | | | Rock2 | | | | | Rasa1 |
| | Itsn2 | 2.75E-13 | 0.00672498 | 0.285 | 0.345 | 7.40E-09 | Dyrk1a | AC149090.1 | | Gm16337 | 1.45E-15 | 0.1631722 | 0.404 | 0.294 | 3.91E-11 | Zfp638 | Jak1 | | | | | Lrmda | | | | Akap10 | | | | | Nufip2 |
| | Tpp2 | 3.82E-13 | -0.0170369 | 0.194 | 0.252 | 1.03E-08 | Asph | Cmtm7 | | Ptpra | 1.82E-15 | -0.0036503 | 0.434 | 0.355 | 4.91E-11 | Tpm3 | Wapl | | | | | Klf3 | | | | Cdkal1 | | | | | Ap3b1 |
| | Nrg3 | 4.44E-13 | 0.58896353 | 0.301 | 0.178 | 1.19E-08 | Laptm5 | Dmtf1 | | Apoe | 2.05E-15 | -0.6017445 | 0.306 | 0.362 | 5.52E-11 | Mtmr3 | Plekho1 | | | | | Tnrc6c | | | | Gab1 | | | | | Peli1 |
| | Nfkb1 | 4.52E-13 | 0.09577606 | 0.331 | 0.376 | 1.22E-08 | Cdkal1 | Larp4b | | Ptgs1 | 2.23E-15 | 0.2243809 | 0.379 | 0.266 | 6.01E-11 | Cttnbp2nl | Clk1 | | | | | Blnk | | | | Kcnk13 | | | | | Zmym4 |
| | Tnpo1 | 5.24E-13 | 0.17219887 | 0.23 | 0.252 | 1.41E-08 | Hnrnpu | Syk | | Dido1 | 2.77E-15 | 0.27842741 | 0.282 | 0.179 | 7.45E-11 | Pds5a | Dmxl1 | | | | | Kcnip4 | | | | Frmd4a | | | | | Ext1 |
| | Lsamp | 5.53E-13 | 0.37399086 | 0.428 | 0.29 | 1.49E-08 | Hpgd | Itgb5 | | Ankhd1 | 3.24E-15 | 0.00489139 | 0.392 | 0.319 | 8.72E-11 | Itga6 | Asxl1 | | | | | Rbm5 | | | | Gtf2i | | | | | Rbm26 |
| | Tlr7 | 1.04E-10 | -0.3675896 | 0.24 | 0.354 | 2.79E-06 | Plek | Ccnd3 | | Tbxas1 | 3.42E-15 | 0.18142498 | 0.578 | 0.447 | 9.19E-11 | Mvb12b | Mitf | | | | | Tmsb4x | | | | Ankrd17 | | | | | Comt |
| | Wsb1 | 6.91E-13 | 0.26217423 | 0.298 | 0.305 | 1.86E-08 | Herc1 | 1-Mar | | Atp9b | 3.63E-15 | -0.1049135 | 0.325 | 0.276 | 9.77E-11 | Nos1ap | B4galt1 | | | | | Arhgap25 | | | | Rhoq | | | | | Dock7 |
| | Atg16l2 | 8.31E-09 | -0.3669834 | 0.192 | 0.287 | 0.0002236 | Trem2 | Ccnl1 | | Il10ra | 4.05E-15 | 0.04937214 | 0.372 | 0.292 | 1.09E-10 | Cyfip1 | Trip12 | | | | | Zeb2 | | | | Cmip | | | | | Dennd4a |
| | Tet3 | 1.03E-12 | -0.0388188 | 0.236 | 0.301 | 2.76E-08 | Magi1 | Arpc2 | | Gsap | 4.89E-15 | -0.3376746 | 0.275 | 0.283 | 1.32E-10 | Basp1 | Mbd5 | | | | | AU020206 | | | | Atp2c1 | | | | | Atp9b |
| | Snhg11 | 1.07E-12 | 0.49281293 | 0.621 | 0.499 | 2.87E-08 | Atp8a1 | Tjp1 | | Tpr | 5.45E-15 | 0.02144421 | 0.277 | 0.213 | 1.47E-10 | Dnm2 | Zfp292 | | | | | Lhfpl2 | | | | Rabep1 | | | | | Cdk14 |
| | Mlph | 6.90E-11 | -0.3632035 | 0.159 | 0.256 | 1.86E-06 | Ppcdc | Mbd5 | | Atp6v0a1 | 6.03E-15 | 0.11624071 | 0.394 | 0.294 | 1.62E-10 | Mgat4a | Zmynd11 | | | | | Gtdc1 | | | | Bin1 | | | | | Rb1 |
| | Pou2f2 | 1.29E-12 | 0.42410099 | 0.366 | 0.317 | 3.47E-08 | Map3k5 | Vrk2 | | Tnpo1 | 6.49E-15 | -0.0811217 | 0.275 | 0.23 | 1.74E-10 | Stag2 | Srsf5 | | | | | Gm42418 | | | | Ankhd1 | | | | | Kansl1l |
| | Clk1 | 1.44E-12 | -0.0838258 | 0.317 | 0.397 | 3.87E-08 | Cd84 | Sh3pxd2a | | Ogt | 7.83E-15 | 0.10498145 | 0.346 | 0.259 | 2.11E-10 | Tbl1xr1 | Vps13b | | | | | Nsmce2 | | | | Clasp2 | | | | | Mafb |
| | Tpm3 | 1.52E-12 | -0.0524286 | 0.255 | 0.326 | 4.08E-08 | Git2 | Atp6v0a1 | | Brd4 | 8.11E-15 | 0.25162776 | 0.371 | 0.257 | 2.18E-10 | Snrnp70 | Insr | | | | | Mbnl2 | | | | Hpgd | | | | | Sipa1l2 |
| | Myo1f | 1.57E-12 | 0.22660766 | 0.518 | 0.529 | 4.21E-08 | Evl | Herc2 | | Fam13b | 8.71E-15 | -0.00395 | 0.308 | 0.251 | 2.34E-10 | Gm26542 | Zfp407 | | | | | Klf12 | | | | Diaph2 | | | | | Mctp1 |
| | Sfpq | 1.71E-12 | -0.0840961 | 0.31 | 0.396 | 4.60E-08 | Snx13 | Wdfy2 | | Hps4 | 8.72E-15 | 0.19068388 | 0.299 | 0.207 | 2.35E-10 | Ptbp3 | Trim44 | | | | | Tns3 | | | | Prkcb | | | | | Zfand6 |
| | Ash1l | 1.82E-12 | 0.17541883 | 0.424 | 0.448 | 4.89E-08 | Nek7 | Kdm6a | | Gfm2 | 9.41E-15 | 0.25830217 | 0.302 | 0.197 | 2.53E-10 | Med12l | Cacna1d | | | | | Akt3 | | | | Sall1 | | | | | Sipa1 |
| | Rapgef1 | 1.82E-12 | -0.0510235 | 0.222 | 0.287 | 4.89E-08 | Eif4g3 | Elmo1 | | Dhx9 | 9.73E-15 | -0.0783827 | 0.276 | 0.231 | 2.62E-10 | Ncoa3 | Plekhm3 | | | | | Mdm4 | | | | Gpr183 | | | | | Hexb |
| | Ncor1 | 1.83E-12 | 0.18032133 | 0.384 | 0.403 | 4.91E-08 | Spi1 | Mef2a | | Slc29a3 | 9.89E-15 | 0.19678124 | 0.261 | 0.174 | 2.66E-10 | Akap10 | Nktr | | | | | Plek | | | | Sdk1 | | | | | Gm48678 |
| | Capza2 | 2.10E-12 | -0.0371261 | 0.264 | 0.335 | 5.64E-08 | Med13l | Fer | | Kmt2e | 1.05E-14 | -0.059907 | 0.386 | 0.32 | 2.83E-10 | Acvr1 | Abca1 | | | | | Gpatch8 | | | | Myh9 | | | | | Sat1 |
| | Dnajc7 | 2.45E-12 | -0.0253688 | 0.243 | 0.306 | 6.59E-08 | Gtf2i | Snx24 | | Wdr33 | 1.10E-14 | -0.0926086 | 0.283 | 0.237 | 2.95E-10 | Rapgef5 | Sfpq | | | | | Pvt1 | | | | Dgkd | | | | | Il6ra |
| | Pten | 2.01E-11 | -0.3555226 | 0.386 | 0.523 | 5.40E-07 | Sh3kbp1 | Lyn | | Clint1 | 1.18E-14 | -0.0114759 | 0.378 | 0.31 | 3.18E-10 | 8030442B05Ri | Clta | | | | | Csmd1 | | | | Gm35188 | | | | | Stag1 |
| | Nlgn1 | 2.99E-12 | 0.58741908 | 0.264 | 0.177 | 8.05E-08 | Mia2 | 8030442B05Rik | | Smad7 | 1.18E-14 | 0.08338352 | 0.294 | 0.214 | 3.18E-10 | Irak2 | Zfand6 | | | | | Peak1 | | | | Pnn | | | | | Ralgapa1 |
| | Rabgap1 | 3.44E-12 | 0.10299763 | 0.23 | 0.267 | 9.25E-08 | Dmxl1 | Acin1 | | Siglech | 1.18E-14 | 0.36261531 | 0.716 | 0.565 | 3.19E-10 | Sptlc2 | Rasa1 | | | | | Smap2 | | | | Abi3 | | | | | Dnajc1 |
| | Atxn7 | 3.93E-12 | 0.09997865 | 0.252 | 0.291 | 1.06E-07 | Phkb | Hnrnpa3 | | Ap3b1 | 1.47E-14 | -0.033646 | 0.354 | 0.295 | 3.96E-10 | Rsrc2 | Nptn | | | | | Fgfr2 | | | | Cd180 | | | | | Tgfbr1 |
| | Fam13b | 4.03E-12 | 0.11995355 | 0.251 | 0.278 | 1.08E-07 | Arid4b | Tulp4 | | C1qa | 1.89E-14 | -0.3542861 | 0.597 | 0.594 | 5.09E-10 | Apc | Havcr2 | | | | | Asxl1 | | | | Frmd4b | | | | | Thoc2 |
| | Cables1 | 2.52E-10 | -0.3554579 | 0.228 | 0.338 | 6.77E-06 | Mllt10 | Brd4 | | Cnot2 | 2.10E-14 | 0.0016272 | 0.4 | 0.329 | 5.65E-10 | Cmtm7 | Itm2b | | | | | Man2b1 | | | | Fam172a | | | | | Fto |
| | Bin2 | 4.14E-12 | -0.0432972 | 0.294 | 0.371 | 1.12E-07 | Usp15 | Fbxl20 | | Hnrnpa3 | 2.13E-14 | 0.24787669 | 0.271 | 0.175 | 5.74E-10 | Numb | Rbfox1 | | | | | Fam105a | | | | Tnrc6b | | | | | Pde4d |
| | Cacna1d | 5.56E-12 | 0.35277536 | 0.383 | 0.352 | 1.50E-07 | Ncoa2 | Dennd5a | | Taok1 | 2.41E-14 | 0.03198472 | 0.338 | 0.267 | 6.47E-10 | Prex1 | Kpna4 | | | | | Itm2b | | | | Smg6 | | | | | Ifngr2 |
| | Anxa3 | 5.67E-12 | 0.35118416 | 0.394 | 0.368 | 1.52E-07 | Glg1 | Spred1 | | Cerk | 2.51E-14 | 0.05564741 | 0.441 | 0.349 | 6.75E-10 | Itsn2 | Atp6v0b | | | | | Parp8 | | | | Rps6ka1 | | | | | Stx7 |
| | Nr6a1 | 3.57E-10 | -0.3543652 | 0.408 | 0.538 | 9.59E-06 | Pnn | Cradd | | Selplg | 2.68E-14 | 0.28440908 | 0.714 | 0.566 | 7.20E-10 | Akap8l | Rnf169 | | | | | Ash1l | | | | Oxr1 | | | | | Dcakd |
| | Nckap1l | 6.65E-12 | -0.083809 | 0.36 | 0.446 | 1.79E-07 | Asxl1 | Tbxas1 | | Tsc22d4 | 2.74E-14 | 0.23664897 | 0.368 | 0.256 | 7.37E-10 | Scaf8 | Anxa3 | | | | | Plgrkt | | | | Mertk | | | | | Prrc2c |
| | Arid2 | 6.87E-12 | -0.0113911 | 0.196 | 0.252 | 1.85E-07 | Rnf180 | Zfp608 | | Zfhx3 | 2.82E-14 | 0.2888454 | 0.92 | 0.844 | 7.59E-10 | Smad3 | Stx7 | | | | | Rbm25 | | | | Mon2 | | | | | Ccnd3 |
| | Slmap | 7.06E-12 | 0.17962425 | 0.257 | 0.271 | 1.90E-07 | Rapgef6 | Zfp950 | | Fth1 | 2.93E-14 | -0.5643802 | 0.2 | 0.273 | 7.88E-10 | Acap2 | Rbm25 | | | | | Cadm1 | | | | Ptprm | | | | | Psme4 |
| | Lrch3 | 8.17E-12 | 0.08543742 | 0.241 | 0.278 | 2.20E-07 | Rock2 | Zdhhc14 | | Zfand3 | 3.28E-14 | 0.1104961 | 0.493 | 0.392 | 8.82E-10 | Rock2 | Stx16 | | | | | Wipf1 | | | | Grb2 | | | | | Mycbp2 |
| | Pag1 | 2.25E-14 | -0.3449295 | 0.75 | 0.864 | 6.06E-10 | Fkbp15 | Fry | | Cd33 | 3.36E-14 | 0.21872179 | 0.347 | 0.243 | 9.05E-10 | Map3k3 | Ppp1r21 | | | | | Cst3 | | | | Arhgap31 | | | | | Inpp4b |
| | Ankrd17 | 8.85E-12 | 0.08277701 | 0.324 | 0.367 | 2.38E-07 | Vps54 | Gm2245 | | Bin2 | 3.42E-14 | 0.23826945 | 0.411 | 0.294 | 9.20E-10 | Chd7 | Park2 | | | | | Gnas | | | | Spi1 | | | | | Pum1 |
| | Gnb1 | 9.01E-12 | 0.05134414 | 0.228 | 0.274 | 2.42E-07 | Ubl3 | Eef2k | | Pkn1 | 4.06E-14 | 0.09770398 | 0.323 | 0.241 | 1.09E-09 | Dapp1 | Hps3 | | | | | Stard9 | | | | Eml4 | | | | | Btbd1 |
| | Fars2 | 1.25E-11 | 0.08508333 | 0.239 | 0.282 | 3.36E-07 | Cltc | Thoc2 | | Tubgcp5 | 4.16E-14 | 0.27892761 | 0.393 | 0.273 | 1.12E-09 | Ski | Fbxo11 | | | | | Fbxo11 | | | | Ints6l | | | | | Clk1 |
| | Khdrbs1 | 1.28E-11 | 0.11595255 | 0.254 | 0.29 | 3.43E-07 | Prex1 | Ubash3b | | Hnrnpm | 4.19E-14 | -0.0456529 | 0.255 | 0.206 | 1.13E-09 | Actr2 | Mdm4 | | | | | Atp8a1 | | | | Nfat5 | | | | | Zfp292 |
| | Mtmr3 | 1.29E-11 | 0.08136337 | 0.3 | 0.343 | 3.47E-07 | Ube2k | Unc93b1 | | Arsb | 4.36E-14 | 0.47027472 | 0.396 | 0.257 | 1.17E-09 | Rtn1 | Ptk2b | | | | | Celf2 | | | | Scamp2 | | | | | Paxbp1 |
| | Pacsin2 | 1.29E-11 | -0.0365275 | 0.23 | 0.291 | 3.48E-07 | Rin2 | Cd300c2 | | Ubash3b | 4.91E-14 | 0.42788622 | 0.641 | 0.491 | 1.32E-09 | Kif21b | Pum1 | | | | | Skap2 | | | | Mylip | | | | | Mlxip |
| | Irf2 | 1.04E-09 | -0.3380388 | 0.322 | 0.441 | 2.79E-05 | Nrros | Man1a | | Tet3 | 5.26E-14 | 0.15874469 | 0.325 | 0.236 | 1.41E-09 | Cep170 | Papola | | | | | Herc1 | | | | Rgs10 | | | | | Sltm |
| | Cyfip1 | 1.40E-11 | -0.0455365 | 0.445 | 0.534 | 3.77E-07 | Stard9 | Tab2 | | Myo1f | 5.41E-14 | -0.2436109 | 0.522 | 0.518 | 1.46E-09 | Lcp2 | Clip1 | | | | | Sh3glb1 | | | | Rasgrp3 | | | | | Cd86 |
| | Rassf2 | 4.39E-17 | -0.3362004 | 0.213 | 0.34 | 1.18E-12 | Pkig | Cerk | | Trem2 | 5.50E-14 | -0.3116771 | 0.332 | 0.343 | 1.48E-09 | Canx | Lpp | | | | | Camk1 | | | | Pros1 | | | | | Nipbl |
| | Gsap | 1.53E-11 | 0.28755416 | 0.283 | 0.278 | 4.12E-07 | Celf1 | Pla2g4a | | Pitpna | 5.57E-14 | -0.1088557 | 0.28 | 0.245 | 1.50E-09 | Map4 | Taok3 | | | | | Huwe1 | | | | Mknk1 | | | | | Nrd1 |
| | B4galt1 | 1.14E-08 | -0.3335281 | 0.286 | 0.391 | 0.00030548 | Fbxo11 | Dennd1a | | Zfp638 | 5.81E-14 | 0.04580896 | 0.411 | 0.328 | 1.56E-09 | Usp34 | Rbm47 | | | | | Abhd12 | | | | Dock1 | | | | | Rbm47 |
| | Dyrk1a | 1.55E-11 | 0.26402973 | 0.266 | 0.263 | 4.18E-07 | Tbl1xr1 | Ralgapa1 | | Tpm3 | 5.97E-14 | 0.06606211 | 0.331 | 0.255 | 1.61E-09 | Chd9 | Arid4a | | | | | Trip12 | | | | Rcbtb2 | | | | | Fnip2 |
| | Sipa1 | 5.21E-09 | -0.320715 | 0.248 | 0.35 | 0.00014014 | Scamp2 | B430010I23Rik | | Wdfy3 | 6.56E-14 | -0.036007 | 0.518 | 0.45 | 1.76E-09 | Spi1 | Larp4b | | | | | Gria2 | | | | Pnisr | | | | | Usp50 |
| | Lpcat2 | 1.27E-14 | -0.3192098 | 0.513 | 0.67 | 3.41E-10 | Bmpr2 | Golm1 | | Mtmr3 | 7.61E-14 | 0.04422083 | 0.378 | 0.3 | 2.05E-09 | AC149090.1 | AW554918 | | | | | Sik2 | | | | Tanc2 | | | | | Atp6v0b |
| | AC149090.1 | 3.68E-15 | -0.3162183 | 0.535 | 0.69 | 9.91E-11 | Rptor | Vsir | | Cttnbp2nl | 8.15E-14 | 0.1219359 | 0.569 | 0.457 | 2.19E-09 | Mtss1 | Psap | | | | | Lcorl | | | | Prex1 | | | | | Dpy19l4 |
| | Cmtm7 | 1.33E-11 | -0.313688 | 0.195 | 0.296 | 3.57E-07 | Dgkd | Rhobtb1 | | Pds5a | 8.63E-14 | 0.18495808 | 0.325 | 0.231 | 2.32E-09 | Abi3 | Gls | | | | | Fam13b | | | | Slc24a3 | | | | | Pik3r1 |
| | Dmtf1 | 2.08E-11 | -0.034512 | 0.205 | 0.265 | 5.60E-07 | Abi1 | Hps4 | | Tns3 | 8.95E-14 | -0.1156288 | 0.517 | 0.467 | 2.41E-09 | Uvrag | Herc1 | | | | | Tbc1d16 | | | | Snx13 | | | | | Gns |
| | Larp4b | 2.14E-11 | -0.0342744 | 0.243 | 0.301 | 5.77E-07 | Zfr | Cryl1 | | Peli1 | 9.16E-14 | -0.1454268 | 0.313 | 0.28 | 2.47E-09 | Fbxw4 | Ttc14 | | | | | C1qc | | | | Kat6a | | | | | Osbpl9 |
| | Syk | 5.58E-09 | -0.3119715 | 0.217 | 0.313 | 0.00015001 | Rps6ka3 | Ktn1 | | Itga6 | 1.02E-13 | 0.28283612 | 0.436 | 0.309 | 2.73E-09 | Gtf2i | Ctsz | | | | | Sesn1 | | | | Irak2 | | | | | Wdfy3 |
| | Asph | 2.29E-11 | 0.33059955 | 0.331 | 0.312 | 6.16E-07 | Ptpra | Snx5 | | Jak1 | 1.03E-13 | -0.0076463 | 0.324 | 0.261 | 2.78E-09 | Dennd5a | St6gal1 | | | | | Fkbp15 | | | | Retreg1 | | | | | Asah1 |
| | Itgb5 | 2.26E-09 | -0.3060159 | 0.595 | 0.714 | 6.08E-05 | Fcho2 | Pik3r5 | | Wapl | 1.13E-13 | -0.0478059 | 0.35 | 0.296 | 3.04E-09 | Bank1 | Man2b1 | | | | | Ncf1 | | | | Mtmr3 | | | | | Skil |
| | Ccnd3 | 2.41E-11 | -0.0444915 | 0.699 | 0.807 | 6.49E-07 | Ankrd12 | Cul1 | | Mvb12b | 1.16E-13 | 0.16171447 | 0.367 | 0.27 | 3.13E-09 | Gsk3b | Vps54 | | | | | Asxl2 | | | | Ehd4 | | | | | St6gal1 |
| | Laptm5 | 2.58E-11 | 0.11182185 | 0.325 | 0.357 | 6.93E-07 | Sh3glb1 | Itga9 | | Nos1ap | 1.16E-13 | 0.23008277 | 0.359 | 0.25 | 3.13E-09 | Lysmd4 | Capza2 | | | | | Tbl1x | | | | Usp15 | | | | | Fryl |
| | Cdkal1 | 2.79E-11 | 0.11333435 | 0.28 | 0.313 | 7.51E-07 | Irak2 | Ppp1r21 | | Plekho1 | 1.18E-13 | -0.0330454 | 0.446 | 0.386 | 3.17E-09 | Extl3 | Acer3 | | | | | P2rx7 | | | | Snx29 | | | | | Ptk2b |
| | Hnrnpu | 2.89E-11 | 0.18798985 | 0.306 | 0.315 | 7.76E-07 | Acer3 | Mlxip | | Cyfip1 | 1.21E-13 | 0.18973497 | 0.57 | 0.445 | 3.26E-09 | Gnb1 | Bclaf1 | | | | | Ctss | | | | Phc2 | | | | | Prpf4b |
| | 1-Mar | 8.71E-12 | -0.2979314 | 0.483 | 0.618 | 2.34E-07 | Cpsf6 | Taok1 | | Basp1 | 1.26E-13 | 0.09659608 | 0.5 | 0.392 | 3.38E-09 | Znrf2 | Rps6ka3 | | | | | Meg3 | | | | Ikzf1 | | | | | Plekho1 |
| | Hpgd | 3.01E-11 | 0.43299022 | 0.306 | 0.268 | 8.10E-07 | Ptp4a2 | Ccdc93 | | Clk1 | 1.46E-13 | -0.1168962 | 0.35 | 0.317 | 3.92E-09 | Prkcd | Asph | | | | | Gna12 | | | | Gatad2b | | | | | Zfp644 |
| | Ccnl1 | 3.36E-11 | -0.0335532 | 0.214 | 0.27 | 9.04E-07 | Pnpla7 | Cdc42 | | Dnm2 | 1.48E-13 | 0.23369217 | 0.424 | 0.313 | 3.97E-09 | Nfkb1 | Fnip1 | | | | | Rnf111 | | | | Fam193a | | | | | Kmt2c |
| | Plek | 3.49E-11 | 0.31985073 | 0.26 | 0.246 | 9.39E-07 | Pros1 | Galnt1 | | Mgat4a | 1.48E-13 | 0.3210603 | 0.575 | 0.43 | 3.98E-09 | Hk2 | Cd84 | | | | | Fth1 | | | | Btbd9 | | | | | Gm26848 |
| | Arpc2 | 3.62E-11 | -0.0606645 | 0.242 | 0.308 | 9.73E-07 | Vps13b | Pld1 | | Dmxl1 | 1.58E-13 | -0.2047043 | 0.311 | 0.29 | 4.25E-09 | Pak2 | Atrx | | | | | Kdm7a | | | | Fnbp1 | | | | | |
| | Tjp1 | 2.32E-07 | -0.2918709 | 0.296 | 0.392 | 0.00625507 | Zmiz1 | Gng2 | | Stag2 | 1.58E-13 | 0.15144513 | 0.359 | 0.267 | 4.26E-09 | Nuak1 | Ash1l | | | | | Ctsz | | | | Rsf1 | | | | | |
| | Mbd5 | 4.31E-11 | -0.0567468 | 0.327 | 0.403 | 1.16E-06 | Map4k3 | Ppp2r5c | | Asxl1 | 1.65E-13 | -0.0286356 | 0.335 | 0.273 | 4.44E-09 | Rap1a | Mlxip | | | | | Asph | | | | Igf1r | | | | | |
| | Herc1 | 4.50E-11 | 0.19735956 | 0.319 | 0.325 | 1.21E-06 | Trip12 | Cbl | | Mitf | 1.82E-13 | -0.273822 | 0.456 | 0.459 | 4.89E-09 | Exoc6b | Setd2 | | | | | Setd2 | | | | Eif4g3 | | | | | |
| | Vrk2 | 1.92E-09 | -0.2897402 | 0.288 | 0.393 | 5.16E-05 | Mylip | Rasa1 | | Tbl1xr1 | 1.88E-13 | 0.077305 | 0.407 | 0.318 | 5.06E-09 | Git2 | Thoc2 | | | | | Apbb2 | | | | Cacnb2 | | | | | |
| | Sh3pxd2a | 4.74E-11 | -0.0660737 | 0.254 | 0.326 | 1.28E-06 | Lhfpl2 | Zfhx3 | | Snrnp70 | 1.94E-13 | 0.16356251 | 0.571 | 0.447 | 5.22E-09 | Nf1 | Ctsh | | | | | R3hdm1 | | | | Sirpa | | | | | |
| | Trem2 | 5.01E-11 | 0.28089164 | 0.343 | 0.322 | 1.35E-06 | Mknk1 | Washc4 | | Gm26542 | 2.12E-13 | 0.38275618 | 0.475 | 0.335 | 5.70E-09 | Kdm3b | Tbc1d16 | | | | | Arid4a | | | | Cux1 | | | | | |
| | Magi1 | 5.12E-11 | 0.31990899 | 0.484 | 0.461 | 1.38E-06 | Smap2 | Sdccag8 | | Ptbp3 | 2.27E-13 | 0.05478694 | 0.422 | 0.339 | 6.11E-09 | Cdk19 | Hnrnpu | | | | | Ank2 | | | | Slmap | | | | | |
| | Atp6v0a1 | 5.14E-11 | -0.0490324 | 0.294 | 0.369 | 1.38E-06 | Tbc1d16 | Akap8l | | B4galt1 | 2.29E-13 | -0.111172 | 0.322 | 0.286 | 6.16E-09 | Rabgap1 | Asah1 | | | | | Pou2f2 | | | | Ncor1 | | | | | |
| | Atp8a1 | 5.48E-11 | 0.07085064 | 0.471 | 0.527 | 1.48E-06 | Chd6 | Rnf169 | | Med12l | 2.33E-13 | 0.2599231 | 0.559 | 0.426 | 6.27E-09 | Rnpep | P3h2 | | | | | Cd81 | | | | Lrch1 | | | | | |
| | Herc2 | 7.42E-09 | -0.2862093 | 0.389 | 0.506 | 0.00019956 | Zmynd11 | Dip2b | | Ncoa3 | 2.43E-13 | 0.2126069 | 0.383 | 0.279 | 6.53E-09 | Ppt1 | Tbl1x | | | | | Cacna1a | | | | Fcho2 | | | | | |
| | Ppcdc | 6.99E-11 | 0.02513938 | 0.323 | 0.374 | 1.88E-06 | Rbfox1 | Lcor | | Akap10 | 2.72E-13 | 0.06928677 | 0.33 | 0.255 | 7.33E-09 | Dpysl2 | Fars2 | | | | | Rnf180 | | | | Maml2 | | | | | |
| | Wdfy2 | 1.13E-12 | -0.2809601 | 0.294 | 0.416 | 3.03E-08 | Ddx17 | Pard3b | | Acvr1 | 2.90E-13 | 0.10617066 | 0.258 | 0.187 | 7.80E-09 | Arhgap30 | Smap2 | | | | | Lamp2 | | | | Rptor | | | | | |
| | Map3k5 | 7.38E-11 | 0.02460644 | 0.301 | 0.356 | 1.98E-06 | Gnas | Pum2 | | Rapgef5 | 3.16E-13 | 0.34632202 | 0.758 | 0.631 | 8.51E-09 | Nr6a1 | Lnpep | | | | | Nsd3 | | | | Map4k3 | | | | | |
| | Kdm6a | 7.67E-11 | -0.0804813 | 0.2 | 0.261 | 2.06E-06 | Wdfy4 | Cd86 | | Trip12 | 3.27E-13 | -0.1392706 | 0.353 | 0.323 | 8.79E-09 | Lrch3 | Ppp1r12a | | | | | Fars2 | | | | Basp1 | | | | | |
| | Cd84 | 7.72E-11 | 0.20195965 | 0.386 | 0.398 | 2.08E-06 | Rnf111 | Fam168a | | Mbd5 | 3.31E-13 | -0.115848 | 0.371 | 0.327 | 8.90E-09 | Gna15 | R3hdm1 | | | | | Laptm5 | | | | Arhgap39 | | | | | |
| | Elmo1 | 4.09E-16 | -0.2802598 | 0.91 | 0.954 | 1.10E-11 | Fam105a | Rab31 | | 8030442B05 | 3.32E-13 | 0.2568975 | 0.495 | 0.365 | 8.92E-09 | Dip2b | Rapgef6 | | | | | Cadm2 | | | | F11r | | | | | |
| | Git2 | 7.85E-11 | 0.02211861 | 0.388 | 0.44 | 2.11E-06 | Kmt2e | Col27a1 | | Irak2 | 3.32E-13 | 0.04984299 | 0.434 | 0.349 | 8.94E-09 | Picalm | Srek1 | | | | | Zfp407 | | | | Chn2 | | | | | |
| | Mef2a | 1.70E-11 | -0.2783916 | 0.866 | 0.916 | 4.58E-07 | Srsf5 | P2ry6 | | Sptlc2 | 3.45E-13 | 0.16463005 | 0.26 | 0.178 | 9.28E-09 | Khdrbs1 | Cnot4 | | | | | Myo1f | | | | Khdrbs1 | | | | | |
| | Evl | 9.38E-11 | 0.16833768 | 0.319 | 0.333 | 2.52E-06 | Lrp6 | Med13 | | Rsrc2 | 3.60E-13 | 0.0452214 | 0.298 | 0.236 | 9.68E-09 | Adap2os | Klf3 | | | | | Wsb1 | | | | Rabgap1 | | | | | |
| | Snx13 | 9.80E-11 | 0.01778305 | 0.279 | 0.336 | 2.64E-06 | Snd1 | Epc1 | | Apc | 3.63E-13 | 0.12220087 | 0.417 | 0.319 | 9.77E-09 | Col27a1 | Gnas | | | | | Mitf | | | | Rb1cc1 | | | | | |
| | Nek7 | 1.02E-10 | 0.17047733 | 0.409 | 0.426 | 2.75E-06 | Hps3 | Ophn1 | | Cmtm7 | 3.69E-13 | 0.39764826 | 0.308 | 0.195 | 9.94E-09 | Rab14 | Akap13 | | | | | Xist | | | | Cttnbp2nl | | | | | |
| | Fer | 2.52E-08 | -0.2709529 | 0.212 | 0.297 | 0.00067756 | Gria2 | Adap2os | | Numb | 4.18E-13 | 0.32142325 | 0.821 | 0.717 | 1.13E-08 | Pik3ap1 | Nipbl | | | | | Ube3a | | | | Nrros | | | | | |
| | Snx24 | 4.34E-10 | -0.2681286 | 0.331 | 0.444 | 1.17E-05 | Cd81 | Fndc3b | | Prex1 | 4.30E-13 | 0.1903034 | 0.468 | 0.356 | 1.16E-08 | Abcc5 | Stard9 | | | | | Osbpl11 | | | | Cfh | | | | | |
| | Lyn | 1.30E-09 | -0.2641459 | 0.73 | 0.824 | 3.51E-05 | Kat6a | Sf3b1 | | Itsn2 | 4.39E-13 | 0.13225845 | 0.375 | 0.285 | 1.18E-08 | Sema4d | Sh3glb1 | | | | | Gm26740 | | | | Fam102b | | | | | |
| | Eif4g3 | 1.09E-10 | 0.10086585 | 0.382 | 0.428 | 2.93E-06 | Spop | Ski | | Akap8l | 4.54E-13 | 0.16673818 | 0.421 | 0.316 | 1.22E-08 | Usp24 | Pid1 | | | | | Nrxn1 | | | | Pias1 | | | | | |
| | Spi1 | 1.11E-10 | 0.31722952 | 0.296 | 0.271 | 2.98E-06 | Akap13 | Rab8b | | Scaf8 | 4.68E-13 | 0.12574594 | 0.273 | 0.197 | 1.26E-08 | Fam193a | Mbtd1 | | | | | Fhit | | | | Gdi2 | | | | | |
| | Med13l | 1.21E-10 | 0.07595734 | 0.307 | 0.348 | 3.27E-06 | Senp2 | Plekho1 | | Zfp292 | 4.82E-13 | -0.0954089 | 0.342 | 0.301 | 1.30E-08 | Gm17231 | Kmt2c | | | | | Spop | | | | Mia2 | | | | | |
| | Gtf2i | 1.28E-10 | 0.02492735 | 0.241 | 0.292 | 3.43E-06 | Abhd12 | Zfc3h1 | | Zmynd11 | 4.98E-13 | -0.0186448 | 0.316 | 0.259 | 1.34E-08 | Slc7a8 | Cog5 | | | | | Nptn | | | | Gapvd1 | | | | | |
| | 8030442B05 | 1.08E-09 | -0.2632615 | 0.365 | 0.482 | 2.91E-05 | Osbpl11 | Lcp2 | | Smad3 | 5.04E-13 | 0.25194908 | 0.29 | 0.193 | 1.36E-08 | Tm6sf1 | Lcorl | | | | | Cmss1 | | | | Senp2 | | | | | |
| | Sh3kbp1 | 1.44E-10 | 0.27533412 | 0.498 | 0.474 | 3.87E-06 | Clta | Itpr2 | | Acap2 | 5.06E-13 | 0.16851071 | 0.513 | 0.398 | 1.36E-08 | Atf7ip | Ptp4a2 | | | | | Snhg11 | | | | Wdr44 | | | | | |
| | Acin1 | 1.47E-10 | -0.0345389 | 0.246 | 0.307 | 3.94E-06 | Rbm5 | Mysm1 | | Rock2 | 5.60E-13 | 0.03595972 | 0.508 | 0.423 | 1.51E-08 | Tab2 | P2rx7 | | | | | Ppp3ca | | | | Nlrp1b | | | | | |
| | Hnrnpa3 | 6.26E-13 | -0.2611525 | 0.175 | 0.271 | 1.68E-08 | Fgd2 | Rab14 | | Srsf5 | 5.72E-13 | -0.0104278 | 0.358 | 0.292 | 1.54E-08 | Whrn | Ctsb | | | | | Arhgap17 | | | | Abi1 | | | | | |
| | Tulp4 | 3.37E-07 | -0.2607071 | 0.202 | 0.281 | 0.00905792 | Lcorl | Ntpcr | | Vps13b | 6.03E-13 | -0.2003091 | 0.367 | 0.353 | 1.62E-08 | Wdfy4 | Zfp950 | | | | | Gm26804 | | | | Phactr2 | | | | | |
| | Mia2 | 1.58E-10 | 0.01840267 | 0.21 | 0.254 | 4.25E-06 | Sirpa | Zfp638 | | Map3k3 | 6.20E-13 | 0.05745348 | 0.442 | 0.354 | 1.67E-08 | Lrrk1 | Rnf111 | | | | | Srrm2 | | | | Son | | | | | |
| | Dmxl1 | 1.73E-10 | 0.12249965 | 0.29 | 0.319 | 4.67E-06 | Srek1 | Havcr2 | | Chd7 | 6.43E-13 | 0.12177571 | 0.449 | 0.347 | 1.73E-08 | Gdi2 | Sltm | | | | | Nkain2 | | | | Tnrc18 | | | | | |
| | Phkb | 1.74E-10 | 0.03894674 | 0.255 | 0.299 | 4.69E-06 | Sik2 | Cd2ap | | Dapp1 | 6.45E-13 | 0.14271596 | 0.418 | 0.319 | 1.74E-08 | Gtf2h2 | Huwe1 | | | | | Pitpna | | | | Ubn2 | | | | | |
| | Brd4 | 1.83E-10 | -0.0142734 | 0.257 | 0.31 | 4.93E-06 | Man1a2 | Cflar | | Ski | 7.12E-13 | 0.18926142 | 0.426 | 0.322 | 1.92E-08 | Man1c1 | Apbb2 | | | | | Dyrk1a | | | | Fcer1g | | | | | |
| | Fbxl20 | 1.85E-10 | -0.0263844 | 0.26 | 0.317 | 4.97E-06 | Spag9 | Itgav | | Insr | 7.73E-13 | -0.2299462 | 0.254 | 0.245 | 2.08E-08 | Csnk1g1 | Rbm5 | | | | | Akap13 | | | | Lair1 | | | | | |
| | Dennd5a | 1.87E-10 | -0.059354 | 0.217 | 0.278 | 5.04E-06 | Thada | Adam10 | | Actr2 | 8.09E-13 | 0.03439882 | 0.405 | 0.33 | 2.18E-08 | Cdkal1 | Gtdc1 | | | | | Sbf2 | | | | Arhgap5 | | | | | |
| | Spred1 | 1.90E-10 | -0.0798556 | 0.196 | 0.258 | 5.12E-06 | Cdk13 | Ifngr1 | | Rtn1 | 8.09E-13 | 0.2551402 | 0.438 | 0.321 | 2.18E-08 | Atxn7l1 | Fyb | | | | | Kmt2e | | | | Sfmbt1 | | | | | |
| | Cradd | 2.52E-09 | -0.2604608 | 0.207 | 0.299 | 6.77E-05 | Pik3cd | Rnf216 | | Kif21b | 8.18E-13 | 0.10849202 | 0.275 | 0.203 | 2.20E-08 | Snd1 | Ctss | | | | | Lsamp | | | | Lims1 | | | | | |
| | Arid4b | 2.12E-10 | 0.17953868 | 0.246 | 0.26 | 5.70E-06 | Ctsz | Rtn1 | | Zfp407 | 8.27E-13 | -0.1596297 | 0.267 | 0.239 | 2.23E-08 | Rcsd1 | Rbm26 | | | | | Glg1 | | | | Ncoa2 | | | | | |
| | Tbxas1 | 9.28E-08 | -0.2563087 | 0.447 | 0.559 | 0.00249781 | Man1c1 | Stambpl1 | | Cep170 | 8.71E-13 | 0.1171047 | 0.332 | 0.25 | 2.34E-08 | Grb2 | Oxct1 | | | | | Atrx | | | | Fgd2 | | | | | |
| | Zfp608 | 2.20E-10 | -0.0140108 | 0.21 | 0.261 | 5.91E-06 | Oxct1 | Trappc9 | | Trim44 | 9.33E-13 | -0.0774086 | 0.308 | 0.266 | 2.51E-08 | Ppcdc | Stag1 | | | | | Zfr | | | | Ankrd11 | | | | | |
| | Zfp950 | 2.23E-10 | -0.0412554 | 0.228 | 0.289 | 6.00E-06 | Rsrc1 | Tbc1d23 | | Lcp2 | 1.01E-12 | 0.18736444 | 0.34 | 0.247 | 2.72E-08 | Ube2k | Bach2 | | | | | Rps6ka3 | | | | Jmjd1c | | | | | |
| | Zdhhc14 | 2.47E-09 | -0.2562924 | 0.275 | 0.375 | 6.66E-05 | Tra2a | Camk2d | | Cacna1d | 1.10E-12 | -0.2898658 | 0.375 | 0.383 | 2.96E-08 | Clasp2 | Trappc8 | | | | | Rab2a | | | | Numb | | | | | |
| | Mllt10 | 2.39E-10 | 0.09918728 | 0.373 | 0.407 | 6.43E-06 | Myh9 | Selplg | | Plekhm3 | 1.24E-12 | -0.0661844 | 0.279 | 0.236 | 3.34E-08 | Tbc1d9 | Map3k5 | | | | | Herc4 | | | | P2ry12 | | | | | |
| | Fry | 4.58E-07 | -0.2546705 | 0.205 | 0.287 | 0.01232512 | Phf20l1 | Ssh2 | | Nktr | 1.31E-12 | -0.1743337 | 0.312 | 0.293 | 3.53E-08 | Fam117b | Bmpr2 | | | | | Zcchc11 | | | | Wdfy4 | | | | | |
| | Gm2245 | 2.90E-11 | -0.2544962 | 0.35 | 0.468 | 7.79E-07 | Scaper | St3gal5 | | Canx | 1.32E-12 | 0.0523964 | 0.283 | 0.217 | 3.55E-08 | Tnrc18 | Heatr5a | | | | | Gm20388 | | | | Ctsc | | | | | |
| | Usp15 | 2.59E-10 | 0.04420995 | 0.219 | 0.26 | 6.98E-06 | Mbnl2 | Zmym4 | | Abca1 | 1.34E-12 | -0.2513724 | 0.365 | 0.367 | 3.62E-08 | Elf1 | Gria2 | | | | | Man1a2 | | | | Nf1 | | | | | |
| | Ncoa2 | 2.65E-10 | 0.08393805 | 0.321 | 0.359 | 7.12E-06 | Srsf11 | Dhx9 | | Map4 | 1.38E-12 | 0.13524751 | 0.469 | 0.364 | 3.71E-08 | Vti1a | Tcf4 | | | | | Tbc1d22a | | | | Il10ra | | | | | |
| | Eef2k | 1.07E-10 | -0.2532548 | 0.231 | 0.327 | 2.88E-06 | Gnaq | Mbnl1 | | Sfpq | 1.44E-12 | -0.1036946 | 0.349 | 0.31 | 3.86E-08 | Ankrd17 | Paxbp1 | | | | | Ptp4a2 | | | | Cpsf6 | | | | | |
| | Thoc2 | 2.80E-10 | -0.0121526 | 0.228 | 0.28 | 7.52E-06 | Abca1 | Kmt2c | | Clta | 1.46E-12 | -0.0020611 | 0.317 | 0.261 | 3.93E-08 | Scaper | Kansl1l | | | | | Ttc14 | | | | Epb41l2 | | | | | |
| | Glg1 | 2.81E-10 | 0.1148743 | 0.272 | 0.304 | 7.55E-06 | Luc7l2 | Ptk2b | | Usp34 | 1.49E-12 | 0.02507689 | 0.452 | 0.377 | 4.01E-08 | Fer | Ifngr2 | | | | | Rbfox1 | | | | Ambra1 | | | | | |
| | Pnn | 3.20E-10 | 0.04384976 | 0.238 | 0.28 | 8.60E-06 | Clasp2 | Inpp5d | | Chd9 | 1.51E-12 | 0.26082243 | 0.876 | 0.773 | 4.07E-08 | Nrf1 | Ralgapa1 | | | | | Arhgap22 | | | | Thada | | | | | |
| | Asxl1 | 3.24E-10 | 0.23321893 | 0.273 | 0.277 | 8.71E-06 | Pitpna | Tacc1 | | Spi1 | 1.57E-12 | 0.05722661 | 0.371 | 0.296 | 4.23E-08 | Wdr44 | Top1 | | | | | Oxct1 | | | | Ube2h | | | | | |
| | Ubash3b | 6.71E-07 | -0.2522715 | 0.491 | 0.597 | 0.01804146 | Scmh1 | Larp1 | | AC149090.1 | 1.69E-12 | 0.2307577 | 0.668 | 0.535 | 4.54E-08 | Bin1 | Sbf2 | | | | | Bclaf1 | | | | Hook3 | | | | | |
| | Rnf180 | 3.28E-10 | 0.15592545 | 0.413 | 0.44 | 8.82E-06 | Pak2 | Tra2b | | Mtss1 | 1.90E-12 | 0.19643779 | 0.267 | 0.182 | 5.11E-08 | Ewsr1 | Phf20l1 | | | | | Abca1 | | | | Tmcc3 | | | | | |
| | Rapgef6 | 3.56E-10 | 0.24844518 | 0.3 | 0.3 | 9.59E-06 | Erbin | Abca9 | | Abi3 | 2.28E-12 | 0.15242366 | 0.382 | 0.291 | 6.13E-08 | Stambpl1 | Cltc | | | | | Rtn3 | | | | Eml6 | | | | | |
| | Unc93b1 | 2.37E-11 | -0.2505901 | 0.533 | 0.665 | 6.38E-07 | Tnrc18 | Usp24 | | Zfand6 | 2.28E-12 | -0.2429035 | 0.26 | 0.26 | 6.14E-08 | Nedd4l | Wsb1 | | | | | Arid1b | | | | Apbb1ip | | | | | |
| | Cd300c2 | 1.59E-06 | -0.2505535 | 0.196 | 0.268 | 0.04272167 | Zfp407 | Myo9a | | Uvrag | 2.39E-12 | 0.16150166 | 0.487 | 0.382 | 6.43E-08 | Chd6 | Akt3 | | | | | Phf20l1 | | | | Lrp6 | | | | | |
| | Man1a | 5.94E-09 | -0.2469156 | 0.375 | 0.486 | 0.00015979 | Kdm2a | Dnajc13 | | Rasa1 | 2.57E-12 | -0.0560387 | 0.303 | 0.251 | 6.91E-08 | Ndfip1 | Ccnd3 | | | | | Srsf5 | | | | Gnb1 | | | | | |
| | Tab2 | 5.72E-10 | -0.245749 | 0.231 | 0.326 | 1.54E-05 | Nf1 | Tbc1d14 | | Fbxw4 | 2.59E-12 | 0.30358771 | 0.269 | 0.175 | 6.97E-08 | Syk | Sipa1l2 | | | | | Exoc4 | | | | Pik3cd | | | | | |
| | Cerk | 4.39E-10 | -0.0747172 | 0.349 | 0.432 | 1.18E-05 | Marcks | Lrp1 | | Gtf2i | 2.81E-12 | 0.05383421 | 0.31 | 0.241 | 7.56E-08 | Myh9 | Rtn3 | | | | | Taok3 | | | | Pak2 | | | | | |
| | Rock2 | 4.57E-10 | 0.00480814 | 0.423 | 0.487 | 1.23E-05 | Pwwp2a | Vav1 | | Dennd5a | 2.82E-12 | 0.10342562 | 0.293 | 0.217 | 7.59E-08 | Plcl2 | Camk1 | | | | | Tenm2 | | | | Lrrk1 | | | | | |
| | Pla2g4a | 4.61E-10 | -0.0921718 | 0.221 | 0.288 | 1.24E-05 | Frrs1 | Nos1ap | | Nptn | 2.90E-12 | -0.0023964 | 0.392 | 0.322 | 7.80E-08 | Washc4 | Comt | | | | | Wapl | | | | Dnmt3a | | | | | |
| | Fkbp15 | 4.66E-10 | 0.0845376 | 0.226 | 0.251 | 1.25E-05 | Phc2 | Soga1 | | Bank1 | 2.93E-12 | 0.32447177 | 0.342 | 0.227 | 7.89E-08 | Nav2 | Ptprc | | | | | Ddx17 | | | | Itsn2 | | | | | |
| | Vps54 | 4.87E-10 | 0.00833766 | 0.236 | 0.284 | 1.31E-05 | Arid1a | Chd7 | | Gsk3b | 3.02E-12 | 0.02138873 | 0.352 | 0.287 | 8.13E-08 | Eif4g3 | Btbd1 | | | | | Cd9 | | | | Lrch3 | | | | | |
| | Dennd1a | 5.04E-10 | -0.0597211 | 0.383 | 0.466 | 1.36E-05 | Rhoq | Mgat5 | | Lysmd4 | 3.24E-12 | 0.23434179 | 0.819 | 0.697 | 8.71E-08 | Agps | Nr3c1 | | | | | Hnrnpu | | | | Rsrc1 | | | | | |
| | Ubl3 | 5.10E-10 | 0.05889922 | 0.217 | 0.255 | 1.37E-05 | Ppp1r12a | Syngr1 | | Extl3 | 3.37E-12 | 0.09573126 | 0.419 | 0.33 | 9.06E-08 | Wdr26 | Ube3a | | | | | Gm26733 | | | | Cdk13 | | | | | |
| | Ralgapa1 | 5.32E-10 | -0.043803 | 0.23 | 0.292 | 1.43E-05 | Fam193a | Zfp644 | | Havcr2 | 3.61E-12 | -0.1125793 | 0.297 | 0.27 | 9.72E-08 | Disc1 | Spop | | | | | Phkb | | | | Arid1a | | | | | |
| | B430010I23R | 3.70E-07 | -0.2440041 | 0.222 | 0.297 | 0.00994835 | Fam172a | Cask | | Gnb1 | 3.64E-12 | 0.21649733 | 0.325 | 0.228 | 9.81E-08 | Pcm1 | Stk3 | | | | | Tgfbr2 | | | | | | | | | |
| | Golm1 | 1.25E-06 | -0.2423233 | 0.288 | 0.372 | 0.03364358 | Apc | Sipa1l2 | | Itm2b | 3.66E-12 | -0.3090265 | 0.514 | 0.529 | 9.85E-08 | Celf1 | Rnf180 | | | | | Gab2 | | | | | | | | | |
| | Cltc | 5.85E-10 | 0.05109323 | 0.315 | 0.351 | 1.57E-05 | Ints6l | Jak1 | | Rbfox1 | 3.83E-12 | -0.549678 | 0.282 | 0.343 | 1.03E-07 | Snx24 | Fnip2 | | | | | Fgf13 | | | | | | | | | |
| | Prex1 | 6.12E-10 | 0.01008651 | 0.356 | 0.412 | 1.65E-05 | Plgrkt | Prpf4b | | Kpna4 | 3.95E-12 | -0.0849857 | 0.342 | 0.297 | 1.06E-07 | Rnf13 | Arid1b | | | | | Plekhm3 | | | | | | | | | |
| | Vsir | 4.28E-07 | -0.2394115 | 0.391 | 0.488 | 0.01152815 | Huwe1 | Usp9x | | Znrf2 | 4.02E-12 | 0.07075983 | 0.275 | 0.21 | 1.08E-07 | Ralgps1 | Arhgap22 | | | | | Srsf11 | | | | | | | | | |
| | Ube2k | 6.17E-10 | 0.0069152 | 0.219 | 0.266 | 1.66E-05 | Ctss | Gm26542 | | Atp6v0b | 4.26E-12 | -0.2166306 | 0.498 | 0.489 | 1.15E-07 | Zdhhc14 | Wipf1 | | | | | Arglu1 | | | | | | | | | |
| | Rin2 | 6.26E-10 | 0.12088022 | 0.276 | 0.304 | 1.69E-05 | Grb2 | Ankfy1 | | Rnf169 | 4.29E-12 | -0.1679023 | 0.438 | 0.414 | 1.15E-07 | Fermt3 | Atp8a1 | | | | | Ttc3 | | | | | | | | | |
| | Rhobtb1 | 9.74E-10 | -0.2386108 | 0.284 | 0.382 | 2.62E-05 | Ubr3 | Gfm2 | | Anxa3 | 4.30E-12 | -0.031637 | 0.461 | 0.394 | 1.16E-07 | Adam10 | Klf12 | | | | | Dlg2 | | | | | | | | | |
| | Hps4 | 2.28E-13 | -0.2374133 | 0.207 | 0.306 | 6.14E-09 | Gphn | Ythdf3 | | Stx7 | 4.32E-12 | -0.0146842 | 0.257 | 0.208 | 1.16E-07 | Adam17 | Mbnl2 | | | | | Ptprd | | | | | | | | | |
| | Cryl1 | 7.77E-10 | -0.07301 | 0.212 | 0.27 | 2.09E-05 | Ankhd1 | Med12l | | Prkcd | 4.36E-12 | 0.19674199 | 0.339 | 0.247 | 1.17E-07 | Spred1 | Med13l | | | | | Psap | | | | | | | | | |
| | Ktn1 | 6.44E-12 | -0.2372553 | 0.209 | 0.304 | 1.73E-07 | Wipf1 | Btbd1 | | Nfkb1 | 4.39E-12 | 0.06428086 | 0.412 | 0.331 | 1.18E-07 | Soga1 | Osbpl8 | | | | | Wwox | | | | | | | | | |
| | Snx5 | 8.13E-10 | -0.0341259 | 0.279 | 0.334 | 2.19E-05 | Fndc3a | Xrn1 | | Rbm25 | 4.40E-12 | -0.1349085 | 0.419 | 0.384 | 1.18E-07 | Mapk14 | Magi1 | | | | | Zmynd11 | | | | | | | | | |
| | Pik3r5 | 8.84E-11 | -0.2346519 | 0.189 | 0.278 | 2.38E-06 | Fnbp1 | Atad2b | | Hk2 | 4.56E-12 | 0.09756906 | 0.294 | 0.228 | 1.23E-07 | Rnf130 | Gpatch8 | | | | | Pid1 | | | | | | | | | |
| | Nrros | 8.60E-10 | 0.03761253 | 0.334 | 0.381 | 2.31E-05 | Fgf13 | Mvb12b | | Stx16 | 4.79E-12 | -0.0640879 | 0.293 | 0.25 | 1.29E-07 | Tgfb1 | Lrmda | | | | | Tcf12 | | | | | | | | | |
| | Cul1 | 9.12E-10 | -0.009738 | 0.286 | 0.343 | 2.45E-05 | Gatad2b | Spire1 | | Ppp1r21 | 4.81E-12 | -0.1159379 | 0.302 | 0.269 | 1.29E-07 | Herc2 | Kdm2a | | | | | Slc38a9 | | | | | | | | | |
| | Itga9 | 1.04E-08 | -0.230548 | 0.288 | 0.383 | 0.00027923 | Ambra1 | Zscan26 | | Park2 | 4.95E-12 | -0.3843037 | 0.423 | 0.564 | 1.33E-07 | Anks1 | Man1a2 | | | | | Nktr | | | | | | | | | |
| | Stard9 | 9.35E-10 | 0.12466198 | 0.364 | 0.391 | 2.52E-05 | Basp1 | Sft2d1 | | Pak2 | 5.05E-12 | 0.02813739 | 0.378 | 0.306 | 1.36E-07 | Fchsd2 | Ddx17 | | | | | Kcnma1 | | | | | | | | | |
| | Pkig | 9.67E-10 | 0.14458006 | 0.282 | 0.308 | 2.60E-05 | Ehd4 | Disc1 | | Nuak1 | 5.14E-12 | 0.28510878 | 0.339 | 0.231 | 1.38E-07 | Cul1 | Smg1 | | | | | Kpna4 | | | | | | | | | |
| | Ppp1r21 | 1.32E-09 | -0.2276897 | 0.269 | 0.362 | 3.56E-05 | Zeb2 | Wdfy3 | | Hps3 | 5.25E-12 | -0.0423583 | 0.308 | 0.257 | 1.41E-07 | Zmiz1 | Atp2b1 | | | | | P3h2 | | | | | | | | | |
| | Mlxip | 3.73E-09 | -0.2273293 | 0.253 | 0.344 | 0.00010036 | Nfat5 | Phf14 | | Fbxo11 | 5.45E-12 | -0.1540813 | 0.328 | 0.297 | 1.47E-07 | Vrk2 | Abhd12 | | | | | Olfr56 | | | | | | | | | |
| | Celf1 | 1.07E-09 | 0.17093852 | 0.354 | 0.371 | 2.87E-05 | Ddx6 | Abl1 | | Rap1a | 5.52E-12 | 0.06966584 | 0.409 | 0.331 | 1.49E-07 | Arid1a | Nrp1 | | | | | Cltc | | | | | | | | | |
| | Taok1 | 1.07E-09 | -0.0179877 | 0.267 | 0.319 | 2.89E-05 | Asxl2 | Zfp652 | | Exoc6b | 5.55E-12 | 0.12160374 | 0.416 | 0.322 | 1.49E-07 | Ube3c | Pde4d | | | | | Lcp1 | | | | | | | | | |
| | Ccdc93 | 1.54E-11 | -0.225121 | 0.165 | 0.25 | 4.14E-07 | Arhgap39 | Ptgs1 | | Mdm4 | 5.63E-12 | -0.0320593 | 0.301 | 0.251 | 1.51E-07 | Ncor1 | Mark3 | | | | | Smg1 | | | | | | | | | |
| | Cdc42 | 1.11E-09 | -0.0316308 | 0.346 | 0.412 | 2.99E-05 | Atp2b1 | Trim44 | | Ptk2b | 5.79E-12 | -0.0297556 | 0.292 | 0.243 | 1.56E-07 | Cryl1 | Gphn | | | | | Arid4b | | | | | | | | | |
| | Fbxo11 | 1.19E-09 | 0.16022631 | 0.297 | 0.319 | 3.20E-05 | Cog5 | Cdk19 | | Git2 | 5.96E-12 | 0.02945348 | 0.457 | 0.388 | 1.60E-07 | Acss1 | Zfp148 | | | | | Gsap | | | | | | | | | |
| | Tbl1xr1 | 1.26E-09 | 0.00894163 | 0.318 | 0.375 | 3.40E-05 | Lrrk1 | Morc3 | | Nf1 | 6.01E-12 | 0.06236009 | 0.519 | 0.429 | 1.62E-07 | Cdc42 | Phip | | | | | Ptprc | | | | | | | | | |
| | Scamp2 | 1.30E-09 | 0.09968392 | 0.263 | 0.289 | 3.50E-05 | Retreg1 | Hectd1 | | Kdm3b | 6.12E-12 | 0.04929818 | 0.362 | 0.294 | 1.65E-07 | Pacsin2 | Sik2 | | | | | Evl | | | | | | | | | |
| | Galnt1 | 3.55E-08 | -0.2239173 | 0.214 | 0.294 | 0.00095536 | Cst3 | Luc7l3 | | Pum1 | 6.13E-12 | -0.0206331 | 0.275 | 0.233 | 1.65E-07 | Spire1 | Usp50 | | | | | Apoe | | | | | | | | | |
| | Pld1 | 4.07E-11 | -0.2218394 | 0.358 | 0.474 | 1.09E-06 | Mitf | Aftph | | Papola | 6.31E-12 | -0.0683046 | 0.327 | 0.282 | 1.70E-07 | Arih1 | Creb1 | | | | | Anxa3 | | | | | | | | | |
| | Gng2 | 6.95E-08 | -0.2217517 | 0.307 | 0.402 | 0.00186987 | Rb1cc1 | Gtf2h2 | | Cdk19 | 6.32E-12 | 0.16378113 | 0.36 | 0.27 | 1.70E-07 | Stab1 | Pik3r1 | | | | | Fmnl2 | | | | | | | | | |
| | Ppp2r5c | 1.47E-09 | -0.0006606 | 0.206 | 0.253 | 3.94E-05 | Rab2a | Sparc | | Rabgap1 | 6.46E-12 | 0.02058805 | 0.291 | 0.23 | 1.74E-07 | Cflar | Ep300 | | | | | Ctsb | | | | | | | | | |
| | Bmpr2 | 1.54E-09 | 0.13731842 | 0.241 | 0.263 | 4.15E-05 | Lamp2 | Pibf1 | | Rnpep | 6.67E-12 | 0.08180066 | 0.257 | 0.191 | 1.80E-07 | Lpcat2 | Zcchc11 | | | | | Pkig | | | | | | | | | |
| | Rptor | 1.55E-09 | 0.05186397 | 0.436 | 0.485 | 4.16E-05 | Smg1 | Kdm3b | | Ppt1 | 6.77E-12 | 0.05894478 | 0.263 | 0.201 | 1.82E-07 | Ccr5 | Tnrc6c | | | | | Pnpla7 | | | | | | | | | |
| | Cbl | 1.75E-09 | -0.0146172 | 0.285 | 0.342 | 4.71E-05 | P2rx7 | Birc6 | | Clip1 | 7.02E-12 | -0.0235989 | 0.264 | 0.215 | 1.89E-07 | Cyth4 | Tgfbr2 | | | | | Fnip1 | | | | | | | | | |
| | Dgkd | 1.76E-09 | 0.18354769 | 0.332 | 0.343 | 4.75E-05 | Osbpl8 | Senp6 | | Dpysl2 | 7.26E-12 | 0.2597408 | 0.251 | 0.161 | 1.95E-07 | Fnbp1 | Nufip2 | | | | | Fndc3a | | | | | | | | | |
| | Abi1 | 1.84E-09 | 0.04129778 | 0.365 | 0.415 | 4.95E-05 | Wdr44 | Ppm1h | | Arhgap30 | 7.34E-12 | 0.10667294 | 0.25 | 0.187 | 1.98E-07 | Pmepa1 | Nrd1 | | | | | Kdm2a | | | | | | | | | |
| | Rasa1 | 1.86E-09 | -0.061123 | 0.251 | 0.316 | 5.00E-05 | Arhgap25 | Acap2 | | Nr6a1 | 7.39E-12 | 0.25832588 | 0.534 | 0.408 | 1.99E-07 | Ube2d3 | Nsd3 | | | | | Rin2 | | | | | | | | | |
| | Zfhx3 | 2.81E-08 | -0.2207252 | 0.844 | 0.901 | 0.00075536 | Wapl | Kif13b | | Lrch3 | 7.53E-12 | 0.09926473 | 0.317 | 0.241 | 2.03E-07 | Thada | Skap2 | | | | | Trem2 | | | | | | | | | |
| | Washc4 | 2.02E-09 | -0.0775016 | 0.202 | 0.26 | 5.45E-05 | Ttc3 | Rock1 | | Gna15 | 7.64E-12 | 0.15598981 | 0.34 | 0.253 | 2.06E-07 | Fmnl3 | Sipa1 | | | | | Cd84 | | | | | | | | | |
| | Sdccag8 | 2.23E-09 | -0.0180975 | 0.251 | 0.307 | 6.01E-05 | Klf3 | Hnrnph1 | | Lpp | 8.39E-12 | -0.0760908 | 0.258 | 0.212 | 2.26E-07 | Tmem243 | Senp6 | | | | | | | | | | | | | | |
| | Zfr | 2.25E-09 | 0.18565769 | 0.339 | 0.341 | 6.06E-05 | Rbm25 | Ccdc50 | | Dip2b | 8.41E-12 | 0.22658055 | 0.786 | 0.66 | 2.26E-07 | Ankfy1 | Peak1 | | | | | | | | | | | | | | |
| | Akap8l | 2.04E-10 | -0.2108061 | 0.316 | 0.419 | 5.48E-06 | Herc4 | Lrba | | Picalm | 8.50E-12 | 0.01921323 | 0.676 | 0.583 | 2.29E-07 | Smurf2 | Zfr | | | | | | | | | | | | | | |
| | Rps6ka3 | 2.32E-09 | 0.04480043 | 0.278 | 0.313 | 6.26E-05 | Ncoa3 | Dnm2 | | Taok3 | 8.79E-12 | -0.0296815 | 0.311 | 0.258 | 2.36E-07 | Ccdc93 | Rab2a | | | | | | | | | | | | | | |
| | Rnf169 | 1.47E-06 | -0.2089886 | 0.414 | 0.504 | 0.03968075 | Cd180 | Arfgef1 | | Khdrbs1 | 9.23E-12 | 0.07076821 | 0.329 | 0.254 | 2.48E-07 | Fcho2 | Dcakd | | | | | | | | | | | | | | |
| | Dip2b | 5.35E-08 | -0.2087049 | 0.66 | 0.765 | 0.00144008 | Pacs1 | Ccnt2 | | Adap2os | 9.53E-12 | 0.03684144 | 0.458 | 0.381 | 2.56E-07 | Mlxipl | Fto | | | | | | | | | | | | | | |
| | Lcor | 2.53E-09 | -0.0413505 | 0.285 | 0.347 | 6.80E-05 | Rsf1 | Mark3 | | Rbm47 | 9.69E-12 | -0.0681543 | 0.399 | 0.346 | 2.61E-07 | Baz2b | Herc4 | | | | | | | | | | | | | | |
| | Pard3b | 2.53E-09 | -0.0162739 | 0.363 | 0.434 | 6.82E-05 | Fth1 | Csf3r | | Col27a1 | 1.02E-11 | 0.20180315 | 0.339 | 0.249 | 2.73E-07 | Cask | Itch | | | | | | | | | | | | | | |

| | | | | | | | |
|---|---|---|---|---|---|---|---|
| Ptpra | 2.54E-09 | 0.10781192 | 0.355 | 0.389 | 6.84E-05 | Nsd3 | Cep68 |
| Fcho2 | 2.58E-09 | 0.0622776 | 0.412 | 0.459 | 6.95E-05 | Mdm4 | Tmem164 |
| Ankrd12 | 2.66E-09 | 0.3289528 | 0.292 | 0.268 | 7.16E-05 | Nsmce2 | Rfx3 |
| Pum2 | 2.68E-09 | -0.0470949 | 0.258 | 0.317 | 7.20E-05 | Mapk14 | Extl3 |
| Cd86 | 6.87E-09 | -0.2075785 | 0.285 | 0.383 | 0.00018474 | Il10ra | Smyd3 |
| Fam168a | 4.80E-09 | -0.2056849 | 0.278 | 0.37 | 0.00012902 | Nptn | Tspan14 |
| Rab31 | 2.76E-09 | -0.0880698 | 0.29 | 0.36 | 7.42E-05 | Setd2 | Atp9b |
| Col27a1 | 2.79E-09 | -0.0951666 | 0.249 | 0.315 | 7.51E-05 | Ssbp2 | Fyb |
| Sh3glb1 | 2.89E-09 | 0.0398345 | 0.272 | 0.307 | 7.78E-05 | Itm2b | Rb1 |
| P2ry6 | 2.92E-09 | -0.093759 | 0.321 | 0.39 | 7.86E-05 | P3h2 | Zmynd8 |
| Med13 | 2.98E-09 | -0.0056294 | 0.211 | 0.263 | 8.01E-05 | Stk3 | Rbm47 |
| Irak2 | 3.06E-09 | 0.01242351 | 0.349 | 0.399 | 8.24E-05 | Ctsh | Rap1b |
| Acer3 | 3.16E-09 | 0.20411489 | 0.367 | 0.366 | 8.51E-05 | Psap | Smad7 |
| Cpsf6 | 3.54E-09 | 0.00927718 | 0.244 | 0.288 | 9.52E-05 | Nrp1 | Ndfip1 |
| Ptp4a2 | 3.54E-09 | 0.04358656 | 0.269 | 0.307 | 9.53E-05 | Smg6 | St3gal6 |
| Pnpla7 | 3.55E-09 | 0.12381072 | 0.401 | 0.431 | 9.56E-05 | Gdi2 | Snrnp70 |
| Pros1 | 3.61E-09 | 0.03020002 | 0.303 | 0.351 | 9.73E-05 | Tnrc6c | Ep300 |
| Epc1 | 5.24E-08 | -0.2043119 | 0.211 | 0.287 | 0.00140988 | Prkcb | Usp50 |
| Vps13b | 3.72E-09 | 0.15230227 | 0.353 | 0.372 | 0.00010011 | Bin1 | Ptbp3 |
| Ophn1 | 2.36E-10 | -0.2036971 | 0.663 | 0.781 | 6.35E-06 | Ube3a | Atf7ip |
| Adap2os | 4.68E-11 | -0.2028092 | 0.381 | 0.491 | 1.26E-06 | Sall1 | Anks1 |
| Fndc3b | 1.53E-13 | -0.2016733 | 0.21 | 0.305 | 4.11E-09 | Gna12 | Atp6v0b |
| Sf3b1 | 4.44E-09 | -0.0424498 | 0.36 | 0.427 | 0.00011955 | Ctsc | Pcm1 |
| Ski | 4.54E-09 | -0.0120417 | 0.322 | 0.378 | 0.00012227 | Skap2 | Stag2 |
| Rab8b | 2.58E-07 | -0.1990547 | 0.328 | 0.415 | 0.0069505 | Sesn1 | Map3k3 |
| Zmiz1 | 4.58E-09 | 0.04365642 | 0.362 | 0.402 | 0.00012331 | Capza1 | Hck |
| Plekho1 | 4.71E-09 | -0.0237987 | 0.386 | 0.447 | 0.00012685 | Dnmt3a | Gm16599 |
| Zfc3h1 | 4.75E-09 | -0.0919631 | 0.21 | 0.271 | 0.0001277 | Rps6ka1 | Ptpn1 |
| Lcp2 | 2.24E-07 | -0.1964402 | 0.247 | 0.325 | 0.00601915 | Pik3ap1 | Serinc3 |
| Map4k3 | 4.86E-09 | 0.16387427 | 0.245 | 0.257 | 0.00013087 | Rtn4 | R3hdm2 |
| Trip12 | 4.91E-09 | 0.13626035 | 0.323 | 0.341 | 0.00013198 | Plekhm3 | Nlk |
| Mylip | 5.07E-09 | 0.00354804 | 0.353 | 0.413 | 0.00013642 | Cux1 | Daglb |
| Lhfpl2 | 5.16E-09 | 0.34842266 | 0.373 | 0.339 | 0.00013878 | Dync1i2 | Plcg2 |
| Itpr2 | 6.93E-07 | -0.1964327 | 0.405 | 0.498 | 0.01863696 | Rtn3 | Vgll4 |
| Mknk1 | 5.25E-09 | 0.2152628 | 0.285 | 0.285 | 0.00014128 | Atrx | Fbxl17 |
| Mysm1 | 2.84E-07 | -0.1962242 | 0.185 | 0.253 | 0.00764623 | AW554918 | Mgat4a |
| Smap2 | 5.59E-09 | 0.11817171 | 0.522 | 0.551 | 0.00015045 | Gtdc1 | Rnf13 |
| Rab14 | 1.85E-06 | -0.1961528 | 0.218 | 0.287 | 0.04969954 | Gpatch8 | Papola |
| Tbc1d16 | 6.06E-09 | 0.1944981 | 0.282 | 0.283 | 0.00016317 | Lims1 | Usp34 |
| Chd6 | 6.27E-09 | 0.07706885 | 0.251 | 0.286 | 0.00016871 | Abi3 | Msi2 |
| Zmynd11 | 6.61E-09 | 0.06808332 | 0.259 | 0.293 | 0.00017789 | Nsd1 | Mafb |
| Ntpcr | 9.25E-10 | -0.1959595 | 0.237 | 0.322 | 2.49E-05 | Rgs10 | Top1 |
| Zfp638 | 9.38E-09 | -0.1958068 | 0.328 | 0.425 | 0.00025228 | Insr | Abcc5 |
| Havcr2 | 6.13E-10 | -0.1928107 | 0.27 | 0.357 | 1.65E-05 | Lrmda | Tsc22d4 |
| Rbfox1 | 7.53E-09 | 0.45580814 | 0.343 | 0.272 | 0.00020266 | Tcf12 | Arih1 |
| Cd2ap | 7.85E-09 | -0.0797533 | 0.272 | 0.337 | 0.00021132 | Slc38a9 | Cnot4 |
| Cflar | 3.63E-07 | -0.1832866 | 0.321 | 0.401 | 0.00977631 | Sfmbt1 | Phf21a |
| Ddx17 | 7.88E-09 | 0.17041908 | 0.336 | 0.346 | 0.00021211 | Fnip1 | Rcsd1 |
| Gnas | 8.18E-09 | 0.40646807 | 0.276 | 0.226 | 0.00022003 | Fus | Nr3c1 |
| Wdfy4 | 8.20E-09 | 0.09220935 | 0.246 | 0.267 | 0.00022066 | Fcer1g | 2610020C07Rik |
| Itgav | 7.37E-10 | -0.183103 | 0.269 | 0.36 | 1.98E-05 | Gab1 | Map4 |
| Rnf111 | 9.27E-09 | 0.03298168 | 0.285 | 0.325 | 0.00024953 | Blnk | Canx |
| Adam10 | 6.54E-10 | -0.182663 | 0.276 | 0.366 | 1.76E-05 | Tnrc6b | Notch2 |
| Fam105a | 9.54E-09 | 0.29808145 | 0.508 | 0.389 | 0.00025661 | Nbea | Csnk1a1 |
| Ifngr1 | 1.92E-11 | -0.1825977 | 0.242 | 0.329 | 5.17E-07 | Gapvd1 | Nipbl |
| Kmt2e | 1.07E-08 | 0.08475626 | 0.32 | 0.353 | 0.00028662 | Etv6 | Ccny |
| Rnf216 | 4.41E-08 | -0.1820704 | 0.501 | 0.607 | 0.00118707 | Fgfr2 | Cdk12 |
| Rtn1 | 3.91E-08 | -0.1817519 | 0.321 | 0.412 | 0.00105319 | Dock11 | Wasf2 |
| Srsf5 | 1.11E-08 | 0.03748667 | 0.292 | 0.336 | 0.00029865 | Ttc14 | Ap3b1 |
| Lrp6 | 1.12E-08 | 0.01904671 | 0.208 | 0.252 | 0.00030066 | Igf1r | Wdr26 |
| Stambpl1 | 3.17E-07 | -0.1809016 | 0.198 | 0.264 | 0.00853089 | Ncf1 | Ppp6r3 |
| Snd1 | 1.13E-08 | 0.06433723 | 0.26 | 0.293 | 0.00030324 | Son | Ext1 |
| Trappc9 | 2.82E-07 | -0.1800545 | 0.199 | 0.268 | 0.00757921 | Zcchc11 | Dapp1 |
| Tbc1d23 | 1.15E-08 | -0.0278272 | 0.221 | 0.269 | 0.00030921 | Kcnk13 | Atxn7l1 |
| Camk2d | 1.18E-08 | -0.0241354 | 0.462 | 0.543 | 0.00031862 | Mbtd1 | Ewsr1 |
| Hps3 | 1.21E-08 | 0.12936071 | 0.257 | 0.277 | 0.00032557 | Fli1 | Peli2 |
| Selplg | 1.40E-08 | -0.1798935 | 0.566 | 0.675 | 0.00037557 | Wac | Nuak1 |
| Ssh2 | 9.17E-07 | -0.1766967 | 0.825 | 0.894 | 0.02466204 | R3hdm1 | Pafah1b1 |
| St3gal5 | 3.10E-08 | -0.1747534 | 0.246 | 0.328 | 0.0008331 | Oxr1 | Gm26848 |
| Zmym4 | 1.35E-08 | -0.0092395 | 0.288 | 0.342 | 0.00036388 | Camk1 | C1qa |
| Gria2 | 1.37E-08 | 0.39696774 | 0.294 | 0.254 | 0.00036946 | Tbl1x | Cd33 |
| Dhx9 | 1.33E-07 | -0.172573 | 0.231 | 0.303 | 0.00358674 | Klf12 | Nisch |
| Mbnl1 | 1.38E-06 | -0.1716221 | 0.791 | 0.868 | 0.03709173 | Lnpep | Cep170 |
| Cd81 | 1.46E-08 | 0.06346175 | 0.349 | 0.4 | 0.00039323 | Kpna4 | Il6ra |
| Kat6a | 1.52E-08 | 0.01627865 | 0.246 | 0.286 | 0.00041011 | Akt3 | Sat1 |
| Kmt2c | 1.59E-08 | -0.0478303 | 0.388 | 0.456 | 0.0004287 | Pias1 | Fermt3 |
| Spop | 1.60E-08 | 0.12999982 | 0.253 | 0.273 | 0.00043032 | Ubn2 | Rbm6 |
| Akap13 | 1.62E-08 | 0.08691338 | 0.472 | 0.513 | 0.00043544 | Gls | Agps |
| Ptk2b | 1.63E-08 | -0.0055393 | 0.243 | 0.29 | 0.00043754 | Strn3 | Pkn1 |
| Inpp5d | 1.41E-06 | -0.1693642 | 0.874 | 0.927 | 0.03786131 | Tcf4 | Setd5 |
| Tacc1 | 1.55E-07 | -0.1683206 | 0.469 | 0.567 | 0.00416633 | Bclaf1 | Bmp2k |
| Larp1 | 1.69E-08 | -0.0948602 | 0.212 | 0.275 | 0.00045604 | Cttnbp2nl | Kansl1l |
| Tra2b | 1.76E-08 | -0.0240418 | 0.209 | 0.252 | 0.00047365 | Lrch1 | Specc1 |
| Senp2 | 1.77E-08 | 0.12536049 | 0.266 | 0.287 | 0.00047613 | C1qc | Rbm26 |
| Abhd12 | 1.77E-08 | 0.11287441 | 0.544 | 0.583 | 0.00047668 | Mon2 | Tpr |
| Osbpl11 | 1.79E-08 | 0.10235569 | 0.313 | 0.337 | 0.00048121 | Itga6 | Gns |
| Clta | 1.81E-08 | 0.10531848 | 0.261 | 0.282 | 0.00048816 | Hdac8 | Casp8 |
| Rbm5 | 1.88E-08 | 0.00558676 | 0.324 | 0.372 | 0.0005046 | Clint1 | Uvrag |
| Abca9 | 4.55E-09 | -0.166903 | 0.434 | 0.538 | 0.00012256 | Pvt1 | Zfp292 |
| Usp24 | 1.64E-08 | -0.166017 | 0.322 | 0.406 | 0.00044092 | Wwox | Ogt |
| Fgd2 | 1.99E-08 | 0.04902819 | 0.478 | 0.529 | 0.00053521 | Tbc1d22a | Mkln1 |
| Lcorl | 2.02E-08 | 0.18781441 | 0.267 | 0.275 | 0.00054259 | Sdk1 | Prrc2c |
| Sirpa | 2.08E-08 | 0.08793527 | 0.541 | 0.579 | 0.00055841 | Hook3 | Rere |
| Srek1 | 2.12E-08 | 0.09692915 | 0.255 | 0.278 | 0.00057031 | Btbd9 | Baz2b |
| Myo9a | 2.32E-08 | -0.0192422 | 0.325 | 0.381 | 0.00062436 | Arid4a | Pmepa1 |
| Dnajc13 | 2.36E-08 | -0.0723512 | 0.202 | 0.256 | 0.00063595 | Lcp1 | Zfand3 |
| Sik2 | 2.47E-08 | 0.16415718 | 0.276 | 0.285 | 0.00066402 | Nktr | Rab3gap1 |
| Tbc1d14 | 2.49E-08 | -0.0133381 | 0.228 | 0.277 | 0.00067033 | Tmsb4x | Prkce |
| Man1a2 | 2.50E-08 | 0.07382076 | 0.429 | 0.463 | 0.00067174 | Rsrc2 | Tcf20 |
| Lrp1 | 3.94E-09 | -0.1631416 | 0.292 | 0.38 | 0.000106 | Ube2h | Slc16a6 |
| Vav1 | 1.94E-08 | -0.1619599 | 0.213 | 0.281 | 0.00052158 | Arhgap5 | Fubp1 |
| Spag9 | 2.65E-08 | 0.10706887 | 0.355 | 0.38 | 0.00071173 | Plxdc2 | Nedd4l |
| Thada | 2.65E-08 | 0.0045467 | 0.217 | 0.262 | 0.0007128 | Eml4 | Arrb2 |
| Nos1ap | 1.70E-06 | -0.1608331 | 0.25 | 0.322 | 0.04576393 | Atp2c1 | Vti1a |
| Soga1 | 2.70E-08 | -0.0043219 | 0.385 | 0.438 | 0.00072624 | Cfh | Cd37 |
| Chd7 | 2.72E-08 | -0.0157065 | 0.347 | 0.403 | 0.00073102 | Pid1 | Smchd1 |
| Cdk13 | 2.74E-08 | 0.00562468 | 0.289 | 0.334 | 0.00073631 | Kdm7a | Comt |
| Pik3cd | 2.78E-08 | 0.05241329 | 0.257 | 0.29 | 0.00074761 | Large1 | Tubgcp5 |
| Mgat5 | 3.45E-08 | -0.1606408 | 0.41 | 0.505 | 0.0009273 | Bach2 | Bbs9 |
| Syngr1 | 2.83E-08 | -0.0928769 | 0.214 | 0.278 | 0.00076136 | Parp8 | Dock10 |
| Zfp644 | 2.86E-08 | -0.06121 | 0.279 | 0.339 | 0.00076989 | F11r | 4930444A19Rik |
| Ctsz | 3.00E-08 | 0.18878236 | 0.257 | 0.265 | 0.00080642 | Camk1d | Actr2 |
| Cask | 1.53E-10 | -0.1601161 | 0.391 | 0.495 | 4.13E-06 | Rabep1 | Exoc6b |
| Man1c1 | 3.04E-08 | 0.02641069 | 0.27 | 0.314 | 0.00081768 | Map4k4 | Synj1 |
| Sipa1l2 | 3.05E-08 | -0.0847925 | 0.331 | 0.402 | 0.00082107 | Fmnl2 | Itch |
| Jak1 | 2.16E-15 | -0.1594553 | 0.261 | 0.362 | 5.82E-11 | Cmip | Sltm |
| Prpf4b | 1.24E-06 | -0.1585725 | 0.391 | 0.475 | 0.03324419 | Celf2 | Rrbp1 |
| Oxct1 | 3.12E-08 | 0.22038075 | 0.292 | 0.286 | 0.00083976 | Dock1 | Slc1a3 |
| Usp9x | 3.15E-08 | -0.0129787 | 0.265 | 0.315 | 0.00084883 | Nav3 | Nrip1 |
| Gm26542 | 3.16E-08 | -0.0567417 | 0.335 | 0.407 | 0.00085006 | Cacnb2 | Phf3 |
| Rsrc1 | 3.18E-08 | 0.0693328 | 0.285 | 0.316 | 0.00085434 | Peak1 | Dock4 |
| Tra2a | 3.33E-08 | 0.00324118 | 0.339 | 0.384 | 0.000895 | Zbtb20 | Aim2 |
| Ankfy1 | 3.38E-08 | -0.0994598 | 0.202 | 0.258 | 0.00090892 | Akap10 | Tmem135 |
| Myh9 | 3.40E-08 | 0.02015688 | 0.244 | 0.284 | 0.00091354 | Heatr5a | Pde4d |
| Phf20l1 | 3.41E-08 | 0.01564138 | 0.261 | 0.304 | 0.00091733 | Mtus1 | Pds5a |
| Gfm2 | 2.03E-11 | -0.1578842 | 0.197 | 0.276 | 5.45E-07 | Ptprm | Rapgef2 |
| Scaper | 3.55E-08 | 0.03428055 | 0.23 | 0.274 | 0.00095492 | Runx1 | Smurf2 |
| Ythdf3 | 1.36E-10 | -0.1572553 | 0.2 | 0.279 | 3.66E-06 | Taok3 | Csmd3 |
| Med12l | 3.56E-08 | -0.0383931 | 0.426 | 0.493 | 0.00095729 | Arhgap22 | Zdhhc20 |
| Mbnl2 | 3.59E-08 | 0.01546824 | 0.464 | 0.526 | 0.00096557 | Gpr183 | Arhgap12 |
| Srsf11 | 3.60E-08 | 0.09958141 | 0.324 | 0.352 | 0.00096937 | Lyst | Gna15 |
| Btbd1 | 3.66E-08 | -0.0758115 | 0.217 | 0.27 | 0.00098561 | Apoe | Cnot2 |
| Gnaq | 3.69E-08 | 0.17621142 | 0.589 | 0.595 | 0.00099285 | Ikzf1 | Lrrc4 |
| Abca1 | 3.72E-08 | 0.0628968 | 0.367 | 0.401 | 0.00100052 | Sbf2 | Arl15 |
| Xrn1 | 3.82E-08 | -0.0395017 | 0.208 | 0.256 | 0.0010277 | Nlrp1b | Mark2 |
| Atad2b | 2.34E-07 | -0.1572416 | 0.255 | 0.324 | 0.00629178 | Jmjd1c | Pik3r1 |
| Luc7l2 | 3.93E-08 | 0.21585638 | 0.574 | 0.571 | 0.00105831 | Phactr2 | Rffl |
| Clasp2 | 3.96E-08 | 0.17640476 | 0.459 | 0.472 | 0.00106547 | Ank2 | Nrd1 |
| Mvb12b | 6.15E-08 | -0.1566221 | 0.27 | 0.345 | 0.00165347 | Ankrd11 | C1qb |
| Pitpna | 4.05E-08 | 0.04384795 | 0.245 | 0.281 | 0.00109032 | Ppp3ca | Hipk2 |
| Scmh1 | 4.28E-08 | 0.03675262 | 0.245 | 0.288 | 0.00115098 | Cmss1 | Gm5086 |
| Spire1 | 4.28E-08 | -0.0357582 | 0.327 | 0.388 | 0.00115161 | Tmcc1 | Chd9 |
| Pak2 | 4.31E-08 | 0.13780086 | 0.306 | 0.318 | 0.00115897 | Rcbtb2 | Stx16 |
| Erbin | 4.35E-08 | 0.05478032 | 0.307 | 0.343 | 0.00117157 | Rasgrp3 | Ulk2 |
| Tnrc18 | 4.36E-08 | 0.12276756 | 0.246 | 0.257 | 0.00117253 | P2ry12 | Taf15 |
| Zscan26 | 7.78E-10 | -0.1559554 | 0.181 | 0.252 | 2.09E-05 | Gm26804 | Bptf |
| Zfp407 | 4.67E-08 | 0.08255733 | 0.239 | 0.269 | 0.00125787 | Pip4k2a | Mier1 |
| Kdm2a | 4.70E-08 | 0.03540972 | 0.318 | 0.352 | 0.0012647 | Maml2 | Slc7a7 |
| Nf1 | 4.71E-08 | 0.13715283 | 0.429 | 0.443 | 0.00126703 | Slc24a3 | Phip |
| Sft2d1 | 4.75E-08 | -0.0347726 | 0.331 | 0.377 | 0.00127689 | Eml6 | Xiap |
| Marcks | 4.79E-08 | 0.05726825 | 0.305 | 0.336 | 0.00128995 | Pnisr | Trio |
| Pwwp2a | 4.86E-08 | 0.03436052 | 0.361 | 0.397 | 0.0013083 | Chn2 | Srgap2 |
| Frrs1 | 4.95E-08 | 0.08173825 | 0.237 | 0.264 | 0.00133121 | Kcnma1 | Pde3b |
| Phc2 | 4.96E-08 | 0.0351457 | 0.266 | 0.302 | 0.00133408 | Frmd4b | Ago3 |
| Disc1 | 5.74E-08 | -0.1552555 | 0.209 | 0.275 | 0.00154423 | Hdac9 | Kansl1 |
| Wdfy3 | 5.22E-08 | -0.0426513 | 0.45 | 0.52 | 0.00140508 | Arglu1 | Hivep3 |
| Phf14 | 1.38E-07 | -0.1514587 | 0.471 | 0.568 | 0.00371377 | Epb41l2 | Tsix |
| Abl1 | 2.77E-12 | -0.1513329 | 0.219 | 0.302 | 7.46E-08 | Diaph2 | Zranb2 |
| Zfp652 | 4.80E-07 | -0.1492922 | 0.447 | 0.537 | 0.01290595 | Dip2c | Slco2b1 |
| Arid1a | 5.65E-08 | 0.07337217 | 0.287 | 0.319 | 0.00151981 | Fam102b | Nrf1 |
| Ptgs1 | 7.03E-09 | -0.1467874 | 0.266 | 0.341 | 0.00018924 | Asap1 | Fam117b |
| Trim44 | 5.79E-08 | -0.0690396 | 0.266 | 0.325 | 0.00155667 | Srrm2 | Nufip2 |
| Cdk19 | 2.39E-10 | -0.1462457 | 0.27 | 0.353 | 6.43E-06 | Snx29 | Rsbn1l |
| Rhoq | 5.83E-08 | 0.02752167 | 0.215 | 0.251 | 0.00156783 | Gab2 | Slc7a8 |
| Ppp1r12a | 5.95E-08 | 0.16955917 | 0.446 | 0.45 | 0.00160169 | Gm26733 | Ubac2 |
| Fam193a | 5.96E-08 | 0.12133437 | 0.297 | 0.315 | 0.00160308 | Ctsb | Srsf7 |
| Fam172a | 6.04E-08 | 0.05802139 | 0.471 | 0.52 | 0.00162461 | Apbb1ip | Nr3c2 |
| Morc3 | 1.09E-08 | -0.1458565 | 0.218 | 0.292 | 0.00029213 | Gm26740 | Dock7 |
| Apc | 6.09E-08 | 0.01128268 | 0.319 | 0.367 | 0.00163739 | Mertk | Ttc28 |
| Hectd1 | 1.06E-06 | -0.1444014 | 0.205 | 0.264 | 0.02848391 | Ptprc | Rap1gds1 |
| Luc7l3 | 1.56E-06 | -0.1438553 | 0.197 | 0.26 | 0.04202322 | Tmcc3 | Zc3h7a |
| Ints6l | 6.20E-08 | 0.06493846 | 0.405 | 0.449 | 0.0016675 | Fhit | Sipa1l1 |
| Plgrkt | 6.44E-08 | 0.09553109 | 0.267 | 0.287 | 0.00173346 | Rbm39 | Tnfrsf11a |
| Huwe1 | 6.45E-08 | 0.07239407 | 0.272 | 0.303 | 0.00173546 | Arhgap31 | Mtdh |
| Ctss | 6.57E-08 | 0.17353456 | 0.649 | 0.667 | 0.00176812 | Tanc2 | Gsk3b |
| Aftph | 1.64E-06 | -0.1436967 | 0.378 | 0.455 | 0.04424701 | Olfr56 | Tmem119 |
| Grb2 | 6.78E-08 | 0.10994785 | 0.312 | 0.333 | 0.00182346 | Immp2l | Trpm7 |
| Gtf2h2 | 6.19E-08 | -0.1415075 | 0.334 | 0.415 | 0.00166561 | Exoc4 | Elk3 |
| Ubr3 | 6.96E-08 | 0.00958259 | 0.254 | 0.295 | 0.00187339 | Numb | Gm48678 |
| Sparc | 1.96E-08 | -0.1411531 | 0.343 | 0.432 | 0.00052787 | Gm42418 | Dst |
| Pibf1 | 7.11E-08 | -0.0411539 | 0.216 | 0.264 | 0.00191247 | Xist | Zfand6 |
| Gphn | 7.32E-08 | 0.22953051 | 0.416 | 0.414 | 0.00197072 | Gm35188 | Ascc3 |
| Kdm3b | 7.37E-08 | -0.0844687 | 0.294 | 0.357 | 0.0019835 | Frmd4a | Creb1 |
| Birc6 | 7.99E-08 | -0.0526358 | 0.299 | 0.361 | 0.0021509 | | |
| Senp6 | 8.14E-08 | -0.0500584 | 0.239 | 0.293 | 0.00219139 | | |
| Ankhd1 | 8.32E-08 | 0.01742673 | 0.319 | 0.367 | 0.00223858 | | |
| Wipf1 | 8.43E-08 | 0.24044985 | 0.348 | 0.334 | 0.00226695 | | |
| Ppm1h | 8.63E-08 | -0.0627519 | 0.499 | 0.579 | 0.00232151 | | |
| Acap2 | 7.00E-08 | -0.1402078 | 0.398 | 0.484 | 0.00188375 | | |
| Kif13b | 8.76E-08 | -0.0655388 | 0.232 | 0.285 | 0.00235754 | | |
| Rock1 | 8.88E-08 | -0.049461 | 0.317 | 0.377 | 0.00238862 | | |
| Hnrnph1 | 9.21E-08 | -0.0402521 | 0.29 | 0.346 | 0.00247832 | | |
| Fndc3a | 9.24E-08 | 0.19830579 | 0.254 | 0.252 | 0.00248542 | | |
| Ccdc50 | 9.24E-08 | -0.0429538 | 0.248 | 0.299 | 0.00248593 | | |
| Lrba | 1.12E-08 | -0.140171 | 0.331 | 0.415 | 0.00030193 | | |
| Fnbp1 | 9.29E-08 | 0.11735164 | 0.389 | 0.407 | 0.00250076 | | |
| Dnm2 | 6.59E-08 | -0.1394472 | 0.313 | 0.389 | 0.00177236 | | |
| Arfgef1 | 9.80E-08 | -0.032725 | 0.203 | 0.252 | 0.00263699 | | |
| Fgf13 | 9.82E-08 | 0.49412785 | 0.401 | 0.344 | 0.00264175 | | |
| Gatad2b | 1.00E-07 | 0.05428648 | 0.227 | 0.256 | 0.00270078 | | |
| Ccnt2 | 1.02E-07 | -0.0594167 | 0.214 | 0.262 | 0.00273923 | | |
| Mark3 | 1.03E-07 | -0.0123242 | 0.3 | 0.347 | 0.00278464 | | |
| Ambra1 | 1.05E-07 | 0.02244324 | 0.337 | 0.381 | 0.00281787 | | |
| Basp1 | 1.07E-07 | 0.13156443 | 0.392 | 0.418 | 0.00287636 | | |
| Csf3r | 1.08E-07 | -0.0822892 | 0.458 | 0.539 | 0.00289716 | | |
| Cep68 | 1.08E-07 | -0.0964571 | 0.223 | 0.28 | 0.00291556 | | |
| Tmem164 | 1.09E-07 | -0.070678 | 0.286 | 0.34 | 0.00294074 | | |
| Rfx3 | 1.12E-07 | -0.0066875 | 0.205 | 0.251 | 0.00301475 | | |
| Ehd4 | 1.12E-07 | 0.009206 | 0.214 | 0.251 | 0.00302115 | | |
| Zeb2 | 1.16E-07 | 0.27546217 | 0.831 | 0.783 | 0.00311702 | | |
| Nfat5 | 1.23E-07 | 0.01989632 | 0.304 | 0.346 | 0.00331522 | | |
| Extl3 | 4.01E-08 | -0.138223 | 0.33 | 0.408 | 0.00108008 | | |
| Ddx6 | 1.24E-07 | 0.07633005 | 0.236 | 0.262 | 0.00334556 | | |
| Asxl2 | 1.26E-07 | 0.14981221 | 0.265 | 0.276 | 0.00338961 | | |
| Arhgap39 | 1.27E-07 | 0.09019037 | 0.422 | 0.457 | 0.00341994 | | |
| Atp2b1 | 1.28E-07 | 0.03919477 | 0.388 | 0.438 | 0.0034557 | | |
| Cog5 | 1.28E-07 | 0.12302008 | 0.29 | 0.309 | 0.00345746 | | |
| Smyd3 | 1.29E-07 | -0.0040763 | 0.437 | 0.501 | 0.00346615 | | |
| Lrrk1 | 1.32E-07 | 0.01564716 | 0.313 | 0.356 | 0.00354504 | | |
| Retreg1 | 1.33E-07 | 0.01037489 | 0.313 | 0.358 | 0.00356992 | | |
| Tspan14 | 1.33E-08 | -0.1380566 | 0.218 | 0.283 | 0.00035845 | | |
| Cst3 | 1.35E-07 | 0.14364449 | 0.86 | 0.914 | 0.00362076 | | |
| Mitf | 1.35E-07 | 0.18603772 | 0.459 | 0.46 | 0.00363177 | | |
| Atp9b | 1.37E-07 | -0.0089074 | 0.276 | 0.321 | 0.00368088 | | |
| Fyb | 1.23E-06 | -0.138028 | 0.67 | 0.759 | 0.03311197 | | |
| Rb1 | 1.39E-07 | -0.0112308 | 0.303 | 0.355 | 0.00373983 | | |
| Zmynd8 | 1.45E-07 | -0.0375054 | 0.222 | 0.273 | 0.00389365 | | |
| Rb1cc1 | 1.48E-07 | 0.0006866 | 0.218 | 0.259 | 0.00397588 | | |
| Rab2a | 1.48E-07 | 0.06373543 | 0.271 | 0.297 | 0.00397981 | | |
| Lamp2 | 1.50E-07 | 0.10874924 | 0.286 | 0.306 | 0.00403664 | | |
| Smg1 | 1.53E-07 | 0.1858745 | 0.377 | 0.383 | 0.00410525 | | |
| P2rx7 | 1.53E-07 | 0.22987778 | 0.322 | 0.346 | 0.00412328 | | |
| Osbpl8 | 1.54E-07 | 0.31425885 | 0.273 | 0.238 | 0.00413273 | | |
| Rbm47 | 2.13E-07 | -0.1373186 | 0.346 | 0.42 | 0.00573884 | | |
| Wdr44 | 1.57E-07 | 0.16935285 | 0.269 | 0.275 | 0.00423709 | | |
| Arhgap25 | 1.63E-07 | 0.01410085 | 0.257 | 0.298 | 0.00437332 | | |
| Wapl | 1.67E-07 | 0.06977482 | 0.296 | 0.322 | 0.00448614 | | |
| Rap1b | 1.68E-07 | -0.0397454 | 0.273 | 0.326 | 0.00452978 | | |
| Ttc3 | 1.77E-07 | 0.22505695 | 0.275 | 0.268 | 0.00475402 | | |

| | | | | | | | |
|---|---|---|---|---|---|---|---|
| Rab14 | 1.02E-11 | 0.02676844 | 0.274 | 0.218 | 2.73E-07 | Sparc | Zfp644 |
| Pik3ap1 | 1.05E-11 | 0.20077354 | 0.434 | 0.33 | 2.82E-07 | Ncoa2 | Laptm5 |
| Arid4a | 1.13E-11 | -0.040953 | 0.316 | 0.267 | 3.05E-07 | Syngr1 | Slc38a9 |
| Larp4b | 1.18E-11 | -0.025751 | 0.288 | 0.243 | 3.18E-07 | Birc6 | Sat1 |
| AW554918 | 1.22E-11 | -0.0235639 | 0.288 | 0.239 | 3.29E-07 | Mllt10 | Cacna1a |
| Psap | 1.22E-11 | -0.176327 | 0.455 | 0.43 | 3.29E-07 | Mapkap1 | Gm26848 |
| Gls | 1.23E-11 | -0.2593927 | 0.283 | 0.283 | 3.30E-07 | Mknk1 | Dpy19l4 |
| Abcc5 | 1.23E-11 | 0.23304253 | 0.291 | 0.199 | 3.30E-07 | Rtn4 | Pkig |
| Sema4d | 1.26E-11 | 0.1645026 | 0.269 | 0.191 | 3.40E-07 | Gm26827 | Phkb |
| Usp24 | 1.26E-11 | 0.09017276 | 0.404 | 0.322 | 3.40E-07 | Hnrnph1 | Psme4 |
| Fam193a | 1.27E-11 | 0.04629836 | 0.365 | 0.297 | 3.42E-07 | Snx2 | Gnaq |
| Herc1 | 1.28E-11 | -0.1639182 | 0.337 | 0.319 | 3.44E-07 | Srsf7 | Fryl |
| Gm17231 | 1.33E-11 | 0.2549882 | 0.31 | 0.215 | 3.59E-07 | Snx5 | Parp8 |
| Ttc14 | 1.35E-11 | -0.0033509 | 0.369 | 0.315 | 3.62E-07 | Ppp6r3 | Gm2245 |
| Ctsz | 1.42E-11 | -0.366088 | 0.237 | 0.257 | 3.81E-07 | Zfp608 | Zmym4 |
| St6gal1 | 1.47E-11 | -0.0741787 | 0.28 | 0.243 | 3.96E-07 | Fndc3b | Arhgap17 |
| Slc7a8 | 1.49E-11 | 0.25477336 | 0.434 | 0.32 | 4.01E-07 | Cfh | Rock1 |
| Tm6sf1 | 1.53E-11 | 0.27854678 | 0.365 | 0.257 | 4.12E-07 | Nckap1l | Arhgap25 |
| Atf7ip | 1.55E-11 | 0.02058206 | 0.316 | 0.252 | 4.16E-07 | Supt20 | Lamp2 |
| Tab2 | 1.60E-11 | 0.18162181 | 0.319 | 0.231 | 4.29E-07 | Eml4 | Rb1 |
| Whrn | 1.60E-11 | 0.13839712 | 0.279 | 0.205 | 4.30E-07 | Slc7a7 | Skil |
| Wdfy4 | 1.73E-11 | 0.15646194 | 0.325 | 0.246 | 4.66E-07 | Hipk2 | Kdm7a |
| Lrrk1 | 1.79E-11 | 0.03165371 | 0.382 | 0.313 | 4.82E-07 | Pag1 | Celf2 |
| Gdi2 | 1.81E-11 | 0.01942578 | 0.412 | 0.347 | 4.88E-07 | Sf3b1 | Pnpla7 |
| Gtf2h2 | 1.83E-11 | 0.17130078 | 0.435 | 0.334 | 4.93E-07 | Nisch | Fubp1 |
| Man1c1 | 1.84E-11 | 0.09486816 | 0.35 | 0.27 | 4.95E-07 | Hmg20a | Marcks |
| Csnk1g1 | 1.96E-11 | 0.00493252 | 0.252 | 0.199 | 5.28E-07 | Pik3r5 | Ncf1 |
| Cdkal1 | 2.02E-11 | 0.03534785 | 0.348 | 0.28 | 5.44E-07 | Abca9 | Lhfpl2 |
| Man2b1 | 2.06E-11 | -0.0881627 | 0.401 | 0.359 | 5.54E-07 | Pwwp2a | Dock7 |
| Atxn7l1 | 2.12E-11 | 0.1026707 | 0.382 | 0.3 | 5.69E-07 | Fry | Cdk14 |
| Snd1 | 2.13E-11 | 0.02523954 | 0.322 | 0.26 | 5.73E-07 | Taf15 | Gna12 |
| Rcsd1 | 2.13E-11 | 0.09785483 | 0.371 | 0.289 | 5.74E-07 | Camk2d | Mctp1 |
| Grb2 | 2.17E-11 | 0.05531228 | 0.385 | 0.312 | 5.83E-07 | Sft2d1 | Mafb |
| Ppcdc | 2.18E-11 | 0.06094433 | 0.396 | 0.323 | 5.86E-07 | Tbc1d23 | Blnk |
| Ube2k | 2.19E-11 | 0.0197111 | 0.272 | 0.219 | 5.89E-07 | Map2k1 | Prrc2c |
| Vps54 | 2.25E-11 | -0.0151275 | 0.286 | 0.236 | 6.05E-07 | Slc16a6 | Pcmtd1 |
| Capza2 | 2.25E-11 | -0.1416526 | 0.29 | 0.264 | 6.06E-07 | Ddx6 | Dyrk1a |
| Clasp2 | 2.36E-11 | 0.02707969 | 0.542 | 0.459 | 6.34E-07 | Map4k4 | Sh3kbp1 |
| Tbc1d9 | 2.36E-11 | 0.03666377 | 0.263 | 0.205 | 6.34E-07 | Mkln1 | Plgrkt |
| Acer3 | 2.38E-11 | -0.0341848 | 0.425 | 0.367 | 6.42E-07 | Sall1 | Fndc3a |
| Fam117b | 2.42E-11 | 0.12009921 | 0.311 | 0.235 | 6.52E-07 | Dennd1a | Rin2 |
| Tnrc18 | 2.57E-11 | 0.22710385 | 0.339 | 0.246 | 6.93E-07 | Csnk1a1 | Tmcc1 |
| Bclaf1 | 2.79E-11 | -0.1524302 | 0.25 | 0.23 | 7.50E-07 | Cables1 | Cd86 |
| Elf1 | 2.87E-11 | 0.07276518 | 0.301 | 0.235 | 7.72E-07 | Tpp2 | Rasal2 |
| Rps6ka3 | 3.02E-11 | -0.0476981 | 0.323 | 0.278 | 8.13E-07 | Peli2 | Lncpint |
| Vti1a | 3.05E-11 | 0.08972549 | 0.423 | 0.339 | 8.22E-07 | Rb1cc1 | Evl |
| Asph | 3.09E-11 | -0.1349081 | 0.363 | 0.331 | 8.32E-07 | Ccdc50 | Kcnma1 |
| Fnip1 | 3.31E-11 | -0.1750506 | 0.325 | 0.311 | 8.91E-07 | Sppl3 | Myo9a |
| Cd84 | 3.38E-11 | -0.06373 | 0.439 | 0.386 | 9.10E-07 | Ccnl1 | Elk3 |
| Ankrd17 | 3.41E-11 | 0.008071 | 0.385 | 0.324 | 9.18E-07 | Ptpn1 | Luc7l2 |
| Scaper | 3.64E-11 | 0.09716416 | 0.309 | 0.23 | 9.79E-07 | Cd2ap | AU020206 |
| Fer | 3.69E-11 | 0.19193195 | 0.293 | 0.212 | 9.93E-07 | Trio | Dock11 |
| Nrf1 | 3.71E-11 | 0.03040052 | 0.268 | 0.215 | 9.97E-07 | Cep68 | Prpf4b |
| Wdr44 | 3.76E-11 | 0.14432916 | 0.351 | 0.269 | 1.01E-06 | Casp8 | Man1a |
| Bin1 | 3.77E-11 | 0.0974107 | 0.622 | 0.523 | 1.01E-06 | Fus | Fkbp15 |
| Ewsr1 | 3.79E-11 | 0.08260513 | 0.306 | 0.236 | 1.02E-06 | Pnisr | Asxl2 |
| Stambpl1 | 3.80E-11 | 0.05208054 | 0.255 | 0.198 | 1.02E-06 | Atxn7 | Ttc3 |
| Atrx | 3.84E-11 | -0.0565178 | 0.408 | 0.36 | 1.03E-06 | Rhobtb1 | Gm48678 |
| Nedd4l | 3.85E-11 | 0.0508144 | 0.261 | 0.2 | 1.03E-06 | Cdk6 | Nbea |
| Chd6 | 3.95E-11 | 0.04003075 | 0.315 | 0.251 | 1.06E-06 | Epc1 | Osbpl9 |
| Ndfip1 | 4.10E-11 | 0.02375148 | 0.269 | 0.213 | 1.10E-06 | Sh3pxd2a | Arglu1 |
| Ash1l | 4.12E-11 | -0.1868423 | 0.432 | 0.424 | 1.11E-06 | Etv6 | Tbc1d22a |
| Syk | 4.31E-11 | 0.11177809 | 0.293 | 0.217 | 1.16E-06 | Slco2b1 | Srsf11 |
| Myh9 | 4.51E-11 | 0.0282172 | 0.301 | 0.244 | 1.21E-06 | Helz | Tcf12 |
| Plcl2 | 4.54E-11 | 0.31620152 | 0.725 | 0.6 | 1.22E-06 | Snx13 | Sesn1 |
| Washc4 | 4.64E-11 | 0.03324791 | 0.259 | 0.202 | 1.25E-06 | Msi2 | Arid4b |
| Nav2 | 4.73E-11 | 0.26667096 | 0.919 | 0.846 | 1.27E-06 | Slmap | Spag9 |
| Mlxip | 4.83E-11 | -0.1470529 | 0.276 | 0.253 | 1.30E-06 | Ints6l | Pvt1 |
| Eif4g3 | 4.87E-11 | 0.00306435 | 0.455 | 0.382 | 1.31E-06 | Srpk2 | Rbm39 |
| Agps | 4.90E-11 | 0.10422714 | 0.344 | 0.268 | 1.32E-06 | Dgkd | Gm44686 |
| Setd2 | 4.93E-11 | -0.0738099 | 0.325 | 0.291 | 1.33E-06 | Golm1 | Fgfr2 |
| Wdr26 | 5.19E-11 | 0.01361218 | 0.332 | 0.276 | 1.40E-06 | Retreg1 | Btbd7 |
| Disc1 | 5.26E-11 | 0.35111279 | 0.313 | 0.209 | 1.42E-06 | Ttc28 | Fam105a |
| Pcm1 | 5.27E-11 | 0.01485308 | 0.31 | 0.255 | 1.42E-06 | Dennd1b | Il6ra |
| Thoc2 | 5.41E-11 | -0.0250323 | 0.273 | 0.228 | 1.46E-06 | Luc7l3 | Ankrd12 |
| Celf1 | 5.43E-11 | 0.0327594 | 0.427 | 0.354 | 1.46E-06 | Pnn | Fmnl2 |
| Snx24 | 5.48E-11 | 0.12198344 | 0.423 | 0.331 | 1.47E-06 | Ktn1 | Ssbp2 |
| Ctsh | 6.38E-11 | -0.1212533 | 0.302 | 0.28 | 1.72E-06 | Vsir | Daglb |
| Rnf13 | 6.44E-11 | 0.08897989 | 0.368 | 0.295 | 1.73E-06 | Crebbp | Lcp1 |
| Ralgps1 | 6.54E-11 | 0.29397807 | 0.257 | 0.166 | 1.76E-06 | Srgap2 | Nsmce2 |
| Zdhhc14 | 6.78E-11 | 0.17183206 | 0.369 | 0.275 | 1.83E-06 | Senp2 | Fgf13 |
| Tbc1d16 | 6.79E-11 | -0.1092722 | 0.314 | 0.282 | 1.83E-06 | Sort1 | Exoc4 |
| Fermt3 | 6.89E-11 | 0.08193297 | 0.368 | 0.294 | 1.85E-06 | Ubl3 | Strbp |
| Adam10 | 6.92E-11 | 0.09952799 | 0.353 | 0.276 | 1.86E-06 | Rab8b | Srrm2 |
| Adam17 | 6.96E-11 | 0.24558357 | 0.262 | 0.177 | 1.87E-06 | Cdk12 | Runx1 |
| Hnrnpu | 7.01E-11 | -0.1823821 | 0.315 | 0.306 | 1.89E-06 | Csad | Ly86 |
| Spred1 | 7.27E-11 | 0.21455672 | 0.281 | 0.196 | 1.96E-06 | Cmip | Tbc1d5 |
| Asah1 | 7.28E-11 | -0.0140175 | 0.264 | 0.218 | 1.96E-06 | Maf | Camk1d |
| Soga1 | 7.36E-11 | 0.32515196 | 0.512 | 0.385 | 1.98E-06 | Hook3 | Dnajc1 |
| P3h2 | 7.38E-11 | -0.2621308 | 0.448 | 0.455 | 1.98E-06 | Aftph | Gab2 |
| Mapk14 | 7.42E-11 | 0.19089252 | 0.414 | 0.317 | 2.00E-06 | Sfmbt1 | Zbtb20 |
| Rnf130 | 7.75E-11 | 0.01506196 | 0.267 | 0.215 | 2.09E-06 | Aff4 | Fat3 |
| Tgfb1 | 7.90E-11 | 0.09863925 | 0.25 | 0.185 | 2.13E-06 | Itga9 | Tgfbr1 |
| Herc2 | 7.91E-11 | 0.25680114 | 0.51 | 0.389 | 2.13E-06 | Wasf2 | Mycbp2 |
| Anks1 | 7.97E-11 | 0.21486727 | 0.315 | 0.228 | 2.14E-06 | Foxn3 | Ank2 |
| Fchsd2 | 8.21E-11 | 0.21254286 | 0.714 | 0.59 | 2.21E-06 | 3110082I17Rik | Glg1 |
| Cul1 | 8.53E-11 | 0.1261583 | 0.37 | 0.286 | 2.29E-06 | Lrba | Olfr56 |
| Tbl1x | 8.72E-11 | -0.0238447 | 0.291 | 0.239 | 2.35E-06 | Lrrc4 | Ext1 |
| Fars2 | 8.75E-11 | -0.1220373 | 0.268 | 0.239 | 2.35E-06 | Smchd1 | Dennd4a |
| Zmiz1 | 9.27E-11 | 0.00051473 | 0.42 | 0.362 | 2.49E-06 | Dock10 | Gm26740 |
| Smap2 | 9.37E-11 | -0.0459688 | 0.582 | 0.522 | 2.52E-06 | Gng2 | Fhit |
| Vrk2 | 9.51E-11 | 0.29114362 | 0.395 | 0.288 | 2.56E-06 | Tor1aip1 | Ppp3ca |
| Arid1a | 9.58E-11 | 0.08025387 | 0.364 | 0.287 | 2.58E-06 | Notch2 | Immp2l |
| Lnpep | 9.74E-11 | -0.0192233 | 0.317 | 0.267 | 2.62E-06 | Garnl3 | Gm26804 |
| Ube3c | 1.00E-10 | 0.06314431 | 0.275 | 0.213 | 2.70E-06 | Pum2 | Zeb2 |
| Ppp1r12a | 1.02E-10 | -0.1676882 | 0.462 | 0.446 | 2.74E-06 | Rptor | Dst |
| R3hdm1 | 1.04E-10 | -0.072338 | 0.343 | 0.302 | 2.81E-06 | Capn3 | Wwox |
| Rapgef6 | 1.05E-10 | -0.026842 | 0.354 | 0.3 | 2.83E-06 | Tlr7 | Hexb |
| Ncor1 | 1.14E-10 | 0.02173954 | 0.449 | 0.384 | 3.05E-06 | Rap1b | Gm26733 |
| Srek1 | 1.16E-10 | -0.2107801 | 0.259 | 0.255 | 3.13E-06 | Ldlrad4 | Inpp4b |
| Cryl1 | 1.25E-10 | 0.16161598 | 0.288 | 0.212 | 3.37E-06 | Tmem135 | Xist |
| Acss1 | 1.28E-10 | 0.13158405 | 0.256 | 0.19 | 3.45E-06 | Sgms1 | |
| Cdc42 | 1.34E-10 | 0.05128645 | 0.422 | 0.346 | 3.60E-06 | Setd5 | |
| Pacsin2 | 1.34E-10 | 0.01721426 | 0.282 | 0.23 | 3.61E-06 | Pip4k2a | |
| Spire1 | 1.36E-10 | 0.02524549 | 0.393 | 0.327 | 3.67E-06 | Itgav | |
| Arih1 | 1.39E-10 | 0.10204806 | 0.423 | 0.339 | 3.74E-06 | Slc24a3 | |
| Cnot4 | 1.40E-10 | -0.0983657 | 0.385 | 0.35 | 3.76E-06 | Il6st | |
| Stab1 | 1.41E-10 | 0.01196117 | 0.317 | 0.241 | 3.79E-06 | Capzb | |
| Cflar | 1.41E-10 | 0.11910973 | 0.403 | 0.321 | 3.79E-06 | Tacc1 | |
| Lpcat2 | 1.45E-10 | 0.08177331 | 0.612 | 0.513 | 3.89E-06 | Mgat5 | |
| Ccr5 | 1.46E-10 | 0.20421594 | 0.339 | 0.248 | 3.94E-06 | Cradd | |
| Klf3 | 1.47E-10 | -0.0148002 | 0.266 | 0.221 | 3.96E-06 | Gapvd1 | |
| Cyth4 | 1.49E-10 | 0.25846551 | 0.611 | 0.489 | 4.02E-06 | Arhgef6 | |
| Fnbp1 | 1.52E-10 | 0.212449 | 0.499 | 0.389 | 4.10E-06 | Plcg2 | |
| Pmepa1 | 1.55E-10 | 0.15167892 | 0.44 | 0.343 | 4.16E-06 | Tec | |
| Ube2d3 | 1.66E-10 | 0.03809067 | 0.265 | 0.209 | 4.48E-06 | Lrch1 | |
| Thada | 1.68E-10 | 0.05198961 | 0.279 | 0.217 | 4.51E-06 | Zfp652 | |
| Fmnl3 | 1.68E-10 | 0.14167485 | 0.387 | 0.3 | 4.52E-06 | Gm37240 | |
| Tmem243 | 1.68E-10 | 0.1899688 | 0.388 | 0.294 | 4.53E-06 | Nr2c2 | |
| Ankfy1 | 1.71E-10 | 0.24304709 | 0.289 | 0.202 | 4.59E-06 | St3gal3 | |
| Smurf2 | 1.72E-10 | 0.19811112 | 0.34 | 0.256 | 4.64E-06 | Rbm6 | |
| Ccdc93 | 1.73E-10 | 0.28811282 | 0.252 | 0.165 | 4.66E-06 | F11r | |
| Fcho2 | 1.77E-10 | 0.02892941 | 0.487 | 0.412 | 4.77E-06 | Ubr3 | |
| Mlxipl | 1.85E-10 | 0.17917339 | 0.253 | 0.182 | 4.97E-06 | Adap2 | |
| Gnas | 1.85E-10 | -0.3949055 | 0.232 | 0.276 | 4.99E-06 | Prkca | |
| Akap13 | 1.89E-10 | -0.0268776 | 0.538 | 0.472 | 5.08E-06 | Fam172a | |
| Nipbl | 1.96E-10 | -0.0137817 | 0.525 | 0.459 | 5.27E-06 | Pten | |
| Stard9 | 2.01E-10 | -0.0357762 | 0.414 | 0.364 | 5.42E-06 | Erbin | |
| Sh3glb1 | 2.13E-10 | -0.0153872 | 0.316 | 0.272 | 5.72E-06 | P2ry6 | |
| Baz2b | 2.25E-10 | 0.07917709 | 0.495 | 0.413 | 6.05E-06 | Arhgap12 | |
| Pid1 | 2.43E-10 | -0.2636288 | 0.543 | 0.532 | 6.55E-06 | Mtus1 | |
| Cask | 2.54E-10 | 0.07430084 | 0.475 | 0.391 | 6.84E-06 | Vgll4 | |
| Sparc | 2.68E-10 | 0.00830857 | 0.409 | 0.343 | 7.20E-06 | Snx6 | |
| Mbtd1 | 2.71E-10 | -0.0528023 | 0.305 | 0.264 | 7.30E-06 | Aim2 | |
| Kmt2c | 2.77E-10 | -0.0539323 | 0.434 | 0.388 | 7.46E-06 | Atxn1 | |
| Cog5 | 2.83E-10 | -0.0341452 | 0.339 | 0.29 | 7.63E-06 | Rffl | |
| Ncoa2 | 2.91E-10 | 0.00480187 | 0.376 | 0.321 | 7.82E-06 | Mon2 | |
| Syngr1 | 3.01E-10 | 0.00269846 | 0.267 | 0.214 | 8.10E-06 | Qk | |
| Birc6 | 3.21E-10 | 0.03473833 | 0.358 | 0.299 | 8.63E-06 | Gatad2b | |
| Lcorl | 3.25E-10 | -0.1606209 | 0.286 | 0.267 | 8.75E-06 | Dock1 | |
| Mllt10 | 3.27E-10 | 0.00969206 | 0.435 | 0.373 | 8.79E-06 | Pan3 | |
| Mapkap1 | 3.31E-10 | 0.00547529 | 0.253 | 0.207 | 8.92E-06 | Ccnt2 | |
| Mknk1 | 3.32E-10 | 0.15627516 | 0.368 | 0.285 | 8.94E-06 | Usp9x | |
| Ptp4a2 | 3.38E-10 | -0.0997834 | 0.296 | 0.269 | 9.08E-06 | Rreb1 | |
| Rtn4 | 3.44E-10 | 0.05061446 | 0.434 | 0.358 | 9.26E-06 | Rtn4rl1 | |
| P2rx7 | 3.51E-10 | -0.2711231 | 0.346 | 0.322 | 9.45E-06 | Rhoq | |
| Ctsb | 3.60E-10 | -0.4279967 | 0.25 | 0.3 | 9.70E-06 | Rsbn1l | |
| Gm26827 | 3.68E-10 | 0.25687492 | 0.303 | 0.211 | 9.90E-06 | Phf3 | |
| Hnrnph1 | 3.70E-10 | 0.03814813 | 0.354 | 0.29 | 9.96E-06 | Eml6 | |
| Snx2 | 3.77E-10 | 0.10624803 | 0.271 | 0.202 | 1.01E-05 | Cdk13 | |
| Srsf7 | 3.82E-10 | 0.00932628 | 0.429 | 0.365 | 1.03E-05 | Acin1 | |
| Snx5 | 3.83E-10 | 0.01648751 | 0.335 | 0.279 | 1.03E-05 | A630001G21Rik | |
| Ppp6r3 | 3.85E-10 | 0.09904843 | 0.272 | 0.211 | 1.04E-05 | Tbc1d14 | |
| Zfp608 | 3.90E-10 | 0.02711554 | 0.259 | 0.21 | 1.05E-05 | Rrbp1 | |
| Fndc3b | 4.03E-10 | 0.12869729 | 0.281 | 0.21 | 1.09E-05 | Ddx5 | |
| Cfh | 4.05E-10 | 0.07746849 | 0.667 | 0.57 | 1.09E-05 | Ikzf1 | |
| Zfp950 | 4.09E-10 | -0.0028261 | 0.276 | 0.228 | 1.10E-05 | Pros1 | |
| Nckap1l | 4.11E-10 | 0.05592381 | 0.43 | 0.36 | 1.11E-05 | Pik3cd | |
| Supt20 | 4.21E-10 | 0.10738657 | 0.254 | 0.189 | 1.13E-05 | Strn3 | |
| Eml4 | 4.23E-10 | 0.04508691 | 0.424 | 0.35 | 1.14E-05 | Bptf | |
| Slc7a7 | 4.53E-10 | 0.13951111 | 0.302 | 0.229 | 1.22E-05 | Nfat5 | |
| Rnf111 | 4.66E-10 | -0.0109802 | 0.336 | 0.285 | 1.25E-05 | Pafah1b1 | |
| Hipk2 | 4.71E-10 | 0.20974066 | 0.314 | 0.228 | 1.27E-05 | Gm16599 | |
| Sltm | 4.74E-10 | -0.1050056 | 0.311 | 0.285 | 1.27E-05 | Vav1 | |
| Huwe1 | 4.77E-10 | -0.0684852 | 0.311 | 0.272 | 1.28E-05 | Xiap | |
| Pag1 | 4.81E-10 | 0.18542692 | 0.853 | 0.75 | 1.30E-05 | Apba1 | |
| Sf3b1 | 5.12E-10 | 0.03194672 | 0.425 | 0.36 | 1.38E-05 | Hdac9 | |
| Nisch | 5.24E-10 | 0.09401234 | 0.29 | 0.222 | 1.41E-05 | Pde3b | |
| Apbb2 | 5.40E-10 | -0.3136357 | 0.289 | 0.305 | 1.45E-05 | Hivep3 | |
| Hmg20a | 5.40E-10 | 0.02509741 | 0.25 | 0.198 | 1.45E-05 | Zc3h7a | |
| Rbm5 | 5.43E-10 | -0.0599485 | 0.359 | 0.324 | 1.46E-05 | Usp15 | |
| Pik3r5 | 5.65E-10 | 0.25262618 | 0.276 | 0.189 | 1.52E-05 | Abi1 | |
| Abca9 | 5.76E-10 | 0.01098995 | 0.508 | 0.434 | 1.55E-05 | Capza1 | |
| Gtdc1 | 5.78E-10 | -0.0180198 | 0.403 | 0.341 | 1.55E-05 | Ifngr1 | |
| Pwwp2a | 5.84E-10 | 0.1086974 | 0.443 | 0.361 | 1.57E-05 | Ulk2 | |
| Fry | 6.13E-10 | 0.27338073 | 0.298 | 0.205 | 1.65E-05 | Rbm33 | |
| Taf15 | 6.61E-10 | 0.022401 | 0.325 | 0.273 | 1.78E-05 | Mark2 | |
| Fyb | 6.71E-10 | -0.092627 | 0.727 | 0.67 | 1.80E-05 | Ivns1abp | |
| Camk2d | 6.78E-10 | 0.13348095 | 0.567 | 0.462 | 1.82E-05 | Ubac2 | |
| Ctss | 6.82E-10 | -0.1886206 | 0.681 | 0.649 | 1.83E-05 | Pias2 | |
| Sft2d1 | 6.87E-10 | 0.129115 | 0.408 | 0.331 | 1.85E-05 | Zeb1 | |
| Rbm26 | 7.01E-10 | -0.1278314 | 0.471 | 0.45 | 1.89E-05 | Itpr2 | |
| Tbc1d23 | 7.13E-10 | 0.11114187 | 0.284 | 0.221 | 1.92E-05 | Cd180 | |
| Oxct1 | 7.20E-10 | -0.1817575 | 0.299 | 0.292 | 1.94E-05 | Cacnb2 | |
| Map2k1 | 7.42E-10 | 0.17710156 | 0.262 | 0.187 | 2.00E-05 | Tjp1 | |
| Stag1 | 7.42E-10 | -0.0635582 | 0.529 | 0.479 | 2.00E-05 | Ankrd11 | |
| Slc16a6 | 7.50E-10 | 0.20074833 | 0.333 | 0.246 | 2.02E-05 | Rcbtb2 | |
| Ddx6 | 7.55E-10 | 0.01921799 | 0.289 | 0.236 | 2.03E-05 | Gpr34 | |
| Bach2 | 7.65E-10 | -0.1114674 | 0.439 | 0.391 | 2.06E-05 | Phc2 | |
| Map4k4 | 7.67E-10 | 0.13629827 | 0.534 | 0.434 | 2.06E-05 | Rab31 | |
| Mkln1 | 7.72E-10 | 0.05535038 | 0.407 | 0.34 | 2.08E-05 | Iffo1 | |
| Trappc8 | 7.92E-10 | -0.0370653 | 0.271 | 0.228 | 2.13E-05 | 1-Mar | |
| Sall1 | 8.14E-10 | 0.10147132 | 0.348 | 0.275 | 2.19E-05 | Pias1 | |
| Map3k5 | 8.22E-10 | -0.0710396 | 0.342 | 0.301 | 2.21E-05 | Fbxl17 | |
| Bmpr2 | 8.53E-10 | -0.1020219 | 0.272 | 0.241 | 2.30E-05 | Larp1 | |
| Dennd1a | 8.82E-10 | 0.01493831 | 0.451 | 0.383 | 2.37E-05 | Chd2 | |
| Heatr5a | 9.81E-10 | -0.0662881 | 0.342 | 0.304 | 2.64E-05 | E2f3 | |
| Csnk1a1 | 1.00E-09 | 0.01342401 | 0.282 | 0.234 | 2.70E-05 | Nudcd3 | |
| Cables1 | 1.00E-09 | 0.12610732 | 0.3 | 0.228 | 2.70E-05 | Tnfrsf11a | |
| Tpp2 | 1.02E-09 | 0.09719982 | 0.255 | 0.194 | 2.74E-05 | Mia2 | |
| Peli2 | 1.04E-09 | 0.04610526 | 0.282 | 0.225 | 2.79E-05 | Ubn2 | |
| Rb1cc1 | 1.07E-09 | 0.01898579 | 0.27 | 0.218 | 2.88E-05 | Fbxl20 | |
| Ccdc50 | 1.11E-09 | 0.11969656 | 0.321 | 0.248 | 2.99E-05 | Plxna4 | |
| Gria2 | 1.13E-09 | -0.4127169 | 0.258 | 0.294 | 3.04E-05 | Arl15 | |
| Sppl3 | 1.18E-09 | 0.16368678 | 0.405 | 0.317 | 3.17E-05 | Rere | |
| Tcf4 | 1.20E-09 | -0.0964387 | 0.594 | 0.548 | 3.24E-05 | Unc93b1 | |
| Paxbp1 | 1.26E-09 | -0.0896152 | 0.258 | 0.23 | 3.39E-05 | Cx3cr1 | |
| Ccnl1 | 1.26E-09 | 0.00647788 | 0.258 | 0.214 | 3.40E-05 | Pld1 | |
| Kansl1l | 1.37E-09 | -0.0910772 | 0.344 | 0.307 | 3.69E-05 | Cpsf6 | |
| Ifngr2 | 1.38E-09 | -0.0174526 | 0.258 | 0.221 | 3.71E-05 | Zcchc7 | |
| Ptpn1 | 1.39E-09 | 0.05672089 | 0.351 | 0.288 | 3.74E-05 | Eef2k | |
| Ralgapa1 | 1.45E-09 | -0.1082393 | 0.26 | 0.23 | 3.91E-05 | Maml3 | |
| Cd2ap | 1.50E-09 | 0.02936032 | 0.329 | 0.272 | 4.02E-05 | Actr3 | |
| Trio | 1.52E-09 | 0.19592369 | 0.401 | 0.309 | 4.08E-05 | Igf1r | |
| Top1 | 1.55E-09 | -0.0629174 | 0.275 | 0.242 | 4.17E-05 | Nrip1 | |
| Sbf2 | 1.56E-09 | -0.0132464 | 0.477 | 0.421 | 4.20E-05 | Arpc2 | |
| Cep68 | 1.77E-09 | 0.08186478 | 0.282 | 0.223 | 4.76E-05 | Map4k3 | |
| Phf20l1 | 1.87E-09 | -0.078979 | 0.294 | 0.261 | 5.02E-05 | Tmcc3 | |
| Casp8 | 1.89E-09 | 0.13200743 | 0.28 | 0.211 | 5.07E-05 | Ago3 | |
| Cltc | 1.90E-09 | -0.0937124 | 0.339 | 0.315 | 5.11E-05 | Ophn1 | |
| Fus | 1.91E-09 | 0.04471491 | 0.523 | 0.45 | 5.14E-05 | Lims1 | |
| Pnisr | 1.93E-09 | 0.04313645 | 0.502 | 0.429 | 5.20E-05 | Mef2a | |
| Atxn7 | 1.96E-09 | 0.00583741 | 0.303 | 0.252 | 5.27E-05 | Rps6ka1 | |
| Rhobtb1 | 1.97E-09 | 0.08888037 | 0.353 | 0.284 | 5.31E-05 | Gm5086 | |
| Cdk6 | 1.99E-09 | 0.25354578 | 0.31 | 0.216 | 5.36E-05 | Hpgds | |
| Wsb1 | 2.00E-09 | -0.0155226 | 0.354 | 0.298 | 5.38E-05 | Tmem164 | |
| Akt3 | 2.05E-09 | -0.4239941 | 0.249 | 0.3 | 5.52E-05 | Trpm7 | |
| Epc1 | 2.10E-09 | 0.11646497 | 0.278 | 0.211 | 5.64E-05 | Bcas3 | |
| Sh3pxd2a | 2.13E-09 | 0.14418966 | 0.33 | 0.254 | 5.73E-05 | Csf1r | |

| | | | | | |
|---|---|---|---|---|---|
| Smad7 | 1.78E-07 | -0.0247258 | 0.214 | 0.264 | 0.00480171 |
| Klf3 | 1.83E-07 | 0.04224557 | 0.221 | 0.252 | 0.00492196 |
| Ndfip1 | 1.87E-07 | -0.056531 | 0.213 | 0.261 | 0.00502999 |
| St3gal6 | 1.89E-07 | -0.0094695 | 0.518 | 0.587 | 0.00507571 |
| Rbm25 | 1.97E-07 | 0.0880982 | 0.384 | 0.42 | 0.00530309 |
| Snrnp70 | 2.03E-07 | -0.0252161 | 0.447 | 0.512 | 0.00545107 |
| Herc4 | 2.10E-07 | 0.11920486 | 0.254 | 0.273 | 0.00565096 |
| Ep300 | 2.11E-07 | -0.0910552 | 0.258 | 0.318 | 0.00567671 |
| Usp50 | 7.88E-09 | -0.1368424 | 0.278 | 0.357 | 0.00021193 |
| Ncoa3 | 2.18E-07 | 0.04717822 | 0.279 | 0.309 | 0.0058654 |
| Ptbp3 | 2.24E-07 | -0.0410745 | 0.339 | 0.397 | 0.0060165 |
| Atf7ip | 1.72E-06 | -0.1356557 | 0.252 | 0.319 | 0.04639886 |
| Anks1 | 2.24E-07 | -0.0091474 | 0.228 | 0.27 | 0.00603797 |
| Cd180 | 2.25E-07 | 0.15437413 | 0.25 | 0.26 | 0.00606158 |
| Pacs1 | 2.31E-07 | 0.09262067 | 0.273 | 0.299 | 0.00620768 |
| Atp6v0b | 1.76E-06 | -0.1344904 | 0.489 | 0.576 | 0.04722185 |
| Pcm1 | 2.68E-08 | -0.1342895 | 0.255 | 0.325 | 0.00072208 |
| Stag2 | 2.46E-07 | -0.0177687 | 0.267 | 0.31 | 0.00661543 |
| Map3k3 | 2.49E-07 | -0.0573568 | 0.354 | 0.425 | 0.00669341 |
| Rsf1 | 2.49E-07 | 0.00641851 | 0.24 | 0.278 | 0.00669676 |
| Fth1 | 2.51E-07 | 0.4292431 | 0.273 | 0.208 | 0.00676701 |
| Hck | 2.54E-07 | -0.0399358 | 0.282 | 0.337 | 0.00682205 |
| Gm16599 | 9.55E-08 | -0.1318771 | 0.253 | 0.32 | 0.00257051 |
| Ptpn1 | 5.32E-08 | -0.1308342 | 0.288 | 0.362 | 0.00143059 |
| Serinc3 | 2.75E-07 | -0.0754763 | 0.59 | 0.675 | 0.00738978 |
| R3hdm2 | 2.77E-07 | -0.0297711 | 0.294 | 0.345 | 0.00744745 |
| Nlk | 7.31E-11 | -0.1294311 | 0.233 | 0.306 | 1.97E-06 |
| Daglb | 5.81E-08 | -0.128101 | 0.247 | 0.313 | 0.00156415 |
| Nsd3 | 3.09E-07 | 0.1771319 | 0.353 | 0.357 | 0.00830248 |
| Plcg2 | 3.83E-09 | -0.1279164 | 0.338 | 0.417 | 0.00010315 |
| Vgll4 | 2.23E-11 | -0.1256547 | 0.23 | 0.302 | 6.00E-07 |
| Fbxl17 | 3.21E-07 | -0.0806233 | 0.489 | 0.569 | 0.00862924 |
| Mdm4 | 3.23E-07 | 0.19481811 | 0.251 | 0.244 | 0.00869651 |
| Mgat4a | 3.24E-07 | -0.0157995 | 0.43 | 0.49 | 0.00870919 |
| Rnf13 | 6.05E-08 | -0.1247747 | 0.295 | 0.367 | 0.0016273 |
| Papola | 3.26E-07 | -0.0861839 | 0.282 | 0.344 | 0.00877326 |
| Usp34 | 4.19E-10 | -0.1246574 | 0.377 | 0.468 | 1.13E-05 |
| Msi2 | 3.37E-07 | -0.0673668 | 0.253 | 0.309 | 0.00906255 |
| Nsmce2 | 3.49E-07 | 0.15618052 | 0.245 | 0.253 | 0.00937969 |
| Mafb | 3.62E-07 | -0.0003981 | 0.286 | 0.331 | 0.00973394 |
| Top1 | 3.63E-07 | -0.0471047 | 0.242 | 0.291 | 0.00976739 |
| Abcc5 | 2.68E-09 | -0.1227217 | 0.199 | 0.267 | 7.22E-05 |
| Mapk14 | 3.64E-07 | 0.03836417 | 0.317 | 0.352 | 0.00979129 |
| Tsc22d4 | 3.08E-08 | -0.1216641 | 0.256 | 0.328 | 0.00082837 |
| Arih1 | 3.90E-07 | -0.0020428 | 0.339 | 0.388 | 0.01048046 |
| Cnot4 | 4.01E-07 | -0.0196092 | 0.35 | 0.403 | 0.01079136 |
| Phf21a | 4.03E-07 | -0.0313766 | 0.392 | 0.458 | 0.01083548 |
| Il10ra | 4.10E-07 | 0.05413176 | 0.292 | 0.324 | 0.01103978 |
| Rcsd1 | 1.41E-16 | -0.1205906 | 0.289 | 0.395 | 3.80E-12 |
| Nr3c1 | 4.56E-07 | -0.1202522 | 0.283 | 0.349 | 0.01227641 |
| 2610020C07 | 1.41E-09 | -0.1189819 | 0.191 | 0.259 | 3.80E-05 |
| Nptn | 4.58E-07 | 0.23022665 | 0.322 | 0.314 | 0.01233137 |
| Setd2 | 4.71E-07 | 0.00877662 | 0.291 | 0.332 | 0.0126773 |
| Map4 | 3.69E-09 | -0.1187002 | 0.364 | 0.451 | 9.93E-05 |
| Ssbp2 | 4.93E-07 | 0.06408719 | 0.297 | 0.33 | 0.0132586 |
| Canx | 5.05E-07 | -0.0670915 | 0.217 | 0.268 | 0.01358643 |
| Notch2 | 1.08E-08 | -0.1181636 | 0.202 | 0.263 | 0.00029152 |
| Csnk1a1 | 5.14E-07 | -0.0425506 | 0.234 | 0.279 | 0.01382548 |
| Itm2b | 5.25E-07 | 0.12051581 | 0.529 | 0.565 | 0.01411485 |
| Nipbl | 5.47E-07 | -0.0794624 | 0.459 | 0.535 | 0.01470549 |
| P3h2 | 5.62E-07 | 0.10803055 | 0.455 | 0.483 | 0.01512247 |
| Stk3 | 5.65E-07 | 0.00296398 | 0.227 | 0.265 | 0.0152153 |
| Ccny | 5.72E-07 | -0.052909 | 0.234 | 0.285 | 0.01538773 |
| Cdk12 | 6.01E-07 | -0.0507117 | 0.274 | 0.323 | 0.01615998 |
| Ctsh | 6.16E-07 | 0.18097509 | 0.28 | 0.272 | 0.01658655 |
| Wasf2 | 6.33E-07 | -0.0574642 | 0.508 | 0.578 | 0.01702172 |
| Ap3b1 | 6.57E-07 | -0.0261125 | 0.295 | 0.338 | 0.01767595 |
| Psap | 6.64E-07 | 0.06494518 | 0.43 | 0.468 | 0.01787454 |
| Nrp1 | 6.64E-07 | 0.23130894 | 0.251 | 0.252 | 0.01787538 |
| Wdr26 | 2.73E-09 | -0.1176309 | 0.276 | 0.347 | 7.35E-05 |
| Ppp6r3 | 3.16E-07 | -0.1169711 | 0.211 | 0.267 | 0.00851186 |
| Ext1 | 6.76E-07 | -0.0518997 | 0.307 | 0.372 | 0.01819404 |
| Dapp1 | 2.17E-10 | -0.1161091 | 0.319 | 0.407 | 5.83E-06 |
| Atxn7l1 | 7.06E-07 | -0.0148829 | 0.3 | 0.348 | 0.01898619 |
| Smg6 | 7.21E-07 | 0.06487396 | 0.368 | 0.395 | 0.01941195 |
| Ewsr1 | 7.54E-07 | -0.087593 | 0.236 | 0.288 | 0.0202912 |
| Peli2 | 7.56E-07 | -0.0990191 | 0.225 | 0.281 | 0.02033159 |
| Gdi2 | 7.57E-07 | 0.09728403 | 0.347 | 0.367 | 0.02036907 |
| Tnrc6c | 7.87E-07 | 0.06324985 | 0.322 | 0.35 | 0.02118019 |
| Prkcb | 7.91E-07 | 0.2136758 | 0.447 | 0.432 | 0.02128382 |
| Nuak1 | 8.05E-07 | -0.0166922 | 0.231 | 0.278 | 0.02165194 |
| Bin1 | 8.24E-07 | 0.1142539 | 0.523 | 0.547 | 0.02215995 |
| Ube3a | 8.51E-07 | 0.01067567 | 0.328 | 0.371 | 0.02289714 |
| Pafah1b1 | 8.56E-07 | -0.0503428 | 0.342 | 0.4 | 0.0230458 |
| Sall1 | 8.64E-07 | 0.14419971 | 0.275 | 0.284 | 0.02325355 |
| Gna12 | 8.84E-07 | 0.1605476 | 0.344 | 0.344 | 0.02377554 |
| Gm26848 | 8.95E-07 | -0.0041939 | 0.306 | 0.251 | 0.02407116 |
| Ctsc | 9.14E-07 | 0.11562656 | 0.232 | 0.254 | 0.02458187 |
| C1qa | 1.23E-07 | -0.1142173 | 0.594 | 0.694 | 0.00332022 |
| Cd33 | 9.48E-07 | -0.0202997 | 0.243 | 0.287 | 0.02549631 |
| Skap2 | 9.76E-07 | 0.20231951 | 0.504 | 0.494 | 0.02625826 |
| Sesn1 | 1.02E-06 | 0.03642736 | 0.242 | 0.267 | 0.02754578 |
| Nisch | 1.04E-06 | -0.0975426 | 0.222 | 0.279 | 0.02803522 |
| Cep170 | 8.41E-13 | -0.1118368 | 0.25 | 0.335 | 2.26E-08 |
| Capza1 | 1.06E-06 | 0.00756818 | 0.233 | 0.268 | 0.02850643 |
| Il6ra | 1.06E-06 | -0.0579553 | 0.385 | 0.448 | 0.02862118 |
| Dnmt3a | 1.07E-06 | 0.06302296 | 0.358 | 0.391 | 0.0287373 |
| Sat1 | 1.11E-06 | -0.047399 | 0.316 | 0.368 | 0.02992773 |
| Rps6ka1 | 1.17E-06 | 0.07196253 | 0.236 | 0.253 | 0.03149571 |
| Fermt3 | 1.17E-06 | -0.0364595 | 0.294 | 0.345 | 0.03155532 |
| Pik3ap1 | 1.17E-06 | 0.01458726 | 0.33 | 0.367 | 0.03161237 |
| Rbm6 | 1.21E-06 | -0.0096937 | 0.328 | 0.371 | 0.03251449 |
| Rtn4 | 1.21E-06 | 0.02037735 | 0.358 | 0.404 | 0.03262596 |
| Agps | 1.22E-06 | -0.0219538 | 0.268 | 0.313 | 0.03290273 |
| Pkn1 | 1.43E-11 | -0.1116591 | 0.241 | 0.32 | 3.86E-07 |
| Plekhm3 | 1.23E-06 | 0.09114054 | 0.236 | 0.256 | 0.03322936 |
| Setd5 | 6.75E-07 | -0.1115548 | 0.32 | 0.391 | 0.01815049 |
| Bmp2k | 5.13E-08 | -0.1108059 | 0.536 | 0.63 | 0.00137959 |
| Cux1 | 1.32E-06 | 0.12882929 | 0.446 | 0.461 | 0.03550222 |
| Dync1i2 | 1.35E-06 | 0.05379456 | 0.224 | 0.253 | 0.03639552 |
| Kansl1l | 1.38E-06 | -0.0508278 | 0.307 | 0.363 | 0.03709086 |
| Specc1 | 8.69E-08 | -0.1103094 | 0.502 | 0.591 | 0.00233703 |
| Rbm26 | 1.41E-06 | -0.053536 | 0.45 | 0.511 | 0.03784834 |
| Tpr | 2.57E-08 | -0.1092523 | 0.213 | 0.276 | 0.00069171 |
| Gns | 1.42E-06 | -0.0453084 | 0.383 | 0.438 | 0.03813239 |
| Casp8 | 1.43E-06 | -0.0875712 | 0.211 | 0.261 | 0.03854768 |
| Uvrag | 3.26E-07 | -0.1086551 | 0.382 | 0.45 | 0.00875832 |
| Rtn3 | 1.49E-06 | 0.13645556 | 0.424 | 0.441 | 0.03999398 |
| Atrx | 1.52E-06 | 0.09234512 | 0.36 | 0.377 | 0.04094255 |
| AW554918 | 1.55E-06 | 0.05735403 | 0.239 | 0.266 | 0.0416352 |
| Zfp292 | 8.17E-10 | -0.1085014 | 0.301 | 0.379 | 2.20E-05 |
| Ogt | 1.43E-08 | -0.1064504 | 0.259 | 0.327 | 0.00038562 |
| Mkln1 | 4.35E-10 | -0.1048694 | 0.34 | 0.421 | 1.17E-05 |
| Gtdc1 | 1.65E-06 | 0.0661774 | 0.341 | 0.378 | 0.04429605 |
| Prrc2c | 2.72E-07 | -0.1046514 | 0.278 | 0.34 | 0.00731156 |
| Rere | 5.12E-07 | -0.1028494 | 0.516 | 0.603 | 0.01378912 |
| Gpatch8 | 1.73E-06 | 0.03254383 | 0.311 | 0.35 | 0.04646213 |
| Baz2b | 1.68E-08 | -0.1018746 | 0.413 | 0.497 | 0.000451 |
| Lims1 | 1.77E-06 | 0.09549739 | 0.414 | 0.44 | 0.04759134 |
| Abi3 | 1.79E-06 | 0.09214334 | 0.291 | 0.307 | 0.04815841 |
| Nsd1 | 1.80E-06 | 0.03670046 | 0.535 | 0.58 | 0.04832673 |
| Rgs10 | 1.81E-06 | 0.03115644 | 0.412 | 0.456 | 0.04877898 |
| Pmepa1 | 1.83E-06 | -0.0482312 | 0.343 | 0.397 | 0.0492948 |
| Zfand3 | 5.71E-10 | -0.1013421 | 0.392 | 0.476 | 1.54E-05 |
| Insr | 1.87E-06 | 0.04510284 | 0.245 | 0.276 | 0.05018203 |
| Lrmda | 1.87E-06 | 0.27597025 | 0.788 | 0.731 | 0.05024057 |
| Tcf12 | 1.91E-06 | 0.0834465 | 0.403 | 0.425 | 0.05135313 |
| Rab3gap1 | 2.08E-06 | -0.1915454 | 0.198 | 0.258 | 0.055952 |
| Prkce | 2.10E-06 | -0.1806233 | 0.391 | 0.479 | 0.05643899 |
| Tcf20 | 2.11E-06 | -0.0577472 | 0.273 | 0.327 | 0.05666926 |
| Slc38a9 | 2.18E-06 | 0.06253937 | 0.239 | 0.264 | 0.05858135 |
| Sfmbt1 | 2.20E-06 | 0.15810269 | 0.265 | 0.273 | 0.05925044 |
| Fnip1 | 2.20E-06 | 0.05910198 | 0.311 | 0.338 | 0.05925299 |
| Slc16a6 | 2.29E-06 | -0.0944377 | 0.246 | 0.3 | 0.06156184 |
| Fubp1 | 2.30E-06 | -0.0262872 | 0.266 | 0.307 | 0.06186629 |
| Nedd4l | 2.32E-06 | -0.0998451 | 0.2 | 0.257 | 0.06230475 |
| Arrb2 | 2.50E-06 | -0.173656 | 0.194 | 0.254 | 0.06719989 |
| Fus | 2.64E-06 | 0.00976051 | 0.45 | 0.494 | 0.07102061 |
| Vti1a | 2.73E-06 | -0.0751973 | 0.339 | 0.401 | 0.07347755 |
| Fcer1g | 2.80E-06 | 0.01109701 | 0.248 | 0.285 | 0.0754644 |
| Gab1 | 2.94E-06 | 0.11661143 | 0.255 | 0.272 | 0.07907263 |
| Cd37 | 2.95E-06 | -0.0904131 | 0.382 | 0.445 | 0.07950523 |
| Smchd1 | 2.97E-06 | -0.1287534 | 0.197 | 0.253 | 0.07995205 |
| Comt | 2.98E-06 | -0.1159291 | 0.317 | 0.381 | 0.08013762 |
| Blnk | 2.98E-06 | 0.17599016 | 0.456 | 0.454 | 0.08023845 |
| Tubgcp5 | 3.00E-06 | -0.0213796 | 0.273 | 0.318 | 0.08067941 |
| Tnrc6b | 3.02E-06 | 0.01577746 | 0.439 | 0.484 | 0.08125594 |
| Bbs9 | 3.07E-06 | -0.2717989 | 0.428 | 0.527 | 0.08258858 |
| Dock10 | 3.09E-06 | -0.1701808 | 0.675 | 0.765 | 0.08326604 |
| Nbea | 3.17E-06 | 0.19590642 | 0.285 | 0.287 | 0.08517488 |
| 4930444A19 | 3.26E-06 | -0.2762362 | 0.323 | 0.412 | 0.08760263 |
| Actr2 | 3.26E-06 | -0.026269 | 0.33 | 0.376 | 0.08764544 |
| Exoc6b | 3.29E-06 | -0.2048595 | 0.322 | 0.406 | 0.08848559 |
| Gapvd1 | 3.37E-06 | 0.07471245 | 0.263 | 0.283 | 0.09068485 |
| Synj1 | 3.38E-06 | -0.0220475 | 0.221 | 0.256 | 0.09085508 |
| Itch | 3.42E-06 | -0.0807119 | 0.373 | 0.437 | 0.09200589 |
| Etv6 | 3.42E-06 | 0.03927057 | 0.499 | 0.544 | 0.09205036 |
| Sltm | 3.44E-06 | -0.0762979 | 0.285 | 0.34 | 0.09259325 |
| Rrbp1 | 3.55E-06 | -0.2397878 | 0.495 | 0.592 | 0.09539377 |
| Slc1a3 | 3.59E-06 | -0.3385296 | 0.242 | 0.33 | 0.09668249 |
| Fgfr2 | 3.62E-06 | 0.14708123 | 0.422 | 0.436 | 0.09745927 |
| Nrip1 | 3.76E-06 | -0.2022399 | 0.496 | 0.593 | 0.1012817 |
| Phf3 | 3.78E-06 | -0.0061872 | 0.249 | 0.285 | 0.10158174 |
| Dock11 | 3.91E-06 | 0.06749942 | 0.333 | 0.359 | 0.1050999 |
| Ttc14 | 4.05E-06 | 0.02381263 | 0.315 | 0.342 | 0.10897042 |
| Igf1r | 4.16E-06 | 0.08427583 | 0.262 | 0.286 | 0.11185454 |
| Dock4 | 4.16E-06 | -0.1882728 | 0.88 | 0.91 | 0.11205084 |
| Aim2 | 4.18E-06 | -0.0841918 | 0.282 | 0.335 | 0.11255775 |
| Tmem135 | 4.23E-06 | -0.0012122 | 0.408 | 0.455 | 0.11370333 |
| Ncf1 | 4.42E-06 | 0.05627194 | 0.264 | 0.29 | 0.11893558 |
| Pde4d | 4.57E-06 | -0.0576254 | 0.343 | 0.403 | 0.12289137 |
| Pds5a | 4.74E-06 | -0.0088026 | 0.231 | 0.271 | 0.12747864 |
| Rapgef2 | 4.76E-06 | -0.0714203 | 0.236 | 0.289 | 0.1280397 |
| Smurf2 | 4.78E-06 | -0.1240972 | 0.256 | 0.31 | 0.12861433 |
| Csmd3 | 4.84E-06 | -0.1985781 | 0.87 | 0.893 | 0.13022947 |
| Zdhhc20 | 4.97E-06 | -0.108114 | 0.241 | 0.298 | 0.13359435 |
| Arhgap12 | 5.00E-06 | -0.0847039 | 0.362 | 0.422 | 0.1346166 |
| Gna15 | 5.03E-06 | -0.1354154 | 0.253 | 0.314 | 0.1354075 |
| Son | 5.08E-06 | 0.01563093 | 0.508 | 0.55 | 0.13668858 |
| Cnot2 | 5.27E-06 | -0.0301968 | 0.329 | 0.376 | 0.14185808 |
| Lrrc4 | 5.40E-06 | -0.1109896 | 0.231 | 0.296 | 0.14536402 |
| Arl15 | 5.41E-06 | -0.1776458 | 0.39 | 0.475 | 0.14545829 |
| Zcchc11 | 5.48E-06 | 0.12004734 | 0.321 | 0.249 | 0.1474583 |
| Kcnk13 | 5.60E-06 | 0.25609247 | 0.28 | 0.259 | 0.15073151 |
| Mark2 | 5.61E-06 | -0.0553825 | 0.239 | 0.285 | 0.15100478 |
| Pik3r1 | 5.65E-06 | -0.1320527 | 0.438 | 0.519 | 0.1519247 |
| Rffl | 5.84E-06 | -0.041805 | 0.238 | 0.283 | 0.15700793 |
| Nrd1 | 6.12E-06 | -0.0499592 | 0.228 | 0.273 | 0.16464858 |
| C1qb | 6.14E-06 | -0.0057428 | 0.563 | 0.628 | 0.16522044 |
| Mbtd1 | 6.70E-06 | 0.07386157 | 0.264 | 0.281 | 0.18014571 |
| Fli1 | 6.72E-06 | 0.05478272 | 0.613 | 0.66 | 0.18084732 |
| Wac | 6.90E-06 | 0.01983571 | 0.281 | 0.311 | 0.18555756 |
| R3hdm1 | 6.90E-06 | 0.00760713 | 0.302 | 0.339 | 0.18559087 |
| Hipk2 | 6.92E-06 | -0.1631177 | 0.228 | 0.293 | 0.18632481 |
| Oxr1 | 7.02E-06 | 0.1327591 | 0.41 | 0.423 | 0.18901097 |
| Gm5086 | 7.22E-06 | -0.0810566 | 0.318 | 0.374 | 0.19436697 |
| Camk1 | 7.68E-06 | 0.12751472 | 0.294 | 0.301 | 0.20670007 |
| Chd9 | 7.91E-06 | -0.1671991 | 0.773 | 0.847 | 0.21295335 |
| Stx16 | 7.92E-06 | -0.018066 | 0.25 | 0.289 | 0.21316349 |
| Ulk2 | 8.12E-06 | -0.0602693 | 0.468 | 0.533 | 0.2184819 |
| Taf15 | 8.24E-06 | -0.0600757 | 0.273 | 0.319 | 0.22159888 |
| Bptf | 8.65E-06 | -0.0036496 | 0.288 | 0.324 | 0.23286262 |
| Tbl1x | 8.84E-06 | 0.05569161 | 0.239 | 0.263 | 0.23776333 |
| Mier1 | 8.85E-06 | -0.1409421 | 0.196 | 0.252 | 0.23821337 |
| Klf12 | 9.15E-06 | 0.19658541 | 0.429 | 0.423 | 0.24607997 |
| Lnpep | 9.65E-06 | 0.02126885 | 0.267 | 0.302 | 0.25962009 |
| Slc7a7 | 9.85E-06 | -0.1113202 | 0.229 | 0.283 | 0.26493575 |
| Phip | 1.05E-05 | -0.0236085 | 0.268 | 0.313 | 0.28222161 |
| Xiap | 1.06E-05 | -0.1737123 | 0.273 | 0.338 | 0.2850409 |
| Kpna4 | 1.11E-05 | 0.07177408 | 0.297 | 0.319 | 0.29913331 |
| Trio | 1.12E-05 | -0.0048515 | 0.309 | 0.347 | 0.30154918 |
| Srgap2 | 1.17E-05 | -0.1722557 | 0.911 | 0.943 | 0.31408975 |
| Pde3b | 1.18E-05 | -0.195258 | 0.724 | 0.8 | 0.31728027 |
| Ago3 | 1.20E-05 | -0.0534458 | 0.248 | 0.291 | 0.32283407 |
| Kansl1 | 1.22E-05 | -0.1092118 | 0.548 | 0.629 | 0.3285325 |
| Akt3 | 1.23E-05 | 0.25262559 | 0.3 | 0.286 | 0.32967044 |
| Pias1 | 1.23E-05 | 0.03337186 | 0.359 | 0.391 | 0.33013846 |
| Ubn2 | 1.23E-05 | 0.05630304 | 0.399 | 0.426 | 0.33065774 |
| Gls | 1.30E-05 | 0.08992342 | 0.283 | 0.298 | 0.34849178 |
| Hivep3 | 1.33E-05 | -0.0621256 | 0.432 | 0.495 | 0.35681112 |
| Strn3 | 1.36E-05 | 0.12715243 | 0.375 | 0.384 | 0.36520808 |
| Tcf4 | 1.37E-05 | 0.02940827 | 0.548 | 0.593 | 0.36875039 |
| Tsix | 1.38E-05 | -0.2147471 | 0.248 | 0.32 | 0.37129359 |
| Zranb2 | 1.40E-05 | -0.1172168 | 0.204 | 0.254 | 0.37633028 |
| Bclaf1 | 1.40E-05 | 0.02122666 | 0.23 | 0.258 | 0.37795347 |
| Slco2b1 | 1.42E-05 | -0.2265109 | 0.596 | 0.687 | 0.38240398 |
| Nrf1 | 1.49E-05 | -0.1165134 | 0.215 | 0.266 | 0.40086158 |
| Cttnbp2nl | 1.51E-05 | 0.00771461 | 0.457 | 0.498 | 0.40533475 |
| Fam117b | 1.51E-05 | -0.0429208 | 0.235 | 0.278 | 0.40693498 |

| | | | | | | |
|---|---|---|---|---|---|---|
| Ccnd3 | 2.19E-09 | -0.2511369 | 0.714 | 0.699 | 5.89E-05 | Zmynd8 |
| Etv6 | 2.24E-09 | 0.08534192 | 0.59 | 0.499 | 6.02E-05 | Susd6 |
| Slco2b1 | 2.28E-09 | 0.28568631 | 0.714 | 0.596 | 6.13E-05 | Pacs1 |
| Helz | 2.33E-09 | 0.09335534 | 0.282 | 0.22 | 6.26E-05 | Slc9a9 |
| Sipa1l2 | 2.33E-09 | -0.0050705 | 0.386 | 0.331 | 6.28E-05 | Zfp36l1 |
| Snx13 | 2.40E-09 | 0.12279746 | 0.356 | 0.279 | 6.47E-05 | Irf2 |
| Msi2 | 2.41E-09 | 0.04652423 | 0.312 | 0.253 | 6.49E-05 | Med13 |
| Slmap | 2.43E-09 | 0.01181649 | 0.306 | 0.257 | 6.54E-05 | Lyn |
| Ints6l | 2.49E-09 | 0.02754223 | 0.477 | 0.405 | 6.71E-05 | Rsf1 |
| Rtn3 | 2.50E-09 | -0.0609994 | 0.474 | 0.424 | 6.74E-05 | Tnrc6b |
| Camk1 | 2.57E-09 | -0.0769086 | 0.323 | 0.294 | 6.90E-05 | Large1 |
| Srpk2 | 2.67E-09 | 0.01881614 | 0.265 | 0.217 | 7.19E-05 | Entpd1 |
| Comt | 2.74E-09 | -0.0934609 | 0.347 | 0.317 | 7.36E-05 | Dock8 |
| Dgkd | 2.74E-09 | 0.11479537 | 0.414 | 0.332 | 7.37E-05 | Hck |
| Ptprc | 2.75E-09 | -0.1825645 | 0.539 | 0.521 | 7.40E-05 | Lgmn |
| Golm1 | 2.76E-09 | 0.26523618 | 0.387 | 0.288 | 7.43E-05 | B430010I23Rik |
| Retreg1 | 2.80E-09 | 0.26370834 | 0.416 | 0.313 | 7.54E-05 | Fgd2 |
| Btbd1 | 2.85E-09 | -0.0170304 | 0.256 | 0.217 | 7.68E-05 | Arhgap39 |
| Nr3c1 | 2.86E-09 | -0.0185469 | 0.331 | 0.283 | 7.70E-05 | Ppm1h |
| Ttc28 | 3.00E-09 | 0.18027017 | 0.501 | 0.401 | 8.06E-05 | Ntpcr |
| Ube3a | 3.03E-09 | -0.115842 | 0.358 | 0.328 | 8.16E-05 | Scmh1 |
| Dennd1b | 3.09E-09 | 0.02614743 | 0.258 | 0.21 | 8.30E-05 | Wnk1 |
| Spop | 3.30E-09 | -0.0406175 | 0.292 | 0.253 | 8.87E-05 | Csf3r |
| Luc7l3 | 3.32E-09 | 0.09021254 | 0.258 | 0.197 | 8.94E-05 | 4930444A19Rik |
| Pnn | 3.39E-09 | 0.10444645 | 0.303 | 0.238 | 9.13E-05 | Nek7 |
| Stk3 | 3.41E-09 | -0.0857407 | 0.251 | 0.227 | 9.18E-05 | Sirpa |
| Rnf180 | 3.64E-09 | -0.0679489 | 0.458 | 0.413 | 9.79E-05 | Ambra1 |
| Ktn1 | 3.64E-09 | 0.21649036 | 0.29 | 0.209 | 9.81E-05 | Fam168a |
| Vsir | 3.72E-09 | 0.11496712 | 0.469 | 0.391 | 0.00010006 | Hdac8 |
| Fnip2 | 3.72E-09 | -0.2023852 | 0.258 | 0.248 | 0.00010013 | Tra2b |
| Crebbp | 3.73E-09 | 0.08626444 | 0.464 | 0.384 | 0.00010025 | Mir142hg |
| Arid1b | 3.75E-09 | -0.1562575 | 0.412 | 0.398 | 0.00010077 | Mylip |
| Arhgap22 | 3.82E-09 | -0.0433074 | 0.528 | 0.469 | 0.00010284 | Smarca2 |
| Srgap2 | 3.84E-09 | 0.20056248 | 0.962 | 0.911 | 0.00010328 | Ube2h |
| Senp2 | 3.91E-09 | 0.01414735 | 0.319 | 0.266 | 0.0001052 | Tcf20 |
| Wipf1 | 3.92E-09 | -0.0464272 | 0.392 | 0.348 | 0.00010539 | Smg6 |
| Sort1 | 3.93E-09 | 0.08235448 | 0.257 | 0.197 | 0.00010562 | Rap1gds1 |
| Ubl3 | 3.98E-09 | 0.02232955 | 0.267 | 0.217 | 0.00010718 | Rgs10 |
| Rab8b | 4.08E-09 | 0.18448253 | 0.421 | 0.328 | 0.00010973 | Frmd4b |
| Cdk12 | 4.12E-09 | 0.1882355 | 0.358 | 0.274 | 0.00011092 | Cd37 |
| Csad | 4.17E-09 | 0.28529399 | 0.271 | 0.185 | 0.00011219 | Btk |
| Cmip | 4.21E-09 | 0.12002525 | 0.421 | 0.334 | 0.00011337 | Kat6a |
| Atp8a1 | 4.24E-09 | -0.0805417 | 0.514 | 0.471 | 0.00011413 | Ehmt1 |
| Maf | 4.28E-09 | 0.16945041 | 0.484 | 0.388 | 0.00011521 | Fam102b |
| Hook3 | 4.32E-09 | 0.1159911 | 0.308 | 0.242 | 0.00011633 | Ascc3 |
| Aftph | 4.46E-09 | 0.03175974 | 0.44 | 0.378 | 0.00012007 | Tra2a |
| Sfmbt1 | 4.59E-09 | 0.08358029 | 0.335 | 0.265 | 0.00012355 | Tcf7l2 |
| Aff4 | 4.70E-09 | 0.05279541 | 0.274 | 0.218 | 0.00012653 | Nav3 |
| Klf12 | 4.73E-09 | -0.0421489 | 0.486 | 0.429 | 0.00012737 | Dnmt3a |
| Mbnl2 | 4.88E-09 | -0.0419171 | 0.517 | 0.464 | 0.00013118 | Ankrd44 |
| Itga9 | 5.21E-09 | 0.2702376 | 0.388 | 0.288 | 0.00014028 | 4933406I18Rik |
| Wasf2 | 5.23E-09 | 0.08273267 | 0.595 | 0.508 | 0.0001407 | Tsix |
| Foxn3 | 5.32E-09 | 0.23168385 | 0.856 | 0.765 | 0.00014307 | Dip2c |
| Med13l | 5.36E-09 | -0.0647546 | 0.342 | 0.307 | 0.00014421 | Serinc3 |
| 3110082I17R | 5.60E-09 | 0.11186741 | 0.251 | 0.191 | 0.00015079 | Inpp5d |
| Lrba | 5.70E-09 | 0.17908628 | 0.423 | 0.331 | 0.00015335 | Zdhhc20 |
| Lrrc4 | 5.73E-09 | 0.0473904 | 0.294 | 0.231 | 0.00015407 | Hectd1 |
| Smchd1 | 6.11E-09 | 0.07781402 | 0.253 | 0.197 | 0.00016449 | Dock4 |
| Dock10 | 6.15E-09 | 0.18114375 | 0.782 | 0.675 | 0.00016549 | Fam49b |
| Gng2 | 6.21E-09 | 0.13307255 | 0.387 | 0.307 | 0.00016704 | Babam2 |
| Tor1aip1 | 6.23E-09 | 0.18322381 | 0.256 | 0.184 | 0.00016771 | Prkce |
| Notch2 | 6.35E-09 | 0.08443718 | 0.258 | 0.202 | 0.0001708 | Pard3b |
| Garnl3 | 6.67E-09 | 0.10794127 | 0.25 | 0.191 | 0.00017958 | Kcnk13 |
| Osbpl8 | 6.68E-09 | -0.195115 | 0.275 | 0.273 | 0.00017984 | Lcor |
| Pum2 | 6.79E-09 | 0.09468953 | 0.325 | 0.258 | 0.00018262 | Rasgrp3 |
| Magi1 | 6.84E-09 | -0.0149307 | 0.551 | 0.484 | 0.00018408 | Btbd9 |
| Rptor | 6.90E-09 | 0.1963584 | 0.539 | 0.436 | 0.00018565 | Scamp2 |
| Capn3 | 7.10E-09 | 0.04747874 | 0.335 | 0.276 | 0.000191 | Calm2 |
| Tlr7 | 7.23E-09 | 0.0114769 | 0.287 | 0.24 | 0.00019455 | Slc1a3 |
| Gpatch8 | 7.35E-09 | -0.046546 | 0.356 | 0.311 | 0.0001977 | Atad2b |
| Rap1b | 7.57E-09 | 0.09434087 | 0.34 | 0.273 | 0.0002038 | Nlk |
| Lrmda | 7.63E-09 | -0.3215072 | 0.733 | 0.788 | 0.00020525 | P4ha1 |
| Kdm2a | 7.75E-09 | -0.05197 | 0.353 | 0.318 | 0.00020856 | Prkcb |
| Ldlrad4 | 7.88E-09 | 0.17689782 | 0.885 | 0.801 | 0.0002119 | St3gal5 |
| Tmem135 | 8.06E-09 | 0.11246621 | 0.496 | 0.408 | 0.00021688 | Sipa1l1 |
| Man1a2 | 8.12E-09 | -0.0838728 | 0.466 | 0.429 | 0.00021856 | Sdccag8 |
| Sgms1 | 8.32E-09 | 0.0266358 | 0.259 | 0.21 | 0.00022385 | Crlf3 |
| Ddx17 | 8.35E-09 | -0.0215909 | 0.382 | 0.336 | 0.00022475 | Jmjd1c |
| Setd5 | 8.43E-09 | 0.00450159 | 0.374 | 0.32 | 0.00022687 | Tulp4 |
| Pip4k2a | 8.43E-09 | 0.09939555 | 0.76 | 0.661 | 0.00022696 | Fli1 |
| Itgav | 8.64E-09 | 0.14197593 | 0.345 | 0.269 | 0.00023248 | P2ry12 |
| Slc24a3 | 9.17E-09 | 0.13701346 | 0.507 | 0.41 | 0.00024673 | Phactr2 |
| Il6st | 9.32E-09 | 0.13001509 | 0.271 | 0.209 | 0.00025087 | Rabep1 |
| Capzb | 9.45E-09 | 0.08572548 | 0.269 | 0.206 | 0.00025419 | Bmp2k |
| Smg1 | 9.76E-09 | -0.0455065 | 0.427 | 0.377 | 0.00026272 | Foxp1 |
| Tacc1 | 9.81E-09 | 0.07336527 | 0.549 | 0.469 | 0.00026396 | Phf21a |
| Mgat5 | 9.90E-09 | 0.19852098 | 0.511 | 0.41 | 0.00026634 | St3gal6 |
| Cradd | 9.97E-09 | 0.30163623 | 0.298 | 0.207 | 0.00026837 | Plxdc2 |
| Atp2b1 | 1.00E-08 | -0.0714196 | 0.436 | 0.388 | 0.00026925 | Rab10 |
| Gapvd1 | 1.01E-08 | 0.03592692 | 0.314 | 0.263 | 0.00027119 | Adgre1 |
| Abhd12 | 1.01E-08 | -0.1418028 | 0.574 | 0.544 | 0.00027218 | Dock2 |
| Nrp1 | 1.02E-08 | -0.237207 | 0.266 | 0.251 | 0.00027545 | Itgb5 |
| Arhgef6 | 1.04E-08 | 0.04152159 | 0.265 | 0.222 | 0.00027882 | Trappc9 |
| Pde4d | 1.12E-08 | -0.0560309 | 0.399 | 0.343 | 0.00030007 | Rsrp1 |
| Plcg2 | 1.12E-08 | 0.15450406 | 0.423 | 0.338 | 0.00030225 | Nsd1 |
| Tec | 1.14E-08 | 0.01621719 | 0.261 | 0.218 | 0.00030664 | Hpgd |
| Lrch1 | 1.17E-08 | 0.10710329 | 0.7 | 0.601 | 0.00031491 | AY036118 |
| Mark3 | 1.18E-08 | -0.055607 | 0.334 | 0.3 | 0.0003187 | A830008E24Rik |
| Zfp652 | 1.28E-08 | 0.0857541 | 0.529 | 0.447 | 0.00034326 | 0610040J01Rik |
| Gm37240 | 1.29E-08 | 0.06544917 | 0.461 | 0.382 | 0.00034768 | Mtdh |
| Nr2c2 | 1.31E-08 | 0.07800409 | 0.26 | 0.205 | 0.00035163 | Arhgap5 |
| St3gal3 | 1.31E-08 | 0.05907435 | 0.263 | 0.215 | 0.00035296 | Cux1 |
| Gphn | 1.32E-08 | -0.2785197 | 0.401 | 0.416 | 0.00035439 | Agmo |
| Rbm6 | 1.35E-08 | 0.12259461 | 0.402 | 0.328 | 0.000362 | Diaph2 |
| F11r | 1.35E-08 | 0.08115775 | 0.319 | 0.26 | 0.00036228 | Sdk1 |
| Ubr3 | 1.36E-08 | 0.03804124 | 0.305 | 0.254 | 0.00036563 | R3hdm2 |
| Adap2 | 1.37E-08 | 0.02258497 | 0.681 | 0.607 | 0.00036845 | Ptprj |
| Zfp148 | 1.39E-08 | -0.0451754 | 0.346 | 0.304 | 0.00037528 | Gab1 |
| Prkca | 1.41E-08 | 0.2098566 | 0.572 | 0.462 | 0.00037816 | Smyd3 |
| Fam172a | 1.43E-08 | 0.00111215 | 0.53 | 0.471 | 0.00038515 | Rapgef2 |
| Pten | 1.46E-08 | 0.02590201 | 0.449 | 0.386 | 0.00039291 | Phf14 |
| Erbin | 1.48E-08 | 0.01080569 | 0.358 | 0.307 | 0.00039717 | Hnrnpa2b1 |
| Phip | 1.55E-08 | -0.072068 | 0.303 | 0.268 | 0.00041769 | Kansl1 |
| P2ry6 | 1.61E-08 | 0.13374612 | 0.394 | 0.321 | 0.00043213 | Specc1 |
| Arhgap12 | 1.67E-08 | 0.00914637 | 0.416 | 0.362 | 0.0004484 | Pitpnc1 |
| Sik2 | 1.68E-08 | -0.0528485 | 0.31 | 0.276 | 0.00045319 | Nfia |
| Usp50 | 1.70E-08 | -0.0999676 | 0.3 | 0.278 | 0.00045789 | Fcrls |
| Creb1 | 1.71E-08 | -0.021536 | 0.297 | 0.256 | 0.00046145 | 4930517O19Rik |
| Mtus1 | 1.75E-08 | 0.02603174 | 0.372 | 0.316 | 0.00047192 | Oxr1 |
| Pik3r1 | 1.81E-08 | -0.0283292 | 0.49 | 0.438 | 0.00048778 | Tanc2 |
| Vgll4 | 1.82E-08 | 0.04263726 | 0.277 | 0.23 | 0.00048979 | Asap1 |
| Snx6 | 1.84E-08 | 0.10010236 | 0.258 | 0.2 | 0.00049401 | Frmd4a |
| Aim2 | 1.86E-08 | 0.00227639 | 0.328 | 0.282 | 0.000501 | Ppp1r9a |
| Atxn1 | 1.87E-08 | 0.15723214 | 0.445 | 0.355 | 0.0005022 | Tg |
| Rffl | 1.88E-08 | 0.21127379 | 0.321 | 0.238 | 0.00050587 | Rnf216 |
| Mon2 | 1.90E-08 | 0.01741615 | 0.299 | 0.253 | 0.0005116 | Chn2 |
| Qk | 1.96E-08 | 0.21714145 | 0.905 | 0.843 | 0.00052674 | Epb41l2 |
| Gatad2b | 2.04E-08 | 0.03735015 | 0.273 | 0.227 | 0.00054882 | Arhgap31 |
| Dock1 | 2.12E-08 | 0.02438255 | 0.325 | 0.275 | 0.0005694 | Rsrc1 |
| Pan3 | 2.16E-08 | 0.03657997 | 0.6 | 0.519 | 0.00058216 | Atp8a2 |
| Ep300 | 2.18E-08 | -0.0187036 | 0.296 | 0.258 | 0.00058566 | Gm35188 |
| Zcchc11 | 2.20E-08 | -0.3472086 | 0.22 | 0.321 | 0.00059136 | Galnt1 |
| Ccnt2 | 2.21E-08 | 0.01586695 | 0.256 | 0.214 | 0.00059386 | Nlrp1b |
| Tnrc6c | 2.27E-08 | -0.0077773 | 0.37 | 0.322 | 0.00061159 | Maml2 |
| Usp9x | 2.31E-08 | 0.02481192 | 0.317 | 0.265 | 0.00062133 | Arhgap45 |
| Tgfbr2 | 2.39E-08 | -0.0422791 | 0.529 | 0.477 | 0.00064307 | App |
| Nufip2 | 2.40E-08 | -0.0124781 | 0.279 | 0.239 | 0.00064696 | Apbb1ip |
| Nrd1 | 2.41E-08 | -0.005176 | 0.271 | 0.228 | 0.00064769 | Atp2c1 |
| Rreb1 | 2.53E-08 | 0.19532072 | 0.795 | 0.694 | 0.00068065 | Lair1 |
| Nsd3 | 2.58E-08 | -0.0593546 | 0.389 | 0.353 | 0.00069363 | Ptprm |
| Rtn4rl1 | 2.59E-08 | 0.24784723 | 0.305 | 0.219 | 0.00069579 | Znrf1 |
| Rhoq | 2.69E-08 | 0.0622334 | 0.268 | 0.215 | 0.0007237 | 9930021J03Rik |
| Rsbn1l | 2.72E-08 | 0.09659561 | 0.307 | 0.246 | 0.00073182 | Ssh2 |
| Phf3 | 2.73E-08 | 0.03226269 | 0.298 | 0.249 | 0.00073544 | Mbnl1 |
| Eml6 | 2.75E-08 | 0.12916769 | 0.314 | 0.245 | 0.00074092 | Bbs9 |
| Cdk13 | 2.76E-08 | 0.04450602 | 0.342 | 0.289 | 0.00074176 | Gm20275 |
| Skap2 | 2.78E-08 | -0.0770211 | 0.546 | 0.504 | 0.00074742 | Plcl1 |
| Acin1 | 2.78E-08 | 0.10042092 | 0.311 | 0.246 | 0.00074912 | Ccny |
| A630001G21 | 2.97E-08 | 0.15218216 | 0.268 | 0.203 | 0.00079882 | Mertk |
| Tbc1d14 | 3.01E-08 | 0.03325566 | 0.279 | 0.228 | 0.00080865 | Wac |
| Rrbp1 | 3.03E-08 | 0.06449543 | 0.578 | 0.495 | 0.00081414 | Mef2c |
| Ddx5 | 3.03E-08 | 0.27056068 | 0.734 | 0.628 | 0.0008158 | Slc8a1 |
| Sipa1 | 3.05E-08 | -0.0208515 | 0.283 | 0.248 | 0.00082115 | Snx29 |
| Ikzf1 | 3.08E-08 | 0.05890942 | 0.743 | 0.653 | 0.00082815 | Macf1 |
| Senp6 | 3.14E-08 | -0.0114634 | 0.28 | 0.239 | 0.0008442 | |
| Pros1 | 3.18E-08 | 0.02714372 | 0.358 | 0.303 | 0.0008549 | |
| Pik3cd | 3.18E-08 | 0.15311074 | 0.33 | 0.257 | 0.00085663 | |
| Strn3 | 3.32E-08 | 0.05353021 | 0.441 | 0.375 | 0.00089285 | |
| Bptf | 3.33E-08 | 0.17733541 | 0.366 | 0.288 | 0.00089711 | |
| Peak1 | 3.39E-08 | -0.305419 | 0.272 | 0.303 | 0.00091165 | |
| Nfat5 | 3.40E-08 | 0.05018969 | 0.368 | 0.304 | 0.00091616 | |
| Zfr | 3.49E-08 | -0.0027934 | 0.384 | 0.339 | 0.00093852 | |
| Pafah1b1 | 3.50E-08 | 0.0086381 | 0.396 | 0.342 | 0.00094251 | |
| Gm16599 | 3.50E-08 | 0.10576616 | 0.315 | 0.253 | 0.00094301 | |
| Rab2a | 3.53E-08 | -0.0622401 | 0.3 | 0.271 | 0.00094987 | |
| Vav1 | 3.68E-08 | 0.10623699 | 0.269 | 0.213 | 0.00099019 | |
| Dcakd | 3.73E-08 | -0.024685 | 0.266 | 0.23 | 0.00100296 | |
| Fto | 3.81E-08 | -0.0028184 | 0.31 | 0.26 | 0.00102639 | |
| Xiap | 3.83E-08 | 0.06644894 | 0.33 | 0.273 | 0.00102957 | |
| Apba1 | 3.96E-08 | 0.06521689 | 0.443 | 0.377 | 0.00106665 | |
| Hdac9 | 4.07E-08 | 0.0444472 | 0.537 | 0.457 | 0.00109406 | |
| Pde3b | 4.08E-08 | 0.2574706 | 0.809 | 0.724 | 0.00109679 | |
| Hivep3 | 4.12E-08 | 0.012082 | 0.502 | 0.432 | 0.00110729 | |
| Herc4 | 4.24E-08 | -0.1310507 | 0.271 | 0.254 | 0.00114002 | |
| Itch | 4.24E-08 | -0.0198595 | 0.425 | 0.373 | 0.00114216 | |
| Zfp644 | 4.38E-08 | -0.0155198 | 0.315 | 0.279 | 0.00117837 | |
| Zc3h7a | 4.45E-08 | 0.08425717 | 0.295 | 0.239 | 0.00119632 | |
| Usp15 | 4.47E-08 | 0.07003071 | 0.271 | 0.219 | 0.00120316 | |
| Laptm5 | 4.47E-08 | -0.1055328 | 0.345 | 0.325 | 0.00120374 | |
| Abi1 | 4.67E-08 | 0.08501317 | 0.435 | 0.365 | 0.00125758 | |
| Capza1 | 4.79E-08 | 0.10201594 | 0.292 | 0.233 | 0.00128958 | |
| Slc38a9 | 4.87E-08 | -0.1006839 | 0.26 | 0.239 | 0.00131091 | |
| Sat1 | 4.95E-08 | -0.1299396 | 0.33 | 0.316 | 0.00133247 | |
| Ifngr1 | 5.09E-08 | 0.09895647 | 0.3 | 0.242 | 0.00136827 | |
| Ulk2 | 5.19E-08 | 0.02097307 | 0.534 | 0.468 | 0.00139615 | |
| Rbm33 | 5.30E-08 | 0.0934465 | 0.262 | 0.208 | 0.0014268 | |
| Cacna1a | 5.34E-08 | -0.1301071 | 0.642 | 0.61 | 0.00143799 | |
| Gm26848 | 5.51E-08 | -0.2165498 | 0.212 | 0.306 | 0.00148155 | |
| Mark2 | 5.86E-08 | 0.05519195 | 0.295 | 0.239 | 0.00157661 | |
| Dpy19l4 | 5.90E-08 | -0.0323574 | 0.267 | 0.235 | 0.00158769 | |
| Ivns1abp | 5.92E-08 | 0.16458615 | 0.707 | 0.602 | 0.00159265 | |
| Ubac2 | 5.95E-08 | 0.07926595 | 0.322 | 0.261 | 0.00160198 | |
| Pkig | 6.18E-08 | -0.0555838 | 0.321 | 0.282 | 0.00166213 | |
| Pias2 | 6.25E-08 | 0.00148321 | 0.252 | 0.212 | 0.00168136 | |
| Phkb | 6.42E-08 | -0.0895861 | 0.277 | 0.255 | 0.001728 | |
| Zeb1 | 6.64E-08 | 0.17474498 | 0.52 | 0.425 | 0.00178719 | |
| Itpr2 | 7.08E-08 | 0.1067134 | 0.486 | 0.405 | 0.00190412 | |
| Cd180 | 7.51E-08 | 0.0323347 | 0.302 | 0.25 | 0.00202021 | |
| Psme4 | 7.67E-08 | -0.019206 | 0.27 | 0.235 | 0.00206259 | |
| Cacnb2 | 7.71E-08 | 0.18798702 | 0.552 | 0.452 | 0.00207448 | |
| Tjp1 | 7.86E-08 | 0.0894136 | 0.358 | 0.296 | 0.00211429 | |
| Gnaq | 8.17E-08 | -0.1004596 | 0.625 | 0.589 | 0.00219954 | |
| Ankrd11 | 8.45E-08 | 0.04580279 | 0.542 | 0.474 | 0.00227389 | |
| Rcbtb2 | 8.60E-08 | 0.0239668 | 0.358 | 0.309 | 0.00231317 | |
| Gpr34 | 8.63E-08 | 0.04839441 | 0.565 | 0.489 | 0.00232236 | |
| Phc2 | 8.86E-08 | 0.01972413 | 0.308 | 0.266 | 0.00238474 | |
| Rab31 | 9.01E-08 | 0.20601366 | 0.373 | 0.29 | 0.00242444 | |
| Iffo1 | 9.14E-08 | 0.11587262 | 0.26 | 0.203 | 0.00245939 | |
| 1-Mar | 9.22E-08 | 0.02264173 | 0.552 | 0.483 | 0.00248152 | |
| Fryl | 9.83E-08 | -0.0064103 | 0.256 | 0.218 | 0.00264485 | |
| Pias1 | 1.01E-07 | 0.02716739 | 0.415 | 0.359 | 0.00270678 | |
| Parp8 | 1.01E-07 | -0.0747574 | 0.416 | 0.377 | 0.00271118 | |
| Gm2245 | 1.01E-07 | -0.064735 | 0.39 | 0.35 | 0.00272905 | |
| Zmym4 | 1.06E-07 | -0.034363 | 0.326 | 0.288 | 0.00284378 | |
| Arhgap17 | 1.13E-07 | -0.0445248 | 0.429 | 0.389 | 0.00303132 | |
| Rock1 | 1.13E-07 | -0.0663254 | 0.348 | 0.317 | 0.00303969 | |
| Arhgap25 | 1.13E-07 | -0.0682276 | 0.285 | 0.257 | 0.00305066 | |
| Fbxl17 | 1.14E-07 | 0.00336768 | 0.554 | 0.489 | 0.00306547 | |
| Larp1 | 1.25E-07 | 0.11204242 | 0.271 | 0.212 | 0.00336046 | |
| Chd2 | 1.25E-07 | 0.08134422 | 0.41 | 0.342 | 0.00336138 | |
| E2f3 | 1.25E-07 | 0.26856507 | 0.268 | 0.19 | 0.00336154 | |
| Nudcd3 | 1.25E-07 | 0.13211964 | 0.257 | 0.197 | 0.0033643 | |
| Tnfrsf11a | 1.27E-07 | 0.2099687 | 0.358 | 0.277 | 0.00341302 | |
| Lamp2 | 1.27E-07 | -0.1157544 | 0.3 | 0.286 | 0.00342674 | |
| Rb1 | 1.28E-07 | -0.115016 | 0.329 | 0.303 | 0.00345371 | |
| Mia2 | 1.30E-07 | 0.1074621 | 0.268 | 0.21 | 0.00351052 | |
| Ubn2 | 1.34E-07 | 0.00366705 | 0.453 | 0.399 | 0.00361889 | |
| Fbxl20 | 1.35E-07 | 0.12748361 | 0.325 | 0.26 | 0.00363157 | |
| Skil | 1.36E-07 | -0.0201374 | 0.256 | 0.218 | 0.00366146 | |
| Plxna4 | 1.37E-07 | 0.23535121 | 0.567 | 0.466 | 0.0036945 | |
| Arl15 | 1.39E-07 | 0.11790294 | 0.467 | 0.39 | 0.0037445 | |
| Rere | 1.48E-07 | 0.18812149 | 0.613 | 0.516 | 0.00398458 | |
| Unc93b1 | 1.52E-07 | 0.19680161 | 0.635 | 0.533 | 0.00408239 | |
| Cx3cr1 | 1.52E-07 | 0.19028486 | 0.778 | 0.681 | 0.0040931 | |
| Kdm7a | 1.53E-07 | -0.1516556 | 0.25 | 0.241 | 0.00411291 | |
| Pld1 | 1.53E-07 | 0.03500619 | 0.419 | 0.358 | 0.00411512 | |
| Cpsf6 | 1.54E-07 | 0.02158995 | 0.287 | 0.244 | 0.00415227 | |
| Celf2 | 1.55E-07 | -0.2275798 | 0.75 | 0.751 | 0.00417157 | |
| Pnpla7 | 1.62E-07 | -0.0477252 | 0.442 | 0.401 | 0.00434788 | |

| | | | | | |
|---|---|---|---|---|---|
| Lrch1 | 1.56E-05 | 0.0890242 | 0.601 | 0.635 | 0.41988647 |
| C1qc | 1.64E-05 | 0.03165755 | 0.578 | 0.628 | 0.44065477 |
| Nufip2 | 1.74E-05 | -0.0201853 | 0.239 | 0.276 | 0.46731205 |
| Rsbn1l | 1.77E-05 | -0.0895088 | 0.246 | 0.296 | 0.47605902 |
| Mon2 | 1.82E-05 | 0.00158937 | 0.253 | 0.286 | 0.48948427 |
| Itga6 | 1.83E-05 | 0.01687372 | 0.309 | 0.346 | 0.49108576 |
| Slc7a8 | 1.94E-05 | -0.1738158 | 0.32 | 0.392 | 0.52171316 |
| Ubac2 | 1.97E-05 | -0.10452 | 0.261 | 0.318 | 0.52986395 |
| Srsf7 | 2.03E-05 | -0.1962546 | 0.365 | 0.447 | 0.54686156 |
| Hdac8 | 2.05E-05 | 0.0632499 | 0.233 | 0.251 | 0.55126017 |
| Clint1 | 2.09E-05 | 0.04594639 | 0.31 | 0.333 | 0.56239099 |
| Nr3c2 | 2.15E-05 | -0.0300132 | 0.209 | 0.25 | 0.57771053 |
| Dock7 | 2.15E-05 | -0.0643029 | 0.219 | 0.268 | 0.5780932 |
| Pvt1 | 2.17E-05 | 0.06295512 | 0.276 | 0.294 | 0.58409466 |
| Wwox | 2.18E-05 | 0.15976772 | 0.446 | 0.449 | 0.58692324 |
| Ttc28 | 2.24E-05 | -0.0968689 | 0.401 | 0.472 | 0.60342546 |
| Tbc1d22a | 2.25E-05 | 0.03873249 | 0.333 | 0.358 | 0.60472477 |
| Sdk1 | 2.28E-05 | 0.12930728 | 0.237 | 0.255 | 0.61349774 |
| Hook3 | 2.43E-05 | 0.05446682 | 0.242 | 0.259 | 0.65331894 |
| Btbd9 | 2.43E-05 | 0.04646173 | 0.455 | 0.487 | 0.65384217 |
| Rap1gds1 | 2.47E-05 | -0.0894288 | 0.594 | 0.671 | 0.66399364 |
| Arid4a | 2.49E-05 | 0.01127472 | 0.267 | 0.302 | 0.66885064 |
| Lcp1 | 2.49E-05 | 0.10749142 | 0.257 | 0.27 | 0.66927029 |
| Zc3h7a | 2.52E-05 | -0.0139595 | 0.239 | 0.27 | 0.67758935 |
| Nktr | 2.56E-05 | 0.09404222 | 0.293 | 0.308 | 0.68950333 |
| Sipa1l1 | 2.57E-05 | -0.200099 | 0.288 | 0.361 | 0.69241866 |
| Tmsb4x | 2.71E-05 | 0.15004128 | 0.389 | 0.398 | 0.72988009 |
| Tnfrsf11a | 2.75E-05 | -0.0970452 | 0.277 | 0.325 | 0.73878777 |
| Rsrc2 | 2.77E-05 | 0.00043008 | 0.236 | 0.262 | 0.7461818 |
| Ube2h | 2.80E-05 | 0.09040936 | 0.31 | 0.326 | 0.75447372 |
| Mtdh | 2.95E-05 | -0.0063826 | 0.503 | 0.546 | 0.79426671 |
| Gsk3b | 2.95E-05 | -0.0629383 | 0.287 | 0.336 | 0.79494522 |
| Tmem119 | 2.99E-05 | -0.2250716 | 0.191 | 0.255 | 0.8057654 |
| Trpm7 | 3.26E-05 | -0.0188837 | 0.257 | 0.289 | 0.87803489 |
| Arhgap5 | 3.38E-05 | 0.06090273 | 0.652 | 0.691 | 0.90864327 |
| Plxdc2 | 3.46E-05 | 0.18232828 | 0.885 | 0.864 | 0.93107094 |
| Eml4 | 3.49E-05 | 0.09717345 | 0.35 | 0.361 | 0.93988492 |
| Elk3 | 3.52E-05 | -0.005262 | 0.296 | 0.333 | 0.94760146 |
| Gm48678 | 3.62E-05 | -0.120239 | 0.297 | 0.358 | 0.97410575 |
| Dst | 3.78E-05 | -0.1810742 | 0.708 | 0.785 | 1 |
| Zfand6 | 3.82E-05 | -0.003899 | 0.26 | 0.296 | 1 |
| Ascc3 | 3.87E-05 | -0.0159901 | 0.257 | 0.297 | 1 |
| Creb1 | 3.96E-05 | -0.0841796 | 0.256 | 0.304 | 1 |
| Atp2c1 | 3.96E-05 | 0.08675583 | 0.364 | 0.384 | 1 |
| Cfh | 4.06E-05 | 0.09268661 | 0.57 | 0.591 | 1 |
| Pid1 | 4.22E-05 | 0.15730647 | 0.532 | 0.531 | 1 |
| Kdm7a | 4.24E-05 | 0.04139115 | 0.241 | 0.265 | 1 |
| Large1 | 4.26E-05 | 0.05332784 | 0.576 | 0.613 | 1 |
| Gbf1 | 4.28E-05 | -0.0400809 | 0.214 | 0.256 | 1 |
| A830008E24 | 4.39E-05 | -0.081603 | 0.217 | 0.268 | 1 |
| Mapkap1 | 4.58E-05 | -0.1157361 | 0.207 | 0.257 | 1 |
| Bach2 | 4.86E-05 | 0.08639287 | 0.391 | 0.42 | 1 |
| Strbp | 4.96E-05 | -0.0374474 | 0.336 | 0.389 | 1 |
| Smarca2 | 5.00E-05 | -0.0862834 | 0.27 | 0.322 | 1 |
| Parp8 | 5.12E-05 | 0.07286164 | 0.377 | 0.4 | 1 |
| Zcchc7 | 5.15E-05 | -0.1304097 | 0.49 | 0.568 | 1 |
| Tg | 5.37E-05 | -0.102948 | 0.221 | 0.268 | 1 |
| F11r | 5.60E-05 | 0.08008082 | 0.26 | 0.269 | 1 |
| 4930517O19 | 5.65E-05 | -0.1001869 | 0.208 | 0.257 | 1 |
| Camk1d | 5.74E-05 | 0.14146048 | 0.708 | 0.724 | 1 |
| Arhgap45 | 5.79E-05 | -0.0973726 | 0.68 | 0.749 | 1 |
| Rabep1 | 5.85E-05 | 0.02173437 | 0.312 | 0.347 | 1 |
| Rsrp1 | 6.13E-05 | -0.0643448 | 0.67 | 0.737 | 1 |
| Ankrd44 | 6.16E-05 | -0.1110524 | 0.645 | 0.723 | 1 |
| Map4k4 | 6.27E-05 | 0.01146326 | 0.434 | 0.468 | 1 |
| Fmnl2 | 6.39E-05 | 0.21395154 | 0.255 | 0.238 | 1 |
| Capn3 | 6.53E-05 | -0.1177414 | 0.276 | 0.333 | 1 |
| Fchsd2 | 6.60E-05 | -0.1140963 | 0.59 | 0.671 | 1 |
| Babam2 | 6.67E-05 | -0.0338178 | 0.288 | 0.328 | 1 |
| Plcl2 | 6.68E-05 | -0.2022776 | 0.6 | 0.684 | 1 |
| Cyth4 | 6.71E-05 | -0.1193362 | 0.489 | 0.558 | 1 |
| Cmip | 6.92E-05 | 0.01590774 | 0.334 | 0.375 | 1 |
| Dock8 | 6.94E-05 | -0.0001507 | 0.726 | 0.781 | 1 |
| Celf2 | 7.30E-05 | 0.21218096 | 0.751 | 0.744 | 1 |
| Fam49b | 7.93E-05 | -0.1461548 | 0.676 | 0.754 | 1 |
| Foxp1 | 8.30E-05 | -0.1105163 | 0.329 | 0.392 | 1 |
| Crlf3 | 8.50E-05 | -0.0506012 | 0.371 | 0.414 | 1 |
| Ptprj | 8.52E-05 | -0.0021572 | 0.656 | 0.712 | 1 |
| Dock1 | 8.70E-05 | 0.17201428 | 0.275 | 0.263 | 1 |
| Nav3 | 8.78E-05 | 0.21833392 | 0.812 | 0.785 | 1 |
| Zfp710 | 8.84E-05 | -0.1902351 | 0.569 | 0.653 | 1 |
| Adap2 | 9.70E-05 | -0.0100657 | 0.607 | 0.661 | 1 |
| Cacnb2 | 9.94E-05 | 0.16904566 | 0.452 | 0.457 | 1 |
| Wdr33 | 0.00010028 | -0.0483033 | 0.237 | 0.28 | 1 |
| Fmnl3 | 0.00010035 | -0.0029227 | 0.3 | 0.335 | 1 |
| Ccr5 | 0.00010405 | -0.0800111 | 0.248 | 0.292 | 1 |
| Elf1 | 0.00010664 | -0.0225741 | 0.235 | 0.266 | 1 |
| Peak1 | 0.00011165 | 0.05455369 | 0.303 | 0.326 | 1 |
| Zbtb20 | 0.00011506 | 0.04494555 | 0.693 | 0.736 | 1 |
| Akap10 | 0.00011575 | 0.01735316 | 0.255 | 0.283 | 1 |
| 0610040J01R | 0.00011587 | -0.0147144 | 0.445 | 0.49 | 1 |
| Heatr5a | 0.00011782 | 0.01164515 | 0.304 | 0.338 | 1 |
| Mtus1 | 0.00012285 | 0.01635338 | 0.316 | 0.343 | 1 |
| Gpr34 | 0.00012418 | -0.1500667 | 0.489 | 0.567 | 1 |
| Pitpnc1 | 0.00012488 | -0.1006402 | 0.477 | 0.544 | 1 |
| Crebbp | 0.00012867 | -0.0498829 | 0.384 | 0.429 | 1 |
| Pbx1 | 0.00012997 | -0.1983621 | 0.201 | 0.26 | 1 |
| Ptprm | 0.00013698 | 0.03546003 | 0.441 | 0.471 | 1 |
| Runx1 | 0.00014851 | 0.06282909 | 0.629 | 0.667 | 1 |
| Taok3 | 0.00015211 | 0.0847757 | 0.258 | 0.271 | 1 |
| Arhgap22 | 0.00015268 | 0.00687707 | 0.469 | 0.506 | 1 |
| Gpr183 | 0.00016241 | 0.02577918 | 0.227 | 0.25 | 1 |
| Zfp148 | 0.00017197 | -0.0753403 | 0.304 | 0.352 | 1 |
| Ivns1abp | 0.00018243 | -0.1171966 | 0.602 | 0.675 | 1 |
| Lyst | 0.00019262 | 0.01430154 | 0.228 | 0.258 | 1 |
| Slc9a9 | 0.00019801 | -0.0783035 | 0.689 | 0.758 | 1 |
| Dnajc1 | 0.00019844 | -0.1507755 | 0.299 | 0.358 | 1 |
| Snx2 | 0.00020193 | -0.1909429 | 0.202 | 0.261 | 1 |
| Ldlrad4 | 0.0002031 | -0.155745 | 0.801 | 0.857 | 1 |
| Zeb1 | 0.00020548 | -0.1081333 | 0.425 | 0.486 | 1 |
| Btbd7 | 0.00021544 | -0.0188097 | 0.23 | 0.257 | 1 |
| Ppp1r9a | 0.00022717 | -0.0300682 | 0.505 | 0.557 | 1 |
| Apoe | 0.0002344 | 0.17981687 | 0.362 | 0.38 | 1 |
| Ikzf1 | 0.0002423 | 0.00665611 | 0.653 | 0.701 | 1 |
| Lncpint | 0.00024426 | -0.0033926 | 0.384 | 0.422 | 1 |
| Sbf2 | 0.00026411 | 0.05262385 | 0.421 | 0.441 | 1 |
| Pan3 | 0.00026577 | -0.0121804 | 0.519 | 0.569 | 1 |
| Hnrnpa2b1 | 0.0002666 | -0.1842005 | 0.545 | 0.625 | 1 |
| Nlrp1b | 0.0002741 | 0.02548292 | 0.228 | 0.254 | 1 |
| Jmjd1c | 0.0002921 | 0.01101548 | 0.602 | 0.642 | 1 |
| Siglech | 0.00029284 | -0.0903409 | 0.565 | 0.632 | 1 |
| Phactr2 | 0.00031076 | 0.02855031 | 0.253 | 0.276 | 1 |
| Ank2 | 0.00031119 | 0.1853114 | 0.535 | 0.518 | 1 |
| Ankrd11 | 0.00032707 | 0.10541784 | 0.474 | 0.476 | 1 |
| Hpgds | 0.00034014 | -0.0746279 | 0.592 | 0.658 | 1 |
| Bcas3 | 0.00037495 | -0.1383515 | 0.279 | 0.331 | 1 |
| Agmo | 0.00039712 | -0.1818848 | 0.54 | 0.618 | 1 |
| Hk2 | 0.00042183 | -0.019149 | 0.228 | 0.252 | 1 |
| Ppp3ca | 0.00048431 | 0.10964338 | 0.639 | 0.646 | 1 |
| Slc8a1 | 0.00049328 | -0.1430593 | 0.858 | 0.882 | 1 |
| Atxn1 | 0.00051303 | -0.1329534 | 0.355 | 0.417 | 1 |
| Stag1 | 0.00056202 | -0.0172748 | 0.479 | 0.519 | 1 |
| Cmss1 | 0.00058489 | 0.07790632 | 0.62 | 0.629 | 1 |
| Fcrls | 0.00060452 | -0.0951327 | 0.438 | 0.498 | 1 |
| Tmcc1 | 0.00062258 | 0.12837516 | 0.415 | 0.417 | 1 |
| Rcbtb2 | 0.00062917 | 0.02338514 | 0.309 | 0.329 | 1 |
| Chd2 | 0.00066481 | -0.0820614 | 0.342 | 0.391 | 1 |
| Rasgrp3 | 0.00067228 | 0.07311599 | 0.433 | 0.452 | 1 |
| Cdk14 | 0.00070195 | -0.0468865 | 0.346 | 0.39 | 1 |
| P2ry12 | 0.00071405 | 0.11594529 | 0.69 | 0.706 | 1 |
| Gm26804 | 0.00076841 | 0.11442187 | 0.383 | 0.321 | 1 |
| Tmem243 | 0.00078225 | -0.195193 | 0.294 | 0.357 | 1 |
| Pcmtd1 | 0.00079758 | -0.0697332 | 0.218 | 0.255 | 1 |
| Fto | 0.00080279 | -0.0451796 | 0.26 | 0.3 | 1 |
| Pip4k2a | 0.0008138 | 0.02999596 | 0.661 | 0.702 | 1 |
| Maml2 | 0.00082694 | 0.1491976 | 0.574 | 0.564 | 1 |
| Slc24a3 | 0.0008741 | 0.06392028 | 0.41 | 0.434 | 1 |
| Eml6 | 0.00087616 | 0.00869823 | 0.245 | 0.273 | 1 |
| Pnisr | 0.00088413 | 0.05102054 | 0.429 | 0.451 | 1 |
| Dock2 | 0.0008921 | -0.0049822 | 0.638 | 0.684 | 1 |
| Nfia | 0.00090451 | -0.120482 | 0.536 | 0.606 | 1 |
| Prkca | 0.00099703 | -0.1111095 | 0.462 | 0.524 | 1 |
| Chn2 | 0.00106787 | 0.06349602 | 0.573 | 0.597 | 1 |
| Kcnma1 | 0.00108531 | 0.20960243 | 0.355 | 0.339 | 1 |
| Csf1r | 0.00117117 | -0.071429 | 0.687 | 0.748 | 1 |
| Frmd4b | 0.00121992 | 0.07429713 | 0.646 | 0.658 | 1 |
| Hdac9 | 0.00131719 | 0.19652143 | 0.457 | 0.424 | 1 |
| Dennd4a | 0.00134945 | -0.0723996 | 0.645 | 0.702 | 1 |
| Arglu1 | 0.00137205 | 0.05718109 | 0.429 | 0.442 | 1 |
| Epb41l2 | 0.00139936 | 0.09362371 | 0.775 | 0.794 | 1 |
| Diaph2 | 0.00145527 | 0.00017283 | 0.656 | 0.7 | 1 |
| Apba1 | 0.00152844 | -0.0068131 | 0.377 | 0.406 | 1 |
| Maf | 0.00163835 | -0.178418 | 0.388 | 0.453 | 1 |
| Rhoh | 0.00186638 | -0.1499257 | 0.295 | 0.347 | 1 |
| Arsb | 0.00189593 | -0.0964095 | 0.257 | 0.299 | 1 |
| Cx3cr1 | 0.00195869 | -0.1141198 | 0.681 | 0.741 | 1 |
| Rreb1 | 0.00196893 | -0.1556873 | 0.694 | 0.756 | 1 |
| Plxna4 | 0.00207947 | -0.1817023 | 0.466 | 0.531 | 1 |
| Rapgef5 | 0.00258981 | -0.1310836 | 0.631 | 0.696 | 1 |
| Dip2c | 0.00277262 | 0.01981497 | 0.3 | 0.326 | 1 |
| Fam102b | 0.00389533 | 0.05742447 | 0.425 | 0.443 | 1 |
| Qk | 0.00401381 | -0.1009998 | 0.843 | 0.888 | 1 |
| Asap1 | 0.00432535 | 0.0567676 | 0.561 | 0.58 | 1 |
| Foxn3 | 0.00506293 | -0.0584026 | 0.765 | 0.815 | 1 |
| Rasal2 | 0.00508279 | -0.0243802 | 0.264 | 0.296 | 1 |
| Srrm2 | 0.00536488 | 0.05112343 | 0.619 | 0.635 | 1 |
| Snx29 | 0.00560596 | 0.04979419 | 0.69 | 0.717 | 1 |
| Mir142hg | 0.00599805 | -0.017477 | 0.255 | 0.279 | 1 |
| Gab2 | 0.00650121 | 0.09310055 | 0.724 | 0.733 | 1 |
| Tgfbr1 | 0.0067821 | -0.0806831 | 0.865 | 0.906 | 1 |
| Osbpl9 | 0.00818818 | -0.119622 | 0.282 | 0.31 | 1 |
| Gm26733 | 0.00840917 | 0.11106507 | 0.256 | 0.211 | 1 |
| Mycbp2 | 0.00884822 | -0.1089308 | 0.702 | 0.754 | 1 |
| Ctsb | 0.00902303 | 0.0359524 | 0.3 | 0.311 | 1 |
| Apbb1ip | 0.01017876 | 0.1116219 | 0.723 | 0.719 | 1 |
| Gm26740 | 0.01102107 | 0.07752901 | 0.249 | 0.251 | 1 |
| Lgmn | 0.01672215 | -0.0292809 | 0.643 | 0.68 | 1 |
| Gm20275 | 0.01836857 | -0.1463153 | 0.264 | 0.306 | 1 |
| App | 0.0184624 | -0.0627836 | 0.248 | 0.276 | 1 |
| Mertk | 0.01977492 | 0.09362722 | 0.824 | 0.841 | 1 |
| Ptprc | 0.02007121 | 0.04347782 | 0.521 | 0.535 | 1 |
| Tmcc3 | 0.0203673 | 0.09017572 | 0.797 | 0.779 | 1 |
| Fhit | 0.02168917 | 0.09843435 | 0.458 | 0.45 | 1 |
| Rbm39 | 0.02483024 | 0.03948902 | 0.713 | 0.729 | 1 |
| Atp8a2 | 0.02563727 | -0.0444533 | 0.536 | 0.573 | 1 |
| Arhgap31 | 0.0296175 | 0.01806513 | 0.265 | 0.279 | 1 |
| Tanc2 | 0.03083279 | 0.09885705 | 0.925 | 0.926 | 1 |
| Olfr56 | 0.03225204 | 0.18848521 | 0.254 | 0.219 | 1 |
| Wnk1 | 0.03265533 | -0.0702047 | 0.639 | 0.68 | 1 |
| Immp2l | 0.03542417 | 0.19101691 | 0.303 | 0.279 | 1 |
| Exoc4 | 0.03954104 | 0.02795055 | 0.509 | 0.518 | 1 |
| Numb | 0.04458575 | 0.01376306 | 0.717 | 0.743 | 1 |
| Mctp1 | 0.04614139 | -0.110217 | 0.521 | 0.566 | 1 |
| Gm37240 | 0.05119335 | -0.0836279 | 0.382 | 0.418 | 1 |
| Plcl1 | 0.10536402 | -0.0307908 | 0.698 | 0.733 | 1 |
| Macf1 | 0.14584153 | -0.0877663 | 0.788 | 0.814 | 1 |
| 4933406I18R | 0.15004463 | -0.0704659 | 0.641 | 0.676 | 1 |
| Maml3 | 0.21622512 | -0.0291019 | 0.76 | 0.787 | 1 |
| Gm42418 | 0.24963047 | 0.10427756 | 0.904 | 0.922 | 1 |
| Xist | 0.31112398 | 0.02725556 | 0.818 | 0.831 | 1 |
| Mef2c | 0.37096279 | -0.0017289 | 0.775 | 0.789 | 1 |
| Gm35188 | 0.4105148 | 0.06041336 | 0.729 | 0.713 | 1 |
| Inpp4b | 0.4233431 | -0.0531823 | 0.711 | 0.734 | 1 |
| Tbc1d5 | 0.43066342 | -0.0205434 | 0.622 | 0.643 | 1 |
| Frmd4a | 0.44904095 | 0.03271465 | 0.776 | 0.784 | 1 |
| Nav2 | 0.54541822 | -0.035369 | 0.846 | 0.861 | 1 |
| Hexb | 0.59405232 | -0.0224753 | 0.844 | 0.846 | 1 |
| Malat1 | 0.88893629 | -0.0081096 | 1 | 1 | 1 |

| | | | | | |
|---|---|---|---|---|---|
| Zcchc7 | 1.64E-07 | 0.08765882 | 0.569 | 0.49 | 0.00441074 |
| Fubp1 | 1.66E-07 | -0.0342473 | 0.298 | 0.266 | 0.00446351 |
| Eef2k | 1.67E-07 | 0.12079359 | 0.292 | 0.231 | 0.00449074 |
| Marcks | 1.67E-07 | -0.1601689 | 0.307 | 0.305 | 0.00449383 |
| Ncf1 | 1.75E-07 | -0.1195673 | 0.275 | 0.264 | 0.00471904 |
| Lhfpl2 | 1.83E-07 | -0.0391731 | 0.419 | 0.373 | 0.00491177 |
| Dock7 | 1.84E-07 | -0.0281991 | 0.258 | 0.219 | 0.00495114 |
| Cdk14 | 1.85E-07 | -0.131637 | 0.374 | 0.346 | 0.00497103 |
| Maml3 | 1.87E-07 | 0.20811298 | 0.846 | 0.76 | 0.00503669 |
| Actr3 | 1.90E-07 | 0.063529 | 0.258 | 0.209 | 0.0051091 |
| Igf1r | 1.90E-07 | 0.00522283 | 0.308 | 0.262 | 0.00511103 |
| Nrip1 | 1.91E-07 | 0.00733862 | 0.56 | 0.496 | 0.00513358 |
| Gna12 | 1.92E-07 | -0.0682184 | 0.368 | 0.344 | 0.00517928 |
| Arpc2 | 1.93E-07 | 0.10545216 | 0.299 | 0.242 | 0.00518113 |
| Mctp1 | 2.04E-07 | -0.0742715 | 0.576 | 0.521 | 0.00548268 |
| Map4k3 | 2.04E-07 | 0.0038616 | 0.287 | 0.245 | 0.00549333 |
| Tmcc3 | 2.10E-07 | 0.20276867 | 0.876 | 0.797 | 0.00564551 |
| Ago3 | 2.10E-07 | 0.05655237 | 0.302 | 0.248 | 0.00564598 |
| Ophn1 | 2.16E-07 | 0.18627165 | 0.761 | 0.663 | 0.00580371 |
| Mafb | 2.18E-07 | -0.2526577 | 0.272 | 0.286 | 0.00586697 |
| Lims1 | 2.21E-07 | 0.20901791 | 0.511 | 0.414 | 0.00595215 |
| Mef2a | 2.25E-07 | 0.19745708 | 0.915 | 0.866 | 0.00604516 |
| Rps6ka1 | 2.26E-07 | 0.11782357 | 0.294 | 0.236 | 0.00607343 |
| Blnk | 2.40E-07 | -0.0763705 | 0.489 | 0.456 | 0.00645239 |
| Gm5086 | 2.56E-07 | 0.04670777 | 0.375 | 0.318 | 0.00688865 |
| Hpgds | 2.61E-07 | 0.15135301 | 0.689 | 0.592 | 0.00702975 |
| Prrc2c | 2.66E-07 | -0.0116797 | 0.316 | 0.278 | 0.00714547 |
| Tmem164 | 2.67E-07 | 0.05667451 | 0.34 | 0.286 | 0.00719293 |
| Trpm7 | 2.76E-07 | 0.00265939 | 0.294 | 0.257 | 0.00741719 |
| Bcas3 | 2.77E-07 | 0.13783555 | 0.35 | 0.279 | 0.00745248 |
| Csf1r | 2.81E-07 | 0.08800038 | 0.775 | 0.687 | 0.00755645 |
| Pcmtd1 | 2.84E-07 | -0.0045765 | 0.254 | 0.218 | 0.00763775 |
| Dyrk1a | 2.85E-07 | -0.1118149 | 0.278 | 0.266 | 0.00767416 |
| Sh3kbp1 | 2.94E-07 | -0.1250816 | 0.509 | 0.498 | 0.00791812 |
| Plgrkt | 2.98E-07 | -0.0906535 | 0.285 | 0.267 | 0.00802611 |
| Zmynd8 | 3.01E-07 | 0.04513967 | 0.269 | 0.222 | 0.00810606 |
| Susd6 | 3.03E-07 | 0.00180364 | 0.291 | 0.247 | 0.00816337 |
| Fndc3a | 3.10E-07 | -0.163723 | 0.258 | 0.254 | 0.00832793 |
| Pacs1 | 3.15E-07 | 0.0184258 | 0.321 | 0.273 | 0.00848751 |
| Slc9a9 | 3.23E-07 | 0.07555383 | 0.78 | 0.689 | 0.00868351 |
| Zfp36l1 | 3.40E-07 | 0.10264789 | 0.253 | 0.198 | 0.00915859 |
| Irf2 | 3.43E-07 | 0.08645372 | 0.39 | 0.322 | 0.00922473 |
| Med13 | 3.51E-07 | 0.07447464 | 0.264 | 0.211 | 0.00945752 |
| Rin2 | 3.58E-07 | -0.1317857 | 0.288 | 0.276 | 0.0096285 |
| Tmcc1 | 4.00E-07 | -0.205211 | 0.413 | 0.415 | 0.01075409 |
| Lyn | 4.01E-07 | 0.10786052 | 0.816 | 0.73 | 0.01077626 |
| Rsf1 | 4.06E-07 | 0.04480645 | 0.285 | 0.24 | 0.01091474 |
| Cd86 | 4.15E-07 | -0.0163985 | 0.33 | 0.285 | 0.01116619 |
| Rasal2 | 4.45E-07 | -0.1712588 | 0.275 | 0.264 | 0.01196936 |
| Tnrc6b | 4.51E-07 | 0.03217787 | 0.499 | 0.439 | 0.01214328 |
| Large1 | 4.59E-07 | 0.07248136 | 0.656 | 0.576 | 0.01235566 |
| Entpd1 | 4.70E-07 | 0.1148185 | 0.646 | 0.559 | 0.01265155 |
| Dock8 | 4.72E-07 | 0.14994475 | 0.815 | 0.726 | 0.01269019 |
| Hck | 5.01E-07 | 0.13004456 | 0.351 | 0.282 | 0.01346935 |
| Lncpint | 5.16E-07 | -0.0651983 | 0.423 | 0.384 | 0.01387516 |
| Lgmn | 5.25E-07 | 0.17943101 | 0.74 | 0.643 | 0.01412043 |
| B430010I23R | 5.42E-07 | 0.23224464 | 0.297 | 0.222 | 0.01457064 |
| Fgd2 | 5.45E-07 | 0.12323006 | 0.563 | 0.478 | 0.01465692 |
| Arhgap39 | 5.55E-07 | 0.0410829 | 0.483 | 0.422 | 0.01492535 |
| Evl | 5.64E-07 | -0.0414924 | 0.349 | 0.319 | 0.01516585 |
| Ppm1h | 6.21E-07 | 0.06046478 | 0.573 | 0.499 | 0.01672122 |
| Kcnma1 | 6.22E-07 | -0.3534056 | 0.316 | 0.355 | 0.0167473 |
| Myo9a | 6.37E-07 | -0.098387 | 0.345 | 0.325 | 0.01715295 |
| Ntpcr | 6.39E-07 | 0.03718462 | 0.281 | 0.237 | 0.01718094 |
| Scmh1 | 6.69E-07 | 0.03652743 | 0.295 | 0.245 | 0.0180038 |
| Wnk1 | 6.80E-07 | 0.13867462 | 0.729 | 0.639 | 0.01830893 |
| Elk3 | 6.92E-07 | -0.0197658 | 0.333 | 0.296 | 0.01863309 |
| Luc7l2 | 7.13E-07 | -0.0855018 | 0.61 | 0.574 | 0.01918856 |
| AU020206 | 7.26E-07 | -0.1644906 | 0.262 | 0.263 | 0.0195417 |
| Csf3r | 7.27E-07 | 0.1367343 | 0.542 | 0.458 | 0.01954976 |
| 4930444A19 | 7.27E-07 | 0.17700425 | 0.405 | 0.323 | 0.01957019 |
| Dock11 | 7.34E-07 | -0.074383 | 0.357 | 0.333 | 0.01975282 |
| Prpf4b | 7.47E-07 | -0.0570375 | 0.428 | 0.391 | 0.02011153 |
| Man1a | 7.65E-07 | -0.0338919 | 0.417 | 0.375 | 0.02059623 |
| Fkbp15 | 8.13E-07 | -0.0437677 | 0.25 | 0.226 | 0.02186971 |
| Nek7 | 8.31E-07 | 0.04024536 | 0.468 | 0.409 | 0.02236193 |
| Asxl2 | 8.33E-07 | -0.0213824 | 0.299 | 0.265 | 0.02242303 |
| Sirpa | 8.65E-07 | 0.05413567 | 0.61 | 0.541 | 0.02328065 |
| Ambra1 | 8.87E-07 | 0.06339898 | 0.396 | 0.337 | 0.02386939 |
| Fam168a | 9.22E-07 | 0.12666877 | 0.344 | 0.278 | 0.02481975 |
| Hdac8 | 9.40E-07 | 0.05525017 | 0.28 | 0.233 | 0.02529616 |
| Tra2b | 9.43E-07 | 0.22470593 | 0.28 | 0.209 | 0.02537506 |
| Mir142hg | 9.90E-07 | 0.03344641 | 0.301 | 0.255 | 0.02663032 |
| Mylip | 1.00E-06 | 0.09558466 | 0.422 | 0.353 | 0.02691794 |
| Smarca2 | 1.01E-06 | 0.08299835 | 0.328 | 0.27 | 0.02712014 |
| Ttc3 | 1.01E-06 | -0.1538835 | 0.283 | 0.275 | 0.02715172 |
| Ube2h | 1.01E-06 | 0.01479639 | 0.357 | 0.31 | 0.02722752 |
| Tcf20 | 1.08E-06 | 0.10124827 | 0.335 | 0.273 | 0.02917508 |
| Smg6 | 1.11E-06 | 0.16239658 | 0.444 | 0.368 | 0.02977736 |
| Rap1gds1 | 1.13E-06 | 0.09369909 | 0.679 | 0.594 | 0.03041636 |
| Gm48678 | 1.16E-06 | -0.0136285 | 0.337 | 0.297 | 0.03111586 |
| Rgs10 | 1.17E-06 | 0.1156458 | 0.483 | 0.412 | 0.03160684 |
| Frmd4b | 1.19E-06 | 0.03792044 | 0.725 | 0.646 | 0.03191252 |
| Cd37 | 1.21E-06 | 0.08457915 | 0.447 | 0.382 | 0.03244669 |
| Nbea | 1.24E-06 | -0.084812 | 0.317 | 0.285 | 0.03339475 |
| Btk | 1.30E-06 | 0.06895513 | 0.257 | 0.21 | 0.03504856 |
| Osbpl9 | 1.30E-06 | -0.1407156 | 0.299 | 0.282 | 0.03505599 |
| Kat6a | 1.35E-06 | 0.01258691 | 0.286 | 0.246 | 0.03636466 |
| Ehmt1 | 1.38E-06 | 0.07691548 | 0.269 | 0.218 | 0.03705648 |
| Arglu1 | 1.42E-06 | -0.0066153 | 0.48 | 0.429 | 0.03828004 |
| Fam102b | 1.43E-06 | 0.17012904 | 0.514 | 0.425 | 0.03859442 |
| Ascc3 | 1.44E-06 | 0.09064434 | 0.313 | 0.257 | 0.03880852 |
| Tra2a | 1.44E-06 | 0.03848167 | 0.389 | 0.339 | 0.03882183 |
| Tcf7l2 | 1.45E-06 | 0.12439712 | 0.3 | 0.239 | 0.03908634 |
| Nav3 | 1.49E-06 | 0.16931588 | 0.878 | 0.812 | 0.04006997 |
| Dnmt3a | 1.51E-06 | 0.02947213 | 0.414 | 0.358 | 0.04050188 |
| Ankrd44 | 1.61E-06 | 0.14886601 | 0.738 | 0.645 | 0.0432751 |
| 4933406I18R | 1.61E-06 | 0.23873503 | 0.734 | 0.641 | 0.04344512 |
| Tbc1d22a | 1.66E-06 | -0.0906235 | 0.349 | 0.333 | 0.04454709 |
| Tsix | 1.71E-06 | 0.09623321 | 0.302 | 0.248 | 0.04598635 |
| Dip2c | 1.81E-06 | 0.02303468 | 0.35 | 0.3 | 0.04865401 |
| Serinc3 | 1.84E-06 | 0.09528689 | 0.672 | 0.59 | 0.04949644 |
| Srsf11 | 1.85E-06 | -0.1015426 | 0.34 | 0.324 | 0.04965216 |
| Inpp5d | 1.89E-06 | 0.12612007 | 0.933 | 0.874 | 0.05081054 |
| Tcf12 | 1.94E-06 | -0.0041163 | 0.442 | 0.403 | 0.05228586 |
| Zdhhc20 | 1.95E-06 | 0.0561622 | 0.288 | 0.241 | 0.05249459 |
| Hectd1 | 1.98E-06 | 0.12471581 | 0.26 | 0.205 | 0.05319363 |
| Dock4 | 1.99E-06 | 0.15990417 | 0.92 | 0.88 | 0.05345918 |
| Fam49b | 2.06E-06 | 0.16828675 | 0.766 | 0.676 | 0.05530478 |
| Sesn1 | 2.11E-06 | -0.0677798 | 0.261 | 0.242 | 0.05665571 |
| Babam2 | 2.20E-06 | 0.08845328 | 0.347 | 0.288 | 0.05917778 |
| Prkce | 2.33E-06 | 0.00909699 | 0.448 | 0.391 | 0.0627433 |
| Pard3b | 2.39E-06 | 0.08568904 | 0.432 | 0.363 | 0.06420288 |
| Kcnk13 | 2.50E-06 | 0.10923378 | 0.344 | 0.28 | 0.06728284 |
| Arid4b | 2.59E-06 | -0.0270183 | 0.279 | 0.246 | 0.0695732 |
| Lcor | 2.77E-06 | 0.07997364 | 0.337 | 0.285 | 0.07462882 |
| Spag9 | 2.80E-06 | -0.1295493 | 0.364 | 0.355 | 0.07536033 |
| Rasgrp3 | 2.80E-06 | 0.00547924 | 0.487 | 0.433 | 0.07539792 |
| Btbd9 | 2.81E-06 | 0.0920914 | 0.529 | 0.455 | 0.07549435 |
| Scamp2 | 2.92E-06 | 0.0659022 | 0.31 | 0.263 | 0.07862789 |
| Calm2 | 2.93E-06 | 0.07804636 | 0.306 | 0.248 | 0.07894299 |
| Slc1a3 | 3.02E-06 | 0.00249445 | 0.282 | 0.242 | 0.08113874 |
| Atad2b | 3.13E-06 | 0.18407293 | 0.323 | 0.255 | 0.08434972 |
| Nlk | 3.18E-06 | 0.26707722 | 0.31 | 0.233 | 0.08563996 |
| P4ha1 | 3.19E-06 | 0.05530667 | 0.254 | 0.217 | 0.08591431 |
| Pvt1 | 3.31E-06 | -0.0469668 | 0.301 | 0.276 | 0.08893419 |
| Prkcb | 3.36E-06 | 0.20153538 | 0.54 | 0.447 | 0.09028891 |
| St3gal5 | 3.37E-06 | 0.18379364 | 0.317 | 0.246 | 0.09066115 |
| Sipa1l1 | 3.39E-06 | 0.11619351 | 0.35 | 0.288 | 0.09125707 |
| Rbm39 | 3.48E-06 | -0.1268978 | 0.73 | 0.713 | 0.09374929 |
| Sdccag8 | 3.49E-06 | 0.05529641 | 0.3 | 0.251 | 0.09392366 |
| Crlf3 | 3.54E-06 | 0.08042437 | 0.428 | 0.371 | 0.09535584 |
| Jmjd1c | 3.63E-06 | 0.02901759 | 0.668 | 0.602 | 0.09767157 |
| Tulp4 | 3.76E-06 | 0.11490719 | 0.251 | 0.202 | 0.10124244 |
| Fli1 | 3.77E-06 | 0.06790332 | 0.692 | 0.613 | 0.10132188 |
| Gm44686 | 3.85E-06 | -0.1338327 | 0.25 | 0.248 | 0.103558 |
| P2ry12 | 4.07E-06 | 0.05679711 | 0.762 | 0.69 | 0.10948158 |
| Phactr2 | 4.10E-06 | 0.05454697 | 0.3 | 0.253 | 0.11037317 |
| Rabep1 | 4.90E-06 | 0.03253201 | 0.355 | 0.312 | 0.13176288 |
| Bmp2k | 4.91E-06 | 0.08611177 | 0.614 | 0.536 | 0.13206874 |
| Foxp1 | 4.99E-06 | 0.19811053 | 0.411 | 0.329 | 0.13435246 |
| Phf21a | 5.10E-06 | 0.09466786 | 0.465 | 0.392 | 0.13729568 |
| Fgfr2 | 5.29E-06 | -0.0404273 | 0.468 | 0.422 | 0.14227134 |
| St3gal6 | 5.58E-06 | 0.04314455 | 0.584 | 0.518 | 0.15007983 |
| Btbd7 | 5.99E-06 | -0.0131407 | 0.258 | 0.23 | 0.16119455 |
| Fam105a | 6.14E-06 | -0.2320971 | 0.413 | 0.508 | 0.16521856 |
| Plxdc2 | 6.15E-06 | 0.1364656 | 0.926 | 0.885 | 0.16541864 |
| Il6ra | 7.65E-06 | -0.0278196 | 0.42 | 0.385 | 0.20593095 |
| Rab10 | 7.79E-06 | 0.08787895 | 0.272 | 0.225 | 0.2095228 |
| Adgre1 | 7.83E-06 | 0.09664646 | 0.26 | 0.211 | 0.21062698 |
| Dock2 | 7.96E-06 | 0.11279238 | 0.723 | 0.638 | 0.21429116 |
| Ankrd12 | 8.38E-06 | -0.1978182 | 0.288 | 0.292 | 0.22557761 |
| Itgb5 | 9.34E-06 | 0.14789745 | 0.681 | 0.595 | 0.25126568 |
| Fmnl2 | 9.39E-06 | -0.2831945 | 0.229 | 0.255 | 0.25266299 |
| Trappc9 | 9.64E-06 | 0.11935446 | 0.253 | 0.199 | 0.25930755 |
| Ssbp2 | 1.05E-05 | -0.1382481 | 0.3 | 0.297 | 0.28255195 |
| Rsrp1 | 1.13E-05 | 0.05417679 | 0.742 | 0.67 | 0.3053165 |
| Nsd1 | 1.28E-05 | 0.03036476 | 0.59 | 0.535 | 0.34326587 |
| Hpgd | 1.43E-05 | 0.0980164 | 0.37 | 0.306 | 0.38393252 |
| AY036118 | 1.49E-05 | 0.29343443 | 0.541 | 0.452 | 0.40118788 |
| Daglb | 1.51E-05 | -0.0093922 | 0.275 | 0.247 | 0.40708595 |
| Lcp1 | 1.55E-05 | -0.0633907 | 0.276 | 0.257 | 0.41834994 |
| Nsmce2 | 1.74E-05 | -0.0147823 | 0.273 | 0.245 | 0.46737095 |
| A830008E24 | 1.81E-05 | 0.22009756 | 0.287 | 0.217 | 0.48713137 |
| 0610040J01R | 1.86E-05 | 0.06106025 | 0.508 | 0.445 | 0.49950812 |
| Mtdh | 1.89E-05 | 0.03436762 | 0.558 | 0.503 | 0.50899973 |
| Arhgap5 | 1.96E-05 | 0.065725 | 0.724 | 0.652 | 0.52613206 |
| Fgf13 | 1.99E-05 | -0.2077638 | 0.423 | 0.401 | 0.53646588 |
| Cux1 | 2.00E-05 | 0.01423747 | 0.498 | 0.446 | 0.53899271 |
| Agmo | 2.13E-05 | 0.16047652 | 0.626 | 0.54 | 0.57396596 |
| Diaph2 | 2.16E-05 | 0.07502504 | 0.728 | 0.656 | 0.58098538 |
| Sdk1 | 2.22E-05 | 0.12575524 | 0.295 | 0.237 | 0.59739836 |
| R3hdm2 | 2.54E-05 | 0.08383793 | 0.345 | 0.294 | 0.68396082 |
| Ptprj | 2.64E-05 | 0.15073577 | 0.739 | 0.656 | 0.71003682 |
| Gab1 | 2.70E-05 | 0.03340215 | 0.292 | 0.255 | 0.72669661 |
| Smyd3 | 2.72E-05 | 0.004495 | 0.486 | 0.437 | 0.73125358 |
| Exoc4 | 2.86E-05 | -0.0575975 | 0.53 | 0.509 | 0.76909508 |
| Strbp | 3.35E-05 | -0.0590804 | 0.367 | 0.336 | 0.90261915 |
| Rapgef2 | 3.90E-05 | 0.01198999 | 0.268 | 0.236 | 1 |
| Srrm2 | 3.91E-05 | -0.095736 | 0.646 | 0.619 | 1 |
| Phf14 | 3.91E-05 | 0.09638253 | 0.537 | 0.471 | 1 |
| Hnrnpa2b1 | 3.99E-05 | 0.018088 | 0.601 | 0.545 | 1 |
| Runx1 | 4.17E-05 | -0.0415914 | 0.672 | 0.629 | 1 |
| Kansl1 | 4.45E-05 | 0.19644056 | 0.634 | 0.548 | 1 |
| Specc1 | 5.28E-05 | 0.02724958 | 0.554 | 0.502 | 1 |
| Ly86 | 5.36E-05 | -0.1399961 | 0.72 | 0.705 | 1 |
| Pitpnc1 | 5.48E-05 | 0.00918742 | 0.527 | 0.477 | 1 |
| Nfia | 6.36E-05 | 0.20837799 | 0.622 | 0.536 | 1 |
| Tbc1d5 | 6.79E-05 | -0.0137747 | 0.672 | 0.622 | 1 |
| Fcrls | 7.54E-05 | 0.14206399 | 0.509 | 0.438 | 1 |
| 4930517O19 | 7.56E-05 | 0.10689262 | 0.258 | 0.208 | 1 |
| Camk1d | 0.00010227 | -0.1216142 | 0.725 | 0.708 | 1 |
| Oxr1 | 0.00010554 | 0.0355773 | 0.461 | 0.41 | 1 |
| Dnajc1 | 0.00010724 | -0.0213173 | 0.329 | 0.299 | 1 |
| Gab2 | 0.00013119 | -0.0357401 | 0.771 | 0.724 | 1 |
| Tanc2 | 0.00014298 | 0.08254019 | 0.958 | 0.925 | 1 |
| Zbtb20 | 0.00014794 | -0.1599376 | 0.692 | 0.693 | 1 |
| Asap1 | 0.00015404 | 0.15512799 | 0.638 | 0.561 | 1 |
| Frmd4a | 0.00015496 | 0.10587856 | 0.841 | 0.776 | 1 |
| Ppp1r9a | 0.00017 | 0.1513515 | 0.58 | 0.505 | 1 |
| Tg | 0.00018501 | 0.13713096 | 0.272 | 0.221 | 1 |
| Rnf216 | 0.00019195 | 0.13675952 | 0.574 | 0.501 | 1 |
| Chn2 | 0.00019254 | 0.09316423 | 0.642 | 0.573 | 1 |
| Epb41l2 | 0.00023145 | 0.12036551 | 0.839 | 0.775 | 1 |
| Arhgap31 | 0.00023626 | 0.11338593 | 0.315 | 0.265 | 1 |
| Rsrc1 | 0.00024844 | 0.00155291 | 0.309 | 0.285 | 1 |
| Fat3 | 0.00026167 | -0.0720223 | 0.346 | 0.327 | 1 |
| Atp8a2 | 0.00026946 | 0.05435888 | 0.593 | 0.536 | 1 |
| Gm35188 | 0.00028114 | 0.14592308 | 0.794 | 0.729 | 1 |
| Tgfbr1 | 0.0002821 | -0.0425198 | 0.907 | 0.865 | 1 |
| Galnt1 | 0.0002865 | 0.11402719 | 0.258 | 0.214 | 1 |
| Nlrp1b | 0.00029805 | 0.08400967 | 0.271 | 0.228 | 1 |
| Maml2 | 0.00029892 | 0.11587309 | 0.644 | 0.574 | 1 |
| Arhgap45 | 0.00030432 | 0.04365118 | 0.737 | 0.68 | 1 |
| App | 0.00032798 | 0.06209947 | 0.292 | 0.248 | 1 |
| Apbb1ip | 0.00035946 | 0.1549875 | 0.788 | 0.723 | 1 |
| Atp2c1 | 0.00036339 | 0.02473427 | 0.4 | 0.364 | 1 |
| Lair1 | 0.00039555 | 0.00619965 | 0.589 | 0.539 | 1 |
| Mycbp2 | 0.0004013 | -0.0336264 | 0.739 | 0.702 | 1 |
| Ptprm | 0.00042745 | 0.17204921 | 0.511 | 0.441 | 1 |
| Ank2 | 0.00046073 | -0.0465713 | 0.559 | 0.535 | 1 |
| Glg1 | 0.000546 | -0.1389742 | 0.27 | 0.272 | 1 |
| Znrf1 | 0.0006608 | 0.27004297 | 0.254 | 0.193 | 1 |
| 9930021J03R | 0.00068689 | 0.0351331 | 0.263 | 0.228 | 1 |
| Ssh2 | 0.0007672 | 0.06502348 | 0.878 | 0.825 | 1 |
| Mbnl1 | 0.00083022 | 0.13128767 | 0.84 | 0.791 | 1 |
| Olfr56 | 0.00086208 | -0.3028095 | 0.199 | 0.254 | 1 |
| Ext1 | 0.00087848 | -0.0517269 | 0.328 | 0.307 | 1 |
| Dennd4a | 0.00093221 | -0.0399754 | 0.68 | 0.645 | 1 |

| | | | | | |
|---|---|---|---|---|---|
| Bbs9 | 0.00120261 | 0.01959875 | 0.471 | 0.428 | 1 |
| Gm26740 | 0.00134726 | -0.0908497 | 0.262 | 0.249 | 1 |
| Fhit | 0.00157488 | -0.0736261 | 0.468 | 0.458 | 1 |
| Gm20275 | 0.00183791 | 0.03787627 | 0.295 | 0.264 | 1 |
| Plcl1 | 0.00195798 | 0.07690794 | 0.758 | 0.698 | 1 |
| Ccny | 0.00198533 | 0.02941002 | 0.261 | 0.234 | 1 |
| Ppp3ca | 0.00232252 | -0.0518516 | 0.666 | 0.639 | 1 |
| Mertk | 0.00232522 | 0.00905499 | 0.867 | 0.824 | 1 |
| Immp2l | 0.0027551 | -0.0290148 | 0.338 | 0.303 | 1 |
| Wac | 0.00341166 | 0.01493417 | 0.304 | 0.281 | 1 |
| Gm26804 | 0.00724237 | -0.1642852 | 0.325 | 0.383 | 1 |
| Mef2c | 0.00964062 | 0.12188244 | 0.822 | 0.775 | 1 |
| Zeb2 | 0.01192212 | -0.1033403 | 0.832 | 0.831 | 1 |
| Dst | 0.01552849 | -0.1222223 | 0.7 | 0.708 | 1 |
| Wwox | 0.01560046 | -0.0659895 | 0.447 | 0.446 | 1 |
| Slc8a1 | 0.01730765 | 0.09647703 | 0.892 | 0.858 | 1 |
| Snx29 | 0.02331953 | 0.09572162 | 0.737 | 0.69 | 1 |
| Hexb | 0.02737739 | -0.0015645 | 0.876 | 0.844 | 1 |
| Macf1 | 0.03459161 | 0.11288373 | 0.822 | 0.788 | 1 |
| Gm26733 | 0.14493117 | -0.1100699 | 0.225 | 0.256 | 1 |
| Inpp4b | 0.14819142 | -0.0445002 | 0.725 | 0.711 | 1 |
| Xist | 0.1868302 | -0.0478719 | 0.83 | 0.818 | 1 |
