## Supplementary table 4 for "DAP12 deficiency alters microglia-oligodendrocyte communication and enhances resilience against tau toxicity"

| | | | | | | | | | | | |
|---|---|---|---|---|---|---|---|---|---|---|---|
| Figure4E_psd95_intensity | | | | | | | | | | | |
| | Dap12+/+ Tau+ | | | | | Dap12-/- Tau+ | | | | | |
| | 0.034274 | 0.092052 | | | | 0.02925 | 0.152633 | 0.567346 | 0.186022 | | |
| | 0.219633 | 0.323166 | 0.457117 | 0.206354 | 0.140781 | 0.388148 | 0.364399 | 0.636669 | 0.139586 | | |
| | 0.077577 | 0.108017 | | | | 0.319351 | 0.011118 | 0.158217 | 0.266614 | | |
| | 0.123175 | 0.180992 | | | | 0.492695 | 0.141608 | 0.478791 | 0.39367 | | |
| | 0.152205 | 0.146154 | 0.254025 | | | 0.539357 | 0.660255 | 1.094244 | 0.030332 | | |
| | 0.170612 | 0.188452 | | | | 0.452265 | 0.013639 | 0.141608 | 0.484137 | | |
| | 0.388541 | 0.138188 | 0.365948 | | | 0.381862 | 0.196102 | 0.484137 | 0.374539 | | |
| | | | | | | 0.347766 | | | | | |
| average | 0.198277 | | | | | 0.342288276 | | | | | |
| Figure4A_DEGs_of_different_exitatory_neurons | | | | | | | | | | | |
| | Dap12+/+ Tau+ | | | | | Dap12-/- Tau+ | | | | | |
| | p_val | avg_log2FC | pct.1 | pct.2 | p_val_adj | | p_val | avg_log2FC | pct.1 | pct.2 | p_val_adj |
| AC149090.1 | 0 | -0.776193402 | 0.765 | 0.932 | 0 | Gm20388 | 0 | 0.041070907 | 0.859 | 0.977 | 0 |
| Gm20388 | 0 | 0.024995127 | 0.977 | 0.81 | 0 | Gm42418 | 1.54E-296 | -0.515146779 | 0.86 | 0.933 | 4.13E-292 |
| Gm26848 | 0 | 0.040915671 | 0.943 | 0.792 | 0 | Gm26848 | 4.12E-199 | -0.071045208 | 0.827 | 0.943 | 1.11E-194 |
| 1-Mar | 0 | -2.932554817 | 0.172 | 0.862 | 0 | Park2 | 1.81E-190 | -0.334077086 | 0.766 | 0.895 | 4.88E-186 |
| Park2 | 9.75E-262 | 0.487604061 | 0.895 | 0.708 | 2.62E-258 | Cmss1 | 1.78E-142 | -0.381467205 | 0.582 | 0.673 | 4.78E-138 |
| Gm47283 | 9.75E-262 | 0.521871319 | 0.358 | 0.199 | 2.62E-257 | Gm26835 | 3.93E-106 | 0.10043744 | 0.784 | 0.853 | 1.06E-101 |
| Gm26835 | 4.68E-208 | -0.114884315 | 0.853 | 0.744 | 1.26E-203 | Gm26804 | 2.60E-90 | -0.113194605 | 0.794 | 0.889 | 6.98E-86 |
| Kcnb2 | 9.22E-199 | 0.423006185 | 0.892 | 0.82 | 2.48E-194 | C230004F18Rik | 2.68E-84 | -0.133877533 | 0.756 | 0.856 | 7.21E-80 |
| Gm26804 | 3.16E-196 | 0.010018677 | 0.889 | 0.764 | 8.52E-192 | Dscaml1 | 7.24E-77 | -0.256396282 | 0.794 | 0.823 | 1.95E-72 |
| Fam19a1 | 2.32E-189 | 0.525630969 | 0.769 | 0.607 | 6.25E-185 | Fam19a2 | 3.81E-76 | -0.268600361 | 0.562 | 0.679 | 1.03E-71 |
| Gm2163 | 1.19E-186 | 0.397404544 | 0.745 | 0.574 | 3.20E-182 | Fam19a1 | 1.14E-73 | -0.246208118 | 0.662 | 0.769 | 3.08E-69 |
| Prkar2a | 3.06E-175 | 0.375178574 | 0.343 | 0.201 | 8.24E-171 | Fam19a5 | 1.74E-73 | -0.273074216 | 0.493 | 0.601 | 4.68E-69 |
| Hspa5 | 1.13E-160 | -0.297169581 | 0.177 | 0.328 | 3.05E-156 | Mkl2 | 1.50E-66 | -0.135211618 | 0.733 | 0.826 | 4.03E-62 |
| Mkl2 | 3.93E-156 | 0.254833625 | 0.826 | 0.679 | 1.06E-151 | Kctd16 | 8.52E-61 | 0.22890818 | 0.901 | 0.878 | 2.29E-56 |
| C230004F18Rik | 3.64E-155 | 0.075567637 | 0.856 | 0.728 | 9.81E-151 | Mlip | 1.34E-55 | 0.217811219 | 0.526 | 0.423 | 3.61E-51 |
| Hsp90b1 | 1.91E-147 | -0.299864389 | 0.24 | 0.395 | 5.14E-143 | Csmd1 | 1.20E-53 | -0.116395834 | 0.994 | 0.997 | 3.22E-49 |
| 2210408I21Rik | 2.59E-145 | 0.371577067 | 0.616 | 0.529 | 6.97E-141 | Galnt1 | 7.61E-53 | 0.174797272 | 0.601 | 0.501 | 2.05E-48 |
| Hectd2 | 5.36E-141 | -0.487895566 | 0.347 | 0.485 | 1.44E-136 | Kcnd3 | 7.92E-52 | -0.234273623 | 0.697 | 0.732 | 2.13E-47 |
| Ptprn | 1.90E-133 | -0.391138094 | 0.581 | 0.697 | 5.12E-129 | Kcnq1ot1 | 2.14E-50 | -0.187845711 | 0.853 | 0.877 | 5.75E-46 |
| Fam19a2 | 4.22E-133 | 0.465056127 | 0.679 | 0.531 | 1.14E-128 | Kcnq3 | 3.27E-50 | -0.194985417 | 0.923 | 0.938 | 8.81E-46 |
| 2410089E03Rik | 4.23E-132 | 0.251791235 | 0.759 | 0.613 | 1.14E-127 | Gm2163 | 1.26E-49 | -0.10693634 | 0.657 | 0.745 | 3.38E-45 |
| Nrg3os | 1.35E-126 | -0.297174755 | 0.921 | 0.953 | 3.63E-122 | Frmd4a | 4.27E-49 | -0.174520457 | 0.961 | 0.972 | 1.15E-44 |
| Prpf4b | 1.85E-126 | -0.3153162 | 0.744 | 0.844 | 4.98E-122 | Zfp507 | 1.69E-48 | 0.151076969 | 0.266 | 0.186 | 4.54E-44 |
| Ddx5 | 8.28E-126 | -0.294216975 | 0.699 | 0.811 | 2.23E-121 | 2010111I01Rik | 5.80E-47 | -0.152729238 | 0.658 | 0.744 | 1.56E-42 |
| Clstn2 | 1.21E-122 | -0.390760949 | 0.902 | 0.929 | 3.27E-118 | 2410089E03Rik | 1.02E-46 | -0.113071311 | 0.674 | 0.759 | 2.74E-42 |
| 2010111I01Rik | 3.48E-122 | 0.284107644 | 0.744 | 0.606 | 9.37E-118 | Shank2 | 1.31E-46 | -0.170601398 | 0.755 | 0.772 | 3.52E-42 |
| Gm20642 | 9.78E-112 | -0.302864541 | 0.89 | 0.936 | 2.63E-107 | 2210408I21Rik | 3.72E-46 | -0.187197222 | 0.58 | 0.616 | 1.00E-41 |
| Srsf7 | 6.88E-111 | -0.280306876 | 0.515 | 0.652 | 1.85E-106 | 4930567K20Rik | 1.03E-45 | 0.205516236 | 0.347 | 0.261 | 2.77E-41 |
| 5031425E22Rik | 2.27E-110 | -0.299991731 | 0.417 | 0.556 | 6.11E-106 | Mkl1 | 3.73E-45 | -0.160087612 | 0.579 | 0.667 | 1.00E-40 |
| Frmd4a | 9.96E-109 | 0.234450393 | 0.972 | 0.955 | 2.68E-104 | Serinc1 | 5.96E-45 | 0.034080115 | 0.337 | 0.277 | 1.60E-40 |
| Ank2 | 1.68E-108 | 0.191730082 | 0.989 | 0.98 | 4.52E-104 | Gm47283 | 5.25E-44 | 0.118748344 | 0.44 | 0.358 | 1.41E-39 |
| Ube3a | 3.25E-105 | -0.258124579 | 0.947 | 0.963 | 8.74E-101 | Acp1 | 6.20E-44 | 0.183708408 | 0.396 | 0.31 | 1.67E-39 |
| Rora | 1.58E-104 | 0.317100866 | 0.901 | 0.855 | 4.24E-100 | 5031415H12Rik | 2.49E-43 | 0.162860828 | 0.327 | 0.245 | 6.69E-39 |
| Hdac9 | 1.26E-103 | 0.360734062 | 0.762 | 0.699 | 3.40E-99 | Ywhag | 2.03E-42 | 0.160030061 | 0.498 | 0.409 | 5.47E-38 |
| Fgf12 | 6.72E-101 | 0.258162213 | 0.932 | 0.905 | 1.81E-96 | Camk1d | 2.39E-40 | -0.18275461 | 0.869 | 0.885 | 6.42E-36 |
| AY036118 | 2.35E-100 | -0.347546669 | 0.618 | 0.724 | 6.32E-96 | Rheb | 3.34E-40 | 0.1455939 | 0.269 | 0.196 | 8.99E-36 |
| Plxna4os3 | 2.37E-97 | -0.30279744 | 0.303 | 0.432 | 6.37E-93 | Spred1 | 3.38E-40 | 0.145346609 | 0.417 | 0.333 | 9.09E-36 |
| Rbm28 | 4.32E-97 | -0.248609119 | 0.37 | 0.505 | 1.16E-92 | AI314180 | 5.40E-40 | -0.141494378 | 0.593 | 0.676 | 1.45E-35 |
| Atp6v0b | 1.07E-96 | -0.295159485 | 0.624 | 0.735 | 2.87E-92 | Jazf1 | 2.78E-39 | -0.156479048 | 0.694 | 0.712 | 7.47E-35 |
| Rbfox1 | 5.03E-96 | 0.21574817 | 0.995 | 0.989 | 1.35E-91 | Robo1 | 1.59E-38 | -0.298191358 | 0.515 | 0.572 | 4.27E-34 |
| AI314180 | 3.98E-94 | 0.232126855 | 0.676 | 0.548 | 1.07E-89 | Mat2a | 2.34E-38 | 0.108819328 | 0.418 | 0.339 | 6.30E-34 |
| Chst9 | 1.04E-93 | -0.506549316 | 0.329 | 0.434 | 2.79E-89 | Otud7a | 7.17E-38 | -0.151088702 | 0.907 | 0.916 | 1.93E-33 |
| Mtus2 | 1.01E-92 | 0.294848285 | 0.688 | 0.603 | 2.73E-88 | Palm2 | 1.02E-37 | -0.18391747 | 0.503 | 0.584 | 2.75E-33 |
| Galnt13 | 1.18E-91 | 0.317382035 | 0.71 | 0.615 | 3.18E-87 | Ywhae | 1.54E-37 | 0.132246985 | 0.426 | 0.345 | 4.15E-33 |
| Thra | 2.83E-90 | 0.272025315 | 0.535 | 0.458 | 7.63E-86 | Itm2c | 3.93E-37 | 0.013734641 | 0.258 | 0.215 | 1.06E-32 |
| Cadps | 5.48E-90 | 0.242741372 | 0.949 | 0.932 | 1.47E-85 | Sipa1l3 | 4.69E-37 | -0.197551088 | 0.591 | 0.624 | 1.26E-32 |
| Grin1 | 7.04E-90 | 0.231723422 | 0.878 | 0.849 | 1.89E-85 | Kcnq2 | 8.92E-37 | -0.116751269 | 0.569 | 0.574 | 2.40E-32 |
| Pdia6 | 1.14E-86 | -0.212415262 | 0.17 | 0.276 | 3.07E-82 | Prickle1 | 1.22E-36 | 0.137561088 | 0.942 | 0.924 | 3.28E-32 |
| Fth1 | 3.94E-85 | -0.128758491 | 0.171 | 0.274 | 1.06E-80 | Zdhhc14 | 1.34E-36 | -0.158557556 | 0.647 | 0.662 | 3.61E-32 |
| Phactr1 | 1.40E-84 | 0.217140226 | 0.971 | 0.948 | 3.78E-80 | Snx3 | 2.40E-36 | 0.10986146 | 0.339 | 0.266 | 6.44E-32 |
| Fam19a5 | 7.84E-84 | 0.256435772 | 0.601 | 0.477 | 2.11E-79 | Slc9a9 | 2.70E-36 | -0.187827376 | 0.521 | 0.539 | 7.28E-32 |
| Dync1i1 | 1.06E-83 | 0.241841991 | 0.852 | 0.8 | 2.86E-79 | Ip6k1 | 4.80E-36 | 0.147846278 | 0.293 | 0.221 | 1.29E-31 |
| Sdk1 | 3.80E-83 | -0.409595013 | 0.214 | 0.317 | 1.02E-78 | Kmt5b | 1.34E-35 | 0.124223304 | 0.398 | 0.321 | 3.59E-31 |
| 4930567K20Rik | 8.50E-83 | -0.311477772 | 0.261 | 0.376 | 2.29E-78 | Srsf7 | 2.01E-35 | 0.127688912 | 0.596 | 0.515 | 5.41E-31 |
| Gabra2 | 2.50E-81 | 0.262136585 | 0.804 | 0.755 | 6.73E-77 | Rhobtb2 | 2.29E-35 | 0.115229342 | 0.292 | 0.224 | 6.16E-31 |
| Anks1b | 1.15E-80 | 0.158037732 | 0.993 | 0.989 | 3.10E-76 | Zfp973 | 3.21E-35 | -0.170002286 | 0.337 | 0.405 | 8.65E-31 |
| Syt7 | 5.74E-80 | 0.281287748 | 0.744 | 0.696 | 1.55E-75 | Gm28928 | 4.38E-35 | -0.209246892 | 0.234 | 0.29 | 1.18E-30 |
| Robo2 | 1.24E-79 | 0.293190846 | 0.829 | 0.812 | 3.34E-75 | 2900011O08Rik | 2.09E-34 | -0.157092329 | 0.465 | 0.544 | 5.61E-30 |
| Prrt1 | 8.98E-79 | -0.188060449 | 0.166 | 0.266 | 2.42E-74 | Pnisr | 3.72E-34 | 0.133059898 | 0.881 | 0.832 | 1.00E-29 |
| Zfp804b | 2.10E-78 | 0.577855992 | 0.37 | 0.282 | 5.66E-74 | Cacna1b | 8.41E-34 | -0.121548779 | 0.662 | 0.666 | 2.26E-29 |
| 4930578G10Rik | 2.30E-78 | 0.275284922 | 0.593 | 0.477 | 6.18E-74 | Crebzf | 9.31E-34 | 0.058897809 | 0.391 | 0.327 | 2.50E-29 |
| 1700085D07Rik | 1.94E-76 | -0.223503219 | 0.31 | 0.426 | 5.21E-72 | Zfp697 | 1.65E-33 | -0.166081474 | 0.39 | 0.426 | 4.45E-29 |
| Cacna1a | 1.23E-75 | -0.283453376 | 0.878 | 0.912 | 3.30E-71 | Spred2 | 2.26E-33 | 0.133094009 | 0.691 | 0.616 | 6.09E-29 |
| Ablim2 | 6.30E-75 | 0.248637607 | 0.67 | 0.616 | 1.69E-70 | Tecr | 2.40E-33 | 0.068381445 | 0.448 | 0.381 | 6.46E-29 |
| Zbtb20 | 9.10E-74 | -0.320271639 | 0.749 | 0.741 | 2.45E-69 | Ppp1r1a | 2.85E-33 | 0.125549978 | 0.289 | 0.222 | 7.66E-29 |
| Gm44151 | 4.28E-70 | -0.287537337 | 0.689 | 0.771 | 1.15E-65 | Aplp1 | 4.54E-33 | 0.008504498 | 0.458 | 0.407 | 1.22E-28 |
| Lrrc4 | 4.63E-70 | -0.268276572 | 0.622 | 0.716 | 1.25E-65 | Rab6a | 5.18E-33 | 0.14606865 | 0.42 | 0.344 | 1.39E-28 |
| Csmd1 | 8.50E-70 | 0.130923827 | 0.997 | 0.994 | 2.29E-65 | Rgl1 | 8.50E-33 | -0.136303596 | 0.63 | 0.644 | 2.29E-28 |
| Smim14 | 1.78E-69 | -0.209086156 | 0.23 | 0.333 | 4.78E-65 | Rbfox1 | 8.51E-33 | -0.128883786 | 0.987 | 0.995 | 2.29E-28 |
| Jakmip1 | 3.93E-69 | 0.230740344 | 0.684 | 0.599 | 1.06E-64 | Atp1a1 | 1.74E-32 | 0.078443768 | 0.457 | 0.387 | 4.69E-28 |
| 2310035C23Rik | 4.01E-69 | 0.231746487 | 0.524 | 0.414 | 1.08E-64 | Slco3a1 | 2.01E-32 | -0.137322476 | 0.496 | 0.51 | 5.42E-28 |
| Ppp2r2b | 4.66E-68 | 0.298871802 | 0.748 | 0.694 | 1.25E-63 | Ube3a | 2.05E-32 | 0.12732753 | 0.96 | 0.947 | 5.51E-28 |
| Hnrnpa2b1 | 2.30E-67 | -0.211656355 | 0.64 | 0.733 | 6.19E-63 | Rnf150 | 3.04E-32 | -0.154149337 | 0.663 | 0.684 | 8.19E-28 |
| Myo16 | 2.37E-67 | 0.279989354 | 0.272 | 0.195 | 6.39E-63 | Caly | 4.66E-32 | 0.050498776 | 0.273 | 0.22 | 1.25E-27 |
| Cntn1 | 2.50E-67 | 0.200902179 | 0.901 | 0.869 | 6.74E-63 | Scn1b | 5.57E-32 | 0.096857679 | 0.272 | 0.208 | 1.50E-27 |
| Zfp697 | 7.01E-67 | 0.233218173 | 0.426 | 0.381 | 1.89E-62 | 2310035C23Rik | 6.14E-32 | -0.151804815 | 0.451 | 0.524 | 1.65E-27 |
| Nrn1 | 1.53E-66 | -0.242728983 | 0.237 | 0.337 | 4.12E-62 | Calm2 | 6.21E-32 | 0.140376716 | 0.621 | 0.544 | 1.67E-27 |
| Ccsap | 2.07E-66 | -0.190707728 | 0.21 | 0.308 | 5.57E-62 | Slc24a5 | 1.07E-31 | 0.132812397 | 0.576 | 0.498 | 2.87E-27 |
| Mapt | 3.25E-66 | 0.201592126 | 0.851 | 0.826 | 8.73E-62 | Amph | 5.76E-31 | -0.133961317 | 0.731 | 0.746 | 1.55E-26 |
| Ksr2 | 3.59E-66 | 0.210787819 | 0.806 | 0.773 | 9.65E-62 | Fam131a | 6.08E-31 | 0.110397511 | 0.505 | 0.43 | 1.64E-26 |

| | | | | | | | | | | | |
|---|---|---|---|---|---|---|---|---|---|---|---|
| P4ha1 | 6.87E-66 | -0.204982056 | 0.382 | 0.493 | 1.85E-61 | Lncpint | 7.54E-31 | -0.134645441 | 0.922 | 0.924 | 2.03E-26 |
| Lrpap1 | 2.51E-65 | -0.232633686 | 0.27 | 0.373 | 6.75E-61 | Galntl6 | 1.18E-30 | 0.002789374 | 0.422 | 0.485 | 3.17E-26 |
| Tmem243 | 3.17E-65 | -0.223992617 | 0.399 | 0.509 | 8.54E-61 | Rchy1 | 1.25E-30 | 0.082650095 | 0.316 | 0.255 | 3.36E-26 |
| Slc9a9 | 7.71E-65 | 0.257675439 | 0.539 | 0.508 | 2.07E-60 | Cx3cl1 | 2.07E-30 | 0.088589418 | 0.313 | 0.252 | 5.58E-26 |
| Nav1 | 2.39E-64 | 0.216817528 | 0.626 | 0.581 | 6.43E-60 | Slc3a2 | 2.90E-30 | 0.08013396 | 0.297 | 0.236 | 7.81E-26 |
| Esrrg | 6.25E-64 | 0.283085563 | 0.489 | 0.41 | 1.68E-59 | Saraf | 4.82E-30 | 0.044237923 | 0.259 | 0.209 | 1.30E-25 |
| Pdia3 | 6.79E-64 | -0.185457128 | 0.203 | 0.298 | 1.83E-59 | Ftx | 7.18E-30 | 0.140598298 | 0.802 | 0.747 | 1.93E-25 |
| Sparcl1 | 7.53E-64 | -0.265587757 | 0.314 | 0.416 | 2.02E-59 | Tra2b | 7.22E-30 | 0.113425865 | 0.417 | 0.346 | 1.94E-25 |
| Ctdspl | 5.00E-63 | 0.22702313 | 0.534 | 0.479 | 1.35E-58 | Gm44257 | 7.57E-30 | 0.159050535 | 0.303 | 0.237 | 2.04E-25 |
| Kcnh7 | 2.49E-62 | 0.318181897 | 0.745 | 0.667 | 6.71E-58 | Smim14 | 1.39E-29 | 0.127566976 | 0.295 | 0.23 | 3.73E-25 |
| Atxn1 | 3.08E-61 | 0.181375477 | 0.938 | 0.91 | 8.28E-57 | Prr14l | 1.82E-29 | 0.128646468 | 0.354 | 0.284 | 4.90E-25 |
| Trim2 | 4.77E-61 | -0.283143611 | 0.837 | 0.874 | 1.28E-56 | Tmem59 | 1.92E-29 | 0.075092032 | 0.314 | 0.255 | 5.17E-25 |
| Dpyd | 9.30E-61 | 0.284132393 | 0.6 | 0.564 | 2.50E-56 | Morf4l1 | 2.02E-29 | 0.083692145 | 0.367 | 0.303 | 5.43E-25 |
| Gm20319 | 1.16E-60 | -0.324858435 | 0.278 | 0.37 | 3.13E-56 | Dync1i1 | 2.30E-29 | -0.129906651 | 0.843 | 0.852 | 6.19E-25 |
| 4930402H24Rik | 5.68E-60 | 0.181030997 | 0.865 | 0.844 | 1.53E-55 | Snhg11 | 3.51E-29 | 0.052252695 | 0.891 | 0.931 | 9.45E-25 |
| Ptprn2 | 6.85E-60 | 0.197593224 | 0.919 | 0.906 | 1.84E-55 | Atp6ap2 | 4.85E-29 | 0.037123976 | 0.279 | 0.233 | 1.31E-24 |
| Ip6k1 | 1.16E-59 | -0.188090455 | 0.221 | 0.314 | 3.11E-55 | Emc10 | 5.64E-29 | 0.056555008 | 0.272 | 0.22 | 1.52E-24 |
| Nrp1 | 2.21E-59 | -0.217982558 | 0.685 | 0.776 | 5.94E-55 | Gm28198 | 5.91E-29 | 0.117249588 | 0.306 | 0.24 | 1.59E-24 |
| Slc35f3 | 4.63E-59 | -0.299768552 | 0.531 | 0.585 | 1.24E-54 | Esco1 | 5.97E-29 | 0.091028788 | 0.408 | 0.341 | 1.61E-24 |
| Rnf112 | 1.38E-58 | -0.265536143 | 0.547 | 0.636 | 3.71E-54 | Atxn7l1 | 8.91E-29 | -0.135074339 | 0.584 | 0.607 | 2.40E-24 |
| AC165953.2 | 2.26E-58 | 0.216078649 | 0.418 | 0.32 | 6.07E-54 | Thrap3 | 1.78E-28 | 0.087494856 | 0.349 | 0.286 | 4.79E-24 |
| Prkag2 | 5.64E-58 | -0.235661033 | 0.811 | 0.856 | 1.52E-53 | Jakmip1 | 3.06E-28 | -0.143301531 | 0.635 | 0.684 | 8.22E-24 |
| Peg3 | 7.92E-58 | 0.240297274 | 0.542 | 0.486 | 2.13E-53 | Srgap3 | 3.22E-28 | -0.117527566 | 0.885 | 0.886 | 8.66E-24 |
| Shank2 | 9.34E-58 | 0.185151034 | 0.772 | 0.758 | 2.51E-53 | Dtx3 | 3.28E-28 | 0.075265083 | 0.354 | 0.292 | 8.83E-24 |
| Nrxn1 | 2.92E-57 | 0.185640139 | 0.995 | 0.989 | 7.85E-53 | Zfp207 | 3.61E-28 | 0.05037004 | 0.446 | 0.387 | 9.71E-24 |
| Gm44257 | 4.10E-57 | -0.223491941 | 0.237 | 0.33 | 1.10E-52 | Syt11 | 4.05E-28 | 0.052252294 | 0.556 | 0.495 | 1.09E-23 |
| Tsix | 1.28E-56 | -0.236933512 | 0.565 | 0.658 | 3.44E-52 | Slc22a17 | 5.03E-28 | 0.026762067 | 0.547 | 0.492 | 1.35E-23 |
| Tmem178b | 6.09E-56 | 0.244461925 | 0.765 | 0.726 | 1.64E-51 | Zfp266 | 8.26E-28 | 0.066520038 | 0.356 | 0.297 | 2.22E-23 |
| Fxr1 | 1.04E-55 | -0.165617896 | 0.426 | 0.528 | 2.80E-51 | Hspa4 | 8.34E-28 | 0.130918547 | 0.458 | 0.386 | 2.24E-23 |
| Gm13269 | 3.27E-55 | -0.261407478 | 0.513 | 0.609 | 8.79E-51 | Lrfn2 | 8.39E-28 | -0.126459996 | 0.676 | 0.689 | 2.26E-23 |
| Pknox2 | 5.31E-55 | 0.250753746 | 0.454 | 0.39 | 1.43E-50 | Lmo4 | 1.11E-27 | 0.082670442 | 0.397 | 0.333 | 3.00E-23 |
| Thy1 | 5.78E-55 | 0.190477812 | 0.272 | 0.199 | 1.56E-50 | Impact | 1.28E-27 | 0.094176648 | 0.298 | 0.237 | 3.45E-23 |
| Dopey2 | 2.11E-54 | 0.199556866 | 0.444 | 0.345 | 5.67E-50 | Ptprn2 | 1.33E-27 | -0.134905431 | 0.912 | 0.919 | 3.58E-23 |
| Mkl1 | 2.42E-54 | 0.096115642 | 0.667 | 0.574 | 6.51E-50 | Cbx5 | 1.37E-27 | 0.069932131 | 0.299 | 0.243 | 3.68E-23 |
| Ccnt2 | 3.12E-54 | -0.158999921 | 0.507 | 0.608 | 8.39E-50 | Rsrc2 | 1.37E-27 | 0.085699091 | 0.47 | 0.402 | 3.68E-23 |
| Tjp1 | 7.20E-54 | 0.178654534 | 0.862 | 0.848 | 1.94E-49 | P4ha1 | 1.55E-27 | 0.140248514 | 0.453 | 0.382 | 4.17E-23 |
| AC129186.1 | 9.29E-54 | 0.345255575 | 0.403 | 0.308 | 2.50E-49 | Psap | 2.06E-27 | -0.008295806 | 0.451 | 0.408 | 5.55E-23 |
| P2ry12 | 1.23E-53 | -0.210926017 | 0.306 | 0.402 | 3.31E-49 | Syp | 2.55E-27 | 0.020251884 | 0.449 | 0.4 | 6.86E-23 |
| 9130011E15Rik | 1.39E-53 | 0.199961815 | 0.447 | 0.35 | 3.75E-49 | Gabra1 | 3.93E-27 | 0.04430078 | 0.393 | 0.34 | 1.06E-22 |
| Ranbp17 | 1.53E-53 | 0.220117311 | 0.346 | 0.282 | 4.12E-49 | Fbxl17 | 4.45E-27 | -0.125991197 | 0.837 | 0.85 | 1.20E-22 |
| Gm43376 | 1.70E-53 | -0.201057678 | 0.569 | 0.665 | 4.56E-49 | Aff4 | 5.09E-27 | 0.120611075 | 0.629 | 0.559 | 1.37E-22 |
| Kcnt2 | 6.59E-53 | 0.252111395 | 0.641 | 0.586 | 1.77E-48 | Hnrnpr | 5.40E-27 | 0.08569293 | 0.501 | 0.434 | 1.45E-22 |
| Vwa8 | 6.69E-53 | 0.209163055 | 0.561 | 0.509 | 1.80E-48 | Nob1 | 5.68E-27 | 0.056542012 | 0.262 | 0.214 | 1.53E-22 |
| Fmn1 | 1.83E-52 | 0.225325586 | 0.443 | 0.355 | 4.92E-48 | Arhgap44 | 7.61E-27 | -0.088345744 | 0.571 | 0.564 | 2.05E-22 |
| Napepld | 2.96E-52 | -0.197885654 | 0.188 | 0.272 | 7.97E-48 | Gm3764 | 9.70E-27 | 0.127561917 | 0.506 | 0.435 | 2.61E-22 |
| Prdm5 | 4.24E-52 | -0.25707193 | 0.185 | 0.267 | 1.14E-47 | 4933427D14Rik | 9.98E-27 | -0.144405446 | 0.246 | 0.303 | 2.69E-22 |
| Gm49003 | 4.84E-52 | -0.196093281 | 0.311 | 0.406 | 1.30E-47 | 1600010M07Rik | 1.15E-26 | 0.092156663 | 0.442 | 0.375 | 3.09E-22 |
| Camk4 | 5.56E-52 | 0.257748668 | 0.565 | 0.49 | 1.50E-47 | Tmed9 | 1.80E-26 | 0.004604203 | 0.308 | 0.272 | 4.85E-22 |
| Usp50 | 5.79E-52 | -0.176950098 | 0.368 | 0.466 | 1.56E-47 | Nrxn2 | 2.65E-26 | -0.089115472 | 0.408 | 0.407 | 7.12E-22 |
| Gtdc1 | 7.70E-52 | 0.197197844 | 0.657 | 0.618 | 2.07E-47 | Nrn1 | 3.50E-26 | 0.098485944 | 0.297 | 0.237 | 9.41E-22 |
| Tra2a | 9.19E-52 | -0.190833404 | 0.663 | 0.745 | 2.47E-47 | Mdga2 | 3.53E-26 | -0.103706355 | 0.97 | 0.978 | 9.50E-22 |
| Dmd | 1.69E-51 | 0.217003141 | 0.921 | 0.886 | 4.55E-47 | Cyfip2 | 3.96E-26 | -0.111207924 | 0.639 | 0.648 | 1.07E-21 |
| Kcnma1 | 1.74E-51 | 0.150637952 | 0.983 | 0.969 | 4.69E-47 | Chst11 | 5.00E-26 | -0.140640293 | 0.416 | 0.443 | 1.34E-21 |
| Sel1l | 2.11E-51 | -0.178177018 | 0.372 | 0.469 | 5.68E-47 | Abi2 | 5.09E-26 | 0.123477398 | 0.678 | 0.613 | 1.37E-21 |
| Hecw2 | 2.53E-51 | 0.217914059 | 0.675 | 0.61 | 6.80E-47 | Pank2 | 5.86E-26 | 0.06117969 | 0.307 | 0.254 | 1.58E-21 |
| Adk | 3.91E-51 | 0.233177006 | 0.488 | 0.434 | 1.05E-46 | Ddx50 | 6.86E-26 | 0.081444064 | 0.567 | 0.501 | 1.85E-21 |
| 3110021N24Rik | 7.83E-51 | -0.191821504 | 0.43 | 0.528 | 2.11E-46 | Hnrnpc | 8.14E-26 | 0.085357353 | 0.348 | 0.287 | 2.19E-21 |
| Nol4l | 7.88E-51 | 0.195208426 | 0.48 | 0.434 | 2.12E-46 | H3f3b | 1.11E-25 | -0.085882971 | 0.269 | 0.268 | 2.97E-21 |
| 9630028H03Rik | 9.61E-51 | -0.302130478 | 0.619 | 0.694 | 2.59E-46 | Brinp2 | 1.46E-25 | -0.171164028 | 0.404 | 0.44 | 3.94E-21 |
| 9330159F19Rik | 1.10E-50 | -0.175111796 | 0.243 | 0.331 | 2.97E-46 | Apbb2 | 1.49E-25 | -0.102777252 | 0.636 | 0.639 | 4.01E-21 |
| Slc22a17 | 1.18E-50 | -0.180729181 | 0.492 | 0.586 | 3.17E-46 | Dcaf7 | 1.52E-25 | 0.031261795 | 0.303 | 0.258 | 4.10E-21 |
| Gm20696 | 3.46E-50 | 0.2078036 | 0.442 | 0.349 | 9.31E-46 | Kcnj3 | 1.72E-25 | 0.142744589 | 0.875 | 0.844 | 4.62E-21 |
| Sobp | 9.48E-50 | 0.192078192 | 0.88 | 0.855 | 2.55E-45 | Ksr1 | 3.15E-25 | -0.109081489 | 0.582 | 0.582 | 8.49E-21 |
| Map2 | 1.32E-49 | -0.176283544 | 0.915 | 0.941 | 3.56E-45 | Cul5 | 3.29E-25 | 0.090721185 | 0.445 | 0.379 | 8.85E-21 |
| Ncoa2 | 1.40E-49 | 0.18376039 | 0.711 | 0.682 | 3.76E-45 | Ankrd12 | 3.76E-25 | -0.122422986 | 0.696 | 0.713 | 1.01E-20 |
| Cpne6 | 1.81E-49 | -0.2059523 | 0.435 | 0.526 | 4.86E-45 | Ube2d2a | 3.94E-25 | 0.100332506 | 0.412 | 0.347 | 1.06E-20 |
| Nav2 | 1.87E-49 | 0.175937573 | 0.929 | 0.909 | 5.03E-45 | Hsp90ab1 | 4.66E-25 | 0.02923022 | 0.342 | 0.298 | 1.25E-20 |
| Klhl2 | 6.07E-49 | -0.169861956 | 0.648 | 0.734 | 1.63E-44 | Scai | 4.75E-25 | 0.126525097 | 0.682 | 0.617 | 1.28E-20 |
| Nudt5 | 8.13E-49 | -0.168921428 | 0.304 | 0.396 | 2.19E-44 | Gm48512 | 5.07E-25 | 0.12523109 | 0.465 | 0.397 | 1.36E-20 |
| 4921534H16Rik | 9.76E-49 | -0.189591497 | 0.508 | 0.604 | 2.62E-44 | Stx12 | 5.19E-25 | 0.083988735 | 0.27 | 0.215 | 1.40E-20 |
| Brinp1 | 1.38E-48 | -0.2471325 | 0.908 | 0.929 | 3.72E-44 | Tia1 | 6.24E-25 | 0.095328032 | 0.523 | 0.455 | 1.68E-20 |
| Rnf19a | 1.78E-48 | -0.19927627 | 0.326 | 0.418 | 4.78E-44 | Canx | 6.60E-25 | 0.071571497 | 0.33 | 0.274 | 1.78E-20 |
| Snrnp48 | 2.22E-48 | -0.160410027 | 0.289 | 0.378 | 5.97E-44 | Cpsf6 | 7.56E-25 | 0.117055818 | 0.587 | 0.519 | 2.03E-20 |
| Palm2 | 2.30E-48 | 0.075433514 | 0.584 | 0.495 | 6.19E-44 | Zc3h15 | 7.80E-25 | 0.088123831 | 0.364 | 0.303 | 2.10E-20 |
| Ralgapa2 | 2.47E-48 | 0.213282315 | 0.691 | 0.645 | 6.64E-44 | Gabbr2 | 8.68E-25 | -0.126952465 | 0.869 | 0.871 | 2.34E-20 |
| Epo | 3.51E-48 | 0.181183537 | 0.485 | 0.39 | 9.45E-44 | Wnk2 | 9.87E-25 | -0.103025085 | 0.578 | 0.585 | 2.66E-20 |
| Rnft1 | 5.28E-48 | -0.129124565 | 0.182 | 0.258 | 1.42E-43 | Yme1l1 | 1.03E-24 | 0.093149917 | 0.418 | 0.355 | 2.76E-20 |
| Hsp90ab1 | 8.59E-48 | -0.138591804 | 0.298 | 0.387 | 2.31E-43 | Syncrip | 1.04E-24 | 0.088020186 | 0.282 | 0.227 | 2.80E-20 |
| Zcchc11 | 9.50E-48 | 0.138798168 | 0.633 | 0.54 | 2.56E-43 | Kbtbd11 | 1.12E-24 | -0.000648233 | 0.306 | 0.272 | 3.02E-20 |
| Gm36975 | 9.56E-48 | -0.266794199 | 0.771 | 0.825 | 2.57E-43 | Exd2 | 1.27E-24 | 0.059631618 | 0.342 | 0.289 | 3.42E-20 |
| Mblac2 | 1.03E-47 | 0.185680541 | 0.353 | 0.304 | 2.76E-43 | Gnptg | 1.30E-24 | 0.042502041 | 0.271 | 0.225 | 3.48E-20 |
| Ankrd12 | 1.36E-47 | 0.163632404 | 0.713 | 0.698 | 3.65E-43 | Spop | 1.37E-24 | 0.093942954 | 0.55 | 0.484 | 3.68E-20 |
| Pcca | 1.39E-47 | 0.166991585 | 0.61 | 0.58 | 3.74E-43 | Vps13a | 1.41E-24 | -0.123936563 | 0.721 | 0.739 | 3.81E-20 |
| Rnpc3 | 1.61E-47 | -0.092106495 | 0.288 | 0.366 | 4.33E-43 | 9330159F19Rik | 1.42E-24 | 0.112194246 | 0.303 | 0.243 | 3.81E-20 |
| 9030624J02Rik | 2.52E-47 | 0.185760551 | 0.334 | 0.248 | 6.77E-43 | Ptprs | 1.66E-24 | -0.091509317 | 0.719 | 0.712 | 4.47E-20 |
| Dnajc3 | 3.24E-47 | -0.155755471 | 0.235 | 0.319 | 8.72E-43 | Paip2 | 2.04E-24 | 0.099399034 | 0.529 | 0.462 | 5.48E-20 |
| Epb41l4b | 4.94E-47 | 0.191961161 | 0.366 | 0.322 | 1.33E-42 | Ccni | 2.24E-24 | 0.072356019 | 0.259 | 0.208 | 6.03E-20 |
| Apc | 5.78E-47 | 0.168347134 | 0.805 | 0.786 | 1.56E-42 | Ccdc47 | 2.52E-24 | 0.076375189 | 0.263 | 0.211 | 6.79E-20 |
| Wipf3 | 1.36E-46 | -0.190881352 | 0.626 | 0.712 | 3.67E-42 | Pkp4 | 2.67E-24 | -0.104867712 | 0.698 | 0.701 | 7.19E-20 |
| Prr14l | 1.79E-46 | -0.16398193 | 0.284 | 0.372 | 4.83E-42 | Cluap1 | 2.75E-24 | 0.061194929 | 0.255 | 0.207 | 7.41E-20 |
| Slc4a3 | 1.92E-46 | 0.189566659 | 0.382 | 0.327 | 5.17E-42 | Slc24a4 | 2.76E-24 | -0.14443721 | 0.414 | 0.435 | 7.41E-20 |
| Dusp11 | 2.07E-46 | -0.141895191 | 0.208 | 0.288 | 5.56E-42 | Specc1 | 2.77E-24 | -0.115100394 | 0.723 | 0.727 | 7.44E-20 |
| Pla2g3 | 2.27E-46 | -0.19031975 | 0.201 | 0.279 | 6.10E-42 | Ndfip1 | 2.96E-24 | 0.103203874 | 0.642 | 0.576 | 7.97E-20 |
| Fchsd2 | 4.00E-46 | 0.19521866 | 0.426 | 0.372 | 1.08E-41 | Dopey1 | 3.18E-24 | -0.135130807 | 0.358 | 0.42 | 8.55E-20 |
| Map2k5 | 5.10E-46 | 0.143152128 | 0.491 | 0.477 | 1.37E-41 | Sgms1 | 3.25E-24 | -0.099056281 | 0.585 | 0.591 | 8.74E-20 |
| Gm12709 | 5.44E-46 | -0.163960783 | 0.195 | 0.274 | 1.46E-41 | Itm2b | 3.44E-24 | 0.012160859 | 0.488 | 0.441 | 9.25E-20 |
| Ddx50 | 6.27E-46 | -0.150364827 | 0.501 | 0.594 | 1.69E-41 | Rbm39 | 3.47E-24 | 0.107441903 | 0.916 | 0.888 | 9.33E-20 |
| 1500035N22Rik | 7.41E-46 | -0.202338985 | 0.291 | 0.378 | 1.99E-41 | Sec62 | 4.17E-24 | 0.093489164 | 0.297 | 0.24 | 1.12E-19 |

| | | | | | | | | | | | |
|---|---|---|---|---|---|---|---|---|---|---|---|
| Macf1 | 8.04E-46 | 0.161006205 | 0.91 | 0.898 | 2.16E-41 | Ifrd1 | 4.52E-24 | 0.099872754 | 0.261 | 0.205 | 1.22E-19 |
| Asic2 | 8.33E-46 | 0.259651885 | 0.907 | 0.876 | 2.24E-41 | Ddx5 | 4.82E-24 | 0.13906172 | 0.75 | 0.699 | 1.30E-19 |
| Wwc1 | 8.68E-46 | 0.197929827 | 0.435 | 0.385 | 2.34E-41 | Ogt | 4.98E-24 | 0.117461624 | 0.674 | 0.61 | 1.34E-19 |
| Hnrnph1 | 8.79E-46 | -0.173615358 | 0.405 | 0.495 | 2.37E-41 | Wwc1 | 5.28E-24 | -0.142902413 | 0.399 | 0.435 | 1.42E-19 |
| Pde4a | 1.40E-45 | -0.194209739 | 0.488 | 0.579 | 3.78E-41 | Clstn2 | 5.62E-24 | 0.141756097 | 0.924 | 0.902 | 1.51E-19 |
| Lrp1b | 1.43E-45 | -0.271998381 | 0.952 | 0.945 | 3.84E-41 | Chgb | 6.76E-24 | 0.082457387 | 0.356 | 0.302 | 1.82E-19 |
| Atp5o | 1.61E-45 | 0.123025562 | 0.667 | 0.577 | 4.33E-41 | Gatad2b | 7.31E-24 | 0.082459595 | 0.602 | 0.537 | 1.97E-19 |
| Clk1 | 5.44E-45 | -0.173927344 | 0.469 | 0.56 | 1.46E-40 | Ndufs1 | 8.48E-24 | 0.062335161 | 0.357 | 0.303 | 2.28E-19 |
| Npepps | 6.88E-45 | 0.165798701 | 0.582 | 0.557 | 1.85E-40 | Slc17a7 | 8.67E-24 | -0.094124472 | 0.7 | 0.702 | 2.33E-19 |
| Zfp207 | 9.74E-45 | -0.09716037 | 0.387 | 0.471 | 2.62E-40 | Pclo | 9.73E-24 | -0.116357259 | 0.878 | 0.888 | 2.62E-19 |
| Chchd6 | 1.59E-44 | 0.168033612 | 0.42 | 0.383 | 4.29E-40 | App | 9.78E-24 | -0.078139681 | 0.859 | 0.852 | 2.63E-19 |
| 4930570B17Rik | 2.43E-44 | -0.171897677 | 0.342 | 0.431 | 6.53E-40 | Srsf10 | 1.08E-23 | 0.033195245 | 0.348 | 0.302 | 2.92E-19 |
| Tmem57 | 2.47E-44 | 0.18901835 | 0.461 | 0.376 | 6.64E-40 | Luzp1 | 1.08E-23 | 0.13318069 | 0.526 | 0.459 | 2.92E-19 |
| Farp1 | 5.62E-44 | 0.241754459 | 0.416 | 0.343 | 1.51E-39 | Rab5b | 1.25E-23 | 0.068814424 | 0.263 | 0.214 | 3.36E-19 |
| Gnptg | 8.42E-44 | -0.13335536 | 0.225 | 0.304 | 2.27E-39 | Bicral | 1.41E-23 | 0.074464062 | 0.342 | 0.287 | 3.79E-19 |
| Canx | 8.84E-44 | -0.136034936 | 0.274 | 0.358 | 2.38E-39 | Camk2d | 1.42E-23 | -0.194853677 | 0.252 | 0.271 | 3.81E-19 |
| Ntng1 | 1.23E-43 | 0.299067985 | 0.563 | 0.514 | 3.32E-39 | Tmem243 | 1.48E-23 | 0.120699383 | 0.465 | 0.399 | 3.97E-19 |
| Scaper | 1.71E-43 | 0.181858612 | 0.648 | 0.602 | 4.60E-39 | Atp6v0d1 | 2.00E-23 | 0.091115667 | 0.296 | 0.239 | 5.38E-19 |
| Gm21992 | 1.78E-43 | 0.15317719 | 0.45 | 0.361 | 4.80E-39 | Atp2a2 | 2.06E-23 | 0.038421744 | 0.628 | 0.575 | 5.54E-19 |
| Slc17a7 | 2.02E-43 | -0.192212845 | 0.702 | 0.771 | 5.44E-39 | Ppp2r2b | 3.10E-23 | -0.163789702 | 0.726 | 0.748 | 8.35E-19 |
| 4833422C13Rik | 3.04E-43 | -0.187042316 | 0.217 | 0.296 | 8.18E-39 | Smarca5 | 3.29E-23 | 0.080039296 | 0.349 | 0.292 | 8.86E-19 |
| Mir124a-1hg | 3.32E-43 | -0.195327321 | 0.535 | 0.62 | 8.94E-39 | Hsp90b1 | 3.48E-23 | 0.033714522 | 0.282 | 0.24 | 9.36E-19 |
| Adam23 | 4.56E-43 | 0.17532211 | 0.567 | 0.527 | 1.23E-38 | Aig1 | 3.55E-23 | -0.079674563 | 0.634 | 0.624 | 9.54E-19 |
| Gm10848 | 8.27E-43 | -0.254820874 | 0.449 | 0.528 | 2.23E-38 | Klf9 | 3.87E-23 | 0.070822185 | 0.319 | 0.266 | 1.04E-18 |
| Ccnjl | 1.16E-42 | -0.266467421 | 0.198 | 0.267 | 3.12E-38 | Atp6v0b | 3.91E-23 | -0.023434875 | 0.652 | 0.624 | 1.05E-18 |
| Hacd4 | 1.17E-42 | -0.173577291 | 0.224 | 0.303 | 3.16E-38 | Cd2ap | 4.06E-23 | 0.081304166 | 0.411 | 0.351 | 1.09E-18 |
| Rtl4 | 1.39E-42 | 0.226349424 | 0.63 | 0.591 | 3.75E-38 | Mysm1 | 4.25E-23 | 0.093096535 | 0.507 | 0.443 | 1.14E-18 |
| Rpl6 | 1.61E-42 | -0.126199526 | 0.314 | 0.397 | 4.34E-38 | Zbtb20 | 4.26E-23 | 0.031419369 | 0.722 | 0.749 | 1.15E-18 |
| Gnao1 | 1.86E-42 | 0.159440344 | 0.714 | 0.702 | 5.02E-38 | C1qtnf4 | 5.02E-23 | 0.026504862 | 0.385 | 0.339 | 1.35E-18 |
| Qk | 3.48E-42 | -0.169719628 | 0.478 | 0.564 | 9.36E-38 | Gnao1 | 6.45E-23 | -0.11134321 | 0.705 | 0.714 | 1.73E-18 |
| Zfyve28 | 3.67E-42 | 0.181785101 | 0.486 | 0.446 | 9.88E-38 | Vdac1 | 6.82E-23 | 0.062682939 | 0.287 | 0.236 | 1.84E-18 |
| Pcsk1n | 5.09E-42 | -0.163084127 | 0.214 | 0.286 | 1.37E-37 | Gm15563 | 6.96E-23 | 0.106007215 | 0.302 | 0.244 | 1.87E-18 |
| Spink10 | 7.19E-42 | -0.179366882 | 0.242 | 0.321 | 1.94E-37 | Lrrfip1 | 7.76E-23 | -0.099491508 | 0.695 | 0.698 | 2.09E-18 |
| 2900011O08Rik | 7.57E-42 | 0.161881125 | 0.544 | 0.455 | 2.04E-37 | 9530059O14Rik | 9.14E-23 | -0.136005166 | 0.743 | 0.755 | 2.46E-18 |
| Kdm7a | 8.08E-42 | -0.210729866 | 0.413 | 0.497 | 2.17E-37 | Rad23b | 1.00E-22 | 0.004354345 | 0.346 | 0.311 | 2.70E-18 |
| Cpne4 | 8.08E-42 | 0.226737289 | 0.517 | 0.451 | 2.17E-37 | Cmip | 1.06E-22 | -0.104426369 | 0.713 | 0.714 | 2.86E-18 |
| Rap1gap2 | 1.27E-41 | 0.208033346 | 0.521 | 0.46 | 3.42E-37 | Ncoa2 | 1.07E-22 | -0.098848317 | 0.707 | 0.711 | 2.89E-18 |
| 2900055J20Rik | 1.83E-41 | 0.192709428 | 0.413 | 0.329 | 4.92E-37 | Vapa | 1.11E-22 | 0.037144238 | 0.321 | 0.276 | 2.99E-18 |
| Acaca | 2.14E-41 | 0.175453328 | 0.525 | 0.485 | 5.76E-37 | Nav1 | 1.17E-22 | -0.103510999 | 0.619 | 0.626 | 3.16E-18 |
| Hmgcll1 | 2.82E-41 | 0.20348292 | 0.366 | 0.299 | 7.59E-37 | Psmd7 | 1.18E-22 | 0.056258259 | 0.32 | 0.27 | 3.19E-18 |
| Plec | 4.63E-41 | 0.159667178 | 0.312 | 0.27 | 1.25E-36 | Sumo1 | 1.33E-22 | 0.098881134 | 0.421 | 0.358 | 3.57E-18 |
| Gm15414 | 5.13E-41 | -0.199058124 | 0.228 | 0.303 | 1.38E-36 | Gsg1l | 1.37E-22 | -0.139050995 | 0.271 | 0.282 | 3.69E-18 |
| Tef | 7.06E-41 | 0.175108544 | 0.413 | 0.368 | 1.90E-36 | Csnk2a1 | 1.42E-22 | 0.061619945 | 0.31 | 0.259 | 3.81E-18 |
| Slc25a42 | 9.02E-41 | -0.152026941 | 0.189 | 0.262 | 2.43E-36 | Dnajb6 | 1.45E-22 | 0.067024495 | 0.365 | 0.309 | 3.91E-18 |
| Cacna1b | 1.05E-40 | 0.138039083 | 0.666 | 0.657 | 2.82E-36 | Ccnt2 | 1.50E-22 | 0.058197253 | 0.566 | 0.507 | 4.04E-18 |
| Sdk2 | 1.16E-40 | -0.118669987 | 0.298 | 0.377 | 3.13E-36 | 5031425E22Rik | 1.65E-22 | 0.088320153 | 0.479 | 0.417 | 4.43E-18 |
| P2ry14 | 1.43E-40 | -0.197637531 | 0.639 | 0.714 | 3.86E-36 | Myo9a | 1.67E-22 | -0.099353676 | 0.757 | 0.762 | 4.49E-18 |
| Igf2bp3 | 1.47E-40 | -0.173449727 | 0.405 | 0.491 | 3.96E-36 | Ccnl2 | 1.70E-22 | 0.065525213 | 0.51 | 0.45 | 4.58E-18 |
| Cmip | 1.47E-40 | 0.138407175 | 0.714 | 0.714 | 3.97E-36 | D830024N08Rik | 1.72E-22 | 0.113619537 | 0.404 | 0.341 | 4.63E-18 |
| Gm26904 | 1.59E-40 | 0.09411617 | 0.626 | 0.544 | 4.28E-36 | Rprd2 | 2.10E-22 | 0.07772132 | 0.431 | 0.371 | 5.64E-18 |
| Kcnq5 | 2.22E-40 | 0.297759939 | 0.623 | 0.555 | 5.97E-36 | Fgf12 | 2.10E-22 | -0.115950765 | 0.922 | 0.932 | 5.64E-18 |
| 4921511C10Rik | 2.61E-40 | -0.197973014 | 0.355 | 0.44 | 7.03E-36 | Galnt18 | 2.10E-22 | -0.135302598 | 0.797 | 0.817 | 5.65E-18 |
| Hcfc2 | 2.93E-40 | -0.142256302 | 0.179 | 0.25 | 7.88E-36 | Usp50 | 2.14E-22 | 0.099913173 | 0.431 | 0.368 | 5.75E-18 |
| Srsf5 | 3.15E-40 | -0.157906715 | 0.395 | 0.477 | 8.46E-36 | Thy1 | 2.19E-22 | 0.01445405 | 0.308 | 0.272 | 5.90E-18 |
| Hpca | 3.95E-40 | -0.184141344 | 0.202 | 0.272 | 1.06E-35 | Psma3 | 2.20E-22 | 0.092073205 | 0.427 | 0.366 | 5.93E-18 |
| Arhgap44 | 4.37E-40 | 0.14346829 | 0.564 | 0.545 | 1.18E-35 | Pafah1b1 | 2.71E-22 | 0.119419577 | 0.743 | 0.689 | 7.30E-18 |
| Nvl | 4.52E-40 | -0.130304925 | 0.613 | 0.696 | 1.22E-35 | Nin | 2.89E-22 | -0.104657613 | 0.591 | 0.602 | 7.77E-18 |
| Ssbp4 | 7.14E-40 | -0.149203661 | 0.328 | 0.412 | 1.92E-35 | Epb41l4b | 3.06E-22 | -0.128723381 | 0.34 | 0.366 | 8.23E-18 |
| Slc25a3 | 9.24E-40 | -0.184985018 | 0.278 | 0.351 | 2.49E-35 | Rap2a | 3.24E-22 | 0.069742399 | 0.27 | 0.221 | 8.73E-18 |
| Nr6a1 | 1.06E-39 | 0.162844639 | 0.57 | 0.541 | 2.86E-35 | Agrn | 3.26E-22 | -0.065850985 | 0.345 | 0.34 | 8.78E-18 |
| Tsga10 | 1.10E-39 | -0.155218095 | 0.32 | 0.403 | 2.95E-35 | Kidins220 | 3.37E-22 | -0.072860993 | 0.652 | 0.644 | 9.08E-18 |
| Srpr | 1.31E-39 | -0.133752201 | 0.269 | 0.347 | 3.54E-35 | Ggt7 | 3.39E-22 | -0.035238327 | 0.388 | 0.366 | 9.12E-18 |
| Ccnl2 | 1.47E-39 | -0.137415846 | 0.45 | 0.537 | 3.96E-35 | Kcnma1 | 3.40E-22 | -0.108107767 | 0.977 | 0.983 | 9.14E-18 |
| Nebl | 1.52E-39 | -0.181767483 | 0.876 | 0.904 | 4.08E-35 | Rtn1 | 3.60E-22 | -0.077897921 | 0.892 | 0.885 | 9.69E-18 |
| Ccdc148 | 1.75E-39 | 0.176362623 | 0.475 | 0.427 | 4.70E-35 | Cdc42 | 3.64E-22 | 0.075722782 | 0.459 | 0.399 | 9.80E-18 |
| Zfp933 | 2.11E-39 | -0.093267411 | 0.241 | 0.311 | 5.69E-35 | Plxna4os3 | 3.94E-22 | 0.092976697 | 0.36 | 0.303 | 1.06E-17 |
| Sh3bp5 | 2.34E-39 | -0.157604011 | 0.493 | 0.579 | 6.30E-35 | Cadps | 4.16E-22 | -0.130029838 | 0.944 | 0.949 | 1.12E-17 |
| Pwwp2a | 3.00E-39 | -0.152697764 | 0.324 | 0.407 | 8.08E-35 | Zfand5 | 4.18E-22 | 0.005693484 | 0.33 | 0.295 | 1.13E-17 |
| Babam2 | 3.72E-39 | 0.183024339 | 0.561 | 0.513 | 1.00E-34 | Sec61a2 | 4.27E-22 | 0.037206242 | 0.486 | 0.433 | 1.15E-17 |
| Csnk1g1 | 4.03E-39 | 0.150313022 | 0.647 | 0.627 | 1.08E-34 | Flnb | 4.31E-22 | -0.111464797 | 0.559 | 0.578 | 1.16E-17 |
| Hnrnpa3 | 4.84E-39 | -0.131023697 | 0.248 | 0.325 | 1.30E-34 | Vmp1 | 4.38E-22 | 0.105796341 | 0.505 | 0.441 | 1.18E-17 |
| Psip1 | 4.89E-39 | -0.153253241 | 0.506 | 0.592 | 1.32E-34 | Cd47 | 4.56E-22 | 0.072329722 | 0.429 | 0.372 | 1.23E-17 |
| Mysm1 | 4.93E-39 | -0.15178207 | 0.443 | 0.529 | 1.33E-34 | Plxna2 | 4.88E-22 | -0.075147398 | 0.616 | 0.603 | 1.31E-17 |
| Soga3 | 5.17E-39 | -0.159437434 | 0.638 | 0.713 | 1.39E-34 | Ablim2 | 5.22E-22 | -0.121056766 | 0.659 | 0.67 | 1.41E-17 |
| Oxct1 | 1.02E-38 | 0.146675721 | 0.625 | 0.608 | 2.75E-34 | Dpp10 | 5.43E-22 | 0.484990865 | 0.383 | 0.423 | 1.46E-17 |
| Rasgef1b | 1.50E-38 | 0.26369802 | 0.453 | 0.387 | 4.04E-34 | Pip5k1a | 6.09E-22 | 0.112552722 | 0.586 | 0.522 | 1.64E-17 |
| Nxpe4 | 1.50E-38 | 0.188205047 | 0.356 | 0.305 | 4.04E-34 | Cox7c | 6.27E-22 | -0.00112941 | 0.265 | 0.239 | 1.69E-17 |
| Sh3pxd2a | 1.84E-38 | 0.157891569 | 0.297 | 0.26 | 4.95E-34 | Synpo | 7.33E-22 | 0.100484552 | 0.382 | 0.322 | 1.97E-17 |
| Myo9a | 1.92E-38 | 0.147811322 | 0.762 | 0.742 | 5.16E-34 | Gm15478 | 7.95E-22 | 0.108029507 | 0.384 | 0.323 | 2.14E-17 |
| Bcas3 | 2.05E-38 | 0.157809825 | 0.665 | 0.634 | 5.52E-34 | Sh3rf3 | 9.37E-22 | -0.12869037 | 0.636 | 0.66 | 2.52E-17 |
| Cox7c | 2.23E-38 | -0.110202832 | 0.239 | 0.311 | 5.99E-34 | Tlk1 | 1.19E-21 | 0.061137395 | 0.418 | 0.365 | 3.20E-17 |
| Cnot3 | 2.37E-38 | -0.131250847 | 0.182 | 0.252 | 6.37E-34 | Ctdspl2 | 1.19E-21 | 0.067241675 | 0.4 | 0.345 | 3.20E-17 |
| Gm44511 | 3.16E-38 | -0.167179043 | 0.436 | 0.473 | 8.51E-34 | Slc25a25 | 1.36E-21 | 0.081927592 | 0.261 | 0.21 | 3.65E-17 |
| Zmiz1 | 3.69E-38 | -0.200261056 | 0.308 | 0.388 | 9.93E-34 | Peg3 | 1.45E-21 | -0.116954833 | 0.535 | 0.542 | 3.91E-17 |
| Sfxn5 | 4.24E-38 | 0.186456764 | 0.309 | 0.256 | 1.14E-33 | Arih2 | 1.50E-21 | 0.051673868 | 0.332 | 0.284 | 4.04E-17 |
| St6galnac5 | 5.03E-38 | 0.161728516 | 0.884 | 0.87 | 1.35E-33 | Ptp4a2 | 1.57E-21 | 0.08838674 | 0.304 | 0.249 | 4.21E-17 |
| Chd2 | 6.41E-38 | -0.168623643 | 0.492 | 0.575 | 1.72E-33 | Gm26691 | 1.62E-21 | -0.164595886 | 0.281 | 0.323 | 4.36E-17 |
| Dock3 | 6.87E-38 | 0.168248772 | 0.887 | 0.865 | 1.85E-33 | Srpr | 1.66E-21 | 0.056588542 | 0.317 | 0.269 | 4.47E-17 |
| Csgalnact1 | 8.12E-38 | 0.216815238 | 0.345 | 0.291 | 2.19E-33 | Rmdn1 | 1.76E-21 | 0.091304487 | 0.586 | 0.524 | 4.73E-17 |
| Lpp | 8.98E-38 | -0.188177697 | 0.36 | 0.443 | 2.42E-33 | Tspan3 | 1.82E-21 | 0.0343628 | 0.27 | 0.229 | 4.91E-17 |
| Tia1 | 1.17E-37 | -0.112823517 | 0.455 | 0.538 | 3.14E-33 | Rnf138 | 1.85E-21 | 0.100046004 | 0.279 | 0.225 | 4.99E-17 |
| Fam133b | 1.29E-37 | -0.103253462 | 0.257 | 0.328 | 3.48E-33 | Trpm7 | 1.94E-21 | 0.103151029 | 0.693 | 0.634 | 5.21E-17 |
| Kcnd3 | 2.26E-37 | 0.183503078 | 0.732 | 0.719 | 6.09E-33 | Rtn4 | 2.25E-21 | 0.073939465 | 0.621 | 0.561 | 6.06E-17 |
| Caskin1 | 2.27E-37 | 0.143743379 | 0.298 | 0.263 | 6.12E-33 | Rpl6 | 2.41E-21 | -0.022653892 | 0.339 | 0.314 | 6.49E-17 |
| Ralyl | 2.31E-37 | 0.170955506 | 0.939 | 0.921 | 6.22E-33 | Galnt16 | 2.44E-21 | -0.091586187 | 0.535 | 0.536 | 6.56E-17 |
| Ccdc66 | 2.81E-37 | -0.118170555 | 0.27 | 0.345 | 7.56E-33 | Xrn2 | 2.60E-21 | 0.031654758 | 0.385 | 0.339 | 7.00E-17 |
| Mdga2 | 3.83E-37 | 0.120523992 | 0.978 | 0.968 | 1.03E-32 | Arpc1a | 2.66E-21 | 0.048238628 | 0.357 | 0.308 | 7.15E-17 |
| Psmc6 | 4.57E-37 | -0.132509773 | 0.358 | 0.44 | 1.23E-32 | Tial1 | 2.88E-21 | 0.074708719 | 0.438 | 0.379 | 7.75E-17 |

| | | | | | | | | | | | |
|---|---|---|---|---|---|---|---|---|---|---|---|
| Nova2 | 4.80E-37 | 0.135692474 | 0.383 | 0.362 | 1.29E-32 | Sumo2 | 2.98E-21 | 0.072918006 | 0.351 | 0.296 | 8.01E-17 |
| Gm3448 | 5.42E-37 | -0.142359819 | 0.233 | 0.307 | 1.46E-32 | Cpeb1 | 3.02E-21 | -0.084502571 | 0.459 | 0.462 | 8.12E-17 |
| Ubash3b | 7.65E-37 | -0.21397165 | 0.231 | 0.302 | 2.06E-32 | Cdk12 | 3.02E-21 | 0.087073613 | 0.649 | 0.588 | 8.13E-17 |
| 1600010M07Rik | 1.02E-36 | -0.155264061 | 0.375 | 0.457 | 2.74E-32 | Traf3ip2 | 3.79E-21 | 0.112228832 | 0.32 | 0.263 | 1.02E-16 |
| Orc4 | 1.53E-36 | -0.110013674 | 0.364 | 0.442 | 4.13E-32 | Wwp1 | 3.94E-21 | 0.052194457 | 0.363 | 0.313 | 1.06E-16 |
| Phactr3 | 1.88E-36 | 0.096230337 | 0.525 | 0.537 | 5.06E-32 | Cstf3 | 4.09E-21 | 0.085651265 | 0.555 | 0.494 | 1.10E-16 |
| 6530409C15Rik | 2.23E-36 | -0.143708726 | 0.272 | 0.349 | 6.01E-32 | Matk | 4.12E-21 | 0.043961522 | 0.602 | 0.549 | 1.11E-16 |
| Odf2 | 3.71E-36 | 0.125343152 | 0.492 | 0.483 | 9.99E-32 | Camkv | 4.39E-21 | 0.075171445 | 0.58 | 0.522 | 1.18E-16 |
| Cpeb1 | 3.87E-36 | 0.13718434 | 0.462 | 0.441 | 1.04E-31 | Enox1 | 4.79E-21 | -0.119963342 | 0.875 | 0.889 | 1.29E-16 |
| Crim1 | 3.97E-36 | 0.181130502 | 0.467 | 0.428 | 1.07E-31 | Homer1 | 5.04E-21 | 0.248542423 | 0.545 | 0.495 | 1.36E-16 |
| Atp9a | 4.59E-36 | 0.137047387 | 0.521 | 0.504 | 1.23E-31 | Ina | 5.40E-21 | 0.122043536 | 0.254 | 0.202 | 1.45E-16 |
| Sycp2 | 5.08E-36 | -0.15998757 | 0.238 | 0.311 | 1.37E-31 | Dym | 5.43E-21 | -0.0850997 | 0.54 | 0.545 | 1.46E-16 |
| Galnt1 | 6.98E-36 | -0.088956653 | 0.501 | 0.58 | 1.88E-31 | Araf | 5.48E-21 | 0.090572109 | 0.447 | 0.387 | 1.47E-16 |
| Spred1 | 7.85E-36 | -0.162846436 | 0.333 | 0.412 | 2.11E-31 | Brd4 | 5.59E-21 | 0.097568152 | 0.569 | 0.507 | 1.50E-16 |
| A930015D03Rik | 8.27E-36 | -0.128259895 | 0.558 | 0.638 | 2.23E-31 | Hspa9 | 6.10E-21 | 0.049633458 | 0.283 | 0.237 | 1.64E-16 |
| Slit3 | 8.35E-36 | 0.179663762 | 0.932 | 0.93 | 2.25E-31 | Hivep3 | 6.61E-21 | -0.102329873 | 0.826 | 0.827 | 1.78E-16 |
| Taok3 | 1.47E-35 | 0.098062047 | 0.432 | 0.431 | 3.95E-31 | Tbce | 6.82E-21 | 0.062522261 | 0.456 | 0.399 | 1.83E-16 |
| Wwp1 | 1.47E-35 | -0.1210506 | 0.313 | 0.389 | 3.96E-31 | Rtn4r | 7.04E-21 | 0.049995221 | 0.255 | 0.212 | 1.89E-16 |
| Smarcad1 | 1.56E-35 | -0.067330933 | 0.359 | 0.428 | 4.19E-31 | Rnf165 | 7.25E-21 | -0.080724682 | 0.486 | 0.488 | 1.95E-16 |
| Krit1 | 1.58E-35 | -0.132917158 | 0.472 | 0.554 | 4.26E-31 | Zfp397 | 7.74E-21 | 0.013292566 | 0.272 | 0.239 | 2.08E-16 |
| Mat2a | 1.90E-35 | -0.124172282 | 0.339 | 0.418 | 5.11E-31 | Ptpn11 | 8.29E-21 | 0.045517999 | 0.332 | 0.285 | 2.23E-16 |
| Gm26749 | 2.05E-35 | -0.108069857 | 0.396 | 0.473 | 5.51E-31 | 2610020C07Rik | 8.63E-21 | 0.091117951 | 0.446 | 0.386 | 2.32E-16 |
| Zfp397 | 2.25E-35 | -0.090821699 | 0.239 | 0.304 | 6.04E-31 | Immp1l | 9.14E-21 | 0.069409415 | 0.537 | 0.478 | 2.46E-16 |
| Thrap3 | 2.33E-35 | -0.062189442 | 0.286 | 0.348 | 6.27E-31 | Btaf1 | 1.00E-20 | 0.048912793 | 0.515 | 0.462 | 2.70E-16 |
| Cntln | 3.03E-35 | 0.139864347 | 0.376 | 0.351 | 8.16E-31 | Lcorl | 1.09E-20 | -0.141832474 | 0.402 | 0.434 | 2.94E-16 |
| Gm28198 | 3.18E-35 | -0.108373071 | 0.24 | 0.311 | 8.54E-31 | Ptpn9 | 1.18E-20 | 0.105246582 | 0.418 | 0.357 | 3.18E-16 |
| Ubl3 | 3.20E-35 | -0.088496912 | 0.258 | 0.325 | 8.60E-31 | Stk39 | 1.36E-20 | -0.08484186 | 0.568 | 0.568 | 3.67E-16 |
| Atp8a2 | 3.53E-35 | 0.166829604 | 0.615 | 0.586 | 9.50E-31 | Hipk3 | 1.45E-20 | 0.075898088 | 0.293 | 0.242 | 3.91E-16 |
| Preb | 3.65E-35 | -0.116933789 | 0.281 | 0.354 | 9.82E-31 | Pcf11 | 1.45E-20 | 0.041498545 | 0.266 | 0.224 | 3.91E-16 |
| 4930545L23Rik | 6.21E-35 | -0.202405108 | 0.282 | 0.356 | 1.67E-30 | Zfp280c | 1.48E-20 | 0.037217308 | 0.386 | 0.341 | 3.98E-16 |
| Kif1b | 6.33E-35 | 0.133221538 | 0.812 | 0.799 | 1.70E-30 | Scg5 | 1.50E-20 | -0.005270925 | 0.546 | 0.51 | 4.03E-16 |
| Shank3 | 6.86E-35 | 0.142856984 | 0.381 | 0.345 | 1.84E-30 | Mcph1 | 1.50E-20 | -0.102905721 | 0.363 | 0.383 | 4.04E-16 |
| Rapgef4 | 7.57E-35 | 0.153489698 | 0.826 | 0.795 | 2.04E-30 | Stx5a | 1.52E-20 | 0.069182562 | 0.301 | 0.251 | 4.08E-16 |
| Chd5 | 8.22E-35 | 0.175493216 | 0.397 | 0.344 | 2.21E-30 | Far1 | 1.62E-20 | 0.03543758 | 0.499 | 0.449 | 4.35E-16 |
| Brd9 | 8.31E-35 | -0.220007578 | 0.518 | 0.586 | 2.24E-30 | Spock1 | 1.73E-20 | -0.22118856 | 0.386 | 0.426 | 4.65E-16 |
| Apbb2 | 8.63E-35 | 0.133730589 | 0.639 | 0.63 | 2.32E-30 | Ndfip2 | 1.84E-20 | 0.092450913 | 0.365 | 0.309 | 4.95E-16 |
| Gm14330 | 8.79E-35 | -0.12794919 | 0.259 | 0.331 | 2.37E-30 | Ak5 | 1.86E-20 | -0.12717826 | 0.804 | 0.817 | 5.01E-16 |
| Disp2 | 9.10E-35 | -0.129469044 | 0.343 | 0.421 | 2.45E-30 | Ppib | 1.99E-20 | -0.03141292 | 0.332 | 0.316 | 5.35E-16 |
| Garem1 | 9.61E-35 | -0.195798609 | 0.547 | 0.622 | 2.59E-30 | Rapgef6 | 2.00E-20 | 0.095338757 | 0.723 | 0.667 | 5.37E-16 |
| Sh3gl1 | 9.70E-35 | -0.10256661 | 0.318 | 0.391 | 2.61E-30 | Ctif | 2.13E-20 | -0.097647888 | 0.509 | 0.518 | 5.73E-16 |
| Snx3 | 1.00E-34 | -0.09120012 | 0.266 | 0.333 | 2.70E-30 | Pigk | 2.14E-20 | 0.127610648 | 0.904 | 0.884 | 5.75E-16 |
| Aak1 | 1.02E-34 | 0.133781674 | 0.841 | 0.831 | 2.75E-30 | Pwwp2a | 2.17E-20 | 0.039684471 | 0.371 | 0.324 | 5.85E-16 |
| Dnmt3a | 1.28E-34 | 0.135916279 | 0.676 | 0.666 | 3.43E-30 | Usp45 | 2.53E-20 | 0.038695978 | 0.443 | 0.397 | 6.82E-16 |
| Dopey1 | 1.28E-34 | 0.157015289 | 0.42 | 0.342 | 3.45E-30 | Snph | 2.58E-20 | -0.087605745 | 0.606 | 0.611 | 6.94E-16 |
| Gm48747 | 1.43E-34 | -0.164729212 | 0.569 | 0.647 | 3.84E-30 | Ddx24 | 2.70E-20 | -0.008170153 | 0.5 | 0.467 | 7.27E-16 |
| Slc6a6 | 1.61E-34 | 0.155752846 | 0.565 | 0.535 | 4.32E-30 | Sdcbp | 2.73E-20 | 0.045835383 | 0.295 | 0.252 | 7.33E-16 |
| Il1rapl1 | 1.74E-34 | 0.197442629 | 0.955 | 0.947 | 4.68E-30 | Lcor | 2.99E-20 | 0.09707336 | 0.62 | 0.559 | 8.03E-16 |
| Slc4a7 | 1.80E-34 | -0.155947189 | 0.42 | 0.501 | 4.84E-30 | Metap1d | 3.00E-20 | 0.054186111 | 0.282 | 0.236 | 8.07E-16 |
| Cntnap2 | 1.86E-34 | 0.162895095 | 0.852 | 0.793 | 5.00E-30 | Mmp17 | 3.16E-20 | 0.021315745 | 0.297 | 0.26 | 8.52E-16 |
| Phf24 | 2.21E-34 | 0.128297963 | 0.605 | 0.594 | 5.96E-30 | Nol4l | 3.60E-20 | -0.10743741 | 0.467 | 0.48 | 9.70E-16 |
| Ttc14 | 2.76E-34 | -0.129272546 | 0.735 | 0.8 | 7.43E-30 | Glt8d1 | 3.71E-20 | 0.043150698 | 0.378 | 0.331 | 9.99E-16 |
| Kcnq1 | 2.93E-34 | -0.025928172 | 0.663 | 0.616 | 7.89E-30 | Gm20275 | 3.74E-20 | -0.106573348 | 0.726 | 0.726 | 1.01E-15 |
| Far1os | 3.03E-34 | -0.153740395 | 0.441 | 0.521 | 8.15E-30 | Shisa4 | 3.82E-20 | 0.036575422 | 0.251 | 0.212 | 1.03E-15 |
| 4732471J01Rik | 3.20E-34 | 0.175891627 | 0.482 | 0.44 | 8.60E-30 | Psmd11 | 3.83E-20 | 0.060478021 | 0.361 | 0.311 | 1.03E-15 |
| Klf12 | 3.51E-34 | 0.182398541 | 0.608 | 0.557 | 9.44E-30 | Eef1a1 | 4.10E-20 | 0.03538125 | 0.276 | 0.237 | 1.10E-15 |
| 4930415C11Rik | 5.99E-34 | -0.243996393 | 0.272 | 0.332 | 1.61E-29 | Ubl3 | 4.21E-20 | 0.071268273 | 0.309 | 0.258 | 1.13E-15 |
| Gm20754 | 6.49E-34 | 0.207116305 | 0.759 | 0.752 | 1.75E-29 | Txlng | 4.52E-20 | 0.086578501 | 0.397 | 0.34 | 1.22E-15 |
| Cnksr2 | 7.35E-34 | -0.156957617 | 0.924 | 0.938 | 1.98E-29 | Snap91 | 4.83E-20 | -0.086034611 | 0.724 | 0.722 | 1.30E-15 |
| Brinp3 | 9.03E-34 | 0.319563906 | 0.444 | 0.385 | 2.43E-29 | Nrf1 | 5.14E-20 | 0.082693531 | 0.396 | 0.34 | 1.38E-15 |
| Myt1l | 1.08E-33 | -0.127401668 | 0.953 | 0.957 | 2.92E-29 | Ttc7b | 5.52E-20 | -0.083493153 | 0.644 | 0.64 | 1.48E-15 |
| Gm15594 | 1.35E-33 | -0.152896385 | 0.381 | 0.459 | 3.64E-29 | Prkca | 5.80E-20 | -0.140966908 | 0.766 | 0.768 | 1.56E-15 |
| Pdlim7 | 1.39E-33 | -0.145807951 | 0.268 | 0.34 | 3.75E-29 | Atad1 | 6.13E-20 | 0.058603678 | 0.33 | 0.281 | 1.65E-15 |
| Pcbp2 | 1.41E-33 | -0.105085521 | 0.364 | 0.439 | 3.79E-29 | Chchd6 | 6.26E-20 | -0.101375951 | 0.403 | 0.42 | 1.69E-15 |
| Car10 | 1.66E-33 | 0.278081063 | 0.612 | 0.566 | 4.47E-29 | Zbtb16 | 6.93E-20 | 0.091750173 | 0.743 | 0.688 | 1.87E-15 |
| Nav3 | 1.86E-33 | 0.132651636 | 0.984 | 0.978 | 5.00E-29 | Gucy1a2 | 7.06E-20 | -0.091308432 | 0.754 | 0.756 | 1.90E-15 |
| Lmbr1 | 2.14E-33 | 0.142409533 | 0.375 | 0.344 | 5.76E-29 | Kifc2 | 7.45E-20 | 0.028032054 | 0.358 | 0.313 | 2.01E-15 |
| Exoc6b | 2.42E-33 | -0.199742205 | 0.8 | 0.846 | 6.51E-29 | Fez1 | 7.60E-20 | 0.049672593 | 0.32 | 0.274 | 2.04E-15 |
| Nos1ap | 2.93E-33 | -0.177303386 | 0.731 | 0.792 | 7.89E-29 | Tmem126a | 7.67E-20 | 0.088586999 | 0.307 | 0.253 | 2.06E-15 |
| Ywhag | 2.99E-33 | -0.158258835 | 0.409 | 0.487 | 8.04E-29 | Rnpc3 | 7.93E-20 | 0.010944591 | 0.322 | 0.288 | 2.13E-15 |
| St7 | 3.18E-33 | -0.138572724 | 0.659 | 0.731 | 8.56E-29 | Dlgap2 | 8.36E-20 | -0.108146648 | 0.97 | 0.98 | 2.25E-15 |
| Bhlhb9 | 3.67E-33 | -0.079597478 | 0.221 | 0.282 | 9.87E-29 | Slc39a10 | 8.42E-20 | 0.044040336 | 0.283 | 0.24 | 2.26E-15 |
| Aig1 | 3.70E-33 | 0.140675614 | 0.624 | 0.608 | 9.96E-29 | Igsf8 | 8.90E-20 | 0.041318527 | 0.283 | 0.241 | 2.40E-15 |
| Arhgef25 | 3.84E-33 | -0.148873335 | 0.446 | 0.525 | 1.03E-28 | Khdrbs1 | 9.03E-20 | 0.065115154 | 0.388 | 0.336 | 2.43E-15 |
| Arpc2 | 3.85E-33 | -0.162449483 | 0.427 | 0.505 | 1.04E-28 | Purg | 1.00E-19 | -0.058939813 | 0.378 | 0.372 | 2.69E-15 |
| Xrn2 | 4.40E-33 | -0.074835079 | 0.339 | 0.406 | 1.18E-28 | Gm26917 | 1.08E-19 | -0.112795805 | 0.425 | 0.443 | 2.92E-15 |
| Gm42303 | 4.98E-33 | -0.179216148 | 0.333 | 0.409 | 1.34E-28 | Hnrnpl | 1.09E-19 | -0.071132074 | 0.469 | 0.463 | 2.93E-15 |
| Gcn1l1 | 5.01E-33 | 0.154480108 | 0.269 | 0.204 | 1.35E-28 | Phf3 | 1.31E-19 | 0.039646556 | 0.639 | 0.587 | 3.51E-15 |
| Ephb2 | 5.95E-33 | 0.152977357 | 0.371 | 0.342 | 1.60E-28 | Kmt2a | 1.35E-19 | -0.087143662 | 0.723 | 0.721 | 3.64E-15 |
| Emc3 | 1.08E-32 | -0.080734571 | 0.193 | 0.251 | 2.91E-28 | Kcnab2 | 1.38E-19 | -0.097168193 | 0.408 | 0.421 | 3.71E-15 |
| Acp1 | 1.10E-32 | -0.15993609 | 0.31 | 0.384 | 2.95E-28 | Tpr | 1.43E-19 | 0.071927765 | 0.486 | 0.43 | 3.85E-15 |
| Ago3 | 1.18E-32 | -0.149162735 | 0.563 | 0.639 | 3.17E-28 | Ip6k2 | 1.49E-19 | 0.04748494 | 0.312 | 0.268 | 4.02E-15 |
| Wdr17 | 1.29E-32 | 0.182324468 | 0.522 | 0.501 | 3.47E-28 | Ep300 | 1.56E-19 | 0.072621214 | 0.427 | 0.371 | 4.19E-15 |
| Fam189a1 | 1.66E-32 | 0.166787748 | 0.798 | 0.765 | 4.47E-28 | Vti1a | 1.65E-19 | -0.095269659 | 0.739 | 0.747 | 4.45E-15 |
| Plekha6 | 1.76E-32 | 0.160801509 | 0.434 | 0.395 | 4.73E-28 | Atl3 | 2.00E-19 | 0.072222297 | 0.288 | 0.238 | 5.39E-15 |
| Timp3 | 1.95E-32 | -0.099248572 | 0.203 | 0.263 | 5.23E-28 | Gm20319 | 2.13E-19 | 0.127550675 | 0.334 | 0.278 | 5.74E-15 |
| Atp2a2 | 2.23E-32 | -0.137091421 | 0.575 | 0.65 | 5.99E-28 | Ryk | 2.25E-19 | -0.007522278 | 0.364 | 0.335 | 6.05E-15 |
| Scn2a | 2.31E-32 | 0.141716674 | 0.835 | 0.812 | 6.23E-28 | Arpc2 | 2.28E-19 | 0.10281985 | 0.487 | 0.427 | 6.14E-15 |
| Cul9 | 2.39E-32 | 0.141630545 | 0.355 | 0.322 | 6.43E-28 | Clec16a | 2.58E-19 | -0.054639232 | 0.506 | 0.494 | 6.95E-15 |
| St3gal3 | 2.40E-32 | 0.128257137 | 0.483 | 0.463 | 6.46E-28 | Snx25 | 2.73E-19 | -0.064581895 | 0.274 | 0.276 | 7.36E-15 |
| Clk2 | 2.42E-32 | -0.113219639 | 0.211 | 0.276 | 6.51E-28 | AY036118 | 2.89E-19 | 0.175905709 | 0.655 | 0.618 | 7.78E-15 |
| Timmdc1 | 2.61E-32 | -0.110053741 | 0.198 | 0.262 | 7.03E-28 | Adgrl1 | 3.02E-19 | -0.026335665 | 0.525 | 0.499 | 8.13E-15 |
| Ptprr | 3.04E-32 | 0.156470593 | 0.478 | 0.442 | 8.19E-28 | Timm23 | 3.08E-19 | 0.018487346 | 0.263 | 0.229 | 8.28E-15 |
| Cap1 | 4.05E-32 | -0.119663665 | 0.264 | 0.332 | 1.09E-27 | Dnal1 | 3.16E-19 | -0.013982581 | 0.269 | 0.248 | 8.51E-15 |
| Eif4g1 | 4.07E-32 | 0.146879899 | 0.405 | 0.372 | 1.10E-27 | Cpe | 3.17E-19 | -0.083597618 | 0.68 | 0.675 | 8.54E-15 |
| Gm15398 | 4.09E-32 | -0.37032065 | 0.237 | 0.254 | 1.10E-27 | Hdac9 | 3.19E-19 | -0.131393633 | 0.757 | 0.762 | 8.59E-15 |
| Pdss2 | 6.25E-32 | 0.106026394 | 0.57 | 0.575 | 1.68E-27 | Stx16 | 3.21E-19 | 0.033122063 | 0.439 | 0.392 | 8.64E-15 |
| Kcnb1 | 6.98E-32 | 0.135067889 | 0.543 | 0.528 | 1.88E-27 | Tro | 3.23E-19 | -0.061009694 | 0.444 | 0.436 | 8.69E-15 |
| Hnrnpdl | 6.99E-32 | -0.119775592 | 0.506 | 0.583 | 1.88E-27 | Qrich1 | 3.36E-19 | 0.078450441 | 0.507 | 0.449 | 9.05E-15 |

| | | | | | | | | | | | |
|---|---|---|---|---|---|---|---|---|---|---|---|
| Birc2 | 9.56E-32 | -0.080375333 | 0.252 | 0.313 | 2.57E-27 | Srr | 3.55E-19 | 0.070055758 | 0.371 | 0.319 | 9.55E-15 |
| Tada2a | 1.16E-31 | 0.099576868 | 0.274 | 0.264 | 3.12E-27 | Srebf2 | 3.78E-19 | 0.028414322 | 0.363 | 0.322 | 1.02E-14 |
| Stxbp1 | 1.17E-31 | 0.126126205 | 0.567 | 0.55 | 3.15E-27 | Krit1 | 4.03E-19 | 0.077972735 | 0.53 | 0.472 | 1.09E-14 |
| Gm26691 | 1.26E-31 | 0.202015717 | 0.323 | 0.273 | 3.38E-27 | Srrt | 4.24E-19 | -0.001750173 | 0.326 | 0.298 | 1.14E-14 |
| Elmod1 | 1.28E-31 | 0.143811056 | 0.705 | 0.69 | 3.45E-27 | Dnaja2 | 4.44E-19 | -0.016771626 | 0.525 | 0.496 | 1.19E-14 |
| Gm35188 | 1.51E-31 | -0.186118126 | 0.523 | 0.593 | 4.06E-27 | AC165953.2 | 4.57E-19 | -0.12240403 | 0.362 | 0.418 | 1.23E-14 |
| Ddx1 | 1.56E-31 | -0.069181789 | 0.288 | 0.349 | 4.21E-27 | A330015K06Rik | 4.66E-19 | 0.092561602 | 0.691 | 0.634 | 1.25E-14 |
| Neurl1a | 1.57E-31 | 0.06702777 | 0.338 | 0.353 | 4.23E-27 | Dennd4a | 4.80E-19 | 0.052023627 | 0.631 | 0.578 | 1.29E-14 |
| Epb41l3 | 1.80E-31 | 0.175327201 | 0.473 | 0.431 | 4.85E-27 | Sh3gl1 | 4.84E-19 | 0.069671134 | 0.37 | 0.318 | 1.30E-14 |
| Kbtbd11 | 1.82E-31 | -0.101276468 | 0.272 | 0.337 | 4.91E-27 | Celf3 | 4.84E-19 | 0.073861164 | 0.538 | 0.482 | 1.30E-14 |
| Galnt11 | 1.96E-31 | -0.119268936 | 0.436 | 0.512 | 5.28E-27 | Camk2a | 4.87E-19 | -0.112081053 | 0.957 | 0.966 | 1.31E-14 |
| Riok3 | 2.11E-31 | -0.09257439 | 0.39 | 0.461 | 5.69E-27 | Phyhipl | 4.88E-19 | 0.08809902 | 0.315 | 0.263 | 1.31E-14 |
| Brip1os | 2.24E-31 | -0.120718195 | 0.378 | 0.453 | 6.04E-27 | Mamld1 | 5.23E-19 | -0.075893406 | 0.336 | 0.339 | 1.41E-14 |
| Ptprd | 2.85E-31 | 0.191248888 | 0.987 | 0.983 | 7.68E-27 | Kcnq1 | 5.36E-19 | -0.104699564 | 0.606 | 0.663 | 1.44E-14 |
| Asb3 | 3.16E-31 | -0.064312673 | 0.527 | 0.595 | 8.50E-27 | Rad17 | 5.63E-19 | 0.029841299 | 0.254 | 0.217 | 1.51E-14 |
| Manf | 3.16E-31 | -0.129500663 | 0.347 | 0.42 | 8.50E-27 | Zfp329 | 5.64E-19 | 0.058681456 | 0.31 | 0.263 | 1.52E-14 |
| Ntm | 3.30E-31 | 0.16636696 | 0.666 | 0.592 | 8.89E-27 | Myo16 | 5.81E-19 | -0.147242392 | 0.236 | 0.272 | 1.56E-14 |
| Pacsin1 | 4.33E-31 | 0.138913318 | 0.736 | 0.727 | 1.16E-26 | Gabrg2 | 6.27E-19 | 0.071075214 | 0.623 | 0.567 | 1.69E-14 |
| Fam69a | 5.51E-31 | 0.152007174 | 0.373 | 0.302 | 1.48E-26 | Ube2v2 | 7.11E-19 | 0.071801515 | 0.284 | 0.236 | 1.91E-14 |
| Sec63 | 5.57E-31 | -0.098165388 | 0.34 | 0.409 | 1.50E-26 | Nrgn | 7.33E-19 | 0.040051633 | 0.481 | 0.431 | 1.97E-14 |
| Dpf3 | 5.64E-31 | -0.175857922 | 0.239 | 0.306 | 1.52E-26 | Tada2a | 7.35E-19 | 0.024491797 | 0.311 | 0.274 | 1.98E-14 |
| Tmem181a | 6.27E-31 | 0.155749822 | 0.524 | 0.47 | 1.69E-26 | Fam168b | 7.36E-19 | 0.050032134 | 0.307 | 0.264 | 1.98E-14 |
| Gm5441 | 6.36E-31 | -0.107593234 | 0.349 | 0.42 | 1.71E-26 | Gm12064 | 7.55E-19 | 0.087212489 | 0.372 | 0.317 | 2.03E-14 |
| Plk2 | 6.70E-31 | -0.156089513 | 0.227 | 0.292 | 1.80E-26 | Trio | 7.63E-19 | -0.096955608 | 0.825 | 0.831 | 2.05E-14 |
| Taf1d | 6.80E-31 | -0.083717334 | 0.226 | 0.286 | 1.83E-26 | Napg | 7.65E-19 | -0.013168666 | 0.299 | 0.276 | 2.06E-14 |
| Slc25a23 | 6.87E-31 | 0.138073751 | 0.463 | 0.437 | 1.85E-26 | Atp1a3 | 7.71E-19 | 0.061592764 | 0.735 | 0.683 | 2.07E-14 |
| Lrrc40 | 8.02E-31 | -0.107647535 | 0.224 | 0.288 | 2.16E-26 | Aplp2 | 7.74E-19 | 0.057155203 | 0.499 | 0.447 | 2.08E-14 |
| Dtx3 | 8.03E-31 | -0.111417151 | 0.292 | 0.362 | 2.16E-26 | Ssbp4 | 7.92E-19 | 0.052213664 | 0.378 | 0.328 | 2.13E-14 |
| Lysmd4 | 8.41E-31 | -0.146239075 | 0.507 | 0.583 | 2.26E-26 | Fam110b | 9.62E-19 | -0.07643298 | 0.418 | 0.417 | 2.59E-14 |
| Ewsr1 | 9.25E-31 | -0.109924737 | 0.575 | 0.649 | 2.49E-26 | Dennd5a | 1.01E-18 | 0.025502131 | 0.371 | 0.332 | 2.72E-14 |
| Rmdn1 | 9.36E-31 | -0.125537537 | 0.524 | 0.599 | 2.52E-26 | Ccdc66 | 1.01E-18 | 0.04766375 | 0.314 | 0.27 | 2.73E-14 |
| Grk3 | 9.63E-31 | 0.153461359 | 0.351 | 0.315 | 2.59E-26 | Tor1aip2 | 1.08E-18 | 0.040487538 | 0.286 | 0.246 | 2.91E-14 |
| Zbtb16 | 1.03E-30 | -0.138101857 | 0.688 | 0.754 | 2.76E-26 | Gm26904 | 1.09E-18 | 0.051471697 | 0.601 | 0.626 | 2.94E-14 |
| Gria4 | 1.08E-30 | 0.205921659 | 0.648 | 0.592 | 2.90E-26 | Cpsf7 | 1.14E-18 | 0.023208971 | 0.293 | 0.257 | 3.06E-14 |
| 4930455C13Rik | 1.12E-30 | -0.094436128 | 0.28 | 0.347 | 3.02E-26 | Pak3 | 1.19E-18 | -0.093479529 | 0.762 | 0.764 | 3.21E-14 |
| Kcna1 | 1.14E-30 | -0.099009523 | 0.251 | 0.315 | 3.07E-26 | Actl6b | 1.24E-18 | 0.004441626 | 0.277 | 0.248 | 3.35E-14 |
| Tpr | 1.29E-30 | -0.075450165 | 0.43 | 0.499 | 3.47E-26 | Klf13 | 1.30E-18 | 0.047944504 | 0.348 | 0.303 | 3.49E-14 |
| Mcu | 1.36E-30 | 0.145986764 | 0.545 | 0.526 | 3.67E-26 | Clstn3 | 1.34E-18 | 0.072606129 | 0.339 | 0.289 | 3.60E-14 |
| Gm47271 | 1.37E-30 | -0.167144751 | 0.291 | 0.362 | 3.69E-26 | N4bp2l2 | 1.40E-18 | 0.065466598 | 0.554 | 0.499 | 3.78E-14 |
| Csnk1a1 | 1.64E-30 | -0.127242374 | 0.634 | 0.705 | 4.41E-26 | Sar1b | 1.41E-18 | 0.043544432 | 0.254 | 0.215 | 3.80E-14 |
| Snhg11 | 1.84E-30 | 0.010428934 | 0.931 | 0.89 | 4.95E-26 | Fat3 | 1.47E-18 | -0.131377297 | 0.765 | 0.773 | 3.95E-14 |
| Top1 | 1.89E-30 | -0.117832767 | 0.584 | 0.658 | 5.07E-26 | Ensa | 1.48E-18 | -0.025862353 | 0.277 | 0.26 | 3.99E-14 |
| Abi1 | 2.28E-30 | -0.128630159 | 0.577 | 0.65 | 6.13E-26 | BC005561 | 1.52E-18 | -0.047063941 | 0.467 | 0.454 | 4.09E-14 |
| Myo18a | 2.76E-30 | 0.113652801 | 0.521 | 0.511 | 7.43E-26 | Chd2 | 1.54E-18 | 0.116119604 | 0.551 | 0.492 | 4.14E-14 |
| Terf1 | 2.90E-30 | -0.09702145 | 0.272 | 0.336 | 7.80E-26 | Dopey2 | 1.57E-18 | -0.112796956 | 0.386 | 0.444 | 4.22E-14 |
| Pde4b | 2.98E-30 | 0.198674081 | 0.529 | 0.457 | 8.03E-26 | Zcchc11 | 1.77E-18 | -0.098863251 | 0.577 | 0.633 | 4.77E-14 |
| Creb1 | 3.03E-30 | -0.07194508 | 0.254 | 0.314 | 8.15E-26 | Mir124a-1hg | 1.79E-18 | 0.035282772 | 0.581 | 0.535 | 4.81E-14 |
| Crebzf | 3.50E-30 | -0.10782976 | 0.327 | 0.398 | 9.41E-26 | Aff3 | 2.07E-18 | -0.098674394 | 0.79 | 0.793 | 5.57E-14 |
| Kmt5b | 3.97E-30 | -0.045403061 | 0.321 | 0.376 | 1.07E-25 | Pcmtd2 | 2.16E-18 | 0.04502494 | 0.293 | 0.25 | 5.81E-14 |
| Ergic1 | 5.63E-30 | 0.111049399 | 0.344 | 0.329 | 1.51E-25 | Fubp1 | 2.18E-18 | 0.086380772 | 0.685 | 0.629 | 5.85E-14 |
| Egfem1 | 7.31E-30 | 0.228550143 | 0.826 | 0.815 | 1.97E-25 | Map2k5 | 2.19E-18 | -0.032089586 | 0.512 | 0.491 | 5.91E-14 |
| C1ql3 | 9.40E-30 | -0.189995759 | 0.299 | 0.369 | 2.53E-25 | Agfg1 | 2.36E-18 | -0.019709447 | 0.387 | 0.364 | 6.35E-14 |
| Tle1 | 9.49E-30 | -0.134580874 | 0.324 | 0.396 | 2.55E-25 | Atg3 | 2.41E-18 | 0.069297643 | 0.285 | 0.238 | 6.50E-14 |
| Trmt1 | 1.06E-29 | -0.081555586 | 0.242 | 0.302 | 2.85E-25 | Ccser2 | 2.44E-18 | -0.036730502 | 0.353 | 0.338 | 6.57E-14 |
| Sptbn4 | 1.07E-29 | 0.117889682 | 0.669 | 0.662 | 2.87E-25 | 2610037D02Rik | 2.45E-18 | 0.059243386 | 0.412 | 0.363 | 6.59E-14 |
| Esf1 | 1.15E-29 | -0.080370841 | 0.251 | 0.312 | 3.09E-25 | Snrnp48 | 2.46E-18 | 0.041769317 | 0.332 | 0.289 | 6.61E-14 |
| Tsc1 | 1.17E-29 | 0.11153243 | 0.4 | 0.388 | 3.16E-25 | Ddx42 | 2.48E-18 | 0.040540792 | 0.343 | 0.3 | 6.67E-14 |
| Ndfip2 | 1.48E-29 | -0.138350399 | 0.309 | 0.38 | 3.97E-25 | Peli1 | 2.54E-18 | 0.041257059 | 0.366 | 0.321 | 6.83E-14 |
| Mark2 | 1.62E-29 | 0.126196806 | 0.583 | 0.565 | 4.37E-25 | Cpne6 | 2.55E-18 | 0.069968029 | 0.489 | 0.435 | 6.86E-14 |
| Wnk1 | 1.63E-29 | 0.116298072 | 0.544 | 0.532 | 4.39E-25 | Pbrm1 | 2.57E-18 | 0.077789785 | 0.445 | 0.39 | 6.92E-14 |
| Slc8a3 | 1.85E-29 | 0.188652678 | 0.324 | 0.281 | 4.98E-25 | Luc7l3 | 2.76E-18 | 0.047742749 | 0.601 | 0.548 | 7.42E-14 |
| Glt8d1 | 2.02E-29 | -0.092004124 | 0.331 | 0.398 | 5.44E-25 | Sars | 2.80E-18 | 0.000339052 | 0.271 | 0.245 | 7.53E-14 |
| Ddx17 | 2.08E-29 | 0.144960232 | 0.549 | 0.518 | 5.58E-25 | Ptprg | 2.85E-18 | -0.117620257 | 0.729 | 0.748 | 7.66E-14 |
| Parg | 2.39E-29 | -0.071442777 | 0.377 | 0.442 | 6.42E-25 | Mau2 | 3.05E-18 | 0.027318539 | 0.323 | 0.286 | 8.20E-14 |
| Araf | 2.42E-29 | -0.105786699 | 0.387 | 0.459 | 6.50E-25 | Slc38a2 | 3.12E-18 | 0.055975212 | 0.363 | 0.316 | 8.39E-14 |
| Caly | 2.43E-29 | -0.075777773 | 0.22 | 0.279 | 6.55E-25 | Ddx1 | 3.12E-18 | -0.013650968 | 0.311 | 0.288 | 8.40E-14 |
| Esco1 | 2.51E-29 | -0.064770555 | 0.341 | 0.402 | 6.76E-25 | Kpna3 | 3.27E-18 | 0.0619069 | 0.445 | 0.394 | 8.79E-14 |
| Kif1a | 3.00E-29 | 0.101980859 | 0.539 | 0.535 | 8.07E-25 | Nova1 | 3.30E-18 | 0.095899426 | 0.254 | 0.206 | 8.89E-14 |
| C630031E19Rik | 3.00E-29 | -0.102141347 | 0.199 | 0.258 | 8.08E-25 | Ap3m1 | 3.37E-18 | 0.045007794 | 0.284 | 0.241 | 9.07E-14 |
| Arhgap32 | 3.01E-29 | 0.113928818 | 0.572 | 0.566 | 8.11E-25 | Atp1b1 | 3.40E-18 | 0.026878738 | 0.787 | 0.746 | 9.14E-14 |
| Zfp266 | 3.33E-29 | -0.086563675 | 0.297 | 0.361 | 8.95E-25 | Tsc22d1 | 3.40E-18 | -0.053966875 | 0.596 | 0.582 | 9.16E-14 |
| Btaf1 | 3.39E-29 | -0.137054209 | 0.462 | 0.536 | 9.12E-25 | Qk | 3.47E-18 | 0.091912307 | 0.536 | 0.478 | 9.33E-14 |
| Ulk2 | 3.48E-29 | 0.122538163 | 0.563 | 0.551 | 9.37E-25 | Palm | 3.52E-18 | 0.056112864 | 0.398 | 0.348 | 9.46E-14 |
| Grid2 | 3.57E-29 | 0.157201895 | 0.691 | 0.637 | 9.62E-25 | Caskin1 | 3.59E-18 | -0.074569945 | 0.291 | 0.298 | 9.65E-14 |
| Actr3 | 4.30E-29 | -0.124878869 | 0.369 | 0.441 | 1.16E-24 | Inpp5f | 3.68E-18 | 0.03998254 | 0.276 | 0.236 | 9.90E-14 |
| Srek1 | 4.38E-29 | -0.107223673 | 0.624 | 0.694 | 1.18E-24 | Appl1 | 3.80E-18 | 0.059040965 | 0.465 | 0.414 | 1.02E-13 |
| Cmss1 | 4.71E-29 | 0.056701334 | 0.673 | 0.701 | 1.27E-24 | Mosmo | 3.88E-18 | 0.050879006 | 0.29 | 0.248 | 1.04E-13 |
| Gm26733 | 4.90E-29 | -0.043052494 | 0.624 | 0.587 | 1.32E-24 | Prkag2 | 4.12E-18 | -0.105382085 | 0.812 | 0.811 | 1.11E-13 |
| Gm16351 | 4.93E-29 | -0.2073771 | 0.317 | 0.385 | 1.33E-24 | Map3k3 | 4.24E-18 | 0.046496504 | 0.539 | 0.49 | 1.14E-13 |
| Eda | 5.38E-29 | 0.190473114 | 0.28 | 0.233 | 1.45E-24 | Ncapd3 | 4.28E-18 | -0.044154494 | 0.368 | 0.356 | 1.15E-13 |
| Gabrg3 | 5.64E-29 | 0.208870525 | 0.584 | 0.531 | 1.52E-24 | Ppme1 | 4.60E-18 | 0.059126932 | 0.393 | 0.343 | 1.24E-13 |
| Gm12353 | 6.13E-29 | -0.083718012 | 0.347 | 0.412 | 1.65E-24 | Dcaf8 | 4.70E-18 | 0.030745079 | 0.409 | 0.366 | 1.27E-13 |
| Grm5 | 6.31E-29 | 0.114907541 | 0.99 | 0.985 | 1.70E-24 | Mob4 | 4.83E-18 | 0.066451976 | 0.279 | 0.233 | 1.30E-13 |
| Cluap1 | 6.42E-29 | -0.065131927 | 0.207 | 0.26 | 1.73E-24 | Pcnp | 4.85E-18 | 0.060904192 | 0.26 | 0.215 | 1.30E-13 |
| Naa16 | 6.69E-29 | -0.087783593 | 0.241 | 0.302 | 1.80E-24 | Nufip2 | 5.04E-18 | 0.077364706 | 0.395 | 0.343 | 1.36E-13 |
| AW554918 | 7.05E-29 | 0.130480937 | 0.428 | 0.404 | 1.90E-24 | Map4k3 | 5.09E-18 | -0.074774805 | 0.659 | 0.652 | 1.37E-13 |
| Dcaf5 | 7.07E-29 | 0.107476979 | 0.438 | 0.428 | 1.90E-24 | Ppp3cb | 5.25E-18 | 0.031305179 | 0.385 | 0.344 | 1.41E-13 |
| Ube2v2 | 7.75E-29 | -0.099104303 | 0.236 | 0.298 | 2.09E-24 | Stau2 | 5.49E-18 | -0.042120357 | 0.692 | 0.671 | 1.48E-13 |
| Ip6k2 | 7.96E-29 | -0.07013035 | 0.268 | 0.325 | 2.14E-24 | Retreg3 | 5.73E-18 | 0.064813619 | 0.254 | 0.209 | 1.54E-13 |
| Mau2 | 8.28E-29 | 0.026472931 | 0.286 | 0.311 | 2.23E-24 | Wac | 5.92E-18 | 0.069270764 | 0.549 | 0.493 | 1.59E-13 |
| Adarb2 | 8.33E-29 | 0.284086381 | 0.392 | 0.411 | 2.24E-24 | Nkain2 | 5.96E-18 | -0.079868454 | 0.985 | 0.991 | 1.60E-13 |
| Ifrd1 | 8.34E-29 | -0.105957277 | 0.205 | 0.265 | 2.24E-24 | Hnrnpdl | 6.25E-18 | 0.030182973 | 0.551 | 0.506 | 1.68E-13 |
| Dnajc19 | 8.41E-29 | -0.086255572 | 0.321 | 0.386 | 2.26E-24 | Ppil2 | 6.35E-18 | 0.008026265 | 0.313 | 0.28 | 1.71E-13 |
| Nudcd3 | 9.50E-29 | -0.072528807 | 0.449 | 0.515 | 2.56E-24 | Psip1 | 6.39E-18 | 0.098088482 | 0.564 | 0.506 | 1.72E-13 |
| Dagla | 9.62E-29 | 0.130726394 | 0.476 | 0.46 | 2.59E-24 | Dhx9 | 6.69E-18 | 0.049908084 | 0.284 | 0.242 | 1.80E-13 |
| Grik4 | 9.89E-29 | -0.190982614 | 0.566 | 0.636 | 2.66E-24 | Map4 | 6.70E-18 | 0.099976557 | 0.557 | 0.499 | 1.80E-13 |
| Dlg2 | 1.00E-28 | 0.079768875 | 0.998 | 0.997 | 2.69E-24 | Pcm1 | 7.69E-18 | 0.017605132 | 0.602 | 0.559 | 2.07E-13 |
| 1700109K24Rik | 1.09E-28 | -0.111234422 | 0.424 | 0.496 | 2.92E-24 | 3110021N24Rik | 7.75E-18 | 0.089815874 | 0.487 | 0.43 | 2.09E-13 |

| | | | | | | | | | | | |
|---|---|---|---|---|---|---|---|---|---|---|---|
| Dhx36 | 1.15E-28 | -0.109456004 | 0.372 | 0.442 | 3.10E-24 | Gm36198 | 7.79E-18 | 0.065724565 | 0.25 | 0.205 | 2.10E-13 |
| Trpc4 | 1.18E-28 | -0.182337159 | 0.503 | 0.575 | 3.19E-24 | Pnn | 7.88E-18 | 0.029229668 | 0.601 | 0.555 | 2.12E-13 |
| Rsrc2 | 1.26E-28 | -0.104986713 | 0.402 | 0.473 | 3.40E-24 | Dnajc5 | 8.08E-18 | 0.064661034 | 0.328 | 0.279 | 2.17E-13 |
| Sil1 | 1.42E-28 | 0.134280773 | 0.263 | 0.235 | 3.83E-24 | Stmn3 | 8.84E-18 | 0.012516879 | 0.391 | 0.354 | 2.38E-13 |
| Sergef | 1.47E-28 | 0.119966313 | 0.477 | 0.47 | 3.94E-24 | Ctcf | 8.84E-18 | 0.028180406 | 0.327 | 0.289 | 2.38E-13 |
| Chrd | 1.51E-28 | -0.135680602 | 0.268 | 0.335 | 4.06E-24 | Chic1 | 8.97E-18 | 0.066685141 | 0.367 | 0.318 | 2.41E-13 |
| Gm42477 | 1.59E-28 | -0.129685109 | 0.286 | 0.355 | 4.29E-24 | Virma | 9.41E-18 | 0.015360938 | 0.37 | 0.334 | 2.53E-13 |
| Gprasp1 | 1.78E-28 | -0.103622606 | 0.532 | 0.604 | 4.80E-24 | Ptpn2 | 9.47E-18 | 0.045828967 | 0.264 | 0.224 | 2.55E-13 |
| Tenm2 | 2.13E-28 | 0.115034527 | 0.992 | 0.983 | 5.73E-24 | Mbtps2 | 9.63E-18 | 0.039576588 | 0.254 | 0.216 | 2.59E-13 |
| Rogdi | 2.19E-28 | -0.084218127 | 0.286 | 0.35 | 5.90E-24 | Actn4 | 1.04E-17 | 0.071772394 | 0.407 | 0.355 | 2.79E-13 |
| Meg3 | 2.32E-28 | 0.137617496 | 0.99 | 0.988 | 6.23E-24 | Lrrc28 | 1.10E-17 | -0.046555064 | 0.323 | 0.312 | 2.97E-13 |
| Dnajb6 | 2.44E-28 | -0.090372243 | 0.309 | 0.374 | 6.58E-24 | Ppp4r2 | 1.11E-17 | 0.033708919 | 0.345 | 0.305 | 2.98E-13 |
| Ccdc47 | 2.46E-28 | -0.071052304 | 0.211 | 0.266 | 6.62E-24 | Arf3 | 1.11E-17 | 0.054237975 | 0.406 | 0.357 | 2.98E-13 |
| Polb | 2.46E-28 | -0.104347552 | 0.486 | 0.558 | 6.63E-24 | Thoc7 | 1.13E-17 | 0.045131336 | 0.257 | 0.218 | 3.03E-13 |
| Kcnk10 | 2.68E-28 | 0.153281606 | 0.364 | 0.323 | 7.21E-24 | Top1 | 1.14E-17 | 0.074878413 | 0.639 | 0.584 | 3.07E-13 |
| Snx29 | 3.44E-28 | 0.090108901 | 0.444 | 0.444 | 9.25E-24 | Pcgf5 | 1.15E-17 | 0.084768993 | 0.321 | 0.272 | 3.08E-13 |
| Hs6st3 | 4.38E-28 | 0.197991991 | 0.803 | 0.748 | 1.18E-23 | Anapc5 | 1.16E-17 | 0.062689843 | 0.409 | 0.359 | 3.11E-13 |
| Gtf2i | 4.44E-28 | 0.106324187 | 0.597 | 0.594 | 1.19E-23 | Tmem178b | 1.17E-17 | -0.101790396 | 0.761 | 0.765 | 3.15E-13 |
| Cry2 | 4.46E-28 | 0.132160151 | 0.318 | 0.285 | 1.20E-23 | Ago3 | 1.19E-17 | 0.069450559 | 0.618 | 0.563 | 3.20E-13 |
| Dennd2a | 4.62E-28 | -0.035119267 | 0.31 | 0.361 | 1.24E-23 | Fyttd1 | 1.23E-17 | 0.033230937 | 0.329 | 0.29 | 3.31E-13 |
| Eif5b | 4.78E-28 | -0.080732638 | 0.372 | 0.437 | 1.29E-23 | Rsbn1 | 1.30E-17 | -0.001526654 | 0.316 | 0.288 | 3.50E-13 |
| Slc25a25 | 4.79E-28 | -0.098128164 | 0.21 | 0.268 | 1.29E-23 | Prkd1 | 1.35E-17 | 0.04173501 | 0.454 | 0.407 | 3.63E-13 |
| Cstf3 | 4.91E-28 | -0.103205725 | 0.494 | 0.565 | 1.32E-23 | Grk3 | 1.36E-17 | -0.114333806 | 0.324 | 0.351 | 3.67E-13 |
| Kcnip4 | 5.75E-28 | 0.199159225 | 0.952 | 0.955 | 1.55E-23 | Fer | 1.38E-17 | -0.061570785 | 0.503 | 0.499 | 3.72E-13 |
| Timm23 | 6.04E-28 | -0.034147711 | 0.229 | 0.273 | 1.63E-23 | Grb2 | 1.40E-17 | 0.067251154 | 0.329 | 0.281 | 3.76E-13 |
| Camk2d | 7.64E-28 | 0.244110906 | 0.271 | 0.22 | 2.06E-23 | Celf4 | 1.58E-17 | -0.123786574 | 0.883 | 0.888 | 4.26E-13 |
| Zkscan3 | 9.41E-28 | 0.12983156 | 0.315 | 0.283 | 2.53E-23 | Calm1 | 1.65E-17 | 0.020023181 | 0.438 | 0.396 | 4.43E-13 |
| Cacna1h | 1.07E-27 | -0.17740946 | 0.268 | 0.332 | 2.88E-23 | Celf5 | 1.66E-17 | 0.067983265 | 0.529 | 0.476 | 4.45E-13 |
| N4bp2l2 | 1.47E-27 | -0.070197271 | 0.499 | 0.564 | 3.95E-23 | Nipsnap2 | 1.71E-17 | 0.045141424 | 0.297 | 0.255 | 4.60E-13 |
| Srcin1 | 1.53E-27 | 0.133223039 | 0.503 | 0.476 | 4.13E-23 | Zfp532 | 1.72E-17 | 0.05709971 | 0.495 | 0.445 | 4.63E-13 |
| Shisa7 | 1.57E-27 | -0.083321799 | 0.284 | 0.346 | 4.23E-23 | Ubxn7 | 1.85E-17 | 0.062809497 | 0.261 | 0.217 | 4.97E-13 |
| Trim37 | 1.58E-27 | 0.098065673 | 0.413 | 0.41 | 4.24E-23 | Mindy2 | 1.86E-17 | 0.009394546 | 0.275 | 0.245 | 5.01E-13 |
| Rufy1 | 1.72E-27 | 0.108475046 | 0.298 | 0.28 | 4.62E-23 | Faim2 | 1.94E-17 | 0.051978605 | 0.304 | 0.263 | 5.22E-13 |
| Chd4 | 2.08E-27 | 0.115037885 | 0.426 | 0.409 | 5.61E-23 | Nr6a1 | 2.00E-17 | -0.107653609 | 0.547 | 0.57 | 5.39E-13 |
| Wasl | 2.17E-27 | -0.104900525 | 0.425 | 0.494 | 5.85E-23 | Sf3b1 | 2.02E-17 | 0.04912347 | 0.653 | 0.602 | 5.43E-13 |
| Rev1 | 2.19E-27 | -0.038402149 | 0.446 | 0.501 | 5.89E-23 | Zfp799 | 2.09E-17 | 0.074083108 | 0.329 | 0.281 | 5.63E-13 |
| Rsf1os1 | 2.22E-27 | -0.08015171 | 0.329 | 0.392 | 5.97E-23 | Crk | 2.09E-17 | 0.055677913 | 0.266 | 0.223 | 5.64E-13 |
| Man2a2 | 2.22E-27 | 0.1108151 | 0.327 | 0.311 | 5.98E-23 | Mgll | 2.10E-17 | 0.022939871 | 0.4 | 0.36 | 5.65E-13 |
| Ppp2r5c | 2.25E-27 | 0.119475525 | 0.553 | 0.541 | 6.06E-23 | Zfp280d | 2.10E-17 | 0.10280927 | 0.644 | 0.589 | 5.65E-13 |
| Edem3 | 2.37E-27 | -0.007514737 | 0.269 | 0.307 | 6.37E-23 | Gabra5 | 2.15E-17 | 0.055469705 | 0.495 | 0.446 | 5.78E-13 |
| Snrnp70 | 2.43E-27 | 0.137235938 | 0.844 | 0.831 | 6.55E-23 | Lancl1 | 2.21E-17 | -0.010068663 | 0.3 | 0.277 | 5.95E-13 |
| Lrrn1 | 2.70E-27 | -0.098925713 | 0.249 | 0.309 | 7.26E-23 | Grina | 2.23E-17 | 0.004434778 | 0.331 | 0.299 | 5.99E-13 |
| Robo1 | 3.13E-27 | 0.232031985 | 0.572 | 0.563 | 8.42E-23 | Clk1 | 2.46E-17 | 0.017384347 | 0.51 | 0.469 | 6.61E-13 |
| Lrrc28 | 3.29E-27 | 0.053023741 | 0.312 | 0.327 | 8.86E-23 | Birc2 | 2.52E-17 | 0.036139447 | 0.29 | 0.252 | 6.77E-13 |
| Trappc8 | 3.41E-27 | -0.02104436 | 0.406 | 0.454 | 9.17E-23 | Cbx3 | 2.55E-17 | 0.016683825 | 0.251 | 0.22 | 6.85E-13 |
| Klf9 | 3.42E-27 | -0.069514487 | 0.266 | 0.322 | 9.20E-23 | Slc6a17 | 2.61E-17 | -0.032819754 | 0.424 | 0.407 | 7.02E-13 |
| Abhd12 | 3.68E-27 | -0.076749163 | 0.403 | 0.467 | 9.91E-23 | Astn1 | 2.66E-17 | -0.070924269 | 0.786 | 0.779 | 7.15E-13 |
| Bbip1 | 3.87E-27 | -0.097951069 | 0.259 | 0.322 | 1.04E-22 | Dmtn | 2.67E-17 | 0.013173941 | 0.434 | 0.396 | 7.19E-13 |
| Kif16b | 4.07E-27 | 0.112927761 | 0.316 | 0.297 | 1.09E-22 | Clta | 2.78E-17 | 0.037741278 | 0.271 | 0.232 | 7.48E-13 |
| Ercc6 | 4.12E-27 | -0.108729014 | 0.245 | 0.307 | 1.11E-22 | Clstn1 | 2.79E-17 | 0.06214031 | 0.537 | 0.486 | 7.52E-13 |
| Prkcz | 4.47E-27 | 0.110274853 | 0.498 | 0.486 | 1.20E-22 | Galnt11 | 2.81E-17 | -0.009845203 | 0.467 | 0.436 | 7.56E-13 |
| Magi3 | 4.52E-27 | 0.132843903 | 0.442 | 0.426 | 1.22E-22 | Dzip3 | 2.93E-17 | -0.010934271 | 0.492 | 0.459 | 7.90E-13 |
| Csnk2a1 | 4.64E-27 | -0.074039166 | 0.259 | 0.316 | 1.25E-22 | Lrrtm4 | 2.95E-17 | -0.057852586 | 0.772 | 0.816 | 7.95E-13 |
| Chst11 | 5.34E-27 | 0.087425496 | 0.443 | 0.449 | 1.44E-22 | Mad1l1 | 2.97E-17 | -0.050698756 | 0.255 | 0.251 | 7.99E-13 |
| Baz1b | 5.52E-27 | 0.118450959 | 0.474 | 0.459 | 1.48E-22 | Setd7 | 3.06E-17 | 0.062560174 | 0.381 | 0.332 | 8.24E-13 |
| Git2 | 5.74E-27 | 0.077031468 | 0.257 | 0.255 | 1.55E-22 | Bbip1 | 3.07E-17 | 0.066348823 | 0.307 | 0.259 | 8.26E-13 |
| Cnih3 | 6.52E-27 | -0.217058586 | 0.301 | 0.363 | 1.76E-22 | Pkd2 | 3.15E-17 | 0.032555622 | 0.28 | 0.242 | 8.48E-13 |
| Ppargc1b | 7.25E-27 | 0.113328837 | 0.349 | 0.329 | 1.95E-22 | Glg1 | 3.23E-17 | -0.079236236 | 0.732 | 0.736 | 8.70E-13 |
| Emc10 | 8.46E-27 | -0.066653386 | 0.22 | 0.273 | 2.28E-22 | Abhd12 | 3.27E-17 | 0.009603289 | 0.438 | 0.403 | 8.80E-13 |
| Ensa | 8.97E-27 | -0.067448117 | 0.26 | 0.316 | 2.41E-22 | R3hdm2 | 3.31E-17 | 0.120775453 | 0.72 | 0.672 | 8.92E-13 |
| Tcf20 | 9.28E-27 | 0.092259141 | 0.741 | 0.747 | 2.50E-22 | Clvs1 | 3.38E-17 | -0.070436224 | 0.619 | 0.612 | 9.10E-13 |
| Ankrd11 | 9.40E-27 | 0.111079528 | 0.772 | 0.769 | 2.53E-22 | Rbm27 | 3.40E-17 | 0.04022958 | 0.298 | 0.259 | 9.15E-13 |
| Pdzrn4 | 9.54E-27 | 0.228257946 | 0.355 | 0.299 | 2.57E-22 | Mdm4 | 3.46E-17 | 0.038168592 | 0.484 | 0.439 | 9.32E-13 |
| Clec16a | 1.01E-26 | -0.06143468 | 0.494 | 0.556 | 2.72E-22 | Cdc27 | 3.50E-17 | 0.059517051 | 0.33 | 0.284 | 9.40E-13 |
| Dync1li1 | 1.02E-26 | 0.04908695 | 0.294 | 0.307 | 2.75E-22 | Xpo7 | 3.78E-17 | 0.045477537 | 0.519 | 0.47 | 1.02E-12 |
| Glt1d1 | 1.22E-26 | 0.094723367 | 0.259 | 0.25 | 3.27E-22 | Gm15594 | 3.87E-17 | 0.099802475 | 0.436 | 0.381 | 1.04E-12 |
| Rab11fip4 | 1.34E-26 | 0.09995129 | 0.415 | 0.41 | 3.60E-22 | Appbp2 | 3.90E-17 | 0.060816762 | 0.308 | 0.262 | 1.05E-12 |
| 4930473A02Rik | 1.36E-26 | -0.132913686 | 0.221 | 0.282 | 3.66E-22 | Ppp1cb | 4.02E-17 | 0.055287757 | 0.422 | 0.373 | 1.08E-12 |
| Mbd2 | 1.42E-26 | -0.049191592 | 0.289 | 0.34 | 3.83E-22 | Nme7 | 4.09E-17 | -0.034145912 | 0.515 | 0.498 | 1.10E-12 |
| Morf4l1 | 1.45E-26 | -0.013003425 | 0.303 | 0.343 | 3.91E-22 | Pcbp2 | 4.30E-17 | 0.046974369 | 0.41 | 0.364 | 1.16E-12 |
| Opa1 | 1.60E-26 | -0.027651907 | 0.342 | 0.388 | 4.31E-22 | Ube2d1 | 4.39E-17 | 0.067389107 | 0.259 | 0.214 | 1.18E-12 |
| Gm26936 | 1.61E-26 | -0.130467921 | 0.194 | 0.253 | 4.33E-22 | Prr16 | 4.44E-17 | 0.182384615 | 0.444 | 0.391 | 1.20E-12 |
| Mia2 | 1.61E-26 | -0.058355004 | 0.305 | 0.361 | 4.34E-22 | Cap1 | 4.49E-17 | 0.018364652 | 0.298 | 0.264 | 1.21E-12 |
| Crebrf | 1.78E-26 | 0.125788251 | 0.322 | 0.291 | 4.78E-22 | Ddx6 | 4.89E-17 | 0.019633809 | 0.325 | 0.29 | 1.32E-12 |
| Hnrnpd | 1.85E-26 | -0.06903588 | 0.233 | 0.287 | 4.98E-22 | Neto1 | 4.97E-17 | -0.063641228 | 0.576 | 0.558 | 1.34E-12 |
| Ly6h | 1.89E-26 | -0.181704531 | 0.496 | 0.558 | 5.09E-22 | Sergef | 5.17E-17 | -0.083152643 | 0.475 | 0.477 | 1.39E-12 |
| Luzp1 | 1.91E-26 | -0.097980922 | 0.459 | 0.527 | 5.14E-22 | Sec63 | 5.23E-17 | 0.088916228 | 0.392 | 0.34 | 1.41E-12 |
| Kdm5a | 1.96E-26 | 0.014064761 | 0.365 | 0.398 | 5.28E-22 | Srek1 | 5.29E-17 | 0.073581886 | 0.677 | 0.624 | 1.42E-12 |
| Spata7 | 1.98E-26 | 0.121702573 | 0.408 | 0.378 | 5.32E-22 | Actr3 | 5.31E-17 | 0.10078337 | 0.424 | 0.369 | 1.43E-12 |
| Lcorl | 2.25E-26 | 0.165058782 | 0.434 | 0.395 | 6.06E-22 | Vgll4 | 5.32E-17 | 0.078627904 | 0.38 | 0.329 | 1.43E-12 |
| Srsf4 | 2.27E-26 | -0.033876117 | 0.246 | 0.289 | 6.11E-22 | Kifap3 | 5.32E-17 | -0.060108022 | 0.53 | 0.525 | 1.43E-12 |
| Srr | 2.35E-26 | -0.072636599 | 0.319 | 0.379 | 6.33E-22 | Mycbp2 | 5.39E-17 | -0.090666836 | 0.97 | 0.973 | 1.45E-12 |
| Fbxl20 | 2.36E-26 | 0.115035335 | 0.53 | 0.518 | 6.34E-22 | Serbp1 | 5.53E-17 | 0.032701655 | 0.289 | 0.252 | 1.49E-12 |
| Sucla2 | 2.36E-26 | -0.084929963 | 0.215 | 0.272 | 6.35E-22 | C1ql3 | 5.64E-17 | 0.147514165 | 0.352 | 0.299 | 1.52E-12 |
| Large1 | 2.41E-26 | -0.158432213 | 0.942 | 0.947 | 6.49E-22 | Ttyh1 | 5.67E-17 | -0.001624495 | 0.4 | 0.369 | 1.53E-12 |
| Gm26565 | 2.70E-26 | 0.141303022 | 0.279 | 0.219 | 7.25E-22 | Nrxn3 | 5.77E-17 | -0.126743869 | 0.933 | 0.954 | 1.55E-12 |
| Med13l | 2.90E-26 | 0.088033296 | 0.622 | 0.628 | 7.81E-22 | Gtf2h2 | 5.83E-17 | 0.012133003 | 0.27 | 0.24 | 1.57E-12 |
| Hecw1 | 2.93E-26 | 0.109102534 | 0.777 | 0.722 | 7.88E-22 | Man1a2 | 5.97E-17 | -0.081986862 | 0.558 | 0.556 | 1.61E-12 |
| Luzp2 | 2.96E-26 | 0.252154868 | 0.256 | 0.213 | 7.96E-22 | Arhgef3 | 6.19E-17 | 0.071693441 | 0.403 | 0.353 | 1.67E-12 |
| Erlec1 | 2.98E-26 | -0.022691599 | 0.271 | 0.312 | 8.01E-22 | Matn2 | 6.27E-17 | -0.02163097 | 0.31 | 0.29 | 1.69E-12 |
| Naa40 | 3.44E-26 | 0.079397239 | 0.31 | 0.31 | 9.26E-22 | B230209E15Rik | 6.56E-17 | 0.067678934 | 0.479 | 0.426 | 1.76E-12 |
| Swt1 | 3.44E-26 | -0.067648185 | 0.264 | 0.319 | 9.26E-22 | Zc3h6 | 6.64E-17 | 0.091612327 | 0.358 | 0.306 | 1.79E-12 |
| Nr3c2 | 3.57E-26 | -0.20702695 | 0.85 | 0.847 | 9.61E-22 | Ggps1 | 6.65E-17 | 0.0325243 | 0.331 | 0.29 | 1.79E-12 |
| Ccnl1 | 3.74E-26 | -0.069012691 | 0.322 | 0.382 | 1.01E-21 | Gm1976 | 6.95E-17 | 0.052759711 | 0.38 | 0.333 | 1.87E-12 |
| Cnot7 | 3.79E-26 | -0.07904356 | 0.203 | 0.256 | 1.02E-21 | Adam10 | 7.09E-17 | 0.040380756 | 0.358 | 0.317 | 1.91E-12 |
| Tnrc6a | 3.99E-26 | -0.12351308 | 0.217 | 0.277 | 1.07E-21 | A930015D03Rik | 7.40E-17 | 0.051599925 | 0.607 | 0.558 | 1.99E-12 |
| Plppr1 | 4.84E-26 | 0.158528569 | 0.304 | 0.274 | 1.30E-21 | Large1 | 7.57E-17 | -0.102891957 | 0.933 | 0.942 | 2.04E-12 |
| Ppp4r2 | 5.06E-26 | -0.083925365 | 0.305 | 0.366 | 1.36E-21 | Shisa7 | 7.58E-17 | 0.021026459 | 0.319 | 0.284 | 2.04E-12 |

| | | | | | | | | | | | |
|---|---|---|---|---|---|---|---|---|---|---|---|
| Polr3a | 5.19E-26 | -0.048563789 | 0.222 | 0.269 | 1.40E-21 | Rbm28 | 8.00E-17 | 0.058597481 | 0.419 | 0.37 | 2.15E-12 |
| Papd4 | 5.20E-26 | 0.139635948 | 0.328 | 0.267 | 1.40E-21 | Pcca | 8.01E-17 | -0.065153137 | 0.614 | 0.61 | 2.15E-12 |
| Gm28750 | 5.56E-26 | -0.121594018 | 0.326 | 0.391 | 1.50E-21 | Gabbr1 | 8.12E-17 | 0.015513201 | 0.467 | 0.428 | 2.19E-12 |
| Galnt17 | 6.04E-26 | -0.191463047 | 0.877 | 0.892 | 1.62E-21 | Abi1 | 8.36E-17 | 0.058969499 | 0.627 | 0.577 | 2.25E-12 |
| Rchy1 | 6.30E-26 | -0.096335302 | 0.255 | 0.315 | 1.70E-21 | Zscan26 | 8.54E-17 | 0.035175455 | 0.355 | 0.316 | 2.30E-12 |
| Supt20 | 6.82E-26 | -0.068912707 | 0.316 | 0.373 | 1.84E-21 | Atf7ip | 8.70E-17 | 0.08213243 | 0.424 | 0.371 | 2.34E-12 |
| Rsbn1l | 7.18E-26 | -0.0912272 | 0.403 | 0.469 | 1.93E-21 | Nudt3 | 8.79E-17 | 0.023008538 | 0.32 | 0.284 | 2.37E-12 |
| Stxbp6 | 7.45E-26 | 0.144571579 | 0.375 | 0.39 | 2.01E-21 | Arel1 | 8.91E-17 | 0.011591716 | 0.265 | 0.236 | 2.40E-12 |
| Acadsb | 7.86E-26 | 0.132904735 | 0.31 | 0.272 | 2.11E-21 | Ppt1 | 9.35E-17 | 0.045458004 | 0.402 | 0.356 | 2.52E-12 |
| Tra2b | 8.04E-26 | -0.059537667 | 0.346 | 0.404 | 2.16E-21 | Chst1 | 9.49E-17 | -0.031623076 | 0.265 | 0.252 | 2.55E-12 |
| Usp29 | 8.14E-26 | 0.195981013 | 0.467 | 0.43 | 2.19E-21 | Lmbrd1 | 9.60E-17 | 0.030995481 | 0.467 | 0.425 | 2.58E-12 |
| Rapgef6 | 8.27E-26 | -0.099286007 | 0.667 | 0.731 | 2.22E-21 | Taf1d | 9.64E-17 | 0.031147187 | 0.261 | 0.226 | 2.59E-12 |
| St18 | 8.63E-26 | -0.153364932 | 0.216 | 0.275 | 2.32E-21 | Grm3 | 9.72E-17 | 0.189439478 | 0.411 | 0.365 | 2.62E-12 |
| Mapk8ip3 | 8.69E-26 | -0.071994425 | 0.466 | 0.53 | 2.34E-21 | Nrg3 | 1.00E-16 | -0.068238337 | 0.995 | 0.997 | 2.69E-12 |
| Hbs1l | 8.91E-26 | -0.058963263 | 0.324 | 0.38 | 2.40E-21 | Dnajc7 | 1.00E-16 | -0.002193604 | 0.424 | 0.392 | 2.70E-12 |
| Zcchc7 | 9.61E-26 | -0.114684195 | 0.841 | 0.882 | 2.58E-21 | Creg2 | 1.09E-16 | 0.023793658 | 0.256 | 0.225 | 2.94E-12 |
| Gsg1l | 9.67E-26 | 0.171653937 | 0.282 | 0.257 | 2.60E-21 | Fbxo41 | 1.10E-16 | -0.048825962 | 0.356 | 0.349 | 2.97E-12 |
| Tmem132d | 1.08E-25 | 0.236896974 | 0.321 | 0.273 | 2.90E-21 | Mapkap1 | 1.10E-16 | -0.020451367 | 0.54 | 0.514 | 2.97E-12 |
| Sema4d | 1.12E-25 | 0.118135222 | 0.329 | 0.309 | 3.02E-21 | Tead1 | 1.11E-16 | -0.039996706 | 0.484 | 0.456 | 2.98E-12 |
| Slc16a2 | 1.17E-25 | 0.137530163 | 0.438 | 0.414 | 3.15E-21 | Rabl6 | 1.13E-16 | 0.052035222 | 0.536 | 0.486 | 3.04E-12 |
| D5Ertd579e | 1.36E-25 | -0.084477752 | 0.419 | 0.484 | 3.67E-21 | Ino80dos | 1.19E-16 | 0.081974442 | 0.356 | 0.306 | 3.20E-12 |
| Capn15 | 1.37E-25 | -0.049095667 | 0.351 | 0.405 | 3.68E-21 | Tspan7 | 1.19E-16 | 0.031817692 | 0.597 | 0.55 | 3.20E-12 |
| Ube4b | 1.41E-25 | 0.120595887 | 0.491 | 0.471 | 3.79E-21 | Gm35188 | 1.23E-16 | 0.118639828 | 0.577 | 0.523 | 3.31E-12 |
| Atp1b1 | 1.41E-25 | -0.110224556 | 0.746 | 0.8 | 3.80E-21 | Psma1 | 1.27E-16 | 0.021239338 | 0.285 | 0.252 | 3.41E-12 |
| Wdr37 | 1.47E-25 | -0.078683611 | 0.378 | 0.441 | 3.94E-21 | Dstyk | 1.27E-16 | 0.044389659 | 0.367 | 0.324 | 3.41E-12 |
| Dym | 1.54E-25 | 0.082123929 | 0.545 | 0.546 | 4.14E-21 | Nfyc | 1.28E-16 | 0.079977611 | 0.366 | 0.315 | 3.43E-12 |
| Smap2 | 1.59E-25 | -0.019279565 | 0.326 | 0.37 | 4.28E-21 | Wdr37 | 1.28E-16 | 0.065324457 | 0.429 | 0.378 | 3.45E-12 |
| Cdc42bpb | 1.65E-25 | 0.108473974 | 0.316 | 0.296 | 4.44E-21 | Ywhaq | 1.33E-16 | 0.052381447 | 0.272 | 0.231 | 3.57E-12 |
| Kdm3b | 1.82E-25 | 0.084042349 | 0.309 | 0.305 | 4.88E-21 | D630045J12Rik | 1.36E-16 | -0.059059677 | 0.49 | 0.481 | 3.65E-12 |
| Txlng | 2.00E-25 | -0.070234135 | 0.34 | 0.4 | 5.39E-21 | Taz | 1.38E-16 | 0.035847615 | 0.269 | 0.232 | 3.70E-12 |
| Clvs2 | 2.02E-25 | -0.144445634 | 0.519 | 0.587 | 5.43E-21 | Gm13269 | 1.42E-16 | 0.03858882 | 0.561 | 0.513 | 3.82E-12 |
| A330008L17Rik | 2.05E-25 | 0.182673674 | 0.343 | 0.332 | 5.52E-21 | Ano6 | 1.45E-16 | -0.081445679 | 0.422 | 0.429 | 3.89E-12 |
| Dcaf7 | 2.14E-25 | 0.030844348 | 0.258 | 0.28 | 5.76E-21 | Ckap5 | 1.45E-16 | 0.042196609 | 0.506 | 0.458 | 3.91E-12 |
| Appl1 | 2.16E-25 | -0.0485002 | 0.414 | 0.47 | 5.82E-21 | Dhx15 | 1.51E-16 | 0.004198171 | 0.285 | 0.258 | 4.06E-12 |
| Adamts3 | 2.20E-25 | 0.120296055 | 0.324 | 0.308 | 5.93E-21 | Nup98 | 1.56E-16 | 0.063296543 | 0.443 | 0.392 | 4.19E-12 |
| Pcnx2 | 2.24E-25 | 0.120402949 | 0.482 | 0.465 | 6.02E-21 | Zfp933 | 1.56E-16 | 0.020277902 | 0.274 | 0.241 | 4.21E-12 |
| Ppp1cb | 2.36E-25 | -0.066687039 | 0.373 | 0.433 | 6.36E-21 | Zfp804a | 1.61E-16 | 0.091416404 | 0.645 | 0.591 | 4.33E-12 |
| Kif5a | 2.40E-25 | 0.134829045 | 0.39 | 0.354 | 6.45E-21 | 1700025G04Rik | 1.64E-16 | -0.052458924 | 0.538 | 0.527 | 4.40E-12 |
| Dnm1 | 2.46E-25 | 0.117616051 | 0.623 | 0.611 | 6.62E-21 | Orai2 | 1.64E-16 | 0.021101619 | 0.257 | 0.224 | 4.42E-12 |
| Hspa4 | 2.49E-25 | -0.102278503 | 0.386 | 0.452 | 6.70E-21 | B3gat1 | 1.70E-16 | -0.052604826 | 0.354 | 0.348 | 4.56E-12 |
| Mkln1os | 2.58E-25 | -0.078731517 | 0.212 | 0.265 | 6.95E-21 | Riok3 | 1.71E-16 | 0.02442784 | 0.431 | 0.39 | 4.60E-12 |
| Glis3 | 2.65E-25 | -0.282894078 | 0.326 | 0.368 | 7.14E-21 | Cep70 | 1.71E-16 | 0.051538573 | 0.513 | 0.463 | 4.61E-12 |
| Pkd1 | 2.66E-25 | 0.102184544 | 0.557 | 0.548 | 7.15E-21 | Atp13a3 | 1.71E-16 | -0.025984268 | 0.346 | 0.328 | 4.61E-12 |
| Lonp2 | 2.71E-25 | -0.064946429 | 0.421 | 0.481 | 7.29E-21 | Zfp106 | 1.73E-16 | 0.003015981 | 0.389 | 0.356 | 4.65E-12 |
| Fras1 | 2.78E-25 | 0.195321825 | 0.288 | 0.233 | 7.48E-21 | Zdhhc6 | 1.77E-16 | 0.090633385 | 0.291 | 0.243 | 4.76E-12 |
| Mical2 | 2.82E-25 | -0.124703326 | 0.815 | 0.857 | 7.58E-21 | Phactr3 | 1.81E-16 | -0.020347871 | 0.552 | 0.525 | 4.87E-12 |
| Snap23 | 3.03E-25 | -0.085583263 | 0.202 | 0.257 | 8.15E-21 | Slc2a3 | 1.83E-16 | 0.078131117 | 0.429 | 0.377 | 4.92E-12 |
| Fkbp5 | 3.24E-25 | -0.117976572 | 0.336 | 0.401 | 8.73E-21 | Zzef1 | 2.01E-16 | -0.009025954 | 0.508 | 0.479 | 5.41E-12 |
| Hnrnpu | 3.45E-25 | 0.111011597 | 0.73 | 0.725 | 9.27E-21 | Hectd2 | 2.14E-16 | 0.04592721 | 0.393 | 0.347 | 5.76E-12 |
| Rsbn1 | 3.54E-25 | -0.034516132 | 0.288 | 0.334 | 9.54E-21 | Vps13d | 2.18E-16 | -0.066967088 | 0.512 | 0.512 | 5.86E-12 |
| D630045J12Rik | 3.71E-25 | 0.058317336 | 0.481 | 0.497 | 9.99E-21 | Setdb1 | 2.18E-16 | 0.050878794 | 0.402 | 0.357 | 5.88E-12 |
| B230334C09Rik | 3.73E-25 | 0.066114051 | 0.446 | 0.457 | 1.00E-20 | Bclaf1 | 2.36E-16 | 0.066551175 | 0.549 | 0.496 | 6.35E-12 |
| Golga1 | 3.80E-25 | -0.061482652 | 0.294 | 0.347 | 1.02E-20 | Scn3a | 2.42E-16 | 0.050369549 | 0.507 | 0.458 | 6.51E-12 |
| Pten | 3.97E-25 | -0.09750741 | 0.547 | 0.614 | 1.07E-20 | Idh3b | 2.46E-16 | -0.004126516 | 0.258 | 0.236 | 6.63E-12 |
| Pkp2 | 4.04E-25 | -0.152109733 | 0.35 | 0.416 | 1.09E-20 | Nsmce2 | 2.55E-16 | -0.035591726 | 0.537 | 0.516 | 6.87E-12 |
| Dido1 | 4.11E-25 | -0.00395931 | 0.322 | 0.358 | 1.11E-20 | Itsn2 | 2.58E-16 | -0.059866384 | 0.583 | 0.577 | 6.93E-12 |
| Atp5c1 | 4.33E-25 | -0.028999031 | 0.22 | 0.261 | 1.17E-20 | Rps6kb1 | 2.59E-16 | 0.037611152 | 0.443 | 0.399 | 6.98E-12 |
| Immt | 4.37E-25 | -0.048437159 | 0.215 | 0.261 | 1.18E-20 | Mapk1ip1 | 2.63E-16 | 0.039373669 | 0.288 | 0.249 | 7.07E-12 |
| Nkain3 | 4.37E-25 | -0.231947459 | 0.664 | 0.705 | 1.18E-20 | Btbd1 | 2.66E-16 | 0.074891658 | 0.331 | 0.283 | 7.15E-12 |
| Ipo11 | 4.43E-25 | 0.060260934 | 0.424 | 0.435 | 1.19E-20 | Reps1 | 2.73E-16 | -0.025311281 | 0.331 | 0.315 | 7.34E-12 |
| Ndufs4 | 4.50E-25 | -0.050391316 | 0.409 | 0.465 | 1.21E-20 | Usp8 | 2.74E-16 | 0.014612374 | 0.298 | 0.266 | 7.37E-12 |
| Thsd7b | 4.71E-25 | -0.047214358 | 0.471 | 0.533 | 1.27E-20 | Foxk1 | 2.77E-16 | 0.046355402 | 0.345 | 0.302 | 7.45E-12 |
| Sptb | 4.81E-25 | 0.066127855 | 0.393 | 0.406 | 1.30E-20 | Dnajb4 | 2.82E-16 | 0.044081872 | 0.25 | 0.213 | 7.58E-12 |
| B3gat1 | 4.89E-25 | 0.095531932 | 0.348 | 0.341 | 1.32E-20 | Fam193b | 2.86E-16 | -0.044223811 | 0.334 | 0.329 | 7.69E-12 |
| Exoc3 | 5.31E-25 | -0.08014035 | 0.27 | 0.327 | 1.43E-20 | Paxbp1 | 2.99E-16 | 0.042227734 | 0.633 | 0.585 | 8.05E-12 |
| Pdzd2 | 5.35E-25 | -0.166435977 | 0.508 | 0.575 | 1.44E-20 | Caprin1 | 3.15E-16 | 0.05992463 | 0.404 | 0.356 | 8.47E-12 |
| Tspoap1 | 5.39E-25 | 0.122603851 | 0.484 | 0.461 | 1.45E-20 | Ift81 | 3.28E-16 | 0.067234693 | 0.295 | 0.25 | 8.82E-12 |
| Cpt1c | 5.59E-25 | -0.109644478 | 0.346 | 0.412 | 1.50E-20 | Zfp692 | 3.29E-16 | 0.026994234 | 0.369 | 0.331 | 8.86E-12 |
| Clk4 | 5.69E-25 | -0.123688225 | 0.422 | 0.49 | 1.53E-20 | Peak1 | 3.33E-16 | -0.127112378 | 0.427 | 0.458 | 8.96E-12 |
| Ino80 | 6.09E-25 | 0.051118615 | 0.479 | 0.497 | 1.64E-20 | Stx6 | 3.36E-16 | 0.059350325 | 0.384 | 0.337 | 9.03E-12 |
| Csnk1g3 | 6.17E-25 | -0.090853971 | 0.469 | 0.534 | 1.66E-20 | Bcr | 3.50E-16 | -0.090839035 | 0.368 | 0.383 | 9.43E-12 |
| Nrbp2 | 6.32E-25 | 0.134619009 | 0.252 | 0.208 | 1.70E-20 | Slc12a5 | 3.54E-16 | 0.04268393 | 0.55 | 0.504 | 9.52E-12 |
| Dctn1 | 6.35E-25 | -0.067945573 | 0.238 | 0.291 | 1.71E-20 | Nrp1 | 3.60E-16 | 0.054054826 | 0.732 | 0.685 | 9.68E-12 |
| Gm47167 | 6.50E-25 | -0.102377977 | 0.274 | 0.335 | 1.75E-20 | Speg | 3.74E-16 | -0.01956708 | 0.367 | 0.346 | 1.01E-11 |
| Phf3 | 6.65E-25 | -0.06163181 | 0.587 | 0.649 | 1.79E-20 | Rabep1 | 3.74E-16 | 0.055347167 | 0.695 | 0.647 | 1.01E-11 |
| Paxbp1 | 6.72E-25 | -0.110377451 | 0.585 | 0.651 | 1.81E-20 | Gps2 | 3.76E-16 | 0.005854927 | 0.281 | 0.254 | 1.01E-11 |
| Xpa | 7.15E-25 | -0.067671725 | 0.204 | 0.253 | 1.92E-20 | Pja2 | 3.81E-16 | 0.029486301 | 0.545 | 0.502 | 1.02E-11 |
| Cadm2 | 7.18E-25 | 0.168646343 | 0.974 | 0.974 | 1.93E-20 | Gtdc1 | 3.82E-16 | -0.075435068 | 0.655 | 0.657 | 1.03E-11 |
| Sorcs2 | 7.20E-25 | -0.114091981 | 0.376 | 0.441 | 1.94E-20 | Gm20696 | 3.82E-16 | -0.116239602 | 0.39 | 0.442 | 1.03E-11 |
| Sec24d | 7.77E-25 | -0.138189816 | 0.222 | 0.281 | 2.09E-20 | Fbxo11 | 4.01E-16 | 0.071649968 | 0.612 | 0.56 | 1.08E-11 |
| Prox1 | 8.22E-25 | -0.133612439 | 0.233 | 0.29 | 2.21E-20 | Taf1 | 4.06E-16 | -0.004264511 | 0.313 | 0.288 | 1.09E-11 |
| Ano6 | 8.61E-25 | 0.100531775 | 0.429 | 0.422 | 2.32E-20 | Gm26724 | 4.15E-16 | 0.067827063 | 0.33 | 0.282 | 1.12E-11 |
| Ccdc58 | 8.93E-25 | -0.077442482 | 0.237 | 0.291 | 2.40E-20 | Rev1 | 4.21E-16 | 0.026881527 | 0.487 | 0.446 | 1.13E-11 |
| Chuk | 9.27E-25 | -0.082027319 | 0.314 | 0.375 | 2.49E-20 | Cdk11b | 4.26E-16 | 0.031555152 | 0.328 | 0.29 | 1.15E-11 |
| Apobec4 | 9.33E-25 | -0.105269758 | 0.264 | 0.323 | 2.51E-20 | Mapk8 | 4.28E-16 | 0.056489167 | 0.426 | 0.378 | 1.15E-11 |
| Rlf | 1.02E-24 | -0.084868908 | 0.478 | 0.542 | 2.75E-20 | Osbpl8 | 4.31E-16 | 0.022839821 | 0.48 | 0.44 | 1.16E-11 |
| Grin2a | 1.04E-24 | -0.152482996 | 0.987 | 0.986 | 2.79E-20 | Zfp1 | 4.41E-16 | 0.028100091 | 0.329 | 0.291 | 1.19E-11 |
| Hacd3 | 1.06E-24 | -0.029634382 | 0.344 | 0.389 | 2.86E-20 | Rnf19a | 4.42E-16 | 0.04184059 | 0.368 | 0.326 | 1.19E-11 |
| Sorl1 | 1.09E-24 | -0.101298053 | 0.521 | 0.588 | 2.94E-20 | Spast | 4.62E-16 | 0.03049814 | 0.373 | 0.335 | 1.24E-11 |
| Rsf1 | 1.19E-24 | 0.078619336 | 0.512 | 0.518 | 3.19E-20 | Pum2 | 4.67E-16 | 0.012905523 | 0.394 | 0.361 | 1.26E-11 |
| 1700025G04Rik | 1.20E-24 | 0.098330887 | 0.527 | 0.527 | 3.23E-20 | Epha10 | 4.84E-16 | -0.118765716 | 0.296 | 0.317 | 1.30E-11 |
| Prickle1 | 1.21E-24 | -0.119213084 | 0.924 | 0.941 | 3.27E-20 | Bod1l | 5.06E-16 | -0.034653151 | 0.466 | 0.449 | 1.36E-11 |
| Usp19 | 1.22E-24 | -0.080909748 | 0.246 | 0.304 | 3.29E-20 | Uhrf2 | 5.12E-16 | 0.064016013 | 0.449 | 0.4 | 1.38E-11 |
| Ankrd13c | 1.24E-24 | -0.081161068 | 0.316 | 0.375 | 3.33E-20 | Creb1 | 5.12E-16 | 0.03351664 | 0.291 | 0.254 | 1.38E-11 |
| Hnrnpc | 1.24E-24 | -0.058264981 | 0.287 | 0.34 | 3.34E-20 | Rnf214 | 5.13E-16 | 0.028498233 | 0.296 | 0.261 | 1.38E-11 |
| Trip11 | 1.25E-24 | -0.011265612 | 0.395 | 0.438 | 3.36E-20 | Kcnc1 | 5.17E-16 | -5.74E-05 | 0.296 | 0.269 | 1.39E-11 |
| Scn1a | 1.31E-24 | 0.109164677 | 0.593 | 0.584 | 3.52E-20 | Zmiz1 | 5.26E-16 | -0.078062281 | 0.305 | 0.308 | 1.42E-11 |
| Hmgcr | 1.33E-24 | -0.117619634 | 0.197 | 0.254 | 3.57E-20 | Ptprk | 5.26E-16 | -0.12204687 | 0.704 | 0.733 | 1.42E-11 |

| | | | | | | | | | | | |
|---|---|---|---|---|---|---|---|---|---|---|---|
| Hlf | 1.36E-24 | 0.123775053 | 0.307 | 0.287 | 3.66E-20 | Nudcd3 | 5.35E-16 | -0.007689412 | 0.478 | 0.449 | 1.44E-11 |
| Phf12 | 1.39E-24 | 0.061941617 | 0.344 | 0.351 | 3.73E-20 | Rc3h2 | 5.40E-16 | 0.042073955 | 0.469 | 0.424 | 1.45E-11 |
| Sipa1l2 | 1.39E-24 | -0.111385258 | 0.223 | 0.279 | 3.74E-20 | Smarca1 | 5.53E-16 | -0.004852472 | 0.289 | 0.265 | 1.49E-11 |
| Yeats2 | 1.45E-24 | -0.032363097 | 0.4 | 0.451 | 3.91E-20 | Hnrnpa3 | 5.71E-16 | 0.057317527 | 0.292 | 0.248 | 1.54E-11 |
| Stt3b | 1.48E-24 | -0.068716822 | 0.304 | 0.36 | 3.98E-20 | Mia3 | 5.93E-16 | 0.062084691 | 0.307 | 0.263 | 1.60E-11 |
| Ccdc57 | 1.49E-24 | 0.112822973 | 0.291 | 0.271 | 4.02E-20 | Suz12 | 5.94E-16 | 0.089917219 | 0.3 | 0.252 | 1.60E-11 |
| Rnf216 | 1.49E-24 | 0.026124785 | 0.45 | 0.479 | 4.02E-20 | Faf2 | 5.95E-16 | 0.051162633 | 0.309 | 0.268 | 1.60E-11 |
| Inpp4b | 1.51E-24 | -0.117957315 | 0.466 | 0.533 | 4.07E-20 | Gmcl1 | 5.97E-16 | 0.026029738 | 0.459 | 0.417 | 1.61E-11 |
| Aste1 | 1.51E-24 | -0.071361198 | 0.201 | 0.25 | 4.08E-20 | Kdm5a | 6.16E-16 | 0.018274705 | 0.402 | 0.365 | 1.66E-11 |
| Camk2g | 1.59E-24 | 0.121724289 | 0.339 | 0.314 | 4.27E-20 | Ccnl1 | 6.39E-16 | 0.013939996 | 0.356 | 0.322 | 1.72E-11 |
| Bicral | 1.59E-24 | -0.029388047 | 0.287 | 0.33 | 4.29E-20 | Immt | 6.68E-16 | 0.035106371 | 0.25 | 0.215 | 1.80E-11 |
| Appbp2 | 1.63E-24 | 0.013477877 | 0.262 | 0.289 | 4.38E-20 | Exoc5 | 6.69E-16 | -0.029956388 | 0.353 | 0.338 | 1.80E-11 |
| Jazf1 | 1.65E-24 | 0.093549337 | 0.712 | 0.715 | 4.45E-20 | Nudt5 | 6.69E-16 | 0.048659682 | 0.347 | 0.304 | 1.80E-11 |
| Tnpo3 | 1.72E-24 | 0.029882406 | 0.399 | 0.425 | 4.63E-20 | Otud7b | 6.77E-16 | 0.05877227 | 0.315 | 0.272 | 1.82E-11 |
| Fam126b | 1.84E-24 | -0.07099503 | 0.446 | 0.508 | 4.96E-20 | Trmt1l | 6.80E-16 | 0.057147701 | 0.299 | 0.256 | 1.83E-11 |
| Echdc1 | 2.04E-24 | -0.09541739 | 0.211 | 0.267 | 5.48E-20 | Uggt1 | 7.03E-16 | -0.012497693 | 0.284 | 0.266 | 1.89E-11 |
| Sdcbp | 2.07E-24 | -0.040908205 | 0.252 | 0.296 | 5.57E-20 | Anp32a | 7.22E-16 | 0.044406499 | 0.288 | 0.249 | 1.94E-11 |
| Osbpl10 | 2.09E-24 | 0.156466525 | 0.36 | 0.326 | 5.62E-20 | Smc3 | 7.50E-16 | 0.072265975 | 0.36 | 0.312 | 2.02E-11 |
| Arfgef2 | 2.10E-24 | 0.125811154 | 0.362 | 0.331 | 5.66E-20 | Fmnl2 | 7.52E-16 | -0.095387327 | 0.645 | 0.649 | 2.02E-11 |
| Fhit | 2.16E-24 | 0.169618467 | 0.481 | 0.437 | 5.82E-20 | Crtc3 | 7.69E-16 | -0.00191694 | 0.504 | 0.471 | 2.07E-11 |
| Rufy3 | 2.43E-24 | -0.076212913 | 0.504 | 0.568 | 6.53E-20 | Thoc1 | 7.73E-16 | 0.049017763 | 0.387 | 0.341 | 2.08E-11 |
| Dync1h1 | 2.50E-24 | 0.074986067 | 0.484 | 0.489 | 6.74E-20 | Nr3c1 | 7.80E-16 | 0.065601114 | 0.405 | 0.356 | 2.10E-11 |
| Exoc6 | 2.64E-24 | 0.004144193 | 0.368 | 0.402 | 7.11E-20 | Sv2a | 7.86E-16 | -0.029914813 | 0.309 | 0.298 | 2.12E-11 |
| Pcmtd1 | 2.73E-24 | -0.080393753 | 0.475 | 0.538 | 7.36E-20 | Ntrk2 | 7.87E-16 | -0.117131317 | 0.825 | 0.808 | 2.12E-11 |
| Ppp1r14c | 2.74E-24 | 0.114355597 | 0.298 | 0.273 | 7.37E-20 | Fam13b | 8.22E-16 | 0.054873561 | 0.452 | 0.404 | 2.21E-11 |
| Gripap1 | 2.75E-24 | 0.046232447 | 0.367 | 0.385 | 7.41E-20 | 4932438A13Rik | 8.31E-16 | -0.077256999 | 0.578 | 0.582 | 2.24E-11 |
| Camkk1 | 2.81E-24 | -0.096423112 | 0.258 | 0.316 | 7.57E-20 | Lmtk3 | 8.42E-16 | -0.016885265 | 0.415 | 0.393 | 2.26E-11 |
| 4930599N23Rik | 2.87E-24 | -0.152200822 | 0.239 | 0.298 | 7.72E-20 | Frs2 | 8.58E-16 | 0.04002304 | 0.395 | 0.352 | 2.31E-11 |
| Virma | 2.88E-24 | -0.009933396 | 0.334 | 0.373 | 7.74E-20 | Kpna1 | 8.97E-16 | 0.043247285 | 0.352 | 0.31 | 2.41E-11 |
| Mob4 | 2.92E-24 | -0.081407509 | 0.233 | 0.288 | 7.85E-20 | Tox | 8.98E-16 | 0.275010245 | 0.324 | 0.286 | 2.42E-11 |
| Mfap3 | 3.01E-24 | -0.07785936 | 0.306 | 0.363 | 8.09E-20 | Slitrk1 | 9.00E-16 | 0.037312956 | 0.253 | 0.218 | 2.42E-11 |
| Rbm26 | 3.02E-24 | 0.097851751 | 0.737 | 0.736 | 8.12E-20 | Disp2 | 9.22E-16 | -0.000814212 | 0.371 | 0.343 | 2.48E-11 |
| Rab6b | 3.12E-24 | 0.070264571 | 0.478 | 0.483 | 8.40E-20 | Cdh10 | 9.43E-16 | 0.170855728 | 0.545 | 0.497 | 2.54E-11 |
| Mapre3 | 3.12E-24 | 0.013219057 | 0.428 | 0.465 | 8.40E-20 | Snd1 | 9.51E-16 | -0.026815756 | 0.579 | 0.555 | 2.56E-11 |
| Tbce | 3.23E-24 | -0.05764294 | 0.399 | 0.458 | 8.69E-20 | Igf1r | 9.74E-16 | -0.074281369 | 0.578 | 0.573 | 2.62E-11 |
| Ispd | 3.31E-24 | 0.128512347 | 0.27 | 0.212 | 8.90E-20 | Col4a2 | 1.00E-15 | -0.075684837 | 0.328 | 0.333 | 2.69E-11 |
| Arel1 | 3.65E-24 | -0.067747589 | 0.236 | 0.288 | 9.81E-20 | U2surp | 1.01E-15 | 0.069561311 | 0.511 | 0.459 | 2.71E-11 |
| Capn7 | 3.89E-24 | -0.082263366 | 0.267 | 0.325 | 1.05E-19 | Ncoa5 | 1.01E-15 | -0.042199031 | 0.407 | 0.394 | 2.72E-11 |
| Prpf39 | 3.95E-24 | -0.048504069 | 0.363 | 0.418 | 1.06E-19 | Fam133b | 1.01E-15 | 0.042744937 | 0.296 | 0.257 | 2.72E-11 |
| Agap3 | 4.00E-24 | 0.011043264 | 0.334 | 0.364 | 1.08E-19 | Fam126b | 1.02E-15 | 0.042383338 | 0.491 | 0.446 | 2.75E-11 |
| Elmo2 | 4.03E-24 | -0.084690196 | 0.25 | 0.306 | 1.09E-19 | Exoc3 | 1.05E-15 | 0.033789452 | 0.306 | 0.27 | 2.82E-11 |
| Gm26694 | 4.06E-24 | 0.016653808 | 0.565 | 0.512 | 1.09E-19 | Mboat2 | 1.06E-15 | -0.045426839 | 0.54 | 0.526 | 2.85E-11 |
| Cacng8 | 4.15E-24 | -0.106970455 | 0.486 | 0.552 | 1.12E-19 | Gpr107 | 1.07E-15 | 0.011106897 | 0.334 | 0.303 | 2.89E-11 |
| Ankrd10 | 4.44E-24 | -0.101855086 | 0.311 | 0.374 | 1.20E-19 | Nsmaf | 1.14E-15 | 0.040632024 | 0.347 | 0.306 | 3.07E-11 |
| Calm1 | 4.59E-24 | -0.076262832 | 0.396 | 0.461 | 1.24E-19 | Rrnad1 | 1.15E-15 | -0.001404566 | 0.284 | 0.259 | 3.08E-11 |
| Usp45 | 4.66E-24 | -0.044334825 | 0.397 | 0.449 | 1.25E-19 | Cacng3 | 1.15E-15 | 0.032642042 | 0.536 | 0.492 | 3.09E-11 |
| Kpna4 | 4.70E-24 | -0.087141263 | 0.431 | 0.495 | 1.27E-19 | Syt13 | 1.16E-15 | 0.012850114 | 0.34 | 0.309 | 3.11E-11 |
| Paip2 | 4.81E-24 | -0.092279396 | 0.462 | 0.527 | 1.29E-19 | Srrm2 | 1.17E-15 | -0.095835257 | 0.921 | 0.93 | 3.16E-11 |
| Pank2 | 4.81E-24 | -0.064419283 | 0.254 | 0.305 | 1.30E-19 | Arhgap32 | 1.18E-15 | -0.046101085 | 0.588 | 0.572 | 3.18E-11 |
| Sumo1 | 4.84E-24 | -0.032043942 | 0.358 | 0.406 | 1.30E-19 | Grik4 | 1.18E-15 | 0.035820287 | 0.613 | 0.566 | 3.19E-11 |
| Zc3h15 | 4.90E-24 | -0.062463883 | 0.303 | 0.357 | 1.32E-19 | Chuk | 1.20E-15 | 0.022580423 | 0.351 | 0.314 | 3.22E-11 |
| Evi5l | 5.23E-24 | 0.106276879 | 0.268 | 0.243 | 1.41E-19 | Nptn | 1.20E-15 | 0.092088753 | 0.897 | 0.874 | 3.22E-11 |
| Fus | 5.75E-24 | -0.097079219 | 0.689 | 0.749 | 1.55E-19 | Nlk | 1.20E-15 | 0.088485967 | 0.693 | 0.642 | 3.23E-11 |
| Lancl1 | 5.88E-24 | 0.031387853 | 0.277 | 0.297 | 1.58E-19 | Sh3bp5 | 1.21E-15 | -0.00849553 | 0.524 | 0.493 | 3.27E-11 |
| Rnf157 | 6.01E-24 | 0.070606453 | 0.586 | 0.597 | 1.62E-19 | Hnrnpd | 1.22E-15 | -0.008314172 | 0.253 | 0.233 | 3.29E-11 |
| Cd46 | 6.18E-24 | 0.070572558 | 0.366 | 0.37 | 1.66E-19 | Rab2a | 1.26E-15 | 0.037349725 | 0.557 | 0.514 | 3.40E-11 |
| Med27 | 6.78E-24 | 0.027507635 | 0.379 | 0.404 | 1.82E-19 | Srsf4 | 1.29E-15 | 0.023514581 | 0.277 | 0.246 | 3.47E-11 |
| Bckdhb | 6.86E-24 | 0.101563508 | 0.338 | 0.32 | 1.85E-19 | Glt1d1 | 1.30E-15 | 0.007245172 | 0.286 | 0.259 | 3.50E-11 |
| Rabl6 | 6.95E-24 | -0.025126657 | 0.486 | 0.537 | 1.87E-19 | Rnf10 | 1.30E-15 | 0.055629255 | 0.275 | 0.233 | 3.50E-11 |
| Snx27 | 7.05E-24 | 0.075215713 | 0.339 | 0.341 | 1.90E-19 | Sqstm1 | 1.31E-15 | 0.055240266 | 0.334 | 0.291 | 3.51E-11 |
| Rasgrf1 | 7.15E-24 | 0.127693686 | 0.899 | 0.884 | 1.92E-19 | Dapk1 | 1.31E-15 | -0.114431863 | 0.787 | 0.798 | 3.54E-11 |
| Snap47 | 7.16E-24 | 0.044263245 | 0.256 | 0.267 | 1.93E-19 | Trank1 | 1.32E-15 | -0.099979537 | 0.717 | 0.724 | 3.55E-11 |
| Gm20404 | 9.19E-24 | -0.052741805 | 0.284 | 0.334 | 2.47E-19 | Nav3 | 1.37E-15 | -0.089995549 | 0.977 | 0.984 | 3.69E-11 |
| Gm37240 | 9.31E-24 | -0.112661496 | 0.76 | 0.813 | 2.50E-19 | Tardbp | 1.38E-15 | 0.003356367 | 0.323 | 0.296 | 3.70E-11 |
| Pcf11 | 1.02E-23 | -0.022615036 | 0.224 | 0.261 | 2.73E-19 | Dpyd | 1.40E-15 | -0.154922953 | 0.575 | 0.6 | 3.76E-11 |
| Bicdl1 | 1.05E-23 | 0.09432281 | 0.518 | 0.512 | 2.82E-19 | Trip11 | 1.40E-15 | -0.02256434 | 0.417 | 0.395 | 3.76E-11 |
| Cul3 | 1.17E-23 | -0.046686184 | 0.342 | 0.393 | 3.15E-19 | Dnajc24 | 1.40E-15 | 0.021703911 | 0.259 | 0.227 | 3.77E-11 |
| Cog5 | 1.24E-23 | 0.094631098 | 0.607 | 0.606 | 3.34E-19 | D130040H23Rik | 1.43E-15 | 0.021443898 | 0.272 | 0.241 | 3.84E-11 |
| Rps6kb2 | 1.28E-23 | -0.074970857 | 0.198 | 0.25 | 3.44E-19 | Eprs | 1.46E-15 | 0.013747953 | 0.298 | 0.267 | 3.92E-11 |
| A330015K06Rik | 1.32E-23 | -0.221138936 | 0.634 | 0.678 | 3.55E-19 | Synrg | 1.48E-15 | 0.075146866 | 0.356 | 0.308 | 3.98E-11 |
| Pacs1 | 1.36E-23 | 0.123941945 | 0.522 | 0.5 | 3.65E-19 | Trip4 | 1.55E-15 | 0.054336527 | 0.269 | 0.229 | 4.17E-11 |
| Dhx15 | 1.39E-23 | -0.069327112 | 0.258 | 0.312 | 3.73E-19 | Ipo11 | 1.56E-15 | -0.060875528 | 0.427 | 0.424 | 4.19E-11 |
| Sppl3 | 1.44E-23 | 0.085860233 | 0.466 | 0.466 | 3.88E-19 | Smap2 | 1.56E-15 | 0.059618104 | 0.372 | 0.326 | 4.20E-11 |
| Nfasc | 1.48E-23 | -0.037504096 | 0.674 | 0.729 | 3.97E-19 | Fam171b | 1.65E-15 | 0.041703588 | 0.543 | 0.497 | 4.44E-11 |
| Akt3 | 1.48E-23 | 0.104744813 | 0.883 | 0.877 | 3.98E-19 | Zfp704 | 1.70E-15 | -0.095922225 | 0.477 | 0.489 | 4.57E-11 |
| Dhdds | 1.50E-23 | 0.090352998 | 0.373 | 0.367 | 4.04E-19 | Rsbn1l | 1.72E-15 | 0.048190471 | 0.449 | 0.403 | 4.64E-11 |
| Otud4 | 1.52E-23 | 0.093013713 | 0.282 | 0.27 | 4.10E-19 | Nova2 | 1.78E-15 | 0.040972081 | 0.427 | 0.383 | 4.80E-11 |
| Fbxo34 | 1.55E-23 | -0.026605128 | 0.438 | 0.488 | 4.18E-19 | 4833420G17Rik | 1.83E-15 | 0.063672474 | 0.348 | 0.302 | 4.93E-11 |
| Dtd1 | 1.66E-23 | 0.117495475 | 0.476 | 0.454 | 4.46E-19 | Cpd | 1.85E-15 | -0.014475963 | 0.335 | 0.314 | 4.97E-11 |
| Spock1 | 1.74E-23 | 0.239924894 | 0.426 | 0.37 | 4.68E-19 | Cep350 | 1.98E-15 | -0.040823886 | 0.479 | 0.463 | 5.33E-11 |
| Arfgef3 | 1.87E-23 | 0.104708275 | 0.621 | 0.615 | 5.04E-19 | Zcchc7 | 1.99E-15 | 0.079072763 | 0.876 | 0.841 | 5.37E-11 |
| Fto | 1.92E-23 | 0.101597058 | 0.725 | 0.719 | 5.17E-19 | Tshz2 | 2.05E-15 | 0.171533162 | 0.28 | 0.273 | 5.51E-11 |
| Gmeb1 | 2.01E-23 | -0.022060228 | 0.227 | 0.264 | 5.40E-19 | Mfap3 | 2.08E-15 | 0.013936026 | 0.337 | 0.306 | 5.61E-11 |
| Ubap2l | 2.02E-23 | -0.004012952 | 0.392 | 0.43 | 5.43E-19 | Fyn | 2.09E-15 | -0.066589147 | 0.584 | 0.585 | 5.61E-11 |
| Celf5 | 2.11E-23 | -0.096626289 | 0.476 | 0.541 | 5.67E-19 | Mtf2 | 2.11E-15 | 0.070758172 | 0.53 | 0.48 | 5.68E-11 |
| Map3k3 | 2.18E-23 | 0.069688912 | 0.49 | 0.498 | 5.87E-19 | Nfix | 2.13E-15 | 0.023582324 | 0.528 | 0.487 | 5.74E-11 |
| Immp1l | 2.21E-23 | -0.076560666 | 0.478 | 0.541 | 5.95E-19 | Gcc2 | 2.15E-15 | 0.013658944 | 0.393 | 0.359 | 5.80E-11 |
| Gnas | 2.21E-23 | 0.117560373 | 0.473 | 0.462 | 5.96E-19 | Gm11149 | 2.16E-15 | 0.008097629 | 0.498 | 0.459 | 5.82E-11 |
| Cx3cl1 | 2.24E-23 | -0.091995745 | 0.252 | 0.308 | 6.02E-19 | Olfm1 | 2.18E-15 | 0.053108534 | 0.642 | 0.592 | 5.87E-11 |
| Zfp280c | 2.33E-23 | 0.100737492 | 0.341 | 0.327 | 6.26E-19 | Smarca4 | 2.20E-15 | -0.00855487 | 0.464 | 0.436 | 5.91E-11 |
| 4930567K12Rik | 2.38E-23 | -0.117955359 | 0.267 | 0.327 | 6.40E-19 | Slc23a2 | 2.20E-15 | 0.037628108 | 0.51 | 0.466 | 5.92E-11 |
| Trappc9 | 2.42E-23 | 0.084692989 | 0.536 | 0.535 | 6.50E-19 | Dner | 2.21E-15 | -0.128628001 | 0.282 | 0.309 | 5.95E-11 |
| Yme1l1 | 2.61E-23 | -0.048394091 | 0.355 | 0.407 | 7.02E-19 | Gabra4 | 2.22E-15 | 0.060973307 | 0.516 | 0.466 | 5.98E-11 |
| Arhgap23 | 2.65E-23 | 0.118205207 | 0.288 | 0.263 | 7.12E-19 | Lonrf2 | 2.26E-15 | 0.018778339 | 0.308 | 0.276 | 6.09E-11 |
| Myrip | 2.71E-23 | 0.096124267 | 0.472 | 0.463 | 7.30E-19 | Etnk1 | 2.29E-15 | 0.047465859 | 0.674 | 0.627 | 6.17E-11 |
| Cdc37l1 | 2.83E-23 | -0.098619622 | 0.524 | 0.589 | 7.61E-19 | Kcnk2 | 2.30E-15 | 0.07289662 | 0.294 | 0.25 | 6.18E-11 |
| Rprd1a | 2.94E-23 | -0.001368632 | 0.398 | 0.434 | 7.90E-19 | Zfp652 | 2.42E-15 | 0.070612763 | 0.484 | 0.433 | 6.52E-11 |
| Zfp1 | 2.99E-23 | -0.044836114 | 0.291 | 0.34 | 8.05E-19 | Tax1bp1 | 2.51E-15 | 0.024895305 | 0.352 | 0.317 | 6.76E-11 |

| | | | | | | | | | | | |
|---|---|---|---|---|---|---|---|---|---|---|---|
| Map3k7 | 3.28E-23 | -0.004657047 | 0.273 | 0.305 | 8.84E-19 | Vti1b | 2.52E-15 | 0.028553353 | 0.334 | 0.297 | 6.78E-11 |
| Rps6kb1 | 3.61E-23 | 0.021731307 | 0.399 | 0.429 | 9.72E-19 | Med1 | 2.71E-15 | 0.061188645 | 0.255 | 0.213 | 7.28E-11 |
| Gm31763 | 3.78E-23 | -0.115978312 | 0.212 | 0.268 | 1.02E-18 | Akap10 | 2.91E-15 | 0.081710446 | 0.44 | 0.389 | 7.83E-11 |
| Vps50 | 3.90E-23 | 0.037607001 | 0.362 | 0.381 | 1.05E-18 | Ppp6r2 | 2.92E-15 | 0.02108898 | 0.557 | 0.518 | 7.86E-11 |
| Cacna2d1 | 3.91E-23 | 0.121266297 | 0.936 | 0.929 | 1.05E-18 | Slc8a2 | 2.94E-15 | 0.03948873 | 0.434 | 0.392 | 7.90E-11 |
| Myl12b | 3.91E-23 | -0.037352963 | 0.217 | 0.259 | 1.05E-18 | Fbxo34 | 2.95E-15 | 0.052060738 | 0.485 | 0.438 | 7.93E-11 |
| Usp48 | 3.93E-23 | 0.075944667 | 0.388 | 0.389 | 1.06E-18 | Ncs1 | 3.00E-15 | -0.004213901 | 0.31 | 0.285 | 8.07E-11 |
| Dgki | 4.00E-23 | 0.138760268 | 0.92 | 0.909 | 1.08E-18 | Trp53bp1 | 3.03E-15 | -0.04123602 | 0.413 | 0.401 | 8.15E-11 |
| Fktn | 4.07E-23 | -0.042337766 | 0.26 | 0.304 | 1.10E-18 | Pld3 | 3.04E-15 | 0.004672555 | 0.267 | 0.242 | 8.19E-11 |
| Ppme1 | 4.12E-23 | -0.086834057 | 0.343 | 0.403 | 1.11E-18 | Sf3b2 | 3.05E-15 | 0.041213665 | 0.318 | 0.278 | 8.22E-11 |
| Rapgef1 | 4.25E-23 | 0.069028469 | 0.41 | 0.417 | 1.14E-18 | Ice1 | 3.06E-15 | -0.0334643 | 0.46 | 0.443 | 8.24E-11 |
| Ncald | 4.28E-23 | 0.162890844 | 0.61 | 0.562 | 1.15E-18 | Terf1 | 3.10E-15 | 0.064689736 | 0.316 | 0.272 | 8.35E-11 |
| Scaf4 | 4.45E-23 | 0.046611094 | 0.349 | 0.362 | 1.20E-18 | Sp4 | 3.14E-15 | 0.000754067 | 0.265 | 0.24 | 8.44E-11 |
| Ppil2 | 4.52E-23 | -0.01601297 | 0.28 | 0.32 | 1.22E-18 | Ttc37 | 3.22E-15 | -0.037788793 | 0.328 | 0.319 | 8.67E-11 |
| Ncor2 | 4.57E-23 | -0.017582828 | 0.413 | 0.46 | 1.23E-18 | Arfgap1 | 3.28E-15 | 0.056361434 | 0.318 | 0.275 | 8.83E-11 |
| Tulp4 | 4.57E-23 | -0.137925739 | 0.528 | 0.593 | 1.23E-18 | Ncor2 | 3.30E-15 | 0.001985457 | 0.446 | 0.413 | 8.87E-11 |
| Gm15489 | 4.79E-23 | -0.132220258 | 0.233 | 0.288 | 1.29E-18 | Chd4 | 3.35E-15 | -0.038800367 | 0.439 | 0.426 | 9.02E-11 |
| Fbxo41 | 4.93E-23 | 0.090437877 | 0.349 | 0.339 | 1.33E-18 | Erlec1 | 3.41E-15 | -0.00945812 | 0.292 | 0.271 | 9.18E-11 |
| Clip1 | 5.16E-23 | 0.063000287 | 0.559 | 0.573 | 1.39E-18 | Plxnc1 | 3.48E-15 | -0.089674596 | 0.239 | 0.254 | 9.35E-11 |
| Atp2b1 | 5.22E-23 | -0.122606214 | 0.933 | 0.946 | 1.40E-18 | Ubac2 | 3.52E-15 | -0.048045338 | 0.43 | 0.422 | 9.46E-11 |
| Vps8 | 5.29E-23 | 0.066305086 | 0.345 | 0.348 | 1.42E-18 | Thoc2 | 3.52E-15 | 0.017813526 | 0.529 | 0.488 | 9.48E-11 |
| Lrch1 | 6.02E-23 | 0.05588436 | 0.494 | 0.512 | 1.62E-18 | Dido1 | 3.58E-15 | 0.033744336 | 0.36 | 0.322 | 9.63E-11 |
| Dlgap4 | 6.02E-23 | 0.091132103 | 0.589 | 0.59 | 1.62E-18 | Fam49a | 3.59E-15 | 0.012318881 | 0.507 | 0.47 | 9.67E-11 |
| Fndc3b | 6.86E-23 | 0.093665687 | 0.403 | 0.4 | 1.85E-18 | Sobp | 3.65E-15 | -0.095224619 | 0.87 | 0.88 | 9.82E-11 |
| Cspp1 | 6.90E-23 | -0.081236285 | 0.551 | 0.613 | 1.86E-18 | Rad54l2 | 3.76E-15 | 0.022301074 | 0.299 | 0.266 | 1.01E-10 |
| Stx16 | 7.04E-23 | -0.084512675 | 0.392 | 0.454 | 1.90E-18 | Xpo1 | 3.76E-15 | 0.029899187 | 0.329 | 0.293 | 1.01E-10 |
| Tmem63b | 7.12E-23 | 0.096683369 | 0.378 | 0.366 | 1.92E-18 | Fus | 3.82E-15 | 0.07925711 | 0.737 | 0.689 | 1.03E-10 |
| Slc38a1 | 7.19E-23 | 0.0820731 | 0.272 | 0.267 | 1.94E-18 | Btbd10 | 3.83E-15 | 0.053504217 | 0.431 | 0.384 | 1.03E-10 |
| Zfp950 | 7.23E-23 | -0.083110293 | 0.535 | 0.598 | 1.95E-18 | Cds2 | 3.95E-15 | 0.026710034 | 0.359 | 0.323 | 1.06E-10 |
| Tex2 | 7.38E-23 | 0.039537664 | 0.351 | 0.368 | 1.98E-18 | Dot1l | 3.96E-15 | -0.014203115 | 0.371 | 0.347 | 1.07E-10 |
| Susd4 | 7.94E-23 | 0.118434298 | 0.662 | 0.65 | 2.14E-18 | Sec14l1 | 3.97E-15 | 0.079433634 | 0.45 | 0.399 | 1.07E-10 |
| Rad17 | 8.27E-23 | -0.019974361 | 0.217 | 0.253 | 2.22E-18 | Lrpap1 | 4.01E-15 | 0.04492768 | 0.309 | 0.27 | 1.08E-10 |
| Nsmce2 | 8.30E-23 | 0.098465306 | 0.516 | 0.512 | 2.23E-18 | Ybx1 | 4.14E-15 | 0.023192733 | 0.277 | 0.245 | 1.11E-10 |
| Gm15577 | 8.33E-23 | -0.092237538 | 0.338 | 0.398 | 2.24E-18 | Actr1b | 4.15E-15 | -0.001801326 | 0.263 | 0.241 | 1.12E-10 |
| Dbn1 | 8.87E-23 | -0.095747297 | 0.521 | 0.586 | 2.39E-18 | Casc4 | 4.16E-15 | 0.068814721 | 0.65 | 0.6 | 1.12E-10 |
| Tmem63c | 8.92E-23 | 0.092968578 | 0.348 | 0.333 | 2.40E-18 | Hdac7 | 4.20E-15 | -0.080097896 | 0.267 | 0.279 | 1.13E-10 |
| Nob1 | 8.94E-23 | -0.053529582 | 0.214 | 0.259 | 2.41E-18 | Ankfy1 | 4.20E-15 | 0.021045739 | 0.359 | 0.324 | 1.13E-10 |
| Atp1a1 | 9.12E-23 | -0.101038518 | 0.387 | 0.451 | 2.45E-18 | Arid1a | 4.27E-15 | 0.031739634 | 0.364 | 0.327 | 1.15E-10 |
| Dock7 | 9.48E-23 | 0.062379554 | 0.547 | 0.56 | 2.55E-18 | Ddx55 | 4.27E-15 | 0.030081503 | 0.349 | 0.313 | 1.15E-10 |
| Eps15l1 | 9.58E-23 | 0.077986231 | 0.452 | 0.455 | 2.58E-18 | Casc3 | 4.28E-15 | 0.052867215 | 0.311 | 0.269 | 1.15E-10 |
| Phyhipl | 1.05E-22 | -0.094100797 | 0.263 | 0.319 | 2.84E-18 | Zfp950 | 4.32E-15 | 0.059828334 | 0.585 | 0.535 | 1.16E-10 |
| Gm12064 | 1.06E-22 | -0.066740413 | 0.317 | 0.372 | 2.85E-18 | Zfp318 | 4.37E-15 | -0.028842751 | 0.277 | 0.264 | 1.18E-10 |
| Fam117b | 1.07E-22 | -0.020190843 | 0.33 | 0.372 | 2.89E-18 | Stk38 | 4.52E-15 | 0.058630723 | 0.456 | 0.408 | 1.22E-10 |
| Dnajc13 | 1.10E-22 | 0.10673789 | 0.385 | 0.364 | 2.97E-18 | Baz1b | 4.52E-15 | -0.082370239 | 0.463 | 0.474 | 1.22E-10 |
| Exd2 | 1.11E-22 | -0.023554161 | 0.289 | 0.33 | 2.98E-18 | Psmd14 | 4.76E-15 | -0.039805836 | 0.501 | 0.487 | 1.28E-10 |
| 4933427D14Rik | 1.13E-22 | 0.1327314 | 0.303 | 0.248 | 3.04E-18 | Nsun7 | 4.84E-15 | -0.007882269 | 0.307 | 0.284 | 1.30E-10 |
| Map7 | 1.16E-22 | 0.121625605 | 0.665 | 0.639 | 3.12E-18 | Gm13963 | 4.91E-15 | 0.110229087 | 0.275 | 0.23 | 1.32E-10 |
| Arpc1a | 1.18E-22 | -0.003432938 | 0.308 | 0.342 | 3.16E-18 | Klf7 | 4.94E-15 | 0.083006667 | 0.354 | 0.306 | 1.33E-10 |
| Crtc1 | 1.19E-22 | 0.086725182 | 0.421 | 0.419 | 3.20E-18 | Smim13 | 5.03E-15 | 0.069527882 | 0.279 | 0.237 | 1.35E-10 |
| Tardbp | 1.21E-22 | 0.024777535 | 0.296 | 0.319 | 3.25E-18 | Gspt1 | 5.06E-15 | 0.065275941 | 0.348 | 0.303 | 1.36E-10 |
| D17Wsu92e | 1.25E-22 | 0.117215548 | 0.25 | 0.197 | 3.36E-18 | Cul3 | 5.07E-15 | 0.038291526 | 0.382 | 0.342 | 1.37E-10 |
| Kdm1a | 1.31E-22 | -0.066843454 | 0.426 | 0.483 | 3.53E-18 | Pip5k1c | 5.13E-15 | 0.005933817 | 0.267 | 0.241 | 1.38E-10 |
| Mef2d | 1.36E-22 | 0.060509348 | 0.378 | 0.386 | 3.65E-18 | Iqsec1 | 5.20E-15 | -0.083401286 | 0.726 | 0.719 | 1.40E-10 |
| Syt16 | 1.38E-22 | 0.116868967 | 0.702 | 0.68 | 3.73E-18 | Abca8b | 5.26E-15 | 0.052366655 | 0.32 | 0.279 | 1.41E-10 |
| Naxd | 1.43E-22 | -0.074367413 | 0.242 | 0.294 | 3.86E-18 | Asph | 5.29E-15 | 0.045229707 | 0.377 | 0.335 | 1.42E-10 |
| Arid1a | 1.46E-22 | 0.023891598 | 0.327 | 0.35 | 3.92E-18 | Myh10 | 5.30E-15 | -0.059525298 | 0.402 | 0.4 | 1.43E-10 |
| Insr | 1.46E-22 | 0.091849889 | 0.471 | 0.464 | 3.93E-18 | Pacsin1 | 5.31E-15 | 0.086607657 | 0.78 | 0.736 | 1.43E-10 |
| Ddhd2 | 1.50E-22 | 0.056541525 | 0.541 | 0.56 | 4.05E-18 | Dlst | 5.47E-15 | 0.01877469 | 0.291 | 0.259 | 1.47E-10 |
| Pcsk5 | 1.51E-22 | -0.20632244 | 0.418 | 0.476 | 4.05E-18 | Pura | 5.69E-15 | -0.023646634 | 0.459 | 0.44 | 1.53E-10 |
| Gm32250 | 1.59E-22 | -0.097486252 | 0.584 | 0.647 | 4.27E-18 | Arid5b | 5.69E-15 | 0.073919395 | 0.452 | 0.401 | 1.53E-10 |
| Dcc | 1.59E-22 | 0.24932262 | 0.754 | 0.727 | 4.28E-18 | Wapl | 5.74E-15 | 0.05388215 | 0.504 | 0.457 | 1.55E-10 |
| Agpat4 | 1.61E-22 | -0.09645304 | 0.399 | 0.461 | 4.33E-18 | Atp5a1 | 5.76E-15 | 0.047548385 | 0.38 | 0.337 | 1.55E-10 |
| D830024N08Rik | 1.61E-22 | -0.043657918 | 0.341 | 0.39 | 4.34E-18 | Neurl1a | 5.88E-15 | -0.016264294 | 0.36 | 0.338 | 1.58E-10 |
| Gcc2 | 1.65E-22 | -0.068434818 | 0.359 | 0.416 | 4.44E-18 | Phf6 | 5.96E-15 | 0.017554846 | 0.328 | 0.295 | 1.60E-10 |
| Bcl11a | 1.65E-22 | 0.053939464 | 0.726 | 0.753 | 4.45E-18 | Capn7 | 5.98E-15 | 0.028520038 | 0.302 | 0.267 | 1.61E-10 |
| Psd3 | 1.67E-22 | -0.150564804 | 0.673 | 0.73 | 4.50E-18 | Wipf2 | 6.28E-15 | 0.042778273 | 0.336 | 0.297 | 1.69E-10 |
| Dzip3 | 1.71E-22 | -0.060792015 | 0.459 | 0.517 | 4.59E-18 | Bin1 | 6.30E-15 | -0.059106085 | 0.493 | 0.49 | 1.69E-10 |
| Pnkd | 1.71E-22 | 0.09447815 | 0.311 | 0.298 | 4.61E-18 | Tnpo3 | 6.51E-15 | -0.011790567 | 0.422 | 0.399 | 1.75E-10 |
| Dcaf6 | 1.81E-22 | 0.105766509 | 0.67 | 0.663 | 4.87E-18 | Psen1 | 6.55E-15 | 0.012648669 | 0.314 | 0.283 | 1.76E-10 |
| Lrrtm4 | 1.82E-22 | 0.300414247 | 0.816 | 0.771 | 4.89E-18 | Frrs1l | 6.63E-15 | -0.002581976 | 0.382 | 0.355 | 1.78E-10 |
| Abca5 | 1.83E-22 | -0.012990881 | 0.323 | 0.362 | 4.94E-18 | Mapk14 | 6.69E-15 | 0.06010757 | 0.476 | 0.427 | 1.80E-10 |
| Rilpl1 | 2.02E-22 | 0.095802816 | 0.328 | 0.322 | 5.44E-18 | Mtch2 | 6.88E-15 | 0.038476952 | 0.321 | 0.282 | 1.85E-10 |
| Pou2f2 | 2.35E-22 | 0.063732773 | 0.286 | 0.294 | 6.32E-18 | Caln1 | 6.91E-15 | -0.102903337 | 0.797 | 0.812 | 1.86E-10 |
| Add1 | 2.36E-22 | 0.058678018 | 0.443 | 0.456 | 6.35E-18 | Prkar1a | 7.01E-15 | 0.034064956 | 0.28 | 0.243 | 1.89E-10 |
| Ahcyl1 | 2.39E-22 | -0.072946541 | 0.43 | 0.49 | 6.44E-18 | Tmem63b | 7.07E-15 | -0.049604721 | 0.386 | 0.378 | 1.90E-10 |
| Peli1 | 2.40E-22 | -0.038555772 | 0.321 | 0.368 | 6.45E-18 | Chordc1 | 7.11E-15 | -0.001006231 | 0.255 | 0.231 | 1.91E-10 |
| U2surp | 2.40E-22 | -0.068494761 | 0.459 | 0.519 | 6.46E-18 | Rab40b | 7.38E-15 | 0.064595921 | 0.411 | 0.363 | 1.99E-10 |
| Abr | 2.40E-22 | 0.11521477 | 0.769 | 0.757 | 6.46E-18 | Ajap1 | 7.43E-15 | -0.072412521 | 0.257 | 0.266 | 2.00E-10 |
| Smarca5 | 2.45E-22 | -0.075590494 | 0.292 | 0.347 | 6.58E-18 | Gnas | 7.43E-15 | -0.044112121 | 0.492 | 0.473 | 2.00E-10 |
| Ndfip1 | 2.52E-22 | -0.101569786 | 0.576 | 0.639 | 6.79E-18 | Smarcad1 | 7.54E-15 | 0.040658014 | 0.402 | 0.359 | 2.03E-10 |
| Hnrnpr | 2.82E-22 | -0.007202602 | 0.434 | 0.476 | 7.60E-18 | Celf1 | 7.78E-15 | 0.086841611 | 0.783 | 0.739 | 2.09E-10 |
| Vgll4 | 2.94E-22 | 0.118462897 | 0.329 | 0.318 | 7.91E-18 | Srgap2 | 7.94E-15 | -0.091282207 | 0.605 | 0.619 | 2.14E-10 |
| Ext2 | 2.99E-22 | 0.042733885 | 0.338 | 0.352 | 8.05E-18 | Ago1 | 8.07E-15 | 0.028974193 | 0.273 | 0.239 | 2.17E-10 |
| Stx5a | 3.02E-22 | -0.091781681 | 0.251 | 0.307 | 8.13E-18 | Papola | 8.17E-15 | 0.052168472 | 0.639 | 0.591 | 2.20E-10 |
| Zc3h14 | 3.08E-22 | -0.040175565 | 0.359 | 0.407 | 8.29E-18 | Bbx | 8.34E-15 | -0.078926181 | 0.335 | 0.344 | 2.24E-10 |
| Tcte2 | 3.12E-22 | 0.134589907 | 0.386 | 0.345 | 8.39E-18 | Rlf | 8.48E-15 | 0.022217063 | 0.518 | 0.478 | 2.28E-10 |
| Ube2d2a | 3.19E-22 | -0.074952284 | 0.347 | 0.404 | 8.58E-18 | Zmym5 | 8.70E-15 | 0.050181527 | 0.255 | 0.216 | 2.34E-10 |
| Inpp4a | 3.30E-22 | 0.093076203 | 0.482 | 0.479 | 8.88E-18 | Fam135a | 8.77E-15 | 0.060284408 | 0.423 | 0.376 | 2.36E-10 |
| Tom1l2 | 3.32E-22 | 0.104436451 | 0.649 | 0.642 | 8.93E-18 | Timm44 | 8.85E-15 | -0.024609566 | 0.293 | 0.279 | 2.38E-10 |
| Itfg1 | 3.38E-22 | 0.111139393 | 0.401 | 0.38 | 9.08E-18 | Nipa1 | 9.00E-15 | 0.030539195 | 0.396 | 0.357 | 2.42E-10 |
| Spata2 | 3.43E-22 | -0.078154464 | 0.209 | 0.259 | 9.23E-18 | Stag2 | 9.21E-15 | 0.049844691 | 0.363 | 0.321 | 2.48E-10 |
| Pdzd4 | 3.56E-22 | 0.085829058 | 0.538 | 0.539 | 9.58E-18 | Tlk2 | 9.22E-15 | 0.071659985 | 0.444 | 0.395 | 2.48E-10 |
| Smyd3 | 3.80E-22 | 0.14851598 | 0.727 | 0.707 | 1.02E-17 | Ahctf1 | 9.24E-15 | 0.04613552 | 0.349 | 0.308 | 2.49E-10 |
| Tmem59 | 3.82E-22 | -0.077722354 | 0.255 | 0.309 | 1.03E-17 | Fndc3a | 9.37E-15 | -0.002032333 | 0.387 | 0.358 | 2.52E-10 |
| Setd5 | 4.18E-22 | 0.072966581 | 0.594 | 0.603 | 1.12E-17 | Cab39l | 9.37E-15 | 0.011095792 | 0.32 | 0.291 | 2.52E-10 |
| Nup98 | 4.30E-22 | -0.049812102 | 0.392 | 0.444 | 1.16E-17 | Nfatc3 | 9.39E-15 | 0.083512531 | 0.285 | 0.24 | 2.53E-10 |
| Sgms1 | 4.35E-22 | 0.073594549 | 0.591 | 0.597 | 1.17E-17 | Pxdn | 9.44E-15 | -0.08494331 | 0.356 | 0.368 | 2.54E-10 |
| Ap2b1 | 4.40E-22 | -0.142170468 | 0.534 | 0.597 | 1.18E-17 | 6-Mar | 9.67E-15 | 0.033730512 | 0.479 | 0.438 | 2.60E-10 |

| | | | | | | | | | | | |
|---|---|---|---|---|---|---|---|---|---|---|---|
| Vapa | 4.51E-22 | -0.055745433 | 0.276 | 0.326 | 1.21E-17 | Sdha | 9.68E-15 | 0.010689391 | 0.301 | 0.271 | 2.60E-10 |
| Zranb1 | 4.66E-22 | -0.031648058 | 0.391 | 0.439 | 1.25E-17 | Phtf1 | 9.77E-15 | -0.043308873 | 0.393 | 0.386 | 2.63E-10 |
| Sun1 | 4.66E-22 | 0.109753217 | 0.393 | 0.37 | 1.25E-17 | Synj1 | 1.02E-14 | -0.079245377 | 0.585 | 0.594 | 2.75E-10 |
| Trip4 | 4.67E-22 | -0.050587367 | 0.229 | 0.274 | 1.26E-17 | Gm48321 | 1.03E-14 | -0.126479083 | 0.256 | 0.286 | 2.76E-10 |
| Spata5 | 4.74E-22 | -0.05795181 | 0.333 | 0.385 | 1.27E-17 | Naxd | 1.03E-14 | 0.02196042 | 0.272 | 0.242 | 2.77E-10 |
| Adcy8 | 4.80E-22 | 0.1513917 | 0.42 | 0.363 | 1.29E-17 | Ube4b | 1.05E-14 | -0.045839025 | 0.503 | 0.491 | 2.82E-10 |
| Zfand5 | 4.85E-22 | -0.038302756 | 0.295 | 0.341 | 1.30E-17 | Abca5 | 1.05E-14 | 0.007497162 | 0.355 | 0.323 | 2.84E-10 |
| Kif3a | 4.95E-22 | -0.096649843 | 0.396 | 0.458 | 1.33E-17 | Pten | 1.06E-14 | 0.037947224 | 0.591 | 0.547 | 2.86E-10 |
| Nin | 5.05E-22 | 0.098335403 | 0.602 | 0.595 | 1.36E-17 | Hmgn3 | 1.06E-14 | 0.003050258 | 0.295 | 0.268 | 2.86E-10 |
| Ggt7 | 5.15E-22 | -0.048739543 | 0.366 | 0.42 | 1.38E-17 | Usp48 | 1.07E-14 | -0.003716105 | 0.416 | 0.388 | 2.87E-10 |
| Rnf20 | 5.65E-22 | -0.054437529 | 0.24 | 0.288 | 1.52E-17 | Exoc2 | 1.07E-14 | -0.063455945 | 0.334 | 0.339 | 2.88E-10 |
| Sf3b6 | 6.50E-22 | -0.035287265 | 0.215 | 0.254 | 1.75E-17 | Ndrg3 | 1.07E-14 | 0.041552197 | 0.577 | 0.532 | 2.88E-10 |
| Nktr | 6.73E-22 | -0.062925877 | 0.659 | 0.716 | 1.81E-17 | Zfp346 | 1.07E-14 | 0.018308737 | 0.272 | 0.241 | 2.88E-10 |
| Gucy1a2 | 6.80E-22 | 0.093456401 | 0.756 | 0.753 | 1.83E-17 | Hnrnpu | 1.08E-14 | -0.057459108 | 0.741 | 0.73 | 2.90E-10 |
| Lrfn2 | 6.98E-22 | 0.096881001 | 0.689 | 0.688 | 1.88E-17 | Nr2c2 | 1.08E-14 | 0.037319995 | 0.507 | 0.463 | 2.92E-10 |
| Suds3 | 7.28E-22 | -0.078727318 | 0.324 | 0.381 | 1.96E-17 | Dgkb | 1.09E-14 | -0.140248136 | 0.726 | 0.753 | 2.93E-10 |
| Ralgps1 | 7.41E-22 | 0.082379276 | 0.69 | 0.692 | 1.99E-17 | Gm13402 | 1.11E-14 | 0.0846834 | 0.301 | 0.255 | 2.98E-10 |
| Wapl | 7.61E-22 | -0.069785805 | 0.457 | 0.516 | 2.05E-17 | Ube2d3 | 1.11E-14 | 0.040544029 | 0.342 | 0.301 | 2.98E-10 |
| Gm48765 | 7.72E-22 | -0.106602507 | 0.273 | 0.33 | 2.08E-17 | H13 | 1.12E-14 | -0.019478536 | 0.269 | 0.253 | 3.02E-10 |
| Grik5 | 7.78E-22 | 0.107417102 | 0.535 | 0.52 | 2.09E-17 | Rasal1 | 1.15E-14 | -0.085070875 | 0.275 | 0.29 | 3.08E-10 |
| Thrb | 7.95E-22 | 0.131976831 | 0.649 | 0.606 | 2.14E-17 | Tmem38a | 1.18E-14 | -0.01357853 | 0.317 | 0.297 | 3.16E-10 |
| Depdc5 | 8.24E-22 | 0.059632844 | 0.389 | 0.396 | 2.22E-17 | Smg7 | 1.18E-14 | 0.003322864 | 0.448 | 0.419 | 3.17E-10 |
| Smchd1 | 8.37E-22 | -0.025031966 | 0.38 | 0.424 | 2.25E-17 | Abhd18 | 1.23E-14 | 0.055512336 | 0.556 | 0.509 | 3.30E-10 |
| Map6 | 8.39E-22 | -0.024846223 | 0.417 | 0.464 | 2.26E-17 | Ddhd2 | 1.23E-14 | -0.002781571 | 0.573 | 0.541 | 3.32E-10 |
| Xrcc4 | 8.54E-22 | -0.0761317 | 0.329 | 0.384 | 2.30E-17 | Slc30a9 | 1.25E-14 | 0.009776133 | 0.28 | 0.253 | 3.36E-10 |
| Fyco1 | 8.91E-22 | -0.040047604 | 0.327 | 0.372 | 2.40E-17 | Fhit | 1.25E-14 | -0.120259077 | 0.451 | 0.481 | 3.37E-10 |
| Larp4b | 9.13E-22 | 0.047227345 | 0.445 | 0.461 | 2.46E-17 | Prpf40b | 1.27E-14 | -0.044437439 | 0.26 | 0.26 | 3.43E-10 |
| Xpo1 | 1.01E-21 | -0.084047518 | 0.293 | 0.349 | 2.72E-17 | Akap8 | 1.29E-14 | 0.021487108 | 0.392 | 0.356 | 3.47E-10 |
| Rabep1 | 1.03E-21 | 0.069757189 | 0.647 | 0.659 | 2.77E-17 | Ptbp3 | 1.30E-14 | 0.0636169 | 0.279 | 0.239 | 3.50E-10 |
| Epc1 | 1.04E-21 | -0.080603191 | 0.487 | 0.548 | 2.80E-17 | A330008L17Rik | 1.32E-14 | -0.127129763 | 0.334 | 0.343 | 3.55E-10 |
| Mapk1ip1 | 1.09E-21 | -0.106167881 | 0.249 | 0.306 | 2.92E-17 | Dgkg | 1.33E-14 | -0.113448934 | 0.815 | 0.824 | 3.57E-10 |
| Setdb1 | 1.09E-21 | 0.051179691 | 0.357 | 0.37 | 2.92E-17 | Cacng7 | 1.38E-14 | 0.057376696 | 0.276 | 0.236 | 3.72E-10 |
| Atp2b4 | 1.15E-21 | 0.13030503 | 0.266 | 0.247 | 3.08E-17 | Gm49353 | 1.40E-14 | -0.109081769 | 0.288 | 0.331 | 3.77E-10 |
| Fubp1 | 1.24E-21 | -0.09503618 | 0.629 | 0.69 | 3.34E-17 | Kdm3b | 1.40E-14 | 0.03973972 | 0.347 | 0.309 | 3.77E-10 |
| Ppm1b | 1.28E-21 | -0.024529154 | 0.26 | 0.298 | 3.45E-17 | Naa15 | 1.40E-14 | 0.058493668 | 0.404 | 0.359 | 3.78E-10 |
| Zcchc14 | 1.29E-21 | 0.085885771 | 0.328 | 0.32 | 3.46E-17 | Simc1 | 1.41E-14 | 0.055483313 | 0.256 | 0.218 | 3.78E-10 |
| Kcnab2 | 1.29E-21 | 0.062578711 | 0.421 | 0.431 | 3.47E-17 | Ap2a1 | 1.44E-14 | 0.055920199 | 0.28 | 0.24 | 3.86E-10 |
| Sptlc1 | 1.30E-21 | -0.068758203 | 0.232 | 0.282 | 3.49E-17 | Fbxw2 | 1.46E-14 | 0.040281055 | 0.313 | 0.276 | 3.94E-10 |
| Sf3b1 | 1.31E-21 | -0.088055669 | 0.602 | 0.663 | 3.51E-17 | Sppl2a | 1.46E-14 | 0.044883967 | 0.334 | 0.295 | 3.94E-10 |
| Sap130 | 1.41E-21 | 0.085908558 | 0.285 | 0.273 | 3.79E-17 | Ddx39b | 1.47E-14 | 0.030085038 | 0.354 | 0.318 | 3.94E-10 |
| Impact | 1.47E-21 | -0.070045495 | 0.237 | 0.289 | 3.96E-17 | 4921534H16Rik | 1.48E-14 | 0.039185459 | 0.555 | 0.508 | 3.99E-10 |
| Trak1 | 1.47E-21 | 0.063170756 | 0.426 | 0.431 | 3.96E-17 | Basp1 | 1.48E-14 | 0.070350332 | 0.598 | 0.548 | 3.99E-10 |
| Slmap | 1.47E-21 | 0.09975728 | 0.423 | 0.407 | 3.97E-17 | Mia2 | 1.49E-14 | 0.010256182 | 0.335 | 0.305 | 4.00E-10 |
| Cep350 | 1.48E-21 | 0.058696379 | 0.463 | 0.474 | 3.98E-17 | Wdfy3 | 1.50E-14 | -0.077887523 | 0.701 | 0.706 | 4.03E-10 |
| Naaladl2 | 1.48E-21 | 0.152908689 | 0.511 | 0.467 | 3.98E-17 | Dnajc13 | 1.50E-14 | -0.035195901 | 0.397 | 0.385 | 4.04E-10 |
| Arl15 | 1.50E-21 | -0.144242905 | 0.764 | 0.809 | 4.02E-17 | Uxs1 | 1.52E-14 | -0.064703154 | 0.323 | 0.327 | 4.08E-10 |
| Tmem175 | 1.58E-21 | -0.073258706 | 0.421 | 0.479 | 4.26E-17 | Eif4e | 1.52E-14 | 0.062052702 | 0.302 | 0.261 | 4.09E-10 |
| Psmd7 | 1.62E-21 | -0.084272188 | 0.27 | 0.324 | 4.35E-17 | Phf24 | 1.52E-14 | -0.045180486 | 0.62 | 0.605 | 4.09E-10 |
| Pura | 1.80E-21 | 0.084332442 | 0.44 | 0.435 | 4.83E-17 | Gm40841 | 1.54E-14 | 0.023895719 | 0.321 | 0.285 | 4.15E-10 |
| B3galt2 | 1.86E-21 | -0.048907928 | 0.331 | 0.381 | 5.00E-17 | Dnm1 | 1.56E-14 | -0.019679224 | 0.652 | 0.623 | 4.21E-10 |
| Pias1 | 1.87E-21 | 0.037398798 | 0.553 | 0.578 | 5.03E-17 | Supt6 | 1.61E-14 | -0.014914404 | 0.341 | 0.32 | 4.34E-10 |
| Mkx | 1.97E-21 | 0.071003975 | 0.246 | 0.256 | 5.29E-17 | Mtif2 | 1.64E-14 | 0.050794852 | 0.424 | 0.379 | 4.41E-10 |
| Susd6 | 2.01E-21 | -0.054820918 | 0.448 | 0.502 | 5.41E-17 | Gnl3l | 1.65E-14 | 0.02336511 | 0.286 | 0.253 | 4.43E-10 |
| Tmem87a | 2.06E-21 | -0.016030412 | 0.221 | 0.254 | 5.53E-17 | Gm10419 | 1.67E-14 | 0.016222041 | 0.316 | 0.284 | 4.50E-10 |
| Ubp1 | 2.09E-21 | 0.107280934 | 0.401 | 0.387 | 5.62E-17 | Vkorc1l1 | 1.72E-14 | 0.034514921 | 0.262 | 0.228 | 4.62E-10 |
| Stk39 | 2.18E-21 | 0.094640344 | 0.568 | 0.561 | 5.86E-17 | Atp5o | 1.74E-14 | -0.062939051 | 0.618 | 0.667 | 4.67E-10 |
| Ryk | 2.19E-21 | -0.02168307 | 0.335 | 0.375 | 5.88E-17 | Sorl1 | 1.74E-14 | -0.00861949 | 0.55 | 0.521 | 4.70E-10 |
| Togaram1 | 2.19E-21 | 0.014469969 | 0.285 | 0.31 | 5.89E-17 | Atp2b4 | 1.75E-14 | -0.091220054 | 0.257 | 0.266 | 4.71E-10 |
| Herc4 | 2.21E-21 | 0.042813811 | 0.36 | 0.375 | 5.93E-17 | Rasa1 | 1.76E-14 | 0.08710832 | 0.525 | 0.474 | 4.73E-10 |
| Yaf2 | 2.31E-21 | -0.003060342 | 0.323 | 0.357 | 6.21E-17 | Eno2 | 1.78E-14 | -0.027780235 | 0.298 | 0.287 | 4.79E-10 |
| Arhgap21 | 2.33E-21 | 0.121178283 | 0.718 | 0.706 | 6.28E-17 | Ndufs4 | 1.79E-14 | 0.038314612 | 0.451 | 0.409 | 4.83E-10 |
| Brwd1 | 2.40E-21 | 0.066010657 | 0.583 | 0.596 | 6.47E-17 | Xkr4 | 1.81E-14 | -0.102886738 | 0.946 | 0.951 | 4.86E-10 |
| Psma3 | 2.50E-21 | -0.083191651 | 0.366 | 0.424 | 6.74E-17 | Rhot1 | 1.83E-14 | 0.022029098 | 0.38 | 0.345 | 4.92E-10 |
| Dcun1d1 | 2.53E-21 | -0.060501901 | 0.211 | 0.257 | 6.80E-17 | Csde1 | 1.87E-14 | 0.02017296 | 0.333 | 0.3 | 5.03E-10 |
| Gm1976 | 2.54E-21 | -0.053497521 | 0.333 | 0.384 | 6.84E-17 | Sptbn1 | 1.87E-14 | -0.079997673 | 0.854 | 0.856 | 5.03E-10 |
| Kmt2a | 2.56E-21 | 0.092933532 | 0.721 | 0.72 | 6.88E-17 | Uvrag | 1.96E-14 | -0.060367848 | 0.6 | 0.593 | 5.28E-10 |
| Ppfia1 | 2.57E-21 | -0.043416895 | 0.337 | 0.384 | 6.91E-17 | Vps50 | 1.99E-14 | 0.0008314 | 0.39 | 0.362 | 5.35E-10 |
| Fam155a | 2.57E-21 | -0.108072184 | 0.989 | 0.987 | 6.93E-17 | Cipc | 1.99E-14 | -0.023383703 | 0.263 | 0.251 | 5.36E-10 |
| Ntng2 | 2.60E-21 | 0.123872342 | 0.396 | 0.379 | 6.99E-17 | Vsnl1 | 2.08E-14 | 0.104845677 | 0.582 | 0.531 | 5.60E-10 |
| Pde4d | 2.61E-21 | 0.155724717 | 0.796 | 0.766 | 7.01E-17 | Tcf20 | 2.10E-14 | -0.058091499 | 0.751 | 0.741 | 5.64E-10 |
| Atp2b3 | 2.65E-21 | 0.056387915 | 0.503 | 0.516 | 7.12E-17 | Glis3 | 2.10E-14 | 0.151733575 | 0.34 | 0.326 | 5.66E-10 |
| Pcmtd2 | 2.68E-21 | -0.043721476 | 0.25 | 0.295 | 7.21E-17 | Trim33 | 2.13E-14 | 0.021419015 | 0.484 | 0.446 | 5.72E-10 |
| Tacc1 | 2.85E-21 | -0.094014787 | 0.555 | 0.617 | 7.67E-17 | Tmem175 | 2.13E-14 | 0.03131845 | 0.461 | 0.421 | 5.72E-10 |
| Hdac7 | 2.88E-21 | 0.123639649 | 0.279 | 0.246 | 7.74E-17 | Ncoa3 | 2.15E-14 | 0.020622568 | 0.268 | 0.236 | 5.78E-10 |
| Cdc42 | 2.93E-21 | -0.090764488 | 0.399 | 0.459 | 7.88E-17 | Map4k5 | 2.15E-14 | -0.048036037 | 0.497 | 0.486 | 5.78E-10 |
| Nfix | 3.10E-21 | 0.089042428 | 0.487 | 0.485 | 8.34E-17 | Dync1li1 | 2.18E-14 | 0.046060551 | 0.333 | 0.294 | 5.85E-10 |
| Tmem117 | 3.15E-21 | 0.113558711 | 0.455 | 0.44 | 8.48E-17 | Ube2q2 | 2.24E-14 | 0.019942741 | 0.282 | 0.251 | 6.03E-10 |
| Rnf115 | 3.23E-21 | -0.057936643 | 0.303 | 0.353 | 8.68E-17 | Usp9x | 2.31E-14 | 0.063977631 | 0.614 | 0.565 | 6.21E-10 |
| Ttc4 | 3.23E-21 | -0.089040373 | 0.204 | 0.255 | 8.70E-17 | Mettl16 | 2.33E-14 | 0.057132972 | 0.311 | 0.268 | 6.26E-10 |
| Lztfl1 | 3.23E-21 | -0.100080543 | 0.35 | 0.409 | 8.70E-17 | Ctdspl | 2.38E-14 | -0.06779114 | 0.536 | 0.534 | 6.41E-10 |
| Trim24 | 3.24E-21 | -0.046433976 | 0.238 | 0.28 | 8.72E-17 | Grik5 | 2.40E-14 | -0.076310351 | 0.529 | 0.535 | 6.46E-10 |
| Rasal1 | 3.32E-21 | 0.116814281 | 0.29 | 0.262 | 8.94E-17 | Mast3 | 2.40E-14 | 0.09329751 | 0.62 | 0.57 | 6.46E-10 |
| Maml3 | 3.49E-21 | -0.176354202 | 0.39 | 0.451 | 9.39E-17 | Mycbp | 2.41E-14 | 0.053623808 | 0.309 | 0.267 | 6.47E-10 |
| Rab27b | 3.55E-21 | -0.099667056 | 0.304 | 0.361 | 9.55E-17 | Srsf1 | 2.44E-14 | 0.025203443 | 0.298 | 0.264 | 6.56E-10 |
| Xpnpep1 | 3.66E-21 | -0.059125693 | 0.222 | 0.269 | 9.85E-17 | Slc8a1 | 2.44E-14 | -0.104351789 | 0.746 | 0.766 | 6.58E-10 |
| Fam49a | 3.81E-21 | 0.06015893 | 0.47 | 0.484 | 1.03E-16 | Trpc1 | 2.45E-14 | 0.025201645 | 0.395 | 0.358 | 6.58E-10 |
| Gsdme | 3.87E-21 | -0.132161955 | 0.452 | 0.514 | 1.04E-16 | Dcun1d5 | 2.48E-14 | 0.030976757 | 0.303 | 0.268 | 6.68E-10 |
| Celf2 | 4.19E-21 | 0.070979997 | 0.998 | 0.997 | 1.13E-16 | Mfsd6 | 2.49E-14 | -0.006641245 | 0.295 | 0.272 | 6.71E-10 |
| Bsn | 4.26E-21 | 0.032200065 | 0.634 | 0.665 | 1.15E-16 | Gm21798 | 2.52E-14 | 0.099511124 | 0.335 | 0.287 | 6.79E-10 |
| Bach2 | 4.35E-21 | 0.154175001 | 0.421 | 0.385 | 1.17E-16 | Mpc1 | 2.52E-14 | 0.035339188 | 0.383 | 0.343 | 6.79E-10 |
| Ero1lb | 4.44E-21 | -0.085348099 | 0.202 | 0.251 | 1.19E-16 | Ranbp2 | 2.53E-14 | 0.060817204 | 0.45 | 0.405 | 6.82E-10 |
| Zzz3 | 4.51E-21 | -0.012287422 | 0.315 | 0.35 | 1.21E-16 | Capn15 | 2.55E-14 | 0.026496065 | 0.386 | 0.351 | 6.87E-10 |
| Sbno1 | 4.59E-21 | 0.021513239 | 0.481 | 0.511 | 1.23E-16 | Eps15 | 2.59E-14 | 0.025907852 | 0.273 | 0.241 | 6.97E-10 |
| Ywhae | 4.62E-21 | -0.01871037 | 0.345 | 0.387 | 1.24E-16 | Cdk13 | 2.65E-14 | -0.014294673 | 0.579 | 0.553 | 7.13E-10 |
| Macrod2os1 | 4.93E-21 | -0.118943329 | 0.392 | 0.454 | 1.33E-16 | Golgb1 | 2.68E-14 | -0.016927326 | 0.508 | 0.486 | 7.21E-10 |
| Efr3b | 5.27E-21 | 0.150421026 | 0.375 | 0.336 | 1.42E-16 | Rbm4b | 2.69E-14 | 0.08814575 | 0.4 | 0.35 | 7.24E-10 |
| Rab2a | 5.48E-21 | -0.044323983 | 0.514 | 0.566 | 1.48E-16 | Prrc2c | 2.69E-14 | -0.068693056 | 0.614 | 0.616 | 7.25E-10 |
| Ube2q2 | 5.62E-21 | 0.02947893 | 0.251 | 0.269 | 1.51E-16 | Acin1 | 2.74E-14 | 0.045415036 | 0.581 | 0.535 | 7.38E-10 |

| | | | | | | | | | | | |
|---|---|---|---|---|---|---|---|---|---|---|---|
| Gtf2h2 | 5.62E-21 | -0.031753782 | 0.24 | 0.279 | 1.51E-16 | Acbd5 | 2.75E-14 | 0.049495861 | 0.298 | 0.259 | 7.39E-10 |
| Sec61a2 | 5.65E-21 | -0.027537634 | 0.433 | 0.481 | 1.52E-16 | Orc3 | 2.77E-14 | 0.023911132 | 0.338 | 0.303 | 7.45E-10 |
| Dlg4 | 5.83E-21 | -0.033471137 | 0.431 | 0.48 | 1.57E-16 | Nfasc | 2.80E-14 | -0.033779786 | 0.694 | 0.674 | 7.53E-10 |
| Fndc9 | 5.84E-21 | -0.137406481 | 0.56 | 0.619 | 1.57E-16 | Nemf | 2.88E-14 | 0.020793559 | 0.405 | 0.37 | 7.75E-10 |
| Rmnd5a | 5.85E-21 | 0.022322371 | 0.35 | 0.376 | 1.57E-16 | Polr3a | 2.88E-14 | 0.02324761 | 0.251 | 0.222 | 7.75E-10 |
| Pkn2 | 5.91E-21 | -0.060732323 | 0.46 | 0.517 | 1.59E-16 | Uhrf1bp1l | 2.88E-14 | -0.001132817 | 0.352 | 0.326 | 7.76E-10 |
| Ppp2r2a | 5.97E-21 | -0.055417526 | 0.361 | 0.412 | 1.61E-16 | Sirpa | 2.90E-14 | -0.001735402 | 0.301 | 0.277 | 7.81E-10 |
| Phf20l1 | 6.08E-21 | -0.112474146 | 0.685 | 0.74 | 1.63E-16 | Mef2d | 2.94E-14 | 0.049942468 | 0.423 | 0.378 | 7.92E-10 |
| Celf3 | 6.26E-21 | -0.095534586 | 0.482 | 0.544 | 1.68E-16 | Pxk | 2.99E-14 | -0.061481773 | 0.297 | 0.299 | 8.03E-10 |
| Nme7 | 6.33E-21 | -0.074278063 | 0.498 | 0.556 | 1.70E-16 | Pdpk1 | 3.00E-14 | 0.040092241 | 0.378 | 0.337 | 8.06E-10 |
| Rad23b | 6.38E-21 | -0.014132701 | 0.311 | 0.348 | 1.72E-16 | Cdc37l1 | 3.04E-14 | -0.012239004 | 0.55 | 0.524 | 8.18E-10 |
| Erp44 | 6.38E-21 | -0.073091263 | 0.265 | 0.317 | 1.72E-16 | Slk | 3.06E-14 | 0.007948474 | 0.324 | 0.297 | 8.25E-10 |
| Nfx1 | 6.42E-21 | -0.030540689 | 0.493 | 0.544 | 1.73E-16 | Tfdp2 | 3.08E-14 | -0.035486269 | 0.487 | 0.468 | 8.30E-10 |
| Eya3 | 6.47E-21 | 0.048963097 | 0.304 | 0.314 | 1.74E-16 | Eps15l1 | 3.20E-14 | -0.025629539 | 0.472 | 0.452 | 8.60E-10 |
| Upf2 | 6.49E-21 | -0.031777531 | 0.402 | 0.449 | 1.75E-16 | Cdadc1 | 3.20E-14 | -0.039223352 | 0.309 | 0.301 | 8.62E-10 |
| Shc3 | 6.86E-21 | 0.139781602 | 0.291 | 0.252 | 1.85E-16 | Atcay | 3.22E-14 | 0.023357043 | 0.404 | 0.37 | 8.66E-10 |
| Acer3 | 7.17E-21 | 0.039898258 | 0.386 | 0.407 | 1.93E-16 | Dennd2a | 3.27E-14 | 0.024095853 | 0.345 | 0.31 | 8.79E-10 |
| Tmem178 | 7.19E-21 | -0.111977119 | 0.396 | 0.454 | 1.93E-16 | Fktn | 3.29E-14 | 0.000479339 | 0.282 | 0.26 | 8.84E-10 |
| Fry | 7.51E-21 | 0.096317377 | 0.89 | 0.885 | 2.02E-16 | Dync1h1 | 3.36E-14 | -0.044704441 | 0.494 | 0.484 | 9.04E-10 |
| Ntrk3 | 7.65E-21 | -0.137785095 | 0.911 | 0.917 | 2.06E-16 | Gmds | 3.38E-14 | -0.069983064 | 0.545 | 0.546 | 9.09E-10 |
| Rasgef1a | 7.71E-21 | 0.101273364 | 0.624 | 0.616 | 2.07E-16 | Fyco1 | 3.40E-14 | 0.004144205 | 0.353 | 0.327 | 9.15E-10 |
| 2610037D02Rik | 8.15E-21 | -0.006496724 | 0.363 | 0.4 | 2.19E-16 | Tet2 | 3.46E-14 | 0.002289381 | 0.352 | 0.324 | 9.30E-10 |
| Ap1b1 | 8.18E-21 | 0.086507103 | 0.281 | 0.267 | 2.20E-16 | Hacd3 | 3.50E-14 | 0.03881186 | 0.383 | 0.344 | 9.42E-10 |
| Clcn3 | 8.20E-21 | -0.074586969 | 0.417 | 0.474 | 2.21E-16 | Rnf20 | 3.55E-14 | -0.023241776 | 0.255 | 0.24 | 9.55E-10 |
| Safb | 8.39E-21 | 0.01641218 | 0.333 | 0.359 | 2.26E-16 | Bmt2 | 3.63E-14 | 0.053508042 | 0.408 | 0.365 | 9.78E-10 |
| Xpo7 | 8.44E-21 | 0.029578571 | 0.47 | 0.496 | 2.27E-16 | Cep170 | 3.68E-14 | -0.035943233 | 0.394 | 0.384 | 9.90E-10 |
| Fnbp1 | 8.50E-21 | 0.094448955 | 0.594 | 0.591 | 2.29E-16 | Cnot2 | 3.71E-14 | -0.010935533 | 0.523 | 0.493 | 9.99E-10 |
| Ptpn2 | 8.65E-21 | -0.048123979 | 0.224 | 0.267 | 2.33E-16 | Dctn4 | 3.72E-14 | 0.046191018 | 0.442 | 0.399 | 1.00E-09 |
| Sec62 | 8.84E-21 | -0.065906359 | 0.24 | 0.289 | 2.38E-16 | Depdc5 | 3.77E-14 | -0.010531472 | 0.413 | 0.389 | 1.02E-09 |
| Dnm1l | 9.18E-21 | -0.046916617 | 0.46 | 0.512 | 2.47E-16 | Yaf2 | 3.84E-14 | 0.050035464 | 0.364 | 0.323 | 1.03E-09 |
| Ccdc88c | 9.25E-21 | 0.134285952 | 0.314 | 0.282 | 2.49E-16 | Alkbh1 | 3.88E-14 | 0.067559125 | 0.262 | 0.221 | 1.04E-09 |
| Lrrtm1 | 9.54E-21 | -0.029311902 | 0.216 | 0.253 | 2.57E-16 | Tmx4 | 3.92E-14 | 0.028175404 | 0.424 | 0.388 | 1.06E-09 |
| Spon1 | 9.77E-21 | -0.146833883 | 0.46 | 0.521 | 2.63E-16 | Nmnat2 | 3.99E-14 | -0.091003871 | 0.658 | 0.67 | 1.07E-09 |
| Brinp2 | 1.03E-20 | 0.137362643 | 0.44 | 0.422 | 2.78E-16 | Trim2 | 4.04E-14 | -0.106005681 | 0.834 | 0.837 | 1.09E-09 |
| Fam118b | 1.04E-20 | -0.076581402 | 0.231 | 0.281 | 2.80E-16 | Ppp4r3a | 4.08E-14 | 0.010841476 | 0.335 | 0.305 | 1.10E-09 |
| Bicra | 1.08E-20 | 0.070731454 | 0.292 | 0.292 | 2.90E-16 | Ncl | 4.12E-14 | -0.014357136 | 0.261 | 0.243 | 1.11E-09 |
| Pbrm1 | 1.09E-20 | -0.040644672 | 0.39 | 0.439 | 2.93E-16 | 2610035D17Rik | 4.14E-14 | -0.007953651 | 0.493 | 0.465 | 1.11E-09 |
| Ppp3cc | 1.10E-20 | 0.060783619 | 0.26 | 0.265 | 2.95E-16 | Hnrnpk | 4.16E-14 | 0.053424183 | 0.27 | 0.23 | 1.12E-09 |
| Larp1 | 1.10E-20 | 0.030417518 | 0.473 | 0.494 | 2.96E-16 | Tpm3 | 4.18E-14 | 0.064178125 | 0.335 | 0.291 | 1.12E-09 |
| Synpo | 1.18E-20 | -0.002310725 | 0.322 | 0.355 | 3.18E-16 | Dnajc18 | 4.37E-14 | 0.042355153 | 0.286 | 0.249 | 1.18E-09 |
| Tmem135 | 1.22E-20 | 0.065907811 | 0.565 | 0.577 | 3.29E-16 | Matr3 | 4.38E-14 | 0.016740409 | 0.27 | 0.239 | 1.18E-09 |
| Zfp609 | 1.28E-20 | 0.085756082 | 0.547 | 0.546 | 3.43E-16 | Ppp2r2a | 4.41E-14 | 0.056016655 | 0.405 | 0.361 | 1.19E-09 |
| Srsf11 | 1.28E-20 | -0.084304435 | 0.7 | 0.755 | 3.46E-16 | Dagla | 4.49E-14 | -0.029936166 | 0.498 | 0.476 | 1.21E-09 |
| 9930021J03Rik | 1.29E-20 | 0.074238159 | 0.564 | 0.57 | 3.46E-16 | Helz | 4.52E-14 | -0.027831373 | 0.494 | 0.475 | 1.22E-09 |
| Zzef1 | 1.29E-20 | 0.02538318 | 0.479 | 0.507 | 3.47E-16 | Gm15398 | 4.60E-14 | 0.21106094 | 0.25 | 0.237 | 1.24E-09 |
| A530046M15Rik | 1.35E-20 | 0.124689926 | 0.289 | 0.258 | 3.63E-16 | Aco2 | 4.62E-14 | 0.028996777 | 0.449 | 0.409 | 1.24E-09 |
| Gm17227 | 1.37E-20 | -0.019154584 | 0.297 | 0.334 | 3.68E-16 | G3bp1 | 4.65E-14 | 0.00463811 | 0.277 | 0.252 | 1.25E-09 |
| Med14 | 1.39E-20 | -0.082032155 | 0.29 | 0.344 | 3.74E-16 | Crtac1 | 4.69E-14 | -0.045131864 | 0.464 | 0.451 | 1.26E-09 |
| Iqgap2 | 1.44E-20 | -0.167002196 | 0.635 | 0.684 | 3.86E-16 | 7-Mar | 4.74E-14 | 0.019272444 | 0.354 | 0.32 | 1.27E-09 |
| Tnks2 | 1.45E-20 | -0.049013575 | 0.36 | 0.41 | 3.91E-16 | Hp1bp3 | 4.79E-14 | -0.017679055 | 0.267 | 0.25 | 1.29E-09 |
| Pak1 | 1.52E-20 | -0.102711351 | 0.338 | 0.397 | 4.08E-16 | Zfp512 | 4.85E-14 | 0.071982872 | 0.251 | 0.21 | 1.31E-09 |
| Airn | 1.53E-20 | -0.164691091 | 0.271 | 0.326 | 4.12E-16 | Agpat4 | 4.85E-14 | 0.027616001 | 0.436 | 0.399 | 1.31E-09 |
| Ube2w | 1.53E-20 | -0.036969711 | 0.478 | 0.527 | 4.13E-16 | Gnl3 | 4.95E-14 | -0.015658399 | 0.274 | 0.257 | 1.33E-09 |
| Xrcc6 | 1.55E-20 | -0.106332541 | 0.472 | 0.534 | 4.16E-16 | Zdhhc20 | 5.07E-14 | 0.064913514 | 0.505 | 0.457 | 1.36E-09 |
| Vps37a | 1.58E-20 | -0.045849004 | 0.236 | 0.279 | 4.24E-16 | Ccdc58 | 5.07E-14 | 0.05596841 | 0.276 | 0.237 | 1.36E-09 |
| Gm12394 | 1.61E-20 | 0.070851789 | 0.558 | 0.497 | 4.34E-16 | Ube2e3 | 5.14E-14 | 0.047485025 | 0.311 | 0.272 | 1.38E-09 |
| Wdr70 | 1.66E-20 | -0.00030873 | 0.344 | 0.377 | 4.47E-16 | Ntan1 | 5.20E-14 | 0.068605917 | 0.262 | 0.22 | 1.40E-09 |
| Mycbp | 1.70E-20 | -0.094082151 | 0.267 | 0.322 | 4.57E-16 | Arhgef7 | 5.32E-14 | -0.040239533 | 0.517 | 0.504 | 1.43E-09 |
| Glp2r | 1.72E-20 | -0.148015842 | 0.53 | 0.581 | 4.62E-16 | Pdlim7 | 5.39E-14 | 0.040688823 | 0.307 | 0.268 | 1.45E-09 |
| Grik3 | 1.76E-20 | 0.123216173 | 0.302 | 0.299 | 4.73E-16 | Plec | 5.44E-14 | -0.006003939 | 0.335 | 0.312 | 1.46E-09 |
| Lamc1 | 1.79E-20 | -0.108488668 | 0.324 | 0.383 | 4.81E-16 | Raf1 | 5.54E-14 | 0.063756176 | 0.455 | 0.407 | 1.49E-09 |
| Drosha | 1.83E-20 | 0.060156931 | 0.25 | 0.249 | 4.93E-16 | Pgbd5 | 5.56E-14 | 0.067044337 | 0.553 | 0.503 | 1.50E-09 |
| BC005561 | 1.86E-20 | -0.097789671 | 0.454 | 0.514 | 5.00E-16 | Usp7 | 5.66E-14 | -0.030327867 | 0.363 | 0.347 | 1.52E-09 |
| Pard3 | 1.87E-20 | -0.149159359 | 0.503 | 0.563 | 5.04E-16 | Chrna7 | 5.71E-14 | -0.045932197 | 0.409 | 0.393 | 1.54E-09 |
| Acin1 | 1.89E-20 | -0.06053193 | 0.535 | 0.592 | 5.09E-16 | Dcakd | 5.75E-14 | 0.04598809 | 0.256 | 0.219 | 1.55E-09 |
| Gdpd1 | 1.96E-20 | -0.002326522 | 0.299 | 0.33 | 5.27E-16 | Ccdc148 | 6.27E-14 | -0.087102068 | 0.461 | 0.475 | 1.69E-09 |
| Parp6 | 1.98E-20 | -0.042398655 | 0.295 | 0.341 | 5.32E-16 | Slc1a1 | 6.30E-14 | -0.006493631 | 0.551 | 0.524 | 1.69E-09 |
| Ppt1 | 1.98E-20 | -0.056507184 | 0.356 | 0.408 | 5.34E-16 | Gnb1 | 6.34E-14 | 0.050348849 | 0.536 | 0.49 | 1.71E-09 |
| Socs7 | 2.05E-20 | 0.048746655 | 0.259 | 0.267 | 5.51E-16 | Nvl | 6.35E-14 | 0.012539332 | 0.65 | 0.613 | 1.71E-09 |
| Scmh1 | 2.06E-20 | 0.069749798 | 0.609 | 0.619 | 5.54E-16 | B3galt2 | 6.39E-14 | 0.017237742 | 0.364 | 0.331 | 1.72E-09 |
| Gtf2f2 | 2.06E-20 | -0.034760203 | 0.288 | 0.329 | 5.54E-16 | Daam1 | 6.51E-14 | -0.077762136 | 0.46 | 0.47 | 1.75E-09 |
| Numa1 | 2.06E-20 | 0.021525433 | 0.346 | 0.372 | 5.55E-16 | Pkig | 6.83E-14 | 0.066507488 | 0.489 | 0.441 | 1.84E-09 |
| Tecpr2 | 2.08E-20 | 0.085834147 | 0.352 | 0.345 | 5.60E-16 | Phf12 | 7.15E-14 | 0.044893241 | 0.384 | 0.344 | 1.92E-09 |
| D130040H23Rik | 2.09E-20 | -0.023390265 | 0.241 | 0.276 | 5.62E-16 | Nrip1 | 7.38E-14 | -0.017451385 | 0.304 | 0.286 | 1.99E-09 |
| Rab40b | 2.15E-20 | 0.026182477 | 0.363 | 0.388 | 5.78E-16 | Zbtb11 | 7.46E-14 | 0.070200242 | 0.413 | 0.366 | 2.01E-09 |
| Sptbn1 | 2.19E-20 | 0.099634321 | 0.856 | 0.85 | 5.90E-16 | Usp47 | 7.47E-14 | 0.028174465 | 0.451 | 0.413 | 2.01E-09 |
| Sfpq | 2.22E-20 | -0.079452577 | 0.696 | 0.751 | 5.98E-16 | Ubxn4 | 7.48E-14 | 0.057381783 | 0.446 | 0.402 | 2.01E-09 |
| Sumo2 | 2.33E-20 | -0.054666599 | 0.296 | 0.346 | 6.27E-16 | Mms19 | 7.50E-14 | 0.020842808 | 0.258 | 0.229 | 2.02E-09 |
| Tspan3 | 2.35E-20 | -0.053451763 | 0.229 | 0.274 | 6.33E-16 | Ric3 | 7.66E-14 | -0.008477395 | 0.292 | 0.272 | 2.06E-09 |
| Dhx30 | 2.42E-20 | 0.061559206 | 0.52 | 0.532 | 6.52E-16 | Usp25 | 7.71E-14 | -0.029530364 | 0.346 | 0.33 | 2.07E-09 |
| Ddx24 | 2.50E-20 | -0.067952036 | 0.467 | 0.523 | 6.72E-16 | Ube2h | 7.81E-14 | 0.007018715 | 0.498 | 0.464 | 2.10E-09 |
| Pias2 | 2.52E-20 | -0.052630635 | 0.524 | 0.577 | 6.78E-16 | Tnks2 | 7.90E-14 | 0.074125169 | 0.408 | 0.36 | 2.13E-09 |
| Dnajc5 | 2.56E-20 | -0.014866487 | 0.279 | 0.314 | 6.89E-16 | Nek1 | 7.98E-14 | -0.054867468 | 0.395 | 0.393 | 2.15E-09 |
| Acss2 | 2.58E-20 | -0.059043151 | 0.286 | 0.334 | 6.94E-16 | Hdgfl3 | 8.02E-14 | 0.034977313 | 0.484 | 0.443 | 2.16E-09 |
| Fnbp4 | 2.60E-20 | 0.067085766 | 0.364 | 0.369 | 7.00E-16 | Lnpep | 8.08E-14 | 0.00673423 | 0.428 | 0.399 | 2.17E-09 |
| Bcr | 2.71E-20 | 0.067094786 | 0.383 | 0.386 | 7.29E-16 | Chd1 | 8.19E-14 | 0.0294804 | 0.275 | 0.242 | 2.20E-09 |
| Qrich1 | 2.77E-20 | 0.027379856 | 0.449 | 0.475 | 7.46E-16 | Ube2w | 8.27E-14 | 0.022748563 | 0.517 | 0.478 | 2.23E-09 |
| Wdr20 | 2.78E-20 | 0.045295811 | 0.29 | 0.3 | 7.48E-16 | Mapkbp1 | 8.75E-14 | 0.013125787 | 0.262 | 0.236 | 2.35E-09 |
| Fbxw11 | 2.84E-20 | -0.077978019 | 0.438 | 0.495 | 7.64E-16 | Chfr | 8.83E-14 | 0.018767506 | 0.531 | 0.494 | 2.38E-09 |
| Acbd5 | 2.91E-20 | -0.020040213 | 0.259 | 0.295 | 7.83E-16 | Pcmt1 | 8.98E-14 | 0.041675202 | 0.422 | 0.381 | 2.42E-09 |
| Dtnb | 2.93E-20 | 0.109192516 | 0.771 | 0.763 | 7.89E-16 | Txndc11 | 9.32E-14 | 0.058832903 | 0.353 | 0.31 | 2.51E-09 |
| Pkd2 | 3.02E-20 | 0.038032198 | 0.242 | 0.255 | 8.11E-16 | Pign | 9.35E-14 | -0.088749893 | 0.28 | 0.305 | 2.51E-09 |
| Dnajc24 | 3.12E-20 | -0.025618774 | 0.227 | 0.263 | 8.40E-16 | Specc1l | 9.52E-14 | -0.013342977 | 0.386 | 0.365 | 2.56E-09 |
| Csmd2 | 3.15E-20 | -0.171179616 | 0.81 | 0.84 | 8.48E-16 | Foxp1 | 9.62E-14 | 0.10585053 | 0.681 | 0.634 | 2.59E-09 |
| Dnajc7 | 3.18E-20 | 0.041205298 | 0.392 | 0.41 | 8.55E-16 | Cul2 | 9.82E-14 | -0.013064182 | 0.283 | 0.265 | 2.64E-09 |
| Rab3ip | 3.31E-20 | -0.014064348 | 0.262 | 0.296 | 8.90E-16 | Ppfia1 | 1.01E-13 | 0.018010296 | 0.369 | 0.337 | 2.72E-09 |
| Cul2 | 3.34E-20 | -0.036187196 | 0.265 | 0.306 | 8.99E-16 | Shprh | 1.03E-13 | -0.004071188 | 0.433 | 0.408 | 2.76E-09 |
| Pcbp3 | 3.37E-20 | 0.100642996 | 0.374 | 0.358 | 9.08E-16 | Elavl3 | 1.03E-13 | 0.073477257 | 0.483 | 0.433 | 2.78E-09 |

| | | | | | | | | | | | |
|---|---|---|---|---|---|---|---|---|---|---|---|
| Dcun1d5 | 3.42E-20 | -0.043828852 | 0.268 | 0.313 | 9.20E-16 | Rsrp1 | 1.04E-13 | -0.04751564 | 0.67 | 0.716 | 2.80E-09 |
| Ckap5 | 3.45E-20 | 0.047749475 | 0.458 | 0.476 | 9.27E-16 | Gpc4 | 1.04E-13 | 0.060846781 | 0.263 | 0.225 | 2.81E-09 |
| Nub1 | 3.52E-20 | 0.082864538 | 0.304 | 0.293 | 9.48E-16 | Mark1 | 1.06E-13 | -0.057558842 | 0.39 | 0.389 | 2.86E-09 |
| Parp1 | 3.52E-20 | -0.091497895 | 0.236 | 0.289 | 9.48E-16 | Actr1a | 1.07E-13 | 0.03866043 | 0.259 | 0.225 | 2.87E-09 |
| Ube2g1 | 3.54E-20 | 0.071571517 | 0.454 | 0.461 | 9.53E-16 | Ogfod1 | 1.07E-13 | 0.012500994 | 0.3 | 0.27 | 2.87E-09 |
| Ssb | 3.63E-20 | -0.03863502 | 0.248 | 0.29 | 9.76E-16 | Exoc6 | 1.08E-13 | -0.003499117 | 0.393 | 0.368 | 2.90E-09 |
| Chp1 | 3.66E-20 | -0.05158505 | 0.245 | 0.289 | 9.84E-16 | Stk3 | 1.08E-13 | -0.049311719 | 0.458 | 0.45 | 2.91E-09 |
| Snx10 | 3.78E-20 | 0.111058548 | 0.381 | 0.35 | 1.02E-15 | Usp19 | 1.08E-13 | 0.047526687 | 0.285 | 0.246 | 2.91E-09 |
| Cenpc1 | 3.82E-20 | 0.025599267 | 0.246 | 0.264 | 1.03E-15 | Kcnq5 | 1.09E-13 | -0.033214471 | 0.583 | 0.623 | 2.93E-09 |
| Eri3 | 4.04E-20 | 0.075540793 | 0.463 | 0.463 | 1.09E-15 | Sel1l | 1.10E-13 | 0.001409201 | 0.399 | 0.372 | 2.95E-09 |
| Uhrf1bp1l | 4.22E-20 | -0.037319069 | 0.326 | 0.369 | 1.13E-15 | Rmnd5a | 1.14E-13 | 0.026859542 | 0.386 | 0.35 | 3.07E-09 |
| Lncpint | 4.26E-20 | 0.093273453 | 0.924 | 0.916 | 1.15E-15 | Kdm5b | 1.15E-13 | 0.034007686 | 0.455 | 0.414 | 3.09E-09 |
| Ift81 | 4.27E-20 | 0.063594735 | 0.25 | 0.248 | 1.15E-15 | Fam155a | 1.16E-13 | -0.070630276 | 0.986 | 0.989 | 3.11E-09 |
| 6530403H02Rik | 4.31E-20 | -0.198246352 | 0.326 | 0.384 | 1.16E-15 | Wnk1 | 1.16E-13 | -0.046070781 | 0.552 | 0.544 | 3.13E-09 |
| Cnot1 | 4.37E-20 | -0.006983947 | 0.433 | 0.471 | 1.18E-15 | Orc4 | 1.17E-13 | 0.033298094 | 0.402 | 0.364 | 3.15E-09 |
| Vps53 | 4.38E-20 | 0.039968518 | 0.267 | 0.275 | 1.18E-15 | Prelid3a | 1.18E-13 | 0.036988495 | 0.425 | 0.384 | 3.16E-09 |
| Xkr4 | 4.54E-20 | 0.127175801 | 0.951 | 0.943 | 1.22E-15 | Tle4 | 1.18E-13 | 0.075027818 | 0.428 | 0.381 | 3.18E-09 |
| Arhgef4 | 4.57E-20 | -0.112122552 | 0.28 | 0.336 | 1.23E-15 | AW554918 | 1.21E-13 | -0.064065309 | 0.427 | 0.428 | 3.27E-09 |
| Dennd5a | 4.70E-20 | 0.012432788 | 0.332 | 0.358 | 1.27E-15 | Rab11fip4 | 1.23E-13 | -0.017779395 | 0.437 | 0.415 | 3.30E-09 |
| 2610020C07Rik | 4.80E-20 | -0.069665739 | 0.386 | 0.442 | 1.29E-15 | Fxr1 | 1.24E-13 | 0.013038719 | 0.46 | 0.426 | 3.35E-09 |
| Ppfia2 | 4.82E-20 | 0.157319838 | 0.951 | 0.941 | 1.30E-15 | Gria1 | 1.27E-13 | -0.101949589 | 0.915 | 0.933 | 3.41E-09 |
| Glmn | 4.98E-20 | -0.008136841 | 0.265 | 0.297 | 1.34E-15 | Heatr3 | 1.27E-13 | 0.019304132 | 0.271 | 0.241 | 3.41E-09 |
| Wdfy3 | 5.06E-20 | 0.072465787 | 0.706 | 0.713 | 1.36E-15 | Ube4a | 1.27E-13 | 0.019180352 | 0.341 | 0.31 | 3.41E-09 |
| Psmd1 | 5.08E-20 | -0.013295213 | 0.354 | 0.392 | 1.37E-15 | Wasf3 | 1.27E-13 | 0.062475611 | 0.415 | 0.369 | 3.41E-09 |
| Usp8 | 5.21E-20 | -0.031848368 | 0.266 | 0.306 | 1.40E-15 | Eml6 | 1.29E-13 | -0.068689078 | 0.848 | 0.842 | 3.46E-09 |
| Ric8b | 5.23E-20 | 0.031809321 | 0.298 | 0.314 | 1.41E-15 | Galnt14 | 1.29E-13 | -0.044167459 | 0.375 | 0.361 | 3.47E-09 |
| Apba1 | 5.51E-20 | 0.100311263 | 0.733 | 0.717 | 1.48E-15 | Ptpa | 1.32E-13 | 5.20E-05 | 0.28 | 0.255 | 3.56E-09 |
| Tbc1d22a | 5.56E-20 | 0.050399145 | 0.318 | 0.327 | 1.50E-15 | Catspere2 | 1.33E-13 | -0.096412907 | 0.409 | 0.43 | 3.57E-09 |
| Arhgef3 | 5.75E-20 | -0.095201659 | 0.353 | 0.41 | 1.55E-15 | Polr3h | 1.35E-13 | 0.076521838 | 0.32 | 0.275 | 3.63E-09 |
| Olfm1 | 5.77E-20 | -0.11697377 | 0.592 | 0.651 | 1.55E-15 | Map3k13 | 1.36E-13 | 0.018440751 | 0.306 | 0.275 | 3.65E-09 |
| Cdk13 | 5.80E-20 | -0.00882629 | 0.553 | 0.594 | 1.56E-15 | Eif2ak4 | 1.36E-13 | 0.018969962 | 0.291 | 0.262 | 3.65E-09 |
| Tax1bp1 | 6.04E-20 | -0.056148078 | 0.317 | 0.366 | 1.62E-15 | Sf1 | 1.36E-13 | 0.016551262 | 0.255 | 0.227 | 3.66E-09 |
| Pus10 | 6.11E-20 | 0.000573582 | 0.279 | 0.307 | 1.64E-15 | Dcaf17 | 1.37E-13 | 0.019228466 | 0.322 | 0.29 | 3.68E-09 |
| Stx8 | 6.28E-20 | 0.025236453 | 0.385 | 0.41 | 1.69E-15 | Wdr47 | 1.39E-13 | 0.043091528 | 0.367 | 0.328 | 3.74E-09 |
| Prpf40a | 6.39E-20 | -0.05116744 | 0.446 | 0.498 | 1.72E-15 | Sik2 | 1.45E-13 | -0.057545513 | 0.43 | 0.426 | 3.91E-09 |
| Uri1 | 6.39E-20 | 0.091354736 | 0.336 | 0.324 | 1.72E-15 | Man2a2 | 1.48E-13 | -0.035430172 | 0.339 | 0.327 | 3.97E-09 |
| Prkd1 | 6.56E-20 | 0.019876443 | 0.407 | 0.445 | 1.77E-15 | Auh | 1.48E-13 | 0.011777177 | 0.346 | 0.317 | 3.98E-09 |
| Ndufs1 | 6.57E-20 | -0.013895973 | 0.303 | 0.34 | 1.77E-15 | Angel2 | 1.50E-13 | 0.049884126 | 0.304 | 0.265 | 4.04E-09 |
| Kif13b | 6.61E-20 | -0.017977339 | 0.358 | 0.398 | 1.78E-15 | Dock7 | 1.57E-13 | -0.043190609 | 0.557 | 0.547 | 4.22E-09 |
| Ccser2 | 7.06E-20 | 0.035566217 | 0.338 | 0.354 | 1.90E-15 | Drg1 | 1.58E-13 | 0.036775714 | 0.616 | 0.571 | 4.25E-09 |
| Etl4 | 7.28E-20 | 0.243414276 | 0.582 | 0.527 | 1.96E-15 | Itgb3bp | 1.60E-13 | 0.064238518 | 0.34 | 0.297 | 4.31E-09 |
| Cfdp1 | 7.39E-20 | -0.005322485 | 0.279 | 0.31 | 1.99E-15 | Wasl | 1.60E-13 | 0.051756175 | 0.469 | 0.425 | 4.31E-09 |
| Bbof1 | 7.48E-20 | -0.087802778 | 0.337 | 0.394 | 2.01E-15 | Sh3gl2 | 1.60E-13 | -0.08700439 | 0.775 | 0.777 | 4.32E-09 |
| B230307C23Rik | 7.50E-20 | 0.005626901 | 0.251 | 0.279 | 2.02E-15 | Rap1gap | 1.63E-13 | 0.036084433 | 0.386 | 0.348 | 4.39E-09 |
| Tpm1 | 7.67E-20 | -0.0537404 | 0.254 | 0.3 | 2.06E-15 | Prpf39 | 1.65E-13 | 0.006380566 | 0.394 | 0.363 | 4.44E-09 |
| Nup214 | 7.83E-20 | 0.027127229 | 0.273 | 0.293 | 2.11E-15 | Tnpo1 | 1.65E-13 | 0.0452524 | 0.571 | 0.527 | 4.44E-09 |
| Tnr | 7.90E-20 | -0.145862411 | 0.863 | 0.882 | 2.12E-15 | Dock9 | 1.66E-13 | -0.07562951 | 0.729 | 0.728 | 4.46E-09 |
| Ints10 | 7.90E-20 | 0.001351577 | 0.327 | 0.358 | 2.13E-15 | Pnpla8 | 1.68E-13 | -0.014331392 | 0.333 | 0.313 | 4.51E-09 |
| Sh3d19 | 8.00E-20 | 0.127264924 | 0.438 | 0.398 | 2.15E-15 | D430042O09Rik | 1.68E-13 | -0.033412247 | 0.468 | 0.452 | 4.52E-09 |
| Atg16l1 | 8.04E-20 | 0.002926868 | 0.353 | 0.382 | 2.16E-15 | Dcaf10 | 1.70E-13 | 0.017719526 | 0.253 | 0.225 | 4.56E-09 |
| Taf1 | 8.07E-20 | 0.012878145 | 0.288 | 0.311 | 2.17E-15 | AC129186.1 | 1.71E-13 | -0.122266253 | 0.355 | 0.403 | 4.61E-09 |
| Ctdspl2 | 8.45E-20 | -0.021384795 | 0.345 | 0.384 | 2.27E-15 | Arhgef28 | 1.73E-13 | -0.058836489 | 0.426 | 0.418 | 4.66E-09 |
| Chrm3 | 8.45E-20 | 0.024380319 | 0.516 | 0.467 | 2.27E-15 | Ergic1 | 1.73E-13 | -0.032872673 | 0.356 | 0.344 | 4.67E-09 |
| Nudt3 | 8.83E-20 | 0.024400386 | 0.284 | 0.303 | 2.38E-15 | Nrbp2 | 1.77E-13 | -0.046651353 | 0.254 | 0.252 | 4.78E-09 |
| Mtmr7 | 8.87E-20 | 0.048740925 | 0.313 | 0.324 | 2.39E-15 | Ccnt1 | 1.78E-13 | 0.063928758 | 0.31 | 0.267 | 4.79E-09 |
| Xpo6 | 9.25E-20 | 0.058536011 | 0.304 | 0.307 | 2.49E-15 | Plk2 | 1.78E-13 | 0.07153955 | 0.268 | 0.227 | 4.79E-09 |
| Arih2 | 9.35E-20 | -0.039710125 | 0.284 | 0.326 | 2.52E-15 | Ints8 | 1.80E-13 | 0.022940114 | 0.287 | 0.256 | 4.84E-09 |
| Zfr2 | 9.43E-20 | 0.022470689 | 0.326 | 0.349 | 2.54E-15 | Lrp11 | 1.82E-13 | 0.005874754 | 0.312 | 0.286 | 4.91E-09 |
| Abcb7 | 9.46E-20 | -0.00293056 | 0.234 | 0.262 | 2.55E-15 | Tmem131 | 1.83E-13 | -0.041937446 | 0.515 | 0.504 | 4.93E-09 |
| Lrch3 | 9.56E-20 | 0.016586656 | 0.447 | 0.477 | 2.57E-15 | Camsap1 | 1.83E-13 | -0.016552664 | 0.251 | 0.237 | 4.94E-09 |
| Ccdc73 | 9.64E-20 | -0.039679515 | 0.252 | 0.294 | 2.59E-15 | Gm28376 | 1.84E-13 | 0.141404658 | 0.782 | 0.766 | 4.95E-09 |
| Ptpn11 | 9.92E-20 | 0.004564052 | 0.285 | 0.312 | 2.67E-15 | Eya3 | 1.84E-13 | 0.025150122 | 0.338 | 0.304 | 4.95E-09 |
| Clip2 | 1.04E-19 | 0.070437444 | 0.268 | 0.262 | 2.79E-15 | Tmem150c | 1.85E-13 | -0.035540287 | 0.427 | 0.413 | 4.98E-09 |
| Catspere2 | 1.04E-19 | 0.111628421 | 0.43 | 0.409 | 2.79E-15 | Cc2d2a | 1.91E-13 | 0.002284703 | 0.257 | 0.234 | 5.13E-09 |
| Gm27153 | 1.04E-19 | -0.088520841 | 0.255 | 0.306 | 2.81E-15 | Caap1 | 1.95E-13 | 0.045362492 | 0.266 | 0.229 | 5.24E-09 |
| Ltn1 | 1.06E-19 | -0.056203639 | 0.206 | 0.25 | 2.86E-15 | Zfand3 | 1.96E-13 | -0.036783396 | 0.685 | 0.668 | 5.27E-09 |
| Sf3b3 | 1.08E-19 | -0.083143412 | 0.319 | 0.374 | 2.90E-15 | Add3 | 1.97E-13 | 0.057101097 | 0.448 | 0.404 | 5.29E-09 |
| Lmbrd1 | 1.09E-19 | 0.060385501 | 0.425 | 0.432 | 2.92E-15 | Asxl2 | 2.04E-13 | 0.010253305 | 0.495 | 0.46 | 5.48E-09 |
| Nrf1 | 1.09E-19 | -0.059822622 | 0.34 | 0.391 | 2.93E-15 | Snx29 | 2.05E-13 | -0.061825327 | 0.442 | 0.444 | 5.53E-09 |
| Unc13b | 1.10E-19 | 0.0993799 | 0.462 | 0.454 | 2.95E-15 | Nsg2 | 2.06E-13 | 0.057968109 | 0.526 | 0.479 | 5.56E-09 |
| Lurap1l | 1.13E-19 | -0.119283191 | 0.297 | 0.353 | 3.03E-15 | Med27 | 2.08E-13 | -0.001121594 | 0.406 | 0.379 | 5.59E-09 |
| Ybx1 | 1.13E-19 | -0.046306415 | 0.245 | 0.288 | 3.05E-15 | Tspan5 | 2.09E-13 | -0.090699479 | 0.794 | 0.791 | 5.62E-09 |
| Zfp973 | 1.17E-19 | 0.120372577 | 0.405 | 0.348 | 3.14E-15 | Mecr | 2.10E-13 | -0.051582952 | 0.441 | 0.437 | 5.64E-09 |
| Rabgap1 | 1.20E-19 | 0.055092482 | 0.639 | 0.651 | 3.22E-15 | Golga1 | 2.12E-13 | 0.031427753 | 0.328 | 0.294 | 5.71E-09 |
| Myh10 | 1.21E-19 | 0.07311434 | 0.4 | 0.399 | 3.24E-15 | Cdh9 | 2.14E-13 | 0.181577639 | 0.514 | 0.491 | 5.75E-09 |
| Rfx7 | 1.23E-19 | 0.087182194 | 0.567 | 0.563 | 3.30E-15 | Kcnb2 | 2.21E-13 | -0.102180851 | 0.884 | 0.892 | 5.93E-09 |
| Usp33 | 1.23E-19 | -0.024503517 | 0.416 | 0.46 | 3.30E-15 | Zwint | 2.23E-13 | -0.007794124 | 0.368 | 0.344 | 6.00E-09 |
| Rab6a | 1.28E-19 | -0.091422846 | 0.344 | 0.401 | 3.44E-15 | Rufy3 | 2.25E-13 | 0.020734469 | 0.542 | 0.504 | 6.05E-09 |
| Smap1 | 1.31E-19 | -0.08960254 | 0.416 | 0.474 | 3.53E-15 | Cdk8 | 2.26E-13 | -0.029435023 | 0.62 | 0.599 | 6.09E-09 |
| Atp6v0d1 | 1.35E-19 | -0.02069468 | 0.239 | 0.274 | 3.63E-15 | Ttc14 | 2.30E-13 | 0.056638503 | 0.777 | 0.735 | 6.19E-09 |
| Palm | 1.40E-19 | 0.061346327 | 0.348 | 0.354 | 3.77E-15 | Ddb2 | 2.30E-13 | 0.063837978 | 0.265 | 0.225 | 6.20E-09 |
| Afdn | 1.40E-19 | -0.084457825 | 0.675 | 0.73 | 3.78E-15 | Elmod1 | 2.34E-13 | 0.069855222 | 0.749 | 0.705 | 6.31E-09 |
| Mras | 1.44E-19 | 0.047689892 | 0.324 | 0.336 | 3.88E-15 | Chd8 | 2.35E-13 | -0.017224214 | 0.419 | 0.401 | 6.32E-09 |
| Helz | 1.45E-19 | 0.062102053 | 0.475 | 0.484 | 3.90E-15 | Rnf216 | 2.35E-13 | -0.023893052 | 0.468 | 0.45 | 6.33E-09 |
| Ahctf1 | 1.47E-19 | -0.018520491 | 0.308 | 0.344 | 3.94E-15 | Rab40c | 2.36E-13 | 0.016130673 | 0.425 | 0.392 | 6.35E-09 |
| Txndc11 | 1.50E-19 | -0.043310564 | 0.31 | 0.355 | 4.03E-15 | Zranb1 | 2.38E-13 | 0.049423259 | 0.435 | 0.391 | 6.39E-09 |
| Smad4 | 1.50E-19 | -0.026217154 | 0.244 | 0.279 | 4.04E-15 | Phlpp1 | 2.38E-13 | -0.067758164 | 0.607 | 0.605 | 6.40E-09 |
| Kcnq3 | 1.52E-19 | 0.104374169 | 0.938 | 0.921 | 4.10E-15 | Rsu1 | 2.39E-13 | -0.083558707 | 0.245 | 0.266 | 6.43E-09 |
| Atcay | 1.54E-19 | 0.070818046 | 0.37 | 0.368 | 4.14E-15 | Ube2r2 | 2.42E-13 | 0.066374643 | 0.412 | 0.366 | 6.50E-09 |
| Apba2 | 1.55E-19 | 0.101310134 | 0.808 | 0.799 | 4.18E-15 | Eri3 | 2.43E-13 | -0.049105847 | 0.47 | 0.463 | 6.53E-09 |
| Ppp4r3a | 1.56E-19 | -0.005293881 | 0.305 | 0.339 | 4.21E-15 | Rap1gds1 | 2.45E-13 | -0.069938997 | 0.574 | 0.58 | 6.60E-09 |
| Pbx1 | 1.59E-19 | 0.11729463 | 0.875 | 0.871 | 4.28E-15 | Pdxdc1 | 2.49E-13 | 0.034728391 | 0.33 | 0.292 | 6.71E-09 |
| Unc13a | 1.63E-19 | 0.0806716 | 0.565 | 0.567 | 4.39E-15 | Mbd2 | 2.51E-13 | 0.0341414 | 0.324 | 0.289 | 6.77E-09 |
| Sacm1l | 1.63E-19 | -0.042854771 | 0.399 | 0.447 | 4.40E-15 | Grik3 | 2.57E-13 | -0.044227945 | 0.322 | 0.302 | 6.90E-09 |
| Thoc2 | 1.65E-19 | -0.047243404 | 0.488 | 0.541 | 4.43E-15 | Dennd6a | 2.62E-13 | 0.051299884 | 0.407 | 0.363 | 7.05E-09 |
| Ascc3 | 1.65E-19 | -0.047819274 | 0.605 | 0.657 | 4.45E-15 | Slc6a15 | 2.67E-13 | 0.0214312 | 0.287 | 0.258 | 7.17E-09 |
| Rap2a | 1.66E-19 | -0.049283787 | 0.221 | 0.262 | 4.48E-15 | Tspoap1 | 2.67E-13 | -0.062613513 | 0.483 | 0.484 | 7.18E-09 |
| Zfand3 | 1.69E-19 | 0.045285011 | 0.668 | 0.689 | 4.56E-15 | Slc25a36 | 2.72E-13 | 0.049492906 | 0.395 | 0.352 | 7.31E-09 |

| | | | | | | | | | | | |
|---|---|---|---|---|---|---|---|---|---|---|---|
| Pitpna | 1.72E-19 | 0.091864284 | 0.513 | 0.501 | 4.62E-15 | Fmn2 | 2.77E-13 | -0.06472047 | 0.745 | 0.745 | 7.46E-09 |
| Anks3 | 1.72E-19 | -0.029096041 | 0.319 | 0.361 | 4.62E-15 | Mapk8ip3 | 2.80E-13 | 0.003090763 | 0.498 | 0.466 | 7.54E-09 |
| Gm28928 | 1.76E-19 | 0.34953533 | 0.29 | 0.237 | 4.74E-15 | Zc3h14 | 2.83E-13 | 0.042920725 | 0.399 | 0.359 | 7.60E-09 |
| Cnrip1 | 1.78E-19 | -0.073021829 | 0.371 | 0.426 | 4.78E-15 | AC149090.1 | 2.83E-13 | 0.083986787 | 0.805 | 0.765 | 7.61E-09 |
| Kcnc3 | 1.81E-19 | -0.017143536 | 0.36 | 0.4 | 4.88E-15 | Esf1 | 2.83E-13 | 0.036080219 | 0.287 | 0.251 | 7.62E-09 |
| Pikfyve | 1.82E-19 | -0.030804111 | 0.243 | 0.281 | 4.89E-15 | Gas7 | 2.84E-13 | -0.027773426 | 0.605 | 0.578 | 7.65E-09 |
| Cnnm1 | 1.83E-19 | 0.071951498 | 0.367 | 0.366 | 4.92E-15 | Lrrc4b | 2.86E-13 | 0.01957761 | 0.269 | 0.24 | 7.69E-09 |
| Zfp644 | 1.89E-19 | -0.025565506 | 0.549 | 0.597 | 5.08E-15 | Rragb | 2.93E-13 | 0.011967933 | 0.321 | 0.292 | 7.89E-09 |
| Mms19 | 1.92E-19 | 0.011377523 | 0.229 | 0.251 | 5.17E-15 | Ube2g1 | 2.95E-13 | -0.006428699 | 0.482 | 0.454 | 7.93E-09 |
| Hectd4 | 1.94E-19 | 0.103356927 | 0.669 | 0.654 | 5.23E-15 | Rcor3 | 2.96E-13 | -0.038971295 | 0.273 | 0.268 | 7.95E-09 |
| Mecr | 1.97E-19 | 0.078665701 | 0.437 | 0.433 | 5.31E-15 | Adcy9 | 2.96E-13 | -0.076681283 | 0.793 | 0.791 | 7.97E-09 |
| Mtif2 | 2.12E-19 | -0.045416356 | 0.379 | 0.428 | 5.71E-15 | Selenoi | 2.97E-13 | 0.041323308 | 0.253 | 0.218 | 7.99E-09 |
| Rad54l2 | 2.14E-19 | -0.008915967 | 0.266 | 0.298 | 5.75E-15 | Vezt | 2.97E-13 | -0.055266754 | 0.435 | 0.432 | 8.00E-09 |
| Fbxw2 | 2.15E-19 | -0.037249711 | 0.276 | 0.316 | 5.78E-15 | Nceh1 | 3.03E-13 | 0.003322945 | 0.305 | 0.281 | 8.16E-09 |
| Ube2o | 2.16E-19 | 0.105178299 | 0.316 | 0.29 | 5.80E-15 | Cfl2 | 3.05E-13 | 0.018728867 | 0.255 | 0.228 | 8.21E-09 |
| Pou2f1 | 2.20E-19 | 0.056678689 | 0.403 | 0.413 | 5.93E-15 | Sh3kbp1 | 3.13E-13 | -0.066631971 | 0.337 | 0.333 | 8.43E-09 |
| Zfp692 | 2.23E-19 | -0.095801695 | 0.331 | 0.388 | 5.99E-15 | Zc3h13 | 3.15E-13 | -0.027386028 | 0.505 | 0.489 | 8.47E-09 |
| Rspry1 | 2.29E-19 | -0.073917138 | 0.265 | 0.315 | 6.15E-15 | Snx1 | 3.17E-13 | -0.004119901 | 0.251 | 0.231 | 8.53E-09 |
| Eml2 | 2.36E-19 | 0.099862818 | 0.32 | 0.298 | 6.36E-15 | Supt20 | 3.18E-13 | 0.02429856 | 0.348 | 0.316 | 8.55E-09 |
| Mast3 | 2.41E-19 | -0.098090452 | 0.57 | 0.628 | 6.50E-15 | Cyth3 | 3.20E-13 | 0.031301424 | 0.311 | 0.278 | 8.60E-09 |
| Atf6 | 2.45E-19 | -0.090050486 | 0.495 | 0.554 | 6.58E-15 | Tle1 | 3.26E-13 | 0.049898994 | 0.366 | 0.324 | 8.78E-09 |
| Wnk2 | 2.61E-19 | 0.078146088 | 0.585 | 0.586 | 7.02E-15 | Gm9801 | 3.35E-13 | 0.001920808 | 0.391 | 0.364 | 9.00E-09 |
| Vezt | 2.62E-19 | 0.063582882 | 0.432 | 0.438 | 7.04E-15 | Itpka | 3.43E-13 | 0.019184545 | 0.399 | 0.366 | 9.24E-09 |
| Gnb1 | 2.67E-19 | 0.055592008 | 0.49 | 0.501 | 7.18E-15 | Gabpb2 | 3.45E-13 | 0.040287292 | 0.276 | 0.242 | 9.28E-09 |
| Grm1 | 2.70E-19 | -0.174001722 | 0.752 | 0.776 | 7.26E-15 | Gtf3c1 | 3.49E-13 | -0.030794432 | 0.357 | 0.346 | 9.39E-09 |
| Ric1 | 2.70E-19 | -0.017193428 | 0.326 | 0.365 | 7.27E-15 | Senp6 | 3.50E-13 | 0.052452907 | 0.612 | 0.566 | 9.43E-09 |
| Dhx9 | 2.79E-19 | -0.021983806 | 0.242 | 0.277 | 7.51E-15 | Scamp5 | 3.55E-13 | 0.033681847 | 0.316 | 0.282 | 9.54E-09 |
| 5031415H12Rik | 2.80E-19 | -0.06678405 | 0.245 | 0.293 | 7.52E-15 | Ext2 | 3.56E-13 | -0.018657623 | 0.354 | 0.338 | 9.58E-09 |
| Phc3 | 2.84E-19 | -0.036463006 | 0.434 | 0.48 | 7.64E-15 | Ndrg4 | 3.57E-13 | -0.035261877 | 0.563 | 0.546 | 9.62E-09 |
| Dcakd | 2.85E-19 | -0.064977019 | 0.219 | 0.265 | 7.67E-15 | Bcl11a | 3.58E-13 | 0.0160915 | 0.763 | 0.726 | 9.63E-09 |
| Zkscan1 | 2.87E-19 | -0.020028183 | 0.217 | 0.25 | 7.73E-15 | Gm15283 | 3.59E-13 | 0.05760693 | 0.268 | 0.228 | 9.65E-09 |
| Ap3b1 | 2.88E-19 | 0.061108767 | 0.482 | 0.494 | 7.74E-15 | Zfhx2 | 3.67E-13 | -0.014516708 | 0.363 | 0.343 | 9.88E-09 |
| Atxn7l1 | 2.98E-19 | 0.094694868 | 0.607 | 0.599 | 8.02E-15 | Zmym4 | 3.79E-13 | 0.068414242 | 0.702 | 0.656 | 1.02E-08 |
| Itga8 | 3.10E-19 | 0.121077651 | 0.589 | 0.58 | 8.35E-15 | Nbr1 | 3.80E-13 | 0.01603102 | 0.307 | 0.279 | 1.02E-08 |
| Exoc2 | 3.20E-19 | 0.022396463 | 0.339 | 0.36 | 8.61E-15 | Dhdds | 3.84E-13 | 0.006760147 | 0.403 | 0.373 | 1.03E-08 |
| Anks1 | 3.34E-19 | 0.069033478 | 0.259 | 0.257 | 8.98E-15 | Adar | 3.92E-13 | -0.033686942 | 0.435 | 0.42 | 1.06E-08 |
| Gps2 | 3.36E-19 | -0.076606862 | 0.254 | 0.304 | 9.05E-15 | Spats2 | 3.93E-13 | 0.052322141 | 0.298 | 0.26 | 1.06E-08 |
| Nbr1 | 3.44E-19 | 0.038333148 | 0.279 | 0.289 | 9.25E-15 | Pnpt1 | 3.94E-13 | -0.001547897 | 0.31 | 0.289 | 1.06E-08 |
| Lrrfip2 | 3.49E-19 | 0.033608521 | 0.322 | 0.34 | 9.38E-15 | Abl1 | 3.97E-13 | 0.054813856 | 0.31 | 0.271 | 1.07E-08 |
| Ptpa | 3.55E-19 | 0.069043407 | 0.255 | 0.252 | 9.54E-15 | Slc44a1 | 4.04E-13 | 0.04072481 | 0.653 | 0.61 | 1.09E-08 |
| Eprs | 3.59E-19 | -0.013966227 | 0.267 | 0.301 | 9.65E-15 | Hdac4 | 4.10E-13 | -0.038593031 | 0.429 | 0.419 | 1.10E-08 |
| Csad | 3.65E-19 | -0.032924964 | 0.222 | 0.26 | 9.82E-15 | Taok1 | 4.19E-13 | 0.045743264 | 0.587 | 0.543 | 1.13E-08 |
| Dennd6a | 3.73E-19 | -0.004536254 | 0.363 | 0.397 | 1.00E-14 | Rgs7 | 4.25E-13 | -0.07959551 | 0.937 | 0.938 | 1.14E-08 |
| Snap91 | 3.83E-19 | 0.079271478 | 0.722 | 0.725 | 1.03E-14 | Usp3 | 4.29E-13 | 0.065155571 | 0.355 | 0.311 | 1.16E-08 |
| Mmd | 3.86E-19 | -0.023518495 | 0.261 | 0.298 | 1.04E-14 | Rprd1a | 4.39E-13 | 0.021036435 | 0.432 | 0.398 | 1.18E-08 |
| Crocc | 3.88E-19 | 0.05738802 | 0.267 | 0.269 | 1.04E-14 | Mdn1 | 4.49E-13 | -0.052435544 | 0.392 | 0.392 | 1.21E-08 |
| Chfr | 3.93E-19 | -0.031941218 | 0.494 | 0.54 | 1.06E-14 | 4930570B17Rik | 4.53E-13 | 0.082933113 | 0.388 | 0.342 | 1.22E-08 |
| Shisa9 | 4.17E-19 | 0.144847836 | 0.685 | 0.683 | 1.12E-14 | Dctn1 | 4.54E-13 | 0.023912326 | 0.269 | 0.238 | 1.22E-08 |
| Usp47 | 4.30E-19 | -0.008064262 | 0.413 | 0.451 | 1.16E-14 | Ago2 | 4.57E-13 | -0.010411775 | 0.391 | 0.37 | 1.23E-08 |
| Gm11149 | 4.30E-19 | -0.079390148 | 0.459 | 0.516 | 1.16E-14 | Acox1 | 4.60E-13 | 0.011119741 | 0.278 | 0.252 | 1.24E-08 |
| Ambra1 | 4.32E-19 | 0.080282298 | 0.715 | 0.72 | 1.16E-14 | Nsmf | 4.67E-13 | 0.011369882 | 0.437 | 0.406 | 1.26E-08 |
| Nrd1 | 4.41E-19 | -0.116144679 | 0.579 | 0.637 | 1.19E-14 | Ints10 | 4.74E-13 | -0.023081596 | 0.342 | 0.327 | 1.28E-08 |
| Scamp5 | 4.43E-19 | 0.007426329 | 0.282 | 0.307 | 1.19E-14 | Phc3 | 4.79E-13 | 0.014501603 | 0.466 | 0.434 | 1.29E-08 |
| Pacrg | 4.44E-19 | 0.109752132 | 0.393 | 0.383 | 1.19E-14 | Lrch1 | 4.79E-13 | -0.032737741 | 0.511 | 0.494 | 1.29E-08 |
| Ipo9 | 4.44E-19 | 0.075190451 | 0.265 | 0.255 | 1.20E-14 | Fam13a | 4.81E-13 | 0.042764791 | 0.29 | 0.255 | 1.29E-08 |
| Micu2 | 4.58E-19 | 0.065580432 | 0.367 | 0.368 | 1.23E-14 | Ski | 4.86E-13 | -0.040755815 | 0.387 | 0.378 | 1.31E-08 |
| Acvr2a | 4.93E-19 | 0.001885723 | 0.452 | 0.493 | 1.33E-14 | Ogdh | 4.87E-13 | 0.03986305 | 0.443 | 0.404 | 1.31E-08 |
| Fam110b | 4.94E-19 | 0.043990806 | 0.417 | 0.434 | 1.33E-14 | Mapre2 | 5.07E-13 | 0.06556883 | 0.545 | 0.499 | 1.37E-08 |
| 9330182L06Rik | 4.98E-19 | 0.105628765 | 0.272 | 0.252 | 1.34E-14 | Stt3b | 5.08E-13 | 0.042871207 | 0.341 | 0.304 | 1.37E-08 |
| Uso1 | 5.10E-19 | -0.052631168 | 0.274 | 0.32 | 1.37E-14 | 4933406I18Rik | 5.17E-13 | -0.038014094 | 0.505 | 0.492 | 1.39E-08 |
| Pja2 | 5.20E-19 | -0.038860743 | 0.502 | 0.551 | 1.40E-14 | Taf15 | 5.47E-13 | 0.01871388 | 0.502 | 0.467 | 1.47E-08 |
| Tfdp2 | 5.22E-19 | 0.073138737 | 0.468 | 0.475 | 1.40E-14 | Slc24a3 | 5.48E-13 | -0.127784476 | 0.655 | 0.67 | 1.47E-08 |
| Gas7 | 5.23E-19 | 0.047389879 | 0.578 | 0.604 | 1.41E-14 | Kdm1a | 5.61E-13 | 0.033194974 | 0.464 | 0.426 | 1.51E-08 |
| Psmb2 | 5.23E-19 | -0.070312333 | 0.295 | 0.345 | 1.41E-14 | Adss | 5.63E-13 | -0.002928221 | 0.279 | 0.257 | 1.52E-08 |
| Homer1 | 5.24E-19 | -0.276138949 | 0.495 | 0.54 | 1.41E-14 | Srrm1 | 5.71E-13 | 0.037221257 | 0.528 | 0.487 | 1.54E-08 |
| Dab1 | 5.28E-19 | 0.103394462 | 0.987 | 0.982 | 1.42E-14 | Tmem234 | 5.72E-13 | 0.03991686 | 0.295 | 0.259 | 1.54E-08 |
| Prmt8 | 5.50E-19 | -0.034256785 | 0.392 | 0.438 | 1.48E-14 | Atxn2l | 5.73E-13 | 0.028757586 | 0.332 | 0.298 | 1.54E-08 |
| Nipa1 | 5.52E-19 | 0.005245894 | 0.357 | 0.389 | 1.49E-14 | Chrm1 | 5.75E-13 | 0.052088292 | 0.263 | 0.225 | 1.55E-08 |
| Tmem132b | 5.71E-19 | 0.104372572 | 0.817 | 0.813 | 1.54E-14 | Uggt2 | 5.95E-13 | -0.063311563 | 0.509 | 0.508 | 1.60E-08 |
| Synrg | 5.73E-19 | -0.073692786 | 0.308 | 0.36 | 1.54E-14 | Crebrf | 6.02E-13 | 0.036089836 | 0.359 | 0.322 | 1.62E-08 |
| Mpc1 | 5.78E-19 | 0.093737393 | 0.343 | 0.331 | 1.55E-14 | Ezh1 | 6.05E-13 | 0.006245944 | 0.258 | 0.235 | 1.63E-08 |
| Ctnna3 | 5.79E-19 | -0.250595119 | 0.554 | 0.597 | 1.56E-14 | Rogdi | 6.06E-13 | 0.041834198 | 0.323 | 0.286 | 1.63E-08 |
| Ccnh | 6.10E-19 | -0.027454222 | 0.328 | 0.368 | 1.64E-14 | Tmem178 | 6.17E-13 | 0.012806837 | 0.431 | 0.396 | 1.66E-08 |
| Fam208a | 6.17E-19 | 0.110107223 | 0.368 | 0.312 | 1.66E-14 | Csrnp3 | 6.19E-13 | 0.091709899 | 0.752 | 0.71 | 1.66E-08 |
| Fyttd1 | 6.27E-19 | -0.002905394 | 0.29 | 0.32 | 1.69E-14 | Scfd1 | 6.26E-13 | 0.041473886 | 0.36 | 0.322 | 1.68E-08 |
| Rab3gap1 | 6.28E-19 | -0.05085637 | 0.479 | 0.531 | 1.69E-14 | Slc16a2 | 6.51E-13 | -0.028937939 | 0.457 | 0.438 | 1.75E-08 |
| Trp53bp1 | 6.43E-19 | 0.048964439 | 0.401 | 0.414 | 1.73E-14 | Xkr6 | 6.55E-13 | -0.063950988 | 0.646 | 0.63 | 1.76E-08 |
| Smoc2 | 6.72E-19 | -0.136902718 | 0.244 | 0.296 | 1.81E-14 | Ireb2 | 6.58E-13 | 0.007616068 | 0.296 | 0.271 | 1.77E-08 |
| Exoc5 | 6.73E-19 | -0.049199102 | 0.338 | 0.384 | 1.81E-14 | Klhl12 | 6.62E-13 | 0.049827148 | 0.266 | 0.23 | 1.78E-08 |
| Map3k13 | 6.83E-19 | 0.013858526 | 0.275 | 0.299 | 1.84E-14 | Rap1gap2 | 6.62E-13 | -0.078145345 | 0.52 | 0.521 | 1.78E-08 |
| Syncrip | 7.00E-19 | -0.048513328 | 0.227 | 0.269 | 1.88E-14 | Akap11 | 6.63E-13 | 0.039383849 | 0.474 | 0.433 | 1.79E-08 |
| Adcy1 | 7.08E-19 | 0.114267659 | 0.579 | 0.57 | 1.90E-14 | Pdzd4 | 6.64E-13 | 0.006876575 | 0.571 | 0.538 | 1.79E-08 |
| Ube2k | 7.24E-19 | 0.02454242 | 0.551 | 0.576 | 1.95E-14 | Clasp1 | 6.65E-13 | -0.061356824 | 0.565 | 0.566 | 1.79E-08 |
| Trim33 | 7.70E-19 | 0.009919183 | 0.446 | 0.476 | 2.07E-14 | Slf2 | 6.69E-13 | 0.036373631 | 0.53 | 0.488 | 1.80E-08 |
| Pcdh9 | 7.80E-19 | 0.150398005 | 0.987 | 0.985 | 2.10E-14 | Cep78 | 7.03E-13 | 0.002664263 | 0.348 | 0.322 | 1.89E-08 |
| Fermt2 | 7.93E-19 | 0.015495823 | 0.371 | 0.4 | 2.13E-14 | Dazap1 | 7.07E-13 | 0.063020031 | 0.309 | 0.268 | 1.90E-08 |
| Acsl6 | 7.95E-19 | -0.067687919 | 0.255 | 0.303 | 2.14E-14 | Gm21992 | 7.11E-13 | -0.090925497 | 0.403 | 0.45 | 1.91E-08 |
| Cacnb4 | 7.98E-19 | 0.103364196 | 0.785 | 0.759 | 2.15E-14 | Phtf2 | 7.28E-13 | 0.004434885 | 0.317 | 0.292 | 1.96E-08 |
| Gm26883 | 8.00E-19 | 0.027270434 | 0.253 | 0.221 | 2.15E-14 | Scn1a | 7.30E-13 | -0.027117196 | 0.614 | 0.593 | 1.96E-08 |
| Ank | 8.43E-19 | 0.091686609 | 0.624 | 0.618 | 2.27E-14 | Ubp1 | 7.31E-13 | -0.026523802 | 0.419 | 0.401 | 1.97E-08 |
| Adarb1 | 8.68E-19 | 0.04867894 | 0.467 | 0.484 | 2.34E-14 | Mta3 | 7.38E-13 | 0.010423082 | 0.407 | 0.375 | 1.98E-08 |
| Unc5d | 8.69E-19 | 0.24947817 | 0.508 | 0.465 | 2.34E-14 | Pcmtd1 | 7.47E-13 | 0.052944253 | 0.52 | 0.475 | 2.01E-08 |
| Kcmf1 | 8.84E-19 | 0.015631216 | 0.374 | 0.4 | 2.38E-14 | Adam22 | 7.69E-13 | -0.064562372 | 0.765 | 0.763 | 2.07E-08 |
| Cul5 | 9.30E-19 | -0.020350004 | 0.379 | 0.419 | 2.50E-14 | Med15 | 7.75E-13 | -0.004095687 | 0.388 | 0.365 | 2.09E-08 |
| Atp11b | 9.41E-19 | 0.084014489 | 0.446 | 0.437 | 2.53E-14 | Kcna1 | 7.78E-13 | 0.025548629 | 0.282 | 0.251 | 2.09E-08 |
| Kat6a | 9.44E-19 | -0.002350883 | 0.265 | 0.294 | 2.54E-14 | Ralgps2 | 7.91E-13 | -0.04245574 | 0.492 | 0.482 | 2.13E-08 |
| Ubxn4 | 9.63E-19 | -0.047488327 | 0.402 | 0.45 | 2.59E-14 | Rab12 | 8.13E-13 | -0.002161228 | 0.266 | 0.246 | 2.19E-08 |
| Gdap1 | 9.68E-19 | -0.041539984 | 0.216 | 0.255 | 2.60E-14 | Otud4 | 8.13E-13 | -0.032042664 | 0.293 | 0.282 | 2.19E-08 |

| | | | | | | | | | | | |
|---|---|---|---|---|---|---|---|---|---|---|---|
| Gm48678 | 9.76E-19 | -0.111187123 | 0.646 | 0.701 | 2.63E-14 | Setd3 | 8.24E-13 | -0.043345469 | 0.294 | 0.289 | 2.22E-08 |
| Rap1gap | 9.86E-19 | -0.013053154 | 0.348 | 0.385 | 2.65E-14 | Ascc3 | 8.30E-13 | 0.021341112 | 0.641 | 0.605 | 2.23E-08 |
| Brd8 | 9.93E-19 | -0.03849423 | 0.263 | 0.303 | 2.67E-14 | Ppig | 8.35E-13 | 0.025906196 | 0.382 | 0.349 | 2.25E-08 |
| Brwd3 | 1.10E-18 | -0.042802191 | 0.24 | 0.28 | 2.95E-14 | Safb2 | 8.36E-13 | -0.016613644 | 0.362 | 0.342 | 2.25E-08 |
| Setx | 1.10E-18 | 0.064201405 | 0.384 | 0.383 | 2.97E-14 | Gm43376 | 8.40E-13 | 0.085707745 | 0.617 | 0.569 | 2.26E-08 |
| Usp12 | 1.11E-18 | 0.006785607 | 0.267 | 0.292 | 3.00E-14 | Hook1 | 8.44E-13 | -0.049830525 | 0.472 | 0.466 | 2.27E-08 |
| Etnk1 | 1.13E-18 | -0.105463617 | 0.627 | 0.683 | 3.03E-14 | Gm11659 | 8.74E-13 | 0.057305643 | 0.284 | 0.245 | 2.35E-08 |
| Ramp1 | 1.13E-18 | -0.01183628 | 0.234 | 0.264 | 3.05E-14 | Ccnc | 8.82E-13 | 0.056027664 | 0.311 | 0.271 | 2.37E-08 |
| Cbx3 | 1.15E-18 | -0.011753881 | 0.22 | 0.25 | 3.08E-14 | Mkx | 8.84E-13 | 0.073476044 | 0.285 | 0.246 | 2.38E-08 |
| Smarca4 | 1.16E-18 | 1.50E-05 | 0.436 | 0.469 | 3.11E-14 | Ccdc30 | 8.90E-13 | -0.039067651 | 0.486 | 0.475 | 2.40E-08 |
| Ep400 | 1.19E-18 | 0.011802129 | 0.476 | 0.506 | 3.21E-14 | Cltb | 9.01E-13 | 0.012614634 | 0.268 | 0.242 | 2.42E-08 |
| Ttc7b | 1.21E-18 | 0.057574738 | 0.64 | 0.654 | 3.25E-14 | Tpm1 | 9.02E-13 | -0.009532103 | 0.271 | 0.254 | 2.43E-08 |
| Tasp1 | 1.21E-18 | 0.034011358 | 0.48 | 0.501 | 3.27E-14 | Asb3 | 9.39E-13 | -0.003733106 | 0.556 | 0.527 | 2.53E-08 |
| Mdn1 | 1.22E-18 | 0.057511172 | 0.392 | 0.393 | 3.27E-14 | Ino80 | 9.44E-13 | 0.020723562 | 0.515 | 0.479 | 2.54E-08 |
| Ythdc2 | 1.28E-18 | -0.012758647 | 0.338 | 0.375 | 3.44E-14 | Fam118b | 9.46E-13 | 0.03328293 | 0.264 | 0.231 | 2.54E-08 |
| Slc35e2 | 1.28E-18 | -0.068696423 | 0.222 | 0.267 | 3.45E-14 | Stx3 | 9.56E-13 | -0.034394333 | 0.295 | 0.287 | 2.57E-08 |
| Srrt | 1.29E-18 | 0.023432587 | 0.298 | 0.316 | 3.48E-14 | Pebp1 | 9.63E-13 | 0.061667313 | 0.347 | 0.303 | 2.59E-08 |
| Itm2b | 1.30E-18 | -0.084333502 | 0.441 | 0.5 | 3.49E-14 | Dennd5b | 9.88E-13 | 0.05856107 | 0.583 | 0.538 | 2.66E-08 |
| Psmd11 | 1.33E-18 | -0.036579173 | 0.311 | 0.354 | 3.57E-14 | Dnajc11 | 1.02E-12 | 0.003918042 | 0.255 | 0.232 | 2.74E-08 |
| Cacng2 | 1.34E-18 | 0.05593456 | 0.307 | 0.316 | 3.62E-14 | Med13l | 1.03E-12 | -0.062744031 | 0.625 | 0.622 | 2.76E-08 |
| Rc3h2 | 1.35E-18 | -0.001091904 | 0.424 | 0.459 | 3.64E-14 | Lrp6 | 1.03E-12 | -0.010996425 | 0.447 | 0.424 | 2.76E-08 |
| Slc25a12 | 1.36E-18 | 0.037347445 | 0.536 | 0.556 | 3.65E-14 | Chchd3 | 1.03E-12 | -0.034735825 | 0.613 | 0.597 | 2.76E-08 |
| Arsb | 1.36E-18 | 0.113073465 | 0.573 | 0.555 | 3.65E-14 | Rufy1 | 1.03E-12 | 0.00640445 | 0.324 | 0.298 | 2.77E-08 |
| 4-Mar | 1.43E-18 | -0.123393277 | 0.235 | 0.287 | 3.84E-14 | Prr14 | 1.05E-12 | 0.031166629 | 0.255 | 0.224 | 2.83E-08 |
| Eftud2 | 1.44E-18 | 0.05333283 | 0.352 | 0.359 | 3.87E-14 | Lrp8 | 1.05E-12 | -0.071988385 | 0.309 | 0.318 | 2.84E-08 |
| Rnf14 | 1.54E-18 | -0.02001978 | 0.296 | 0.332 | 4.14E-14 | Vps37a | 1.06E-12 | -0.006749124 | 0.254 | 0.236 | 2.84E-08 |
| Neo1 | 1.54E-18 | 0.001211283 | 0.478 | 0.514 | 4.14E-14 | Trpc4 | 1.08E-12 | 0.029925868 | 0.539 | 0.503 | 2.90E-08 |
| Fam171a1 | 1.56E-18 | 0.041174938 | 0.386 | 0.398 | 4.20E-14 | Tnrc6a | 1.08E-12 | 0.091424109 | 0.258 | 0.217 | 2.91E-08 |
| Mapk14 | 1.57E-18 | 0.07866434 | 0.427 | 0.424 | 4.22E-14 | BC052040 | 1.08E-12 | -0.076628176 | 0.248 | 0.263 | 2.92E-08 |
| R3hdm1 | 1.58E-18 | -0.215177484 | 0.922 | 0.935 | 4.25E-14 | Zbtb38 | 1.08E-12 | 0.025791831 | 0.431 | 0.395 | 2.92E-08 |
| Asxl2 | 1.59E-18 | 0.020160846 | 0.46 | 0.49 | 4.29E-14 | Eftud2 | 1.10E-12 | 0.004364983 | 0.379 | 0.352 | 2.96E-08 |
| Pkp4 | 1.62E-18 | 0.089532619 | 0.701 | 0.694 | 4.35E-14 | Mbtd1 | 1.10E-12 | -0.016963235 | 0.566 | 0.545 | 2.96E-08 |
| 8030462N17Rik | 1.67E-18 | -0.06861275 | 0.292 | 0.341 | 4.48E-14 | Ispd | 1.11E-12 | -0.094904848 | 0.229 | 0.27 | 2.98E-08 |
| Zfp704 | 1.73E-18 | 0.058815221 | 0.489 | 0.502 | 4.65E-14 | Tspan13 | 1.11E-12 | 0.063028402 | 0.417 | 0.373 | 2.99E-08 |
| Hk1 | 1.76E-18 | 0.081091743 | 0.336 | 0.325 | 4.73E-14 | Ncoa6 | 1.12E-12 | 0.005023057 | 0.429 | 0.398 | 3.01E-08 |
| Rbm39 | 1.76E-18 | -0.077467387 | 0.888 | 0.916 | 4.73E-14 | Rnf2 | 1.13E-12 | 0.040676931 | 0.25 | 0.216 | 3.03E-08 |
| Tmem234 | 1.83E-18 | -0.079890612 | 0.259 | 0.31 | 4.92E-14 | Ankrd13c | 1.13E-12 | 0.031638376 | 0.349 | 0.316 | 3.04E-08 |
| Arhgef28 | 1.84E-18 | -0.109795943 | 0.418 | 0.475 | 4.94E-14 | Trim24 | 1.13E-12 | 0.024068624 | 0.267 | 0.238 | 3.04E-08 |
| Chordc1 | 1.85E-18 | -0.049290569 | 0.231 | 0.275 | 4.97E-14 | Atp6v1b2 | 1.13E-12 | -0.016870498 | 0.403 | 0.385 | 3.05E-08 |
| Map3k2 | 1.88E-18 | 0.021463647 | 0.393 | 0.417 | 5.06E-14 | Dlgap3 | 1.16E-12 | -0.005092886 | 0.437 | 0.411 | 3.12E-08 |
| Heatr3 | 1.95E-18 | 0.040686859 | 0.241 | 0.251 | 5.24E-14 | Herc3 | 1.17E-12 | 0.009010822 | 0.51 | 0.477 | 3.15E-08 |
| Ireb2 | 2.05E-18 | 0.025090724 | 0.271 | 0.287 | 5.50E-14 | Pik3ca | 1.17E-12 | -0.004622245 | 0.322 | 0.3 | 3.16E-08 |
| Ap3b2 | 2.07E-18 | 0.031982829 | 0.343 | 0.359 | 5.56E-14 | Lurap1l | 1.18E-12 | -0.043950468 | 0.302 | 0.297 | 3.16E-08 |
| Pum1 | 2.08E-18 | -0.027995315 | 0.527 | 0.573 | 5.59E-14 | Plcb1 | 1.18E-12 | -0.093468563 | 0.959 | 0.965 | 3.17E-08 |
| Sppl2a | 2.10E-18 | 0.010518829 | 0.295 | 0.321 | 5.64E-14 | Trappc9 | 1.18E-12 | -0.076978828 | 0.523 | 0.536 | 3.17E-08 |
| P4htm | 2.11E-18 | 0.059401886 | 0.283 | 0.284 | 5.67E-14 | Srrm3 | 1.19E-12 | -0.013604653 | 0.618 | 0.592 | 3.21E-08 |
| Chd8 | 2.11E-18 | 0.026807161 | 0.401 | 0.421 | 5.69E-14 | Capza1 | 1.20E-12 | 0.01168171 | 0.317 | 0.289 | 3.22E-08 |
| Kcnq2 | 2.16E-18 | 0.02340859 | 0.574 | 0.601 | 5.81E-14 | Prmt8 | 1.20E-12 | 0.008239234 | 0.423 | 0.392 | 3.24E-08 |
| Nbeal1 | 2.23E-18 | 0.077907228 | 0.359 | 0.356 | 6.00E-14 | Nt5c2 | 1.21E-12 | 0.045102254 | 0.633 | 0.589 | 3.26E-08 |
| Atg3 | 2.23E-18 | -0.061736611 | 0.238 | 0.282 | 6.00E-14 | Mtmr1 | 1.26E-12 | 0.030895857 | 0.35 | 0.314 | 3.40E-08 |
| Rbm25 | 2.24E-18 | 0.030730325 | 0.646 | 0.676 | 6.03E-14 | Uso1 | 1.28E-12 | 0.017205037 | 0.303 | 0.274 | 3.44E-08 |
| Ttc37 | 2.27E-18 | -0.020078276 | 0.319 | 0.355 | 6.11E-14 | Ap2a2 | 1.30E-12 | -0.050744583 | 0.391 | 0.389 | 3.50E-08 |
| Rnf182 | 2.28E-18 | -0.076426642 | 0.285 | 0.333 | 6.14E-14 | Dag1 | 1.32E-12 | 0.060351021 | 0.357 | 0.314 | 3.55E-08 |
| Usp9x | 2.35E-18 | -0.0486107 | 0.565 | 0.616 | 6.33E-14 | Gm42477 | 1.35E-12 | 0.060283113 | 0.328 | 0.286 | 3.63E-08 |
| Gm13402 | 2.38E-18 | -0.081485775 | 0.255 | 0.305 | 6.40E-14 | Pkia | 1.38E-12 | 0.044296353 | 0.454 | 0.414 | 3.71E-08 |
| Magi2 | 2.38E-18 | -0.120101585 | 0.987 | 0.984 | 6.40E-14 | Mier1 | 1.38E-12 | 0.033322579 | 0.381 | 0.343 | 3.72E-08 |
| Fyn | 2.40E-18 | 0.066680248 | 0.585 | 0.588 | 6.45E-14 | Scfd2 | 1.39E-12 | -0.050130226 | 0.467 | 0.464 | 3.74E-08 |
| Psma1 | 2.43E-18 | -0.070268613 | 0.252 | 0.3 | 6.53E-14 | Srsf5 | 1.40E-12 | 0.024714365 | 0.432 | 0.395 | 3.76E-08 |
| Rgl1 | 2.47E-18 | 0.025085403 | 0.644 | 0.675 | 6.64E-14 | Fam117b | 1.44E-12 | 0.041973423 | 0.369 | 0.33 | 3.86E-08 |
| Psen1 | 2.58E-18 | 0.02360139 | 0.283 | 0.303 | 6.94E-14 | Yeats2 | 1.45E-12 | -0.009701248 | 0.423 | 0.4 | 3.89E-08 |
| Rab11fip2 | 2.63E-18 | -0.037160467 | 0.425 | 0.474 | 7.09E-14 | Rab27b | 1.46E-12 | 0.06297447 | 0.346 | 0.304 | 3.94E-08 |
| Nufip2 | 2.77E-18 | -0.042542878 | 0.343 | 0.388 | 7.47E-14 | Gm17227 | 1.46E-12 | 0.022211856 | 0.328 | 0.297 | 3.94E-08 |
| Fgf13 | 2.78E-18 | 0.10674815 | 0.381 | 0.377 | 7.48E-14 | Bbs9 | 1.48E-12 | -0.021990763 | 0.355 | 0.337 | 3.98E-08 |
| Clta | 2.78E-18 | -0.034143195 | 0.232 | 0.269 | 7.48E-14 | Thra | 1.48E-12 | -0.040638011 | 0.547 | 0.535 | 3.99E-08 |
| Tanc1 | 2.79E-18 | -0.139364052 | 0.628 | 0.679 | 7.50E-14 | D430041D05Rik | 1.49E-12 | -0.054871042 | 0.719 | 0.71 | 4.01E-08 |
| Gm44686 | 2.79E-18 | -0.104365434 | 0.34 | 0.396 | 7.52E-14 | Faxc | 1.50E-12 | 0.030413839 | 0.252 | 0.223 | 4.02E-08 |
| Nf1 | 2.82E-18 | 0.071746537 | 0.64 | 0.648 | 7.58E-14 | Coro2b | 1.52E-12 | -0.006916706 | 0.429 | 0.404 | 4.08E-08 |
| Supt6 | 2.83E-18 | 0.012227606 | 0.32 | 0.344 | 7.62E-14 | Gm37240 | 1.53E-12 | 0.060129379 | 0.8 | 0.76 | 4.11E-08 |
| Lmo3 | 2.83E-18 | -0.084501225 | 0.415 | 0.471 | 7.62E-14 | Slc6a6 | 1.56E-12 | -0.045457712 | 0.579 | 0.565 | 4.20E-08 |
| Tlk1 | 2.87E-18 | -0.030016797 | 0.365 | 0.407 | 7.72E-14 | Rab7 | 1.57E-12 | 0.034892439 | 0.281 | 0.247 | 4.22E-08 |
| Camta2 | 2.89E-18 | -0.066348483 | 0.284 | 0.333 | 7.79E-14 | Polb | 1.58E-12 | 0.024944054 | 0.524 | 0.486 | 4.26E-08 |
| Hnrnpll | 2.93E-18 | -0.107067122 | 0.353 | 0.408 | 7.87E-14 | Tpst1 | 1.59E-12 | 0.040547072 | 0.253 | 0.22 | 4.27E-08 |
| Phf20 | 2.94E-18 | -0.06758235 | 0.417 | 0.47 | 7.91E-14 | Inpp4a | 1.60E-12 | -0.044720954 | 0.493 | 0.482 | 4.32E-08 |
| Bcl2l1 | 3.00E-18 | 0.050296444 | 0.252 | 0.26 | 8.06E-14 | Safb | 1.63E-12 | -0.008518983 | 0.355 | 0.333 | 4.39E-08 |
| Vps41 | 3.03E-18 | -0.004448767 | 0.472 | 0.509 | 8.16E-14 | Ncoa7 | 1.64E-12 | -0.050110872 | 0.638 | 0.631 | 4.41E-08 |
| Grb10 | 3.09E-18 | 0.117565263 | 0.265 | 0.237 | 8.31E-14 | Mapre3 | 1.65E-12 | -0.029862299 | 0.444 | 0.428 | 4.44E-08 |
| Scfd2 | 3.09E-18 | 0.061729605 | 0.464 | 0.469 | 8.31E-14 | Ubr5 | 1.65E-12 | -0.072381506 | 0.64 | 0.644 | 4.45E-08 |
| Adss | 3.10E-18 | 0.086295216 | 0.257 | 0.241 | 8.33E-14 | Dab1 | 1.67E-12 | -0.077245552 | 0.978 | 0.987 | 4.49E-08 |
| Ptbp3 | 3.14E-18 | 0.009770261 | 0.239 | 0.263 | 8.45E-14 | Ktn1 | 1.68E-12 | -0.009647081 | 0.584 | 0.559 | 4.52E-08 |
| Cdadc1 | 3.27E-18 | -0.017713784 | 0.301 | 0.337 | 8.79E-14 | Dcun1d2 | 1.68E-12 | 0.011233206 | 0.297 | 0.27 | 4.53E-08 |
| Mllt3 | 3.32E-18 | -0.076438899 | 0.748 | 0.797 | 8.93E-14 | Zfp608 | 1.71E-12 | 0.10645296 | 0.51 | 0.463 | 4.59E-08 |
| Eif4g3 | 3.41E-18 | 0.085593498 | 0.773 | 0.772 | 9.17E-14 | Rab6b | 1.72E-12 | -0.051097093 | 0.482 | 0.478 | 4.62E-08 |
| Cyb5b | 3.48E-18 | -0.019941973 | 0.222 | 0.256 | 9.36E-14 | Manf | 1.72E-12 | 0.036673243 | 0.384 | 0.347 | 4.62E-08 |
| Caprin1 | 3.49E-18 | -0.016225538 | 0.356 | 0.393 | 9.40E-14 | Lrrn2 | 1.74E-12 | 0.022073764 | 0.564 | 0.527 | 4.67E-08 |
| Atp1a3 | 3.50E-18 | 0.078181207 | 0.683 | 0.69 | 9.42E-14 | Kif13b | 1.74E-12 | -0.029701916 | 0.372 | 0.358 | 4.68E-08 |
| Uxs1 | 3.61E-18 | 0.037212736 | 0.327 | 0.34 | 9.72E-14 | Tmem135 | 1.76E-12 | -0.036761256 | 0.579 | 0.565 | 4.75E-08 |
| Mapk8 | 3.62E-18 | -0.046978349 | 0.378 | 0.425 | 9.75E-14 | Akap9 | 1.79E-12 | -0.073917694 | 0.46 | 0.466 | 4.82E-08 |
| Fgfr2 | 3.68E-18 | -0.110556073 | 0.897 | 0.919 | 9.90E-14 | Osbp2 | 1.80E-12 | -0.079644955 | 0.647 | 0.65 | 4.85E-08 |
| Hacd2 | 3.70E-18 | 0.022855831 | 0.308 | 0.327 | 9.95E-14 | Eif5b | 1.85E-12 | 0.04630613 | 0.412 | 0.372 | 4.98E-08 |
| Cacng3 | 3.73E-18 | 0.123458899 | 0.492 | 0.472 | 1.00E-13 | Fgfr1op2 | 1.85E-12 | 0.046334529 | 0.285 | 0.248 | 4.99E-08 |
| Adgrl1 | 3.90E-18 | 0.067387382 | 0.499 | 0.504 | 1.05E-13 | Ano3 | 1.89E-12 | 0.118250037 | 0.763 | 0.737 | 5.09E-08 |
| Camkk2 | 3.95E-18 | 0.09804128 | 0.342 | 0.32 | 1.06E-13 | Farsb | 1.90E-12 | -0.017134158 | 0.31 | 0.293 | 5.10E-08 |
| Rps6kc1 | 3.96E-18 | 0.055487745 | 0.334 | 0.337 | 1.07E-13 | Rab3gap1 | 1.94E-12 | 0.050762435 | 0.523 | 0.479 | 5.21E-08 |
| Kif5b | 3.97E-18 | -0.028626008 | 0.434 | 0.478 | 1.07E-13 | Ap3b2 | 1.98E-12 | 0.002803652 | 0.369 | 0.343 | 5.32E-08 |
| Fam172a | 4.00E-18 | 0.076231977 | 0.728 | 0.733 | 1.08E-13 | Klhl3 | 2.01E-12 | -0.045100341 | 0.6 | 0.59 | 5.42E-08 |
| Mbtd1 | 4.27E-18 | 0.064402421 | 0.545 | 0.552 | 1.15E-13 | Sarnp | 2.05E-12 | 0.036904616 | 0.393 | 0.356 | 5.52E-08 |
| Ndrg3 | 4.41E-18 | 0.061695129 | 0.532 | 0.542 | 1.19E-13 | Atp6v1h | 2.06E-12 | -0.021714027 | 0.449 | 0.431 | 5.53E-08 |
| Ivd | 4.43E-18 | 0.0777225 | 0.263 | 0.252 | 1.19E-13 | Fbxl20 | 2.08E-12 | 0.016177861 | 0.563 | 0.53 | 5.60E-08 |

| | | | | | | | | | | | |
|---|---|---|---|---|---|---|---|---|---|---|---|
| Tbcd | 4.46E-18 | 0.076808067 | 0.372 | 0.364 | 1.20E-13 | Pcyt2 | 2.08E-12 | 0.019691149 | 0.388 | 0.353 | 5.60E-08 |
| Gabra1 | 4.48E-18 | 0.033511962 | 0.34 | 0.358 | 1.21E-13 | Kif2a | 2.11E-12 | 0.03757963 | 0.465 | 0.426 | 5.68E-08 |
| Fkbp1a | 4.55E-18 | -0.091133633 | 0.26 | 0.311 | 1.23E-13 | Taok3 | 2.12E-12 | -0.030195055 | 0.445 | 0.432 | 5.70E-08 |
| Mgea5 | 4.68E-18 | 0.092994077 | 0.433 | 0.377 | 1.26E-13 | Ube2l3 | 2.14E-12 | 0.034444308 | 0.262 | 0.23 | 5.77E-08 |
| Ptpra | 4.71E-18 | -0.124259721 | 0.595 | 0.65 | 1.27E-13 | Sacm1l | 2.17E-12 | 0.033866031 | 0.437 | 0.399 | 5.83E-08 |
| Uimc1 | 4.75E-18 | -0.032274809 | 0.299 | 0.339 | 1.28E-13 | Zyg11b | 2.17E-12 | 0.035019427 | 0.383 | 0.346 | 5.84E-08 |
| Ift57 | 4.83E-18 | 0.003311073 | 0.23 | 0.254 | 1.30E-13 | Zmym2 | 2.22E-12 | 0.015323703 | 0.518 | 0.483 | 5.96E-08 |
| Trpc4ap | 4.87E-18 | 0.056627547 | 0.319 | 0.323 | 1.31E-13 | Emsy | 2.25E-12 | -0.011977354 | 0.423 | 0.402 | 6.05E-08 |
| Zcchc17 | 4.87E-18 | -0.045128487 | 0.214 | 0.253 | 1.31E-13 | Ilf3 | 2.25E-12 | 0.010109325 | 0.329 | 0.301 | 6.05E-08 |
| Cacng7 | 5.04E-18 | 0.005594381 | 0.236 | 0.258 | 1.36E-13 | Add1 | 2.25E-12 | 0.038519608 | 0.483 | 0.443 | 6.06E-08 |
| Stx6 | 5.06E-18 | 0.028978241 | 0.337 | 0.353 | 1.36E-13 | P4htm | 2.30E-12 | 0.02676462 | 0.313 | 0.283 | 6.19E-08 |
| Senp2 | 5.07E-18 | -0.000370693 | 0.317 | 0.347 | 1.37E-13 | Itgb1 | 2.37E-12 | 0.037907138 | 0.264 | 0.23 | 6.37E-08 |
| 0610010F05Rik | 5.09E-18 | 0.034722633 | 0.364 | 0.381 | 1.37E-13 | Gmeb1 | 2.40E-12 | 0.049519543 | 0.262 | 0.227 | 6.46E-08 |
| Gon4l | 5.24E-18 | 0.065413447 | 0.582 | 0.588 | 1.41E-13 | Lpgat1 | 2.41E-12 | -0.033229459 | 0.387 | 0.376 | 6.49E-08 |
| Snca | 5.41E-18 | -0.126592545 | 0.517 | 0.574 | 1.45E-13 | Dlg4 | 2.42E-12 | 0.026255343 | 0.466 | 0.431 | 6.52E-08 |
| Sf3b2 | 5.48E-18 | 0.015830597 | 0.278 | 0.299 | 1.47E-13 | Sos1 | 2.44E-12 | -0.01430615 | 0.388 | 0.367 | 6.56E-08 |
| Stx18 | 5.99E-18 | -0.033018067 | 0.328 | 0.368 | 1.61E-13 | Rab11fip2 | 2.48E-12 | 0.022480158 | 0.462 | 0.425 | 6.68E-08 |
| Tnks | 6.09E-18 | 0.027465376 | 0.424 | 0.447 | 1.64E-13 | Zmynd8 | 2.52E-12 | 0.034910227 | 0.717 | 0.677 | 6.78E-08 |
| Ubr4 | 6.16E-18 | 0.03693363 | 0.248 | 0.257 | 1.66E-13 | Gdi2 | 2.56E-12 | 0.018483467 | 0.275 | 0.247 | 6.88E-08 |
| Safb2 | 6.34E-18 | 0.025487011 | 0.342 | 0.363 | 1.71E-13 | Dlgap4 | 2.58E-12 | -0.038416932 | 0.605 | 0.589 | 6.93E-08 |
| Rps6ka2 | 6.50E-18 | 0.122314145 | 0.368 | 0.335 | 1.75E-13 | Tcp11l1 | 2.60E-12 | 0.039147467 | 0.331 | 0.295 | 7.00E-08 |
| Mib1 | 6.71E-18 | -0.067510689 | 0.461 | 0.515 | 1.81E-13 | Ppp2r5c | 2.63E-12 | -0.032186872 | 0.57 | 0.553 | 7.07E-08 |
| Eef2k | 6.78E-18 | 0.097945732 | 0.282 | 0.262 | 1.82E-13 | Pcsk7 | 2.63E-12 | 0.016228106 | 0.418 | 0.385 | 7.09E-08 |
| Pex14 | 6.81E-18 | 0.0628197 | 0.262 | 0.261 | 1.83E-13 | Tbc1d23 | 2.74E-12 | 0.036756826 | 0.348 | 0.313 | 7.36E-08 |
| Frs2 | 6.85E-18 | 0.003347417 | 0.352 | 0.381 | 1.84E-13 | Ccnh | 2.77E-12 | 0.037085937 | 0.364 | 0.328 | 7.45E-08 |
| Ccdc6 | 7.01E-18 | -0.026236122 | 0.278 | 0.313 | 1.89E-13 | Pdss2 | 2.80E-12 | 0.017501338 | 0.605 | 0.57 | 7.52E-08 |
| Gm26724 | 7.09E-18 | -0.076991167 | 0.282 | 0.333 | 1.91E-13 | Hagh | 2.87E-12 | 0.01721161 | 0.255 | 0.229 | 7.71E-08 |
| Anp32a | 7.15E-18 | 0.008227933 | 0.249 | 0.271 | 1.92E-13 | R3hdm1 | 2.87E-12 | 0.085998326 | 0.939 | 0.922 | 7.73E-08 |
| Ankfy1 | 7.20E-18 | 0.043583308 | 0.324 | 0.334 | 1.94E-13 | Osbpl9 | 2.95E-12 | 0.042731374 | 0.517 | 0.475 | 7.94E-08 |
| Ktn1 | 7.60E-18 | -0.090393056 | 0.559 | 0.615 | 2.05E-13 | Ywhaz | 3.03E-12 | 0.046996936 | 0.527 | 0.484 | 8.15E-08 |
| Phf6 | 7.82E-18 | -0.048230521 | 0.295 | 0.339 | 2.10E-13 | Ophn1 | 3.06E-12 | -0.04700702 | 0.415 | 0.407 | 8.23E-08 |
| Pip5k1a | 7.90E-18 | -0.057432558 | 0.522 | 0.575 | 2.13E-13 | Rab28 | 3.09E-12 | -0.014048999 | 0.376 | 0.358 | 8.33E-08 |
| Trpm3 | 7.98E-18 | -0.151220383 | 0.617 | 0.612 | 2.15E-13 | Klhl2 | 3.11E-12 | 0.012072525 | 0.685 | 0.648 | 8.38E-08 |
| Dhx57 | 8.12E-18 | 0.042452934 | 0.371 | 0.382 | 2.19E-13 | Nmt2 | 3.14E-12 | 0.001571116 | 0.433 | 0.406 | 8.44E-08 |
| Camsap2 | 8.22E-18 | -0.006211784 | 0.372 | 0.404 | 2.21E-13 | Dlg1 | 3.16E-12 | 0.057688721 | 0.688 | 0.645 | 8.50E-08 |
| Fam184a | 8.25E-18 | 0.050633573 | 0.297 | 0.303 | 2.22E-13 | Eml4 | 3.21E-12 | -0.017156607 | 0.559 | 0.535 | 8.63E-08 |
| Sec24a | 8.36E-18 | 0.035795865 | 0.354 | 0.369 | 2.25E-13 | Adcy1 | 3.22E-12 | -0.044505619 | 0.596 | 0.579 | 8.68E-08 |
| Ranbp2 | 8.49E-18 | -0.055945076 | 0.405 | 0.454 | 2.28E-13 | Dhx57 | 3.30E-12 | -0.013687622 | 0.392 | 0.371 | 8.89E-08 |
| Ap3m1 | 8.51E-18 | -0.078774913 | 0.241 | 0.29 | 2.29E-13 | Vamp4 | 3.34E-12 | 0.02829922 | 0.271 | 0.24 | 8.98E-08 |
| Mnat1 | 8.54E-18 | -0.011377743 | 0.411 | 0.448 | 2.30E-13 | Zfp398 | 3.39E-12 | 0.006723517 | 0.46 | 0.43 | 9.12E-08 |
| Desi2 | 8.88E-18 | -0.016452262 | 0.438 | 0.478 | 2.39E-13 | Rnf13 | 3.43E-12 | 0.001416608 | 0.311 | 0.286 | 9.23E-08 |
| Ice1 | 9.03E-18 | 0.013063155 | 0.443 | 0.47 | 2.43E-13 | Map3k7 | 3.46E-12 | -0.007930967 | 0.292 | 0.273 | 9.30E-08 |
| Spen | 9.15E-18 | 0.037731784 | 0.31 | 0.323 | 2.46E-13 | Med12l | 3.46E-12 | -0.072868936 | 0.715 | 0.718 | 9.30E-08 |
| Cd2ap | 9.15E-18 | 0.035406414 | 0.351 | 0.366 | 2.46E-13 | Wdr26 | 3.47E-12 | 0.021935852 | 0.373 | 0.341 | 9.34E-08 |
| Gsk3b | 9.19E-18 | 0.062106097 | 0.623 | 0.63 | 2.47E-13 | Arhgap35 | 3.50E-12 | 0.004415895 | 0.476 | 0.445 | 9.41E-08 |
| Chpt1 | 9.26E-18 | -0.05924816 | 0.309 | 0.355 | 2.49E-13 | Fam76a | 3.52E-12 | 0.049918371 | 0.256 | 0.222 | 9.46E-08 |
| Plaa | 9.38E-18 | 0.002969254 | 0.24 | 0.264 | 2.52E-13 | Satb2 | 3.60E-12 | 0.128553198 | 0.31 | 0.267 | 9.69E-08 |
| Cdk8 | 9.44E-18 | 0.028901543 | 0.599 | 0.624 | 2.54E-13 | Rasgef1a | 3.64E-12 | -0.054178152 | 0.631 | 0.624 | 9.80E-08 |
| Plekhm3 | 9.51E-18 | 0.058808058 | 0.42 | 0.425 | 2.56E-13 | Gm15577 | 3.68E-12 | -0.063268352 | 0.344 | 0.338 | 9.91E-08 |
| Kpna3 | 9.79E-18 | 0.024917254 | 0.394 | 0.415 | 2.63E-13 | Ubr1 | 3.70E-12 | 0.031279926 | 0.437 | 0.401 | 9.96E-08 |
| Trim46 | 9.85E-18 | 0.097055005 | 0.331 | 0.311 | 2.65E-13 | Nrp2 | 3.72E-12 | -0.063214408 | 0.403 | 0.399 | 1.00E-07 |
| Ddx46 | 9.87E-18 | -0.064885424 | 0.438 | 0.49 | 2.66E-13 | Fubp3 | 3.81E-12 | 0.019210846 | 0.287 | 0.259 | 1.03E-07 |
| Cdon | 1.02E-17 | 0.072687488 | 0.291 | 0.293 | 2.75E-13 | Mphosph9 | 3.84E-12 | -0.007431081 | 0.404 | 0.382 | 1.03E-07 |
| Gmds | 1.03E-17 | 0.096933591 | 0.546 | 0.534 | 2.78E-13 | Kdm5c | 3.86E-12 | 0.033834954 | 0.413 | 0.376 | 1.04E-07 |
| Tox4 | 1.07E-17 | -0.073069444 | 0.214 | 0.26 | 2.88E-13 | Plekha1 | 3.87E-12 | 0.034512479 | 0.372 | 0.335 | 1.04E-07 |
| Npas2 | 1.07E-17 | 0.133517206 | 0.439 | 0.392 | 2.88E-13 | Smad3 | 3.88E-12 | -0.02369434 | 0.432 | 0.407 | 1.04E-07 |
| Dscaml1 | 1.08E-17 | 0.097830708 | 0.823 | 0.825 | 2.90E-13 | Faah | 3.88E-12 | 0.005324628 | 0.52 | 0.488 | 1.04E-07 |
| Mosmo | 1.08E-17 | -0.046180754 | 0.248 | 0.289 | 2.90E-13 | Mnat1 | 3.91E-12 | 0.03365418 | 0.45 | 0.411 | 1.05E-07 |
| Mmd2 | 1.11E-17 | -0.042572519 | 0.303 | 0.346 | 2.99E-13 | Parp6 | 3.93E-12 | -0.010372207 | 0.313 | 0.295 | 1.06E-07 |
| Sv2b | 1.11E-17 | 0.117827449 | 0.549 | 0.492 | 2.99E-13 | Nup214 | 3.93E-12 | 0.019513707 | 0.303 | 0.273 | 1.06E-07 |
| Tial1 | 1.13E-17 | -0.053502143 | 0.379 | 0.43 | 3.03E-13 | Ppp1r14c | 3.96E-12 | 0.071663121 | 0.34 | 0.298 | 1.06E-07 |
| Arpp21 | 1.14E-17 | 0.085062754 | 0.951 | 0.949 | 3.06E-13 | Larp4b | 4.11E-12 | -0.026108822 | 0.461 | 0.445 | 1.11E-07 |
| Ap4s1 | 1.14E-17 | 0.069039179 | 0.255 | 0.25 | 3.07E-13 | 2-Mar | 4.12E-12 | 0.007021621 | 0.259 | 0.235 | 1.11E-07 |
| Ppp3cb | 1.15E-17 | -0.025093207 | 0.344 | 0.384 | 3.10E-13 | Plaa | 4.25E-12 | 0.003458561 | 0.26 | 0.24 | 1.14E-07 |
| Nrg2 | 1.16E-17 | 0.143575634 | 0.536 | 0.521 | 3.11E-13 | Neo1 | 4.28E-12 | -0.012697057 | 0.501 | 0.478 | 1.15E-07 |
| Agk | 1.16E-17 | 0.073059422 | 0.284 | 0.278 | 3.12E-13 | Pgm2l1 | 4.32E-12 | 0.046960528 | 0.358 | 0.32 | 1.16E-07 |
| Lnpk | 1.25E-17 | 0.071158842 | 0.358 | 0.358 | 3.36E-13 | Leng8 | 4.36E-12 | 0.032370309 | 0.427 | 0.389 | 1.17E-07 |
| Commd10 | 1.26E-17 | -0.007073741 | 0.286 | 0.316 | 3.38E-13 | Itch | 4.37E-12 | 0.029852809 | 0.578 | 0.54 | 1.17E-07 |
| Mindy3 | 1.26E-17 | -0.002604853 | 0.293 | 0.323 | 3.40E-13 | Stxbp1 | 4.39E-12 | 0.020364986 | 0.603 | 0.567 | 1.18E-07 |
| Uggt1 | 1.27E-17 | -0.052274382 | 0.266 | 0.31 | 3.42E-13 | Tmem191c | 4.41E-12 | 0.042071936 | 0.586 | 0.544 | 1.19E-07 |
| Ppp1r12b | 1.31E-17 | 0.094062367 | 0.699 | 0.693 | 3.52E-13 | Hnrnpll | 4.42E-12 | 0.059339194 | 0.395 | 0.353 | 1.19E-07 |
| Jak2 | 1.35E-17 | -0.085118792 | 0.247 | 0.296 | 3.65E-13 | Sorcs2 | 4.46E-12 | -0.050606104 | 0.387 | 0.376 | 1.20E-07 |
| Prdm2 | 1.36E-17 | -0.01706137 | 0.331 | 0.366 | 3.66E-13 | Pdzrn4 | 4.53E-12 | -0.137890116 | 0.358 | 0.355 | 1.22E-07 |
| Ddx10 | 1.37E-17 | 0.043838787 | 0.359 | 0.369 | 3.68E-13 | Kif3a | 4.54E-12 | 0.032615134 | 0.432 | 0.396 | 1.22E-07 |
| Fndc3a | 1.42E-17 | -0.087270138 | 0.358 | 0.411 | 3.82E-13 | Slc17a5 | 4.59E-12 | 0.053434816 | 0.328 | 0.29 | 1.23E-07 |
| Grm7 | 1.45E-17 | 0.06761068 | 0.99 | 0.982 | 3.90E-13 | Kcnip2 | 4.64E-12 | 0.022356867 | 0.374 | 0.341 | 1.25E-07 |
| Gnl3 | 1.48E-17 | -0.014675928 | 0.257 | 0.289 | 3.97E-13 | Smg5 | 4.69E-12 | 0.038628478 | 0.28 | 0.247 | 1.26E-07 |
| Pkia | 1.50E-17 | -0.049770893 | 0.414 | 0.46 | 4.04E-13 | St13 | 4.70E-12 | 0.053311822 | 0.311 | 0.272 | 1.26E-07 |
| Eaf2 | 1.51E-17 | -0.064917919 | 0.214 | 0.258 | 4.05E-13 | Ppp2r2c | 4.76E-12 | -0.034811707 | 0.522 | 0.507 | 1.28E-07 |
| Pdpk1 | 1.53E-17 | -0.014990779 | 0.337 | 0.373 | 4.12E-13 | Car10 | 4.77E-12 | -0.157997149 | 0.604 | 0.612 | 1.28E-07 |
| Tro | 1.54E-17 | 0.004772739 | 0.436 | 0.466 | 4.14E-13 | Csnk1g3 | 4.80E-12 | 0.033619314 | 0.508 | 0.469 | 1.29E-07 |
| Golgb1 | 1.57E-17 | 0.073104025 | 0.486 | 0.484 | 4.23E-13 | Ap4s1 | 4.88E-12 | 0.042117312 | 0.289 | 0.255 | 1.31E-07 |
| Camta1 | 1.62E-17 | 0.100215669 | 0.833 | 0.828 | 4.35E-13 | Strbp | 4.89E-12 | 0.076220913 | 0.958 | 0.951 | 1.32E-07 |
| Col4a2 | 1.69E-17 | 0.081542418 | 0.333 | 0.327 | 4.55E-13 | Iws1 | 4.91E-12 | 0.050036217 | 0.312 | 0.275 | 1.32E-07 |
| Gatad2b | 1.73E-17 | -0.010672918 | 0.537 | 0.577 | 4.66E-13 | Fkbp1a | 5.04E-12 | 0.04129329 | 0.294 | 0.26 | 1.36E-07 |
| Usp13 | 1.74E-17 | 0.004031966 | 0.232 | 0.255 | 4.68E-13 | Msra | 5.04E-12 | 0.082722221 | 0.897 | 0.874 | 1.36E-07 |
| Pcm1 | 1.76E-17 | 0.010181524 | 0.559 | 0.591 | 4.74E-13 | Zfr | 5.06E-12 | 0.058348068 | 0.77 | 0.73 | 1.36E-07 |
| Trpc1 | 1.80E-17 | -0.006786104 | 0.358 | 0.392 | 4.86E-13 | Klhl32 | 5.11E-12 | 0.011630563 | 0.492 | 0.458 | 1.37E-07 |
| Zfp532 | 1.85E-17 | -0.031540243 | 0.445 | 0.49 | 4.98E-13 | Ccbe1 | 5.21E-12 | -0.088747476 | 0.325 | 0.32 | 1.40E-07 |
| Fam13b | 1.86E-17 | 0.020773877 | 0.404 | 0.427 | 5.00E-13 | Igf2bp3 | 5.30E-12 | 0.079028201 | 0.451 | 0.405 | 1.43E-07 |
| Map7d2 | 1.92E-17 | 0.083921224 | 0.477 | 0.471 | 5.15E-13 | Tbcel | 5.32E-12 | -0.012727001 | 0.293 | 0.277 | 1.43E-07 |
| Smarca1 | 1.92E-17 | -0.02670384 | 0.265 | 0.3 | 5.16E-13 | Mtch1 | 5.36E-12 | 0.016610101 | 0.259 | 0.233 | 1.44E-07 |
| Cyfip2 | 1.93E-17 | 0.049284107 | 0.648 | 0.666 | 5.20E-13 | Cpt1c | 5.40E-12 | 0.013226462 | 0.376 | 0.346 | 1.45E-07 |
| Zfp831 | 1.95E-17 | 0.053354728 | 0.331 | 0.336 | 5.25E-13 | Snap47 | 5.48E-12 | -0.011545075 | 0.271 | 0.256 | 1.47E-07 |
| Ptcd3 | 1.97E-17 | -0.037797603 | 0.217 | 0.254 | 5.30E-13 | Ankrd28 | 5.57E-12 | 0.043016701 | 0.473 | 0.433 | 1.50E-07 |
| Kifap3 | 1.97E-17 | 0.028829127 | 0.525 | 0.548 | 5.30E-13 | Icam5 | 5.60E-12 | -0.010828526 | 0.508 | 0.486 | 1.51E-07 |
| Strada | 1.98E-17 | 0.075442887 | 0.35 | 0.344 | 5.31E-13 | Dab2ip | 5.66E-12 | -0.014371737 | 0.291 | 0.273 | 1.52E-07 |
| Sqstm1 | 2.02E-17 | 0.008837213 | 0.291 | 0.317 | 5.43E-13 | Ppm1a | 5.70E-12 | 0.039594532 | 0.3 | 0.265 | 1.53E-07 |

| | | | | | | | | | | | |
|---|---|---|---|---|---|---|---|---|---|---|---|
| Snx1 | 2.04E-17 | -0.033771096 | 0.231 | 0.266 | 5.48E-13 | Clock | 5.70E-12 | -0.031511086 | 0.526 | 0.513 | 1.53E-07 |
| Cab39 | 2.11E-17 | -0.015738022 | 0.328 | 0.362 | 5.68E-13 | Foxk2 | 5.72E-12 | 0.019487612 | 0.32 | 0.291 | 1.54E-07 |
| Aftph | 2.16E-17 | 0.011861116 | 0.401 | 0.427 | 5.81E-13 | Apba2 | 5.84E-12 | -0.043039996 | 0.821 | 0.808 | 1.57E-07 |
| Snapc3 | 2.17E-17 | -0.037708077 | 0.366 | 0.409 | 5.85E-13 | Kpna4 | 5.84E-12 | 0.039903857 | 0.471 | 0.431 | 1.57E-07 |
| Chd3 | 2.20E-17 | -0.010375922 | 0.384 | 0.42 | 5.91E-13 | Gak | 5.88E-12 | -0.00437664 | 0.313 | 0.292 | 1.58E-07 |
| Casc4 | 2.23E-17 | -0.100455837 | 0.6 | 0.655 | 6.01E-13 | Trappc8 | 5.89E-12 | 0.007233052 | 0.434 | 0.406 | 1.58E-07 |
| Ypel2 | 2.25E-17 | 0.06343635 | 0.256 | 0.258 | 6.04E-13 | Megf9 | 5.93E-12 | -0.034926369 | 0.488 | 0.476 | 1.60E-07 |
| Rhot1 | 2.25E-17 | 0.008185528 | 0.345 | 0.37 | 6.06E-13 | Sgsm3 | 6.01E-12 | 0.032372844 | 0.261 | 0.23 | 1.62E-07 |
| Sugp1 | 2.25E-17 | 0.049346977 | 0.312 | 0.317 | 6.07E-13 | Phf14 | 6.04E-12 | -0.001240965 | 0.634 | 0.604 | 1.63E-07 |
| Ube2d3 | 2.26E-17 | -0.033461761 | 0.301 | 0.342 | 6.09E-13 | Pfkp | 6.29E-12 | -0.035052961 | 0.409 | 0.398 | 1.69E-07 |
| Acot7 | 2.29E-17 | 0.053722809 | 0.31 | 0.311 | 6.15E-13 | Pogz | 6.32E-12 | 0.034150741 | 0.334 | 0.3 | 1.70E-07 |
| Ppp4r3b | 2.31E-17 | -0.008954846 | 0.354 | 0.388 | 6.20E-13 | Negr1 | 6.43E-12 | -0.131728341 | 0.949 | 0.956 | 1.73E-07 |
| St13 | 2.35E-17 | -0.05904175 | 0.272 | 0.317 | 6.32E-13 | Lmbr1 | 6.51E-12 | -0.067078682 | 0.367 | 0.375 | 1.75E-07 |
| E130308A19Rik | 2.38E-17 | -0.005642399 | 0.234 | 0.26 | 6.40E-13 | Nfx1 | 6.52E-12 | -0.018131267 | 0.515 | 0.493 | 1.76E-07 |
| Ep300 | 2.39E-17 | -0.037099182 | 0.371 | 0.415 | 6.42E-13 | Slc7a8 | 6.53E-12 | 0.036665019 | 0.369 | 0.333 | 1.76E-07 |
| Dot1l | 2.39E-17 | -0.034426345 | 0.347 | 0.389 | 6.44E-13 | Dis3l2 | 6.67E-12 | -0.016972119 | 0.656 | 0.633 | 1.80E-07 |
| Pgbd5 | 2.42E-17 | -0.121895709 | 0.503 | 0.56 | 6.52E-13 | Tmem161b | 6.68E-12 | -0.01801306 | 0.444 | 0.422 | 1.80E-07 |
| Unc5a | 2.56E-17 | 0.056255264 | 0.322 | 0.326 | 6.88E-13 | Zfyve1 | 6.69E-12 | 0.05651306 | 0.327 | 0.287 | 1.80E-07 |
| Atp2c1 | 2.60E-17 | -0.082674336 | 0.662 | 0.714 | 7.01E-13 | Prepl | 6.70E-12 | -0.033929348 | 0.339 | 0.331 | 1.80E-07 |
| 3110082I17Rik | 2.77E-17 | 0.060576913 | 0.284 | 0.286 | 7.44E-13 | Anks3 | 6.70E-12 | 0.036620993 | 0.356 | 0.319 | 1.80E-07 |
| Ophn1 | 2.80E-17 | -0.041877943 | 0.407 | 0.453 | 7.53E-13 | Ep400 | 6.78E-12 | -0.012084793 | 0.5 | 0.476 | 1.82E-07 |
| Dis3l2 | 2.84E-17 | 0.069402367 | 0.633 | 0.64 | 7.63E-13 | Acsl6 | 6.83E-12 | 0.00326887 | 0.275 | 0.255 | 1.84E-07 |
| Uhrf2 | 2.88E-17 | -0.012551099 | 0.4 | 0.437 | 7.76E-13 | Cdkl5 | 6.87E-12 | 0.062224048 | 0.534 | 0.491 | 1.85E-07 |
| Srrm3 | 2.91E-17 | 0.061722452 | 0.592 | 0.6 | 7.82E-13 | Ric1 | 6.89E-12 | -0.008501204 | 0.345 | 0.326 | 1.85E-07 |
| Arhgef7 | 2.91E-17 | 0.025108602 | 0.504 | 0.526 | 7.82E-13 | Ap1b1 | 7.06E-12 | -0.04781441 | 0.281 | 0.281 | 1.90E-07 |
| Tmeff1 | 2.92E-17 | 0.072922571 | 0.26 | 0.253 | 7.86E-13 | Atp2b3 | 7.46E-12 | 0.034497849 | 0.542 | 0.503 | 2.01E-07 |
| Chrm1 | 2.92E-17 | -0.005705544 | 0.225 | 0.252 | 7.87E-13 | Tmem131l | 7.47E-12 | -0.04172777 | 0.259 | 0.255 | 2.01E-07 |
| 11-Sep | 2.95E-17 | 0.086768355 | 0.442 | 0.434 | 7.93E-13 | Klhl24 | 7.53E-12 | 0.019790306 | 0.28 | 0.252 | 2.03E-07 |
| Fntb | 2.97E-17 | -0.02651891 | 0.301 | 0.338 | 7.99E-13 | Ndufa10 | 7.53E-12 | 0.009755659 | 0.276 | 0.252 | 2.03E-07 |
| Cstf2 | 2.99E-17 | 0.090670611 | 0.251 | 0.229 | 8.05E-13 | Chpt1 | 7.55E-12 | 0.060140472 | 0.348 | 0.309 | 2.03E-07 |
| Tecpr1 | 3.00E-17 | 0.018286655 | 0.291 | 0.312 | 8.08E-13 | Irf2 | 7.61E-12 | -0.014544807 | 0.414 | 0.394 | 2.05E-07 |
| Tead1 | 3.04E-17 | 0.059495395 | 0.456 | 0.476 | 8.18E-13 | Adam23 | 7.63E-12 | -0.049570657 | 0.573 | 0.567 | 2.05E-07 |
| Flnb | 3.07E-17 | 0.029445195 | 0.578 | 0.6 | 8.25E-13 | Unc5a | 7.67E-12 | -0.001706183 | 0.344 | 0.322 | 2.06E-07 |
| Mmp17 | 3.08E-17 | -0.015788689 | 0.26 | 0.292 | 8.29E-13 | Wdr20 | 7.68E-12 | 0.030523829 | 0.323 | 0.29 | 2.07E-07 |
| Phf14 | 3.13E-17 | -0.081566186 | 0.604 | 0.658 | 8.43E-13 | Mlxip | 7.68E-12 | 0.040630024 | 0.287 | 0.252 | 2.07E-07 |
| Ankrd27 | 3.18E-17 | 0.100913247 | 0.251 | 0.217 | 8.55E-13 | Eif4g1 | 7.88E-12 | -0.003865188 | 0.431 | 0.405 | 2.12E-07 |
| Prrc2c | 3.20E-17 | 0.001029666 | 0.616 | 0.653 | 8.60E-13 | Usp14 | 7.93E-12 | 0.042331141 | 0.327 | 0.291 | 2.13E-07 |
| Cdc27 | 3.25E-17 | -0.012814137 | 0.284 | 0.316 | 8.74E-13 | Ranbp17 | 8.00E-12 | -0.067240456 | 0.341 | 0.346 | 2.15E-07 |
| Lrp11 | 3.28E-17 | -0.014777432 | 0.286 | 0.32 | 8.82E-13 | Kif1a | 8.05E-12 | -0.0159187 | 0.559 | 0.539 | 2.16E-07 |
| Ogfod1 | 3.32E-17 | -0.007465796 | 0.27 | 0.3 | 8.95E-13 | Lnpk | 8.17E-12 | 0.017928593 | 0.39 | 0.358 | 2.20E-07 |
| Ubr1 | 3.34E-17 | -0.010135532 | 0.401 | 0.435 | 9.00E-13 | Lrrc40 | 8.19E-12 | 0.042321259 | 0.257 | 0.224 | 2.20E-07 |
| Arhgap26 | 3.35E-17 | 0.11100714 | 0.71 | 0.669 | 9.02E-13 | Gm47271 | 8.21E-12 | 0.078093093 | 0.334 | 0.291 | 2.21E-07 |
| Lrrc4c | 3.40E-17 | 0.079759853 | 0.918 | 0.896 | 9.16E-13 | Nisch | 8.27E-12 | -0.04061311 | 0.515 | 0.504 | 2.23E-07 |
| Kifc2 | 3.41E-17 | 0.013072019 | 0.313 | 0.341 | 9.16E-13 | Rtf1 | 8.29E-12 | 0.019051522 | 0.356 | 0.325 | 2.23E-07 |
| Stim2 | 3.47E-17 | -0.095709509 | 0.511 | 0.567 | 9.32E-13 | Cers5 | 8.40E-12 | -0.001426339 | 0.308 | 0.288 | 2.26E-07 |
| Ppig | 3.54E-17 | 0.013845128 | 0.349 | 0.373 | 9.53E-13 | Upf2 | 8.47E-12 | -0.015934674 | 0.422 | 0.402 | 2.28E-07 |
| Arfgap1 | 3.55E-17 | -0.013215106 | 0.275 | 0.306 | 9.55E-13 | Sema4d | 8.51E-12 | -0.049871736 | 0.335 | 0.329 | 2.29E-07 |
| Mllt10 | 3.56E-17 | 0.074820926 | 0.768 | 0.776 | 9.57E-13 | Trrap | 8.67E-12 | -0.019470054 | 0.419 | 0.4 | 2.33E-07 |
| Dixdc1 | 3.58E-17 | 0.038452533 | 0.315 | 0.329 | 9.64E-13 | Rb1cc1 | 8.72E-12 | 0.041809221 | 0.6 | 0.559 | 2.35E-07 |
| Tpp2 | 3.59E-17 | -0.04390312 | 0.325 | 0.368 | 9.66E-13 | Ccar1 | 8.73E-12 | 0.049927607 | 0.498 | 0.456 | 2.35E-07 |
| Cd47 | 3.61E-17 | 0.04380003 | 0.372 | 0.386 | 9.70E-13 | Napb | 8.77E-12 | 0.015101736 | 0.316 | 0.288 | 2.36E-07 |
| Ube3c | 3.61E-17 | 0.030102855 | 0.583 | 0.608 | 9.72E-13 | Zfc3h1 | 8.85E-12 | -0.02148759 | 0.407 | 0.392 | 2.38E-07 |
| Pum2 | 3.72E-17 | -0.020985551 | 0.361 | 0.398 | 1.00E-12 | Cabin1 | 8.87E-12 | -0.033412285 | 0.508 | 0.494 | 2.39E-07 |
| St7l | 3.78E-17 | -0.007109168 | 0.371 | 0.404 | 1.02E-12 | Scmh1 | 8.91E-12 | 0.027856883 | 0.648 | 0.609 | 2.40E-07 |
| Shprh | 3.79E-17 | 0.018072186 | 0.408 | 0.432 | 1.02E-12 | Pdpr | 9.04E-12 | 0.046008241 | 0.262 | 0.227 | 2.43E-07 |
| Bod1l | 3.85E-17 | -0.002276504 | 0.449 | 0.482 | 1.04E-12 | Rapgef1 | 9.11E-12 | -0.01252839 | 0.432 | 0.41 | 2.45E-07 |
| Btbd3 | 3.87E-17 | -0.005311027 | 0.325 | 0.359 | 1.04E-12 | Phc2 | 9.22E-12 | -0.015008099 | 0.537 | 0.515 | 2.48E-07 |
| Ogdh | 3.93E-17 | 0.022527617 | 0.404 | 0.425 | 1.06E-12 | Arnt | 9.25E-12 | 0.059101095 | 0.454 | 0.411 | 2.49E-07 |
| Usp24 | 3.95E-17 | 0.017339485 | 0.449 | 0.476 | 1.06E-12 | Armc9 | 9.27E-12 | 0.007371532 | 0.564 | 0.532 | 2.50E-07 |
| Cc2d2a | 4.03E-17 | 0.00191414 | 0.234 | 0.258 | 1.08E-12 | Esrrg | 9.31E-12 | -0.10176461 | 0.47 | 0.489 | 2.50E-07 |
| Dstyk | 4.05E-17 | -0.018776908 | 0.324 | 0.36 | 1.09E-12 | Pik3c2a | 9.36E-12 | 0.039256569 | 0.379 | 0.342 | 2.52E-07 |
| Pik3c2a | 4.06E-17 | -0.057496691 | 0.342 | 0.39 | 1.09E-12 | 6530403H02Rik | 9.48E-12 | -0.143086554 | 0.311 | 0.326 | 2.55E-07 |
| Ccl25 | 4.08E-17 | 0.001450341 | 0.282 | 0.308 | 1.10E-12 | Prpf40a | 9.64E-12 | 0.026070739 | 0.481 | 0.446 | 2.59E-07 |
| Lrp1 | 4.13E-17 | 0.073034101 | 0.388 | 0.384 | 1.11E-12 | Zfp157 | 9.64E-12 | 0.000553558 | 0.268 | 0.248 | 2.60E-07 |
| Cemip | 4.19E-17 | 0.057068348 | 0.354 | 0.371 | 1.13E-12 | Rnf182 | 9.76E-12 | 0.047296295 | 0.321 | 0.285 | 2.63E-07 |
| Lrp6 | 4.22E-17 | -0.014275294 | 0.424 | 0.462 | 1.14E-12 | Fam171a1 | 9.82E-12 | -0.059671265 | 0.381 | 0.386 | 2.64E-07 |
| Plekhm2 | 4.23E-17 | 0.045044881 | 0.308 | 0.317 | 1.14E-12 | Gfm2 | 9.84E-12 | 0.042502596 | 0.329 | 0.293 | 2.65E-07 |
| Fnip2 | 4.27E-17 | 0.096751537 | 0.315 | 0.302 | 1.15E-12 | Mitf | 9.88E-12 | -0.091175289 | 0.262 | 0.28 | 2.66E-07 |
| Slc9a7 | 4.38E-17 | 0.071832406 | 0.423 | 0.419 | 1.18E-12 | Arhgef4 | 1.00E-11 | -0.028700879 | 0.295 | 0.28 | 2.69E-07 |
| Zfp280d | 4.40E-17 | -0.070985528 | 0.589 | 0.643 | 1.18E-12 | Camk2g | 1.01E-11 | 0.034034135 | 0.374 | 0.339 | 2.71E-07 |
| Rcor3 | 4.44E-17 | 0.062650366 | 0.268 | 0.265 | 1.20E-12 | Smarcc2 | 1.01E-11 | 0.024183563 | 0.396 | 0.365 | 2.72E-07 |
| Trak2 | 4.57E-17 | -0.059749148 | 0.228 | 0.27 | 1.23E-12 | Zzz3 | 1.02E-11 | 0.045339567 | 0.352 | 0.315 | 2.76E-07 |
| Smurf1 | 4.58E-17 | 0.082872383 | 0.311 | 0.3 | 1.23E-12 | Gle1 | 1.03E-11 | 0.055581132 | 0.26 | 0.224 | 2.77E-07 |
| Rptor | 4.59E-17 | 0.047425502 | 0.584 | 0.6 | 1.23E-12 | Ptpn5 | 1.04E-11 | -0.016596622 | 0.371 | 0.352 | 2.79E-07 |
| Atl3 | 4.64E-17 | -0.039365616 | 0.238 | 0.277 | 1.25E-12 | Rnf14 | 1.04E-11 | 0.01781961 | 0.324 | 0.296 | 2.79E-07 |
| Gm42439 | 4.70E-17 | -0.120359327 | 0.651 | 0.702 | 1.27E-12 | Abca3 | 1.04E-11 | 0.005963525 | 0.306 | 0.282 | 2.81E-07 |
| Neto1 | 4.73E-17 | 0.10606394 | 0.558 | 0.554 | 1.27E-12 | Pip4k2a | 1.06E-11 | -0.068562819 | 0.327 | 0.338 | 2.86E-07 |
| Gm6994 | 4.74E-17 | -0.115883021 | 0.307 | 0.36 | 1.27E-12 | Epc1 | 1.10E-11 | 0.051590135 | 0.529 | 0.487 | 2.95E-07 |
| Fubp3 | 4.74E-17 | 0.032775826 | 0.259 | 0.27 | 1.28E-12 | Fbxl5 | 1.10E-11 | 0.039594167 | 0.417 | 0.378 | 2.97E-07 |
| Ttc21b | 4.74E-17 | -0.01723549 | 0.255 | 0.287 | 1.28E-12 | Fam120b | 1.12E-11 | -0.008349312 | 0.391 | 0.371 | 3.02E-07 |
| Rab3gap2 | 4.78E-17 | 0.052463854 | 0.377 | 0.382 | 1.29E-12 | Ankrd10 | 1.12E-11 | 0.002478754 | 0.336 | 0.311 | 3.02E-07 |
| Faf2 | 4.87E-17 | 0.004987352 | 0.268 | 0.292 | 1.31E-12 | Spindoc | 1.12E-11 | 0.035558077 | 0.278 | 0.246 | 3.03E-07 |
| Itsn2 | 4.93E-17 | 0.06821421 | 0.577 | 0.578 | 1.33E-12 | Fbxw7 | 1.13E-11 | -0.008741861 | 0.538 | 0.514 | 3.03E-07 |
| Lmtk3 | 5.06E-17 | 0.066393647 | 0.393 | 0.392 | 1.36E-12 | Ascc2 | 1.13E-11 | -0.021325237 | 0.294 | 0.281 | 3.03E-07 |
| Setd2 | 5.21E-17 | 0.005104122 | 0.449 | 0.482 | 1.40E-12 | Kif5b | 1.13E-11 | 0.049460983 | 0.476 | 0.434 | 3.04E-07 |
| Trpm7 | 5.22E-17 | -0.088226668 | 0.634 | 0.687 | 1.40E-12 | Aatk | 1.13E-11 | -0.028265649 | 0.269 | 0.26 | 3.05E-07 |
| Vps13d | 5.24E-17 | -5.84E-05 | 0.512 | 0.544 | 1.41E-12 | Snrnp70 | 1.14E-11 | -0.033660724 | 0.86 | 0.844 | 3.08E-07 |
| Ccdc30 | 5.30E-17 | -0.04265726 | 0.475 | 0.521 | 1.43E-12 | Uri1 | 1.16E-11 | 0.003544159 | 0.361 | 0.336 | 3.11E-07 |
| Pak2 | 5.35E-17 | -0.027008438 | 0.252 | 0.288 | 1.44E-12 | Cdc40 | 1.17E-11 | 0.025456805 | 0.493 | 0.458 | 3.14E-07 |
| Thada | 5.68E-17 | 0.028214732 | 0.444 | 0.462 | 1.53E-12 | Msantd2 | 1.18E-11 | 0.025823265 | 0.264 | 0.234 | 3.17E-07 |
| Gm26905 | 5.68E-17 | -0.08754693 | 0.334 | 0.297 | 1.53E-12 | Ssb | 1.20E-11 | 0.005127288 | 0.269 | 0.248 | 3.24E-07 |
| Rnf38 | 5.75E-17 | 0.053275464 | 0.262 | 0.265 | 1.55E-12 | Unc5d | 1.20E-11 | -0.135999981 | 0.467 | 0.508 | 3.24E-07 |
| Wwox | 5.82E-17 | 0.112840457 | 0.519 | 0.516 | 1.57E-12 | Clcn3 | 1.21E-11 | 0.039006738 | 0.453 | 0.417 | 3.26E-07 |
| Rnf13 | 5.84E-17 | 0.070835509 | 0.286 | 0.281 | 1.57E-12 | Zkscan3 | 1.21E-11 | -0.012966211 | 0.332 | 0.315 | 3.27E-07 |
| Lnpep | 5.85E-17 | 0.009221902 | 0.399 | 0.426 | 1.57E-12 | Fam214a | 1.22E-11 | -0.046438663 | 0.427 | 0.422 | 3.27E-07 |
| Clasp1 | 5.89E-17 | 0.070372618 | 0.566 | 0.57 | 1.58E-12 | Map4k4 | 1.22E-11 | 0.021348618 | 0.46 | 0.427 | 3.29E-07 |
| Reep1 | 5.95E-17 | 0.075172114 | 0.547 | 0.544 | 1.60E-12 | Cnot1 | 1.22E-11 | -0.003954193 | 0.458 | 0.433 | 3.29E-07 |
| Calm2 | 6.06E-17 | -0.067219273 | 0.544 | 0.598 | 1.63E-12 | Rnf157 | 1.24E-11 | 0.007434263 | 0.617 | 0.586 | 3.34E-07 |

| | | | | | | | | | | | |
|---|---|---|---|---|---|---|---|---|---|---|---|
| Snx25 | 6.15E-17 | 0.009703204 | 0.276 | 0.296 | 1.65E-12 | Trim35 | 1.25E-11 | 0.018071088 | 0.555 | 0.52 | 3.36E-07 |
| Lrig2 | 6.24E-17 | 0.015771413 | 0.44 | 0.465 | 1.68E-12 | Map3k2 | 1.26E-11 | -0.024416038 | 0.408 | 0.393 | 3.38E-07 |
| Ap2a1 | 6.25E-17 | -0.069881767 | 0.24 | 0.286 | 1.68E-12 | Parg | 1.28E-11 | -0.009143368 | 0.397 | 0.377 | 3.45E-07 |
| Mia3 | 6.28E-17 | -0.06027483 | 0.263 | 0.308 | 1.69E-12 | Rtn3 | 1.29E-11 | 0.073865177 | 0.909 | 0.887 | 3.48E-07 |
| Scfd1 | 6.28E-17 | -0.002913961 | 0.322 | 0.352 | 1.69E-12 | Zfp644 | 1.30E-11 | 0.042757272 | 0.589 | 0.549 | 3.50E-07 |
| Slc39a10 | 6.34E-17 | -0.058305191 | 0.24 | 0.284 | 1.71E-12 | Hnrnph3 | 1.30E-11 | 0.0172732 | 0.412 | 0.38 | 3.51E-07 |
| Fam219a | 6.35E-17 | 0.078029009 | 0.513 | 0.514 | 1.71E-12 | Asxl1 | 1.31E-11 | 0.038289877 | 0.502 | 0.463 | 3.52E-07 |
| Thsd7a | 6.39E-17 | 0.204823312 | 0.444 | 0.412 | 1.72E-12 | Sfswap | 1.33E-11 | -0.068316886 | 0.524 | 0.531 | 3.58E-07 |
| Agfg2 | 6.42E-17 | 0.083124168 | 0.327 | 0.316 | 1.73E-12 | Tmem260 | 1.35E-11 | 0.009790725 | 0.296 | 0.271 | 3.63E-07 |
| Prkar1a | 6.66E-17 | -0.064920158 | 0.243 | 0.289 | 1.79E-12 | Scaf4 | 1.35E-11 | -0.013095359 | 0.366 | 0.349 | 3.64E-07 |
| Mcph1 | 6.67E-17 | 0.060640122 | 0.383 | 0.385 | 1.80E-12 | Sbno1 | 1.36E-11 | 0.031490771 | 0.519 | 0.481 | 3.65E-07 |
| Capza1 | 6.81E-17 | 0.009672657 | 0.289 | 0.31 | 1.83E-12 | Slc7a6 | 1.38E-11 | 0.038460736 | 0.366 | 0.33 | 3.70E-07 |
| Rprd2 | 6.90E-17 | -0.017375008 | 0.371 | 0.408 | 1.86E-12 | Kcnc3 | 1.39E-11 | -0.005797535 | 0.384 | 0.36 | 3.73E-07 |
| Dnm2 | 6.96E-17 | 0.04005098 | 0.454 | 0.471 | 1.87E-12 | Gtf2a1 | 1.39E-11 | 0.04774212 | 0.262 | 0.227 | 3.73E-07 |
| Vopp1 | 7.04E-17 | -0.009651408 | 0.231 | 0.26 | 1.90E-12 | Ppp4r3b | 1.40E-11 | 0.023804946 | 0.386 | 0.354 | 3.77E-07 |
| Prrc2b | 7.14E-17 | 0.083918486 | 0.516 | 0.504 | 1.92E-12 | Sestd1 | 1.41E-11 | -0.016253734 | 0.342 | 0.325 | 3.80E-07 |
| Serbp1 | 7.17E-17 | -0.050454933 | 0.252 | 0.294 | 1.93E-12 | Lztr1 | 1.43E-11 | -0.036516409 | 0.25 | 0.246 | 3.85E-07 |
| Cnot6l | 7.27E-17 | -0.035249067 | 0.362 | 0.403 | 1.96E-12 | Myo9b | 1.45E-11 | -0.031816573 | 0.396 | 0.386 | 3.91E-07 |
| Kmt2e | 7.46E-17 | 0.038053802 | 0.628 | 0.654 | 2.01E-12 | Slc16a10 | 1.46E-11 | 0.007738306 | 0.256 | 0.233 | 3.92E-07 |
| Pxk | 7.53E-17 | -0.031824442 | 0.299 | 0.339 | 2.03E-12 | Rps6kc1 | 1.47E-11 | -0.043424564 | 0.337 | 0.334 | 3.95E-07 |
| Atg4c | 7.74E-17 | 0.025771168 | 0.329 | 0.347 | 2.08E-12 | Focad | 1.47E-11 | -0.040972289 | 0.654 | 0.645 | 3.95E-07 |
| Ubn1 | 7.84E-17 | -0.010753314 | 0.297 | 0.327 | 2.11E-12 | Odf2 | 1.49E-11 | -0.040999659 | 0.503 | 0.492 | 4.00E-07 |
| Mtmr3 | 7.87E-17 | 0.047621011 | 0.528 | 0.542 | 2.12E-12 | Fermt2 | 1.53E-11 | 0.012407815 | 0.401 | 0.371 | 4.10E-07 |
| Srpk1 | 7.96E-17 | 0.057690464 | 0.349 | 0.35 | 2.14E-12 | Comt | 1.53E-11 | -0.011709801 | 0.293 | 0.277 | 4.13E-07 |
| Elavl1 | 8.07E-17 | -8.20E-05 | 0.293 | 0.319 | 2.17E-12 | Psmb2 | 1.54E-11 | 0.015652803 | 0.323 | 0.295 | 4.15E-07 |
| Ski | 8.14E-17 | 0.049565302 | 0.378 | 0.387 | 2.19E-12 | Clpx | 1.56E-11 | 0.028292109 | 0.281 | 0.251 | 4.20E-07 |
| Fam131a | 8.36E-17 | -0.002346262 | 0.43 | 0.465 | 2.25E-12 | Ankhd1 | 1.59E-11 | 0.046552372 | 0.63 | 0.588 | 4.27E-07 |
| Ints6 | 8.45E-17 | -0.058284242 | 0.398 | 0.447 | 2.27E-12 | Metap2 | 1.59E-11 | -0.009187502 | 0.362 | 0.341 | 4.29E-07 |
| Kdm5b | 8.70E-17 | 0.017256975 | 0.414 | 0.441 | 2.34E-12 | Kat6a | 1.65E-11 | 0.02940193 | 0.297 | 0.265 | 4.45E-07 |
| Ppip5k1 | 8.83E-17 | 0.043303287 | 0.448 | 0.463 | 2.38E-12 | Tsnax | 1.66E-11 | 0.000607212 | 0.276 | 0.254 | 4.45E-07 |
| Nphp4 | 9.07E-17 | 0.037861653 | 0.301 | 0.31 | 2.44E-12 | Ubr2 | 1.66E-11 | -0.009654764 | 0.506 | 0.482 | 4.47E-07 |
| Col4a3bp | 9.43E-17 | 0.003104917 | 0.333 | 0.359 | 2.54E-12 | Cyld | 1.67E-11 | 0.031670727 | 0.43 | 0.394 | 4.51E-07 |
| Nrxn3 | 9.56E-17 | 0.14646596 | 0.954 | 0.934 | 2.57E-12 | Grb14 | 1.68E-11 | 0.020430492 | 0.34 | 0.311 | 4.51E-07 |
| Cyld | 9.59E-17 | -0.024616266 | 0.394 | 0.434 | 2.58E-12 | N4bp1 | 1.68E-11 | 0.009168785 | 0.278 | 0.253 | 4.52E-07 |
| Zfhx2 | 9.72E-17 | 0.028681297 | 0.343 | 0.358 | 2.62E-12 | Trim44 | 1.68E-11 | -0.024916368 | 0.611 | 0.591 | 4.53E-07 |
| Ago2 | 9.86E-17 | -0.02545375 | 0.37 | 0.407 | 2.65E-12 | Setx | 1.72E-11 | 0.036478122 | 0.42 | 0.384 | 4.64E-07 |
| Miga1 | 9.91E-17 | 0.061925738 | 0.323 | 0.322 | 2.67E-12 | Micu2 | 1.74E-11 | 0.010058451 | 0.394 | 0.367 | 4.69E-07 |
| Mfsd4a | 9.96E-17 | 0.090121409 | 0.375 | 0.362 | 2.68E-12 | Tnk2 | 1.76E-11 | 0.010887231 | 0.275 | 0.249 | 4.74E-07 |
| Wac | 1.01E-16 | -0.042103025 | 0.493 | 0.542 | 2.71E-12 | Gda | 1.76E-11 | 0.042896338 | 0.43 | 0.392 | 4.74E-07 |
| Crk | 1.03E-16 | -0.032482725 | 0.223 | 0.258 | 2.78E-12 | Mindy3 | 1.80E-11 | 0.028492124 | 0.326 | 0.293 | 4.84E-07 |
| Arid2 | 1.05E-16 | 0.002808613 | 0.298 | 0.326 | 2.84E-12 | Tnrc18 | 1.80E-11 | 0.022949168 | 0.411 | 0.379 | 4.84E-07 |
| Dmtn | 1.06E-16 | 0.028441603 | 0.396 | 0.418 | 2.86E-12 | Rapgef5 | 1.83E-11 | 0.072982297 | 0.722 | 0.68 | 4.92E-07 |
| Arfgef1 | 1.07E-16 | -0.011287519 | 0.524 | 0.562 | 2.87E-12 | Nhsl2 | 1.83E-11 | -0.067081411 | 0.489 | 0.485 | 4.92E-07 |
| Ap1g1 | 1.08E-16 | -0.035790614 | 0.325 | 0.365 | 2.91E-12 | Fryl | 1.83E-11 | -0.055075328 | 0.498 | 0.499 | 4.93E-07 |
| Iws1 | 1.10E-16 | -0.035192058 | 0.275 | 0.313 | 2.95E-12 | Trmt1 | 1.85E-11 | 0.030784784 | 0.272 | 0.242 | 4.97E-07 |
| Rbm27 | 1.10E-16 | -0.013488082 | 0.259 | 0.288 | 2.95E-12 | Ilf2 | 1.86E-11 | 0.021839085 | 0.26 | 0.232 | 4.99E-07 |
| Morn1 | 1.10E-16 | 0.076824298 | 0.251 | 0.238 | 2.95E-12 | Rp9 | 1.86E-11 | 0.035550293 | 0.367 | 0.332 | 5.01E-07 |
| Fgfr1 | 1.10E-16 | -0.064015544 | 0.242 | 0.285 | 2.96E-12 | Prkar2a | 1.87E-11 | 0.064541107 | 0.384 | 0.343 | 5.03E-07 |
| Wasf1 | 1.10E-16 | -0.033571767 | 0.673 | 0.718 | 2.97E-12 | Ythdf3 | 1.95E-11 | -0.01038862 | 0.348 | 0.328 | 5.25E-07 |
| Syngap1 | 1.11E-16 | -0.015415467 | 0.272 | 0.304 | 2.99E-12 | Gm26749 | 1.96E-11 | 0.04712162 | 0.436 | 0.396 | 5.28E-07 |
| Cpd | 1.12E-16 | 0.078691963 | 0.314 | 0.304 | 3.01E-12 | Malat1 | 2.01E-11 | 0.080745698 | 0.998 | 1 | 5.40E-07 |
| Khdrbs1 | 1.12E-16 | -0.005383013 | 0.336 | 0.366 | 3.01E-12 | Vps53 | 2.01E-11 | 0.001381337 | 0.287 | 0.267 | 5.41E-07 |
| Tmem260 | 1.15E-16 | 0.004788268 | 0.271 | 0.294 | 3.10E-12 | Lrrn1 | 2.05E-11 | 0.022258522 | 0.277 | 0.249 | 5.52E-07 |
| 4833420G17Rik | 1.20E-16 | -0.026255258 | 0.302 | 0.34 | 3.23E-12 | Scamp1 | 2.11E-11 | -0.015310699 | 0.359 | 0.341 | 5.69E-07 |
| Rasa2 | 1.24E-16 | 0.093061268 | 0.32 | 0.298 | 3.33E-12 | Jak1 | 2.12E-11 | -0.00896596 | 0.393 | 0.374 | 5.70E-07 |
| Gng7 | 1.33E-16 | 0.054492574 | 0.251 | 0.256 | 3.58E-12 | Oxct1 | 2.13E-11 | -0.044864098 | 0.635 | 0.625 | 5.74E-07 |
| 2610035D17Rik | 1.33E-16 | -0.059079743 | 0.465 | 0.514 | 3.59E-12 | Luc7l | 2.14E-11 | 0.002468091 | 0.335 | 0.312 | 5.77E-07 |
| Clpb | 1.42E-16 | 0.020608114 | 0.263 | 0.278 | 3.81E-12 | Rbbp6 | 2.15E-11 | -0.011156697 | 0.364 | 0.346 | 5.79E-07 |
| Gabrg2 | 1.47E-16 | 0.045454426 | 0.567 | 0.586 | 3.94E-12 | Emc2 | 2.22E-11 | 0.023864937 | 0.253 | 0.225 | 5.98E-07 |
| Srrm1 | 1.49E-16 | -0.054858832 | 0.487 | 0.538 | 4.02E-12 | Lrrfip2 | 2.23E-11 | -0.001245957 | 0.345 | 0.322 | 5.99E-07 |
| Gm15563 | 1.49E-16 | -0.098688918 | 0.244 | 0.292 | 4.02E-12 | Crocc | 2.23E-11 | 0.008344191 | 0.291 | 0.267 | 6.01E-07 |
| Cul4a | 1.51E-16 | 0.033066965 | 0.25 | 0.26 | 4.07E-12 | Gm14330 | 2.24E-11 | 0.057419998 | 0.296 | 0.259 | 6.03E-07 |
| Setd3 | 1.53E-16 | 0.056828744 | 0.289 | 0.287 | 4.11E-12 | Kdm4c | 2.26E-11 | 0.049586309 | 0.601 | 0.559 | 6.09E-07 |
| Lztr1 | 1.53E-16 | -0.081929721 | 0.246 | 0.294 | 4.12E-12 | Fto | 2.30E-11 | -0.065964932 | 0.719 | 0.725 | 6.18E-07 |
| Map4k5 | 1.53E-16 | 0.045098856 | 0.486 | 0.5 | 4.12E-12 | Brd8 | 2.31E-11 | -0.005392093 | 0.281 | 0.263 | 6.23E-07 |
| Hmg20a | 1.53E-16 | -0.004561057 | 0.424 | 0.456 | 4.13E-12 | Abhd2 | 2.33E-11 | 0.024188771 | 0.322 | 0.292 | 6.26E-07 |
| Zfat | 1.58E-16 | 0.003435844 | 0.245 | 0.268 | 4.26E-12 | St3gal3 | 2.34E-11 | -0.037775229 | 0.491 | 0.483 | 6.31E-07 |
| Naa15 | 1.59E-16 | -0.020405271 | 0.359 | 0.396 | 4.27E-12 | 4732471J01Rik | 2.36E-11 | -0.084046085 | 0.47 | 0.482 | 6.34E-07 |
| Dyrk1a | 1.62E-16 | 0.005150403 | 0.495 | 0.528 | 4.35E-12 | Axin1 | 2.39E-11 | 0.062633332 | 0.275 | 0.238 | 6.44E-07 |
| BC052040 | 1.63E-16 | 0.070038961 | 0.263 | 0.254 | 4.38E-12 | Preb | 2.42E-11 | 0.031117429 | 0.313 | 0.281 | 6.51E-07 |
| Nek1 | 1.63E-16 | 0.019569646 | 0.393 | 0.415 | 4.39E-12 | Kcmf1 | 2.43E-11 | -0.020528864 | 0.388 | 0.374 | 6.53E-07 |
| Cfap36 | 1.65E-16 | 0.006535438 | 0.321 | 0.349 | 4.43E-12 | Sec23a | 2.43E-11 | 0.029468715 | 0.321 | 0.289 | 6.54E-07 |
| Dmxl2 | 1.67E-16 | 0.073667127 | 0.659 | 0.664 | 4.50E-12 | 9130011E15Rik | 2.51E-11 | -0.079460793 | 0.402 | 0.447 | 6.76E-07 |
| Dexi | 1.71E-16 | -0.034693776 | 0.218 | 0.252 | 4.60E-12 | Gucy1b1 | 2.53E-11 | 0.028218525 | 0.309 | 0.277 | 6.82E-07 |
| Ncoa5 | 1.73E-16 | 0.024412573 | 0.394 | 0.416 | 4.65E-12 | Fcho2 | 2.56E-11 | 0.012241075 | 0.49 | 0.46 | 6.90E-07 |
| Usp7 | 1.78E-16 | 0.031771588 | 0.347 | 0.364 | 4.79E-12 | Glmn | 2.58E-11 | -0.008177548 | 0.281 | 0.265 | 6.95E-07 |
| Sesn1 | 1.79E-16 | 0.071522951 | 0.454 | 0.453 | 4.83E-12 | Crtc1 | 2.59E-11 | -0.033626241 | 0.434 | 0.421 | 6.96E-07 |
| Pvt1 | 1.84E-16 | 0.055396157 | 0.505 | 0.517 | 4.96E-12 | Cdip1 | 2.61E-11 | 0.004172373 | 0.423 | 0.397 | 7.02E-07 |
| Lrrc49 | 1.85E-16 | 0.019791308 | 0.375 | 0.395 | 4.99E-12 | Ank | 2.62E-11 | -0.026042971 | 0.643 | 0.624 | 7.06E-07 |
| Sirpa | 1.86E-16 | 0.019982746 | 0.277 | 0.296 | 4.99E-12 | Pde4a | 2.65E-11 | -0.019865218 | 0.508 | 0.488 | 7.13E-07 |
| H13 | 1.87E-16 | 0.043135246 | 0.253 | 0.26 | 5.04E-12 | Tsc22d2 | 2.67E-11 | -0.018222267 | 0.33 | 0.315 | 7.18E-07 |
| Hdac4 | 1.91E-16 | 0.041996711 | 0.419 | 0.43 | 5.15E-12 | Plppr4 | 2.67E-11 | 0.077004122 | 0.623 | 0.579 | 7.20E-07 |
| Dennd4a | 1.93E-16 | 0.060171993 | 0.578 | 0.586 | 5.19E-12 | Lemd3 | 2.71E-11 | -0.001157706 | 0.257 | 0.237 | 7.30E-07 |
| Eno2 | 1.95E-16 | -0.017715343 | 0.287 | 0.321 | 5.26E-12 | Kansl3 | 2.74E-11 | -0.004277974 | 0.274 | 0.255 | 7.37E-07 |
| Dlgap3 | 1.97E-16 | 0.068545134 | 0.411 | 0.412 | 5.30E-12 | Sorbs2os | 2.74E-11 | -0.081534203 | 0.758 | 0.747 | 7.38E-07 |
| Slc30a9 | 2.01E-16 | -0.019080385 | 0.253 | 0.287 | 5.42E-12 | Arfgef2 | 2.77E-11 | -0.011007278 | 0.382 | 0.362 | 7.44E-07 |
| Dennd4c | 2.04E-16 | 0.070240418 | 0.295 | 0.289 | 5.48E-12 | Ssbp3 | 2.85E-11 | -0.069727427 | 0.403 | 0.411 | 7.67E-07 |
| Mta3 | 2.08E-16 | 0.041212554 | 0.375 | 0.389 | 5.60E-12 | Edem3 | 2.87E-11 | -0.01697853 | 0.283 | 0.269 | 7.73E-07 |
| Cdip1 | 2.08E-16 | -0.020088118 | 0.397 | 0.435 | 5.60E-12 | Socs7 | 2.87E-11 | 0.030666373 | 0.29 | 0.259 | 7.73E-07 |
| Btbd10 | 2.08E-16 | -0.001761606 | 0.384 | 0.416 | 5.60E-12 | Usp29 | 2.89E-11 | -0.099530233 | 0.469 | 0.467 | 7.77E-07 |
| Foxk2 | 2.09E-16 | -0.019038326 | 0.291 | 0.323 | 5.61E-12 | Bicra | 2.89E-11 | 0.023096599 | 0.32 | 0.292 | 7.79E-07 |
| Slc6a15 | 2.11E-16 | 0.099680656 | 0.258 | 0.236 | 5.67E-12 | Map3k12 | 2.96E-11 | 0.010235824 | 0.255 | 0.231 | 7.97E-07 |
| Lrpprc | 2.13E-16 | 0.056763638 | 0.386 | 0.39 | 5.74E-12 | Slc7a14 | 2.97E-11 | -0.107541172 | 0.641 | 0.652 | 7.99E-07 |
| Dok6 | 2.15E-16 | -0.098665994 | 0.875 | 0.904 | 5.78E-12 | P2ry12 | 2.97E-11 | 0.040306527 | 0.342 | 0.306 | 8.00E-07 |
| Msi2 | 2.15E-16 | 0.008836326 | 0.478 | 0.51 | 5.80E-12 | Kctd1 | 3.02E-11 | -0.050630594 | 0.454 | 0.447 | 8.12E-07 |
| Brd1 | 2.17E-16 | 0.017090289 | 0.347 | 0.37 | 5.84E-12 | Kcnc2 | 3.07E-11 | -0.16848167 | 0.307 | 0.335 | 8.26E-07 |
| Ptpro | 2.24E-16 | 0.023437874 | 0.234 | 0.254 | 6.04E-12 | Jph4 | 3.11E-11 | -0.009858017 | 0.326 | 0.308 | 8.37E-07 |
| Klf13 | 2.25E-16 | -0.064518237 | 0.303 | 0.35 | 6.06E-12 | Pde8b | 3.12E-11 | 0.022984586 | 0.652 | 0.615 | 8.39E-07 |

| | | | | | | | | | | | |
|---|---|---|---|---|---|---|---|---|---|---|---|
| Papola | 2.26E-16 | 0.032877783 | 0.591 | 0.615 | 6.08E-12 | Snx24 | 3.13E-11 | -0.002145958 | 0.423 | 0.4 | 8.43E-07 |
| Btbd1 | 2.33E-16 | -0.042377494 | 0.283 | 0.324 | 6.28E-12 | Reep3 | 3.21E-11 | 0.072703864 | 0.262 | 0.224 | 8.63E-07 |
| Ulk4 | 2.37E-16 | 0.072190393 | 0.49 | 0.486 | 6.37E-12 | Cep295 | 3.33E-11 | -0.009938504 | 0.26 | 0.245 | 8.96E-07 |
| Micu3 | 2.40E-16 | 0.048036595 | 0.431 | 0.443 | 6.47E-12 | Tbcd | 3.36E-11 | -0.048306454 | 0.374 | 0.372 | 9.04E-07 |
| Fip1l1 | 2.55E-16 | 0.006649617 | 0.305 | 0.33 | 6.85E-12 | Sppl3 | 3.42E-11 | 0.041653029 | 0.506 | 0.466 | 9.21E-07 |
| St8sia1 | 2.56E-16 | 0.093709554 | 0.273 | 0.251 | 6.89E-12 | Tub | 3.45E-11 | -0.020396665 | 0.497 | 0.48 | 9.27E-07 |
| Ldah | 2.58E-16 | 0.006560247 | 0.276 | 0.299 | 6.94E-12 | Bmpr1a | 3.48E-11 | 0.049596768 | 0.439 | 0.399 | 9.36E-07 |
| Phtf1 | 2.66E-16 | -0.032033416 | 0.386 | 0.426 | 7.15E-12 | Pspc1 | 3.53E-11 | 0.042041742 | 0.461 | 0.423 | 9.50E-07 |
| Dph6 | 2.66E-16 | 0.060034942 | 0.285 | 0.285 | 7.16E-12 | Esyt2 | 3.56E-11 | 0.007580683 | 0.65 | 0.621 | 9.59E-07 |
| H3f3b | 2.71E-16 | -0.07703438 | 0.268 | 0.318 | 7.30E-12 | Xrcc6 | 3.57E-11 | 0.030547136 | 0.511 | 0.472 | 9.61E-07 |
| Rtf1 | 2.77E-16 | 0.005466425 | 0.325 | 0.351 | 7.46E-12 | Xylt1 | 3.58E-11 | -0.097219896 | 0.579 | 0.593 | 9.63E-07 |
| A630089N07Rik | 2.78E-16 | 0.055917593 | 0.347 | 0.354 | 7.47E-12 | Cfap36 | 3.61E-11 | -0.004985505 | 0.344 | 0.321 | 9.70E-07 |
| Rai1 | 2.79E-16 | 0.059113294 | 0.47 | 0.476 | 7.49E-12 | Slc25a3 | 3.65E-11 | -0.010361634 | 0.296 | 0.278 | 9.82E-07 |
| Trhde | 2.81E-16 | 0.150418852 | 0.414 | 0.387 | 7.56E-12 | Ptpra | 3.66E-11 | 0.013197173 | 0.629 | 0.595 | 9.84E-07 |
| Cltb | 2.83E-16 | -0.024290608 | 0.242 | 0.274 | 7.61E-12 | Trit1 | 3.69E-11 | 0.003145416 | 0.25 | 0.23 | 9.93E-07 |
| Smg6 | 2.83E-16 | 0.080889759 | 0.751 | 0.753 | 7.62E-12 | 5031439G07Rik | 3.80E-11 | -0.0145348 | 0.375 | 0.357 | 1.02E-06 |
| Mtmr1 | 2.92E-16 | -0.046566788 | 0.314 | 0.357 | 7.86E-12 | Prpf4b | 3.81E-11 | 0.06123699 | 0.783 | 0.744 | 1.03E-06 |
| Slc44a1 | 2.92E-16 | -0.064238478 | 0.61 | 0.662 | 7.87E-12 | Frmd6 | 3.82E-11 | 0.084491138 | 0.305 | 0.265 | 1.03E-06 |
| Kcnip2 | 2.94E-16 | -0.025660337 | 0.341 | 0.38 | 7.92E-12 | Agfg2 | 3.94E-11 | 0.029713321 | 0.36 | 0.327 | 1.06E-06 |
| Tm9sf4 | 2.96E-16 | 0.011114859 | 0.284 | 0.307 | 7.96E-12 | Copa | 3.95E-11 | 0.024849802 | 0.382 | 0.35 | 1.06E-06 |
| Slc22a23 | 2.97E-16 | 0.097120192 | 0.406 | 0.389 | 7.99E-12 | Gpatch2 | 3.97E-11 | -0.046149087 | 0.403 | 0.396 | 1.07E-06 |
| Gpatch2 | 3.00E-16 | 0.062040344 | 0.396 | 0.397 | 8.08E-12 | Hcn1 | 4.00E-11 | 0.016778501 | 0.573 | 0.608 | 1.08E-06 |
| Clpx | 3.05E-16 | -0.010462669 | 0.251 | 0.28 | 8.19E-12 | Dixdc1 | 4.04E-11 | 0.019853584 | 0.343 | 0.315 | 1.09E-06 |
| Fbxw7 | 3.07E-16 | -0.027012788 | 0.514 | 0.555 | 8.25E-12 | Smchd1 | 4.11E-11 | 0.013202545 | 0.409 | 0.38 | 1.11E-06 |
| Etv1 | 3.13E-16 | -0.151823338 | 0.242 | 0.289 | 8.43E-12 | Hectd1 | 4.21E-11 | 0.044608449 | 0.413 | 0.374 | 1.13E-06 |
| Ccser1 | 3.14E-16 | 0.094456718 | 0.961 | 0.954 | 8.46E-12 | Nrd1 | 4.31E-11 | 0.038954132 | 0.618 | 0.579 | 1.16E-06 |
| Zdhhc6 | 3.15E-16 | -0.066244021 | 0.243 | 0.288 | 8.47E-12 | Fndc3b | 4.40E-11 | -0.068163605 | 0.398 | 0.403 | 1.18E-06 |
| Mdh1 | 3.19E-16 | -0.058373805 | 0.215 | 0.257 | 8.60E-12 | Spire1 | 4.44E-11 | 0.022037264 | 0.59 | 0.554 | 1.19E-06 |
| Tango2 | 3.22E-16 | 0.003829386 | 0.392 | 0.422 | 8.65E-12 | Cacng2 | 4.63E-11 | -0.016922706 | 0.325 | 0.307 | 1.25E-06 |
| Otulin | 3.42E-16 | -0.071059259 | 0.42 | 0.471 | 9.21E-12 | Cdc73 | 4.63E-11 | 0.023335528 | 0.397 | 0.364 | 1.25E-06 |
| Kdm2b | 3.49E-16 | -0.00648331 | 0.334 | 0.366 | 9.40E-12 | Tex2 | 4.72E-11 | 0.004284915 | 0.375 | 0.351 | 1.27E-06 |
| Phc2 | 3.52E-16 | 0.058039747 | 0.515 | 0.521 | 9.48E-12 | Tacc2 | 4.75E-11 | -0.000109399 | 0.294 | 0.272 | 1.28E-06 |
| Capza2 | 3.54E-16 | -0.037236043 | 0.218 | 0.255 | 9.52E-12 | Spaca6 | 4.78E-11 | 0.002649876 | 0.252 | 0.232 | 1.29E-06 |
| Slk | 3.61E-16 | -0.021596129 | 0.297 | 0.331 | 9.72E-12 | Osbpl6 | 4.89E-11 | -0.07533284 | 0.774 | 0.773 | 1.31E-06 |
| Irf2 | 3.75E-16 | -0.036288468 | 0.394 | 0.435 | 1.01E-11 | Ppm1b | 4.94E-11 | 0.0150737 | 0.285 | 0.26 | 1.33E-06 |
| Srebf2 | 3.77E-16 | 0.080182032 | 0.322 | 0.314 | 1.02E-11 | Ddx46 | 4.97E-11 | 0.050489542 | 0.479 | 0.438 | 1.34E-06 |
| Pdxdc1 | 3.87E-16 | -0.012096509 | 0.292 | 0.326 | 1.04E-11 | Syngap1 | 4.99E-11 | 0.026082935 | 0.302 | 0.272 | 1.34E-06 |
| Eif4e | 3.91E-16 | -0.018874067 | 0.261 | 0.293 | 1.05E-11 | Sap130 | 5.00E-11 | -0.024083051 | 0.295 | 0.285 | 1.34E-06 |
| Osbpl9 | 3.91E-16 | -0.011350883 | 0.475 | 0.515 | 1.05E-11 | Ncoa1 | 5.03E-11 | -0.058198952 | 0.817 | 0.809 | 1.35E-06 |
| Smg7 | 3.98E-16 | -0.008453168 | 0.419 | 0.454 | 1.07E-11 | Ptprn | 5.04E-11 | 0.018069315 | 0.613 | 0.581 | 1.36E-06 |
| Rnf10 | 3.99E-16 | -0.007647495 | 0.233 | 0.261 | 1.07E-11 | Xrn1 | 5.07E-11 | 0.044321785 | 0.559 | 0.519 | 1.36E-06 |
| Prkg1 | 4.01E-16 | 0.112423922 | 0.891 | 0.883 | 1.08E-11 | P2ry14 | 5.10E-11 | 0.067205847 | 0.682 | 0.639 | 1.37E-06 |
| Nbas | 4.03E-16 | -0.010416055 | 0.414 | 0.45 | 1.08E-11 | Aftph | 5.17E-11 | 0.008694233 | 0.429 | 0.401 | 1.39E-06 |
| Fbxl2 | 4.11E-16 | 0.050579766 | 0.531 | 0.541 | 1.11E-11 | Ppp6r3 | 5.19E-11 | 0.006902839 | 0.499 | 0.469 | 1.40E-06 |
| Alkbh8 | 4.14E-16 | -0.000165947 | 0.226 | 0.251 | 1.11E-11 | Mmd | 5.27E-11 | 0.021557442 | 0.288 | 0.261 | 1.42E-06 |
| Wdr7 | 4.15E-16 | 0.073240557 | 0.635 | 0.635 | 1.12E-11 | Lrig2 | 5.28E-11 | 0.044685075 | 0.478 | 0.44 | 1.42E-06 |
| Strn4 | 4.20E-16 | 0.009402173 | 0.345 | 0.369 | 1.13E-11 | Arhgap26 | 5.29E-11 | -0.086274297 | 0.699 | 0.71 | 1.42E-06 |
| Fnip1 | 4.30E-16 | 0.028023532 | 0.416 | 0.436 | 1.16E-11 | D10Wsu102e | 5.38E-11 | -0.06277232 | 0.38 | 0.381 | 1.45E-06 |
| Iqsec2 | 4.40E-16 | 0.095463147 | 0.662 | 0.65 | 1.18E-11 | Fam222b | 5.47E-11 | 0.046705736 | 0.463 | 0.423 | 1.47E-06 |
| Cdk11b | 4.46E-16 | -0.036276553 | 0.29 | 0.328 | 1.20E-11 | Zfp91 | 5.51E-11 | -0.000911968 | 0.271 | 0.251 | 1.48E-06 |
| Iqsec3 | 4.53E-16 | -0.033807156 | 0.386 | 0.427 | 1.22E-11 | Stum | 5.52E-11 | -0.048467219 | 0.343 | 0.339 | 1.48E-06 |
| Hook3 | 4.56E-16 | -0.037850477 | 0.475 | 0.52 | 1.23E-11 | Arfgef1 | 5.56E-11 | 0.00706588 | 0.554 | 0.524 | 1.50E-06 |
| Rb1cc1 | 4.61E-16 | -0.033190909 | 0.559 | 0.605 | 1.24E-11 | Susd6 | 5.59E-11 | -0.010008012 | 0.471 | 0.448 | 1.50E-06 |
| Jph4 | 4.62E-16 | 0.05102417 | 0.308 | 0.311 | 1.24E-11 | Cdon | 5.60E-11 | 0.036130431 | 0.324 | 0.291 | 1.51E-06 |
| Gpr107 | 4.70E-16 | 0.043745815 | 0.303 | 0.311 | 1.27E-11 | Arhgef11 | 5.62E-11 | -0.03137799 | 0.426 | 0.42 | 1.51E-06 |
| Mef2a | 4.76E-16 | -0.070597597 | 0.568 | 0.62 | 1.28E-11 | Srcin1 | 5.63E-11 | -0.013544143 | 0.524 | 0.503 | 1.51E-06 |
| Agfg1 | 4.78E-16 | 0.030250195 | 0.364 | 0.38 | 1.29E-11 | Chd3 | 5.87E-11 | 0.012557358 | 0.413 | 0.384 | 1.58E-06 |
| Gtf3c1 | 4.87E-16 | 0.021083604 | 0.346 | 0.365 | 1.31E-11 | Nfib | 5.89E-11 | -0.088624431 | 0.797 | 0.807 | 1.59E-06 |
| Svop | 4.96E-16 | 0.0668612 | 0.563 | 0.568 | 1.33E-11 | Atrx | 5.96E-11 | -0.028027154 | 0.751 | 0.731 | 1.60E-06 |
| Inhba | 5.01E-16 | -0.094773162 | 0.275 | 0.321 | 1.35E-11 | Ttc8 | 6.14E-11 | 0.003208704 | 0.521 | 0.494 | 1.65E-06 |
| Miga2 | 5.08E-16 | 0.06194687 | 0.279 | 0.275 | 1.37E-11 | Hacd2 | 6.16E-11 | -0.022946124 | 0.321 | 0.308 | 1.66E-06 |
| Syt13 | 5.11E-16 | -0.072871457 | 0.309 | 0.357 | 1.38E-11 | Nap1l1 | 6.23E-11 | -0.002475336 | 0.251 | 0.233 | 1.68E-06 |
| Zdhhc20 | 5.13E-16 | 0.029967662 | 0.457 | 0.477 | 1.38E-11 | Iqce | 6.28E-11 | -0.017581516 | 0.337 | 0.322 | 1.69E-06 |
| Top2b | 5.15E-16 | -0.033218444 | 0.359 | 0.399 | 1.38E-11 | Gm10785 | 6.30E-11 | 0.017268325 | 0.428 | 0.395 | 1.70E-06 |
| Tdrd3 | 5.20E-16 | 0.030465804 | 0.245 | 0.258 | 1.40E-11 | Ewsr1 | 6.35E-11 | 0.00190837 | 0.606 | 0.575 | 1.71E-06 |
| Ppp6r2 | 5.25E-16 | 0.022895089 | 0.518 | 0.542 | 1.41E-11 | Naa16 | 6.37E-11 | 0.012650687 | 0.267 | 0.241 | 1.72E-06 |
| Dnal1 | 5.26E-16 | 0.029189482 | 0.248 | 0.26 | 1.42E-11 | Cadm1 | 6.41E-11 | -0.077808947 | 0.932 | 0.93 | 1.72E-06 |
| Fmnl2 | 5.33E-16 | 0.086727277 | 0.649 | 0.65 | 1.43E-11 | Hmbox1 | 6.43E-11 | 0.014355692 | 0.518 | 0.485 | 1.73E-06 |
| Cenpp | 5.47E-16 | -0.072800299 | 0.296 | 0.343 | 1.47E-11 | Dgkd | 6.47E-11 | -0.018689123 | 0.53 | 0.51 | 1.74E-06 |
| Grin2b | 5.59E-16 | -0.079508338 | 0.988 | 0.986 | 1.50E-11 | Gtpbp2 | 6.47E-11 | -0.018807494 | 0.275 | 0.263 | 1.74E-06 |
| Flywch1 | 5.59E-16 | 0.066696701 | 0.263 | 0.258 | 1.50E-11 | Gm26906 | 6.51E-11 | -0.098257187 | 0.37 | 0.407 | 1.75E-06 |
| Lars2 | 5.66E-16 | 0.022536128 | 0.234 | 0.25 | 1.52E-11 | Abhd17b | 6.73E-11 | -0.016008819 | 0.328 | 0.313 | 1.81E-06 |
| Slc38a9 | 5.94E-16 | 0.039214042 | 0.458 | 0.474 | 1.60E-11 | Rph3a | 6.86E-11 | 0.075459837 | 0.317 | 0.278 | 1.85E-06 |
| Abl2 | 6.08E-16 | 0.050080973 | 0.573 | 0.585 | 1.63E-11 | Npepps | 6.86E-11 | 0.009832032 | 0.614 | 0.582 | 1.85E-06 |
| Diaph1 | 6.09E-16 | 0.038271905 | 0.249 | 0.257 | 1.64E-11 | Arl3 | 7.17E-11 | 0.054795666 | 0.407 | 0.368 | 1.93E-06 |
| Zfp654 | 6.21E-16 | -0.024243763 | 0.31 | 0.345 | 1.67E-11 | Zcchc18 | 7.22E-11 | -0.007550512 | 0.403 | 0.385 | 1.94E-06 |
| Syt11 | 6.32E-16 | 0.05541697 | 0.495 | 0.504 | 1.70E-11 | Tbc1d19 | 7.31E-11 | -0.039108448 | 0.352 | 0.35 | 1.97E-06 |
| Chgb | 6.39E-16 | -0.048462383 | 0.302 | 0.347 | 1.72E-11 | Rictor | 7.33E-11 | -0.029784596 | 0.417 | 0.406 | 1.97E-06 |
| Mprip | 6.49E-16 | -0.019185498 | 0.673 | 0.713 | 1.75E-11 | Dhx30 | 7.35E-11 | -0.002193247 | 0.548 | 0.52 | 1.98E-06 |
| Fam168b | 6.61E-16 | -0.00585502 | 0.264 | 0.29 | 1.78E-11 | Ddx17 | 7.36E-11 | -0.0465338 | 0.552 | 0.549 | 1.98E-06 |
| Suz12 | 6.63E-16 | -0.025596891 | 0.252 | 0.287 | 1.79E-11 | Prox1 | 7.48E-11 | 0.051643035 | 0.265 | 0.233 | 2.01E-06 |
| Copa | 6.76E-16 | 0.017924007 | 0.35 | 0.372 | 1.82E-11 | Macf1 | 7.52E-11 | -0.06920881 | 0.912 | 0.91 | 2.02E-06 |
| Sos1 | 6.85E-16 | 0.006403918 | 0.367 | 0.394 | 1.84E-11 | Cntnap5c | 7.61E-11 | -0.176937031 | 0.316 | 0.342 | 2.05E-06 |
| Iqsec1 | 7.05E-16 | 0.0667559 | 0.719 | 0.725 | 1.90E-11 | Ccdc57 | 7.66E-11 | -0.011224889 | 0.31 | 0.291 | 2.06E-06 |
| Ubap2 | 7.15E-16 | -0.008556124 | 0.273 | 0.302 | 1.93E-11 | Ube2k | 7.68E-11 | 0.057612379 | 0.593 | 0.551 | 2.07E-06 |
| Inpp5f | 7.16E-16 | -0.01165127 | 0.236 | 0.265 | 1.93E-11 | 8030462N17Rik | 7.70E-11 | 0.030291728 | 0.323 | 0.292 | 2.07E-06 |
| Wdr47 | 7.21E-16 | -0.018284481 | 0.328 | 0.362 | 1.94E-11 | Epb41l1 | 7.70E-11 | -0.076965278 | 0.581 | 0.586 | 2.07E-06 |
| Ube2e3 | 7.38E-16 | -0.019473079 | 0.272 | 0.305 | 1.99E-11 | Trip12 | 7.83E-11 | 0.010706321 | 0.66 | 0.629 | 2.11E-06 |
| Actl6b | 7.53E-16 | -0.005918936 | 0.248 | 0.275 | 2.03E-11 | Rreb1 | 7.87E-11 | -0.077120197 | 0.29 | 0.296 | 2.12E-06 |
| Zc3h13 | 7.58E-16 | -0.043994077 | 0.489 | 0.535 | 2.04E-11 | Senp2 | 7.87E-11 | 9.46E-05 | 0.339 | 0.317 | 2.12E-06 |
| Trps1 | 7.59E-16 | 0.115976987 | 0.324 | 0.285 | 2.04E-11 | Slc38a1 | 7.93E-11 | 0.003496078 | 0.294 | 0.272 | 2.13E-06 |
| Ccdc88a | 7.59E-16 | 0.028388182 | 0.729 | 0.756 | 2.04E-11 | Mprip | 8.01E-11 | -0.027513015 | 0.692 | 0.673 | 2.15E-06 |
| Myo9b | 7.81E-16 | 0.006375987 | 0.386 | 0.412 | 2.10E-11 | Slc25a12 | 8.01E-11 | 0.029544537 | 0.572 | 0.536 | 2.16E-06 |
| Bmt2 | 7.91E-16 | -0.012804365 | 0.365 | 0.398 | 2.13E-11 | Sptbn2 | 8.09E-11 | 0.025447398 | 0.521 | 0.486 | 2.18E-06 |
| Rbm5 | 8.10E-16 | 0.071001634 | 0.653 | 0.657 | 2.18E-11 | Slc7a6os | 8.14E-11 | 0.020223368 | 0.304 | 0.274 | 2.19E-06 |
| Mecp2 | 8.11E-16 | -0.020610504 | 0.392 | 0.429 | 2.18E-11 | Ttc17 | 8.26E-11 | -0.002623433 | 0.523 | 0.499 | 2.22E-06 |
| Itch | 8.31E-16 | -0.022283915 | 0.54 | 0.58 | 2.24E-11 | Map6 | 8.29E-11 | -0.021185343 | 0.434 | 0.417 | 2.23E-06 |
| 7-Mar | 8.40E-16 | -0.003544198 | 0.32 | 0.349 | 2.26E-11 | Spen | 8.37E-11 | 0.016948608 | 0.338 | 0.31 | 2.25E-06 |

| | | | | | | | | | | | |
|---|---|---|---|---|---|---|---|---|---|---|---|
| Ctps2 | 8.41E-16 | 0.09903018 | 0.279 | 0.253 | 2.26E-11 | Zc3h12b | 8.60E-11 | -0.054351628 | 0.563 | 0.559 | 2.31E-06 |
| Far1 | 8.54E-16 | -0.04998479 | 0.449 | 0.497 | 2.30E-11 | Sez6l2 | 8.64E-11 | 0.008977357 | 0.49 | 0.463 | 2.32E-06 |
| Glyr1 | 8.65E-16 | 0.053942246 | 0.312 | 0.314 | 2.33E-11 | Adgrb2 | 8.74E-11 | 0.011846977 | 0.488 | 0.456 | 2.35E-06 |
| Abl1 | 8.66E-16 | -0.002043627 | 0.271 | 0.296 | 2.33E-11 | Limk2 | 8.75E-11 | 0.013972401 | 0.295 | 0.27 | 2.35E-06 |
| Vdac1 | 8.71E-16 | -0.037262649 | 0.236 | 0.274 | 2.34E-11 | Thap3 | 8.82E-11 | 0.025889742 | 0.276 | 0.247 | 2.37E-06 |
| Gabrb1 | 8.73E-16 | -0.113292379 | 0.955 | 0.961 | 2.35E-11 | Cnrip1 | 8.90E-11 | 0.038250538 | 0.407 | 0.371 | 2.39E-06 |
| Ddhd1 | 8.79E-16 | -0.010485796 | 0.414 | 0.45 | 2.37E-11 | Prdm10 | 8.93E-11 | 0.044531646 | 0.353 | 0.317 | 2.40E-06 |
| Baz2a | 8.87E-16 | 0.058669452 | 0.31 | 0.308 | 2.39E-11 | Csnk2a2 | 9.71E-11 | 0.001516273 | 0.315 | 0.295 | 2.61E-06 |
| Gnptab | 8.89E-16 | -0.047130799 | 0.3 | 0.342 | 2.39E-11 | Gm26836 | 9.81E-11 | 0.099570086 | 0.36 | 0.332 | 2.64E-06 |
| Ggps1 | 9.12E-16 | -0.027828967 | 0.29 | 0.328 | 2.45E-11 | Insr | 9.84E-11 | -0.060303087 | 0.467 | 0.471 | 2.65E-06 |
| Tor1aip2 | 9.13E-16 | -0.052240896 | 0.246 | 0.288 | 2.46E-11 | Spata5 | 1.01E-10 | -0.022643843 | 0.346 | 0.333 | 2.71E-06 |
| Pde3b | 9.20E-16 | 0.047988352 | 0.334 | 0.342 | 2.48E-11 | Adgrb1 | 1.01E-10 | -0.043294743 | 0.575 | 0.564 | 2.71E-06 |
| Kcnh1 | 9.25E-16 | 0.068647783 | 0.498 | 0.507 | 2.49E-11 | Rab11fip3 | 1.03E-10 | -0.009690953 | 0.62 | 0.593 | 2.76E-06 |
| Alkbh1 | 9.36E-16 | -0.07523114 | 0.221 | 0.265 | 2.52E-11 | Slc16a7 | 1.03E-10 | 0.051193417 | 0.582 | 0.541 | 2.78E-06 |
| Pds5a | 9.39E-16 | -0.013483401 | 0.44 | 0.476 | 2.53E-11 | Rcor1 | 1.03E-10 | 0.055456693 | 0.407 | 0.368 | 2.78E-06 |
| Cep295 | 9.41E-16 | -0.007286572 | 0.245 | 0.271 | 2.53E-11 | Cyth1 | 1.04E-10 | -0.034653593 | 0.414 | 0.406 | 2.79E-06 |
| Casc3 | 9.41E-16 | 0.007633856 | 0.269 | 0.292 | 2.53E-11 | Trim37 | 1.04E-10 | 0.021201003 | 0.446 | 0.413 | 2.79E-06 |
| Dclk1 | 9.42E-16 | 0.069338435 | 0.947 | 0.938 | 2.53E-11 | Rnps1 | 1.06E-10 | -0.021416929 | 0.318 | 0.305 | 2.84E-06 |
| Slc1a1 | 9.53E-16 | 0.013374451 | 0.524 | 0.549 | 2.56E-11 | Fbxo42 | 1.06E-10 | 0.003917351 | 0.293 | 0.271 | 2.85E-06 |
| Mapk10 | 9.58E-16 | 0.079973138 | 0.917 | 0.92 | 2.58E-11 | Nek9 | 1.06E-10 | 0.028878977 | 0.295 | 0.265 | 2.86E-06 |
| Emsy | 9.62E-16 | -0.016738626 | 0.402 | 0.439 | 2.59E-11 | Pak2 | 1.07E-10 | 0.041558878 | 0.286 | 0.252 | 2.89E-06 |
| Ric3 | 9.68E-16 | 0.000804645 | 0.272 | 0.297 | 2.61E-11 | Gm38393 | 1.08E-10 | -0.039975461 | 0.304 | 0.302 | 2.90E-06 |
| Mtor | 9.78E-16 | 0.05253768 | 0.318 | 0.321 | 2.63E-11 | Arntl | 1.08E-10 | 0.039673796 | 0.333 | 0.3 | 2.90E-06 |
| Rcor1 | 9.95E-16 | -0.012585166 | 0.368 | 0.402 | 2.68E-11 | Zfp827 | 1.09E-10 | -0.051689984 | 0.317 | 0.319 | 2.92E-06 |
| Zfp236 | 1.04E-15 | -0.030784756 | 0.299 | 0.335 | 2.79E-11 | Hk1 | 1.11E-10 | 0.040036907 | 0.372 | 0.336 | 2.99E-06 |
| Smc3 | 1.07E-15 | -0.048186481 | 0.312 | 0.354 | 2.87E-11 | 1700109K24Rik | 1.14E-10 | -0.031197543 | 0.44 | 0.424 | 3.06E-06 |
| Fam193b | 1.09E-15 | 0.048322039 | 0.329 | 0.334 | 2.94E-11 | Atg16l1 | 1.14E-10 | -0.006442942 | 0.372 | 0.353 | 3.06E-06 |
| Dync1i2 | 1.10E-15 | -0.052936173 | 0.738 | 0.782 | 2.95E-11 | Ift57 | 1.14E-10 | 0.015980992 | 0.256 | 0.23 | 3.06E-06 |
| Rundc3b | 1.10E-15 | 0.016139779 | 0.464 | 0.489 | 2.97E-11 | Tbk1 | 1.14E-10 | 0.008950007 | 0.292 | 0.268 | 3.06E-06 |
| Ppp2r2c | 1.11E-15 | 0.030289388 | 0.507 | 0.53 | 2.99E-11 | Mcf2l | 1.14E-10 | -0.026774159 | 0.477 | 0.463 | 3.08E-06 |
| Ccnt1 | 1.12E-15 | -0.030166126 | 0.267 | 0.302 | 3.02E-11 | Atp9a | 1.14E-10 | 0.007987983 | 0.549 | 0.521 | 3.08E-06 |
| Cwc27 | 1.12E-15 | 0.006308426 | 0.424 | 0.453 | 3.02E-11 | Fgf14 | 1.15E-10 | 0.049470439 | 0.988 | 0.993 | 3.09E-06 |
| Bclaf1 | 1.13E-15 | -0.024592871 | 0.496 | 0.538 | 3.04E-11 | Mtmr12 | 1.15E-10 | 0.043591775 | 0.487 | 0.448 | 3.10E-06 |
| Dnajc18 | 1.14E-15 | 0.010392353 | 0.249 | 0.271 | 3.05E-11 | Mfsd4a | 1.15E-10 | 0.003576044 | 0.399 | 0.375 | 3.11E-06 |
| Pnisr | 1.14E-15 | -0.039110431 | 0.832 | 0.87 | 3.06E-11 | Sec24b | 1.17E-10 | -0.043881102 | 0.464 | 0.461 | 3.14E-06 |
| Abhd18 | 1.16E-15 | -0.020832824 | 0.509 | 0.549 | 3.12E-11 | Crim1 | 1.17E-10 | -0.051409444 | 0.476 | 0.467 | 3.16E-06 |
| Scai | 1.16E-15 | -0.073832522 | 0.617 | 0.668 | 3.12E-11 | Commd10 | 1.18E-10 | 0.039283682 | 0.317 | 0.286 | 3.17E-06 |
| Hdlbp | 1.17E-15 | -0.0239731 | 0.312 | 0.349 | 3.14E-11 | Snx2 | 1.20E-10 | 0.015012141 | 0.253 | 0.231 | 3.22E-06 |
| Agpat3 | 1.17E-15 | 0.033293304 | 0.289 | 0.301 | 3.15E-11 | Dync1li2 | 1.20E-10 | 0.002250997 | 0.364 | 0.34 | 3.23E-06 |
| Ncs1 | 1.19E-15 | -0.007905843 | 0.285 | 0.314 | 3.21E-11 | Smad4 | 1.21E-10 | -0.013156558 | 0.257 | 0.244 | 3.25E-06 |
| 9530059O14Rik | 1.22E-15 | 0.070460036 | 0.755 | 0.758 | 3.28E-11 | Sos2 | 1.21E-10 | 0.021307929 | 0.302 | 0.275 | 3.25E-06 |
| Rbm10 | 1.22E-15 | -0.005778527 | 0.259 | 0.285 | 3.29E-11 | Actr2 | 1.21E-10 | 0.048077592 | 0.457 | 0.418 | 3.27E-06 |
| Vti1b | 1.24E-15 | -0.002306939 | 0.297 | 0.326 | 3.35E-11 | Copg2 | 1.23E-10 | 0.026894762 | 0.461 | 0.427 | 3.30E-06 |
| Ube2h | 1.28E-15 | 0.007487865 | 0.464 | 0.492 | 3.43E-11 | Nktr | 1.23E-10 | 0.00802905 | 0.69 | 0.659 | 3.30E-06 |
| Usp32 | 1.29E-15 | 0.066103969 | 0.609 | 0.615 | 3.47E-11 | Fgfr2 | 1.23E-10 | 0.079405011 | 0.916 | 0.897 | 3.31E-06 |
| Dcaf8 | 1.32E-15 | 0.031639133 | 0.366 | 0.382 | 3.54E-11 | Nf2 | 1.25E-10 | 0.04693969 | 0.32 | 0.285 | 3.35E-06 |
| Ccnc | 1.32E-15 | 0.00581016 | 0.271 | 0.295 | 3.55E-11 | Fam184a | 1.25E-10 | -0.005632382 | 0.315 | 0.297 | 3.37E-06 |
| Slc7a6os | 1.32E-15 | -0.067356455 | 0.274 | 0.32 | 3.56E-11 | Lrrc4 | 1.25E-10 | 0.03506256 | 0.659 | 0.622 | 3.37E-06 |
| Nfyc | 1.33E-15 | -0.031526008 | 0.315 | 0.353 | 3.57E-11 | Mmp24 | 1.27E-10 | 0.016541782 | 0.262 | 0.237 | 3.40E-06 |
| Far2 | 1.39E-15 | -0.035975496 | 0.327 | 0.367 | 3.73E-11 | Atp8a2 | 1.28E-10 | -0.062781501 | 0.613 | 0.615 | 3.44E-06 |
| Mast4 | 1.40E-15 | -0.083470714 | 0.695 | 0.743 | 3.77E-11 | Sv2b | 1.28E-10 | -0.107502037 | 0.512 | 0.549 | 3.45E-06 |
| Csnk2a2 | 1.46E-15 | 0.024625771 | 0.295 | 0.308 | 3.93E-11 | Adarb1 | 1.28E-10 | -0.041733925 | 0.475 | 0.467 | 3.45E-06 |
| Pcgf5 | 1.48E-15 | -0.003454864 | 0.272 | 0.299 | 3.98E-11 | Miat | 1.28E-10 | 0.067217335 | 0.754 | 0.715 | 3.46E-06 |
| Cep170 | 1.48E-15 | 0.015627504 | 0.384 | 0.405 | 3.99E-11 | Cnot4 | 1.29E-10 | -0.000670354 | 0.705 | 0.677 | 3.47E-06 |
| Arhgap39 | 1.49E-15 | -0.080892788 | 0.755 | 0.8 | 4.01E-11 | Sec11a | 1.29E-10 | 0.006362798 | 0.27 | 0.248 | 3.47E-06 |
| Mamld1 | 1.52E-15 | -0.062719593 | 0.339 | 0.382 | 4.10E-11 | Zdhhc8 | 1.35E-10 | 0.017722636 | 0.264 | 0.238 | 3.63E-06 |
| Zfp385b | 1.53E-15 | 0.114965988 | 0.622 | 0.577 | 4.11E-11 | Elp2 | 1.35E-10 | 0.007625078 | 0.254 | 0.233 | 3.64E-06 |
| Arhgap35 | 1.53E-15 | 0.043199452 | 0.445 | 0.459 | 4.11E-11 | Sparcl1 | 1.39E-10 | 0.047176436 | 0.35 | 0.314 | 3.73E-06 |
| Prkar1b | 1.54E-15 | -0.006847751 | 0.334 | 0.366 | 4.14E-11 | Slc9a1 | 1.40E-10 | 0.028986155 | 0.257 | 0.228 | 3.78E-06 |
| D130043K22Rik | 1.54E-15 | 0.101326861 | 0.406 | 0.383 | 4.15E-11 | Rab3gap2 | 1.41E-10 | -0.038643328 | 0.383 | 0.377 | 3.79E-06 |
| Clip4 | 1.58E-15 | 0.0604812 | 0.273 | 0.269 | 4.25E-11 | Ubap2l | 1.42E-10 | 0.023930723 | 0.424 | 0.392 | 3.81E-06 |
| Clint1 | 1.59E-15 | -0.017363707 | 0.406 | 0.442 | 4.29E-11 | Dph6 | 1.43E-10 | 0.030673753 | 0.315 | 0.285 | 3.84E-06 |
| Nkain2 | 1.63E-15 | 0.118944835 | 0.991 | 0.986 | 4.39E-11 | Ttll11 | 1.44E-10 | -0.048682687 | 0.645 | 0.637 | 3.87E-06 |
| Speg | 1.64E-15 | 0.082455701 | 0.346 | 0.334 | 4.41E-11 | Trim9 | 1.44E-10 | -0.062397281 | 0.843 | 0.836 | 3.88E-06 |
| Cdh11 | 1.68E-15 | -0.020623897 | 0.683 | 0.728 | 4.52E-11 | Acot7 | 1.45E-10 | -0.050250115 | 0.307 | 0.31 | 3.90E-06 |
| Kdm2a | 1.70E-15 | 0.033679287 | 0.517 | 0.537 | 4.57E-11 | Hlf | 1.46E-10 | -0.060206041 | 0.306 | 0.307 | 3.93E-06 |
| Purg | 1.70E-15 | 0.007951508 | 0.372 | 0.398 | 4.57E-11 | Atp6v1a | 1.46E-10 | 0.012390143 | 0.365 | 0.338 | 3.94E-06 |
| Gapvd1 | 1.72E-15 | 0.036875785 | 0.527 | 0.543 | 4.63E-11 | Arhgap5 | 1.48E-10 | 0.048209281 | 0.431 | 0.392 | 3.97E-06 |
| Actn4 | 1.73E-15 | 0.001959474 | 0.355 | 0.382 | 4.66E-11 | Map2k2 | 1.48E-10 | 0.030351327 | 0.262 | 0.233 | 3.99E-06 |
| B4galt6 | 1.74E-15 | -0.019096645 | 0.219 | 0.25 | 4.67E-11 | Lclat1 | 1.50E-10 | 0.000675348 | 0.349 | 0.328 | 4.03E-06 |
| Cbl | 1.75E-15 | -0.026490001 | 0.402 | 0.441 | 4.70E-11 | Zfyve28 | 1.50E-10 | -0.068267598 | 0.482 | 0.486 | 4.04E-06 |
| Lpgat1 | 1.75E-15 | 0.04005962 | 0.376 | 0.387 | 4.70E-11 | Pou2f1 | 1.52E-10 | -0.013682971 | 0.422 | 0.403 | 4.10E-06 |
| Sbf2 | 1.77E-15 | 0.099900264 | 0.764 | 0.754 | 4.76E-11 | Zc2hc1a | 1.53E-10 | -0.014369041 | 0.273 | 0.259 | 4.11E-06 |
| Taf1b | 1.83E-15 | 0.017014048 | 0.37 | 0.391 | 4.93E-11 | Map2k6 | 1.54E-10 | 0.045158838 | 0.441 | 0.403 | 4.15E-06 |
| Ncl | 1.84E-15 | -0.031159959 | 0.243 | 0.278 | 4.95E-11 | Slc24a2 | 1.56E-10 | -0.103372907 | 0.921 | 0.922 | 4.19E-06 |
| Tab2 | 1.84E-15 | 0.038490371 | 0.296 | 0.307 | 4.96E-11 | Mgrn1 | 1.59E-10 | -0.025292273 | 0.39 | 0.378 | 4.28E-06 |
| Rims1 | 1.88E-15 | 0.08947427 | 0.972 | 0.968 | 5.07E-11 | Zfp654 | 1.60E-10 | 0.027897939 | 0.341 | 0.31 | 4.30E-06 |
| 4930419G24Rik | 1.92E-15 | -0.217607696 | 0.531 | 0.573 | 5.17E-11 | Psmd1 | 1.61E-10 | 0.015271841 | 0.382 | 0.354 | 4.33E-06 |
| Plekha1 | 2.00E-15 | 0.009286629 | 0.335 | 0.361 | 5.39E-11 | Rap1a | 1.62E-10 | 0.027238365 | 0.365 | 0.334 | 4.35E-06 |
| Cep83 | 2.01E-15 | -0.03585505 | 0.361 | 0.401 | 5.41E-11 | Wdr70 | 1.65E-10 | -0.001730593 | 0.366 | 0.344 | 4.44E-06 |
| Ext1 | 2.04E-15 | 0.083620178 | 0.772 | 0.777 | 5.50E-11 | Zc3h7a | 1.68E-10 | 0.01943398 | 0.47 | 0.437 | 4.51E-06 |
| Btrc | 2.14E-15 | 0.060533136 | 0.641 | 0.646 | 5.75E-11 | Drosha | 1.69E-10 | -0.061375272 | 0.238 | 0.25 | 4.54E-06 |
| Rab7 | 2.15E-15 | -0.007678544 | 0.247 | 0.274 | 5.77E-11 | Epc2 | 1.70E-10 | 0.021811484 | 0.298 | 0.27 | 4.57E-06 |
| Usf3 | 2.15E-15 | -0.022312745 | 0.243 | 0.274 | 5.78E-11 | Ppp2r5e | 1.72E-10 | -0.006832007 | 0.522 | 0.5 | 4.62E-06 |
| Gucy1b1 | 2.16E-15 | 0.068311175 | 0.277 | 0.272 | 5.81E-11 | Anks1 | 1.73E-10 | -0.066764994 | 0.25 | 0.259 | 4.64E-06 |
| Dlst | 2.18E-15 | -0.002396363 | 0.259 | 0.284 | 5.85E-11 | Pias1 | 1.75E-10 | 0.025459587 | 0.588 | 0.553 | 4.70E-06 |
| Agap2 | 2.18E-15 | 0.053665716 | 0.254 | 0.253 | 5.87E-11 | 5330417C22Rik | 1.75E-10 | 0.008349526 | 0.513 | 0.483 | 4.71E-06 |
| Plxna2 | 2.19E-15 | 0.066693882 | 0.603 | 0.61 | 5.89E-11 | 11-Sep | 1.78E-10 | 0.01558945 | 0.473 | 0.442 | 4.80E-06 |
| Ino80d | 2.19E-15 | -0.005110469 | 0.299 | 0.326 | 5.89E-11 | Desi1 | 1.84E-10 | 0.015254016 | 0.361 | 0.333 | 4.96E-06 |
| Ralgapb | 2.19E-15 | 0.052295917 | 0.408 | 0.415 | 5.90E-11 | Shoc2 | 1.87E-10 | -0.014174604 | 0.25 | 0.238 | 5.03E-06 |
| Smg1 | 2.20E-15 | 0.006794023 | 0.622 | 0.654 | 5.93E-11 | Rab10 | 1.88E-10 | 0.062574623 | 0.527 | 0.486 | 5.05E-06 |
| Mtch1 | 2.21E-15 | -0.04507453 | 0.233 | 0.271 | 5.94E-11 | Pikfyve | 1.88E-10 | 0.022258307 | 0.27 | 0.243 | 5.06E-06 |
| Ppp2r3a | 2.27E-15 | -0.083553704 | 0.492 | 0.544 | 6.10E-11 | Acer3 | 1.90E-10 | 0.002038533 | 0.412 | 0.386 | 5.11E-06 |
| Cnot2 | 2.27E-15 | -0.009469311 | 0.493 | 0.53 | 6.12E-11 | Vps54 | 1.90E-10 | 0.024177827 | 0.404 | 0.372 | 5.11E-06 |
| Slc4a4 | 2.31E-15 | -0.20968644 | 0.45 | 0.489 | 6.22E-11 | Klf12 | 1.90E-10 | -0.097262455 | 0.594 | 0.608 | 5.11E-06 |
| Prpf40b | 2.33E-15 | 0.061072798 | 0.26 | 0.253 | 6.26E-11 | Smap1 | 1.90E-10 | 0.028731618 | 0.449 | 0.416 | 5.12E-06 |
| Gm11659 | 2.36E-15 | -0.009528687 | 0.245 | 0.271 | 6.34E-11 | Ddhd1 | 1.92E-10 | -0.071740813 | 0.401 | 0.414 | 5.16E-06 |
| Hectd1 | 2.37E-15 | -0.050306148 | 0.374 | 0.42 | 6.38E-11 | Aagab | 1.93E-10 | 0.026814969 | 0.381 | 0.35 | 5.19E-06 |

| | | | | | | | | | | | |
|---|---|---|---|---|---|---|---|---|---|---|---|
| Nsun7 | 2.41E-15 | -0.030518059 | 0.284 | 0.319 | 6.48E-11 | Lztfl1 | 1.94E-10 | 0.04374755 | 0.386 | 0.35 | 5.22E-06 |
| Atp6v1b2 | 2.41E-15 | 0.004129729 | 0.385 | 0.412 | 6.49E-11 | Ulk2 | 1.97E-10 | 0.040910948 | 0.603 | 0.563 | 5.30E-06 |
| Ash1l | 2.48E-15 | 0.055268235 | 0.721 | 0.734 | 6.68E-11 | 9330182L06Rik | 1.98E-10 | -0.047304419 | 0.272 | 0.272 | 5.32E-06 |
| Celf4 | 2.56E-15 | 0.112025028 | 0.888 | 0.891 | 6.88E-11 | Vwa8 | 2.02E-10 | -0.053483166 | 0.562 | 0.561 | 5.42E-06 |
| Syp | 2.57E-15 | 0.024765974 | 0.4 | 0.423 | 6.90E-11 | Eea1 | 2.03E-10 | -0.026212133 | 0.296 | 0.287 | 5.45E-06 |
| Ptpn4 | 2.57E-15 | 0.027186404 | 0.46 | 0.481 | 6.91E-11 | Gpr137c | 2.06E-10 | -0.001563038 | 0.317 | 0.297 | 5.55E-06 |
| Ppm1h | 2.61E-15 | -0.091541213 | 0.631 | 0.682 | 7.04E-11 | Klhl7 | 2.08E-10 | 0.036348926 | 0.536 | 0.498 | 5.61E-06 |
| Arnt | 2.62E-15 | 0.002119911 | 0.411 | 0.442 | 7.04E-11 | Cul1 | 2.09E-10 | 0.043971864 | 0.48 | 0.442 | 5.62E-06 |
| Serinc1 | 2.64E-15 | -0.006673246 | 0.277 | 0.31 | 7.09E-11 | Kdm6a | 2.09E-10 | 0.03073462 | 0.406 | 0.372 | 5.62E-06 |
| Arhgap5 | 2.65E-15 | -0.065302366 | 0.392 | 0.442 | 7.13E-11 | Wdfy2 | 2.09E-10 | -0.042167334 | 0.264 | 0.265 | 5.64E-06 |
| Slc25a26 | 2.65E-15 | 0.039563884 | 0.258 | 0.265 | 7.14E-11 | Nxpe4 | 2.17E-10 | -0.002193436 | 0.379 | 0.356 | 5.85E-06 |
| Fbxo38 | 2.66E-15 | -0.054903405 | 0.3 | 0.344 | 7.17E-11 | 3-Sep | 2.19E-10 | -0.035959487 | 0.336 | 0.333 | 5.90E-06 |
| Cpsf6 | 2.67E-15 | -0.025221281 | 0.519 | 0.563 | 7.17E-11 | Rnf217 | 2.21E-10 | -0.031981537 | 0.295 | 0.286 | 5.94E-06 |
| Cbfa2t2 | 2.76E-15 | -0.038390346 | 0.525 | 0.571 | 7.41E-11 | Actn1 | 2.25E-10 | -0.052760815 | 0.345 | 0.348 | 6.04E-06 |
| Nemf | 2.84E-15 | 0.010756452 | 0.37 | 0.394 | 7.65E-11 | Bcl2l1 | 2.26E-10 | -0.021037164 | 0.265 | 0.252 | 6.08E-06 |
| Malat1 | 2.85E-15 | -0.1984629 | 1 | 0.999 | 7.67E-11 | Soga3 | 2.31E-10 | 0.010892602 | 0.671 | 0.638 | 6.21E-06 |
| Arf3 | 2.86E-15 | 0.053285085 | 0.357 | 0.359 | 7.70E-11 | Erc2 | 2.36E-10 | -0.050691324 | 0.974 | 0.982 | 6.35E-06 |
| Snx6 | 2.87E-15 | -0.014916712 | 0.223 | 0.251 | 7.72E-11 | Nploc4 | 2.37E-10 | -0.011003167 | 0.33 | 0.315 | 6.37E-06 |
| Ndrg4 | 2.87E-15 | -0.089064788 | 0.546 | 0.598 | 7.74E-11 | Gnptab | 2.39E-10 | -0.013972632 | 0.314 | 0.3 | 6.44E-06 |
| Rapgef5 | 2.88E-15 | -0.136257863 | 0.68 | 0.726 | 7.75E-11 | Rab3ip | 2.40E-10 | -0.013214302 | 0.275 | 0.262 | 6.46E-06 |
| Ttll11 | 2.88E-15 | 0.028161344 | 0.637 | 0.661 | 7.76E-11 | Pi4kb | 2.41E-10 | 0.023680253 | 0.256 | 0.231 | 6.47E-06 |
| Cdc73 | 2.89E-15 | 0.006087345 | 0.364 | 0.391 | 7.78E-11 | Rbm33 | 2.41E-10 | -0.018478308 | 0.451 | 0.436 | 6.47E-06 |
| Reep3 | 2.99E-15 | -0.068472844 | 0.224 | 0.266 | 8.05E-11 | Setd2 | 2.41E-10 | 0.006342278 | 0.478 | 0.449 | 6.49E-06 |
| Sgsm3 | 3.02E-15 | -0.026768469 | 0.23 | 0.264 | 8.11E-11 | Lrpprc | 2.43E-10 | -0.034737749 | 0.393 | 0.386 | 6.54E-06 |
| Garnl3 | 3.02E-15 | 0.138560994 | 0.338 | 0.299 | 8.13E-11 | Dgkz | 2.49E-10 | 0.07079788 | 0.468 | 0.426 | 6.69E-06 |
| Aagab | 3.04E-15 | -0.056728268 | 0.35 | 0.395 | 8.17E-11 | Hbs1l | 2.50E-10 | 0.00202573 | 0.346 | 0.324 | 6.73E-06 |
| Prdm10 | 3.11E-15 | 0.013064892 | 0.317 | 0.338 | 8.36E-11 | Psmc6 | 2.51E-10 | -0.007942135 | 0.379 | 0.358 | 6.76E-06 |
| Pls3 | 3.11E-15 | -0.091193394 | 0.34 | 0.389 | 8.38E-11 | Adamts3 | 2.53E-10 | -0.00600945 | 0.344 | 0.324 | 6.80E-06 |
| Fer | 3.15E-15 | 0.03695861 | 0.499 | 0.514 | 8.49E-11 | Gm16351 | 2.55E-10 | 0.044366716 | 0.352 | 0.317 | 6.87E-06 |
| Rnf111 | 3.18E-15 | -0.025407527 | 0.413 | 0.452 | 8.56E-11 | St3gal5 | 2.57E-10 | 0.030870888 | 0.428 | 0.394 | 6.92E-06 |
| Mfhas1 | 3.20E-15 | 0.048566866 | 0.325 | 0.33 | 8.61E-11 | Rora | 2.60E-10 | -0.092539239 | 0.901 | 0.901 | 7.01E-06 |
| Chst1 | 3.20E-15 | -0.082697985 | 0.252 | 0.298 | 8.62E-11 | Myef2 | 2.66E-10 | 0.035383476 | 0.491 | 0.455 | 7.17E-06 |
| Nploc4 | 3.24E-15 | 0.041658129 | 0.315 | 0.324 | 8.71E-11 | Alcam | 2.67E-10 | 0.102997929 | 0.523 | 0.48 | 7.19E-06 |
| Arnt2 | 3.26E-15 | 0.030065515 | 0.419 | 0.439 | 8.78E-11 | Sdccag8 | 2.69E-10 | -0.024545093 | 0.548 | 0.534 | 7.25E-06 |
| Edrf1 | 3.28E-15 | -0.006090609 | 0.228 | 0.252 | 8.83E-11 | Pde3b | 2.71E-10 | -0.052183885 | 0.334 | 0.334 | 7.30E-06 |
| Rgs7 | 3.39E-15 | 0.067900593 | 0.938 | 0.929 | 9.13E-11 | Cadm3 | 2.72E-10 | -0.009279491 | 0.266 | 0.25 | 7.33E-06 |
| Rab40c | 3.43E-15 | 0.024461256 | 0.392 | 0.411 | 9.22E-11 | Chic2 | 2.73E-10 | 0.041225126 | 0.256 | 0.225 | 7.33E-06 |
| Gspt1 | 3.49E-15 | -0.013275761 | 0.303 | 0.333 | 9.39E-11 | Jak2 | 2.75E-10 | 0.009514477 | 0.27 | 0.247 | 7.39E-06 |
| Tmem245 | 3.50E-15 | 0.072671531 | 0.324 | 0.317 | 9.41E-11 | Ipo9 | 2.75E-10 | 0.000859364 | 0.283 | 0.265 | 7.41E-06 |
| Trip12 | 3.51E-15 | -0.009279007 | 0.629 | 0.667 | 9.45E-11 | Tecpr2 | 2.76E-10 | 0.018832258 | 0.381 | 0.352 | 7.43E-06 |
| Pigk | 3.57E-15 | -0.118332333 | 0.884 | 0.902 | 9.61E-11 | Gstz1 | 2.76E-10 | -0.003989851 | 0.262 | 0.243 | 7.44E-06 |
| Dhx32 | 3.57E-15 | 0.022573378 | 0.253 | 0.266 | 9.61E-11 | Pds5a | 2.77E-10 | 0.0382346 | 0.478 | 0.44 | 7.45E-06 |
| Dennd1a | 3.62E-15 | 0.083785196 | 0.878 | 0.877 | 9.74E-11 | Epha3 | 2.78E-10 | -0.000181175 | 0.488 | 0.458 | 7.48E-06 |
| Mpdz | 3.66E-15 | -0.031750684 | 0.406 | 0.446 | 9.84E-11 | Csnk1g1 | 2.80E-10 | -0.028453229 | 0.663 | 0.647 | 7.55E-06 |
| Ppp2r5e | 3.66E-15 | 0.028249821 | 0.5 | 0.518 | 9.85E-11 | Far1os | 2.83E-10 | 0.019752472 | 0.47 | 0.441 | 7.61E-06 |
| Actr3b | 3.69E-15 | 0.075263763 | 0.477 | 0.474 | 9.92E-11 | Timp4 | 2.83E-10 | -0.04773501 | 0.296 | 0.298 | 7.62E-06 |
| 6-Mar | 3.72E-15 | 0.05292445 | 0.438 | 0.443 | 1.00E-10 | Kif5c | 2.83E-10 | -0.045479107 | 0.662 | 0.657 | 7.63E-06 |
| Foxk1 | 3.73E-15 | -0.011334908 | 0.302 | 0.333 | 1.00E-10 | Dennd1b | 2.90E-10 | -0.026388633 | 0.514 | 0.5 | 7.80E-06 |
| Zyg11b | 3.76E-15 | -0.003135566 | 0.346 | 0.375 | 1.01E-10 | Rars2 | 2.92E-10 | -0.010746232 | 0.375 | 0.356 | 7.85E-06 |
| Gm48383 | 3.76E-15 | -0.06132894 | 0.337 | 0.382 | 1.01E-10 | 7-Sep | 2.92E-10 | 0.04237781 | 0.363 | 0.328 | 7.86E-06 |
| Btbd7 | 3.79E-15 | 0.037079458 | 0.416 | 0.431 | 1.02E-10 | Nrxn1 | 2.93E-10 | -0.049972188 | 0.989 | 0.995 | 7.88E-06 |
| Smc5 | 3.90E-15 | -0.050864173 | 0.336 | 0.379 | 1.05E-10 | Fam219a | 2.93E-10 | -0.006448415 | 0.539 | 0.513 | 7.90E-06 |
| Dctn4 | 3.91E-15 | -0.047740732 | 0.399 | 0.443 | 1.05E-10 | Aak1 | 2.94E-10 | -0.051287688 | 0.848 | 0.841 | 7.92E-06 |
| Aco2 | 3.94E-15 | 0.025370533 | 0.409 | 0.428 | 1.06E-10 | Pcdh7 | 2.95E-10 | -0.123698625 | 0.513 | 0.552 | 7.94E-06 |
| Anapc1 | 3.94E-15 | 0.034169752 | 0.268 | 0.277 | 1.06E-10 | Tbc1d22a | 2.96E-10 | -0.029077901 | 0.325 | 0.318 | 7.97E-06 |
| Memo1 | 4.01E-15 | 0.015229884 | 0.414 | 0.438 | 1.08E-10 | Foxj3 | 2.97E-10 | 0.045540088 | 0.358 | 0.322 | 8.00E-06 |
| Farsb | 4.02E-15 | 0.023208494 | 0.293 | 0.31 | 1.08E-10 | Zfp445 | 2.98E-10 | -0.021846066 | 0.489 | 0.474 | 8.01E-06 |
| Brd4 | 4.07E-15 | 0.01053417 | 0.507 | 0.536 | 1.09E-10 | Tgfbr3 | 2.99E-10 | -0.049502675 | 0.416 | 0.414 | 8.03E-06 |
| Hspa9 | 4.11E-15 | -0.043874009 | 0.237 | 0.275 | 1.11E-10 | Arhgap21 | 3.02E-10 | -0.016002008 | 0.744 | 0.718 | 8.14E-06 |
| Tmx4 | 4.18E-15 | -0.028714082 | 0.388 | 0.427 | 1.12E-10 | Tra2a | 3.03E-10 | 0.074839337 | 0.703 | 0.663 | 8.14E-06 |
| Fam13c | 4.19E-15 | -0.006533966 | 0.494 | 0.527 | 1.13E-10 | Cers4 | 3.06E-10 | -0.008310643 | 0.401 | 0.379 | 8.22E-06 |
| Snd1 | 4.22E-15 | -0.077255789 | 0.555 | 0.606 | 1.14E-10 | Htt | 3.06E-10 | -0.00383039 | 0.505 | 0.48 | 8.23E-06 |
| 2810403A07Rik | 4.28E-15 | 0.088986019 | 0.278 | 0.232 | 1.15E-10 | Cdkal1 | 3.07E-10 | 0.014598654 | 0.544 | 0.514 | 8.25E-06 |
| Sarnp | 4.37E-15 | -0.03323079 | 0.356 | 0.396 | 1.18E-10 | 4930578G10Rik | 3.09E-10 | -0.009637898 | 0.555 | 0.593 | 8.32E-06 |
| Tmem38a | 4.44E-15 | -0.023128052 | 0.297 | 0.331 | 1.20E-10 | Dmxl1 | 3.10E-10 | -0.023067906 | 0.677 | 0.659 | 8.34E-06 |
| Tbcel | 4.46E-15 | 0.024806833 | 0.277 | 0.292 | 1.20E-10 | Rbm26 | 3.12E-10 | -0.026873774 | 0.753 | 0.737 | 8.39E-06 |
| Rnf214 | 4.50E-15 | 0.030161102 | 0.261 | 0.272 | 1.21E-10 | Ttll7 | 3.14E-10 | -0.024499926 | 0.427 | 0.416 | 8.46E-06 |
| Acox1 | 4.52E-15 | -0.02197504 | 0.252 | 0.284 | 1.22E-10 | Dpf3 | 3.17E-10 | -0.002201333 | 0.26 | 0.239 | 8.52E-06 |
| Mvb12b | 4.53E-15 | 0.025330666 | 0.383 | 0.4 | 1.22E-10 | Rapgef4 | 3.18E-10 | -0.063070215 | 0.829 | 0.826 | 8.56E-06 |
| Cab39l | 4.63E-15 | -0.01313724 | 0.291 | 0.322 | 1.25E-10 | Snx32 | 3.20E-10 | 0.018700842 | 0.306 | 0.279 | 8.62E-06 |
| 1700084C06Rik | 4.68E-15 | -0.056126872 | 0.314 | 0.358 | 1.26E-10 | Cdh12 | 3.22E-10 | 0.134020328 | 0.718 | 0.685 | 8.68E-06 |
| Nsun2 | 4.70E-15 | -0.018712293 | 0.349 | 0.384 | 1.26E-10 | Ap2b1 | 3.27E-10 | -0.0068116 | 0.556 | 0.534 | 8.79E-06 |
| Gpatch8 | 4.73E-15 | -0.06326232 | 0.807 | 0.847 | 1.27E-10 | Gprasp1 | 3.27E-10 | 0.026338125 | 0.566 | 0.532 | 8.79E-06 |
| Taf15 | 4.83E-15 | 0.011789094 | 0.467 | 0.494 | 1.30E-10 | Dyrk1a | 3.30E-10 | -0.023683207 | 0.51 | 0.495 | 8.88E-06 |
| Dpp8 | 4.93E-15 | 0.028188385 | 0.311 | 0.325 | 1.33E-10 | Btbd7 | 3.33E-10 | -0.036428628 | 0.425 | 0.416 | 8.95E-06 |
| Ankrd28 | 4.95E-15 | 0.024634434 | 0.433 | 0.452 | 1.33E-10 | Syn1 | 3.34E-10 | 0.060107115 | 0.549 | 0.509 | 8.98E-06 |
| Matn2 | 5.03E-15 | -0.02224429 | 0.29 | 0.322 | 1.35E-10 | Ap3d1 | 3.35E-10 | 0.015078086 | 0.269 | 0.244 | 9.02E-06 |
| Zbtb38 | 5.07E-15 | 0.050642667 | 0.395 | 0.402 | 1.36E-10 | Tmeff1 | 3.38E-10 | 0.031913074 | 0.289 | 0.26 | 9.08E-06 |
| Pogz | 5.12E-15 | 0.001487237 | 0.3 | 0.326 | 1.38E-10 | AU040320 | 3.40E-10 | -0.020384901 | 0.365 | 0.352 | 9.14E-06 |
| Vps9d1 | 5.18E-15 | -0.006917648 | 0.279 | 0.306 | 1.39E-10 | Prickle2 | 3.45E-10 | -0.067047251 | 0.893 | 0.891 | 9.27E-06 |
| Nrp2 | 5.19E-15 | 0.058540662 | 0.399 | 0.41 | 1.40E-10 | Swt1 | 3.48E-10 | 0.011589972 | 0.288 | 0.264 | 9.37E-06 |
| Gm48512 | 5.26E-15 | -0.085792858 | 0.397 | 0.447 | 1.41E-10 | Ptpn1 | 3.49E-10 | 0.036665396 | 0.474 | 0.44 | 9.40E-06 |
| Ap2a2 | 5.29E-15 | 0.002357923 | 0.389 | 0.416 | 1.42E-10 | Pus7 | 3.50E-10 | -0.038640617 | 0.271 | 0.268 | 9.42E-06 |
| Cnnm2 | 5.35E-15 | 0.044545228 | 0.395 | 0.404 | 1.44E-10 | Ivd | 3.50E-10 | -0.002533342 | 0.282 | 0.263 | 9.43E-06 |
| Ptp4a2 | 5.48E-15 | -0.034500081 | 0.249 | 0.284 | 1.47E-10 | Ddx10 | 3.51E-10 | -0.035761989 | 0.365 | 0.359 | 9.45E-06 |
| Fbxo11 | 5.55E-15 | -0.042632759 | 0.56 | 0.605 | 1.49E-10 | Dnmt3a | 3.68E-10 | -0.057153804 | 0.681 | 0.676 | 9.90E-06 |
| Cpsf7 | 5.59E-15 | -0.025156262 | 0.257 | 0.289 | 1.50E-10 | Osbpl10 | 3.79E-10 | -0.079152581 | 0.354 | 0.36 | 1.02E-05 |
| Stim1 | 5.90E-15 | 0.044013012 | 0.346 | 0.355 | 1.59E-10 | Tm9sf4 | 3.85E-10 | 0.007343801 | 0.307 | 0.284 | 1.04E-05 |
| Zhx3 | 5.95E-15 | 0.034625878 | 0.359 | 0.371 | 1.60E-10 | Gm28379 | 3.87E-10 | -0.050576972 | 0.304 | 0.304 | 1.04E-05 |
| Sec23a | 6.01E-15 | 0.050964256 | 0.289 | 0.291 | 1.62E-10 | Pik3cb | 3.89E-10 | 0.029666602 | 0.265 | 0.237 | 1.05E-05 |
| Lonrf2 | 6.10E-15 | -0.0256902 | 0.276 | 0.309 | 1.64E-10 | Tdrd3 | 3.92E-10 | -0.018752027 | 0.256 | 0.245 | 1.05E-05 |
| Fmn2 | 6.11E-15 | 0.079663391 | 0.745 | 0.741 | 1.64E-10 | Mapt | 4.07E-10 | -0.057594835 | 0.859 | 0.851 | 1.10E-05 |
| Fam13a | 6.12E-15 | 0.056688786 | 0.255 | 0.254 | 1.65E-10 | Golga4 | 4.08E-10 | -0.06228058 | 0.407 | 0.409 | 1.10E-05 |
| Evi5 | 6.18E-15 | 0.054617288 | 0.395 | 0.399 | 1.66E-10 | Gtf2i | 4.09E-10 | 0.022733618 | 0.631 | 0.597 | 1.10E-05 |
| Fbrsl1 | 6.19E-15 | 0.025106032 | 0.354 | 0.373 | 1.67E-10 | Strn4 | 4.13E-10 | 0.013738936 | 0.371 | 0.345 | 1.11E-05 |
| Wdr60 | 6.28E-15 | -0.063194957 | 0.356 | 0.402 | 1.69E-10 | Gm15952 | 4.13E-10 | 0.055443107 | 0.301 | 0.264 | 1.11E-05 |
| Map4k4 | 6.35E-15 | 0.01302978 | 0.427 | 0.451 | 1.71E-10 | Spns2 | 4.16E-10 | -0.040860395 | 0.278 | 0.275 | 1.12E-05 |
| Cramp1l | 6.52E-15 | 0.020359459 | 0.269 | 0.283 | 1.75E-10 | Dnm1l | 4.18E-10 | 0.009978399 | 0.49 | 0.46 | 1.13E-05 |

| | | | | | | | | | | | |
|---|---|---|---|---|---|---|---|---|---|---|---|
| Myh9 | 6.61E-15 | -0.077649525 | 0.271 | 0.316 | 1.78E-10 | Chka | 4.19E-10 | -0.011007398 | 0.626 | 0.603 | 1.13E-05 |
| Idh3b | 6.62E-15 | -0.067826785 | 0.236 | 0.279 | 1.78E-10 | Ephb2 | 4.28E-10 | -0.053055283 | 0.374 | 0.371 | 1.15E-05 |
| Kansl3 | 6.67E-15 | 0.032085716 | 0.255 | 0.265 | 1.80E-10 | Ap1g1 | 4.29E-10 | 0.038784424 | 0.358 | 0.325 | 1.15E-05 |
| Lars | 6.85E-15 | -0.001959952 | 0.347 | 0.375 | 1.84E-10 | Suds3 | 4.34E-10 | 0.036861074 | 0.357 | 0.324 | 1.17E-05 |
| Atl1 | 6.90E-15 | 0.046037677 | 0.361 | 0.367 | 1.86E-10 | Micu1 | 4.37E-10 | -0.016456161 | 0.446 | 0.427 | 1.18E-05 |
| Mga | 7.00E-15 | 0.024232181 | 0.388 | 0.407 | 1.88E-10 | Dcun1d4 | 4.46E-10 | -0.031829763 | 0.25 | 0.246 | 1.20E-05 |
| Lman2l | 7.08E-15 | -0.05297339 | 0.317 | 0.36 | 1.91E-10 | Gm26905 | 4.53E-10 | -0.002032316 | 0.302 | 0.334 | 1.22E-05 |
| Kdm4c | 7.10E-15 | -0.043225674 | 0.559 | 0.604 | 1.91E-10 | Cbfa2t2 | 4.54E-10 | 0.030066838 | 0.561 | 0.525 | 1.22E-05 |
| Kidins220 | 7.11E-15 | 0.035327166 | 0.644 | 0.664 | 1.91E-10 | Stxbp5 | 4.66E-10 | -0.044861236 | 0.683 | 0.671 | 1.25E-05 |
| Mapre2 | 7.24E-15 | -0.000669758 | 0.499 | 0.529 | 1.95E-10 | Glrb | 4.69E-10 | -0.030532882 | 0.459 | 0.445 | 1.26E-05 |
| Gm37679 | 7.25E-15 | -0.081283424 | 0.497 | 0.548 | 1.95E-10 | Tnik | 4.69E-10 | -0.066618534 | 0.933 | 0.938 | 1.26E-05 |
| Wipf2 | 7.39E-15 | 0.027974956 | 0.297 | 0.309 | 1.99E-10 | Atxn7 | 4.71E-10 | -0.031708087 | 0.405 | 0.398 | 1.27E-05 |
| Atp5a1 | 7.44E-15 | -0.047646501 | 0.337 | 0.38 | 2.00E-10 | Usf3 | 4.73E-10 | 0.032151052 | 0.271 | 0.243 | 1.27E-05 |
| Itsn1 | 7.50E-15 | 0.086189747 | 0.626 | 0.619 | 2.02E-10 | Ppp3cc | 4.75E-10 | -0.000546774 | 0.281 | 0.26 | 1.28E-05 |
| Xpr1 | 7.53E-15 | -0.127397975 | 0.482 | 0.531 | 2.03E-10 | Dync1i2 | 4.81E-10 | -0.029669672 | 0.754 | 0.738 | 1.29E-05 |
| Csde1 | 7.56E-15 | -0.037890756 | 0.3 | 0.339 | 2.03E-10 | 1700085D07Rik | 4.83E-10 | 0.011557471 | 0.337 | 0.31 | 1.30E-05 |
| Epb41 | 7.61E-15 | 0.121853181 | 0.28 | 0.252 | 2.05E-10 | Srpk1 | 4.86E-10 | 0.026745501 | 0.38 | 0.349 | 1.31E-05 |
| Cdc42se2 | 7.66E-15 | -0.048771557 | 0.518 | 0.565 | 2.06E-10 | Epha4 | 4.88E-10 | 0.086471957 | 0.773 | 0.739 | 1.31E-05 |
| Pcmt1 | 7.91E-15 | -0.025304116 | 0.381 | 0.418 | 2.13E-10 | Gm32250 | 4.90E-10 | 0.01964376 | 0.619 | 0.584 | 1.32E-05 |
| Klhl22 | 7.92E-15 | 0.027804552 | 0.312 | 0.325 | 2.13E-10 | Usp32 | 4.90E-10 | -0.002470286 | 0.635 | 0.609 | 1.32E-05 |
| Chd6 | 7.95E-15 | 0.085156611 | 0.623 | 0.612 | 2.14E-10 | Ino80d | 5.03E-10 | 0.049656678 | 0.334 | 0.299 | 1.35E-05 |
| Trim44 | 7.98E-15 | -0.03531802 | 0.591 | 0.634 | 2.15E-10 | 9030624J02Rik | 5.03E-10 | -0.084768923 | 0.294 | 0.334 | 1.35E-05 |
| Mtmr12 | 8.06E-15 | -0.07526043 | 0.448 | 0.498 | 2.17E-10 | Stk25 | 5.10E-10 | -0.002569165 | 0.254 | 0.235 | 1.37E-05 |
| Scaf8 | 8.06E-15 | 0.035234474 | 0.547 | 0.564 | 2.17E-10 | Fntb | 5.11E-10 | 0.055135061 | 0.338 | 0.301 | 1.37E-05 |
| Smarcc1 | 8.07E-15 | 0.008238286 | 0.379 | 0.404 | 2.17E-10 | Mapk1 | 5.15E-10 | 0.045367476 | 0.682 | 0.643 | 1.39E-05 |
| Herc3 | 8.11E-15 | -0.04767774 | 0.477 | 0.524 | 2.18E-10 | Ppargc1b | 5.23E-10 | -0.036101681 | 0.355 | 0.349 | 1.41E-05 |
| Prr14 | 8.13E-15 | -0.024990489 | 0.224 | 0.256 | 2.19E-10 | Arglu1 | 5.29E-10 | -0.083726312 | 0.666 | 0.677 | 1.42E-05 |
| Mier1 | 8.24E-15 | -0.025378249 | 0.343 | 0.38 | 2.22E-10 | Arhgap20 | 5.44E-10 | 0.049181975 | 0.499 | 0.46 | 1.46E-05 |
| Epb41l1 | 8.43E-15 | 0.096264385 | 0.586 | 0.576 | 2.27E-10 | Rassf8 | 5.45E-10 | 0.06421747 | 0.398 | 0.359 | 1.47E-05 |
| Itgb3bp | 8.44E-15 | -0.051575331 | 0.297 | 0.339 | 2.27E-10 | Naa40 | 5.47E-10 | 0.008544575 | 0.335 | 0.31 | 1.47E-05 |
| Prickle2 | 8.45E-15 | 0.091268926 | 0.891 | 0.89 | 2.27E-10 | Epb41l3 | 5.52E-10 | -0.047566833 | 0.481 | 0.473 | 1.49E-05 |
| Htt | 8.49E-15 | -0.022586522 | 0.48 | 0.519 | 2.29E-10 | Carmil1 | 5.53E-10 | -0.035118283 | 0.466 | 0.451 | 1.49E-05 |
| Prepl | 8.64E-15 | 0.040916441 | 0.331 | 0.339 | 2.33E-10 | Fip1l1 | 5.59E-10 | -0.018180157 | 0.317 | 0.305 | 1.50E-05 |
| Fam193a | 8.80E-15 | 0.055523311 | 0.602 | 0.608 | 2.37E-10 | Ankib1 | 5.59E-10 | -0.059543801 | 0.452 | 0.455 | 1.50E-05 |
| Vamp4 | 8.81E-15 | -0.013441662 | 0.24 | 0.269 | 2.37E-10 | Zswim8 | 5.67E-10 | -0.020836205 | 0.348 | 0.336 | 1.53E-05 |
| Arhgef11 | 8.94E-15 | 0.006363218 | 0.42 | 0.443 | 2.41E-10 | Jakmip3 | 5.67E-10 | 0.022684844 | 0.507 | 0.475 | 1.53E-05 |
| Map2k1 | 8.97E-15 | 0.035190292 | 0.353 | 0.368 | 2.41E-10 | Hdlbp | 5.72E-10 | 0.033427038 | 0.345 | 0.312 | 1.54E-05 |
| Trrap | 9.06E-15 | 0.031221832 | 0.4 | 0.415 | 2.44E-10 | Tef | 5.78E-10 | -0.044237689 | 0.418 | 0.413 | 1.55E-05 |
| Gm19710 | 9.17E-15 | -0.047476514 | 0.543 | 0.591 | 2.47E-10 | Slmap | 5.93E-10 | -0.062783636 | 0.415 | 0.423 | 1.60E-05 |
| Smim13 | 9.70E-15 | -0.001757836 | 0.237 | 0.26 | 2.61E-10 | Parp8 | 5.95E-10 | -0.072391201 | 0.416 | 0.425 | 1.60E-05 |
| Ttc8 | 9.72E-15 | 0.035365252 | 0.494 | 0.51 | 2.62E-10 | Tcerg1 | 6.11E-10 | 0.006756992 | 0.315 | 0.294 | 1.64E-05 |
| Wasf3 | 9.74E-15 | -0.011673287 | 0.369 | 0.401 | 2.62E-10 | B230307C23Rik | 6.12E-10 | 0.0361624 | 0.282 | 0.251 | 1.65E-05 |
| AU040320 | 9.89E-15 | 0.039839006 | 0.352 | 0.36 | 2.66E-10 | Svop | 6.15E-10 | 0.008368473 | 0.592 | 0.563 | 1.65E-05 |
| Gtpbp2 | 9.92E-15 | -0.042684658 | 0.263 | 0.3 | 2.67E-10 | 3110082I17Rik | 6.15E-10 | -0.014460357 | 0.3 | 0.284 | 1.66E-05 |
| Rev3l | 1.02E-14 | -0.066339997 | 0.643 | 0.691 | 2.75E-10 | Cstf2 | 6.38E-10 | -0.046858158 | 0.245 | 0.251 | 1.72E-05 |
| Taok1 | 1.04E-14 | 0.010822636 | 0.543 | 0.571 | 2.79E-10 | Ambra1 | 6.49E-10 | 0.025887658 | 0.749 | 0.715 | 1.75E-05 |
| Kansl1l | 1.04E-14 | -0.096453399 | 0.637 | 0.686 | 2.81E-10 | Thada | 6.60E-10 | 0.011325248 | 0.471 | 0.444 | 1.77E-05 |
| Stag2 | 1.05E-14 | -0.011558094 | 0.321 | 0.351 | 2.82E-10 | Atf6 | 6.61E-10 | 0.008341968 | 0.523 | 0.495 | 1.78E-05 |
| Ctcf | 1.06E-14 | -0.007917081 | 0.289 | 0.317 | 2.85E-10 | Ipo8 | 6.70E-10 | 0.001943966 | 0.269 | 0.249 | 1.80E-05 |
| Dync1li2 | 1.06E-14 | 0.074606435 | 0.34 | 0.331 | 2.86E-10 | Bsn | 6.70E-10 | -0.021740864 | 0.653 | 0.634 | 1.80E-05 |
| Scn3a | 1.08E-14 | -0.041321536 | 0.458 | 0.503 | 2.90E-10 | Mgat4c | 6.70E-10 | 0.016673988 | 0.473 | 0.503 | 1.80E-05 |
| Baz2b | 1.08E-14 | -0.076246902 | 0.659 | 0.707 | 2.91E-10 | Snapc3 | 6.84E-10 | 0.010050873 | 0.391 | 0.366 | 1.84E-05 |
| Nr2c2 | 1.10E-14 | 0.054443704 | 0.463 | 0.471 | 2.95E-10 | Brip1os | 6.84E-10 | 0.000199907 | 0.403 | 0.378 | 1.84E-05 |
| Rars2 | 1.10E-14 | 0.01796632 | 0.356 | 0.377 | 2.96E-10 | Ttc39c | 6.98E-10 | -0.012934446 | 0.306 | 0.293 | 1.88E-05 |
| Tlk2 | 1.10E-14 | 0.004896159 | 0.395 | 0.421 | 2.97E-10 | Rai1 | 7.06E-10 | -0.025977912 | 0.486 | 0.47 | 1.90E-05 |
| Stk25 | 1.12E-14 | -0.051129236 | 0.235 | 0.275 | 3.02E-10 | Pag1 | 7.13E-10 | -0.011099832 | 0.251 | 0.236 | 1.92E-05 |
| Tbc1d19 | 1.19E-14 | 0.028625575 | 0.35 | 0.361 | 3.20E-10 | Ttll5 | 7.19E-10 | -0.042865644 | 0.451 | 0.448 | 1.93E-05 |
| Rbl2 | 1.19E-14 | 0.016573185 | 0.241 | 0.257 | 3.20E-10 | Acss2 | 7.31E-10 | 0.011813247 | 0.31 | 0.286 | 1.97E-05 |
| Nmt2 | 1.20E-14 | 0.033572417 | 0.406 | 0.418 | 3.24E-10 | Med14 | 7.36E-10 | 0.014667212 | 0.316 | 0.29 | 1.98E-05 |
| Iqce | 1.21E-14 | 0.037470514 | 0.322 | 0.33 | 3.26E-10 | Pak1 | 7.48E-10 | 0.042880991 | 0.372 | 0.338 | 2.01E-05 |
| Tmed9 | 1.22E-14 | -0.049338383 | 0.272 | 0.313 | 3.27E-10 | Shtn1 | 7.53E-10 | 0.012290684 | 0.619 | 0.589 | 2.02E-05 |
| Zwint | 1.25E-14 | -0.071917234 | 0.344 | 0.391 | 3.37E-10 | Dennd4c | 7.64E-10 | 0.002341656 | 0.314 | 0.295 | 2.06E-05 |
| Epha3 | 1.29E-14 | -0.064545447 | 0.458 | 0.506 | 3.48E-10 | Zfp236 | 7.65E-10 | 0.006297108 | 0.32 | 0.299 | 2.06E-05 |
| Ptprz1 | 1.29E-14 | 0.021107436 | 0.299 | 0.318 | 3.48E-10 | Tab2 | 7.65E-10 | -0.05039418 | 0.291 | 0.296 | 2.06E-05 |
| Taf3 | 1.30E-14 | 0.01682397 | 0.287 | 0.305 | 3.50E-10 | Snta1 | 7.73E-10 | 0.012993678 | 0.287 | 0.264 | 2.08E-05 |
| Ocrl | 1.31E-14 | 0.004256358 | 0.399 | 0.424 | 3.52E-10 | Arih1 | 7.77E-10 | 0.049074493 | 0.707 | 0.668 | 2.09E-05 |
| Pnpla8 | 1.32E-14 | -0.005514031 | 0.313 | 0.341 | 3.56E-10 | Zfp638 | 7.93E-10 | 0.064682921 | 0.693 | 0.654 | 2.13E-05 |
| Slc17a5 | 1.36E-14 | 0.009162947 | 0.29 | 0.31 | 3.66E-10 | Pitpna | 8.07E-10 | -0.023414856 | 0.528 | 0.513 | 2.17E-05 |
| Ncoa6 | 1.39E-14 | 0.00664031 | 0.398 | 0.426 | 3.75E-10 | Sptlc1 | 8.26E-10 | 0.02568525 | 0.259 | 0.232 | 2.22E-05 |
| Arid4a | 1.41E-14 | -0.022746195 | 0.396 | 0.432 | 3.80E-10 | Pcnx2 | 8.45E-10 | -0.059391969 | 0.479 | 0.482 | 2.27E-05 |
| Aff2 | 1.41E-14 | -0.136568675 | 0.479 | 0.53 | 3.81E-10 | Srrm4 | 8.54E-10 | -0.074784242 | 0.649 | 0.651 | 2.30E-05 |
| Med13 | 1.44E-14 | -0.029494702 | 0.459 | 0.499 | 3.87E-10 | Setd5 | 8.61E-10 | -0.049535848 | 0.599 | 0.594 | 2.32E-05 |
| Epn2 | 1.47E-14 | -0.038369646 | 0.547 | 0.591 | 3.94E-10 | Sun1 | 8.66E-10 | -0.030631426 | 0.4 | 0.393 | 2.33E-05 |
| Tcerg1 | 1.47E-14 | 0.005008577 | 0.294 | 0.316 | 3.97E-10 | Clpb | 8.67E-10 | -0.035348328 | 0.265 | 0.263 | 2.33E-05 |
| Eef1a1 | 1.48E-14 | -0.06844664 | 0.237 | 0.28 | 3.99E-10 | Mgea5 | 8.67E-10 | -0.07918609 | 0.393 | 0.433 | 2.33E-05 |
| Fam49b | 1.49E-14 | 0.084945278 | 0.562 | 0.557 | 4.00E-10 | Tecpr1 | 8.75E-10 | 0.015961995 | 0.317 | 0.291 | 2.36E-05 |
| Usp37 | 1.49E-14 | -0.014946237 | 0.421 | 0.46 | 4.01E-10 | Clk4 | 8.80E-10 | -0.00546631 | 0.444 | 0.422 | 2.37E-05 |
| Cntn4 | 1.50E-14 | 0.189075327 | 0.497 | 0.459 | 4.04E-10 | Nipsnap1 | 8.81E-10 | 0.020468628 | 0.303 | 0.275 | 2.37E-05 |
| Kif2a | 1.56E-14 | 0.019397337 | 0.426 | 0.448 | 4.19E-10 | Pam | 8.85E-10 | 0.069329127 | 0.7 | 0.662 | 2.38E-05 |
| Brdt | 1.56E-14 | -0.018960443 | 0.258 | 0.288 | 4.20E-10 | Acadsb | 8.86E-10 | 0.045175014 | 0.345 | 0.31 | 2.38E-05 |
| Ccdc39 | 1.58E-14 | -0.041865457 | 0.23 | 0.265 | 4.25E-10 | Fam13c | 8.94E-10 | 0.047686222 | 0.532 | 0.494 | 2.41E-05 |
| Ube4a | 1.58E-14 | 0.021969191 | 0.31 | 0.326 | 4.26E-10 | C78859 | 9.01E-10 | -0.037783435 | 0.328 | 0.325 | 2.43E-05 |
| Hipk3 | 1.60E-14 | -0.05570366 | 0.242 | 0.282 | 4.31E-10 | 4931406P16Rik | 9.09E-10 | 0.041465598 | 0.265 | 0.234 | 2.45E-05 |
| Ints8 | 1.60E-14 | -0.034122379 | 0.256 | 0.29 | 4.31E-10 | Usp15 | 9.10E-10 | -0.036444305 | 0.57 | 0.56 | 2.45E-05 |
| Reps1 | 1.64E-14 | 0.018372219 | 0.315 | 0.331 | 4.40E-10 | Brd9 | 9.28E-10 | 0.06524516 | 0.559 | 0.518 | 2.50E-05 |
| Cep290 | 1.65E-14 | -0.015735486 | 0.237 | 0.265 | 4.43E-10 | Dnm2 | 9.35E-10 | -0.005785576 | 0.478 | 0.454 | 2.52E-05 |
| Epc2 | 1.66E-14 | 0.002174475 | 0.27 | 0.295 | 4.46E-10 | Rbm10 | 9.38E-10 | 0.036065523 | 0.29 | 0.259 | 2.52E-05 |
| Tmcc3 | 1.67E-14 | 0.060217294 | 0.297 | 0.299 | 4.50E-10 | Flywch1 | 9.44E-10 | -0.00393888 | 0.28 | 0.263 | 2.54E-05 |
| Sltm | 1.68E-14 | 0.041711491 | 0.695 | 0.715 | 4.51E-10 | Ankrd33b | 9.67E-10 | 0.027965618 | 0.53 | 0.495 | 2.60E-05 |
| Myo6 | 1.69E-14 | 0.078387764 | 0.302 | 0.29 | 4.54E-10 | Mapk10 | 9.82E-10 | -0.069142719 | 0.92 | 0.917 | 2.64E-05 |
| Zfp106 | 1.72E-14 | -0.018395874 | 0.356 | 0.39 | 4.62E-10 | Ythdc2 | 9.91E-10 | -0.006724184 | 0.357 | 0.338 | 2.67E-05 |
| Mok | 1.72E-14 | 0.005698285 | 0.235 | 0.256 | 4.62E-10 | Mtdh | 1.01E-09 | -0.013104856 | 0.606 | 0.583 | 2.72E-05 |
| Tbc1d23 | 1.78E-14 | -0.030981227 | 0.313 | 0.349 | 4.78E-10 | Ube2o | 1.03E-09 | -0.027907141 | 0.324 | 0.316 | 2.78E-05 |
| Dmxl1 | 1.78E-14 | -0.100290682 | 0.659 | 0.707 | 4.78E-10 | Mvb12b | 1.03E-09 | -0.038929918 | 0.39 | 0.383 | 2.78E-05 |
| Eif2ak4 | 1.80E-14 | 0.037914678 | 0.262 | 0.268 | 4.85E-10 | Scaper | 1.03E-09 | -0.041485769 | 0.657 | 0.648 | 2.78E-05 |
| Sdha | 1.88E-14 | 0.011435242 | 0.271 | 0.292 | 5.06E-10 | Arpp21 | 1.04E-09 | 0.075857282 | 0.96 | 0.951 | 2.79E-05 |
| Atrn | 1.89E-14 | 0.040689886 | 0.578 | 0.594 | 5.08E-10 | 4930473A02Rik | 1.04E-09 | 0.056897098 | 0.255 | 0.221 | 2.80E-05 |
| Rnps1 | 1.90E-14 | 0.079200114 | 0.305 | 0.291 | 5.10E-10 | Ric8b | 1.06E-09 | -0.033280892 | 0.303 | 0.298 | 2.84E-05 |

| | | | | | | | | | | | |
|---|---|---|---|---|---|---|---|---|---|---|---|
| Ubr2 | 1.91E-14 | 0.024482943 | 0.482 | 0.501 | 5.14E-10 | Rc3h1 | 1.07E-09 | 0.058121817 | 0.386 | 0.349 | 2.87E-05 |
| Papd5 | 1.92E-14 | 0.105087032 | 0.305 | 0.261 | 5.17E-10 | Zufsp | 1.08E-09 | -0.081875508 | 0.219 | 0.254 | 2.90E-05 |
| Psmd14 | 1.94E-14 | 0.005608174 | 0.487 | 0.514 | 5.23E-10 | Abcb7 | 1.08E-09 | 2.48E-05 | 0.253 | 0.234 | 2.92E-05 |
| Prelid3a | 1.99E-14 | -0.033077816 | 0.384 | 0.424 | 5.36E-10 | Abcc5 | 1.09E-09 | 0.024645663 | 0.628 | 0.594 | 2.93E-05 |
| Fcho2 | 2.07E-14 | 0.00267355 | 0.46 | 0.489 | 5.56E-10 | Dhx36 | 1.10E-09 | 0.007315391 | 0.396 | 0.372 | 2.97E-05 |
| Nsmf | 2.07E-14 | -0.065158188 | 0.406 | 0.452 | 5.57E-10 | Gm48239 | 1.11E-09 | -0.051846609 | 0.283 | 0.284 | 2.97E-05 |
| Cep85l | 2.10E-14 | -0.036010427 | 0.298 | 0.336 | 5.65E-10 | Sltm | 1.11E-09 | -0.004419971 | 0.721 | 0.695 | 2.98E-05 |
| Macrod1 | 2.10E-14 | 0.097245583 | 0.28 | 0.262 | 5.65E-10 | Mical3 | 1.11E-09 | -0.040950283 | 0.719 | 0.704 | 2.99E-05 |
| Atp6v0a1 | 2.13E-14 | -0.070851729 | 0.758 | 0.801 | 5.72E-10 | Armc8 | 1.11E-09 | 0.001825441 | 0.397 | 0.373 | 2.99E-05 |
| Cacul1 | 2.15E-14 | 0.006187893 | 0.25 | 0.271 | 5.77E-10 | Fbxo38 | 1.11E-09 | 0.002174231 | 0.32 | 0.3 | 3.00E-05 |
| Copg2 | 2.18E-14 | 0.065720954 | 0.427 | 0.425 | 5.86E-10 | Cemip | 1.12E-09 | 0.012779408 | 0.382 | 0.354 | 3.01E-05 |
| Cadps2 | 2.23E-14 | 0.125939469 | 0.464 | 0.454 | 6.00E-10 | Shisa6 | 1.12E-09 | -0.109137086 | 0.733 | 0.742 | 3.01E-05 |
| D10Wsu102e | 2.29E-14 | 0.038290304 | 0.381 | 0.395 | 6.15E-10 | St7l | 1.13E-09 | 0.017918426 | 0.4 | 0.371 | 3.03E-05 |
| Kirrel3 | 2.32E-14 | -0.166538377 | 0.726 | 0.721 | 6.24E-10 | Clint1 | 1.14E-09 | -0.026105312 | 0.418 | 0.406 | 3.06E-05 |
| Cdk12 | 2.37E-14 | -0.011437025 | 0.588 | 0.625 | 6.39E-10 | Hacd4 | 1.18E-09 | 0.048941178 | 0.256 | 0.224 | 3.18E-05 |
| Ola1 | 2.38E-14 | -0.000308504 | 0.444 | 0.474 | 6.40E-10 | Stx18 | 1.20E-09 | 0.027314717 | 0.356 | 0.328 | 3.23E-05 |
| Frrs1l | 2.40E-14 | 0.03776125 | 0.355 | 0.368 | 6.45E-10 | Rbm25 | 1.20E-09 | 0.003719212 | 0.675 | 0.646 | 3.23E-05 |
| Ubac2 | 2.40E-14 | -0.003888068 | 0.422 | 0.452 | 6.46E-10 | Lin7a | 1.21E-09 | 0.029828816 | 0.373 | 0.341 | 3.27E-05 |
| Timp4 | 2.42E-14 | 0.00389996 | 0.298 | 0.321 | 6.51E-10 | Sntb2 | 1.22E-09 | 0.087609762 | 0.603 | 0.562 | 3.27E-05 |
| Rab11fip3 | 2.44E-14 | 0.039432116 | 0.593 | 0.611 | 6.55E-10 | Reep1 | 1.24E-09 | 0.030000354 | 0.582 | 0.547 | 3.34E-05 |
| Spop | 2.51E-14 | 0.009709129 | 0.484 | 0.513 | 6.76E-10 | Kdm7a | 1.24E-09 | 0.07346997 | 0.454 | 0.413 | 3.35E-05 |
| Ttc17 | 2.51E-14 | 0.047216514 | 0.499 | 0.507 | 6.76E-10 | Pnkd | 1.25E-09 | 0.008490465 | 0.334 | 0.311 | 3.37E-05 |
| Gabpb2 | 2.53E-14 | 0.006225551 | 0.242 | 0.263 | 6.80E-10 | Clvs2 | 1.25E-09 | 0.067369884 | 0.559 | 0.519 | 3.37E-05 |
| C2cd5 | 2.55E-14 | -0.056193669 | 0.66 | 0.707 | 6.86E-10 | Unc13b | 1.27E-09 | -0.031229685 | 0.474 | 0.462 | 3.43E-05 |
| Efr3a | 2.55E-14 | -0.125741284 | 0.46 | 0.511 | 6.87E-10 | Iqsec2 | 1.28E-09 | -0.04191229 | 0.672 | 0.662 | 3.46E-05 |
| Mfsd6 | 2.56E-14 | 0.027129865 | 0.272 | 0.286 | 6.89E-10 | Camsap2 | 1.29E-09 | -0.024318547 | 0.383 | 0.372 | 3.48E-05 |
| Grb2 | 2.56E-14 | -0.033650452 | 0.281 | 0.315 | 6.89E-10 | Prkce | 1.29E-09 | -0.071014457 | 0.962 | 0.967 | 3.48E-05 |
| Magi1 | 2.57E-14 | 0.084050091 | 0.59 | 0.598 | 6.92E-10 | Scn2a | 1.32E-09 | -0.05636915 | 0.834 | 0.835 | 3.55E-05 |
| Tsc22d1 | 2.58E-14 | -0.023925212 | 0.582 | 0.621 | 6.93E-10 | Chd6 | 1.32E-09 | 0.036438428 | 0.66 | 0.623 | 3.56E-05 |
| Ncapd3 | 2.60E-14 | -0.019096792 | 0.356 | 0.389 | 6.98E-10 | Sgcz | 1.32E-09 | 0.002219555 | 0.442 | 0.479 | 3.56E-05 |
| 5330417C22Rik | 2.62E-14 | 0.072629679 | 0.483 | 0.481 | 7.04E-10 | Dpysl2 | 1.32E-09 | 0.02324279 | 0.531 | 0.499 | 3.56E-05 |
| Rasa1 | 2.64E-14 | -0.008725153 | 0.474 | 0.507 | 7.10E-10 | Cry2 | 1.33E-09 | 0.000768339 | 0.339 | 0.318 | 3.57E-05 |
| Vav3 | 2.67E-14 | 0.025149744 | 0.307 | 0.33 | 7.18E-10 | Cltc | 1.33E-09 | 0.025298227 | 0.378 | 0.348 | 3.58E-05 |
| Ppp6r3 | 2.67E-14 | 0.044032584 | 0.469 | 0.482 | 7.18E-10 | Dlat | 1.34E-09 | -0.002291622 | 0.267 | 0.25 | 3.60E-05 |
| Sec11a | 2.72E-14 | -0.034694253 | 0.248 | 0.283 | 7.33E-10 | Brwd1 | 1.34E-09 | -0.020890862 | 0.6 | 0.583 | 3.61E-05 |
| Fbxl17 | 2.73E-14 | 0.075191455 | 0.85 | 0.851 | 7.35E-10 | Git2 | 1.34E-09 | -0.021457695 | 0.266 | 0.257 | 3.61E-05 |
| Tet2 | 2.76E-14 | -0.031617567 | 0.324 | 0.36 | 7.42E-10 | Cdh8 | 1.35E-09 | 0.00299962 | 0.786 | 0.757 | 3.62E-05 |
| Jakmip3 | 2.81E-14 | 0.008430614 | 0.475 | 0.501 | 7.57E-10 | Ndufaf7 | 1.35E-09 | -0.00392791 | 0.305 | 0.287 | 3.62E-05 |
| Tgfbr3 | 2.81E-14 | 0.069158506 | 0.414 | 0.411 | 7.57E-10 | Gria3 | 1.35E-09 | -0.077698024 | 0.92 | 0.921 | 3.64E-05 |
| Larp4 | 2.88E-14 | 0.002285998 | 0.26 | 0.282 | 7.74E-10 | Pum1 | 1.35E-09 | 0.020101585 | 0.559 | 0.527 | 3.65E-05 |
| Abca3 | 2.95E-14 | 0.053613043 | 0.282 | 0.281 | 7.93E-10 | Lingo1 | 1.37E-09 | -0.036367322 | 0.539 | 0.53 | 3.67E-05 |
| Armc8 | 2.95E-14 | -0.018712778 | 0.373 | 0.407 | 7.93E-10 | Prkcz | 1.37E-09 | 0.022517039 | 0.528 | 0.498 | 3.68E-05 |
| Drg1 | 2.96E-14 | -0.066436986 | 0.571 | 0.62 | 7.97E-10 | Zfp292 | 1.38E-09 | -0.024037966 | 0.513 | 0.497 | 3.71E-05 |
| Dgkd | 3.02E-14 | -0.027069398 | 0.51 | 0.55 | 8.14E-10 | Hdac8 | 1.39E-09 | -0.011904792 | 0.59 | 0.568 | 3.75E-05 |
| Nr3c1 | 3.11E-14 | 0.048253439 | 0.356 | 0.363 | 8.36E-10 | Tmcc3 | 1.39E-09 | 0.023967805 | 0.326 | 0.297 | 3.75E-05 |
| Plxna4 | 3.14E-14 | -0.135763858 | 0.908 | 0.912 | 8.45E-10 | Ppp4r4 | 1.40E-09 | -0.062391024 | 0.404 | 0.405 | 3.77E-05 |
| Tmem131 | 3.16E-14 | 0.008857377 | 0.504 | 0.531 | 8.52E-10 | Agpat3 | 1.42E-09 | 0.010566064 | 0.312 | 0.289 | 3.83E-05 |
| Tnrc6c | 3.25E-14 | 0.082123696 | 0.939 | 0.931 | 8.75E-10 | Oprm1 | 1.42E-09 | 0.025470527 | 0.324 | 0.294 | 3.83E-05 |
| Napg | 3.27E-14 | -0.014967187 | 0.276 | 0.305 | 8.79E-10 | Cask | 1.45E-09 | -0.006334208 | 0.581 | 0.558 | 3.90E-05 |
| Ajap1 | 3.31E-14 | 0.057222874 | 0.266 | 0.266 | 8.92E-10 | Gm44151 | 1.46E-09 | 0.089656737 | 0.727 | 0.689 | 3.93E-05 |
| Daam1 | 3.32E-14 | 0.068791511 | 0.47 | 0.465 | 8.93E-10 | Lin52 | 1.47E-09 | 0.018759733 | 0.298 | 0.273 | 3.97E-05 |
| Kdm6a | 3.39E-14 | -0.032263343 | 0.372 | 0.41 | 9.13E-10 | Pde10a | 1.48E-09 | -0.102258754 | 0.873 | 0.882 | 3.98E-05 |
| Gm15283 | 3.40E-14 | -0.027710132 | 0.228 | 0.261 | 9.14E-10 | 4930415C11Rik | 1.48E-09 | 0.079952126 | 0.309 | 0.272 | 3.98E-05 |
| Akap8 | 3.46E-14 | -0.041208584 | 0.356 | 0.398 | 9.31E-10 | Eml2 | 1.50E-09 | -0.017177538 | 0.333 | 0.32 | 4.03E-05 |
| Gabbr1 | 3.51E-14 | -0.022754406 | 0.428 | 0.465 | 9.45E-10 | Chrd | 1.50E-09 | 0.026790662 | 0.297 | 0.268 | 4.04E-05 |
| 2610507B11Rik | 3.56E-14 | 0.037825719 | 0.294 | 0.304 | 9.57E-10 | Elp4 | 1.51E-09 | -0.042889691 | 0.489 | 0.48 | 4.06E-05 |
| Pcyt2 | 3.72E-14 | -0.068909654 | 0.353 | 0.401 | 1.00E-09 | Slit1 | 1.51E-09 | -0.065248543 | 0.703 | 0.701 | 4.07E-05 |
| Gak | 3.82E-14 | 0.020616201 | 0.292 | 0.308 | 1.03E-09 | Asxl3 | 1.52E-09 | -0.005431887 | 0.363 | 0.344 | 4.09E-05 |
| Atp8a1 | 3.84E-14 | 0.095027589 | 0.789 | 0.78 | 1.03E-09 | Rabgap1 | 1.52E-09 | 0.005341532 | 0.665 | 0.639 | 4.09E-05 |
| Ggnbp2 | 3.87E-14 | 0.006980261 | 0.469 | 0.497 | 1.04E-09 | Zdhhc2 | 1.52E-09 | -0.00867899 | 0.355 | 0.337 | 4.09E-05 |
| Cul1 | 4.01E-14 | -0.034615916 | 0.442 | 0.482 | 1.08E-09 | Slc4a3 | 1.52E-09 | -0.029341192 | 0.391 | 0.382 | 4.10E-05 |
| 5730522E02Rik | 4.09E-14 | 0.107071956 | 0.89 | 0.872 | 1.10E-09 | B230334C09Rik | 1.53E-09 | -0.038997989 | 0.452 | 0.446 | 4.11E-05 |
| Ptpn1 | 4.13E-14 | -0.052324609 | 0.44 | 0.484 | 1.11E-09 | Ccny | 1.53E-09 | -0.043589517 | 0.597 | 0.594 | 4.12E-05 |
| Nwd2 | 4.24E-14 | 0.122097177 | 0.435 | 0.41 | 1.14E-09 | Miga2 | 1.55E-09 | 0.014105985 | 0.302 | 0.279 | 4.17E-05 |
| Mphosph9 | 4.24E-14 | 0.045431289 | 0.382 | 0.39 | 1.14E-09 | Larp4 | 1.55E-09 | 0.018406794 | 0.285 | 0.26 | 4.17E-05 |
| Zfyve1 | 4.25E-14 | 0.013038345 | 0.287 | 0.304 | 1.14E-09 | Lrp1 | 1.55E-09 | -0.027192568 | 0.399 | 0.388 | 4.18E-05 |
| Mon2 | 4.31E-14 | 0.011859578 | 0.434 | 0.456 | 1.16E-09 | Cplx2 | 1.56E-09 | 0.006907534 | 0.338 | 0.315 | 4.20E-05 |
| Vps13c | 4.37E-14 | -0.122135513 | 0.603 | 0.648 | 1.18E-09 | Klhl22 | 1.57E-09 | -0.015579503 | 0.323 | 0.312 | 4.23E-05 |
| Mapkbp1 | 4.39E-14 | 0.002578893 | 0.236 | 0.257 | 1.18E-09 | Atrn | 1.58E-09 | 0.005614274 | 0.606 | 0.578 | 4.25E-05 |
| Ehmt1 | 4.42E-14 | -0.007960981 | 0.561 | 0.596 | 1.19E-09 | Dbn1 | 1.58E-09 | 0.030399431 | 0.556 | 0.521 | 4.26E-05 |
| Ralgps2 | 4.51E-14 | -0.00108184 | 0.482 | 0.512 | 1.21E-09 | Bzw2 | 1.59E-09 | 0.025905207 | 0.369 | 0.338 | 4.29E-05 |
| Ctnna2 | 4.60E-14 | 0.090572603 | 0.967 | 0.958 | 1.24E-09 | Fchsd2 | 1.62E-09 | -0.056960652 | 0.427 | 0.426 | 4.35E-05 |
| Usp25 | 4.63E-14 | 0.008901494 | 0.33 | 0.354 | 1.25E-09 | Xpo6 | 1.68E-09 | -0.002379248 | 0.322 | 0.304 | 4.52E-05 |
| Pik3ca | 4.71E-14 | 0.008058623 | 0.3 | 0.32 | 1.27E-09 | Hipk2 | 1.68E-09 | -0.04085148 | 0.33 | 0.331 | 4.53E-05 |
| Ankhd1 | 4.78E-14 | 0.019229904 | 0.588 | 0.614 | 1.29E-09 | Scn3b | 1.69E-09 | 0.025803114 | 0.424 | 0.391 | 4.55E-05 |
| Akap10 | 4.82E-14 | 0.029746377 | 0.389 | 0.404 | 1.30E-09 | Dcaf1 | 1.69E-09 | 0.033476776 | 0.351 | 0.32 | 4.56E-05 |
| Trit1 | 4.85E-14 | -0.006012428 | 0.23 | 0.254 | 1.30E-09 | Gm15489 | 1.70E-09 | 0.074011984 | 0.269 | 0.233 | 4.57E-05 |
| Asxl3 | 4.88E-14 | 0.041026506 | 0.344 | 0.355 | 1.31E-09 | Cwf19l2 | 1.74E-09 | 0.030975259 | 0.273 | 0.245 | 4.69E-05 |
| Usp46 | 4.99E-14 | -0.040407161 | 0.374 | 0.413 | 1.34E-09 | Cwc27 | 1.75E-09 | -0.031926995 | 0.434 | 0.424 | 4.70E-05 |
| Srsf10 | 5.00E-14 | -0.032175348 | 0.302 | 0.338 | 1.35E-09 | Sort1 | 1.76E-09 | 0.029285108 | 0.649 | 0.615 | 4.74E-05 |
| Ywhaz | 5.00E-14 | -0.046426499 | 0.484 | 0.528 | 1.35E-09 | Atg4c | 1.77E-09 | -0.009284928 | 0.345 | 0.329 | 4.75E-05 |
| Rragb | 5.09E-14 | 0.00905797 | 0.292 | 0.314 | 1.37E-09 | Dicer1 | 1.77E-09 | 0.022713211 | 0.36 | 0.331 | 4.77E-05 |
| Zmym6 | 5.12E-14 | 0.042487301 | 0.28 | 0.286 | 1.38E-09 | Camkk1 | 1.77E-09 | -0.047573681 | 0.258 | 0.258 | 4.77E-05 |
| Pibf1 | 5.23E-14 | -0.00020371 | 0.465 | 0.495 | 1.41E-09 | Zdhhc17 | 1.78E-09 | 0.025267887 | 0.396 | 0.368 | 4.78E-05 |
| Metap1d | 5.24E-14 | -0.055113693 | 0.236 | 0.276 | 1.41E-09 | Bbof1 | 1.80E-09 | 0.001176765 | 0.359 | 0.337 | 4.84E-05 |
| Gpr45 | 5.33E-14 | 0.077971418 | 0.448 | 0.44 | 1.43E-09 | Ptprz1 | 1.82E-09 | -0.073567193 | 0.293 | 0.299 | 4.90E-05 |
| Atxn7 | 5.37E-14 | 0.006114221 | 0.398 | 0.42 | 1.44E-09 | Pbx1 | 1.82E-09 | -0.059765965 | 0.88 | 0.875 | 4.91E-05 |
| Ddx42 | 5.42E-14 | 0.009555974 | 0.3 | 0.319 | 1.46E-09 | Rimbp2 | 1.85E-09 | -0.061734223 | 0.752 | 0.742 | 4.98E-05 |
| Cabp1 | 5.43E-14 | 0.086419286 | 0.332 | 0.312 | 1.46E-09 | Ankrd6 | 1.85E-09 | -0.049055234 | 0.306 | 0.304 | 4.98E-05 |
| Tnk2 | 5.51E-14 | 0.040496426 | 0.249 | 0.256 | 1.48E-09 | C530008M17Rik | 1.86E-09 | -0.051620833 | 0.459 | 0.45 | 4.99E-05 |
| Shank1 | 5.57E-14 | 0.044128755 | 0.583 | 0.594 | 1.50E-09 | Ppp1r16b | 1.86E-09 | 0.007142105 | 0.373 | 0.35 | 5.01E-05 |
| Abhd17b | 5.78E-14 | -0.024888265 | 0.313 | 0.347 | 1.55E-09 | Ttc21b | 1.90E-09 | -0.023780061 | 0.26 | 0.255 | 5.12E-05 |
| Slc24a5 | 5.81E-14 | -0.040736316 | 0.498 | 0.544 | 1.56E-09 | Kif16b | 1.94E-09 | -0.052542284 | 0.314 | 0.316 | 5.23E-05 |
| Sdccag8 | 5.81E-14 | -0.026209293 | 0.534 | 0.574 | 1.56E-09 | Ralgps1 | 1.94E-09 | -0.037842972 | 0.7 | 0.69 | 5.23E-05 |
| Spast | 5.81E-14 | 0.010414629 | 0.335 | 0.354 | 1.56E-09 | Gpc5 | 1.99E-09 | -0.144055006 | 0.242 | 0.276 | 5.35E-05 |
| Orc2 | 5.87E-14 | 0.054017352 | 0.302 | 0.301 | 1.58E-09 | Evi5l | 1.99E-09 | -0.031045698 | 0.272 | 0.268 | 5.36E-05 |
| Plpbp | 5.96E-14 | -0.070089659 | 0.214 | 0.255 | 1.60E-09 | Fam81a | 2.03E-09 | 0.04525666 | 0.279 | 0.247 | 5.47E-05 |
| Scamp1 | 6.02E-14 | 0.055434992 | 0.341 | 0.343 | 1.62E-09 | Dcaf5 | 2.05E-09 | -0.008083311 | 0.458 | 0.438 | 5.53E-05 |

| | | | | | | | | | | | |
|---|---|---|---|---|---|---|---|---|---|---|---|
| Ylpm1 | 6.11E-14 | 0.042914046 | 0.643 | 0.657 | 1.64E-09 | Snx30 | 2.07E-09 | 0.013668366 | 0.353 | 0.328 | 5.58E-05 |
| Efl1 | 6.24E-14 | 0.010002573 | 0.233 | 0.251 | 1.68E-09 | Kdm2a | 2.09E-09 | 0.057570723 | 0.558 | 0.517 | 5.62E-05 |
| Auh | 6.36E-14 | -0.006805528 | 0.317 | 0.344 | 1.71E-09 | Garem1 | 2.12E-09 | -0.037745363 | 0.554 | 0.547 | 5.70E-05 |
| Nipsnap2 | 6.36E-14 | 0.051964579 | 0.255 | 0.256 | 1.71E-09 | C130071C03Rik | 2.13E-09 | -0.009387338 | 0.7 | 0.675 | 5.73E-05 |
| Cabin1 | 6.49E-14 | 0.04329485 | 0.494 | 0.503 | 1.75E-09 | Slc25a27 | 2.14E-09 | 0.046253045 | 0.429 | 0.393 | 5.76E-05 |
| Chic1 | 6.51E-14 | -0.057938712 | 0.318 | 0.36 | 1.75E-09 | Cacna1d | 2.14E-09 | -0.076579362 | 0.863 | 0.874 | 5.77E-05 |
| Dcaf10 | 6.53E-14 | -0.006682273 | 0.225 | 0.25 | 1.76E-09 | Gm26565 | 2.16E-09 | -0.0832119 | 0.243 | 0.279 | 5.80E-05 |
| Klhl7 | 6.55E-14 | -0.042360517 | 0.498 | 0.541 | 1.76E-09 | Dnttip1 | 2.16E-09 | 0.008855803 | 0.251 | 0.231 | 5.82E-05 |
| Stmn3 | 6.59E-14 | 0.053062878 | 0.354 | 0.359 | 1.77E-09 | Urgcp | 2.18E-09 | 0.041437688 | 0.401 | 0.367 | 5.87E-05 |
| Nt5c2 | 6.67E-14 | 0.078172971 | 0.589 | 0.588 | 1.79E-09 | Pkn2 | 2.19E-09 | -0.007636969 | 0.479 | 0.46 | 5.88E-05 |
| Greb1l | 6.75E-14 | -0.049785024 | 0.312 | 0.352 | 1.82E-09 | Mfhas1 | 2.20E-09 | -0.036074347 | 0.33 | 0.325 | 5.92E-05 |
| Ncoa7 | 6.78E-14 | 0.002453742 | 0.631 | 0.661 | 1.82E-09 | Jarid2 | 2.20E-09 | 0.041971385 | 0.478 | 0.443 | 5.93E-05 |
| Snrk | 6.87E-14 | -0.012380249 | 0.226 | 0.252 | 1.85E-09 | Slc43a2 | 2.21E-09 | -0.030448016 | 0.336 | 0.329 | 5.94E-05 |
| Eea1 | 7.02E-14 | 0.057745458 | 0.287 | 0.284 | 1.89E-09 | Zeb1 | 2.22E-09 | -0.046118282 | 0.614 | 0.605 | 5.98E-05 |
| B230209E15Rik | 7.11E-14 | 0.00693563 | 0.426 | 0.454 | 1.91E-09 | Zfp821 | 2.25E-09 | 0.027735743 | 0.264 | 0.237 | 6.05E-05 |
| Thoc7 | 7.18E-14 | -0.039125022 | 0.218 | 0.252 | 1.93E-09 | Mbnl1 | 2.27E-09 | 0.006614778 | 0.428 | 0.404 | 6.11E-05 |
| Gm28375 | 7.31E-14 | -0.041838087 | 0.412 | 0.455 | 1.97E-09 | Cacna1h | 2.27E-09 | -0.036427651 | 0.274 | 0.268 | 6.11E-05 |
| Zfp318 | 7.45E-14 | -0.010440636 | 0.264 | 0.292 | 2.00E-09 | Fam120a | 2.30E-09 | -0.024721724 | 0.43 | 0.418 | 6.19E-05 |
| Kif3c | 7.52E-14 | 0.044810942 | 0.321 | 0.326 | 2.02E-09 | Ehmt1 | 2.33E-09 | 0.043257897 | 0.599 | 0.561 | 6.26E-05 |
| Foxj3 | 7.53E-14 | 0.015990787 | 0.322 | 0.341 | 2.03E-09 | Csmd2 | 2.33E-09 | -0.049566545 | 0.817 | 0.81 | 6.26E-05 |
| Rc3h1 | 7.57E-14 | 0.021025123 | 0.349 | 0.365 | 2.04E-09 | Fbxw11 | 2.41E-09 | 0.025883401 | 0.469 | 0.438 | 6.48E-05 |
| Prpf6 | 7.62E-14 | 0.000123878 | 0.274 | 0.299 | 2.05E-09 | Mkln1 | 2.48E-09 | -0.035378149 | 0.542 | 0.532 | 6.67E-05 |
| Tpm3 | 7.83E-14 | -0.032448171 | 0.291 | 0.326 | 2.11E-09 | Gm47167 | 2.50E-09 | 0.061463489 | 0.31 | 0.274 | 6.72E-05 |
| Pds5b | 7.90E-14 | 0.009759265 | 0.508 | 0.536 | 2.13E-09 | Slc22a23 | 2.50E-09 | -0.062793819 | 0.398 | 0.406 | 6.74E-05 |
| Cyb5r4 | 8.26E-14 | -0.035195527 | 0.22 | 0.254 | 2.22E-09 | Cttn | 2.51E-09 | -0.009591818 | 0.298 | 0.283 | 6.75E-05 |
| Eml1 | 8.27E-14 | 0.055169952 | 0.328 | 0.332 | 2.23E-09 | Dtnbp1 | 2.53E-09 | 0.015683048 | 0.26 | 0.236 | 6.81E-05 |
| Anapc5 | 8.38E-14 | 0.056151944 | 0.359 | 0.358 | 2.25E-09 | Uimc1 | 2.55E-09 | 0.017131753 | 0.325 | 0.299 | 6.86E-05 |
| Wdr33 | 8.55E-14 | -0.00551675 | 0.588 | 0.623 | 2.30E-09 | Rras2 | 2.56E-09 | 0.033347806 | 0.452 | 0.418 | 6.88E-05 |
| Mtch2 | 8.57E-14 | 0.034324162 | 0.282 | 0.292 | 2.31E-09 | Dhx32 | 2.59E-09 | -0.007068301 | 0.267 | 0.253 | 6.96E-05 |
| Fam120b | 8.59E-14 | 0.001833032 | 0.371 | 0.395 | 2.31E-09 | Anks1b | 2.63E-09 | -0.046579336 | 0.991 | 0.993 | 7.07E-05 |
| Ubr3 | 8.94E-14 | 0.070379126 | 0.689 | 0.689 | 2.40E-09 | Atl1 | 2.65E-09 | -0.004820214 | 0.38 | 0.361 | 7.12E-05 |
| Kcnn3 | 9.06E-14 | 0.024459279 | 0.249 | 0.264 | 2.44E-09 | Il1rapl1 | 2.68E-09 | -0.077962118 | 0.943 | 0.955 | 7.21E-05 |
| Aplp2 | 9.14E-14 | -0.049906799 | 0.447 | 0.492 | 2.46E-09 | Gigyf2 | 2.81E-09 | 0.014487053 | 0.563 | 0.533 | 7.57E-05 |
| Inpp5b | 9.26E-14 | 0.038684134 | 0.263 | 0.267 | 2.49E-09 | Sptb | 2.84E-09 | -0.014386343 | 0.411 | 0.393 | 7.65E-05 |
| Gbf1 | 9.61E-14 | 0.041414968 | 0.58 | 0.591 | 2.59E-09 | Dclk2 | 2.89E-09 | -0.044154472 | 0.465 | 0.459 | 7.78E-05 |
| Stxbp4 | 9.62E-14 | -0.008659337 | 0.242 | 0.267 | 2.59E-09 | Pard3 | 2.91E-09 | 0.037940246 | 0.54 | 0.503 | 7.83E-05 |
| Slc16a7 | 9.65E-14 | 0.064368821 | 0.541 | 0.546 | 2.60E-09 | Map7 | 2.91E-09 | -0.031961936 | 0.68 | 0.665 | 7.83E-05 |
| Pcsk7 | 9.76E-14 | -0.01606361 | 0.385 | 0.42 | 2.63E-09 | Rbm6 | 2.94E-09 | 0.049758878 | 0.758 | 0.723 | 7.91E-05 |
| Arhgef2 | 9.82E-14 | 0.041702862 | 0.369 | 0.379 | 2.64E-09 | Bmpr2 | 2.96E-09 | 0.068800372 | 0.521 | 0.482 | 7.98E-05 |
| Zfp652 | 9.87E-14 | 0.00164706 | 0.433 | 0.461 | 2.66E-09 | Wdr33 | 2.99E-09 | 0.051559678 | 0.626 | 0.588 | 8.05E-05 |
| Raf1 | 9.99E-14 | -0.008990595 | 0.407 | 0.44 | 2.69E-09 | Wdr7 | 3.00E-09 | -0.057414575 | 0.634 | 0.635 | 8.06E-05 |
| Vps54 | 1.03E-13 | -0.020001836 | 0.372 | 0.407 | 2.78E-09 | Camta2 | 3.00E-09 | 0.009063941 | 0.307 | 0.284 | 8.07E-05 |
| Abi2 | 1.03E-13 | -0.015634084 | 0.613 | 0.649 | 2.78E-09 | Gm27153 | 3.01E-09 | -0.059510384 | 0.246 | 0.255 | 8.09E-05 |
| Rab10 | 1.05E-13 | -0.016437525 | 0.486 | 0.521 | 2.83E-09 | Cep112 | 3.02E-09 | -0.109993747 | 0.494 | 0.503 | 8.13E-05 |
| Mgll | 1.07E-13 | 0.063305406 | 0.36 | 0.364 | 2.89E-09 | Atp2c1 | 3.02E-09 | 0.029614784 | 0.697 | 0.662 | 8.13E-05 |
| Med15 | 1.07E-13 | -0.015671193 | 0.365 | 0.397 | 2.89E-09 | Selenow | 3.03E-09 | -0.025496858 | 0.254 | 0.247 | 8.15E-05 |
| Otud7b | 1.08E-13 | 0.057567112 | 0.272 | 0.271 | 2.90E-09 | Hspa12a | 3.03E-09 | 0.02461155 | 0.37 | 0.341 | 8.15E-05 |
| Bmpr2 | 1.09E-13 | -0.018005133 | 0.482 | 0.516 | 2.92E-09 | Rngtt | 3.07E-09 | 0.001287366 | 0.487 | 0.464 | 8.27E-05 |
| Dnaja2 | 1.09E-13 | -0.010417995 | 0.496 | 0.529 | 2.94E-09 | Acvr2a | 3.11E-09 | -0.043804672 | 0.462 | 0.452 | 8.35E-05 |
| Ube2l3 | 1.12E-13 | -0.009922156 | 0.23 | 0.254 | 3.01E-09 | Mcu | 3.12E-09 | -0.017951236 | 0.565 | 0.545 | 8.39E-05 |
| Acvr1 | 1.12E-13 | -0.060114875 | 0.47 | 0.516 | 3.02E-09 | Exoc1 | 3.15E-09 | 0.007708386 | 0.251 | 0.231 | 8.48E-05 |
| Gm28379 | 1.14E-13 | -0.028400828 | 0.304 | 0.337 | 3.05E-09 | Dnajc19 | 3.16E-09 | -0.035007226 | 0.328 | 0.321 | 8.50E-05 |
| Adap1 | 1.14E-13 | 0.078640513 | 0.265 | 0.246 | 3.07E-09 | Pex14 | 3.17E-09 | 0.006626017 | 0.284 | 0.262 | 8.53E-05 |
| Macrod2 | 1.16E-13 | 0.077958567 | 0.976 | 0.971 | 3.13E-09 | Upp2 | 3.18E-09 | -0.02636184 | 0.758 | 0.742 | 8.56E-05 |
| Flrt2 | 1.17E-13 | -0.148310965 | 0.392 | 0.44 | 3.14E-09 | Tnrc6c | 3.19E-09 | -0.063907482 | 0.937 | 0.939 | 8.59E-05 |
| Anapc10 | 1.17E-13 | -0.031425027 | 0.227 | 0.258 | 3.15E-09 | Zfp182 | 3.25E-09 | -0.017954345 | 0.266 | 0.255 | 8.76E-05 |
| Fbxo42 | 1.17E-13 | 0.048322598 | 0.271 | 0.271 | 3.16E-09 | Arnt2 | 3.28E-09 | -0.031678188 | 0.43 | 0.419 | 8.82E-05 |
| Camkmt | 1.18E-13 | 0.0470142 | 0.527 | 0.539 | 3.18E-09 | Rnf112 | 3.28E-09 | -0.020083336 | 0.565 | 0.547 | 8.84E-05 |
| Atp2b2 | 1.19E-13 | 0.097465093 | 0.845 | 0.826 | 3.19E-09 | Lrch3 | 3.31E-09 | 0.005094597 | 0.472 | 0.447 | 8.89E-05 |
| Nrxn2 | 1.19E-13 | 0.049946719 | 0.407 | 0.413 | 3.21E-09 | Pou2f2 | 3.32E-09 | 0.014613967 | 0.311 | 0.286 | 8.93E-05 |
| Usp15 | 1.24E-13 | 0.012012124 | 0.56 | 0.587 | 3.33E-09 | Atxn2 | 3.36E-09 | 0.010868099 | 0.597 | 0.568 | 9.05E-05 |
| Polr3h | 1.24E-13 | -0.049653805 | 0.275 | 0.314 | 3.33E-09 | Gbf1 | 3.37E-09 | 0.01155063 | 0.608 | 0.58 | 9.06E-05 |
| Gm9801 | 1.27E-13 | 0.001255881 | 0.364 | 0.391 | 3.41E-09 | Trim46 | 3.41E-09 | 0.001925879 | 0.353 | 0.331 | 9.17E-05 |
| Hnrnpk | 1.27E-13 | -0.067243978 | 0.23 | 0.272 | 3.42E-09 | Stx8 | 3.45E-09 | -0.005102527 | 0.405 | 0.385 | 9.29E-05 |
| Elp4 | 1.28E-13 | 0.013183322 | 0.48 | 0.507 | 3.45E-09 | Kcnk10 | 3.46E-09 | 0.051386678 | 0.4 | 0.364 | 9.32E-05 |
| Ascc2 | 1.31E-13 | 0.007295867 | 0.281 | 0.302 | 3.51E-09 | Dnah9 | 3.47E-09 | -0.075145495 | 0.399 | 0.406 | 9.34E-05 |
| Apex2 | 1.32E-13 | 0.042414725 | 0.444 | 0.453 | 3.55E-09 | Micu3 | 3.51E-09 | -0.007196315 | 0.451 | 0.431 | 9.45E-05 |
| Fam214a | 1.33E-13 | 0.018001659 | 0.422 | 0.442 | 3.57E-09 | Slc44a5 | 3.62E-09 | -0.055111727 | 0.622 | 0.615 | 9.75E-05 |
| Chm | 1.33E-13 | -0.017035961 | 0.4 | 0.434 | 3.59E-09 | Atp9b | 3.67E-09 | -0.048431334 | 0.72 | 0.718 | 9.88E-05 |
| Rap1a | 1.34E-13 | -0.019262967 | 0.334 | 0.367 | 3.61E-09 | Fam193a | 3.73E-09 | -0.003602085 | 0.626 | 0.602 | 0.000100436 |
| Spns2 | 1.35E-13 | -0.040478438 | 0.275 | 0.312 | 3.63E-09 | Snx10 | 3.78E-09 | -0.01347341 | 0.396 | 0.381 | 0.000101827 |
| Shtn1 | 1.38E-13 | 0.053669593 | 0.589 | 0.596 | 3.72E-09 | Mga | 3.79E-09 | 0.0024107 | 0.411 | 0.388 | 0.000101865 |
| Rsrc1 | 1.38E-13 | 0.04588587 | 0.51 | 0.522 | 3.72E-09 | Atp11b | 3.79E-09 | -0.000635961 | 0.467 | 0.446 | 0.000101873 |
| Ociad2 | 1.40E-13 | -0.105317812 | 0.258 | 0.302 | 3.77E-09 | Kdm2b | 3.82E-09 | 0.023457026 | 0.364 | 0.334 | 0.000102859 |
| Gdi2 | 1.42E-13 | 0.004340119 | 0.247 | 0.269 | 3.83E-09 | Bcl9 | 3.85E-09 | 0.004776273 | 0.358 | 0.337 | 0.000103479 |
| Iqcb1 | 1.44E-13 | -0.025532699 | 0.269 | 0.301 | 3.88E-09 | Glp2r | 3.89E-09 | 0.09820403 | 0.567 | 0.53 | 0.000104782 |
| Gda | 1.45E-13 | 0.020589866 | 0.392 | 0.41 | 3.91E-09 | Gm12394 | 3.95E-09 | -0.057621411 | 0.517 | 0.558 | 0.000106411 |
| Orc3 | 1.51E-13 | -0.033730769 | 0.303 | 0.339 | 4.05E-09 | Mmaa | 3.96E-09 | -0.007612392 | 0.271 | 0.257 | 0.000106551 |
| Hspa4l | 1.51E-13 | -0.016061638 | 0.278 | 0.308 | 4.06E-09 | Timm9 | 3.98E-09 | -0.015070284 | 0.325 | 0.311 | 0.000106978 |
| Unc5c | 1.53E-13 | 0.066468595 | 0.722 | 0.737 | 4.11E-09 | Asah1 | 4.03E-09 | -0.010075798 | 0.251 | 0.238 | 0.000108347 |
| Foxp1 | 1.53E-13 | -0.137300795 | 0.634 | 0.675 | 4.12E-09 | Nlgn1 | 4.03E-09 | -0.055039181 | 0.983 | 0.99 | 0.000108534 |
| Igf1r | 1.56E-13 | 0.039119414 | 0.573 | 0.59 | 4.19E-09 | Nalcn | 4.06E-09 | -0.027659492 | 0.766 | 0.749 | 0.000109265 |
| Rbbp6 | 1.56E-13 | -0.021960974 | 0.346 | 0.38 | 4.20E-09 | Mpp6 | 4.06E-09 | 0.064181142 | 0.474 | 0.435 | 0.000109346 |
| Sez6 | 1.56E-13 | 0.095232832 | 0.278 | 0.26 | 4.20E-09 | Chp1 | 4.10E-09 | 0.045484639 | 0.276 | 0.245 | 0.000110313 |
| Sf1 | 1.60E-13 | -0.003806152 | 0.227 | 0.251 | 4.30E-09 | Sh3glb1 | 4.11E-09 | -0.002790189 | 0.257 | 0.24 | 0.000110504 |
| Ezh1 | 1.60E-13 | -0.002843144 | 0.235 | 0.256 | 4.31E-09 | Zmat4 | 4.18E-09 | 0.120502996 | 0.727 | 0.696 | 0.000112505 |
| Gm14372 | 1.71E-13 | -0.020841583 | 0.223 | 0.253 | 4.59E-09 | 6530409C15Rik | 4.19E-09 | 0.030976668 | 0.302 | 0.272 | 0.00011261 |
| Rngtt | 1.76E-13 | 0.00533602 | 0.464 | 0.489 | 4.73E-09 | Lyn | 4.19E-09 | 0.055610115 | 0.286 | 0.252 | 0.000112666 |
| Prkacb | 1.77E-13 | 0.019973186 | 0.3 | 0.317 | 4.77E-09 | Mecp2 | 4.20E-09 | 0.007620199 | 0.419 | 0.392 | 0.000113115 |
| Ccar1 | 1.78E-13 | -0.012378155 | 0.456 | 0.49 | 4.78E-09 | Cspp1 | 4.23E-09 | -0.001932981 | 0.575 | 0.551 | 0.000113891 |
| Agrp | 1.79E-13 | -0.0690717 | 0.223 | 0.264 | 4.83E-09 | Nbea | 4.24E-09 | -0.059480046 | 0.976 | 0.982 | 0.000114082 |
| Sos2 | 1.82E-13 | 0.041336194 | 0.275 | 0.281 | 4.90E-09 | Rasa2 | 4.32E-09 | 0.009530754 | 0.344 | 0.32 | 0.000116174 |
| Ubr5 | 1.90E-13 | 0.066435231 | 0.644 | 0.646 | 5.10E-09 | Gdpd1 | 4.37E-09 | 0.003305808 | 0.319 | 0.299 | 0.000117669 |
| Mrps9 | 1.90E-13 | 0.00684514 | 0.244 | 0.264 | 5.13E-09 | Fbxl2 | 4.39E-09 | 0.00819117 | 0.557 | 0.531 | 0.000118157 |
| Mink1 | 1.93E-13 | 0.022047105 | 0.469 | 0.491 | 5.20E-09 | Ubn1 | 4.44E-09 | 0.006531214 | 0.318 | 0.297 | 0.000119456 |
| Cyth1 | 1.95E-13 | 0.068284979 | 0.406 | 0.401 | 5.25E-09 | Cdh2 | 4.44E-09 | -0.072838155 | 0.746 | 0.757 | 0.000119585 |
| Adar | 1.98E-13 | 0.068618556 | 0.42 | 0.416 | 5.32E-09 | Marf1 | 4.46E-09 | 0.000957847 | 0.294 | 0.275 | 0.000120051 |
| Kcnmb4 | 2.00E-13 | -0.037799245 | 0.241 | 0.275 | 5.38E-09 | Ankrd11 | 4.48E-09 | -0.036091552 | 0.784 | 0.772 | 0.000120627 |

| | | | | | | | | | | | |
|---|---|---|---|---|---|---|---|---|---|---|---|
| Pus7 | 2.02E-13 | 0.053280987 | 0.268 | 0.268 | 5.44E-09 | Magi1 | 4.48E-09 | -0.021154833 | 0.609 | 0.59 | 0.000120658 |
| Tub | 2.07E-13 | -0.004279948 | 0.48 | 0.51 | 5.57E-09 | S100pbp | 4.51E-09 | 0.020217502 | 0.256 | 0.231 | 0.000121368 |
| Zfp827 | 2.12E-13 | 0.045436963 | 0.319 | 0.323 | 5.71E-09 | Sik3 | 4.56E-09 | -0.024143706 | 0.733 | 0.712 | 0.000122713 |
| Zdhhc17 | 2.16E-13 | -0.023068791 | 0.368 | 0.401 | 5.82E-09 | Cfdp1 | 4.59E-09 | 0.036462681 | 0.308 | 0.279 | 0.000123536 |
| Sar1b | 2.17E-13 | -0.048050065 | 0.215 | 0.25 | 5.83E-09 | Cenpc1 | 4.60E-09 | -0.000735987 | 0.264 | 0.246 | 0.000123842 |
| Slc3a2 | 2.19E-13 | -0.041624959 | 0.236 | 0.272 | 5.90E-09 | Ola1 | 4.61E-09 | 0.011951391 | 0.472 | 0.444 | 0.000123988 |
| Pam | 2.21E-13 | 0.047230071 | 0.662 | 0.625 | 5.95E-09 | Ahcyl1 | 4.65E-09 | 0.024513171 | 0.46 | 0.43 | 0.000125014 |
| Pde7a | 2.21E-13 | -0.007074751 | 0.403 | 0.431 | 5.95E-09 | Nae1 | 4.69E-09 | 2.48E-05 | 0.295 | 0.275 | 0.000126115 |
| Slc35f1 | 2.23E-13 | 0.137079432 | 0.498 | 0.452 | 6.01E-09 | Ccl25 | 4.80E-09 | -0.003882808 | 0.298 | 0.282 | 0.000129092 |
| Tnpo1 | 2.25E-13 | 0.006346199 | 0.527 | 0.554 | 6.06E-09 | Sptbn4 | 4.88E-09 | 0.03097439 | 0.704 | 0.669 | 0.000131344 |
| Chd1 | 2.26E-13 | -0.039184052 | 0.242 | 0.276 | 6.09E-09 | Picalm | 4.96E-09 | 0.013000157 | 0.374 | 0.349 | 0.00013351 |
| Gpr137c | 2.28E-13 | 0.040874824 | 0.297 | 0.309 | 6.14E-09 | Prkdc | 4.97E-09 | -0.013465669 | 0.305 | 0.291 | 0.000133714 |
| Lrfn5 | 2.31E-13 | 0.089841263 | 0.955 | 0.947 | 6.22E-09 | Tnr | 4.99E-09 | -0.082867232 | 0.867 | 0.863 | 0.000134267 |
| Tsc22d2 | 2.32E-13 | -0.010243685 | 0.315 | 0.343 | 6.24E-09 | Strada | 5.02E-09 | 0.002900245 | 0.372 | 0.35 | 0.000135191 |
| Eefsec | 2.33E-13 | 0.018827186 | 0.399 | 0.421 | 6.26E-09 | Glyr1 | 5.26E-09 | 0.013603118 | 0.337 | 0.312 | 0.000141605 |
| Mgat5 | 2.33E-13 | 0.015181568 | 0.505 | 0.527 | 6.27E-09 | Fras1 | 5.27E-09 | -0.123508682 | 0.265 | 0.288 | 0.000141702 |
| Mycbp2 | 2.36E-13 | -0.07838739 | 0.973 | 0.975 | 6.34E-09 | St7 | 5.27E-09 | -0.033975891 | 0.671 | 0.659 | 0.000141884 |
| Bdp1 | 2.40E-13 | -0.043813883 | 0.387 | 0.428 | 6.44E-09 | Rab14 | 5.39E-09 | -0.009030999 | 0.286 | 0.271 | 0.000145057 |
| Mphosph8 | 2.40E-13 | -0.061765108 | 0.266 | 0.307 | 6.46E-09 | Arid4a | 5.40E-09 | 0.025832412 | 0.426 | 0.396 | 0.000145265 |
| Zfp182 | 2.41E-13 | -0.014904138 | 0.255 | 0.282 | 6.47E-09 | Brwd3 | 5.42E-09 | -0.00772956 | 0.254 | 0.24 | 0.000145707 |
| Atf7ip | 2.41E-13 | 0.004872406 | 0.371 | 0.397 | 6.49E-09 | Tnrc6b | 5.42E-09 | 0.060299979 | 0.828 | 0.798 | 0.000145921 |
| Ddi2 | 2.46E-13 | -0.015064747 | 0.282 | 0.31 | 6.62E-09 | Nfkb1 | 5.48E-09 | 0.008963621 | 0.368 | 0.345 | 0.000147577 |
| Lzts1 | 2.50E-13 | -0.029407827 | 0.319 | 0.353 | 6.72E-09 | Eipr1 | 5.53E-09 | -0.020281496 | 0.337 | 0.326 | 0.000148776 |
| Akap9 | 2.55E-13 | 0.037062387 | 0.466 | 0.482 | 6.86E-09 | Limch1 | 5.63E-09 | -0.044367037 | 0.659 | 0.649 | 0.000151563 |
| Ptprk | 2.56E-13 | 0.098294855 | 0.733 | 0.731 | 6.89E-09 | Pkd1 | 5.64E-09 | 0.016104434 | 0.588 | 0.557 | 0.00015188 |
| Bin1 | 2.57E-13 | 0.048622578 | 0.49 | 0.499 | 6.91E-09 | Cdc42se2 | 5.81E-09 | 0.007022005 | 0.547 | 0.518 | 0.000156211 |
| G3bp1 | 2.61E-13 | 0.021814244 | 0.252 | 0.266 | 7.02E-09 | Bckdhb | 5.86E-09 | -0.041019835 | 0.338 | 0.338 | 0.000157543 |
| Atad1 | 2.66E-13 | -0.020094169 | 0.281 | 0.312 | 7.17E-09 | Mirg | 5.89E-09 | -0.045767081 | 0.353 | 0.354 | 0.000158529 |
| Snx13 | 2.69E-13 | 0.027481364 | 0.354 | 0.367 | 7.23E-09 | Mllt3 | 5.95E-09 | -0.044251943 | 0.755 | 0.748 | 0.000160028 |
| Ttc19 | 2.80E-13 | -0.050702163 | 0.648 | 0.693 | 7.53E-09 | Ccdc88a | 6.02E-09 | -0.037705705 | 0.74 | 0.729 | 0.000161914 |
| Mapkap1 | 2.82E-13 | 0.004629357 | 0.514 | 0.542 | 7.60E-09 | Plekhm2 | 6.03E-09 | 0.010186221 | 0.332 | 0.308 | 0.000162366 |
| Mtf2 | 2.86E-13 | -0.017895596 | 0.48 | 0.515 | 7.70E-09 | Gm48678 | 6.04E-09 | 0.038304449 | 0.682 | 0.646 | 0.000162572 |
| Armc9 | 2.99E-13 | 0.040241507 | 0.532 | 0.546 | 8.05E-09 | Rapgef4os1 | 6.10E-09 | 0.050353058 | 0.405 | 0.367 | 0.000164104 |
| Pign | 3.05E-13 | 0.079791012 | 0.305 | 0.288 | 8.20E-09 | Sh3pxd2a | 6.12E-09 | -0.039670481 | 0.3 | 0.297 | 0.00016462 |
| Pgm2l1 | 3.15E-13 | -0.01322908 | 0.32 | 0.349 | 8.47E-09 | Ptpn12 | 6.15E-09 | 0.00299788 | 0.44 | 0.416 | 0.000165534 |
| Pak7 | 3.15E-13 | 0.089656075 | 0.665 | 0.648 | 8.49E-09 | Ramp1 | 6.23E-09 | 0.019440099 | 0.257 | 0.234 | 0.000167518 |
| Prkdc | 3.18E-13 | -0.005047788 | 0.291 | 0.313 | 8.56E-09 | Mark2 | 6.37E-09 | -0.010484547 | 0.603 | 0.583 | 0.000171436 |
| Cntnap5a | 3.19E-13 | 0.164198495 | 0.647 | 0.62 | 8.59E-09 | Nsd1 | 6.43E-09 | 0.044307488 | 0.823 | 0.791 | 0.000173066 |
| Picalm | 3.21E-13 | 0.015101908 | 0.349 | 0.366 | 8.64E-09 | Plekhm3 | 6.45E-09 | -0.005355434 | 0.439 | 0.42 | 0.000173548 |
| Man1a2 | 3.25E-13 | 0.032365722 | 0.556 | 0.578 | 8.74E-09 | Agap3 | 6.48E-09 | 0.007181474 | 0.357 | 0.334 | 0.000174328 |
| Chrna7 | 3.27E-13 | 0.058531237 | 0.393 | 0.4 | 8.80E-09 | Gsdme | 6.68E-09 | 0.07732569 | 0.491 | 0.452 | 0.000179621 |
| Ptpn12 | 3.30E-13 | -0.008900099 | 0.416 | 0.448 | 8.87E-09 | Reps2 | 6.74E-09 | 0.025319226 | 0.666 | 0.635 | 0.000181398 |
| Abhd2 | 3.30E-13 | 0.000781626 | 0.292 | 0.315 | 8.88E-09 | Pds5b | 6.76E-09 | 0.001578723 | 0.533 | 0.508 | 0.000181839 |
| Clstn1 | 3.37E-13 | 0.084574267 | 0.486 | 0.473 | 9.08E-09 | Prpf6 | 6.78E-09 | 0.035124076 | 0.304 | 0.274 | 0.000182412 |
| Sacs | 3.48E-13 | -0.018264367 | 0.278 | 0.309 | 9.37E-09 | Acat1 | 6.81E-09 | 0.01491778 | 0.412 | 0.387 | 0.00018336 |
| Sestd1 | 3.51E-13 | 0.012759903 | 0.325 | 0.345 | 9.44E-09 | Gpi1 | 7.07E-09 | -0.041299642 | 0.355 | 0.356 | 0.000190218 |
| Urgcp | 3.52E-13 | -0.009505467 | 0.367 | 0.397 | 9.47E-09 | Serpini1 | 7.09E-09 | 0.026079207 | 0.365 | 0.335 | 0.000190798 |
| Nsmaf | 3.55E-13 | -0.033349735 | 0.306 | 0.341 | 9.54E-09 | Fars2 | 7.14E-09 | -0.003698124 | 0.602 | 0.579 | 0.000192199 |
| Atp6v1h | 3.56E-13 | -0.004066974 | 0.431 | 0.458 | 9.58E-09 | Trpc4ap | 7.21E-09 | -0.027177082 | 0.328 | 0.319 | 0.000193969 |
| Arhgap20 | 3.60E-13 | -0.091978165 | 0.46 | 0.509 | 9.69E-09 | Susd1 | 7.22E-09 | 0.028816294 | 0.265 | 0.241 | 0.000194195 |
| Ddx6 | 3.64E-13 | 0.009581816 | 0.29 | 0.311 | 9.78E-09 | Plppr5 | 7.35E-09 | 0.052004437 | 0.471 | 0.434 | 0.000197763 |
| Mindy2 | 3.69E-13 | 0.010380376 | 0.245 | 0.262 | 9.93E-09 | Cadps2 | 7.36E-09 | -0.094319054 | 0.448 | 0.464 | 0.000198059 |
| Cdkal1 | 3.74E-13 | 0.009942039 | 0.514 | 0.538 | 1.01E-08 | Cnnm1 | 7.48E-09 | -0.036163855 | 0.37 | 0.367 | 0.000201387 |
| Rreb1 | 3.81E-13 | 0.094221239 | 0.296 | 0.285 | 1.03E-08 | Gm48765 | 7.70E-09 | 0.057729114 | 0.307 | 0.273 | 0.000207083 |
| Gmcl1 | 3.88E-13 | 0.059814983 | 0.417 | 0.418 | 1.04E-08 | Rps6ka5 | 7.70E-09 | -0.06839992 | 0.279 | 0.289 | 0.000207211 |
| Luc7l3 | 3.91E-13 | 0.026570447 | 0.548 | 0.57 | 1.05E-08 | Ccdc73 | 7.80E-09 | 0.028041414 | 0.279 | 0.252 | 0.000209778 |
| Fez1 | 3.94E-13 | -0.03375871 | 0.274 | 0.308 | 1.06E-08 | Tango2 | 7.82E-09 | -0.004393122 | 0.412 | 0.392 | 0.000210325 |
| Napb | 3.98E-13 | -0.006362875 | 0.288 | 0.315 | 1.07E-08 | Ttc19 | 7.94E-09 | 0.047553277 | 0.684 | 0.648 | 0.000213606 |
| Fgf14 | 3.99E-13 | 0.036860286 | 0.993 | 0.989 | 1.07E-08 | Phf20l1 | 7.99E-09 | 0.008161346 | 0.714 | 0.685 | 0.000214925 |
| Zc3h12b | 4.04E-13 | 0.06000799 | 0.559 | 0.562 | 1.09E-08 | Rab30 | 8.08E-09 | -0.042179697 | 0.369 | 0.367 | 0.000217373 |
| Ino80dos | 4.05E-13 | -0.030675823 | 0.306 | 0.34 | 1.09E-08 | Fam172a | 8.23E-09 | -0.037816351 | 0.74 | 0.728 | 0.000221343 |
| Rnf169 | 4.13E-13 | -0.035206049 | 0.619 | 0.66 | 1.11E-08 | Pitpnc1 | 8.27E-09 | -0.095225563 | 0.362 | 0.368 | 0.000222588 |
| Atp13a3 | 4.16E-13 | 0.056415076 | 0.328 | 0.325 | 1.12E-08 | Mast4 | 8.41E-09 | -0.0520317 | 0.716 | 0.695 | 0.000226297 |
| Iqgap1 | 4.17E-13 | 0.063215416 | 0.309 | 0.306 | 1.12E-08 | Rgs6 | 8.46E-09 | -0.172463097 | 0.263 | 0.282 | 0.000227569 |
| Gm28376 | 4.25E-13 | -0.095501296 | 0.766 | 0.791 | 1.14E-08 | Sil1 | 8.60E-09 | 0.038765026 | 0.292 | 0.263 | 0.00023133 |
| Clock | 4.25E-13 | 0.001174093 | 0.513 | 0.541 | 1.14E-08 | Cabp1 | 8.67E-09 | -0.011205778 | 0.348 | 0.332 | 0.000233159 |
| Sec24b | 4.29E-13 | -0.000820315 | 0.461 | 0.488 | 1.15E-08 | Cul9 | 8.67E-09 | -0.021150874 | 0.366 | 0.355 | 0.000233303 |
| Pi4ka | 4.37E-13 | 0.055978264 | 0.746 | 0.75 | 1.18E-08 | Dip2a | 8.71E-09 | -0.076317829 | 0.27 | 0.286 | 0.000234468 |
| Adamts17 | 4.45E-13 | -0.080077229 | 0.43 | 0.478 | 1.20E-08 | Hnrnph1 | 8.93E-09 | 0.021905723 | 0.436 | 0.405 | 0.000240265 |
| Arhgap15 | 4.47E-13 | -0.111234614 | 0.227 | 0.269 | 1.20E-08 | Larp1 | 8.94E-09 | 0.016303426 | 0.501 | 0.473 | 0.000240517 |
| Zswim5 | 4.48E-13 | 0.002481938 | 0.343 | 0.367 | 1.20E-08 | Arhgap17 | 9.11E-09 | -0.034553456 | 0.435 | 0.427 | 0.000245134 |
| Luc7l | 4.52E-13 | 0.052737461 | 0.312 | 0.313 | 1.22E-08 | Dnajc10 | 9.17E-09 | -0.023479539 | 0.264 | 0.258 | 0.000246645 |
| Rrnad1 | 4.54E-13 | -0.002386729 | 0.259 | 0.283 | 1.22E-08 | Lmo7 | 9.24E-09 | 0.000760914 | 0.583 | 0.555 | 0.000248604 |
| Prkcg | 4.64E-13 | 0.084051485 | 0.613 | 0.585 | 1.25E-08 | Inhba | 9.27E-09 | -0.011715059 | 0.289 | 0.275 | 0.000249447 |
| Trim35 | 4.71E-13 | -0.044237906 | 0.52 | 0.563 | 1.27E-08 | Rasal2 | 9.55E-09 | 0.079565344 | 0.884 | 0.862 | 0.000257031 |
| Lrba | 4.72E-13 | 0.063468572 | 0.671 | 0.674 | 1.27E-08 | Fbrsl1 | 9.67E-09 | -0.035696688 | 0.359 | 0.354 | 0.000260227 |
| Specc1l | 4.87E-13 | 0.041794735 | 0.365 | 0.371 | 1.31E-08 | Luzp2 | 9.73E-09 | 0.03369038 | 0.232 | 0.256 | 0.000261862 |
| Tox | 4.97E-13 | -0.133394697 | 0.286 | 0.271 | 1.34E-08 | Gm42439 | 9.76E-09 | 0.073944716 | 0.688 | 0.651 | 0.000262742 |
| Gm26954 | 4.99E-13 | -0.041202795 | 0.231 | 0.264 | 1.34E-08 | 4921511C10Rik | 9.77E-09 | 0.033082467 | 0.388 | 0.355 | 0.000262765 |
| Gm48003 | 5.01E-13 | -0.099713791 | 0.223 | 0.265 | 1.35E-08 | Galnt13 | 9.79E-09 | -0.082586042 | 0.695 | 0.71 | 0.000263375 |
| Wdpcp | 5.09E-13 | -0.030188435 | 0.273 | 0.307 | 1.37E-08 | Apc | 9.81E-09 | -0.044457616 | 0.814 | 0.805 | 0.000264053 |
| Gm26906 | 5.10E-13 | 0.065055341 | 0.407 | 0.362 | 1.37E-08 | Znrf1 | 9.83E-09 | 0.04074548 | 0.464 | 0.43 | 0.000264438 |
| Stk38 | 5.13E-13 | -0.020474869 | 0.408 | 0.443 | 1.38E-08 | Spock2 | 9.96E-09 | 0.029502928 | 0.427 | 0.397 | 0.000267941 |
| Igsf8 | 5.13E-13 | -0.010283842 | 0.241 | 0.268 | 1.38E-08 | Camkk2 | 9.99E-09 | -0.01755329 | 0.355 | 0.342 | 0.000268739 |
| Znrf1 | 5.46E-13 | 0.014733496 | 0.43 | 0.451 | 1.47E-08 | Pcnx | 1.00E-08 | -0.03492438 | 0.604 | 0.594 | 0.000269005 |
| Ctif | 5.53E-13 | 0.015279343 | 0.518 | 0.54 | 1.49E-08 | Dmtf1 | 1.02E-08 | -0.001756567 | 0.286 | 0.268 | 0.000274769 |
| Rapgefl1 | 5.75E-13 | 0.033706207 | 0.359 | 0.37 | 1.55E-08 | Xrcc4 | 1.05E-08 | 0.002646215 | 0.349 | 0.329 | 0.000281929 |
| Sec14l1 | 5.86E-13 | 0.00266489 | 0.399 | 0.426 | 1.58E-08 | Senp7 | 1.05E-08 | -0.015425383 | 0.562 | 0.543 | 0.000283091 |
| Pdss1 | 5.89E-13 | 0.002980673 | 0.251 | 0.271 | 1.59E-08 | Myrip | 1.06E-08 | -0.044034817 | 0.474 | 0.472 | 0.000283997 |
| Afg1l | 6.02E-13 | 0.027528354 | 0.284 | 0.295 | 1.62E-08 | Tnks | 1.06E-08 | -0.032002355 | 0.433 | 0.424 | 0.000286082 |
| Ythdf3 | 6.07E-13 | -0.019250774 | 0.328 | 0.36 | 1.63E-08 | Gm37679 | 1.08E-08 | -0.027999518 | 0.511 | 0.497 | 0.000289352 |
| Secisbp2l | 6.19E-13 | -0.033450594 | 0.235 | 0.266 | 1.67E-08 | Vps9d1 | 1.08E-08 | -0.030836624 | 0.284 | 0.279 | 0.000289812 |
| Lrtm1 | 6.31E-13 | -0.037799324 | 0.331 | 0.369 | 1.70E-08 | Camkmt | 1.08E-08 | -0.03988856 | 0.533 | 0.527 | 0.000289899 |
| 5031439G07Rik | 6.32E-13 | 0.005922934 | 0.357 | 0.382 | 1.70E-08 | Ldah | 1.08E-08 | -0.001008534 | 0.293 | 0.276 | 0.000290905 |
| Axin1 | 6.35E-13 | 0.015297777 | 0.238 | 0.254 | 1.71E-08 | Actr3b | 1.10E-08 | -0.017047734 | 0.492 | 0.477 | 0.000294802 |
| Rab14 | 6.46E-13 | -0.01020892 | 0.271 | 0.299 | 1.74E-08 | Rnf111 | 1.10E-08 | 0.010918903 | 0.438 | 0.413 | 0.000296925 |
| Tspan13 | 6.47E-13 | -0.038794204 | 0.373 | 0.412 | 1.74E-08 | Cop1 | 1.11E-08 | -0.034066065 | 0.411 | 0.406 | 0.000297833 |
| Zfp799 | 6.86E-13 | -0.034094443 | 0.281 | 0.313 | 1.85E-08 | Mib1 | 1.11E-08 | 0.038747954 | 0.495 | 0.461 | 0.00029809 |

| | | | | | | | | | | | |
|---|---|---|---|---|---|---|---|---|---|---|---|
| Syne1 | 6.87E-13 | 0.106907288 | 0.879 | 0.869 | 1.85E-08 | Gm15414 | 1.11E-08 | 0.064055463 | 0.262 | 0.228 | 0.000298844 |
| Zmym2 | 6.91E-13 | -0.029948997 | 0.483 | 0.523 | 1.86E-08 | Usp24 | 1.13E-08 | -0.01280961 | 0.466 | 0.449 | 0.000304116 |
| Srrm2 | 6.91E-13 | 0.087301537 | 0.93 | 0.927 | 1.86E-08 | Senp5 | 1.15E-08 | 0.054502698 | 0.448 | 0.411 | 0.000309142 |
| Mark1 | 7.00E-13 | -0.033607933 | 0.389 | 0.426 | 1.88E-08 | Meg3 | 1.15E-08 | -0.062617707 | 0.984 | 0.99 | 0.000310132 |
| Rab28 | 7.06E-13 | 0.004321187 | 0.358 | 0.38 | 1.90E-08 | Zfyve9 | 1.18E-08 | 0.019709685 | 0.638 | 0.607 | 0.000317971 |
| Tmem161b | 7.11E-13 | -0.050327848 | 0.422 | 0.466 | 1.91E-08 | Fkbp5 | 1.19E-08 | -0.065502814 | 0.329 | 0.336 | 0.000320438 |
| Setd7 | 7.11E-13 | 0.031640372 | 0.332 | 0.345 | 1.91E-08 | Gpr155 | 1.23E-08 | -0.01999209 | 0.259 | 0.248 | 0.000332083 |
| Foxn3 | 7.13E-13 | 0.121820866 | 0.535 | 0.511 | 1.92E-08 | Anapc1 | 1.24E-08 | -0.020558124 | 0.277 | 0.268 | 0.000332609 |
| Uvrag | 7.13E-13 | 0.004801585 | 0.593 | 0.622 | 1.92E-08 | Zeb2 | 1.25E-08 | 0.06301235 | 0.747 | 0.754 | 0.000336645 |
| Ssbp3 | 7.25E-13 | 0.050823418 | 0.411 | 0.415 | 1.95E-08 | Tmem120b | 1.25E-08 | 0.037209953 | 0.261 | 0.233 | 0.00033758 |
| Specc1 | 7.37E-13 | -0.080068371 | 0.727 | 0.769 | 1.98E-08 | E130307A14Rik | 1.27E-08 | 0.012822546 | 0.599 | 0.571 | 0.000340786 |
| Spock3 | 7.50E-13 | 0.120577699 | 0.456 | 0.423 | 2.02E-08 | Baz2b | 1.27E-08 | 0.034599115 | 0.694 | 0.659 | 0.000341978 |
| Ttyh1 | 7.55E-13 | -0.036057286 | 0.369 | 0.407 | 2.03E-08 | Tmem167 | 1.32E-08 | -0.00569608 | 0.319 | 0.301 | 0.000355336 |
| Ddx55 | 7.59E-13 | -0.018156414 | 0.313 | 0.342 | 2.04E-08 | Top2b | 1.33E-08 | -0.00835396 | 0.377 | 0.359 | 0.000358338 |
| Ppfia3 | 7.64E-13 | 0.02677008 | 0.28 | 0.291 | 2.06E-08 | Crmp1 | 1.34E-08 | 0.023642538 | 0.33 | 0.305 | 0.000359364 |
| Cnot4 | 7.74E-13 | -0.04898352 | 0.677 | 0.719 | 2.08E-08 | Ildr2 | 1.36E-08 | 0.007274862 | 0.504 | 0.479 | 0.000365102 |
| Slc24a4 | 7.88E-13 | 0.00852165 | 0.435 | 0.463 | 2.12E-08 | Elovl6 | 1.36E-08 | 0.007172836 | 0.357 | 0.335 | 0.000366417 |
| Zfp398 | 7.92E-13 | -0.011189717 | 0.43 | 0.462 | 2.13E-08 | Smc6 | 1.37E-08 | 0.016550516 | 0.292 | 0.267 | 0.000367473 |
| Ror1 | 7.96E-13 | -0.130292739 | 0.338 | 0.384 | 2.14E-08 | Baz2a | 1.38E-08 | 0.009582523 | 0.331 | 0.31 | 0.000370801 |
| Gabra5 | 8.17E-13 | 0.014212098 | 0.446 | 0.466 | 2.20E-08 | Smarca2 | 1.40E-08 | -0.005895653 | 0.523 | 0.498 | 0.000376809 |
| Fbxl5 | 8.38E-13 | -0.033402823 | 0.378 | 0.416 | 2.25E-08 | Msi2 | 1.41E-08 | -0.006903654 | 0.497 | 0.478 | 0.000380131 |
| Zhx2 | 8.46E-13 | -0.025219928 | 0.265 | 0.296 | 2.28E-08 | Sacs | 1.43E-08 | -0.018509374 | 0.29 | 0.278 | 0.000383749 |
| Esyt2 | 8.57E-13 | 0.044068606 | 0.621 | 0.631 | 2.31E-08 | Kansl1l | 1.45E-08 | -0.010191603 | 0.658 | 0.637 | 0.000389553 |
| Stk4 | 8.76E-13 | -0.021755299 | 0.274 | 0.304 | 2.36E-08 | Cbl | 1.46E-08 | 0.009849148 | 0.426 | 0.402 | 0.00039346 |
| Faah | 8.93E-13 | 0.030572589 | 0.488 | 0.505 | 2.40E-08 | Itfg1 | 1.47E-08 | -0.010096632 | 0.418 | 0.401 | 0.000396569 |
| Hnrnpl | 8.95E-13 | 0.032928174 | 0.463 | 0.477 | 2.41E-08 | Tcaf1 | 1.48E-08 | -0.010815127 | 0.534 | 0.52 | 0.000398722 |
| Ddx39b | 9.00E-13 | -0.059914136 | 0.318 | 0.361 | 2.42E-08 | Mtus2 | 1.48E-08 | 0.013105997 | 0.717 | 0.688 | 0.000398854 |
| N4bp1 | 9.13E-13 | -0.006417073 | 0.253 | 0.277 | 2.46E-08 | Nsd3 | 1.49E-08 | -0.020395344 | 0.758 | 0.739 | 0.00040139 |
| Bend6 | 9.24E-13 | 0.067146618 | 0.312 | 0.306 | 2.49E-08 | Txndc16 | 1.49E-08 | 0.036975643 | 0.333 | 0.303 | 0.000401737 |
| Tmtc1 | 9.42E-13 | -0.157237519 | 0.452 | 0.495 | 2.53E-08 | Hnrnpa2b1 | 1.50E-08 | 0.005257414 | 0.667 | 0.64 | 0.000402302 |
| Rfx3 | 9.59E-13 | -0.23304325 | 0.737 | 0.755 | 2.58E-08 | Phka2 | 1.50E-08 | -0.02859116 | 0.32 | 0.314 | 0.000404551 |
| Myef2 | 9.60E-13 | 0.050250485 | 0.455 | 0.461 | 2.58E-08 | Gm26733 | 1.50E-08 | -0.011704834 | 0.593 | 0.624 | 0.00040463 |
| Slf1 | 9.63E-13 | 0.049398993 | 0.35 | 0.354 | 2.59E-08 | Grip1 | 1.51E-08 | -0.06575261 | 0.676 | 0.675 | 0.000405521 |
| Zfp148 | 9.65E-13 | 0.020322513 | 0.616 | 0.639 | 2.60E-08 | Mink1 | 1.61E-08 | -0.038518776 | 0.476 | 0.469 | 0.000432083 |
| Thoc1 | 9.76E-13 | -0.021977701 | 0.341 | 0.375 | 2.63E-08 | Dzip1 | 1.61E-08 | -0.031010516 | 0.306 | 0.298 | 0.000432786 |
| Smarcc2 | 9.76E-13 | 0.039344441 | 0.365 | 0.371 | 2.63E-08 | Brd1 | 1.62E-08 | -0.005624543 | 0.365 | 0.347 | 0.000435608 |
| Lrrfip1 | 9.80E-13 | 0.029809974 | 0.698 | 0.718 | 2.64E-08 | Pde7a | 1.64E-08 | -0.023060699 | 0.412 | 0.403 | 0.000442332 |
| Smad3 | 9.83E-13 | 0.083672351 | 0.407 | 0.409 | 2.64E-08 | Tulp3 | 1.65E-08 | 0.025066226 | 0.276 | 0.25 | 0.000443667 |
| Wdfy2 | 1.00E-12 | 0.051819036 | 0.265 | 0.263 | 2.70E-08 | Rptor | 1.66E-08 | -0.000480901 | 0.609 | 0.584 | 0.000445767 |
| Mkln1 | 1.04E-12 | 0.023166118 | 0.532 | 0.553 | 2.79E-08 | Slc38a9 | 1.66E-08 | -0.021751092 | 0.471 | 0.458 | 0.000446753 |
| Pip4k2a | 1.04E-12 | 0.059561516 | 0.338 | 0.337 | 2.81E-08 | Tmtc1 | 1.68E-08 | -0.0141191 | 0.469 | 0.452 | 0.000452782 |
| Lcor | 1.05E-12 | -0.023697523 | 0.559 | 0.597 | 2.81E-08 | Orc2 | 1.68E-08 | 0.017275173 | 0.327 | 0.302 | 0.000452908 |
| Bcl11b | 1.08E-12 | -0.090277876 | 0.536 | 0.584 | 2.91E-08 | Mtor | 1.70E-08 | -0.034821617 | 0.322 | 0.318 | 0.000456846 |
| Eipr1 | 1.08E-12 | 0.005760585 | 0.326 | 0.345 | 2.91E-08 | Kansl1 | 1.70E-08 | 0.061538776 | 0.842 | 0.816 | 0.000456973 |
| Vmp1 | 1.09E-12 | -0.060089117 | 0.441 | 0.486 | 2.92E-08 | Fgfr1 | 1.72E-08 | 0.005879554 | 0.262 | 0.242 | 0.000464018 |
| Ranbp10 | 1.12E-12 | -0.020549573 | 0.292 | 0.321 | 3.02E-08 | Sugp2 | 1.75E-08 | -0.016900272 | 0.321 | 0.311 | 0.000470009 |
| Arhgef18 | 1.14E-12 | 0.027238826 | 0.273 | 0.282 | 3.08E-08 | Map3k5 | 1.75E-08 | -0.061753223 | 0.31 | 0.315 | 0.000471061 |
| Aplp1 | 1.17E-12 | -0.043928157 | 0.407 | 0.451 | 3.15E-08 | Avl9 | 1.76E-08 | -0.002743368 | 0.26 | 0.246 | 0.00047427 |
| Slc25a36 | 1.18E-12 | 0.018194953 | 0.352 | 0.371 | 3.18E-08 | Mon2 | 1.77E-08 | 0.005257319 | 0.456 | 0.434 | 0.000476312 |
| Senp6 | 1.19E-12 | 0.005465909 | 0.566 | 0.595 | 3.21E-08 | Gm12353 | 1.78E-08 | 0.014826664 | 0.371 | 0.347 | 0.000479237 |
| Tet3 | 1.20E-12 | -0.064296875 | 0.424 | 0.466 | 3.22E-08 | Cntln | 1.80E-08 | -0.055083562 | 0.369 | 0.376 | 0.000483125 |
| Dcaf1 | 1.24E-12 | 0.011222437 | 0.32 | 0.337 | 3.33E-08 | Zfat | 1.82E-08 | 0.004926771 | 0.264 | 0.245 | 0.00048852 |
| Snx2 | 1.25E-12 | -0.035714318 | 0.231 | 0.263 | 3.37E-08 | Gpcpd1 | 1.82E-08 | 0.027779374 | 0.394 | 0.365 | 0.000490632 |
| Ergic2 | 1.28E-12 | -0.012243021 | 0.258 | 0.284 | 3.45E-08 | Adam11 | 1.83E-08 | -0.01421233 | 0.269 | 0.258 | 0.000492215 |
| Prkca | 1.32E-12 | 0.06528772 | 0.768 | 0.774 | 3.55E-08 | Sez6l | 1.84E-08 | -0.058288692 | 0.796 | 0.795 | 0.00049493 |
| Nexmif | 1.34E-12 | 0.046529719 | 0.484 | 0.495 | 3.60E-08 | Tln2 | 1.86E-08 | -0.020591034 | 0.532 | 0.517 | 0.000500204 |
| Cnot10 | 1.36E-12 | 0.01973312 | 0.361 | 0.378 | 3.65E-08 | Vps8 | 1.89E-08 | -0.002583332 | 0.364 | 0.345 | 0.000508349 |
| Kcnj6 | 1.38E-12 | -0.113498914 | 0.847 | 0.867 | 3.72E-08 | Smc5 | 1.89E-08 | 0.03653092 | 0.367 | 0.336 | 0.000508549 |
| Tcp11l1 | 1.39E-12 | 0.004686951 | 0.295 | 0.317 | 3.73E-08 | Gsk3b | 1.90E-08 | -0.033179719 | 0.631 | 0.623 | 0.000511607 |
| Fbxw8 | 1.43E-12 | 0.041509138 | 0.26 | 0.262 | 3.84E-08 | Dync2h1 | 1.91E-08 | -0.039355779 | 0.518 | 0.512 | 0.000512635 |
| Pdcd4 | 1.48E-12 | 0.066759588 | 0.438 | 0.438 | 3.97E-08 | 1700084C06Rik | 1.91E-08 | 0.026372554 | 0.342 | 0.314 | 0.00051336 |
| Nisch | 1.51E-12 | 0.053017865 | 0.504 | 0.511 | 4.06E-08 | Rilpl1 | 1.91E-08 | -0.026810616 | 0.338 | 0.328 | 0.000513748 |
| Zdhhc2 | 1.53E-12 | -0.087680454 | 0.337 | 0.382 | 4.12E-08 | Cacna2d3 | 1.92E-08 | -0.11349834 | 0.83 | 0.838 | 0.000515366 |
| Map4k3 | 1.54E-12 | 0.0235534 | 0.652 | 0.671 | 4.14E-08 | Ptpn4 | 1.95E-08 | -0.022308681 | 0.474 | 0.46 | 0.000525947 |
| Sorcs3 | 1.57E-12 | -0.24369175 | 0.475 | 0.465 | 4.23E-08 | Eda | 1.97E-08 | -0.097776089 | 0.264 | 0.28 | 0.000530074 |
| Hdgfl3 | 1.58E-12 | -0.011428133 | 0.443 | 0.475 | 4.26E-08 | Slf1 | 1.99E-08 | -0.018346815 | 0.363 | 0.35 | 0.000536574 |
| Nck2 | 1.63E-12 | -0.014240224 | 0.367 | 0.396 | 4.38E-08 | Kif3c | 2.00E-08 | 0.049065897 | 0.355 | 0.321 | 0.000537559 |
| Atp6v1a | 1.65E-12 | -0.051558955 | 0.338 | 0.378 | 4.44E-08 | Stk4 | 2.00E-08 | -0.031020595 | 0.277 | 0.274 | 0.00053798 |
| Atf2 | 1.66E-12 | 0.027120623 | 0.513 | 0.529 | 4.46E-08 | Mblac2 | 2.00E-08 | -0.049011661 | 0.354 | 0.353 | 0.000538038 |
| Gpr158 | 1.73E-12 | 0.05538403 | 0.864 | 0.858 | 4.64E-08 | Rbfox2 | 2.10E-08 | -0.053492764 | 0.85 | 0.846 | 0.000565146 |
| Atp6ap2 | 1.73E-12 | -0.010726574 | 0.233 | 0.259 | 4.65E-08 | Ptk2b | 2.11E-08 | -0.056330917 | 0.704 | 0.702 | 0.000567709 |
| Snx30 | 1.75E-12 | -0.032649393 | 0.328 | 0.363 | 4.71E-08 | Gapvd1 | 2.14E-08 | 0.001592965 | 0.55 | 0.527 | 0.000575012 |
| Gm3764 | 1.76E-12 | -0.025278983 | 0.435 | 0.473 | 4.73E-08 | Clip4 | 2.14E-08 | -0.019765491 | 0.283 | 0.273 | 0.000576847 |
| Ubxn11 | 1.77E-12 | 0.015714619 | 0.318 | 0.336 | 4.76E-08 | Tmcc1 | 2.16E-08 | 0.0128419 | 0.841 | 0.815 | 0.000580642 |
| Mettl16 | 1.79E-12 | -0.00593641 | 0.268 | 0.293 | 4.81E-08 | 4930455C13Rik | 2.16E-08 | 0.048965727 | 0.313 | 0.28 | 0.000582518 |
| Aatf | 1.81E-12 | 0.03104823 | 0.311 | 0.322 | 4.88E-08 | Farp1 | 2.18E-08 | -0.10241778 | 0.386 | 0.416 | 0.000587866 |
| Fam234b | 1.82E-12 | -0.043600128 | 0.221 | 0.256 | 4.89E-08 | Taf1b | 2.19E-08 | -0.006381743 | 0.387 | 0.37 | 0.000589093 |
| Gnaq | 1.86E-12 | 0.05876526 | 0.83 | 0.836 | 5.01E-08 | Plekhg5 | 2.22E-08 | 0.032058615 | 0.662 | 0.627 | 0.000596858 |
| Slco3a1 | 1.89E-12 | -0.054054744 | 0.51 | 0.553 | 5.10E-08 | Nub1 | 2.23E-08 | 0.033466237 | 0.334 | 0.304 | 0.000600271 |
| Hagh | 1.89E-12 | -0.036722491 | 0.229 | 0.261 | 5.10E-08 | Gng7 | 2.26E-08 | 0.05342578 | 0.283 | 0.251 | 0.000607138 |
| Gm16183 | 1.96E-12 | 0.09070422 | 0.39 | 0.366 | 5.27E-08 | Thrb | 2.26E-08 | -0.075691728 | 0.634 | 0.649 | 0.000608118 |
| Ptpn9 | 1.96E-12 | -0.038506836 | 0.357 | 0.395 | 5.28E-08 | Usp33 | 2.27E-08 | 0.03394268 | 0.448 | 0.416 | 0.000609899 |
| Tbl1xr1 | 1.96E-12 | -0.012771842 | 0.401 | 0.432 | 5.28E-08 | Rapgefl1 | 2.31E-08 | -0.011121018 | 0.374 | 0.359 | 0.000620356 |
| Uggt2 | 1.97E-12 | 0.069536362 | 0.508 | 0.503 | 5.29E-08 | Cramp1l | 2.32E-08 | 0.003111854 | 0.287 | 0.269 | 0.000623517 |
| Tmem131l | 1.97E-12 | 0.043218274 | 0.255 | 0.257 | 5.29E-08 | Slain1 | 2.34E-08 | 0.022377855 | 0.267 | 0.244 | 0.000629501 |
| Fat3 | 2.03E-12 | 0.096936752 | 0.773 | 0.766 | 5.46E-08 | Kcnip3 | 2.37E-08 | -0.058606912 | 0.531 | 0.532 | 0.000637663 |
| Lin52 | 2.03E-12 | 0.00785286 | 0.273 | 0.293 | 5.47E-08 | Gm11417 | 2.38E-08 | -0.054476163 | 0.276 | 0.278 | 0.0006402 |
| Acbd6 | 2.12E-12 | 0.017971309 | 0.483 | 0.504 | 5.70E-08 | Plppr2 | 2.38E-08 | -0.006956351 | 0.367 | 0.349 | 0.000641247 |
| Parp8 | 2.14E-12 | 0.066290021 | 0.425 | 0.426 | 5.77E-08 | Iqsec3 | 2.40E-08 | -0.015296441 | 0.4 | 0.386 | 0.000645855 |
| Tiam1 | 2.15E-12 | 0.126153094 | 0.652 | 0.645 | 5.79E-08 | Gng2 | 2.42E-08 | 0.032035834 | 0.531 | 0.497 | 0.000651988 |
| Elovl6 | 2.18E-12 | -0.049522772 | 0.335 | 0.373 | 5.87E-08 | Tiam2 | 2.46E-08 | 0.021561244 | 0.28 | 0.255 | 0.000660588 |
| Smg5 | 2.19E-12 | 0.020577222 | 0.247 | 0.258 | 5.90E-08 | Kcnab1 | 2.46E-08 | 0.032671744 | 0.778 | 0.745 | 0.000661588 |
| Sugp2 | 2.27E-12 | 0.014587076 | 0.311 | 0.329 | 6.11E-08 | Nsun2 | 2.50E-08 | -0.014726374 | 0.362 | 0.349 | 0.000671338 |
| Micu1 | 2.29E-12 | 0.024234941 | 0.427 | 0.447 | 6.15E-08 | Phf8 | 2.51E-08 | -0.021533525 | 0.26 | 0.252 | 0.000676468 |
| Fam222b | 2.30E-12 | -0.035503072 | 0.423 | 0.46 | 6.18E-08 | Lekr1 | 2.55E-08 | 0.040389205 | 0.28 | 0.253 | 0.000685666 |
| Gm37459 | 2.30E-12 | -0.126199114 | 0.394 | 0.44 | 6.19E-08 | Zcchc14 | 2.56E-08 | -0.021153749 | 0.34 | 0.328 | 0.000689759 |
| Trpc6 | 2.38E-12 | 0.100506562 | 0.374 | 0.377 | 6.40E-08 | Fbxw8 | 2.57E-08 | -0.043641794 | 0.259 | 0.26 | 0.000691077 |
| Usp4 | 2.38E-12 | 0.010973915 | 0.282 | 0.3 | 6.40E-08 | Lpp | 2.57E-08 | -0.014365573 | 0.375 | 0.36 | 0.000691175 |

| | | | | | | | | | | | |
|---|---|---|---|---|---|---|---|---|---|---|---|
| Zswim8 | 2.48E-12 | 0.027453744 | 0.336 | 0.348 | 6.68E-08 | Atg16l2 | 2.59E-08 | -0.050323107 | 0.251 | 0.258 | 0.00069668 |
| Sorbs1 | 2.50E-12 | 0.072704746 | 0.838 | 0.833 | 6.74E-08 | Usp12 | 2.63E-08 | 0.027030935 | 0.295 | 0.267 | 0.000706558 |
| Abcc5 | 2.53E-12 | -0.028724221 | 0.594 | 0.634 | 6.81E-08 | Shc3 | 2.65E-08 | 0.023799457 | 0.319 | 0.291 | 0.000711948 |
| Zfyve9 | 2.60E-12 | -0.019149785 | 0.607 | 0.642 | 6.99E-08 | Pias2 | 2.66E-08 | -0.025927532 | 0.535 | 0.524 | 0.000715137 |
| Ppm1a | 2.61E-12 | 0.023992724 | 0.265 | 0.277 | 7.02E-08 | Shisa9 | 2.67E-08 | -0.0878357 | 0.675 | 0.685 | 0.000717209 |
| Cers5 | 2.64E-12 | 0.032118339 | 0.288 | 0.297 | 7.10E-08 | Raph1 | 2.67E-08 | 0.044168792 | 0.474 | 0.44 | 0.000718157 |
| Dennd1b | 2.67E-12 | 0.00783632 | 0.5 | 0.525 | 7.18E-08 | Mtmr3 | 2.68E-08 | 0.005125023 | 0.553 | 0.528 | 0.000720273 |
| Setd4 | 2.67E-12 | -0.069198832 | 0.3 | 0.342 | 7.18E-08 | Gripap1 | 2.69E-08 | 0.017547535 | 0.394 | 0.367 | 0.000724927 |
| Nipbl | 2.71E-12 | -0.052557133 | 0.612 | 0.656 | 7.29E-08 | Nsf | 2.74E-08 | 0.065944459 | 0.894 | 0.88 | 0.000736314 |
| Astn1 | 2.78E-12 | 0.05209669 | 0.779 | 0.79 | 7.49E-08 | Hnrnpm | 2.80E-08 | 0.004558869 | 0.436 | 0.413 | 0.000752266 |
| Cdc40 | 2.80E-12 | -0.01090087 | 0.458 | 0.49 | 7.53E-08 | Zfp148 | 2.83E-08 | 0.000417254 | 0.642 | 0.616 | 0.000760699 |
| Rnf165 | 2.81E-12 | 0.009519406 | 0.488 | 0.51 | 7.55E-08 | Gns | 2.85E-08 | -0.005348949 | 0.267 | 0.253 | 0.000767779 |
| Lrrn3 | 2.85E-12 | -0.045041821 | 0.27 | 0.305 | 7.66E-08 | Wasf1 | 2.91E-08 | 0.003612547 | 0.699 | 0.673 | 0.000783448 |
| Phtf2 | 2.89E-12 | -0.00101172 | 0.292 | 0.314 | 7.78E-08 | Nipbl | 2.91E-08 | 0.030425557 | 0.645 | 0.612 | 0.000783705 |
| Snx24 | 2.92E-12 | -0.004447009 | 0.4 | 0.427 | 7.87E-08 | Mast2 | 2.96E-08 | -0.017435631 | 0.532 | 0.517 | 0.000796674 |
| Fam135a | 2.93E-12 | 0.040344648 | 0.376 | 0.383 | 7.87E-08 | Rufy2 | 3.02E-08 | -0.013550823 | 0.516 | 0.498 | 0.000812159 |
| Coro2b | 2.95E-12 | -0.014622538 | 0.404 | 0.436 | 7.94E-08 | Nrg3os | 3.05E-08 | 0.042725778 | 0.938 | 0.921 | 0.000820463 |
| Slf2 | 2.96E-12 | -0.00282523 | 0.488 | 0.519 | 7.95E-08 | Agtpbp1 | 3.10E-08 | -0.045961797 | 0.571 | 0.568 | 0.000833005 |
| D430042O09Rik | 2.98E-12 | 0.018885412 | 0.452 | 0.472 | 8.01E-08 | Lars | 3.10E-08 | 0.019017475 | 0.372 | 0.347 | 0.000835024 |
| Spats2 | 2.98E-12 | 0.023195065 | 0.26 | 0.271 | 8.03E-08 | Cdh11 | 3.12E-08 | 0.025677143 | 0.715 | 0.683 | 0.000839885 |
| Spock2 | 3.08E-12 | 0.043865971 | 0.397 | 0.401 | 8.29E-08 | Cnot6l | 3.12E-08 | -0.004487874 | 0.38 | 0.362 | 0.000840676 |
| Cers4 | 3.13E-12 | -0.003826518 | 0.379 | 0.408 | 8.43E-08 | Clip1 | 3.16E-08 | -0.02343009 | 0.573 | 0.559 | 0.000849162 |
| Ppp1r16b | 3.15E-12 | -0.051450905 | 0.35 | 0.39 | 8.48E-08 | Mgat5 | 3.20E-08 | -0.057815899 | 0.495 | 0.505 | 0.000860835 |
| Dicer1 | 3.23E-12 | -0.011788388 | 0.331 | 0.359 | 8.69E-08 | Unc13a | 3.25E-08 | 0.018217667 | 0.595 | 0.565 | 0.000874391 |
| Zdhhc21 | 3.25E-12 | -0.008381111 | 0.305 | 0.331 | 8.75E-08 | Prkacb | 3.34E-08 | 0.048851198 | 0.332 | 0.3 | 0.000897663 |
| Fam171b | 3.26E-12 | -0.002956534 | 0.497 | 0.526 | 8.78E-08 | Bdp1 | 3.41E-08 | -0.004391826 | 0.405 | 0.387 | 0.000917017 |
| Mitf | 3.29E-12 | -0.088579726 | 0.28 | 0.323 | 8.85E-08 | Syndig1 | 3.43E-08 | -0.061190532 | 0.346 | 0.352 | 0.000923866 |
| Trmt1l | 3.38E-12 | -0.035492758 | 0.256 | 0.289 | 9.09E-08 | Mphosph8 | 3.44E-08 | 0.034971201 | 0.294 | 0.266 | 0.000925023 |
| Zfc3h1 | 3.50E-12 | -0.016096379 | 0.392 | 0.425 | 9.41E-08 | Scaf8 | 3.44E-08 | -0.029155245 | 0.558 | 0.547 | 0.000925848 |
| Fbxo10 | 3.52E-12 | -0.073651375 | 0.283 | 0.325 | 9.47E-08 | Tcte2 | 3.48E-08 | -0.072030901 | 0.368 | 0.386 | 0.000936227 |
| Raver2 | 3.54E-12 | 0.017947142 | 0.364 | 0.385 | 9.54E-08 | Pygb | 3.50E-08 | -0.036281454 | 0.324 | 0.32 | 0.000941728 |
| Fut8 | 3.58E-12 | -0.048745776 | 0.664 | 0.707 | 9.63E-08 | Agk | 3.56E-08 | -0.016524511 | 0.295 | 0.284 | 0.000957462 |
| Snph | 3.59E-12 | -0.028243587 | 0.611 | 0.649 | 9.65E-08 | Gm38604 | 3.66E-08 | -0.005596193 | 0.278 | 0.261 | 0.000984945 |
| Rb1 | 3.66E-12 | -0.02137059 | 0.476 | 0.512 | 9.86E-08 | Trak1 | 3.67E-08 | 0.009611266 | 0.451 | 0.426 | 0.000986506 |
| Angptl2 | 3.71E-12 | -0.07613684 | 0.29 | 0.333 | 9.99E-08 | Opa1 | 3.69E-08 | 0.009936009 | 0.364 | 0.342 | 0.000993333 |
| Gm26917 | 3.94E-12 | -0.001352025 | 0.443 | 0.476 | 1.06E-07 | Cep83 | 3.70E-08 | -0.01735027 | 0.372 | 0.361 | 0.000996043 |
| Rufy2 | 3.96E-12 | -0.01302437 | 0.498 | 0.531 | 1.06E-07 | Rubcn | 3.79E-08 | 0.011458775 | 0.252 | 0.232 | 0.001020356 |
| Lrrc8d | 4.04E-12 | 0.059958877 | 0.326 | 0.323 | 1.09E-07 | Numa1 | 3.79E-08 | 0.014424306 | 0.371 | 0.346 | 0.001020557 |
| Fryl | 4.04E-12 | 0.03184633 | 0.499 | 0.512 | 1.09E-07 | Clip2 | 3.79E-08 | -0.013165005 | 0.28 | 0.268 | 0.001020862 |
| Hnrnph3 | 4.11E-12 | 0.035638407 | 0.38 | 0.389 | 1.11E-07 | Rnf169 | 3.81E-08 | -0.005193833 | 0.643 | 0.619 | 0.001024952 |
| Ttll7 | 4.17E-12 | 0.044586046 | 0.416 | 0.42 | 1.12E-07 | Secisbp2l | 3.81E-08 | 0.033106418 | 0.26 | 0.235 | 0.001025225 |
| Nup93 | 4.23E-12 | -0.079509343 | 0.441 | 0.488 | 1.14E-07 | Erp44 | 3.83E-08 | 0.006658082 | 0.284 | 0.265 | 0.001029504 |
| Stk3 | 4.41E-12 | 0.006985523 | 0.45 | 0.474 | 1.19E-07 | Lmtk2 | 3.91E-08 | 0.05250129 | 0.541 | 0.506 | 0.001051569 |
| Basp1 | 4.42E-12 | -0.056700495 | 0.548 | 0.593 | 1.19E-07 | Herc2 | 3.96E-08 | -0.028372031 | 0.643 | 0.634 | 0.001065408 |
| Pnpt1 | 4.44E-12 | 0.001034995 | 0.289 | 0.31 | 1.20E-07 | Pdcd4 | 3.97E-08 | -0.00982081 | 0.458 | 0.438 | 0.001068187 |
| Gm48239 | 4.55E-12 | 0.075032738 | 0.284 | 0.271 | 1.23E-07 | Rnf38 | 4.00E-08 | 0.015145539 | 0.286 | 0.262 | 0.001075136 |
| Gfod1 | 4.84E-12 | 0.077379212 | 0.647 | 0.646 | 1.30E-07 | Satb1 | 4.09E-08 | 0.063466882 | 0.273 | 0.241 | 0.001101469 |
| 7-Sep | 4.90E-12 | -0.067761826 | 0.328 | 0.37 | 1.32E-07 | Rsf1 | 4.11E-08 | -0.019377587 | 0.525 | 0.512 | 0.001106179 |
| Adam10 | 4.91E-12 | -0.024811906 | 0.317 | 0.348 | 1.32E-07 | Ncor1 | 4.11E-08 | 0.050806154 | 0.768 | 0.735 | 0.001106318 |
| Sfmbt1 | 5.00E-12 | 0.013131128 | 0.338 | 0.358 | 1.35E-07 | Ehbp1 | 4.12E-08 | -0.043001054 | 0.68 | 0.675 | 0.001109325 |
| Anapc4 | 5.01E-12 | -0.031447967 | 0.251 | 0.284 | 1.35E-07 | Zranb2 | 4.31E-08 | -0.009906853 | 0.736 | 0.713 | 0.001161025 |
| Dab2ip | 5.03E-12 | 0.028423578 | 0.273 | 0.284 | 1.35E-07 | Cacul1 | 4.39E-08 | 0.003874225 | 0.268 | 0.25 | 0.00118162 |
| Rtn4 | 5.25E-12 | -0.012945607 | 0.561 | 0.598 | 1.41E-07 | Nuak1 | 4.41E-08 | -0.040536373 | 0.25 | 0.249 | 0.001187483 |
| Tnrc18 | 5.30E-12 | 0.052407382 | 0.379 | 0.378 | 1.43E-07 | Sgsm2 | 4.51E-08 | 0.021975637 | 0.45 | 0.421 | 0.001214681 |
| Crebbp | 5.42E-12 | -0.03800936 | 0.688 | 0.728 | 1.46E-07 | Myo6 | 4.55E-08 | 0.008053446 | 0.323 | 0.302 | 0.001225467 |
| Gria3 | 5.45E-12 | 0.076756492 | 0.921 | 0.92 | 1.47E-07 | Ust | 4.56E-08 | -0.03796551 | 0.555 | 0.546 | 0.00122691 |
| Dag1 | 5.75E-12 | 0.014803154 | 0.314 | 0.332 | 1.55E-07 | Rnf115 | 4.56E-08 | 0.025503599 | 0.33 | 0.303 | 0.001227552 |
| Mdm4 | 5.83E-12 | 0.00878599 | 0.439 | 0.462 | 1.57E-07 | Elavl1 | 4.60E-08 | 0.003431776 | 0.311 | 0.293 | 0.001237577 |
| Mark3 | 5.85E-12 | 0.039499579 | 0.727 | 0.742 | 1.57E-07 | Abcc8 | 4.65E-08 | -0.002329316 | 0.251 | 0.235 | 0.001250538 |
| Tsnax | 5.85E-12 | -0.013688348 | 0.254 | 0.28 | 1.57E-07 | Fnbp4 | 4.66E-08 | -0.015286263 | 0.377 | 0.364 | 0.001253819 |
| Atxn2 | 5.88E-12 | 0.041749234 | 0.568 | 0.577 | 1.58E-07 | Zdhhc21 | 4.70E-08 | 0.034942042 | 0.335 | 0.305 | 0.001263443 |
| Capzb | 5.91E-12 | 0.030114469 | 0.293 | 0.3 | 1.59E-07 | Klhl29 | 4.73E-08 | -0.077984946 | 0.788 | 0.794 | 0.001272431 |
| Zer1 | 6.03E-12 | 0.01905121 | 0.367 | 0.381 | 1.62E-07 | Magi3 | 4.73E-08 | -0.06514116 | 0.433 | 0.442 | 0.001273587 |
| Ankib1 | 6.19E-12 | 0.037551736 | 0.455 | 0.465 | 1.66E-07 | Gm44686 | 4.74E-08 | 0.055151309 | 0.374 | 0.34 | 0.001274332 |
| 4930532I03Rik | 6.42E-12 | -0.031289264 | 0.399 | 0.437 | 1.73E-07 | Map2k1 | 4.82E-08 | 0.023825788 | 0.38 | 0.353 | 0.001296064 |
| Epg5 | 6.43E-12 | 0.015670019 | 0.255 | 0.269 | 1.73E-07 | Fam69a | 4.85E-08 | -0.085676545 | 0.343 | 0.373 | 0.001305508 |
| Nae1 | 6.45E-12 | -0.022491583 | 0.275 | 0.304 | 1.73E-07 | Ccdc6 | 4.91E-08 | -0.011117136 | 0.288 | 0.278 | 0.001321329 |
| Chd9 | 6.48E-12 | -0.075038518 | 0.801 | 0.836 | 1.74E-07 | Herc4 | 5.06E-08 | 0.023146333 | 0.385 | 0.36 | 0.001361484 |
| C78859 | 6.65E-12 | -0.030058536 | 0.325 | 0.358 | 1.79E-07 | Ctnnd2 | 5.13E-08 | -0.076126111 | 0.963 | 0.967 | 0.001381653 |
| Kif21a | 6.88E-12 | -0.061269531 | 0.764 | 0.802 | 1.85E-07 | Ncam1 | 5.19E-08 | -0.059095153 | 0.889 | 0.884 | 0.001395136 |
| Zscan26 | 6.94E-12 | 0.01601157 | 0.316 | 0.334 | 1.87E-07 | Chm | 5.23E-08 | 0.018198334 | 0.428 | 0.4 | 0.001407323 |
| Pced1b | 7.09E-12 | 0.063655369 | 0.307 | 0.298 | 1.91E-07 | 5330434G04Rik | 5.23E-08 | -0.013340878 | 0.669 | 0.651 | 0.001408291 |
| Tmem167 | 7.12E-12 | 0.025494694 | 0.301 | 0.314 | 1.92E-07 | Nf1 | 5.27E-08 | -0.021540154 | 0.657 | 0.64 | 0.00141909 |
| Rictor | 7.17E-12 | 0.032750114 | 0.406 | 0.416 | 1.93E-07 | Chd5 | 5.36E-08 | -0.068516791 | 0.386 | 0.397 | 0.001441453 |
| Tmem29 | 7.29E-12 | -0.026031293 | 0.222 | 0.25 | 1.96E-07 | Rnf25 | 5.36E-08 | -0.052605405 | 0.27 | 0.276 | 0.001443361 |
| Slc8a2 | 7.30E-12 | 0.026070878 | 0.392 | 0.407 | 1.96E-07 | Gm16105 | 5.46E-08 | -0.042440197 | 0.297 | 0.297 | 0.001470018 |
| Tcaf1 | 7.33E-12 | -0.021748568 | 0.52 | 0.554 | 1.97E-07 | Otulin | 5.47E-08 | 0.051281039 | 0.456 | 0.42 | 0.001470535 |
| Rbm33 | 7.34E-12 | 0.016607543 | 0.436 | 0.456 | 1.97E-07 | Pus10 | 5.53E-08 | 0.014692253 | 0.3 | 0.279 | 0.001488966 |
| Mir9-3hg | 7.35E-12 | -0.023576181 | 0.43 | 0.464 | 1.98E-07 | Cnnm2 | 5.54E-08 | 0.031978163 | 0.424 | 0.395 | 0.001489746 |
| Ppp1r2 | 7.37E-12 | -0.049180429 | 0.224 | 0.258 | 1.98E-07 | Tom1l2 | 5.54E-08 | 0.004860749 | 0.674 | 0.649 | 0.001491724 |
| Dmtf1 | 7.45E-12 | -0.02866918 | 0.268 | 0.299 | 2.01E-07 | Arhgef2 | 5.68E-08 | 0.016133488 | 0.395 | 0.369 | 0.001528727 |
| Il1rap | 7.52E-12 | -0.16082886 | 0.595 | 0.627 | 2.02E-07 | Prdm2 | 5.69E-08 | -0.014824003 | 0.343 | 0.331 | 0.001531932 |
| Tecr | 7.56E-12 | -0.000676033 | 0.381 | 0.409 | 2.03E-07 | Prrc2b | 5.72E-08 | -0.011636243 | 0.532 | 0.516 | 0.001537812 |
| Jak1 | 7.91E-12 | -0.035579774 | 0.374 | 0.409 | 2.13E-07 | 4930590L20Rik | 5.84E-08 | 0.0433262 | 0.277 | 0.248 | 0.001571133 |
| Ift88 | 8.06E-12 | 0.018665051 | 0.3 | 0.315 | 2.17E-07 | Mrps9 | 5.84E-08 | -0.018223894 | 0.254 | 0.244 | 0.001571659 |
| Zc3h6 | 8.15E-12 | 0.013199492 | 0.306 | 0.322 | 2.19E-07 | Brdt | 5.88E-08 | 0.024520216 | 0.283 | 0.258 | 0.001583168 |
| Smurf2 | 8.19E-12 | 0.02258265 | 0.374 | 0.389 | 2.20E-07 | A230057D06Rik | 5.92E-08 | -0.046753428 | 0.757 | 0.755 | 0.001593049 |
| Mpp6 | 8.46E-12 | -0.002582443 | 0.435 | 0.461 | 2.28E-07 | Sidt1 | 5.94E-08 | -0.026670686 | 0.669 | 0.654 | 0.001598097 |
| Tbc1d9b | 8.59E-12 | -0.005114562 | 0.249 | 0.271 | 2.31E-07 | Fut8 | 6.06E-08 | 0.059579409 | 0.699 | 0.664 | 0.001631871 |
| Pygb | 8.83E-12 | 0.029142213 | 0.32 | 0.329 | 2.38E-07 | Ypel2 | 6.11E-08 | 0.007283571 | 0.277 | 0.256 | 0.001645329 |
| Tenm1 | 8.85E-12 | -0.135742602 | 0.683 | 0.719 | 2.38E-07 | Map2 | 6.13E-08 | -0.051517428 | 0.919 | 0.915 | 0.001649584 |
| Sv2a | 9.07E-12 | 0.037324783 | 0.298 | 0.305 | 2.44E-07 | Ift88 | 6.17E-08 | -0.005546161 | 0.317 | 0.3 | 0.001658975 |
| Arid4b | 9.16E-12 | 0.026512245 | 0.61 | 0.628 | 2.46E-07 | Ppp2r5a | 6.20E-08 | 0.026463236 | 0.264 | 0.24 | 0.001667149 |
| Cenpv | 9.16E-12 | 0.062263065 | 0.314 | 0.309 | 2.47E-07 | Usp46 | 6.20E-08 | 0.063140789 | 0.409 | 0.374 | 0.001667779 |
| Zfp407 | 9.18E-12 | 0.001831424 | 0.491 | 0.518 | 2.47E-07 | Tmem181a | 6.22E-08 | -0.029268677 | 0.528 | 0.524 | 0.001673175 |
| Dpysl2 | 9.37E-12 | 0.062115631 | 0.499 | 0.498 | 2.52E-07 | Sez6 | 6.22E-08 | -0.06227884 | 0.268 | 0.278 | 0.001673538 |
| Dnajc6 | 9.48E-12 | 0.035698319 | 0.722 | 0.735 | 2.55E-07 | Agap2 | 6.40E-08 | -0.008624245 | 0.266 | 0.254 | 0.001721284 |
| Srpk2 | 9.49E-12 | 0.030013045 | 0.717 | 0.735 | 2.55E-07 | Gm42303 | 6.54E-08 | 0.068843919 | 0.368 | 0.333 | 0.001759779 |

| | | | | | | | | | | | |
|---|---|---|---|---|---|---|---|---|---|---|---|
| Taf2 | 9.92E-12 | -0.001984363 | 0.229 | 0.25 | 2.67E-07 | 1110051M20Rik | 6.59E-08 | 0.017042001 | 0.733 | 0.706 | 0.001774339 |
| Hook1 | 1.01E-11 | 0.035709174 | 0.466 | 0.479 | 2.71E-07 | Tbc1d9b | 6.88E-08 | 0.040768615 | 0.278 | 0.249 | 0.001852279 |
| Zfp277 | 1.02E-11 | 0.015718238 | 0.29 | 0.305 | 2.74E-07 | Kcnb1 | 6.92E-08 | -0.020297467 | 0.56 | 0.543 | 0.001862069 |
| Gm40841 | 1.07E-11 | 0.075286382 | 0.285 | 0.281 | 2.89E-07 | Sf3b3 | 6.93E-08 | 0.037182905 | 0.349 | 0.319 | 0.001865163 |
| Fgfr1op2 | 1.09E-11 | -0.034156156 | 0.248 | 0.281 | 2.94E-07 | Sycp2 | 6.97E-08 | 0.053444438 | 0.269 | 0.238 | 0.001874754 |
| Nek9 | 1.10E-11 | 0.025322154 | 0.265 | 0.275 | 2.95E-07 | Klhdc10 | 6.98E-08 | 0.019045699 | 0.724 | 0.695 | 0.001877999 |
| Asxl1 | 1.17E-11 | 0.033385894 | 0.463 | 0.476 | 3.15E-07 | Fam149b | 7.00E-08 | 0.01022619 | 0.28 | 0.261 | 0.001883627 |
| Chka | 1.17E-11 | -0.067505951 | 0.603 | 0.647 | 3.15E-07 | Gm20687 | 7.07E-08 | 0.076649568 | 0.389 | 0.371 | 0.001901565 |
| Tulp3 | 1.23E-11 | 0.022063561 | 0.25 | 0.262 | 3.30E-07 | Macrod2 | 7.07E-08 | -0.060027598 | 0.974 | 0.976 | 0.001902128 |
| Lgi1 | 1.25E-11 | -0.118193889 | 0.602 | 0.644 | 3.37E-07 | Efr3a | 7.12E-08 | 0.028734886 | 0.49 | 0.46 | 0.001915688 |
| Gigyf2 | 1.26E-11 | 0.024664514 | 0.533 | 0.553 | 3.38E-07 | Gtf2f2 | 7.16E-08 | -0.003343463 | 0.303 | 0.288 | 0.001926856 |
| Scg5 | 1.30E-11 | 0.035397532 | 0.51 | 0.524 | 3.51E-07 | Med13 | 7.19E-08 | 0.024763123 | 0.49 | 0.459 | 0.001935309 |
| Cacna1e | 1.31E-11 | -0.08201797 | 0.952 | 0.96 | 3.52E-07 | Wdr60 | 7.27E-08 | -0.012583783 | 0.369 | 0.356 | 0.001956685 |
| Pspc1 | 1.35E-11 | -0.018366645 | 0.423 | 0.455 | 3.63E-07 | Psd3 | 7.27E-08 | -0.005942305 | 0.697 | 0.673 | 0.001957234 |
| Phka2 | 1.36E-11 | 0.048184681 | 0.314 | 0.315 | 3.65E-07 | Phip | 7.32E-08 | 0.00472761 | 0.553 | 0.528 | 0.001969966 |
| Zfr | 1.38E-11 | -0.013949442 | 0.73 | 0.761 | 3.71E-07 | Stxbp4 | 7.32E-08 | 0.004727475 | 0.259 | 0.242 | 0.00197074 |
| Trappc10 | 1.42E-11 | -0.009365913 | 0.246 | 0.271 | 3.81E-07 | Pvt1 | 7.34E-08 | 0.026117629 | 0.534 | 0.505 | 0.001974696 |
| 1700024B18Rik | 1.45E-11 | -0.066838136 | 0.367 | 0.409 | 3.89E-07 | Fam168a | 7.35E-08 | -0.009168338 | 0.748 | 0.727 | 0.001978614 |
| Uba5 | 1.46E-11 | -0.057217471 | 0.234 | 0.271 | 3.94E-07 | Adap1 | 7.44E-08 | 0.001739419 | 0.282 | 0.265 | 0.002001499 |
| Gm11417 | 1.47E-11 | 0.066773444 | 0.278 | 0.272 | 3.96E-07 | Vav3 | 7.44E-08 | 0.062564639 | 0.339 | 0.307 | 0.002002211 |
| Gm28905 | 1.49E-11 | 0.127634357 | 0.28 | 0.24 | 4.00E-07 | Ccdc50 | 7.62E-08 | 0.009936714 | 0.293 | 0.273 | 0.002049331 |
| Ccnd2 | 1.52E-11 | 0.055816283 | 0.25 | 0.245 | 4.10E-07 | Rab8b | 7.87E-08 | 0.048462562 | 0.325 | 0.292 | 0.002117227 |
| Sort1 | 1.54E-11 | 0.045382249 | 0.615 | 0.626 | 4.14E-07 | Nwd2 | 7.93E-08 | 0.01919555 | 0.462 | 0.435 | 0.002132613 |
| Rnf130 | 1.55E-11 | 0.052534825 | 0.361 | 0.364 | 4.16E-07 | Gria2 | 8.03E-08 | 0.050637182 | 0.992 | 0.992 | 0.002161045 |
| Strn | 1.56E-11 | -0.022719596 | 0.301 | 0.331 | 4.21E-07 | Tsc1 | 8.11E-08 | 0.004849702 | 0.421 | 0.4 | 0.002182755 |
| Ap3s1 | 1.57E-11 | 0.008619847 | 0.249 | 0.267 | 4.22E-07 | Lrrn3 | 8.13E-08 | 0.038509754 | 0.299 | 0.27 | 0.002187551 |
| Zfp329 | 1.57E-11 | -0.016508205 | 0.263 | 0.29 | 4.22E-07 | Mras | 8.21E-08 | 0.013657113 | 0.347 | 0.324 | 0.002208602 |
| Nrgn | 1.58E-11 | -0.025915121 | 0.431 | 0.468 | 4.26E-07 | Rps6ka2 | 8.22E-08 | -0.060065579 | 0.36 | 0.368 | 0.002211626 |
| Zmym4 | 1.62E-11 | 0.048622017 | 0.656 | 0.669 | 4.36E-07 | Rsrc1 | 8.38E-08 | -0.004296719 | 0.532 | 0.51 | 0.002255847 |
| Zfp292 | 1.68E-11 | 0.005867991 | 0.497 | 0.521 | 4.52E-07 | Arhgef25 | 8.47E-08 | 0.005862756 | 0.469 | 0.446 | 0.002279041 |
| Gnl3l | 1.74E-11 | -0.023355515 | 0.253 | 0.282 | 4.69E-07 | Arhgef18 | 8.48E-08 | -0.021862096 | 0.281 | 0.273 | 0.002281664 |
| Cbx5 | 1.74E-11 | -0.050242717 | 0.243 | 0.279 | 4.69E-07 | Klc1 | 8.69E-08 | -0.012665628 | 0.463 | 0.446 | 0.002338824 |
| Sptbn2 | 1.76E-11 | -0.053552192 | 0.486 | 0.528 | 4.73E-07 | Tbc1d1 | 8.71E-08 | -0.04734545 | 0.433 | 0.428 | 0.002343204 |
| Traf3 | 1.78E-11 | 0.059389533 | 0.447 | 0.445 | 4.78E-07 | Ripor2 | 8.75E-08 | -0.023569683 | 0.552 | 0.54 | 0.002354565 |
| Kpna1 | 1.82E-11 | -0.035533222 | 0.31 | 0.344 | 4.89E-07 | Setd4 | 8.86E-08 | 0.029353148 | 0.328 | 0.3 | 0.002385206 |
| Atad2b | 1.82E-11 | 0.012783228 | 0.51 | 0.533 | 4.90E-07 | 4930517O19Rik | 8.88E-08 | 0.040918127 | 0.333 | 0.302 | 0.002388267 |
| Slc1a2 | 1.85E-11 | 0.07265546 | 0.545 | 0.524 | 4.97E-07 | Tulp4 | 8.99E-08 | 0.009568759 | 0.554 | 0.528 | 0.002419974 |
| Lmtk2 | 1.86E-11 | 0.035137177 | 0.506 | 0.514 | 5.00E-07 | Lmo3 | 9.12E-08 | 0.042594549 | 0.448 | 0.415 | 0.002453589 |
| Smc6 | 1.87E-11 | -0.001959319 | 0.267 | 0.29 | 5.03E-07 | Mmd2 | 9.12E-08 | 0.000455735 | 0.321 | 0.303 | 0.002453675 |
| Rab12 | 1.89E-11 | 0.000516274 | 0.246 | 0.265 | 5.08E-07 | Cenpv | 9.18E-08 | -0.016625927 | 0.326 | 0.314 | 0.002469245 |
| Fam120c | 1.90E-11 | 0.046085414 | 0.442 | 0.449 | 5.12E-07 | 2610507B11Rik | 9.26E-08 | -0.016527835 | 0.304 | 0.294 | 0.002490529 |
| Limk2 | 1.91E-11 | 0.03412443 | 0.27 | 0.278 | 5.14E-07 | Ubxn11 | 9.37E-08 | 0.023312023 | 0.344 | 0.318 | 0.002522265 |
| Dip2b | 1.92E-11 | -0.046276477 | 0.659 | 0.7 | 5.16E-07 | Map3k4 | 9.57E-08 | -0.03685964 | 0.358 | 0.352 | 0.002575307 |
| Mmaa | 1.94E-11 | -0.005720411 | 0.257 | 0.279 | 5.23E-07 | Tbck | 9.70E-08 | -0.035850559 | 0.393 | 0.393 | 0.002609147 |
| Fam120a | 2.01E-11 | 0.00173924 | 0.418 | 0.443 | 5.41E-07 | Ercc6 | 9.73E-08 | 0.015148487 | 0.265 | 0.245 | 0.002619216 |
| Dnajc1 | 2.02E-11 | -0.093158722 | 0.775 | 0.806 | 5.42E-07 | Iqcb1 | 9.98E-08 | 0.040550085 | 0.297 | 0.269 | 0.002685719 |
| Zc2hc1a | 2.08E-11 | 0.0213174 | 0.259 | 0.27 | 5.60E-07 | Pcdh19 | 9.99E-08 | 0.03509256 | 0.327 | 0.301 | 0.002689287 |
| 1110051M20Rik | 2.10E-11 | 0.03981506 | 0.706 | 0.718 | 5.65E-07 | Tmem63c | 1.01E-07 | -0.039027596 | 0.348 | 0.348 | 0.002706953 |
| Zranb2 | 2.16E-11 | -0.01813717 | 0.713 | 0.75 | 5.82E-07 | Slc25a23 | 1.03E-07 | -0.012226631 | 0.482 | 0.463 | 0.002766234 |
| Dzank1 | 2.18E-11 | 0.04303039 | 0.35 | 0.356 | 5.86E-07 | Ranbp9 | 1.03E-07 | 0.040746854 | 0.605 | 0.572 | 0.002782524 |
| Pcdh15 | 2.21E-11 | -0.209279743 | 0.447 | 0.42 | 5.94E-07 | Spon1 | 1.04E-07 | 0.030384626 | 0.493 | 0.46 | 0.002788505 |
| Arl3 | 2.25E-11 | 0.017279266 | 0.368 | 0.383 | 6.06E-07 | Dnm3 | 1.04E-07 | -0.035623515 | 0.547 | 0.538 | 0.002798813 |
| Ube2r2 | 2.34E-11 | 0.003545388 | 0.366 | 0.388 | 6.30E-07 | Arsb | 1.04E-07 | 0.006988295 | 0.599 | 0.573 | 0.002799338 |
| Ccdc149 | 2.36E-11 | 0.026034842 | 0.273 | 0.283 | 6.34E-07 | Flrt2 | 1.04E-07 | 0.055742651 | 0.426 | 0.392 | 0.002800284 |
| N4bp2l1 | 2.41E-11 | 0.010388216 | 0.312 | 0.329 | 6.47E-07 | Fsd1l | 1.04E-07 | -0.026379189 | 0.26 | 0.256 | 0.002804444 |
| Rgs17 | 2.41E-11 | -0.051299951 | 0.377 | 0.415 | 6.48E-07 | Pcdh15 | 1.05E-07 | 0.08892126 | 0.422 | 0.447 | 0.00281907 |
| Plppr4 | 2.42E-11 | -0.053712286 | 0.579 | 0.621 | 6.52E-07 | Sybu | 1.05E-07 | 0.058851939 | 0.826 | 0.799 | 0.002822829 |
| Rbfox3 | 2.44E-11 | 0.091394998 | 0.906 | 0.894 | 6.56E-07 | Sgip1 | 1.05E-07 | -0.058090698 | 0.894 | 0.889 | 0.002838277 |
| Stx3 | 2.46E-11 | -0.043201162 | 0.287 | 0.321 | 6.61E-07 | 1500035N22Rik | 1.06E-07 | -0.016814101 | 0.303 | 0.291 | 0.002850433 |
| Lekr1 | 2.51E-11 | 0.018251746 | 0.253 | 0.266 | 6.75E-07 | Cttnbp2 | 1.06E-07 | 0.071790949 | 0.957 | 0.954 | 0.002854812 |
| Nptn | 2.53E-11 | -0.059142104 | 0.874 | 0.899 | 6.82E-07 | Cul4a | 1.07E-07 | 0.01079542 | 0.27 | 0.25 | 0.00288287 |
| Rock1 | 2.61E-11 | -0.011227991 | 0.655 | 0.688 | 7.01E-07 | Iqgap1 | 1.08E-07 | -0.006621624 | 0.325 | 0.309 | 0.002893422 |
| Tnik | 2.63E-11 | 0.067187498 | 0.938 | 0.93 | 7.07E-07 | Ccnd2 | 1.09E-07 | -0.00698859 | 0.263 | 0.25 | 0.002944037 |
| Pxdn | 2.65E-11 | 0.048810866 | 0.368 | 0.371 | 7.14E-07 | Sugp1 | 1.10E-07 | -0.003768548 | 0.327 | 0.312 | 0.002949844 |
| Phkb | 2.75E-11 | 0.060029714 | 0.297 | 0.291 | 7.40E-07 | Asic2 | 1.12E-07 | -0.089380925 | 0.9 | 0.907 | 0.003021058 |
| Klhl32 | 2.82E-11 | 0.03976062 | 0.458 | 0.469 | 7.60E-07 | Ddi2 | 1.14E-07 | -0.009501846 | 0.295 | 0.282 | 0.003070953 |
| Rnf138 | 2.83E-11 | -0.023558246 | 0.225 | 0.252 | 7.61E-07 | Mir9-3hg | 1.15E-07 | 0.006271218 | 0.452 | 0.43 | 0.003097891 |
| Tbl1x | 2.96E-11 | 0.089742106 | 0.286 | 0.26 | 7.97E-07 | Ergic2 | 1.17E-07 | 0.001885597 | 0.276 | 0.258 | 0.003156426 |
| Scn3b | 2.99E-11 | -0.065004855 | 0.391 | 0.434 | 8.05E-07 | Col4a3bp | 1.19E-07 | 0.013630555 | 0.354 | 0.333 | 0.003200771 |
| Dclk2 | 3.14E-11 | 0.034808443 | 0.459 | 0.472 | 8.44E-07 | Elmo2 | 1.23E-07 | 0.032252927 | 0.276 | 0.25 | 0.003313268 |
| Msantd2 | 3.21E-11 | 0.004789928 | 0.234 | 0.252 | 8.65E-07 | Wdpcp | 1.24E-07 | -0.002817655 | 0.291 | 0.273 | 0.003324617 |
| Ablim1 | 3.23E-11 | 0.091578054 | 0.6 | 0.59 | 8.70E-07 | Cep85l | 1.26E-07 | -0.001930094 | 0.317 | 0.298 | 0.003380288 |
| Gpi1 | 3.34E-11 | -0.019622998 | 0.356 | 0.385 | 8.98E-07 | Pcbp3 | 1.26E-07 | -0.044168359 | 0.372 | 0.374 | 0.003395095 |
| Fam78b | 3.35E-11 | 0.037813884 | 0.261 | 0.266 | 9.00E-07 | D5Ertd579e | 1.28E-07 | 0.030467498 | 0.45 | 0.419 | 0.00343481 |
| Bcl9 | 3.42E-11 | 0.049654988 | 0.337 | 0.337 | 9.21E-07 | Nol4 | 1.28E-07 | -0.033969044 | 0.718 | 0.706 | 0.003441952 |
| Pnn | 3.44E-11 | 0.008683184 | 0.555 | 0.583 | 9.25E-07 | Galc | 1.29E-07 | 0.031313725 | 0.291 | 0.263 | 0.003464749 |
| Slc2a3 | 3.45E-11 | -0.009781227 | 0.377 | 0.405 | 9.28E-07 | Ints6 | 1.30E-07 | 0.018435258 | 0.424 | 0.398 | 0.003498263 |
| Xylt1 | 3.45E-11 | 0.089886465 | 0.593 | 0.59 | 9.28E-07 | Fam78b | 1.30E-07 | -0.032356599 | 0.263 | 0.261 | 0.003508652 |
| Sbf1 | 3.46E-11 | -0.050517164 | 0.227 | 0.261 | 9.32E-07 | Shank3 | 1.31E-07 | -0.043134102 | 0.379 | 0.381 | 0.003525089 |
| Adcy9 | 3.52E-11 | 0.055015484 | 0.791 | 0.796 | 9.48E-07 | Cacng8 | 1.33E-07 | 0.056267706 | 0.521 | 0.486 | 0.003565726 |
| Actr2 | 3.65E-11 | -0.038265633 | 0.418 | 0.455 | 9.81E-07 | Desi2 | 1.33E-07 | 0.037597849 | 0.47 | 0.438 | 0.003568505 |
| Ablim3 | 3.65E-11 | 0.066906182 | 0.276 | 0.274 | 9.83E-07 | Asap2 | 1.34E-07 | -0.052763212 | 0.405 | 0.401 | 0.003596231 |
| Cpeb4 | 3.66E-11 | 0.033666836 | 0.546 | 0.563 | 9.86E-07 | Edrf1 | 1.38E-07 | 0.027064082 | 0.252 | 0.228 | 0.003705057 |
| Cask | 3.67E-11 | 0.009556464 | 0.558 | 0.579 | 9.88E-07 | Myo18a | 1.38E-07 | -0.030676382 | 0.529 | 0.521 | 0.003716374 |
| Mgat4a | 3.71E-11 | 0.028871736 | 0.281 | 0.289 | 9.99E-07 | Phkb | 1.38E-07 | -0.044279131 | 0.297 | 0.297 | 0.003721753 |
| Spire1 | 3.76E-11 | -0.004018227 | 0.554 | 0.584 | 1.01E-06 | 4932443L11Rik | 1.39E-07 | 0.048178141 | 0.28 | 0.25 | 0.003734857 |
| Cltc | 3.78E-11 | -0.051065009 | 0.348 | 0.384 | 1.02E-06 | Samd12 | 1.42E-07 | -0.07815804 | 0.884 | 0.893 | 0.003809181 |
| Cep120 | 3.87E-11 | 0.019426984 | 0.259 | 0.268 | 1.04E-06 | Apex2 | 1.43E-07 | -0.024798201 | 0.455 | 0.444 | 0.003835407 |
| Cacna1d | 3.93E-11 | 0.085679634 | 0.874 | 0.866 | 1.06E-06 | Vps41 | 1.43E-07 | -0.011999505 | 0.489 | 0.472 | 0.003837116 |
| Lmo4 | 3.95E-11 | 0.033804999 | 0.333 | 0.346 | 1.06E-06 | Memo1 | 1.49E-07 | 0.015187968 | 0.438 | 0.414 | 0.004017654 |
| Nfkb1 | 4.08E-11 | 0.027883518 | 0.345 | 0.356 | 1.10E-06 | Son | 1.50E-07 | 0.044217197 | 0.765 | 0.734 | 0.004032872 |
| Ppp2r5a | 4.15E-11 | -0.016041269 | 0.24 | 0.266 | 1.12E-06 | Lysmd4 | 1.50E-07 | -0.038595791 | 0.515 | 0.507 | 0.004047038 |
| Bzw2 | 4.18E-11 | -0.001390062 | 0.338 | 0.361 | 1.13E-06 | Clmn | 1.53E-07 | -0.052285623 | 0.251 | 0.257 | 0.004130203 |
| Acat1 | 4.21E-11 | -0.000639693 | 0.387 | 0.409 | 1.13E-06 | Mef2a | 1.58E-07 | -0.006977282 | 0.588 | 0.568 | 0.004259759 |
| Stau2 | 4.27E-11 | -0.026888268 | 0.671 | 0.707 | 1.15E-06 | Rasgrp1 | 1.58E-07 | 0.078357716 | 0.428 | 0.393 | 0.004263851 |
| Sh3rf3 | 4.64E-11 | 0.045345327 | 0.66 | 0.67 | 1.25E-06 | Gm19710 | 1.59E-07 | 0.047902882 | 0.579 | 0.543 | 0.004281519 |
| Asph | 4.68E-11 | -0.019603125 | 0.335 | 0.362 | 1.26E-06 | Cpeb4 | 1.62E-07 | 0.025080699 | 0.578 | 0.546 | 0.00434727 |
| Angpt2 | 4.70E-11 | -0.037336462 | 0.226 | 0.257 | 1.27E-06 | Lonp2 | 1.63E-07 | 0.007096422 | 0.444 | 0.421 | 0.004381636 |

| | | | | | | | | | | | |
|---|---|---|---|---|---|---|---|---|---|---|---|
| Tspan5 | 4.70E-11 | -0.114815688 | 0.791 | 0.822 | 1.27E-06 | Akap8l | 1.67E-07 | 0.016560066 | 0.798 | 0.772 | 0.004485836 |
| Klf7 | 4.73E-11 | 0.026102835 | 0.306 | 0.318 | 1.27E-06 | Tnfrsf21 | 1.68E-07 | -0.017113021 | 0.289 | 0.278 | 0.004525314 |
| Tiam2 | 4.76E-11 | -0.018079198 | 0.255 | 0.282 | 1.28E-06 | St5 | 1.72E-07 | 0.010858183 | 0.291 | 0.27 | 0.004617945 |
| Dnajc10 | 4.87E-11 | -0.014971197 | 0.258 | 0.281 | 1.31E-06 | Birc6 | 1.74E-07 | 0.02426712 | 0.752 | 0.723 | 0.004693764 |
| Pag1 | 4.88E-11 | -0.019212222 | 0.236 | 0.26 | 1.31E-06 | 4430402I18Rik | 1.76E-07 | 0.045139277 | 0.542 | 0.506 | 0.004730294 |
| Cyfip1 | 5.06E-11 | 0.016327365 | 0.554 | 0.579 | 1.36E-06 | Acvr1 | 1.79E-07 | 0.023167077 | 0.499 | 0.47 | 0.004805882 |
| Mast2 | 5.08E-11 | 0.049721654 | 0.517 | 0.52 | 1.37E-06 | Ica1l | 1.80E-07 | 0.030020797 | 0.327 | 0.3 | 0.004835253 |
| Mgrn1 | 5.09E-11 | -0.000255835 | 0.378 | 0.401 | 1.37E-06 | Nckap1 | 1.82E-07 | 0.025865721 | 0.738 | 0.708 | 0.004909146 |
| Zbtb11 | 5.23E-11 | 0.0148913 | 0.366 | 0.38 | 1.41E-06 | Ppp1r21 | 1.82E-07 | 0.014130727 | 0.274 | 0.254 | 0.004909959 |
| C1qtnf4 | 5.26E-11 | 0.043244233 | 0.339 | 0.35 | 1.42E-06 | Psmd9 | 1.83E-07 | 0.013711002 | 0.27 | 0.25 | 0.004913814 |
| Sec22a | 5.28E-11 | 0.004535659 | 0.258 | 0.276 | 1.42E-06 | Rb1 | 1.85E-07 | -0.017117799 | 0.491 | 0.476 | 0.004977559 |
| Nedd4l | 5.32E-11 | -0.121040941 | 0.838 | 0.853 | 1.43E-06 | Ror1 | 1.88E-07 | -0.027691071 | 0.352 | 0.338 | 0.005056277 |
| Adgrb2 | 5.35E-11 | 0.025427375 | 0.456 | 0.476 | 1.44E-06 | Tbc1d5 | 1.92E-07 | -0.038058125 | 0.63 | 0.626 | 0.00515569 |
| Agbl4 | 5.52E-11 | 0.114580357 | 0.742 | 0.711 | 1.49E-06 | Rabgap1l | 1.94E-07 | -0.043603984 | 0.902 | 0.894 | 0.005208458 |
| Lclat1 | 5.57E-11 | 0.027179125 | 0.328 | 0.337 | 1.50E-06 | Ints6l | 1.97E-07 | 0.008537244 | 0.318 | 0.299 | 0.005304797 |
| Tmem150c | 5.57E-11 | -0.046238199 | 0.413 | 0.449 | 1.50E-06 | Hook3 | 2.00E-07 | 0.035903267 | 0.507 | 0.475 | 0.005379328 |
| Rab30 | 5.59E-11 | 0.05003498 | 0.367 | 0.368 | 1.50E-06 | Tasp1 | 2.01E-07 | -0.017026628 | 0.494 | 0.48 | 0.005402213 |
| Numb | 5.65E-11 | -0.018429374 | 0.45 | 0.481 | 1.52E-06 | Pan3 | 2.08E-07 | -0.002291717 | 0.683 | 0.66 | 0.00560894 |
| St3gal5 | 5.89E-11 | -0.041859182 | 0.394 | 0.431 | 1.58E-06 | Lman2l | 2.09E-07 | -0.000584091 | 0.335 | 0.317 | 0.0056271 |
| Slc23a2 | 6.12E-11 | 0.01365252 | 0.466 | 0.485 | 1.65E-06 | Smarcc1 | 2.10E-07 | 0.017888925 | 0.404 | 0.379 | 0.005653513 |
| Cntn5 | 6.19E-11 | 0.140982741 | 0.721 | 0.7 | 1.66E-06 | Smpd4 | 2.10E-07 | -0.008738957 | 0.288 | 0.276 | 0.005657703 |
| Cplx2 | 6.19E-11 | 0.002097096 | 0.315 | 0.339 | 1.67E-06 | Fam49b | 2.11E-07 | -0.04975966 | 0.563 | 0.562 | 0.00568445 |
| Fam168a | 6.41E-11 | -0.008049259 | 0.727 | 0.758 | 1.72E-06 | Slc4a7 | 2.11E-07 | 0.02903545 | 0.45 | 0.42 | 0.005688325 |
| Erc1 | 6.55E-11 | 0.012158525 | 0.651 | 0.67 | 1.76E-06 | Fbxo10 | 2.12E-07 | -0.039245446 | 0.283 | 0.283 | 0.005708292 |
| Orai2 | 6.59E-11 | -0.07231311 | 0.224 | 0.26 | 1.77E-06 | Hmg20a | 2.14E-07 | 0.034250422 | 0.454 | 0.424 | 0.005747576 |
| Hmbox1 | 6.86E-11 | 0.013482256 | 0.485 | 0.508 | 1.84E-06 | Gm28375 | 2.14E-07 | 0.044543052 | 0.445 | 0.412 | 0.0057477 |
| Gm10785 | 6.87E-11 | -0.047629078 | 0.395 | 0.435 | 1.85E-06 | Greb1l | 2.14E-07 | -0.02336594 | 0.322 | 0.312 | 0.005752397 |
| Ankrd33b | 6.90E-11 | -0.128497331 | 0.495 | 0.538 | 1.86E-06 | Agap1 | 2.16E-07 | -0.051772767 | 0.682 | 0.679 | 0.005820897 |
| Nsd2 | 7.06E-11 | 0.039919349 | 0.481 | 0.491 | 1.90E-06 | Sgk1 | 2.20E-07 | 0.042575903 | 0.343 | 0.313 | 0.005913297 |
| Osbpl8 | 7.06E-11 | -0.006130967 | 0.44 | 0.464 | 1.90E-06 | Eefsec | 2.23E-07 | -0.007209972 | 0.416 | 0.399 | 0.00601238 |
| Slc24a3 | 7.20E-11 | -0.112051295 | 0.67 | 0.706 | 1.94E-06 | Pdss1 | 2.24E-07 | 0.012050539 | 0.27 | 0.251 | 0.006032004 |
| Sclt1 | 7.23E-11 | -0.011953597 | 0.533 | 0.565 | 1.95E-06 | Bptf | 2.28E-07 | -0.001730386 | 0.616 | 0.595 | 0.0061295 |
| Grm8 | 7.29E-11 | 0.159978796 | 0.403 | 0.37 | 1.96E-06 | Sbf1 | 2.28E-07 | 0.031092179 | 0.252 | 0.227 | 0.006130366 |
| Sh3gl2 | 7.51E-11 | 0.073953622 | 0.777 | 0.776 | 2.02E-06 | Sfmbt1 | 2.28E-07 | 0.008530368 | 0.36 | 0.338 | 0.006132663 |
| Metap2 | 7.52E-11 | -0.023104592 | 0.341 | 0.372 | 2.02E-06 | Clasp2 | 2.32E-07 | 0.001541924 | 0.772 | 0.749 | 0.006255604 |
| Fbxw4 | 7.54E-11 | -0.00330523 | 0.413 | 0.437 | 2.03E-06 | Phlpp2 | 2.32E-07 | -0.014629573 | 0.341 | 0.329 | 0.006255826 |
| Gls | 7.97E-11 | -0.056478242 | 0.83 | 0.859 | 2.15E-06 | Cnot10 | 2.33E-07 | 0.017383522 | 0.384 | 0.361 | 0.006259362 |
| Hp1bp3 | 8.01E-11 | 0.013198325 | 0.25 | 0.265 | 2.16E-06 | Zswim5 | 2.36E-07 | 0.034593752 | 0.371 | 0.343 | 0.006363458 |
| Fars2 | 8.10E-11 | 0.007740145 | 0.579 | 0.605 | 2.18E-06 | Fnip2 | 2.37E-07 | -0.017652381 | 0.327 | 0.315 | 0.006372132 |
| Phlpp2 | 8.10E-11 | 0.038008566 | 0.329 | 0.334 | 2.18E-06 | Ccdc149 | 2.39E-07 | -0.000809452 | 0.291 | 0.273 | 0.006440616 |
| Slc39a11 | 8.15E-11 | 0.046611136 | 0.27 | 0.272 | 2.19E-06 | Kif3b | 2.43E-07 | -0.012659665 | 0.285 | 0.274 | 0.006531221 |
| Atg10 | 8.25E-11 | 0.080147518 | 0.303 | 0.285 | 2.22E-06 | Kif26b | 2.44E-07 | 0.016343154 | 0.376 | 0.351 | 0.006555141 |
| Kdm5c | 8.42E-11 | -0.005661673 | 0.376 | 0.402 | 2.26E-06 | Acsl1 | 2.47E-07 | 0.003226377 | 0.257 | 0.24 | 0.006648026 |
| Nipsnap1 | 8.44E-11 | 0.045214227 | 0.275 | 0.275 | 2.27E-06 | Sec24a | 2.51E-07 | 0.01557956 | 0.377 | 0.354 | 0.006742167 |
| Adgrb1 | 8.63E-11 | -0.009579516 | 0.564 | 0.597 | 2.32E-06 | Fbxw4 | 2.53E-07 | 0.010504905 | 0.436 | 0.413 | 0.0068056 |
| Eml4 | 8.68E-11 | 0.011756008 | 0.535 | 0.559 | 2.33E-06 | Cd46 | 2.57E-07 | -0.012188855 | 0.378 | 0.366 | 0.006904641 |
| Ehbp1 | 8.77E-11 | 0.071375202 | 0.675 | 0.671 | 2.36E-06 | Tacc1 | 2.62E-07 | 0.047913662 | 0.59 | 0.555 | 0.007042836 |
| Upp2 | 9.06E-11 | 0.071173427 | 0.742 | 0.733 | 2.44E-06 | St8sia1 | 2.63E-07 | -0.021424849 | 0.281 | 0.273 | 0.007072267 |
| Supt3 | 9.08E-11 | -0.018536659 | 0.393 | 0.423 | 2.44E-06 | Robo2 | 2.65E-07 | -0.082808433 | 0.831 | 0.829 | 0.007132736 |
| Kif5c | 9.38E-11 | 0.050188998 | 0.657 | 0.657 | 2.52E-06 | Csad | 2.67E-07 | 0.042388191 | 0.25 | 0.222 | 0.007172222 |
| Ndufaf2 | 9.39E-11 | -0.037361978 | 0.233 | 0.265 | 2.53E-06 | Zer1 | 2.67E-07 | 0.02575875 | 0.394 | 0.367 | 0.00719659 |
| Nf2 | 9.64E-11 | -0.043450819 | 0.285 | 0.319 | 2.59E-06 | Pdzd2 | 2.68E-07 | -0.114106194 | 0.497 | 0.508 | 0.007200809 |
| Galntl6 | 1.01E-10 | 0.009553185 | 0.485 | 0.448 | 2.71E-06 | Cyfip1 | 2.69E-07 | 0.013271792 | 0.581 | 0.554 | 0.007242973 |
| Cop1 | 1.05E-10 | -0.02788605 | 0.406 | 0.438 | 2.81E-06 | Stam2 | 2.70E-07 | 0.018598283 | 0.257 | 0.237 | 0.007267546 |
| Tmeff2 | 1.05E-10 | 0.076137437 | 0.861 | 0.842 | 2.82E-06 | Bicdl1 | 2.72E-07 | 0.006219337 | 0.541 | 0.518 | 0.007315254 |
| Klhdc10 | 1.05E-10 | -0.044691019 | 0.695 | 0.733 | 2.83E-06 | Srsf11 | 2.75E-07 | 0.005380701 | 0.728 | 0.7 | 0.00739886 |
| Naa35 | 1.06E-10 | 0.008945259 | 0.298 | 0.314 | 2.85E-06 | 0610010F05Rik | 2.82E-07 | 0.01190933 | 0.388 | 0.364 | 0.007593927 |
| Txndc16 | 1.08E-10 | 0.023710793 | 0.303 | 0.314 | 2.91E-06 | Ptbp2 | 2.84E-07 | 0.018563682 | 0.467 | 0.44 | 0.007643327 |
| Frmpd4 | 1.10E-10 | 0.130796738 | 0.923 | 0.918 | 2.95E-06 | Lims1 | 2.85E-07 | 0.019165099 | 0.33 | 0.307 | 0.007667897 |
| Mbnl1 | 1.11E-10 | -0.012381514 | 0.404 | 0.431 | 2.98E-06 | Slc8a3 | 2.85E-07 | -0.06069852 | 0.322 | 0.324 | 0.007677887 |
| Kctd16 | 1.11E-10 | 0.102141732 | 0.878 | 0.859 | 2.98E-06 | Mark3 | 2.87E-07 | -0.02591588 | 0.741 | 0.727 | 0.007721993 |
| Zc3h7a | 1.25E-10 | 0.022126191 | 0.437 | 0.454 | 3.35E-06 | Cap2 | 2.88E-07 | 0.067203639 | 0.779 | 0.752 | 0.007761381 |
| Becn1 | 1.25E-10 | -0.044146649 | 0.222 | 0.255 | 3.35E-06 | Cdk17 | 2.93E-07 | 0.041896992 | 0.651 | 0.617 | 0.007872862 |
| Tbck | 1.29E-10 | -0.0110295 | 0.393 | 0.418 | 3.47E-06 | Evl | 2.96E-07 | -0.013965354 | 0.371 | 0.359 | 0.007966046 |
| Kdm4b | 1.35E-10 | 0.016990648 | 0.329 | 0.342 | 3.62E-06 | Slc39a11 | 2.97E-07 | -0.033373397 | 0.275 | 0.27 | 0.007986705 |
| Bptf | 1.36E-10 | 0.040057136 | 0.595 | 0.603 | 3.65E-06 | Lrba | 2.97E-07 | -0.024043365 | 0.684 | 0.671 | 0.007992061 |
| Cwf19l2 | 1.41E-10 | -0.006537967 | 0.245 | 0.267 | 3.79E-06 | Afdn | 2.97E-07 | 0.010784279 | 0.702 | 0.675 | 0.007997346 |
| Lrrc8b | 1.43E-10 | -0.009712078 | 0.32 | 0.344 | 3.84E-06 | Lrrtm3 | 2.98E-07 | -0.033949889 | 0.461 | 0.447 | 0.008026113 |
| Mcf2l | 1.43E-10 | 0.003470875 | 0.463 | 0.486 | 3.86E-06 | Gm48747 | 3.00E-07 | -0.048001733 | 0.569 | 0.569 | 0.008061433 |
| Tbc1d32 | 1.46E-10 | 0.037949228 | 0.27 | 0.274 | 3.93E-06 | Arid4b | 3.00E-07 | 0.027603161 | 0.641 | 0.61 | 0.008066055 |
| Ldlrad4 | 1.48E-10 | 0.004392075 | 0.712 | 0.74 | 3.98E-06 | Usp4 | 3.06E-07 | 0.037589537 | 0.31 | 0.282 | 0.008226328 |
| Cadm3 | 1.51E-10 | -0.014487996 | 0.25 | 0.276 | 4.07E-06 | Rgs17 | 3.12E-07 | -0.048136833 | 0.376 | 0.377 | 0.008395121 |
| Eps15 | 1.54E-10 | -0.02050369 | 0.241 | 0.267 | 4.16E-06 | Gnaq | 3.14E-07 | -0.044372825 | 0.835 | 0.83 | 0.008449935 |
| Sphkap | 1.56E-10 | -0.114328757 | 0.762 | 0.788 | 4.20E-06 | Stim2 | 3.16E-07 | -0.016846646 | 0.528 | 0.511 | 0.008491192 |
| Ift74 | 1.62E-10 | -0.014278447 | 0.235 | 0.259 | 4.37E-06 | Mef2c | 3.28E-07 | 0.023331076 | 0.578 | 0.546 | 0.008830653 |
| Glrb | 1.64E-10 | 0.001007998 | 0.445 | 0.471 | 4.41E-06 | Gpm6b | 3.29E-07 | -0.035319668 | 0.834 | 0.827 | 0.008840911 |
| Ica1l | 1.66E-10 | -0.020177943 | 0.3 | 0.328 | 4.48E-06 | Cdc42bpb | 3.31E-07 | -0.028090198 | 0.32 | 0.316 | 0.008897616 |
| Enah | 1.70E-10 | 0.030859033 | 0.761 | 0.778 | 4.56E-06 | Fnip1 | 3.35E-07 | 0.022040623 | 0.444 | 0.416 | 0.009003573 |
| 2900026A02Rik | 1.70E-10 | 0.065992644 | 0.338 | 0.326 | 4.57E-06 | Lingo2 | 3.35E-07 | 0.02651557 | 0.85 | 0.869 | 0.009009535 |
| Cux1 | 1.71E-10 | 0.031489772 | 0.321 | 0.334 | 4.59E-06 | Hectd4 | 3.37E-07 | -0.029397948 | 0.682 | 0.669 | 0.00907777 |
| Emc2 | 1.71E-10 | -0.031297971 | 0.225 | 0.254 | 4.61E-06 | Kmt2e | 3.46E-07 | 0.005013857 | 0.654 | 0.628 | 0.009311956 |
| Snta1 | 1.72E-10 | -0.073861466 | 0.264 | 0.303 | 4.62E-06 | Ryr3 | 3.51E-07 | -0.073028183 | 0.846 | 0.846 | 0.009430962 |
| Nrip1 | 1.75E-10 | 0.004066564 | 0.286 | 0.304 | 4.70E-06 | N4bp2l1 | 3.51E-07 | -0.027276381 | 0.318 | 0.312 | 0.009432354 |
| Dazap1 | 1.75E-10 | -0.02774745 | 0.268 | 0.297 | 4.71E-06 | Tmem117 | 3.51E-07 | -0.047059617 | 0.456 | 0.455 | 0.009433487 |
| Lrrn2 | 1.75E-10 | 0.009330648 | 0.527 | 0.551 | 4.72E-06 | Jakmip2 | 3.57E-07 | -0.021685741 | 0.709 | 0.694 | 0.009614878 |
| Focad | 1.76E-10 | -0.046821555 | 0.645 | 0.683 | 4.73E-06 | Acaca | 3.63E-07 | -0.041646549 | 0.526 | 0.525 | 0.009770414 |
| Rps6ka5 | 1.76E-10 | 0.076978673 | 0.289 | 0.276 | 4.74E-06 | Ap3b1 | 3.70E-07 | -0.007966184 | 0.5 | 0.482 | 0.009949597 |
| Pde8b | 1.79E-10 | 0.055986259 | 0.615 | 0.623 | 4.80E-06 | Pknox2 | 3.77E-07 | -0.005031028 | 0.474 | 0.454 | 0.010138435 |
| Tbk1 | 1.81E-10 | -0.030214765 | 0.268 | 0.298 | 4.86E-06 | Abhd6 | 3.84E-07 | -0.010353832 | 0.391 | 0.378 | 0.010331333 |
| Akap11 | 1.81E-10 | -0.022799959 | 0.433 | 0.466 | 4.86E-06 | Susd4 | 3.97E-07 | -0.01638509 | 0.679 | 0.662 | 0.010677042 |
| Usp31 | 1.81E-10 | -0.01809133 | 0.253 | 0.278 | 4.88E-06 | Nrg1 | 3.97E-07 | -0.093998247 | 0.767 | 0.794 | 0.010693853 |
| Marf1 | 1.81E-10 | 0.026886312 | 0.275 | 0.283 | 4.88E-06 | Ralgapb | 3.99E-07 | -0.024090072 | 0.419 | 0.408 | 0.010736736 |
| Abhd6 | 1.88E-10 | 0.008112877 | 0.378 | 0.397 | 5.06E-06 | Ralgapa2 | 4.10E-07 | -0.061558634 | 0.689 | 0.691 | 0.011024824 |
| Slc7a6 | 1.95E-10 | 0.009899211 | 0.33 | 0.348 | 5.24E-06 | Gon4l | 4.16E-07 | 0.002761219 | 0.602 | 0.582 | 0.011184583 |
| Map4 | 1.98E-10 | 0.012692935 | 0.499 | 0.519 | 5.34E-06 | Arid2 | 4.21E-07 | -0.014443998 | 0.311 | 0.298 | 0.011319023 |
| Comt | 2.00E-10 | 0.021105615 | 0.277 | 0.289 | 5.39E-06 | Tmem57 | 4.23E-07 | -0.070933546 | 0.431 | 0.461 | 0.011369412 |
| Tspan7 | 2.02E-10 | -0.037015919 | 0.55 | 0.588 | 5.43E-06 | 2010300C02Rik | 4.39E-07 | 0.031060386 | 0.734 | 0.704 | 0.011805218 |
| Clmn | 2.06E-10 | 0.072486146 | 0.257 | 0.246 | 5.54E-06 | Grin2a | 4.42E-07 | -0.068406428 | 0.982 | 0.987 | 0.011905446 |

| | | | | | | | | | | |
|---|---|---|---|---|---|---|---|---|---|---|
| Cipc | 2.06E-10 | 0.002745408 | 0.251 | 0.268 | 5.55E-06 | Dst | 4.47E-07 | -0.051306538 | 0.857 | 0.857 | 0.012037942 |
| Wdr26 | 2.09E-10 | -0.024813326 | 0.341 | 0.371 | 5.62E-06 | Oxr1 | 4.57E-07 | -0.053739764 | 0.896 | 0.891 | 0.012306033 |
| Itpka | 2.11E-10 | 0.064322824 | 0.366 | 0.358 | 5.67E-06 | Sclt1 | 4.59E-07 | -0.003714306 | 0.554 | 0.533 | 0.012351184 |
| Galnt9 | 2.19E-10 | -0.035561969 | 0.409 | 0.446 | 5.90E-06 | Usp40 | 4.60E-07 | 0.050818406 | 0.369 | 0.338 | 0.012385385 |
| Nlgn1 | 2.20E-10 | 0.071529857 | 0.99 | 0.982 | 5.92E-06 | Gls | 4.74E-07 | 0.015916397 | 0.854 | 0.83 | 0.012760678 |
| Cfl2 | 2.32E-10 | -0.020101406 | 0.228 | 0.254 | 6.24E-06 | Fat4 | 4.75E-07 | -0.008480306 | 0.26 | 0.248 | 0.012785643 |
| Psmd9 | 2.35E-10 | 0.011334161 | 0.25 | 0.263 | 6.32E-06 | Iars2 | 4.76E-07 | 0.021445474 | 0.26 | 0.237 | 0.012813567 |
| Srsf1 | 2.36E-10 | -0.053548802 | 0.264 | 0.302 | 6.34E-06 | Pkp2 | 4.77E-07 | -0.028762034 | 0.358 | 0.35 | 0.012847206 |
| Xist | 2.36E-10 | 0.031321437 | 0.967 | 0.951 | 6.35E-06 | Ulk4 | 4.79E-07 | -0.02446233 | 0.498 | 0.49 | 0.01288711 |
| Gpm6b | 2.41E-10 | 0.074238163 | 0.827 | 0.823 | 6.48E-06 | Gpatch8 | 5.12E-07 | 0.044297584 | 0.833 | 0.807 | 0.013767033 |
| 2-Mar | 2.41E-10 | -0.027807976 | 0.235 | 0.263 | 6.50E-06 | Ablim3 | 5.16E-07 | -0.040570474 | 0.277 | 0.276 | 0.013895801 |
| Rapgef2 | 2.42E-10 | -0.086341093 | 0.824 | 0.852 | 6.50E-06 | Rbl2 | 5.17E-07 | 0.025981265 | 0.265 | 0.241 | 0.013914939 |
| Slc8a1 | 2.45E-10 | 0.04053334 | 0.766 | 0.755 | 6.59E-06 | Syt7 | 5.21E-07 | -0.052490481 | 0.75 | 0.744 | 0.014031574 |
| Kcnc1 | 2.46E-10 | 0.060637259 | 0.269 | 0.262 | 6.61E-06 | Gm26694 | 5.23E-07 | -0.011885595 | 0.538 | 0.565 | 0.014082969 |
| Nckap1 | 2.55E-10 | -0.017229186 | 0.708 | 0.743 | 6.86E-06 | Grb10 | 5.32E-07 | -0.025019782 | 0.271 | 0.265 | 0.014302032 |
| Ppargc1a | 2.63E-10 | 0.031418368 | 0.333 | 0.342 | 7.09E-06 | Btbd3 | 5.33E-07 | 0.013816986 | 0.347 | 0.325 | 0.01433637 |
| Ptprs | 2.64E-10 | 0.05513676 | 0.712 | 0.72 | 7.10E-06 | Naa35 | 5.47E-07 | -0.033950681 | 0.299 | 0.298 | 0.014725129 |
| Hnrnpm | 2.68E-10 | -0.003271003 | 0.413 | 0.438 | 7.22E-06 | Zmynd11 | 5.58E-07 | 0.025303253 | 0.702 | 0.673 | 0.015018759 |
| Raph1 | 2.77E-10 | 0.02689517 | 0.44 | 0.451 | 7.46E-06 | Miga1 | 5.59E-07 | -0.037098578 | 0.32 | 0.323 | 0.015051358 |
| 1700111E14Rik | 2.77E-10 | -0.084304917 | 0.614 | 0.655 | 7.46E-06 | Zhx3 | 5.62E-07 | -0.002478021 | 0.375 | 0.359 | 0.015120738 |
| Slc12a5 | 2.87E-10 | 0.031067696 | 0.504 | 0.519 | 7.72E-06 | Nbeal1 | 5.66E-07 | -0.020686257 | 0.368 | 0.359 | 0.015229467 |
| Gm16599 | 2.95E-10 | -0.034955037 | 0.255 | 0.286 | 7.92E-06 | Abl2 | 5.69E-07 | -0.026128644 | 0.582 | 0.573 | 0.015306935 |
| Sox5 | 2.99E-10 | 0.071026421 | 0.432 | 0.392 | 8.05E-06 | Atad2b | 5.71E-07 | 0.032935696 | 0.54 | 0.51 | 0.01536568 |
| Herc2 | 3.05E-10 | -0.02819418 | 0.634 | 0.668 | 8.20E-06 | Zfp277 | 5.74E-07 | -0.023003197 | 0.295 | 0.29 | 0.015452761 |
| C530008M17Rik | 3.06E-10 | 0.068534997 | 0.45 | 0.448 | 8.23E-06 | Tet3 | 5.80E-07 | -0.000497293 | 0.442 | 0.424 | 0.015593521 |
| Plppr2 | 3.10E-10 | 0.038559715 | 0.349 | 0.357 | 8.33E-06 | Nck2 | 5.85E-07 | -0.00402703 | 0.382 | 0.367 | 0.01574522 |
| Ttc39c | 3.14E-10 | -0.029845717 | 0.293 | 0.322 | 8.46E-06 | Cenpp | 5.85E-07 | 0.068262792 | 0.328 | 0.296 | 0.01575158 |
| Ccny | 3.28E-10 | -0.0343716 | 0.594 | 0.631 | 8.84E-06 | Nphp4 | 5.87E-07 | -0.022016328 | 0.306 | 0.301 | 0.0157932 |
| 8-Sep | 3.29E-10 | 0.049663844 | 0.28 | 0.277 | 8.84E-06 | Nedd4 | 5.97E-07 | 0.046183932 | 0.619 | 0.585 | 0.016053337 |
| Zfp445 | 3.30E-10 | 0.007924001 | 0.474 | 0.494 | 8.87E-06 | Frmd5 | 6.15E-07 | 0.07868832 | 0.681 | 0.649 | 0.016535592 |
| Matr3 | 3.34E-10 | 0.005131111 | 0.239 | 0.257 | 9.00E-06 | Epg5 | 6.16E-07 | -0.008691404 | 0.268 | 0.255 | 0.016581979 |
| Sema5a | 3.36E-10 | -0.052012438 | 0.373 | 0.412 | 9.04E-06 | Cab39 | 6.23E-07 | 0.011541299 | 0.346 | 0.328 | 0.016755169 |
| Ints3 | 3.42E-10 | -0.029280663 | 0.249 | 0.278 | 9.21E-06 | Unc5c | 6.25E-07 | -0.050375413 | 0.718 | 0.722 | 0.016817259 |
| Plxnc1 | 3.48E-10 | 0.029063132 | 0.254 | 0.262 | 9.37E-06 | Tmod2 | 6.30E-07 | 0.011272884 | 0.504 | 0.481 | 0.016947174 |
| Nuak1 | 3.49E-10 | 0.038611294 | 0.249 | 0.252 | 9.38E-06 | Ppp2r3a | 6.37E-07 | 0.025978825 | 0.52 | 0.492 | 0.017126404 |
| Pclo | 3.51E-10 | 0.06705955 | 0.888 | 0.886 | 9.45E-06 | Nnat | 6.48E-07 | -0.02443363 | 0.326 | 0.322 | 0.017425132 |
| Cblb | 3.56E-10 | -0.105548034 | 0.459 | 0.501 | 9.59E-06 | Lamp2 | 6.61E-07 | 0.041924156 | 0.281 | 0.253 | 0.017787997 |
| Me3 | 3.64E-10 | 0.068312866 | 0.684 | 0.677 | 9.78E-06 | Parp1 | 6.79E-07 | 0.008401527 | 0.255 | 0.236 | 0.018269369 |
| E130307A14Rik | 3.66E-10 | 0.04369357 | 0.571 | 0.578 | 9.86E-06 | Sesn1 | 6.83E-07 | -0.022934079 | 0.466 | 0.454 | 0.018387233 |
| Klhl3 | 3.69E-10 | -0.012561264 | 0.59 | 0.621 | 9.93E-06 | Dtd1 | 6.84E-07 | 0.007716427 | 0.496 | 0.476 | 0.018413357 |
| Traip | 3.74E-10 | -0.040479387 | 0.238 | 0.27 | 1.01E-05 | Tomm40 | 6.92E-07 | 0.033634282 | 0.271 | 0.245 | 0.018609036 |
| Gfm2 | 3.76E-10 | 0.003727194 | 0.293 | 0.311 | 1.01E-05 | Itsn1 | 6.97E-07 | -0.043717257 | 0.629 | 0.626 | 0.018754497 |
| Spindoc | 3.81E-10 | 0.022872438 | 0.246 | 0.255 | 1.02E-05 | Gria4 | 6.99E-07 | -0.069767306 | 0.631 | 0.648 | 0.018797767 |
| Rnf217 | 3.84E-10 | 0.034285353 | 0.286 | 0.29 | 1.03E-05 | Map2k4 | 7.06E-07 | 0.022960874 | 0.712 | 0.684 | 0.018988847 |
| Ildr2 | 3.89E-10 | 0.032099345 | 0.479 | 0.49 | 1.05E-05 | Ptchd4 | 7.10E-07 | 0.068909711 | 0.471 | 0.437 | 0.019102761 |
| Nsd1 | 3.93E-10 | 0.066957483 | 0.791 | 0.79 | 1.06E-05 | Hspa4l | 7.18E-07 | -0.034262875 | 0.279 | 0.278 | 0.019325573 |
| Adck1 | 4.04E-10 | 0.03768728 | 0.251 | 0.253 | 1.09E-05 | Fnbp1 | 7.19E-07 | 0.009559348 | 0.618 | 0.594 | 0.019348484 |
| Bbx | 4.13E-10 | -0.040636313 | 0.344 | 0.377 | 1.11E-05 | Lrrc8b | 7.24E-07 | 0.015142085 | 0.339 | 0.32 | 0.019481667 |
| Golga4 | 4.21E-10 | 0.035203597 | 0.409 | 0.419 | 1.13E-05 | 1700111E14Rik | 7.25E-07 | 0.016322703 | 0.645 | 0.614 | 0.019519735 |
| Klhl12 | 4.39E-10 | -0.011882019 | 0.23 | 0.252 | 1.18E-05 | Wipf3 | 7.32E-07 | -0.058812776 | 0.633 | 0.626 | 0.019688446 |
| Xrn1 | 4.43E-10 | -0.019777248 | 0.519 | 0.552 | 1.19E-05 | Bcas3 | 7.41E-07 | -0.004320068 | 0.684 | 0.665 | 0.019944548 |
| Senp7 | 4.44E-10 | -0.004125843 | 0.543 | 0.572 | 1.19E-05 | Atf2 | 7.58E-07 | 0.037111051 | 0.544 | 0.513 | 0.020395375 |
| Akap13 | 4.53E-10 | -0.052327137 | 0.277 | 0.312 | 1.22E-05 | Ube3c | 7.74E-07 | 0.015065793 | 0.608 | 0.583 | 0.020832137 |
| Mgat4c | 4.55E-10 | 0.181174298 | 0.503 | 0.461 | 1.22E-05 | Kif1b | 7.91E-07 | 8.68E-05 | 0.832 | 0.812 | 0.02127254 |
| Epha5 | 4.68E-10 | -0.05972161 | 0.894 | 0.919 | 1.26E-05 | Dpp6 | 7.96E-07 | -0.055755851 | 0.95 | 0.956 | 0.021426406 |
| Cep70 | 4.74E-10 | -0.026821881 | 0.463 | 0.497 | 1.27E-05 | Sympk | 8.01E-07 | -0.00173623 | 0.356 | 0.338 | 0.021546378 |
| Gabra4 | 4.79E-10 | 0.014035362 | 0.466 | 0.487 | 1.29E-05 | Filip1l | 8.03E-07 | -9.89E-05 | 0.259 | 0.242 | 0.021617327 |
| Ccbe1 | 4.89E-10 | 0.09857989 | 0.32 | 0.308 | 1.32E-05 | Dnajc6 | 8.07E-07 | -0.031846293 | 0.729 | 0.722 | 0.021726746 |
| Aff4 | 4.91E-10 | -0.016713073 | 0.559 | 0.589 | 1.32E-05 | Gramd1b | 8.16E-07 | 0.004984831 | 0.681 | 0.657 | 0.021958831 |
| Cds2 | 4.91E-10 | 0.037522623 | 0.323 | 0.326 | 1.32E-05 | Arhgef12 | 8.17E-07 | 0.031985104 | 0.613 | 0.583 | 0.021969962 |
| Reps2 | 5.02E-10 | -0.063269643 | 0.635 | 0.675 | 1.35E-05 | 4930587E11Rik | 8.26E-07 | 0.087613409 | 0.382 | 0.349 | 0.022211763 |
| Hlcs | 5.07E-10 | -0.030427237 | 0.227 | 0.254 | 1.36E-05 | Trappc10 | 8.60E-07 | -0.002297387 | 0.261 | 0.246 | 0.023148678 |
| Rpgrip1l | 5.19E-10 | 0.012952765 | 0.239 | 0.252 | 1.40E-05 | Csgalnact1 | 8.63E-07 | -0.010095378 | 0.36 | 0.345 | 0.023209289 |
| Zdhhc14 | 5.29E-10 | -0.05130163 | 0.662 | 0.697 | 1.42E-05 | Akap7 | 8.80E-07 | 0.02584331 | 0.256 | 0.235 | 0.023666745 |
| Pcdh19 | 5.39E-10 | 0.000154119 | 0.301 | 0.322 | 1.45E-05 | Ssh2 | 9.08E-07 | -0.043980775 | 0.782 | 0.779 | 0.024437546 |
| Nfia | 5.43E-10 | -0.109643268 | 0.593 | 0.632 | 1.46E-05 | Gpr45 | 9.11E-07 | -0.016424559 | 0.458 | 0.448 | 0.024500405 |
| Clstn3 | 5.47E-10 | -0.040359443 | 0.289 | 0.321 | 1.47E-05 | Dmxl2 | 9.50E-07 | 0.022143768 | 0.688 | 0.659 | 0.025559721 |
| Hs2st1 | 5.47E-10 | 0.035135158 | 0.579 | 0.592 | 1.47E-05 | Rfx7 | 9.58E-07 | 0.035056045 | 0.598 | 0.567 | 0.025781456 |
| Faim2 | 5.87E-10 | 0.001396493 | 0.263 | 0.28 | 1.58E-05 | Ubn2 | 9.59E-07 | 0.049501848 | 0.8 | 0.772 | 0.025794276 |
| Sez6l2 | 5.93E-10 | -0.023758767 | 0.463 | 0.495 | 1.60E-05 | Ift74 | 9.66E-07 | 0.006837433 | 0.252 | 0.235 | 0.025992785 |
| Tacc2 | 5.94E-10 | -0.014180785 | 0.272 | 0.298 | 1.60E-05 | Zfp385b | 9.78E-07 | -0.054172768 | 0.621 | 0.622 | 0.026310737 |
| E2f3 | 5.98E-10 | -0.087033133 | 0.259 | 0.296 | 1.61E-05 | Aifm3 | 9.87E-07 | 0.022496136 | 0.314 | 0.289 | 0.026567233 |
| Cdk17 | 6.14E-10 | -0.049046535 | 0.617 | 0.656 | 1.65E-05 | Acbd6 | 9.93E-07 | -0.005499884 | 0.496 | 0.483 | 0.026712221 |
| Nmnat2 | 6.35E-10 | 0.043214867 | 0.67 | 0.67 | 1.71E-05 | Cacnb4 | 1.02E-06 | 0.094324856 | 0.811 | 0.785 | 0.027315062 |
| Cog4 | 6.83E-10 | 0.001391928 | 0.261 | 0.279 | 1.84E-05 | Gm49003 | 1.04E-06 | 0.030368347 | 0.337 | 0.311 | 0.028108487 |
| Atxn2l | 6.86E-10 | 0.008682949 | 0.298 | 0.315 | 1.85E-05 | Zhx2 | 1.05E-06 | 0.00067785 | 0.283 | 0.265 | 0.028217848 |
| Son | 6.89E-10 | -0.078753063 | 0.734 | 0.768 | 1.85E-05 | Lrtm1 | 1.07E-06 | 0.00375395 | 0.351 | 0.331 | 0.028841509 |
| Tusc3 | 6.94E-10 | 0.059201486 | 0.565 | 0.563 | 1.87E-05 | Smg1 | 1.07E-06 | 0.000706993 | 0.643 | 0.622 | 0.028870648 |
| Usp40 | 6.96E-10 | -0.026216884 | 0.338 | 0.366 | 1.87E-05 | Plekha6 | 1.09E-06 | -0.050906607 | 0.429 | 0.434 | 0.029264673 |
| Slc7a8 | 7.03E-10 | 0.00949809 | 0.333 | 0.349 | 1.89E-05 | Sms | 1.09E-06 | 0.022178829 | 0.312 | 0.289 | 0.029313123 |
| Tln2 | 7.03E-10 | 0.047033988 | 0.517 | 0.521 | 1.89E-05 | Evi5 | 1.09E-06 | 0.026725557 | 0.421 | 0.395 | 0.029341743 |
| Pld3 | 7.13E-10 | 0.002546925 | 0.242 | 0.262 | 1.92E-05 | Ptprd | 1.11E-06 | -0.048866943 | 0.98 | 0.987 | 0.029925435 |
| Syt17 | 7.17E-10 | 0.034073112 | 0.377 | 0.389 | 1.93E-05 | E2f3 | 1.11E-06 | -0.033832575 | 0.259 | 0.259 | 0.029971299 |
| Trio | 7.18E-10 | 0.036706161 | 0.831 | 0.844 | 1.93E-05 | Rbfox3 | 1.11E-06 | 0.045434195 | 0.918 | 0.906 | 0.029979335 |
| Psme4 | 7.29E-10 | -0.010492208 | 0.617 | 0.645 | 1.96E-05 | Ggnbp2 | 1.13E-06 | 0.00845207 | 0.492 | 0.469 | 0.030323875 |
| Jarid2 | 7.32E-10 | -0.014859606 | 0.443 | 0.469 | 1.97E-05 | Mical2 | 1.15E-06 | 0.038998209 | 0.841 | 0.815 | 0.031034516 |
| Cdh18 | 7.34E-10 | 0.245344101 | 0.458 | 0.434 | 1.97E-05 | Ubr4 | 1.16E-06 | -0.008425775 | 0.26 | 0.248 | 0.031089658 |
| Ptbp2 | 7.41E-10 | 0.009161379 | 0.44 | 0.459 | 1.99E-05 | Ubap2 | 1.18E-06 | 0.001750114 | 0.29 | 0.273 | 0.031859525 |
| Jph1 | 7.43E-10 | 0.090737006 | 0.596 | 0.602 | 2.00E-05 | Fgf13 | 1.19E-06 | 0.009799643 | 0.404 | 0.381 | 0.032036965 |
| Chchd3 | 7.68E-10 | 0.014873606 | 0.597 | 0.618 | 2.07E-05 | Zfr2 | 1.20E-06 | -0.004403958 | 0.342 | 0.326 | 0.032231849 |
| Sfswap | 7.75E-10 | 0.052514825 | 0.531 | 0.531 | 2.09E-05 | Ctnna2 | 1.21E-06 | -0.045398221 | 0.959 | 0.967 | 0.032596368 |
| Clvs1 | 8.21E-10 | -0.012712279 | 0.612 | 0.642 | 2.21E-05 | Snca | 1.24E-06 | 0.027026247 | 0.544 | 0.517 | 0.033403147 |
| Dpp10 | 8.37E-10 | -0.380125705 | 0.423 | 0.384 | 2.25E-05 | Camk2b | 1.24E-06 | 0.022695352 | 0.785 | 0.758 | 0.033488708 |
| Hivep3 | 8.43E-10 | 0.050119337 | 0.827 | 0.834 | 2.27E-05 | Rgs11 | 1.27E-06 | 0.014749325 | 0.262 | 0.243 | 0.034123662 |
| Dlat | 8.47E-10 | -0.013656739 | 0.25 | 0.272 | 2.28E-05 | Xist | 1.27E-06 | -0.035653185 | 0.955 | 0.967 | 0.034257604 |
| Dync2h1 | 8.50E-10 | -0.055377068 | 0.512 | 0.553 | 2.29E-05 | Gm15738 | 1.27E-06 | 0.018210694 | 0.844 | 0.819 | 0.034280463 |
| Madd | 8.76E-10 | -0.017563881 | 0.48 | 0.51 | 2.36E-05 | 9930021J03Rik | 1.28E-06 | -0.002364589 | 0.584 | 0.564 | 0.034518986 |
| AC165271.1 | 9.07E-10 | -0.104158988 | 0.316 | 0.34 | 2.44E-05 | Ranbp10 | 1.30E-06 | 0.014842809 | 0.314 | 0.292 | 0.0350986 |

| | | | | | | | | | | | |
|---|---|---|---|---|---|---|---|---|---|---|---|
| Dnah9 | 9.53E-10 | -0.046312123 | 0.406 | 0.443 | 2.56E-05 | Tbc1d12 | 1.31E-06 | -0.002806142 | 0.308 | 0.292 | 0.035352397 |
| Nedd4 | 9.71E-10 | 0.041517788 | 0.585 | 0.598 | 2.61E-05 | Plcl2 | 1.32E-06 | 0.02860263 | 0.57 | 0.54 | 0.035490276 |
| Aff3 | 9.81E-10 | -0.101107293 | 0.793 | 0.824 | 2.64E-05 | Supt3 | 1.34E-06 | 0.003331956 | 0.413 | 0.393 | 0.036033777 |
| Gpcpd1 | 9.92E-10 | -0.018940184 | 0.365 | 0.393 | 2.67E-05 | Traf3 | 1.34E-06 | -0.019128498 | 0.459 | 0.447 | 0.036037763 |
| Gm16054 | 9.99E-10 | -0.069867832 | 0.216 | 0.251 | 2.69E-05 | Wsb1 | 1.34E-06 | 0.019323449 | 0.291 | 0.268 | 0.036107545 |
| Tle4 | 1.01E-09 | -0.063595709 | 0.381 | 0.419 | 2.72E-05 | Kcnh1 | 1.35E-06 | -0.028370682 | 0.506 | 0.498 | 0.036267999 |
| Smarca2 | 1.03E-09 | 0.082738482 | 0.498 | 0.483 | 2.78E-05 | Ablim1 | 1.36E-06 | -0.054782328 | 0.604 | 0.6 | 0.03646279 |
| Zeb2 | 1.05E-09 | -0.104395048 | 0.754 | 0.766 | 2.82E-05 | Csnk1a1 | 1.36E-06 | 0.036290239 | 0.664 | 0.634 | 0.036507322 |
| Slc43a2 | 1.06E-09 | 0.024746842 | 0.329 | 0.338 | 2.86E-05 | 4933413L06Rik | 1.41E-06 | 0.101852373 | 0.305 | 0.278 | 0.037972721 |
| Gstz1 | 1.12E-09 | 0.00751872 | 0.243 | 0.258 | 3.02E-05 | Aatf | 1.43E-06 | -0.013244781 | 0.322 | 0.311 | 0.038363782 |
| Tbc1d12 | 1.13E-09 | 0.047014446 | 0.292 | 0.289 | 3.05E-05 | Nbas | 1.43E-06 | -0.007907011 | 0.428 | 0.414 | 0.038556988 |
| Zcchc18 | 1.16E-09 | -0.065134455 | 0.385 | 0.425 | 3.13E-05 | Tusc3 | 1.43E-06 | 0.0479005 | 0.598 | 0.565 | 0.038608595 |
| Rras2 | 1.17E-09 | 0.006175013 | 0.418 | 0.442 | 3.15E-05 | Syt14 | 1.44E-06 | 0.008060405 | 0.568 | 0.546 | 0.038788214 |
| Gm20275 | 1.18E-09 | 0.006850882 | 0.726 | 0.752 | 3.19E-05 | Atp6v0a1 | 1.47E-06 | 0.034985416 | 0.785 | 0.758 | 0.039676195 |
| Psmd4 | 1.20E-09 | -0.0311386 | 0.227 | 0.255 | 3.23E-05 | Prkar1b | 1.54E-06 | -0.013053449 | 0.347 | 0.334 | 0.041303127 |
| Dzip1 | 1.24E-09 | 0.05508251 | 0.298 | 0.295 | 3.33E-05 | Lgi1 | 1.54E-06 | -0.023444968 | 0.617 | 0.602 | 0.041524556 |
| Lin7a | 1.24E-09 | 0.074020023 | 0.341 | 0.326 | 3.34E-05 | Sec22a | 1.55E-06 | 0.004237121 | 0.274 | 0.258 | 0.041713269 |
| 3-Sep | 1.26E-09 | -0.011416439 | 0.333 | 0.356 | 3.39E-05 | 8-Sep | 1.55E-06 | 0.005277648 | 0.299 | 0.28 | 0.041715053 |
| Klc1 | 1.29E-09 | -0.014619827 | 0.446 | 0.477 | 3.46E-05 | Cnih3 | 1.56E-06 | -0.029134066 | 0.308 | 0.301 | 0.042024727 |
| Crtc3 | 1.34E-09 | -0.025579918 | 0.471 | 0.503 | 3.59E-05 | Nhlrc2 | 1.59E-06 | 0.032185931 | 0.251 | 0.228 | 0.042748072 |
| Otud7a | 1.37E-09 | 0.065913856 | 0.916 | 0.913 | 3.69E-05 | Fgd4 | 1.61E-06 | -0.028789853 | 0.541 | 0.534 | 0.043236005 |
| Rims2 | 1.48E-09 | 0.039952886 | 0.868 | 0.87 | 3.99E-05 | Bend6 | 1.66E-06 | 0.02024279 | 0.334 | 0.312 | 0.044787563 |
| Mapk1 | 1.48E-09 | 0.01862178 | 0.643 | 0.665 | 3.99E-05 | Gabrb2 | 1.67E-06 | -0.049504116 | 0.76 | 0.752 | 0.044801161 |
| Gm48091 | 1.50E-09 | -0.083111309 | 0.62 | 0.659 | 4.05E-05 | Arhgap23 | 1.67E-06 | -0.029001812 | 0.293 | 0.288 | 0.044820664 |
| Sema6d | 1.60E-09 | 0.094093717 | 0.593 | 0.598 | 4.29E-05 | Plxdc2 | 1.67E-06 | -0.048351107 | 0.324 | 0.32 | 0.044827987 |
| Ndufaf7 | 1.60E-09 | 0.029902472 | 0.287 | 0.294 | 4.30E-05 | Dcaf6 | 1.68E-06 | -0.013172074 | 0.686 | 0.67 | 0.045149012 |
| Actr1b | 1.61E-09 | -0.004493412 | 0.241 | 0.262 | 4.33E-05 | Ash1l | 1.68E-06 | 0.014429537 | 0.746 | 0.721 | 0.045168701 |
| Angel2 | 1.69E-09 | -0.012937166 | 0.265 | 0.289 | 4.56E-05 | Dtnb | 1.73E-06 | -0.056630051 | 0.765 | 0.771 | 0.046426785 |
| Actn1 | 1.73E-09 | 0.020865131 | 0.348 | 0.359 | 4.65E-05 | Rundc3b | 1.74E-06 | 0.007359641 | 0.487 | 0.464 | 0.046946038 |
| Dnttip1 | 1.83E-09 | -0.037888527 | 0.231 | 0.261 | 4.93E-05 | Faf1 | 1.83E-06 | -0.02560725 | 0.747 | 0.736 | 0.049325047 |
| Map2k4 | 1.85E-09 | -0.048329753 | 0.684 | 0.721 | 4.98E-05 | Lzts1 | 1.86E-06 | -0.027455124 | 0.328 | 0.319 | 0.050115377 |
| Timm44 | 1.88E-09 | 0.050035909 | 0.279 | 0.274 | 5.05E-05 | Cntnap2 | 1.90E-06 | -0.029502694 | 0.828 | 0.852 | 0.051228257 |
| Sms | 1.92E-09 | -0.058947001 | 0.289 | 0.324 | 5.15E-05 | Myo1d | 1.94E-06 | -0.024848151 | 0.267 | 0.263 | 0.052333684 |
| Sipa1l3 | 1.93E-09 | -0.096661659 | 0.624 | 0.662 | 5.19E-05 | Myo5a | 1.96E-06 | -0.016882246 | 0.667 | 0.651 | 0.052657473 |
| Pafah1b1 | 1.93E-09 | -0.001215042 | 0.689 | 0.718 | 5.19E-05 | Vps13b | 1.96E-06 | -0.040536673 | 0.818 | 0.814 | 0.052701627 |
| Tnfrsf21 | 1.93E-09 | 0.005835711 | 0.278 | 0.294 | 5.19E-05 | Ints3 | 1.97E-06 | 0.004735605 | 0.267 | 0.249 | 0.052928534 |
| Slc4a8 | 1.98E-09 | -0.002975915 | 0.275 | 0.295 | 5.33E-05 | Map7d2 | 1.99E-06 | -0.000475422 | 0.497 | 0.477 | 0.053511605 |
| Galnt16 | 1.99E-09 | 0.029931758 | 0.536 | 0.546 | 5.36E-05 | Cep120 | 2.00E-06 | -0.004035587 | 0.271 | 0.259 | 0.053776322 |
| Fam149b | 2.08E-09 | -0.017196096 | 0.261 | 0.285 | 5.61E-05 | Papd5 | 2.01E-06 | -0.065024911 | 0.276 | 0.305 | 0.05401942 |
| Zfp91 | 2.11E-09 | -0.001958321 | 0.251 | 0.27 | 5.68E-05 | Syt16 | 2.04E-06 | 0.021374539 | 0.727 | 0.702 | 0.05492197 |
| Dcun1d2 | 2.17E-09 | 0.008620671 | 0.27 | 0.286 | 5.84E-05 | Pacs1 | 2.09E-06 | 0.005001386 | 0.545 | 0.522 | 0.056130367 |
| Phip | 2.18E-09 | -0.015791479 | 0.528 | 0.559 | 5.86E-05 | Galnt9 | 2.12E-06 | 0.004926028 | 0.432 | 0.409 | 0.057141665 |
| Rph3a | 2.19E-09 | 0.047022795 | 0.278 | 0.276 | 5.89E-05 | Raver2 | 2.12E-06 | 0.050726354 | 0.393 | 0.364 | 0.057155734 |
| Slc25a27 | 2.23E-09 | -0.021474794 | 0.393 | 0.421 | 5.99E-05 | Mtmr7 | 2.13E-06 | -0.02518752 | 0.32 | 0.313 | 0.057244876 |
| Huwe1 | 2.47E-09 | 0.034431348 | 0.667 | 0.681 | 6.65E-05 | Epb41 | 2.15E-06 | 0.028177566 | 0.303 | 0.28 | 0.057750157 |
| Tbc1d5 | 2.47E-09 | 0.061917605 | 0.626 | 0.622 | 6.66E-05 | Numb | 2.17E-06 | 0.045424539 | 0.48 | 0.45 | 0.05849059 |
| Ahi1 | 2.64E-09 | -0.056376853 | 0.982 | 0.985 | 7.09E-05 | Pls3 | 2.19E-06 | -0.000134369 | 0.356 | 0.34 | 0.059056149 |
| Rassf8 | 2.69E-09 | -0.001509529 | 0.359 | 0.382 | 7.23E-05 | Ankrd27 | 2.23E-06 | -0.036026829 | 0.247 | 0.251 | 0.060086452 |
| Prkaa2 | 2.73E-09 | -0.003747884 | 0.238 | 0.257 | 7.34E-05 | Msh3 | 2.30E-06 | -0.002060747 | 0.254 | 0.24 | 0.061827549 |
| Ap3d1 | 2.78E-09 | -0.003351635 | 0.244 | 0.261 | 7.49E-05 | Rtl4 | 2.31E-06 | -0.056839371 | 0.623 | 0.63 | 0.062064681 |
| Hs6st2 | 2.82E-09 | 0.082498465 | 0.305 | 0.272 | 7.59E-05 | Airn | 2.31E-06 | 0.026365893 | 0.294 | 0.271 | 0.062102203 |
| Zfp638 | 2.88E-09 | -0.051466764 | 0.654 | 0.691 | 7.74E-05 | Strn | 2.32E-06 | 0.01836711 | 0.323 | 0.301 | 0.062473374 |
| Osbpl1a | 2.92E-09 | 0.088812952 | 0.356 | 0.324 | 7.87E-05 | Ptpre | 2.37E-06 | -0.017934868 | 0.536 | 0.524 | 0.063863011 |
| Oxr1 | 2.96E-09 | 0.045365294 | 0.891 | 0.895 | 7.97E-05 | Nfat5 | 2.38E-06 | 0.020135735 | 0.652 | 0.626 | 0.064036292 |
| Icam5 | 2.97E-09 | 0.036060322 | 0.486 | 0.497 | 8.00E-05 | Fam208a | 2.38E-06 | -0.063763183 | 0.338 | 0.368 | 0.064049805 |
| Pcdh7 | 3.04E-09 | 0.124467939 | 0.552 | 0.543 | 8.18E-05 | Adcy8 | 2.54E-06 | 0.03322687 | 0.446 | 0.42 | 0.068338263 |
| Braf | 3.05E-09 | 0.029261043 | 0.691 | 0.704 | 8.22E-05 | Aff2 | 2.62E-06 | 0.003322628 | 0.498 | 0.479 | 0.070498348 |
| Mpped2 | 3.08E-09 | 0.065297065 | 0.571 | 0.567 | 8.29E-05 | Gm26854 | 2.65E-06 | -0.050289353 | 0.27 | 0.278 | 0.071280089 |
| Evl | 3.16E-09 | 0.029177234 | 0.359 | 0.363 | 8.50E-05 | Usp20 | 2.71E-06 | -0.002364662 | 0.261 | 0.248 | 0.073049092 |
| Bmpr1a | 3.17E-09 | -0.007202121 | 0.399 | 0.423 | 8.52E-05 | Fstl4 | 2.76E-06 | -0.094117281 | 0.588 | 0.586 | 0.074259647 |
| Stum | 3.24E-09 | -0.042928851 | 0.339 | 0.372 | 8.73E-05 | Plppr1 | 2.80E-06 | -0.026147304 | 0.312 | 0.304 | 0.075289414 |
| Nceh1 | 3.29E-09 | 0.021977361 | 0.281 | 0.29 | 8.86E-05 | Mcc | 2.83E-06 | 0.026442199 | 0.312 | 0.288 | 0.076223471 |
| Nsg2 | 3.31E-09 | -0.039539056 | 0.479 | 0.516 | 8.91E-05 | Vps13c | 2.85E-06 | 0.07746741 | 0.633 | 0.603 | 0.076659026 |
| Larp1b | 3.34E-09 | 0.066678222 | 0.253 | 0.242 | 8.98E-05 | Sptan1 | 2.87E-06 | 0.049324419 | 0.758 | 0.729 | 0.077218632 |
| Nhsl2 | 3.34E-09 | -0.047014209 | 0.485 | 0.521 | 8.98E-05 | Slc9a7 | 2.90E-06 | -0.037978352 | 0.422 | 0.423 | 0.077897839 |
| Eml5 | 3.41E-09 | -0.046133986 | 0.774 | 0.807 | 9.19E-05 | Ptprr | 2.99E-06 | -0.016253374 | 0.491 | 0.478 | 0.080470525 |
| Caap1 | 3.63E-09 | -0.030352108 | 0.229 | 0.256 | 9.76E-05 | Shank1 | 3.02E-06 | -0.000820855 | 0.6 | 0.583 | 0.081333846 |
| Lingo2 | 3.75E-09 | 0.102666283 | 0.869 | 0.842 | 0.000101008 | Gm20404 | 3.03E-06 | 0.00688741 | 0.302 | 0.284 | 0.081490181 |
| Dcaf17 | 3.78E-09 | 0.021199345 | 0.29 | 0.3 | 0.000101813 | Smurf2 | 3.03E-06 | 0.028418533 | 0.4 | 0.374 | 0.081577674 |
| Tmod2 | 3.82E-09 | 0.04828863 | 0.481 | 0.481 | 0.000102718 | Ppp1r13b | 3.06E-06 | 0.04506249 | 0.644 | 0.613 | 0.082281874 |
| Arhgap12 | 3.85E-09 | 0.048747449 | 0.385 | 0.392 | 0.000103567 | Parm1 | 3.21E-06 | -0.064946844 | 0.281 | 0.287 | 0.086251075 |
| Faf1 | 3.94E-09 | 0.056197119 | 0.736 | 0.735 | 0.000106128 | Sfpq | 3.21E-06 | -0.018988485 | 0.711 | 0.696 | 0.086304358 |
| Immp2l | 3.97E-09 | 0.068661368 | 0.712 | 0.709 | 0.00010688 | Sorbs1 | 3.21E-06 | -0.049795195 | 0.84 | 0.838 | 0.08642378 |
| Epha10 | 4.05E-09 | -0.112076431 | 0.317 | 0.355 | 0.000108982 | Ppfia3 | 3.40E-06 | -0.011893281 | 0.289 | 0.28 | 0.091390378 |
| Ptk2 | 4.10E-09 | -0.07584129 | 0.846 | 0.865 | 0.00011033 | Tbc1d32 | 3.40E-06 | -0.031973571 | 0.27 | 0.27 | 0.091464589 |
| Camk1d | 4.17E-09 | 0.054966858 | 0.885 | 0.892 | 0.000112284 | Ylpm1 | 3.43E-06 | -0.033458838 | 0.649 | 0.643 | 0.092370425 |
| Ilf3 | 4.27E-09 | 0.020104471 | 0.301 | 0.309 | 0.000114824 | Xpr1 | 3.47E-06 | 0.007374151 | 0.504 | 0.482 | 0.093479377 |
| Ndufa10 | 4.40E-09 | -0.053241717 | 0.252 | 0.285 | 0.000118451 | Pibf1 | 3.53E-06 | -0.040637551 | 0.468 | 0.465 | 0.095045856 |
| Gm26699 | 4.51E-09 | -0.026603855 | 0.415 | 0.398 | 0.000121285 | Kcnmb4 | 3.62E-06 | 0.024391596 | 0.262 | 0.241 | 0.09733773 |
| 4931406P16Rik | 4.58E-09 | -0.005049247 | 0.234 | 0.252 | 0.000123218 | Cobl | 3.67E-06 | -0.071239071 | 0.285 | 0.289 | 0.098712749 |
| Snap25 | 4.64E-09 | -0.093874489 | 0.929 | 0.942 | 0.000124791 | Apobec4 | 3.69E-06 | 0.019925954 | 0.285 | 0.264 | 0.099171819 |
| Ppib | 4.72E-09 | -0.035553236 | 0.316 | 0.346 | 0.00012692 | Unc79 | 3.70E-06 | -0.040519882 | 0.851 | 0.844 | 0.099679287 |
| Lrrc8c | 4.75E-09 | 0.019325993 | 0.261 | 0.272 | 0.000127718 | Snx27 | 3.72E-06 | -0.011011681 | 0.352 | 0.339 | 0.099968325 |
| Gm45645 | 4.80E-09 | -0.048989421 | 0.23 | 0.261 | 0.000129173 | Usp31 | 3.83E-06 | 0.023040519 | 0.274 | 0.253 | 0.103141813 |
| Ttll5 | 4.95E-09 | -0.00751713 | 0.448 | 0.473 | 0.000133148 | Mgat4a | 3.85E-06 | -0.010970696 | 0.291 | 0.281 | 0.103590228 |
| Cit | 5.20E-09 | 0.108765842 | 0.261 | 0.228 | 0.000140026 | Rit2 | 3.92E-06 | 0.030636817 | 0.659 | 0.631 | 0.105465483 |
| Gm10419 | 5.30E-09 | 0.010316184 | 0.284 | 0.3 | 0.000142626 | Braf | 3.97E-06 | 0.010085105 | 0.714 | 0.691 | 0.106796273 |
| Pan3 | 5.39E-09 | 0.032245519 | 0.66 | 0.674 | 0.000144974 | Ppp3ca | 3.97E-06 | 0.054855452 | 0.982 | 0.985 | 0.106837739 |
| Ranbp9 | 5.52E-09 | -0.043642072 | 0.572 | 0.608 | 0.000148497 | Gm48091 | 4.02E-06 | 0.027757602 | 0.649 | 0.62 | 0.10803905 |
| Srgap2 | 5.64E-09 | -0.005198826 | 0.619 | 0.646 | 0.000151715 | Zmym6 | 4.09E-06 | -0.010481722 | 0.289 | 0.28 | 0.110039737 |
| Hipk2 | 5.69E-09 | 0.004744841 | 0.331 | 0.348 | 0.000153121 | Ubr3 | 4.14E-06 | -0.008216205 | 0.707 | 0.689 | 0.111329744 |
| Pkig | 5.70E-09 | 0.028783688 | 0.441 | 0.457 | 0.000153392 | Grid2 | 4.27E-06 | -0.065467016 | 0.682 | 0.691 | 0.114777659 |
| Cttn | 5.90E-09 | -0.000142767 | 0.283 | 0.301 | 0.000158886 | Adck1 | 4.40E-06 | -0.004948498 | 0.262 | 0.251 | 0.11836738 |
| Znrf3 | 5.94E-09 | 0.034588186 | 0.34 | 0.347 | 0.000159755 | Uck2 | 4.55E-06 | 0.030049528 | 0.262 | 0.239 | 0.12230941 |
| Ksr1 | 6.03E-09 | 0.003866106 | 0.582 | 0.606 | 0.000162217 | Phf20 | 4.70E-06 | 0.021201501 | 0.44 | 0.417 | 0.126430586 |
| Wscd2 | 6.06E-09 | 0.017310084 | 0.465 | 0.479 | 0.000163076 | Ppp1r12a | 4.75E-06 | 0.015320181 | 0.75 | 0.726 | 0.127895724 |
| Atxn10 | 6.19E-09 | -0.021047653 | 0.61 | 0.641 | 0.000166634 | Rock1 | 4.78E-06 | -0.035691488 | 0.659 | 0.655 | 0.128687216 |
| Hacd1 | 6.30E-09 | 0.0317902 | 0.256 | 0.259 | 0.000169596 | Kcnn2 | 4.79E-06 | 0.001334765 | 0.663 | 0.644 | 0.128874415 |

| | | | | | | | | | | | |
|---|---|---|---|---|---|---|---|---|---|---|---|
| Ogt | 6.37E-09 | -0.02518532 | 0.61 | 0.644 | 0.000171301 | Nsd2 | 4.85E-06 | -0.00890841 | 0.495 | 0.481 | 0.130573906 |
| Pcdh17 | 6.79E-09 | -0.088281039 | 0.425 | 0.464 | 0.000182755 | Ptpn3 | 4.88E-06 | 0.022523344 | 0.443 | 0.419 | 0.131425291 |
| Myo5a | 6.80E-09 | 0.058558291 | 0.651 | 0.649 | 0.000182848 | Trafd1 | 4.90E-06 | 0.028186989 | 0.298 | 0.274 | 0.131888083 |
| Rgs6 | 6.86E-09 | 0.131534562 | 0.282 | 0.281 | 0.000184712 | Gm5441 | 5.11E-06 | -0.011135127 | 0.363 | 0.349 | 0.137444233 |
| Adam11 | 7.30E-09 | 0.048947333 | 0.258 | 0.253 | 0.000196388 | A830082K12Rik | 5.27E-06 | -0.002552884 | 0.382 | 0.367 | 0.141751832 |
| Ppp1r21 | 7.35E-09 | -0.002414493 | 0.254 | 0.271 | 0.000197649 | Btrc | 5.38E-06 | 0.011708331 | 0.665 | 0.641 | 0.144678633 |
| Kif3b | 7.46E-09 | 0.022490695 | 0.274 | 0.282 | 0.000200812 | Akap6 | 5.57E-06 | -0.034093458 | 0.908 | 0.902 | 0.149740034 |
| Hace1 | 7.50E-09 | 0.044290369 | 0.506 | 0.51 | 0.00020186 | Bicd1 | 5.82E-06 | 0.035498887 | 0.746 | 0.718 | 0.156692441 |
| Mboat2 | 7.69E-09 | -0.00843523 | 0.526 | 0.553 | 0.000206785 | Pcsk2os2 | 5.92E-06 | 0.01716315 | 0.263 | 0.241 | 0.159405629 |
| Atp9b | 7.69E-09 | -0.019646217 | 0.718 | 0.749 | 0.000207002 | Gpr158 | 5.98E-06 | -0.047774323 | 0.862 | 0.864 | 0.160985339 |
| Sgsm2 | 7.83E-09 | 0.038061789 | 0.421 | 0.43 | 0.000210751 | Me3 | 6.03E-06 | -0.04764344 | 0.685 | 0.684 | 0.16218334 |
| Lrp8 | 8.06E-09 | -0.019995204 | 0.318 | 0.344 | 0.000216795 | Arid1b | 6.08E-06 | -0.041445062 | 0.797 | 0.793 | 0.163505099 |
| Cpeb2 | 8.68E-09 | -0.055108304 | 0.228 | 0.259 | 0.000233633 | Gabrg3 | 6.11E-06 | -0.057085355 | 0.593 | 0.584 | 0.164490815 |
| Ankrd6 | 8.68E-09 | 0.02936047 | 0.304 | 0.314 | 0.000233635 | Tsga10 | 6.19E-06 | 0.027199213 | 0.344 | 0.32 | 0.16665158 |
| Banf2 | 8.76E-09 | 0.081944065 | 0.325 | 0.292 | 0.000235795 | 4930545L23Rik | 6.20E-06 | -0.038303732 | 0.284 | 0.282 | 0.166942621 |
| Akap8l | 8.78E-09 | -0.035664685 | 0.772 | 0.804 | 0.000236292 | Kdm4b | 6.22E-06 | 0.020294521 | 0.35 | 0.329 | 0.167332781 |
| Ncor1 | 8.79E-09 | -0.044983475 | 0.735 | 0.769 | 0.000236587 | Larp1b | 6.25E-06 | -0.025414788 | 0.257 | 0.253 | 0.168090488 |
| Hdac8 | 8.94E-09 | 0.002666106 | 0.568 | 0.591 | 0.000240536 | Rtn4rl1 | 6.33E-06 | -0.011391214 | 0.379 | 0.366 | 0.170357094 |
| Gm38393 | 9.10E-09 | 0.02019245 | 0.302 | 0.312 | 0.000244931 | Ptprt | 6.43E-06 | -0.051647267 | 0.294 | 0.322 | 0.173144388 |
| Hspa12a | 9.11E-09 | -0.001371656 | 0.341 | 0.361 | 0.000245065 | Ldlrad4 | 6.44E-06 | 0.004812307 | 0.734 | 0.712 | 0.173412249 |
| Stag1 | 9.54E-09 | 0.076520894 | 0.828 | 0.821 | 0.000256729 | Ntng2 | 6.60E-06 | 0.003395035 | 0.414 | 0.396 | 0.177469901 |
| Cpeb3 | 9.58E-09 | -0.002458691 | 0.727 | 0.738 | 0.000257645 | Adgrl3 | 6.77E-06 | -0.067601705 | 0.949 | 0.954 | 0.182234636 |
| Agrn | 9.63E-09 | 0.0321252 | 0.34 | 0.346 | 0.000258982 | Rbm5 | 6.98E-06 | 0.026579337 | 0.68 | 0.653 | 0.187736258 |
| Itgbl1 | 9.76E-09 | -0.008879829 | 0.272 | 0.296 | 0.000262676 | Myh9 | 6.98E-06 | 0.03471451 | 0.295 | 0.271 | 0.18776295 |
| Fgd4 | 1.01E-08 | 0.003232736 | 0.534 | 0.556 | 0.000270649 | Khdrbs3 | 7.15E-06 | -0.10438607 | 0.64 | 0.666 | 0.192445797 |
| Rnf150 | 1.02E-08 | 0.048709403 | 0.684 | 0.691 | 0.000273371 | Agbl4 | 7.20E-06 | -0.085879846 | 0.724 | 0.742 | 0.193663639 |
| Bbs9 | 1.02E-08 | 0.023508412 | 0.337 | 0.347 | 0.000273576 | Rsf1os1 | 7.24E-06 | -0.000191342 | 0.344 | 0.329 | 0.194677344 |
| Dennd5b | 1.02E-08 | -0.006683252 | 0.538 | 0.563 | 0.000274356 | Srpk2 | 7.44E-06 | 0.047870881 | 0.745 | 0.717 | 0.200112817 |
| Crtac1 | 1.05E-08 | 0.016223608 | 0.451 | 0.467 | 0.000281679 | Huwe1 | 7.44E-06 | -0.008121941 | 0.685 | 0.667 | 0.200250649 |
| Trim9 | 1.09E-08 | -0.072215225 | 0.836 | 0.859 | 0.000293579 | Far2 | 7.46E-06 | 0.020049562 | 0.349 | 0.327 | 0.200669112 |
| Ptpn5 | 1.14E-08 | 0.036843801 | 0.352 | 0.36 | 0.000307625 | Chl1 | 7.46E-06 | 0.015878156 | 0.691 | 0.666 | 0.200695118 |
| Tomm40 | 1.16E-08 | -0.014591592 | 0.245 | 0.269 | 0.000313314 | Gm26883 | 7.46E-06 | -0.002500083 | 0.238 | 0.253 | 0.20073665 |
| Trank1 | 1.22E-08 | 0.036548031 | 0.724 | 0.737 | 0.000327817 | Ephb1 | 7.75E-06 | -0.028396321 | 0.321 | 0.316 | 0.208521846 |
| Sidt1 | 1.35E-08 | 0.050577824 | 0.654 | 0.648 | 0.000364163 | Etv1 | 7.81E-06 | 0.089342836 | 0.269 | 0.242 | 0.210040012 |
| C130071C03Rik | 1.37E-08 | 0.017310688 | 0.675 | 0.694 | 0.00036811 | Tmem132d | 8.04E-06 | -0.113766249 | 0.303 | 0.321 | 0.216309 |
| Zeb1 | 1.38E-08 | 0.031983885 | 0.605 | 0.613 | 0.000370752 | Zfp462 | 8.07E-06 | -0.07830187 | 0.333 | 0.352 | 0.217257203 |
| Spag9 | 1.39E-08 | -0.043906629 | 0.737 | 0.77 | 0.000374787 | Ankrd17 | 8.13E-06 | 0.008884041 | 0.687 | 0.664 | 0.218718902 |
| Mtcl1 | 1.46E-08 | 0.068626153 | 0.308 | 0.297 | 0.000394136 | Pi4ka | 8.20E-06 | 0.010982004 | 0.767 | 0.746 | 0.220558102 |
| Rtn4rl1 | 1.47E-08 | 0.006421623 | 0.366 | 0.385 | 0.000395514 | Pacrg | 8.24E-06 | -0.048330153 | 0.389 | 0.393 | 0.221712789 |
| Kcnq1ot1 | 1.47E-08 | -0.077154984 | 0.877 | 0.885 | 0.00039587 | Prkaa2 | 8.31E-06 | 0.03709566 | 0.263 | 0.238 | 0.223643022 |
| Fsd1l | 1.49E-08 | 0.013811873 | 0.256 | 0.267 | 0.000402253 | Spata7 | 8.34E-06 | 0.015731302 | 0.428 | 0.408 | 0.224316868 |
| Gm38604 | 1.50E-08 | -0.007317881 | 0.261 | 0.281 | 0.000404169 | Tsix | 8.34E-06 | -0.007692317 | 0.582 | 0.565 | 0.22448766 |
| 4933413L06Rik | 1.51E-08 | -0.115450851 | 0.278 | 0.309 | 0.000405614 | Plekha5 | 8.39E-06 | -0.055989233 | 0.847 | 0.856 | 0.225654707 |
| Arntl | 1.58E-08 | 0.001290201 | 0.3 | 0.316 | 0.000425364 | Gm28750 | 8.56E-06 | 0.029965428 | 0.349 | 0.326 | 0.230219401 |
| Arhgef9 | 1.60E-08 | -0.046799195 | 0.619 | 0.655 | 0.00042927 | Cntn3 | 9.06E-06 | 0.094232709 | 0.683 | 0.654 | 0.243667435 |
| Kctd1 | 1.66E-08 | 0.042815713 | 0.447 | 0.448 | 0.000445472 | Diaph2 | 9.08E-06 | -0.059975106 | 0.732 | 0.737 | 0.244208237 |
| Syn1 | 1.66E-08 | -0.010190196 | 0.509 | 0.533 | 0.000446653 | Chrm3 | 9.11E-06 | -0.046501235 | 0.488 | 0.516 | 0.245248687 |
| Pcnx | 1.70E-08 | -0.034756882 | 0.594 | 0.626 | 0.000457867 | Wdfy1 | 9.15E-06 | -0.003990375 | 0.252 | 0.241 | 0.24606754 |
| Gab2 | 1.76E-08 | 0.080526855 | 0.313 | 0.301 | 0.000472693 | Spag9 | 9.39E-06 | 0.019723015 | 0.761 | 0.737 | 0.252673811 |
| Mef2c | 1.76E-08 | 0.009537505 | 0.546 | 0.572 | 0.000474675 | Cds1 | 9.47E-06 | -0.00819366 | 0.256 | 0.247 | 0.25469025 |
| Epha7 | 1.80E-08 | -0.09603677 | 0.689 | 0.719 | 0.000483397 | Lrrc49 | 9.55E-06 | 0.013709691 | 0.395 | 0.375 | 0.257014609 |
| Stxbp5 | 1.82E-08 | -0.053612632 | 0.671 | 0.706 | 0.000490787 | Slc1a2 | 9.62E-06 | 0.001976894 | 0.563 | 0.545 | 0.258710919 |
| Gnai1 | 1.85E-08 | 0.017287589 | 0.33 | 0.339 | 0.000496852 | Chst9 | 9.76E-06 | 0.011725982 | 0.348 | 0.329 | 0.262485488 |
| Timm9 | 2.02E-08 | -0.000698281 | 0.311 | 0.328 | 0.000544454 | Exoc6b | 9.81E-06 | -0.035575473 | 0.803 | 0.8 | 0.263858658 |
| Rbm6 | 2.08E-08 | -0.005237506 | 0.723 | 0.751 | 0.000560086 | Ldb2 | 9.90E-06 | 0.095186946 | 0.503 | 0.49 | 0.266341346 |
| Syt14 | 2.09E-08 | 0.018721932 | 0.546 | 0.563 | 0.000561315 | Cog5 | 9.94E-06 | -0.016531053 | 0.617 | 0.607 | 0.267561995 |
| Cyth3 | 2.13E-08 | -0.006480715 | 0.278 | 0.296 | 0.000574136 | Sema6d | 1.01E-05 | -0.034835223 | 0.614 | 0.593 | 0.272945137 |
| Lyn | 2.18E-08 | -0.037527587 | 0.252 | 0.282 | 0.000586472 | A630089N07Rik | 1.03E-05 | -0.030846014 | 0.348 | 0.347 | 0.276815816 |
| Galnt14 | 2.19E-08 | -0.023148503 | 0.361 | 0.388 | 0.00058975 | Banf2 | 1.07E-05 | -0.070731828 | 0.3 | 0.325 | 0.287954198 |
| Nell1 | 2.28E-08 | 0.145272528 | 0.384 | 0.35 | 0.000613104 | Pitpnm2 | 1.08E-05 | -0.017240021 | 0.582 | 0.564 | 0.290877239 |
| Dip2a | 2.40E-08 | 0.057426955 | 0.286 | 0.28 | 0.000646318 | Pkn1 | 1.09E-05 | -0.007854748 | 0.256 | 0.244 | 0.293131238 |
| Zfp157 | 2.41E-08 | -0.024119304 | 0.248 | 0.272 | 0.000647803 | Anapc4 | 1.10E-05 | 0.004746525 | 0.265 | 0.251 | 0.295481553 |
| Crym | 2.53E-08 | -0.037770737 | 0.257 | 0.285 | 0.000681614 | Stim1 | 1.11E-05 | 0.024713284 | 0.368 | 0.346 | 0.298578397 |
| Msh3 | 2.60E-08 | 0.017326529 | 0.24 | 0.25 | 0.000700237 | Sgsm1 | 1.13E-05 | -0.03612272 | 0.338 | 0.341 | 0.305023437 |
| Rgs7bp | 2.69E-08 | -0.110239181 | 0.809 | 0.824 | 0.000722723 | Thsd7b | 1.18E-05 | -0.07810053 | 0.474 | 0.471 | 0.317706167 |
| Traf3ip2 | 2.71E-08 | 0.010525315 | 0.263 | 0.277 | 0.000730039 | Lrrc8c | 1.20E-05 | -0.047383926 | 0.253 | 0.261 | 0.321913974 |
| Cdk14 | 2.72E-08 | 0.060482004 | 0.901 | 0.893 | 0.00073069 | Eml1 | 1.21E-05 | 6.26E-05 | 0.342 | 0.328 | 0.324739722 |
| Ralgapa1 | 2.73E-08 | -0.091290832 | 0.788 | 0.817 | 0.000733545 | Snx13 | 1.21E-05 | 0.015929052 | 0.373 | 0.354 | 0.32485137 |
| Abca8b | 2.75E-08 | 0.039100943 | 0.279 | 0.279 | 0.000738638 | Epn2 | 1.23E-05 | 0.0186584 | 0.573 | 0.547 | 0.331581692 |
| 2010300C02Rik | 2.81E-08 | -0.077684556 | 0.704 | 0.738 | 0.000756378 | Tpp2 | 1.27E-05 | 0.031786338 | 0.349 | 0.325 | 0.340501723 |
| Zfp451 | 2.82E-08 | -0.025326196 | 0.282 | 0.308 | 0.000757547 | Dzank1 | 1.27E-05 | -0.035539856 | 0.35 | 0.35 | 0.340924928 |
| Arglu1 | 2.90E-08 | 0.095074577 | 0.677 | 0.656 | 0.000781361 | Pex5l | 1.31E-05 | -0.024910965 | 0.735 | 0.756 | 0.351633746 |
| Flrt3 | 2.94E-08 | -0.028564086 | 0.414 | 0.445 | 0.000790366 | Atp8a1 | 1.31E-05 | -0.03389978 | 0.795 | 0.789 | 0.351700394 |
| Filip1l | 2.96E-08 | -0.052159204 | 0.242 | 0.274 | 0.000796919 | Zfp609 | 1.31E-05 | 0.011434119 | 0.571 | 0.547 | 0.353568558 |
| Ldb2 | 2.98E-08 | 0.051687219 | 0.49 | 0.454 | 0.00080291 | Mllt10 | 1.31E-05 | -0.005296641 | 0.788 | 0.768 | 0.353790627 |
| Dtna | 3.09E-08 | -0.077373794 | 0.636 | 0.672 | 0.000830279 | Flrt3 | 1.34E-05 | 0.021740832 | 0.439 | 0.414 | 0.359631515 |
| Klhl29 | 3.10E-08 | 0.012099599 | 0.794 | 0.817 | 0.000835153 | Galnt7 | 1.34E-05 | -0.002592261 | 0.252 | 0.241 | 0.360688096 |
| Rp9 | 3.13E-08 | 0.013490295 | 0.332 | 0.345 | 0.000841006 | Cux1 | 1.37E-05 | -0.015656136 | 0.329 | 0.321 | 0.368907147 |
| Rnf25 | 3.19E-08 | -0.031667015 | 0.276 | 0.305 | 0.000856995 | Atxn10 | 1.45E-05 | 0.020941379 | 0.635 | 0.61 | 0.390130846 |
| Phf8 | 3.19E-08 | -0.016109526 | 0.252 | 0.274 | 0.000858899 | C2cd5 | 1.46E-05 | 0.013233464 | 0.685 | 0.66 | 0.392097913 |
| Ryr3 | 3.37E-08 | 0.084404152 | 0.846 | 0.836 | 0.000907734 | Tmem245 | 1.48E-05 | -0.03825626 | 0.321 | 0.324 | 0.397060802 |
| Nyap2 | 3.45E-08 | 0.071417757 | 0.426 | 0.415 | 0.000928584 | Taf3 | 1.49E-05 | -0.030347242 | 0.289 | 0.287 | 0.400556005 |
| Elavl3 | 3.51E-08 | 0.027375672 | 0.433 | 0.444 | 0.000944595 | Madd | 1.52E-05 | 0.033736002 | 0.508 | 0.48 | 0.407868575 |
| A230057D06Rik | 3.57E-08 | 0.055479234 | 0.755 | 0.752 | 0.000959782 | Mapk4 | 1.52E-05 | 0.043922001 | 0.434 | 0.408 | 0.408113437 |
| Frmd4b | 3.58E-08 | -0.081186198 | 0.466 | 0.503 | 0.000962597 | Gm20642 | 1.53E-05 | 0.021688936 | 0.908 | 0.89 | 0.412862767 |
| Zmynd8 | 3.68E-08 | -0.043023516 | 0.677 | 0.711 | 0.000990163 | 2900026A02Rik | 1.56E-05 | -0.010613512 | 0.348 | 0.338 | 0.420986845 |
| Rap1gds1 | 3.77E-08 | -0.036633539 | 0.58 | 0.611 | 0.001015552 | Inpp5b | 1.59E-05 | 0.005762518 | 0.279 | 0.263 | 0.428037725 |
| Kmt2c | 3.80E-08 | 0.025103452 | 0.685 | 0.697 | 0.00102119 | Scn8a | 1.66E-05 | -0.029425708 | 0.873 | 0.867 | 0.446156076 |
| Birc6 | 3.90E-08 | 0.030138183 | 0.723 | 0.736 | 0.001048538 | Rims1 | 1.70E-05 | -0.050993725 | 0.97 | 0.972 | 0.457437507 |
| Zmynd11 | 3.91E-08 | -0.023394449 | 0.673 | 0.705 | 0.001053274 | Kif21a | 1.76E-05 | 0.019258453 | 0.787 | 0.764 | 0.474339322 |
| Gm20687 | 4.02E-08 | 0.041948849 | 0.371 | 0.338 | 0.001080864 | Ppm1l | 1.78E-05 | 0.039127529 | 0.603 | 0.577 | 0.47909525 |
| Slc44a5 | 4.03E-08 | 0.083371722 | 0.615 | 0.604 | 0.001084656 | Strn3 | 1.79E-05 | 0.014467091 | 0.74 | 0.717 | 0.482929261 |
| Hap1 | 4.05E-08 | 0.016891612 | 0.244 | 0.258 | 0.001089736 | Thsd4 | 1.80E-05 | -0.033739036 | 0.267 | 0.263 | 0.484217797 |
| Gria2 | 4.06E-08 | -0.034146636 | 0.992 | 0.993 | 0.001092115 | Hace1 | 1.86E-05 | 0.006685308 | 0.526 | 0.506 | 0.499205173 |
| Trafd1 | 4.09E-08 | 0.053354791 | 0.274 | 0.266 | 0.001101057 | Wscd2 | 1.92E-05 | -0.007248761 | 0.479 | 0.465 | 0.515790601 |
| Cdh2 | 4.25E-08 | -0.041036847 | 0.757 | 0.789 | 0.001142759 | Gfod1 | 1.94E-05 | 0.000344148 | 0.668 | 0.647 | 0.521408406 |
| C2cd2 | 4.30E-08 | -0.079771216 | 0.306 | 0.341 | 0.001156864 | 9630028H03Rik | 1.96E-05 | -0.012530439 | 0.636 | 0.619 | 0.528720967 |
| Rapgef4os1 | 4.34E-08 | 0.002822601 | 0.367 | 0.389 | 0.001169067 | Smurf1 | 1.97E-05 | 0.000767953 | 0.326 | 0.311 | 0.530833845 |

| | | | | | | | | | | | |
|---|---|---|---|---|---|---|---|---|---|---|---|
| Sptan1 | 4.47E-08 | 0.033457364 | 0.729 | 0.737 | 0.001203739 | Eef2k | 1.98E-05 | 0.009061535 | 0.301 | 0.282 | 0.531437075 |
| Astn2 | 4.66E-08 | 0.139596846 | 0.561 | 0.542 | 0.001252778 | Sbf2 | 2.00E-05 | -0.028967369 | 0.775 | 0.764 | 0.537704128 |
| Slc6a17 | 4.69E-08 | -0.019708476 | 0.407 | 0.433 | 0.001261324 | Apba1 | 2.04E-05 | -0.024543703 | 0.743 | 0.733 | 0.547859628 |
| St5 | 4.71E-08 | -0.047959614 | 0.27 | 0.299 | 0.00126744 | Dtnbos | 2.05E-05 | -0.012526388 | 0.405 | 0.393 | 0.550432814 |
| Zfp608 | 4.92E-08 | -0.078058369 | 0.463 | 0.5 | 0.001324696 | Usp34 | 2.08E-05 | -0.000189982 | 0.829 | 0.811 | 0.559987661 |
| Vcl | 4.95E-08 | -0.078765115 | 0.248 | 0.282 | 0.001330555 | D130043K22Rik | 2.16E-05 | -0.006119868 | 0.419 | 0.406 | 0.580348639 |
| Gm15478 | 4.95E-08 | 0.026652765 | 0.323 | 0.337 | 0.001333018 | Tmed5 | 2.16E-05 | 0.033626909 | 0.253 | 0.231 | 0.581951568 |
| Vps13a | 5.15E-08 | 0.002255048 | 0.739 | 0.762 | 0.001384563 | Trpm3 | 2.19E-05 | -0.014110332 | 0.598 | 0.617 | 0.58933842 |
| Pfkp | 5.17E-08 | 0.019394554 | 0.398 | 0.408 | 0.001390615 | Cers6 | 2.20E-05 | -0.046779166 | 0.753 | 0.754 | 0.590917594 |
| Map3k4 | 5.33E-08 | -0.032326774 | 0.352 | 0.381 | 0.001435423 | Itpr1 | 2.21E-05 | 0.019782618 | 0.705 | 0.681 | 0.59536063 |
| Adam22 | 5.46E-08 | 0.036091302 | 0.763 | 0.773 | 0.001467795 | Sorbs2 | 2.25E-05 | -0.074957979 | 0.873 | 0.873 | 0.605132029 |
| Strn3 | 5.59E-08 | -0.011422187 | 0.717 | 0.745 | 0.001505033 | Slc35f4 | 2.26E-05 | -0.01970696 | 0.546 | 0.537 | 0.608243229 |
| Col4a1 | 5.61E-08 | -0.006027164 | 0.247 | 0.264 | 0.001509854 | Usp37 | 2.26E-05 | 0.036498077 | 0.449 | 0.421 | 0.608893302 |
| 4430402I18Rik | 5.65E-08 | 0.038034699 | 0.506 | 0.519 | 0.00152079 | Tjp1 | 2.29E-05 | -0.043148754 | 0.862 | 0.862 | 0.616562224 |
| Usp14 | 5.73E-08 | -0.014375502 | 0.291 | 0.313 | 0.00154083 | Ap3s1 | 2.32E-05 | 0.006704747 | 0.265 | 0.249 | 0.623010503 |
| Nsd3 | 5.81E-08 | -0.011447057 | 0.739 | 0.766 | 0.001563924 | Ppm1h | 2.36E-05 | 0.002413298 | 0.651 | 0.631 | 0.634735452 |
| Rtn1 | 6.12E-08 | 0.075360116 | 0.885 | 0.882 | 0.001647301 | Chd9 | 2.42E-05 | -0.002030835 | 0.818 | 0.801 | 0.651883874 |
| Arih1 | 6.18E-08 | 0.00072253 | 0.668 | 0.691 | 0.001663711 | Efr3b | 2.43E-05 | -0.044074934 | 0.376 | 0.375 | 0.653064208 |
| Megf9 | 6.52E-08 | 0.001908028 | 0.476 | 0.495 | 0.001754718 | Rev3l | 2.57E-05 | 0.040574473 | 0.67 | 0.643 | 0.691362882 |
| Ankrd17 | 6.77E-08 | -0.056031361 | 0.664 | 0.699 | 0.001820574 | Kmt2c | 2.64E-05 | -0.022505532 | 0.694 | 0.685 | 0.711396808 |
| Agtpbp1 | 7.04E-08 | -0.013310985 | 0.568 | 0.594 | 0.001893139 | Crym | 2.71E-05 | -0.029579612 | 0.259 | 0.257 | 0.729693807 |
| Desi1 | 7.04E-08 | -0.022707258 | 0.333 | 0.359 | 0.001893922 | Fut9 | 2.76E-05 | 0.070956865 | 0.651 | 0.623 | 0.743031909 |
| Arhgef12 | 7.04E-08 | 0.00734789 | 0.583 | 0.605 | 0.001894159 | Smg6 | 2.81E-05 | -0.01134811 | 0.766 | 0.751 | 0.755617674 |
| Diaph2 | 7.29E-08 | 0.088315657 | 0.737 | 0.726 | 0.001962342 | Pde7b | 2.84E-05 | 0.091948687 | 0.362 | 0.334 | 0.764179801 |
| Crmp1 | 7.44E-08 | -0.030892301 | 0.305 | 0.333 | 0.002001306 | Slc25a26 | 2.99E-05 | -0.020487561 | 0.262 | 0.258 | 0.80502133 |
| Ptpre | 7.55E-08 | 0.04843858 | 0.524 | 0.527 | 0.002030616 | Prkcb | 3.09E-05 | -0.03303254 | 0.783 | 0.776 | 0.830629447 |
| App | 7.78E-08 | 0.059339175 | 0.852 | 0.849 | 0.002093582 | 1700024B18Rik | 3.26E-05 | -0.021938989 | 0.376 | 0.367 | 0.878139231 |
| Arid1b | 8.13E-08 | -0.005103004 | 0.793 | 0.817 | 0.002187986 | Mpped2 | 3.30E-05 | -0.004385531 | 0.588 | 0.571 | 0.887859694 |
| Srrm4 | 8.17E-08 | 0.045912526 | 0.651 | 0.659 | 0.002197616 | Dpp8 | 3.39E-05 | 0.001868432 | 0.324 | 0.311 | 0.912652991 |
| Cttnbp2 | 8.62E-08 | -0.072006966 | 0.954 | 0.954 | 0.002318606 | Tcf12 | 3.54E-05 | 0.001130866 | 0.729 | 0.71 | 0.951661057 |
| Map2k6 | 8.70E-08 | 0.035494381 | 0.403 | 0.408 | 0.002340877 | Zfp831 | 3.56E-05 | -0.013394131 | 0.339 | 0.331 | 0.957717866 |
| Plppr5 | 8.84E-08 | 0.03056317 | 0.434 | 0.442 | 0.002378536 | Slit3 | 3.60E-05 | -0.06811638 | 0.928 | 0.932 | 0.968964485 |
| Glg1 | 9.48E-08 | -0.020780314 | 0.736 | 0.763 | 0.002552101 | Ksr2 | 3.61E-05 | 0.012095957 | 0.826 | 0.806 | 0.970739516 |
| R3hdm2 | 9.84E-08 | -0.101250018 | 0.672 | 0.703 | 0.002646823 | 4930419G24Rik | 3.62E-05 | 0.096895339 | 0.523 | 0.531 | 0.975283378 |
| Arhgap17 | 9.95E-08 | 0.001056269 | 0.427 | 0.447 | 0.002677534 | Arhgap39 | 3.78E-05 | -0.013378409 | 0.773 | 0.755 | 1 |
| Camkv | 1.02E-07 | -0.001965794 | 0.522 | 0.546 | 0.002738461 | Rps6ka3 | 3.79E-05 | -0.022909249 | 0.427 | 0.421 | 1 |
| Gm45740 | 1.12E-07 | -0.072304331 | 0.236 | 0.268 | 0.003012621 | Kcnip4 | 3.81E-05 | -0.068105953 | 0.94 | 0.952 | 1 |
| Pebp1 | 1.15E-07 | -0.02361739 | 0.303 | 0.331 | 0.003100343 | Dok6 | 3.85E-05 | -0.033359294 | 0.881 | 0.875 | 1 |
| A830082K12Rik | 1.17E-07 | 0.003020456 | 0.367 | 0.384 | 0.003160135 | Hs2st1 | 3.86E-05 | -0.006964998 | 0.592 | 0.579 | 1 |
| Hmgn3 | 1.18E-07 | -0.050938718 | 0.268 | 0.301 | 0.003184787 | Nup93 | 4.02E-05 | -0.013011439 | 0.45 | 0.441 | 1 |
| Sh3glb1 | 1.26E-07 | 0.005300984 | 0.24 | 0.255 | 0.003401668 | Dtna | 4.09E-05 | -0.01369993 | 0.654 | 0.636 | 1 |
| Fam208b | 1.27E-07 | 0.046835561 | 0.344 | 0.311 | 0.003418946 | Sfi1 | 4.10E-05 | 0.003167066 | 0.271 | 0.258 | 1 |
| Aifm3 | 1.27E-07 | 0.043423345 | 0.289 | 0.286 | 0.003425722 | Sdk2 | 4.21E-05 | -0.021795202 | 0.312 | 0.298 | 1 |
| Tnrc6b | 1.36E-07 | 0.044510269 | 0.798 | 0.805 | 0.003660842 | Ctps2 | 4.27E-05 | -0.023612138 | 0.282 | 0.279 | 1 |
| Ncam2 | 1.48E-07 | 0.1027489 | 0.781 | 0.766 | 0.003983042 | Ccdc88c | 4.27E-05 | -0.041494295 | 0.31 | 0.314 | 1 |
| Sntg1 | 1.54E-07 | -0.06566103 | 0.937 | 0.938 | 0.004155503 | Celf2 | 4.37E-05 | -0.022734316 | 0.997 | 0.998 | 1 |
| Plxdc2 | 1.57E-07 | 0.073651924 | 0.32 | 0.306 | 0.004225062 | Atg10 | 4.41E-05 | -0.024491931 | 0.305 | 0.303 | 1 |
| 4932438A13Rik | 1.60E-07 | -0.039838792 | 0.582 | 0.615 | 0.004291751 | Zfp451 | 4.52E-05 | 0.015879596 | 0.3 | 0.282 | 1 |
| Phlpp1 | 1.60E-07 | -0.011650759 | 0.605 | 0.633 | 0.00429889 | 4930402H24Rik | 4.57E-05 | -0.013674712 | 0.875 | 0.865 | 1 |
| Sfi1 | 1.65E-07 | 0.042602269 | 0.258 | 0.253 | 0.004440917 | A330102I10Rik | 4.62E-05 | 0.02990627 | 0.332 | 0.31 | 1 |
| Nnat | 1.66E-07 | -0.069690204 | 0.322 | 0.353 | 0.004462922 | Slc4a8 | 4.86E-05 | 0.002776513 | 0.289 | 0.275 | 1 |
| Prkcb | 1.66E-07 | 0.051676425 | 0.776 | 0.76 | 0.004479612 | Grin1 | 4.94E-05 | -0.009060474 | 0.89 | 0.878 | 1 |
| Xkr6 | 1.67E-07 | 0.030871282 | 0.63 | 0.645 | 0.004489057 | Cdh13 | 5.27E-05 | -0.136759792 | 0.379 | 0.396 | 1 |
| Homer2 | 1.69E-07 | 0.017340717 | 0.316 | 0.325 | 0.004534044 | Cntnap5b | 5.51E-05 | -0.070606427 | 0.517 | 0.533 | 1 |
| Gm42418 | 1.69E-07 | 0.116893921 | 0.933 | 0.924 | 0.004537404 | Lamc1 | 5.53E-05 | 0.010977108 | 0.342 | 0.324 | 1 |
| Kat6b | 1.74E-07 | 0.034512702 | 0.649 | 0.658 | 0.004679726 | Dock4 | 5.73E-05 | -0.047068479 | 0.903 | 0.908 | 1 |
| Htr4 | 1.75E-07 | -0.046256068 | 0.475 | 0.508 | 0.004714551 | Lrrc8d | 5.84E-05 | -0.020688588 | 0.332 | 0.326 | 1 |
| Mical3 | 1.78E-07 | 0.013815942 | 0.704 | 0.725 | 0.00478104 | Dnajc1 | 5.87E-05 | -0.036607971 | 0.78 | 0.775 | 1 |
| Ica1 | 1.82E-07 | -0.02144738 | 0.731 | 0.758 | 0.004904538 | Acyp2 | 6.21E-05 | -0.025339352 | 0.444 | 0.441 | 1 |
| Gm13883 | 1.85E-07 | -0.060490124 | 0.23 | 0.26 | 0.004983288 | Cnksr2 | 6.27E-05 | 0.057653128 | 0.93 | 0.924 | 1 |
| Avl9 | 1.88E-07 | -0.008444495 | 0.246 | 0.263 | 0.005065013 | Grid1 | 6.37E-05 | -0.045789175 | 0.883 | 0.881 | 1 |
| Tcf12 | 1.89E-07 | 0.050447241 | 0.71 | 0.712 | 0.005085572 | Gab2 | 6.37E-05 | -0.065998017 | 0.303 | 0.313 | 1 |
| Pcsk2os2 | 1.90E-07 | -0.036437022 | 0.241 | 0.267 | 0.005099728 | Ppip5k1 | 6.46E-05 | 0.032418625 | 0.473 | 0.448 | 1 |
| Cdkl5 | 1.92E-07 | 0.012362395 | 0.491 | 0.504 | 0.005159128 | Rgs7bp | 6.55E-05 | 0.017250403 | 0.83 | 0.809 | 1 |
| Nalcn | 2.08E-07 | 0.044563546 | 0.749 | 0.748 | 0.005587658 | Crebbp | 6.58E-05 | 0.029560328 | 0.713 | 0.688 | 1 |
| Dgkz | 2.12E-07 | -0.038612864 | 0.426 | 0.457 | 0.005695576 | Afg1l | 6.60E-05 | -0.005004233 | 0.295 | 0.284 | 1 |
| Ubn2 | 2.15E-07 | -0.023985541 | 0.772 | 0.799 | 0.005797628 | D17Wsu92e | 6.65E-05 | -0.049902875 | 0.225 | 0.25 | 1 |
| Map3k5 | 2.19E-07 | 0.0334872 | 0.315 | 0.314 | 0.005880514 | Tanc1 | 7.07E-05 | -0.070039547 | 0.623 | 0.628 | 1 |
| Snrpn | 2.19E-07 | -0.005105692 | 0.273 | 0.289 | 0.005901058 | Dennd1a | 7.26E-05 | -0.033334549 | 0.886 | 0.878 | 1 |
| Lingo1 | 2.26E-07 | 0.032175009 | 0.53 | 0.531 | 0.006079332 | Fbxl7 | 7.36E-05 | 0.053864867 | 0.32 | 0.295 | 1 |
| Cep78 | 2.26E-07 | -0.016496278 | 0.322 | 0.345 | 0.006091978 | Itgbl1 | 7.62E-05 | -0.005670269 | 0.285 | 0.272 | 1 |
| Luc7l2 | 2.38E-07 | 0.060176101 | 0.918 | 0.921 | 0.006402581 | Itga8 | 7.87E-05 | 0.050258235 | 0.616 | 0.589 | 1 |
| 4933406I18Rik | 2.42E-07 | -0.004995297 | 0.492 | 0.515 | 0.006507232 | Rpgr | 7.93E-05 | 0.014012629 | 0.26 | 0.243 | 1 |
| Miat | 2.43E-07 | 0.052189103 | 0.715 | 0.719 | 0.006527494 | Gm48383 | 8.43E-05 | -0.00279795 | 0.349 | 0.337 | 1 |
| Mad1l1 | 2.43E-07 | -0.068050293 | 0.251 | 0.283 | 0.00654614 | Adgrb3 | 8.45E-05 | 0.043089509 | 0.989 | 0.99 | 1 |
| Baiap2 | 2.65E-07 | -0.003130799 | 0.643 | 0.668 | 0.007119601 | Akap13 | 8.45E-05 | -0.044046067 | 0.275 | 0.277 | 1 |
| Ncoa1 | 2.69E-07 | 0.044937379 | 0.809 | 0.818 | 0.007244337 | Ocrl | 8.48E-05 | -0.00612249 | 0.409 | 0.399 | 1 |
| Bicd1 | 2.73E-07 | -0.006189657 | 0.718 | 0.744 | 0.007343478 | A530046M15Rik | 8.49E-05 | -0.040634459 | 0.287 | 0.289 | 1 |
| Rsu1 | 2.75E-07 | 0.032520987 | 0.266 | 0.266 | 0.007407204 | Lrp1b | 8.50E-05 | 0.006289775 | 0.942 | 0.952 | 1 |
| Clasp2 | 2.91E-07 | 0.049322064 | 0.749 | 0.751 | 0.007824348 | Zfp407 | 8.64E-05 | -0.026850214 | 0.497 | 0.491 | 1 |
| Amph | 2.92E-07 | -0.048191548 | 0.746 | 0.776 | 0.007860749 | Gm30054 | 8.71E-05 | 0.027894081 | 0.306 | 0.284 | 1 |
| Klhl24 | 2.95E-07 | 0.030867636 | 0.252 | 0.253 | 0.007934159 | Fam208b | 8.95E-05 | -0.049127074 | 0.318 | 0.344 | 1 |
| Dtnbos | 2.97E-07 | -0.03202071 | 0.393 | 0.422 | 0.007980947 | Kalrn | 9.00E-05 | -0.010075742 | 0.99 | 0.995 | 1 |
| Slc38a2 | 3.02E-07 | 0.032764497 | 0.316 | 0.321 | 0.008132156 | Dgki | 9.35E-05 | -0.062091921 | 0.921 | 0.92 | 1 |
| Synpr | 3.05E-07 | -0.106019967 | 0.452 | 0.465 | 0.008216352 | Garnl3 | 9.65E-05 | -0.064298629 | 0.324 | 0.338 | 1 |
| Grm3 | 3.19E-07 | 0.088694418 | 0.365 | 0.334 | 0.008575334 | Gm36975 | 0.00010091 | 0.013689653 | 0.793 | 0.771 | 1 |
| Ano4 | 3.21E-07 | 0.091031825 | 0.44 | 0.424 | 0.008634086 | Vcl | 0.000105267 | 0.030538572 | 0.27 | 0.248 | 1 |
| Ptpn3 | 3.21E-07 | -0.025166989 | 0.419 | 0.443 | 0.008634878 | Runx2 | 0.000105498 | -0.019379409 | 0.66 | 0.648 | 1 |
| Wsb1 | 3.30E-07 | -0.043190156 | 0.268 | 0.298 | 0.008889921 | Tmeff2 | 0.000105765 | -0.047146862 | 0.861 | 0.861 | 1 |
| Gng2 | 3.39E-07 | -0.032303293 | 0.497 | 0.527 | 0.00913072 | Rasgef1b | 0.000107012 | -0.074326319 | 0.43 | 0.453 | 1 |
| Sympk | 3.64E-07 | 0.019343511 | 0.338 | 0.346 | 0.009797101 | Trps1 | 0.000110263 | -0.064544794 | 0.316 | 0.324 | 1 |
| Btbd9 | 3.70E-07 | -0.057258296 | 0.856 | 0.876 | 0.009968431 | Arhgef9 | 0.000110824 | 0.023582913 | 0.643 | 0.619 | 1 |
| Atrx | 3.72E-07 | 0.020488101 | 0.731 | 0.748 | 0.010010247 | Sorcs3 | 0.000110851 | -0.073060763 | 0.46 | 0.475 | 1 |
| Satb1 | 3.92E-07 | -0.011166113 | 0.241 | 0.261 | 0.010535676 | Rspry1 | 0.000112887 | 0.012817092 | 0.281 | 0.265 | 1 |
| Csmd3 | 4.06E-07 | 0.056167189 | 0.968 | 0.966 | 0.010917076 | Angptl2 | 0.000131694 | 0.018137141 | 0.309 | 0.29 | 1 |
| Synj1 | 4.13E-07 | -0.00912202 | 0.594 | 0.619 | 0.011100599 | Syt17 | 0.000134549 | 0.003271619 | 0.392 | 0.377 | 1 |
| Csrnp3 | 4.13E-07 | -0.025375506 | 0.71 | 0.739 | 0.011116823 | Tenm4 | 0.000136041 | -0.071738892 | 0.736 | 0.745 | 1 |
| Fam81a | 4.39E-07 | -0.013323459 | 0.247 | 0.266 | 0.011813924 | Mtcl1 | 0.000138616 | -0.034079926 | 0.308 | 0.308 | 1 |

| | | | | | | | | | | | |
|---|---|---|---|---|---|---|---|---|---|---|---|
| Osbp2 | 4.48E-07 | 0.020991211 | 0.65 | 0.665 | 0.012060206 | Rasgrf2 | 0.00013998 | -0.056935421 | 0.349 | 0.355 | 1 |
| Sgsm1 | 4.53E-07 | 0.038773124 | 0.341 | 0.34 | 0.012196627 | Camta1 | 0.000140716 | -0.028737461 | 0.839 | 0.833 | 1 |
| Lyst | 4.62E-07 | -0.068524932 | 0.657 | 0.689 | 0.012435718 | Fam120c | 0.000143704 | 0.001527203 | 0.457 | 0.442 | 1 |
| Erc2 | 4.88E-07 | -0.145042123 | 0.982 | 0.976 | 0.013138612 | Slc4a10 | 0.000151231 | 0.00515908 | 0.825 | 0.806 | 1 |
| Vti1a | 5.06E-07 | 0.027063482 | 0.747 | 0.759 | 0.013627282 | Edil3 | 0.000156479 | 0.100750764 | 0.605 | 0.584 | 1 |
| Ccdc50 | 5.29E-07 | -0.021949242 | 0.273 | 0.296 | 0.014227591 | Maml3 | 0.000158598 | 0.004954978 | 0.409 | 0.39 | 1 |
| Tbc1d1 | 5.46E-07 | 0.05968873 | 0.428 | 0.408 | 0.014696146 | Capzb | 0.000159271 | -0.018387254 | 0.296 | 0.293 | 1 |
| Gramd1b | 5.46E-07 | 0.060112683 | 0.657 | 0.653 | 0.014700679 | Mpped1 | 0.000167736 | 0.039448904 | 0.336 | 0.312 | 1 |
| Man1a | 5.60E-07 | -0.129558503 | 0.341 | 0.348 | 0.015058653 | Ahi1 | 0.000169811 | 0.03075524 | 0.983 | 0.982 | 1 |
| Acyp2 | 5.65E-07 | 0.020853618 | 0.441 | 0.45 | 0.015202379 | Gm16599 | 0.000169966 | -0.017286135 | 0.263 | 0.255 | 1 |
| Shisa6 | 5.71E-07 | -0.090834961 | 0.742 | 0.766 | 0.015355621 | Ano4 | 0.00017239 | -0.071992999 | 0.425 | 0.44 | 1 |
| Mlxip | 5.79E-07 | 0.034868743 | 0.252 | 0.252 | 0.015580717 | Erc1 | 0.000174552 | 0.012220004 | 0.67 | 0.651 | 1 |
| Cdyl2 | 5.87E-07 | -0.029978707 | 0.229 | 0.252 | 0.015784643 | Cyp7b1 | 0.000174822 | -0.019365108 | 0.474 | 0.459 | 1 |
| Rasgrp1 | 5.92E-07 | -0.018092084 | 0.393 | 0.417 | 0.015917491 | Abr | 0.000177287 | -0.028802903 | 0.776 | 0.769 | 1 |
| Camk2a | 6.10E-07 | -0.077812052 | 0.966 | 0.969 | 0.016407621 | Sfxn5 | 0.000178302 | -0.03202564 | 0.309 | 0.309 | 1 |
| Serpini1 | 6.20E-07 | -0.014362386 | 0.335 | 0.358 | 0.016687141 | Frmd4b | 0.000183375 | -0.032869726 | 0.479 | 0.466 | 1 |
| Mcc | 6.29E-07 | 0.060368374 | 0.288 | 0.287 | 0.016916379 | Pcdh17 | 0.000183516 | 0.003714148 | 0.442 | 0.425 | 1 |
| Myo1d | 6.89E-07 | 0.028302993 | 0.263 | 0.266 | 0.018552317 | Sgcd | 0.00018756 | -0.069229711 | 0.574 | 0.592 | 1 |
| Cacna1c | 6.99E-07 | -0.072775367 | 0.922 | 0.923 | 0.01880053 | Cblb | 0.000190545 | -0.034914724 | 0.46 | 0.459 | 1 |
| 4932443L11Rik | 7.37E-07 | 0.009978369 | 0.25 | 0.262 | 0.019825675 | Herc1 | 0.000195104 | -0.016929488 | 0.784 | 0.773 | 1 |
| Adgra1 | 8.01E-07 | 0.002541319 | 0.256 | 0.272 | 0.021539186 | Gm1992 | 0.000200738 | 0.005108182 | 0.499 | 0.482 | 1 |
| Ust | 8.14E-07 | -0.040879236 | 0.546 | 0.575 | 0.02189155 | Lsamp | 0.000202041 | -0.082852948 | 0.988 | 0.99 | 1 |
| Ripor2 | 8.22E-07 | -0.006093008 | 0.54 | 0.558 | 0.02211822 | Tenm2 | 0.000206896 | 0.031375534 | 0.993 | 0.992 | 1 |
| Herc1 | 8.24E-07 | 0.038195321 | 0.773 | 0.777 | 0.022175086 | Sertad2 | 0.000210605 | 0.041801464 | 0.315 | 0.291 | 1 |
| Cntn3 | 8.84E-07 | 0.039284429 | 0.654 | 0.631 | 0.023793655 | Psme4 | 0.000216051 | -0.0100184 | 0.63 | 0.617 | 1 |
| Adcy2 | 9.77E-07 | 0.066656257 | 0.825 | 0.816 | 0.026286864 | Sel1l3 | 0.000216469 | 0.01608004 | 0.39 | 0.372 | 1 |
| Add3 | 9.81E-07 | 0.030305019 | 0.404 | 0.408 | 0.026396737 | Mctp1 | 0.000223048 | -0.076818081 | 0.763 | 0.776 | 1 |
| Asap2 | 9.97E-07 | -0.066489844 | 0.401 | 0.433 | 0.026832252 | Slc4a4 | 0.000223105 | 0.024563105 | 0.476 | 0.45 | 1 |
| Ppip5k2 | 1.01E-06 | -0.014177436 | 0.236 | 0.254 | 0.02728572 | 2900055J20Rik | 0.000226672 | 0.059525898 | 0.428 | 0.413 | 1 |
| Ppp4r4 | 1.01E-06 | -0.037484856 | 0.405 | 0.433 | 0.027303471 | Tbl1xr1 | 0.000233813 | 0.001587272 | 0.415 | 0.401 | 1 |
| Gm1992 | 1.19E-06 | -0.014793845 | 0.482 | 0.505 | 0.032139295 | Phf21a | 0.000238526 | 0.023222241 | 0.773 | 0.751 | 1 |
| Grip1 | 1.21E-06 | -0.040767608 | 0.675 | 0.702 | 0.032553805 | Gabrb1 | 0.000251187 | -0.040447696 | 0.959 | 0.955 | 1 |
| Kcnk2 | 1.22E-06 | -0.084650624 | 0.25 | 0.279 | 0.032844774 | Mpdz | 0.00025293 | -0.012413519 | 0.414 | 0.406 | 1 |
| Igsf9b | 1.23E-06 | -0.08271026 | 0.277 | 0.308 | 0.033157176 | Pde1a | 0.000269649 | -0.039293339 | 0.619 | 0.642 | 1 |
| Rimbp2 | 1.27E-06 | -0.027850855 | 0.742 | 0.769 | 0.034153583 | Macrod2os1 | 0.000271901 | 0.033432858 | 0.415 | 0.392 | 1 |
| Dner | 1.37E-06 | 0.077933668 | 0.309 | 0.291 | 0.036874403 | 4930567K12Rik | 0.000282286 | -0.002362206 | 0.279 | 0.267 | 1 |
| Spred2 | 1.38E-06 | -0.053350351 | 0.616 | 0.649 | 0.037234813 | A230006K03Rik | 0.000288455 | -0.057278647 | 0.584 | 0.598 | 1 |
| Cacna2d3 | 1.41E-06 | 0.012823231 | 0.838 | 0.86 | 0.037945108 | Nexmif | 0.000292133 | -0.016021355 | 0.494 | 0.484 | 1 |
| Tanc2 | 1.56E-06 | 0.050732589 | 0.931 | 0.924 | 0.04200972 | Foxn3 | 0.000293197 | 0.042705233 | 0.559 | 0.535 | 1 |
| Senp5 | 1.61E-06 | 0.01613117 | 0.411 | 0.42 | 0.043276537 | Unc80 | 0.000295334 | -0.027428741 | 0.891 | 0.884 | 1 |
| Sgk1 | 1.64E-06 | 0.032540453 | 0.313 | 0.318 | 0.04415705 | Ralgapa1 | 0.000299331 | -0.010036819 | 0.801 | 0.788 | 1 |
| Chn1 | 1.65E-06 | -0.007748064 | 0.845 | 0.868 | 0.044367171 | Sphkap | 0.000301251 | 0.023246443 | 0.784 | 0.762 | 1 |
| Sgip1 | 1.73E-06 | 0.051303216 | 0.889 | 0.89 | 0.046447455 | Ppp1r12b | 0.00030622 | -0.02513685 | 0.707 | 0.699 | 1 |
| Ftx | 1.78E-06 | -0.000781564 | 0.747 | 0.772 | 0.047909936 | Spink10 | 0.000309864 | 0.037699729 | 0.262 | 0.242 | 1 |
| Kalrn | 1.80E-06 | 0.040122003 | 0.995 | 0.991 | 0.048380451 | Znrf3 | 0.000310095 | -0.025807597 | 0.339 | 0.34 | 1 |
| Lrrc7 | 1.82E-06 | -0.053125895 | 0.988 | 0.986 | 0.049015416 | Ank3 | 0.000320817 | -0.016123138 | 0.984 | 0.99 | 1 |
| A330102I10Rik | 2.02E-06 | 0.048797121 | 0.31 | 0.303 | 0.054247962 | Cntn5 | 0.00032302 | -0.113763306 | 0.7 | 0.721 | 1 |
| Galc | 2.05E-06 | 0.030413192 | 0.263 | 0.264 | 0.055165167 | Sh3rf1 | 0.000329104 | -0.073042495 | 0.438 | 0.451 | 1 |
| Mtdh | 2.24E-06 | -0.01716076 | 0.583 | 0.61 | 0.060303459 | Gm26699 | 0.000336767 | 0.054479381 | 0.428 | 0.415 | 1 |
| Zfp804a | 2.48E-06 | -0.130609118 | 0.591 | 0.622 | 0.06676138 | Mbnl2 | 0.000339709 | 0.040767623 | 0.87 | 0.854 | 1 |
| Cadm1 | 2.73E-06 | 0.041331486 | 0.93 | 0.921 | 0.073532725 | Cacna1e | 0.000346075 | -0.040461627 | 0.948 | 0.952 | 1 |
| Leng8 | 2.74E-06 | 0.004986869 | 0.389 | 0.408 | 0.073674709 | Macrod1 | 0.000348678 | 0.046942635 | 0.303 | 0.28 | 1 |
| Mctp1 | 2.80E-06 | 0.049890847 | 0.776 | 0.751 | 0.075260661 | Hmgcll1 | 0.000349047 | -0.032647501 | 0.364 | 0.366 | 1 |
| Smpd4 | 3.20E-06 | -0.014837622 | 0.276 | 0.296 | 0.086213317 | Ptk2 | 0.000350053 | -0.021353926 | 0.856 | 0.846 | 1 |
| Lamp2 | 3.29E-06 | -0.003493861 | 0.253 | 0.268 | 0.088438141 | Brinp1 | 0.000351481 | 0.039504535 | 0.918 | 0.908 | 1 |
| Pde10a | 3.30E-06 | -0.101864181 | 0.882 | 0.866 | 0.088887438 | Ssbp2 | 0.000352265 | 0.000886425 | 0.779 | 0.763 | 1 |
| Zfp462 | 3.51E-06 | 0.075841838 | 0.352 | 0.338 | 0.094409717 | Exoc4 | 0.000353969 | -0.010459444 | 0.878 | 0.866 | 1 |
| Gm49353 | 3.85E-06 | 0.05544151 | 0.331 | 0.301 | 0.103657016 | Cog4 | 0.000354195 | 0.005199056 | 0.274 | 0.261 | 1 |
| Usp34 | 4.06E-06 | 0.030827819 | 0.811 | 0.82 | 0.10933849 | Gnai1 | 0.000356736 | 0.02038549 | 0.347 | 0.33 | 1 |
| Frmd5 | 4.21E-06 | -0.06757626 | 0.649 | 0.679 | 0.113323395 | Jam3 | 0.000362805 | 0.044916295 | 0.32 | 0.297 | 1 |
| Sel1l3 | 4.27E-06 | 0.032028705 | 0.372 | 0.374 | 0.114918268 | Gm16183 | 0.000375427 | -0.001947142 | 0.403 | 0.39 | 1 |
| Prex2 | 4.28E-06 | -0.065487247 | 0.242 | 0.269 | 0.115220907 | Tshz3 | 0.000379675 | -0.048736279 | 0.334 | 0.329 | 1 |
| Stmn2 | 4.49E-06 | 0.067221841 | 0.31 | 0.291 | 0.120746821 | Gm6994 | 0.000395852 | 0.008948694 | 0.322 | 0.307 | 1 |
| Ints6l | 4.75E-06 | -0.010209651 | 0.299 | 0.318 | 0.127833458 | Togaram1 | 0.000399593 | -0.004585848 | 0.295 | 0.285 | 1 |
| Ak5 | 4.79E-06 | 0.070868546 | 0.817 | 0.812 | 0.12883254 | Nfia | 0.000408022 | -0.047462202 | 0.599 | 0.593 | 1 |
| Dpp6 | 4.93E-06 | -0.062163649 | 0.956 | 0.95 | 0.132760064 | Arhgap12 | 0.000412574 | -0.012023674 | 0.395 | 0.385 | 1 |
| Gm15738 | 5.00E-06 | -0.048706538 | 0.819 | 0.843 | 0.134416617 | Camk4 | 0.00045475 | -0.0506664 | 0.557 | 0.565 | 1 |
| Snx32 | 5.23E-06 | 0.0054209 | 0.279 | 0.291 | 0.140684156 | Htr4 | 0.000456613 | 0.031842221 | 0.498 | 0.475 | 1 |
| Ncam1 | 5.45E-06 | 0.064427177 | 0.884 | 0.883 | 0.146666435 | Gpm6a | 0.000471703 | -0.000389453 | 0.871 | 0.859 | 1 |
| Agap1 | 6.51E-06 | 0.024311959 | 0.679 | 0.695 | 0.175229495 | Gm20754 | 0.00050351 | 0.028451912 | 0.747 | 0.759 | 1 |
| Gm16105 | 7.03E-06 | -0.042247412 | 0.297 | 0.325 | 0.189130335 | Gm10848 | 0.000520716 | 0.007226896 | 0.469 | 0.449 | 1 |
| Kansl1 | 7.54E-06 | 0.024876721 | 0.816 | 0.828 | 0.202791546 | Dlg2 | 0.000522494 | -0.01340321 | 0.995 | 0.998 | 1 |
| Epha4 | 7.70E-06 | -0.029063246 | 0.739 | 0.766 | 0.207263902 | Epha6 | 0.000548035 | -0.023670875 | 0.9 | 0.914 | 1 |
| Ctnnd2 | 7.74E-06 | -0.036172479 | 0.967 | 0.972 | 0.208140968 | Ank2 | 0.000581844 | -0.036418909 | 0.987 | 0.989 | 1 |
| Ephb1 | 7.88E-06 | -0.023003099 | 0.316 | 0.339 | 0.211898419 | Man1a | 0.000626646 | 0.109069758 | 0.36 | 0.341 | 1 |
| Aatk | 7.92E-06 | 0.00682996 | 0.26 | 0.272 | 0.21298587 | Add2 | 0.000659594 | -0.045424733 | 0.787 | 0.783 | 1 |
| Jakmip2 | 9.20E-06 | -0.02226914 | 0.694 | 0.718 | 0.247580497 | Tmem132b | 0.000683978 | -0.031533685 | 0.821 | 0.817 | 1 |
| Cobl | 1.01E-05 | 0.062416716 | 0.289 | 0.285 | 0.272535821 | Gm44511 | 0.000692478 | -0.028276472 | 0.414 | 0.436 | 1 |
| Selenow | 1.05E-05 | -0.011705717 | 0.247 | 0.265 | 0.281943883 | Hap1 | 0.000695666 | -0.008057567 | 0.253 | 0.244 | 1 |
| Lmo7 | 1.06E-05 | -0.035716362 | 0.555 | 0.582 | 0.285835873 | Col25a1 | 0.000707221 | -0.051355145 | 0.356 | 0.357 | 1 |
| Rps6ka3 | 1.11E-05 | 0.028088739 | 0.421 | 0.423 | 0.297513331 | Rock2 | 0.000716482 | -0.004690433 | 0.7 | 0.685 | 1 |
| Rmnd1 | 1.12E-05 | -0.030276553 | 0.231 | 0.253 | 0.30031611 | Mbp | 0.000721461 | 0.036197169 | 0.33 | 0.308 | 1 |
| Nlk | 1.12E-05 | 0.010853021 | 0.642 | 0.656 | 0.300412478 | Ppargc1a | 0.000746967 | 0.004263718 | 0.346 | 0.333 | 1 |
| Grina | 1.12E-05 | -0.04158666 | 0.299 | 0.328 | 0.301876178 | Fam189a1 | 0.000752836 | -0.039444906 | 0.792 | 0.798 | 1 |
| Arid5b | 1.43E-05 | 0.01261344 | 0.401 | 0.412 | 0.383966679 | Rnf130 | 0.000756125 | 0.010864285 | 0.377 | 0.361 | 1 |
| Peak1 | 1.50E-05 | -0.053337694 | 0.458 | 0.483 | 0.402950495 | Stox2 | 0.000823972 | -0.038809532 | 0.723 | 0.721 | 1 |
| Scn8a | 1.58E-05 | 0.046751081 | 0.867 | 0.867 | 0.425541262 | Kcnt2 | 0.000847036 | 0.033371855 | 0.663 | 0.641 | 1 |
| Mpped1 | 1.63E-05 | 0.048679131 | 0.312 | 0.309 | 0.437613742 | Ndst4 | 0.000897863 | 0.064571719 | 0.32 | 0.298 | 1 |
| Ptprg | 1.66E-05 | -0.034496278 | 0.748 | 0.77 | 0.446114262 | Gm2164 | 0.000913779 | 0.05776605 | 0.333 | 0.31 | 1 |
| Kcnc2 | 1.68E-05 | 0.024055142 | 0.335 | 0.308 | 0.451173179 | Jmjd1c | 0.00093815 | 0.016790014 | 0.797 | 0.778 | 1 |
| Unc79 | 1.69E-05 | -0.001321073 | 0.844 | 0.864 | 0.453427888 | Adarb2 | 0.000942999 | 0.026461382 | 0.396 | 0.392 | 1 |
| Pitpnc1 | 1.72E-05 | -0.031172281 | 0.368 | 0.393 | 0.462446453 | Pak7 | 0.000963125 | 0.010539669 | 0.685 | 0.665 | 1 |
| Gpm6a | 1.77E-05 | -0.057078103 | 0.859 | 0.874 | 0.476988485 | Nebl | 0.000984187 | -0.054607634 | 0.874 | 0.876 | 1 |
| Msra | 1.90E-05 | -0.049281423 | 0.874 | 0.893 | 0.511671423 | Dkk3 | 0.000984963 | 0.034721203 | 0.398 | 0.376 | 1 |
| Rsf1os2 | 2.10E-05 | 0.008937975 | 0.242 | 0.251 | 0.565052916 | Adk | 0.000985981 | -0.037846174 | 0.49 | 0.488 | 1 |
| Lims1 | 2.14E-05 | 0.051959279 | 0.307 | 0.296 | 0.574489635 | 4930532I03Rik | 0.001006082 | 0.043093754 | 0.423 | 0.399 | 1 |
| Plcb1 | 2.23E-05 | -0.098063151 | 0.965 | 0.963 | 0.600681985 | Smyd3 | 0.001006735 | -0.001635412 | 0.74 | 0.727 | 1 |
| Fstl4 | 2.27E-05 | 0.022948072 | 0.586 | 0.603 | 0.609736787 | Gabra2 | 0.001049472 | 0.036731339 | 0.822 | 0.804 | 1 |

| | | | | | | | | | | | |
|---|---|---|---|---|---|---|---|---|---|---|---|
| Syndig1 | 2.35E-05 | 0.008482675 | 0.352 | 0.361 | 0.632860889 | Kif5a | 0.001097735 | 0.001977661 | 0.402 | 0.39 | 1 |
| Srgap3 | 2.41E-05 | 0.04415055 | 0.886 | 0.885 | 0.64907487 | Bicc1 | 0.001099827 | 0.029868054 | 0.259 | 0.24 | 1 |
| Psap | 2.45E-05 | -0.009614316 | 0.408 | 0.434 | 0.660461599 | Wdr17 | 0.001149237 | -0.025187011 | 0.536 | 0.522 | 1 |
| Matk | 2.66E-05 | 0.010150023 | 0.549 | 0.567 | 0.715219234 | Adgrl2 | 0.001155122 | -0.051086246 | 0.282 | 0.304 | 1 |
| Sntb2 | 2.87E-05 | -0.079967153 | 0.562 | 0.591 | 0.771186404 | Arfgef3 | 0.001184786 | 0.014754765 | 0.641 | 0.621 | 1 |
| Pid1 | 2.98E-05 | 0.086031426 | 0.405 | 0.378 | 0.80251941 | Adcy2 | 0.001208596 | -0.022456721 | 0.835 | 0.825 | 1 |
| Ptprj | 3.06E-05 | -0.085475111 | 0.795 | 0.816 | 0.82290716 | Baiap2 | 0.001244447 | -0.001226701 | 0.659 | 0.643 | 1 |
| Itpr1 | 3.37E-05 | -0.080402165 | 0.681 | 0.706 | 0.906418661 | Chn1 | 0.001274102 | 0.029827627 | 0.861 | 0.845 | 1 |
| Asap1 | 3.55E-05 | 0.053104921 | 0.606 | 0.582 | 0.954223478 | Il1rap | 0.00131027 | 0.031576999 | 0.618 | 0.595 | 1 |
| Gm48749 | 3.66E-05 | 0.084095593 | 0.283 | 0.264 | 0.983460625 | Cntn4 | 0.001400449 | -0.058028001 | 0.474 | 0.497 | 1 |
| Parm1 | 4.05E-05 | -0.030831535 | 0.287 | 0.308 | 1 | Meis2 | 0.001401147 | -0.066986835 | 0.325 | 0.327 | 1 |
| Pde1a | 4.29E-05 | -0.122039566 | 0.642 | 0.639 | 1 | Naaladl2 | 0.00143503 | -0.047337706 | 0.504 | 0.511 | 1 |
| Plekhg5 | 4.33E-05 | -0.009313331 | 0.627 | 0.651 | 1 | Hs6st2 | 0.001440005 | 0.015636105 | 0.32 | 0.305 | 1 |
| Tmem108 | 4.42E-05 | 0.096011792 | 0.699 | 0.69 | 1 | Cpeb3 | 0.001453196 | 0.009173041 | 0.743 | 0.727 | 1 |
| Arap2 | 4.57E-05 | -0.086079711 | 0.311 | 0.309 | 1 | Cntnap5a | 0.001465233 | -0.015892489 | 0.625 | 0.647 | 1 |
| Khdrbs3 | 4.57E-05 | -0.011327977 | 0.666 | 0.641 | 1 | Nyap2 | 0.001488416 | 0.025174661 | 0.444 | 0.426 | 1 |
| Cdh13 | 4.76E-05 | -0.022717163 | 0.396 | 0.423 | 1 | Immp2l | 0.001494872 | -0.00014027 | 0.725 | 0.712 | 1 |
| Ptchd4 | 5.04E-05 | -0.000513075 | 0.437 | 0.458 | 1 | Nedd4l | 0.00151493 | 0.00796212 | 0.854 | 0.838 | 1 |
| Gm15952 | 5.29E-05 | 0.006314239 | 0.264 | 0.275 | 1 | Wwox | 0.001523892 | -0.018568427 | 0.53 | 0.519 | 1 |
| Kcnn2 | 5.35E-05 | -0.129388818 | 0.644 | 0.669 | 1 | Tiam1 | 0.001534679 | -0.061791741 | 0.646 | 0.652 | 1 |
| Sertad2 | 5.42E-05 | -0.017429919 | 0.291 | 0.31 | 1 | Magi2 | 0.001547719 | -0.039041413 | 0.984 | 0.987 | 1 |
| Frmd6 | 5.91E-05 | -0.091543432 | 0.265 | 0.288 | 1 | Elavl4 | 0.001553929 | 0.047068498 | 0.319 | 0.297 | 1 |
| Fnbp1l | 6.09E-05 | -0.089592004 | 0.252 | 0.274 | 1 | Man2a1 | 0.00161676 | 0.055327757 | 0.331 | 0.309 | 1 |
| Rab8b | 6.11E-05 | 0.037108165 | 0.292 | 0.292 | 1 | Maml2 | 0.001635783 | -0.122170489 | 0.478 | 0.478 | 1 |
| Usp3 | 6.13E-05 | -0.005314675 | 0.311 | 0.329 | 1 | Ica1 | 0.001686532 | 0.001223224 | 0.746 | 0.731 | 1 |
| Grik2 | 6.29E-05 | 0.065166362 | 0.959 | 0.958 | 1 | Atp2b2 | 0.001709285 | 0.020879847 | 0.862 | 0.845 | 1 |
| Runx2 | 6.63E-05 | 0.018996618 | 0.648 | 0.661 | 1 | Fry | 0.001716115 | 0.026103883 | 0.903 | 0.89 | 1 |
| Runx1t1 | 6.69E-05 | 0.029159792 | 0.515 | 0.489 | 1 | Kirrel3 | 0.001725722 | -0.050252375 | 0.706 | 0.726 | 1 |
| Slc2a13 | 7.38E-05 | -0.044368656 | 0.679 | 0.704 | 1 | Kcnj6 | 0.001748904 | -0.040229336 | 0.842 | 0.847 | 1 |
| Syn3 | 7.91E-05 | -0.004179266 | 0.559 | 0.58 | 1 | Dcc | 0.001749059 | 0.013953386 | 0.744 | 0.754 | 1 |
| Mirg | 8.30E-05 | -0.00988947 | 0.354 | 0.371 | 1 | Cdc42bpa | 0.001765368 | -0.027171289 | 0.862 | 0.859 | 1 |
| Prr16 | 9.19E-05 | 0.057323055 | 0.391 | 0.376 | 1 | Gcn1l1 | 0.001800362 | -0.039025919 | 0.249 | 0.269 | 1 |
| Pitpnm2 | 9.21E-05 | 0.032616547 | 0.564 | 0.575 | 1 | Npas2 | 0.00199759 | -0.003045445 | 0.447 | 0.439 | 1 |
| Ssh2 | 9.67E-05 | 0.007065167 | 0.779 | 0.794 | 1 | Lrfn5 | 0.002050617 | 0.044362051 | 0.955 | 0.955 | 1 |
| Slc7a14 | 0.000103205 | -0.022375307 | 0.652 | 0.675 | 1 | C2cd2 | 0.002089817 | -0.001138671 | 0.317 | 0.306 | 1 |
| Nrg3 | 0.000104773 | -0.046414703 | 0.997 | 0.997 | 1 | Ptprj | 0.002139295 | -0.02044318 | 0.805 | 0.795 | 1 |
| Sik3 | 0.000111714 | -0.038446234 | 0.712 | 0.733 | 1 | Kat6b | 0.002153016 | 0.01570591 | 0.669 | 0.649 | 1 |
| Nfat5 | 0.000112576 | 0.010093095 | 0.626 | 0.64 | 1 | Papd4 | 0.002259818 | -0.046326488 | 0.308 | 0.328 | 1 |
| Ank3 | 0.000115774 | -0.022185239 | 0.99 | 0.985 | 1 | Setbp1 | 0.002274254 | -0.033898396 | 0.913 | 0.915 | 1 |
| Kcnip3 | 0.000116283 | -0.043953776 | 0.532 | 0.557 | 1 | Epha5 | 0.002327422 | 0.060913137 | 0.906 | 0.894 | 1 |
| Nol4 | 0.000119493 | -0.003614571 | 0.706 | 0.72 | 1 | Ntm | 0.00234511 | 0.016921593 | 0.65 | 0.666 | 1 |
| Lsamp | 0.00012541 | 0.087857595 | 0.99 | 0.989 | 1 | Babam2 | 0.00239004 | -0.027229617 | 0.562 | 0.561 | 1 |
| Nsf | 0.00013429 | 0.047294261 | 0.88 | 0.883 | 1 | Hacd1 | 0.002390891 | -0.000916567 | 0.262 | 0.256 | 1 |
| Kif26b | 0.000137243 | -0.02618888 | 0.351 | 0.373 | 1 | Cdk14 | 0.002484352 | -0.007525932 | 0.912 | 0.901 | 1 |
| Vps13b | 0.000138265 | 0.018298475 | 0.814 | 0.825 | 1 | Dlgap1 | 0.00264483 | -0.027602258 | 0.986 | 0.991 | 1 |
| Gnal | 0.000140039 | 0.00131489 | 0.347 | 0.359 | 1 | Zfp804b | 0.002686608 | -0.122356055 | 0.358 | 0.37 | 1 |
| Negr1 | 0.000140883 | 0.088412029 | 0.956 | 0.954 | 1 | 2810403A07Rik | 0.002697598 | -0.04080221 | 0.26 | 0.278 | 1 |
| Plcl1 | 0.0001432 | -0.012297184 | 0.493 | 0.519 | 1 | Sh3d19 | 0.00275654 | 0.044526952 | 0.459 | 0.438 | 1 |
| 5330434G04Rik | 0.000147165 | -0.015930152 | 0.651 | 0.668 | 1 | Diaph1 | 0.002882848 | -0.001324445 | 0.258 | 0.249 | 1 |
| Jam3 | 0.000158455 | -0.022525175 | 0.297 | 0.317 | 1 | Rab3c | 0.002895321 | -0.036861202 | 0.474 | 0.476 | 1 |
| Kcnd2 | 0.000159992 | 0.087613733 | 0.985 | 0.983 | 1 | Ctnna3 | 0.003007882 | 0.048505133 | 0.576 | 0.554 | 1 |
| Dnm3 | 0.000161808 | -0.030552835 | 0.538 | 0.56 | 1 | Chst8 | 0.003102123 | 0.011703394 | 0.253 | 0.239 | 1 |
| Kcnab1 | 0.0001639 | 0.04406218 | 0.745 | 0.733 | 1 | Adamts17 | 0.00315116 | 0.010139432 | 0.447 | 0.43 | 1 |
| Nrg1 | 0.000166476 | 0.10599422 | 0.794 | 0.793 | 1 | Rims2 | 0.003203235 | -0.007813492 | 0.873 | 0.868 | 1 |
| Sik2 | 0.000167068 | -0.109503028 | 0.426 | 0.452 | 1 | Dmd | 0.003231929 | -0.047432829 | 0.916 | 0.921 | 1 |
| Ndst4 | 0.000167526 | 0.057460599 | 0.298 | 0.274 | 1 | Auts2 | 0.003234146 | -0.02945232 | 0.972 | 0.977 | 1 |
| Celf1 | 0.000169047 | -0.005157566 | 0.739 | 0.758 | 1 | Trpc5 | 0.003387572 | 0.011222334 | 0.541 | 0.525 | 1 |
| Tmem191c | 0.000170488 | 0.071873324 | 0.544 | 0.528 | 1 | St6galnac3 | 0.003431494 | 0.018782009 | 0.768 | 0.751 | 1 |
| Gm21798 | 0.000172289 | -0.100326429 | 0.287 | 0.308 | 1 | Dip2b | 0.003452541 | 0.033062586 | 0.679 | 0.659 | 1 |
| A230006K03Rik | 0.000174371 | 0.057730884 | 0.598 | 0.597 | 1 | Ttc3 | 0.00366324 | -0.042324609 | 0.933 | 0.937 | 1 |
| Zmat4 | 0.000184768 | -0.028591013 | 0.696 | 0.72 | 1 | Kcnh7 | 0.003771805 | -0.039699846 | 0.736 | 0.745 | 1 |
| Rbm4b | 0.000185547 | -0.015897125 | 0.35 | 0.372 | 1 | Nckap5 | 0.003853746 | 0.025611084 | 0.363 | 0.348 | 1 |
| Alcam | 0.000194126 | 0.046985572 | 0.48 | 0.471 | 1 | Ccser1 | 0.004088622 | -0.033477965 | 0.963 | 0.961 | 1 |
| Mettl23 | 0.000194643 | 0.042394278 | 0.304 | 0.28 | 1 | Khdrbs2 | 0.004123187 | 0.063042791 | 0.787 | 0.772 | 1 |
| Rab3c | 0.000217329 | 0.001676096 | 0.476 | 0.485 | 1 | Fam135b | 0.004197842 | -0.025858706 | 0.877 | 0.867 | 1 |
| Cyp7b1 | 0.000225331 | -0.091415519 | 0.459 | 0.483 | 1 | Akt3 | 0.004356792 | -0.005528692 | 0.892 | 0.883 | 1 |
| Sh3kbp1 | 0.000252371 | -0.085864678 | 0.333 | 0.358 | 1 | Pced1b | 0.004365899 | 0.014470434 | 0.319 | 0.307 | 1 |
| Sorbs2os | 0.000255949 | -0.013032387 | 0.747 | 0.766 | 1 | Prkcg | 0.004411141 | 0.007025604 | 0.623 | 0.613 | 1 |
| Limch1 | 0.000259221 | -0.031969739 | 0.649 | 0.672 | 1 | Arl15 | 0.004420022 | 0.024494751 | 0.782 | 0.764 | 1 |
| Sgcz | 0.000260843 | 0.08952315 | 0.479 | 0.452 | 1 | Grm7 | 0.004652907 | -0.010738533 | 0.986 | 0.99 | 1 |
| Cacnb2 | 0.000262331 | -0.101240413 | 0.896 | 0.901 | 1 | Hecw1 | 0.004711865 | 0.054404858 | 0.791 | 0.777 | 1 |
| Gm30054 | 0.000268464 | -0.039007443 | 0.284 | 0.306 | 1 | Slc2a13 | 0.004830011 | 0.021243799 | 0.697 | 0.679 | 1 |
| Ppp1r12a | 0.000268913 | -0.019887972 | 0.726 | 0.746 | 1 | Dip2c | 0.005422307 | -0.038340971 | 0.929 | 0.93 | 1 |
| Atg16l2 | 0.000271941 | 0.04075229 | 0.258 | 0.25 | 1 | Sorcs1 | 0.005438788 | -0.035755764 | 0.243 | 0.261 | 1 |
| Dscam | 0.000275254 | -0.04793666 | 0.901 | 0.899 | 1 | Snap25 | 0.005583464 | 0.040709463 | 0.938 | 0.929 | 1 |
| Hs3st4 | 0.000279969 | -0.091721697 | 0.591 | 0.616 | 1 | Eif4g3 | 0.005629739 | 0.01169989 | 0.789 | 0.773 | 1 |
| Nfib | 0.000299702 | -0.043631085 | 0.807 | 0.827 | 1 | Cntn1 | 0.005672662 | -0.029945241 | 0.902 | 0.901 | 1 |
| Rasgrf2 | 0.000304139 | 0.060882447 | 0.355 | 0.347 | 1 | A830018L16Rik | 0.005675843 | -0.04769909 | 0.742 | 0.747 | 1 |
| Dock9 | 0.000317066 | 0.007248287 | 0.728 | 0.739 | 1 | Tenm1 | 0.005699955 | -0.050419639 | 0.677 | 0.683 | 1 |
| Samd5 | 0.000348586 | 0.07344178 | 0.352 | 0.331 | 1 | Eml5 | 0.006063167 | 0.008546971 | 0.79 | 0.774 | 1 |
| Slc4a10 | 0.000388815 | -0.012571878 | 0.806 | 0.824 | 1 | Sdccag3 | 0.006141415 | -0.043765263 | 0.266 | 0.282 | 1 |
| Sgcd | 0.000398046 | 0.109246233 | 0.592 | 0.58 | 1 | Stxbp6 | 0.006166784 | 0.064701276 | 0.395 | 0.375 | 1 |
| Ppp1r13b | 0.000410295 | -0.02128376 | 0.613 | 0.634 | 1 | Atrnl1 | 0.006200368 | -0.043464644 | 0.846 | 0.85 | 1 |
| Gm26518 | 0.000412995 | -0.000912461 | 0.323 | 0.309 | 1 | Cdh18 | 0.006731987 | -0.07709164 | 0.462 | 0.458 | 1 |
| Sema3e | 0.000422756 | -0.071301838 | 0.305 | 0.328 | 1 | Enah | 0.006732256 | 0.031951949 | 0.778 | 0.761 | 1 |
| Map1b | 0.000446507 | -0.065862856 | 0.907 | 0.908 | 1 | Morn1 | 0.006793109 | 0.006241402 | 0.26 | 0.251 | 1 |
| Dlgap2 | 0.000459033 | 0.029878341 | 0.98 | 0.975 | 1 | Snrpn | 0.007207807 | 0.003887019 | 0.279 | 0.273 | 1 |
| Ptk2b | 0.000466881 | -0.027505348 | 0.702 | 0.723 | 1 | Ppp1r9a | 0.007237873 | -0.039016038 | 0.818 | 0.817 | 1 |
| Gm11099 | 0.000469728 | 0.057920077 | 0.302 | 0.29 | 1 | Frmpd4 | 0.007369168 | -0.03430301 | 0.933 | 0.923 | 1 |
| Gns | 0.000483347 | 0.030310203 | 0.253 | 0.249 | 1 | Adgra1 | 0.007501567 | -0.007516379 | 0.263 | 0.256 | 1 |
| Maml2 | 0.000483739 | -0.012851579 | 0.478 | 0.503 | 1 | Nkain3 | 0.00785149 | -0.052803939 | 0.654 | 0.664 | 1 |
| Rbfox2 | 0.000485425 | 0.00946181 | 0.846 | 0.86 | 1 | Grm8 | 0.008029185 | -0.06426186 | 0.388 | 0.403 | 1 |
| Gm26854 | 0.000486906 | 0.022945157 | 0.278 | 0.278 | 1 | Nrg2 | 0.008082579 | -0.031717967 | 0.541 | 0.536 | 1 |
| Dock4 | 0.000496351 | -0.071370288 | 0.908 | 0.914 | 1 | Sema5a | 0.008566407 | 0.022074972 | 0.391 | 0.373 | 1 |
| Carmil1 | 0.000540975 | -0.073571866 | 0.451 | 0.476 | 1 | Hecw2 | 0.008793683 | 0.086701261 | 0.69 | 0.675 | 1 |
| Gria1 | 0.000551925 | -0.052190892 | 0.933 | 0.926 | 1 | Osbpl1a | 0.009073393 | 0.036027803 | 0.374 | 0.356 | 1 |
| Cdc42bpa | 0.000627151 | -0.000814655 | 0.859 | 0.873 | 1 | Fnbp1l | 0.009546944 | 0.041857355 | 0.268 | 0.252 | 1 |
| Tmem200a | 0.000663714 | 0.061224175 | 0.295 | 0.278 | 1 | Trhde | 0.009801541 | -0.029866325 | 0.403 | 0.414 | 1 |

| | | | | | | | | | | | |
|---|---|---|---|---|---|---|---|---|---|---|---|
| Caln1 | 0.000670547 | 0.024149744 | 0.812 | 0.798 | 1 | Samd5 | 0.009819259 | 0.059954656 | 0.368 | 0.352 | 1 |
| Dst | 0.000683973 | -0.001195446 | 0.857 | 0.871 | 1 | Bcl11b | 0.009884192 | 0.013980553 | 0.552 | 0.536 | 1 |
| Adgrl2 | 0.000714656 | 0.056054334 | 0.304 | 0.282 | 1 | Gnal | 0.011225862 | -0.005561099 | 0.354 | 0.347 | 1 |
| Prkce | 0.000729762 | -0.055257585 | 0.967 | 0.965 | 1 | Gm21954 | 0.011658833 | 0.004931819 | 0.485 | 0.474 | 1 |
| Ppm1l | 0.000739432 | -0.051759442 | 0.577 | 0.597 | 1 | Dgkh | 0.011704595 | 0.067301716 | 0.684 | 0.67 | 1 |
| Ryr2 | 0.000769169 | -0.06222781 | 0.975 | 0.974 | 1 | Gphn | 0.013052475 | -0.034900183 | 0.928 | 0.931 | 1 |
| Chsy3 | 0.000818762 | 0.053881578 | 0.603 | 0.606 | 1 | Syn2 | 0.01321426 | 0.003137449 | 0.892 | 0.881 | 1 |
| Pde4dip | 0.000836955 | 0.024980535 | 0.849 | 0.854 | 1 | Mettl23 | 0.013290334 | 0.007951289 | 0.296 | 0.304 | 1 |
| Lrrtm3 | 0.000837766 | -0.048426699 | 0.447 | 0.47 | 1 | Lrrc7 | 0.013710088 | -0.015484272 | 0.985 | 0.988 | 1 |
| Med12l | 0.000858662 | 0.003487039 | 0.718 | 0.73 | 1 | Rapgef2 | 0.013792903 | 0.007959344 | 0.838 | 0.824 | 1 |
| Rsrp1 | 0.000876871 | 0.072140214 | 0.716 | 0.707 | 1 | Sox5 | 0.014775342 | -0.020309439 | 0.414 | 0.432 | 1 |
| Meis2 | 0.000880317 | 0.09237557 | 0.327 | 0.309 | 1 | Syne1 | 0.014859936 | -0.002069373 | 0.89 | 0.879 | 1 |
| Sh3rf1 | 0.000882383 | 0.051092321 | 0.451 | 0.446 | 1 | Grin2b | 0.015281042 | 0.01901689 | 0.985 | 0.988 | 1 |
| Rit2 | 0.000902546 | -0.038500694 | 0.631 | 0.654 | 1 | Iqgap2 | 0.015407901 | -0.053023939 | 0.634 | 0.635 | 1 |
| Slit1 | 0.000905004 | -0.056108475 | 0.701 | 0.723 | 1 | Nell2 | 0.015646641 | -0.035108133 | 0.923 | 0.922 | 1 |
| Cap2 | 0.000944121 | 0.046342681 | 0.752 | 0.748 | 1 | Tbl1x | 0.016075253 | -0.011203644 | 0.289 | 0.286 | 1 |
| Gm26836 | 0.001036858 | -0.044832492 | 0.332 | 0.338 | 1 | Sipa1l1 | 0.016173319 | -0.039037162 | 0.864 | 0.866 | 1 |
| Tshz2 | 0.001095969 | 0.000890704 | 0.273 | 0.264 | 1 | Pid1 | 0.017991458 | -0.058887592 | 0.391 | 0.405 | 1 |
| Dock10 | 0.001164685 | -0.065393686 | 0.441 | 0.459 | 1 | 5730522E02Rik | 0.018120903 | 0.028073868 | 0.901 | 0.89 | 1 |
| Gabrb3 | 0.001298304 | 0.02972212 | 0.98 | 0.977 | 1 | Cpne8 | 0.018136811 | 0.057040752 | 0.287 | 0.273 | 1 |
| Phf21a | 0.001329274 | 0.013559267 | 0.751 | 0.763 | 1 | Cacnb2 | 0.018562979 | -0.043452266 | 0.905 | 0.896 | 1 |
| Sez6l | 0.001405145 | -0.020612446 | 0.795 | 0.812 | 1 | Pde4dip | 0.019949053 | 0.003853313 | 0.861 | 0.849 | 1 |
| Gabrb2 | 0.001437127 | 0.022911324 | 0.752 | 0.759 | 1 | Cacna1a | 0.021638477 | -0.016327268 | 0.885 | 0.878 | 1 |
| 4930517O19Rik | 0.001526667 | 0.017981748 | 0.302 | 0.306 | 1 | Stmn2 | 0.022007478 | -0.015545602 | 0.313 | 0.31 | 1 |
| Cdh8 | 0.001602187 | -0.022057008 | 0.757 | 0.773 | 1 | Slc35f3 | 0.022437367 | 0.073801247 | 0.535 | 0.531 | 1 |
| Syt1 | 0.001630473 | 0.004405508 | 0.993 | 0.989 | 1 | Csmd3 | 0.02315235 | -0.014333236 | 0.962 | 0.968 | 1 |
| Acap2 | 0.00181673 | -0.025589067 | 0.786 | 0.805 | 1 | Stag1 | 0.02329524 | 0.012737233 | 0.841 | 0.828 | 1 |
| Gm10754 | 0.00190907 | 0.0411668 | 0.266 | 0.246 | 1 | Ntng1 | 0.023508835 | 0.05772632 | 0.58 | 0.563 | 1 |
| Pde7b | 0.001923792 | -0.021896277 | 0.334 | 0.349 | 1 | Kcnd2 | 0.023559012 | -0.023461783 | 0.983 | 0.985 | 1 |
| Pip5k1b | 0.002093226 | 0.020889626 | 0.678 | 0.689 | 1 | Ryr2 | 0.024022494 | -0.024661808 | 0.97 | 0.975 | 1 |
| Galnt18 | 0.002152748 | 0.00962026 | 0.817 | 0.824 | 1 | Hs3st4 | 0.025071855 | -0.048423331 | 0.581 | 0.591 | 1 |
| Zufsp | 0.002223149 | 0.032141244 | 0.254 | 0.235 | 1 | Thsd7a | 0.025684426 | 0.06863356 | 0.449 | 0.444 | 1 |
| Nell2 | 0.002313965 | 0.033914801 | 0.922 | 0.921 | 1 | Atp2b1 | 0.025748128 | 0.055462785 | 0.935 | 0.933 | 1 |
| Chl1 | 0.002518425 | -0.034738114 | 0.666 | 0.687 | 1 | Dock10 | 0.026114685 | 0.030541026 | 0.446 | 0.441 | 1 |
| Stox2 | 0.002752499 | 0.037557593 | 0.721 | 0.729 | 1 | St6galnac5 | 0.027504895 | -0.015550222 | 0.889 | 0.884 | 1 |
| Ahcyl2 | 0.00280482 | -0.075708709 | 0.839 | 0.849 | 1 | Ext1 | 0.031248634 | 0.069216192 | 0.78 | 0.772 | 1 |
| Sipa1l1 | 0.002899983 | 0.011522426 | 0.866 | 0.877 | 1 | Grm5 | 0.031416913 | -0.002260453 | 0.986 | 0.99 | 1 |
| Tmem126a | 0.002983952 | -0.015493666 | 0.253 | 0.272 | 1 | Syn3 | 0.031421684 | -0.00566928 | 0.568 | 0.559 | 1 |
| Ptprt | 0.003057953 | -0.009805377 | 0.322 | 0.307 | 1 | Plxna4 | 0.033247549 | -0.039999182 | 0.902 | 0.908 | 1 |
| Cpe | 0.003085264 | -0.058058981 | 0.675 | 0.695 | 1 | AC165271.1 | 0.037923755 | -0.030934107 | 0.306 | 0.316 | 1 |
| Exoc4 | 0.003216398 | -0.031334291 | 0.866 | 0.879 | 1 | Igsf9b | 0.039531833 | 0.006152242 | 0.285 | 0.277 | 1 |
| Nrcam | 0.003240626 | -0.052739508 | 0.94 | 0.942 | 1 | 4930509J09Rik | 0.040400887 | 0.040145979 | 0.668 | 0.657 | 1 |
| Jmjd1c | 0.003425799 | -0.028571703 | 0.778 | 0.796 | 1 | St8sia5 | 0.044144249 | 0.002991842 | 0.266 | 0.259 | 1 |
| Fbxl7 | 0.003598628 | -0.088738557 | 0.295 | 0.315 | 1 | Ube2e2 | 0.044253568 | -0.029818614 | 0.954 | 0.958 | 1 |
| Cep112 | 0.003874456 | 0.009898058 | 0.503 | 0.51 | 1 | Ggact | 0.04455189 | 0.028831137 | 0.448 | 0.432 | 1 |
| Gm48321 | 0.003976401 | 0.046421648 | 0.286 | 0.274 | 1 | Gm10754 | 0.044656212 | -0.031554886 | 0.252 | 0.266 | 1 |
| A830018L16Rik | 0.004267708 | -0.053894044 | 0.747 | 0.744 | 1 | Phactr1 | 0.045611433 | -0.021140159 | 0.969 | 0.971 | 1 |
| Dlc1 | 0.004361516 | 0.094635007 | 0.325 | 0.308 | 1 | Gm26561 | 0.046996778 | -0.022399241 | 0.269 | 0.282 | 1 |
| Dgkh | 0.00446484 | -0.00347716 | 0.67 | 0.69 | 1 | Fndc9 | 0.047153008 | 0.015864478 | 0.576 | 0.56 | 1 |
| Gabbr2 | 0.004775334 | 0.015449427 | 0.871 | 0.877 | 1 | Gabrb3 | 0.048862926 | -0.018055034 | 0.979 | 0.98 | 1 |
| Thsd4 | 0.005689229 | -0.011293323 | 0.263 | 0.274 | 1 | Pcsk2 | 0.048980953 | 0.009596957 | 0.894 | 0.885 | 1 |
| 4930509J09Rik | 0.00571845 | 0.019857595 | 0.657 | 0.67 | 1 | Atxn1 | 0.049445684 | -0.031993479 | 0.934 | 0.938 | 1 |
| Pex5l | 0.005820541 | 0.010005969 | 0.756 | 0.768 | 1 | Pcsk5 | 0.049782269 | -0.005001221 | 0.431 | 0.418 | 1 |
| Tcf4 | 0.00584564 | -0.026788039 | 0.966 | 0.971 | 1 | Abhd15 | 0.051722357 | -0.033632595 | 0.256 | 0.267 | 1 |
| St8sia5 | 0.006009955 | 0.020440819 | 0.259 | 0.258 | 1 | Gm29237 | 0.057171204 | -0.034430936 | 0.405 | 0.415 | 1 |
| Rtn3 | 0.006469559 | 0.002982834 | 0.887 | 0.898 | 1 | Sema3e | 0.057699292 | 0.026150903 | 0.317 | 0.305 | 1 |
| Grb14 | 0.007017493 | -0.017239268 | 0.311 | 0.323 | 1 | Gm26518 | 0.060246577 | 0.028213779 | 0.327 | 0.323 | 1 |
| Sdccag3 | 0.007452258 | -0.022635884 | 0.282 | 0.282 | 1 | Ndst3 | 0.06024918 | 0.016924341 | 0.377 | 0.368 | 1 |
| Auts2 | 0.00825075 | 0.014197929 | 0.977 | 0.972 | 1 | Cadm2 | 0.060966501 | 0.003387813 | 0.97 | 0.974 | 1 |
| Pak3 | 0.008367239 | -0.002462955 | 0.764 | 0.774 | 1 | Gm37459 | 0.062300635 | 0.069956434 | 0.395 | 0.394 | 1 |
| Mlip | 0.008414769 | -0.016910579 | 0.423 | 0.404 | 1 | Etl4 | 0.065555804 | -0.032524518 | 0.57 | 0.582 | 1 |
| Oprm1 | 0.008420027 | -0.00620645 | 0.294 | 0.301 | 1 | Homer2 | 0.069174249 | 0.029216669 | 0.329 | 0.316 | 1 |
| Gm26561 | 0.008680548 | -0.01702821 | 0.282 | 0.278 | 1 | Arap2 | 0.070130018 | 0.052772744 | 0.325 | 0.311 | 1 |
| Hcn1 | 0.008700544 | -0.021815166 | 0.608 | 0.589 | 1 | Mbd5 | 0.073301927 | -0.014359978 | 0.874 | 0.871 | 1 |
| Rgs11 | 0.008764893 | -0.012945906 | 0.243 | 0.256 | 1 | Lyst | 0.078142641 | -0.035650594 | 0.654 | 0.657 | 1 |
| Sorbs2 | 0.009258329 | 0.032642622 | 0.873 | 0.863 | 1 | Il1rapl2 | 0.078359993 | -0.063384633 | 0.251 | 0.26 | 1 |
| Tmcc1 | 0.009619418 | -0.000427966 | 0.815 | 0.827 | 1 | Nos1ap | 0.078817654 | 0.003153945 | 0.74 | 0.731 | 1 |
| Hivep2 | 0.010827363 | -0.040228196 | 0.954 | 0.96 | 1 | Dclk1 | 0.079209267 | 0.017644934 | 0.951 | 0.947 | 1 |
| Sybu | 0.010897948 | 0.015309429 | 0.799 | 0.806 | 1 | Cacna2d1 | 0.079852794 | 0.007333848 | 0.943 | 0.936 | 1 |
| Gm30382 | 0.011127222 | -0.075149336 | 0.249 | 0.257 | 1 | Pde4b | 0.08199064 | -0.053016734 | 0.515 | 0.529 | 1 |
| Slc24a2 | 0.011462964 | -0.034966425 | 0.922 | 0.923 | 1 | Adamtsl1 | 0.083787078 | -0.046061147 | 0.264 | 0.266 | 1 |
| Adamtsl1 | 0.011906732 | 0.059698531 | 0.266 | 0.258 | 1 | Fmn1 | 0.084800466 | 0.016569028 | 0.453 | 0.443 | 1 |
| Gm29237 | 0.012184108 | 0.035654833 | 0.415 | 0.397 | 1 | Rasgrf1 | 0.093441948 | -0.004355368 | 0.904 | 0.899 | 1 |
| Ppp3ca | 0.014804767 | -0.013470205 | 0.985 | 0.983 | 1 | Dock3 | 0.099324779 | 0.004732762 | 0.895 | 0.887 | 1 |
| Dapk1 | 0.014852073 | 0.021365936 | 0.798 | 0.804 | 1 | Lrrc4c | 0.102033074 | 0.067164314 | 0.921 | 0.918 | 1 |
| Zfpm2 | 0.015054112 | 0.026456147 | 0.675 | 0.684 | 1 | Gm10563 | 0.108503383 | -0.005987263 | 0.367 | 0.362 | 1 |
| Rock2 | 0.015082102 | -0.03549446 | 0.685 | 0.702 | 1 | Rere | 0.112031263 | 0.01487906 | 0.926 | 0.921 | 1 |
| Adgrb3 | 0.016530977 | 0.010176737 | 0.99 | 0.988 | 1 | Jph1 | 0.125746413 | 0.043222602 | 0.608 | 0.596 | 1 |
| Opcml | 0.017075842 | 0.024477837 | 0.983 | 0.985 | 1 | Ntrk3 | 0.128226892 | -0.020665836 | 0.909 | 0.911 | 1 |
| Cers6 | 0.017098367 | -0.004518712 | 0.754 | 0.766 | 1 | Luc7l2 | 0.13166802 | 0.011050566 | 0.925 | 0.918 | 1 |
| Camk2b | 0.017552763 | -0.004093894 | 0.758 | 0.771 | 1 | Elavl2 | 0.134518717 | 0.038663246 | 0.3 | 0.288 | 1 |
| Ntrk2 | 0.017773272 | -0.044336502 | 0.808 | 0.809 | 1 | B3galt1 | 0.138130241 | 0.016516933 | 0.918 | 0.911 | 1 |
| Col25a1 | 0.01829128 | 0.027547171 | 0.357 | 0.353 | 1 | Slit2 | 0.148815662 | -0.017383421 | 0.324 | 0.326 | 1 |
| Man2a1 | 0.018525268 | -0.052191663 | 0.309 | 0.326 | 1 | Cpne4 | 0.15104488 | -0.017425325 | 0.513 | 0.517 | 1 |
| Cntnap5b | 0.018864958 | -0.067736875 | 0.533 | 0.552 | 1 | Ahcyl2 | 0.154374287 | -0.000403012 | 0.829 | 0.839 | 1 |
| Elavl4 | 0.019631611 | -0.061895463 | 0.297 | 0.313 | 1 | Gm11099 | 0.155018838 | -0.024856997 | 0.3 | 0.302 | 1 |
| Dlg1 | 0.021054168 | -0.025013756 | 0.645 | 0.658 | 1 | Trpc6 | 0.157032449 | 0.050049808 | 0.384 | 0.374 | 1 |
| Gm10563 | 0.023707619 | -0.010744354 | 0.362 | 0.376 | 1 | Nav2 | 0.171580283 | -0.011959677 | 0.934 | 0.929 | 1 |
| Plcl2 | 0.024724418 | 0.013763319 | 0.54 | 0.54 | 1 | Gm26551 | 0.179362062 | -0.020833131 | 0.343 | 0.354 | 1 |
| Slc35f4 | 0.024989218 | -0.033024675 | 0.537 | 0.553 | 1 | Runx1t1 | 0.181940424 | 0.025320934 | 0.517 | 0.515 | 1 |
| Rere | 0.025639914 | 0.031413546 | 0.921 | 0.92 | 1 | Nr3c2 | 0.184519402 | 0.021021632 | 0.844 | 0.85 | 1 |
| Gm2164 | 0.028646891 | 0.054010739 | 0.31 | 0.311 | 1 | Gap43 | 0.187747059 | 0.027872392 | 0.25 | 0.24 | 1 |
| Add2 | 0.03213963 | 0.017863275 | 0.783 | 0.79 | 1 | Dscam | 0.188888051 | -0.024824999 | 0.904 | 0.901 | 1 |
| Rasal2 | 0.032644839 | -0.033502835 | 0.862 | 0.873 | 1 | Map1b | 0.1925528 | -0.026688748 | 0.906 | 0.907 | 1 |
| Slit2 | 0.032793994 | 0.019420858 | 0.326 | 0.312 | 1 | Asap1 | 0.199809607 | 0.05271299 | 0.613 | 0.606 | 1 |
| Ndst3 | 0.033127976 | 0.016455308 | 0.368 | 0.369 | 1 | Ociad2 | 0.226245035 | 0.00574168 | 0.264 | 0.258 | 1 |
| Ggact | 0.035757904 | -0.02391101 | 0.432 | 0.446 | 1 | Nrcam | 0.239990638 | 0.012613772 | 0.945 | 0.94 | 1 |
| Fam135b | 0.037729509 | -0.017631124 | 0.867 | 0.877 | 1 | Synpr | 0.242537735 | -0.075598585 | 0.445 | 0.452 | 1 |

| | | | | | |
|---|---|---|---|---|---|
| Samd12 | 0.038080246 | 0.039058505 | 0.893 | 0.888 | 1 |
| Enox1 | 0.041387135 | -0.007968684 | 0.889 | 0.897 | 1 |
| Nckap5 | 0.043970544 | 0.00696565 | 0.348 | 0.349 | 1 |
| B3galt1 | 0.044139555 | 0.010232052 | 0.911 | 0.91 | 1 |
| Dip2c | 0.045731485 | -0.036603136 | 0.93 | 0.938 | 1 |
| Pcsk2 | 0.046020859 | 0.008905147 | 0.885 | 0.877 | 1 |
| Grid1 | 0.046069221 | -0.010861263 | 0.881 | 0.891 | 1 |
| Gpc5 | 0.051221506 | -0.024566623 | 0.276 | 0.27 | 1 |
| Dkk3 | 0.051567931 | -0.030822828 | 0.376 | 0.39 | 1 |
| Cdh12 | 0.069533871 | -0.053608121 | 0.685 | 0.698 | 1 |
| Dlgap1 | 0.069617828 | 0.001270228 | 0.991 | 0.989 | 1 |
| Abhd15 | 0.071116576 | -0.019940905 | 0.267 | 0.268 | 1 |
| Kcnj3 | 0.072614812 | 0.006490077 | 0.844 | 0.853 | 1 |
| Osbpl6 | 0.073426006 | -0.078529584 | 0.773 | 0.783 | 1 |
| Ccdc85a | 0.074470745 | 0.019958891 | 0.709 | 0.7 | 1 |
| Ssbp2 | 0.074967233 | -0.045684366 | 0.763 | 0.774 | 1 |
| Epha6 | 0.078770543 | -0.024409601 | 0.914 | 0.91 | 1 |
| Ppp1r9a | 0.080685933 | 0.015708361 | 0.817 | 0.824 | 1 |
| Ube2e2 | 0.084007305 | -0.02081206 | 0.958 | 0.953 | 1 |
| Cdh9 | 0.08448686 | -0.039368088 | 0.491 | 0.506 | 1 |
| Gm26551 | 0.089540469 | 0.005382577 | 0.354 | 0.345 | 1 |
| D430041D05Rik | 0.089738376 | 0.007667608 | 0.71 | 0.709 | 1 |
| Cntnap5c | 0.09889825 | 0.042460304 | 0.342 | 0.336 | 1 |
| Trpc5 | 0.120635218 | -0.042926325 | 0.525 | 0.538 | 1 |
| Dgkg | 0.123382103 | -0.018248195 | 0.824 | 0.817 | 1 |
| Elavl2 | 0.146275265 | 0.017466717 | 0.288 | 0.282 | 1 |
| Mbp | 0.153612957 | -0.011026813 | 0.308 | 0.319 | 1 |
| Mapk4 | 0.169819007 | -0.031663542 | 0.408 | 0.397 | 1 |
| Ano3 | 0.171086825 | -0.030723027 | 0.737 | 0.733 | 1 |
| Syn2 | 0.172900416 | 0.0013857 | 0.881 | 0.886 | 1 |
| Mbnl2 | 0.177279223 | 0.005189023 | 0.854 | 0.858 | 1 |
| St6galnac3 | 0.199862701 | -0.037464322 | 0.751 | 0.757 | 1 |
| Ttc3 | 0.216038077 | 0.015337241 | 0.937 | 0.941 | 1 |
| Nbea | 0.224817846 | 0.009945476 | 0.982 | 0.979 | 1 |
| Khdrbs2 | 0.224844233 | -0.015103931 | 0.772 | 0.777 | 1 |
| Ppm1e | 0.225044637 | -0.022177767 | 0.924 | 0.93 | 1 |
| Unc80 | 0.232585916 | 0.01445423 | 0.884 | 0.887 | 1 |
| Gm43713 | 0.2387791 | -0.036768084 | 0.624 | 0.634 | 1 |
| Atrnl1 | 0.266194544 | 0.004727283 | 0.85 | 0.855 | 1 |
| Gphn | 0.296171529 | 0.011498742 | 0.931 | 0.931 | 1 |
| Setbp1 | 0.316413895 | -0.019620511 | 0.915 | 0.919 | 1 |
| Cpne8 | 0.326328616 | 0.018376704 | 0.273 | 0.265 | 1 |
| Dgkb | 0.332683449 | 0.015381969 | 0.753 | 0.754 | 1 |
| Il1rapl2 | 0.334312807 | 0.082902862 | 0.26 | 0.253 | 1 |
| Mbd5 | 0.348951502 | -0.015107318 | 0.871 | 0.878 | 1 |
| Tenm3 | 0.36110403 | -0.050480266 | 0.564 | 0.561 | 1 |
| Strbp | 0.384223905 | -0.000788705 | 0.951 | 0.955 | 1 |
| Akap6 | 0.393390576 | 0.008357089 | 0.902 | 0.901 | 1 |
| Tenm4 | 0.399567416 | -0.043511698 | 0.745 | 0.737 | 1 |
| Stxbp5l | 0.499904746 | -0.000801498 | 0.909 | 0.913 | 1 |
| Satb2 | 0.502284496 | -0.004696866 | 0.267 | 0.268 | 1 |
| Eml6 | 0.569727258 | -0.026308359 | 0.842 | 0.847 | 1 |
| Srsf2 | 0.571719426 | 0.02268057 | 0.278 | 0.274 | 1 |
| Edil3 | 0.640130005 | -0.039911873 | 0.584 | 0.586 | 1 |
| Rabgap1l | 0.649731274 | -0.020895731 | 0.894 | 0.898 | 1 |
| Sorcs1 | 0.664722391 | -0.003852088 | 0.261 | 0.256 | 1 |
| Gm21954 | 0.70899218 | -0.026879376 | 0.474 | 0.478 | 1 |
| Fut9 | 0.74775248 | -0.033686242 | 0.623 | 0.625 | 1 |
| Htr1f | 0.748752055 | -0.004989491 | 0.299 | 0.3 | 1 |
| Cdh10 | 0.759927001 | -0.003969089 | 0.497 | 0.496 | 1 |
| Vsnl1 | 0.760320268 | -0.034728585 | 0.531 | 0.535 | 1 |
| Tshz3 | 0.769625078 | -0.018096646 | 0.329 | 0.325 | 1 |
| Mmp16 | 0.895645464 | 0.023425183 | 0.864 | 0.864 | 1 |
| Plekha5 | 0.901569395 | 0.005354607 | 0.856 | 0.857 | 1 |
| 4930587E11Rik | 0.925114109 | -0.032854126 | 0.349 | 0.35 | 1 |
| Adgrl3 | 0.938642058 | -0.018688215 | 0.954 | 0.955 | 1 |

| | | | | | |
|---|---|---|---|---|---|
| Syt1 | 0.245056213 | 0.015325158 | 0.992 | 0.993 | 1 |
| Spock3 | 0.267110604 | 0.013505673 | 0.464 | 0.456 | 1 |
| Epo | 0.275414747 | -0.003639061 | 0.477 | 0.485 | 1 |
| Bach2 | 0.281006156 | 0.017131273 | 0.431 | 0.421 | 1 |
| Cacna1c | 0.290480797 | -0.041498145 | 0.921 | 0.922 | 1 |
| Cit | 0.292244371 | -0.021631626 | 0.253 | 0.261 | 1 |
| Btbd9 | 0.293886353 | 0.036295329 | 0.859 | 0.856 | 1 |
| Grik2 | 0.344268308 | 0.025207488 | 0.957 | 0.959 | 1 |
| Ppfia2 | 0.346201864 | 0.043359448 | 0.95 | 0.951 | 1 |
| Tmem108 | 0.369455758 | 0.029705988 | 0.704 | 0.699 | 1 |
| Zfpm2 | 0.376562069 | -0.037210617 | 0.673 | 0.675 | 1 |
| Prkg1 | 0.385145104 | -0.035993414 | 0.89 | 0.891 | 1 |
| Tmem200a | 0.393201886 | -0.013928981 | 0.289 | 0.295 | 1 |
| Gm48749 | 0.410216902 | 0.00474025 | 0.288 | 0.283 | 1 |
| Rfx3 | 0.423709285 | 0.049772149 | 0.741 | 0.737 | 1 |
| Htr1f | 0.436028584 | -0.017415228 | 0.296 | 0.299 | 1 |
| Ncam2 | 0.441421059 | 0.037507656 | 0.783 | 0.781 | 1 |
| Ralyl | 0.451333626 | 0.025846322 | 0.943 | 0.939 | 1 |
| Stxbp5l | 0.463052786 | 0.012776637 | 0.913 | 0.909 | 1 |
| Acap2 | 0.464988145 | 0.008148598 | 0.793 | 0.786 | 1 |
| Inpp4b | 0.468364515 | -0.018096314 | 0.468 | 0.466 | 1 |
| Gm43713 | 0.475356076 | 0.004793384 | 0.627 | 0.624 | 1 |
| Pip5k1b | 0.475934773 | -7.40E-07 | 0.684 | 0.678 | 1 |
| Slc35f1 | 0.502964282 | -0.002045177 | 0.498 | 0.498 | 1 |
| Ly6h | 0.518960724 | -0.013576196 | 0.491 | 0.496 | 1 |
| Ppm1e | 0.56486757 | 0.010704849 | 0.927 | 0.924 | 1 |
| Egfem1 | 0.575069501 | -0.047739634 | 0.821 | 0.826 | 1 |
| Plcl1 | 0.575338319 | -0.037451036 | 0.487 | 0.493 | 1 |
| Pde4d | 0.595666039 | -0.001173281 | 0.797 | 0.796 | 1 |
| Chsy3 | 0.605316477 | 0.004204128 | 0.605 | 0.603 | 1 |
| Epha7 | 0.611670276 | 0.012352318 | 0.695 | 0.689 | 1 |
| Tcf4 | 0.620222694 | -0.002797435 | 0.968 | 0.966 | 1 |
| Tanc2 | 0.620415101 | 0.000373822 | 0.934 | 0.931 | 1 |
| Galnt17 | 0.62240843 | 0.001636116 | 0.878 | 0.877 | 1 |
| Tenm3 | 0.62517048 | 0.063891342 | 0.568 | 0.564 | 1 |
| Mmp16 | 0.706212585 | 0.013797339 | 0.867 | 0.864 | 1 |
| Gm28905 | 0.730061385 | 0.033420745 | 0.282 | 0.28 | 1 |
| Grm1 | 0.769579097 | -0.001718065 | 0.753 | 0.752 | 1 |
| Hs6st3 | 0.776382651 | -0.015659099 | 0.805 | 0.803 | 1 |
| Nell1 | 0.777989155 | -0.003100367 | 0.38 | 0.384 | 1 |
| Pcdh9 | 0.789111845 | -0.007636814 | 0.986 | 0.987 | 1 |
| Brinp3 | 0.794131464 | 0.060974832 | 0.447 | 0.444 | 1 |
| Ncald | 0.803748795 | 0.011702657 | 0.615 | 0.61 | 1 |
| Myt1l | 0.813400539 | 0.004319822 | 0.953 | 0.953 | 1 |
| Sntg1 | 0.863435899 | -0.010090172 | 0.937 | 0.937 | 1 |
| Hivep2 | 0.867434459 | -0.008200308 | 0.953 | 0.954 | 1 |
| Dlc1 | 0.943173286 | 0.008057076 | 0.327 | 0.325 | 1 |
| Opcml | 0.964457701 | -0.027554105 | 0.983 | 0.983 | 1 |
| Ccdc85a | 0.966123707 | 0.023825821 | 0.711 | 0.709 | 1 |
| Astn2 | 0.971165054 | 0.021953082 | 0.562 | 0.561 | 1 |
| Srsf2 | 0.986028157 | 0.004210321 | 0.279 | 0.278 | 1 |

Figure4E_psd95_intensity

| | Dap12+/+ Tau+ | | | | |
|---|---|---|---|---|---|
| | 0.034274 | 0.092052 | | | |
| | 0.219633 | 0.323166 | 0.457117 | 0.206354 | 0.140781 |
| | 0.077577 | 0.108017 | | | |
| | 0.123175 | 0.180992 | | | |
| | 0.152205 | 0.146154 | 0.254025 | | |
| | 0.170612 | 0.188452 | | | |
| | 0.388541 | 0.138188 | 0.365948 | | |
| average | 0.198277 | | | | |

| Dap12-/- Tau+ | | | |
|---|---|---|---|
| 0.02925 | 0.152633 | 0.567346 | 0.186022 |
| 0.388148 | 0.364399 | 0.636669 | 0.139586 |
| 0.319351 | 0.011118 | 0.158217 | 0.266614 |
| 0.492695 | 0.141608 | 0.478791 | 0.39367 |
| 0.539357 | 0.660255 | 1.094244 | 0.030332 |
| 0.452265 | 0.013639 | 0.141608 | 0.484137 |
| 0.381862 | 0.196102 | 0.484137 | 0.374539 |
| 0.347766 | | | |
| 0.34228828 | | | |
