## Supplementary table 5 for "DAP12 deficiency alters microglia-oligodendrocyte communication and enhances resilience against tau toxicity"

Figure5C_Hubgene_in_different_modules

| gene_name | module | color | kME | gene_name | module | color | kME | gene_name | module | color | kME | gene_name | module | color | kME |
|---|---|---|---|---|---|---|---|---|---|---|---|---|---|---|---|
| total gene number | 151 | | | total gene number | 449 | | | total gene number | 74 | | | total gene number | 150 | | |
| Odr4 | Module1 | Blue | 0.211815984 | Kcnip4 | Module2 | turquoise | 0.555778623 | Shisa3 | Module3 | yellow | 0.006427636 | Lypd6 | Module4 | Brown | 0.124203628 |
| Pip4p2 | Module1 | Blue | 0.143801795 | Rbfox1 | Module2 | turquoise | 0.494951526 | Serp1 | Module3 | yellow | 0.008805332 | Wasf3 | Module4 | Brown | 0.125015053 |
| Emc8 | Module1 | Blue | 0.094562566 | Csmd1 | Module2 | turquoise | 0.485471031 | AC161607.1 | Module3 | yellow | 0.012387072 | Mcam | Module4 | Brown | 0.125867496 |
| Gm26672 | Module1 | Blue | 0.259611874 | Nrg3 | Module2 | turquoise | 0.475727394 | Letm2 | Module3 | yellow | 0.013345837 | Xrcc6 | Module4 | Brown | 0.128243913 |
| Gcn1 | Module1 | Blue | 0.11032844 | Celf2 | Module2 | turquoise | 0.450852671 | Aatk | Module3 | yellow | 0.033376665 | Kcnab1 | Module4 | Brown | 0.128469298 |
| Ro60 | Module1 | Blue | 0.114616486 | Meg3 | Module2 | turquoise | 0.431865044 | Elmod2 | Module3 | yellow | 0.073968101 | Tmem163 | Module4 | Brown | 0.130702178 |
| Pik3r3 | Module1 | Blue | 0.100462811 | Nrxn3 | Module2 | turquoise | 0.427477518 | Ank2 | Module3 | yellow | 0.081405609 | Ephb2 | Module4 | Brown | 0.130705139 |
| Pwwp3a | Module1 | Blue | 0.103604601 | Lrrtm4 | Module2 | turquoise | 0.416149754 | Fnbp1 | Module3 | yellow | 0.093057371 | Cntfr | Module4 | Brown | 0.131255738 |
| Rab4b | Module1 | Blue | 0.112687522 | Kcnd2 | Module2 | turquoise | 0.378615468 | Xist | Module3 | yellow | 0.101015496 | Qk | Module4 | Brown | 0.133193955 |
| 3110056K07Rik | Module1 | Blue | 0.138871324 | Fgf14 | Module2 | turquoise | 0.375898097 | Cdc37l1 | Module3 | yellow | 0.103831688 | Lrp4 | Module4 | Brown | 0.133527165 |
| Ndor1 | Module1 | Blue | 0.084396779 | Dlgap1 | Module2 | turquoise | 0.362709104 | Mbp | Module3 | yellow | 0.107207401 | Kank1 | Module4 | Brown | 0.134477348 |
| Afg3l1 | Module1 | Blue | 0.131523706 | Fam155a | Module2 | turquoise | 0.354442179 | Limch1 | Module3 | yellow | 0.110649886 | Cdc42se2 | Module4 | Brown | 0.136084929 |
| Rpl13 | Module1 | Blue | 0.083220099 | Snhg11 | Module2 | turquoise | 0.344643439 | Rtn4 | Module3 | yellow | 0.111406056 | Aebp1 | Module4 | Brown | 0.136479108 |
| Gm20342 | Module1 | Blue | 0.25882323 | Cntnap2 | Module2 | turquoise | 0.341975732 | Cdk19 | Module3 | yellow | 0.118225414 | Gm4221 | Module4 | Brown | 0.136889538 |
| Pip4p1 | Module1 | Blue | 0.089956974 | Lsamp | Module2 | turquoise | 0.338391303 | St18 | Module3 | yellow | 0.121788021 | Ncoa7 | Module4 | Brown | 0.137828691 |
| Cct6a | Module1 | Blue | 0.207118825 | Dab1 | Module2 | turquoise | 0.335824837 | Plp1 | Module3 | yellow | 0.136663672 | Vangl1 | Module4 | Brown | 0.138827288 |
| Lypla1 | Module1 | Blue | 0.141160569 | Syt1 | Module2 | turquoise | 0.335723995 | Pcdh9 | Module3 | yellow | 0.148890826 | Zfhx2 | Module4 | Brown | 0.140490357 |
| Gm49336 | Module1 | Blue | 0.099618229 | Epha6 | Module2 | turquoise | 0.332607399 | Gm29455 | Module3 | yellow | 0.175231478 | Pde1c | Module4 | Brown | 0.144001212 |
| Gm47664 | Module1 | Blue | 0.141450726 | Grin2a | Module2 | turquoise | 0.330190917 | Map7 | Module3 | yellow | 0.178149248 | Stox2 | Module4 | Brown | 0.144477429 |
| E330020D12Rik | Module1 | Blue | 0.158598281 | Hs6st3 | Module2 | turquoise | 0.329231236 | Gm32005 | Module3 | yellow | 0.178938478 | Cdh6 | Module4 | Brown | 0.144953692 |
| BC065397 | Module1 | Blue | 0.132542772 | Gria1 | Module2 | turquoise | 0.320482434 | Mboat1 | Module3 | yellow | 0.1932768 | Csmd2 | Module4 | Brown | 0.145829457 |
| Ctnnd1 | Module1 | Blue | 0.153040402 | Lrrc7 | Module2 | turquoise | 0.316908326 | Ccdc152 | Module3 | yellow | 0.193962652 | Mast4 | Module4 | Brown | 0.147292492 |
| Entr1 | Module1 | Blue | 0.223882118 | Frmpd4 | Module2 | turquoise | 0.309271538 | Slc5a11 | Module3 | yellow | 0.217401396 | 4-Sep | Module4 | Brown | 0.147863417 |
| Dgcr2 | Module1 | Blue | 0.118291439 | Nrxn1 | Module2 | turquoise | 0.309181183 | Slc24a2 | Module3 | yellow | 0.218320606 | Arhgap24 | Module4 | Brown | 0.148454807 |
| Zc3h18 | Module1 | Blue | 0.145380764 | Erc2 | Module2 | turquoise | 0.302427982 | Plekhh2 | Module3 | yellow | 0.219562439 | Slc4a10 | Module4 | Brown | 0.149193795 |
| Tex52 | Module1 | Blue | 0.088526678 | Kalrn | Module2 | turquoise | 0.301171194 | Lama2 | Module3 | yellow | 0.223749671 | Dock4 | Module4 | Brown | 0.149712132 |
| Sys1 | Module1 | Blue | 0.364225246 | Dlgap2 | Module2 | turquoise | 0.291986867 | Plin3 | Module3 | yellow | 0.224645509 | Fign | Module4 | Brown | 0.149861039 |
| Gm13684 | Module1 | Blue | 0.123363471 | Fam19a1 | Module2 | turquoise | 0.283197874 | Sh3kbp1 | Module3 | yellow | 0.228477835 | Tom1l2 | Module4 | Brown | 0.149968759 |
| Rnf8 | Module1 | Blue | 0.306349073 | Lrfn5 | Module2 | turquoise | 0.276404195 | Gab2 | Module3 | yellow | 0.230211537 | Rapgef3 | Module4 | Brown | 0.150391012 |
| Nmt1 | Module1 | Blue | 0.112786701 | Galntl6 | Module2 | turquoise | 0.272835444 | Shroom3 | Module3 | yellow | 0.230494575 | Kcna6 | Module4 | Brown | 0.150645441 |
| 3830406C13Rik | Module1 | Blue | 0.117856561 | Egfem1 | Module2 | turquoise | 0.272260315 | Sgk3 | Module3 | yellow | 0.23477251 | 5031439G07Rik | Module4 | Brown | 0.150966184 |
| Srcap | Module1 | Blue | 0.222187899 | Kcnq5 | Module2 | turquoise | 0.268880849 | Mtss1 | Module3 | yellow | 0.236094617 | Rasal1 | Module4 | Brown | 0.151061288 |
| Ilrun | Module1 | Blue | 0.274621342 | Grm5 | Module2 | turquoise | 0.268601651 | 4930587E11Rik | Module3 | yellow | 0.24026368 | Hepacam | Module4 | Brown | 0.152070095 |
| Xndc1 | Module1 | Blue | 0.189053525 | Slit3 | Module2 | turquoise | 0.265013317 | Rasgrp3 | Module3 | yellow | 0.24127717 | Snx30 | Module4 | Brown | 0.152125318 |
| Slc49a4 | Module1 | Blue | 0.085780322 | Tenm2 | Module2 | turquoise | 0.263038854 | Galnt7 | Module3 | yellow | 0.247770762 | Cttnbp2 | Module4 | Brown | 0.152864167 |
| Gm12940 | Module1 | Blue | 0.060656979 | Gabrb1 | Module2 | turquoise | 0.259636313 | Ephb1 | Module3 | yellow | 0.260074606 | Lgi3 | Module4 | Brown | 0.152893421 |
| Per3 | Module1 | Blue | 0.138382621 | Rims1 | Module2 | turquoise | 0.25681722 | 1700111E14Rik | Module3 | yellow | 0.263340393 | Edil3 | Module4 | Brown | 0.153641034 |
| Synj2bp | Module1 | Blue | 0.05834142 | Lingo2 | Module2 | turquoise | 0.255765548 | Gm34408 | Module3 | yellow | 0.2664648 | Adamtsl1 | Module4 | Brown | 0.154931714 |
| Rxylt1 | Module1 | Blue | 0.157572188 | Sntg1 | Module2 | turquoise | 0.253354854 | St6galnac3 | Module3 | yellow | 0.270842306 | Kif21b | Module4 | Brown | 0.155070204 |
| Stimate | Module1 | Blue | 0.158669865 | Cacnb2 | Module2 | turquoise | 0.252120175 | Carns1 | Module3 | yellow | 0.274861841 | Amph | Module4 | Brown | 0.156490753 |
| Mrtfa | Module1 | Blue | 0.052482636 | Gria2 | Module2 | turquoise | 0.248833636 | Kctd3 | Module3 | yellow | 0.275057363 | Phldb1 | Module4 | Brown | 0.156607292 |
| Cox4i1 | Module1 | Blue | 0.093132993 | Grin2b | Module2 | turquoise | 0.24803054 | St3gal4 | Module3 | yellow | 0.276785696 | Fbn2 | Module4 | Brown | 0.157535513 |
| Banp | Module1 | Blue | 0.148165321 | Ryr2 | Module2 | turquoise | 0.247046547 | Lrrtm3 | Module3 | yellow | 0.27781339 | Trim2 | Module4 | Brown | 0.158326006 |
| Tnrc6a | Module1 | Blue | 0.069366757 | Gabrb3 | Module2 | turquoise | 0.2419487 | Nkain2 | Module3 | yellow | 0.289349188 | Bcas1 | Module4 | Brown | 0.158756549 |
| Tubb3 | Module1 | Blue | 0.083044719 | Prkg1 | Module2 | turquoise | 0.228040316 | Bcat1 | Module3 | yellow | 0.290106037 | Mgat5 | Module4 | Brown | 0.158823944 |
| Gm35438.1 | Module1 | Blue | 0.085799561 | Chrm3 | Module2 | turquoise | 0.227500078 | Rftn1 | Module3 | yellow | 0.291528503 | Cacnb4 | Module4 | Brown | 0.15909962 |
| Ccpg1 | Module1 | Blue | 0.056117137 | Dgkb | Module2 | turquoise | 0.226974244 | Myo1e | Module3 | yellow | 0.294321048 | Meis2 | Module4 | Brown | 0.159227234 |
| Oip5os1 | Module1 | Blue | 0.05647064 | Arpp21 | Module2 | turquoise | 0.226394516 | Slc25a27 | Module3 | yellow | 0.295685279 | Abca8a | Module4 | Brown | 0.160595216 |
| 2810468N07Rik | Module1 | Blue | 0.126645259 | Ryr3 | Module2 | turquoise | 0.226078036 | Piezo2 | Module3 | yellow | 0.295726102 | Syne1 | Module4 | Brown | 0.161841256 |
| C730002L08Rik | Module1 | Blue | 0.100494862 | Hs3st4 | Module2 | turquoise | 0.225856582 | Tspan9 | Module3 | yellow | 0.301572849 | Ptprj | Module4 | Brown | 0.161926599 |
| Dbndd2 | Module1 | Blue | 0.058530139 | Kcnb2 | Module2 | turquoise | 0.224619429 | Cdh20 | Module3 | yellow | 0.303738696 | Sh3gl3 | Module4 | Brown | 0.162130736 |
| 1700012D14Rik | Module1 | Blue | 0.163564924 | Nrg1 | Module2 | turquoise | 0.224233628 | Rapgef5 | Module3 | yellow | 0.306355393 | Hmgcll1 | Module4 | Brown | 0.162480442 |
| Commd1 | Module1 | Blue | 0.102837843 | Ntm | Module2 | turquoise | 0.222847904 | Spock3 | Module3 | yellow | 0.306642357 | Desi1 | Module4 | Brown | 0.162844474 |
| AY036118 | Module1 | Blue | 0.216368559 | Grip1 | Module2 | turquoise | 0.221618304 | Efr3b | Module3 | yellow | 0.307387459 | Nbas | Module4 | Brown | 0.164138556 |
| Resf1 | Module1 | Blue | 0.104270611 | Trpm3 | Module2 | turquoise | 0.22034692 | Crybg3 | Module3 | yellow | 0.311932595 | Pcdh15 | Module4 | Brown | 0.164445684 |
| 4930581F22Rik | Module1 | Blue | 0.052579061 | Asic2 | Module2 | turquoise | 0.216826917 | Hapln2 | Module3 | yellow | 0.316051844 | Cd9 | Module4 | Brown | 0.164668525 |
| Mtrex | Module1 | Blue | 0.138592411 | Gm42418 | Module2 | turquoise | 0.216390173 | Grik1 | Module3 | yellow | 0.317984455 | Slc12a2 | Module4 | Brown | 0.164953211 |
| Gpbp1 | Module1 | Blue | 0.125892347 | Gm28928 | Module2 | turquoise | 0.216020893 | Mapre2 | Module3 | yellow | 0.33355574 | Gabrr2 | Module4 | Brown | 0.164992007 |
| Cryzl1 | Module1 | Blue | 0.553141678 | Sgcz | Module2 | turquoise | 0.212535372 | Zdhhc20 | Module3 | yellow | 0.339627626 | Grid2 | Module4 | Brown | 0.165503316 |
| Gm9895 | Module1 | Blue | 0.141344758 | Cacna1e | Module2 | turquoise | 0.209087324 | Usp31 | Module3 | yellow | 0.359841133 | Phlpp1 | Module4 | Brown | 0.166322993 |
| Tuba1a | Module1 | Blue | 0.105344636 | Epha5 | Module2 | turquoise | 0.20854226 | Selenop | Module3 | yellow | 0.364294577 | Cadm1 | Module4 | Brown | 0.166672862 |
| Snhg14 | Module1 | Blue | 0.140587459 | Fam19a2 | Module2 | turquoise | 0.2075473 | A330049N07Rik | Module3 | yellow | 0.366994418 | Gfra1 | Module4 | Brown | 0.171261045 |
| Tut7 | Module1 | Blue | 0.160402475 | Nrcam | Module2 | turquoise | 0.205461522 | Ctnna3 | Module3 | yellow | 0.383453175 | Dscam | Module4 | Brown | 0.175950411 |
| Khdc4 | Module1 | Blue | 0.198200611 | Dok6 | Module2 | turquoise | 0.205302974 | Robo1 | Module3 | yellow | 0.38619678 | App | Module4 | Brown | 0.176720754 |
| Zup1 | Module1 | Blue | 0.10160633 | Cntn5 | Module2 | turquoise | 0.204049889 | Tln2 | Module3 | yellow | 0.388143268 | Mapt | Module4 | Brown | 0.177111154 |
| Gm50323 | Module1 | Blue | 0.099418901 | Ptprn2 | Module2 | turquoise | 0.200642581 | Apod | Module3 | yellow | 0.396738343 | Hook3 | Module4 | Brown | 0.177789921 |
| Zfhx2os | Module1 | Blue | 0.089406566 | Kcnh7 | Module2 | turquoise | 0.199865256 | Dgki | Module3 | yellow | 0.420780819 | Ick | Module4 | Brown | 0.178348579 |
| Ate1 | Module1 | Blue | 0.123804613 | Clstn2 | Module2 | turquoise | 0.197562111 | Agmo | Module3 | yellow | 0.430847158 | Fnip2 | Module4 | Brown | 0.180825786 |
| Maco1 | Module1 | Blue | 0.129733002 | Etl4 | Module2 | turquoise | 0.195818575 | Cntn3 | Module3 | yellow | 0.443680454 | Cerk | Module4 | Brown | 0.181901065 |
| Evi2a | Module1 | Blue | 0.09061597 | Shisa6 | Module2 | turquoise | 0.192584752 | Slco3a1 | Module3 | yellow | 0.44670459 | Chsy3 | Module4 | Brown | 0.182639899 |
| Gas5 | Module1 | Blue | 0.066962744 | Tenm3 | Module2 | turquoise | 0.188430355 | Ctnna2 | Module3 | yellow | 0.446951842 | Kif13a | Module4 | Brown | 0.182865245 |
| 4933400F21Rik | Module1 | Blue | 0.217441238 | Gria3 | Module2 | turquoise | 0.185882526 | Tmem117 | Module3 | yellow | 0.481138917 | Bfsp2 | Module4 | Brown | 0.184025911 |
| Gm49797 | Module1 | Blue | 0.161004835 | Nebl | Module2 | turquoise | 0.185218075 | Enpp6 | Module3 | yellow | 0.494905732 | Inpp5a | Module4 | Brown | 0.184669089 |
| Gm28320 | Module1 | Blue | 0.08681026 | Unc80 | Module2 | turquoise | 0.184194509 | Anln | Module3 | yellow | 0.513005019 | Bin1 | Module4 | Brown | 0.184720526 |
| Atp6v0c | Module1 | Blue | 0.057590859 | Snap25 | Module2 | turquoise | 0.182407452 | | | | | Npas3 | Module4 | Brown | 0.184722002 |
| Cpne1 | Module1 | Blue | 0.101294029 | Myt1l | Module2 | turquoise | 0.18229506 | | | | | Dlg1 | Module4 | Brown | 0.185700908 |
| Tasor2 | Module1 | Blue | 0.295838444 | Opcml | Module2 | turquoise | 0.182087317 | | | | | Fbxw15 | Module4 | Brown | 0.186933655 |
| Galnt2 | Module1 | Blue | 0.141371901 | Rgs7 | Module2 | turquoise | 0.180286761 | | | | | Pcdh7 | Module4 | Brown | 0.186967525 |
| Gm33255 | Module1 | Blue | 0.143519315 | Kcnj3 | Module2 | turquoise | 0.17997255 | | | | | Mal | Module4 | Brown | 0.187257145 |
| B230206H07Rik | Module1 | Blue | 0.100962356 | 5730522E02Rik | Module2 | turquoise | 0.17995622 | | | | | Zfpm2 | Module4 | Brown | 0.187257994 |
| Gse1 | Module1 | Blue | 0.091273201 | Robo2 | Module2 | turquoise | 0.178604253 | | | | | Foxn3 | Module4 | Brown | 0.188803935 |
| Fam53b | Module1 | Blue | 0.183747458 | Epha7 | Module2 | turquoise | 0.178478627 | | | | | Hivep2 | Module4 | Brown | 0.189449449 |
| Tent4b | Module1 | Blue | 0.119316658 | Ntrk3 | Module2 | turquoise | 0.178144757 | | | | | Dpyd | Module4 | Brown | 0.190115227 |
| Adnp | Module1 | Blue | 0.198638351 | Dgkg | Module2 | turquoise | 0.177797955 | | | | | Zfp638 | Module4 | Brown | 0.190631789 |
| 6430590A07Rik | Module1 | Blue | 0.142341799 | Plcb1 | Module2 | turquoise | 0.177080802 | | | | | Oxr1 | Module4 | Brown | 0.190766694 |
| Zfp276 | Module1 | Blue | 0.123403192 | St6galnac5 | Module2 | turquoise | 0.176743276 | | | | | Dusp10 | Module4 | Brown | 0.19227149 |
| Lncppara | Module1 | Blue | 0.043115165 | Sorbs2 | Module2 | turquoise | 0.175423536 | | | | | Nfia | Module4 | Brown | 0.192890069 |
| Gm46367 | Module1 | Blue | 0.149054852 | Celf4 | Module2 | turquoise | 0.175132578 | | | | | Tnik | Module4 | Brown | 0.193997405 |
| A330076H08Rik | Module1 | Blue | 0.075465788 | Gabra2 | Module2 | turquoise | 0.174492561 | | | | | Pde8a | Module4 | Brown | 0.195782521 |
| Tent2 | Module1 | Blue | 0.16437705 | Mgat4c | Module2 | turquoise | 0.174109815 | | | | | Csmd3 | Module4 | Brown | 0.196233801 |
| Dpm1 | Module1 | Blue | 0.164702961 | Gabrg3 | Module2 | turquoise | 0.173661106 | | | | | Fgfr2 | Module4 | Brown | 0.196563325 |
| Gm16759 | Module1 | Blue | 0.064881335 | Cntnap5a | Module2 | turquoise | 0.172944605 | | | | | Pvt1 | Module4 | Brown | 0.197581954 |
| D7Ertd443e | Module1 | Blue | 0.436433997 | Nav3 | Module2 | turquoise | 0.17245516 | | | | | Ttll5 | Module4 | Brown | 0.199410351 |
| Gtf3c2 | Module1 | Blue | 0.363224696 | Sgcd | Module2 | turquoise | 0.171957119 | | | | | Unc5c | Module4 | Brown | 0.203315769 |
| Prnp | Module1 | Blue | 0.315040974 | Kcnc2 | Module2 | turquoise | 0.171830854 | | | | | Ppp1r16b | Module4 | Brown | 0.204520095 |
| 2900052L18Rik | Module1 | Blue | 0.077364129 | Mctp1 | Module2 | turquoise | 0.171329532 | | | | | Serinc5 | Module4 | Brown | 0.205397722 |
| Firre | Module1 | Blue | 0.061885298 | Mapk10 | Module2 | turquoise | 0.170465692 | | | | | Man1a | Module4 | Brown | 0.205750023 |
| Vps35l | Module1 | Blue | 0.162710151 | Stxbp5l | Module2 | turquoise | 0.167320138 | | | | | Tbc1d5 | Module4 | Brown | 0.207854344 |
| Armh3 | Module1 | Blue | 0.064875046 | Tnr | Module2 | turquoise | 0.1665851 | | | | | Cadps | Module4 | Brown | 0.208751012 |
| Gm10863 | Module1 | Blue | 0.108133727 | Dpp10 | Module2 | turquoise | 0.165916282 | | | | | B3galt1 | Module4 | Brown | 0.208821461 |
| Glcci1 | Module1 | Blue | 0.111990138 | Brinp1 | Module2 | turquoise | 0.165616694 | | | | | Spock1 | Module4 | Brown | 0.208992838 |
| Gm17106 | Module1 | Blue | 0.047870877 | Kcnt2 | Module2 | turquoise | 0.165563099 | | | | | Maml2 | Module4 | Brown | 0.20921823 |
| Oga | Module1 | Blue | 0.240257191 | Ptprg | Module2 | turquoise | 0.165155554 | | | | | Ust | Module4 | Brown | 0.210090802 |
| 1810026B05Rik | Module1 | Blue | 0.041948331 | Car10 | Module2 | turquoise | 0.16491606 | | | | | Nav2 | Module4 | Brown | 0.211720255 |
| Insyn2a | Module1 | Blue | 0.16948306 | Plxna4 | Module2 | turquoise | 0.163280264 | | | | | Kit | Module4 | Brown | 0.213512954 |
| Cplane1 | Module1 | Blue | 0.177655024 | C230004F18Rik | Module2 | turquoise | 0.163108248 | | | | | Tmod1 | Module4 | Brown | 0.216544399 |
| 4930512B01Rik | Module1 | Blue | 0.048448502 | Dpp6 | Module2 | turquoise | 0.162120104 | | | | | Add3 | Module4 | Brown | 0.217233152 |
| Mrtfb | Module1 | Blue | 0.155648629 | Cntnap5c | Module2 | turquoise | 0.161992994 | | | | | Creb5 | Module4 | Brown | 0.220916179 |
| Pabpn1 | Module1 | Blue | 0.152852017 | Ppp2r2b | Module2 | turquoise | 0.161843709 | | | | | Dock10 | Module4 | Brown | 0.22139863 |
| A330023F24Rik | Module1 | Blue | 0.211881784 | Nkain3 | Module2 | turquoise | 0.160515534 | | | | | Il33 | Module4 | Brown | 0.2252577 |
| Gm19522 | Module1 | Blue | 0.102803614 | Zfp385b | Module2 | turquoise | 0.159658659 | | | | | Ccp110 | Module4 | Brown | 0.229626005 |
| Dop1a | Module1 | Blue | 0.069628831 | A830018L16Rik | Module2 | turquoise | 0.159148032 | | | | | C030029H02Rik | Module4 | Brown | 0.230627769 |
| Btbd8 | Module1 | Blue | 0.253317644 | Ntng1 | Module2 | turquoise | 0.158335504 | | | | | Nckap5 | Module4 | Brown | 0.2320776 |
| Zswim6 | Module1 | Blue | 0.104509529 | Hcn1 | Module2 | turquoise | 0.157897284 | | | | | Pak1 | Module4 | Brown | 0.232598356 |
| Umad1 | Module1 | Blue | 0.113130581 | Pde1a | Module2 | turquoise | 0.15737135 | | | | | Arap2 | Module4 | Brown | 0.233136622 |
| 9330117O12Rik | Module1 | Blue | 0.008716251 | Camk2a | Module2 | turquoise | 0.157070891 | | | | | Myo18a | Module4 | Brown | 0.233480389 |
| Slc12a6 | Module1 | Blue | 0.03432616 | Khdrbs2 | Module2 | turquoise | 0.155350452 | | | | | Rhoj | Module4 | Brown | 0.233908755 |
| 4632427E13Rik | Module1 | Blue | 0.135347703 | Cnksr2 | Module2 | turquoise | 0.154161848 | | | | | 4930419G24Rik | Module4 | Brown | 0.234872927 |
| Tut4 | Module1 | Blue | 0.127579649 | Adcy2 | Module2 | turquoise | 0.152520265 | | | | | Gm42756 | Module4 | Brown | 0.237702085 |
| 9130019P16Rik | Module1 | Blue | 0.116727027 | Park2 | Module2 | turquoise | 0.152080913 | | | | | Fbxl5 | Module4 | Brown | 0.241204707 |
| Dop1b | Module1 | Blue | 0.099858639 | Cdh12 | Module2 | turquoise | 0.149929409 | | | | | Phactr3 | Module4 | Brown | 0.24160962 |
| Ecpas | Module1 | Blue | 0.2355146 | Tenm1 | Module2 | turquoise | 0.149528984 | | | | | Auts2 | Module4 | Brown | 0.245593961 |
| Itprid2 | Module1 | Blue | 0.127630107 | Cacna1d | Module2 | turquoise | 0.148491856 | | | | | Opalin | Module4 | Brown | 0.245737168 |
| Tcf25 | Module1 | Blue | 0.171677879 | Atrnl1 | Module2 | turquoise | 0.148268689 | | | | | Add1 | Module4 | Brown | 0.250884677 |
| Crppa | Module1 | Blue | 0.061415425 | Nol4 | Module2 | turquoise | 0.14819007 | | | | | Ank3 | Module4 | Brown | 0.251970968 |
| Relch | Module1 | Blue | 0.284764713 | Gabrb2 | Module2 | turquoise | 0.14809804 | | | | | Grik2 | Module4 | Brown | 0.255547786 |
| Atf7 | Module1 | Blue | 0.080519442 | Agbl4 | Module2 | turquoise | 0.147239685 | | | | | Tmod2 | Module4 | Brown | 0.256283203 |
| Ube2b | Module1 | Blue | 0.247785837 | Cmss1 | Module2 | turquoise | 0.146949046 | | | | | Tspan2 | Module4 | Brown | 0.258376466 |
| Cox16 | Module1 | Blue | 0.195575244 | Zfp804b | Module2 | turquoise | 0.146841172 | | | | | Erbin | Module4 | Brown | 0.26354107 |
| Plaat3 | Module1 | Blue | 0.366844469 | Cap2 | Module2 | turquoise | 0.146702903 | | | | | Exoc6b | Module4 | Brown | 0.265240228 |
| Stard13 | Module1 | Blue | 0.078872826 | Fat3 | Module2 | turquoise | 0.146496455 | | | | | Fam13c | Module4 | Brown | 0.269170468 |
| Tafa5 | Module1 | Blue | 0.083089728 | Gm20754 | Module2 | turquoise | 0.146370031 | | | | | Slc13a1 | Module4 | Brown | 0.269570714 |
| Mir99ahg | Module1 | Blue | 0.113422853 | Syn2 | Module2 | turquoise | 0.146001023 | | | | | Daam1 | Module4 | Brown | 0.270628621 |
| 5330438D12Rik | Module1 | Blue | 0.118221703 | Ppm1e | Module2 | turquoise | 0.145804496 | | | | | Dock1 | Module4 | Brown | 0.271580734 |
| Rhou | Module1 | Blue | 0.141528357 | Samd12 | Module2 | turquoise | 0.144676528 | | | | | Prickle1 | Module4 | Brown | 0.271835237 |
| 4930511M06Rik | Module1 | Blue | 0.235262337 | Hecw1 | Module2 | turquoise | 0.144500152 | | | | | Ablim2 | Module4 | Brown | 0.274764006 |
| Pakap.1 | Module1 | Blue | 0.112914192 | Neto1 | Module2 | turquoise | 0.144347513 | | | | | Zfp536 | Module4 | Brown | 0.278290361 |
| 1500004A13Rik | Module1 | Blue | 0.074868232 | Ak5 | Module2 | turquoise | 0.141681012 | | | | | 7630403G23Rik | Module4 | Brown | 0.302303066 |
| Tasor | Module1 | Blue | 0.109812108 | Khdrbs3 | Module2 | turquoise | 0.141291555 | | | | | Nfasc | Module4 | Brown | 0.302851628 |
| Aopep | Module1 | Blue | 0.074192496 | Zmat4 | Module2 | turquoise | 0.139973623 | | | | | Srgap1 | Module4 | Brown | 0.30538409 |
| Gm19500 | Module1 | Blue | 0.109858365 | Trim9 | Module2 | turquoise | 0.139958836 | | | | | Clmn | Module4 | Brown | 0.312203918 |
| Schip1 | Module1 | Blue | 0.11739501 | Unc79 | Module2 | turquoise | 0.13936627 | | | | | Pde4b | Module4 | Brown | 0.316352807 |
| 4930420G21Rik | Module1 | Blue | 0.14900919 | Rgs7bp | Module2 | turquoise | 0.138900932 | | | | | Frmd5 | Module4 | Brown | 0.327382549 |
| Dleu2 | Module1 | Blue | 0.108013339 | Rasgrf1 | Module2 | turquoise | 0.135683712 | | | | | Plekha1 | Module4 | Brown | 0.329910326 |
| Prkn | Module1 | Blue | 0.117613417 | Camk4 | Module2 | turquoise | 0.135192885 | | | | | Pex5l | Module4 | Brown | 0.330695861 |
| Pakap | Module1 | Blue | 0.141398516 | Fam135b | Module2 | turquoise | 0.134920669 | | | | | Ano4 | Module4 | Brown | 0.339904619 |
| Gm4258 | Module1 | Blue | 0.091527662 | Nlgn1 | Module2 | turquoise | 0.133640373 | | | | | Synpr | Module4 | Brown | 0.381246437 |
| Mir100hg | Module1 | Blue | 0.079221695 | Kcnn2 | Module2 | turquoise | 0.13288755 | | | | | Ptgds | Module4 | Brown | 0.4034237 |
| Trf | Module1 | Blue | 0.111549024 | Sox5 | Module2 | turquoise | 0.132523953 | | | | | Erbb4 | Module4 | Brown | 0.40817164 |
| Sox2ot | Module1 | Blue | 0.079752558 | Gpr158 | Module2 | turquoise | 0.131991103 | | | | | Kirrel3 | Module4 | Brown | 0.542997048 |

Figure5DE_Dap12_deletion_in_tauopathy_affects_different_modules

Dap12+/+ tau+ vs tau-

| p_val | avg_log2FC | pct.1 | pct.2 | p_val_adj | module | Color |
|---|---|---|---|---|---|---|
| 1.83E-109 | 3.59677084 | 0.611 | 0.193 | 7.34E-109 | Module2 | Turquoise |
| 4.79E-85 | -5 | 0 | 0.238 | 1.92E-84 | Module1 | Blue |
| 6.78E-14 | -0.7312236 | 0.343 | 0.486 | 2.71E-01 | Module3 | Yellow |
| 0.241930429 | -0.3251143 | 0.459 | 0.457 | 0.967722 | Module4 | Brown |

Dap12-/- tau+ vs Dap12+/+ tau+

| p_val | avg_log2FC | pct.1 | pct.2 | p_val_adj | module | |
|---|---|---|---|---|---|---|
| 1.09E-170 | -5 | 0.093 | 0.611 | 4.34E-170 | Module2 | Turquoise |
| 5.26E-80 | >5 | 0.222 | 0 | 2.10E-79 | Module1 | Blue |
| 4.34E-12 | 0.641217007 | 0.561 | 0.459 | 7.30E-01 | Module4 | Yellow |
| 1.37E-06 | 0.427194874 | 0.431 | 0.343 | 5.48E-02 | Module3 | Brown |

sFigure5B_trajectory_analysis_of_OPC_and_OL

Samples Dimensions

3265 2

lineages: 1

Lineage1: 5 2 1 3

curves: 1

Curve1: Length: 17.384 | Samples: 3265

sFigure5CD_OPC+Oligodendrocyte integrated cell ratio of different clusters

| | WT_1 | WT_2 | WT_3 | DAP12WT_ | DAP12WT_2 | DAP12WT_ | DAP12KO_1 | DAP12KO_2 | DAP12KO_3 | | By Genotype | DAP12KO | DAP12WT | WT |
|---|---|---|---|---|---|---|---|---|---|---|---|---|---|---|
| 1 | 537 | 613 | 455 | 588 | 665 | 347 | 606 | 845 | 498 | | 1 | 1949 | 1600 | 1605 |
| 2 | 327 | 214 | 152 | 654 | 857 | 1208 | 453 | 418 | 274 | | 2 | 1445 | 2719 | 793 |
| 3 | 315 | 290 | 194 | 326 | 381 | 355 | 307 | 439 | 221 | | 3 | 967 | 1062 | 799 |
| 4 | 258 | 191 | 118 | 229 | 342 | 180 | 234 | 238 | 148 | | 4 | 620 | 751 | 567 |
| 5 | 238 | 181 | 130 | 197 | 321 | 222 | 162 | 228 | 107 | | 5 | 497 | 740 | 549 |
| 6 | 21 | 15 | 4 | 5 | 14 | 10 | 10 | 8 | 3 | | 6 | 21 | 29 | 40 |
| Sum | 1696 | 1504 | 1053 | 1999 | 2580 | 2322 | 1772 | 2176 | 1251 | | | | | |

| | WT_1 | WT_2 | WT_3 | DAP12WT_ | DAP12WT_2 | DAP12WT_ | DAP12KO_1 | DAP12KO_2 | DAP12KO_3 |
|---|---|---|---|---|---|---|---|---|---|
| 1 | 0.31662736 | 0.40757979 | 0.43209877 | 0.294147 | 0.25775194 | 0.14944 | 0.34198646 | 0.38832721 | 0.39808154 |
| 2 | 0.1928066 | 0.14228723 | 0.14434948 | 0.327164 | 0.33217054 | 0.52024 | 0.25564334 | 0.19209559 | 0.21902478 |
| 3 | 0.18573113 | 0.19281915 | 0.18423552 | 0.163082 | 0.14767442 | 0.15289 | 0.17325056 | 0.20174632 | 0.17665867 |
| 4 | 0.15212264 | 0.12699468 | 0.11206078 | 0.114557 | 0.13255814 | 0.07752 | 0.13205418 | 0.109375 | 0.11830536 |
| 5 | 0.14033019 | 0.12034575 | 0.12345679 | 0.098549 | 0.12441861 | 0.09561 | 0.09142212 | 0.10477941 | 0.08553158 |
| 6 | 0.01238208 | 0.0099734 | 0.00379867 | 0.002501 | 0.00542636 | 0.00431 | 0.00564334 | 0.00367647 | 0.00239808 |

sFigure5E_GO_Biological_Process_2021 of Turquoise module

| | Term | Overlap | P.value | usted.P.va | Old.Adjusted.P.value | Genes | db | module | Combined.Score | db | module | Genes |
|---|---|---|---|---|---|---|---|---|---|---|---|---|
| 713 | tic transmissior | 24/306 | 3 | 6.90E-10 | 8.08E-07 | 21 | 9.68645E+12 | 1088 | 22814261555 | GO_Biological_Proces | Module2 | GRIA1;GABRB3;GABRA2;CHRM3;GRIA2;SNAP25;GABRB1; PTPRN2;SYT1;KCND2;NRXN1;GABRG3;CACNA1E;GRIN2B;SHISA6;RIMS1; CACNB2;GRIN2A;GRM5;DLGAP1;ASIC2;ERC2;DLGAP2;GRIA3 |
| 714 | -synaptic signal | 19/244 | 1 | 1.88E-04 | 2.83E-03 | 20 | 5.1166E+12 | 799 | 3.68811E+11 | GO_Biological_Proces | Module2 | GRIA1;GABRB3;GABRA2;CHRM3;GRIA2;SNAP25;GABRB1;SYT1;KCND2;NRXN1;GABRG3; CACNA1E;GRIN2B;CACNB2;GRIN2A;GRM5;DLGAP1;ASIC2;GRIA3 |
| 715 | smitter recepto | 9/58 | 1 | 1.76E+02 | 6.57E+04 | 40 | 67055393586 | 1008 | 34635688586 | GO_Biological_Proces | Module2 | GRIA1;GRIA2;GRIN2A;LRRC7;DLGAP1;DLGAP2;GRIN2B;GRIA3;SHISA6 |
| 716 | on channel acti | 9/89 | 6 | 1.80E+03 | 4.75E+05 | 24 | 5.02747E+12 | 519 | 5.68788E+11 | GO_Biological_Proces | Module2 | GRIA1;CACNB2;GRIA2;GRIN2A;FGF14;LRRC7;GRIN2B;GRIA3;SHISA6 |
| 717 | A receptor acti | 6/32 | 1 | 9.48E+04 | 4.85E+07 | 48 | 7.90507E+12 | 894 | 3.33362E+11 | GO_Biological_Proces | Module2 | GRIA1;GRIA2;GRIN2A;LRRC7;GRIN2B;GRIA3 |
| 718 | r signaling path | 6/37 | 2 | 7.53E+05 | 1.11E+07 | 40 | 9.10776E+11 | 712 | 1.78389E+11 | GO_Biological_Proces | Module2 | GRIA2;GRIN2A;GRM5;PLCB1;GRIN2B;GRIA3 |
| 719 | genesis (GO:00 | 11/240 | 3 | 2.72E+05 | 1.69E+07 | 10 | 6.168E+12 | 182 | 9.84249E+11 | GO_Biological_Proces | Module2 | EPHA5;ROBO2;EPHA7;EPHA6;DAB1;DOK6;NRXN1;NRXN3;TNR;NRCAM;SLIT3 |
| 720 | uidance (GO:00 | 10/203 | 7 | 2.77E+05 | 5.53E+07 | 11 | 3.45423E+12 | 186 | 4.72397E+11 | GO_Biological_Proces | Module2 | EPHA5;ROBO2;EPHA7;EPHA6;DAB1;DOK6;NRXN1;NRXN3;TNR;SLIT3 |
| 721 | al synaptic trans | 8/109 | 7 | 3.71E+05 | 5.53E+07 | 17 | 4.60611E+11 | 279 | 9.50473E+11 | GO_Biological_Proces | Module2 | RIMS1;GRM5;MCTP1;CLSTN2;SYT1;NRG3;DLGAP1;DLGAP2 |
| 722 | itter transport ( | 6/73 | 1 | 7.44E+06 | 1.07468E+14 | 18 | 8.9457E+12 | 250 | 5.33811E+11 | GO_Biological_Proces | Module2 | GABRA2;SNAP25;RIMS1;PTPRN2;SYT1;NRXN1 |

| sFigure5F_Overlap of mOL3 and OPC+OL2 | OPC+OL2 | mOL3 | DEGs of mOL3 | DEGs of OPC+OL2 |
|---|---|---|---|---|

| | gene | p_val.x | avg_log2FC.x | pct.1.x | pct.2.x | p_val_adj.x | p_val.y | avg_log2FC.y | pct.1.y | pct.2.y | p_val_adj.y |
|---|---|---|---|---|---|---|---|---|---|---|---|
| 96 | Slc13a1 | 6.47E-21 | 0.658497365 | 0.439 | 0.471 | 9.39E-18 | 3.32E-24 | 0.006431522 | 0.207 | 0.258 | 8.94E-20 |
| 99 | Clic4 | 1.28E-15 | 0.640749277 | 0.608 | 0.613 | 1.86E-12 | 2.36E-55 | -0.074533903 | 0.257 | 0.35 | 6.37E-51 |
| 79 | Kirrel3 | 4.53E-35 | 0.56271785 | 0.787 | 0.8 | 6.58E-32 | 1.95E-22 | 0.364477488 | 0.594 | 0.51 | 5.25E-18 |
| 21 | Apoe | 4.53E-111 | 0.543674256 | 0.476 | 0.552 | ####### | 4.26E-68 | 0.14474695 | 0.448 | 0.539 | 1.15E-63 |
| 42 | Lrrtm4 | 5.80E-66 | 0.533444283 | 0.733 | 0.845 | 8.42E-63 | 7.60E-148 | 1.213324077 | 0.274 | 0.095 | 2.05E-143 |
| 92 | Gatm | 1.52E-23 | 0.468692818 | 0.777 | 0.837 | 2.21E-20 | 1.17E-61 | 0.036253955 | 0.518 | 0.62 | 3.15E-57 |
| 43 | Nrxn3 | 7.25E-66 | 0.463728583 | 0.749 | 0.867 | 1.05E-62 | 4.20E-149 | 1.21760068 | 0.31 | 0.129 | 1.13E-144 |
| 77 | C030029H02Rik | 6.33E-36 | 0.407169119 | 0.775 | 0.842 | 9.19E-33 | 7.53E-43 | 0.265767531 | 0.632 | 0.632 | 2.03E-38 |
| 23 | Ptgds | 4.68E-108 | 0.387140544 | 0.54 | 0.643 | ####### | 3.39E-20 | -0.075074059 | 0.49 | 0.584 | 9.13E-16 |
| 60 | Slc38a2 | 4.22E-52 | 0.383773988 | 0.514 | 0.633 | 6.12E-49 | 9.36E-50 | -0.040938569 | 0.366 | 0.465 | 2.52E-45 |
| 101 | Unc5c | 2.31E-14 | 0.350653974 | 0.857 | 0.895 | 3.35E-11 | 4.12E-22 | -0.054662462 | 0.728 | 0.807 | 1.11E-17 |
| 53 | Cldn11 | 3.12E-55 | 0.324736309 | 0.575 | 0.723 | 4.52E-52 | 3.39E-63 | 0.008361785 | 0.365 | 0.461 | 9.13E-59 |
| 78 | Mag | 9.71E-36 | 0.263061185 | 0.743 | 0.855 | 1.41E-32 | 3.98E-66 | -0.173422971 | 0.553 | 0.711 | 1.07E-61 |
| 40 | Gm42756 | 1.28E-68 | 0.248090821 | 0.382 | 0.562 | 1.86E-65 | 3.50E-31 | -0.255706205 | 0.177 | 0.263 | 9.43E-27 |
| 100 | Sema6a | 1.66E-14 | 0.240180555 | 0.733 | 0.808 | 2.41E-11 | 1.65E-51 | 0.155647847 | 0.327 | 0.378 | 4.45E-47 |
| 93 | 9330111N05Rik | 1.87E-21 | 0.234001331 | 0.53 | 0.633 | 2.71E-18 | 3.72E-09 | -0.189740956 | 0.208 | 0.257 | 0.000100185 |
| 55 | Mobp | 9.49E-55 | 0.216389096 | 0.758 | 0.889 | 1.38E-51 | 7.03E-63 | -0.292421403 | 0.642 | 0.803 | 1.89E-58 |
| 84 | Tspan2 | 6.71E-30 | 0.207602275 | 0.685 | 0.794 | 9.74E-27 | 2.62E-42 | 0.022669257 | 0.482 | 0.565 | 7.04E-38 |
| 97 | Chn2 | 1.12E-20 | 0.204343687 | 0.605 | 0.699 | 1.62E-17 | 1.01E-46 | -0.067539244 | 0.206 | 0.281 | 2.73E-42 |
| 48 | Mal | 2.52E-59 | 0.193369376 | 0.572 | 0.738 | 3.66E-56 | 1.46E-61 | -0.168890813 | 0.363 | 0.501 | 3.93E-57 |
| 94 | Cobll1 | 3.52E-21 | 0.188336154 | 0.662 | 0.76 | 5.11E-18 | 1.10E-60 | 0.0411556 | 0.304 | 0.382 | 2.96E-56 |
| 70 | Pcdh15 | 7.43E-43 | 0.188160044 | 0.627 | 0.771 | 1.08E-39 | 1.89E-29 | 0.243633702 | 0.303 | 0.311 | 5.08E-25 |
| 108 | Fign | 0.00416497 | 0.168654733 | 0.603 | 0.642 | 1 | 2.41E-43 | -0.015051956 | 0.259 | 0.332 | 6.49E-39 |
| 83 | Sox10 | 1.34E-31 | 0.164953178 | 0.42 | 0.54 | 1.95E-28 | 5.71E-58 | -0.000233343 | 0.277 | 0.362 | 1.54E-53 |
| 56 | Plekhh1 | 6.23E-54 | 0.161763167 | 0.612 | 0.771 | 9.04E-51 | 1.71E-74 | -0.104216935 | 0.439 | 0.58 | 4.61E-70 |
| 102 | Erbb4 | 4.25E-13 | 0.1617239 | 0.899 | 0.941 | 6.16E-10 | 8.02E-13 | -0.015700624 | 0.761 | 0.807 | 2.16E-08 |
| 104 | Abca8a | 3.64E-12 | 0.160155061 | 0.608 | 0.693 | 5.28E-09 | 2.99E-40 | -0.03007555 | 0.226 | 0.296 | 8.06E-36 |
| 95 | Creb5 | 5.31E-21 | 0.156957622 | 0.673 | 0.774 | 7.70E-18 | 3.58E-56 | -0.012703156 | 0.314 | 0.404 | 9.63E-52 |
| 75 | Ninj2 | 1.42E-36 | 0.14192221 | 0.698 | 0.824 | 2.06E-33 | 2.08E-40 | 0.027303619 | 0.38 | 0.457 | 5.59E-36 |
| 80 | Dcc | 1.28E-34 | 0.137246297 | 0.74 | 0.859 | 1.85E-31 | 4.72E-25 | 0.153265829 | 0.37 | 0.4 | 1.27E-20 |
| 69 | Elmo1 | 1.06E-43 | 0.133469415 | 0.817 | 0.924 | 1.54E-40 | 2.75E-53 | -0.300951194 | 0.673 | 0.818 | 7.40E-49 |
| 106 | Cobl | 7.44E-07 | 0.128914521 | 0.604 | 0.665 | 0.00108 | 6.85E-46 | -0.000358497 | 0.217 | 0.283 | 1.84E-41 |
| 13 | Il33 | 3.65E-141 | 0.121066961 | 0.483 | 0.705 | ####### | 5.22E-30 | 0.042881108 | 0.416 | 0.476 | 1.41E-25 |
| 71 | Aspa | 8.57E-43 | 0.114917807 | 0.789 | 0.905 | 1.24E-39 | 7.96E-49 | -0.029196053 | 0.65 | 0.766 | 2.14E-44 |
| 44 | Car2 | 4.49E-65 | 0.113106427 | 0.698 | 0.852 | 6.51E-62 | 3.88E-42 | -0.02667842 | 0.588 | 0.693 | 1.04E-37 |
| 29 | Adamtsl1 | 2.19E-91 | 0.111252709 | 0.539 | 0.746 | 3.17E-88 | 1.09E-48 | -0.020583843 | 0.368 | 0.461 | 2.93E-44 |
| 89 | Tenm2 | 1.50E-24 | 0.107993608 | 0.82 | 0.906 | 2.18E-21 | 2.53E-88 | 0.787871799 | 0.491 | 0.383 | 6.82E-84 |
| 103 | Slc6a1 | 1.91E-12 | 0.099209329 | 0.314 | 0.395 | 2.77E-09 | 4.66E-59 | -0.089111539 | 0.189 | 0.277 | 1.26E-54 |
| 72 | Adipor2 | 1.51E-42 | 0.097683824 | 0.698 | 0.831 | 2.19E-39 | 7.56E-55 | -0.286470428 | 0.535 | 0.695 | 2.04E-50 |
| 109 | Prkcq | 0.00430686 | 0.09638253 | 0.548 | 0.576 | 1 | 5.44E-46 | -0.051963839 | 0.192 | 0.263 | 1.46E-41 |
| 88 | Sh3d19 | 1.35E-24 | 0.092765687 | 0.644 | 0.759 | 1.96E-21 | 1.94E-55 | -0.062486788 | 0.433 | 0.552 | 5.23E-51 |
| 74 | Mog | 1.58E-39 | 0.089400173 | 0.782 | 0.897 | 2.29E-36 | 2.84E-48 | -0.085081438 | 0.658 | 0.782 | 7.65E-44 |
| 35 | 1700047M11Rik | 2.46E-80 | 0.07538195 | 0.479 | 0.682 | 3.56E-77 | 1.64E-63 | -0.122162684 | 0.327 | 0.447 | 4.41E-59 |
| 86 | Lama2 | 1.35E-28 | 0.068509068 | 0.647 | 0.772 | 1.96E-25 | 1.28E-28 | -0.122154653 | 0.277 | 0.358 | 3.45E-24 |
| 61 | Pde8a | 7.23E-52 | 0.060932695 | 0.72 | 0.866 | 1.05E-48 | 1.03E-54 | -0.038996623 | 0.524 | 0.64 | 2.78E-50 |
| 27 | Glul | 1.60E-97 | 0.052760106 | 0.341 | 0.571 | 2.32E-94 | 2.03E-50 | -0.199838866 | 0.186 | 0.288 | 5.47E-46 |
| 76 | Grid2 | 1.44E-36 | 0.049976941 | 0.78 | 0.895 | 2.10E-33 | 2.44E-45 | -0.040937949 | 0.565 | 0.681 | 6.56E-41 |
| 107 | Stk39 | 1.87E-06 | 0.040512945 | 0.573 | 0.625 | 0.00272 | 6.10E-52 | -0.047978947 | 0.219 | 0.298 | 1.64E-47 |
| 105 | Plp1 | 1.34E-10 | 0.030297446 | 0.977 | 0.995 | 1.94E-07 | 2.41E-38 | -0.18244021 | 0.965 | 0.988 | 6.50E-34 |
| 36 | Sytl2 | 4.82E-76 | 0.025382117 | 0.716 | 0.888 | 6.99E-73 | 4.31E-48 | 0.182561396 | 0.347 | 0.382 | 1.16E-43 |
| 73 | Plxdc2 | 3.60E-41 | 0.023549269 | 0.856 | 0.951 | 5.22E-38 | 2.27E-40 | -0.076390901 | 0.696 | 0.803 | 6.11E-36 |
| 67 | Zfp536 | 2.82E-46 | 0.023185598 | 0.83 | 0.932 | 4.10E-43 | 3.82E-30 | -0.207716632 | 0.765 | 0.86 | 1.03E-25 |
| 90 | Prkca | 3.76E-24 | 0.001831846 | 0.709 | 0.816 | 5.46E-21 | 3.28E-55 | -0.111153313 | 0.329 | 0.448 | 8.83E-51 |
| 54 | Pstpip2 | 9.31E-55 | -0.00473266 | 0.604 | 0.773 | 1.35E-51 | 8.98E-59 | -0.182567643 | 0.418 | 0.562 | 2.42E-54 |
| 31 | Fa2h | 9.35E-87 | -0.011067991 | 0.751 | 0.923 | 1.36E-83 | 1.05E-61 | -0.103097908 | 0.477 | 0.613 | 2.82E-57 |
| 8 | D7Ertd443e | 1.40E-154 | -0.012923997 | 0.382 | 0.673 | ####### | 7.05E-63 | -0.571415538 | 0.245 | 0.418 | 1.90E-58 |
| 47 | Lrrc4c | 2.43E-60 | -0.015273615 | 0.75 | 0.899 | 3.52E-57 | 2.98E-25 | -0.156624403 | 0.644 | 0.745 | 8.02E-21 |
| 91 | Tmtc2 | 4.23E-24 | -0.016107756 | 0.601 | 0.718 | 6.14E-21 | 3.52E-57 | -0.147528554 | 0.325 | 0.454 | 9.47E-53 |
| 64 | Heg1 | 1.81E-49 | -0.030217751 | 0.476 | 0.657 | 2.63E-46 | 4.99E-47 | -0.229041145 | 0.16 | 0.256 | 1.34E-42 |
| 22 | Apod | 3.61E-110 | -0.031837546 | 0.465 | 0.706 | ####### | 1.86E-66 | -0.50370207 | 0.351 | 0.537 | 5.00E-62 |
| 66 | Samd4 | 8.14E-47 | -0.033295659 | 0.577 | 0.744 | 1.18E-43 | 8.22E-41 | 0.077430243 | 0.24 | 0.289 | 2.21E-36 |
| 12 | Sec14l5 | 6.03E-144 | -0.037801965 | 0.593 | 0.844 | ####### | 2.49E-68 | -0.220395921 | 0.496 | 0.662 | 6.70E-64 |
| 68 | Sema6d | 6.36E-44 | -0.039261583 | 0.716 | 0.856 | 9.23E-41 | 8.80E-42 | 0.070669812 | 0.316 | 0.373 | 2.37E-37 |
| 45 | Neat1 | 1.28E-64 | -0.041760634 | 0.735 | 0.893 | 1.86E-61 | 4.90E-39 | -0.206758197 | 0.665 | 0.791 | 1.32E-34 |
| 25 | Oxr1 | 3.44E-103 | -0.046727252 | 0.57 | 0.778 | ####### | 8.40E-30 | -0.080580193 | 0.524 | 0.617 | 2.26E-25 |
| 2 | Hapln2 | 1.20E-238 | -0.04779294 | 0.283 | 0.593 | ####### | 7.79E-100 | -0.329747972 | 0.246 | 0.427 | 2.10E-95 |
| 34 | Spock1 | 2.94E-81 | -0.053589158 | 0.582 | 0.788 | 4.27E-78 | 1.05E-24 | -0.261821714 | 0.39 | 0.499 | 2.84E-20 |
| 37 | Pam | 1.55E-73 | -0.062102007 | 0.578 | 0.785 | 2.24E-70 | 1.97E-38 | 0.070369235 | 0.207 | 0.255 | 5.31E-34 |
| 28 | Gjc3 | 8.15E-92 | -0.062667942 | 0.653 | 0.856 | 1.18E-88 | 4.81E-81 | -0.307498356 | 0.492 | 0.682 | 1.30E-76 |
| 46 | Ugt8a | 3.81E-62 | -0.07312668 | 0.791 | 0.93 | 5.52E-59 | 8.17E-60 | -0.134583411 | 0.477 | 0.616 | 2.20E-55 |
| 51 | Gm16168 | 1.93E-55 | -0.075830835 | 0.766 | 0.907 | 2.80E-52 | 7.66E-46 | -0.216888429 | 0.5 | 0.644 | 2.06E-41 |
| 33 | Cdh19 | 1.85E-84 | -0.077179449 | 0.547 | 0.764 | 2.69E-81 | 1.65E-45 | -0.308131852 | 0.39 | 0.535 | 4.45E-41 |
| 7 | Alcam | 3.91E-161 | -0.078872105 | 0.221 | 0.492 | ####### | 3.08E-35 | -0.114205993 | 0.183 | 0.259 | 8.30E-31 |
| 6 | Rcan2 | 4.37E-162 | -0.084931966 | 0.339 | 0.601 | ####### | 2.84E-81 | -0.398482062 | 0.288 | 0.471 | 7.64E-77 |
| 24 | Pde1c | 1.44E-103 | -0.089516159 | 0.534 | 0.778 | ####### | 9.22E-34 | 0.024033934 | 0.256 | 0.312 | 2.48E-29 |
| 4 | Gm37459 | 2.43E-188 | -0.092124603 | 0.36 | 0.652 | ####### | 1.49E-65 | -0.49599198 | 0.305 | 0.485 | 4.00E-61 |
| 52 | Bcas1 | 1.96E-55 | -0.096562418 | 0.711 | 0.867 | 2.85E-52 | 3.13E-69 | -0.251316165 | 0.455 | 0.625 | 8.43E-65 |
| 38 | Gab1 | 6.58E-72 | -0.096836819 | 0.744 | 0.908 | 9.55E-69 | 5.73E-56 | -0.161714415 | 0.455 | 0.595 | 1.54E-51 |
| 85 | Sorcs1 | 1.56E-29 | -0.105664417 | 0.737 | 0.842 | 2.26E-26 | 1.37E-47 | -0.096439451 | 0.536 | 0.659 | 3.69E-43 |
| 50 | Stxbp6 | 4.25E-56 | -0.106766632 | 0.656 | 0.825 | 6.17E-53 | 2.30E-49 | -0.06691668 | 0.408 | 0.519 | 6.18E-45 |
| 5 | 4930419G24Rik | 5.92E-186 | -0.108622733 | 0.409 | 0.686 | ####### | 9.55E-95 | -0.541144846 | 0.395 | 0.62 | 2.57E-90 |
| 3 | Gria4 | 2.01E-216 | -0.111196407 | 0.434 | 0.769 | ####### | 2.71E-31 | 0.225431348 | 0.32 | 0.334 | 7.30E-27 |
| 32 | Lrrtm3 | 3.46E-86 | -0.114343234 | 0.364 | 0.591 | 5.02E-83 | 3.29E-64 | -0.391215226 | 0.265 | 0.428 | 8.84E-60 |
| 87 | Pex5l | 3.10E-28 | -0.117287706 | 0.903 | 0.966 | 4.50E-25 | 1.20E-48 | -0.331977909 | 0.774 | 0.887 | 3.23E-44 |
| 98 | Mbp | 5.94E-20 | -0.123822665 | 0.978 | 0.995 | 8.61E-17 | 1.03E-60 | -0.226609831 | 0.967 | 0.991 | 2.76E-56 |
| 81 | St18 | 2.20E-34 | -0.129393824 | 0.97 | 0.996 | 3.19E-31 | 2.09E-26 | -0.160812766 | 0.926 | 0.97 | 5.63E-22 |
| 57 | Elovl7 | 6.97E-53 | -0.132448157 | 0.602 | 0.776 | 1.01E-49 | 3.97E-54 | -0.258305567 | 0.186 | 0.298 | 1.07E-49 |
| 82 | Ephb1 | 1.08E-31 | -0.13755315 | 0.656 | 0.768 | 1.56E-28 | 1.44E-65 | -0.316724132 | 0.265 | 0.416 | 3.87E-61 |
| 10 | Dock5 | 3.95E-146 | -0.137553278 | 0.525 | 0.817 | ####### | 1.89E-62 | -0.198578599 | 0.232 | 0.352 | 5.09E-58 |
| 49 | Npas3 | 2.83E-58 | -0.139280318 | 0.717 | 0.874 | 4.11E-55 | 1.24E-32 | -0.061669261 | 0.465 | 0.559 | 3.33E-28 |
| 58 | Ctnna3 | 1.10E-52 | -0.140986412 | 0.822 | 0.938 | 1.60E-49 | 2.64E-46 | -0.302940347 | 0.653 | 0.792 | 7.10E-42 |
| 30 | Trps1 | 1.04E-90 | -0.14318268 | 0.389 | 0.628 | 1.51E-87 | 3.99E-38 | 0.046853391 | 0.245 | 0.301 | 1.08E-33 |
| 65 | Prr5l | 5.20E-49 | -0.148000284 | 0.918 | 0.98 | 7.54E-46 | 4.17E-56 | -0.301193518 | 0.811 | 0.916 | 1.12E-51 |
| 18 | Daam2 | 1.67E-121 | -0.153148036 | 0.593 | 0.839 | ####### | 3.45E-70 | -0.327123278 | 0.463 | 0.646 | 9.30E-66 |
| 41 | Ptn | 4.41E-66 | -0.156788314 | 0.607 | 0.799 | 6.40E-63 | 3.60E-58 | -0.203623008 | 0.394 | 0.538 | 9.69E-54 |
| 19 | Rasgrp3 | 7.34E-115 | -0.165794075 | 0.44 | 0.714 | ####### | 3.79E-49 | -0.225692176 | 0.161 | 0.258 | 1.02E-44 |
| 63 | Enpp2 | 5.83E-51 | -0.177119898 | 0.857 | 0.96 | 8.46E-48 | 2.34E-27 | -0.07082672 | 0.758 | 0.846 | 6.31E-23 |
| 11 | Prox1 | 2.80E-145 | -0.180624053 | 0.498 | 0.795 | ####### | 2.80E-50 | -0.090871847 | 0.177 | 0.257 | 7.54E-46 |
| 26 | Ubash3b | 1.75E-98 | -0.199314429 | 0.464 | 0.718 | 2.53E-95 | 1.13E-55 | -0.307578627 | 0.199 | 0.327 | 3.03E-51 |
| 62 | Cdh20 | 3.59E-51 | -0.23862983 | 0.771 | 0.898 | 5.21E-48 | 1.05E-53 | -0.321716737 | 0.465 | 0.627 | 2.82E-49 |
| 9 | Pcdh11x | 7.18E-149 | -0.243083666 | 0.245 | 0.526 | ####### | 9.27E-32 | -0.308245913 | 0.196 | 0.296 | 2.49E-27 |
| 15 | Plekhg1 | 5.47E-130 | -0.291686178 | 0.543 | 0.814 | ####### | 4.34E-61 | -0.087422241 | 0.395 | 0.519 | 1.17E-56 |
| 16 | Spock3 | 1.46E-129 | -0.339608453 | 0.58 | 0.844 | ####### | 2.38E-39 | -0.265351769 | 0.413 | 0.546 | 6.40E-35 |
| 39 | St3gal4 | 5.81E-71 | -0.362694285 | 0.655 | 0.837 | 8.43E-68 | 7.24E-32 | -0.183483337 | 0.198 | 0.281 | 1.95E-27 |
| 20 | A330049N07Rik | 5.63E-112 | -0.398302276 | 0.595 | 0.832 | ####### | 1.76E-63 | -0.393581653 | 0.25 | 0.408 | 4.75E-59 |
| 1 | A230001M10Rik | 0 | -0.410634548 | 0.388 | 0.831 | 0 | 1.71E-52 | -0.167880555 | 0.338 | 0.461 | 4.61E-48 |
| 17 | Rnf220 | 4.79E-127 | -0.449203963 | 0.834 | 0.978 | ####### | 1.39E-49 | -0.244831492 | 0.687 | 0.824 | 3.74E-45 |
| 14 | Anln | 3.61E-133 | -0.480559351 | 0.363 | 0.663 | ####### | 4.89E-112 | -0.720921735 | 0.237 | 0.471 | 1.32E-107 |
| 59 | Robo1 | 3.51E-52 | -0.48679591 | 0.579 | 0.728 | 5.10E-49 | 2.63E-15 | -0.338671531 | 0.194 | 0.266 | 7.08E-11 |

| | p_val | avg_log2FC | pct.1 | pct.2 | p_val_adj |
|---|---|---|---|---|---|
| Kcnip4 | 3.50E-256 | 1.638148626 | 0.485 | 0.189 | 9.43E-252 |
| Meg3 | 1.15E-209 | 1.191499385 | 0.735 | 0.46 | 3.11E-205 |
| Rbfox1 | 1.97E-202 | 1.416472422 | 0.352 | 0.124 | 5.30E-198 |
| Csmd1 | 6.45E-188 | 1.384201597 | 0.37 | 0.142 | 1.74E-183 |
| Nrg3 | 2.27E-184 | 1.362780868 | 0.367 | 0.152 | 6.10E-180 |
| Celf2 | 1.49E-156 | 1.129681296 | 0.401 | 0.197 | 4.00E-152 |
| Nrxn3 | 4.20E-149 | 1.21760068 | 0.31 | 0.129 | 1.13E-144 |
| Lrrtm4 | 7.60E-148 | 1.213324077 | 0.274 | 0.095 | 2.05E-143 |
| Snhg11 | 7.12E-142 | 0.925375742 | 0.667 | 0.482 | 1.92E-137 |
| Syt1 | 1.82E-125 | 1.010849493 | 0.298 | 0.181 | 4.91E-121 |
| Nrxn1 | 1.53E-117 | 0.938643336 | 0.397 | 0.286 | 4.11E-113 |
| Gria2 | 2.18E-115 | 0.791775315 | 0.353 | 0.276 | 5.87E-111 |
| Dlgap1 | 6.19E-114 | 1.005356356 | 0.273 | 0.127 | 1.67E-109 |
| Anln | 4.89E-112 | -0.720921735 | 0.237 | 0.471 | 1.32E-107 |
| Lsamp | 8.06E-104 | 0.770785934 | 0.632 | 0.514 | 2.17E-99 |
| Hapln2 | 7.79E-100 | -0.329747972 | 0.246 | 0.427 | 2.10E-95 |
| 4930419G24Rik | 9.55E-95 | -0.541144846 | 0.395 | 0.62 | 2.57E-90 |
| Tenm2 | 2.53E-88 | 0.787871799 | 0.491 | 0.383 | 6.82E-84 |
| Gm11149 | 2.91E-88 | -0.376701074 | 0.29 | 0.477 | 7.84E-84 |
| 1-Mar | 9.02E-84 | -1.076608992 | 0.131 | 0.312 | 2.43E-79 |
| Jam3 | 4.15E-82 | -0.011313289 | 0.282 | 0.388 | 1.12E-77 |
| Arhgef10 | 1.19E-81 | -0.167342592 | 0.354 | 0.505 | 3.19E-77 |
| Rcan2 | 2.84E-81 | -0.398482062 | 0.288 | 0.471 | 7.64E-77 |
| Gjc3 | 4.81E-81 | -0.307498356 | 0.492 | 0.682 | 1.30E-76 |
| Prrc2c | 5.51E-81 | -0.06275944 | 0.246 | 0.359 | 1.48E-76 |
| Rbm25 | 2.08E-80 | 0.070215854 | 0.298 | 0.388 | 5.61E-76 |
| Srsf7 | 5.73E-79 | -0.158320817 | 0.229 | 0.356 | 1.54E-74 |
| Acap2 | 1.29E-78 | 0.049094926 | 0.32 | 0.417 | 3.46E-74 |
| Atp2b1 | 2.89E-78 | 0.297258623 | 0.261 | 0.294 | 7.78E-74 |
| Itpk1 | 3.11E-78 | -0.243795793 | 0.268 | 0.419 | 8.38E-74 |
| Aplp2 | 2.99E-77 | -0.094544578 | 0.191 | 0.294 | 8.04E-73 |
| Rab31 | 2.74E-76 | -0.316985405 | 0.486 | 0.675 | 7.38E-72 |
| Ptpra | 1.03E-75 | -0.024196427 | 0.263 | 0.363 | 2.78E-71 |
| Rcbtb1 | 1.15E-75 | -0.136824309 | 0.364 | 0.507 | 3.10E-71 |
| Cmss1 | 2.20E-75 | 0.324688519 | 0.546 | 0.566 | 5.93E-71 |
| Paip2 | 2.78E-75 | -0.204084316 | 0.239 | 0.37 | 7.49E-71 |
| Mettl7a1 | 6.37E-75 | -0.296372495 | 0.169 | 0.301 | 1.72E-70 |
| Mat2a | 6.92E-75 | -0.181384461 | 0.313 | 0.457 | 1.86E-70 |
| Rab40c | 1.54E-74 | -0.025694402 | 0.257 | 0.361 | 4.14E-70 |
| Plekhh1 | 1.71E-74 | -0.104216935 | 0.439 | 0.58 | 4.61E-70 |
| Klf13 | 1.93E-74 | -0.103011591 | 0.288 | 0.411 | 5.19E-70 |
| Itm2b | 4.46E-74 | -0.037880103 | 0.222 | 0.322 | 1.20E-69 |
| Nipbl | 4.86E-74 | 0.034493432 | 0.288 | 0.378 | 1.31E-69 |
| Ppp2r2a | 6.38E-74 | -0.094076173 | 0.259 | 0.373 | 1.72E-69 |
| Camk1d | 1.19E-73 | 0.295624341 | 0.286 | 0.308 | 3.20E-69 |
| Otud7b | 1.59E-73 | -0.158895537 | 0.421 | 0.572 | 4.28E-69 |
| Tra2a | 7.93E-73 | -0.009236161 | 0.323 | 0.428 | 2.13E-68 |
| Gm44866 | 7.96E-73 | -0.601876914 | 0.109 | 0.255 | 2.14E-68 |
| Cpeb2 | 1.19E-72 | -0.006874226 | 0.244 | 0.335 | 3.20E-68 |
| Ptpn11 | 2.04E-72 | -0.185290974 | 0.211 | 0.333 | 5.50E-68 |
| Dpysl2 | 2.32E-72 | -0.078769997 | 0.336 | 0.456 | 6.24E-68 |
| Rffl | 2.50E-72 | -0.169278847 | 0.275 | 0.407 | 6.72E-68 |
| Cyth1 | 3.49E-72 | -0.09944906 | 0.298 | 0.42 | 9.39E-68 |
| Slc44a1 | 6.57E-72 | -0.212627339 | 0.453 | 0.62 | 1.77E-67 |
| Kcna1 | 1.12E-71 | -0.169203759 | 0.251 | 0.377 | 3.02E-67 |
| Cntn2 | 1.19E-71 | 0.031804852 | 0.359 | 0.452 | 3.21E-67 |
| Ddx17 | 1.72E-71 | 0.084589107 | 0.378 | 0.46 | 4.62E-67 |
| Zfp280d | 2.45E-71 | -0.054446742 | 0.234 | 0.335 | 6.59E-67 |
| Trpm7 | 2.66E-71 | 0.022181956 | 0.265 | 0.352 | 7.16E-67 |
| Wdfy3 | 3.60E-71 | 0.044247477 | 0.26 | 0.339 | 9.69E-67 |
| Grb14 | 1.28E-70 | -0.016061381 | 0.4 | 0.517 | 3.44E-66 |
| Stmn4 | 1.84E-70 | -0.044043377 | 0.208 | 0.299 | 4.94E-66 |
| Hnrnpa2b1 | 1.90E-70 | -0.18441906 | 0.388 | 0.539 | 5.11E-66 |
| Abca2 | 2.23E-70 | -0.184232258 | 0.199 | 0.316 | 6.01E-66 |
| Pnisr | 3.26E-70 | -0.062921096 | 0.418 | 0.544 | 8.78E-66 |
| Daam2 | 3.45E-70 | -0.327123278 | 0.463 | 0.646 | 9.30E-66 |
| Ermn | 3.91E-70 | -0.215340575 | 0.256 | 0.392 | 1.05E-65 |
| Litaf | 4.14E-70 | -0.195327394 | 0.276 | 0.41 | 1.11E-65 |
| Ogt | 4.40E-70 | -0.065293385 | 0.221 | 0.321 | 1.19E-65 |
| Clk1 | 4.98E-70 | -0.088219783 | 0.23 | 0.336 | 1.34E-65 |
| Zbtb16 | 6.41E-70 | -0.268392011 | 0.269 | 0.42 | 1.73E-65 |
| Pard3 | 6.86E-70 | -0.198745199 | 0.35 | 0.501 | 1.85E-65 |
| Kmt2c | 7.54E-70 | -0.01142178 | 0.307 | 0.408 | 2.03E-65 |
| Spag9 | 1.04E-69 | -0.024341369 | 0.362 | 0.47 | 2.80E-65 |
| Usp34 | 1.06E-69 | 0.107863081 | 0.343 | 0.415 | 2.85E-65 |
| Epc1 | 2.28E-69 | -0.036983051 | 0.204 | 0.293 | 6.12E-65 |
| Bcas1 | 3.13E-69 | -0.251316165 | 0.455 | 0.625 | 8.43E-65 |
| Ddr1 | 5.00E-69 | -0.067815114 | 0.231 | 0.33 | 1.35E-64 |
| Arid4b | 5.01E-69 | -0.027670587 | 0.331 | 0.435 | 1.35E-64 |
| Srek1 | 8.71E-69 | 0.047054436 | 0.207 | 0.28 | 2.34E-64 |
| Srrm1 | 8.91E-69 | -0.025673957 | 0.203 | 0.288 | 2.40E-64 |
| Rsrp1 | 1.01E-68 | -0.105390074 | 0.488 | 0.636 | 2.72E-64 |
| Sez6l2 | 1.86E-68 | -0.245568496 | 0.271 | 0.414 | 5.02E-64 |
| Sec14l5 | 2.49E-68 | -0.220395921 | 0.496 | 0.662 | 6.70E-64 |
| Sptbn1 | 2.63E-68 | 0.077114712 | 0.38 | 0.465 | 7.07E-64 |
| Thra | 2.81E-68 | 0.178181094 | 0.284 | 0.332 | 7.55E-64 |
| Atrx | 3.57E-68 | 0.035563403 | 0.295 | 0.384 | 9.62E-64 |
| Cd81 | 3.88E-68 | -0.010297152 | 0.222 | 0.31 | 1.05E-63 |
| Rbm39 | 4.18E-68 | -0.035653514 | 0.51 | 0.639 | 1.13E-63 |
| Zfp532 | 4.20E-68 | -0.143255803 | 0.235 | 0.351 | 1.13E-63 |
| Apoe | 4.26E-68 | 0.14474695 | 0.448 | 0.539 | 1.15E-63 |
| Pafah1b1 | 4.71E-68 | -0.163294813 | 0.408 | 0.553 | 1.27E-63 |
| Itch | 4.80E-68 | -0.092909106 | 0.413 | 0.543 | 1.29E-63 |
| Myo1d | 6.52E-68 | 0.013754494 | 0.406 | 0.509 | 1.75E-63 |
| Ube2d1 | 9.49E-68 | -0.08806118 | 0.279 | 0.389 | 2.55E-63 |
| Pacrg | 1.28E-67 | -0.401301236 | 0.502 | 0.687 | 3.45E-63 |
| Cnot4 | 1.48E-67 | 0.029122616 | 0.223 | 0.301 | 3.98E-63 |
| Rock1 | 1.48E-67 | 0.060878941 | 0.226 | 0.294 | 3.98E-63 |
| Nkain1 | 1.64E-67 | -0.083324756 | 0.267 | 0.373 | 4.43E-63 |
| Megf10 | 2.59E-67 | -0.149642409 | 0.205 | 0.316 | 6.96E-63 |
| Usp54 | 3.26E-67 | -0.164655965 | 0.432 | 0.579 | 8.79E-63 |
| Bcas3 | 4.49E-67 | -0.143730282 | 0.327 | 0.463 | 1.21E-62 |
| Atrn | 5.25E-67 | 0.003633573 | 0.259 | 0.345 | 1.41E-62 |
| Dip2a | 7.85E-67 | -0.098543358 | 0.399 | 0.527 | 2.11E-62 |
| Rnf13 | 9.75E-67 | -0.041120275 | 0.386 | 0.498 | 2.63E-62 |
| Pnn | 1.05E-66 | -0.015556188 | 0.216 | 0.304 | 2.83E-62 |
| Ttyh2 | 1.11E-66 | -0.007145442 | 0.354 | 0.456 | 2.99E-62 |
| Crebbp | 1.46E-66 | 0.109910713 | 0.27 | 0.333 | 3.94E-62 |
| Ppp1r12a | 1.48E-66 | 0.067420746 | 0.277 | 0.35 | 4.00E-62 |
| Srrm2 | 1.58E-66 | -0.055999838 | 0.448 | 0.575 | 4.26E-62 |
| Apod | 1.86E-66 | -0.50370207 | 0.351 | 0.537 | 5.00E-62 |
| Csnk1a1 | 1.95E-66 | 0.013786541 | 0.204 | 0.28 | 5.26E-62 |
| Aatk | 2.54E-66 | 0.110535576 | 0.467 | 0.545 | 6.83E-62 |
| Dixdc1 | 2.85E-66 | -0.059588256 | 0.282 | 0.388 | 7.66E-62 |
| Dpy19l1 | 3.54E-66 | -0.03724027 | 0.202 | 0.286 | 9.52E-62 |
| Rb1 | 3.78E-66 | -0.07521711 | 0.209 | 0.306 | 1.02E-61 |
| Sf3b1 | 3.91E-66 | 0.000873009 | 0.2 | 0.278 | 1.05E-61 |
| Mag | 3.98E-66 | -0.173422971 | 0.553 | 0.711 | 1.07E-61 |
| Dync1li2 | 4.15E-66 | 0.008777599 | 0.208 | 0.284 | 1.12E-61 |
| Tmbim1 | 5.11E-66 | 0.045163708 | 0.201 | 0.267 | 1.38E-61 |
| Epb41l3 | 5.24E-66 | 0.067926659 | 0.339 | 0.422 | 1.41E-61 |
| Clk4 | 6.86E-66 | 0.061693842 | 0.234 | 0.305 | 1.85E-61 |
| Neo1 | 1.22E-65 | 0.031557051 | 0.349 | 0.439 | 3.27E-61 |
| Ephb1 | 1.44E-65 | -0.316724132 | 0.265 | 0.416 | 3.87E-61 |
| Scd2 | 1.49E-65 | -0.172082841 | 0.46 | 0.613 | 4.00E-61 |
| Gm37459 | 1.49E-65 | -0.49599198 | 0.305 | 0.485 | 4.00E-61 |
| Exoc4 | 1.78E-65 | 0.019796403 | 0.38 | 0.479 | 4.80E-61 |
| Arhgap39 | 2.14E-65 | -0.113213883 | 0.299 | 0.414 | 5.75E-61 |
| Hook3 | 2.25E-65 | -0.005396973 | 0.31 | 0.403 | 6.05E-61 |
| Dnm2 | 2.33E-65 | -0.081646424 | 0.189 | 0.282 | 6.28E-61 |
| Bcl2l1 | 2.48E-65 | -0.161571461 | 0.339 | 0.473 | 6.67E-61 |
| Dmxl1 | 2.97E-65 | 0.151664324 | 0.295 | 0.348 | 7.99E-61 |
| Aig1 | 3.95E-65 | 0.010894025 | 0.307 | 0.397 | 1.06E-60 |
| Inpp5f | 4.26E-65 | -0.153286063 | 0.222 | 0.332 | 1.15E-60 |
| 2610035D17Rik | 4.46E-65 | -0.296833968 | 0.251 | 0.396 | 1.20E-60 |
| Tmcc3 | 4.64E-65 | -0.273627081 | 0.354 | 0.511 | 1.25E-60 |
| Rbm26 | 4.79E-65 | 0.072472305 | 0.278 | 0.349 | 1.29E-60 |
| Atp6v0b | 5.42E-65 | 0.092134753 | 0.27 | 0.341 | 1.46E-60 |
| Prpf4b | 5.57E-65 | -0.051995763 | 0.327 | 0.438 | 1.50E-60 |
| Gm26936 | 5.86E-65 | -0.248572945 | 0.21 | 0.339 | 1.58E-60 |
| Zcchc7 | 6.87E-65 | -0.139341929 | 0.402 | 0.542 | 1.85E-60 |
| Son | 8.16E-65 | 0.007437758 | 0.403 | 0.506 | 2.20E-60 |
| Chd6 | 8.56E-65 | 0.123314969 | 0.28 | 0.336 | 2.30E-60 |
| Dennd5a | 1.88E-64 | -0.105286585 | 0.234 | 0.338 | 5.06E-60 |
| Hnrnpdl | 2.03E-64 | -0.018706054 | 0.268 | 0.357 | 5.47E-60 |
| Dennd1a | 2.40E-64 | -0.094926135 | 0.301 | 0.414 | 6.46E-60 |
| Rab10 | 2.59E-64 | -0.122121448 | 0.23 | 0.337 | 6.96E-60 |
| Ankrd28 | 2.71E-64 | -0.125217232 | 0.449 | 0.587 | 7.29E-60 |
| Ptk2 | 3.02E-64 | -0.082036486 | 0.473 | 0.606 | 8.13E-60 |
| Nrd1 | 3.10E-64 | -0.062659513 | 0.217 | 0.308 | 8.35E-60 |
| Lrrtm3 | 3.29E-64 | -0.391215226 | 0.265 | 0.428 | 8.84E-60 |
| Fus | 3.41E-64 | -0.129706368 | 0.255 | 0.366 | 9.19E-60 |
| Zranb2 | 3.46E-64 | 0.126046557 | 0.209 | 0.26 | 9.30E-60 |
| Dnajc1 | 3.88E-64 | 0.020233626 | 0.222 | 0.299 | 1.04E-59 |
| Atp2c1 | 4.63E-64 | 0.105722063 | 0.234 | 0.294 | 1.25E-59 |
| Rbm5 | 4.64E-64 | 0.023006738 | 0.263 | 0.343 | 1.25E-59 |
| Pik3c2b | 4.65E-64 | -0.008417588 | 0.274 | 0.363 | 1.25E-59 |
| Picalm | 4.90E-64 | -0.165155194 | 0.33 | 0.462 | 1.32E-59 |
| AY036118 | 6.13E-64 | -0.250237234 | 0.28 | 0.424 | 1.65E-59 |
| Agpat4 | 6.53E-64 | -0.186542518 | 0.225 | 0.341 | 1.76E-59 |
| Abcc5 | 6.72E-64 | -0.018859098 | 0.238 | 0.323 | 1.81E-59 |
| Sgk1 | 6.77E-64 | -0.28974472 | 0.257 | 0.403 | 1.82E-59 |
| St3gal3 | 7.18E-64 | -0.143253767 | 0.257 | 0.371 | 1.93E-59 |
| Plekhg3 | 8.54E-64 | -0.012344874 | 0.222 | 0.301 | 2.30E-59 |
| Pik3r1 | 9.10E-64 | -0.090621571 | 0.288 | 0.399 | 2.45E-59 |
| Kctd13 | 9.14E-64 | -0.163361385 | 0.255 | 0.371 | 2.46E-59 |
| Ankib1 | 1.16E-63 | 0.077343204 | 0.352 | 0.429 | 3.13E-59 |
| Sltm | 1.19E-63 | 0.014324184 | 0.296 | 0.38 | 3.21E-59 |
| Cttnbp2 | 1.23E-63 | 0.169991741 | 0.458 | 0.521 | 3.30E-59 |
| Ywhaq | 1.25E-63 | -0.083263485 | 0.313 | 0.424 | 3.37E-59 |
| Ttc14 | 1.33E-63 | 0.041470208 | 0.265 | 0.343 | 3.58E-59 |
| Ccdc50 | 1.38E-63 | -0.076491658 | 0.33 | 0.445 | 3.70E-59 |
| Ncor1 | 1.40E-63 | -0.003838776 | 0.289 | 0.377 | 3.76E-59 |
| 1700047M11Rik | 1.64E-63 | -0.122162684 | 0.327 | 0.447 | 4.41E-59 |
| A330049N07Rik | 1.76E-63 | -0.393581653 | 0.25 | 0.408 | 4.75E-59 |
| Secisbp2l | 1.79E-63 | -0.182182844 | 0.185 | 0.289 | 4.81E-59 |
| Nrbp2 | 1.94E-63 | 0.143016363 | 0.305 | 0.357 | 5.23E-59 |
| Cipc | 2.30E-63 | -0.076802349 | 0.366 | 0.48 | 6.19E-59 |
| Fto | 2.56E-63 | -0.046823685 | 0.286 | 0.384 | 6.90E-59 |
| Arih1 | 3.02E-63 | -0.002778048 | 0.272 | 0.357 | 8.14E-59 |
| Cldn11 | 3.39E-63 | 0.008361785 | 0.365 | 0.461 | 9.13E-59 |
| Retreg1 | 3.90E-63 | -0.04065905 | 0.277 | 0.376 | 1.05E-58 |
| Ncoa2 | 3.95E-63 | 0.043123605 | 0.329 | 0.411 | 1.06E-58 |
| Mobp | 7.03E-63 | -0.292421403 | 0.642 | 0.803 | 1.89E-58 |
| D7Ertd443e | 7.05E-63 | -0.571415538 | 0.245 | 0.418 | 1.90E-58 |
| Lrch3 | 7.40E-63 | -0.159746743 | 0.275 | 0.394 | 1.99E-58 |
| Osbpl1a | 7.41E-63 | -0.04965754 | 0.311 | 0.413 | 1.99E-58 |
| Arhgap21 | 8.69E-63 | -0.179227061 | 0.307 | 0.438 | 2.34E-58 |
| Foxp1 | 9.27E-63 | -0.168464092 | 0.175 | 0.279 | 2.50E-58 |
| Ash1l | 1.04E-62 | 0.023431129 | 0.256 | 0.335 | 2.80E-58 |
| Pacs2 | 1.13E-62 | -0.053350339 | 0.238 | 0.326 | 3.05E-58 |
| Bmp2k | 1.19E-62 | -0.041916404 | 0.181 | 0.261 | 3.21E-58 |
| Csrp1 | 1.21E-62 | -0.127748677 | 0.308 | 0.427 | 3.25E-58 |
| Gna12 | 1.36E-62 | -0.137586585 | 0.252 | 0.364 | 3.65E-58 |
| Ppp3ca | 1.67E-62 | 0.25749511 | 0.438 | 0.475 | 4.50E-58 |
| Tnrc6c | 1.76E-62 | 0.092295376 | 0.335 | 0.404 | 4.72E-58 |
| Opcml | 1.84E-62 | 0.465088116 | 0.469 | 0.44 | 4.97E-58 |
| Hnrnpu | 1.85E-62 | 0.048641822 | 0.293 | 0.368 | 4.98E-58 |
| Eps15 | 1.87E-62 | 0.026190366 | 0.257 | 0.335 | 5.03E-58 |
| Dock5 | 1.89E-62 | -0.198578599 | 0.232 | 0.352 | 5.09E-58 |
| Plekha5 | 2.27E-62 | 0.182852998 | 0.272 | 0.317 | 6.11E-58 |
| Map4k4 | 2.38E-62 | -0.096688451 | 0.352 | 0.469 | 6.41E-58 |
| Scamp5 | 3.26E-62 | -0.080736583 | 0.184 | 0.272 | 8.78E-58 |
| Fubp1 | 5.70E-62 | 0.053163157 | 0.206 | 0.271 | 1.53E-57 |
| Srcin1 | 5.99E-62 | -0.060308227 | 0.321 | 0.425 | 1.61E-57 |
| Herc1 | 6.63E-62 | 0.133145581 | 0.225 | 0.275 | 1.78E-57 |
| Cdh2 | 8.07E-62 | -0.054417462 | 0.344 | 0.452 | 2.17E-57 |
| Ccdc88a | 8.19E-62 | 0.055295532 | 0.407 | 0.493 | 2.20E-57 |
| Igsf11 | 8.31E-62 | -0.155870005 | 0.175 | 0.273 | 2.24E-57 |
| Lgi3 | 8.58E-62 | 0.070743022 | 0.234 | 0.298 | 2.31E-57 |
| 4930402H24Rik | 8.62E-62 | 0.058020843 | 0.519 | 0.617 | 2.32E-57 |
| Epn2 | 8.62E-62 | -0.105876273 | 0.337 | 0.453 | 2.32E-57 |
| Trim35 | 9.62E-62 | 0.022875732 | 0.316 | 0.402 | 2.59E-57 |

| | p_val | avg_log2FC | pct.1 | pct.2 | p_val_adj |
|---|---|---|---|---|---|
| Slc13a1 | 6.47E-21 | 0.65849736 | 0.44 | 0.47 | 9.39E-18 |
| Clic4 | 1.28E-15 | 0.64074928 | 0.61 | 0.61 | 1.86E-12 |
| Kirrel3 | 4.53E-35 | 0.56271785 | 0.79 | 0.8 | 6.58E-32 |
| Apoe | 4.53E-111 | 0.54367426 | 0.48 | 0.55 | 6.57E-108 |
| Lrrtm4 | 5.80E-66 | 0.53344428 | 0.73 | 0.85 | 8.42E-63 |
| Epha6 | 4.92E-94 | 0.51697823 | 0.69 | 0.85 | 7.14E-91 |
| Ttr | 9.72E-121 | 0.48802331 | 0.51 | 0.73 | 1.41E-117 |
| Gatm | 1.52E-23 | 0.46869282 | 0.78 | 0.84 | 2.21E-20 |
| Nrxn3 | 7.25E-66 | 0.46372858 | 0.75 | 0.87 | 1.05E-62 |
| C030029H0 | 6.33E-36 | 0.40716912 | 0.78 | 0.84 | 9.19E-33 |
| Galntl6 | 2.14E-44 | 0.39942152 | 0.64 | 0.76 | 3.11E-41 |
| 7630403G2 | 1.06E-182 | 0.39689787 | 0.27 | 0.5 | 1.54E-179 |
| Ptgds | 4.68E-108 | 0.38714054 | 0.54 | 0.64 | 6.79E-105 |
| Crtac1 | 7.93E-18 | 0.38643449 | 0.56 | 0.55 | 1.15E-14 |
| Slc38a2 | 4.22E-52 | 0.38377399 | 0.51 | 0.63 | 6.12E-49 |
| Cntn4 | 1.90E-06 | 0.37100068 | 0.46 | 0.5 | 0.00275261 |
| Hs3st4 | 4.65E-47 | 0.36731699 | 0.59 | 0.69 | 6.74E-44 |
| Unc5c | 2.31E-14 | 0.35065397 | 0.86 | 0.9 | 3.35E-11 |
| Hs6st3 | 3.94E-32 | 0.34913383 | 0.66 | 0.77 | 5.71E-29 |
| Maml2 | 1.85E-10 | 0.32580391 | 0.41 | 0.48 | 2.68E-07 |
| Cldn11 | 3.12E-55 | 0.32473631 | 0.58 | 0.72 | 4.52E-52 |
| Cst3 | 1.89E-58 | 0.30767474 | 0.42 | 0.58 | 2.74E-55 |
| Cntnap2 | 8.04E-61 | 0.30634743 | 0.71 | 0.84 | 1.17E-57 |
| Ahcyl2 | 3.76E-09 | 0.27735941 | 0.59 | 0.65 | 5.45E-06 |
| Sphkap | 1.39E-65 | 0.27221142 | 0.55 | 0.72 | 2.01E-62 |
| Mag | 9.71E-36 | 0.26306119 | 0.74 | 0.86 | 1.41E-32 |
| Ntm | 5.55E-73 | 0.26155752 | 0.51 | 0.72 | 8.05E-70 |
| Trpm3 | 8.74E-31 | 0.24984343 | 0.52 | 0.66 | 1.27E-27 |
| Gm4275 | 1.28E-68 | 0.24809082 | 0.38 | 0.56 | 1.86E-65 |
| Trhde | 1.53E-06 | 0.24400779 | 0.47 | 0.42 | 0.00221359 |
| Bicc1 | 1.80E-51 | 0.24282089 | 0.54 | 0.71 | 2.61E-48 |
| Sema6a | 1.66E-14 | 0.24018056 | 0.73 | 0.81 | 2.41E-11 |
| Ryr3 | 1.01E-15 | 0.23591439 | 0.62 | 0.7 | 1.47E-12 |
| Cntn5 | 9.14E-54 | 0.23534501 | 0.58 | 0.74 | 1.33E-50 |
| 9330111N0 | 1.87E-21 | 0.23400133 | 0.53 | 0.63 | 2.71E-18 |
| A230006K0 | 2.75E-12 | 0.23100844 | 0.42 | 0.48 | 3.99E-09 |
| Etl4 | 2.21E-69 | 0.2307426 | 0.6 | 0.79 | 3.21E-66 |
| Gpc5 | 7.17E-32 | 0.22877716 | 0.52 | 0.65 | 1.04E-28 |
| Egfem1 | 3.22E-30 | 0.22701652 | 0.58 | 0.72 | 4.67E-27 |
| Kcnt2 | 7.27E-20 | 0.2242976 | 0.59 | 0.64 | 1.05E-16 |
| Tmem16 | 4.95E-18 | 0.22344317 | 0.52 | 0.43 | 7.19E-15 |
| Nrg1 | 8.85E-16 | 0.2223584 | 0.52 | 0.62 | 1.28E-12 |
| Rhoj | 0.29943105 | 0.21807796 | 0.29 | 0.3 | 1 |
| Fbn2 | 1.41E-19 | 0.21674217 | 0.59 | 0.68 | 2.05E-16 |
| Mobp | 9.49E-55 | 0.2163891 | 0.76 | 0.89 | 1.38E-51 |
| Cacnb2 | 1.84E-13 | 0.21331135 | 0.61 | 0.69 | 2.67E-10 |
| Thrb | 2.09E-06 | 0.20945207 | 0.57 | 0.51 | 0.00303729 |
| Tspan2 | 6.71E-30 | 0.20760228 | 0.69 | 0.79 | 9.74E-27 |
| Grip1 | 3.93E-13 | 0.20721219 | 0.59 | 0.66 | 5.71E-10 |
| Il1rapl2 | 7.73E-17 | 0.2045162 | 0.47 | 0.57 | 1.12E-13 |
| Chn2 | 1.12E-20 | 0.20434369 | 0.61 | 0.7 | 1.62E-17 |
| Asic2 | 3.49E-14 | 0.20111291 | 0.6 | 0.68 | 5.06E-11 |
| Prkcg | 1.88E-11 | 0.19955198 | 0.52 | 0.57 | 2.73E-08 |
| Kcnc2 | 7.79E-10 | 0.19830826 | 0.55 | 0.61 | 1.13E-06 |
| Epha7 | 1.69E-84 | 0.19721821 | 0.61 | 0.81 | 2.45E-81 |
| Tenm4 | 8.32E-66 | 0.19626489 | 0.57 | 0.75 | 1.21E-62 |
| Slc35f1 | 1.52E-12 | 0.19529701 | 0.45 | 0.54 | 2.20E-09 |
| Htr2c | 1.68E-13 | 0.19461217 | 0.54 | 0.49 | 2.44E-10 |
| Mal | 2.52E-59 | 0.19336938 | 0.57 | 0.74 | 3.66E-56 |
| 5033421B0 | 8.24E-30 | 0.19212559 | 0.56 | 0.68 | 1.20E-26 |
| Brinp3 | 8.23E-16 | 0.19061983 | 0.49 | 0.59 | 1.19E-12 |
| Cobll1 | 3.52E-21 | 0.18833615 | 0.66 | 0.76 | 5.11E-18 |
| Pcdh15 | 7.43E-43 | 0.18816004 | 0.63 | 0.77 | 1.08E-39 |
| Synpr | 8.53E-54 | 0.18533351 | 0.61 | 0.76 | 1.24E-50 |
| Fbxw15 | 1.42E-167 | 0.18128348 | 0.2 | 0.42 | 2.06E-164 |
| Kcnq5 | 2.31E-23 | 0.18032718 | 0.63 | 0.74 | 3.35E-20 |
| Chrm3 | 3.22E-50 | 0.17835954 | 0.58 | 0.74 | 4.67E-47 |
| Stmn2 | 2.66E-14 | 0.17689113 | 0.37 | 0.28 | 3.86E-11 |
| Ano3 | 7.04E-16 | 0.17600257 | 0.4 | 0.49 | 1.02E-12 |
| Ccnd3 | 7.36E-41 | 0.17553756 | 0.27 | 0.42 | 1.07E-37 |
| Tenm3 | 5.39E-11 | 0.17145198 | 0.6 | 0.68 | 7.82E-08 |
| Fign | 0.00416497 | 0.16865473 | 0.6 | 0.64 | 1 |
| Lcp1 | 0.00080855 | 0.16665612 | 0.22 | 0.25 | 1 |
| Lingo2 | 4.45E-84 | 0.16566564 | 0.69 | 0.86 | 6.46E-81 |
| Astn2 | 4.33E-48 | 0.1649715 | 0.39 | 0.56 | 6.28E-45 |
| Sox10 | 1.34E-31 | 0.16495318 | 0.42 | 0.54 | 1.95E-28 |
| Plekhh1 | 6.23E-54 | 0.16176317 | 0.61 | 0.77 | 9.04E-51 |
| Erbb4 | 4.25E-13 | 0.1617239 | 0.9 | 0.94 | 6.16E-10 |
| Kcnh7 | 9.12E-15 | 0.16076762 | 0.62 | 0.71 | 1.32E-11 |
| Abca8a | 3.64E-12 | 0.16015506 | 0.61 | 0.69 | 5.28E-09 |
| Adarb2 | 6.03E-43 | 0.15791123 | 0.55 | 0.72 | 8.75E-40 |
| 1700042O1 | 5.27E-05 | 0.15713052 | 0.24 | 0.28 | 0.07640878 |
| Creb5 | 5.31E-21 | 0.15695762 | 0.67 | 0.77 | 7.70E-18 |
| Dner | 0.04386467 | 0.15224288 | 0.4 | 0.39 | 1 |
| Nxph1 | 2.06E-24 | 0.15018958 | 0.42 | 0.55 | 2.99E-21 |
| Cntnap5 | 1.88E-166 | 0.14776446 | 0.45 | 0.77 | 2.72E-163 |
| Hcn1 | 4.77E-25 | 0.14574644 | 0.57 | 0.69 | 6.92E-22 |
| Sv2b | 3.53E-41 | 0.14349119 | 0.42 | 0.57 | 5.12E-38 |
| Ninj2 | 1.42E-36 | 0.14192221 | 0.7 | 0.82 | 2.06E-33 |
| Hhip | 9.62E-36 | 0.14092265 | 0.6 | 0.74 | 1.40E-32 |
| Pde3a | 9.54E-06 | 0.14089721 | 0.41 | 0.47 | 0.01384757 |
| Grm1 | 2.69E-98 | 0.14046493 | 0.53 | 0.76 | 3.90E-95 |
| Sgcz | 4.91E-12 | 0.1394017 | 0.58 | 0.66 | 7.12E-09 |
| Dcc | 1.28E-34 | 0.1372463 | 0.74 | 0.86 | 1.85E-31 |
| Dpp10 | 2.01E-91 | 0.13654432 | 0.39 | 0.63 | 2.91E-88 |
| Cnr1 | 9.81E-44 | 0.13490968 | 0.5 | 0.33 | 1.42E-40 |
| Cdh9 | 6.23E-31 | 0.13399877 | 0.52 | 0.65 | 9.04E-28 |
| Khdrbs2 | 3.41E-07 | 0.13398087 | 0.53 | 0.6 | 0.00049456 |
| Elmo1 | 1.06E-43 | 0.13346941 | 0.82 | 0.92 | 1.54E-40 |
| Camk4 | 0.35947986 | 0.13285231 | 0.52 | 0.53 | 1 |
| Cobl | 7.44E-07 | 0.12891452 | 0.6 | 0.67 | 0.00107963 |
| Slc44a5 | 7.33E-17 | 0.12865027 | 0.54 | 0.44 | 1.06E-13 |
| Lypd6 | 0.00617315 | 0.12732911 | 0.57 | 0.6 | 1 |
| Man1a | 2.54E-59 | 0.12697499 | 0.59 | 0.77 | 3.69E-56 |
| Sugct | 6.79E-08 | 0.12668146 | 0.52 | 0.45 | 9.85E-05 |
| Dkk3 | 3.65E-21 | 0.12654627 | 0.21 | 0.29 | 5.30E-18 |
| Ntng1 | 3.46E-80 | 0.12525482 | 0.42 | 0.64 | 5.03E-77 |
| Nckap5 | 3.51E-22 | 0.12375143 | 0.57 | 0.68 | 5.09E-19 |
| Pde4d | 2.15E-48 | 0.12334966 | 0.61 | 0.78 | 3.11E-45 |
| Cdh12 | 6.02E-11 | 0.12196786 | 0.49 | 0.57 | 8.74E-08 |
| Ptprg | 1.79E-11 | 0.12148298 | 0.57 | 0.66 | 2.60E-08 |
| 2010300C0 | 8.99E-25 | 0.12119775 | 0.55 | 0.66 | 1.30E-21 |
| Il33 | 3.65E-141 | 0.12106696 | 0.48 | 0.71 | 5.30E-138 |
| Sox5 | 1.64E-21 | 0.12073484 | 0.54 | 0.66 | 2.38E-18 |
| Epha3 | 1.25E-05 | 0.12042657 | 0.46 | 0.52 | 0.01811505 |
| Clstn2 | 5.59E-82 | 0.11917851 | 0.43 | 0.66 | 8.10E-79 |
| Bcas1os | 0.05702983 | 0.11844841 | 0.49 | 0.5 | 1 |
| Gm3264 | 1.17E-24 | 0.11619843 | 0.56 | 0.67 | 1.70E-21 |
| Col23a1 | 3.17E-90 | 0.11608173 | 0.25 | 0.48 | 4.61E-87 |
| Aspa | 8.57E-43 | 0.11491781 | 0.79 | 0.91 | 1.24E-39 |
| Gfra1 | 1.96E-12 | 0.11350068 | 0.41 | 0.5 | 2.85E-09 |
| Car2 | 4.49E-65 | 0.11310643 | 0.7 | 0.85 | 6.51E-62 |
| Cntnap5 | 6.33E-153 | 0.11184255 | 0.38 | 0.69 | 9.19E-150 |
| Adamtsl | 2.19E-91 | 0.11125271 | 0.54 | 0.75 | 3.17E-88 |
| Tenm2 | 1.50E-24 | 0.10799361 | 0.82 | 0.91 | 2.18E-21 |
| Thsd7b | 0.00122444 | 0.10777259 | 0.57 | 0.61 | 1 |
| Pcdh7 | 4.96E-08 | 0.10603361 | 0.46 | 0.53 | 7.20E-05 |
| Chgb | 0.00946433 | 0.10478765 | 0.41 | 0.44 | 1 |
| Galnt17 | 1.75E-26 | 0.10478691 | 0.48 | 0.61 | 2.54E-23 |
| Zfp385b | 1.47E-76 | 0.10476632 | 0.58 | 0.79 | 2.14E-73 |
| Ralyl | 1.06E-40 | 0.10178056 | 0.68 | 0.82 | 1.54E-37 |
| Meis2 | 2.28E-28 | 0.10156213 | 0.38 | 0.51 | 3.30E-25 |
| Col19a1 | 2.01E-09 | 0.09939748 | 0.47 | 0.41 | 2.91E-06 |
| Mctp1 | 4.57E-43 | 0.09928899 | 0.51 | 0.68 | 6.64E-40 |
| Slc6a1 | 1.91E-12 | 0.09920933 | 0.31 | 0.4 | 2.77E-09 |
| Htr4 | 9.15E-167 | 0.09833291 | 0.23 | 0.55 | 1.33E-163 |
| A330008L1 | 1.32E-76 | 0.09814422 | 0.51 | 0.72 | 1.92E-73 |
| Ankrd33 | 4.15E-22 | 0.09795803 | 0.53 | 0.63 | 6.03E-19 |
| Adipor2 | 1.51E-42 | 0.09768382 | 0.7 | 0.83 | 2.19E-39 |
| Ptprz1 | 5.03E-43 | 0.09745341 | 0.4 | 0.57 | 7.30E-40 |
| Slit2 | 7.04E-05 | 0.09660943 | 0.54 | 0.59 | 0.10219588 |
| Svep1 | 2.29E-09 | 0.09648023 | 0.4 | 0.46 | 3.32E-06 |
| Hectd2 | 0.00216727 | 0.09644828 | 0.39 | 0.43 | 1 |
| Prkcq | 0.00430686 | 0.09638253 | 0.55 | 0.58 | 1 |
| Arhgap2 | 3.74E-07 | 0.09505697 | 0.43 | 0.4 | 0.00054255 |
| Sh3d19 | 1.35E-24 | 0.09276569 | 0.64 | 0.76 | 1.96E-21 |
| Hexb | 1.29E-28 | 0.09247235 | 0.4 | 0.54 | 1.87E-25 |
| Dlx1as | 3.59E-25 | 0.09198435 | 0.35 | 0.47 | 5.21E-22 |
| Tgfbr1 | 0.00231422 | 0.09178401 | 0.45 | 0.49 | 1 |
| Ndrg2 | 0.00589455 | 0.09123174 | 0.32 | 0.3 | 1 |
| Mog | 1.58E-39 | 0.08940017 | 0.78 | 0.9 | 2.29E-36 |
| A830018L1 | 1.72E-80 | 0.08704598 | 0.45 | 0.68 | 2.50E-77 |
| Efr3a | 6.94E-14 | 0.08623415 | 0.39 | 0.48 | 1.01E-10 |
| Gm2075 | 4.83E-50 | 0.08603716 | 0.53 | 0.71 | 7.00E-47 |
| Pde10a | 0.00278489 | 0.08419294 | 0.54 | 0.57 | 1 |
| Runx2 | 2.56E-07 | 0.0836829 | 0.48 | 0.42 | 0.00037216 |
| Slc16a2 | 0.12788309 | 0.08347328 | 0.54 | 0.52 | 1 |
| Ptprt | 4.34E-123 | 0.08102744 | 0.31 | 0.59 | 6.30E-120 |
| Prex2 | 5.47E-10 | 0.08030653 | 0.47 | 0.39 | 7.93E-07 |
| Wnt5b | 2.57E-46 | 0.0799183 | 0.39 | 0.56 | 3.73E-43 |
| Mpp7 | 0.04796418 | 0.07969741 | 0.46 | 0.49 | 1 |
| Cpne4 | 4.65E-91 | 0.0792312 | 0.42 | 0.66 | 6.75E-88 |
| Neto1 | 1.28E-11 | 0.07880313 | 0.57 | 0.65 | 1.86E-08 |
| Gm1458 | 5.64E-18 | 0.07880039 | 0.49 | 0.4 | 8.19E-15 |
| Ptchd4 | 1.33E-13 | 0.07809661 | 0.49 | 0.58 | 1.93E-10 |
| Cacna2d | 2.00E-11 | 0.07773702 | 0.54 | 0.62 | 2.91E-08 |
| Spag6l | 4.63E-149 | 0.07760871 | 0.28 | 0.59 | 6.72E-146 |
| Atp10b | 1.70E-67 | 0.07665219 | 0.49 | 0.69 | 2.46E-64 |
| Adgrl2 | 1.62E-61 | 0.07627844 | 0.29 | 0.48 | 2.35E-58 |
| 1700047M1 | 2.46E-80 | 0.07538195 | 0.48 | 0.68 | 3.56E-77 |
| Nr4a2 | 1.30E-32 | 0.07495886 | 0.55 | 0.65 | 1.88E-29 |
| Chsy3 | 4.35E-32 | 0.07476277 | 0.53 | 0.67 | 6.31E-29 |
| Acvr1c | 1.95E-14 | 0.07461359 | 0.43 | 0.52 | 2.82E-11 |
| Sned1 | 3.47E-160 | 0.07367965 | 0.37 | 0.69 | 5.03E-157 |
| Sorcs3 | 7.76E-16 | 0.07324492 | 0.53 | 0.63 | 1.13E-12 |
| Nwd2 | 8.56E-16 | 0.0730161 | 0.44 | 0.54 | 1.24E-12 |
| Htra1 | 1.30E-44 | 0.07271749 | 0.27 | 0.42 | 1.88E-41 |
| Gm3038 | 3.73E-06 | 0.07147778 | 0.44 | 0.39 | 0.00540925 |
| Gm4837 | 1.13E-118 | 0.07075187 | 0.27 | 0.51 | 1.64E-115 |
| Cacng3 | 3.12E-20 | 0.07074833 | 0.46 | 0.57 | 4.53E-17 |
| Sst | 5.32E-30 | 0.07071591 | 0.42 | 0.56 | 7.72E-27 |
| Clu | 7.26E-21 | 0.07002892 | 0.25 | 0.35 | 1.05E-17 |
| Rfx3 | 8.15E-38 | 0.06894727 | 0.45 | 0.6 | 1.18E-34 |
| Ccdc162 | 3.06E-17 | 0.06869698 | 0.44 | 0.34 | 4.45E-14 |
| Lama2 | 1.35E-28 | 0.06850907 | 0.65 | 0.77 | 1.96E-25 |
| Nell1 | 1.24E-22 | 0.06847681 | 0.51 | 0.63 | 1.79E-19 |
| Fgf12 | 2.17E-131 | 0.06836007 | 0.43 | 0.72 | 3.15E-128 |
| Rasgrf2 | 2.17E-12 | 0.06835789 | 0.51 | 0.42 | 3.15E-09 |
| Tshz3 | 0.00453567 | 0.06727834 | 0.49 | 0.47 | 1 |
| Mcc | 2.78E-07 | 0.06710094 | 0.5 | 0.44 | 0.00040393 |
| Slc2a13 | 0.01002757 | 0.0669946 | 0.55 | 0.56 | 1 |
| Eps8l2 | 1.98E-36 | 0.0662306 | 0.42 | 0.58 | 2.88E-33 |
| Flrt2 | 6.51E-05 | 0.06608423 | 0.51 | 0.56 | 0.0944688 |
| Caln1 | 3.88E-179 | 0.06606707 | 0.39 | 0.73 | 5.63E-176 |
| Galnt9 | 3.52E-37 | 0.06602126 | 0.42 | 0.57 | 5.11E-34 |
| Cadps2 | 0.00650435 | 0.06537883 | 0.55 | 0.59 | 1 |
| Slc7a11 | 8.49E-132 | 0.06459741 | 0.09 | 0.32 | 1.23E-128 |
| Dnah14 | 1.03E-06 | 0.06436004 | 0.32 | 0.38 | 0.00149024 |
| Camk2d | 1.73E-17 | 0.06338457 | 0.45 | 0.56 | 2.51E-14 |
| Pcsk1 | 9.89E-07 | 0.06329594 | 0.45 | 0.39 | 0.00143492 |
| Raver2 | 0.00015285 | 0.0632687 | 0.51 | 0.46 | 0.22179205 |
| Efna5 | 1.57E-49 | 0.06321503 | 0.34 | 0.52 | 2.28E-46 |
| Sgcd | 2.06E-172 | 0.06213614 | 0.31 | 0.65 | 2.99E-169 |
| Iqgap2 | 0.54350714 | 0.06189951 | 0.54 | 0.55 | 1 |
| Pde8a | 7.23E-52 | 0.0609327 | 0.72 | 0.87 | 1.05E-48 |
| Slco1c1 | 3.27E-115 | 0.06076327 | 0.14 | 0.37 | 4.75E-112 |
| Gli2 | 8.08E-07 | 0.06045358 | 0.29 | 0.35 | 0.0011722 |
| Reln | 5.68E-118 | 0.06025623 | 0.19 | 0.45 | 8.25E-115 |
| Teddm2 | 2.44E-11 | 0.06019929 | 0.33 | 0.4 | 3.53E-08 |
| Prss12 | 5.02E-22 | 0.05981404 | 0.28 | 0.4 | 7.29E-19 |
| Hs3st2 | 2.06E-44 | 0.05960752 | 0.27 | 0.13 | 2.99E-41 |
| Rxfp1 | 4.55E-79 | 0.05960532 | 0.45 | 0.68 | 6.60E-76 |
| Inpp4b | 2.00E-18 | 0.05926537 | 0.54 | 0.43 | 2.90E-15 |

| | | | | | | | | | | | |
|---|---|---|---|---|---|---|---|---|---|---|---|
| Nedd4 | 1.00E-61 | 0.048747647 | 0.239 | 0.307 | 2.70E-57 | 4930447C0 | 1.73E-25 | 0.05912211 | 0.38 | 0.5 | 2.51E-22 |
| Fa2h | 1.05E-61 | -0.103097908 | 0.477 | 0.613 | 2.82E-57 | Col25a1 | 1.47E-09 | 0.05884271 | 0.46 | 0.54 | 2.14E-06 |
| Gatm | 1.17E-61 | 0.036253955 | 0.518 | 0.62 | 3.15E-57 | Rgs6 | 1.40E-37 | 0.05884249 | 0.37 | 0.53 | 2.03E-34 |
| Mal | 1.46E-61 | -0.168890813 | 0.363 | 0.501 | 3.93E-57 | Itgad | 1.22E-13 | 0.0579301 | 0.33 | 0.42 | 1.78E-10 |
| Sfpq | 1.55E-61 | -0.094824179 | 0.266 | 0.368 | 4.17E-57 | Npr3 | 2.46E-21 | 0.05752732 | 0.55 | 0.63 | 3.56E-18 |
| Nktr | 1.70E-61 | 0.006624398 | 0.287 | 0.37 | 4.59E-57 | Rdh5 | 0.32328288 | 0.05645728 | 0.39 | 0.4 | 1 |
| Ubr3 | 1.93E-61 | 0.052327325 | 0.245 | 0.313 | 5.19E-57 | Hck | 2.04E-29 | 0.05582145 | 0.47 | 0.34 | 2.96E-26 |
| Akap9 | 2.00E-61 | -0.018871412 | 0.231 | 0.315 | 5.38E-57 | Lin28b | 0.08208738 | 0.05547587 | 0.42 | 0.44 | 1 |
| 4932438A13Rik | 2.05E-61 | 0.080173865 | 0.211 | 0.268 | 5.52E-57 | Kcnj13 | 3.80E-67 | 0.05506314 | 0.26 | 0.47 | 5.51E-64 |
| Ubr2 | 2.34E-61 | 0.092394986 | 0.255 | 0.318 | 6.29E-57 | Clmp | 1.39E-123 | 0.05472932 | 0.13 | 0.38 | 2.02E-120 |
| Gm35188 | 2.50E-61 | -0.31431863 | 0.235 | 0.379 | 6.74E-57 | Gm4832 | 2.52E-111 | 0.05443683 | 0.45 | 0.21 | 3.66E-108 |
| Ralgapa1 | 3.02E-61 | 0.090587013 | 0.3 | 0.368 | 8.13E-57 | Vstm2a | 0.00118689 | 0.05406421 | 0.41 | 0.46 | 1 |
| Aplp1 | 3.05E-61 | -0.079626199 | 0.381 | 0.501 | 8.21E-57 | Mgat4c | 5.99E-129 | 0.05402404 | 0.35 | 0.64 | 8.69E-126 |
| Plekhg1 | 4.34E-61 | -0.087422241 | 0.395 | 0.519 | 1.17E-56 | Dcdc2a | 0.00330078 | 0.05319433 | 0.52 | 0.54 | 1 |
| Ttll7 | 4.55E-61 | -0.029894105 | 0.556 | 0.679 | 1.23E-56 | Tmem10 | 0.00013801 | 0.05318507 | 0.54 | 0.58 | 0.20025696 |
| Chd9 | 4.94E-61 | 0.039232456 | 0.224 | 0.296 | 1.33E-56 | Cdh6 | 0.00013982 | 0.05282656 | 0.52 | 0.57 | 0.20288537 |
| Zmynd8 | 5.63E-61 | 0.026155076 | 0.245 | 0.318 | 1.51E-56 | Glul | 1.60E-97 | 0.05276011 | 0.34 | 0.57 | 2.32E-94 |
| Ablim2 | 6.20E-61 | 0.358612456 | 0.323 | 0.315 | 1.67E-56 | Zfpm2 | 9.63E-56 | 0.0526659 | 0.52 | 0.71 | 1.40E-52 |
| 5031439G07Rik | 6.25E-61 | -0.099737109 | 0.359 | 0.475 | 1.68E-56 | Col5a2 | 4.53E-11 | 0.0524802 | 0.39 | 0.31 | 6.58E-08 |
| Actn4 | 6.30E-61 | -0.227845036 | 0.151 | 0.256 | 1.70E-56 | Pdzrn3 | 1.60E-06 | 0.05129824 | 0.41 | 0.47 | 0.0023219 |
| Syt11 | 6.32E-61 | -0.005417105 | 0.315 | 0.406 | 1.70E-56 | Grid2 | 1.44E-36 | 0.04997694 | 0.78 | 0.9 | 2.10E-33 |
| Krit1 | 6.47E-61 | 0.005732273 | 0.214 | 0.289 | 1.74E-56 | Dlx6os1 | 2.36E-14 | 0.04972171 | 0.34 | 0.43 | 3.43E-11 |
| Atp11a | 6.68E-61 | -0.062692783 | 0.181 | 0.264 | 1.80E-56 | Frmpd4 | 3.14E-50 | 0.04959095 | 0.6 | 0.77 | 4.56E-47 |
| Tubb4a | 7.58E-61 | 0.028980955 | 0.591 | 0.702 | 2.04E-56 | Nell1os | 0.02690318 | 0.04958542 | 0.39 | 0.42 | 1 |
| Atf7ip | 1.02E-60 | -0.094395659 | 0.228 | 0.325 | 2.75E-56 | Notch2 | 5.42E-87 | 0.04914657 | 0.38 | 0.63 | 7.86E-84 |
| Mbp | 1.03E-60 | -0.226609831 | 0.967 | 0.991 | 2.76E-56 | Gap43 | 0.0460337 | 0.0485527 | 0.33 | 0.35 | 1 |
| Nkain2 | 1.09E-60 | -0.208873463 | 0.998 | 0.999 | 2.93E-56 | Shisa6 | 3.85E-07 | 0.04783875 | 0.56 | 0.62 | 0.00055895 |
| Cobll1 | 1.10E-60 | 0.0411556 | 0.304 | 0.382 | 2.96E-56 | Angpt1 | 0.00024225 | 0.04753375 | 0.51 | 0.46 | 0.3514994 |
| Mllt10 | 1.15E-60 | 0.076693959 | 0.271 | 0.336 | 3.10E-56 | Lgmn | 1.86E-116 | 0.04730745 | 0.19 | 0.43 | 2.70E-113 |
| Sbf2 | 1.24E-60 | 0.051879148 | 0.276 | 0.349 | 3.34E-56 | Hunk | 0.00024429 | 0.04645964 | 0.54 | 0.57 | 0.35446115 |
| Ddx5 | 1.24E-60 | -0.157039511 | 0.41 | 0.546 | 3.34E-56 | Aldh1a1 | 7.74E-12 | 0.04640457 | 0.39 | 0.32 | 1.12E-08 |
| Mast3 | 1.29E-60 | -0.131627153 | 0.549 | 0.697 | 3.48E-56 | Il31ra | 0.80364386 | 0.0460861 | 0.42 | 0.43 | 1 |
| Phldb1 | 1.38E-60 | 0.088016057 | 0.526 | 0.606 | 3.71E-56 | Nhsl2 | 3.35E-197 | 0.04499418 | 0.14 | 0.47 | 4.86E-194 |
| Srsf11 | 1.41E-60 | -0.015111519 | 0.258 | 0.343 | 3.80E-56 | L3mbtl3 | 4.77E-23 | 0.0447077 | 0.36 | 0.48 | 6.91E-20 |
| Stxbp3 | 1.66E-60 | -0.15059943 | 0.226 | 0.334 | 4.47E-56 | Trpc4 | 2.51E-166 | 0.04444081 | 0.47 | 0.79 | 3.65E-163 |
| St6gal1 | 1.80E-60 | -0.234567648 | 0.34 | 0.484 | 4.84E-56 | Mkx | 6.93E-36 | 0.04443588 | 0.43 | 0.29 | 1.01E-32 |
| 1810062O18Rik | 2.56E-60 | -0.21955348 | 0.163 | 0.269 | 6.90E-56 | Satb2 | 2.62E-50 | 0.04427483 | 0.38 | 0.56 | 3.80E-47 |
| Srsf2 | 2.78E-60 | 0.01291978 | 0.211 | 0.285 | 7.49E-56 | Htr1f | 3.22E-111 | 0.04407118 | 0.24 | 0.5 | 4.68E-108 |
| Ccnl2 | 2.96E-60 | 0.017213025 | 0.19 | 0.259 | 7.97E-56 | Zmat4 | 1.41E-52 | 0.04386934 | 0.56 | 0.74 | 2.05E-49 |
| Ppp1r21 | 3.13E-60 | 0.005943372 | 0.215 | 0.288 | 8.42E-56 | Mndal | 9.33E-34 | 0.04312266 | 0.41 | 0.55 | 1.35E-30 |
| Arid1b | 3.21E-60 | -0.077215905 | 0.257 | 0.358 | 8.63E-56 | 4930509J09 | 1.87E-22 | 0.04299122 | 0.51 | 0.63 | 2.72E-19 |
| Birc6 | 3.89E-60 | 0.032134472 | 0.273 | 0.347 | 1.05E-55 | Lhfpl3 | 3.30E-171 | 0.04298497 | 0.24 | 0.57 | 4.79E-168 |
| Usp31 | 3.93E-60 | -0.37001066 | 0.158 | 0.286 | 1.06E-55 | Gm2815 | 1.79E-74 | 0.04293622 | 0.36 | 0.58 | 2.60E-71 |
| Vps13b | 6.03E-60 | 0.095880134 | 0.286 | 0.346 | 1.62E-55 | Aebp1 | 1.25E-71 | 0.04282654 | 0.32 | 0.53 | 1.82E-68 |
| Nsd1 | 7.13E-60 | 0.028308063 | 0.36 | 0.445 | 1.92E-55 | Dnah9 | 1.75E-144 | 0.04246889 | 0.36 | 0.67 | 2.54E-141 |
| Dock1 | 7.20E-60 | 0.006103506 | 0.464 | 0.566 | 1.94E-55 | Stac | 0.00380902 | 0.04175774 | 0.52 | 0.51 | 1 |
| Ambra1 | 7.88E-60 | -0.014554258 | 0.257 | 0.343 | 2.12E-55 | Arhgap3 | 4.33E-13 | 0.04159348 | 0.34 | 0.43 | 6.28E-10 |
| Ugt8a | 8.17E-60 | -0.134583411 | 0.477 | 0.616 | 2.20E-55 | Neurod6 | 3.99E-88 | 0.04147895 | 0.4 | 0.64 | 5.78E-85 |
| Kat2b | 1.05E-59 | -2.57E-05 | 0.268 | 0.352 | 2.84E-55 | Dach2 | 1.60E-06 | 0.0413357 | 0.39 | 0.46 | 0.00232124 |
| Ppfibp2 | 1.09E-59 | -0.086359222 | 0.245 | 0.343 | 2.92E-55 | Xylt1 | 5.14E-26 | 0.04051337 | 0.55 | 0.68 | 7.46E-23 |
| Herc4 | 1.18E-59 | -0.115044844 | 0.185 | 0.276 | 3.17E-55 | Stk39 | 1.87E-06 | 0.04051295 | 0.57 | 0.63 | 0.00271632 |
| Myo6 | 1.49E-59 | -0.0544741 | 0.486 | 0.607 | 4.01E-55 | Atp1a2 | 7.00E-17 | 0.03981379 | 0.26 | 0.36 | 1.02E-13 |
| Phf3 | 1.50E-59 | 0.028840125 | 0.261 | 0.335 | 4.03E-55 | Adam12 | 2.48E-42 | 0.03952125 | 0.28 | 0.44 | 3.60E-39 |
| 4-Sep | 1.79E-59 | 0.08108312 | 0.494 | 0.576 | 4.82E-55 | Ranbp3 | 0.00020286 | 0.03942095 | 0.46 | 0.49 | 0.29434938 |
| Kazn | 2.03E-59 | -0.045305894 | 0.216 | 0.298 | 5.46E-55 | Cntnap4 | 4.39E-07 | 0.03889007 | 0.48 | 0.55 | 0.00063725 |
| Ankhd1 | 3.06E-59 | -0.004172579 | 0.227 | 0.306 | 8.25E-55 | Gm3169 | 4.02E-10 | 0.03878375 | 0.39 | 0.31 | 5.84E-07 |
| Slc6a1 | 4.66E-59 | -0.089111539 | 0.189 | 0.277 | 1.26E-54 | Lrp4 | 7.40E-26 | 0.03871062 | 0.15 | 0.25 | 1.07E-22 |
| Zfp280c | 4.91E-59 | -0.079212064 | 0.271 | 0.371 | 1.32E-54 | Zfp804a | 4.37E-13 | 0.0385768 | 0.55 | 0.63 | 6.35E-10 |
| Tmeff1 | 5.71E-59 | -0.154131689 | 0.275 | 0.395 | 1.54E-54 | Sntg2 | 0.24248515 | 0.03832566 | 0.53 | 0.52 | 1 |
| Dock9 | 5.98E-59 | -0.196897109 | 0.397 | 0.541 | 1.61E-54 | Lrrk2 | 0.17570924 | 0.03799165 | 0.55 | 0.54 | 1 |
| Bptf | 6.18E-59 | 0.023460091 | 0.228 | 0.301 | 1.66E-54 | Htr7 | 1.45E-07 | 0.03795041 | 0.5 | 0.51 | 0.00020976 |
| Myo9a | 6.49E-59 | 0.135215572 | 0.319 | 0.375 | 1.75E-54 | Gna14 | 1.28E-08 | 0.03785338 | 0.25 | 0.19 | 1.86E-05 |
| Dst | 6.93E-59 | 0.009111783 | 0.589 | 0.7 | 1.87E-54 | Necab1 | 2.00E-17 | 0.03777541 | 0.43 | 0.54 | 2.90E-14 |
| Tnfaip6 | 7.62E-59 | -0.115042751 | 0.217 | 0.315 | 2.05E-54 | Trpc6 | 8.40E-08 | 0.03728664 | 0.49 | 0.55 | 0.00012189 |
| Shtn1 | 7.96E-59 | -0.047968743 | 0.472 | 0.589 | 2.14E-54 | Adcy8 | 0.00070566 | 0.03706015 | 0.51 | 0.54 | 1 |
| Pstpip2 | 8.98E-59 | -0.182567643 | 0.418 | 0.562 | 2.42E-54 | Prlr | 0.00200062 | 0.03687908 | 0.41 | 0.46 | 1 |
| Apc | 1.03E-58 | 0.072534733 | 0.38 | 0.453 | 2.77E-54 | Man2a1 | 5.80E-10 | 0.03634331 | 0.55 | 0.63 | 8.42E-07 |
| Hipk2 | 1.04E-58 | 0.005136589 | 0.478 | 0.584 | 2.80E-54 | Kcnc1 | 0.28792073 | 0.03555849 | 0.34 | 0.34 | 1 |
| R3hdm1 | 1.15E-58 | 0.143346905 | 0.376 | 0.432 | 3.09E-54 | C130073E2 | 4.37E-63 | 0.03499937 | 0.41 | 0.25 | 6.34E-60 |
| Lpgat1 | 1.30E-58 | 0.008436028 | 0.403 | 0.5 | 3.49E-54 | Zdbf2 | 0.21904646 | 0.03450673 | 0.5 | 0.49 | 1 |
| Dtna | 1.32E-58 | 0.146149987 | 0.341 | 0.395 | 3.56E-54 | Grm8 | 3.41E-18 | 0.03443187 | 0.37 | 0.48 | 4.94E-15 |
| Ttll5 | 1.44E-58 | -0.077716984 | 0.293 | 0.395 | 3.89E-54 | Gja1 | 4.61E-11 | 0.03382266 | 0.26 | 0.34 | 6.69E-08 |
| Tnrc6b | 1.46E-58 | -0.047507254 | 0.332 | 0.435 | 3.93E-54 | L3mbtl4 | 0.0005594 | 0.03373758 | 0.26 | 0.22 | 0.81169463 |
| Arhgef28 | 1.92E-58 | -0.001636607 | 0.326 | 0.412 | 5.18E-54 | Tenm1 | 3.06E-36 | 0.03307329 | 0.47 | 0.62 | 4.43E-33 |
| Sh3kbp1 | 2.23E-58 | -0.208033321 | 0.222 | 0.339 | 6.01E-54 | Spef2 | 1.15E-112 | 0.03303287 | 0.31 | 0.59 | 1.68E-109 |
| Snx29 | 2.46E-58 | -0.090959272 | 0.238 | 0.336 | 6.61E-54 | Car10 | 2.80E-13 | 0.03301582 | 0.53 | 0.61 | 4.06E-10 |
| Dock3 | 2.90E-58 | 0.008770263 | 0.332 | 0.421 | 7.80E-54 | Npas2 | 8.77E-129 | 0.03288083 | 0.41 | 0.69 | 1.27E-125 |
| Ankrd12 | 3.08E-58 | 0.104195332 | 0.314 | 0.377 | 8.29E-54 | Cemip | 1.82E-88 | 0.03283223 | 0.46 | 0.7 | 2.64E-85 |
| Ncoa1 | 3.47E-58 | 0.148776062 | 0.274 | 0.325 | 9.34E-54 | Arhgap2 | 1.04E-55 | 0.03195895 | 0.35 | 0.54 | 1.51E-52 |
| Ptn | 3.60E-58 | -0.203623008 | 0.394 | 0.538 | 9.69E-54 | Mid1 | 5.63E-19 | 0.03184161 | 0.2 | 0.3 | 8.17E-16 |
| Dennd5b | 3.98E-58 | -0.149082553 | 0.236 | 0.344 | 1.07E-53 | Rab3b | 9.82E-15 | 0.03158729 | 0.54 | 0.64 | 1.42E-11 |
| Mef2a | 4.11E-58 | 0.042781779 | 0.202 | 0.267 | 1.11E-53 | Serpini1 | 2.49E-14 | 0.03155031 | 0.54 | 0.63 | 3.61E-11 |
| Plxnb3 | 4.63E-58 | -0.16197201 | 0.16 | 0.252 | 1.25E-53 | Galnt13 | 3.71E-09 | 0.03129572 | 0.45 | 0.5 | 5.38E-06 |
| Tacc1 | 5.54E-58 | -0.240328358 | 0.19 | 0.306 | 1.49E-53 | Cntnap5 | 1.16E-46 | 0.03120192 | 0.33 | 0.51 | 1.68E-43 |
| Slco3a1 | 5.60E-58 | -0.577318784 | 0.141 | 0.282 | 1.51E-53 | Slc35f4 | 4.76E-47 | 0.03102461 | 0.46 | 0.64 | 6.90E-44 |
| Sox10 | 5.71E-58 | -0.000233343 | 0.277 | 0.362 | 1.54E-53 | Fyb | 2.04E-95 | 0.03090823 | 0.14 | 0.36 | 2.96E-92 |
| Wdr33 | 5.87E-58 | -0.132885114 | 0.171 | 0.262 | 1.58E-53 | Ntsr2 | 8.08E-31 | 0.03080854 | 0.43 | 0.3 | 1.17E-27 |
| Kdm2a | 6.53E-58 | -0.015575488 | 0.18 | 0.253 | 1.76E-53 | Gabra5 | 1.67E-34 | 0.03044282 | 0.45 | 0.6 | 2.42E-31 |
| Arnt2 | 7.33E-58 | -0.025446011 | 0.255 | 0.337 | 1.97E-53 | Plp1 | 1.34E-10 | 0.03029745 | 0.98 | 1 | 1.94E-07 |
| Kdm6a | 7.78E-58 | -0.031844588 | 0.276 | 0.369 | 2.09E-53 | Bmp6 | 8.04E-58 | 0.03017736 | 0.47 | 0.66 | 1.17E-54 |
| Wnk1 | 9.45E-58 | 0.02217013 | 0.524 | 0.624 | 2.54E-53 | Tbc1d1 | 0.08804896 | 0.0300763 | 0.53 | 0.53 | 1 |
| Rtn3 | 1.06E-57 | 0.023392362 | 0.269 | 0.347 | 2.84E-53 | Smoc2 | 5.62E-74 | 0.02985271 | 0.29 | 0.52 | 8.15E-71 |
| Pip4k2a | 1.06E-57 | -0.070937091 | 0.506 | 0.629 | 2.85E-53 | Fras1 | 1.31E-11 | 0.02979535 | 0.52 | 0.61 | 1.90E-08 |
| Kansl1 | 1.11E-57 | 0.022347309 | 0.342 | 0.429 | 2.99E-53 | Crispld1 | 5.97E-211 | 0.02962499 | 0.07 | 0.38 | 8.66E-208 |
| Kmt2e | 1.30E-57 | -0.008764885 | 0.278 | 0.361 | 3.49E-53 | Samd5 | 8.01E-05 | 0.02943417 | 0.51 | 0.57 | 0.11619644 |
| Pcm1 | 1.53E-57 | -0.032186799 | 0.212 | 0.293 | 4.12E-53 | Kcnn2 | 1.68E-41 | 0.02914575 | 0.42 | 0.59 | 2.43E-38 |
| Lrba | 1.77E-57 | 0.073810914 | 0.247 | 0.309 | 4.77E-53 | Apbb1ip | 7.26E-75 | 0.0291367 | 0.2 | 0.4 | 1.05E-71 |
| Cdkal1 | 1.80E-57 | -0.155549726 | 0.166 | 0.26 | 4.86E-53 | Emcn | 8.00E-12 | 0.02907885 | 0.41 | 0.33 | 1.16E-08 |
| Faf1 | 1.81E-57 | 0.000283038 | 0.213 | 0.286 | 4.87E-53 | Vtn | 0.00171426 | 0.02907568 | 0.53 | 0.56 | 1 |
| Atg4c | 1.82E-57 | -0.116626577 | 0.181 | 0.273 | 4.91E-53 | Gm1394 | 7.60E-23 | 0.02891706 | 0.34 | 0.46 | 1.10E-19 |
| Mink1 | 2.21E-57 | -0.126834309 | 0.234 | 0.335 | 5.96E-53 | Fgf1 | 6.21E-06 | 0.02870896 | 0.49 | 0.46 | 0.00901651 |
| Gtf2i | 2.77E-57 | 0.036971284 | 0.221 | 0.285 | 7.45E-53 | Tmtc1 | 0.08252043 | 0.02844636 | 0.53 | 0.53 | 1 |
| Huwe1 | 3.41E-57 | 0.030789843 | 0.265 | 0.336 | 9.17E-53 | Hgf | 0.27054597 | 0.02812223 | 0.35 | 0.36 | 1 |
| Baz2b | 3.50E-57 | -0.194020511 | 0.509 | 0.66 | 9.42E-53 | Fyb2 | 2.58E-13 | 0.02712086 | 0.41 | 0.32 | 3.75E-10 |
| Tmtc2 | 3.52E-57 | -0.147528554 | 0.325 | 0.454 | 9.47E-53 | Car12 | 1.45E-12 | 0.02688877 | 0.39 | 0.47 | 2.11E-09 |
| Jakmip3 | 4.00E-57 | -0.148622032 | 0.212 | 0.313 | 1.08E-52 | D5Ertd61 | 7.66E-13 | 0.02669757 | 0.48 | 0.57 | 1.11E-09 |
| Ikzf2 | 4.00E-57 | -0.135241075 | 0.196 | 0.299 | 1.08E-52 | Cdh13 | 1.75E-11 | 0.02654818 | 0.44 | 0.52 | 2.53E-08 |
| Luc7l3 | 4.83E-57 | -0.000842019 | 0.195 | 0.264 | 1.30E-52 | Lpar1 | 1.12E-208 | 0.02612491 | 0.43 | 0.78 | 1.63E-205 |
| Slc25a13 | 4.94E-57 | -0.186430224 | 0.252 | 0.371 | 1.33E-52 | Tll2 | 4.71E-14 | 0.02605816 | 0.35 | 0.44 | 6.83E-11 |
| Ccnt2 | 5.15E-57 | 0.091738333 | 0.202 | 0.251 | 1.39E-52 | Tshz2 | 1.42E-10 | 0.02599895 | 0.46 | 0.53 | 2.06E-07 |
| Ank | 5.40E-57 | -0.06410116 | 0.249 | 0.344 | 1.45E-52 | Itga8 | 2.62E-42 | 0.02577112 | 0.56 | 0.72 | 3.80E-39 |
| Gm48742 | 5.53E-57 | -0.306228225 | 0.158 | 0.275 | 1.49E-52 | Sytl2 | 4.82E-76 | 0.02538212 | 0.72 | 0.89 | 6.99E-73 |
| Setd5 | 5.73E-57 | 0.010195687 | 0.245 | 0.318 | 1.54E-52 | 9530059O1 | 6.48E-144 | 0.02530737 | 0.43 | 0.72 | 9.40E-141 |
| Nsmce2 | 5.80E-57 | -0.013259586 | 0.176 | 0.25 | 1.56E-52 | Hpse2 | 1.52E-25 | 0.02502264 | 0.37 | 0.27 | 2.20E-22 |
| Mdm4 | 6.41E-57 | -0.006794486 | 0.197 | 0.268 | 1.73E-52 | Arhgap1 | 2.42E-22 | 0.02498886 | 0.35 | 0.47 | 3.51E-19 |
| Fam172a | 6.84E-57 | 0.012032018 | 0.318 | 0.403 | 1.84E-52 | Mgp | 5.97E-13 | 0.02476803 | 0.34 | 0.43 | 8.66E-10 |
| Magi1 | 7.40E-57 | -0.020889878 | 0.33 | 0.43 | 1.99E-52 | Gprin3 | 5.25E-79 | 0.02446525 | 0.37 | 0.17 | 7.62E-76 |
| Mkln1 | 8.31E-57 | -0.019650524 | 0.22 | 0.294 | 2.24E-52 | Syt17 | 1.30E-99 | 0.02384603 | 0.11 | 0.32 | 1.89E-96 |
| Apba1 | 1.08E-56 | 0.147305745 | 0.358 | 0.412 | 2.90E-52 | Plxdc2 | 3.60E-41 | 0.02354927 | 0.86 | 0.95 | 5.22E-38 |
| Snrnp70 | 1.23E-56 | 0.049406279 | 0.46 | 0.546 | 3.30E-52 | Cacng2 | 2.34E-61 | 0.02354855 | 0.27 | 0.46 | 3.39E-58 |
| Ckap5 | 1.23E-56 | -0.051582472 | 0.189 | 0.265 | 3.30E-52 | Pdzd2 | 5.77E-13 | 0.02338353 | 0.36 | 0.35 | 8.37E-10 |
| Arrdc3 | 1.36E-56 | -0.155490539 | 0.221 | 0.329 | 3.67E-52 | Zfp536 | 2.82E-46 | 0.0231856 | 0.83 | 0.93 | 4.10E-43 |
| Fam168a | 1.45E-56 | -0.043182464 | 0.221 | 0.305 | 3.91E-52 | Sv2c | 1.54E-41 | 0.02270751 | 0.41 | 0.25 | 2.24E-38 |
| Cpeb3 | 1.53E-56 | -0.030641379 | 0.247 | 0.331 | 4.13E-52 | Gm1075 | 2.23E-32 | 0.02225416 | 0.37 | 0.52 | 3.24E-29 |
| Kansl1l | 1.72E-56 | -0.051740044 | 0.306 | 0.409 | 4.62E-52 | Coro6 | 4.05E-235 | 0.02193622 | 0.12 | 0.48 | 5.88E-232 |
| Rbm6 | 2.04E-56 | 0.044924416 | 0.31 | 0.384 | 5.49E-52 | Gm1623 | 3.37E-101 | 0.02185819 | 0.18 | 0.43 | 4.89E-98 |
| Vmp1 | 2.36E-56 | -0.187472936 | 0.642 | 0.789 | 6.35E-52 | Cntnap3 | 8.13E-47 | 0.02168945 | 0.23 | 0.39 | 1.18E-43 |
| Macf1 | 2.46E-56 | 0.03840031 | 0.377 | 0.464 | 6.63E-52 | Slc6a20a | 0.05104387 | 0.02143562 | 0.31 | 0.29 | 1 |
| Creb5 | 3.58E-56 | -0.012703156 | 0.314 | 0.404 | 9.63E-52 | Tead1 | 2.69E-148 | 0.02136576 | 0.21 | 0.51 | 3.90E-145 |
| Add3 | 3.67E-56 | -0.289846886 | 0.241 | 0.372 | 9.88E-52 | Npas4 | 2.16E-40 | 0.02127456 | 0.33 | 0.49 | 3.14E-37 |
| Prr5l | 4.17E-56 | -0.301193518 | 0.811 | 0.916 | 1.12E-51 | Arhgap1 | 4.33E-14 | 0.02113916 | 0.45 | 0.54 | 6.29E-11 |
| Abl1 | 4.75E-56 | -0.087307998 | 0.172 | 0.253 | 1.28E-51 | Cpa6 | 2.91E-129 | 0.0209448 | 0.09 | 0.33 | 4.23E-126 |
| Gab1 | 5.73E-56 | -0.161714415 | 0.455 | 0.595 | 1.54E-51 | Lyz2 | 5.90E-63 | 0.02089924 | 0.26 | 0.11 | 8.57E-60 |
| Nbas | 6.60E-56 | 0.056342632 | 0.522 | 0.611 | 1.78E-51 | Myo1f | 2.47E-125 | 0.02069706 | 0.24 | 0.53 | 3.58E-122 |
| Nfix | 8.01E-56 | -0.042269595 | 0.194 | 0.271 | 2.16E-51 | C1ql3 | 1.49E-57 | 0.02034722 | 0.31 | 0.51 | 2.17E-54 |
| Nsd3 | 1.07E-55 | -0.008872912 | 0.233 | 0.306 | 2.87E-51 | Sptlc3 | 2.79E-11 | 0.0202015 | 0.43 | 0.35 | 4.06E-08 |
| Pcmtd1 | 1.09E-55 | -0.00736507 | 0.181 | 0.25 | 2.94E-51 | Adamts1 | 4.05E-21 | 0.01979474 | 0.46 | 0.58 | 5.88E-18 |
| Ubash3b | 1.13E-55 | -0.307578627 | 0.199 | 0.327 | 3.03E-51 | Ak9 | 6.65E-84 | 0.01978655 | 0.26 | 0.08 | 9.64E-81 |
| Hepacam | 1.13E-55 | -0.116085527 | 0.301 | 0.407 | 3.05E-51 | Slc13a4 | 1.55E-10 | 0.0195008 | 0.47 | 0.38 | 2.24E-07 |
| Vps8 | 1.21E-55 | -0.128227867 | 0.268 | 0.374 | 3.26E-51 | Afap1 | 4.57E-34 | 0.01949263 | 0.48 | 0.63 | 6.63E-31 |
| Klf7 | 1.65E-55 | -0.113370464 | 0.217 | 0.31 | 4.45E-51 | Cftr | 1.36E-07 | 0.01933725 | 0.23 | 0.29 | 0.00019665 |
| Jmjd1c | 1.66E-55 | 0.043037873 | 0.266 | 0.338 | 4.46E-51 | Mlip | 2.32E-67 | 0.01932535 | 0.48 | 0.69 | 3.36E-64 |
| Slain1 | 1.74E-55 | -0.100968353 | 0.427 | 0.55 | 4.68E-51 | Atp13a5 | 0.0022392 | 0.01922342 | 0.44 | 0.48 | 1 |
| Pde4dip | 1.88E-55 | 0.101401627 | 0.198 | 0.25 | 5.05E-51 | Adgrv1 | 1.13E-99 | 0.01910703 | 0.26 | 0.5 | 1.64E-96 |
| Rps6ka5 | 1.90E-55 | -0.080047159 | 0.221 | 0.31 | 5.12E-51 | Pip5k1b | 0.00033725 | 0.01875724 | 0.51 | 0.5 | 0.48935263 |
| Arglu1 | 1.92E-55 | 0.037053855 | 0.331 | 0.41 | 5.17E-51 | 4921539H0 | 7.93E-26 | 0.01851472 | 0.41 | 0.54 | 1.15E-22 |
| Sh3d19 | 1.94E-55 | -0.062486788 | 0.433 | 0.552 | 5.23E-51 | Tbx15 | 0.18720111 | 0.01850649 | 0.32 | 0.3 | 1 |
| Clic4 | 2.36E-55 | -0.074533903 | 0.257 | 0.35 | 6.37E-51 | Baiap2l1 | 0.24242281 | 0.01847099 | 0.5 | 0.51 | 1 |
| Rsf1 | 2.37E-55 | -0.028972185 | 0.181 | 0.254 | 6.37E-51 | Rab38 | 4.53E-06 | 0.01821084 | 0.36 | 0.42 | 0.00657674 |
| Glrb | 2.39E-55 | -0.054619993 | 0.181 | 0.259 | 6.44E-51 | Abca9 | 3.41E-71 | 0.01819751 | 0.35 | 0.16 | 4.94E-68 |
| Arhgap5 | 2.51E-55 | -0.093160079 | 0.234 | 0.329 | 6.74E-51 | Mbnl3 | 0.00228674 | 0.01807286 | 0.27 | 0.31 | 1 |
| Mtdh | 2.54E-55 | 0.058400799 | 0.224 | 0.286 | 6.82E-51 | Lrtm1 | 1.55E-15 | 0.01783258 | 0.18 | 0.27 | 2.25E-12 |
| Srpk2 | 2.63E-55 | -0.01142143 | 0.19 | 0.261 | 7.08E-51 | Cald1 | 1.43E-23 | 0.01766279 | 0.43 | 0.56 | 2.07E-20 |
| Ptma | 2.80E-55 | -0.010310828 | 0.197 | 0.268 | 7.52E-51 | Adamts2 | 1.04E-85 | 0.01736792 | 0.53 | 0.76 | 1.51E-82 |
| Ankrd11 | 2.87E-55 | 0.014940459 | 0.214 | 0.282 | 7.71E-51 | Plcxd3 | 1.11E-05 | 0.01733512 | 0.4 | 0.37 | 0.0160998 |
| Prkca | 3.28E-55 | -0.111153313 | 0.329 | 0.448 | 8.83E-51 | Rasgrp1 | 2.26E-93 | 0.0171177 | 0.31 | 0.56 | 3.27E-90 |
| Il1rap | 3.66E-55 | -0.096965328 | 0.453 | 0.576 | 9.85E-51 | Trpv4 | 8.00E-14 | 0.01705406 | 0.4 | 0.31 | 1.16E-10 |
| Phip | 3.75E-55 | 0.185141667 | 0.247 | 0.282 | 1.01E-50 | Vsnl1 | 1.02E-14 | 0.01670044 | 0.46 | 0.56 | 1.48E-11 |
| Nf1 | 3.76E-55 | -0.009269398 | 0.325 | 0.411 | 1.01E-50 | Sox6 | 2.10E-64 | 0.01664875 | 0.38 | 0.59 | 3.04E-61 |
| Kidins220 | 3.78E-55 | 0.006119996 | 0.19 | 0.255 | 1.02E-50 | Rbpms | 0.37342468 | 0.0166049 | 0.41 | 0.43 | 1 |
| Polr3e | 4.07E-55 | -0.355947566 | 0.181 | 0.307 | 1.10E-50 | Tspan18 | 5.42E-07 | 0.01580765 | 0.41 | 0.48 | 0.00078644 |
| Qdpr | 4.10E-55 | -0.013939319 | 0.219 | 0.293 | 1.10E-50 | Mdga1 | 2.15E-07 | 0.01576683 | 0.45 | 0.39 | 0.00031135 |
| Rabgap1l | 5.61E-55 | 0.015124527 | 0.394 | 0.484 | 1.51E-50 | Lmo3 | 6.11E-19 | 0.01562267 | 0.18 | 0.27 | 8.86E-16 |
| Galnt7 | 5.65E-55 | -0.203770686 | 0.184 | 0.29 | 1.52E-50 | Gm1084 | 3.94E-05 | 0.01557281 | 0.35 | 0.4 | 0.0571934 |
| Mbnl1 | 6.44E-55 | -0.048430314 | 0.28 | 0.367 | 1.73E-50 | Gm1719 | 7.81E-14 | 0.01543337 | 0.41 | 0.5 | 1.13E-10 |
| Adipor2 | 7.56E-55 | -0.286470428 | 0.535 | 0.695 | 2.04E-50 | 4930473D1 | 1.23E-15 | 0.0151913 | 0.26 | 0.35 | 1.79E-12 |
| Cog5 | 8.54E-55 | 0.054971397 | 0.249 | 0.312 | 2.30E-50 | Sema3a | 3.23E-05 | 0.01499331 | 0.44 | 0.49 | 0.04687929 |
| Inf2 | 8.55E-55 | -0.049849411 | 0.182 | 0.258 | 2.30E-50 | Junb | 1.58E-191 | 0.0148982 | 0.18 | 0.52 | 2.30E-188 |
| Smad7 | 9.62E-55 | -0.025867754 | 0.318 | 0.411 | 2.59E-50 | Lurap1l | 3.33E-14 | 0.01422618 | 0.46 | 0.56 | 4.83E-11 |
| Snx30 | 9.70E-55 | -0.102520637 | 0.237 | 0.332 | 2.61E-50 | Met | 4.84E-12 | 0.01405061 | 0.28 | 0.2 | 7.03E-09 |
| Slc25a27 | 9.93E-55 | -0.26177104 | 0.218 | 0.338 | 2.67E-50 | Adamts1 | 0.69712078 | 0.01396786 | 0.45 | 0.46 | 1 |
| Tmem63a | 9.98E-55 | -0.036531631 | 0.215 | 0.292 | 2.69E-50 | Gm2926 | 8.95E-11 | 0.01392638 | 0.25 | 0.32 | 1.30E-07 |
| Gnao1 | 1.01E-54 | -0.224494451 | 0.56 | 0.713 | 2.71E-50 | A330093E2 | 8.42E-15 | 0.01378085 | 0.46 | 0.37 | 1.22E-11 |
| Pde8a | 1.03E-54 | -0.038996623 | 0.524 | 0.64 | 2.78E-50 | Rarb | 6.29E-67 | 0.01371171 | 0.5 | 0.3 | 9.12E-64 |
| 8-Mar | 1.04E-54 | -0.100274095 | 0.171 | 0.255 | 2.79E-50 | Rab3c | 5.45E-08 | 0.01355794 | 0.56 | 0.62 | 7.91E-05 |
| Zdhhc20 | 1.13E-54 | -0.271360951 | 0.455 | 0.612 | 3.03E-50 | Scml4 | 1.23E-64 | 0.01351782 | 0.17 | 0.36 | 1.79E-61 |
| Abhd17b | 1.46E-54 | -0.043008416 | 0.47 | 0.582 | 3.93E-50 | Fam163a | 0.00135732 | 0.01308862 | 0.37 | 0.42 | 1 |
| Nckap1 | 1.51E-54 | -0.001696462 | 0.265 | 0.342 | 4.07E-50 | Cd37 | 1.92E-06 | 0.01305769 | 0.49 | 0.46 | 0.0027812 |
| Zfp445 | 1.59E-54 | 0.058931105 | 0.197 | 0.252 | 4.29E-50 | Otx2os1 | 1.95E-13 | 0.01305588 | 0.36 | 0.45 | 2.82E-10 |
| Npepps | 1.70E-54 | 0.015127179 | 0.311 | 0.389 | 4.57E-50 | Spag16 | 0.14605747 | 0.01304618 | 0.33 | 0.35 | 1 |
| Pls1 | 2.21E-54 | -0.062276554 | 0.297 | 0.395 | 5.94E-50 | Gm2817 | 9.00E-30 | 0.01282494 | 0.47 | 0.33 | 1.31E-26 |
| Akap8l | 2.31E-54 | -0.062628131 | 0.225 | 0.312 | 6.21E-50 | Ndnf | 3.16E-18 | 0.0127835 | 0.45 | 0.35 | 4.59E-15 |
| Txndc16 | 2.88E-54 | -0.030893199 | 0.279 | 0.362 | 7.74E-50 | Fnbp1l | 1.69E-14 | 0.01241588 | 0.32 | 0.42 | 2.45E-11 |
| 4931406P16Rik | 2.88E-54 | -0.005215211 | 0.186 | 0.253 | 7.75E-50 | Rbm47 | 0.00157156 | 0.01241474 | 0.4 | 0.36 | 1 |
| Stag1 | 2.93E-54 | -0.051627188 | 0.29 | 0.382 | 7.89E-50 | Fam129a | 8.64E-10 | 0.01221391 | 0.23 | 0.3 | 1.25E-06 |
| Asxl2 | 2.99E-54 | -0.148649211 | 0.181 | 0.274 | 8.04E-50 | Efcab6 | 8.46E-09 | 0.01221289 | 0.47 | 0.53 | 1.23E-05 |
| Rap1a | 3.13E-54 | -0.049784354 | 0.27 | 0.358 | 8.43E-50 | Aoah | 0.0981987 | 0.01211709 | 0.38 | 0.36 | 1 |
| Taf15 | 3.53E-54 | -0.032248081 | 0.187 | 0.257 | 9.50E-50 | Tll1 | 0.00184452 | 0.01180123 | 0.42 | 0.47 | 1 |
| Mycbp2 | 3.56E-54 | 0.055831204 | 0.579 | 0.67 | 9.60E-50 | Slc26a4 | 2.66E-228 | 0.01172197 | 0.31 | 0.7 | 3.86E-225 |
| Agap1 | 3.71E-54 | -0.130007107 | 0.584 | 0.724 | 9.98E-50 | Sema3d | 9.34E-254 | 0.01161274 | 0.08 | 0.43 | 1.36E-250 |
| Crebrf | 3.81E-54 | 0.116121057 | 0.212 | 0.258 | 1.02E-49 | Epha10 | 1.77E-36 | 0.01120967 | 0.25 | 0.39 | 2.57E-33 |
| Elovl7 | 3.97E-54 | -0.258305567 | 0.186 | 0.298 | 1.07E-49 | P3h2 | 0.10183959 | 0.01100975 | 0.45 | 0.44 | 1 |
| Mta3 | 4.15E-54 | -0.059202683 | 0.211 | 0.291 | 1.12E-49 | Pcsk5 | 9.68E-19 | 0.0108523 | 0.47 | 0.58 | 1.41E-15 |
| Trip12 | 4.31E-54 | 0.042817725 | 0.226 | 0.289 | 1.16E-49 | Slc4a5 | 2.58E-39 | 0.01082434 | 0.38 | 0.54 | 3.74E-36 |
| Fxr1 | 4.59E-54 | -0.007116593 | 0.211 | 0.279 | 1.24E-49 | Zbbx | 7.09E-12 | 0.01079158 | 0.32 | 0.4 | 1.03E-08 |
| Foxn3 | 4.67E-54 | -0.002239587 | 0.464 | 0.564 | 1.26E-49 | Ndst3 | 5.54E-111 | 0.01070623 | 0.21 | 0.47 | 8.04E-108 |
| Arhgef12 | 4.92E-54 | -0.010928803 | 0.218 | 0.293 | 1.32E-49 | Arhgap2 | 1.12E-110 | 0.01068621 | 0.27 | 0.54 | 1.62E-107 |
| Pkp4 | 5.28E-54 | -0.007108554 | 0.479 | 0.584 | 1.42E-49 | Matn2 | 4.00E-29 | 0.01056722 | 0.42 | 0.56 | 5.81E-26 |
| Map4 | 5.33E-54 | -0.104206477 | 0.375 | 0.493 | 1.44E-49 | Pappa | 7.31E-40 | 0.01037113 | 0.27 | 0.14 | 1.06E-36 |
| Shisa4 | 6.01E-54 | -0.149229113 | 0.22 | 0.321 | 1.62E-49 | Trem2 | 1.14E-87 | 0.01036102 | 0.17 | 0.39 | 1.66E-84 |
| Hip1 | 6.06E-54 | -0.112619039 | 0.232 | 0.329 | 1.63E-49 | Ccdc114 | 0.14754612 | 0.01035915 | 0.39 | 0.4 | 1 |
| Mbd5 | 6.09E-54 | 0.100767487 | 0.291 | 0.352 | 1.64E-49 | Arhgap2 | 0.00826847 | 0.01012492 | 0.46 | 0.5 | 1 |
| Phf21a | 7.27E-54 | 0.00528235 | 0.273 | 0.352 | 1.96E-49 | Gm3850 | 3.04E-142 | 0.01000702 | 0.11 | 0.32 | 4.42E-139 |
| Dlg1 | 7.89E-54 | 0.064085698 | 0.538 | 0.619 | 2.12E-49 | Cfap65 | 9.43E-81 | 0.00983916 | 0.43 | 0.66 | 1.37E-77 |
| Dnajb2 | 9.01E-54 | 0.134496262 | 0.232 | 0.274 | 2.43E-49 | Moxd1 | 2.96E-54 | 0.0098175 | 0.3 | 0.49 | 4.29E-51 |
| Cdh20 | 1.05E-53 | -0.321716737 | 0.465 | 0.627 | 2.82E-49 | Npsr1 | 0.00031703 | 0.00976626 | 0.47 | 0.52 | 0.46001655 |
| Fam193a | 1.05E-53 | 0.016641066 | 0.205 | 0.272 | 2.82E-49 | Dsp | 7.48E-72 | 0.00950342 | 0.33 | 0.55 | 1.09E-68 |
| Tanc2 | 1.08E-53 | -0.021867943 | 0.265 | 0.351 | 2.92E-49 | Gpc4 | 2.18E-06 | 0.00946213 | 0.44 | 0.51 | 0.00316581 |
| Rsrc1 | 1.15E-53 | 0.039750031 | 0.229 | 0.294 | 3.09E-49 | Gm4270 | 1.76E-119 | 0.0092592 | 0.13 | 0.39 | 2.56E-116 |
| Cdk12 | 1.55E-53 | -0.06019426 | 0.221 | 0.3 | 4.18E-49 | Ahnak | 0.65901496 | 0.00914322 | 0.33 | 0.32 | 1 |
| Gm42418 | 1.63E-53 | 0.356292006 | 0.85 | 0.837 | 4.40E-49 | Adgrl4 | 1.59E-30 | 0.00912937 | 0.36 | 0.5 | 2.30E-27 |
| Insr | 1.71E-53 | -0.066195861 | 0.186 | 0.264 | 4.60E-49 | Fam20a | 0.05411631 | 0.00891694 | 0.5 | 0.52 | 1 |
| Cpm | 1.71E-53 | -0.092898135 | 0.187 | 0.271 | 4.60E-49 | Gm3555 | 3.80E-55 | 0.00872945 | 0.12 | 0.27 | 5.51E-52 |
| Brd4 | 2.02E-53 | -0.079949016 | 0.172 | 0.25 | 5.45E-49 | Pparg | 5.37E-06 | 0.00869757 | 0.42 | 0.48 | 0.00779372 |

| | | | | | | | | | | | | |
|---|---|---|---|---|---|---|---|---|---|---|---|---|
| Gpatch8 | 2.14E-53 | -0.068898842 | 0.395 | 0.503 | 5.77E-49 | | Gad2 | 4.11E-30 | 0.0085587 | 0.3 | 0.44 | 5.96E-27 |
| Elmo1 | 2.75E-53 | -0.300951194 | 0.673 | 0.818 | 7.40E-49 | | Myo3b | 0.83763572 | 0.00850001 | 0.4 | 0.39 | 1 |
| Ttc3 | 3.15E-53 | 0.32753307 | 0.261 | 0.264 | 8.48E-49 | | Igfbp2 | 4.34E-53 | 0.0082717 | 0.12 | 0.27 | 6.30E-50 |
| Fbxl5 | 3.17E-53 | 0.08209296 | 0.329 | 0.387 | 8.53E-49 | | Rnf182 | 2.68E-22 | 0.00825958 | 0.43 | 0.55 | 3.89E-19 |
| Scaper | 3.25E-53 | 0.107569467 | 0.226 | 0.274 | 8.74E-49 | | Gm2674 | 1.45E-170 | 0.00820758 | 0.15 | 0.47 | 2.11E-167 |
| Gm42413 | 3.28E-53 | -0.0227909 | 0.335 | 0.43 | 8.84E-49 | | Mamdc | 1.96E-64 | 0.0081386 | 0.35 | 0.56 | 2.84E-61 |
| Gnaq | 3.30E-53 | 0.03103939 | 0.314 | 0.389 | 8.88E-49 | | Glra3 | 2.82E-96 | 0.00809965 | 0.07 | 0.25 | 4.10E-93 |
| Fryl | 3.38E-53 | -0.017499649 | 0.436 | 0.537 | 9.11E-49 | | Kcne2 | 1.23E-97 | 0.00800326 | 0.19 | 0.42 | 1.78E-94 |
| Efnb3 | 3.53E-53 | 0.037653735 | 0.222 | 0.289 | 9.50E-49 | | Tmem20 | 0.03433951 | 0.00793802 | 0.47 | 0.48 | 1 |
| Mtmr2 | 3.61E-53 | -0.074081513 | 0.239 | 0.326 | 9.71E-49 | | Plch2 | 2.11E-21 | 0.00791262 | 0.33 | 0.45 | 3.07E-18 |
| Daam1 | 3.71E-53 | 0.071830356 | 0.473 | 0.548 | 9.98E-49 | | Fn1 | 9.07E-34 | 0.0078921 | 0.35 | 0.5 | 1.32E-30 |
| Bicd1 | 4.11E-53 | -0.074915431 | 0.27 | 0.365 | 1.11E-48 | | Tbxas1 | 7.93E-11 | 0.00777152 | 0.47 | 0.55 | 1.15E-07 |
| Fbxo7 | 4.27E-53 | -0.031012717 | 0.191 | 0.262 | 1.15E-48 | | Calb1 | 4.95E-140 | 0.00750807 | 0.1 | 0.35 | 7.19E-137 |
| Rb1cc1 | 5.32E-53 | 0.05724459 | 0.198 | 0.253 | 1.43E-48 | | Mylk | 2.49E-63 | 0.00747094 | 0.38 | 0.2 | 3.62E-60 |
| Srsf5 | 6.22E-53 | 0.067869708 | 0.227 | 0.284 | 1.68E-48 | | Gm2063 | 2.96E-208 | 0.00743858 | 0.16 | 0.51 | 4.30E-205 |
| Ago3 | 6.34E-53 | -0.064652676 | 0.222 | 0.304 | 1.71E-48 | | Dchs2 | 9.36E-44 | 0.00728989 | 0.36 | 0.53 | 1.36E-40 |
| Wac | 6.93E-53 | -0.024943221 | 0.204 | 0.275 | 1.87E-48 | | Blnk | 5.91E-57 | 0.00726238 | 0.27 | 0.46 | 8.58E-54 |
| Spop | 7.96E-53 | -0.080315323 | 0.175 | 0.256 | 2.14E-48 | | Rph3a | 1.86E-10 | 0.00719284 | 0.53 | 0.46 | 2.70E-07 |
| Map4k5 | 1.22E-52 | 0.04240623 | 0.349 | 0.426 | 3.29E-48 | | 1700007F1 | 0.20203691 | 0.0071821 | 0.42 | 0.44 | 1 |
| Gnai1 | 1.28E-52 | -0.005021242 | 0.281 | 0.361 | 3.43E-48 | | Id1 | 3.22E-06 | 0.00714359 | 0.39 | 0.46 | 0.00467436 |
| Lrrc8b | 1.69E-52 | -0.024980182 | 0.211 | 0.284 | 4.56E-48 | | Pkp2 | 4.71E-150 | 0.00666182 | 0.38 | 0.69 | 6.83E-147 |
| A230001M10Ri | 1.71E-52 | -0.167880555 | 0.338 | 0.461 | 4.61E-48 | | Phkg1 | 2.39E-17 | 0.00662271 | 0.4 | 0.5 | 3.47E-14 |
| Amph | 2.26E-52 | 0.121445688 | 0.291 | 0.338 | 6.09E-48 | | Scube1 | 2.62E-61 | 0.00650305 | 0.37 | 0.58 | 3.81E-58 |
| Chd7 | 2.34E-52 | 0.082183038 | 0.282 | 0.341 | 6.30E-48 | | Dcn | 2.12E-37 | 0.00645167 | 0.42 | 0.58 | 3.08E-34 |
| Elf2 | 2.61E-52 | -0.105507871 | 0.169 | 0.25 | 7.03E-48 | | Ar | 9.96E-06 | 0.00632849 | 0.31 | 0.36 | 0.01445863 |
| Atad2b | 2.73E-52 | -0.104977207 | 0.241 | 0.335 | 7.34E-48 | | Gm5149 | 2.40E-187 | 0.00618042 | 0.16 | 0.5 | 3.49E-184 |
| Tom1l2 | 2.84E-52 | -0.083418284 | 0.28 | 0.378 | 7.65E-48 | | Ptger3 | 4.87E-32 | 0.00614051 | 0.36 | 0.5 | 7.06E-29 |
| Immp1l | 3.72E-52 | -0.049115766 | 0.23 | 0.312 | 1.00E-47 | | Siglech | 4.36E-25 | 0.00605146 | 0.16 | 0.26 | 6.33E-22 |
| Zfr | 5.16E-52 | 0.109176451 | 0.268 | 0.319 | 1.39E-47 | | Vwa3a | 7.32E-05 | 0.00599739 | 0.21 | 0.25 | 0.10618373 |
| Slc20a2 | 5.18E-52 | -0.114197481 | 0.226 | 0.318 | 1.40E-47 | | Lrriq1 | 1.09E-05 | 0.0058503 | 0.28 | 0.33 | 0.01588769 |
| Gatad2b | 5.85E-52 | -0.081365722 | 0.189 | 0.271 | 1.58E-47 | | Nhs | 6.03E-13 | 0.00575596 | 0.35 | 0.44 | 8.75E-10 |
| Stk39 | 6.10E-52 | -0.047978947 | 0.219 | 0.298 | 1.64E-47 | | Klhl1 | 2.25E-97 | 0.00551591 | 0.13 | 0.36 | 3.26E-94 |
| Tmcc1 | 6.23E-52 | -0.129418693 | 0.579 | 0.715 | 1.68E-47 | | Phka1 | 9.49E-13 | 0.00550508 | 0.45 | 0.42 | 1.38E-09 |
| Cdc42se2 | 8.13E-52 | -0.009973342 | 0.229 | 0.3 | 2.19E-47 | | Gm5860 | 9.14E-07 | 0.00541298 | 0.42 | 0.47 | 0.00132579 |
| Ccp110 | 8.80E-52 | -0.047943574 | 0.35 | 0.446 | 2.37E-47 | | Id4 | 0.00933765 | 0.00535761 | 0.43 | 0.4 | 1 |
| Specc1 | 1.11E-51 | 0.016000042 | 0.285 | 0.361 | 3.00E-47 | | Cdh7 | 5.95E-44 | 0.00533531 | 0.25 | 0.41 | 8.63E-41 |
| Cul1 | 1.39E-51 | -0.08500341 | 0.175 | 0.255 | 3.75E-47 | | Gm1941 | 3.53E-214 | 0.00531211 | 0.12 | 0.46 | 5.12E-211 |
| Gm48678 | 1.42E-51 | -0.055858954 | 0.458 | 0.573 | 3.82E-47 | | Ly6c1 | 7.99E-95 | 0.00522688 | 0.26 | 0.51 | 1.16E-91 |
| Ubn2 | 1.45E-51 | -0.055898229 | 0.29 | 0.382 | 3.90E-47 | | Atp10a | 7.53E-17 | 0.0052229 | 0.48 | 0.59 | 1.09E-13 |
| Sema6a | 1.65E-51 | 0.155647847 | 0.327 | 0.378 | 4.45E-47 | | Coch | 3.34E-11 | 0.00518085 | 0.31 | 0.39 | 4.84E-08 |
| Add1 | 1.86E-51 | -0.115221527 | 0.291 | 0.393 | 5.02E-47 | | Galnt14 | 3.08E-15 | 0.00506869 | 0.32 | 0.41 | 4.47E-12 |
| Fam13b | 2.04E-51 | -0.031501338 | 0.198 | 0.269 | 5.49E-47 | | Frem1 | 0.17090045 | 0.00491362 | 0.36 | 0.36 | 1 |
| Wdr20 | 2.14E-51 | -0.081189626 | 0.194 | 0.273 | 5.75E-47 | | Igsf9b | 4.30E-118 | 0.00485954 | 0.14 | 0.39 | 6.24E-115 |
| Ypel2 | 2.35E-51 | -0.079289873 | 0.541 | 0.662 | 6.34E-47 | | Gm4874 | 0.09330596 | 0.00456148 | 0.34 | 0.36 | 1 |
| Trim59 | 3.28E-51 | -0.018852359 | 0.202 | 0.275 | 8.83E-47 | | Kitl | 1.02E-88 | 0.00453179 | 0.26 | 0.5 | 1.47E-85 |
| Phf14 | 3.32E-51 | -0.024098605 | 0.192 | 0.262 | 8.95E-47 | | Fcrls | 0.08038947 | 0.0045221 | 0.46 | 0.44 | 1 |
| App | 3.57E-51 | -0.021744337 | 0.664 | 0.776 | 9.61E-47 | | Igfbp7 | 1.48E-38 | 0.00438076 | 0.38 | 0.23 | 2.15E-35 |
| Dnmt3a | 3.81E-51 | 0.067203683 | 0.217 | 0.273 | 1.02E-46 | | Il16 | 2.82E-112 | 0.00415106 | 0.22 | 0.49 | 4.09E-109 |
| Xrcc6 | 3.98E-51 | -0.006745244 | 0.322 | 0.409 | 1.07E-46 | | Chrm2 | 0.05318409 | 0.00413562 | 0.49 | 0.52 | 1 |
| Glg1 | 4.16E-51 | -0.009958779 | 0.203 | 0.272 | 1.12E-46 | | Ptpru | 4.78E-57 | 0.00412437 | 0.48 | 0.3 | 6.94E-54 |
| Vti1a | 4.43E-51 | -0.00596328 | 0.247 | 0.325 | 1.19E-46 | | Slco2b1 | 1.41E-162 | 0.00410643 | 0.12 | 0.41 | 2.05E-159 |
| E130308A19Rik | 4.52E-51 | -0.04501108 | 0.258 | 0.341 | 1.22E-46 | | Gm3625 | 1.25E-203 | 0.00406421 | 0.14 | 0.48 | 1.81E-200 |
| Adam22 | 6.21E-51 | 0.017532446 | 0.221 | 0.289 | 1.67E-46 | | Slc4a4 | 3.51E-85 | 0.00400702 | 0.29 | 0.53 | 5.09E-82 |
| Desi1 | 6.80E-51 | 0.008763532 | 0.234 | 0.304 | 1.83E-46 | | Maob | 9.84E-54 | 0.00395286 | 0.14 | 0.3 | 1.43E-50 |
| Kctd3 | 7.36E-51 | -0.253156758 | 0.201 | 0.31 | 1.98E-46 | | Slc9a2 | 0.00298941 | 0.0039328 | 0.5 | 0.54 | 1 |
| Pan3 | 7.47E-51 | 0.012121579 | 0.21 | 0.275 | 2.01E-46 | | Neb | 5.46E-08 | 0.00378224 | 0.42 | 0.36 | 7.92E-05 |
| Cadm1 | 7.73E-51 | 0.108153077 | 0.328 | 0.392 | 2.08E-46 | | Grin2c | 0.08375426 | 0.00341393 | 0.46 | 0.44 | 1 |
| Tspan5 | 8.37E-51 | -0.14385047 | 0.324 | 0.441 | 2.25E-46 | | A830008E2 | 0.07719147 | 0.00334243 | 0.31 | 0.29 | 1 |
| Lncpint | 8.52E-51 | 0.108707756 | 0.267 | 0.32 | 2.29E-46 | | Ano1 | 0.00017184 | 0.00319166 | 0.4 | 0.45 | 0.24934577 |
| Ehbp1 | 9.34E-51 | 0.099795186 | 0.229 | 0.281 | 2.51E-46 | | Syndig1 | 7.34E-06 | 0.00297209 | 0.39 | 0.45 | 0.01065535 |
| Tjp1 | 1.39E-50 | 0.086728665 | 0.32 | 0.383 | 3.73E-46 | | Gm1717 | 1.74E-05 | 0.00295795 | 0.43 | 0.49 | 0.02531084 |
| Arhgef2 | 1.40E-50 | 0.020599407 | 0.195 | 0.254 | 3.76E-46 | | Drc7 | 0.81686403 | 0.00282886 | 0.4 | 0.39 | 1 |
| Babam2 | 1.46E-50 | -0.036669734 | 0.204 | 0.279 | 3.93E-46 | | Epsti1 | 0.01301029 | 0.00280006 | 0.34 | 0.31 | 1 |
| Ralgps1 | 1.49E-50 | -0.082619555 | 0.327 | 0.427 | 4.02E-46 | | Nr4a3 | 2.13E-40 | 0.00271481 | 0.36 | 0.53 | 3.10E-37 |
| Glul | 2.03E-50 | -0.199838866 | 0.186 | 0.288 | 5.47E-46 | | Igsf5 | 1.59E-67 | 0.00262473 | 0.09 | 0.25 | 2.31E-64 |
| Iqsec1 | 2.10E-50 | -0.136753035 | 0.291 | 0.398 | 5.65E-46 | | Alkal2 | 2.71E-06 | 0.00247379 | 0.25 | 0.31 | 0.00393444 |
| Eml1 | 2.59E-50 | -0.42872703 | 0.138 | 0.258 | 6.98E-46 | | Bank1 | 8.96E-20 | 0.00230549 | 0.36 | 0.26 | 1.30E-16 |
| Prox1 | 2.80E-50 | -0.090871847 | 0.177 | 0.257 | 7.54E-46 | | Fgf10 | 4.44E-34 | 0.00225208 | 0.3 | 0.44 | 6.45E-31 |
| Fth1 | 3.52E-50 | -0.01650048 | 0.419 | 0.522 | 9.49E-46 | | Mageb1 | 0.39768458 | 0.00219897 | 0.24 | 0.25 | 1 |
| Sort1 | 3.98E-50 | -0.031638606 | 0.255 | 0.337 | 1.07E-45 | | Slc2a1 | 1.51E-82 | 0.00211808 | 0.41 | 0.64 | 2.20E-79 |
| Luc7l2 | 5.88E-50 | 0.130125279 | 0.579 | 0.642 | 1.58E-45 | | Sh3rf1 | 8.53E-40 | 0.00205861 | 0.17 | 0.31 | 1.24E-36 |
| Slc38a2 | 9.36E-50 | -0.040938569 | 0.366 | 0.465 | 2.52E-45 | | Rcsd1 | 0.07208963 | 0.00200689 | 0.24 | 0.26 | 1 |
| Frmd4b | 9.43E-50 | -0.061192317 | 0.545 | 0.661 | 2.54E-45 | | Myo1b | 3.25E-05 | 0.00191685 | 0.43 | 0.37 | 0.04711984 |
| Rnf220 | 1.39E-49 | -0.244831492 | 0.687 | 0.824 | 3.74E-45 | | Prkca | 3.76E-24 | 0.00183185 | 0.71 | 0.82 | 5.46E-21 |
| Acaca | 1.45E-49 | -0.128437236 | 0.348 | 0.463 | 3.90E-45 | | Rtl4 | 3.91E-22 | 0.00182593 | 0.48 | 0.59 | 5.68E-19 |
| Tjp2 | 1.52E-49 | -0.115311561 | 0.172 | 0.253 | 4.09E-45 | | Scn9a | 2.33E-176 | 0.00169528 | 0.18 | 0.51 | 3.38E-173 |
| Kif13b | 1.74E-49 | -0.148073892 | 0.532 | 0.666 | 4.69E-45 | | Tmem17 | 3.88E-12 | 0.00166102 | 0.48 | 0.57 | 5.62E-09 |
| Zmym4 | 1.97E-49 | 0.066413411 | 0.22 | 0.274 | 5.30E-45 | | Hapln1 | 0.19746991 | 0.00160453 | 0.42 | 0.43 | 1 |
| Phactr3 | 2.08E-49 | 0.01769354 | 0.242 | 0.309 | 5.59E-45 | | 3-Mar | 9.05E-100 | 0.00157306 | 0.22 | 0.47 | 1.31E-96 |
| Wipf1 | 2.10E-49 | -0.112478499 | 0.255 | 0.349 | 5.64E-45 | | 9330185C1 | 1.04E-283 | 0.00156594 | 0.15 | 0.57 | 1.52E-280 |
| Rabgap1 | 2.10E-49 | 0.026035428 | 0.259 | 0.324 | 5.65E-45 | | Ptchd1 | 3.07E-06 | 0.00152668 | 0.3 | 0.36 | 0.00445928 |
| Stxbp6 | 2.30E-49 | -0.06691668 | 0.408 | 0.519 | 6.18E-45 | | Kcnmb2 | 5.14E-31 | 0.00136027 | 0.43 | 0.58 | 7.46E-28 |
| Strn3 | 2.36E-49 | 0.002878784 | 0.278 | 0.353 | 6.35E-45 | | Penk | 2.24E-28 | 0.00121129 | 0.28 | 0.41 | 3.25E-25 |
| Cerk | 2.59E-49 | 0.028290767 | 0.267 | 0.336 | 6.96E-45 | | Skap1 | 9.15673121502032e-311 | 0.00117659 | 0.07 | 0.47 | 1.33E-307 |
| Olfml1 | 3.38E-49 | -0.001184281 | 0.217 | 0.285 | 9.10E-45 | | Arl15 | 7.49E-35 | 0.00117524 | 0.48 | 0.63 | 1.09E-31 |
| Usp9x | 3.61E-49 | -0.043841169 | 0.195 | 0.264 | 9.72E-45 | | Glis3 | 5.16E-05 | 0.00117184 | 0.47 | 0.53 | 0.07483255 |
| Rasgrp3 | 3.79E-49 | -0.225692176 | 0.161 | 0.258 | 1.02E-44 | | Tmtc4 | 7.25E-51 | 0.00107368 | 0.12 | 0.27 | 1.05E-47 |
| Akap6 | 5.35E-49 | 0.034355592 | 0.38 | 0.46 | 1.44E-44 | | Cfap52 | 1.67E-110 | 0.00070857 | 0.22 | 0.48 | 2.42E-107 |
| A330015K06Rik | 5.42E-49 | -0.130112406 | 0.418 | 0.541 | 1.46E-44 | | Kcns3 | 4.84E-93 | 0.00068713 | 0.23 | 0.48 | 7.03E-90 |
| Rere | 5.59E-49 | -0.142764977 | 0.504 | 0.637 | 1.50E-44 | | Otof | 2.04E-09 | 0.00064151 | 0.28 | 0.21 | 2.96E-06 |
| Pclo | 6.27E-49 | 0.234399949 | 0.23 | 0.254 | 1.69E-44 | | Gm1215 | 1.57E-43 | 0.00048639 | 0.34 | 0.52 | 2.28E-40 |
| Tanc1 | 7.28E-49 | -0.012289563 | 0.204 | 0.271 | 1.96E-44 | | Slc25a2 | 4.54E-82 | 0.00040887 | 0.35 | 0.59 | 6.59E-79 |
| Tcf12 | 7.60E-49 | -0.100512082 | 0.615 | 0.742 | 2.05E-44 | | Tox2 | 1.23E-15 | 0.00037836 | 0.26 | 0.36 | 1.78E-12 |
| Aspa | 7.96E-49 | -0.029196053 | 0.65 | 0.766 | 2.14E-44 | | Kcnj8 | 2.31E-77 | 0.00036839 | 0.17 | 0.38 | 3.34E-74 |
| Phf20l1 | 8.66E-49 | 0.042545508 | 0.211 | 0.267 | 2.33E-44 | | Col4a4 | 1.74E-41 | 0.00030294 | 0.12 | 0.25 | 2.53E-38 |
| Mical3 | 9.18E-49 | -0.037692669 | 0.185 | 0.253 | 2.47E-44 | | Pltp | 3.15E-265 | 0.00021993 | 0.07 | 0.44 | 4.57E-262 |
| Kif13a | 9.34E-49 | 0.059841271 | 0.495 | 0.577 | 2.51E-44 | | Gm4744 | 4.14E-124 | 0.00016457 | 0.21 | 0.48 | 6.01E-121 |
| Adamtsl1 | 1.09E-48 | -0.020583843 | 0.368 | 0.461 | 2.93E-44 | | Ccdc3 | 6.67E-53 | 8.79E-05 | 0.16 | 0.32 | 9.68E-50 |
| Cep97 | 1.13E-48 | 0.05813916 | 0.213 | 0.265 | 3.05E-44 | | Lef1 | 5.73E-88 | -2.66E-05 | 0.22 | 0.45 | 8.32E-85 |
| Fut8 | 1.19E-48 | -0.060051642 | 0.199 | 0.277 | 3.20E-44 | | Gm1515 | 0.00620353 | -7.90E-05 | 0.35 | 0.38 | 1 |
| Pex5l | 1.20E-48 | -0.331977909 | 0.774 | 0.887 | 3.23E-44 | | Rspo2 | 1.89E-35 | -0.0002062 | 0.37 | 0.53 | 2.75E-32 |
| Bin1 | 1.34E-48 | -0.031620576 | 0.609 | 0.72 | 3.62E-44 | | Gm6994 | 1.12E-71 | -0.0002292 | 0.13 | 0.3 | 1.62E-68 |
| Prkcz | 2.11E-48 | 0.011705364 | 0.237 | 0.302 | 5.69E-44 | | Pde5a | 2.40E-32 | -0.0002739 | 0.41 | 0.27 | 3.49E-29 |
| Trim44 | 2.53E-48 | -0.198405923 | 0.175 | 0.27 | 6.81E-44 | | Il10ra | 4.89E-287 | -0.0002937 | 0.06 | 0.44 | 7.10E-284 |
| Mog | 2.84E-48 | -0.085081438 | 0.658 | 0.782 | 7.65E-44 | | Pde11a | 1.43E-74 | -0.0005698 | 0.31 | 0.53 | 2.08E-71 |
| Sytl2 | 4.31E-48 | 0.182561396 | 0.347 | 0.382 | 1.16E-43 | | Gm1223 | 6.03E-07 | -0.000573 | 0.23 | 0.29 | 0.00087557 |
| Arid5b | 4.83E-48 | -0.034942284 | 0.215 | 0.291 | 1.30E-43 | | Glra2 | 4.61E-11 | -0.0009076 | 0.32 | 0.4 | 6.69E-08 |
| Slc12a2 | 5.48E-48 | -0.08650474 | 0.499 | 0.616 | 1.47E-43 | | Ifitm3 | 8.02E-28 | -0.0011478 | 0.45 | 0.59 | 1.16E-24 |
| Fbxl17 | 5.52E-48 | 0.042883228 | 0.257 | 0.32 | 1.49E-43 | | Maf | 0.00018757 | -0.0011561 | 0.44 | 0.4 | 0.27216117 |
| Kndc1 | 9.02E-48 | -0.104318987 | 0.252 | 0.347 | 2.43E-43 | | Abca12 | 3.59E-42 | -0.0013316 | 0.2 | 0.35 | 5.21E-39 |
| Nfib | 9.64E-48 | 0.014173799 | 0.224 | 0.292 | 2.59E-43 | | Egln3 | 3.56E-182 | -0.0013343 | 0.1 | 0.4 | 5.16E-179 |
| Sorcs1 | 1.37E-47 | -0.096439451 | 0.536 | 0.659 | 3.69E-43 | | Edaradd | 0.02895794 | -0.0013894 | 0.24 | 0.27 | 1 |
| Klhl2 | 2.82E-47 | -0.095118319 | 0.567 | 0.687 | 7.59E-43 | | C230072F1 | 4.23E-185 | -0.0013894 | 0.05 | 0.32 | 6.13E-182 |
| Tmod2 | 2.95E-47 | -0.167049975 | 0.529 | 0.666 | 7.94E-43 | | D630024D0 | 2.55E-78 | -0.0016872 | 0.36 | 0.59 | 3.70E-75 |
| Ppp2r3a | 3.50E-47 | -0.099659511 | 0.53 | 0.652 | 9.43E-43 | | Cit | 1.91E-15 | -0.0017617 | 0.35 | 0.44 | 2.77E-12 |
| Ssh2 | 3.95E-47 | -0.10743109 | 0.537 | 0.663 | 1.06E-42 | | Elavl2 | 3.44E-39 | -0.0018487 | 0.45 | 0.61 | 4.99E-36 |
| Heg1 | 4.99E-47 | -0.229041145 | 0.16 | 0.256 | 1.34E-42 | | Col14a1 | 2.79E-07 | -0.0020741 | 0.3 | 0.25 | 0.00040542 |
| Apbb2 | 7.04E-47 | -0.126400774 | 0.348 | 0.463 | 1.89E-42 | | B4galnt3 | 2.03E-07 | -0.0023845 | 0.32 | 0.39 | 0.00029384 |
| Mapre2 | 8.44E-47 | -0.229865714 | 0.205 | 0.312 | 2.27E-42 | | Adamts1 | 2.95E-17 | -0.0024788 | 0.39 | 0.49 | 4.28E-14 |
| Tcf4 | 8.49E-47 | 0.060129573 | 0.565 | 0.65 | 2.28E-42 | | Ly6a | 0.07774871 | -0.0025468 | 0.49 | 0.47 | 1 |
| Pcyt2 | 8.76E-47 | 0.003310724 | 0.202 | 0.269 | 2.36E-42 | | Foxd1 | 3.11E-224 | -0.0025831 | 0.04 | 0.35 | 4.52E-221 |
| Slk | 9.63E-47 | -0.032070999 | 0.2 | 0.268 | 2.59E-42 | | Dpp4 | 5.61E-191 | -0.0026179 | 0.11 | 0.42 | 8.14E-188 |
| Phactr1 | 9.76E-47 | 0.078107112 | 0.545 | 0.628 | 2.63E-42 | | Rai14 | 1.89E-80 | -0.0027073 | 0.26 | 0.49 | 2.74E-77 |
| Chn2 | 1.01E-46 | -0.067539244 | 0.206 | 0.281 | 2.73E-42 | | Aqp4 | 1.54E-09 | -0.0027081 | 0.45 | 0.53 | 2.24E-06 |
| Itsn2 | 1.08E-46 | 0.000396093 | 0.449 | 0.541 | 2.91E-42 | | Il1r1 | 7.56E-34 | -0.002796 | 0.48 | 0.63 | 1.10E-30 |
| Dip2b | 1.41E-46 | -0.099045854 | 0.498 | 0.615 | 3.80E-42 | | Hcrtr2 | 2.23E-64 | -0.0030784 | 0.19 | 0.38 | 3.23E-61 |
| Synj2 | 1.73E-46 | -0.060749691 | 0.27 | 0.35 | 4.65E-42 | | Slc16a10 | 1.67E-24 | -0.0031124 | 0.44 | 0.57 | 2.42E-21 |
| Dip2c | 1.96E-46 | 0.122439415 | 0.263 | 0.314 | 5.27E-42 | | Hmcn1 | 0.00022542 | -0.0031286 | 0.38 | 0.42 | 0.32708467 |
| Ctnna3 | 2.64E-46 | -0.302940347 | 0.653 | 0.792 | 7.10E-42 | | St8sia6 | 4.51E-08 | -0.0032354 | 0.37 | 0.43 | 6.55E-05 |
| Plekha1 | 2.67E-46 | -0.007491868 | 0.403 | 0.493 | 7.19E-42 | | Mettl24 | 1.65E-209 | -0.0032774 | 0.06 | 0.37 | 2.39E-206 |
| Myo18a | 2.84E-46 | 0.020934574 | 0.28 | 0.344 | 7.64E-42 | | Lair1 | 0.00130573 | -0.0033445 | 0.4 | 0.36 | 1 |
| Lcorl | 5.21E-46 | 0.090788518 | 0.242 | 0.29 | 1.40E-41 | | Pde1a | 3.38E-16 | -0.0034904 | 0.58 | 0.68 | 4.90E-13 |
| Prkcq | 5.44E-46 | -0.051963839 | 0.192 | 0.263 | 1.46E-41 | | Oprm1 | 1.10E-05 | -0.0035323 | 0.41 | 0.47 | 0.0160024 |
| Nfat5 | 5.69E-46 | 0.083911234 | 0.235 | 0.288 | 1.53E-41 | | Tfcp2l1 | 3.40E-228 | -0.0035555 | 0.01 | 0.28 | 4.94E-225 |
| Cobl | 6.85E-46 | -0.000358497 | 0.217 | 0.283 | 1.84E-41 | | Ngf | 0.27908539 | -0.0035859 | 0.47 | 0.47 | 1 |
| Gm16168 | 7.66E-46 | -0.216888429 | 0.5 | 0.644 | 2.06E-41 | | Gm1604 | 2.67E-56 | -0.0035937 | 0.17 | 0.34 | 3.88E-53 |
| Slc9a9 | 7.77E-46 | 0.013295925 | 0.237 | 0.301 | 2.09E-41 | | Syt2 | 1.16E-28 | -0.004008 | 0.17 | 0.29 | 1.68E-25 |
| Ext1 | 1.22E-45 | -0.026763441 | 0.212 | 0.285 | 3.29E-41 | | Klf2 | 2.13E-08 | -0.0040797 | 0.36 | 0.43 | 3.09E-05 |
| Dis3l2 | 1.62E-45 | -0.1095222 | 0.256 | 0.35 | 4.35E-41 | | Cyyr1 | 2.28E-38 | -0.0041482 | 0.34 | 0.5 | 3.31E-35 |
| Cdh19 | 1.65E-45 | -0.308131852 | 0.39 | 0.535 | 4.45E-41 | | Syt10 | 2.19E-27 | -0.0043557 | 0.31 | 0.2 | 3.17E-24 |
| Arhgap23 | 1.97E-45 | -0.080891796 | 0.657 | 0.775 | 5.29E-41 | | Khdrbs3 | 3.64E-56 | -0.0044759 | 0.39 | 0.59 | 5.28E-53 |
| Dnm3 | 2.33E-45 | -0.137664067 | 0.739 | 0.857 | 6.27E-41 | | Col4a5 | 0.00017477 | -0.0045074 | 0.46 | 0.51 | 0.25358932 |
| Grid2 | 2.44E-45 | -0.040937949 | 0.565 | 0.681 | 6.56E-41 | | Gm1622 | 0.0174929 | -0.0045088 | 0.39 | 0.38 | 1 |
| Clmn | 2.88E-45 | 0.04019419 | 0.498 | 0.577 | 7.75E-41 | | Frmpd3 | 1.59E-14 | -0.0045243 | 0.39 | 0.49 | 2.30E-11 |
| Gm20275 | 3.00E-45 | -0.032287879 | 0.216 | 0.29 | 8.07E-41 | | Pstpip2 | 9.31E-55 | -0.0047327 | 0.6 | 0.77 | 1.35E-51 |
| Zfp638 | 3.13E-45 | -0.003643976 | 0.475 | 0.566 | 8.42E-41 | | Clic6 | 5.09E-305 | -0.0047633 | 0.06 | 0.45 | 7.38E-302 |
| Hecw2 | 3.18E-45 | -0.209186327 | 0.708 | 0.836 | 8.56E-41 | | Abcb5 | 2.43E-25 | -0.0049286 | 0.35 | 0.48 | 3.53E-22 |
| Arap2 | 3.41E-45 | 0.020073885 | 0.325 | 0.396 | 9.18E-41 | | Adgra1 | 7.73E-19 | -0.0049864 | 0.33 | 0.44 | 1.12E-15 |
| Dock4 | 3.73E-45 | 0.157759514 | 0.328 | 0.37 | 1.00E-40 | | Nid1 | 4.75E-133 | -0.0050055 | 0.06 | 0.29 | 6.89E-130 |
| Kif21a | 6.91E-45 | 0.04122927 | 0.558 | 0.64 | 1.86E-40 | | Id3 | 2.49E-10 | -0.0053819 | 0.41 | 0.49 | 3.61E-07 |
| Ppp1r9a | 7.39E-45 | 0.048815111 | 0.218 | 0.274 | 1.99E-40 | | Col6a1 | 0.00033566 | -0.0053843 | 0.24 | 0.28 | 0.4870384 |
| Nfia | 2.15E-44 | -0.069042187 | 0.482 | 0.587 | 5.78E-40 | | Wdr72 | 3.19E-89 | -0.0058163 | 0.36 | 0.61 | 4.63E-86 |
| Igf1r | 2.37E-44 | -0.043187289 | 0.247 | 0.32 | 6.37E-40 | | Trpc3 | 9.28E-21 | -0.0059265 | 0.3 | 0.22 | 1.35E-17 |
| Kcnab1 | 2.91E-44 | -0.145705868 | 0.411 | 0.534 | 7.82E-40 | | Prdm6 | 0 | -0.00615 | 0.06 | 0.5 | 0 |
| Jph1 | 3.90E-44 | -0.009998704 | 0.255 | 0.327 | 1.05E-39 | | Paqr5 | 1.34E-85 | -0.006244 | 0.38 | 0.62 | 1.95E-82 |
| Trim2 | 7.02E-44 | 0.004913448 | 0.539 | 0.632 | 1.89E-39 | | Crhbp | 1.39E-33 | -0.006377 | 0.4 | 0.55 | 2.02E-30 |
| Dscaml1 | 1.20E-43 | -0.176142793 | 0.596 | 0.731 | 3.23E-39 | | Dnah6 | 0.00048028 | -0.0064731 | 0.24 | 0.28 | 0.69689254 |
| Gpm6b | 1.23E-43 | -0.220463221 | 0.686 | 0.816 | 3.32E-39 | | Cdh8 | 5.90E-167 | -0.0064742 | 0.4 | 0.73 | 8.56E-164 |
| Kif1b | 1.26E-43 | -0.072671613 | 0.645 | 0.759 | 3.38E-39 | | Col1a2 | 2.50E-15 | -0.0064985 | 0.32 | 0.41 | 3.62E-12 |
| Erbin | 1.34E-43 | 0.063689045 | 0.653 | 0.731 | 3.60E-39 | | Rxfp2 | 1.11E-186 | -0.0066042 | 0.03 | 0.28 | 1.61E-183 |
| Psd3 | 1.56E-43 | 0.097590602 | 0.264 | 0.312 | 4.21E-39 | | Scn5a | 2.75E-35 | -0.0066563 | 0.35 | 0.5 | 3.99E-32 |
| Map7d2 | 1.97E-43 | -0.210719462 | 0.163 | 0.256 | 5.31E-39 | | 8030442B0 | 1.20E-99 | -0.0068609 | 0.34 | 0.6 | 1.74E-96 |
| Anks1b | 2.04E-43 | 0.309832404 | 0.55 | 0.55 | 5.48E-39 | | Fbln1 | 9.88131291682493e-324 | -0.0068652 | 0.06 | 0.47 | 1.4337785042313e-320 |
| Fign | 2.41E-43 | -0.015051956 | 0.259 | 0.332 | 6.49E-39 | | St8sia2 | 9.50E-264 | -0.0068676 | 0.09 | 0.47 | 1.38E-260 |
| C030029H02Rik | 7.53E-43 | 0.265767531 | 0.632 | 0.632 | 2.03E-38 | | Brinp2 | 5.97E-13 | -0.0070225 | 0.38 | 0.32 | 8.67E-10 |
| Zdhhc14 | 8.28E-43 | -0.071397951 | 0.459 | 0.561 | 2.23E-38 | | Plpp3 | 1.45E-10 | -0.0070239 | 0.44 | 0.51 | 2.10E-07 |
| Rasal2 | 9.17E-43 | -0.017700239 | 0.206 | 0.275 | 2.47E-38 | | Ccdc141 | 0.10047012 | -0.0070826 | 0.45 | 0.46 | 1 |
| Opalin | 1.52E-42 | 0.173443599 | 0.297 | 0.325 | 4.08E-38 | | Usp18 | 1.46E-07 | -0.0072198 | 0.34 | 0.3 | 0.00021163 |
| Tulp4 | 1.55E-42 | -0.064918041 | 0.535 | 0.641 | 4.16E-38 | | Nyap2 | 4.72E-06 | -0.0074228 | 0.55 | 0.61 | 0.00684303 |
| Gm28376 | 2.38E-42 | -0.079605646 | 0.369 | 0.476 | 6.40E-38 | | Gm1526 | 0.00048508 | -0.007521 | 0.5 | 0.54 | 0.70384637 |
| Kcnk13 | 2.61E-42 | -0.177923962 | 0.22 | 0.315 | 7.04E-38 | | Ddr2 | 0.00022941 | -0.0076259 | 0.27 | 0.24 | 0.33287927 |
| Tspan2 | 2.62E-42 | 0.022669257 | 0.482 | 0.565 | 7.04E-38 | | A830019P0 | 2.90E-55 | -0.0077619 | 0.48 | 0.67 | 4.21E-52 |
| Dnajc6 | 3.23E-42 | 0.109713949 | 0.251 | 0.295 | 8.69E-38 | | Rgs12 | 4.60E-117 | -0.0077845 | 0.23 | 0.51 | 6.67E-114 |
| Car2 | 3.88E-42 | -0.02667842 | 0.588 | 0.693 | 1.04E-37 | | Fstl4 | 2.01E-14 | -0.0078929 | 0.44 | 0.54 | 2.92E-11 |
| Nbea | 4.57E-42 | 0.004907225 | 0.623 | 0.721 | 1.23E-37 | | Cd74 | 1.01E-39 | -0.0079285 | 0.33 | 0.5 | 1.46E-36 |
| Otud7a | 5.78E-42 | -0.249568724 | 0.44 | 0.581 | 1.56E-37 | | Cabcoco | 2.70E-155 | -0.0079476 | 0.15 | 0.45 | 3.92E-152 |
| Ube3a | 5.84E-42 | 0.184821906 | 0.243 | 0.275 | 1.57E-37 | | Mrvi1 | 5.67E-50 | -0.0079503 | 0.26 | 0.44 | 8.23E-47 |
| Sik2 | 6.77E-42 | -0.066434947 | 0.187 | 0.255 | 1.82E-37 | | Mapk4 | 1.74E-11 | -0.0079818 | 0.33 | 0.4 | 2.53E-08 |
| Rap1gds1 | 8.03E-42 | -0.14800998 | 0.178 | 0.26 | 2.16E-37 | | Cfap44 | 4.40E-59 | -0.0080947 | 0.36 | 0.56 | 6.38E-56 |
| Sema6d | 8.80E-42 | 0.070669812 | 0.316 | 0.373 | 2.37E-37 | | Lrrc3b | 3.09E-122 | -0.0081291 | 0.07 | 0.29 | 4.49E-119 |
| Rev3l | 1.51E-41 | 0.036555818 | 0.223 | 0.276 | 4.07E-37 | | Cdh18 | 7.26E-07 | -0.008231 | 0.53 | 0.56 | 0.00105316 |
| Zbtb20 | 1.72E-41 | 0.057810928 | 0.747 | 0.831 | 4.62E-37 | | Pecam1 | 1.93E-194 | -0.0083193 | 0.02 | 0.26 | 2.80E-191 |
| Zfp704 | 2.61E-41 | -0.016931307 | 0.214 | 0.278 | 7.03E-37 | | Tmem44 | 3.04E-59 | -0.0083325 | 0.36 | 0.19 | 4.41E-56 |
| Nlgn1 | 2.70E-41 | 0.304478686 | 0.568 | 0.581 | 7.26E-37 | | Neto2 | 3.12E-12 | -0.0083339 | 0.52 | 0.47 | 4.53E-09 |
| Tmem243 | 2.85E-41 | -0.030590764 | 0.19 | 0.253 | 7.68E-37 | | Myo5b | 4.05E-114 | -0.0083997 | 0.29 | 0.56 | 5.88E-111 |
| Enox1 | 3.53E-41 | -0.106192929 | 0.575 | 0.694 | 9.51E-37 | | Slc26a7 | 2.36E-154 | -0.008524 | 0.14 | 0.44 | 3.43E-151 |
| Srgap1 | 3.77E-41 | 0.224111921 | 0.311 | 0.33 | 1.01E-36 | | Npnt | 3.56E-79 | -0.0087686 | 0.22 | 0.44 | 5.17E-76 |
| Fmnl2 | 7.46E-41 | -0.168033103 | 0.467 | 0.593 | 2.01E-36 | | Gpr34 | 9.00E-05 | -0.0089712 | 0.27 | 0.23 | 0.13061182 |
| Samd4 | 8.22E-41 | 0.077430243 | 0.24 | 0.289 | 2.21E-36 | | Etv1 | 2.96E-07 | -0.0091269 | 0.36 | 0.37 | 0.00042881 |
| Gphn | 1.20E-40 | -0.060606199 | 0.674 | 0.782 | 3.23E-36 | | Pamr1 | 0.01589698 | -0.009269 | 0.36 | 0.35 | 1 |
| Ninj2 | 2.08E-40 | 0.027303619 | 0.38 | 0.457 | 5.59E-36 | | Postn | 5.41E-241 | -0.0095602 | 0.03 | 0.34 | 7.84E-238 |
| Plxdc2 | 2.27E-40 | -0.076390901 | 0.696 | 0.803 | 6.11E-36 | | Bnc2 | 1.18E-285 | -0.0096712 | 0.09 | 0.48 | 1.72E-282 |
| Abca8a | 2.99E-40 | -0.03007555 | 0.226 | 0.296 | 8.06E-36 | | Vgll3 | 3.55E-213 | -0.009885 | 0.1 | 0.44 | 5.15E-210 |
| Limch1 | 3.88E-40 | 0.011915733 | 0.595 | 0.684 | 1.04E-35 | | 4933413L0 | 2.06E-215 | -0.009887 | 0.21 | 0.59 | 2.99E-212 |
| Gm47271 | 4.45E-40 | -0.175877631 | 0.187 | 0.277 | 1.20E-35 | | Rspo3 | 9.49E-52 | -0.010136 | 0.41 | 0.6 | 1.38E-48 |
| Dgki | 5.13E-40 | -0.200118828 | 0.518 | 0.65 | 1.38E-35 | | Pdzph1 | 4.84E-184 | -0.0106116 | 0.16 | 0.49 | 7.03E-181 |
| Sik3 | 1.00E-39 | -0.116576875 | 0.641 | 0.759 | 2.71E-35 | | Cnih3 | 1.26E-08 | -0.0107103 | 0.43 | 0.5 | 1.83E-05 |
| Stox2 | 1.08E-39 | 0.042361266 | 0.237 | 0.294 | 2.92E-35 | | Lrrk1 | 3.45E-82 | -0.0107331 | 0.23 | 0.45 | 5.00E-79 |
| Rftn1 | 1.09E-39 | -0.213373303 | 0.182 | 0.273 | 2.94E-35 | | Cyth4 | 1.14E-76 | -0.0107617 | 0.27 | 0.49 | 1.65E-73 |
| Pak1 | 1.97E-39 | -0.055804664 | 0.601 | 0.706 | 5.29E-35 | | Olfm3 | 7.46E-10 | -0.0107852 | 0.54 | 0.61 | 1.08E-06 |
| Ppp1r16b | 2.09E-39 | -0.05011914 | 0.628 | 0.735 | 5.64E-35 | | Fa2h | 9.35E-87 | -0.011068 | 0.75 | 0.92 | 1.36E-83 |
| Spock3 | 2.38E-39 | -0.265351769 | 0.413 | 0.546 | 6.40E-35 | | Ccdc170 | 8.74E-59 | -0.0111569 | 0.36 | 0.55 | 1.27E-55 |
| Neat1 | 4.90E-39 | -0.206758197 | 0.665 | 0.791 | 1.32E-34 | | Rnf152 | 4.60E-06 | -0.0112417 | 0.45 | 0.51 | 0.00667184 |
| Naaladl2 | 5.08E-39 | -0.074363831 | 0.438 | 0.546 | 1.37E-34 | | Ankrd55 | 5.05E-291 | -0.0112593 | 0.09 | 0.49 | 7.33E-288 |
| Gas7 | 8.01E-39 | 0.03477835 | 0.233 | 0.289 | 2.16E-34 | | Scara5 | 1.94E-152 | -0.0113589 | 0.06 | 0.31 | 2.81E-149 |
| Mbnl2 | 1.10E-38 | -0.089859411 | 0.683 | 0.794 | 2.97E-34 | | Gm3061 | 3.42E-10 | -0.0113631 | 0.48 | 0.56 | 4.96E-07 |
| Rtn4 | 1.15E-38 | -0.035531483 | 0.717 | 0.816 | 3.09E-34 | | Tgfbr2 | 3.96E-90 | -0.0113912 | 0.24 | 0.47 | 5.75E-87 |
| Pam | 1.97E-38 | 0.070369235 | 0.207 | 0.255 | 5.31E-34 | | Tbx18 | 1.46E-189 | -0.0116283 | 0.23 | 0.59 | 2.12E-186 |
| Plp1 | 2.41E-38 | -0.18244021 | 0.965 | 0.988 | 6.50E-34 | | B230110G1 | 7.51E-58 | -0.0118562 | 0.32 | 0.52 | 1.09E-54 |
| Cacnb4 | 2.85E-38 | 0.078089262 | 0.211 | 0.26 | 7.68E-34 | | Fap | 7.02E-12 | -0.0118667 | 0.32 | 0.4 | 1.02E-08 |
| Smg6 | 3.18E-38 | -0.050961924 | 0.236 | 0.305 | 8.56E-34 | | Cdh1 | 9.20E-124 | -0.0118998 | 0.36 | 0.64 | 1.33E-120 |
| Trps1 | 3.99E-38 | 0.046853391 | 0.245 | 0.301 | 1.08E-33 | | Slc1a2 | 1.43E-216 | -0.0122822 | 0.2 | 0.54 | 2.07E-213 |
| Ncam2 | 1.39E-37 | -0.168771393 | 0.63 | 0.75 | 3.73E-33 | | Gm2890 | 4.86E-138 | -0.012364 | 0.35 | 0.65 | 7.05E-135 |
| Tmem117 | 1.73E-37 | -0.240153099 | 0.171 | 0.264 | 4.65E-33 | | Pgm5 | 0.20447587 | -0.0124539 | 0.34 | 0.33 | 1 |

| | | | | | | | | | | | |
|---|---|---|---|---|---|---|---|---|---|---|---|
| Ncam1 | 6.66E-37 | -0.15436643 | 0.769 | 0.874 | 1.79E-32 | Nos1 | 1.66E-80 | -0.0125448 | 0.43 | 0.66 | 2.40E-77 |
| Kif6 | 1.12E-36 | -0.078561257 | 0.182 | 0.251 | 3.03E-32 | Fhad1 | 4.36E-71 | -0.0125584 | 0.13 | 0.32 | 6.32E-68 |
| Camkmt | 2.17E-36 | -0.049277501 | 0.201 | 0.268 | 5.85E-32 | Esr1 | 2.96E-101 | -0.0127898 | 0.44 | 0.7 | 4.29E-98 |
| Exoc6b | 3.34E-36 | -0.109424458 | 0.628 | 0.739 | 8.99E-32 | D7Ertd44 | 1.40E-154 | -0.012924 | 0.38 | 0.67 | 2.03E-151 |
| Negr1 | 3.59E-36 | 0.182235595 | 0.227 | 0.261 | 9.67E-32 | Col4a6 | 0.38719768 | -0.0130802 | 0.33 | 0.34 | 1 |
| Slc22a23 | 1.30E-35 | 0.01395908 | 0.232 | 0.286 | 3.49E-31 | Pth2r | 2.63E-21 | -0.0132163 | 0.36 | 0.48 | 3.81E-18 |
| Adgrb3 | 1.34E-35 | 0.4230345 | 0.331 | 0.305 | 3.59E-31 | Grpr | 2.78E-110 | -0.0134246 | 0.31 | 0.57 | 4.03E-107 |
| Strbp | 1.41E-35 | -0.037674717 | 0.387 | 0.474 | 3.79E-31 | Ezr | 4.52E-42 | -0.0134485 | 0.28 | 0.44 | 6.56E-39 |
| Eml6 | 1.88E-35 | 0.102022092 | 0.249 | 0.283 | 5.07E-31 | Slc13a3 | 1.72E-18 | -0.0135144 | 0.49 | 0.6 | 2.49E-15 |
| Dmd | 2.20E-35 | 0.206427226 | 0.263 | 0.286 | 5.94E-31 | Osbpl10 | 2.51E-06 | -0.0135785 | 0.27 | 0.31 | 0.00364135 |
| Fgfr2 | 2.70E-35 | -0.076732415 | 0.825 | 0.912 | 7.27E-31 | Lepr | 6.55E-151 | -0.0137929 | 0.35 | 0.67 | 9.50E-148 |
| Cdh10 | 2.91E-35 | -0.017306724 | 0.405 | 0.493 | 7.82E-31 | Gm2977 | 6.13E-85 | -0.0141586 | 0.38 | 0.62 | 8.90E-82 |
| Alcam | 3.08E-35 | -0.114205993 | 0.183 | 0.259 | 8.30E-31 | Tmem23 | 5.87E-117 | -0.0142667 | 0.06 | 0.27 | 8.51E-114 |
| Ptprk | 4.32E-35 | -0.187920354 | 0.653 | 0.773 | 1.16E-30 | Osbpl3 | 2.50E-06 | -0.014362 | 0.45 | 0.51 | 0.00363352 |
| Nfasc | 5.91E-35 | -0.053560583 | 0.576 | 0.672 | 1.59E-30 | Gm4853 | 0.04110396 | -0.0143931 | 0.28 | 0.29 | 1 |
| Atp8a1 | 5.97E-35 | -0.078848 | 0.665 | 0.768 | 1.61E-30 | Lhfp | 4.34E-31 | -0.014435 | 0.29 | 0.42 | 6.30E-28 |
| Cdc37l1 | 6.52E-34 | -0.105153221 | 0.72 | 0.824 | 1.76E-29 | Il34 | 4.44E-75 | -0.0145517 | 0.42 | 0.64 | 6.44E-72 |
| Pde1c | 9.22E-34 | 0.024033934 | 0.256 | 0.312 | 2.48E-29 | Fli1 | 1.10E-75 | -0.0148656 | 0.23 | 0.44 | 1.59E-72 |
| Grik2 | 1.13E-33 | 0.078248088 | 0.64 | 0.702 | 3.03E-29 | Gm3055 | 1.26E-49 | -0.0148779 | 0.29 | 0.47 | 1.83E-46 |
| 7-Sep | 1.28E-33 | -0.154325751 | 0.677 | 0.79 | 3.44E-29 | Bcar3 | 5.13E-20 | -0.015162 | 0.48 | 0.59 | 7.44E-17 |
| Pcdh9 | 1.74E-33 | -0.170995008 | 0.997 | 0.999 | 4.68E-29 | Lrrc4c | 2.43E-60 | -0.0152736 | 0.75 | 0.9 | 3.52E-57 |
| Gm38304 | 2.56E-33 | -0.235746762 | 0.182 | 0.269 | 6.89E-29 | Slc39a1 | 5.31E-88 | -0.0153994 | 0.19 | 0.42 | 7.71E-85 |
| Kcnq1ot1 | 4.27E-33 | -0.01395831 | 0.213 | 0.269 | 1.15E-28 | Fbln2 | 6.02E-06 | -0.0155723 | 0.33 | 0.32 | 0.00873439 |
| Pak7 | 1.14E-32 | -0.023902077 | 0.642 | 0.734 | 3.08E-28 | Bdnf | 2.58E-150 | -0.0156782 | 0.19 | 0.49 | 3.74E-147 |
| Npas3 | 1.24E-32 | -0.061669261 | 0.465 | 0.559 | 3.33E-28 | Tmtc2 | 4.23E-24 | -0.0161078 | 0.6 | 0.72 | 6.14E-21 |
| St6galnac3 | 1.87E-32 | -0.196212886 | 0.816 | 0.903 | 5.05E-28 | Lrig1 | 2.77E-22 | -0.0161436 | 0.22 | 0.33 | 4.02E-19 |
| St3gal4 | 7.24E-32 | -0.183483337 | 0.198 | 0.281 | 1.95E-27 | Itgbl1 | 6.55E-63 | -0.0161691 | 0.41 | 0.62 | 9.50E-60 |
| Pcdh11x | 9.27E-32 | -0.308245913 | 0.196 | 0.296 | 2.49E-27 | Sorcs2 | 0.00050159 | -0.0162073 | 0.37 | 0.35 | 0.72780395 |
| Cdk19 | 1.20E-31 | -0.088155983 | 0.671 | 0.772 | 3.22E-27 | Mpped1 | 1.71E-27 | -0.0163238 | 0.28 | 0.41 | 2.49E-24 |
| Gria4 | 2.71E-31 | 0.225431348 | 0.32 | 0.334 | 7.30E-27 | 9330158H0 | 9.36E-210 | -0.0163835 | 0.18 | 0.54 | 1.36E-206 |
| Xkr4 | 3.35E-31 | 0.233896841 | 0.391 | 0.402 | 9.01E-27 | Gm2951 | 8.05E-42 | -0.0167831 | 0.32 | 0.47 | 1.17E-38 |
| Gm42756 | 3.50E-31 | -0.255706205 | 0.177 | 0.263 | 9.43E-27 | Gpr176 | 0.01131707 | -0.0170541 | 0.49 | 0.47 | 1 |
| Nav3 | 3.58E-31 | 0.41960387 | 0.257 | 0.223 | 9.63E-27 | Slc16a9 | 3.59E-51 | -0.0171164 | 0.38 | 0.56 | 5.21E-48 |
| Plcl1 | 2.56E-30 | -0.205034784 | 0.95 | 0.98 | 6.88E-26 | Grip1os | 6.46E-07 | -0.018271 | 0.33 | 0.38 | 0.0009372 |
| Prickle1 | 3.16E-30 | 0.225239363 | 0.359 | 0.367 | 8.50E-26 | Sel1l3 | 8.23E-34 | -0.018316 | 0.52 | 0.67 | 1.19E-30 |
| Zfp536 | 3.82E-30 | -0.207716632 | 0.765 | 0.86 | 1.03E-25 | Gpc6 | 6.01E-69 | -0.0190018 | 0.41 | 0.63 | 8.71E-66 |
| B3galt1 | 4.16E-30 | -0.012778927 | 0.631 | 0.712 | 1.12E-25 | Tmem7 | 8.51E-260 | -0.0190725 | 0.02 | 0.32 | 1.24E-256 |
| Wwox | 4.43E-30 | -0.165644509 | 0.26 | 0.348 | 1.19E-25 | Gm1396 | 2.78E-21 | -0.0191347 | 0.38 | 0.5 | 4.03E-18 |
| Il33 | 5.22E-30 | 0.042881108 | 0.416 | 0.476 | 1.41E-25 | Mrc1 | 3.06E-258 | -0.0191425 | 0.05 | 0.38 | 4.45E-255 |
| Tbc1d5 | 7.63E-30 | -0.150571678 | 0.702 | 0.807 | 2.05E-25 | Socs2 | 5.41E-97 | -0.01934 | 0.43 | 0.68 | 7.85E-94 |
| Oxr1 | 8.40E-30 | -0.080580193 | 0.524 | 0.617 | 2.26E-25 | Cyp7b1 | 2.63E-203 | -0.0194447 | 0.12 | 0.45 | 3.82E-200 |
| Tmem178b | 1.62E-29 | -0.08613169 | 0.658 | 0.756 | 4.37E-25 | Gnal | 2.03E-20 | -0.019551 | 0.46 | 0.37 | 2.94E-17 |
| Cadps | 1.65E-29 | 0.364670143 | 0.253 | 0.237 | 4.44E-25 | Pcdh19 | 0.01194099 | -0.0198102 | 0.51 | 0.54 | 1 |
| Pcdh15 | 1.89E-29 | 0.243633702 | 0.303 | 0.311 | 5.08E-25 | Ly86 | 6.00E-151 | -0.0200198 | 0.22 | 0.54 | 8.71E-148 |
| Xist | 2.87E-29 | 0.001967738 | 0.679 | 0.762 | 7.72E-25 | Gm1383 | 4.36E-25 | -0.020155 | 0.33 | 0.45 | 6.33E-22 |
| Map7 | 6.99E-29 | -0.159624536 | 0.842 | 0.918 | 1.88E-24 | Gm4610 | 1.11E-14 | -0.0201582 | 0.3 | 0.39 | 1.61E-11 |
| Lama2 | 1.28E-28 | -0.122154653 | 0.277 | 0.358 | 3.45E-24 | Atp13a4 | 4.89E-14 | -0.0203782 | 0.36 | 0.45 | 7.10E-11 |
| Dlg2 | 3.72E-28 | 0.093066715 | 0.81 | 0.87 | 1.00E-23 | Fstl5 | 2.49E-05 | -0.0205805 | 0.54 | 0.53 | 0.03613284 |
| Enpp2 | 2.34E-27 | -0.07082672 | 0.758 | 0.846 | 6.31E-23 | Wdr95 | 7.63E-65 | -0.0205902 | 0.14 | 0.32 | 1.11E-61 |
| Auts2 | 6.68E-27 | 0.162389902 | 0.683 | 0.707 | 1.80E-22 | Gm1362 | 1.25E-69 | -0.0209278 | 0.24 | 0.45 | 1.81E-66 |
| Ccser1 | 1.31E-26 | 0.205100264 | 0.35 | 0.366 | 3.52E-22 | Gm1622 | 1.44E-160 | -0.0209644 | 0.05 | 0.31 | 2.09E-157 |
| Il1rapl1 | 1.58E-26 | 0.131792157 | 0.591 | 0.637 | 4.25E-22 | Rbms1 | 1.73E-35 | -0.0211121 | 0.24 | 0.38 | 2.51E-32 |
| St18 | 2.09E-26 | -0.160812766 | 0.926 | 0.97 | 5.63E-22 | Cdh23 | 1.64E-148 | -0.0211408 | 0.2 | 0.5 | 2.38E-145 |
| Acyp2 | 4.69E-26 | 0.001402355 | 0.255 | 0.308 | 1.26E-21 | Adgrg6 | 1.17E-10 | -0.0219883 | 0.41 | 0.49 | 1.69E-07 |
| Lrrc4c | 2.98E-25 | -0.156624403 | 0.644 | 0.745 | 8.02E-21 | Nipal2 | 2.64E-05 | -0.0221461 | 0.38 | 0.4 | 0.03835946 |
| Cdc42bpa | 3.01E-25 | -0.173116346 | 0.755 | 0.846 | 8.10E-21 | Gm2184 | 7.91E-198 | -0.0221723 | 0.02 | 0.27 | 1.15E-194 |
| Cadm2 | 3.81E-25 | 0.151705035 | 0.714 | 0.751 | 1.03E-20 | 6530403H0 | 1.27E-164 | -0.0224166 | 0.33 | 0.67 | 1.84E-161 |
| Clasp2 | 4.32E-25 | 0.030738918 | 0.736 | 0.8 | 1.16E-20 | Prdm16 | 1.47E-255 | -0.022431 | 0.23 | 0.64 | 2.14E-252 |
| Slc8a1 | 4.33E-25 | -0.115601454 | 0.675 | 0.77 | 1.17E-20 | Col9a3 | 2.56E-09 | -0.0225069 | 0.24 | 0.3 | 3.71E-06 |
| Dcc | 4.72E-25 | 0.153265829 | 0.37 | 0.4 | 1.27E-20 | Chrna7 | 1.61E-12 | -0.0226297 | 0.47 | 0.56 | 2.33E-09 |
| Mdga2 | 5.01E-25 | 0.073648601 | 0.722 | 0.782 | 1.35E-20 | Stk33 | 1.36E-07 | -0.0235165 | 0.46 | 0.45 | 0.00019665 |
| Csmd3 | 5.40E-25 | -0.116106546 | 0.774 | 0.86 | 1.45E-20 | 4930544I0 | 1.22E-25 | -0.0237666 | 0.44 | 0.57 | 1.77E-22 |
| Spock1 | 1.05E-24 | -0.261821714 | 0.39 | 0.499 | 2.84E-20 | Adamts1 | 4.33E-80 | -0.0238101 | 0.33 | 0.55 | 6.29E-77 |
| Malat1 | 1.49E-24 | 0.106557666 | 1 | 1 | 4.01E-20 | Trpc5 | 4.32E-37 | -0.0238237 | 0.46 | 0.62 | 6.27E-34 |
| Slc13a1 | 3.32E-24 | 0.006431522 | 0.207 | 0.258 | 8.94E-20 | Nrp2 | 1.71E-88 | -0.0240461 | 0.32 | 0.56 | 2.49E-85 |
| Prickle2 | 3.84E-24 | -0.114860582 | 0.712 | 0.805 | 1.03E-19 | Gm1348 | 4.07E-08 | -0.0242456 | 0.33 | 0.39 | 5.90E-05 |
| Slc24a2 | 4.66E-24 | -0.141335598 | 0.979 | 0.989 | 1.25E-19 | Chst8 | 4.29E-55 | -0.024457 | 0.24 | 0.42 | 6.22E-52 |
| Ctnna2 | 5.38E-24 | -0.302075265 | 0.492 | 0.604 | 1.45E-19 | Fgfr3 | 4.66E-264 | -0.0244735 | 0.06 | 0.4 | 6.76E-261 |
| Ube2e2 | 8.38E-24 | 0.097089704 | 0.677 | 0.719 | 2.26E-19 | Gpr149 | 7.58E-16 | -0.024799 | 0.36 | 0.45 | 1.10E-12 |
| Kirrel3 | 1.95E-22 | 0.364477488 | 0.594 | 0.51 | 5.25E-18 | Sdk2 | 1.34E-10 | -0.025128 | 0.51 | 0.45 | 1.94E-07 |
| Unc5c | 4.12E-22 | -0.054662462 | 0.728 | 0.807 | 1.11E-17 | Nnat | 2.30E-74 | -0.0252696 | 0.21 | 0.42 | 3.33E-71 |
| Ano4 | 3.56E-21 | 0.118352977 | 0.821 | 0.851 | 9.59E-17 | Gm211 | 7.24E-20 | -0.0255186 | 0.54 | 0.66 | 1.05E-16 |
| Ank3 | 1.61E-20 | 0.01957283 | 0.817 | 0.875 | 4.35E-16 | Tprkb | 3.55E-58 | -0.0258151 | 0.21 | 0.38 | 5.15E-55 |
| Fam208a | 2.23E-20 | 0.116398869 | 0.241 | 0.254 | 6.00E-16 | Sgms2 | 1.39E-181 | -0.0260756 | 0.06 | 0.34 | 2.02E-178 |
| Macrod2 | 2.72E-20 | 0.173850793 | 0.37 | 0.377 | 7.31E-16 | Whrn | 0.00075911 | -0.0266294 | 0.35 | 0.35 | 1 |
| Ptgds | 3.39E-20 | -0.075074059 | 0.49 | 0.584 | 9.13E-16 | Gm1326 | 2.40E-78 | -0.0274371 | 0.28 | 0.51 | 3.49E-75 |
| Park2 | 9.97E-20 | 0.290648949 | 0.492 | 0.442 | 2.68E-15 | Plcb4 | 2.04E-46 | -0.0274917 | 0.39 | 0.57 | 2.96E-43 |
| Kcnq3 | 3.56E-19 | -0.007678502 | 0.633 | 0.7 | 9.58E-15 | Stard5 | 4.79E-26 | -0.0275843 | 0.32 | 0.45 | 6.96E-23 |
| Gm26733 | 9.02E-19 | 0.109469803 | 0.238 | 0.258 | 2.43E-14 | Ccbe1 | 3.60E-16 | -0.0276367 | 0.39 | 0.48 | 5.22E-13 |
| Fnbp1 | 1.43E-18 | -0.045977818 | 0.819 | 0.883 | 3.84E-14 | Ndst4 | 1.12E-105 | -0.0278149 | 0.31 | 0.57 | 1.62E-102 |
| Dock10 | 3.41E-18 | -0.014541861 | 0.897 | 0.947 | 9.19E-14 | 4930467D2 | 7.77E-101 | -0.0281229 | 0.16 | 0.39 | 1.13E-97 |
| Mettl23 | 4.20E-17 | -0.081014537 | 0.202 | 0.252 | 1.13E-12 | Hdac9 | 1.32E-16 | -0.0284595 | 0.57 | 0.66 | 1.92E-13 |
| 2010111I01Rik | 7.63E-17 | 0.107485925 | 0.295 | 0.31 | 2.05E-12 | Gm2968 | 3.26E-06 | -0.0285131 | 0.31 | 0.34 | 0.00473647 |
| Mapt | 2.68E-16 | -0.046481112 | 0.876 | 0.929 | 7.22E-12 | Bmpr1b | 5.35E-220 | -0.0286659 | 0.2 | 0.58 | 7.76E-217 |
| Edil3 | 2.32E-15 | -0.097997152 | 0.952 | 0.978 | 6.25E-11 | Rreb1 | 8.46E-38 | -0.0289846 | 0.41 | 0.57 | 1.23E-34 |
| Robo1 | 2.63E-15 | -0.338671531 | 0.194 | 0.266 | 7.08E-11 | Tbc1d9 | 1.08E-09 | -0.0290106 | 0.38 | 0.44 | 1.57E-06 |
| Zeb2 | 5.98E-15 | -0.024575318 | 0.839 | 0.893 | 1.61E-10 | Cpne2 | 1.13E-19 | -0.0291437 | 0.35 | 0.45 | 1.64E-16 |
| Grm3 | 6.53E-14 | -0.041825243 | 0.788 | 0.848 | 1.76E-09 | Kcnk5 | 3.88E-118 | -0.0291829 | 0.31 | 0.59 | 5.63E-115 |
| Erbb4 | 8.02E-13 | -0.015700624 | 0.761 | 0.807 | 2.16E-08 | Ros1 | 4.63E-81 | -0.0292137 | 0.42 | 0.64 | 6.72E-78 |
| Gm26694 | 2.28E-12 | 0.016975804 | 0.271 | 0.3 | 6.13E-08 | Cntn6 | 1.24E-16 | -0.0292836 | 0.3 | 0.36 | 1.79E-13 |
| Ank2 | 2.44E-12 | -0.077448473 | 0.932 | 0.963 | 6.57E-08 | Adra1b | 4.29E-46 | -0.0295543 | 0.35 | 0.52 | 6.22E-43 |
| Qk | 2.85E-12 | -0.067298439 | 0.956 | 0.979 | 7.66E-08 | Gsg1l | 1.64E-69 | -0.0299332 | 0.11 | 0.28 | 2.38E-66 |
| Phlpp1 | 3.60E-12 | 0.004029196 | 0.907 | 0.944 | 9.69E-08 | Heg1 | 1.81E-49 | -0.0302178 | 0.48 | 0.66 | 2.63E-46 |
| Gm26804 | 4.04E-12 | 0.107280873 | 0.401 | 0.413 | 1.09E-07 | Adamts | 1.77E-56 | -0.0302358 | 0.25 | 0.14 | 2.57E-53 |
| Frmd5 | 7.97E-11 | -0.087797031 | 0.923 | 0.956 | 2.14E-06 | Ipcef1 | 2.53E-135 | -0.0306888 | 0.06 | 0.29 | 3.68E-132 |
| Gm4593 | 2.41E-10 | 0.138523525 | 0.444 | 0.436 | 6.50E-06 | Cp | 6.13E-25 | -0.0308403 | 0.27 | 0.39 | 8.90E-22 |
| Mast4 | 3.16E-10 | -0.078919597 | 0.919 | 0.952 | 8.50E-06 | Pdgfd | 6.08E-199 | -0.0309924 | 0.07 | 0.37 | 8.83E-196 |
| 9330111N05Rik | 3.72E-09 | -0.189740956 | 0.208 | 0.257 | 0.000100185 | 2900026A0 | 2.08E-136 | -0.0316164 | 0.21 | 0.5 | 3.02E-133 |
| Lrp1b | 1.07E-08 | -0.057234933 | 0.961 | 0.979 | 0.000289259 | Vcan | 8.15E-208 | -0.0316699 | 0.21 | 0.58 | 1.18E-204 |
| Pde4b | 1.48E-08 | -0.066226879 | 0.996 | 0.998 | 0.000399651 | Apod | 3.61E-110 | -0.0318375 | 0.47 | 0.71 | 5.24E-107 |
| Grm7 | 1.15E-07 | 0.005172175 | 0.457 | 0.477 | 0.00308767 | Cx3cr1 | 5.04E-62 | -0.031966 | 0.24 | 0.43 | 7.32E-59 |
| Tmeff2 | 2.57E-07 | -0.0425151 | 0.963 | 0.981 | 0.00692046 | Eda | 2.15E-25 | -0.0324283 | 0.19 | 0.31 | 3.12E-22 |
| Gm20388 | 1.36E-06 | 0.15305098 | 0.586 | 0.558 | 0.036491459 | Vwc2l | 3.08E-173 | -0.0324353 | 0.41 | 0.74 | 4.48E-170 |
| AC110241.2 | 3.34E-05 | 0.131075582 | 0.389 | 0.373 | 0.898775149 | Gulp1 | 6.63E-14 | -0.0328063 | 0.46 | 0.4 | 9.62E-11 |
| Palm2 | 3.96E-05 | 0.088336847 | 0.505 | 0.496 | 1 | Adamts | 2.70E-10 | -0.032949 | 0.47 | 0.52 | 3.92E-07 |
| Ptprd | 0.02131165 | 0.021793605 | 0.983 | 0.99 | 1 | Ptprc | 4.47E-107 | -0.0330967 | 0.42 | 0.68 | 6.48E-104 |
| Magi2 | 0.036742626 | -0.009574877 | 0.992 | 0.995 | 1 | Samd4 | 8.14E-47 | -0.0332957 | 0.58 | 0.74 | 1.18E-43 |
| | | | | | | Crim1 | 6.68E-72 | -0.0338011 | 0.4 | 0.62 | 9.70E-69 |
| | | | | | | Nr2f2 | 1.41E-09 | -0.0341372 | 0.27 | 0.33 | 2.05E-06 |
| | | | | | | Adamtsl | 5.20E-08 | -0.034282 | 0.49 | 0.45 | 7.54E-05 |
| | | | | | | Zbtb7c | 1.78E-07 | -0.0346746 | 0.4 | 0.41 | 0.0002581 |
| | | | | | | Mei4 | 8.79E-11 | -0.0347446 | 0.52 | 0.61 | 1.28E-07 |
| | | | | | | Atp2b4 | 1.37E-34 | -0.0348978 | 0.25 | 0.4 | 1.99E-31 |
| | | | | | | Agbl4 | 9.15E-21 | -0.0349501 | 0.56 | 0.67 | 1.33E-17 |
| | | | | | | Mir670h | 2.28E-07 | -0.0351354 | 0.44 | 0.51 | 0.00033133 |
| | | | | | | Steap2 | 1.34E-18 | -0.0356065 | 0.39 | 0.48 | 1.94E-15 |
| | | | | | | Cped1 | 5.31E-51 | -0.0358533 | 0.29 | 0.47 | 7.70E-48 |
| | | | | | | Gpr63 | 7.09E-133 | -0.0359287 | 0.16 | 0.44 | 1.03E-129 |
| | | | | | | Unc5d | 1.40E-121 | -0.0364717 | 0.39 | 0.68 | 2.03E-118 |
| | | | | | | Nhsl1 | 6.39E-53 | -0.0364871 | 0.15 | 0.31 | 9.27E-50 |
| | | | | | | Gm487 | 7.03E-129 | -0.0372475 | 0.18 | 0.46 | 1.02E-125 |
| | | | | | | Epas1 | 6.13E-09 | -0.03759 | 0.52 | 0.47 | 8.89E-06 |
| | | | | | | Sec14l5 | 6.03E-144 | -0.037802 | 0.59 | 0.84 | 8.75E-141 |
| | | | | | | D030068K2 | 1.66E-92 | -0.0378265 | 0.18 | 0.41 | 2.41E-89 |
| | | | | | | Bace2 | 1.13E-49 | -0.0378433 | 0.35 | 0.53 | 1.65E-46 |
| | | | | | | Kit | 6.66E-42 | -0.038602 | 0.39 | 0.56 | 9.67E-39 |
| | | | | | | Luzp2 | 8.35E-31 | -0.0387172 | 0.52 | 0.66 | 1.21E-27 |
| | | | | | | Sema6d | 6.36E-44 | -0.0392616 | 0.72 | 0.86 | 9.23E-41 |
| | | | | | | Rerg | 1.56E-20 | -0.0393055 | 0.52 | 0.63 | 2.26E-17 |
| | | | | | | Irak2 | 2.49E-17 | -0.0394433 | 0.42 | 0.52 | 3.61E-14 |
| | | | | | | Gpc3 | 4.07E-248 | -0.0396777 | 0.18 | 0.58 | 5.90E-245 |
| | | | | | | Wwtr1 | 1.19E-225 | -0.0398695 | 0.09 | 0.44 | 1.73E-222 |
| | | | | | | Strip2 | 1.47E-175 | -0.040031 | 0.37 | 0.71 | 2.14E-172 |
| | | | | | | Col11a1 | 6.97E-14 | -0.0402085 | 0.28 | 0.28 | 1.01E-10 |
| | | | | | | Gm3381 | 1.68E-119 | -0.0403527 | 0.3 | 0.59 | 2.44E-116 |
| | | | | | | Cacna2d | 9.70E-108 | -0.0407084 | 0.11 | 0.33 | 1.41E-104 |
| | | | | | | Dgkh | 2.50E-44 | -0.0409372 | 0.46 | 0.63 | 3.62E-41 |
| | | | | | | Syt6 | 6.58E-76 | -0.0410511 | 0.14 | 0.33 | 9.55E-73 |
| | | | | | | C530008M1 | 7.66E-11 | -0.041562 | 0.33 | 0.36 | 1.11E-07 |
| | | | | | | Neat1 | 1.28E-64 | -0.0417606 | 0.74 | 0.89 | 1.86E-61 |
| | | | | | | Runx1t1 | 1.77E-11 | -0.0418357 | 0.56 | 0.62 | 2.56E-08 |
| | | | | | | Zfp804b | 1.48E-150 | -0.0424033 | 0.34 | 0.66 | 2.15E-147 |
| | | | | | | Lmntd1 | 2.44E-17 | -0.0425502 | 0.35 | 0.44 | 3.55E-14 |
| | | | | | | Kif26b | 1.79E-26 | -0.0427974 | 0.31 | 0.43 | 2.59E-23 |
| | | | | | | Btbd11 | 4.70E-60 | -0.0431546 | 0.44 | 0.28 | 6.82E-57 |
| | | | | | | Esrrg | 1.54E-07 | -0.0432085 | 0.52 | 0.53 | 0.00022316 |
| | | | | | | Slc7a2 | 1.49E-59 | -0.0432583 | 0.32 | 0.51 | 2.16E-56 |
| | | | | | | Pag1 | 1.23E-08 | -0.0434831 | 0.48 | 0.52 | 1.79E-05 |
| | | | | | | Nkain3 | 7.12E-27 | -0.0436375 | 0.57 | 0.69 | 1.03E-23 |
| | | | | | | Gm2669 | 2.62E-31 | -0.0436431 | 0.39 | 0.53 | 3.80E-28 |
| | | | | | | Zfp521 | 5.65E-11 | -0.0436514 | 0.41 | 0.41 | 8.20E-08 |
| | | | | | | Ank1 | 3.10E-08 | -0.0441198 | 0.54 | 0.53 | 4.50E-05 |
| | | | | | | Rbpj | 6.98E-20 | -0.0446753 | 0.46 | 0.57 | 1.01E-16 |
| | | | | | | Garnl3 | 2.68E-54 | -0.0450869 | 0.5 | 0.69 | 3.89E-51 |
| | | | | | | Hs3st5 | 1.51E-19 | -0.045501 | 0.38 | 0.31 | 2.19E-16 |
| | | | | | | Gm4459 | 1.14E-43 | -0.0457158 | 0.38 | 0.25 | 1.66E-40 |
| | | | | | | Myof | 5.59E-124 | -0.0461981 | 0.31 | 0.6 | 8.11E-121 |
| | | | | | | Frmpd1 | 2.39E-173 | -0.0463398 | 0.48 | 0.19 | 3.47E-170 |
| | | | | | | Oxr1 | 3.44E-103 | -0.0467273 | 0.57 | 0.78 | 5.00E-100 |
| | | | | | | Satb1 | 1.08E-23 | -0.0473112 | 0.39 | 0.51 | 1.57E-20 |
| | | | | | | Ptpn13 | 4.38E-17 | -0.0476886 | 0.52 | 0.62 | 6.36E-14 |
| | | | | | | Hapln2 | 1.20E-238 | -0.0477929 | 0.28 | 0.59 | 1.74E-235 |
| | | | | | | Spats2l | 9.24E-97 | -0.0481023 | 0.17 | 0.4 | 1.34E-93 |
| | | | | | | Gm4850 | 3.74E-22 | -0.0481879 | 0.52 | 0.64 | 5.43E-19 |
| | | | | | | Grk5 | 6.49E-12 | -0.0482062 | 0.49 | 0.48 | 9.42E-09 |
| | | | | | | Gabra3 | 1.70E-16 | -0.0485443 | 0.26 | 0.34 | 2.47E-13 |
| | | | | | | Elfn1 | 1.02E-14 | -0.0492093 | 0.29 | 0.36 | 1.48E-11 |
| | | | | | | Prrg1 | 4.32E-40 | -0.0497579 | 0.52 | 0.68 | 6.27E-37 |
| | | | | | | F3 | 1.32E-139 | -0.0502033 | 0.2 | 0.5 | 1.92E-136 |
| | | | | | | Dgkb | 4.53E-82 | -0.0511169 | 0.51 | 0.74 | 6.57E-79 |
| | | | | | | Gpnmb | 3.85E-11 | -0.051628 | 0.42 | 0.38 | 5.58E-08 |
| | | | | | | Bcl2 | 2.21E-15 | -0.0521217 | 0.54 | 0.46 | 3.21E-12 |
| | | | | | | Cdc14a | 4.75E-14 | -0.0521243 | 0.35 | 0.42 | 6.89E-11 |
| | | | | | | Gad1 | 1.79E-86 | -0.0521429 | 0.13 | 0.33 | 2.59E-83 |
| | | | | | | 0610040J0 | 1.39E-14 | -0.0525623 | 0.51 | 0.54 | 2.01E-11 |
| | | | | | | Spock1 | 2.94E-81 | -0.0535892 | 0.58 | 0.79 | 4.27E-78 |
| | | | | | | Pard3bos | 4.87E-158 | -0.0537046 | 0.12 | 0.4 | 7.06E-155 |
| | | | | | | Arsj | 2.05E-07 | -0.054341 | 0.4 | 0.43 | 0.00029767 |
| | | | | | | Car13 | 2.21E-260 | -0.0544897 | 0.11 | 0.5 | 3.21E-257 |
| | | | | | | Prom1 | 3.68E-18 | -0.054718 | 0.48 | 0.59 | 5.34E-15 |
| | | | | | | Lin7a | 5.00E-13 | -0.054759 | 0.48 | 0.42 | 7.25E-10 |
| | | | | | | Nwd1 | 4.24E-110 | -0.0551417 | 0.06 | 0.26 | 6.15E-107 |
| | | | | | | Sh3rf3 | 4.50E-35 | -0.0552456 | 0.4 | 0.53 | 6.53E-32 |
| | | | | | | Ptpro | 2.89E-33 | -0.0558341 | 0.34 | 0.29 | 4.19E-30 |
| | | | | | | Gm4532 | 1.04E-17 | -0.0563453 | 0.35 | 0.45 | 1.52E-14 |
| | | | | | | Smoc1 | 9.40E-92 | -0.0566396 | 0.15 | 0.37 | 1.36E-88 |
| | | | | | | Cpq | 1.51E-12 | -0.0573564 | 0.5 | 0.52 | 2.18E-09 |
| | | | | | | 2600014E2 | 4.92E-06 | -0.0576476 | 0.42 | 0.4 | 0.00714342 |
| | | | | | | Fat3 | 2.34E-29 | -0.0585026 | 0.53 | 0.66 | 3.40E-26 |
| | | | | | | Spon1 | 1.16E-54 | -0.0587859 | 0.4 | 0.57 | 1.69E-51 |
| | | | | | | Ntn4 | 7.25E-21 | -0.0589257 | 0.31 | 0.4 | 1.05E-17 |
| | | | | | | Utrn | 2.27E-27 | -0.0590492 | 0.38 | 0.33 | 3.30E-24 |
| | | | | | | Fam189a | 9.56E-63 | -0.0592478 | 0.3 | 0.5 | 1.39E-59 |
| | | | | | | Rnf144b | 2.55E-132 | -0.0592612 | 0.09 | 0.34 | 3.70E-129 |
| | | | | | | Sema3e | 7.36E-76 | -0.0597924 | 0.34 | 0.57 | 1.07E-72 |
| | | | | | | Siah3 | 5.46E-54 | -0.0601525 | 0.23 | 0.4 | 7.92E-51 |
| | | | | | | Unc93b1 | 9.14E-298 | -0.0610377 | 0.16 | 0.59 | 1.33E-294 |
| | | | | | | Grin3a | 2.05E-124 | -0.0610667 | 0.24 | 0.53 | 2.98E-121 |
| | | | | | | Rgs20 | 4.64E-54 | -0.0611366 | 0.27 | 0.45 | 6.74E-51 |
| | | | | | | Pla2g7 | 1.67E-99 | -0.0614906 | 0.15 | 0.38 | 2.43E-96 |
| | | | | | | Pam | 1.55E-73 | -0.062102 | 0.58 | 0.79 | 2.24E-70 |
| | | | | | | Gjc3 | 8.15E-92 | -0.0626679 | 0.65 | 0.86 | 1.18E-88 |
| | | | | | | Airn | 2.64E-38 | -0.0626691 | 0.4 | 0.28 | 3.83E-35 |
| | | | | | | Tgfb2 | 3.49E-12 | -0.0630109 | 0.47 | 0.43 | 5.07E-09 |
| | | | | | | Dpf3 | 2.06E-29 | -0.063641 | 0.42 | 0.55 | 2.99E-26 |
| | | | | | | Dlc1 | 1.59E-13 | -0.0639722 | 0.48 | 0.51 | 2.31E-10 |
| | | | | | | Wdr17 | 2.99E-24 | -0.0642501 | 0.46 | 0.57 | 4.34E-21 |
| | | | | | | Rgcc | 1.71E-119 | -0.0650346 | 0.28 | 0.14 | 2.48E-116 |
| | | | | | | Gm1539 | 2.50E-59 | -0.066327 | 0.17 | 0.34 | 3.63E-56 |
| | | | | | | Fhod3 | 5.30E-21 | -0.0663347 | 0.32 | 0.42 | 7.69E-18 |
| | | | | | | Pou6f2 | 4.47E-45 | -0.0667214 | 0.23 | 0.39 | 6.49E-42 |
| | | | | | | Hs6st2 | 1.97E-20 | -0.0676006 | 0.41 | 0.51 | 2.85E-17 |
| | | | | | | Eps8 | 3.08E-22 | -0.0677497 | 0.27 | 0.3 | 4.46E-19 |
| | | | | | | Arhgap6 | 9.28E-47 | -0.0686912 | 0.32 | 0.49 | 1.35E-43 |

| | | | | | |
|---|---|---|---|---|---|
| Tcf7l1 | 1.28E-09 | -0.068808 | 0.3 | 0.35 | 1.85E-06 |
| Arhgef2 | 3.83E-13 | -0.0696477 | 0.45 | 0.5 | 5.56E-10 |
| Fmn1 | 6.75E-59 | -0.0699144 | 0.44 | 0.64 | 9.80E-56 |
| 9430041J1 | 1.17E-42 | -0.0699583 | 0.29 | 0.45 | 1.70E-39 |
| Dach1 | 2.41E-76 | -0.0701975 | 0.24 | 0.46 | 3.50E-73 |
| Pde7b | 6.34E-127 | -0.0703224 | 0.39 | 0.67 | 9.19E-124 |
| Acsl3 | 2.20E-131 | -0.0704685 | 0.39 | 0.68 | 3.20E-128 |
| Vwc2 | 4.64E-85 | -0.0721687 | 0.41 | 0.24 | 6.73E-82 |
| 9530026P0 | 6.36E-46 | -0.0722591 | 0.33 | 0.51 | 9.23E-43 |
| Cnksr3 | 4.73E-46 | -0.0729281 | 0.27 | 0.44 | 6.86E-43 |
| Ugt8a | 3.81E-62 | -0.0731267 | 0.79 | 0.93 | 5.52E-59 |
| Igsf21 | 9.27E-19 | -0.0738485 | 0.4 | 0.45 | 1.34E-15 |
| Adra1a | 4.81E-23 | -0.0747833 | 0.27 | 0.36 | 6.97E-20 |
| Efemp1 | 3.01E-189 | -0.074844 | 0.29 | 0.64 | 4.37E-186 |
| Snrpn | 8.43E-29 | -0.0753372 | 0.48 | 0.61 | 1.22E-25 |
| Sdk1 | 4.89E-21 | -0.0755206 | 0.32 | 0.38 | 7.09E-18 |
| Rfx4 | 1.63E-21 | -0.0756408 | 0.4 | 0.47 | 2.37E-18 |
| Gm1616 | 1.93E-55 | -0.0758308 | 0.77 | 0.91 | 2.80E-52 |
| Cdh19 | 1.85E-84 | -0.0771794 | 0.55 | 0.76 | 2.69E-81 |
| Alcam | 3.91E-161 | -0.0788721 | 0.22 | 0.49 | 5.67E-158 |
| Sema3d | 1.02E-25 | -0.0797717 | 0.24 | 0.33 | 1.47E-22 |
| Enpp1 | 8.75E-66 | -0.0801015 | 0.52 | 0.72 | 1.27E-62 |
| Lama4 | 2.22E-20 | -0.0804565 | 0.33 | 0.37 | 3.22E-17 |
| Emid1 | 1.16E-35 | -0.0808265 | 0.43 | 0.42 | 1.68E-32 |
| Foxp2 | 1.03E-31 | -0.0821114 | 0.44 | 0.57 | 1.49E-28 |
| Bcan | 3.73E-23 | -0.0830143 | 0.49 | 0.52 | 5.41E-20 |
| Grik3 | 1.87E-68 | -0.0835847 | 0.1 | 0.26 | 2.71E-65 |
| Rorb | 2.67E-30 | -0.083614 | 0.52 | 0.43 | 3.87E-27 |
| Rcan2 | 4.37E-162 | -0.084932 | 0.34 | 0.6 | 6.34E-159 |
| Parm1 | 1.01E-96 | -0.0857733 | 0.41 | 0.66 | 1.46E-93 |
| Rasgef1 | 2.48E-09 | -0.0860131 | 0.52 | 0.55 | 3.60E-06 |
| Nectin3 | 2.23E-26 | -0.0860183 | 0.41 | 0.54 | 3.23E-23 |
| Prr16 | 3.52E-14 | -0.0861259 | 0.54 | 0.5 | 5.11E-11 |
| Pdgfra | 1.74E-240 | -0.0863452 | 0.07 | 0.42 | 2.52E-237 |
| Pbx3 | 1.69E-84 | -0.086704 | 0.41 | 0.65 | 2.46E-81 |
| Eya1 | 1.52E-185 | -0.0876062 | 0.16 | 0.49 | 2.21E-182 |
| Ror1 | 1.45E-43 | -0.0884177 | 0.22 | 0.38 | 2.11E-40 |
| P2rx7 | 5.28E-10 | -0.0891528 | 0.47 | 0.47 | 7.66E-07 |
| Gabrg3 | 5.75E-55 | -0.0892586 | 0.51 | 0.69 | 8.34E-52 |
| Pde1c | 1.44E-103 | -0.0895162 | 0.53 | 0.78 | 2.09E-100 |
| Vav3 | 5.08E-51 | -0.0895478 | 0.45 | 0.62 | 7.37E-48 |
| Tox | 1.12E-36 | -0.0899886 | 0.5 | 0.63 | 1.63E-33 |
| Gm2179 | 1.30E-100 | -0.0900265 | 0.18 | 0.41 | 1.88E-97 |
| Plppr1 | 2.07E-158 | -0.0906867 | 0.19 | 0.5 | 3.01E-155 |
| Gm3745 | 2.43E-188 | -0.0921246 | 0.36 | 0.65 | 3.53E-185 |
| Kcnk2 | 3.75E-16 | -0.0921286 | 0.4 | 0.4 | 5.44E-13 |
| Tle4 | 1.84E-195 | -0.0922473 | 0.2 | 0.54 | 2.67E-192 |
| Mertk | 4.37E-39 | -0.0927307 | 0.29 | 0.41 | 6.34E-36 |
| Plpp4 | 2.14E-47 | -0.0929796 | 0.46 | 0.4 | 3.10E-44 |
| Cpne8 | 1.29E-46 | -0.0941757 | 0.44 | 0.6 | 1.88E-43 |
| Pdzrn4 | 7.10E-63 | -0.0948351 | 0.3 | 0.49 | 1.03E-59 |
| Phldb2 | 1.42E-40 | -0.0958641 | 0.4 | 0.56 | 2.06E-37 |
| Bcas1 | 1.96E-55 | -0.0965624 | 0.71 | 0.87 | 2.85E-52 |
| Gab1 | 6.58E-72 | -0.0968368 | 0.74 | 0.91 | 9.55E-69 |
| Bmper | 7.59E-96 | -0.0977326 | 0.16 | 0.38 | 1.10E-92 |
| Chst9 | 6.45E-107 | -0.0980088 | 0.3 | 0.56 | 9.36E-104 |
| Shc4 | 1.04E-28 | -0.0983252 | 0.52 | 0.6 | 1.51E-25 |
| Gm3368 | 3.46E-35 | -0.09871 | 0.39 | 0.35 | 5.03E-32 |
| Lypd6b | 1.58E-37 | -0.0987858 | 0.48 | 0.64 | 2.29E-34 |
| Cux2 | 6.28E-51 | -0.0991935 | 0.27 | 0.45 | 9.11E-48 |
| Ntng2 | 1.76E-13 | -0.1009135 | 0.5 | 0.58 | 2.55E-10 |
| 4930555F0 | 8.47E-16 | -0.1012379 | 0.46 | 0.44 | 1.23E-12 |
| Ppfibp1 | 2.21E-34 | -0.101344 | 0.28 | 0.38 | 3.21E-31 |
| Zfp462 | 8.23E-47 | -0.1014418 | 0.56 | 0.73 | 1.19E-43 |
| Rab27b | 3.90E-67 | -0.1029026 | 0.39 | 0.59 | 5.66E-64 |
| Ghr | 3.36E-32 | -0.1032837 | 0.4 | 0.32 | 4.87E-29 |
| Ltbp1 | 1.25E-22 | -0.1042205 | 0.44 | 0.45 | 1.81E-19 |
| Grik1 | 2.64E-14 | -0.105386 | 0.62 | 0.7 | 3.83E-11 |
| Sorcs1 | 1.56E-29 | -0.1056644 | 0.74 | 0.84 | 2.26E-26 |
| Gm3685 | 2.20E-50 | -0.1058409 | 0.32 | 0.5 | 3.19E-47 |
| Shisa9 | 2.15E-37 | -0.1064912 | 0.43 | 0.57 | 3.11E-34 |
| Itgb5 | 1.40E-128 | -0.1065812 | 0.1 | 0.35 | 2.03E-125 |
| Stxbp6 | 4.25E-56 | -0.1067666 | 0.66 | 0.83 | 6.17E-53 |
| Tox3 | 2.78E-52 | -0.1071421 | 0.15 | 0.29 | 4.03E-49 |
| Etv5 | 1.69E-13 | -0.1080241 | 0.55 | 0.57 | 2.45E-10 |
| 4930419G2 | 5.92E-186 | -0.1086227 | 0.41 | 0.69 | 8.60E-183 |
| Serpine | 3.31E-139 | -0.1086392 | 0.08 | 0.32 | 4.80E-136 |
| Grik4 | 1.27E-40 | -0.1093213 | 0.48 | 0.64 | 1.84E-37 |
| Gm2164 | 1.80E-109 | -0.1094924 | 0.25 | 0.52 | 2.61E-106 |
| Cspg4 | 1.35E-68 | -0.1101959 | 0.15 | 0.33 | 1.96E-65 |
| Sulf1 | 1.77E-31 | -0.1111167 | 0.18 | 0.29 | 2.57E-28 |
| Gria4 | 2.01E-216 | -0.1111964 | 0.43 | 0.77 | 2.92E-213 |
| Greb1l | 4.18E-173 | -0.1115074 | 0.19 | 0.52 | 6.07E-170 |
| Lpcat2 | 1.18E-20 | -0.1121125 | 0.4 | 0.49 | 1.71E-17 |
| Tes | 5.79E-94 | -0.1123011 | 0.18 | 0.4 | 8.40E-91 |
| Mef2c | 3.70E-29 | -0.1129419 | 0.39 | 0.52 | 5.37E-26 |
| Rlbp1 | 4.12E-45 | -0.1131385 | 0.25 | 0.38 | 5.98E-42 |
| Tmem19 | 2.16E-32 | -0.1135971 | 0.28 | 0.42 | 3.13E-29 |
| Gm1305 | 3.19E-19 | -0.1138593 | 0.37 | 0.38 | 4.63E-16 |
| Fbxl7 | 6.30E-21 | -0.1139175 | 0.37 | 0.38 | 9.14E-18 |
| Galnt10 | 7.69E-15 | -0.1141084 | 0.54 | 0.54 | 1.12E-11 |
| Rbms3 | 8.85E-32 | -0.1141949 | 0.33 | 0.46 | 1.28E-28 |
| Lrrtm3 | 3.46E-86 | -0.1143432 | 0.36 | 0.59 | 5.02E-83 |
| Usp29 | 1.25E-15 | -0.1147281 | 0.51 | 0.56 | 1.82E-12 |
| Unc13c | 7.42E-38 | -0.1153442 | 0.4 | 0.41 | 1.08E-34 |
| Shroom | 3.87E-13 | -0.1158538 | 0.47 | 0.56 | 5.61E-10 |
| Homer1 | 5.35E-46 | -0.1164803 | 0.48 | 0.65 | 7.76E-43 |
| Pex5l | 3.10E-28 | -0.1172877 | 0.9 | 0.97 | 4.50E-25 |
| Myo16 | 5.24E-47 | -0.1174549 | 0.48 | 0.63 | 7.60E-44 |
| 4930578G1 | 7.17E-27 | -0.1176069 | 0.39 | 0.5 | 1.04E-23 |
| Olfm2 | 2.15E-24 | -0.1184968 | 0.5 | 0.52 | 3.12E-21 |
| Cacng4 | 1.73E-54 | -0.1190439 | 0.32 | 0.42 | 2.52E-51 |
| Phactr2 | 6.68E-69 | -0.1210248 | 0.24 | 0.45 | 9.70E-66 |
| Kcnip1 | 7.20E-52 | -0.1217922 | 0.38 | 0.53 | 1.05E-48 |
| Epb41l4 | 1.40E-26 | -0.1225929 | 0.55 | 0.67 | 2.03E-23 |
| Mbp | 5.94E-20 | -0.1238227 | 0.98 | 1 | 8.61E-17 |
| Itga9 | 5.55E-168 | -0.1252266 | 0.05 | 0.3 | 8.05E-165 |
| Cachd1 | 2.05E-117 | -0.1253783 | 0.22 | 0.49 | 2.98E-114 |
| Cep112 | 3.02E-44 | -0.1262149 | 0.49 | 0.66 | 4.39E-41 |
| Ldb2 | 3.23E-15 | -0.1277809 | 0.42 | 0.47 | 4.68E-12 |
| St18 | 2.20E-34 | -0.1293938 | 0.97 | 1 | 3.19E-31 |
| Sncaip | 9.29E-20 | -0.1311023 | 0.32 | 0.36 | 1.35E-16 |
| Elovl7 | 6.97E-53 | -0.1324482 | 0.6 | 0.78 | 1.01E-49 |
| Kcnh8 | 1.65E-43 | -0.1335229 | 0.38 | 0.34 | 2.39E-40 |
| Slc1a3 | 5.92E-43 | -0.1355045 | 0.47 | 0.64 | 8.59E-40 |
| E130114P1 | 7.30E-69 | -0.1359737 | 0.15 | 0.32 | 1.06E-65 |
| Rad51b | 3.09E-14 | -0.1365837 | 0.33 | 0.29 | 4.49E-11 |
| Zfhx4 | 8.73E-252 | -0.1370546 | 0.16 | 0.52 | 1.27E-248 |
| Ephb1 | 1.08E-31 | -0.1375532 | 0.66 | 0.77 | 1.56E-28 |
| Dock5 | 3.95E-146 | -0.1375533 | 0.53 | 0.82 | 5.73E-143 |
| Sema5b | 2.09E-141 | -0.137688 | 0.07 | 0.31 | 3.03E-138 |
| Ptprr | 1.92E-15 | -0.1384564 | 0.47 | 0.52 | 2.78E-12 |
| Npas3 | 2.83E-58 | -0.1392803 | 0.72 | 0.87 | 4.11E-55 |
| Ctnna3 | 1.10E-52 | -0.1409864 | 0.82 | 0.94 | 1.60E-49 |
| Stk32a | 4.28E-40 | -0.1411087 | 0.51 | 0.53 | 6.21E-37 |
| Akap13 | 3.59E-21 | -0.1420334 | 0.43 | 0.52 | 5.21E-18 |
| Elavl4 | 4.58E-40 | -0.1430608 | 0.43 | 0.57 | 6.65E-37 |
| Itga4 | 2.82E-201 | -0.1431406 | 0.15 | 0.5 | 4.09E-198 |
| Trps1 | 1.04E-90 | -0.1431827 | 0.39 | 0.63 | 1.51E-87 |
| Gm3212 | 9.31E-25 | -0.1459649 | 0.23 | 0.31 | 1.35E-21 |
| Megf11 | 1.91E-103 | -0.1473547 | 0.27 | 0.51 | 2.77E-100 |
| Prr5l | 5.20E-49 | -0.1480003 | 0.92 | 0.98 | 7.54E-46 |
| Gm2031 | 5.25E-182 | -0.1487472 | 0.18 | 0.52 | 7.62E-179 |
| Paqr8 | 2.43E-12 | -0.1510236 | 0.47 | 0.49 | 3.52E-09 |
| Daam2 | 1.67E-121 | -0.153148 | 0.59 | 0.84 | 2.42E-118 |
| Errfi1 | 5.26E-40 | -0.153545 | 0.27 | 0.43 | 7.63E-37 |
| Zfhx3 | 7.02E-119 | -0.1545713 | 0.19 | 0.45 | 1.02E-115 |
| Plce1 | 1.91E-152 | -0.1558525 | 0.09 | 0.36 | 2.77E-149 |
| Ptn | 4.41E-66 | -0.1567883 | 0.61 | 0.8 | 6.40E-63 |
| Syne2 | 5.12E-80 | -0.1569251 | 0.34 | 0.57 | 7.44E-77 |
| Calcrl | 2.69E-61 | -0.1631585 | 0.41 | 0.57 | 3.90E-58 |
| Svil | 1.92E-37 | -0.1644381 | 0.31 | 0.44 | 2.79E-34 |
| Rasgrp3 | 7.34E-115 | -0.1657941 | 0.44 | 0.71 | 1.07E-111 |
| Dpyd | 5.59E-50 | -0.1689296 | 0.43 | 0.61 | 8.12E-47 |
| Epb41l2 | 2.23E-65 | -0.1715609 | 0.41 | 0.62 | 3.24E-62 |
| Enpp2 | 5.83E-51 | -0.1771199 | 0.86 | 0.96 | 8.46E-48 |
| Kctd8 | 1.15E-96 | -0.1771247 | 0.16 | 0.39 | 1.67E-93 |
| Fgf13 | 1.52E-27 | -0.1789898 | 0.48 | 0.58 | 2.20E-24 |
| Prox1 | 2.80E-145 | -0.1806241 | 0.5 | 0.8 | 4.07E-142 |
| 4930587E1 | 9.56E-41 | -0.186659 | 0.38 | 0.54 | 1.39E-37 |
| 4930445B1 | 5.09E-52 | -0.1881157 | 0.28 | 0.45 | 7.38E-49 |
| Maml3 | 1.18E-34 | -0.1903028 | 0.43 | 0.57 | 1.71E-31 |
| Alk | 4.46E-24 | -0.1905268 | 0.39 | 0.42 | 6.47E-21 |
| Thsd4 | 3.40E-62 | -0.1926853 | 0.4 | 0.6 | 4.93E-59 |
| Sema5a | 1.53E-21 | -0.1972765 | 0.47 | 0.54 | 2.22E-18 |
| Pla2g4a | 9.34E-56 | -0.1975705 | 0.49 | 0.68 | 1.36E-52 |
| Itpr2 | 1.33E-95 | -0.1985984 | 0.3 | 0.55 | 1.93E-92 |
| Ubash3 | 1.75E-98 | -0.1993144 | 0.46 | 0.72 | 2.53E-95 |
| Pard3b | 4.32E-21 | -0.2037506 | 0.45 | 0.52 | 6.26E-18 |
| Tcf7l2 | 7.38E-61 | -0.2041938 | 0.35 | 0.56 | 1.07E-57 |
| Kank1 | 6.00E-42 | -0.2049941 | 0.47 | 0.52 | 8.70E-39 |
| Nkx2-2o | 1.68E-38 | -0.2059628 | 0.46 | 0.62 | 2.44E-35 |
| Arhgap1 | 4.51E-249 | -0.2110108 | 0.34 | 0.75 | 6.54E-246 |
| Eya4 | 8.48E-93 | -0.2147041 | 0.19 | 0.42 | 1.23E-89 |
| Lrmda | 1.48E-99 | -0.2174709 | 0.31 | 0.56 | 2.15E-96 |
| Cldn14 | 1.26E-96 | -0.2175898 | 0.36 | 0.61 | 1.83E-93 |
| Tnr | 3.34E-62 | -0.2212163 | 0.53 | 0.72 | 4.85E-59 |
| 6030407O0 | 8.89E-48 | -0.223347 | 0.37 | 0.49 | 1.29E-44 |
| Prkcb | 3.68E-230 | -0.2265684 | 0.31 | 0.68 | 5.34E-227 |
| Efr3b | 5.85E-78 | -0.2323212 | 0.39 | 0.62 | 8.49E-75 |
| Cdh20 | 3.59E-51 | -0.2386298 | 0.77 | 0.9 | 5.21E-48 |
| Pcdh11 | 7.18E-149 | -0.2430837 | 0.25 | 0.53 | 1.04E-145 |
| Ptprm | 2.43E-64 | -0.2499235 | 0.38 | 0.59 | 3.52E-61 |
| Piezo2 | 3.13E-89 | -0.2565741 | 0.43 | 0.66 | 4.54E-86 |
| Gm816 | 6.41E-124 | -0.2778923 | 0.46 | 0.74 | 9.30E-121 |
| 9630013A2 | 4.24E-30 | -0.2871281 | 0.25 | 0.38 | 6.16E-27 |
| Tmem13 | 2.42E-74 | -0.291516 | 0.48 | 0.69 | 3.51E-71 |
| Plekhg1 | 5.47E-130 | -0.2916862 | 0.54 | 0.81 | 7.93E-127 |
| Mob3b | 8.81E-90 | -0.2940206 | 0.57 | 0.79 | 1.28E-86 |
| Abtb2 | 1.48E-82 | -0.316433 | 0.44 | 0.67 | 2.15E-79 |
| Spock3 | 1.46E-129 | -0.3396085 | 0.58 | 0.84 | 2.13E-126 |
| Myo1e | 1.05E-91 | -0.3517414 | 0.57 | 0.79 | 1.53E-88 |
| St3gal4 | 5.81E-71 | -0.3626943 | 0.66 | 0.84 | 8.43E-68 |
| Thsd7a | 2.36E-59 | -0.3658756 | 0.49 | 0.67 | 3.43E-56 |
| A330049N0 | 5.63E-112 | -0.3983023 | 0.6 | 0.83 | 8.16E-109 |
| A230001M1 | 0 | -0.4106345 | 0.39 | 0.83 | 0 |
| Rnf220 | 4.79E-127 | -0.449204 | 0.83 | 0.98 | 6.95E-124 |
| Anln | 3.61E-133 | -0.4805594 | 0.36 | 0.66 | 5.24E-130 |
| Robo1 | 3.51E-52 | -0.4867959 | 0.58 | 0.73 | 5.10E-49 |
| Enpp6 | 2.74E-285 | -0.6438328 | 0.25 | 0.69 | 3.97E-282 |
| Cntn3 | 1.43E-193 | -0.7296133 | 0.39 | 0.75 | 2.08E-190 |
| mt-Nd1 | 3.16E-48 | -0.7438684 | 0.52 | 0.51 | 4.58E-45 |
| mt-Nd2 | 1.26E-114 | -0.7760417 | 0.16 | 0.4 | 1.83E-111 |
| Agmo | 7.59E-109 | -0.7769718 | 0.41 | 0.64 | 1.10E-105 |
| mt-Nd4 | 8.87E-154 | -0.9262412 | 0.25 | 0.57 | 1.29E-150 |
| mt-Cyt | 4.46E-56 | -1.0252253 | 0.45 | 0.56 | 6.48E-53 |
| mt-Co2 | 1.44E-86 | -1.0743948 | 0.2 | 0.41 | 2.09E-83 |
| mt-Co1 | 3.82E-57 | -1.2139153 | 0.3 | 0.45 | 5.54E-54 |
| mt-Co3 | 7.20E-66 | -1.3081391 | 0.31 | 0.35 | 1.04E-62 |
| mt-Atp | 2.23E-59 | -1.333445 | 0.26 | 0.34 | 3.23E-56 |
