## Supplementary table 6 for "DAP12 deficiency alters microglia-oligodendrocyte communication and enhances resilience against tau toxicity"

Figure6A_Oligodendroocyte_cell_ratio_of_different_clusters

| | WT_1 | WT_2 | WT_3 | DAP12WT_1 | DAP12WT_2 | DAP12WT_3 | DAP12KO_1 | DAP12KO_2 | DAP12KO_3 |
|---|---|---|---|---|---|---|---|---|---|
| 1 | 640 | 610 | 416 | 646 | 693 | 465 | 667 | 882 | 476 |
| 2 | 562 | 439 | 323 | 445 | 522 | 146 | 719 | 930 | 482 |
| 3 | 26 | 11 | 9 | 460 | 656 | 1290 | 38 | 20 | 4 |
| 4 | 225 | 160 | 86 | 208 | 293 | 127 | 215 | 226 | 127 |
| 5 | 19 | 8 | 8 | 6 | 11 | 13 | 12 | 14 | 7 |
| sum | 1472 | 1228 | 842 | 1765 | 2175 | 2041 | 1651 | 2072 | 1096 |

| | WT_1 | WT_2 | WT_3 | DAP12WT_1 | DAP12WT_2 | DAP12WT_3 | DAP12KO_1 | DAP12KO_2 | DAP12KO_3 |
|---|---|---|---|---|---|---|---|---|---|
| 1 | 0.43478261 | 0.49674267 | 0.49406176 | 0.36600567 | 0.31862069 | 0.2278295 | 0.40399758 | 0.42567568 | 0.43430657 |
| 2 | 0.38179348 | 0.35749186 | 0.38361045 | 0.25212465 | 0.24 | 0.07153356 | 0.43549364 | 0.4488417 | 0.43978102 |
| 3 | 0.01766304 | 0.00895765 | 0.01068884 | 0.26062323 | 0.3016092 | 0.63204312 | 0.02301635 | 0.00965251 | 0.00364964 |
| 4 | 0.15285326 | 0.13029316 | 0.10213777 | 0.11784703 | 0.13471264 | 0.0622244 | 0.13022411 | 0.10907336 | 0.11587591 |
| 5 | 0.01290761 | 0.00651466 | 0.00950119 | 0.00339943 | 0.00505747 | 0.00636943 | 0.00726832 | 0.00675676 | 0.00638686 |

Figure6C_GSEA_correlation_between_mouse_oligodendrocytes_and_previous_reported_oligodendrocyte_population

| Cluster_num | NAME | GS<br> follow link to MSigDB | SIZE | ES | NES | NOM p-val | FDR q-val | FWER p-val | RANK AT MAX | LEADING EDGE |
|---|---|---|---|---|---|---|---|---|---|---|
| mOL1 | IMMATURE OLI | IMMATURE OLI | 17 | -0.5416986 | -2.3176298 | 0 | 0 | 0 | 86 | tags=35%, list=10%, signal=38% |
| mOL1 | MATURE OLI | MATURE OLI | 142 | 0.26242098 | 1.4879508 | 0.031948883 | 0.10643972 | 0.124 | 256 | tags=38%, list=29%, signal=45% |
| mOL1 | OPC | OPC | 58 | 0.18265876 | 0.8715811 | 0.6477684 | 0.64914936 | 0.885 | 181 | tags=22%, list=20%, signal=26% |
| mOL2 | OPC | OPC | 23 | -0.5700282 | -1.3320397 | 0.116987176 | 0.0884434 | 0.1 | 90 | tags=52%, list=32%, signal=70% |
| mOL2 | MATURE OLI | MATURE OLI | 61 | 0.34545216 | 0.98409534 | 0.5035461 | 0.5092736 | 0.371 | 41 | tags=23%, list=15%, signal=21% |
| mOL3 | MATURE OLI | MATURE OLI | 128 | -0.17657064 | -1.369618 | 0.076771654 | 0.112843364 | 0.173 | 469 | tags=76%, list=63%, signal=170% |
| mOL3 | IMMATURE OLI | IMMATURE OLI | 18 | 0.66874576 | 2.8143473 | 0 | 0 | 0 | 48 | tags=33%, list=6%, signal=35% |
| mOL3 | OPC | OPC | 55 | 0.37983254 | 2.363158 | 0 | 0 | 0 | 252 | tags=58%, list=34%, signal=81% |
| mOL5 | MATURE OLI | MATURE OLI | 104 | -0.51075405 | -2.6691675 | 0 | 0 | 0 | 144 | tags=37%, list=9%, signal=38% |
| mOL5 | OPC | OPC | 74 | 0.7299593 | 2.208004 | 0 | 0 | 0 | 128 | tags=38%, list=8%, signal=39% |
| mOL5 | IMMATURE OLI | IMMATURE OLI | 15 | 0.6346251 | 1.5568635 | 0.052369077 | 0.028862212 | 0.08 | 44 | tags=33%, list=3%, signal=34% |

Figure6F_Western_validation_of_Tenm2_and_Nrg3

Tenm2/GAPDH

| Dap12+/+ tau+ vs tau- | Dap12+/+ tau+ vs tau- | Dap12-/- tau+ vs Dap12+/+ tau+ |
|---|---|---|
| 0.653445174 | 0.833875169 | 1.192320581 |
| 0.602435533 | 1.204758688 | 0.591657505 |
| 0.91740705 | 1.719650047 | 0.68869032 |
| 1.092437671 | 1.240391841 | 0.572513889 |
| 0.555184938 | 1.148379824 | 1.17996848 |
| 0.670170036 | 1.602722822 | 0.980323554 |

Nrg3/GAPDH

| Dap12+/+ tau+ vs tau- | Dap12+/+ tau+ vs tau- | Dap12-/- tau+ vs Dap12+/+ tau+ |
|---|---|---|
| 0.115755422 | 0.256504795 | 0.336640456 |
| 0.061754959 | 0.441702002 | 0.246953051 |
| 0.021574175 | 0.465124913 | 0.216917031 |
| 0.035801939 | 0.260136635 | 0.16279985 |
| 0.038138544 | 0.303294606 | 0.284595826 |
| 0.074926505 | 0.386012442 | 0.126167001 |

Figure6LM_IHC_staining_of_OLIG2_MBP

Olig2 IHC

| Dap12+/+ tau+ vs tau- | Dap12-/- tau+ vs Dap12+/+ tau+ |
|---|---|
| 0.633569482 | 1.560220497 |
| 0.947961209 | 1.48201867 |
| 0.850528431 | 1.445483631 |
| 1.161107393 | 1.121898339 |
| 1.311807465 | 1.42808615 |
| 0.989259337 | 1.098802191 |
| 1.105162808 | 1.108906756 |
| 0.964621371 | 1.020007231 |
| 1.035982504 | 0.868710749 |

Mbp IHC

| Dap12+/+ tau+ vs tau- | Dap12-/- tau+ vs Dap12+/+ tau+ |
|---|---|
| 1.09632568 | 1.250090848 |
| 0.831444495 | 1.137905036 |
| 1.076855414 | 1.227014003 |
| 1.060665854 | 1.188755947 |
| 0.773639918 | 1.064731074 |
| 1.101924272 | 1.022937758 |
| 1.059144368 | 1.072397937 |
| | 1.407279358 |
| | 1.481166518 |

sFigure7C_IPA pathways mOL3_VS_Others

| Ingenuity Canonical Pathways | -log(p-value) | Ratio | z-score | Molecules |
|---|---|---|---|---|
| Calcium Signaling | 1.45 | 0.0335 | 2.449 | ATP2B1,CAMK1D,CREBBP,GRIA2,GRIA4,PPP3CA,RCAN2 |
| RHOGDI Signaling | 1.41 | 0.0327 | 2.236 | ARHGEF10,CDH19,CDH20,CREBBP,GNA12,GNAO1,PRKCA |
| Dilated Cardiomyopathy Signaling Pathway | 0.775 | 0.0274 | 2 | CAMK1D,DMD,GAB1,PTPN11 |
| Protein Kinase A Signaling | 1.22 | 0.0258 | 1.897 | ADD1,ADD3,CREBBP,DCC,PLCL1,PPP3CA,PRKCA,PTPN11,PTPRK,UBASH3B |
| Synaptogenesis Signaling Pathway | 4.33 | 0.0493 | 1.291 | APOE,CADM1,CDH19,CDH20,CREBBP,EPHB1,GRIA2,GRIA4,NLGN1,NRXN1,Nrxn3,PAFAH1B1,PIK3R1,STXBP3,SYT1 |
| Role of Osteoclasts in Rheumatoid Arthritis Signaling Pathway | 0.823 | 0.0235 | 1.134 | CREBBP,CTNNA2,CTNNA3,IL1RAP,IL1RAPL1,PIK3R1,PPP3CA |
| Chaperone Mediated Autophagy Signaling Pathway | 0 | 0.00913 | 1 | CREBBP,PIK3R1,PPP3CA,PRR5L |
| RAR Activation | 0 | 0.00978 | 1 | CREBBP,NCOA1,PIK3R1,ZBTB16 |
| AMPK Signaling | 0.863 | 0.0253 | 1 | ACACA,CREBBP,GNA12,GNAO1,PIK3R1,PPP2R3A |
| PI3K/AKT Signaling | 0.777 | 0.0253 | 1 | BCL2L1,GAB1,INPP5F,PIK3R1,PPP2R3A |
| PPAR Signaling | 1.16 | 0.0381 | 1 | CREBBP,IL1RAP,IL1RAPL1,NCOA1 |
| cAMP-mediated signaling | 0.372 | 0.0177 | 1 | CAMK1D,CREBBP,GNAO1,PPP3CA |
| Autophagy | 0.43 | 0.019 | 1 | CREBBP,PIK3R1,PPP2R3A,PPP3CA |
| Xenobiotic Metabolism PXR Signaling Pathway | 0.597 | 0.023 | 1 | CREBBP,NCOA1,PRKCA,UGT8 |
| Synaptic Long Term Potentiation | 1.9 | 0.0462 | 0.816 | CREBBP,GRIA2,GRIA4,PLCL1,PPP3CA,PRKCA |
| HIF1α Signaling | 1.12 | 0.0299 | 0.816 | CAMK1D,CREBBP,NCOA1,PIK3R1,PPP3CA,PRKCA |
| Amyotrophic Lateral Sclerosis Signaling | 2.19 | 0.0531 | 0.447 | BCL2L1,GLUL,GRIA2,GRIA4,PIK3R1,PPP3CA |
| GABAergic Receptor Signaling Pathway (Enhanced) | 1.33 | 0.0376 | 0.447 | GLUL,GNAO1,NLGN1,NRXN1,Nrxn3 |
| Cachexia Signaling Pathway | 0 | 0.0142 | 0.447 | IL1RAP,IL1RAPL1,MALAT1,PIK3R1,PRKCA |
| IL-12 Signaling and Production in Macrophages | 0.728 | 0.0243 | 0.447 | APOD,APOE,NCOA1,PIK3R1,PRKCA |
| Opioid Signaling Pathway | 0.684 | 0.0222 | 0.447 | CAMK1D,CREBBP,GNA12,GNAO1,PPP3CA,PRKCA |
| Endocannabinoid Neuronal Synapse Pathway | 1.17 | 0.0338 | 0.447 | GNAO1,GRIA2,GRIA4,PLCL1,PPP3CA |
| Senescence Pathway | 0.393 | 0.0176 | 0.447 | ASXL2,CREBBP,PIK3R1,PPP2R3A,PPP3CA |
| Xenobiotic Metabolism CAR Signaling Pathway | 0.959 | 0.0291 | 0.447 | CREBBP,NCOA1,PPP2R3A,PRKCA,UGT8 |
| Sertoli Cell Germ Cell Junction Signaling Pathway (Enhanced) | 1.27 | 0.0304 | 0.378 | CTNNA2,IL1RAP,IL1RAPL1,LAMA2,MTMR2,PIK3R1,PPP2R3A |
| CLEAR Signaling Pathway | 0.628 | 0.0213 | 0 | CREBBP,FGFR2,PPP2R3A,PPP3CA,PRKCA,VPS8 |
| IL-10 Signaling | 0.836 | 0.029 | 0 | BCL2L1,CREBBP,IL1RAP,IL1RAPL1 |
| Activin Inhibin Signaling Pathway | 0.471 | 0.02 | 0 | CREBBP,IL1RAP,IL1RAPL1,PIK3R1 |
| DHCR24 Signaling Pathway | 1.92 | 0.0465 | 0 | APOD,APOE,NLGN1,PIK3R1,PRKCA,THRA |
| LXR/RXR Activation | 1.61 | 0.0446 | 0 | ACACA,APOD,APOE,IL1RAP,IL1RAPL1 |
| CTLA4 Signaling in Cytotoxic T Lymphocytes | 0 | 0.0102 | 0 | BCL2L1,PIK3R1,PPP2R3A,PTPN11 |
| Neuropathic Pain Signaling in Dorsal Horn Neurons | 2.47 | 0.0606 | 0 | CAMK1D,GRIA2,GRIA4,PIK3R1,PLCL1,PRKCA |
| CREB Signaling in Neurons | 0.451 | 0.0172 | 0 | ADGRB3,CREBBP,FGFR2,GNA12,GNAO1,GRIA2,GRIA4,PIK3R1,PLCL1,PRKCA |
| roduction of Nitric Oxide and Reactive Oxygen Species in Macrophage | 1.32 | 0.0337 | 0 | APOD,APOE,CREBBP,PIK3R1,PPP2R3A,PRKCA |
| Gα12/13 Signaling | 0.904 | 0.0308 | 0 | CDH19,CDH20,GNA12,PIK3R1 |
| ILK Signaling | 0.511 | 0.0209 | 0 | ACTN4,CREBBP,PIK3R1,PPP2R3A |
| Phospholipase C Signaling | 0 | 0.00548 | 0 | ARHGEF10,CREBBP,PPP3CA,PRKCA |
| Role of NFAT in Cardiac Hypertrophy | 0.981 | 0.0274 | 0 | CAMK1D,PIK3R1,PLCL1,PPP3CA,PRKCA,RCAN2 |
| WNT/Ca+ pathway | 1.79 | 0.0606 | 0 | CREBBP,PLCL1,PPP3CA,PRKCA |
| ERK/MAPK Signaling | 0.438 | 0.0192 | 0 | CREBBP,PIK3R1,PPP2R3A,PRKCA |
| Synaptic Long Term Depression | 1.64 | 0.0366 | -0.378 | GNA12,GNAO1,GRIA2,GRIA4,PLCL1,PPP2R3A,PRKCA |
| B Cell Receptor Signaling | 0.325 | 0.0158 | -0.378 | BCL2L1,CREBBP,GAB1,INPP5F,PIK3R1,PPP3CA,PTPN11 |
| G-Protein Coupled Receptor Signaling | 0 | 0.0104 | -0.378 | ADGRB3,CREBBP,GNA12,GNAO1,PIK3R1,PPP3CA,PRKCA |
| Wound Healing Signaling Pathway | 0.561 | 0.0209 | -0.447 | FGFR2,IL1RAP,IL1RAPL1,LAMA2,PRKCA |
| Role of NFAT in Regulation of the Immune Response | 0 | 0.00775 | -0.447 | GNA12,GNAO1,PIK3R1,PPP3CA,RCAN2 |
| Dopamine-DARPP32 Feedback in cAMP Signaling | 0.891 | 0.0276 | -0.447 | CREBBP,PLCL1,PPP2R3A,PPP3CA,PRKCA |
| VEGF Signaling | 1.87 | 0.0521 | -0.447 | ACTN4,BCL2L1,PIK3R1,PRKCA,PTPN11 |
| Glutaminergic Receptor Signaling Pathway (Enhanced) | 1.74 | 0.0318 | -0.632 | CREBBP,DGKI,GLUL,GNAO1,GRIA2,GRIA4,PIK3R1,PLCL1,PPP3CA,PRKCA |
| Systemic Lupus Erythematosus in B Cell Signaling Pathway | 0.323 | 0.0157 | -0.707 | BCL2L1,GAB1,INPP5F,PIK3R1,PPP3CA,PRKCA,PTPN11,RASGRP3 |
| S100 Family Signaling Pathway | 0 | 0.00844 | -0.816 | ADGRB3,CREBBP,FGFR2,PIK3R1,PLCL1,PRKCA |
| TR/RXR Activation | 2.01 | 0.0488 | -0.816 | ACACA,MBP,NCOA1,PIK3R1,RCAN2,THRA |
| Role of Tissue Factor in Cancer | 1.12 | 0.03 | -0.816 | BCL2L1,GNA12,PIK3R1,PPP2R3A,PRKCA,PTPN11 |
| Oxytocin in Brain Signaling Pathway | 0.53 | 0.0214 | -1 | CREBBP,GNAO1,PIK3R1,PRKCA |
| Pulmonary Fibrosis Idiopathic Signaling Pathway | 0 | 0.0126 | -1 | CREBBP,FGFR2,GNA12,PIK3R1 |
| ID1 Signaling Pathway | 0.488 | 0.0204 | -1 | BCL2L1,CREBBP,FGFR2,PIK3R1 |
| Neutrophil Extracellular Trap Signaling Pathway | 0 | 0.0114 | -1 | PIK3R1,PLCL1,PPP3CA,PRKCA |
| Neuregulin Signaling | 1.04 | 0.0345 | -1 | NRG3,PIK3R1,PRKCA,PTPN11 |
| Actin Cytoskeleton Signaling | 0.34 | 0.0169 | -1 | ACTN4,GNA12,PIK3R1,SSH2 |
| IL-33 Signaling Pathway | 0.625 | 0.0237 | -1 | BCL2L1,CREBBP,IL1RAP,PIK3R1 |
| IL-15 Production | 1.03 | 0.0342 | -1 | AATK,CLK1,EPHB1,FGFR2 |
| p70S6K Signaling | 0 | 0.0102 | -1 | PIK3R1,PLCL1,PPP2R3A,PRKCA |
| P2Y Purigenic Receptor Signaling Pathway | 0.895 | 0.0305 | -1 | CREBBP,PIK3R1,PLCL1,PRKCA |
| NGF Signaling | 1.02 | 0.0339 | -1 | CREBBP,GAB1,PIK3R1,PTPN11 |
| Neurotrophin/TRK Signaling | 2.26 | 0.0649 | -1 | CREBBP,GAB1,PIK3R1,PTPN11,SORCS1 |
| Osteoarthritis Pathway | 0.616 | 0.022 | -1 | CREBBP,CTNNA2,IL1RAP,IL1RAPL1,SIK3 |
| Insulin Secretion Signaling Pathway | 0.259 | 0.015 | -1 | CREBBP,PIK3R1,PLCL1,PRKCA |
| Oxytocin Signaling Pathway | 0.964 | 0.0256 | -1.134 | ATP2B1,CREBBP,GNA12,GNAO1,PIK3R1,PPP3CA,PRKCA |
| Breast Cancer Regulation by Stathmin1 | 0 | 0.0123 | -1.134 | ADGRB3,ARHGEF10,CAMK1D,CREBBP,PIK3R1,PPP2R3A,PRKCA |
| Multiple Sclerosis Signaling Pathway | 0.789 | 0.0255 | -1.342 | CTNNA3,MBP,MOG,PLP1,UBR2 |
| Fc Epsilon RI Signaling | 1.59 | 0.0442 | -1.342 | GAB1,INPP5F,PIK3R1,PRKCA,PTPN11 |
| Huntington's Disease Signaling | 0.97 | 0.0257 | -1.342 | BCL2L1,CREBBP,DNM3,HIP1,PIK3R1,PRKCA,SGK1 |
| Acetylcholine Receptor Signaling Pathway | 0.835 | 0.0265 | -1.342 | CREBBP,GNAO1,PIK3R1,PLCL1,PRKCA |
| PPARα/RXRα Activation | 1.24 | 0.0323 | -1.342 | ADIPOR2,CREBBP,IL1RAP,IL1RAPL1,PLCL1,PRKCA |
| Chronic Myeloid Leukemia Signaling | 0.428 | 0.0183 | -1.342 | BCL2L1,PIK3R1,PLCL1,PPP3CA,PTPN11 |
| ERK5 Signaling | 2.36 | 0.0685 | -1.342 | CREBBP,GAB1,GNA12,PTPN11,SGK1 |
| Signaling by Rho Family GTPases | 1.41 | 0.0308 | -1.342 | ARHGEF10,CDH19,CDH20,GNA12,GNAO1,PARD3,PIK3R1,WIPF1 |
| IL-4 Signaling | 0 | 0.0134 | -1.342 | BCL2L1,CREBBP,GAB1,PIK3R1,PRR5L |
| Insulin Receptor Signaling | 1.28 | 0.0362 | -1.342 | GAB1,INPP5F,PIK3R1,PTPN11,SGK1 |
| IL-6 Signaling | 1.43 | 0.04 | -1.342 | IL1RAP,IL1RAPL1,PIK3R1,PTPN11,TNFAIP6 |
| Phagosome Formation | 0 | 0.00774 | -1.342 | ADGRB3,ELMO1,PIK3R1,PRKCA,WIPF1 |
| Systemic Lupus Erythematosus in T Cell Signaling Pathway | 0 | 0.0117 | -1.342 | CREBBP,GNA12,PIK3R1,PPP2R3A,PPP3CA |
| Hepatic Fibrosis Signaling Pathway | 0 | 0.0123 | -1.342 | CREBBP,IL1RAP,IL1RAPL1,PIK3R1,PRKCA |
| Xenobiotic Metabolism General Signaling Pathway | 1.34 | 0.0379 | -1.342 | ABCA2,CREBBP,PIK3R1,PRKCA,UGT8 |
| Ephrin Receptor Signaling | 1.12 | 0.03 | -1.342 | CREBBP,EPHB1,GNA12,GNAO1,PTPN11,WIPF1 |
| Estrogen Receptor Signaling | 0.627 | 0.02 | -1.414 | CREBBP,DDX5,GNA12,GNAO1,NCOA1,PIK3R1,PLCL1,PRKCA |
| Sertoli Cell-Sertoli Cell Junction Signaling | 3.08 | 0.0462 | -1.508 | ACTN4,CDH19,CDH20,CREBBP,CTNNA2,EPN2,IL1RAP,IL1RAPL1,LAMA2,PIK3R1,TJP2 |
| Thrombin Signaling | 1.37 | 0.0321 | -1.633 | ARHGEF10,CAMK1D,GNA12,GNAO1,PIK3R1,PLCL1,PRKCA |
| FGF Signaling | 2.78 | 0.0698 | -1.633 | CREBBP,FGFR2,GAB1,PIK3R1,PRKCA,PTPN11 |
| Serotonin Receptor Signaling | 0 | 0.0135 | -1.633 | CREBBP,GNA12,GNAO1,PIK3R1,PLCL1,PRKCA |
| D-myo-inositol-5-phosphate Metabolism | 2.18 | 0.0428 | -1.89 | INPP5F,ITPK1,MTMR2,PPFIBP2,PPP2R3A,PPP3CA,PTPN11,SSH2 |
| D-myo-inositol (1,4,5,6)-Tetrakisphosphate Biosynthesis | 2.42 | 0.0468 | -1.89 | INPP5F,ITPK1,MTMR2,PPFIBP2,PPP2R3A,PPP3CA,PTPN11,SSH2 |
| D-myo-inositol (3,4,5,6)-tetrakisphosphate Biosynthesis | 2.42 | 0.0468 | -1.89 | INPP5F,ITPK1,MTMR2,PPFIBP2,PPP2R3A,PPP3CA,PTPN11,SSH2 |
| 3-phosphoinositide Degradation | 2.25 | 0.044 | -1.89 | INPP5F,ITPK1,MTMR2,PPFIBP2,PPP2R3A,PPP3CA,PTPN11,SSH2 |
| Cardiac Hypertrophy Signaling (Enhanced) | 0 | 0.0134 | -1.89 | FGFR2,GNA12,INPP5F,PIK3R1,PLCL1,PPP3CA,PRKCA |
| Myelination Signaling Pathway | 1.72 | 0.0315 | -1.897 | CREBBP,CYTH1,FGFR2,LAMA2,MAG,MBP,PIK3R1,PLP1,PPP3CA,PTPN11 |
| MicroRNA Biogenesis Signaling Pathway | 0.586 | 0.0227 | -2 | DDX5,Fus,HNRNPA2B1,PIK3R1 |
| Orexin Signaling Pathway | 0.352 | 0.0172 | -2 | PIK3R1,PLCL1,PRKCA,PTPN11 |
| IL-8 Signaling | 0.758 | 0.0249 | -2 | BCL2L1,GNA12,GNAO1,PIK3R1,PRKCA |
| HGF Signaling | 0.913 | 0.031 | -2 | GAB1,PIK3R1,PRKCA,PTPN11 |
| Aldosterone Signaling in Epithelial Cells | 0.991 | 0.0298 | -2 | DNAJB2,PIK3R1,PLCL1,PRKCA,SGK1 |
| SAPK/JNK Signaling | 0 | 0.0128 | -2 | GAB1,GNA12,MINK1,PIK3R1 |
| ation of the Epithelial Mesenchymal Transition by Growth Factors Pat | 0.525 | 0.0213 | -2 | FGFR2,GAB1,PIK3R1,PTPN11 |
| Leukocyte Extravasation Signaling | 2.18 | 0.0428 | -2.121 | ACTN4,CTNNA2,CTNNA3,EDIL3,PIK3R1,PRKCA,PTPN11,WIPF1 |
| Superpathway of Inositol Phosphate Compounds | 2.22 | 0.0402 | -2.121 | INPP5F,ITPK1,MTMR2,PIK3R1,PPFIBP2,PPP2R3A,PPP3CA,PTPN11,SSH2 |
| 3-phosphoinositide Biosynthesis | 2.58 | 0.0457 | -2.121 | INPP5F,ITPK1,MTMR2,PIK3R1,PPFIBP2,PPP2R3A,PPP3CA,PTPN11,SSH2 |
| Endothelin-1 Signaling | 1.24 | 0.0323 | -2.449 | GAB1,GNA12,GNAO1,PIK3R1,PLCL1,PRKCA |

sFigure7D_IPA pathways mOL3_VS_Others Upstream Regulators

| Upstream Regulator | Molecule Type | Predicted Activation State | Activation z-score | Flags | p-value of overlap | Target Molecules in Dataset | Mechanistic Network |
|---|---|---|---|---|---|---|---|
| Alpha catenin | group | Activated | 2.2 | bias | 0.0212 | NL2,LAMA2,SGK1,TNFAIP6,WIF | |
| mir-802 | microRNA | Activated | 2.646 | bias | 0.00481 | RRDC3,CDH19,GNAO1,HECW2, | |
| SOX2 | transcription regulator | Inhibited | -3.159 | | 3.84E-09 | 1700047M11Rik,ARHGEF10,BCAS1,FA2H,Foxp1,GAB1,GJC3,MAG,MBP,MOBP,MOG,OPALIN,PCDH15,PIK3R1,PLP1,PLXNB3,SEMA6A,SLC44A1,UGT8,WIPF1 | |

| | | | | | | |
|---|---|---|---|---|---|---|
| MEF2C | transcription regulator | Inhibited | -2.646 | bias | 0.0000541 | EPHB1,HAPLN2,MAG,MBP,MOE |
| HOXC6 | transcription regulator | Inhibited | -2.236 | | 0.0000546 | 2,MEGF10,PDE4DIP,SLC44A1,ZN |
| ARID1A | transcription regulator | Inhibited | -2.236 | | 0.128 | RP1,NCAM1,NEGR1,PLCL1,SPOC |
| TCF7L2 | transcription regulator | Inhibited | -5.379 | bias | 1.9E-37 | 1700047M11Rik,AATK,ADD1,ADIPOR2,ANKIB1,ANLN,APOD,ARHGEF10,CSRP1,DOCK9,EPHB1,FA2H,FGFR2,FMNL2,GAB1,GLUL,HAPLN2,IL1RAP,KLHL2,LGI3,MAG,MAL,MAP7,MBP,MOBP,MOG,MTMR2,NKAIN1,NKAIN2,OPALIN,OTUD7B,PEX5L,PICALM,PLEKHH1,PLP1,PRICKLE1,PTPN11,RAB31,RCBTB1,RFFL,SECISBP2L,SLC44A1,SNX30,ST18,STXBP3,SYTL2,TBC1D5,UGT8,YPEL2,ZNF536 |

All molecules regulated by SOX2 and TCF7L2

| Symbol | Entrez Gene Name | Gene Symbol - mouse (Entrez Gene) | Expr p-value | Expr Log Ratio | xpr False Discovery Rate (q-valu | Location | Family | Drugs | Entrez Gene ID for Human | Entrez Gene ID for Mouse | Gene ID f |
|---|---|---|---|---|---|---|---|---|---|---|---|
| 1700047M11Rik | RIKEN cDNA 1700047M11 gene | 1700047M11Rik | 8.73E-69 | -0.138811456 | 2.35E-64 | Other | other | | | 67330 | |
| ADD1 | adducin 1 | Add1 | 5.49E-55 | -0.10800732 | 1.48E-50 | Cytoplasm | other | | 118 | 11518 | 24170 |
| ARHGEF10 | guanine nucleotide exchange facto | Arhgef10 | 1.25E-79 | -0.170490809 | 3.37E-75 | Cytoplasm | enzyme | | 9639 | 234094 | 306618 |
| BCAS1 | in enriched myelin associated prote | Bcas1 | 1.06E-70 | -0.242033183 | 2.86E-66 | Plasma Membrane | other | | 8537 | 76960 | 246755 |
| FA2H | fatty acid 2-hydroxylase | Fa2h | 2.28E-62 | -0.102695023 | 6.14E-58 | Cytoplasm | enzyme | | 79152 | 338521 | 307855 |
| Foxp1 | forkhead box P1 | Foxp1 | 5.57E-66 | -0.185934577 | 1.50E-61 | Nucleus | transcription regulator | | | 108655 | 297480 |
| GAB1 | GRB2 associated binding protein 1 | Gab1 | 1.19E-58 | -0.155276071 | 3.21E-54 | Cytoplasm | other | | 2549 | 14388 | 361388 |
| GJC3 | gap junction protein gamma 3 | Gjc3 | 3.35E-83 | -0.304689252 | 9.01E-79 | Extracellular Space | transporter | | 349149 | 118446 | 288529 |
| MAG | myelin associated glycoprotein | Mag | 6.50E-71 | -0.194706879 | 1.75E-66 | Plasma Membrane | other | | 4099 | 17136 | 29409 |
| MBP | myelin basic protein | Mbp | 2.67E-62 | -0.2224608 | 7.18E-58 | Extracellular Space | other | | 4155 | 17196 | |
| MOBP | associated oligodendrocyte basic p | Mobp | 1.55E-67 | -0.296821282 | 4.17E-63 | Cytoplasm | other | | 4336 | 17433 | 25037 |
| MOG | nyelin oligodendrocyte glycoprotei | Mog | 3.41E-52 | -0.110262248 | 9.17E-48 | Extracellular Space | other | | 4340 | 17441 | 24558 |
| PIK3R1 | hoinositide-3-kinase regulatory sub | Pik3r1 | 5.67E-66 | -0.122732421 | 1.53E-61 | Cytoplasm | kinase | | 5295 | 18708 | 25513 |
| PLP1 | proteolipid protein 1 | Plp1 | 1.18E-39 | -0.17969041 | 3.17E-35 | Plasma Membrane | other | | 5354 | 18823 | 24943 |
| PLXNB3 | plexin B3 | Plxnb3 | 4.00E-59 | -0.161994973 | 1.08E-54 | Plasma Membrane | transmembrane receptor | | 5365 | 140571 | 363517 |
| SECISBP2L | SECIS binding protein 2 like | Secisbp2l | 3.04E-68 | -0.200972127 | 8.18E-64 | Other | other | | 9728 | 70354 | 296115 |
| SLC44A1 | solute carrier family 44 member 1 | Slc44a1 | 1.91E-76 | -0.225308676 | 5.13E-72 | Plasma Membrane | transporter | | 23446 | 100434 | 85254 |
| SOX2 | SRY-box transcription factor 2 | | | | | Nucleus | transcription regulator | | 6657 | 20674 | 499593 |
| TCF7L2 | transcription factor 7 like 2 | | | | | Nucleus | transcription regulator | | 6934 | 21416 | 679869 |
| UGT8 | UDP glycosyltransferase 8 | Ugt8a | 6.74E-64 | -0.147082446 | 1.81E-59 | Cytoplasm | enzyme | | 7368 | 22239 | 50555 |
| WIPF1 | WASL interacting protein family men | Wipf1 | 3.77E-52 | -0.123371353 | 1.02E-47 | Cytoplasm | other | | 7456 | 215280 | 117538 |

All relationships to SOX2 and TCF7L2

| From Molecule(s) | To Molecule(s) | Catalyst(s) |
|---|---|---|
| SOX2 | 1700047M11Rik | |
| SOX2 | ARHGEF10 | |
| SOX2 | BCAS1 | |
| SOX2 | FA2H | |
| SOX2 | Foxp1 | |
| SOX2 | GAB1 | |
| SOX2 | GJC3 | |
| SOX2 | MAG | |
| SOX2 | MBP | |
| SOX2 | MOBP | |
| SOX2 | MOG | |
| SOX2 | PIK3R1 | |
| SOX2 | PLP1 | |
| SOX2 | PLXNB3 | |
| SOX2 | SLC44A1 | |
| SOX2 | UGT8 | |
| SOX2 | WIPF1 | |
| SOX2 | SOX2 | |
| SOX2 | SOX2 | |
| TCF7L2 | 1700047M11Rik | |
| TCF7L2 | ADD1 | |
| TCF7L2 | ARHGEF10 | |
| TCF7L2 | FA2H | |
| TCF7L2 | GAB1 | |
| TCF7L2 | MAG | |
| TCF7L2 | MBP | |
| TCF7L2 | MOBP | |
| TCF7L2 | MOG | |
| TCF7L2 | PLP1 | |
| TCF7L2 | SLC44A1 | |
| TCF7L2 | SOX2 | |
| TCF7L2 | UGT8 | |
